## Supplementary material for "Conserved water molecules as structural ligands modulating pathogenic variation in human protein binding sites": Dataset S1

| source_id | classification | binding_site | binding_site_single_water | protein_binding_site |
|---|---|---|---|---|
| rs121918343 | pathogenic | 1 | 1 | 0 |
| rs267606649 | pathogenic | 1 | 1 | 1 |
| rs121918342 | pathogenic | 0 | 1 | 0 |
| rs267606650 | pathogenic | 1 | 1 | 1 |
| rs1130335 | benign | 1 | 1 | 1 |
| rs199476087 | pathogenic | 1 | 1 | 1 |
| rs772163112 | benign | 0 | 0 | 0 |
| rs755793 | benign | 0 | 0 | 0 |
| rs121918155 | pathogenic | 1 | 1 | 0 |
| rs121918156 | pathogenic | 1 | 1 | 0 |
| rs121918154 | pathogenic | 1 | 1 | 1 |
| rs121918151 | pathogenic | 1 | 1 | 1 |
| rs121918144 | pathogenic | 0 | 1 | 0 |
| rs121918146 | pathogenic | 1 | 1 | 1 |
| rs121918160 | pathogenic | 1 | 1 | 1 |
| rs121918150 | pathogenic | 1 | 1 | 1 |
| rs121918141 | pathogenic | 1 | 1 | 1 |
| rs142742242 | pathogenic | 1 | 1 | 1 |
| rs121918142 | pathogenic | 1 | 1 | 1 |
| rs121918157 | pathogenic | 1 | 1 | 1 |
| rs121918101 | pathogenic | 1 | 1 | 1 |
| rs121908260 | pathogenic | 1 | 1 | 1 |
| rs80356667 | pathogenic | 1 | 1 | 1 |
| rs80356668 | pathogenic | 1 | 1 | 1 |
| rs397514540 | pathogenic | 1 | 1 | 1 |
| rs104894685 | pathogenic | 0 | 1 | 0 |
| rs199869995 | pathogenic | 1 | 1 | 1 |
| rs121909072 | pathogenic | 0 | 0 | 0 |
| rs143669310 | benign | 1 | 1 | 1 |
| rs747347498 | benign | 0 | 0 | 0 |
| rs138283222 | benign | 0 | 0 | 0 |
| rs118203931 | pathogenic | 1 | 1 | 1 |
| rs118203933 | pathogenic | 1 | 1 | 0 |
| rs118203932 | pathogenic | 1 | 1 | 0 |
| rs2228063 | benign | 1 | 1 | 1 |
| rs377767360 | pathogenic | 0 | 0 | 0 |
| rs397518413 | pathogenic | 1 | 1 | 1 |
| rs1800458 | benign | 1 | 1 | 0 |
| rs121918083 | pathogenic | 1 | 1 | 1 |
| rs121918094 | pathogenic | 1 | 1 | 1 |
| rs121918098 | pathogenic | 1 | 1 | 1 |
| rs121918093 | pathogenic | 1 | 1 | 1 |
| rs79977247 | pathogenic | 1 | 1 | 1 |
| rs28933979 | pathogenic | 1 | 1 | 1 |
| rs121918068 | pathogenic | 1 | 1 | 1 |
| rs121918077 | pathogenic | 1 | 1 | 1 |
| rs11541796 | pathogenic | 1 | 1 | 1 |
| rs104894665 | pathogenic | 1 | 1 | 1 |
| rs121918090 | pathogenic | 1 | 1 | 1 |
| rs121918081 | pathogenic | 1 | 1 | 1 |
| rs121918080 | pathogenic | 1 | 1 | 1 |
| rs121918076 | pathogenic | 1 | 1 | 1 |
| rs121918097 | pathogenic | 1 | 1 | 1 |

| | | | | |
|---|---|---|---|---|
| rs121918079 | pathogenic | 1 | 1 | 1 |
| rs121918069 | pathogenic | 1 | 1 | 1 |
| rs121918086 | pathogenic | 1 | 1 | 1 |
| rs121918091 | pathogenic | 1 | 1 | 1 |
| rs121918085 | pathogenic | 1 | 1 | 1 |
| rs121918100 | pathogenic | 1 | 1 | 1 |
| rs121918084 | pathogenic | 1 | 1 | 1 |
| rs121918071 | pathogenic | 1 | 1 | 1 |
| rs121918072 | pathogenic | 1 | 1 | 1 |
| rs121918082 | pathogenic | 1 | 1 | 1 |
| rs121918087 | pathogenic | 1 | 1 | 1 |
| rs76410435 | benign | 1 | 1 | 1 |
| rs121918089 | pathogenic | 1 | 1 | 1 |
| rs121918073 | pathogenic | 1 | 1 | 1 |
| rs121918075 | pathogenic | 1 | 1 | 1 |
| rs121918088 | pathogenic | 1 | 1 | 1 |
| rs76992529 | pathogenic | 1 | 1 | 1 |
| rs536294863 | benign | 1 | 1 | 1 |
| rs730881162 | benign | 1 | 1 | 1 |
| rs113994087 | pathogenic | 1 | 1 | 1 |
| rs74716434 | benign | 1 | 1 | 1 |
| rs121918694 | pathogenic | 1 | 1 | 1 |
| rs121918707 | pathogenic | 1 | 1 | 1 |
| rs121918695 | pathogenic | 1 | 1 | 0 |
| rs121918690 | pathogenic | 1 | 1 | 0 |
| rs121918696 | pathogenic | 1 | 1 | 0 |
| rs121918693 | pathogenic | 1 | 1 | 0 |
| rs28999969 | pathogenic | 1 | 1 | 1 |
| rs121918697 | pathogenic | 1 | 1 | 1 |
| rs121918688 | pathogenic | 1 | 1 | 0 |
| rs28999970 | pathogenic | 1 | 1 | 0 |
| rs121918686 | pathogenic | 1 | 1 | 0 |
| rs28999971 | pathogenic | 1 | 1 | 1 |
| rs121918698 | pathogenic | 1 | 1 | 1 |
| rs121918691 | pathogenic | 1 | 1 | 0 |
| rs121918692 | pathogenic | 1 | 1 | 1 |
| rs121918703 | pathogenic | 1 | 1 | 1 |
| rs121918687 | pathogenic | 1 | 1 | 1 |
| rs28933408 | pathogenic | 1 | 1 | 1 |
| rs121918702 | pathogenic | 1 | 1 | 1 |
| rs17875899 | benign | 1 | 1 | 1 |
| rs111033620 | pathogenic | 1 | 1 | 1 |
| rs111033622 | pathogenic | 1 | 1 | 1 |
| rs111033621 | pathogenic | 1 | 1 | 1 |
| rs111033618 | pathogenic | 1 | 1 | 1 |
| rs5918 | benign | 0 | 0 | 0 |
| rs121918452 | pathogenic | 0 | 0 | 0 |
| rs121918445 | pathogenic | 1 | 1 | 1 |
| rs5917 | benign | 1 | 1 | 1 |
| rs121918444 | pathogenic | 1 | 1 | 1 |
| rs121918446 | pathogenic | 1 | 1 | 1 |
| rs121918449 | pathogenic | 0 | 0 | 0 |
| rs121918448 | benign | 0 | 0 | 0 |
| rs13306487 | benign | 1 | 1 | 1 |

| | | | | |
|---|---|---|---|---|
| rs1805044 | benign | 0 | 0 | 0 |
| rs117078377 | benign | 0 | 0 | 0 |
| rs138132026 | benign | 1 | 1 | 1 |
| rs75762896 | benign | 1 | 1 | 0 |
| rs121918043 | pathogenic | 1 | 1 | 0 |
| rs143581690 | benign | 1 | 1 | 0 |
| rs147654263 | benign | 0 | 0 | 0 |
| rs587782529 | pathogenic | 1 | 1 | 1 |
| rs121912664 | pathogenic | 1 | 1 | 1 |
| rs150293825 | benign | 1 | 1 | 1 |
| rs17882252 | benign | 1 | 1 | 1 |
| rs121912662 | pathogenic | 1 | 1 | 1 |
| rs35993958 | benign | 1 | 1 | 1 |
| rs606231253 | pathogenic | 1 | 1 | 1 |
| rs2227983 | benign | 1 | 1 | 1 |
| rs587779351 | pathogenic | 0 | 0 | 0 |
| rs779773463 | pathogenic | 1 | 1 | 1 |
| rs713040 | benign | 1 | 1 | 1 |
| rs334 | pathogenic | 1 | 1 | 1 |
| rs33930165 | pathogenic | 1 | 1 | 1 |
| rs33918131 | benign | 1 | 1 | 1 |
| rs63750783 | pathogenic | 1 | 1 | 1 |
| rs33986703 | pathogenic | 1 | 1 | 1 |
| rs33972047 | pathogenic | 1 | 1 | 1 |
| rs33936254 | pathogenic | 1 | 1 | 1 |
| rs33959855 | benign | 1 | 1 | 1 |
| rs33950507 | pathogenic | 1 | 1 | 1 |
| rs33916412 | pathogenic | 1 | 1 | 0 |
| rs1135071 | pathogenic | 1 | 1 | 1 |
| rs33960103 | pathogenic | 1 | 1 | 1 |
| rs11549407 | pathogenic | 1 | 1 | 1 |
| rs33922842 | pathogenic | 1 | 1 | 1 |
| rs35278874 | benign | 1 | 1 | 1 |
| rs34621955 | benign | 1 | 1 | 1 |
| rs33995148 | pathogenic | 1 | 1 | 1 |
| rs33945705 | benign | 1 | 1 | 1 |
| rs33993568 | benign | 1 | 1 | 1 |
| rs35553496 | benign | 1 | 1 | 1 |
| rs33913712 | pathogenic | 1 | 1 | 1 |
| rs33917785 | pathogenic | 1 | 1 | 1 |
| rs34515413 | benign | 1 | 1 | 1 |
| rs35256489 | pathogenic | 1 | 1 | 0 |
| rs34484056 | pathogenic | 1 | 1 | 1 |
| rs33947020 | benign | 1 | 1 | 1 |
| rs33946267 | pathogenic | 1 | 1 | 1 |
| rs33966761 | pathogenic | 1 | 1 | 1 |
| rs36020563 | benign | 1 | 1 | 1 |
| rs122454124 | pathogenic | 1 | 1 | 1 |
| rs122454126 | pathogenic | 1 | 1 | 0 |
| rs122454127 | pathogenic | 1 | 1 | 1 |
| rs122454130 | pathogenic | 1 | 1 | 1 |
| rs122454125 | pathogenic | 1 | 1 | 1 |
| rs122454131 | pathogenic | 1 | 1 | 0 |
| rs121908302 | pathogenic | 0 | 1 | 0 |

| | | | | |
|---|---|---|---|---|
| rs77829017 | pathogenic | 0 | 1 | 0 |
| rs1141814 | pathogenic | 0 | 1 | 0 |
| rs121908312 | pathogenic | 0 | 1 | 0 |
| rs79653797 | pathogenic | 1 | 1 | 0 |
| rs439898 | pathogenic | 1 | 1 | 0 |
| rs121908299 | pathogenic | 0 | 1 | 0 |
| rs80356763 | pathogenic | 1 | 1 | 1 |
| rs781152868 | pathogenic | 0 | 1 | 0 |
| rs121908297 | pathogenic | 0 | 1 | 0 |
| rs398123531 | benign | 0 | 1 | 0 |
| rs398123532 | pathogenic | 0 | 1 | 0 |
| rs364897 | pathogenic | 1 | 1 | 1 |
| rs1064644 | pathogenic | 0 | 1 | 0 |
| rs409652 | pathogenic | 1 | 1 | 1 |
| rs121908300 | pathogenic | 0 | 1 | 0 |
| rs381737 | pathogenic | 0 | 1 | 0 |
| rs74500255 | pathogenic | 0 | 1 | 0 |
| rs121908313 | pathogenic | 0 | 1 | 0 |
| rs367968666 | pathogenic | 0 | 1 | 0 |
| rs78973108 | pathogenic | 1 | 1 | 1 |
| rs121908298 | pathogenic | 1 | 1 | 1 |
| rs78396650 | pathogenic | 0 | 0 | 0 |
| rs78198234 | pathogenic | 1 | 1 | 0 |
| rs121908304 | pathogenic | 1 | 1 | 1 |
| rs76539814 | pathogenic | 1 | 1 | 1 |
| rs121908305 | pathogenic | 0 | 1 | 0 |
| rs121908306 | pathogenic | 1 | 1 | 0 |
| rs121908308 | pathogenic | 1 | 1 | 1 |
| rs121908307 | pathogenic | 0 | 1 | 0 |
| rs75548401 | benign | 0 | 1 | 0 |
| rs76763715 | pathogenic | 0 | 0 | 0 |
| rs121908314 | pathogenic | 0 | 1 | 0 |
| rs398123528 | pathogenic | 0 | 1 | 0 |
| rs121908311 | pathogenic | 0 | 0 | 0 |
| rs121908310 | pathogenic | 1 | 1 | 0 |
| rs74598136 | pathogenic | 0 | 1 | 0 |
| rs1064651 | pathogenic | 1 | 1 | 1 |
| rs77369218 | pathogenic | 1 | 1 | 1 |
| rs121908295 | pathogenic | 0 | 1 | 0 |
| rs421016 | pathogenic | 0 | 1 | 0 |
| rs368060 | pathogenic | 0 | 1 | 0 |
| rs80356771 | pathogenic | 1 | 1 | 1 |
| rs80356772 | pathogenic | 1 | 1 | 1 |
| rs121908301 | pathogenic | 0 | 0 | 0 |
| rs747506979 | pathogenic | 0 | 1 | 0 |
| rs75822236 | pathogenic | 0 | 1 | 0 |
| rs1990760 | benign | 1 | 1 | 0 |
| rs28934568 | pathogenic | 1 | 1 | 0 |
| rs34833812 | benign | 1 | 1 | 0 |
| rs193922666 | benign | 1 | 1 | 0 |
| rs104893812 | pathogenic | 1 | 1 | 0 |
| rs144701411 | benign | 0 | 0 | 0 |
| rs104893813 | pathogenic | 0 | 1 | 0 |
| rs199660234 | pathogenic | 1 | 1 | 0 |

| | | | | |
|---|---|---|---|---|
| rs104893814 | pathogenic | 1 | 1 | 0 |
| rs143095746 | benign | 1 | 1 | 1 |
| rs104893807 | pathogenic | 1 | 1 | 0 |
| rs104893811 | pathogenic | 0 | 0 | 0 |
| rs104893816 | pathogenic | 0 | 0 | 0 |
| rs121918714 | pathogenic | 0 | 1 | 0 |
| rs104893810 | pathogenic | 1 | 1 | 0 |
| rs104893815 | pathogenic | 1 | 1 | 0 |
| rs4303 | benign | 0 | 1 | 0 |
| rs121912704 | pathogenic | 0 | 1 | 0 |
| rs56394458 | benign | 1 | 1 | 0 |
| rs397514688 | pathogenic | 0 | 0 | 0 |
| rs367797185 | pathogenic | 1 | 1 | 0 |
| rs12709426 | benign | 0 | 1 | 0 |
| rs2228547 | benign | 0 | 0 | 0 |
| rs111033546 | pathogenic | 0 | 0 | 0 |
| rs111033553 | pathogenic | 0 | 0 | 0 |
| rs111033554 | pathogenic | 1 | 1 | 1 |
| rs111033544 | pathogenic | 1 | 1 | 1 |
| rs111033555 | pathogenic | 1 | 1 | 1 |
| rs111033545 | pathogenic | 1 | 1 | 1 |
| rs111033549 | pathogenic | 0 | 1 | 0 |
| rs111033552 | pathogenic | 1 | 1 | 1 |
| rs752204035 | benign | 0 | 0 | 0 |
| rs11569534 | benign | 0 | 0 | 0 |
| rs28937571 | pathogenic | 1 | 1 | 1 |
| rs104894215 | pathogenic | 1 | 1 | 1 |
| rs3888798 | benign | 1 | 1 | 1 |
| rs764263034 | benign | 1 | 1 | 1 |
| rs1800973 | benign | 1 | 1 | 1 |
| rs367623154 | benign | 1 | 1 | 1 |
| rs201230446 | pathogenic | 1 | 1 | 1 |
| rs267606766 | pathogenic | 0 | 1 | 0 |
| rs267606769 | pathogenic | 0 | 1 | 0 |
| rs267606767 | pathogenic | 1 | 1 | 0 |
| rs267606768 | pathogenic | 0 | 1 | 0 |
| rs61733129 | benign | 0 | 1 | 0 |
| rs201947120 | pathogenic | 1 | 1 | 1 |
| rs283413 | benign | 0 | 1 | 0 |
| rs36119840 | benign | 0 | 0 | 0 |
| rs281865419 | pathogenic | 1 | 1 | 1 |
| rs137852937 | pathogenic | 0 | 1 | 0 |
| rs2230709 | benign | 0 | 1 | 0 |
| rs6695033 | benign | 0 | 1 | 0 |
| rs180177032 | pathogenic | 1 | 1 | 1 |
| rs180177033 | pathogenic | 1 | 1 | 1 |
| rs121913351 | pathogenic | 1 | 1 | 1 |
| rs397507473 | pathogenic | 1 | 1 | 1 |
| rs121913355 | pathogenic | 1 | 1 | 1 |
| rs121913357 | pathogenic | 1 | 1 | 1 |
| rs180177036 | pathogenic | 1 | 1 | 1 |
| rs180177039 | pathogenic | 1 | 1 | 1 |
| rs180177038 | pathogenic | 1 | 1 | 1 |
| rs180177040 | pathogenic | 1 | 1 | 1 |

| | | | | |
|---|---|---|---|---|
| rs121913338 | pathogenic | 1 | 1 | 1 |
| rs121913366 | pathogenic | 1 | 1 | 1 |
| rs121913369 | pathogenic | 1 | 1 | 1 |
| rs113488022 | pathogenic | 1 | 1 | 1 |
| rs121913364 | pathogenic | 1 | 1 | 1 |
| rs180177042 | pathogenic | 1 | 1 | 1 |
| rs200461617 | benign | 0 | 0 | 0 |
| rs141609745 | benign | 0 | 0 | 0 |
| rs104894383 | pathogenic | 1 | 1 | 1 |
| rs104894384 | pathogenic | 1 | 1 | 1 |
| rs104894381 | pathogenic | 1 | 1 | 0 |
| rs147710408 | benign | 1 | 1 | 1 |
| rs77357563 | benign | 1 | 1 | 1 |
| rs104894378 | pathogenic | 0 | 0 | 0 |
| rs104894382 | pathogenic | 0 | 0 | 0 |
| rs137853292 | pathogenic | 1 | 1 | 1 |
| rs587778834 | pathogenic | 0 | 0 | 0 |
| rs137853294 | pathogenic | 1 | 1 | 1 |
| rs551747882 | benign | 0 | 0 | 0 |
| rs3092903 | benign | 1 | 1 | 1 |
| rs137853296 | pathogenic | 0 | 0 | 0 |
| rs398122917 | pathogenic | 0 | 1 | 0 |
| rs41304992 | benign | 0 | 0 | 0 |
| rs121912708 | pathogenic | 1 | 1 | 1 |
| rs140102105 | pathogenic | 1 | 1 | 1 |
| rs376917645 | pathogenic | 1 | 1 | 0 |
| rs372660425 | pathogenic | 1 | 1 | 0 |
| rs555896752 | pathogenic | 1 | 1 | 1 |
| rs777829351 | pathogenic | 1 | 1 | 0 |
| rs121912711 | pathogenic | 1 | 1 | 0 |
| rs754449549 | pathogenic | 0 | 1 | 0 |
| rs141701364 | benign | 0 | 1 | 0 |
| rs761011399 | benign | 1 | 1 | 1 |
| rs121912710 | benign | 1 | 1 | 0 |
| rs121912707 | pathogenic | 1 | 1 | 0 |
| rs747597620 | pathogenic | 0 | 1 | 0 |
| rs186558364 | pathogenic | 1 | 1 | 1 |
| rs556400964 | pathogenic | 1 | 1 | 1 |
| rs61757684 | benign | 1 | 1 | 1 |
| rs121917766 | pathogenic | 1 | 1 | 1 |
| rs121918248 | pathogenic | 0 | 0 | 0 |
| rs398123278 | pathogenic | 0 | 0 | 0 |
| rs121918251 | pathogenic | 1 | 1 | 1 |
| rs727504022 | pathogenic | 0 | 0 | 0 |
| rs190834116 | pathogenic | 1 | 1 | 1 |
| rs121918249 | pathogenic | 1 | 1 | 0 |
| rs121918257 | pathogenic | 1 | 1 | 0 |
| rs142068898 | benign | 0 | 0 | 0 |
| rs757000253 | pathogenic | 0 | 0 | 0 |
| rs200908035 | pathogenic | 1 | 1 | 0 |
| rs760782399 | pathogenic | 0 | 0 | 0 |
| rs778702777 | pathogenic | 0 | 0 | 0 |
| rs121918258 | pathogenic | 1 | 1 | 0 |
| rs121918256 | pathogenic | 0 | 0 | 0 |

| | | | | |
|---|---|---|---|---|
| rs200596762 | pathogenic | 1 | 1 | 0 |
| rs547709692 | pathogenic | 0 | 0 | 0 |
| rs758577372 | pathogenic | 0 | 1 | 0 |
| rs564069299 | pathogenic | 1 | 1 | 1 |
| rs772552898 | pathogenic | 1 | 1 | 1 |
| rs368790885 | pathogenic | 1 | 1 | 1 |
| rs121918250 | pathogenic | 0 | 0 | 0 |
| rs533755473 | pathogenic | 1 | 1 | 1 |
| rs753288303 | pathogenic | 1 | 1 | 1 |
| rs2229385 | benign | 0 | 0 | 0 |
| rs1141321 | benign | 0 | 0 | 0 |
| rs753564352 | pathogenic | 0 | 0 | 0 |
| rs775593146 | pathogenic | 0 | 0 | 0 |
| rs121918254 | pathogenic | 0 | 0 | 0 |
| rs8589 | benign | 1 | 1 | 0 |
| rs777758903 | pathogenic | 0 | 0 | 0 |
| rs140600746 | pathogenic | 1 | 1 | 0 |
| rs121918255 | pathogenic | 1 | 1 | 0 |
| rs121918252 | pathogenic | 0 | 0 | 0 |
| rs779990936 | pathogenic | 1 | 1 | 0 |
| rs121912455 | pathogenic | 1 | 1 | 1 |
| rs121912459 | pathogenic | 1 | 1 | 1 |
| rs74315452 | pathogenic | 1 | 1 | 1 |
| rs121912451 | pathogenic | 1 | 1 | 1 |
| rs79006549 | pathogenic | 0 | 0 | 0 |
| rs28934889 | benign | 0 | 1 | 0 |
| rs111033557 | benign | 1 | 1 | 1 |
| rs28934597 | pathogenic | 1 | 1 | 1 |
| rs28934596 | pathogenic | 1 | 1 | 1 |
| rs28934595 | pathogenic | 1 | 1 | 1 |
| rs146519482 | pathogenic | 1 | 1 | 1 |
| rs111033563 | pathogenic | 1 | 1 | 1 |
| rs75385605 | benign | 1 | 1 | 1 |
| rs121908569 | pathogenic | 1 | 1 | 1 |
| rs35719940 | benign | 0 | 0 | 0 |
| rs121918664 | pathogenic | 0 | 0 | 0 |
| rs104893665 | pathogenic | 1 | 1 | 1 |
| rs104893666 | pathogenic | 1 | 1 | 1 |
| rs200845457 | benign | 1 | 1 | 1 |
| rs779204655 | pathogenic | 1 | 1 | 1 |
| rs121917747 | pathogenic | 1 | 1 | 1 |
| rs6042 | benign | 1 | 1 | 1 |
| rs104894380 | pathogenic | 0 | 0 | 0 |
| rs369083541 | pathogenic | 1 | 1 | 1 |
| rs145243532 | benign | 1 | 1 | 1 |
| rs121909500 | pathogenic | 1 | 1 | 1 |
| rs140993310 | benign | 1 | 1 | 1 |
| rs200837270 | pathogenic | 1 | 1 | 1 |
| rs121918484 | pathogenic | 1 | 1 | 1 |
| rs121918306 | pathogenic | 0 | 0 | 0 |
| rs118204089 | pathogenic | 1 | 1 | 0 |
| rs118204083 | pathogenic | 1 | 1 | 0 |
| rs201953584 | pathogenic | 1 | 1 | 1 |
| rs74315362 | pathogenic | 1 | 1 | 1 |

| | | | | |
|---|---|---|---|---|
| rs118204084 | pathogenic | 1 | 1 | 0 |
| rs118204085 | pathogenic | 1 | 1 | 1 |
| rs113403872 | pathogenic | 1 | 1 | 0 |
| rs1052176 | benign | 1 | 1 | 1 |
| rs2031640 | benign | 0 | 0 | 0 |
| rs104894230 | pathogenic | 1 | 1 | 1 |
| rs104894229 | pathogenic | 1 | 1 | 1 |
| rs104894228 | pathogenic | 1 | 1 | 1 |
| rs104894226 | pathogenic | 1 | 1 | 1 |
| rs121917757 | pathogenic | 1 | 1 | 1 |
| rs121917758 | pathogenic | 1 | 1 | 1 |
| rs28933406 | pathogenic | 1 | 1 | 1 |
| rs121917756 | pathogenic | 1 | 1 | 1 |
| rs104894227 | pathogenic | 1 | 1 | 1 |
| rs369106578 | pathogenic | 1 | 1 | 1 |
| rs397517142 | benign | 1 | 1 | 1 |
| rs104894231 | pathogenic | 1 | 1 | 1 |
| rs121917759 | pathogenic | 1 | 1 | 1 |
| rs104893827 | pathogenic | 0 | 0 | 0 |
| rs104893826 | pathogenic | 1 | 1 | 1 |
| rs5030802 | pathogenic | 1 | 1 | 1 |
| rs5030827 | pathogenic | 1 | 1 | 0 |
| rs119103277 | pathogenic | 1 | 1 | 1 |
| rs5030808 | pathogenic | 0 | 1 | 0 |
| rs5030809 | pathogenic | 1 | 1 | 1 |
| rs765978945 | pathogenic | 1 | 1 | 1 |
| rs104893824 | pathogenic | 1 | 1 | 1 |
| rs104893831 | pathogenic | 0 | 1 | 0 |
| rs104893830 | pathogenic | 0 | 1 | 0 |
| rs119103278 | pathogenic | 0 | 1 | 0 |
| rs5030818 | pathogenic | 1 | 1 | 1 |
| rs397516444 | pathogenic | 1 | 1 | 1 |
| rs28940297 | pathogenic | 1 | 1 | 1 |
| rs267607170 | pathogenic | 1 | 1 | 1 |
| rs104893825 | pathogenic | 1 | 1 | 1 |
| rs5030820 | pathogenic | 0 | 1 | 0 |
| rs5030821 | pathogenic | 0 | 1 | 0 |
| rs5030823 | pathogenic | 1 | 1 | 1 |
| rs5030824 | pathogenic | 1 | 1 | 1 |
| rs28940301 | pathogenic | 0 | 1 | 0 |
| rs28940300 | pathogenic | 0 | 1 | 0 |
| rs28940298 | pathogenic | 0 | 1 | 0 |
| rs200019083 | benign | 0 | 0 | 0 |
| rs759748655 | pathogenic | 0 | 0 | 0 |
| rs121917739 | pathogenic | 1 | 1 | 1 |
| rs75353611 | pathogenic | 1 | 1 | 1 |
| rs79744198 | benign | 1 | 1 | 1 |
| rs79228041 | pathogenic | 1 | 1 | 1 |
| rs80106970 | benign | 1 | 1 | 1 |
| rs375353223 | benign | 0 | 0 | 0 |
| rs56311081 | benign | 0 | 0 | 0 |
| rs55985569 | benign | 1 | 1 | 1 |
| rs45483396 | benign | 0 | 0 | 0 |
| rs35284565 | benign | 1 | 1 | 1 |

| | | | | |
|---|---|---|---|---|
| rs34349517 | benign | 0 | 0 | 0 |
| rs35776110 | benign | 0 | 0 | 0 |
| rs33917957 | benign | 1 | 1 | 1 |
| rs587778445 | benign | 0 | 0 | 0 |
| rs34624984 | benign | 0 | 0 | 0 |
| rs701753 | benign | 0 | 0 | 0 |
| rs855314 | benign | 0 | 0 | 0 |
| rs121918371 | pathogenic | 1 | 1 | 0 |
| rs398122912 | pathogenic | 0 | 1 | 0 |
| rs1126728 | benign | 0 | 0 | 0 |
| rs587777404 | pathogenic | 1 | 1 | 0 |
| rs11208257 | benign | 0 | 0 | 0 |
| rs6676290 | benign | 0 | 0 | 0 |
| rs104894118 | pathogenic | 0 | 0 | 0 |
| rs121918655 | pathogenic | 1 | 1 | 1 |
| rs104894120 | pathogenic | 1 | 1 | 1 |
| rs137852238 | pathogenic | 0 | 0 | 0 |
| rs137852237 | pathogenic | 0 | 0 | 0 |
| rs137852499 | pathogenic | 1 | 1 | 1 |
| rs137852480 | pathogenic | 1 | 1 | 1 |
| rs137852494 | pathogenic | 1 | 1 | 1 |
| rs137852502 | pathogenic | 1 | 1 | 1 |
| rs137852500 | pathogenic | 1 | 1 | 1 |
| rs137852487 | pathogenic | 1 | 1 | 1 |
| rs137852481 | pathogenic | 1 | 1 | 1 |
| rs137852501 | pathogenic | 0 | 0 | 0 |
| rs137852478 | pathogenic | 1 | 1 | 1 |
| rs137852485 | pathogenic | 1 | 1 | 1 |
| rs137852482 | pathogenic | 1 | 1 | 0 |
| rs137852483 | pathogenic | 0 | 1 | 0 |
| rs137852477 | pathogenic | 1 | 1 | 0 |
| rs137852496 | pathogenic | 1 | 1 | 0 |
| rs137852484 | pathogenic | 0 | 1 | 0 |
| rs137852498 | pathogenic | 1 | 1 | 0 |
| rs137852497 | pathogenic | 1 | 1 | 0 |
| rs137852493 | pathogenic | 1 | 1 | 0 |
| rs137852504 | pathogenic | 1 | 1 | 1 |
| rs267606863 | pathogenic | 1 | 1 | 1 |
| rs137852486 | pathogenic | 1 | 1 | 1 |
| rs137852479 | pathogenic | 1 | 1 | 1 |
| rs112122950 | benign | 0 | 0 | 0 |
| rs72546648 | benign | 1 | 1 | 1 |
| rs72552291 | pathogenic | 0 | 0 | 0 |
| rs79184941 | pathogenic | 1 | 1 | 1 |
| rs77543610 | pathogenic | 1 | 1 | 1 |
| rs121918505 | pathogenic | 1 | 1 | 1 |
| rs776587763 | pathogenic | 1 | 1 | 1 |
| rs121918497 | pathogenic | 1 | 1 | 1 |
| rs121918499 | pathogenic | 1 | 1 | 1 |
| rs121918501 | pathogenic | 1 | 1 | 1 |
| rs121918500 | pathogenic | 0 | 0 | 0 |
| rs121918504 | pathogenic | 1 | 1 | 1 |
| rs121918510 | pathogenic | 1 | 1 | 1 |
| rs121918493 | pathogenic | 1 | 1 | 1 |

| | | | | |
|---|---|---|---|---|
| rs121918489 | pathogenic | 0 | 0 | 0 |
| rs121918495 | pathogenic | 1 | 1 | 1 |
| rs121918488 | pathogenic | 0 | 0 | 0 |
| rs121918496 | pathogenic | 0 | 0 | 0 |
| rs121918492 | pathogenic | 1 | 1 | 1 |
| rs121918494 | pathogenic | 1 | 1 | 1 |
| rs121918502 | pathogenic | 1 | 1 | 1 |
| rs121918490 | pathogenic | 1 | 1 | 1 |
| rs72466451 | pathogenic | 1 | 1 | 1 |
| rs66468541 | pathogenic | 0 | 0 | 0 |
| rs137852662 | pathogenic | 0 | 0 | 0 |
| rs137852663 | pathogenic | 0 | 0 | 0 |
| rs606231407 | pathogenic | 0 | 0 | 0 |
| rs137852233 | pathogenic | 1 | 1 | 1 |
| rs112306225 | benign | 0 | 0 | 0 |
| rs387906808 | pathogenic | 1 | 1 | 0 |
| rs730882094 | pathogenic | 1 | 1 | 1 |
| rs373869746 | pathogenic | 1 | 1 | 1 |
| rs777524402 | pathogenic | 1 | 1 | 1 |
| rs769737896 | pathogenic | 0 | 1 | 0 |
| rs13306515 | benign | 1 | 1 | 0 |
| rs746982741 | pathogenic | 0 | 0 | 0 |
| rs2230200 | benign | 1 | 1 | 1 |
| rs1047286 | benign | 0 | 0 | 0 |
| rs141737564 | benign | 0 | 0 | 0 |
| rs774826179 | benign | 0 | 0 | 0 |
| rs11569422 | benign | 1 | 1 | 1 |
| rs146613648 | benign | 0 | 0 | 0 |
| rs144432231 | benign | 0 | 0 | 0 |
| rs140655115 | benign | 0 | 0 | 0 |
| rs387906837 | pathogenic | 1 | 1 | 0 |
| rs121909076 | pathogenic | 1 | 1 | 0 |
| rs121909077 | pathogenic | 0 | 0 | 0 |
| rs121909074 | pathogenic | 0 | 0 | 0 |
| rs137853330 | pathogenic | 1 | 1 | 1 |
| rs137853331 | pathogenic | 1 | 1 | 1 |
| rs137853332 | pathogenic | 1 | 1 | 1 |
| rs28931586 | pathogenic | 1 | 1 | 0 |
| rs11569321 | benign | 1 | 1 | 1 |
| rs119486097 | benign | 0 | 0 | 0 |
| rs1799807 | pathogenic | 1 | 1 | 1 |
| rs201820739 | pathogenic | 1 | 1 | 0 |
| rs114706984 | pathogenic | 1 | 1 | 0 |
| rs140080572 | pathogenic | 1 | 1 | 0 |
| rs28933389 | pathogenic | 0 | 1 | 0 |
| rs16849700 | benign | 1 | 1 | 0 |
| rs115624085 | pathogenic | 1 | 1 | 1 |
| rs747196387 | pathogenic | 1 | 1 | 1 |
| rs104893684 | pathogenic | 1 | 1 | 0 |
| rs121918557 | pathogenic | 1 | 1 | 0 |
| rs121918556 | pathogenic | 1 | 1 | 1 |
| rs398124571 | pathogenic | 1 | 1 | 1 |
| rs776631396 | pathogenic | 0 | 1 | 0 |
| rs398124576 | pathogenic | 1 | 1 | 1 |

| | | | | |
|---|---|---|---|---|
| rs398124581 | pathogenic | 1 | 1 | 1 |
| rs398124582 | pathogenic | 1 | 1 | 1 |
| rs149766077 | pathogenic | 0 | 1 | 0 |
| rs79761867 | pathogenic | 0 | 1 | 0 |
| rs148905512 | pathogenic | 0 | 1 | 0 |
| rs121965004 | pathogenic | 1 | 1 | 1 |
| rs386834233 | pathogenic | 0 | 1 | 0 |
| rs398124598 | pathogenic | 0 | 1 | 0 |
| rs398124603 | pathogenic | 1 | 1 | 1 |
| rs546716073 | benign | 1 | 1 | 1 |
| rs200310662 | benign | 1 | 1 | 1 |
| rs115494727 | benign | 1 | 1 | 1 |
| rs781513008 | benign | 0 | 0 | 0 |
| rs587776398 | benign | 1 | 1 | 0 |
| rs35288908 | benign | 1 | 1 | 1 |
| rs150634562 | pathogenic | 1 | 1 | 0 |
| rs142948132 | pathogenic | 1 | 1 | 1 |
| rs132630321 | pathogenic | 1 | 1 | 1 |
| rs132630317 | pathogenic | 1 | 1 | 1 |
| rs132630320 | pathogenic | 0 | 0 | 0 |
| rs781394318 | pathogenic | 1 | 1 | 1 |
| rs111033624 | pathogenic | 1 | 1 | 1 |
| rs111033630 | pathogenic | 1 | 1 | 1 |
| rs111033627 | pathogenic | 1 | 1 | 1 |
| rs193922287 | pathogenic | 1 | 1 | 1 |
| rs759072800 | pathogenic | 0 | 1 | 0 |
| rs104894010 | pathogenic | 0 | 0 | 0 |
| rs193922296 | pathogenic | 0 | 0 | 0 |
| rs104894006 | pathogenic | 1 | 1 | 1 |
| rs193922311 | pathogenic | 1 | 1 | 0 |
| rs80356654 | pathogenic | 1 | 1 | 0 |
| rs144723656 | benign | 1 | 1 | 1 |
| rs148311934 | pathogenic | 1 | 1 | 0 |
| rs80356655 | pathogenic | 1 | 1 | 1 |
| rs769268803 | pathogenic | 1 | 1 | 0 |
| rs104894008 | pathogenic | 0 | 1 | 0 |
| rs104894005 | pathogenic | 1 | 1 | 1 |
| rs104894009 | pathogenic | 1 | 1 | 0 |
| rs193922340 | pathogenic | 0 | 0 | 0 |
| rs104894016 | pathogenic | 0 | 1 | 0 |
| rs193921338 | pathogenic | 0 | 0 | 0 |
| rs193922272 | pathogenic | 1 | 1 | 1 |
| rs104894012 | pathogenic | 1 | 1 | 0 |
| rs193922285 | pathogenic | 1 | 1 | 1 |
| rs104894336 | pathogenic | 1 | 1 | 1 |
| rs28931580 | pathogenic | 1 | 1 | 1 |
| rs104894326 | pathogenic | 1 | 1 | 1 |
| rs104894331 | pathogenic | 0 | 0 | 0 |
| rs104894338 | pathogenic | 0 | 0 | 0 |
| rs104894333 | pathogenic | 1 | 1 | 1 |
| rs104894330 | pathogenic | 1 | 1 | 1 |
| rs143886391 | benign | 1 | 1 | 1 |
| rs104894335 | pathogenic | 1 | 1 | 1 |
| rs104894337 | pathogenic | 0 | 0 | 0 |

| | | | | |
|---|---|---|---|---|
| rs104894328 | pathogenic | 0 | 0 | 0 |
| rs104894341 | pathogenic | 1 | 1 | 0 |
| rs104894329 | pathogenic | 0 | 0 | 0 |
| rs137852850 | pathogenic | 1 | 1 | 1 |
| rs137852854 | pathogenic | 0 | 1 | 0 |
| rs137852849 | pathogenic | 0 | 1 | 0 |
| rs137852848 | pathogenic | 1 | 1 | 1 |
| rs137852852 | pathogenic | 0 | 1 | 0 |
| rs137852853 | pathogenic | 0 | 1 | 0 |
| rs137852847 | pathogenic | 0 | 1 | 0 |
| rs137852846 | pathogenic | 0 | 1 | 0 |
| rs137852844 | pathogenic | 0 | 1 | 0 |
| rs144411158 | pathogenic | 0 | 0 | 0 |
| rs2302234 | benign | 0 | 0 | 0 |
| rs139620139 | pathogenic | 0 | 0 | 0 |
| rs146689929 | benign | 0 | 0 | 0 |
| rs373328681 | pathogenic | 1 | 1 | 0 |
| rs387906620 | pathogenic | 1 | 1 | 1 |
| rs1799983 | benign | 0 | 1 | 0 |
| rs28999111 | pathogenic | 1 | 1 | 1 |
| rs104894544 | pathogenic | 1 | 1 | 1 |
| rs104894546 | pathogenic | 1 | 1 | 1 |
| rs28999112 | pathogenic | 0 | 1 | 0 |
| rs104894545 | pathogenic | 1 | 1 | 1 |
| rs104894547 | pathogenic | 1 | 1 | 1 |
| rs74315310 | pathogenic | 0 | 0 | 0 |
| rs267606852 | pathogenic | 0 | 1 | 0 |
| rs137853586 | pathogenic | 0 | 1 | 0 |
| rs137853582 | pathogenic | 1 | 1 | 1 |
| rs61754634 | pathogenic | 1 | 1 | 1 |
| rs137853587 | pathogenic | 0 | 1 | 0 |
| rs267606851 | pathogenic | 1 | 1 | 1 |
| rs267606853 | pathogenic | 1 | 1 | 1 |
| rs137853584 | pathogenic | 1 | 1 | 1 |
| rs137853585 | pathogenic | 1 | 1 | 1 |
| rs460184 | pathogenic | 1 | 1 | 1 |
| rs104893751 | pathogenic | 0 | 1 | 0 |
| rs77358162 | benign | 0 | 0 | 0 |
| rs387906936 | pathogenic | 0 | 0 | 0 |
| rs769525399 | pathogenic | 0 | 0 | 0 |
| rs371271054 | pathogenic | 1 | 1 | 0 |
| rs118204000 | pathogenic | 0 | 0 | 0 |
| rs138568622 | pathogenic | 1 | 1 | 1 |
| rs34509359 | benign | 0 | 0 | 0 |
| rs104894130 | pathogenic | 1 | 1 | 0 |
| rs281865424 | pathogenic | 0 | 0 | 0 |
| rs121912778 | pathogenic | 1 | 1 | 0 |
| rs771055145 | pathogenic | 1 | 1 | 0 |
| rs776174514 | pathogenic | 1 | 1 | 0 |
| rs545986367 | pathogenic | 1 | 1 | 1 |
| rs570768038 | benign | 1 | 1 | 1 |
| rs74315442 | pathogenic | 1 | 1 | 1 |
| rs79716074 | benign | 1 | 1 | 1 |
| rs748486078 | pathogenic | 1 | 1 | 1 |

| | | | | |
|---|---|---|---|---|
| rs2397084 | benign | 1 | 1 | 1 |
| rs11465553 | benign | 1 | 1 | 1 |
| rs144576902 | benign | 1 | 1 | 1 |
| rs763780 | benign | 1 | 1 | 1 |
| rs267606870 | pathogenic | 1 | 1 | 0 |
| rs121913502 | pathogenic | 1 | 1 | 0 |
| rs118101777 | benign | 1 | 1 | 1 |
| rs118053940 | benign | 0 | 0 | 0 |
| rs368771578 | benign | 1 | 1 | 1 |
| rs11540654 | pathogenic | 1 | 1 | 1 |
| rs121912658 | pathogenic | 1 | 1 | 1 |
| rs28934873 | pathogenic | 0 | 1 | 0 |
| rs780442292 | pathogenic | 1 | 1 | 1 |
| rs758781593 | benign | 1 | 1 | 1 |
| rs28934875 | pathogenic | 1 | 1 | 1 |
| rs28934874 | pathogenic | 1 | 1 | 0 |
| rs587782705 | pathogenic | 1 | 1 | 1 |
| rs587780068 | pathogenic | 1 | 1 | 1 |
| rs587782144 | pathogenic | 1 | 1 | 1 |
| rs138729528 | pathogenic | 1 | 1 | 1 |
| rs28934578 | pathogenic | 1 | 1 | 1 |
| rs397514495 | pathogenic | 1 | 1 | 1 |
| rs121912665 | pathogenic | 1 | 1 | 1 |
| rs397516435 | pathogenic | 1 | 1 | 1 |
| rs587778720 | pathogenic | 1 | 1 | 1 |
| rs397516436 | pathogenic | 1 | 1 | 1 |
| rs587781386 | benign | 1 | 1 | 1 |
| rs121912666 | pathogenic | 1 | 1 | 0 |
| rs587780073 | pathogenic | 1 | 1 | 1 |
| rs587782664 | pathogenic | 1 | 1 | 1 |
| rs730882005 | pathogenic | 1 | 1 | 1 |
| rs28934573 | pathogenic | 1 | 1 | 1 |
| rs121912655 | pathogenic | 1 | 1 | 1 |
| rs28934575 | pathogenic | 1 | 1 | 1 |
| rs121912656 | pathogenic | 1 | 1 | 1 |
| rs11540652 | pathogenic | 1 | 1 | 1 |
| rs121912651 | pathogenic | 1 | 1 | 1 |
| rs730882027 | pathogenic | 1 | 1 | 1 |
| rs121912653 | pathogenic | 1 | 1 | 1 |
| rs121912652 | pathogenic | 1 | 1 | 1 |
| rs121912657 | pathogenic | 0 | 1 | 0 |
| rs121913343 | pathogenic | 1 | 1 | 1 |
| rs28934576 | pathogenic | 1 | 1 | 1 |
| rs763098116 | pathogenic | 1 | 1 | 1 |
| rs121912660 | pathogenic | 1 | 1 | 1 |
| rs28934574 | pathogenic | 1 | 1 | 1 |
| rs371409680 | pathogenic | 1 | 1 | 1 |
| rs121912667 | pathogenic | 1 | 1 | 1 |
| rs121912663 | pathogenic | 1 | 1 | 1 |
| rs201744589 | pathogenic | 0 | 0 | 0 |
| rs33926449 | benign | 0 | 0 | 0 |
| rs74315302 | pathogenic | 0 | 1 | 0 |
| rs202040269 | pathogenic | 1 | 1 | 1 |
| rs375958814 | pathogenic | 1 | 1 | 1 |

| | | | | |
|---|---|---|---|---|
| rs121909624 | pathogenic | 1 | 1 | 1 |
| rs121909621 | pathogenic | 1 | 1 | 1 |
| rs121909622 | pathogenic | 1 | 1 | 0 |
| rs4220 | benign | 1 | 1 | 1 |
| rs147559466 | pathogenic | 1 | 1 | 1 |
| rs757434857 | pathogenic | 1 | 1 | 1 |
| rs398123072 | pathogenic | 1 | 1 | 1 |
| rs121434280 | pathogenic | 0 | 1 | 0 |
| rs398123074 | pathogenic | 0 | 1 | 0 |
| rs762114560 | pathogenic | 1 | 1 | 1 |
| rs121434283 | pathogenic | 1 | 1 | 0 |
| rs121434277 | pathogenic | 0 | 1 | 0 |
| rs745844469 | pathogenic | 1 | 1 | 1 |
| rs121434279 | pathogenic | 1 | 1 | 0 |
| rs121434278 | pathogenic | 1 | 1 | 1 |
| rs373715782 | pathogenic | 1 | 1 | 1 |
| rs200724875 | pathogenic | 1 | 1 | 1 |
| rs149678400 | pathogenic | 0 | 1 | 0 |
| rs770273135 | pathogenic | 1 | 1 | 1 |
| rs121434276 | pathogenic | 0 | 1 | 0 |
| rs121434281 | pathogenic | 1 | 1 | 0 |
| rs768884003 | pathogenic | 1 | 1 | 1 |
| rs201375579 | pathogenic | 0 | 0 | 0 |
| rs121434274 | pathogenic | 1 | 1 | 0 |
| rs121434282 | pathogenic | 1 | 1 | 0 |
| rs779759347 | pathogenic | 0 | 1 | 0 |
| rs77931234 | pathogenic | 1 | 1 | 1 |
| rs148207467 | pathogenic | 1 | 1 | 1 |
| rs766140986 | pathogenic | 1 | 1 | 1 |
| rs121434275 | pathogenic | 1 | 1 | 1 |
| rs748110745 | pathogenic | 1 | 1 | 1 |
| rs104894274 | pathogenic | 1 | 1 | 1 |
| rs104894273 | pathogenic | 0 | 1 | 0 |
| rs104894278 | pathogenic | 1 | 1 | 1 |
| rs104894275 | pathogenic | 1 | 1 | 1 |
| rs104894277 | pathogenic | 1 | 1 | 1 |
| rs104894276 | pathogenic | 0 | 0 | 0 |
| rs104894280 | pathogenic | 1 | 1 | 1 |
| rs145882709 | pathogenic | 1 | 1 | 1 |
| rs104894279 | pathogenic | 1 | 1 | 1 |
| rs80356670 | pathogenic | 1 | 1 | 1 |
| rs121918102 | pathogenic | 1 | 1 | 1 |
| rs80356671 | pathogenic | 1 | 1 | 1 |
| rs80356672 | pathogenic | 1 | 1 | 1 |
| rs758609113 | pathogenic | 1 | 1 | 0 |
| rs121918188 | pathogenic | 0 | 1 | 0 |
| rs121918189 | pathogenic | 1 | 1 | 0 |
| rs121918191 | pathogenic | 1 | 1 | 1 |
| rs1769259 | benign | 1 | 1 | 1 |
| rs140222720 | benign | 1 | 1 | 1 |
| rs2297085 | benign | 1 | 1 | 1 |
| rs200679026 | pathogenic | 1 | 1 | 1 |
| rs121918192 | pathogenic | 0 | 1 | 0 |
| rs145771944 | benign | 0 | 1 | 0 |

| | | | | |
|---|---|---|---|---|
| rs201508880 | pathogenic | 0 | 1 | 0 |
| rs17221854 | pathogenic | 0 | 0 | 0 |
| rs120074189 | pathogenic | 1 | 1 | 1 |
| rs120074190 | pathogenic | 1 | 1 | 1 |
| rs758600283 | benign | 1 | 1 | 1 |
| rs200305064 | pathogenic | 1 | 1 | 1 |
| rs121917891 | pathogenic | 1 | 1 | 1 |
| rs3772809 | benign | 0 | 1 | 0 |
| rs387906619 | pathogenic | 1 | 1 | 0 |
| rs2304681 | benign | 0 | 1 | 0 |
| rs121917849 | pathogenic | 0 | 0 | 0 |
| rs143010236 | pathogenic | 0 | 0 | 0 |
| rs121917850 | pathogenic | 1 | 1 | 0 |
| rs35916840 | pathogenic | 1 | 1 | 0 |
| rs28939688 | pathogenic | 1 | 1 | 1 |
| rs11258194 | benign | 1 | 1 | 1 |
| rs199840952 | pathogenic | 1 | 1 | 1 |
| rs121918159 | pathogenic | 1 | 1 | 1 |
| rs200396166 | pathogenic | 0 | 0 | 0 |
| rs757966746 | pathogenic | 0 | 1 | 0 |
| rs121918608 | pathogenic | 0 | 1 | 0 |
| rs74315401 | pathogenic | 1 | 1 | 1 |
| rs11538758 | pathogenic | 0 | 0 | 0 |
| rs74315402 | pathogenic | 1 | 1 | 1 |
| rs1799990 | pathogenic | 1 | 1 | 1 |
| rs74315410 | pathogenic | 1 | 1 | 1 |
| rs150351644 | benign | 1 | 1 | 1 |
| rs16990018 | benign | 1 | 1 | 1 |
| rs74315403 | pathogenic | 1 | 1 | 1 |
| rs74315408 | pathogenic | 1 | 1 | 1 |
| rs74315411 | pathogenic | 1 | 1 | 1 |
| rs74315413 | pathogenic | 1 | 1 | 1 |
| rs74315405 | pathogenic | 1 | 1 | 1 |
| rs28933385 | pathogenic | 1 | 1 | 1 |
| rs74315412 | pathogenic | 1 | 1 | 1 |
| rs74315407 | pathogenic | 1 | 1 | 1 |
| rs74315406 | pathogenic | 1 | 1 | 1 |
| rs11571117 | benign | 1 | 1 | 1 |
| rs121917742 | pathogenic | 1 | 1 | 1 |
| rs149782619 | pathogenic | 1 | 1 | 1 |
| rs397514492 | pathogenic | 1 | 1 | 1 |
| rs397514489 | pathogenic | 0 | 1 | 0 |
| rs397514493 | pathogenic | 1 | 1 | 1 |
| rs373849532 | pathogenic | 1 | 1 | 1 |
| rs121965048 | pathogenic | 1 | 1 | 1 |
| rs121965037 | pathogenic | 1 | 1 | 1 |
| rs121965060 | pathogenic | 0 | 1 | 0 |
| rs386833603 | pathogenic | 1 | 1 | 1 |
| rs121965038 | pathogenic | 1 | 1 | 1 |
| rs121965039 | pathogenic | 1 | 1 | 1 |
| rs121965040 | pathogenic | 1 | 1 | 1 |
| rs267606925 | pathogenic | 1 | 1 | 1 |
| rs121965056 | pathogenic | 0 | 1 | 0 |
| rs121965059 | pathogenic | 0 | 0 | 0 |

| | | | | |
|---|---|---|---|---|
| rs386833615 | pathogenic | 1 | 1 | 0 |
| rs121965051 | pathogenic | 1 | 1 | 0 |
| rs121965046 | pathogenic | 0 | 1 | 0 |
| rs121965052 | pathogenic | 0 | 1 | 0 |
| rs386833617 | pathogenic | 0 | 1 | 0 |
| rs386833618 | pathogenic | 1 | 1 | 0 |
| rs121965041 | pathogenic | 0 | 1 | 0 |
| rs121965042 | pathogenic | 0 | 1 | 0 |
| rs121965057 | pathogenic | 1 | 1 | 1 |
| rs386833621 | pathogenic | 1 | 1 | 1 |
| rs121965049 | pathogenic | 1 | 1 | 1 |
| rs386833623 | pathogenic | 1 | 1 | 1 |
| rs121965047 | pathogenic | 1 | 1 | 1 |
| rs121965053 | pathogenic | 0 | 1 | 0 |
| rs121965045 | pathogenic | 0 | 1 | 0 |
| rs121965054 | pathogenic | 0 | 1 | 0 |
| rs121965043 | pathogenic | 0 | 1 | 0 |
| rs386833598 | pathogenic | 0 | 0 | 0 |
| rs137854545 | pathogenic | 1 | 1 | 1 |
| rs137854544 | pathogenic | 1 | 1 | 1 |
| rs104894455 | pathogenic | 0 | 1 | 0 |
| rs1049564 | benign | 0 | 1 | 0 |
| rs104894460 | pathogenic | 1 | 1 | 1 |
| rs104894453 | pathogenic | 1 | 1 | 1 |
| rs104894450 | pathogenic | 0 | 1 | 0 |
| rs104894454 | pathogenic | 0 | 1 | 0 |
| rs104894452 | pathogenic | 1 | 1 | 1 |
| rs104894451 | pathogenic | 0 | 1 | 0 |
| rs104894368 | pathogenic | 1 | 1 | 1 |
| rs730880950 | pathogenic | 1 | 1 | 1 |
| rs121909535 | pathogenic | 1 | 1 | 1 |
| rs121909537 | pathogenic | 1 | 1 | 1 |
| rs121909536 | pathogenic | 1 | 1 | 1 |
| rs121909542 | pathogenic | 1 | 1 | 1 |
| rs121909538 | pathogenic | 1 | 1 | 1 |
| rs121909539 | pathogenic | 1 | 1 | 1 |
| rs121909540 | pathogenic | 1 | 1 | 1 |
| rs17560 | benign | 1 | 1 | 1 |
| rs121909543 | pathogenic | 0 | 1 | 0 |
| rs121909544 | pathogenic | 1 | 1 | 1 |
| rs2307456 | benign | 0 | 1 | 0 |
| rs745778317 | pathogenic | 0 | 1 | 0 |
| rs759607901 | pathogenic | 0 | 1 | 0 |
| rs201188114 | benign | 0 | 0 | 0 |
| rs1800472 | benign | 0 | 0 | 0 |
| rs587780728 | benign | 1 | 1 | 1 |
| rs144386518 | benign | 1 | 1 | 1 |
| rs281865439 | pathogenic | 1 | 1 | 1 |
| rs140175796 | pathogenic | 1 | 1 | 1 |
| rs5030843 | pathogenic | 1 | 1 | 0 |
| rs75166491 | pathogenic | 1 | 1 | 0 |
| rs199475679 | pathogenic | 1 | 1 | 1 |
| rs281865440 | pathogenic | 1 | 1 | 1 |
| rs74486803 | pathogenic | 0 | 1 | 0 |

| | | | | |
|---|---|---|---|---|
| rs199475602 | pathogenic | 0 | 1 | 0 |
| rs5030844 | pathogenic | 1 | 1 | 0 |
| rs62514927 | pathogenic | 1 | 1 | 1 |
| rs62514928 | pathogenic | 1 | 1 | 1 |
| rs62516152 | pathogenic | 0 | 1 | 0 |
| rs62507348 | pathogenic | 1 | 1 | 1 |
| rs76687508 | pathogenic | 1 | 1 | 1 |
| rs62508730 | pathogenic | 1 | 1 | 1 |
| rs62508588 | pathogenic | 0 | 1 | 0 |
| rs5030846 | pathogenic | 0 | 1 | 0 |
| rs74503222 | pathogenic | 1 | 1 | 0 |
| rs5030847 | pathogenic | 1 | 1 | 0 |
| rs5030850 | pathogenic | 1 | 1 | 1 |
| rs778154939 | pathogenic | 0 | 1 | 0 |
| rs62514952 | pathogenic | 1 | 1 | 1 |
| rs78655458 | pathogenic | 1 | 1 | 1 |
| rs62508698 | pathogenic | 1 | 1 | 0 |
| rs5030851 | pathogenic | 1 | 1 | 0 |
| rs62642933 | pathogenic | 1 | 1 | 1 |
| rs5030853 | pathogenic | 1 | 1 | 1 |
| rs62642935 | pathogenic | 1 | 1 | 1 |
| rs199475615 | pathogenic | 0 | 0 | 0 |
| rs62516062 | pathogenic | 1 | 1 | 0 |
| rs62516092 | pathogenic | 0 | 0 | 0 |
| rs62516095 | pathogenic | 1 | 1 | 1 |
| rs62642937 | pathogenic | 1 | 1 | 1 |
| rs62516141 | pathogenic | 0 | 1 | 0 |
| rs5030856 | pathogenic | 1 | 1 | 1 |
| rs5030857 | pathogenic | 1 | 1 | 1 |
| rs5030858 | pathogenic | 1 | 1 | 1 |
| rs79931499 | pathogenic | 1 | 1 | 1 |
| rs5030860 | pathogenic | 1 | 1 | 1 |
| rs281865437 | pathogenic | 1 | 1 | 1 |
| rs62644499 | pathogenic | 1 | 1 | 1 |
| rs147689487 | benign | 0 | 0 | 0 |
| rs397515417 | pathogenic | 1 | 1 | 1 |
| rs121909332 | pathogenic | 1 | 1 | 1 |
| rs121909330 | pathogenic | 1 | 1 | 0 |
| rs121909329 | pathogenic | 1 | 1 | 0 |
| rs121909335 | pathogenic | 1 | 1 | 1 |
| rs387906789 | pathogenic | 1 | 1 | 1 |
| rs121909334 | pathogenic | 1 | 1 | 1 |
| rs121909331 | pathogenic | 1 | 1 | 1 |
| rs372487178 | pathogenic | 1 | 1 | 1 |
| rs387906937 | pathogenic | 1 | 1 | 0 |
| rs16845 | benign | 1 | 1 | 1 |
| rs672601313 | pathogenic | 1 | 1 | 1 |
| rs117805308 | benign | 0 | 0 | 0 |
| rs121434261 | pathogenic | 1 | 1 | 1 |
| rs141629512 | benign | 0 | 0 | 0 |
| rs199957176 | benign | 1 | 1 | 1 |
| rs2230461 | benign | 1 | 1 | 1 |
| rs764618040 | pathogenic | 1 | 1 | 1 |
| rs267606949 | pathogenic | 1 | 1 | 1 |

| | | | | |
|---|---|---|---|---|
| rs587777483 | pathogenic | 1 | 1 | 1 |
| rs267606947 | pathogenic | 1 | 1 | 1 |
| rs6941583 | benign | 0 | 0 | 0 |
| rs749597959 | benign | 1 | 1 | 1 |
| rs182340753 | benign | 1 | 1 | 1 |
| rs146678221 | benign | 0 | 0 | 0 |
| rs11539202 | benign | 0 | 0 | 0 |
| rs148723565 | benign | 0 | 0 | 0 |
| rs61752925 | benign | 1 | 1 | 1 |
| rs113309941 | pathogenic | 0 | 0 | 0 |
| rs121434520 | pathogenic | 1 | 1 | 1 |
| rs121434517 | pathogenic | 1 | 1 | 1 |
| rs267607027 | pathogenic | 1 | 1 | 1 |
| rs138603088 | pathogenic | 0 | 0 | 0 |
| rs121434516 | pathogenic | 1 | 1 | 0 |
| rs121434521 | pathogenic | 0 | 0 | 0 |
| rs145735112 | benign | 1 | 1 | 1 |
| rs121434519 | pathogenic | 1 | 1 | 1 |
| rs75326924 | pathogenic | 1 | 1 | 0 |
| rs142186404 | pathogenic | 0 | 0 | 0 |
| rs121918035 | pathogenic | 0 | 0 | 0 |
| rs374385878 | pathogenic | 1 | 1 | 0 |
| rs11557652 | benign | 0 | 0 | 0 |
| rs118203986 | pathogenic | 1 | 1 | 1 |
| rs141675330 | benign | 1 | 1 | 1 |
| rs104893927 | pathogenic | 0 | 0 | 0 |
| rs104893935 | pathogenic | 0 | 0 | 0 |
| rs104893933 | pathogenic | 1 | 1 | 0 |
| rs104893934 | pathogenic | 0 | 0 | 0 |
| rs145183203 | benign | 1 | 1 | 0 |
| rs2230806 | benign | 0 | 0 | 0 |
| rs9282541 | benign | 0 | 0 | 0 |
| rs559110055 | benign | 0 | 0 | 0 |
| rs200030513 | benign | 0 | 0 | 0 |
| rs9282543 | benign | 0 | 0 | 0 |
| rs147675550 | benign | 0 | 0 | 0 |
| rs137854496 | pathogenic | 0 | 0 | 0 |
| rs374190304 | benign | 0 | 0 | 0 |
| rs114620717 | benign | 0 | 0 | 0 |
| rs2066718 | benign | 0 | 0 | 0 |
| rs35819696 | benign | 0 | 0 | 0 |
| rs138880920 | benign | 0 | 0 | 0 |
| rs2066715 | benign | 0 | 0 | 0 |
| rs35207495 | benign | 0 | 0 | 0 |
| rs2066714 | benign | 0 | 0 | 0 |
| rs28937313 | pathogenic | 0 | 0 | 0 |
| rs28937314 | pathogenic | 0 | 0 | 0 |
| rs137854495 | pathogenic | 0 | 0 | 0 |
| rs140365800 | benign | 0 | 0 | 0 |
| rs13306073 | benign | 0 | 0 | 0 |
| rs28933692 | pathogenic | 0 | 0 | 0 |
| rs33918808 | benign | 0 | 0 | 0 |
| rs76881554 | benign | 0 | 0 | 0 |
| rs143180998 | benign | 0 | 0 | 0 |

| | | | | |
|---|---|---|---|---|
| rs137854500 | pathogenic | 0 | 0 | 0 |
| rs538614702 | benign | 0 | 0 | 0 |
| rs201879964 | benign | 0 | 0 | 0 |
| rs2230808 | benign | 0 | 0 | 0 |
| rs138422574 | benign | 0 | 0 | 0 |
| rs137854498 | pathogenic | 0 | 0 | 0 |
| rs150125857 | benign | 0 | 0 | 0 |
| rs146292819 | pathogenic | 0 | 0 | 0 |
| rs142688906 | benign | 0 | 0 | 0 |
| rs137854499 | pathogenic | 0 | 0 | 0 |
| rs34879708 | benign | 0 | 0 | 0 |
| rs121913028 | pathogenic | 1 | 1 | 1 |
| rs1041951 | benign | 1 | 1 | 0 |
| rs138714094 | benign | 1 | 1 | 0 |
| rs200650502 | benign | 0 | 1 | 0 |
| rs370708663 | pathogenic | 1 | 1 | 1 |
| rs118204040 | pathogenic | 0 | 1 | 0 |
| rs146899669 | pathogenic | 1 | 1 | 0 |
| rs118204039 | pathogenic | 0 | 1 | 0 |
| rs121909233 | pathogenic | 0 | 0 | 0 |
| rs121909225 | pathogenic | 1 | 1 | 0 |
| rs121909236 | pathogenic | 0 | 0 | 0 |
| rs398123317 | pathogenic | 0 | 0 | 0 |
| rs121909226 | pathogenic | 1 | 1 | 0 |
| rs121909238 | pathogenic | 1 | 1 | 0 |
| rs121909230 | pathogenic | 0 | 0 | 0 |
| rs121909237 | pathogenic | 0 | 0 | 0 |
| rs121909222 | pathogenic | 1 | 1 | 0 |
| rs121909223 | pathogenic | 1 | 1 | 0 |
| rs121909218 | pathogenic | 1 | 1 | 0 |
| rs121909229 | pathogenic | 1 | 1 | 0 |
| rs121909224 | pathogenic | 1 | 1 | 0 |
| rs121909241 | pathogenic | 0 | 0 | 0 |
| rs121909221 | pathogenic | 0 | 0 | 0 |
| rs121913294 | pathogenic | 0 | 0 | 0 |
| rs121909234 | pathogenic | 0 | 0 | 0 |
| rs121909240 | pathogenic | 0 | 0 | 0 |
| rs767623493 | benign | 0 | 0 | 0 |
| rs121909239 | pathogenic | 0 | 0 | 0 |
| rs120074145 | pathogenic | 1 | 1 | 1 |
| rs120074148 | pathogenic | 1 | 1 | 1 |
| rs762991875 | pathogenic | 0 | 0 | 0 |
| rs148639841 | pathogenic | 1 | 1 | 1 |
| rs199524907 | pathogenic | 1 | 1 | 1 |
| rs120074141 | pathogenic | 1 | 1 | 1 |
| rs120074144 | pathogenic | 0 | 1 | 0 |
| rs120074146 | pathogenic | 1 | 1 | 1 |
| rs120074147 | pathogenic | 0 | 1 | 0 |
| rs120074143 | pathogenic | 0 | 1 | 0 |
| rs120074140 | pathogenic | 1 | 1 | 0 |
| rs28937315 | pathogenic | 1 | 1 | 1 |
| rs76066357 | benign | 1 | 1 | 1 |
| rs137852911 | pathogenic | 0 | 1 | 0 |
| rs137852907 | pathogenic | 1 | 1 | 0 |

| | | | | |
|---|---|---|---|---|
| rs137852910 | pathogenic | 1 | 1 | 1 |
| rs537585211 | benign | 0 | 0 | 0 |
| rs202134424 | pathogenic | 0 | 0 | 0 |
| rs200930463 | pathogenic | 0 | 0 | 0 |
| rs587777241 | pathogenic | 1 | 1 | 0 |
| rs77563738 | pathogenic | 0 | 0 | 0 |
| rs750868279 | pathogenic | 1 | 1 | 0 |
| rs148936893 | pathogenic | 0 | 0 | 0 |
| rs146597836 | pathogenic | 0 | 0 | 0 |
| rs775440641 | pathogenic | 1 | 1 | 1 |
| rs770689762 | pathogenic | 0 | 1 | 0 |
| rs376785840 | pathogenic | 0 | 0 | 0 |
| rs137852531 | pathogenic | 0 | 0 | 0 |
| rs137852538 | pathogenic | 1 | 1 | 0 |
| rs137852529 | pathogenic | 0 | 1 | 0 |
| rs137852528 | pathogenic | 0 | 1 | 0 |
| rs137852535 | pathogenic | 1 | 1 | 0 |
| rs137853045 | pathogenic | 0 | 1 | 0 |
| rs137853046 | pathogenic | 1 | 1 | 1 |
| rs137853043 | pathogenic | 1 | 1 | 1 |
| rs137853044 | pathogenic | 1 | 1 | 1 |
| rs137853047 | pathogenic | 1 | 1 | 1 |
| rs267606990 | pathogenic | 1 | 1 | 1 |
| rs751437780 | pathogenic | 1 | 1 | 0 |
| rs121918461 | pathogenic | 1 | 1 | 1 |
| rs121918460 | pathogenic | 1 | 1 | 1 |
| rs121918459 | pathogenic | 1 | 1 | 0 |
| rs397507511 | pathogenic | 1 | 1 | 1 |
| rs121918454 | pathogenic | 1 | 1 | 1 |
| rs121918453 | pathogenic | 1 | 1 | 1 |
| rs121918462 | pathogenic | 1 | 1 | 0 |
| rs121918465 | pathogenic | 1 | 1 | 0 |
| rs121918464 | pathogenic | 1 | 1 | 0 |
| rs121918466 | pathogenic | 1 | 1 | 1 |
| rs397507521 | pathogenic | 1 | 1 | 1 |
| rs397507527 | pathogenic | 1 | 1 | 1 |
| rs777603059 | pathogenic | 0 | 1 | 0 |
| rs121918456 | pathogenic | 1 | 1 | 1 |
| rs121918463 | pathogenic | 1 | 1 | 1 |
| rs121918455 | pathogenic | 0 | 1 | 0 |
| rs28933386 | pathogenic | 0 | 1 | 0 |
| rs774356443 | benign | 1 | 1 | 1 |
| rs121918467 | pathogenic | 1 | 1 | 1 |
| rs121918468 | pathogenic | 1 | 1 | 1 |
| rs121918469 | pathogenic | 1 | 1 | 1 |
| rs397507539 | pathogenic | 1 | 1 | 0 |
| rs121918458 | pathogenic | 1 | 1 | 1 |
| rs121918470 | pathogenic | 1 | 1 | 1 |
| rs35289681 | benign | 0 | 1 | 0 |
| rs4302331 | benign | 0 | 1 | 0 |
| rs11544803 | pathogenic | 1 | 1 | 1 |
| rs116254156 | benign | 0 | 0 | 0 |
| rs104893909 | pathogenic | 1 | 1 | 1 |
| rs104893911 | pathogenic | 1 | 1 | 1 |

| | | | | |
|---|---|---|---|---|
| rs104893908 | pathogenic | 1 | 1 | 0 |
| rs258751 | benign | 1 | 1 | 1 |
| rs104893914 | pathogenic | 1 | 1 | 0 |
| rs121909727 | pathogenic | 1 | 1 | 1 |
| rs104893910 | pathogenic | 1 | 1 | 1 |
| rs121909726 | pathogenic | 1 | 1 | 1 |
| rs104893912 | pathogenic | 1 | 1 | 1 |
| rs138407155 | benign | 1 | 1 | 1 |
| rs574061789 | benign | 1 | 1 | 1 |
| rs121912718 | pathogenic | 1 | 1 | 1 |
| rs121912719 | pathogenic | 1 | 1 | 1 |
| rs140770089 | benign | 1 | 1 | 1 |
| rs28931573 | pathogenic | 1 | 1 | 1 |
| rs121912717 | pathogenic | 1 | 1 | 1 |
| rs376456598 | pathogenic | 0 | 0 | 0 |
| rs398123499 | pathogenic | 1 | 1 | 1 |
| rs773048903 | pathogenic | 1 | 1 | 0 |
| rs398123503 | pathogenic | 1 | 1 | 1 |
| rs375785084 | pathogenic | 0 | 1 | 0 |
| rs137852874 | pathogenic | 1 | 1 | 1 |
| rs373713279 | pathogenic | 1 | 1 | 1 |
| rs137852873 | pathogenic | 1 | 1 | 0 |
| rs367823977 | pathogenic | 1 | 1 | 1 |
| rs398123508 | pathogenic | 1 | 1 | 1 |
| rs137852871 | pathogenic | 1 | 1 | 1 |
| rs753698250 | pathogenic | 0 | 1 | 0 |
| rs182923857 | pathogenic | 1 | 1 | 1 |
| rs148090804 | benign | 1 | 1 | 1 |
| rs137852870 | pathogenic | 1 | 1 | 1 |
| rs267607131 | pathogenic | 1 | 1 | 0 |
| rs137852902 | pathogenic | 0 | 0 | 0 |
| rs61741646 | benign | 1 | 1 | 1 |
| rs137852905 | pathogenic | 0 | 0 | 0 |
| rs121918681 | pathogenic | 0 | 1 | 0 |
| rs8177238 | benign | 1 | 1 | 0 |
| rs5030776 | pathogenic | 1 | 1 | 1 |
| rs104894572 | pathogenic | 0 | 0 | 0 |
| rs104894576 | pathogenic | 0 | 0 | 0 |
| rs80084721 | pathogenic | 0 | 0 | 0 |
| rs104894146 | pathogenic | 0 | 1 | 0 |
| rs104894138 | pathogenic | 1 | 1 | 0 |
| rs104894153 | pathogenic | 1 | 1 | 0 |
| rs104894135 | pathogenic | 1 | 1 | 0 |
| rs104894147 | pathogenic | 1 | 1 | 0 |
| rs104894148 | pathogenic | 1 | 1 | 0 |
| rs104894154 | pathogenic | 1 | 1 | 0 |
| rs104894150 | pathogenic | 1 | 1 | 0 |
| rs104894136 | pathogenic | 1 | 1 | 0 |
| rs104894144 | pathogenic | 0 | 0 | 0 |
| rs104894137 | pathogenic | 0 | 0 | 0 |
| rs61754278 | pathogenic | 1 | 1 | 0 |
| rs104894149 | pathogenic | 1 | 1 | 0 |
| rs104894139 | pathogenic | 1 | 1 | 0 |
| rs104894142 | pathogenic | 0 | 0 | 0 |

| | | | | |
|---|---|---|---|---|
| rs104894143 | pathogenic | 0 | 0 | 0 |
| rs104894145 | pathogenic | 0 | 0 | 0 |
| rs121909545 | benign | 0 | 1 | 0 |
| rs782308462 | pathogenic | 1 | 1 | 1 |
| rs375002796 | pathogenic | 0 | 1 | 0 |
| rs201727685 | pathogenic | 0 | 0 | 0 |
| rs768050261 | pathogenic | 1 | 1 | 1 |
| rs780851340 | benign | 0 | 0 | 0 |
| rs139824241 | benign | 1 | 1 | 0 |
| rs757486575 | pathogenic | 1 | 1 | 0 |
| rs750831299 | pathogenic | 1 | 1 | 1 |
| rs767602961 | pathogenic | 1 | 1 | 1 |
| rs121913661 | pathogenic | 1 | 1 | 1 |
| rs777692271 | pathogenic | 1 | 1 | 1 |
| rs121913659 | pathogenic | 1 | 1 | 1 |
| rs587777533 | pathogenic | 1 | 1 | 1 |
| rs104894210 | pathogenic | 0 | 0 | 0 |
| rs104894216 | pathogenic | 1 | 1 | 1 |
| rs104894207 | pathogenic | 1 | 1 | 1 |
| rs104894208 | pathogenic | 0 | 1 | 0 |
| rs104894214 | pathogenic | 1 | 1 | 1 |
| rs104894211 | pathogenic | 1 | 1 | 1 |
| rs61749384 | pathogenic | 0 | 0 | 0 |
| rs61750071 | pathogenic | 1 | 1 | 1 |
| rs61750074 | pathogenic | 0 | 0 | 0 |
| rs11063988 | benign | 0 | 0 | 0 |
| rs216311 | benign | 0 | 0 | 0 |
| rs1800382 | benign | 1 | 1 | 1 |
| rs11063987 | benign | 1 | 1 | 1 |
| rs387906847 | pathogenic | 1 | 1 | 1 |
| rs73003466 | benign | 0 | 0 | 0 |
| rs149787233 | benign | 1 | 1 | 1 |
| rs144451841 | pathogenic | 1 | 1 | 1 |
| rs141638421 | pathogenic | 1 | 1 | 1 |
| rs121434497 | pathogenic | 0 | 0 | 0 |
| rs121434498 | pathogenic | 0 | 0 | 0 |
| rs267607230 | pathogenic | 1 | 1 | 0 |
| rs121434499 | pathogenic | 1 | 1 | 0 |
| rs546749336 | benign | 1 | 1 | 1 |
| rs138873805 | benign | 0 | 0 | 0 |
| rs730880522 | pathogenic | 0 | 0 | 0 |
| rs200371894 | benign | 0 | 0 | 0 |
| rs730880509 | benign | 0 | 0 | 0 |
| rs267606704 | pathogenic | 1 | 1 | 1 |
| rs267606706 | pathogenic | 0 | 0 | 0 |
| rs267606705 | pathogenic | 1 | 1 | 1 |
| rs267606707 | pathogenic | 0 | 0 | 0 |
| rs387906665 | pathogenic | 1 | 1 | 1 |
| rs267606708 | pathogenic | 1 | 1 | 1 |
| rs148368481 | benign | 0 | 0 | 0 |
| rs13204 | benign | 0 | 0 | 0 |
| rs140547520 | pathogenic | 1 | 1 | 1 |
| rs755234990 | benign | 1 | 1 | 0 |
| rs151184052 | benign | 0 | 0 | 0 |

| | | | | |
|---|---|---|---|---|
| rs3736358 | benign | 0 | 0 | 0 |
| rs3736360 | benign | 0 | 0 | 0 |
| rs1138469 | benign | 0 | 0 | 0 |
| rs74315351 | pathogenic | 0 | 1 | 0 |
| rs137853051 | pathogenic | 1 | 1 | 0 |
| rs74315353 | pathogenic | 0 | 1 | 0 |
| rs74315352 | pathogenic | 0 | 1 | 0 |
| rs74315354 | pathogenic | 1 | 1 | 1 |
| rs28938172 | pathogenic | 1 | 1 | 1 |
| rs52836744 | pathogenic | 0 | 0 | 0 |
| rs121913159 | pathogenic | 1 | 1 | 1 |
| rs121913145 | pathogenic | 1 | 1 | 1 |
| rs891087 | benign | 1 | 1 | 1 |
| rs9282757 | benign | 1 | 1 | 1 |
| rs397517404 | pathogenic | 1 | 1 | 1 |
| rs199918720 | benign | 1 | 1 | 0 |
| rs368299411 | benign | 1 | 1 | 0 |
| rs139399951 | benign | 1 | 1 | 0 |
| rs763981974 | benign | 1 | 1 | 0 |
| rs144242114 | benign | 0 | 0 | 0 |
| rs750965801 | benign | 0 | 0 | 0 |
| rs561310777 | benign | 0 | 0 | 0 |
| rs148185335 | benign | 0 | 0 | 0 |
| rs147742157 | benign | 0 | 0 | 0 |
| rs375664373 | pathogenic | 1 | 1 | 1 |
| rs777327250 | pathogenic | 0 | 0 | 0 |
| rs121908471 | pathogenic | 1 | 1 | 0 |
| rs2301612 | pathogenic | 1 | 1 | 0 |
| rs121908476 | pathogenic | 0 | 0 | 0 |
| rs36220239 | benign | 0 | 0 | 0 |
| rs121908473 | pathogenic | 0 | 0 | 0 |
| rs28647808 | benign | 0 | 0 | 0 |
| rs36090624 | benign | 0 | 0 | 0 |
| rs121908002 | pathogenic | 1 | 1 | 1 |
| rs4251545 | benign | 1 | 1 | 1 |
| rs78705382 | pathogenic | 0 | 0 | 0 |
| rs75184679 | pathogenic | 1 | 1 | 1 |
| rs74555752 | pathogenic | 0 | 0 | 0 |
| rs56320207 | benign | 1 | 1 | 1 |
| rs28730837 | pathogenic | 1 | 1 | 1 |
| rs119469014 | pathogenic | 1 | 1 | 1 |
| rs119468010 | pathogenic | 1 | 1 | 1 |
| rs35220041 | benign | 0 | 0 | 0 |
| rs762321137 | pathogenic | 1 | 1 | 1 |
| rs28931570 | pathogenic | 1 | 1 | 1 |
| rs709932 | benign | 1 | 1 | 1 |
| rs746531546 | benign | 1 | 1 | 1 |
| rs6647 | benign | 1 | 1 | 1 |
| rs141095970 | benign | 1 | 1 | 1 |
| rs143370956 | benign | 1 | 1 | 1 |
| rs28929474 | pathogenic | 1 | 1 | 1 |
| rs61761869 | pathogenic | 1 | 1 | 1 |
| rs1303 | benign | 1 | 1 | 1 |
| rs149447502 | benign | 0 | 0 | 0 |

| | | | | |
|---|---|---|---|---|
| rs267607180 | pathogenic | 1 | 1 | 1 |
| rs150304121 | benign | 0 | 0 | 0 |
| rs147630375 | benign | 1 | 1 | 1 |
| rs11539445 | benign | 1 | 1 | 1 |
| rs372098364 | pathogenic | 1 | 1 | 0 |
| rs760072286 | benign | 0 | 0 | 0 |
| rs745595833 | pathogenic | 1 | 1 | 1 |
| rs587777214 | pathogenic | 0 | 0 | 0 |
| rs150318528 | benign | 0 | 0 | 0 |
| rs137853254 | pathogenic | 1 | 1 | 1 |
| rs137853255 | pathogenic | 0 | 0 | 0 |
| rs137853253 | pathogenic | 0 | 0 | 0 |
| rs202166915 | benign | 0 | 0 | 0 |
| rs137853259 | pathogenic | 0 | 0 | 0 |
| rs2229137 | benign | 0 | 0 | 0 |
| rs137853258 | pathogenic | 1 | 1 | 0 |
| rs137853252 | pathogenic | 1 | 1 | 1 |
| rs137853256 | pathogenic | 0 | 0 | 0 |
| rs137853250 | pathogenic | 1 | 1 | 1 |
| rs121434615 | pathogenic | 0 | 0 | 0 |
| rs115014558 | benign | 1 | 1 | 1 |
| rs34116584 | benign | 1 | 1 | 1 |
| rs34885252 | benign | 1 | 1 | 1 |
| rs180177262 | pathogenic | 1 | 1 | 1 |
| rs180177162 | pathogenic | 1 | 1 | 1 |
| rs121908523 | pathogenic | 1 | 1 | 1 |
| rs180177170 | pathogenic | 1 | 1 | 1 |
| rs180177180 | pathogenic | 1 | 1 | 1 |
| rs767586362 | pathogenic | 1 | 1 | 1 |
| rs121908521 | pathogenic | 1 | 1 | 1 |
| rs180177203 | pathogenic | 1 | 1 | 1 |
| rs138025751 | pathogenic | 1 | 1 | 1 |
| rs180177210 | pathogenic | 0 | 1 | 0 |
| rs180177211 | pathogenic | 0 | 1 | 0 |
| rs121908524 | pathogenic | 1 | 1 | 0 |
| rs121908530 | pathogenic | 1 | 1 | 0 |
| rs180177227 | pathogenic | 0 | 1 | 0 |
| rs180177230 | pathogenic | 1 | 1 | 1 |
| rs121908529 | pathogenic | 0 | 1 | 0 |
| rs117195882 | benign | 1 | 1 | 0 |
| rs180177238 | pathogenic | 1 | 1 | 0 |
| rs34664134 | benign | 1 | 1 | 1 |
| rs180177246 | pathogenic | 0 | 1 | 0 |
| rs536352238 | pathogenic | 0 | 0 | 0 |
| rs121908520 | pathogenic | 1 | 1 | 0 |
| rs180177248 | pathogenic | 1 | 1 | 0 |
| rs180177252 | pathogenic | 1 | 1 | 1 |
| rs121908526 | pathogenic | 1 | 1 | 1 |
| rs121908525 | pathogenic | 1 | 1 | 1 |
| rs147106773 | benign | 1 | 1 | 1 |
| rs180177264 | pathogenic | 0 | 1 | 0 |
| rs146525143 | pathogenic | 1 | 1 | 1 |
| rs140992177 | benign | 1 | 1 | 1 |
| rs13408961 | benign | 0 | 1 | 0 |

| | | | | |
|---|---|---|---|---|
| rs4426527 | benign | 1 | 1 | 1 |
| rs3088308 | benign | 0 | 1 | 0 |
| rs4469829 | benign | 1 | 1 | 0 |
| rs104886003 | pathogenic | 1 | 1 | 1 |
| rs121913286 | pathogenic | 1 | 1 | 1 |
| rs121913279 | pathogenic | 1 | 1 | 1 |
| rs121434636 | pathogenic | 1 | 1 | 1 |
| rs755299132 | pathogenic | 1 | 1 | 1 |
| rs748318386 | pathogenic | 1 | 1 | 1 |
| rs387906757 | pathogenic | 1 | 1 | 1 |
| rs121965064 | pathogenic | 1 | 1 | 1 |
| rs28934608 | pathogenic | 0 | 0 | 0 |
| rs5967 | benign | 1 | 1 | 1 |
| rs773905328 | pathogenic | 0 | 0 | 0 |
| rs104894256 | pathogenic | 0 | 1 | 0 |
| rs28931612 | pathogenic | 0 | 1 | 0 |
| rs104894263 | pathogenic | 1 | 1 | 1 |
| rs607969 | benign | 1 | 1 | 0 |
| rs104894262 | pathogenic | 0 | 1 | 0 |
| rs104894268 | pathogenic | 1 | 1 | 1 |
| rs104894264 | pathogenic | 0 | 1 | 0 |
| rs104894259 | pathogenic | 1 | 1 | 0 |
| rs2959656 | benign | 0 | 0 | 0 |
| rs121913035 | pathogenic | 1 | 1 | 0 |
| rs139605474 | benign | 0 | 0 | 0 |
| rs3817552 | benign | 0 | 0 | 0 |
| rs150541115 | benign | 0 | 0 | 0 |
| rs562763010 | benign | 1 | 1 | 1 |
| rs200053119 | pathogenic | 0 | 0 | 0 |
| rs73826386 | benign | 0 | 0 | 0 |
| rs143655258 | benign | 0 | 0 | 0 |
| rs758857853 | pathogenic | 1 | 1 | 0 |
| rs2228380 | benign | 0 | 0 | 0 |
| rs141078733 | benign | 1 | 1 | 1 |
| rs55824172 | pathogenic | 1 | 1 | 1 |
| rs756751089 | pathogenic | 1 | 1 | 1 |
| rs121964990 | pathogenic | 0 | 0 | 0 |
| rs143555186 | benign | 0 | 0 | 0 |
| rs121964993 | pathogenic | 1 | 1 | 0 |
| rs121964992 | pathogenic | 1 | 1 | 1 |
| rs121964991 | pathogenic | 1 | 1 | 0 |
| rs121964988 | pathogenic | 1 | 1 | 1 |
| rs4687 | benign | 1 | 1 | 1 |
| rs267607125 | pathogenic | 1 | 1 | 1 |
| rs751682878 | benign | 1 | 1 | 1 |
| rs267607126 | pathogenic | 1 | 1 | 1 |
| rs768713596 | pathogenic | 0 | 0 | 0 |
| rs147288996 | benign | 0 | 0 | 0 |
| rs387906639 | pathogenic | 1 | 1 | 1 |
| rs730881069 | pathogenic | 1 | 1 | 1 |
| rs730881071 | pathogenic | 1 | 1 | 1 |
| rs397516348 | pathogenic | 0 | 0 | 0 |
| rs121917760 | pathogenic | 0 | 0 | 0 |
| rs104894724 | pathogenic | 1 | 1 | 1 |

| | | | | |
|---|---|---|---|---|
| rs397516349 | pathogenic | 1 | 1 | 1 |
| rs12075 | benign | 0 | 0 | 0 |
| rs147549987 | benign | 0 | 0 | 0 |
| rs56079734 | benign | 0 | 0 | 0 |
| rs1017842 | benign | 1 | 1 | 1 |
| rs137852593 | pathogenic | 1 | 1 | 1 |
| rs137852571 | pathogenic | 1 | 1 | 1 |
| rs137852591 | pathogenic | 1 | 1 | 1 |
| rs9332971 | pathogenic | 1 | 1 | 1 |
| rs755226547 | pathogenic | 1 | 1 | 1 |
| rs79206939 | benign | 0 | 0 | 0 |
| rs121918214 | pathogenic | 1 | 1 | 0 |
| rs757075712 | pathogenic | 1 | 1 | 0 |
| rs150483902 | benign | 1 | 1 | 1 |
| rs113993973 | pathogenic | 0 | 1 | 0 |
| rs946616 | benign | 0 | 0 | 0 |
| rs113993976 | pathogenic | 1 | 1 | 0 |
| rs151022264 | benign | 0 | 1 | 0 |
| rs113993977 | pathogenic | 1 | 1 | 0 |
| rs113993978 | pathogenic | 1 | 1 | 0 |
| rs2228499 | benign | 0 | 1 | 0 |
| rs113993979 | pathogenic | 1 | 1 | 0 |
| rs113993980 | pathogenic | 1 | 1 | 0 |
| rs113993983 | pathogenic | 0 | 1 | 0 |
| rs150547274 | pathogenic | 1 | 1 | 0 |
| rs113993984 | pathogenic | 1 | 1 | 0 |
| rs113993986 | pathogenic | 1 | 1 | 0 |
| rs34313873 | benign | 0 | 1 | 0 |
| rs113993988 | pathogenic | 1 | 1 | 0 |
| rs199638097 | pathogenic | 1 | 1 | 1 |
| rs118204447 | pathogenic | 1 | 1 | 1 |
| rs11862754 | benign | 0 | 0 | 0 |
| rs118204445 | pathogenic | 0 | 0 | 0 |
| rs118204441 | pathogenic | 0 | 0 | 0 |
| rs118204438 | pathogenic | 0 | 0 | 0 |
| rs118204436 | pathogenic | 0 | 0 | 0 |
| rs146093755 | pathogenic | 0 | 0 | 0 |
| rs781439830 | pathogenic | 0 | 0 | 0 |
| rs398123438 | pathogenic | 0 | 0 | 0 |
| rs398123439 | pathogenic | 0 | 0 | 0 |
| rs118204444 | pathogenic | 0 | 0 | 0 |
| rs7187889 | benign | 0 | 0 | 0 |
| rs118204435 | pathogenic | 1 | 1 | 0 |
| rs34745339 | benign | 0 | 0 | 0 |
| rs398123440 | pathogenic | 0 | 0 | 0 |
| rs118204442 | pathogenic | 0 | 0 | 0 |
| rs770053354 | pathogenic | 0 | 0 | 0 |
| rs118204448 | pathogenic | 0 | 0 | 0 |
| rs118204443 | pathogenic | 0 | 0 | 0 |
| rs118204446 | pathogenic | 0 | 0 | 0 |
| rs746756997 | pathogenic | 0 | 0 | 0 |
| rs267606838 | pathogenic | 0 | 0 | 0 |
| rs118204437 | pathogenic | 0 | 0 | 0 |
| rs398123429 | pathogenic | 0 | 0 | 0 |

| | | | | |
|---|---|---|---|---|
| rs398123430 | pathogenic | 0 | 0 | 0 |
| rs2303269 | benign | 0 | 0 | 0 |
| rs114703967 | benign | 0 | 0 | 0 |
| rs118204439 | pathogenic | 0 | 0 | 0 |
| rs151296605 | benign | 0 | 0 | 0 |
| rs118204440 | pathogenic | 0 | 0 | 0 |
| rs78127134 | benign | 0 | 0 | 0 |
| rs372893383 | pathogenic | 0 | 0 | 0 |
| rs61753266 | pathogenic | 1 | 1 | 1 |
| rs139579928 | benign | 1 | 1 | 1 |
| rs56275071 | pathogenic | 1 | 1 | 1 |
| rs121909736 | pathogenic | 1 | 1 | 0 |
| rs121909737 | pathogenic | 1 | 1 | 0 |
| rs121909735 | pathogenic | 1 | 1 | 0 |
| rs121909731 | pathogenic | 1 | 1 | 1 |
| rs121909734 | pathogenic | 1 | 1 | 1 |
| rs121909733 | pathogenic | 1 | 1 | 1 |
| rs121909732 | pathogenic | 1 | 1 | 1 |
| rs121909730 | pathogenic | 1 | 1 | 1 |
| rs202198360 | benign | 1 | 1 | 1 |
| rs587777294 | pathogenic | 1 | 1 | 0 |
| rs587777297 | pathogenic | 1 | 1 | 0 |
| rs104893913 | pathogenic | 1 | 1 | 0 |
| rs17860424 | pathogenic | 1 | 1 | 1 |
| rs1045485 | benign | 1 | 1 | 1 |
| rs13926 | benign | 0 | 0 | 0 |
| rs79343837 | benign | 0 | 0 | 0 |
| rs45516306 | benign | 1 | 1 | 1 |
| rs139849355 | benign | 1 | 1 | 1 |
| rs104894874 | pathogenic | 1 | 1 | 1 |
| rs104894870 | pathogenic | 1 | 1 | 1 |
| rs137852221 | pathogenic | 1 | 1 | 1 |
| rs104894869 | pathogenic | 1 | 1 | 1 |
| rs104894880 | pathogenic | 1 | 1 | 1 |
| rs104894872 | pathogenic | 1 | 1 | 1 |
| rs104894868 | pathogenic | 1 | 1 | 1 |
| rs104894867 | pathogenic | 1 | 1 | 1 |
| rs104894871 | pathogenic | 1 | 1 | 1 |
| rs104894877 | pathogenic | 1 | 1 | 1 |
| rs104894883 | pathogenic | 1 | 1 | 1 |
| rs104894875 | pathogenic | 1 | 1 | 1 |
| rs104894876 | pathogenic | 1 | 1 | 1 |
| rs104894878 | pathogenic | 1 | 1 | 1 |
| rs137852220 | pathogenic | 1 | 1 | 1 |
| rs28933684 | pathogenic | 1 | 1 | 1 |
| rs104894873 | pathogenic | 1 | 1 | 1 |
| rs138409264 | benign | 0 | 0 | 0 |
| rs2236722 | benign | 0 | 0 | 0 |
| rs28757184 | benign | 0 | 0 | 0 |
| rs121434538 | pathogenic | 0 | 0 | 0 |
| rs700519 | benign | 0 | 0 | 0 |
| rs80051519 | pathogenic | 0 | 0 | 0 |
| rs121434536 | pathogenic | 1 | 1 | 0 |
| rs121434534 | pathogenic | 1 | 1 | 0 |

| | | | | |
|---|---|---|---|---|
| rs78310315 | pathogenic | 1 | 1 | 0 |
| rs540839115 | pathogenic | 1 | 1 | 1 |
| rs80338731 | pathogenic | 1 | 1 | 1 |
| rs180177151 | pathogenic | 1 | 1 | 1 |
| rs180177152 | pathogenic | 0 | 0 | 0 |
| rs80338732 | pathogenic | 1 | 1 | 1 |
| rs80338675 | pathogenic | 1 | 1 | 1 |
| rs80338676 | pathogenic | 1 | 1 | 1 |
| rs180177153 | pathogenic | 0 | 0 | 0 |
| rs180177154 | pathogenic | 1 | 1 | 1 |
| rs5896 | benign | 1 | 1 | 1 |
| rs121918477 | pathogenic | 0 | 0 | 0 |
| rs121918428 | pathogenic | 1 | 1 | 1 |
| rs121918427 | pathogenic | 1 | 1 | 1 |
| rs544300820 | benign | 1 | 1 | 1 |
| rs121918110 | pathogenic | 1 | 1 | 1 |
| rs1057141 | benign | 0 | 0 | 0 |
| rs1135216 | benign | 1 | 1 | 1 |
| rs121917702 | pathogenic | 1 | 1 | 1 |
| rs28941474 | pathogenic | 0 | 0 | 0 |
| rs104893944 | pathogenic | 1 | 1 | 1 |
| rs104893947 | pathogenic | 1 | 1 | 0 |
| rs104893943 | pathogenic | 1 | 1 | 1 |
| rs104893942 | pathogenic | 1 | 1 | 1 |
| rs104893940 | pathogenic | 1 | 1 | 1 |
| rs34045013 | benign | 0 | 0 | 0 |
| rs143083947 | pathogenic | 0 | 0 | 0 |
| rs145786986 | benign | 0 | 0 | 0 |
| rs119450942 | pathogenic | 0 | 0 | 0 |
| rs374259530 | pathogenic | 0 | 0 | 0 |
| rs756210458 | pathogenic | 0 | 0 | 0 |
| rs11089991 | benign | 0 | 0 | 0 |
| rs28941471 | pathogenic | 0 | 1 | 0 |
| rs750614500 | pathogenic | 0 | 1 | 0 |
| rs199761158 | benign | 1 | 1 | 1 |
| rs119450944 | pathogenic | 1 | 1 | 1 |
| rs373458753 | pathogenic | 1 | 1 | 1 |
| rs202064195 | pathogenic | 1 | 1 | 1 |
| rs776496275 | pathogenic | 1 | 1 | 1 |
| rs761493155 | pathogenic | 1 | 1 | 1 |
| rs763542069 | pathogenic | 0 | 0 | 0 |
| rs119450941 | pathogenic | 0 | 0 | 0 |
| rs119450940 | pathogenic | 0 | 0 | 0 |
| rs777821034 | pathogenic | 1 | 1 | 1 |
| rs771121666 | pathogenic | 1 | 1 | 1 |
| rs372895468 | pathogenic | 1 | 1 | 1 |
| rs61750603 | pathogenic | 0 | 0 | 0 |
| rs76863441 | pathogenic | 0 | 0 | 0 |
| rs142601143 | benign | 1 | 1 | 1 |
| rs185832993 | benign | 1 | 1 | 0 |
| rs17860568 | benign | 0 | 0 | 0 |
| rs142064825 | benign | 0 | 0 | 0 |
| rs564076183 | benign | 0 | 0 | 0 |
| rs386834034 | pathogenic | 0 | 0 | 0 |

| | | | | |
|---|---|---|---|---|
| rs147663769 | benign | 0 | 0 | 0 |
| rs776478974 | benign | 0 | 0 | 0 |
| rs111840783 | benign | 0 | 0 | 0 |
| rs1516446 | benign | 1 | 1 | 1 |
| rs143749154 | benign | 1 | 1 | 1 |
| rs80257412 | benign | 0 | 0 | 0 |
| rs142943120 | benign | 0 | 0 | 0 |
| rs144553482 | benign | 0 | 0 | 0 |
| rs35997569 | benign | 0 | 0 | 0 |
| rs772909106 | pathogenic | 0 | 0 | 0 |
| rs149433837 | benign | 1 | 1 | 1 |
| rs532178791 | pathogenic | 0 | 0 | 0 |
| rs145588689 | pathogenic | 1 | 1 | 0 |
| rs17843776 | benign | 1 | 1 | 0 |
| rs121917890 | pathogenic | 1 | 1 | 1 |
| rs121917892 | pathogenic | 0 | 0 | 0 |
| rs606231289 | pathogenic | 0 | 0 | 0 |
| rs370011798 | pathogenic | 0 | 0 | 0 |
| rs34689457 | benign | 0 | 0 | 0 |
| rs118204451 | pathogenic | 0 | 0 | 0 |
| rs118204452 | pathogenic | 1 | 1 | 1 |
| rs652785 | benign | 0 | 0 | 0 |
| rs35378126 | benign | 0 | 0 | 0 |
| rs80358214 | pathogenic | 1 | 1 | 1 |
| rs80358215 | pathogenic | 0 | 1 | 0 |
| rs267606858 | pathogenic | 1 | 1 | 1 |
| rs143137713 | pathogenic | 1 | 1 | 1 |
| rs782168634 | benign | 0 | 0 | 0 |
| rs143873938 | benign | 0 | 0 | 0 |
| rs782776511 | benign | 0 | 0 | 0 |
| rs201093148 | benign | 0 | 0 | 0 |
| rs267606815 | pathogenic | 0 | 0 | 0 |
| rs137853311 | pathogenic | 0 | 0 | 0 |
| rs267606817 | pathogenic | 0 | 0 | 0 |
| rs72650006 | benign | 0 | 0 | 0 |
| rs727503638 | benign | 1 | 1 | 0 |
| rs189286381 | benign | 0 | 0 | 0 |
| rs376644553 | benign | 0 | 0 | 0 |
| rs121434610 | pathogenic | 1 | 1 | 1 |
| rs267607076 | pathogenic | 1 | 1 | 1 |
| rs201275730 | benign | 0 | 0 | 0 |
| rs121909587 | pathogenic | 0 | 0 | 0 |
| rs121909558 | pathogenic | 1 | 1 | 0 |
| rs28929469 | pathogenic | 1 | 1 | 0 |
| rs121909547 | pathogenic | 1 | 1 | 0 |
| rs121909552 | pathogenic | 1 | 1 | 0 |
| rs121909573 | pathogenic | 1 | 1 | 1 |
| rs121909567 | pathogenic | 1 | 1 | 0 |
| rs121909569 | pathogenic | 0 | 0 | 0 |
| rs121909563 | pathogenic | 1 | 1 | 0 |
| rs121909570 | pathogenic | 1 | 1 | 1 |
| rs121909571 | pathogenic | 1 | 1 | 0 |
| rs121909572 | pathogenic | 1 | 1 | 1 |
| rs121909565 | pathogenic | 1 | 1 | 1 |

| | | | | |
|---|---|---|---|---|
| rs121909557 | pathogenic | 1 | 1 | 1 |
| rs121909568 | pathogenic | 1 | 1 | 1 |
| rs121909566 | pathogenic | 1 | 1 | 1 |
| rs121909549 | pathogenic | 1 | 1 | 1 |
| rs121909554 | pathogenic | 1 | 1 | 1 |
| rs121909550 | pathogenic | 1 | 1 | 1 |
| rs121909546 | pathogenic | 1 | 1 | 0 |
| rs121909555 | pathogenic | 1 | 1 | 1 |
| rs121909564 | pathogenic | 1 | 1 | 1 |
| rs548046212 | pathogenic | 0 | 0 | 0 |
| rs147353781 | benign | 0 | 0 | 0 |
| rs104894678 | pathogenic | 1 | 1 | 1 |
| rs1130426 | benign | 0 | 0 | 0 |
| rs104894677 | pathogenic | 1 | 1 | 1 |
| rs141917423 | benign | 1 | 1 | 1 |
| rs574777883 | benign | 0 | 0 | 0 |
| rs150199226 | benign | 1 | 1 | 1 |
| rs199618701 | benign | 0 | 1 | 0 |
| rs121964923 | pathogenic | 0 | 0 | 0 |
| rs121964924 | pathogenic | 0 | 0 | 0 |
| rs201258823 | pathogenic | 0 | 1 | 0 |
| rs267606773 | pathogenic | 0 | 0 | 0 |
| rs35638832 | benign | 0 | 0 | 0 |
| rs121908632 | pathogenic | 1 | 1 | 1 |
| rs121908625 | pathogenic | 1 | 1 | 1 |
| rs748949603 | pathogenic | 0 | 0 | 0 |
| rs757523840 | pathogenic | 0 | 0 | 0 |
| rs62541771 | pathogenic | 0 | 0 | 0 |
| rs776384541 | pathogenic | 1 | 1 | 0 |
| rs121908627 | pathogenic | 1 | 1 | 0 |
| rs28937594 | pathogenic | 0 | 0 | 0 |
| rs121907888 | pathogenic | 0 | 0 | 0 |
| rs121907889 | pathogenic | 0 | 1 | 0 |
| rs28941785 | pathogenic | 1 | 1 | 1 |
| rs28941786 | pathogenic | 1 | 1 | 1 |
| rs1021737 | benign | 0 | 0 | 0 |
| rs121908370 | pathogenic | 1 | 1 | 1 |
| rs63750083 | pathogenic | 0 | 0 | 0 |
| rs17883862 | benign | 0 | 0 | 0 |
| rs587782070 | pathogenic | 1 | 1 | 1 |
| rs587781269 | pathogenic | 1 | 1 | 1 |
| rs28909982 | pathogenic | 1 | 1 | 1 |
| rs373648967 | benign | 1 | 1 | 1 |
| rs368570187 | benign | 1 | 1 | 1 |
| rs730881701 | pathogenic | 1 | 1 | 1 |
| rs137853007 | pathogenic | 0 | 0 | 0 |
| rs575910805 | benign | 0 | 0 | 0 |
| rs104893875 | pathogenic | 1 | 1 | 1 |
| rs104893877 | pathogenic | 1 | 1 | 1 |
| rs119485093 | pathogenic | 1 | 1 | 1 |
| rs119485092 | pathogenic | 0 | 0 | 0 |
| rs119485090 | pathogenic | 0 | 0 | 0 |
| rs28942099 | pathogenic | 1 | 1 | 1 |
| rs373730800 | pathogenic | 1 | 1 | 1 |

| | | | | |
|---|---|---|---|---|
| rs113993995 | pathogenic | 0 | 0 | 0 |
| rs201070132 | pathogenic | 0 | 0 | 0 |
| rs113993996 | pathogenic | 0 | 0 | 0 |
| rs113993998 | pathogenic | 0 | 0 | 0 |
| rs78950939 | pathogenic | 1 | 1 | 1 |
| rs56378716 | pathogenic | 1 | 1 | 1 |
| rs754921704 | benign | 0 | 1 | 0 |
| rs766379103 | pathogenic | 1 | 1 | 0 |
| rs121912515 | pathogenic | 0 | 1 | 0 |
| rs764831888 | benign | 1 | 1 | 0 |
| rs572527340 | benign | 0 | 0 | 0 |
| rs35969327 | benign | 0 | 0 | 0 |
| rs35814893 | benign | 0 | 0 | 0 |
| rs34032300 | benign | 0 | 0 | 0 |
| rs80358260 | pathogenic | 0 | 0 | 0 |
| rs151220873 | benign | 0 | 0 | 0 |
| rs80358261 | pathogenic | 1 | 1 | 0 |
| rs11694 | pathogenic | 0 | 0 | 0 |
| rs80358264 | pathogenic | 0 | 0 | 0 |
| rs80358266 | pathogenic | 1 | 1 | 0 |
| rs104894458 | pathogenic | 1 | 1 | 1 |
| rs121918028 | pathogenic | 0 | 0 | 0 |
| rs11583680 | pathogenic | 0 | 0 | 0 |
| rs4715 | benign | 1 | 1 | 1 |
| rs34957318 | benign | 0 | 0 | 0 |
| rs1124 | benign | 1 | 1 | 1 |
| rs104894778 | pathogenic | 1 | 1 | 1 |
| rs104894777 | pathogenic | 1 | 1 | 1 |
| rs104894769 | pathogenic | 1 | 1 | 1 |
| rs104894768 | pathogenic | 1 | 1 | 1 |
| rs104894771 | pathogenic | 1 | 1 | 1 |
| rs575372249 | benign | 1 | 1 | 1 |
| rs118203914 | pathogenic | 0 | 0 | 0 |
| rs118203916 | pathogenic | 1 | 1 | 0 |
| rs140154690 | benign | 0 | 0 | 0 |
| rs34382405 | benign | 0 | 0 | 0 |
| rs73718082 | benign | 0 | 0 | 0 |
| rs764932724 | benign | 0 | 0 | 0 |
| rs554931092 | pathogenic | 1 | 1 | 0 |
| rs202060864 | pathogenic | 0 | 0 | 0 |
| rs116567033 | benign | 0 | 0 | 0 |
| rs11243011 | benign | 0 | 0 | 0 |
| rs142073519 | pathogenic | 0 | 0 | 0 |
| rs397514611 | pathogenic | 1 | 1 | 0 |
| rs397514612 | pathogenic | 0 | 0 | 0 |
| rs372054960 | pathogenic | 0 | 0 | 0 |
| rs775690041 | pathogenic | 0 | 0 | 0 |
| rs148921184 | pathogenic | 0 | 0 | 0 |
| rs12711521 | benign | 0 | 0 | 0 |
| rs2273346 | benign | 0 | 0 | 0 |
| rs12085877 | benign | 1 | 1 | 1 |
| rs144471433 | benign | 1 | 1 | 1 |
| rs397514671 | pathogenic | 1 | 1 | 1 |
| rs76394680 | benign | 0 | 0 | 0 |

| | | | | |
|---|---|---|---|---|
| rs78289603 | pathogenic | 1 | 1 | 1 |
| rs137852913 | pathogenic | 1 | 1 | 1 |
| rs137852914 | pathogenic | 1 | 1 | 1 |
| rs121918653 | pathogenic | 1 | 1 | 1 |
| rs35515200 | benign | 1 | 1 | 0 |
| rs17876030 | benign | 1 | 1 | 1 |
| rs556623443 | benign | 1 | 1 | 1 |
| rs121434365 | pathogenic | 1 | 1 | 1 |
| rs121434364 | pathogenic | 0 | 0 | 0 |
| rs41400249 | benign | 0 | 0 | 0 |
| rs149440376 | benign | 1 | 1 | 1 |
| rs181590179 | benign | 0 | 0 | 0 |
| rs113993968 | pathogenic | 0 | 0 | 0 |
| rs587776445 | pathogenic | 1 | 1 | 1 |
| rs113993969 | pathogenic | 1 | 1 | 1 |
| rs113993970 | pathogenic | 1 | 1 | 1 |
| rs587776449 | pathogenic | 1 | 1 | 1 |
| rs113993971 | pathogenic | 1 | 1 | 1 |
| rs267607194 | pathogenic | 1 | 1 | 1 |
| rs397514745 | pathogenic | 0 | 0 | 0 |
| rs201650281 | pathogenic | 0 | 0 | 0 |
| rs150529876 | benign | 1 | 1 | 1 |
| rs77573084 | benign | 0 | 0 | 0 |
| rs398122379 | pathogenic | 1 | 1 | 0 |
| rs765478968 | benign | 0 | 0 | 0 |
| rs201399681 | benign | 0 | 0 | 0 |
| rs17159287 | benign | 0 | 0 | 0 |
| rs538571144 | benign | 0 | 0 | 0 |
| rs137852647 | pathogenic | 0 | 0 | 0 |
| rs201358272 | pathogenic | 1 | 1 | 1 |
| rs181251337 | benign | 0 | 0 | 0 |
| rs3213119 | benign | 0 | 0 | 0 |
| rs137853102 | pathogenic | 1 | 1 | 0 |
| rs4956145 | benign | 0 | 0 | 0 |
| rs146036912 | pathogenic | 1 | 1 | 1 |
| rs375717077 | pathogenic | 1 | 1 | 1 |
| rs137853103 | pathogenic | 1 | 1 | 1 |
| rs776184830 | pathogenic | 1 | 1 | 1 |
| rs200273673 | pathogenic | 1 | 1 | 1 |
| rs267607025 | pathogenic | 1 | 1 | 1 |
| rs267607024 | pathogenic | 1 | 1 | 1 |
| rs515726188 | pathogenic | 1 | 1 | 1 |
| rs147315735 | benign | 0 | 0 | 0 |
| rs121918308 | pathogenic | 1 | 1 | 0 |
| rs515726195 | pathogenic | 0 | 0 | 0 |
| rs515726196 | pathogenic | 1 | 1 | 0 |
| rs121918311 | pathogenic | 0 | 1 | 0 |
| rs121918309 | pathogenic | 0 | 0 | 0 |
| rs387906891 | pathogenic | 0 | 0 | 0 |
| rs182614164 | pathogenic | 1 | 1 | 0 |
| rs121918307 | pathogenic | 0 | 0 | 0 |
| rs201440849 | benign | 0 | 0 | 0 |
| rs118203957 | pathogenic | 0 | 0 | 0 |
| rs4986790 | benign | 0 | 0 | 0 |

| | | | | |
|---|---|---|---|---|
| rs281864928 | pathogenic | 1 | 1 | 0 |
| rs267607156 | pathogenic | 0 | 0 | 0 |
| rs55916807 | benign | 0 | 0 | 0 |
| rs749750052 | benign | 0 | 0 | 0 |
| rs74382477 | benign | 0 | 0 | 0 |
| rs201016638 | benign | 0 | 0 | 0 |
| rs376683989 | benign | 0 | 0 | 0 |
| rs144439703 | pathogenic | 0 | 0 | 0 |
| rs779122022 | benign | 0 | 0 | 0 |
| rs121918056 | pathogenic | 0 | 0 | 0 |
| rs113994093 | pathogenic | 1 | 1 | 1 |
| rs141367015 | pathogenic | 0 | 0 | 0 |
| rs775538075 | pathogenic | 0 | 0 | 0 |
| rs121918055 | pathogenic | 0 | 0 | 0 |
| rs754246294 | benign | 1 | 1 | 1 |
| rs2307447 | benign | 1 | 1 | 1 |
| rs113994096 | pathogenic | 1 | 1 | 1 |
| rs367610201 | pathogenic | 1 | 1 | 1 |
| rs758438414 | pathogenic | 1 | 1 | 0 |
| rs121918046 | pathogenic | 0 | 0 | 0 |
| rs201871736 | benign | 0 | 0 | 0 |
| rs773994204 | benign | 0 | 0 | 0 |
| rs2307450 | benign | 1 | 1 | 1 |
| rs200257554 | benign | 0 | 0 | 0 |
| rs185645212 | benign | 0 | 0 | 0 |
| rs121918054 | pathogenic | 0 | 0 | 0 |
| rs113994097 | pathogenic | 0 | 0 | 0 |
| rs202037973 | pathogenic | 0 | 0 | 0 |
| rs769827124 | pathogenic | 1 | 1 | 0 |
| rs41549716 | benign | 1 | 1 | 1 |
| rs113994098 | pathogenic | 1 | 1 | 0 |
| rs144500145 | pathogenic | 1 | 1 | 0 |
| rs121918053 | pathogenic | 1 | 1 | 0 |
| rs121918050 | pathogenic | 1 | 1 | 0 |
| rs368587966 | pathogenic | 0 | 0 | 0 |
| rs139590686 | pathogenic | 0 | 0 | 0 |
| rs121918048 | pathogenic | 1 | 1 | 0 |
| rs768653086 | pathogenic | 1 | 1 | 0 |
| rs113994099 | pathogenic | 1 | 1 | 0 |
| rs121918051 | pathogenic | 1 | 1 | 0 |
| rs753160398 | pathogenic | 1 | 1 | 0 |
| rs201477273 | pathogenic | 1 | 1 | 0 |
| rs142347031 | pathogenic | 0 | 0 | 0 |
| rs121918049 | pathogenic | 0 | 0 | 0 |
| rs267606959 | pathogenic | 1 | 1 | 0 |
| rs201732356 | pathogenic | 0 | 0 | 0 |
| rs368435864 | pathogenic | 0 | 0 | 0 |
| rs764036283 | pathogenic | 0 | 0 | 0 |
| rs2307442 | benign | 0 | 0 | 0 |
| rs2307440 | benign | 0 | 0 | 0 |
| rs548076633 | pathogenic | 0 | 1 | 0 |
| rs750792237 | benign | 0 | 0 | 0 |
| rs3087374 | benign | 0 | 0 | 0 |
| rs121918060 | pathogenic | 0 | 1 | 0 |

| | | | | |
|---|---|---|---|---|
| rs397514765 | pathogenic | 1 | 1 | 1 |
| rs121918063 | pathogenic | 0 | 0 | 0 |
| rs121918058 | pathogenic | 1 | 1 | 1 |
| rs121918064 | pathogenic | 0 | 0 | 0 |
| rs36033115 | benign | 0 | 1 | 0 |
| rs121918057 | pathogenic | 0 | 1 | 0 |
| rs121918065 | pathogenic | 1 | 1 | 1 |
| rs121918061 | pathogenic | 1 | 1 | 1 |
| rs121918066 | pathogenic | 0 | 1 | 0 |
| rs707555 | benign | 0 | 1 | 0 |
| rs8192708 | benign | 0 | 1 | 0 |
| rs11552145 | benign | 0 | 0 | 0 |
| rs1804160 | benign | 0 | 1 | 0 |
| rs34103538 | benign | 0 | 1 | 0 |
| rs751393950 | pathogenic | 1 | 1 | 1 |
| rs121907979 | pathogenic | 1 | 1 | 1 |
| rs121907975 | pathogenic | 1 | 1 | 1 |
| rs121907962 | pathogenic | 1 | 1 | 1 |
| rs121907957 | pathogenic | 0 | 0 | 0 |
| rs121907953 | pathogenic | 1 | 1 | 1 |
| rs28941770 | pathogenic | 1 | 1 | 1 |
| rs28941771 | pathogenic | 1 | 1 | 1 |
| rs121907969 | pathogenic | 1 | 1 | 1 |
| rs121907973 | pathogenic | 1 | 1 | 1 |
| rs1800429 | pathogenic | 0 | 0 | 0 |
| rs121907976 | pathogenic | 1 | 1 | 0 |
| rs121907961 | pathogenic | 0 | 0 | 0 |
| rs121907974 | pathogenic | 0 | 0 | 0 |
| rs121907959 | pathogenic | 0 | 0 | 0 |
| rs121907971 | pathogenic | 1 | 1 | 0 |
| rs121907954 | pathogenic | 0 | 0 | 0 |
| rs121907977 | pathogenic | 0 | 0 | 0 |
| rs121907963 | pathogenic | 1 | 1 | 0 |
| rs370266293 | pathogenic | 1 | 1 | 0 |
| rs1800430 | benign | 0 | 0 | 0 |
| rs121907958 | pathogenic | 0 | 0 | 0 |
| rs202173526 | pathogenic | 0 | 0 | 0 |
| rs121907978 | pathogenic | 0 | 0 | 0 |
| rs121907981 | pathogenic | 0 | 1 | 0 |
| rs145012038 | benign | 1 | 1 | 1 |
| rs121907968 | pathogenic | 1 | 1 | 1 |
| rs121907966 | pathogenic | 1 | 1 | 1 |
| rs121907956 | pathogenic | 1 | 1 | 1 |
| rs28942071 | pathogenic | 1 | 1 | 1 |
| rs121907955 | pathogenic | 1 | 1 | 1 |
| rs4727853 | benign | 0 | 0 | 0 |
| rs201880593 | benign | 0 | 0 | 0 |
| rs201434579 | benign | 1 | 1 | 1 |
| rs74551128 | pathogenic | 1 | 1 | 0 |
| rs193922500 | pathogenic | 0 | 0 | 0 |
| rs121909009 | pathogenic | 1 | 1 | 0 |
| rs121908805 | pathogenic | 1 | 1 | 0 |
| rs139573311 | pathogenic | 0 | 0 | 0 |
| rs213950 | benign | 0 | 0 | 0 |

| | | | | |
|---|---|---|---|---|
| rs79282516 | pathogenic | 0 | 0 | 0 |
| rs121909017 | pathogenic | 1 | 1 | 1 |
| rs77101217 | pathogenic | 1 | 1 | 1 |
| rs1800092 | pathogenic | 0 | 1 | 0 |
| rs77646904 | pathogenic | 0 | 0 | 0 |
| rs1800095 | benign | 1 | 1 | 1 |
| rs387906368 | pathogenic | 1 | 1 | 1 |
| rs113993959 | pathogenic | 1 | 1 | 1 |
| rs121908755 | pathogenic | 1 | 1 | 1 |
| rs121909005 | pathogenic | 1 | 1 | 1 |
| rs121908757 | pathogenic | 1 | 1 | 1 |
| rs75527207 | pathogenic | 1 | 1 | 0 |
| rs121909013 | pathogenic | 1 | 1 | 0 |
| rs121909044 | pathogenic | 1 | 1 | 0 |
| rs74597325 | pathogenic | 1 | 1 | 0 |
| rs75549581 | pathogenic | 0 | 0 | 0 |
| rs80055610 | pathogenic | 1 | 1 | 1 |
| rs1800097 | pathogenic | 0 | 0 | 0 |
| rs121909006 | pathogenic | 0 | 1 | 0 |
| rs397508276 | pathogenic | 0 | 0 | 0 |
| rs397508296 | pathogenic | 1 | 1 | 1 |
| rs139468767 | pathogenic | 0 | 0 | 0 |
| rs121908759 | pathogenic | 1 | 1 | 0 |
| rs121909033 | pathogenic | 1 | 1 | 0 |
| rs397508331 | pathogenic | 0 | 0 | 0 |
| rs387907279 | pathogenic | 1 | 1 | 1 |
| rs546530618 | benign | 0 | 0 | 0 |
| rs558259739 | benign | 0 | 0 | 0 |
| rs201827340 | benign | 0 | 0 | 0 |
| rs148865119 | pathogenic | 1 | 1 | 0 |
| rs2229413 | benign | 1 | 1 | 1 |
| rs34040148 | benign | 1 | 1 | 1 |
| rs121964964 | pathogenic | 1 | 1 | 1 |
| rs760214620 | pathogenic | 1 | 1 | 1 |
| rs775992753 | pathogenic | 1 | 1 | 1 |
| rs770095972 | pathogenic | 1 | 1 | 1 |
| rs781444670 | pathogenic | 1 | 1 | 1 |
| rs147474549 | pathogenic | 0 | 0 | 0 |
| rs121964965 | pathogenic | 0 | 1 | 0 |
| rs121964963 | pathogenic | 1 | 1 | 1 |
| rs755952006 | pathogenic | 1 | 1 | 1 |
| rs121964970 | pathogenic | 1 | 1 | 1 |
| rs121964973 | pathogenic | 1 | 1 | 1 |
| rs773734233 | pathogenic | 1 | 1 | 1 |
| rs758236584 | pathogenic | 1 | 1 | 1 |
| rs149119723 | pathogenic | 1 | 1 | 0 |
| rs5742905 | pathogenic | 0 | 0 | 0 |
| rs121964962 | pathogenic | 1 | 1 | 1 |
| rs781567152 | pathogenic | 1 | 1 | 0 |
| rs398123151 | pathogenic | 1 | 1 | 1 |
| rs760417941 | pathogenic | 1 | 1 | 1 |
| rs774926464 | pathogenic | 0 | 0 | 0 |
| rs121964972 | pathogenic | 1 | 1 | 0 |
| rs372010465 | pathogenic | 1 | 1 | 1 |

| | | | | |
|---|---|---|---|---|
| rs763036586 | pathogenic | 1 | 1 | 1 |
| rs769080151 | pathogenic | 1 | 1 | 1 |
| rs121964967 | pathogenic | 1 | 1 | 1 |
| rs28934892 | pathogenic | 1 | 1 | 0 |
| rs28934891 | pathogenic | 1 | 1 | 1 |
| rs121964971 | pathogenic | 1 | 1 | 1 |
| rs121964968 | pathogenic | 1 | 1 | 1 |
| rs150828989 | benign | 1 | 1 | 1 |
| rs3775291 | benign | 1 | 1 | 0 |
| rs104894419 | pathogenic | 1 | 1 | 1 |
| rs2232642 | benign | 0 | 0 | 0 |
| rs104894551 | pathogenic | 1 | 1 | 0 |
| rs766328537 | pathogenic | 0 | 1 | 0 |
| rs104894553 | pathogenic | 1 | 1 | 0 |
| rs80099330 | benign | 0 | 0 | 0 |
| rs199565861 | pathogenic | 0 | 0 | 0 |
| rs104894548 | pathogenic | 0 | 0 | 0 |
| rs104894549 | pathogenic | 0 | 0 | 0 |
| rs104894550 | pathogenic | 1 | 1 | 1 |
| rs12948217 | pathogenic | 1 | 1 | 1 |
| rs761064915 | pathogenic | 0 | 0 | 0 |
| rs28940279 | pathogenic | 1 | 1 | 1 |
| rs774323189 | pathogenic | 1 | 1 | 1 |
| rs28940574 | pathogenic | 0 | 0 | 0 |
| rs515299 | benign | 1 | 1 | 1 |
| rs10127939 | benign | 1 | 1 | 1 |
| rs396991 | benign | 1 | 1 | 1 |
| rs145956175 | benign | 0 | 1 | 0 |
| rs747838255 | pathogenic | 1 | 1 | 1 |
| rs762659685 | pathogenic | 1 | 1 | 0 |
| rs104894831 | pathogenic | 0 | 1 | 0 |
| rs104894848 | pathogenic | 1 | 1 | 0 |
| rs104894833 | pathogenic | 0 | 1 | 0 |
| rs781838005 | pathogenic | 0 | 1 | 0 |
| rs104894834 | pathogenic | 1 | 1 | 0 |
| rs372966991 | pathogenic | 1 | 1 | 0 |
| rs138886989 | benign | 1 | 1 | 1 |
| rs28935195 | pathogenic | 0 | 1 | 0 |
| rs28935196 | pathogenic | 0 | 1 | 0 |
| rs372416832 | pathogenic | 1 | 1 | 1 |
| rs28935197 | pathogenic | 1 | 1 | 0 |
| rs28935486 | pathogenic | 1 | 1 | 0 |
| rs28935487 | pathogenic | 1 | 1 | 0 |
| rs28935488 | pathogenic | 1 | 1 | 1 |
| rs28935495 | pathogenic | 1 | 1 | 1 |
| rs28935485 | pathogenic | 1 | 1 | 1 |
| rs140329381 | pathogenic | 0 | 1 | 0 |
| rs104894830 | pathogenic | 0 | 1 | 0 |
| rs104894846 | pathogenic | 0 | 1 | 0 |
| rs28935489 | pathogenic | 0 | 1 | 0 |
| rs104894828 | pathogenic | 1 | 1 | 0 |
| rs28935491 | pathogenic | 1 | 1 | 0 |
| rs104894832 | pathogenic | 1 | 1 | 0 |
| rs28935492 | pathogenic | 1 | 1 | 0 |

| | | | | |
|---|---|---|---|---|
| rs730880437 | benign | 1 | 1 | 0 |
| rs28935493 | pathogenic | 0 | 1 | 0 |
| rs28935494 | pathogenic | 1 | 1 | 1 |
| rs111422676 | pathogenic | 1 | 1 | 1 |
| rs397515869 | benign | 1 | 1 | 0 |
| rs782449839 | pathogenic | 0 | 1 | 0 |
| rs104894852 | pathogenic | 1 | 1 | 1 |
| rs116929575 | pathogenic | 1 | 1 | 1 |
| rs7023954 | benign | 0 | 0 | 0 |
| rs184520335 | benign | 1 | 1 | 1 |
| rs143080527 | benign | 0 | 0 | 0 |
| rs1050565 | benign | 0 | 0 | 0 |
| rs11545035 | benign | 0 | 0 | 0 |
| rs267607190 | pathogenic | 1 | 1 | 1 |
| rs200283987 | benign | 0 | 0 | 0 |
| rs372556807 | benign | 1 | 1 | 0 |
| rs563073392 | benign | 0 | 0 | 0 |
| rs4539 | benign | 0 | 0 | 0 |
| rs28931609 | pathogenic | 1 | 1 | 1 |
| rs121912978 | pathogenic | 1 | 1 | 1 |
| rs104894072 | pathogenic | 1 | 1 | 1 |
| rs551933154 | benign | 1 | 1 | 1 |
| rs752962897 | benign | 1 | 1 | 0 |
| rs121912977 | pathogenic | 1 | 1 | 0 |
| rs4537 | benign | 1 | 1 | 1 |
| rs201830462 | benign | 0 | 0 | 0 |
| rs4544 | benign | 0 | 0 | 0 |
| rs562670189 | benign | 0 | 0 | 0 |
| rs4538 | benign | 0 | 1 | 0 |
| rs61757294 | pathogenic | 0 | 0 | 0 |
| rs5313 | benign | 0 | 0 | 0 |
| rs4545 | benign | 0 | 0 | 0 |
| rs72554627 | pathogenic | 1 | 1 | 1 |
| rs72554626 | pathogenic | 1 | 1 | 1 |
| rs387906861 | pathogenic | 1 | 1 | 1 |
| rs749655461 | pathogenic | 1 | 1 | 1 |
| rs121909606 | pathogenic | 1 | 1 | 1 |
| rs121909607 | pathogenic | 1 | 1 | 1 |
| rs200203992 | benign | 0 | 0 | 0 |
| rs6050 | benign | 0 | 0 | 0 |
| rs184635235 | benign | 0 | 0 | 0 |
| rs139146037 | benign | 0 | 0 | 0 |
| rs121909612 | pathogenic | 0 | 0 | 0 |
| rs78506343 | pathogenic | 0 | 0 | 0 |
| rs121918138 | pathogenic | 1 | 1 | 1 |
| rs121918139 | pathogenic | 1 | 1 | 1 |
| rs121918140 | pathogenic | 0 | 0 | 0 |
| rs36063028 | benign | 1 | 1 | 1 |
| rs45495192 | benign | 1 | 1 | 1 |
| rs45458802 | benign | 1 | 1 | 1 |
| rs552865793 | pathogenic | 1 | 1 | 1 |
| rs753744791 | pathogenic | 1 | 1 | 1 |
| rs777518512 | benign | 1 | 1 | 1 |
| rs142708991 | benign | 1 | 1 | 0 |

| | | | | |
|---|---|---|---|---|
| rs746069957 | benign | 0 | 0 | 0 |
| rs1334811 | benign | 0 | 0 | 0 |
| rs138894008 | benign | 1 | 1 | 0 |
| rs35030851 | benign | 1 | 1 | 0 |
| rs35378598 | benign | 0 | 0 | 0 |
| rs35818764 | benign | 1 | 1 | 1 |
| rs4631561 | benign | 1 | 1 | 1 |
| rs121918660 | pathogenic | 0 | 0 | 0 |
| rs373175587 | benign | 0 | 0 | 0 |
| rs61731956 | pathogenic | 1 | 1 | 1 |
| rs180177305 | pathogenic | 0 | 0 | 0 |
| rs119490108 | pathogenic | 0 | 0 | 0 |
| rs180177312 | pathogenic | 1 | 1 | 0 |
| rs180177314 | pathogenic | 1 | 1 | 0 |
| rs180177322 | pathogenic | 1 | 1 | 1 |
| rs180177323 | pathogenic | 1 | 1 | 1 |
| rs180177325 | pathogenic | 0 | 0 | 0 |
| rs35196441 | benign | 0 | 0 | 0 |
| rs28763904 | benign | 0 | 0 | 0 |
| rs2229519 | benign | 0 | 0 | 0 |
| rs137852886 | pathogenic | 0 | 0 | 0 |
| rs137852892 | pathogenic | 0 | 0 | 0 |
| rs770427750 | pathogenic | 0 | 0 | 0 |
| rs137852887 | pathogenic | 0 | 0 | 0 |
| rs137852893 | pathogenic | 0 | 0 | 0 |
| rs80338671 | pathogenic | 1 | 1 | 0 |
| rs2228389 | benign | 0 | 0 | 0 |
| rs80338672 | pathogenic | 0 | 0 | 0 |
| rs201958741 | pathogenic | 0 | 0 | 0 |
| rs80338673 | pathogenic | 0 | 0 | 0 |
| rs137852888 | pathogenic | 0 | 0 | 0 |
| rs137852889 | pathogenic | 0 | 0 | 0 |
| rs137852894 | pathogenic | 0 | 0 | 0 |
| rs137852890 | pathogenic | 0 | 0 | 0 |
| rs137852891 | pathogenic | 0 | 0 | 0 |
| rs766935302 | pathogenic | 0 | 0 | 0 |
| rs193074572 | benign | 0 | 0 | 0 |
| rs555210692 | benign | 1 | 1 | 0 |
| rs119103242 | pathogenic | 0 | 1 | 0 |
| rs143725101 | benign | 0 | 1 | 0 |
| rs28940589 | pathogenic | 1 | 1 | 1 |
| rs113994145 | pathogenic | 0 | 0 | 0 |
| rs773872994 | benign | 0 | 0 | 0 |
| rs28940590 | pathogenic | 1 | 1 | 0 |
| rs145400751 | benign | 1 | 1 | 1 |
| rs587782331 | benign | 0 | 0 | 0 |
| rs730881787 | benign | 0 | 0 | 0 |
| rs587778531 | benign | 0 | 0 | 0 |
| rs781652274 | benign | 0 | 0 | 0 |
| rs55882234 | benign | 0 | 0 | 0 |
| rs2020908 | benign | 0 | 0 | 0 |
| rs63750741 | pathogenic | 1 | 1 | 1 |
| rs41295268 | benign | 1 | 1 | 1 |
| rs63750897 | benign | 1 | 1 | 0 |

| | | | | |
|---|---|---|---|---|
| rs63751005 | benign | 1 | 1 | 0 |
| rs587779223 | benign | 0 | 0 | 0 |
| rs202127474 | benign | 0 | 0 | 0 |
| rs374230313 | benign | 0 | 0 | 0 |
| rs2020912 | benign | 0 | 0 | 0 |
| rs63751017 | pathogenic | 0 | 0 | 0 |
| rs587779246 | pathogenic | 0 | 0 | 0 |
| rs63749919 | pathogenic | 0 | 0 | 0 |
| rs63750563 | pathogenic | 1 | 1 | 1 |
| rs63749999 | pathogenic | 0 | 0 | 0 |
| rs200492211 | pathogenic | 0 | 0 | 0 |
| rs576269342 | benign | 0 | 0 | 0 |
| rs267608075 | benign | 0 | 0 | 0 |
| rs149605979 | benign | 0 | 0 | 0 |
| rs63750252 | benign | 0 | 0 | 0 |
| rs754289472 | benign | 0 | 0 | 0 |
| rs63750119 | pathogenic | 0 | 0 | 0 |
| rs587782109 | benign | 0 | 0 | 0 |
| rs34625968 | benign | 1 | 1 | 1 |
| rs764786814 | benign | 1 | 1 | 1 |
| rs199594809 | benign | 1 | 1 | 1 |
| rs74315364 | pathogenic | 1 | 1 | 1 |
| rs201253919 | pathogenic | 0 | 0 | 0 |
| rs387906828 | pathogenic | 1 | 1 | 1 |
| rs34521609 | benign | 1 | 1 | 1 |
| rs199472894 | pathogenic | 0 | 0 | 0 |
| rs199472902 | pathogenic | 0 | 0 | 0 |
| rs121912505 | pathogenic | 0 | 0 | 0 |
| rs28928905 | pathogenic | 0 | 0 | 0 |
| rs374376640 | benign | 0 | 0 | 0 |
| rs202102276 | benign | 0 | 0 | 0 |
| rs199472918 | pathogenic | 0 | 0 | 0 |
| rs121912516 | pathogenic | 1 | 1 | 1 |
| rs121912504 | pathogenic | 0 | 0 | 0 |
| rs9333649 | pathogenic | 0 | 0 | 0 |
| rs121912508 | pathogenic | 0 | 0 | 0 |
| rs199473428 | pathogenic | 1 | 1 | 1 |
| rs104894021 | pathogenic | 0 | 0 | 0 |
| rs28928904 | pathogenic | 1 | 1 | 1 |
| rs199473039 | benign | 1 | 1 | 1 |
| rs199472958 | pathogenic | 1 | 1 | 1 |
| rs1137617 | pathogenic | 1 | 1 | 1 |
| rs199473532 | pathogenic | 1 | 1 | 1 |
| rs199472988 | pathogenic | 0 | 0 | 0 |
| rs189014161 | pathogenic | 0 | 0 | 0 |
| rs12720441 | pathogenic | 0 | 0 | 0 |
| rs199473001 | pathogenic | 1 | 1 | 0 |
| rs121912506 | pathogenic | 1 | 1 | 0 |
| rs773724817 | pathogenic | 1 | 1 | 1 |
| rs118204002 | pathogenic | 1 | 1 | 1 |
| rs138556525 | pathogenic | 1 | 1 | 1 |
| rs72558181 | pathogenic | 1 | 1 | 1 |
| rs118204005 | pathogenic | 1 | 1 | 1 |
| rs118204004 | pathogenic | 0 | 0 | 0 |

| | | | | |
|---|---|---|---|---|
| rs118204001 | pathogenic | 1 | 1 | 1 |
| rs118204006 | pathogenic | 1 | 1 | 1 |
| rs118204003 | pathogenic | 0 | 0 | 0 |
| rs121913147 | pathogenic | 1 | 1 | 0 |
| rs2229429 | benign | 1 | 1 | 0 |
| rs182552223 | benign | 0 | 0 | 0 |
| rs146588336 | benign | 0 | 0 | 0 |
| rs121964983 | pathogenic | 0 | 1 | 0 |
| rs121964982 | pathogenic | 0 | 0 | 0 |
| rs386833679 | pathogenic | 0 | 0 | 0 |
| rs386833680 | pathogenic | 0 | 0 | 0 |
| rs121964981 | pathogenic | 1 | 1 | 0 |
| rs121964984 | pathogenic | 0 | 0 | 0 |
| rs144971200 | benign | 0 | 0 | 0 |
| rs121964985 | pathogenic | 1 | 1 | 0 |
| rs141246107 | benign | 0 | 0 | 0 |
| rs104894561 | pathogenic | 1 | 1 | 0 |
| rs377023736 | pathogenic | 0 | 0 | 0 |
| rs121964907 | pathogenic | 1 | 1 | 1 |
| rs121964909 | pathogenic | 1 | 1 | 1 |
| rs765070743 | pathogenic | 0 | 0 | 0 |
| rs2228119 | benign | 0 | 0 | 0 |
| rs745976989 | pathogenic | 1 | 1 | 1 |
| rs121964904 | pathogenic | 0 | 0 | 0 |
| rs386833430 | pathogenic | 1 | 1 | 1 |
| rs267606992 | pathogenic | 0 | 0 | 0 |
| rs61752063 | pathogenic | 0 | 0 | 0 |
| rs61752067 | pathogenic | 1 | 1 | 1 |
| rs104894934 | pathogenic | 0 | 0 | 0 |
| rs104894929 | pathogenic | 1 | 1 | 1 |
| rs80189640 | benign | 0 | 0 | 0 |
| rs137854547 | pathogenic | 1 | 1 | 1 |
| rs137854540 | pathogenic | 1 | 1 | 1 |
| rs782092363 | pathogenic | 1 | 1 | 1 |
| rs147783915 | benign | 0 | 0 | 0 |
| rs28933687 | pathogenic | 1 | 1 | 1 |
| rs149503319 | benign | 0 | 0 | 0 |
| rs1805147 | benign | 0 | 0 | 0 |
| rs587777218 | pathogenic | 1 | 1 | 1 |
| rs200786705 | benign | 0 | 0 | 0 |
| rs56219475 | benign | 0 | 0 | 0 |
| rs587781172 | benign | 0 | 0 | 0 |
| rs757094189 | pathogenic | 0 | 0 | 0 |
| rs140811086 | benign | 1 | 1 | 1 |
| rs574088829 | pathogenic | 1 | 1 | 1 |
| rs121918525 | pathogenic | 0 | 0 | 0 |
| rs121918527 | pathogenic | 0 | 0 | 0 |
| rs121964848 | pathogenic | 0 | 1 | 0 |
| rs121964845 | pathogenic | 1 | 1 | 1 |
| rs121964846 | pathogenic | 0 | 1 | 0 |
| rs17217716 | benign | 1 | 1 | 1 |
| rs141711342 | benign | 0 | 0 | 0 |
| rs372619120 | benign | 0 | 0 | 0 |
| rs63750002 | benign | 1 | 1 | 1 |

| | | | | |
|---|---|---|---|---|
| rs193922373 | benign | 0 | 0 | 0 |
| rs41295286 | benign | 0 | 0 | 0 |
| rs17217772 | benign | 0 | 0 | 0 |
| rs145649774 | benign | 0 | 0 | 0 |
| rs63750214 | pathogenic | 0 | 0 | 0 |
| rs63750716 | benign | 0 | 0 | 0 |
| rs202026056 | benign | 0 | 0 | 0 |
| rs4987188 | benign | 1 | 1 | 0 |
| rs267607938 | benign | 0 | 0 | 0 |
| rs17224367 | benign | 0 | 0 | 0 |
| rs63750006 | benign | 0 | 0 | 0 |
| rs587781373 | benign | 0 | 0 | 0 |
| rs63751207 | pathogenic | 1 | 1 | 0 |
| rs730881759 | benign | 0 | 0 | 0 |
| rs372350768 | benign | 1 | 1 | 1 |
| rs55778204 | benign | 1 | 1 | 0 |
| rs63749910 | benign | 0 | 0 | 0 |
| rs63750844 | benign | 0 | 0 | 0 |
| rs28929483 | pathogenic | 0 | 0 | 0 |
| rs61756468 | benign | 0 | 0 | 0 |
| rs63750875 | pathogenic | 1 | 1 | 0 |
| rs776528054 | benign | 1 | 1 | 0 |
| rs751939698 | benign | 0 | 0 | 0 |
| rs587780685 | benign | 0 | 0 | 0 |
| rs730881763 | benign | 1 | 1 | 0 |
| rs373226409 | benign | 0 | 0 | 0 |
| rs202145681 | benign | 1 | 1 | 1 |
| rs63749841 | benign | 1 | 1 | 1 |
| rs63750757 | benign | 1 | 1 | 1 |
| rs587778527 | benign | 1 | 1 | 1 |
| rs63750849 | benign | 0 | 0 | 0 |
| rs587779969 | benign | 0 | 0 | 0 |
| rs121908046 | pathogenic | 1 | 1 | 1 |
| rs28940882 | pathogenic | 1 | 1 | 0 |
| rs28940883 | pathogenic | 1 | 1 | 1 |
| rs727503943 | pathogenic | 1 | 1 | 1 |
| rs137853859 | pathogenic | 1 | 1 | 1 |
| rs121908045 | pathogenic | 1 | 1 | 0 |
| rs137853861 | pathogenic | 1 | 1 | 0 |
| rs3180383 | pathogenic | 0 | 1 | 0 |
| rs144431814 | benign | 0 | 0 | 0 |
| rs139502866 | benign | 0 | 0 | 0 |
| rs17524437 | benign | 0 | 0 | 0 |
| rs760960779 | benign | 0 | 0 | 0 |
| rs148960463 | pathogenic | 0 | 0 | 0 |
| rs111033647 | pathogenic | 1 | 1 | 1 |
| rs111033648 | pathogenic | 0 | 0 | 0 |
| rs1800461 | benign | 0 | 0 | 0 |
| rs111033656 | pathogenic | 0 | 0 | 0 |
| rs111033663 | pathogenic | 1 | 1 | 0 |
| rs111033669 | pathogenic | 1 | 1 | 1 |
| rs111033690 | pathogenic | 0 | 0 | 0 |
| rs111033686 | pathogenic | 0 | 0 | 0 |
| rs111033687 | pathogenic | 0 | 0 | 0 |

| | | | | |
|---|---|---|---|---|
| rs111033695 | pathogenic | 0 | 0 | 0 |
| rs111033697 | pathogenic | 0 | 0 | 0 |
| rs111033693 | pathogenic | 0 | 0 | 0 |
| rs111033694 | pathogenic | 0 | 0 | 0 |
| rs111033721 | pathogenic | 1 | 1 | 1 |
| rs111033722 | pathogenic | 0 | 0 | 0 |
| rs111033725 | pathogenic | 1 | 1 | 0 |
| rs75391579 | pathogenic | 1 | 1 | 1 |
| rs111033728 | pathogenic | 1 | 1 | 1 |
| rs111033735 | pathogenic | 1 | 1 | 1 |
| rs111033739 | pathogenic | 1 | 1 | 1 |
| rs111033736 | pathogenic | 0 | 0 | 0 |
| rs111033744 | pathogenic | 1 | 1 | 1 |
| rs111033750 | pathogenic | 1 | 1 | 1 |
| rs111033752 | pathogenic | 0 | 0 | 0 |
| rs111033751 | pathogenic | 0 | 0 | 0 |
| rs111033846 | pathogenic | 0 | 0 | 0 |
| rs368166217 | pathogenic | 0 | 0 | 0 |
| rs111033764 | pathogenic | 0 | 0 | 0 |
| rs111033847 | benign | 0 | 0 | 0 |
| rs111033778 | pathogenic | 0 | 0 | 0 |
| rs111033773 | pathogenic | 0 | 0 | 0 |
| rs111033792 | pathogenic | 0 | 0 | 0 |
| rs367543266 | pathogenic | 1 | 1 | 1 |
| rs111033794 | pathogenic | 0 | 0 | 0 |
| rs111033802 | pathogenic | 0 | 0 | 0 |
| rs111033803 | pathogenic | 0 | 0 | 0 |
| rs111033809 | pathogenic | 1 | 1 | 0 |
| rs121918376 | pathogenic | 0 | 1 | 0 |
| rs121918377 | pathogenic | 0 | 1 | 0 |
| rs121918378 | pathogenic | 1 | 1 | 1 |
| rs121918374 | pathogenic | 1 | 1 | 1 |
| rs137852379 | pathogenic | 0 | 0 | 0 |
| rs137852388 | pathogenic | 1 | 1 | 1 |
| rs137852368 | pathogenic | 0 | 0 | 0 |
| rs28935499 | pathogenic | 0 | 0 | 0 |
| rs28933668 | pathogenic | 0 | 0 | 0 |
| rs28933669 | pathogenic | 0 | 0 | 0 |
| rs28933670 | pathogenic | 0 | 0 | 0 |
| rs28933671 | pathogenic | 0 | 0 | 0 |
| rs28933672 | pathogenic | 0 | 0 | 0 |
| rs28937272 | pathogenic | 0 | 0 | 0 |
| rs35383156 | benign | 0 | 0 | 0 |
| rs137852418 | pathogenic | 0 | 0 | 0 |
| rs137852419 | pathogenic | 0 | 0 | 0 |
| rs137852428 | pathogenic | 0 | 0 | 0 |
| rs137852436 | pathogenic | 0 | 0 | 0 |
| rs28937285 | pathogenic | 0 | 0 | 0 |
| rs35213384 | benign | 1 | 1 | 1 |
| rs3810950 | benign | 0 | 0 | 0 |
| rs201479289 | pathogenic | 0 | 0 | 0 |
| rs121912820 | pathogenic | 0 | 1 | 0 |
| rs121912815 | pathogenic | 0 | 1 | 0 |
| rs115510708 | benign | 0 | 0 | 0 |

| | | | | |
|---|---|---|---|---|
| rs868749 | benign | 0 | 0 | 0 |
| rs121912823 | pathogenic | 0 | 0 | 0 |
| rs115212829 | benign | 0 | 0 | 0 |
| rs121912822 | pathogenic | 0 | 0 | 0 |
| rs121912816 | pathogenic | 0 | 1 | 0 |
| rs4838544 | benign | 0 | 0 | 0 |
| rs121912818 | pathogenic | 0 | 0 | 0 |
| rs121912821 | pathogenic | 0 | 0 | 0 |
| rs121912817 | pathogenic | 0 | 0 | 0 |
| rs8178992 | benign | 0 | 0 | 0 |
| rs121912819 | pathogenic | 1 | 1 | 0 |
| rs80097077 | benign | 1 | 1 | 0 |
| rs121909359 | pathogenic | 1 | 1 | 0 |
| rs121909358 | pathogenic | 1 | 1 | 1 |
| rs121909361 | pathogenic | 1 | 1 | 1 |
| rs121909357 | pathogenic | 0 | 1 | 0 |
| rs121909365 | pathogenic | 1 | 1 | 1 |
| rs6413484 | pathogenic | 1 | 1 | 1 |
| rs121909373 | pathogenic | 1 | 1 | 1 |
| rs121909366 | pathogenic | 1 | 1 | 1 |
| rs121909367 | pathogenic | 1 | 1 | 1 |
| rs121909368 | pathogenic | 1 | 1 | 1 |
| rs121909369 | pathogenic | 0 | 0 | 0 |
| rs121909363 | pathogenic | 1 | 1 | 1 |
| rs45588036 | pathogenic | 1 | 1 | 1 |
| rs121909364 | pathogenic | 1 | 1 | 1 |
| rs80358282 | benign | 0 | 0 | 0 |
| rs80358284 | pathogenic | 1 | 1 | 1 |
| rs61735303 | benign | 0 | 0 | 0 |
| rs2229274 | benign | 0 | 0 | 0 |
| rs145006491 | benign | 0 | 0 | 0 |
| rs121913580 | pathogenic | 1 | 1 | 0 |
| rs730882196 | pathogenic | 0 | 0 | 0 |
| rs153477 | benign | 0 | 0 | 0 |
| rs153478 | benign | 1 | 1 | 0 |
| rs61740602 | benign | 1 | 1 | 0 |
| rs104893892 | pathogenic | 1 | 1 | 0 |
| rs121909595 | pathogenic | 1 | 1 | 1 |
| rs28931605 | pathogenic | 1 | 1 | 1 |
| rs121909597 | pathogenic | 0 | 1 | 0 |
| rs202233735 | pathogenic | 0 | 1 | 0 |
| rs121909596 | pathogenic | 1 | 1 | 1 |
| rs75156162 | benign | 0 | 1 | 0 |
| rs2305430 | benign | 1 | 1 | 1 |
| rs150318966 | benign | 0 | 0 | 0 |
| rs121909598 | pathogenic | 0 | 1 | 0 |
| rs5744175 | benign | 0 | 0 | 0 |
| rs28935476 | pathogenic | 1 | 1 | 1 |
| rs28935475 | pathogenic | 1 | 1 | 1 |
| rs122461163 | pathogenic | 1 | 1 | 1 |
| rs62626305 | pathogenic | 1 | 1 | 1 |
| rs587777157 | pathogenic | 0 | 0 | 0 |
| rs137854478 | pathogenic | 1 | 1 | 0 |
| rs137854465 | pathogenic | 0 | 0 | 0 |

| | | | | |
|---|---|---|---|---|
| rs137854470 | pathogenic | 0 | 0 | 0 |
| rs137854482 | pathogenic | 0 | 0 | 0 |
| rs137854473 | pathogenic | 0 | 0 | 0 |
| rs137854456 | pathogenic | 0 | 0 | 0 |
| rs140598 | benign | 0 | 0 | 0 |
| rs104894105 | pathogenic | 1 | 1 | 1 |
| rs104894107 | pathogenic | 1 | 1 | 1 |
| rs104894106 | pathogenic | 1 | 1 | 1 |
| rs74621026 | benign | 1 | 1 | 1 |
| rs2290332 | benign | 0 | 0 | 0 |
| rs780202604 | pathogenic | 1 | 1 | 1 |
| rs74315305 | pathogenic | 1 | 1 | 0 |
| rs766751878 | pathogenic | 1 | 1 | 1 |
| rs137854603 | pathogenic | 1 | 1 | 1 |
| rs139592595 | benign | 1 | 1 | 0 |
| rs150565592 | benign | 1 | 1 | 1 |
| rs763520126 | benign | 0 | 0 | 0 |
| rs55980502 | benign | 0 | 0 | 0 |
| rs137852812 | pathogenic | 0 | 0 | 0 |
| rs137852813 | pathogenic | 0 | 0 | 0 |
| rs190222208 | benign | 0 | 0 | 0 |
| rs138459502 | pathogenic | 0 | 0 | 0 |
| rs730881041 | pathogenic | 0 | 0 | 0 |
| rs267607080 | pathogenic | 0 | 0 | 0 |
| rs137852814 | pathogenic | 0 | 0 | 0 |
| rs267607079 | pathogenic | 0 | 0 | 0 |
| rs192378417 | benign | 0 | 0 | 0 |
| rs1288775 | benign | 0 | 0 | 0 |
| rs80338737 | pathogenic | 1 | 1 | 1 |
| rs397514708 | pathogenic | 0 | 0 | 0 |
| rs775933965 | pathogenic | 0 | 0 | 0 |
| rs747608698 | benign | 0 | 1 | 0 |
| rs1145086 | benign | 0 | 1 | 0 |
| rs121912953 | pathogenic | 1 | 1 | 0 |
| rs243849 | benign | 0 | 0 | 0 |
| rs121912955 | pathogenic | 1 | 1 | 1 |
| rs17859943 | benign | 0 | 1 | 0 |
| rs755557988 | benign | 1 | 1 | 1 |
| rs10195 | benign | 0 | 0 | 0 |
| rs61735357 | benign | 1 | 1 | 1 |
| rs137882374 | benign | 0 | 0 | 0 |
| rs5388 | benign | 0 | 0 | 0 |
| rs6352 | benign | 0 | 0 | 0 |
| rs76645461 | pathogenic | 1 | 1 | 0 |
| rs1050828 | pathogenic | 0 | 0 | 0 |
| rs1050829 | pathogenic | 0 | 0 | 0 |
| rs137852314 | pathogenic | 0 | 0 | 0 |
| rs137852331 | pathogenic | 0 | 0 | 0 |
| rs5030872 | pathogenic | 0 | 0 | 0 |
| rs267606836 | pathogenic | 0 | 0 | 0 |
| rs5030868 | pathogenic | 0 | 0 | 0 |
| rs137852330 | pathogenic | 0 | 1 | 0 |
| rs137852326 | pathogenic | 1 | 1 | 1 |
| rs137852318 | pathogenic | 1 | 1 | 1 |

| | | | | |
|---|---|---|---|---|
| rs74575103 | benign | 0 | 0 | 0 |
| rs137852327 | pathogenic | 1 | 1 | 1 |
| rs76723693 | pathogenic | 0 | 0 | 0 |
| rs5030869 | pathogenic | 0 | 0 | 0 |
| rs34193178 | pathogenic | 1 | 1 | 1 |
| rs137852316 | pathogenic | 1 | 1 | 0 |
| rs398123546 | pathogenic | 1 | 1 | 1 |
| rs72554665 | pathogenic | 0 | 0 | 0 |
| rs72554664 | pathogenic | 0 | 0 | 0 |
| rs28939068 | pathogenic | 1 | 1 | 1 |
| rs59301204 | pathogenic | 1 | 1 | 1 |
| rs143715750 | benign | 1 | 1 | 1 |
| rs35263902 | pathogenic | 1 | 1 | 1 |
| rs202214518 | benign | 1 | 1 | 1 |
| rs34076756 | benign | 0 | 0 | 0 |
| rs199592341 | pathogenic | 1 | 1 | 0 |
| rs121908833 | pathogenic | 1 | 1 | 1 |
| rs121908834 | pathogenic | 1 | 1 | 0 |
| rs138454151 | benign | 0 | 0 | 0 |
| rs141323599 | benign | 0 | 0 | 0 |
| rs137853200 | pathogenic | 0 | 0 | 0 |
| rs137853199 | pathogenic | 0 | 0 | 0 |
| rs139787163 | benign | 0 | 0 | 0 |
| rs61731097 | benign | 1 | 1 | 1 |
| rs61729795 | benign | 0 | 0 | 0 |
| rs149242330 | benign | 0 | 0 | 0 |
| rs587777490 | pathogenic | 1 | 1 | 1 |
| rs398123011 | pathogenic | 1 | 1 | 0 |
| rs775186117 | pathogenic | 0 | 0 | 0 |
| rs72677237 | benign | 0 | 0 | 0 |
| rs104894163 | pathogenic | 1 | 1 | 1 |
| rs121909588 | pathogenic | 0 | 0 | 0 |
| rs1042579 | benign | 0 | 0 | 0 |
| rs121909273 | pathogenic | 0 | 0 | 0 |
| rs121909274 | pathogenic | 0 | 0 | 0 |
| rs119473034 | pathogenic | 1 | 1 | 1 |
| rs398123065 | pathogenic | 0 | 0 | 0 |
| rs587777042 | pathogenic | 1 | 1 | 0 |
| rs761413864 | pathogenic | 1 | 1 | 1 |
| rs118203965 | pathogenic | 0 | 0 | 0 |
| rs118203966 | pathogenic | 0 | 0 | 0 |
| rs1801201 | benign | 1 | 1 | 0 |
| rs1136201 | benign | 1 | 1 | 1 |
| rs144148008 | benign | 0 | 0 | 0 |
| rs200118378 | benign | 1 | 1 | 1 |
| rs112984118 | benign | 1 | 1 | 1 |
| rs149446102 | benign | 1 | 1 | 1 |
| rs1427463 | benign | 0 | 0 | 0 |
| rs201936720 | benign | 1 | 1 | 1 |
| rs61751983 | benign | 1 | 1 | 1 |
| rs104894632 | pathogenic | 1 | 1 | 1 |
| rs149202905 | benign | 0 | 0 | 0 |
| rs2230210 | benign | 1 | 1 | 1 |
| rs111631988 | benign | 0 | 0 | 0 |

| | | | | |
|---|---|---|---|---|
| rs137852550 | pathogenic | 0 | 0 | 0 |
| rs5963403 | benign | 0 | 0 | 0 |
| rs138018739 | benign | 0 | 0 | 0 |
| rs62640590 | benign | 0 | 0 | 0 |
| rs121909581 | pathogenic | 1 | 1 | 1 |
| rs202017913 | benign | 1 | 1 | 1 |
| rs143163513 | benign | 0 | 0 | 0 |
| rs238238 | benign | 0 | 1 | 0 |
| rs143945974 | benign | 0 | 0 | 0 |
| rs238239 | benign | 1 | 1 | 1 |
| rs121918403 | pathogenic | 0 | 1 | 0 |
| rs61735456 | benign | 0 | 1 | 0 |
| rs121918404 | pathogenic | 1 | 1 | 0 |
| rs12674488 | benign | 0 | 0 | 0 |
| rs201599657 | pathogenic | 0 | 0 | 0 |
| rs1057090 | benign | 0 | 0 | 0 |
| rs1057091 | benign | 0 | 0 | 0 |
| rs587783740 | benign | 0 | 0 | 0 |
| rs75956769 | benign | 0 | 0 | 0 |
| rs201691299 | benign | 0 | 0 | 0 |
| rs62621087 | benign | 0 | 0 | 0 |
| rs201308454 | benign | 0 | 0 | 0 |
| rs145657360 | benign | 0 | 0 | 0 |
| rs121434592 | pathogenic | 1 | 1 | 0 |
| rs375650413 | benign | 0 | 0 | 0 |
| rs121434437 | pathogenic | 1 | 1 | 0 |
| rs763494292 | pathogenic | 0 | 0 | 0 |
| rs121964997 | pathogenic | 1 | 1 | 0 |
| rs121964996 | pathogenic | 0 | 0 | 0 |
| rs41283425 | benign | 1 | 1 | 1 |
| rs376289528 | benign | 1 | 1 | 1 |
| rs144171604 | benign | 1 | 1 | 1 |
| rs782440692 | pathogenic | 1 | 1 | 1 |
| rs782058451 | pathogenic | 1 | 1 | 1 |
| rs78437817 | benign | 0 | 0 | 0 |
| rs200327969 | benign | 1 | 1 | 1 |
| rs143043662 | benign | 0 | 0 | 0 |
| rs111966833 | benign | 1 | 1 | 1 |
| rs35523678 | benign | 0 | 0 | 0 |
| rs515726208 | pathogenic | 0 | 0 | 0 |
| rs397517239 | pathogenic | 1 | 1 | 1 |
| rs148966602 | benign | 1 | 1 | 1 |
| rs76371972 | benign | 1 | 1 | 1 |
| rs2229968 | benign | 1 | 1 | 1 |
| rs61751538 | benign | 1 | 1 | 1 |
| rs200800978 | pathogenic | 1 | 1 | 1 |
| rs148695964 | benign | 1 | 1 | 1 |
| rs140042418 | benign | 1 | 1 | 1 |
| rs116185403 | benign | 0 | 0 | 0 |
| rs142828669 | benign | 1 | 1 | 1 |
| rs139832048 | benign | 0 | 1 | 0 |
| rs58750568 | benign | 0 | 1 | 0 |
| rs77681311 | benign | 0 | 0 | 0 |
| rs151208452 | benign | 0 | 0 | 0 |

| | | | | |
|---|---|---|---|---|
| rs76463933 | benign | 0 | 0 | 0 |
| rs144434647 | benign | 0 | 0 | 0 |
| rs117361485 | benign | 0 | 0 | 0 |
| rs587777588 | pathogenic | 0 | 0 | 0 |
| rs80338796 | pathogenic | 0 | 0 | 0 |
| rs121434594 | pathogenic | 0 | 0 | 0 |
| rs146766138 | pathogenic | 1 | 1 | 1 |
| rs28939679 | pathogenic | 1 | 1 | 1 |
| rs748754134 | pathogenic | 1 | 1 | 0 |
| rs764276946 | pathogenic | 1 | 1 | 1 |
| rs111033588 | pathogenic | 1 | 1 | 1 |
| rs201633414 | pathogenic | 1 | 1 | 1 |
| rs121909262 | pathogenic | 1 | 1 | 1 |
| rs104893702 | pathogenic | 1 | 1 | 1 |
| rs121909265 | pathogenic | 1 | 1 | 1 |
| rs121909266 | pathogenic | 1 | 1 | 1 |
| rs193922439 | pathogenic | 1 | 1 | 0 |
| rs104893691 | pathogenic | 1 | 1 | 1 |
| rs104893695 | pathogenic | 0 | 0 | 0 |
| rs104893708 | pathogenic | 1 | 1 | 1 |
| rs121909260 | pathogenic | 0 | 0 | 0 |
| rs104893696 | pathogenic | 0 | 0 | 0 |
| rs121909267 | pathogenic | 0 | 0 | 0 |
| rs121909263 | pathogenic | 0 | 0 | 0 |
| rs121909264 | pathogenic | 0 | 0 | 0 |
| rs104893694 | pathogenic | 1 | 1 | 1 |
| rs121909268 | pathogenic | 1 | 1 | 1 |
| rs104893689 | pathogenic | 1 | 1 | 0 |
| rs104893707 | pathogenic | 1 | 1 | 0 |
| rs104893697 | pathogenic | 1 | 1 | 1 |
| rs28936684 | pathogenic | 1 | 1 | 1 |
| rs62269092 | benign | 1 | 1 | 0 |
| rs121909259 | pathogenic | 1 | 1 | 0 |
| rs548403340 | benign | 1 | 1 | 0 |
| rs142818334 | benign | 0 | 0 | 0 |
| rs12493789 | benign | 1 | 1 | 1 |
| rs193922423 | pathogenic | 1 | 1 | 0 |
| rs104893719 | pathogenic | 1 | 1 | 1 |
| rs117375173 | benign | 1 | 1 | 0 |
| rs104893712 | pathogenic | 0 | 0 | 0 |
| rs121917776 | pathogenic | 1 | 1 | 1 |
| rs767325003 | benign | 1 | 1 | 1 |
| rs35824453 | benign | 0 | 0 | 0 |
| rs141641266 | benign | 0 | 0 | 0 |
| rs1050239 | benign | 0 | 0 | 0 |
| rs141303384 | benign | 0 | 0 | 0 |
| rs12720458 | pathogenic | 0 | 0 | 0 |
| rs368507376 | pathogenic | 1 | 1 | 1 |
| rs17215500 | pathogenic | 1 | 1 | 1 |
| rs120074188 | pathogenic | 1 | 1 | 1 |
| rs397508097 | pathogenic | 1 | 1 | 1 |
| rs138551008 | pathogenic | 0 | 0 | 0 |
| rs387907139 | pathogenic | 1 | 1 | 1 |
| rs765487627 | pathogenic | 0 | 0 | 0 |

| | | | | |
|---|---|---|---|---|
| rs587778587 | benign | 0 | 0 | 0 |
| rs45568339 | benign | 0 | 0 | 0 |
| rs78179744 | benign | 0 | 0 | 0 |
| rs45624036 | benign | 0 | 0 | 0 |
| rs61755166 | benign | 0 | 0 | 0 |
| rs180177132 | pathogenic | 1 | 1 | 1 |
| rs587776527 | pathogenic | 0 | 0 | 0 |
| rs373783514 | benign | 0 | 0 | 0 |
| rs730881896 | benign | 0 | 0 | 0 |
| rs118203998 | pathogenic | 0 | 0 | 0 |
| rs41281039 | pathogenic | 1 | 1 | 1 |
| rs118204430 | pathogenic | 1 | 1 | 1 |
| rs1800546 | pathogenic | 1 | 1 | 0 |
| rs76917243 | pathogenic | 0 | 1 | 0 |
| rs78340951 | pathogenic | 0 | 1 | 0 |
| rs77718928 | pathogenic | 0 | 0 | 0 |
| rs369586696 | pathogenic | 0 | 1 | 0 |
| rs745517517 | pathogenic | 1 | 1 | 0 |
| rs372872777 | pathogenic | 0 | 0 | 0 |
| rs748704459 | pathogenic | 0 | 0 | 0 |
| rs769215411 | pathogenic | 0 | 0 | 0 |
| rs139425890 | pathogenic | 0 | 0 | 0 |
| rs779694939 | pathogenic | 1 | 1 | 0 |
| rs773729410 | pathogenic | 0 | 0 | 0 |
| rs121908623 | pathogenic | 1 | 1 | 1 |
| rs121908622 | pathogenic | 1 | 1 | 1 |
| rs121908621 | pathogenic | 1 | 1 | 1 |
| rs762106720 | pathogenic | 1 | 1 | 0 |
| rs199877522 | pathogenic | 0 | 0 | 0 |
| rs74315291 | pathogenic | 1 | 1 | 1 |
| rs79137415 | benign | 0 | 0 | 0 |
| rs121913045 | pathogenic | 0 | 0 | 0 |
| rs34295337 | pathogenic | 0 | 0 | 0 |
| rs121913046 | pathogenic | 0 | 0 | 0 |
| rs121913047 | pathogenic | 0 | 0 | 0 |
| rs121913048 | pathogenic | 0 | 0 | 0 |
| rs267607166 | pathogenic | 0 | 0 | 0 |
| rs17850684 | benign | 1 | 1 | 1 |
| rs119476051 | pathogenic | 1 | 1 | 1 |
| rs17850683 | benign | 1 | 1 | 1 |
| rs119476049 | pathogenic | 1 | 1 | 1 |
| rs119476046 | pathogenic | 1 | 1 | 1 |
| rs119476048 | pathogenic | 0 | 0 | 0 |
| rs119476047 | pathogenic | 1 | 1 | 1 |
| rs28939094 | pathogenic | 1 | 1 | 1 |
| rs119476050 | pathogenic | 0 | 0 | 0 |
| rs387906941 | pathogenic | 1 | 1 | 1 |
| rs41511344 | pathogenic | 1 | 1 | 0 |
| rs748112833 | pathogenic | 1 | 1 | 1 |
| rs74342080 | benign | 1 | 1 | 0 |
| rs116482870 | benign | 0 | 0 | 0 |
| rs104893743 | pathogenic | 0 | 0 | 0 |
| rs117179004 | benign | 0 | 0 | 0 |
| rs587779409 | pathogenic | 1 | 1 | 1 |

| | | | | |
|---|---|---|---|---|
| rs3745406 | benign | 1 | 1 | 0 |
| rs387906696 | pathogenic | 0 | 0 | 0 |
| rs72549369 | pathogenic | 1 | 1 | 0 |
| rs45523133 | benign | 0 | 0 | 0 |
| rs119460972 | pathogenic | 0 | 1 | 0 |
| rs34929837 | benign | 0 | 1 | 0 |
| rs17323225 | benign | 0 | 1 | 0 |
| rs17011368 | benign | 0 | 0 | 0 |
| rs748902766 | pathogenic | 1 | 1 | 1 |
| rs387907292 | pathogenic | 0 | 0 | 0 |
| rs387907293 | pathogenic | 1 | 1 | 0 |
| rs371526758 | pathogenic | 1 | 1 | 1 |
| rs142968179 | pathogenic | 1 | 1 | 1 |
| rs387907291 | pathogenic | 1 | 1 | 0 |
| rs121907949 | pathogenic | 0 | 0 | 0 |
| rs121907947 | pathogenic | 0 | 0 | 0 |
| rs121907950 | pathogenic | 1 | 1 | 1 |
| rs121907948 | pathogenic | 1 | 1 | 1 |
| rs4926 | benign | 0 | 0 | 0 |
| rs748809996 | pathogenic | 0 | 0 | 0 |
| rs1799883 | benign | 1 | 1 | 0 |
| rs104893974 | pathogenic | 0 | 1 | 0 |
| rs104893973 | pathogenic | 0 | 1 | 0 |
| rs34452707 | benign | 0 | 0 | 0 |
| rs119467003 | pathogenic | 1 | 1 | 1 |
| rs121908261 | pathogenic | 0 | 1 | 0 |
| rs28933985 | pathogenic | 1 | 1 | 1 |
| rs80356669 | pathogenic | 1 | 1 | 1 |
| rs28936677 | pathogenic | 1 | 1 | 0 |
| rs121918326 | pathogenic | 0 | 0 | 0 |
| rs121918324 | pathogenic | 1 | 1 | 0 |
| rs121918325 | pathogenic | 1 | 1 | 0 |
| rs41270025 | pathogenic | 1 | 1 | 0 |
| rs121918323 | pathogenic | 0 | 0 | 0 |
| rs12735723 | pathogenic | 1 | 1 | 0 |
| rs36013429 | benign | 0 | 0 | 0 |
| rs28936676 | pathogenic | 1 | 1 | 0 |
| rs771809901 | pathogenic | 1 | 1 | 1 |
| rs758350837 | benign | 1 | 1 | 0 |
| rs121908192 | pathogenic | 1 | 1 | 1 |
| rs147568500 | pathogenic | 1 | 1 | 1 |
| rs3729734 | benign | 0 | 0 | 0 |
| rs371536347 | benign | 1 | 1 | 1 |
| rs80358193 | pathogenic | 1 | 1 | 1 |
| rs119476044 | pathogenic | 1 | 1 | 1 |
| rs111292798 | benign | 1 | 1 | 1 |
| rs121908063 | pathogenic | 1 | 1 | 0 |
| rs121908065 | pathogenic | 1 | 1 | 1 |
| rs121908062 | pathogenic | 1 | 1 | 1 |
| rs121908064 | pathogenic | 1 | 1 | 1 |
| rs61746421 | benign | 1 | 1 | 1 |
| rs34886916 | benign | 1 | 1 | 1 |
| rs118204045 | pathogenic | 0 | 0 | 0 |
| rs138607412 | benign | 1 | 1 | 1 |

| | | | | |
|---|---|---|---|---|
| rs118204046 | pathogenic | 0 | 0 | 0 |
| rs118204048 | pathogenic | 0 | 0 | 0 |
| rs118204044 | pathogenic | 0 | 0 | 0 |
| rs149664056 | benign | 0 | 0 | 0 |
| rs121909285 | pathogenic | 0 | 0 | 0 |
| rs1802498 | benign | 0 | 0 | 0 |
| rs1065761 | benign | 0 | 0 | 0 |
| rs119477054 | pathogenic | 0 | 0 | 0 |
| rs12720097 | benign | 0 | 0 | 0 |
| rs747281324 | pathogenic | 1 | 1 | 0 |
| rs367543028 | pathogenic | 1 | 1 | 0 |
| rs146077918 | benign | 0 | 0 | 0 |
| rs142551229 | benign | 0 | 0 | 0 |
| rs367543039 | pathogenic | 0 | 0 | 0 |
| rs587779884 | pathogenic | 1 | 1 | 0 |
| rs137853153 | pathogenic | 0 | 0 | 0 |
| rs367543029 | pathogenic | 0 | 0 | 0 |
| rs587783037 | pathogenic | 1 | 1 | 0 |
| rs142928725 | benign | 0 | 0 | 0 |
| rs28385141 | benign | 0 | 0 | 0 |
| rs587779888 | pathogenic | 0 | 0 | 0 |
| rs121909801 | pathogenic | 1 | 1 | 1 |
| rs121909790 | pathogenic | 1 | 1 | 0 |
| rs121909797 | pathogenic | 1 | 1 | 0 |
| rs121909794 | pathogenic | 1 | 1 | 1 |
| rs121909791 | pathogenic | 1 | 1 | 1 |
| rs121909793 | pathogenic | 1 | 1 | 1 |
| rs147977279 | benign | 1 | 1 | 1 |
| rs147352564 | benign | 0 | 0 | 0 |
| rs368758045 | benign | 1 | 1 | 1 |
| rs114498261 | benign | 1 | 1 | 0 |
| rs35484156 | benign | 0 | 0 | 0 |
| rs139176878 | benign | 0 | 0 | 0 |
| rs11543934 | benign | 0 | 0 | 0 |
| rs34192549 | benign | 0 | 0 | 0 |
| rs568736295 | benign | 0 | 0 | 0 |
| rs55747232 | benign | 0 | 0 | 0 |
| rs781887272 | benign | 0 | 0 | 0 |
| rs146273341 | benign | 0 | 0 | 0 |
| rs41473544 | benign | 0 | 0 | 0 |
| rs137852680 | pathogenic | 0 | 0 | 0 |
| rs55891000 | benign | 0 | 0 | 0 |
| rs140351189 | benign | 0 | 0 | 0 |
| rs137852679 | pathogenic | 1 | 1 | 0 |
| rs137852678 | pathogenic | 0 | 0 | 0 |
| rs137852677 | pathogenic | 1 | 1 | 1 |
| rs138620818 | benign | 0 | 0 | 0 |
| rs121908531 | pathogenic | 0 | 0 | 0 |
| rs72554307 | pathogenic | 0 | 0 | 0 |
| rs72554308 | pathogenic | 0 | 0 | 0 |
| rs72554312 | pathogenic | 1 | 1 | 0 |
| rs67939655 | pathogenic | 0 | 0 | 0 |
| rs67486158 | pathogenic | 0 | 0 | 0 |
| rs72554338 | pathogenic | 1 | 1 | 1 |

| | | | | |
|---|---|---|---|---|
| rs1800324 | pathogenic | 1 | 1 | 1 |
| rs72554356 | pathogenic | 1 | 1 | 1 |
| rs66656800 | pathogenic | 1 | 1 | 1 |
| rs68026851 | pathogenic | 1 | 1 | 1 |
| rs72558412 | pathogenic | 0 | 0 | 0 |
| rs72558415 | pathogenic | 0 | 0 | 0 |
| rs72558423 | pathogenic | 0 | 0 | 0 |
| rs67120076 | pathogenic | 1 | 1 | 1 |
| rs1800328 | pathogenic | 1 | 1 | 1 |
| rs72558451 | pathogenic | 1 | 1 | 1 |
| rs72558454 | pathogenic | 1 | 1 | 1 |
| rs672601372 | pathogenic | 0 | 0 | 0 |
| rs139644798 | pathogenic | 0 | 0 | 0 |
| rs369398935 | pathogenic | 0 | 0 | 0 |
| rs267607035 | pathogenic | 0 | 0 | 0 |
| rs267607036 | pathogenic | 1 | 1 | 1 |
| rs2302566 | benign | 1 | 1 | 0 |
| rs115959591 | benign | 0 | 0 | 0 |
| rs773876739 | pathogenic | 0 | 0 | 0 |
| rs1799967 | benign | 0 | 0 | 0 |
| rs80356968 | benign | 0 | 0 | 0 |
| rs28897695 | benign | 1 | 1 | 1 |
| rs80357314 | benign | 0 | 0 | 0 |
| rs80357169 | benign | 0 | 0 | 0 |
| rs397507239 | benign | 1 | 1 | 1 |
| rs80187739 | pathogenic | 1 | 1 | 1 |
| rs41293459 | pathogenic | 1 | 1 | 1 |
| rs55770810 | pathogenic | 1 | 1 | 1 |
| rs80356860 | pathogenic | 0 | 0 | 0 |
| rs56195342 | benign | 0 | 0 | 0 |
| rs80357442 | benign | 0 | 0 | 0 |
| rs41293463 | pathogenic | 1 | 1 | 1 |
| rs80357069 | pathogenic | 0 | 0 | 0 |
| rs397509275 | benign | 0 | 0 | 0 |
| rs80356920 | benign | 0 | 0 | 0 |
| rs397509284 | pathogenic | 0 | 0 | 0 |
| rs80357286 | benign | 0 | 0 | 0 |
| rs80357274 | benign | 0 | 0 | 0 |
| rs765656957 | benign | 0 | 0 | 0 |
| rs755427809 | benign | 0 | 0 | 0 |
| rs80357322 | benign | 0 | 0 | 0 |
| rs121434430 | pathogenic | 0 | 0 | 0 |
| rs121918450 | pathogenic | 1 | 1 | 1 |
| rs1532268 | benign | 0 | 0 | 0 |
| rs2303080 | benign | 0 | 0 | 0 |
| rs10064631 | benign | 0 | 0 | 0 |
| rs162036 | benign | 0 | 0 | 0 |
| rs2287780 | benign | 0 | 0 | 0 |
| rs16879334 | benign | 0 | 0 | 0 |
| rs137853061 | pathogenic | 1 | 1 | 0 |
| rs10380 | benign | 0 | 0 | 0 |
| rs1494558 | benign | 0 | 0 | 0 |
| rs1494555 | benign | 0 | 0 | 0 |
| rs192994060 | benign | 0 | 0 | 0 |

| | | | | |
|---|---|---|---|---|
| rs147128920 | benign | 1 | 1 | 0 |
| rs201803986 | pathogenic | 0 | 0 | 0 |
| rs746419489 | pathogenic | 0 | 1 | 0 |
| rs764396564 | pathogenic | 0 | 0 | 0 |
| rs758304537 | pathogenic | 0 | 0 | 0 |
| rs267606763 | pathogenic | 0 | 0 | 0 |
| rs267606762 | pathogenic | 0 | 0 | 0 |
| rs150702945 | pathogenic | 1 | 1 | 1 |
| rs202047589 | pathogenic | 0 | 0 | 0 |
| rs755562733 | pathogenic | 0 | 0 | 0 |
| rs267606764 | pathogenic | 1 | 1 | 1 |
| rs138207257 | pathogenic | 1 | 1 | 0 |
| rs149150736 | pathogenic | 1 | 1 | 1 |
| rs777046879 | pathogenic | 0 | 0 | 0 |
| rs104894408 | pathogenic | 1 | 1 | 1 |
| rs1801002 | pathogenic | 1 | 1 | 1 |
| rs28929485 | pathogenic | 1 | 1 | 1 |
| rs104894396 | pathogenic | 1 | 1 | 1 |
| rs371024165 | pathogenic | 0 | 0 | 0 |
| rs35887622 | pathogenic | 1 | 1 | 1 |
| rs587783644 | pathogenic | 1 | 1 | 1 |
| rs72474224 | pathogenic | 1 | 1 | 1 |
| rs111033296 | pathogenic | 1 | 1 | 1 |
| rs104894407 | pathogenic | 1 | 1 | 1 |
| rs72561723 | pathogenic | 1 | 1 | 1 |
| rs104894398 | pathogenic | 1 | 1 | 1 |
| rs28931594 | pathogenic | 1 | 1 | 1 |
| rs104894412 | pathogenic | 1 | 1 | 1 |
| rs111033297 | pathogenic | 1 | 1 | 1 |
| rs104894404 | pathogenic | 1 | 1 | 1 |
| rs104894410 | pathogenic | 1 | 1 | 1 |
| rs104894403 | pathogenic | 1 | 1 | 1 |
| rs397516870 | benign | 0 | 0 | 0 |
| rs200023879 | pathogenic | 1 | 1 | 1 |
| rs121912968 | pathogenic | 0 | 0 | 0 |
| rs104894402 | pathogenic | 1 | 1 | 1 |
| rs28931593 | pathogenic | 1 | 1 | 1 |
| rs104894397 | pathogenic | 0 | 0 | 0 |
| rs80338944 | pathogenic | 0 | 0 | 0 |
| rs199883710 | pathogenic | 0 | 0 | 0 |
| rs727504302 | pathogenic | 0 | 0 | 0 |
| rs145216882 | benign | 1 | 1 | 1 |
| rs781534323 | pathogenic | 1 | 1 | 1 |
| rs111033218 | benign | 1 | 1 | 1 |
| rs104894409 | pathogenic | 0 | 0 | 0 |
| rs80338945 | pathogenic | 1 | 1 | 1 |
| rs111033299 | pathogenic | 0 | 0 | 0 |
| rs143343083 | pathogenic | 1 | 1 | 1 |
| rs80338946 | pathogenic | 0 | 0 | 0 |
| rs150529554 | pathogenic | 0 | 0 | 0 |
| rs111033295 | pathogenic | 0 | 0 | 0 |
| rs111033188 | benign | 0 | 0 | 0 |
| rs397516874 | pathogenic | 0 | 0 | 0 |
| rs76434661 | pathogenic | 0 | 0 | 0 |

| | | | | |
|---|---|---|---|---|
| rs80338948 | pathogenic | 0 | 0 | 0 |
| rs104894401 | pathogenic | 0 | 0 | 0 |
| rs767178508 | pathogenic | 0 | 0 | 0 |
| rs772264564 | pathogenic | 0 | 0 | 0 |
| rs28931592 | pathogenic | 0 | 0 | 0 |
| rs80338949 | pathogenic | 1 | 1 | 1 |
| rs376898963 | benign | 1 | 1 | 1 |
| rs111033360 | benign | 0 | 0 | 0 |
| rs774518779 | pathogenic | 1 | 1 | 1 |
| rs28931595 | pathogenic | 1 | 1 | 1 |
| rs771748289 | pathogenic | 0 | 0 | 0 |
| rs104894406 | pathogenic | 0 | 0 | 0 |
| rs76838169 | benign | 0 | 0 | 0 |
| rs111033294 | pathogenic | 0 | 0 | 0 |
| rs200214298 | pathogenic | 0 | 0 | 0 |
| rs28940283 | pathogenic | 0 | 1 | 0 |
| rs1047883 | benign | 0 | 0 | 0 |
| rs138254425 | benign | 0 | 0 | 0 |
| rs121912592 | pathogenic | 1 | 1 | 0 |
| rs369061090 | pathogenic | 1 | 1 | 0 |
| rs758724746 | pathogenic | 0 | 1 | 0 |
| rs121912595 | pathogenic | 0 | 0 | 0 |
| rs76340296 | benign | 0 | 0 | 0 |
| rs140578009 | benign | 1 | 1 | 1 |
| rs74445297 | benign | 0 | 0 | 0 |
| rs78599516 | benign | 1 | 1 | 0 |
| rs104894035 | pathogenic | 1 | 1 | 0 |
| rs79451216 | benign | 0 | 0 | 0 |
| rs104894036 | pathogenic | 0 | 0 | 0 |
| rs73343757 | benign | 0 | 0 | 0 |
| rs121917722 | pathogenic | 1 | 1 | 1 |
| rs747700126 | pathogenic | 0 | 1 | 0 |
| rs121917725 | pathogenic | 1 | 1 | 1 |
| rs121917721 | pathogenic | 1 | 1 | 1 |
| rs121917723 | pathogenic | 1 | 1 | 0 |
| rs17569 | benign | 1 | 1 | 1 |
| rs2230062 | benign | 1 | 1 | 1 |
| rs17570 | benign | 0 | 0 | 0 |
| rs121917724 | pathogenic | 0 | 1 | 0 |
| rs199422235 | pathogenic | 0 | 0 | 0 |
| rs199422237 | pathogenic | 1 | 1 | 1 |
| rs199422234 | pathogenic | 1 | 1 | 0 |
| rs140015315 | pathogenic | 1 | 1 | 1 |
| rs730880298 | pathogenic | 1 | 1 | 1 |
| rs779692470 | pathogenic | 0 | 0 | 0 |
| rs369152939 | pathogenic | 0 | 0 | 0 |
| rs527236040 | pathogenic | 0 | 0 | 0 |
| rs587776985 | pathogenic | 1 | 1 | 1 |
| rs147077598 | pathogenic | 1 | 1 | 1 |
| rs128620185 | pathogenic | 1 | 1 | 1 |
| rs128620189 | pathogenic | 0 | 0 | 0 |
| rs128621190 | pathogenic | 0 | 0 | 0 |
| rs61754648 | benign | 0 | 0 | 0 |
| rs137852906 | pathogenic | 0 | 0 | 0 |

| | | | | |
|---|---|---|---|---|
| rs76811038 | pathogenic | 0 | 0 | 0 |
| rs149468422 | benign | 0 | 0 | 0 |
| rs5911 | benign | 0 | 0 | 0 |
| rs28942112 | pathogenic | 1 | 1 | 1 |
| rs143275858 | pathogenic | 1 | 1 | 0 |
| rs28362261 | benign | 0 | 0 | 0 |
| rs28362263 | benign | 1 | 1 | 1 |
| rs558707786 | pathogenic | 0 | 0 | 0 |
| rs138775799 | pathogenic | 0 | 0 | 0 |
| rs748170941 | pathogenic | 0 | 0 | 0 |
| rs121908380 | pathogenic | 0 | 0 | 0 |
| rs765123255 | pathogenic | 1 | 1 | 1 |
| rs730881832 | pathogenic | 0 | 1 | 0 |
| rs587782165 | benign | 1 | 1 | 1 |
| rs762307622 | pathogenic | 0 | 0 | 0 |
| rs34612342 | pathogenic | 1 | 1 | 0 |
| rs143353451 | pathogenic | 0 | 1 | 0 |
| rs747993448 | pathogenic | 0 | 1 | 0 |
| rs376561094 | pathogenic | 0 | 0 | 0 |
| rs776487884 | pathogenic | 1 | 1 | 0 |
| rs34126013 | pathogenic | 1 | 1 | 0 |
| rs200495564 | pathogenic | 1 | 1 | 0 |
| rs140342925 | pathogenic | 1 | 1 | 0 |
| rs587780749 | pathogenic | 1 | 1 | 0 |
| rs773087549 | pathogenic | 1 | 1 | 0 |
| rs730881833 | pathogenic | 0 | 0 | 0 |
| rs374950566 | pathogenic | 1 | 1 | 0 |
| rs587780752 | benign | 0 | 0 | 0 |
| rs3219489 | benign | 0 | 0 | 0 |
| rs199742231 | benign | 0 | 0 | 0 |
| rs587780082 | pathogenic | 0 | 0 | 0 |
| rs35881803 | benign | 0 | 0 | 0 |
| rs41305623 | benign | 1 | 1 | 1 |
| rs121917856 | pathogenic | 1 | 1 | 1 |
| rs576284122 | benign | 0 | 0 | 0 |
| rs142036701 | benign | 0 | 0 | 0 |
| rs121917852 | pathogenic | 1 | 1 | 1 |
| rs121917854 | pathogenic | 1 | 1 | 1 |
| rs121917853 | pathogenic | 1 | 1 | 1 |
| rs121917857 | pathogenic | 0 | 1 | 0 |
| rs193922740 | pathogenic | 1 | 1 | 1 |
| rs565779970 | pathogenic | 1 | 1 | 1 |
| rs6723526 | benign | 0 | 0 | 0 |
| rs141094096 | pathogenic | 1 | 1 | 1 |
| rs104894637 | pathogenic | 0 | 0 | 0 |
| rs104894636 | pathogenic | 0 | 0 | 0 |
| rs104894642 | pathogenic | 0 | 0 | 0 |
| rs104894638 | pathogenic | 0 | 0 | 0 |
| rs753666460 | pathogenic | 0 | 0 | 0 |
| rs104894643 | pathogenic | 1 | 1 | 0 |
| rs34520362 | pathogenic | 0 | 0 | 0 |
| rs374621913 | pathogenic | 0 | 0 | 0 |
| rs113641837 | benign | 0 | 0 | 0 |
| rs104894635 | pathogenic | 0 | 0 | 0 |

| | | | | |
|---|---|---|---|---|
| rs143947056 | pathogenic | 0 | 0 | 0 |
| rs138504221 | pathogenic | 0 | 0 | 0 |
| rs9894254 | benign | 0 | 0 | 0 |
| rs104894640 | pathogenic | 1 | 1 | 1 |
| rs58786455 | benign | 0 | 0 | 0 |
| rs144143780 | pathogenic | 0 | 0 | 0 |
| rs62620232 | benign | 0 | 0 | 0 |
| rs34297805 | benign | 1 | 1 | 1 |
| rs104894641 | pathogenic | 0 | 0 | 0 |
| rs148876719 | benign | 0 | 0 | 0 |
| rs104894639 | pathogenic | 0 | 0 | 0 |
| rs7503034 | benign | 0 | 0 | 0 |
| rs201631570 | benign | 0 | 0 | 0 |
| rs150181226 | pathogenic | 0 | 0 | 0 |
| rs377546036 | pathogenic | 0 | 0 | 0 |
| rs267606700 | pathogenic | 0 | 0 | 0 |
| rs267606701 | pathogenic | 1 | 1 | 0 |
| rs267606699 | pathogenic | 1 | 1 | 0 |
| rs9903215 | benign | 0 | 0 | 0 |
| rs139486406 | benign | 0 | 0 | 0 |
| rs143791614 | benign | 0 | 0 | 0 |
| rs573483474 | benign | 0 | 0 | 0 |
| rs2032892 | benign | 0 | 0 | 0 |
| rs137852619 | pathogenic | 0 | 0 | 0 |
| rs200243548 | benign | 0 | 0 | 0 |
| rs115702388 | benign | 0 | 0 | 0 |
| rs138085358 | pathogenic | 0 | 0 | 0 |
| rs121912789 | pathogenic | 1 | 1 | 1 |
| rs763407972 | pathogenic | 1 | 1 | 1 |
| rs147278302 | benign | 0 | 0 | 0 |
| rs374540411 | benign | 0 | 0 | 0 |
| rs121912790 | pathogenic | 1 | 1 | 0 |
| rs752049917 | benign | 0 | 0 | 0 |
| rs139640491 | benign | 1 | 1 | 1 |
| rs138010738 | benign | 1 | 1 | 0 |
| rs201328768 | pathogenic | 1 | 1 | 0 |
| rs2135720 | benign | 1 | 1 | 0 |
| rs201816080 | benign | 1 | 1 | 0 |
| rs754391973 | pathogenic | 0 | 0 | 0 |
| rs139087859 | benign | 0 | 0 | 0 |
| rs6805421 | benign | 0 | 0 | 0 |
| rs146266900 | benign | 0 | 0 | 0 |
| rs397509419 | pathogenic | 0 | 0 | 0 |
| rs149980411 | benign | 0 | 0 | 0 |
| rs747711488 | pathogenic | 1 | 1 | 1 |
| rs149474339 | pathogenic | 1 | 1 | 1 |
| rs758252808 | pathogenic | 1 | 1 | 0 |
| rs745756308 | pathogenic | 0 | 0 | 0 |
| rs4986958 | benign | 0 | 0 | 0 |
| rs74315444 | pathogenic | 0 | 0 | 0 |
| rs61752137 | pathogenic | 0 | 0 | 0 |
| rs8179065 | benign | 1 | 1 | 1 |
| rs758361736 | pathogenic | 1 | 1 | 0 |
| rs139794067 | pathogenic | 1 | 1 | 1 |

| | | | | |
|---|---|---|---|---|
| rs730880954 | pathogenic | 1 | 1 | 1 |
| rs730880955 | pathogenic | 1 | 1 | 1 |
| rs150634297 | pathogenic | 0 | 0 | 0 |
| rs104893750 | pathogenic | 1 | 1 | 1 |
| rs104893748 | pathogenic | 1 | 1 | 1 |
| rs104893749 | pathogenic | 1 | 1 | 1 |
| rs753520553 | pathogenic | 1 | 1 | 0 |
| rs61737294 | benign | 0 | 0 | 0 |
| rs104894601 | pathogenic | 0 | 0 | 0 |
| rs376090795 | pathogenic | 0 | 0 | 0 |
| rs104894596 | pathogenic | 0 | 0 | 0 |
| rs104894595 | pathogenic | 0 | 0 | 0 |
| rs104894598 | pathogenic | 0 | 0 | 0 |
| rs104894597 | pathogenic | 0 | 0 | 0 |
| rs104894591 | pathogenic | 0 | 0 | 0 |
| rs104894593 | pathogenic | 0 | 0 | 0 |
| rs104894594 | pathogenic | 0 | 0 | 0 |
| rs104894590 | pathogenic | 0 | 0 | 0 |
| rs86312 | benign | 0 | 0 | 0 |
| rs77874543 | benign | 0 | 0 | 0 |
| rs121912590 | pathogenic | 1 | 1 | 1 |
| rs121912589 | pathogenic | 1 | 1 | 1 |
| rs121912586 | pathogenic | 1 | 1 | 1 |
| rs121912585 | pathogenic | 1 | 1 | 1 |
| rs121912587 | pathogenic | 1 | 1 | 1 |
| rs762909359 | pathogenic | 0 | 0 | 0 |
| rs28931610 | pathogenic | 0 | 0 | 0 |
| rs104894097 | pathogenic | 1 | 1 | 1 |
| rs775176191 | benign | 0 | 0 | 0 |
| rs199907548 | pathogenic | 0 | 0 | 0 |
| rs104894095 | pathogenic | 1 | 1 | 1 |
| rs104894109 | pathogenic | 1 | 1 | 1 |
| rs372266620 | benign | 1 | 1 | 1 |
| rs104894099 | pathogenic | 0 | 0 | 0 |
| rs137854599 | pathogenic | 1 | 1 | 1 |
| rs104894094 | pathogenic | 0 | 0 | 0 |
| rs104894104 | pathogenic | 0 | 0 | 0 |
| rs6413463 | benign | 1 | 1 | 1 |
| rs104894098 | pathogenic | 0 | 0 | 0 |
| rs6413464 | benign | 0 | 0 | 0 |
| rs3731249 | benign | 0 | 0 | 0 |
| rs16931179 | benign | 0 | 0 | 0 |
| rs2734715 | benign | 0 | 0 | 0 |
| rs759080581 | pathogenic | 0 | 0 | 0 |
| rs1800746 | benign | 0 | 0 | 0 |
| rs202189057 | pathogenic | 0 | 0 | 0 |
| rs121908204 | pathogenic | 0 | 0 | 0 |
| rs756564767 | pathogenic | 0 | 0 | 0 |
| rs34758634 | benign | 1 | 1 | 1 |
| rs28940573 | pathogenic | 0 | 0 | 0 |
| rs119455955 | pathogenic | 1 | 1 | 0 |
| rs752164603 | pathogenic | 1 | 1 | 0 |
| rs763162812 | pathogenic | 0 | 0 | 0 |
| rs119455957 | pathogenic | 0 | 0 | 0 |

| | | | | |
|---|---|---|---|---|
| rs119455958 | pathogenic | 1 | 1 | 0 |
| rs750428882 | pathogenic | 0 | 0 | 0 |
| rs141482368 | benign | 0 | 0 | 0 |
| rs119455953 | pathogenic | 0 | 0 | 0 |
| rs119455954 | pathogenic | 0 | 0 | 0 |
| rs112812685 | benign | 1 | 1 | 0 |
| rs138744051 | benign | 1 | 1 | 0 |
| rs121908200 | pathogenic | 0 | 0 | 0 |
| rs119455956 | pathogenic | 0 | 0 | 0 |
| rs78970155 | benign | 0 | 0 | 0 |
| rs2834167 | benign | 0 | 0 | 0 |
| rs387907326 | pathogenic | 1 | 1 | 1 |
| rs2107732 | benign | 0 | 0 | 0 |
| rs11552377 | benign | 0 | 0 | 0 |
| rs137852843 | pathogenic | 0 | 0 | 0 |
| rs104894783 | pathogenic | 0 | 0 | 0 |
| rs122457137 | pathogenic | 1 | 1 | 1 |
| rs104894779 | pathogenic | 1 | 1 | 1 |
| rs104894786 | pathogenic | 0 | 0 | 0 |
| rs104894784 | pathogenic | 0 | 0 | 0 |
| rs104894785 | pathogenic | 0 | 0 | 0 |
| rs201870761 | pathogenic | 0 | 0 | 0 |
| rs104894781 | pathogenic | 0 | 0 | 0 |
| rs28897728 | benign | 0 | 0 | 0 |
| rs370273690 | pathogenic | 1 | 1 | 0 |
| rs9332739 | benign | 0 | 0 | 0 |
| rs150878060 | benign | 0 | 0 | 0 |
| rs573509224 | benign | 0 | 0 | 0 |
| rs36221133 | benign | 0 | 0 | 0 |
| rs2230009 | benign | 0 | 0 | 0 |
| rs267607008 | pathogenic | 0 | 0 | 0 |
| rs5985 | benign | 1 | 1 | 1 |
| rs5982 | benign | 1 | 1 | 1 |
| rs5987 | benign | 0 | 0 | 0 |
| rs5988 | benign | 0 | 0 | 0 |
| rs121918680 | pathogenic | 1 | 1 | 0 |
| rs121918679 | pathogenic | 1 | 1 | 0 |
| rs7845 | benign | 1 | 1 | 0 |
| rs1049296 | benign | 1 | 1 | 1 |
| rs121918677 | pathogenic | 1 | 1 | 0 |
| rs200086262 | pathogenic | 1 | 1 | 0 |
| rs746930990 | pathogenic | 1 | 1 | 0 |
| rs104893878 | pathogenic | 1 | 1 | 1 |
| rs121912468 | pathogenic | 1 | 1 | 1 |
| rs121912467 | pathogenic | 1 | 1 | 1 |
| rs145351926 | benign | 1 | 1 | 1 |
| rs141212743 | pathogenic | 1 | 1 | 1 |
| rs397515473 | pathogenic | 1 | 1 | 0 |
| rs121918313 | pathogenic | 0 | 1 | 0 |
| rs73015965 | pathogenic | 0 | 0 | 0 |
| rs121918030 | pathogenic | 0 | 1 | 0 |
| rs77782321 | benign | 0 | 0 | 0 |
| rs374661051 | pathogenic | 1 | 1 | 1 |
| rs267606854 | pathogenic | 0 | 0 | 0 |

| | | | | |
|---|---|---|---|---|
| rs41279678 | benign | 0 | 0 | 0 |
| rs61754640 | benign | 0 | 0 | 0 |
| rs199473314 | pathogenic | 0 | 0 | 0 |
| rs137854601 | pathogenic | 0 | 0 | 0 |
| rs137854614 | pathogenic | 0 | 0 | 0 |
| rs137854615 | pathogenic | 0 | 0 | 0 |
| rs199473635 | pathogenic | 0 | 0 | 0 |
| rs121434478 | pathogenic | 1 | 1 | 1 |
| rs775407864 | pathogenic | 1 | 1 | 1 |
| rs12706927 | benign | 0 | 1 | 0 |
| rs775614261 | benign | 0 | 0 | 0 |
| rs768926261 | benign | 0 | 0 | 0 |
| rs114057640 | benign | 0 | 0 | 0 |
| rs387906886 | pathogenic | 1 | 1 | 0 |
| rs138658585 | benign | 0 | 0 | 0 |
| rs4880091 | benign | 0 | 0 | 0 |
| rs121918552 | pathogenic | 1 | 1 | 1 |
| rs10794537 | benign | 0 | 0 | 0 |
| rs121965022 | pathogenic | 1 | 1 | 0 |
| rs121965020 | pathogenic | 0 | 0 | 0 |
| rs758452450 | pathogenic | 0 | 0 | 0 |
| rs747981483 | benign | 0 | 0 | 0 |
| rs58037052 | benign | 0 | 0 | 0 |
| rs121965029 | pathogenic | 1 | 1 | 0 |
| rs147490060 | benign | 0 | 0 | 0 |
| rs3755955 | benign | 0 | 0 | 0 |
| rs148789453 | pathogenic | 0 | 0 | 0 |
| rs121965030 | pathogenic | 0 | 0 | 0 |
| rs76722191 | benign | 0 | 0 | 0 |
| rs199801029 | pathogenic | 0 | 0 | 0 |
| rs121965033 | pathogenic | 0 | 0 | 0 |
| rs368454909 | pathogenic | 1 | 1 | 0 |
| rs6831280 | benign | 0 | 0 | 0 |
| rs121965024 | pathogenic | 0 | 0 | 0 |
| rs121965019 | pathogenic | 0 | 0 | 0 |
| rs11934801 | benign | 0 | 0 | 0 |
| rs73066479 | benign | 0 | 0 | 0 |
| rs121965027 | pathogenic | 0 | 0 | 0 |
| rs121965026 | pathogenic | 1 | 1 | 0 |
| rs121965021 | pathogenic | 0 | 0 | 0 |
| rs398123257 | benign | 0 | 0 | 0 |
| rs121965031 | pathogenic | 0 | 0 | 0 |
| rs202003795 | pathogenic | 0 | 0 | 0 |
| rs559510809 | pathogenic | 0 | 0 | 0 |
| rs762936223 | benign | 0 | 0 | 0 |
| rs137854580 | pathogenic | 1 | 1 | 1 |
| rs112263157 | benign | 0 | 0 | 0 |
| rs2289181 | benign | 0 | 0 | 0 |
| rs138034915 | benign | 0 | 1 | 0 |
| rs528367092 | pathogenic | 0 | 1 | 0 |
| rs1042393 | benign | 0 | 0 | 0 |
| rs370950728 | pathogenic | 0 | 1 | 0 |
| rs530478036 | benign | 0 | 1 | 0 |
| rs374569672 | benign | 0 | 1 | 0 |

| | | | | |
|---|---|---|---|---|
| rs121907944 | pathogenic | 0 | 1 | 0 |
| rs200856561 | pathogenic | 0 | 1 | 0 |
| rs577915581 | pathogenic | 0 | 1 | 0 |
| rs201896815 | pathogenic | 0 | 1 | 0 |
| rs121907945 | pathogenic | 0 | 1 | 0 |
| rs121907940 | pathogenic | 0 | 1 | 0 |
| rs543300039 | pathogenic | 0 | 1 | 0 |
| rs121907936 | pathogenic | 0 | 1 | 0 |
| rs766074609 | pathogenic | 0 | 1 | 0 |
| rs755253527 | pathogenic | 0 | 1 | 0 |
| rs560575383 | pathogenic | 0 | 1 | 0 |
| rs528369909 | benign | 0 | 1 | 0 |
| rs770610356 | pathogenic | 0 | 1 | 0 |
| rs747610090 | pathogenic | 0 | 1 | 0 |
| rs145712232 | benign | 0 | 1 | 0 |
| rs7215458 | benign | 1 | 1 | 0 |
| rs535644999 | pathogenic | 0 | 1 | 0 |
| rs772883420 | pathogenic | 1 | 1 | 0 |
| rs398123169 | pathogenic | 0 | 1 | 0 |
| rs121907937 | pathogenic | 0 | 1 | 0 |
| rs121907941 | pathogenic | 1 | 1 | 0 |
| rs121907942 | pathogenic | 0 | 1 | 0 |
| rs779556619 | pathogenic | 0 | 1 | 0 |
| rs1800307 | benign | 0 | 1 | 0 |
| rs377544304 | pathogenic | 1 | 1 | 0 |
| rs549029029 | pathogenic | 0 | 1 | 0 |
| rs753269119 | pathogenic | 1 | 1 | 1 |
| rs757617999 | pathogenic | 0 | 1 | 0 |
| rs28937909 | pathogenic | 0 | 1 | 0 |
| rs28940868 | pathogenic | 0 | 1 | 0 |
| rs368438393 | pathogenic | 0 | 1 | 0 |
| rs536906561 | pathogenic | 0 | 1 | 0 |
| rs374143224 | pathogenic | 0 | 1 | 0 |
| rs778418246 | pathogenic | 0 | 1 | 0 |
| rs398123172 | pathogenic | 0 | 1 | 0 |
| rs121907938 | pathogenic | 0 | 1 | 0 |
| rs1800312 | pathogenic | 0 | 1 | 0 |
| rs1126690 | benign | 0 | 1 | 0 |
| rs1800314 | benign | 0 | 1 | 0 |
| rs369532274 | pathogenic | 0 | 1 | 0 |
| rs121907943 | pathogenic | 0 | 1 | 0 |
| rs780321415 | pathogenic | 0 | 1 | 0 |
| rs377286472 | benign | 0 | 1 | 0 |
| rs121918406 | pathogenic | 1 | 1 | 1 |
| rs147484870 | benign | 0 | 0 | 0 |
| rs193922671 | pathogenic | 0 | 0 | 0 |
| rs201397978 | benign | 1 | 1 | 1 |
| rs34884895 | benign | 1 | 1 | 1 |
| rs138329459 | benign | 0 | 0 | 0 |
| rs2289366 | benign | 1 | 1 | 1 |
| rs141858334 | benign | 1 | 1 | 1 |
| rs59907886 | benign | 1 | 1 | 1 |
| rs137852653 | pathogenic | 0 | 0 | 0 |
| rs137852652 | pathogenic | 0 | 0 | 0 |

| | | | | |
|---|---|---|---|---|
| rs137852656 | pathogenic | 0 | 0 | 0 |
| rs61739916 | benign | 0 | 0 | 0 |
| rs28936668 | pathogenic | 0 | 0 | 0 |
| rs137852651 | pathogenic | 0 | 0 | 0 |
| rs137852650 | pathogenic | 0 | 0 | 0 |
| rs28936669 | pathogenic | 0 | 0 | 0 |
| rs137852654 | pathogenic | 0 | 0 | 0 |
| rs28936368 | pathogenic | 1 | 1 | 0 |
| rs137852655 | pathogenic | 0 | 0 | 0 |
| rs61752496 | benign | 0 | 0 | 0 |
| rs750098055 | benign | 0 | 0 | 0 |
| rs764149793 | benign | 0 | 0 | 0 |
| rs724160017 | pathogenic | 0 | 1 | 0 |
| rs724160019 | pathogenic | 0 | 0 | 0 |
| rs202219398 | pathogenic | 1 | 1 | 0 |
| rs724160023 | pathogenic | 1 | 1 | 0 |
| rs761953453 | pathogenic | 0 | 0 | 0 |
| rs80338835 | pathogenic | 1 | 1 | 1 |
| rs56284940 | benign | 0 | 0 | 0 |
| rs74575749 | benign | 0 | 0 | 0 |
| rs587777075 | pathogenic | 1 | 1 | 1 |
| rs641153 | benign | 0 | 0 | 0 |
| rs12614 | benign | 0 | 0 | 0 |
| rs117314762 | benign | 0 | 0 | 0 |
| rs537478097 | benign | 1 | 1 | 1 |
| rs144812066 | benign | 0 | 0 | 0 |
| rs4151651 | benign | 0 | 0 | 0 |
| rs138207668 | benign | 1 | 1 | 1 |
| rs149101394 | benign | 1 | 1 | 1 |
| rs4151659 | benign | 1 | 1 | 1 |
| rs45484591 | benign | 1 | 1 | 1 |
| rs761991070 | pathogenic | 0 | 0 | 0 |
| rs755948940 | pathogenic | 0 | 0 | 0 |
| rs137853097 | pathogenic | 0 | 0 | 0 |
| rs775297352 | benign | 0 | 0 | 0 |
| rs28943591 | benign | 0 | 0 | 0 |
| rs191468413 | pathogenic | 1 | 1 | 0 |
| rs11539471 | benign | 1 | 1 | 1 |
| rs11205 | benign | 0 | 0 | 0 |
| rs768409400 | pathogenic | 0 | 0 | 0 |
| rs762013077 | pathogenic | 0 | 0 | 0 |
| rs768474112 | pathogenic | 0 | 0 | 0 |
| rs121965063 | pathogenic | 1 | 1 | 1 |
| rs143648758 | pathogenic | 0 | 0 | 0 |
| rs121965066 | pathogenic | 0 | 0 | 0 |
| rs752907087 | pathogenic | 0 | 0 | 0 |
| rs747702749 | pathogenic | 0 | 0 | 0 |
| rs754087775 | pathogenic | 0 | 0 | 0 |
| rs121918641 | pathogenic | 1 | 1 | 1 |
| rs121918642 | pathogenic | 1 | 1 | 1 |
| rs121918640 | pathogenic | 1 | 1 | 1 |
| rs121918637 | pathogenic | 1 | 1 | 1 |
| rs121918644 | pathogenic | 1 | 1 | 1 |
| rs16840544 | benign | 0 | 0 | 0 |

| | | | | |
|---|---|---|---|---|
| rs1042391 | benign | 1 | 1 | 1 |
| rs104893744 | pathogenic | 1 | 1 | 1 |
| rs120074136 | pathogenic | 0 | 0 | 0 |
| rs1805081 | benign | 1 | 1 | 0 |
| rs80358251 | benign | 0 | 0 | 0 |
| rs774588125 | benign | 0 | 0 | 0 |
| rs120074134 | pathogenic | 0 | 0 | 0 |
| rs139751448 | pathogenic | 0 | 0 | 0 |
| rs77080672 | benign | 0 | 0 | 0 |
| rs372445155 | pathogenic | 1 | 1 | 0 |
| rs13381670 | benign | 0 | 0 | 0 |
| rs1788799 | benign | 0 | 0 | 0 |
| rs376213990 | pathogenic | 0 | 0 | 0 |
| rs757475924 | pathogenic | 0 | 0 | 0 |
| rs80358253 | pathogenic | 0 | 0 | 0 |
| rs1805082 | benign | 0 | 0 | 0 |
| rs120074130 | pathogenic | 0 | 0 | 0 |
| rs768999208 | pathogenic | 0 | 0 | 0 |
| rs34302553 | benign | 0 | 0 | 0 |
| rs28940897 | pathogenic | 0 | 0 | 0 |
| rs1140458 | benign | 0 | 0 | 0 |
| rs143124972 | pathogenic | 0 | 0 | 0 |
| rs120074135 | pathogenic | 0 | 0 | 0 |
| rs543206298 | pathogenic | 0 | 0 | 0 |
| rs120074132 | pathogenic | 0 | 0 | 0 |
| rs759826138 | pathogenic | 0 | 0 | 0 |
| rs28942108 | pathogenic | 0 | 0 | 0 |
| rs80358254 | pathogenic | 0 | 0 | 0 |
| rs80358257 | pathogenic | 0 | 0 | 0 |
| rs781261962 | pathogenic | 0 | 0 | 0 |
| rs28942107 | pathogenic | 0 | 0 | 0 |
| rs80358258 | pathogenic | 0 | 0 | 0 |
| rs80358259 | pathogenic | 1 | 1 | 0 |
| rs141440861 | benign | 0 | 0 | 0 |
| rs28942106 | pathogenic | 0 | 0 | 0 |
| rs778878523 | pathogenic | 0 | 0 | 0 |
| rs28942105 | pathogenic | 0 | 0 | 0 |
| rs61731969 | benign | 0 | 0 | 0 |
| rs200444084 | pathogenic | 0 | 0 | 0 |
| rs113371321 | pathogenic | 0 | 0 | 0 |
| rs369098773 | pathogenic | 0 | 0 | 0 |
| rs35248744 | benign | 0 | 0 | 0 |
| rs758902805 | pathogenic | 0 | 0 | 0 |
| rs120074131 | pathogenic | 0 | 0 | 0 |
| rs1805084 | benign | 0 | 0 | 0 |
| rs117793540 | benign | 0 | 0 | 0 |
| rs530058063 | benign | 0 | 0 | 0 |
| rs1064422 | pathogenic | 1 | 1 | 1 |
| rs104894506 | pathogenic | 1 | 1 | 1 |
| rs104894507 | pathogenic | 1 | 1 | 1 |
| rs104894508 | pathogenic | 1 | 1 | 0 |
| rs28999113 | pathogenic | 1 | 1 | 0 |
| rs281860266 | pathogenic | 0 | 0 | 0 |
| rs104894961 | pathogenic | 1 | 1 | 0 |

| | | | | |
|---|---|---|---|---|
| rs122463168 | pathogenic | 0 | 0 | 0 |
| rs199469624 | pathogenic | 1 | 1 | 1 |
| rs150107747 | benign | 0 | 0 | 0 |
| rs144984628 | pathogenic | 0 | 0 | 0 |
| rs28935171 | pathogenic | 0 | 0 | 0 |
| rs137852443 | pathogenic | 1 | 1 | 0 |
| rs28933674 | pathogenic | 1 | 1 | 1 |
| rs28933675 | pathogenic | 1 | 1 | 1 |
| rs28933676 | pathogenic | 1 | 1 | 1 |
| rs28933677 | pathogenic | 1 | 1 | 1 |
| rs28933678 | pathogenic | 0 | 0 | 0 |
| rs28933679 | pathogenic | 1 | 1 | 1 |
| rs28933680 | pathogenic | 0 | 0 | 0 |
| rs28933681 | pathogenic | 0 | 0 | 0 |
| rs28933682 | pathogenic | 0 | 0 | 0 |
| rs28937294 | pathogenic | 1 | 1 | 0 |
| rs137852459 | pathogenic | 0 | 0 | 0 |
| rs137852461 | pathogenic | 0 | 0 | 0 |
| rs137852464 | pathogenic | 1 | 1 | 1 |
| rs137852465 | pathogenic | 1 | 1 | 1 |
| rs1800297 | benign | 1 | 1 | 0 |
| rs137852374 | pathogenic | 1 | 1 | 1 |
| rs672601312 | pathogenic | 1 | 1 | 1 |
| rs387907196 | pathogenic | 1 | 1 | 1 |
| rs148348866 | benign | 0 | 0 | 0 |
| rs141138948 | pathogenic | 1 | 1 | 1 |
| rs137852685 | pathogenic | 1 | 1 | 0 |
| rs104893900 | pathogenic | 1 | 1 | 0 |
| rs137852686 | pathogenic | 1 | 1 | 1 |
| rs104893906 | pathogenic | 1 | 1 | 0 |
| rs587782207 | pathogenic | 1 | 1 | 1 |
| rs75086406 | pathogenic | 0 | 0 | 0 |
| rs199822819 | pathogenic | 0 | 0 | 0 |
| rs398123166 | pathogenic | 1 | 1 | 1 |
| rs121913123 | pathogenic | 1 | 1 | 1 |
| rs772190176 | pathogenic | 0 | 1 | 0 |
| rs587782216 | pathogenic | 1 | 1 | 1 |
| rs398123159 | pathogenic | 1 | 1 | 1 |
| rs121913122 | pathogenic | 1 | 1 | 1 |
| rs755436052 | pathogenic | 1 | 1 | 1 |
| rs200796606 | pathogenic | 1 | 1 | 1 |
| rs2070080 | pathogenic | 0 | 1 | 0 |
| rs149616320 | benign | 0 | 0 | 0 |
| rs143529458 | benign | 0 | 0 | 0 |
| rs267607077 | pathogenic | 1 | 1 | 1 |
| rs143898031 | benign | 1 | 1 | 0 |
| rs148304857 | benign | 1 | 1 | 1 |
| rs137852809 | pathogenic | 0 | 0 | 0 |
| rs3765623 | benign | 0 | 0 | 0 |
| rs16944397 | benign | 0 | 0 | 0 |
| rs730880167 | benign | 0 | 0 | 0 |
| rs187108957 | benign | 0 | 0 | 0 |
| rs117342470 | benign | 0 | 0 | 0 |
| rs751200138 | pathogenic | 1 | 1 | 1 |

| | | | | |
|---|---|---|---|---|
| rs755799528 | pathogenic | 0 | 0 | 0 |
| rs537484504 | benign | 0 | 0 | 0 |
| rs748944640 | pathogenic | 0 | 0 | 0 |
| rs199774121 | pathogenic | 0 | 0 | 0 |
| rs766094434 | pathogenic | 1 | 1 | 0 |
| rs777321035 | pathogenic | 1 | 1 | 1 |
| rs146269774 | benign | 0 | 1 | 0 |
| rs144085805 | benign | 0 | 1 | 0 |
| rs367643028 | benign | 0 | 0 | 0 |
| rs42524 | benign | 0 | 0 | 0 |
| rs28940586 | pathogenic | 0 | 0 | 0 |
| rs74315327 | pathogenic | 1 | 1 | 1 |
| rs199676861 | pathogenic | 0 | 0 | 0 |
| rs115802744 | benign | 1 | 1 | 0 |
| rs752866643 | pathogenic | 1 | 1 | 1 |
| rs28941784 | pathogenic | 1 | 1 | 1 |
| rs398124434 | pathogenic | 1 | 1 | 1 |
| rs756414548 | pathogenic | 1 | 1 | 1 |
| rs376128990 | pathogenic | 1 | 1 | 1 |
| rs746219370 | pathogenic | 1 | 1 | 1 |
| rs756195708 | pathogenic | 1 | 1 | 1 |
| rs369296618 | pathogenic | 1 | 1 | 1 |
| rs9593 | benign | 1 | 1 | 1 |
| rs139510844 | benign | 0 | 0 | 0 |
| rs28382158 | benign | 0 | 0 | 0 |
| rs28919570 | pathogenic | 0 | 0 | 0 |
| rs587777331 | pathogenic | 0 | 0 | 0 |
| rs587777333 | pathogenic | 0 | 0 | 0 |
| rs587777332 | pathogenic | 0 | 0 | 0 |
| rs774980346 | pathogenic | 0 | 0 | 0 |
| rs370934093 | pathogenic | 0 | 0 | 0 |
| rs121913479 | pathogenic | 1 | 1 | 1 |
| rs121913484 | pathogenic | 0 | 0 | 0 |
| rs121913485 | pathogenic | 1 | 1 | 1 |
| rs28931614 | pathogenic | 1 | 1 | 1 |
| rs17881656 | benign | 1 | 1 | 1 |
| rs28931615 | pathogenic | 1 | 1 | 1 |
| rs147846832 | benign | 0 | 0 | 0 |
| rs80356506 | pathogenic | 0 | 0 | 0 |
| rs80356494 | pathogenic | 0 | 0 | 0 |
| rs121908894 | pathogenic | 0 | 0 | 0 |
| rs121908895 | pathogenic | 0 | 0 | 0 |
| rs80356508 | pathogenic | 0 | 0 | 0 |
| rs80356507 | pathogenic | 0 | 0 | 0 |
| rs121908896 | pathogenic | 0 | 0 | 0 |
| rs745372589 | pathogenic | 1 | 1 | 0 |
| rs183898408 | benign | 0 | 0 | 0 |
| rs200844292 | benign | 0 | 0 | 0 |
| rs146897002 | benign | 0 | 0 | 0 |
| rs121964869 | pathogenic | 1 | 1 | 1 |
| rs201768800 | benign | 1 | 1 | 1 |
| rs9807556 | benign | 1 | 1 | 1 |
| rs184721031 | benign | 0 | 0 | 0 |
| rs201104206 | benign | 0 | 0 | 0 |

| | | | | |
|---|---|---|---|---|
| rs188677538 | benign | 0 | 0 | 0 |
| rs149528866 | benign | 0 | 0 | 0 |
| rs115382168 | benign | 0 | 0 | 0 |
| rs78846023 | benign | 1 | 1 | 1 |
| rs61730662 | benign | 0 | 0 | 0 |
| rs121918368 | pathogenic | 0 | 0 | 0 |
| rs566433112 | pathogenic | 1 | 1 | 1 |
| rs121909342 | pathogenic | 0 | 0 | 0 |
| rs67586389 | pathogenic | 0 | 0 | 0 |
| rs72466485 | pathogenic | 0 | 0 | 0 |
| rs72466486 | pathogenic | 1 | 1 | 1 |
| rs72466487 | pathogenic | 1 | 1 | 1 |
| rs267607112 | pathogenic | 0 | 0 | 0 |
| rs118204013 | pathogenic | 0 | 0 | 0 |
| rs121913495 | pathogenic | 0 | 0 | 0 |
| rs11554273 | pathogenic | 0 | 0 | 0 |
| rs193922477 | pathogenic | 1 | 1 | 1 |
| rs193922479 | pathogenic | 1 | 1 | 1 |
| rs72547554 | pathogenic | 0 | 0 | 0 |
| rs72547561 | pathogenic | 1 | 1 | 1 |
| rs72547562 | pathogenic | 1 | 1 | 0 |
| rs115977487 | benign | 1 | 1 | 0 |
| rs72547564 | pathogenic | 0 | 0 | 0 |
| rs72547568 | pathogenic | 1 | 1 | 0 |
| rs72547569 | pathogenic | 1 | 1 | 1 |
| rs72547571 | pathogenic | 0 | 0 | 0 |
| rs72547575 | pathogenic | 1 | 1 | 1 |
| rs544994462 | benign | 0 | 0 | 0 |
| rs190470017 | benign | 0 | 0 | 0 |
| rs199473266 | pathogenic | 1 | 1 | 1 |
| rs137854602 | pathogenic | 0 | 0 | 0 |
| rs143739249 | pathogenic | 0 | 0 | 0 |
| rs397514697 | pathogenic | 1 | 1 | 1 |
| rs373764886 | pathogenic | 0 | 0 | 0 |
| rs370242930 | pathogenic | 0 | 0 | 0 |
| rs377145777 | pathogenic | 1 | 1 | 1 |
| rs191256186 | benign | 0 | 0 | 0 |
| rs150070901 | benign | 1 | 1 | 1 |
| rs138098032 | benign | 0 | 0 | 0 |
| rs117731813 | benign | 1 | 1 | 1 |
| rs104894488 | pathogenic | 1 | 1 | 1 |
| rs11302 | benign | 0 | 1 | 0 |
| rs121918447 | pathogenic | 1 | 1 | 1 |
| rs138205828 | benign | 1 | 1 | 0 |
| rs149858290 | benign | 0 | 0 | 0 |
| rs750764003 | pathogenic | 0 | 1 | 0 |
| rs141381896 | benign | 0 | 0 | 0 |
| rs149027682 | benign | 0 | 0 | 0 |
| rs121908415 | pathogenic | 0 | 0 | 0 |
| rs121908416 | pathogenic | 0 | 0 | 0 |
| rs121908417 | pathogenic | 0 | 0 | 0 |
| rs587780306 | benign | 0 | 0 | 0 |
| rs3218725 | benign | 0 | 0 | 0 |
| rs3218729 | benign | 0 | 0 | 0 |

| | | | | |
|---|---|---|---|---|
| rs387906917 | pathogenic | 0 | 0 | 0 |
| rs200652608 | pathogenic | 0 | 0 | 0 |
| rs3218721 | benign | 0 | 0 | 0 |
| rs63750869 | pathogenic | 1 | 1 | 1 |
| rs118203910 | pathogenic | 0 | 1 | 0 |
| rs9332701 | benign | 1 | 1 | 0 |
| rs6027 | benign | 0 | 0 | 0 |
| rs121909110 | pathogenic | 0 | 1 | 0 |
| rs762550967 | pathogenic | 1 | 1 | 0 |
| rs763615602 | pathogenic | 1 | 1 | 0 |
| rs144907403 | benign | 0 | 0 | 0 |
| rs104893631 | pathogenic | 1 | 1 | 0 |
| rs144181978 | pathogenic | 1 | 1 | 0 |
| rs74874677 | benign | 1 | 1 | 1 |
| rs104893632 | pathogenic | 0 | 1 | 0 |
| rs371342464 | benign | 1 | 1 | 1 |
| rs104893633 | pathogenic | 1 | 1 | 0 |
| rs201088712 | pathogenic | 0 | 0 | 0 |
| rs201857541 | benign | 1 | 1 | 1 |
| rs139517732 | pathogenic | 1 | 1 | 0 |
| rs191057824 | benign | 1 | 1 | 0 |
| rs35683768 | benign | 1 | 1 | 0 |
| rs104895271 | pathogenic | 0 | 0 | 0 |
| rs104895217 | pathogenic | 1 | 1 | 1 |
| rs104895223 | pathogenic | 1 | 1 | 1 |
| rs104895218 | pathogenic | 1 | 1 | 1 |
| rs104895225 | pathogenic | 1 | 1 | 1 |
| rs4149637 | benign | 1 | 1 | 1 |
| rs104895219 | pathogenic | 1 | 1 | 1 |
| rs104895220 | pathogenic | 1 | 1 | 1 |
| rs104895228 | pathogenic | 1 | 1 | 1 |
| rs104895221 | pathogenic | 1 | 1 | 1 |
| rs104895222 | pathogenic | 1 | 1 | 1 |
| rs104895247 | pathogenic | 0 | 0 | 0 |
| rs28940585 | pathogenic | 0 | 0 | 0 |
| rs561367199 | pathogenic | 1 | 1 | 1 |
| rs143793815 | benign | 0 | 0 | 0 |
| rs28941473 | pathogenic | 1 | 1 | 1 |
| rs751590073 | pathogenic | 1 | 1 | 1 |
| rs111407265 | benign | 1 | 1 | 1 |
| rs373697663 | pathogenic | 1 | 1 | 1 |
| rs369879957 | pathogenic | 0 | 0 | 0 |
| rs28941472 | pathogenic | 1 | 1 | 1 |
| rs28940287 | pathogenic | 1 | 1 | 1 |
| rs77625410 | benign | 0 | 0 | 0 |
| rs111033477 | benign | 0 | 0 | 0 |
| rs115123584 | benign | 0 | 0 | 0 |
| rs374576916 | benign | 0 | 0 | 0 |
| rs397516317 | pathogenic | 0 | 0 | 0 |
| rs368657015 | pathogenic | 0 | 0 | 0 |
| rs397516321 | pathogenic | 0 | 0 | 0 |
| rs397516322 | pathogenic | 0 | 0 | 0 |
| rs111033215 | pathogenic | 0 | 0 | 0 |
| rs199606180 | pathogenic | 0 | 0 | 0 |

| | | | | |
|---|---|---|---|---|
| rs111033192 | pathogenic | 0 | 0 | 0 |
| rs948962 | benign | 1 | 1 | 1 |
| rs142293185 | benign | 0 | 0 | 0 |
| rs376764423 | pathogenic | 0 | 0 | 0 |
| rs773844428 | pathogenic | 0 | 0 | 0 |
| rs186644871 | benign | 0 | 0 | 0 |
| rs200313391 | benign | 0 | 0 | 0 |
| rs377670513 | pathogenic | 0 | 0 | 0 |
| rs1132036 | benign | 0 | 0 | 0 |
| rs111033231 | benign | 0 | 0 | 0 |
| rs145945630 | pathogenic | 0 | 0 | 0 |
| rs121912984 | pathogenic | 1 | 1 | 1 |
| rs1800435 | benign | 0 | 1 | 0 |
| rs121912980 | pathogenic | 0 | 0 | 0 |
| rs121912982 | pathogenic | 1 | 1 | 1 |
| rs121912983 | pathogenic | 0 | 0 | 0 |
| rs121912981 | pathogenic | 0 | 0 | 0 |
| rs1057868 | benign | 0 | 0 | 0 |
| rs56048668 | benign | 0 | 0 | 0 |
| rs55865465 | benign | 0 | 0 | 0 |
| rs748096324 | benign | 1 | 1 | 0 |
| rs150475750 | benign | 0 | 0 | 0 |
| rs143025739 | benign | 1 | 1 | 0 |
| rs79702016 | benign | 1 | 1 | 0 |
| rs41338945 | benign | 0 | 0 | 0 |
| rs146406037 | benign | 1 | 1 | 1 |
| rs143274491 | benign | 0 | 0 | 0 |
| rs3829986 | benign | 0 | 0 | 0 |
| rs149168939 | benign | 0 | 0 | 0 |
| rs1800385 | benign | 0 | 0 | 0 |
| rs1800386 | pathogenic | 0 | 0 | 0 |
| rs121918524 | pathogenic | 1 | 1 | 1 |
| rs121912700 | pathogenic | 0 | 0 | 0 |
| rs104893930 | pathogenic | 1 | 1 | 1 |
| rs104893931 | pathogenic | 0 | 0 | 0 |
| rs116125936 | benign | 0 | 0 | 0 |
| rs627441 | benign | 0 | 0 | 0 |
| rs104894481 | pathogenic | 0 | 0 | 0 |
| rs137853129 | pathogenic | 0 | 0 | 0 |
| rs137853130 | pathogenic | 1 | 1 | 0 |
| rs28940282 | pathogenic | 0 | 0 | 0 |
| rs2229989 | benign | 0 | 0 | 0 |
| rs104894647 | pathogenic | 0 | 0 | 0 |
| rs1061170 | benign | 1 | 1 | 1 |
| rs576402053 | benign | 0 | 0 | 0 |
| rs34972246 | benign | 0 | 0 | 0 |
| rs192725607 | benign | 0 | 0 | 0 |
| rs137852615 | pathogenic | 1 | 1 | 0 |
| rs137852617 | pathogenic | 1 | 1 | 1 |
| rs137852611 | pathogenic | 0 | 0 | 0 |
| rs137852612 | pathogenic | 0 | 0 | 0 |
| rs137852614 | pathogenic | 1 | 1 | 1 |
| rs137852610 | pathogenic | 1 | 1 | 1 |
| rs137852618 | pathogenic | 1 | 1 | 1 |

| | | | | |
|---|---|---|---|---|
| rs137852613 | pathogenic | 1 | 1 | 0 |
| rs137852609 | pathogenic | 0 | 0 | 0 |
| rs63750306 | pathogenic | 1 | 1 | 1 |
| rs63750265 | pathogenic | 1 | 1 | 1 |
| rs63751287 | pathogenic | 1 | 1 | 1 |
| rs199723282 | benign | 0 | 0 | 0 |
| rs121917807 | pathogenic | 0 | 0 | 0 |
| rs121917809 | pathogenic | 0 | 0 | 0 |
| rs75134564 | pathogenic | 0 | 0 | 0 |
| rs267606930 | pathogenic | 1 | 1 | 0 |
| rs121909302 | pathogenic | 0 | 0 | 0 |
| rs121909303 | pathogenic | 0 | 0 | 0 |
| rs121909299 | pathogenic | 0 | 0 | 0 |
| rs121909301 | pathogenic | 0 | 0 | 0 |
| rs121909300 | pathogenic | 1 | 1 | 0 |
| rs111033589 | pathogenic | 0 | 0 | 0 |
| rs137853059 | pathogenic | 0 | 0 | 0 |
| rs137853057 | pathogenic | 1 | 1 | 1 |
| rs1801474 | benign | 0 | 0 | 0 |
| rs137853060 | pathogenic | 1 | 1 | 1 |
| rs137853058 | pathogenic | 1 | 1 | 1 |
| rs34424986 | pathogenic | 0 | 0 | 0 |
| rs199657839 | benign | 1 | 1 | 1 |
| rs56092260 | benign | 0 | 0 | 0 |
| rs1801582 | benign | 1 | 1 | 1 |
| rs1801334 | benign | 1 | 1 | 1 |
| rs191486604 | pathogenic | 0 | 0 | 0 |
| rs121908051 | pathogenic | 0 | 0 | 0 |
| rs121908055 | pathogenic | 1 | 1 | 0 |
| rs28940886 | pathogenic | 0 | 0 | 0 |
| rs121908050 | pathogenic | 0 | 0 | 0 |
| rs28940887 | pathogenic | 0 | 0 | 0 |
| rs387906300 | pathogenic | 0 | 0 | 0 |
| rs28942087 | pathogenic | 0 | 0 | 0 |
| rs121908049 | pathogenic | 1 | 1 | 0 |
| rs121908054 | pathogenic | 1 | 1 | 0 |
| rs121908048 | pathogenic | 0 | 0 | 0 |
| rs28940888 | pathogenic | 1 | 1 | 1 |
| rs121908053 | pathogenic | 1 | 1 | 0 |
| rs122467169 | pathogenic | 1 | 1 | 1 |
| rs122467172 | pathogenic | 1 | 1 | 1 |
| rs122467170 | pathogenic | 1 | 1 | 1 |
| rs28935477 | pathogenic | 1 | 1 | 0 |
| rs28942088 | pathogenic | 0 | 0 | 0 |
| rs751478142 | pathogenic | 1 | 1 | 0 |
| rs765368797 | pathogenic | 0 | 0 | 0 |
| rs147810437 | benign | 0 | 0 | 0 |
| rs121918209 | pathogenic | 1 | 1 | 1 |
| rs121918208 | pathogenic | 1 | 1 | 1 |
| rs121918210 | pathogenic | 1 | 1 | 1 |
| rs35515638 | benign | 1 | 1 | 1 |
| rs121918207 | pathogenic | 1 | 1 | 1 |
| rs141298312 | benign | 0 | 0 | 0 |
| rs121918211 | pathogenic | 0 | 0 | 0 |

| | | | | |
|---|---|---|---|---|
| rs149660059 | benign | 0 | 0 | 0 |
| rs200670286 | pathogenic | 1 | 1 | 1 |
| rs121918212 | pathogenic | 1 | 1 | 1 |
| rs121918213 | pathogenic | 1 | 1 | 1 |
| rs121918205 | pathogenic | 1 | 1 | 1 |
| rs104894025 | pathogenic | 1 | 1 | 1 |
| rs104894026 | pathogenic | 1 | 1 | 1 |
| rs104894028 | pathogenic | 0 | 1 | 0 |
| rs104894029 | pathogenic | 0 | 1 | 0 |
| rs119458971 | pathogenic | 0 | 0 | 0 |
| rs119458969 | pathogenic | 1 | 1 | 1 |
| rs1801591 | benign | 1 | 1 | 1 |
| rs140169311 | benign | 0 | 0 | 0 |
| rs753363278 | benign | 1 | 1 | 1 |
| rs199763682 | pathogenic | 0 | 0 | 0 |
| rs119458970 | pathogenic | 1 | 1 | 1 |
| rs763722868 | pathogenic | 0 | 0 | 0 |
| rs56404770 | benign | 0 | 0 | 0 |
| rs34221241 | benign | 0 | 0 | 0 |
| rs121909625 | pathogenic | 0 | 0 | 0 |
| rs550921485 | pathogenic | 0 | 0 | 0 |
| rs1800454 | benign | 0 | 0 | 0 |
| rs241447 | benign | 0 | 0 | 0 |
| rs2230606 | benign | 1 | 1 | 0 |
| rs5036 | pathogenic | 1 | 1 | 1 |
| rs13306788 | benign | 0 | 0 | 0 |
| rs28929480 | pathogenic | 0 | 0 | 0 |
| rs121912749 | pathogenic | 0 | 0 | 0 |
| rs56361140 | pathogenic | 0 | 0 | 0 |
| rs28931583 | pathogenic | 1 | 1 | 1 |
| rs36211715 | pathogenic | 1 | 1 | 1 |
| rs138049878 | pathogenic | 1 | 1 | 1 |
| rs727503253 | pathogenic | 1 | 1 | 1 |
| rs267606908 | pathogenic | 0 | 0 | 0 |
| rs121913631 | pathogenic | 1 | 1 | 1 |
| rs36211716 | benign | 1 | 1 | 1 |
| rs121913639 | pathogenic | 1 | 1 | 1 |
| rs730880761 | pathogenic | 1 | 1 | 1 |
| rs121913629 | pathogenic | 1 | 1 | 1 |
| rs754368658 | pathogenic | 0 | 0 | 0 |
| rs777090017 | pathogenic | 0 | 0 | 0 |
| rs267606913 | pathogenic | 1 | 1 | 1 |
| rs773138218 | pathogenic | 0 | 0 | 0 |
| rs371295780 | pathogenic | 0 | 0 | 0 |
| rs387906830 | pathogenic | 1 | 1 | 0 |
| rs771364038 | pathogenic | 0 | 1 | 0 |
| rs772731615 | pathogenic | 0 | 0 | 0 |
| rs137853063 | pathogenic | 0 | 0 | 0 |
| rs387907076 | pathogenic | 0 | 1 | 0 |
| rs35460768 | benign | 0 | 0 | 0 |
| rs2278196 | benign | 1 | 1 | 1 |
| rs119103275 | pathogenic | 0 | 0 | 0 |
| rs119103276 | pathogenic | 1 | 1 | 1 |
| rs374402066 | pathogenic | 1 | 1 | 1 |

| | | | | |
|---|---|---|---|---|
| rs119103274 | pathogenic | 0 | 0 | 0 |
| rs2274064 | benign | 0 | 0 | 0 |
| rs267607023 | pathogenic | 1 | 1 | 0 |
| rs142117489 | benign | 1 | 1 | 0 |
| rs371258270 | benign | 0 | 0 | 0 |
| rs34716057 | benign | 0 | 0 | 0 |
| rs201778606 | benign | 0 | 0 | 0 |
| rs201438792 | benign | 0 | 0 | 0 |
| rs140547417 | benign | 0 | 0 | 0 |
| rs5881 | benign | 0 | 0 | 0 |
| rs752298084 | benign | 0 | 0 | 0 |
| rs34855278 | benign | 0 | 0 | 0 |
| rs5880 | benign | 1 | 1 | 0 |
| rs5882 | benign | 0 | 0 | 0 |
| rs1800777 | benign | 0 | 0 | 0 |
| rs577301331 | benign | 0 | 0 | 0 |
| rs137853313 | pathogenic | 0 | 0 | 0 |
| rs28935169 | pathogenic | 0 | 0 | 0 |
| rs137853315 | pathogenic | 0 | 0 | 0 |
| rs137853314 | pathogenic | 0 | 0 | 0 |
| rs28935469 | pathogenic | 0 | 0 | 0 |
| rs137853318 | pathogenic | 0 | 0 | 0 |
| rs28935470 | pathogenic | 0 | 0 | 0 |
| rs3729986 | benign | 0 | 0 | 0 |
| rs730880619 | pathogenic | 0 | 0 | 0 |
| rs201012766 | benign | 0 | 0 | 0 |
| rs727503218 | pathogenic | 0 | 0 | 0 |
| rs11570052 | benign | 0 | 0 | 0 |
| rs730880623 | pathogenic | 0 | 0 | 0 |
| rs138753870 | benign | 0 | 0 | 0 |
| rs397516068 | pathogenic | 0 | 0 | 0 |
| rs3729989 | benign | 1 | 1 | 1 |
| rs397516074 | pathogenic | 0 | 0 | 0 |
| rs561449819 | pathogenic | 0 | 0 | 0 |
| rs121917779 | benign | 0 | 0 | 0 |
| rs142881576 | pathogenic | 0 | 0 | 0 |
| rs77384282 | benign | 0 | 0 | 0 |
| rs201865787 | benign | 0 | 0 | 0 |
| rs142244640 | benign | 0 | 0 | 0 |
| rs200488706 | benign | 0 | 0 | 0 |
| rs137853291 | pathogenic | 1 | 1 | 0 |
| rs28933990 | pathogenic | 1 | 1 | 0 |
| rs140570189 | benign | 0 | 0 | 0 |
| rs201588178 | benign | 0 | 0 | 0 |
| rs104893846 | pathogenic | 0 | 0 | 0 |
| rs754545360 | pathogenic | 0 | 1 | 0 |
| rs104893851 | pathogenic | 0 | 1 | 0 |
| rs104893849 | pathogenic | 0 | 0 | 0 |
| rs571038432 | pathogenic | 1 | 1 | 0 |
| rs2270655 | benign | 1 | 1 | 1 |
| rs315952 | benign | 0 | 0 | 0 |
| rs151176548 | benign | 0 | 0 | 0 |
| rs122459147 | pathogenic | 1 | 1 | 0 |
| rs122458140 | pathogenic | 0 | 0 | 0 |

| | | | | |
|---|---|---|---|---|
| rs122458142 | pathogenic | 1 | 1 | 0 |
| rs267606812 | pathogenic | 1 | 1 | 0 |
| rs267606813 | pathogenic | 1 | 1 | 0 |
| rs122458143 | pathogenic | 0 | 0 | 0 |
| rs122459146 | pathogenic | 0 | 0 | 0 |
| rs122458144 | pathogenic | 0 | 0 | 0 |
| rs122458145 | pathogenic | 0 | 0 | 0 |
| rs113994007 | pathogenic | 1 | 1 | 0 |
| rs758746181 | pathogenic | 1 | 1 | 1 |
| rs145034527 | pathogenic | 1 | 1 | 0 |
| rs119468006 | pathogenic | 1 | 1 | 0 |
| rs201908721 | pathogenic | 0 | 0 | 0 |
| rs755933881 | pathogenic | 1 | 1 | 0 |
| rs771578775 | pathogenic | 0 | 0 | 0 |
| rs17849927 | benign | 0 | 0 | 0 |
| rs139133094 | benign | 0 | 0 | 0 |
| rs12593 | benign | 0 | 0 | 0 |
| rs578189699 | pathogenic | 1 | 1 | 0 |
| rs119468008 | pathogenic | 0 | 0 | 0 |
| rs768524626 | benign | 0 | 0 | 0 |
| rs119468009 | pathogenic | 0 | 0 | 0 |
| rs119468004 | pathogenic | 0 | 0 | 0 |
| rs199874519 | pathogenic | 0 | 0 | 0 |
| rs376478331 | pathogenic | 0 | 0 | 0 |
| rs61995958 | pathogenic | 0 | 0 | 0 |
| rs199714731 | benign | 0 | 0 | 0 |
| rs149185954 | benign | 0 | 0 | 0 |
| rs121918353 | pathogenic | 0 | 0 | 0 |
| rs1131695 | benign | 0 | 0 | 0 |
| rs28939668 | pathogenic | 0 | 0 | 0 |
| rs758432471 | pathogenic | 1 | 1 | 1 |
| rs752746786 | pathogenic | 1 | 1 | 1 |
| rs28935471 | pathogenic | 0 | 0 | 0 |
| rs80338837 | pathogenic | 0 | 0 | 0 |
| rs137853312 | pathogenic | 0 | 0 | 0 |
| rs28935472 | pathogenic | 0 | 0 | 0 |
| rs28935473 | pathogenic | 0 | 0 | 0 |
| rs267606752 | pathogenic | 0 | 0 | 0 |
| rs3917973 | benign | 0 | 0 | 0 |
| rs3918018 | benign | 0 | 0 | 0 |
| rs757976755 | pathogenic | 0 | 0 | 0 |
| rs121918329 | pathogenic | 1 | 1 | 0 |
| rs770114459 | pathogenic | 0 | 0 | 0 |
| rs6662272 | benign | 0 | 0 | 0 |
| rs121918242 | pathogenic | 1 | 1 | 0 |
| rs121918240 | pathogenic | 1 | 1 | 0 |
| rs200094982 | pathogenic | 0 | 0 | 0 |
| rs121918241 | pathogenic | 0 | 0 | 0 |
| rs140522266 | pathogenic | 1 | 1 | 0 |
| rs121918243 | pathogenic | 1 | 1 | 0 |
| rs370596113 | pathogenic | 1 | 1 | 0 |
| rs398124295 | pathogenic | 1 | 1 | 0 |
| rs587776889 | pathogenic | 1 | 1 | 0 |
| rs747527726 | pathogenic | 1 | 1 | 0 |

| | | | | |
|---|---|---|---|---|
| rs112631212 | benign | 0 | 0 | 0 |
| rs104894605 | pathogenic | 0 | 0 | 0 |
| rs104894607 | pathogenic | 0 | 0 | 0 |
| rs104894606 | pathogenic | 0 | 0 | 0 |
| rs143873873 | benign | 0 | 0 | 0 |
| rs104894388 | pathogenic | 1 | 1 | 0 |
| rs104894391 | pathogenic | 1 | 1 | 0 |
| rs5350 | benign | 1 | 1 | 1 |
| rs104894390 | pathogenic | 1 | 1 | 1 |
| rs5346 | benign | 1 | 1 | 1 |
| rs121912500 | pathogenic | 0 | 0 | 0 |
| rs74315450 | pathogenic | 1 | 1 | 0 |
| rs121913396 | pathogenic | 1 | 1 | 1 |
| rs28931588 | pathogenic | 1 | 1 | 1 |
| rs121913400 | pathogenic | 0 | 0 | 0 |
| rs28931589 | pathogenic | 1 | 1 | 1 |
| rs121913403 | pathogenic | 0 | 0 | 0 |
| rs121913413 | pathogenic | 0 | 0 | 0 |
| rs760107623 | pathogenic | 1 | 1 | 0 |
| rs16866378 | benign | 1 | 1 | 0 |
| rs201025290 | pathogenic | 1 | 1 | 0 |
| rs12720356 | benign | 1 | 1 | 0 |
| rs55882956 | benign | 0 | 1 | 0 |
| rs4986933 | benign | 1 | 1 | 0 |
| rs2020957 | benign | 1 | 1 | 1 |
| rs1800124 | benign | 1 | 1 | 1 |
| rs150077735 | benign | 1 | 1 | 1 |
| rs201419000 | benign | 0 | 0 | 0 |
| rs772044176 | pathogenic | 0 | 0 | 0 |
| rs138346388 | benign | 0 | 0 | 0 |
| rs121964916 | pathogenic | 0 | 0 | 0 |
| rs112534524 | pathogenic | 0 | 0 | 0 |
| rs182078921 | benign | 1 | 1 | 1 |
| rs113273712 | benign | 0 | 0 | 0 |
| rs74817407 | benign | 1 | 1 | 1 |
| rs61733901 | benign | 1 | 1 | 1 |
| rs121964912 | pathogenic | 1 | 1 | 1 |
| rs41278047 | benign | 1 | 1 | 1 |
| rs7437875 | benign | 1 | 1 | 1 |
| rs113460688 | benign | 1 | 1 | 1 |
| rs754572081 | benign | 1 | 1 | 1 |
| rs201082692 | benign | 1 | 1 | 1 |
| rs202085803 | benign | 1 | 1 | 1 |
| rs141320511 | benign | 0 | 0 | 0 |
| rs78501403 | benign | 0 | 0 | 0 |
| rs28939675 | pathogenic | 0 | 0 | 0 |
| rs74315522 | pathogenic | 0 | 0 | 0 |
| rs121434639 | pathogenic | 1 | 1 | 0 |
| rs121434640 | pathogenic | 1 | 1 | 1 |
| rs757282628 | benign | 1 | 1 | 1 |
| rs386834010 | pathogenic | 0 | 0 | 0 |
| rs200863680 | pathogenic | 0 | 0 | 0 |
| rs193919336 | pathogenic | 1 | 1 | 0 |
| rs386834011 | pathogenic | 0 | 0 | 0 |

| | | | | |
|---|---|---|---|---|
| rs28940869 | pathogenic | 0 | 0 | 0 |
| rs386834014 | pathogenic | 1 | 1 | 0 |
| rs267606960 | pathogenic | 0 | 0 | 0 |
| rs28942068 | pathogenic | 0 | 0 | 0 |
| rs193919335 | pathogenic | 0 | 0 | 0 |
| rs74374973 | pathogenic | 0 | 0 | 0 |
| rs386834018 | pathogenic | 0 | 0 | 0 |
| rs267606962 | pathogenic | 0 | 0 | 0 |
| rs6659553 | benign | 0 | 0 | 0 |
| rs29281 | benign | 0 | 0 | 0 |
| rs782265090 | benign | 0 | 0 | 0 |
| rs35910969 | pathogenic | 0 | 0 | 0 |
| rs121909113 | pathogenic | 1 | 1 | 1 |
| rs29001571 | pathogenic | 1 | 1 | 1 |
| rs587781250 | pathogenic | 1 | 1 | 1 |
| rs28939680 | pathogenic | 1 | 1 | 1 |
| rs28939681 | pathogenic | 1 | 1 | 1 |
| rs121909112 | pathogenic | 1 | 1 | 1 |
| rs28937568 | pathogenic | 1 | 1 | 1 |
| rs1799895 | pathogenic | 0 | 0 | 0 |
| rs386833431 | pathogenic | 1 | 1 | 1 |
| rs121964905 | pathogenic | 1 | 1 | 1 |
| rs121964906 | pathogenic | 1 | 1 | 1 |
| rs11551664 | pathogenic | 1 | 1 | 1 |
| rs104894705 | pathogenic | 1 | 1 | 1 |
| rs104893805 | pathogenic | 0 | 0 | 0 |
| rs104894205 | pathogenic | 0 | 0 | 0 |
| rs104894204 | pathogenic | 1 | 1 | 0 |
| rs62642926 | pathogenic | 0 | 0 | 0 |
| rs5030841 | pathogenic | 1 | 1 | 1 |
| rs75193786 | pathogenic | 1 | 1 | 1 |
| rs62508682 | pathogenic | 0 | 0 | 0 |
| rs3850625 | benign | 1 | 1 | 1 |
| rs141908793 | benign | 0 | 0 | 0 |
| rs187864727 | benign | 0 | 0 | 0 |
| rs548909101 | benign | 0 | 0 | 0 |
| rs3825394 | benign | 0 | 1 | 0 |
| rs143548402 | benign | 0 | 0 | 0 |
| rs267607146 | pathogenic | 0 | 0 | 0 |
| rs267607144 | pathogenic | 0 | 0 | 0 |
| rs267607148 | pathogenic | 1 | 1 | 1 |
| rs267607143 | pathogenic | 0 | 0 | 0 |
| rs267607145 | pathogenic | 0 | 0 | 0 |
| rs121912636 | pathogenic | 1 | 1 | 1 |
| rs121912634 | pathogenic | 1 | 1 | 1 |
| rs570675468 | benign | 0 | 0 | 0 |
| rs368861241 | pathogenic | 1 | 1 | 1 |
| rs104894528 | pathogenic | 1 | 1 | 1 |
| rs77493891 | benign | 0 | 0 | 0 |
| rs62640903 | benign | 0 | 0 | 0 |
| rs149926091 | benign | 1 | 1 | 1 |
| rs641081 | benign | 0 | 0 | 0 |
| rs4930199 | benign | 0 | 0 | 0 |
| rs121913020 | pathogenic | 0 | 0 | 0 |

| | | | | |
|---|---|---|---|---|
| rs1799793 | benign | 0 | 0 | 0 |
| rs121913016 | pathogenic | 0 | 0 | 0 |
| rs121913025 | pathogenic | 0 | 0 | 0 |
| rs121913019 | pathogenic | 0 | 0 | 0 |
| rs376556895 | pathogenic | 1 | 1 | 1 |
| rs121913024 | pathogenic | 1 | 1 | 1 |
| rs121913021 | pathogenic | 0 | 0 | 0 |
| rs121913023 | pathogenic | 0 | 0 | 0 |
| rs41556519 | pathogenic | 0 | 0 | 0 |
| rs121913022 | pathogenic | 0 | 0 | 0 |
| rs144564120 | pathogenic | 0 | 0 | 0 |
| rs121913026 | pathogenic | 1 | 1 | 1 |
| rs121913018 | pathogenic | 1 | 1 | 1 |
| rs142954643 | benign | 0 | 0 | 0 |
| rs76655496 | benign | 0 | 0 | 0 |
| rs104894371 | pathogenic | 1 | 1 | 0 |
| rs137964936 | benign | 0 | 0 | 0 |
| rs759736132 | pathogenic | 0 | 0 | 0 |
| rs387906595 | pathogenic | 1 | 1 | 0 |
| rs768248216 | pathogenic | 0 | 0 | 0 |
| rs548231008 | benign | 0 | 0 | 0 |
| rs757094384 | benign | 0 | 0 | 0 |
| rs867529 | benign | 0 | 0 | 0 |
| rs13045 | benign | 1 | 1 | 1 |
| rs121908570 | pathogenic | 0 | 0 | 0 |
| rs201872586 | benign | 1 | 1 | 0 |
| rs149092990 | benign | 0 | 0 | 0 |
| rs115585711 | benign | 0 | 0 | 0 |
| rs137909416 | benign | 1 | 1 | 0 |
| rs143388949 | benign | 1 | 1 | 1 |
| rs73137716 | benign | 1 | 1 | 1 |
| rs121908947 | pathogenic | 1 | 1 | 1 |
| rs398122904 | pathogenic | 1 | 1 | 1 |
| rs35715497 | benign | 1 | 1 | 0 |
| rs35462421 | benign | 0 | 0 | 0 |
| rs119475043 | pathogenic | 0 | 0 | 0 |
| rs119475042 | pathogenic | 0 | 0 | 0 |
| rs1076669 | benign | 1 | 1 | 0 |
| rs141146885 | benign | 0 | 0 | 0 |
| rs3026906 | pathogenic | 0 | 0 | 0 |
| rs75362221 | benign | 1 | 1 | 1 |
| rs138472116 | benign | 1 | 1 | 1 |
| rs371040282 | pathogenic | 1 | 1 | 1 |
| rs4988496 | benign | 0 | 0 | 0 |
| rs121918117 | pathogenic | 1 | 1 | 1 |
| rs4988498 | benign | 0 | 0 | 0 |
| rs5035 | benign | 0 | 0 | 0 |
| rs121918322 | pathogenic | 1 | 1 | 1 |
| rs370359540 | benign | 0 | 0 | 0 |
| rs758972589 | pathogenic | 0 | 0 | 0 |
| rs144856889 | benign | 0 | 0 | 0 |
| rs121434623 | pathogenic | 1 | 1 | 1 |
| rs61735600 | benign | 0 | 0 | 0 |
| rs72550870 | benign | 1 | 1 | 1 |

| | | | | |
|---|---|---|---|---|
| rs56392418 | benign | 0 | 0 | 0 |
| rs41307788 | benign | 1 | 1 | 1 |
| rs387907239 | pathogenic | 1 | 1 | 0 |
| rs28931584 | pathogenic | 0 | 0 | 0 |
| rs121912745 | pathogenic | 0 | 0 | 0 |
| rs121912744 | pathogenic | 0 | 0 | 0 |
| rs121912754 | pathogenic | 1 | 1 | 1 |
| rs121912746 | pathogenic | 0 | 0 | 0 |
| rs121912757 | pathogenic | 0 | 0 | 0 |
| rs121912758 | pathogenic | 0 | 0 | 0 |
| rs121912748 | pathogenic | 0 | 0 | 0 |
| rs121912755 | pathogenic | 0 | 0 | 0 |
| rs121912741 | pathogenic | 0 | 0 | 0 |
| rs121912753 | pathogenic | 0 | 0 | 0 |
| rs121912750 | pathogenic | 0 | 0 | 0 |
| rs2285644 | pathogenic | 0 | 0 | 0 |
| rs121912751 | pathogenic | 0 | 0 | 0 |
| rs5026 | benign | 0 | 0 | 0 |
| rs121912759 | pathogenic | 0 | 0 | 0 |
| rs28931585 | pathogenic | 0 | 0 | 0 |
| rs17122769 | benign | 0 | 0 | 0 |
| rs28362322 | benign | 1 | 1 | 1 |
| rs3824874 | benign | 0 | 0 | 0 |
| rs146004831 | benign | 0 | 0 | 0 |
| rs142155860 | benign | 0 | 0 | 0 |
| rs121434404 | pathogenic | 0 | 0 | 0 |
| rs61735578 | benign | 1 | 1 | 1 |
| rs558018 | benign | 0 | 0 | 0 |
| rs61735577 | benign | 1 | 1 | 1 |
| rs76784113 | benign | 0 | 0 | 0 |
| rs199469640 | pathogenic | 0 | 0 | 0 |
| rs199469628 | pathogenic | 1 | 1 | 0 |
| rs199469629 | pathogenic | 0 | 1 | 0 |
| rs387907156 | pathogenic | 0 | 1 | 0 |
| rs199469643 | pathogenic | 0 | 0 | 0 |
| rs199469646 | pathogenic | 0 | 0 | 0 |
| rs766294940 | pathogenic | 0 | 0 | 0 |
| rs387907347 | pathogenic | 0 | 0 | 0 |
| rs137853590 | pathogenic | 1 | 1 | 1 |
| rs137853589 | pathogenic | 1 | 1 | 1 |
| rs137853588 | pathogenic | 1 | 1 | 1 |
| rs55909005 | benign | 0 | 0 | 0 |
| rs4581 | benign | 0 | 0 | 0 |
| rs63750123 | benign | 0 | 0 | 0 |
| rs10254120 | benign | 0 | 0 | 0 |
| rs121434629 | pathogenic | 1 | 1 | 0 |
| rs200313585 | benign | 0 | 0 | 0 |
| rs143162541 | benign | 1 | 1 | 0 |
| rs760615315 | benign | 0 | 0 | 0 |
| rs768488890 | benign | 1 | 1 | 1 |
| rs116373169 | benign | 1 | 1 | 0 |
| rs63750871 | pathogenic | 1 | 1 | 0 |
| rs116349687 | benign | 0 | 0 | 0 |
| rs375289386 | benign | 0 | 0 | 0 |

| | | | | |
|---|---|---|---|---|
| rs760228510 | pathogenic | 1 | 1 | 1 |
| rs587779343 | pathogenic | 1 | 1 | 1 |
| rs573125799 | benign | 0 | 0 | 0 |
| rs1805322 | benign | 0 | 0 | 0 |
| rs267608153 | pathogenic | 0 | 0 | 0 |
| rs139194813 | benign | 0 | 0 | 0 |
| rs530021751 | benign | 0 | 0 | 0 |
| rs200640585 | pathogenic | 0 | 0 | 0 |
| rs139438201 | benign | 0 | 0 | 0 |
| rs200513014 | benign | 0 | 0 | 0 |
| rs567102013 | benign | 0 | 0 | 0 |
| rs121908608 | pathogenic | 0 | 0 | 0 |
| rs121908606 | pathogenic | 0 | 0 | 0 |
| rs17103671 | benign | 0 | 0 | 0 |
| rs34725281 | benign | 0 | 0 | 0 |
| rs104894421 | pathogenic | 1 | 1 | 0 |
| rs121918145 | pathogenic | 1 | 1 | 1 |
| rs121918152 | pathogenic | 1 | 1 | 1 |
| rs56379106 | benign | 0 | 0 | 0 |
| rs4987076 | benign | 0 | 0 | 0 |
| rs370243877 | pathogenic | 0 | 0 | 0 |
| rs17679445 | benign | 1 | 1 | 1 |
| rs550423482 | pathogenic | 1 | 1 | 1 |
| rs104894629 | pathogenic | 1 | 1 | 1 |
| rs773450573 | pathogenic | 1 | 1 | 1 |
| rs116679717 | benign | 0 | 0 | 0 |
| rs372000848 | pathogenic | 0 | 0 | 0 |
| rs138595973 | benign | 0 | 0 | 0 |
| rs115244417 | benign | 0 | 0 | 0 |
| rs11575292 | benign | 0 | 0 | 0 |
| rs137853211 | pathogenic | 1 | 1 | 1 |
| rs137853207 | pathogenic | 0 | 0 | 0 |
| rs137853210 | pathogenic | 1 | 1 | 1 |
| rs6262 | benign | 0 | 0 | 0 |
| rs6263 | benign | 0 | 1 | 0 |
| rs137853208 | pathogenic | 0 | 1 | 0 |
| rs137853212 | pathogenic | 1 | 1 | 0 |
| rs137853209 | pathogenic | 1 | 1 | 1 |
| rs201951824 | pathogenic | 0 | 0 | 0 |
| rs11575542 | benign | 0 | 0 | 0 |
| rs199681788 | benign | 0 | 1 | 0 |
| rs748835902 | benign | 1 | 1 | 1 |
| rs753719949 | benign | 0 | 0 | 0 |
| rs267606647 | pathogenic | 0 | 0 | 0 |
| rs267606648 | pathogenic | 1 | 1 | 1 |
| rs267606643 | pathogenic | 0 | 1 | 0 |
| rs559947967 | pathogenic | 1 | 1 | 0 |
| rs267606645 | pathogenic | 1 | 1 | 1 |
| rs12116440 | benign | 0 | 0 | 0 |
| rs121908539 | pathogenic | 0 | 0 | 0 |
| rs137853096 | pathogenic | 1 | 1 | 0 |
| rs148363262 | benign | 1 | 1 | 0 |
| rs587777442 | pathogenic | 0 | 0 | 0 |
| rs25640 | benign | 1 | 1 | 1 |

| | | | | |
|---|---|---|---|---|
| rs28943589 | benign | 0 | 0 | 0 |
| rs1143650 | benign | 1 | 1 | 1 |
| rs150886026 | benign | 0 | 1 | 0 |
| rs121908133 | pathogenic | 1 | 1 | 0 |
| rs121908131 | pathogenic | 0 | 0 | 0 |
| rs754477154 | pathogenic | 0 | 0 | 0 |
| rs747334007 | benign | 0 | 0 | 0 |
| rs121908132 | pathogenic | 0 | 0 | 0 |
| rs104894103 | pathogenic | 1 | 1 | 0 |
| rs140088149 | benign | 0 | 0 | 0 |
| rs149803148 | benign | 0 | 0 | 0 |
| rs11064480 | benign | 0 | 0 | 0 |
| rs74435397 | pathogenic | 1 | 1 | 1 |
| rs587783626 | pathogenic | 1 | 1 | 1 |
| rs148211042 | pathogenic | 1 | 1 | 1 |
| rs17849502 | benign | 0 | 0 | 0 |
| rs13306575 | pathogenic | 1 | 1 | 1 |
| rs201426733 | benign | 0 | 0 | 0 |
| rs182278784 | benign | 1 | 1 | 1 |
| rs77786415 | benign | 1 | 1 | 1 |
| rs200192119 | benign | 1 | 1 | 1 |
| rs12877501 | benign | 1 | 1 | 1 |
| rs77985244 | benign | 1 | 1 | 1 |
| rs5907 | pathogenic | 1 | 1 | 0 |
| rs2305178 | benign | 0 | 0 | 0 |
| rs3135867 | benign | 0 | 0 | 0 |
| rs121913482 | pathogenic | 1 | 1 | 1 |
| rs121913483 | pathogenic | 1 | 1 | 1 |
| rs4647924 | pathogenic | 1 | 1 | 1 |
| rs121913111 | pathogenic | 1 | 1 | 1 |
| rs104894002 | pathogenic | 0 | 0 | 0 |
| rs104894001 | pathogenic | 0 | 0 | 0 |
| rs75932628 | benign | 0 | 0 | 0 |
| rs201258663 | pathogenic | 0 | 0 | 0 |
| rs104893998 | pathogenic | 0 | 0 | 0 |
| rs2234253 | benign | 1 | 1 | 0 |
| rs121908402 | pathogenic | 1 | 1 | 0 |
| rs1805389 | benign | 0 | 0 | 0 |
| rs1805388 | benign | 0 | 0 | 0 |
| rs748385144 | pathogenic | 0 | 0 | 0 |
| rs104894420 | pathogenic | 0 | 0 | 0 |
| rs104894418 | pathogenic | 0 | 0 | 0 |
| rs61731910 | benign | 0 | 0 | 0 |
| rs758723288 | pathogenic | 1 | 1 | 0 |
| rs201865375 | pathogenic | 1 | 1 | 0 |
| rs565090080 | pathogenic | 1 | 1 | 1 |
| rs113042166 | benign | 0 | 0 | 0 |
| rs61736326 | benign | 1 | 1 | 1 |
| rs12030699 | benign | 1 | 1 | 1 |
| rs121913578 | pathogenic | 1 | 1 | 1 |
| rs61739582 | benign | 0 | 0 | 0 |
| rs745983207 | pathogenic | 0 | 0 | 0 |
| rs763524380 | benign | 0 | 0 | 0 |
| rs121912792 | pathogenic | 0 | 0 | 0 |

| | | | | |
|---|---|---|---|---|
| rs121912793 | pathogenic | 0 | 0 | 0 |
| rs387906599 | pathogenic | 0 | 0 | 0 |
| rs121912794 | pathogenic | 0 | 1 | 0 |
| rs121912795 | pathogenic | 0 | 0 | 0 |
| rs121912796 | pathogenic | 1 | 1 | 0 |
| rs121912797 | pathogenic | 0 | 1 | 0 |
| rs121912798 | pathogenic | 1 | 1 | 1 |
| rs80359812 | pathogenic | 1 | 1 | 0 |
| rs776583130 | benign | 1 | 1 | 0 |
| rs80359814 | pathogenic | 1 | 1 | 0 |
| rs202060209 | pathogenic | 1 | 1 | 0 |
| rs267607061 | pathogenic | 1 | 1 | 0 |
| rs267607060 | pathogenic | 0 | 0 | 0 |
| rs80359816 | pathogenic | 1 | 1 | 0 |
| rs80359818 | pathogenic | 1 | 1 | 0 |
| rs387907313 | pathogenic | 0 | 0 | 0 |
| rs587784395 | benign | 0 | 0 | 0 |
| rs5811 | benign | 0 | 0 | 0 |
| rs121909740 | pathogenic | 1 | 1 | 0 |
| rs148518827 | benign | 1 | 1 | 0 |
| rs121909739 | pathogenic | 1 | 1 | 0 |
| rs780528770 | benign | 1 | 1 | 0 |
| rs13306758 | pathogenic | 0 | 0 | 0 |
| rs267607059 | pathogenic | 0 | 0 | 0 |
| rs61730638 | benign | 0 | 0 | 0 |
| rs104894533 | pathogenic | 0 | 0 | 0 |
| rs104894527 | pathogenic | 0 | 0 | 0 |
| rs200503569 | pathogenic | 1 | 1 | 1 |
| rs80338700 | pathogenic | 1 | 1 | 1 |
| rs765433263 | pathogenic | 1 | 1 | 1 |
| rs104894530 | pathogenic | 1 | 1 | 1 |
| rs80338701 | pathogenic | 1 | 1 | 1 |
| rs141498002 | pathogenic | 1 | 1 | 1 |
| rs104894525 | pathogenic | 0 | 0 | 0 |
| rs80338703 | pathogenic | 0 | 0 | 0 |
| rs28936415 | pathogenic | 1 | 1 | 0 |
| rs150719105 | pathogenic | 0 | 0 | 0 |
| rs148032587 | pathogenic | 0 | 0 | 0 |
| rs150577656 | pathogenic | 0 | 0 | 0 |
| rs190521996 | pathogenic | 0 | 0 | 0 |
| rs104894526 | pathogenic | 1 | 1 | 1 |
| rs80338704 | pathogenic | 0 | 0 | 0 |
| rs199562225 | pathogenic | 1 | 1 | 1 |
| rs532870929 | pathogenic | 0 | 0 | 0 |
| rs398123309 | pathogenic | 0 | 0 | 0 |
| rs78290141 | pathogenic | 0 | 0 | 0 |
| rs80338705 | pathogenic | 0 | 0 | 0 |
| rs104894531 | pathogenic | 1 | 1 | 1 |
| rs80338706 | pathogenic | 0 | 0 | 0 |
| rs80338707 | pathogenic | 0 | 0 | 0 |
| rs80338708 | pathogenic | 0 | 0 | 0 |
| rs80338709 | pathogenic | 0 | 0 | 0 |
| rs104893833 | benign | 0 | 0 | 0 |
| rs61744585 | benign | 0 | 0 | 0 |

| | | | | |
|---|---|---|---|---|
| rs73569515 | benign | 0 | 0 | 0 |
| rs398123373 | pathogenic | 0 | 0 | 0 |
| rs756854513 | pathogenic | 0 | 0 | 0 |
| rs58232698 | benign | 0 | 0 | 0 |
| rs9600455 | benign | 0 | 0 | 0 |
| rs201016593 | pathogenic | 0 | 0 | 0 |
| rs756039188 | pathogenic | 0 | 0 | 0 |
| rs121908024 | pathogenic | 0 | 0 | 0 |
| rs373144619 | benign | 0 | 0 | 0 |
| rs138078086 | benign | 0 | 0 | 0 |
| rs376207800 | pathogenic | 0 | 0 | 0 |
| rs749038326 | pathogenic | 1 | 1 | 0 |
| rs372845091 | benign | 0 | 0 | 0 |
| rs377271627 | pathogenic | 0 | 0 | 0 |
| rs563390335 | pathogenic | 0 | 0 | 0 |
| rs13306510 | benign | 0 | 0 | 0 |
| rs764042910 | pathogenic | 0 | 0 | 0 |
| rs201384282 | benign | 1 | 1 | 1 |
| rs730882086 | pathogenic | 1 | 1 | 1 |
| rs756613387 | pathogenic | 1 | 1 | 1 |
| rs121908028 | benign | 1 | 1 | 0 |
| rs773328511 | benign | 0 | 0 | 0 |
| rs749220643 | pathogenic | 1 | 1 | 0 |
| rs730882089 | pathogenic | 0 | 0 | 0 |
| rs146651743 | benign | 0 | 0 | 0 |
| rs148698650 | pathogenic | 0 | 0 | 0 |
| rs140241383 | benign | 0 | 0 | 0 |
| rs757252110 | pathogenic | 0 | 0 | 0 |
| rs13306512 | pathogenic | 1 | 1 | 1 |
| rs121908043 | pathogenic | 1 | 1 | 1 |
| rs773658037 | pathogenic | 0 | 0 | 0 |
| rs774439908 | pathogenic | 0 | 0 | 0 |
| rs779732323 | pathogenic | 1 | 1 | 0 |
| rs749780672 | pathogenic | 0 | 0 | 0 |
| rs375651668 | benign | 1 | 1 | 1 |
| rs193922568 | pathogenic | 1 | 1 | 0 |
| rs750363970 | benign | 1 | 1 | 0 |
| rs5932 | benign | 0 | 0 | 0 |
| rs755154048 | pathogenic | 1 | 1 | 0 |
| rs759876319 | pathogenic | 0 | 0 | 0 |
| rs764550980 | pathogenic | 1 | 1 | 0 |
| rs779940524 | pathogenic | 0 | 0 | 0 |
| rs563382937 | pathogenic | 1 | 1 | 1 |
| rs28942083 | pathogenic | 0 | 1 | 0 |
| rs770744861 | pathogenic | 0 | 1 | 0 |
| rs121908031 | pathogenic | 0 | 1 | 0 |
| rs776217028 | pathogenic | 0 | 1 | 0 |
| rs763625913 | pathogenic | 0 | 0 | 0 |
| rs3736772 | benign | 0 | 0 | 0 |
| rs104894696 | benign | 1 | 1 | 1 |
| rs121918177 | pathogenic | 0 | 1 | 0 |
| rs121918181 | pathogenic | 1 | 1 | 0 |
| rs121918174 | pathogenic | 0 | 1 | 0 |
| rs121918182 | pathogenic | 0 | 0 | 0 |

| | | | | |
|---|---|---|---|---|
| rs121918175 | pathogenic | 1 | 1 | 1 |
| rs121918185 | pathogenic | 1 | 1 | 1 |
| rs121918173 | pathogenic | 1 | 1 | 0 |
| rs121918178 | pathogenic | 1 | 1 | 1 |
| rs121918179 | pathogenic | 1 | 1 | 1 |
| rs121918183 | pathogenic | 0 | 0 | 0 |
| rs121918176 | pathogenic | 1 | 1 | 1 |
| rs121918172 | pathogenic | 0 | 1 | 0 |
| rs121918184 | pathogenic | 0 | 0 | 0 |
| rs121434560 | pathogenic | 1 | 1 | 0 |
| rs121434561 | pathogenic | 1 | 1 | 0 |
| rs3218716 | pathogenic | 1 | 1 | 1 |
| rs145532615 | benign | 0 | 0 | 0 |
| rs149840927 | pathogenic | 0 | 0 | 0 |
| rs730880762 | pathogenic | 0 | 0 | 0 |
| rs730880764 | pathogenic | 0 | 0 | 0 |
| rs730880765 | pathogenic | 0 | 0 | 0 |
| rs587782962 | pathogenic | 0 | 0 | 0 |
| rs141735183 | pathogenic | 0 | 0 | 0 |
| rs397516187 | pathogenic | 0 | 0 | 0 |
| rs727505026 | pathogenic | 0 | 0 | 0 |
| rs529700838 | pathogenic | 0 | 0 | 0 |
| rs540263945 | pathogenic | 0 | 0 | 0 |
| rs149103761 | benign | 0 | 0 | 0 |
| rs730880906 | pathogenic | 0 | 0 | 0 |
| rs730880785 | pathogenic | 0 | 0 | 0 |
| rs397516199 | pathogenic | 0 | 0 | 0 |
| rs730880725 | benign | 0 | 0 | 0 |
| rs730880796 | pathogenic | 0 | 0 | 0 |
| rs201307101 | benign | 0 | 0 | 0 |
| rs397516214 | pathogenic | 0 | 0 | 0 |
| rs3729823 | benign | 0 | 0 | 0 |
| rs397516218 | pathogenic | 0 | 0 | 0 |
| rs121913647 | pathogenic | 0 | 0 | 0 |
| rs373514686 | pathogenic | 0 | 0 | 0 |
| rs763538103 | pathogenic | 0 | 0 | 0 |
| rs121913650 | pathogenic | 0 | 0 | 0 |
| rs139222507 | pathogenic | 0 | 0 | 0 |
| rs369940645 | pathogenic | 0 | 0 | 0 |
| rs763073072 | pathogenic | 0 | 0 | 0 |
| rs145734640 | pathogenic | 0 | 0 | 0 |
| rs730880821 | pathogenic | 0 | 0 | 0 |
| rs28933098 | pathogenic | 0 | 0 | 0 |
| rs372381770 | pathogenic | 0 | 0 | 0 |
| rs146796870 | benign | 0 | 0 | 0 |
| rs104894485 | pathogenic | 1 | 1 | 1 |
| rs35407712 | benign | 0 | 0 | 0 |
| rs35842750 | benign | 0 | 0 | 0 |
| rs2721191 | benign | 0 | 0 | 0 |
| rs35098923 | benign | 0 | 0 | 0 |
| rs34293591 | benign | 1 | 1 | 0 |
| rs36023964 | benign | 0 | 0 | 0 |
| rs4251691 | benign | 0 | 0 | 0 |
| rs34666647 | benign | 0 | 0 | 0 |

| | | | | |
|---|---|---|---|---|
| rs4925828 | benign | 0 | 0 | 0 |
| rs35348691 | benign | 0 | 0 | 0 |
| rs36078464 | benign | 0 | 0 | 0 |
| rs34915097 | benign | 0 | 0 | 0 |
| rs35101495 | benign | 0 | 0 | 0 |
| rs141568342 | pathogenic | 0 | 0 | 0 |
| rs1800571 | pathogenic | 0 | 0 | 0 |
| rs140566084 | benign | 1 | 1 | 1 |
| rs727503791 | pathogenic | 1 | 1 | 0 |
| rs545215807 | pathogenic | 0 | 0 | 0 |
| rs113994168 | pathogenic | 0 | 0 | 0 |
| rs113994167 | pathogenic | 0 | 0 | 0 |
| rs200788251 | pathogenic | 0 | 0 | 0 |
| rs200573371 | pathogenic | 0 | 0 | 0 |
| rs771874163 | pathogenic | 1 | 1 | 1 |
| rs112406105 | pathogenic | 1 | 1 | 1 |
| rs118204015 | pathogenic | 1 | 1 | 1 |
| rs113994169 | pathogenic | 0 | 0 | 0 |
| rs118204018 | pathogenic | 0 | 1 | 0 |
| rs138834083 | pathogenic | 0 | 0 | 0 |
| rs533055438 | pathogenic | 1 | 1 | 1 |
| rs2309689 | pathogenic | 1 | 1 | 1 |
| rs138058572 | pathogenic | 1 | 1 | 1 |
| rs118204017 | pathogenic | 1 | 1 | 0 |
| rs766742117 | pathogenic | 1 | 1 | 1 |
| rs751995154 | pathogenic | 1 | 1 | 1 |
| rs200771970 | pathogenic | 0 | 0 | 0 |
| rs118204014 | pathogenic | 0 | 0 | 0 |
| rs148584617 | pathogenic | 1 | 1 | 1 |
| rs121912720 | pathogenic | 0 | 0 | 0 |
| rs121912721 | pathogenic | 0 | 0 | 0 |
| rs28929476 | pathogenic | 0 | 0 | 0 |
| rs28931574 | pathogenic | 1 | 1 | 1 |
| rs12718465 | benign | 0 | 0 | 0 |
| rs144689354 | pathogenic | 0 | 0 | 0 |
| rs149095705 | pathogenic | 0 | 0 | 0 |
| rs572839648 | benign | 1 | 1 | 1 |
| rs121909715 | pathogenic | 1 | 1 | 0 |
| rs34077456 | benign | 1 | 1 | 1 |
| rs397517876 | benign | 0 | 0 | 0 |
| rs1064074 | benign | 0 | 0 | 0 |
| rs142751309 | benign | 0 | 0 | 0 |
| rs116275562 | benign | 0 | 0 | 0 |
| rs148095899 | benign | 0 | 0 | 0 |
| rs751838040 | pathogenic | 0 | 0 | 0 |
| rs121434324 | pathogenic | 1 | 1 | 1 |
| rs143367518 | pathogenic | 0 | 0 | 0 |
| rs121434325 | pathogenic | 0 | 0 | 0 |
| rs397515328 | pathogenic | 0 | 0 | 0 |
| rs376039938 | pathogenic | 0 | 0 | 0 |
| rs387907243 | pathogenic | 0 | 0 | 0 |
| rs28934876 | pathogenic | 0 | 0 | 0 |
| rs387907245 | pathogenic | 0 | 0 | 0 |
| rs138544311 | benign | 0 | 0 | 0 |

| | | | | |
|---|---|---|---|---|
| rs643788 | benign | 0 | 0 | 0 |
| rs749977756 | benign | 0 | 0 | 0 |
| rs1800574 | benign | 0 | 0 | 0 |
| rs137853336 | pathogenic | 1 | 1 | 1 |
| rs111916034 | benign | 0 | 0 | 0 |
| rs307826 | benign | 0 | 0 | 0 |
| rs35874891 | benign | 0 | 0 | 0 |
| rs111033580 | pathogenic | 1 | 1 | 1 |
| rs111033581 | pathogenic | 1 | 1 | 1 |
| rs121908373 | pathogenic | 0 | 0 | 0 |
| rs767122713 | pathogenic | 0 | 0 | 0 |
| rs2233213 | benign | 0 | 0 | 0 |
| rs121908605 | pathogenic | 1 | 1 | 0 |
| rs11887534 | benign | 0 | 0 | 0 |
| rs4148211 | benign | 0 | 0 | 0 |
| rs80025980 | benign | 0 | 0 | 0 |
| rs137854891 | pathogenic | 1 | 1 | 0 |
| rs74358901 | benign | 0 | 0 | 0 |
| rs9282574 | benign | 0 | 0 | 0 |
| rs137852993 | pathogenic | 0 | 0 | 0 |
| rs137852990 | pathogenic | 0 | 0 | 0 |
| rs137852987 | pathogenic | 0 | 0 | 0 |
| rs4148217 | benign | 0 | 0 | 0 |
| rs140690030 | benign | 0 | 0 | 0 |
| rs137852992 | pathogenic | 0 | 0 | 0 |
| rs6544718 | benign | 0 | 0 | 0 |
| rs137852989 | pathogenic | 0 | 0 | 0 |
| rs150893072 | benign | 1 | 1 | 1 |
| rs28931613 | pathogenic | 1 | 1 | 1 |
| rs143789597 | benign | 1 | 1 | 1 |
| rs121913037 | pathogenic | 1 | 1 | 0 |
| rs188802138 | pathogenic | 1 | 1 | 0 |
| rs121913038 | pathogenic | 1 | 1 | 0 |
| rs146835206 | benign | 0 | 0 | 0 |
| rs121913041 | pathogenic | 1 | 1 | 1 |
| rs121913039 | pathogenic | 1 | 1 | 1 |
| rs149977726 | pathogenic | 1 | 1 | 0 |
| rs121913036 | pathogenic | 0 | 0 | 0 |
| rs121913040 | pathogenic | 0 | 0 | 0 |
| rs765007158 | benign | 0 | 0 | 0 |
| rs112723255 | benign | 0 | 0 | 0 |
| rs574164748 | benign | 0 | 0 | 0 |
| rs80338734 | pathogenic | 1 | 1 | 0 |
| rs104894694 | pathogenic | 1 | 1 | 0 |
| rs753198836 | pathogenic | 0 | 0 | 0 |
| rs760101382 | pathogenic | 0 | 0 | 0 |
| rs121909272 | pathogenic | 1 | 1 | 0 |
| rs370421531 | pathogenic | 0 | 0 | 0 |
| rs565109128 | benign | 0 | 0 | 0 |
| rs17851582 | benign | 0 | 0 | 0 |
| rs753228876 | benign | 0 | 0 | 0 |
| rs141471799 | benign | 0 | 0 | 0 |
| rs281864916 | pathogenic | 1 | 1 | 0 |
| rs121918409 | pathogenic | 1 | 1 | 1 |

| | | | | |
|---|---|---|---|---|
| rs121918413 | pathogenic | 1 | 1 | 1 |
| rs121918411 | pathogenic | 1 | 1 | 1 |
| rs121918408 | pathogenic | 1 | 1 | 1 |
| rs121918412 | pathogenic | 1 | 1 | 1 |
| rs121918410 | pathogenic | 1 | 1 | 1 |
| rs116474260 | benign | 1 | 1 | 1 |
| rs281864919 | pathogenic | 1 | 1 | 1 |
| rs397514452 | pathogenic | 1 | 1 | 0 |
| rs747843638 | benign | 0 | 0 | 0 |
| rs774357609 | pathogenic | 1 | 1 | 1 |
| rs201804005 | benign | 0 | 0 | 0 |
| rs746485916 | pathogenic | 0 | 0 | 0 |
| rs764131178 | pathogenic | 0 | 0 | 0 |
| rs374094065 | pathogenic | 0 | 0 | 0 |
| rs76151636 | pathogenic | 1 | 1 | 0 |
| rs753250853 | pathogenic | 0 | 0 | 0 |
| rs761632029 | pathogenic | 0 | 0 | 0 |
| rs775541743 | pathogenic | 0 | 0 | 0 |
| rs778749563 | pathogenic | 0 | 0 | 0 |
| rs60431989 | pathogenic | 1 | 1 | 0 |
| rs755554442 | pathogenic | 1 | 1 | 0 |
| rs749085322 | pathogenic | 0 | 0 | 0 |
| rs756029120 | pathogenic | 1 | 1 | 0 |
| rs137853279 | pathogenic | 1 | 1 | 0 |
| rs777354267 | pathogenic | 1 | 1 | 1 |
| rs142940486 | benign | 1 | 1 | 1 |
| rs140167393 | benign | 1 | 1 | 1 |
| rs2066827 | benign | 0 | 0 | 0 |
| rs765681672 | benign | 0 | 0 | 0 |
| rs137985549 | benign | 0 | 0 | 0 |
| rs74315340 | pathogenic | 1 | 1 | 1 |
| rs74315341 | pathogenic | 0 | 0 | 0 |
| rs74315335 | pathogenic | 0 | 0 | 0 |
| rs61730974 | benign | 0 | 0 | 0 |
| rs121909193 | pathogenic | 0 | 0 | 0 |
| rs74315334 | pathogenic | 0 | 0 | 0 |
| rs74315329 | pathogenic | 0 | 0 | 0 |
| rs74315330 | pathogenic | 0 | 0 | 0 |
| rs121909194 | pathogenic | 0 | 0 | 0 |
| rs28936694 | pathogenic | 0 | 0 | 0 |
| rs74315336 | pathogenic | 0 | 0 | 0 |
| rs74315338 | pathogenic | 1 | 1 | 1 |
| rs74315328 | pathogenic | 0 | 0 | 0 |
| rs74315331 | pathogenic | 0 | 0 | 0 |
| rs74315332 | pathogenic | 0 | 0 | 0 |
| rs141696651 | benign | 0 | 0 | 0 |
| rs7041 | benign | 0 | 0 | 0 |
| rs4588 | benign | 0 | 0 | 0 |
| rs121908981 | pathogenic | 1 | 1 | 1 |
| rs606231459 | pathogenic | 0 | 0 | 0 |
| rs398123053 | pathogenic | 0 | 0 | 0 |
| rs774259802 | benign | 0 | 0 | 0 |
| rs121909582 | pathogenic | 1 | 1 | 1 |
| rs104893680 | pathogenic | 1 | 1 | 1 |

| | | | | |
|---|---|---|---|---|
| rs137854907 | pathogenic | 0 | 0 | 0 |
| rs765715798 | pathogenic | 0 | 0 | 0 |
| rs104893678 | pathogenic | 0 | 0 | 0 |
| rs104893679 | pathogenic | 0 | 1 | 0 |
| rs104893681 | pathogenic | 1 | 1 | 0 |
| rs121909640 | pathogenic | 1 | 1 | 0 |
| rs780705654 | pathogenic | 0 | 0 | 0 |
| rs201672146 | benign | 1 | 1 | 0 |
| rs12136658 | benign | 0 | 0 | 0 |
| rs56127440 | benign | 0 | 0 | 0 |
| rs104894433 | pathogenic | 1 | 1 | 1 |
| rs104894435 | pathogenic | 0 | 1 | 0 |
| rs104894437 | pathogenic | 1 | 1 | 1 |
| rs104894441 | pathogenic | 1 | 1 | 1 |
| rs104894440 | pathogenic | 1 | 1 | 1 |
| rs104894445 | pathogenic | 1 | 1 | 1 |
| rs104894436 | pathogenic | 0 | 0 | 0 |
| rs104894438 | pathogenic | 0 | 0 | 0 |
| rs200891969 | pathogenic | 1 | 1 | 0 |
| rs104894443 | pathogenic | 1 | 1 | 1 |
| rs104894434 | pathogenic | 1 | 1 | 1 |
| rs41298442 | pathogenic | 1 | 1 | 1 |
| rs104894442 | pathogenic | 1 | 1 | 1 |
| rs35453854 | benign | 0 | 0 | 0 |
| rs121912898 | pathogenic | 0 | 0 | 0 |
| rs121912897 | pathogenic | 1 | 1 | 0 |
| rs138260625 | benign | 0 | 0 | 0 |
| rs147802438 | benign | 1 | 1 | 0 |
| rs146443310 | benign | 1 | 1 | 0 |
| rs747319628 | pathogenic | 1 | 1 | 0 |
| rs1801195 | benign | 0 | 0 | 0 |
| rs145045341 | benign | 1 | 1 | 0 |
| rs116001173 | benign | 1 | 1 | 0 |
| rs587784175 | pathogenic | 0 | 0 | 0 |
| rs104893857 | pathogenic | 0 | 0 | 0 |
| rs104893862 | pathogenic | 1 | 1 | 0 |
| rs104893858 | pathogenic | 1 | 1 | 1 |
| rs121909249 | pathogenic | 0 | 1 | 0 |
| rs121909248 | pathogenic | 1 | 1 | 1 |
| rs104893859 | pathogenic | 1 | 1 | 0 |
| rs187063864 | pathogenic | 1 | 1 | 0 |
| rs121917817 | pathogenic | 1 | 1 | 0 |
| rs121917818 | pathogenic | 0 | 0 | 0 |
| rs753594601 | pathogenic | 0 | 0 | 0 |
| rs28937869 | pathogenic | 1 | 1 | 0 |
| rs6165 | benign | 0 | 0 | 0 |
| rs200089714 | pathogenic | 1 | 1 | 0 |
| rs372000714 | pathogenic | 0 | 0 | 0 |
| rs2228206 | benign | 0 | 0 | 0 |
| rs560096 | benign | 0 | 0 | 0 |
| rs137852666 | pathogenic | 0 | 0 | 0 |
| rs10896380 | benign | 0 | 0 | 0 |
| rs773690764 | pathogenic | 0 | 0 | 0 |
| rs201060167 | pathogenic | 0 | 0 | 0 |

| | | | | |
|---|---|---|---|---|
| rs137852670 | pathogenic | 0 | 0 | 0 |
| rs780594709 | pathogenic | 0 | 0 | 0 |
| rs145226920 | pathogenic | 0 | 0 | 0 |
| rs137852665 | pathogenic | 0 | 0 | 0 |
| rs724159960 | pathogenic | 0 | 0 | 0 |
| rs756985703 | pathogenic | 0 | 0 | 0 |
| rs137852667 | pathogenic | 0 | 0 | 0 |
| rs151079750 | pathogenic | 1 | 1 | 0 |
| rs2306636 | benign | 0 | 0 | 0 |
| rs72647885 | benign | 1 | 1 | 0 |
| rs377584435 | pathogenic | 1 | 1 | 1 |
| rs757583846 | pathogenic | 1 | 1 | 0 |
| rs121918148 | pathogenic | 0 | 0 | 0 |
| rs149201802 | benign | 0 | 0 | 0 |
| rs587781904 | pathogenic | 1 | 1 | 1 |
| rs143421325 | benign | 0 | 0 | 0 |
| rs587781253 | pathogenic | 0 | 0 | 0 |
| rs587780331 | benign | 0 | 0 | 0 |
| rs151001016 | benign | 0 | 0 | 0 |
| rs150888094 | benign | 0 | 0 | 0 |
| rs149581331 | benign | 0 | 0 | 0 |
| rs774840535 | benign | 0 | 0 | 0 |
| rs138428684 | benign | 0 | 0 | 0 |
| rs149300055 | benign | 0 | 0 | 0 |
| rs571005578 | benign | 0 | 0 | 0 |
| rs200731839 | benign | 0 | 0 | 0 |
| rs121965007 | pathogenic | 0 | 0 | 0 |
| rs121965013 | pathogenic | 0 | 1 | 0 |
| rs121965009 | pathogenic | 0 | 0 | 0 |
| rs1800457 | benign | 1 | 1 | 0 |
| rs121965006 | pathogenic | 1 | 1 | 0 |
| rs121965008 | pathogenic | 0 | 0 | 0 |
| rs61732609 | pathogenic | 0 | 0 | 0 |
| rs121965011 | pathogenic | 0 | 0 | 0 |
| rs121965015 | pathogenic | 0 | 0 | 0 |
| rs121965018 | pathogenic | 1 | 1 | 0 |
| rs121965016 | pathogenic | 0 | 0 | 0 |
| rs77806 | benign | 0 | 0 | 0 |
| rs149731356 | benign | 0 | 0 | 0 |
| rs72549407 | pathogenic | 0 | 0 | 0 |
| rs200363929 | benign | 0 | 0 | 0 |
| rs376403708 | benign | 1 | 1 | 0 |
| rs56376197 | benign | 1 | 1 | 1 |
| rs200237973 | benign | 1 | 1 | 1 |
| rs548330075 | benign | 0 | 0 | 0 |
| rs148291350 | benign | 0 | 0 | 0 |
| rs71579353 | pathogenic | 0 | 0 | 0 |
| rs146278697 | benign | 0 | 0 | 0 |
| rs141033098 | benign | 0 | 0 | 0 |
| rs397517234 | pathogenic | 0 | 0 | 0 |
| rs71579374 | benign | 0 | 0 | 0 |
| rs140381835 | benign | 0 | 0 | 0 |
| rs137852702 | pathogenic | 0 | 0 | 0 |
| rs137852696 | pathogenic | 1 | 1 | 1 |

| | | | | |
|---|---|---|---|---|
| rs137852697 | pathogenic | 0 | 0 | 0 |
| rs386833639 | pathogenic | 0 | 0 | 0 |
| rs386833640 | pathogenic | 0 | 0 | 0 |
| rs386833641 | benign | 0 | 0 | 0 |
| rs137852701 | pathogenic | 0 | 0 | 0 |
| rs386833642 | pathogenic | 0 | 0 | 0 |
| rs137852695 | pathogenic | 0 | 0 | 0 |
| rs386833646 | pathogenic | 0 | 0 | 0 |
| rs137852700 | pathogenic | 1 | 1 | 0 |
| rs386833649 | pathogenic | 1 | 1 | 0 |
| rs386833650 | pathogenic | 0 | 0 | 0 |
| rs148412181 | pathogenic | 0 | 0 | 0 |
| rs386833655 | pathogenic | 0 | 0 | 0 |
| rs386833656 | pathogenic | 1 | 1 | 0 |
| rs137852698 | pathogenic | 0 | 0 | 0 |
| rs386833661 | pathogenic | 0 | 0 | 0 |
| rs386833670 | pathogenic | 0 | 0 | 0 |
| rs111033632 | pathogenic | 1 | 1 | 1 |
| rs34308573 | benign | 1 | 1 | 1 |
| rs376048533 | pathogenic | 1 | 1 | 0 |
| rs3747517 | benign | 1 | 1 | 0 |
| rs137853328 | pathogenic | 0 | 0 | 0 |
| rs77273740 | benign | 0 | 0 | 0 |
| rs267606760 | pathogenic | 0 | 0 | 0 |
| rs77576840 | pathogenic | 0 | 0 | 0 |
| rs45465204 | benign | 1 | 1 | 0 |
| rs5320 | benign | 0 | 0 | 0 |
| rs5324 | benign | 0 | 0 | 0 |
| rs4531 | benign | 0 | 0 | 0 |
| rs267606761 | pathogenic | 0 | 0 | 0 |
| rs75215331 | pathogenic | 0 | 0 | 0 |
| rs6271 | benign | 0 | 0 | 0 |
| rs201643277 | pathogenic | 0 | 0 | 0 |
| rs4252250 | benign | 0 | 0 | 0 |
| rs137853580 | pathogenic | 0 | 0 | 0 |
| rs368287711 | pathogenic | 0 | 0 | 0 |
| rs137853579 | pathogenic | 0 | 0 | 0 |
| rs3135932 | benign | 0 | 0 | 0 |
| rs2228055 | benign | 0 | 0 | 0 |
| rs140322345 | benign | 1 | 1 | 1 |
| rs1154510 | pathogenic | 0 | 0 | 0 |
| rs11833399 | benign | 0 | 0 | 0 |
| rs36023382 | benign | 1 | 1 | 1 |
| rs61754444 | benign | 0 | 0 | 0 |
| rs28935168 | pathogenic | 0 | 0 | 0 |
| rs28934907 | pathogenic | 0 | 0 | 0 |
| rs28934904 | pathogenic | 1 | 1 | 0 |
| rs61748392 | pathogenic | 1 | 1 | 0 |
| rs28934908 | pathogenic | 0 | 0 | 0 |
| rs28934905 | pathogenic | 0 | 0 | 0 |
| rs28934906 | pathogenic | 1 | 1 | 0 |
| rs606231307 | pathogenic | 0 | 0 | 0 |
| rs9333296 | benign | 0 | 0 | 0 |
| rs3218614 | benign | 0 | 0 | 0 |

| | | | | |
|---|---|---|---|---|
| rs2230792 | benign | 0 | 0 | 0 |
| rs2230793 | benign | 0 | 0 | 0 |
| rs2230794 | benign | 0 | 0 | 0 |
| rs749052963 | pathogenic | 0 | 0 | 0 |
| rs2230798 | benign | 0 | 0 | 0 |
| rs2230795 | benign | 0 | 0 | 0 |
| rs3204145 | benign | 0 | 0 | 0 |
| rs1538660 | benign | 0 | 0 | 0 |
| rs9878928 | benign | 0 | 0 | 0 |
| rs104893742 | pathogenic | 0 | 0 | 0 |
| rs28936702 | pathogenic | 0 | 0 | 0 |
| rs28936703 | pathogenic | 0 | 0 | 0 |
| rs769392 | benign | 0 | 0 | 0 |
| rs769402 | benign | 1 | 1 | 1 |
| rs104894884 | pathogenic | 1 | 1 | 1 |
| rs1801316 | benign | 1 | 1 | 1 |
| rs104894885 | pathogenic | 1 | 1 | 1 |
| rs782572 | benign | 0 | 0 | 0 |
| rs151166046 | pathogenic | 0 | 0 | 0 |
| rs772153760 | pathogenic | 1 | 1 | 0 |
| rs35405862 | benign | 0 | 0 | 0 |
| rs146571352 | pathogenic | 0 | 1 | 0 |
| rs374698153 | pathogenic | 0 | 0 | 0 |
| rs7594497 | benign | 0 | 0 | 0 |
| rs80338716 | pathogenic | 0 | 0 | 0 |
| rs80338719 | pathogenic | 1 | 1 | 1 |
| rs80338721 | pathogenic | 0 | 0 | 0 |
| rs730882249 | pathogenic | 0 | 0 | 0 |
| rs146262009 | pathogenic | 0 | 0 | 0 |
| rs104895083 | pathogenic | 1 | 1 | 1 |
| rs104895100 | benign | 1 | 1 | 1 |
| rs104895085 | pathogenic | 0 | 0 | 0 |
| rs28940580 | pathogenic | 0 | 0 | 0 |
| rs104895098 | pathogenic | 0 | 0 | 0 |
| rs28940578 | pathogenic | 0 | 0 | 0 |
| rs61752717 | pathogenic | 0 | 0 | 0 |
| rs28940579 | pathogenic | 0 | 0 | 0 |
| rs104895097 | pathogenic | 0 | 0 | 0 |
| rs140425651 | benign | 1 | 1 | 0 |
| rs768090444 | pathogenic | 0 | 0 | 0 |
| rs764459544 | pathogenic | 0 | 0 | 0 |
| rs80338802 | pathogenic | 1 | 1 | 1 |
| rs149969786 | pathogenic | 1 | 1 | 0 |
| rs115311625 | pathogenic | 1 | 1 | 1 |
| rs778768583 | pathogenic | 1 | 1 | 1 |
| rs892090 | benign | 0 | 0 | 0 |
| rs28929498 | pathogenic | 1 | 1 | 1 |
| rs137854467 | pathogenic | 0 | 0 | 0 |
| rs137854480 | pathogenic | 0 | 0 | 0 |
| rs148682262 | benign | 0 | 0 | 0 |
| rs137852744 | pathogenic | 0 | 0 | 0 |
| rs137852745 | pathogenic | 0 | 1 | 0 |
| rs137852749 | pathogenic | 0 | 1 | 0 |
| rs137852746 | pathogenic | 0 | 1 | 0 |

| | | | | |
|---|---|---|---|---|
| rs145633958 | benign | 0 | 0 | 0 |
| rs141679950 | benign | 0 | 0 | 0 |
| rs76397662 | benign | 1 | 1 | 1 |
| rs370736139 | benign | 0 | 0 | 0 |
| rs781750106 | benign | 0 | 0 | 0 |
| rs546866208 | benign | 0 | 0 | 0 |
| rs142338976 | benign | 0 | 0 | 0 |
| rs115272158 | benign | 0 | 0 | 0 |
| rs115423919 | benign | 0 | 0 | 0 |
| rs537874538 | benign | 0 | 0 | 0 |
| rs9282834 | benign | 0 | 0 | 0 |
| rs201745826 | benign | 0 | 0 | 0 |
| rs75873440 | pathogenic | 0 | 0 | 0 |
| rs147219360 | benign | 0 | 0 | 0 |
| rs77939446 | pathogenic | 0 | 0 | 0 |
| rs77558292 | pathogenic | 0 | 0 | 0 |
| rs80069458 | pathogenic | 0 | 0 | 0 |
| rs76262710 | pathogenic | 0 | 0 | 0 |
| rs79781594 | pathogenic | 0 | 0 | 0 |
| rs77316810 | pathogenic | 0 | 0 | 0 |
| rs77503355 | pathogenic | 0 | 0 | 0 |
| rs79890926 | pathogenic | 0 | 0 | 0 |
| rs377767402 | benign | 0 | 0 | 0 |
| rs121913308 | pathogenic | 0 | 0 | 0 |
| rs55846256 | benign | 0 | 0 | 0 |
| rs75076352 | pathogenic | 0 | 0 | 0 |
| rs75996173 | pathogenic | 0 | 0 | 0 |
| rs77709286 | pathogenic | 0 | 0 | 0 |
| rs151058706 | benign | 0 | 0 | 0 |
| rs138964358 | benign | 0 | 0 | 0 |
| rs137852353 | pathogenic | 1 | 1 | 0 |
| rs201196020 | benign | 0 | 0 | 0 |
| rs137853201 | pathogenic | 1 | 1 | 1 |
| rs113994174 | pathogenic | 1 | 1 | 1 |
| rs104893674 | pathogenic | 1 | 1 | 1 |
| rs779519249 | benign | 1 | 1 | 1 |
| rs9606756 | benign | 0 | 0 | 0 |
| rs75680863 | benign | 1 | 1 | 1 |
| rs35838082 | benign | 0 | 0 | 0 |
| rs1801198 | benign | 0 | 0 | 0 |
| rs117458738 | benign | 1 | 1 | 0 |
| rs9621049 | benign | 1 | 1 | 0 |
| rs1131603 | benign | 1 | 1 | 0 |
| rs4820889 | benign | 0 | 0 | 0 |
| rs777767978 | pathogenic | 1 | 1 | 0 |
| rs147399860 | benign | 1 | 1 | 1 |
| rs104893950 | pathogenic | 0 | 0 | 0 |
| rs777308612 | benign | 0 | 0 | 0 |
| rs151091410 | benign | 0 | 0 | 0 |
| rs11537667 | benign | 0 | 0 | 0 |
| rs267606621 | pathogenic | 1 | 1 | 0 |
| rs143058902 | benign | 0 | 0 | 0 |
| rs370261904 | pathogenic | 0 | 0 | 0 |
| rs141169746 | benign | 0 | 0 | 0 |

| | | | | |
|---|---|---|---|---|
| rs111033436 | benign | 0 | 0 | 0 |
| rs148533341 | benign | 0 | 0 | 0 |
| rs143827620 | benign | 0 | 0 | 0 |
| rs4935502 | benign | 0 | 0 | 0 |
| rs61735473 | benign | 0 | 0 | 0 |
| rs267606932 | pathogenic | 1 | 1 | 0 |
| rs61730754 | benign | 0 | 0 | 0 |
| rs374491359 | pathogenic | 1 | 1 | 1 |
| rs761800077 | benign | 1 | 1 | 1 |
| rs760391381 | benign | 1 | 1 | 1 |
| rs104893885 | pathogenic | 0 | 0 | 0 |
| rs104893886 | pathogenic | 1 | 1 | 1 |
| rs104893884 | pathogenic | 0 | 0 | 0 |
| rs147715509 | benign | 0 | 0 | 0 |
| rs115533243 | benign | 0 | 0 | 0 |
| rs397509390 | pathogenic | 0 | 1 | 0 |
| rs80357134 | pathogenic | 0 | 0 | 0 |
| rs766004110 | benign | 1 | 1 | 1 |
| rs769650474 | benign | 0 | 0 | 0 |
| rs398122635 | benign | 1 | 1 | 0 |
| rs199522616 | pathogenic | 0 | 0 | 0 |
| rs28897672 | pathogenic | 1 | 1 | 0 |
| rs80357086 | pathogenic | 1 | 1 | 1 |
| rs80357064 | pathogenic | 1 | 1 | 0 |
| rs80356936 | benign | 0 | 0 | 0 |
| rs28897673 | benign | 0 | 0 | 0 |
| rs146747324 | benign | 0 | 0 | 0 |
| rs78635798 | pathogenic | 0 | 0 | 0 |
| rs75146158 | pathogenic | 1 | 1 | 1 |
| rs587782886 | benign | 0 | 0 | 0 |
| rs121434634 | pathogenic | 0 | 0 | 0 |
| rs121434635 | pathogenic | 0 | 0 | 0 |
| rs587777778 | pathogenic | 0 | 0 | 0 |
| rs118203967 | pathogenic | 0 | 0 | 0 |
| rs751375244 | pathogenic | 1 | 1 | 1 |
| rs104893863 | pathogenic | 1 | 1 | 0 |
| rs104893865 | pathogenic | 0 | 0 | 0 |
| rs104893864 | pathogenic | 1 | 1 | 1 |
| rs104893866 | pathogenic | 1 | 1 | 0 |
| rs561738497 | benign | 0 | 0 | 0 |
| rs57599352 | pathogenic | 1 | 1 | 1 |
| rs267607448 | pathogenic | 1 | 1 | 1 |
| rs63751596 | pathogenic | 0 | 0 | 0 |
| rs63751612 | benign | 0 | 0 | 0 |
| rs63751684 | benign | 0 | 0 | 0 |
| rs267607861 | benign | 0 | 0 | 0 |
| rs35831931 | benign | 0 | 0 | 0 |
| rs63750876 | benign | 0 | 0 | 0 |
| rs41295284 | benign | 0 | 0 | 0 |
| rs35502531 | benign | 0 | 0 | 0 |
| rs63750449 | benign | 0 | 0 | 0 |
| rs63750899 | pathogenic | 0 | 0 | 0 |
| rs55907433 | benign | 0 | 0 | 0 |
| rs63750217 | pathogenic | 0 | 0 | 0 |

| | | | | |
|---|---|---|---|---|
| rs63751275 | pathogenic | 0 | 0 | 0 |
| rs63750702 | benign | 1 | 1 | 1 |
| rs63750114 | benign | 0 | 0 | 0 |
| rs2020873 | benign | 0 | 0 | 0 |
| rs140195825 | benign | 1 | 1 | 1 |
| rs182763805 | benign | 0 | 0 | 0 |
| rs533696377 | benign | 0 | 0 | 0 |
| rs41278788 | benign | 0 | 0 | 0 |
| rs104894193 | pathogenic | 1 | 1 | 0 |
| rs3824915 | benign | 1 | 1 | 1 |
| rs104894196 | pathogenic | 0 | 0 | 0 |
| rs78681604 | benign | 1 | 1 | 1 |
| rs77321498 | benign | 1 | 1 | 1 |
| rs137852965 | pathogenic | 0 | 0 | 0 |
| rs137852961 | pathogenic | 0 | 0 | 0 |
| rs137852962 | pathogenic | 1 | 1 | 0 |
| rs753376100 | pathogenic | 0 | 0 | 0 |
| rs137852968 | pathogenic | 1 | 1 | 1 |
| rs137852959 | pathogenic | 1 | 1 | 0 |
| rs137852967 | pathogenic | 0 | 0 | 0 |
| rs113437301 | benign | 1 | 1 | 1 |
| rs61747482 | benign | 1 | 1 | 1 |
| rs2234919 | benign | 1 | 1 | 0 |
| rs121908863 | pathogenic | 0 | 0 | 0 |
| rs140455771 | pathogenic | 0 | 0 | 0 |
| rs121908810 | pathogenic | 1 | 1 | 1 |
| rs143879853 | benign | 0 | 0 | 0 |
| rs188319622 | benign | 0 | 0 | 0 |
| rs121918645 | pathogenic | 0 | 0 | 0 |
| rs115352435 | pathogenic | 0 | 0 | 0 |
| rs121434427 | pathogenic | 1 | 1 | 1 |
| rs121434428 | pathogenic | 0 | 0 | 0 |
| rs35086265 | benign | 1 | 1 | 1 |
| rs11576415 | benign | 0 | 0 | 0 |
| rs121434429 | pathogenic | 1 | 1 | 1 |
| rs4074536 | benign | 1 | 1 | 1 |
| rs10801999 | benign | 0 | 0 | 0 |
| rs121434550 | pathogenic | 0 | 0 | 0 |
| rs148057999 | pathogenic | 0 | 0 | 0 |
| rs28730716 | benign | 0 | 0 | 0 |
| rs142036299 | benign | 0 | 0 | 0 |
| rs762381137 | pathogenic | 0 | 0 | 0 |
| rs121434549 | pathogenic | 0 | 0 | 0 |
| rs139228801 | pathogenic | 0 | 0 | 0 |
| rs147219463 | benign | 0 | 0 | 0 |
| rs1131857 | benign | 0 | 0 | 0 |
| rs121917874 | pathogenic | 0 | 1 | 0 |
| rs2228056 | benign | 0 | 0 | 0 |
| rs121917870 | pathogenic | 0 | 0 | 0 |
| rs121917866 | pathogenic | 0 | 0 | 0 |
| rs11921054 | benign | 0 | 0 | 0 |
| rs121917868 | pathogenic | 0 | 0 | 0 |
| rs28931603 | pathogenic | 0 | 0 | 0 |
| rs104895295 | pathogenic | 1 | 1 | 0 |

| | | | | |
|---|---|---|---|---|
| rs104895382 | pathogenic | 0 | 0 | 0 |
| rs104895298 | pathogenic | 0 | 0 | 0 |
| rs104895300 | pathogenic | 0 | 0 | 0 |
| rs104895332 | pathogenic | 0 | 0 | 0 |
| rs758026399 | pathogenic | 1 | 1 | 1 |
| rs104895366 | pathogenic | 0 | 0 | 0 |
| rs104895304 | pathogenic | 0 | 0 | 0 |
| rs759997079 | pathogenic | 0 | 0 | 0 |
| rs121917789 | pathogenic | 0 | 0 | 0 |
| rs104895319 | pathogenic | 0 | 0 | 0 |
| rs104895317 | pathogenic | 1 | 1 | 0 |
| rs28934897 | pathogenic | 0 | 0 | 0 |
| rs104895380 | benign | 0 | 0 | 0 |
| rs104895360 | pathogenic | 0 | 0 | 0 |
| rs28936396 | pathogenic | 1 | 1 | 0 |
| rs121909307 | pathogenic | 1 | 1 | 1 |
| rs28938472 | pathogenic | 1 | 1 | 0 |
| rs121909308 | pathogenic | 1 | 1 | 0 |
| rs121909309 | pathogenic | 0 | 0 | 0 |
| rs121913013 | benign | 1 | 1 | 1 |
| rs201855245 | benign | 1 | 1 | 1 |
| rs2230232 | benign | 1 | 1 | 0 |
| rs113451409 | benign | 0 | 0 | 0 |
| rs121913011 | pathogenic | 1 | 1 | 0 |
| rs185821167 | pathogenic | 1 | 1 | 0 |
| rs2230234 | benign | 0 | 0 | 0 |
| rs193922639 | benign | 1 | 1 | 0 |
| rs774863785 | pathogenic | 1 | 1 | 0 |
| rs576404380 | pathogenic | 1 | 1 | 0 |
| rs2230235 | benign | 1 | 1 | 0 |
| rs200484060 | pathogenic | 1 | 1 | 0 |
| rs199761749 | benign | 0 | 0 | 0 |
| rs199681901 | benign | 0 | 0 | 0 |
| rs121908951 | pathogenic | 0 | 0 | 0 |
| rs769251460 | benign | 0 | 0 | 0 |
| rs28937320 | pathogenic | 0 | 0 | 0 |
| rs200607527 | pathogenic | 0 | 0 | 0 |
| rs137852640 | pathogenic | 1 | 1 | 1 |
| rs137852636 | pathogenic | 1 | 1 | 1 |
| rs137852638 | pathogenic | 0 | 0 | 0 |
| rs142637231 | pathogenic | 0 | 0 | 0 |
| rs151328418 | benign | 0 | 1 | 0 |
| rs137852639 | pathogenic | 1 | 1 | 1 |
| rs34605667 | benign | 0 | 0 | 0 |
| rs104894073 | pathogenic | 1 | 1 | 0 |
| rs56298569 | pathogenic | 0 | 0 | 0 |
| rs757600616 | pathogenic | 0 | 0 | 0 |
| rs139194636 | pathogenic | 0 | 0 | 0 |
| rs369517993 | pathogenic | 1 | 1 | 1 |
| rs138846036 | benign | 1 | 1 | 1 |
| rs200808744 | pathogenic | 1 | 1 | 1 |
| rs2255543 | benign | 0 | 0 | 0 |
| rs149165166 | pathogenic | 0 | 0 | 0 |
| rs544956641 | pathogenic | 1 | 1 | 1 |

| | | | | |
|---|---|---|---|---|
| rs28941783 | pathogenic | 0 | 0 | 0 |
| rs562853291 | pathogenic | 1 | 1 | 1 |
| rs28942100 | pathogenic | 0 | 0 | 0 |
| rs120074174 | pathogenic | 1 | 1 | 1 |
| rs120074171 | pathogenic | 1 | 1 | 1 |
| rs120074173 | pathogenic | 0 | 0 | 0 |
| rs120074172 | pathogenic | 1 | 1 | 1 |
| rs767159114 | pathogenic | 1 | 1 | 1 |
| rs104893657 | pathogenic | 0 | 0 | 0 |
| rs104893656 | pathogenic | 0 | 0 | 0 |
| rs104893659 | pathogenic | 0 | 0 | 0 |
| rs104893658 | pathogenic | 0 | 0 | 0 |
| rs200386310 | pathogenic | 0 | 0 | 0 |
| rs2072712 | benign | 0 | 0 | 0 |
| rs35274867 | benign | 0 | 0 | 0 |
| rs35343172 | benign | 0 | 0 | 0 |
| rs10079250 | benign | 0 | 0 | 0 |
| rs34951517 | benign | 0 | 0 | 0 |
| rs267606695 | pathogenic | 0 | 0 | 0 |
| rs149391728 | benign | 0 | 0 | 0 |
| rs387906598 | pathogenic | 1 | 1 | 0 |
| rs137854578 | pathogenic | 0 | 0 | 0 |
| rs141519952 | benign | 0 | 0 | 0 |
| rs146572883 | benign | 0 | 0 | 0 |
| rs111866410 | benign | 0 | 0 | 0 |
| rs139387758 | benign | 1 | 1 | 1 |
| rs35672074 | benign | 1 | 1 | 1 |
| rs121434322 | pathogenic | 0 | 0 | 0 |
| rs150884181 | benign | 0 | 0 | 0 |
| rs35867471 | benign | 0 | 0 | 0 |
| rs2867802 | benign | 0 | 0 | 0 |
| rs137902889 | benign | 0 | 0 | 0 |
| rs369804864 | pathogenic | 0 | 0 | 0 |
| rs781050795 | pathogenic | 0 | 0 | 0 |
| rs2305799 | benign | 0 | 0 | 0 |
| rs2229531 | benign | 0 | 0 | 0 |
| rs147025508 | benign | 0 | 0 | 0 |
| rs76857106 | pathogenic | 0 | 0 | 0 |
| rs7247284 | benign | 0 | 0 | 0 |
| rs62619782 | benign | 0 | 0 | 0 |
| rs377244188 | pathogenic | 0 | 0 | 0 |
| rs75718910 | pathogenic | 0 | 0 | 0 |
| rs79843600 | pathogenic | 0 | 0 | 0 |
| rs75037667 | pathogenic | 1 | 1 | 1 |
| rs121913006 | pathogenic | 0 | 0 | 0 |
| rs397517925 | pathogenic | 1 | 1 | 1 |
| rs267606831 | pathogenic | 1 | 1 | 1 |
| rs121909667 | pathogenic | 1 | 1 | 1 |
| rs121909669 | pathogenic | 1 | 1 | 1 |
| rs121909668 | pathogenic | 1 | 1 | 1 |
| rs121909671 | pathogenic | 1 | 1 | 1 |
| rs534596036 | benign | 0 | 0 | 0 |
| rs144037629 | pathogenic | 0 | 0 | 0 |
| rs11546406 | benign | 0 | 0 | 0 |

| | | | | |
|---|---|---|---|---|
| rs143697995 | pathogenic | 0 | 0 | 0 |
| rs121908331 | pathogenic | 0 | 0 | 0 |
| rs149377346 | benign | 0 | 0 | 0 |
| rs104894086 | pathogenic | 1 | 1 | 0 |
| rs104894089 | pathogenic | 0 | 0 | 0 |
| rs104894090 | pathogenic | 1 | 1 | 0 |
| rs387907235 | pathogenic | 0 | 0 | 0 |
| rs137852689 | pathogenic | 0 | 0 | 0 |
| rs137852690 | pathogenic | 1 | 1 | 0 |
| rs104894087 | pathogenic | 0 | 0 | 0 |
| rs104894085 | pathogenic | 0 | 0 | 0 |
| rs79383654 | benign | 0 | 0 | 0 |
| rs121434239 | pathogenic | 0 | 0 | 0 |
| rs121434240 | pathogenic | 0 | 0 | 0 |
| rs121434236 | pathogenic | 0 | 0 | 0 |
| rs121434238 | pathogenic | 1 | 1 | 1 |
| rs121918107 | pathogenic | 1 | 1 | 1 |
| rs121918103 | pathogenic | 1 | 1 | 1 |
| rs121918104 | pathogenic | 1 | 1 | 1 |
| rs61747614 | benign | 1 | 1 | 1 |
| rs555329870 | benign | 0 | 0 | 0 |
| rs186094931 | benign | 0 | 0 | 0 |
| rs267606785 | pathogenic | 0 | 0 | 0 |
| rs34474955 | benign | 0 | 0 | 0 |
| rs1126497 | benign | 1 | 1 | 1 |
| rs397514661 | pathogenic | 0 | 0 | 0 |
| rs189732445 | benign | 0 | 0 | 0 |
| rs146685071 | benign | 0 | 0 | 0 |
| rs121908636 | pathogenic | 1 | 1 | 0 |
| rs121908643 | pathogenic | 0 | 0 | 0 |
| rs121908644 | pathogenic | 1 | 1 | 1 |
| rs575001023 | pathogenic | 1 | 1 | 1 |
| rs121908646 | pathogenic | 1 | 1 | 0 |
| rs121908638 | pathogenic | 1 | 1 | 1 |
| rs777828000 | pathogenic | 1 | 1 | 1 |
| rs74923032 | benign | 0 | 0 | 0 |
| rs192838388 | pathogenic | 1 | 1 | 1 |
| rs148918985 | pathogenic | 1 | 1 | 1 |
| rs773540764 | benign | 1 | 1 | 1 |
| rs370595480 | pathogenic | 1 | 1 | 1 |
| rs762387914 | pathogenic | 1 | 1 | 1 |
| rs121908642 | pathogenic | 1 | 1 | 1 |
| rs183276875 | pathogenic | 1 | 1 | 1 |
| rs121908648 | pathogenic | 1 | 1 | 1 |
| rs121908647 | pathogenic | 1 | 1 | 1 |
| rs121908640 | pathogenic | 1 | 1 | 1 |
| rs771937610 | pathogenic | 1 | 1 | 1 |
| rs121908641 | pathogenic | 1 | 1 | 1 |
| rs5745068 | benign | 0 | 0 | 0 |
| rs73155056 | benign | 1 | 1 | 1 |
| rs148788180 | benign | 0 | 0 | 0 |
| rs121918137 | pathogenic | 0 | 0 | 0 |
| rs41269593 | benign | 0 | 0 | 0 |
| rs11159 | benign | 1 | 1 | 0 |

| | | | | |
|---|---|---|---|---|
| rs114733570 | pathogenic | 1 | 1 | 0 |
| rs3211783 | benign | 0 | 0 | 0 |
| rs4762 | benign | 0 | 0 | 0 |
| rs74315283 | pathogenic | 0 | 0 | 0 |
| rs111033559 | pathogenic | 0 | 0 | 0 |
| rs111033560 | pathogenic | 1 | 1 | 1 |
| rs749962743 | benign | 0 | 0 | 0 |
| rs267606986 | pathogenic | 0 | 0 | 0 |
| rs34516635 | benign | 0 | 0 | 0 |
| rs117440569 | benign | 0 | 0 | 0 |
| rs61740877 | benign | 0 | 0 | 0 |
| rs121918352 | pathogenic | 0 | 0 | 0 |
| rs183974372 | benign | 0 | 0 | 0 |
| rs140511204 | benign | 1 | 1 | 0 |
| rs6040067 | benign | 0 | 0 | 0 |
| rs199674138 | benign | 0 | 0 | 0 |
| rs121918350 | pathogenic | 0 | 0 | 0 |
| rs138292988 | benign | 1 | 1 | 1 |
| rs145288947 | benign | 0 | 0 | 0 |
| rs369714529 | benign | 0 | 0 | 0 |
| rs200945419 | benign | 0 | 0 | 0 |
| rs116959874 | benign | 0 | 0 | 0 |
| rs141659620 | pathogenic | 1 | 1 | 0 |
| rs111475461 | benign | 0 | 0 | 0 |
| rs61755320 | pathogenic | 0 | 0 | 0 |
| rs748600162 | pathogenic | 1 | 1 | 1 |
| rs369227537 | pathogenic | 1 | 1 | 1 |
| rs372981030 | pathogenic | 1 | 1 | 1 |
| rs137854486 | pathogenic | 0 | 0 | 0 |
| rs137854487 | pathogenic | 0 | 0 | 0 |
| rs199905054 | pathogenic | 0 | 0 | 0 |
| rs771438170 | pathogenic | 0 | 0 | 0 |
| rs774949844 | pathogenic | 0 | 0 | 0 |
| rs4899145 | benign | 0 | 0 | 0 |
| rs200642525 | benign | 0 | 0 | 0 |
| rs190140598 | pathogenic | 0 | 0 | 0 |
| rs2250889 | benign | 0 | 0 | 0 |
| rs17577 | benign | 0 | 0 | 0 |
| rs142032681 | benign | 0 | 0 | 0 |
| rs144577963 | benign | 0 | 0 | 0 |
| rs61754623 | benign | 0 | 0 | 0 |
| rs139873200 | benign | 0 | 0 | 0 |
| rs532530906 | benign | 0 | 0 | 0 |
| rs145762716 | benign | 0 | 0 | 0 |
| rs34062020 | benign | 0 | 0 | 0 |
| rs35258161 | benign | 0 | 0 | 0 |
| rs149792649 | benign | 0 | 0 | 0 |
| rs78101774 | benign | 0 | 0 | 0 |
| rs766466297 | pathogenic | 0 | 1 | 0 |
| rs140467171 | pathogenic | 0 | 0 | 0 |
| rs141340466 | pathogenic | 0 | 0 | 0 |
| rs201555303 | pathogenic | 0 | 0 | 0 |
| rs143592405 | pathogenic | 0 | 0 | 0 |
| rs781249411 | pathogenic | 0 | 0 | 0 |

| | | | | |
|---|---|---|---|---|
| rs116644127 | benign | 0 | 0 | 0 |
| rs151146863 | benign | 1 | 1 | 1 |
| rs116440799 | benign | 0 | 0 | 0 |
| rs138958351 | benign | 0 | 0 | 0 |
| rs143319354 | benign | 0 | 0 | 0 |
| rs768669208 | pathogenic | 1 | 1 | 1 |
| rs28940289 | pathogenic | 1 | 1 | 0 |
| rs745656120 | pathogenic | 1 | 1 | 0 |
| rs387906987 | pathogenic | 0 | 0 | 0 |
| rs147240777 | benign | 0 | 0 | 0 |
| rs140945833 | benign | 0 | 0 | 0 |
| rs146614981 | benign | 0 | 0 | 0 |
| rs149878562 | benign | 0 | 0 | 0 |
| rs28938168 | pathogenic | 0 | 0 | 0 |
| rs33939927 | pathogenic | 1 | 1 | 1 |
| rs34995376 | pathogenic | 1 | 1 | 1 |
| rs201034481 | pathogenic | 1 | 1 | 1 |
| rs774069983 | benign | 0 | 0 | 0 |
| rs199422226 | pathogenic | 1 | 1 | 0 |
| rs199422225 | pathogenic | 1 | 1 | 1 |
| rs199422224 | pathogenic | 1 | 1 | 0 |
| rs149271416 | pathogenic | 1 | 1 | 0 |
| rs1801318 | benign | 1 | 1 | 1 |
| rs372691318 | pathogenic | 1 | 1 | 1 |
| rs147685849 | benign | 0 | 0 | 0 |
| rs387907199 | pathogenic | 0 | 0 | 0 |
| rs767122069 | pathogenic | 0 | 0 | 0 |
| rs2302427 | benign | 0 | 0 | 0 |
| rs201135441 | benign | 0 | 0 | 0 |
| rs201861847 | pathogenic | 0 | 0 | 0 |
| rs137852952 | pathogenic | 0 | 0 | 0 |
| rs16986560 | benign | 0 | 0 | 0 |
| rs147344332 | pathogenic | 1 | 1 | 1 |
| rs36027220 | pathogenic | 0 | 0 | 0 |
| rs192401229 | benign | 0 | 0 | 0 |
| rs121909106 | pathogenic | 1 | 1 | 0 |
| rs118204103 | pathogenic | 1 | 1 | 0 |
| rs118204104 | pathogenic | 1 | 1 | 0 |
| rs118204105 | pathogenic | 1 | 1 | 0 |
| rs118204106 | pathogenic | 0 | 0 | 0 |
| rs118204119 | pathogenic | 0 | 0 | 0 |
| rs118204107 | pathogenic | 0 | 0 | 0 |
| rs118204094 | pathogenic | 0 | 0 | 0 |
| rs118204098 | pathogenic | 1 | 1 | 0 |
| rs118204101 | pathogenic | 1 | 1 | 0 |
| rs118204096 | pathogenic | 1 | 1 | 0 |
| rs118204108 | pathogenic | 0 | 0 | 0 |
| rs536814318 | pathogenic | 0 | 0 | 0 |
| rs34413634 | pathogenic | 1 | 1 | 0 |
| rs118204100 | pathogenic | 0 | 0 | 0 |
| rs118204109 | pathogenic | 0 | 0 | 0 |
| rs118204116 | pathogenic | 1 | 1 | 0 |
| rs118204110 | pathogenic | 0 | 0 | 0 |
| rs118204099 | pathogenic | 0 | 0 | 0 |

| | | | | |
|---|---|---|---|---|
| rs118204111 | pathogenic | 0 | 0 | 0 |
| rs118204112 | pathogenic | 0 | 0 | 0 |
| rs118204113 | pathogenic | 0 | 0 | 0 |
| rs118204114 | pathogenic | 0 | 0 | 0 |
| rs118204115 | pathogenic | 0 | 0 | 0 |
| rs150428209 | benign | 0 | 0 | 0 |
| rs144949995 | benign | 0 | 0 | 0 |
| rs190023981 | benign | 0 | 0 | 0 |
| rs104894611 | pathogenic | 1 | 1 | 1 |
| rs28937580 | pathogenic | 1 | 1 | 1 |
| rs121908949 | pathogenic | 0 | 0 | 0 |
| rs104894612 | pathogenic | 1 | 1 | 1 |
| rs104894609 | pathogenic | 1 | 1 | 1 |
| rs104894610 | pathogenic | 1 | 1 | 1 |
| rs104894615 | pathogenic | 0 | 0 | 0 |
| rs104894603 | pathogenic | 0 | 0 | 0 |
| rs104894602 | pathogenic | 1 | 1 | 1 |
| rs121908948 | pathogenic | 1 | 1 | 1 |
| rs104894608 | pathogenic | 1 | 1 | 1 |
| rs371545673 | benign | 0 | 0 | 0 |
| rs199925238 | benign | 0 | 0 | 0 |
| rs146313451 | benign | 0 | 0 | 0 |
| rs121918193 | pathogenic | 1 | 1 | 0 |
| rs2228500 | benign | 0 | 0 | 0 |
| rs121918194 | pathogenic | 1 | 1 | 0 |
| rs121918196 | pathogenic | 0 | 0 | 0 |
| rs41291971 | benign | 0 | 0 | 0 |
| rs199473690 | pathogenic | 0 | 0 | 0 |
| rs2228613 | benign | 0 | 0 | 0 |
| rs267607130 | pathogenic | 1 | 1 | 1 |
| rs77615401 | benign | 1 | 1 | 1 |
| rs374618872 | benign | 1 | 1 | 1 |
| rs146165014 | benign | 0 | 0 | 0 |
| rs144586070 | benign | 0 | 0 | 0 |
| rs121918541 | pathogenic | 0 | 0 | 0 |
| rs142105943 | benign | 0 | 0 | 0 |
| rs121918540 | pathogenic | 0 | 0 | 0 |
| rs6791 | benign | 0 | 0 | 0 |
| rs61736587 | pathogenic | 0 | 0 | 0 |
| rs61736586 | benign | 0 | 0 | 0 |
| rs121917741 | pathogenic | 1 | 1 | 1 |
| rs41302885 | benign | 0 | 0 | 0 |
| rs141026028 | benign | 0 | 0 | 0 |
| rs193922493 | pathogenic | 0 | 0 | 0 |
| rs73598374 | benign | 0 | 0 | 0 |
| rs121908725 | pathogenic | 1 | 1 | 0 |
| rs199422328 | pathogenic | 0 | 1 | 0 |
| rs11555566 | benign | 1 | 1 | 1 |
| rs267606634 | pathogenic | 0 | 0 | 0 |
| rs121908714 | pathogenic | 1 | 1 | 0 |
| rs121908717 | pathogenic | 1 | 1 | 0 |
| rs267606635 | pathogenic | 1 | 1 | 0 |
| rs121908739 | pathogenic | 0 | 0 | 0 |
| rs121908735 | pathogenic | 0 | 1 | 0 |

| | | | | |
|---|---|---|---|---|
| rs121908716 | pathogenic | 0 | 1 | 0 |
| rs121908723 | pathogenic | 0 | 1 | 0 |
| rs121908721 | pathogenic | 0 | 0 | 0 |
| rs199422327 | pathogenic | 0 | 0 | 0 |
| rs121908715 | pathogenic | 0 | 0 | 0 |
| rs587783563 | pathogenic | 0 | 0 | 0 |
| rs104894780 | pathogenic | 0 | 0 | 0 |
| rs56030372 | pathogenic | 1 | 1 | 1 |
| rs104894782 | pathogenic | 1 | 1 | 1 |
| rs149495971 | benign | 0 | 0 | 0 |
| rs118203942 | pathogenic | 1 | 1 | 0 |
| rs118203939 | pathogenic | 0 | 0 | 0 |
| rs118203938 | pathogenic | 0 | 0 | 0 |
| rs118203943 | pathogenic | 0 | 0 | 0 |
| rs118203940 | pathogenic | 0 | 0 | 0 |
| rs727503809 | pathogenic | 0 | 0 | 0 |
| rs1065757 | benign | 0 | 0 | 0 |
| rs1071598 | benign | 0 | 0 | 0 |
| rs25414 | benign | 0 | 0 | 0 |
| rs118203944 | pathogenic | 0 | 0 | 0 |
| rs118203941 | pathogenic | 0 | 0 | 0 |
| rs144222167 | benign | 0 | 0 | 0 |
| rs201101343 | pathogenic | 0 | 0 | 0 |
| rs144601717 | benign | 0 | 0 | 0 |
| rs267607129 | pathogenic | 0 | 0 | 0 |
| rs104894729 | pathogenic | 0 | 0 | 0 |
| rs104894727 | pathogenic | 0 | 0 | 0 |
| rs104894725 | pathogenic | 0 | 0 | 0 |
| rs104894515 | pathogenic | 1 | 1 | 1 |
| rs74688377 | benign | 0 | 0 | 0 |
| rs121908513 | pathogenic | 1 | 1 | 0 |
| rs121908509 | pathogenic | 0 | 0 | 0 |
| rs121908514 | pathogenic | 1 | 1 | 0 |
| rs121908512 | pathogenic | 1 | 1 | 0 |
| rs121908510 | pathogenic | 0 | 0 | 0 |
| rs121908511 | pathogenic | 1 | 1 | 1 |
| rs121908518 | pathogenic | 1 | 1 | 1 |
| rs34358665 | benign | 0 | 0 | 0 |
| rs147568834 | benign | 0 | 0 | 0 |
| rs201717599 | benign | 0 | 0 | 0 |
| rs61744487 | benign | 0 | 0 | 0 |
| rs370627877 | pathogenic | 1 | 1 | 1 |
| rs368593151 | pathogenic | 1 | 1 | 1 |
| rs114363936 | benign | 0 | 0 | 0 |
| rs137853333 | pathogenic | 0 | 0 | 0 |
| rs137853080 | pathogenic | 1 | 1 | 1 |
| rs137853077 | pathogenic | 1 | 1 | 1 |
| rs780749732 | benign | 1 | 1 | 1 |
| rs137853081 | pathogenic | 1 | 1 | 0 |
| rs137853078 | pathogenic | 0 | 0 | 0 |
| rs121913315 | pathogenic | 1 | 1 | 1 |
| rs752513525 | pathogenic | 1 | 1 | 1 |
| rs104894918 | pathogenic | 0 | 0 | 0 |
| rs104894917 | pathogenic | 0 | 0 | 0 |

| | | | | |
|---|---|---|---|---|
| rs104894919 | pathogenic | 0 | 0 | 0 |
| rs28939672 | pathogenic | 0 | 0 | 0 |
| rs104894173 | pathogenic | 0 | 0 | 0 |
| rs104894179 | pathogenic | 0 | 0 | 0 |
| rs104894174 | pathogenic | 1 | 1 | 0 |
| rs104894178 | pathogenic | 1 | 1 | 0 |
| rs28939714 | pathogenic | 1 | 1 | 1 |
| rs104894270 | pathogenic | 1 | 1 | 1 |
| rs3765571 | benign | 0 | 0 | 0 |
| rs752669339 | pathogenic | 1 | 1 | 1 |
| rs537043237 | pathogenic | 1 | 1 | 1 |
| rs774047299 | pathogenic | 1 | 1 | 1 |
| rs121434583 | pathogenic | 1 | 1 | 1 |
| rs121907927 | pathogenic | 0 | 0 | 0 |
| rs121907925 | pathogenic | 0 | 1 | 0 |
| rs746212325 | pathogenic | 0 | 0 | 0 |
| rs754637179 | pathogenic | 0 | 0 | 0 |
| rs780882740 | pathogenic | 0 | 0 | 0 |
| rs121918130 | pathogenic | 0 | 0 | 0 |
| rs121918129 | pathogenic | 0 | 0 | 0 |
| rs13297509 | pathogenic | 0 | 0 | 0 |
| rs746867724 | pathogenic | 0 | 0 | 0 |
| rs121918128 | pathogenic | 0 | 0 | 0 |
| rs752300607 | pathogenic | 0 | 0 | 0 |
| rs121964860 | pathogenic | 0 | 0 | 0 |
| rs45466197 | pathogenic | 1 | 1 | 1 |
| rs3730238 | benign | 1 | 1 | 1 |
| rs730881125 | pathogenic | 1 | 1 | 1 |
| rs121964861 | pathogenic | 1 | 1 | 1 |
| rs397516484 | pathogenic | 0 | 0 | 0 |
| rs367785431 | pathogenic | 0 | 0 | 0 |
| rs141121678 | pathogenic | 0 | 0 | 0 |
| rs12720263 | benign | 0 | 0 | 0 |
| rs2304256 | benign | 0 | 0 | 0 |
| rs2304255 | benign | 0 | 0 | 0 |
| rs121909045 | pathogenic | 0 | 0 | 0 |
| rs397508796 | pathogenic | 0 | 0 | 0 |
| rs151020603 | pathogenic | 0 | 0 | 0 |
| rs77284892 | pathogenic | 0 | 0 | 0 |
| rs368505753 | pathogenic | 0 | 0 | 0 |
| rs115545701 | pathogenic | 0 | 0 | 0 |
| rs121908749 | pathogenic | 0 | 0 | 0 |
| rs75961395 | pathogenic | 0 | 0 | 0 |
| rs121908750 | pathogenic | 0 | 0 | 0 |
| rs121908751 | pathogenic | 0 | 0 | 0 |
| rs397508464 | pathogenic | 0 | 0 | 0 |
| rs397508461 | pathogenic | 0 | 0 | 0 |
| rs121909031 | pathogenic | 0 | 0 | 0 |
| rs113993958 | pathogenic | 0 | 0 | 0 |
| rs77834169 | pathogenic | 0 | 0 | 0 |
| rs78655421 | pathogenic | 0 | 0 | 0 |
| rs141723617 | benign | 0 | 0 | 0 |
| rs35516286 | benign | 0 | 0 | 0 |
| rs1800079 | pathogenic | 1 | 1 | 0 |

| | | | | |
|---|---|---|---|---|
| rs80282562 | pathogenic | 0 | 0 | 0 |
| rs121908752 | pathogenic | 0 | 0 | 0 |
| rs556880586 | pathogenic | 0 | 0 | 0 |
| rs121908760 | pathogenic | 0 | 0 | 0 |
| rs397508350 | pathogenic | 0 | 0 | 0 |
| rs374946172 | pathogenic | 0 | 0 | 0 |
| rs397508387 | pathogenic | 0 | 0 | 0 |
| rs201386642 | pathogenic | 0 | 0 | 0 |
| rs121909012 | pathogenic | 0 | 0 | 0 |
| rs121909008 | pathogenic | 0 | 0 | 0 |
| rs149790377 | pathogenic | 0 | 0 | 0 |
| rs397508442 | pathogenic | 0 | 0 | 0 |
| rs121909035 | pathogenic | 0 | 0 | 0 |
| rs386134230 | pathogenic | 0 | 0 | 0 |
| rs1800112 | benign | 0 | 0 | 0 |
| rs144055758 | pathogenic | 0 | 0 | 0 |
| rs142394380 | pathogenic | 0 | 0 | 0 |
| rs121909036 | pathogenic | 0 | 0 | 0 |
| rs78194216 | pathogenic | 0 | 0 | 0 |
| rs121909019 | pathogenic | 0 | 0 | 0 |
| rs121909020 | pathogenic | 0 | 0 | 0 |
| rs200321110 | pathogenic | 0 | 0 | 0 |
| rs202179988 | pathogenic | 0 | 0 | 0 |
| rs121909037 | pathogenic | 0 | 0 | 0 |
| rs186045772 | pathogenic | 0 | 0 | 0 |
| rs139304906 | pathogenic | 0 | 0 | 0 |
| rs78802634 | pathogenic | 0 | 0 | 0 |
| rs121908761 | pathogenic | 0 | 0 | 0 |
| rs36210737 | pathogenic | 0 | 0 | 0 |
| rs139729994 | pathogenic | 0 | 0 | 0 |
| rs79850223 | pathogenic | 0 | 0 | 0 |
| rs1800120 | benign | 0 | 0 | 0 |
| rs74767530 | pathogenic | 0 | 0 | 0 |
| rs374705585 | pathogenic | 0 | 0 | 0 |
| rs1136287 | benign | 0 | 0 | 0 |
| rs6828 | benign | 0 | 0 | 0 |
| rs104894856 | pathogenic | 0 | 0 | 0 |
| rs104894860 | pathogenic | 0 | 0 | 0 |
| rs61736892 | pathogenic | 0 | 0 | 0 |
| rs145807417 | benign | 0 | 0 | 0 |
| rs104894853 | pathogenic | 0 | 0 | 0 |
| rs199422227 | pathogenic | 0 | 0 | 0 |
| rs113993946 | pathogenic | 0 | 0 | 0 |
| rs199422231 | pathogenic | 0 | 0 | 0 |
| rs199422228 | pathogenic | 0 | 0 | 0 |
| rs142195196 | benign | 0 | 1 | 0 |
| rs148656104 | benign | 1 | 1 | 1 |
| rs141944844 | benign | 0 | 0 | 0 |
| rs104894342 | pathogenic | 0 | 0 | 0 |
| rs104894343 | pathogenic | 0 | 0 | 0 |
| rs104894344 | pathogenic | 0 | 0 | 0 |
| rs573322878 | benign | 0 | 0 | 0 |
| rs104894347 | pathogenic | 0 | 0 | 0 |
| rs28937882 | pathogenic | 0 | 0 | 0 |

| | | | | |
|---|---|---|---|---|
| rs193922702 | pathogenic | 0 | 0 | 0 |
| rs200786104 | benign | 0 | 0 | 0 |
| rs690016544 | pathogenic | 1 | 1 | 1 |
| rs28943592 | benign | 0 | 0 | 0 |
| rs28943594 | benign | 0 | 0 | 0 |
| rs121908957 | pathogenic | 0 | 0 | 0 |
| rs115408010 | benign | 1 | 1 | 1 |
| rs77591659 | benign | 0 | 0 | 0 |
| rs121918511 | pathogenic | 0 | 0 | 0 |
| rs538359948 | benign | 0 | 0 | 0 |
| rs76708142 | benign | 0 | 0 | 0 |
| rs121907913 | pathogenic | 1 | 1 | 0 |
| rs121907920 | pathogenic | 0 | 0 | 0 |
| rs121907923 | pathogenic | 1 | 1 | 0 |
| rs121907928 | pathogenic | 1 | 1 | 0 |
| rs121907919 | pathogenic | 1 | 1 | 0 |
| rs121907918 | pathogenic | 1 | 1 | 0 |
| rs74315493 | pathogenic | 1 | 1 | 1 |
| rs74315494 | pathogenic | 1 | 1 | 1 |
| rs145446060 | benign | 1 | 1 | 1 |
| rs550409143 | benign | 1 | 1 | 0 |

| compound_binding_site | cofactor_binding_site | nucleic_binding_site | ion_binding_site |
|---|---|---|---|
| 1 | 0 | 0 | 0 |
| 1 | 0 | 0 | 0 |
| 0 | 0 | 0 | 0 |
| 1 | 0 | 0 | 0 |
| 0 | 0 | 0 | 0 |
| 0 | 0 | 0 | 0 |
| 0 | 0 | 0 | 0 |
| 0 | 0 | 0 | 0 |
| 1 | 0 | 0 | 1 |
| 1 | 0 | 0 | 0 |
| 0 | 0 | 0 | 1 |
| 1 | 0 | 1 | 0 |
| 0 | 0 | 0 | 0 |
| 0 | 0 | 0 | 0 |
| 0 | 0 | 0 | 0 |
| 0 | 0 | 0 | 0 |
| 0 | 0 | 0 | 0 |
| 1 | 0 | 0 | 1 |
| 1 | 0 | 1 | 0 |
| 0 | 0 | 0 | 0 |
| 1 | 0 | 0 | 1 |
| 1 | 0 | 0 | 0 |
| 1 | 0 | 0 | 0 |
| 1 | 0 | 0 | 0 |
| 0 | 0 | 0 | 0 |
| 0 | 0 | 0 | 0 |
| 1 | 0 | 0 | 1 |
| 0 | 0 | 0 | 0 |
| 0 | 0 | 0 | 0 |
| 0 | 0 | 0 | 0 |
| 0 | 0 | 0 | 0 |
| 1 | 0 | 0 | 0 |
| 1 | 0 | 0 | 1 |
| 1 | 0 | 0 | 0 |
| 0 | 0 | 0 | 0 |
| 0 | 0 | 0 | 0 |
| 0 | 0 | 0 | 0 |
| 1 | 0 | 0 | 1 |
| 1 | 0 | 0 | 1 |
| 1 | 0 | 0 | 1 |
| 1 | 0 | 0 | 0 |
| 1 | 0 | 0 | 0 |
| 1 | 0 | 0 | 0 |
| 1 | 0 | 0 | 0 |
| 1 | 0 | 0 | 0 |
| 1 | 0 | 0 | 0 |
| 1 | 0 | 0 | 0 |
| 1 | 0 | 0 | 0 |
| 1 | 0 | 0 | 0 |
| 1 | 0 | 0 | 0 |
| 1 | 0 | 0 | 0 |
| 1 | 0 | 0 | 0 |
| 1 | 0 | 0 | 0 |

| | | | |
|---|---|---|---|
| 1 | 0 | 0 | 1 |
| 1 | 0 | 0 | 0 |
| 1 | 0 | 0 | 0 |
| 1 | 0 | 0 | 0 |
| 0 | 0 | 0 | 0 |
| 1 | 0 | 0 | 0 |
| 0 | 0 | 0 | 0 |
| 1 | 0 | 0 | 0 |
| 1 | 0 | 0 | 0 |
| 1 | 0 | 0 | 1 |
| 1 | 0 | 0 | 0 |
| 1 | 0 | 0 | 0 |
| 1 | 0 | 0 | 0 |
| 1 | 0 | 0 | 0 |
| 1 | 0 | 0 | 0 |
| 1 | 0 | 0 | 0 |
| 1 | 0 | 0 | 0 |
| 1 | 0 | 0 | 1 |
| 1 | 0 | 0 | 1 |
| 1 | 1 | 0 | 1 |
| 1 | 0 | 0 | 0 |
| 1 | 0 | 0 | 0 |
| 1 | 0 | 0 | 0 |
| 1 | 0 | 0 | 0 |
| 1 | 1 | 0 | 0 |
| 1 | 0 | 0 | 0 |
| 1 | 0 | 0 | 0 |
| 1 | 0 | 0 | 0 |
| 1 | 0 | 0 | 0 |
| 1 | 0 | 0 | 0 |
| 1 | 1 | 0 | 0 |
| 1 | 1 | 0 | 0 |
| 1 | 1 | 0 | 0 |
| 1 | 1 | 0 | 0 |
| 1 | 1 | 0 | 0 |
| 1 | 0 | 0 | 0 |
| 1 | 1 | 0 | 0 |
| 1 | 0 | 0 | 0 |
| 1 | 0 | 0 | 0 |
| 1 | 1 | 0 | 0 |
| 0 | 0 | 0 | 0 |
| 0 | 0 | 0 | 0 |
| 0 | 0 | 0 | 0 |
| 0 | 0 | 0 | 0 |
| 0 | 0 | 0 | 0 |
| 0 | 0 | 0 | 0 |
| 0 | 0 | 0 | 0 |
| 1 | 0 | 0 | 1 |
| 0 | 0 | 0 | 0 |
| 1 | 0 | 0 | 0 |
| 1 | 0 | 0 | 0 |
| 0 | 0 | 0 | 0 |
| 0 | 0 | 0 | 0 |
| 0 | 0 | 0 | 0 |

| | | | |
|---|---|---|---|
| 0 | 0 | 0 | 0 |
| 0 | 0 | 0 | 0 |
| 0 | 0 | 0 | 0 |
| 1 | 0 | 0 | 0 |
| 1 | 0 | 0 | 0 |
| 1 | 0 | 0 | 0 |
| 0 | 0 | 0 | 0 |
| 0 | 0 | 1 | 0 |
| 0 | 0 | 1 | 0 |
| 0 | 0 | 1 | 0 |
| 0 | 0 | 1 | 0 |
| 0 | 0 | 1 | 0 |
| 0 | 0 | 0 | 0 |
| 0 | 0 | 0 | 0 |
| 1 | 0 | 0 | 0 |
| 0 | 0 | 0 | 0 |
| 0 | 0 | 0 | 0 |
| 1 | 0 | 0 | 0 |
| 1 | 0 | 0 | 0 |
| 1 | 0 | 0 | 0 |
| 1 | 0 | 0 | 0 |
| 1 | 1 | 0 | 0 |
| 1 | 0 | 0 | 0 |
| 1 | 0 | 0 | 0 |
| 0 | 0 | 0 | 0 |
| 0 | 0 | 0 | 0 |
| 1 | 0 | 0 | 0 |
| 1 | 1 | 0 | 0 |
| 1 | 0 | 0 | 0 |
| 1 | 0 | 0 | 0 |
| 1 | 1 | 0 | 0 |
| 1 | 1 | 0 | 0 |
| 1 | 1 | 0 | 0 |
| 1 | 1 | 0 | 0 |
| 1 | 1 | 0 | 0 |
| 1 | 1 | 0 | 0 |
| 1 | 1 | 0 | 0 |
| 1 | 1 | 0 | 0 |
| 1 | 1 | 0 | 0 |
| 1 | 1 | 0 | 0 |
| 1 | 1 | 0 | 0 |
| 1 | 1 | 0 | 0 |
| 1 | 0 | 0 | 0 |
| 1 | 0 | 0 | 0 |
| 1 | 0 | 0 | 0 |
| 1 | 1 | 0 | 0 |
| 1 | 0 | 0 | 0 |
| 1 | 1 | 1 | 0 |
| 1 | 1 | 1 | 0 |
| 1 | 1 | 0 | 0 |
| 1 | 1 | 0 | 0 |
| 1 | 0 | 0 | 0 |
| 1 | 0 | 0 | 0 |
| 0 | 0 | 0 | 0 |

| | | | |
|---|---|---|---|
| 0 | 0 | 0 | 0 |
| 0 | 0 | 0 | 0 |
| 0 | 0 | 0 | 0 |
| 1 | 0 | 0 | 0 |
| 1 | 0 | 0 | 0 |
| 0 | 0 | 0 | 0 |
| 0 | 0 | 0 | 0 |
| 0 | 0 | 0 | 0 |
| 0 | 0 | 0 | 0 |
| 0 | 0 | 0 | 0 |
| 0 | 0 | 0 | 0 |
| 0 | 0 | 0 | 0 |
| 0 | 0 | 0 | 0 |
| 0 | 0 | 0 | 0 |
| 0 | 0 | 0 | 0 |
| 0 | 0 | 0 | 0 |
| 0 | 0 | 0 | 0 |
| 0 | 0 | 0 | 0 |
| 0 | 0 | 0 | 0 |
| 0 | 0 | 0 | 0 |
| 0 | 0 | 0 | 0 |
| 0 | 0 | 0 | 0 |
| 1 | 0 | 0 | 0 |
| 1 | 0 | 0 | 0 |
| 0 | 0 | 0 | 0 |
| 0 | 0 | 0 | 0 |
| 1 | 0 | 0 | 0 |
| 0 | 0 | 0 | 0 |
| 0 | 0 | 0 | 0 |
| 0 | 0 | 0 | 0 |
| 0 | 0 | 0 | 0 |
| 0 | 0 | 0 | 0 |
| 0 | 0 | 0 | 0 |
| 0 | 0 | 0 | 0 |
| 1 | 0 | 0 | 0 |
| 0 | 0 | 0 | 0 |
| 0 | 0 | 0 | 0 |
| 0 | 0 | 0 | 0 |
| 0 | 0 | 0 | 0 |
| 0 | 0 | 0 | 0 |
| 0 | 0 | 0 | 0 |
| 0 | 0 | 0 | 0 |
| 0 | 0 | 0 | 0 |
| 0 | 0 | 0 | 0 |
| 0 | 0 | 0 | 0 |
| 0 | 0 | 0 | 0 |
| 0 | 0 | 1 | 0 |
| 1 | 0 | 0 | 0 |
| 1 | 0 | 0 | 0 |
| 1 | 1 | 0 | 0 |
| 1 | 0 | 0 | 0 |
| 0 | 0 | 0 | 0 |
| 0 | 0 | 0 | 0 |
| 1 | 0 | 0 | 0 |

| | | | |
|---|---|---|---|
| 1 | 0 | 0 | 0 |
| 1 | 0 | 0 | 0 |
| 1 | 0 | 0 | 0 |
| 0 | 0 | 0 | 0 |
| 0 | 0 | 0 | 0 |
| 0 | 0 | 0 | 0 |
| 1 | 0 | 0 | 0 |
| 1 | 0 | 0 | 0 |
| 0 | 0 | 0 | 0 |
| 0 | 0 | 0 | 0 |
| 1 | 0 | 0 | 0 |
| 0 | 0 | 0 | 0 |
| 1 | 0 | 0 | 0 |
| 0 | 0 | 0 | 0 |
| 0 | 0 | 0 | 0 |
| 0 | 0 | 0 | 0 |
| 0 | 0 | 0 | 0 |
| 0 | 0 | 0 | 0 |
| 0 | 0 | 0 | 0 |
| 0 | 0 | 0 | 0 |
| 0 | 0 | 0 | 0 |
| 0 | 0 | 0 | 0 |
| 0 | 0 | 0 | 0 |
| 0 | 0 | 0 | 0 |
| 0 | 0 | 0 | 0 |
| 0 | 0 | 0 | 0 |
| 1 | 0 | 0 | 0 |
| 1 | 0 | 0 | 0 |
| 1 | 0 | 0 | 0 |
| 1 | 0 | 0 | 0 |
| 1 | 0 | 0 | 0 |
| 1 | 0 | 0 | 0 |
| 0 | 0 | 0 | 0 |
| 0 | 0 | 0 | 0 |
| 1 | 0 | 0 | 0 |
| 0 | 0 | 0 | 0 |
| 0 | 0 | 0 | 0 |
| 0 | 0 | 0 | 0 |
| 0 | 0 | 0 | 0 |
| 0 | 0 | 0 | 0 |
| 0 | 0 | 0 | 0 |
| 0 | 0 | 0 | 0 |
| 0 | 0 | 0 | 0 |
| 0 | 0 | 0 | 0 |
| 1 | 1 | 0 | 0 |
| 1 | 1 | 0 | 0 |
| 1 | 1 | 0 | 1 |
| 1 | 1 | 0 | 1 |
| 1 | 1 | 0 | 1 |
| 1 | 1 | 0 | 1 |
| 1 | 1 | 0 | 0 |
| 1 | 1 | 0 | 1 |
| 1 | 1 | 0 | 1 |
| 1 | 1 | 0 | 1 |

| | | | |
|---|---|---|---|
| 1 | 1 | 0 | 1 |
| 1 | 1 | 0 | 1 |
| 1 | 1 | 0 | 1 |
| 1 | 1 | 0 | 1 |
| 1 | 1 | 0 | 1 |
| 1 | 0 | 0 | 0 |
| 0 | 0 | 0 | 0 |
| 0 | 0 | 0 | 0 |
| 0 | 0 | 0 | 0 |
| 1 | 0 | 0 | 0 |
| 0 | 0 | 1 | 0 |
| 0 | 0 | 0 | 0 |
| 0 | 0 | 0 | 0 |
| 0 | 0 | 0 | 0 |
| 0 | 0 | 0 | 0 |
| 0 | 0 | 0 | 0 |
| 0 | 0 | 0 | 0 |
| 0 | 0 | 0 | 0 |
| 0 | 0 | 0 | 0 |
| 0 | 0 | 0 | 0 |
| 0 | 0 | 0 | 0 |
| 0 | 0 | 0 | 0 |
| 0 | 0 | 0 | 0 |
| 0 | 0 | 0 | 0 |
| 0 | 0 | 0 | 0 |
| 1 | 1 | 0 | 0 |
| 1 | 1 | 0 | 0 |
| 1 | 1 | 0 | 0 |
| 0 | 1 | 0 | 0 |
| 1 | 0 | 0 | 0 |
| 0 | 0 | 0 | 0 |
| 0 | 0 | 0 | 0 |
| 0 | 0 | 0 | 0 |
| 1 | 0 | 0 | 0 |
| 1 | 1 | 0 | 0 |
| 0 | 0 | 0 | 0 |
| 0 | 0 | 0 | 0 |
| 0 | 0 | 0 | 0 |
| 0 | 0 | 0 | 0 |
| 1 | 0 | 0 | 0 |
| 0 | 0 | 0 | 0 |
| 0 | 0 | 0 | 0 |
| 0 | 0 | 0 | 0 |
| 0 | 0 | 0 | 0 |
| 0 | 0 | 0 | 0 |
| 0 | 1 | 0 | 0 |
| 0 | 1 | 0 | 0 |
| 0 | 0 | 0 | 0 |
| 0 | 0 | 0 | 0 |
| 0 | 1 | 0 | 0 |
| 0 | 0 | 0 | 0 |
| 0 | 0 | 0 | 0 |
| 0 | 1 | 0 | 0 |
| 0 | 0 | 0 | 0 |

| | | | |
|---|---|---|---|
| 1 | 1 | 0 | 0 |
| 0 | 0 | 0 | 0 |
| 0 | 0 | 0 | 0 |
| 0 | 0 | 0 | 0 |
| 0 | 0 | 0 | 0 |
| 0 | 0 | 0 | 0 |
| 0 | 0 | 0 | 0 |
| 0 | 0 | 0 | 0 |
| 0 | 0 | 0 | 0 |
| 0 | 0 | 0 | 0 |
| 0 | 0 | 0 | 0 |
| 0 | 0 | 0 | 0 |
| 0 | 0 | 0 | 0 |
| 0 | 0 | 0 | 0 |
| 0 | 1 | 0 | 0 |
| 0 | 0 | 0 | 0 |
| 0 | 1 | 0 | 0 |
| 0 | 1 | 0 | 0 |
| 0 | 0 | 0 | 0 |
| 0 | 1 | 0 | 0 |
| 1 | 0 | 0 | 1 |
| 1 | 1 | 0 | 0 |
| 1 | 0 | 0 | 0 |
| 1 | 0 | 0 | 1 |
| 0 | 0 | 0 | 0 |
| 0 | 0 | 0 | 0 |
| 0 | 0 | 0 | 0 |
| 0 | 0 | 0 | 0 |
| 1 | 0 | 0 | 0 |
| 0 | 0 | 0 | 0 |
| 1 | 0 | 0 | 0 |
| 0 | 0 | 0 | 0 |
| 0 | 0 | 0 | 0 |
| 0 | 0 | 0 | 0 |
| 0 | 0 | 0 | 0 |
| 0 | 0 | 0 | 0 |
| 0 | 0 | 0 | 0 |
| 1 | 1 | 0 | 0 |
| 1 | 1 | 0 | 0 |
| 1 | 1 | 0 | 0 |
| 0 | 0 | 0 | 0 |
| 0 | 0 | 0 | 0 |
| 0 | 0 | 0 | 0 |
| 1 | 0 | 0 | 0 |
| 1 | 0 | 0 | 1 |
| 1 | 0 | 0 | 1 |
| 1 | 0 | 0 | 0 |
| 1 | 0 | 0 | 1 |
| 1 | 0 | 0 | 0 |
| 0 | 0 | 0 | 0 |
| 1 | 1 | 0 | 0 |
| 1 | 1 | 0 | 0 |
| 0 | 0 | 0 | 0 |
| 0 | 0 | 0 | 0 |

| | | | |
|---|---|---|---|
| 1 | 0 | 0 | 0 |
| 1 | 0 | 0 | 0 |
| 1 | 0 | 0 | 0 |
| 0 | 0 | 0 | 0 |
| 0 | 0 | 0 | 0 |
| 1 | 1 | 0 | 1 |
| 1 | 1 | 0 | 1 |
| 1 | 1 | 0 | 1 |
| 1 | 1 | 0 | 1 |
| 1 | 1 | 0 | 1 |
| 1 | 1 | 0 | 1 |
| 1 | 1 | 0 | 1 |
| 1 | 1 | 0 | 0 |
| 1 | 1 | 0 | 0 |
| 1 | 1 | 1 | 1 |
| 0 | 0 | 0 | 0 |
| 1 | 1 | 0 | 0 |
| 1 | 1 | 0 | 0 |
| 0 | 0 | 0 | 0 |
| 0 | 0 | 0 | 0 |
| 0 | 0 | 0 | 0 |
| 1 | 0 | 0 | 0 |
| 1 | 0 | 0 | 0 |
| 0 | 0 | 0 | 0 |
| 1 | 0 | 0 | 0 |
| 1 | 0 | 0 | 0 |
| 1 | 0 | 0 | 0 |
| 0 | 0 | 0 | 0 |
| 0 | 0 | 0 | 0 |
| 0 | 0 | 0 | 0 |
| 0 | 0 | 0 | 0 |
| 0 | 0 | 0 | 0 |
| 0 | 0 | 0 | 0 |
| 0 | 0 | 0 | 0 |
| 0 | 0 | 0 | 0 |
| 0 | 0 | 0 | 0 |
| 0 | 0 | 0 | 0 |
| 0 | 0 | 0 | 0 |
| 0 | 0 | 0 | 0 |
| 0 | 0 | 0 | 0 |
| 0 | 0 | 0 | 0 |
| 0 | 0 | 0 | 0 |
| 0 | 0 | 0 | 0 |
| 0 | 0 | 0 | 0 |
| 0 | 0 | 0 | 0 |
| 1 | 0 | 0 | 0 |
| 1 | 0 | 0 | 0 |
| 1 | 0 | 0 | 0 |
| 1 | 0 | 0 | 0 |
| 0 | 0 | 0 | 0 |
| 0 | 0 | 0 | 0 |
| 0 | 0 | 0 | 0 |
| 0 | 0 | 0 | 0 |
| 0 | 0 | 0 | 0 |

| | | | |
|---|---|---|---|
| 0 | 0 | 0 | 0 |
| 0 | 0 | 0 | 0 |
| 0 | 0 | 0 | 0 |
| 0 | 0 | 0 | 0 |
| 0 | 0 | 0 | 0 |
| 0 | 0 | 0 | 0 |
| 0 | 0 | 0 | 0 |
| 1 | 0 | 0 | 1 |
| 0 | 0 | 0 | 0 |
| 0 | 0 | 0 | 0 |
| 1 | 0 | 0 | 0 |
| 0 | 0 | 0 | 0 |
| 0 | 0 | 0 | 0 |
| 0 | 0 | 0 | 0 |
| 0 | 0 | 0 | 0 |
| 1 | 0 | 0 | 0 |
| 0 | 0 | 0 | 0 |
| 0 | 0 | 0 | 0 |
| 0 | 0 | 0 | 0 |
| 0 | 0 | 0 | 0 |
| 0 | 0 | 0 | 0 |
| 0 | 0 | 0 | 0 |
| 0 | 0 | 0 | 0 |
| 1 | 1 | 0 | 1 |
| 0 | 0 | 0 | 0 |
| 0 | 0 | 0 | 0 |
| 0 | 0 | 0 | 0 |
| 1 | 1 | 0 | 1 |
| 1 | 1 | 0 | 0 |
| 0 | 0 | 0 | 0 |
| 1 | 0 | 0 | 0 |
| 1 | 1 | 0 | 0 |
| 0 | 0 | 0 | 0 |
| 1 | 0 | 0 | 0 |
| 1 | 1 | 0 | 0 |
| 1 | 0 | 0 | 0 |
| 1 | 1 | 0 | 1 |
| 1 | 1 | 0 | 1 |
| 1 | 0 | 0 | 0 |
| 0 | 0 | 0 | 0 |
| 0 | 0 | 0 | 0 |
| 0 | 0 | 0 | 0 |
| 0 | 0 | 0 | 0 |
| 0 | 0 | 0 | 0 |
| 0 | 0 | 0 | 0 |
| 0 | 0 | 0 | 0 |
| 0 | 0 | 0 | 0 |
| 0 | 0 | 0 | 0 |
| 0 | 0 | 0 | 0 |
| 0 | 0 | 0 | 0 |
| 0 | 0 | 0 | 0 |
| 0 | 0 | 0 | 0 |
| 0 | 0 | 0 | 0 |
| 0 | 0 | 0 | 0 |

| | | | |
|---|---|---|---|
| 0 | 0 | 0 | 0 |
| 0 | 0 | 0 | 0 |
| 0 | 0 | 0 | 0 |
| 0 | 0 | 0 | 0 |
| 0 | 0 | 0 | 0 |
| 0 | 0 | 0 | 0 |
| 0 | 0 | 0 | 0 |
| 0 | 0 | 0 | 0 |
| 0 | 0 | 0 | 0 |
| 0 | 0 | 0 | 0 |
| 0 | 0 | 0 | 0 |
| 0 | 0 | 0 | 0 |
| 0 | 0 | 0 | 0 |
| 1 | 1 | 0 | 0 |
| 0 | 0 | 0 | 0 |
| 1 | 0 | 0 | 0 |
| 0 | 0 | 0 | 1 |
| 0 | 0 | 0 | 0 |
| 0 | 0 | 0 | 0 |
| 0 | 0 | 0 | 0 |
| 0 | 0 | 0 | 0 |
| 0 | 0 | 0 | 0 |
| 0 | 0 | 0 | 0 |
| 0 | 0 | 0 | 0 |
| 0 | 0 | 0 | 0 |
| 0 | 0 | 0 | 0 |
| 0 | 0 | 0 | 0 |
| 0 | 0 | 0 | 0 |
| 0 | 0 | 0 | 0 |
| 0 | 0 | 0 | 0 |
| 1 | 0 | 0 | 0 |
| 1 | 0 | 0 | 0 |
| 0 | 0 | 0 | 0 |
| 0 | 0 | 0 | 0 |
| 0 | 0 | 0 | 0 |
| 0 | 0 | 0 | 0 |
| 0 | 0 | 0 | 0 |
| 0 | 0 | 0 | 0 |
| 0 | 0 | 0 | 0 |
| 0 | 0 | 0 | 0 |
| 1 | 0 | 0 | 0 |
| 1 | 0 | 0 | 0 |
| 1 | 0 | 0 | 0 |
| 1 | 0 | 0 | 0 |
| 0 | 0 | 0 | 0 |
| 1 | 0 | 0 | 0 |
| 0 | 0 | 0 | 0 |
| 0 | 0 | 0 | 0 |
| 1 | 0 | 0 | 0 |
| 1 | 0 | 0 | 0 |
| 0 | 0 | 0 | 0 |
| 0 | 0 | 0 | 0 |
| 0 | 0 | 0 | 0 |
| 0 | 0 | 0 | 0 |

| | | | |
|---|---|---|---|
| 0 | 0 | 0 | 0 |
| 0 | 0 | 0 | 0 |
| 0 | 0 | 0 | 0 |
| 0 | 0 | 0 | 0 |
| 0 | 0 | 0 | 0 |
| 0 | 0 | 0 | 0 |
| 0 | 0 | 0 | 0 |
| 0 | 0 | 0 | 0 |
| 0 | 0 | 0 | 0 |
| 0 | 0 | 0 | 0 |
| 0 | 0 | 0 | 0 |
| 0 | 0 | 0 | 0 |
| 0 | 0 | 0 | 0 |
| 0 | 0 | 0 | 0 |
| 0 | 0 | 0 | 0 |
| 1 | 0 | 0 | 1 |
| 0 | 0 | 0 | 0 |
| 0 | 0 | 0 | 0 |
| 0 | 0 | 0 | 0 |
| 0 | 0 | 0 | 0 |
| 0 | 0 | 0 | 0 |
| 1 | 1 | 0 | 0 |
| 1 | 1 | 0 | 0 |
| 1 | 0 | 0 | 0 |
| 1 | 0 | 0 | 0 |
| 0 | 0 | 0 | 0 |
| 0 | 0 | 0 | 0 |
| 0 | 0 | 0 | 0 |
| 0 | 0 | 0 | 0 |
| 1 | 0 | 0 | 0 |
| 1 | 0 | 0 | 0 |
| 1 | 0 | 0 | 0 |
| 1 | 1 | 0 | 0 |
| 1 | 1 | 0 | 0 |
| 1 | 0 | 0 | 0 |
| 0 | 0 | 0 | 0 |
| 0 | 0 | 0 | 0 |
| 0 | 1 | 0 | 0 |
| 0 | 0 | 0 | 0 |
| 0 | 0 | 0 | 0 |
| 0 | 0 | 0 | 0 |
| 1 | 1 | 0 | 0 |
| 1 | 0 | 0 | 0 |
| 1 | 0 | 0 | 0 |
| 0 | 0 | 0 | 0 |
| 0 | 0 | 0 | 0 |
| 0 | 0 | 0 | 0 |
| 0 | 0 | 0 | 0 |
| 0 | 0 | 0 | 0 |
| 0 | 0 | 0 | 0 |
| 0 | 0 | 0 | 0 |
| 0 | 0 | 0 | 0 |
| 0 | 0 | 0 | 0 |
| 0 | 0 | 0 | 0 |

| | | | |
|---|---|---|---|
| 0 | 0 | 0 | 0 |
| 1 | 0 | 0 | 0 |
| 0 | 0 | 0 | 0 |
| 0 | 0 | 0 | 0 |
| 0 | 0 | 0 | 0 |
| 0 | 0 | 0 | 0 |
| 1 | 0 | 0 | 0 |
| 0 | 0 | 0 | 0 |
| 0 | 0 | 0 | 0 |
| 0 | 0 | 0 | 0 |
| 0 | 0 | 0 | 0 |
| 0 | 0 | 0 | 0 |
| 0 | 0 | 0 | 0 |
| 0 | 0 | 0 | 0 |
| 0 | 0 | 0 | 0 |
| 0 | 0 | 0 | 0 |
| 0 | 1 | 0 | 0 |
| 0 | 0 | 0 | 0 |
| 0 | 0 | 0 | 0 |
| 0 | 0 | 0 | 0 |
| 1 | 0 | 0 | 0 |
| 0 | 0 | 0 | 0 |
| 0 | 0 | 0 | 0 |
| 1 | 1 | 0 | 0 |
| 0 | 0 | 0 | 0 |
| 0 | 0 | 0 | 0 |
| 0 | 0 | 0 | 0 |
| 0 | 0 | 0 | 0 |
| 1 | 0 | 0 | 0 |
| 0 | 0 | 0 | 0 |
| 0 | 0 | 0 | 0 |
| 0 | 0 | 0 | 0 |
| 0 | 0 | 0 | 0 |
| 0 | 0 | 0 | 0 |
| 0 | 0 | 0 | 0 |
| 1 | 0 | 0 | 0 |
| 0 | 0 | 0 | 0 |
| 0 | 0 | 0 | 0 |
| 0 | 0 | 0 | 0 |
| 0 | 0 | 0 | 0 |
| 1 | 1 | 0 | 0 |
| 0 | 0 | 0 | 0 |
| 0 | 0 | 0 | 0 |
| 0 | 0 | 0 | 0 |
| 0 | 0 | 0 | 0 |
| 0 | 0 | 0 | 0 |
| 1 | 0 | 0 | 0 |
| 1 | 0 | 0 | 1 |
| 1 | 0 | 0 | 0 |
| 0 | 0 | 0 | 0 |
| 0 | 0 | 0 | 0 |
| 0 | 0 | 0 | 0 |
| 1 | 0 | 0 | 0 |
| 1 | 0 | 0 | 0 |

| | | | |
|---|---|---|---|
| 1 | 0 | 0 | 0 |
| 0 | 0 | 0 | 0 |
| 0 | 0 | 0 | 0 |
| 0 | 0 | 0 | 0 |
| 1 | 1 | 0 | 0 |
| 1 | 1 | 0 | 0 |
| 0 | 0 | 0 | 0 |
| 0 | 0 | 0 | 0 |
| 1 | 0 | 0 | 0 |
| 1 | 0 | 0 | 0 |
| 1 | 0 | 1 | 0 |
| 0 | 0 | 0 | 0 |
| 0 | 0 | 1 | 0 |
| 0 | 0 | 1 | 0 |
| 0 | 0 | 1 | 0 |
| 1 | 0 | 0 | 0 |
| 1 | 0 | 0 | 0 |
| 0 | 0 | 0 | 0 |
| 0 | 0 | 0 | 0 |
| 1 | 0 | 0 | 1 |
| 1 | 0 | 0 | 1 |
| 1 | 0 | 0 | 0 |
| 0 | 0 | 0 | 0 |
| 0 | 0 | 0 | 0 |
| 0 | 0 | 0 | 0 |
| 0 | 0 | 0 | 0 |
| 0 | 0 | 0 | 0 |
| 1 | 0 | 0 | 0 |
| 0 | 0 | 0 | 0 |
| 0 | 0 | 0 | 1 |
| 0 | 0 | 0 | 1 |
| 0 | 0 | 1 | 1 |
| 0 | 0 | 1 | 1 |
| 0 | 0 | 1 | 1 |
| 0 | 0 | 1 | 1 |
| 0 | 0 | 1 | 0 |
| 0 | 0 | 1 | 0 |
| 0 | 0 | 0 | 0 |
| 0 | 0 | 0 | 0 |
| 0 | 0 | 0 | 0 |
| 0 | 0 | 0 | 0 |
| 0 | 0 | 1 | 0 |
| 0 | 0 | 1 | 0 |
| 0 | 0 | 1 | 0 |
| 0 | 0 | 1 | 0 |
| 0 | 0 | 1 | 0 |
| 0 | 0 | 1 | 0 |
| 0 | 0 | 1 | 0 |
| 0 | 0 | 1 | 0 |
| 0 | 0 | 0 | 0 |
| 0 | 0 | 0 | 0 |
| 0 | 0 | 0 | 0 |
| 0 | 0 | 0 | 0 |
| 1 | 0 | 0 | 0 |

| | | | |
|---|---|---|---|
| 0 | 0 | 0 | 0 |
| 0 | 0 | 0 | 0 |
| 0 | 0 | 0 | 0 |
| 1 | 0 | 0 | 0 |
| 0 | 0 | 0 | 0 |
| 0 | 0 | 0 | 0 |
| 0 | 0 | 0 | 0 |
| 0 | 0 | 0 | 0 |
| 0 | 0 | 0 | 0 |
| 0 | 0 | 0 | 0 |
| 1 | 1 | 0 | 0 |
| 0 | 0 | 0 | 0 |
| 1 | 1 | 0 | 0 |
| 1 | 1 | 0 | 0 |
| 0 | 1 | 0 | 0 |
| 0 | 0 | 0 | 0 |
| 0 | 0 | 0 | 0 |
| 0 | 0 | 0 | 0 |
| 0 | 0 | 0 | 0 |
| 0 | 0 | 0 | 0 |
| 1 | 1 | 0 | 0 |
| 0 | 1 | 0 | 0 |
| 0 | 0 | 0 | 0 |
| 0 | 1 | 0 | 0 |
| 1 | 1 | 0 | 0 |
| 0 | 0 | 0 | 0 |
| 0 | 0 | 0 | 0 |
| 1 | 1 | 0 | 0 |
| 1 | 1 | 0 | 0 |
| 0 | 1 | 0 | 0 |
| 0 | 0 | 0 | 0 |
| 0 | 0 | 0 | 0 |
| 0 | 0 | 0 | 0 |
| 0 | 0 | 0 | 0 |
| 0 | 0 | 0 | 0 |
| 0 | 0 | 0 | 0 |
| 0 | 0 | 0 | 0 |
| 0 | 0 | 0 | 0 |
| 0 | 0 | 0 | 0 |
| 0 | 0 | 0 | 0 |
| 1 | 0 | 0 | 0 |
| 1 | 0 | 0 | 0 |
| 1 | 0 | 0 | 0 |
| 1 | 0 | 0 | 0 |
| 1 | 1 | 0 | 1 |
| 0 | 0 | 0 | 0 |
| 1 | 0 | 0 | 0 |
| 0 | 0 | 0 | 0 |
| 1 | 0 | 0 | 0 |
| 0 | 0 | 0 | 0 |
| 1 | 0 | 0 | 0 |
| 1 | 0 | 0 | 0 |
| 0 | 0 | 0 | 0 |
| 0 | 0 | 0 | 0 |

| | | | |
|---|---|---|---|
| 0 | 0 | 0 | 0 |
| 0 | 0 | 0 | 0 |
| 0 | 0 | 0 | 0 |
| 0 | 0 | 0 | 0 |
| 1 | 0 | 0 | 0 |
| 1 | 0 | 0 | 0 |
| 1 | 0 | 0 | 0 |
| 0 | 0 | 0 | 0 |
| 1 | 1 | 0 | 0 |
| 0 | 0 | 0 | 0 |
| 0 | 0 | 0 | 0 |
| 0 | 0 | 0 | 0 |
| 1 | 1 | 0 | 0 |
| 1 | 0 | 0 | 0 |
| 0 | 0 | 0 | 0 |
| 0 | 0 | 0 | 0 |
| 1 | 1 | 0 | 0 |
| 0 | 0 | 0 | 0 |
| 0 | 0 | 0 | 0 |
| 0 | 0 | 0 | 0 |
| 0 | 0 | 0 | 0 |
| 0 | 0 | 0 | 0 |
| 0 | 0 | 0 | 0 |
| 0 | 0 | 0 | 0 |
| 1 | 0 | 0 | 0 |
| 0 | 0 | 0 | 0 |
| 1 | 0 | 0 | 0 |
| 0 | 0 | 0 | 0 |
| 0 | 0 | 0 | 0 |
| 0 | 0 | 0 | 0 |
| 1 | 0 | 0 | 0 |
| 1 | 0 | 0 | 0 |
| 0 | 0 | 0 | 0 |
| 0 | 0 | 0 | 0 |
| 1 | 0 | 0 | 0 |
| 0 | 0 | 0 | 0 |
| 0 | 0 | 0 | 0 |
| 1 | 0 | 0 | 0 |
| 0 | 0 | 0 | 0 |
| 1 | 0 | 0 | 0 |
| 1 | 0 | 0 | 0 |
| 0 | 0 | 0 | 0 |
| 0 | 0 | 0 | 0 |
| 1 | 1 | 0 | 0 |
| 1 | 0 | 0 | 0 |
| 1 | 1 | 0 | 0 |
| 0 | 0 | 0 | 0 |
| 0 | 0 | 0 | 0 |
| 0 | 0 | 0 | 0 |
| 0 | 0 | 0 | 0 |
| 1 | 1 | 0 | 0 |
| 0 | 0 | 0 | 0 |
| 0 | 0 | 0 | 0 |
| 0 | 0 | 0 | 0 |

| | | | |
|---|---|---|---|
| 0 | 1 | 0 | 0 |
| 1 | 0 | 0 | 0 |
| 0 | 0 | 0 | 0 |
| 0 | 0 | 0 | 0 |
| 0 | 0 | 0 | 0 |
| 1 | 1 | 0 | 0 |
| 0 | 0 | 0 | 0 |
| 0 | 0 | 0 | 0 |
| 0 | 0 | 0 | 0 |
| 1 | 0 | 0 | 0 |
| 1 | 0 | 0 | 0 |
| 0 | 0 | 0 | 0 |
| 0 | 0 | 0 | 0 |
| 0 | 0 | 0 | 0 |
| 0 | 0 | 0 | 0 |
| 0 | 0 | 0 | 0 |
| 0 | 0 | 0 | 0 |
| 0 | 0 | 0 | 0 |
| 0 | 0 | 0 | 0 |
| 0 | 0 | 0 | 0 |
| 0 | 0 | 0 | 0 |
| 0 | 0 | 0 | 0 |
| 0 | 0 | 0 | 0 |
| 1 | 1 | 0 | 0 |
| 0 | 0 | 0 | 0 |
| 0 | 0 | 0 | 0 |
| 1 | 1 | 1 | 0 |
| 0 | 0 | 0 | 0 |
| 0 | 0 | 0 | 0 |
| 0 | 0 | 0 | 0 |
| 1 | 1 | 1 | 0 |
| 1 | 0 | 0 | 0 |
| 1 | 0 | 0 | 0 |
| 1 | 0 | 0 | 0 |
| 1 | 0 | 0 | 0 |
| 1 | 1 | 1 | 0 |
| 1 | 1 | 1 | 0 |
| 0 | 0 | 0 | 0 |
| 0 | 0 | 0 | 0 |
| 1 | 1 | 1 | 0 |
| 0 | 0 | 0 | 0 |
| 0 | 0 | 0 | 0 |
| 0 | 0 | 0 | 0 |
| 0 | 0 | 0 | 0 |
| 0 | 0 | 0 | 0 |
| 0 | 0 | 0 | 0 |
| 0 | 0 | 0 | 0 |
| 0 | 0 | 0 | 0 |
| 1 | 0 | 0 | 0 |
| 1 | 0 | 0 | 0 |
| 1 | 0 | 0 | 0 |
| 0 | 0 | 0 | 0 |
| 0 | 0 | 0 | 0 |
| 0 | 0 | 0 | 0 |

| | | | |
|---|---|---|---|
| 0 | 0 | 0 | 0 |
| 1 | 0 | 0 | 0 |
| 0 | 0 | 0 | 0 |
| 0 | 0 | 0 | 0 |
| 0 | 0 | 0 | 0 |
| 0 | 0 | 0 | 0 |
| 0 | 0 | 0 | 0 |
| 0 | 0 | 0 | 0 |
| 0 | 0 | 0 | 0 |
| 0 | 0 | 0 | 0 |
| 1 | 1 | 0 | 0 |
| 1 | 1 | 0 | 0 |
| 0 | 0 | 0 | 0 |
| 0 | 0 | 0 | 0 |
| 0 | 0 | 0 | 0 |
| 1 | 0 | 0 | 0 |
| 1 | 0 | 0 | 0 |
| 1 | 1 | 0 | 1 |
| 0 | 0 | 0 | 0 |
| 0 | 0 | 0 | 0 |
| 0 | 0 | 0 | 0 |
| 0 | 0 | 0 | 0 |
| 1 | 0 | 0 | 1 |
| 0 | 0 | 0 | 0 |
| 0 | 0 | 0 | 0 |
| 1 | 0 | 0 | 0 |
| 0 | 0 | 0 | 0 |
| 0 | 0 | 0 | 0 |
| 0 | 0 | 0 | 0 |
| 0 | 0 | 0 | 0 |
| 0 | 0 | 0 | 0 |
| 0 | 0 | 0 | 0 |
| 0 | 0 | 0 | 0 |
| 0 | 0 | 0 | 0 |
| 0 | 0 | 0 | 0 |
| 1 | 0 | 0 | 0 |
| 1 | 0 | 0 | 0 |
| 1 | 1 | 0 | 0 |
| 1 | 1 | 0 | 0 |
| 0 | 0 | 0 | 0 |
| 0 | 0 | 0 | 0 |
| 1 | 0 | 0 | 0 |
| 0 | 0 | 0 | 0 |
| 1 | 0 | 0 | 0 |
| 1 | 1 | 0 | 0 |
| 0 | 0 | 0 | 0 |
| 0 | 0 | 0 | 0 |
| 0 | 0 | 0 | 0 |
| 0 | 0 | 0 | 0 |
| 0 | 0 | 0 | 0 |
| 0 | 0 | 0 | 0 |
| 1 | 0 | 0 | 0 |
| 0 | 0 | 0 | 0 |
| 0 | 0 | 0 | 0 |

| | | | |
|---|---|---|---|
| 0 | 0 | 0 | 0 |
| 1 | 1 | 0 | 0 |
| 0 | 0 | 0 | 0 |
| 1 | 0 | 0 | 0 |
| 1 | 0 | 0 | 0 |
| 0 | 0 | 0 | 0 |
| 0 | 0 | 0 | 0 |
| 0 | 0 | 0 | 0 |
| 0 | 0 | 0 | 0 |
| 0 | 0 | 0 | 0 |
| 0 | 0 | 0 | 0 |
| 1 | 1 | 0 | 0 |
| 1 | 1 | 0 | 1 |
| 0 | 0 | 0 | 0 |
| 0 | 1 | 0 | 0 |
| 0 | 0 | 0 | 0 |
| 0 | 0 | 0 | 0 |
| 0 | 0 | 0 | 0 |
| 0 | 0 | 0 | 0 |
| 0 | 0 | 0 | 0 |
| 0 | 0 | 0 | 0 |
| 1 | 0 | 0 | 0 |
| 0 | 0 | 0 | 0 |
| 0 | 0 | 0 | 0 |
| 0 | 0 | 0 | 0 |
| 0 | 0 | 0 | 0 |
| 0 | 0 | 0 | 0 |
| 1 | 0 | 0 | 0 |
| 0 | 0 | 0 | 0 |
| 0 | 0 | 0 | 0 |
| 0 | 0 | 0 | 0 |
| 0 | 0 | 0 | 0 |
| 0 | 0 | 0 | 0 |
| 0 | 0 | 0 | 0 |
| 0 | 0 | 0 | 0 |
| 0 | 0 | 0 | 0 |
| 0 | 0 | 0 | 0 |
| 0 | 0 | 0 | 0 |
| 0 | 0 | 0 | 0 |
| 0 | 0 | 0 | 0 |
| 0 | 0 | 0 | 0 |
| 0 | 0 | 0 | 0 |
| 0 | 0 | 0 | 0 |
| 0 | 0 | 0 | 0 |
| 0 | 0 | 0 | 0 |
| 0 | 0 | 0 | 0 |
| 0 | 0 | 0 | 0 |
| 0 | 0 | 0 | 0 |
| 0 | 0 | 0 | 0 |
| 0 | 0 | 0 | 0 |
| 0 | 0 | 0 | 0 |
| 0 | 0 | 0 | 0 |
| 0 | 0 | 0 | 0 |
| 0 | 0 | 0 | 0 |

| | | | |
|---|---|---|---|
| 0 | 0 | 0 | 0 |
| 0 | 0 | 0 | 0 |
| 0 | 0 | 0 | 0 |
| 0 | 0 | 0 | 0 |
| 0 | 0 | 0 | 0 |
| 0 | 0 | 0 | 0 |
| 0 | 0 | 0 | 0 |
| 0 | 0 | 0 | 0 |
| 0 | 0 | 0 | 0 |
| 0 | 0 | 0 | 0 |
| 0 | 0 | 0 | 0 |
| 0 | 0 | 0 | 0 |
| 1 | 0 | 0 | 0 |
| 1 | 1 | 0 | 0 |
| 0 | 0 | 0 | 0 |
| 1 | 0 | 0 | 0 |
| 0 | 0 | 0 | 0 |
| 1 | 0 | 0 | 0 |
| 0 | 0 | 0 | 0 |
| 0 | 0 | 0 | 0 |
| 1 | 0 | 0 | 0 |
| 0 | 0 | 0 | 0 |
| 0 | 0 | 0 | 0 |
| 1 | 0 | 0 | 0 |
| 1 | 0 | 0 | 0 |
| 0 | 0 | 0 | 0 |
| 0 | 0 | 0 | 0 |
| 1 | 0 | 0 | 0 |
| 1 | 0 | 0 | 0 |
| 1 | 0 | 0 | 0 |
| 1 | 0 | 0 | 0 |
| 1 | 0 | 0 | 0 |
| 0 | 0 | 0 | 0 |
| 0 | 0 | 0 | 0 |
| 0 | 0 | 0 | 0 |
| 0 | 0 | 0 | 0 |
| 0 | 0 | 0 | 0 |
| 0 | 0 | 0 | 0 |
| 0 | 0 | 0 | 0 |
| 0 | 0 | 0 | 0 |
| 0 | 0 | 0 | 0 |
| 0 | 0 | 0 | 0 |
| 0 | 0 | 0 | 0 |
| 0 | 0 | 0 | 0 |
| 1 | 1 | 0 | 0 |
| 0 | 0 | 0 | 0 |
| 0 | 0 | 0 | 0 |
| 0 | 0 | 0 | 0 |
| 0 | 0 | 0 | 0 |
| 1 | 1 | 0 | 0 |
| 1 | 0 | 0 | 0 |
| 0 | 0 | 0 | 0 |
| 0 | 0 | 0 | 0 |
| 0 | 0 | 0 | 1 |

| | | | |
|---|---|---|---|
| 0 | 0 | 0 | 0 |
| 0 | 0 | 0 | 0 |
| 0 | 0 | 0 | 0 |
| 0 | 0 | 0 | 0 |
| 1 | 0 | 0 | 1 |
| 0 | 0 | 0 | 0 |
| 1 | 0 | 0 | 0 |
| 0 | 0 | 0 | 0 |
| 0 | 0 | 0 | 0 |
| 0 | 0 | 0 | 0 |
| 0 | 0 | 0 | 0 |
| 0 | 0 | 0 | 0 |
| 0 | 0 | 0 | 0 |
| 1 | 0 | 0 | 0 |
| 0 | 0 | 0 | 0 |
| 0 | 0 | 0 | 0 |
| 1 | 1 | 0 | 0 |
| 0 | 0 | 0 | 0 |
| 1 | 0 | 0 | 0 |
| 1 | 0 | 0 | 0 |
| 0 | 0 | 0 | 0 |
| 0 | 0 | 0 | 0 |
| 0 | 0 | 0 | 0 |
| 1 | 0 | 0 | 0 |
| 1 | 0 | 0 | 0 |
| 1 | 0 | 0 | 0 |
| 1 | 0 | 0 | 0 |
| 1 | 0 | 0 | 0 |
| 1 | 0 | 0 | 0 |
| 1 | 0 | 0 | 0 |
| 1 | 0 | 0 | 0 |
| 1 | 0 | 0 | 0 |
| 1 | 0 | 0 | 0 |
| 1 | 0 | 0 | 0 |
| 0 | 0 | 0 | 0 |
| 0 | 0 | 0 | 0 |
| 0 | 0 | 0 | 0 |
| 1 | 0 | 0 | 0 |
| 1 | 0 | 0 | 0 |
| 0 | 0 | 0 | 0 |
| 0 | 0 | 0 | 0 |
| 1 | 0 | 0 | 0 |
| 0 | 0 | 0 | 0 |
| 1 | 0 | 0 | 0 |
| 1 | 0 | 0 | 0 |
| 1 | 0 | 0 | 0 |
| 1 | 0 | 0 | 0 |
| 1 | 0 | 0 | 0 |
| 0 | 0 | 0 | 0 |
| 0 | 0 | 0 | 0 |
| 1 | 0 | 0 | 0 |
| 0 | 0 | 0 | 0 |
| 1 | 0 | 0 | 0 |
| 1 | 0 | 0 | 0 |

| | | | |
|---|---|---|---|
| 1 | 0 | 0 | 0 |
| 1 | 0 | 0 | 0 |
| 1 | 0 | 0 | 0 |
| 1 | 0 | 0 | 0 |
| 1 | 0 | 0 | 0 |
| 1 | 0 | 0 | 0 |
| 0 | 0 | 0 | 0 |
| 1 | 0 | 0 | 0 |
| 1 | 0 | 0 | 0 |
| 1 | 0 | 0 | 0 |
| 1 | 0 | 0 | 0 |
| 1 | 0 | 0 | 0 |
| 1 | 0 | 0 | 0 |
| 1 | 0 | 0 | 0 |
| 0 | 0 | 0 | 0 |
| 0 | 0 | 0 | 0 |
| 1 | 1 | 0 | 0 |
| 1 | 1 | 0 | 0 |
| 0 | 0 | 0 | 0 |
| 0 | 0 | 0 | 0 |
| 0 | 0 | 0 | 0 |
| 1 | 1 | 0 | 1 |
| 0 | 0 | 0 | 0 |
| 0 | 0 | 0 | 0 |
| 0 | 0 | 0 | 0 |
| 0 | 0 | 0 | 0 |
| 0 | 0 | 0 | 0 |
| 0 | 0 | 0 | 0 |
| 0 | 0 | 0 | 0 |
| 1 | 0 | 1 | 0 |
| 0 | 0 | 0 | 0 |
| 0 | 0 | 0 | 0 |
| 0 | 0 | 0 | 0 |
| 0 | 0 | 0 | 0 |
| 0 | 0 | 0 | 0 |
| 0 | 0 | 0 | 0 |
| 0 | 0 | 0 | 0 |
| 0 | 0 | 0 | 0 |
| 0 | 0 | 0 | 0 |
| 0 | 0 | 0 | 0 |
| 1 | 1 | 0 | 0 |
| 1 | 1 | 0 | 0 |
| 1 | 0 | 0 | 0 |
| 1 | 1 | 0 | 0 |
| 1 | 0 | 0 | 0 |
| 1 | 1 | 0 | 0 |
| 1 | 0 | 0 | 0 |
| 1 | 0 | 0 | 0 |
| 0 | 0 | 0 | 0 |
| 0 | 0 | 0 | 0 |
| 1 | 0 | 0 | 0 |
| 1 | 0 | 0 | 0 |
| 1 | 0 | 0 | 0 |
| 0 | 0 | 0 | 0 |

| | | | |
|---|---|---|---|
| 0 | 0 | 0 | 0 |
| 0 | 0 | 0 | 0 |
| 0 | 0 | 0 | 0 |
| 1 | 1 | 1 | 0 |
| 0 | 0 | 0 | 0 |
| 0 | 0 | 0 | 0 |
| 0 | 0 | 0 | 0 |
| 0 | 0 | 0 | 0 |
| 0 | 1 | 0 | 0 |
| 0 | 1 | 0 | 0 |
| 0 | 1 | 0 | 0 |
| 0 | 0 | 0 | 0 |
| 0 | 0 | 0 | 0 |
| 0 | 0 | 0 | 0 |
| 1 | 0 | 0 | 0 |
| 1 | 0 | 0 | 0 |
| 0 | 0 | 0 | 0 |
| 0 | 0 | 0 | 0 |
| 1 | 0 | 0 | 0 |
| 0 | 0 | 0 | 0 |
| 1 | 0 | 0 | 0 |
| 1 | 0 | 0 | 0 |
| 0 | 0 | 0 | 0 |
| 0 | 0 | 0 | 0 |
| 0 | 0 | 0 | 0 |
| 0 | 0 | 0 | 0 |
| 0 | 0 | 0 | 0 |
| 0 | 0 | 1 | 0 |
| 0 | 0 | 0 | 0 |
| 0 | 1 | 0 | 0 |
| 0 | 0 | 0 | 0 |
| 0 | 0 | 0 | 0 |
| 0 | 0 | 0 | 0 |
| 0 | 0 | 0 | 0 |
| 0 | 0 | 0 | 0 |
| 0 | 0 | 0 | 0 |
| 1 | 0 | 0 | 0 |
| 1 | 0 | 0 | 0 |
| 1 | 0 | 0 | 0 |
| 0 | 0 | 0 | 0 |
| 0 | 0 | 0 | 0 |
| 0 | 0 | 0 | 0 |
| 0 | 0 | 0 | 0 |
| 0 | 0 | 0 | 0 |
| 0 | 0 | 0 | 0 |
| 0 | 0 | 0 | 0 |
| 0 | 0 | 0 | 0 |
| 0 | 0 | 0 | 1 |
| 0 | 0 | 0 | 0 |
| 0 | 0 | 0 | 0 |
| 0 | 0 | 0 | 0 |
| 0 | 0 | 0 | 0 |
| 1 | 1 | 0 | 0 |
| 0 | 0 | 0 | 0 |

| | | | |
|---|---|---|---|
| 0 | 0 | 0 | 0 |
| 0 | 0 | 0 | 0 |
| 0 | 0 | 0 | 0 |
| 0 | 0 | 0 | 0 |
| 1 | 0 | 0 | 0 |
| 0 | 0 | 0 | 0 |
| 0 | 0 | 0 | 0 |
| 0 | 0 | 0 | 0 |
| 0 | 0 | 0 | 0 |
| 0 | 0 | 0 | 0 |
| 0 | 0 | 0 | 0 |
| 0 | 0 | 0 | 0 |
| 0 | 0 | 0 | 0 |
| 0 | 0 | 0 | 0 |
| 0 | 0 | 0 | 0 |
| 0 | 0 | 0 | 0 |
| 0 | 0 | 0 | 0 |
| 0 | 0 | 0 | 0 |
| 0 | 0 | 0 | 0 |
| 0 | 0 | 0 | 0 |
| 0 | 0 | 0 | 0 |
| 0 | 0 | 0 | 0 |
| 0 | 0 | 0 | 0 |
| 0 | 0 | 0 | 0 |
| 0 | 0 | 0 | 0 |
| 0 | 0 | 0 | 0 |
| 0 | 0 | 0 | 0 |
| 0 | 0 | 0 | 0 |
| 0 | 0 | 0 | 0 |
| 0 | 0 | 0 | 0 |
| 0 | 0 | 0 | 0 |
| 0 | 0 | 0 | 0 |
| 0 | 0 | 0 | 0 |
| 0 | 0 | 0 | 0 |
| 0 | 0 | 0 | 0 |
| 0 | 0 | 0 | 0 |
| 0 | 0 | 0 | 0 |
| 0 | 0 | 0 | 0 |
| 0 | 0 | 0 | 0 |
| 0 | 0 | 0 | 0 |
| 1 | 1 | 0 | 0 |
| 0 | 0 | 0 | 0 |
| 0 | 0 | 0 | 0 |
| 0 | 0 | 0 | 0 |
| 0 | 0 | 0 | 0 |
| 1 | 0 | 0 | 0 |
| 1 | 0 | 0 | 0 |
| 0 | 0 | 0 | 0 |
| 1 | 0 | 0 | 0 |
| 0 | 0 | 0 | 0 |
| 0 | 0 | 0 | 0 |
| 1 | 0 | 0 | 0 |
| 1 | 0 | 0 | 0 |
| 0 | 0 | 0 | 0 |

| | | | |
|---|---|---|---|
| 0 | 0 | 0 | 0 |
| 0 | 0 | 0 | 0 |
| 0 | 0 | 0 | 0 |
| 0 | 0 | 0 | 0 |
| 1 | 0 | 0 | 0 |
| 0 | 0 | 0 | 0 |
| 0 | 0 | 0 | 0 |
| 0 | 0 | 0 | 0 |
| 0 | 0 | 0 | 0 |
| 0 | 0 | 0 | 0 |
| 0 | 0 | 0 | 0 |
| 0 | 0 | 0 | 0 |
| 0 | 0 | 0 | 0 |
| 0 | 0 | 0 | 0 |
| 0 | 0 | 0 | 0 |
| 1 | 1 | 0 | 1 |
| 0 | 0 | 0 | 0 |
| 0 | 0 | 0 | 0 |
| 0 | 0 | 0 | 0 |
| 0 | 0 | 0 | 0 |
| 0 | 0 | 0 | 0 |
| 0 | 0 | 0 | 0 |
| 0 | 0 | 0 | 0 |
| 0 | 0 | 0 | 0 |
| 0 | 0 | 0 | 0 |
| 0 | 0 | 0 | 0 |
| 0 | 0 | 0 | 0 |
| 1 | 0 | 0 | 0 |
| 1 | 0 | 0 | 0 |
| 0 | 0 | 0 | 0 |
| 1 | 1 | 0 | 0 |
| 0 | 0 | 0 | 0 |
| 0 | 0 | 0 | 0 |
| 0 | 0 | 0 | 0 |
| 1 | 1 | 0 | 0 |
| 1 | 1 | 0 | 0 |
| 0 | 0 | 0 | 0 |
| 1 | 0 | 0 | 0 |
| 0 | 0 | 0 | 0 |
| 1 | 1 | 0 | 0 |
| 0 | 1 | 0 | 0 |
| 0 | 0 | 0 | 0 |
| 0 | 0 | 0 | 0 |
| 0 | 0 | 0 | 0 |
| 1 | 1 | 0 | 0 |
| 1 | 1 | 0 | 0 |
| 1 | 1 | 0 | 0 |
| 0 | 0 | 0 | 0 |
| 0 | 0 | 0 | 0 |
| 0 | 0 | 0 | 0 |
| 0 | 0 | 0 | 0 |
| 0 | 0 | 0 | 0 |
| 0 | 0 | 0 | 0 |
| 0 | 0 | 0 | 0 |

| | | | |
|---|---|---|---|
| 0 | 0 | 0 | 0 |
| 0 | 0 | 0 | 0 |
| 1 | 0 | 0 | 0 |
| 1 | 0 | 0 | 0 |
| 0 | 0 | 0 | 0 |
| 1 | 0 | 0 | 0 |
| 0 | 0 | 0 | 0 |
| 0 | 0 | 0 | 0 |
| 0 | 0 | 0 | 0 |
| 0 | 1 | 0 | 0 |
| 0 | 0 | 0 | 0 |
| 0 | 0 | 0 | 0 |
| 0 | 0 | 0 | 0 |
| 0 | 0 | 0 | 0 |
| 0 | 0 | 0 | 0 |
| 0 | 0 | 0 | 0 |
| 0 | 0 | 0 | 0 |
| 1 | 0 | 0 | 0 |
| 0 | 0 | 0 | 0 |
| 1 | 0 | 0 | 0 |
| 0 | 0 | 0 | 0 |
| 1 | 0 | 0 | 0 |
| 0 | 0 | 0 | 0 |
| 1 | 0 | 0 | 0 |
| 0 | 0 | 0 | 0 |
| 0 | 0 | 0 | 0 |
| 0 | 0 | 0 | 0 |
| 0 | 0 | 0 | 0 |
| 0 | 0 | 0 | 0 |
| 0 | 0 | 0 | 0 |
| 0 | 0 | 0 | 0 |
| 1 | 0 | 0 | 0 |
| 0 | 0 | 0 | 0 |
| 0 | 0 | 0 | 0 |
| 0 | 0 | 0 | 0 |
| 0 | 0 | 0 | 0 |
| 0 | 0 | 0 | 0 |
| 0 | 0 | 0 | 0 |
| 1 | 1 | 0 | 0 |
| 1 | 1 | 0 | 0 |
| 1 | 1 | 0 | 0 |
| 1 | 1 | 0 | 0 |
| 1 | 1 | 0 | 0 |
| 1 | 0 | 0 | 0 |
| 1 | 0 | 0 | 0 |
| 1 | 0 | 0 | 0 |
| 0 | 0 | 0 | 0 |
| 0 | 0 | 0 | 0 |
| 0 | 0 | 0 | 0 |
| 0 | 0 | 0 | 0 |
| 0 | 0 | 0 | 0 |
| 0 | 0 | 0 | 0 |
| 0 | 0 | 0 | 0 |
| 0 | 0 | 0 | 0 |

| | | | |
|---|---|---|---|
| 0 | 0 | 0 | 0 |
| 0 | 0 | 0 | 0 |
| 0 | 0 | 0 | 0 |
| 0 | 0 | 0 | 0 |
| 0 | 0 | 0 | 0 |
| 1 | 0 | 0 | 0 |
| 1 | 0 | 0 | 0 |
| 0 | 0 | 0 | 0 |
| 0 | 0 | 0 | 0 |
| 0 | 0 | 0 | 0 |
| 0 | 0 | 0 | 0 |
| 1 | 0 | 0 | 0 |
| 0 | 0 | 1 | 0 |
| 0 | 0 | 0 | 0 |
| 0 | 0 | 0 | 0 |
| 0 | 0 | 0 | 0 |
| 1 | 0 | 0 | 0 |
| 0 | 0 | 0 | 0 |
| 1 | 0 | 0 | 0 |
| 1 | 0 | 0 | 0 |
| 0 | 0 | 0 | 0 |
| 1 | 0 | 0 | 0 |
| 1 | 1 | 0 | 0 |
| 0 | 0 | 0 | 0 |
| 1 | 1 | 0 | 0 |
| 1 | 1 | 0 | 0 |
| 1 | 1 | 0 | 0 |
| 0 | 0 | 0 | 0 |
| 1 | 0 | 0 | 0 |
| 0 | 0 | 0 | 0 |
| 0 | 0 | 0 | 0 |
| 0 | 0 | 0 | 0 |
| 0 | 0 | 0 | 0 |
| 0 | 0 | 0 | 0 |
| 0 | 0 | 0 | 0 |
| 0 | 0 | 0 | 0 |
| 0 | 0 | 0 | 0 |
| 0 | 0 | 0 | 0 |
| 0 | 0 | 0 | 0 |
| 0 | 0 | 0 | 0 |
| 0 | 0 | 0 | 0 |
| 0 | 0 | 0 | 0 |
| 0 | 0 | 0 | 0 |
| 0 | 0 | 0 | 0 |
| 0 | 0 | 0 | 0 |
| 0 | 0 | 0 | 0 |
| 0 | 0 | 0 | 0 |
| 0 | 0 | 0 | 0 |
| 0 | 0 | 0 | 0 |
| 0 | 0 | 0 | 0 |
| 0 | 0 | 0 | 0 |
| 0 | 0 | 0 | 0 |
| 0 | 0 | 0 | 0 |
| 0 | 0 | 0 | 0 |

| | | | |
|---|---|---|---|
| 0 | 0 | 0 | 0 |
| 0 | 0 | 0 | 0 |
| 0 | 0 | 0 | 0 |
| 0 | 0 | 0 | 0 |
| 0 | 0 | 0 | 0 |
| 0 | 0 | 0 | 0 |
| 0 | 0 | 0 | 0 |
| 0 | 0 | 0 | 0 |
| 0 | 0 | 0 | 0 |
| 0 | 0 | 0 | 0 |
| 0 | 1 | 0 | 0 |
| 0 | 1 | 0 | 0 |
| 0 | 1 | 0 | 0 |
| 0 | 1 | 0 | 0 |
| 0 | 0 | 0 | 0 |
| 0 | 1 | 0 | 0 |
| 0 | 1 | 0 | 0 |
| 1 | 1 | 0 | 0 |
| 0 | 1 | 0 | 0 |
| 1 | 0 | 0 | 0 |
| 1 | 1 | 0 | 0 |
| 1 | 1 | 0 | 0 |
| 0 | 0 | 1 | 1 |
| 0 | 0 | 0 | 0 |
| 1 | 0 | 0 | 0 |
| 0 | 0 | 0 | 0 |
| 0 | 0 | 0 | 0 |
| 0 | 0 | 0 | 0 |
| 0 | 0 | 0 | 0 |
| 0 | 0 | 0 | 0 |
| 0 | 0 | 0 | 0 |
| 0 | 0 | 0 | 0 |
| 0 | 0 | 0 | 0 |
| 0 | 0 | 0 | 0 |
| 0 | 0 | 0 | 0 |
| 0 | 0 | 0 | 0 |
| 0 | 0 | 0 | 0 |
| 0 | 0 | 0 | 0 |
| 0 | 0 | 0 | 0 |
| 0 | 0 | 0 | 0 |
| 1 | 0 | 0 | 0 |
| 0 | 0 | 0 | 0 |
| 1 | 0 | 0 | 0 |
| 1 | 0 | 0 | 0 |
| 0 | 0 | 0 | 0 |
| 1 | 0 | 0 | 0 |
| 0 | 0 | 0 | 0 |
| 0 | 0 | 0 | 0 |
| 0 | 0 | 0 | 0 |
| 0 | 0 | 0 | 0 |
| 0 | 0 | 0 | 0 |
| 0 | 0 | 0 | 0 |
| 1 | 1 | 0 | 0 |
| 1 | 1 | 0 | 0 |

| | | | |
|---|---|---|---|
| 1 | 1 | 0 | 0 |
| 0 | 0 | 0 | 0 |
| 0 | 0 | 0 | 0 |
| 0 | 0 | 0 | 0 |
| 0 | 0 | 0 | 0 |
| 0 | 0 | 0 | 0 |
| 0 | 1 | 0 | 0 |
| 0 | 0 | 0 | 0 |
| 0 | 0 | 0 | 0 |
| 0 | 1 | 0 | 0 |
| 1 | 0 | 0 | 0 |
| 0 | 0 | 0 | 0 |
| 0 | 0 | 1 | 0 |
| 0 | 0 | 1 | 0 |
| 0 | 0 | 0 | 0 |
| 0 | 0 | 0 | 0 |
| 0 | 0 | 0 | 0 |
| 0 | 1 | 0 | 0 |
| 0 | 0 | 0 | 0 |
| 0 | 0 | 0 | 0 |
| 1 | 0 | 0 | 0 |
| 0 | 0 | 0 | 1 |
| 0 | 0 | 0 | 0 |
| 0 | 0 | 0 | 0 |
| 0 | 0 | 0 | 0 |
| 0 | 0 | 0 | 0 |
| 0 | 0 | 0 | 0 |
| 0 | 0 | 0 | 0 |
| 0 | 0 | 0 | 0 |
| 0 | 0 | 0 | 0 |
| 0 | 0 | 0 | 0 |
| 0 | 0 | 0 | 0 |
| 0 | 0 | 0 | 0 |
| 0 | 0 | 0 | 0 |
| 0 | 0 | 0 | 0 |
| 0 | 0 | 0 | 0 |
| 1 | 0 | 0 | 0 |
| 0 | 0 | 0 | 0 |
| 1 | 0 | 0 | 0 |
| 0 | 0 | 0 | 0 |
| 0 | 0 | 0 | 0 |
| 0 | 0 | 0 | 0 |
| 0 | 0 | 0 | 0 |
| 0 | 0 | 0 | 0 |
| 0 | 0 | 0 | 0 |
| 0 | 0 | 0 | 0 |
| 0 | 0 | 0 | 0 |
| 0 | 0 | 0 | 0 |
| 0 | 0 | 0 | 0 |
| 1 | 0 | 0 | 0 |
| 0 | 0 | 0 | 0 |
| 0 | 0 | 0 | 0 |
| 0 | 0 | 0 | 0 |
| 0 | 0 | 0 | 0 |

| | | | |
|---|---|---|---|
| 0 | 0 | 0 | 0 |
| 0 | 0 | 0 | 0 |
| 0 | 0 | 0 | 0 |
| 0 | 0 | 0 | 0 |
| 0 | 0 | 0 | 0 |
| 0 | 0 | 0 | 0 |
| 0 | 0 | 0 | 0 |
| 0 | 0 | 0 | 0 |
| 0 | 0 | 0 | 0 |
| 0 | 0 | 0 | 0 |
| 0 | 0 | 0 | 0 |
| 0 | 0 | 0 | 0 |
| 0 | 0 | 1 | 0 |
| 1 | 0 | 0 | 0 |
| 1 | 0 | 0 | 0 |
| 0 | 0 | 0 | 0 |
| 0 | 0 | 0 | 0 |
| 0 | 0 | 0 | 0 |
| 0 | 0 | 0 | 0 |
| 0 | 0 | 0 | 0 |
| 0 | 0 | 0 | 0 |
| 0 | 0 | 0 | 0 |
| 0 | 0 | 0 | 0 |
| 1 | 1 | 0 | 1 |
| 0 | 0 | 0 | 0 |
| 1 | 1 | 0 | 0 |
| 1 | 1 | 0 | 1 |
| 0 | 0 | 0 | 0 |
| 0 | 0 | 0 | 0 |
| 0 | 0 | 0 | 0 |
| 0 | 0 | 0 | 0 |
| 0 | 0 | 0 | 0 |
| 0 | 0 | 0 | 0 |
| 0 | 0 | 0 | 0 |
| 0 | 0 | 0 | 0 |
| 1 | 0 | 0 | 0 |
| 0 | 0 | 0 | 0 |
| 0 | 0 | 0 | 0 |
| 0 | 0 | 0 | 0 |
| 0 | 0 | 0 | 0 |
| 0 | 0 | 0 | 0 |
| 0 | 0 | 0 | 0 |
| 1 | 0 | 0 | 0 |
| 1 | 0 | 0 | 0 |
| 1 | 0 | 0 | 0 |
| 1 | 0 | 0 | 0 |
| 1 | 0 | 0 | 0 |
| 0 | 0 | 0 | 0 |
| 0 | 0 | 0 | 0 |
| 1 | 0 | 0 | 0 |
| 1 | 0 | 0 | 0 |
| 0 | 0 | 0 | 0 |
| 1 | 0 | 0 | 0 |
| 1 | 0 | 0 | 0 |

| | | | |
|---|---|---|---|
| 0 | 0 | 0 | 0 |
| 1 | 0 | 0 | 0 |
| 0 | 0 | 0 | 0 |
| 0 | 0 | 0 | 0 |
| 0 | 0 | 0 | 0 |
| 0 | 0 | 0 | 0 |
| 1 | 0 | 0 | 0 |
| 0 | 0 | 0 | 0 |
| 0 | 0 | 0 | 0 |
| 0 | 0 | 0 | 0 |
| 0 | 0 | 0 | 0 |
| 1 | 1 | 0 | 0 |
| 0 | 0 | 0 | 0 |
| 0 | 0 | 0 | 0 |
| 0 | 0 | 0 | 0 |
| 0 | 0 | 0 | 0 |
| 0 | 0 | 0 | 0 |
| 0 | 0 | 0 | 0 |
| 0 | 0 | 0 | 0 |
| 0 | 0 | 0 | 0 |
| 0 | 0 | 0 | 0 |
| 0 | 0 | 0 | 0 |
| 0 | 0 | 0 | 0 |
| 0 | 0 | 0 | 0 |
| 0 | 0 | 0 | 0 |
| 0 | 0 | 0 | 0 |
| 0 | 0 | 0 | 0 |
| 0 | 0 | 0 | 0 |
| 1 | 1 | 0 | 0 |
| 1 | 0 | 0 | 0 |
| 0 | 0 | 0 | 0 |
| 0 | 0 | 0 | 0 |
| 0 | 0 | 0 | 0 |
| 0 | 0 | 0 | 0 |
| 0 | 0 | 0 | 0 |
| 0 | 0 | 0 | 0 |
| 0 | 0 | 0 | 0 |
| 0 | 0 | 0 | 0 |
| 0 | 0 | 0 | 0 |
| 0 | 0 | 0 | 0 |
| 0 | 0 | 0 | 0 |
| 0 | 0 | 0 | 0 |
| 0 | 0 | 0 | 0 |
| 0 | 0 | 0 | 0 |
| 0 | 0 | 0 | 0 |
| 0 | 0 | 0 | 0 |
| 0 | 0 | 0 | 0 |
| 0 | 0 | 0 | 0 |
| 0 | 0 | 0 | 0 |
| 0 | 0 | 0 | 0 |
| 0 | 0 | 0 | 0 |
| 0 | 0 | 0 | 0 |
| 0 | 0 | 1 | 0 |
| 0 | 0 | 0 | 0 |

| | | | |
|---|---|---|---|
| 0 | 0 | 0 | 0 |
| 0 | 0 | 0 | 0 |
| 0 | 0 | 0 | 0 |
| 0 | 0 | 0 | 0 |
| 1 | 0 | 0 | 0 |
| 0 | 0 | 0 | 0 |
| 0 | 0 | 0 | 0 |
| 1 | 1 | 0 | 0 |
| 0 | 0 | 0 | 0 |
| 1 | 1 | 0 | 0 |
| 0 | 0 | 0 | 0 |
| 0 | 0 | 0 | 0 |
| 0 | 0 | 0 | 0 |
| 0 | 0 | 0 | 0 |
| 0 | 0 | 0 | 0 |
| 0 | 0 | 0 | 0 |
| 1 | 0 | 0 | 0 |
| 0 | 0 | 0 | 0 |
| 0 | 0 | 0 | 0 |
| 1 | 0 | 0 | 0 |
| 1 | 0 | 0 | 0 |
| 0 | 0 | 0 | 0 |
| 0 | 0 | 0 | 0 |
| 0 | 0 | 0 | 0 |
| 0 | 0 | 0 | 0 |
| 0 | 0 | 0 | 0 |
| 1 | 0 | 0 | 0 |
| 0 | 0 | 0 | 0 |
| 0 | 0 | 0 | 0 |
| 1 | 0 | 0 | 0 |
| 0 | 0 | 0 | 0 |
| 0 | 0 | 0 | 0 |
| 0 | 0 | 0 | 0 |
| 1 | 1 | 0 | 0 |
| 0 | 0 | 0 | 0 |
| 0 | 0 | 0 | 0 |
| 0 | 0 | 0 | 0 |
| 0 | 0 | 0 | 0 |
| 0 | 1 | 0 | 0 |
| 0 | 0 | 0 | 0 |
| 0 | 0 | 0 | 0 |
| 0 | 0 | 0 | 0 |
| 0 | 0 | 0 | 0 |
| 1 | 1 | 0 | 0 |
| 0 | 0 | 0 | 0 |
| 0 | 0 | 0 | 0 |
| 0 | 0 | 0 | 0 |
| 0 | 0 | 0 | 0 |
| 0 | 0 | 0 | 0 |
| 0 | 0 | 0 | 0 |
| 0 | 0 | 0 | 0 |
| 1 | 0 | 0 | 0 |
| 0 | 0 | 0 | 0 |
| 0 | 0 | 0 | 0 |

| | | | |
|---|---|---|---|
| 1 | 0 | 0 | 1 |
| 1 | 0 | 0 | 1 |
| 1 | 0 | 0 | 1 |
| 0 | 0 | 0 | 0 |
| 0 | 0 | 0 | 0 |
| 0 | 0 | 0 | 0 |
| 1 | 0 | 0 | 0 |
| 0 | 0 | 0 | 0 |
| 0 | 0 | 0 | 0 |
| 0 | 0 | 0 | 0 |
| 0 | 0 | 1 | 0 |
| 0 | 0 | 0 | 0 |
| 0 | 0 | 0 | 0 |
| 0 | 0 | 0 | 0 |
| 0 | 0 | 0 | 0 |
| 0 | 0 | 0 | 0 |
| 1 | 0 | 0 | 0 |
| 0 | 0 | 0 | 0 |
| 0 | 0 | 0 | 0 |
| 0 | 0 | 0 | 0 |
| 0 | 0 | 0 | 0 |
| 0 | 0 | 0 | 0 |
| 0 | 0 | 0 | 0 |
| 1 | 1 | 0 | 0 |
| 0 | 0 | 0 | 0 |
| 0 | 0 | 0 | 0 |
| 0 | 0 | 0 | 0 |
| 0 | 0 | 0 | 0 |
| 0 | 0 | 0 | 0 |
| 0 | 0 | 0 | 0 |
| 0 | 0 | 0 | 0 |
| 0 | 0 | 0 | 0 |
| 0 | 1 | 0 | 0 |
| 0 | 0 | 0 | 0 |
| 1 | 1 | 0 | 0 |
| 0 | 0 | 0 | 0 |
| 0 | 0 | 0 | 0 |
| 0 | 0 | 0 | 0 |
| 0 | 0 | 0 | 0 |
| 0 | 0 | 0 | 0 |
| 0 | 0 | 0 | 0 |
| 1 | 0 | 0 | 1 |
| 0 | 0 | 0 | 0 |
| 1 | 0 | 0 | 1 |
| 0 | 0 | 0 | 0 |
| 1 | 0 | 0 | 1 |
| 0 | 0 | 0 | 0 |
| 0 | 0 | 0 | 0 |
| 0 | 0 | 0 | 0 |
| 1 | 0 | 0 | 0 |
| 0 | 0 | 0 | 0 |
| 0 | 0 | 0 | 0 |
| 0 | 0 | 0 | 0 |
| 0 | 0 | 0 | 0 |

| | | | |
|---|---|---|---|
| 0 | 0 | 0 | 0 |
| 0 | 0 | 0 | 0 |
| 0 | 0 | 0 | 0 |
| 0 | 0 | 0 | 0 |
| 0 | 0 | 0 | 0 |
| 0 | 0 | 0 | 0 |
| 0 | 0 | 0 | 0 |
| 0 | 0 | 0 | 0 |
| 0 | 0 | 0 | 0 |
| 0 | 0 | 0 | 0 |
| 0 | 0 | 0 | 0 |
| 0 | 0 | 0 | 0 |
| 0 | 0 | 0 | 0 |
| 0 | 0 | 0 | 0 |
| 0 | 0 | 0 | 0 |
| 0 | 0 | 0 | 0 |
| 0 | 0 | 0 | 0 |
| 0 | 0 | 0 | 0 |
| 0 | 0 | 1 | 0 |
| 0 | 0 | 0 | 0 |
| 0 | 0 | 0 | 0 |
| 0 | 0 | 0 | 0 |
| 0 | 0 | 0 | 0 |
| 0 | 0 | 0 | 0 |
| 0 | 0 | 0 | 0 |
| 0 | 0 | 0 | 0 |
| 0 | 0 | 0 | 0 |
| 0 | 0 | 0 | 0 |
| 0 | 0 | 1 | 0 |
| 0 | 0 | 0 | 0 |
| 0 | 0 | 1 | 0 |
| 1 | 0 | 0 | 0 |
| 1 | 1 | 1 | 0 |
| 1 | 1 | 1 | 0 |
| 0 | 0 | 0 | 0 |
| 0 | 0 | 0 | 0 |
| 0 | 1 | 0 | 0 |
| 1 | 1 | 1 | 0 |
| 1 | 1 | 1 | 0 |
| 0 | 0 | 1 | 0 |
| 0 | 0 | 1 | 0 |
| 1 | 0 | 0 | 0 |
| 0 | 0 | 0 | 0 |
| 0 | 0 | 0 | 0 |
| 1 | 1 | 0 | 0 |
| 0 | 0 | 0 | 0 |
| 0 | 0 | 0 | 0 |
| 0 | 0 | 0 | 0 |
| 0 | 0 | 0 | 0 |
| 0 | 0 | 0 | 0 |
| 0 | 0 | 0 | 0 |
| 0 | 0 | 0 | 0 |
| 0 | 0 | 0 | 0 |
| 0 | 0 | 0 | 0 |

| | | | |
|---|---|---|---|
| 0 | 0 | 0 | 0 |
| 0 | 0 | 0 | 0 |
| 0 | 0 | 0 | 0 |
| 0 | 0 | 0 | 0 |
| 0 | 0 | 0 | 0 |
| 0 | 0 | 0 | 0 |
| 0 | 0 | 0 | 0 |
| 0 | 0 | 0 | 0 |
| 0 | 0 | 0 | 0 |
| 0 | 0 | 0 | 0 |
| 0 | 0 | 0 | 0 |
| 0 | 0 | 0 | 0 |
| 0 | 0 | 0 | 0 |
| 0 | 0 | 0 | 0 |
| 0 | 0 | 0 | 0 |
| 0 | 0 | 0 | 0 |
| 0 | 0 | 0 | 0 |
| 0 | 0 | 0 | 0 |
| 0 | 0 | 0 | 0 |
| 1 | 0 | 0 | 0 |
| 1 | 0 | 0 | 0 |
| 0 | 0 | 0 | 0 |
| 0 | 0 | 0 | 0 |
| 0 | 0 | 0 | 0 |
| 0 | 0 | 0 | 0 |
| 1 | 0 | 0 | 0 |
| 0 | 0 | 0 | 0 |
| 0 | 0 | 0 | 0 |
| 0 | 0 | 0 | 0 |
| 1 | 0 | 0 | 0 |
| 0 | 0 | 0 | 0 |
| 0 | 0 | 0 | 0 |
| 1 | 0 | 0 | 0 |
| 1 | 0 | 0 | 0 |
| 0 | 0 | 0 | 0 |
| 0 | 0 | 0 | 0 |
| 0 | 0 | 0 | 0 |
| 0 | 0 | 0 | 0 |
| 0 | 0 | 0 | 0 |
| 0 | 0 | 0 | 0 |
| 0 | 0 | 0 | 0 |
| 0 | 0 | 0 | 0 |
| 0 | 0 | 0 | 0 |
| 0 | 0 | 0 | 0 |
| 0 | 0 | 0 | 0 |
| 0 | 0 | 0 | 0 |
| 0 | 0 | 0 | 0 |
| 0 | 1 | 0 | 0 |
| 0 | 1 | 0 | 0 |
| 0 | 0 | 0 | 0 |
| 0 | 1 | 0 | 0 |
| 0 | 1 | 0 | 0 |
| 0 | 0 | 0 | 0 |
| 0 | 0 | 0 | 0 |

| | | | |
|---|---|---|---|
| 0 | 0 | 0 | 0 |
| 0 | 0 | 0 | 0 |
| 1 | 1 | 0 | 1 |
| 0 | 0 | 0 | 0 |
| 0 | 0 | 0 | 0 |
| 0 | 0 | 0 | 0 |
| 0 | 0 | 0 | 0 |
| 0 | 1 | 0 | 0 |
| 1 | 1 | 0 | 0 |
| 1 | 1 | 0 | 0 |
| 1 | 1 | 0 | 0 |
| 1 | 1 | 0 | 0 |
| 1 | 1 | 0 | 0 |
| 0 | 1 | 0 | 0 |
| 0 | 1 | 0 | 0 |
| 0 | 0 | 0 | 0 |
| 0 | 0 | 0 | 0 |
| 0 | 0 | 0 | 0 |
| 0 | 0 | 0 | 0 |
| 0 | 0 | 0 | 0 |
| 0 | 0 | 0 | 0 |
| 0 | 0 | 0 | 0 |
| 0 | 1 | 0 | 0 |
| 0 | 1 | 0 | 0 |
| 0 | 0 | 0 | 0 |
| 0 | 0 | 1 | 0 |
| 0 | 0 | 0 | 0 |
| 0 | 0 | 0 | 0 |
| 0 | 0 | 0 | 0 |
| 0 | 1 | 0 | 0 |
| 0 | 0 | 0 | 0 |
| 0 | 0 | 0 | 0 |
| 0 | 0 | 0 | 0 |
| 0 | 0 | 0 | 0 |
| 0 | 0 | 0 | 0 |
| 0 | 0 | 0 | 0 |
| 0 | 0 | 0 | 0 |
| 0 | 0 | 0 | 0 |
| 0 | 0 | 0 | 0 |
| 1 | 0 | 0 | 0 |
| 1 | 1 | 0 | 0 |
| 1 | 0 | 0 | 0 |
| 0 | 0 | 0 | 0 |
| 0 | 1 | 0 | 0 |
| 1 | 1 | 0 | 0 |
| 0 | 1 | 0 | 0 |
| 0 | 0 | 0 | 0 |
| 1 | 0 | 0 | 0 |
| 0 | 1 | 0 | 0 |
| 0 | 0 | 0 | 0 |
| 0 | 0 | 0 | 0 |
| 0 | 0 | 0 | 0 |
| 1 | 1 | 0 | 0 |
| 0 | 0 | 0 | 0 |

| | | | |
|---|---|---|---|
| 0 | 0 | 0 | 0 |
| 0 | 0 | 0 | 0 |
| 1 | 0 | 0 | 0 |
| 1 | 1 | 0 | 0 |
| 1 | 1 | 0 | 0 |
| 0 | 0 | 0 | 0 |
| 0 | 0 | 0 | 0 |
| 0 | 0 | 0 | 0 |
| 1 | 0 | 0 | 0 |
| 0 | 0 | 0 | 0 |
| 0 | 0 | 0 | 0 |
| 1 | 0 | 0 | 1 |
| 0 | 0 | 0 | 0 |
| 1 | 0 | 0 | 0 |
| 0 | 0 | 0 | 0 |
| 0 | 0 | 0 | 0 |
| 0 | 0 | 0 | 0 |
| 0 | 0 | 0 | 0 |
| 0 | 0 | 0 | 0 |
| 0 | 0 | 0 | 0 |
| 0 | 0 | 0 | 0 |
| 1 | 0 | 0 | 1 |
| 1 | 0 | 0 | 0 |
| 0 | 0 | 0 | 0 |
| 0 | 0 | 0 | 0 |
| 0 | 0 | 0 | 0 |
| 0 | 0 | 0 | 0 |
| 0 | 0 | 0 | 0 |
| 1 | 0 | 0 | 0 |
| 1 | 0 | 0 | 0 |
| 0 | 0 | 0 | 0 |
| 0 | 0 | 0 | 0 |
| 0 | 0 | 0 | 0 |
| 0 | 0 | 0 | 0 |
| 0 | 0 | 0 | 0 |
| 0 | 0 | 0 | 0 |
| 1 | 0 | 0 | 0 |
| 0 | 0 | 0 | 0 |
| 0 | 0 | 0 | 0 |
| 1 | 0 | 0 | 0 |
| 1 | 0 | 0 | 0 |
| 1 | 0 | 0 | 0 |
| 1 | 0 | 0 | 0 |
| 0 | 0 | 0 | 0 |
| 0 | 0 | 0 | 0 |
| 1 | 0 | 0 | 0 |
| 0 | 0 | 0 | 0 |
| 0 | 0 | 0 | 0 |
| 0 | 0 | 0 | 0 |
| 0 | 0 | 0 | 0 |
| 0 | 0 | 0 | 0 |
| 1 | 0 | 0 | 0 |
| 1 | 0 | 0 | 0 |
| 1 | 0 | 0 | 0 |

| | | | |
|---|---|---|---|
| 1 | 0 | 0 | 0 |
| 0 | 0 | 0 | 0 |
| 0 | 0 | 0 | 0 |
| 0 | 0 | 0 | 0 |
| 1 | 0 | 0 | 0 |
| 0 | 0 | 0 | 0 |
| 1 | 0 | 0 | 0 |
| 0 | 0 | 0 | 0 |
| 0 | 0 | 0 | 0 |
| 0 | 0 | 0 | 0 |
| 0 | 0 | 0 | 0 |
| 0 | 0 | 0 | 0 |
| 0 | 0 | 0 | 0 |
| 0 | 0 | 0 | 0 |
| 0 | 0 | 0 | 0 |
| 1 | 0 | 0 | 0 |
| 0 | 0 | 0 | 0 |
| 0 | 0 | 0 | 0 |
| 0 | 0 | 0 | 0 |
| 0 | 0 | 0 | 0 |
| 0 | 0 | 0 | 0 |
| 0 | 0 | 0 | 0 |
| 1 | 0 | 0 | 0 |
| 1 | 0 | 0 | 0 |
| 0 | 0 | 0 | 0 |
| 0 | 0 | 0 | 0 |
| 0 | 0 | 0 | 0 |
| 0 | 0 | 0 | 0 |
| 0 | 0 | 0 | 0 |
| 0 | 0 | 0 | 0 |
| 0 | 0 | 0 | 0 |
| 0 | 0 | 0 | 0 |
| 1 | 0 | 0 | 0 |
| 0 | 0 | 0 | 0 |
| 0 | 0 | 0 | 0 |
| 0 | 0 | 0 | 0 |
| 0 | 0 | 0 | 0 |
| 0 | 0 | 0 | 0 |
| 0 | 0 | 0 | 0 |
| 0 | 0 | 0 | 0 |
| 0 | 0 | 0 | 0 |
| 0 | 0 | 0 | 0 |
| 0 | 0 | 0 | 0 |
| 0 | 0 | 0 | 0 |
| 0 | 1 | 0 | 0 |
| 0 | 0 | 0 | 0 |
| 0 | 0 | 0 | 0 |
| 0 | 0 | 0 | 0 |
| 0 | 0 | 0 | 0 |
| 0 | 0 | 0 | 0 |
| 0 | 0 | 0 | 0 |
| 0 | 0 | 0 | 0 |
| 0 | 0 | 0 | 0 |
| 1 | 0 | 0 | 0 |

| | | | |
|---|---|---|---|
| 0 | 0 | 0 | 0 |
| 0 | 0 | 0 | 0 |
| 0 | 0 | 0 | 0 |
| 0 | 0 | 0 | 0 |
| 0 | 0 | 0 | 0 |
| 0 | 0 | 0 | 0 |
| 0 | 0 | 0 | 0 |
| 0 | 0 | 0 | 0 |
| 0 | 0 | 0 | 0 |
| 1 | 0 | 0 | 0 |
| 0 | 0 | 0 | 0 |
| 0 | 0 | 0 | 0 |
| 1 | 1 | 0 | 0 |
| 1 | 1 | 0 | 0 |
| 1 | 1 | 0 | 0 |
| 1 | 1 | 0 | 0 |
| 0 | 0 | 0 | 0 |
| 0 | 0 | 0 | 0 |
| 0 | 0 | 0 | 0 |
| 0 | 0 | 0 | 0 |
| 0 | 0 | 0 | 0 |
| 0 | 0 | 0 | 0 |
| 0 | 0 | 0 | 0 |
| 0 | 0 | 0 | 0 |
| 0 | 0 | 0 | 0 |
| 0 | 0 | 0 | 1 |
| 0 | 0 | 0 | 0 |
| 0 | 0 | 0 | 0 |
| 0 | 0 | 0 | 0 |
| 0 | 0 | 0 | 0 |
| 0 | 0 | 0 | 0 |
| 0 | 0 | 0 | 0 |
| 0 | 0 | 0 | 0 |
| 0 | 0 | 0 | 0 |
| 0 | 0 | 0 | 0 |
| 0 | 0 | 0 | 0 |
| 0 | 0 | 0 | 0 |
| 1 | 1 | 0 | 0 |
| 0 | 0 | 0 | 0 |
| 0 | 0 | 0 | 0 |
| 1 | 1 | 0 | 0 |
| 0 | 0 | 0 | 0 |
| 0 | 0 | 0 | 0 |
| 1 | 0 | 0 | 1 |
| 0 | 0 | 0 | 0 |
| 0 | 0 | 0 | 0 |
| 0 | 0 | 0 | 0 |
| 0 | 0 | 0 | 0 |
| 0 | 0 | 0 | 0 |
| 0 | 0 | 0 | 0 |
| 0 | 0 | 0 | 0 |
| 0 | 0 | 1 | 0 |
| 0 | 0 | 1 | 0 |
| 0 | 0 | 1 | 0 |

| | | | |
|---|---|---|---|
| 0 | 0 | 1 | 0 |
| 0 | 0 | 0 | 0 |
| 0 | 0 | 0 | 0 |
| 0 | 0 | 0 | 0 |
| 0 | 0 | 0 | 0 |
| 0 | 0 | 0 | 0 |
| 0 | 0 | 0 | 0 |
| 0 | 0 | 0 | 0 |
| 0 | 0 | 0 | 0 |
| 0 | 0 | 0 | 0 |
| 0 | 0 | 0 | 0 |
| 0 | 0 | 0 | 0 |
| 0 | 0 | 0 | 0 |
| 0 | 0 | 0 | 0 |
| 0 | 0 | 0 | 0 |
| 0 | 0 | 0 | 0 |
| 0 | 0 | 0 | 0 |
| 0 | 0 | 0 | 0 |
| 0 | 0 | 0 | 0 |
| 0 | 0 | 0 | 0 |
| 0 | 0 | 0 | 0 |
| 0 | 0 | 0 | 0 |
| 0 | 0 | 0 | 0 |
| 0 | 1 | 0 | 1 |
| 0 | 0 | 0 | 0 |
| 0 | 0 | 0 | 0 |
| 0 | 0 | 0 | 0 |
| 0 | 0 | 0 | 0 |
| 0 | 0 | 0 | 0 |
| 0 | 0 | 0 | 0 |
| 0 | 0 | 0 | 0 |
| 0 | 0 | 0 | 0 |
| 0 | 0 | 0 | 0 |
| 0 | 0 | 0 | 0 |
| 0 | 0 | 0 | 0 |
| 0 | 0 | 0 | 0 |
| 0 | 0 | 0 | 0 |
| 0 | 0 | 0 | 0 |
| 0 | 0 | 0 | 0 |
| 0 | 0 | 0 | 0 |
| 0 | 0 | 0 | 0 |
| 0 | 0 | 0 | 0 |
| 0 | 0 | 0 | 0 |
| 0 | 0 | 0 | 0 |
| 0 | 0 | 0 | 0 |
| 0 | 0 | 0 | 0 |
| 1 | 0 | 0 | 0 |
| 1 | 0 | 0 | 0 |
| 0 | 0 | 0 | 0 |
| 0 | 1 | 0 | 0 |
| 1 | 1 | 0 | 1 |
| 1 | 1 | 0 | 1 |
| 1 | 1 | 0 | 1 |
| 0 | 0 | 0 | 0 |

| | | | |
|---|---|---|---|
| 1 | 1 | 0 | 0 |
| 0 | 0 | 0 | 0 |
| 0 | 0 | 0 | 0 |
| 0 | 0 | 0 | 0 |
| 0 | 0 | 0 | 0 |
| 0 | 0 | 0 | 0 |
| 0 | 0 | 0 | 0 |
| 0 | 0 | 0 | 0 |
| 0 | 0 | 0 | 0 |
| 0 | 0 | 0 | 0 |
| 0 | 0 | 0 | 0 |
| 0 | 1 | 0 | 0 |
| 0 | 0 | 0 | 0 |
| 0 | 0 | 0 | 0 |
| 0 | 1 | 0 | 0 |
| 0 | 0 | 0 | 0 |
| 1 | 1 | 0 | 0 |
| 0 | 0 | 0 | 0 |
| 0 | 0 | 0 | 0 |
| 1 | 0 | 0 | 0 |
| 0 | 0 | 0 | 0 |
| 0 | 0 | 0 | 0 |
| 0 | 0 | 0 | 0 |
| 0 | 0 | 0 | 0 |
| 0 | 0 | 0 | 0 |
| 0 | 0 | 0 | 0 |
| 0 | 0 | 0 | 0 |
| 0 | 0 | 0 | 0 |
| 0 | 0 | 0 | 0 |
| 0 | 0 | 0 | 0 |
| 0 | 0 | 0 | 0 |
| 0 | 0 | 0 | 0 |
| 0 | 0 | 0 | 0 |
| 1 | 0 | 0 | 0 |
| 0 | 0 | 0 | 0 |
| 0 | 1 | 0 | 0 |
| 0 | 0 | 0 | 0 |
| 0 | 0 | 0 | 0 |
| 0 | 0 | 0 | 0 |
| 0 | 0 | 0 | 0 |
| 0 | 0 | 0 | 0 |
| 0 | 0 | 0 | 0 |
| 0 | 0 | 0 | 0 |
| 0 | 0 | 0 | 0 |
| 0 | 0 | 0 | 0 |
| 0 | 0 | 0 | 0 |
| 0 | 0 | 0 | 0 |
| 0 | 0 | 0 | 0 |
| 1 | 1 | 0 | 0 |
| 0 | 0 | 0 | 0 |
| 0 | 0 | 0 | 0 |
| 0 | 0 | 0 | 0 |
| 0 | 0 | 0 | 0 |
| 0 | 0 | 0 | 0 |

| | | | |
|---|---|---|---|
| 0 | 0 | 0 | 0 |
| 0 | 0 | 0 | 0 |
| 0 | 0 | 0 | 0 |
| 0 | 0 | 0 | 0 |
| 0 | 0 | 0 | 0 |
| 0 | 0 | 0 | 0 |
| 0 | 0 | 0 | 0 |
| 0 | 1 | 0 | 0 |
| 0 | 0 | 0 | 0 |
| 0 | 0 | 0 | 0 |
| 0 | 0 | 0 | 0 |
| 0 | 0 | 0 | 0 |
| 0 | 0 | 1 | 0 |
| 0 | 0 | 0 | 0 |
| 0 | 0 | 1 | 0 |
| 0 | 0 | 1 | 0 |
| 0 | 0 | 0 | 0 |
| 0 | 0 | 0 | 0 |
| 0 | 0 | 0 | 0 |
| 0 | 0 | 0 | 0 |
| 0 | 1 | 0 | 0 |
| 0 | 1 | 0 | 0 |
| 0 | 0 | 0 | 0 |
| 0 | 0 | 0 | 0 |
| 0 | 1 | 0 | 0 |
| 0 | 0 | 0 | 0 |
| 0 | 0 | 0 | 0 |
| 0 | 1 | 0 | 0 |
| 0 | 0 | 0 | 0 |
| 0 | 0 | 0 | 0 |
| 0 | 0 | 0 | 0 |
| 0 | 0 | 0 | 0 |
| 1 | 1 | 0 | 0 |
| 1 | 1 | 0 | 0 |
| 1 | 1 | 0 | 0 |
| 0 | 1 | 0 | 0 |
| 1 | 0 | 0 | 0 |
| 0 | 1 | 0 | 0 |
| 1 | 1 | 0 | 0 |
| 0 | 0 | 0 | 0 |
| 0 | 0 | 0 | 0 |
| 0 | 0 | 0 | 0 |
| 0 | 0 | 0 | 0 |
| 0 | 0 | 0 | 0 |
| 0 | 0 | 0 | 0 |
| 0 | 0 | 0 | 0 |
| 0 | 0 | 0 | 0 |
| 0 | 0 | 0 | 0 |
| 0 | 0 | 0 | 0 |
| 1 | 1 | 0 | 0 |
| 1 | 1 | 0 | 0 |
| 0 | 0 | 0 | 0 |
| 0 | 0 | 0 | 0 |
| 0 | 0 | 0 | 0 |

| | | | |
|---|---|---|---|
| 0 | 0 | 0 | 0 |
| 0 | 0 | 0 | 0 |
| 0 | 0 | 0 | 0 |
| 0 | 0 | 0 | 0 |
| 0 | 0 | 0 | 0 |
| 0 | 0 | 0 | 0 |
| 1 | 1 | 0 | 0 |
| 1 | 1 | 0 | 0 |
| 0 | 0 | 0 | 0 |
| 0 | 0 | 0 | 0 |
| 0 | 0 | 0 | 0 |
| 0 | 0 | 0 | 0 |
| 0 | 0 | 0 | 0 |
| 0 | 0 | 0 | 0 |
| 0 | 0 | 0 | 0 |
| 0 | 0 | 0 | 0 |
| 0 | 0 | 0 | 0 |
| 0 | 0 | 0 | 0 |
| 0 | 0 | 0 | 0 |
| 0 | 0 | 0 | 0 |
| 0 | 0 | 0 | 0 |
| 0 | 0 | 0 | 0 |
| 0 | 0 | 0 | 0 |
| 0 | 0 | 0 | 0 |
| 0 | 0 | 0 | 0 |
| 0 | 0 | 0 | 0 |
| 0 | 0 | 0 | 0 |
| 1 | 1 | 0 | 0 |
| 0 | 0 | 0 | 0 |
| 0 | 0 | 0 | 0 |
| 0 | 0 | 0 | 0 |
| 0 | 0 | 0 | 0 |
| 0 | 0 | 0 | 0 |
| 0 | 0 | 0 | 0 |
| 0 | 0 | 0 | 0 |
| 0 | 0 | 0 | 0 |
| 0 | 0 | 0 | 0 |
| 0 | 0 | 0 | 0 |
| 0 | 0 | 0 | 0 |
| 0 | 0 | 0 | 0 |
| 0 | 0 | 0 | 0 |
| 0 | 0 | 0 | 0 |
| 0 | 0 | 0 | 0 |
| 0 | 0 | 0 | 0 |
| 0 | 0 | 0 | 0 |
| 0 | 0 | 0 | 0 |
| 0 | 0 | 0 | 0 |
| 0 | 0 | 0 | 0 |
| 0 | 0 | 0 | 0 |
| 0 | 0 | 0 | 0 |
| 0 | 0 | 0 | 0 |
| 0 | 0 | 0 | 0 |
| 0 | 0 | 0 | 0 |
| 0 | 0 | 0 | 0 |

| | | | |
|---|---|---|---|
| 0 | 0 | 0 | 0 |
| 0 | 0 | 0 | 0 |
| 0 | 0 | 0 | 0 |
| 0 | 0 | 0 | 0 |
| 0 | 0 | 0 | 0 |
| 0 | 0 | 0 | 0 |
| 0 | 0 | 0 | 0 |
| 0 | 0 | 0 | 0 |
| 0 | 0 | 0 | 0 |
| 0 | 0 | 0 | 0 |
| 1 | 0 | 0 | 0 |
| 1 | 1 | 0 | 0 |
| 1 | 0 | 0 | 0 |
| 0 | 0 | 0 | 0 |
| 0 | 0 | 0 | 0 |
| 0 | 0 | 0 | 0 |
| 0 | 0 | 0 | 0 |
| 0 | 0 | 0 | 0 |
| 0 | 0 | 0 | 0 |
| 0 | 0 | 0 | 0 |
| 0 | 0 | 0 | 0 |
| 0 | 0 | 0 | 0 |
| 0 | 0 | 0 | 0 |
| 0 | 0 | 0 | 0 |
| 0 | 0 | 0 | 0 |
| 0 | 0 | 0 | 0 |
| 0 | 0 | 0 | 0 |
| 0 | 0 | 0 | 0 |
| 0 | 0 | 0 | 0 |
| 0 | 0 | 0 | 0 |
| 0 | 0 | 0 | 0 |
| 0 | 1 | 0 | 0 |
| 0 | 0 | 0 | 0 |
| 0 | 0 | 0 | 0 |
| 1 | 0 | 0 | 0 |
| 1 | 0 | 0 | 0 |
| 1 | 0 | 0 | 0 |
| 0 | 0 | 0 | 0 |
| 0 | 0 | 0 | 0 |
| 0 | 0 | 0 | 0 |
| 0 | 0 | 0 | 0 |
| 0 | 0 | 0 | 0 |
| 0 | 0 | 0 | 0 |
| 0 | 0 | 0 | 0 |
| 0 | 0 | 0 | 0 |
| 0 | 0 | 0 | 0 |
| 0 | 0 | 0 | 0 |
| 0 | 0 | 0 | 0 |
| 0 | 0 | 0 | 0 |
| 0 | 0 | 0 | 0 |
| 0 | 0 | 0 | 0 |
| 0 | 0 | 0 | 0 |
| 0 | 0 | 0 | 1 |
| 0 | 0 | 0 | 0 |

| | | | |
|---|---|---|---|
| 0 | 0 | 0 | 0 |
| 0 | 0 | 0 | 0 |
| 0 | 0 | 0 | 0 |
| 0 | 0 | 0 | 0 |
| 0 | 0 | 0 | 0 |
| 0 | 0 | 0 | 0 |
| 0 | 0 | 0 | 0 |
| 0 | 0 | 0 | 0 |
| 0 | 0 | 0 | 0 |
| 0 | 0 | 0 | 0 |
| 0 | 0 | 0 | 0 |
| 1 | 0 | 0 | 0 |
| 0 | 0 | 0 | 0 |
| 0 | 0 | 0 | 0 |
| 0 | 0 | 1 | 0 |
| 0 | 0 | 0 | 0 |
| 0 | 0 | 0 | 0 |
| 0 | 0 | 0 | 0 |
| 0 | 0 | 0 | 0 |
| 0 | 0 | 0 | 0 |
| 0 | 0 | 0 | 0 |
| 0 | 0 | 0 | 0 |
| 0 | 0 | 0 | 0 |
| 0 | 0 | 0 | 0 |
| 0 | 0 | 0 | 0 |
| 0 | 0 | 0 | 0 |
| 0 | 0 | 0 | 0 |
| 0 | 0 | 0 | 0 |
| 0 | 0 | 0 | 0 |
| 0 | 0 | 0 | 0 |
| 0 | 0 | 0 | 0 |
| 0 | 0 | 0 | 0 |
| 0 | 0 | 0 | 0 |
| 1 | 0 | 0 | 0 |
| 0 | 0 | 0 | 0 |
| 1 | 0 | 0 | 1 |
| 0 | 0 | 0 | 0 |
| 0 | 0 | 0 | 0 |
| 0 | 0 | 0 | 0 |
| 0 | 0 | 0 | 0 |
| 0 | 0 | 0 | 0 |
| 0 | 0 | 0 | 0 |
| 0 | 0 | 0 | 0 |
| 0 | 1 | 0 | 0 |
| 0 | 0 | 0 | 0 |
| 0 | 0 | 0 | 0 |
| 0 | 0 | 0 | 0 |
| 0 | 0 | 0 | 0 |
| 0 | 0 | 0 | 0 |
| 0 | 0 | 0 | 0 |
| 0 | 0 | 0 | 0 |
| 0 | 0 | 0 | 0 |
| 0 | 0 | 0 | 0 |
| 0 | 0 | 0 | 0 |

| | | | |
|---|---|---|---|
| 0 | 0 | 0 | 0 |
| 0 | 0 | 0 | 0 |
| 0 | 0 | 0 | 0 |
| 0 | 0 | 0 | 0 |
| 0 | 0 | 0 | 0 |
| 0 | 1 | 0 | 0 |
| 0 | 0 | 0 | 0 |
| 0 | 0 | 0 | 0 |
| 0 | 0 | 0 | 0 |
| 0 | 0 | 0 | 0 |
| 0 | 0 | 0 | 0 |
| 0 | 0 | 0 | 0 |
| 0 | 0 | 0 | 0 |
| 0 | 0 | 0 | 0 |
| 0 | 0 | 0 | 0 |
| 0 | 1 | 0 | 0 |
| 1 | 1 | 0 | 0 |
| 0 | 0 | 1 | 0 |
| 0 | 0 | 0 | 0 |
| 0 | 0 | 0 | 0 |
| 0 | 0 | 0 | 0 |
| 0 | 0 | 0 | 0 |
| 0 | 0 | 0 | 0 |
| 0 | 0 | 0 | 0 |
| 0 | 0 | 0 | 0 |
| 0 | 0 | 0 | 0 |
| 0 | 0 | 0 | 0 |
| 1 | 0 | 0 | 0 |
| 0 | 0 | 0 | 0 |
| 0 | 0 | 0 | 0 |
| 0 | 0 | 1 | 0 |
| 0 | 0 | 0 | 0 |
| 0 | 0 | 0 | 0 |
| 0 | 0 | 0 | 0 |
| 0 | 0 | 0 | 0 |
| 0 | 0 | 0 | 0 |
| 0 | 0 | 0 | 0 |
| 1 | 1 | 0 | 0 |
| 0 | 0 | 0 | 0 |
| 0 | 0 | 0 | 0 |
| 0 | 0 | 0 | 0 |
| 1 | 0 | 0 | 0 |
| 1 | 0 | 0 | 0 |
| 0 | 0 | 0 | 0 |
| 0 | 0 | 0 | 0 |
| 0 | 0 | 0 | 0 |
| 0 | 0 | 0 | 0 |
| 0 | 0 | 0 | 0 |
| 0 | 0 | 0 | 0 |
| 0 | 0 | 0 | 0 |
| 0 | 0 | 0 | 0 |
| 0 | 0 | 0 | 0 |
| 0 | 0 | 0 | 0 |
| 0 | 0 | 0 | 0 |

| | | | |
|---|---|---|---|
| 0 | 0 | 0 | 0 |
| 0 | 0 | 0 | 0 |
| 0 | 0 | 0 | 0 |
| 0 | 0 | 0 | 0 |
| 0 | 0 | 0 | 0 |
| 0 | 0 | 0 | 0 |
| 0 | 0 | 0 | 0 |
| 0 | 0 | 0 | 0 |
| 0 | 0 | 0 | 0 |
| 0 | 0 | 0 | 0 |
| 0 | 0 | 0 | 0 |
| 0 | 0 | 0 | 0 |
| 1 | 0 | 0 | 0 |
| 0 | 0 | 0 | 0 |
| 0 | 0 | 0 | 0 |
| 0 | 0 | 0 | 0 |
| 0 | 0 | 0 | 0 |
| 0 | 0 | 0 | 0 |
| 0 | 0 | 0 | 0 |
| 0 | 0 | 0 | 0 |
| 0 | 0 | 0 | 0 |
| 0 | 0 | 0 | 0 |
| 0 | 0 | 0 | 0 |
| 1 | 0 | 0 | 0 |
| 0 | 0 | 0 | 0 |
| 0 | 0 | 0 | 0 |
| 0 | 0 | 0 | 0 |
| 1 | 1 | 0 | 1 |
| 0 | 0 | 0 | 0 |
| 0 | 0 | 0 | 0 |
| 0 | 0 | 0 | 0 |
| 0 | 0 | 0 | 0 |
| 0 | 0 | 0 | 0 |
| 0 | 0 | 0 | 0 |
| 0 | 0 | 0 | 0 |
| 0 | 0 | 0 | 0 |
| 0 | 0 | 0 | 0 |
| 0 | 0 | 0 | 0 |
| 0 | 0 | 0 | 0 |
| 0 | 0 | 0 | 0 |
| 0 | 0 | 0 | 0 |
| 0 | 0 | 0 | 0 |
| 0 | 0 | 0 | 0 |
| 0 | 0 | 0 | 0 |
| 0 | 0 | 0 | 0 |
| 0 | 0 | 0 | 0 |
| 0 | 0 | 0 | 0 |
| 0 | 0 | 0 | 0 |
| 0 | 0 | 0 | 0 |
| 0 | 0 | 0 | 0 |
| 0 | 0 | 0 | 0 |
| 0 | 0 | 0 | 0 |
| 0 | 0 | 0 | 0 |
| 0 | 0 | 0 | 0 |

| | | | |
|---|---|---|---|
| 0 | 0 | 0 | 0 |
| 0 | 0 | 0 | 0 |
| 0 | 0 | 0 | 0 |
| 0 | 0 | 0 | 0 |
| 0 | 0 | 0 | 0 |
| 0 | 0 | 0 | 0 |
| 0 | 0 | 0 | 0 |
| 0 | 0 | 0 | 0 |
| 1 | 0 | 0 | 0 |
| 1 | 0 | 0 | 0 |
| 1 | 0 | 0 | 0 |
| 0 | 0 | 0 | 0 |
| 1 | 0 | 0 | 0 |
| 0 | 0 | 0 | 0 |
| 0 | 0 | 0 | 0 |
| 1 | 0 | 0 | 0 |
| 0 | 0 | 0 | 0 |
| 0 | 0 | 0 | 0 |
| 0 | 0 | 0 | 0 |
| 0 | 0 | 0 | 0 |
| 0 | 0 | 0 | 0 |
| 0 | 0 | 0 | 0 |
| 0 | 0 | 0 | 0 |
| 0 | 0 | 0 | 0 |
| 0 | 0 | 0 | 0 |
| 0 | 0 | 0 | 0 |
| 0 | 0 | 0 | 0 |
| 1 | 0 | 0 | 0 |
| 1 | 0 | 0 | 0 |
| 0 | 0 | 0 | 0 |
| 0 | 0 | 0 | 0 |
| 0 | 0 | 0 | 0 |
| 1 | 0 | 0 | 0 |
| 0 | 0 | 0 | 0 |
| 0 | 0 | 0 | 0 |
| 0 | 0 | 0 | 0 |
| 0 | 0 | 0 | 0 |
| 0 | 0 | 0 | 0 |
| 0 | 0 | 0 | 0 |
| 0 | 0 | 0 | 0 |
| 1 | 0 | 0 | 0 |
| 1 | 0 | 0 | 0 |
| 0 | 0 | 0 | 0 |
| 0 | 0 | 0 | 0 |
| 0 | 0 | 0 | 0 |
| 0 | 0 | 0 | 0 |
| 0 | 0 | 0 | 0 |
| 0 | 0 | 0 | 0 |
| 0 | 0 | 0 | 0 |
| 0 | 0 | 0 | 0 |
| 0 | 0 | 0 | 0 |
| 0 | 0 | 0 | 0 |
| 0 | 0 | 0 | 0 |
| 0 | 0 | 0 | 0 |

| | | | |
|---|---|---|---|
| 0 | 0 | 0 | 0 |
| 0 | 0 | 0 | 0 |
| 0 | 0 | 0 | 0 |
| 0 | 0 | 0 | 0 |
| 0 | 0 | 0 | 0 |
| 0 | 0 | 0 | 0 |
| 0 | 0 | 0 | 0 |
| 0 | 0 | 0 | 0 |
| 0 | 0 | 0 | 0 |
| 0 | 0 | 0 | 0 |
| 1 | 0 | 0 | 0 |
| 0 | 0 | 0 | 0 |
| 1 | 0 | 0 | 0 |
| 0 | 0 | 0 | 0 |
| 0 | 0 | 0 | 0 |
| 0 | 0 | 0 | 0 |
| 0 | 0 | 0 | 0 |
| 0 | 1 | 0 | 0 |
| 0 | 0 | 0 | 0 |
| 0 | 0 | 0 | 0 |
| 0 | 0 | 0 | 0 |
| 0 | 0 | 0 | 0 |
| 1 | 0 | 0 | 0 |
| 0 | 0 | 0 | 0 |
| 0 | 0 | 0 | 0 |
| 1 | 0 | 0 | 0 |
| 1 | 0 | 0 | 0 |
| 1 | 0 | 0 | 0 |
| 0 | 0 | 0 | 0 |
| 0 | 0 | 0 | 0 |
| 0 | 0 | 0 | 0 |
| 0 | 0 | 0 | 0 |
| 0 | 0 | 0 | 0 |
| 0 | 0 | 0 | 0 |
| 0 | 0 | 0 | 0 |
| 0 | 0 | 0 | 0 |
| 0 | 0 | 0 | 0 |
| 0 | 0 | 0 | 0 |
| 0 | 0 | 0 | 0 |
| 0 | 0 | 0 | 0 |
| 1 | 1 | 0 | 0 |
| 0 | 0 | 0 | 0 |
| 0 | 0 | 0 | 0 |
| 0 | 0 | 0 | 0 |
| 0 | 0 | 0 | 0 |
| 0 | 0 | 0 | 0 |
| 0 | 0 | 0 | 0 |
| 1 | 0 | 0 | 0 |
| 0 | 0 | 0 | 0 |
| 0 | 0 | 0 | 0 |
| 0 | 0 | 0 | 0 |
| 0 | 0 | 0 | 0 |
| 0 | 0 | 0 | 0 |
| 0 | 0 | 0 | 0 |

| | | | |
|---|---|---|---|
| 1 | 0 | 0 | 0 |
| 0 | 0 | 0 | 0 |
| 1 | 0 | 0 | 0 |
| 0 | 0 | 0 | 0 |
| 0 | 0 | 0 | 0 |
| 0 | 0 | 0 | 0 |
| 0 | 0 | 0 | 0 |
| 0 | 0 | 0 | 0 |
| 1 | 1 | 0 | 0 |
| 0 | 0 | 0 | 0 |
| 1 | 1 | 0 | 0 |
| 1 | 1 | 0 | 0 |
| 0 | 0 | 0 | 0 |
| 1 | 1 | 0 | 0 |
| 0 | 0 | 0 | 0 |
| 0 | 0 | 0 | 0 |
| 0 | 0 | 0 | 0 |
| 0 | 0 | 0 | 0 |
| 0 | 0 | 0 | 0 |
| 0 | 0 | 0 | 0 |
| 1 | 0 | 0 | 0 |
| 0 | 0 | 0 | 0 |
| 0 | 0 | 0 | 0 |
| 0 | 0 | 0 | 0 |
| 1 | 0 | 0 | 0 |
| 0 | 0 | 0 | 0 |
| 1 | 0 | 0 | 0 |
| 1 | 0 | 0 | 0 |
| 1 | 1 | 0 | 0 |
| 0 | 0 | 0 | 0 |
| 1 | 1 | 0 | 0 |
| 1 | 1 | 0 | 0 |
| 1 | 1 | 0 | 0 |
| 0 | 0 | 0 | 0 |
| 1 | 1 | 0 | 0 |
| 0 | 0 | 0 | 0 |
| 1 | 0 | 0 | 0 |
| 0 | 0 | 0 | 0 |
| 1 | 1 | 0 | 0 |
| 1 | 1 | 0 | 0 |
| 1 | 0 | 0 | 0 |
| 0 | 0 | 0 | 0 |
| 0 | 0 | 0 | 0 |
| 1 | 0 | 0 | 0 |
| 0 | 0 | 0 | 0 |
| 0 | 0 | 0 | 0 |
| 1 | 0 | 0 | 0 |
| 0 | 0 | 0 | 0 |
| 1 | 0 | 0 | 0 |
| 0 | 0 | 0 | 0 |
| 0 | 0 | 0 | 0 |
| 0 | 0 | 0 | 0 |
| 0 | 0 | 0 | 0 |
| 0 | 0 | 0 | 0 |

| | | | |
|---|---|---|---|
| 0 | 0 | 0 | 0 |
| 0 | 0 | 0 | 0 |
| 0 | 0 | 0 | 0 |
| 0 | 0 | 0 | 0 |
| 0 | 0 | 0 | 0 |
| 0 | 0 | 0 | 0 |
| 0 | 0 | 0 | 0 |
| 0 | 0 | 0 | 0 |
| 0 | 0 | 0 | 0 |
| 1 | 1 | 0 | 0 |
| 1 | 1 | 0 | 0 |
| 0 | 0 | 0 | 0 |
| 0 | 0 | 0 | 0 |
| 0 | 0 | 0 | 0 |
| 1 | 0 | 1 | 0 |
| 0 | 0 | 0 | 0 |
| 0 | 0 | 0 | 0 |
| 1 | 0 | 1 | 0 |
| 0 | 0 | 0 | 0 |
| 0 | 0 | 0 | 0 |
| 0 | 0 | 0 | 0 |
| 0 | 0 | 1 | 0 |
| 1 | 0 | 1 | 0 |
| 0 | 0 | 1 | 0 |
| 0 | 0 | 1 | 0 |
| 0 | 0 | 1 | 1 |
| 0 | 0 | 1 | 1 |
| 0 | 0 | 0 | 0 |
| 0 | 0 | 0 | 0 |
| 0 | 0 | 0 | 0 |
| 0 | 0 | 0 | 0 |
| 0 | 0 | 0 | 0 |
| 0 | 0 | 0 | 0 |
| 0 | 0 | 0 | 0 |
| 0 | 0 | 0 | 0 |
| 0 | 0 | 0 | 0 |
| 0 | 0 | 0 | 0 |
| 0 | 0 | 0 | 0 |
| 0 | 0 | 0 | 0 |
| 0 | 0 | 0 | 0 |
| 0 | 0 | 0 | 0 |
| 0 | 0 | 0 | 0 |
| 0 | 0 | 0 | 0 |
| 0 | 0 | 1 | 0 |
| 0 | 0 | 0 | 0 |
| 0 | 0 | 0 | 0 |
| 0 | 0 | 0 | 0 |
| 0 | 0 | 0 | 0 |
| 0 | 0 | 0 | 0 |
| 0 | 0 | 0 | 0 |
| 1 | 0 | 0 | 0 |
| 0 | 0 | 0 | 0 |
| 0 | 0 | 0 | 0 |
| 0 | 0 | 0 | 0 |

| | | | |
|---|---|---|---|
| 0 | 0 | 0 | 0 |
| 0 | 0 | 0 | 0 |
| 0 | 0 | 0 | 0 |
| 1 | 0 | 0 | 0 |
| 0 | 0 | 0 | 0 |
| 0 | 0 | 0 | 0 |
| 0 | 0 | 0 | 0 |
| 0 | 0 | 0 | 0 |
| 1 | 0 | 0 | 0 |
| 1 | 0 | 0 | 0 |
| 0 | 0 | 0 | 0 |
| 0 | 0 | 0 | 0 |
| 0 | 0 | 0 | 0 |
| 0 | 0 | 0 | 0 |
| 0 | 0 | 0 | 0 |
| 0 | 0 | 0 | 0 |
| 0 | 0 | 1 | 0 |
| 0 | 0 | 0 | 0 |
| 0 | 0 | 0 | 0 |
| 0 | 0 | 0 | 0 |
| 0 | 0 | 0 | 0 |
| 0 | 0 | 0 | 0 |
| 0 | 0 | 0 | 0 |
| 0 | 0 | 0 | 0 |
| 0 | 0 | 0 | 0 |
| 0 | 0 | 0 | 0 |
| 0 | 0 | 0 | 0 |
| 0 | 0 | 0 | 0 |
| 0 | 0 | 0 | 0 |
| 0 | 0 | 0 | 0 |
| 0 | 0 | 0 | 0 |
| 0 | 0 | 0 | 0 |
| 0 | 0 | 0 | 0 |
| 0 | 0 | 0 | 0 |
| 0 | 0 | 0 | 0 |
| 0 | 0 | 0 | 0 |
| 0 | 0 | 0 | 0 |
| 0 | 0 | 0 | 0 |
| 0 | 0 | 0 | 0 |
| 0 | 0 | 0 | 0 |
| 0 | 0 | 0 | 0 |
| 0 | 0 | 0 | 0 |
| 0 | 0 | 0 | 0 |
| 0 | 0 | 0 | 0 |
| 0 | 0 | 0 | 0 |
| 0 | 0 | 0 | 0 |
| 0 | 0 | 0 | 0 |
| 0 | 0 | 0 | 0 |
| 0 | 0 | 0 | 0 |
| 0 | 1 | 0 | 0 |
| 0 | 0 | 0 | 0 |
| 0 | 0 | 0 | 0 |
| 0 | 0 | 0 | 0 |
| 0 | 0 | 0 | 0 |

| | | | |
|---|---|---|---|
| 0 | 0 | 1 | 0 |
| 0 | 0 | 0 | 0 |
| 0 | 0 | 0 | 0 |
| 0 | 0 | 0 | 0 |
| 0 | 0 | 0 | 0 |
| 0 | 0 | 0 | 0 |
| 0 | 0 | 0 | 0 |
| 1 | 0 | 0 | 0 |
| 0 | 0 | 0 | 0 |
| 0 | 0 | 0 | 0 |
| 0 | 0 | 0 | 0 |
| 1 | 0 | 0 | 0 |
| 0 | 0 | 0 | 0 |
| 0 | 0 | 0 | 0 |
| 0 | 0 | 0 | 0 |
| 0 | 0 | 0 | 0 |
| 0 | 0 | 0 | 0 |
| 0 | 0 | 0 | 0 |
| 0 | 0 | 0 | 0 |
| 0 | 0 | 0 | 0 |
| 0 | 0 | 0 | 0 |
| 0 | 0 | 0 | 0 |
| 0 | 0 | 0 | 0 |
| 0 | 0 | 0 | 0 |
| 0 | 0 | 0 | 0 |
| 0 | 0 | 0 | 0 |
| 0 | 0 | 0 | 0 |
| 0 | 0 | 0 | 0 |
| 0 | 0 | 0 | 0 |
| 0 | 0 | 0 | 0 |
| 0 | 0 | 0 | 0 |
| 0 | 0 | 0 | 0 |
| 0 | 0 | 0 | 0 |
| 0 | 0 | 0 | 0 |
| 0 | 0 | 0 | 0 |
| 0 | 0 | 0 | 0 |
| 0 | 0 | 0 | 0 |
| 0 | 0 | 0 | 0 |
| 0 | 0 | 0 | 0 |
| 0 | 0 | 0 | 0 |
| 0 | 0 | 0 | 0 |
| 0 | 0 | 0 | 0 |
| 0 | 0 | 0 | 0 |
| 0 | 0 | 0 | 0 |
| 0 | 0 | 0 | 0 |
| 0 | 0 | 0 | 0 |
| 0 | 0 | 0 | 0 |
| 0 | 0 | 0 | 0 |
| 0 | 0 | 0 | 0 |
| 0 | 0 | 0 | 0 |
| 0 | 0 | 0 | 0 |
| 0 | 0 | 0 | 0 |
| 0 | 0 | 0 | 0 |
| 0 | 0 | 0 | 0 |

| | | | |
|---|---|---|---|
| 0 | 0 | 0 | 0 |
| 0 | 0 | 0 | 0 |
| 0 | 0 | 0 | 0 |
| 0 | 0 | 0 | 0 |
| 0 | 0 | 0 | 0 |
| 0 | 0 | 0 | 0 |
| 0 | 0 | 0 | 0 |
| 0 | 0 | 0 | 0 |
| 0 | 0 | 0 | 0 |
| 0 | 0 | 0 | 0 |
| 0 | 0 | 0 | 0 |
| 0 | 0 | 0 | 0 |
| 0 | 0 | 0 | 0 |
| 0 | 0 | 0 | 0 |
| 0 | 0 | 0 | 0 |
| 0 | 0 | 0 | 0 |
| 0 | 0 | 0 | 0 |
| 0 | 0 | 0 | 0 |
| 0 | 0 | 0 | 1 |
| 0 | 1 | 0 | 1 |
| 0 | 0 | 0 | 0 |
| 0 | 0 | 0 | 0 |
| 0 | 0 | 0 | 0 |
| 0 | 0 | 0 | 0 |
| 0 | 0 | 0 | 0 |
| 1 | 0 | 0 | 0 |
| 1 | 0 | 0 | 0 |
| 0 | 0 | 0 | 0 |
| 0 | 0 | 0 | 0 |
| 0 | 0 | 0 | 0 |
| 0 | 0 | 0 | 0 |
| 0 | 0 | 0 | 0 |
| 0 | 0 | 0 | 0 |
| 1 | 0 | 0 | 1 |
| 1 | 0 | 0 | 0 |
| 1 | 0 | 0 | 1 |
| 1 | 0 | 0 | 0 |
| 0 | 0 | 0 | 0 |
| 0 | 0 | 0 | 0 |
| 0 | 0 | 0 | 0 |
| 0 | 0 | 0 | 0 |
| 1 | 0 | 0 | 0 |
| 1 | 0 | 0 | 1 |
| 1 | 0 | 0 | 0 |
| 0 | 0 | 0 | 0 |
| 0 | 0 | 0 | 0 |
| 0 | 0 | 0 | 0 |
| 0 | 0 | 0 | 0 |
| 0 | 0 | 0 | 0 |
| 1 | 0 | 0 | 0 |
| 0 | 0 | 0 | 0 |
| 0 | 0 | 0 | 0 |
| 0 | 0 | 0 | 0 |
| 0 | 0 | 0 | 0 |

| | | | |
|---|---|---|---|
| 0 | 0 | 0 | 0 |
| 0 | 0 | 0 | 0 |
| 0 | 0 | 0 | 0 |
| 1 | 0 | 0 | 0 |
| 1 | 0 | 0 | 0 |
| 0 | 0 | 0 | 0 |
| 1 | 0 | 0 | 0 |
| 0 | 0 | 0 | 0 |
| 0 | 0 | 0 | 0 |
| 0 | 0 | 0 | 0 |
| 0 | 0 | 0 | 0 |
| 1 | 0 | 1 | 0 |
| 0 | 0 | 0 | 0 |
| 0 | 0 | 0 | 0 |
| 0 | 0 | 0 | 0 |
| 0 | 0 | 1 | 0 |
| 0 | 0 | 0 | 0 |
| 0 | 0 | 0 | 0 |
| 0 | 0 | 0 | 0 |
| 0 | 0 | 1 | 0 |
| 1 | 0 | 1 | 0 |
| 1 | 0 | 0 | 0 |
| 1 | 0 | 0 | 0 |
| 1 | 0 | 0 | 0 |
| 0 | 0 | 1 | 0 |
| 0 | 0 | 0 | 0 |
| 1 | 0 | 0 | 0 |
| 0 | 0 | 0 | 0 |
| 0 | 0 | 0 | 0 |
| 0 | 0 | 0 | 0 |
| 0 | 0 | 0 | 0 |
| 0 | 0 | 0 | 0 |
| 0 | 0 | 0 | 0 |
| 0 | 0 | 0 | 0 |
| 0 | 0 | 0 | 0 |
| 0 | 0 | 0 | 0 |
| 0 | 0 | 0 | 1 |
| 0 | 0 | 0 | 1 |
| 0 | 0 | 0 | 1 |
| 0 | 0 | 0 | 0 |
| 0 | 0 | 0 | 0 |
| 0 | 0 | 0 | 0 |
| 0 | 0 | 0 | 0 |
| 0 | 0 | 0 | 0 |
| 0 | 0 | 0 | 0 |
| 0 | 0 | 0 | 0 |
| 0 | 0 | 0 | 0 |
| 0 | 0 | 0 | 0 |
| 0 | 0 | 0 | 0 |
| 0 | 0 | 0 | 0 |
| 0 | 0 | 0 | 0 |
| 0 | 0 | 0 | 0 |
| 0 | 0 | 0 | 0 |
| 0 | 0 | 0 | 0 |

| | | | |
|---|---|---|---|
| 0 | 0 | 0 | 0 |
| 0 | 0 | 0 | 0 |
| 0 | 0 | 0 | 0 |
| 0 | 0 | 0 | 0 |
| 0 | 0 | 0 | 0 |
| 0 | 0 | 0 | 0 |
| 0 | 0 | 0 | 0 |
| 0 | 0 | 0 | 0 |
| 0 | 0 | 0 | 0 |
| 0 | 0 | 0 | 0 |
| 0 | 0 | 0 | 0 |
| 0 | 0 | 0 | 0 |
| 0 | 0 | 0 | 0 |
| 0 | 0 | 0 | 0 |
| 0 | 0 | 0 | 0 |
| 0 | 0 | 0 | 0 |
| 0 | 1 | 0 | 0 |
| 0 | 1 | 0 | 0 |
| 0 | 0 | 0 | 0 |
| 0 | 0 | 0 | 0 |
| 0 | 0 | 0 | 0 |
| 0 | 0 | 0 | 0 |
| 0 | 0 | 0 | 0 |
| 0 | 0 | 0 | 0 |
| 0 | 0 | 0 | 0 |
| 0 | 0 | 0 | 0 |
| 0 | 0 | 0 | 0 |
| 0 | 0 | 0 | 0 |
| 0 | 0 | 0 | 0 |
| 0 | 0 | 0 | 0 |
| 0 | 0 | 0 | 0 |
| 1 | 0 | 0 | 0 |
| 0 | 0 | 0 | 0 |
| 0 | 0 | 0 | 0 |
| 0 | 0 | 0 | 0 |
| 0 | 0 | 0 | 0 |
| 0 | 0 | 0 | 0 |
| 0 | 0 | 0 | 0 |
| 0 | 0 | 0 | 0 |
| 0 | 0 | 0 | 0 |
| 0 | 0 | 0 | 0 |
| 0 | 0 | 0 | 0 |
| 0 | 0 | 0 | 0 |
| 0 | 0 | 0 | 0 |
| 0 | 0 | 0 | 0 |
| 0 | 0 | 0 | 0 |
| 1 | 1 | 0 | 0 |
| 0 | 0 | 0 | 0 |
| 0 | 0 | 0 | 0 |
| 0 | 0 | 0 | 0 |
| 0 | 0 | 0 | 0 |
| 0 | 0 | 0 | 0 |
| 1 | 0 | 0 | 0 |
| 0 | 0 | 0 | 0 |

| | | | |
|---|---|---|---|
| 0 | 0 | 0 | 0 |
| 0 | 0 | 0 | 0 |
| 0 | 0 | 0 | 0 |
| 0 | 0 | 0 | 0 |
| 0 | 0 | 0 | 0 |
| 0 | 0 | 0 | 0 |
| 1 | 0 | 0 | 0 |
| 0 | 0 | 0 | 0 |
| 0 | 0 | 0 | 0 |
| 0 | 0 | 0 | 0 |
| 0 | 0 | 0 | 0 |
| 0 | 0 | 0 | 0 |
| 0 | 0 | 0 | 0 |
| 0 | 0 | 0 | 0 |
| 0 | 0 | 0 | 0 |
| 0 | 0 | 0 | 0 |
| 0 | 0 | 0 | 0 |
| 0 | 0 | 0 | 0 |
| 0 | 0 | 0 | 0 |
| 0 | 0 | 0 | 0 |
| 0 | 0 | 0 | 0 |
| 0 | 0 | 0 | 0 |
| 0 | 0 | 0 | 0 |
| 0 | 0 | 0 | 0 |
| 0 | 0 | 0 | 0 |
| 0 | 0 | 0 | 0 |
| 0 | 0 | 0 | 0 |
| 0 | 0 | 0 | 0 |
| 0 | 0 | 0 | 0 |
| 0 | 0 | 0 | 0 |
| 0 | 0 | 0 | 0 |
| 0 | 0 | 0 | 0 |
| 0 | 0 | 0 | 0 |
| 0 | 0 | 0 | 0 |
| 0 | 0 | 0 | 0 |
| 0 | 0 | 0 | 0 |
| 0 | 0 | 0 | 0 |
| 0 | 0 | 0 | 0 |
| 0 | 0 | 0 | 0 |
| 0 | 0 | 0 | 0 |
| 0 | 0 | 0 | 0 |
| 0 | 0 | 0 | 0 |
| 0 | 0 | 0 | 0 |
| 0 | 0 | 0 | 0 |
| 0 | 0 | 0 | 0 |
| 0 | 0 | 0 | 0 |
| 0 | 0 | 0 | 0 |
| 0 | 0 | 0 | 0 |
| 0 | 0 | 0 | 0 |
| 0 | 0 | 0 | 0 |
| 0 | 0 | 0 | 0 |
| 0 | 0 | 0 | 0 |
| 0 | 0 | 0 | 0 |
| 0 | 0 | 0 | 0 |

| | | | |
|---|---|---|---|
| 0 | 0 | 0 | 0 |
| 0 | 0 | 0 | 0 |
| 0 | 0 | 0 | 0 |
| 0 | 0 | 0 | 0 |
| 0 | 0 | 0 | 0 |
| 0 | 0 | 0 | 0 |
| 0 | 0 | 0 | 0 |
| 0 | 0 | 0 | 0 |
| 0 | 0 | 0 | 0 |
| 0 | 0 | 0 | 0 |
| 0 | 0 | 0 | 0 |
| 0 | 0 | 0 | 0 |
| 0 | 0 | 0 | 0 |
| 0 | 0 | 0 | 0 |
| 0 | 0 | 0 | 0 |
| 0 | 0 | 0 | 0 |
| 0 | 0 | 0 | 0 |
| 0 | 0 | 0 | 0 |
| 0 | 0 | 0 | 0 |
| 0 | 0 | 0 | 0 |
| 0 | 0 | 0 | 0 |
| 0 | 0 | 0 | 0 |
| 0 | 0 | 0 | 0 |
| 0 | 0 | 0 | 0 |
| 1 | 1 | 0 | 0 |
| 0 | 0 | 0 | 0 |
| 0 | 0 | 0 | 0 |
| 0 | 0 | 0 | 0 |
| 0 | 0 | 0 | 0 |
| 0 | 0 | 0 | 0 |
| 0 | 0 | 0 | 0 |
| 0 | 0 | 0 | 0 |
| 0 | 0 | 0 | 0 |
| 0 | 0 | 0 | 0 |
| 0 | 0 | 0 | 0 |
| 1 | 0 | 0 | 0 |
| 1 | 0 | 0 | 1 |
| 1 | 0 | 0 | 0 |
| 0 | 0 | 0 | 0 |
| 1 | 0 | 0 | 0 |
| 1 | 1 | 0 | 0 |
| 1 | 1 | 0 | 0 |
| 0 | 0 | 0 | 0 |
| 0 | 0 | 0 | 0 |
| 0 | 0 | 0 | 0 |
| 0 | 0 | 0 | 0 |
| 0 | 0 | 0 | 0 |
| 0 | 0 | 0 | 0 |
| 0 | 0 | 0 | 0 |
| 0 | 0 | 0 | 0 |
| 0 | 0 | 0 | 0 |
| 0 | 0 | 0 | 0 |
| 0 | 0 | 0 | 0 |
| 0 | 0 | 0 | 0 |

| | | | |
|---|---|---|---|
| 0 | 0 | 0 | 0 |
| 0 | 0 | 0 | 0 |
| 0 | 0 | 0 | 0 |
| 0 | 0 | 0 | 0 |
| 0 | 0 | 0 | 0 |
| 0 | 0 | 0 | 0 |
| 0 | 0 | 0 | 0 |
| 0 | 0 | 0 | 0 |
| 0 | 0 | 0 | 0 |
| 0 | 0 | 0 | 0 |
| 0 | 0 | 0 | 0 |
| 0 | 0 | 0 | 0 |
| 0 | 0 | 0 | 0 |
| 1 | 0 | 0 | 0 |
| 0 | 0 | 0 | 0 |
| 0 | 0 | 0 | 0 |
| 1 | 1 | 0 | 0 |
| 0 | 0 | 0 | 0 |
| 0 | 0 | 0 | 0 |
| 0 | 0 | 0 | 0 |
| 0 | 0 | 0 | 0 |
| 0 | 0 | 0 | 0 |
| 0 | 0 | 0 | 0 |
| 1 | 0 | 0 | 0 |
| 0 | 0 | 0 | 0 |
| 0 | 0 | 0 | 0 |
| 0 | 0 | 0 | 0 |
| 0 | 0 | 0 | 0 |
| 0 | 0 | 0 | 0 |
| 0 | 0 | 0 | 0 |
| 0 | 0 | 0 | 0 |
| 1 | 0 | 0 | 0 |
| 0 | 0 | 0 | 0 |
| 0 | 0 | 0 | 0 |
| 0 | 0 | 0 | 0 |
| 0 | 0 | 0 | 0 |
| 0 | 0 | 0 | 0 |
| 0 | 0 | 0 | 0 |
| 0 | 0 | 0 | 0 |
| 0 | 0 | 0 | 0 |
| 0 | 0 | 0 | 0 |
| 0 | 0 | 0 | 0 |
| 0 | 0 | 0 | 0 |
| 0 | 0 | 0 | 0 |
| 0 | 0 | 0 | 0 |
| 0 | 0 | 0 | 0 |
| 0 | 0 | 0 | 0 |
| 0 | 0 | 0 | 0 |
| 0 | 0 | 0 | 0 |
| 0 | 0 | 0 | 0 |
| 0 | 0 | 0 | 0 |
| 0 | 0 | 0 | 0 |
| 0 | 0 | 0 | 0 |
| 0 | 0 | 0 | 0 |

| | | | |
|---|---|---|---|
| 0 | 0 | 0 | 0 |
| 0 | 0 | 0 | 0 |
| 0 | 0 | 0 | 0 |
| 0 | 0 | 0 | 0 |
| 0 | 0 | 0 | 0 |
| 0 | 0 | 0 | 0 |
| 0 | 0 | 0 | 0 |
| 0 | 0 | 0 | 0 |
| 0 | 0 | 0 | 0 |
| 0 | 0 | 0 | 0 |
| 0 | 0 | 0 | 0 |
| 0 | 0 | 0 | 0 |
| 0 | 0 | 0 | 0 |
| 0 | 0 | 0 | 0 |
| 0 | 0 | 0 | 0 |
| 0 | 0 | 0 | 0 |
| 0 | 0 | 0 | 0 |
| 1 | 0 | 0 | 0 |
| 0 | 0 | 0 | 0 |
| 0 | 0 | 0 | 0 |
| 0 | 0 | 0 | 0 |
| 0 | 0 | 0 | 0 |
| 0 | 0 | 0 | 0 |
| 0 | 0 | 0 | 0 |
| 1 | 0 | 0 | 0 |
| 0 | 0 | 0 | 0 |
| 0 | 0 | 0 | 0 |
| 0 | 0 | 0 | 0 |
| 0 | 0 | 0 | 0 |
| 0 | 0 | 0 | 0 |
| 0 | 0 | 0 | 0 |
| 0 | 0 | 0 | 0 |
| 0 | 0 | 0 | 0 |
| 0 | 0 | 0 | 0 |
| 0 | 0 | 0 | 0 |
| 0 | 0 | 0 | 0 |
| 0 | 0 | 0 | 0 |
| 0 | 0 | 0 | 0 |
| 0 | 0 | 0 | 0 |
| 0 | 0 | 0 | 0 |
| 0 | 0 | 0 | 0 |
| 0 | 0 | 0 | 0 |
| 0 | 0 | 0 | 0 |
| 0 | 0 | 0 | 0 |
| 0 | 0 | 0 | 0 |
| 0 | 0 | 0 | 0 |
| 0 | 0 | 0 | 0 |
| 0 | 0 | 0 | 0 |
| 0 | 0 | 0 | 0 |
| 0 | 0 | 0 | 0 |
| 0 | 0 | 0 | 0 |
| 0 | 0 | 0 | 0 |
| 0 | 0 | 0 | 0 |
| 0 | 0 | 0 | 0 |

| | | | |
|---|---|---|---|
| 0 | 0 | 0 | 0 |
| 0 | 0 | 0 | 0 |
| 0 | 0 | 0 | 0 |
| 0 | 0 | 0 | 0 |
| 0 | 0 | 0 | 0 |
| 0 | 0 | 0 | 0 |
| 0 | 0 | 0 | 0 |
| 0 | 0 | 0 | 0 |
| 0 | 0 | 0 | 0 |
| 0 | 0 | 0 | 0 |
| 0 | 0 | 0 | 0 |
| 0 | 0 | 0 | 0 |
| 0 | 0 | 0 | 0 |
| 0 | 0 | 0 | 0 |
| 1 | 1 | 0 | 0 |
| 1 | 1 | 0 | 0 |
| 0 | 0 | 0 | 0 |
| 0 | 0 | 0 | 0 |
| 0 | 0 | 0 | 0 |
| 0 | 0 | 0 | 0 |
| 0 | 0 | 0 | 0 |
| 0 | 0 | 0 | 0 |
| 0 | 0 | 0 | 0 |
| 0 | 0 | 0 | 0 |
| 0 | 0 | 0 | 0 |
| 0 | 0 | 0 | 0 |
| 0 | 0 | 0 | 0 |
| 1 | 0 | 0 | 0 |
| 0 | 0 | 0 | 0 |
| 1 | 0 | 0 | 0 |
| 0 | 0 | 0 | 0 |
| 0 | 0 | 0 | 0 |
| 0 | 0 | 0 | 0 |
| 0 | 0 | 0 | 0 |
| 0 | 0 | 0 | 0 |
| 0 | 0 | 0 | 0 |
| 1 | 0 | 0 | 0 |
| 0 | 0 | 0 | 0 |
| 0 | 0 | 0 | 0 |
| 0 | 0 | 0 | 0 |
| 0 | 0 | 0 | 0 |
| 0 | 0 | 0 | 0 |
| 0 | 0 | 0 | 0 |
| 0 | 0 | 0 | 0 |
| 0 | 0 | 0 | 0 |
| 0 | 0 | 0 | 0 |
| 0 | 0 | 0 | 0 |
| 0 | 0 | 0 | 0 |
| 0 | 0 | 0 | 0 |
| 0 | 0 | 0 | 0 |
| 0 | 0 | 0 | 0 |
| 0 | 0 | 0 | 0 |
| 0 | 0 | 0 | 0 |
| 0 | 0 | 0 | 0 |

| | | | |
|---|---|---|---|
| 0 | 0 | 0 | 0 |
| 0 | 0 | 0 | 0 |
| 0 | 0 | 0 | 0 |
| 0 | 0 | 0 | 0 |
| 0 | 0 | 0 | 0 |
| 0 | 0 | 0 | 0 |
| 0 | 0 | 0 | 0 |
| 0 | 0 | 0 | 0 |
| 0 | 0 | 0 | 0 |
| 0 | 0 | 0 | 0 |
| 0 | 0 | 0 | 0 |
| 0 | 0 | 0 | 0 |
| 0 | 0 | 0 | 0 |
| 0 | 0 | 0 | 0 |
| 0 | 0 | 0 | 0 |
| 0 | 0 | 0 | 0 |
| 0 | 0 | 0 | 0 |
| 0 | 0 | 0 | 0 |
| 0 | 0 | 0 | 0 |
| 0 | 0 | 0 | 0 |
| 0 | 0 | 0 | 0 |
| 0 | 0 | 0 | 0 |
| 0 | 0 | 0 | 0 |
| 0 | 0 | 0 | 0 |
| 0 | 0 | 0 | 0 |
| 0 | 0 | 0 | 0 |
| 0 | 0 | 0 | 0 |
| 0 | 0 | 0 | 0 |
| 0 | 0 | 0 | 0 |
| 0 | 0 | 0 | 0 |
| 0 | 0 | 0 | 0 |
| 0 | 0 | 0 | 0 |
| 0 | 0 | 0 | 0 |
| 0 | 0 | 0 | 0 |
| 0 | 0 | 0 | 0 |
| 0 | 0 | 0 | 0 |
| 0 | 0 | 0 | 0 |
| 0 | 0 | 0 | 0 |
| 0 | 0 | 0 | 0 |
| 0 | 0 | 0 | 0 |
| 0 | 0 | 0 | 0 |
| 0 | 0 | 0 | 0 |
| 0 | 0 | 0 | 0 |
| 0 | 0 | 0 | 0 |
| 0 | 0 | 0 | 0 |
| 0 | 0 | 0 | 0 |
| 0 | 0 | 0 | 0 |
| 0 | 0 | 0 | 0 |
| 1 | 0 | 0 | 1 |
| 0 | 0 | 0 | 0 |
| 1 | 0 | 0 | 0 |
| 1 | 0 | 0 | 0 |
| 0 | 0 | 0 | 0 |
| 0 | 0 | 0 | 1 |

| | | | |
|---|---|---|---|
| 0 | 0 | 0 | 0 |
| 0 | 0 | 0 | 0 |
| 0 | 0 | 0 | 0 |
| 0 | 0 | 0 | 0 |
| 0 | 0 | 0 | 0 |
| 0 | 0 | 0 | 0 |
| 0 | 0 | 0 | 0 |
| 0 | 0 | 0 | 0 |
| 0 | 0 | 0 | 0 |
| 0 | 0 | 0 | 0 |
| 0 | 0 | 0 | 0 |
| 0 | 0 | 0 | 0 |
| 0 | 0 | 0 | 0 |
| 0 | 0 | 0 | 0 |
| 0 | 0 | 0 | 0 |
| 0 | 0 | 0 | 0 |
| 0 | 0 | 0 | 0 |
| 0 | 0 | 0 | 0 |
| 0 | 0 | 0 | 0 |
| 0 | 0 | 0 | 0 |
| 0 | 0 | 0 | 0 |
| 0 | 0 | 0 | 0 |
| 0 | 0 | 0 | 0 |
| 0 | 0 | 0 | 0 |
| 0 | 0 | 0 | 0 |
| 0 | 0 | 0 | 0 |
| 0 | 0 | 1 | 0 |
| 0 | 0 | 1 | 0 |
| 0 | 0 | 1 | 0 |
| 0 | 0 | 1 | 0 |
| 0 | 0 | 0 | 0 |
| 0 | 0 | 0 | 0 |
| 0 | 0 | 0 | 0 |
| 1 | 0 | 0 | 0 |
| 1 | 0 | 0 | 0 |
| 0 | 0 | 0 | 0 |
| 0 | 0 | 0 | 0 |
| 0 | 0 | 0 | 0 |
| 0 | 0 | 0 | 0 |
| 1 | 0 | 0 | 0 |
| 0 | 0 | 0 | 0 |
| 0 | 0 | 0 | 0 |
| 0 | 0 | 0 | 0 |
| 0 | 0 | 0 | 0 |
| 0 | 0 | 0 | 0 |
| 0 | 1 | 0 | 0 |
| 0 | 0 | 0 | 0 |
| 0 | 0 | 0 | 0 |
| 0 | 0 | 0 | 0 |
| 0 | 0 | 0 | 0 |
| 0 | 0 | 0 | 0 |
| 0 | 0 | 0 | 0 |
| 0 | 0 | 0 | 0 |
| 0 | 0 | 0 | 0 |

| | | | |
|---|---|---|---|
| 0 | 0 | 0 | 0 |
| 0 | 0 | 0 | 0 |
| 0 | 0 | 0 | 0 |
| 0 | 0 | 0 | 0 |
| 0 | 0 | 0 | 0 |
| 0 | 0 | 0 | 1 |
| 0 | 0 | 0 | 0 |
| 0 | 0 | 0 | 0 |
| 0 | 0 | 0 | 0 |
| 0 | 0 | 0 | 0 |
| 0 | 0 | 0 | 0 |
| 0 | 0 | 0 | 0 |
| 0 | 0 | 0 | 0 |
| 0 | 1 | 0 | 0 |
| 0 | 1 | 0 | 0 |
| 0 | 1 | 0 | 0 |
| 0 | 1 | 0 | 0 |
| 0 | 1 | 0 | 0 |
| 0 | 0 | 0 | 0 |
| 0 | 0 | 0 | 0 |
| 0 | 0 | 0 | 0 |
| 0 | 0 | 0 | 0 |
| 0 | 0 | 0 | 0 |
| 0 | 0 | 0 | 0 |
| 0 | 0 | 0 | 0 |
| 0 | 0 | 0 | 0 |
| 0 | 0 | 0 | 0 |
| 0 | 0 | 0 | 0 |
| 0 | 0 | 0 | 0 |
| 0 | 0 | 0 | 0 |
| 0 | 0 | 0 | 0 |
| 0 | 0 | 0 | 0 |
| 0 | 0 | 0 | 0 |
| 0 | 0 | 0 | 0 |
| 0 | 0 | 0 | 0 |
| 0 | 0 | 0 | 0 |
| 0 | 0 | 0 | 0 |
| 0 | 0 | 0 | 0 |
| 0 | 0 | 0 | 0 |
| 0 | 0 | 0 | 0 |
| 0 | 0 | 0 | 0 |
| 0 | 0 | 0 | 0 |
| 0 | 0 | 0 | 0 |
| 0 | 0 | 0 | 0 |
| 0 | 0 | 0 | 0 |
| 0 | 0 | 0 | 0 |
| 0 | 0 | 0 | 0 |
| 0 | 0 | 0 | 0 |
| 0 | 0 | 0 | 0 |
| 0 | 0 | 0 | 0 |
| 0 | 0 | 0 | 0 |
| 0 | 0 | 0 | 0 |
| 0 | 0 | 0 | 0 |
| 0 | 0 | 0 | 0 |

| | | | |
|---|---|---|---|
| 0 | 0 | 0 | 0 |
| 0 | 0 | 0 | 0 |
| 0 | 0 | 0 | 0 |
| 1 | 0 | 0 | 0 |
| 0 | 0 | 0 | 0 |
| 0 | 0 | 0 | 0 |
| 0 | 0 | 0 | 0 |
| 0 | 0 | 0 | 0 |
| 0 | 0 | 0 | 0 |
| 0 | 0 | 0 | 0 |
| 0 | 0 | 0 | 0 |
| 0 | 0 | 0 | 0 |
| 0 | 0 | 0 | 0 |
| 0 | 0 | 0 | 0 |
| 0 | 0 | 0 | 0 |
| 0 | 0 | 0 | 0 |
| 0 | 0 | 0 | 0 |
| 0 | 0 | 0 | 0 |
| 0 | 0 | 0 | 0 |
| 0 | 0 | 0 | 0 |
| 1 | 1 | 0 | 0 |
| 1 | 1 | 0 | 0 |
| 0 | 0 | 0 | 0 |
| 0 | 1 | 0 | 0 |
| 0 | 0 | 0 | 0 |
| 0 | 0 | 0 | 0 |
| 0 | 0 | 0 | 0 |
| 0 | 0 | 0 | 0 |
| 0 | 0 | 0 | 0 |
| 0 | 0 | 0 | 0 |
| 0 | 0 | 0 | 0 |
| 0 | 0 | 0 | 0 |
| 0 | 0 | 0 | 0 |
| 0 | 0 | 0 | 0 |
| 0 | 0 | 0 | 0 |
| 0 | 0 | 0 | 0 |
| 0 | 0 | 0 | 0 |
| 0 | 0 | 0 | 0 |
| 0 | 0 | 0 | 0 |
| 1 | 0 | 0 | 0 |
| 1 | 0 | 0 | 0 |
| 0 | 0 | 0 | 0 |
| 0 | 0 | 0 | 0 |
| 1 | 0 | 0 | 0 |
| 0 | 0 | 0 | 0 |
| 0 | 0 | 0 | 0 |
| 0 | 0 | 0 | 0 |
| 0 | 0 | 0 | 0 |
| 0 | 0 | 0 | 0 |
| 0 | 0 | 0 | 0 |
| 0 | 0 | 0 | 0 |
| 0 | 0 | 0 | 0 |
| 0 | 0 | 0 | 0 |
| 0 | 0 | 0 | 0 |

| | | | |
|---|---|---|---|
| 0 | 0 | 0 | 0 |
| 0 | 0 | 0 | 0 |
| 0 | 0 | 0 | 0 |
| 0 | 0 | 0 | 0 |
| 0 | 0 | 0 | 0 |
| 1 | 0 | 0 | 0 |
| 0 | 0 | 0 | 0 |
| 0 | 0 | 0 | 0 |
| 0 | 1 | 0 | 0 |
| 1 | 1 | 0 | 0 |
| 0 | 0 | 0 | 0 |
| 1 | 1 | 0 | 0 |
| 1 | 0 | 0 | 0 |
| 0 | 0 | 0 | 0 |
| 0 | 0 | 0 | 0 |
| 0 | 0 | 0 | 0 |
| 1 | 1 | 0 | 0 |
| 0 | 0 | 0 | 0 |
| 0 | 0 | 0 | 0 |
| 1 | 0 | 0 | 0 |
| 0 | 0 | 0 | 0 |
| 0 | 0 | 0 | 0 |
| 0 | 0 | 0 | 0 |
| 0 | 0 | 0 | 0 |
| 0 | 0 | 0 | 0 |
| 0 | 0 | 0 | 0 |
| 0 | 0 | 0 | 0 |
| 0 | 0 | 0 | 0 |
| 0 | 0 | 0 | 0 |
| 0 | 0 | 0 | 0 |
| 1 | 0 | 0 | 0 |
| 0 | 0 | 0 | 0 |
| 0 | 0 | 0 | 0 |
| 0 | 0 | 0 | 0 |
| 0 | 0 | 0 | 0 |
| 1 | 0 | 0 | 0 |
| 0 | 0 | 0 | 0 |
| 0 | 0 | 0 | 0 |
| 0 | 0 | 0 | 0 |
| 0 | 0 | 0 | 0 |
| 0 | 0 | 0 | 0 |
| 0 | 0 | 0 | 0 |
| 0 | 0 | 0 | 0 |
| 0 | 0 | 0 | 0 |
| 0 | 0 | 0 | 0 |
| 0 | 0 | 0 | 0 |
| 0 | 0 | 0 | 0 |
| 0 | 0 | 0 | 0 |
| 0 | 0 | 0 | 0 |
| 0 | 0 | 0 | 0 |
| 0 | 0 | 0 | 0 |
| 0 | 0 | 0 | 0 |
| 0 | 0 | 0 | 0 |
| 0 | 0 | 0 | 0 |

| | | | |
|---|---|---|---|
| 0 | 0 | 0 | 0 |
| 0 | 0 | 0 | 0 |
| 0 | 0 | 0 | 0 |
| 0 | 0 | 0 | 0 |
| 0 | 0 | 0 | 0 |
| 0 | 0 | 0 | 0 |
| 0 | 0 | 0 | 0 |
| 0 | 0 | 0 | 0 |
| 0 | 0 | 0 | 0 |
| 0 | 0 | 0 | 0 |
| 0 | 0 | 0 | 0 |
| 0 | 0 | 0 | 0 |
| 0 | 0 | 0 | 0 |
| 0 | 0 | 0 | 0 |
| 0 | 0 | 0 | 1 |
| 0 | 0 | 0 | 0 |
| 0 | 0 | 0 | 0 |
| 0 | 0 | 0 | 0 |
| 0 | 0 | 0 | 0 |
| 0 | 0 | 0 | 0 |
| 0 | 0 | 0 | 0 |
| 0 | 0 | 0 | 0 |
| 0 | 0 | 0 | 0 |
| 0 | 0 | 0 | 0 |
| 0 | 0 | 0 | 0 |
| 0 | 0 | 0 | 0 |
| 0 | 0 | 0 | 0 |
| 0 | 0 | 0 | 0 |
| 0 | 0 | 0 | 0 |
| 0 | 0 | 0 | 0 |
| 0 | 0 | 0 | 0 |
| 0 | 0 | 0 | 0 |
| 0 | 0 | 0 | 0 |
| 0 | 0 | 0 | 0 |
| 0 | 0 | 0 | 0 |
| 0 | 0 | 0 | 0 |
| 0 | 0 | 0 | 0 |
| 0 | 0 | 0 | 0 |
| 0 | 0 | 0 | 0 |
| 0 | 0 | 1 | 0 |
| 0 | 0 | 0 | 0 |
| 0 | 0 | 0 | 0 |
| 0 | 0 | 0 | 0 |
| 1 | 0 | 0 | 0 |
| 0 | 0 | 0 | 0 |
| 0 | 0 | 0 | 0 |
| 0 | 0 | 0 | 0 |
| 0 | 0 | 0 | 0 |
| 1 | 0 | 0 | 1 |
| 0 | 0 | 0 | 0 |
| 0 | 0 | 0 | 0 |
| 0 | 0 | 0 | 0 |
| 0 | 0 | 0 | 0 |
| 0 | 0 | 0 | 0 |

| | | | |
|---|---|---|---|
| 0 | 0 | 0 | 0 |
| 0 | 0 | 0 | 0 |
| 0 | 0 | 0 | 0 |
| 0 | 0 | 0 | 0 |
| 0 | 0 | 0 | 0 |
| 0 | 0 | 0 | 0 |
| 0 | 0 | 0 | 0 |
| 0 | 0 | 0 | 0 |
| 0 | 0 | 0 | 0 |
| 1 | 0 | 0 | 0 |
| 0 | 0 | 0 | 0 |
| 0 | 0 | 0 | 0 |
| 0 | 0 | 0 | 0 |
| 0 | 0 | 0 | 0 |
| 1 | 0 | 0 | 0 |
| 0 | 0 | 0 | 0 |
| 0 | 0 | 0 | 0 |
| 0 | 0 | 0 | 0 |
| 0 | 0 | 0 | 0 |
| 0 | 0 | 0 | 0 |
| 0 | 0 | 0 | 0 |
| 0 | 0 | 0 | 0 |
| 0 | 0 | 0 | 0 |
| 0 | 0 | 0 | 0 |
| 1 | 0 | 0 | 0 |
| 0 | 0 | 0 | 0 |
| 0 | 0 | 0 | 0 |
| 0 | 0 | 0 | 0 |
| 0 | 0 | 0 | 0 |
| 0 | 0 | 0 | 0 |
| 0 | 0 | 0 | 0 |
| 0 | 0 | 0 | 0 |
| 0 | 0 | 0 | 0 |
| 0 | 0 | 0 | 0 |
| 1 | 0 | 0 | 0 |
| 1 | 0 | 0 | 0 |
| 0 | 0 | 0 | 0 |
| 0 | 0 | 0 | 0 |
| 1 | 0 | 0 | 0 |
| 0 | 0 | 1 | 0 |
| 0 | 0 | 1 | 0 |
| 0 | 0 | 1 | 0 |
| 0 | 0 | 1 | 0 |
| 0 | 0 | 0 | 0 |
| 1 | 0 | 0 | 0 |
| 0 | 0 | 0 | 0 |
| 0 | 0 | 0 | 0 |
| 0 | 0 | 0 | 0 |
| 0 | 0 | 1 | 0 |
| 0 | 0 | 0 | 0 |
| 0 | 0 | 0 | 0 |
| 0 | 0 | 0 | 0 |
| 0 | 0 | 0 | 0 |
| 0 | 0 | 0 | 0 |

| | | | |
|---|---|---|---|
| 0 | 0 | 0 | 0 |
| 0 | 0 | 0 | 0 |
| 0 | 0 | 0 | 0 |
| 0 | 0 | 0 | 0 |
| 0 | 0 | 0 | 0 |
| 0 | 0 | 0 | 0 |
| 0 | 0 | 0 | 0 |
| 0 | 0 | 0 | 0 |
| 0 | 0 | 0 | 0 |
| 0 | 0 | 0 | 0 |
| 0 | 1 | 0 | 0 |
| 0 | 0 | 0 | 0 |
| 0 | 0 | 0 | 0 |
| 0 | 0 | 0 | 0 |
| 0 | 0 | 0 | 0 |
| 0 | 1 | 0 | 0 |
| 0 | 0 | 0 | 0 |
| 0 | 0 | 0 | 0 |
| 0 | 0 | 0 | 0 |
| 0 | 0 | 0 | 0 |
| 0 | 0 | 0 | 0 |
| 0 | 0 | 0 | 0 |
| 0 | 0 | 0 | 0 |
| 0 | 0 | 0 | 0 |
| 0 | 0 | 0 | 0 |
| 0 | 0 | 0 | 0 |
| 0 | 0 | 0 | 0 |
| 0 | 0 | 0 | 0 |
| 0 | 0 | 0 | 0 |
| 0 | 0 | 0 | 0 |
| 0 | 0 | 0 | 0 |
| 0 | 0 | 0 | 0 |
| 0 | 0 | 0 | 0 |
| 0 | 0 | 0 | 0 |
| 0 | 0 | 0 | 0 |
| 0 | 0 | 0 | 0 |
| 0 | 0 | 0 | 0 |
| 0 | 0 | 0 | 0 |
| 0 | 0 | 0 | 0 |
| 0 | 0 | 0 | 0 |
| 0 | 0 | 0 | 0 |
| 0 | 0 | 0 | 0 |
| 0 | 0 | 0 | 0 |
| 0 | 0 | 0 | 0 |
| 1 | 1 | 0 | 0 |
| 0 | 0 | 0 | 0 |
| 0 | 0 | 0 | 0 |
| 0 | 0 | 0 | 0 |
| 0 | 0 | 0 | 0 |
| 0 | 0 | 0 | 0 |
| 0 | 0 | 0 | 0 |
| 0 | 0 | 0 | 0 |
| 0 | 0 | 0 | 0 |
| 0 | 0 | 0 | 0 |

| | | | |
|---|---|---|---|
| 0 | 0 | 0 | 0 |
| 0 | 0 | 0 | 0 |
| 0 | 0 | 1 | 0 |
| 1 | 0 | 0 | 0 |
| 0 | 0 | 0 | 0 |
| 0 | 0 | 0 | 0 |
| 0 | 0 | 0 | 0 |
| 0 | 0 | 0 | 0 |
| 0 | 0 | 0 | 0 |
| 0 | 0 | 0 | 0 |
| 0 | 0 | 0 | 0 |
| 0 | 0 | 0 | 0 |
| 1 | 0 | 0 | 0 |
| 0 | 0 | 0 | 0 |
| 0 | 0 | 0 | 0 |
| 0 | 0 | 0 | 0 |
| 0 | 0 | 0 | 0 |
| 0 | 0 | 0 | 0 |
| 0 | 0 | 0 | 0 |
| 0 | 0 | 0 | 0 |
| 0 | 0 | 0 | 0 |
| 0 | 0 | 0 | 0 |
| 0 | 0 | 0 | 0 |
| 0 | 0 | 0 | 0 |
| 0 | 0 | 0 | 0 |
| 0 | 0 | 0 | 0 |
| 0 | 0 | 0 | 0 |
| 0 | 0 | 0 | 0 |
| 0 | 0 | 0 | 0 |
| 0 | 0 | 0 | 0 |
| 0 | 0 | 0 | 0 |
| 0 | 0 | 0 | 0 |
| 0 | 0 | 0 | 0 |
| 0 | 0 | 0 | 0 |
| 0 | 0 | 0 | 0 |
| 0 | 0 | 0 | 0 |
| 0 | 0 | 0 | 0 |
| 0 | 0 | 0 | 0 |
| 0 | 0 | 0 | 0 |
| 0 | 0 | 0 | 0 |
| 0 | 1 | 0 | 0 |
| 1 | 1 | 0 | 0 |
| 0 | 0 | 0 | 0 |
| 0 | 0 | 0 | 0 |
| 0 | 0 | 0 | 0 |
| 0 | 0 | 0 | 0 |
| 0 | 0 | 0 | 0 |
| 0 | 0 | 0 | 0 |
| 0 | 1 | 0 | 0 |
| 0 | 0 | 0 | 0 |
| 0 | 0 | 0 | 0 |
| 0 | 0 | 0 | 0 |
| 0 | 0 | 0 | 1 |
| 0 | 0 | 0 | 0 |

| | | | |
|---|---|---|---|
| 0 | 0 | 0 | 1 |
| 0 | 0 | 0 | 1 |
| 0 | 0 | 0 | 1 |
| 0 | 0 | 0 | 0 |
| 0 | 0 | 0 | 0 |
| 0 | 0 | 0 | 0 |
| 0 | 0 | 0 | 0 |
| 1 | 0 | 0 | 0 |
| 0 | 0 | 0 | 0 |
| 1 | 1 | 0 | 0 |
| 0 | 1 | 0 | 0 |
| 0 | 0 | 0 | 0 |
| 1 | 1 | 0 | 0 |
| 0 | 0 | 0 | 0 |
| 0 | 0 | 0 | 0 |
| 0 | 0 | 0 | 0 |
| 0 | 0 | 0 | 0 |
| 1 | 0 | 0 | 0 |
| 0 | 0 | 0 | 0 |
| 0 | 0 | 0 | 0 |
| 0 | 0 | 0 | 0 |
| 0 | 0 | 0 | 0 |
| 0 | 0 | 0 | 0 |
| 0 | 0 | 0 | 0 |
| 0 | 0 | 0 | 0 |
| 0 | 0 | 0 | 0 |
| 0 | 0 | 0 | 0 |
| 0 | 0 | 0 | 0 |
| 0 | 0 | 0 | 0 |
| 0 | 0 | 0 | 0 |
| 0 | 0 | 0 | 0 |
| 1 | 0 | 0 | 0 |
| 0 | 0 | 0 | 0 |
| 0 | 0 | 0 | 0 |
| 0 | 0 | 0 | 0 |
| 0 | 0 | 0 | 0 |
| 0 | 0 | 0 | 0 |
| 0 | 0 | 0 | 0 |
| 0 | 0 | 0 | 0 |
| 0 | 0 | 0 | 0 |
| 0 | 0 | 0 | 0 |
| 1 | 1 | 0 | 0 |
| 0 | 0 | 0 | 0 |
| 0 | 0 | 0 | 0 |
| 1 | 0 | 0 | 0 |
| 0 | 1 | 0 | 0 |
| 0 | 0 | 0 | 0 |
| 0 | 0 | 0 | 0 |
| 0 | 1 | 0 | 0 |
| 1 | 1 | 0 | 0 |
| 1 | 1 | 0 | 0 |
| 1 | 1 | 0 | 0 |
| 1 | 1 | 0 | 0 |
| 1 | 1 | 0 | 0 |

| | | | |
|---|---|---|---|
| 0 | 0 | 0 | 0 |
| 0 | 0 | 0 | 0 |
| 0 | 0 | 0 | 0 |
| 0 | 0 | 0 | 0 |
| 0 | 0 | 0 | 0 |
| 1 | 0 | 0 | 0 |
| 1 | 0 | 0 | 0 |
| 0 | 0 | 0 | 0 |
| 0 | 0 | 0 | 0 |
| 1 | 0 | 0 | 0 |
| 0 | 0 | 0 | 0 |
| 0 | 0 | 1 | 0 |
| 0 | 0 | 0 | 0 |
| 0 | 0 | 0 | 0 |
| 0 | 0 | 0 | 0 |
| 0 | 0 | 0 | 0 |
| 0 | 0 | 0 | 0 |
| 0 | 0 | 0 | 0 |
| 0 | 0 | 0 | 0 |
| 0 | 0 | 0 | 0 |
| 1 | 1 | 0 | 0 |
| 1 | 0 | 0 | 0 |
| 0 | 0 | 0 | 0 |
| 0 | 0 | 1 | 0 |
| 0 | 0 | 0 | 0 |
| 0 | 0 | 1 | 0 |
| 0 | 0 | 0 | 0 |
| 0 | 0 | 0 | 0 |
| 0 | 0 | 0 | 0 |
| 0 | 0 | 0 | 0 |
| 0 | 0 | 0 | 0 |
| 0 | 0 | 0 | 0 |
| 0 | 0 | 0 | 0 |
| 0 | 0 | 0 | 0 |
| 0 | 0 | 0 | 0 |
| 0 | 0 | 0 | 0 |
| 1 | 0 | 0 | 0 |
| 0 | 0 | 0 | 0 |
| 1 | 0 | 0 | 0 |
| 1 | 0 | 0 | 0 |
| 1 | 0 | 0 | 0 |
| 0 | 0 | 0 | 0 |
| 0 | 0 | 0 | 0 |
| 0 | 0 | 0 | 0 |
| 0 | 0 | 0 | 0 |
| 0 | 0 | 0 | 0 |
| 0 | 0 | 0 | 0 |
| 1 | 0 | 1 | 0 |
| 0 | 0 | 0 | 0 |
| 0 | 0 | 0 | 0 |
| 0 | 0 | 0 | 0 |
| 0 | 0 | 0 | 0 |
| 1 | 1 | 0 | 0 |
| 0 | 0 | 0 | 0 |

| | | | |
|---|---|---|---|
| 0 | 0 | 0 | 0 |
| 0 | 1 | 0 | 0 |
| 0 | 0 | 0 | 0 |
| 0 | 0 | 0 | 0 |
| 0 | 0 | 0 | 0 |
| 0 | 0 | 0 | 0 |
| 0 | 0 | 0 | 0 |
| 0 | 0 | 0 | 0 |
| 0 | 0 | 0 | 0 |
| 0 | 0 | 0 | 0 |
| 0 | 0 | 0 | 0 |
| 0 | 0 | 0 | 0 |
| 0 | 0 | 0 | 0 |
| 0 | 0 | 0 | 0 |
| 0 | 0 | 0 | 0 |
| 0 | 0 | 0 | 0 |
| 0 | 0 | 0 | 0 |
| 0 | 0 | 0 | 0 |
| 0 | 0 | 0 | 0 |
| 0 | 0 | 0 | 0 |
| 1 | 0 | 0 | 0 |
| 0 | 0 | 0 | 0 |
| 0 | 0 | 0 | 0 |
| 0 | 0 | 0 | 0 |
| 1 | 0 | 0 | 0 |
| 0 | 0 | 0 | 0 |
| 0 | 0 | 0 | 0 |
| 0 | 0 | 0 | 1 |
| 0 | 0 | 0 | 0 |
| 0 | 0 | 0 | 0 |
| 0 | 0 | 0 | 0 |
| 0 | 0 | 0 | 0 |
| 0 | 0 | 0 | 0 |
| 0 | 0 | 0 | 0 |
| 0 | 0 | 0 | 0 |
| 0 | 0 | 0 | 0 |
| 0 | 0 | 0 | 0 |
| 0 | 0 | 0 | 0 |
| 0 | 0 | 0 | 0 |
| 0 | 0 | 0 | 0 |
| 0 | 0 | 0 | 0 |
| 0 | 0 | 0 | 0 |
| 0 | 0 | 0 | 0 |
| 0 | 0 | 0 | 0 |
| 0 | 0 | 0 | 0 |
| 0 | 0 | 0 | 0 |
| 0 | 0 | 0 | 0 |
| 0 | 0 | 0 | 0 |
| 0 | 0 | 0 | 0 |
| 0 | 0 | 0 | 0 |
| 0 | 0 | 0 | 0 |
| 0 | 0 | 0 | 0 |
| 0 | 0 | 0 | 0 |
| 0 | 0 | 0 | 0 |

| | | | |
|---|---|---|---|
| 0 | 0 | 0 | 0 |
| 0 | 0 | 0 | 0 |
| 0 | 0 | 0 | 0 |
| 0 | 0 | 0 | 0 |
| 0 | 0 | 0 | 0 |
| 0 | 0 | 0 | 0 |
| 0 | 0 | 0 | 0 |
| 0 | 0 | 0 | 0 |
| 0 | 0 | 0 | 0 |
| 0 | 0 | 0 | 0 |
| 0 | 0 | 0 | 0 |
| 0 | 0 | 0 | 0 |
| 0 | 0 | 0 | 0 |
| 0 | 0 | 0 | 0 |
| 0 | 0 | 0 | 0 |
| 1 | 0 | 0 | 0 |
| 0 | 0 | 0 | 0 |
| 0 | 0 | 0 | 0 |
| 0 | 1 | 0 | 0 |
| 0 | 0 | 0 | 0 |
| 0 | 0 | 0 | 0 |
| 0 | 0 | 0 | 0 |
| 0 | 0 | 0 | 0 |
| 0 | 0 | 0 | 0 |
| 0 | 0 | 0 | 0 |
| 1 | 0 | 0 | 0 |
| 0 | 0 | 0 | 0 |
| 0 | 0 | 0 | 0 |
| 0 | 0 | 0 | 0 |
| 0 | 0 | 0 | 0 |
| 0 | 0 | 0 | 0 |
| 0 | 0 | 0 | 0 |
| 0 | 0 | 0 | 0 |
| 1 | 1 | 0 | 0 |
| 0 | 0 | 0 | 0 |
| 0 | 0 | 0 | 0 |
| 0 | 0 | 0 | 0 |
| 0 | 0 | 0 | 0 |
| 0 | 0 | 0 | 0 |
| 0 | 0 | 0 | 0 |
| 0 | 0 | 0 | 0 |
| 0 | 0 | 0 | 0 |
| 0 | 0 | 0 | 0 |
| 0 | 0 | 0 | 0 |
| 0 | 0 | 0 | 0 |
| 0 | 0 | 0 | 0 |
| 0 | 0 | 0 | 0 |
| 1 | 0 | 0 | 0 |
| 0 | 0 | 0 | 0 |
| 0 | 0 | 0 | 0 |
| 0 | 0 | 0 | 0 |
| 0 | 0 | 0 | 0 |
| 0 | 0 | 0 | 0 |
| 1 | 0 | 0 | 1 |

| | | | |
|---|---|---|---|
| 0 | 0 | 0 | 0 |
| 0 | 0 | 0 | 0 |
| 0 | 0 | 1 | 0 |
| 0 | 0 | 0 | 0 |
| 0 | 0 | 0 | 0 |
| 0 | 0 | 0 | 0 |
| 0 | 0 | 0 | 0 |
| 0 | 0 | 0 | 0 |
| 0 | 0 | 0 | 0 |
| 0 | 0 | 0 | 0 |
| 0 | 0 | 0 | 0 |
| 0 | 0 | 0 | 0 |
| 0 | 0 | 0 | 0 |
| 0 | 0 | 0 | 0 |
| 0 | 0 | 0 | 0 |
| 0 | 0 | 0 | 0 |
| 0 | 0 | 0 | 0 |
| 0 | 0 | 0 | 0 |
| 0 | 0 | 0 | 0 |
| 0 | 0 | 0 | 0 |
| 0 | 0 | 0 | 0 |
| 0 | 0 | 0 | 0 |
| 0 | 0 | 0 | 0 |
| 0 | 0 | 0 | 0 |
| 0 | 0 | 0 | 0 |
| 0 | 0 | 0 | 0 |
| 0 | 0 | 0 | 0 |
| 0 | 0 | 0 | 0 |
| 0 | 0 | 0 | 0 |
| 0 | 0 | 0 | 0 |
| 0 | 0 | 0 | 0 |
| 1 | 0 | 0 | 0 |
| 0 | 0 | 0 | 0 |
| 0 | 0 | 0 | 0 |
| 0 | 0 | 0 | 0 |
| 0 | 0 | 0 | 0 |
| 0 | 0 | 0 | 0 |
| 0 | 0 | 0 | 0 |
| 0 | 0 | 0 | 0 |
| 0 | 0 | 0 | 0 |
| 1 | 1 | 0 | 0 |
| 0 | 0 | 0 | 0 |
| 0 | 0 | 0 | 0 |
| 0 | 0 | 0 | 0 |
| 0 | 0 | 0 | 0 |
| 1 | 1 | 0 | 0 |
| 0 | 0 | 0 | 0 |
| 1 | 1 | 0 | 0 |
| 0 | 0 | 0 | 0 |
| 1 | 0 | 0 | 0 |
| 1 | 0 | 0 | 0 |
| 1 | 0 | 0 | 0 |
| 0 | 0 | 0 | 0 |
| 0 | 0 | 0 | 0 |

| | | | |
|---|---|---|---|
| 1 | 0 | 0 | 0 |
| 0 | 0 | 0 | 0 |
| 0 | 0 | 0 | 0 |
| 0 | 0 | 0 | 0 |
| 0 | 0 | 0 | 0 |
| 0 | 0 | 0 | 0 |
| 0 | 0 | 0 | 0 |
| 0 | 0 | 0 | 0 |
| 0 | 0 | 0 | 0 |
| 0 | 0 | 0 | 0 |
| 0 | 0 | 0 | 0 |
| 0 | 0 | 0 | 0 |
| 0 | 0 | 0 | 0 |
| 0 | 0 | 0 | 0 |
| 0 | 0 | 0 | 0 |
| 1 | 1 | 1 | 0 |
| 0 | 0 | 0 | 0 |
| 1 | 0 | 0 | 0 |
| 0 | 0 | 0 | 0 |
| 0 | 0 | 0 | 0 |
| 0 | 0 | 0 | 0 |
| 0 | 1 | 0 | 0 |
| 0 | 0 | 0 | 0 |
| 0 | 1 | 0 | 0 |
| 0 | 1 | 0 | 0 |
| 0 | 0 | 0 | 0 |
| 0 | 0 | 0 | 0 |
| 0 | 0 | 0 | 0 |
| 0 | 0 | 0 | 0 |
| 0 | 0 | 0 | 0 |
| 0 | 0 | 0 | 0 |
| 0 | 0 | 0 | 0 |
| 1 | 1 | 0 | 0 |
| 0 | 0 | 0 | 0 |
| 0 | 0 | 0 | 0 |
| 0 | 0 | 0 | 0 |
| 0 | 1 | 0 | 0 |
| 1 | 1 | 0 | 0 |
| 0 | 0 | 0 | 0 |
| 0 | 0 | 0 | 0 |
| 0 | 0 | 0 | 0 |
| 0 | 0 | 0 | 0 |
| 0 | 0 | 0 | 0 |
| 0 | 0 | 0 | 0 |
| 1 | 1 | 0 | 0 |
| 0 | 0 | 0 | 0 |
| 1 | 0 | 0 | 0 |
| 1 | 1 | 0 | 0 |
| 0 | 0 | 0 | 0 |
| 0 | 0 | 0 | 0 |
| 1 | 1 | 0 | 0 |
| 1 | 1 | 0 | 0 |
| 0 | 0 | 0 | 0 |
| 1 | 1 | 0 | 0 |

| | | | |
|---|---|---|---|
| 0 | 0 | 0 | 0 |
| 0 | 0 | 0 | 0 |
| 0 | 0 | 0 | 0 |
| 1 | 1 | 0 | 0 |
| 0 | 0 | 0 | 0 |
| 0 | 0 | 0 | 0 |
| 0 | 0 | 0 | 0 |
| 0 | 0 | 0 | 0 |
| 1 | 0 | 0 | 0 |
| 0 | 0 | 0 | 0 |
| 0 | 0 | 0 | 0 |
| 0 | 0 | 0 | 0 |
| 1 | 0 | 1 | 0 |
| 1 | 0 | 0 | 0 |
| 0 | 0 | 0 | 0 |
| 0 | 0 | 0 | 0 |
| 0 | 0 | 0 | 0 |
| 0 | 0 | 0 | 0 |
| 0 | 0 | 0 | 0 |
| 0 | 0 | 0 | 0 |
| 0 | 0 | 0 | 0 |
| 0 | 0 | 0 | 0 |
| 0 | 0 | 0 | 0 |
| 1 | 0 | 0 | 0 |
| 0 | 0 | 0 | 0 |
| 0 | 0 | 0 | 0 |
| 0 | 0 | 0 | 0 |
| 0 | 0 | 0 | 0 |
| 0 | 0 | 0 | 0 |
| 0 | 0 | 0 | 0 |
| 0 | 0 | 0 | 0 |
| 0 | 0 | 0 | 0 |
| 0 | 0 | 0 | 0 |
| 0 | 0 | 0 | 0 |
| 0 | 0 | 0 | 0 |
| 0 | 0 | 0 | 0 |
| 0 | 0 | 0 | 0 |
| 0 | 0 | 0 | 0 |
| 0 | 0 | 0 | 0 |
| 0 | 0 | 0 | 0 |
| 0 | 0 | 0 | 0 |
| 0 | 0 | 0 | 0 |
| 0 | 0 | 0 | 0 |
| 1 | 1 | 0 | 0 |
| 1 | 1 | 0 | 0 |
| 0 | 0 | 0 | 0 |
| 0 | 0 | 0 | 0 |
| 0 | 1 | 0 | 0 |
| 0 | 1 | 0 | 0 |
| 0 | 1 | 0 | 0 |
| 0 | 0 | 0 | 0 |
| 0 | 0 | 0 | 0 |
| 0 | 0 | 0 | 0 |
| 0 | 0 | 0 | 0 |

| | | | |
|---|---|---|---|
| 0 | 0 | 0 | 0 |
| 0 | 0 | 0 | 0 |
| 0 | 0 | 0 | 0 |
| 0 | 0 | 0 | 0 |
| 1 | 0 | 0 | 0 |
| 0 | 0 | 0 | 0 |
| 0 | 0 | 0 | 0 |
| 1 | 0 | 0 | 0 |
| 1 | 0 | 0 | 0 |
| 1 | 0 | 0 | 0 |
| 1 | 0 | 0 | 0 |
| 1 | 0 | 0 | 0 |
| 0 | 0 | 0 | 0 |
| 1 | 0 | 0 | 0 |
| 1 | 0 | 0 | 0 |
| 0 | 0 | 0 | 0 |
| 0 | 0 | 0 | 0 |
| 0 | 0 | 0 | 0 |
| 1 | 0 | 0 | 0 |
| 1 | 0 | 0 | 0 |
| 1 | 0 | 0 | 0 |
| 1 | 0 | 0 | 0 |
| 0 | 0 | 0 | 0 |
| 0 | 0 | 0 | 0 |
| 0 | 0 | 0 | 0 |
| 0 | 0 | 0 | 0 |
| 0 | 0 | 0 | 0 |
| 0 | 0 | 0 | 0 |
| 0 | 0 | 0 | 0 |
| 0 | 0 | 0 | 0 |
| 0 | 0 | 0 | 0 |
| 0 | 0 | 0 | 0 |
| 1 | 0 | 0 | 0 |
| 0 | 0 | 0 | 0 |
| 0 | 0 | 0 | 0 |
| 1 | 0 | 0 | 0 |
| 0 | 0 | 0 | 0 |
| 0 | 0 | 0 | 0 |
| 0 | 0 | 0 | 0 |
| 0 | 0 | 0 | 0 |
| 0 | 0 | 0 | 0 |
| 0 | 0 | 0 | 0 |
| 0 | 0 | 0 | 0 |
| 0 | 0 | 0 | 0 |
| 0 | 0 | 0 | 0 |
| 0 | 0 | 0 | 0 |
| 0 | 0 | 0 | 0 |
| 0 | 0 | 0 | 0 |
| 0 | 0 | 0 | 0 |
| 0 | 0 | 0 | 0 |
| 0 | 0 | 0 | 0 |
| 0 | 0 | 0 | 0 |
| 0 | 0 | 0 | 0 |
| 0 | 0 | 0 | 0 |

| | | | |
|---|---|---|---|
| 0 | 0 | 0 | 0 |
| 0 | 0 | 0 | 0 |
| 0 | 0 | 0 | 0 |
| 0 | 0 | 0 | 0 |
| 0 | 0 | 0 | 0 |
| 0 | 0 | 0 | 0 |
| 0 | 0 | 0 | 0 |
| 0 | 0 | 0 | 0 |
| 0 | 0 | 0 | 0 |
| 0 | 0 | 0 | 0 |
| 0 | 0 | 0 | 0 |
| 0 | 0 | 0 | 1 |
| 0 | 0 | 0 | 0 |
| 0 | 0 | 0 | 0 |
| 0 | 0 | 0 | 0 |
| 0 | 0 | 0 | 0 |
| 0 | 0 | 0 | 0 |
| 0 | 0 | 0 | 1 |
| 0 | 0 | 0 | 0 |
| 0 | 0 | 0 | 0 |
| 0 | 0 | 0 | 1 |
| 0 | 0 | 0 | 0 |
| 0 | 0 | 0 | 0 |
| 0 | 0 | 0 | 0 |
| 0 | 0 | 0 | 0 |
| 0 | 0 | 0 | 0 |
| 0 | 0 | 0 | 0 |
| 0 | 0 | 0 | 0 |
| 0 | 0 | 0 | 1 |
| 0 | 0 | 0 | 0 |
| 0 | 0 | 0 | 0 |
| 0 | 0 | 0 | 0 |
| 0 | 0 | 0 | 0 |
| 0 | 0 | 0 | 0 |
| 0 | 0 | 0 | 0 |
| 0 | 0 | 0 | 0 |
| 0 | 0 | 0 | 0 |
| 0 | 0 | 0 | 0 |
| 0 | 0 | 0 | 0 |
| 0 | 0 | 0 | 0 |
| 0 | 0 | 0 | 0 |
| 0 | 0 | 0 | 0 |
| 0 | 0 | 0 | 0 |
| 0 | 0 | 0 | 0 |
| 0 | 0 | 0 | 0 |
| 0 | 0 | 0 | 0 |
| 0 | 0 | 0 | 0 |
| 0 | 0 | 0 | 0 |
| 0 | 0 | 0 | 0 |
| 0 | 0 | 0 | 0 |
| 0 | 0 | 0 | 0 |
| 0 | 0 | 0 | 0 |
| 0 | 0 | 0 | 0 |
| 0 | 0 | 0 | 0 |

| | | | |
|---|---|---|---|
| 0 | 0 | 0 | 0 |
| 0 | 0 | 0 | 0 |
| 1 | 0 | 0 | 0 |
| 0 | 0 | 0 | 0 |
| 1 | 0 | 0 | 0 |
| 0 | 0 | 0 | 0 |
| 0 | 0 | 0 | 0 |
| 0 | 0 | 0 | 0 |
| 0 | 0 | 0 | 0 |
| 1 | 1 | 0 | 0 |
| 0 | 0 | 1 | 0 |
| 0 | 0 | 0 | 0 |
| 0 | 0 | 0 | 0 |
| 0 | 0 | 0 | 0 |
| 0 | 0 | 0 | 0 |
| 0 | 0 | 0 | 0 |
| 0 | 0 | 0 | 0 |
| 0 | 0 | 0 | 0 |
| 0 | 0 | 0 | 0 |
| 0 | 0 | 0 | 0 |
| 0 | 0 | 0 | 0 |
| 0 | 0 | 0 | 0 |
| 0 | 0 | 0 | 0 |
| 0 | 0 | 0 | 0 |
| 0 | 0 | 0 | 0 |
| 0 | 0 | 0 | 0 |
| 0 | 0 | 0 | 0 |
| 0 | 0 | 0 | 0 |
| 0 | 0 | 0 | 0 |
| 0 | 0 | 0 | 0 |
| 0 | 0 | 0 | 0 |
| 0 | 0 | 0 | 0 |
| 0 | 0 | 0 | 0 |
| 0 | 0 | 0 | 0 |
| 0 | 0 | 0 | 0 |
| 0 | 0 | 0 | 0 |
| 0 | 0 | 0 | 0 |
| 0 | 0 | 0 | 0 |
| 0 | 0 | 0 | 0 |
| 0 | 0 | 0 | 0 |
| 0 | 0 | 0 | 0 |
| 0 | 0 | 0 | 0 |
| 0 | 0 | 0 | 0 |
| 0 | 0 | 0 | 0 |
| 0 | 0 | 0 | 0 |
| 1 | 0 | 0 | 0 |
| 0 | 0 | 0 | 0 |
| 0 | 0 | 0 | 0 |
| 0 | 0 | 0 | 0 |
| 0 | 0 | 0 | 0 |
| 1 | 0 | 0 | 0 |
| 0 | 0 | 0 | 0 |
| 0 | 0 | 0 | 0 |
| 0 | 0 | 0 | 0 |

| | | | |
|---|---|---|---|
| 0 | 0 | 0 | 0 |
| 0 | 0 | 0 | 0 |
| 0 | 0 | 0 | 0 |
| 0 | 0 | 0 | 0 |
| 0 | 0 | 0 | 0 |
| 0 | 0 | 0 | 0 |
| 0 | 0 | 0 | 0 |
| 0 | 0 | 0 | 0 |
| 1 | 1 | 0 | 0 |
| 0 | 0 | 0 | 0 |
| 0 | 0 | 0 | 0 |
| 0 | 0 | 0 | 0 |
| 0 | 0 | 0 | 0 |
| 0 | 0 | 0 | 0 |
| 0 | 1 | 0 | 0 |
| 0 | 1 | 0 | 0 |
| 0 | 1 | 0 | 0 |
| 0 | 0 | 0 | 0 |
| 0 | 0 | 0 | 0 |
| 0 | 0 | 0 | 0 |
| 0 | 1 | 0 | 0 |
| 0 | 0 | 0 | 0 |
| 0 | 0 | 0 | 0 |
| 1 | 1 | 0 | 0 |
| 0 | 1 | 0 | 0 |
| 0 | 1 | 0 | 0 |
| 0 | 0 | 0 | 0 |
| 0 | 0 | 0 | 0 |
| 0 | 0 | 0 | 0 |
| 0 | 0 | 0 | 0 |
| 0 | 0 | 0 | 0 |
| 0 | 0 | 0 | 0 |
| 0 | 0 | 0 | 0 |
| 0 | 0 | 0 | 0 |
| 0 | 0 | 0 | 0 |
| 0 | 0 | 0 | 0 |
| 0 | 0 | 0 | 0 |
| 1 | 0 | 0 | 0 |
| 0 | 0 | 0 | 0 |
| 0 | 0 | 0 | 0 |
| 0 | 0 | 0 | 0 |
| 0 | 0 | 0 | 0 |
| 0 | 0 | 0 | 0 |
| 0 | 0 | 0 | 0 |
| 0 | 0 | 0 | 0 |
| 0 | 0 | 0 | 0 |
| 0 | 0 | 0 | 0 |
| 0 | 0 | 0 | 0 |
| 0 | 0 | 0 | 0 |
| 0 | 0 | 0 | 0 |
| 0 | 0 | 0 | 0 |
| 0 | 0 | 0 | 0 |
| 0 | 0 | 0 | 0 |
| 0 | 0 | 0 | 0 |

| | | | |
|---|---|---|---|
| 0 | 0 | 0 | 0 |
| 0 | 0 | 0 | 0 |
| 0 | 0 | 0 | 0 |
| 0 | 0 | 1 | 0 |
| 0 | 0 | 0 | 0 |
| 0 | 0 | 0 | 0 |
| 0 | 0 | 0 | 0 |
| 0 | 0 | 0 | 0 |
| 0 | 0 | 0 | 0 |
| 0 | 0 | 0 | 0 |
| 0 | 0 | 0 | 0 |
| 0 | 0 | 0 | 0 |
| 1 | 0 | 0 | 0 |
| 0 | 0 | 0 | 0 |
| 0 | 0 | 0 | 0 |
| 0 | 0 | 0 | 0 |
| 0 | 1 | 0 | 0 |
| 0 | 0 | 0 | 0 |
| 0 | 0 | 0 | 0 |
| 0 | 0 | 0 | 0 |
| 0 | 0 | 0 | 0 |
| 0 | 0 | 0 | 0 |
| 0 | 0 | 0 | 0 |
| 0 | 0 | 0 | 0 |
| 0 | 0 | 0 | 0 |
| 0 | 0 | 0 | 0 |
| 0 | 0 | 0 | 0 |
| 0 | 0 | 0 | 0 |
| 0 | 0 | 0 | 0 |
| 0 | 0 | 0 | 0 |
| 1 | 0 | 0 | 0 |
| 1 | 0 | 0 | 0 |
| 1 | 0 | 0 | 0 |
| 0 | 0 | 0 | 0 |
| 1 | 0 | 0 | 0 |
| 1 | 0 | 0 | 0 |
| 1 | 0 | 0 | 0 |
| 0 | 0 | 0 | 0 |
| 0 | 0 | 0 | 0 |
| 0 | 0 | 0 | 0 |
| 0 | 0 | 0 | 0 |
| 0 | 0 | 0 | 0 |
| 0 | 1 | 0 | 0 |
| 0 | 1 | 0 | 0 |
| 0 | 0 | 0 | 0 |
| 0 | 0 | 0 | 0 |
| 1 | 0 | 0 | 0 |
| 0 | 0 | 0 | 0 |
| 0 | 0 | 0 | 0 |
| 0 | 0 | 0 | 0 |
| 0 | 0 | 0 | 0 |
| 0 | 0 | 0 | 0 |
| 1 | 0 | 0 | 0 |
| 0 | 0 | 0 | 0 |

| | | | |
|---|---|---|---|
| 0 | 0 | 0 | 0 |
| 1 | 0 | 0 | 0 |
| 1 | 0 | 0 | 0 |
| 0 | 0 | 0 | 0 |
| 0 | 0 | 0 | 0 |
| 0 | 0 | 0 | 0 |
| 0 | 0 | 0 | 0 |
| 0 | 1 | 0 | 0 |
| 0 | 0 | 0 | 0 |
| 0 | 0 | 0 | 0 |
| 0 | 0 | 0 | 0 |
| 0 | 0 | 0 | 0 |
| 0 | 0 | 0 | 0 |
| 0 | 0 | 0 | 0 |
| 1 | 0 | 0 | 0 |
| 0 | 0 | 0 | 0 |
| 0 | 0 | 0 | 0 |
| 0 | 0 | 0 | 0 |
| 0 | 0 | 0 | 0 |
| 1 | 0 | 0 | 0 |
| 1 | 0 | 0 | 0 |
| 0 | 0 | 0 | 0 |
| 1 | 0 | 0 | 0 |
| 1 | 0 | 0 | 0 |
| 0 | 0 | 0 | 0 |
| 0 | 0 | 0 | 0 |
| 0 | 0 | 0 | 0 |
| 0 | 0 | 0 | 0 |
| 0 | 0 | 0 | 0 |
| 0 | 0 | 0 | 0 |
| 0 | 0 | 0 | 0 |
| 0 | 0 | 0 | 0 |
| 0 | 0 | 0 | 0 |
| 0 | 0 | 0 | 0 |
| 0 | 0 | 0 | 0 |
| 0 | 0 | 0 | 0 |
| 0 | 0 | 0 | 0 |
| 0 | 0 | 0 | 0 |
| 0 | 0 | 0 | 0 |
| 0 | 0 | 0 | 0 |
| 0 | 0 | 0 | 0 |
| 0 | 0 | 0 | 0 |
| 0 | 0 | 0 | 0 |
| 0 | 0 | 0 | 0 |
| 0 | 0 | 0 | 0 |
| 0 | 0 | 0 | 0 |
| 0 | 0 | 0 | 0 |
| 0 | 0 | 0 | 0 |
| 0 | 0 | 0 | 0 |
| 0 | 0 | 0 | 0 |
| 0 | 0 | 0 | 0 |
| 0 | 0 | 0 | 0 |
| 0 | 0 | 0 | 0 |
| 1 | 1 | 0 | 1 |

| | | | |
|---|---|---|---|
| 0 | 0 | 0 | 0 |
| 0 | 0 | 0 | 0 |
| 0 | 0 | 0 | 0 |
| 0 | 0 | 0 | 0 |
| 0 | 1 | 0 | 0 |
| 0 | 0 | 0 | 0 |
| 0 | 0 | 0 | 0 |
| 0 | 1 | 0 | 0 |
| 0 | 0 | 0 | 0 |
| 0 | 0 | 0 | 0 |
| 0 | 0 | 0 | 0 |
| 0 | 0 | 0 | 0 |
| 0 | 0 | 0 | 0 |
| 0 | 0 | 0 | 0 |
| 0 | 1 | 0 | 1 |
| 0 | 1 | 0 | 0 |
| 0 | 0 | 0 | 0 |
| 0 | 0 | 0 | 0 |
| 0 | 1 | 0 | 0 |
| 0 | 1 | 0 | 1 |
| 0 | 0 | 0 | 0 |
| 0 | 0 | 0 | 0 |
| 0 | 0 | 0 | 0 |
| 0 | 0 | 0 | 0 |
| 0 | 0 | 0 | 0 |
| 1 | 0 | 0 | 0 |
| 0 | 0 | 0 | 0 |
| 0 | 0 | 1 | 0 |
| 0 | 0 | 1 | 0 |
| 0 | 0 | 1 | 0 |
| 0 | 0 | 0 | 0 |
| 0 | 0 | 0 | 0 |
| 0 | 0 | 0 | 0 |
| 0 | 0 | 0 | 0 |
| 0 | 0 | 0 | 0 |
| 0 | 0 | 1 | 0 |
| 0 | 0 | 0 | 0 |
| 0 | 0 | 0 | 0 |
| 1 | 0 | 1 | 0 |
| 0 | 0 | 1 | 0 |
| 1 | 1 | 0 | 0 |
| 0 | 1 | 0 | 0 |
| 0 | 0 | 0 | 0 |
| 0 | 0 | 0 | 0 |
| 1 | 0 | 0 | 0 |
| 0 | 0 | 0 | 0 |
| 0 | 0 | 1 | 0 |
| 0 | 0 | 0 | 0 |
| 0 | 0 | 0 | 0 |
| 0 | 0 | 0 | 0 |
| 0 | 0 | 0 | 0 |
| 0 | 0 | 0 | 0 |
| 0 | 0 | 0 | 0 |
| 0 | 0 | 0 | 0 |

| | | | |
|---|---|---|---|
| 0 | 0 | 0 | 0 |
| 0 | 0 | 0 | 0 |
| 0 | 0 | 0 | 0 |
| 0 | 0 | 0 | 0 |
| 0 | 0 | 0 | 0 |
| 0 | 0 | 0 | 0 |
| 0 | 0 | 0 | 0 |
| 1 | 1 | 0 | 0 |
| 0 | 0 | 0 | 0 |
| 0 | 0 | 0 | 0 |
| 0 | 0 | 0 | 0 |
| 0 | 0 | 0 | 1 |
| 0 | 0 | 0 | 0 |
| 0 | 0 | 0 | 0 |
| 0 | 0 | 0 | 0 |
| 0 | 0 | 0 | 0 |
| 0 | 0 | 0 | 0 |
| 0 | 0 | 0 | 0 |
| 0 | 0 | 0 | 0 |
| 0 | 0 | 0 | 0 |
| 0 | 0 | 0 | 0 |
| 0 | 0 | 0 | 0 |
| 0 | 0 | 0 | 0 |
| 0 | 0 | 0 | 0 |
| 0 | 0 | 0 | 0 |
| 0 | 0 | 0 | 0 |
| 0 | 0 | 0 | 0 |
| 0 | 0 | 0 | 0 |
| 0 | 0 | 0 | 0 |
| 0 | 1 | 0 | 0 |
| 0 | 1 | 0 | 0 |
| 0 | 0 | 0 | 0 |
| 0 | 0 | 0 | 0 |
| 0 | 0 | 0 | 0 |
| 0 | 0 | 0 | 0 |
| 0 | 1 | 0 | 0 |
| 0 | 0 | 0 | 0 |
| 0 | 0 | 0 | 0 |
| 0 | 0 | 0 | 0 |
| 0 | 0 | 0 | 0 |
| 0 | 0 | 0 | 0 |
| 1 | 1 | 0 | 0 |
| 1 | 0 | 0 | 0 |
| 0 | 0 | 0 | 0 |
| 0 | 0 | 0 | 0 |
| 0 | 0 | 0 | 0 |
| 0 | 0 | 0 | 0 |
| 0 | 0 | 0 | 0 |
| 0 | 0 | 0 | 0 |
| 0 | 0 | 0 | 0 |
| 0 | 0 | 0 | 0 |
| 0 | 0 | 0 | 0 |
| 0 | 0 | 0 | 0 |
| 0 | 0 | 0 | 0 |

| | | | |
|---|---|---|---|
| 0 | 0 | 0 | 0 |
| 0 | 0 | 0 | 0 |
| 0 | 0 | 0 | 0 |
| 0 | 0 | 0 | 0 |
| 0 | 0 | 0 | 0 |
| 0 | 0 | 0 | 0 |
| 0 | 0 | 0 | 0 |
| 0 | 0 | 0 | 0 |
| 1 | 0 | 0 | 0 |
| 1 | 0 | 0 | 0 |
| 0 | 0 | 0 | 0 |
| 0 | 0 | 0 | 0 |
| 0 | 0 | 0 | 0 |
| 1 | 0 | 0 | 0 |
| 0 | 0 | 0 | 0 |
| 0 | 0 | 0 | 0 |
| 0 | 0 | 0 | 0 |
| 0 | 0 | 0 | 0 |
| 0 | 0 | 0 | 0 |
| 1 | 1 | 0 | 0 |
| 0 | 0 | 1 | 0 |
| 0 | 0 | 0 | 0 |
| 0 | 0 | 0 | 0 |
| 0 | 0 | 0 | 0 |
| 0 | 0 | 0 | 0 |
| 0 | 0 | 0 | 0 |
| 0 | 0 | 0 | 0 |
| 0 | 0 | 0 | 0 |
| 0 | 0 | 0 | 0 |
| 0 | 0 | 0 | 0 |
| 0 | 0 | 0 | 0 |
| 0 | 0 | 0 | 0 |
| 0 | 0 | 0 | 0 |
| 0 | 0 | 0 | 0 |
| 0 | 0 | 0 | 0 |
| 0 | 0 | 0 | 0 |
| 0 | 0 | 0 | 0 |
| 0 | 0 | 0 | 0 |
| 0 | 0 | 0 | 0 |
| 0 | 0 | 0 | 0 |
| 0 | 0 | 0 | 0 |
| 0 | 0 | 0 | 0 |
| 0 | 0 | 0 | 0 |
| 0 | 0 | 0 | 0 |
| 0 | 0 | 0 | 0 |
| 0 | 0 | 0 | 0 |
| 1 | 0 | 1 | 0 |
| 1 | 0 | 0 | 0 |
| 0 | 0 | 0 | 0 |
| 0 | 0 | 0 | 0 |
| 0 | 0 | 1 | 0 |
| 0 | 0 | 0 | 0 |
| 0 | 0 | 0 | 0 |
| 0 | 0 | 0 | 0 |

| | | | |
|---|---|---|---|
| 0 | 0 | 0 | 0 |
| 0 | 0 | 0 | 0 |
| 0 | 0 | 0 | 0 |
| 0 | 0 | 0 | 0 |
| 0 | 0 | 0 | 0 |
| 0 | 0 | 0 | 0 |
| 0 | 0 | 0 | 0 |
| 0 | 0 | 0 | 0 |
| 0 | 0 | 0 | 0 |
| 0 | 0 | 0 | 0 |
| 0 | 0 | 0 | 0 |
| 0 | 0 | 0 | 0 |
| 0 | 0 | 0 | 0 |
| 0 | 0 | 0 | 0 |
| 0 | 0 | 0 | 0 |
| 0 | 0 | 0 | 0 |
| 0 | 0 | 0 | 0 |
| 0 | 0 | 0 | 0 |
| 0 | 0 | 0 | 0 |
| 1 | 0 | 0 | 0 |
| 0 | 0 | 0 | 0 |
| 0 | 0 | 0 | 0 |
| 0 | 0 | 0 | 0 |
| 0 | 0 | 0 | 0 |
| 0 | 0 | 0 | 0 |
| 0 | 0 | 0 | 0 |
| 0 | 0 | 0 | 0 |
| 0 | 0 | 0 | 0 |
| 0 | 0 | 0 | 0 |
| 0 | 0 | 0 | 0 |
| 0 | 0 | 0 | 0 |
| 0 | 0 | 0 | 0 |
| 0 | 0 | 0 | 0 |
| 0 | 0 | 0 | 0 |
| 0 | 0 | 0 | 0 |
| 0 | 0 | 0 | 0 |
| 0 | 0 | 0 | 0 |
| 0 | 0 | 0 | 0 |
| 1 | 0 | 0 | 1 |
| 0 | 0 | 0 | 0 |
| 0 | 0 | 0 | 0 |
| 0 | 0 | 0 | 0 |
| 0 | 0 | 0 | 1 |
| 0 | 0 | 0 | 1 |
| 0 | 0 | 0 | 1 |
| 0 | 0 | 0 | 0 |
| 0 | 0 | 0 | 0 |
| 0 | 0 | 0 | 0 |
| 0 | 0 | 0 | 0 |
| 0 | 0 | 0 | 0 |
| 0 | 0 | 0 | 0 |
| 0 | 0 | 0 | 0 |
| 0 | 0 | 0 | 0 |
| 0 | 0 | 0 | 0 |

| | | | |
|---|---|---|---|
| 0 | 0 | 0 | 0 |
| 0 | 0 | 0 | 0 |
| 0 | 0 | 0 | 0 |
| 0 | 0 | 0 | 0 |
| 0 | 0 | 0 | 0 |
| 0 | 0 | 0 | 0 |
| 0 | 0 | 0 | 0 |
| 0 | 0 | 0 | 0 |
| 0 | 0 | 0 | 0 |
| 0 | 0 | 0 | 0 |
| 0 | 0 | 0 | 0 |
| 0 | 0 | 0 | 0 |
| 0 | 0 | 0 | 0 |
| 0 | 0 | 0 | 0 |
| 0 | 0 | 0 | 0 |
| 0 | 0 | 0 | 0 |
| 0 | 0 | 0 | 0 |
| 0 | 0 | 0 | 0 |
| 0 | 0 | 0 | 0 |
| 0 | 0 | 0 | 0 |
| 0 | 0 | 0 | 0 |
| 0 | 0 | 0 | 0 |
| 0 | 0 | 0 | 0 |
| 0 | 0 | 0 | 0 |
| 0 | 0 | 0 | 0 |
| 0 | 0 | 0 | 0 |
| 0 | 0 | 0 | 0 |
| 0 | 0 | 0 | 0 |
| 0 | 0 | 0 | 0 |
| 0 | 0 | 0 | 0 |
| 0 | 0 | 0 | 0 |
| 0 | 0 | 0 | 0 |
| 1 | 1 | 0 | 1 |
| 1 | 1 | 0 | 1 |
| 1 | 0 | 0 | 0 |
| 0 | 0 | 0 | 0 |
| 0 | 0 | 0 | 0 |
| 0 | 0 | 0 | 0 |
| 0 | 0 | 0 | 0 |
| 0 | 0 | 0 | 0 |
| 0 | 0 | 0 | 0 |
| 0 | 0 | 0 | 0 |
| 1 | 1 | 0 | 0 |
| 0 | 0 | 0 | 0 |
| 1 | 0 | 0 | 0 |
| 0 | 0 | 0 | 0 |
| 0 | 0 | 0 | 0 |
| 0 | 0 | 0 | 0 |
| 0 | 0 | 0 | 0 |
| 0 | 0 | 0 | 0 |
| 1 | 0 | 0 | 0 |
| 0 | 0 | 0 | 0 |
| 0 | 0 | 0 | 0 |
| 0 | 0 | 0 | 0 |

| | | | |
|---|---|---|---|
| 0 | 0 | 0 | 0 |
| 0 | 0 | 0 | 0 |
| 0 | 0 | 0 | 0 |
| 0 | 0 | 0 | 0 |
| 0 | 0 | 0 | 0 |
| 0 | 0 | 0 | 0 |
| 0 | 0 | 0 | 0 |
| 0 | 0 | 0 | 0 |
| 0 | 0 | 0 | 0 |
| 0 | 0 | 0 | 0 |
| 0 | 0 | 0 | 0 |
| 0 | 0 | 0 | 0 |
| 0 | 0 | 0 | 0 |
| 0 | 0 | 0 | 0 |
| 0 | 0 | 0 | 0 |
| 0 | 0 | 0 | 0 |
| 0 | 0 | 0 | 0 |
| 0 | 0 | 0 | 0 |
| 0 | 0 | 0 | 0 |
| 0 | 0 | 0 | 1 |
| 0 | 0 | 0 | 0 |
| 0 | 0 | 0 | 1 |
| 0 | 0 | 0 | 1 |
| 0 | 0 | 0 | 1 |
| 0 | 0 | 0 | 0 |
| 0 | 0 | 0 | 0 |
| 0 | 0 | 0 | 0 |
| 0 | 0 | 0 | 0 |
| 0 | 0 | 0 | 0 |
| 0 | 0 | 0 | 0 |
| 0 | 0 | 0 | 0 |
| 0 | 0 | 0 | 0 |
| 0 | 0 | 0 | 0 |
| 0 | 0 | 0 | 0 |
| 0 | 0 | 0 | 0 |
| 0 | 1 | 0 | 0 |
| 0 | 0 | 0 | 0 |
| 0 | 0 | 0 | 0 |
| 0 | 1 | 0 | 0 |
| 0 | 0 | 0 | 0 |
| 0 | 0 | 0 | 0 |
| 0 | 0 | 0 | 0 |
| 0 | 0 | 0 | 0 |
| 0 | 0 | 0 | 0 |
| 0 | 0 | 0 | 0 |
| 0 | 0 | 0 | 0 |
| 0 | 0 | 0 | 0 |
| 0 | 0 | 0 | 0 |
| 0 | 0 | 0 | 0 |
| 0 | 0 | 0 | 0 |
| 0 | 0 | 0 | 0 |
| 0 | 0 | 0 | 0 |
| 0 | 0 | 0 | 0 |
| 0 | 0 | 0 | 0 |

| | | | |
|---|---|---|---|
| 0 | 0 | 0 | 0 |
| 0 | 0 | 0 | 0 |
| 0 | 0 | 0 | 0 |
| 0 | 0 | 0 | 0 |
| 0 | 0 | 0 | 0 |
| 0 | 0 | 0 | 0 |
| 0 | 0 | 0 | 0 |
| 0 | 0 | 0 | 0 |
| 0 | 0 | 1 | 0 |
| 0 | 0 | 1 | 0 |
| 0 | 0 | 0 | 0 |
| 0 | 0 | 0 | 0 |
| 0 | 0 | 0 | 0 |
| 0 | 0 | 0 | 0 |
| 0 | 0 | 0 | 0 |
| 0 | 1 | 0 | 0 |
| 0 | 0 | 0 | 0 |
| 0 | 0 | 0 | 0 |
| 1 | 1 | 0 | 0 |
| 0 | 0 | 0 | 0 |
| 0 | 0 | 0 | 0 |
| 0 | 0 | 0 | 0 |
| 0 | 0 | 0 | 0 |
| 0 | 0 | 0 | 0 |
| 0 | 0 | 0 | 0 |
| 0 | 0 | 0 | 0 |
| 0 | 0 | 0 | 0 |
| 0 | 0 | 0 | 0 |
| 0 | 0 | 0 | 0 |
| 0 | 0 | 0 | 0 |
| 0 | 0 | 0 | 0 |
| 0 | 0 | 0 | 0 |
| 0 | 0 | 0 | 0 |
| 0 | 0 | 0 | 0 |
| 0 | 0 | 0 | 0 |
| 0 | 0 | 0 | 0 |
| 0 | 0 | 0 | 0 |
| 0 | 0 | 0 | 0 |
| 0 | 0 | 0 | 0 |
| 0 | 0 | 0 | 0 |
| 0 | 0 | 0 | 0 |
| 0 | 0 | 0 | 0 |
| 0 | 0 | 0 | 0 |
| 0 | 0 | 0 | 0 |
| 0 | 0 | 0 | 0 |
| 0 | 0 | 0 | 0 |
| 0 | 0 | 0 | 0 |
| 0 | 0 | 0 | 0 |
| 0 | 0 | 0 | 0 |
| 0 | 0 | 0 | 0 |
| 0 | 0 | 0 | 0 |
| 0 | 0 | 0 | 0 |
| 0 | 0 | 0 | 0 |
| 1 | 0 | 0 | 0 |

| | | | |
|---|---|---|---|
| 0 | 0 | 0 | 0 |
| 0 | 0 | 0 | 0 |
| 0 | 0 | 0 | 0 |
| 0 | 0 | 0 | 0 |
| 0 | 0 | 0 | 0 |
| 0 | 0 | 0 | 0 |
| 0 | 0 | 0 | 0 |
| 0 | 0 | 0 | 0 |
| 0 | 0 | 0 | 0 |
| 0 | 0 | 0 | 0 |
| 1 | 0 | 0 | 0 |
| 0 | 0 | 0 | 0 |
| 0 | 0 | 0 | 0 |
| 0 | 0 | 0 | 0 |
| 1 | 1 | 0 | 1 |
| 0 | 0 | 0 | 0 |
| 1 | 0 | 0 | 0 |
| 1 | 1 | 0 | 0 |
| 0 | 0 | 0 | 0 |
| 0 | 0 | 0 | 0 |
| 0 | 0 | 0 | 0 |
| 0 | 0 | 0 | 0 |
| 0 | 0 | 0 | 0 |
| 0 | 0 | 0 | 1 |
| 0 | 0 | 0 | 0 |
| 0 | 0 | 0 | 0 |
| 0 | 0 | 0 | 0 |
| 0 | 0 | 0 | 1 |
| 0 | 0 | 0 | 0 |
| 0 | 0 | 0 | 0 |
| 0 | 0 | 0 | 0 |
| 0 | 0 | 0 | 0 |
| 0 | 0 | 0 | 0 |
| 0 | 0 | 0 | 0 |
| 0 | 0 | 0 | 0 |
| 0 | 0 | 0 | 0 |
| 0 | 0 | 0 | 0 |
| 0 | 0 | 0 | 0 |
| 0 | 0 | 0 | 0 |
| 0 | 0 | 0 | 0 |
| 0 | 0 | 0 | 0 |
| 0 | 0 | 0 | 0 |
| 0 | 0 | 0 | 0 |
| 0 | 0 | 0 | 0 |
| 0 | 0 | 1 | 0 |
| 0 | 0 | 0 | 0 |
| 0 | 0 | 0 | 0 |
| 0 | 0 | 0 | 0 |
| 0 | 0 | 0 | 0 |
| 0 | 0 | 0 | 0 |
| 0 | 0 | 0 | 0 |
| 0 | 0 | 0 | 0 |
| 0 | 0 | 0 | 0 |
| 0 | 0 | 0 | 0 |

| | | | |
|---|---|---|---|
| 0 | 0 | 0 | 0 |
| 0 | 0 | 0 | 0 |
| 0 | 0 | 0 | 0 |
| 0 | 0 | 0 | 0 |
| 0 | 0 | 0 | 0 |
| 0 | 0 | 0 | 0 |
| 1 | 0 | 0 | 1 |
| 1 | 0 | 0 | 0 |
| 0 | 0 | 0 | 0 |
| 0 | 0 | 0 | 0 |
| 0 | 0 | 0 | 0 |
| 0 | 0 | 0 | 0 |
| 0 | 0 | 0 | 0 |
| 0 | 0 | 0 | 0 |
| 0 | 0 | 0 | 0 |
| 0 | 0 | 0 | 0 |
| 0 | 0 | 0 | 0 |
| 0 | 0 | 0 | 0 |
| 0 | 0 | 0 | 0 |
| 0 | 0 | 0 | 0 |
| 1 | 0 | 0 | 0 |
| 0 | 0 | 0 | 0 |
| 0 | 0 | 0 | 0 |
| 0 | 0 | 0 | 0 |
| 0 | 0 | 0 | 0 |
| 0 | 0 | 0 | 0 |
| 0 | 0 | 0 | 0 |
| 0 | 0 | 0 | 0 |
| 0 | 0 | 0 | 0 |
| 0 | 0 | 0 | 0 |
| 0 | 0 | 0 | 0 |
| 0 | 0 | 0 | 0 |
| 0 | 0 | 0 | 0 |
| 0 | 0 | 0 | 0 |
| 0 | 0 | 0 | 0 |
| 0 | 0 | 0 | 0 |
| 0 | 0 | 0 | 0 |
| 0 | 0 | 0 | 0 |
| 0 | 0 | 0 | 0 |
| 0 | 0 | 0 | 0 |
| 0 | 0 | 0 | 0 |
| 0 | 0 | 0 | 0 |
| 0 | 0 | 0 | 0 |
| 0 | 0 | 0 | 0 |
| 0 | 0 | 0 | 0 |
| 0 | 0 | 0 | 0 |
| 0 | 0 | 0 | 0 |
| 0 | 0 | 0 | 0 |
| 0 | 0 | 0 | 0 |
| 0 | 0 | 0 | 0 |
| 0 | 0 | 0 | 0 |
| 0 | 0 | 0 | 0 |
| 0 | 0 | 0 | 0 |
| 0 | 0 | 0 | 0 |

| | | | |
|---|---|---|---|
| 0 | 0 | 0 | 0 |
| 0 | 0 | 0 | 0 |
| 0 | 0 | 0 | 0 |
| 1 | 0 | 0 | 0 |
| 0 | 0 | 0 | 0 |
| 1 | 0 | 0 | 0 |
| 0 | 0 | 0 | 0 |
| 0 | 0 | 0 | 0 |
| 1 | 0 | 0 | 0 |
| 0 | 0 | 0 | 0 |
| 0 | 0 | 0 | 0 |
| 0 | 0 | 0 | 0 |
| 0 | 0 | 0 | 0 |
| 0 | 0 | 0 | 0 |
| 0 | 0 | 0 | 0 |
| 0 | 0 | 0 | 0 |
| 0 | 0 | 0 | 0 |
| 0 | 0 | 0 | 0 |
| 1 | 0 | 0 | 0 |
| 0 | 0 | 0 | 0 |
| 0 | 0 | 0 | 0 |
| 0 | 0 | 0 | 0 |
| 0 | 0 | 0 | 0 |
| 0 | 0 | 0 | 0 |
| 0 | 0 | 0 | 0 |
| 0 | 0 | 0 | 0 |
| 0 | 0 | 0 | 0 |
| 0 | 0 | 0 | 0 |
| 0 | 1 | 0 | 0 |
| 0 | 0 | 0 | 0 |
| 0 | 0 | 0 | 0 |
| 0 | 0 | 0 | 0 |
| 0 | 1 | 0 | 0 |
| 1 | 1 | 0 | 0 |
| 1 | 0 | 0 | 0 |
| 0 | 0 | 0 | 0 |
| 0 | 0 | 0 | 0 |
| 0 | 0 | 0 | 0 |
| 0 | 0 | 0 | 0 |
| 0 | 0 | 0 | 0 |
| 1 | 0 | 0 | 0 |
| 0 | 0 | 0 | 0 |
| 0 | 0 | 0 | 0 |
| 0 | 0 | 0 | 0 |
| 0 | 0 | 0 | 0 |
| 0 | 0 | 0 | 0 |
| 0 | 0 | 0 | 0 |
| 0 | 0 | 0 | 0 |
| 0 | 0 | 0 | 0 |
| 0 | 0 | 0 | 0 |
| 0 | 0 | 0 | 0 |
| 0 | 0 | 0 | 0 |
| 0 | 0 | 0 | 0 |
| 1 | 0 | 0 | 0 |

| | | | |
|---|---|---|---|
| 1 | 0 | 0 | 1 |
| 0 | 0 | 0 | 0 |
| 0 | 0 | 0 | 0 |
| 0 | 0 | 0 | 0 |
| 0 | 0 | 0 | 0 |
| 0 | 0 | 0 | 0 |
| 0 | 0 | 0 | 0 |
| 0 | 0 | 0 | 0 |
| 0 | 0 | 0 | 0 |
| 0 | 0 | 0 | 0 |
| 0 | 0 | 0 | 0 |
| 0 | 0 | 0 | 0 |
| 0 | 0 | 0 | 0 |
| 0 | 0 | 0 | 0 |
| 0 | 0 | 0 | 0 |
| 0 | 0 | 0 | 0 |
| 0 | 0 | 0 | 0 |
| 0 | 0 | 0 | 0 |
| 0 | 0 | 0 | 0 |
| 0 | 0 | 0 | 0 |
| 0 | 0 | 0 | 0 |
| 0 | 0 | 0 | 0 |
| 0 | 1 | 0 | 0 |
| 0 | 0 | 0 | 0 |
| 0 | 0 | 0 | 0 |
| 0 | 0 | 0 | 0 |
| 0 | 0 | 0 | 0 |
| 0 | 0 | 0 | 0 |
| 0 | 0 | 0 | 0 |
| 0 | 0 | 0 | 0 |
| 0 | 0 | 0 | 0 |
| 0 | 0 | 0 | 0 |
| 0 | 0 | 0 | 0 |
| 0 | 0 | 0 | 0 |
| 0 | 0 | 0 | 0 |
| 0 | 0 | 0 | 0 |
| 0 | 0 | 0 | 0 |
| 0 | 0 | 0 | 0 |
| 0 | 0 | 0 | 0 |
| 0 | 0 | 0 | 0 |
| 0 | 0 | 0 | 0 |
| 0 | 0 | 0 | 0 |
| 0 | 0 | 0 | 0 |
| 0 | 0 | 0 | 0 |
| 0 | 0 | 0 | 0 |
| 0 | 0 | 0 | 0 |
| 0 | 0 | 0 | 0 |
| 0 | 0 | 0 | 0 |
| 0 | 0 | 0 | 0 |
| 0 | 0 | 0 | 0 |
| 0 | 0 | 0 | 0 |
| 0 | 0 | 0 | 0 |
| 0 | 0 | 0 | 0 |
| 0 | 0 | 0 | 0 |

| | | | |
|---|---|---|---|
| 0 | 0 | 0 | 0 |
| 0 | 0 | 0 | 0 |
| 0 | 0 | 0 | 0 |
| 0 | 0 | 0 | 0 |
| 0 | 0 | 0 | 0 |
| 0 | 0 | 0 | 0 |
| 0 | 1 | 0 | 0 |
| 0 | 1 | 0 | 0 |
| 0 | 0 | 0 | 0 |
| 0 | 0 | 0 | 0 |
| 0 | 0 | 0 | 0 |
| 0 | 0 | 0 | 0 |
| 0 | 0 | 0 | 0 |
| 0 | 0 | 0 | 0 |
| 0 | 0 | 0 | 0 |
| 0 | 0 | 0 | 0 |
| 1 | 0 | 0 | 0 |
| 0 | 0 | 0 | 0 |
| 1 | 0 | 0 | 0 |
| 1 | 0 | 0 | 0 |
| 1 | 0 | 0 | 0 |
| 1 | 0 | 0 | 0 |
| 0 | 0 | 0 | 0 |
| 0 | 0 | 0 | 0 |
| 0 | 0 | 0 | 0 |
| 0 | 0 | 0 | 0 |
| 0 | 0 | 0 | 0 |
| 0 | 0 | 0 | 0 |
| 0 | 0 | 0 | 0 |
| 0 | 0 | 0 | 0 |
| 0 | 0 | 0 | 0 |
| 0 | 0 | 0 | 0 |
| 0 | 0 | 0 | 0 |
| 0 | 0 | 0 | 0 |
| 0 | 0 | 0 | 0 |
| 0 | 0 | 1 | 0 |
| 1 | 0 | 0 | 0 |
| 1 | 0 | 0 | 0 |
| 1 | 0 | 0 | 0 |
| 0 | 0 | 0 | 0 |
| 0 | 0 | 0 | 0 |
| 0 | 0 | 0 | 0 |
| 0 | 0 | 0 | 0 |
| 1 | 0 | 0 | 0 |
| 1 | 0 | 0 | 0 |
| 1 | 0 | 0 | 0 |
| 0 | 0 | 0 | 0 |
| 0 | 0 | 0 | 0 |
| 1 | 0 | 0 | 0 |
| 0 | 0 | 0 | 0 |
| 0 | 0 | 0 | 0 |
| 1 | 0 | 0 | 0 |
| 0 | 0 | 0 | 0 |
| 0 | 0 | 0 | 0 |

| | | | |
|---|---|---|---|
| 0 | 0 | 0 | 0 |
| 0 | 0 | 0 | 0 |
| 0 | 0 | 0 | 0 |
| 0 | 0 | 0 | 0 |
| 0 | 0 | 0 | 0 |
| 0 | 0 | 0 | 0 |
| 0 | 0 | 0 | 0 |
| 0 | 0 | 0 | 0 |
| 0 | 0 | 0 | 0 |
| 0 | 0 | 0 | 0 |
| 0 | 0 | 0 | 0 |
| 0 | 0 | 0 | 0 |
| 0 | 0 | 0 | 0 |
| 0 | 0 | 0 | 0 |
| 0 | 0 | 0 | 0 |
| 0 | 0 | 0 | 0 |
| 0 | 0 | 0 | 0 |
| 0 | 0 | 0 | 0 |
| 0 | 0 | 0 | 0 |
| 0 | 0 | 0 | 0 |
| 0 | 0 | 0 | 0 |
| 0 | 0 | 0 | 0 |
| 1 | 1 | 0 | 0 |
| 0 | 0 | 0 | 0 |
| 0 | 1 | 0 | 0 |
| 0 | 0 | 0 | 0 |
| 0 | 0 | 0 | 0 |
| 0 | 0 | 0 | 0 |
| 0 | 0 | 0 | 0 |
| 0 | 0 | 0 | 0 |
| 0 | 0 | 0 | 0 |
| 0 | 0 | 0 | 0 |
| 0 | 0 | 0 | 0 |
| 0 | 0 | 0 | 0 |
| 0 | 0 | 0 | 0 |
| 0 | 0 | 0 | 0 |
| 0 | 0 | 0 | 0 |
| 0 | 0 | 0 | 0 |
| 0 | 0 | 0 | 0 |
| 0 | 0 | 0 | 0 |
| 0 | 0 | 0 | 0 |
| 0 | 0 | 0 | 0 |
| 0 | 0 | 0 | 0 |
| 0 | 0 | 0 | 0 |
| 0 | 0 | 0 | 0 |
| 1 | 0 | 0 | 1 |
| 0 | 0 | 0 | 0 |
| 0 | 0 | 0 | 0 |
| 0 | 0 | 0 | 0 |
| 1 | 0 | 0 | 0 |
| 1 | 0 | 0 | 0 |
| 1 | 0 | 0 | 0 |
| 0 | 0 | 0 | 0 |
| 0 | 0 | 0 | 0 |

| | | | |
|---|---|---|---|
| 0 | 0 | 0 | 0 |
| 0 | 0 | 0 | 0 |
| 0 | 0 | 0 | 0 |
| 0 | 0 | 0 | 0 |
| 0 | 0 | 0 | 0 |
| 0 | 0 | 0 | 0 |
| 0 | 0 | 0 | 0 |
| 0 | 0 | 0 | 0 |
| 0 | 0 | 0 | 0 |
| 0 | 0 | 0 | 0 |
| 0 | 0 | 0 | 1 |
| 0 | 0 | 0 | 0 |
| 0 | 0 | 0 | 0 |
| 0 | 0 | 0 | 0 |
| 0 | 0 | 0 | 0 |
| 0 | 0 | 0 | 0 |
| 0 | 0 | 0 | 0 |
| 0 | 0 | 0 | 0 |
| 0 | 0 | 0 | 0 |
| 0 | 0 | 0 | 0 |
| 0 | 0 | 0 | 0 |
| 0 | 0 | 0 | 0 |
| 0 | 0 | 0 | 0 |
| 0 | 0 | 0 | 0 |
| 0 | 0 | 0 | 0 |
| 0 | 0 | 0 | 0 |
| 0 | 0 | 0 | 0 |
| 0 | 0 | 0 | 0 |
| 0 | 0 | 0 | 0 |
| 0 | 0 | 0 | 0 |
| 0 | 1 | 0 | 0 |
| 0 | 0 | 0 | 0 |
| 1 | 1 | 0 | 0 |
| 1 | 1 | 0 | 0 |
| 0 | 0 | 0 | 0 |
| 0 | 0 | 0 | 0 |
| 0 | 0 | 0 | 0 |
| 0 | 0 | 0 | 0 |
| 0 | 0 | 0 | 0 |
| 0 | 0 | 0 | 0 |
| 0 | 0 | 0 | 0 |
| 0 | 1 | 0 | 0 |
| 0 | 0 | 0 | 0 |
| 0 | 0 | 0 | 0 |
| 0 | 0 | 0 | 0 |
| 1 | 0 | 0 | 0 |
| 1 | 0 | 0 | 0 |
| 1 | 0 | 0 | 0 |
| 1 | 1 | 1 | 0 |
| 0 | 0 | 0 | 0 |
| 1 | 1 | 1 | 1 |
| 0 | 0 | 0 | 0 |
| 0 | 0 | 0 | 0 |
| 0 | 0 | 0 | 0 |

| | | | |
|---|---|---|---|
| 0 | 0 | 0 | 0 |
| 0 | 0 | 0 | 0 |
| 0 | 0 | 0 | 0 |
| 0 | 0 | 0 | 0 |
| 1 | 0 | 0 | 0 |
| 1 | 0 | 0 | 0 |
| 0 | 0 | 0 | 0 |
| 0 | 0 | 0 | 0 |
| 0 | 0 | 0 | 0 |
| 0 | 0 | 0 | 0 |
| 0 | 0 | 0 | 0 |
| 0 | 0 | 0 | 0 |
| 0 | 0 | 0 | 0 |
| 0 | 0 | 0 | 0 |
| 0 | 0 | 0 | 0 |
| 0 | 0 | 0 | 0 |
| 0 | 0 | 0 | 0 |
| 0 | 0 | 0 | 0 |
| 0 | 0 | 0 | 0 |
| 0 | 0 | 0 | 0 |
| 0 | 0 | 0 | 0 |
| 0 | 0 | 0 | 0 |
| 0 | 0 | 0 | 0 |
| 0 | 0 | 0 | 0 |
| 0 | 0 | 0 | 0 |
| 0 | 0 | 0 | 0 |
| 0 | 0 | 0 | 0 |
| 0 | 0 | 0 | 0 |
| 0 | 0 | 0 | 0 |
| 0 | 0 | 0 | 0 |
| 0 | 0 | 0 | 0 |
| 0 | 0 | 0 | 0 |
| 0 | 0 | 0 | 0 |
| 0 | 0 | 0 | 0 |
| 0 | 0 | 0 | 0 |
| 0 | 0 | 0 | 0 |
| 0 | 0 | 0 | 0 |
| 0 | 0 | 0 | 0 |
| 0 | 0 | 0 | 0 |
| 0 | 0 | 0 | 0 |
| 0 | 0 | 0 | 0 |
| 0 | 0 | 0 | 0 |
| 0 | 0 | 0 | 0 |
| 0 | 0 | 0 | 0 |
| 0 | 0 | 0 | 0 |
| 0 | 0 | 0 | 0 |
| 0 | 0 | 0 | 0 |
| 0 | 0 | 0 | 0 |
| 0 | 0 | 0 | 0 |
| 0 | 0 | 0 | 0 |
| 0 | 0 | 0 | 0 |
| 0 | 0 | 0 | 0 |
| 0 | 0 | 0 | 0 |
| 0 | 1 | 0 | 0 |

| | | | |
|---|---|---|---|
| 0 | 0 | 0 | 0 |
| 0 | 0 | 0 | 0 |
| 0 | 0 | 0 | 0 |
| 0 | 0 | 0 | 0 |
| 0 | 0 | 0 | 0 |
| 0 | 0 | 0 | 0 |
| 0 | 0 | 0 | 0 |
| 0 | 0 | 0 | 0 |
| 0 | 0 | 0 | 0 |
| 0 | 0 | 0 | 0 |
| 0 | 0 | 0 | 0 |
| 0 | 0 | 0 | 0 |
| 0 | 0 | 0 | 0 |
| 0 | 0 | 0 | 0 |
| 0 | 0 | 0 | 0 |
| 0 | 0 | 0 | 0 |
| 0 | 0 | 0 | 0 |
| 0 | 0 | 0 | 0 |
| 0 | 0 | 0 | 0 |
| 0 | 0 | 0 | 0 |
| 0 | 0 | 0 | 0 |
| 0 | 0 | 0 | 0 |
| 0 | 0 | 0 | 0 |
| 0 | 0 | 0 | 0 |
| 0 | 0 | 0 | 0 |
| 0 | 0 | 0 | 0 |
| 0 | 0 | 0 | 0 |
| 0 | 0 | 0 | 0 |
| 0 | 0 | 0 | 0 |
| 0 | 0 | 0 | 0 |
| 0 | 0 | 0 | 0 |
| 0 | 0 | 0 | 0 |
| 0 | 0 | 0 | 0 |
| 0 | 0 | 0 | 0 |
| 0 | 0 | 0 | 0 |
| 0 | 0 | 0 | 0 |
| 0 | 0 | 0 | 0 |
| 0 | 0 | 0 | 0 |
| 0 | 0 | 0 | 0 |
| 0 | 0 | 0 | 0 |
| 0 | 0 | 0 | 0 |
| 0 | 0 | 0 | 0 |
| 0 | 0 | 0 | 0 |
| 0 | 0 | 0 | 0 |
| 0 | 0 | 0 | 0 |
| 0 | 0 | 0 | 0 |
| 0 | 0 | 0 | 0 |
| 0 | 0 | 0 | 0 |
| 0 | 0 | 0 | 0 |
| 0 | 0 | 0 | 0 |
| 0 | 0 | 0 | 0 |
| 0 | 0 | 0 | 0 |
| 0 | 0 | 0 | 0 |
| 0 | 0 | 0 | 0 |

| | | | |
|---|---|---|---|
| 0 | 0 | 0 | 0 |
| 0 | 0 | 0 | 0 |
| 0 | 0 | 0 | 0 |
| 0 | 0 | 0 | 0 |
| 0 | 0 | 0 | 0 |
| 0 | 0 | 0 | 0 |
| 0 | 0 | 0 | 0 |
| 0 | 0 | 0 | 0 |
| 0 | 0 | 0 | 0 |
| 0 | 0 | 0 | 0 |
| 0 | 0 | 0 | 0 |
| 0 | 0 | 1 | 0 |
| 0 | 0 | 0 | 0 |
| 0 | 0 | 1 | 0 |
| 0 | 0 | 1 | 0 |
| 0 | 0 | 1 | 0 |
| 0 | 0 | 1 | 0 |
| 0 | 0 | 0 | 0 |
| 0 | 0 | 0 | 0 |
| 0 | 0 | 0 | 0 |
| 1 | 0 | 0 | 0 |

WaterConsLevel0.0

| glycan_binding_site | water_binding_site | water_only_binding_site | non_binding_site |
|---|---|---|---|
| 0 | 1 | 0 | 0 |
| 0 | 1 | 0 | 0 |
| 0 | 1 | 1 | 1 |
| 0 | 1 | 0 | 0 |
| 0 | 0 | 0 | 0 |
| 0 | 0 | 0 | 0 |
| 0 | 0 | 0 | 1 |
| 0 | 0 | 0 | 1 |
| 0 | 1 | 0 | 0 |
| 0 | 1 | 0 | 0 |
| 0 | 1 | 0 | 0 |
| 0 | 1 | 0 | 0 |
| 0 | 1 | 1 | 1 |
| 1 | 1 | 0 | 0 |
| 0 | 1 | 0 | 0 |
| 0 | 1 | 0 | 0 |
| 0 | 1 | 0 | 0 |
| 0 | 1 | 0 | 0 |
| 0 | 1 | 0 | 0 |
| 0 | 1 | 0 | 0 |
| 0 | 1 | 0 | 0 |
| 1 | 1 | 0 | 0 |
| 1 | 1 | 0 | 0 |
| 0 | 1 | 0 | 0 |
| 0 | 1 | 0 | 0 |
| 0 | 1 | 1 | 1 |
| 0 | 1 | 0 | 0 |
| 0 | 0 | 0 | 1 |
| 0 | 0 | 0 | 0 |
| 0 | 0 | 0 | 1 |
| 0 | 0 | 0 | 1 |
| 0 | 1 | 0 | 0 |
| 0 | 1 | 0 | 0 |
| 0 | 1 | 0 | 0 |
| 0 | 1 | 0 | 0 |
| 0 | 0 | 0 | 1 |
| 0 | 0 | 0 | 0 |
| 0 | 1 | 0 | 0 |
| 0 | 1 | 0 | 0 |
| 0 | 1 | 0 | 0 |
| 0 | 1 | 0 | 0 |
| 0 | 1 | 0 | 0 |
| 0 | 1 | 0 | 0 |
| 0 | 1 | 0 | 0 |
| 0 | 1 | 0 | 0 |
| 0 | 1 | 0 | 0 |
| 0 | 1 | 0 | 0 |
| 0 | 1 | 0 | 0 |
| 0 | 1 | 0 | 0 |
| 0 | 1 | 0 | 0 |
| 0 | 1 | 0 | 0 |
| 0 | 1 | 0 | 0 |
| 0 | 1 | 0 | 0 |

| | | | |
|---|---|---|---|
| 0 | 1 | 0 | 0 |
| 0 | 1 | 0 | 0 |
| 0 | 1 | 0 | 0 |
| 0 | 1 | 0 | 0 |
| 0 | 1 | 0 | 0 |
| 0 | 1 | 0 | 0 |
| 0 | 1 | 0 | 0 |
| 0 | 1 | 0 | 0 |
| 0 | 1 | 0 | 0 |
| 0 | 1 | 0 | 0 |
| 0 | 1 | 0 | 0 |
| 0 | 1 | 0 | 0 |
| 0 | 1 | 0 | 0 |
| 0 | 1 | 0 | 0 |
| 0 | 1 | 0 | 0 |
| 0 | 1 | 0 | 0 |
| 0 | 1 | 0 | 0 |
| 0 | 1 | 0 | 0 |
| 0 | 1 | 0 | 0 |
| 0 | 1 | 0 | 0 |
| 0 | 1 | 0 | 0 |
| 0 | 1 | 0 | 0 |
| 0 | 1 | 0 | 0 |
| 0 | 1 | 0 | 0 |
| 0 | 1 | 0 | 0 |
| 0 | 1 | 0 | 0 |
| 0 | 1 | 0 | 0 |
| 0 | 1 | 0 | 0 |
| 0 | 0 | 0 | 0 |
| 0 | 0 | 0 | 0 |
| 0 | 0 | 0 | 0 |
| 0 | 0 | 0 | 0 |
| 0 | 0 | 0 | 0 |
| 0 | 0 | 0 | 0 |
| 0 | 1 | 0 | 0 |
| 0 | 1 | 0 | 0 |
| 0 | 0 | 0 | 0 |
| 0 | 1 | 0 | 0 |
| 0 | 1 | 0 | 0 |
| 0 | 1 | 0 | 0 |
| 1 | 0 | 0 | 0 |
| 0 | 0 | 0 | 0 |
| 1 | 0 | 0 | 0 |
| 1 | 1 | 0 | 0 |
| 0 | 0 | 0 | 0 |
| 0 | 0 | 0 | 1 |
| 0 | 0 | 0 | 1 |
| 0 | 1 | 0 | 0 |
| 1 | 0 | 0 | 0 |
| 0 | 0 | 0 | 0 |
| 0 | 0 | 0 | 0 |
| 0 | 0 | 0 | 1 |
| 0 | 0 | 0 | 1 |
| 0 | 0 | 0 | 0 |

| | | | |
|---|---|---|---|
| 0 | 0 | 0 | 1 |
| 0 | 0 | 0 | 1 |
| 0 | 0 | 0 | 0 |
| 0 | 0 | 0 | 0 |
| 0 | 0 | 0 | 0 |
| 0 | 0 | 0 | 0 |
| 0 | 0 | 0 | 1 |
| 0 | 1 | 0 | 0 |
| 0 | 1 | 0 | 0 |
| 0 | 0 | 0 | 0 |
| 0 | 1 | 0 | 0 |
| 0 | 1 | 0 | 0 |
| 0 | 0 | 0 | 0 |
| 1 | 0 | 0 | 0 |
| 1 | 0 | 0 | 0 |
| 0 | 0 | 0 | 1 |
| 0 | 0 | 0 | 0 |
| 0 | 1 | 0 | 0 |
| 0 | 1 | 0 | 0 |
| 0 | 1 | 0 | 0 |
| 0 | 1 | 0 | 0 |
| 0 | 1 | 0 | 0 |
| 0 | 1 | 0 | 0 |
| 0 | 1 | 0 | 0 |
| 0 | 1 | 0 | 0 |
| 0 | 1 | 0 | 0 |
| 0 | 1 | 0 | 0 |
| 0 | 1 | 0 | 0 |
| 0 | 1 | 0 | 0 |
| 0 | 1 | 0 | 0 |
| 0 | 1 | 0 | 0 |
| 0 | 1 | 0 | 0 |
| 0 | 1 | 0 | 0 |
| 0 | 1 | 0 | 0 |
| 0 | 1 | 0 | 0 |
| 0 | 1 | 0 | 0 |
| 0 | 1 | 0 | 0 |
| 0 | 1 | 0 | 0 |
| 0 | 1 | 0 | 0 |
| 0 | 1 | 0 | 0 |
| 0 | 1 | 0 | 0 |
| 0 | 1 | 0 | 0 |
| 0 | 1 | 0 | 0 |
| 0 | 1 | 0 | 0 |
| 0 | 1 | 0 | 0 |
| 1 | 1 | 0 | 0 |
| 0 | 1 | 0 | 0 |
| 0 | 1 | 0 | 0 |
| 0 | 1 | 0 | 0 |
| 0 | 1 | 0 | 0 |
| 0 | 1 | 0 | 0 |
| 0 | 1 | 0 | 0 |
| 0 | 1 | 1 | 1 |

| | | | |
|---|---|---|---|
| 0 | 1 | 1 | 1 |
| 0 | 1 | 1 | 1 |
| 0 | 1 | 1 | 1 |
| 0 | 1 | 0 | 0 |
| 0 | 1 | 0 | 0 |
| 0 | 1 | 1 | 1 |
| 0 | 1 | 0 | 0 |
| 0 | 1 | 1 | 1 |
| 0 | 1 | 1 | 1 |
| 0 | 1 | 1 | 1 |
| 0 | 1 | 1 | 1 |
| 0 | 1 | 0 | 0 |
| 0 | 1 | 1 | 1 |
| 0 | 0 | 0 | 0 |
| 0 | 1 | 1 | 1 |
| 0 | 1 | 1 | 1 |
| 0 | 1 | 1 | 1 |
| 0 | 1 | 1 | 1 |
| 0 | 1 | 1 | 1 |
| 0 | 1 | 0 | 0 |
| 0 | 1 | 0 | 0 |
| 0 | 0 | 0 | 1 |
| 0 | 1 | 0 | 0 |
| 0 | 1 | 0 | 0 |
| 0 | 1 | 0 | 0 |
| 0 | 1 | 1 | 1 |
| 0 | 1 | 0 | 0 |
| 0 | 1 | 0 | 0 |
| 0 | 1 | 1 | 1 |
| 0 | 1 | 1 | 1 |
| 0 | 0 | 0 | 1 |
| 0 | 1 | 1 | 1 |
| 0 | 1 | 1 | 1 |
| 0 | 0 | 0 | 1 |
| 0 | 1 | 0 | 0 |
| 0 | 1 | 1 | 1 |
| 1 | 1 | 0 | 0 |
| 1 | 1 | 0 | 0 |
| 0 | 1 | 1 | 1 |
| 0 | 1 | 1 | 1 |
| 0 | 1 | 1 | 1 |
| 0 | 1 | 0 | 0 |
| 0 | 1 | 0 | 0 |
| 0 | 0 | 0 | 1 |
| 0 | 1 | 1 | 1 |
| 0 | 1 | 1 | 1 |
| 0 | 0 | 0 | 0 |
| 0 | 1 | 0 | 0 |
| 0 | 0 | 0 | 0 |
| 0 | 1 | 0 | 0 |
| 0 | 0 | 0 | 0 |
| 0 | 0 | 0 | 1 |
| 0 | 1 | 1 | 1 |
| 0 | 1 | 0 | 0 |

| | | | |
|---|---|---|---|
| 0 | 0 | 0 | 0 |
| 0 | 1 | 0 | 0 |
| 0 | 1 | 0 | 0 |
| 0 | 0 | 0 | 1 |
| 0 | 0 | 0 | 1 |
| 0 | 1 | 1 | 1 |
| 0 | 1 | 0 | 0 |
| 0 | 1 | 0 | 0 |
| 0 | 1 | 1 | 1 |
| 0 | 1 | 1 | 1 |
| 1 | 1 | 0 | 0 |
| 0 | 0 | 0 | 1 |
| 1 | 1 | 0 | 0 |
| 0 | 1 | 1 | 1 |
| 0 | 0 | 0 | 1 |
| 0 | 0 | 0 | 1 |
| 0 | 0 | 0 | 1 |
| 0 | 1 | 0 | 0 |
| 0 | 0 | 0 | 0 |
| 0 | 1 | 0 | 0 |
| 0 | 0 | 0 | 0 |
| 0 | 1 | 1 | 1 |
| 0 | 1 | 0 | 0 |
| 0 | 0 | 0 | 1 |
| 0 | 0 | 0 | 1 |
| 0 | 1 | 0 | 0 |
| 0 | 1 | 0 | 0 |
| 0 | 0 | 0 | 0 |
| 0 | 1 | 0 | 0 |
| 0 | 1 | 0 | 0 |
| 0 | 1 | 0 | 0 |
| 0 | 1 | 0 | 0 |
| 0 | 1 | 1 | 1 |
| 0 | 1 | 1 | 1 |
| 0 | 1 | 0 | 0 |
| 0 | 1 | 1 | 1 |
| 0 | 1 | 1 | 1 |
| 0 | 1 | 0 | 0 |
| 0 | 1 | 1 | 1 |
| 0 | 0 | 0 | 1 |
| 0 | 1 | 0 | 0 |
| 0 | 1 | 1 | 1 |
| 0 | 1 | 1 | 1 |
| 0 | 1 | 1 | 1 |
| 0 | 1 | 0 | 0 |
| 0 | 1 | 0 | 0 |
| 0 | 1 | 0 | 0 |
| 0 | 1 | 0 | 0 |
| 0 | 1 | 0 | 0 |
| 0 | 1 | 0 | 0 |
| 0 | 1 | 0 | 0 |
| 0 | 1 | 0 | 0 |
| 0 | 1 | 0 | 0 |
| 0 | 1 | 0 | 0 |

| | | | |
|---|---|---|---|
| 0 | 1 | 0 | 0 |
| 0 | 1 | 0 | 0 |
| 0 | 1 | 0 | 0 |
| 0 | 1 | 0 | 0 |
| 0 | 1 | 0 | 0 |
| 0 | 1 | 0 | 0 |
| 0 | 0 | 0 | 1 |
| 0 | 0 | 0 | 1 |
| 0 | 0 | 0 | 0 |
| 0 | 0 | 0 | 0 |
| 0 | 0 | 0 | 0 |
| 0 | 0 | 0 | 0 |
| 0 | 0 | 0 | 0 |
| 0 | 0 | 0 | 1 |
| 0 | 0 | 0 | 1 |
| 0 | 0 | 0 | 0 |
| 0 | 0 | 0 | 1 |
| 0 | 0 | 0 | 0 |
| 0 | 0 | 0 | 1 |
| 0 | 0 | 0 | 0 |
| 0 | 0 | 0 | 1 |
| 0 | 1 | 1 | 1 |
| 0 | 0 | 0 | 1 |
| 0 | 1 | 0 | 0 |
| 0 | 1 | 0 | 0 |
| 0 | 1 | 0 | 0 |
| 0 | 1 | 0 | 0 |
| 0 | 1 | 0 | 0 |
| 0 | 1 | 0 | 0 |
| 0 | 1 | 0 | 0 |
| 0 | 1 | 1 | 1 |
| 0 | 1 | 1 | 1 |
| 0 | 1 | 0 | 0 |
| 0 | 1 | 0 | 0 |
| 0 | 1 | 0 | 0 |
| 0 | 1 | 1 | 1 |
| 0 | 0 | 0 | 0 |
| 0 | 1 | 0 | 0 |
| 0 | 1 | 0 | 0 |
| 0 | 1 | 0 | 0 |
| 0 | 0 | 0 | 1 |
| 0 | 0 | 0 | 1 |
| 0 | 0 | 0 | 0 |
| 0 | 0 | 0 | 1 |
| 0 | 0 | 0 | 0 |
| 0 | 0 | 0 | 0 |
| 0 | 0 | 0 | 0 |
| 0 | 0 | 0 | 1 |
| 0 | 0 | 0 | 1 |
| 0 | 0 | 0 | 0 |
| 0 | 0 | 0 | 1 |
| 0 | 0 | 0 | 1 |
| 0 | 0 | 0 | 0 |
| 0 | 0 | 0 | 1 |

| | | | |
|---|---|---|---|
| 0 | 0 | 0 | 0 |
| 0 | 0 | 0 | 1 |
| 0 | 1 | 1 | 1 |
| 0 | 1 | 0 | 0 |
| 0 | 1 | 0 | 0 |
| 0 | 0 | 0 | 0 |
| 0 | 0 | 0 | 1 |
| 0 | 1 | 0 | 0 |
| 0 | 0 | 0 | 0 |
| 0 | 0 | 0 | 1 |
| 0 | 0 | 0 | 1 |
| 0 | 0 | 0 | 1 |
| 0 | 0 | 0 | 1 |
| 0 | 0 | 0 | 1 |
| 0 | 0 | 0 | 0 |
| 0 | 0 | 0 | 1 |
| 0 | 0 | 0 | 0 |
| 0 | 0 | 0 | 0 |
| 0 | 0 | 0 | 1 |
| 0 | 0 | 0 | 0 |
| 0 | 1 | 0 | 0 |
| 0 | 1 | 0 | 0 |
| 0 | 1 | 0 | 0 |
| 0 | 1 | 0 | 0 |
| 0 | 0 | 0 | 1 |
| 0 | 1 | 1 | 1 |
| 0 | 1 | 0 | 0 |
| 0 | 1 | 0 | 0 |
| 0 | 1 | 0 | 0 |
| 0 | 1 | 0 | 0 |
| 0 | 1 | 0 | 0 |
| 0 | 1 | 0 | 0 |
| 0 | 0 | 0 | 0 |
| 0 | 0 | 0 | 0 |
| 0 | 0 | 0 | 1 |
| 0 | 0 | 0 | 1 |
| 0 | 0 | 0 | 0 |
| 0 | 1 | 0 | 0 |
| 0 | 1 | 0 | 0 |
| 0 | 0 | 0 | 0 |
| 0 | 0 | 0 | 0 |
| 0 | 1 | 0 | 0 |
| 0 | 0 | 0 | 1 |
| 0 | 1 | 0 | 0 |
| 0 | 1 | 0 | 0 |
| 0 | 1 | 0 | 0 |
| 0 | 1 | 0 | 0 |
| 0 | 1 | 0 | 0 |
| 1 | 1 | 0 | 0 |
| 0 | 0 | 0 | 1 |
| 0 | 1 | 0 | 0 |
| 0 | 1 | 0 | 0 |
| 0 | 0 | 0 | 0 |
| 0 | 1 | 0 | 0 |

| | | | |
|---|---|---|---|
| 0 | 1 | 0 | 0 |
| 0 | 1 | 0 | 0 |
| 0 | 1 | 0 | 0 |
| 0 | 1 | 0 | 0 |
| 0 | 0 | 0 | 1 |
| 0 | 1 | 0 | 0 |
| 0 | 1 | 0 | 0 |
| 0 | 1 | 0 | 0 |
| 0 | 1 | 0 | 0 |
| 0 | 1 | 0 | 0 |
| 0 | 1 | 0 | 0 |
| 0 | 1 | 0 | 0 |
| 0 | 1 | 0 | 0 |
| 0 | 1 | 0 | 0 |
| 0 | 1 | 0 | 0 |
| 0 | 1 | 0 | 0 |
| 0 | 1 | 0 | 0 |
| 0 | 1 | 0 | 0 |
| 0 | 0 | 0 | 1 |
| 0 | 0 | 0 | 0 |
| 0 | 0 | 0 | 0 |
| 0 | 0 | 0 | 0 |
| 0 | 1 | 0 | 0 |
| 0 | 1 | 1 | 1 |
| 0 | 0 | 0 | 0 |
| 0 | 0 | 0 | 0 |
| 0 | 0 | 0 | 0 |
| 0 | 1 | 1 | 1 |
| 0 | 1 | 1 | 1 |
| 0 | 1 | 1 | 1 |
| 0 | 1 | 0 | 0 |
| 0 | 0 | 0 | 0 |
| 0 | 1 | 0 | 0 |
| 0 | 1 | 0 | 0 |
| 0 | 1 | 0 | 0 |
| 0 | 1 | 1 | 1 |
| 0 | 1 | 1 | 1 |
| 0 | 0 | 0 | 0 |
| 0 | 1 | 0 | 0 |
| 0 | 1 | 1 | 1 |
| 0 | 1 | 1 | 1 |
| 0 | 1 | 1 | 1 |
| 0 | 0 | 0 | 1 |
| 0 | 0 | 0 | 1 |
| 0 | 1 | 0 | 0 |
| 0 | 0 | 0 | 0 |
| 0 | 1 | 0 | 0 |
| 0 | 0 | 0 | 0 |
| 0 | 0 | 0 | 0 |
| 0 | 0 | 0 | 1 |
| 0 | 0 | 0 | 1 |
| 0 | 0 | 0 | 0 |
| 0 | 0 | 0 | 1 |
| 0 | 0 | 0 | 0 |

| | | | |
|---|---|---|---|
| 0 | 0 | 0 | 1 |
| 0 | 0 | 0 | 1 |
| 0 | 0 | 0 | 0 |
| 0 | 0 | 0 | 1 |
| 0 | 0 | 0 | 1 |
| 0 | 0 | 0 | 1 |
| 0 | 0 | 0 | 1 |
| 0 | 1 | 0 | 0 |
| 0 | 1 | 1 | 1 |
| 0 | 0 | 0 | 1 |
| 0 | 0 | 0 | 0 |
| 0 | 0 | 0 | 1 |
| 0 | 0 | 0 | 1 |
| 0 | 0 | 0 | 1 |
| 0 | 1 | 0 | 0 |
| 0 | 0 | 0 | 0 |
| 0 | 0 | 0 | 1 |
| 0 | 0 | 0 | 1 |
| 0 | 1 | 0 | 0 |
| 0 | 1 | 0 | 0 |
| 0 | 1 | 0 | 0 |
| 0 | 1 | 0 | 0 |
| 0 | 1 | 0 | 0 |
| 0 | 1 | 0 | 0 |
| 0 | 1 | 0 | 0 |
| 0 | 0 | 0 | 1 |
| 0 | 1 | 0 | 0 |
| 0 | 1 | 0 | 0 |
| 0 | 1 | 0 | 0 |
| 0 | 1 | 1 | 1 |
| 0 | 1 | 0 | 0 |
| 0 | 1 | 0 | 0 |
| 0 | 1 | 1 | 1 |
| 0 | 1 | 0 | 0 |
| 0 | 1 | 0 | 0 |
| 0 | 1 | 0 | 0 |
| 0 | 1 | 0 | 0 |
| 0 | 1 | 0 | 0 |
| 0 | 1 | 0 | 0 |
| 0 | 1 | 0 | 0 |
| 0 | 0 | 0 | 1 |
| 0 | 0 | 0 | 0 |
| 0 | 0 | 0 | 1 |
| 0 | 0 | 0 | 0 |
| 0 | 0 | 0 | 0 |
| 1 | 0 | 0 | 0 |
| 0 | 0 | 0 | 0 |
| 1 | 0 | 0 | 0 |
| 0 | 0 | 0 | 0 |
| 0 | 0 | 0 | 0 |
| 0 | 0 | 0 | 1 |
| 0 | 0 | 0 | 0 |
| 0 | 0 | 0 | 0 |
| 0 | 0 | 0 | 0 |

| | | | |
|---|---|---|---|
| 0 | 0 | 0 | 1 |
| 0 | 0 | 0 | 0 |
| 0 | 0 | 0 | 1 |
| 0 | 0 | 0 | 1 |
| 1 | 0 | 0 | 0 |
| 0 | 0 | 0 | 0 |
| 1 | 0 | 0 | 0 |
| 1 | 0 | 0 | 0 |
| 0 | 0 | 0 | 0 |
| 0 | 0 | 0 | 1 |
| 0 | 0 | 0 | 1 |
| 0 | 0 | 0 | 1 |
| 0 | 0 | 0 | 1 |
| 1 | 1 | 0 | 0 |
| 0 | 0 | 0 | 1 |
| 0 | 0 | 0 | 0 |
| 0 | 0 | 0 | 0 |
| 0 | 0 | 0 | 0 |
| 1 | 1 | 0 | 0 |
| 0 | 1 | 1 | 1 |
| 1 | 1 | 0 | 0 |
| 0 | 0 | 0 | 1 |
| 0 | 0 | 0 | 0 |
| 0 | 0 | 0 | 1 |
| 0 | 0 | 0 | 1 |
| 0 | 0 | 0 | 1 |
| 0 | 0 | 0 | 0 |
| 0 | 0 | 0 | 1 |
| 0 | 0 | 0 | 1 |
| 0 | 0 | 0 | 1 |
| 0 | 0 | 0 | 0 |
| 0 | 0 | 0 | 0 |
| 0 | 0 | 0 | 1 |
| 0 | 0 | 0 | 1 |
| 0 | 0 | 0 | 0 |
| 0 | 0 | 0 | 0 |
| 0 | 0 | 0 | 0 |
| 1 | 0 | 0 | 0 |
| 0 | 0 | 0 | 0 |
| 0 | 0 | 0 | 1 |
| 0 | 1 | 0 | 0 |
| 0 | 1 | 0 | 0 |
| 0 | 1 | 0 | 0 |
| 0 | 1 | 0 | 0 |
| 0 | 1 | 1 | 1 |
| 1 | 1 | 0 | 0 |
| 0 | 1 | 0 | 0 |
| 0 | 0 | 0 | 0 |
| 0 | 1 | 0 | 0 |
| 1 | 1 | 0 | 0 |
| 0 | 1 | 0 | 0 |
| 0 | 1 | 0 | 0 |
| 0 | 1 | 1 | 1 |
| 0 | 0 | 0 | 0 |

| | | | |
|---|---|---|---|
| 0 | 1 | 0 | 0 |
| 0 | 1 | 0 | 0 |
| 0 | 1 | 1 | 1 |
| 0 | 1 | 1 | 1 |
| 0 | 1 | 1 | 1 |
| 0 | 1 | 0 | 0 |
| 0 | 1 | 1 | 1 |
| 0 | 1 | 1 | 1 |
| 0 | 1 | 0 | 0 |
| 0 | 1 | 0 | 0 |
| 1 | 1 | 0 | 0 |
| 1 | 1 | 0 | 0 |
| 0 | 0 | 0 | 1 |
| 1 | 1 | 0 | 0 |
| 0 | 0 | 0 | 0 |
| 0 | 0 | 0 | 0 |
| 0 | 1 | 0 | 0 |
| 0 | 0 | 0 | 0 |
| 0 | 0 | 0 | 0 |
| 0 | 0 | 0 | 1 |
| 0 | 0 | 0 | 0 |
| 0 | 1 | 0 | 0 |
| 0 | 0 | 0 | 0 |
| 0 | 0 | 0 | 0 |
| 0 | 1 | 0 | 0 |
| 0 | 1 | 1 | 1 |
| 0 | 0 | 0 | 1 |
| 0 | 0 | 0 | 1 |
| 0 | 1 | 0 | 0 |
| 0 | 1 | 0 | 0 |
| 0 | 0 | 0 | 0 |
| 0 | 0 | 0 | 0 |
| 0 | 0 | 0 | 0 |
| 0 | 1 | 0 | 0 |
| 0 | 1 | 0 | 0 |
| 0 | 1 | 1 | 1 |
| 0 | 1 | 0 | 0 |
| 0 | 1 | 0 | 0 |
| 0 | 0 | 0 | 1 |
| 0 | 1 | 1 | 1 |
| 0 | 0 | 0 | 1 |
| 0 | 1 | 0 | 0 |
| 0 | 0 | 0 | 0 |
| 0 | 0 | 0 | 0 |
| 0 | 0 | 0 | 0 |
| 0 | 0 | 0 | 0 |
| 0 | 0 | 0 | 0 |
| 0 | 0 | 0 | 1 |
| 0 | 0 | 0 | 1 |
| 0 | 0 | 0 | 0 |
| 0 | 0 | 0 | 0 |
| 0 | 0 | 0 | 0 |
| 0 | 0 | 0 | 0 |
| 0 | 0 | 0 | 1 |

| | | | |
|---|---|---|---|
| 0 | 0 | 0 | 1 |
| 0 | 0 | 0 | 0 |
| 0 | 0 | 0 | 1 |
| 0 | 1 | 0 | 0 |
| 0 | 1 | 1 | 1 |
| 0 | 1 | 1 | 1 |
| 0 | 1 | 0 | 0 |
| 0 | 1 | 1 | 1 |
| 0 | 1 | 1 | 1 |
| 0 | 1 | 1 | 1 |
| 0 | 1 | 1 | 1 |
| 0 | 1 | 1 | 1 |
| 0 | 0 | 0 | 1 |
| 0 | 0 | 0 | 1 |
| 0 | 0 | 0 | 1 |
| 0 | 0 | 0 | 1 |
| 0 | 0 | 0 | 0 |
| 0 | 0 | 0 | 0 |
| 0 | 1 | 1 | 1 |
| 0 | 0 | 0 | 0 |
| 0 | 0 | 0 | 0 |
| 0 | 1 | 0 | 0 |
| 0 | 1 | 1 | 1 |
| 0 | 1 | 0 | 0 |
| 0 | 0 | 0 | 0 |
| 0 | 0 | 0 | 1 |
| 0 | 1 | 1 | 1 |
| 0 | 1 | 1 | 1 |
| 0 | 1 | 0 | 0 |
| 0 | 0 | 0 | 0 |
| 0 | 1 | 1 | 1 |
| 0 | 1 | 0 | 0 |
| 0 | 1 | 0 | 0 |
| 0 | 1 | 0 | 0 |
| 0 | 1 | 0 | 0 |
| 0 | 0 | 0 | 0 |
| 0 | 1 | 1 | 1 |
| 0 | 0 | 0 | 1 |
| 0 | 0 | 0 | 1 |
| 0 | 0 | 0 | 1 |
| 0 | 0 | 0 | 0 |
| 0 | 0 | 0 | 1 |
| 0 | 0 | 0 | 0 |
| 0 | 0 | 0 | 1 |
| 1 | 0 | 0 | 0 |
| 0 | 0 | 0 | 1 |
| 0 | 1 | 0 | 0 |
| 0 | 1 | 0 | 0 |
| 1 | 1 | 0 | 0 |
| 0 | 0 | 0 | 0 |
| 0 | 0 | 0 | 0 |
| 0 | 0 | 0 | 0 |
| 0 | 0 | 0 | 0 |
| 0 | 0 | 0 | 0 |

| | | | |
|---|---|---|---|
| 0 | 0 | 0 | 0 |
| 0 | 0 | 0 | 0 |
| 0 | 0 | 0 | 0 |
| 0 | 0 | 0 | 0 |
| 0 | 1 | 0 | 0 |
| 0 | 1 | 0 | 0 |
| 0 | 1 | 0 | 0 |
| 0 | 0 | 0 | 1 |
| 0 | 1 | 0 | 0 |
| 0 | 1 | 0 | 0 |
| 0 | 1 | 0 | 0 |
| 0 | 1 | 1 | 1 |
| 0 | 1 | 0 | 0 |
| 0 | 1 | 0 | 0 |
| 0 | 1 | 0 | 0 |
| 0 | 1 | 0 | 0 |
| 0 | 1 | 0 | 0 |
| 0 | 1 | 0 | 0 |
| 0 | 1 | 0 | 0 |
| 0 | 1 | 0 | 0 |
| 0 | 1 | 0 | 0 |
| 0 | 1 | 0 | 0 |
| 0 | 1 | 0 | 0 |
| 0 | 1 | 0 | 0 |
| 0 | 1 | 0 | 0 |
| 0 | 1 | 0 | 0 |
| 0 | 1 | 0 | 0 |
| 0 | 1 | 0 | 0 |
| 0 | 1 | 0 | 0 |
| 0 | 1 | 0 | 0 |
| 0 | 1 | 0 | 0 |
| 0 | 1 | 0 | 0 |
| 0 | 1 | 0 | 0 |
| 0 | 1 | 0 | 0 |
| 0 | 1 | 0 | 0 |
| 0 | 1 | 0 | 0 |
| 0 | 1 | 0 | 0 |
| 0 | 1 | 0 | 0 |
| 0 | 1 | 0 | 0 |
| 0 | 1 | 0 | 0 |
| 0 | 1 | 1 | 1 |
| 0 | 1 | 0 | 0 |
| 0 | 1 | 0 | 0 |
| 0 | 1 | 0 | 0 |
| 0 | 1 | 0 | 0 |
| 0 | 1 | 0 | 0 |
| 0 | 1 | 0 | 0 |
| 0 | 1 | 0 | 0 |
| 0 | 1 | 0 | 0 |
| 0 | 0 | 0 | 1 |
| 0 | 0 | 0 | 1 |
| 0 | 1 | 1 | 1 |
| 1 | 1 | 0 | 0 |
| 0 | 1 | 0 | 0 |

| | | | |
|---|---|---|---|
| 0 | 0 | 0 | 0 |
| 0 | 0 | 0 | 0 |
| 1 | 1 | 0 | 0 |
| 1 | 1 | 0 | 0 |
| 0 | 0 | 0 | 0 |
| 0 | 0 | 0 | 0 |
| 0 | 1 | 0 | 0 |
| 0 | 1 | 1 | 1 |
| 0 | 1 | 1 | 1 |
| 0 | 0 | 0 | 0 |
| 0 | 1 | 0 | 0 |
| 0 | 1 | 1 | 1 |
| 0 | 1 | 0 | 0 |
| 0 | 1 | 0 | 0 |
| 0 | 1 | 0 | 0 |
| 0 | 0 | 0 | 0 |
| 0 | 0 | 0 | 0 |
| 0 | 1 | 1 | 1 |
| 0 | 1 | 0 | 0 |
| 0 | 1 | 1 | 1 |
| 0 | 1 | 0 | 0 |
| 0 | 1 | 0 | 0 |
| 0 | 0 | 0 | 1 |
| 0 | 1 | 0 | 0 |
| 0 | 1 | 0 | 0 |
| 0 | 1 | 1 | 1 |
| 0 | 1 | 0 | 0 |
| 0 | 0 | 0 | 0 |
| 0 | 1 | 0 | 0 |
| 0 | 1 | 0 | 0 |
| 0 | 1 | 0 | 0 |
| 0 | 0 | 0 | 0 |
| 0 | 1 | 1 | 1 |
| 0 | 0 | 0 | 0 |
| 0 | 1 | 0 | 0 |
| 0 | 0 | 0 | 0 |
| 0 | 0 | 0 | 1 |
| 0 | 0 | 0 | 0 |
| 0 | 0 | 0 | 0 |
| 0 | 0 | 0 | 0 |
| 0 | 1 | 0 | 0 |
| 0 | 1 | 0 | 0 |
| 0 | 1 | 0 | 0 |
| 0 | 1 | 0 | 0 |
| 0 | 1 | 0 | 0 |
| 0 | 1 | 1 | 1 |
| 0 | 1 | 0 | 0 |
| 0 | 1 | 0 | 0 |
| 0 | 1 | 0 | 0 |
| 0 | 1 | 0 | 0 |
| 0 | 1 | 0 | 0 |
| 0 | 1 | 0 | 0 |
| 0 | 1 | 1 | 1 |
| 0 | 1 | 1 | 1 |

| | | | |
|---|---|---|---|
| 0 | 1 | 1 | 1 |
| 0 | 0 | 0 | 1 |
| 0 | 0 | 0 | 0 |
| 0 | 0 | 0 | 0 |
| 0 | 1 | 0 | 0 |
| 0 | 1 | 0 | 0 |
| 0 | 1 | 0 | 0 |
| 0 | 1 | 1 | 1 |
| 0 | 1 | 0 | 0 |
| 0 | 1 | 1 | 1 |
| 0 | 0 | 0 | 1 |
| 0 | 0 | 0 | 1 |
| 0 | 1 | 0 | 0 |
| 0 | 0 | 0 | 0 |
| 0 | 1 | 0 | 0 |
| 0 | 0 | 0 | 0 |
| 0 | 1 | 0 | 0 |
| 1 | 1 | 0 | 0 |
| 0 | 0 | 0 | 1 |
| 0 | 1 | 1 | 1 |
| 0 | 1 | 1 | 1 |
| 0 | 0 | 0 | 0 |
| 0 | 0 | 0 | 1 |
| 0 | 0 | 0 | 0 |
| 0 | 0 | 0 | 0 |
| 0 | 1 | 0 | 0 |
| 0 | 0 | 0 | 0 |
| 0 | 0 | 0 | 0 |
| 0 | 1 | 0 | 0 |
| 0 | 0 | 0 | 0 |
| 0 | 1 | 0 | 0 |
| 0 | 1 | 0 | 0 |
| 0 | 0 | 0 | 0 |
| 0 | 0 | 0 | 0 |
| 0 | 1 | 0 | 0 |
| 0 | 0 | 0 | 0 |
| 0 | 1 | 0 | 0 |
| 0 | 1 | 0 | 0 |
| 1 | 1 | 0 | 0 |
| 0 | 1 | 0 | 0 |
| 0 | 1 | 0 | 0 |
| 0 | 1 | 1 | 1 |
| 0 | 1 | 0 | 0 |
| 0 | 1 | 0 | 0 |
| 0 | 1 | 0 | 0 |
| 0 | 1 | 0 | 0 |
| 0 | 1 | 1 | 1 |
| 0 | 1 | 0 | 0 |
| 0 | 1 | 0 | 0 |
| 0 | 1 | 0 | 0 |
| 0 | 1 | 0 | 0 |
| 0 | 0 | 0 | 0 |
| 0 | 1 | 1 | 1 |
| 0 | 0 | 0 | 1 |

| | | | |
|---|---|---|---|
| 0 | 1 | 0 | 0 |
| 0 | 1 | 0 | 0 |
| 0 | 1 | 1 | 1 |
| 0 | 1 | 1 | 1 |
| 0 | 1 | 1 | 1 |
| 0 | 1 | 0 | 0 |
| 0 | 1 | 1 | 1 |
| 0 | 1 | 1 | 1 |
| 0 | 1 | 0 | 0 |
| 0 | 0 | 0 | 0 |
| 0 | 1 | 0 | 0 |
| 0 | 1 | 0 | 0 |
| 0 | 1 | 0 | 0 |
| 0 | 1 | 1 | 1 |
| 0 | 1 | 1 | 1 |
| 0 | 1 | 1 | 1 |
| 0 | 1 | 1 | 1 |
| 0 | 0 | 0 | 1 |
| 0 | 0 | 0 | 0 |
| 0 | 0 | 0 | 0 |
| 0 | 1 | 1 | 1 |
| 0 | 1 | 1 | 1 |
| 0 | 1 | 0 | 0 |
| 0 | 1 | 0 | 0 |
| 0 | 1 | 1 | 1 |
| 0 | 1 | 1 | 1 |
| 0 | 1 | 0 | 0 |
| 0 | 1 | 1 | 1 |
| 0 | 0 | 0 | 0 |
| 0 | 0 | 0 | 0 |
| 0 | 1 | 0 | 0 |
| 0 | 1 | 0 | 0 |
| 0 | 1 | 0 | 0 |
| 1 | 1 | 0 | 0 |
| 1 | 1 | 0 | 0 |
| 0 | 1 | 0 | 0 |
| 0 | 1 | 0 | 0 |
| 0 | 1 | 0 | 0 |
| 0 | 1 | 1 | 1 |
| 0 | 1 | 0 | 0 |
| 0 | 1 | 1 | 1 |
| 0 | 1 | 1 | 1 |
| 0 | 1 | 1 | 1 |
| 0 | 0 | 0 | 1 |
| 0 | 0 | 0 | 1 |
| 0 | 0 | 0 | 0 |
| 0 | 0 | 0 | 0 |
| 0 | 0 | 0 | 0 |
| 0 | 0 | 0 | 0 |
| 0 | 1 | 0 | 0 |
| 0 | 1 | 0 | 0 |
| 0 | 1 | 0 | 0 |
| 0 | 1 | 0 | 0 |
| 0 | 1 | 1 | 1 |

| | | | |
|---|---|---|---|
| 0 | 1 | 1 | 1 |
| 0 | 1 | 0 | 0 |
| 0 | 1 | 0 | 0 |
| 0 | 1 | 0 | 0 |
| 0 | 1 | 1 | 1 |
| 0 | 1 | 0 | 0 |
| 0 | 1 | 0 | 0 |
| 0 | 1 | 0 | 0 |
| 0 | 1 | 1 | 1 |
| 0 | 1 | 1 | 1 |
| 0 | 1 | 0 | 0 |
| 0 | 1 | 0 | 0 |
| 0 | 1 | 0 | 0 |
| 0 | 1 | 1 | 1 |
| 0 | 1 | 0 | 0 |
| 0 | 1 | 0 | 0 |
| 0 | 1 | 0 | 0 |
| 0 | 1 | 0 | 0 |
| 0 | 1 | 0 | 0 |
| 0 | 1 | 0 | 0 |
| 0 | 1 | 0 | 0 |
| 0 | 0 | 0 | 1 |
| 0 | 1 | 0 | 0 |
| 0 | 0 | 0 | 1 |
| 0 | 1 | 0 | 0 |
| 0 | 1 | 0 | 0 |
| 0 | 1 | 1 | 1 |
| 0 | 1 | 0 | 0 |
| 0 | 1 | 0 | 0 |
| 0 | 1 | 0 | 0 |
| 0 | 1 | 0 | 0 |
| 0 | 1 | 0 | 0 |
| 0 | 1 | 0 | 0 |
| 0 | 1 | 0 | 0 |
| 0 | 0 | 0 | 1 |
| 0 | 1 | 0 | 0 |
| 0 | 1 | 0 | 0 |
| 0 | 1 | 0 | 0 |
| 0 | 1 | 0 | 0 |
| 0 | 0 | 0 | 0 |
| 0 | 0 | 0 | 0 |
| 0 | 0 | 0 | 0 |
| 0 | 0 | 0 | 0 |
| 0 | 0 | 0 | 0 |
| 0 | 0 | 0 | 0 |
| 0 | 0 | 0 | 0 |
| 0 | 0 | 0 | 0 |
| 0 | 0 | 0 | 1 |
| 0 | 1 | 0 | 0 |
| 0 | 0 | 0 | 1 |
| 0 | 0 | 0 | 0 |
| 0 | 0 | 0 | 0 |
| 0 | 1 | 0 | 0 |
| 0 | 0 | 0 | 0 |

| | | | |
|---|---|---|---|
| 0 | 1 | 0 | 0 |
| 0 | 1 | 0 | 0 |
| 0 | 0 | 0 | 1 |
| 0 | 1 | 0 | 0 |
| 0 | 0 | 0 | 0 |
| 0 | 0 | 0 | 1 |
| 0 | 0 | 0 | 1 |
| 0 | 0 | 0 | 1 |
| 0 | 0 | 0 | 0 |
| 0 | 0 | 0 | 1 |
| 0 | 1 | 0 | 0 |
| 0 | 1 | 0 | 0 |
| 0 | 1 | 0 | 0 |
| 0 | 0 | 0 | 1 |
| 0 | 1 | 0 | 0 |
| 0 | 0 | 0 | 1 |
| 0 | 1 | 0 | 0 |
| 0 | 1 | 0 | 0 |
| 1 | 0 | 0 | 0 |
| 0 | 0 | 0 | 1 |
| 0 | 0 | 0 | 1 |
| 0 | 0 | 0 | 0 |
| 0 | 0 | 0 | 1 |
| 0 | 0 | 0 | 0 |
| 0 | 0 | 0 | 0 |
| 0 | 0 | 0 | 1 |
| 0 | 0 | 0 | 1 |
| 0 | 0 | 0 | 0 |
| 0 | 0 | 0 | 1 |
| 1 | 0 | 0 | 0 |
| 0 | 0 | 0 | 1 |
| 0 | 0 | 0 | 1 |
| 0 | 0 | 0 | 1 |
| 0 | 0 | 0 | 1 |
| 0 | 0 | 0 | 1 |
| 0 | 0 | 0 | 1 |
| 0 | 0 | 0 | 1 |
| 0 | 0 | 0 | 1 |
| 0 | 0 | 0 | 1 |
| 0 | 0 | 0 | 1 |
| 0 | 0 | 0 | 1 |
| 0 | 0 | 0 | 1 |
| 0 | 0 | 0 | 1 |
| 0 | 0 | 0 | 1 |
| 0 | 0 | 0 | 1 |
| 0 | 0 | 0 | 1 |
| 0 | 0 | 0 | 1 |
| 0 | 0 | 0 | 1 |
| 0 | 0 | 0 | 1 |
| 0 | 0 | 0 | 1 |
| 0 | 0 | 0 | 1 |
| 0 | 0 | 0 | 1 |
| 0 | 0 | 0 | 1 |
| 0 | 0 | 0 | 1 |

| | | | |
|---|---|---|---|
| 0 | 0 | 0 | 1 |
| 0 | 0 | 0 | 1 |
| 0 | 0 | 0 | 1 |
| 0 | 0 | 0 | 1 |
| 0 | 0 | 0 | 1 |
| 0 | 0 | 0 | 1 |
| 0 | 0 | 0 | 1 |
| 0 | 0 | 0 | 1 |
| 0 | 0 | 0 | 1 |
| 0 | 0 | 0 | 1 |
| 0 | 0 | 0 | 1 |
| 0 | 0 | 0 | 0 |
| 0 | 1 | 0 | 0 |
| 0 | 1 | 0 | 0 |
| 0 | 1 | 1 | 1 |
| 0 | 1 | 0 | 0 |
| 0 | 1 | 1 | 1 |
| 0 | 1 | 0 | 0 |
| 0 | 1 | 1 | 1 |
| 0 | 0 | 0 | 1 |
| 0 | 0 | 0 | 0 |
| 0 | 0 | 0 | 1 |
| 0 | 0 | 0 | 1 |
| 0 | 0 | 0 | 0 |
| 0 | 0 | 0 | 0 |
| 0 | 0 | 0 | 1 |
| 0 | 0 | 0 | 1 |
| 0 | 0 | 0 | 0 |
| 0 | 0 | 0 | 0 |
| 0 | 0 | 0 | 0 |
| 0 | 0 | 0 | 0 |
| 0 | 0 | 0 | 0 |
| 0 | 0 | 0 | 1 |
| 0 | 0 | 0 | 1 |
| 0 | 0 | 0 | 1 |
| 0 | 0 | 0 | 1 |
| 0 | 0 | 0 | 1 |
| 0 | 0 | 0 | 1 |
| 0 | 0 | 0 | 1 |
| 0 | 1 | 0 | 0 |
| 0 | 1 | 0 | 0 |
| 0 | 0 | 0 | 1 |
| 0 | 1 | 0 | 0 |
| 0 | 1 | 0 | 0 |
| 0 | 1 | 0 | 0 |
| 0 | 1 | 1 | 1 |
| 0 | 1 | 0 | 0 |
| 0 | 1 | 1 | 1 |
| 0 | 1 | 1 | 1 |
| 0 | 1 | 0 | 0 |
| 0 | 1 | 0 | 0 |
| 0 | 1 | 0 | 0 |
| 0 | 1 | 1 | 1 |
| 0 | 1 | 0 | 0 |

| | | | |
|---|---|---|---|
| 0 | 1 | 0 | 0 |
| 0 | 0 | 0 | 1 |
| 0 | 0 | 0 | 1 |
| 0 | 0 | 0 | 1 |
| 0 | 1 | 0 | 0 |
| 0 | 0 | 0 | 1 |
| 0 | 0 | 0 | 0 |
| 0 | 0 | 0 | 1 |
| 0 | 0 | 0 | 1 |
| 0 | 0 | 0 | 0 |
| 0 | 1 | 1 | 1 |
| 0 | 0 | 0 | 1 |
| 0 | 0 | 0 | 1 |
| 0 | 1 | 0 | 0 |
| 0 | 1 | 1 | 1 |
| 0 | 1 | 1 | 1 |
| 0 | 1 | 0 | 0 |
| 0 | 1 | 1 | 1 |
| 0 | 1 | 0 | 0 |
| 0 | 1 | 0 | 0 |
| 0 | 1 | 0 | 0 |
| 0 | 1 | 0 | 0 |
| 0 | 0 | 0 | 0 |
| 0 | 1 | 0 | 0 |
| 0 | 1 | 0 | 0 |
| 0 | 1 | 0 | 0 |
| 0 | 1 | 0 | 0 |
| 0 | 1 | 0 | 0 |
| 0 | 1 | 0 | 0 |
| 0 | 1 | 0 | 0 |
| 0 | 1 | 0 | 0 |
| 0 | 1 | 0 | 0 |
| 0 | 1 | 0 | 0 |
| 0 | 1 | 0 | 0 |
| 0 | 1 | 0 | 0 |
| 0 | 1 | 0 | 0 |
| 0 | 1 | 1 | 1 |
| 0 | 1 | 0 | 0 |
| 0 | 1 | 0 | 0 |
| 0 | 1 | 1 | 1 |
| 0 | 1 | 1 | 1 |
| 0 | 1 | 0 | 0 |
| 0 | 0 | 0 | 0 |
| 0 | 1 | 0 | 0 |
| 0 | 1 | 0 | 0 |
| 0 | 1 | 0 | 0 |
| 0 | 1 | 0 | 0 |
| 0 | 1 | 0 | 0 |
| 0 | 1 | 1 | 1 |
| 0 | 1 | 1 | 1 |
| 0 | 0 | 0 | 0 |
| 0 | 0 | 0 | 1 |
| 0 | 1 | 0 | 0 |
| 0 | 1 | 0 | 0 |

| | | | |
|---|---|---|---|
| 0 | 0 | 0 | 0 |
| 0 | 1 | 0 | 0 |
| 0 | 1 | 0 | 0 |
| 0 | 1 | 0 | 0 |
| 0 | 1 | 0 | 0 |
| 0 | 1 | 0 | 0 |
| 0 | 1 | 0 | 0 |
| 0 | 0 | 0 | 0 |
| 0 | 0 | 0 | 0 |
| 0 | 0 | 0 | 0 |
| 0 | 0 | 0 | 0 |
| 0 | 0 | 0 | 0 |
| 0 | 0 | 0 | 0 |
| 0 | 0 | 0 | 0 |
| 0 | 0 | 0 | 1 |
| 0 | 1 | 0 | 0 |
| 0 | 1 | 0 | 0 |
| 0 | 1 | 0 | 0 |
| 0 | 1 | 1 | 1 |
| 0 | 0 | 0 | 0 |
| 0 | 1 | 0 | 0 |
| 0 | 1 | 0 | 0 |
| 0 | 1 | 0 | 0 |
| 0 | 1 | 0 | 0 |
| 0 | 1 | 0 | 0 |
| 0 | 1 | 1 | 1 |
| 0 | 1 | 0 | 0 |
| 0 | 1 | 0 | 0 |
| 0 | 1 | 0 | 0 |
| 0 | 0 | 0 | 0 |
| 0 | 0 | 0 | 1 |
| 0 | 0 | 0 | 0 |
| 0 | 0 | 0 | 1 |
| 0 | 1 | 1 | 1 |
| 1 | 1 | 0 | 0 |
| 0 | 0 | 0 | 0 |
| 0 | 0 | 0 | 1 |
| 0 | 0 | 0 | 1 |
| 0 | 0 | 0 | 1 |
| 0 | 1 | 1 | 1 |
| 0 | 1 | 0 | 0 |
| 0 | 1 | 0 | 0 |
| 0 | 1 | 0 | 0 |
| 0 | 1 | 0 | 0 |
| 0 | 1 | 0 | 0 |
| 0 | 1 | 0 | 0 |
| 0 | 0 | 0 | 0 |
| 0 | 0 | 0 | 0 |
| 0 | 0 | 0 | 1 |
| 0 | 0 | 0 | 1 |
| 0 | 1 | 0 | 0 |
| 0 | 1 | 0 | 0 |
| 0 | 1 | 0 | 0 |
| 0 | 0 | 0 | 1 |

| | | | |
|---|---|---|---|
| 0 | 0 | 0 | 1 |
| 0 | 0 | 0 | 1 |
| 0 | 1 | 1 | 1 |
| 0 | 1 | 0 | 0 |
| 0 | 1 | 1 | 1 |
| 0 | 0 | 0 | 1 |
| 0 | 0 | 0 | 0 |
| 0 | 0 | 0 | 1 |
| 0 | 0 | 0 | 0 |
| 0 | 0 | 0 | 0 |
| 0 | 0 | 0 | 0 |
| 0 | 0 | 0 | 0 |
| 0 | 0 | 0 | 0 |
| 0 | 0 | 0 | 0 |
| 0 | 0 | 0 | 0 |
| 0 | 0 | 0 | 0 |
| 0 | 0 | 0 | 1 |
| 0 | 1 | 0 | 0 |
| 0 | 1 | 0 | 0 |
| 0 | 1 | 1 | 1 |
| 0 | 1 | 0 | 0 |
| 0 | 0 | 0 | 0 |
| 0 | 0 | 0 | 1 |
| 0 | 0 | 0 | 0 |
| 0 | 0 | 0 | 1 |
| 0 | 0 | 0 | 1 |
| 0 | 0 | 0 | 1 |
| 0 | 0 | 0 | 0 |
| 0 | 0 | 0 | 0 |
| 0 | 0 | 0 | 0 |
| 0 | 0 | 0 | 1 |
| 0 | 0 | 0 | 0 |
| 0 | 0 | 0 | 0 |
| 0 | 0 | 0 | 0 |
| 0 | 0 | 0 | 1 |
| 0 | 0 | 0 | 1 |
| 0 | 1 | 0 | 0 |
| 0 | 1 | 0 | 0 |
| 0 | 1 | 0 | 0 |
| 0 | 0 | 0 | 1 |
| 0 | 0 | 0 | 1 |
| 0 | 0 | 0 | 1 |
| 0 | 0 | 0 | 1 |
| 0 | 0 | 0 | 0 |
| 0 | 0 | 0 | 1 |
| 0 | 0 | 0 | 0 |
| 0 | 0 | 0 | 1 |
| 0 | 0 | 0 | 0 |
| 0 | 0 | 0 | 0 |
| 0 | 0 | 0 | 1 |
| 0 | 0 | 0 | 1 |
| 0 | 1 | 0 | 0 |
| 0 | 1 | 0 | 0 |
| 0 | 0 | 0 | 1 |

| | | | |
|---|---|---|---|
| 0 | 0 | 0 | 1 |
| 0 | 0 | 0 | 1 |
| 0 | 0 | 0 | 1 |
| 0 | 1 | 1 | 1 |
| 0 | 1 | 0 | 0 |
| 0 | 1 | 1 | 1 |
| 0 | 1 | 1 | 1 |
| 0 | 1 | 0 | 0 |
| 0 | 1 | 0 | 0 |
| 0 | 0 | 0 | 1 |
| 0 | 0 | 0 | 0 |
| 0 | 0 | 0 | 0 |
| 1 | 0 | 0 | 0 |
| 1 | 0 | 0 | 0 |
| 0 | 0 | 0 | 0 |
| 1 | 0 | 0 | 0 |
| 1 | 0 | 0 | 0 |
| 1 | 0 | 0 | 0 |
| 1 | 0 | 0 | 0 |
| 0 | 0 | 0 | 1 |
| 0 | 0 | 0 | 1 |
| 0 | 0 | 0 | 1 |
| 0 | 0 | 0 | 1 |
| 0 | 0 | 0 | 1 |
| 0 | 0 | 0 | 0 |
| 0 | 0 | 0 | 1 |
| 1 | 0 | 0 | 0 |
| 1 | 0 | 0 | 0 |
| 0 | 0 | 0 | 1 |
| 0 | 0 | 0 | 1 |
| 0 | 0 | 0 | 1 |
| 0 | 0 | 0 | 1 |
| 0 | 0 | 0 | 1 |
| 0 | 1 | 0 | 0 |
| 0 | 1 | 0 | 0 |
| 0 | 0 | 0 | 1 |
| 0 | 0 | 0 | 0 |
| 0 | 0 | 0 | 1 |
| 0 | 0 | 0 | 0 |
| 0 | 1 | 0 | 0 |
| 0 | 1 | 0 | 0 |
| 0 | 1 | 0 | 0 |
| 0 | 0 | 0 | 1 |
| 0 | 0 | 0 | 0 |
| 0 | 0 | 0 | 0 |
| 1 | 0 | 0 | 0 |
| 1 | 1 | 0 | 0 |
| 1 | 0 | 0 | 0 |
| 0 | 0 | 0 | 0 |
| 0 | 1 | 0 | 0 |
| 0 | 1 | 0 | 0 |
| 1 | 1 | 0 | 0 |
| 1 | 0 | 0 | 0 |
| 0 | 0 | 0 | 1 |

| | | | |
|---|---|---|---|
| 0 | 0 | 0 | 0 |
| 0 | 0 | 0 | 1 |
| 0 | 0 | 0 | 0 |
| 0 | 0 | 0 | 0 |
| 0 | 1 | 0 | 0 |
| 0 | 0 | 0 | 1 |
| 0 | 0 | 0 | 0 |
| 0 | 0 | 0 | 1 |
| 0 | 0 | 0 | 1 |
| 0 | 0 | 0 | 0 |
| 0 | 0 | 0 | 1 |
| 0 | 0 | 0 | 1 |
| 0 | 0 | 0 | 1 |
| 0 | 0 | 0 | 1 |
| 0 | 0 | 0 | 1 |
| 0 | 1 | 0 | 0 |
| 0 | 1 | 0 | 0 |
| 0 | 0 | 0 | 1 |
| 0 | 0 | 0 | 0 |
| 0 | 0 | 0 | 1 |
| 0 | 0 | 0 | 0 |
| 0 | 0 | 0 | 0 |
| 0 | 0 | 0 | 0 |
| 0 | 1 | 0 | 0 |
| 0 | 1 | 0 | 0 |
| 0 | 1 | 0 | 0 |
| 0 | 1 | 0 | 0 |
| 0 | 0 | 0 | 0 |
| 0 | 0 | 0 | 0 |
| 0 | 1 | 0 | 0 |
| 0 | 1 | 0 | 0 |
| 0 | 1 | 0 | 0 |
| 0 | 1 | 1 | 1 |
| 0 | 1 | 1 | 1 |
| 0 | 1 | 0 | 0 |
| 0 | 1 | 0 | 0 |
| 0 | 1 | 1 | 1 |
| 0 | 1 | 0 | 0 |
| 0 | 1 | 1 | 1 |
| 0 | 1 | 0 | 0 |
| 0 | 1 | 0 | 0 |
| 0 | 1 | 0 | 0 |
| 0 | 1 | 1 | 1 |
| 0 | 0 | 0 | 1 |
| 0 | 1 | 0 | 0 |
| 0 | 1 | 0 | 0 |
| 0 | 1 | 0 | 0 |
| 0 | 1 | 0 | 0 |
| 0 | 1 | 0 | 0 |
| 0 | 1 | 0 | 0 |
| 0 | 1 | 1 | 1 |
| 0 | 1 | 0 | 0 |
| 0 | 1 | 0 | 0 |
| 0 | 1 | 1 | 1 |

| | | | |
|---|---|---|---|
| 0 | 1 | 0 | 0 |
| 0 | 1 | 1 | 1 |
| 0 | 1 | 0 | 0 |
| 0 | 0 | 0 | 0 |
| 0 | 0 | 0 | 0 |
| 0 | 0 | 0 | 0 |
| 0 | 1 | 0 | 0 |
| 0 | 0 | 0 | 0 |
| 0 | 0 | 0 | 0 |
| 0 | 1 | 0 | 0 |
| 0 | 0 | 0 | 0 |
| 0 | 0 | 0 | 1 |
| 0 | 0 | 0 | 0 |
| 0 | 0 | 0 | 1 |
| 0 | 1 | 1 | 1 |
| 0 | 1 | 1 | 1 |
| 0 | 1 | 0 | 0 |
| 0 | 1 | 0 | 0 |
| 0 | 1 | 1 | 1 |
| 0 | 1 | 0 | 0 |
| 0 | 1 | 1 | 1 |
| 0 | 1 | 0 | 0 |
| 0 | 0 | 0 | 1 |
| 0 | 1 | 0 | 0 |
| 0 | 0 | 0 | 1 |
| 0 | 0 | 0 | 1 |
| 0 | 0 | 0 | 1 |
| 0 | 0 | 0 | 0 |
| 0 | 0 | 0 | 1 |
| 0 | 0 | 0 | 1 |
| 0 | 0 | 0 | 1 |
| 1 | 0 | 0 | 0 |
| 0 | 0 | 0 | 1 |
| 0 | 0 | 0 | 0 |
| 0 | 1 | 0 | 0 |
| 0 | 0 | 0 | 0 |
| 0 | 0 | 0 | 1 |
| 0 | 0 | 0 | 1 |
| 0 | 1 | 0 | 0 |
| 0 | 0 | 0 | 0 |
| 0 | 1 | 0 | 0 |
| 0 | 1 | 0 | 0 |
| 0 | 1 | 0 | 0 |
| 0 | 0 | 0 | 0 |
| 0 | 1 | 0 | 0 |
| 0 | 1 | 0 | 0 |
| 0 | 0 | 0 | 1 |
| 0 | 0 | 0 | 1 |
| 0 | 0 | 0 | 0 |
| 0 | 0 | 0 | 0 |
| 0 | 0 | 0 | 0 |
| 0 | 0 | 0 | 1 |
| 0 | 0 | 0 | 1 |
| 0 | 0 | 0 | 0 |

| | | | |
|---|---|---|---|
| 0 | 0 | 0 | 0 |
| 0 | 0 | 0 | 1 |
| 0 | 0 | 0 | 1 |
| 0 | 0 | 0 | 1 |
| 0 | 0 | 0 | 0 |
| 0 | 1 | 0 | 0 |
| 0 | 1 | 0 | 0 |
| 0 | 1 | 0 | 0 |
| 0 | 1 | 0 | 0 |
| 0 | 1 | 0 | 0 |
| 0 | 0 | 0 | 1 |
| 0 | 1 | 0 | 0 |
| 0 | 0 | 0 | 0 |
| 0 | 0 | 0 | 0 |
| 0 | 1 | 1 | 1 |
| 0 | 0 | 0 | 1 |
| 0 | 1 | 0 | 0 |
| 0 | 1 | 1 | 1 |
| 0 | 1 | 0 | 0 |
| 0 | 1 | 0 | 0 |
| 0 | 1 | 1 | 1 |
| 0 | 1 | 0 | 0 |
| 0 | 1 | 0 | 0 |
| 0 | 1 | 1 | 1 |
| 0 | 1 | 0 | 0 |
| 0 | 1 | 0 | 0 |
| 0 | 1 | 0 | 0 |
| 0 | 1 | 1 | 1 |
| 0 | 1 | 0 | 0 |
| 0 | 0 | 0 | 0 |
| 0 | 0 | 0 | 0 |
| 0 | 0 | 0 | 1 |
| 0 | 0 | 0 | 1 |
| 0 | 0 | 0 | 1 |
| 0 | 0 | 0 | 1 |
| 0 | 0 | 0 | 1 |
| 0 | 0 | 0 | 1 |
| 0 | 0 | 0 | 1 |
| 0 | 0 | 0 | 1 |
| 0 | 0 | 0 | 1 |
| 0 | 0 | 0 | 1 |
| 0 | 0 | 0 | 1 |
| 1 | 0 | 0 | 0 |
| 0 | 0 | 0 | 1 |
| 0 | 0 | 0 | 1 |
| 0 | 0 | 0 | 1 |
| 0 | 0 | 0 | 1 |
| 0 | 0 | 0 | 1 |
| 0 | 0 | 0 | 1 |
| 0 | 0 | 0 | 1 |
| 0 | 0 | 0 | 1 |
| 0 | 0 | 0 | 1 |
| 0 | 0 | 0 | 1 |
| 0 | 0 | 0 | 1 |

| | | | |
|---|---|---|---|
| 0 | 0 | 0 | 1 |
| 0 | 0 | 0 | 1 |
| 0 | 0 | 0 | 1 |
| 0 | 0 | 0 | 1 |
| 0 | 0 | 0 | 1 |
| 0 | 0 | 0 | 1 |
| 0 | 0 | 0 | 1 |
| 0 | 0 | 0 | 1 |
| 0 | 1 | 0 | 0 |
| 0 | 0 | 0 | 0 |
| 0 | 0 | 0 | 0 |
| 0 | 0 | 0 | 0 |
| 0 | 0 | 0 | 0 |
| 0 | 0 | 0 | 0 |
| 0 | 0 | 0 | 0 |
| 0 | 0 | 0 | 0 |
| 0 | 0 | 0 | 0 |
| 0 | 0 | 0 | 0 |
| 0 | 1 | 0 | 0 |
| 0 | 1 | 0 | 0 |
| 1 | 0 | 0 | 0 |
| 1 | 0 | 0 | 0 |
| 0 | 1 | 0 | 0 |
| 0 | 1 | 0 | 0 |
| 0 | 1 | 0 | 0 |
| 0 | 0 | 0 | 1 |
| 0 | 0 | 0 | 1 |
| 0 | 0 | 0 | 0 |
| 0 | 0 | 0 | 0 |
| 1 | 0 | 0 | 0 |
| 1 | 0 | 0 | 0 |
| 0 | 0 | 0 | 0 |
| 1 | 0 | 0 | 0 |
| 1 | 0 | 0 | 0 |
| 0 | 0 | 0 | 0 |
| 0 | 0 | 0 | 0 |
| 0 | 0 | 0 | 0 |
| 0 | 1 | 0 | 0 |
| 0 | 1 | 0 | 0 |
| 0 | 0 | 0 | 0 |
| 0 | 0 | 0 | 0 |
| 0 | 0 | 0 | 0 |
| 1 | 0 | 0 | 0 |
| 1 | 0 | 0 | 0 |
| 1 | 0 | 0 | 0 |
| 0 | 0 | 0 | 0 |
| 0 | 0 | 0 | 1 |
| 0 | 0 | 0 | 1 |
| 0 | 0 | 0 | 1 |
| 0 | 0 | 0 | 1 |
| 0 | 0 | 0 | 1 |
| 0 | 0 | 0 | 1 |
| 0 | 1 | 0 | 0 |
| 0 | 1 | 0 | 0 |

| | | | |
|---|---|---|---|
| 0 | 1 | 0 | 0 |
| 0 | 0 | 0 | 0 |
| 0 | 0 | 0 | 0 |
| 0 | 1 | 0 | 0 |
| 0 | 0 | 0 | 1 |
| 0 | 0 | 0 | 0 |
| 0 | 0 | 0 | 0 |
| 0 | 0 | 0 | 0 |
| 0 | 0 | 0 | 1 |
| 0 | 0 | 0 | 0 |
| 1 | 0 | 0 | 0 |
| 0 | 0 | 0 | 1 |
| 0 | 0 | 0 | 0 |
| 0 | 0 | 0 | 0 |
| 0 | 0 | 0 | 0 |
| 0 | 0 | 0 | 0 |
| 0 | 0 | 0 | 1 |
| 0 | 0 | 0 | 0 |
| 0 | 0 | 0 | 0 |
| 0 | 0 | 0 | 1 |
| 0 | 1 | 0 | 0 |
| 0 | 0 | 0 | 0 |
| 0 | 1 | 0 | 0 |
| 0 | 1 | 0 | 0 |
| 0 | 1 | 0 | 0 |
| 0 | 0 | 0 | 1 |
| 0 | 0 | 0 | 1 |
| 0 | 0 | 0 | 1 |
| 0 | 0 | 0 | 1 |
| 0 | 0 | 0 | 1 |
| 0 | 0 | 0 | 1 |
| 0 | 0 | 0 | 1 |
| 0 | 1 | 1 | 1 |
| 0 | 1 | 1 | 1 |
| 0 | 0 | 0 | 0 |
| 0 | 1 | 0 | 0 |
| 0 | 0 | 0 | 0 |
| 0 | 1 | 0 | 0 |
| 0 | 1 | 0 | 0 |
| 0 | 1 | 0 | 0 |
| 0 | 0 | 0 | 1 |
| 0 | 0 | 0 | 1 |
| 0 | 0 | 0 | 1 |
| 0 | 0 | 0 | 0 |
| 0 | 0 | 0 | 0 |
| 0 | 0 | 0 | 0 |
| 0 | 0 | 0 | 1 |
| 0 | 0 | 0 | 1 |
| 0 | 0 | 0 | 0 |
| 0 | 0 | 0 | 0 |
| 0 | 0 | 0 | 1 |
| 0 | 0 | 0 | 1 |
| 0 | 0 | 0 | 1 |
| 0 | 0 | 0 | 1 |

| | | | |
|---|---|---|---|
| 0 | 0 | 0 | 1 |
| 0 | 0 | 0 | 1 |
| 0 | 0 | 0 | 1 |
| 0 | 0 | 0 | 0 |
| 0 | 0 | 0 | 0 |
| 0 | 0 | 0 | 1 |
| 0 | 0 | 0 | 1 |
| 0 | 0 | 0 | 1 |
| 0 | 0 | 0 | 1 |
| 0 | 0 | 0 | 1 |
| 0 | 0 | 0 | 0 |
| 0 | 0 | 0 | 1 |
| 0 | 1 | 0 | 0 |
| 0 | 0 | 0 | 0 |
| 0 | 0 | 0 | 0 |
| 0 | 0 | 0 | 1 |
| 0 | 0 | 0 | 1 |
| 0 | 0 | 0 | 1 |
| 0 | 0 | 0 | 1 |
| 0 | 0 | 0 | 1 |
| 0 | 0 | 0 | 0 |
| 0 | 0 | 0 | 1 |
| 0 | 0 | 0 | 1 |
| 0 | 1 | 0 | 0 |
| 0 | 1 | 1 | 1 |
| 0 | 1 | 0 | 0 |
| 0 | 1 | 0 | 0 |
| 0 | 0 | 0 | 1 |
| 0 | 0 | 0 | 1 |
| 0 | 0 | 0 | 1 |
| 0 | 0 | 0 | 1 |
| 0 | 0 | 0 | 1 |
| 0 | 0 | 0 | 1 |
| 0 | 0 | 0 | 1 |
| 0 | 0 | 0 | 1 |
| 0 | 0 | 0 | 0 |
| 0 | 0 | 0 | 1 |
| 0 | 0 | 0 | 1 |
| 0 | 0 | 0 | 0 |
| 0 | 0 | 0 | 0 |
| 0 | 0 | 0 | 1 |
| 0 | 0 | 0 | 1 |
| 1 | 0 | 0 | 0 |
| 1 | 0 | 0 | 0 |
| 0 | 0 | 0 | 0 |
| 0 | 0 | 0 | 0 |
| 1 | 0 | 0 | 0 |
| 1 | 0 | 0 | 0 |
| 0 | 0 | 0 | 1 |
| 1 | 0 | 0 | 0 |
| 1 | 0 | 0 | 0 |
| 1 | 1 | 0 | 0 |
| 1 | 1 | 0 | 0 |
| 1 | 0 | 0 | 0 |

| | | | |
|---|---|---|---|
| 0 | 1 | 0 | 0 |
| 0 | 1 | 0 | 0 |
| 1 | 0 | 0 | 0 |
| 1 | 0 | 0 | 0 |
| 1 | 0 | 0 | 0 |
| 1 | 0 | 0 | 0 |
| 0 | 0 | 0 | 0 |
| 0 | 0 | 0 | 0 |
| 0 | 0 | 0 | 0 |
| 0 | 0 | 0 | 1 |
| 0 | 0 | 0 | 1 |
| 0 | 0 | 0 | 0 |
| 0 | 0 | 0 | 1 |
| 0 | 0 | 0 | 0 |
| 0 | 0 | 0 | 0 |
| 0 | 0 | 0 | 1 |
| 0 | 0 | 0 | 0 |
| 0 | 1 | 1 | 1 |
| 0 | 0 | 0 | 1 |
| 0 | 0 | 0 | 1 |
| 0 | 1 | 1 | 1 |
| 0 | 0 | 0 | 1 |
| 0 | 0 | 0 | 1 |
| 0 | 0 | 0 | 0 |
| 0 | 0 | 0 | 0 |
| 0 | 0 | 0 | 1 |
| 0 | 0 | 0 | 1 |
| 0 | 0 | 0 | 1 |
| 0 | 0 | 0 | 0 |
| 0 | 0 | 0 | 0 |
| 0 | 0 | 0 | 1 |
| 0 | 0 | 0 | 1 |
| 0 | 1 | 1 | 1 |
| 0 | 1 | 0 | 0 |
| 0 | 1 | 0 | 0 |
| 0 | 0 | 0 | 1 |
| 0 | 0 | 0 | 0 |
| 0 | 0 | 0 | 1 |
| 0 | 0 | 0 | 1 |
| 0 | 0 | 0 | 0 |
| 0 | 0 | 0 | 0 |
| 0 | 0 | 0 | 0 |
| 0 | 0 | 0 | 0 |
| 0 | 0 | 0 | 0 |
| 0 | 0 | 0 | 0 |
| 0 | 0 | 0 | 1 |
| 0 | 0 | 0 | 1 |
| 0 | 0 | 0 | 0 |
| 0 | 0 | 0 | 0 |
| 0 | 0 | 0 | 0 |
| 0 | 0 | 0 | 1 |
| 0 | 0 | 0 | 1 |
| 0 | 0 | 0 | 0 |
| 0 | 0 | 0 | 0 |

| | | | |
|---|---|---|---|
| 0 | 0 | 0 | 1 |
| 0 | 0 | 0 | 1 |
| 0 | 0 | 0 | 1 |
| 0 | 0 | 0 | 1 |
| 0 | 1 | 0 | 0 |
| 0 | 1 | 0 | 0 |
| 0 | 1 | 1 | 1 |
| 0 | 0 | 0 | 0 |
| 0 | 1 | 1 | 1 |
| 0 | 0 | 0 | 0 |
| 0 | 0 | 0 | 1 |
| 0 | 0 | 0 | 1 |
| 0 | 0 | 0 | 1 |
| 0 | 0 | 0 | 1 |
| 0 | 0 | 0 | 1 |
| 0 | 0 | 0 | 1 |
| 0 | 0 | 0 | 0 |
| 0 | 0 | 0 | 1 |
| 0 | 0 | 0 | 1 |
| 0 | 0 | 0 | 0 |
| 0 | 0 | 0 | 0 |
| 0 | 0 | 0 | 1 |
| 0 | 0 | 0 | 1 |
| 0 | 0 | 0 | 0 |
| 0 | 0 | 0 | 1 |
| 0 | 0 | 0 | 0 |
| 0 | 0 | 0 | 0 |
| 0 | 0 | 0 | 0 |
| 0 | 0 | 0 | 0 |
| 0 | 0 | 0 | 0 |
| 0 | 0 | 0 | 0 |
| 0 | 0 | 0 | 0 |
| 0 | 0 | 0 | 1 |
| 0 | 0 | 0 | 0 |
| 0 | 0 | 0 | 1 |
| 0 | 0 | 0 | 1 |
| 0 | 0 | 0 | 1 |
| 0 | 0 | 0 | 1 |
| 0 | 0 | 0 | 0 |
| 0 | 0 | 0 | 1 |
| 0 | 0 | 0 | 1 |
| 0 | 0 | 0 | 1 |
| 0 | 0 | 0 | 1 |
| 0 | 1 | 0 | 0 |
| 0 | 0 | 0 | 1 |
| 0 | 0 | 0 | 1 |
| 0 | 0 | 0 | 1 |
| 0 | 0 | 0 | 1 |
| 0 | 0 | 0 | 1 |
| 0 | 0 | 0 | 1 |
| 0 | 0 | 0 | 0 |
| 0 | 1 | 0 | 0 |
| 0 | 0 | 0 | 0 |
| 0 | 0 | 0 | 1 |

| | | | |
|---|---|---|---|
| 0 | 1 | 0 | 0 |
| 0 | 1 | 0 | 0 |
| 0 | 1 | 0 | 0 |
| 0 | 0 | 0 | 0 |
| 1 | 1 | 0 | 0 |
| 1 | 1 | 0 | 0 |
| 0 | 1 | 0 | 0 |
| 0 | 0 | 0 | 0 |
| 0 | 0 | 0 | 1 |
| 0 | 0 | 0 | 1 |
| 0 | 0 | 0 | 0 |
| 0 | 0 | 0 | 1 |
| 0 | 0 | 0 | 1 |
| 0 | 0 | 0 | 0 |
| 0 | 0 | 0 | 0 |
| 0 | 0 | 0 | 0 |
| 0 | 1 | 0 | 0 |
| 0 | 0 | 0 | 0 |
| 0 | 0 | 0 | 0 |
| 0 | 0 | 0 | 1 |
| 0 | 0 | 0 | 1 |
| 0 | 0 | 0 | 0 |
| 0 | 0 | 0 | 1 |
| 0 | 1 | 0 | 0 |
| 0 | 0 | 0 | 1 |
| 0 | 0 | 0 | 1 |
| 0 | 0 | 0 | 1 |
| 0 | 0 | 0 | 1 |
| 0 | 0 | 0 | 1 |
| 0 | 0 | 0 | 0 |
| 0 | 0 | 0 | 1 |
| 0 | 0 | 0 | 1 |
| 0 | 0 | 0 | 0 |
| 0 | 0 | 0 | 1 |
| 0 | 1 | 0 | 0 |
| 0 | 1 | 0 | 0 |
| 0 | 1 | 0 | 0 |
| 0 | 0 | 0 | 0 |
| 0 | 0 | 0 | 0 |
| 0 | 0 | 0 | 0 |
| 0 | 0 | 0 | 0 |
| 0 | 1 | 0 | 0 |
| 0 | 0 | 0 | 1 |
| 0 | 1 | 0 | 0 |
| 0 | 0 | 0 | 1 |
| 0 | 0 | 0 | 0 |
| 0 | 1 | 1 | 1 |
| 0 | 0 | 0 | 1 |
| 0 | 0 | 0 | 1 |
| 0 | 0 | 0 | 0 |
| 0 | 0 | 0 | 1 |
| 0 | 0 | 0 | 1 |
| 0 | 0 | 0 | 1 |
| 0 | 0 | 0 | 1 |

| | | | |
|---|---|---|---|
| 1 | 1 | 0 | 0 |
| 0 | 0 | 0 | 1 |
| 0 | 0 | 0 | 1 |
| 0 | 0 | 0 | 1 |
| 0 | 0 | 0 | 1 |
| 0 | 0 | 0 | 1 |
| 0 | 0 | 0 | 1 |
| 0 | 0 | 0 | 1 |
| 0 | 0 | 0 | 1 |
| 0 | 0 | 0 | 1 |
| 0 | 0 | 0 | 0 |
| 0 | 0 | 0 | 1 |
| 0 | 0 | 0 | 1 |
| 0 | 0 | 0 | 1 |
| 0 | 0 | 0 | 0 |
| 0 | 0 | 0 | 0 |
| 0 | 0 | 0 | 0 |
| 0 | 0 | 0 | 0 |
| 0 | 0 | 0 | 0 |
| 0 | 0 | 0 | 1 |
| 0 | 0 | 0 | 1 |
| 0 | 0 | 0 | 1 |
| 0 | 0 | 0 | 0 |
| 0 | 0 | 0 | 1 |
| 0 | 0 | 0 | 1 |
| 0 | 0 | 0 | 1 |
| 0 | 0 | 0 | 1 |
| 0 | 0 | 0 | 1 |
| 0 | 0 | 0 | 0 |
| 0 | 0 | 0 | 0 |
| 0 | 0 | 0 | 0 |
| 0 | 1 | 0 | 0 |
| 0 | 1 | 0 | 0 |
| 0 | 0 | 0 | 0 |
| 0 | 0 | 0 | 1 |
| 0 | 0 | 0 | 1 |
| 0 | 0 | 0 | 0 |
| 0 | 0 | 0 | 0 |
| 0 | 1 | 0 | 0 |
| 0 | 1 | 0 | 0 |
| 0 | 1 | 0 | 0 |
| 0 | 0 | 0 | 0 |
| 0 | 0 | 0 | 1 |
| 0 | 0 | 0 | 1 |
| 0 | 0 | 0 | 0 |
| 0 | 0 | 0 | 1 |
| 0 | 0 | 0 | 1 |
| 0 | 0 | 0 | 1 |
| 0 | 0 | 0 | 1 |
| 0 | 0 | 0 | 1 |
| 0 | 1 | 1 | 1 |
| 0 | 0 | 0 | 1 |
| 0 | 0 | 0 | 1 |
| 0 | 1 | 1 | 1 |

| | | | |
|---|---|---|---|
| 0 | 1 | 0 | 0 |
| 0 | 0 | 0 | 1 |
| 0 | 1 | 0 | 0 |
| 0 | 0 | 0 | 1 |
| 0 | 1 | 1 | 1 |
| 0 | 1 | 1 | 1 |
| 0 | 1 | 0 | 0 |
| 0 | 1 | 0 | 0 |
| 0 | 1 | 1 | 1 |
| 0 | 1 | 1 | 1 |
| 0 | 1 | 1 | 1 |
| 0 | 0 | 0 | 1 |
| 0 | 1 | 1 | 1 |
| 0 | 1 | 1 | 1 |
| 0 | 0 | 0 | 0 |
| 0 | 0 | 0 | 0 |
| 0 | 0 | 0 | 0 |
| 0 | 0 | 0 | 0 |
| 0 | 0 | 0 | 1 |
| 0 | 1 | 0 | 0 |
| 0 | 1 | 0 | 0 |
| 0 | 0 | 0 | 0 |
| 0 | 0 | 0 | 0 |
| 0 | 0 | 0 | 0 |
| 0 | 0 | 0 | 1 |
| 0 | 1 | 0 | 0 |
| 0 | 0 | 0 | 1 |
| 0 | 0 | 0 | 1 |
| 0 | 0 | 0 | 1 |
| 0 | 1 | 0 | 0 |
| 0 | 0 | 0 | 1 |
| 0 | 0 | 0 | 1 |
| 0 | 0 | 0 | 0 |
| 0 | 0 | 0 | 0 |
| 0 | 0 | 0 | 1 |
| 0 | 0 | 0 | 1 |
| 0 | 0 | 0 | 1 |
| 0 | 0 | 0 | 1 |
| 0 | 1 | 1 | 1 |
| 0 | 0 | 0 | 0 |
| 0 | 0 | 0 | 0 |
| 0 | 1 | 0 | 0 |
| 0 | 1 | 0 | 0 |
| 0 | 0 | 0 | 0 |
| 0 | 0 | 0 | 0 |
| 0 | 0 | 0 | 1 |
| 0 | 0 | 0 | 1 |
| 0 | 0 | 0 | 0 |
| 0 | 0 | 0 | 0 |
| 0 | 0 | 0 | 1 |
| 0 | 0 | 0 | 0 |
| 0 | 0 | 0 | 0 |
| 0 | 0 | 0 | 1 |
| 0 | 0 | 0 | 1 |

| | | | |
|---|---|---|---|
| 0 | 0 | 0 | 1 |
| 0 | 1 | 0 | 0 |
| 0 | 1 | 0 | 0 |
| 0 | 1 | 1 | 1 |
| 0 | 0 | 0 | 1 |
| 0 | 0 | 0 | 0 |
| 0 | 0 | 0 | 0 |
| 0 | 0 | 0 | 0 |
| 0 | 0 | 0 | 0 |
| 0 | 0 | 0 | 0 |
| 0 | 0 | 0 | 0 |
| 0 | 0 | 0 | 0 |
| 0 | 0 | 0 | 0 |
| 0 | 0 | 0 | 0 |
| 0 | 0 | 0 | 0 |
| 0 | 0 | 0 | 1 |
| 0 | 1 | 0 | 0 |
| 0 | 0 | 0 | 1 |
| 0 | 1 | 1 | 1 |
| 0 | 0 | 0 | 1 |
| 0 | 0 | 0 | 0 |
| 0 | 0 | 0 | 1 |
| 0 | 1 | 0 | 0 |
| 0 | 0 | 0 | 0 |
| 0 | 0 | 0 | 1 |
| 0 | 0 | 0 | 0 |
| 0 | 0 | 0 | 1 |
| 0 | 0 | 0 | 1 |
| 0 | 0 | 0 | 1 |
| 0 | 1 | 0 | 0 |
| 0 | 1 | 0 | 0 |
| 0 | 0 | 0 | 0 |
| 0 | 1 | 0 | 0 |
| 0 | 1 | 0 | 0 |
| 0 | 1 | 0 | 0 |
| 0 | 1 | 0 | 0 |
| 0 | 1 | 0 | 0 |
| 0 | 0 | 0 | 1 |
| 0 | 1 | 1 | 1 |
| 0 | 1 | 0 | 0 |
| 0 | 1 | 0 | 0 |
| 0 | 0 | 0 | 0 |
| 0 | 0 | 0 | 0 |
| 0 | 1 | 0 | 0 |
| 0 | 1 | 0 | 0 |
| 0 | 1 | 0 | 0 |
| 0 | 0 | 0 | 1 |
| 0 | 1 | 0 | 0 |
| 0 | 0 | 0 | 0 |
| 0 | 0 | 0 | 0 |
| 0 | 0 | 0 | 0 |
| 0 | 0 | 0 | 1 |
| 0 | 1 | 0 | 0 |
| 0 | 1 | 0 | 0 |

| | | | |
|---|---|---|---|
| 0 | 1 | 0 | 0 |
| 0 | 1 | 0 | 0 |
| 0 | 0 | 0 | 0 |
| 0 | 0 | 0 | 0 |
| 0 | 0 | 0 | 0 |
| 0 | 0 | 0 | 0 |
| 0 | 0 | 0 | 0 |
| 0 | 0 | 0 | 0 |
| 1 | 0 | 0 | 0 |
| 0 | 0 | 0 | 0 |
| 0 | 0 | 0 | 1 |
| 0 | 1 | 0 | 0 |
| 0 | 1 | 1 | 1 |
| 0 | 0 | 0 | 0 |
| 0 | 0 | 0 | 1 |
| 0 | 0 | 0 | 1 |
| 0 | 0 | 0 | 1 |
| 0 | 0 | 0 | 1 |
| 0 | 0 | 0 | 0 |
| 0 | 0 | 0 | 0 |
| 0 | 0 | 0 | 1 |
| 0 | 1 | 0 | 0 |
| 0 | 1 | 0 | 0 |
| 0 | 0 | 0 | 1 |
| 0 | 0 | 0 | 0 |
| 0 | 0 | 0 | 0 |
| 0 | 0 | 0 | 0 |
| 0 | 1 | 1 | 1 |
| 0 | 1 | 0 | 0 |
| 0 | 0 | 0 | 0 |
| 0 | 1 | 1 | 1 |
| 1 | 1 | 0 | 0 |
| 0 | 1 | 1 | 1 |
| 0 | 1 | 1 | 1 |
| 1 | 1 | 0 | 0 |
| 1 | 1 | 0 | 0 |
| 1 | 1 | 0 | 0 |
| 0 | 1 | 1 | 1 |
| 0 | 1 | 1 | 1 |
| 0 | 1 | 0 | 0 |
| 1 | 0 | 0 | 0 |
| 0 | 1 | 0 | 0 |
| 0 | 1 | 0 | 0 |
| 1 | 1 | 0 | 0 |
| 1 | 1 | 0 | 0 |
| 0 | 1 | 0 | 0 |
| 0 | 1 | 1 | 1 |
| 0 | 1 | 1 | 1 |
| 0 | 1 | 1 | 1 |
| 0 | 1 | 1 | 1 |
| 1 | 1 | 0 | 0 |
| 0 | 1 | 0 | 0 |
| 0 | 1 | 0 | 0 |
| 0 | 1 | 0 | 0 |

| | | | |
|---|---|---|---|
| 0 | 1 | 0 | 0 |
| 0 | 1 | 1 | 1 |
| 0 | 1 | 0 | 0 |
| 0 | 1 | 0 | 0 |
| 0 | 1 | 0 | 0 |
| 0 | 1 | 1 | 1 |
| 0 | 1 | 0 | 0 |
| 0 | 0 | 0 | 0 |
| 0 | 0 | 0 | 1 |
| 0 | 1 | 0 | 0 |
| 0 | 0 | 0 | 1 |
| 0 | 0 | 0 | 1 |
| 0 | 0 | 0 | 1 |
| 0 | 0 | 0 | 0 |
| 0 | 0 | 0 | 1 |
| 0 | 0 | 0 | 0 |
| 0 | 0 | 0 | 1 |
| 0 | 0 | 0 | 1 |
| 0 | 0 | 0 | 0 |
| 0 | 0 | 0 | 0 |
| 0 | 0 | 0 | 0 |
| 0 | 0 | 0 | 0 |
| 0 | 0 | 0 | 0 |
| 0 | 0 | 0 | 0 |
| 0 | 0 | 0 | 0 |
| 0 | 0 | 0 | 1 |
| 0 | 0 | 0 | 1 |
| 0 | 0 | 0 | 1 |
| 0 | 1 | 1 | 1 |
| 0 | 0 | 0 | 1 |
| 0 | 0 | 0 | 1 |
| 0 | 0 | 0 | 1 |
| 0 | 0 | 0 | 0 |
| 0 | 0 | 0 | 0 |
| 0 | 0 | 0 | 0 |
| 0 | 0 | 0 | 0 |
| 0 | 0 | 0 | 0 |
| 0 | 0 | 0 | 0 |
| 0 | 0 | 0 | 1 |
| 0 | 0 | 0 | 1 |
| 0 | 0 | 0 | 1 |
| 0 | 0 | 0 | 1 |
| 0 | 0 | 0 | 1 |
| 0 | 0 | 0 | 1 |
| 0 | 1 | 0 | 0 |
| 0 | 0 | 0 | 0 |
| 0 | 0 | 0 | 1 |
| 0 | 0 | 0 | 0 |
| 0 | 0 | 0 | 0 |
| 0 | 0 | 0 | 0 |
| 0 | 0 | 0 | 0 |
| 0 | 0 | 0 | 0 |
| 0 | 0 | 0 | 0 |
| 0 | 0 | 0 | 0 |

| | | | |
|---|---|---|---|
| 0 | 0 | 0 | 1 |
| 0 | 0 | 0 | 1 |
| 1 | 0 | 0 | 0 |
| 1 | 0 | 0 | 0 |
| 0 | 0 | 0 | 1 |
| 0 | 0 | 0 | 0 |
| 0 | 0 | 0 | 0 |
| 0 | 0 | 0 | 1 |
| 0 | 0 | 0 | 1 |
| 0 | 1 | 0 | 0 |
| 0 | 0 | 0 | 1 |
| 0 | 0 | 0 | 1 |
| 0 | 1 | 0 | 0 |
| 0 | 1 | 0 | 0 |
| 0 | 1 | 0 | 0 |
| 0 | 1 | 0 | 0 |
| 0 | 0 | 0 | 1 |
| 0 | 0 | 0 | 1 |
| 0 | 0 | 0 | 1 |
| 0 | 0 | 0 | 1 |
| 0 | 0 | 0 | 1 |
| 0 | 0 | 0 | 1 |
| 0 | 0 | 0 | 1 |
| 0 | 0 | 0 | 1 |
| 0 | 0 | 0 | 1 |
| 0 | 0 | 0 | 0 |
| 0 | 0 | 0 | 1 |
| 0 | 0 | 0 | 1 |
| 0 | 0 | 0 | 1 |
| 0 | 0 | 0 | 1 |
| 0 | 0 | 0 | 1 |
| 0 | 0 | 0 | 1 |
| 0 | 0 | 0 | 1 |
| 0 | 0 | 0 | 1 |
| 0 | 0 | 0 | 1 |
| 0 | 0 | 0 | 1 |
| 0 | 0 | 0 | 1 |
| 0 | 0 | 0 | 0 |
| 0 | 1 | 1 | 1 |
| 0 | 1 | 1 | 1 |
| 0 | 1 | 0 | 0 |
| 0 | 0 | 0 | 1 |
| 0 | 0 | 0 | 1 |
| 0 | 1 | 0 | 0 |
| 0 | 0 | 0 | 0 |
| 0 | 0 | 0 | 1 |
| 0 | 0 | 0 | 1 |
| 0 | 0 | 0 | 1 |
| 0 | 0 | 0 | 1 |
| 0 | 0 | 0 | 1 |
| 0 | 0 | 0 | 1 |
| 0 | 0 | 0 | 0 |
| 0 | 0 | 0 | 0 |
| 0 | 0 | 0 | 0 |

| | | | |
|---|---|---|---|
| 0 | 0 | 0 | 0 |
| 0 | 0 | 0 | 1 |
| 0 | 0 | 0 | 1 |
| 0 | 0 | 0 | 1 |
| 0 | 0 | 0 | 1 |
| 0 | 0 | 0 | 1 |
| 0 | 0 | 0 | 1 |
| 0 | 0 | 0 | 1 |
| 0 | 0 | 0 | 0 |
| 0 | 0 | 0 | 1 |
| 0 | 0 | 0 | 1 |
| 0 | 0 | 0 | 1 |
| 0 | 0 | 0 | 1 |
| 0 | 0 | 0 | 1 |
| 0 | 0 | 0 | 1 |
| 0 | 0 | 0 | 1 |
| 0 | 0 | 0 | 1 |
| 0 | 0 | 0 | 1 |
| 0 | 0 | 0 | 0 |
| 0 | 0 | 0 | 0 |
| 0 | 0 | 0 | 0 |
| 0 | 1 | 0 | 0 |
| 0 | 0 | 0 | 1 |
| 0 | 0 | 0 | 0 |
| 0 | 0 | 0 | 0 |
| 0 | 0 | 0 | 1 |
| 0 | 0 | 0 | 1 |
| 0 | 0 | 0 | 1 |
| 0 | 0 | 0 | 1 |
| 0 | 0 | 0 | 1 |
| 0 | 0 | 0 | 1 |
| 0 | 0 | 0 | 1 |
| 0 | 0 | 0 | 0 |
| 0 | 0 | 0 | 1 |
| 0 | 0 | 0 | 1 |
| 0 | 0 | 0 | 1 |
| 0 | 0 | 0 | 0 |
| 0 | 0 | 0 | 1 |
| 0 | 0 | 0 | 0 |
| 0 | 0 | 0 | 0 |
| 0 | 0 | 0 | 0 |
| 0 | 0 | 0 | 0 |
| 0 | 0 | 0 | 0 |
| 0 | 0 | 0 | 1 |
| 0 | 0 | 0 | 1 |
| 0 | 0 | 0 | 1 |
| 0 | 0 | 0 | 0 |
| 0 | 0 | 0 | 0 |
| 0 | 0 | 0 | 0 |
| 0 | 1 | 0 | 0 |
| 0 | 1 | 0 | 0 |
| 0 | 1 | 0 | 0 |
| 0 | 1 | 0 | 0 |
| 0 | 0 | 0 | 1 |

| | | | |
|---|---|---|---|
| 0 | 0 | 0 | 0 |
| 0 | 0 | 0 | 0 |
| 0 | 0 | 0 | 1 |
| 1 | 0 | 0 | 0 |
| 1 | 0 | 0 | 0 |
| 0 | 0 | 0 | 1 |
| 0 | 0 | 0 | 1 |
| 0 | 1 | 1 | 1 |
| 0 | 0 | 0 | 1 |
| 0 | 0 | 0 | 1 |
| 0 | 0 | 0 | 1 |
| 0 | 0 | 0 | 0 |
| 0 | 0 | 0 | 1 |
| 0 | 0 | 0 | 1 |
| 0 | 0 | 0 | 0 |
| 0 | 0 | 0 | 1 |
| 0 | 0 | 0 | 0 |
| 0 | 0 | 0 | 1 |
| 0 | 0 | 0 | 0 |
| 1 | 0 | 0 | 0 |
| 0 | 0 | 0 | 1 |
| 0 | 0 | 0 | 1 |
| 0 | 0 | 0 | 0 |
| 0 | 0 | 0 | 1 |
| 0 | 0 | 0 | 0 |
| 0 | 0 | 0 | 1 |
| 0 | 0 | 0 | 1 |
| 0 | 0 | 0 | 0 |
| 0 | 0 | 0 | 1 |
| 0 | 0 | 0 | 0 |
| 0 | 0 | 0 | 1 |
| 0 | 0 | 0 | 0 |
| 0 | 0 | 0 | 0 |
| 0 | 1 | 0 | 0 |
| 0 | 0 | 0 | 1 |
| 0 | 0 | 0 | 0 |
| 0 | 0 | 0 | 1 |
| 0 | 0 | 0 | 1 |
| 0 | 0 | 0 | 0 |
| 0 | 0 | 0 | 1 |
| 0 | 0 | 0 | 1 |
| 0 | 0 | 0 | 1 |
| 0 | 0 | 0 | 1 |
| 0 | 0 | 0 | 0 |
| 0 | 0 | 0 | 0 |
| 0 | 0 | 0 | 1 |
| 0 | 0 | 0 | 1 |
| 0 | 1 | 1 | 1 |
| 0 | 1 | 0 | 0 |
| 0 | 1 | 1 | 1 |
| 0 | 0 | 0 | 0 |
| 0 | 0 | 0 | 1 |
| 0 | 0 | 0 | 1 |
| 0 | 0 | 0 | 0 |

| | | | |
|---|---|---|---|
| 0 | 0 | 0 | 1 |
| 0 | 0 | 0 | 1 |
| 0 | 0 | 0 | 1 |
| 0 | 0 | 0 | 1 |
| 0 | 0 | 0 | 1 |
| 0 | 0 | 0 | 1 |
| 0 | 0 | 0 | 1 |
| 0 | 0 | 0 | 0 |
| 0 | 0 | 0 | 1 |
| 0 | 0 | 0 | 1 |
| 0 | 0 | 0 | 1 |
| 0 | 0 | 0 | 1 |
| 0 | 0 | 0 | 0 |
| 0 | 0 | 0 | 1 |
| 0 | 0 | 0 | 0 |
| 0 | 0 | 0 | 0 |
| 0 | 0 | 0 | 1 |
| 0 | 0 | 0 | 1 |
| 0 | 0 | 0 | 1 |
| 0 | 0 | 0 | 1 |
| 0 | 0 | 0 | 0 |
| 0 | 0 | 0 | 0 |
| 0 | 0 | 0 | 1 |
| 0 | 0 | 0 | 1 |
| 0 | 0 | 0 | 0 |
| 0 | 0 | 0 | 1 |
| 0 | 0 | 0 | 0 |
| 0 | 0 | 0 | 0 |
| 0 | 0 | 0 | 0 |
| 0 | 0 | 0 | 0 |
| 0 | 0 | 0 | 1 |
| 0 | 0 | 0 | 1 |
| 0 | 1 | 0 | 0 |
| 0 | 1 | 0 | 0 |
| 0 | 1 | 0 | 0 |
| 0 | 1 | 0 | 0 |
| 0 | 1 | 0 | 0 |
| 0 | 0 | 0 | 0 |
| 0 | 1 | 0 | 0 |
| 0 | 1 | 1 | 1 |
| 0 | 0 | 0 | 1 |
| 0 | 0 | 0 | 1 |
| 0 | 0 | 0 | 1 |
| 0 | 0 | 0 | 1 |
| 0 | 0 | 0 | 1 |
| 0 | 0 | 0 | 0 |
| 0 | 0 | 0 | 1 |
| 0 | 0 | 0 | 1 |
| 0 | 0 | 0 | 1 |
| 0 | 0 | 0 | 0 |
| 0 | 0 | 0 | 0 |
| 0 | 0 | 0 | 1 |
| 0 | 0 | 0 | 1 |
| 0 | 0 | 0 | 1 |

| | | | |
|---|---|---|---|
| 0 | 0 | 0 | 1 |
| 0 | 0 | 0 | 1 |
| 0 | 0 | 0 | 1 |
| 0 | 0 | 0 | 1 |
| 0 | 0 | 0 | 0 |
| 0 | 0 | 0 | 1 |
| 0 | 0 | 0 | 0 |
| 0 | 0 | 0 | 0 |
| 0 | 0 | 0 | 0 |
| 0 | 0 | 0 | 0 |
| 0 | 0 | 0 | 0 |
| 0 | 0 | 0 | 1 |
| 0 | 0 | 0 | 0 |
| 0 | 0 | 0 | 0 |
| 0 | 0 | 0 | 1 |
| 0 | 0 | 0 | 1 |
| 0 | 0 | 0 | 1 |
| 0 | 0 | 0 | 1 |
| 0 | 0 | 0 | 1 |
| 0 | 0 | 0 | 1 |
| 0 | 0 | 0 | 1 |
| 0 | 0 | 0 | 1 |
| 0 | 0 | 0 | 1 |
| 0 | 0 | 0 | 0 |
| 0 | 0 | 0 | 1 |
| 0 | 0 | 0 | 1 |
| 0 | 0 | 0 | 1 |
| 0 | 0 | 0 | 0 |
| 0 | 1 | 1 | 1 |
| 0 | 1 | 1 | 1 |
| 0 | 1 | 0 | 0 |
| 0 | 1 | 0 | 0 |
| 0 | 0 | 0 | 1 |
| 0 | 0 | 0 | 0 |
| 0 | 0 | 0 | 1 |
| 0 | 0 | 0 | 1 |
| 0 | 0 | 0 | 1 |
| 0 | 0 | 0 | 1 |
| 0 | 0 | 0 | 1 |
| 0 | 0 | 0 | 1 |
| 0 | 0 | 0 | 1 |
| 0 | 0 | 0 | 1 |
| 0 | 0 | 0 | 1 |
| 0 | 0 | 0 | 1 |
| 0 | 0 | 0 | 1 |
| 0 | 0 | 0 | 1 |
| 0 | 0 | 0 | 1 |
| 0 | 0 | 0 | 1 |
| 0 | 0 | 0 | 0 |
| 0 | 0 | 0 | 1 |
| 0 | 0 | 0 | 1 |
| 0 | 1 | 1 | 1 |
| 0 | 1 | 1 | 1 |
| 0 | 0 | 0 | 1 |

| | | | |
|---|---|---|---|
| 0 | 0 | 0 | 1 |
| 0 | 0 | 0 | 1 |
| 0 | 0 | 0 | 1 |
| 0 | 0 | 0 | 1 |
| 0 | 1 | 1 | 1 |
| 0 | 0 | 0 | 1 |
| 0 | 0 | 0 | 1 |
| 0 | 0 | 0 | 1 |
| 0 | 0 | 0 | 1 |
| 0 | 0 | 0 | 1 |
| 0 | 0 | 0 | 0 |
| 0 | 0 | 0 | 0 |
| 0 | 0 | 0 | 0 |
| 1 | 1 | 0 | 0 |
| 1 | 0 | 0 | 0 |
| 0 | 1 | 1 | 1 |
| 1 | 0 | 0 | 0 |
| 0 | 0 | 0 | 0 |
| 0 | 0 | 0 | 0 |
| 0 | 0 | 0 | 0 |
| 0 | 0 | 0 | 0 |
| 1 | 0 | 0 | 0 |
| 0 | 0 | 0 | 1 |
| 0 | 0 | 0 | 0 |
| 1 | 0 | 0 | 0 |
| 1 | 0 | 0 | 0 |
| 0 | 0 | 0 | 1 |
| 0 | 0 | 0 | 0 |
| 0 | 0 | 0 | 1 |
| 0 | 0 | 0 | 1 |
| 0 | 0 | 0 | 1 |
| 0 | 0 | 0 | 0 |
| 0 | 0 | 0 | 1 |
| 0 | 0 | 0 | 1 |
| 0 | 0 | 0 | 0 |
| 0 | 0 | 0 | 0 |
| 0 | 0 | 0 | 0 |
| 0 | 1 | 0 | 0 |
| 0 | 0 | 0 | 0 |
| 0 | 1 | 1 | 1 |
| 0 | 1 | 1 | 1 |
| 0 | 1 | 0 | 0 |
| 0 | 1 | 1 | 1 |
| 0 | 0 | 0 | 0 |
| 0 | 0 | 0 | 1 |
| 0 | 1 | 1 | 1 |
| 0 | 0 | 0 | 1 |
| 0 | 1 | 0 | 0 |
| 0 | 1 | 0 | 0 |
| 0 | 1 | 0 | 0 |
| 0 | 1 | 0 | 0 |
| 0 | 0 | 0 | 1 |
| 0 | 0 | 0 | 0 |
| 0 | 0 | 0 | 1 |

| | | | |
|---|---|---|---|
| 0 | 0 | 0 | 1 |
| 0 | 0 | 0 | 1 |
| 0 | 0 | 0 | 1 |
| 0 | 0 | 0 | 1 |
| 0 | 0 | 0 | 1 |
| 0 | 0 | 0 | 0 |
| 0 | 0 | 0 | 0 |
| 0 | 0 | 0 | 0 |
| 0 | 0 | 0 | 0 |
| 0 | 0 | 0 | 1 |
| 1 | 1 | 0 | 0 |
| 1 | 0 | 0 | 0 |
| 0 | 0 | 0 | 0 |
| 0 | 0 | 0 | 0 |
| 0 | 0 | 0 | 0 |
| 0 | 0 | 0 | 0 |
| 0 | 0 | 0 | 1 |
| 0 | 0 | 0 | 1 |
| 0 | 0 | 0 | 1 |
| 0 | 0 | 0 | 1 |
| 0 | 0 | 0 | 1 |
| 0 | 0 | 0 | 1 |
| 0 | 0 | 0 | 1 |
| 0 | 0 | 0 | 1 |
| 0 | 0 | 0 | 1 |
| 0 | 0 | 0 | 1 |
| 0 | 0 | 0 | 1 |
| 0 | 0 | 0 | 1 |
| 0 | 1 | 0 | 0 |
| 0 | 0 | 0 | 1 |
| 0 | 0 | 0 | 1 |
| 0 | 1 | 1 | 1 |
| 0 | 1 | 1 | 1 |
| 0 | 1 | 0 | 0 |
| 0 | 0 | 0 | 1 |
| 0 | 1 | 0 | 0 |
| 0 | 1 | 1 | 1 |
| 0 | 0 | 0 | 0 |
| 0 | 0 | 0 | 1 |
| 0 | 0 | 0 | 0 |
| 0 | 0 | 0 | 1 |
| 0 | 0 | 0 | 1 |
| 0 | 0 | 0 | 1 |
| 0 | 0 | 0 | 0 |
| 0 | 0 | 0 | 1 |
| 0 | 0 | 0 | 1 |
| 0 | 0 | 0 | 1 |
| 0 | 0 | 0 | 1 |
| 0 | 0 | 0 | 1 |
| 0 | 0 | 0 | 1 |
| 0 | 0 | 0 | 1 |
| 0 | 1 | 1 | 1 |
| 0 | 0 | 0 | 0 |
| 0 | 0 | 0 | 0 |

| | | | |
|---|---|---|---|
| 0 | 0 | 0 | 1 |
| 0 | 0 | 0 | 0 |
| 0 | 0 | 0 | 1 |
| 0 | 0 | 0 | 1 |
| 0 | 1 | 0 | 0 |
| 0 | 0 | 0 | 0 |
| 0 | 0 | 0 | 0 |
| 0 | 0 | 0 | 1 |
| 0 | 0 | 0 | 1 |
| 0 | 0 | 0 | 0 |
| 0 | 0 | 0 | 0 |
| 0 | 0 | 0 | 0 |
| 0 | 0 | 0 | 0 |
| 0 | 0 | 0 | 0 |
| 0 | 0 | 0 | 1 |
| 0 | 0 | 0 | 0 |
| 0 | 1 | 0 | 0 |
| 0 | 0 | 0 | 0 |
| 0 | 0 | 0 | 1 |
| 0 | 0 | 0 | 1 |
| 0 | 0 | 0 | 1 |
| 0 | 0 | 0 | 1 |
| 0 | 0 | 0 | 1 |
| 0 | 0 | 0 | 0 |
| 0 | 0 | 0 | 1 |
| 0 | 0 | 0 | 1 |
| 0 | 0 | 0 | 0 |
| 0 | 0 | 0 | 0 |
| 0 | 0 | 0 | 1 |
| 0 | 0 | 0 | 1 |
| 0 | 0 | 0 | 0 |
| 0 | 0 | 0 | 1 |
| 0 | 0 | 0 | 1 |
| 0 | 0 | 0 | 1 |
| 0 | 0 | 0 | 1 |
| 0 | 0 | 0 | 0 |
| 0 | 0 | 0 | 1 |
| 0 | 0 | 0 | 0 |
| 0 | 1 | 0 | 0 |
| 0 | 0 | 0 | 1 |
| 0 | 0 | 0 | 1 |
| 0 | 0 | 0 | 0 |
| 0 | 0 | 0 | 0 |
| 0 | 0 | 0 | 1 |
| 0 | 0 | 0 | 0 |
| 0 | 0 | 0 | 0 |
| 0 | 0 | 0 | 0 |
| 0 | 0 | 0 | 1 |
| 0 | 0 | 0 | 0 |
| 0 | 0 | 0 | 0 |
| 0 | 0 | 0 | 0 |
| 0 | 0 | 0 | 1 |
| 0 | 0 | 0 | 0 |
| 0 | 0 | 0 | 1 |

| | | | |
|---|---|---|---|
| 0 | 0 | 0 | 1 |
| 0 | 0 | 0 | 1 |
| 0 | 0 | 0 | 1 |
| 0 | 0 | 0 | 1 |
| 0 | 0 | 0 | 0 |
| 0 | 0 | 0 | 0 |
| 0 | 0 | 0 | 1 |
| 0 | 1 | 1 | 1 |
| 0 | 0 | 0 | 1 |
| 0 | 1 | 0 | 0 |
| 0 | 1 | 1 | 1 |
| 0 | 1 | 1 | 1 |
| 0 | 1 | 0 | 0 |
| 0 | 0 | 0 | 1 |
| 0 | 0 | 0 | 1 |
| 0 | 0 | 0 | 1 |
| 0 | 0 | 0 | 1 |
| 0 | 0 | 0 | 1 |
| 0 | 0 | 0 | 1 |
| 0 | 0 | 0 | 1 |
| 0 | 0 | 0 | 1 |
| 0 | 0 | 0 | 1 |
| 0 | 0 | 0 | 1 |
| 0 | 1 | 0 | 0 |
| 0 | 0 | 0 | 1 |
| 1 | 0 | 0 | 0 |
| 0 | 0 | 0 | 1 |
| 0 | 0 | 0 | 0 |
| 0 | 0 | 0 | 1 |
| 0 | 0 | 0 | 0 |
| 0 | 0 | 0 | 0 |
| 0 | 0 | 0 | 0 |
| 0 | 0 | 0 | 0 |
| 0 | 0 | 0 | 0 |
| 0 | 0 | 0 | 1 |
| 0 | 0 | 0 | 0 |
| 0 | 0 | 0 | 1 |
| 0 | 0 | 0 | 0 |
| 0 | 0 | 0 | 1 |
| 0 | 0 | 0 | 1 |
| 0 | 0 | 0 | 0 |
| 0 | 0 | 0 | 0 |
| 0 | 0 | 0 | 0 |
| 0 | 0 | 0 | 0 |
| 0 | 0 | 0 | 0 |
| 0 | 0 | 0 | 0 |
| 0 | 0 | 0 | 0 |
| 0 | 1 | 0 | 0 |
| 0 | 0 | 0 | 1 |
| 0 | 0 | 0 | 0 |
| 0 | 1 | 1 | 1 |
| 0 | 1 | 1 | 1 |
| 0 | 0 | 0 | 1 |
| 0 | 0 | 0 | 1 |

| | | | |
|---|---|---|---|
| 0 | 0 | 0 | 1 |
| 0 | 0 | 0 | 1 |
| 0 | 0 | 0 | 1 |
| 0 | 0 | 0 | 1 |
| 0 | 0 | 0 | 1 |
| 0 | 0 | 0 | 1 |
| 0 | 0 | 0 | 0 |
| 0 | 0 | 0 | 0 |
| 0 | 0 | 0 | 0 |
| 0 | 0 | 0 | 0 |
| 0 | 0 | 0 | 0 |
| 0 | 0 | 0 | 0 |
| 0 | 0 | 0 | 0 |
| 0 | 0 | 0 | 0 |
| 0 | 0 | 0 | 0 |
| 0 | 0 | 0 | 0 |
| 1 | 0 | 0 | 0 |
| 0 | 0 | 0 | 0 |
| 0 | 0 | 0 | 1 |
| 0 | 0 | 0 | 0 |
| 0 | 0 | 0 | 1 |
| 0 | 0 | 0 | 1 |
| 0 | 0 | 0 | 1 |
| 0 | 0 | 0 | 1 |
| 0 | 0 | 0 | 1 |
| 0 | 0 | 0 | 0 |
| 0 | 0 | 0 | 0 |
| 0 | 0 | 0 | 0 |
| 0 | 0 | 0 | 0 |
| 0 | 0 | 0 | 0 |
| 0 | 0 | 0 | 0 |
| 1 | 0 | 0 | 0 |
| 0 | 0 | 0 | 0 |
| 1 | 0 | 0 | 0 |
| 0 | 0 | 0 | 1 |
| 1 | 0 | 0 | 0 |
| 1 | 0 | 0 | 0 |
| 0 | 0 | 0 | 0 |
| 1 | 0 | 0 | 0 |
| 0 | 0 | 0 | 1 |
| 0 | 0 | 0 | 0 |
| 0 | 0 | 0 | 0 |
| 0 | 0 | 0 | 1 |
| 0 | 0 | 0 | 1 |
| 0 | 0 | 0 | 1 |
| 0 | 0 | 0 | 1 |
| 0 | 0 | 0 | 1 |
| 0 | 0 | 0 | 0 |
| 0 | 0 | 0 | 0 |
| 0 | 0 | 0 | 0 |
| 0 | 0 | 0 | 0 |
| 0 | 0 | 0 | 1 |
| 0 | 0 | 0 | 0 |
| 0 | 0 | 0 | 1 |

| | | | |
|---|---|---|---|
| 0 | 0 | 0 | 1 |
| 0 | 0 | 0 | 1 |
| 0 | 0 | 0 | 1 |
| 0 | 0 | 0 | 1 |
| 0 | 0 | 0 | 1 |
| 0 | 0 | 0 | 0 |
| 0 | 0 | 0 | 1 |
| 0 | 0 | 0 | 1 |
| 0 | 0 | 0 | 1 |
| 0 | 0 | 0 | 1 |
| 0 | 1 | 0 | 0 |
| 0 | 1 | 0 | 0 |
| 0 | 1 | 0 | 0 |
| 0 | 1 | 1 | 1 |
| 0 | 1 | 1 | 1 |
| 0 | 0 | 0 | 1 |
| 0 | 1 | 1 | 1 |
| 0 | 0 | 0 | 0 |
| 0 | 0 | 0 | 1 |
| 0 | 0 | 0 | 1 |
| 0 | 0 | 0 | 1 |
| 0 | 0 | 0 | 1 |
| 0 | 0 | 0 | 0 |
| 0 | 0 | 0 | 1 |
| 0 | 0 | 0 | 0 |
| 0 | 0 | 0 | 0 |
| 0 | 0 | 0 | 0 |
| 0 | 0 | 0 | 0 |
| 0 | 0 | 0 | 1 |
| 0 | 0 | 0 | 0 |
| 0 | 0 | 0 | 1 |
| 0 | 0 | 0 | 1 |
| 0 | 0 | 0 | 1 |
| 0 | 0 | 0 | 1 |
| 0 | 0 | 0 | 1 |
| 0 | 0 | 0 | 1 |
| 0 | 0 | 0 | 1 |
| 0 | 0 | 0 | 0 |
| 0 | 0 | 0 | 0 |
| 0 | 0 | 0 | 0 |
| 0 | 1 | 0 | 0 |
| 0 | 0 | 0 | 0 |
| 0 | 0 | 0 | 1 |
| 0 | 0 | 0 | 0 |
| 0 | 0 | 0 | 0 |
| 0 | 0 | 0 | 1 |
| 0 | 0 | 0 | 0 |
| 0 | 1 | 0 | 0 |
| 0 | 0 | 0 | 0 |
| 1 | 0 | 0 | 0 |
| 0 | 0 | 0 | 1 |
| 0 | 0 | 0 | 1 |
| 0 | 0 | 0 | 1 |
| 0 | 1 | 0 | 0 |

| | | | |
|---|---|---|---|
| 0 | 0 | 0 | 0 |
| 0 | 0 | 0 | 1 |
| 0 | 1 | 0 | 0 |
| 0 | 0 | 0 | 1 |
| 0 | 1 | 1 | 1 |
| 0 | 1 | 1 | 1 |
| 0 | 1 | 1 | 1 |
| 0 | 0 | 0 | 1 |
| 0 | 1 | 0 | 0 |
| 0 | 0 | 0 | 1 |
| 0 | 1 | 0 | 0 |
| 0 | 1 | 0 | 0 |
| 0 | 1 | 0 | 0 |
| 0 | 1 | 0 | 0 |
| 0 | 0 | 0 | 1 |
| 0 | 0 | 0 | 1 |
| 0 | 0 | 0 | 0 |
| 0 | 0 | 0 | 0 |
| 0 | 0 | 0 | 1 |
| 0 | 0 | 0 | 1 |
| 0 | 0 | 0 | 0 |
| 0 | 1 | 1 | 1 |
| 0 | 1 | 1 | 1 |
| 0 | 0 | 0 | 1 |
| 0 | 1 | 0 | 0 |
| 0 | 1 | 1 | 1 |
| 0 | 1 | 0 | 0 |
| 0 | 1 | 0 | 0 |
| 0 | 1 | 0 | 0 |
| 0 | 0 | 0 | 1 |
| 0 | 0 | 0 | 0 |
| 0 | 0 | 0 | 0 |
| 0 | 0 | 0 | 0 |
| 0 | 0 | 0 | 1 |
| 0 | 0 | 0 | 0 |
| 0 | 0 | 0 | 1 |
| 0 | 0 | 0 | 0 |
| 0 | 0 | 0 | 0 |
| 0 | 1 | 0 | 0 |
| 0 | 0 | 0 | 0 |
| 0 | 0 | 0 | 0 |
| 0 | 0 | 0 | 1 |
| 0 | 1 | 0 | 0 |
| 0 | 1 | 0 | 0 |
| 0 | 1 | 0 | 0 |
| 1 | 0 | 0 | 0 |
| 0 | 0 | 0 | 0 |
| 1 | 0 | 0 | 0 |
| 1 | 0 | 0 | 0 |
| 1 | 0 | 0 | 0 |
| 0 | 0 | 0 | 0 |
| 0 | 0 | 0 | 0 |
| 0 | 0 | 0 | 1 |
| 0 | 0 | 0 | 0 |

| | | | |
|---|---|---|---|
| 0 | 0 | 0 | 1 |
| 0 | 0 | 0 | 1 |
| 0 | 0 | 0 | 1 |
| 0 | 0 | 0 | 1 |
| 0 | 0 | 0 | 1 |
| 0 | 0 | 0 | 1 |
| 0 | 0 | 0 | 1 |
| 0 | 0 | 0 | 1 |
| 0 | 0 | 0 | 1 |
| 0 | 0 | 0 | 0 |
| 0 | 0 | 0 | 0 |
| 0 | 0 | 0 | 1 |
| 0 | 0 | 0 | 1 |
| 0 | 0 | 0 | 1 |
| 0 | 0 | 0 | 0 |
| 0 | 0 | 0 | 1 |
| 0 | 0 | 0 | 1 |
| 0 | 0 | 0 | 0 |
| 0 | 0 | 0 | 1 |
| 0 | 0 | 0 | 1 |
| 0 | 0 | 0 | 1 |
| 0 | 1 | 0 | 0 |
| 0 | 0 | 0 | 0 |
| 0 | 1 | 0 | 0 |
| 0 | 1 | 0 | 0 |
| 0 | 1 | 0 | 0 |
| 0 | 1 | 0 | 0 |
| 0 | 0 | 0 | 0 |
| 0 | 0 | 0 | 1 |
| 0 | 0 | 0 | 0 |
| 1 | 0 | 0 | 0 |
| 0 | 0 | 0 | 1 |
| 0 | 0 | 0 | 1 |
| 0 | 0 | 0 | 1 |
| 0 | 0 | 0 | 1 |
| 0 | 0 | 0 | 1 |
| 0 | 0 | 0 | 1 |
| 0 | 0 | 0 | 1 |
| 0 | 0 | 0 | 1 |
| 0 | 0 | 0 | 1 |
| 0 | 0 | 0 | 1 |
| 0 | 0 | 0 | 1 |
| 0 | 0 | 0 | 1 |
| 0 | 0 | 0 | 0 |
| 0 | 0 | 0 | 1 |
| 0 | 0 | 0 | 0 |
| 0 | 0 | 0 | 1 |
| 0 | 0 | 0 | 1 |
| 0 | 0 | 0 | 1 |
| 0 | 0 | 0 | 1 |
| 0 | 1 | 0 | 0 |
| 0 | 0 | 0 | 1 |
| 0 | 0 | 0 | 1 |
| 0 | 0 | 0 | 0 |

| | | | |
|---|---|---|---|
| 0 | 1 | 0 | 0 |
| 0 | 0 | 0 | 0 |
| 0 | 0 | 0 | 0 |
| 0 | 1 | 0 | 0 |
| 0 | 0 | 0 | 1 |
| 0 | 0 | 0 | 1 |
| 0 | 0 | 0 | 1 |
| 0 | 0 | 0 | 0 |
| 0 | 1 | 0 | 0 |
| 0 | 1 | 0 | 0 |
| 0 | 1 | 0 | 0 |
| 0 | 0 | 0 | 1 |
| 0 | 0 | 0 | 1 |
| 0 | 0 | 0 | 1 |
| 0 | 0 | 0 | 1 |
| 0 | 0 | 0 | 0 |
| 0 | 0 | 0 | 0 |
| 0 | 0 | 0 | 1 |
| 0 | 0 | 0 | 1 |
| 0 | 0 | 0 | 1 |
| 0 | 0 | 0 | 1 |
| 0 | 1 | 0 | 0 |
| 0 | 0 | 0 | 1 |
| 0 | 0 | 0 | 1 |
| 0 | 1 | 0 | 0 |
| 0 | 0 | 0 | 0 |
| 0 | 0 | 0 | 0 |
| 0 | 0 | 0 | 0 |
| 0 | 0 | 0 | 1 |
| 0 | 0 | 0 | 1 |
| 0 | 0 | 0 | 1 |
| 0 | 0 | 0 | 0 |
| 0 | 0 | 0 | 1 |
| 0 | 0 | 0 | 1 |
| 0 | 0 | 0 | 1 |
| 0 | 0 | 0 | 1 |
| 0 | 0 | 0 | 1 |
| 0 | 0 | 0 | 1 |
| 0 | 0 | 0 | 1 |
| 0 | 0 | 0 | 1 |
| 0 | 0 | 0 | 1 |
| 0 | 0 | 0 | 1 |
| 0 | 0 | 0 | 0 |
| 0 | 0 | 0 | 1 |
| 0 | 0 | 0 | 1 |
| 0 | 0 | 0 | 1 |
| 0 | 0 | 0 | 1 |
| 0 | 0 | 0 | 1 |
| 0 | 0 | 0 | 1 |
| 0 | 0 | 0 | 0 |
| 0 | 0 | 0 | 1 |
| 0 | 0 | 0 | 1 |
| 0 | 0 | 0 | 1 |
| 0 | 0 | 0 | 1 |

| | | | |
|---|---|---|---|
| 0 | 0 | 0 | 0 |
| 0 | 0 | 0 | 1 |
| 0 | 1 | 1 | 1 |
| 0 | 0 | 0 | 1 |
| 0 | 0 | 0 | 1 |
| 0 | 0 | 0 | 1 |
| 0 | 0 | 0 | 1 |
| 0 | 1 | 0 | 0 |
| 0 | 0 | 0 | 1 |
| 0 | 0 | 0 | 1 |
| 0 | 0 | 0 | 0 |
| 0 | 0 | 0 | 0 |
| 0 | 1 | 0 | 0 |
| 0 | 0 | 0 | 1 |
| 0 | 0 | 0 | 0 |
| 0 | 0 | 0 | 0 |
| 0 | 0 | 0 | 0 |
| 0 | 0 | 0 | 0 |
| 0 | 0 | 0 | 1 |
| 0 | 0 | 0 | 0 |
| 0 | 0 | 0 | 0 |
| 0 | 0 | 0 | 0 |
| 0 | 0 | 0 | 0 |
| 0 | 0 | 0 | 0 |
| 0 | 0 | 0 | 0 |
| 0 | 0 | 0 | 0 |
| 0 | 0 | 0 | 0 |
| 0 | 0 | 0 | 0 |
| 0 | 0 | 0 | 0 |
| 0 | 0 | 0 | 0 |
| 0 | 0 | 0 | 0 |
| 0 | 0 | 0 | 0 |
| 0 | 0 | 0 | 1 |
| 0 | 0 | 0 | 0 |
| 0 | 0 | 0 | 1 |
| 0 | 0 | 0 | 0 |
| 0 | 0 | 0 | 0 |
| 0 | 0 | 0 | 1 |
| 0 | 0 | 0 | 1 |
| 0 | 0 | 0 | 1 |
| 0 | 0 | 0 | 1 |
| 0 | 0 | 0 | 0 |
| 0 | 0 | 0 | 0 |
| 0 | 0 | 0 | 0 |
| 0 | 0 | 0 | 1 |
| 0 | 0 | 0 | 0 |
| 0 | 0 | 0 | 1 |
| 0 | 0 | 0 | 0 |
| 0 | 0 | 0 | 1 |
| 0 | 0 | 0 | 1 |
| 0 | 0 | 0 | 1 |
| 0 | 0 | 0 | 1 |
| 0 | 0 | 0 | 1 |
| 0 | 0 | 0 | 1 |

| | | | |
|---|---|---|---|
| 0 | 0 | 0 | 1 |
| 0 | 0 | 0 | 1 |
| 0 | 0 | 0 | 1 |
| 0 | 0 | 0 | 1 |
| 0 | 0 | 0 | 1 |
| 0 | 0 | 0 | 0 |
| 0 | 0 | 0 | 0 |
| 0 | 0 | 0 | 1 |
| 0 | 0 | 0 | 0 |
| 0 | 0 | 0 | 0 |
| 0 | 0 | 0 | 1 |
| 0 | 0 | 0 | 1 |
| 0 | 0 | 0 | 1 |
| 0 | 0 | 0 | 1 |
| 0 | 0 | 0 | 1 |
| 0 | 1 | 1 | 1 |
| 0 | 0 | 0 | 1 |
| 0 | 0 | 0 | 1 |
| 0 | 1 | 0 | 0 |
| 0 | 1 | 0 | 0 |
| 0 | 1 | 1 | 1 |
| 0 | 0 | 0 | 1 |
| 0 | 0 | 0 | 1 |
| 0 | 0 | 0 | 0 |
| 0 | 0 | 0 | 1 |
| 0 | 0 | 0 | 0 |
| 0 | 0 | 0 | 0 |
| 0 | 0 | 0 | 1 |
| 0 | 0 | 0 | 1 |
| 0 | 0 | 0 | 1 |
| 0 | 1 | 0 | 0 |
| 0 | 1 | 1 | 1 |
| 0 | 1 | 0 | 0 |
| 0 | 1 | 0 | 0 |
| 0 | 1 | 0 | 0 |
| 0 | 1 | 0 | 0 |
| 0 | 1 | 0 | 0 |
| 0 | 0 | 0 | 1 |
| 0 | 1 | 1 | 1 |
| 0 | 0 | 0 | 1 |
| 0 | 1 | 0 | 0 |
| 0 | 1 | 0 | 0 |
| 0 | 1 | 0 | 0 |
| 0 | 0 | 0 | 0 |
| 0 | 0 | 0 | 1 |
| 0 | 0 | 0 | 1 |
| 0 | 0 | 0 | 1 |
| 0 | 0 | 0 | 0 |
| 0 | 0 | 0 | 0 |
| 0 | 0 | 0 | 0 |
| 0 | 0 | 0 | 1 |
| 0 | 0 | 0 | 1 |
| 0 | 0 | 0 | 1 |
| 0 | 0 | 0 | 1 |

| | | | |
|---|---|---|---|
| 0 | 0 | 0 | 1 |
| 0 | 0 | 0 | 1 |
| 0 | 0 | 0 | 1 |
| 0 | 0 | 0 | 0 |
| 0 | 0 | 0 | 0 |
| 0 | 0 | 0 | 1 |
| 0 | 1 | 0 | 0 |
| 0 | 0 | 0 | 1 |
| 0 | 0 | 0 | 1 |
| 0 | 0 | 0 | 1 |
| 0 | 0 | 0 | 1 |
| 0 | 0 | 0 | 0 |
| 0 | 1 | 1 | 1 |
| 0 | 0 | 0 | 0 |
| 0 | 0 | 0 | 1 |
| 0 | 0 | 0 | 0 |
| 0 | 1 | 1 | 1 |
| 0 | 1 | 1 | 1 |
| 0 | 0 | 0 | 1 |
| 0 | 1 | 0 | 0 |
| 0 | 0 | 0 | 0 |
| 0 | 0 | 0 | 0 |
| 0 | 0 | 0 | 0 |
| 0 | 0 | 0 | 0 |
| 0 | 0 | 0 | 0 |
| 0 | 0 | 0 | 1 |
| 0 | 0 | 0 | 0 |
| 0 | 0 | 0 | 1 |
| 0 | 0 | 0 | 1 |
| 0 | 0 | 0 | 1 |
| 0 | 0 | 0 | 1 |
| 0 | 0 | 0 | 1 |
| 0 | 0 | 0 | 0 |
| 0 | 1 | 0 | 0 |
| 0 | 0 | 0 | 1 |
| 0 | 0 | 0 | 1 |
| 0 | 1 | 0 | 0 |
| 0 | 1 | 0 | 0 |
| 0 | 1 | 0 | 0 |
| 0 | 1 | 1 | 1 |
| 0 | 0 | 0 | 0 |
| 0 | 1 | 0 | 0 |
| 0 | 0 | 0 | 1 |
| 0 | 0 | 0 | 0 |
| 0 | 0 | 0 | 1 |
| 0 | 0 | 0 | 1 |
| 0 | 0 | 0 | 1 |
| 0 | 0 | 0 | 1 |
| 0 | 0 | 0 | 1 |
| 1 | 0 | 0 | 0 |
| 0 | 0 | 0 | 1 |
| 0 | 0 | 0 | 1 |
| 0 | 0 | 0 | 1 |
| 0 | 0 | 0 | 1 |

| | | | |
|---|---|---|---|
| 0 | 0 | 0 | 1 |
| 0 | 0 | 0 | 1 |
| 0 | 0 | 0 | 1 |
| 0 | 0 | 0 | 0 |
| 0 | 0 | 0 | 1 |
| 0 | 0 | 0 | 1 |
| 0 | 0 | 0 | 1 |
| 0 | 0 | 0 | 0 |
| 0 | 0 | 0 | 1 |
| 0 | 0 | 0 | 1 |
| 0 | 0 | 0 | 1 |
| 0 | 0 | 0 | 1 |
| 0 | 0 | 0 | 1 |
| 0 | 0 | 0 | 1 |
| 0 | 0 | 0 | 1 |
| 0 | 0 | 0 | 1 |
| 0 | 0 | 0 | 0 |
| 0 | 0 | 0 | 0 |
| 0 | 0 | 0 | 1 |
| 0 | 0 | 0 | 1 |
| 0 | 0 | 0 | 1 |
| 0 | 0 | 0 | 1 |
| 0 | 0 | 0 | 1 |
| 0 | 0 | 0 | 1 |
| 0 | 0 | 0 | 1 |
| 0 | 0 | 0 | 1 |
| 0 | 0 | 0 | 1 |
| 0 | 1 | 0 | 0 |
| 0 | 1 | 0 | 0 |
| 0 | 0 | 0 | 1 |
| 0 | 0 | 0 | 1 |
| 0 | 0 | 0 | 0 |
| 0 | 0 | 0 | 1 |
| 0 | 0 | 0 | 0 |
| 1 | 0 | 0 | 0 |
| 1 | 0 | 0 | 0 |
| 1 | 0 | 0 | 0 |
| 1 | 0 | 0 | 0 |
| 0 | 0 | 0 | 1 |
| 0 | 0 | 0 | 1 |
| 0 | 0 | 0 | 1 |
| 0 | 0 | 0 | 1 |
| 0 | 0 | 0 | 1 |
| 0 | 0 | 0 | 1 |
| 0 | 0 | 0 | 0 |
| 0 | 0 | 0 | 0 |
| 0 | 0 | 0 | 0 |
| 0 | 0 | 0 | 1 |
| 0 | 0 | 0 | 1 |
| 0 | 0 | 0 | 1 |
| 0 | 0 | 0 | 1 |
| 0 | 0 | 0 | 0 |
| 0 | 1 | 0 | 0 |
| 0 | 0 | 0 | 0 |

| | | | |
|---|---|---|---|
| 0 | 0 | 0 | 0 |
| 0 | 0 | 0 | 0 |
| 0 | 0 | 0 | 1 |
| 0 | 0 | 0 | 0 |
| 0 | 0 | 0 | 0 |
| 0 | 0 | 0 | 0 |
| 0 | 0 | 0 | 0 |
| 0 | 0 | 0 | 1 |
| 0 | 0 | 0 | 1 |
| 0 | 0 | 0 | 1 |
| 0 | 0 | 0 | 1 |
| 0 | 0 | 0 | 1 |
| 0 | 0 | 0 | 1 |
| 0 | 0 | 0 | 1 |
| 0 | 0 | 0 | 1 |
| 0 | 0 | 0 | 1 |
| 0 | 0 | 0 | 1 |
| 0 | 0 | 0 | 1 |
| 0 | 0 | 0 | 1 |
| 0 | 0 | 0 | 1 |
| 0 | 0 | 0 | 0 |
| 0 | 0 | 0 | 0 |
| 0 | 0 | 0 | 0 |
| 0 | 0 | 0 | 0 |
| 0 | 0 | 0 | 0 |
| 0 | 0 | 0 | 1 |
| 0 | 0 | 0 | 1 |
| 0 | 0 | 0 | 0 |
| 0 | 0 | 0 | 1 |
| 0 | 0 | 0 | 1 |
| 0 | 0 | 0 | 0 |
| 0 | 0 | 0 | 0 |
| 0 | 0 | 0 | 0 |
| 0 | 0 | 0 | 1 |
| 0 | 0 | 0 | 0 |
| 0 | 0 | 0 | 1 |
| 0 | 0 | 0 | 1 |
| 0 | 0 | 0 | 0 |
| 0 | 0 | 0 | 1 |
| 0 | 0 | 0 | 1 |
| 0 | 0 | 0 | 1 |
| 0 | 0 | 0 | 1 |
| 0 | 0 | 0 | 1 |
| 0 | 0 | 0 | 1 |
| 0 | 0 | 0 | 1 |
| 0 | 0 | 0 | 1 |
| 0 | 0 | 0 | 1 |
| 0 | 0 | 0 | 1 |
| 0 | 0 | 0 | 0 |
| 0 | 0 | 0 | 1 |
| 1 | 0 | 0 | 0 |
| 1 | 0 | 0 | 0 |
| 0 | 0 | 0 | 1 |
| 0 | 0 | 0 | 1 |

| | | | |
|---|---|---|---|
| 1 | 0 | 0 | 0 |
| 0 | 0 | 0 | 1 |
| 0 | 0 | 0 | 1 |
| 0 | 0 | 0 | 1 |
| 0 | 0 | 0 | 1 |
| 1 | 0 | 0 | 0 |
| 1 | 0 | 0 | 0 |
| 0 | 0 | 0 | 1 |
| 0 | 0 | 0 | 1 |
| 0 | 0 | 0 | 1 |
| 0 | 0 | 0 | 1 |
| 0 | 0 | 0 | 0 |
| 0 | 0 | 0 | 1 |
| 0 | 0 | 0 | 1 |
| 0 | 0 | 0 | 1 |
| 0 | 0 | 0 | 1 |
| 0 | 0 | 0 | 0 |
| 0 | 0 | 0 | 0 |
| 0 | 0 | 0 | 1 |
| 0 | 0 | 0 | 1 |
| 0 | 0 | 0 | 1 |
| 0 | 0 | 0 | 1 |
| 0 | 0 | 0 | 1 |
| 0 | 0 | 0 | 1 |
| 0 | 0 | 0 | 0 |
| 0 | 0 | 0 | 1 |
| 0 | 0 | 0 | 1 |
| 0 | 0 | 0 | 1 |
| 0 | 0 | 0 | 1 |
| 0 | 0 | 0 | 1 |
| 0 | 0 | 0 | 1 |
| 0 | 0 | 0 | 0 |
| 0 | 0 | 0 | 0 |
| 0 | 0 | 0 | 1 |
| 0 | 0 | 0 | 1 |
| 0 | 1 | 0 | 0 |
| 1 | 0 | 0 | 0 |
| 0 | 1 | 0 | 0 |
| 1 | 1 | 0 | 0 |
| 0 | 1 | 0 | 0 |
| 0 | 0 | 0 | 0 |
| 0 | 0 | 0 | 0 |
| 0 | 0 | 0 | 0 |
| 0 | 0 | 0 | 0 |
| 0 | 0 | 0 | 0 |
| 0 | 0 | 0 | 0 |
| 0 | 1 | 0 | 0 |
| 1 | 0 | 0 | 0 |
| 0 | 1 | 1 | 1 |
| 0 | 0 | 0 | 1 |
| 0 | 1 | 1 | 1 |
| 0 | 0 | 0 | 1 |
| 0 | 0 | 0 | 0 |
| 0 | 0 | 0 | 1 |

| | | | |
|---|---|---|---|
| 0 | 0 | 0 | 1 |
| 0 | 0 | 0 | 1 |
| 0 | 0 | 0 | 1 |
| 0 | 0 | 0 | 1 |
| 0 | 0 | 0 | 1 |
| 0 | 0 | 0 | 1 |
| 0 | 0 | 0 | 1 |
| 0 | 1 | 0 | 0 |
| 0 | 0 | 0 | 0 |
| 0 | 1 | 1 | 1 |
| 0 | 0 | 0 | 1 |
| 0 | 0 | 0 | 1 |
| 0 | 0 | 0 | 1 |
| 0 | 1 | 0 | 0 |
| 0 | 0 | 0 | 1 |
| 0 | 0 | 0 | 1 |
| 0 | 1 | 0 | 0 |
| 0 | 0 | 0 | 1 |
| 1 | 0 | 0 | 0 |
| 0 | 0 | 0 | 1 |
| 0 | 0 | 0 | 1 |
| 0 | 0 | 0 | 1 |
| 0 | 0 | 0 | 1 |
| 0 | 0 | 0 | 0 |
| 0 | 0 | 0 | 1 |
| 0 | 0 | 0 | 1 |
| 0 | 0 | 0 | 1 |
| 0 | 0 | 0 | 1 |
| 0 | 0 | 0 | 1 |
| 0 | 0 | 0 | 1 |
| 0 | 0 | 0 | 1 |
| 1 | 0 | 0 | 0 |
| 0 | 0 | 0 | 1 |
| 0 | 0 | 0 | 1 |
| 0 | 0 | 0 | 1 |
| 0 | 0 | 0 | 1 |
| 0 | 0 | 0 | 1 |
| 0 | 0 | 0 | 1 |
| 1 | 0 | 0 | 0 |
| 0 | 0 | 0 | 1 |
| 0 | 0 | 0 | 1 |
| 0 | 0 | 0 | 1 |
| 0 | 0 | 0 | 1 |
| 0 | 0 | 0 | 1 |
| 0 | 0 | 0 | 1 |
| 0 | 0 | 0 | 0 |
| 0 | 0 | 0 | 1 |
| 0 | 0 | 0 | 1 |
| 0 | 1 | 1 | 1 |
| 0 | 1 | 1 | 1 |
| 0 | 0 | 0 | 1 |
| 0 | 1 | 1 | 1 |
| 0 | 1 | 1 | 1 |
| 0 | 1 | 1 | 1 |

| | | | |
|---|---|---|---|
| 0 | 1 | 1 | 1 |
| 0 | 1 | 1 | 1 |
| 0 | 1 | 1 | 1 |
| 0 | 1 | 1 | 1 |
| 0 | 1 | 1 | 1 |
| 0 | 1 | 1 | 1 |
| 0 | 1 | 1 | 1 |
| 0 | 1 | 1 | 1 |
| 0 | 1 | 1 | 1 |
| 0 | 1 | 1 | 1 |
| 0 | 1 | 1 | 1 |
| 0 | 1 | 1 | 1 |
| 0 | 1 | 1 | 1 |
| 0 | 1 | 1 | 1 |
| 0 | 1 | 1 | 1 |
| 1 | 1 | 0 | 0 |
| 0 | 1 | 1 | 1 |
| 0 | 1 | 0 | 0 |
| 0 | 1 | 1 | 1 |
| 0 | 1 | 1 | 1 |
| 1 | 1 | 0 | 0 |
| 0 | 1 | 1 | 1 |
| 0 | 1 | 1 | 1 |
| 0 | 1 | 1 | 1 |
| 0 | 1 | 0 | 0 |
| 0 | 1 | 1 | 1 |
| 0 | 1 | 0 | 0 |
| 0 | 1 | 1 | 1 |
| 0 | 1 | 1 | 1 |
| 0 | 1 | 1 | 1 |
| 0 | 1 | 1 | 1 |
| 0 | 1 | 1 | 1 |
| 0 | 1 | 1 | 1 |
| 0 | 1 | 1 | 1 |
| 0 | 1 | 1 | 1 |
| 0 | 1 | 1 | 1 |
| 0 | 1 | 1 | 1 |
| 0 | 1 | 1 | 1 |
| 0 | 1 | 1 | 1 |
| 0 | 1 | 1 | 1 |
| 0 | 1 | 1 | 1 |
| 0 | 1 | 1 | 1 |
| 0 | 1 | 1 | 1 |
| 0 | 0 | 0 | 0 |
| 0 | 0 | 0 | 1 |
| 0 | 0 | 0 | 1 |
| 0 | 0 | 0 | 0 |
| 0 | 0 | 0 | 0 |
| 0 | 0 | 0 | 1 |
| 0 | 0 | 0 | 0 |
| 0 | 0 | 0 | 0 |
| 0 | 0 | 0 | 0 |
| 0 | 0 | 0 | 1 |
| 0 | 0 | 0 | 1 |

| | | | |
|---|---|---|---|
| 0 | 0 | 0 | 1 |
| 0 | 0 | 0 | 1 |
| 0 | 0 | 0 | 1 |
| 0 | 0 | 0 | 1 |
| 0 | 0 | 0 | 1 |
| 0 | 0 | 0 | 1 |
| 0 | 0 | 0 | 1 |
| 1 | 0 | 0 | 0 |
| 0 | 0 | 0 | 1 |
| 0 | 0 | 0 | 1 |
| 0 | 0 | 0 | 1 |
| 0 | 0 | 0 | 1 |
| 0 | 1 | 1 | 1 |
| 0 | 0 | 0 | 1 |
| 0 | 1 | 0 | 0 |
| 0 | 0 | 0 | 0 |
| 0 | 0 | 0 | 1 |
| 0 | 0 | 0 | 0 |
| 0 | 0 | 0 | 1 |
| 0 | 0 | 0 | 1 |
| 0 | 0 | 0 | 0 |
| 0 | 0 | 0 | 1 |
| 0 | 0 | 0 | 1 |
| 0 | 0 | 0 | 1 |
| 0 | 0 | 0 | 0 |
| 0 | 0 | 0 | 1 |
| 0 | 0 | 0 | 1 |
| 1 | 1 | 0 | 0 |
| 0 | 0 | 0 | 0 |
| 1 | 0 | 0 | 0 |
| 1 | 0 | 0 | 0 |
| 0 | 0 | 0 | 1 |
| 0 | 0 | 0 | 1 |
| 0 | 0 | 0 | 1 |
| 0 | 0 | 0 | 1 |
| 0 | 0 | 0 | 1 |
| 0 | 1 | 0 | 0 |
| 0 | 0 | 0 | 0 |
| 0 | 0 | 0 | 1 |
| 0 | 0 | 0 | 1 |
| 0 | 0 | 0 | 1 |
| 0 | 0 | 0 | 1 |
| 0 | 0 | 0 | 0 |
| 0 | 0 | 0 | 1 |
| 0 | 0 | 0 | 1 |
| 0 | 0 | 0 | 1 |
| 0 | 0 | 0 | 1 |
| 0 | 0 | 0 | 1 |
| 0 | 0 | 0 | 0 |
| 0 | 0 | 0 | 0 |
| 0 | 0 | 0 | 0 |
| 0 | 0 | 0 | 0 |
| 0 | 0 | 0 | 0 |
| 0 | 0 | 0 | 1 |

| | | | |
|---|---|---|---|
| 0 | 0 | 0 | 0 |
| 0 | 0 | 0 | 0 |
| 0 | 0 | 0 | 1 |
| 1 | 0 | 0 | 0 |
| 0 | 0 | 0 | 1 |
| 0 | 0 | 0 | 1 |
| 0 | 0 | 0 | 1 |
| 0 | 0 | 0 | 1 |
| 0 | 0 | 0 | 1 |
| 1 | 0 | 0 | 0 |
| 0 | 0 | 0 | 1 |
| 0 | 0 | 0 | 1 |
| 0 | 0 | 0 | 1 |
| 0 | 0 | 0 | 1 |
| 0 | 0 | 0 | 1 |
| 0 | 0 | 0 | 1 |
| 0 | 0 | 0 | 1 |
| 0 | 0 | 0 | 1 |
| 0 | 0 | 0 | 1 |
| 0 | 0 | 0 | 1 |
| 0 | 0 | 0 | 1 |
| 0 | 0 | 0 | 1 |
| 0 | 0 | 0 | 1 |
| 0 | 0 | 0 | 1 |
| 0 | 0 | 0 | 1 |
| 0 | 0 | 0 | 1 |
| 0 | 0 | 0 | 1 |
| 0 | 0 | 0 | 1 |
| 0 | 0 | 0 | 1 |
| 0 | 0 | 0 | 1 |
| 0 | 0 | 0 | 1 |
| 0 | 0 | 0 | 1 |
| 1 | 0 | 0 | 0 |
| 0 | 0 | 0 | 1 |
| 0 | 0 | 0 | 1 |
| 0 | 0 | 0 | 1 |
| 0 | 0 | 0 | 1 |
| 0 | 0 | 0 | 1 |
| 0 | 0 | 0 | 1 |
| 0 | 0 | 0 | 1 |
| 0 | 0 | 0 | 1 |
| 0 | 0 | 0 | 1 |
| 0 | 0 | 0 | 1 |
| 0 | 0 | 0 | 1 |
| 0 | 0 | 0 | 1 |
| 0 | 0 | 0 | 1 |
| 0 | 0 | 0 | 1 |
| 0 | 1 | 0 | 0 |
| 0 | 1 | 0 | 0 |
| 0 | 1 | 0 | 0 |
| 0 | 0 | 0 | 0 |
| 0 | 1 | 0 | 0 |
| 0 | 0 | 0 | 1 |
| 0 | 0 | 0 | 0 |

| | | | |
|---|---|---|---|
| 0 | 0 | 0 | 1 |
| 0 | 0 | 0 | 0 |
| 0 | 0 | 0 | 1 |
| 0 | 0 | 0 | 1 |
| 0 | 0 | 0 | 1 |
| 1 | 0 | 0 | 0 |
| 0 | 0 | 0 | 0 |
| 0 | 0 | 0 | 0 |
| 1 | 0 | 0 | 0 |
| 0 | 0 | 0 | 0 |
| 0 | 0 | 0 | 1 |
| 0 | 0 | 0 | 0 |
| 0 | 0 | 0 | 1 |
| 0 | 0 | 0 | 1 |
| 0 | 0 | 0 | 1 |
| 1 | 0 | 0 | 0 |
| 0 | 0 | 0 | 1 |
| 0 | 0 | 0 | 1 |
| 0 | 0 | 0 | 0 |
| 0 | 0 | 0 | 0 |
| 1 | 0 | 0 | 0 |
| 1 | 0 | 0 | 0 |
| 0 | 0 | 0 | 0 |
| 0 | 0 | 0 | 0 |
| 0 | 0 | 0 | 1 |
| 0 | 0 | 0 | 0 |
| 0 | 1 | 0 | 0 |
| 0 | 0 | 0 | 0 |
| 0 | 1 | 0 | 0 |
| 0 | 1 | 0 | 0 |
| 0 | 1 | 0 | 0 |
| 0 | 0 | 0 | 1 |
| 0 | 0 | 0 | 1 |
| 0 | 1 | 0 | 0 |
| 0 | 1 | 0 | 0 |
| 0 | 1 | 1 | 1 |
| 0 | 0 | 0 | 0 |
| 0 | 0 | 0 | 0 |
| 0 | 1 | 0 | 0 |
| 0 | 1 | 0 | 0 |
| 0 | 1 | 0 | 0 |
| 0 | 1 | 1 | 1 |
| 0 | 0 | 0 | 1 |
| 0 | 0 | 0 | 1 |
| 0 | 0 | 0 | 0 |
| 0 | 0 | 0 | 0 |
| 0 | 0 | 0 | 0 |
| 0 | 0 | 0 | 1 |
| 0 | 0 | 0 | 1 |
| 0 | 0 | 0 | 1 |
| 0 | 0 | 0 | 1 |
| 0 | 0 | 0 | 1 |
| 0 | 0 | 0 | 1 |
| 0 | 0 | 0 | 0 |

| | | | |
|---|---|---|---|
| 0 | 0 | 0 | 1 |
| 0 | 0 | 0 | 1 |
| 0 | 0 | 0 | 1 |
| 0 | 0 | 0 | 1 |
| 1 | 0 | 0 | 0 |
| 0 | 0 | 0 | 0 |
| 0 | 1 | 1 | 1 |
| 0 | 1 | 1 | 1 |
| 0 | 0 | 0 | 1 |
| 0 | 0 | 0 | 1 |
| 0 | 0 | 0 | 1 |
| 0 | 0 | 0 | 0 |
| 0 | 0 | 0 | 1 |
| 0 | 0 | 0 | 0 |
| 0 | 1 | 0 | 0 |
| 0 | 1 | 0 | 0 |
| 0 | 1 | 0 | 0 |
| 0 | 1 | 0 | 0 |
| 0 | 1 | 0 | 0 |
| 0 | 1 | 0 | 0 |
| 0 | 1 | 0 | 0 |
| 0 | 1 | 0 | 0 |
| 0 | 0 | 0 | 0 |
| 0 | 0 | 0 | 1 |
| 0 | 0 | 0 | 1 |
| 0 | 0 | 0 | 1 |
| 0 | 0 | 0 | 1 |
| 0 | 0 | 0 | 1 |
| 0 | 0 | 0 | 1 |
| 0 | 0 | 0 | 1 |
| 0 | 0 | 0 | 1 |
| 0 | 0 | 0 | 0 |
| 0 | 0 | 0 | 1 |
| 0 | 0 | 0 | 0 |
| 0 | 0 | 0 | 0 |
| 0 | 0 | 0 | 0 |
| 0 | 0 | 0 | 0 |
| 0 | 0 | 0 | 1 |
| 0 | 0 | 0 | 1 |
| 0 | 0 | 0 | 1 |
| 0 | 0 | 0 | 1 |
| 0 | 0 | 0 | 1 |
| 0 | 0 | 0 | 1 |
| 0 | 0 | 0 | 1 |
| 0 | 0 | 0 | 1 |
| 1 | 0 | 0 | 0 |
| 0 | 0 | 0 | 1 |
| 0 | 0 | 0 | 1 |
| 0 | 0 | 0 | 1 |
| 1 | 1 | 0 | 0 |
| 1 | 1 | 0 | 0 |
| 0 | 0 | 0 | 0 |
| 0 | 0 | 0 | 1 |
| 0 | 0 | 0 | 1 |

| | | | |
|---|---|---|---|
| 0 | 0 | 0 | 1 |
| 0 | 0 | 0 | 1 |
| 0 | 0 | 0 | 1 |
| 0 | 0 | 0 | 0 |
| 0 | 0 | 0 | 1 |
| 0 | 0 | 0 | 1 |
| 0 | 0 | 0 | 0 |
| 0 | 0 | 0 | 1 |
| 0 | 0 | 0 | 1 |
| 0 | 0 | 0 | 1 |
| 0 | 0 | 0 | 0 |
| 0 | 1 | 0 | 0 |
| 0 | 0 | 0 | 1 |
| 0 | 0 | 0 | 1 |
| 0 | 0 | 0 | 1 |
| 0 | 0 | 0 | 1 |
| 0 | 1 | 0 | 0 |
| 0 | 1 | 0 | 0 |
| 0 | 0 | 0 | 1 |
| 0 | 1 | 0 | 0 |
| 0 | 1 | 0 | 0 |
| 0 | 1 | 0 | 0 |
| 0 | 0 | 0 | 1 |
| 0 | 1 | 0 | 0 |
| 0 | 0 | 0 | 0 |
| 0 | 0 | 0 | 1 |
| 0 | 0 | 0 | 0 |
| 0 | 0 | 0 | 1 |
| 0 | 0 | 0 | 1 |
| 0 | 0 | 0 | 0 |
| 0 | 0 | 0 | 1 |
| 0 | 0 | 0 | 1 |
| 0 | 0 | 0 | 0 |
| 0 | 0 | 0 | 1 |
| 0 | 0 | 0 | 1 |
| 0 | 0 | 0 | 0 |
| 0 | 0 | 0 | 1 |
| 0 | 0 | 0 | 0 |
| 0 | 0 | 0 | 1 |
| 0 | 1 | 0 | 0 |
| 0 | 1 | 0 | 0 |
| 0 | 1 | 1 | 1 |
| 0 | 0 | 0 | 0 |
| 0 | 0 | 0 | 0 |
| 0 | 0 | 0 | 1 |
| 0 | 1 | 1 | 1 |
| 0 | 0 | 0 | 1 |
| 0 | 0 | 0 | 1 |
| 0 | 0 | 0 | 1 |
| 0 | 0 | 0 | 1 |
| 0 | 0 | 0 | 1 |
| 0 | 0 | 0 | 1 |
| 0 | 0 | 0 | 1 |
| 0 | 0 | 0 | 1 |

| | | | |
|---|---|---|---|
| 0 | 0 | 0 | 1 |
| 0 | 0 | 0 | 1 |
| 0 | 0 | 0 | 1 |
| 0 | 0 | 0 | 0 |
| 0 | 1 | 1 | 1 |
| 0 | 0 | 0 | 0 |
| 0 | 0 | 0 | 1 |
| 0 | 1 | 1 | 1 |
| 0 | 1 | 0 | 0 |
| 0 | 1 | 0 | 0 |
| 0 | 0 | 0 | 1 |
| 0 | 1 | 0 | 0 |
| 0 | 1 | 0 | 0 |
| 0 | 1 | 0 | 0 |
| 0 | 1 | 1 | 1 |
| 0 | 0 | 0 | 0 |
| 0 | 0 | 0 | 0 |
| 0 | 0 | 0 | 1 |
| 1 | 1 | 0 | 0 |
| 1 | 0 | 0 | 0 |
| 1 | 0 | 0 | 0 |
| 1 | 0 | 0 | 0 |
| 0 | 0 | 0 | 1 |
| 0 | 0 | 0 | 0 |
| 0 | 0 | 0 | 0 |
| 0 | 0 | 0 | 0 |
| 0 | 0 | 0 | 0 |
| 0 | 0 | 0 | 0 |
| 0 | 0 | 0 | 0 |
| 0 | 0 | 0 | 0 |
| 0 | 0 | 0 | 0 |
| 0 | 0 | 0 | 0 |
| 0 | 0 | 0 | 0 |
| 0 | 0 | 0 | 1 |
| 0 | 0 | 0 | 1 |
| 0 | 1 | 0 | 0 |
| 0 | 0 | 0 | 1 |
| 0 | 1 | 0 | 0 |
| 0 | 1 | 0 | 0 |
| 0 | 1 | 0 | 0 |
| 0 | 1 | 0 | 0 |
| 0 | 0 | 0 | 1 |
| 0 | 0 | 0 | 0 |
| 0 | 0 | 0 | 0 |
| 0 | 0 | 0 | 1 |
| 0 | 0 | 0 | 1 |
| 0 | 0 | 0 | 1 |
| 0 | 0 | 0 | 1 |
| 0 | 0 | 0 | 1 |
| 0 | 0 | 0 | 1 |
| 0 | 0 | 0 | 1 |
| 0 | 0 | 0 | 1 |
| 0 | 0 | 0 | 1 |
| 0 | 0 | 0 | 1 |

| | | | |
|---|---|---|---|
| 0 | 0 | 0 | 1 |
| 0 | 0 | 0 | 0 |
| 0 | 0 | 0 | 1 |
| 0 | 0 | 0 | 1 |
| 0 | 0 | 0 | 1 |
| 0 | 0 | 0 | 1 |
| 0 | 0 | 0 | 1 |
| 0 | 0 | 0 | 1 |
| 0 | 0 | 0 | 1 |
| 0 | 0 | 0 | 1 |
| 0 | 0 | 0 | 1 |
| 0 | 0 | 0 | 0 |
| 0 | 1 | 1 | 1 |
| 0 | 0 | 0 | 1 |
| 0 | 1 | 0 | 0 |
| 0 | 0 | 0 | 1 |
| 0 | 0 | 0 | 1 |
| 0 | 0 | 0 | 1 |
| 0 | 0 | 0 | 1 |
| 0 | 0 | 0 | 1 |
| 1 | 0 | 0 | 0 |
| 0 | 0 | 0 | 1 |
| 1 | 0 | 0 | 0 |
| 1 | 0 | 0 | 0 |
| 0 | 0 | 0 | 1 |
| 0 | 0 | 0 | 0 |
| 0 | 0 | 0 | 1 |
| 0 | 0 | 0 | 1 |
| 0 | 0 | 0 | 1 |
| 0 | 0 | 0 | 1 |
| 0 | 0 | 0 | 1 |
| 0 | 1 | 0 | 0 |
| 0 | 0 | 0 | 1 |
| 0 | 0 | 0 | 0 |
| 0 | 0 | 0 | 1 |
| 0 | 0 | 0 | 1 |
| 0 | 0 | 0 | 1 |
| 0 | 0 | 0 | 1 |
| 0 | 0 | 0 | 1 |
| 0 | 0 | 0 | 0 |
| 0 | 0 | 0 | 1 |
| 0 | 0 | 0 | 1 |
| 0 | 0 | 0 | 1 |
| 0 | 0 | 0 | 0 |
| 0 | 0 | 0 | 1 |
| 0 | 0 | 0 | 1 |
| 0 | 0 | 0 | 1 |
| 1 | 0 | 0 | 0 |
| 0 | 1 | 0 | 0 |
| 0 | 0 | 0 | 1 |
| 0 | 0 | 0 | 1 |
| 0 | 0 | 0 | 0 |
| 1 | 0 | 0 | 0 |
| 0 | 1 | 0 | 0 |

| | | | |
|---|---|---|---|
| 1 | 0 | 0 | 0 |
| 0 | 0 | 0 | 1 |
| 0 | 0 | 0 | 0 |
| 0 | 0 | 0 | 0 |
| 0 | 0 | 0 | 0 |
| 0 | 0 | 0 | 1 |
| 0 | 0 | 0 | 1 |
| 0 | 0 | 0 | 1 |
| 0 | 0 | 0 | 1 |
| 0 | 1 | 0 | 0 |
| 0 | 0 | 0 | 1 |
| 0 | 0 | 0 | 1 |
| 0 | 0 | 0 | 1 |
| 0 | 0 | 0 | 1 |
| 0 | 0 | 0 | 0 |
| 0 | 0 | 0 | 1 |
| 0 | 0 | 0 | 1 |
| 0 | 0 | 0 | 0 |
| 0 | 0 | 0 | 1 |
| 0 | 0 | 0 | 0 |
| 0 | 0 | 0 | 0 |
| 0 | 0 | 0 | 1 |
| 0 | 0 | 0 | 0 |
| 0 | 0 | 0 | 1 |
| 0 | 0 | 0 | 0 |
| 0 | 0 | 0 | 0 |
| 0 | 0 | 0 | 1 |
| 0 | 0 | 0 | 1 |
| 1 | 0 | 0 | 0 |
| 0 | 0 | 0 | 1 |
| 0 | 0 | 0 | 1 |
| 0 | 0 | 0 | 1 |
| 0 | 0 | 0 | 1 |
| 0 | 0 | 0 | 1 |
| 0 | 0 | 0 | 0 |
| 0 | 0 | 0 | 0 |
| 0 | 0 | 0 | 1 |
| 0 | 0 | 0 | 0 |
| 0 | 0 | 0 | 0 |
| 0 | 0 | 0 | 0 |
| 0 | 0 | 0 | 0 |
| 0 | 0 | 0 | 0 |
| 0 | 0 | 0 | 0 |
| 0 | 0 | 0 | 1 |
| 0 | 1 | 0 | 0 |
| 0 | 0 | 0 | 1 |
| 0 | 0 | 0 | 1 |
| 0 | 0 | 0 | 0 |
| 0 | 0 | 0 | 0 |
| 0 | 0 | 0 | 0 |
| 0 | 0 | 0 | 0 |
| 0 | 1 | 0 | 0 |
| 0 | 0 | 0 | 1 |
| 0 | 0 | 0 | 1 |

| | | | |
|---|---|---|---|
| 0 | 0 | 0 | 1 |
| 0 | 0 | 0 | 0 |
| 0 | 0 | 0 | 0 |
| 0 | 0 | 0 | 0 |
| 0 | 0 | 0 | 0 |
| 0 | 0 | 0 | 0 |
| 0 | 1 | 0 | 0 |
| 0 | 1 | 1 | 1 |
| 0 | 1 | 1 | 1 |
| 0 | 0 | 0 | 1 |
| 0 | 0 | 0 | 0 |
| 0 | 0 | 0 | 0 |
| 0 | 0 | 0 | 1 |
| 0 | 0 | 0 | 0 |
| 0 | 0 | 0 | 1 |
| 0 | 0 | 0 | 0 |
| 0 | 0 | 0 | 1 |
| 0 | 0 | 0 | 1 |
| 0 | 0 | 0 | 1 |
| 0 | 0 | 0 | 1 |
| 0 | 0 | 0 | 1 |
| 0 | 0 | 0 | 1 |
| 0 | 0 | 0 | 1 |
| 1 | 0 | 0 | 0 |
| 0 | 0 | 0 | 0 |
| 0 | 0 | 0 | 1 |
| 0 | 0 | 0 | 1 |
| 0 | 0 | 0 | 1 |
| 0 | 0 | 0 | 1 |
| 0 | 0 | 0 | 0 |
| 0 | 0 | 0 | 0 |
| 0 | 0 | 0 | 0 |
| 0 | 0 | 0 | 0 |
| 0 | 0 | 0 | 1 |
| 0 | 0 | 0 | 0 |
| 0 | 0 | 0 | 0 |
| 0 | 0 | 0 | 0 |
| 0 | 0 | 0 | 0 |
| 0 | 0 | 0 | 0 |
| 0 | 0 | 0 | 1 |
| 0 | 0 | 0 | 1 |
| 0 | 0 | 0 | 0 |
| 0 | 0 | 0 | 1 |
| 0 | 0 | 0 | 1 |
| 0 | 1 | 0 | 0 |
| 0 | 1 | 1 | 1 |
| 0 | 0 | 0 | 1 |
| 0 | 0 | 0 | 1 |
| 0 | 1 | 1 | 1 |
| 0 | 0 | 0 | 1 |
| 0 | 0 | 0 | 0 |
| 0 | 0 | 0 | 1 |
| 0 | 0 | 0 | 0 |
| 0 | 0 | 0 | 0 |

| | | | |
|---|---|---|---|
| 0 | 0 | 0 | 1 |
| 0 | 0 | 0 | 1 |
| 0 | 0 | 0 | 0 |
| 0 | 0 | 0 | 0 |
| 0 | 0 | 0 | 1 |
| 0 | 0 | 0 | 1 |
| 0 | 0 | 0 | 1 |
| 0 | 0 | 0 | 1 |
| 0 | 0 | 0 | 1 |
| 0 | 0 | 0 | 1 |
| 0 | 0 | 0 | 1 |
| 0 | 0 | 0 | 1 |
| 0 | 0 | 0 | 0 |
| 0 | 0 | 0 | 1 |
| 0 | 0 | 0 | 1 |
| 0 | 0 | 0 | 1 |
| 0 | 0 | 0 | 1 |
| 0 | 0 | 0 | 1 |
| 0 | 0 | 0 | 1 |
| 0 | 0 | 0 | 1 |
| 0 | 0 | 0 | 1 |
| 0 | 0 | 0 | 1 |
| 0 | 0 | 0 | 1 |
| 0 | 0 | 0 | 1 |
| 0 | 0 | 0 | 1 |
| 0 | 0 | 0 | 1 |
| 0 | 0 | 0 | 1 |
| 0 | 0 | 0 | 1 |
| 0 | 0 | 0 | 1 |
| 0 | 0 | 0 | 1 |
| 0 | 0 | 0 | 1 |
| 0 | 0 | 0 | 0 |
| 0 | 0 | 0 | 1 |
| 0 | 0 | 0 | 1 |
| 0 | 0 | 0 | 1 |
| 0 | 0 | 0 | 1 |
| 0 | 0 | 0 | 1 |
| 0 | 0 | 0 | 1 |
| 0 | 0 | 0 | 1 |
| 0 | 0 | 0 | 1 |
| 0 | 0 | 0 | 0 |
| 0 | 0 | 0 | 0 |
| 0 | 0 | 0 | 1 |
| 0 | 0 | 0 | 1 |
| 0 | 0 | 0 | 1 |
| 0 | 1 | 1 | 1 |
| 0 | 1 | 1 | 1 |
| 0 | 0 | 0 | 1 |
| 0 | 0 | 0 | 0 |
| 0 | 0 | 0 | 0 |
| 0 | 0 | 0 | 1 |
| 0 | 0 | 0 | 1 |
| 0 | 0 | 0 | 0 |
| 0 | 0 | 0 | 1 |

| | | | |
|---|---|---|---|
| 0 | 0 | 0 | 0 |
| 0 | 0 | 0 | 0 |
| 0 | 0 | 0 | 0 |
| 0 | 0 | 0 | 1 |
| 0 | 0 | 0 | 1 |
| 0 | 0 | 0 | 1 |
| 0 | 0 | 0 | 1 |
| 0 | 1 | 0 | 0 |
| 0 | 1 | 0 | 0 |
| 0 | 0 | 0 | 0 |
| 0 | 0 | 0 | 0 |
| 0 | 0 | 0 | 1 |
| 0 | 0 | 0 | 0 |
| 0 | 0 | 0 | 1 |
| 0 | 0 | 0 | 1 |
| 0 | 0 | 0 | 1 |
| 0 | 0 | 0 | 1 |
| 0 | 0 | 0 | 0 |
| 0 | 0 | 0 | 1 |
| 0 | 0 | 0 | 1 |
| 0 | 0 | 0 | 1 |
| 0 | 0 | 0 | 1 |
| 0 | 0 | 0 | 1 |
| 0 | 0 | 0 | 1 |
| 0 | 0 | 0 | 1 |
| 0 | 0 | 0 | 1 |
| 0 | 0 | 0 | 1 |
| 0 | 0 | 0 | 1 |
| 0 | 0 | 0 | 1 |
| 0 | 0 | 0 | 1 |
| 0 | 0 | 0 | 0 |
| 0 | 1 | 0 | 0 |
| 0 | 0 | 0 | 1 |
| 0 | 0 | 0 | 1 |
| 0 | 0 | 0 | 1 |
| 0 | 0 | 0 | 1 |
| 0 | 0 | 0 | 1 |
| 0 | 0 | 0 | 1 |
| 0 | 0 | 0 | 1 |
| 0 | 0 | 0 | 1 |
| 0 | 0 | 0 | 1 |
| 0 | 0 | 0 | 0 |
| 0 | 0 | 0 | 1 |
| 0 | 0 | 0 | 1 |
| 0 | 0 | 0 | 0 |
| 0 | 0 | 0 | 0 |
| 0 | 0 | 0 | 1 |
| 0 | 0 | 0 | 1 |
| 0 | 0 | 0 | 0 |
| 0 | 0 | 0 | 0 |
| 0 | 0 | 0 | 0 |
| 0 | 0 | 0 | 0 |
| 0 | 0 | 0 | 0 |
| 0 | 0 | 0 | 0 |

| | | | |
|---|---|---|---|
| 0 | 0 | 0 | 1 |
| 0 | 0 | 0 | 1 |
| 0 | 0 | 0 | 1 |
| 0 | 0 | 0 | 1 |
| 0 | 0 | 0 | 1 |
| 0 | 0 | 0 | 0 |
| 0 | 0 | 0 | 0 |
| 0 | 0 | 0 | 0 |
| 0 | 0 | 0 | 0 |
| 0 | 1 | 0 | 0 |
| 0 | 0 | 0 | 1 |
| 0 | 0 | 0 | 0 |
| 0 | 0 | 0 | 0 |
| 0 | 0 | 0 | 0 |
| 0 | 0 | 0 | 1 |
| 0 | 0 | 0 | 0 |
| 0 | 0 | 0 | 1 |
| 0 | 0 | 0 | 1 |
| 1 | 0 | 0 | 0 |
| 1 | 0 | 0 | 0 |
| 0 | 1 | 0 | 0 |
| 0 | 0 | 0 | 0 |
| 0 | 1 | 1 | 1 |
| 0 | 0 | 0 | 0 |
| 0 | 0 | 0 | 0 |
| 0 | 0 | 0 | 0 |
| 0 | 0 | 0 | 0 |
| 0 | 0 | 0 | 1 |
| 0 | 0 | 0 | 1 |
| 0 | 0 | 0 | 1 |
| 0 | 0 | 0 | 1 |
| 0 | 0 | 0 | 1 |
| 0 | 0 | 0 | 0 |
| 0 | 0 | 0 | 1 |
| 0 | 0 | 0 | 0 |
| 0 | 1 | 0 | 0 |
| 0 | 1 | 0 | 0 |
| 0 | 0 | 0 | 0 |
| 0 | 1 | 0 | 0 |
| 0 | 1 | 0 | 0 |
| 0 | 1 | 0 | 0 |
| 1 | 0 | 0 | 0 |
| 0 | 0 | 0 | 0 |
| 0 | 0 | 0 | 1 |
| 0 | 0 | 0 | 1 |
| 0 | 0 | 0 | 1 |
| 0 | 0 | 0 | 1 |
| 0 | 0 | 0 | 0 |
| 0 | 0 | 0 | 0 |
| 0 | 0 | 0 | 0 |
| 0 | 0 | 0 | 1 |
| 0 | 0 | 0 | 1 |
| 0 | 0 | 0 | 0 |
| 0 | 0 | 0 | 1 |

| | | | |
|---|---|---|---|
| 0 | 0 | 0 | 1 |
| 0 | 0 | 0 | 0 |
| 0 | 0 | 0 | 1 |
| 0 | 0 | 0 | 1 |
| 0 | 0 | 0 | 1 |
| 0 | 0 | 0 | 1 |
| 0 | 0 | 0 | 1 |
| 0 | 0 | 0 | 1 |
| 0 | 0 | 0 | 1 |
| 0 | 0 | 0 | 1 |
| 0 | 0 | 0 | 1 |
| 0 | 0 | 0 | 1 |
| 0 | 0 | 0 | 0 |
| 0 | 0 | 0 | 0 |
| 0 | 0 | 0 | 0 |
| 0 | 0 | 0 | 0 |
| 0 | 0 | 0 | 0 |
| 0 | 0 | 0 | 0 |
| 0 | 0 | 0 | 0 |
| 0 | 0 | 0 | 1 |
| 1 | 1 | 0 | 0 |
| 0 | 0 | 0 | 0 |
| 0 | 0 | 0 | 0 |
| 0 | 0 | 0 | 0 |
| 0 | 0 | 0 | 0 |
| 0 | 0 | 0 | 1 |
| 0 | 0 | 0 | 1 |
| 0 | 0 | 0 | 0 |
| 0 | 0 | 0 | 1 |
| 0 | 0 | 0 | 0 |
| 0 | 0 | 0 | 0 |
| 0 | 0 | 0 | 1 |
| 0 | 0 | 0 | 0 |
| 0 | 0 | 0 | 1 |
| 0 | 0 | 0 | 1 |
| 0 | 0 | 0 | 1 |
| 0 | 1 | 1 | 1 |
| 0 | 0 | 0 | 1 |
| 0 | 0 | 0 | 1 |
| 0 | 0 | 0 | 1 |
| 0 | 0 | 0 | 0 |
| 0 | 0 | 0 | 1 |
| 0 | 0 | 0 | 1 |
| 0 | 0 | 0 | 0 |
| 0 | 0 | 0 | 0 |
| 0 | 0 | 0 | 1 |
| 0 | 0 | 0 | 0 |
| 0 | 0 | 0 | 0 |
| 0 | 0 | 0 | 1 |
| 0 | 0 | 0 | 1 |
| 0 | 0 | 0 | 0 |
| 0 | 0 | 0 | 1 |
| 0 | 0 | 0 | 1 |
| 0 | 0 | 0 | 1 |

| | | | |
|---|---|---|---|
| 0 | 0 | 0 | 1 |
| 0 | 0 | 0 | 1 |
| 0 | 0 | 0 | 1 |
| 0 | 0 | 0 | 1 |
| 0 | 0 | 0 | 0 |
| 0 | 0 | 0 | 0 |
| 0 | 0 | 0 | 1 |
| 0 | 0 | 0 | 1 |
| 0 | 0 | 0 | 1 |
| 0 | 0 | 0 | 1 |
| 0 | 0 | 0 | 1 |
| 0 | 0 | 0 | 0 |
| 0 | 0 | 0 | 0 |
| 0 | 0 | 0 | 1 |
| 0 | 0 | 0 | 1 |
| 0 | 0 | 0 | 0 |
| 0 | 0 | 0 | 1 |
| 0 | 0 | 0 | 1 |
| 0 | 0 | 0 | 0 |
| 0 | 0 | 0 | 1 |
| 0 | 0 | 0 | 1 |
| 0 | 0 | 0 | 1 |
| 0 | 0 | 0 | 1 |
| 0 | 0 | 0 | 0 |
| 0 | 0 | 0 | 1 |
| 0 | 0 | 0 | 0 |
| 0 | 0 | 0 | 1 |
| 0 | 0 | 0 | 1 |
| 1 | 0 | 0 | 0 |
| 1 | 0 | 0 | 0 |
| 0 | 0 | 0 | 0 |
| 0 | 0 | 0 | 0 |
| 0 | 0 | 0 | 0 |
| 0 | 0 | 0 | 0 |
| 0 | 0 | 0 | 1 |
| 0 | 0 | 0 | 1 |
| 0 | 0 | 0 | 1 |
| 1 | 0 | 0 | 0 |
| 0 | 0 | 0 | 1 |
| 0 | 0 | 0 | 1 |
| 0 | 0 | 0 | 0 |
| 0 | 0 | 0 | 0 |
| 0 | 0 | 0 | 0 |
| 0 | 0 | 0 | 1 |
| 0 | 0 | 0 | 0 |
| 0 | 0 | 0 | 1 |
| 0 | 0 | 0 | 1 |
| 0 | 0 | 0 | 0 |
| 0 | 0 | 0 | 1 |
| 0 | 0 | 0 | 1 |
| 0 | 0 | 0 | 1 |
| 0 | 1 | 0 | 0 |
| 0 | 0 | 0 | 1 |
| 0 | 0 | 0 | 0 |

| | | | |
|---|---|---|---|
| 0 | 0 | 0 | 1 |
| 0 | 1 | 0 | 0 |
| 0 | 0 | 0 | 0 |
| 0 | 0 | 0 | 1 |
| 0 | 0 | 0 | 1 |
| 0 | 0 | 0 | 1 |
| 0 | 0 | 0 | 0 |
| 0 | 0 | 0 | 1 |
| 0 | 0 | 0 | 1 |
| 0 | 0 | 0 | 1 |
| 0 | 0 | 0 | 1 |
| 0 | 0 | 0 | 1 |
| 0 | 0 | 0 | 1 |
| 0 | 0 | 0 | 1 |
| 0 | 0 | 0 | 1 |
| 0 | 0 | 0 | 1 |
| 0 | 0 | 0 | 1 |
| 0 | 0 | 0 | 1 |
| 0 | 0 | 0 | 1 |
| 0 | 0 | 0 | 1 |
| 0 | 0 | 0 | 1 |
| 0 | 0 | 0 | 0 |
| 0 | 0 | 0 | 1 |
| 0 | 0 | 0 | 1 |
| 0 | 0 | 0 | 1 |
| 0 | 0 | 0 | 1 |
| 0 | 0 | 0 | 0 |
| 0 | 0 | 0 | 1 |
| 0 | 0 | 0 | 0 |
| 0 | 0 | 0 | 1 |
| 0 | 0 | 0 | 1 |
| 0 | 0 | 0 | 0 |
| 0 | 1 | 1 | 1 |
| 0 | 1 | 1 | 1 |
| 0 | 0 | 0 | 1 |
| 0 | 0 | 0 | 1 |
| 0 | 0 | 0 | 1 |
| 0 | 0 | 0 | 1 |
| 0 | 1 | 0 | 0 |
| 0 | 1 | 0 | 0 |
| 0 | 1 | 0 | 0 |
| 0 | 0 | 0 | 1 |
| 0 | 0 | 0 | 1 |
| 0 | 0 | 0 | 1 |
| 0 | 0 | 0 | 1 |
| 0 | 1 | 0 | 0 |
| 0 | 0 | 0 | 1 |
| 0 | 0 | 0 | 0 |
| 0 | 0 | 0 | 1 |
| 0 | 1 | 0 | 0 |
| 0 | 0 | 0 | 0 |
| 0 | 1 | 0 | 0 |
| 0 | 0 | 0 | 1 |
| 0 | 0 | 0 | 1 |

| | | | |
|---|---|---|---|
| 0 | 0 | 0 | 0 |
| 0 | 0 | 0 | 0 |
| 0 | 0 | 0 | 1 |
| 0 | 0 | 0 | 1 |
| 0 | 0 | 0 | 1 |
| 0 | 0 | 0 | 1 |
| 0 | 0 | 0 | 1 |
| 0 | 0 | 0 | 1 |
| 0 | 0 | 0 | 1 |
| 0 | 0 | 0 | 1 |
| 0 | 0 | 0 | 1 |
| 0 | 0 | 0 | 1 |
| 0 | 0 | 0 | 1 |
| 0 | 0 | 0 | 1 |
| 0 | 0 | 0 | 1 |
| 0 | 0 | 0 | 0 |
| 0 | 0 | 0 | 0 |
| 0 | 1 | 0 | 0 |
| 0 | 0 | 0 | 1 |
| 0 | 0 | 0 | 1 |
| 0 | 0 | 0 | 1 |
| 0 | 1 | 0 | 0 |
| 0 | 0 | 0 | 0 |
| 0 | 0 | 0 | 0 |
| 0 | 0 | 0 | 0 |
| 0 | 0 | 0 | 1 |
| 0 | 0 | 0 | 1 |
| 0 | 0 | 0 | 1 |
| 0 | 0 | 0 | 1 |
| 0 | 0 | 0 | 1 |
| 0 | 0 | 0 | 0 |
| 0 | 0 | 0 | 1 |
| 0 | 0 | 0 | 0 |
| 0 | 0 | 0 | 1 |
| 0 | 1 | 1 | 1 |
| 0 | 1 | 1 | 1 |
| 0 | 1 | 0 | 0 |
| 0 | 0 | 0 | 0 |
| 0 | 0 | 0 | 1 |
| 0 | 0 | 0 | 1 |
| 0 | 1 | 1 | 1 |
| 0 | 1 | 0 | 0 |
| 0 | 0 | 0 | 1 |
| 0 | 0 | 0 | 1 |
| 0 | 1 | 0 | 0 |
| 0 | 1 | 1 | 1 |
| 0 | 0 | 0 | 0 |
| 0 | 1 | 0 | 0 |
| 0 | 0 | 0 | 1 |
| 0 | 0 | 0 | 1 |
| 0 | 1 | 0 | 0 |
| 0 | 1 | 0 | 0 |
| 0 | 0 | 0 | 1 |
| 0 | 1 | 0 | 0 |

| | | | |
|---|---|---|---|
| 0 | 0 | 0 | 1 |
| 0 | 0 | 0 | 0 |
| 0 | 1 | 1 | 1 |
| 0 | 1 | 0 | 0 |
| 0 | 0 | 0 | 1 |
| 0 | 0 | 0 | 1 |
| 0 | 0 | 0 | 1 |
| 0 | 0 | 0 | 1 |
| 0 | 0 | 0 | 0 |
| 0 | 0 | 0 | 1 |
| 0 | 0 | 0 | 1 |
| 0 | 0 | 0 | 1 |
| 0 | 0 | 0 | 0 |
| 0 | 1 | 0 | 0 |
| 0 | 0 | 0 | 0 |
| 0 | 0 | 0 | 1 |
| 0 | 0 | 0 | 0 |
| 0 | 0 | 0 | 1 |
| 0 | 1 | 0 | 0 |
| 0 | 0 | 0 | 0 |
| 0 | 0 | 0 | 0 |
| 0 | 1 | 0 | 0 |
| 0 | 0 | 0 | 0 |
| 1 | 0 | 0 | 0 |
| 0 | 0 | 0 | 1 |
| 0 | 0 | 0 | 1 |
| 0 | 0 | 0 | 0 |
| 0 | 0 | 0 | 0 |
| 0 | 0 | 0 | 0 |
| 0 | 0 | 0 | 0 |
| 0 | 0 | 0 | 1 |
| 0 | 0 | 0 | 1 |
| 0 | 0 | 0 | 1 |
| 0 | 0 | 0 | 1 |
| 0 | 0 | 0 | 1 |
| 1 | 0 | 0 | 0 |
| 1 | 0 | 0 | 0 |
| 0 | 0 | 0 | 1 |
| 0 | 0 | 0 | 1 |
| 0 | 0 | 0 | 1 |
| 0 | 0 | 0 | 1 |
| 0 | 0 | 0 | 1 |
| 0 | 0 | 0 | 1 |
| 0 | 1 | 0 | 0 |
| 0 | 1 | 0 | 0 |
| 0 | 0 | 0 | 0 |
| 0 | 0 | 0 | 1 |
| 0 | 0 | 0 | 0 |
| 0 | 0 | 0 | 0 |
| 0 | 0 | 0 | 0 |
| 0 | 0 | 0 | 1 |
| 0 | 0 | 0 | 1 |
| 0 | 0 | 0 | 1 |
| 0 | 0 | 0 | 1 |

| | | | |
|---|---|---|---|
| 0 | 0 | 0 | 1 |
| 0 | 0 | 0 | 1 |
| 0 | 1 | 1 | 1 |
| 0 | 0 | 0 | 1 |
| 0 | 0 | 0 | 0 |
| 0 | 1 | 1 | 1 |
| 0 | 0 | 0 | 0 |
| 0 | 0 | 0 | 0 |
| 0 | 0 | 0 | 0 |
| 0 | 0 | 0 | 0 |
| 0 | 0 | 0 | 0 |
| 0 | 0 | 0 | 0 |
| 0 | 0 | 0 | 1 |
| 0 | 0 | 0 | 0 |
| 0 | 0 | 0 | 0 |
| 0 | 0 | 0 | 1 |
| 0 | 0 | 0 | 1 |
| 0 | 0 | 0 | 1 |
| 0 | 0 | 0 | 0 |
| 0 | 0 | 0 | 0 |
| 0 | 0 | 0 | 0 |
| 0 | 0 | 0 | 0 |
| 0 | 0 | 0 | 1 |
| 0 | 0 | 0 | 1 |
| 0 | 0 | 0 | 1 |
| 0 | 0 | 0 | 1 |
| 0 | 0 | 0 | 1 |
| 0 | 0 | 0 | 0 |
| 0 | 0 | 0 | 0 |
| 0 | 0 | 0 | 0 |
| 0 | 0 | 0 | 0 |
| 0 | 0 | 0 | 0 |
| 0 | 0 | 0 | 0 |
| 0 | 0 | 0 | 1 |
| 0 | 0 | 0 | 1 |
| 0 | 0 | 0 | 0 |
| 0 | 0 | 0 | 1 |
| 0 | 0 | 0 | 1 |
| 0 | 0 | 0 | 1 |
| 0 | 0 | 0 | 1 |
| 0 | 0 | 0 | 0 |
| 0 | 0 | 0 | 1 |
| 0 | 0 | 0 | 0 |
| 0 | 0 | 0 | 1 |
| 0 | 0 | 0 | 1 |
| 0 | 0 | 0 | 1 |
| 0 | 0 | 0 | 1 |
| 0 | 0 | 0 | 0 |
| 0 | 0 | 0 | 1 |
| 0 | 0 | 0 | 1 |
| 0 | 0 | 0 | 1 |
| 0 | 0 | 0 | 1 |
| 0 | 0 | 0 | 1 |
| 0 | 0 | 0 | 1 |

| | | | |
|---|---|---|---|
| 0 | 0 | 0 | 1 |
| 0 | 0 | 0 | 1 |
| 0 | 0 | 0 | 1 |
| 0 | 0 | 0 | 1 |
| 0 | 0 | 0 | 1 |
| 0 | 0 | 0 | 1 |
| 0 | 0 | 0 | 1 |
| 0 | 0 | 0 | 1 |
| 0 | 0 | 0 | 1 |
| 0 | 0 | 0 | 1 |
| 0 | 0 | 0 | 1 |
| 0 | 0 | 0 | 0 |
| 0 | 0 | 0 | 1 |
| 0 | 0 | 0 | 1 |
| 0 | 0 | 0 | 1 |
| 0 | 0 | 0 | 1 |
| 0 | 0 | 0 | 1 |
| 0 | 0 | 0 | 0 |
| 0 | 0 | 0 | 0 |
| 0 | 0 | 0 | 0 |
| 0 | 0 | 0 | 0 |
| 0 | 0 | 0 | 1 |
| 1 | 0 | 0 | 0 |
| 0 | 0 | 0 | 1 |
| 0 | 0 | 0 | 1 |
| 0 | 0 | 0 | 1 |
| 0 | 0 | 0 | 1 |
| 0 | 0 | 0 | 1 |
| 0 | 0 | 0 | 0 |
| 0 | 0 | 0 | 0 |
| 0 | 0 | 0 | 1 |
| 0 | 0 | 0 | 1 |
| 1 | 0 | 0 | 0 |
| 0 | 0 | 0 | 1 |
| 0 | 0 | 0 | 0 |
| 1 | 0 | 0 | 0 |
| 1 | 0 | 0 | 0 |
| 0 | 0 | 0 | 1 |
| 1 | 0 | 0 | 0 |
| 0 | 0 | 0 | 1 |
| 1 | 0 | 0 | 0 |
| 0 | 0 | 0 | 1 |
| 0 | 0 | 0 | 0 |
| 0 | 1 | 1 | 1 |
| 0 | 1 | 1 | 1 |
| 0 | 1 | 1 | 1 |
| 0 | 1 | 1 | 1 |
| 0 | 0 | 0 | 1 |
| 0 | 0 | 0 | 1 |
| 0 | 0 | 0 | 0 |
| 0 | 1 | 1 | 1 |
| 1 | 0 | 0 | 0 |
| 0 | 1 | 1 | 1 |
| 0 | 0 | 0 | 1 |

| | | | |
|---|---|---|---|
| 0 | 1 | 0 | 0 |
| 0 | 1 | 0 | 0 |
| 0 | 0 | 0 | 0 |
| 0 | 0 | 0 | 0 |
| 0 | 1 | 0 | 0 |
| 0 | 0 | 0 | 1 |
| 0 | 1 | 0 | 0 |
| 0 | 1 | 1 | 1 |
| 0 | 0 | 0 | 1 |
| 0 | 0 | 0 | 0 |
| 0 | 0 | 0 | 0 |
| 0 | 0 | 0 | 0 |
| 0 | 0 | 0 | 1 |
| 0 | 0 | 0 | 1 |
| 0 | 0 | 0 | 1 |
| 0 | 0 | 0 | 1 |
| 0 | 0 | 0 | 1 |
| 0 | 0 | 0 | 1 |
| 0 | 0 | 0 | 1 |
| 0 | 0 | 0 | 1 |
| 0 | 0 | 0 | 1 |
| 0 | 0 | 0 | 1 |
| 0 | 0 | 0 | 1 |
| 0 | 0 | 0 | 1 |
| 0 | 0 | 0 | 1 |
| 0 | 0 | 0 | 1 |
| 0 | 0 | 0 | 1 |
| 0 | 0 | 0 | 1 |
| 0 | 0 | 0 | 1 |
| 0 | 0 | 0 | 1 |
| 0 | 0 | 0 | 1 |
| 0 | 0 | 0 | 1 |
| 0 | 0 | 0 | 1 |
| 0 | 0 | 0 | 1 |
| 0 | 0 | 0 | 1 |
| 0 | 0 | 0 | 1 |
| 0 | 0 | 0 | 1 |
| 0 | 0 | 0 | 1 |
| 0 | 0 | 0 | 1 |
| 0 | 0 | 0 | 1 |
| 0 | 0 | 0 | 1 |
| 0 | 0 | 0 | 1 |
| 0 | 0 | 0 | 1 |
| 0 | 0 | 0 | 1 |
| 0 | 0 | 0 | 1 |
| 0 | 0 | 0 | 0 |
| 0 | 0 | 0 | 1 |
| 0 | 0 | 0 | 1 |
| 0 | 0 | 0 | 1 |
| 0 | 0 | 0 | 1 |
| 0 | 0 | 0 | 0 |
| 0 | 0 | 0 | 1 |
| 0 | 0 | 0 | 1 |
| 0 | 0 | 0 | 1 |

| | | | |
|---|---|---|---|
| 0 | 0 | 0 | 1 |
| 0 | 0 | 0 | 1 |
| 0 | 0 | 0 | 1 |
| 0 | 0 | 0 | 1 |
| 0 | 0 | 0 | 1 |
| 0 | 0 | 0 | 1 |
| 0 | 0 | 0 | 1 |
| 0 | 0 | 0 | 0 |
| 0 | 1 | 0 | 0 |
| 0 | 0 | 0 | 1 |
| 0 | 0 | 0 | 1 |
| 0 | 0 | 0 | 1 |
| 0 | 0 | 0 | 1 |
| 0 | 0 | 0 | 1 |
| 0 | 1 | 0 | 0 |
| 0 | 1 | 0 | 0 |
| 0 | 0 | 0 | 0 |
| 0 | 0 | 0 | 1 |
| 0 | 1 | 1 | 1 |
| 0 | 0 | 0 | 1 |
| 0 | 1 | 0 | 0 |
| 0 | 1 | 0 | 0 |
| 0 | 1 | 0 | 0 |
| 0 | 0 | 0 | 0 |
| 0 | 1 | 0 | 0 |
| 0 | 1 | 0 | 0 |
| 0 | 0 | 0 | 1 |
| 0 | 0 | 0 | 1 |
| 0 | 0 | 0 | 0 |
| 0 | 0 | 0 | 1 |
| 0 | 0 | 0 | 1 |
| 0 | 0 | 0 | 1 |
| 0 | 0 | 0 | 0 |
| 0 | 0 | 0 | 1 |
| 0 | 0 | 0 | 1 |
| 0 | 0 | 0 | 1 |
| 0 | 0 | 0 | 0 |
| 0 | 0 | 0 | 0 |
| 0 | 0 | 0 | 0 |
| 0 | 0 | 0 | 1 |
| 0 | 0 | 0 | 1 |
| 0 | 0 | 0 | 1 |
| 0 | 0 | 0 | 1 |
| 0 | 0 | 0 | 1 |
| 0 | 0 | 0 | 1 |
| 0 | 0 | 0 | 0 |
| 0 | 0 | 0 | 1 |
| 0 | 0 | 0 | 1 |
| 0 | 0 | 0 | 1 |
| 0 | 0 | 0 | 1 |
| 0 | 0 | 0 | 1 |
| 0 | 0 | 0 | 1 |
| 0 | 0 | 0 | 1 |
| 0 | 0 | 0 | 1 |

| | | | |
|---|---|---|---|
| 0 | 0 | 0 | 1 |
| 0 | 0 | 0 | 1 |
| 0 | 0 | 0 | 1 |
| 0 | 1 | 0 | 0 |
| 0 | 0 | 0 | 1 |
| 0 | 0 | 0 | 1 |
| 0 | 0 | 0 | 1 |
| 0 | 0 | 0 | 0 |
| 0 | 0 | 0 | 0 |
| 0 | 0 | 0 | 1 |
| 0 | 0 | 0 | 1 |
| 0 | 0 | 0 | 1 |
| 0 | 0 | 0 | 0 |
| 0 | 0 | 0 | 1 |
| 0 | 0 | 0 | 1 |
| 0 | 0 | 0 | 1 |
| 0 | 0 | 0 | 0 |
| 0 | 0 | 0 | 1 |
| 0 | 0 | 0 | 1 |
| 0 | 0 | 0 | 1 |
| 0 | 0 | 0 | 1 |
| 0 | 0 | 0 | 1 |
| 0 | 0 | 0 | 1 |
| 0 | 0 | 0 | 1 |
| 0 | 0 | 0 | 1 |
| 0 | 0 | 0 | 1 |
| 0 | 0 | 0 | 1 |
| 0 | 0 | 0 | 0 |
| 0 | 0 | 0 | 0 |
| 0 | 0 | 0 | 0 |
| 0 | 0 | 0 | 0 |
| 0 | 0 | 0 | 0 |
| 0 | 0 | 0 | 0 |
| 0 | 0 | 0 | 1 |
| 0 | 0 | 0 | 0 |
| 0 | 0 | 0 | 0 |
| 0 | 0 | 0 | 0 |
| 0 | 0 | 0 | 1 |
| 0 | 0 | 0 | 1 |
| 0 | 0 | 0 | 1 |
| 0 | 0 | 0 | 1 |
| 0 | 0 | 0 | 1 |
| 0 | 0 | 0 | 0 |
| 0 | 0 | 0 | 0 |
| 0 | 0 | 0 | 1 |
| 0 | 0 | 0 | 1 |
| 0 | 0 | 0 | 0 |
| 0 | 0 | 0 | 1 |
| 0 | 0 | 0 | 1 |
| 0 | 0 | 0 | 1 |
| 0 | 0 | 0 | 1 |
| 0 | 0 | 0 | 1 |
| 0 | 0 | 0 | 0 |
| 0 | 0 | 0 | 0 |

| | | | |
|---|---|---|---|
| 0 | 0 | 0 | 0 |
| 0 | 0 | 0 | 0 |
| 0 | 0 | 0 | 0 |
| 0 | 0 | 0 | 0 |
| 0 | 0 | 0 | 0 |
| 0 | 0 | 0 | 0 |
| 0 | 0 | 0 | 0 |
| 0 | 0 | 0 | 0 |
| 0 | 0 | 0 | 1 |
| 0 | 1 | 0 | 0 |
| 0 | 0 | 0 | 1 |
| 0 | 0 | 0 | 1 |
| 0 | 0 | 0 | 1 |
| 0 | 0 | 0 | 1 |
| 0 | 0 | 0 | 0 |
| 0 | 0 | 0 | 1 |
| 0 | 0 | 0 | 1 |
| 0 | 0 | 0 | 1 |
| 0 | 0 | 0 | 1 |
| 0 | 0 | 0 | 0 |
| 0 | 0 | 0 | 0 |
| 0 | 0 | 0 | 1 |
| 0 | 0 | 0 | 0 |
| 0 | 0 | 0 | 0 |
| 0 | 0 | 0 | 0 |
| 0 | 0 | 0 | 0 |
| 0 | 0 | 0 | 0 |
| 0 | 0 | 0 | 1 |
| 0 | 0 | 0 | 1 |
| 0 | 0 | 0 | 1 |
| 0 | 0 | 0 | 0 |
| 0 | 0 | 0 | 1 |
| 0 | 0 | 0 | 1 |
| 0 | 0 | 0 | 1 |
| 0 | 0 | 0 | 1 |
| 0 | 0 | 0 | 1 |
| 0 | 0 | 0 | 1 |
| 0 | 0 | 0 | 1 |
| 0 | 0 | 0 | 1 |
| 0 | 0 | 0 | 1 |
| 0 | 0 | 0 | 1 |
| 0 | 0 | 0 | 0 |
| 0 | 0 | 0 | 1 |
| 0 | 0 | 0 | 1 |
| 0 | 0 | 0 | 1 |
| 0 | 0 | 0 | 1 |
| 0 | 0 | 0 | 1 |
| 0 | 0 | 0 | 1 |
| 0 | 0 | 0 | 0 |
| 0 | 0 | 0 | 1 |
| 0 | 0 | 0 | 1 |
| 0 | 0 | 0 | 1 |
| 0 | 1 | 0 | 0 |
| 0 | 1 | 0 | 0 |

| | | | |
|---|---|---|---|
| 0 | 0 | 0 | 1 |
| 0 | 0 | 0 | 1 |
| 0 | 0 | 0 | 1 |
| 0 | 1 | 1 | 1 |
| 0 | 1 | 0 | 0 |
| 1 | 0 | 0 | 0 |
| 0 | 0 | 0 | 1 |
| 0 | 0 | 0 | 0 |
| 0 | 0 | 0 | 1 |
| 0 | 0 | 0 | 1 |
| 0 | 0 | 0 | 0 |
| 0 | 1 | 1 | 1 |
| 0 | 1 | 0 | 0 |
| 0 | 0 | 0 | 0 |
| 0 | 1 | 0 | 0 |
| 0 | 1 | 0 | 0 |
| 0 | 0 | 0 | 1 |
| 0 | 0 | 0 | 1 |
| 0 | 0 | 0 | 0 |
| 0 | 1 | 0 | 0 |
| 0 | 1 | 0 | 0 |
| 0 | 1 | 0 | 0 |
| 0 | 1 | 0 | 0 |
| 0 | 0 | 0 | 1 |
| 0 | 0 | 0 | 1 |
| 0 | 0 | 0 | 0 |
| 0 | 0 | 0 | 1 |
| 0 | 0 | 0 | 0 |
| 0 | 0 | 0 | 0 |
| 0 | 0 | 0 | 0 |
| 0 | 0 | 0 | 1 |
| 1 | 0 | 0 | 0 |
| 1 | 0 | 0 | 0 |
| 0 | 0 | 0 | 1 |
| 0 | 0 | 0 | 1 |
| 0 | 0 | 0 | 0 |
| 0 | 0 | 0 | 0 |
| 0 | 1 | 1 | 1 |
| 0 | 1 | 0 | 0 |
| 0 | 1 | 0 | 0 |
| 0 | 1 | 0 | 0 |
| 0 | 0 | 0 | 0 |
| 0 | 0 | 0 | 1 |
| 0 | 0 | 0 | 1 |
| 0 | 0 | 0 | 0 |
| 0 | 0 | 0 | 1 |
| 0 | 0 | 0 | 0 |
| 0 | 0 | 0 | 1 |
| 0 | 0 | 0 | 1 |
| 0 | 0 | 0 | 1 |
| 0 | 0 | 0 | 1 |
| 0 | 0 | 0 | 1 |
| 0 | 0 | 0 | 1 |
| 0 | 0 | 0 | 1 |

| | | | |
|---|---|---|---|
| 0 | 0 | 0 | 1 |
| 0 | 0 | 0 | 1 |
| 0 | 0 | 0 | 1 |
| 0 | 0 | 0 | 1 |
| 0 | 0 | 0 | 1 |
| 0 | 0 | 0 | 1 |
| 0 | 0 | 0 | 1 |
| 0 | 0 | 0 | 0 |
| 0 | 0 | 0 | 1 |
| 1 | 0 | 0 | 0 |
| 0 | 0 | 0 | 0 |
| 0 | 0 | 0 | 0 |
| 0 | 0 | 0 | 1 |
| 0 | 0 | 0 | 1 |
| 0 | 0 | 0 | 0 |
| 0 | 0 | 0 | 1 |
| 0 | 0 | 0 | 1 |
| 0 | 0 | 0 | 1 |
| 0 | 0 | 0 | 1 |
| 0 | 0 | 0 | 1 |
| 0 | 0 | 0 | 1 |
| 0 | 0 | 0 | 1 |
| 0 | 0 | 0 | 1 |
| 0 | 0 | 0 | 1 |
| 0 | 0 | 0 | 1 |
| 0 | 0 | 0 | 1 |
| 0 | 0 | 0 | 1 |
| 0 | 1 | 1 | 1 |
| 0 | 0 | 0 | 1 |
| 0 | 0 | 0 | 0 |
| 0 | 0 | 0 | 0 |
| 0 | 0 | 0 | 1 |
| 0 | 0 | 0 | 1 |
| 0 | 0 | 0 | 1 |
| 0 | 0 | 0 | 1 |
| 0 | 0 | 0 | 0 |
| 0 | 0 | 0 | 1 |
| 0 | 0 | 0 | 1 |
| 0 | 0 | 0 | 1 |
| 0 | 0 | 0 | 1 |
| 0 | 0 | 0 | 1 |
| 0 | 0 | 0 | 0 |
| 0 | 0 | 0 | 0 |
| 0 | 1 | 0 | 0 |
| 0 | 0 | 0 | 1 |
| 0 | 0 | 0 | 1 |
| 0 | 0 | 0 | 1 |
| 0 | 0 | 0 | 1 |
| 0 | 0 | 0 | 1 |
| 0 | 0 | 0 | 1 |
| 0 | 0 | 0 | 1 |
| 0 | 0 | 0 | 1 |
| 0 | 0 | 0 | 1 |
| 0 | 0 | 0 | 0 |

| | | | |
|---|---|---|---|
| 0 | 0 | 0 | 1 |
| 0 | 0 | 0 | 1 |
| 0 | 0 | 0 | 1 |
| 0 | 0 | 0 | 1 |
| 0 | 0 | 0 | 1 |
| 0 | 0 | 0 | 1 |
| 0 | 0 | 0 | 1 |
| 0 | 0 | 0 | 1 |
| 0 | 0 | 0 | 0 |
| 0 | 0 | 0 | 0 |
| 0 | 0 | 0 | 1 |
| 0 | 0 | 0 | 1 |
| 0 | 0 | 0 | 1 |
| 0 | 0 | 0 | 0 |
| 0 | 0 | 0 | 1 |
| 0 | 0 | 0 | 1 |
| 0 | 0 | 0 | 1 |
| 1 | 0 | 0 | 0 |
| 1 | 0 | 0 | 0 |
| 0 | 0 | 0 | 0 |
| 0 | 0 | 0 | 0 |
| 0 | 0 | 0 | 1 |
| 0 | 0 | 0 | 1 |
| 0 | 0 | 0 | 1 |
| 0 | 0 | 0 | 1 |
| 1 | 0 | 0 | 0 |
| 0 | 0 | 0 | 1 |
| 0 | 0 | 0 | 1 |
| 0 | 0 | 0 | 1 |
| 0 | 0 | 0 | 1 |
| 0 | 0 | 0 | 1 |
| 0 | 0 | 0 | 1 |
| 0 | 0 | 0 | 1 |
| 0 | 0 | 0 | 1 |
| 0 | 0 | 0 | 1 |
| 0 | 0 | 0 | 1 |
| 0 | 0 | 0 | 1 |
| 0 | 0 | 0 | 1 |
| 0 | 0 | 0 | 1 |
| 0 | 0 | 0 | 0 |
| 0 | 0 | 0 | 1 |
| 0 | 0 | 0 | 1 |
| 0 | 0 | 0 | 0 |
| 0 | 0 | 0 | 1 |
| 0 | 0 | 0 | 1 |
| 0 | 0 | 0 | 1 |
| 0 | 0 | 0 | 0 |
| 0 | 0 | 0 | 0 |
| 0 | 0 | 0 | 1 |
| 0 | 0 | 0 | 1 |
| 0 | 0 | 0 | 0 |
| 0 | 0 | 0 | 1 |
| 0 | 0 | 0 | 1 |
| 0 | 0 | 0 | 1 |

| | | | |
|---|---|---|---|
| 0 | 0 | 0 | 1 |
| 0 | 0 | 0 | 1 |
| 0 | 0 | 0 | 1 |
| 0 | 0 | 0 | 1 |
| 0 | 0 | 0 | 1 |
| 0 | 0 | 0 | 1 |
| 0 | 0 | 0 | 1 |
| 0 | 0 | 0 | 1 |
| 0 | 0 | 0 | 1 |
| 0 | 0 | 0 | 1 |
| 0 | 0 | 0 | 1 |
| 0 | 0 | 0 | 1 |
| 0 | 0 | 0 | 1 |
| 0 | 0 | 0 | 0 |
| 0 | 0 | 0 | 0 |
| 0 | 0 | 0 | 0 |
| 0 | 0 | 0 | 0 |
| 0 | 0 | 0 | 1 |
| 0 | 0 | 0 | 1 |
| 0 | 0 | 0 | 0 |
| 0 | 0 | 0 | 1 |
| 0 | 1 | 1 | 1 |
| 0 | 0 | 0 | 1 |
| 0 | 0 | 0 | 1 |
| 0 | 0 | 0 | 1 |
| 0 | 0 | 0 | 0 |
| 0 | 0 | 0 | 1 |
| 0 | 0 | 0 | 1 |
| 0 | 0 | 0 | 1 |
| 0 | 0 | 0 | 0 |
| 0 | 0 | 0 | 0 |
| 0 | 0 | 0 | 1 |
| 0 | 0 | 0 | 1 |
| 0 | 0 | 0 | 1 |
| 0 | 0 | 0 | 1 |
| 0 | 0 | 0 | 1 |
| 0 | 0 | 0 | 1 |
| 0 | 0 | 0 | 1 |
| 0 | 0 | 0 | 0 |
| 0 | 0 | 0 | 1 |
| 0 | 0 | 0 | 1 |
| 0 | 1 | 0 | 0 |
| 0 | 1 | 0 | 0 |
| 0 | 1 | 0 | 0 |
| 0 | 0 | 0 | 0 |
| 0 | 0 | 0 | 1 |
| 0 | 0 | 0 | 0 |
| 0 | 0 | 0 | 1 |
| 0 | 0 | 0 | 1 |
| 0 | 0 | 0 | 1 |
| 0 | 0 | 0 | 1 |
| 0 | 1 | 1 | 1 |
| 0 | 1 | 1 | 1 |
| 0 | 1 | 1 | 1 |

| | | | |
|---|---|---|---|
| 0 | 0 | 0 | 1 |
| 0 | 0 | 0 | 1 |
| 0 | 0 | 0 | 0 |
| 0 | 0 | 0 | 1 |
| 0 | 0 | 0 | 1 |
| 0 | 0 | 0 | 1 |
| 0 | 0 | 0 | 1 |
| 0 | 0 | 0 | 1 |
| 0 | 0 | 0 | 1 |
| 0 | 0 | 0 | 1 |
| 0 | 0 | 0 | 1 |
| 0 | 0 | 0 | 1 |
| 0 | 0 | 0 | 1 |
| 0 | 0 | 0 | 1 |
| 0 | 0 | 0 | 1 |
| 0 | 0 | 0 | 1 |
| 0 | 0 | 0 | 1 |
| 0 | 0 | 0 | 1 |
| 0 | 0 | 0 | 1 |
| 0 | 0 | 0 | 1 |
| 0 | 0 | 0 | 1 |
| 0 | 0 | 0 | 1 |
| 0 | 0 | 0 | 1 |
| 0 | 0 | 0 | 1 |
| 0 | 0 | 0 | 1 |
| 0 | 0 | 0 | 1 |
| 0 | 0 | 0 | 1 |
| 0 | 0 | 0 | 1 |
| 0 | 0 | 0 | 1 |
| 0 | 0 | 0 | 1 |
| 1 | 0 | 0 | 0 |
| 0 | 0 | 0 | 1 |
| 0 | 1 | 0 | 0 |
| 0 | 1 | 0 | 0 |
| 0 | 1 | 0 | 0 |
| 0 | 0 | 0 | 0 |
| 0 | 0 | 0 | 1 |
| 0 | 0 | 0 | 0 |
| 0 | 0 | 0 | 1 |
| 0 | 0 | 0 | 1 |
| 1 | 0 | 0 | 0 |
| 1 | 0 | 0 | 0 |
| 0 | 0 | 0 | 0 |
| 0 | 0 | 0 | 1 |
| 0 | 0 | 0 | 0 |
| 0 | 0 | 0 | 0 |
| 0 | 0 | 0 | 1 |
| 0 | 0 | 0 | 1 |
| 0 | 0 | 0 | 1 |
| 0 | 0 | 0 | 1 |
| 0 | 0 | 0 | 0 |
| 0 | 0 | 0 | 1 |
| 0 | 0 | 0 | 1 |
| 0 | 0 | 0 | 1 |

| | | | |
|---|---|---|---|
| 0 | 0 | 0 | 1 |
| 0 | 0 | 0 | 1 |
| 0 | 0 | 0 | 1 |
| 0 | 0 | 0 | 1 |
| 0 | 0 | 0 | 1 |
| 1 | 0 | 0 | 0 |
| 0 | 0 | 0 | 1 |
| 0 | 0 | 0 | 0 |
| 0 | 0 | 0 | 0 |
| 0 | 0 | 0 | 0 |
| 0 | 0 | 0 | 1 |
| 0 | 1 | 0 | 0 |
| 0 | 0 | 0 | 1 |
| 0 | 0 | 0 | 1 |
| 0 | 0 | 0 | 1 |
| 0 | 1 | 1 | 1 |
| 0 | 0 | 0 | 1 |
| 0 | 0 | 0 | 0 |
| 0 | 0 | 0 | 1 |
| 0 | 0 | 0 | 0 |
| 0 | 0 | 0 | 1 |
| 0 | 0 | 0 | 0 |
| 0 | 0 | 0 | 0 |
| 0 | 0 | 0 | 0 |
| 0 | 0 | 0 | 1 |
| 0 | 0 | 0 | 1 |
| 0 | 0 | 0 | 1 |
| 0 | 0 | 0 | 1 |
| 0 | 0 | 0 | 0 |
| 0 | 0 | 0 | 1 |
| 0 | 0 | 0 | 1 |
| 0 | 0 | 0 | 1 |
| 0 | 0 | 0 | 1 |
| 0 | 0 | 0 | 1 |
| 0 | 0 | 0 | 0 |
| 0 | 0 | 0 | 0 |
| 0 | 0 | 0 | 1 |
| 0 | 0 | 0 | 0 |
| 0 | 0 | 0 | 0 |
| 0 | 0 | 0 | 1 |
| 0 | 0 | 0 | 0 |
| 0 | 0 | 0 | 0 |
| 0 | 0 | 0 | 1 |
| 0 | 0 | 0 | 1 |
| 0 | 0 | 0 | 1 |
| 0 | 0 | 0 | 1 |
| 0 | 0 | 0 | 1 |
| 0 | 0 | 0 | 1 |
| 0 | 0 | 0 | 1 |
| 0 | 0 | 0 | 1 |
| 0 | 0 | 0 | 1 |
| 0 | 0 | 0 | 1 |
| 0 | 0 | 0 | 1 |
| 0 | 0 | 0 | 1 |

| | | | |
|---|---|---|---|
| 0 | 0 | 0 | 1 |
| 0 | 0 | 0 | 0 |
| 0 | 0 | 0 | 1 |
| 0 | 0 | 0 | 1 |
| 0 | 0 | 0 | 0 |
| 0 | 0 | 0 | 1 |
| 0 | 0 | 0 | 1 |
| 0 | 0 | 0 | 1 |
| 0 | 0 | 0 | 0 |
| 0 | 0 | 0 | 0 |
| 0 | 0 | 0 | 1 |
| 0 | 0 | 0 | 0 |
| 0 | 0 | 0 | 0 |
| 0 | 0 | 0 | 1 |
| 0 | 0 | 0 | 1 |
| 0 | 0 | 0 | 0 |
| 0 | 0 | 0 | 1 |
| 0 | 0 | 0 | 0 |
| 0 | 1 | 0 | 0 |
| 0 | 0 | 0 | 1 |
| 0 | 0 | 0 | 0 |
| 0 | 0 | 0 | 0 |
| 1 | 0 | 0 | 0 |
| 0 | 0 | 0 | 1 |
| 0 | 0 | 0 | 1 |
| 0 | 0 | 0 | 0 |
| 0 | 0 | 0 | 1 |
| 0 | 0 | 0 | 1 |
| 0 | 0 | 0 | 1 |
| 0 | 0 | 0 | 1 |
| 0 | 0 | 0 | 0 |
| 0 | 0 | 0 | 1 |
| 0 | 0 | 0 | 0 |
| 0 | 0 | 0 | 1 |
| 0 | 0 | 0 | 0 |
| 0 | 0 | 0 | 0 |
| 0 | 0 | 0 | 1 |
| 0 | 0 | 0 | 1 |
| 0 | 0 | 0 | 1 |
| 0 | 0 | 0 | 1 |
| 0 | 0 | 0 | 1 |
| 0 | 0 | 0 | 1 |
| 0 | 0 | 0 | 1 |
| 0 | 0 | 0 | 1 |
| 0 | 0 | 0 | 1 |
| 0 | 0 | 0 | 1 |
| 0 | 1 | 1 | 1 |
| 0 | 0 | 0 | 1 |
| 0 | 0 | 0 | 1 |
| 0 | 0 | 0 | 1 |
| 0 | 0 | 0 | 1 |
| 0 | 0 | 0 | 1 |
| 0 | 0 | 0 | 1 |
| 0 | 0 | 0 | 0 |

| | | | |
|---|---|---|---|
| 0 | 0 | 0 | 1 |
| 0 | 0 | 0 | 1 |
| 0 | 0 | 0 | 1 |
| 0 | 0 | 0 | 1 |
| 0 | 0 | 0 | 0 |
| 0 | 0 | 0 | 1 |
| 0 | 0 | 0 | 1 |
| 0 | 0 | 0 | 1 |
| 0 | 0 | 0 | 1 |
| 0 | 0 | 0 | 1 |
| 0 | 0 | 0 | 0 |
| 0 | 0 | 0 | 1 |
| 0 | 0 | 0 | 1 |
| 0 | 0 | 0 | 1 |
| 0 | 0 | 0 | 0 |
| 0 | 0 | 0 | 0 |
| 0 | 0 | 0 | 0 |
| 0 | 0 | 0 | 0 |
| 0 | 0 | 0 | 1 |
| 0 | 0 | 0 | 0 |
| 0 | 0 | 0 | 0 |
| 1 | 0 | 0 | 0 |
| 0 | 0 | 0 | 1 |
| 0 | 0 | 0 | 0 |
| 1 | 0 | 0 | 0 |
| 0 | 0 | 0 | 1 |
| 1 | 0 | 0 | 0 |
| 0 | 0 | 0 | 0 |
| 1 | 0 | 0 | 0 |
| 1 | 0 | 0 | 0 |
| 1 | 0 | 0 | 0 |
| 0 | 0 | 0 | 1 |
| 0 | 0 | 0 | 1 |
| 0 | 0 | 0 | 1 |
| 0 | 0 | 0 | 1 |
| 0 | 0 | 0 | 1 |
| 0 | 0 | 0 | 1 |
| 0 | 1 | 0 | 0 |
| 0 | 0 | 0 | 0 |
| 0 | 0 | 0 | 1 |
| 0 | 0 | 0 | 1 |
| 0 | 1 | 1 | 1 |
| 0 | 0 | 0 | 0 |
| 0 | 0 | 0 | 1 |
| 0 | 0 | 0 | 0 |
| 0 | 0 | 0 | 1 |
| 0 | 0 | 0 | 1 |
| 0 | 0 | 0 | 1 |
| 0 | 1 | 0 | 0 |
| 0 | 1 | 0 | 0 |
| 0 | 1 | 0 | 0 |
| 0 | 0 | 0 | 1 |
| 0 | 0 | 0 | 1 |
| 0 | 0 | 0 | 0 |

| | | | |
|---|---|---|---|
| 0 | 0 | 0 | 1 |
| 0 | 1 | 0 | 0 |
| 0 | 0 | 0 | 1 |
| 0 | 1 | 0 | 0 |
| 0 | 1 | 0 | 0 |
| 0 | 0 | 0 | 1 |
| 0 | 1 | 0 | 0 |
| 0 | 1 | 0 | 0 |
| 0 | 0 | 0 | 1 |
| 0 | 0 | 0 | 1 |
| 0 | 0 | 0 | 1 |
| 0 | 0 | 0 | 1 |
| 0 | 0 | 0 | 1 |
| 0 | 0 | 0 | 1 |
| 0 | 0 | 0 | 1 |
| 0 | 0 | 0 | 1 |
| 0 | 0 | 0 | 1 |
| 0 | 0 | 0 | 1 |
| 0 | 0 | 0 | 1 |
| 0 | 0 | 0 | 1 |
| 1 | 1 | 0 | 0 |
| 0 | 0 | 0 | 1 |
| 0 | 0 | 0 | 1 |
| 0 | 0 | 0 | 1 |
| 0 | 0 | 0 | 1 |
| 0 | 0 | 0 | 0 |
| 0 | 0 | 0 | 0 |
| 0 | 0 | 0 | 1 |
| 0 | 0 | 0 | 1 |
| 0 | 0 | 0 | 1 |
| 0 | 0 | 0 | 1 |
| 0 | 0 | 0 | 1 |
| 0 | 0 | 0 | 1 |
| 0 | 0 | 0 | 1 |
| 0 | 0 | 0 | 1 |
| 0 | 0 | 0 | 1 |
| 0 | 0 | 0 | 1 |
| 0 | 0 | 0 | 1 |
| 0 | 0 | 0 | 1 |
| 0 | 0 | 0 | 1 |
| 0 | 0 | 0 | 1 |
| 0 | 0 | 0 | 1 |
| 0 | 0 | 0 | 1 |
| 0 | 0 | 0 | 0 |
| 0 | 0 | 0 | 1 |
| 0 | 0 | 0 | 0 |
| 0 | 0 | 0 | 0 |
| 0 | 0 | 0 | 0 |
| 0 | 0 | 0 | 0 |
| 0 | 0 | 0 | 0 |
| 0 | 0 | 0 | 0 |
| 0 | 0 | 0 | 1 |
| 0 | 0 | 0 | 1 |
| 0 | 0 | 0 | 1 |

| | | | |
|---|---|---|---|
| 0 | 0 | 0 | 1 |
| 0 | 0 | 0 | 1 |
| 0 | 0 | 0 | 1 |
| 0 | 0 | 0 | 0 |
| 0 | 0 | 0 | 1 |
| 0 | 0 | 0 | 0 |
| 0 | 0 | 0 | 1 |
| 0 | 0 | 0 | 1 |
| 0 | 0 | 0 | 0 |
| 0 | 0 | 0 | 1 |
| 0 | 0 | 0 | 1 |
| 0 | 0 | 0 | 1 |
| 0 | 0 | 0 | 1 |
| 0 | 0 | 0 | 1 |
| 0 | 0 | 0 | 1 |
| 0 | 0 | 0 | 0 |
| 0 | 0 | 0 | 0 |
| 0 | 0 | 0 | 0 |
| 0 | 0 | 0 | 0 |
| 0 | 0 | 0 | 0 |
| 0 | 0 | 0 | 1 |
| 0 | 0 | 0 | 1 |
| 0 | 0 | 0 | 1 |
| 0 | 0 | 0 | 1 |
| 0 | 0 | 0 | 0 |
| 0 | 0 | 0 | 1 |
| 0 | 0 | 0 | 1 |
| 0 | 0 | 0 | 1 |
| 0 | 0 | 0 | 0 |
| 0 | 0 | 0 | 1 |
| 0 | 1 | 0 | 0 |
| 0 | 1 | 0 | 0 |
| 0 | 0 | 0 | 0 |
| 0 | 1 | 0 | 0 |
| 0 | 1 | 0 | 0 |
| 0 | 0 | 0 | 1 |
| 0 | 1 | 0 | 0 |
| 0 | 1 | 0 | 0 |
| 0 | 1 | 0 | 0 |
| 0 | 1 | 0 | 0 |
| 0 | 1 | 0 | 0 |
| 0 | 0 | 0 | 0 |
| 0 | 0 | 0 | 0 |
| 0 | 0 | 0 | 0 |
| 0 | 0 | 0 | 0 |
| 0 | 0 | 0 | 0 |
| 0 | 0 | 0 | 0 |
| 0 | 0 | 0 | 0 |
| 0 | 0 | 0 | 1 |
| 0 | 0 | 0 | 0 |
| 0 | 0 | 0 | 1 |
| 0 | 0 | 0 | 1 |
| 0 | 0 | 0 | 1 |
| 0 | 0 | 0 | 0 |

| | | | |
|---|---|---|---|
| 0 | 1 | 0 | 0 |
| 0 | 0 | 0 | 1 |
| 0 | 0 | 0 | 1 |
| 0 | 0 | 0 | 1 |
| 0 | 0 | 0 | 1 |
| 0 | 0 | 0 | 0 |
| 0 | 0 | 0 | 1 |
| 0 | 0 | 0 | 1 |
| 0 | 0 | 0 | 1 |
| 0 | 0 | 0 | 1 |
| 0 | 0 | 0 | 1 |
| 0 | 0 | 0 | 1 |
| 0 | 0 | 0 | 1 |
| 1 | 0 | 0 | 0 |
| 0 | 0 | 0 | 1 |
| 0 | 0 | 0 | 1 |
| 0 | 0 | 0 | 1 |
| 0 | 0 | 0 | 0 |
| 0 | 0 | 0 | 1 |
| 0 | 0 | 0 | 1 |
| 0 | 0 | 0 | 1 |
| 0 | 0 | 0 | 1 |
| 0 | 0 | 0 | 0 |
| 0 | 0 | 0 | 1 |
| 0 | 0 | 0 | 1 |
| 0 | 0 | 0 | 0 |
| 0 | 0 | 0 | 0 |
| 0 | 0 | 0 | 0 |
| 0 | 0 | 0 | 1 |
| 0 | 0 | 0 | 1 |
| 0 | 0 | 0 | 1 |
| 0 | 0 | 0 | 1 |
| 0 | 0 | 0 | 1 |
| 0 | 0 | 0 | 1 |
| 0 | 0 | 0 | 1 |
| 0 | 0 | 0 | 1 |
| 0 | 0 | 0 | 1 |
| 0 | 0 | 0 | 1 |
| 0 | 0 | 0 | 1 |
| 0 | 0 | 0 | 1 |
| 0 | 0 | 0 | 1 |
| 0 | 0 | 0 | 1 |
| 0 | 0 | 0 | 1 |
| 0 | 0 | 0 | 1 |
| 0 | 0 | 0 | 1 |
| 0 | 0 | 0 | 1 |
| 0 | 0 | 0 | 1 |
| 0 | 0 | 0 | 1 |
| 0 | 1 | 1 | 1 |
| 0 | 0 | 0 | 1 |
| 0 | 0 | 0 | 1 |
| 0 | 0 | 0 | 1 |
| 0 | 0 | 0 | 1 |
| 0 | 0 | 0 | 1 |

| | | | |
|---|---|---|---|
| 0 | 0 | 0 | 1 |
| 0 | 0 | 0 | 0 |
| 0 | 0 | 0 | 1 |
| 0 | 0 | 0 | 1 |
| 0 | 0 | 0 | 1 |
| 0 | 0 | 0 | 0 |
| 0 | 0 | 0 | 0 |
| 0 | 0 | 0 | 0 |
| 0 | 0 | 0 | 1 |
| 0 | 0 | 0 | 1 |
| 0 | 0 | 0 | 1 |
| 0 | 0 | 0 | 1 |
| 0 | 0 | 0 | 1 |
| 0 | 0 | 0 | 1 |
| 0 | 0 | 0 | 0 |
| 0 | 0 | 0 | 0 |
| 0 | 0 | 0 | 0 |
| 0 | 0 | 0 | 1 |
| 0 | 0 | 0 | 0 |
| 0 | 0 | 0 | 0 |
| 0 | 0 | 0 | 0 |
| 0 | 0 | 0 | 0 |
| 0 | 0 | 0 | 0 |
| 0 | 0 | 0 | 0 |
| 0 | 0 | 0 | 1 |
| 0 | 0 | 0 | 1 |
| 0 | 0 | 0 | 1 |
| 0 | 0 | 0 | 1 |
| 0 | 0 | 0 | 1 |
| 0 | 0 | 0 | 1 |
| 0 | 0 | 0 | 1 |
| 0 | 0 | 0 | 1 |
| 0 | 0 | 0 | 0 |
| 0 | 0 | 0 | 1 |
| 0 | 0 | 0 | 1 |
| 0 | 0 | 0 | 0 |
| 0 | 0 | 0 | 0 |
| 0 | 0 | 0 | 0 |
| 0 | 0 | 0 | 0 |
| 0 | 0 | 0 | 1 |
| 0 | 0 | 0 | 1 |
| 0 | 0 | 0 | 1 |
| 0 | 0 | 0 | 1 |
| 0 | 0 | 0 | 0 |
| 0 | 0 | 0 | 0 |
| 0 | 0 | 0 | 0 |
| 0 | 0 | 0 | 1 |
| 0 | 0 | 0 | 1 |
| 0 | 0 | 0 | 0 |
| 0 | 0 | 0 | 1 |
| 0 | 0 | 0 | 1 |
| 0 | 0 | 0 | 0 |
| 0 | 0 | 0 | 1 |
| 0 | 0 | 0 | 1 |

| | | | |
|---|---|---|---|
| 0 | 0 | 0 | 1 |
| 0 | 0 | 0 | 1 |
| 0 | 0 | 0 | 1 |
| 0 | 0 | 0 | 1 |
| 0 | 0 | 0 | 1 |
| 0 | 0 | 0 | 1 |
| 0 | 0 | 0 | 1 |
| 0 | 0 | 0 | 1 |
| 0 | 0 | 0 | 0 |
| 0 | 0 | 0 | 0 |
| 0 | 0 | 0 | 1 |
| 0 | 0 | 0 | 0 |
| 0 | 0 | 0 | 0 |
| 0 | 0 | 0 | 0 |
| 0 | 0 | 0 | 1 |
| 0 | 0 | 0 | 1 |
| 0 | 0 | 0 | 0 |
| 0 | 0 | 0 | 0 |
| 0 | 0 | 0 | 0 |
| 0 | 0 | 0 | 1 |
| 0 | 0 | 0 | 1 |
| 0 | 0 | 0 | 1 |
| 0 | 0 | 0 | 0 |
| 0 | 0 | 0 | 1 |
| 0 | 0 | 0 | 0 |
| 0 | 0 | 0 | 1 |
| 0 | 0 | 0 | 1 |
| 0 | 0 | 0 | 1 |
| 0 | 0 | 0 | 1 |
| 0 | 0 | 0 | 0 |
| 0 | 0 | 0 | 0 |
| 0 | 0 | 0 | 0 |
| 0 | 0 | 0 | 1 |
| 0 | 0 | 0 | 1 |
| 0 | 0 | 0 | 1 |
| 0 | 0 | 0 | 1 |
| 0 | 0 | 0 | 1 |
| 0 | 0 | 0 | 1 |
| 0 | 0 | 0 | 1 |
| 0 | 0 | 0 | 1 |
| 0 | 1 | 0 | 0 |
| 0 | 0 | 0 | 1 |
| 0 | 0 | 0 | 1 |
| 0 | 0 | 0 | 1 |
| 0 | 0 | 0 | 1 |
| 0 | 0 | 0 | 0 |
| 0 | 1 | 1 | 1 |
| 0 | 0 | 0 | 0 |
| 0 | 0 | 0 | 1 |
| 0 | 1 | 0 | 0 |
| 0 | 1 | 0 | 0 |
| 0 | 0 | 0 | 0 |
| 0 | 0 | 0 | 1 |
| 0 | 1 | 1 | 1 |

| | | | |
|---|---|---|---|
| 0 | 1 | 1 | 1 |
| 0 | 1 | 1 | 1 |
| 0 | 0 | 0 | 1 |
| 0 | 0 | 0 | 1 |
| 0 | 0 | 0 | 1 |
| 0 | 0 | 0 | 1 |
| 0 | 0 | 0 | 1 |
| 0 | 0 | 0 | 0 |
| 0 | 0 | 0 | 0 |
| 0 | 0 | 0 | 1 |
| 0 | 0 | 0 | 0 |
| 0 | 0 | 0 | 1 |
| 0 | 0 | 0 | 1 |
| 0 | 0 | 0 | 1 |
| 0 | 0 | 0 | 1 |
| 0 | 0 | 0 | 1 |
| 0 | 0 | 0 | 1 |
| 0 | 0 | 0 | 1 |
| 0 | 0 | 0 | 1 |
| 0 | 0 | 0 | 1 |
| 0 | 0 | 0 | 1 |
| 0 | 0 | 0 | 1 |
| 0 | 0 | 0 | 1 |
| 0 | 0 | 0 | 1 |
| 0 | 0 | 0 | 1 |
| 0 | 0 | 0 | 1 |
| 0 | 0 | 0 | 1 |
| 0 | 0 | 0 | 1 |
| 0 | 0 | 0 | 0 |
| 0 | 0 | 0 | 1 |
| 0 | 0 | 0 | 0 |
| 0 | 0 | 0 | 1 |
| 0 | 0 | 0 | 0 |
| 0 | 1 | 0 | 0 |
| 0 | 0 | 0 | 1 |
| 0 | 0 | 0 | 0 |
| 0 | 0 | 0 | 0 |
| 0 | 0 | 0 | 1 |
| 0 | 0 | 0 | 1 |
| 0 | 0 | 0 | 1 |
| 0 | 0 | 0 | 1 |
| 0 | 0 | 0 | 0 |
| 0 | 0 | 0 | 0 |
| 0 | 0 | 0 | 1 |
| 0 | 0 | 0 | 1 |
| 0 | 1 | 0 | 0 |
| 0 | 0 | 0 | 0 |
| 0 | 1 | 0 | 0 |
| 0 | 0 | 0 | 0 |
| 0 | 0 | 0 | 1 |
| 0 | 1 | 0 | 0 |
| 0 | 0 | 0 | 0 |
| 0 | 0 | 0 | 1 |
| 0 | 0 | 0 | 1 |

| | | | |
|---|---|---|---|
| 0 | 0 | 0 | 1 |
| 0 | 0 | 0 | 1 |
| 0 | 0 | 0 | 1 |
| 0 | 0 | 0 | 1 |
| 0 | 0 | 0 | 0 |
| 0 | 0 | 0 | 0 |
| 0 | 0 | 0 | 0 |
| 0 | 0 | 0 | 0 |
| 0 | 0 | 0 | 1 |
| 0 | 0 | 0 | 0 |
| 0 | 0 | 0 | 0 |
| 0 | 0 | 0 | 0 |
| 0 | 0 | 0 | 0 |
| 0 | 0 | 0 | 1 |
| 0 | 1 | 1 | 1 |
| 0 | 0 | 0 | 1 |
| 0 | 0 | 0 | 1 |
| 0 | 0 | 0 | 1 |
| 0 | 0 | 0 | 1 |
| 0 | 0 | 0 | 1 |
| 0 | 0 | 0 | 1 |
| 0 | 0 | 0 | 1 |
| 0 | 0 | 0 | 1 |
| 0 | 0 | 0 | 1 |
| 0 | 0 | 0 | 1 |
| 0 | 0 | 0 | 0 |
| 0 | 0 | 0 | 0 |
| 0 | 0 | 0 | 0 |
| 0 | 0 | 0 | 0 |
| 0 | 0 | 0 | 1 |
| 0 | 0 | 0 | 1 |
| 0 | 0 | 0 | 1 |
| 0 | 0 | 0 | 1 |
| 0 | 0 | 0 | 1 |
| 0 | 0 | 0 | 1 |
| 0 | 0 | 0 | 1 |
| 0 | 0 | 0 | 1 |
| 0 | 0 | 0 | 1 |
| 0 | 0 | 0 | 1 |
| 0 | 0 | 0 | 1 |
| 0 | 0 | 0 | 1 |
| 0 | 0 | 0 | 1 |
| 0 | 0 | 0 | 1 |
| 0 | 0 | 0 | 1 |
| 0 | 0 | 0 | 1 |
| 0 | 0 | 0 | 1 |
| 0 | 0 | 0 | 1 |
| 0 | 0 | 0 | 1 |
| 0 | 0 | 0 | 1 |
| 0 | 0 | 0 | 1 |
| 0 | 0 | 0 | 1 |
| 0 | 0 | 0 | 1 |
| 0 | 0 | 0 | 1 |
| 0 | 0 | 0 | 0 |

| | | | |
|---|---|---|---|
| 0 | 0 | 0 | 1 |
| 0 | 0 | 0 | 1 |
| 0 | 0 | 0 | 1 |
| 0 | 0 | 0 | 1 |
| 0 | 0 | 0 | 1 |
| 0 | 0 | 0 | 1 |
| 0 | 0 | 0 | 1 |
| 0 | 0 | 0 | 1 |
| 0 | 0 | 0 | 1 |
| 0 | 0 | 0 | 1 |
| 0 | 0 | 0 | 1 |
| 0 | 0 | 0 | 1 |
| 0 | 0 | 0 | 1 |
| 0 | 0 | 0 | 1 |
| 0 | 0 | 0 | 1 |
| 0 | 0 | 0 | 1 |
| 0 | 0 | 0 | 1 |
| 0 | 0 | 0 | 1 |
| 0 | 0 | 0 | 1 |
| 0 | 0 | 0 | 1 |
| 0 | 0 | 0 | 1 |
| 0 | 0 | 0 | 1 |
| 0 | 0 | 0 | 1 |
| 0 | 0 | 0 | 1 |
| 0 | 0 | 0 | 1 |
| 0 | 0 | 0 | 1 |
| 0 | 0 | 0 | 1 |
| 0 | 0 | 0 | 1 |
| 0 | 0 | 0 | 1 |
| 0 | 0 | 0 | 1 |
| 0 | 0 | 0 | 1 |
| 0 | 0 | 0 | 1 |
| 0 | 0 | 0 | 1 |
| 0 | 0 | 0 | 1 |
| 0 | 0 | 0 | 1 |
| 0 | 0 | 0 | 1 |
| 0 | 0 | 0 | 1 |
| 0 | 0 | 0 | 1 |
| 0 | 0 | 0 | 1 |
| 0 | 0 | 0 | 1 |
| 0 | 0 | 0 | 1 |
| 0 | 0 | 0 | 1 |
| 0 | 0 | 0 | 1 |
| 0 | 0 | 0 | 1 |
| 0 | 0 | 0 | 1 |
| 0 | 0 | 0 | 1 |
| 0 | 1 | 1 | 1 |
| 0 | 0 | 0 | 0 |
| 0 | 0 | 0 | 1 |
| 0 | 0 | 0 | 1 |
| 0 | 0 | 0 | 1 |
| 0 | 0 | 0 | 1 |
| 0 | 0 | 0 | 1 |
| 0 | 0 | 0 | 1 |

| | | | |
|---|---|---|---|
| 0 | 0 | 0 | 1 |
| 0 | 0 | 0 | 1 |
| 0 | 0 | 0 | 0 |
| 0 | 0 | 0 | 1 |
| 0 | 0 | 0 | 1 |
| 0 | 0 | 0 | 1 |
| 0 | 0 | 0 | 0 |
| 0 | 0 | 0 | 1 |
| 0 | 0 | 0 | 1 |
| 0 | 0 | 0 | 1 |
| 0 | 0 | 0 | 1 |
| 0 | 0 | 0 | 0 |
| 0 | 0 | 0 | 1 |
| 0 | 0 | 0 | 0 |
| 0 | 0 | 0 | 0 |
| 0 | 0 | 0 | 0 |
| 0 | 0 | 0 | 0 |
| 0 | 0 | 0 | 0 |
| 0 | 0 | 0 | 0 |
| 0 | 0 | 0 | 0 |
| 0 | 0 | 0 | 0 |

| surface_non_binding_site | non_binding_site_single_water | surface_non_binding_site_single_water |
|---|---|---|
| 0 | 0 | 0 |
| 0 | 0 | 0 |
| 0 | 0 | 0 |
| 0 | 0 | 0 |
| 0 | 0 | 0 |
| 0 | 0 | 0 |
| 1 | 1 | 1 |
| 1 | 1 | 1 |
| 0 | 0 | 0 |
| 0 | 0 | 0 |
| 0 | 0 | 0 |
| 0 | 0 | 0 |
| 0 | 0 | 0 |
| 0 | 0 | 0 |
| 0 | 0 | 0 |
| 0 | 0 | 0 |
| 0 | 0 | 0 |
| 0 | 0 | 0 |
| 0 | 0 | 0 |
| 0 | 0 | 0 |
| 0 | 0 | 0 |
| 0 | 0 | 0 |
| 0 | 0 | 0 |
| 0 | 0 | 0 |
| 0 | 0 | 0 |
| 1 | 0 | 0 |
| 0 | 0 | 0 |
| 1 | 1 | 1 |
| 0 | 0 | 0 |
| 0 | 1 | 0 |
| 0 | 1 | 0 |
| 0 | 0 | 0 |
| 0 | 0 | 0 |
| 0 | 0 | 0 |
| 0 | 0 | 0 |
| 1 | 1 | 1 |
| 0 | 0 | 0 |
| 0 | 0 | 0 |
| 0 | 0 | 0 |
| 0 | 0 | 0 |
| 0 | 0 | 0 |
| 0 | 0 | 0 |
| 0 | 0 | 0 |
| 0 | 0 | 0 |
| 0 | 0 | 0 |
| 0 | 0 | 0 |
| 0 | 0 | 0 |
| 0 | 0 | 0 |
| 0 | 0 | 0 |
| 0 | 0 | 0 |
| 0 | 0 | 0 |
| 0 | 0 | 0 |
| 0 | 0 | 0 |

| | | |
|---|---|---|
| 0 | 0 | 0 |
| 0 | 0 | 0 |
| 0 | 0 | 0 |
| 0 | 0 | 0 |
| 0 | 0 | 0 |
| 0 | 0 | 0 |
| 0 | 0 | 0 |
| 0 | 0 | 0 |
| 0 | 0 | 0 |
| 0 | 0 | 0 |
| 0 | 0 | 0 |
| 0 | 0 | 0 |
| 0 | 0 | 0 |
| 0 | 0 | 0 |
| 0 | 0 | 0 |
| 0 | 0 | 0 |
| 0 | 0 | 0 |
| 0 | 0 | 0 |
| 0 | 0 | 0 |
| 0 | 0 | 0 |
| 0 | 0 | 0 |
| 0 | 0 | 0 |
| 0 | 0 | 0 |
| 0 | 0 | 0 |
| 0 | 0 | 0 |
| 0 | 0 | 0 |
| 0 | 0 | 0 |
| 0 | 0 | 0 |
| 0 | 0 | 0 |
| 0 | 0 | 0 |
| 0 | 0 | 0 |
| 0 | 0 | 0 |
| 0 | 0 | 0 |
| 0 | 0 | 0 |
| 0 | 0 | 0 |
| 0 | 0 | 0 |
| 0 | 0 | 0 |
| 0 | 0 | 0 |
| 0 | 0 | 0 |
| 0 | 0 | 0 |
| 0 | 0 | 0 |
| 0 | 0 | 0 |
| 0 | 0 | 0 |
| 0 | 0 | 0 |
| 0 | 0 | 0 |
| 1 | 1 | 1 |
| 0 | 1 | 0 |
| 0 | 0 | 0 |
| 0 | 0 | 0 |
| 0 | 0 | 0 |
| 0 | 0 | 0 |
| 1 | 1 | 1 |
| 1 | 1 | 1 |
| 0 | 0 | 0 |

| | | |
|---|---|---|
| 0 | 1 | 0 |
| 0 | 1 | 0 |
| 0 | 0 | 0 |
| 0 | 0 | 0 |
| 0 | 0 | 0 |
| 0 | 0 | 0 |
| 0 | 1 | 0 |
| 0 | 0 | 0 |
| 0 | 0 | 0 |
| 0 | 0 | 0 |
| 0 | 0 | 0 |
| 0 | 0 | 0 |
| 0 | 0 | 0 |
| 0 | 0 | 0 |
| 0 | 0 | 0 |
| 1 | 1 | 1 |
| 0 | 0 | 0 |
| 0 | 0 | 0 |
| 0 | 0 | 0 |
| 0 | 0 | 0 |
| 0 | 0 | 0 |
| 0 | 0 | 0 |
| 0 | 0 | 0 |
| 0 | 0 | 0 |
| 0 | 0 | 0 |
| 0 | 0 | 0 |
| 0 | 0 | 0 |
| 0 | 0 | 0 |
| 0 | 0 | 0 |
| 0 | 0 | 0 |
| 0 | 0 | 0 |
| 0 | 0 | 0 |
| 0 | 0 | 0 |
| 0 | 0 | 0 |
| 0 | 0 | 0 |
| 0 | 0 | 0 |
| 0 | 0 | 0 |
| 0 | 0 | 0 |
| 0 | 0 | 0 |
| 0 | 0 | 0 |
| 0 | 0 | 0 |
| 0 | 0 | 0 |
| 0 | 0 | 0 |
| 0 | 0 | 0 |
| 0 | 0 | 0 |
| 0 | 0 | 0 |
| 0 | 0 | 0 |
| 0 | 0 | 0 |
| 0 | 0 | 0 |
| 0 | 0 | 0 |
| 0 | 0 | 0 |
| 0 | 0 | 0 |
| 0 | 0 | 0 |
| 0 | 0 | 0 |

| | | |
|---|---|---|
| 1 | 0 | 0 |
| 1 | 0 | 0 |
| 1 | 0 | 0 |
| 0 | 0 | 0 |
| 0 | 0 | 0 |
| 0 | 0 | 0 |
| 0 | 0 | 0 |
| 1 | 0 | 0 |
| 0 | 0 | 0 |
| 1 | 0 | 0 |
| 1 | 0 | 0 |
| 0 | 0 | 0 |
| 1 | 0 | 0 |
| 0 | 0 | 0 |
| 0 | 0 | 0 |
| 1 | 0 | 0 |
| 1 | 0 | 0 |
| 1 | 0 | 0 |
| 1 | 0 | 0 |
| 0 | 0 | 0 |
| 0 | 0 | 0 |
| 0 | 1 | 0 |
| 0 | 0 | 0 |
| 0 | 0 | 0 |
| 0 | 0 | 0 |
| 1 | 0 | 0 |
| 0 | 0 | 0 |
| 0 | 0 | 0 |
| 1 | 0 | 0 |
| 1 | 0 | 0 |
| 1 | 1 | 1 |
| 0 | 0 | 0 |
| 0 | 0 | 0 |
| 0 | 1 | 0 |
| 0 | 0 | 0 |
| 0 | 0 | 0 |
| 0 | 0 | 0 |
| 0 | 0 | 0 |
| 0 | 0 | 0 |
| 1 | 0 | 0 |
| 1 | 0 | 0 |
| 0 | 0 | 0 |
| 0 | 0 | 0 |
| 1 | 1 | 1 |
| 1 | 0 | 0 |
| 1 | 0 | 0 |
| 0 | 0 | 0 |
| 0 | 0 | 0 |
| 0 | 0 | 0 |
| 0 | 0 | 0 |
| 0 | 0 | 0 |
| 0 | 1 | 0 |
| 1 | 0 | 0 |
| 0 | 0 | 0 |

| | | |
|---|---|---|
| 0 | 0 | 0 |
| 0 | 0 | 0 |
| 0 | 0 | 0 |
| 1 | 1 | 1 |
| 1 | 1 | 1 |
| 1 | 0 | 0 |
| 0 | 0 | 0 |
| 0 | 0 | 0 |
| 1 | 0 | 0 |
| 1 | 0 | 0 |
| 0 | 0 | 0 |
| 0 | 1 | 0 |
| 0 | 0 | 0 |
| 1 | 0 | 0 |
| 0 | 1 | 0 |
| 1 | 1 | 1 |
| 1 | 1 | 1 |
| 0 | 0 | 0 |
| 0 | 0 | 0 |
| 0 | 0 | 0 |
| 0 | 0 | 0 |
| 1 | 0 | 0 |
| 0 | 0 | 0 |
| 0 | 1 | 0 |
| 1 | 1 | 1 |
| 0 | 0 | 0 |
| 0 | 0 | 0 |
| 0 | 0 | 0 |
| 0 | 0 | 0 |
| 0 | 0 | 0 |
| 0 | 0 | 0 |
| 0 | 0 | 0 |
| 1 | 0 | 0 |
| 1 | 0 | 0 |
| 0 | 0 | 0 |
| 1 | 0 | 0 |
| 0 | 0 | 0 |
| 0 | 0 | 0 |
| 1 | 0 | 0 |
| 0 | 1 | 0 |
| 0 | 0 | 0 |
| 1 | 0 | 0 |
| 1 | 0 | 0 |
| 1 | 0 | 0 |
| 0 | 0 | 0 |
| 0 | 0 | 0 |
| 0 | 0 | 0 |
| 0 | 0 | 0 |
| 0 | 0 | 0 |
| 0 | 0 | 0 |
| 0 | 0 | 0 |
| 0 | 0 | 0 |
| 0 | 0 | 0 |
| 0 | 0 | 0 |

| | | |
|---|---|---|
| 0 | 0 | 0 |
| 0 | 0 | 0 |
| 0 | 0 | 0 |
| 0 | 0 | 0 |
| 0 | 0 | 0 |
| 0 | 0 | 0 |
| 0 | 1 | 0 |
| 0 | 1 | 0 |
| 0 | 0 | 0 |
| 0 | 0 | 0 |
| 0 | 0 | 0 |
| 0 | 0 | 0 |
| 0 | 0 | 0 |
| 0 | 1 | 0 |
| 0 | 1 | 0 |
| 0 | 0 | 0 |
| 0 | 1 | 0 |
| 0 | 0 | 0 |
| 1 | 1 | 1 |
| 0 | 0 | 0 |
| 1 | 1 | 1 |
| 1 | 0 | 0 |
| 1 | 1 | 1 |
| 0 | 0 | 0 |
| 0 | 0 | 0 |
| 0 | 0 | 0 |
| 0 | 0 | 0 |
| 0 | 0 | 0 |
| 0 | 0 | 0 |
| 0 | 0 | 0 |
| 0 | 0 | 0 |
| 1 | 0 | 0 |
| 0 | 0 | 0 |
| 0 | 0 | 0 |
| 0 | 0 | 0 |
| 0 | 0 | 0 |
| 0 | 0 | 0 |
| 0 | 0 | 0 |
| 0 | 0 | 0 |
| 0 | 0 | 0 |
| 0 | 1 | 0 |
| 0 | 1 | 0 |
| 0 | 0 | 0 |
| 0 | 1 | 0 |
| 0 | 0 | 0 |
| 0 | 0 | 0 |
| 0 | 0 | 0 |
| 1 | 1 | 1 |
| 1 | 1 | 1 |
| 0 | 0 | 0 |
| 0 | 1 | 0 |
| 1 | 1 | 1 |
| 0 | 0 | 0 |
| 0 | 1 | 0 |

| | | |
|---|---|---|
| 0 | 0 | 0 |
| 1 | 1 | 1 |
| 0 | 0 | 0 |
| 0 | 0 | 0 |
| 0 | 0 | 0 |
| 0 | 0 | 0 |
| 0 | 1 | 0 |
| 0 | 0 | 0 |
| 0 | 0 | 0 |
| 1 | 1 | 1 |
| 1 | 1 | 1 |
| 1 | 1 | 1 |
| 1 | 1 | 1 |
| 1 | 1 | 1 |
| 0 | 0 | 0 |
| 1 | 1 | 1 |
| 0 | 0 | 0 |
| 0 | 0 | 0 |
| 1 | 1 | 1 |
| 0 | 0 | 0 |
| 0 | 0 | 0 |
| 0 | 0 | 0 |
| 0 | 0 | 0 |
| 0 | 0 | 0 |
| 1 | 1 | 1 |
| 1 | 0 | 0 |
| 0 | 0 | 0 |
| 0 | 0 | 0 |
| 0 | 0 | 0 |
| 0 | 0 | 0 |
| 0 | 0 | 0 |
| 0 | 0 | 0 |
| 0 | 0 | 0 |
| 0 | 0 | 0 |
| 1 | 1 | 1 |
| 1 | 1 | 1 |
| 0 | 0 | 0 |
| 0 | 0 | 0 |
| 0 | 0 | 0 |
| 0 | 0 | 0 |
| 0 | 0 | 0 |
| 0 | 0 | 0 |
| 0 | 1 | 0 |
| 0 | 0 | 0 |
| 0 | 0 | 0 |
| 0 | 0 | 0 |
| 0 | 0 | 0 |
| 0 | 0 | 0 |
| 0 | 0 | 0 |
| 1 | 1 | 1 |
| 0 | 0 | 0 |
| 0 | 0 | 0 |
| 0 | 0 | 0 |
| 0 | 0 | 0 |

| | | |
|---|---|---|
| 0 | 0 | 0 |
| 0 | 0 | 0 |
| 0 | 0 | 0 |
| 0 | 0 | 0 |
| 1 | 1 | 1 |
| 0 | 0 | 0 |
| 0 | 0 | 0 |
| 0 | 0 | 0 |
| 0 | 0 | 0 |
| 0 | 0 | 0 |
| 0 | 0 | 0 |
| 0 | 0 | 0 |
| 0 | 0 | 0 |
| 0 | 0 | 0 |
| 0 | 0 | 0 |
| 0 | 0 | 0 |
| 0 | 0 | 0 |
| 0 | 0 | 0 |
| 1 | 1 | 1 |
| 0 | 0 | 0 |
| 0 | 0 | 0 |
| 0 | 0 | 0 |
| 0 | 0 | 0 |
| 1 | 0 | 0 |
| 0 | 0 | 0 |
| 0 | 0 | 0 |
| 0 | 0 | 0 |
| 1 | 0 | 0 |
| 1 | 0 | 0 |
| 1 | 0 | 0 |
| 0 | 0 | 0 |
| 0 | 0 | 0 |
| 0 | 0 | 0 |
| 0 | 0 | 0 |
| 0 | 0 | 0 |
| 1 | 0 | 0 |
| 1 | 0 | 0 |
| 0 | 0 | 0 |
| 0 | 0 | 0 |
| 1 | 0 | 0 |
| 1 | 0 | 0 |
| 1 | 0 | 0 |
| 0 | 1 | 0 |
| 1 | 1 | 1 |
| 0 | 0 | 0 |
| 0 | 0 | 0 |
| 0 | 0 | 0 |
| 0 | 0 | 0 |
| 0 | 0 | 0 |
| 0 | 1 | 0 |
| 1 | 1 | 1 |
| 0 | 0 | 0 |
| 1 | 1 | 1 |
| 0 | 0 | 0 |

| | | |
|---|---|---|
| 1 | 1 | 1 |
| 1 | 1 | 1 |
| 0 | 0 | 0 |
| 1 | 1 | 1 |
| 1 | 1 | 1 |
| 1 | 1 | 1 |
| 1 | 1 | 1 |
| 0 | 0 | 0 |
| 1 | 0 | 0 |
| 1 | 1 | 1 |
| 0 | 0 | 0 |
| 1 | 1 | 1 |
| 1 | 1 | 1 |
| 0 | 1 | 0 |
| 0 | 0 | 0 |
| 0 | 0 | 0 |
| 1 | 1 | 1 |
| 1 | 1 | 1 |
| 0 | 0 | 0 |
| 0 | 0 | 0 |
| 0 | 0 | 0 |
| 0 | 0 | 0 |
| 0 | 0 | 0 |
| 0 | 0 | 0 |
| 0 | 0 | 0 |
| 1 | 1 | 1 |
| 0 | 0 | 0 |
| 0 | 0 | 0 |
| 0 | 0 | 0 |
| 1 | 0 | 0 |
| 0 | 0 | 0 |
| 0 | 0 | 0 |
| 0 | 0 | 0 |
| 0 | 0 | 0 |
| 0 | 0 | 0 |
| 0 | 0 | 0 |
| 0 | 0 | 0 |
| 0 | 0 | 0 |
| 0 | 0 | 0 |
| 0 | 0 | 0 |
| 0 | 1 | 0 |
| 0 | 0 | 0 |
| 1 | 1 | 1 |
| 0 | 0 | 0 |
| 0 | 0 | 0 |
| 0 | 0 | 0 |
| 0 | 0 | 0 |
| 0 | 0 | 0 |
| 0 | 0 | 0 |
| 0 | 0 | 0 |
| 0 | 1 | 0 |
| 0 | 0 | 0 |
| 0 | 0 | 0 |
| 0 | 0 | 0 |

| | | |
|---|---|---|
| 1 | 1 | 1 |
| 0 | 0 | 0 |
| 0 | 1 | 0 |
| 0 | 1 | 0 |
| 0 | 0 | 0 |
| 0 | 0 | 0 |
| 0 | 0 | 0 |
| 0 | 0 | 0 |
| 0 | 0 | 0 |
| 0 | 1 | 0 |
| 1 | 1 | 1 |
| 1 | 1 | 1 |
| 1 | 1 | 1 |
| 0 | 0 | 0 |
| 1 | 1 | 1 |
| 0 | 0 | 0 |
| 0 | 0 | 0 |
| 0 | 0 | 0 |
| 0 | 0 | 0 |
| 1 | 0 | 0 |
| 0 | 0 | 0 |
| 1 | 1 | 1 |
| 0 | 0 | 0 |
| 0 | 1 | 0 |
| 1 | 1 | 1 |
| 1 | 1 | 1 |
| 0 | 0 | 0 |
| 1 | 1 | 1 |
| 1 | 1 | 1 |
| 1 | 1 | 1 |
| 0 | 0 | 0 |
| 0 | 0 | 0 |
| 1 | 1 | 1 |
| 1 | 1 | 1 |
| 0 | 0 | 0 |
| 0 | 0 | 0 |
| 0 | 0 | 0 |
| 0 | 0 | 0 |
| 0 | 0 | 0 |
| 1 | 1 | 1 |
| 0 | 0 | 0 |
| 0 | 0 | 0 |
| 0 | 0 | 0 |
| 0 | 0 | 0 |
| 1 | 0 | 0 |
| 0 | 0 | 0 |
| 0 | 0 | 0 |
| 0 | 0 | 0 |
| 0 | 0 | 0 |
| 0 | 0 | 0 |
| 0 | 0 | 0 |
| 0 | 0 | 0 |
| 1 | 0 | 0 |
| 0 | 0 | 0 |

| | | |
|---|---|---|
| 0 | 0 | 0 |
| 0 | 0 | 0 |
| 0 | 0 | 0 |
| 0 | 0 | 0 |
| 0 | 0 | 0 |
| 0 | 0 | 0 |
| 0 | 0 | 0 |
| 1 | 0 | 0 |
| 0 | 0 | 0 |
| 0 | 0 | 0 |
| 0 | 0 | 0 |
| 0 | 0 | 0 |
| 1 | 1 | 1 |
| 0 | 0 | 0 |
| 0 | 0 | 0 |
| 0 | 0 | 0 |
| 0 | 0 | 0 |
| 0 | 0 | 0 |
| 0 | 0 | 0 |
| 1 | 1 | 1 |
| 0 | 0 | 0 |
| 0 | 0 | 0 |
| 0 | 0 | 0 |
| 0 | 0 | 0 |
| 0 | 0 | 0 |
| 1 | 0 | 0 |
| 1 | 1 | 1 |
| 0 | 1 | 0 |
| 0 | 0 | 0 |
| 0 | 0 | 0 |
| 0 | 0 | 0 |
| 0 | 0 | 0 |
| 0 | 0 | 0 |
| 0 | 0 | 0 |
| 0 | 0 | 0 |
| 1 | 0 | 0 |
| 0 | 0 | 0 |
| 0 | 0 | 0 |
| 1 | 1 | 1 |
| 1 | 0 | 0 |
| 0 | 1 | 0 |
| 0 | 0 | 0 |
| 0 | 0 | 0 |
| 0 | 0 | 0 |
| 0 | 0 | 0 |
| 0 | 0 | 0 |
| 0 | 0 | 0 |
| 1 | 1 | 1 |
| 1 | 1 | 1 |
| 0 | 0 | 0 |
| 0 | 0 | 0 |
| 0 | 0 | 0 |
| 0 | 0 | 0 |
| 1 | 1 | 1 |

| | | |
|---|---|---|
| 1 | 1 | 1 |
| 0 | 0 | 0 |
| 0 | 1 | 0 |
| 0 | 0 | 0 |
| 1 | 0 | 0 |
| 1 | 0 | 0 |
| 0 | 0 | 0 |
| 1 | 0 | 0 |
| 1 | 0 | 0 |
| 1 | 0 | 0 |
| 1 | 0 | 0 |
| 1 | 0 | 0 |
| 1 | 1 | 1 |
| 1 | 1 | 1 |
| 1 | 1 | 1 |
| 1 | 1 | 1 |
| 0 | 0 | 0 |
| 0 | 0 | 0 |
| 1 | 0 | 0 |
| 0 | 0 | 0 |
| 0 | 0 | 0 |
| 0 | 0 | 0 |
| 1 | 0 | 0 |
| 0 | 0 | 0 |
| 0 | 0 | 0 |
| 1 | 1 | 1 |
| 1 | 0 | 0 |
| 1 | 0 | 0 |
| 0 | 0 | 0 |
| 0 | 0 | 0 |
| 0 | 0 | 0 |
| 0 | 0 | 0 |
| 0 | 0 | 0 |
| 0 | 0 | 0 |
| 0 | 0 | 0 |
| 0 | 0 | 0 |
| 1 | 0 | 0 |
| 1 | 1 | 1 |
| 1 | 1 | 1 |
| 1 | 1 | 1 |
| 0 | 0 | 0 |
| 1 | 1 | 1 |
| 0 | 0 | 0 |
| 1 | 1 | 1 |
| 0 | 0 | 0 |
| 0 | 1 | 0 |
| 0 | 0 | 0 |
| 0 | 0 | 0 |
| 0 | 0 | 0 |
| 0 | 0 | 0 |
| 0 | 0 | 0 |
| 0 | 0 | 0 |
| 0 | 0 | 0 |
| 0 | 0 | 0 |

| | | |
|---|---|---|
| 0 | 0 | 0 |
| 0 | 0 | 0 |
| 0 | 0 | 0 |
| 0 | 0 | 0 |
| 0 | 0 | 0 |
| 0 | 0 | 0 |
| 0 | 0 | 0 |
| 1 | 1 | 1 |
| 0 | 0 | 0 |
| 0 | 0 | 0 |
| 0 | 0 | 0 |
| 1 | 0 | 0 |
| 0 | 0 | 0 |
| 0 | 0 | 0 |
| 0 | 0 | 0 |
| 0 | 0 | 0 |
| 0 | 0 | 0 |
| 0 | 0 | 0 |
| 0 | 0 | 0 |
| 0 | 0 | 0 |
| 0 | 0 | 0 |
| 0 | 0 | 0 |
| 0 | 0 | 0 |
| 0 | 0 | 0 |
| 0 | 0 | 0 |
| 0 | 0 | 0 |
| 0 | 0 | 0 |
| 0 | 0 | 0 |
| 0 | 0 | 0 |
| 0 | 0 | 0 |
| 0 | 0 | 0 |
| 0 | 0 | 0 |
| 0 | 0 | 0 |
| 0 | 0 | 0 |
| 0 | 0 | 0 |
| 0 | 0 | 0 |
| 0 | 0 | 0 |
| 0 | 0 | 0 |
| 0 | 0 | 0 |
| 0 | 0 | 0 |
| 1 | 0 | 0 |
| 0 | 0 | 0 |
| 0 | 0 | 0 |
| 0 | 0 | 0 |
| 0 | 0 | 0 |
| 0 | 0 | 0 |
| 0 | 0 | 0 |
| 0 | 0 | 0 |
| 0 | 0 | 0 |
| 0 | 1 | 0 |
| 0 | 1 | 0 |
| 1 | 0 | 0 |
| 0 | 0 | 0 |
| 0 | 0 | 0 |

| | | |
|---|---|---|
| 0 | 0 | 0 |
| 0 | 0 | 0 |
| 0 | 0 | 0 |
| 0 | 0 | 0 |
| 0 | 0 | 0 |
| 0 | 0 | 0 |
| 0 | 0 | 0 |
| 1 | 0 | 0 |
| 1 | 0 | 0 |
| 0 | 0 | 0 |
| 0 | 0 | 0 |
| 1 | 0 | 0 |
| 0 | 0 | 0 |
| 0 | 0 | 0 |
| 0 | 0 | 0 |
| 0 | 0 | 0 |
| 0 | 0 | 0 |
| 1 | 0 | 0 |
| 0 | 0 | 0 |
| 1 | 0 | 0 |
| 0 | 0 | 0 |
| 0 | 0 | 0 |
| 1 | 1 | 1 |
| 0 | 0 | 0 |
| 0 | 0 | 0 |
| 1 | 0 | 0 |
| 0 | 0 | 0 |
| 0 | 0 | 0 |
| 0 | 0 | 0 |
| 0 | 0 | 0 |
| 0 | 0 | 0 |
| 0 | 0 | 0 |
| 1 | 0 | 0 |
| 0 | 0 | 0 |
| 0 | 0 | 0 |
| 0 | 0 | 0 |
| 1 | 1 | 1 |
| 0 | 0 | 0 |
| 0 | 0 | 0 |
| 0 | 0 | 0 |
| 0 | 0 | 0 |
| 0 | 0 | 0 |
| 0 | 0 | 0 |
| 0 | 0 | 0 |
| 0 | 0 | 0 |
| 0 | 0 | 0 |
| 0 | 0 | 0 |
| 0 | 0 | 0 |
| 0 | 0 | 0 |
| 0 | 0 | 0 |
| 0 | 0 | 0 |
| 0 | 0 | 0 |
| 0 | 0 | 0 |
| 0 | 0 | 0 |

| | | |
|---|---|---|
| 1 | 0 | 0 |
| 0 | 1 | 0 |
| 0 | 0 | 0 |
| 0 | 0 | 0 |
| 0 | 0 | 0 |
| 0 | 0 | 0 |
| 0 | 0 | 0 |
| 0 | 0 | 0 |
| 0 | 0 | 0 |
| 1 | 0 | 0 |
| 0 | 1 | 0 |
| 1 | 1 | 1 |
| 0 | 0 | 0 |
| 0 | 0 | 0 |
| 0 | 0 | 0 |
| 0 | 0 | 0 |
| 0 | 0 | 0 |
| 0 | 0 | 0 |
| 1 | 1 | 1 |
| 1 | 0 | 0 |
| 1 | 0 | 0 |
| 0 | 0 | 0 |
| 0 | 1 | 0 |
| 0 | 0 | 0 |
| 0 | 0 | 0 |
| 0 | 0 | 0 |
| 0 | 0 | 0 |
| 0 | 0 | 0 |
| 0 | 0 | 0 |
| 0 | 0 | 0 |
| 0 | 0 | 0 |
| 0 | 0 | 0 |
| 0 | 0 | 0 |
| 0 | 0 | 0 |
| 0 | 0 | 0 |
| 0 | 0 | 0 |
| 0 | 0 | 0 |
| 0 | 0 | 0 |
| 0 | 0 | 0 |
| 0 | 0 | 0 |
| 0 | 0 | 0 |
| 1 | 0 | 0 |
| 0 | 0 | 0 |
| 0 | 0 | 0 |
| 0 | 0 | 0 |
| 0 | 0 | 0 |
| 1 | 0 | 0 |
| 0 | 0 | 0 |
| 0 | 0 | 0 |
| 0 | 0 | 0 |
| 0 | 0 | 0 |
| 0 | 0 | 0 |
| 1 | 0 | 0 |
| 0 | 1 | 0 |

| | | |
|---|---|---|
| 0 | 0 | 0 |
| 0 | 0 | 0 |
| 1 | 0 | 0 |
| 1 | 0 | 0 |
| 1 | 0 | 0 |
| 0 | 0 | 0 |
| 1 | 0 | 0 |
| 0 | 0 | 0 |
| 0 | 0 | 0 |
| 0 | 0 | 0 |
| 0 | 0 | 0 |
| 0 | 0 | 0 |
| 0 | 0 | 0 |
| 1 | 0 | 0 |
| 1 | 0 | 0 |
| 1 | 0 | 0 |
| 0 | 0 | 0 |
| 1 | 1 | 1 |
| 0 | 0 | 0 |
| 0 | 0 | 0 |
| 1 | 0 | 0 |
| 1 | 0 | 0 |
| 0 | 0 | 0 |
| 0 | 0 | 0 |
| 1 | 0 | 0 |
| 0 | 0 | 0 |
| 0 | 0 | 0 |
| 1 | 0 | 0 |
| 0 | 0 | 0 |
| 0 | 0 | 0 |
| 0 | 0 | 0 |
| 0 | 0 | 0 |
| 0 | 0 | 0 |
| 0 | 0 | 0 |
| 0 | 0 | 0 |
| 0 | 0 | 0 |
| 0 | 0 | 0 |
| 0 | 0 | 0 |
| 0 | 0 | 0 |
| 0 | 0 | 0 |
| 1 | 0 | 0 |
| 1 | 0 | 0 |
| 1 | 0 | 0 |
| 1 | 1 | 1 |
| 1 | 1 | 1 |
| 0 | 0 | 0 |
| 0 | 0 | 0 |
| 0 | 0 | 0 |
| 0 | 0 | 0 |
| 0 | 0 | 0 |
| 0 | 0 | 0 |
| 0 | 0 | 0 |
| 0 | 0 | 0 |
| 1 | 0 | 0 |

| | | |
|---|---|---|
| 1 | 0 | 0 |
| 0 | 0 | 0 |
| 0 | 0 | 0 |
| 0 | 0 | 0 |
| 1 | 0 | 0 |
| 0 | 0 | 0 |
| 0 | 0 | 0 |
| 0 | 0 | 0 |
| 1 | 0 | 0 |
| 1 | 0 | 0 |
| 0 | 0 | 0 |
| 0 | 0 | 0 |
| 0 | 0 | 0 |
| 1 | 0 | 0 |
| 0 | 0 | 0 |
| 0 | 0 | 0 |
| 0 | 0 | 0 |
| 0 | 0 | 0 |
| 0 | 0 | 0 |
| 0 | 0 | 0 |
| 0 | 0 | 0 |
| 1 | 1 | 1 |
| 0 | 0 | 0 |
| 0 | 1 | 0 |
| 0 | 0 | 0 |
| 0 | 0 | 0 |
| 0 | 0 | 0 |
| 0 | 0 | 0 |
| 0 | 0 | 0 |
| 0 | 0 | 0 |
| 0 | 0 | 0 |
| 0 | 0 | 0 |
| 0 | 0 | 0 |
| 0 | 0 | 0 |
| 0 | 1 | 0 |
| 0 | 0 | 0 |
| 0 | 0 | 0 |
| 0 | 0 | 0 |
| 0 | 0 | 0 |
| 0 | 0 | 0 |
| 0 | 0 | 0 |
| 0 | 0 | 0 |
| 0 | 0 | 0 |
| 0 | 0 | 0 |
| 0 | 0 | 0 |
| 0 | 0 | 0 |
| 0 | 0 | 0 |
| 1 | 1 | 1 |
| 0 | 0 | 0 |
| 1 | 1 | 1 |
| 0 | 0 | 0 |
| 0 | 0 | 0 |
| 0 | 0 | 0 |
| 0 | 0 | 0 |

| | | |
|---|---|---|
| 0 | 0 | 0 |
| 0 | 0 | 0 |
| 1 | 1 | 1 |
| 0 | 0 | 0 |
| 0 | 0 | 0 |
| 0 | 1 | 0 |
| 0 | 1 | 0 |
| 0 | 1 | 0 |
| 0 | 0 | 0 |
| 0 | 1 | 0 |
| 0 | 0 | 0 |
| 0 | 0 | 0 |
| 0 | 0 | 0 |
| 1 | 1 | 1 |
| 0 | 0 | 0 |
| 0 | 1 | 0 |
| 0 | 0 | 0 |
| 0 | 0 | 0 |
| 0 | 0 | 0 |
| 1 | 1 | 1 |
| 1 | 1 | 1 |
| 0 | 0 | 0 |
| 1 | 1 | 1 |
| 0 | 0 | 0 |
| 0 | 0 | 0 |
| 1 | 1 | 1 |
| 1 | 1 | 1 |
| 0 | 0 | 0 |
| 1 | 1 | 1 |
| 0 | 0 | 0 |
| 0 | 1 | 0 |
| 0 | 1 | 0 |
| 1 | 1 | 1 |
| 0 | 1 | 0 |
| 1 | 1 | 1 |
| 1 | 1 | 1 |
| 1 | 1 | 1 |
| 1 | 1 | 1 |
| 1 | 1 | 1 |
| 1 | 1 | 1 |
| 1 | 1 | 1 |
| 1 | 1 | 1 |
| 0 | 1 | 0 |
| 0 | 1 | 0 |
| 0 | 1 | 0 |
| 0 | 1 | 0 |
| 0 | 1 | 0 |
| 0 | 1 | 0 |
| 0 | 1 | 0 |
| 0 | 1 | 0 |
| 0 | 1 | 0 |
| 0 | 1 | 0 |
| 0 | 1 | 0 |
| 0 | 1 | 0 |

| | | |
|---|---|---|
| 0 | 1 | 0 |
| 1 | 1 | 1 |
| 1 | 1 | 1 |
| 1 | 1 | 1 |
| 1 | 1 | 1 |
| 1 | 1 | 1 |
| 1 | 1 | 1 |
| 0 | 1 | 0 |
| 0 | 1 | 0 |
| 0 | 1 | 0 |
| 0 | 1 | 0 |
| 0 | 0 | 0 |
| 0 | 0 | 0 |
| 0 | 0 | 0 |
| 1 | 0 | 0 |
| 0 | 0 | 0 |
| 1 | 0 | 0 |
| 0 | 0 | 0 |
| 1 | 0 | 0 |
| 1 | 1 | 1 |
| 0 | 0 | 0 |
| 1 | 1 | 1 |
| 1 | 1 | 1 |
| 0 | 0 | 0 |
| 0 | 0 | 0 |
| 1 | 1 | 1 |
| 1 | 1 | 1 |
| 0 | 0 | 0 |
| 0 | 0 | 0 |
| 0 | 0 | 0 |
| 0 | 0 | 0 |
| 0 | 0 | 0 |
| 0 | 1 | 0 |
| 0 | 1 | 0 |
| 1 | 1 | 1 |
| 1 | 1 | 1 |
| 1 | 1 | 1 |
| 1 | 1 | 1 |
| 1 | 1 | 1 |
| 0 | 0 | 0 |
| 0 | 0 | 0 |
| 0 | 1 | 0 |
| 0 | 0 | 0 |
| 0 | 0 | 0 |
| 0 | 0 | 0 |
| 1 | 0 | 0 |
| 0 | 0 | 0 |
| 1 | 0 | 0 |
| 1 | 0 | 0 |
| 0 | 0 | 0 |
| 0 | 0 | 0 |
| 0 | 0 | 0 |
| 0 | 0 | 0 |
| 0 | 0 | 0 |

| | | |
|---|---|---|
| 0 | 0 | 0 |
| 0 | 1 | 0 |
| 1 | 1 | 1 |
| 1 | 1 | 1 |
| 0 | 0 | 0 |
| 1 | 1 | 1 |
| 0 | 0 | 0 |
| 1 | 1 | 1 |
| 1 | 1 | 1 |
| 0 | 0 | 0 |
| 1 | 0 | 0 |
| 1 | 1 | 1 |
| 1 | 1 | 1 |
| 0 | 0 | 0 |
| 1 | 0 | 0 |
| 1 | 0 | 0 |
| 0 | 0 | 0 |
| 1 | 0 | 0 |
| 0 | 0 | 0 |
| 0 | 0 | 0 |
| 0 | 0 | 0 |
| 0 | 0 | 0 |
| 0 | 0 | 0 |
| 0 | 0 | 0 |
| 0 | 0 | 0 |
| 0 | 0 | 0 |
| 0 | 0 | 0 |
| 0 | 0 | 0 |
| 0 | 0 | 0 |
| 0 | 0 | 0 |
| 0 | 0 | 0 |
| 0 | 0 | 0 |
| 0 | 0 | 0 |
| 0 | 0 | 0 |
| 0 | 0 | 0 |
| 0 | 0 | 0 |
| 1 | 0 | 0 |
| 0 | 0 | 0 |
| 0 | 0 | 0 |
| 0 | 0 | 0 |
| 0 | 0 | 0 |
| 0 | 0 | 0 |
| 0 | 0 | 0 |
| 0 | 0 | 0 |
| 0 | 0 | 0 |
| 0 | 0 | 0 |
| 0 | 0 | 0 |
| 0 | 0 | 0 |
| 1 | 0 | 0 |
| 1 | 0 | 0 |
| 0 | 0 | 0 |
| 1 | 1 | 1 |
| 0 | 0 | 0 |
| 0 | 0 | 0 |

| | | |
|---|---|---|
| 0 | 0 | 0 |
| 0 | 0 | 0 |
| 0 | 0 | 0 |
| 0 | 0 | 0 |
| 0 | 0 | 0 |
| 0 | 0 | 0 |
| 0 | 0 | 0 |
| 0 | 0 | 0 |
| 0 | 0 | 0 |
| 0 | 0 | 0 |
| 0 | 0 | 0 |
| 0 | 0 | 0 |
| 0 | 0 | 0 |
| 0 | 0 | 0 |
| 0 | 1 | 0 |
| 0 | 0 | 0 |
| 0 | 0 | 0 |
| 0 | 0 | 0 |
| 1 | 0 | 0 |
| 0 | 0 | 0 |
| 0 | 0 | 0 |
| 0 | 0 | 0 |
| 0 | 0 | 0 |
| 0 | 0 | 0 |
| 0 | 0 | 0 |
| 1 | 0 | 0 |
| 0 | 0 | 0 |
| 0 | 0 | 0 |
| 0 | 0 | 0 |
| 0 | 0 | 0 |
| 1 | 1 | 1 |
| 0 | 0 | 0 |
| 1 | 1 | 1 |
| 1 | 0 | 0 |
| 0 | 0 | 0 |
| 0 | 0 | 0 |
| 1 | 1 | 1 |
| 1 | 1 | 1 |
| 1 | 1 | 1 |
| 1 | 0 | 0 |
| 0 | 0 | 0 |
| 0 | 0 | 0 |
| 0 | 0 | 0 |
| 0 | 0 | 0 |
| 0 | 0 | 0 |
| 0 | 0 | 0 |
| 0 | 0 | 0 |
| 0 | 0 | 0 |
| 1 | 1 | 1 |
| 1 | 1 | 1 |
| 0 | 0 | 0 |
| 0 | 0 | 0 |
| 0 | 0 | 0 |
| 0 | 1 | 0 |

| | | |
|---|---|---|
| 1 | 1 | 1 |
| 1 | 1 | 1 |
| 1 | 0 | 0 |
| 0 | 0 | 0 |
| 1 | 0 | 0 |
| 1 | 1 | 1 |
| 0 | 0 | 0 |
| 1 | 1 | 1 |
| 0 | 0 | 0 |
| 0 | 0 | 0 |
| 0 | 0 | 0 |
| 0 | 0 | 0 |
| 0 | 0 | 0 |
| 0 | 0 | 0 |
| 0 | 0 | 0 |
| 0 | 0 | 0 |
| 0 | 1 | 0 |
| 0 | 0 | 0 |
| 0 | 0 | 0 |
| 1 | 0 | 0 |
| 0 | 0 | 0 |
| 0 | 0 | 0 |
| 1 | 1 | 1 |
| 0 | 0 | 0 |
| 1 | 1 | 1 |
| 1 | 1 | 1 |
| 0 | 1 | 0 |
| 0 | 0 | 0 |
| 0 | 0 | 0 |
| 0 | 0 | 0 |
| 1 | 1 | 1 |
| 0 | 0 | 0 |
| 0 | 0 | 0 |
| 0 | 0 | 0 |
| 0 | 1 | 0 |
| 0 | 1 | 0 |
| 0 | 0 | 0 |
| 0 | 0 | 0 |
| 0 | 0 | 0 |
| 1 | 1 | 1 |
| 1 | 1 | 1 |
| 0 | 1 | 0 |
| 0 | 1 | 0 |
| 0 | 0 | 0 |
| 1 | 1 | 1 |
| 0 | 0 | 0 |
| 1 | 1 | 1 |
| 0 | 0 | 0 |
| 0 | 0 | 0 |
| 1 | 1 | 1 |
| 0 | 1 | 0 |
| 0 | 0 | 0 |
| 0 | 0 | 0 |
| 1 | 1 | 1 |

| | | |
|---|---|---|
| 1 | 1 | 1 |
| 1 | 1 | 1 |
| 0 | 1 | 0 |
| 0 | 0 | 0 |
| 0 | 0 | 0 |
| 1 | 0 | 0 |
| 1 | 0 | 0 |
| 0 | 0 | 0 |
| 0 | 0 | 0 |
| 1 | 1 | 1 |
| 0 | 0 | 0 |
| 0 | 0 | 0 |
| 0 | 0 | 0 |
| 0 | 0 | 0 |
| 0 | 0 | 0 |
| 0 | 0 | 0 |
| 0 | 0 | 0 |
| 0 | 0 | 0 |
| 0 | 0 | 0 |
| 0 | 1 | 0 |
| 1 | 1 | 1 |
| 1 | 1 | 1 |
| 1 | 1 | 1 |
| 1 | 1 | 1 |
| 0 | 0 | 0 |
| 0 | 1 | 0 |
| 0 | 0 | 0 |
| 0 | 0 | 0 |
| 1 | 1 | 1 |
| 1 | 1 | 1 |
| 1 | 1 | 1 |
| 1 | 1 | 1 |
| 1 | 1 | 1 |
| 0 | 0 | 0 |
| 0 | 0 | 0 |
| 1 | 1 | 1 |
| 0 | 0 | 0 |
| 0 | 1 | 0 |
| 0 | 0 | 0 |
| 0 | 0 | 0 |
| 0 | 0 | 0 |
| 0 | 0 | 0 |
| 0 | 1 | 0 |
| 0 | 0 | 0 |
| 0 | 0 | 0 |
| 0 | 0 | 0 |
| 0 | 0 | 0 |
| 0 | 0 | 0 |
| 0 | 0 | 0 |
| 0 | 0 | 0 |
| 0 | 0 | 0 |
| 0 | 0 | 0 |
| 0 | 0 | 0 |
| 0 | 1 | 0 |

| | | |
|---|---|---|
| 0 | 0 | 0 |
| 0 | 1 | 0 |
| 0 | 0 | 0 |
| 0 | 0 | 0 |
| 0 | 0 | 0 |
| 1 | 1 | 1 |
| 0 | 0 | 0 |
| 1 | 1 | 1 |
| 0 | 1 | 0 |
| 0 | 0 | 0 |
| 0 | 1 | 0 |
| 0 | 1 | 0 |
| 0 | 1 | 0 |
| 0 | 1 | 0 |
| 0 | 1 | 0 |
| 0 | 0 | 0 |
| 0 | 0 | 0 |
| 0 | 1 | 0 |
| 0 | 0 | 0 |
| 1 | 1 | 1 |
| 0 | 0 | 0 |
| 0 | 0 | 0 |
| 0 | 0 | 0 |
| 0 | 0 | 0 |
| 0 | 0 | 0 |
| 0 | 0 | 0 |
| 0 | 0 | 0 |
| 0 | 0 | 0 |
| 0 | 0 | 0 |
| 0 | 0 | 0 |
| 0 | 0 | 0 |
| 0 | 0 | 0 |
| 1 | 0 | 0 |
| 1 | 0 | 0 |
| 0 | 0 | 0 |
| 0 | 0 | 0 |
| 0 | 0 | 0 |
| 0 | 0 | 0 |
| 1 | 0 | 0 |
| 0 | 0 | 0 |
| 0 | 0 | 0 |
| 0 | 0 | 0 |
| 1 | 0 | 0 |
| 0 | 1 | 0 |
| 0 | 0 | 0 |
| 0 | 0 | 0 |
| 0 | 0 | 0 |
| 0 | 0 | 0 |
| 0 | 0 | 0 |
| 0 | 0 | 0 |
| 1 | 0 | 0 |
| 0 | 0 | 0 |
| 0 | 0 | 0 |
| 1 | 0 | 0 |

| | | |
|---|---|---|
| 0 | 0 | 0 |
| 1 | 0 | 0 |
| 0 | 0 | 0 |
| 0 | 0 | 0 |
| 0 | 0 | 0 |
| 0 | 0 | 0 |
| 0 | 0 | 0 |
| 0 | 0 | 0 |
| 0 | 0 | 0 |
| 0 | 0 | 0 |
| 0 | 0 | 0 |
| 1 | 1 | 1 |
| 0 | 0 | 0 |
| 1 | 1 | 1 |
| 1 | 0 | 0 |
| 1 | 0 | 0 |
| 0 | 0 | 0 |
| 0 | 0 | 0 |
| 0 | 0 | 0 |
| 0 | 0 | 0 |
| 1 | 0 | 0 |
| 0 | 0 | 0 |
| 0 | 1 | 0 |
| 0 | 0 | 0 |
| 1 | 1 | 1 |
| 1 | 1 | 1 |
| 1 | 1 | 1 |
| 0 | 0 | 0 |
| 1 | 1 | 1 |
| 1 | 1 | 1 |
| 1 | 1 | 1 |
| 0 | 0 | 0 |
| 1 | 1 | 1 |
| 0 | 0 | 0 |
| 0 | 0 | 0 |
| 0 | 0 | 0 |
| 0 | 1 | 0 |
| 1 | 1 | 1 |
| 0 | 0 | 0 |
| 0 | 0 | 0 |
| 0 | 0 | 0 |
| 0 | 0 | 0 |
| 0 | 0 | 0 |
| 0 | 0 | 0 |
| 0 | 0 | 0 |
| 0 | 0 | 0 |
| 1 | 1 | 1 |
| 1 | 1 | 1 |
| 0 | 0 | 0 |
| 0 | 0 | 0 |
| 0 | 0 | 0 |
| 1 | 1 | 1 |
| 1 | 1 | 1 |
| 0 | 0 | 0 |

| | | |
|---|---|---|
| 0 | 0 | 0 |
| 0 | 1 | 0 |
| 0 | 1 | 0 |
| 1 | 1 | 1 |
| 0 | 0 | 0 |
| 0 | 0 | 0 |
| 0 | 0 | 0 |
| 0 | 0 | 0 |
| 0 | 0 | 0 |
| 0 | 0 | 0 |
| 1 | 1 | 1 |
| 0 | 0 | 0 |
| 0 | 0 | 0 |
| 0 | 0 | 0 |
| 1 | 0 | 0 |
| 1 | 1 | 1 |
| 0 | 0 | 0 |
| 1 | 0 | 0 |
| 0 | 0 | 0 |
| 0 | 0 | 0 |
| 1 | 0 | 0 |
| 0 | 0 | 0 |
| 0 | 0 | 0 |
| 1 | 0 | 0 |
| 0 | 0 | 0 |
| 0 | 0 | 0 |
| 0 | 0 | 0 |
| 1 | 0 | 0 |
| 0 | 0 | 0 |
| 0 | 0 | 0 |
| 0 | 0 | 0 |
| 1 | 1 | 1 |
| 1 | 1 | 1 |
| 1 | 1 | 1 |
| 0 | 1 | 0 |
| 0 | 1 | 0 |
| 0 | 1 | 0 |
| 1 | 1 | 1 |
| 1 | 1 | 1 |
| 1 | 1 | 1 |
| 0 | 1 | 0 |
| 1 | 1 | 1 |
| 0 | 0 | 0 |
| 0 | 1 | 0 |
| 1 | 1 | 1 |
| 1 | 1 | 1 |
| 0 | 1 | 0 |
| 1 | 1 | 1 |
| 1 | 1 | 1 |
| 1 | 1 | 1 |
| 1 | 1 | 1 |
| 1 | 1 | 1 |
| 1 | 1 | 1 |
| 1 | 1 | 1 |

| | | |
|---|---|---|
| 0 | 1 | 0 |
| 0 | 1 | 0 |
| 1 | 1 | 1 |
| 1 | 1 | 1 |
| 1 | 1 | 1 |
| 1 | 1 | 1 |
| 1 | 1 | 1 |
| 1 | 1 | 1 |
| 0 | 0 | 0 |
| 0 | 0 | 0 |
| 0 | 0 | 0 |
| 0 | 0 | 0 |
| 0 | 0 | 0 |
| 0 | 0 | 0 |
| 0 | 0 | 0 |
| 0 | 0 | 0 |
| 0 | 0 | 0 |
| 0 | 0 | 0 |
| 0 | 0 | 0 |
| 0 | 0 | 0 |
| 0 | 0 | 0 |
| 0 | 0 | 0 |
| 0 | 0 | 0 |
| 0 | 0 | 0 |
| 0 | 0 | 0 |
| 1 | 1 | 1 |
| 1 | 1 | 1 |
| 0 | 0 | 0 |
| 0 | 0 | 0 |
| 0 | 0 | 0 |
| 0 | 0 | 0 |
| 0 | 0 | 0 |
| 0 | 0 | 0 |
| 0 | 0 | 0 |
| 0 | 0 | 0 |
| 0 | 0 | 0 |
| 0 | 0 | 0 |
| 0 | 0 | 0 |
| 0 | 0 | 0 |
| 0 | 0 | 0 |
| 0 | 0 | 0 |
| 0 | 0 | 0 |
| 0 | 0 | 0 |
| 0 | 0 | 0 |
| 0 | 0 | 0 |
| 0 | 0 | 0 |
| 1 | 1 | 1 |
| 0 | 1 | 0 |
| 1 | 1 | 1 |
| 1 | 1 | 1 |
| 1 | 1 | 1 |
| 1 | 1 | 1 |
| 0 | 0 | 0 |
| 0 | 0 | 0 |

| | | |
|---|---|---|
| 0 | 0 | 0 |
| 0 | 0 | 0 |
| 0 | 0 | 0 |
| 0 | 0 | 0 |
| 0 | 1 | 0 |
| 0 | 0 | 0 |
| 0 | 0 | 0 |
| 0 | 0 | 0 |
| 1 | 1 | 1 |
| 0 | 0 | 0 |
| 0 | 0 | 0 |
| 0 | 1 | 0 |
| 0 | 0 | 0 |
| 0 | 0 | 0 |
| 0 | 0 | 0 |
| 0 | 0 | 0 |
| 1 | 1 | 1 |
| 0 | 0 | 0 |
| 0 | 0 | 0 |
| 1 | 1 | 1 |
| 0 | 0 | 0 |
| 0 | 0 | 0 |
| 0 | 0 | 0 |
| 0 | 0 | 0 |
| 0 | 0 | 0 |
| 0 | 1 | 0 |
| 0 | 1 | 0 |
| 1 | 1 | 1 |
| 1 | 1 | 1 |
| 0 | 1 | 0 |
| 1 | 1 | 1 |
| 1 | 1 | 1 |
| 1 | 0 | 0 |
| 1 | 0 | 0 |
| 0 | 0 | 0 |
| 0 | 0 | 0 |
| 0 | 0 | 0 |
| 0 | 0 | 0 |
| 0 | 0 | 0 |
| 0 | 0 | 0 |
| 1 | 1 | 1 |
| 1 | 1 | 1 |
| 1 | 1 | 1 |
| 0 | 0 | 0 |
| 0 | 0 | 0 |
| 0 | 0 | 0 |
| 1 | 1 | 1 |
| 0 | 1 | 0 |
| 0 | 0 | 0 |
| 0 | 0 | 0 |
| 1 | 1 | 1 |
| 1 | 1 | 1 |
| 1 | 1 | 1 |
| 1 | 1 | 1 |

| | | |
|---|---|---|
| 0 | 1 | 0 |
| 1 | 1 | 1 |
| 0 | 1 | 0 |
| 0 | 0 | 0 |
| 0 | 0 | 0 |
| 1 | 1 | 1 |
| 1 | 1 | 1 |
| 1 | 1 | 1 |
| 1 | 1 | 1 |
| 1 | 1 | 1 |
| 0 | 0 | 0 |
| 0 | 1 | 0 |
| 0 | 0 | 0 |
| 0 | 0 | 0 |
| 0 | 0 | 0 |
| 1 | 1 | 1 |
| 1 | 1 | 1 |
| 0 | 1 | 0 |
| 1 | 1 | 1 |
| 1 | 1 | 1 |
| 0 | 0 | 0 |
| 1 | 1 | 1 |
| 1 | 1 | 1 |
| 0 | 0 | 0 |
| 1 | 0 | 0 |
| 0 | 0 | 0 |
| 0 | 0 | 0 |
| 1 | 1 | 1 |
| 1 | 1 | 1 |
| 1 | 1 | 1 |
| 1 | 1 | 1 |
| 1 | 1 | 1 |
| 1 | 1 | 1 |
| 1 | 1 | 1 |
| 1 | 1 | 1 |
| 0 | 0 | 0 |
| 1 | 1 | 1 |
| 1 | 1 | 1 |
| 0 | 0 | 0 |
| 0 | 0 | 0 |
| 1 | 1 | 1 |
| 0 | 1 | 0 |
| 0 | 0 | 0 |
| 0 | 0 | 0 |
| 0 | 0 | 0 |
| 0 | 0 | 0 |
| 0 | 0 | 0 |
| 0 | 0 | 0 |
| 1 | 1 | 1 |
| 0 | 0 | 0 |
| 0 | 0 | 0 |
| 0 | 0 | 0 |
| 0 | 0 | 0 |
| 0 | 0 | 0 |

| | | |
|---|---|---|
| 0 | 0 | 0 |
| 0 | 0 | 0 |
| 0 | 0 | 0 |
| 0 | 0 | 0 |
| 0 | 0 | 0 |
| 0 | 0 | 0 |
| 0 | 0 | 0 |
| 0 | 0 | 0 |
| 0 | 0 | 0 |
| 0 | 1 | 0 |
| 0 | 1 | 0 |
| 0 | 0 | 0 |
| 1 | 1 | 1 |
| 0 | 0 | 0 |
| 0 | 0 | 0 |
| 0 | 1 | 0 |
| 0 | 0 | 0 |
| 1 | 0 | 0 |
| 1 | 1 | 1 |
| 1 | 1 | 1 |
| 0 | 0 | 0 |
| 0 | 1 | 0 |
| 0 | 1 | 0 |
| 0 | 0 | 0 |
| 0 | 0 | 0 |
| 0 | 1 | 0 |
| 0 | 1 | 0 |
| 1 | 1 | 1 |
| 0 | 0 | 0 |
| 0 | 0 | 0 |
| 1 | 1 | 1 |
| 1 | 1 | 1 |
| 0 | 0 | 0 |
| 0 | 0 | 0 |
| 0 | 0 | 0 |
| 0 | 1 | 0 |
| 0 | 0 | 0 |
| 0 | 1 | 0 |
| 0 | 1 | 0 |
| 0 | 0 | 0 |
| 0 | 0 | 0 |
| 0 | 0 | 0 |
| 0 | 0 | 0 |
| 0 | 0 | 0 |
| 0 | 0 | 0 |
| 1 | 1 | 1 |
| 0 | 1 | 0 |
| 0 | 0 | 0 |
| 0 | 0 | 0 |
| 0 | 0 | 0 |
| 1 | 1 | 1 |
| 1 | 1 | 1 |
| 0 | 0 | 0 |
| 0 | 0 | 0 |

| | | |
|---|---|---|
| 1 | 1 | 1 |
| 1 | 1 | 1 |
| 1 | 1 | 1 |
| 1 | 1 | 1 |
| 0 | 0 | 0 |
| 0 | 0 | 0 |
| 1 | 0 | 0 |
| 0 | 0 | 0 |
| 1 | 0 | 0 |
| 0 | 0 | 0 |
| 1 | 1 | 1 |
| 1 | 1 | 1 |
| 1 | 1 | 1 |
| 1 | 1 | 1 |
| 1 | 1 | 1 |
| 1 | 1 | 1 |
| 0 | 0 | 0 |
| 1 | 1 | 1 |
| 1 | 1 | 1 |
| 0 | 0 | 0 |
| 0 | 0 | 0 |
| 0 | 1 | 0 |
| 0 | 1 | 0 |
| 0 | 0 | 0 |
| 0 | 1 | 0 |
| 0 | 0 | 0 |
| 0 | 0 | 0 |
| 0 | 0 | 0 |
| 0 | 0 | 0 |
| 0 | 0 | 0 |
| 0 | 0 | 0 |
| 0 | 0 | 0 |
| 0 | 1 | 0 |
| 0 | 0 | 0 |
| 1 | 1 | 1 |
| 1 | 1 | 1 |
| 1 | 1 | 1 |
| 1 | 1 | 1 |
| 0 | 0 | 0 |
| 1 | 1 | 1 |
| 1 | 1 | 1 |
| 1 | 1 | 1 |
| 1 | 1 | 1 |
| 0 | 0 | 0 |
| 1 | 1 | 1 |
| 1 | 1 | 1 |
| 1 | 1 | 1 |
| 1 | 1 | 1 |
| 1 | 1 | 1 |
| 1 | 1 | 1 |
| 0 | 0 | 0 |
| 0 | 0 | 0 |
| 0 | 0 | 0 |
| 0 | 1 | 0 |

| | | |
|---|---|---|
| 0 | 0 | 0 |
| 0 | 0 | 0 |
| 0 | 0 | 0 |
| 0 | 0 | 0 |
| 0 | 0 | 0 |
| 0 | 0 | 0 |
| 0 | 0 | 0 |
| 0 | 0 | 0 |
| 1 | 1 | 1 |
| 1 | 1 | 1 |
| 0 | 0 | 0 |
| 1 | 1 | 1 |
| 1 | 1 | 1 |
| 0 | 0 | 0 |
| 0 | 0 | 0 |
| 0 | 0 | 0 |
| 0 | 0 | 0 |
| 0 | 0 | 0 |
| 0 | 0 | 0 |
| 1 | 1 | 1 |
| 1 | 1 | 1 |
| 0 | 0 | 0 |
| 1 | 1 | 1 |
| 0 | 0 | 0 |
| 1 | 1 | 1 |
| 1 | 1 | 1 |
| 1 | 1 | 1 |
| 0 | 1 | 0 |
| 0 | 1 | 0 |
| 0 | 0 | 0 |
| 0 | 1 | 0 |
| 1 | 1 | 1 |
| 0 | 0 | 0 |
| 1 | 1 | 1 |
| 0 | 0 | 0 |
| 0 | 0 | 0 |
| 0 | 0 | 0 |
| 0 | 0 | 0 |
| 0 | 0 | 0 |
| 0 | 0 | 0 |
| 0 | 0 | 0 |
| 0 | 0 | 0 |
| 1 | 1 | 1 |
| 0 | 0 | 0 |
| 1 | 1 | 1 |
| 0 | 0 | 0 |
| 1 | 0 | 0 |
| 1 | 1 | 1 |
| 1 | 1 | 1 |
| 0 | 0 | 0 |
| 1 | 1 | 1 |
| 1 | 1 | 1 |
| 0 | 1 | 0 |
| 1 | 1 | 1 |

| | | |
|---|---|---|
| 0 | 0 | 0 |
| 0 | 1 | 0 |
| 1 | 1 | 1 |
| 0 | 1 | 0 |
| 0 | 1 | 0 |
| 0 | 1 | 0 |
| 0 | 1 | 0 |
| 1 | 1 | 1 |
| 1 | 1 | 1 |
| 1 | 1 | 1 |
| 0 | 0 | 0 |
| 1 | 1 | 1 |
| 1 | 1 | 1 |
| 0 | 1 | 0 |
| 0 | 0 | 0 |
| 0 | 0 | 0 |
| 0 | 0 | 0 |
| 0 | 0 | 0 |
| 0 | 0 | 0 |
| 0 | 1 | 0 |
| 0 | 1 | 0 |
| 1 | 1 | 1 |
| 0 | 0 | 0 |
| 0 | 1 | 0 |
| 0 | 1 | 0 |
| 0 | 1 | 0 |
| 1 | 1 | 1 |
| 1 | 1 | 1 |
| 0 | 0 | 0 |
| 0 | 0 | 0 |
| 0 | 0 | 0 |
| 0 | 0 | 0 |
| 0 | 0 | 0 |
| 0 | 0 | 0 |
| 1 | 1 | 1 |
| 1 | 1 | 1 |
| 0 | 0 | 0 |
| 0 | 0 | 0 |
| 0 | 0 | 0 |
| 0 | 0 | 0 |
| 0 | 0 | 0 |
| 0 | 0 | 0 |
| 1 | 1 | 1 |
| 1 | 1 | 1 |
| 0 | 0 | 0 |
| 1 | 1 | 1 |
| 1 | 1 | 1 |
| 1 | 1 | 1 |
| 1 | 1 | 1 |
| 1 | 1 | 1 |
| 1 | 0 | 0 |
| 0 | 1 | 0 |
| 0 | 1 | 0 |
| 1 | 0 | 0 |

| | | |
|---|---|---|
| 0 | 0 | 0 |
| 1 | 1 | 1 |
| 0 | 0 | 0 |
| 0 | 1 | 0 |
| 1 | 0 | 0 |
| 0 | 0 | 0 |
| 0 | 0 | 0 |
| 0 | 0 | 0 |
| 1 | 0 | 0 |
| 1 | 0 | 0 |
| 0 | 0 | 0 |
| 1 | 1 | 1 |
| 1 | 0 | 0 |
| 1 | 0 | 0 |
| 0 | 0 | 0 |
| 0 | 0 | 0 |
| 0 | 0 | 0 |
| 0 | 0 | 0 |
| 1 | 1 | 1 |
| 0 | 0 | 0 |
| 0 | 0 | 0 |
| 0 | 0 | 0 |
| 0 | 0 | 0 |
| 0 | 0 | 0 |
| 1 | 1 | 1 |
| 0 | 0 | 0 |
| 1 | 1 | 1 |
| 0 | 1 | 0 |
| 1 | 1 | 1 |
| 0 | 0 | 0 |
| 1 | 1 | 1 |
| 1 | 1 | 1 |
| 0 | 0 | 0 |
| 0 | 0 | 0 |
| 1 | 1 | 1 |
| 1 | 1 | 1 |
| 1 | 1 | 1 |
| 0 | 1 | 0 |
| 1 | 0 | 0 |
| 0 | 0 | 0 |
| 0 | 0 | 0 |
| 0 | 0 | 0 |
| 0 | 0 | 0 |
| 0 | 0 | 0 |
| 0 | 0 | 0 |
| 0 | 1 | 0 |
| 0 | 1 | 0 |
| 0 | 0 | 0 |
| 0 | 0 | 0 |
| 1 | 1 | 1 |
| 0 | 0 | 0 |
| 0 | 0 | 0 |
| 0 | 1 | 0 |
| 1 | 1 | 1 |

| | | |
|---|---|---|
| 1 | 1 | 1 |
| 0 | 0 | 0 |
| 0 | 0 | 0 |
| 1 | 0 | 0 |
| 1 | 1 | 1 |
| 0 | 0 | 0 |
| 0 | 0 | 0 |
| 0 | 0 | 0 |
| 0 | 0 | 0 |
| 0 | 0 | 0 |
| 0 | 0 | 0 |
| 0 | 0 | 0 |
| 0 | 0 | 0 |
| 0 | 0 | 0 |
| 0 | 0 | 0 |
| 0 | 1 | 0 |
| 0 | 0 | 0 |
| 1 | 1 | 1 |
| 1 | 0 | 0 |
| 1 | 1 | 1 |
| 0 | 0 | 0 |
| 1 | 1 | 1 |
| 0 | 0 | 0 |
| 0 | 0 | 0 |
| 1 | 1 | 1 |
| 0 | 0 | 0 |
| 0 | 1 | 0 |
| 0 | 1 | 0 |
| 0 | 1 | 0 |
| 0 | 0 | 0 |
| 0 | 0 | 0 |
| 0 | 0 | 0 |
| 0 | 0 | 0 |
| 0 | 0 | 0 |
| 0 | 0 | 0 |
| 0 | 0 | 0 |
| 0 | 0 | 0 |
| 1 | 1 | 1 |
| 1 | 0 | 0 |
| 0 | 0 | 0 |
| 0 | 0 | 0 |
| 0 | 0 | 0 |
| 0 | 0 | 0 |
| 0 | 0 | 0 |
| 0 | 0 | 0 |
| 0 | 0 | 0 |
| 1 | 1 | 1 |
| 0 | 0 | 0 |
| 0 | 0 | 0 |
| 0 | 0 | 0 |
| 0 | 0 | 0 |
| 0 | 1 | 0 |
| 0 | 0 | 0 |
| 0 | 0 | 0 |

| | | |
|---|---|---|
| 0 | 0 | 0 |
| 0 | 0 | 0 |
| 0 | 0 | 0 |
| 0 | 0 | 0 |
| 0 | 0 | 0 |
| 0 | 0 | 0 |
| 0 | 0 | 0 |
| 0 | 0 | 0 |
| 0 | 0 | 0 |
| 0 | 0 | 0 |
| 1 | 1 | 1 |
| 0 | 0 | 0 |
| 1 | 0 | 0 |
| 0 | 0 | 0 |
| 0 | 1 | 0 |
| 1 | 1 | 1 |
| 1 | 1 | 1 |
| 1 | 1 | 1 |
| 0 | 0 | 0 |
| 0 | 0 | 0 |
| 1 | 1 | 1 |
| 0 | 0 | 0 |
| 0 | 0 | 0 |
| 1 | 1 | 1 |
| 0 | 0 | 0 |
| 0 | 0 | 0 |
| 0 | 0 | 0 |
| 1 | 0 | 0 |
| 0 | 0 | 0 |
| 0 | 0 | 0 |
| 1 | 0 | 0 |
| 0 | 0 | 0 |
| 1 | 0 | 0 |
| 1 | 0 | 0 |
| 0 | 0 | 0 |
| 0 | 0 | 0 |
| 0 | 0 | 0 |
| 1 | 0 | 0 |
| 1 | 0 | 0 |
| 0 | 0 | 0 |
| 0 | 0 | 0 |
| 0 | 0 | 0 |
| 0 | 0 | 0 |
| 0 | 0 | 0 |
| 0 | 0 | 0 |
| 0 | 0 | 0 |
| 0 | 0 | 0 |
| 0 | 0 | 0 |
| 0 | 0 | 0 |
| 0 | 0 | 0 |
| 0 | 0 | 0 |
| 0 | 0 | 0 |
| 0 | 0 | 0 |
| 0 | 0 | 0 |

| | | |
|---|---|---|
| 0 | 0 | 0 |
| 1 | 0 | 0 |
| 0 | 0 | 0 |
| 0 | 0 | 0 |
| 0 | 0 | 0 |
| 1 | 0 | 0 |
| 0 | 0 | 0 |
| 0 | 0 | 0 |
| 1 | 1 | 1 |
| 0 | 0 | 0 |
| 0 | 1 | 0 |
| 1 | 1 | 1 |
| 1 | 1 | 1 |
| 0 | 0 | 0 |
| 1 | 1 | 1 |
| 0 | 0 | 0 |
| 1 | 1 | 1 |
| 1 | 1 | 1 |
| 0 | 0 | 0 |
| 0 | 0 | 0 |
| 0 | 0 | 0 |
| 0 | 0 | 0 |
| 0 | 0 | 0 |
| 0 | 0 | 0 |
| 0 | 0 | 0 |
| 0 | 1 | 0 |
| 1 | 1 | 1 |
| 1 | 1 | 1 |
| 1 | 0 | 0 |
| 1 | 1 | 1 |
| 1 | 1 | 1 |
| 0 | 1 | 0 |
| 0 | 0 | 0 |
| 0 | 0 | 0 |
| 0 | 0 | 0 |
| 0 | 0 | 0 |
| 0 | 0 | 0 |
| 0 | 0 | 0 |
| 0 | 1 | 0 |
| 0 | 1 | 0 |
| 0 | 1 | 0 |
| 0 | 1 | 0 |
| 0 | 1 | 0 |
| 0 | 1 | 0 |
| 0 | 0 | 0 |
| 0 | 0 | 0 |
| 0 | 1 | 0 |
| 0 | 0 | 0 |
| 0 | 0 | 0 |
| 0 | 0 | 0 |
| 0 | 0 | 0 |
| 0 | 0 | 0 |
| 0 | 0 | 0 |
| 0 | 0 | 0 |

| | | |
|---|---|---|
| 1 | 1 | 1 |
| 1 | 1 | 1 |
| 0 | 0 | 0 |
| 0 | 0 | 0 |
| 1 | 1 | 1 |
| 0 | 0 | 0 |
| 0 | 0 | 0 |
| 1 | 1 | 1 |
| 1 | 1 | 1 |
| 0 | 0 | 0 |
| 1 | 1 | 1 |
| 1 | 1 | 1 |
| 0 | 0 | 0 |
| 0 | 0 | 0 |
| 0 | 0 | 0 |
| 0 | 0 | 0 |
| 0 | 1 | 0 |
| 0 | 1 | 0 |
| 1 | 1 | 1 |
| 1 | 1 | 1 |
| 1 | 1 | 1 |
| 0 | 1 | 0 |
| 1 | 1 | 1 |
| 1 | 1 | 1 |
| 1 | 1 | 1 |
| 0 | 0 | 0 |
| 1 | 1 | 1 |
| 1 | 1 | 1 |
| 1 | 1 | 1 |
| 0 | 1 | 0 |
| 0 | 1 | 0 |
| 1 | 1 | 1 |
| 1 | 1 | 1 |
| 1 | 1 | 1 |
| 1 | 1 | 1 |
| 1 | 1 | 1 |
| 1 | 1 | 1 |
| 0 | 0 | 0 |
| 0 | 0 | 0 |
| 1 | 0 | 0 |
| 0 | 0 | 0 |
| 1 | 1 | 1 |
| 1 | 1 | 1 |
| 0 | 0 | 0 |
| 0 | 0 | 0 |
| 0 | 1 | 0 |
| 0 | 1 | 0 |
| 0 | 1 | 0 |
| 1 | 1 | 1 |
| 1 | 1 | 1 |
| 1 | 1 | 1 |
| 0 | 0 | 0 |
| 0 | 0 | 0 |
| 0 | 0 | 0 |

| | | |
|---|---|---|
| 0 | 0 | 0 |
| 1 | 1 | 1 |
| 1 | 1 | 1 |
| 1 | 1 | 1 |
| 1 | 1 | 1 |
| 1 | 1 | 1 |
| 1 | 1 | 1 |
| 1 | 1 | 1 |
| 0 | 0 | 0 |
| 1 | 1 | 1 |
| 1 | 1 | 1 |
| 1 | 1 | 1 |
| 0 | 1 | 0 |
| 0 | 1 | 0 |
| 1 | 1 | 1 |
| 1 | 1 | 1 |
| 0 | 1 | 0 |
| 0 | 1 | 0 |
| 0 | 0 | 0 |
| 0 | 0 | 0 |
| 0 | 0 | 0 |
| 0 | 0 | 0 |
| 0 | 1 | 0 |
| 0 | 0 | 0 |
| 0 | 0 | 0 |
| 1 | 1 | 1 |
| 1 | 1 | 1 |
| 1 | 1 | 1 |
| 1 | 1 | 1 |
| 1 | 1 | 1 |
| 1 | 1 | 1 |
| 1 | 1 | 1 |
| 0 | 0 | 0 |
| 1 | 1 | 1 |
| 1 | 1 | 1 |
| 0 | 1 | 0 |
| 0 | 0 | 0 |
| 1 | 1 | 1 |
| 0 | 0 | 0 |
| 0 | 0 | 0 |
| 0 | 0 | 0 |
| 0 | 0 | 0 |
| 0 | 0 | 0 |
| 1 | 1 | 1 |
| 1 | 1 | 1 |
| 1 | 1 | 1 |
| 0 | 0 | 0 |
| 0 | 0 | 0 |
| 0 | 0 | 0 |
| 0 | 0 | 0 |
| 0 | 0 | 0 |
| 0 | 0 | 0 |
| 0 | 0 | 0 |
| 0 | 1 | 0 |

| | | |
|---|---|---|
| 0 | 0 | 0 |
| 0 | 0 | 0 |
| 0 | 1 | 0 |
| 0 | 0 | 0 |
| 0 | 0 | 0 |
| 1 | 1 | 1 |
| 1 | 1 | 1 |
| 1 | 0 | 0 |
| 1 | 1 | 1 |
| 1 | 1 | 1 |
| 1 | 1 | 1 |
| 0 | 0 | 0 |
| 1 | 1 | 1 |
| 1 | 1 | 1 |
| 0 | 0 | 0 |
| 1 | 1 | 1 |
| 0 | 0 | 0 |
| 1 | 1 | 1 |
| 0 | 0 | 0 |
| 0 | 0 | 0 |
| 1 | 1 | 1 |
| 1 | 1 | 1 |
| 0 | 0 | 0 |
| 1 | 1 | 1 |
| 0 | 0 | 0 |
| 1 | 1 | 1 |
| 1 | 1 | 1 |
| 0 | 0 | 0 |
| 1 | 1 | 1 |
| 0 | 0 | 0 |
| 0 | 1 | 0 |
| 0 | 0 | 0 |
| 0 | 0 | 0 |
| 0 | 0 | 0 |
| 1 | 1 | 1 |
| 0 | 0 | 0 |
| 1 | 1 | 1 |
| 1 | 1 | 1 |
| 0 | 0 | 0 |
| 1 | 1 | 1 |
| 1 | 1 | 1 |
| 1 | 1 | 1 |
| 1 | 1 | 1 |
| 0 | 0 | 0 |
| 0 | 0 | 0 |
| 1 | 1 | 1 |
| 1 | 1 | 1 |
| 1 | 0 | 0 |
| 0 | 0 | 0 |
| 1 | 0 | 0 |
| 0 | 0 | 0 |
| 1 | 1 | 1 |
| 1 | 1 | 1 |
| 0 | 0 | 0 |

| | | |
|---|---|---|
| 1 | 1 | 1 |
| 1 | 1 | 1 |
| 1 | 1 | 1 |
| 1 | 1 | 1 |
| 0 | 1 | 0 |
| 1 | 1 | 1 |
| 1 | 1 | 1 |
| 0 | 0 | 0 |
| 1 | 1 | 1 |
| 1 | 1 | 1 |
| 1 | 1 | 1 |
| 1 | 1 | 1 |
| 0 | 0 | 0 |
| 1 | 1 | 1 |
| 0 | 0 | 0 |
| 0 | 0 | 0 |
| 1 | 1 | 1 |
| 1 | 1 | 1 |
| 1 | 1 | 1 |
| 1 | 1 | 1 |
| 0 | 0 | 0 |
| 0 | 0 | 0 |
| 1 | 1 | 1 |
| 1 | 1 | 1 |
| 0 | 0 | 0 |
| 1 | 1 | 1 |
| 0 | 0 | 0 |
| 0 | 0 | 0 |
| 0 | 0 | 0 |
| 0 | 0 | 0 |
| 0 | 1 | 0 |
| 0 | 1 | 0 |
| 0 | 0 | 0 |
| 0 | 0 | 0 |
| 0 | 0 | 0 |
| 0 | 0 | 0 |
| 0 | 0 | 0 |
| 0 | 0 | 0 |
| 0 | 0 | 0 |
| 1 | 0 | 0 |
| 1 | 1 | 1 |
| 0 | 1 | 0 |
| 0 | 1 | 0 |
| 0 | 1 | 0 |
| 0 | 1 | 0 |
| 0 | 0 | 0 |
| 0 | 1 | 0 |
| 0 | 1 | 0 |
| 1 | 1 | 1 |
| 0 | 0 | 0 |
| 0 | 0 | 0 |
| 1 | 1 | 1 |
| 1 | 1 | 1 |
| 1 | 1 | 1 |

| | | |
|---|---|---|
| 1 | 1 | 1 |
| 1 | 1 | 1 |
| 1 | 1 | 1 |
| 1 | 1 | 1 |
| 0 | 0 | 0 |
| 1 | 1 | 1 |
| 0 | 0 | 0 |
| 0 | 0 | 0 |
| 0 | 0 | 0 |
| 0 | 0 | 0 |
| 0 | 0 | 0 |
| 1 | 1 | 1 |
| 0 | 0 | 0 |
| 0 | 0 | 0 |
| 1 | 1 | 1 |
| 1 | 1 | 1 |
| 1 | 1 | 1 |
| 1 | 1 | 1 |
| 1 | 1 | 1 |
| 1 | 1 | 1 |
| 1 | 1 | 1 |
| 1 | 1 | 1 |
| 1 | 1 | 1 |
| 0 | 0 | 0 |
| 1 | 1 | 1 |
| 1 | 1 | 1 |
| 1 | 1 | 1 |
| 0 | 0 | 0 |
| 1 | 0 | 0 |
| 1 | 0 | 0 |
| 0 | 0 | 0 |
| 0 | 0 | 0 |
| 1 | 1 | 1 |
| 0 | 0 | 0 |
| 0 | 1 | 0 |
| 1 | 1 | 1 |
| 1 | 1 | 1 |
| 1 | 1 | 1 |
| 1 | 1 | 1 |
| 1 | 1 | 1 |
| 1 | 1 | 1 |
| 1 | 1 | 1 |
| 1 | 1 | 1 |
| 1 | 1 | 1 |
| 1 | 1 | 1 |
| 1 | 1 | 1 |
| 1 | 1 | 1 |
| 1 | 1 | 1 |
| 0 | 0 | 0 |
| 0 | 1 | 0 |
| 1 | 1 | 1 |
| 1 | 0 | 0 |
| 1 | 0 | 0 |
| 1 | 1 | 1 |

| | | |
|---|---|---|
| 1 | 1 | 1 |
| 0 | 1 | 0 |
| 1 | 1 | 1 |
| 0 | 1 | 0 |
| 1 | 0 | 0 |
| 1 | 1 | 1 |
| 1 | 1 | 1 |
| 1 | 1 | 1 |
| 1 | 1 | 1 |
| 1 | 1 | 1 |
| 0 | 0 | 0 |
| 0 | 0 | 0 |
| 0 | 0 | 0 |
| 0 | 0 | 0 |
| 0 | 0 | 0 |
| 1 | 0 | 0 |
| 0 | 0 | 0 |
| 0 | 0 | 0 |
| 0 | 0 | 0 |
| 0 | 0 | 0 |
| 0 | 0 | 0 |
| 0 | 0 | 0 |
| 0 | 1 | 0 |
| 0 | 0 | 0 |
| 0 | 0 | 0 |
| 0 | 0 | 0 |
| 1 | 1 | 1 |
| 0 | 0 | 0 |
| 0 | 1 | 0 |
| 0 | 1 | 0 |
| 0 | 1 | 0 |
| 0 | 0 | 0 |
| 1 | 1 | 1 |
| 1 | 1 | 1 |
| 0 | 0 | 0 |
| 0 | 0 | 0 |
| 0 | 0 | 0 |
| 0 | 0 | 0 |
| 0 | 0 | 0 |
| 1 | 0 | 0 |
| 1 | 0 | 0 |
| 0 | 0 | 0 |
| 1 | 0 | 0 |
| 0 | 0 | 0 |
| 1 | 1 | 1 |
| 1 | 0 | 0 |
| 1 | 1 | 1 |
| 0 | 0 | 0 |
| 0 | 0 | 0 |
| 0 | 0 | 0 |
| 0 | 0 | 0 |
| 1 | 1 | 1 |
| 0 | 0 | 0 |
| 1 | 1 | 1 |

| | | |
|---|---|---|
| 1 | 1 | 1 |
| 1 | 1 | 1 |
| 1 | 1 | 1 |
| 1 | 1 | 1 |
| 1 | 1 | 1 |
| 0 | 0 | 0 |
| 0 | 0 | 0 |
| 0 | 0 | 0 |
| 0 | 0 | 0 |
| 0 | 1 | 0 |
| 0 | 0 | 0 |
| 0 | 0 | 0 |
| 0 | 0 | 0 |
| 0 | 0 | 0 |
| 0 | 0 | 0 |
| 0 | 0 | 0 |
| 1 | 1 | 1 |
| 0 | 1 | 0 |
| 1 | 1 | 1 |
| 1 | 1 | 1 |
| 1 | 1 | 1 |
| 1 | 1 | 1 |
| 1 | 1 | 1 |
| 1 | 1 | 1 |
| 1 | 1 | 1 |
| 1 | 1 | 1 |
| 1 | 1 | 1 |
| 1 | 1 | 1 |
| 0 | 0 | 0 |
| 0 | 1 | 0 |
| 0 | 1 | 0 |
| 1 | 0 | 0 |
| 1 | 0 | 0 |
| 0 | 0 | 0 |
| 1 | 1 | 1 |
| 0 | 0 | 0 |
| 1 | 0 | 0 |
| 0 | 0 | 0 |
| 1 | 1 | 1 |
| 0 | 0 | 0 |
| 1 | 1 | 1 |
| 1 | 1 | 1 |
| 0 | 1 | 0 |
| 0 | 0 | 0 |
| 1 | 1 | 1 |
| 1 | 1 | 1 |
| 1 | 1 | 1 |
| 1 | 1 | 1 |
| 1 | 1 | 1 |
| 1 | 1 | 1 |
| 1 | 1 | 1 |
| 1 | 0 | 0 |
| 0 | 0 | 0 |
| 0 | 0 | 0 |

| | | |
|---|---|---|
| 1 | 1 | 1 |
| 0 | 0 | 0 |
| 1 | 1 | 1 |
| 1 | 1 | 1 |
| 0 | 0 | 0 |
| 0 | 0 | 0 |
| 0 | 0 | 0 |
| 1 | 1 | 1 |
| 1 | 1 | 1 |
| 0 | 0 | 0 |
| 0 | 0 | 0 |
| 0 | 0 | 0 |
| 0 | 0 | 0 |
| 0 | 0 | 0 |
| 1 | 1 | 1 |
| 0 | 0 | 0 |
| 0 | 0 | 0 |
| 0 | 0 | 0 |
| 1 | 1 | 1 |
| 1 | 1 | 1 |
| 1 | 1 | 1 |
| 1 | 1 | 1 |
| 1 | 1 | 1 |
| 0 | 0 | 0 |
| 0 | 1 | 0 |
| 0 | 1 | 0 |
| 0 | 0 | 0 |
| 0 | 0 | 0 |
| 1 | 1 | 1 |
| 1 | 1 | 1 |
| 0 | 0 | 0 |
| 1 | 1 | 1 |
| 1 | 1 | 1 |
| 0 | 1 | 0 |
| 1 | 1 | 1 |
| 0 | 0 | 0 |
| 0 | 1 | 0 |
| 0 | 0 | 0 |
| 0 | 0 | 0 |
| 0 | 1 | 0 |
| 0 | 1 | 0 |
| 0 | 0 | 0 |
| 0 | 0 | 0 |
| 0 | 1 | 0 |
| 0 | 0 | 0 |
| 0 | 0 | 0 |
| 0 | 0 | 0 |
| 0 | 1 | 0 |
| 0 | 0 | 0 |
| 0 | 0 | 0 |
| 0 | 0 | 0 |
| 1 | 1 | 1 |
| 0 | 0 | 0 |
| 1 | 1 | 1 |

| | | |
|---|---|---|
| 1 | 1 | 1 |
| 1 | 1 | 1 |
| 1 | 1 | 1 |
| 1 | 1 | 1 |
| 0 | 0 | 0 |
| 0 | 0 | 0 |
| 1 | 1 | 1 |
| 1 | 0 | 0 |
| 1 | 1 | 1 |
| 0 | 0 | 0 |
| 1 | 0 | 0 |
| 1 | 0 | 0 |
| 0 | 0 | 0 |
| 1 | 1 | 1 |
| 1 | 1 | 1 |
| 1 | 1 | 1 |
| 1 | 1 | 1 |
| 1 | 1 | 1 |
| 1 | 1 | 1 |
| 1 | 1 | 1 |
| 1 | 1 | 1 |
| 0 | 1 | 0 |
| 1 | 1 | 1 |
| 0 | 0 | 0 |
| 1 | 1 | 1 |
| 0 | 0 | 0 |
| 1 | 1 | 1 |
| 0 | 0 | 0 |
| 1 | 1 | 1 |
| 0 | 0 | 0 |
| 0 | 0 | 0 |
| 0 | 0 | 0 |
| 0 | 0 | 0 |
| 0 | 0 | 0 |
| 0 | 1 | 0 |
| 0 | 0 | 0 |
| 1 | 1 | 1 |
| 0 | 0 | 0 |
| 1 | 1 | 1 |
| 1 | 1 | 1 |
| 0 | 0 | 0 |
| 0 | 0 | 0 |
| 0 | 0 | 0 |
| 0 | 0 | 0 |
| 0 | 0 | 0 |
| 0 | 0 | 0 |
| 0 | 0 | 0 |
| 0 | 0 | 0 |
| 1 | 1 | 1 |
| 0 | 0 | 0 |
| 1 | 0 | 0 |
| 1 | 0 | 0 |
| 1 | 1 | 1 |
| 1 | 1 | 1 |

| | | |
|---|---|---|
| 1 | 1 | 1 |
| 1 | 1 | 1 |
| 1 | 1 | 1 |
| 0 | 1 | 0 |
| 0 | 1 | 0 |
| 0 | 1 | 0 |
| 0 | 0 | 0 |
| 0 | 0 | 0 |
| 0 | 0 | 0 |
| 0 | 0 | 0 |
| 0 | 0 | 0 |
| 0 | 0 | 0 |
| 0 | 0 | 0 |
| 0 | 0 | 0 |
| 0 | 0 | 0 |
| 0 | 0 | 0 |
| 0 | 0 | 0 |
| 0 | 0 | 0 |
| 1 | 1 | 1 |
| 0 | 0 | 0 |
| 0 | 1 | 0 |
| 0 | 1 | 0 |
| 0 | 1 | 0 |
| 1 | 1 | 1 |
| 0 | 1 | 0 |
| 0 | 0 | 0 |
| 0 | 0 | 0 |
| 0 | 0 | 0 |
| 0 | 0 | 0 |
| 0 | 0 | 0 |
| 0 | 0 | 0 |
| 0 | 0 | 0 |
| 0 | 0 | 0 |
| 0 | 0 | 0 |
| 1 | 1 | 1 |
| 0 | 0 | 0 |
| 0 | 0 | 0 |
| 0 | 0 | 0 |
| 0 | 0 | 0 |
| 0 | 1 | 0 |
| 0 | 0 | 0 |
| 0 | 0 | 0 |
| 1 | 1 | 1 |
| 0 | 1 | 0 |
| 1 | 1 | 1 |
| 0 | 1 | 0 |
| 0 | 1 | 0 |
| 0 | 0 | 0 |
| 0 | 0 | 0 |
| 0 | 0 | 0 |
| 0 | 0 | 0 |
| 0 | 1 | 0 |
| 0 | 0 | 0 |
| 1 | 1 | 1 |

| | | |
|---|---|---|
| 0 | 1 | 0 |
| 1 | 1 | 1 |
| 1 | 1 | 1 |
| 1 | 1 | 1 |
| 1 | 1 | 1 |
| 0 | 0 | 0 |
| 0 | 1 | 0 |
| 1 | 1 | 1 |
| 1 | 1 | 1 |
| 1 | 1 | 1 |
| 0 | 0 | 0 |
| 0 | 0 | 0 |
| 0 | 0 | 0 |
| 0 | 0 | 0 |
| 0 | 0 | 0 |
| 0 | 1 | 0 |
| 1 | 0 | 0 |
| 0 | 0 | 0 |
| 1 | 1 | 1 |
| 1 | 1 | 1 |
| 1 | 1 | 1 |
| 1 | 1 | 1 |
| 0 | 0 | 0 |
| 1 | 1 | 1 |
| 0 | 0 | 0 |
| 0 | 0 | 0 |
| 0 | 0 | 0 |
| 0 | 0 | 0 |
| 1 | 1 | 1 |
| 0 | 0 | 0 |
| 1 | 1 | 1 |
| 0 | 1 | 0 |
| 0 | 1 | 0 |
| 0 | 1 | 0 |
| 1 | 1 | 1 |
| 1 | 1 | 1 |
| 0 | 1 | 0 |
| 0 | 0 | 0 |
| 0 | 0 | 0 |
| 0 | 0 | 0 |
| 0 | 0 | 0 |
| 0 | 0 | 0 |
| 1 | 1 | 1 |
| 0 | 0 | 0 |
| 0 | 0 | 0 |
| 1 | 1 | 1 |
| 0 | 0 | 0 |
| 0 | 0 | 0 |
| 0 | 0 | 0 |
| 0 | 0 | 0 |
| 1 | 1 | 1 |
| 1 | 1 | 1 |
| 1 | 1 | 1 |
| 0 | 0 | 0 |

| | | |
|---|---|---|
| 0 | 0 | 0 |
| 1 | 1 | 1 |
| 0 | 0 | 0 |
| 0 | 1 | 0 |
| 1 | 0 | 0 |
| 1 | 0 | 0 |
| 1 | 0 | 0 |
| 1 | 1 | 1 |
| 0 | 0 | 0 |
| 1 | 1 | 1 |
| 0 | 0 | 0 |
| 0 | 0 | 0 |
| 0 | 0 | 0 |
| 0 | 0 | 0 |
| 1 | 1 | 1 |
| 1 | 1 | 1 |
| 0 | 0 | 0 |
| 0 | 0 | 0 |
| 0 | 1 | 0 |
| 1 | 1 | 1 |
| 0 | 0 | 0 |
| 1 | 0 | 0 |
| 1 | 0 | 0 |
| 1 | 1 | 1 |
| 0 | 0 | 0 |
| 1 | 0 | 0 |
| 0 | 0 | 0 |
| 0 | 0 | 0 |
| 0 | 0 | 0 |
| 1 | 1 | 1 |
| 0 | 0 | 0 |
| 0 | 0 | 0 |
| 0 | 0 | 0 |
| 1 | 1 | 1 |
| 0 | 0 | 0 |
| 1 | 1 | 1 |
| 0 | 0 | 0 |
| 0 | 0 | 0 |
| 0 | 0 | 0 |
| 0 | 0 | 0 |
| 0 | 0 | 0 |
| 1 | 1 | 1 |
| 0 | 0 | 0 |
| 0 | 0 | 0 |
| 0 | 0 | 0 |
| 0 | 0 | 0 |
| 0 | 0 | 0 |
| 0 | 0 | 0 |
| 0 | 0 | 0 |
| 0 | 0 | 0 |
| 0 | 0 | 0 |
| 0 | 0 | 0 |
| 1 | 1 | 1 |
| 0 | 0 | 0 |

| | | |
|---|---|---|
| 0 | 1 | 0 |
| 1 | 1 | 1 |
| 1 | 1 | 1 |
| 1 | 1 | 1 |
| 1 | 1 | 1 |
| 1 | 1 | 1 |
| 1 | 1 | 1 |
| 1 | 1 | 1 |
| 1 | 1 | 1 |
| 0 | 0 | 0 |
| 0 | 0 | 0 |
| 1 | 1 | 1 |
| 1 | 1 | 1 |
| 1 | 1 | 1 |
| 0 | 0 | 0 |
| 1 | 1 | 1 |
| 1 | 1 | 1 |
| 0 | 0 | 0 |
| 0 | 1 | 0 |
| 0 | 1 | 0 |
| 0 | 1 | 0 |
| 0 | 0 | 0 |
| 0 | 0 | 0 |
| 0 | 0 | 0 |
| 0 | 0 | 0 |
| 0 | 0 | 0 |
| 0 | 0 | 0 |
| 0 | 0 | 0 |
| 1 | 1 | 1 |
| 0 | 0 | 0 |
| 0 | 0 | 0 |
| 1 | 1 | 1 |
| 1 | 1 | 1 |
| 1 | 1 | 1 |
| 1 | 1 | 1 |
| 1 | 1 | 1 |
| 1 | 1 | 1 |
| 1 | 1 | 1 |
| 1 | 1 | 1 |
| 1 | 1 | 1 |
| 1 | 1 | 1 |
| 1 | 1 | 1 |
| 1 | 1 | 1 |
| 0 | 0 | 0 |
| 1 | 1 | 1 |
| 0 | 0 | 0 |
| 1 | 1 | 1 |
| 1 | 1 | 1 |
| 1 | 1 | 1 |
| 1 | 1 | 1 |
| 0 | 0 | 0 |
| 1 | 1 | 1 |
| 1 | 1 | 1 |
| 0 | 0 | 0 |

| | | |
|---|---|---|
| 0 | 0 | 0 |
| 0 | 0 | 0 |
| 0 | 0 | 0 |
| 0 | 0 | 0 |
| 1 | 1 | 1 |
| 0 | 1 | 0 |
| 1 | 1 | 1 |
| 0 | 0 | 0 |
| 0 | 0 | 0 |
| 0 | 0 | 0 |
| 0 | 0 | 0 |
| 1 | 1 | 1 |
| 1 | 1 | 1 |
| 1 | 1 | 1 |
| 1 | 1 | 1 |
| 0 | 0 | 0 |
| 0 | 0 | 0 |
| 0 | 1 | 0 |
| 0 | 1 | 0 |
| 1 | 1 | 1 |
| 1 | 1 | 1 |
| 0 | 0 | 0 |
| 1 | 1 | 1 |
| 0 | 1 | 0 |
| 0 | 0 | 0 |
| 0 | 0 | 0 |
| 0 | 0 | 0 |
| 0 | 0 | 0 |
| 1 | 1 | 1 |
| 1 | 1 | 1 |
| 1 | 1 | 1 |
| 0 | 0 | 0 |
| 1 | 1 | 1 |
| 1 | 1 | 1 |
| 1 | 1 | 1 |
| 1 | 1 | 1 |
| 1 | 1 | 1 |
| 1 | 1 | 1 |
| 1 | 1 | 1 |
| 1 | 1 | 1 |
| 1 | 1 | 1 |
| 1 | 1 | 1 |
| 0 | 0 | 0 |
| 0 | 1 | 0 |
| 0 | 1 | 0 |
| 1 | 1 | 1 |
| 0 | 1 | 0 |
| 1 | 1 | 1 |
| 1 | 1 | 1 |
| 0 | 0 | 0 |
| 1 | 1 | 1 |
| 1 | 1 | 1 |
| 1 | 1 | 1 |
| 1 | 1 | 1 |

| | | |
|---|---|---|
| 0 | 0 | 0 |
| 1 | 1 | 1 |
| 0 | 0 | 0 |
| 0 | 1 | 0 |
| 1 | 1 | 1 |
| 1 | 1 | 1 |
| 1 | 1 | 1 |
| 0 | 0 | 0 |
| 1 | 1 | 1 |
| 1 | 1 | 1 |
| 0 | 0 | 0 |
| 0 | 0 | 0 |
| 0 | 0 | 0 |
| 1 | 1 | 1 |
| 0 | 0 | 0 |
| 0 | 0 | 0 |
| 0 | 0 | 0 |
| 0 | 0 | 0 |
| 0 | 1 | 0 |
| 0 | 0 | 0 |
| 0 | 0 | 0 |
| 0 | 0 | 0 |
| 0 | 0 | 0 |
| 0 | 0 | 0 |
| 0 | 0 | 0 |
| 0 | 0 | 0 |
| 0 | 0 | 0 |
| 0 | 0 | 0 |
| 0 | 0 | 0 |
| 0 | 0 | 0 |
| 0 | 0 | 0 |
| 0 | 0 | 0 |
| 1 | 1 | 1 |
| 0 | 0 | 0 |
| 1 | 1 | 1 |
| 0 | 0 | 0 |
| 0 | 0 | 0 |
| 1 | 1 | 1 |
| 1 | 1 | 1 |
| 0 | 1 | 0 |
| 0 | 1 | 0 |
| 0 | 0 | 0 |
| 0 | 0 | 0 |
| 0 | 0 | 0 |
| 1 | 1 | 1 |
| 0 | 0 | 0 |
| 1 | 1 | 1 |
| 0 | 0 | 0 |
| 0 | 1 | 0 |
| 0 | 1 | 0 |
| 0 | 1 | 0 |
| 0 | 1 | 0 |
| 0 | 1 | 0 |
| 1 | 1 | 1 |

| | | |
|---|---|---|
| 1 | 1 | 1 |
| 1 | 1 | 1 |
| 0 | 1 | 0 |
| 1 | 1 | 1 |
| 1 | 1 | 1 |
| 0 | 0 | 0 |
| 0 | 0 | 0 |
| 1 | 1 | 1 |
| 0 | 0 | 0 |
| 0 | 0 | 0 |
| 1 | 1 | 1 |
| 0 | 1 | 0 |
| 1 | 1 | 1 |
| 1 | 1 | 1 |
| 1 | 1 | 1 |
| 0 | 0 | 0 |
| 1 | 1 | 1 |
| 1 | 1 | 1 |
| 0 | 0 | 0 |
| 0 | 0 | 0 |
| 0 | 0 | 0 |
| 1 | 1 | 1 |
| 1 | 1 | 1 |
| 0 | 0 | 0 |
| 1 | 1 | 1 |
| 0 | 0 | 0 |
| 0 | 0 | 0 |
| 1 | 1 | 1 |
| 1 | 1 | 1 |
| 1 | 1 | 1 |
| 0 | 0 | 0 |
| 1 | 0 | 0 |
| 0 | 0 | 0 |
| 0 | 0 | 0 |
| 0 | 0 | 0 |
| 0 | 0 | 0 |
| 0 | 0 | 0 |
| 1 | 1 | 1 |
| 0 | 0 | 0 |
| 0 | 1 | 0 |
| 0 | 0 | 0 |
| 0 | 0 | 0 |
| 0 | 0 | 0 |
| 0 | 0 | 0 |
| 1 | 1 | 1 |
| 0 | 1 | 0 |
| 0 | 1 | 0 |
| 0 | 0 | 0 |
| 0 | 0 | 0 |
| 0 | 0 | 0 |
| 1 | 1 | 1 |
| 0 | 1 | 0 |
| 1 | 1 | 1 |
| 1 | 1 | 1 |

| | | |
|---|---|---|
| 1 | 1 | 1 |
| 0 | 1 | 0 |
| 0 | 1 | 0 |
| 0 | 0 | 0 |
| 0 | 0 | 0 |
| 1 | 1 | 1 |
| 0 | 0 | 0 |
| 1 | 1 | 1 |
| 1 | 1 | 1 |
| 1 | 1 | 1 |
| 1 | 1 | 1 |
| 0 | 0 | 0 |
| 1 | 0 | 0 |
| 0 | 0 | 0 |
| 1 | 1 | 1 |
| 0 | 0 | 0 |
| 0 | 0 | 0 |
| 0 | 0 | 0 |
| 1 | 1 | 1 |
| 0 | 0 | 0 |
| 0 | 0 | 0 |
| 0 | 0 | 0 |
| 0 | 0 | 0 |
| 0 | 0 | 0 |
| 0 | 0 | 0 |
| 1 | 1 | 1 |
| 0 | 0 | 0 |
| 0 | 1 | 0 |
| 0 | 1 | 0 |
| 0 | 1 | 0 |
| 0 | 1 | 0 |
| 0 | 1 | 0 |
| 0 | 0 | 0 |
| 0 | 0 | 0 |
| 1 | 1 | 1 |
| 1 | 1 | 1 |
| 0 | 0 | 0 |
| 0 | 0 | 0 |
| 0 | 0 | 0 |
| 1 | 0 | 0 |
| 0 | 0 | 0 |
| 0 | 0 | 0 |
| 1 | 1 | 1 |
| 0 | 0 | 0 |
| 1 | 1 | 1 |
| 1 | 1 | 1 |
| 1 | 1 | 1 |
| 1 | 1 | 1 |
| 1 | 1 | 1 |
| 0 | 0 | 0 |
| 1 | 1 | 1 |
| 1 | 1 | 1 |
| 1 | 1 | 1 |
| 0 | 1 | 0 |

| | | |
|---|---|---|
| 1 | 1 | 1 |
| 1 | 1 | 1 |
| 1 | 1 | 1 |
| 0 | 0 | 0 |
| 1 | 1 | 1 |
| 1 | 1 | 1 |
| 0 | 1 | 0 |
| 0 | 0 | 0 |
| 1 | 1 | 1 |
| 1 | 1 | 1 |
| 1 | 1 | 1 |
| 1 | 1 | 1 |
| 1 | 1 | 1 |
| 0 | 1 | 0 |
| 0 | 1 | 0 |
| 0 | 1 | 0 |
| 0 | 0 | 0 |
| 0 | 0 | 0 |
| 0 | 1 | 0 |
| 1 | 1 | 1 |
| 0 | 1 | 0 |
| 1 | 1 | 1 |
| 0 | 1 | 0 |
| 1 | 1 | 1 |
| 1 | 1 | 1 |
| 1 | 1 | 1 |
| 1 | 1 | 1 |
| 0 | 0 | 0 |
| 0 | 0 | 0 |
| 1 | 1 | 1 |
| 1 | 1 | 1 |
| 0 | 0 | 0 |
| 0 | 1 | 0 |
| 0 | 0 | 0 |
| 0 | 0 | 0 |
| 0 | 0 | 0 |
| 0 | 0 | 0 |
| 0 | 0 | 0 |
| 1 | 1 | 1 |
| 1 | 1 | 1 |
| 1 | 1 | 1 |
| 0 | 1 | 0 |
| 1 | 1 | 1 |
| 1 | 1 | 1 |
| 0 | 0 | 0 |
| 0 | 0 | 0 |
| 0 | 0 | 0 |
| 1 | 1 | 1 |
| 1 | 1 | 1 |
| 1 | 1 | 1 |
| 1 | 1 | 1 |
| 0 | 0 | 0 |
| 0 | 0 | 0 |
| 0 | 0 | 0 |

| | | |
|---|---|---|
| 0 | 0 | 0 |
| 0 | 0 | 0 |
| 1 | 1 | 1 |
| 0 | 0 | 0 |
| 0 | 0 | 0 |
| 0 | 0 | 0 |
| 0 | 0 | 0 |
| 1 | 1 | 1 |
| 1 | 1 | 1 |
| 1 | 1 | 1 |
| 1 | 1 | 1 |
| 1 | 1 | 1 |
| 0 | 1 | 0 |
| 0 | 1 | 0 |
| 1 | 1 | 1 |
| 1 | 1 | 1 |
| 1 | 1 | 1 |
| 1 | 1 | 1 |
| 1 | 1 | 1 |
| 1 | 1 | 1 |
| 0 | 0 | 0 |
| 0 | 0 | 0 |
| 0 | 0 | 0 |
| 0 | 0 | 0 |
| 0 | 0 | 0 |
| 1 | 1 | 1 |
| 1 | 1 | 1 |
| 0 | 0 | 0 |
| 0 | 1 | 0 |
| 0 | 1 | 0 |
| 0 | 0 | 0 |
| 0 | 0 | 0 |
| 0 | 0 | 0 |
| 1 | 1 | 1 |
| 0 | 0 | 0 |
| 1 | 1 | 1 |
| 0 | 1 | 0 |
| 0 | 0 | 0 |
| 1 | 1 | 1 |
| 1 | 1 | 1 |
| 0 | 1 | 0 |
| 0 | 1 | 0 |
| 1 | 1 | 1 |
| 1 | 1 | 1 |
| 1 | 1 | 1 |
| 1 | 1 | 1 |
| 1 | 1 | 1 |
| 1 | 1 | 1 |
| 0 | 0 | 0 |
| 1 | 1 | 1 |
| 0 | 0 | 0 |
| 0 | 0 | 0 |
| 1 | 1 | 1 |
| 0 | 1 | 0 |

| | | |
|---|---|---|
| 0 | 0 | 0 |
| 1 | 1 | 1 |
| 1 | 1 | 1 |
| 1 | 1 | 1 |
| 1 | 1 | 1 |
| 0 | 0 | 0 |
| 0 | 0 | 0 |
| 1 | 1 | 1 |
| 0 | 1 | 0 |
| 1 | 1 | 1 |
| 1 | 1 | 1 |
| 0 | 0 | 0 |
| 0 | 1 | 0 |
| 1 | 1 | 1 |
| 0 | 1 | 0 |
| 1 | 1 | 1 |
| 0 | 0 | 0 |
| 0 | 0 | 0 |
| 1 | 1 | 1 |
| 1 | 1 | 1 |
| 1 | 1 | 1 |
| 1 | 1 | 1 |
| 1 | 1 | 1 |
| 1 | 1 | 1 |
| 0 | 0 | 0 |
| 1 | 1 | 1 |
| 1 | 1 | 1 |
| 1 | 1 | 1 |
| 1 | 1 | 1 |
| 1 | 1 | 1 |
| 0 | 1 | 0 |
| 0 | 0 | 0 |
| 0 | 0 | 0 |
| 0 | 1 | 0 |
| 1 | 1 | 1 |
| 0 | 0 | 0 |
| 0 | 0 | 0 |
| 0 | 0 | 0 |
| 0 | 0 | 0 |
| 0 | 0 | 0 |
| 0 | 0 | 0 |
| 0 | 0 | 0 |
| 0 | 0 | 0 |
| 0 | 0 | 0 |
| 0 | 0 | 0 |
| 0 | 0 | 0 |
| 0 | 0 | 0 |
| 0 | 0 | 0 |
| 1 | 0 | 0 |
| 1 | 1 | 1 |
| 1 | 0 | 0 |
| 1 | 1 | 1 |
| 0 | 0 | 0 |
| 1 | 1 | 1 |

| | | |
|---|---|---|
| 1 | 1 | 1 |
| 1 | 1 | 1 |
| 1 | 1 | 1 |
| 1 | 1 | 1 |
| 1 | 1 | 1 |
| 1 | 1 | 1 |
| 1 | 1 | 1 |
| 0 | 0 | 0 |
| 0 | 0 | 0 |
| 1 | 0 | 0 |
| 1 | 1 | 1 |
| 1 | 1 | 1 |
| 1 | 1 | 1 |
| 0 | 0 | 0 |
| 1 | 1 | 1 |
| 1 | 1 | 1 |
| 0 | 0 | 0 |
| 1 | 1 | 1 |
| 0 | 0 | 0 |
| 0 | 1 | 0 |
| 1 | 1 | 1 |
| 1 | 1 | 1 |
| 1 | 1 | 1 |
| 0 | 0 | 0 |
| 1 | 1 | 1 |
| 1 | 1 | 1 |
| 0 | 1 | 0 |
| 0 | 1 | 0 |
| 0 | 1 | 0 |
| 0 | 1 | 0 |
| 1 | 1 | 1 |
| 0 | 0 | 0 |
| 1 | 1 | 1 |
| 0 | 1 | 0 |
| 1 | 1 | 1 |
| 1 | 1 | 1 |
| 1 | 1 | 1 |
| 1 | 1 | 1 |
| 0 | 0 | 0 |
| 1 | 1 | 1 |
| 1 | 1 | 1 |
| 1 | 1 | 1 |
| 0 | 1 | 0 |
| 1 | 1 | 1 |
| 1 | 1 | 1 |
| 0 | 0 | 0 |
| 0 | 1 | 0 |
| 0 | 1 | 0 |
| 1 | 0 | 0 |
| 1 | 0 | 0 |
| 0 | 1 | 0 |
| 1 | 0 | 0 |
| 0 | 0 | 0 |
| 1 | 0 | 0 |

| | | |
|---|---|---|
| 1 | 0 | 0 |
| 1 | 0 | 0 |
| 1 | 0 | 0 |
| 1 | 0 | 0 |
| 0 | 0 | 0 |
| 0 | 0 | 0 |
| 0 | 0 | 0 |
| 0 | 0 | 0 |
| 1 | 0 | 0 |
| 1 | 0 | 0 |
| 1 | 0 | 0 |
| 1 | 0 | 0 |
| 1 | 0 | 0 |
| 0 | 0 | 0 |
| 1 | 0 | 0 |
| 0 | 0 | 0 |
| 1 | 0 | 0 |
| 0 | 0 | 0 |
| 1 | 0 | 0 |
| 0 | 0 | 0 |
| 0 | 0 | 0 |
| 1 | 0 | 0 |
| 1 | 0 | 0 |
| 0 | 0 | 0 |
| 0 | 0 | 0 |
| 1 | 0 | 0 |
| 0 | 0 | 0 |
| 0 | 0 | 0 |
| 0 | 0 | 0 |
| 1 | 0 | 0 |
| 1 | 0 | 0 |
| 0 | 0 | 0 |
| 1 | 0 | 0 |
| 1 | 0 | 0 |
| 1 | 0 | 0 |
| 1 | 0 | 0 |
| 1 | 0 | 0 |
| 1 | 0 | 0 |
| 1 | 0 | 0 |
| 1 | 0 | 0 |
| 1 | 0 | 0 |
| 1 | 0 | 0 |
| 1 | 0 | 0 |
| 0 | 0 | 0 |
| 1 | 1 | 1 |
| 1 | 1 | 1 |
| 0 | 0 | 0 |
| 0 | 0 | 0 |
| 0 | 1 | 0 |
| 0 | 0 | 0 |
| 0 | 0 | 0 |
| 0 | 0 | 0 |
| 1 | 1 | 1 |
| 1 | 1 | 1 |

| | | |
|---|---|---|
| 1 | 1 | 1 |
| 1 | 1 | 1 |
| 1 | 1 | 1 |
| 1 | 1 | 1 |
| 1 | 1 | 1 |
| 1 | 1 | 1 |
| 1 | 1 | 1 |
| 0 | 0 | 0 |
| 1 | 1 | 1 |
| 1 | 1 | 1 |
| 0 | 1 | 0 |
| 0 | 1 | 0 |
| 1 | 0 | 0 |
| 1 | 1 | 1 |
| 0 | 0 | 0 |
| 0 | 0 | 0 |
| 0 | 1 | 0 |
| 0 | 0 | 0 |
| 1 | 1 | 1 |
| 1 | 1 | 1 |
| 0 | 0 | 0 |
| 0 | 1 | 0 |
| 0 | 1 | 0 |
| 1 | 1 | 1 |
| 0 | 0 | 0 |
| 1 | 1 | 1 |
| 0 | 1 | 0 |
| 0 | 0 | 0 |
| 0 | 0 | 0 |
| 0 | 0 | 0 |
| 0 | 0 | 0 |
| 1 | 1 | 1 |
| 1 | 1 | 1 |
| 0 | 1 | 0 |
| 1 | 1 | 1 |
| 1 | 1 | 1 |
| 0 | 0 | 0 |
| 0 | 0 | 0 |
| 1 | 1 | 1 |
| 1 | 1 | 1 |
| 1 | 1 | 1 |
| 1 | 1 | 1 |
| 0 | 0 | 0 |
| 0 | 1 | 0 |
| 1 | 1 | 1 |
| 1 | 1 | 1 |
| 1 | 1 | 1 |
| 1 | 1 | 1 |
| 0 | 0 | 0 |
| 0 | 0 | 0 |
| 0 | 0 | 0 |
| 0 | 0 | 0 |
| 0 | 0 | 0 |
| 1 | 1 | 1 |

| | | |
|---|---|---|
| 0 | 0 | 0 |
| 0 | 0 | 0 |
| 0 | 1 | 0 |
| 0 | 0 | 0 |
| 1 | 1 | 1 |
| 0 | 1 | 0 |
| 0 | 1 | 0 |
| 1 | 1 | 1 |
| 1 | 1 | 1 |
| 0 | 0 | 0 |
| 1 | 1 | 1 |
| 0 | 1 | 0 |
| 1 | 1 | 1 |
| 0 | 1 | 0 |
| 0 | 1 | 0 |
| 0 | 1 | 0 |
| 0 | 1 | 0 |
| 1 | 1 | 1 |
| 1 | 1 | 1 |
| 0 | 1 | 0 |
| 0 | 1 | 0 |
| 0 | 1 | 0 |
| 0 | 1 | 0 |
| 0 | 1 | 0 |
| 0 | 1 | 0 |
| 0 | 1 | 0 |
| 0 | 1 | 0 |
| 0 | 1 | 0 |
| 0 | 1 | 0 |
| 0 | 1 | 0 |
| 1 | 1 | 1 |
| 1 | 1 | 1 |
| 0 | 0 | 0 |
| 1 | 1 | 1 |
| 1 | 1 | 1 |
| 1 | 1 | 1 |
| 1 | 1 | 1 |
| 0 | 1 | 0 |
| 1 | 1 | 1 |
| 1 | 1 | 1 |
| 1 | 1 | 1 |
| 1 | 1 | 1 |
| 1 | 1 | 1 |
| 0 | 1 | 0 |
| 0 | 1 | 0 |
| 1 | 1 | 1 |
| 1 | 1 | 1 |
| 0 | 0 | 0 |
| 0 | 0 | 0 |
| 0 | 0 | 0 |
| 0 | 0 | 0 |
| 0 | 0 | 0 |
| 1 | 1 | 1 |
| 0 | 0 | 0 |

| | | |
|---|---|---|
| 1 | 1 | 1 |
| 0 | 0 | 0 |
| 1 | 1 | 1 |
| 0 | 1 | 0 |
| 0 | 1 | 0 |
| 0 | 0 | 0 |
| 0 | 0 | 0 |
| 0 | 0 | 0 |
| 0 | 0 | 0 |
| 0 | 0 | 0 |
| 0 | 1 | 0 |
| 0 | 0 | 0 |
| 0 | 1 | 0 |
| 1 | 1 | 1 |
| 0 | 1 | 0 |
| 0 | 0 | 0 |
| 1 | 1 | 1 |
| 1 | 1 | 1 |
| 0 | 0 | 0 |
| 0 | 0 | 0 |
| 0 | 0 | 0 |
| 0 | 0 | 0 |
| 0 | 0 | 0 |
| 0 | 0 | 0 |
| 0 | 1 | 0 |
| 0 | 0 | 0 |
| 0 | 0 | 0 |
| 0 | 0 | 0 |
| 0 | 0 | 0 |
| 0 | 0 | 0 |
| 0 | 0 | 0 |
| 1 | 1 | 1 |
| 1 | 1 | 1 |
| 0 | 0 | 0 |
| 0 | 0 | 0 |
| 1 | 0 | 0 |
| 0 | 0 | 0 |
| 0 | 0 | 0 |
| 0 | 0 | 0 |
| 0 | 0 | 0 |
| 0 | 0 | 0 |
| 1 | 0 | 0 |
| 1 | 1 | 1 |
| 1 | 1 | 1 |
| 0 | 0 | 0 |
| 0 | 0 | 0 |
| 0 | 0 | 0 |
| 1 | 1 | 1 |
| 1 | 1 | 1 |
| 1 | 1 | 1 |
| 1 | 1 | 1 |
| 1 | 1 | 1 |
| 1 | 1 | 1 |
| 0 | 0 | 0 |

| | | |
|---|---|---|
| 1 | 1 | 1 |
| 1 | 1 | 1 |
| 1 | 1 | 1 |
| 1 | 1 | 1 |
| 0 | 0 | 0 |
| 0 | 0 | 0 |
| 1 | 0 | 0 |
| 1 | 0 | 0 |
| 0 | 1 | 0 |
| 1 | 1 | 1 |
| 1 | 1 | 1 |
| 0 | 0 | 0 |
| 1 | 1 | 1 |
| 0 | 0 | 0 |
| 0 | 0 | 0 |
| 0 | 0 | 0 |
| 0 | 0 | 0 |
| 0 | 0 | 0 |
| 0 | 0 | 0 |
| 0 | 0 | 0 |
| 0 | 0 | 0 |
| 0 | 0 | 0 |
| 0 | 0 | 0 |
| 0 | 1 | 0 |
| 0 | 1 | 0 |
| 0 | 1 | 0 |
| 1 | 1 | 1 |
| 1 | 1 | 1 |
| 1 | 1 | 1 |
| 1 | 1 | 1 |
| 1 | 1 | 1 |
| 0 | 0 | 0 |
| 1 | 1 | 1 |
| 0 | 0 | 0 |
| 0 | 0 | 0 |
| 0 | 0 | 0 |
| 0 | 0 | 0 |
| 1 | 1 | 1 |
| 1 | 1 | 1 |
| 0 | 1 | 0 |
| 1 | 1 | 1 |
| 1 | 1 | 1 |
| 1 | 1 | 1 |
| 0 | 1 | 0 |
| 1 | 1 | 1 |
| 0 | 0 | 0 |
| 1 | 1 | 1 |
| 1 | 1 | 1 |
| 1 | 1 | 1 |
| 0 | 0 | 0 |
| 0 | 0 | 0 |
| 0 | 0 | 0 |
| 0 | 1 | 0 |
| 0 | 1 | 0 |

| | | |
|---|---|---|
| 0 | 1 | 0 |
| 0 | 1 | 0 |
| 0 | 1 | 0 |
| 0 | 0 | 0 |
| 0 | 1 | 0 |
| 1 | 1 | 1 |
| 0 | 0 | 0 |
| 0 | 1 | 0 |
| 1 | 1 | 1 |
| 1 | 1 | 1 |
| 0 | 0 | 0 |
| 0 | 0 | 0 |
| 1 | 1 | 1 |
| 1 | 1 | 1 |
| 0 | 1 | 0 |
| 0 | 1 | 0 |
| 0 | 0 | 0 |
| 0 | 0 | 0 |
| 1 | 1 | 1 |
| 0 | 0 | 0 |
| 0 | 0 | 0 |
| 0 | 0 | 0 |
| 0 | 1 | 0 |
| 0 | 0 | 0 |
| 0 | 0 | 0 |
| 1 | 1 | 1 |
| 0 | 0 | 0 |
| 0 | 1 | 0 |
| 1 | 1 | 1 |
| 0 | 0 | 0 |
| 0 | 1 | 0 |
| 1 | 1 | 1 |
| 0 | 0 | 0 |
| 0 | 1 | 0 |
| 0 | 1 | 0 |
| 0 | 0 | 0 |
| 1 | 1 | 1 |
| 0 | 0 | 0 |
| 1 | 1 | 1 |
| 0 | 0 | 0 |
| 0 | 0 | 0 |
| 1 | 0 | 0 |
| 0 | 0 | 0 |
| 0 | 0 | 0 |
| 1 | 1 | 1 |
| 1 | 0 | 0 |
| 0 | 1 | 0 |
| 1 | 1 | 1 |
| 1 | 1 | 1 |
| 1 | 1 | 1 |
| 1 | 1 | 1 |
| 0 | 1 | 0 |
| 0 | 1 | 0 |
| 0 | 1 | 0 |

| | | |
|---|---|---|
| 0 | 1 | 0 |
| 0 | 1 | 0 |
| 0 | 1 | 0 |
| 0 | 0 | 0 |
| 1 | 0 | 0 |
| 0 | 0 | 0 |
| 1 | 1 | 1 |
| 1 | 0 | 0 |
| 0 | 0 | 0 |
| 0 | 0 | 0 |
| 1 | 1 | 1 |
| 0 | 0 | 0 |
| 0 | 0 | 0 |
| 0 | 0 | 0 |
| 1 | 0 | 0 |
| 0 | 0 | 0 |
| 0 | 0 | 0 |
| 0 | 1 | 0 |
| 0 | 0 | 0 |
| 0 | 0 | 0 |
| 0 | 0 | 0 |
| 0 | 0 | 0 |
| 1 | 1 | 1 |
| 0 | 0 | 0 |
| 0 | 0 | 0 |
| 0 | 0 | 0 |
| 0 | 0 | 0 |
| 0 | 0 | 0 |
| 0 | 0 | 0 |
| 0 | 0 | 0 |
| 0 | 0 | 0 |
| 0 | 0 | 0 |
| 0 | 0 | 0 |
| 0 | 1 | 0 |
| 0 | 1 | 0 |
| 0 | 0 | 0 |
| 1 | 1 | 1 |
| 0 | 0 | 0 |
| 0 | 0 | 0 |
| 0 | 0 | 0 |
| 0 | 0 | 0 |
| 1 | 1 | 1 |
| 0 | 0 | 0 |
| 0 | 0 | 0 |
| 1 | 1 | 1 |
| 0 | 1 | 0 |
| 1 | 1 | 1 |
| 1 | 1 | 1 |
| 1 | 1 | 1 |
| 0 | 1 | 0 |
| 1 | 1 | 1 |
| 1 | 1 | 1 |
| 1 | 1 | 1 |
| 1 | 1 | 1 |

| | | |
|---|---|---|
| 1 | 1 | 1 |
| 0 | 0 | 0 |
| 1 | 1 | 1 |
| 1 | 1 | 1 |
| 1 | 1 | 1 |
| 1 | 1 | 1 |
| 1 | 1 | 1 |
| 1 | 1 | 1 |
| 1 | 1 | 1 |
| 0 | 1 | 0 |
| 1 | 1 | 1 |
| 0 | 0 | 0 |
| 1 | 0 | 0 |
| 0 | 1 | 0 |
| 0 | 0 | 0 |
| 0 | 1 | 0 |
| 0 | 1 | 0 |
| 1 | 1 | 1 |
| 1 | 1 | 1 |
| 1 | 1 | 1 |
| 0 | 0 | 0 |
| 0 | 1 | 0 |
| 0 | 0 | 0 |
| 0 | 0 | 0 |
| 1 | 1 | 1 |
| 0 | 0 | 0 |
| 1 | 1 | 1 |
| 1 | 1 | 1 |
| 1 | 1 | 1 |
| 1 | 1 | 1 |
| 1 | 1 | 1 |
| 0 | 0 | 0 |
| 1 | 1 | 1 |
| 0 | 0 | 0 |
| 1 | 1 | 1 |
| 0 | 1 | 0 |
| 0 | 1 | 0 |
| 0 | 1 | 0 |
| 1 | 1 | 1 |
| 0 | 0 | 0 |
| 1 | 1 | 1 |
| 1 | 1 | 1 |
| 0 | 1 | 0 |
| 0 | 0 | 0 |
| 1 | 1 | 1 |
| 1 | 1 | 1 |
| 1 | 1 | 1 |
| 0 | 0 | 0 |
| 0 | 0 | 0 |
| 1 | 1 | 1 |
| 1 | 1 | 1 |
| 0 | 0 | 0 |
| 0 | 0 | 0 |
| 0 | 0 | 0 |

| | | |
|---|---|---|
| 0 | 0 | 0 |
| 1 | 1 | 1 |
| 0 | 0 | 0 |
| 0 | 0 | 0 |
| 0 | 0 | 0 |
| 1 | 1 | 1 |
| 1 | 1 | 1 |
| 0 | 1 | 0 |
| 1 | 1 | 1 |
| 0 | 0 | 0 |
| 1 | 1 | 1 |
| 0 | 1 | 0 |
| 1 | 1 | 1 |
| 1 | 1 | 1 |
| 0 | 0 | 0 |
| 0 | 1 | 0 |
| 1 | 1 | 1 |
| 0 | 0 | 0 |
| 1 | 1 | 1 |
| 0 | 0 | 0 |
| 0 | 0 | 0 |
| 1 | 1 | 1 |
| 0 | 0 | 0 |
| 1 | 1 | 1 |
| 0 | 0 | 0 |
| 0 | 0 | 0 |
| 1 | 1 | 1 |
| 0 | 1 | 0 |
| 0 | 0 | 0 |
| 1 | 1 | 1 |
| 1 | 1 | 1 |
| 1 | 1 | 1 |
| 1 | 1 | 1 |
| 0 | 1 | 0 |
| 0 | 0 | 0 |
| 0 | 0 | 0 |
| 1 | 1 | 1 |
| 0 | 0 | 0 |
| 0 | 0 | 0 |
| 0 | 0 | 0 |
| 0 | 0 | 0 |
| 0 | 0 | 0 |
| 0 | 0 | 0 |
| 0 | 1 | 0 |
| 0 | 0 | 0 |
| 1 | 1 | 1 |
| 0 | 1 | 0 |
| 0 | 0 | 0 |
| 0 | 0 | 0 |
| 0 | 0 | 0 |
| 0 | 0 | 0 |
| 0 | 0 | 0 |
| 1 | 1 | 1 |
| 1 | 1 | 1 |

| | | |
|---|---|---|
| 1 | 1 | 1 |
| 0 | 0 | 0 |
| 0 | 0 | 0 |
| 0 | 0 | 0 |
| 0 | 0 | 0 |
| 0 | 0 | 0 |
| 0 | 0 | 0 |
| 1 | 0 | 0 |
| 1 | 0 | 0 |
| 1 | 1 | 1 |
| 0 | 0 | 0 |
| 0 | 0 | 0 |
| 1 | 1 | 1 |
| 0 | 0 | 0 |
| 1 | 1 | 1 |
| 0 | 0 | 0 |
| 1 | 1 | 1 |
| 1 | 1 | 1 |
| 0 | 1 | 0 |
| 0 | 1 | 0 |
| 0 | 1 | 0 |
| 1 | 1 | 1 |
| 0 | 1 | 0 |
| 0 | 0 | 0 |
| 0 | 0 | 0 |
| 1 | 1 | 1 |
| 1 | 1 | 1 |
| 1 | 1 | 1 |
| 1 | 1 | 1 |
| 0 | 0 | 0 |
| 0 | 0 | 0 |
| 0 | 0 | 0 |
| 0 | 0 | 0 |
| 1 | 1 | 1 |
| 0 | 0 | 0 |
| 0 | 0 | 0 |
| 0 | 0 | 0 |
| 0 | 0 | 0 |
| 0 | 0 | 0 |
| 1 | 1 | 1 |
| 1 | 1 | 1 |
| 0 | 0 | 0 |
| 1 | 1 | 1 |
| 1 | 1 | 1 |
| 0 | 0 | 0 |
| 1 | 0 | 0 |
| 1 | 1 | 1 |
| 0 | 1 | 0 |
| 1 | 0 | 0 |
| 0 | 1 | 0 |
| 0 | 0 | 0 |
| 1 | 1 | 1 |
| 0 | 0 | 0 |
| 0 | 0 | 0 |

| | | |
|---|---|---|
| 0 | 1 | 0 |
| 1 | 1 | 1 |
| 0 | 0 | 0 |
| 0 | 0 | 0 |
| 1 | 1 | 1 |
| 1 | 1 | 1 |
| 1 | 1 | 1 |
| 1 | 1 | 1 |
| 1 | 1 | 1 |
| 1 | 1 | 1 |
| 1 | 1 | 1 |
| 1 | 1 | 1 |
| 0 | 0 | 0 |
| 1 | 1 | 1 |
| 1 | 1 | 1 |
| 1 | 1 | 1 |
| 1 | 1 | 1 |
| 1 | 1 | 1 |
| 1 | 1 | 1 |
| 1 | 1 | 1 |
| 1 | 1 | 1 |
| 1 | 1 | 1 |
| 1 | 1 | 1 |
| 1 | 1 | 1 |
| 1 | 1 | 1 |
| 1 | 1 | 1 |
| 0 | 1 | 0 |
| 1 | 1 | 1 |
| 1 | 1 | 1 |
| 1 | 1 | 1 |
| 1 | 1 | 1 |
| 0 | 0 | 0 |
| 1 | 1 | 1 |
| 0 | 1 | 0 |
| 1 | 1 | 1 |
| 1 | 1 | 1 |
| 0 | 1 | 0 |
| 1 | 1 | 1 |
| 1 | 1 | 1 |
| 1 | 1 | 1 |
| 0 | 0 | 0 |
| 0 | 0 | 0 |
| 1 | 1 | 1 |
| 1 | 1 | 1 |
| 1 | 1 | 1 |
| 1 | 0 | 0 |
| 1 | 0 | 0 |
| 1 | 1 | 1 |
| 0 | 0 | 0 |
| 0 | 0 | 0 |
| 1 | 1 | 1 |
| 0 | 1 | 0 |
| 0 | 0 | 0 |
| 1 | 1 | 1 |

| | | |
|---|---|---|
| 0 | 0 | 0 |
| 0 | 0 | 0 |
| 0 | 0 | 0 |
| 1 | 1 | 1 |
| 1 | 1 | 1 |
| 1 | 1 | 1 |
| 1 | 1 | 1 |
| 0 | 0 | 0 |
| 0 | 0 | 0 |
| 0 | 0 | 0 |
| 0 | 0 | 0 |
| 1 | 1 | 1 |
| 0 | 0 | 0 |
| 1 | 1 | 1 |
| 1 | 1 | 1 |
| 1 | 1 | 1 |
| 1 | 1 | 1 |
| 0 | 0 | 0 |
| 1 | 1 | 1 |
| 1 | 1 | 1 |
| 1 | 1 | 1 |
| 1 | 1 | 1 |
| 0 | 1 | 0 |
| 1 | 1 | 1 |
| 1 | 1 | 1 |
| 0 | 1 | 0 |
| 1 | 1 | 1 |
| 1 | 1 | 1 |
| 1 | 1 | 1 |
| 1 | 1 | 1 |
| 0 | 0 | 0 |
| 0 | 0 | 0 |
| 1 | 1 | 1 |
| 1 | 1 | 1 |
| 1 | 1 | 1 |
| 1 | 1 | 1 |
| 1 | 1 | 1 |
| 1 | 1 | 1 |
| 1 | 1 | 1 |
| 1 | 1 | 1 |
| 1 | 1 | 1 |
| 0 | 0 | 0 |
| 1 | 1 | 1 |
| 1 | 1 | 1 |
| 0 | 0 | 0 |
| 0 | 0 | 0 |
| 1 | 1 | 1 |
| 1 | 1 | 1 |
| 0 | 0 | 0 |
| 0 | 0 | 0 |
| 0 | 0 | 0 |
| 0 | 0 | 0 |
| 0 | 0 | 0 |
| 0 | 0 | 0 |

| | | |
|---|---|---|
| 1 | 1 | 1 |
| 1 | 1 | 1 |
| 1 | 1 | 1 |
| 1 | 1 | 1 |
| 1 | 1 | 1 |
| 0 | 0 | 0 |
| 0 | 0 | 0 |
| 0 | 0 | 0 |
| 0 | 0 | 0 |
| 0 | 0 | 0 |
| 0 | 1 | 0 |
| 0 | 0 | 0 |
| 0 | 0 | 0 |
| 0 | 0 | 0 |
| 0 | 1 | 0 |
| 0 | 0 | 0 |
| 0 | 1 | 0 |
| 0 | 1 | 0 |
| 0 | 0 | 0 |
| 0 | 0 | 0 |
| 0 | 0 | 0 |
| 0 | 0 | 0 |
| 1 | 0 | 0 |
| 0 | 0 | 0 |
| 0 | 0 | 0 |
| 0 | 0 | 0 |
| 0 | 0 | 0 |
| 1 | 1 | 1 |
| 1 | 1 | 1 |
| 1 | 1 | 1 |
| 1 | 1 | 1 |
| 1 | 1 | 1 |
| 0 | 0 | 0 |
| 0 | 1 | 0 |
| 0 | 0 | 0 |
| 0 | 0 | 0 |
| 0 | 0 | 0 |
| 0 | 0 | 0 |
| 0 | 0 | 0 |
| 0 | 0 | 0 |
| 0 | 0 | 0 |
| 0 | 0 | 0 |
| 0 | 0 | 0 |
| 1 | 1 | 1 |
| 1 | 1 | 1 |
| 1 | 1 | 1 |
| 1 | 1 | 1 |
| 0 | 0 | 0 |
| 0 | 0 | 0 |
| 0 | 0 | 0 |
| 1 | 1 | 1 |
| 1 | 1 | 1 |
| 0 | 0 | 0 |
| 1 | 1 | 1 |

| | | |
|---|---|---|
| 1 | 1 | 1 |
| 0 | 0 | 0 |
| 0 | 1 | 0 |
| 0 | 1 | 0 |
| 1 | 1 | 1 |
| 1 | 1 | 1 |
| 0 | 1 | 0 |
| 1 | 1 | 1 |
| 0 | 1 | 0 |
| 1 | 1 | 1 |
| 1 | 1 | 1 |
| 1 | 1 | 1 |
| 0 | 0 | 0 |
| 0 | 0 | 0 |
| 0 | 0 | 0 |
| 0 | 0 | 0 |
| 0 | 0 | 0 |
| 0 | 0 | 0 |
| 0 | 0 | 0 |
| 0 | 1 | 0 |
| 0 | 0 | 0 |
| 0 | 0 | 0 |
| 0 | 0 | 0 |
| 0 | 0 | 0 |
| 0 | 0 | 0 |
| 1 | 1 | 1 |
| 1 | 1 | 1 |
| 0 | 0 | 0 |
| 0 | 1 | 0 |
| 0 | 0 | 0 |
| 0 | 0 | 0 |
| 1 | 1 | 1 |
| 0 | 0 | 0 |
| 1 | 1 | 1 |
| 1 | 1 | 1 |
| 1 | 1 | 1 |
| 1 | 0 | 0 |
| 1 | 1 | 1 |
| 1 | 1 | 1 |
| 1 | 1 | 1 |
| 0 | 0 | 0 |
| 1 | 1 | 1 |
| 1 | 1 | 1 |
| 0 | 0 | 0 |
| 0 | 0 | 0 |
| 1 | 1 | 1 |
| 0 | 0 | 0 |
| 0 | 0 | 0 |
| 1 | 1 | 1 |
| 1 | 1 | 1 |
| 0 | 0 | 0 |
| 1 | 1 | 1 |
| 1 | 1 | 1 |
| 1 | 1 | 1 |

| | | |
|---|---|---|
| 1 | 1 | 1 |
| 1 | 1 | 1 |
| 1 | 1 | 1 |
| 1 | 1 | 1 |
| 0 | 0 | 0 |
| 0 | 0 | 0 |
| 1 | 1 | 1 |
| 1 | 1 | 1 |
| 1 | 1 | 1 |
| 0 | 1 | 0 |
| 1 | 1 | 1 |
| 0 | 0 | 0 |
| 0 | 0 | 0 |
| 1 | 1 | 1 |
| 1 | 1 | 1 |
| 0 | 0 | 0 |
| 0 | 1 | 0 |
| 1 | 1 | 1 |
| 0 | 0 | 0 |
| 1 | 1 | 1 |
| 1 | 1 | 1 |
| 1 | 1 | 1 |
| 1 | 1 | 1 |
| 0 | 0 | 0 |
| 1 | 1 | 1 |
| 0 | 0 | 0 |
| 1 | 1 | 1 |
| 1 | 1 | 1 |
| 0 | 0 | 0 |
| 0 | 0 | 0 |
| 0 | 0 | 0 |
| 0 | 0 | 0 |
| 0 | 0 | 0 |
| 0 | 0 | 0 |
| 1 | 1 | 1 |
| 1 | 1 | 1 |
| 1 | 1 | 1 |
| 0 | 0 | 0 |
| 1 | 1 | 1 |
| 1 | 1 | 1 |
| 0 | 0 | 0 |
| 0 | 0 | 0 |
| 0 | 0 | 0 |
| 1 | 1 | 1 |
| 0 | 0 | 0 |
| 0 | 1 | 0 |
| 0 | 1 | 0 |
| 0 | 0 | 0 |
| 1 | 1 | 1 |
| 1 | 1 | 1 |
| 1 | 1 | 1 |
| 0 | 0 | 0 |
| 1 | 1 | 1 |
| 0 | 0 | 0 |

| | | |
|---|---|---|
| 1 | 1 | 1 |
| 0 | 0 | 0 |
| 0 | 0 | 0 |
| 0 | 1 | 0 |
| 1 | 1 | 1 |
| 1 | 1 | 1 |
| 0 | 0 | 0 |
| 1 | 1 | 1 |
| 0 | 1 | 0 |
| 0 | 1 | 0 |
| 1 | 1 | 1 |
| 1 | 1 | 1 |
| 1 | 1 | 1 |
| 1 | 1 | 1 |
| 1 | 1 | 1 |
| 1 | 1 | 1 |
| 1 | 1 | 1 |
| 1 | 1 | 1 |
| 1 | 1 | 1 |
| 1 | 1 | 1 |
| 1 | 1 | 1 |
| 0 | 0 | 0 |
| 0 | 1 | 0 |
| 0 | 1 | 0 |
| 1 | 1 | 1 |
| 1 | 1 | 1 |
| 0 | 0 | 0 |
| 1 | 1 | 1 |
| 0 | 0 | 0 |
| 0 | 1 | 0 |
| 1 | 1 | 1 |
| 0 | 0 | 0 |
| 0 | 0 | 0 |
| 1 | 0 | 0 |
| 1 | 1 | 1 |
| 1 | 1 | 1 |
| 1 | 1 | 1 |
| 1 | 1 | 1 |
| 0 | 0 | 0 |
| 0 | 0 | 0 |
| 0 | 0 | 0 |
| 1 | 1 | 1 |
| 1 | 1 | 1 |
| 0 | 1 | 0 |
| 0 | 1 | 0 |
| 0 | 0 | 0 |
| 1 | 1 | 1 |
| 0 | 0 | 0 |
| 1 | 1 | 1 |
| 0 | 0 | 0 |
| 0 | 0 | 0 |
| 0 | 0 | 0 |
| 1 | 1 | 1 |
| 1 | 1 | 1 |

| | | |
|---|---|---|
| 0 | 0 | 0 |
| 0 | 0 | 0 |
| 1 | 1 | 1 |
| 1 | 1 | 1 |
| 1 | 1 | 1 |
| 1 | 1 | 1 |
| 1 | 1 | 1 |
| 1 | 1 | 1 |
| 1 | 1 | 1 |
| 1 | 1 | 1 |
| 1 | 1 | 1 |
| 1 | 1 | 1 |
| 1 | 1 | 1 |
| 1 | 1 | 1 |
| 1 | 1 | 1 |
| 0 | 0 | 0 |
| 0 | 0 | 0 |
| 0 | 0 | 0 |
| 1 | 1 | 1 |
| 1 | 1 | 1 |
| 0 | 1 | 0 |
| 0 | 0 | 0 |
| 0 | 0 | 0 |
| 0 | 0 | 0 |
| 0 | 0 | 0 |
| 1 | 1 | 1 |
| 1 | 1 | 1 |
| 1 | 1 | 1 |
| 1 | 1 | 1 |
| 1 | 1 | 1 |
| 0 | 0 | 0 |
| 0 | 1 | 0 |
| 0 | 0 | 0 |
| 1 | 1 | 1 |
| 1 | 0 | 0 |
| 1 | 0 | 0 |
| 0 | 0 | 0 |
| 0 | 0 | 0 |
| 0 | 1 | 0 |
| 1 | 1 | 1 |
| 1 | 0 | 0 |
| 0 | 0 | 0 |
| 1 | 1 | 1 |
| 0 | 1 | 0 |
| 0 | 0 | 0 |
| 1 | 0 | 0 |
| 0 | 0 | 0 |
| 0 | 0 | 0 |
| 1 | 1 | 1 |
| 0 | 1 | 0 |
| 0 | 0 | 0 |
| 0 | 0 | 0 |
| 1 | 1 | 1 |
| 0 | 0 | 0 |

| | | |
|---|---|---|
| 0 | 1 | 0 |
| 0 | 0 | 0 |
| 1 | 0 | 0 |
| 0 | 0 | 0 |
| 1 | 1 | 1 |
| 1 | 1 | 1 |
| 1 | 1 | 1 |
| 0 | 1 | 0 |
| 0 | 0 | 0 |
| 1 | 1 | 1 |
| 0 | 1 | 0 |
| 0 | 1 | 0 |
| 0 | 0 | 0 |
| 0 | 0 | 0 |
| 0 | 0 | 0 |
| 1 | 1 | 1 |
| 0 | 0 | 0 |
| 0 | 1 | 0 |
| 0 | 0 | 0 |
| 0 | 0 | 0 |
| 0 | 0 | 0 |
| 0 | 0 | 0 |
| 0 | 0 | 0 |
| 0 | 0 | 0 |
| 0 | 1 | 0 |
| 0 | 1 | 0 |
| 0 | 0 | 0 |
| 0 | 0 | 0 |
| 0 | 0 | 0 |
| 0 | 0 | 0 |
| 1 | 1 | 1 |
| 1 | 1 | 1 |
| 1 | 1 | 1 |
| 1 | 1 | 1 |
| 1 | 1 | 1 |
| 0 | 0 | 0 |
| 0 | 0 | 0 |
| 0 | 1 | 0 |
| 1 | 1 | 1 |
| 1 | 1 | 1 |
| 0 | 1 | 0 |
| 1 | 1 | 1 |
| 1 | 1 | 1 |
| 0 | 0 | 0 |
| 0 | 0 | 0 |
| 0 | 0 | 0 |
| 1 | 1 | 1 |
| 0 | 0 | 0 |
| 0 | 0 | 0 |
| 0 | 0 | 0 |
| 1 | 1 | 1 |
| 1 | 1 | 1 |
| 0 | 1 | 0 |
| 0 | 1 | 0 |

| | | |
|---|---|---|
| 0 | 1 | 0 |
| 1 | 1 | 1 |
| 1 | 0 | 0 |
| 1 | 1 | 1 |
| 0 | 0 | 0 |
| 1 | 0 | 0 |
| 0 | 0 | 0 |
| 0 | 0 | 0 |
| 0 | 0 | 0 |
| 0 | 0 | 0 |
| 0 | 0 | 0 |
| 0 | 0 | 0 |
| 0 | 1 | 0 |
| 0 | 0 | 0 |
| 0 | 0 | 0 |
| 1 | 1 | 1 |
| 1 | 1 | 1 |
| 1 | 1 | 1 |
| 0 | 0 | 0 |
| 0 | 0 | 0 |
| 0 | 0 | 0 |
| 0 | 0 | 0 |
| 0 | 1 | 0 |
| 0 | 1 | 0 |
| 1 | 1 | 1 |
| 1 | 1 | 1 |
| 1 | 1 | 1 |
| 0 | 0 | 0 |
| 0 | 0 | 0 |
| 0 | 0 | 0 |
| 0 | 0 | 0 |
| 0 | 0 | 0 |
| 0 | 0 | 0 |
| 0 | 1 | 0 |
| 1 | 1 | 1 |
| 0 | 0 | 0 |
| 1 | 1 | 1 |
| 1 | 1 | 1 |
| 1 | 1 | 1 |
| 1 | 1 | 1 |
| 0 | 0 | 0 |
| 1 | 1 | 1 |
| 0 | 0 | 0 |
| 0 | 1 | 0 |
| 1 | 1 | 1 |
| 1 | 1 | 1 |
| 1 | 1 | 1 |
| 0 | 0 | 0 |
| 1 | 1 | 1 |
| 1 | 1 | 1 |
| 1 | 1 | 1 |
| 1 | 1 | 1 |
| 1 | 1 | 1 |
| 0 | 1 | 0 |

| | | |
|---|---|---|
| 0 | 1 | 0 |
| 1 | 1 | 1 |
| 1 | 1 | 1 |
| 0 | 1 | 0 |
| 1 | 1 | 1 |
| 0 | 1 | 0 |
| 0 | 1 | 0 |
| 0 | 1 | 0 |
| 0 | 1 | 0 |
| 0 | 1 | 0 |
| 0 | 1 | 0 |
| 0 | 0 | 0 |
| 0 | 1 | 0 |
| 0 | 1 | 0 |
| 0 | 1 | 0 |
| 0 | 1 | 0 |
| 0 | 1 | 0 |
| 0 | 0 | 0 |
| 0 | 0 | 0 |
| 0 | 0 | 0 |
| 0 | 0 | 0 |
| 0 | 1 | 0 |
| 0 | 0 | 0 |
| 1 | 1 | 1 |
| 1 | 1 | 1 |
| 1 | 1 | 1 |
| 1 | 1 | 1 |
| 0 | 1 | 0 |
| 0 | 0 | 0 |
| 0 | 0 | 0 |
| 1 | 1 | 1 |
| 1 | 1 | 1 |
| 0 | 0 | 0 |
| 1 | 1 | 1 |
| 0 | 0 | 0 |
| 0 | 0 | 0 |
| 0 | 0 | 0 |
| 1 | 1 | 1 |
| 0 | 0 | 0 |
| 1 | 1 | 1 |
| 0 | 0 | 0 |
| 1 | 1 | 1 |
| 0 | 0 | 0 |
| 1 | 0 | 0 |
| 1 | 0 | 0 |
| 1 | 0 | 0 |
| 1 | 0 | 0 |
| 0 | 1 | 0 |
| 0 | 1 | 0 |
| 0 | 0 | 0 |
| 1 | 0 | 0 |
| 0 | 0 | 0 |
| 1 | 0 | 0 |
| 1 | 1 | 1 |

| | | |
|---|---|---|
| 0 | 0 | 0 |
| 0 | 0 | 0 |
| 0 | 0 | 0 |
| 0 | 0 | 0 |
| 0 | 0 | 0 |
| 1 | 1 | 1 |
| 0 | 0 | 0 |
| 0 | 0 | 0 |
| 1 | 1 | 1 |
| 0 | 0 | 0 |
| 0 | 0 | 0 |
| 0 | 0 | 0 |
| 0 | 1 | 0 |
| 0 | 1 | 0 |
| 0 | 1 | 0 |
| 0 | 1 | 0 |
| 0 | 1 | 0 |
| 0 | 1 | 0 |
| 0 | 1 | 0 |
| 0 | 1 | 0 |
| 0 | 1 | 0 |
| 0 | 1 | 0 |
| 0 | 1 | 0 |
| 0 | 1 | 0 |
| 0 | 1 | 0 |
| 0 | 1 | 0 |
| 0 | 1 | 0 |
| 0 | 1 | 0 |
| 0 | 1 | 0 |
| 0 | 1 | 0 |
| 0 | 1 | 0 |
| 0 | 1 | 0 |
| 0 | 1 | 0 |
| 0 | 1 | 0 |
| 0 | 1 | 0 |
| 0 | 1 | 0 |
| 0 | 1 | 0 |
| 0 | 1 | 0 |
| 0 | 1 | 0 |
| 0 | 1 | 0 |
| 0 | 1 | 0 |
| 0 | 1 | 0 |
| 0 | 1 | 0 |
| 0 | 1 | 0 |
| 0 | 1 | 0 |
| 0 | 0 | 0 |
| 1 | 1 | 1 |
| 1 | 1 | 1 |
| 1 | 1 | 1 |
| 1 | 1 | 1 |
| 0 | 0 | 0 |
| 1 | 1 | 1 |
| 1 | 1 | 1 |
| 0 | 1 | 0 |

| | | |
|---|---|---|
| 1 | 1 | 1 |
| 1 | 1 | 1 |
| 1 | 1 | 1 |
| 1 | 1 | 1 |
| 0 | 1 | 0 |
| 0 | 1 | 0 |
| 0 | 1 | 0 |
| 0 | 0 | 0 |
| 0 | 0 | 0 |
| 1 | 1 | 1 |
| 0 | 1 | 0 |
| 1 | 1 | 1 |
| 1 | 1 | 1 |
| 1 | 1 | 1 |
| 0 | 0 | 0 |
| 0 | 0 | 0 |
| 0 | 0 | 0 |
| 1 | 1 | 1 |
| 1 | 0 | 0 |
| 1 | 1 | 1 |
| 0 | 0 | 0 |
| 0 | 0 | 0 |
| 0 | 0 | 0 |
| 0 | 0 | 0 |
| 0 | 0 | 0 |
| 0 | 0 | 0 |
| 0 | 1 | 0 |
| 1 | 1 | 1 |
| 0 | 0 | 0 |
| 1 | 1 | 1 |
| 1 | 1 | 1 |
| 1 | 1 | 1 |
| 0 | 0 | 0 |
| 1 | 1 | 1 |
| 1 | 1 | 1 |
| 1 | 1 | 1 |
| 0 | 0 | 0 |
| 0 | 0 | 0 |
| 0 | 0 | 0 |
| 1 | 1 | 1 |
| 1 | 1 | 1 |
| 1 | 1 | 1 |
| 1 | 1 | 1 |
| 1 | 1 | 1 |
| 0 | 1 | 0 |
| 0 | 0 | 0 |
| 1 | 1 | 1 |
| 0 | 1 | 0 |
| 1 | 1 | 1 |
| 0 | 1 | 0 |
| 1 | 1 | 1 |
| 1 | 1 | 1 |
| 0 | 1 | 0 |
| 1 | 1 | 1 |

| | | |
|---|---|---|
| 1 | 1 | 1 |
| 0 | 1 | 0 |
| 1 | 1 | 1 |
| 0 | 0 | 0 |
| 1 | 1 | 1 |
| 1 | 1 | 1 |
| 1 | 1 | 1 |
| 0 | 0 | 0 |
| 0 | 0 | 0 |
| 1 | 1 | 1 |
| 1 | 1 | 1 |
| 1 | 1 | 1 |
| 0 | 0 | 0 |
| 0 | 1 | 0 |
| 0 | 1 | 0 |
| 0 | 1 | 0 |
| 0 | 0 | 0 |
| 1 | 1 | 1 |
| 1 | 1 | 1 |
| 1 | 1 | 1 |
| 1 | 1 | 1 |
| 1 | 1 | 1 |
| 1 | 1 | 1 |
| 1 | 1 | 1 |
| 0 | 1 | 0 |
| 1 | 1 | 1 |
| 0 | 1 | 0 |
| 0 | 0 | 0 |
| 0 | 0 | 0 |
| 0 | 0 | 0 |
| 0 | 0 | 0 |
| 0 | 0 | 0 |
| 0 | 0 | 0 |
| 0 | 1 | 0 |
| 0 | 0 | 0 |
| 0 | 0 | 0 |
| 0 | 0 | 0 |
| 1 | 1 | 1 |
| 1 | 1 | 1 |
| 1 | 1 | 1 |
| 1 | 1 | 1 |
| 0 | 1 | 0 |
| 0 | 0 | 0 |
| 0 | 0 | 0 |
| 1 | 1 | 1 |
| 1 | 1 | 1 |
| 0 | 0 | 0 |
| 1 | 1 | 1 |
| 1 | 1 | 1 |
| 1 | 1 | 1 |
| 1 | 1 | 1 |
| 1 | 1 | 1 |
| 0 | 0 | 0 |
| 0 | 0 | 0 |

| | | |
|---|---|---|
| 0 | 0 | 0 |
| 0 | 0 | 0 |
| 0 | 0 | 0 |
| 0 | 0 | 0 |
| 0 | 0 | 0 |
| 0 | 0 | 0 |
| 0 | 0 | 0 |
| 0 | 0 | 0 |
| 1 | 1 | 1 |
| 0 | 0 | 0 |
| 1 | 1 | 1 |
| 1 | 1 | 1 |
| 1 | 1 | 1 |
| 1 | 1 | 1 |
| 0 | 0 | 0 |
| 1 | 1 | 1 |
| 1 | 1 | 1 |
| 1 | 1 | 1 |
| 1 | 1 | 1 |
| 0 | 0 | 0 |
| 0 | 0 | 0 |
| 1 | 1 | 1 |
| 0 | 0 | 0 |
| 0 | 0 | 0 |
| 0 | 0 | 0 |
| 0 | 0 | 0 |
| 0 | 0 | 0 |
| 0 | 1 | 0 |
| 0 | 1 | 0 |
| 0 | 1 | 0 |
| 0 | 0 | 0 |
| 1 | 1 | 1 |
| 1 | 1 | 1 |
| 1 | 1 | 1 |
| 1 | 1 | 1 |
| 1 | 1 | 1 |
| 1 | 1 | 1 |
| 1 | 1 | 1 |
| 1 | 1 | 1 |
| 1 | 1 | 1 |
| 0 | 1 | 0 |
| 0 | 0 | 0 |
| 1 | 1 | 1 |
| 1 | 1 | 1 |
| 1 | 1 | 1 |
| 0 | 1 | 0 |
| 1 | 1 | 1 |
| 1 | 1 | 1 |
| 0 | 0 | 0 |
| 0 | 1 | 0 |
| 1 | 1 | 1 |
| 1 | 1 | 1 |
| 0 | 0 | 0 |
| 0 | 0 | 0 |

| | | |
|---|---|---|
| 1 | 1 | 1 |
| 1 | 1 | 1 |
| 0 | 1 | 0 |
| 1 | 0 | 0 |
| 0 | 0 | 0 |
| 0 | 0 | 0 |
| 0 | 1 | 0 |
| 0 | 0 | 0 |
| 1 | 1 | 1 |
| 1 | 1 | 1 |
| 0 | 0 | 0 |
| 1 | 0 | 0 |
| 0 | 0 | 0 |
| 0 | 0 | 0 |
| 0 | 0 | 0 |
| 0 | 0 | 0 |
| 1 | 1 | 1 |
| 1 | 1 | 1 |
| 0 | 0 | 0 |
| 0 | 0 | 0 |
| 0 | 0 | 0 |
| 0 | 0 | 0 |
| 0 | 0 | 0 |
| 1 | 1 | 1 |
| 1 | 1 | 1 |
| 0 | 0 | 0 |
| 1 | 1 | 1 |
| 0 | 0 | 0 |
| 0 | 0 | 0 |
| 0 | 0 | 0 |
| 0 | 1 | 0 |
| 0 | 0 | 0 |
| 0 | 0 | 0 |
| 1 | 1 | 1 |
| 1 | 1 | 1 |
| 0 | 0 | 0 |
| 0 | 0 | 0 |
| 1 | 0 | 0 |
| 0 | 0 | 0 |
| 0 | 0 | 0 |
| 0 | 0 | 0 |
| 0 | 0 | 0 |
| 0 | 1 | 0 |
| 1 | 1 | 1 |
| 0 | 0 | 0 |
| 0 | 1 | 0 |
| 0 | 0 | 0 |
| 1 | 1 | 1 |
| 1 | 1 | 1 |
| 1 | 1 | 1 |
| 1 | 1 | 1 |
| 1 | 1 | 1 |
| 0 | 1 | 0 |
| 1 | 1 | 1 |

| | | |
|---|---|---|
| 1 | 1 | 1 |
| 1 | 1 | 1 |
| 1 | 1 | 1 |
| 0 | 1 | 0 |
| 1 | 1 | 1 |
| 1 | 1 | 1 |
| 0 | 1 | 0 |
| 0 | 0 | 0 |
| 1 | 1 | 1 |
| 0 | 0 | 0 |
| 0 | 0 | 0 |
| 0 | 0 | 0 |
| 0 | 1 | 0 |
| 1 | 1 | 1 |
| 0 | 0 | 0 |
| 0 | 1 | 0 |
| 0 | 1 | 0 |
| 0 | 1 | 0 |
| 0 | 1 | 0 |
| 0 | 1 | 0 |
| 1 | 1 | 1 |
| 1 | 1 | 1 |
| 1 | 1 | 1 |
| 0 | 1 | 0 |
| 1 | 1 | 1 |
| 1 | 1 | 1 |
| 1 | 1 | 1 |
| 1 | 0 | 0 |
| 1 | 1 | 1 |
| 0 | 0 | 0 |
| 0 | 0 | 0 |
| 1 | 1 | 1 |
| 1 | 1 | 1 |
| 0 | 1 | 0 |
| 0 | 1 | 0 |
| 0 | 0 | 0 |
| 1 | 1 | 1 |
| 1 | 1 | 1 |
| 1 | 1 | 1 |
| 1 | 1 | 1 |
| 1 | 1 | 1 |
| 0 | 0 | 0 |
| 0 | 0 | 0 |
| 0 | 0 | 0 |
| 1 | 1 | 1 |
| 1 | 1 | 1 |
| 1 | 1 | 1 |
| 1 | 1 | 1 |
| 1 | 1 | 1 |
| 1 | 1 | 1 |
| 1 | 1 | 1 |
| 1 | 1 | 1 |
| 1 | 1 | 1 |
| 0 | 0 | 0 |

| | | |
|---|---|---|
| 1 | 1 | 1 |
| 1 | 1 | 1 |
| 1 | 1 | 1 |
| 1 | 1 | 1 |
| 1 | 1 | 1 |
| 1 | 1 | 1 |
| 1 | 1 | 1 |
| 0 | 1 | 0 |
| 0 | 0 | 0 |
| 0 | 0 | 0 |
| 1 | 1 | 1 |
| 0 | 1 | 0 |
| 0 | 1 | 0 |
| 0 | 0 | 0 |
| 1 | 1 | 1 |
| 1 | 1 | 1 |
| 1 | 1 | 1 |
| 0 | 0 | 0 |
| 0 | 0 | 0 |
| 0 | 0 | 0 |
| 0 | 0 | 0 |
| 0 | 1 | 0 |
| 1 | 1 | 1 |
| 1 | 1 | 1 |
| 1 | 1 | 1 |
| 0 | 0 | 0 |
| 1 | 1 | 1 |
| 0 | 1 | 0 |
| 1 | 1 | 1 |
| 1 | 1 | 1 |
| 1 | 1 | 1 |
| 1 | 1 | 1 |
| 1 | 1 | 1 |
| 1 | 1 | 1 |
| 1 | 1 | 1 |
| 1 | 1 | 1 |
| 1 | 1 | 1 |
| 1 | 1 | 1 |
| 1 | 1 | 1 |
| 0 | 0 | 0 |
| 1 | 1 | 1 |
| 1 | 1 | 1 |
| 0 | 0 | 0 |
| 1 | 1 | 1 |
| 1 | 1 | 1 |
| 1 | 1 | 1 |
| 0 | 0 | 0 |
| 0 | 0 | 0 |
| 1 | 1 | 1 |
| 1 | 1 | 1 |
| 0 | 0 | 0 |
| 1 | 1 | 1 |
| 0 | 1 | 0 |
| 1 | 1 | 1 |

| | | |
|---|---|---|
| 1 | 1 | 1 |
| 1 | 1 | 1 |
| 1 | 1 | 1 |
| 1 | 1 | 1 |
| 1 | 1 | 1 |
| 1 | 1 | 1 |
| 1 | 1 | 1 |
| 0 | 1 | 0 |
| 1 | 1 | 1 |
| 1 | 1 | 1 |
| 1 | 1 | 1 |
| 1 | 1 | 1 |
| 1 | 1 | 1 |
| 0 | 0 | 0 |
| 0 | 0 | 0 |
| 0 | 0 | 0 |
| 0 | 0 | 0 |
| 0 | 1 | 0 |
| 1 | 1 | 1 |
| 0 | 0 | 0 |
| 1 | 1 | 1 |
| 1 | 0 | 0 |
| 1 | 1 | 1 |
| 1 | 1 | 1 |
| 1 | 1 | 1 |
| 0 | 0 | 0 |
| 0 | 1 | 0 |
| 1 | 1 | 1 |
| 1 | 1 | 1 |
| 0 | 0 | 0 |
| 0 | 0 | 0 |
| 1 | 1 | 1 |
| 1 | 1 | 1 |
| 1 | 1 | 1 |
| 1 | 1 | 1 |
| 1 | 1 | 1 |
| 1 | 1 | 1 |
| 1 | 1 | 1 |
| 0 | 0 | 0 |
| 1 | 1 | 1 |
| 1 | 1 | 1 |
| 0 | 0 | 0 |
| 0 | 0 | 0 |
| 0 | 0 | 0 |
| 0 | 0 | 0 |
| 1 | 1 | 1 |
| 0 | 0 | 0 |
| 1 | 1 | 1 |
| 1 | 1 | 1 |
| 1 | 1 | 1 |
| 0 | 1 | 0 |
| 1 | 0 | 0 |
| 1 | 0 | 0 |
| 1 | 0 | 0 |

| | | |
|---|---|---|
| 1 | 1 | 1 |
| 1 | 1 | 1 |
| 0 | 0 | 0 |
| 1 | 1 | 1 |
| 1 | 1 | 1 |
| 1 | 1 | 1 |
| 1 | 1 | 1 |
| 0 | 1 | 0 |
| 1 | 1 | 1 |
| 1 | 1 | 1 |
| 1 | 1 | 1 |
| 0 | 1 | 0 |
| 0 | 1 | 0 |
| 0 | 1 | 0 |
| 0 | 1 | 0 |
| 0 | 1 | 0 |
| 0 | 1 | 0 |
| 0 | 1 | 0 |
| 0 | 1 | 0 |
| 0 | 1 | 0 |
| 0 | 1 | 0 |
| 0 | 1 | 0 |
| 0 | 1 | 0 |
| 0 | 1 | 0 |
| 0 | 1 | 0 |
| 0 | 1 | 0 |
| 0 | 1 | 0 |
| 0 | 1 | 0 |
| 1 | 1 | 1 |
| 1 | 1 | 1 |
| 0 | 0 | 0 |
| 1 | 1 | 1 |
| 0 | 0 | 0 |
| 0 | 0 | 0 |
| 0 | 0 | 0 |
| 0 | 0 | 0 |
| 1 | 1 | 1 |
| 0 | 0 | 0 |
| 1 | 1 | 1 |
| 1 | 1 | 1 |
| 0 | 0 | 0 |
| 0 | 0 | 0 |
| 0 | 0 | 0 |
| 1 | 1 | 1 |
| 0 | 0 | 0 |
| 0 | 0 | 0 |
| 1 | 1 | 1 |
| 1 | 1 | 1 |
| 1 | 1 | 1 |
| 1 | 1 | 1 |
| 0 | 0 | 0 |
| 1 | 1 | 1 |
| 1 | 1 | 1 |
| 1 | 1 | 1 |

| | | |
|---|---|---|
| 1 | 1 | 1 |
| 1 | 1 | 1 |
| 1 | 1 | 1 |
| 1 | 1 | 1 |
| 1 | 1 | 1 |
| 0 | 0 | 0 |
| 1 | 1 | 1 |
| 0 | 0 | 0 |
| 0 | 0 | 0 |
| 0 | 0 | 0 |
| 1 | 1 | 1 |
| 0 | 0 | 0 |
| 1 | 1 | 1 |
| 1 | 1 | 1 |
| 1 | 1 | 1 |
| 1 | 0 | 0 |
| 1 | 1 | 1 |
| 0 | 0 | 0 |
| 1 | 1 | 1 |
| 0 | 0 | 0 |
| 1 | 1 | 1 |
| 0 | 0 | 0 |
| 0 | 0 | 0 |
| 0 | 0 | 0 |
| 1 | 1 | 1 |
| 0 | 1 | 0 |
| 1 | 1 | 1 |
| 1 | 1 | 1 |
| 0 | 0 | 0 |
| 1 | 1 | 1 |
| 1 | 1 | 1 |
| 1 | 1 | 1 |
| 1 | 1 | 1 |
| 1 | 1 | 1 |
| 0 | 0 | 0 |
| 0 | 0 | 0 |
| 1 | 1 | 1 |
| 0 | 0 | 0 |
| 0 | 0 | 0 |
| 0 | 1 | 0 |
| 0 | 0 | 0 |
| 0 | 0 | 0 |
| 0 | 1 | 0 |
| 0 | 1 | 0 |
| 1 | 1 | 1 |
| 0 | 1 | 0 |
| 1 | 1 | 1 |
| 1 | 1 | 1 |
| 1 | 1 | 1 |
| 1 | 1 | 1 |
| 1 | 1 | 1 |
| 1 | 1 | 1 |
| 1 | 1 | 1 |
| 1 | 1 | 1 |

| | | |
|---|---|---|
| 1 | 1 | 1 |
| 0 | 0 | 0 |
| 0 | 1 | 0 |
| 1 | 1 | 1 |
| 0 | 0 | 0 |
| 1 | 1 | 1 |
| 1 | 1 | 1 |
| 1 | 1 | 1 |
| 0 | 0 | 0 |
| 0 | 0 | 0 |
| 1 | 1 | 1 |
| 0 | 0 | 0 |
| 0 | 0 | 0 |
| 1 | 1 | 1 |
| 1 | 1 | 1 |
| 0 | 0 | 0 |
| 0 | 1 | 0 |
| 0 | 0 | 0 |
| 0 | 0 | 0 |
| 1 | 1 | 1 |
| 0 | 0 | 0 |
| 0 | 0 | 0 |
| 0 | 0 | 0 |
| 1 | 1 | 1 |
| 0 | 1 | 0 |
| 0 | 0 | 0 |
| 1 | 1 | 1 |
| 1 | 1 | 1 |
| 1 | 1 | 1 |
| 1 | 1 | 1 |
| 0 | 0 | 0 |
| 1 | 1 | 1 |
| 0 | 0 | 0 |
| 1 | 1 | 1 |
| 0 | 0 | 0 |
| 0 | 0 | 0 |
| 1 | 1 | 1 |
| 1 | 1 | 1 |
| 1 | 1 | 1 |
| 1 | 1 | 1 |
| 1 | 1 | 1 |
| 1 | 1 | 1 |
| 1 | 1 | 1 |
| 1 | 1 | 1 |
| 1 | 1 | 1 |
| 1 | 1 | 1 |
| 1 | 0 | 0 |
| 1 | 1 | 1 |
| 1 | 1 | 1 |
| 1 | 1 | 1 |
| 1 | 1 | 1 |
| 1 | 1 | 1 |
| 1 | 1 | 1 |
| 0 | 0 | 0 |

| | | |
|---|---|---|
| 1 | 1 | 1 |
| 1 | 1 | 1 |
| 1 | 1 | 1 |
| 1 | 1 | 1 |
| 0 | 0 | 0 |
| 1 | 1 | 1 |
| 0 | 1 | 0 |
| 0 | 1 | 0 |
| 0 | 1 | 0 |
| 1 | 1 | 1 |
| 0 | 0 | 0 |
| 1 | 1 | 1 |
| 1 | 1 | 1 |
| 1 | 1 | 1 |
| 0 | 0 | 0 |
| 0 | 0 | 0 |
| 0 | 0 | 0 |
| 0 | 0 | 0 |
| 0 | 1 | 0 |
| 0 | 0 | 0 |
| 0 | 0 | 0 |
| 0 | 0 | 0 |
| 1 | 1 | 1 |
| 0 | 0 | 0 |
| 0 | 0 | 0 |
| 1 | 1 | 1 |
| 0 | 0 | 0 |
| 0 | 0 | 0 |
| 0 | 0 | 0 |
| 0 | 0 | 0 |
| 0 | 0 | 0 |
| 1 | 1 | 1 |
| 1 | 1 | 1 |
| 0 | 1 | 0 |
| 1 | 1 | 1 |
| 1 | 1 | 1 |
| 1 | 1 | 1 |
| 0 | 0 | 0 |
| 0 | 0 | 0 |
| 0 | 1 | 0 |
| 1 | 1 | 1 |
| 0 | 0 | 0 |
| 0 | 0 | 0 |
| 0 | 1 | 0 |
| 0 | 0 | 0 |
| 1 | 1 | 1 |
| 1 | 1 | 1 |
| 1 | 1 | 1 |
| 0 | 0 | 0 |
| 0 | 0 | 0 |
| 0 | 0 | 0 |
| 1 | 1 | 1 |
| 0 | 1 | 0 |
| 0 | 0 | 0 |

| | | |
|---|---|---|
| 1 | 1 | 1 |
| 0 | 0 | 0 |
| 1 | 1 | 1 |
| 0 | 0 | 0 |
| 0 | 0 | 0 |
| 0 | 1 | 0 |
| 0 | 0 | 0 |
| 0 | 0 | 0 |
| 1 | 1 | 1 |
| 1 | 1 | 1 |
| 1 | 1 | 1 |
| 1 | 1 | 1 |
| 1 | 1 | 1 |
| 1 | 1 | 1 |
| 1 | 1 | 1 |
| 1 | 1 | 1 |
| 1 | 1 | 1 |
| 0 | 1 | 0 |
| 1 | 1 | 1 |
| 1 | 1 | 1 |
| 0 | 0 | 0 |
| 0 | 1 | 0 |
| 0 | 1 | 0 |
| 0 | 1 | 0 |
| 0 | 1 | 0 |
| 0 | 0 | 0 |
| 0 | 0 | 0 |
| 1 | 1 | 1 |
| 1 | 1 | 1 |
| 1 | 1 | 1 |
| 1 | 1 | 1 |
| 1 | 1 | 1 |
| 1 | 1 | 1 |
| 0 | 1 | 0 |
| 1 | 1 | 1 |
| 1 | 1 | 1 |
| 1 | 1 | 1 |
| 1 | 1 | 1 |
| 0 | 1 | 0 |
| 1 | 1 | 1 |
| 1 | 1 | 1 |
| 0 | 1 | 0 |
| 1 | 1 | 1 |
| 0 | 0 | 0 |
| 1 | 1 | 1 |
| 0 | 0 | 0 |
| 0 | 0 | 0 |
| 0 | 0 | 0 |
| 0 | 0 | 0 |
| 0 | 0 | 0 |
| 0 | 0 | 0 |
| 0 | 1 | 0 |
| 1 | 1 | 1 |
| 1 | 1 | 1 |

| | | |
|---|---|---|
| 1 | 1 | 1 |
| 1 | 1 | 1 |
| 1 | 1 | 1 |
| 0 | 0 | 0 |
| 1 | 1 | 1 |
| 0 | 0 | 0 |
| 1 | 1 | 1 |
| 1 | 1 | 1 |
| 0 | 0 | 0 |
| 1 | 1 | 1 |
| 1 | 1 | 1 |
| 0 | 1 | 0 |
| 1 | 1 | 1 |
| 1 | 1 | 1 |
| 1 | 1 | 1 |
| 0 | 0 | 0 |
| 0 | 0 | 0 |
| 0 | 0 | 0 |
| 0 | 0 | 0 |
| 0 | 0 | 0 |
| 1 | 1 | 1 |
| 1 | 1 | 1 |
| 1 | 1 | 1 |
| 1 | 1 | 1 |
| 0 | 0 | 0 |
| 1 | 1 | 1 |
| 1 | 1 | 1 |
| 1 | 1 | 1 |
| 0 | 0 | 0 |
| 1 | 1 | 1 |
| 0 | 0 | 0 |
| 0 | 0 | 0 |
| 0 | 0 | 0 |
| 0 | 0 | 0 |
| 0 | 0 | 0 |
| 1 | 1 | 1 |
| 0 | 0 | 0 |
| 0 | 0 | 0 |
| 0 | 0 | 0 |
| 0 | 0 | 0 |
| 0 | 0 | 0 |
| 0 | 0 | 0 |
| 0 | 0 | 0 |
| 0 | 0 | 0 |
| 0 | 0 | 0 |
| 0 | 0 | 0 |
| 0 | 0 | 0 |
| 0 | 0 | 0 |
| 1 | 1 | 1 |
| 0 | 0 | 0 |
| 1 | 1 | 1 |
| 1 | 1 | 1 |
| 0 | 1 | 0 |
| 0 | 0 | 0 |

| | | |
|---|---|---|
| 0 | 0 | 0 |
| 0 | 1 | 0 |
| 1 | 1 | 1 |
| 1 | 1 | 1 |
| 1 | 1 | 1 |
| 0 | 0 | 0 |
| 1 | 1 | 1 |
| 0 | 1 | 0 |
| 1 | 1 | 1 |
| 0 | 1 | 0 |
| 0 | 1 | 0 |
| 0 | 1 | 0 |
| 1 | 1 | 1 |
| 0 | 0 | 0 |
| 0 | 1 | 0 |
| 1 | 1 | 1 |
| 1 | 1 | 1 |
| 0 | 0 | 0 |
| 1 | 1 | 1 |
| 1 | 1 | 1 |
| 1 | 1 | 1 |
| 1 | 1 | 1 |
| 0 | 0 | 0 |
| 1 | 1 | 1 |
| 1 | 1 | 1 |
| 0 | 0 | 0 |
| 0 | 0 | 0 |
| 0 | 0 | 0 |
| 1 | 1 | 1 |
| 0 | 1 | 0 |
| 1 | 1 | 1 |
| 1 | 1 | 1 |
| 1 | 1 | 1 |
| 1 | 1 | 1 |
| 1 | 1 | 1 |
| 1 | 1 | 1 |
| 1 | 1 | 1 |
| 1 | 1 | 1 |
| 1 | 1 | 1 |
| 1 | 1 | 1 |
| 1 | 1 | 1 |
| 1 | 1 | 1 |
| 1 | 1 | 1 |
| 1 | 1 | 1 |
| 1 | 1 | 1 |
| 1 | 1 | 1 |
| 1 | 1 | 1 |
| 1 | 1 | 1 |
| 1 | 0 | 0 |
| 1 | 1 | 1 |
| 1 | 1 | 1 |
| 1 | 1 | 1 |
| 1 | 1 | 1 |
| 1 | 1 | 1 |

| | | |
|---|---|---|
| 1 | 1 | 1 |
| 0 | 0 | 0 |
| 1 | 1 | 1 |
| 1 | 1 | 1 |
| 1 | 1 | 1 |
| 0 | 0 | 0 |
| 0 | 0 | 0 |
| 0 | 0 | 0 |
| 1 | 1 | 1 |
| 1 | 1 | 1 |
| 0 | 1 | 0 |
| 1 | 1 | 1 |
| 1 | 1 | 1 |
| 1 | 1 | 1 |
| 0 | 0 | 0 |
| 0 | 0 | 0 |
| 0 | 0 | 0 |
| 1 | 1 | 1 |
| 0 | 0 | 0 |
| 0 | 0 | 0 |
| 0 | 0 | 0 |
| 0 | 0 | 0 |
| 0 | 0 | 0 |
| 0 | 0 | 0 |
| 0 | 1 | 0 |
| 1 | 1 | 1 |
| 1 | 1 | 1 |
| 0 | 1 | 0 |
| 0 | 1 | 0 |
| 1 | 1 | 1 |
| 1 | 1 | 1 |
| 1 | 1 | 1 |
| 0 | 0 | 0 |
| 1 | 1 | 1 |
| 0 | 1 | 0 |
| 0 | 0 | 0 |
| 0 | 0 | 0 |
| 0 | 0 | 0 |
| 0 | 0 | 0 |
| 1 | 1 | 1 |
| 0 | 1 | 0 |
| 1 | 1 | 1 |
| 1 | 1 | 1 |
| 0 | 0 | 0 |
| 0 | 0 | 0 |
| 0 | 0 | 0 |
| 0 | 1 | 0 |
| 1 | 1 | 1 |
| 0 | 0 | 0 |
| 1 | 1 | 1 |
| 1 | 1 | 1 |
| 0 | 0 | 0 |
| 1 | 1 | 1 |
| 0 | 1 | 0 |

| | | |
|---|---|---|
| 1 | 1 | 1 |
| 0 | 1 | 0 |
| 1 | 1 | 1 |
| 1 | 1 | 1 |
| 1 | 1 | 1 |
| 1 | 1 | 1 |
| 0 | 1 | 0 |
| 1 | 1 | 1 |
| 0 | 0 | 0 |
| 0 | 0 | 0 |
| 1 | 1 | 1 |
| 0 | 0 | 0 |
| 0 | 0 | 0 |
| 0 | 0 | 0 |
| 1 | 1 | 1 |
| 1 | 1 | 1 |
| 0 | 0 | 0 |
| 0 | 0 | 0 |
| 0 | 0 | 0 |
| 1 | 1 | 1 |
| 1 | 1 | 1 |
| 1 | 1 | 1 |
| 0 | 0 | 0 |
| 1 | 1 | 1 |
| 0 | 0 | 0 |
| 1 | 1 | 1 |
| 1 | 1 | 1 |
| 1 | 1 | 1 |
| 1 | 1 | 1 |
| 0 | 0 | 0 |
| 0 | 0 | 0 |
| 0 | 0 | 0 |
| 1 | 1 | 1 |
| 0 | 1 | 0 |
| 1 | 1 | 1 |
| 1 | 1 | 1 |
| 1 | 1 | 1 |
| 0 | 1 | 0 |
| 0 | 1 | 0 |
| 0 | 1 | 0 |
| 0 | 0 | 0 |
| 1 | 1 | 1 |
| 1 | 1 | 1 |
| 1 | 1 | 1 |
| 1 | 1 | 1 |
| 0 | 0 | 0 |
| 1 | 0 | 0 |
| 0 | 0 | 0 |
| 1 | 1 | 1 |
| 0 | 0 | 0 |
| 0 | 0 | 0 |
| 0 | 0 | 0 |
| 0 | 1 | 0 |
| 1 | 0 | 0 |

| | | |
|---|---|---|
| 1 | 0 | 0 |
| 0 | 0 | 0 |
| 0 | 1 | 0 |
| 1 | 1 | 1 |
| 0 | 1 | 0 |
| 0 | 1 | 0 |
| 0 | 1 | 0 |
| 0 | 0 | 0 |
| 0 | 0 | 0 |
| 0 | 1 | 0 |
| 0 | 0 | 0 |
| 1 | 1 | 1 |
| 1 | 1 | 1 |
| 1 | 1 | 1 |
| 0 | 1 | 0 |
| 1 | 1 | 1 |
| 1 | 1 | 1 |
| 1 | 1 | 1 |
| 1 | 1 | 1 |
| 0 | 1 | 0 |
| 1 | 1 | 1 |
| 1 | 1 | 1 |
| 1 | 1 | 1 |
| 1 | 1 | 1 |
| 0 | 1 | 0 |
| 0 | 1 | 0 |
| 0 | 1 | 0 |
| 0 | 1 | 0 |
| 0 | 0 | 0 |
| 0 | 1 | 0 |
| 0 | 0 | 0 |
| 1 | 1 | 1 |
| 0 | 0 | 0 |
| 0 | 0 | 0 |
| 0 | 1 | 0 |
| 0 | 0 | 0 |
| 0 | 0 | 0 |
| 1 | 1 | 1 |
| 0 | 1 | 0 |
| 0 | 1 | 0 |
| 1 | 1 | 1 |
| 0 | 0 | 0 |
| 0 | 0 | 0 |
| 1 | 1 | 1 |
| 1 | 1 | 1 |
| 0 | 0 | 0 |
| 0 | 0 | 0 |
| 0 | 0 | 0 |
| 0 | 0 | 0 |
| 0 | 1 | 0 |
| 0 | 0 | 0 |
| 0 | 0 | 0 |
| 1 | 1 | 1 |
| 1 | 1 | 1 |

| | | |
|---|---|---|
| 1 | 1 | 1 |
| 1 | 1 | 1 |
| 1 | 1 | 1 |
| 1 | 1 | 1 |
| 0 | 0 | 0 |
| 0 | 0 | 0 |
| 0 | 0 | 0 |
| 0 | 0 | 0 |
| 1 | 1 | 1 |
| 0 | 0 | 0 |
| 0 | 0 | 0 |
| 0 | 0 | 0 |
| 0 | 0 | 0 |
| 1 | 1 | 1 |
| 1 | 0 | 0 |
| 0 | 1 | 0 |
| 1 | 1 | 1 |
| 1 | 1 | 1 |
| 1 | 1 | 1 |
| 0 | 1 | 0 |
| 1 | 1 | 1 |
| 1 | 1 | 1 |
| 1 | 1 | 1 |
| 1 | 1 | 1 |
| 1 | 1 | 1 |
| 0 | 0 | 0 |
| 0 | 0 | 0 |
| 0 | 0 | 0 |
| 0 | 0 | 0 |
| 0 | 1 | 0 |
| 0 | 1 | 0 |
| 0 | 1 | 0 |
| 1 | 1 | 1 |
| 0 | 1 | 0 |
| 0 | 1 | 0 |
| 1 | 1 | 1 |
| 1 | 1 | 1 |
| 1 | 1 | 1 |
| 1 | 1 | 1 |
| 1 | 1 | 1 |
| 1 | 1 | 1 |
| 1 | 1 | 1 |
| 1 | 1 | 1 |
| 1 | 1 | 1 |
| 1 | 1 | 1 |
| 1 | 1 | 1 |
| 1 | 1 | 1 |
| 1 | 1 | 1 |
| 1 | 1 | 1 |
| 1 | 1 | 1 |
| 1 | 1 | 1 |
| 1 | 1 | 1 |
| 1 | 1 | 1 |
| 0 | 0 | 0 |

| | | |
|---|---|---|
| 1 | 1 | 1 |
| 1 | 1 | 1 |
| 1 | 1 | 1 |
| 0 | 1 | 0 |
| 0 | 1 | 0 |
| 0 | 1 | 0 |
| 0 | 1 | 0 |
| 0 | 1 | 0 |
| 1 | 1 | 1 |
| 1 | 1 | 1 |
| 1 | 1 | 1 |
| 1 | 1 | 1 |
| 1 | 1 | 1 |
| 1 | 1 | 1 |
| 1 | 1 | 1 |
| 1 | 1 | 1 |
| 1 | 1 | 1 |
| 1 | 1 | 1 |
| 1 | 1 | 1 |
| 1 | 1 | 1 |
| 1 | 1 | 1 |
| 1 | 1 | 1 |
| 1 | 1 | 1 |
| 1 | 1 | 1 |
| 1 | 1 | 1 |
| 1 | 1 | 1 |
| 1 | 1 | 1 |
| 1 | 1 | 1 |
| 1 | 1 | 1 |
| 1 | 1 | 1 |
| 1 | 1 | 1 |
| 1 | 1 | 1 |
| 1 | 1 | 1 |
| 0 | 1 | 0 |
| 1 | 1 | 1 |
| 1 | 1 | 1 |
| 1 | 1 | 1 |
| 1 | 1 | 1 |
| 1 | 1 | 1 |
| 1 | 1 | 1 |
| 0 | 1 | 0 |
| 1 | 1 | 1 |
| 1 | 1 | 1 |
| 1 | 1 | 1 |
| 0 | 1 | 0 |
| 1 | 0 | 0 |
| 0 | 0 | 0 |
| 1 | 1 | 1 |
| 0 | 1 | 0 |
| 1 | 1 | 1 |
| 1 | 1 | 1 |
| 0 | 1 | 0 |
| 0 | 1 | 0 |
| 0 | 1 | 0 |

| | | |
|---|---|---|
| 0 | 1 | 0 |
| 1 | 1 | 1 |
| 0 | 0 | 0 |
| 1 | 1 | 1 |
| 1 | 1 | 1 |
| 1 | 1 | 1 |
| 0 | 0 | 0 |
| 1 | 1 | 1 |
| 1 | 1 | 1 |
| 0 | 1 | 0 |
| 0 | 1 | 0 |
| 0 | 0 | 0 |
| 1 | 1 | 1 |
| 0 | 0 | 0 |
| 0 | 0 | 0 |
| 0 | 0 | 0 |
| 0 | 0 | 0 |
| 0 | 0 | 0 |
| 0 | 0 | 0 |
| 0 | 0 | 0 |
| 0 | 0 | 0 |

| organism_scientific | evo_cons | cluster_id | pdb_id | chain_id | resi | resn | change | uniprot_id |
|---|---|---|---|---|---|---|---|---|
| HOMO SAPIENS | 5 | 87 | 3uzw | A | 106 | LEU | p.Leu106Phe | P51857 |
| HOMO SAPIENS | 3 | 87 | 3uzw | A | 133 | PRO | p.Pro133Arg | P51857 |
| HOMO SAPIENS | 0 | 87 | 3uzw | A | 198 | PRO | p.Pro198Leu | P51857 |
| HOMO SAPIENS | 7 | 87 | 3uzw | A | 261 | ARG | p.Arg261Cys | P51857 |
| HOMO SAPIENS | 0 | 4693 | 3mk1 | A | 3 | PRO | p.Pro25Leu | P05187 |
| HOMO SAPIENS | 8 | 4830 | 2h64 | B | 101 | CYS | p.Cys124Arg | P36894 |
| HOMO SAPIENS | 2 | 4830 | 2h64 | B | 116 | PRO | p.Pro139Ser | P36894 |
| HOMO SAPIENS | 0 | 6937 | 3dar | A | 186 | MET | p.Met186Thr | P21802 |
| HOMO SAPIENS | 3 | 5 | 1aut | C | 30 | GLN | p.Gln226His | P04070 |
| HOMO SAPIENS | 1 | 5 | 1aut | C | 68 | LEU | p.Leu265Phe | P04070 |
| HOMO SAPIENS | 0 | 5 | 1aut | C | 75 | ARG | p.Arg272Cys | P04070 |
| HOMO SAPIENS | 6 | 5 | 1aut | C | 92 | PRO | p.Pro289Leu | P04070 |
| HOMO SAPIENS | 0 | 5 | 1aut | C | 104 | ALA | p.Ala301Val | P04070 |
| HOMO SAPIENS | 1 | 5 | 1aut | C | 112 | ALA | p.Ala309Thr | P04070 |
| HOMO SAPIENS | 2 | 5 | 1aut | C | 115 | SER | p.Ser312Leu | P04070 |
| HOMO SAPIENS | 1 | 5 | 1aut | C | 133 | GLY | p.Gly334Ser | P04070 |
| HOMO SAPIENS | 0 | 5 | 1aut | C | 147 | ARG | p.Arg348Ter | P04070 |
| HOMO SAPIENS | 8 | 5 | 1aut | C | 194 | ASP | p.Asp401Asn | P04070 |
| HOMO SAPIENS | 7 | 5 | 1aut | C | 237 | TRP | p.Trp444Cys | P04070 |
| HOMO SAPIENS | 0 | 5 | 1aut | C | 238 | ILE | p.Ile445Met | P04070 |
| HOMO SAPIENS | 8 | 43 | 1aiy | F | 10 | HIS | p.His34Asp | P01308 |
| HOMO SAPIENS | 2 | 43 | 1aiy | F | 22 | ARG | p.Arg46Gln | P01308 |
| HOMO SAPIENS | 4 | 43 | 1aiy | F | 23 | GLY | p.Gly47Val | P01308 |
| HOMO SAPIENS | 7 | 43 | 1aiy | F | 24 | PHE | p.Phe48Ser | P01308 |
| HOMO SAPIENS | 5 | 24 | 2fg8 | A | 33 | THR | p.Thr30Ile | P02792 |
| HOMO SAPIENS | 4 | 24 | 2fg8 | A | 99 | ALA | p.Ala96Thr | P02792 |
| HOMO SAPIENS | 5 | 24 | 2fg8 | A | 107 | GLU | p.Glu104Ter | P02792 |
| HOMO SAPIENS | 3 | 5409 | 5bnq | A | 199 | MET | p.Met199Lys | O14788 |
| HOMO SAPIENS | -1 | 839 | 3swc | A | 959 | GLN | p.Gln990His | Q9H9B1 |
| HOMO SAPIENS | -1 | 839 | 3swc | A | 968 | ALA | p.Ala999Val | Q9H9B1 |
| HOMO SAPIENS | -1 | 839 | 3swc | A | 973 | SER | p.Ser1004Ile | Q9H9B1 |
| HOMO SAPIENS | 8 | 17 | 1eou | A | 18 | LYS | p.Lys18Glu | P00918 |
| HOMO SAPIENS | 2 | 17 | 1eou | A | 107 | HIS | p.His107Tyr | P00918 |
| HOMO SAPIENS | 1 | 17 | 1eou | A | 236 | PRO | p.Pro236His | P00918 |
| HOMO SAPIENS | 6 | 17 | 1eou | A | 252 | ASN | p.Asn252Asp | P00918 |
| HOMO SAPIENS | 2 | 9140 | 5c4v | A | 445 | ARG | p.Arg445Ter | Q13485 |
| HOMO SAPIENS | 7 | 9140 | 5c4v | A | 496 | ARG | p.Arg496Cys | Q13485 |
| HOMO SAPIENS | -1 | 42 | 4pwi | A | 6 | GLY | p.Gly26Ser | P02766 |
| HOMO SAPIENS | 6 | 42 | 4pwi | A | 10 | CYS | p.Cys30Arg | P02766 |
| HOMO SAPIENS | 8 | 42 | 4pwi | A | 12 | LEU | p.Leu32Pro | P02766 |
| HOMO SAPIENS | 9 | 42 | 4pwi | A | 18 | ASP | p.Asp38Gly | P02766 |
| HOMO SAPIENS | 8 | 42 | 4pwi | A | 20 | VAL | p.Val40Ile | P02766 |
| HOMO SAPIENS | 4 | 42 | 4pwi | A | 30 | MET | p.Val50Ala | P02766 |
| HOMO SAPIENS | 4 | 42 | 4pwi | A | 30 | MET | p.Val50Leu | P02766 |
| HOMO SAPIENS | 6 | 42 | 4pwi | A | 33 | PHE | p.Phe53Ile | P02766 |
| HOMO SAPIENS | 7 | 42 | 4pwi | A | 36 | ALA | p.Ala56Pro | P02766 |
| HOMO SAPIENS | 7 | 42 | 4pwi | A | 42 | GLU | p.Glu62Gly | P02766 |
| HOMO SAPIENS | 8 | 42 | 4pwi | A | 44 | PHE | p.Phe64Ser | P02766 |
| HOMO SAPIENS | 4 | 42 | 4pwi | A | 47 | GLY | p.Gly67Ala | P02766 |
| HOMO SAPIENS | 6 | 42 | 4pwi | A | 49 | THR | p.Thr69Ala | P02766 |
| HOMO SAPIENS | 6 | 42 | 4pwi | A | 50 | SER | p.Ser70Ile | P02766 |
| HOMO SAPIENS | 6 | 42 | 4pwi | A | 50 | SER | p.Ser70Arg | P02766 |
| HOMO SAPIENS | 0 | 42 | 4pwi | A | 53 | GLY | p.Gly73Glu | P02766 |

| | | | | | | | | |
|---|---|---|---|---|---|---|---|---|
| HOMO SAPIENS | 7 | 42 | 4pwi | A | 55 | LEU | p.Leu75Pro | P02766 |
| HOMO SAPIENS | 6 | 42 | 4pwi | A | 58 | LEU | p.Leu78His | P02766 |
| HOMO SAPIENS | 7 | 42 | 4pwi | A | 61 | GLU | p.Glu81Lys | P02766 |
| HOMO SAPIENS | 8 | 42 | 4pwi | A | 64 | PHE | p.Phe84Leu | P02766 |
| HOMO SAPIENS | 8 | 42 | 4pwi | A | 68 | ILE | p.Ile88Leu | P02766 |
| HOMO SAPIENS | 8 | 42 | 4pwi | A | 69 | TYR | p.Tyr89His | P02766 |
| HOMO SAPIENS | 4 | 42 | 4pwi | A | 71 | VAL | p.Val91Ala | P02766 |
| HOMO SAPIENS | 6 | 42 | 4pwi | A | 77 | SER | p.Ser97Tyr | P02766 |
| HOMO SAPIENS | 4 | 42 | 4pwi | A | 84 | ILE | p.Ile104Ser | P02766 |
| HOMO SAPIENS | 8 | 42 | 4pwi | A | 89 | GLU | p.Glu109Gln | P02766 |
| HOMO SAPIENS | 8 | 42 | 4pwi | A | 97 | ALA | p.Ala117Gly | P02766 |
| HOMO SAPIENS | 7 | 42 | 4pwi | A | 99 | ASP | p.Asp119Asn | P02766 |
| HOMO SAPIENS | 9 | 42 | 4pwi | A | 107 | ILE | p.Ile127Val | P02766 |
| HOMO SAPIENS | 9 | 42 | 4pwi | A | 111 | LEU | p.Leu131Met | P02766 |
| HOMO SAPIENS | 7 | 42 | 4pwi | A | 114 | TYR | p.Tyr134Cys | P02766 |
| HOMO SAPIENS | 7 | 42 | 4pwi | A | 114 | TYR | p.Tyr134His | P02766 |
| HOMO SAPIENS | 7 | 42 | 4pwi | A | 122 | VAL | p.Val142Ile | P02766 |
| HOMO SAPIENS | -1 | 42 | 4pwi | A | 126 | LYS | p.Lys146Arg | P02766 |
| HOMO SAPIENS | -1 | 42 | 4pwi | A | 127 | GLU | p.Glu147Ter | P02766 |
| HOMO SAPIENS | -1 | 1312 | 4dce | A | 1275 | ARG | p.Arg1275Gln | Q9UM73 |
| HOMO SAPIENS | -1 | 1312 | 4dce | A | 1280 | ALA | p.Ala1280Val | Q9UM73 |
| HOMO SAPIENS | 1 | 20 | 1y0x | X | 234 | ALA | p.Ala234Thr | P10828 |
| HOMO SAPIENS | 0 | 20 | 1y0x | X | 243 | ARG | p.Arg243Trp | P10828 |
| HOMO SAPIENS | 3 | 20 | 1y0x | X | 316 | ARG | p.Arg316His | P10828 |
| HOMO SAPIENS | 1 | 20 | 1y0x | X | 317 | ALA | p.Ala317Thr | P10828 |
| HOMO SAPIENS | 2 | 20 | 1y0x | X | 320 | ARG | p.Arg320Cys | P10828 |
| HOMO SAPIENS | 2 | 20 | 1y0x | X | 320 | ARG | p.Arg320His | P10828 |
| HOMO SAPIENS | 0 | 20 | 1y0x | X | 332 | GLY | p.Gly332Arg | P10828 |
| HOMO SAPIENS | 1 | 20 | 1y0x | X | 338 | ARG | p.Arg338Trp | P10828 |
| HOMO SAPIENS | 0 | 20 | 1y0x | X | 340 | GLN | p.Gln340His | P10828 |
| HOMO SAPIENS | 0 | 20 | 1y0x | X | 345 | GLY | p.Gly345Val | P10828 |
| HOMO SAPIENS | 0 | 20 | 1y0x | X | 345 | GLY | p.Gly345Ser | P10828 |
| HOMO SAPIENS | 0 | 20 | 1y0x | X | 347 | GLY | p.Gly347Glu | P10828 |
| HOMO SAPIENS | 1 | 20 | 1y0x | X | 438 | ARG | p.Arg438His | P10828 |
| HOMO SAPIENS | 0 | 20 | 1y0x | X | 442 | MET | p.Met442Val | P10828 |
| HOMO SAPIENS | 0 | 20 | 1y0x | X | 443 | LYS | p.Lys443Glu | P10828 |
| HOMO SAPIENS | 0 | 20 | 1y0x | X | 446 | CYS | p.Cys446Arg | P10828 |
| HOMO SAPIENS | 2 | 20 | 1y0x | X | 453 | PRO | p.Pro453His | P10828 |
| HOMO SAPIENS | 2 | 20 | 1y0x | X | 453 | PRO | p.Pro453Thr | P10828 |
| HOMO SAPIENS | 3 | 20 | 1y0x | X | 459 | PHE | p.Phe459Cys | P10828 |
| HOMO SAPIENS | 0 | 4409 | 3qb7 | D | 87 | GLU | p.Glu109Lys | P31785 |
| HOMO SAPIENS | 1 | 4409 | 3qb7 | D | 92 | GLY | p.Gly114Asp | P31785 |
| HOMO SAPIENS | 1 | 4409 | 3qb7 | D | 93 | CYS | p.Cys115Arg | P31785 |
| HOMO SAPIENS | 0 | 4409 | 3qb7 | D | 131 | ILE | p.Ile153Asn | P31785 |
| HOMO SAPIENS | 9 | 4409 | 3qb7 | D | 200 | ARG | p.Arg222Cys | P31785 |
| HOMO SAPIENS | 0 | 3891 | 3ije | B | 33 | LEU | p.Leu59Pro | P05106 |
| HOMO SAPIENS | 0 | 3891 | 3ije | B | 117 | LEU | p.Leu143Trp | P05106 |
| HOMO SAPIENS | 3 | 3891 | 3ije | B | 119 | ASP | p.Asp145Tyr | P05106 |
| HOMO SAPIENS | 2 | 3891 | 3ije | B | 143 | ARG | p.Arg169Gln | P05106 |
| HOMO SAPIENS | 2 | 3891 | 3ije | B | 214 | ARG | p.Arg240Gln | P05106 |
| HOMO SAPIENS | 2 | 3891 | 3ije | B | 214 | ARG | p.Arg240Trp | P05106 |
| HOMO SAPIENS | 2 | 3891 | 3ije | B | 374 | CYS | p.Cys400Tyr | P05106 |
| HOMO SAPIENS | 1 | 3891 | 3ije | B | 407 | PRO | p.Pro433Ala | P05106 |
| HOMO SAPIENS | 0 | 3891 | 3ije | B | 489 | ARG | p.Arg515Gln | P05106 |

| | | | | | | | | |
|---|---|---|---|---|---|---|---|---|
| HOMO SAPIENS | -1 | 9613 | 5mke | B | 28 | ARG | p.Arg28Pro | Q13563 |
| HOMO SAPIENS | -1 | 9613 | 5mke | B | 190 | ALA | p.Ala190Thr | Q13563 |
| HOMO SAPIENS | 4 | 9613 | 5mke | B | 262 | VAL | p.Val262Met | Q13563 |
| HOMO SAPIENS | 0 | 9613 | 5mke | B | 482 | PHE | p.Phe482Cys | Q13563 |
| HOMO SAPIENS | 4 | 9613 | 5mke | B | 511 | ASP | p.Asp511Val | Q13563 |
| HOMO SAPIENS | 5 | 9613 | 5mke | B | 516 | VAL | p.Val516Leu | Q13563 |
| HOMO SAPIENS | -1 | 9613 | 5mke | B | 807 | ARG | p.Arg807Gln | Q13563 |
| HOMO SAPIENS | 4 | 2428 | 1olh | C | 337 | ARG | p.Arg337Cys | P04637 |
| HOMO SAPIENS | 4 | 2428 | 1olh | C | 337 | ARG | p.Arg337His | P04637 |
| HOMO SAPIENS | 8 | 2428 | 1olh | C | 338 | PHE | p.Phe338Leu | P04637 |
| HOMO SAPIENS | 8 | 2428 | 1olh | C | 339 | GLU | p.Glu339Lys | P04637 |
| HOMO SAPIENS | 7 | 2428 | 1olh | C | 344 | LEU | p.Leu344Pro | P04637 |
| HOMO SAPIENS | 0 | 2428 | 1olh | C | 360 | GLY | p.Gly360Ala | P04637 |
| HOMO SAPIENS | 7 | 4867 | 1mox | B | 404 | GLY | p.Gly428Asp | P00533 |
| HOMO SAPIENS | 5 | 4867 | 1mox | B | 497 | ARG | p.Arg521Lys | P00533 |
| HOMO SAPIENS | 1 | 4630 | 1fu1 | A | 43 | TRP | p.Trp43Arg | Q13426 |
| HOMO SAPIENS | 2 | 4630 | 1fu1 | A | 161 | ARG | p.Arg161Ter | Q13426 |
| HOMO SAPIENS | 2 | 15 | 1lfy | B | 2 | HIS | p.His3Gln | P68871 |
| HOMO SAPIENS | 2 | 15 | 1lfy | B | 6 | GLU | p.Glu7Ala | P68871 |
| HOMO SAPIENS | 2 | 15 | 1lfy | B | 6 | GLU | p.Glu7Lys | P68871 |
| HOMO SAPIENS | 4 | 15 | 1lfy | B | 9 | SER | p.Ser10Cys | P68871 |
| HOMO SAPIENS | 6 | 15 | 1lfy | B | 15 | TRP | p.Trp16Ter | P68871 |
| HOMO SAPIENS | 3 | 15 | 1lfy | B | 17 | LYS | p.Lys18Glu | P68871 |
| HOMO SAPIENS | 6 | 15 | 1lfy | B | 19 | ASN | p.Asn20Ser | P68871 |
| HOMO SAPIENS | 6 | 15 | 1lfy | B | 22 | GLU | p.Glu23Ala | P68871 |
| HOMO SAPIENS | 6 | 15 | 1lfy | B | 22 | GLU | p.Glu23Lys | P68871 |
| HOMO SAPIENS | 8 | 15 | 1lfy | B | 26 | GLU | p.Glu27Lys | P68871 |
| HOMO SAPIENS | 8 | 15 | 1lfy | B | 28 | LEU | p.Leu29Pro | P68871 |
| HOMO SAPIENS | 8 | 15 | 1lfy | B | 30 | ARG | p.Arg31Ser | P68871 |
| HOMO SAPIENS | 8 | 15 | 1lfy | B | 30 | ARG | p.Arg31Lys | P68871 |
| HOMO SAPIENS | 6 | 15 | 1lfy | B | 39 | GLN | p.Gln40Ter | P68871 |
| HOMO SAPIENS | 2 | 15 | 1lfy | B | 43 | GLU | p.Glu44Gln | P68871 |
| HOMO SAPIENS | 6 | 15 | 1lfy | B | 57 | ASN | p.Asn58Lys | P68871 |
| HOMO SAPIENS | 6 | 15 | 1lfy | B | 59 | LYS | p.Lys60Asn | P68871 |
| HOMO SAPIENS | 8 | 15 | 1lfy | B | 61 | LYS | p.Lys62Glu | P68871 |
| HOMO SAPIENS | 4 | 15 | 1lfy | B | 73 | ASP | p.Asp74Tyr | P68871 |
| HOMO SAPIENS | 4 | 15 | 1lfy | B | 87 | THR | p.Thr88Ile | P68871 |
| HOMO SAPIENS | 4 | 15 | 1lfy | B | 87 | THR | p.Thr88Pro | P68871 |
| HOMO SAPIENS | 5 | 15 | 1lfy | B | 90 | GLU | p.Glu91Lys | P68871 |
| HOMO SAPIENS | 6 | 15 | 1lfy | B | 91 | LEU | p.Leu92Pro | P68871 |
| HOMO SAPIENS | 5 | 15 | 1lfy | B | 97 | HIS | p.His98Gln | P68871 |
| HOMO SAPIENS | 6 | 15 | 1lfy | B | 110 | LEU | p.Leu111Pro | P68871 |
| HOMO SAPIENS | 6 | 15 | 1lfy | B | 113 | VAL | p.Val114Glu | P68871 |
| HOMO SAPIENS | 2 | 15 | 1lfy | B | 119 | GLY | p.Gly120Ala | P68871 |
| HOMO SAPIENS | 4 | 15 | 1lfy | B | 121 | GLU | p.Glu122Gln | P68871 |
| HOMO SAPIENS | 5 | 15 | 1lfy | B | 134 | VAL | p.Val135Glu | P68871 |
| HOMO SAPIENS | 3 | 15 | 1lfy | B | 143 | HIS | p.His144Gln | P68871 |
| HOMO SAPIENS | 4 | 91 | 5d9k | A | 75 | GLY | p.Gly75Val | P51812 |
| HOMO SAPIENS | 5 | 91 | 5d9k | A | 82 | VAL | p.Val82Phe | P51812 |
| HOMO SAPIENS | 0 | 91 | 5d9k | A | 114 | ARG | p.Arg114Trp | P51812 |
| HOMO SAPIENS | 4 | 91 | 5d9k | A | 189 | ILE | p.Ile189Lys | P51812 |
| HOMO SAPIENS | -1 | 91 | 5d9k | A | 227 | SER | p.Ser227Ala | P51812 |
| HOMO SAPIENS | 1 | 91 | 5d9k | A | 268 | PHE | p.Phe268Ser | P51812 |
| HOMO SAPIENS | 0 | 954 | 2vt0 | A | 15 | VAL | p.Val54Leu | P04062 |

| | | | | | | | | |
|---|---|---|---|---|---|---|---|---|
| HOMO SAPIENS | 5 | 954 | 2vt0 | A | 46 | GLY | p.Gly85Glu | P04062 |
| HOMO SAPIENS | 5 | 954 | 2vt0 | A | 48 | ARG | p.Arg87Trp | P04062 |
| HOMO SAPIENS | 6 | 954 | 2vt0 | A | 79 | LYS | p.Lys118Asn | P04062 |
| HOMO SAPIENS | 0 | 954 | 2vt0 | A | 120 | ARG | p.Arg159Gln | P04062 |
| HOMO SAPIENS | 0 | 954 | 2vt0 | A | 120 | ARG | p.Arg159Trp | P04062 |
| HOMO SAPIENS | 0 | 954 | 2vt0 | A | 122 | PRO | p.Pro161Ser | P04062 |
| HOMO SAPIENS | 5 | 954 | 2vt0 | A | 131 | ARG | p.Arg170Leu | P04062 |
| HOMO SAPIENS | 5 | 954 | 2vt0 | A | 135 | TYR | p.Tyr174Cys | P04062 |
| HOMO SAPIENS | 0 | 954 | 2vt0 | A | 157 | LYS | p.Lys196Gln | P04062 |
| HOMO SAPIENS | 5 | 954 | 2vt0 | A | 163 | ARG | p.Arg202Gln | P04062 |
| HOMO SAPIENS | 5 | 954 | 2vt0 | A | 170 | ARG | p.Arg209Cys | P04062 |
| HOMO SAPIENS | 5 | 954 | 2vt0 | A | 188 | ASN | p.Asn227Ser | P04062 |
| HOMO SAPIENS | 3 | 954 | 2vt0 | A | 196 | SER | p.Ser235Pro | P04062 |
| HOMO SAPIENS | 5 | 954 | 2vt0 | A | 202 | GLY | p.Gly241Arg | P04062 |
| HOMO SAPIENS | 0 | 954 | 2vt0 | A | 212 | TYR | p.Tyr251His | P04062 |
| HOMO SAPIENS | 0 | 954 | 2vt0 | A | 213 | PHE | p.Phe252Ile | P04062 |
| HOMO SAPIENS | 5 | 954 | 2vt0 | A | 216 | PHE | p.Phe255Tyr | P04062 |
| HOMO SAPIENS | 6 | 954 | 2vt0 | A | 251 | PHE | p.Phe290Leu | P04062 |
| HOMO SAPIENS | 5 | 954 | 2vt0 | A | 255 | HIS | p.His294Gln | P04062 |
| HOMO SAPIENS | 6 | 954 | 2vt0 | A | 257 | ARG | p.Arg296Gln | P04062 |
| HOMO SAPIENS | 5 | 954 | 2vt0 | A | 289 | PRO | p.Pro328Leu | P04062 |
| HOMO SAPIENS | 0 | 954 | 2vt0 | A | 309 | ALA | p.Ala348Val | P04062 |
| HOMO SAPIENS | 0 | 954 | 2vt0 | A | 311 | HIS | p.His350Arg | P04062 |
| HOMO SAPIENS | 6 | 954 | 2vt0 | A | 312 | TRP | p.Trp351Cys | P04062 |
| HOMO SAPIENS | 5 | 954 | 2vt0 | A | 323 | THR | p.Thr362Ile | P04062 |
| HOMO SAPIENS | 5 | 954 | 2vt0 | A | 325 | GLY | p.Gly364Arg | P04062 |
| HOMO SAPIENS | 6 | 954 | 2vt0 | A | 342 | CYS | p.Cys381Gly | P04062 |
| HOMO SAPIENS | 5 | 954 | 2vt0 | A | 353 | ARG | p.Arg392Gly | P04062 |
| HOMO SAPIENS | 3 | 954 | 2vt0 | A | 364 | SER | p.Ser403Thr | P04062 |
| HOMO SAPIENS | 5 | 954 | 2vt0 | A | 369 | THR | p.Thr408Met | P04062 |
| HOMO SAPIENS | 3 | 954 | 2vt0 | A | 370 | ASN | p.Asn409Ser | P04062 |
| HOMO SAPIENS | 0 | 954 | 2vt0 | A | 371 | LEU | p.Leu410Val | P04062 |
| HOMO SAPIENS | 0 | 954 | 2vt0 | A | 375 | VAL | p.Val414Leu | P04062 |
| HOMO SAPIENS | 0 | 954 | 2vt0 | A | 377 | GLY | p.Gly416Ser | P04062 |
| HOMO SAPIENS | 8 | 954 | 2vt0 | A | 398 | VAL | p.Val437Phe | P04062 |
| HOMO SAPIENS | 0 | 954 | 2vt0 | A | 401 | PRO | p.Pro440Leu | P04062 |
| HOMO SAPIENS | 3 | 954 | 2vt0 | A | 409 | ASP | p.Asp448His | P04062 |
| HOMO SAPIENS | 3 | 954 | 2vt0 | A | 409 | ASP | p.Asp448Val | P04062 |
| HOMO SAPIENS | 0 | 954 | 2vt0 | A | 415 | PRO | p.Pro454Arg | P04062 |
| HOMO SAPIENS | 5 | 954 | 2vt0 | A | 444 | LEU | p.Leu483Arg | P04062 |
| HOMO SAPIENS | 5 | 954 | 2vt0 | A | 456 | ALA | p.Ala495Pro | P04062 |
| HOMO SAPIENS | 5 | 954 | 2vt0 | A | 463 | ARG | p.Arg502Cys | P04062 |
| HOMO SAPIENS | 5 | 954 | 2vt0 | A | 463 | ARG | p.Arg502His | P04062 |
| HOMO SAPIENS | 5 | 954 | 2vt0 | A | 478 | GLY | p.Gly517Ser | P04062 |
| HOMO SAPIENS | 5 | 954 | 2vt0 | A | 496 | ARG | p.Arg535Cys | P04062 |
| HOMO SAPIENS | 5 | 954 | 2vt0 | A | 496 | ARG | p.Arg535His | P04062 |
| HOMO SAPIENS | 1 | 9037 | 2rqb | A | 946 | ALA | p.Ala946Thr | Q9BYX4 |
| HOMO SAPIENS | 0 | 253 | 5E+092 | A | 308 | LEU | p.Leu308Pro | P37173 |
| HOMO SAPIENS | 0 | 253 | 5E+092 | A | 315 | THR | p.Thr315Met | P37173 |
| HOMO SAPIENS | 0 | 253 | 5E+092 | A | 328 | HIS | p.His328Gln | P37173 |
| HOMO SAPIENS | 0 | 253 | 5E+092 | A | 336 | TYR | p.Tyr336Asn | P37173 |
| HOMO SAPIENS | 0 | 253 | 5E+092 | A | 348 | ARG | p.Arg348Cys | P37173 |
| HOMO SAPIENS | 0 | 253 | 5E+092 | A | 355 | ALA | p.Ala355Pro | P37173 |
| HOMO SAPIENS | 1 | 253 | 5E+092 | A | 356 | ARG | p.Arg356Gly | P37173 |

| | | | | | | | | |
|---|---|---|---|---|---|---|---|---|
| HOMO SAPIENS | 0 | 253 | 5E+092 | A | 357 | GLY | p.Gly357Trp | P37173 |
| HOMO SAPIENS | 1 | 253 | 5E+092 | A | 403 | ARG | p.Arg403His | P37173 |
| HOMO SAPIENS | 5 | 253 | 5E+092 | A | 449 | SER | p.Ser449Phe | P37173 |
| HOMO SAPIENS | 0 | 253 | 5E+092 | A | 460 | ARG | p.Arg460Cys | P37173 |
| HOMO SAPIENS | 0 | 253 | 5E+092 | A | 460 | ARG | p.Arg460His | P37173 |
| HOMO SAPIENS | 2 | 253 | 5E+092 | A | 526 | GLU | p.Glu526Gln | P37173 |
| HOMO SAPIENS | 5 | 253 | 5E+092 | A | 528 | ARG | p.Arg528Cys | P37173 |
| HOMO SAPIENS | 5 | 253 | 5E+092 | A | 528 | ARG | p.Arg528His | P37173 |
| HOMO SAPIENS | 6 | 158 | 5amc | A | 231 | ARG | p.Arg260Leu | P12821 |
| HOMO SAPIENS | 7 | 158 | 5amc | A | 237 | TYR | p.Tyr266Ter | P12821 |
| HOMO SAPIENS | 6 | 158 | 5amc | A | 325 | GLY | p.Gly354Arg | P12821 |
| HOMO SAPIENS | 0 | 158 | 5amc | A | 467 | ARG | p.Arg496Ter | P12821 |
| HOMO SAPIENS | 4 | 158 | 5amc | A | 479 | ARG | p.Arg508Ter | P12821 |
| HOMO SAPIENS | 0 | 158 | 5amc | A | 563 | ASP | p.Asp592Gly | P12821 |
| HOMO SAPIENS | -1 | 33062 | 1gr3 | A | 545 | GLY | p.Gly545Arg | Q03692 |
| HOMO SAPIENS | 5 | 33062 | 1gr3 | A | 591 | CYS | p.Cys591Arg | Q03692 |
| HOMO SAPIENS | 2 | 33062 | 1gr3 | A | 595 | GLY | p.Gly595Glu | Q03692 |
| HOMO SAPIENS | 0 | 33062 | 1gr3 | A | 597 | TYR | p.Tyr597Cys | Q03692 |
| HOMO SAPIENS | 5 | 33062 | 1gr3 | A | 598 | TYR | p.Tyr598Asp | Q03692 |
| HOMO SAPIENS | 7 | 33062 | 1gr3 | A | 600 | SER | p.Ser600Pro | Q03692 |
| HOMO SAPIENS | 1 | 33062 | 1gr3 | A | 614 | LEU | p.Leu614Pro | Q03692 |
| HOMO SAPIENS | 6 | 33062 | 1gr3 | A | 651 | TRP | p.Trp651Arg | Q03692 |
| HOMO SAPIENS | 7 | 33062 | 1gr3 | A | 671 | SER | p.Ser671Pro | Q03692 |
| HOMO SAPIENS | -1 | 38257 | 2l2i | B | 87 | PRO | p.Pro87Ala | Q13351 |
| HOMO SAPIENS | 2 | 2036 | 1ghq | A | 231 | GLY | p.Gly1224Asp | P01024 |
| HOMO SAPIENS | 5 | 3594 | 4cde | F | 388 | TYR | p.Tyr412Cys | P53634 |
| HOMO SAPIENS | 9 | 3594 | 4cde | F | 405 | TRP | p.Trp429Cys | P53634 |
| HOMO SAPIENS | 0 | 3594 | 4cde | F | 429 | ILE | p.Ile453Val | P53634 |
| HOMO SAPIENS | 7 | 18 | 1jkb | A | 34 | TRP | p.Trp52Ter | P61626 |
| HOMO SAPIENS | 9 | 18 | 1jkb | A | 70 | THR | p.Thr88Asn | P61626 |
| HOMO SAPIENS | 8 | 18 | 1jkb | A | 121 | VAL | p.Val139Ile | P61626 |
| HOMO SAPIENS | 2 | 414 | 3fj6 | A | 136 | ARG | p.Arg135Cys | Q02127 |
| HOMO SAPIENS | 1 | 414 | 3fj6 | A | 153 | GLY | p.Gly152Arg | Q02127 |
| HOMO SAPIENS | 0 | 414 | 3fj6 | A | 200 | ARG | p.Arg199Cys | Q02127 |
| HOMO SAPIENS | 0 | 414 | 3fj6 | A | 203 | GLY | p.Gly202Ala | Q02127 |
| HOMO SAPIENS | 0 | 414 | 3fj6 | A | 245 | ARG | p.Arg244Trp | Q02127 |
| HOMO SAPIENS | 0 | 414 | 3fj6 | A | 342 | ALA | p.Ala341Val | Q02127 |
| HOMO SAPIENS | 1 | 414 | 3fj6 | A | 347 | ARG | p.Arg346Trp | Q02127 |
| HOMO SAPIENS | 6 | 103 | 1u3w | A | 77 | GLY | p.Gly78Ter | P00326 |
| HOMO SAPIENS | -1 | 8369 | 4ux8 | F | 16 | ARG | p.Arg93Trp | P39905 |
| HOMO SAPIENS | 0 | 366 | 2zjc | A | 32 | GLY | p.Arg108Trp | P01375 |
| HOMO SAPIENS | 0 | 1457 | 3v9i | A | 352 | LEU | p.Ser352Leu | P30038 |
| HOMO SAPIENS | 0 | 1457 | 3v9i | A | 470 | VAL | p.Val470Ile | P30038 |
| HOMO SAPIENS | 0 | 1457 | 3v9i | A | 473 | THR | p.Thr473Ala | P30038 |
| HOMO SAPIENS | 1 | 2 | 4wo5 | A | 462 | ARG | p.Arg462Ile | P15056 |
| HOMO SAPIENS | 3 | 2 | 4wo5 | A | 463 | ILE | p.Ile463Ser | P15056 |
| HOMO SAPIENS | 1 | 2 | 4wo5 | A | 466 | GLY | p.Gly466Val | P15056 |
| HOMO SAPIENS | 1 | 2 | 4wo5 | A | 468 | PHE | p.Phe468Ser | P15056 |
| HOMO SAPIENS | 2 | 2 | 4wo5 | A | 469 | GLY | p.Gly469Ala | P15056 |
| HOMO SAPIENS | 2 | 2 | 4wo5 | A | 469 | GLY | p.Gly469Arg | P15056 |
| HOMO SAPIENS | 2 | 2 | 4wo5 | A | 485 | LEU | p.Leu485Phe | P15056 |
| HOMO SAPIENS | 0 | 2 | 4wo5 | A | 501 | GLU | p.Glu501Gly | P15056 |
| HOMO SAPIENS | 0 | 2 | 4wo5 | A | 501 | GLU | p.Glu501Lys | P15056 |
| HOMO SAPIENS | 7 | 2 | 4wo5 | A | 581 | ASN | p.Asn581Asp | P15056 |

| | | | | | | | | |
|---|---|---|---|---|---|---|---|---|
| HOMO SAPIENS | 8 | 2 | 4wo5 | A | 594 | ASP | p.Asp594Gly | P15056 |
| HOMO SAPIENS | 0 | 2 | 4wo5 | A | 597 | LEU | p.Leu597Arg | P15056 |
| HOMO SAPIENS | 0 | 2 | 4wo5 | A | 597 | LEU | p.Leu597Val | P15056 |
| HOMO SAPIENS | 0 | 2 | 4wo5 | A | 600 | VAL | p.Val600Glu | P15056 |
| HOMO SAPIENS | 2 | 2 | 4wo5 | A | 601 | LYS | p.Lys601Glu | P15056 |
| HOMO SAPIENS | 8 | 2 | 4wo5 | A | 638 | ASP | p.Asp638Glu | P15056 |
| HOMO SAPIENS | -1 | 30366 | 5bqd | B | 19 | ALA | p.Ala19Ser | Q99593 |
| HOMO SAPIENS | -1 | 30366 | 5bqd | B | 23 | PRO | p.Pro23His | Q99593 |
| HOMO SAPIENS | 1 | 30366 | 5bqd | B | 49 | GLN | p.Gln49Lys | Q99593 |
| HOMO SAPIENS | 2 | 30366 | 5bqd | B | 54 | ILE | p.Ile54Thr | Q99593 |
| HOMO SAPIENS | 8 | 30366 | 5bqd | B | 80 | GLY | p.Gly80Arg | Q99593 |
| HOMO SAPIENS | 6 | 30366 | 5bqd | B | 106 | ILE | p.Ile106Val | Q99593 |
| HOMO SAPIENS | 7 | 30366 | 5bqd | B | 111 | ASP | p.Asp111Tyr | Q99593 |
| HOMO SAPIENS | -1 | 30366 | 5bqd | B | 237 | ARG | p.Arg237Gln | Q99593 |
| HOMO SAPIENS | -1 | 30366 | 5bqd | B | 237 | ARG | p.Arg237Trp | Q99593 |
| HOMO SAPIENS | 5 | 4789 | 1n4m | A | 567 | SER | p.Ser567Leu | P06400 |
| HOMO SAPIENS | 0 | 4789 | 1n4m | A | 658 | ALA | p.Ala658Asp | P06400 |
| HOMO SAPIENS | 7 | 4789 | 1n4m | A | 661 | ARG | p.Arg661Trp | P06400 |
| HOMO SAPIENS | 6 | 4789 | 1n4m | A | 668 | ARG | p.Arg668His | P06400 |
| HOMO SAPIENS | 8 | 4789 | 1n4m | A | 697 | ASP | p.Asp697Glu | P06400 |
| HOMO SAPIENS | 6 | 4789 | 1n4m | A | 712 | CYS | p.Cys712Arg | P06400 |
| HOMO SAPIENS | 1 | 1798 | 2r5g | A | 101 | HIS | p.His101Gln | O15247 |
| HOMO SAPIENS | 0 | 1798 | 2r5g | A | 160 | PRO | p.Pro160Ala | O15247 |
| HOMO SAPIENS | 2 | 22 | 4x0t | A | 82 | ARG | p.Arg110Ter | P49419 |
| HOMO SAPIENS | 0 | 22 | 4x0t | A | 149 | ALA | p.Ala177Pro | P49419 |
| HOMO SAPIENS | 4 | 22 | 4x0t | A | 164 | THR | p.Thr192Met | P49419 |
| HOMO SAPIENS | 9 | 22 | 4x0t | A | 167 | ASN | p.Asn195Ser | P49419 |
| HOMO SAPIENS | 0 | 22 | 4x0t | A | 175 | TRP | p.Trp203Gly | P49419 |
| HOMO SAPIENS | 0 | 22 | 4x0t | A | 194 | THR | p.Thr222Ala | P49419 |
| HOMO SAPIENS | 0 | 22 | 4x0t | A | 273 | ASN | p.Asn301Ile | P49419 |
| HOMO SAPIENS | 0 | 22 | 4x0t | A | 307 | ARG | p.Arg335Gln | P49419 |
| HOMO SAPIENS | 3 | 22 | 4x0t | A | 313 | SER | p.Ser341Asn | P49419 |
| HOMO SAPIENS | 0 | 22 | 4x0t | A | 329 | ILE | p.Ile357Val | P49419 |
| HOMO SAPIENS | 1 | 22 | 4x0t | A | 380 | TYR | p.Tyr408Ter | P49419 |
| HOMO SAPIENS | 9 | 22 | 4x0t | A | 399 | GLU | p.Glu427Gln | P49419 |
| HOMO SAPIENS | 0 | 22 | 4x0t | A | 406 | TYR | p.Tyr434Cys | P49419 |
| HOMO SAPIENS | 0 | 22 | 4x0t | A | 431 | ILE | p.Ile459Phe | P49419 |
| HOMO SAPIENS | 4 | 22 | 4x0t | A | 477 | GLY | p.Gly505Arg | P49419 |
| HOMO SAPIENS | 0 | 22 | 4x0t | A | 495 | THR | p.Thr523Ala | P49419 |
| HOMO SAPIENS | 8 | 11042 | 1y6b | A | 1145 | PRO | p.Pro1147Ser | P35968 |
| HOMO SAPIENS | -1 | 3867 | 2xij | A | 18 | GLN | p.Gln18Ter | P22033 |
| HOMO SAPIENS | -1 | 3867 | 2xij | A | 31 | ARG | p.Arg31Ter | P22033 |
| HOMO SAPIENS | 2 | 3867 | 2xij | A | 93 | ARG | p.Arg93His | P22033 |
| HOMO SAPIENS | 0 | 3867 | 2xij | A | 94 | GLY | p.Gly94Arg | P22033 |
| HOMO SAPIENS | 2 | 3867 | 2xij | A | 95 | PRO | p.Pro95Leu | P22033 |
| HOMO SAPIENS | 2 | 3867 | 2xij | A | 105 | TRP | p.Trp105Arg | P22033 |
| HOMO SAPIENS | 2 | 3867 | 2xij | A | 108 | ARG | p.Arg108Cys | P22033 |
| HOMO SAPIENS | 3 | 3867 | 2xij | A | 154 | ARG | p.Arg154His | P22033 |
| HOMO SAPIENS | 3 | 3867 | 2xij | A | 156 | ASP | p.Asp156Val | P22033 |
| HOMO SAPIENS | 0 | 3867 | 2xij | A | 189 | ASN | p.Asn189Ile | P22033 |
| HOMO SAPIENS | 0 | 3867 | 2xij | A | 191 | ALA | p.Ala191Glu | P22033 |
| HOMO SAPIENS | 0 | 3867 | 2xij | A | 203 | GLY | p.Gly203Arg | P22033 |
| HOMO SAPIENS | 2 | 3867 | 2xij | A | 215 | GLY | p.Gly215Ser | P22033 |
| HOMO SAPIENS | 0 | 3867 | 2xij | A | 219 | ASN | p.Asn219Tyr | P22033 |

| | | | | | | | | |
|---|---|---|---|---|---|---|---|---|
| HOMO SAPIENS | 3 | 3867 | 2xij | A | 228 | ARG | p.Arg228Ter | P22033 |
| HOMO SAPIENS | 1 | 3867 | 2xij | A | 297 | THR | p.Thr297Ile | P22033 |
| HOMO SAPIENS | 0 | 3867 | 2xij | A | 326 | ARG | p.Arg326Lys | P22033 |
| HOMO SAPIENS | 3 | 3867 | 2xij | A | 369 | ARG | p.Arg369His | P22033 |
| HOMO SAPIENS | 3 | 3867 | 2xij | A | 369 | ARG | p.Arg369Cys | P22033 |
| HOMO SAPIENS | 0 | 3867 | 2xij | A | 370 | THR | p.Thr370Pro | P22033 |
| HOMO SAPIENS | 0 | 3867 | 2xij | A | 377 | ALA | p.Ala377Glu | P22033 |
| HOMO SAPIENS | 2 | 3867 | 2xij | A | 426 | GLY | p.Gly426Ala | P22033 |
| HOMO SAPIENS | 2 | 3867 | 2xij | A | 427 | GLY | p.Gly427Asp | P22033 |
| HOMO SAPIENS | 1 | 3867 | 2xij | A | 499 | THR | p.Ala499Thr | P22033 |
| HOMO SAPIENS | 2 | 3867 | 2xij | A | 532 | ARG | p.Arg532His | P22033 |
| HOMO SAPIENS | 3 | 3867 | 2xij | A | 555 | ALA | p.Ala555Thr | P22033 |
| HOMO SAPIENS | 0 | 3867 | 2xij | A | 573 | PHE | p.Phe573Ser | P22033 |
| HOMO SAPIENS | 3 | 3867 | 2xij | A | 623 | GLY | p.Gly623Arg | P22033 |
| HOMO SAPIENS | 5 | 3867 | 2xij | A | 671 | VAL | p.Ile671Val | P22033 |
| HOMO SAPIENS | 4 | 3867 | 2xij | A | 694 | ARG | p.Arg694Trp | P22033 |
| HOMO SAPIENS | 1 | 3867 | 2xij | A | 700 | MET | p.Met700Lys | P22033 |
| HOMO SAPIENS | 8 | 3867 | 2xij | A | 703 | GLY | p.Gly703Arg | P22033 |
| HOMO SAPIENS | 0 | 3867 | 2xij | A | 717 | GLY | p.Gly717Val | P22033 |
| HOMO SAPIENS | 1 | 3867 | 2xij | A | 727 | ARG | p.Arg727Ter | P22033 |
| HOMO SAPIENS | -1 | 52 | 3k91 | B | 72 | GLY | p.Gly73Ser | P00441 |
| HOMO SAPIENS | 6 | 52 | 3k91 | B | 96 | ASP | p.Asp97Asn | P00441 |
| HOMO SAPIENS | 5 | 52 | 3k91 | B | 112 | ILE | p.Ile113Thr | P00441 |
| HOMO SAPIENS | -1 | 52 | 3k91 | B | 134 | SER | p.Ser135Asn | P00441 |
| HOMO SAPIENS | 0 | 3950 | 4fom | A | 296 | ASN | p.Asn296His | Q9NQS3 |
| HOMO SAPIENS | 1 | 10 | 1de4 | A | 31 | VAL | p.Val53Met | Q30201 |
| HOMO SAPIENS | 0 | 10 | 1de4 | A | 37 | VAL | p.Val59Met | Q30201 |
| HOMO SAPIENS | 0 | 10 | 1de4 | A | 71 | GLY | p.Gly93Arg | Q30201 |
| HOMO SAPIENS | 0 | 10 | 1de4 | A | 83 | ILE | p.Ile105Thr | Q30201 |
| HOMO SAPIENS | 0 | 10 | 1de4 | A | 105 | GLN | p.Gln127His | Q30201 |
| HOMO SAPIENS | 1 | 10 | 1de4 | A | 146 | GLU | p.Glu168Ter | Q30201 |
| HOMO SAPIENS | 2 | 10 | 1de4 | A | 261 | GLN | p.Gln283Pro | Q30201 |
| HOMO SAPIENS | 0 | 8038 | 4x7n | A | 586 | ILE | p.Ile586Leu | Q9NZJ5 |
| HOMO SAPIENS | 0 | 8038 | 4x7n | A | 588 | ARG | p.Arg588Gln | Q9NZJ5 |
| HOMO SAPIENS | 0 | 35175 | 5ugw | A | 1062 | ALA | p.Ala1062Thr | O14746 |
| HOMO SAPIENS | 2 | 35175 | 5ugw | A | 1090 | VAL | p.Val1090Met | O14746 |
| HOMO SAPIENS | 0 | 1722 | 4z3k | C | 147 | ARG | p.Arg150Gly | P35270 |
| HOMO SAPIENS | 0 | 1722 | 4z3k | C | 160 | PRO | p.Pro163Leu | P35270 |
| HOMO SAPIENS | 0 | 1722 | 4z3k | C | 207 | ARG | p.Arg210Trp | P35270 |
| HOMO SAPIENS | 0 | 1722 | 4z3k | C | 216 | ARG | p.Arg219Ter | P35270 |
| HOMO SAPIENS | 0 | 1722 | 4z3k | C | 248 | LYS | p.Lys251Ter | P35270 |
| HOMO SAPIENS | 1 | 2832 | 4x8v | L | 115 | HIS | p.His175Gln | P08709 |
| HOMO SAPIENS | 0 | 825 | 1emh | A | 242 | PHE | p.Phe251Ser | P13051 |
| HOMO SAPIENS | 0 | 122 | 5uwk | A | 109 | ARG | p.Arg109Ter | P45452 |
| HOMO SAPIENS | 1 | 122 | 5uwk | A | 170 | LYS | p.Lys170Thr | P45452 |
| HOMO SAPIENS | 9 | 122 | 5uwk | A | 232 | HIS | p.His232Asn | P45452 |
| HOMO SAPIENS | 5 | 122 | 5uwk | A | 257 | ASP | p.Asp257Val | P45452 |
| HOMO SAPIENS | 9 | 1074 | 3p5i | A | 264 | TRP | p.Trp264Arg | Q9UJ71 |
| HOMO SAPIENS | 0 | 146 | 3r3g | A | 14C | GLU | p.Glu352Lys | P00734 |
| HOMO SAPIENS | 3 | 9522 | 1wyq | A | 83 | LEU | p.Leu253Pro | O15020 |
| HOMO SAPIENS | 8 | 67 | 2vgb | A | 130 | SER | p.Ser130Tyr | P30613 |
| HOMO SAPIENS | 8 | 67 | 2vgb | A | 163 | ARG | p.Arg163Cys | P30613 |
| HOMO SAPIENS | 4 | 67 | 2vgb | A | 241 | GLU | p.Glu241Ter | P30613 |
| HOMO SAPIENS | 9 | 67 | 2vgb | A | 384 | THR | p.Thr384Met | P30613 |

| | | | | | | | | |
|---|---|---|---|---|---|---|---|---|
| HOMO SAPIENS | 0 | 67 | 2vgb | A | 421 | GLN | p.Gln421Lys | P30613 |
| HOMO SAPIENS | 6 | 67 | 2vgb | A | 479 | ARG | p.Arg479His | P30613 |
| HOMO SAPIENS | 8 | 67 | 2vgb | A | 510 | ARG | p.Arg510Gln | P30613 |
| HOMO SAPIENS | 4 | 67 | 2vgb | A | 569 | ARG | p.Arg569Trp | P30613 |
| HOMO SAPIENS | 0 | 9088 | 5v44 | A | 232 | ASN | p.Asn232Lys | Q9NZJ4 |
| HOMO SAPIENS | 4 | 16 | 3lbn | A | 12 | GLY | p.Gly12Val | P01112 |
| HOMO SAPIENS | 4 | 16 | 3lbn | A | 12 | GLY | p.Gly12Ser | P01112 |
| HOMO SAPIENS | 4 | 16 | 3lbn | A | 13 | GLY | p.Gly13Cys | P01112 |
| HOMO SAPIENS | 4 | 16 | 3lbn | A | 13 | GLY | p.Gly13Asp | P01112 |
| HOMO SAPIENS | 5 | 16 | 3lbn | A | 22 | GLN | p.Gln22Lys | P01112 |
| HOMO SAPIENS | 5 | 16 | 3lbn | A | 58 | THR | p.Thr58Ile | P01112 |
| HOMO SAPIENS | 3 | 16 | 3lbn | A | 61 | GLN | p.Gln61Lys | P01112 |
| HOMO SAPIENS | 1 | 16 | 3lbn | A | 63 | GLU | p.Glu63Lys | P01112 |
| HOMO SAPIENS | 9 | 16 | 3lbn | A | 117 | LYS | p.Lys117Arg | P01112 |
| HOMO SAPIENS | 6 | 16 | 3lbn | A | 123 | ARG | p.Arg123Cys | P01112 |
| HOMO SAPIENS | 1 | 16 | 3lbn | A | 138 | GLY | p.Gly138Ser | P01112 |
| HOMO SAPIENS | 9 | 16 | 3lbn | A | 146 | ALA | p.Ala146Thr | P01112 |
| HOMO SAPIENS | 9 | 16 | 3lbn | A | 146 | ALA | p.Ala146Val | P01112 |
| HOMO SAPIENS | 1 | 359 | 5nvy | C | 63 | LEU | p.Leu63Pro | P40337 |
| HOMO SAPIENS | 1 | 359 | 5nvy | C | 64 | ARG | p.Arg64Pro | P40337 |
| HOMO SAPIENS | 1 | 359 | 5nvy | C | 70 | GLU | p.Glu70Lys | P40337 |
| HOMO SAPIENS | 0 | 359 | 5nvy | C | 84 | VAL | p.Val84Leu | P40337 |
| HOMO SAPIENS | 1 | 359 | 5nvy | C | 88 | TRP | p.Trp88Ser | P40337 |
| HOMO SAPIENS | 1 | 359 | 5nvy | C | 93 | GLY | p.Gly93Cys | P40337 |
| HOMO SAPIENS | 1 | 359 | 5nvy | C | 98 | TYR | p.Tyr98His | P40337 |
| HOMO SAPIENS | 1 | 359 | 5nvy | C | 111 | SER | p.Ser111Arg | P40337 |
| HOMO SAPIENS | 1 | 359 | 5nvy | C | 112 | TYR | p.Tyr112His | P40337 |
| HOMO SAPIENS | 1 | 359 | 5nvy | C | 126 | ASP | p.Asp126Tyr | P40337 |
| HOMO SAPIENS | 1 | 359 | 5nvy | C | 130 | VAL | p.Val130Leu | P40337 |
| HOMO SAPIENS | 1 | 359 | 5nvy | C | 135 | LEU | p.Leu135Phe | P40337 |
| HOMO SAPIENS | 9 | 359 | 5nvy | C | 161 | ARG | p.Arg161Gly | P40337 |
| HOMO SAPIENS | 7 | 359 | 5nvy | C | 162 | CYS | p.Cys162Phe | P40337 |
| HOMO SAPIENS | 4 | 359 | 5nvy | C | 163 | LEU | p.Leu163Pro | P40337 |
| HOMO SAPIENS | 3 | 359 | 5nvy | C | 164 | GLN | p.Gln164Arg | P40337 |
| HOMO SAPIENS | 9 | 359 | 5nvy | C | 166 | VAL | p.Val166Phe | P40337 |
| HOMO SAPIENS | 7 | 359 | 5nvy | C | 167 | ARG | p.Arg167Gly | P40337 |
| HOMO SAPIENS | 7 | 359 | 5nvy | C | 167 | ARG | p.Arg167Gln | P40337 |
| HOMO SAPIENS | 1 | 359 | 5nvy | C | 183 | SER | p.Ser183Ter | P40337 |
| HOMO SAPIENS | 2 | 359 | 5nvy | C | 188 | LEU | p.Leu188Val | P40337 |
| HOMO SAPIENS | 1 | 359 | 5nvy | C | 191 | HIS | p.His191Asp | P40337 |
| HOMO SAPIENS | 1 | 359 | 5nvy | C | 192 | PRO | p.Pro192Ser | P40337 |
| HOMO SAPIENS | 1 | 359 | 5nvy | C | 200 | ARG | p.Arg200Trp | P40337 |
| HOMO SAPIENS | -1 | 359 | 5nvy | C | 211 | MET | p.Met211Leu | P40337 |
| HOMO SAPIENS | 0 | 14099 | 4btm | A | 282 | ARG | p.Arg258Gln | Q5TCY1 |
| HOMO SAPIENS | 0 | 696 | 5h1b | A | 150 | ARG | p.Arg150Gln | Q06609 |
| HOMO SAPIENS | -1 | 208 | 2bxl | A | 1 | ASP | p.Asp25Val | P02768 |
| HOMO SAPIENS | 0 | 208 | 2bxl | A | 269 | ASP | p.Asp293Gly | P02768 |
| HOMO SAPIENS | 0 | 208 | 2bxl | A | 570 | GLU | p.Glu594Lys | P02768 |
| HOMO SAPIENS | 0 | 208 | 2bxl | A | 573 | LYS | p.Lys597Glu | P02768 |
| HOMO SAPIENS | -1 | 12813 | 2uzx | B | 35 | MET | p.Met35Leu | P08581 |
| HOMO SAPIENS | 1 | 12813 | 2uzx | B | 156 | SER | p.Ser156Leu | P08581 |
| HOMO SAPIENS | 0 | 12813 | 2uzx | B | 168 | GLU | p.Glu168Asp | P08581 |
| HOMO SAPIENS | 2 | 12813 | 2uzx | B | 211 | LEU | p.Leu211Trp | P08581 |
| HOMO SAPIENS | 2 | 12813 | 2uzx | B | 218 | ARG | p.Arg218Ser | P08581 |

| | | | | | | | | |
|---|---|---|---|---|---|---|---|---|
| HOMO SAPIENS | 1 | 12813 | 2uzx | B | 238 | LEU | p.Leu238Ser | P08581 |
| HOMO SAPIENS | 4 | 12813 | 2uzx | B | 320 | ALA | p.Ala320Val | P08581 |
| HOMO SAPIENS | 1 | 12813 | 2uzx | B | 375 | ASN | p.Asn375Ser | P08581 |
| HOMO SAPIENS | 0 | 12813 | 2uzx | B | 575 | LEU | p.Leu575Ile | P08581 |
| HOMO SAPIENS | 1 | 10780 | 4ejx | A | 348 | ARG | p.Arg367Cys | P00450 |
| HOMO SAPIENS | 2 | 10780 | 4ejx | A | 525 | ASP | p.Asp544Glu | P00450 |
| HOMO SAPIENS | 1 | 1407 | 5jn5 | A | 88 | ILE | p.Ile88Val | P36871 |
| HOMO SAPIENS | 7 | 1407 | 5jn5 | A | 115 | THR | p.Thr115Ala | P36871 |
| HOMO SAPIENS | 3 | 1407 | 5jn5 | A | 121 | GLY | p.Gly121Arg | P36871 |
| HOMO SAPIENS | 2 | 1407 | 5jn5 | A | 221 | ARG | p.Arg221Cys | P36871 |
| HOMO SAPIENS | 6 | 1407 | 5jn5 | A | 263 | TYR | p.Asp263Tyr | P36871 |
| HOMO SAPIENS | 3 | 1407 | 5jn5 | A | 420 | TYR | p.Tyr420His | P36871 |
| HOMO SAPIENS | 0 | 1407 | 5jn5 | A | 501 | VAL | p.Val501Ile | P36871 |
| HOMO SAPIENS | -1 | 1878 | 1yow | A | 255 | ARG | p.Arg255Leu | Q13285 |
| HOMO SAPIENS | 6 | 1878 | 1yow | A | 293 | ASP | p.Asp293Asn | Q13285 |
| HOMO SAPIENS | 1 | 1878 | 1yow | A | 437 | LEU | p.Leu437Gln | Q13285 |
| HOMO SAPIENS | 0 | 3071 | 2wpm | E | 145 | ARG | p.Arg191His | P00740 |
| HOMO SAPIENS | 0 | 3071 | 2wpm | E | 145 | ARG | p.Arg191Cys | P00740 |
| HOMO SAPIENS | 3 | 589 | 3ggc | B | 15 | GLY | p.Gly16Ser | P00492 |
| HOMO SAPIENS | 4 | 589 | 3ggc | B | 40 | LEU | p.Leu41Pro | P00492 |
| HOMO SAPIENS | 2 | 589 | 3ggc | B | 50 | ARG | p.Arg51Gly | P00492 |
| HOMO SAPIENS | 5 | 589 | 3ggc | B | 51 | ASP | p.Asp52Gly | P00492 |
| HOMO SAPIENS | 0 | 589 | 3ggc | B | 57 | GLY | p.Gly58Arg | P00492 |
| HOMO SAPIENS | 6 | 589 | 3ggc | B | 69 | GLY | p.Gly70Glu | P00492 |
| HOMO SAPIENS | 1 | 589 | 3ggc | B | 73 | PHE | p.Phe74Leu | P00492 |
| HOMO SAPIENS | 0 | 589 | 3ggc | B | 77 | LEU | p.Leu78Val | P00492 |
| HOMO SAPIENS | 3 | 589 | 3ggc | B | 79 | ASP | p.Asp80Val | P00492 |
| HOMO SAPIENS | 0 | 589 | 3ggc | B | 103 | SER | p.Ser104Arg | P00492 |
| HOMO SAPIENS | 0 | 589 | 3ggc | B | 109 | SER | p.Ser110Leu | P00492 |
| HOMO SAPIENS | 1 | 589 | 3ggc | B | 129 | VAL | p.Val130Asp | P00492 |
| HOMO SAPIENS | 4 | 589 | 3ggc | B | 131 | ILE | p.Ile132Met | P00492 |
| HOMO SAPIENS | 7 | 589 | 3ggc | B | 142 | MET | p.Met143Lys | P00492 |
| HOMO SAPIENS | 0 | 589 | 3ggc | B | 160 | ALA | p.Ala161Ser | P00492 |
| HOMO SAPIENS | 6 | 589 | 3ggc | B | 167 | THR | p.Thr168Ile | P00492 |
| HOMO SAPIENS | 0 | 589 | 3ggc | B | 169 | ARG | p.Arg170Ter | P00492 |
| HOMO SAPIENS | 5 | 589 | 3ggc | B | 175 | PRO | p.Pro176Leu | P00492 |
| HOMO SAPIENS | 8 | 589 | 3ggc | B | 193 | ASP | p.Asp194Glu | P00492 |
| HOMO SAPIENS | 8 | 589 | 3ggc | B | 193 | ASP | p.Asp194Asn | P00492 |
| HOMO SAPIENS | 8 | 589 | 3ggc | B | 198 | PHE | p.Phe199Val | P00492 |
| HOMO SAPIENS | 8 | 589 | 3ggc | B | 200 | ASP | p.Asp201Gly | P00492 |
| HOMO SAPIENS | 0 | 4269 | 2pla | B | 174 | GLU | p.Glu174Lys | Q8N335 |
| HOMO SAPIENS | 1 | 4269 | 2pla | B | 220 | ARG | p.Arg220His | Q8N335 |
| HOMO SAPIENS | 4 | 4269 | 2pla | B | 280 | ALA | p.Ala280Val | Q8N335 |
| HOMO SAPIENS | 0 | 2712 | 1ii4 | E | 252 | TRP | p.Ser252Leu | P21802 |
| HOMO SAPIENS | 0 | 2712 | 1ii4 | E | 253 | PRO | p.Pro253Arg | P21802 |
| HOMO SAPIENS | 5 | 2712 | 1ii4 | E | 267 | SER | p.Ser267Pro | P21802 |
| HOMO SAPIENS | 0 | 2712 | 1ii4 | E | 278 | CYS | p.Cys278Phe | P21802 |
| HOMO SAPIENS | 0 | 2712 | 1ii4 | E | 289 | GLN | p.Gln289Pro | P21802 |
| HOMO SAPIENS | 0 | 2712 | 1ii4 | E | 290 | TRP | p.Trp290Cys | P21802 |
| HOMO SAPIENS | 0 | 2712 | 1ii4 | E | 290 | TRP | p.Trp290Arg | P21802 |
| HOMO SAPIENS | 0 | 2712 | 1ii4 | E | 292 | LYS | p.Lys292Glu | P21802 |
| HOMO SAPIENS | 0 | 2712 | 1ii4 | E | 315 | ALA | p.Ala315Ser | P21802 |
| HOMO SAPIENS | 0 | 2712 | 1ii4 | E | 321 | ASP | p.Asp321Ala | P21802 |
| HOMO SAPIENS | 1 | 2712 | 1ii4 | E | 328 | TYR | p.Tyr328Cys | P21802 |

| | | | | | | | | |
|---|---|---|---|---|---|---|---|---|
| HOMO SAPIENS | 9 | 2712 | 1ii4 | E | 340 | TYR | p.Tyr340His | P21802 |
| HOMO SAPIENS | 1 | 2712 | 1ii4 | E | 341 | THR | p.Thr341Pro | P21802 |
| HOMO SAPIENS | 0 | 2712 | 1ii4 | E | 342 | CYS | p.Cys342Arg | P21802 |
| HOMO SAPIENS | 0 | 2712 | 1ii4 | E | 342 | CYS | p.Cys342Trp | P21802 |
| HOMO SAPIENS | 0 | 2712 | 1ii4 | E | 344 | ALA | p.Ala344Gly | P21802 |
| HOMO SAPIENS | 0 | 2712 | 1ii4 | E | 347 | SER | p.Ser347Cys | P21802 |
| HOMO SAPIENS | 0 | 2712 | 1ii4 | E | 351 | SER | p.Ser351Cys | P21802 |
| HOMO SAPIENS | 1 | 2712 | 1ii4 | E | 354 | SER | p.Ser354Cys | P21802 |
| HOMO SAPIENS | 2 | 53 | 4pj1 | H | 5 | ASP | p.Asp29Gly | P10809 |
| HOMO SAPIENS | 0 | 53 | 4pj1 | H | 74 | VAL | p.Val98Ile | P10809 |
| HOMO SAPIENS | 0 | 29968 | 2fdb | M | 71 | LYS | p.Lys89Glu | P55075 |
| HOMO SAPIENS | 2 | 29968 | 2fdb | M | 98 | ARG | p.Arg116Gly | P55075 |
| HOMO SAPIENS | 2 | 29968 | 2fdb | M | 122 | GLY | p.Gly140Ser | P55075 |
| HOMO SAPIENS | 8 | 5061 | 5f85 | B | 60 | GLY | p.Gly106Ser | P00740 |
| HOMO SAPIENS | 0 | 35944 | 1h6h | A | 85 | THR | p.Thr85Asn | Q15080 |
| HOMO SAPIENS | 8 | 35944 | 1h6h | A | 105 | ARG | p.Arg105Gln | Q15080 |
| HOMO SAPIENS | 7 | 12016 | 1hj7 | A | 295 | ASN | p.Asn316Ser | P01130 |
| HOMO SAPIENS | 5 | 12016 | 1hj7 | A | 302 | GLY | p.Gly323Ser | P01130 |
| HOMO SAPIENS | 6 | 12016 | 1hj7 | A | 319 | CYS | p.Cys340Trp | P01130 |
| HOMO SAPIENS | 7 | 12016 | 1hj7 | A | 329 | ARG | p.Arg350Ter | P01130 |
| HOMO SAPIENS | 5 | 12016 | 1hj7 | A | 331 | CYS | p.Cys352Ter | P01130 |
| HOMO SAPIENS | 2 | 12016 | 1hj7 | A | 345 | GLN | p.Gln366Arg | P01130 |
| HOMO SAPIENS | 2 | 1481 | 2i07 | A | 239 | TYR | p.Tyr261Ter | P01024 |
| HOMO SAPIENS | -1 | 1481 | 2i07 | A | 292 | LEU | p.Pro314Leu | P01024 |
| HOMO SAPIENS | 2 | 1481 | 2i07 | A | 326 | ILE | p.Ile348Val | P01024 |
| HOMO SAPIENS | 2 | 1481 | 2i07 | A | 413 | GLU | p.Glu435Lys | P01024 |
| HOMO SAPIENS | 4 | 1481 | 2i07 | A | 447 | GLU | p.Glu469Asp | P01024 |
| HOMO SAPIENS | 2 | 1481 | 2i07 | A | 597 | VAL | p.Val619Met | P01024 |
| HOMO SAPIENS | 5 | 1481 | 2i07 | A | 603 | ILE | p.Ile625Phe | P01024 |
| HOMO SAPIENS | 0 | 1481 | 2i07 | A | 611 | LYS | p.Lys633Arg | P01024 |
| HOMO SAPIENS | 0 | 11123 | 3c5n | A | 315 | GLY | p.Gly368Trp | O00294 |
| HOMO SAPIENS | 8 | 11123 | 3c5n | A | 329 | PHE | p.Phe382Ser | O00294 |
| HOMO SAPIENS | 5 | 11123 | 3c5n | A | 429 | ARG | p.Arg482Trp | O00294 |
| HOMO SAPIENS | 8 | 11123 | 3c5n | A | 438 | PHE | p.Phe491Leu | O00294 |
| HOMO SAPIENS | 3 | 4398 | 3fx0 | B | 1288 | ALA | p.Ala288Gly | Q9Y6K9 |
| HOMO SAPIENS | 7 | 4398 | 3fx0 | B | 1315 | GLU | p.Glu315Ala | Q9Y6K9 |
| HOMO SAPIENS | 8 | 4398 | 3fx0 | B | 1319 | ARG | p.Arg319Gln | Q9Y6K9 |
| HOMO SAPIENS | 6 | 3989 | 3qd6 | S | 83 | CYS | p.Cys83Arg | P25942 |
| HOMO SAPIENS | 1 | 3989 | 3qd6 | S | 124 | SER | p.Ser124Leu | P25942 |
| HOMO SAPIENS | 2 | 24003 | 2m0v | A | 225 | GLU | p.Glu225Lys | O14745 |
| HOMO SAPIENS | 2 | 46 | 4aqd | A | 70 | ASP | p.Asp98Gly | P06276 |
| HOMO SAPIENS | 4 | 46 | 4aqd | A | 115 | GLY | p.Gly143Asp | P06276 |
| HOMO SAPIENS | 3 | 46 | 4aqd | A | 184 | ALA | p.Ala212Val | P06276 |
| HOMO SAPIENS | 0 | 46 | 4aqd | A | 225 | GLY | p.Gly253Ter | P06276 |
| HOMO SAPIENS | 2 | 46 | 4aqd | A | 243 | THR | p.Thr271Met | P06276 |
| HOMO SAPIENS | 2 | 46 | 4aqd | A | 255 | GLU | p.Glu283Asp | P06276 |
| HOMO SAPIENS | 5 | 46 | 4aqd | A | 267 | LYS | p.Lys295Arg | P06276 |
| HOMO SAPIENS | 3 | 46 | 4aqd | A | 271 | GLU | p.Glu299Ter | P06276 |
| HOMO SAPIENS | 2 | 46 | 4aqd | A | 307 | LEU | p.Leu335Pro | P06276 |
| HOMO SAPIENS | 1 | 46 | 4aqd | A | 330 | LEU | p.Leu358Ile | P06276 |
| HOMO SAPIENS | 0 | 46 | 4aqd | A | 497 | GLU | p.Glu525Val | P06276 |
| HOMO SAPIENS | 3 | 786 | 1ols | B | 51 | GLY | p.Gly101Asp | P21953 |
| HOMO SAPIENS | 2 | 786 | 1ols | B | 87 | ALA | p.Ala137Val | P21953 |
| HOMO SAPIENS | 0 | 786 | 1ols | B | 110 | ILE | p.Ile160Ser | P21953 |

| | | | | | | | | |
|---|---|---|---|---|---|---|---|---|
| HOMO SAPIENS | 4 | 786 | 1ols | B | 120 | ARG | p.Arg170Cys | P21953 |
| HOMO SAPIENS | 2 | 786 | 1ols | B | 126 | ASN | p.Asn176Tyr | P21953 |
| HOMO SAPIENS | 0 | 786 | 1ols | B | 133 | ARG | p.Arg183Gly | P21953 |
| HOMO SAPIENS | 0 | 786 | 1ols | B | 133 | ARG | p.Arg183Pro | P21953 |
| HOMO SAPIENS | 0 | 786 | 1ols | B | 135 | PRO | p.Pro185Leu | P21953 |
| HOMO SAPIENS | 3 | 786 | 1ols | B | 156 | HIS | p.His206Tyr | P21953 |
| HOMO SAPIENS | 0 | 786 | 1ols | B | 228 | GLY | p.Gly278Ser | P21953 |
| HOMO SAPIENS | 1 | 786 | 1ols | B | 235 | ARG | p.Arg285Ter | P21953 |
| HOMO SAPIENS | 6 | 786 | 1ols | B | 274 | ARG | p.Arg324Ter | P21953 |
| HOMO SAPIENS | 1 | 831 | 4zt1 | B | 48 | VAL | p.Val202Ile | P12830 |
| HOMO SAPIENS | 6 | 831 | 4zt1 | B | 70 | ARG | p.Arg224Cys | P12830 |
| HOMO SAPIENS | 5 | 831 | 4zt1 | B | 79 | HIS | p.His233Gln | P12830 |
| HOMO SAPIENS | 4 | 831 | 4zt1 | B | 120 | GLY | p.Gly274Ser | P12830 |
| HOMO SAPIENS | 7 | 831 | 4zt1 | B | 187 | TYR | p.Tyr341Ter | P12830 |
| HOMO SAPIENS | 0 | 8497 | 3sla | A | 287 | ASN | p.Asn287Ser | P35222 |
| HOMO SAPIENS | 7 | 23986 | 2ke1 | A | 301 | VAL | p.Val301Met | O43918 |
| HOMO SAPIENS | 0 | 4967 | 1rj7 | B | 334 | ARG | p.Arg334His | Q92838 |
| HOMO SAPIENS | 0 | 4967 | 1rj7 | B | 338 | THR | p.Thr338Met | Q92838 |
| HOMO SAPIENS | 3 | 4967 | 1rj7 | B | 349 | ALA | p.Ala349Thr | Q92838 |
| HOMO SAPIENS | 2 | 4967 | 1rj7 | B | 358 | GLN | p.Gln358Glu | Q92838 |
| HOMO SAPIENS | 1 | 4967 | 1rj7 | B | 375 | LYS | p.Lys375Glu | Q92838 |
| HOMO SAPIENS | 9 | 7623 | 1d4t | A | 32 | ARG | p.Arg32Thr | O60880 |
| HOMO SAPIENS | 6 | 7623 | 1d4t | A | 55 | ARG | p.Arg55Leu | O60880 |
| HOMO SAPIENS | 0 | 7623 | 1d4t | A | 68 | THR | p.Thr68Ile | O60880 |
| HOMO SAPIENS | 6 | 705 | 3idh | A | 59 | PRO | p.Pro59Ser | P35557 |
| HOMO SAPIENS | 5 | 705 | 3idh | A | 124 | ASP | p.Asp124Asn | P35557 |
| HOMO SAPIENS | 4 | 705 | 3idh | A | 131 | SER | p.Ser131Pro | P35557 |
| HOMO SAPIENS | 0 | 705 | 3idh | A | 147 | GLY | p.Gly147Asp | P35557 |
| HOMO SAPIENS | 5 | 705 | 3idh | A | 186 | ARG | p.Arg186Ter | P35557 |
| HOMO SAPIENS | 0 | 705 | 3idh | A | 202 | MET | p.Met202Thr | P35557 |
| HOMO SAPIENS | 2 | 705 | 3idh | A | 210 | MET | p.Met210Lys | P35557 |
| HOMO SAPIENS | 1 | 705 | 3idh | A | 215 | TYR | p.Tyr215Ter | P35557 |
| HOMO SAPIENS | 0 | 705 | 3idh | A | 226 | VAL | p.Val226Met | P35557 |
| HOMO SAPIENS | 9 | 705 | 3idh | A | 228 | THR | p.Thr228Met | P35557 |
| HOMO SAPIENS | 9 | 705 | 3idh | A | 256 | GLU | p.Glu256Lys | P35557 |
| HOMO SAPIENS | 2 | 705 | 3idh | A | 261 | GLY | p.Gly261Arg | P35557 |
| HOMO SAPIENS | 6 | 705 | 3idh | A | 279 | GLU | p.Glu279Gln | P35557 |
| HOMO SAPIENS | 7 | 705 | 3idh | A | 299 | GLY | p.Gly299Arg | P35557 |
| HOMO SAPIENS | 2 | 705 | 3idh | A | 318 | GLY | p.Gly318Trp | P35557 |
| HOMO SAPIENS | 4 | 705 | 3idh | A | 378 | ALA | p.Ala378Thr | P35557 |
| HOMO SAPIENS | 0 | 705 | 3idh | A | 387 | ALA | p.Ala387Glu | P35557 |
| HOMO SAPIENS | 5 | 705 | 3idh | A | 414 | LYS | p.Lys414Glu | P35557 |
| HOMO SAPIENS | 0 | 705 | 3idh | A | 455 | VAL | p.Val455Met | P35557 |
| HOMO SAPIENS | -1 | 705 | 3idh | A | 462 | MET | p.Met462Ile | P35557 |
| HOMO SAPIENS | 1 | 509 | 4nef | A | 22 | LEU | p.Leu22Val | P41181 |
| HOMO SAPIENS | 4 | 509 | 4nef | A | 57 | GLN | p.Gln57Pro | P41181 |
| HOMO SAPIENS | 8 | 509 | 4nef | A | 64 | GLY | p.Gly64Arg | P41181 |
| HOMO SAPIENS | 7 | 509 | 4nef | A | 68 | ASN | p.Asn68Ser | P41181 |
| HOMO SAPIENS | 1 | 509 | 4nef | A | 100 | GLY | p.Gly100Val | P41181 |
| HOMO SAPIENS | 0 | 509 | 4nef | A | 125 | THR | p.Thr125Met | P41181 |
| HOMO SAPIENS | 0 | 509 | 4nef | A | 126 | THR | p.Thr126Met | P41181 |
| HOMO SAPIENS | 4 | 509 | 4nef | A | 146 | PHE | p.Phe146Leu | P41181 |
| HOMO SAPIENS | 0 | 509 | 4nef | A | 175 | GLY | p.Gly175Arg | P41181 |
| HOMO SAPIENS | 5 | 509 | 4nef | A | 181 | CYS | p.Cys181Trp | P41181 |

| | | | | | | | | | |
|---|---|---|---|---|---|---|---|---|---|
| HOMO SAPIENS | 9 | 509 | 4nef | A | | 187 | ARG | p.Arg187Cys | P41181 |
| HOMO SAPIENS | 0 | 509 | 4nef | A | | 190 | ALA | p.Ala190Thr | P41181 |
| HOMO SAPIENS | 0 | 509 | 4nef | A | | 216 | SER | p.Ser216Pro | P41181 |
| HOMO SAPIENS | 4 | 2242 | 1z70 | X | | 1155 | SER | p.Ser155Pro | Q8NBK3 |
| HOMO SAPIENS | 3 | 2242 | 1z70 | X | | 1218 | CYS | p.Cys218Tyr | Q8NBK3 |
| HOMO SAPIENS | 0 | 2242 | 1z70 | X | | 1279 | ALA | p.Ala279Val | Q8NBK3 |
| HOMO SAPIENS | -1 | 2242 | 1z70 | X | | 1336 | OCS | p.Cys336Arg | Q8NBK3 |
| HOMO SAPIENS | 4 | 2242 | 1z70 | X | | 1345 | ARG | p.Arg345Cys | Q8NBK3 |
| HOMO SAPIENS | 2 | 2242 | 1z70 | X | | 1348 | ALA | p.Ala348Pro | Q8NBK3 |
| HOMO SAPIENS | 5 | 2242 | 1z70 | X | | 1349 | ARG | p.Arg349Gln | Q8NBK3 |
| HOMO SAPIENS | 5 | 2242 | 1z70 | X | | 1349 | ARG | p.Arg349Trp | Q8NBK3 |
| HOMO SAPIENS | 3 | 2242 | 1z70 | X | | 1359 | SER | p.Ser359Ter | Q8NBK3 |
| HOMO SAPIENS | 5 | 8269 | 5wrr | B | | 136 | ARG | p.Arg136Ter | Q96MK3 |
| HOMO SAPIENS | 1 | 8269 | 5wrr | B | | 332 | LYS | p.Asn332Lys | Q96MK3 |
| HOMO SAPIENS | 3 | 8269 | 5wrr | B | | 478 | ARG | p.Arg478Ter | Q96MK3 |
| HOMO SAPIENS | 5 | 8269 | 5wrr | B | | 504 | THR | p.Thr504Ile | Q96MK3 |
| HOMO SAPIENS | 5 | 1411 | 4h1s | B | | 221 | SER | p.Ser221Ter | P21589 |
| HOMO SAPIENS | 1 | 1411 | 4h1s | B | | 358 | CYS | p.Cys358Tyr | P21589 |
| HOMO SAPIENS | 2 | 23 | 5vvd | B | | 298 | GLU | p.Glu298Asp | P29474 |
| HOMO SAPIENS | 0 | 19 | 5jlh | E | | 88 | THR | p.Thr89Ile | P63261 |
| HOMO SAPIENS | 6 | 19 | 5jlh | E | | 117 | LYS | p.Lys118Met | P63261 |
| HOMO SAPIENS | 5 | 19 | 5jlh | E | | 263 | PRO | p.Pro264Leu | P63261 |
| HOMO SAPIENS | 1 | 19 | 5jlh | E | | 277 | THR | p.Thr278Ile | P63261 |
| HOMO SAPIENS | 6 | 19 | 5jlh | E | | 331 | PRO | p.Pro332Ala | P63261 |
| HOMO SAPIENS | 5 | 19 | 5jlh | E | | 369 | VAL | p.Val370Ala | P63261 |
| HOMO SAPIENS | 0 | 14306 | 4w4o | C | | 92 | ARG | p.Arg92Ter | P12314 |
| HOMO SAPIENS | 3 | 603 | 1nuh | A | | 4 | THR | p.Thr5Ile | P06744 |
| HOMO SAPIENS | 2 | 603 | 1nuh | A | | 19 | HIS | p.His20Pro | P06744 |
| HOMO SAPIENS | 8 | 603 | 1nuh | A | | 158 | GLY | p.Gly159Ser | P06744 |
| HOMO SAPIENS | 6 | 603 | 1nuh | A | | 223 | THR | p.Thr224Met | P06744 |
| HOMO SAPIENS | 0 | 603 | 1nuh | A | | 338 | LEU | p.Leu339Pro | P06744 |
| HOMO SAPIENS | 6 | 603 | 1nuh | A | | 342 | GLN | p.Gln343Arg | P06744 |
| HOMO SAPIENS | 6 | 603 | 1nuh | A | | 374 | THR | p.Thr375Arg | P06744 |
| HOMO SAPIENS | 4 | 603 | 1nuh | A | | 524 | ILE | p.Ile525Thr | P06744 |
| HOMO SAPIENS | 4 | 603 | 1nuh | A | | 538 | ASP | p.Asp539Asn | P06744 |
| HOMO SAPIENS | 1 | 3107 | 3kzj | A | | 1197 | VAL | p.Val1197Ala | P08603 |
| HOMO SAPIENS | 9 | 1748 | 2noh | A | | 46 | ARG | p.Arg46Gln | O15527 |
| HOMO SAPIENS | 2 | 11750 | 3s4y | A | | 33 | ARG | p.Arg33His | Q9H3S4 |
| HOMO SAPIENS | 2 | 11750 | 3s4y | A | | 40 | LEU | p.Leu40Pro | Q9H3S4 |
| HOMO SAPIENS | 4 | 11750 | 3s4y | A | | 142 | LEU | p.Leu142Phe | Q9H3S4 |
| HOMO SAPIENS | 5 | 11750 | 3s4y | A | | 219 | ASN | p.Asn219Ser | Q9H3S4 |
| HOMO SAPIENS | 1 | 2591 | 5vnj | A | | 382 | PHE | p.Phe382Leu | Q15436 |
| HOMO SAPIENS | 2 | 2591 | 5vnj | A | | 702 | MET | p.Met702Val | Q15436 |
| HOMO SAPIENS | 1 | 2426 | 5m8t | A | | 87 | ARG | p.Arg87Trp | P17643 |
| HOMO SAPIENS | 1 | 2426 | 5m8t | A | | 166 | SER | p.Ser166Ter | P17643 |
| HOMO SAPIENS | 0 | 2426 | 5m8t | A | | 356 | ARG | p.Arg356Gln | P17643 |
| HOMO SAPIENS | 2 | 2426 | 5m8t | A | | 374 | SER | p.Arg374Ter | P17643 |
| HOMO SAPIENS | 6 | 2426 | 5m8t | A | | 378 | ASN | p.Asn378Ser | P17643 |
| HOMO SAPIENS | 6 | 2426 | 5m8t | A | | 382 | LEU | p.Leu382Pro | P17643 |
| HOMO SAPIENS | 7 | 10097 | 4n6v | | 1 | 46 | GLN | p.Gln46Ter | P04080 |
| HOMO SAPIENS | 3 | 10097 | 4n6v | | 1 | 65 | VAL | p.Val65Ile | P04080 |
| HOMO SAPIENS | 8 | 10097 | 4n6v | | 1 | 68 | ARG | p.Arg68Ter | P04080 |
| HOMO SAPIENS | 2 | 859 | 4z9a | A | | 105 | GLN | p.Gln106Arg | P24666 |
| HOMO SAPIENS | 8 | 1420 | 5nan | F | | 95 | SER | p.Ser95Leu | Q96PD4 |

| | | | | | | | | |
|---|---|---|---|---|---|---|---|---|
| HOMO SAPIENS | 6 | 1420 | 5nan | F | 126 | GLU | p.Glu126Gly | Q96PD4 |
| HOMO SAPIENS | 8 | 1420 | 5nan | F | 155 | VAL | p.Val155Ile | Q96PD4 |
| HOMO SAPIENS | 6 | 1420 | 5nan | F | 157 | PRO | p.Pro157Ser | Q96PD4 |
| HOMO SAPIENS | -1 | 1420 | 5nan | F | 161 | HIS | p.His161Arg | Q96PD4 |
| HOMO SAPIENS | 3 | 288 | 5i96 | B | 140 | GLN | p.Arg140Gly | P48735 |
| HOMO SAPIENS | 3 | 288 | 5i96 | B | 140 | GLN | p.Arg140Gln | P48735 |
| HOMO SAPIENS | 2 | 288 | 5i96 | B | 261 | ARG | p.Arg261His | P48735 |
| HOMO SAPIENS | 2 | 288 | 5i96 | B | 435 | THR | p.Thr435Met | P48735 |
| HOMO SAPIENS | 9 | 120 | 1tup | B | 107 | TYR | p.Tyr107His | P04637 |
| HOMO SAPIENS | 8 | 120 | 1tup | B | 110 | ARG | p.Arg110Leu | P04637 |
| HOMO SAPIENS | 5 | 120 | 1tup | B | 120 | LYS | p.Lys120Glu | P04637 |
| HOMO SAPIENS | 4 | 120 | 1tup | B | 133 | MET | p.Met133Thr | P04637 |
| HOMO SAPIENS | 3 | 120 | 1tup | B | 134 | PHE | p.Phe134Cys | P04637 |
| HOMO SAPIENS | 8 | 120 | 1tup | B | 136 | GLN | p.Gln136His | P04637 |
| HOMO SAPIENS | 9 | 120 | 1tup | B | 138 | ALA | p.Ala138Pro | P04637 |
| HOMO SAPIENS | 9 | 120 | 1tup | B | 151 | PRO | p.Pro151Ser | P04637 |
| HOMO SAPIENS | 9 | 120 | 1tup | B | 152 | PRO | p.Pro152Leu | P04637 |
| HOMO SAPIENS | 9 | 120 | 1tup | B | 158 | ARG | p.Arg158Cys | P04637 |
| HOMO SAPIENS | 9 | 120 | 1tup | B | 158 | ARG | p.Arg158His | P04637 |
| HOMO SAPIENS | 9 | 120 | 1tup | B | 175 | ARG | p.Arg175Cys | P04637 |
| HOMO SAPIENS | 9 | 120 | 1tup | B | 175 | ARG | p.Arg175Leu | P04637 |
| HOMO SAPIENS | 8 | 120 | 1tup | B | 181 | ARG | p.Arg181Leu | P04637 |
| HOMO SAPIENS | 9 | 120 | 1tup | B | 189 | ALA | p.Ala189Val | P04637 |
| HOMO SAPIENS | 9 | 120 | 1tup | B | 196 | ARG | p.Arg196Gly | P04637 |
| HOMO SAPIENS | 9 | 120 | 1tup | B | 213 | ARG | p.Arg213Gln | P04637 |
| HOMO SAPIENS | 9 | 120 | 1tup | B | 213 | ARG | p.Arg213Ter | P04637 |
| HOMO SAPIENS | 9 | 120 | 1tup | B | 214 | HIS | p.His214Gln | P04637 |
| HOMO SAPIENS | 9 | 120 | 1tup | B | 220 | TYR | p.Tyr220Cys | P04637 |
| HOMO SAPIENS | 0 | 120 | 1tup | B | 234 | TYR | p.Tyr234Cys | P04637 |
| HOMO SAPIENS | 9 | 120 | 1tup | B | 237 | MET | p.Met237Ile | P04637 |
| HOMO SAPIENS | 0 | 120 | 1tup | B | 238 | CYS | p.Cys238Tyr | P04637 |
| HOMO SAPIENS | 9 | 120 | 1tup | B | 241 | SER | p.Ser241Phe | P04637 |
| HOMO SAPIENS | 9 | 120 | 1tup | B | 242 | CYS | p.Cys242Ser | P04637 |
| HOMO SAPIENS | 9 | 120 | 1tup | B | 245 | GLY | p.Gly245Cys | P04637 |
| HOMO SAPIENS | 9 | 120 | 1tup | B | 245 | GLY | p.Gly245Asp | P04637 |
| HOMO SAPIENS | 9 | 120 | 1tup | B | 248 | ARG | p.Arg248Leu | P04637 |
| HOMO SAPIENS | 9 | 120 | 1tup | B | 248 | ARG | p.Arg248Trp | P04637 |
| HOMO SAPIENS | 9 | 120 | 1tup | B | 251 | ILE | p.Ile251Ser | P04637 |
| HOMO SAPIENS | 9 | 120 | 1tup | B | 252 | LEU | p.Leu252Pro | P04637 |
| HOMO SAPIENS | 9 | 120 | 1tup | B | 258 | GLU | p.Glu258Lys | P04637 |
| HOMO SAPIENS | 8 | 120 | 1tup | B | 272 | VAL | p.Val272Leu | P04637 |
| HOMO SAPIENS | 9 | 120 | 1tup | B | 273 | ARG | p.Arg273Cys | P04637 |
| HOMO SAPIENS | 9 | 120 | 1tup | B | 273 | ARG | p.Arg273His | P04637 |
| HOMO SAPIENS | 9 | 120 | 1tup | B | 277 | CYS | p.Cys277Phe | P04637 |
| HOMO SAPIENS | 9 | 120 | 1tup | B | 280 | ARG | p.Arg280Ile | P04637 |
| HOMO SAPIENS | 9 | 120 | 1tup | B | 282 | ARG | p.Arg282Trp | P04637 |
| HOMO SAPIENS | 9 | 120 | 1tup | B | 283 | ARG | p.Arg283His | P04637 |
| HOMO SAPIENS | 9 | 120 | 1tup | B | 285 | GLU | p.Glu285Val | P04637 |
| HOMO SAPIENS | -1 | 120 | 1tup | B | 292 | LYS | p.Lys292Ile | P04637 |
| HOMO SAPIENS | -1 | 120 | 1tup | B | 298 | GLU | p.Glu298Lys | P04637 |
| HOMO SAPIENS | -1 | 2791 | 4ygb | A | 39 | VAL | p.Val39Ala | P49257 |
| HOMO SAPIENS | 2 | 106 | 4x6j | A | 32 | GLY | p.Gly146Arg | P43235 |
| HOMO SAPIENS | 0 | 106 | 4x6j | A | 76 | GLN | p.Gln190Ter | P43235 |
| HOMO SAPIENS | 7 | 106 | 4x6j | A | 198 | ARG | p.Arg312Ter | P43235 |

| | | | | | | | | |
|---|---|---|---|---|---|---|---|---|
| HOMO SAPIENS | 5 | 490 | 2hlo | B | 172 | LEU | p.Leu202Gln | P02675 |
| HOMO SAPIENS | 0 | 490 | 2hlo | B | 353 | LEU | p.Leu383Arg | P02675 |
| HOMO SAPIENS | 4 | 490 | 2hlo | B | 400 | GLY | p.Gly430Asp | P02675 |
| HOMO SAPIENS | 7 | 490 | 2hlo | B | 448 | ARG | p.Arg478Lys | P02675 |
| HOMO SAPIENS | 1 | 289 | 4p13 | B | 18 | GLU | p.Glu43Lys | P11310 |
| HOMO SAPIENS | 2 | 289 | 4p13 | B | 20 | GLN | p.Gln45Arg | P11310 |
| HOMO SAPIENS | 5 | 289 | 4p13 | B | 28 | ARG | p.Arg53Cys | P11310 |
| HOMO SAPIENS | 1 | 289 | 4p13 | B | 42 | TYR | p.Tyr67His | P11310 |
| HOMO SAPIENS | 1 | 289 | 4p13 | B | 53 | ILE | p.Ile78Thr | P11310 |
| HOMO SAPIENS | 1 | 289 | 4p13 | B | 59 | LEU | p.Leu84Phe | P11310 |
| HOMO SAPIENS | 2 | 289 | 4p13 | B | 96 | THR | p.Thr121Ile | P11310 |
| HOMO SAPIENS | 0 | 289 | 4p13 | B | 124 | MET | p.Met149Ile | P11310 |
| HOMO SAPIENS | 5 | 289 | 4p13 | B | 143 | ASP | p.Asp168Ala | P11310 |
| HOMO SAPIENS | 7 | 289 | 4p13 | B | 168 | THR | p.Thr193Ala | P11310 |
| HOMO SAPIENS | 1 | 289 | 4p13 | B | 170 | GLY | p.Gly195Arg | P11310 |
| HOMO SAPIENS | 2 | 289 | 4p13 | B | 181 | ARG | p.Arg206Cys | P11310 |
| HOMO SAPIENS | 2 | 289 | 4p13 | B | 181 | ARG | p.Arg206His | P11310 |
| HOMO SAPIENS | 8 | 289 | 4p13 | B | 203 | THR | p.Thr228Ile | P11310 |
| HOMO SAPIENS | 2 | 289 | 4p13 | B | 208 | ILE | p.Ile233Asn | P11310 |
| HOMO SAPIENS | 1 | 289 | 4p13 | B | 219 | CYS | p.Cys244Arg | P11310 |
| HOMO SAPIENS | 6 | 289 | 4p13 | B | 220 | SER | p.Ser245Leu | P11310 |
| HOMO SAPIENS | 5 | 289 | 4p13 | B | 228 | GLU | p.Glu253Lys | P11310 |
| HOMO SAPIENS | 2 | 289 | 4p13 | B | 241 | ASP | p.Asp266Gly | P11310 |
| HOMO SAPIENS | 1 | 289 | 4p13 | B | 242 | GLY | p.Gly267Arg | P11310 |
| HOMO SAPIENS | 7 | 289 | 4p13 | B | 256 | ARG | p.Arg281Thr | P11310 |
| HOMO SAPIENS | 4 | 289 | 4p13 | B | 269 | ARG | p.Arg294Thr | P11310 |
| HOMO SAPIENS | 4 | 289 | 4p13 | B | 304 | GLU | p.Lys329Glu | P11310 |
| HOMO SAPIENS | 1 | 289 | 4p13 | B | 324 | ARG | p.Arg349Ter | P11310 |
| HOMO SAPIENS | 3 | 289 | 4p13 | B | 326 | THR | p.Thr351Ile | P11310 |
| HOMO SAPIENS | 7 | 289 | 4p13 | B | 350 | ILE | p.Ile375Thr | P11310 |
| HOMO SAPIENS | 3 | 289 | 4p13 | B | 359 | GLU | p.Glu384Ter | P11310 |
| HOMO SAPIENS | 1 | 4563 | 3i2b | A | 16 | ARG | p.Arg16Cys | Q03393 |
| HOMO SAPIENS | 5 | 4563 | 3i2b | A | 25 | ARG | p.Arg25Gln | Q03393 |
| HOMO SAPIENS | 1 | 4563 | 3i2b | A | 47 | ASN | p.Asn47Asp | Q03393 |
| HOMO SAPIENS | 8 | 4563 | 3i2b | A | 52 | ASN | p.Asn52Ser | Q03393 |
| HOMO SAPIENS | 1 | 4563 | 3i2b | A | 56 | VAL | p.Val56Met | Q03393 |
| HOMO SAPIENS | 1 | 4563 | 3i2b | A | 87 | PRO | p.Pro87Ser | Q03393 |
| HOMO SAPIENS | 1 | 4563 | 3i2b | A | 96 | ASP | p.Asp96Asn | Q03393 |
| HOMO SAPIENS | 0 | 4563 | 3i2b | A | 99 | TYR | p.Tyr99Ter | Q03393 |
| HOMO SAPIENS | 2 | 4563 | 3i2b | A | 116 | ASP | p.Asp116Gly | Q03393 |
| HOMO SAPIENS | 2 | 41 | 3exx | A | 1 | GLY | p.Gly90Cys | P01308 |
| HOMO SAPIENS | 5 | 41 | 3exx | A | 3 | VAL | p.Val92Leu | P01308 |
| HOMO SAPIENS | 4 | 41 | 3exx | A | 7 | CYS | p.Cys96Ser | P01308 |
| HOMO SAPIENS | 9 | 41 | 3exx | A | 19 | TYR | p.Tyr108Cys | P01308 |
| HOMO SAPIENS | 9 | 131 | 4mjo | D | 118 | ASP | p.Asp119Asn | P09467 |
| HOMO SAPIENS | 0 | 131 | 4mjo | D | 163 | GLY | p.Gly164Ser | P09467 |
| HOMO SAPIENS | 0 | 131 | 4mjo | D | 176 | ALA | p.Ala177Asp | P09467 |
| HOMO SAPIENS | 5 | 131 | 4mjo | D | 193 | PHE | p.Phe194Ser | P09467 |
| HOMO SAPIENS | 4 | 131 | 4mjo | D | 217 | ARG | p.Arg218Lys | P09467 |
| HOMO SAPIENS | 4 | 131 | 4mjo | D | 223 | VAL | p.Val224Ile | P09467 |
| HOMO SAPIENS | 4 | 131 | 4mjo | D | 232 | PHE | p.Phe233Ile | P09467 |
| HOMO SAPIENS | 5 | 131 | 4mjo | D | 243 | ARG | p.Arg244Trp | P09467 |
| HOMO SAPIENS | 0 | 131 | 4mjo | D | 283 | PRO | p.Pro284Arg | P09467 |
| HOMO SAPIENS | 0 | 131 | 4mjo | D | 288 | MET | p.Met289Leu | P09467 |

| | | | | | | | | |
|---|---|---|---|---|---|---|---|---|
| HOMO SAPIENS | 0 | 131 | 4mjo | D | 323 | ASP | p.Asp324Asn | P09467 |
| HOMO SAPIENS | -1 | 13122 | 3hfc | A | 583 | ARG | p.Arg583Cys | P51787 |
| HOMO SAPIENS | 1 | 13122 | 3hfc | A | 587 | THR | p.Thr587Met | P51787 |
| HOMO SAPIENS | 0 | 13122 | 3hfc | A | 589 | GLY | p.Gly589Asp | P51787 |
| HOMO SAPIENS | 4 | 3002 | 3nbq | B | 94 | ARG | p.Arg94Cys | Q16831 |
| HOMO SAPIENS | 1 | 3616 | 3dbp | A | 97 | ILE | p.Ile286Asn | P11172 |
| HOMO SAPIENS | 0 | 3616 | 3dbp | A | 240 | GLY | p.Gly429Arg | P11172 |
| HOMO SAPIENS | 0 | 3616 | 3dbp | A | 257 | ILE | p.Ile446Val | P11172 |
| HOMO SAPIENS | 1 | 584 | 1drf | A | 79 | LEU | p.Leu80Phe | P00374 |
| HOMO SAPIENS | 3 | 1609 | 3qa2 | B | 49 | VAL | p.Val49Ile | P50053 |
| HOMO SAPIENS | 0 | 20560 | 5mug | A | 101 | HIS | p.His101Gln | P49638 |
| HOMO SAPIENS | 2 | 20560 | 5mug | A | 120 | ALA | p.Ala120Thr | P49638 |
| HOMO SAPIENS | 3 | 20560 | 5mug | A | 192 | ARG | p.Arg192His | P49638 |
| HOMO SAPIENS | 3 | 20560 | 5mug | A | 221 | ARG | p.Arg221Trp | P49638 |
| HOMO SAPIENS | 4 | 256 | 3vtv | A | 65 | THR | p.Glu50Lys | Q9GZQ8 |
| HOMO SAPIENS | 4 | 256 | 3vtv | A | 113 | VAL | p.Met98Lys | Q9GZQ8 |
| HOMO SAPIENS | 4 | 5397 | 1m61 | A | 192 | ARG | p.Arg192Trp | P43403 |
| HOMO SAPIENS | 5 | 39881 | 1aut | L | 107 | HIS | p.His149Pro | P04070 |
| HOMO SAPIENS | 5 | 4010 | 4uf8 | I | 33 | ALA | p.Ala33Val | Q13501 |
| HOMO SAPIENS | 3 | 313 | 5w49 | A | 98 | PRO | p.Pro98Leu | P23526 |
| HOMO SAPIENS | 2 | 313 | 5w49 | A | 143 | TYR | p.Tyr143Cys | P23526 |
| HOMO SAPIENS | -1 | 1643 | 2m8t | A | 102 | PRO | p.Pro102Leu | P04156 |
| HOMO SAPIENS | -1 | 1643 | 2m8t | A | 105 | PRO | p.Pro105Leu | P04156 |
| HOMO SAPIENS | -1 | 1643 | 2m8t | A | 117 | ALA | p.Ala117Val | P04156 |
| HOMO SAPIENS | 5 | 1643 | 2m8t | A | 129 | MET | p.Met129Val | P04156 |
| HOMO SAPIENS | 5 | 1643 | 2m8t | A | 131 | GLY | p.Gly131Val | P04156 |
| HOMO SAPIENS | 2 | 1643 | 2m8t | A | 142 | GLY | p.Gly142Ser | P04156 |
| HOMO SAPIENS | 3 | 1643 | 2m8t | A | 171 | ASN | p.Asn171Ser | P04156 |
| HOMO SAPIENS | 9 | 1643 | 2m8t | A | 178 | ASP | p.Asp178Asn | P04156 |
| HOMO SAPIENS | 9 | 1643 | 2m8t | A | 180 | VAL | p.Val180Ile | P04156 |
| HOMO SAPIENS | 9 | 1643 | 2m8t | A | 183 | THR | p.Thr183Ala | P04156 |
| HOMO SAPIENS | 9 | 1643 | 2m8t | A | 187 | HIS | p.His187Arg | P04156 |
| HOMO SAPIENS | 8 | 1643 | 2m8t | A | 198 | PHE | p.Phe198Ser | P04156 |
| HOMO SAPIENS | 9 | 1643 | 2m8t | A | 200 | GLU | p.Glu200Lys | P04156 |
| HOMO SAPIENS | 9 | 1643 | 2m8t | A | 208 | ARG | p.Arg208His | P04156 |
| HOMO SAPIENS | 3 | 1643 | 2m8t | A | 210 | VAL | p.Val210Ile | P04156 |
| HOMO SAPIENS | 6 | 1643 | 2m8t | A | 217 | GLN | p.Gln217Arg | P04156 |
| HOMO SAPIENS | 1 | 29 | 5kos | A | 151 | GLY | p.Gly217Arg | P00797 |
| HOMO SAPIENS | 1 | 29 | 5kos | A | 164 | ARG | p.Arg230Lys | P00797 |
| HOMO SAPIENS | 3 | 504 | 4zkv | B | 37 | ARG | p.Arg37Pro | P49773 |
| HOMO SAPIENS | 2 | 504 | 4zkv | B | 62 | GLN | p.Gln62Ter | P49773 |
| HOMO SAPIENS | 2 | 504 | 4zkv | B | 84 | CYS | p.Cys84Arg | P49773 |
| HOMO SAPIENS | 4 | 504 | 4zkv | B | 93 | GLY | p.Gly93Asp | P49773 |
| HOMO SAPIENS | 9 | 504 | 4zkv | B | 112 | HIS | p.His112Asn | P49773 |
| HOMO SAPIENS | 0 | 74 | 5vwo | C | 54 | ASN | p.Asn54Lys | P04181 |
| HOMO SAPIENS | 0 | 74 | 5vwo | C | 55 | TYR | p.Tyr55His | P04181 |
| HOMO SAPIENS | 2 | 74 | 5vwo | C | 90 | GLN | p.Gln90Glu | P04181 |
| HOMO SAPIENS | 6 | 74 | 5vwo | C | 91 | GLY | p.Gly91Glu | P04181 |
| HOMO SAPIENS | 2 | 74 | 5vwo | C | 93 | CYS | p.Cys93Phe | P04181 |
| HOMO SAPIENS | 1 | 74 | 5vwo | C | 154 | ARG | p.Arg154Leu | P04181 |
| HOMO SAPIENS | 0 | 74 | 5vwo | C | 180 | ARG | p.Arg180Thr | P04181 |
| HOMO SAPIENS | 0 | 74 | 5vwo | C | 199 | PRO | p.Pro199Gln | P04181 |
| HOMO SAPIENS | 4 | 74 | 5vwo | C | 209 | TYR | p.Tyr209Ter | P04181 |
| HOMO SAPIENS | 0 | 74 | 5vwo | C | 226 | ALA | p.Ala226Val | P04181 |

| | | | | | | | | |
|---|---|---|---|---|---|---|---|---|
| HOMO SAPIENS | 5 | 74 | 5vwo | C | 233 | GLN | p.Gln233Arg | P04181 |
| HOMO SAPIENS | 5 | 74 | 5vwo | C | 241 | PRO | p.Pro241Leu | P04181 |
| HOMO SAPIENS | 2 | 74 | 5vwo | C | 245 | TYR | p.Tyr245Cys | P04181 |
| HOMO SAPIENS | 6 | 74 | 5vwo | C | 250 | ARG | p.Arg250Pro | P04181 |
| HOMO SAPIENS | 6 | 74 | 5vwo | C | 250 | ARG | p.Arg250Ter | P04181 |
| HOMO SAPIENS | 8 | 74 | 5vwo | C | 267 | THR | p.Thr267Ile | P04181 |
| HOMO SAPIENS | 0 | 74 | 5vwo | C | 270 | ALA | p.Ala270Pro | P04181 |
| HOMO SAPIENS | 0 | 74 | 5vwo | C | 271 | ARG | p.Arg271Lys | P04181 |
| HOMO SAPIENS | 1 | 74 | 5vwo | C | 299 | TYR | p.Tyr299Ter | P04181 |
| HOMO SAPIENS | 0 | 74 | 5vwo | C | 318 | GLU | p.Glu318Lys | P04181 |
| HOMO SAPIENS | 1 | 74 | 5vwo | C | 319 | HIS | p.His319Tyr | P04181 |
| HOMO SAPIENS | 1 | 74 | 5vwo | C | 331 | ARG | p.Arg331Ter | P04181 |
| HOMO SAPIENS | 0 | 74 | 5vwo | C | 332 | VAL | p.Val332Met | P04181 |
| HOMO SAPIENS | 1 | 74 | 5vwo | C | 353 | GLY | p.Gly353Asp | P04181 |
| HOMO SAPIENS | 2 | 74 | 5vwo | C | 375 | GLY | p.Gly375Ala | P04181 |
| HOMO SAPIENS | 0 | 74 | 5vwo | C | 394 | CYS | p.Cys394Arg | P04181 |
| HOMO SAPIENS | 0 | 74 | 5vwo | C | 402 | LEU | p.Leu402Pro | P04181 |
| HOMO SAPIENS | 0 | 74 | 5vwo | C | 436 | ILE | p.Ile436Asn | P04181 |
| HOMO SAPIENS | 7 | 24356 | 4az0 | A | 104 | VAL | p.Val132Met | P10619 |
| HOMO SAPIENS | 4 | 24356 | 4az0 | A | 221 | TYR | p.Tyr249Asn | P10619 |
| HOMO SAPIENS | 3 | 245 | 1rsz | A | 24 | ARG | p.Arg24Ter | P00491 |
| HOMO SAPIENS | 3 | 245 | 1rsz | A | 51 | SER | p.Gly51Ser | P00491 |
| HOMO SAPIENS | 1 | 245 | 1rsz | A | 58 | ARG | p.Arg58Ter | P00491 |
| HOMO SAPIENS | 4 | 245 | 1rsz | A | 89 | GLU | p.Glu89Lys | P00491 |
| HOMO SAPIENS | 6 | 245 | 1rsz | A | 128 | ASP | p.Asp128Gly | P00491 |
| HOMO SAPIENS | 0 | 245 | 1rsz | A | 174 | ALA | p.Ala174Pro | P00491 |
| HOMO SAPIENS | 8 | 245 | 1rsz | A | 192 | TYR | p.Tyr192Cys | P00491 |
| HOMO SAPIENS | 8 | 245 | 1rsz | A | 234 | ARG | p.Arg234Pro | P00491 |
| HOMO SAPIENS | 0 | 37 | 5tby | E | 22 | GLU | p.Glu22Lys | P10916 |
| HOMO SAPIENS | 1 | 37 | 5tby | E | 86 | PHE | p.Phe86Ser | P10916 |
| HOMO SAPIENS | 9 | 71 | 5m9j | A | 12 | GLN | p.Gln36Leu | P03950 |
| HOMO SAPIENS | 3 | 71 | 5m9j | A | 17 | LYS | p.Lys41Glu | P03950 |
| HOMO SAPIENS | 3 | 71 | 5m9j | A | 17 | LYS | p.Lys41Ile | P03950 |
| HOMO SAPIENS | 4 | 71 | 5m9j | A | 28 | SER | p.Ser52Asn | P03950 |
| HOMO SAPIENS | 7 | 71 | 5m9j | A | 31 | ARG | p.Arg55Lys | P03950 |
| HOMO SAPIENS | 9 | 71 | 5m9j | A | 39 | CYS | p.Cys63Trp | P03950 |
| HOMO SAPIENS | 9 | 71 | 5m9j | A | 40 | LYS | p.Lys64Ile | P03950 |
| HOMO SAPIENS | 2 | 71 | 5m9j | A | 60 | GLU | p.Lys84Glu | P03950 |
| HOMO SAPIENS | 0 | 71 | 5m9j | A | 112 | PRO | p.Pro136Leu | P03950 |
| HOMO SAPIENS | 9 | 71 | 5m9j | A | 113 | VAL | p.Val137Ile | P03950 |
| HOMO SAPIENS | 3 | 326 | 5dqi | A | 209 | GLY | p.Gly209Val | Q9Y253 |
| HOMO SAPIENS | 1 | 326 | 5dqi | A | 242 | SER | p.Ser242Ter | Q9Y253 |
| HOMO SAPIENS | 1 | 326 | 5dqi | A | 303 | ARG | p.Arg303Ter | Q9Y253 |
| HOMO SAPIENS | 1 | 8203 | 3iu3 | K | 132 | VAL | p.Val153Ile | P01589 |
| HOMO SAPIENS | 0 | 4995 | 5ffo | H | 234 | THR | p.Thr263Ile | P01137 |
| HOMO SAPIENS | 5 | 38609 | 2ruk | A | 49 | ASP | p.Asp49His | P04637 |
| HOMO SAPIENS | 5 | 38609 | 2ruk | A | 58 | PRO | p.Pro58Arg | P04637 |
| HOMO SAPIENS | 0 | 1607 | 6pah | A | 117 | THR | p.Thr117Ile | P00439 |
| HOMO SAPIENS | 3 | 1607 | 6pah | A | 145 | ASP | p.Asp145Val | P00439 |
| HOMO SAPIENS | 5 | 1607 | 6pah | A | 158 | ARG | p.Arg158Gln | P00439 |
| HOMO SAPIENS | 5 | 1607 | 6pah | A | 158 | ARG | p.Arg158Trp | P00439 |
| HOMO SAPIENS | 5 | 1607 | 6pah | A | 169 | ARG | p.Arg169His | P00439 |
| HOMO SAPIENS | 5 | 1607 | 6pah | A | 169 | ARG | p.Arg169Cys | P00439 |
| HOMO SAPIENS | 3 | 1607 | 6pah | A | 176 | ARG | p.Arg176Leu | P00439 |

| | | | | | | | | |
|---|---|---|---|---|---|---|---|---|
| HOMO SAPIENS | 3 | 1607 | 6pah | A | 177 | VAL | p.Val177Leu | P00439 |
| HOMO SAPIENS | 1 | 1607 | 6pah | A | 194 | LEU | p.Leu194Pro | P00439 |
| HOMO SAPIENS | 1 | 1607 | 6pah | A | 204 | TYR | p.Tyr204Cys | P00439 |
| HOMO SAPIENS | 1 | 1607 | 6pah | A | 204 | TYR | p.Tyr204Ter | P00439 |
| HOMO SAPIENS | 2 | 1607 | 6pah | A | 230 | VAL | p.Val230Ile | P00439 |
| HOMO SAPIENS | 4 | 1607 | 6pah | A | 232 | GLN | p.Gln232Ter | P00439 |
| HOMO SAPIENS | 4 | 1607 | 6pah | A | 241 | ARG | p.Arg241Cys | P00439 |
| HOMO SAPIENS | 4 | 1607 | 6pah | A | 241 | ARG | p.Arg241His | P00439 |
| HOMO SAPIENS | 5 | 1607 | 6pah | A | 243 | ARG | p.Arg243Gln | P00439 |
| HOMO SAPIENS | 5 | 1607 | 6pah | A | 243 | ARG | p.Arg243Ter | P00439 |
| HOMO SAPIENS | 7 | 1607 | 6pah | A | 249 | LEU | p.Leu249Phe | P00439 |
| HOMO SAPIENS | 5 | 1607 | 6pah | A | 252 | ARG | p.Arg252Gly | P00439 |
| HOMO SAPIENS | 5 | 1607 | 6pah | A | 261 | ARG | p.Arg261Ter | P00439 |
| HOMO SAPIENS | 4 | 1607 | 6pah | A | 267 | GLN | p.Gln267Leu | P00439 |
| HOMO SAPIENS | 1 | 1607 | 6pah | A | 272 | GLY | p.Gly272Ter | P00439 |
| HOMO SAPIENS | 2 | 1607 | 6pah | A | 277 | TYR | p.Tyr277Asp | P00439 |
| HOMO SAPIENS | 7 | 1607 | 6pah | A | 280 | GLU | p.Glu280Lys | P00439 |
| HOMO SAPIENS | 8 | 1607 | 6pah | A | 281 | PRO | p.Pro281Leu | P00439 |
| HOMO SAPIENS | 0 | 1607 | 6pah | A | 299 | PHE | p.Phe299Cys | P00439 |
| HOMO SAPIENS | 5 | 1607 | 6pah | A | 300 | ALA | p.Ala300Ser | P00439 |
| HOMO SAPIENS | 5 | 1607 | 6pah | A | 309 | ALA | p.Ala309Asp | P00439 |
| HOMO SAPIENS | 4 | 1607 | 6pah | A | 320 | LYS | p.Lys320Asn | P00439 |
| HOMO SAPIENS | 0 | 1607 | 6pah | A | 345 | ALA | p.Ala345Ser | P00439 |
| HOMO SAPIENS | 0 | 1607 | 6pah | A | 348 | LEU | p.Leu348Val | P00439 |
| HOMO SAPIENS | 6 | 1607 | 6pah | A | 356 | TYR | p.Tyr356Ter | P00439 |
| HOMO SAPIENS | 2 | 1607 | 6pah | A | 380 | THR | p.Thr380Met | P00439 |
| HOMO SAPIENS | 0 | 1607 | 6pah | A | 386 | TYR | p.Tyr386Cys | P00439 |
| HOMO SAPIENS | 5 | 1607 | 6pah | A | 390 | GLU | p.Glu390Gly | P00439 |
| HOMO SAPIENS | 5 | 1607 | 6pah | A | 403 | ALA | p.Ala403Val | P00439 |
| HOMO SAPIENS | 4 | 1607 | 6pah | A | 408 | ARG | p.Arg408Trp | P00439 |
| HOMO SAPIENS | 3 | 1607 | 6pah | A | 413 | ARG | p.Arg413Pro | P00439 |
| HOMO SAPIENS | 4 | 1607 | 6pah | A | 414 | TYR | p.Tyr414Cys | P00439 |
| HOMO SAPIENS | 4 | 1607 | 6pah | A | 414 | TYR | p.Tyr414His | P00439 |
| HOMO SAPIENS | 4 | 1607 | 6pah | A | 415 | ASP | p.Asp415Asn | P00439 |
| HOMO SAPIENS | -1 | 676 | 5dc5 | A | 7 | PRO | p.Pro7Leu | Q9BY41 |
| HOMO SAPIENS | 0 | 676 | 5dc5 | A | 311 | THR | p.Thr311Met | Q9BY41 |
| HOMO SAPIENS | 1 | 124 | 5ftk | E | 95 | ARG | p.Arg95Gly | P55072 |
| HOMO SAPIENS | 1 | 124 | 5ftk | E | 155 | ARG | p.Arg155Cys | P55072 |
| HOMO SAPIENS | 1 | 124 | 5ftk | E | 155 | ARG | p.Arg155His | P55072 |
| HOMO SAPIENS | 1 | 124 | 5ftk | E | 159 | ARG | p.Arg159His | P55072 |
| HOMO SAPIENS | 1 | 124 | 5ftk | E | 159 | ARG | p.Arg159Gly | P55072 |
| HOMO SAPIENS | 0 | 124 | 5ftk | E | 191 | ARG | p.Arg191Gln | P55072 |
| HOMO SAPIENS | 0 | 124 | 5ftk | E | 232 | ALA | p.Ala232Glu | P55072 |
| HOMO SAPIENS | 5 | 5442 | 1fgg | B | 223 | GLY | p.Gly223Ser | O94766 |
| HOMO SAPIENS | 0 | 5442 | 1fgg | B | 277 | ARG | p.Arg277Gln | O94766 |
| HOMO SAPIENS | 0 | 9072 | 2gys | B | 225 | GLU | p.Glu249Gln | P32927 |
| HOMO SAPIENS | 0 | 9072 | 2gys | B | 247 | SER | p.Ser271Ter | P32927 |
| HOMO SAPIENS | 0 | 9072 | 2gys | B | 294 | ASP | p.Asp312Asn | P32927 |
| HOMO SAPIENS | 0 | 1484 | 4zrj | A | 62 | PHE | p.Phe62Ser | P35240 |
| HOMO SAPIENS | 3 | 1484 | 4zrj | A | 205 | MET | p.Met205Val | P35240 |
| HOMO SAPIENS | 3 | 1484 | 4zrj | A | 301 | ILE | p.Ile301Met | P35240 |
| HOMO SAPIENS | 4 | 40923 | 2enq | A | 68 | ILE | p.Ile391Met | P42336 |
| HOMO SAPIENS | 1 | 6179 | 5ofv | A | 125 | THR | p.Thr125Met | O43175 |
| HOMO SAPIENS | 0 | 6179 | 5ofv | A | 135 | ARG | p.Arg135Trp | O43175 |

| | | | | | | | | |
|---|---|---|---|---|---|---|---|---|
| HOMO SAPIENS | 1 | 6179 | 5ofv | A | 163 | ARG | p.Arg163Gln | O43175 |
| HOMO SAPIENS | 9 | 6179 | 5ofv | A | 261 | VAL | p.Val261Met | O43175 |
| HOMO SAPIENS | 7 | 34118 | 2i5o | A | 24 | MET | p.Met647Leu | Q9Y253 |
| HOMO SAPIENS | 8 | 3234 | 5c84 | A | 311 | LYS | p.Lys311Asn | P98170 |
| HOMO SAPIENS | 3 | 3234 | 5c84 | A | 321 | ALA | p.Ala321Gly | P98170 |
| HOMO SAPIENS | -1 | 15129 | 1zy8 | K | 1 | GLY | p.Gly54Ser | O00330 |
| HOMO SAPIENS | -1 | 15129 | 1zy8 | K | 48 | THR | p.Thr101Ala | O00330 |
| HOMO SAPIENS | -1 | 15129 | 1zy8 | K | 105 | GLU | p.Glu158Asp | O00330 |
| HOMO SAPIENS | 8 | 15129 | 1zy8 | K | 136 | ARG | p.Arg189His | O00330 |
| HOMO SAPIENS | -1 | 15129 | 1zy8 | K | 195 | GLN | p.Gln248Ter | O00330 |
| HOMO SAPIENS | 8 | 202 | 4bzb | A | 123 | HIS | p.His123Pro | Q9Y3Z3 |
| HOMO SAPIENS | 4 | 202 | 4bzb | A | 145 | ARG | p.Arg145Ter | Q9Y3Z3 |
| HOMO SAPIENS | 6 | 202 | 4bzb | A | 164 | ARG | p.Arg164Ter | Q9Y3Z3 |
| HOMO SAPIENS | 3 | 202 | 4bzb | A | 201 | ILE | p.Ile201Asn | Q9Y3Z3 |
| HOMO SAPIENS | 0 | 202 | 4bzb | A | 209 | GLY | p.Gly209Ser | Q9Y3Z3 |
| HOMO SAPIENS | 0 | 202 | 4bzb | A | 254 | MET | p.Met254Val | Q9Y3Z3 |
| HOMO SAPIENS | 5 | 202 | 4bzb | A | 531 | ARG | p.Arg531Ser | Q9Y3Z3 |
| HOMO SAPIENS | 6 | 202 | 4bzb | A | 548 | GLN | p.Gln548Ter | Q9Y3Z3 |
| HOMO SAPIENS | 3 | 30947 | 5lgd | A | 90 | PRO | p.Pro90Ser | P16671 |
| HOMO SAPIENS | 6 | 30947 | 5lgd | A | 254 | PHE | p.Phe254Leu | P16671 |
| HOMO SAPIENS | 0 | 30947 | 5lgd | A | 413 | ILE | p.Ile413Leu | P16671 |
| HOMO SAPIENS | 2 | 3906 | 5jtc | A | 735 | ARG | p.Arg735Trp | Q12797 |
| HOMO SAPIENS | 1 | 45 | 5uyx | C | 146 | GLU | p.Glu146Val | P48643 |
| HOMO SAPIENS | 1 | 45 | 5uyx | C | 147 | HIS | p.His147Arg | P48643 |
| HOMO SAPIENS | 0 | 45 | 5uyx | C | 362 | ILE | p.Ile362Met | P48643 |
| HOMO SAPIENS | 8 | 39596 | 1mhn | A | 95 | GLY | p.Gly95Arg | Q16637 |
| HOMO SAPIENS | 4 | 39596 | 1mhn | A | 111 | ALA | p.Ala111Gly | Q16637 |
| HOMO SAPIENS | 9 | 39596 | 1mhn | A | 116 | ILE | p.Ile116Phe | Q16637 |
| HOMO SAPIENS | 1 | 39596 | 1mhn | A | 136 | GLN | p.Gln136Glu | Q16637 |
| HOMO SAPIENS | 1 | 6408 | 5xjy | A | 85 | PRO | p.Pro85Leu | O95477 |
| HOMO SAPIENS | 0 | 6408 | 5xjy | A | 219 | ARG | p.Arg219Lys | O95477 |
| HOMO SAPIENS | 0 | 6408 | 5xjy | A | 230 | ARG | p.Arg230Cys | O95477 |
| HOMO SAPIENS | 1 | 6408 | 5xjy | A | 281 | MET | p.Met281Thr | O95477 |
| HOMO SAPIENS | -1 | 6408 | 5xjy | A | 343 | ALA | p.Ala343Gly | O95477 |
| HOMO SAPIENS | 1 | 6408 | 5xjy | A | 399 | VAL | p.Val399Ala | O95477 |
| HOMO SAPIENS | 1 | 6408 | 5xjy | A | 496 | ARG | p.Arg496Trp | O95477 |
| HOMO SAPIENS | 1 | 6408 | 5xjy | A | 590 | TRP | p.Trp590Ser | O95477 |
| HOMO SAPIENS | 1 | 6408 | 5xjy | A | 638 | ARG | p.Arg638Gln | O95477 |
| HOMO SAPIENS | 4 | 6408 | 5xjy | A | 697 | ALA | p.Ala697Thr | O95477 |
| HOMO SAPIENS | 1 | 6408 | 5xjy | A | 771 | VAL | p.Val771Met | O95477 |
| HOMO SAPIENS | 1 | 6408 | 5xjy | A | 774 | THR | p.Thr774Pro | O95477 |
| HOMO SAPIENS | 1 | 6408 | 5xjy | A | 776 | LYS | p.Lys776Asn | O95477 |
| HOMO SAPIENS | 0 | 6408 | 5xjy | A | 825 | VAL | p.Val825Ile | O95477 |
| HOMO SAPIENS | -1 | 6408 | 5xjy | A | 868 | GLU | p.Glu868Lys | O95477 |
| HOMO SAPIENS | -1 | 6408 | 5xjy | A | 883 | ILE | p.Ile883Met | O95477 |
| HOMO SAPIENS | 0 | 6408 | 5xjy | A | 935 | ASN | p.Asn935Ser | O95477 |
| HOMO SAPIENS | 0 | 6408 | 5xjy | A | 935 | ASN | p.Asn935His | O95477 |
| HOMO SAPIENS | 0 | 6408 | 5xjy | A | 937 | ALA | p.Ala937Val | O95477 |
| HOMO SAPIENS | 0 | 6408 | 5xjy | A | 1018 | ASP | p.Asp1018Gly | O95477 |
| HOMO SAPIENS | 0 | 6408 | 5xjy | A | 1096 | VAL | p.Val1096Ile | O95477 |
| HOMO SAPIENS | 0 | 6408 | 5xjy | A | 1099 | ASP | p.Asp1099Tyr | O95477 |
| HOMO SAPIENS | 0 | 6408 | 5xjy | A | 1172 | GLU | p.Glu1172Asp | O95477 |
| HOMO SAPIENS | 0 | 6408 | 5xjy | A | 1181 | SER | p.Ser1181Phe | O95477 |
| HOMO SAPIENS | 0 | 6408 | 5xjy | A | 1182 | ALA | p.Ala1182Thr | O95477 |

| | | | | | | | | |
|---|---|---|---|---|---|---|---|---|
| HOMO SAPIENS | -1 | 6408 | 5xjy | A | 1289 | ASP | p.Asp1289Asn | O95477 |
| HOMO SAPIENS | 0 | 6408 | 5xjy | A | 1399 | THR | p.Thr1399Ser | O95477 |
| HOMO SAPIENS | 1 | 6408 | 5xjy | A | 1408 | LEU | p.Leu1408Phe | O95477 |
| HOMO SAPIENS | 0 | 6408 | 5xjy | A | 1587 | LYS | p.Lys1587Arg | O95477 |
| HOMO SAPIENS | 1 | 6408 | 5xjy | A | 1674 | VAL | p.Val1674Ile | O95477 |
| HOMO SAPIENS | 1 | 6408 | 5xjy | A | 1680 | ARG | p.Arg1680Trp | O95477 |
| HOMO SAPIENS | 1 | 6408 | 5xjy | A | 1680 | ARG | p.Arg1680Gln | O95477 |
| HOMO SAPIENS | 0 | 6408 | 5xjy | A | 1800 | ASN | p.Asn1800His | O95477 |
| HOMO SAPIENS | -1 | 6408 | 5xjy | A | 1925 | ARG | p.Arg1925Gln | O95477 |
| HOMO SAPIENS | 0 | 6408 | 5xjy | A | 2009 | PHE | p.Phe2009Ser | O95477 |
| HOMO SAPIENS | -1 | 6408 | 5xjy | A | 2243 | ASP | p.Asp2243Glu | O95477 |
| HOMO SAPIENS | 4 | 23760 | 2mut | A | 231 | LEU | p.Phe231Leu | P07992 |
| HOMO SAPIENS | 6 | 1460 | 2hre | B | 96 | ARG | p.Arg96Gln | P22830 |
| HOMO SAPIENS | 8 | 1460 | 2hre | B | 129 | GLY | p.Gly129Arg | P22830 |
| HOMO SAPIENS | 9 | 1460 | 2hre | B | 172 | GLU | p.Glu172Lys | P22830 |
| HOMO SAPIENS | 4 | 1460 | 2hre | B | 285 | GLN | p.Gln285Arg | P22830 |
| HOMO SAPIENS | 5 | 1460 | 2hre | B | 362 | VAL | p.Val362Gly | P22830 |
| HOMO SAPIENS | 6 | 1460 | 2hre | B | 411 | CYS | p.Cys411Gly | P22830 |
| HOMO SAPIENS | 6 | 1460 | 2hre | B | 417 | PHE | p.Phe417Ser | P22830 |
| HOMO SAPIENS | 0 | 2210 | 5bug | A | 19 | ASP | p.Asp19Asn | P60484 |
| HOMO SAPIENS | 0 | 2210 | 5bug | A | 35 | MET | p.Met35Arg | P60484 |
| HOMO SAPIENS | 1 | 2210 | 5bug | A | 61 | HIS | p.His61Asp | P60484 |
| HOMO SAPIENS | 0 | 2210 | 5bug | A | 68 | TYR | p.Tyr68His | P60484 |
| HOMO SAPIENS | 8 | 2210 | 5bug | A | 70 | LEU | p.Leu70Pro | P60484 |
| HOMO SAPIENS | 6 | 2210 | 5bug | A | 93 | HIS | p.His93Arg | P60484 |
| HOMO SAPIENS | 1 | 2210 | 5bug | A | 112 | LEU | p.Leu112Pro | P60484 |
| HOMO SAPIENS | 0 | 2210 | 5bug | A | 121 | ALA | p.Ala121Gly | P60484 |
| HOMO SAPIENS | 3 | 2210 | 5bug | A | 123 | HIS | p.His123Arg | P60484 |
| HOMO SAPIENS | 7 | 2210 | 5bug | A | 124 | CYS | p.Cys124Arg | P60484 |
| HOMO SAPIENS | 9 | 2210 | 5bug | A | 129 | GLY | p.Gly129Glu | P60484 |
| HOMO SAPIENS | 9 | 2210 | 5bug | A | 130 | ARG | p.Arg130Gln | P60484 |
| HOMO SAPIENS | 9 | 2210 | 5bug | A | 130 | ARG | p.Arg130Gly | P60484 |
| HOMO SAPIENS | 0 | 2210 | 5bug | A | 132 | GLY | p.Gly132Val | P60484 |
| HOMO SAPIENS | 0 | 2210 | 5bug | A | 170 | SER | p.Ser170Arg | P60484 |
| HOMO SAPIENS | 1 | 2210 | 5bug | A | 173 | ARG | p.Arg173His | P60484 |
| HOMO SAPIENS | 0 | 2210 | 5bug | A | 217 | VAL | p.Val217Ile | P60484 |
| HOMO SAPIENS | 0 | 2210 | 5bug | A | 241 | PHE | p.Phe241Ser | P60484 |
| HOMO SAPIENS | 0 | 2210 | 5bug | A | 250 | CYS | p.Cys250Ter | P60484 |
| HOMO SAPIENS | 1 | 2210 | 5bug | A | 252 | ASP | p.Asp252Gly | P60484 |
| HOMO SAPIENS | 3 | 123 | 2ibu | A | 93 | ASN | p.Asn93Ser | P24752 |
| HOMO SAPIENS | 0 | 123 | 2ibu | A | 145 | GLN | p.Gln145Glu | P24752 |
| HOMO SAPIENS | 0 | 123 | 2ibu | A | 152 | GLY | p.Gly152Ala | P24752 |
| HOMO SAPIENS | 2 | 123 | 2ibu | A | 158 | ASN | p.Asn158Asp | P24752 |
| HOMO SAPIENS | 2 | 123 | 2ibu | A | 158 | ASN | p.Asn158Ser | P24752 |
| HOMO SAPIENS | 5 | 123 | 2ibu | A | 183 | GLY | p.Gly183Arg | P24752 |
| HOMO SAPIENS | 3 | 123 | 2ibu | A | 272 | GLN | p.Gln272Ter | P24752 |
| HOMO SAPIENS | 3 | 123 | 2ibu | A | 312 | ILE | p.Ile312Thr | P24752 |
| HOMO SAPIENS | 1 | 123 | 2ibu | A | 333 | ALA | p.Ala333Pro | P24752 |
| HOMO SAPIENS | 5 | 123 | 2ibu | A | 379 | GLY | p.Gly379Val | P24752 |
| HOMO SAPIENS | 0 | 123 | 2ibu | A | 380 | ALA | p.Ala380Thr | P24752 |
| HOMO SAPIENS | 2 | 369 | 5nrw | A | 1175 | TYR | p.Tyr1175Cys | Q92793 |
| HOMO SAPIENS | 2 | 830 | 2vc2 | A | 116 | LEU | p.Leu147Val | P08514 |
| HOMO SAPIENS | 0 | 830 | 2vc2 | A | 183 | LEU | p.Leu214Pro | P08514 |
| HOMO SAPIENS | 4 | 830 | 2vc2 | A | 242 | GLY | p.Gly273Asp | P08514 |

| | | | | | | | | |
|---|---|---|---|---|---|---|---|---|
| HOMO SAPIENS | 6 | 830 | 2vc2 | A | 324 | GLU | p.Glu355Lys | P08514 |
| HOMO SAPIENS | -1 | 10353 | 5ejc | E | 989 | ALA | p.Ala989Ser | Q92574 |
| HOMO SAPIENS | 0 | 15448 | 3lgg | A | 21 | GLY | p.Gly47Arg | Q9NZK5 |
| HOMO SAPIENS | 0 | 15448 | 3lgg | A | 21 | GLY | p.Gly47Val | Q9NZK5 |
| HOMO SAPIENS | 9 | 15448 | 3lgg | A | 86 | HIS | p.His112Gln | Q9NZK5 |
| HOMO SAPIENS | 0 | 15448 | 3lgg | A | 143 | ARG | p.Arg169Gln | Q9NZK5 |
| HOMO SAPIENS | 3 | 15448 | 3lgg | A | 221 | ALA | p.Ala247Val | Q9NZK5 |
| HOMO SAPIENS | 0 | 15448 | 3lgg | A | 225 | PRO | p.Pro251Leu | Q9NZK5 |
| HOMO SAPIENS | 0 | 15448 | 3lgg | A | 283 | MET | p.Met309Ile | Q9NZK5 |
| HOMO SAPIENS | 2 | 15448 | 3lgg | A | 334 | THR | p.Thr360Ala | Q9NZK5 |
| HOMO SAPIENS | 7 | 15448 | 3lgg | A | 357 | GLY | p.Gly383Ser | Q9NZK5 |
| HOMO SAPIENS | 0 | 15448 | 3lgg | A | 427 | TYR | p.Tyr453Cys | Q9NZK5 |
| HOMO SAPIENS | 0 | 872 | 2y3i | A | 87 | LEU | p.Leu88Pro | P00558 |
| HOMO SAPIENS | 1 | 872 | 2y3i | A | 163 | ASP | p.Asp164Val | P00558 |
| HOMO SAPIENS | 4 | 872 | 2y3i | A | 205 | ARG | p.Arg206Pro | P00558 |
| HOMO SAPIENS | 4 | 872 | 2y3i | A | 267 | ASP | p.Asp268Asn | P00558 |
| HOMO SAPIENS | 8 | 872 | 2y3i | A | 284 | ASP | p.Asp285Val | P00558 |
| HOMO SAPIENS | 4 | 8 | 5jco | B | 188 | ILE | p.Ile188Leu | Q71U36 |
| HOMO SAPIENS | 7 | 8 | 5jco | B | 263 | PRO | p.Pro263Thr | Q71U36 |
| HOMO SAPIENS | 6 | 8 | 5jco | B | 264 | ARG | p.Arg264Cys | Q71U36 |
| HOMO SAPIENS | 7 | 8 | 5jco | B | 402 | ARG | p.Arg402His | Q71U36 |
| HOMO SAPIENS | 7 | 8 | 5jco | B | 419 | SER | p.Ser419Leu | Q71U36 |
| HOMO SAPIENS | -1 | 880 | 4nwf | A | 2 | THR | p.Thr2Ile | Q06124 |
| HOMO SAPIENS | 2 | 880 | 4nwf | A | 58 | ASN | p.Asn58Ser | Q06124 |
| HOMO SAPIENS | 4 | 880 | 4nwf | A | 61 | ASP | p.Asp61Gly | Q06124 |
| HOMO SAPIENS | 2 | 880 | 4nwf | A | 62 | TYR | p.Tyr62Asp | Q06124 |
| HOMO SAPIENS | 3 | 880 | 4nwf | A | 63 | TYR | p.Tyr63Cys | Q06124 |
| HOMO SAPIENS | 2 | 880 | 4nwf | A | 69 | GLU | p.Glu69Gln | Q06124 |
| HOMO SAPIENS | 2 | 880 | 4nwf | A | 72 | ALA | p.Ala72Gly | Q06124 |
| HOMO SAPIENS | 2 | 880 | 4nwf | A | 72 | ALA | p.Ala72Ser | Q06124 |
| HOMO SAPIENS | 3 | 880 | 4nwf | A | 73 | THR | p.Thr73Ile | Q06124 |
| HOMO SAPIENS | 5 | 880 | 4nwf | A | 76 | GLU | p.Glu76Ala | Q06124 |
| HOMO SAPIENS | 5 | 880 | 4nwf | A | 76 | GLU | p.Glu76Lys | Q06124 |
| HOMO SAPIENS | 3 | 880 | 4nwf | A | 79 | GLN | p.Gln79Arg | Q06124 |
| HOMO SAPIENS | 5 | 880 | 4nwf | A | 152 | ARG | p.Arg152His | Q06124 |
| HOMO SAPIENS | 2 | 880 | 4nwf | A | 268 | GLY | p.Gly268Cys | Q06124 |
| HOMO SAPIENS | 9 | 880 | 4nwf | A | 276 | LYS | p.Lys276Arg | Q06124 |
| HOMO SAPIENS | 9 | 880 | 4nwf | A | 279 | TYR | p.Tyr279Cys | Q06124 |
| HOMO SAPIENS | 5 | 880 | 4nwf | A | 285 | PHE | p.Phe285Ser | Q06124 |
| HOMO SAPIENS | 0 | 880 | 4nwf | A | 308 | ASP | p.Asn308Ser | Q06124 |
| HOMO SAPIENS | 0 | 880 | 4nwf | A | 308 | ASP | p.Asn308Asp | Q06124 |
| HOMO SAPIENS | 5 | 880 | 4nwf | A | 392 | ALA | p.Ala392Thr | Q06124 |
| HOMO SAPIENS | 1 | 880 | 4nwf | A | 411 | THR | p.Thr415Met | Q06124 |
| HOMO SAPIENS | 0 | 880 | 4nwf | A | 461 | ALA | p.Ala465Thr | Q06124 |
| HOMO SAPIENS | 0 | 880 | 4nwf | A | 464 | GLY | p.Gly468Ala | Q06124 |
| HOMO SAPIENS | 1 | 880 | 4nwf | A | 491 | PRO | p.Pro495Ser | Q06124 |
| HOMO SAPIENS | 1 | 880 | 4nwf | A | 502 | SER | p.Ser506Thr | Q06124 |
| HOMO SAPIENS | 8 | 880 | 4nwf | A | 510 | GLN | p.Gln514Pro | Q06124 |
| HOMO SAPIENS | 4 | 1872 | 3wf1 | D | 109 | ARG | p.Arg109Trp | P16278 |
| HOMO SAPIENS | 0 | 1872 | 3wf1 | D | 521 | ARG | p.Arg521Cys | P16278 |
| HOMO SAPIENS | 5 | 910 | 5xte | A | 45 | SER | p.Ser45Phe | O14949 |
| HOMO SAPIENS | 2 | 9017 | 5f0k | A | 382 | LYS | p.Lys382Arg | Q96QK1 |
| HOMO SAPIENS | 1 | 710 | 4lsj | A | 559 | ILE | p.Ile559Asn | P04150 |
| HOMO SAPIENS | 5 | 710 | 4lsj | A | 571 | VAL | p.Val571Ala | P04150 |

| | | | | | | | | |
|---|---|---|---|---|---|---|---|---|
| HOMO SAPIENS | 0 | 710 | 4lsj | A | 641 | ASP | p.Asp641Val | P04150 |
| HOMO SAPIENS | 1 | 710 | 4lsj | A | 678 | ASP | p.Asp678Glu | P04150 |
| HOMO SAPIENS | 2 | 710 | 4lsj | A | 679 | GLY | p.Gly679Ser | P04150 |
| HOMO SAPIENS | 1 | 710 | 4lsj | A | 737 | PHE | p.Phe737Leu | P04150 |
| HOMO SAPIENS | 1 | 710 | 4lsj | A | 747 | ILE | p.Ile747Met | P04150 |
| HOMO SAPIENS | 1 | 710 | 4lsj | A | 753 | LEU | p.Leu753Phe | P04150 |
| HOMO SAPIENS | 1 | 710 | 4lsj | A | 773 | LEU | p.Leu773Pro | P04150 |
| HOMO SAPIENS | 1 | 2145 | 1av1 | B | 71 | PHE | p.Phe95Tyr | P02647 |
| HOMO SAPIENS | 1 | 2145 | 1av1 | B | 128 | GLU | p.Glu152Lys | P02647 |
| HOMO SAPIENS | 1 | 2145 | 1av1 | B | 136 | GLU | p.Glu160Lys | P02647 |
| HOMO SAPIENS | 2 | 2145 | 1av1 | B | 143 | PRO | p.Pro167Arg | P02647 |
| HOMO SAPIENS | 6 | 2145 | 1av1 | B | 164 | ALA | p.Ala188Ser | P02647 |
| HOMO SAPIENS | 7 | 2145 | 1av1 | B | 173 | ARG | p.Arg197Cys | P02647 |
| HOMO SAPIENS | 2 | 2145 | 1av1 | B | 198 | GLU | p.Glu222Lys | P02647 |
| HOMO SAPIENS | -1 | 3432 | 2beu | A | 1 | SER | p.Ser46Ter | P12694 |
| HOMO SAPIENS | 5 | 3432 | 2beu | A | 79 | ARG | p.Arg124Trp | P12694 |
| HOMO SAPIENS | 7 | 3432 | 2beu | A | 114 | ARG | p.Arg159Gln | P12694 |
| HOMO SAPIENS | 6 | 3432 | 2beu | A | 166 | THR | p.Thr211Met | P12694 |
| HOMO SAPIENS | 4 | 3432 | 2beu | A | 175 | ALA | p.Ala220Val | P12694 |
| HOMO SAPIENS | 5 | 3432 | 2beu | A | 204 | GLY | p.Gly249Ser | P12694 |
| HOMO SAPIENS | 4 | 3432 | 2beu | A | 209 | ALA | p.Ala254Asp | P12694 |
| HOMO SAPIENS | 6 | 3432 | 2beu | A | 220 | ARG | p.Arg265Trp | P12694 |
| HOMO SAPIENS | 6 | 3432 | 2beu | A | 234 | TYR | p.Tyr279Cys | P12694 |
| HOMO SAPIENS | 5 | 3432 | 2beu | A | 240 | ALA | p.Ala285Pro | P12694 |
| HOMO SAPIENS | 4 | 3432 | 2beu | A | 245 | GLY | p.Gly290Arg | P12694 |
| HOMO SAPIENS | 5 | 3432 | 2beu | A | 269 | ARG | p.Arg314Ter | P12694 |
| HOMO SAPIENS | 3 | 3432 | 2beu | A | 301 | ARG | p.Arg346Cys | P12694 |
| HOMO SAPIENS | 5 | 3432 | 2beu | A | 359 | ASN | p.Asn404Ser | P12694 |
| HOMO SAPIENS | 4 | 3432 | 2beu | A | 393 | TYR | p.Tyr438Asn | P12694 |
| HOMO SAPIENS | 8 | 4541 | 1rr8 | C | 533 | ASP | p.Asp533Gly | P11387 |
| HOMO SAPIENS | 2 | 3247 | 1tzn | d | 1105 | GLY | p.Gly105Asp | P58335 |
| HOMO SAPIENS | 3 | 3247 | 1tzn | d | 1111 | ARG | p.Arg111Cys | P58335 |
| HOMO SAPIENS | 2 | 3247 | 1tzn | d | 1189 | ILE | p.Ile189Thr | P58335 |
| HOMO SAPIENS | 9 | 467 | 1suv | C | 58 | ASP | p.Asp77Asn | P02787 |
| HOMO SAPIENS | 2 | 467 | 1suv | C | 277 | ASP | p.Asp296Gly | P02787 |
| HOMO SAPIENS | 9 | 6485 | 1xwd | B | 51 | CYS | p.Cys69Gly | P01225 |
| HOMO SAPIENS | 1 | 694 | 1wuu | A | 28 | PRO | p.Pro28Thr | P51570 |
| HOMO SAPIENS | 0 | 694 | 1wuu | A | 32 | VAL | p.Val32Met | P51570 |
| HOMO SAPIENS | 0 | 694 | 1wuu | A | 198 | ALA | p.Ala198Val | P51570 |
| HOMO SAPIENS | 0 | 143 | 4nkx | A | 93 | PHE | p.Phe93Cys | P05093 |
| HOMO SAPIENS | 3 | 143 | 4nkx | A | 96 | ARG | p.Arg96Trp | P05093 |
| HOMO SAPIENS | 3 | 143 | 4nkx | A | 96 | ARG | p.Arg96Gln | P05093 |
| HOMO SAPIENS | 0 | 143 | 4nkx | A | 106 | SER | p.Ser106Pro | P05093 |
| HOMO SAPIENS | 1 | 143 | 4nkx | A | 114 | PHE | p.Phe114Val | P05093 |
| HOMO SAPIENS | 1 | 143 | 4nkx | A | 116 | ASP | p.Asp116Val | P05093 |
| HOMO SAPIENS | 3 | 143 | 4nkx | A | 125 | ARG | p.Arg125Gln | P05093 |
| HOMO SAPIENS | 0 | 143 | 4nkx | A | 201 | TYR | p.Tyr201Asp | P05093 |
| HOMO SAPIENS | 0 | 143 | 4nkx | A | 239 | ARG | p.Arg239Ter | P05093 |
| HOMO SAPIENS | 0 | 143 | 4nkx | A | 329 | TYR | p.Tyr329Asp | P05093 |
| HOMO SAPIENS | 1 | 143 | 4nkx | A | 342 | PRO | p.Pro342Thr | P05093 |
| HOMO SAPIENS | 2 | 143 | 4nkx | A | 347 | ARG | p.Arg347His | P05093 |
| HOMO SAPIENS | 2 | 143 | 4nkx | A | 347 | ARG | p.Arg347Cys | P05093 |
| HOMO SAPIENS | 1 | 143 | 4nkx | A | 358 | ARG | p.Arg358Gln | P05093 |
| HOMO SAPIENS | 0 | 143 | 4nkx | A | 362 | ARG | p.Arg362Cys | P05093 |

| | | | | | | | | |
|---|---|---|---|---|---|---|---|---|
| HOMO SAPIENS | 1 | 143 | 4nkx | A | 406 | TRP | p.Trp406Arg | P05093 |
| HOMO SAPIENS | 0 | 143 | 4nkx | A | 428 | PRO | p.Pro428Leu | P05093 |
| HOMO SAPIENS | 6 | 1441 | 1uuh | A | 46 | ARG | p.Arg46Gly | P16070 |
| HOMO SAPIENS | 3 | 82 | 4wlv | D | 37 | GLY | p.Gly37Arg | P40926 |
| HOMO SAPIENS | 0 | 82 | 4wlv | D | 133 | PRO | p.Pro133Leu | P40926 |
| HOMO SAPIENS | 2 | 392 | 5xtd | A | 56 | SER | p.Ser56Pro | P49821 |
| HOMO SAPIENS | 2 | 392 | 5xtd | A | 59 | ARG | p.Arg59Ter | P49821 |
| HOMO SAPIENS | 2 | 392 | 5xtd | A | 74 | ASP | p.Asp74Asn | P49821 |
| HOMO SAPIENS | 4 | 392 | 5xtd | A | 116 | ASN | p.Asn116Lys | P49821 |
| HOMO SAPIENS | 4 | 392 | 5xtd | A | 117 | ALA | p.Ala117Thr | P49821 |
| HOMO SAPIENS | 5 | 392 | 5xtd | A | 122 | PRO | p.Pro122Leu | P49821 |
| HOMO SAPIENS | 3 | 392 | 5xtd | A | 160 | GLY | p.Gly160Glu | P49821 |
| HOMO SAPIENS | 4 | 392 | 5xtd | A | 214 | GLU | p.Glu214Lys | P49821 |
| HOMO SAPIENS | 2 | 392 | 5xtd | A | 357 | MET | p.Met357Thr | P49821 |
| HOMO SAPIENS | 4 | 392 | 5xtd | A | 423 | THR | p.Thr423Met | P49821 |
| HOMO SAPIENS | 0 | 22593 | 3pdf | A | 8 | TYR | p.Tyr32Ter | P53634 |
| HOMO SAPIENS | 0 | 22593 | 3pdf | A | 15 | TRP | p.Trp39Ser | P53634 |
| HOMO SAPIENS | 0 | 22593 | 3pdf | A | 103 | HIS | p.His127Pro | P53634 |
| HOMO SAPIENS | 9 | 22593 | 3pdf | A | 228 | GLN | p.Gln252Leu | P53634 |
| HOMO SAPIENS | 3 | 22593 | 3pdf | A | 262 | GLN | p.Gln286Arg | P53634 |
| HOMO SAPIENS | 4 | 22593 | 3pdf | A | 277 | GLY | p.Gly301Ser | P53634 |
| HOMO SAPIENS | 0 | 22593 | 3pdf | A | 323 | TYR | p.Tyr347Cys | P53634 |
| HOMO SAPIENS | 5 | 3461 | 4c2a | A | 1306 | GLN | p.Arg1306Trp | P04275 |
| HOMO SAPIENS | 5 | 3461 | 4c2a | A | 1374 | ARG | p.Arg1374Cys | P04275 |
| HOMO SAPIENS | 5 | 3461 | 4c2a | A | 1379 | ARG | p.Arg1379Cys | P04275 |
| HOMO SAPIENS | 4 | 3461 | 4c2a | A | 1380 | ILE | p.Ile1380Val | P04275 |
| HOMO SAPIENS | 0 | 3461 | 4c2a | A | 1381 | ALA | p.Thr1381Ala | P04275 |
| HOMO SAPIENS | 5 | 3461 | 4c2a | A | 1399 | ARG | p.Arg1399His | P04275 |
| HOMO SAPIENS | 5 | 3461 | 4c2a | A | 1435 | ASN | p.Asn1435Ser | P04275 |
| HOMO SAPIENS | 0 | 4533 | 5uj7 | C | 174 | TYR | p.Tyr174Cys | O43929 |
| HOMO SAPIENS | 0 | 4533 | 5uj7 | C | 202 | LEU | p.Leu202Val | O43929 |
| HOMO SAPIENS | 0 | 1374 | 3j07 | O | 39 | PRO | p.Pro39Leu | P02511 |
| HOMO SAPIENS | 0 | 1374 | 3j07 | O | 107 | ARG | p.Arg107His | P02511 |
| HOMO SAPIENS | 0 | 1374 | 3j07 | O | 157 | ARG | p.Arg157His | P02511 |
| HOMO SAPIENS | -1 | 942 | 1s9i | B | 57 | PHE | p.Phe57Cys | P36507 |
| HOMO SAPIENS | -1 | 942 | 1s9i | B | 57 | PHE | p.Phe57Val | P36507 |
| HOMO SAPIENS | 2 | 942 | 1s9i | B | 128 | PRO | p.Pro128Gln | P36507 |
| HOMO SAPIENS | 1 | 942 | 1s9i | B | 134 | TYR | p.Tyr134His | P36507 |
| HOMO SAPIENS | 1 | 942 | 1s9i | B | 188 | HIS | p.His188Gln | P36507 |
| HOMO SAPIENS | 0 | 942 | 1s9i | B | 262 | VAL | p.Val262Phe | P36507 |
| HOMO SAPIENS | 0 | 942 | 1s9i | B | 263 | GLY | p.Gly263Arg | P36507 |
| HOMO SAPIENS | -1 | 942 | 1s9i | B | 298 | PRO | p.Pro298Leu | P36507 |
| HOMO SAPIENS | -1 | 942 | 1s9i | B | 305 | VAL | p.Val305Ile | P36507 |
| HOMO SAPIENS | 0 | 3245 | 1fbv | A | 367 | GLN | p.Gln367Pro | P22681 |
| HOMO SAPIENS | 0 | 3245 | 1fbv | A | 371 | TYR | p.Tyr371His | P22681 |
| HOMO SAPIENS | 1 | 3245 | 1fbv | A | 382 | LYS | p.Lys382Glu | P22681 |
| HOMO SAPIENS | 2 | 3245 | 1fbv | A | 390 | ASP | p.Asp390Tyr | P22681 |
| HOMO SAPIENS | 9 | 3245 | 1fbv | A | 396 | CYS | p.Cys396Arg | P22681 |
| HOMO SAPIENS | 7 | 3245 | 1fbv | A | 420 | ARG | p.Arg420Gln | P22681 |
| HOMO SAPIENS | 1 | 3245 | 1fbv | A | 429 | ILE | p.Ile429Met | P22681 |
| HOMO SAPIENS | 0 | 2604 | 2pbd | P | 111 | LEU | p.Leu112Met | P07737 |
| HOMO SAPIENS | 3 | 2604 | 2pbd | P | 116 | GLU | p.Glu117Gly | P07737 |
| HOMO SAPIENS | 4 | 2013 | 3b7w | A | 472 | ARG | p.Arg472Gly | Q08AH3 |
| HOMO SAPIENS | 0 | 24346 | 3sh5 | A | 85 | ARG | p.Arg4281His | P98160 |

| | | | | | | | | |
|---|---|---|---|---|---|---|---|---|
| HOMO SAPIENS | 0 | 24346 | 3sh5 | A | 122 | ARG | p.Arg4318Trp | P98160 |
| HOMO SAPIENS | 0 | 24346 | 3sh5 | A | 135 | SER | p.Ser4331Asn | P98160 |
| HOMO SAPIENS | 0 | 24346 | 3sh5 | A | 136 | VAL | p.Val4332Ile | P98160 |
| HOMO SAPIENS | 0 | 707 | 4p2g | A | 26 | MET | p.Met26Ile | Q99497 |
| HOMO SAPIENS | 5 | 707 | 4p2g | A | 39 | ALA | p.Ala39Ser | Q99497 |
| HOMO SAPIENS | 5 | 707 | 4p2g | A | 64 | GLU | p.Glu64Asp | Q99497 |
| HOMO SAPIENS | 4 | 707 | 4p2g | A | 149 | ASP | p.Asp149Ala | Q99497 |
| HOMO SAPIENS | 6 | 707 | 4p2g | A | 163 | GLU | p.Glu163Lys | Q99497 |
| HOMO SAPIENS | 6 | 707 | 4p2g | A | 166 | LEU | p.Leu166Pro | Q99497 |
| HOMO SAPIENS | 3 | 5121 | 4xss | E | 31 | GLY | p.Gly58Arg | P06213 |
| HOMO SAPIENS | 2 | 5121 | 4xss | E | 119 | ILE | p.Ile146Met | P06213 |
| HOMO SAPIENS | 8 | 5121 | 4xss | E | 209 | HIS | p.His236Arg | P06213 |
| HOMO SAPIENS | 2 | 5121 | 4xss | E | 234 | ASP | p.Asp261Glu | P06213 |
| HOMO SAPIENS | 1 | 5121 | 4xss | E | 276 | GLN | p.Gln303His | P06213 |
| HOMO SAPIENS | 1 | 27790 | 5erp | B | 140 | THR | p.Thr275Met | Q02487 |
| HOMO SAPIENS | 0 | 27790 | 5erp | B | 150 | ILE | p.Ile285Thr | Q02487 |
| HOMO SAPIENS | 1 | 27790 | 5erp | B | 205 | THR | p.Thr340Ala | Q02487 |
| HOMO SAPIENS | 1 | 27790 | 5erp | B | 223 | THR | p.Thr358Ile | Q02487 |
| HOMO SAPIENS | 0 | 27790 | 5erp | B | 301 | GLY | p.Gly436Ala | Q02487 |
| HOMO SAPIENS | -1 | 27790 | 5erp | B | 315 | ARG | p.Arg450Ser | Q02487 |
| HOMO SAPIENS | 0 | 27790 | 5erp | B | 369 | SER | p.Ser504Gly | Q02487 |
| HOMO SAPIENS | 0 | 27790 | 5erp | B | 385 | ILE | p.Ile520Thr | Q02487 |
| HOMO SAPIENS | 0 | 27790 | 5erp | B | 461 | ALA | p.Ala596Val | Q02487 |
| HOMO SAPIENS | 0 | 27790 | 5erp | B | 503 | GLN | p.Gln638His | Q02487 |
| HOMO SAPIENS | 5 | 6599 | 4nsc | A | 129 | ARG | p.Arg129Leu | Q9BPX6 |
| HOMO SAPIENS | -1 | 6599 | 4nsc | A | 183 | GLN | p.Gln183Ter | Q9BPX6 |
| HOMO SAPIENS | 0 | 20420 | 3ghm | A | 398 | ARG | p.Arg398His | Q76LX8 |
| HOMO SAPIENS | 0 | 20420 | 3ghm | A | 448 | GLN | p.Gln448Glu | Q76LX8 |
| HOMO SAPIENS | 1 | 20420 | 3ghm | A | 449 | GLN | p.Gln449Ter | Q76LX8 |
| HOMO SAPIENS | 0 | 20420 | 3ghm | A | 456 | GLN | p.Gln456His | Q76LX8 |
| HOMO SAPIENS | 1 | 20420 | 3ghm | A | 528 | ARG | p.Arg528Gly | Q76LX8 |
| HOMO SAPIENS | 0 | 20420 | 3ghm | A | 618 | PRO | p.Pro618Ala | Q76LX8 |
| HOMO SAPIENS | 0 | 20420 | 3ghm | A | 625 | ARG | p.Arg625His | Q76LX8 |
| HOMO SAPIENS | 0 | 275 | 4rmz | A | 293 | GLN | p.Gln293Ter | Q9NWZ3 |
| HOMO SAPIENS | 0 | 275 | 4rmz | A | 428 | ALA | p.Ala428Thr | Q9NWZ3 |
| HOMO SAPIENS | 2 | 10458 | 3puf | B | 138 | LEU | p.Leu138Phe | Q5TBB1 |
| HOMO SAPIENS | 6 | 10458 | 3puf | B | 177 | ALA | p.Ala177Thr | Q5TBB1 |
| HOMO SAPIENS | -1 | 10458 | 3puf | B | 185 | VAL | p.Val185Gly | Q5TBB1 |
| HOMO SAPIENS | 0 | 26648 | 2n90 | A | 36 | ARG | p.Arg444Gln | P04629 |
| HOMO SAPIENS | 2 | 977 | 1dnw | D | 166 | ALA | p.Ala332Val | P05164 |
| HOMO SAPIENS | 4 | 977 | 1dnw | D | 333 | ARG | p.Arg499Cys | P05164 |
| HOMO SAPIENS | 1 | 977 | 1dnw | D | 403 | ARG | p.Arg569Trp | P05164 |
| HOMO SAPIENS | 0 | 25704 | 3ajb | A | 82 | GLN | p.Gln82Arg | P56589 |
| HOMO SAPIENS | 3 | 255 | 3t1p | A | 38 | TYR | p.Tyr62Ter | P01009 |
| HOMO SAPIENS | 3 | 255 | 3t1p | A | 39 | ARG | p.Arg63Cys | P01009 |
| HOMO SAPIENS | 0 | 255 | 3t1p | A | 101 | ARG | p.Arg125His | P01009 |
| HOMO SAPIENS | 0 | 255 | 3t1p | A | 212 | GLN | p.Gln236Arg | P01009 |
| HOMO SAPIENS | 0 | 255 | 3t1p | A | 213 | VAL | p.Val237Ala | P01009 |
| HOMO SAPIENS | 1 | 255 | 3t1p | A | 269 | HIS | p.His293Gln | P01009 |
| HOMO SAPIENS | 8 | 255 | 3t1p | A | 341 | ASP | p.Asp365Asn | P01009 |
| HOMO SAPIENS | 9 | 255 | 3t1p | A | 342 | GLU | p.Glu366Lys | P01009 |
| HOMO SAPIENS | 0 | 255 | 3t1p | A | 369 | PRO | p.Pro393Ser | P01009 |
| HOMO SAPIENS | 0 | 255 | 3t1p | A | 376 | GLU | p.Glu400Asp | P01009 |
| HOMO SAPIENS | -1 | 5175 | 3zxi | A | 35 | ALA | p.Ala35Asp | Q9Y2Z4 |

| | | | | | | | | | |
|---|---|---|---|---|---|---|---|---|---|
| HOMO SAPIENS | 0 | 5175 | 3zxi | A | | 52 | PHE | p.Phe52Leu | Q9Y2Z4 |
| HOMO SAPIENS | -1 | 5175 | 3zxi | A | | 68 | GLY | p.Gly68Ser | Q9Y2Z4 |
| HOMO SAPIENS | 0 | 5175 | 3zxi | A | | 179 | LYS | p.Lys179Gln | Q9Y2Z4 |
| HOMO SAPIENS | 0 | 5175 | 3zxi | A | | 191 | GLY | p.Gly191Val | Q9Y2Z4 |
| HOMO SAPIENS | 0 | 5175 | 3zxi | A | | 251 | ILE | p.Ile251Val | Q9Y2Z4 |
| HOMO SAPIENS | 2 | 5175 | 3zxi | A | | 295 | ASP | p.Asp295Asn | Q9Y2Z4 |
| HOMO SAPIENS | 0 | 5175 | 3zxi | A | | 311 | ASP | p.Asp311Glu | Q9Y2Z4 |
| HOMO SAPIENS | 0 | 5175 | 3zxi | A | | 360 | ARG | p.Arg360Ter | Q9Y2Z4 |
| HOMO SAPIENS | 1 | 3016 | 3exg | | 3 | 4 | ASP | p.Asp33His | P08559 |
| HOMO SAPIENS | 1 | 3016 | 3exg | | 3 | 176 | PHE | p.Phe205Leu | P08559 |
| HOMO SAPIENS | 1 | 3016 | 3exg | | 3 | 214 | TYR | p.Tyr243Asn | P08559 |
| HOMO SAPIENS | 1 | 3016 | 3exg | | 3 | 229 | ASP | p.Asp258Ala | P08559 |
| HOMO SAPIENS | 0 | 3016 | 3exg | | 3 | 233 | VAL | p.Val262Ile | P08559 |
| HOMO SAPIENS | 1 | 3016 | 3exg | | 3 | 234 | ARG | p.Arg263Gly | P08559 |
| HOMO SAPIENS | 0 | 3016 | 3exg | | 3 | 253 | MET | p.Met282Leu | P08559 |
| HOMO SAPIENS | 7 | 3016 | 3exg | | 3 | 259 | ARG | p.Arg288His | P08559 |
| HOMO SAPIENS | -1 | 3016 | 3exg | | 3 | 273 | ARG | p.Arg302Cys | P08559 |
| HOMO SAPIENS | 2 | 3016 | 3exg | | 3 | 286 | ASP | p.Asp315Asn | P08559 |
| HOMO SAPIENS | 1 | 3016 | 3exg | | 3 | 349 | ARG | p.Arg378His | P08559 |
| HOMO SAPIENS | 5 | 6729 | 4a0l | E | | 572 | ARG | p.Arg572Cys | Q13620 |
| HOMO SAPIENS | 0 | 1067 | 4kyo | A | | 9 | THR | p.Thr9Asn | P21549 |
| HOMO SAPIENS | 0 | 1067 | 4kyo | A | | 11 | PRO | p.Pro11Leu | P21549 |
| HOMO SAPIENS | 0 | 1067 | 4kyo | A | | 22 | ASN | p.Asn22Ser | P21549 |
| HOMO SAPIENS | 4 | 1067 | 4kyo | A | | 25 | LEU | p.Leu25Arg | P21549 |
| HOMO SAPIENS | 7 | 1067 | 4kyo | A | | 36 | ARG | p.Arg36His | P21549 |
| HOMO SAPIENS | 3 | 1067 | 4kyo | A | | 41 | GLY | p.Gly41Arg | P21549 |
| HOMO SAPIENS | 0 | 1067 | 4kyo | A | | 42 | GLY | p.Gly42Glu | P21549 |
| HOMO SAPIENS | 3 | 1067 | 4kyo | A | | 56 | ILE | p.Ile56Asn | P21549 |
| HOMO SAPIENS | 3 | 1067 | 4kyo | A | | 59 | GLU | p.Glu59Lys | P21549 |
| HOMO SAPIENS | 3 | 1067 | 4kyo | A | | 66 | TYR | p.Tyr66Ter | P21549 |
| HOMO SAPIENS | 3 | 1067 | 4kyo | A | | 111 | ARG | p.Arg111Leu | P21549 |
| HOMO SAPIENS | 7 | 1067 | 4kyo | A | | 118 | ARG | p.Arg118His | P21549 |
| HOMO SAPIENS | 3 | 1067 | 4kyo | A | | 122 | ARG | p.Arg122Ter | P21549 |
| HOMO SAPIENS | 3 | 1067 | 4kyo | A | | 124 | HIS | p.His124Pro | P21549 |
| HOMO SAPIENS | 0 | 1067 | 4kyo | A | | 152 | PHE | p.Phe152Ile | P21549 |
| HOMO SAPIENS | 1 | 1067 | 4kyo | A | | 156 | GLY | p.Gly156Arg | P21549 |
| HOMO SAPIENS | 0 | 1067 | 4kyo | A | | 161 | GLY | p.Gly161Arg | P21549 |
| HOMO SAPIENS | 3 | 1067 | 4kyo | A | | 166 | LEU | p.Leu166Pro | P21549 |
| HOMO SAPIENS | 5 | 1067 | 4kyo | A | | 170 | GLY | p.Gly170Arg | P21549 |
| HOMO SAPIENS | 5 | 1067 | 4kyo | A | | 186 | ALA | p.Ala186Val | P21549 |
| HOMO SAPIENS | 3 | 1067 | 4kyo | A | | 187 | SER | p.Ser187Phe | P21549 |
| HOMO SAPIENS | 3 | 1067 | 4kyo | A | | 197 | ARG | p.Arg197Pro | P21549 |
| HOMO SAPIENS | 5 | 1067 | 4kyo | A | | 201 | ASP | p.Asp201Glu | P21549 |
| HOMO SAPIENS | 0 | 1067 | 4kyo | A | | 202 | ILE | p.Ile202Asn | P21549 |
| HOMO SAPIENS | 1 | 1067 | 4kyo | A | | 205 | SER | p.Ser205Pro | P21549 |
| HOMO SAPIENS | 1 | 1067 | 4kyo | A | | 205 | SER | p.Ser205Ter | P21549 |
| HOMO SAPIENS | 2 | 1067 | 4kyo | A | | 216 | GLY | p.Gly216Arg | P21549 |
| HOMO SAPIENS | 5 | 1067 | 4kyo | A | | 233 | ARG | p.Arg233Cys | P21549 |
| HOMO SAPIENS | 5 | 1067 | 4kyo | A | | 244 | ILE | p.Ile244Thr | P21549 |
| HOMO SAPIENS | 5 | 1067 | 4kyo | A | | 244 | ILE | p.Ile244Met | P21549 |
| HOMO SAPIENS | 4 | 1067 | 4kyo | A | | 253 | CYS | p.Cys253Arg | P21549 |
| HOMO SAPIENS | 7 | 1067 | 4kyo | A | | 274 | GLU | p.Glu274Asp | P21549 |
| HOMO SAPIENS | 7 | 1067 | 4kyo | A | | 279 | ILE | p.Ile279Thr | P21549 |
| HOMO SAPIENS | 6 | 1067 | 4kyo | A | | 295 | ALA | p.Ala295Thr | P21549 |

| | | | | | | | | |
|---|---|---|---|---|---|---|---|---|
| HOMO SAPIENS | 5 | 1067 | 4kyo | A | 340 | ILE | p.Ile340Met | P21549 |
| HOMO SAPIENS | 1 | 790 | 2ric | C | 270 | SER | p.Ser290Thr | P35247 |
| HOMO SAPIENS | 0 | 790 | 2ric | C | 289 | GLU | p.Glu309Lys | P35247 |
| HOMO SAPIENS | 1 | 212 | 5sxe | A | 545 | GLU | p.Glu545Lys | P42336 |
| HOMO SAPIENS | 3 | 212 | 5sxe | A | 546 | GLN | p.Gln546Glu | P42336 |
| HOMO SAPIENS | 2 | 212 | 5sxe | A | 1047 | HIS | p.His1047Leu | P42336 |
| HOMO SAPIENS | 7 | 152 | 2zqq | F | 197 | ARG | p.Arg197Ter | Q13825 |
| HOMO SAPIENS | 0 | 152 | 2zqq | F | 248 | ALA | p.Ala248Thr | Q13825 |
| HOMO SAPIENS | 0 | 152 | 2zqq | F | 275 | ALA | p.Ala275Val | Q13825 |
| HOMO SAPIENS | 5 | 152 | 2zqq | F | 331 | LYS | p.Lys331Ter | Q13825 |
| HOMO SAPIENS | 6 | 17801 | 2j8l | A | 283 | PHE | p.Phe301Leu | P03951 |
| HOMO SAPIENS | 3 | 17801 | 2j8l | A | 308 | ARG | p.Arg326Cys | P03951 |
| HOMO SAPIENS | 0 | 17801 | 2j8l | A | 321 | CYS | p.Cys339Phe | P03951 |
| HOMO SAPIENS | 3 | 17801 | 2j8l | A | 351 | TYR | p.Tyr369Ter | P03951 |
| HOMO SAPIENS | 7 | 1802 | 5db0 | A | 22 | LEU | p.Leu22Arg | O00255 |
| HOMO SAPIENS | 7 | 1802 | 5db0 | A | 26 | GLU | p.Glu26Lys | O00255 |
| HOMO SAPIENS | 7 | 1802 | 5db0 | A | 139 | HIS | p.His139Asp | O00255 |
| HOMO SAPIENS | 0 | 1802 | 5db0 | A | 171 | ARG | p.Arg176Gln | O00255 |
| HOMO SAPIENS | 0 | 1802 | 5db0 | A | 184 | VAL | p.Val189Glu | O00255 |
| HOMO SAPIENS | 7 | 1802 | 5db0 | A | 255 | GLU | p.Glu260Lys | O00255 |
| HOMO SAPIENS | 4 | 1802 | 5db0 | A | 418 | ASP | p.Asp423Asn | O00255 |
| HOMO SAPIENS | 0 | 1802 | 5db0 | A | 436 | TRP | p.Trp441Arg | O00255 |
| HOMO SAPIENS | -1 | 1802 | 5db0 | A | 541 | ALA | p.Thr546Ala | O00255 |
| HOMO SAPIENS | 4 | 1802 | 5db0 | A | 552 | THR | p.Thr557Ser | O00255 |
| HOMO SAPIENS | 0 | 3450 | 2yuz | A | 36 | PRO | p.Pro461Gln | Q00872 |
| HOMO SAPIENS | 1 | 3450 | 2yuz | A | 56 | GLN | p.His481Gln | Q00872 |
| HOMO SAPIENS | 2 | 3450 | 2yuz | A | 68 | ALA | p.Ala493Val | Q00872 |
| HOMO SAPIENS | 0 | 3211 | 3rqg | A | 72 | VAL | p.Val72Leu | Q9BUL8 |
| HOMO SAPIENS | 2 | 4168 | 4q4f | A | 121 | ARG | p.Arg121Ter | Q14108 |
| HOMO SAPIENS | 2 | 4168 | 4q4f | A | 121 | ARG | p.Arg121Gln | Q14108 |
| HOMO SAPIENS | 2 | 4168 | 4q4f | A | 159 | MET | p.Met159Val | Q14108 |
| HOMO SAPIENS | 4 | 4168 | 4q4f | A | 363 | HIS | p.His363Asn | Q14108 |
| HOMO SAPIENS | 2 | 4168 | 4q4f | A | 396 | VAL | p.Val396Ile | Q14108 |
| HOMO SAPIENS | 0 | 1448 | 1y97 | A | 87 | ARG | p.Arg130Ter | Q9BQ50 |
| HOMO SAPIENS | 0 | 3444 | 4iwp | A | 151 | SER | p.Ser151Phe | Q9UHD2 |
| HOMO SAPIENS | 0 | 3444 | 4iwp | A | 401 | LYS | p.Lys401Glu | Q9UHD2 |
| HOMO SAPIENS | 0 | 116 | 1zy8 | B | 194 | GLY | p.Gly229Cys | P09622 |
| HOMO SAPIENS | 0 | 116 | 1zy8 | B | 255 | ASP | p.Asp290Gly | P09622 |
| HOMO SAPIENS | 2 | 116 | 1zy8 | B | 326 | MET | p.Met361Val | P09622 |
| HOMO SAPIENS | 0 | 116 | 1zy8 | B | 340 | GLU | p.Glu375Lys | P09622 |
| HOMO SAPIENS | 2 | 116 | 1zy8 | B | 358 | ILE | p.Ile393Thr | P09622 |
| HOMO SAPIENS | 1 | 116 | 1zy8 | B | 453 | PRO | p.Pro488Leu | P09622 |
| HOMO SAPIENS | 7 | 127 | 4qo7 | B | 160 | SER | p.Ser161Arg | P00338 |
| HOMO SAPIENS | 0 | 2025 | 1wrl | A | 8 | ALA | p.Ala8Val | P63316 |
| HOMO SAPIENS | 8 | 2025 | 1wrl | A | 24 | PHE | p.Phe24Leu | P63316 |
| HOMO SAPIENS | 0 | 2025 | 1wrl | A | 84 | SER | p.Cys84Tyr | P63316 |
| HOMO SAPIENS | 8 | 3791 | 1mhd | B | 93 | ARG | p.Arg93Ter | P84022 |
| HOMO SAPIENS | 0 | 1575 | 4phc | A | 205 | GLY | p.Gly205Asp | P12081 |
| HOMO SAPIENS | 0 | 1575 | 4phc | A | 454 | TYR | p.Tyr454Ser | P12081 |
| HOMO SAPIENS | 0 | 39620 | 1ozs | B | 135 | ARG | p.Arg136Gln | P19429 |
| HOMO SAPIENS | 5 | 39620 | 1ozs | B | 140 | ARG | p.Arg141Trp | P19429 |
| HOMO SAPIENS | 5 | 39620 | 1ozs | B | 142 | THR | p.Thr143Asn | P19429 |
| HOMO SAPIENS | 5 | 39620 | 1ozs | B | 143 | LEU | p.Leu144Gln | P19429 |
| HOMO SAPIENS | 5 | 39620 | 1ozs | B | 144 | ARG | p.Arg145Gly | P19429 |

| | | | | | | | | |
|---|---|---|---|---|---|---|---|---|
| HOMO SAPIENS | 5 | 39620 | 1ozs | B | 144 | ARG | p.Arg145Gln | P19429 |
| HOMO SAPIENS | -1 | 20934 | 4nuv | D | 42 | GLY | p.Gly42Asp | Q16570 |
| HOMO SAPIENS | -1 | 40565 | 5lcw | S | 4 | VAL | p.Val4Met | O60566 |
| HOMO SAPIENS | 0 | 40565 | 5lcw | S | 40 | THR | p.Thr40Met | O60566 |
| HOMO SAPIENS | 4 | 40565 | 5lcw | S | 198 | ALA | p.Glu390Asp | O60566 |
| HOMO SAPIENS | 4 | 758 | 4oiu | A | 726 | ARG | p.Arg727Leu | P10275 |
| HOMO SAPIENS | 4 | 758 | 4oiu | A | 730 | VAL | p.Val731Met | P10275 |
| HOMO SAPIENS | 2 | 758 | 4oiu | A | 798 | GLN | p.Gln799Glu | P10275 |
| HOMO SAPIENS | 3 | 758 | 4oiu | A | 855 | ARG | p.Arg856His | P10275 |
| HOMO SAPIENS | 1 | 758 | 4oiu | A | 886 | MET | p.Met887Val | P10275 |
| HOMO SAPIENS | 0 | 3303 | 4ie0 | A | 134 | ALA | p.Ala134Thr | Q9C0B1 |
| HOMO SAPIENS | 9 | 3303 | 4ie0 | A | 316 | ARG | p.Arg316Gln | Q9C0B1 |
| HOMO SAPIENS | 1 | 41502 | 3fgh | A | 136 | PRO | p.Pro178Leu | Q00059 |
| HOMO SAPIENS | 5 | 175 | 3ceh | B | 58 | THR | p.Thr59Lys | P06737 |
| HOMO SAPIENS | 8 | 175 | 3ceh | B | 93 | ARG | p.Arg94Ter | P06737 |
| HOMO SAPIENS | 5 | 175 | 3ceh | B | 221 | VAL | p.Val222Ile | P06737 |
| HOMO SAPIENS | 0 | 175 | 3ceh | B | 338 | ASN | p.Asn339Ser | P06737 |
| HOMO SAPIENS | 1 | 175 | 3ceh | B | 372 | PHE | p.Phe373Leu | P06737 |
| HOMO SAPIENS | 9 | 175 | 3ceh | B | 376 | ASN | p.Asn377Lys | P06737 |
| HOMO SAPIENS | 8 | 175 | 3ceh | B | 398 | ARG | p.Arg399Ter | P06737 |
| HOMO SAPIENS | 5 | 175 | 3ceh | B | 424 | ARG | p.Arg425Pro | P06737 |
| HOMO SAPIENS | 9 | 175 | 3ceh | B | 455 | VAL | p.Val456Met | P06737 |
| HOMO SAPIENS | 8 | 175 | 3ceh | B | 490 | ARG | p.Arg491Cys | P06737 |
| HOMO SAPIENS | 8 | 175 | 3ceh | B | 631 | ASN | p.Asn632Ile | P06737 |
| HOMO SAPIENS | 9 | 175 | 3ceh | B | 648 | TYR | p.Tyr649Ter | P06737 |
| HOMO SAPIENS | 9 | 175 | 3ceh | B | 672 | GLU | p.Glu673Lys | P06737 |
| HOMO SAPIENS | 9 | 175 | 3ceh | B | 674 | SER | p.Ser675Leu | P06737 |
| HOMO SAPIENS | 0 | 175 | 3ceh | B | 805 | ILE | p.Ile806Leu | P06737 |
| HOMO SAPIENS | 0 | 175 | 3ceh | B | 820 | TYR | p.Tyr821His | P06737 |
| HOMO SAPIENS | 1 | 6750 | 4fdj | A | 47 | GLY | p.Gly47Arg | P34059 |
| HOMO SAPIENS | 9 | 6750 | 4fdj | A | 60 | ASP | p.Asp60Asn | P34059 |
| HOMO SAPIENS | 5 | 6750 | 4fdj | A | 67 | LEU | p.Leu67Met | P34059 |
| HOMO SAPIENS | 0 | 6750 | 4fdj | A | 69 | PHE | p.Phe69Val | P34059 |
| HOMO SAPIENS | 3 | 6750 | 4fdj | A | 94 | ARG | p.Arg94Gly | P34059 |
| HOMO SAPIENS | 0 | 6750 | 4fdj | A | 113 | ILE | p.Ile113Phe | P34059 |
| HOMO SAPIENS | 0 | 6750 | 4fdj | A | 138 | VAL | p.Val138Ala | P34059 |
| HOMO SAPIENS | 0 | 6750 | 4fdj | A | 139 | GLY | p.Gly139Ser | P34059 |
| HOMO SAPIENS | 0 | 6750 | 4fdj | A | 151 | PRO | p.Pro151Thr | P34059 |
| HOMO SAPIENS | 0 | 6750 | 4fdj | A | 155 | GLY | p.Gly155Arg | P34059 |
| HOMO SAPIENS | 0 | 6750 | 4fdj | A | 159 | TRP | p.Trp159Ter | P34059 |
| HOMO SAPIENS | 0 | 6750 | 4fdj | A | 162 | SER | p.Ser162Phe | P34059 |
| HOMO SAPIENS | 0 | 6750 | 4fdj | A | 200 | THR | p.Thr200Met | P34059 |
| HOMO SAPIENS | 0 | 6750 | 4fdj | A | 204 | ASN | p.Asn204Lys | P34059 |
| HOMO SAPIENS | 0 | 6750 | 4fdj | A | 231 | ALA | p.Ala231Gly | P34059 |
| HOMO SAPIENS | 2 | 6750 | 4fdj | A | 235 | THR | p.Thr235Lys | P34059 |
| HOMO SAPIENS | 1 | 6750 | 4fdj | A | 259 | ARG | p.Arg259Gln | P34059 |
| HOMO SAPIENS | 0 | 6750 | 4fdj | A | 287 | SER | p.Ser287Leu | P34059 |
| HOMO SAPIENS | 0 | 6750 | 4fdj | A | 291 | ALA | p.Ala291Thr | P34059 |
| HOMO SAPIENS | 3 | 6750 | 4fdj | A | 301 | GLY | p.Gly301Cys | P34059 |
| HOMO SAPIENS | 0 | 6750 | 4fdj | A | 312 | THR | p.Thr312Ser | P34059 |
| HOMO SAPIENS | 0 | 6750 | 4fdj | A | 318 | MET | p.Met318Arg | P34059 |
| HOMO SAPIENS | 0 | 6750 | 4fdj | A | 340 | GLY | p.Gly340Asp | P34059 |
| HOMO SAPIENS | 0 | 6750 | 4fdj | A | 386 | ARG | p.Arg386Cys | P34059 |
| HOMO SAPIENS | 0 | 6750 | 4fdj | A | 391 | MET | p.Met391Val | P34059 |

| | | | | | | | | |
|---|---|---|---|---|---|---|---|---|
| HOMO SAPIENS | 0 | 6750 | 4fdj | A | 392 | ALA | p.Ala392Val | P34059 |
| HOMO SAPIENS | 0 | 6750 | 4fdj | A | 393 | ALA | p.Ala393Ser | P34059 |
| HOMO SAPIENS | 0 | 6750 | 4fdj | A | 459 | ALA | p.Ala459Val | P34059 |
| HOMO SAPIENS | 0 | 6750 | 4fdj | A | 473 | GLN | p.Gln473Ter | P34059 |
| HOMO SAPIENS | 2 | 6750 | 4fdj | A | 480 | VAL | p.Val480Phe | P34059 |
| HOMO SAPIENS | 0 | 6750 | 4fdj | A | 487 | ASN | p.Asn487Ser | P34059 |
| HOMO SAPIENS | 0 | 6750 | 4fdj | A | 488 | VAL | p.Val488Met | P34059 |
| HOMO SAPIENS | 0 | 6750 | 4fdj | A | 520 | TRP | p.Trp520Ter | P34059 |
| HOMO SAPIENS | 4 | 1088 | 2p3u | A | 102 | GLU | p.Glu142Lys | P00742 |
| HOMO SAPIENS | 6 | 370 | 1nr1 | D | 73 | ASP | p.Asp126His | P00367 |
| HOMO SAPIENS | 3 | 370 | 1nr1 | D | 221 | ARG | p.Arg274Cys | P00367 |
| HOMO SAPIENS | 2 | 370 | 1nr1 | D | 265 | ARG | p.Arg318Lys | P00367 |
| HOMO SAPIENS | 1 | 370 | 1nr1 | D | 269 | ARG | p.Arg322His | P00367 |
| HOMO SAPIENS | 1 | 370 | 1nr1 | D | 296 | GLU | p.Glu349Ala | P00367 |
| HOMO SAPIENS | 1 | 370 | 1nr1 | D | 445 | SER | p.Ser498Leu | P00367 |
| HOMO SAPIENS | 1 | 370 | 1nr1 | D | 446 | GLY | p.Gly499Asp | P00367 |
| HOMO SAPIENS | 1 | 370 | 1nr1 | D | 446 | GLY | p.Gly499Ser | P00367 |
| HOMO SAPIENS | 1 | 370 | 1nr1 | D | 448 | SER | p.Ser501Pro | P00367 |
| HOMO SAPIENS | 2 | 370 | 1nr1 | D | 454 | HIS | p.His507Tyr | P00367 |
| HOMO SAPIENS | 1 | 7720 | 2he0 | A | 147 | HIS | p.His2018Gln | P46531 |
| HOMO SAPIENS | 2 | 11844 | 5l0v | A | 218 | ARG | p.Arg218Ter | Q8NBL1 |
| HOMO SAPIENS | 2 | 11844 | 5l0v | A | 279 | ARG | p.Arg279Trp | Q8NBL1 |
| HOMO SAPIENS | 9 | 1485 | 5va0 | A | 477 | ARG | p.Arg477His | P04150 |
| HOMO SAPIENS | 0 | 235 | 2fun | B | 2248 | ARG | p.Arg248Trp | Q14790 |
| HOMO SAPIENS | 4 | 235 | 2fun | B | 2285 | HIS | p.Asp285His | Q14790 |
| HOMO SAPIENS | 0 | 4822 | 5y3n | A | 307 | ARG | p.Arg307Gly | Q12931 |
| HOMO SAPIENS | 8 | 32928 | 4m6w | A | 1886 | HIS | p.His1886Tyr | Q8IYD8 |
| HOMO SAPIENS | 0 | 2711 | 4y5v | F | 22 | GLY | p.Gly46Glu | P19235 |
| HOMO SAPIENS | -1 | 2711 | 4y5v | F | 166 | ALA | p.Ala190Ser | P19235 |
| HOMO SAPIENS | 4 | 5161 | 5bpu | A | 42 | HIS | p.His42Arg | Q00604 |
| HOMO SAPIENS | 0 | 5161 | 5bpu | A | 44 | TYR | p.Tyr44Cys | Q00604 |
| HOMO SAPIENS | 0 | 5161 | 5bpu | A | 45 | VAL | p.Val45Glu | Q00604 |
| HOMO SAPIENS | 0 | 5161 | 5bpu | A | 60 | VAL | p.Val60Glu | Q00604 |
| HOMO SAPIENS | 0 | 5161 | 5bpu | A | 61 | LEU | p.Leu61Phe | Q00604 |
| HOMO SAPIENS | 7 | 5161 | 5bpu | A | 69 | CYS | p.Cys69Ser | Q00604 |
| HOMO SAPIENS | 0 | 5161 | 5bpu | A | 75 | SER | p.Ser75Cys | Q00604 |
| HOMO SAPIENS | 0 | 5161 | 5bpu | A | 90 | ARG | p.Arg90Pro | Q00604 |
| HOMO SAPIENS | 9 | 5161 | 5bpu | A | 96 | CYS | p.Cys96Tyr | Q00604 |
| HOMO SAPIENS | 9 | 5161 | 5bpu | A | 96 | CYS | p.Cys96Trp | Q00604 |
| HOMO SAPIENS | 0 | 5161 | 5bpu | A | 101 | SER | p.Ser101Phe | Q00604 |
| HOMO SAPIENS | 0 | 5161 | 5bpu | A | 105 | ALA | p.Ala105Thr | Q00604 |
| HOMO SAPIENS | 0 | 5161 | 5bpu | A | 110 | CYS | p.Cys110Gly | Q00604 |
| HOMO SAPIENS | 0 | 5161 | 5bpu | A | 121 | ARG | p.Arg121Trp | Q00604 |
| HOMO SAPIENS | 0 | 5161 | 5bpu | A | 121 | ARG | p.Arg121Leu | Q00604 |
| HOMO SAPIENS | 0 | 5161 | 5bpu | A | 124 | LEU | p.Leu124Phe | Q00604 |
| HOMO SAPIENS | 8 | 5161 | 5bpu | A | 128 | CYS | p.Cys128Ter | Q00604 |
| HOMO SAPIENS | 1 | 13944 | 5xqh | A | 238 | ARG | p.Arg238His | Q9GZN7 |
| HOMO SAPIENS | -1 | 7508 | 5jl6 | A | 39 | TRP | p.Trp39Arg | P11511 |
| HOMO SAPIENS | 0 | 7508 | 5jl6 | A | 201 | THR | p.Thr201Met | P11511 |
| HOMO SAPIENS | 0 | 7508 | 5jl6 | A | 210 | GLU | p.Glu210Lys | P11511 |
| HOMO SAPIENS | 1 | 7508 | 5jl6 | A | 264 | ARG | p.Arg264Cys | P11511 |
| HOMO SAPIENS | 2 | 7508 | 5jl6 | A | 365 | ARG | p.Arg365Gln | P11511 |
| HOMO SAPIENS | 7 | 7508 | 5jl6 | A | 375 | ARG | p.Arg375Cys | P11511 |
| HOMO SAPIENS | 8 | 7508 | 5jl6 | A | 435 | ARG | p.Arg435Cys | P11511 |

| | | | | | | | | |
|---|---|---|---|---|---|---|---|---|
| HOMO SAPIENS | 9 | 7508 | 5jl6 | A | 437 | CYS | p.Cys437Tyr | P11511 |
| HOMO SAPIENS | -1 | 30206 | 5l95 | B | 246 | ARG | p.Arg246Ter | Q9GZZ9 |
| HOMO SAPIENS | 4 | 875 | 3efh | B | 43 | GLU | p.Glu43Asp | P60891 |
| HOMO SAPIENS | 5 | 875 | 3efh | B | 65 | ASP | p.Asp65Asn | P60891 |
| HOMO SAPIENS | 0 | 875 | 3efh | B | 87 | ALA | p.Ala87Thr | P60891 |
| HOMO SAPIENS | 2 | 875 | 3efh | B | 115 | MET | p.Met115Thr | P60891 |
| HOMO SAPIENS | 8 | 875 | 3efh | B | 133 | GLN | p.Gln133Pro | P60891 |
| HOMO SAPIENS | 8 | 875 | 3efh | B | 152 | LEU | p.Leu152Pro | P60891 |
| HOMO SAPIENS | 7 | 875 | 3efh | B | 290 | ILE | p.Ile290Thr | P60891 |
| HOMO SAPIENS | 6 | 875 | 3efh | B | 306 | GLY | p.Gly306Arg | P60891 |
| HOMO SAPIENS | 0 | 7978 | 5edm | A | 122 | MET | p.Thr165Met | P00734 |
| HOMO SAPIENS | -1 | 7978 | 5edm | A | 257 | ARG | p.Arg314Cys | P00734 |
| HOMO SAPIENS | 0 | 1716 | 1b98 | M | 126 | ARG | p.Arg206Gln | P34130 |
| HOMO SAPIENS | 0 | 1716 | 1b98 | M | 126 | ARG | p.Arg206Trp | P34130 |
| HOMO SAPIENS | 9 | 3960 | 2qyp | B | 28 | ILE | p.Ile338Val | P07602 |
| HOMO SAPIENS | 1 | 3960 | 2qyp | B | 39 | LEU | p.Leu349Pro | P07602 |
| HOMO SAPIENS | 0 | 31414 | 5u1d | A | 333 | ILE | p.Ile393Val | Q03518 |
| HOMO SAPIENS | 0 | 31414 | 5u1d | A | 637 | ASP | p.Asp697Gly | Q03518 |
| HOMO SAPIENS | 0 | 31414 | 5u1d | A | 659 | ARG | p.Arg719Gln | Q03518 |
| HOMO SAPIENS | 3 | 119 | 2pho | A | 11 | ILE | p.Ile11Thr | P05089 |
| HOMO SAPIENS | 8 | 119 | 2pho | A | 21 | ARG | p.Arg21Ter | P05089 |
| HOMO SAPIENS | 0 | 119 | 2pho | A | 122 | TRP | p.Trp122Ter | P05089 |
| HOMO SAPIENS | 6 | 119 | 2pho | A | 138 | GLY | p.Gly138Val | P05089 |
| HOMO SAPIENS | 5 | 119 | 2pho | A | 290 | THR | p.Thr290Ser | P05089 |
| HOMO SAPIENS | 5 | 119 | 2pho | A | 291 | ARG | p.Arg291Ter | P05089 |
| HOMO SAPIENS | 0 | 3732 | 2i99 | A | 247 | TYR | p.Tyr247Ter | Q14894 |
| HOMO SAPIENS | -1 | 1382 | 4ffx | A | 2 | ALA | p.Ala2Val | P30566 |
| HOMO SAPIENS | 1 | 1382 | 4ffx | A | 72 | ILE | p.Ile72Met | P30566 |
| HOMO SAPIENS | 1 | 1382 | 4ffx | A | 100 | PRO | p.Pro100Ala | P30566 |
| HOMO SAPIENS | 0 | 1382 | 4ffx | A | 114 | TYR | p.Tyr114His | P30566 |
| HOMO SAPIENS | 1 | 1382 | 4ffx | A | 141 | ARG | p.Arg141Trp | P30566 |
| HOMO SAPIENS | 0 | 1382 | 4ffx | A | 147 | LYS | p.Lys147Met | P30566 |
| HOMO SAPIENS | 3 | 1382 | 4ffx | A | 190 | ARG | p.Arg190Gln | P30566 |
| HOMO SAPIENS | 3 | 1382 | 4ffx | A | 190 | ARG | p.Arg190Ter | P30566 |
| HOMO SAPIENS | 0 | 1382 | 4ffx | A | 217 | HIS | p.His217Asp | P30566 |
| HOMO SAPIENS | 2 | 1382 | 4ffx | A | 246 | LYS | p.Lys246Glu | P30566 |
| HOMO SAPIENS | 5 | 1382 | 4ffx | A | 303 | ARG | p.Arg303Cys | P30566 |
| HOMO SAPIENS | 1 | 1382 | 4ffx | A | 318 | PRO | p.Pro318Leu | P30566 |
| HOMO SAPIENS | 6 | 1382 | 4ffx | A | 332 | ASP | p.Asp332His | P30566 |
| HOMO SAPIENS | 6 | 1382 | 4ffx | A | 337 | ARG | p.Arg337Ter | P30566 |
| HOMO SAPIENS | 1 | 1382 | 4ffx | A | 396 | ARG | p.Arg396His | P30566 |
| HOMO SAPIENS | 1 | 1382 | 4ffx | A | 426 | ARG | p.Arg426His | P30566 |
| HOMO SAPIENS | 0 | 1382 | 4ffx | A | 438 | SER | p.Ser438Pro | P30566 |
| HOMO SAPIENS | 1 | 1382 | 4ffx | A | 447 | SER | p.Ser447Pro | P30566 |
| HOMO SAPIENS | 2 | 1382 | 4ffx | A | 448 | SER | p.Ser448Pro | P30566 |
| HOMO SAPIENS | 2 | 1382 | 4ffx | A | 450 | THR | p.Thr450Ser | P30566 |
| HOMO SAPIENS | 0 | 6212 | 1ao3 | A | 46 | SER | p.Ser1731Thr | P04275 |
| HOMO SAPIENS | 0 | 1711 | 3f9c | A | 279 | VAL | p.Val279Phe | Q13093 |
| HOMO SAPIENS | 0 | 6112 | 4fvl | A | 312 | HIS | p.His312Arg | P45452 |
| HOMO SAPIENS | 0 | 6112 | 4fvl | A | 323 | THR | p.Thr323Met | P45452 |
| HOMO SAPIENS | 0 | 6112 | 4fvl | A | 390 | ASP | p.Asp390Gly | P45452 |
| HOMO SAPIENS | 0 | 6112 | 4fvl | A | 458 | ARG | p.Arg458Cys | P45452 |
| HOMO SAPIENS | 2 | 6180 | 5ggn | A | 97 | ARG | p.Arg97Trp | Q8WZA1 |
| HOMO SAPIENS | 2 | 6180 | 5ggn | A | 215 | ARG | p.Arg215Ter | Q8WZA1 |

| | | | | | | | | |
|---|---|---|---|---|---|---|---|---|
| HOMO SAPIENS | -1 | 5526 | 4ak3 | A | 11 | ASP | p.Asp1232Asn | P02461 |
| HOMO SAPIENS | 6 | 5526 | 4ak3 | A | 38 | ALA | p.Ala1259Thr | P02461 |
| HOMO SAPIENS | 0 | 5526 | 4ak3 | A | 92 | LYS | p.Lys1313Arg | P02461 |
| HOMO SAPIENS | 0 | 5526 | 4ak3 | A | 132 | GLN | p.His1353Gln | P02461 |
| HOMO SAPIENS | 4 | 16616 | 1hci | B | 433 | SER | p.Ser433Leu | P35609 |
| HOMO SAPIENS | 1 | 16616 | 1hci | B | 475 | ASP | p.Asp475Asn | P35609 |
| HOMO SAPIENS | 1 | 16616 | 1hci | B | 476 | ALA | p.Ala476Ser | P35609 |
| HOMO SAPIENS | 2 | 16616 | 1hci | B | 507 | MET | p.Met507Val | P35609 |
| HOMO SAPIENS | 1 | 16616 | 1hci | B | 604 | MET | p.Met604Val | P35609 |
| HOMO SAPIENS | 2 | 16616 | 1hci | B | 640 | ARG | p.Arg640His | P35609 |
| HOMO SAPIENS | 1 | 16616 | 1hci | B | 721 | ARG | p.Arg721Cys | P35609 |
| HOMO SAPIENS | -1 | 3402 | 5iaa | B | 57 | MET | p.Met57Val | Q9GZZ9 |
| HOMO SAPIENS | 7 | 3250 | 3irq | B | 193 | PRO | p.Pro193Ala | P55265 |
| HOMO SAPIENS | 2 | 15200 | 2wns | A | 30 | SER | p.Ser30Gly | P11172 |
| HOMO SAPIENS | 0 | 15200 | 2wns | A | 96 | ARG | p.Arg96Gly | P11172 |
| HOMO SAPIENS | 0 | 15200 | 2wns | A | 109 | VAL | p.Val109Gly | P11172 |
| HOMO SAPIENS | 2 | 15759 | 4x4w | B | 140 | LEU | p.Leu166Ser | Q96Q11 |
| HOMO SAPIENS | 0 | 15759 | 4x4w | B | 197 | ILE | p.Ile223Thr | Q96Q11 |
| HOMO SAPIENS | 0 | 3156 | 3w03 | B | 14 | ALA | p.Ala14Thr | Q9H9Q4 |
| HOMO SAPIENS | 0 | 3156 | 3w03 | B | 57 | ARG | p.Arg57Gly | Q9H9Q4 |
| HOMO SAPIENS | 0 | 3156 | 3w03 | B | 123 | CYS | p.Cys123Arg | Q9H9Q4 |
| HOMO SAPIENS | 0 | 11693 | 3ojy | A | 63 | GLN | p.Gln93Lys | P07357 |
| HOMO SAPIENS | 0 | 1421 | 2qc8 | E | 275 | GLU | p.Glu275Asp | P15104 |
| HOMO SAPIENS | 9 | 1421 | 2qc8 | E | 324 | ARG | p.Arg324Cys | P15104 |
| HOMO SAPIENS | 6 | 1421 | 2qc8 | E | 341 | ARG | p.Arg341Cys | P15104 |
| HOMO SAPIENS | 6 | 2850 | 3qvb | A | 83 | THR | p.Thr83Met | P46976 |
| HOMO SAPIENS | 8 | 2850 | 3qvb | A | 102 | ASP | p.Asp102His | P46976 |
| HOMO SAPIENS | 1 | 24314 | 4m9p | A | 480 | PRO | p.Pro480Leu | P21333 |
| HOMO SAPIENS | 0 | 24314 | 4m9p | A | 528 | VAL | p.Val528Met | P21333 |
| HOMO SAPIENS | 0 | 24314 | 4m9p | A | 590 | VAL | p.Val590Ile | P21333 |
| HOMO SAPIENS | 0 | 24314 | 4m9p | A | 605 | ASP | p.Asp605Asn | P21333 |
| HOMO SAPIENS | 1 | 24314 | 4m9p | A | 637 | PRO | p.Pro637Gln | P21333 |
| HOMO SAPIENS | 0 | 24314 | 4m9p | A | 656 | LEU | p.Leu656Phe | P21333 |
| HOMO SAPIENS | 2 | 24314 | 4m9p | A | 711 | VAL | p.Val711Asp | P21333 |
| HOMO SAPIENS | 0 | 4951 | 5joe | A | 40 | THR | p.Thr9621Ser | Q8WZ42 |
| HOMO SAPIENS | 1 | 4951 | 5joe | A | 41 | GLU | p.Glu9622Gln | Q8WZ42 |
| HOMO SAPIENS | 0 | 4951 | 5joe | A | 82 | ALA | p.Ala9663Ser | Q8WZ42 |
| HOMO SAPIENS | 4 | 4951 | 5joe | A | 89 | ILE | p.Ile9670Asn | Q8WZ42 |
| HOMO SAPIENS | 0 | 9664 | 3c6m | A | 56 | GLY | p.Gly56Ser | P52788 |
| HOMO SAPIENS | 0 | 9664 | 3c6m | A | 132 | VAL | p.Val132Gly | P52788 |
| HOMO SAPIENS | 1 | 9664 | 3c6m | A | 190 | THR | p.Thr190Ile | P52788 |
| HOMO SAPIENS | -1 | 12118 | 5hcd | B | 19 | GLN | p.Gln19Ter | P01031 |
| HOMO SAPIENS | 0 | 1583 | 2b4x | L | 7 | ILE | p.Ile39Asn | P01008 |
| HOMO SAPIENS | 0 | 1583 | 2b4x | L | 24 | ARG | p.Arg56Cys | P01008 |
| HOMO SAPIENS | 0 | 1583 | 2b4x | L | 47 | ARG | p.Arg79Cys | P01008 |
| HOMO SAPIENS | 0 | 1583 | 2b4x | L | 47 | ARG | p.Arg79His | P01008 |
| HOMO SAPIENS | 1 | 1583 | 2b4x | L | 95 | CYS | p.Cys127Arg | P01008 |
| HOMO SAPIENS | 1 | 1583 | 2b4x | L | 99 | LEU | p.Leu131Phe | P01008 |
| HOMO SAPIENS | 0 | 1583 | 2b4x | L | 116 | SER | p.Ser148Pro | P01008 |
| HOMO SAPIENS | 0 | 1583 | 2b4x | L | 129 | ARG | p.Arg161Gln | P01008 |
| HOMO SAPIENS | 0 | 1583 | 2b4x | L | 135 | ASN | p.Asn167Thr | P01008 |
| HOMO SAPIENS | 2 | 1583 | 2b4x | L | 187 | ASN | p.Asn219Asp | P01008 |
| HOMO SAPIENS | 0 | 1583 | 2b4x | L | 191 | SER | p.Ser223Pro | P01008 |
| HOMO SAPIENS | 2 | 1583 | 2b4x | L | 349 | SER | p.Ser381Pro | P01008 |

| | | | | | | | | |
|---|---|---|---|---|---|---|---|---|
| HOMO SAPIENS | 0 | 1583 | 2b4x | L | 382 | ALA | p.Ala414Thr | P01008 |
| HOMO SAPIENS | 0 | 1583 | 2b4x | L | 387 | ALA | p.Ala419Val | P01008 |
| HOMO SAPIENS | 0 | 1583 | 2b4x | L | 392 | GLY | p.Gly424Asp | P01008 |
| HOMO SAPIENS | 0 | 1583 | 2b4x | L | 393 | ARG | p.Arg425His | P01008 |
| HOMO SAPIENS | 0 | 1583 | 2b4x | L | 393 | ARG | p.Arg425Cys | P01008 |
| HOMO SAPIENS | 0 | 1583 | 2b4x | L | 394 | SER | p.Ser426Leu | P01008 |
| HOMO SAPIENS | 0 | 1583 | 2b4x | L | 404 | ALA | p.Ala436Thr | P01008 |
| HOMO SAPIENS | 2 | 1583 | 2b4x | L | 407 | PRO | p.Pro439Leu | P01008 |
| HOMO SAPIENS | 2 | 1583 | 2b4x | L | 429 | PRO | p.Pro461Leu | P01008 |
| HOMO SAPIENS | 0 | 11647 | 2a1t | S | 78 | ALA | p.Ala78Ser | P38117 |
| HOMO SAPIENS | 0 | 11647 | 2a1t | S | 98 | ARG | p.Arg98Cys | P38117 |
| HOMO SAPIENS | 8 | 11647 | 2a1t | S | 128 | ASP | p.Asp128Asn | P38117 |
| HOMO SAPIENS | 5 | 11647 | 2a1t | S | 154 | THR | p.Thr154Met | P38117 |
| HOMO SAPIENS | 6 | 11647 | 2a1t | S | 164 | ARG | p.Arg164Gln | P38117 |
| HOMO SAPIENS | 5 | 11647 | 2a1t | S | 174 | ARG | p.Arg174His | P38117 |
| HOMO SAPIENS | -1 | 11200 | 4tx5 | A | 4 | ILE | p.Ile59Val | Q9NR28 |
| HOMO SAPIENS | 0 | 11200 | 4tx5 | A | 169 | GLY | p.Gly224Arg | Q9NR28 |
| HOMO SAPIENS | 3 | 417 | 2vr2 | A | 6 | ARG | p.Arg6Gln | Q14117 |
| HOMO SAPIENS | 1 | 417 | 2vr2 | A | 334 | GLN | p.Gln334Arg | Q14117 |
| HOMO SAPIENS | 2 | 417 | 2vr2 | A | 360 | TRP | p.Trp360Arg | Q14117 |
| HOMO SAPIENS | 0 | 417 | 2vr2 | A | 379 | SER | p.Ser379Arg | Q14117 |
| HOMO SAPIENS | 0 | 417 | 2vr2 | A | 435 | GLY | p.Gly435Arg | Q14117 |
| HOMO SAPIENS | 0 | 12498 | 3eo3 | B | 423 | ILE | p.Ile423Val | Q9Y223 |
| HOMO SAPIENS | 5 | 12498 | 3eo3 | B | 572 | VAL | p.Val572Leu | Q9Y223 |
| HOMO SAPIENS | 0 | 12498 | 3eo3 | B | 576 | GLY | p.Gly576Glu | Q9Y223 |
| HOMO SAPIENS | 0 | 12498 | 3eo3 | B | 587 | ILE | p.Ile587Thr | Q9Y223 |
| HOMO SAPIENS | 0 | 12498 | 3eo3 | B | 615 | SER | p.Ser615Ter | Q9Y223 |
| HOMO SAPIENS | 1 | 12498 | 3eo3 | B | 631 | ALA | p.Ala631Val | Q9Y223 |
| HOMO SAPIENS | 1 | 12498 | 3eo3 | B | 669 | GLY | p.Gly669Arg | Q9Y223 |
| HOMO SAPIENS | 1 | 12498 | 3eo3 | B | 696 | VAL | p.Val696Met | Q9Y223 |
| HOMO SAPIENS | 2 | 12498 | 3eo3 | B | 712 | MET | p.Met712Thr | Q9Y223 |
| HOMO SAPIENS | 0 | 1568 | 1r74 | A | 49 | LEU | p.Leu50Pro | Q14749 |
| HOMO SAPIENS | 0 | 1568 | 1r74 | A | 176 | HIS | p.His177Asn | Q14749 |
| HOMO SAPIENS | 3 | 147 | 5eig | A | 67 | THR | p.Thr67Ile | P32929 |
| HOMO SAPIENS | 3 | 147 | 5eig | A | 240 | GLN | p.Gln240Glu | P32929 |
| HOMO SAPIENS | -1 | 147 | 5eig | A | 403 | SER | p.Ser403Ile | P32929 |
| HOMO SAPIENS | 8 | 20036 | 2lsr | A | 31 | ARG | p.Arg31Ter | Q9Y6N9 |
| HOMO SAPIENS | -1 | 41382 | 4uis | B | 431 | UNK | p.Ala431Glu | P49768 |
| HOMO SAPIENS | -1 | 8536 | 3i6w | B | 85 | PRO | p.Pro85Leu | O96017 |
| HOMO SAPIENS | 0 | 8536 | 3i6w | B | 93 | TRP | p.Trp93Ter | O96017 |
| HOMO SAPIENS | 0 | 8536 | 3i6w | B | 95 | ARG | p.Arg95Ter | O96017 |
| HOMO SAPIENS | 0 | 8536 | 3i6w | B | 117 | ARG | p.Arg117Gly | O96017 |
| HOMO SAPIENS | 0 | 8536 | 3i6w | B | 119 | LYS | p.Lys119Arg | O96017 |
| HOMO SAPIENS | 0 | 8536 | 3i6w | B | 137 | ARG | p.Arg137Gln | O96017 |
| HOMO SAPIENS | 0 | 8536 | 3i6w | B | 137 | ARG | p.Arg137Ter | O96017 |
| HOMO SAPIENS | 0 | 8536 | 3i6w | B | 145 | ARG | p.Arg145Trp | O96017 |
| HOMO SAPIENS | 0 | 8536 | 3i6w | B | 160 | ILE | p.Ile160Met | O96017 |
| HOMO SAPIENS | 0 | 41322 | 4bxl | C | 46 | GLU | p.Glu46Lys | P37840 |
| HOMO SAPIENS | 5 | 41322 | 4bxl | C | 53 | ALA | p.Ala53Thr | P37840 |
| HOMO SAPIENS | 2 | 24966 | 3hve | A | 15 | ARG | p.Arg15Ser | Q9H2C0 |
| HOMO SAPIENS | 1 | 24966 | 3hve | A | 138 | ARG | p.Arg138His | Q9H2C0 |
| HOMO SAPIENS | 0 | 24966 | 3hve | A | 201 | ARG | p.Arg201Ter | Q9H2C0 |
| HOMO SAPIENS | 0 | 11680 | 5an9 | J | 8 | ASN | p.Asn8Lys | Q9Y3A5 |
| HOMO SAPIENS | 1 | 11680 | 5an9 | J | 33 | LYS | p.Lys33Arg | Q9Y3A5 |

| | | | | | | | | |
|---|---|---|---|---|---|---|---|---|
| HOMO SAPIENS | 1 | 11680 | 5an9 | J | 126 | ARG | p.Arg126Thr | Q9Y3A5 |
| HOMO SAPIENS | 1 | 11680 | 5an9 | J | 130 | VAL | p.Val130Met | Q9Y3A5 |
| HOMO SAPIENS | 1 | 11680 | 5an9 | J | 169 | ARG | p.Arg169Cys | Q9Y3A5 |
| HOMO SAPIENS | 1 | 11680 | 5an9 | J | 218 | ARG | p.Arg218Ter | Q9Y3A5 |
| HOMO SAPIENS | 4 | 978 | 3zs0 | B | 7 | TYR | p.Tyr173Cys | P05164 |
| HOMO SAPIENS | 1 | 978 | 3zs0 | B | 85 | MET | p.Met251Thr | P05164 |
| HOMO SAPIENS | 4 | 5140 | 2l4r | A | 52 | CYS | p.Cys52Ter | Q12809 |
| HOMO SAPIENS | 4 | 5140 | 2l4r | A | 63 | PRO | p.Pro63His | Q12809 |
| HOMO SAPIENS | 0 | 5140 | 2l4r | A | 100 | ARG | p.Arg100Gly | Q12809 |
| HOMO SAPIENS | 4 | 5140 | 2l4r | A | 129 | PHE | p.Phe129Leu | Q12809 |
| HOMO SAPIENS | 0 | 25836 | 2gy7 | B | 103 | GLU | p.Glu103Asp | Q02763 |
| HOMO SAPIENS | 0 | 25836 | 2gy7 | B | 148 | ILE | p.Ile148Thr | Q02763 |
| HOMO SAPIENS | 0 | 25836 | 2gy7 | B | 226 | ALA | p.Ala226Val | Q02763 |
| HOMO SAPIENS | 0 | 25836 | 2gy7 | B | 391 | THR | p.Thr391Ile | Q02763 |
| HOMO SAPIENS | 4 | 12563 | 5kwy | D | 20 | GLU | p.Glu20Ter | P61916 |
| HOMO SAPIENS | 2 | 12563 | 5kwy | D | 30 | VAL | p.Val30Met | P61916 |
| HOMO SAPIENS | 4 | 12563 | 5kwy | D | 39 | VAL | p.Val39Met | P61916 |
| HOMO SAPIENS | 8 | 12563 | 5kwy | D | 67 | SER | p.Ser67Pro | P61916 |
| HOMO SAPIENS | 8 | 12563 | 5kwy | D | 99 | CYS | p.Cys99Arg | P61916 |
| HOMO SAPIENS | 2 | 12563 | 5kwy | D | 118 | GLU | p.Glu118Ter | P61916 |
| HOMO SAPIENS | 2 | 12563 | 5kwy | D | 120 | PRO | p.Pro120Ser | P61916 |
| HOMO SAPIENS | -1 | 1679 | 1pmk | A | -2 | VAL | p.Val374Phe | P00747 |
| HOMO SAPIENS | -1 | 5315 | 2w2n | P | 53 | ALA | p.Ala53Val | Q8NBP7 |
| HOMO SAPIENS | 8 | 13496 | 2yad | A | 138 | THR | p.Asn138Thr | P11686 |
| HOMO SAPIENS | -1 | 13496 | 2yad | A | 173 | ARG | p.Arg167Gln | P11686 |
| HOMO SAPIENS | 2 | 13496 | 2yad | A | 186 | SER | p.Asn186Ser | P11686 |
| HOMO SAPIENS | 0 | 4426 | 3qd6 | C | 123 | ALA | p.Ala123Glu | P29965 |
| HOMO SAPIENS | 1 | 4426 | 3qd6 | C | 140 | TRP | p.Trp140Gly | P29965 |
| HOMO SAPIENS | 0 | 4426 | 3qd6 | C | 155 | LEU | p.Leu155Pro | P29965 |
| HOMO SAPIENS | 6 | 4426 | 3qd6 | C | 227 | GLY | p.Gly227Val | P29965 |
| HOMO SAPIENS | 0 | 4426 | 3qd6 | C | 235 | ALA | p.Ala235Pro | P29965 |
| HOMO SAPIENS | 0 | 10734 | 3dyd | B | 45 | PRO | p.Pro45Ser | P17735 |
| HOMO SAPIENS | -1 | 10734 | 3dyd | B | 57 | ARG | p.Arg57Ter | P17735 |
| HOMO SAPIENS | 8 | 10734 | 3dyd | B | 417 | ARG | p.Arg417Ter | P17735 |
| HOMO SAPIENS | 0 | 6956 | 4pzs | A | 1600 | LEU | p.Leu1600Phe | Q09472 |
| HOMO SAPIENS | 0 | 9370 | 3hfv | A | 21 | SER | p.Ser57Cys | O95363 |
| HOMO SAPIENS | 1 | 9370 | 3hfv | A | 25 | ASP | p.Asp61Glu | O95363 |
| HOMO SAPIENS | 1 | 9370 | 3hfv | A | 130 | ALA | p.Ala166Thr | O95363 |
| HOMO SAPIENS | 5 | 9370 | 3hfv | A | 148 | ASP | p.Asp184Asn | O95363 |
| HOMO SAPIENS | 0 | 9370 | 3hfv | A | 187 | ARG | p.Arg223Cys | O95363 |
| HOMO SAPIENS | 0 | 9370 | 3hfv | A | 210 | THR | p.Thr246Met | O95363 |
| HOMO SAPIENS | 1 | 9370 | 3hfv | A | 244 | ASN | p.Asn280Ser | O95363 |
| HOMO SAPIENS | 2 | 9370 | 3hfv | A | 288 | TYR | p.Tyr324Cys | O95363 |
| HOMO SAPIENS | 5 | 9370 | 3hfv | A | 293 | ILE | p.Ile329Thr | O95363 |
| HOMO SAPIENS | 1 | 9370 | 3hfv | A | 355 | ASP | p.Asp391Val | O95363 |
| HOMO SAPIENS | 1 | 9370 | 3hfv | A | 371 | THR | p.Thr407Met | O95363 |
| HOMO SAPIENS | 1 | 9370 | 3hfv | A | 383 | ARG | p.Arg419Cys | O95363 |
| HOMO SAPIENS | 1 | 9370 | 3hfv | A | 387 | ARG | p.Arg423Gln | O95363 |
| HOMO SAPIENS | 0 | 14401 | 1q3x | A | 371 | ASP | p.Asp371Tyr | O00187 |
| HOMO SAPIENS | 0 | 14401 | 1q3x | A | 377 | VAL | p.Val377Ala | O00187 |
| HOMO SAPIENS | 0 | 14401 | 1q3x | A | 439 | ARG | p.Arg439His | O00187 |
| HOMO SAPIENS | 0 | 14401 | 1q3x | A | 577 | GLN | p.Gln577His | O00187 |
| HOMO SAPIENS | 2 | 18270 | 3teq | A | 429 | ARG | p.Arg429Cys | Q13586 |
| HOMO SAPIENS | -1 | 16081 | 4mqt | A | 231 | VAL | p.Val231Ile | P08172 |

| | | | | | | | | |
|---|---|---|---|---|---|---|---|---|
| HOMO SAPIENS | 9 | 6607 | 3whu | B | 81 | ASP | p.Asp81His | Q8NI22 |
| HOMO SAPIENS | 9 | 6607 | 3whu | B | 129 | ASP | p.Asp129Glu | Q8NI22 |
| HOMO SAPIENS | 9 | 6607 | 3whu | B | 136 | ILE | p.Ile136Thr | Q8NI22 |
| HOMO SAPIENS | 0 | 7208 | 2e9w | D | 11 | ASN | p.Asn36Ser | P21583 |
| HOMO SAPIENS | 0 | 24566 | 4bdx | A | 10 | LEU | p.Leu140Val | P00748 |
| HOMO SAPIENS | 5 | 24566 | 4bdx | A | 77 | PRO | p.Ala207Pro | P00748 |
| HOMO SAPIENS | 7 | 606 | 3qnf | A | 319 | MET | p.Met319Val | Q9NZ08 |
| HOMO SAPIENS | 8 | 4097 | 1ydl | A | 21 | LEU | p.Leu21Pro | Q6ZYL4 |
| HOMO SAPIENS | 6 | 4097 | 1ydl | A | 56 | ARG | p.Arg56Ter | Q6ZYL4 |
| HOMO SAPIENS | 8 | 36691 | 1dqb | A | 42 | ARG | p.Arg403Lys | P07204 |
| HOMO SAPIENS | 0 | 5642 | 3j7y | X | 59 | ARG | p.Arg59Pro | Q13084 |
| HOMO SAPIENS | 1 | 5642 | 3j7y | X | 204 | VAL | p.Val204Met | Q13084 |
| HOMO SAPIENS | 1 | 1119 | 3nwu | B | 252 | ALA | p.Ala252Thr | Q92743 |
| HOMO SAPIENS | 3 | 1119 | 3nwu | B | 274 | ARG | p.Arg274Gln | Q92743 |
| HOMO SAPIENS | 9 | 1119 | 3nwu | B | 297 | VAL | p.Val297Met | Q92743 |
| HOMO SAPIENS | -1 | 1119 | 3nwu | B | 302 | ARG | p.Arg302Ter | Q92743 |
| HOMO SAPIENS | 9 | 1119 | 3nwu | B | 321 | ALA | p.Ala321Thr | Q92743 |
| HOMO SAPIENS | -1 | 1119 | 3nwu | B | 370 | ARG | p.Arg370Ter | Q92743 |
| HOMO SAPIENS | 0 | 416 | 4ycu | B | 105 | LEU | p.Leu105His | Q15046 |
| HOMO SAPIENS | 0 | 416 | 4ycu | B | 145 | TYR | p.Tyr145His | Q15046 |
| HOMO SAPIENS | 0 | 416 | 4ycu | B | 200 | PRO | p.Pro200Leu | Q15046 |
| HOMO SAPIENS | 0 | 416 | 4ycu | B | 201 | TYR | p.Tyr201His | Q15046 |
| HOMO SAPIENS | 0 | 416 | 4ycu | B | 420 | ARG | p.Arg420Cys | Q15046 |
| HOMO SAPIENS | 9 | 3297 | 4fpb | A | 680 | THR | p.Thr680Ser | P19367 |
| HOMO SAPIENS | 0 | 6771 | 2zt6 | A | 201 | ALA | p.Ala255Val | P41250 |
| HOMO SAPIENS | 0 | 6771 | 2zt6 | A | 202 | ASP | p.Asp256His | P41250 |
| HOMO SAPIENS | 0 | 6771 | 2zt6 | A | 334 | ARG | p.Arg388Gln | P41250 |
| HOMO SAPIENS | -1 | 6771 | 2zt6 | A | 439 | ASN | p.Asn493Ser | P41250 |
| HOMO SAPIENS | -1 | 6771 | 2zt6 | A | 500 | ASP | p.Asp554Asn | P41250 |
| HOMO SAPIENS | 1 | 6771 | 2zt6 | A | 581 | SER | p.Ser635Leu | P41250 |
| HOMO SAPIENS | -1 | 6771 | 2zt6 | A | 684 | GLU | p.Glu738Lys | P41250 |
| HOMO SAPIENS | 0 | 2687 | 4grw | B | 276 | VAL | p.Val298Phe | P29460 |
| HOMO SAPIENS | 6 | 913 | 1m75 | B | 45 | ASP | p.Asp57Glu | Q16836 |
| HOMO SAPIENS | 2 | 913 | 1m75 | B | 74 | PRO | p.Leu86Pro | Q16836 |
| HOMO SAPIENS | 3 | 913 | 1m75 | B | 214 | TYR | p.Tyr226His | Q16836 |
| HOMO SAPIENS | 3 | 913 | 1m75 | B | 224 | ARG | p.Arg236Ter | Q16836 |
| HOMO SAPIENS | 4 | 913 | 1m75 | B | 246 | PRO | p.Pro258Leu | Q16836 |
| HOMO SAPIENS | 2 | 4713 | 4djn | B | 40 | ARG | p.Arg40Cys | Q7LG56 |
| HOMO SAPIENS | 1 | 4713 | 4djn | B | 41 | ARG | p.Arg41Pro | Q7LG56 |
| HOMO SAPIENS | 2 | 4713 | 4djn | B | 110 | ARG | p.Arg110His | Q7LG56 |
| HOMO SAPIENS | 2 | 4713 | 4djn | B | 121 | ARG | p.Arg121His | Q7LG56 |
| HOMO SAPIENS | 9 | 4713 | 4djn | B | 131 | GLU | p.Glu131Lys | Q7LG56 |
| HOMO SAPIENS | 2 | 4713 | 4djn | B | 182 | THR | p.Thr182Ala | Q7LG56 |
| HOMO SAPIENS | 6 | 4713 | 4djn | B | 194 | GLU | p.Glu194Lys | Q7LG56 |
| HOMO SAPIENS | 2 | 4713 | 4djn | B | 211 | ARG | p.Arg211Lys | Q7LG56 |
| HOMO SAPIENS | 0 | 4713 | 4djn | B | 224 | ILE | p.Ile224Ser | Q7LG56 |
| HOMO SAPIENS | 3 | 4713 | 4djn | B | 229 | GLY | p.Gly229Val | Q7LG56 |
| HOMO SAPIENS | 2 | 4713 | 4djn | B | 236 | CYS | p.Cys236Phe | Q7LG56 |
| HOMO SAPIENS | 0 | 4713 | 4djn | B | 273 | GLY | p.Gly273Ser | Q7LG56 |
| HOMO SAPIENS | -1 | 4713 | 4djn | B | 282 | MSE | p.Met282Ile | Q7LG56 |
| HOMO SAPIENS | 3 | 4713 | 4djn | B | 284 | GLN | p.Gln284Ter | Q7LG56 |
| HOMO SAPIENS | 3 | 4713 | 4djn | B | 305 | ALA | p.Ala305Ser | Q7LG56 |
| HOMO SAPIENS | -1 | 8974 | 5n7i | A | 500 | ARG | p.Arg500Ter | Q8N4S9 |
| HOMO SAPIENS | 0 | 17072 | 3fxi | B | 299 | ASP | p.Asp299Gly | O00206 |

| | | | | | | | | |
|---|---|---|---|---|---|---|---|---|
| HOMO SAPIENS | 1 | 14355 | 2wwk | T | 55 | ILE | p.Ile34306Asn | Q8WZ42 |
| HOMO SAPIENS | 0 | 14355 | 2wwk | T | 64 | LEU | p.Leu34315Pro | Q8WZ42 |
| HOMO SAPIENS | 2 | 17797 | 1xu1 | S | 72 | ARG | p.Arg72His | O14836 |
| HOMO SAPIENS | -1 | 10836 | 4ztu | A | 39 | GLN | p.Gln39Arg | P54098 |
| HOMO SAPIENS | -1 | 10836 | 4ztu | A | 42 | ARG | p.Arg42Gln | P54098 |
| HOMO SAPIENS | -1 | 10836 | 4ztu | A | 45 | GLN | p.Gln45Arg | P54098 |
| HOMO SAPIENS | -1 | 10836 | 4ztu | A | 52 | GLN | p.Gln52Glu | P54098 |
| HOMO SAPIENS | 0 | 10836 | 4ztu | A | 88 | PHE | p.Phe88Leu | P54098 |
| HOMO SAPIENS | 1 | 10836 | 4ztu | A | 194 | ALA | p.Ala194Val | P54098 |
| HOMO SAPIENS | 1 | 10836 | 4ztu | A | 227 | ARG | p.Arg227Trp | P54098 |
| HOMO SAPIENS | 0 | 10836 | 4ztu | A | 232 | ARG | p.Arg232His | P54098 |
| HOMO SAPIENS | 1 | 10836 | 4ztu | A | 285 | GLN | p.Gln285His | P54098 |
| HOMO SAPIENS | 1 | 10836 | 4ztu | A | 434 | TYR | p.Tyr434His | P54098 |
| HOMO SAPIENS | -1 | 10836 | 4ztu | A | 511 | SER | p.Ser511Asn | P54098 |
| HOMO SAPIENS | 0 | 10836 | 4ztu | A | 533 | CYS | p.Cys533Trp | P54098 |
| HOMO SAPIENS | 1 | 10836 | 4ztu | A | 546 | ARG | p.Arg546Cys | P54098 |
| HOMO SAPIENS | 1 | 10836 | 4ztu | A | 587 | PRO | p.Pro587Leu | P54098 |
| HOMO SAPIENS | 0 | 10836 | 4ztu | A | 603 | MET | p.Met603Thr | P54098 |
| HOMO SAPIENS | 0 | 10836 | 4ztu | A | 623 | LEU | p.Leu623Trp | P54098 |
| HOMO SAPIENS | -1 | 10836 | 4ztu | A | 627 | ARG | p.Arg627Trp | P54098 |
| HOMO SAPIENS | -1 | 10836 | 4ztu | A | 628 | ARG | p.Arg628Pro | P54098 |
| HOMO SAPIENS | 0 | 10836 | 4ztu | A | 635 | PRO | p.Pro635Leu | P54098 |
| HOMO SAPIENS | 0 | 10836 | 4ztu | A | 662 | GLU | p.Glu662Lys | P54098 |
| HOMO SAPIENS | -1 | 10836 | 4ztu | A | 674 | GLY | p.Gly674Asp | P54098 |
| HOMO SAPIENS | -1 | 10836 | 4ztu | A | 722 | ARG | p.Arg722His | P54098 |
| HOMO SAPIENS | -1 | 10836 | 4ztu | A | 737 | GLY | p.Gly737Arg | P54098 |
| HOMO SAPIENS | 0 | 10836 | 4ztu | A | 748 | TRP | p.Trp748Ser | P54098 |
| HOMO SAPIENS | 0 | 10836 | 4ztu | A | 749 | PHE | p.Phe749Ser | P54098 |
| HOMO SAPIENS | 1 | 10836 | 4ztu | A | 807 | ARG | p.Arg807Cys | P54098 |
| HOMO SAPIENS | 1 | 10836 | 4ztu | A | 831 | TYR | p.Tyr831Cys | P54098 |
| HOMO SAPIENS | 8 | 10836 | 4ztu | A | 848 | GLY | p.Gly848Ser | P54098 |
| HOMO SAPIENS | 1 | 10836 | 4ztu | A | 852 | ARG | p.Arg852Cys | P54098 |
| HOMO SAPIENS | 9 | 10836 | 4ztu | A | 853 | ARG | p.Arg853Trp | P54098 |
| HOMO SAPIENS | 1 | 10836 | 4ztu | A | 864 | ASN | p.Asn864Ser | P54098 |
| HOMO SAPIENS | 1 | 10836 | 4ztu | A | 879 | GLN | p.Gln879Arg | P54098 |
| HOMO SAPIENS | 1 | 10836 | 4ztu | A | 914 | THR | p.Thr914Pro | P54098 |
| HOMO SAPIENS | 6 | 10836 | 4ztu | A | 932 | HIS | p.His932Tyr | P54098 |
| HOMO SAPIENS | 2 | 10836 | 4ztu | A | 944 | GLU | p.Glu944Lys | P54098 |
| HOMO SAPIENS | 6 | 10836 | 4ztu | A | 955 | TYR | p.Tyr955Cys | P54098 |
| HOMO SAPIENS | 1 | 10836 | 4ztu | A | 957 | ALA | p.Ala957Ser | P54098 |
| HOMO SAPIENS | 1 | 10836 | 4ztu | A | 957 | ALA | p.Ala957Val | P54098 |
| HOMO SAPIENS | 1 | 10836 | 4ztu | A | 964 | ARG | p.Arg964Cys | P54098 |
| HOMO SAPIENS | 1 | 10836 | 4ztu | A | 966 | LEU | p.Leu966Arg | P54098 |
| HOMO SAPIENS | 0 | 10836 | 4ztu | A | 1051 | GLY | p.Gly1051Arg | P54098 |
| HOMO SAPIENS | 1 | 10836 | 4ztu | A | 1073 | PRO | p.Pro1073Leu | P54098 |
| HOMO SAPIENS | 1 | 10836 | 4ztu | A | 1096 | ARG | p.Arg1096Cys | P54098 |
| HOMO SAPIENS | 1 | 10836 | 4ztu | A | 1096 | ARG | p.Arg1096His | P54098 |
| HOMO SAPIENS | 0 | 10836 | 4ztu | A | 1116 | MET | p.Met1116Thr | P54098 |
| HOMO SAPIENS | 1 | 10836 | 4ztu | A | 1142 | ARG | p.Arg1142Trp | P54098 |
| HOMO SAPIENS | 1 | 10836 | 4ztu | A | 1146 | ARG | p.Arg1146Cys | P54098 |
| HOMO SAPIENS | 1 | 10836 | 4ztu | A | 1157 | ASN | p.Asn1157Ser | P54098 |
| HOMO SAPIENS | -1 | 10836 | 4ztu | A | 1234 | ARG | p.Arg1234Gln | P54098 |
| HOMO SAPIENS | -1 | 10836 | 4ztu | A | 1236 | GLN | p.Gln1236His | P54098 |
| HOMO SAPIENS | 5 | 2666 | 1jph | A | 62 | PRO | p.Pro62Leu | P06132 |

| | | | | | | | | |
|---|---|---|---|---|---|---|---|---|
| HOMO SAPIENS | 5 | 2666 | 1jph | A | 116 | GLN | p.Gln116Ter | P06132 |
| HOMO SAPIENS | 0 | 2666 | 1jph | A | 165 | MET | p.Met165Arg | P06132 |
| HOMO SAPIENS | 6 | 2666 | 1jph | A | 167 | GLU | p.Glu167Lys | P06132 |
| HOMO SAPIENS | 0 | 2666 | 1jph | A | 195 | LEU | p.Leu195Phe | P06132 |
| HOMO SAPIENS | 7 | 2666 | 1jph | A | 253 | LEU | p.Leu253Gln | P06132 |
| HOMO SAPIENS | 0 | 2666 | 1jph | A | 281 | GLY | p.Gly281Glu | P06132 |
| HOMO SAPIENS | 9 | 2666 | 1jph | A | 304 | ASN | p.Asn304Lys | P06132 |
| HOMO SAPIENS | 8 | 2666 | 1jph | A | 311 | TYR | p.Tyr311Cys | P06132 |
| HOMO SAPIENS | 5 | 2666 | 1jph | A | 332 | ARG | p.Arg332His | P06132 |
| HOMO SAPIENS | 1 | 889 | 1m51 | A | 184 | LEU | p.Val184Leu | P35558 |
| HOMO SAPIENS | 0 | 889 | 1m51 | A | 267 | VAL | p.Ile267Val | P35558 |
| HOMO SAPIENS | 2 | 889 | 1m51 | A | 276 | GLU | p.Glu276Lys | P35558 |
| HOMO SAPIENS | 6 | 889 | 1m51 | A | 368 | VAL | p.Val368Ile | P35558 |
| HOMO SAPIENS | 0 | 889 | 1m51 | A | 579 | SER | p.Ser579Arg | P35558 |
| HOMO SAPIENS | 1 | 4084 | 2gjx | A | 28 | GLN | p.Gln28Ter | P06865 |
| HOMO SAPIENS | 0 | 4084 | 2gjx | A | 39 | LEU | p.Leu39Arg | P06865 |
| HOMO SAPIENS | 0 | 4084 | 2gjx | A | 127 | LEU | p.Leu127Arg | P06865 |
| HOMO SAPIENS | 6 | 4084 | 2gjx | A | 137 | ARG | p.Arg137Ter | P06865 |
| HOMO SAPIENS | 6 | 4084 | 2gjx | A | 170 | ARG | p.Arg170Gln | P06865 |
| HOMO SAPIENS | 9 | 4084 | 2gjx | A | 178 | ARG | p.Arg178Cys | P06865 |
| HOMO SAPIENS | 9 | 4084 | 2gjx | A | 178 | ARG | p.Arg178His | P06865 |
| HOMO SAPIENS | 5 | 4084 | 2gjx | A | 180 | TYR | p.Tyr180His | P06865 |
| HOMO SAPIENS | 5 | 4084 | 2gjx | A | 180 | TYR | p.Tyr180Ter | P06865 |
| HOMO SAPIENS | 1 | 4084 | 2gjx | A | 197 | LYS | p.Lys197Thr | P06865 |
| HOMO SAPIENS | 0 | 4084 | 2gjx | A | 200 | VAL | p.Val200Met | P06865 |
| HOMO SAPIENS | 9 | 4084 | 2gjx | A | 204 | HIS | p.His204Arg | P06865 |
| HOMO SAPIENS | 5 | 4084 | 2gjx | A | 210 | SER | p.Ser210Phe | P06865 |
| HOMO SAPIENS | 0 | 4084 | 2gjx | A | 211 | PHE | p.Phe211Ser | P06865 |
| HOMO SAPIENS | 2 | 4084 | 2gjx | A | 250 | GLY | p.Gly250Asp | P06865 |
| HOMO SAPIENS | 9 | 4084 | 2gjx | A | 258 | ASP | p.Asp258His | P06865 |
| HOMO SAPIENS | 0 | 4084 | 2gjx | A | 269 | GLY | p.Gly269Ser | P06865 |
| HOMO SAPIENS | 0 | 4084 | 2gjx | A | 301 | MET | p.Met301Arg | P06865 |
| HOMO SAPIENS | 3 | 4084 | 2gjx | A | 393 | ARG | p.Arg393Ter | P06865 |
| HOMO SAPIENS | 3 | 4084 | 2gjx | A | 393 | ARG | p.Arg393Pro | P06865 |
| HOMO SAPIENS | 0 | 4084 | 2gjx | A | 399 | ASN | p.Asn399Asp | P06865 |
| HOMO SAPIENS | 2 | 4084 | 2gjx | A | 420 | TRP | p.Trp420Cys | P06865 |
| HOMO SAPIENS | 1 | 4084 | 2gjx | A | 434 | PHE | p.Phe434Leu | P06865 |
| HOMO SAPIENS | 0 | 4084 | 2gjx | A | 454 | GLY | p.Gly454Ser | P06865 |
| HOMO SAPIENS | 1 | 4084 | 2gjx | A | 474 | TRP | p.Trp474Cys | P06865 |
| HOMO SAPIENS | 0 | 4084 | 2gjx | A | 479 | ALA | p.Ala479Thr | P06865 |
| HOMO SAPIENS | 5 | 4084 | 2gjx | A | 485 | TRP | p.Trp485Arg | P06865 |
| HOMO SAPIENS | 4 | 4084 | 2gjx | A | 499 | ARG | p.Arg499Cys | P06865 |
| HOMO SAPIENS | 4 | 4084 | 2gjx | A | 499 | ARG | p.Arg499His | P06865 |
| HOMO SAPIENS | 2 | 4084 | 2gjx | A | 504 | ARG | p.Arg504Cys | P06865 |
| HOMO SAPIENS | 2 | 4084 | 2gjx | A | 504 | ARG | p.Arg504His | P06865 |
| HOMO SAPIENS | -1 | 1943 | 1xmj | A | 417 | ASN | p.Asn417Lys | P13569 |
| HOMO SAPIENS | -1 | 1943 | 1xmj | A | 422 | SER | p.Ser422Phe | P13569 |
| HOMO SAPIENS | 1 | 1943 | 1xmj | A | 438 | THR | p.Thr438Ala | P13569 |
| HOMO SAPIENS | 0 | 1943 | 1xmj | A | 455 | ALA | p.Ala455Glu | P13569 |
| HOMO SAPIENS | 4 | 1943 | 1xmj | A | 456 | VAL | p.Val456Ala | P13569 |
| HOMO SAPIENS | 9 | 1943 | 1xmj | A | 458 | GLY | p.Gly458Val | P13569 |
| HOMO SAPIENS | 8 | 1943 | 1xmj | A | 466 | SER | p.Ser466Ter | P13569 |
| HOMO SAPIENS | 0 | 1943 | 1xmj | A | 467 | LEU | p.Leu467Pro | P13569 |
| HOMO SAPIENS | 1 | 1943 | 1xmj | A | 470 | MET | p.Val470Met | P13569 |

| | | | | | | | | |
|---|---|---|---|---|---|---|---|---|
| HOMO SAPIENS | 5 | 1943 | 1xmj | A | 480 | GLY | p.Gly480Cys | P13569 |
| HOMO SAPIENS | 3 | 1943 | 1xmj | A | 492 | SER | p.Ser492Phe | P13569 |
| HOMO SAPIENS | 8 | 1943 | 1xmj | A | 493 | GLN | p.Gln493Ter | P13569 |
| HOMO SAPIENS | 5 | 1943 | 1xmj | A | 506 | ILE | p.Ile506Met | P13569 |
| HOMO SAPIENS | 3 | 1943 | 1xmj | A | 520 | VAL | p.Val520Phe | P13569 |
| HOMO SAPIENS | 1 | 1943 | 1xmj | A | 528 | GLU | p.Glu528Asp | P13569 |
| HOMO SAPIENS | 1 | 1943 | 1xmj | A | 534 | ALA | p.Ala534Glu | P13569 |
| HOMO SAPIENS | 4 | 1943 | 1xmj | A | 542 | GLY | p.Gly542Ter | P13569 |
| HOMO SAPIENS | 6 | 1943 | 1xmj | A | 549 | SER | p.Ser549Asn | P13569 |
| HOMO SAPIENS | 6 | 1943 | 1xmj | A | 549 | SER | p.Ser549Arg | P13569 |
| HOMO SAPIENS | 6 | 1943 | 1xmj | A | 549 | SER | p.Ser549Arg | P13569 |
| HOMO SAPIENS | 6 | 1943 | 1xmj | A | 551 | GLY | p.Gly551Asp | P13569 |
| HOMO SAPIENS | 6 | 1943 | 1xmj | A | 551 | GLY | p.Gly551Ser | P13569 |
| HOMO SAPIENS | 6 | 1943 | 1xmj | A | 553 | GLN | p.Arg553Gln | P13569 |
| HOMO SAPIENS | 6 | 1943 | 1xmj | A | 553 | GLN | p.Arg553Ter | P13569 |
| HOMO SAPIENS | 0 | 1943 | 1xmj | A | 559 | ALA | p.Ala559Thr | P13569 |
| HOMO SAPIENS | 7 | 1943 | 1xmj | A | 560 | ARG | p.Arg560Lys | P13569 |
| HOMO SAPIENS | 4 | 1943 | 1xmj | A | 562 | VAL | p.Val562Ile | P13569 |
| HOMO SAPIENS | 3 | 1943 | 1xmj | A | 563 | TYR | p.Tyr563Asn | P13569 |
| HOMO SAPIENS | 3 | 1943 | 1xmj | A | 569 | TYR | p.Tyr569Asp | P13569 |
| HOMO SAPIENS | 3 | 1943 | 1xmj | A | 585 | GLU | p.Glu585Ter | P13569 |
| HOMO SAPIENS | 1 | 1943 | 1xmj | A | 618 | ILE | p.Ile618Thr | P13569 |
| HOMO SAPIENS | 9 | 1943 | 1xmj | A | 622 | GLY | p.Gly622Asp | P13569 |
| HOMO SAPIENS | -1 | 1943 | 1xmj | A | 648 | ASP | p.Asp648Val | P13569 |
| HOMO SAPIENS | 0 | 1943 | 1xmj | A | 673 | GLY | p.Gly673Ter | P13569 |
| HOMO SAPIENS | 5 | 5111 | 4rec | A | 749 | ARG | p.Arg749Ter | Q9Y2M0 |
| HOMO SAPIENS | -1 | 4580 | 4l0d | B | 2 | GLY | p.Pro2Leu | P35520 |
| HOMO SAPIENS | -1 | 4580 | 4l0d | B | 12 | PRO | p.Pro12Ser | P35520 |
| HOMO SAPIENS | -1 | 4580 | 4l0d | B | 18 | ARG | p.Arg18Cys | P35520 |
| HOMO SAPIENS | 1 | 4580 | 4l0d | B | 49 | PRO | p.Pro49Leu | P35520 |
| HOMO SAPIENS | 2 | 4580 | 4l0d | B | 70 | PRO | p.Pro70Leu | P35520 |
| HOMO SAPIENS | 1 | 4580 | 4l0d | B | 102 | LYS | p.Lys102Gln | P35520 |
| HOMO SAPIENS | 7 | 4580 | 4l0d | B | 114 | ALA | p.Ala114Val | P35520 |
| HOMO SAPIENS | 5 | 4580 | 4l0d | B | 116 | GLY | p.Gly116Arg | P35520 |
| HOMO SAPIENS | 8 | 4580 | 4l0d | B | 121 | ARG | p.Arg121Cys | P35520 |
| HOMO SAPIENS | 8 | 4580 | 4l0d | B | 121 | ARG | p.Arg121His | P35520 |
| HOMO SAPIENS | 5 | 4580 | 4l0d | B | 125 | ARG | p.Arg125Gln | P35520 |
| HOMO SAPIENS | 4 | 4580 | 4l0d | B | 134 | GLY | p.Gly134Arg | P35520 |
| HOMO SAPIENS | 4 | 4580 | 4l0d | B | 139 | GLY | p.Gly139Arg | P35520 |
| HOMO SAPIENS | 0 | 4580 | 4l0d | B | 145 | PRO | p.Pro145Leu | P35520 |
| HOMO SAPIENS | 0 | 4580 | 4l0d | B | 148 | GLY | p.Gly148Arg | P35520 |
| HOMO SAPIENS | 0 | 4580 | 4l0d | B | 168 | VAL | p.Val168Met | P35520 |
| HOMO SAPIENS | 2 | 4580 | 4l0d | B | 191 | THR | p.Thr191Met | P35520 |
| HOMO SAPIENS | 5 | 4580 | 4l0d | B | 234 | ASP | p.Asp234Asn | P35520 |
| HOMO SAPIENS | 0 | 4580 | 4l0d | B | 257 | THR | p.Thr257Met | P35520 |
| HOMO SAPIENS | 4 | 4580 | 4l0d | B | 262 | THR | p.Thr262Met | P35520 |
| HOMO SAPIENS | 0 | 4580 | 4l0d | B | 278 | ILE | p.Ile278Thr | P35520 |
| HOMO SAPIENS | 2 | 4580 | 4l0d | B | 307 | GLY | p.Gly307Ser | P35520 |
| HOMO SAPIENS | 4 | 4580 | 4l0d | B | 320 | VAL | p.Val320Ala | P35520 |
| HOMO SAPIENS | 2 | 4580 | 4l0d | B | 336 | ARG | p.Arg336Cys | P35520 |
| HOMO SAPIENS | 2 | 4580 | 4l0d | B | 336 | ARG | p.Arg336His | P35520 |
| HOMO SAPIENS | 0 | 4580 | 4l0d | B | 351 | GLY | p.Gly351Arg | P35520 |
| HOMO SAPIENS | 0 | 4580 | 4l0d | B | 353 | THR | p.Thr353Met | P35520 |
| HOMO SAPIENS | 0 | 4580 | 4l0d | B | 371 | VAL | p.Val371Met | P35520 |

| | | | | | | | | |
|---|---|---|---|---|---|---|---|---|
| HOMO SAPIENS | 2 | 4580 | 4l0d | B | 379 | ARG | p.Arg379Gln | P35520 |
| HOMO SAPIENS | 2 | 4580 | 4l0d | B | 379 | ARG | p.Arg379Trp | P35520 |
| HOMO SAPIENS | 2 | 4580 | 4l0d | B | 384 | LYS | p.Lys384Glu | P35520 |
| HOMO SAPIENS | 0 | 4580 | 4l0d | B | 422 | PRO | p.Pro422Leu | P35520 |
| HOMO SAPIENS | 0 | 4580 | 4l0d | B | 444 | ASP | p.Asp444Asn | P35520 |
| HOMO SAPIENS | 0 | 4580 | 4l0d | B | 466 | SER | p.Ser466Leu | P35520 |
| HOMO SAPIENS | 0 | 4580 | 4l0d | B | 539 | LEU | p.Leu539Ser | P35520 |
| HOMO SAPIENS | 0 | 4580 | 4l0d | B | 548 | ARG | p.Arg548Gln | P35520 |
| HOMO SAPIENS | 1 | 6536 | 3ulu | A | 412 | LEU | p.Leu412Phe | O15455 |
| HOMO SAPIENS | 1 | 28493 | 3ii6 | X | 814 | ARG | p.Arg814Ter | P49917 |
| HOMO SAPIENS | 1 | 28493 | 3ii6 | X | 857 | ALA | p.Ala857Thr | P49917 |
| HOMO SAPIENS | 9 | 3543 | 2q51 | A | 24 | GLU | p.Glu24Gly | P45381 |
| HOMO SAPIENS | 1 | 3543 | 2q51 | A | 27 | GLY | p.Gly27Arg | P45381 |
| HOMO SAPIENS | 9 | 3543 | 2q51 | A | 71 | ARG | p.Arg71His | P45381 |
| HOMO SAPIENS | -1 | 3543 | 2q51 | A | 82 | MSE | p.Met82Thr | P45381 |
| HOMO SAPIENS | 0 | 3543 | 2q51 | A | 143 | ILE | p.Ile143Phe | P45381 |
| HOMO SAPIENS | 1 | 3543 | 2q51 | A | 152 | CYS | p.Cys152Arg | P45381 |
| HOMO SAPIENS | 2 | 3543 | 2q51 | A | 218 | CYS | p.Cys218Ter | P45381 |
| HOMO SAPIENS | 1 | 3543 | 2q51 | A | 231 | TYR | p.Tyr231Cys | P45381 |
| HOMO SAPIENS | 1 | 3543 | 2q51 | A | 231 | TYR | p.Tyr231Ter | P45381 |
| HOMO SAPIENS | 2 | 3543 | 2q51 | A | 274 | GLY | p.Gly274Arg | P45381 |
| HOMO SAPIENS | 6 | 3543 | 2q51 | A | 285 | GLU | p.Glu285Ala | P45381 |
| HOMO SAPIENS | 5 | 3543 | 2q51 | A | 287 | ALA | p.Ala287Thr | P45381 |
| HOMO SAPIENS | 2 | 3543 | 2q51 | A | 305 | ALA | p.Ala305Glu | P45381 |
| HOMO SAPIENS | 2 | 24687 | 1hfi | A | 25 | SER | p.Ser890Ile | P08603 |
| HOMO SAPIENS | 0 | 1508 | 3wn5 | C | 45 | LEU | p.Leu66Arg | P08637 |
| HOMO SAPIENS | 0 | 1508 | 3wn5 | C | 155 | VAL | p.Phe176Val | P08637 |
| HOMO SAPIENS | 0 | 750 | 5tpa | A | 54 | VAL | p.Val452Met | Q12879 |
| HOMO SAPIENS | 9 | 750 | 5tpa | A | 120 | ARG | p.Arg518Cys | Q12879 |
| HOMO SAPIENS | 0 | 750 | 5tpa | A | 199 | ALA | p.Ala716Thr | Q12879 |
| HOMO SAPIENS | 4 | 1363 | 3hg2 | B | 40 | PRO | p.Pro40Ser | P06280 |
| HOMO SAPIENS | 3 | 1363 | 3hg2 | B | 65 | SER | p.Ser65Thr | P06280 |
| HOMO SAPIENS | 4 | 1363 | 3hg2 | B | 66 | GLU | p.Glu66Gln | P06280 |
| HOMO SAPIENS | 2 | 1363 | 3hg2 | B | 80 | GLY | p.Gly80Asp | P06280 |
| HOMO SAPIENS | 3 | 1363 | 3hg2 | B | 112 | ARG | p.Arg112Cys | P06280 |
| HOMO SAPIENS | 3 | 1363 | 3hg2 | B | 112 | ARG | p.Arg112His | P06280 |
| HOMO SAPIENS | 0 | 1363 | 3hg2 | B | 139 | ASN | p.Asn139Ser | P06280 |
| HOMO SAPIENS | 7 | 1363 | 3hg2 | B | 156 | ALA | p.Ala156Thr | P06280 |
| HOMO SAPIENS | 5 | 1363 | 3hg2 | B | 162 | TRP | p.Trp162Arg | P06280 |
| HOMO SAPIENS | 5 | 1363 | 3hg2 | B | 207 | TYR | p.Tyr207His | P06280 |
| HOMO SAPIENS | 0 | 1363 | 3hg2 | B | 215 | ASN | p.Asn215Ser | P06280 |
| HOMO SAPIENS | 0 | 1363 | 3hg2 | B | 264 | ASP | p.Asp264Val | P06280 |
| HOMO SAPIENS | 5 | 1363 | 3hg2 | B | 266 | ASP | p.Asp266Val | P06280 |
| HOMO SAPIENS | 2 | 1363 | 3hg2 | B | 269 | VAL | p.Val269Ala | P06280 |
| HOMO SAPIENS | 3 | 1363 | 3hg2 | B | 272 | ASN | p.Asn272Ser | P06280 |
| HOMO SAPIENS | 3 | 1363 | 3hg2 | B | 279 | GLN | p.Gln279Glu | P06280 |
| HOMO SAPIENS | 0 | 1363 | 3hg2 | B | 289 | ILE | p.Ile289Val | P06280 |
| HOMO SAPIENS | 0 | 1363 | 3hg2 | B | 296 | MET | p.Met296Val | P06280 |
| HOMO SAPIENS | 0 | 1363 | 3hg2 | B | 296 | MET | p.Met296Ile | P06280 |
| HOMO SAPIENS | 0 | 1363 | 3hg2 | B | 297 | SER | p.Ser297Phe | P06280 |
| HOMO SAPIENS | 2 | 1363 | 3hg2 | B | 301 | ARG | p.Arg301Gln | P06280 |
| HOMO SAPIENS | 3 | 1363 | 3hg2 | B | 327 | GLN | p.Gln327Lys | P06280 |
| HOMO SAPIENS | 0 | 1363 | 3hg2 | B | 328 | GLY | p.Gly328Arg | P06280 |
| HOMO SAPIENS | 0 | 1363 | 3hg2 | B | 328 | GLY | p.Gly328Ala | P06280 |

| | | | | | | | | |
|---|---|---|---|---|---|---|---|---|
| HOMO SAPIENS | 2 | 1363 | 3hg2 | B | 331 | LEU | p.Leu331Phe | P06280 |
| HOMO SAPIENS | 0 | 1363 | 3hg2 | B | 342 | ARG | p.Arg342Gln | P06280 |
| HOMO SAPIENS | 1 | 1363 | 3hg2 | B | 361 | GLY | p.Gly361Arg | P06280 |
| HOMO SAPIENS | 1 | 1363 | 3hg2 | B | 363 | ARG | p.Arg363His | P06280 |
| HOMO SAPIENS | 1 | 1363 | 3hg2 | B | 385 | THR | p.Thr385Ala | P06280 |
| HOMO SAPIENS | 1 | 1363 | 3hg2 | B | 399 | TRP | p.Trp399Ser | P06280 |
| HOMO SAPIENS | 1 | 1363 | 3hg2 | B | 410 | THR | p.Thr410Ala | P06280 |
| HOMO SAPIENS | 2 | 3904 | 3caa | B | 385 | MET | p.Met414Val | P01011 |
| HOMO SAPIENS | 0 | 558 | 5eub | A | 56 | VAL | p.Val56Ile | Q13126 |
| HOMO SAPIENS | 0 | 558 | 5eub | A | 189 | TRP | p.Trp189Leu | Q13126 |
| HOMO SAPIENS | 0 | 558 | 5eub | A | 218 | ALA | p.Ala218Gly | Q13126 |
| HOMO SAPIENS | 1 | 3865 | 1cb5 | B | 443 | ILE | p.Ile443Val | Q13867 |
| HOMO SAPIENS | 2 | 14329 | 4m92 | A | 24 | ILE | p.Ile65Val | Q13454 |
| HOMO SAPIENS | 1 | 7381 | 5kz5 | k | 50 | GLY | p.Gly50Glu | Q9H1K1 |
| HOMO SAPIENS | 0 | 1102 | 4fdh | E | 86 | PRO | p.Pro86Ala | P19099 |
| HOMO SAPIENS | 0 | 1102 | 4fdh | E | 118 | ALA | p.Ala118Thr | P19099 |
| HOMO SAPIENS | 0 | 1102 | 4fdh | E | 159 | PRO | p.Pro159Leu | P19099 |
| HOMO SAPIENS | 0 | 1102 | 4fdh | E | 173 | LYS | p.Lys173Arg | P19099 |
| HOMO SAPIENS | 0 | 1102 | 4fdh | E | 181 | ARG | p.Arg181Trp | P19099 |
| HOMO SAPIENS | 0 | 1102 | 4fdh | E | 185 | THR | p.Thr185Ile | P19099 |
| HOMO SAPIENS | 0 | 1102 | 4fdh | E | 198 | GLU | p.Glu198Asp | P19099 |
| HOMO SAPIENS | 0 | 1102 | 4fdh | E | 214 | HIS | p.His214Asp | P19099 |
| HOMO SAPIENS | 0 | 1102 | 4fdh | E | 251 | LYS | p.Lys251Arg | P19099 |
| HOMO SAPIENS | 0 | 1102 | 4fdh | E | 255 | GLU | p.Glu255Ter | P19099 |
| HOMO SAPIENS | 0 | 1102 | 4fdh | E | 281 | ASN | p.Asn281Ser | P19099 |
| HOMO SAPIENS | 0 | 1102 | 4fdh | E | 320 | ALA | p.Ala320Val | P19099 |
| HOMO SAPIENS | 0 | 1102 | 4fdh | E | 339 | ILE | p.Ile339Thr | P19099 |
| HOMO SAPIENS | 0 | 1102 | 4fdh | E | 341 | ARG | p.Arg341Cys | P19099 |
| HOMO SAPIENS | 2 | 1102 | 4fdh | E | 374 | ARG | p.Arg374Trp | P19099 |
| HOMO SAPIENS | 0 | 1102 | 4fdh | E | 386 | VAL | p.Val386Ala | P19099 |
| HOMO SAPIENS | 0 | 1102 | 4fdh | E | 390 | LEU | p.Leu390Phe | P19099 |
| HOMO SAPIENS | -1 | 1102 | 4fdh | E | 435 | GLY | p.Gly435Ser | P19099 |
| HOMO SAPIENS | 0 | 1102 | 4fdh | E | 461 | LEU | p.Leu461Pro | P19099 |
| HOMO SAPIENS | 0 | 1102 | 4fdh | E | 498 | THR | p.Thr498Ala | P19099 |
| HOMO SAPIENS | 6 | 10292 | 2of5 | F | 128 | GLY | p.Gly128Arg | P78560 |
| HOMO SAPIENS | 6 | 10292 | 2of5 | F | 170 | ARG | p.Arg170Cys | P78560 |
| HOMO SAPIENS | -1 | 18965 | 3ghg | A | 16 | ARG | p.Arg35Cys | P02671 |
| HOMO SAPIENS | -1 | 18965 | 3ghg | A | 16 | ARG | p.Arg35His | P02671 |
| HOMO SAPIENS | -1 | 18965 | 3ghg | A | 283 | PRO | p.Pro302Ala | P02671 |
| HOMO SAPIENS | -1 | 18965 | 3ghg | A | 312 | THR | p.Thr331Ala | P02671 |
| HOMO SAPIENS | -1 | 18965 | 3ghg | A | 381 | SER | p.Ser400Phe | P02671 |
| HOMO SAPIENS | -1 | 18965 | 3ghg | A | 463 | VAL | p.Val482Met | P02671 |
| HOMO SAPIENS | -1 | 18965 | 3ghg | A | 526 | GLU | p.Glu545Val | P02671 |
| HOMO SAPIENS | -1 | 18965 | 3ghg | A | 554 | ARG | p.Arg573Leu | P02671 |
| HOMO SAPIENS | 8 | 3023 | 4ttb | A | 101 | ARG | p.Arg101Trp | Q6PHW0 |
| HOMO SAPIENS | 2 | 3023 | 4ttb | A | 116 | ILE | p.Ile116Thr | Q6PHW0 |
| HOMO SAPIENS | 0 | 3023 | 4ttb | A | 220 | ALA | p.Ala220Thr | Q6PHW0 |
| HOMO SAPIENS | 2 | 3023 | 4ttb | A | 271 | GLU | p.Glu271Lys | Q6PHW0 |
| HOMO SAPIENS | 3 | 33226 | 1ya5 | T | 11 | SER | p.Ser11Ter | O15273 |
| HOMO SAPIENS | 6 | 33226 | 1ya5 | T | 64 | SER | p.Ser64Leu | O15273 |
| HOMO SAPIENS | 6 | 33226 | 1ya5 | T | 70 | ARG | p.Arg70Gln | O15273 |
| HOMO SAPIENS | 3 | 33226 | 1ya5 | T | 75 | GLY | p.Gly75Ser | O15273 |
| HOMO SAPIENS | 0 | 33226 | 1ya5 | T | 87 | ARG | p.Arg87Trp | O15273 |
| HOMO SAPIENS | 2 | 5667 | 3dax | A | 398 | PRO | p.Pro398Ala | P22680 |

| | | | | | | | | |
|---|---|---|---|---|---|---|---|---|
| HOMO SAPIENS | 0 | 12861 | 5mya | A | 464 | ASN | p.Asn464His | Q02763 |
| HOMO SAPIENS | 2 | 12861 | 5mya | A | 486 | VAL | p.Val486Ile | Q02763 |
| HOMO SAPIENS | 0 | 12861 | 5mya | A | 522 | ARG | p.Arg522His | Q02763 |
| HOMO SAPIENS | 2 | 12861 | 5mya | A | 600 | VAL | p.Val600Leu | Q02763 |
| HOMO SAPIENS | 0 | 12861 | 5mya | A | 634 | LEU | p.Leu634Phe | Q02763 |
| HOMO SAPIENS | 0 | 12861 | 5mya | A | 684 | HIS | p.His684Gln | Q02763 |
| HOMO SAPIENS | 0 | 12861 | 5mya | A | 724 | ALA | p.Ala724Thr | Q02763 |
| HOMO SAPIENS | 0 | 7743 | 3qzw | G | 90 | GLY | p.Gly111Ser | P01732 |
| HOMO SAPIENS | 2 | 9498 | 3kci | A | 4267 | ASP | p.Asp4267Glu | O95714 |
| HOMO SAPIENS | 5 | 201 | 3ips | B | 415 | ARG | p.Arg415Gln | Q13133 |
| HOMO SAPIENS | 3 | 229 | 2q50 | A | 68 | LEU | p.Leu68Pro | Q9UBQ7 |
| HOMO SAPIENS | 2 | 229 | 2q50 | A | 99 | ARG | p.Arg99Ter | Q9UBQ7 |
| HOMO SAPIENS | 3 | 229 | 2q50 | A | 160 | GLY | p.Gly160Arg | Q9UBQ7 |
| HOMO SAPIENS | 5 | 229 | 2q50 | A | 165 | GLY | p.Gly165Asp | Q9UBQ7 |
| HOMO SAPIENS | 4 | 229 | 2q50 | A | 302 | ARG | p.Arg302Cys | Q9UBQ7 |
| HOMO SAPIENS | 4 | 229 | 2q50 | A | 302 | ARG | p.Arg302His | Q9UBQ7 |
| HOMO SAPIENS | -1 | 229 | 2q50 | A | 322 | MSE | p.Met322Arg | Q9UBQ7 |
| HOMO SAPIENS | -1 | 4519 | 5clw | A | 40 | PRO | p.Pro40Thr | Q04446 |
| HOMO SAPIENS | 1 | 4519 | 5clw | A | 59 | ILE | p.Ile59Thr | Q04446 |
| HOMO SAPIENS | 0 | 4519 | 5clw | A | 190 | ARG | p.Arg190Gly | Q04446 |
| HOMO SAPIENS | 1 | 4519 | 5clw | A | 224 | LEU | p.Leu224Pro | Q04446 |
| HOMO SAPIENS | 0 | 4519 | 5clw | A | 236 | GLN | p.Gln236His | Q04446 |
| HOMO SAPIENS | 1 | 4519 | 5clw | A | 254 | THR | p.Thr254Ala | Q04446 |
| HOMO SAPIENS | 0 | 4519 | 5clw | A | 257 | PHE | p.Phe257Leu | Q04446 |
| HOMO SAPIENS | 1 | 4519 | 5clw | A | 262 | ARG | p.Arg262Cys | Q04446 |
| HOMO SAPIENS | 3 | 4519 | 5clw | A | 329 | TYR | p.Tyr329Ser | Q04446 |
| HOMO SAPIENS | 0 | 4519 | 5clw | A | 507 | THR | p.Thr507Ala | Q04446 |
| HOMO SAPIENS | 0 | 4519 | 5clw | A | 515 | ARG | p.Arg515Cys | Q04446 |
| HOMO SAPIENS | 0 | 4519 | 5clw | A | 515 | ARG | p.Arg515His | Q04446 |
| HOMO SAPIENS | 0 | 4519 | 5clw | A | 524 | ARG | p.Arg524Gln | Q04446 |
| HOMO SAPIENS | 0 | 4519 | 5clw | A | 524 | ARG | p.Arg524Ter | Q04446 |
| HOMO SAPIENS | 0 | 4519 | 5clw | A | 545 | HIS | p.His545Arg | Q04446 |
| HOMO SAPIENS | 2 | 4519 | 5clw | A | 548 | TRP | p.Trp548Ter | Q04446 |
| HOMO SAPIENS | 0 | 4519 | 5clw | A | 592 | GLU | p.Glu592Ter | Q04446 |
| HOMO SAPIENS | 0 | 4519 | 5clw | A | 628 | HIS | p.His628Arg | Q04446 |
| HOMO SAPIENS | 0 | 4519 | 5clw | A | 637 | ARG | p.Arg637Ter | Q04446 |
| HOMO SAPIENS | 0 | 4519 | 5clw | A | 673 | ALA | p.Ala673Thr | Q04446 |
| HOMO SAPIENS | 1 | 10405 | 3bg3 | D | 507 | VAL | p.Val507Ile | P11498 |
| HOMO SAPIENS | 0 | 10405 | 3bg3 | D | 583 | ARG | p.Arg583Leu | P11498 |
| HOMO SAPIENS | 1 | 10405 | 3bg3 | D | 589 | LYS | p.Lys589Asn | P11498 |
| HOMO SAPIENS | 6 | 10405 | 3bg3 | D | 610 | ALA | p.Ala610Thr | P11498 |
| HOMO SAPIENS | 2 | 10405 | 3bg3 | D | 631 | ARG | p.Arg631Gln | P11498 |
| HOMO SAPIENS | 1 | 10405 | 3bg3 | D | 732 | ARG | p.Arg732Gln | P11498 |
| HOMO SAPIENS | 6 | 10405 | 3bg3 | D | 743 | MET | p.Met743Ile | P11498 |
| HOMO SAPIENS | 0 | 10405 | 3bg3 | D | 750 | THR | p.Thr750Met | P11498 |
| HOMO SAPIENS | -1 | 13751 | 2o8d | B | 350 | ALA | p.Ala350Val | P52701 |
| HOMO SAPIENS | -1 | 13751 | 2o8d | B | 352 | VAL | p.Val352Ile | P52701 |
| HOMO SAPIENS | -1 | 13751 | 2o8d | B | 355 | GLY | p.Gly355Ser | P52701 |
| HOMO SAPIENS | 0 | 13751 | 2o8d | B | 385 | ARG | p.Arg385Trp | P52701 |
| HOMO SAPIENS | 0 | 13751 | 2o8d | B | 390 | ASP | p.Asp390Glu | P52701 |
| HOMO SAPIENS | 0 | 13751 | 2o8d | B | 396 | LEU | p.Leu396Val | P52701 |
| HOMO SAPIENS | 0 | 13751 | 2o8d | B | 449 | LEU | p.Leu449Pro | P52701 |
| HOMO SAPIENS | 0 | 13751 | 2o8d | B | 468 | ARG | p.Arg468His | P52701 |
| HOMO SAPIENS | 0 | 13751 | 2o8d | B | 503 | SER | p.Ser503Cys | P52701 |

| | | | | | | | | |
|---|---|---|---|---|---|---|---|---|
| HOMO SAPIENS | 1 | 13751 | 2o8d | B | 509 | VAL | p.Val509Ala | P52701 |
| HOMO SAPIENS | 0 | 13751 | 2o8d | B | 675 | GLU | p.Glu675Asp | P52701 |
| HOMO SAPIENS | 0 | 13751 | 2o8d | B | 795 | ILE | p.Ile795Thr | P52701 |
| HOMO SAPIENS | 0 | 13751 | 2o8d | B | 850 | TYR | p.Tyr850Ter | P52701 |
| HOMO SAPIENS | 0 | 13751 | 2o8d | B | 878 | VAL | p.Val878Ala | P52701 |
| HOMO SAPIENS | 0 | 13751 | 2o8d | B | 911 | ARG | p.Arg911Ter | P52701 |
| HOMO SAPIENS | 0 | 13751 | 2o8d | B | 922 | ARG | p.Arg922Ter | P52701 |
| HOMO SAPIENS | 0 | 13751 | 2o8d | B | 969 | TYR | p.Tyr969Cys | P52701 |
| HOMO SAPIENS | 0 | 13751 | 2o8d | B | 1005 | ARG | p.Arg1005Ter | P52701 |
| HOMO SAPIENS | 0 | 13751 | 2o8d | B | 1035 | ARG | p.Arg1035Ter | P52701 |
| HOMO SAPIENS | 1 | 13751 | 2o8d | B | 1048 | GLN | p.Gln1048Ter | P52701 |
| HOMO SAPIENS | 1 | 13751 | 2o8d | B | 1051 | VAL | p.Val1051Ile | P52701 |
| HOMO SAPIENS | 0 | 13751 | 2o8d | B | 1054 | ILE | p.Ile1054Phe | P52701 |
| HOMO SAPIENS | 0 | 13751 | 2o8d | B | 1054 | ILE | p.Ile1054Met | P52701 |
| HOMO SAPIENS | 0 | 13751 | 2o8d | B | 1163 | GLU | p.Glu1163Val | P52701 |
| HOMO SAPIENS | 0 | 13751 | 2o8d | B | 1237 | GLU | p.Glu1237Asp | P52701 |
| HOMO SAPIENS | 0 | 13751 | 2o8d | B | 1242 | ARG | p.Arg1242His | P52701 |
| HOMO SAPIENS | -1 | 13751 | 2o8d | B | 1278 | PRO | p.Pro1278Thr | P52701 |
| HOMO SAPIENS | 0 | 13751 | 2o8d | B | 1304 | ARG | p.Arg1304Lys | P52701 |
| HOMO SAPIENS | 0 | 13751 | 2o8d | B | 1317 | HIS | p.His1317Gln | P52701 |
| HOMO SAPIENS | 0 | 13751 | 2o8d | B | 1329 | SER | p.Ser1329Leu | P52701 |
| HOMO SAPIENS | 3 | 13273 | 4g8k | A | 265 | GLU | p.Glu265Ter | Q05823 |
| HOMO SAPIENS | -1 | 11088 | 5uj7 | A | 666 | ARG | p.Arg666Trp | Q13415 |
| HOMO SAPIENS | 4 | 11088 | 5uj7 | A | 720 | ARG | p.Arg720Gln | Q13415 |
| HOMO SAPIENS | 0 | 11088 | 5uj7 | A | 816 | MET | p.Met816Thr | Q13415 |
| HOMO SAPIENS | 1 | 4352 | 5va1 | A | 421 | THR | p.Thr421Met | Q12809 |
| HOMO SAPIENS | 0 | 4352 | 5va1 | A | 451 | PRO | p.Pro451Leu | Q12809 |
| HOMO SAPIENS | 0 | 4352 | 5va1 | A | 470 | ASN | p.Asn470Asp | Q12809 |
| HOMO SAPIENS | 0 | 4352 | 5va1 | A | 490 | ALA | p.Ala490Thr | Q12809 |
| HOMO SAPIENS | 0 | 4352 | 5va1 | A | 491 | VAL | p.Val491Ile | Q12809 |
| HOMO SAPIENS | 1 | 4352 | 5va1 | A | 533 | VAL | p.Val533Gly | Q12809 |
| HOMO SAPIENS | 0 | 4352 | 5va1 | A | 552 | LEU | p.Leu552Ser | Q12809 |
| HOMO SAPIENS | 0 | 4352 | 5va1 | A | 558 | ALA | p.Ala558Pro | Q12809 |
| HOMO SAPIENS | 1 | 4352 | 5va1 | A | 561 | ALA | p.Ala561Val | Q12809 |
| HOMO SAPIENS | 0 | 4352 | 5va1 | A | 572 | GLY | p.Gly572Arg | Q12809 |
| HOMO SAPIENS | -1 | 4352 | 5va1 | A | 582 | ARG | p.Arg582Cys | Q12809 |
| HOMO SAPIENS | 0 | 4352 | 5va1 | A | 584 | GLY | p.Gly584Ser | Q12809 |
| HOMO SAPIENS | 0 | 4352 | 5va1 | A | 588 | ASN | p.Asn588Lys | Q12809 |
| HOMO SAPIENS | 0 | 4352 | 5va1 | A | 593 | ILE | p.Ile593Arg | Q12809 |
| HOMO SAPIENS | 1 | 4352 | 5va1 | A | 627 | PHE | p.Phe627Leu | Q12809 |
| HOMO SAPIENS | 0 | 4352 | 5va1 | A | 630 | VAL | p.Val630Leu | Q12809 |
| HOMO SAPIENS | 0 | 4352 | 5va1 | A | 652 | TYR | p.Tyr652Ter | Q12809 |
| HOMO SAPIENS | 0 | 4352 | 5va1 | A | 711 | ILE | p.Ile711Val | Q12809 |
| HOMO SAPIENS | 0 | 4352 | 5va1 | A | 735 | SER | p.Ser735Leu | Q12809 |
| HOMO SAPIENS | 0 | 4352 | 5va1 | A | 744 | ARG | p.Arg744Ter | Q12809 |
| HOMO SAPIENS | 0 | 4352 | 5va1 | A | 784 | ARG | p.Arg784Trp | Q12809 |
| HOMO SAPIENS | 1 | 4352 | 5va1 | A | 820 | GLY | p.Gly820Arg | Q12809 |
| HOMO SAPIENS | 5 | 4352 | 5va1 | A | 822 | VAL | p.Val822Met | Q12809 |
| HOMO SAPIENS | 1 | 4352 | 5va1 | A | 863 | ARG | p.Arg863Ter | Q12809 |
| HOMO SAPIENS | 5 | 303 | 2obv | A | 55 | ALA | p.Ala55Asp | Q00266 |
| HOMO SAPIENS | 7 | 303 | 2obv | A | 259 | ALA | p.Ala259Gly | Q00266 |
| HOMO SAPIENS | 9 | 303 | 2obv | A | 264 | ARG | p.Arg264His | Q00266 |
| HOMO SAPIENS | 9 | 303 | 2obv | A | 264 | ARG | p.Arg264Cys | Q00266 |
| HOMO SAPIENS | 0 | 303 | 2obv | A | 305 | LEU | p.Leu305Pro | Q00266 |

| | | | | | | | | |
|---|---|---|---|---|---|---|---|---|
| HOMO SAPIENS | 7 | 303 | 2obv | A | 322 | ILE | p.Ile322Met | Q00266 |
| HOMO SAPIENS | 0 | 303 | 2obv | A | 336 | GLY | p.Gly336Arg | Q00266 |
| HOMO SAPIENS | 0 | 303 | 2obv | A | 357 | PRO | p.Pro357Leu | Q00266 |
| HOMO SAPIENS | 0 | 2077 | 4zxb | E | 462 | ASN | p.Asn489Ser | P06213 |
| HOMO SAPIENS | -1 | 2077 | 4zxb | E | 519 | ASP | p.Asp546Glu | P06213 |
| HOMO SAPIENS | 2 | 2077 | 4zxb | E | 819 | THR | p.Thr858Ala | P06213 |
| HOMO SAPIENS | 0 | 2077 | 4zxb | E | 907 | ASP | p.Asp946Glu | P06213 |
| HOMO SAPIENS | 7 | 3274 | 1wsv | B | 14 | HIS | p.His42Arg | P48728 |
| HOMO SAPIENS | 0 | 3274 | 1wsv | B | 19 | GLY | p.Gly47Arg | P48728 |
| HOMO SAPIENS | 4 | 3274 | 1wsv | B | 45 | ARG | p.Arg73Cys | P48728 |
| HOMO SAPIENS | 1 | 3274 | 1wsv | B | 49 | SER | p.Ser77Leu | P48728 |
| HOMO SAPIENS | 3 | 3274 | 1wsv | B | 241 | GLY | p.Gly269Asp | P48728 |
| HOMO SAPIENS | 7 | 3274 | 1wsv | B | 248 | ASP | p.Asp276His | P48728 |
| HOMO SAPIENS | 0 | 3274 | 1wsv | B | 272 | MET | p.Met300Val | P48728 |
| HOMO SAPIENS | 4 | 3274 | 1wsv | B | 292 | ARG | p.Arg320His | P48728 |
| HOMO SAPIENS | 0 | 3274 | 1wsv | B | 354 | ARG | p.Arg382Gln | P48728 |
| HOMO SAPIENS | -1 | 435 | 3uyt | A | 44 | THR | p.Thr44Ala | P48730 |
| HOMO SAPIENS | 1 | 4698 | 5ife | C | 666 | LYS | p.Lys666Asn | O75533 |
| HOMO SAPIENS | 1 | 16917 | 1apy | C | 37 | GLY | p.Gly60Asp | P20933 |
| HOMO SAPIENS | 0 | 16917 | 1apy | C | 49 | SER | p.Ser72Pro | P20933 |
| HOMO SAPIENS | 0 | 16917 | 1apy | C | 84 | ARG | p.Arg107Ter | P20933 |
| HOMO SAPIENS | 0 | 16917 | 1apy | C | 126 | SER | p.Thr149Ser | P20933 |
| HOMO SAPIENS | 0 | 16917 | 1apy | C | 135 | TRP | p.Trp158Ter | P20933 |
| HOMO SAPIENS | 0 | 16917 | 1apy | C | 140 | CYS | p.Cys163Ser | P20933 |
| HOMO SAPIENS | 0 | 16917 | 1apy | C | 145 | TRP | p.Trp168Ter | P20933 |
| HOMO SAPIENS | 0 | 10413 | 4frw | A | 185 | THR | p.Thr185Met | Q96NY8 |
| HOMO SAPIENS | 0 | 5217 | 5n6w | G | 96 | TRP | p.Trp96Arg | O15537 |
| HOMO SAPIENS | 1 | 5217 | 5n6w | G | 102 | ARG | p.Arg102Trp | O15537 |
| HOMO SAPIENS | 0 | 5217 | 5n6w | G | 109 | GLY | p.Gly109Arg | O15537 |
| HOMO SAPIENS | -1 | 5217 | 5n6w | G | 223 | CYS | p.Cys223Arg | O15537 |
| HOMO SAPIENS | 0 | 9485 | 4z6y | E | 138 | ALA | p.Ala138Asp | Q9P0N9 |
| HOMO SAPIENS | 3 | 4504 | 4mwt | B | 411 | GLY | p.Gly439Ser | P10619 |
| HOMO SAPIENS | 3 | 4504 | 4mwt | B | 412 | PHE | p.Phe440Val | P10619 |
| HOMO SAPIENS | 6 | 6391 | 4kxy | A | 318 | ARG | p.Arg318Cys | P29401 |
| HOMO SAPIENS | 6 | 19546 | 2bx6 | A | 87 | THR | p.Thr87Ile | O75695 |
| HOMO SAPIENS | 1 | 19546 | 2bx6 | A | 118 | ARG | p.Arg118His | O75695 |
| HOMO SAPIENS | 1 | 19546 | 2bx6 | A | 201 | ILE | p.Ile201Val | O75695 |
| HOMO SAPIENS | 1 | 19546 | 2bx6 | A | 282 | ARG | p.Arg282Trp | O75695 |
| HOMO SAPIENS | 1 | 5265 | 5usr | H | 68 | ARG | p.Arg68Leu | Q9HD34 |
| HOMO SAPIENS | 2 | 1012 | 4nyi | S | 569 | LEU | p.Leu569Val | Q07889 |
| HOMO SAPIENS | 2 | 1012 | 4nyi | S | 655 | PRO | p.Pro655Leu | Q07889 |
| HOMO SAPIENS | 4 | 1012 | 4nyi | S | 663 | ILE | p.Ile663Met | Q07889 |
| HOMO SAPIENS | 2 | 1012 | 4nyi | S | 702 | TYR | p.Tyr702Cys | Q07889 |
| HOMO SAPIENS | 2 | 1012 | 4nyi | S | 708 | ALA | p.Ala708Thr | Q07889 |
| HOMO SAPIENS | 4 | 1012 | 4nyi | S | 733 | ILE | p.Ile733Phe | Q07889 |
| HOMO SAPIENS | 1 | 1848 | 3g1n | A | 4013 | ARG | p.Arg4013Trp | Q7Z6Z7 |
| HOMO SAPIENS | 1 | 1848 | 3g1n | A | 4187 | ARG | p.Arg4187Cys | Q7Z6Z7 |
| HOMO SAPIENS | 1 | 77 | 4pod | A | 41 | CYS | p.Cys79Tyr | P60174 |
| HOMO SAPIENS | 9 | 77 | 4pod | A | 104 | GLU | p.Glu142Asp | P60174 |
| HOMO SAPIENS | 1 | 77 | 4pod | A | 122 | GLY | p.Gly160Arg | P60174 |
| HOMO SAPIENS | -1 | 8672 | 3thz | A | 8 | THR | p.Thr8Met | P43246 |
| HOMO SAPIENS | 0 | 8672 | 3thz | A | 19 | PHE | p.Phe19Leu | P43246 |
| HOMO SAPIENS | 0 | 8672 | 3thz | A | 23 | PHE | p.Phe23Leu | P43246 |
| HOMO SAPIENS | 0 | 8672 | 3thz | A | 96 | ARG | p.Arg96His | P43246 |

| | | | | | | | | |
|---|---|---|---|---|---|---|---|---|
| HOMO SAPIENS | 0 | 8672 | 3thz | A | 102 | VAL | p.Val102Ile | P43246 |
| HOMO SAPIENS | 0 | 8672 | 3thz | A | 106 | ARG | p.Arg106Lys | P43246 |
| HOMO SAPIENS | 0 | 8672 | 3thz | A | 127 | ASN | p.Asn127Ser | P43246 |
| HOMO SAPIENS | 0 | 8672 | 3thz | A | 128 | LEU | p.Leu128Val | P43246 |
| HOMO SAPIENS | 0 | 8672 | 3thz | A | 163 | VAL | p.Val163Gly | P43246 |
| HOMO SAPIENS | 0 | 8672 | 3thz | A | 169 | ILE | p.Ile169Val | P43246 |
| HOMO SAPIENS | 0 | 8672 | 3thz | A | 315 | GLY | p.Gly315Asp | P43246 |
| HOMO SAPIENS | -1 | 8672 | 3thz | A | 322 | GLY | p.Gly322Asp | P43246 |
| HOMO SAPIENS | 0 | 8672 | 3thz | A | 331 | ASN | p.Asn331Asp | P43246 |
| HOMO SAPIENS | 1 | 8672 | 3thz | A | 390 | LEU | p.Leu390Phe | P43246 |
| HOMO SAPIENS | 0 | 8672 | 3thz | A | 419 | GLN | p.Gln419Lys | P43246 |
| HOMO SAPIENS | 0 | 8672 | 3thz | A | 447 | PHE | p.Phe447Leu | P43246 |
| HOMO SAPIENS | 0 | 8672 | 3thz | A | 524 | ARG | p.Arg524Pro | P43246 |
| HOMO SAPIENS | 0 | 8672 | 3thz | A | 539 | PHE | p.Phe539Leu | P43246 |
| HOMO SAPIENS | -1 | 8672 | 3thz | A | 546 | LYS | p.Lys546Asn | P43246 |
| HOMO SAPIENS | 0 | 8672 | 3thz | A | 564 | THR | p.Thr564Ala | P43246 |
| HOMO SAPIENS | 0 | 8672 | 3thz | A | 577 | ILE | p.Ile577Thr | P43246 |
| HOMO SAPIENS | 0 | 8672 | 3thz | A | 588 | TYR | p.Tyr588Ter | P43246 |
| HOMO SAPIENS | 0 | 8672 | 3thz | A | 622 | PRO | p.Pro622Leu | P43246 |
| HOMO SAPIENS | 0 | 8672 | 3thz | A | 629 | GLN | p.Gln629Arg | P43246 |
| HOMO SAPIENS | 0 | 8672 | 3thz | A | 636 | ALA | p.Ala636Pro | P43246 |
| HOMO SAPIENS | 2 | 8672 | 3thz | A | 642 | VAL | p.Val642Ile | P43246 |
| HOMO SAPIENS | 1 | 8672 | 3thz | A | 654 | ASP | p.Asp654Glu | P43246 |
| HOMO SAPIENS | 0 | 8672 | 3thz | A | 662 | GLN | p.Gln662His | P43246 |
| HOMO SAPIENS | 2 | 8672 | 3thz | A | 681 | GLN | p.Gln681His | P43246 |
| HOMO SAPIENS | 0 | 8672 | 3thz | A | 707 | CYS | p.Cys707Ser | P43246 |
| HOMO SAPIENS | 0 | 8672 | 3thz | A | 809 | GLU | p.Glu809Lys | P43246 |
| HOMO SAPIENS | 0 | 8672 | 3thz | A | 813 | MET | p.Met813Val | P43246 |
| HOMO SAPIENS | 1 | 8672 | 3thz | A | 834 | ALA | p.Ala834Thr | P43246 |
| HOMO SAPIENS | 0 | 8672 | 3thz | A | 849 | LEU | p.Leu849Val | P43246 |
| HOMO SAPIENS | -1 | 8672 | 3thz | A | 860 | SER | p.Ser860Leu | P43246 |
| HOMO SAPIENS | -1 | 8672 | 3thz | A | 934 | THR | p.Thr934Lys | P43246 |
| HOMO SAPIENS | 2 | 292 | 1i3k | A | 34 | ASN | p.Asn34Ser | Q14376 |
| HOMO SAPIENS | 7 | 292 | 1i3k | A | 90 | GLY | p.Gly90Glu | Q14376 |
| HOMO SAPIENS | 2 | 292 | 1i3k | A | 103 | ASP | p.Asp103Gly | Q14376 |
| HOMO SAPIENS | 2 | 292 | 1i3k | A | 136 | TYR | p.Tyr136Ter | Q14376 |
| HOMO SAPIENS | 0 | 292 | 1i3k | A | 169 | ARG | p.Arg169Trp | Q14376 |
| HOMO SAPIENS | 0 | 292 | 1i3k | A | 183 | LEU | p.Leu183Pro | Q14376 |
| HOMO SAPIENS | 0 | 292 | 1i3k | A | 302 | GLY | p.Gly302Asp | Q14376 |
| HOMO SAPIENS | 0 | 292 | 1i3k | A | 313 | LEU | p.Leu313Met | Q14376 |
| HOMO SAPIENS | 1 | 2084 | 5vc8 | A | 295 | GLN | p.Gln295Leu | O96028 |
| HOMO SAPIENS | -1 | 10157 | 5wrw | A | 7 | GLY | p.Gly7Ala | O76094 |
| HOMO SAPIENS | -1 | 10157 | 5wrw | A | 7 | GLY | p.Gly7Arg | O76094 |
| HOMO SAPIENS | -1 | 10157 | 5wrw | A | 8 | GLY | p.Gly8Ala | O76094 |
| HOMO SAPIENS | 0 | 3113 | 3in5 | A | 29 | GLU | p.Glu29Lys | Q9UBT6 |
| HOMO SAPIENS | 5 | 4546 | 5in3 | B | 44 | VAL | p.Val44Leu | P07902 |
| HOMO SAPIENS | -1 | 4546 | 5in3 | B | 51 | ARG | p.Arg51Leu | P07902 |
| HOMO SAPIENS | -1 | 4546 | 5in3 | B | 62 | LEU | p.Leu62Met | P07902 |
| HOMO SAPIENS | 4 | 4546 | 5in3 | B | 66 | PRO | p.Pro66His | P07902 |
| HOMO SAPIENS | 2 | 4546 | 5in3 | B | 74 | LEU | p.Leu74Pro | P07902 |
| HOMO SAPIENS | 5 | 4546 | 5in3 | B | 97 | ASN | p.Asn97Ser | P07902 |
| HOMO SAPIENS | 1 | 4546 | 5in3 | B | 135 | SER | p.Ser135Leu | P07902 |
| HOMO SAPIENS | 5 | 4546 | 5in3 | B | 138 | THR | p.Thr138Met | P07902 |
| HOMO SAPIENS | 7 | 4546 | 5in3 | B | 139 | LEU | p.Leu139Pro | P07902 |

| | | | | | | | | |
|---|---|---|---|---|---|---|---|---|
| HOMO SAPIENS | 2 | 4546 | 5in3 | B | 142 | MET | p.Met142Lys | P07902 |
| HOMO SAPIENS | 1 | 4546 | 5in3 | B | 143 | SER | p.Ser143Leu | P07902 |
| HOMO SAPIENS | 2 | 4546 | 5in3 | B | 148 | ARG | p.Arg148Gly | P07902 |
| HOMO SAPIENS | 2 | 4546 | 5in3 | B | 148 | ARG | p.Arg148Gln | P07902 |
| HOMO SAPIENS | 5 | 4546 | 5in3 | B | 183 | PRO | p.Pro183Thr | P07902 |
| HOMO SAPIENS | 5 | 4546 | 5in3 | B | 185 | PRO | p.Pro185His | P07902 |
| HOMO SAPIENS | 5 | 4546 | 5in3 | B | 186 | HIS | p.His186Tyr | P07902 |
| HOMO SAPIENS | 7 | 4546 | 5in3 | B | 188 | GLN | p.Gln188Arg | P07902 |
| HOMO SAPIENS | 5 | 4546 | 5in3 | B | 195 | LEU | p.Leu195Pro | P07902 |
| HOMO SAPIENS | 4 | 4546 | 5in3 | B | 201 | ARG | p.Arg201His | P07902 |
| HOMO SAPIENS | 4 | 4546 | 5in3 | B | 201 | ARG | p.Arg201Cys | P07902 |
| HOMO SAPIENS | 4 | 4546 | 5in3 | B | 203 | GLU | p.Glu203Lys | P07902 |
| HOMO SAPIENS | 2 | 4546 | 5in3 | B | 209 | TYR | p.Tyr209Cys | P07902 |
| HOMO SAPIENS | 5 | 4546 | 5in3 | B | 223 | ARG | p.Arg223Ser | P07902 |
| HOMO SAPIENS | 5 | 4546 | 5in3 | B | 226 | LEU | p.Leu226Pro | P07902 |
| HOMO SAPIENS | 5 | 4546 | 5in3 | B | 226 | LEU | p.Leu226Val | P07902 |
| HOMO SAPIENS | 2 | 4546 | 5in3 | B | 227 | LEU | p.Leu227Pro | P07902 |
| HOMO SAPIENS | 8 | 4546 | 5in3 | B | 258 | ARG | p.Arg258Cys | P07902 |
| HOMO SAPIENS | 1 | 4546 | 5in3 | B | 265 | PRO | p.Pro265Ala | P07902 |
| HOMO SAPIENS | 8 | 4546 | 5in3 | B | 268 | THR | p.Thr268Asn | P07902 |
| HOMO SAPIENS | 7 | 4546 | 5in3 | B | 278 | ILE | p.Ile278Asn | P07902 |
| HOMO SAPIENS | 5 | 4546 | 5in3 | B | 285 | LYS | p.Lys285Asn | P07902 |
| HOMO SAPIENS | 1 | 4546 | 5in3 | B | 319 | HIS | p.His319Gln | P07902 |
| HOMO SAPIENS | 4 | 4546 | 5in3 | B | 321 | HIS | p.His321Tyr | P07902 |
| HOMO SAPIENS | 4 | 4546 | 5in3 | B | 325 | PRO | p.Pro325Leu | P07902 |
| HOMO SAPIENS | 2 | 4546 | 5in3 | B | 328 | ARG | p.Arg328His | P07902 |
| HOMO SAPIENS | 2 | 4546 | 5in3 | B | 329 | SER | p.Ser329Phe | P07902 |
| HOMO SAPIENS | 5 | 4546 | 5in3 | B | 334 | LYS | p.Lys334Arg | P07902 |
| HOMO SAPIENS | 6 | 1290 | 5uav | C | 119 | ARG | p.Arg119Gly | P32322 |
| HOMO SAPIENS | 6 | 1290 | 5uav | C | 119 | ARG | p.Arg119His | P32322 |
| HOMO SAPIENS | 3 | 1290 | 5uav | C | 251 | ARG | p.Arg251His | P32322 |
| HOMO SAPIENS | 7 | 1290 | 5uav | C | 266 | ARG | p.Arg266Gln | P32322 |
| HOMO SAPIENS | 0 | 9618 | 2r7e | A | 22 | GLY | p.Gly41Cys | P00451 |
| HOMO SAPIENS | 2 | 9618 | 2r7e | A | 113 | GLU | p.Glu132Asp | P00451 |
| HOMO SAPIENS | -1 | 9618 | 2r7e | A | 336 | ARG | p.Arg355Ter | P00451 |
| HOMO SAPIENS | 0 | 9618 | 2r7e | A | 372 | ARG | p.Arg391His | P00451 |
| HOMO SAPIENS | 0 | 9618 | 2r7e | A | 373 | SER | p.Ser392Leu | P00451 |
| HOMO SAPIENS | 0 | 9618 | 2r7e | A | 373 | SER | p.Ser392Pro | P00451 |
| HOMO SAPIENS | 3 | 9618 | 2r7e | A | 386 | ILE | p.Ile405Ser | P00451 |
| HOMO SAPIENS | 6 | 9618 | 2r7e | A | 390 | GLU | p.Glu409Gly | P00451 |
| HOMO SAPIENS | 2 | 9618 | 2r7e | A | 412 | LEU | p.Leu431Phe | P00451 |
| HOMO SAPIENS | 5 | 9618 | 2r7e | A | 425 | LYS | p.Lys444Arg | P00451 |
| HOMO SAPIENS | 0 | 9618 | 2r7e | A | 484 | ARG | p.Arg503His | P00451 |
| HOMO SAPIENS | 2 | 9618 | 2r7e | A | 531 | ARG | p.Arg550His | P00451 |
| HOMO SAPIENS | 3 | 9618 | 2r7e | A | 535 | SER | p.Ser554Gly | P00451 |
| HOMO SAPIENS | 2 | 9618 | 2r7e | A | 593 | ARG | p.Arg612Cys | P00451 |
| HOMO SAPIENS | 0 | 9618 | 2r7e | A | 704 | ALA | p.Ala723Thr | P00451 |
| HOMO SAPIENS | 0 | 9618 | 2r7e | A | 720 | GLU | p.Glu739Lys | P00451 |
| HOMO SAPIENS | 1 | 13085 | 3ud1 | A | 1075 | ILE | p.Thr1075Ile | P16157 |
| HOMO SAPIENS | -1 | 3289 | 2fy4 | A | 2 | ALA | p.Ala120Thr | P28329 |
| HOMO SAPIENS | 7 | 3289 | 2fy4 | A | 18 | VAL | p.Val136Met | P28329 |
| HOMO SAPIENS | 5 | 3289 | 2fy4 | A | 92 | LEU | p.Leu210Pro | P28329 |
| HOMO SAPIENS | 2 | 3289 | 2fy4 | A | 93 | PRO | p.Pro211Ala | P28329 |
| HOMO SAPIENS | 4 | 3289 | 2fy4 | A | 131 | LEU | p.Leu249Val | P28329 |

| | | | | | | | | |
|---|---|---|---|---|---|---|---|---|
| HOMO SAPIENS | 2 | 3289 | 2fy4 | A | 181 | PRO | p.Pro299Leu | P28329 |
| HOMO SAPIENS | 0 | 3289 | 2fy4 | A | 218 | ILE | p.Ile336Thr | P28329 |
| HOMO SAPIENS | 2 | 3289 | 2fy4 | A | 261 | ASP | p.Asp379His | P28329 |
| HOMO SAPIENS | 0 | 3289 | 2fy4 | A | 302 | ARG | p.Arg420Cys | P28329 |
| HOMO SAPIENS | 5 | 3289 | 2fy4 | A | 323 | GLU | p.Glu441Lys | P28329 |
| HOMO SAPIENS | 0 | 3289 | 2fy4 | A | 343 | MET | p.Val461Met | P28329 |
| HOMO SAPIENS | 2 | 3289 | 2fy4 | A | 364 | ARG | p.Arg482Gly | P28329 |
| HOMO SAPIENS | 3 | 3289 | 2fy4 | A | 380 | SER | p.Ser498Leu | P28329 |
| HOMO SAPIENS | 6 | 3289 | 2fy4 | A | 388 | VAL | p.Val506Leu | P28329 |
| HOMO SAPIENS | 7 | 3289 | 2fy4 | A | 429 | HIS | p.His547Gln | P28329 |
| HOMO SAPIENS | 8 | 3289 | 2fy4 | A | 442 | ARG | p.Arg560His | P28329 |
| HOMO SAPIENS | 3 | 3289 | 2fy4 | A | 443 | ARG | p.Arg561Gln | P28329 |
| HOMO SAPIENS | 0 | 4137 | 3hhr | B | 38 | CYS | p.Cys56Ter | P10912 |
| HOMO SAPIENS | 0 | 4137 | 3hhr | B | 43 | ARG | p.Arg61Ter | P10912 |
| HOMO SAPIENS | 0 | 4137 | 3hhr | B | 44 | GLU | p.Glu62Lys | P10912 |
| HOMO SAPIENS | 0 | 4137 | 3hhr | B | 96 | PHE | p.Phe114Ser | P10912 |
| HOMO SAPIENS | 5 | 4137 | 3hhr | B | 131 | PRO | p.Pro149Gln | P10912 |
| HOMO SAPIENS | 0 | 4137 | 3hhr | B | 144 | VAL | p.Val162Ile | P10912 |
| HOMO SAPIENS | 0 | 4137 | 3hhr | B | 150 | HIS | p.His168Gln | P10912 |
| HOMO SAPIENS | 0 | 4137 | 3hhr | B | 152 | ASP | p.Asp170His | P10912 |
| HOMO SAPIENS | 0 | 4137 | 3hhr | B | 153 | ILE | p.Ile171Thr | P10912 |
| HOMO SAPIENS | 0 | 4137 | 3hhr | B | 154 | GLN | p.Gln172Pro | P10912 |
| HOMO SAPIENS | 0 | 4137 | 3hhr | B | 155 | VAL | p.Val173Gly | P10912 |
| HOMO SAPIENS | 0 | 4137 | 3hhr | B | 217 | ARG | p.Arg235Ter | P10912 |
| HOMO SAPIENS | 5 | 4137 | 3hhr | B | 224 | GLU | p.Glu242Asp | P10912 |
| HOMO SAPIENS | 5 | 4137 | 3hhr | B | 224 | GLU | p.Glu242Ter | P10912 |
| HOMO SAPIENS | 6 | 2493 | 5bpb | A | 69 | HIS | p.His69Tyr | Q9ULV1 |
| HOMO SAPIENS | 6 | 2493 | 5bpb | A | 105 | MET | p.Met105Val | Q9ULV1 |
| HOMO SAPIENS | -1 | 2493 | 5bpb | A | 168 | PRO | p.Pro168Ser | Q9ULV1 |
| HOMO SAPIENS | 0 | 32123 | 4ccz | A | 314 | ASP | p.Asp314Asn | Q99707 |
| HOMO SAPIENS | 0 | 32123 | 4ccz | A | 345 | VAL | p.Val345Phe | Q99707 |
| HOMO SAPIENS | 1 | 32123 | 4ccz | A | 585 | ARG | p.Arg585Ter | Q99707 |
| HOMO SAPIENS | 4 | 4618 | 2ag4 | A | 26 | PRO | p.Pro55Leu | P17900 |
| HOMO SAPIENS | 4 | 4618 | 2ag4 | A | 30 | VAL | p.Ile59Val | P17900 |
| HOMO SAPIENS | 2 | 4618 | 2ag4 | A | 40 | VAL | p.Met69Val | P17900 |
| HOMO SAPIENS | 2 | 4618 | 2ag4 | A | 124 | VAL | p.Val153Ala | P17900 |
| HOMO SAPIENS | 8 | 4618 | 2ag4 | A | 140 | ARG | p.Arg169Pro | P17900 |
| HOMO SAPIENS | 5 | 1082 | 1hk0 | X | 14 | ARG | p.Arg15Cys | P07320 |
| HOMO SAPIENS | 7 | 1082 | 1hk0 | X | 23 | PRO | p.Pro24Thr | P07320 |
| HOMO SAPIENS | 6 | 1082 | 1hk0 | X | 36 | ARG | p.Arg37Ser | P07320 |
| HOMO SAPIENS | 5 | 1082 | 1hk0 | X | 55 | TYR | p.Tyr56Ter | P07320 |
| HOMO SAPIENS | 3 | 1082 | 1hk0 | X | 58 | ARG | p.Arg59His | P07320 |
| HOMO SAPIENS | 7 | 1082 | 1hk0 | X | 88 | HIS | p.His88Gln | P07320 |
| HOMO SAPIENS | 6 | 1082 | 1hk0 | X | 95 | ARG | p.Arg95Ser | P07320 |
| HOMO SAPIENS | 5 | 1082 | 1hk0 | X | 126 | VAL | p.Val126Met | P07320 |
| HOMO SAPIENS | 8 | 1082 | 1hk0 | X | 157 | TRP | p.Trp157Ter | P07320 |
| HOMO SAPIENS | 0 | 22879 | 3j0a | B | 644 | ILE | p.Ile644Phe | O60602 |
| HOMO SAPIENS | 8 | 6 | 2o23 | B | 122 | LEU | p.Leu122Val | Q99714 |
| HOMO SAPIENS | 1 | 6 | 2o23 | B | 130 | ARG | p.Arg130Cys | Q99714 |
| HOMO SAPIENS | 1 | 6 | 2o23 | B | 247 | ASN | p.Asn247Ser | Q99714 |
| HOMO SAPIENS | 0 | 6 | 2o23 | B | 249 | GLU | p.Glu249Gln | Q99714 |
| HOMO SAPIENS | 1 | 12910 | 4hl4 | A | 67 | ARG | p.Arg67Ter | Q96BZ9 |
| HOMO SAPIENS | 8 | 16105 | 1lmj | A | 7 | GLU | p.Glu1073Lys | P35555 |
| HOMO SAPIENS | 8 | 16105 | 1lmj | A | 8 | CYS | p.Cys1074Arg | P35555 |

| | | | | | | | | |
|---|---|---|---|---|---|---|---|---|
| HOMO SAPIENS | 9 | 16105 | 1lmj | A | 51 | CYS | p.Cys1117Tyr | P35555 |
| HOMO SAPIENS | 9 | 16105 | 1lmj | A | 63 | CYS | p.Cys1129Tyr | P35555 |
| HOMO SAPIENS | 9 | 16105 | 1lmj | A | 65 | ASN | p.Asn1131Tyr | P35555 |
| HOMO SAPIENS | 4 | 16105 | 1lmj | A | 71 | ARG | p.Arg1137Pro | P35555 |
| HOMO SAPIENS | 1 | 16105 | 1lmj | A | 82 | PRO | p.Pro1148Ala | P35555 |
| HOMO SAPIENS | 0 | 4795 | 3s5f | A | 106 | LEU | p.Leu106Ser | Q16595 |
| HOMO SAPIENS | 0 | 4795 | 3s5f | A | 130 | GLY | p.Gly130Val | Q16595 |
| HOMO SAPIENS | 0 | 4795 | 3s5f | A | 154 | ILE | p.Ile154Phe | Q16595 |
| HOMO SAPIENS | -1 | 4795 | 3s5f | A | 209 | ASP | p.Asp209Gly | Q16595 |
| HOMO SAPIENS | -1 | 2884 | 4lli | A | 1655 | ASP | p.Asp1655Glu | Q9Y4I1 |
| HOMO SAPIENS | 0 | 1786 | 7pck | C | 56P | TYR | p.Tyr71Ter | P43235 |
| HOMO SAPIENS | 0 | 1786 | 7pck | C | 64P | GLY | p.Gly79Glu | P43235 |
| HOMO SAPIENS | 7 | 38286 | 2l53 | B | 1914 | ARG | p.Arg1914Gly | Q14524 |
| HOMO SAPIENS | 1 | 38286 | 2l53 | B | 1924 | ALA | p.Ala1924Thr | Q14524 |
| HOMO SAPIENS | 0 | 31747 | 3ksy | A | 25 | PRO | p.Pro25Ser | Q07889 |
| HOMO SAPIENS | 3 | 31747 | 3ksy | A | 37 | THR | p.Thr37Ala | Q07889 |
| HOMO SAPIENS | 0 | 31747 | 3ksy | A | 54 | GLN | p.Gln54His | Q07889 |
| HOMO SAPIENS | -1 | 31747 | 3ksy | A | 190 | ASP | p.Asp190Glu | Q07889 |
| HOMO SAPIENS | 2 | 31747 | 3ksy | A | 266 | THR | p.Thr266Lys | Q07889 |
| HOMO SAPIENS | 1 | 31747 | 3ksy | A | 269 | MET | p.Met269Arg | Q07889 |
| HOMO SAPIENS | 1 | 31747 | 3ksy | A | 340 | PRO | p.Pro340Ser | Q07889 |
| HOMO SAPIENS | 1 | 31747 | 3ksy | A | 423 | ASN | p.Asn423Lys | Q07889 |
| HOMO SAPIENS | 1 | 31747 | 3ksy | A | 424 | GLU | p.Glu424Lys | Q07889 |
| HOMO SAPIENS | 1 | 31747 | 3ksy | A | 432 | TRP | p.Trp432Arg | Q07889 |
| HOMO SAPIENS | 1 | 31747 | 3ksy | A | 552 | ARG | p.Arg552Gly | Q07889 |
| HOMO SAPIENS | 1 | 31747 | 3ksy | A | 552 | ARG | p.Arg552Ser | Q07889 |
| HOMO SAPIENS | 3 | 7529 | 6jdw | A | 93 | ILE | p.Ile93Met | P50440 |
| HOMO SAPIENS | 2 | 7529 | 6jdw | A | 110 | GLN | p.Gln110His | P50440 |
| HOMO SAPIENS | 5 | 7529 | 6jdw | A | 149 | TRP | p.Trp149Ter | P50440 |
| HOMO SAPIENS | 0 | 7529 | 6jdw | A | 169 | ARG | p.Arg169Ter | P50440 |
| HOMO SAPIENS | 0 | 7529 | 6jdw | A | 260 | ARG | p.Arg260Ter | P50440 |
| HOMO SAPIENS | 3 | 7529 | 6jdw | A | 344 | ILE | p.Ile344Val | P50440 |
| HOMO SAPIENS | 5 | 7529 | 6jdw | A | 418 | LEU | p.Leu418Ile | P50440 |
| HOMO SAPIENS | 1 | 14788 | 1eak | A | 101 | ARG | p.Arg101His | P08253 |
| HOMO SAPIENS | 0 | 14788 | 1eak | A | 383 | ASP | p.Asp383Glu | P08253 |
| HOMO SAPIENS | 7 | 14788 | 1eak | A | 404 | GLN | p.Glu404Lys | P08253 |
| HOMO SAPIENS | 2 | 14788 | 1eak | A | 447 | ALA | p.Ala447Val | P08253 |
| HOMO SAPIENS | 0 | 1255 | 5xtc | p | 8 | PRO | p.Pro8His | Q9Y6M9 |
| HOMO SAPIENS | 0 | 1255 | 5xtc | p | 146 | PRO | p.Pro146Ser | Q9Y6M9 |
| HOMO SAPIENS | 3 | 6721 | 1a22 | A | 25 | PHE | p.Phe51Tyr | P01241 |
| HOMO SAPIENS | 2 | 6721 | 1a22 | A | 95 | SER | p.Ser121Arg | P01241 |
| HOMO SAPIENS | 2 | 6721 | 1a22 | A | 110 | VAL | p.Val136Ile | P01241 |
| HOMO SAPIENS | 0 | 1380 | 5i6z | A | 605 | LYS | p.Lys605Asn | P31645 |
| HOMO SAPIENS | 0 | 941 | 1qki | D | 48 | ILE | p.Ile48Thr | P11413 |
| HOMO SAPIENS | 3 | 941 | 1qki | D | 68 | VAL | p.Val68Met | P11413 |
| HOMO SAPIENS | 2 | 941 | 1qki | D | 126 | ASN | p.Asn126Asp | P11413 |
| HOMO SAPIENS | 8 | 941 | 1qki | D | 163 | GLY | p.Gly163Ser | P11413 |
| HOMO SAPIENS | 6 | 941 | 1qki | D | 165 | ASN | p.Asn165Asp | P11413 |
| HOMO SAPIENS | 4 | 941 | 1qki | D | 181 | ASP | p.Asp181Val | P11413 |
| HOMO SAPIENS | 4 | 941 | 1qki | D | 182 | ARG | p.Arg182Trp | P11413 |
| HOMO SAPIENS | 5 | 941 | 1qki | D | 188 | SER | p.Ser188Phe | P11413 |
| HOMO SAPIENS | 9 | 941 | 1qki | D | 198 | ARG | p.Arg198Cys | P11413 |
| HOMO SAPIENS | 8 | 941 | 1qki | D | 213 | VAL | p.Val213Leu | P11413 |
| HOMO SAPIENS | 5 | 941 | 1qki | D | 282 | ASP | p.Asp282His | P11413 |

| | | | | | | | | |
|---|---|---|---|---|---|---|---|---|
| HOMO SAPIENS | 7 | 941 | 1qki | D | 285 | ARG | p.Arg285His | P11413 |
| HOMO SAPIENS | 5 | 941 | 1qki | D | 291 | VAL | p.Val291Met | P11413 |
| HOMO SAPIENS | 8 | 941 | 1qki | D | 323 | LEU | p.Leu323Pro | P11413 |
| HOMO SAPIENS | 3 | 941 | 1qki | D | 335 | ALA | p.Ala335Thr | P11413 |
| HOMO SAPIENS | 6 | 941 | 1qki | D | 350 | ASP | p.Asp350His | P11413 |
| HOMO SAPIENS | 5 | 941 | 1qki | D | 393 | ARG | p.Arg393His | P11413 |
| HOMO SAPIENS | 5 | 941 | 1qki | D | 454 | ARG | p.Arg454Cys | P11413 |
| HOMO SAPIENS | 2 | 941 | 1qki | D | 459 | LEU | p.Arg459Leu | P11413 |
| HOMO SAPIENS | 4 | 941 | 1qki | D | 463 | ARG | p.Arg463His | P11413 |
| HOMO SAPIENS | 4 | 2483 | 1tij | A | 68 | LEU | p.Leu94Gln | P01034 |
| HOMO SAPIENS | 2 | 20632 | 3v5b | A | 331 | ARG | p.Arg331Pro | P02545 |
| HOMO SAPIENS | 2 | 20632 | 3v5b | A | 374 | HIS | p.His374Gln | P02545 |
| HOMO SAPIENS | 4 | 13929 | 5xtd | J | 75 | ARG | p.Arg75His | Q16795 |
| HOMO SAPIENS | 3 | 13929 | 5xtd | J | 243 | VAL | p.Val243Ala | Q16795 |
| HOMO SAPIENS | 1 | 13929 | 5xtd | J | 294 | PRO | p.Pro294Leu | Q16795 |
| HOMO SAPIENS | 2 | 13929 | 5xtd | J | 321 | ARG | p.Arg321Pro | Q16795 |
| HOMO SAPIENS | 0 | 9411 | 5thl | A | 41 | ARG | p.Gly41Arg | P54577 |
| HOMO SAPIENS | 0 | 9411 | 5thl | A | 196 | GLU | p.Glu196Lys | P54577 |
| HOMO SAPIENS | 0 | 9411 | 5thl | A | 316 | ALA | p.Ala316Ser | P54577 |
| HOMO SAPIENS | 0 | 9411 | 5thl | A | 339 | ALA | p.Ala339Thr | P54577 |
| HOMO SAPIENS | 4 | 16273 | 2e8n | A | 46 | THR | p.Thr940Ile | P29317 |
| HOMO SAPIENS | 7 | 16273 | 2e8n | A | 54 | GLY | p.Gly948Trp | P29317 |
| HOMO SAPIENS | 6 | 16273 | 2e8n | A | 65 | ALA | p.Ala959Thr | P29317 |
| HOMO SAPIENS | 7 | 30387 | 2k9y | A | 541 | VAL | p.Val541Met | P29317 |
| HOMO SAPIENS | 0 | 6961 | 4nha | A | 843 | ILE | p.Ile853Val | Q9UPY3 |
| HOMO SAPIENS | -1 | 6961 | 4nha | A | 862 | ALA | p.Ala872Thr | Q9UPY3 |
| HOMO SAPIENS | 3 | 15708 | 4pzg | A | 16 | ARG | p.Arg16Ter | Q9Y5X0 |
| HOMO SAPIENS | 4 | 15708 | 4pzg | A | 51 | ARG | p.Arg51Gln | Q9Y5X0 |
| HOMO SAPIENS | 3 | 30166 | 4o00 | A | 44 | ARG | p.Arg14257Ter | Q8WZ42 |
| HOMO SAPIENS | 2 | 30166 | 4o00 | A | 54 | ARG | p.Arg14267His | Q8WZ42 |
| HOMO SAPIENS | 8 | 11158 | 4hca | A | 275 | TRP | p.Trp274Arg | P23771 |
| HOMO SAPIENS | 2 | 4328 | 4e0s | A | 1476 | ARG | p.Arg1476Ter | P01031 |
| HOMO SAPIENS | 0 | 11462 | 1dx5 | I | 455 | ALA | p.Ala473Val | P07204 |
| HOMO SAPIENS | -1 | 11150 | 2fic | A | 45 | LYS | p.Lys35Asn | O00499 |
| HOMO SAPIENS | 6 | 11150 | 2fic | A | 161 | ASP | p.Asp151Asn | O00499 |
| HOMO SAPIENS | 8 | 27382 | 4mqv | B | 17 | ARG | p.Arg17Ter | Q9NZC9 |
| HOMO SAPIENS | -1 | 37991 | 1q0p | A | 231 | GLN | p.Gln256Ter | P00751 |
| HOMO SAPIENS | 1 | 443 | 5xte | U | 215 | LEU | p.Leu215Phe | P08574 |
| HOMO SAPIENS | 8 | 21471 | 5y4e | A | 1461 | ASN | p.Thr692Ala | Q01484 |
| HOMO SAPIENS | -1 | 35443 | 3um3 | B | 129 | ASP | p.Asp129Val | Q9H444 |
| HOMO SAPIENS | -1 | 35443 | 3um3 | B | 161 | GLU | p.Glu161Lys | Q9H444 |
| HOMO SAPIENS | 6 | 28781 | 2n2a | B | 111 | ILE | p.Ile654Val | P04626 |
| HOMO SAPIENS | 7 | 28781 | 2n2a | B | 112 | ILE | p.Ile655Val | P04626 |
| HOMO SAPIENS | -1 | 3346 | 4ztz | C | 61 | GLY | p.Gly61Arg | Q9UHN1 |
| HOMO SAPIENS | 4 | 3346 | 4ztz | C | 103 | GLY | p.Gly103Ser | Q9UHN1 |
| HOMO SAPIENS | 4 | 3346 | 4ztz | C | 133 | HIS | p.His133Tyr | Q9UHN1 |
| HOMO SAPIENS | -1 | 3346 | 4ztz | C | 153 | LEU | p.Leu153Val | Q9UHN1 |
| HOMO SAPIENS | -1 | 3346 | 4ztz | C | 169 | ALA | p.Ala169Thr | Q9UHN1 |
| HOMO SAPIENS | 7 | 3346 | 4ztz | C | 369 | ARG | p.Arg369Gly | Q9UHN1 |
| HOMO SAPIENS | 4 | 3346 | 4ztz | C | 386 | ASP | p.Asp386Glu | Q9UHN1 |
| HOMO SAPIENS | 4 | 3346 | 4ztz | C | 451 | GLY | p.Gly451Glu | Q9UHN1 |
| HOMO SAPIENS | 4 | 2131 | 5fo9 | E | 1549 | LEU | p.Leu1549Met | P01024 |
| HOMO SAPIENS | 2 | 2131 | 5fo9 | E | 1619 | SER | p.Ser1619Arg | P01024 |
| HOMO SAPIENS | 3 | 12662 | 4jhp | C | 75 | ILE | p.Ile75Val | Q92834 |

| | | | | | | | | |
|---|---|---|---|---|---|---|---|---|
| HOMO SAPIENS | 1 | 12662 | 4jhp | C | 173 | GLY | p.Gly173Arg | Q92834 |
| HOMO SAPIENS | 0 | 12662 | 4jhp | C | 184 | GLN | p.Gln184His | Q92834 |
| HOMO SAPIENS | 3 | 12662 | 4jhp | C | 262 | ALA | p.Ala262Gly | Q92834 |
| HOMO SAPIENS | 3 | 12662 | 4jhp | C | 302 | CYS | p.Cys302Tyr | Q92834 |
| HOMO SAPIENS | 5 | 1093 | 2jg9 | A | 186 | GLN | p.Gln208Ter | P02745 |
| HOMO SAPIENS | 0 | 13181 | 4r4l | A | 12 | MET | p.Met13Ile | Q13976 |
| HOMO SAPIENS | 8 | 25939 | 4iyp | A | 106 | ARG | p.Arg106Gln | P78318 |
| HOMO SAPIENS | 1 | 217 | 2xsx | A | 71 | SER | p.Asn71Ser | P13929 |
| HOMO SAPIENS | 0 | 217 | 2xsx | A | 76 | ALA | p.Ala76Pro | P13929 |
| HOMO SAPIENS | 1 | 217 | 2xsx | A | 85 | ALA | p.Val85Ala | P13929 |
| HOMO SAPIENS | 8 | 217 | 2xsx | A | 156 | GLY | p.Gly156Asp | P13929 |
| HOMO SAPIENS | 5 | 217 | 2xsx | A | 332 | VAL | p.Val332Ile | P13929 |
| HOMO SAPIENS | 7 | 217 | 2xsx | A | 374 | GLY | p.Gly374Glu | P13929 |
| HOMO SAPIENS | 1 | 5508 | 3szm | A | 682 | THR | p.Thr682Asn | Q8NEM0 |
| HOMO SAPIENS | 0 | 5508 | 3szm | A | 715 | TRP | p.Trp715Ter | Q8NEM0 |
| HOMO SAPIENS | 1 | 5508 | 3szm | A | 761 | VAL | p.Ala761Val | Q8NEM0 |
| HOMO SAPIENS | 1 | 5508 | 3szm | A | 828 | PRO | p.Pro828Ser | Q8NEM0 |
| HOMO SAPIENS | 4 | 5508 | 3szm | A | 829 | GLU | p.Glu829Asp | Q8NEM0 |
| HOMO SAPIENS | 2 | 16353 | 2q0z | X | 171 | GLY | p.Gly1978Ala | O75643 |
| HOMO SAPIENS | 2 | 16353 | 2q0z | X | 188 | ALA | p.Ala1995Thr | O75643 |
| HOMO SAPIENS | 0 | 4127 | 3swr | A | 860 | GLY | p.Gly860Arg | P26358 |
| HOMO SAPIENS | -1 | 4127 | 3swr | A | 1122 | SER | p.Ser1122Cys | P26358 |
| HOMO SAPIENS | 0 | 4127 | 3swr | A | 1216 | ASP | p.Asp1216Glu | P26358 |
| HOMO SAPIENS | 0 | 6382 | 3o96 | A | 17 | GLU | p.Glu17Lys | P31749 |
| HOMO SAPIENS | 6 | 14314 | 5xtc | Q | 71 | PRO | p.Pro71Leu | O75306 |
| HOMO SAPIENS | 2 | 5886 | 4fao | E | 22 | ARG | p.Arg40His | Q13705 |
| HOMO SAPIENS | 1 | 2534 | 3mp4 | A | 37 | GLU | p.Glu37Ter | P35914 |
| HOMO SAPIENS | 3 | 2534 | 3mp4 | A | 41 | MET | p.Arg41Gln | P35914 |
| HOMO SAPIENS | 1 | 2534 | 3mp4 | A | 70 | VAL | p.Val70Leu | P35914 |
| HOMO SAPIENS | 2 | 2196 | 3ifq | A | 142 | ARG | p.Arg142His | P14923 |
| HOMO SAPIENS | 2 | 2196 | 3ifq | A | 153 | ASP | p.Asp153Glu | P14923 |
| HOMO SAPIENS | 3 | 2196 | 3ifq | A | 176 | ARG | p.Arg176Gln | P14923 |
| HOMO SAPIENS | 3 | 2196 | 3ifq | A | 265 | ARG | p.Arg265His | P14923 |
| HOMO SAPIENS | 2 | 2196 | 3ifq | A | 301 | GLU | p.Glu301Gly | P14923 |
| HOMO SAPIENS | 0 | 2196 | 3ifq | A | 456 | VAL | p.Val456Ile | P14923 |
| HOMO SAPIENS | 3 | 2196 | 3ifq | A | 603 | VAL | p.Val603Leu | P14923 |
| HOMO SAPIENS | 0 | 2196 | 3ifq | A | 648 | VAL | p.Val648Ile | P14923 |
| HOMO SAPIENS | 1 | 8746 | 1cgj | I | 32 | PRO | p.Pro55Ser | P00995 |
| HOMO SAPIENS | 1 | 8746 | 1cgj | I | 44 | ARG | p.Arg67His | P00995 |
| HOMO SAPIENS | 1 | 8746 | 1cgj | I | 44 | ARG | p.Arg67Cys | P00995 |
| HOMO SAPIENS | 3 | 3212 | 1syq | A | 105 | ARG | p.Arg105Ter | P18206 |
| HOMO SAPIENS | 3 | 3212 | 1syq | A | 126 | PHE | p.Phe126Leu | P18206 |
| HOMO SAPIENS | 4 | 10028 | 3eto | A | 1609 | ARG | p.Arg1608His | P46531 |
| HOMO SAPIENS | 1 | 10028 | 3eto | A | 1672 | VAL | p.Val1671Ile | P46531 |
| HOMO SAPIENS | 5 | 10028 | 3eto | A | 1692 | GLN | p.Gln1691His | P46531 |
| HOMO SAPIENS | 3 | 3749 | 5xtc | Z | 70 | GLY | p.Gly70Ter | O43676 |
| HOMO SAPIENS | 2 | 17039 | 3brv | B | 57 | GLU | p.Glu57Lys | Q9Y6K9 |
| HOMO SAPIENS | 8 | 3589 | 2fh3 | A | 433 | VAL | p.Val460Leu | P06396 |
| HOMO SAPIENS | 1 | 3589 | 2fh3 | A | 454 | ARG | p.Arg481Cys | P06396 |
| HOMO SAPIENS | 4 | 3589 | 2fh3 | A | 498 | ARG | p.Arg525His | P06396 |
| HOMO SAPIENS | 2 | 3589 | 2fh3 | A | 520 | TYR | p.Tyr547Cys | P06396 |
| HOMO SAPIENS | 1 | 3589 | 2fh3 | A | 529 | GLY | p.Gly556Arg | P06396 |
| HOMO SAPIENS | 2 | 3589 | 2fh3 | A | 536 | THR | p.Thr563Ser | P06396 |
| HOMO SAPIENS | 2 | 3589 | 2fh3 | A | 579 | VAL | p.Val606Met | P06396 |

| | | | | | | | | |
|---|---|---|---|---|---|---|---|---|
| HOMO SAPIENS | 1 | 3589 | 2fh3 | A | 589 | THR | p.Thr616Met | P06396 |
| HOMO SAPIENS | 5 | 3589 | 2fh3 | A | 668 | THR | p.Thr695Met | P06396 |
| HOMO SAPIENS | 0 | 148 | 1lcl | A | 130 | ILE | p.Ile130Ser | Q05315 |
| HOMO SAPIENS | -1 | 24727 | 4fj3 | P | 237 | ALA | p.Ala237Thr | P04049 |
| HOMO SAPIENS | 0 | 24727 | 4fj3 | P | 257 | SER | p.Ser257Leu | P04049 |
| HOMO SAPIENS | 0 | 24727 | 4fj3 | P | 261 | PRO | p.Pro261Ala | P04049 |
| HOMO SAPIENS | 0 | 2949 | 5xtb | B | 77 | ARG | p.Arg77Gly | O00217 |
| HOMO SAPIENS | 0 | 2949 | 5xtb | B | 79 | PRO | p.Pro79Leu | O00217 |
| HOMO SAPIENS | 0 | 2949 | 5xtb | B | 100 | ALA | p.Ala100Val | O00217 |
| HOMO SAPIENS | 1 | 2949 | 5xtb | B | 115 | LYS | p.Lys115Glu | O00217 |
| HOMO SAPIENS | 0 | 2949 | 5xtb | B | 138 | ARG | p.Arg138His | O00217 |
| HOMO SAPIENS | 1 | 3959 | 5k5s | B | 25 | ARG | p.Arg25Ter | P41180 |
| HOMO SAPIENS | 4 | 3959 | 5k5s | B | 39 | PRO | p.Pro39Ala | P41180 |
| HOMO SAPIENS | 0 | 3959 | 5k5s | B | 47 | LYS | p.Lys47Asn | P41180 |
| HOMO SAPIENS | 2 | 3959 | 5k5s | B | 62 | ARG | p.Arg62Met | P41180 |
| HOMO SAPIENS | 2 | 3959 | 5k5s | B | 66 | ARG | p.Arg66Cys | P41180 |
| HOMO SAPIENS | 4 | 3959 | 5k5s | B | 90 | ASN | p.Asn90Thr | P41180 |
| HOMO SAPIENS | 1 | 3959 | 5k5s | B | 116 | ALA | p.Ala116Thr | P41180 |
| HOMO SAPIENS | 1 | 3959 | 5k5s | B | 118 | ASN | p.Asn118Lys | P41180 |
| HOMO SAPIENS | -1 | 3959 | 5k5s | B | 125 | LEU | p.Leu125Pro | P41180 |
| HOMO SAPIENS | -1 | 3959 | 5k5s | B | 127 | GLU | p.Glu127Ala | P41180 |
| HOMO SAPIENS | -1 | 3959 | 5k5s | B | 128 | PHE | p.Phe128Leu | P41180 |
| HOMO SAPIENS | -1 | 3959 | 5k5s | B | 131 | CYS | p.Cys131Trp | P41180 |
| HOMO SAPIENS | 1 | 3959 | 5k5s | B | 138 | THR | p.Thr138Met | P41180 |
| HOMO SAPIENS | 0 | 3959 | 5k5s | B | 143 | GLY | p.Gly143Glu | P41180 |
| HOMO SAPIENS | 5 | 3959 | 5k5s | B | 151 | THR | p.Thr151Met | P41180 |
| HOMO SAPIENS | 4 | 3959 | 5k5s | B | 180 | PHE | p.Phe180Cys | P41180 |
| HOMO SAPIENS | 0 | 3959 | 5k5s | B | 185 | ARG | p.Arg185Gln | P41180 |
| HOMO SAPIENS | 0 | 3959 | 5k5s | B | 185 | ARG | p.Arg185Ter | P41180 |
| HOMO SAPIENS | 2 | 3959 | 5k5s | B | 191 | GLU | p.Glu191Lys | P41180 |
| HOMO SAPIENS | 2 | 3959 | 5k5s | B | 227 | ARG | p.Arg227Leu | P41180 |
| HOMO SAPIENS | 2 | 3959 | 5k5s | B | 250 | GLU | p.Glu250Lys | P41180 |
| HOMO SAPIENS | 6 | 3959 | 5k5s | B | 297 | GLU | p.Glu297Lys | P41180 |
| HOMO SAPIENS | 1 | 3959 | 5k5s | B | 336 | LYS | p.Lys336Asn | P41180 |
| HOMO SAPIENS | 6 | 3959 | 5k5s | B | 429 | HIS | p.His429Tyr | P41180 |
| HOMO SAPIENS | 0 | 3959 | 5k5s | B | 445 | THR | p.Thr445Ala | P41180 |
| HOMO SAPIENS | 3 | 3959 | 5k5s | B | 509 | GLY | p.Gly509Arg | P41180 |
| HOMO SAPIENS | 1 | 3959 | 5k5s | B | 553 | GLY | p.Gly553Arg | P41180 |
| HOMO SAPIENS | 3 | 3959 | 5k5s | B | 592 | ASN | p.Asn592Ser | P41180 |
| HOMO SAPIENS | -1 | 3959 | 5k5s | B | 604 | GLU | p.Glu604Lys | P41180 |
| HOMO SAPIENS | 4 | 2564 | 5l0c | C | 975 | ARG | p.Arg975Trp | P18206 |
| HOMO SAPIENS | 4 | 2564 | 5l0c | C | 975 | ARG | p.Arg975Leu | P18206 |
| HOMO SAPIENS | 1 | 5695 | 5i8r | A | 294 | ARG | p.Arg294Gln | P17405 |
| HOMO SAPIENS | 0 | 5695 | 5i8r | A | 485 | ALA | p.Ala485Val | P17405 |
| HOMO SAPIENS | 1 | 5695 | 5i8r | A | 506 | GLY | p.Gly506Arg | P17405 |
| HOMO SAPIENS | -1 | 1978 | 3avs | A | 1060 | HIS | p.His1060Leu | O15550 |
| HOMO SAPIENS | 0 | 11173 | 4v0c | B | 362 | LYS | p.Lys362Arg | P51787 |
| HOMO SAPIENS | 3 | 11173 | 4v0c | B | 381 | CYS | p.Cys381Tyr | P51787 |
| HOMO SAPIENS | 7 | 11173 | 4v0c | B | 518 | ARG | p.Arg518Gly | P51787 |
| HOMO SAPIENS | 2 | 11173 | 4v0c | B | 525 | ALA | p.Ala525Thr | P51787 |
| HOMO SAPIENS | 0 | 11173 | 4v0c | B | 530 | GLN | p.Gln530Ter | P51787 |
| HOMO SAPIENS | -1 | 11173 | 4v0c | B | 536 | TYR | p.Tyr536Ter | P51787 |
| HOMO SAPIENS | 2 | 1113 | 5xtd | N | 60 | ARG | p.Arg60Ter | Q9UI09 |
| HOMO SAPIENS | 3 | 6061 | 2mmz | A | 148 | LEU | p.Leu148Ser | Q86SX6 |

| | | | | | | | | |
|---|---|---|---|---|---|---|---|---|
| HOMO SAPIENS | -1 | 26081 | 3eu7 | A | 837 | GLU | p.Glu837Ter | Q86YC2 |
| HOMO SAPIENS | 0 | 26081 | 3eu7 | A | 864 | PRO | p.Pro864Ser | Q86YC2 |
| HOMO SAPIENS | 5 | 26081 | 3eu7 | A | 891 | CYS | p.Cys891Trp | Q86YC2 |
| HOMO SAPIENS | 0 | 26081 | 3eu7 | A | 932 | VAL | p.Val932Met | Q86YC2 |
| HOMO SAPIENS | 5 | 26081 | 3eu7 | A | 961 | LEU | p.Leu961Met | Q86YC2 |
| HOMO SAPIENS | 5 | 26081 | 3eu7 | A | 1038 | TRP | p.Trp1038Ter | Q86YC2 |
| HOMO SAPIENS | -1 | 26081 | 3eu7 | A | 1086 | ARG | p.Arg1086Ter | Q86YC2 |
| HOMO SAPIENS | 0 | 26081 | 3eu7 | A | 1122 | ASP | p.Asp1122Glu | Q86YC2 |
| HOMO SAPIENS | 5 | 26081 | 3eu7 | A | 1158 | HIS | p.His1158Gln | Q86YC2 |
| HOMO SAPIENS | 5 | 26081 | 3eu7 | A | 1183 | TYR | p.Tyr1183Ter | Q86YC2 |
| HOMO SAPIENS | 4 | 199 | 1qo5 | C | 45 | ARG | p.Arg46Trp | P05062 |
| HOMO SAPIENS | 5 | 199 | 1qo5 | C | 147 | TRP | p.Trp148Arg | P05062 |
| HOMO SAPIENS | 0 | 199 | 1qo5 | C | 149 | ALA | p.Ala150Pro | P05062 |
| HOMO SAPIENS | 0 | 199 | 1qo5 | C | 174 | ALA | p.Ala175Asp | P05062 |
| HOMO SAPIENS | 0 | 199 | 1qo5 | C | 334 | ASN | p.Asn335Lys | P05062 |
| HOMO SAPIENS | 0 | 199 | 1qo5 | C | 337 | ALA | p.Ala338Val | P05062 |
| HOMO SAPIENS | 5 | 199 | 1qo5 | C | 342 | TYR | p.Tyr343His | P05062 |
| HOMO SAPIENS | 0 | 6910 | 4zht | B | 59 | ARG | p.Arg59Ter | Q9Y223 |
| HOMO SAPIENS | 4 | 6910 | 4zht | B | 129 | ARG | p.Arg129Ter | Q9Y223 |
| HOMO SAPIENS | 4 | 6910 | 4zht | B | 129 | ARG | p.Arg129Gln | Q9Y223 |
| HOMO SAPIENS | 1 | 6910 | 4zht | B | 162 | ARG | p.Arg162Cys | Q9Y223 |
| HOMO SAPIENS | 0 | 6910 | 4zht | B | 176 | ASP | p.Asp176Val | Q9Y223 |
| HOMO SAPIENS | 0 | 6910 | 4zht | B | 216 | VAL | p.Val216Ala | Q9Y223 |
| HOMO SAPIENS | 0 | 6910 | 4zht | B | 246 | ARG | p.Arg246Trp | Q9Y223 |
| HOMO SAPIENS | 0 | 6910 | 4zht | B | 263 | ARG | p.Arg263Leu | Q9Y223 |
| HOMO SAPIENS | 0 | 6910 | 4zht | B | 266 | ARG | p.Arg266Gln | Q9Y223 |
| HOMO SAPIENS | 0 | 6910 | 4zht | B | 266 | ARG | p.Arg266Trp | Q9Y223 |
| HOMO SAPIENS | 0 | 6910 | 4zht | B | 277 | ARG | p.Arg277Cys | Q9Y223 |
| HOMO SAPIENS | 0 | 6910 | 4zht | B | 378 | ASP | p.Asp378Tyr | Q9Y223 |
| HOMO SAPIENS | 3 | 211 | 5kxi | C | 262 | VAL | p.Val287Leu | P17787 |
| HOMO SAPIENS | 0 | 211 | 5kxi | C | 369 | PHE | p.Phe478Leu | P17787 |
| HOMO SAPIENS | 0 | 2454 | 5iy9 | V | null | PHE | p.Phe99Ser | P19447 |
| HOMO SAPIENS | 0 | 2454 | 5iy9 | V | null | ARG | p.Arg109Ter | P19447 |
| HOMO SAPIENS | 0 | 2454 | 5iy9 | V | null | THR | p.Thr119Pro | P19447 |
| HOMO SAPIENS | 0 | 2454 | 5iy9 | V | 425 | ARG | p.Arg425Ter | P19447 |
| HOMO SAPIENS | 1 | 2454 | 5iy9 | V | 545 | GLN | p.Gln545Ter | P19447 |
| HOMO SAPIENS | -1 | 3805 | 3rbq | A | 57 | LYS | p.Lys57Ter | Q13432 |
| HOMO SAPIENS | 2 | 2649 | 4ido | A | 43 | ASP | p.Asp43Glu | Q8WXF7 |
| HOMO SAPIENS | 2 | 2649 | 4ido | A | 157 | LEU | p.Leu157Trp | Q8WXF7 |
| HOMO SAPIENS | 2 | 2649 | 4ido | A | 193 | PHE | p.Phe193Cys | Q8WXF7 |
| HOMO SAPIENS | 7 | 2649 | 4ido | A | 217 | ARG | p.Arg217Gln | Q8WXF7 |
| HOMO SAPIENS | 4 | 2649 | 4ido | A | 239 | ARG | p.Arg239Cys | Q8WXF7 |
| HOMO SAPIENS | 3 | 2649 | 4ido | A | 258 | HIS | p.His258Arg | Q8WXF7 |
| HOMO SAPIENS | 4 | 2649 | 4ido | A | 259 | SER | p.Ser259Tyr | Q8WXF7 |
| HOMO SAPIENS | 4 | 2649 | 4ido | A | 408 | MET | p.Met408Val | Q8WXF7 |
| HOMO SAPIENS | 2 | 2649 | 4ido | A | 415 | ARG | p.Arg415Trp | Q8WXF7 |
| HOMO SAPIENS | 3 | 2649 | 4ido | A | 416 | ARG | p.Arg416Cys | Q8WXF7 |
| HOMO SAPIENS | 5 | 2203 | 3wff | A | 810 | LEU | p.Ser810Leu | P08235 |
| HOMO SAPIENS | 4 | 12648 | 5ep6 | B | 696 | GLU | p.Glu696Lys | Q9UHD2 |
| HOMO SAPIENS | 0 | 30622 | 2pe4 | A | 90 | GLU | p.Glu90Asp | Q12794 |
| HOMO SAPIENS | 0 | 30622 | 2pe4 | A | 256 | GLY | p.Gly256Arg | Q12794 |
| HOMO SAPIENS | 1 | 30622 | 2pe4 | A | 268 | GLU | p.Glu268Lys | Q12794 |
| HOMO SAPIENS | 0 | 30622 | 2pe4 | A | 392 | THR | p.Thr392Met | Q12794 |
| HOMO SAPIENS | 5 | 5411 | 1lya | A | 93 | SER | p.Ser157Leu | P07339 |

| | | | | | | | | |
|---|---|---|---|---|---|---|---|---|
| HOMO SAPIENS | 3 | 5743 | 2uzp | A | 189 | ASN | p.Asn189Lys | P05129 |
| HOMO SAPIENS | 1 | 12529 | 3kfd | J | 21 | ASN | p.Asn45Ser | P36897 |
| HOMO SAPIENS | 1 | 1063 | 2e1q | B | 149 | ARG | p.Arg149Cys | P47989 |
| HOMO SAPIENS | -1 | 1063 | 2e1q | B | 172 | GLY | p.Gly172Arg | P47989 |
| HOMO SAPIENS | 4 | 1063 | 2e1q | B | 228 | ARG | p.Arg228Ter | P47989 |
| HOMO SAPIENS | 5 | 1063 | 2e1q | B | 395 | LYS | p.Lys395Met | P47989 |
| HOMO SAPIENS | 2 | 1063 | 2e1q | B | 646 | ILE | p.Ile646Val | P47989 |
| HOMO SAPIENS | 4 | 1063 | 2e1q | B | 703 | ILE | p.Ile703Val | P47989 |
| HOMO SAPIENS | 8 | 2525 | 1kqn | F | 18 | ASN | p.Asn18Ser | Q9HAN9 |
| HOMO SAPIENS | 0 | 2525 | 1kqn | F | 151 | VAL | p.Val151Phe | Q9HAN9 |
| HOMO SAPIENS | 0 | 2525 | 1kqn | F | 153 | LEU | p.Leu153Val | Q9HAN9 |
| HOMO SAPIENS | 0 | 2525 | 1kqn | F | 169 | TRP | p.Trp169Ter | Q9HAN9 |
| HOMO SAPIENS | 0 | 2525 | 1kqn | F | 207 | ARG | p.Arg207Trp | Q9HAN9 |
| HOMO SAPIENS | 0 | 2525 | 1kqn | F | 273 | ASN | p.Asn273Asp | Q9HAN9 |
| HOMO SAPIENS | 0 | 12299 | 5duq | A | 432 | VAL | p.Val454Glu | P05155 |
| HOMO SAPIENS | 0 | 12299 | 5duq | A | 436 | ALA | p.Ala458Thr | P05155 |
| HOMO SAPIENS | 0 | 12299 | 5duq | A | 443 | ALA | p.Ala465Val | P05155 |
| HOMO SAPIENS | 0 | 12299 | 5duq | A | 444 | ARG | p.Arg466His | P05155 |
| HOMO SAPIENS | 0 | 12299 | 5duq | A | 458 | MET | p.Val480Met | P05155 |
| HOMO SAPIENS | 0 | 327 | 1uhp | A | 36 | ASP | p.Asp274Asn | Q9UPQ7 |
| HOMO SAPIENS | 0 | 64 | 2mo5 | A | 54 | ALA | p.Ala55Thr | P12104 |
| HOMO SAPIENS | 2 | 35969 | 1jrh | I | 60 | CYS | p.Cys77Tyr | P15260 |
| HOMO SAPIENS | 4 | 35969 | 1jrh | I | 70 | ILE | p.Ile87Thr | P15260 |
| HOMO SAPIENS | 2 | 1068 | 1mu7 | A | 304 | ARG | p.Arg304Gln | Q9NUW8 |
| HOMO SAPIENS | 8 | 1068 | 1mu7 | A | 493 | HIS | p.His493Arg | Q9NUW8 |
| HOMO SAPIENS | 0 | 32360 | 2kqp | A | 31 | ARG | p.Arg55Cys | P01308 |
| HOMO SAPIENS | 0 | 32360 | 2kqp | A | 65 | ARG | p.Arg89His | P01308 |
| HOMO SAPIENS | 0 | 32360 | 2kqp | A | 65 | ARG | p.Arg89Cys | P01308 |
| HOMO SAPIENS | 6 | 10805 | 3nks | A | 12 | ILE | p.Ile12Thr | P50336 |
| HOMO SAPIENS | 0 | 10805 | 3nks | A | 20 | HIS | p.His20Pro | P50336 |
| HOMO SAPIENS | 0 | 10805 | 3nks | A | 59 | ARG | p.Arg59Trp | P50336 |
| HOMO SAPIENS | 0 | 10805 | 3nks | A | 168 | ARG | p.Arg168Cys | P50336 |
| HOMO SAPIENS | 0 | 10805 | 3nks | A | 168 | ARG | p.Arg168His | P50336 |
| HOMO SAPIENS | 2 | 10805 | 3nks | A | 232 | GLY | p.Gly232Arg | P50336 |
| HOMO SAPIENS | 3 | 10805 | 3nks | A | 256 | PRO | p.Pro256Arg | P50336 |
| HOMO SAPIENS | 0 | 10805 | 3nks | A | 304 | ARG | p.Arg304His | P50336 |
| HOMO SAPIENS | 0 | 10805 | 3nks | A | 349 | ASP | p.Asp349Ala | P50336 |
| HOMO SAPIENS | 9 | 7008 | 3u2l | A | 125 | GLN | p.Gln125Ter | P55789 |
| HOMO SAPIENS | 4 | 7008 | 3u2l | A | 172 | HIS | p.His172Asn | P55789 |
| HOMO SAPIENS | 4 | 7008 | 3u2l | A | 194 | ARG | p.Arg194His | P55789 |
| HOMO SAPIENS | 3 | 7008 | 3u2l | A | 198 | GLY | p.Gly198Ser | P55789 |
| HOMO SAPIENS | 0 | 20821 | 3e0g | A | 65 | HIS | p.His116Tyr | P42702 |
| HOMO SAPIENS | 3 | 1151 | 5l9v | A | 278 | ASP | p.Asp278Asn | Q9GZT9 |
| HOMO SAPIENS | 4 | 1151 | 5l9v | A | 317 | CYS | p.Pro317Arg | Q9GZT9 |
| HOMO SAPIENS | 5 | 1151 | 5l9v | A | 371 | ARG | p.Arg371His | Q9GZT9 |
| HOMO SAPIENS | 2 | 3935 | 1gwb | A | 36 | ARG | p.Arg36Gly | P07359 |
| HOMO SAPIENS | 1 | 3935 | 1gwb | A | 73 | LEU | p.Leu73Phe | P07359 |
| HOMO SAPIENS | 1 | 3935 | 1gwb | A | 172 | ALA | p.Ala172Val | P07359 |
| HOMO SAPIENS | 0 | 3935 | 1gwb | A | 249 | GLY | p.Gly249Val | P07359 |
| HOMO SAPIENS | 0 | 3935 | 1gwb | A | 255 | MET | p.Met255Val | P07359 |
| HOMO SAPIENS | 2 | 14541 | 5cuz | A | 138 | GLU | p.Glu138Lys | Q9H3L0 |
| HOMO SAPIENS | 2 | 14541 | 5cuz | A | 143 | SER | p.Ser143Ile | Q9H3L0 |
| HOMO SAPIENS | 4 | 14541 | 5cuz | A | 182 | THR | p.Thr182Asn | Q9H3L0 |
| HOMO SAPIENS | 6 | 14541 | 5cuz | A | 206 | ASN | p.Asn206Ile | Q9H3L0 |

| | | | | | | | | |
|---|---|---|---|---|---|---|---|---|
| HOMO SAPIENS | 0 | 14541 | 5cuz | A | 249 | TYR | p.Tyr249Cys | Q9H3L0 |
| HOMO SAPIENS | 4 | 14541 | 5cuz | A | 250 | ARG | p.Arg250Ter | Q9H3L0 |
| HOMO SAPIENS | 2 | 14541 | 5cuz | A | 259 | LEU | p.Leu259Pro | Q9H3L0 |
| HOMO SAPIENS | 0 | 4206 | 4fao | D | 30 | PRO | p.Pro30Ser | P37023 |
| HOMO SAPIENS | 1 | 4206 | 4fao | D | 50 | TRP | p.Trp50Cys | P37023 |
| HOMO SAPIENS | 0 | 16553 | 3v30 | A | 251 | GLN | p.Gln251Glu | O14593 |
| HOMO SAPIENS | 0 | 5012 | 5hbf | A | 442 | ALA | p.Ala442Gly | Q13231 |
| HOMO SAPIENS | 5 | 4844 | 5tdc | A | 136 | HIS | p.His136Arg | Q8IWV7 |
| HOMO SAPIENS | 0 | 15435 | 4cgz | A | 643 | ARG | p.Arg643His | P54132 |
| HOMO SAPIENS | 1 | 15435 | 4cgz | A | 672 | GLN | p.Gln672Arg | P54132 |
| HOMO SAPIENS | 1 | 15435 | 4cgz | A | 700 | GLN | p.Gln700Ter | P54132 |
| HOMO SAPIENS | 0 | 15435 | 4cgz | A | 707 | PRO | p.Pro707Ser | P54132 |
| HOMO SAPIENS | 0 | 15435 | 4cgz | A | 755 | LYS | p.Lys755Glu | P54132 |
| HOMO SAPIENS | 0 | 15435 | 4cgz | A | 881 | TRP | p.Trp881Ter | P54132 |
| HOMO SAPIENS | 2 | 15435 | 4cgz | A | 899 | ARG | p.Arg899Ter | P54132 |
| HOMO SAPIENS | 0 | 15435 | 4cgz | A | 1036 | CYS | p.Cys1036Phe | P54132 |
| HOMO SAPIENS | 0 | 15435 | 4cgz | A | 1055 | CYS | p.Cys1055Ser | P54132 |
| HOMO SAPIENS | 0 | 15435 | 4cgz | A | 1139 | ARG | p.Arg1139Ter | P54132 |
| HOMO SAPIENS | -1 | 15435 | 4cgz | A | 1198 | VAL | p.Val1198Met | P54132 |
| HOMO SAPIENS | -1 | 15435 | 4cgz | A | 1205 | VAL | p.Val1205Ile | P54132 |
| HOMO SAPIENS | 0 | 15435 | 4cgz | A | 1251 | GLU | p.Glu1251Gln | P54132 |
| HOMO SAPIENS | 0 | 11185 | 1kb4 | A | 30 | ARG | p.Arg30Ter | P11473 |
| HOMO SAPIENS | 5 | 11185 | 1kb4 | A | 33 | GLY | p.Gly33Asp | P11473 |
| HOMO SAPIENS | 5 | 11185 | 1kb4 | A | 46 | GLY | p.Gly46Asp | P11473 |
| HOMO SAPIENS | 6 | 11185 | 1kb4 | A | 50 | ARG | p.Arg50Gln | P11473 |
| HOMO SAPIENS | 9 | 11185 | 1kb4 | A | 73 | ARG | p.Arg73Gln | P11473 |
| HOMO SAPIENS | 9 | 11185 | 1kb4 | A | 80 | ARG | p.Arg80Gln | P11473 |
| HOMO SAPIENS | 0 | 5382 | 3fl7 | A | 41 | LEU | p.Leu41Val | P29317 |
| HOMO SAPIENS | 2 | 5382 | 3fl7 | A | 175 | ARG | p.Arg175Cys | P29317 |
| HOMO SAPIENS | 1 | 5382 | 3fl7 | A | 195 | ARG | p.Arg195Cys | P29317 |
| HOMO SAPIENS | 0 | 5382 | 3fl7 | A | 232 | ASP | p.Asp232Gly | P29317 |
| HOMO SAPIENS | 0 | 5382 | 3fl7 | A | 277 | SER | p.Ser277Leu | P29317 |
| HOMO SAPIENS | 0 | 5382 | 3fl7 | A | 315 | ARG | p.Arg315Gln | P29317 |
| HOMO SAPIENS | 7 | 5382 | 3fl7 | A | 350 | PRO | p.Pro350Thr | P29317 |
| HOMO SAPIENS | 0 | 5382 | 3fl7 | A | 391 | GLY | p.Gly391Arg | P29317 |
| HOMO SAPIENS | 0 | 5382 | 3fl7 | A | 418 | VAL | p.Val418Ile | P29317 |
| HOMO SAPIENS | 0 | 5382 | 3fl7 | A | 511 | THR | p.Thr511Met | P29317 |
| HOMO SAPIENS | 2 | 29168 | 4p22 | B | 144 | VAL | p.Val144Ile | P22314 |
| HOMO SAPIENS | 5 | 4348 | 1bf5 | A | 241 | ARG | p.Arg241Gln | P42224 |
| HOMO SAPIENS | 2 | 4348 | 1bf5 | A | 266 | VAL | p.Val266Ile | P42224 |
| HOMO SAPIENS | 4 | 4348 | 1bf5 | A | 320 | GLU | p.Glu320Gln | P42224 |
| HOMO SAPIENS | 1 | 4348 | 1bf5 | A | 372 | ASN | p.Asn372Lys | P42224 |
| HOMO SAPIENS | 1 | 4348 | 1bf5 | A | 447 | ASP | p.Asp447Glu | P42224 |
| HOMO SAPIENS | 2 | 4348 | 1bf5 | A | 463 | GLN | p.Gln463His | P42224 |
| HOMO SAPIENS | 4 | 4348 | 1bf5 | A | 600 | LEU | p.Leu600Pro | P42224 |
| HOMO SAPIENS | 0 | 4348 | 1bf5 | A | 706 | LEU | p.Leu706Ser | P42224 |
| HOMO SAPIENS | 4 | 4991 | 4bej | B | 141 | ASN | p.Asn141Ser | O00429 |
| HOMO SAPIENS | 0 | 4991 | 4bej | B | 395 | ALA | p.Ala395Asp | O00429 |
| HOMO SAPIENS | 2 | 11792 | 1fvo | A | 40 | ARG | p.Arg40Cys | P00480 |
| HOMO SAPIENS | 2 | 11792 | 1fvo | A | 40 | ARG | p.Arg40His | P00480 |
| HOMO SAPIENS | 0 | 11792 | 1fvo | A | 45 | LEU | p.Leu45Pro | P00480 |
| HOMO SAPIENS | 1 | 11792 | 1fvo | A | 47 | ASN | p.Asn47Ile | P00480 |
| HOMO SAPIENS | 0 | 11792 | 1fvo | A | 50 | GLY | p.Gly50Arg | P00480 |
| HOMO SAPIENS | 5 | 11792 | 1fvo | A | 87 | GLU | p.Glu87Lys | P00480 |

| | | | | | | | | |
|---|---|---|---|---|---|---|---|---|
| HOMO SAPIENS | 3 | 11792 | 1fvo | A | 111 | LEU | p.Leu111Pro | P00480 |
| HOMO SAPIENS | 1 | 11792 | 1fvo | A | 125 | THR | p.Thr125Met | P00480 |
| HOMO SAPIENS | 3 | 11792 | 1fvo | A | 129 | ARG | p.Arg129His | P00480 |
| HOMO SAPIENS | 9 | 11792 | 1fvo | A | 141 | ARG | p.Arg141Pro | P00480 |
| HOMO SAPIENS | 0 | 11792 | 1fvo | A | 206 | MET | p.Met206Arg | P00480 |
| HOMO SAPIENS | 0 | 11792 | 1fvo | A | 207 | SER | p.Ser207Arg | P00480 |
| HOMO SAPIENS | 0 | 11792 | 1fvo | A | 216 | GLN | p.Gln216Glu | P00480 |
| HOMO SAPIENS | 1 | 11792 | 1fvo | A | 225 | PRO | p.Pro225Leu | P00480 |
| HOMO SAPIENS | 0 | 11792 | 1fvo | A | 270 | ARG | p.Gln270Arg | P00480 |
| HOMO SAPIENS | 0 | 11792 | 1fvo | A | 270 | ARG | p.Gln270Ter | P00480 |
| HOMO SAPIENS | 2 | 11792 | 1fvo | A | 277 | ARG | p.Arg277Trp | P00480 |
| HOMO SAPIENS | 0 | 3721 | 4r3z | B | 2 | ASP | p.Asp2Gly | P54136 |
| HOMO SAPIENS | 0 | 3721 | 4r3z | B | 456 | SER | p.Ser456Leu | P54136 |
| HOMO SAPIENS | 5 | 3721 | 4r3z | B | 512 | ARG | p.Arg512Gln | P54136 |
| HOMO SAPIENS | 3 | 5749 | 3hl2 | A | 239 | ALA | p.Ala239Thr | Q9HD40 |
| HOMO SAPIENS | 1 | 5749 | 3hl2 | A | 334 | TYR | p.Tyr334Cys | Q9HD40 |
| HOMO SAPIENS | 1 | 5749 | 3hl2 | A | 452 | LYS | p.Lys452Asn | Q9HD40 |
| HOMO SAPIENS | -1 | 5749 | 3hl2 | A | 465 | ARG | p.Arg465Leu | Q9HD40 |
| HOMO SAPIENS | -1 | 5749 | 3hl2 | A | 489 | ASP | p.Asp489Gly | Q9HD40 |
| HOMO SAPIENS | 5 | 1721 | 3k15 | A | 1652 | MET | p.Met1652Ile | P38398 |
| HOMO SAPIENS | 5 | 1721 | 3k15 | A | 1652 | MET | p.Met1652Lys | P38398 |
| HOMO SAPIENS | 5 | 1721 | 3k15 | A | 1662 | PHE | p.Phe1662Cys | P38398 |
| HOMO SAPIENS | 5 | 1721 | 3k15 | A | 1664 | LEU | p.Leu1664Pro | P38398 |
| HOMO SAPIENS | 0 | 1721 | 3k15 | A | 1665 | VAL | p.Val1665Met | P38398 |
| HOMO SAPIENS | 5 | 1721 | 3k15 | A | 1690 | LYS | p.Lys1690Gln | P38398 |
| HOMO SAPIENS | 5 | 1721 | 3k15 | A | 1692 | ASP | p.Asp1692Asn | P38398 |
| HOMO SAPIENS | 5 | 1721 | 3k15 | A | 1699 | ARG | p.Arg1699Gln | P38398 |
| HOMO SAPIENS | 5 | 1721 | 3k15 | A | 1699 | ARG | p.Arg1699Trp | P38398 |
| HOMO SAPIENS | 3 | 1721 | 3k15 | A | 1706 | GLY | p.Gly1706Glu | P38398 |
| HOMO SAPIENS | 5 | 1721 | 3k15 | A | 1720 | THR | p.Thr1720Ala | P38398 |
| HOMO SAPIENS | 5 | 1721 | 3k15 | A | 1751 | ARG | p.Arg1751Pro | P38398 |
| HOMO SAPIENS | 4 | 1721 | 3k15 | A | 1775 | MET | p.Met1775Arg | P38398 |
| HOMO SAPIENS | 2 | 1721 | 3k15 | A | 1788 | GLY | p.Gly1788Val | P38398 |
| HOMO SAPIENS | 5 | 1721 | 3k15 | A | 1794 | GLU | p.Glu1794Asp | P38398 |
| HOMO SAPIENS | 5 | 1721 | 3k15 | A | 1804 | VAL | p.Val1804Asp | P38398 |
| HOMO SAPIENS | 3 | 1721 | 3k15 | A | 1815 | TRP | p.Trp1815Ter | P38398 |
| HOMO SAPIENS | 1 | 1721 | 3k15 | A | 1819 | ASN | p.Asn1819Ser | P38398 |
| HOMO SAPIENS | 5 | 1721 | 3k15 | A | 1856 | PRO | p.Pro1856Ser | P38398 |
| HOMO SAPIENS | 5 | 1721 | 3k15 | A | 1858 | ILE | p.Ile1858Leu | P38398 |
| HOMO SAPIENS | 5 | 1721 | 3k15 | A | 1858 | ILE | p.Ile1858Thr | P38398 |
| HOMO SAPIENS | 5 | 1721 | 3k15 | A | 1859 | PRO | p.Pro1859Arg | P38398 |
| HOMO SAPIENS | 2 | 22086 | 3kq4 | B | 1297 | PRO | p.Pro1297Leu | O60494 |
| HOMO SAPIENS | 1 | 12135 | 2k9j | B | 724 | ARG | p.Arg750Ter | P05106 |
| HOMO SAPIENS | -1 | 24027 | 2qtl | A | 175 | LEU | p.Ser202Leu | Q9UBK8 |
| HOMO SAPIENS | 0 | 24027 | 2qtl | A | 257 | SER | p.Ser284Thr | Q9UBK8 |
| HOMO SAPIENS | 0 | 24027 | 2qtl | A | 333 | LEU | p.Leu360Val | Q9UBK8 |
| HOMO SAPIENS | 0 | 24027 | 2qtl | A | 350 | LYS | p.Lys377Arg | Q9UBK8 |
| HOMO SAPIENS | 0 | 24027 | 2qtl | A | 415 | ARG | p.Arg442Cys | Q9UBK8 |
| HOMO SAPIENS | 2 | 24027 | 2qtl | A | 450 | PRO | p.Pro477Arg | Q9UBK8 |
| HOMO SAPIENS | 4 | 24027 | 2qtl | A | 487 | GLY | p.Gly514Arg | Q9UBK8 |
| HOMO SAPIENS | 1 | 24027 | 2qtl | A | 595 | HIS | p.His622Tyr | Q9UBK8 |
| HOMO SAPIENS | 0 | 5758 | 3di2 | B | 46 | THR | p.Thr66Ile | P16871 |
| HOMO SAPIENS | 1 | 5758 | 3di2 | B | 118 | VAL | p.Ile138Val | P16871 |
| HOMO SAPIENS | 4 | 848 | 5oa3 | Y | 79 | LEU | p.Leu79Val | P62847 |

| | | | | | | | | |
|---|---|---|---|---|---|---|---|---|
| HOMO SAPIENS | 2 | 848 | 5oa3 | Y | 124 | ASN | p.Asn124Ser | P62847 |
| HOMO SAPIENS | 0 | 81 | 3s5o | A | 36 | ALA | p.Ala36Glu | Q86XE5 |
| HOMO SAPIENS | 0 | 81 | 3s5o | A | 39 | TYR | p.Tyr39Ter | Q86XE5 |
| HOMO SAPIENS | 0 | 81 | 3s5o | A | 45 | PRO | p.Pro45Leu | Q86XE5 |
| HOMO SAPIENS | 0 | 81 | 3s5o | A | 70 | ARG | p.Arg70Ter | Q86XE5 |
| HOMO SAPIENS | 0 | 81 | 3s5o | A | 70 | ARG | p.Arg70Pro | Q86XE5 |
| HOMO SAPIENS | 1 | 81 | 3s5o | A | 97 | ARG | p.Arg97Cys | Q86XE5 |
| HOMO SAPIENS | 4 | 81 | 3s5o | A | 113 | GLU | p.Glu113Lys | Q86XE5 |
| HOMO SAPIENS | 1 | 81 | 3s5o | A | 190 | PRO | p.Pro190Leu | Q86XE5 |
| HOMO SAPIENS | 0 | 81 | 3s5o | A | 245 | VAL | p.Val245Phe | Q86XE5 |
| HOMO SAPIENS | 0 | 81 | 3s5o | A | 257 | CYS | p.Cys257Gly | Q86XE5 |
| HOMO SAPIENS | 0 | 81 | 3s5o | A | 287 | GLY | p.Gly287Val | Q86XE5 |
| HOMO SAPIENS | 7 | 81 | 3s5o | A | 303 | ARG | p.Arg303Cys | Q86XE5 |
| HOMO SAPIENS | 0 | 81 | 3s5o | A | 325 | GLY | p.Gly325Ser | Q86XE5 |
| HOMO SAPIENS | -1 | 622 | 5era | A | 12 | GLY | p.Gly12Arg | P29033 |
| HOMO SAPIENS | -1 | 622 | 5era | A | 12 | GLY | p.Gly12Asp | P29033 |
| HOMO SAPIENS | -1 | 622 | 5era | A | 17 | SER | p.Ser17Phe | P29033 |
| HOMO SAPIENS | 5 | 622 | 5era | A | 24 | TRP | p.Trp24Ter | P29033 |
| HOMO SAPIENS | 0 | 622 | 5era | A | 32 | ARG | p.Arg32Cys | P29033 |
| HOMO SAPIENS | 5 | 622 | 5era | A | 34 | MET | p.Met34Thr | P29033 |
| HOMO SAPIENS | 5 | 622 | 5era | A | 36 | LEU | p.Leu36Pro | P29033 |
| HOMO SAPIENS | 5 | 622 | 5era | A | 37 | VAL | p.Val37Ile | P29033 |
| HOMO SAPIENS | 5 | 622 | 5era | A | 40 | ALA | p.Ala40Glu | P29033 |
| HOMO SAPIENS | 5 | 622 | 5era | A | 44 | TRP | p.Trp44Cys | P29033 |
| HOMO SAPIENS | 5 | 622 | 5era | A | 45 | GLY | p.Gly45Glu | P29033 |
| HOMO SAPIENS | 5 | 622 | 5era | A | 47 | GLU | p.Glu47Ter | P29033 |
| HOMO SAPIENS | 5 | 622 | 5era | A | 50 | ASP | p.Asp50Asn | P29033 |
| HOMO SAPIENS | 5 | 622 | 5era | A | 54 | ASN | p.Asn54Lys | P29033 |
| HOMO SAPIENS | 5 | 622 | 5era | A | 57 | GLN | p.Gln57Ter | P29033 |
| HOMO SAPIENS | 5 | 622 | 5era | A | 59 | GLY | p.Gly59Ala | P29033 |
| HOMO SAPIENS | 5 | 622 | 5era | A | 59 | GLY | p.Gly59Ser | P29033 |
| HOMO SAPIENS | 5 | 622 | 5era | A | 66 | ASP | p.Asp66His | P29033 |
| HOMO SAPIENS | 5 | 622 | 5era | A | 68 | TYR | p.Tyr68Cys | P29033 |
| HOMO SAPIENS | 5 | 622 | 5era | A | 70 | PRO | p.Pro70Ala | P29033 |
| HOMO SAPIENS | 5 | 622 | 5era | A | 73 | HIS | p.His73Arg | P29033 |
| HOMO SAPIENS | 5 | 622 | 5era | A | 75 | ARG | p.Arg75Trp | P29033 |
| HOMO SAPIENS | 5 | 622 | 5era | A | 75 | ARG | p.Arg75Gln | P29033 |
| HOMO SAPIENS | 2 | 622 | 5era | A | 77 | TRP | p.Trp77Arg | P29033 |
| HOMO SAPIENS | 2 | 622 | 5era | A | 77 | TRP | p.Trp77Ter | P29033 |
| HOMO SAPIENS | 0 | 622 | 5era | A | 80 | GLN | p.Gln80Ter | P29033 |
| HOMO SAPIENS | 0 | 622 | 5era | A | 80 | GLN | p.Gln80Pro | P29033 |
| HOMO SAPIENS | 5 | 622 | 5era | A | 81 | LEU | p.Leu81Val | P29033 |
| HOMO SAPIENS | 5 | 622 | 5era | A | 82 | ILE | p.Ile82Met | P29033 |
| HOMO SAPIENS | 5 | 622 | 5era | A | 83 | PHE | p.Phe83Leu | P29033 |
| HOMO SAPIENS | 5 | 622 | 5era | A | 84 | VAL | p.Val84Leu | P29033 |
| HOMO SAPIENS | 5 | 622 | 5era | A | 90 | LEU | p.Leu90Pro | P29033 |
| HOMO SAPIENS | 5 | 622 | 5era | A | 95 | VAL | p.Val95Met | P29033 |
| HOMO SAPIENS | -1 | 622 | 5era | A | 100 | HIS | p.His100Tyr | P29033 |
| HOMO SAPIENS | -1 | 622 | 5era | A | 113 | SER | p.Ser113Arg | P29033 |
| HOMO SAPIENS | -1 | 622 | 5era | A | 119 | GLU | p.Glu119Lys | P29033 |
| HOMO SAPIENS | -1 | 622 | 5era | A | 122 | LYS | p.Lys122Ile | P29033 |
| HOMO SAPIENS | -1 | 622 | 5era | A | 123 | THR | p.Thr123Asn | P29033 |
| HOMO SAPIENS | -1 | 622 | 5era | A | 124 | GLN | p.Gln124Ter | P29033 |
| HOMO SAPIENS | 5 | 622 | 5era | A | 139 | SER | p.Ser139Asn | P29033 |

| | | | | | | | | |
|---|---|---|---|---|---|---|---|---|
| HOMO SAPIENS | 5 | 622 | 5era | A | 143 | ARG | p.Arg143Trp | P29033 |
| HOMO SAPIENS | 5 | 622 | 5era | A | 143 | ARG | p.Arg143Gln | P29033 |
| HOMO SAPIENS | 0 | 622 | 5era | A | 147 | GLU | p.Glu147Lys | P29033 |
| HOMO SAPIENS | 7 | 622 | 5era | A | 155 | TYR | p.Tyr155Ter | P29033 |
| HOMO SAPIENS | 2 | 622 | 5era | A | 159 | ASP | p.Asp159Val | P29033 |
| HOMO SAPIENS | 5 | 622 | 5era | A | 163 | MET | p.Met163Val | P29033 |
| HOMO SAPIENS | 5 | 622 | 5era | A | 165 | ARG | p.Arg165Trp | P29033 |
| HOMO SAPIENS | 5 | 622 | 5era | A | 167 | VAL | p.Val167Met | P29033 |
| HOMO SAPIENS | 5 | 622 | 5era | A | 169 | CYS | p.Cys169Tyr | P29033 |
| HOMO SAPIENS | 5 | 622 | 5era | A | 179 | ASP | p.Asp179Asn | P29033 |
| HOMO SAPIENS | 0 | 622 | 5era | A | 199 | SER | p.Ser199Phe | P29033 |
| HOMO SAPIENS | 0 | 622 | 5era | A | 202 | CYS | p.Cys202Phe | P29033 |
| HOMO SAPIENS | 5 | 622 | 5era | A | 203 | ILE | p.Ile203Thr | P29033 |
| HOMO SAPIENS | 5 | 622 | 5era | A | 206 | ASN | p.Asn206Ser | P29033 |
| HOMO SAPIENS | 0 | 8103 | 5dot | A | 163 | GLY | p.Gly163Ter | P31327 |
| HOMO SAPIENS | 0 | 8103 | 5dot | A | 337 | HIS | p.His337Arg | P31327 |
| HOMO SAPIENS | 3 | 8103 | 5dot | A | 344 | THR | p.Thr344Ala | P31327 |
| HOMO SAPIENS | 1 | 8103 | 5dot | A | 405 | THR | p.Thr405Asn | P31327 |
| HOMO SAPIENS | 5 | 8103 | 5dot | A | 544 | THR | p.Thr544Met | P31327 |
| HOMO SAPIENS | 5 | 8103 | 5dot | A | 716 | ASN | p.Asn716Lys | P31327 |
| HOMO SAPIENS | -1 | 8103 | 5dot | A | 780 | ARG | p.Arg780His | P31327 |
| HOMO SAPIENS | 0 | 8103 | 5dot | A | 982 | GLY | p.Gly982Asp | P31327 |
| HOMO SAPIENS | 3 | 8103 | 5dot | A | 1119 | ALA | p.Ala1119Ser | P31327 |
| HOMO SAPIENS | 1 | 8103 | 5dot | A | 1376 | GLY | p.Gly1376Ser | P31327 |
| HOMO SAPIENS | 4 | 6833 | 1l8l | B | 27 | ARG | p.Arg27Ser | P78330 |
| HOMO SAPIENS | 3 | 6833 | 1l8l | B | 32 | ASP | p.Asp32Gly | P78330 |
| HOMO SAPIENS | 3 | 6833 | 1l8l | B | 32 | ASP | p.Asp32Asn | P78330 |
| HOMO SAPIENS | 1 | 6833 | 1l8l | B | 49 | ARG | p.Arg49Trp | P78330 |
| HOMO SAPIENS | 0 | 6833 | 1l8l | B | 52 | MET | p.Met52Thr | P78330 |
| HOMO SAPIENS | 0 | 6833 | 1l8l | B | 83 | GLN | p.Gln83His | P78330 |
| HOMO SAPIENS | 6 | 810 | 2iw2 | A | 185 | ARG | p.Arg184Gln | P12955 |
| HOMO SAPIENS | 0 | 810 | 2iw2 | A | 213 | ALA | p.Ala212Pro | P12955 |
| HOMO SAPIENS | 1 | 810 | 2iw2 | A | 266 | ARG | p.Arg265Ter | P12955 |
| HOMO SAPIENS | 9 | 810 | 2iw2 | A | 277 | ASP | p.Asp276Asn | P12955 |
| HOMO SAPIENS | 0 | 810 | 2iw2 | A | 279 | GLY | p.Gly278Asp | P12955 |
| HOMO SAPIENS | 9 | 810 | 2iw2 | A | 378 | HIS | p.His377Gln | P12955 |
| HOMO SAPIENS | 0 | 810 | 2iw2 | A | 389 | ARG | p.Arg388His | P12955 |
| HOMO SAPIENS | 0 | 810 | 2iw2 | A | 436 | PHE | p.Leu435Phe | P12955 |
| HOMO SAPIENS | 0 | 810 | 2iw2 | A | 449 | GLY | p.Gly448Arg | P12955 |
| HOMO SAPIENS | -1 | 1530 | 5fwj | A | 388 | ALA | p.Ala388Pro | P41229 |
| HOMO SAPIENS | 4 | 1530 | 5fwj | A | 451 | SER | p.Ser451Arg | P41229 |
| HOMO SAPIENS | 3 | 1530 | 5fwj | A | 731 | LEU | p.Leu731Phe | P41229 |
| HOMO SAPIENS | 3 | 578 | 5net | B | 123 | ALA | p.Ala143Thr | P18564 |
| HOMO SAPIENS | 6 | 578 | 5net | B | 176 | PRO | p.Pro196Thr | P18564 |
| HOMO SAPIENS | 3 | 578 | 5net | B | 280 | GLU | p.Glu300Lys | P18564 |
| HOMO SAPIENS | -1 | 5247 | 4j15 | A | 274 | ALA | p.Ala274Val | P14868 |
| HOMO SAPIENS | -1 | 5247 | 4j15 | A | 280 | HIS | p.His280Leu | P14868 |
| HOMO SAPIENS | 2 | 5247 | 4j15 | A | 460 | ARG | p.Arg460His | P14868 |
| HOMO SAPIENS | 5 | 5247 | 4j15 | A | 494 | ARG | p.Arg494Cys | P14868 |
| HOMO SAPIENS | 7 | 2566 | 1b55 | A | 28 | ARG | p.Arg28His | Q06187 |
| HOMO SAPIENS | 5 | 2566 | 1b55 | A | 33 | THR | p.Thr33Pro | Q06187 |
| HOMO SAPIENS | 0 | 2566 | 1b55 | A | 113 | VAL | p.Val113Asp | Q06187 |
| HOMO SAPIENS | 0 | 356 | 5a2s | A | 786 | ALA | p.Ala786Thr | P56524 |
| HOMO SAPIENS | 5 | 2960 | 4cak | A | 553 | ARG | p.Arg584Ter | P08514 |

| | | | | | | | | |
|---|---|---|---|---|---|---|---|---|
| HOMO SAPIENS | 5 | 2960 | 4cak | A | 565 | ILE | p.Ile596Thr | P08514 |
| HOMO SAPIENS | -1 | 2960 | 4cak | A | 841 | LEU | p.Leu872Met | P08514 |
| HOMO SAPIENS | -1 | 2960 | 4cak | A | 843 | ILE | p.Ile874Ser | P08514 |
| HOMO SAPIENS | 0 | 5054 | 4k8r | B | 216 | PHE | p.Phe216Leu | Q8NBP7 |
| HOMO SAPIENS | 1 | 5054 | 4k8r | B | 417 | HIS | p.His417Gln | Q8NBP7 |
| HOMO SAPIENS | 1 | 5054 | 4k8r | B | 425 | ASN | p.Asn425Ser | Q8NBP7 |
| HOMO SAPIENS | 1 | 5054 | 4k8r | B | 443 | ALA | p.Ala443Thr | Q8NBP7 |
| HOMO SAPIENS | 0 | 27283 | 3n5n | X | 71 | HIS | p.His82Arg | Q9UIF7 |
| HOMO SAPIENS | 1 | 27283 | 3n5n | X | 83 | ARG | p.Arg94Ter | Q9UIF7 |
| HOMO SAPIENS | 8 | 27283 | 3n5n | X | 89 | TRP | p.Trp100Ter | Q9UIF7 |
| HOMO SAPIENS | 8 | 27283 | 3n5n | X | 90 | TYR | p.Tyr101Ter | Q9UIF7 |
| HOMO SAPIENS | 7 | 27283 | 3n5n | X | 95 | ARG | p.Arg106Trp | Q9UIF7 |
| HOMO SAPIENS | 7 | 27283 | 3n5n | X | 117 | TRP | p.Trp128Arg | Q9UIF7 |
| HOMO SAPIENS | 0 | 27283 | 3n5n | X | 137 | GLY | p.Gly148Arg | Q9UIF7 |
| HOMO SAPIENS | 5 | 27283 | 3n5n | X | 142 | TRP | p.Trp153Ter | Q9UIF7 |
| HOMO SAPIENS | 0 | 27283 | 3n5n | X | 165 | TYR | p.Tyr176Cys | Q9UIF7 |
| HOMO SAPIENS | 0 | 27283 | 3n5n | X | 168 | ARG | p.Arg179His | Q9UIF7 |
| HOMO SAPIENS | 0 | 27283 | 3n5n | X | 168 | ARG | p.Arg179Cys | Q9UIF7 |
| HOMO SAPIENS | 0 | 27283 | 3n5n | X | 196 | GLN | p.Gln207Ter | Q9UIF7 |
| HOMO SAPIENS | 8 | 27283 | 3n5n | X | 201 | VAL | p.Val212Met | Q9UIF7 |
| HOMO SAPIENS | 9 | 27283 | 3n5n | X | 227 | ARG | p.Arg238Trp | Q9UIF7 |
| HOMO SAPIENS | 9 | 27283 | 3n5n | X | 231 | ARG | p.Arg242Cys | Q9UIF7 |
| HOMO SAPIENS | 9 | 27283 | 3n5n | X | 231 | ARG | p.Arg242His | Q9UIF7 |
| HOMO SAPIENS | 3 | 27283 | 3n5n | X | 232 | VAL | p.Val243Phe | Q9UIF7 |
| HOMO SAPIENS | 3 | 27283 | 3n5n | X | 246 | GLN | p.Gln257Ter | Q9UIF7 |
| HOMO SAPIENS | 9 | 27283 | 3n5n | X | 272 | GLY | p.Gly283Glu | Q9UIF7 |
| HOMO SAPIENS | 9 | 27283 | 3n5n | X | 281 | PRO | p.Pro292Leu | Q9UIF7 |
| HOMO SAPIENS | -1 | 27283 | 3n5n | X | 314 | ASP | p.Asp325Glu | Q9UIF7 |
| HOMO SAPIENS | 0 | 27283 | 3n5n | X | 324 | GLN | p.Gln335His | Q9UIF7 |
| HOMO SAPIENS | 0 | 27283 | 3n5n | X | 324 | GLN | p.Gln335Arg | Q9UIF7 |
| HOMO SAPIENS | 0 | 27283 | 3n5n | X | 324 | GLN | p.Gln335Ter | Q9UIF7 |
| HOMO SAPIENS | -1 | 20852 | 4p60 | B | 243 | ARG | p.Arg243Lys | O75746 |
| HOMO SAPIENS | -1 | 1875 | 5dd2 | G | 152 | VAL | p.Val179Leu | P06396 |
| HOMO SAPIENS | 9 | 2126 | 3n1p | B | 46 | PRO | p.Pro46Leu | Q14623 |
| HOMO SAPIENS | 8 | 2126 | 3n1p | B | 76 | GLU | p.Glu76Ala | Q14623 |
| HOMO SAPIENS | 9 | 2126 | 3n1p | B | 77 | ARG | p.Arg77Ser | Q14623 |
| HOMO SAPIENS | 6 | 2126 | 3n1p | B | 95 | GLU | p.Glu95Lys | Q14623 |
| HOMO SAPIENS | 9 | 2126 | 3n1p | B | 100 | ASP | p.Asp100Glu | Q14623 |
| HOMO SAPIENS | 9 | 2126 | 3n1p | B | 131 | GLU | p.Glu131Lys | Q14623 |
| HOMO SAPIENS | 0 | 2126 | 3n1p | B | 190 | VAL | p.Val190Ala | Q14623 |
| HOMO SAPIENS | 5 | 40410 | 3ioz | B | 70 | GLN | p.Gln70Leu | P20963 |
| HOMO SAPIENS | 1 | 5754 | 5lkz | A | 1108 | THR | p.Tyr1191Ter | Q09472 |
| HOMO SAPIENS | 3 | 8159 | 1tiu | A | 3 | GLU | p.Lys12679Glu | Q8WZ42 |
| HOMO SAPIENS | 4 | 3275 | 4ldq | A | 210 | ARG | p.Arg210Gln | Q9BV20 |
| HOMO SAPIENS | 0 | 8143 | 4miv | B | 66 | SER | p.Ser66Trp | P51688 |
| HOMO SAPIENS | 1 | 8143 | 4miv | B | 74 | ARG | p.Arg74Cys | P51688 |
| HOMO SAPIENS | 0 | 8143 | 4miv | B | 128 | PRO | p.Pro128Leu | P51688 |
| HOMO SAPIENS | 1 | 8143 | 4miv | B | 150 | ARG | p.Arg150Gln | P51688 |
| HOMO SAPIENS | 0 | 8143 | 4miv | B | 191 | GLY | p.Gly191Arg | P51688 |
| HOMO SAPIENS | 0 | 8143 | 4miv | B | 206 | ARG | p.Arg206Pro | P51688 |
| HOMO SAPIENS | 0 | 8143 | 4miv | B | 225 | PHE | p.Phe225Leu | P51688 |
| HOMO SAPIENS | 0 | 8143 | 4miv | B | 233 | ARG | p.Arg233Ter | P51688 |
| HOMO SAPIENS | 0 | 8143 | 4miv | B | 234 | ALA | p.Ala234Gly | P51688 |
| HOMO SAPIENS | 0 | 8143 | 4miv | B | 245 | ARG | p.Arg245His | P51688 |

| | | | | | | | | |
|---|---|---|---|---|---|---|---|---|
| HOMO SAPIENS | 0 | 8143 | 4miv | B | 293 | PRO | p.Pro293Ser | P51688 |
| HOMO SAPIENS | 1 | 8143 | 4miv | B | 298 | SER | p.Ser298Pro | P51688 |
| HOMO SAPIENS | 0 | 8143 | 4miv | B | 361 | VAL | p.Val361Ile | P51688 |
| HOMO SAPIENS | 0 | 8143 | 4miv | B | 369 | GLU | p.Glu369Lys | P51688 |
| HOMO SAPIENS | 0 | 8143 | 4miv | B | 372 | MET | p.Met372Ile | P51688 |
| HOMO SAPIENS | 1 | 8143 | 4miv | B | 380 | GLN | p.Gln380Arg | P51688 |
| HOMO SAPIENS | 0 | 8143 | 4miv | B | 387 | VAL | p.Val387Met | P51688 |
| HOMO SAPIENS | 0 | 8143 | 4miv | B | 394 | MET | p.Met394Ile | P51688 |
| HOMO SAPIENS | 0 | 8143 | 4miv | B | 433 | ARG | p.Arg433Gln | P51688 |
| HOMO SAPIENS | 1 | 8143 | 4miv | B | 446 | HIS | p.His446Arg | P51688 |
| HOMO SAPIENS | 5 | 8143 | 4miv | B | 447 | GLU | p.Glu447Lys | P51688 |
| HOMO SAPIENS | 2 | 8143 | 4miv | B | 456 | ARG | p.Arg456His | P51688 |
| HOMO SAPIENS | 1 | 21880 | 2oo9 | B | 857 | LEU | p.Leu857Phe | P22681 |
| HOMO SAPIENS | 0 | 9467 | 1s1d | A | 154 | LEU | p.Leu224Pro | Q8WVQ1 |
| HOMO SAPIENS | 0 | 9467 | 1s1d | A | 156 | VAL | p.Val226Met | Q8WVQ1 |
| HOMO SAPIENS | 0 | 9467 | 1s1d | A | 229 | PRO | p.Pro299Leu | Q8WVQ1 |
| HOMO SAPIENS | 4 | 9467 | 1s1d | A | 230 | ARG | p.Arg300Cys | Q8WVQ1 |
| HOMO SAPIENS | 4 | 9467 | 1s1d | A | 230 | ARG | p.Arg300His | Q8WVQ1 |
| HOMO SAPIENS | 0 | 9467 | 1s1d | A | 253 | ALA | p.Ala323Thr | Q8WVQ1 |
| HOMO SAPIENS | 2 | 9467 | 1s1d | A | 305 | MET | p.Met375Leu | Q8WVQ1 |
| HOMO SAPIENS | -1 | 2361 | 5llm | A | 5 | ASN | p.Asn5Lys | P43003 |
| HOMO SAPIENS | 0 | 2361 | 5llm | A | 71 | TYR | p.Tyr71Cys | P43003 |
| HOMO SAPIENS | -1 | 2361 | 5llm | A | 211 | LYS | p.Glu219Asp | Q15758 |
| HOMO SAPIENS | 8 | 2361 | 5llm | A | 270 | PRO | p.Pro290Arg | P43003 |
| HOMO SAPIENS | 1 | 2361 | 5llm | A | 321 | THR | p.Ala329Thr | Q15758 |
| HOMO SAPIENS | 1 | 2361 | 5llm | A | 365 | ARG | p.Arg385His | P43003 |
| HOMO SAPIENS | 1 | 2361 | 5llm | A | 479 | ARG | p.Arg499Gln | P43003 |
| HOMO SAPIENS | 3 | 5412 | 4obz | D | 165 | PHE | p.Phe229Ile | P07339 |
| HOMO SAPIENS | 2 | 5412 | 4obz | D | 187 | ASP | p.Asp251Asn | P07339 |
| HOMO SAPIENS | 0 | 5412 | 4obz | D | 218 | GLY | p.Gly282Arg | P07339 |
| HOMO SAPIENS | 0 | 5412 | 4obz | D | 245 | ARG | p.Arg309His | P07339 |
| HOMO SAPIENS | 2 | 5412 | 4obz | D | 319 | TRP | p.Trp383Cys | P07339 |
| HOMO SAPIENS | 0 | 3544 | 5h64 | A | 277 | MET | p.Met277Ile | P42345 |
| HOMO SAPIENS | 0 | 14230 | 2kbx | A | 145 | ARG | p.Arg145Ser | Q13418 |
| HOMO SAPIENS | 2 | 34970 | 4xhz | A | 834 | ARG | p.Arg855Trp | Q96QU1 |
| HOMO SAPIENS | 4 | 34970 | 4xhz | A | 854 | SER | p.Ser875Ter | Q96QU1 |
| HOMO SAPIENS | 1 | 34970 | 4xhz | A | 908 | ARG | p.Arg929Gln | Q96QU1 |
| HOMO SAPIENS | 1 | 34970 | 4xhz | A | 941 | ARG | p.Arg962Cys | Q96QU1 |
| HOMO SAPIENS | 0 | 34970 | 4xhz | A | 970 | ARG | p.Arg991Ter | Q96QU1 |
| HOMO SAPIENS | 0 | 34970 | 4xhz | A | 976 | GLU | p.Glu997Gly | Q96QU1 |
| HOMO SAPIENS | 0 | 25734 | 4asc | A | 345 | SER | p.Asn345Ser | Q2TBA0 |
| HOMO SAPIENS | 0 | 25734 | 4asc | A | 514 | VAL | p.Val514Met | Q2TBA0 |
| HOMO SAPIENS | 6 | 25734 | 4asc | A | 528 | GLU | p.Glu528Lys | Q2TBA0 |
| HOMO SAPIENS | 0 | 25734 | 4asc | A | 580 | LEU | p.Leu580Ile | Q2TBA0 |
| HOMO SAPIENS | 8 | 2356 | 3kse | D | 22 | LYS | p.Lys22Ter | P01040 |
| HOMO SAPIENS | 1 | 2356 | 3kse | D | 58 | ARG | p.Arg58Ter | P01040 |
| HOMO SAPIENS | 2 | 6000 | 1jqd | A | 60 | GLY | p.Gly60Asp | P50135 |
| HOMO SAPIENS | 9 | 6000 | 1jqd | A | 208 | LEU | p.Leu208Pro | P50135 |
| HOMO SAPIENS | 0 | 14741 | 5eh1 | A | 58 | THR | p.Thr58Arg | P38484 |
| HOMO SAPIENS | 0 | 14741 | 5eh1 | A | 168 | THR | p.Thr168Asn | P38484 |
| HOMO SAPIENS | 1 | 4814 | 4kxk | D | 427 | ARG | p.Arg427Ter | P50542 |
| HOMO SAPIENS | 0 | 188 | 2xt3 | A | 49 | ASP | p.Asp52Asn | Q2M1P5 |
| HOMO SAPIENS | 3 | 188 | 2xt3 | A | 142 | TYR | p.Tyr145Ser | Q2M1P5 |
| HOMO SAPIENS | 9 | 8681 | 5tby | D | 57 | ALA | p.Ala57Asp | P08590 |

| | | | | | | | | |
|---|---|---|---|---|---|---|---|---|
| HOMO SAPIENS | 3 | 8681 | 5tby | D | 62 | ASP | p.Asp62Asn | P08590 |
| HOMO SAPIENS | 0 | 8681 | 5tby | D | 65 | PRO | p.Pro65Arg | P08590 |
| HOMO SAPIENS | 7 | 8681 | 5tby | D | 79 | VAL | p.Val79Ile | P08590 |
| HOMO SAPIENS | 0 | 8681 | 5tby | D | 143 | GLU | p.Glu143Lys | P08590 |
| HOMO SAPIENS | 0 | 8681 | 5tby | D | 149 | MET | p.Met149Val | P08590 |
| HOMO SAPIENS | 3 | 8681 | 5tby | D | 154 | ARG | p.Arg154His | P08590 |
| HOMO SAPIENS | 6 | 30984 | 4xwh | A | 140 | TYR | p.Tyr140Cys | P54802 |
| HOMO SAPIENS | 4 | 30984 | 4xwh | A | 141 | SER | p.Ser141Thr | P54802 |
| HOMO SAPIENS | 2 | 30984 | 4xwh | A | 234 | ARG | p.Arg234Cys | P54802 |
| HOMO SAPIENS | 2 | 30984 | 4xwh | A | 336 | GLU | p.Glu336Ter | P54802 |
| HOMO SAPIENS | 6 | 30984 | 4xwh | A | 482 | ARG | p.Arg482Trp | P54802 |
| HOMO SAPIENS | 2 | 30984 | 4xwh | A | 521 | PRO | p.Pro521Leu | P54802 |
| HOMO SAPIENS | 0 | 30984 | 4xwh | A | 565 | ARG | p.Arg565Gln | P54802 |
| HOMO SAPIENS | 0 | 30984 | 4xwh | A | 565 | ARG | p.Arg565Trp | P54802 |
| HOMO SAPIENS | 6 | 30984 | 4xwh | A | 626 | ARG | p.Arg626Ter | P54802 |
| HOMO SAPIENS | 6 | 30984 | 4xwh | A | 643 | ARG | p.Arg643His | P54802 |
| HOMO SAPIENS | 6 | 30984 | 4xwh | A | 643 | ARG | p.Arg643Cys | P54802 |
| HOMO SAPIENS | 2 | 30984 | 4xwh | A | 674 | ARG | p.Arg674His | P54802 |
| HOMO SAPIENS | 2 | 30984 | 4xwh | A | 737 | ARG | p.Arg737Gly | P54802 |
| HOMO SAPIENS | 5 | 10545 | 1oqe | N | 6 | PRO | p.Pro21Arg | Q96RJ3 |
| HOMO SAPIENS | 1 | 16995 | 5d3a | B | 12 | MET | p.Met947Arg | Q7Z4S6 |
| HOMO SAPIENS | 1 | 16995 | 5d3a | B | 12 | MET | p.Met947Val | Q7Z4S6 |
| HOMO SAPIENS | 2 | 16995 | 5d3a | B | 19 | ARG | p.Arg954Gln | Q7Z4S6 |
| HOMO SAPIENS | 2 | 16995 | 5d3a | B | 19 | ARG | p.Arg954Trp | Q7Z4S6 |
| HOMO SAPIENS | 0 | 16995 | 5d3a | B | 75 | ILE | p.Ile1010Thr | Q7Z4S6 |
| HOMO SAPIENS | 0 | 17091 | 2yin | B | 1324 | GLN | p.Gln1324Ter | Q92608 |
| HOMO SAPIENS | 5 | 27606 | 1lm7 | B | 2366 | ARG | p.Arg2366Cys | P15924 |
| HOMO SAPIENS | 2 | 15731 | 1bi7 | B | 24 | ARG | p.Arg24Pro | P42771 |
| HOMO SAPIENS | 0 | 15731 | 1bi7 | B | 28 | VAL | p.Val28Gly | P42771 |
| HOMO SAPIENS | 0 | 15731 | 1bi7 | B | 49 | ILE | p.Ile49Thr | P42771 |
| HOMO SAPIENS | 0 | 15731 | 1bi7 | B | 53 | MET | p.Met53Ile | P42771 |
| HOMO SAPIENS | 2 | 15731 | 1bi7 | B | 56 | SER | p.Ser56Ile | P42771 |
| HOMO SAPIENS | 1 | 15731 | 1bi7 | B | 57 | ALA | p.Ala57Val | P42771 |
| HOMO SAPIENS | 4 | 15731 | 1bi7 | B | 59 | VAL | p.Val59Gly | P42771 |
| HOMO SAPIENS | 9 | 15731 | 1bi7 | B | 89 | GLY | p.Gly89Asp | P42771 |
| HOMO SAPIENS | 0 | 15731 | 1bi7 | B | 101 | GLY | p.Gly101Trp | P42771 |
| HOMO SAPIENS | 0 | 15731 | 1bi7 | B | 114 | PRO | p.Pro114Ser | P42771 |
| HOMO SAPIENS | 9 | 15731 | 1bi7 | B | 123 | HIS | p.His123Gln | P42771 |
| HOMO SAPIENS | 8 | 15731 | 1bi7 | B | 126 | VAL | p.Val126Asp | P42771 |
| HOMO SAPIENS | 7 | 15731 | 1bi7 | B | 127 | ALA | p.Ala127Ser | P42771 |
| HOMO SAPIENS | -1 | 15731 | 1bi7 | B | 148 | ALA | p.Ala148Thr | P42771 |
| HOMO SAPIENS | -1 | 19858 | 3jbk | M | 866 | ALA | p.Ala934Val | P18206 |
| HOMO SAPIENS | 0 | 19484 | 3ee6 | B | 62 | SER | p.Ser62Leu | O14773 |
| HOMO SAPIENS | 1 | 19484 | 3ee6 | B | 66 | GLN | p.Gln66Ter | O14773 |
| HOMO SAPIENS | 0 | 19484 | 3ee6 | B | 100 | GLN | p.Gln100Arg | O14773 |
| HOMO SAPIENS | 0 | 19484 | 3ee6 | B | 104 | LEU | p.Leu104Ter | O14773 |
| HOMO SAPIENS | 0 | 19484 | 3ee6 | B | 127 | ARG | p.Arg127Gln | O14773 |
| HOMO SAPIENS | 0 | 19484 | 3ee6 | B | 127 | ARG | p.Arg127Ter | O14773 |
| HOMO SAPIENS | -1 | 19484 | 3ee6 | B | 185 | ARG | p.Arg185Cys | O14773 |
| HOMO SAPIENS | 0 | 19484 | 3ee6 | B | 206 | ARG | p.Arg206Cys | O14773 |
| HOMO SAPIENS | 0 | 19484 | 3ee6 | B | 208 | ARG | p.Arg208Ter | O14773 |
| HOMO SAPIENS | 0 | 19484 | 3ee6 | B | 214 | GLN | p.Gln214Ter | O14773 |
| HOMO SAPIENS | 1 | 19484 | 3ee6 | B | 276 | ASP | p.Asp276Val | O14773 |
| HOMO SAPIENS | 0 | 19484 | 3ee6 | B | 284 | GLY | p.Gly284Val | O14773 |

| | | | | | | | | |
|---|---|---|---|---|---|---|---|---|
| HOMO SAPIENS | 0 | 19484 | 3ee6 | B | 286 | ASN | p.Asn286Ser | O14773 |
| HOMO SAPIENS | 0 | 19484 | 3ee6 | B | 339 | ARG | p.Arg339Gly | O14773 |
| HOMO SAPIENS | 2 | 19484 | 3ee6 | B | 345 | MET | p.Met345Leu | O14773 |
| HOMO SAPIENS | 0 | 19484 | 3ee6 | B | 365 | CYS | p.Cys365Arg | O14773 |
| HOMO SAPIENS | 0 | 19484 | 3ee6 | B | 365 | CYS | p.Cys365Tyr | O14773 |
| HOMO SAPIENS | 0 | 19484 | 3ee6 | B | 373 | GLN | p.Gln373Glu | O14773 |
| HOMO SAPIENS | 6 | 19484 | 3ee6 | B | 418 | ARG | p.Arg418Gln | O14773 |
| HOMO SAPIENS | 7 | 19484 | 3ee6 | B | 422 | GLN | p.Gln422His | O14773 |
| HOMO SAPIENS | 0 | 19484 | 3ee6 | B | 447 | ARG | p.Arg447His | O14773 |
| HOMO SAPIENS | 0 | 19484 | 3ee6 | B | 466 | VAL | p.Val466Met | O14773 |
| HOMO SAPIENS | 0 | 4654 | 5t5w | A | 47 | LYS | p.Lys47Glu | Q08334 |
| HOMO SAPIENS | 0 | 4654 | 5t5w | A | 141 | GLU | p.Glu141Ter | Q08334 |
| HOMO SAPIENS | -1 | 18815 | 4wj7 | A | 53 | VAL | p.Val53Ile | Q9BSQ5 |
| HOMO SAPIENS | 0 | 18815 | 4wj7 | A | 120 | VAL | p.Val120Ile | Q9BSQ5 |
| HOMO SAPIENS | 0 | 18815 | 4wj7 | A | 198 | LEU | p.Leu198Arg | Q9BSQ5 |
| HOMO SAPIENS | 0 | 4602 | 1mjd | A | 47 | SER | p.Ser47Arg | O43602 |
| HOMO SAPIENS | 6 | 4602 | 1mjd | A | 59 | ARG | p.Arg59Leu | O43602 |
| HOMO SAPIENS | 7 | 4602 | 1mjd | A | 62 | ASP | p.Asp62Asn | O43602 |
| HOMO SAPIENS | 7 | 4602 | 1mjd | A | 71 | ALA | p.Ala71Ser | O43602 |
| HOMO SAPIENS | 7 | 4602 | 1mjd | A | 78 | ARG | p.Arg78Leu | O43602 |
| HOMO SAPIENS | 7 | 4602 | 1mjd | A | 89 | ARG | p.Arg89Gly | O43602 |
| HOMO SAPIENS | 8 | 4602 | 1mjd | A | 104 | ILE | p.Ile104Phe | O43602 |
| HOMO SAPIENS | 7 | 4602 | 1mjd | A | 125 | TYR | p.Tyr125His | O43602 |
| HOMO SAPIENS | 5 | 34467 | 1n0w | B | 1529 | GLY | p.Gly1529Arg | P51587 |
| HOMO SAPIENS | 8 | 5738 | 1u31 | A | 130 | ASN | p.Asn1009Lys | Q13423 |
| HOMO SAPIENS | 0 | 5760 | 2i6s | A | 298 | GLU | p.Glu318Asp | P06681 |
| HOMO SAPIENS | 0 | 5760 | 2i6s | A | 573 | ARG | p.Arg593Gln | P06681 |
| HOMO SAPIENS | 0 | 5760 | 2i6s | A | 592 | SER | p.Ser612Asn | P06681 |
| HOMO SAPIENS | 0 | 5760 | 2i6s | A | 621 | VAL | p.Val641Ala | P06681 |
| HOMO SAPIENS | 1 | 10977 | 2fby | A | 114 | VAL | p.Val114Ile | Q14191 |
| HOMO SAPIENS | 0 | 10977 | 2fby | A | 135 | LYS | p.Lys135Glu | Q14191 |
| HOMO SAPIENS | -1 | 1175 | 1qrk | A | 34 | VAL | p.Val35Leu | P00488 |
| HOMO SAPIENS | 4 | 1175 | 1qrk | A | 564 | PRO | p.Pro565Leu | P00488 |
| HOMO SAPIENS | 0 | 1175 | 1qrk | A | 650 | VAL | p.Val651Ile | P00488 |
| HOMO SAPIENS | 0 | 1175 | 1qrk | A | 651 | GLU | p.Gln652Glu | P00488 |
| HOMO SAPIENS | 3 | 909 | 4h0w | A | 375 | GLU | p.Glu394Lys | P02787 |
| HOMO SAPIENS | 0 | 909 | 4h0w | A | 458 | ALA | p.Ala477Pro | P02787 |
| HOMO SAPIENS | 6 | 909 | 4h0w | A | 515 | TYR | p.Tyr534Ter | P02787 |
| HOMO SAPIENS | 7 | 909 | 4h0w | A | 570 | PRO | p.Pro589Ser | P02787 |
| HOMO SAPIENS | 3 | 909 | 4h0w | A | 652 | GLY | p.Gly671Glu | P02787 |
| HOMO SAPIENS | 5 | 1964 | 2car | A | 151 | TRP | p.Trp151Ter | Q9BY32 |
| HOMO SAPIENS | 7 | 1964 | 2car | A | 178 | ARG | p.Arg178Cys | Q9BY32 |
| HOMO SAPIENS | 0 | 144 | 2n0a | J | 30 | ALA | p.Ala30Pro | P37840 |
| HOMO SAPIENS | 1 | 12288 | 4q58 | D | 1225 | ARG | p.Arg1225His | P16144 |
| HOMO SAPIENS | 2 | 12288 | 4q58 | D | 1281 | ARG | p.Arg1281Trp | P16144 |
| HOMO SAPIENS | 2 | 12288 | 4q58 | D | 1287 | ASN | p.Asn1287Lys | P16144 |
| HOMO SAPIENS | 0 | 5085 | 3s94 | A | 360 | ARG | p.Arg360His | O75581 |
| HOMO SAPIENS | 0 | 5085 | 3s94 | A | 433 | ASN | p.Asn433Ser | O75581 |
| HOMO SAPIENS | 2 | 5085 | 3s94 | A | 611 | ARG | p.Arg611Cys | O75581 |
| HOMO SAPIENS | 0 | 15452 | 4dur | A | 19 | LYS | p.Lys38Glu | P00747 |
| HOMO SAPIENS | 2 | 15452 | 4dur | A | 216 | ARG | p.Arg235His | P00747 |
| HOMO SAPIENS | 7 | 8406 | 4wo1 | D | 28 | VAL | p.Val55Leu | O43914 |
| HOMO SAPIENS | 2 | 895 | 5xte | K | 183 | ARG | p.Arg183Trp | P22695 |
| HOMO SAPIENS | 2 | 4191 | 4wnd | A | 120 | ARG | p.Arg127Ter | P81274 |

| | | | | | | | | |
|---|---|---|---|---|---|---|---|---|
| HOMO SAPIENS | 2 | 4191 | 4wnd | A | 120 | ARG | p.Arg127Gln | P81274 |
| HOMO SAPIENS | 1 | 4191 | 4wnd | A | 349 | GLY | p.Gly356Arg | P81274 |
| HOMO SAPIENS | 0 | 41734 | 2kbi | A | 1777 | VAL | p.Val1777Met | Q14524 |
| HOMO SAPIENS | 1 | 41734 | 2kbi | A | 1784 | GLU | p.Glu1784Lys | Q14524 |
| HOMO SAPIENS | 5 | 41734 | 2kbi | A | 1795 | TYR | p.Tyr1795Cys | Q14524 |
| HOMO SAPIENS | 5 | 41734 | 2kbi | A | 1795 | TYR | p.Tyr1795His | Q14524 |
| HOMO SAPIENS | 7 | 41734 | 2kbi | A | 1826 | ARG | p.Arg1826Cys | Q14524 |
| HOMO SAPIENS | 3 | 2119 | 1pl0 | A | 426 | LYS | p.Lys426Arg | P31939 |
| HOMO SAPIENS | 5 | 3475 | 5u62 | C | 50 | LEU | p.Leu50Ser | Q15910 |
| HOMO SAPIENS | 3 | 379 | 4uez | A | 200 | TYR | p.Tyr200Ter | P15085 |
| HOMO SAPIENS | 0 | 8195 | 5xaw | B | 67 | ARG | p.Arg67Trp | Q8IWT1 |
| HOMO SAPIENS | 0 | 8195 | 5xaw | B | 94 | THR | p.Thr94Met | Q8IWT1 |
| HOMO SAPIENS | 1 | 5711 | 1fo3 | A | 265 | ARG | p.Arg265His | Q9UKM7 |
| HOMO SAPIENS | 4 | 5711 | 1fo3 | A | 334 | ARG | p.Arg334Cys | Q9UKM7 |
| HOMO SAPIENS | 1 | 5711 | 1fo3 | A | 478 | LYS | p.Lys478Arg | Q9UKM7 |
| HOMO SAPIENS | 1 | 5711 | 1fo3 | A | 682 | ASP | p.Asp682Glu | Q9UKM7 |
| HOMO SAPIENS | 0 | 170 | 2n3y | A | 41 | GLY | p.Gly42Ser | P99999 |
| HOMO SAPIENS | 0 | 4122 | 3w81 | A | 33 | HIS | p.His33Gln | P35475 |
| HOMO SAPIENS | 0 | 4122 | 3w81 | A | 64 | TYR | p.Tyr64Ter | P35475 |
| HOMO SAPIENS | 0 | 4122 | 3w81 | A | 70 | GLN | p.Gln70Ter | P35475 |
| HOMO SAPIENS | 0 | 4122 | 3w81 | A | 75 | ALA | p.Ala75Thr | P35475 |
| HOMO SAPIENS | 0 | 4122 | 3w81 | A | 79 | ALA | p.Ala79Val | P35475 |
| HOMO SAPIENS | 0 | 4122 | 3w81 | A | 79 | ALA | p.Ala79Thr | P35475 |
| HOMO SAPIENS | 0 | 4122 | 3w81 | A | 89 | ARG | p.Arg89Gln | P35475 |
| HOMO SAPIENS | 0 | 4122 | 3w81 | A | 99 | THR | p.Thr99Ile | P35475 |
| HOMO SAPIENS | 0 | 4122 | 3w81 | A | 105 | ARG | p.Arg105Gln | P35475 |
| HOMO SAPIENS | 0 | 4122 | 3w81 | A | 238 | LEU | p.Leu238Gln | P35475 |
| HOMO SAPIENS | 0 | 4122 | 3w81 | A | 300 | ALA | p.Ala300Thr | P35475 |
| HOMO SAPIENS | 0 | 4122 | 3w81 | A | 322 | VAL | p.Val322Glu | P35475 |
| HOMO SAPIENS | 0 | 4122 | 3w81 | A | 327 | ALA | p.Ala327Pro | P35475 |
| HOMO SAPIENS | 0 | 4122 | 3w81 | A | 346 | LEU | p.Leu346Arg | P35475 |
| HOMO SAPIENS | 0 | 4122 | 3w81 | A | 349 | ASP | p.Asp349Asn | P35475 |
| HOMO SAPIENS | 0 | 4122 | 3w81 | A | 361 | ALA | p.Ala361Thr | P35475 |
| HOMO SAPIENS | 0 | 4122 | 3w81 | A | 366 | THR | p.Thr366Pro | P35475 |
| HOMO SAPIENS | 0 | 4122 | 3w81 | A | 402 | TRP | p.Trp402Ter | P35475 |
| HOMO SAPIENS | 0 | 4122 | 3w81 | A | 409 | GLY | p.Gly409Arg | P35475 |
| HOMO SAPIENS | 0 | 4122 | 3w81 | A | 454 | VAL | p.Val454Ile | P35475 |
| HOMO SAPIENS | 1 | 4122 | 3w81 | A | 490 | LEU | p.Leu490Pro | P35475 |
| HOMO SAPIENS | 1 | 4122 | 3w81 | A | 492 | ARG | p.Arg492Pro | P35475 |
| HOMO SAPIENS | 0 | 4122 | 3w81 | A | 533 | PRO | p.Pro533Arg | P35475 |
| HOMO SAPIENS | 0 | 4122 | 3w81 | A | 591 | ALA | p.Ala591Thr | P35475 |
| HOMO SAPIENS | 7 | 4122 | 3w81 | A | 619 | ARG | p.Arg619Gly | P35475 |
| HOMO SAPIENS | 0 | 15325 | 3wvz | B | 54 | VAL | p.Val54Leu | Q53FT3 |
| HOMO SAPIENS | 3 | 3228 | 5iz8 | A | 421 | TRP | p.Trp421Ter | P25054 |
| HOMO SAPIENS | 2 | 3228 | 5iz8 | A | 437 | PRO | p.Pro437Arg | P25054 |
| HOMO SAPIENS | 3 | 3228 | 5iz8 | A | 499 | ARG | p.Arg499Ter | P25054 |
| HOMO SAPIENS | -1 | 29229 | 5buo | A | 539 | SER | p.Ser614Gly | P05067 |
| HOMO SAPIENS | -1 | 34931 | 4n7v | C | 54 | SER | p.Ser54Leu | O94986 |
| HOMO SAPIENS | 0 | 2450 | 5kzx | A | 123 | GLY | p.Gly123Glu | P10253 |
| HOMO SAPIENS | 1 | 2450 | 5kzx | A | 190 | ARG | p.Arg190His | P10253 |
| HOMO SAPIENS | -1 | 2450 | 5kzx | A | 199 | HIS | p.His199Arg | P10253 |
| HOMO SAPIENS | 0 | 2450 | 5kzx | A | 219 | GLY | p.Gly219Arg | P10253 |
| HOMO SAPIENS | 0 | 2450 | 5kzx | A | 220 | VAL | p.Val220Leu | P10253 |
| HOMO SAPIENS | 0 | 2450 | 5kzx | A | 222 | VAL | p.Val222Met | P10253 |

| | | | | | | | | |
|---|---|---|---|---|---|---|---|---|
| HOMO SAPIENS | 0 | 2450 | 5kzx | A | 237 | ALA | p.Ala237Val | P10253 |
| HOMO SAPIENS | 0 | 2450 | 5kzx | A | 251 | SER | p.Ser251Leu | P10253 |
| HOMO SAPIENS | 0 | 2450 | 5kzx | A | 254 | SER | p.Ser254Leu | P10253 |
| HOMO SAPIENS | 0 | 2450 | 5kzx | A | 262 | GLU | p.Glu262Lys | P10253 |
| HOMO SAPIENS | 0 | 2450 | 5kzx | A | 293 | GLY | p.Gly293Arg | P10253 |
| HOMO SAPIENS | 0 | 2450 | 5kzx | A | 299 | LEU | p.Leu299Arg | P10253 |
| HOMO SAPIENS | 0 | 2450 | 5kzx | A | 309 | GLY | p.Gly309Arg | P10253 |
| HOMO SAPIENS | 0 | 2450 | 5kzx | A | 318 | MET | p.Met318Thr | P10253 |
| HOMO SAPIENS | 0 | 2450 | 5kzx | A | 355 | LEU | p.Leu355Pro | P10253 |
| HOMO SAPIENS | 1 | 2450 | 5kzx | A | 361 | PRO | p.Pro361Leu | P10253 |
| HOMO SAPIENS | 4 | 2450 | 5kzx | A | 408 | MET | p.Met408Val | P10253 |
| HOMO SAPIENS | 1 | 2450 | 5kzx | A | 429 | GLN | p.Gln429Glu | P10253 |
| HOMO SAPIENS | 4 | 2450 | 5kzx | A | 437 | ARG | p.Arg437Cys | P10253 |
| HOMO SAPIENS | 0 | 2450 | 5kzx | A | 439 | MET | p.Met439Lys | P10253 |
| HOMO SAPIENS | 1 | 2450 | 5kzx | A | 448 | SER | p.Ser448Thr | P10253 |
| HOMO SAPIENS | 0 | 2450 | 5kzx | A | 451 | PRO | p.Pro451Arg | P10253 |
| HOMO SAPIENS | 1 | 2450 | 5kzx | A | 459 | ASP | p.Asp459His | P10253 |
| HOMO SAPIENS | 8 | 2450 | 5kzx | A | 481 | TRP | p.Trp481Arg | P10253 |
| HOMO SAPIENS | 1 | 2450 | 5kzx | A | 489 | ASP | p.Asp489Asn | P10253 |
| HOMO SAPIENS | 0 | 2450 | 5kzx | A | 521 | GLU | p.Glu521Lys | P10253 |
| HOMO SAPIENS | 0 | 2450 | 5kzx | A | 529 | SER | p.Ser529Val | P10253 |
| HOMO SAPIENS | 0 | 2450 | 5kzx | A | 545 | PRO | p.Pro545Leu | P10253 |
| HOMO SAPIENS | 0 | 2450 | 5kzx | A | 552 | LEU | p.Leu552Pro | P10253 |
| HOMO SAPIENS | 0 | 2450 | 5kzx | A | 576 | GLY | p.Gly576Ser | P10253 |
| HOMO SAPIENS | 9 | 2450 | 5kzx | A | 600 | ARG | p.Arg600His | P10253 |
| HOMO SAPIENS | 8 | 2450 | 5kzx | A | 615 | GLY | p.Gly615Arg | P10253 |
| HOMO SAPIENS | 3 | 2450 | 5kzx | A | 619 | SER | p.Ser619Asn | P10253 |
| HOMO SAPIENS | 0 | 2450 | 5kzx | A | 638 | GLY | p.Gly638Trp | P10253 |
| HOMO SAPIENS | 0 | 2450 | 5kzx | A | 643 | GLY | p.Gly643Arg | P10253 |
| HOMO SAPIENS | 9 | 2450 | 5kzx | A | 645 | ASP | p.Asp645Glu | P10253 |
| HOMO SAPIENS | 9 | 2450 | 5kzx | A | 645 | ASP | p.Asp645His | P10253 |
| HOMO SAPIENS | 0 | 2450 | 5kzx | A | 648 | GLY | p.Gly648Ser | P10253 |
| HOMO SAPIENS | 5 | 2450 | 5kzx | A | 660 | ARG | p.Arg660His | P10253 |
| HOMO SAPIENS | 8 | 2450 | 5kzx | A | 672 | ARG | p.Arg672Gln | P10253 |
| HOMO SAPIENS | 0 | 2450 | 5kzx | A | 702 | ARG | p.Arg702Leu | P10253 |
| HOMO SAPIENS | 3 | 2450 | 5kzx | A | 725 | ARG | p.Arg725Trp | P10253 |
| HOMO SAPIENS | 0 | 2450 | 5kzx | A | 746 | TRP | p.Trp746Cys | P10253 |
| HOMO SAPIENS | 0 | 2450 | 5kzx | A | 780 | ILE | p.Val780Ile | P10253 |
| HOMO SAPIENS | 1 | 2450 | 5kzx | A | 816 | VAL | p.Val816Ile | P10253 |
| HOMO SAPIENS | 0 | 2450 | 5kzx | A | 838 | GLN | p.Gln838Ter | P10253 |
| HOMO SAPIENS | 2 | 2450 | 5kzx | A | 854 | ARG | p.Arg854Ter | P10253 |
| HOMO SAPIENS | 3 | 2450 | 5kzx | A | 870 | ARG | p.Arg870Ter | P10253 |
| HOMO SAPIENS | 0 | 2450 | 5kzx | A | 890 | VAL | p.Val890Leu | P10253 |
| HOMO SAPIENS | 6 | 1882 | 4km6 | A | 175 | CYS | p.Cys175Ter | P15328 |
| HOMO SAPIENS | 3 | 14767 | 1lm5 | A | 2628 | LEU | p.Leu2628Pro | P15924 |
| HOMO SAPIENS | 5 | 14767 | 1lm5 | A | 2655 | ALA | p.Ala2655Gly | P15924 |
| HOMO SAPIENS | 3 | 14767 | 1lm5 | A | 2731 | TYR | p.Tyr2731His | P15924 |
| HOMO SAPIENS | 1 | 14767 | 1lm5 | A | 2767 | THR | p.Thr2767Asn | P15924 |
| HOMO SAPIENS | -1 | 14767 | 1lm5 | A | 2819 | MET | p.Met2819Leu | P15924 |
| HOMO SAPIENS | -1 | 11246 | 4y5o | A | 289 | SER | p.Ser289Asn | Q9BSQ5 |
| HOMO SAPIENS | 0 | 7380 | 5kz5 | I | 60 | ARG | p.Arg60His | Q16595 |
| HOMO SAPIENS | 0 | 7380 | 5kz5 | I | 76 | MET | p.Met76Val | Q16595 |
| HOMO SAPIENS | 2 | 22187 | 3fby | A | 328 | CYS | p.Cys328Arg | P49747 |
| HOMO SAPIENS | 2 | 22187 | 3fby | A | 342 | ASP | p.Asp342Tyr | P49747 |

| | | | | | | | | |
|---|---|---|---|---|---|---|---|---|
| HOMO SAPIENS | 2 | 22187 | 3fby | A | 348 | CYS | p.Cys348Arg | P49747 |
| HOMO SAPIENS | 2 | 22187 | 3fby | A | 386 | ASN | p.Asn386Asp | P49747 |
| HOMO SAPIENS | 2 | 22187 | 3fby | A | 453 | ASN | p.Asn453Ser | P49747 |
| HOMO SAPIENS | 2 | 22187 | 3fby | A | 468 | CYS | p.Cys468Tyr | P49747 |
| HOMO SAPIENS | 3 | 22187 | 3fby | A | 472 | ASP | p.Asp472Tyr | P49747 |
| HOMO SAPIENS | 3 | 22187 | 3fby | A | 473 | ASP | p.Asp473Gly | P49747 |
| HOMO SAPIENS | 2 | 22187 | 3fby | A | 523 | ASN | p.Asn523Lys | P49747 |
| HOMO SAPIENS | 1 | 22187 | 3fby | A | 718 | ARG | p.Arg718Trp | P49747 |
| HOMO SAPIENS | 1 | 22187 | 3fby | A | 719 | GLY | p.Gly719Asp | P49747 |
| HOMO SAPIENS | 0 | 22187 | 3fby | A | 756 | GLN | p.Gln756Arg | P49747 |
| HOMO SAPIENS | -1 | 2784 | 5kvh | A | 88 | MET | p.Met88Val | O95831 |
| HOMO SAPIENS | -1 | 2784 | 5kvh | A | 96 | ASN | p.Asn96Ser | O95831 |
| HOMO SAPIENS | 0 | 2784 | 5kvh | A | 282 | THR | p.Thr282Met | O95831 |
| HOMO SAPIENS | 1 | 2784 | 5kvh | A | 366 | ASN | p.Asn366Ser | O95831 |
| HOMO SAPIENS | 3 | 2784 | 5kvh | A | 463 | ARG | p.Arg463Ile | O95831 |
| HOMO SAPIENS | 0 | 2784 | 5kvh | A | 498 | VAL | p.Val498Met | O95831 |
| HOMO SAPIENS | -1 | 2784 | 5kvh | A | 549 | ALA | p.Ala549Val | O95831 |
| HOMO SAPIENS | 0 | 38316 | 2lnk | C | 1933 | ARG | p.Arg1933Ter | P35579 |
| HOMO SAPIENS | 0 | 33808 | 2miu | A | 29 | VAL | p.Val29Met | Q14192 |
| HOMO SAPIENS | 1 | 33808 | 2miu | A | 37 | ALA | p.Ala37Ser | Q14192 |
| HOMO SAPIENS | 2 | 20994 | 2hey | R | 65 | ARG | p.Arg65Cys | P43489 |
| HOMO SAPIENS | -1 | 5687 | 2xwj | I | 7 | ARG | p.Arg32Gln | P00751 |
| HOMO SAPIENS | -1 | 5687 | 2xwj | I | 7 | ARG | p.Arg32Trp | P00751 |
| HOMO SAPIENS | 0 | 5687 | 2xwj | I | 49 | ARG | p.Arg74His | P00751 |
| HOMO SAPIENS | 0 | 5687 | 2xwj | I | 177 | ARG | p.Arg202Gly | P00751 |
| HOMO SAPIENS | 0 | 5687 | 2xwj | I | 217 | ILE | p.Ile242Leu | P00751 |
| HOMO SAPIENS | -1 | 5687 | 2xwj | I | 227 | GLY | p.Gly252Ser | P00751 |
| HOMO SAPIENS | 0 | 5687 | 2xwj | I | 483 | HIS | p.His508Gln | P00751 |
| HOMO SAPIENS | 0 | 5687 | 2xwj | I | 508 | LYS | p.Lys533Arg | P00751 |
| HOMO SAPIENS | 0 | 5687 | 2xwj | I | 540 | LYS | p.Lys565Glu | P00751 |
| HOMO SAPIENS | 0 | 5687 | 2xwj | I | 541 | GLU | p.Glu566Ala | P00751 |
| HOMO SAPIENS | 2 | 953 | 3d4j | A | 249 | PHE | p.Phe249Ser | P53602 |
| HOMO SAPIENS | 1 | 953 | 3d4j | A | 292 | ASN | p.Asn292Ser | P53602 |
| HOMO SAPIENS | 0 | 3300 | 1s9c | A | 140 | ASN | p.Asn457Tyr | P51659 |
| HOMO SAPIENS | 2 | 3300 | 1s9c | A | 167 | ILE | p.Ile484Val | P51659 |
| HOMO SAPIENS | 0 | 3300 | 1s9c | A | 174 | ALA | p.Ala491Thr | P51659 |
| HOMO SAPIENS | 8 | 3300 | 1s9c | A | 193 | ASP | p.Asp510Asn | P51659 |
| HOMO SAPIENS | 3 | 3300 | 1s9c | A | 194 | TRP | p.Trp511Arg | P51659 |
| HOMO SAPIENS | 0 | 3300 | 1s9c | A | 242 | VAL | p.Ile559Val | P51659 |
| HOMO SAPIENS | 0 | 10810 | 5i25 | A | 5 | GLN | p.Gln23Ter | P03951 |
| HOMO SAPIENS | 0 | 10810 | 5i25 | A | 55 | TRP | p.Trp73Ter | P03951 |
| HOMO SAPIENS | 0 | 10810 | 5i25 | A | 91 | ALA | p.Ala109Thr | P03951 |
| HOMO SAPIENS | 0 | 10810 | 5i25 | A | 117 | GLU | p.Glu135Ter | P03951 |
| HOMO SAPIENS | 0 | 10810 | 5i25 | A | 118 | CYS | p.Cys136Ter | P03951 |
| HOMO SAPIENS | 0 | 10810 | 5i25 | A | 128 | CYS | p.Cys146Ter | P03951 |
| HOMO SAPIENS | 0 | 10810 | 5i25 | A | 210 | ARG | p.Arg228Ter | P03951 |
| HOMO SAPIENS | 0 | 10810 | 5i25 | A | 226 | GLN | p.Gln244Ter | P03951 |
| HOMO SAPIENS | 0 | 10810 | 5i25 | A | 233 | GLN | p.Gln251Ter | P03951 |
| HOMO SAPIENS | 0 | 16668 | 3lbx | A | 28 | ARG | p.Arg28Leu | P02549 |
| HOMO SAPIENS | 0 | 16668 | 3lbx | A | 28 | ARG | p.Arg28Cys | P02549 |
| HOMO SAPIENS | 1 | 16668 | 3lbx | A | 41 | ARG | p.Arg41Trp | P02549 |
| HOMO SAPIENS | 1 | 16668 | 3lbx | A | 45 | ARG | p.Arg45Ser | P02549 |
| HOMO SAPIENS | 2 | 16668 | 3lbx | A | 48 | LYS | p.Lys48Arg | P02549 |
| HOMO SAPIENS | 1 | 16668 | 3lbx | A | 152 | ASP | p.Asp152Asn | P02549 |

| | | | | | | | | |
|---|---|---|---|---|---|---|---|---|
| HOMO SAPIENS | 0 | 226 | 2ble | A | 256 | ILE | p.Phe256Ile | P36959 |
| HOMO SAPIENS | 2 | 36808 | 4c7n | A | 3 | SER | p.Ser357Pro | O75030 |
| HOMO SAPIENS | 0 | 4092 | 5jnx | A | 113 | CYS | p.Cys113Arg | O15118 |
| HOMO SAPIENS | 4 | 4092 | 5jnx | A | 215 | HIS | p.His215Arg | O15118 |
| HOMO SAPIENS | 6 | 4092 | 5jnx | A | 237 | PRO | p.Pro237Ser | O15118 |
| HOMO SAPIENS | 0 | 4092 | 5jnx | A | 363 | ALA | p.Ala363Val | O15118 |
| HOMO SAPIENS | 0 | 4092 | 5jnx | A | 378 | VAL | p.Val378Ala | O15118 |
| HOMO SAPIENS | 8 | 4092 | 5jnx | A | 404 | ARG | p.Arg404Gln | O15118 |
| HOMO SAPIENS | 8 | 4092 | 5jnx | A | 411 | ARG | p.Arg411Gln | O15118 |
| HOMO SAPIENS | 2 | 4092 | 5jnx | A | 474 | PRO | p.Pro474Leu | O15118 |
| HOMO SAPIENS | 8 | 4092 | 5jnx | A | 511 | THR | p.Thr511Met | O15118 |
| HOMO SAPIENS | 0 | 4092 | 5jnx | A | 642 | MET | p.Met642Ile | O15118 |
| HOMO SAPIENS | 2 | 4092 | 5jnx | A | 664 | VAL | p.Val664Met | O15118 |
| HOMO SAPIENS | 0 | 4092 | 5jnx | A | 734 | SER | p.Ser734Ile | O15118 |
| HOMO SAPIENS | -1 | 4092 | 5jnx | A | 775 | GLN | p.Gln775Pro | O15118 |
| HOMO SAPIENS | 0 | 4092 | 5jnx | A | 858 | ILE | p.Ile858Val | O15118 |
| HOMO SAPIENS | 0 | 4092 | 5jnx | A | 889 | VAL | p.Val889Met | O15118 |
| HOMO SAPIENS | 0 | 4092 | 5jnx | A | 910 | GLY | p.Gly910Ser | O15118 |
| HOMO SAPIENS | 0 | 4092 | 5jnx | A | 911 | GLY | p.Gly911Ser | O15118 |
| HOMO SAPIENS | 0 | 4092 | 5jnx | A | 928 | GLN | p.Gln928Pro | O15118 |
| HOMO SAPIENS | 0 | 4092 | 5jnx | A | 931 | ASN | p.Asn931Lys | O15118 |
| HOMO SAPIENS | 0 | 4092 | 5jnx | A | 940 | SER | p.Ser940Leu | O15118 |
| HOMO SAPIENS | 0 | 4092 | 5jnx | A | 950 | VAL | p.Val950Met | O15118 |
| HOMO SAPIENS | 0 | 4092 | 5jnx | A | 954 | SER | p.Ser954Leu | O15118 |
| HOMO SAPIENS | 0 | 4092 | 5jnx | A | 958 | ARG | p.Arg958Gln | O15118 |
| HOMO SAPIENS | 0 | 4092 | 5jnx | A | 958 | ARG | p.Arg958Ter | O15118 |
| HOMO SAPIENS | -1 | 4092 | 5jnx | A | 978 | ARG | p.Arg978Cys | O15118 |
| HOMO SAPIENS | 0 | 4092 | 5jnx | A | 992 | GLY | p.Gly992Trp | O15118 |
| HOMO SAPIENS | 0 | 4092 | 5jnx | A | 1007 | PRO | p.Pro1007Ala | O15118 |
| HOMO SAPIENS | 0 | 4092 | 5jnx | A | 1019 | TYR | p.Tyr1019Cys | O15118 |
| HOMO SAPIENS | 2 | 4092 | 5jnx | A | 1035 | ALA | p.Ala1035Val | O15118 |
| HOMO SAPIENS | 2 | 4092 | 5jnx | A | 1054 | ALA | p.Ala1054Thr | O15118 |
| HOMO SAPIENS | 2 | 4092 | 5jnx | A | 1061 | ILE | p.Ile1061Thr | O15118 |
| HOMO SAPIENS | 0 | 4092 | 5jnx | A | 1073 | GLY | p.Gly1073Ser | O15118 |
| HOMO SAPIENS | 2 | 4092 | 5jnx | A | 1088 | TYR | p.Tyr1088Cys | O15118 |
| HOMO SAPIENS | 2 | 4092 | 5jnx | A | 1142 | MET | p.Met1142Thr | O15118 |
| HOMO SAPIENS | 2 | 4092 | 5jnx | A | 1156 | ASN | p.Asn1156Ser | O15118 |
| HOMO SAPIENS | 0 | 4092 | 5jnx | A | 1179 | MET | p.Met1179Val | O15118 |
| HOMO SAPIENS | 0 | 4092 | 5jnx | A | 1186 | ARG | p.Arg1186His | O15118 |
| HOMO SAPIENS | 0 | 4092 | 5jnx | A | 1187 | ALA | p.Ala1187Gly | O15118 |
| HOMO SAPIENS | 2 | 4092 | 5jnx | A | 1189 | GLU | p.Glu1189Gly | O15118 |
| HOMO SAPIENS | 2 | 4092 | 5jnx | A | 1200 | SER | p.Ser1200Gly | O15118 |
| HOMO SAPIENS | 2 | 4092 | 5jnx | A | 1205 | THR | p.Thr1205Lys | O15118 |
| HOMO SAPIENS | 0 | 4092 | 5jnx | A | 1213 | LEU | p.Leu1213Phe | O15118 |
| HOMO SAPIENS | -1 | 4092 | 5jnx | A | 1266 | ARG | p.Arg1266Gln | O15118 |
| HOMO SAPIENS | 9 | 16498 | 4i6o | A | 64 | ARG | p.Arg735Trp | P01024 |
| HOMO SAPIENS | 1 | 21902 | 3un9 | C | 751 | ARG | p.Arg751His | Q86UT6 |
| HOMO SAPIENS | 7 | 20391 | 2h32 | B | 105 | PRO | p.Pro142Leu | P15814 |
| HOMO SAPIENS | 6 | 1635 | 4x45 | A | 65 | ASP | p.Asp65Val | P07741 |
| HOMO SAPIENS | 1 | 1635 | 4x45 | A | 98 | TRP | p.Trp98Ter | P07741 |
| HOMO SAPIENS | 0 | 1635 | 4x45 | A | 110 | LEU | p.Leu110Pro | P07741 |
| HOMO SAPIENS | 7 | 1635 | 4x45 | A | 136 | MET | p.Met136Thr | P07741 |
| HOMO SAPIENS | 1 | 1635 | 4x45 | A | 150 | VAL | p.Val150Phe | P07741 |
| HOMO SAPIENS | 4 | 34810 | 2rpc | A | 16 | CYS | p.Cys253Ser | O60481 |

| | | | | | | | | |
|---|---|---|---|---|---|---|---|---|
| HOMO SAPIENS | 1 | 34810 | 2rpc | A | 18 | TRP | p.Trp255Gly | O60481 |
| HOMO SAPIENS | 1 | 14359 | 4hxi | A | 78 | MET | p.Met78Val | Q9UH77 |
| HOMO SAPIENS | 0 | 5342 | 4d9t | A | 628 | GLU | p.Glu628Asp | P51812 |
| HOMO SAPIENS | -1 | 5342 | 4d9t | A | 723 | ARG | p.Arg723His | P51812 |
| HOMO SAPIENS | -1 | 5342 | 4d9t | A | 729 | ARG | p.Arg729Gln | P51812 |
| HOMO SAPIENS | 1 | 9619 | 3j2s | B | 1781 | ARG | p.Arg1800Cys | P00451 |
| HOMO SAPIENS | 0 | 9619 | 3j2s | B | 1823 | MET | p.Met1842Ile | P00451 |
| HOMO SAPIENS | 1 | 9619 | 3j2s | B | 1825 | PRO | p.Pro1844Ser | P00451 |
| HOMO SAPIENS | 0 | 9619 | 3j2s | B | 1826 | THR | p.Thr1845Pro | P00451 |
| HOMO SAPIENS | 0 | 9619 | 3j2s | B | 1834 | ALA | p.Ala1853Val | P00451 |
| HOMO SAPIENS | 0 | 9619 | 3j2s | B | 1846 | ASP | p.Asp1865Asn | P00451 |
| HOMO SAPIENS | 1 | 9619 | 3j2s | B | 1848 | HIS | p.His1867Arg | P00451 |
| HOMO SAPIENS | 0 | 9619 | 3j2s | B | 1854 | PRO | p.Pro1873Arg | P00451 |
| HOMO SAPIENS | 1 | 9619 | 3j2s | B | 1885 | GLU | p.Glu1904Lys | P00451 |
| HOMO SAPIENS | 0 | 9619 | 3j2s | B | 1922 | ASN | p.Asn1941Ser | P00451 |
| HOMO SAPIENS | 1 | 9619 | 3j2s | B | 1941 | ARG | p.Arg1960Gln | P00451 |
| HOMO SAPIENS | 4 | 9619 | 3j2s | B | 2105 | TYR | p.Tyr2124Cys | P00451 |
| HOMO SAPIENS | 6 | 9619 | 3j2s | B | 2150 | ARG | p.Arg2169His | P00451 |
| HOMO SAPIENS | 0 | 9619 | 3j2s | B | 2159 | ARG | p.Arg2178Cys | P00451 |
| HOMO SAPIENS | 0 | 9619 | 3j2s | B | 2159 | ARG | p.Arg2178His | P00451 |
| HOMO SAPIENS | 2 | 9619 | 3j2s | B | 2238 | MET | p.Met2257Val | P00451 |
| HOMO SAPIENS | 2 | 9619 | 3j2s | B | 2300 | PRO | p.Pro2319Ser | P00451 |
| HOMO SAPIENS | 1 | 667 | 1z2m | A | 127 | GLU | p.Glu127Ter | P05161 |
| HOMO SAPIENS | 0 | 15717 | 2nn6 | G | 30 | GLY | p.Gly31Ala | Q9NQT5 |
| HOMO SAPIENS | -1 | 15717 | 2nn6 | G | 55 | ASN | p.Asn56His | Q9NQT5 |
| HOMO SAPIENS | 7 | 15717 | 2nn6 | G | 131 | ASP | p.Asp132Ala | Q9NQT5 |
| HOMO SAPIENS | 7 | 784 | 3rkq | A | 161 | ARG | p.Arg161Pro | P52952 |
| HOMO SAPIENS | 6 | 784 | 3rkq | A | 178 | THR | p.Thr178Met | P52952 |
| HOMO SAPIENS | 7 | 784 | 3rkq | A | 183 | LYS | p.Lys183Glu | P52952 |
| HOMO SAPIENS | 8 | 784 | 3rkq | A | 190 | ARG | p.Arg190Cys | P52952 |
| HOMO SAPIENS | 6 | 385 | 30000 | C | 70 | ALA | p.Ala70Thr | P07954 |
| HOMO SAPIENS | 2 | 385 | 30000 | C | 101 | ARG | p.Arg101Pro | P07954 |
| HOMO SAPIENS | 2 | 385 | 30000 | C | 174 | PRO | p.Pro174Arg | P07954 |
| HOMO SAPIENS | 6 | 385 | 30000 | C | 187 | SER | p.Ser187Ter | P07954 |
| HOMO SAPIENS | 9 | 385 | 30000 | C | 233 | ARG | p.Arg233His | P07954 |
| HOMO SAPIENS | 3 | 385 | 30000 | C | 273 | ALA | p.Ala273Thr | P07954 |
| HOMO SAPIENS | 4 | 385 | 30000 | C | 334 | SER | p.Ser334Arg | P07954 |
| HOMO SAPIENS | 6 | 385 | 30000 | C | 340 | ASN | p.Asn340Lys | P07954 |
| HOMO SAPIENS | 8 | 385 | 30000 | C | 343 | ARG | p.Arg343Ter | P07954 |
| HOMO SAPIENS | 3 | 385 | 30000 | C | 350 | ARG | p.Arg350Trp | P07954 |
| HOMO SAPIENS | 4 | 385 | 30000 | C | 376 | GLN | p.Gln376Pro | P07954 |
| HOMO SAPIENS | 6 | 385 | 30000 | C | 434 | CYS | p.Cys434Ter | P07954 |
| HOMO SAPIENS | 0 | 3316 | 4f91 | B | 934 | THR | p.Thr934Ala | O75643 |
| HOMO SAPIENS | 0 | 3316 | 4f91 | B | 1002 | ASN | p.Asn1002Ser | O75643 |
| HOMO SAPIENS | 0 | 3316 | 4f91 | B | 1087 | SER | p.Ser1087Leu | O75643 |
| HOMO SAPIENS | 4 | 3316 | 4f91 | B | 1389 | VAL | p.Val1389Ile | O75643 |
| HOMO SAPIENS | 0 | 619 | 5hbt | B | 195 | ASP | p.Asp240His | P02708 |
| HOMO SAPIENS | 1 | 619 | 5hbt | B | 209 | ARG | p.Arg254Leu | P02708 |
| HOMO SAPIENS | 0 | 18171 | 2y25 | B | 1408 | ASP | p.Asp1408Asn | P52179 |
| HOMO SAPIENS | 0 | 18171 | 2y25 | B | 1453 | MET | p.Met1453Thr | P52179 |
| HOMO SAPIENS | 0 | 18171 | 2y25 | B | 1459 | ILE | p.Ile1459Lys | P52179 |
| HOMO SAPIENS | 0 | 18171 | 2y25 | B | 1558 | PHE | p.Phe1558Leu | P52179 |
| HOMO SAPIENS | 0 | 18171 | 2y25 | B | 1573 | ARG | p.Arg1573Gln | P52179 |
| HOMO SAPIENS | 7 | 18171 | 2y25 | B | 1663 | VAL | p.Val1663Met | P52179 |

| | | | | | | | | |
|---|---|---|---|---|---|---|---|---|
| HOMO SAPIENS | 8 | 14262 | 2fcw | B | 117 | SER | p.Ser138Ter | P01130 |
| HOMO SAPIENS | 8 | 14262 | 2fcw | B | 118 | ASP | p.Asp139Glu | P01130 |
| HOMO SAPIENS | 9 | 14262 | 2fcw | B | 119 | GLU | p.Glu140Lys | P01130 |
| HOMO SAPIENS | 7 | 14262 | 2fcw | B | 122 | CYS | p.Cys143Ter | P01130 |
| HOMO SAPIENS | 6 | 14262 | 2fcw | B | 134 | CYS | p.Cys155Ter | P01130 |
| HOMO SAPIENS | 7 | 14262 | 2fcw | B | 147 | ASP | p.Asp168Glu | P01130 |
| HOMO SAPIENS | 4 | 2624 | 3b7b | A | 855 | VAL | p.Val886Met | Q9H9B1 |
| HOMO SAPIENS | 7 | 2624 | 3b7b | A | 868 | ILE | p.Ile899Val | Q9H9B1 |
| HOMO SAPIENS | 0 | 2624 | 3b7b | A | 892 | GLU | p.Glu923Lys | Q9H9B1 |
| HOMO SAPIENS | 0 | 4791 | 5ctd | B | 68 | LEU | p.Pro549Ala | Q14055 |
| HOMO SAPIENS | 5 | 460 | 4ui1 | D | 80 | CYS | p.Cys80Arg | Q6ZVN8 |
| HOMO SAPIENS | 3 | 460 | 4ui1 | D | 101 | LEU | p.Leu101Pro | Q6ZVN8 |
| HOMO SAPIENS | 0 | 40782 | 1sy6 | A | 47 | LYS | p.Lys69Ter | P09693 |
| HOMO SAPIENS | -1 | 1092 | 2idx | A | 62 | THR | p.Thr62Met | Q96EY8 |
| HOMO SAPIENS | 6 | 1092 | 2idx | A | 183 | HIS | p.His183Leu | Q96EY8 |
| HOMO SAPIENS | 7 | 1092 | 2idx | A | 186 | ARG | p.Arg186Trp | Q96EY8 |
| HOMO SAPIENS | 6 | 1092 | 2idx | A | 190 | ARG | p.Arg190Cys | Q96EY8 |
| HOMO SAPIENS | 6 | 1092 | 2idx | A | 190 | ARG | p.Arg190His | Q96EY8 |
| HOMO SAPIENS | 7 | 1092 | 2idx | A | 191 | ARG | p.Arg191Trp | Q96EY8 |
| HOMO SAPIENS | 7 | 1092 | 2idx | A | 191 | ARG | p.Arg191Gln | Q96EY8 |
| HOMO SAPIENS | 6 | 1092 | 2idx | A | 195 | ARG | p.Arg195His | Q96EY8 |
| HOMO SAPIENS | 1 | 1092 | 2idx | A | 234 | GLN | p.Gln234Ter | Q96EY8 |
| HOMO SAPIENS | 0 | 1092 | 2idx | A | 239 | MET | p.Met239Lys | Q96EY8 |
| HOMO SAPIENS | -1 | 6418 | 5fur | G | 11 | THR | p.Thr11Ile | P21675 |
| HOMO SAPIENS | -1 | 6418 | 5fur | G | 290 | LEU | p.Leu269Val | P21675 |
| HOMO SAPIENS | 0 | 6100 | 1wiq | A | 240 | ARG | p.Arg265Trp | P01730 |
| HOMO SAPIENS | 0 | 13757 | 4ye6 | A | 45 | GLY | p.Gly45Val | P47897 |
| HOMO SAPIENS | 1 | 13757 | 4ye6 | A | 57 | TYR | p.Tyr57His | P47897 |
| HOMO SAPIENS | 8 | 13757 | 4ye6 | A | 403 | ARG | p.Arg403Trp | P47897 |
| HOMO SAPIENS | 7 | 13757 | 4ye6 | A | 524 | ARG | p.Arg524Trp | P47897 |
| HOMO SAPIENS | 6 | 13757 | 4ye6 | A | 690 | ARG | p.Arg690Cys | P47897 |
| HOMO SAPIENS | 0 | 26958 | 2lzl | A | 370 | GLY | p.Gly370Cys | P22607 |
| HOMO SAPIENS | 0 | 26958 | 2lzl | A | 371 | SER | p.Ser371Cys | P22607 |
| HOMO SAPIENS | 0 | 26958 | 2lzl | A | 373 | TYR | p.Tyr373Cys | P22607 |
| HOMO SAPIENS | 0 | 26958 | 2lzl | A | 380 | GLY | p.Gly380Arg | P22607 |
| HOMO SAPIENS | 0 | 26958 | 2lzl | A | 384 | PHE | p.Phe384Leu | P22607 |
| HOMO SAPIENS | 5 | 26958 | 2lzl | A | 391 | ALA | p.Ala391Glu | P22607 |
| HOMO SAPIENS | 6 | 11325 | 2wa6 | A | 121 | TYR | p.Tyr121Phe | O75369 |
| HOMO SAPIENS | 2 | 11325 | 2wa6 | A | 161 | PHE | p.Phe161Cys | O75369 |
| HOMO SAPIENS | 0 | 11325 | 2wa6 | A | 171 | LEU | p.Leu171Arg | O75369 |
| HOMO SAPIENS | 2 | 11325 | 2wa6 | A | 173 | ALA | p.Ala173Val | O75369 |
| HOMO SAPIENS | 0 | 11325 | 2wa6 | A | 202 | MET | p.Met202Val | O75369 |
| HOMO SAPIENS | 3 | 11325 | 2wa6 | A | 227 | GLU | p.Glu227Lys | O75369 |
| HOMO SAPIENS | 0 | 11325 | 2wa6 | A | 234 | LEU | p.Leu234Val | O75369 |
| HOMO SAPIENS | 2 | 11325 | 2wa6 | A | 235 | SER | p.Ser235Pro | O75369 |
| HOMO SAPIENS | 9 | 7240 | 3tgx | A | 137 | TYR | p.Tyr156Cys | Q9HBE5 |
| HOMO SAPIENS | 0 | 35479 | 4elj | A | 307 | THR | p.Thr307Ile | P06400 |
| HOMO SAPIENS | 0 | 35479 | 4elj | A | 310 | GLY | p.Gly310Glu | P06400 |
| HOMO SAPIENS | 0 | 35479 | 4elj | A | 377 | THR | p.Thr377Ser | P06400 |
| HOMO SAPIENS | 9 | 36706 | 1he7 | A | 359 | TYR | p.Tyr359Cys | P04629 |
| HOMO SAPIENS | 4 | 6538 | 4d0e | A | 504 | ARG | p.Arg504His | P46531 |
| HOMO SAPIENS | 1 | 26518 | 5fm5 | M | 600 | GLU | p.Glu600Val | P52179 |
| HOMO SAPIENS | -1 | 26518 | 5fm5 | M | 651 | ARG | p.Arg651Gln | P52179 |
| HOMO SAPIENS | -1 | 26518 | 5fm5 | M | 660 | PRO | p.Pro660Ala | P52179 |

| | | | | | | | | |
|---|---|---|---|---|---|---|---|---|
| HOMO SAPIENS | -1 | 26518 | 5fm5 | M | 688 | THR | p.Thr688Ser | P52179 |
| HOMO SAPIENS | -1 | 26518 | 5fm5 | M | 704 | GLU | p.Glu704Lys | P52179 |
| HOMO SAPIENS | -1 | 26518 | 5fm5 | M | 727 | THR | p.Thr727Ala | P52179 |
| HOMO SAPIENS | 2 | 27054 | 4rhp | B | 209 | LEU | p.Leu209Val | O75208 |
| HOMO SAPIENS | -1 | 27054 | 4rhp | B | 288 | LYS | p.Lys288Asn | O75208 |
| HOMO SAPIENS | 5 | 2384 | 5fqd | B | 419 | ARG | p.Arg419Ter | Q96SW2 |
| HOMO SAPIENS | 4 | 5154 | 2hl3 | A | 56 | LYS | p.Lys56Arg | Q14203 |
| HOMO SAPIENS | 0 | 5154 | 2hl3 | A | 59 | GLY | p.Gly59Ser | Q14203 |
| HOMO SAPIENS | 0 | 5154 | 2hl3 | A | 71 | GLY | p.Gly71Ala | Q14203 |
| HOMO SAPIENS | 0 | 5154 | 2hl3 | A | 71 | GLY | p.Gly71Arg | Q14203 |
| HOMO SAPIENS | 7 | 5154 | 2hl3 | A | 72 | THR | p.Thr72Pro | Q14203 |
| HOMO SAPIENS | 2 | 5154 | 2hl3 | A | 74 | GLN | p.Gln74Pro | Q14203 |
| HOMO SAPIENS | 2 | 20039 | 2l1g | A | 9 | GLY | p.Gly9Cys | Q9NVV9 |
| HOMO SAPIENS | 5 | 20039 | 2l1g | A | 81 | PHE | p.Phe81Leu | Q9NVV9 |
| HOMO SAPIENS | -1 | 97 | 5uz7 | A | 201 | ARG | p.Arg201His | P63092 |
| HOMO SAPIENS | -1 | 97 | 5uz7 | A | 201 | ARG | p.Arg201Cys | P63092 |
| HOMO SAPIENS | 0 | 3257 | 4b7w | B | 278 | GLU | p.Glu278Asp | P41235 |
| HOMO SAPIENS | 1 | 3257 | 4b7w | B | 331 | ARG | p.Arg331Cys | P41235 |
| HOMO SAPIENS | 0 | 2814 | 4qgk | B | 10 | GLN | p.Gln10Ter | P51648 |
| HOMO SAPIENS | 0 | 2814 | 4qgk | B | 177 | ARG | p.Arg177Ter | P51648 |
| HOMO SAPIENS | 9 | 2814 | 4qgk | B | 184 | THR | p.Thr184Met | P51648 |
| HOMO SAPIENS | 1 | 2814 | 4qgk | B | 188 | ALA | p.Ala188Val | P51648 |
| HOMO SAPIENS | 0 | 2814 | 4qgk | B | 214 | CYS | p.Cys214Tyr | P51648 |
| HOMO SAPIENS | 0 | 2814 | 4qgk | B | 245 | ASP | p.Asp245Asn | P51648 |
| HOMO SAPIENS | 0 | 2814 | 4qgk | B | 266 | LYS | p.Lys266Asn | P51648 |
| HOMO SAPIENS | 0 | 2814 | 4qgk | B | 315 | PRO | p.Pro315Ser | P51648 |
| HOMO SAPIENS | 0 | 2814 | 4qgk | B | 386 | ASN | p.Asn386Ser | P51648 |
| HOMO SAPIENS | -1 | 5486 | 5u6o | B | 47 | GLY | p.Gly47Val | O60741 |
| HOMO SAPIENS | 1 | 21475 | 3pdy | A | 679 | THR | p.Thr789Ile | Q15149 |
| HOMO SAPIENS | 0 | 32890 | 4djc | B | 1501 | LEU | p.Leu1501Val | Q14524 |
| HOMO SAPIENS | -1 | 32890 | 4djc | B | 1512 | ARG | p.Arg1512Trp | Q14524 |
| HOMO SAPIENS | 0 | 26683 | 2xze | B | 38 | ARG | p.Arg38Cys | O95630 |
| HOMO SAPIENS | 0 | 26683 | 2xze | B | 100 | PHE | p.Phe100Tyr | O95630 |
| HOMO SAPIENS | -1 | 23175 | 5dka | A | 163 | SER | p.Ser163Leu | Q96EQ8 |
| HOMO SAPIENS | -1 | 23175 | 5dka | A | 174 | ARG | p.Arg174Cys | Q96EQ8 |
| HOMO SAPIENS | 6 | 21177 | 3k1r | A | 155 | ARG | p.Arg155Ter | Q9Y6N9 |
| HOMO SAPIENS | 0 | 1237 | 2wwm | O | 40 | PRO | p.Pro40Arg | O75147 |
| HOMO SAPIENS | 1 | 2543 | 5ewl | B | 64 | VAL | p.Val64Leu | Q13224 |
| HOMO SAPIENS | 2 | 2543 | 5ewl | B | 271 | ALA | p.Ala271Glu | Q13224 |
| HOMO SAPIENS | 3 | 34262 | 2mng | A | 613 | PHE | p.Phe613Leu | Q9Y3Q4 |
| HOMO SAPIENS | 7 | 34262 | 2mng | A | 672 | SER | p.Ser672Arg | Q9Y3Q4 |
| HOMO SAPIENS | 4 | 1667 | 1f05 | A | 321 | LYS | p.Lys321Met | P37837 |
| HOMO SAPIENS | 0 | 25179 | 2knc | B | 752 | SER | p.Ser778Pro | P05106 |
| HOMO SAPIENS | 0 | 2185 | 2ig7 | A | 50 | TYR | p.Tyr50Cys | Q9Y259 |
| HOMO SAPIENS | 0 | 2185 | 2ig7 | A | 224 | ASN | p.Asn224His | Q9Y259 |
| HOMO SAPIENS | 0 | 2185 | 2ig7 | A | 270 | TYR | p.Tyr270Ter | Q9Y259 |
| HOMO SAPIENS | 0 | 2185 | 2ig7 | A | 328 | GLN | p.Gln328Arg | Q9Y259 |
| HOMO SAPIENS | 3 | 2201 | 2r0o | A | 179 | PRO | p.Pro179Leu | O43707 |
| HOMO SAPIENS | 3 | 2201 | 2r0o | A | 255 | GLU | p.Lys255Glu | O43707 |
| HOMO SAPIENS | 3 | 2201 | 2r0o | A | 259 | THR | p.Thr259Ile | O43707 |
| HOMO SAPIENS | 4 | 2201 | 2r0o | A | 262 | SER | p.Ser262Pro | O43707 |
| HOMO SAPIENS | -1 | 31221 | 2wvr | C | 44 | THR | p.Thr44Pro | Q9H211 |
| HOMO SAPIENS | -1 | 31221 | 2wvr | C | 135 | ALA | p.Ala135Val | Q9H211 |
| HOMO SAPIENS | -1 | 31221 | 2wvr | C | 456 | GLU | p.Glu456Ala | Q9H211 |

| | | | | | | | | |
|---|---|---|---|---|---|---|---|---|
| HOMO SAPIENS | -1 | 31221 | 2wvr | C | 462 | ARG | p.Arg462Gln | Q9H211 |
| HOMO SAPIENS | -1 | 31221 | 2wvr | C | 468 | GLU | p.Glu468Lys | Q9H211 |
| HOMO SAPIENS | -1 | 31221 | 2wvr | C | 537 | ALA | p.Ala537Val | Q9H211 |
| HOMO SAPIENS | 5 | 2576 | 5o3l | C | 363 | VAL | p.Val680Ile | P10636 |
| HOMO SAPIENS | 5 | 1345 | 1czt | A | 37 | ARG | p.Arg2102Cys | P12259 |
| HOMO SAPIENS | 1 | 1345 | 1czt | A | 83 | MET | p.Met2148Thr | P12259 |
| HOMO SAPIENS | 1 | 1345 | 1czt | A | 157 | ASP | p.Asp2222Gly | P12259 |
| HOMO SAPIENS | 8 | 35985 | 1lo1 | A | 110 | ALA | p.Ala110Val | O95718 |
| HOMO SAPIENS | 1 | 617 | 2ocp | B | 44 | GLU | p.Glu44Lys | Q16854 |
| HOMO SAPIENS | 5 | 617 | 2ocp | B | 46 | ASN | p.Asn46Ser | Q16854 |
| HOMO SAPIENS | 0 | 617 | 2ocp | B | 59 | THR | p.Thr59Met | Q16854 |
| HOMO SAPIENS | 7 | 617 | 2ocp | B | 142 | ARG | p.Arg142Lys | Q16854 |
| HOMO SAPIENS | 3 | 617 | 2ocp | B | 154 | ASN | p.Asn154Lys | Q16854 |
| HOMO SAPIENS | 3 | 617 | 2ocp | B | 170 | GLN | p.Gln170Arg | Q16854 |
| HOMO SAPIENS | 3 | 617 | 2ocp | B | 227 | GLU | p.Glu227Lys | Q16854 |
| HOMO SAPIENS | 3 | 617 | 2ocp | B | 233 | LYS | p.Lys233Glu | Q16854 |
| HOMO SAPIENS | 3 | 617 | 2ocp | B | 255 | ASP | p.Asp255Tyr | Q16854 |
| HOMO SAPIENS | -1 | 38975 | 5itz | D | 421 | GLN | p.Gln421His | Q9HC77 |
| HOMO SAPIENS | 2 | 6235 | 2f8v | A | 15 | VAL | p.Val15Ile | Q8WZ42 |
| HOMO SAPIENS | 0 | 6235 | 2f8v | A | 54 | VAL | p.Val54Met | Q8WZ42 |
| HOMO SAPIENS | 0 | 6235 | 2f8v | A | 59 | SER | p.Ser59Arg | Q8WZ42 |
| HOMO SAPIENS | 0 | 6235 | 2f8v | A | 60 | ASP | p.Asp60Tyr | Q8WZ42 |
| HOMO SAPIENS | 0 | 12630 | 1ext | B | 12 | ASP | p.Asp41Glu | P19438 |
| HOMO SAPIENS | 3 | 12630 | 1ext | B | 30 | CYS | p.Cys59Arg | P19438 |
| HOMO SAPIENS | 3 | 12630 | 1ext | B | 30 | CYS | p.Cys59Ser | P19438 |
| HOMO SAPIENS | 5 | 12630 | 1ext | B | 33 | CYS | p.Cys62Tyr | P19438 |
| HOMO SAPIENS | 5 | 12630 | 1ext | B | 33 | CYS | p.Cys62Gly | P19438 |
| HOMO SAPIENS | 1 | 12630 | 1ext | B | 46 | PRO | p.Pro75Leu | P19438 |
| HOMO SAPIENS | 3 | 12630 | 1ext | B | 50 | THR | p.Thr79Met | P19438 |
| HOMO SAPIENS | 6 | 12630 | 1ext | B | 52 | CYS | p.Cys81Phe | P19438 |
| HOMO SAPIENS | 6 | 12630 | 1ext | B | 70 | CYS | p.Cys99Ser | P19438 |
| HOMO SAPIENS | 5 | 12630 | 1ext | B | 88 | CYS | p.Cys117Arg | P19438 |
| HOMO SAPIENS | 5 | 12630 | 1ext | B | 88 | CYS | p.Cys117Tyr | P19438 |
| HOMO SAPIENS | -1 | 12630 | 1ext | B | 170 | ILE | p.Ile199Asn | P19438 |
| HOMO SAPIENS | 0 | 994 | 1aos | B | 95 | ARG | p.Arg95Cys | P04424 |
| HOMO SAPIENS | 0 | 994 | 1aos | B | 111 | ARG | p.Arg111Gln | P04424 |
| HOMO SAPIENS | 0 | 994 | 1aos | B | 131 | THR | p.Thr131Met | P04424 |
| HOMO SAPIENS | 6 | 994 | 1aos | B | 178 | VAL | p.Val178Met | P04424 |
| HOMO SAPIENS | 9 | 994 | 1aos | B | 182 | ARG | p.Arg182Gln | P04424 |
| HOMO SAPIENS | 9 | 994 | 1aos | B | 186 | ARG | p.Arg186Trp | P04424 |
| HOMO SAPIENS | 8 | 994 | 1aos | B | 193 | ARG | p.Arg193Gln | P04424 |
| HOMO SAPIENS | 5 | 994 | 1aos | B | 217 | ARG | p.Arg217Ter | P04424 |
| HOMO SAPIENS | 0 | 994 | 1aos | B | 286 | GLN | p.Gln286Arg | P04424 |
| HOMO SAPIENS | 5 | 994 | 1aos | B | 379 | ARG | p.Arg379Cys | P04424 |
| HOMO SAPIENS | 4 | 16066 | 5mv9 | A | 1719 | TYR | p.Tyr1719Cys | Q13402 |
| HOMO SAPIENS | -1 | 16066 | 5mv9 | A | 1739 | ARG | p.Arg1739Ter | Q13402 |
| HOMO SAPIENS | 4 | 16066 | 5mv9 | A | 1775 | ILE | p.Ile1775Thr | Q13402 |
| HOMO SAPIENS | 2 | 16066 | 5mv9 | A | 1786 | SER | p.Ser1786Thr | Q13402 |
| HOMO SAPIENS | 4 | 16066 | 5mv9 | A | 1798 | GLN | p.Gln1798Ter | Q13402 |
| HOMO SAPIENS | 0 | 16066 | 5mv9 | A | 1858 | LEU | p.Leu1858Pro | Q13402 |
| HOMO SAPIENS | 6 | 16066 | 5mv9 | A | 1873 | ARG | p.Arg1873Trp | Q13402 |
| HOMO SAPIENS | 6 | 16066 | 5mv9 | A | 1873 | ARG | p.Arg1873Leu | Q13402 |
| HOMO SAPIENS | 6 | 16066 | 5mv9 | A | 1883 | ARG | p.Arg1883Gln | Q13402 |
| HOMO SAPIENS | 4 | 16066 | 5mv9 | A | 1887 | PRO | p.Pro1887Leu | Q13402 |

| | | | | | | | | |
|---|---|---|---|---|---|---|---|---|
| HOMO SAPIENS | 5 | 16066 | 5mv9 | A | 1942 | GLY | p.Gly1942Ter | Q13402 |
| HOMO SAPIENS | 5 | 16066 | 5mv9 | A | 1954 | LEU | p.Leu1954Ile | Q13402 |
| HOMO SAPIENS | 5 | 16066 | 5mv9 | A | 1956 | VAL | p.Val1956Ile | Q13402 |
| HOMO SAPIENS | 5 | 16066 | 5mv9 | A | 1967 | ARG | p.Arg1967Ter | Q13402 |
| HOMO SAPIENS | 5 | 16066 | 5mv9 | A | 1990 | GLN | p.Gln1990Ter | Q13402 |
| HOMO SAPIENS | 2 | 16066 | 5mv9 | A | 2005 | LYS | p.Lys2005Glu | Q13402 |
| HOMO SAPIENS | 1 | 16066 | 5mv9 | A | 2072 | VAL | p.Val2072Ile | Q13402 |
| HOMO SAPIENS | 4 | 16066 | 5mv9 | A | 2109 | THR | p.Thr2109Ile | Q13402 |
| HOMO SAPIENS | 4 | 16066 | 5mv9 | A | 2142 | ASP | p.Asp2142Asn | Q13402 |
| HOMO SAPIENS | -1 | 16066 | 5mv9 | A | 2214 | GLY | p.Gly2214Ser | Q13402 |
| HOMO SAPIENS | 7 | 27942 | 1deb | A | 24 | ARG | p.Arg24Ter | P25054 |
| HOMO SAPIENS | 0 | 876 | 5hnr | B | 12 | PHE | p.Phe12Leu | P13716 |
| HOMO SAPIENS | 0 | 876 | 5hnr | B | 59 | LYS | p.Lys59Asn | P13716 |
| HOMO SAPIENS | -1 | 876 | 5hnr | B | 133 | GLY | p.Gly133Arg | P13716 |
| HOMO SAPIENS | 2 | 876 | 5hnr | B | 240 | ARG | p.Arg240Trp | P13716 |
| HOMO SAPIENS | 0 | 876 | 5hnr | B | 274 | ALA | p.Ala274Thr | P13716 |
| HOMO SAPIENS | 0 | 876 | 5hnr | B | 275 | VAL | p.Val275Met | P13716 |
| HOMO SAPIENS | 0 | 1142 | 3qe2 | A | 503 | VAL | p.Ala500Val | P16435 |
| HOMO SAPIENS | 8 | 14358 | 4dkd | C | 32 | VAL | p.Val32Gly | P07333 |
| HOMO SAPIENS | 0 | 14358 | 4dkd | C | 60 | LEU | p.Leu60Val | P07333 |
| HOMO SAPIENS | 7 | 14358 | 4dkd | C | 75 | THR | p.Thr75Ile | P07333 |
| HOMO SAPIENS | -1 | 14358 | 4dkd | C | 90 | PRO | p.Pro90Thr | P07333 |
| HOMO SAPIENS | 0 | 14358 | 4dkd | C | 106 | ARG | p.Arg106Trp | P07333 |
| HOMO SAPIENS | 0 | 14358 | 4dkd | C | 241 | ASN | p.Asn241Asp | P07333 |
| HOMO SAPIENS | 0 | 14358 | 4dkd | C | 245 | ALA | p.Ala245Ser | P07333 |
| HOMO SAPIENS | 0 | 14358 | 4dkd | C | 255 | ASN | p.Asn255Ile | P07333 |
| HOMO SAPIENS | 0 | 14358 | 4dkd | C | 268 | VAL | p.Val268Ile | P07333 |
| HOMO SAPIENS | 0 | 14358 | 4dkd | C | 279 | VAL | p.Val279Met | P07333 |
| HOMO SAPIENS | 0 | 14358 | 4dkd | C | 294 | ARG | p.Arg294Gln | P07333 |
| HOMO SAPIENS | 5 | 8968 | 3ppy | A | 1565 | VAL | p.Val1565Leu | P04275 |
| HOMO SAPIENS | 4 | 8968 | 3ppy | A | 1584 | TYR | p.Tyr1584Cys | P04275 |
| HOMO SAPIENS | 5 | 5341 | 3k3n | A | 279 | PHE | p.Phe315Ser | Q9UPP1 |
| HOMO SAPIENS | 0 | 29463 | 1q7l | A | 197 | ARG | p.Arg197Trp | Q03154 |
| HOMO SAPIENS | 0 | 33761 | 2leh | B | 30 | ASP | p.Asp30Asn | Q16637 |
| HOMO SAPIENS | 6 | 33761 | 2leh | B | 44 | ASP | p.Asp44Val | Q16637 |
| HOMO SAPIENS | -1 | 12249 | 2pnr | C | 224 | GLU | p.Glu310Lys | P10515 |
| HOMO SAPIENS | -1 | 12249 | 2pnr | C | 232 | VAL | p.Val318Ala | P10515 |
| HOMO SAPIENS | -1 | 7 | 5whk | B | -9 | ALA | p.Ala11Pro | P61769 |
| HOMO SAPIENS | 8 | 20298 | 4euw | A | 154 | PHE | p.Phe154Leu | P48436 |
| HOMO SAPIENS | 9 | 20298 | 4euw | A | 158 | ALA | p.Ala158Thr | P48436 |
| HOMO SAPIENS | 5 | 20298 | 4euw | A | 165 | HIS | p.His165Tyr | P48436 |
| HOMO SAPIENS | 3 | 20298 | 4euw | A | 169 | HIS | p.His169Gln | P48436 |
| HOMO SAPIENS | 0 | 20298 | 4euw | A | 173 | LYS | p.Lys173Glu | P48436 |
| HOMO SAPIENS | 1 | 16465 | 2jgx | A | 402 | TYR | p.Tyr402His | P08603 |
| HOMO SAPIENS | 0 | 8993 | 3v8o | A | 659 | LEU | p.Leu659Arg | Q14315 |
| HOMO SAPIENS | 6 | 8993 | 3v8o | A | 693 | ASP | p.Asp693Ala | Q14315 |
| HOMO SAPIENS | 0 | 8993 | 3v8o | A | 709 | ALA | p.Ala709Thr | Q14315 |
| HOMO SAPIENS | 2 | 3852 | 5es4 | B | 106 | ASP | p.Asp128Asn | P05107 |
| HOMO SAPIENS | 2 | 3852 | 5es4 | B | 116 | SER | p.Ser138Pro | P05107 |
| HOMO SAPIENS | 1 | 3852 | 5es4 | B | 127 | LEU | p.Leu149Pro | P05107 |
| HOMO SAPIENS | 0 | 3852 | 5es4 | B | 147 | GLY | p.Gly169Arg | P05107 |
| HOMO SAPIENS | 3 | 3852 | 5es4 | B | 156 | PRO | p.Pro178Leu | P05107 |
| HOMO SAPIENS | 0 | 3852 | 5es4 | B | 174 | LYS | p.Lys196Thr | P05107 |
| HOMO SAPIENS | 2 | 3852 | 5es4 | B | 251 | GLY | p.Gly273Arg | P05107 |

| | | | | | | | | |
|---|---|---|---|---|---|---|---|---|
| HOMO SAPIENS | 2 | 3852 | 5es4 | B | 329 | ASN | p.Asn351Ser | P05107 |
| HOMO SAPIENS | 2 | 3852 | 5es4 | B | 571 | ARG | p.Arg593Cys | P05107 |
| HOMO SAPIENS | 7 | 5962 | 5fn2 | B | 146 | MET | p.Met146Leu | P49768 |
| HOMO SAPIENS | 3 | 5962 | 5fn2 | B | 166 | LEU | p.Leu166Pro | P49768 |
| HOMO SAPIENS | 1 | 5962 | 5fn2 | B | 233 | MET | p.Met233Leu | P49768 |
| HOMO SAPIENS | 1 | 5962 | 5fn2 | B | 261 | VAL | p.Val261Gly | P49768 |
| HOMO SAPIENS | 1 | 5962 | 5fn2 | B | 266 | GLY | p.Gly266Ser | P49768 |
| HOMO SAPIENS | -1 | 5962 | 5fn2 | B | 333 | ASP | p.Asp333Gly | P49768 |
| HOMO SAPIENS | 6 | 1918 | 3dlx | A | 58 | THR | p.Thr58Met | P55809 |
| HOMO SAPIENS | 5 | 1918 | 3dlx | A | 133 | VAL | p.Val133Glu | P55809 |
| HOMO SAPIENS | 5 | 1918 | 3dlx | A | 219 | GLY | p.Gly219Glu | P55809 |
| HOMO SAPIENS | 0 | 1918 | 3dlx | A | 221 | VAL | p.Val221Met | P55809 |
| HOMO SAPIENS | 5 | 1918 | 3dlx | A | 283 | SER | p.Ser283Ter | P55809 |
| HOMO SAPIENS | 4 | 1918 | 3dlx | A | 324 | GLY | p.Gly324Glu | P55809 |
| HOMO SAPIENS | 0 | 1918 | 3dlx | A | 456 | CYS | p.Cys456Phe | P55809 |
| HOMO SAPIENS | 0 | 9010 | 2wfr | A | 162 | LEU | p.Leu162Pro | O43323 |
| HOMO SAPIENS | 5 | 3770 | 5n38 | A | 56 | VAL | p.Val56Glu | O60260 |
| HOMO SAPIENS | 1 | 3770 | 5n38 | A | 161 | LYS | p.Lys161Asn | O60260 |
| HOMO SAPIENS | 1 | 3770 | 5n38 | A | 167 | SER | p.Ser167Asn | O60260 |
| HOMO SAPIENS | 1 | 3770 | 5n38 | A | 211 | LYS | p.Lys211Asn | O60260 |
| HOMO SAPIENS | 1 | 3770 | 5n38 | A | 212 | CYS | p.Cys212Tyr | O60260 |
| HOMO SAPIENS | 1 | 3770 | 5n38 | A | 275 | ARG | p.Arg275Trp | O60260 |
| HOMO SAPIENS | 2 | 3770 | 5n38 | A | 334 | ARG | p.Arg334Cys | O60260 |
| HOMO SAPIENS | 1 | 3770 | 5n38 | A | 366 | ARG | p.Arg366Trp | O60260 |
| HOMO SAPIENS | -1 | 3770 | 5n38 | A | 380 | VAL | p.Val380Leu | O60260 |
| HOMO SAPIENS | 1 | 3770 | 5n38 | A | 394 | ASP | p.Asp394Asn | O60260 |
| HOMO SAPIENS | 1 | 3770 | 5n38 | A | 430 | GLY | p.Gly430Asp | O60260 |
| HOMO SAPIENS | -1 | 2467 | 4xx1 | A | 10 | PRO | p.Pro34Leu | P04180 |
| HOMO SAPIENS | 4 | 2467 | 4xx1 | A | 83 | TYR | p.Tyr107Ter | P04180 |
| HOMO SAPIENS | 4 | 2467 | 4xx1 | A | 93 | ALA | p.Ala117Thr | P04180 |
| HOMO SAPIENS | 4 | 2467 | 4xx1 | A | 123 | THR | p.Thr147Ile | P04180 |
| HOMO SAPIENS | 4 | 2467 | 4xx1 | A | 135 | ARG | p.Arg159Trp | P04180 |
| HOMO SAPIENS | 3 | 2467 | 4xx1 | A | 158 | ARG | p.Arg182Cys | P04180 |
| HOMO SAPIENS | 0 | 2467 | 4xx1 | A | 209 | LEU | p.Leu233Pro | P04180 |
| HOMO SAPIENS | 1 | 2467 | 4xx1 | A | 228 | ASN | p.Asn252Lys | P04180 |
| HOMO SAPIENS | 4 | 2467 | 4xx1 | A | 252 | MET | p.Met276Lys | P04180 |
| HOMO SAPIENS | 3 | 2467 | 4xx1 | A | 293 | MET | p.Met317Ile | P04180 |
| HOMO SAPIENS | 5 | 2467 | 4xx1 | A | 321 | THR | p.Thr345Met | P04180 |
| HOMO SAPIENS | 9 | 2467 | 4xx1 | A | 347 | THR | p.Thr371Met | P04180 |
| HOMO SAPIENS | 1 | 5834 | 3qrf | G | 371 | PHE | p.Phe371Cys | Q9BZS1 |
| HOMO SAPIENS | 1 | 5834 | 3qrf | G | 373 | PHE | p.Phe373Ala | Q9BZS1 |
| HOMO SAPIENS | 9 | 5834 | 3qrf | G | 384 | ALA | p.Ala384Thr | Q9BZS1 |
| HOMO SAPIENS | 6 | 5834 | 3qrf | G | 397 | ARG | p.Arg397Trp | Q9BZS1 |
| HOMO SAPIENS | -1 | 11869 | 4km8 | A | 15 | PRO | p.Pro15Leu | Q9UMX1 |
| HOMO SAPIENS | 5 | 262 | 4kbo | A | 126 | ARG | p.Arg126Trp | P38646 |
| HOMO SAPIENS | 3 | 262 | 4kbo | A | 128 | TYR | p.Tyr128Cys | P38646 |
| HOMO SAPIENS | -1 | 11347 | 4uf9 | D | 117 | ALA | p.Ala117Val | Q13501 |
| HOMO SAPIENS | 0 | 2173 | 4ah6 | A | 45 | SER | p.Ser45Gly | Q6PI48 |
| HOMO SAPIENS | 2 | 2173 | 4ah6 | A | 152 | CYS | p.Cys152Phe | Q6PI48 |
| HOMO SAPIENS | 4 | 2173 | 4ah6 | A | 179 | ARG | p.Arg179His | Q6PI48 |
| HOMO SAPIENS | 0 | 2173 | 4ah6 | A | 196 | LYS | p.Lys196Arg | Q6PI48 |
| HOMO SAPIENS | 8 | 2173 | 4ah6 | A | 263 | ARG | p.Arg263Gln | Q6PI48 |
| HOMO SAPIENS | 1 | 2173 | 4ah6 | A | 338 | GLY | p.Gly338Glu | Q6PI48 |
| HOMO SAPIENS | 0 | 2173 | 4ah6 | A | 425 | GLU | p.Glu425Ter | Q6PI48 |

| | | | | | | | | |
|---|---|---|---|---|---|---|---|---|
| HOMO SAPIENS | 1 | 2173 | 4ah6 | A | 548 | LEU | p.Leu548Met | Q6PI48 |
| HOMO SAPIENS | 0 | 2173 | 4ah6 | A | 609 | ARG | p.Arg609Trp | Q6PI48 |
| HOMO SAPIENS | 2 | 2173 | 4ah6 | A | 613 | LEU | p.Leu613Phe | Q6PI48 |
| HOMO SAPIENS | 0 | 2173 | 4ah6 | A | 626 | LEU | p.Leu626Gln | Q6PI48 |
| HOMO SAPIENS | 0 | 2173 | 4ah6 | A | 626 | LEU | p.Leu626Val | Q6PI48 |
| HOMO SAPIENS | 7 | 3203 | 2cn1 | A | 87 | ASP | p.Asp137Val | Q9H0P0 |
| HOMO SAPIENS | 8 | 3203 | 2cn1 | A | 166 | GLN | p.Gln216Ter | Q9H0P0 |
| HOMO SAPIENS | 8 | 3203 | 2cn1 | A | 179 | ASN | p.Asn229Ser | Q9H0P0 |
| HOMO SAPIENS | 7 | 3203 | 2cn1 | A | 230 | GLY | p.Gly280Arg | Q9H0P0 |
| HOMO SAPIENS | 2 | 3424 | 2a1t | R | 116 | GLY | p.Gly116Arg | P13804 |
| HOMO SAPIENS | 2 | 3424 | 2a1t | R | 157 | VAL | p.Val157Gly | P13804 |
| HOMO SAPIENS | 2 | 3424 | 2a1t | R | 171 | THR | p.Thr171Ile | P13804 |
| HOMO SAPIENS | 0 | 3424 | 2a1t | R | 178 | THR | p.Thr178Arg | P13804 |
| HOMO SAPIENS | 0 | 3424 | 2a1t | R | 194 | VAL | p.Val194Met | P13804 |
| HOMO SAPIENS | 1 | 3424 | 2a1t | R | 209 | ARG | p.Arg209Ter | P13804 |
| HOMO SAPIENS | 6 | 3424 | 2a1t | R | 266 | THR | p.Thr266Met | P13804 |
| HOMO SAPIENS | 0 | 27598 | 1g1c | A | 26 | ARG | p.Arg2098Gly | Q8WZ42 |
| HOMO SAPIENS | 1 | 27598 | 1g1c | A | 46 | ILE | p.Ile2118Thr | Q8WZ42 |
| HOMO SAPIENS | 0 | 32885 | 4bsk | A | 149 | ASN | p.Asn149Asp | P35916 |
| HOMO SAPIENS | -1 | 8074 | 3ghg | B | 17 | ARG | p.Arg47Ter | P02675 |
| HOMO SAPIENS | -1 | 447 | 5g4p | A | 190 | ALA | p.Ala190Thr | P84074 |
| HOMO SAPIENS | 0 | 10761 | 5u1d | B | 379 | VAL | p.Val379Ile | Q03519 |
| HOMO SAPIENS | 0 | 10761 | 5u1d | B | 665 | THR | p.Thr665Ala | Q03519 |
| HOMO SAPIENS | 5 | 29251 | 5fn7 | A | 79 | ILE | p.Ile294Leu | P08575 |
| HOMO SAPIENS | -1 | 27727 | 4ky9 | A | 56 | LYS | p.Lys56Glu | P02730 |
| HOMO SAPIENS | 2 | 27727 | 4ky9 | A | 72 | GLU | p.Glu72Asp | P02730 |
| HOMO SAPIENS | 2 | 27727 | 4ky9 | A | 90 | GLU | p.Glu90Lys | P02730 |
| HOMO SAPIENS | 1 | 27727 | 4ky9 | A | 130 | GLY | p.Gly130Arg | P02730 |
| HOMO SAPIENS | 2 | 27727 | 4ky9 | A | 150 | ARG | p.Arg150Ter | P02730 |
| HOMO SAPIENS | 0 | 27727 | 4ky9 | A | 327 | PRO | p.Pro327Arg | P02730 |
| HOMO SAPIENS | 2 | 13202 | 2fxo | B | 870 | ARG | p.Arg870His | P12883 |
| HOMO SAPIENS | 2 | 13202 | 2fxo | B | 870 | ARG | p.Arg870Cys | P12883 |
| HOMO SAPIENS | 4 | 13202 | 2fxo | B | 904 | ARG | p.Arg904Cys | P12883 |
| HOMO SAPIENS | 4 | 13202 | 2fxo | B | 906 | ASP | p.Asp906Gly | P12883 |
| HOMO SAPIENS | 2 | 13202 | 2fxo | B | 908 | LEU | p.Leu908Val | P12883 |
| HOMO SAPIENS | 2 | 13202 | 2fxo | B | 923 | ASN | p.Asn923Lys | P12883 |
| HOMO SAPIENS | 7 | 13202 | 2fxo | B | 935 | GLU | p.Glu935Lys | P12883 |
| HOMO SAPIENS | 7 | 13202 | 2fxo | B | 935 | GLU | p.Glu935Val | P12883 |
| HOMO SAPIENS | 8 | 13202 | 2fxo | B | 949 | GLU | p.Glu949Lys | P12883 |
| HOMO SAPIENS | 0 | 13200 | 2a1s | A | 349 | ARG | p.Arg349Trp | O95453 |
| HOMO SAPIENS | 0 | 13200 | 2a1s | A | 421 | LYS | p.Lys421Arg | O95453 |
| HOMO SAPIENS | 6 | 3196 | 5xtb | T | 115 | CYS | p.Cys115Tyr | O75380 |
| HOMO SAPIENS | 0 | 1108 | 5ukf | A | 89 | ARG | p.Arg89Gln | Q99986 |
| HOMO SAPIENS | 0 | 1108 | 5ukf | A | 119 | HIS | p.His119Arg | Q99986 |
| HOMO SAPIENS | 0 | 1108 | 5ukf | A | 133 | ARG | p.Arg133Cys | Q99986 |
| HOMO SAPIENS | 0 | 1108 | 5ukf | A | 236 | VAL | p.Val236Leu | Q99986 |
| HOMO SAPIENS | 0 | 1108 | 5ukf | A | 321 | ARG | p.Arg321Cys | Q99986 |
| HOMO SAPIENS | -1 | 1108 | 5ukf | A | 358 | ARG | p.Arg358Ter | Q99986 |
| HOMO SAPIENS | 0 | 5842 | 4yli | C | 204 | GLY | p.Gly204Ser | Q9BWP8 |
| HOMO SAPIENS | -1 | 31362 | 4wqo | A | 25 | PRO | p.Pro25Leu | P40337 |
| HOMO SAPIENS | 2 | 10107 | 2j8h | A | 34 | VAL | p.Val31886Ile | Q8WZ42 |
| HOMO SAPIENS | 8 | 20190 | 1wm5 | A | 77 | ARG | p.Arg77Gln | P19878 |
| HOMO SAPIENS | 1 | 20190 | 1wm5 | A | 100 | GLN | p.Gln100Ter | P19878 |
| HOMO SAPIENS | 0 | 20190 | 1wm5 | A | 102 | ARG | p.Arg102Ter | P19878 |

| | | | | | | | | | |
|---|---|---|---|---|---|---|---|---|---|
| HOMO SAPIENS | 0 | 20190 | 1wm5 | A | | 128 | ALA | p.Ala128Val | P19878 |
| HOMO SAPIENS | 0 | 20190 | 1wm5 | A | | 181 | LYS | p.Lys181Arg | P19878 |
| HOMO SAPIENS | 7 | 1278 | 5a2q | a | | 33 | ASP | p.Asp33Asn | P62854 |
| HOMO SAPIENS | 3 | 19515 | 2obd | A | | 11 | ILE | p.Ile28Met | P11597 |
| HOMO SAPIENS | 3 | 19515 | 2obd | A | | 89 | VAL | p.Val106Met | P11597 |
| HOMO SAPIENS | 3 | 19515 | 2obd | A | | 137 | ARG | p.Arg154Trp | P11597 |
| HOMO SAPIENS | 3 | 19515 | 2obd | A | | 139 | ASP | p.Asp156Asn | P11597 |
| HOMO SAPIENS | 3 | 19515 | 2obd | A | | 204 | SER | p.Ser221Arg | P11597 |
| HOMO SAPIENS | 0 | 19515 | 2obd | A | | 297 | GLU | p.Glu314Lys | P11597 |
| HOMO SAPIENS | 0 | 19515 | 2obd | A | | 314 | GLY | p.Gly331Ser | P11597 |
| HOMO SAPIENS | 0 | 19515 | 2obd | A | | 332 | SER | p.Ser349Tyr | P11597 |
| HOMO SAPIENS | 0 | 19515 | 2obd | A | | 368 | VAL | p.Val385Met | P11597 |
| HOMO SAPIENS | 0 | 19515 | 2obd | A | | 373 | ALA | p.Ala390Pro | P11597 |
| HOMO SAPIENS | 3 | 19515 | 2obd | A | | 405 | ILE | p.Val422Ile | P11597 |
| HOMO SAPIENS | 0 | 19515 | 2obd | A | | 451 | ARG | p.Arg468Gln | P11597 |
| HOMO SAPIENS | 0 | 19515 | 2obd | A | | 451 | ARG | p.Arg468Ter | P11597 |
| HOMO SAPIENS | 0 | 6626 | 3hop | A | | 39 | ALA | p.Ala39Gly | P21333 |
| HOMO SAPIENS | 1 | 6626 | 3hop | A | | 82 | GLU | p.Glu82Val | P21333 |
| HOMO SAPIENS | 1 | 6626 | 3hop | A | | 128 | ALA | p.Ala128Val | P21333 |
| HOMO SAPIENS | 2 | 6626 | 3hop | A | | 203 | ASP | p.Asp203Tyr | P21333 |
| HOMO SAPIENS | 1 | 6626 | 3hop | A | | 207 | PRO | p.Pro207Leu | P21333 |
| HOMO SAPIENS | 2 | 6626 | 3hop | A | | 210 | CYS | p.Cys210Phe | P21333 |
| HOMO SAPIENS | 7 | 6626 | 3hop | A | | 254 | GLU | p.Glu254Lys | P21333 |
| HOMO SAPIENS | 4 | 4210 | 2avg | A | | 8 | VAL | p.Val158Met | Q14896 |
| HOMO SAPIENS | 6 | 4210 | 2avg | A | | 15 | GLU | p.Glu165Asp | Q14896 |
| HOMO SAPIENS | 4 | 4210 | 2avg | A | | 27 | ARG | p.Arg177His | Q14896 |
| HOMO SAPIENS | 0 | 4210 | 2avg | A | | 28 | VAL | p.Val178Met | Q14896 |
| HOMO SAPIENS | 6 | 4210 | 2avg | A | | 39 | VAL | p.Val189Ile | Q14896 |
| HOMO SAPIENS | 2 | 4210 | 2avg | A | | 52 | LYS | p.Lys202Gln | Q14896 |
| HOMO SAPIENS | 4 | 4210 | 2avg | A | | 67 | SER | p.Ser217Gly | Q14896 |
| HOMO SAPIENS | 6 | 4210 | 2avg | A | | 69 | VAL | p.Val219Leu | Q14896 |
| HOMO SAPIENS | 2 | 4210 | 2avg | A | | 86 | GLY | p.Ser236Gly | Q14896 |
| HOMO SAPIENS | 4 | 4210 | 2avg | A | | 108 | GLU | p.Glu258Lys | Q14896 |
| HOMO SAPIENS | 0 | 22237 | 1w6k | A | | 588 | GLY | p.Gly588Ser | P48449 |
| HOMO SAPIENS | 0 | 10765 | 4a5w | B | | 98 | GLU | p.Glu98Ala | P13671 |
| HOMO SAPIENS | 0 | 10765 | 4a5w | B | | 596 | ARG | p.Arg596Ter | P13671 |
| HOMO SAPIENS | -1 | 15515 | 3hx3 | A | | 11 | MSE | p.Met10Lys | P12271 |
| HOMO SAPIENS | 8 | 15515 | 3hx3 | A | | 64 | ARG | p.Arg64Gln | P12271 |
| HOMO SAPIENS | 0 | 15515 | 3hx3 | A | | 182 | PHE | p.Phe182Cys | P12271 |
| HOMO SAPIENS | 0 | 15515 | 3hx3 | A | | 216 | ARG | p.Arg216Gln | P12271 |
| HOMO SAPIENS | -1 | 15515 | 3hx3 | A | | 226 | MSE | p.Met226Lys | P12271 |
| HOMO SAPIENS | 0 | 15515 | 3hx3 | A | | 234 | TRP | p.Arg234Trp | P12271 |
| HOMO SAPIENS | 0 | 15515 | 3hx3 | A | | 273 | LEU | p.Leu273Phe | P12271 |
| HOMO SAPIENS | 0 | 15515 | 3hx3 | A | | 292 | THR | p.Thr292Met | P12271 |
| HOMO SAPIENS | 0 | 5152 | 2www | D | | 95 | GLN | p.Gln95Ter | Q8IVH4 |
| HOMO SAPIENS | 0 | 5152 | 2www | D | | 133 | GLN | p.Gln133Ter | Q8IVH4 |
| HOMO SAPIENS | 0 | 5152 | 2www | D | | 145 | ARG | p.Arg145Ter | Q8IVH4 |
| HOMO SAPIENS | 0 | 5152 | 2www | D | | 207 | TYR | p.Tyr207Cys | Q8IVH4 |
| HOMO SAPIENS | 8 | 5152 | 2www | D | | 330 | ARG | p.Arg330Ter | Q8IVH4 |
| HOMO SAPIENS | 0 | 5152 | 2www | D | | 363 | GLN | p.Gln363His | Q8IVH4 |
| HOMO SAPIENS | 1 | 1626 | 1ilr | | 1 | 105 | SER | p.Ser130Arg | P18510 |
| HOMO SAPIENS | -1 | 929 | 5loh | A | | 12 | GLY | p.Gly12Arg | Q96GX5 |
| HOMO SAPIENS | 9 | 3206 | 1x63 | A | | 21 | CYS | p.Cys104Arg | Q13642 |
| HOMO SAPIENS | 8 | 3206 | 1x63 | A | | 39 | TRP | p.Trp122Ser | Q13642 |

| | | | | | | | | |
|---|---|---|---|---|---|---|---|---|
| HOMO SAPIENS | 9 | 3206 | 1x63 | A | 40 | HIS | p.His123Tyr | Q13642 |
| HOMO SAPIENS | 9 | 3206 | 1x63 | A | 40 | HIS | p.His123Leu | Q13642 |
| HOMO SAPIENS | 9 | 3206 | 1x63 | A | 40 | HIS | p.His123Gln | Q13642 |
| HOMO SAPIENS | 3 | 3206 | 1x63 | A | 49 | CYS | p.Cys132Phe | Q13642 |
| HOMO SAPIENS | 3 | 3206 | 1x63 | A | 67 | CYS | p.Cys150Tyr | Q13642 |
| HOMO SAPIENS | 3 | 3206 | 1x63 | A | 70 | CYS | p.Cys153Arg | Q13642 |
| HOMO SAPIENS | 3 | 3206 | 1x63 | A | 70 | CYS | p.Cys153Tyr | Q13642 |
| HOMO SAPIENS | 5 | 1052 | 3ecs | E | 208 | ASN | p.Asn208Tyr | Q14232 |
| HOMO SAPIENS | 2 | 1052 | 3ecs | E | 275 | TYR | p.Tyr275Cys | Q14232 |
| HOMO SAPIENS | 0 | 24253 | 5i35 | A | 271 | ARG | p.Arg271Cys | Q8NI60 |
| HOMO SAPIENS | 0 | 24253 | 5i35 | A | 272 | GLY | p.Gly272Asp | Q8NI60 |
| HOMO SAPIENS | 0 | 24253 | 5i35 | A | 299 | ARG | p.Arg299Trp | Q8NI60 |
| HOMO SAPIENS | 1 | 24253 | 5i35 | A | 339 | ALA | p.Ala339Thr | Q8NI60 |
| HOMO SAPIENS | 0 | 24253 | 5i35 | A | 348 | ARG | p.Arg348Ter | Q8NI60 |
| HOMO SAPIENS | 0 | 24253 | 5i35 | A | 395 | ILE | p.Ile395Met | Q8NI60 |
| HOMO SAPIENS | 0 | 24253 | 5i35 | A | 396 | ASP | p.Asp396Glu | Q8NI60 |
| HOMO SAPIENS | 0 | 24253 | 5i35 | A | 480 | PHE | p.Phe480Leu | Q8NI60 |
| HOMO SAPIENS | 2 | 24253 | 5i35 | A | 511 | THR | p.Thr511Met | Q8NI60 |
| HOMO SAPIENS | 0 | 24253 | 5i35 | A | 514 | TYR | p.Tyr514Cys | Q8NI60 |
| HOMO SAPIENS | 0 | 24253 | 5i35 | A | 538 | ARG | p.Arg538Gln | Q8NI60 |
| HOMO SAPIENS | 0 | 24253 | 5i35 | A | 549 | GLY | p.Gly549Ser | Q8NI60 |
| HOMO SAPIENS | 0 | 24253 | 5i35 | A | 551 | GLU | p.Glu551Lys | Q8NI60 |
| HOMO SAPIENS | -1 | 24253 | 5i35 | A | 555 | MSE | p.Met555Ile | Q8NI60 |
| HOMO SAPIENS | 0 | 24253 | 5i35 | A | 568 | GLU | p.Glu568Ter | Q8NI60 |
| HOMO SAPIENS | 0 | 24253 | 5i35 | A | 602 | PRO | p.Pro602Arg | Q8NI60 |
| HOMO SAPIENS | -1 | 15505 | 4goj | C | 22 | GLY | p.Gly22Val | Q13432 |
| HOMO SAPIENS | 5 | 23902 | 2lrr | A | 765 | HIS | p.His765Gln | P38935 |
| HOMO SAPIENS | 1 | 29650 | 2vj2 | A | 234 | CYS | p.Cys234Tyr | P78504 |
| HOMO SAPIENS | 0 | 29650 | 2vj2 | A | 255 | TYR | p.Tyr255Ter | P78504 |
| HOMO SAPIENS | 3 | 29650 | 2vj2 | A | 274 | GLY | p.Gly274Asp | P78504 |
| HOMO SAPIENS | 1 | 105 | 5he0 | B | 77 | GLY | p.Gly77Ser | P62873 |
| HOMO SAPIENS | 2 | 105 | 5he0 | B | 80 | ILE | p.Ile80Thr | P62873 |
| HOMO SAPIENS | 3 | 4587 | 3rgh | A | 159 | ASP | p.Asp1159Ala | P21333 |
| HOMO SAPIENS | 3 | 4587 | 3rgh | A | 184 | ASP | p.Asp1184Glu | P21333 |
| HOMO SAPIENS | 2 | 4587 | 3rgh | A | 186 | SER | p.Ser1186Leu | P21333 |
| HOMO SAPIENS | 4 | 4587 | 3rgh | A | 188 | ALA | p.Ala1188Thr | P21333 |
| HOMO SAPIENS | 2 | 4587 | 3rgh | A | 199 | SER | p.Ser1199Leu | P21333 |
| HOMO SAPIENS | 0 | 20283 | 4n3w | A | 1278 | GLU | p.Glu1278Lys | Q92793 |
| HOMO SAPIENS | 1 | 14294 | 2d9q | B | 208 | MET | p.Met231Thr | Q99062 |
| HOMO SAPIENS | 6 | 14294 | 2d9q | B | 297 | ASP | p.Asp320Asn | Q99062 |
| HOMO SAPIENS | 0 | 11642 | 3bpt | A | 66 | ARG | p.Arg66Trp | Q6NVY1 |
| HOMO SAPIENS | 0 | 11642 | 3bpt | A | 122 | TYR | p.Tyr122Cys | Q6NVY1 |
| HOMO SAPIENS | 0 | 11642 | 3bpt | A | 345 | GLY | p.Gly345Ser | Q6NVY1 |
| HOMO SAPIENS | 1 | 3942 | 3sby | A | 60 | ASP | p.Asp60His | Q9Y4U1 |
| HOMO SAPIENS | 0 | 3942 | 3sby | A | 111 | ARG | p.Arg111Ter | Q9Y4U1 |
| HOMO SAPIENS | 1 | 3942 | 3sby | A | 116 | LEU | p.Leu116Pro | Q9Y4U1 |
| HOMO SAPIENS | 1 | 3942 | 3sby | A | 130 | TYR | p.Tyr130Cys | Q9Y4U1 |
| HOMO SAPIENS | 1 | 3942 | 3sby | A | 132 | ARG | p.Arg132Ter | Q9Y4U1 |
| HOMO SAPIENS | 1 | 3942 | 3sby | A | 147 | GLY | p.Gly147Ala | Q9Y4U1 |
| HOMO SAPIENS | 1 | 3942 | 3sby | A | 161 | ARG | p.Arg161Gln | Q9Y4U1 |
| HOMO SAPIENS | 1 | 3942 | 3sby | A | 161 | ARG | p.Arg161Ter | Q9Y4U1 |
| HOMO SAPIENS | 1 | 3942 | 3sby | A | 203 | TRP | p.Trp203Ter | Q9Y4U1 |
| HOMO SAPIENS | 1 | 3942 | 3sby | A | 203 | TRP | p.Trp203Ter | Q9Y4U1 |
| HOMO SAPIENS | 1 | 3942 | 3sby | A | 205 | TYR | p.Tyr205Ter | Q9Y4U1 |

| | | | | | | | | | |
|---|---|---|---|---|---|---|---|---|---|
| HOMO SAPIENS | 0 | 38340 | 2mbh | B | | 39 | MET | p.Met39Leu | Q13351 |
| HOMO SAPIENS | 2 | 7360 | 4k30 | A | | 430 | LEU | p.Leu430Pro | Q8N159 |
| HOMO SAPIENS | 2 | 7360 | 4k30 | A | | 433 | GLU | p.Glu433Asp | Q8N159 |
| HOMO SAPIENS | 7 | 7360 | 4k30 | A | | 484 | TRP | p.Trp484Arg | Q8N159 |
| HOMO SAPIENS | 1 | 41903 | 2xs6 | A | | 151 | PRO | p.Pro151Ser | O00459 |
| HOMO SAPIENS | 0 | 172 | 5glh | A | | 183 | ALA | p.Ala183Gly | P24530 |
| HOMO SAPIENS | 0 | 172 | 5glh | A | | 201 | ARG | p.Arg201Ter | P24530 |
| HOMO SAPIENS | 0 | 172 | 5glh | A | | 244 | THR | p.Thr244Met | P24530 |
| HOMO SAPIENS | 0 | 172 | 5glh | A | | 253 | ARG | p.Arg253Ter | P24530 |
| HOMO SAPIENS | 9 | 172 | 5glh | A | | 1017 | ILE | p.Leu17Phe | P00720 |
| HOMO SAPIENS | -1 | 3342 | 1ljm | A | | 1058 | HIS | p.His58Asn | Q01196 |
| HOMO SAPIENS | 4 | 3342 | 1ljm | A | | 1174 | ARG | p.Arg174Gln | Q01196 |
| HOMO SAPIENS | 5 | 25607 | 1p22 | C | | 32 | ASP | p.Asp32Ala | P35222 |
| HOMO SAPIENS | 5 | 25607 | 1p22 | C | | 32 | ASP | p.Asp32Tyr | P35222 |
| HOMO SAPIENS | -1 | 25607 | 1p22 | C | | 33 | SEP | p.Ser33Phe | P35222 |
| HOMO SAPIENS | 5 | 25607 | 1p22 | C | | 34 | GLY | p.Gly34Glu | P35222 |
| HOMO SAPIENS | -1 | 25607 | 1p22 | C | | 37 | SEP | p.Ser37Cys | P35222 |
| HOMO SAPIENS | -1 | 25607 | 1p22 | C | | 41 | THR | p.Thr41Ile | P35222 |
| HOMO SAPIENS | 1 | 48 | 4uow | | 9 | 60 | GLN | p.Arg34111His | Q8WZ42 |
| HOMO SAPIENS | 0 | 48 | 4uow | | 9 | 64 | LEU | p.Val34115Ala | Q8WZ42 |
| HOMO SAPIENS | 0 | 1127 | 5c01 | A | | 638 | ARG | p.Arg638Ter | P29597 |
| HOMO SAPIENS | 0 | 1127 | 5c01 | A | | 684 | ILE | p.Ile684Ser | P29597 |
| HOMO SAPIENS | 0 | 1127 | 5c01 | A | | 703 | ARG | p.Arg703Trp | P29597 |
| HOMO SAPIENS | 5 | 16389 | 2aq0 | B | | 128 | ALA | p.Ala860Asp | Q92889 |
| HOMO SAPIENS | 2 | 16389 | 2aq0 | B | | 141 | ILE | p.Ile873Val | Q92889 |
| HOMO SAPIENS | 5 | 16389 | 2aq0 | B | | 143 | GLU | p.Glu875Gly | Q92889 |
| HOMO SAPIENS | 0 | 16389 | 2aq0 | B | | 180 | GLY | p.Gly912Arg | Q92889 |
| HOMO SAPIENS | 0 | 15816 | 2xrc | A | | 89 | THR | p.Thr107Ala | P05156 |
| HOMO SAPIENS | 0 | 15816 | 2xrc | A | | 133 | ASN | p.Asn151Ser | P05156 |
| HOMO SAPIENS | 0 | 15816 | 2xrc | A | | 185 | THR | p.Thr203Ile | P05156 |
| HOMO SAPIENS | 0 | 15816 | 2xrc | A | | 225 | GLY | p.Gly243Asp | P05156 |
| HOMO SAPIENS | 0 | 15816 | 2xrc | A | | 243 | GLY | p.Gly261Ala | P05156 |
| HOMO SAPIENS | 0 | 15816 | 2xrc | A | | 269 | GLY | p.Gly287Arg | P05156 |
| HOMO SAPIENS | -1 | 15816 | 2xrc | A | | 288 | ILE | p.Ile306Val | P05156 |
| HOMO SAPIENS | -1 | 15816 | 2xrc | A | | 388 | ARG | p.Arg406His | P05156 |
| HOMO SAPIENS | 7 | 15816 | 2xrc | A | | 398 | ILE | p.Ile416Leu | P05156 |
| HOMO SAPIENS | 9 | 15816 | 2xrc | A | | 400 | HIS | p.His418Leu | P05156 |
| HOMO SAPIENS | 0 | 15816 | 2xrc | A | | 423 | LYS | p.Lys441Arg | P05156 |
| HOMO SAPIENS | -1 | 15816 | 2xrc | A | | 530 | GLU | p.Glu548Gln | P05156 |
| HOMO SAPIENS | 0 | 15816 | 2xrc | A | | 535 | PRO | p.Pro553Ala | P05156 |
| HOMO SAPIENS | 0 | 15816 | 2xrc | A | | 536 | GLU | p.Glu554Gly | P05156 |
| HOMO SAPIENS | 1 | 12385 | 4zlp | B | | 1450 | ALA | p.Ala1450Pro | Q9UM47 |
| HOMO SAPIENS | 7 | 12385 | 4zlp | B | | 1482 | GLY | p.Gly1482Arg | Q9UM47 |
| HOMO SAPIENS | 2 | 12385 | 4zlp | B | | 1518 | LEU | p.Leu1518Met | Q9UM47 |
| HOMO SAPIENS | 0 | 12385 | 4zlp | B | | 1560 | ARG | p.Arg1560Pro | Q9UM47 |
| HOMO SAPIENS | 0 | 1227 | 4a04 | A | | 148 | PHE | p.Phe148Tyr | O43435 |
| HOMO SAPIENS | 4 | 1227 | 4a04 | A | | 194 | HIS | p.His194Gln | O43435 |
| HOMO SAPIENS | 2 | 8017 | 3ei4 | B | | 244 | LYS | p.Lys244Glu | Q92466 |
| HOMO SAPIENS | 1 | 8017 | 3ei4 | B | | 273 | ARG | p.Arg273His | Q92466 |
| HOMO SAPIENS | 3 | 36112 | 2k8f | B | | 9 | SER | p.Ser9Arg | P04637 |
| HOMO SAPIENS | 2 | 12880 | 5ggf | A | | 265 | ARG | p.Arg265His | Q8WZA1 |
| HOMO SAPIENS | 2 | 12880 | 5ggf | A | | 287 | ILE | p.Ile287Ser | Q8WZA1 |
| HOMO SAPIENS | 2 | 12880 | 5ggf | A | | 311 | ARG | p.Arg311Gln | Q8WZA1 |
| HOMO SAPIENS | 2 | 12880 | 5ggf | A | | 425 | TRP | p.Trp425Ser | Q8WZA1 |

| | | | | | | | | |
|---|---|---|---|---|---|---|---|---|
| HOMO SAPIENS | 2 | 12880 | 5ggf | A | 442 | ARG | p.Arg442Cys | Q8WZA1 |
| HOMO SAPIENS | 2 | 12880 | 5ggf | A | 448 | GLY | p.Gly448Arg | Q8WZA1 |
| HOMO SAPIENS | 0 | 12880 | 5ggf | A | 490 | CYS | p.Cys490Tyr | Q8WZA1 |
| HOMO SAPIENS | 0 | 12880 | 5ggf | A | 493 | PRO | p.Pro493Arg | Q8WZA1 |
| HOMO SAPIENS | 3 | 12880 | 5ggf | A | 550 | SER | p.Ser550Asn | Q8WZA1 |
| HOMO SAPIENS | 1 | 12880 | 5ggf | A | 556 | ASP | p.Asp556Asn | Q8WZA1 |
| HOMO SAPIENS | 0 | 12880 | 5ggf | A | 580 | ARG | p.Arg580Ter | Q8WZA1 |
| HOMO SAPIENS | 2 | 12880 | 5ggf | A | 605 | ARG | p.Arg605Pro | Q8WZA1 |
| HOMO SAPIENS | 0 | 12880 | 5ggf | A | 623 | VAL | p.Met623Val | Q8WZA1 |
| HOMO SAPIENS | 1 | 10187 | 4qw2 | A | 145 | ALA | p.Ala145Ser | Q06787 |
| HOMO SAPIENS | 1 | 10187 | 4qw2 | A | 174 | ASN | p.Asn174Lys | Q06787 |
| HOMO SAPIENS | 0 | 24002 | 2m0t | A | 110 | LEU | p.Leu110Val | O14745 |
| HOMO SAPIENS | 9 | 3126 | 4mjh | A | 99 | LEU | p.Leu99Met | P04792 |
| HOMO SAPIENS | 1 | 3126 | 4mjh | A | 127 | ARG | p.Arg127Trp | P04792 |
| HOMO SAPIENS | 1 | 3126 | 4mjh | A | 127 | ARG | p.Arg127Leu | P04792 |
| HOMO SAPIENS | 1 | 3126 | 4mjh | A | 135 | SER | p.Ser135Phe | P04792 |
| HOMO SAPIENS | 2 | 3126 | 4mjh | A | 136 | ARG | p.Arg136Trp | P04792 |
| HOMO SAPIENS | 1 | 3126 | 4mjh | A | 140 | ARG | p.Arg140Gly | P04792 |
| HOMO SAPIENS | 0 | 3126 | 4mjh | A | 151 | THR | p.Thr151Ile | P04792 |
| HOMO SAPIENS | -1 | 15780 | 2jlp | A | 213 | ARG | p.Arg231Gly | P08294 |
| HOMO SAPIENS | 8 | 3578 | 1apz | D | 203 | GLY | p.Gly226Asp | P20933 |
| HOMO SAPIENS | 1 | 3578 | 1apz | D | 279 | GLY | p.Gly302Arg | P20933 |
| HOMO SAPIENS | 0 | 3578 | 1apz | D | 283 | CYS | p.Cys306Arg | P20933 |
| HOMO SAPIENS | 5 | 1335 | 5xtd | C | 115 | ARG | p.Arg115Cys | O75251 |
| HOMO SAPIENS | 0 | 1335 | 5xtd | C | 122 | VAL | p.Val122Met | O75251 |
| HOMO SAPIENS | 2 | 5063 | 2le4 | A | 37 | ARG | p.Arg74Pro | P48431 |
| HOMO SAPIENS | 1 | 25870 | 2o10 | A | 44 | LEU | p.Leu44Pro | P50461 |
| HOMO SAPIENS | 4 | 25870 | 2o10 | A | 58 | CYS | p.Cys58Gly | P50461 |
| HOMO SAPIENS | 0 | 18748 | 5fii | C | 39 | PHE | p.Phe39Leu | P00439 |
| HOMO SAPIENS | 6 | 18748 | 5fii | C | 48 | LEU | p.Leu48Ser | P00439 |
| HOMO SAPIENS | 2 | 18748 | 5fii | C | 65 | ILE | p.Ile65Asn | P00439 |
| HOMO SAPIENS | 3 | 18748 | 5fii | C | 95 | ILE | p.Ile95Phe | P00439 |
| HOMO SAPIENS | 1 | 14261 | 2vay | B | 1539 | ARG | p.Arg1539Cys | Q13698 |
| HOMO SAPIENS | 0 | 2494 | 4dx1 | A | 183 | GLU | p.Glu183Asp | Q9HBA0 |
| HOMO SAPIENS | 0 | 2494 | 4dx1 | A | 217 | ALA | p.Ala217Ser | Q9HBA0 |
| HOMO SAPIENS | 0 | 2494 | 4dx1 | A | 217 | ALA | p.Ala217Val | Q9HBA0 |
| HOMO SAPIENS | 4 | 2494 | 4dx1 | A | 224 | ARG | p.Arg224Trp | Q9HBA0 |
| HOMO SAPIENS | 4 | 2494 | 4dx1 | A | 254 | VAL | p.Val254Leu | Q9HBA0 |
| HOMO SAPIENS | 0 | 2494 | 4dx1 | A | 269 | ARG | p.Arg269Cys | Q9HBA0 |
| HOMO SAPIENS | 0 | 2494 | 4dx1 | A | 269 | ARG | p.Arg269His | Q9HBA0 |
| HOMO SAPIENS | 0 | 2494 | 4dx1 | A | 278 | GLU | p.Glu278Lys | Q9HBA0 |
| HOMO SAPIENS | 0 | 2494 | 4dx1 | A | 315 | ARG | p.Arg315Trp | Q9HBA0 |
| HOMO SAPIENS | 0 | 2494 | 4dx1 | A | 316 | ARG | p.Arg316Cys | Q9HBA0 |
| HOMO SAPIENS | 0 | 2494 | 4dx1 | A | 331 | ILE | p.Ile331Phe | Q9HBA0 |
| HOMO SAPIENS | 0 | 2494 | 4dx1 | A | 333 | ASP | p.Asp333Gly | Q9HBA0 |
| HOMO SAPIENS | 1 | 2494 | 4dx1 | A | 365 | VAL | p.Val365Met | Q9HBA0 |
| HOMO SAPIENS | 0 | 23998 | 2l1r | B | 161 | ARG | p.Arg162Trp | P19429 |
| HOMO SAPIENS | 0 | 8354 | 4f0x | D | 187 | SER | p.Ser187Ter | O95822 |
| HOMO SAPIENS | 2 | 8354 | 4f0x | D | 259 | GLY | p.Gly259Ala | O95822 |
| HOMO SAPIENS | 2 | 8354 | 4f0x | D | 284 | THR | p.Thr284Ala | O95822 |
| HOMO SAPIENS | 3 | 8354 | 4f0x | D | 309 | LYS | p.Lys309Gln | O95822 |
| HOMO SAPIENS | 0 | 21182 | 4apo | A | 228 | GLN | p.Gln228Lys | O00170 |
| HOMO SAPIENS | 1 | 21182 | 4apo | A | 307 | GLN | p.Gln307Arg | O00170 |
| HOMO SAPIENS | 0 | 2287 | 5of4 | B | 112 | ARG | p.Arg112His | P18074 |

| | | | | | | | | |
|---|---|---|---|---|---|---|---|---|
| HOMO SAPIENS | 0 | 2287 | 5of4 | B | 312 | ASP | p.Asp312Asn | P18074 |
| HOMO SAPIENS | 3 | 2287 | 5of4 | B | 461 | LEU | p.Leu461Val | P18074 |
| HOMO SAPIENS | 0 | 2287 | 5of4 | B | 485 | LEU | p.Leu485Pro | P18074 |
| HOMO SAPIENS | 1 | 2287 | 5of4 | B | 541 | SER | p.Ser541Arg | P18074 |
| HOMO SAPIENS | 0 | 2287 | 5of4 | B | 616 | ARG | p.Arg616Pro | P18074 |
| HOMO SAPIENS | 0 | 2287 | 5of4 | B | 616 | ARG | p.Arg616Trp | P18074 |
| HOMO SAPIENS | 1 | 2287 | 5of4 | B | 658 | ARG | p.Arg658Cys | P18074 |
| HOMO SAPIENS | 1 | 2287 | 5of4 | B | 681 | ASP | p.Asp681Asn | P18074 |
| HOMO SAPIENS | 1 | 2287 | 5of4 | B | 683 | ARG | p.Arg683Trp | P18074 |
| HOMO SAPIENS | 0 | 2287 | 5of4 | B | 713 | GLY | p.Gly713Arg | P18074 |
| HOMO SAPIENS | 0 | 2287 | 5of4 | B | 717 | ALA | p.Ala717Gly | P18074 |
| HOMO SAPIENS | 0 | 2287 | 5of4 | B | 722 | ARG | p.Arg722Trp | P18074 |
| HOMO SAPIENS | 0 | 2287 | 5of4 | B | 725 | ALA | p.Ala725Pro | P18074 |
| HOMO SAPIENS | 4 | 12188 | 4its | A | 122 | ARG | p.Arg122Gly | Q9Y606 |
| HOMO SAPIENS | 4 | 12188 | 4its | A | 133 | ASP | p.Asp133Asn | Q9Y606 |
| HOMO SAPIENS | 4 | 12188 | 4its | A | 144 | ARG | p.Arg144Trp | Q9Y606 |
| HOMO SAPIENS | 0 | 31529 | 1kgd | A | 761 | ARG | p.Arg766Gln | O14936 |
| HOMO SAPIENS | 1 | 31529 | 1kgd | A | 824 | GLN | p.Gln829Arg | O14936 |
| HOMO SAPIENS | 0 | 10906 | 2h63 | C | 18 | ARG | p.Arg18Ter | P53004 |
| HOMO SAPIENS | 9 | 27767 | 4wyq | A | 280 | LEU | p.Leu278Phe | Q9UPY3 |
| HOMO SAPIENS | 0 | 27767 | 4wyq | A | 297 | SER | p.Ser295Cys | Q9UPY3 |
| HOMO SAPIENS | 1 | 27767 | 4wyq | A | 334 | TYR | p.Tyr332Ter | Q9UPY3 |
| HOMO SAPIENS | 7 | 7775 | 5sv7 | C | 136 | SER | p.Ser136Cys | Q9NZJ5 |
| HOMO SAPIENS | 2 | 7775 | 5sv7 | C | 166 | GLN | p.Gln166Arg | Q9NZJ5 |
| HOMO SAPIENS | 2 | 7775 | 5sv7 | C | 332 | GLU | p.Glu332Ter | Q9NZJ5 |
| HOMO SAPIENS | 0 | 21449 | 2e9w | B | 84 | THR | p.Thr84Met | P10721 |
| HOMO SAPIENS | 0 | 21449 | 2e9w | B | 168 | ALA | p.Ala168Glu | P10721 |
| HOMO SAPIENS | 0 | 21449 | 2e9w | B | 178 | ALA | p.Ala178Thr | P10721 |
| HOMO SAPIENS | 0 | 21449 | 2e9w | B | 293 | ASN | p.Asn293Ser | P10721 |
| HOMO SAPIENS | -1 | 21449 | 2e9w | B | 318 | MET | p.Met318Leu | P10721 |
| HOMO SAPIENS | 0 | 21449 | 2e9w | B | 374 | VAL | p.Val374Ile | P10721 |
| HOMO SAPIENS | 6 | 3010 | 3flg | A | 66 | SER | p.Ser270Pro | Q9UBC3 |
| HOMO SAPIENS | 1 | 33918 | 2xzz | A | 760 | ARG | p.Arg760Ter | P22735 |
| HOMO SAPIENS | 2 | 1254 | 5xtc | w | 65 | ASN | p.Asn65Ser | O95299 |
| HOMO SAPIENS | 1 | 1254 | 5xtc | w | 238 | GLU | p.Glu238Lys | O95299 |
| HOMO SAPIENS | 3 | 10245 | 3siv | B | 194 | ALA | p.Ala194Glu | Q8WWY3 |
| HOMO SAPIENS | 4 | 10245 | 3siv | B | 216 | ALA | p.Ala216Pro | Q8WWY3 |
| HOMO SAPIENS | 0 | 2670 | 3dwb | A | 341 | THR | p.Thr341Ile | P42892 |
| HOMO SAPIENS | 1 | 2670 | 3dwb | A | 462 | SER | p.Ser462Asn | P42892 |
| HOMO SAPIENS | 1 | 2670 | 3dwb | A | 754 | ARG | p.Arg754Cys | P42892 |
| HOMO SAPIENS | 1 | 2961 | 5xtb | O | 40 | VAL | p.Val40Met | P19404 |
| HOMO SAPIENS | 5 | 2961 | 5xtb | O | 172 | ILE | p.Ile172Thr | P19404 |
| HOMO SAPIENS | 5 | 2961 | 5xtb | O | 190 | ASP | p.Asp190Gly | P19404 |
| HOMO SAPIENS | 5 | 29655 | 2xdg | A | 57 | ALA | p.Ala57Thr | Q02643 |
| HOMO SAPIENS | 4 | 29655 | 2xdg | A | 72 | GLU | p.Glu72Ter | Q02643 |
| HOMO SAPIENS | 0 | 29655 | 2xdg | A | 121 | GLU | p.Glu121Asp | Q02643 |
| HOMO SAPIENS | -1 | 17935 | 1hyn | P | 38 | ASP | p.Asp38Ala | P02730 |
| HOMO SAPIENS | 7 | 12825 | 1g82 | D | 99 | SER | p.Ser99Asn | P31371 |
| HOMO SAPIENS | 6 | 24927 | 1jm7 | B | 85 | VAL | p.Val85Leu | Q99728 |
| HOMO SAPIENS | 1 | 24927 | 1jm7 | B | 112 | ARG | p.Arg112Ter | Q99728 |
| HOMO SAPIENS | 0 | 24927 | 1jm7 | B | 116 | HIS | p.His116Tyr | Q99728 |
| HOMO SAPIENS | 7 | 92 | 5hde | A | 61 | LYS | p.Lys61Arg | Q05209 |
| HOMO SAPIENS | 0 | 17246 | 1szb | A | 84 | ARG | p.Arg99Gln | O00187 |
| HOMO SAPIENS | 1 | 17246 | 1szb | A | 105 | ASP | p.Asp120Gly | O00187 |

| | | | | | | | | |
|---|---|---|---|---|---|---|---|---|
| HOMO SAPIENS | 0 | 17246 | 1szb | A | 111 | PRO | p.Pro126Leu | O00187 |
| HOMO SAPIENS | 9 | 17246 | 1szb | A | 141 | CYS | p.Cys156Tyr | O00187 |
| HOMO SAPIENS | 0 | 32473 | 1b72 | A | 207 | ARG | p.Arg207Cys | P14653 |
| HOMO SAPIENS | 0 | 1920 | 4yzf | B | 488 | VAL | p.Val488Met | P02730 |
| HOMO SAPIENS | 0 | 1920 | 4yzf | B | 589 | ARG | p.Arg589Cys | P02730 |
| HOMO SAPIENS | 0 | 1920 | 4yzf | B | 589 | ARG | p.Arg589His | P02730 |
| HOMO SAPIENS | 0 | 1920 | 4yzf | B | 602 | ARG | p.Arg602Pro | P02730 |
| HOMO SAPIENS | 0 | 1920 | 4yzf | B | 613 | SER | p.Ser613Phe | P02730 |
| HOMO SAPIENS | -1 | 1920 | 4yzf | B | 646 | ARG | p.Arg646Gln | P02730 |
| HOMO SAPIENS | -1 | 1920 | 4yzf | B | 646 | ARG | p.Arg646Trp | P02730 |
| HOMO SAPIENS | 0 | 1920 | 4yzf | B | 701 | GLY | p.Gly701Asp | P02730 |
| HOMO SAPIENS | 0 | 1920 | 4yzf | B | 760 | ARG | p.Arg760Gln | P02730 |
| HOMO SAPIENS | 0 | 1920 | 4yzf | B | 771 | GLY | p.Gly771Asp | P02730 |
| HOMO SAPIENS | 0 | 1920 | 4yzf | B | 773 | SER | p.Ser773Pro | P02730 |
| HOMO SAPIENS | 0 | 1920 | 4yzf | B | 837 | THR | p.Thr837Ala | P02730 |
| HOMO SAPIENS | 0 | 1920 | 4yzf | B | 854 | PRO | p.Pro854Leu | P02730 |
| HOMO SAPIENS | 0 | 1920 | 4yzf | B | 858 | ALA | p.Ala858Asp | P02730 |
| HOMO SAPIENS | 0 | 1920 | 4yzf | B | 862 | VAL | p.Val862Ile | P02730 |
| HOMO SAPIENS | 0 | 1920 | 4yzf | B | 868 | PRO | p.Pro868Leu | P02730 |
| HOMO SAPIENS | 0 | 1920 | 4yzf | B | 870 | ARG | p.Arg870Trp | P02730 |
| HOMO SAPIENS | 0 | 9863 | 2juj | A | 54 | VAL | p.Val904Ile | P22681 |
| HOMO SAPIENS | 0 | 21934 | 2na7 | A | 184 | ILE | p.Ile184Val | P25445 |
| HOMO SAPIENS | -1 | 6746 | 1lw3 | A | 3 | THR | p.Lys3Thr | Q13614 |
| HOMO SAPIENS | -1 | 6746 | 1lw3 | A | 44 | VAL | p.Val44Ile | Q13614 |
| HOMO SAPIENS | 3 | 6746 | 1lw3 | A | 183 | ALA | p.Ala183Ser | Q13614 |
| HOMO SAPIENS | 4 | 6746 | 1lw3 | A | 482 | GLN | p.Gln482Ter | Q13614 |
| HOMO SAPIENS | 4 | 6746 | 1lw3 | A | 502 | GLU | p.Glu502Gln | Q13614 |
| HOMO SAPIENS | 3 | 6746 | 1lw3 | A | 545 | ASN | p.Asn545Ser | Q13614 |
| HOMO SAPIENS | 0 | 6746 | 1lw3 | A | 586 | ARG | p.Arg586Trp | Q13614 |
| HOMO SAPIENS | -1 | 6746 | 1lw3 | A | 602 | ALA | p.Ala602Gly | Q13614 |
| HOMO SAPIENS | 1 | 319 | 4ch9 | B | 336 | ARG | p.Arg336Ile | Q9UH77 |
| HOMO SAPIENS | 3 | 319 | 4ch9 | B | 340 | ALA | p.Ala340Val | Q9UH77 |
| HOMO SAPIENS | 0 | 319 | 4ch9 | B | 384 | ARG | p.Arg384Gln | Q9UH77 |
| HOMO SAPIENS | 5 | 319 | 4ch9 | B | 426 | PRO | p.Pro426Leu | Q9UH77 |
| HOMO SAPIENS | 3 | 319 | 4ch9 | B | 431 | ARG | p.Arg431Gln | Q9UH77 |
| HOMO SAPIENS | 2 | 319 | 4ch9 | B | 575 | ARG | p.Arg575Trp | Q9UH77 |
| HOMO SAPIENS | 0 | 574 | 2qkk | A | 142 | VAL | p.Val142Ile | O60930 |
| HOMO SAPIENS | 0 | 24021 | 2n8y | A | 752 | ARG | p.Arg752Gln | P12814 |
| HOMO SAPIENS | 0 | 18693 | 2y7j | B | 44 | ARG | p.Arg44Ter | P15735 |
| HOMO SAPIENS | 0 | 18693 | 2y7j | B | 106 | VAL | p.Val106Glu | P15735 |
| HOMO SAPIENS | 3 | 18693 | 2y7j | B | 189 | GLY | p.Gly189Glu | P15735 |
| HOMO SAPIENS | 0 | 17087 | 2ifg | A | 237 | THR | p.Thr237Met | P04629 |
| HOMO SAPIENS | 0 | 7980 | 1qub | A | 247 | VAL | p.Val266Leu | P02749 |
| HOMO SAPIENS | -1 | 2205 | 1h7u | A | 18 | ILE | p.Ile18Val | P54278 |
| HOMO SAPIENS | -1 | 2205 | 1h7u | A | 20 | ARG | p.Arg20Gln | P54278 |
| HOMO SAPIENS | 9 | 2205 | 1h7u | A | 46 | SER | p.Ser46Ile | P54278 |
| HOMO SAPIENS | 0 | 2205 | 1h7u | A | 60 | ASP | p.Asp60Glu | P54278 |
| HOMO SAPIENS | 0 | 2205 | 1h7u | A | 80 | PHE | p.Phe80Leu | P54278 |
| HOMO SAPIENS | 0 | 2205 | 1h7u | A | 118 | SER | p.Ser118Arg | P54278 |
| HOMO SAPIENS | 0 | 2205 | 1h7u | A | 126 | HIS | p.His126Gln | P54278 |
| HOMO SAPIENS | 0 | 2205 | 1h7u | A | 128 | SER | p.Ser128Leu | P54278 |
| HOMO SAPIENS | 1 | 2205 | 1h7u | A | 134 | ARG | p.Arg134Ter | P54278 |
| HOMO SAPIENS | 0 | 2205 | 1h7u | A | 166 | LEU | p.Leu166Pro | P54278 |
| HOMO SAPIENS | 1 | 2205 | 1h7u | A | 191 | TYR | p.Tyr191Cys | P54278 |

| | | | | | | | | |
|---|---|---|---|---|---|---|---|---|
| HOMO SAPIENS | 0 | 2205 | 1h7u | A | 211 | ARG | p.Arg211Ter | P54278 |
| HOMO SAPIENS | 0 | 2205 | 1h7u | A | 233 | GLN | p.Gln233Ter | P54278 |
| HOMO SAPIENS | 0 | 2205 | 1h7u | A | 255 | TYR | p.Tyr255Ter | P54278 |
| HOMO SAPIENS | 0 | 2205 | 1h7u | A | 277 | THR | p.Thr277Lys | P54278 |
| HOMO SAPIENS | 0 | 2205 | 1h7u | A | 301 | LYS | p.Lys301Asn | P54278 |
| HOMO SAPIENS | 0 | 2205 | 1h7u | A | 312 | MET | p.Met312Ile | P54278 |
| HOMO SAPIENS | 0 | 2205 | 1h7u | A | 312 | MET | p.Met312Thr | P54278 |
| HOMO SAPIENS | 0 | 2205 | 1h7u | A | 315 | ARG | p.Arg315Ter | P54278 |
| HOMO SAPIENS | 0 | 2205 | 1h7u | A | 318 | TYR | p.Tyr318Cys | P54278 |
| HOMO SAPIENS | 0 | 2205 | 1h7u | A | 335 | ASN | p.Asn335Ile | P54278 |
| HOMO SAPIENS | 0 | 2205 | 1h7u | A | 360 | ILE | p.Ile360Met | P54278 |
| HOMO SAPIENS | 6 | 20754 | 5mpo | B | 7 | VAL | p.Val7Phe | O96033 |
| HOMO SAPIENS | 4 | 20754 | 5mpo | B | 64 | VAL | p.Val64Ile | O96033 |
| HOMO SAPIENS | 5 | 34944 | 4qam | B | 1114 | ASP | p.Asp1114Gly | Q96KN7 |
| HOMO SAPIENS | 5 | 34944 | 4qam | B | 1240 | GLY | p.Gly1240Glu | Q96KN7 |
| HOMO SAPIENS | 9 | 40496 | 3vnn | A | 278 | ARG | p.Arg278His | P49917 |
| HOMO SAPIENS | 6 | 30131 | 4dt7 | E | 40 | PRO | p.Pro210Leu | P04070 |
| HOMO SAPIENS | -1 | 30131 | 4dt7 | E | 50 | ARG | p.Arg220Trp | P04070 |
| HOMO SAPIENS | 7 | 15311 | 2pqt | A | 64 | ARG | p.Arg64Trp | P18440 |
| HOMO SAPIENS | 2 | 15311 | 2pqt | A | 149 | VAL | p.Val149Ile | P18440 |
| HOMO SAPIENS | -1 | 19581 | 3hy8 | A | 33 | ASP | p.Asp33Val | Q9NVS9 |
| HOMO SAPIENS | 5 | 19581 | 3hy8 | A | 116 | ARG | p.Arg116Gln | Q9NVS9 |
| HOMO SAPIENS | 4 | 19581 | 3hy8 | A | 225 | ARG | p.Arg225His | Q9NVS9 |
| HOMO SAPIENS | 2 | 19581 | 3hy8 | A | 229 | TRP | p.Arg229Trp | Q9NVS9 |
| HOMO SAPIENS | 2 | 19581 | 3hy8 | A | 229 | TRP | p.Arg229Gln | Q9NVS9 |
| HOMO SAPIENS | 0 | 10266 | 3t1i | A | 41 | ASP | p.Asp41His | P49959 |
| HOMO SAPIENS | 1 | 10266 | 3t1i | A | 305 | ARG | p.Arg305Trp | P49959 |
| HOMO SAPIENS | 0 | 10266 | 3t1i | A | 308 | PHE | p.Phe308Leu | P49959 |
| HOMO SAPIENS | 0 | 10266 | 3t1i | A | 334 | SER | p.Ser334Arg | P49959 |
| HOMO SAPIENS | 1 | 1600 | 3rbf | A | 61 | GLU | p.Glu61Asp | P20711 |
| HOMO SAPIENS | 0 | 1600 | 3rbf | A | 91 | ALA | p.Ala91Val | P20711 |
| HOMO SAPIENS | -1 | 1600 | 3rbf | A | 102 | GLY | p.Gly102Ser | P20711 |
| HOMO SAPIENS | 3 | 1600 | 3rbf | A | 147 | SER | p.Ser147Arg | P20711 |
| HOMO SAPIENS | 1 | 1600 | 3rbf | A | 210 | PRO | p.Pro210Leu | P20711 |
| HOMO SAPIENS | 0 | 1600 | 3rbf | A | 217 | MET | p.Met217Val | P20711 |
| HOMO SAPIENS | 4 | 1600 | 3rbf | A | 250 | SER | p.Ser250Phe | P20711 |
| HOMO SAPIENS | 2 | 1600 | 3rbf | A | 275 | ALA | p.Ala275Thr | P20711 |
| HOMO SAPIENS | 1 | 1600 | 3rbf | A | 309 | PHE | p.Phe309Leu | P20711 |
| HOMO SAPIENS | -1 | 1600 | 3rbf | A | 347 | ARG | p.Arg347Gln | P20711 |
| HOMO SAPIENS | 0 | 1600 | 3rbf | A | 462 | ARG | p.Arg462Gln | P20711 |
| HOMO SAPIENS | 6 | 20830 | 5y4f | A | 714 | THR | p.Thr715Ile | Q01484 |
| HOMO SAPIENS | 7 | 20830 | 5y4f | A | 719 | ASP | p.Asp720Glu | Q01484 |
| HOMO SAPIENS | 0 | 20830 | 5y4f | A | 814 | VAL | p.Val815Ile | Q01484 |
| HOMO SAPIENS | -1 | 302 | 2c9y | A | 9 | GLU | p.Glu9Ter | P54819 |
| HOMO SAPIENS | 0 | 302 | 2c9y | A | 103 | ARG | p.Arg103Trp | P54819 |
| HOMO SAPIENS | 3 | 302 | 2c9y | A | 165 | ASP | p.Asp165Gly | P54819 |
| HOMO SAPIENS | 0 | 302 | 2c9y | A | 182 | ALA | p.Ala182Asp | P54819 |
| HOMO SAPIENS | 2 | 302 | 2c9y | A | 186 | ARG | p.Arg186Cys | P54819 |
| HOMO SAPIENS | 0 | 302 | 2c9y | A | 209 | ALA | p.Ala209Thr | P54819 |
| HOMO SAPIENS | -1 | 16920 | 1jlj | B | 10 | ASN | p.Asn10Tyr | Q9NQX3 |
| HOMO SAPIENS | 9 | 8462 | 1zbq | B | 16 | GLY | p.Gly16Ser | P51659 |
| HOMO SAPIENS | 5 | 8462 | 1zbq | B | 19 | ALA | p.Ala19Gly | P51659 |
| HOMO SAPIENS | 1 | 8462 | 1zbq | B | 34 | ALA | p.Ala34Val | P51659 |
| HOMO SAPIENS | 1 | 8462 | 1zbq | B | 106 | ARG | p.Arg106His | P51659 |

| | | | | | | | | |
|---|---|---|---|---|---|---|---|---|
| HOMO SAPIENS | 0 | 8462 | 1zbq | B | 140 | LYS | p.Lys140Asn | P51659 |
| HOMO SAPIENS | 0 | 8462 | 1zbq | B | 292 | THR | p.Thr292Ser | P51659 |
| HOMO SAPIENS | 0 | 4375 | 4ndi | A | 199 | ARG | p.Arg213His | Q7Z2E3 |
| HOMO SAPIENS | 3 | 4375 | 4ndi | A | 201 | HIS | p.His215Arg | Q7Z2E3 |
| HOMO SAPIENS | 0 | 4375 | 4ndi | A | 206 | PRO | p.Pro220Leu | Q7Z2E3 |
| HOMO SAPIENS | 2 | 4375 | 4ndi | A | 223 | LEU | p.Leu237Phe | Q7Z2E3 |
| HOMO SAPIENS | 0 | 4375 | 4ndi | A | 245 | ARG | p.Arg259His | Q7Z2E3 |
| HOMO SAPIENS | 0 | 4375 | 4ndi | A | 263 | VAL | p.Val277Gly | Q7Z2E3 |
| HOMO SAPIENS | 0 | 4375 | 4ndi | A | 279 | TRP | p.Trp293Ter | Q7Z2E3 |
| HOMO SAPIENS | 0 | 27004 | 4ccg | Y | 321 | ASP | p.Asp321Glu | Q9NW38 |
| HOMO SAPIENS | -1 | 27004 | 4ccg | Y | 372 | GLY | p.Gly372Ala | Q9NW38 |
| HOMO SAPIENS | -1 | 6383 | 5fai | A | 34 | ALA | p.Ala34Gly | Q92979 |
| HOMO SAPIENS | 8 | 6383 | 5fai | A | 86 | ASP | p.Asp86Gly | Q92979 |
| HOMO SAPIENS | 2 | 24585 | 4mi5 | A | 684 | ARG | p.Arg679Cys | Q15910 |
| HOMO SAPIENS | 0 | 15967 | 3nhb | A | 723 | ARG | p.Arg723Gln | Q9NP58 |
| HOMO SAPIENS | 5 | 18152 | 1oey | A | 389 | HIS | p.His389Gln | P19878 |
| HOMO SAPIENS | 5 | 18152 | 1oey | A | 395 | ARG | p.Arg395Trp | P19878 |
| HOMO SAPIENS | -1 | 1044 | 1li1 | F | 2 | VAL | p.Val1486Ile | P08572 |
| HOMO SAPIENS | 6 | 1044 | 1li1 | F | 37 | GLU | p.Glu1521Lys | P08572 |
| HOMO SAPIENS | 5 | 1044 | 1li1 | F | 157 | SER | p.Ser1639Leu | P08572 |
| HOMO SAPIENS | 5 | 1044 | 1li1 | F | 179 | ARG | p.Arg1662His | P08572 |
| HOMO SAPIENS | 5 | 1044 | 1li1 | F | 180 | GLY | p.Gly1663Arg | P08572 |
| HOMO SAPIENS | 6 | 20076 | 4uyi | A | 671 | LEU | p.Leu671Ser | Q8IY92 |
| HOMO SAPIENS | 0 | 19588 | 1jmj | A | 189 | ARG | p.Arg208His | P05546 |
| HOMO SAPIENS | -1 | 38023 | 1ry7 | B | 65 | GLY | p.Gly65Arg | P22607 |
| HOMO SAPIENS | -1 | 38023 | 1ry7 | B | 139 | ASP | p.Asp139Glu | P22607 |
| HOMO SAPIENS | 0 | 38023 | 1ry7 | B | 248 | ARG | p.Arg248Cys | P22607 |
| HOMO SAPIENS | 0 | 38023 | 1ry7 | B | 249 | SER | p.Ser249Cys | P22607 |
| HOMO SAPIENS | 0 | 38023 | 1ry7 | B | 250 | PRO | p.Pro250Arg | P22607 |
| HOMO SAPIENS | 0 | 38023 | 1ry7 | B | 322 | GLU | p.Glu322Lys | P22607 |
| HOMO SAPIENS | 5 | 16884 | 5eli | A | 33 | GLN | p.Gln33Ter | Q9NZC2 |
| HOMO SAPIENS | 0 | 16884 | 5eli | A | 44 | TRP | p.Trp44Ter | Q9NZC2 |
| HOMO SAPIENS | 0 | 16884 | 5eli | A | 47 | ARG | p.Arg47His | Q9NZC2 |
| HOMO SAPIENS | 0 | 16884 | 5eli | A | 66 | THR | p.Thr66Met | Q9NZC2 |
| HOMO SAPIENS | 0 | 16884 | 5eli | A | 78 | TRP | p.Trp78Ter | Q9NZC2 |
| HOMO SAPIENS | 1 | 16884 | 5eli | A | 96 | THR | p.Thr96Lys | Q9NZC2 |
| HOMO SAPIENS | 9 | 16884 | 5eli | A | 126 | VAL | p.Val126Gly | Q9NZC2 |
| HOMO SAPIENS | -1 | 12267 | 3w5o | A | 3 | ALA | p.Ala3Val | P49917 |
| HOMO SAPIENS | 1 | 12267 | 3w5o | A | 9 | THR | p.Thr9Ile | P49917 |
| HOMO SAPIENS | 2 | 12267 | 3w5o | A | 188 | ARG | p.Arg188Gln | P49917 |
| HOMO SAPIENS | 0 | 12267 | 3w5o | A | 469 | GLY | p.Gly469Glu | P49917 |
| HOMO SAPIENS | 1 | 12267 | 3w5o | A | 580 | ARG | p.Arg580Ter | P49917 |
| HOMO SAPIENS | 0 | 12267 | 3w5o | A | 600 | GLU | p.Glu600Lys | P49917 |
| HOMO SAPIENS | 9 | 2839 | 2hw5 | B | 54 | ARG | p.Arg54Cys | P30084 |
| HOMO SAPIENS | 9 | 2839 | 2hw5 | B | 59 | ASN | p.Asn59Ser | P30084 |
| HOMO SAPIENS | 0 | 2839 | 2hw5 | B | 273 | LYS | p.Lys273Glu | P30084 |
| HOMO SAPIENS | 1 | 12266 | 2o2k | B | 939 | GLY | p.Gly939Cys | Q99707 |
| HOMO SAPIENS | 5 | 12266 | 2o2k | B | 1164 | ARG | p.Arg1164His | Q99707 |
| HOMO SAPIENS | 1 | 12266 | 2o2k | B | 1166 | LEU | p.Leu1166Met | Q99707 |
| HOMO SAPIENS | 5 | 12266 | 2o2k | B | 1173 | PRO | p.Pro1173Leu | Q99707 |
| HOMO SAPIENS | 5 | 12266 | 2o2k | B | 1222 | ASN | p.Asn1222Ser | Q99707 |
| HOMO SAPIENS | 0 | 30891 | 3ch4 | B | 138 | ARG | p.Arg138Ter | Q15126 |
| HOMO SAPIENS | -1 | 4433 | 1qlz | A | 54 | GLY | p.Gly54Ser | P04156 |
| HOMO SAPIENS | 0 | 22568 | 3cxl | A | 20 | LEU | p.Leu20Phe | P15882 |

| | | | | | | | | |
|---|---|---|---|---|---|---|---|---|
| HOMO SAPIENS | 0 | 22568 | 3cxl | A | 126 | ILE | p.Ile126Met | P15882 |
| HOMO SAPIENS | 0 | 22568 | 3cxl | A | 141 | PRO | p.Pro141Leu | P15882 |
| HOMO SAPIENS | 0 | 22568 | 3cxl | A | 143 | TYR | p.Tyr143His | P15882 |
| HOMO SAPIENS | 0 | 22568 | 3cxl | A | 223 | ALA | p.Ala223Val | P15882 |
| HOMO SAPIENS | 0 | 22568 | 3cxl | A | 228 | GLY | p.Gly228Ser | P15882 |
| HOMO SAPIENS | 0 | 22568 | 3cxl | A | 252 | PRO | p.Pro252Gln | P15882 |
| HOMO SAPIENS | 0 | 22568 | 3cxl | A | 313 | GLU | p.Glu313Lys | P15882 |
| HOMO SAPIENS | 3 | 2668 | 5eqg | A | 34 | ASN | p.Asn34Ile | P11166 |
| HOMO SAPIENS | 3 | 2668 | 5eqg | A | 77 | MET | p.Met77Val | P11166 |
| HOMO SAPIENS | 6 | 2668 | 5eqg | A | 91 | GLY | p.Gly91Asp | P11166 |
| HOMO SAPIENS | 6 | 2668 | 5eqg | A | 92 | ARG | p.Arg92Trp | P11166 |
| HOMO SAPIENS | 6 | 2668 | 5eqg | A | 93 | ARG | p.Arg93Trp | P11166 |
| HOMO SAPIENS | 0 | 2668 | 5eqg | A | 95 | SER | p.Ser95Ile | P11166 |
| HOMO SAPIENS | 3 | 2668 | 5eqg | A | 126 | ARG | p.Arg126His | P11166 |
| HOMO SAPIENS | 3 | 2668 | 5eqg | A | 126 | ARG | p.Arg126Cys | P11166 |
| HOMO SAPIENS | 4 | 2668 | 5eqg | A | 232 | ARG | p.Arg232Cys | P11166 |
| HOMO SAPIENS | 3 | 2668 | 5eqg | A | 249 | ARG | p.Arg249Gln | P11166 |
| HOMO SAPIENS | 3 | 2668 | 5eqg | A | 255 | LYS | p.Lys255Arg | P11166 |
| HOMO SAPIENS | 0 | 2668 | 5eqg | A | 275 | ALA | p.Ala275Thr | P11166 |
| HOMO SAPIENS | 4 | 2668 | 5eqg | A | 299 | GLU | p.Glu299Lys | P11166 |
| HOMO SAPIENS | 6 | 2668 | 5eqg | A | 314 | GLY | p.Gly314Ser | P11166 |
| HOMO SAPIENS | 7 | 2668 | 5eqg | A | 315 | ILE | p.Ile315Met | P11166 |
| HOMO SAPIENS | -1 | 2668 | 5eqg | A | 458 | ARG | p.Arg458Trp | P11166 |
| HOMO SAPIENS | -1 | 2668 | 5eqg | A | 468 | ARG | p.Arg468Trp | P11166 |
| HOMO SAPIENS | 3 | 2291 | 2amy | A | 31 | PHE | p.Phe31Leu | O15305 |
| HOMO SAPIENS | 7 | 2291 | 2amy | A | 32 | LEU | p.Leu32Arg | O15305 |
| HOMO SAPIENS | 7 | 2291 | 2amy | A | 65 | ASP | p.Asp65Tyr | O15305 |
| HOMO SAPIENS | 1 | 2291 | 2amy | A | 108 | ALA | p.Ala108Val | O15305 |
| HOMO SAPIENS | 3 | 2291 | 2amy | A | 113 | PRO | p.Pro113Leu | O15305 |
| HOMO SAPIENS | 3 | 2291 | 2amy | A | 113 | PRO | p.Pro113Ser | O15305 |
| HOMO SAPIENS | 6 | 2291 | 2amy | A | 117 | GLY | p.Gly117Arg | O15305 |
| HOMO SAPIENS | 6 | 2291 | 2amy | A | 119 | PHE | p.Phe119Leu | O15305 |
| HOMO SAPIENS | 6 | 2291 | 2amy | A | 123 | ARG | p.Arg123Gln | O15305 |
| HOMO SAPIENS | 0 | 2291 | 2amy | A | 129 | VAL | p.Val129Met | O15305 |
| HOMO SAPIENS | 7 | 2291 | 2amy | A | 139 | GLU | p.Glu139Lys | O15305 |
| HOMO SAPIENS | 5 | 2291 | 2amy | A | 141 | ARG | p.Arg141His | O15305 |
| HOMO SAPIENS | 7 | 2291 | 2amy | A | 144 | PHE | p.Phe144Leu | O15305 |
| HOMO SAPIENS | 6 | 2291 | 2amy | A | 148 | ASP | p.Asp148Asn | O15305 |
| HOMO SAPIENS | 6 | 2291 | 2amy | A | 153 | ILE | p.Ile153Thr | O15305 |
| HOMO SAPIENS | 2 | 2291 | 2amy | A | 157 | PHE | p.Phe157Ser | O15305 |
| HOMO SAPIENS | 1 | 2291 | 2amy | A | 162 | ARG | p.Arg162Trp | O15305 |
| HOMO SAPIENS | 6 | 2291 | 2amy | A | 188 | ASP | p.Asp188Gly | O15305 |
| HOMO SAPIENS | 5 | 2291 | 2amy | A | 194 | ARG | p.Arg194Ter | O15305 |
| HOMO SAPIENS | 0 | 2291 | 2amy | A | 207 | PHE | p.Phe207Ser | O15305 |
| HOMO SAPIENS | 5 | 2291 | 2amy | A | 208 | GLY | p.Gly208Ala | O15305 |
| HOMO SAPIENS | 1 | 2291 | 2amy | A | 216 | ASN | p.Asn216Ile | O15305 |
| HOMO SAPIENS | 5 | 2291 | 2amy | A | 218 | HIS | p.His218Leu | O15305 |
| HOMO SAPIENS | 6 | 2291 | 2amy | A | 223 | ASP | p.Asp223Glu | O15305 |
| HOMO SAPIENS | 6 | 2291 | 2amy | A | 226 | THR | p.Thr226Ser | O15305 |
| HOMO SAPIENS | 5 | 2291 | 2amy | A | 231 | VAL | p.Val231Met | O15305 |
| HOMO SAPIENS | 1 | 2291 | 2amy | A | 237 | THR | p.Thr237Met | O15305 |
| HOMO SAPIENS | 1 | 2291 | 2amy | A | 241 | CYS | p.Cys241Ser | O15305 |
| HOMO SAPIENS | 5 | 35907 | 1d6g | A | 21 | GLY | p.Gly21Arg | P32238 |
| HOMO SAPIENS | -1 | 33209 | 1x9d | A | 197 | PRO | p.Pro197Leu | Q9UKM7 |

| | | | | | | | | |
|---|---|---|---|---|---|---|---|---|
| HOMO SAPIENS | -1 | 33209 | 1x9d | A | 201 | PRO | p.Pro201Leu | Q9UKM7 |
| HOMO SAPIENS | 6 | 21305 | 4yep | B | 1326 | ARG | p.Arg1326Ter | P24043 |
| HOMO SAPIENS | 6 | 21305 | 4yep | B | 1350 | ARG | p.Arg1350Ter | P24043 |
| HOMO SAPIENS | 0 | 21553 | 3qyb | A | 1120 | VAL | p.Val1119Ala | O60343 |
| HOMO SAPIENS | 2 | 21553 | 3qyb | A | 1148 | THR | p.Thr1147Met | O60343 |
| HOMO SAPIENS | -1 | 30645 | 3m0c | C | -17 | TRP | p.Trp4Ter | P01130 |
| HOMO SAPIENS | -1 | 30645 | 3m0c | C | -17 | TRP | p.Trp4Ter | P01130 |
| HOMO SAPIENS | -1 | 30645 | 3m0c | C | 12 | GLN | p.Gln33Ter | P01130 |
| HOMO SAPIENS | -1 | 30645 | 3m0c | C | 15 | ASP | p.Asp36Glu | P01130 |
| HOMO SAPIENS | -1 | 30645 | 3m0c | C | 28 | SER | p.Ser49Arg | P01130 |
| HOMO SAPIENS | -1 | 30645 | 3m0c | C | 41 | THR | p.Thr62Met | P01130 |
| HOMO SAPIENS | -1 | 30645 | 3m0c | C | 69 | ASP | p.Asp90Asn | P01130 |
| HOMO SAPIENS | -1 | 30645 | 3m0c | C | 76 | ASN | p.Asn97Lys | P01130 |
| HOMO SAPIENS | -1 | 30645 | 3m0c | C | 78 | SER | p.Ser99Ter | P01130 |
| HOMO SAPIENS | -1 | 30645 | 3m0c | C | 81 | GLN | p.Gln102Ter | P01130 |
| HOMO SAPIENS | -1 | 30645 | 3m0c | C | 84 | PRO | p.Pro105Ser | P01130 |
| HOMO SAPIENS | -1 | 30645 | 3m0c | C | 194 | ARG | p.Arg215Cys | P01130 |
| HOMO SAPIENS | -1 | 30645 | 3m0c | C | 198 | GLY | p.Gly219Asp | P01130 |
| HOMO SAPIENS | -1 | 30645 | 3m0c | C | 201 | CYS | p.Cys222Tyr | P01130 |
| HOMO SAPIENS | -1 | 30645 | 3m0c | C | 201 | CYS | p.Cys222Ter | P01130 |
| HOMO SAPIENS | -1 | 30645 | 3m0c | C | 206 | ASP | p.Asp227Glu | P01130 |
| HOMO SAPIENS | -1 | 30645 | 3m0c | C | 249 | CYS | p.Cys270Ter | P01130 |
| HOMO SAPIENS | -1 | 30645 | 3m0c | C | 250 | VAL | p.Val271Ile | P01130 |
| HOMO SAPIENS | 0 | 30645 | 3m0c | C | 255 | CYS | p.Cys276Tyr | P01130 |
| HOMO SAPIENS | 0 | 30645 | 3m0c | C | 255 | CYS | p.Cys276Ter | P01130 |
| HOMO SAPIENS | 0 | 30645 | 3m0c | C | 256 | GLU | p.Glu277Lys | P01130 |
| HOMO SAPIENS | 0 | 30645 | 3m0c | C | 265 | SER | p.Ser286Arg | P01130 |
| HOMO SAPIENS | 0 | 30645 | 3m0c | C | 284 | TRP | p.Trp305Ter | P01130 |
| HOMO SAPIENS | 0 | 30645 | 3m0c | C | 292 | CYS | p.Cys313Ter | P01130 |
| HOMO SAPIENS | 1 | 30645 | 3m0c | C | 385 | ARG | p.Arg406Trp | P01130 |
| HOMO SAPIENS | 0 | 30645 | 3m0c | C | 395 | ARG | p.Arg416Gln | P01130 |
| HOMO SAPIENS | 0 | 30645 | 3m0c | C | 398 | TYR | p.Tyr419Ter | P01130 |
| HOMO SAPIENS | 1 | 30645 | 3m0c | C | 415 | VAL | p.Val436Ala | P01130 |
| HOMO SAPIENS | 1 | 30645 | 3m0c | C | 424 | ASP | p.Asp445Glu | P01130 |
| HOMO SAPIENS | 1 | 30645 | 3m0c | C | 425 | LEU | p.Leu446Val | P01130 |
| HOMO SAPIENS | -1 | 30645 | 3m0c | C | 440 | GLY | p.Gly461Cys | P01130 |
| HOMO SAPIENS | -1 | 30645 | 3m0c | C | 441 | VAL | p.Val462Ile | P01130 |
| HOMO SAPIENS | 1 | 30645 | 3m0c | C | 447 | VAL | p.Val468Ile | P01130 |
| HOMO SAPIENS | 1 | 30645 | 3m0c | C | 489 | THR | p.Thr510Met | P01130 |
| HOMO SAPIENS | 0 | 30645 | 3m0c | C | 524 | GLY | p.Gly545Glu | P01130 |
| HOMO SAPIENS | 1 | 30645 | 3m0c | C | 615 | GLY | p.Gly636Ser | P01130 |
| HOMO SAPIENS | 1 | 30645 | 3m0c | C | 625 | LEU | p.Leu646Ile | P01130 |
| HOMO SAPIENS | 1 | 30645 | 3m0c | C | 630 | ASP | p.Asp651Gly | P01130 |
| HOMO SAPIENS | 3 | 30645 | 3m0c | C | 646 | CYS | p.Cys667Tyr | P01130 |
| HOMO SAPIENS | 4 | 30645 | 3m0c | C | 654 | GLY | p.Gly675Ser | P01130 |
| HOMO SAPIENS | 6 | 30645 | 3m0c | C | 660 | CYS | p.Cys681Ter | P01130 |
| HOMO SAPIENS | 5 | 30645 | 3m0c | C | 676 | ALA | p.Ala697Pro | P01130 |
| HOMO SAPIENS | -1 | 30645 | 3m0c | C | 749 | GLN | p.Gln770Ter | P01130 |
| HOMO SAPIENS | -1 | 41323 | 4bxo | A | 1812 | PRO | p.Pro1812Ala | Q8IYD8 |
| HOMO SAPIENS | 5 | 8335 | 3h0t | C | 12 | GLY | p.Gly71Asp | P81172 |
| HOMO SAPIENS | 6 | 609 | 1bhg | B | 148 | PRO | p.Pro148Ser | P08236 |
| HOMO SAPIENS | 1 | 609 | 1bhg | B | 176 | LEU | p.Leu176Phe | P08236 |
| HOMO SAPIENS | 6 | 609 | 1bhg | B | 216 | ARG | p.Arg216Trp | P08236 |
| HOMO SAPIENS | 0 | 609 | 1bhg | B | 350 | LYS | p.Lys350Asn | P08236 |

| | | | | | | | | |
|---|---|---|---|---|---|---|---|---|
| HOMO SAPIENS | 1 | 609 | 1bhg | B | 354 | ALA | p.Ala354Val | P08236 |
| HOMO SAPIENS | 1 | 609 | 1bhg | B | 357 | ARG | p.Arg357Ter | P08236 |
| HOMO SAPIENS | 0 | 609 | 1bhg | B | 382 | ARG | p.Arg382Cys | P08236 |
| HOMO SAPIENS | 0 | 609 | 1bhg | B | 495 | TYR | p.Tyr495Cys | P08236 |
| HOMO SAPIENS | 2 | 609 | 1bhg | B | 507 | TRP | p.Trp507Ter | P08236 |
| HOMO SAPIENS | 0 | 609 | 1bhg | B | 577 | ARG | p.Arg577Leu | P08236 |
| HOMO SAPIENS | 1 | 609 | 1bhg | B | 611 | ARG | p.Arg611Trp | P08236 |
| HOMO SAPIENS | 0 | 609 | 1bhg | B | 619 | ALA | p.Ala619Val | P08236 |
| HOMO SAPIENS | 0 | 609 | 1bhg | B | 627 | TRP | p.Trp627Cys | P08236 |
| HOMO SAPIENS | 5 | 3422 | 1x9n | A | 566 | GLU | p.Glu566Lys | P18858 |
| HOMO SAPIENS | 2 | 3422 | 1x9n | A | 771 | ARG | p.Arg771Trp | P18858 |
| HOMO SAPIENS | 0 | 3279 | 5tby | B | 797 | ALA | p.Ala797Thr | P12883 |
| HOMO SAPIENS | -1 | 3279 | 5tby | B | 982 | MET | p.Met982Thr | P12883 |
| HOMO SAPIENS | -1 | 3279 | 5tby | B | 992 | LEU | p.Leu992Met | P12883 |
| HOMO SAPIENS | -1 | 3279 | 5tby | B | 1004 | GLN | p.Gln1004Glu | P12883 |
| HOMO SAPIENS | -1 | 3279 | 5tby | B | 1013 | GLU | p.Glu1013Lys | P12883 |
| HOMO SAPIENS | -1 | 3279 | 5tby | B | 1032 | ASP | p.Asp1032Asn | P12883 |
| HOMO SAPIENS | -1 | 3279 | 5tby | B | 1053 | ARG | p.Arg1053Gln | P12883 |
| HOMO SAPIENS | -1 | 3279 | 5tby | B | 1117 | GLU | p.Glu1117Ter | P12883 |
| HOMO SAPIENS | -1 | 3279 | 5tby | B | 1193 | ARG | p.Arg1193His | P12883 |
| HOMO SAPIENS | -1 | 3279 | 5tby | B | 1205 | GLU | p.Glu1205Lys | P12883 |
| HOMO SAPIENS | -1 | 3279 | 5tby | B | 1207 | ILE | p.Ile1207Met | P12883 |
| HOMO SAPIENS | -1 | 3279 | 5tby | B | 1250 | ARG | p.Arg1250Pro | P12883 |
| HOMO SAPIENS | -1 | 3279 | 5tby | B | 1259 | HIS | p.His1259Gln | P12883 |
| HOMO SAPIENS | -1 | 3279 | 5tby | B | 1272 | ASP | p.Asp1272Asn | P12883 |
| HOMO SAPIENS | -1 | 3279 | 5tby | B | 1295 | GLU | p.Glu1295Lys | P12883 |
| HOMO SAPIENS | -1 | 3279 | 5tby | B | 1335 | SER | p.Ser1335Leu | P12883 |
| HOMO SAPIENS | -1 | 3279 | 5tby | B | 1399 | GLU | p.Glu1399Asp | P12883 |
| HOMO SAPIENS | -1 | 3279 | 5tby | B | 1413 | SER | p.Ser1413Ter | P12883 |
| HOMO SAPIENS | -1 | 3279 | 5tby | B | 1459 | LYS | p.Lys1459Asn | P12883 |
| HOMO SAPIENS | -1 | 3279 | 5tby | B | 1467 | LEU | p.Leu1467Val | P12883 |
| HOMO SAPIENS | -1 | 3279 | 5tby | B | 1491 | SER | p.Ser1491Cys | P12883 |
| HOMO SAPIENS | -1 | 3279 | 5tby | B | 1496 | GLU | p.Glu1496Ala | P12883 |
| HOMO SAPIENS | -1 | 3279 | 5tby | B | 1500 | ARG | p.Arg1500Pro | P12883 |
| HOMO SAPIENS | -1 | 3279 | 5tby | B | 1606 | ARG | p.Arg1606His | P12883 |
| HOMO SAPIENS | -1 | 3279 | 5tby | B | 1664 | ASN | p.Asn1664Lys | P12883 |
| HOMO SAPIENS | -1 | 3279 | 5tby | B | 1712 | ARG | p.Arg1712Trp | P12883 |
| HOMO SAPIENS | -1 | 3279 | 5tby | B | 1769 | LEU | p.Leu1769Met | P12883 |
| HOMO SAPIENS | -1 | 3279 | 5tby | B | 1808 | GLY | p.Gly1808Ser | P12883 |
| HOMO SAPIENS | -1 | 3279 | 5tby | B | 1818 | ARG | p.Arg1818Trp | P12883 |
| HOMO SAPIENS | -1 | 3279 | 5tby | B | 1820 | ARG | p.Arg1820Trp | P12883 |
| HOMO SAPIENS | -1 | 3279 | 5tby | B | 1844 | GLU | p.Glu1844Lys | P12883 |
| HOMO SAPIENS | -1 | 3279 | 5tby | B | 1845 | ARG | p.Arg1845Trp | P12883 |
| HOMO SAPIENS | -1 | 3279 | 5tby | B | 1854 | THR | p.Thr1854Met | P12883 |
| HOMO SAPIENS | -1 | 3279 | 5tby | B | 1909 | ARG | p.Arg1909Trp | P12883 |
| HOMO SAPIENS | 2 | 2260 | 4kl1 | A | 553 | ASP | p.Asp553Asn | Q9Y3Q4 |
| HOMO SAPIENS | 0 | 40570 | 5lst | A | 522 | ARG | p.Arg522Cys | O94761 |
| HOMO SAPIENS | 0 | 40570 | 5lst | A | 522 | ARG | p.Arg522His | O94761 |
| HOMO SAPIENS | 0 | 40570 | 5lst | A | 591 | PRO | p.Pro591Leu | O94761 |
| HOMO SAPIENS | 0 | 40570 | 5lst | A | 793 | PRO | p.Pro793Ser | O94761 |
| HOMO SAPIENS | 4 | 40570 | 5lst | A | 799 | VAL | p.Val799Met | O94761 |
| HOMO SAPIENS | 0 | 40570 | 5lst | A | 1004 | ARG | p.Arg1004Trp | O94761 |
| HOMO SAPIENS | 0 | 40570 | 5lst | A | 1005 | ARG | p.Arg1005Gln | O94761 |
| HOMO SAPIENS | 0 | 40570 | 5lst | A | 1021 | ARG | p.Arg1021Gln | O94761 |

| | | | | | | | | |
|---|---|---|---|---|---|---|---|---|
| HOMO SAPIENS | 0 | 40570 | 5lst | A | 1043 | LEU | p.Leu1043Pro | O94761 |
| HOMO SAPIENS | 0 | 40570 | 5lst | A | 1045 | ALA | p.Ala1045Thr | O94761 |
| HOMO SAPIENS | 0 | 40570 | 5lst | A | 1105 | GLY | p.Gly1105Asp | O94761 |
| HOMO SAPIENS | 0 | 40570 | 5lst | A | 1105 | GLY | p.Gly1105Ser | O94761 |
| HOMO SAPIENS | -1 | 40570 | 5lst | A | 1113 | GLY | p.Gly1113Arg | O94761 |
| HOMO SAPIENS | -1 | 18865 | 1gxc | A | 64 | GLU | p.Glu64Lys | O96017 |
| HOMO SAPIENS | -1 | 11673 | 3dzu | D | 85 | PRO | p.Pro113Gln | P37231 |
| HOMO SAPIENS | 0 | 19383 | 3b96 | A | 63 | LYS | p.Lys103Arg | P49748 |
| HOMO SAPIENS | 4 | 19383 | 3b96 | A | 140 | ALA | p.Ala180Thr | P49748 |
| HOMO SAPIENS | 1 | 19383 | 3b96 | A | 145 | GLY | p.Gly185Ser | P49748 |
| HOMO SAPIENS | 0 | 19383 | 3b96 | A | 220 | THR | p.Thr260Met | P49748 |
| HOMO SAPIENS | 5 | 19383 | 3b96 | A | 243 | VAL | p.Val283Ala | P49748 |
| HOMO SAPIENS | 0 | 19383 | 3b96 | A | 249 | GLY | p.Gly289Arg | P49748 |
| HOMO SAPIENS | 0 | 19383 | 3b96 | A | 254 | GLY | p.Gly294Glu | P49748 |
| HOMO SAPIENS | 7 | 19383 | 3b96 | A | 326 | ARG | p.Arg366Cys | P49748 |
| HOMO SAPIENS | 7 | 19383 | 3b96 | A | 326 | ARG | p.Arg366His | P49748 |
| HOMO SAPIENS | 5 | 19383 | 3b96 | A | 342 | LYS | p.Lys382Gln | P49748 |
| HOMO SAPIENS | 0 | 19383 | 3b96 | A | 369 | THR | p.Thr409Met | P49748 |
| HOMO SAPIENS | 5 | 19383 | 3b96 | A | 376 | ALA | p.Ala416Ser | P49748 |
| HOMO SAPIENS | 0 | 19383 | 3b96 | A | 385 | ALA | p.Ala425Thr | P49748 |
| HOMO SAPIENS | 5 | 19383 | 3b96 | A | 399 | GLY | p.Gly439Asp | P49748 |
| HOMO SAPIENS | 7 | 19383 | 3b96 | A | 401 | GLY | p.Gly441Asp | P49748 |
| HOMO SAPIENS | 9 | 19383 | 3b96 | A | 413 | ARG | p.Arg453Gln | P49748 |
| HOMO SAPIENS | 2 | 19383 | 3b96 | A | 418 | PHE | p.Phe458Leu | P49748 |
| HOMO SAPIENS | 6 | 19383 | 3b96 | A | 419 | ARG | p.Arg459Trp | P49748 |
| HOMO SAPIENS | 6 | 19383 | 3b96 | A | 419 | ARG | p.Arg459Gln | P49748 |
| HOMO SAPIENS | -1 | 19383 | 3b96 | A | 471 | ARG | p.Arg511Gln | P49748 |
| HOMO SAPIENS | 0 | 19383 | 3b96 | A | 573 | ARG | p.Arg613Trp | P49748 |
| HOMO SAPIENS | 0 | 19383 | 3b96 | A | 575 | ARG | p.Arg615Gln | P49748 |
| HOMO SAPIENS | 0 | 22286 | 3k2s | A | 3 | PRO | p.Pro27His | P02647 |
| HOMO SAPIENS | 0 | 22286 | 3k2s | A | 4 | PRO | p.Pro28Arg | P02647 |
| HOMO SAPIENS | 0 | 22286 | 3k2s | A | 10 | ARG | p.Arg34Leu | P02647 |
| HOMO SAPIENS | 0 | 22286 | 3k2s | A | 26 | GLY | p.Gly50Arg | P02647 |
| HOMO SAPIENS | 0 | 22286 | 3k2s | A | 37 | ALA | p.Ala61Thr | P02647 |
| HOMO SAPIENS | 1 | 1622 | 2qrv | D | 631 | ARG | p.Arg635Trp | Q9Y6K1 |
| HOMO SAPIENS | 1 | 1622 | 2qrv | D | 900 | PRO | p.Pro904Leu | Q9Y6K1 |
| HOMO SAPIENS | 6 | 40796 | 1v18 | B | 1509 | LEU | p.Leu1509Val | P25054 |
| HOMO SAPIENS | 1 | 22375 | 4pki | G | 1214 | LYS | p.Asp214Asn | P28289 |
| HOMO SAPIENS | 1 | 16307 | 5mv8 | B | 483 | ALA | p.Ala483Ser | Q9Y6N9 |
| HOMO SAPIENS | 6 | 16307 | 5mv8 | B | 506 | ASN | p.Asn506Lys | Q9Y6N9 |
| HOMO SAPIENS | 4 | 16307 | 5mv8 | B | 519 | ASP | p.Glu519Asp | Q9Y6N9 |
| HOMO SAPIENS | 0 | 16307 | 5mv8 | B | 530 | GLY | p.Gly530Arg | Q9Y6N9 |
| HOMO SAPIENS | 4 | 36221 | 4cvo | A | 127 | VAL | p.Val127Ile | Q03468 |
| HOMO SAPIENS | 7 | 36221 | 4cvo | A | 134 | ARG | p.Arg134Trp | Q03468 |
| HOMO SAPIENS | 0 | 36221 | 4cvo | A | 156 | GLN | p.Gln156Ter | Q03468 |
| HOMO SAPIENS | 0 | 10829 | 4a11 | B | 13 | GLU | p.Glu13Ter | Q13216 |
| HOMO SAPIENS | 1 | 10829 | 4a11 | B | 100 | TYR | p.Tyr100Ter | Q13216 |
| HOMO SAPIENS | 0 | 10829 | 4a11 | B | 160 | ALA | p.Ala160Val | Q13216 |
| HOMO SAPIENS | 1 | 6759 | 5lev | A | 29 | ILE | p.Ile29Phe | Q9H3H5 |
| HOMO SAPIENS | 0 | 6759 | 5lev | A | 108 | MET | p.Met108Ile | Q9H3H5 |
| HOMO SAPIENS | 2 | 6759 | 5lev | A | 117 | VAL | p.Val117Ile | Q9H3H5 |
| HOMO SAPIENS | 1 | 6759 | 5lev | A | 170 | TYR | p.Tyr170Cys | Q9H3H5 |
| HOMO SAPIENS | 0 | 6759 | 5lev | A | 264 | GLY | p.Val264Gly | Q9H3H5 |
| HOMO SAPIENS | 0 | 6759 | 5lev | A | 332 | PHE | p.Phe332Val | Q9H3H5 |

| | | | | | | | | |
|---|---|---|---|---|---|---|---|---|
| HOMO SAPIENS | 7 | 6759 | 5lev | A | 393 | ILE | p.Ile393Val | Q9H3H5 |
| HOMO SAPIENS | -1 | 33473 | 5m5e | C | 9 | ASN | p.Asn31Ser | P31785 |
| HOMO SAPIENS | 0 | 26155 | 1ic8 | B | 98 | ALA | p.Ala98Val | P20823 |
| HOMO SAPIENS | 0 | 15329 | 4iqr | B | 127 | ARG | p.Arg136Trp | P41235 |
| HOMO SAPIENS | 0 | 20725 | 4bsj | A | 350 | GLU | p.Glu350Val | P35916 |
| HOMO SAPIENS | 0 | 20725 | 4bsj | A | 494 | THR | p.Thr494Ala | P35916 |
| HOMO SAPIENS | 0 | 20725 | 4bsj | A | 527 | ASN | p.Asn527Ser | P35916 |
| HOMO SAPIENS | 6 | 29959 | 1xiw | B | 47 | ARG | p.Arg68Ter | P04234 |
| HOMO SAPIENS | 2 | 29959 | 1xiw | B | 72 | CYS | p.Cys93Ter | P04234 |
| HOMO SAPIENS | 3 | 20242 | 5tjc | A | 102 | ARG | p.Arg102Ter | Q9GZU1 |
| HOMO SAPIENS | 3 | 20242 | 5tjc | A | 232 | THR | p.Thr232Pro | Q9GZU1 |
| HOMO SAPIENS | 7 | 9590 | 4ap8 | A | 50 | THR | p.Thr50Ala | O96007 |
| HOMO SAPIENS | 4 | 9590 | 4ap8 | A | 168 | GLU | p.Glu168Lys | O96007 |
| HOMO SAPIENS | -1 | 4297 | 5do7 | B | 19 | ASP | p.Asp19His | Q9H221 |
| HOMO SAPIENS | -1 | 4297 | 5do7 | B | 54 | TYR | p.Tyr54Cys | Q9H221 |
| HOMO SAPIENS | -1 | 4297 | 5do7 | B | 80 | CYS | p.Cys80Tyr | Q9H221 |
| HOMO SAPIENS | 5 | 4297 | 5do7 | B | 107 | SER | p.Ser107Ter | Q9H221 |
| HOMO SAPIENS | 1 | 4297 | 5do7 | B | 164 | ARG | p.Arg164Pro | Q9H221 |
| HOMO SAPIENS | 0 | 4297 | 5do7 | B | 210 | VAL | p.Val210Met | Q9H221 |
| HOMO SAPIENS | 0 | 4297 | 5do7 | B | 231 | PRO | p.Pro231Thr | Q9H221 |
| HOMO SAPIENS | 0 | 4297 | 5do7 | B | 263 | ARG | p.Arg263Gln | Q9H221 |
| HOMO SAPIENS | 0 | 4297 | 5do7 | B | 361 | TRP | p.Trp361Ter | Q9H221 |
| HOMO SAPIENS | 0 | 4297 | 5do7 | B | 400 | THR | p.Thr400Lys | Q9H221 |
| HOMO SAPIENS | 0 | 4297 | 5do7 | B | 549 | ALA | p.Ala549Pro | Q9H221 |
| HOMO SAPIENS | 0 | 4297 | 5do7 | B | 596 | LEU | p.Leu596Arg | Q9H221 |
| HOMO SAPIENS | 0 | 4297 | 5do7 | B | 632 | VAL | p.Val632Ala | Q9H221 |
| HOMO SAPIENS | 0 | 4297 | 5do7 | B | 658 | TYR | p.Tyr658Ter | Q9H221 |
| HOMO SAPIENS | 2 | 9356 | 4igg | B | 206 | GLN | p.Gln206His | P35221 |
| HOMO SAPIENS | 6 | 2179 | 2wk6 | A | 44 | ARG | p.Arg44Gln | P19971 |
| HOMO SAPIENS | 6 | 2179 | 2wk6 | A | 81 | ARG | p.Arg81Gln | P19971 |
| HOMO SAPIENS | 6 | 2179 | 2wk6 | A | 145 | GLY | p.Gly145Arg | P19971 |
| HOMO SAPIENS | 7 | 2179 | 2wk6 | A | 146 | ARG | p.Arg146His | P19971 |
| HOMO SAPIENS | 7 | 2179 | 2wk6 | A | 153 | GLY | p.Gly153Ser | P19971 |
| HOMO SAPIENS | 0 | 2179 | 2wk6 | A | 173 | MET | p.Met173Ile | P19971 |
| HOMO SAPIENS | 6 | 2179 | 2wk6 | A | 202 | ARG | p.Arg202Lys | P19971 |
| HOMO SAPIENS | 6 | 2179 | 2wk6 | A | 208 | VAL | p.Val208Met | P19971 |
| HOMO SAPIENS | 5 | 2179 | 2wk6 | A | 222 | LYS | p.Lys222Arg | P19971 |
| HOMO SAPIENS | 5 | 2179 | 2wk6 | A | 289 | GLU | p.Glu289Ala | P19971 |
| HOMO SAPIENS | 2 | 2179 | 2wk6 | A | 311 | GLY | p.Gly311Arg | P19971 |
| HOMO SAPIENS | 2 | 2179 | 2wk6 | A | 376 | ARG | p.Arg376Pro | P19971 |
| HOMO SAPIENS | 2 | 2179 | 2wk6 | A | 465 | ALA | p.Ala465Thr | P19971 |
| HOMO SAPIENS | -1 | 11856 | 3orh | A | 3 | ALA | p.Ala3Ser | Q14353 |
| HOMO SAPIENS | 1 | 11856 | 3orh | A | 20 | TRP | p.Trp20Ser | Q14353 |
| HOMO SAPIENS | 1 | 11856 | 3orh | A | 50 | MET | p.Met50Leu | Q14353 |
| HOMO SAPIENS | 1 | 11856 | 3orh | A | 110 | VAL | p.Val110Phe | Q14353 |
| HOMO SAPIENS | 8 | 11856 | 3orh | A | 164 | GLY | p.Gly164Asp | Q14353 |
| HOMO SAPIENS | 1 | 11856 | 3orh | A | 169 | CYS | p.Cys169Tyr | Q14353 |
| HOMO SAPIENS | 1 | 11856 | 3orh | A | 174 | TRP | p.Trp174Ter | Q14353 |
| HOMO SAPIENS | 2 | 11856 | 3orh | A | 196 | ALA | p.Ala196Val | Q14353 |
| HOMO SAPIENS | 1 | 11856 | 3orh | A | 209 | THR | p.Thr209Met | Q14353 |
| HOMO SAPIENS | 1 | 11856 | 3orh | A | 219 | ASP | p.Asp219Asn | Q14353 |
| HOMO SAPIENS | 1 | 11856 | 3orh | A | 224 | ALA | p.Ala224Thr | Q14353 |
| HOMO SAPIENS | 1 | 13244 | 2m6i | D | 218 | ARG | p.Arg246Leu | P23415 |
| HOMO SAPIENS | 1 | 13244 | 2m6i | D | 244 | ILE | p.Ile272Asn | P23415 |

| | | | | | | | | |
|---|---|---|---|---|---|---|---|---|
| HOMO SAPIENS | 5 | 13244 | 2m6i | D | 250 | PRO | p.Pro278Thr | P23415 |
| HOMO SAPIENS | 6 | 13244 | 2m6i | D | 266 | GLN | p.Gln294His | P23415 |
| HOMO SAPIENS | 1 | 13244 | 2m6i | D | 271 | ARG | p.Arg299Leu | P23415 |
| HOMO SAPIENS | 1 | 13244 | 2m6i | D | 276 | LYS | p.Lys304Glu | P23415 |
| HOMO SAPIENS | 1 | 13244 | 2m6i | D | 279 | TYR | p.Tyr307Cys | P23415 |
| HOMO SAPIENS | 0 | 13244 | 2m6i | D | 316 | GLY | p.Gly378Ser | P23415 |
| HOMO SAPIENS | 1 | 13244 | 2m6i | D | 400 | ARG | p.Arg428His | P23415 |
| HOMO SAPIENS | 3 | 1967 | 2i6b | A | 301 | ALA | p.Ala318Glu | P55263 |
| HOMO SAPIENS | 4 | 11096 | 1h8b | A | 61 | ALA | p.Ala882Thr | P35609 |
| HOMO SAPIENS | 0 | 15423 | 5ngz | A | 2 | GLN | p.Gln2Glu | Q9NPD8 |
| HOMO SAPIENS | 3 | 30475 | 3lpw | A | 21 | SER | p.Ser22894Tyr | Q8WZ42 |
| HOMO SAPIENS | 6 | 32687 | 2arf | A | 1041 | ARG | p.Arg1041Trp | P35670 |
| HOMO SAPIENS | 8 | 32687 | 2arf | A | 1061 | GLY | p.Gly1061Glu | P35670 |
| HOMO SAPIENS | 6 | 32687 | 2arf | A | 1064 | GLU | p.Glu1064Ala | P35670 |
| HOMO SAPIENS | 2 | 32687 | 2arf | A | 1069 | HIS | p.His1069Gln | P35670 |
| HOMO SAPIENS | 2 | 32687 | 2arf | A | 1088 | LEU | p.Leu1088Ter | P35670 |
| HOMO SAPIENS | 0 | 32687 | 2arf | A | 1099 | GLY | p.Gly1099Ser | P35670 |
| HOMO SAPIENS | 4 | 32687 | 2arf | A | 1106 | VAL | p.Val1106Asp | P35670 |
| HOMO SAPIENS | 2 | 32687 | 2arf | A | 1142 | GLN | p.Gln1142His | P35670 |
| HOMO SAPIENS | 8 | 32687 | 2arf | A | 1148 | ILE | p.Ile1148Thr | P35670 |
| HOMO SAPIENS | 6 | 32687 | 2arf | A | 1151 | ARG | p.Arg1151Cys | P35670 |
| HOMO SAPIENS | 4 | 32687 | 2arf | A | 1169 | MET | p.Met1169Val | P35670 |
| HOMO SAPIENS | 4 | 32687 | 2arf | A | 1173 | GLU | p.Glu1173Lys | P35670 |
| HOMO SAPIENS | 2 | 32687 | 2arf | A | 1176 | GLY | p.Gly1176Arg | P35670 |
| HOMO SAPIENS | 3 | 15544 | 5uq3 | C | 575 | PRO | p.Pro69Leu | P46527 |
| HOMO SAPIENS | 6 | 15544 | 5uq3 | C | 592 | GLU | p.Glu86Asp | P46527 |
| HOMO SAPIENS | 6 | 15544 | 5uq3 | C | 599 | ARG | p.Arg93Trp | P46527 |
| HOMO SAPIENS | -1 | 15544 | 5uq3 | C | 615 | VAL | p.Val109Gly | P46527 |
| HOMO SAPIENS | -1 | 15544 | 5uq3 | C | 631 | SER | p.Ser125Pro | P46527 |
| HOMO SAPIENS | -1 | 15544 | 5uq3 | C | 639 | PRO | p.Pro133Ser | P46527 |
| HOMO SAPIENS | 2 | 2703 | 4wxq | A | 245 | CYS | p.Cys245Tyr | Q99972 |
| HOMO SAPIENS | 0 | 2703 | 4wxq | A | 252 | GLY | p.Gly252Arg | Q99972 |
| HOMO SAPIENS | 1 | 2703 | 4wxq | A | 337 | GLN | p.Gln337Arg | Q99972 |
| HOMO SAPIENS | 5 | 2703 | 4wxq | A | 347 | TYR | p.Tyr347Ter | Q99972 |
| HOMO SAPIENS | 3 | 2703 | 4wxq | A | 364 | GLY | p.Gly364Val | Q99972 |
| HOMO SAPIENS | 2 | 2703 | 4wxq | A | 367 | GLY | p.Gly367Arg | Q99972 |
| HOMO SAPIENS | 2 | 2703 | 4wxq | A | 368 | GLN | p.Gln368Ter | Q99972 |
| HOMO SAPIENS | 6 | 2703 | 4wxq | A | 370 | PRO | p.Pro370Leu | Q99972 |
| HOMO SAPIENS | 1 | 2703 | 4wxq | A | 380 | ASP | p.Asp380His | Q99972 |
| HOMO SAPIENS | 8 | 2703 | 4wxq | A | 399 | GLY | p.Gly399Val | Q99972 |
| HOMO SAPIENS | 0 | 2703 | 4wxq | A | 423 | LYS | p.Lys423Glu | Q99972 |
| HOMO SAPIENS | 3 | 2703 | 4wxq | A | 433 | CYS | p.Cys433Arg | Q99972 |
| HOMO SAPIENS | 3 | 2703 | 4wxq | A | 437 | TYR | p.Tyr437His | Q99972 |
| HOMO SAPIENS | 2 | 2703 | 4wxq | A | 477 | ILE | p.Ile477Ser | Q99972 |
| HOMO SAPIENS | 3 | 2703 | 4wxq | A | 480 | ASN | p.Asn480Lys | Q99972 |
| HOMO SAPIENS | -1 | 34100 | 2fot | C | 1171 | GLN | p.Gln1173Pro | Q13813 |
| HOMO SAPIENS | 0 | 8773 | 1kw2 | A | 432 | ASP | p.Asp432Glu | P02774 |
| HOMO SAPIENS | 0 | 8773 | 1kw2 | A | 436 | LYS | p.Lys436Thr | P02774 |
| HOMO SAPIENS | 0 | 8402 | 4dzo | B | 649 | TYR | p.Tyr649Ter | Q9Y6D9 |
| HOMO SAPIENS | 0 | 39962 | 1nst | A | 611 | GLY | p.Gly611Ser | P52848 |
| HOMO SAPIENS | 2 | 22578 | 3dof | B | 45 | MET | p.Met45Arg | Q9Y2Y0 |
| HOMO SAPIENS | 1 | 23290 | 5j1s | A | 292 | ILE | p.Ile292Thr | O14656 |
| HOMO SAPIENS | 4 | 9505 | 4j1y | A | 534 | ARG | p.Arg534Trp | P09871 |
| HOMO SAPIENS | 9 | 260 | 2h57 | A | 31 | THR | p.Thr31Met | Q9H0F7 |

| | | | | | | | | |
|---|---|---|---|---|---|---|---|---|
| HOMO SAPIENS | 0 | 260 | 2h57 | A | 91 | ILE | p.Ile91Thr | Q9H0F7 |
| HOMO SAPIENS | 0 | 260 | 2h57 | A | 121 | ARG | p.Arg121His | Q9H0F7 |
| HOMO SAPIENS | 0 | 260 | 2h57 | A | 122 | ARG | p.Arg122Ter | Q9H0F7 |
| HOMO SAPIENS | 2 | 260 | 2h57 | A | 169 | GLY | p.Gly169Ala | Q9H0F7 |
| HOMO SAPIENS | 4 | 260 | 2h57 | A | 170 | LEU | p.Leu170Trp | Q9H0F7 |
| HOMO SAPIENS | 4 | 34023 | 2cr3 | A | 18 | GLY | p.Gly48Ser | P11362 |
| HOMO SAPIENS | 0 | 37667 | 3ii7 | A | 420 | ARG | p.Arg420Cys | Q8IXQ5 |
| HOMO SAPIENS | 0 | 41720 | 2k9u | A | 68 | ARG | p.Arg2364His | Q14315 |
| HOMO SAPIENS | 1 | 29250 | 5fmv | A | 568 | HIS | p.His568Gln | P08575 |
| HOMO SAPIENS | 1 | 1280 | 1fb1 | A | 69 | PRO | p.Pro69Leu | P30793 |
| HOMO SAPIENS | 3 | 1280 | 1fb1 | A | 88 | ARG | p.Arg88Trp | P30793 |
| HOMO SAPIENS | 2 | 1280 | 1fb1 | A | 108 | GLY | p.Gly108Asp | P30793 |
| HOMO SAPIENS | 4 | 1280 | 1fb1 | A | 134 | ASP | p.Asp134Val | P30793 |
| HOMO SAPIENS | 6 | 1280 | 1fb1 | A | 135 | ILE | p.Ile135Lys | P30793 |
| HOMO SAPIENS | 9 | 1280 | 1fb1 | A | 144 | HIS | p.His144Pro | P30793 |
| HOMO SAPIENS | 9 | 1280 | 1fb1 | A | 184 | ARG | p.Arg184His | P30793 |
| HOMO SAPIENS | 1 | 1280 | 1fb1 | A | 196 | ALA | p.Ala196Ser | P30793 |
| HOMO SAPIENS | 3 | 1280 | 1fb1 | A | 201 | GLY | p.Gly201Glu | P30793 |
| HOMO SAPIENS | 2 | 1280 | 1fb1 | A | 204 | VAL | p.Val204Ile | P30793 |
| HOMO SAPIENS | 2 | 1280 | 1fb1 | A | 211 | MET | p.Met211Ile | P30793 |
| HOMO SAPIENS | 0 | 1280 | 1fb1 | A | 221 | MET | p.Met221Thr | P30793 |
| HOMO SAPIENS | 3 | 1280 | 1fb1 | A | 224 | LYS | p.Lys224Arg | P30793 |
| HOMO SAPIENS | 2 | 1280 | 1fb1 | A | 249 | ARG | p.Arg249Ser | P30793 |
| HOMO SAPIENS | 8 | 41352 | 4k12 | A | 47 | ILE | p.Ile551Thr | P08603 |
| HOMO SAPIENS | 9 | 20703 | 5nir | A | 57 | CYS | p.Cys57Tyr | P02458 |
| HOMO SAPIENS | 9 | 20703 | 5nir | A | 64 | CYS | p.Cys64Ter | P02458 |
| HOMO SAPIENS | 0 | 20087 | 4cmn | A | 299 | MET | p.Met299Ile | Q01968 |
| HOMO SAPIENS | 1 | 20819 | 3aaf | A | 979 | ILE | p.Ile979Met | Q14191 |
| HOMO SAPIENS | 1 | 20819 | 3aaf | A | 984 | LEU | p.Leu984Ile | Q14191 |
| HOMO SAPIENS | 3 | 20819 | 3aaf | A | 987 | ARG | p.Arg987Ter | Q14191 |
| HOMO SAPIENS | -1 | 20819 | 3aaf | A | 1074 | PHE | p.Leu1074Phe | Q14191 |
| HOMO SAPIENS | 0 | 3619 | 3dem | A | 206 | ASN | p.Asn225Ser | P48740 |
| HOMO SAPIENS | 1 | 3619 | 3dem | A | 269 | ARG | p.Arg288Leu | P48740 |
| HOMO SAPIENS | 0 | 3105 | 3ooi | A | 164 | TYR | p.Tyr2015His | Q96L73 |
| HOMO SAPIENS | 6 | 15828 | 2lkx | A | 16 | LEU | p.Leu100Gln | Q99697 |
| HOMO SAPIENS | 8 | 15828 | 2lkx | A | 24 | ARG | p.Arg108His | Q99697 |
| HOMO SAPIENS | 2 | 15828 | 2lkx | A | 30 | THR | p.Thr114Pro | Q99697 |
| HOMO SAPIENS | 8 | 15828 | 2lkx | A | 45 | VAL | p.Val129Leu | Q99697 |
| HOMO SAPIENS | 8 | 15828 | 2lkx | A | 46 | ARG | p.Arg130Trp | Q99697 |
| HOMO SAPIENS | 9 | 15828 | 2lkx | A | 53 | ARG | p.Arg137Pro | Q99697 |
| HOMO SAPIENS | 9 | 13017 | 4irq | A | 141 | ARG | p.Arg141Trp | Q9UBV7 |
| HOMO SAPIENS | 0 | 13017 | 4irq | A | 186 | ALA | p.Ala186Asp | Q9UBV7 |
| HOMO SAPIENS | 0 | 13017 | 4irq | A | 206 | LEU | p.Leu206Pro | Q9UBV7 |
| HOMO SAPIENS | 1 | 13017 | 4irq | A | 214 | CYS | p.Cys214Tyr | Q9UBV7 |
| HOMO SAPIENS | 0 | 13017 | 4irq | A | 270 | ARG | p.Arg270Cys | Q9UBV7 |
| HOMO SAPIENS | -1 | 13053 | 4mqw | X | 307 | ALA | p.Ala307Thr | P23945 |
| HOMO SAPIENS | 0 | 19406 | 4b3g | A | 43 | ARG | p.Arg43Ter | P38935 |
| HOMO SAPIENS | 0 | 19406 | 4b3g | A | 46 | CYS | p.Cys46Ter | P38935 |
| HOMO SAPIENS | 0 | 19406 | 4b3g | A | 75 | ALA | p.Ala75Thr | P38935 |
| HOMO SAPIENS | 0 | 19406 | 4b3g | A | 201 | LEU | p.Leu201Ser | P38935 |
| HOMO SAPIENS | 3 | 19406 | 4b3g | A | 213 | HIS | p.His213Arg | P38935 |
| HOMO SAPIENS | 0 | 19406 | 4b3g | A | 275 | ILE | p.Ile275Val | P38935 |
| HOMO SAPIENS | -1 | 19406 | 4b3g | A | 320 | ARG | p.Arg320Ter | P38935 |
| HOMO SAPIENS | 0 | 19406 | 4b3g | A | 361 | LEU | p.Leu361Pro | P38935 |

| | | | | | | | | |
|---|---|---|---|---|---|---|---|---|
| HOMO SAPIENS | 0 | 19406 | 4b3g | A | 369 | PHE | p.Phe369Leu | P38935 |
| HOMO SAPIENS | 0 | 19406 | 4b3g | A | 493 | THR | p.Thr493Ile | P38935 |
| HOMO SAPIENS | 0 | 19406 | 4b3g | A | 496 | CYS | p.Cys496Ter | P38935 |
| HOMO SAPIENS | 0 | 19406 | 4b3g | A | 514 | GLU | p.Glu514Lys | P38935 |
| HOMO SAPIENS | 0 | 19406 | 4b3g | A | 528 | ALA | p.Ala528Thr | P38935 |
| HOMO SAPIENS | 0 | 19406 | 4b3g | A | 531 | PRO | p.Pro531Thr | P38935 |
| HOMO SAPIENS | 0 | 19406 | 4b3g | A | 580 | VAL | p.Val580Ile | P38935 |
| HOMO SAPIENS | 6 | 19406 | 4b3g | A | 603 | ARG | p.Arg603His | P38935 |
| HOMO SAPIENS | 0 | 21501 | 5jde | A | 40 | SER | p.Ser2831Asn | Q8WZ42 |
| HOMO SAPIENS | 5 | 21501 | 5jde | A | 189 | ALA | p.Ala2980Thr | Q8WZ42 |
| HOMO SAPIENS | 6 | 15008 | 3mop | H | 12 | ARG | p.Arg12Cys | Q9NWZ3 |
| HOMO SAPIENS | 8 | 17455 | 3jtc | C | 15 | ARG | p.Arg57Trp | P04070 |
| HOMO SAPIENS | -1 | 17455 | 3jtc | C | 20 | CGU | p.Glu62Ala | P04070 |
| HOMO SAPIENS | 0 | 27795 | 5gox | B | 625 | TYR | p.Tyr625Ter | Q92878 |
| HOMO SAPIENS | 0 | 27795 | 5gox | B | 653 | SER | p.Ser653Ter | Q92878 |
| HOMO SAPIENS | -1 | 6558 | 5nug | A | 137 | VAL | p.Val137Ile | Q14204 |
| HOMO SAPIENS | -1 | 6558 | 5nug | A | 567 | ARG | p.Arg567His | Q14204 |
| HOMO SAPIENS | -1 | 6558 | 5nug | A | 1140 | THR | p.Thr1140Met | Q14204 |
| HOMO SAPIENS | -1 | 6558 | 5nug | A | 1991 | ASP | p.Asp1991Asn | Q14204 |
| HOMO SAPIENS | -1 | 6558 | 5nug | A | 2401 | LYS | p.Lys2401Asn | Q14204 |
| HOMO SAPIENS | 0 | 6558 | 5nug | A | 2488 | ARG | p.Arg2488His | Q14204 |
| HOMO SAPIENS | 0 | 6558 | 5nug | A | 3907 | HIS | p.His3907Gln | Q14204 |
| HOMO SAPIENS | 0 | 6558 | 5nug | A | 3981 | THR | p.Thr3981Arg | Q14204 |
| HOMO SAPIENS | 0 | 6558 | 5nug | A | 4394 | THR | p.Thr4394Met | Q14204 |
| HOMO SAPIENS | 0 | 6558 | 5nug | A | 4573 | ASN | p.Asn4573Ser | Q14204 |
| HOMO SAPIENS | 0 | 6558 | 5nug | A | 4603 | SER | p.Ser4603Ile | Q14204 |
| HOMO SAPIENS | 1 | 4244 | 1umk | A | 57 | ARG | p.Arg58Gln | P00387 |
| HOMO SAPIENS | 1 | 4244 | 1umk | A | 72 | LEU | p.Leu73Pro | P00387 |
| HOMO SAPIENS | 1 | 4244 | 1umk | A | 105 | VAL | p.Val106Met | P00387 |
| HOMO SAPIENS | 0 | 4244 | 1umk | A | 116 | THR | p.Thr117Ser | P00387 |
| HOMO SAPIENS | 8 | 4244 | 1umk | A | 127 | SER | p.Ser128Pro | P00387 |
| HOMO SAPIENS | 0 | 4244 | 1umk | A | 148 | LEU | p.Leu149Pro | P00387 |
| HOMO SAPIENS | 0 | 4244 | 1umk | A | 159 | ARG | p.Arg160Ter | P00387 |
| HOMO SAPIENS | 0 | 4244 | 1umk | A | 203 | CYS | p.Cys204Arg | P00387 |
| HOMO SAPIENS | 0 | 4244 | 1umk | A | 203 | CYS | p.Cys204Tyr | P00387 |
| HOMO SAPIENS | 2 | 4244 | 1umk | A | 239 | ASP | p.Asp240Gly | P00387 |
| HOMO SAPIENS | 0 | 4244 | 1umk | A | 291 | GLY | p.Gly292Asp | P00387 |
| HOMO SAPIENS | 4 | 34495 | 1s35 | A | 1151 | ASP | p.Asn1151Asp | P11277 |
| HOMO SAPIENS | 3 | 30110 | 3zqs | A | 224 | THR | p.Thr224Ala | Q9NW38 |
| HOMO SAPIENS | 3 | 17588 | 5dse | B | 53 | LEU | p.Leu53Pro | Q9BYI3 |
| HOMO SAPIENS | 0 | 1564 | 4yqf | B | 72 | HIS | p.His347Gln | Q9UHD8 |
| HOMO SAPIENS | 0 | 17673 | 4jnw | A | 32 | ARG | p.Arg32186His | Q8WZ42 |
| HOMO SAPIENS | 3 | 17673 | 4jnw | A | 140 | ILE | p.Ile32294Val | Q8WZ42 |
| HOMO SAPIENS | 5 | 17673 | 4jnw | A | 269 | ALA | p.Ala32423Thr | Q8WZ42 |
| HOMO SAPIENS | 3 | 12858 | 3s97 | C | 302 | LEU | p.Leu302Pro | Q12860 |
| HOMO SAPIENS | 0 | 6147 | 4nn0 | A | 114 | ARG | p.Arg114Gln | Q9BXJ0 |
| HOMO SAPIENS | 0 | 7549 | 1tr2 | A | 277 | LEU | p.Leu277Met | P18206 |
| HOMO SAPIENS | 1 | 7549 | 1tr2 | A | 413 | ALA | p.Ala413Thr | P18206 |
| HOMO SAPIENS | 1 | 7549 | 1tr2 | A | 519 | ILE | p.Ile519Leu | P18206 |
| HOMO SAPIENS | 5 | 7549 | 1tr2 | A | 547 | ARG | p.Arg547Ter | P18206 |
| HOMO SAPIENS | 2 | 7549 | 1tr2 | A | 636 | HIS | p.His636Arg | P18206 |
| HOMO SAPIENS | 2 | 7549 | 1tr2 | A | 824 | ILE | p.Ile824Met | P18206 |
| HOMO SAPIENS | 2 | 13900 | 3gro | A | 45 | CYS | p.Cys45Tyr | P50897 |
| HOMO SAPIENS | 2 | 13900 | 3gro | A | 75 | THR | p.Thr75Pro | P50897 |

| | | | | | | | | |
|---|---|---|---|---|---|---|---|---|
| HOMO SAPIENS | 2 | 13900 | 3gro | A | 79 | ASP | p.Asp79Gly | P50897 |
| HOMO SAPIENS | 5 | 13900 | 3gro | A | 91 | GLN | p.Gln91Pro | P50897 |
| HOMO SAPIENS | 5 | 13900 | 3gro | A | 96 | CYS | p.Cys96Tyr | P50897 |
| HOMO SAPIENS | 0 | 13900 | 3gro | A | 104 | LYS | p.Lys104Ter | P50897 |
| HOMO SAPIENS | 2 | 13900 | 3gro | A | 108 | GLY | p.Gly108Arg | P50897 |
| HOMO SAPIENS | 2 | 13900 | 3gro | A | 109 | TYR | p.Tyr109Asp | P50897 |
| HOMO SAPIENS | 0 | 13900 | 3gro | A | 122 | ARG | p.Arg122Trp | P50897 |
| HOMO SAPIENS | 0 | 13900 | 3gro | A | 138 | SER | p.Ser138Leu | P50897 |
| HOMO SAPIENS | 5 | 13900 | 3gro | A | 151 | ARG | p.Arg151Ter | P50897 |
| HOMO SAPIENS | 7 | 13900 | 3gro | A | 164 | ARG | p.Arg164Ter | P50897 |
| HOMO SAPIENS | 0 | 13900 | 3gro | A | 177 | GLN | p.Gln177Glu | P50897 |
| HOMO SAPIENS | 0 | 13900 | 3gro | A | 181 | VAL | p.Val181Leu | P50897 |
| HOMO SAPIENS | 0 | 13900 | 3gro | A | 184 | GLU | p.Glu184Lys | P50897 |
| HOMO SAPIENS | 5 | 13900 | 3gro | A | 186 | TRP | p.Trp186Ter | P50897 |
| HOMO SAPIENS | 0 | 13900 | 3gro | A | 219 | LEU | p.Leu219Gln | P50897 |
| HOMO SAPIENS | 0 | 13900 | 3gro | A | 222 | LEU | p.Leu222Pro | P50897 |
| HOMO SAPIENS | 5 | 13900 | 3gro | A | 296 | TRP | p.Trp296Ter | P50897 |
| HOMO SAPIENS | 6 | 35028 | 5mwb | A | 444 | CYS | p.Cys444Tyr | Q04721 |
| HOMO SAPIENS | 0 | 35028 | 5mwb | A | 514 | ASN | p.Asn514Ser | Q04721 |
| HOMO SAPIENS | 5 | 1483 | 4gl2 | A | 822 | ARG | p.Arg822Gln | Q9BYX4 |
| HOMO SAPIENS | 0 | 1483 | 4gl2 | A | 843 | ARG | p.His843Arg | Q9BYX4 |
| HOMO SAPIENS | -1 | 28838 | 3cl3 | D | 153 | LEU | p.Leu153Arg | Q9Y6K9 |
| HOMO SAPIENS | 2 | 22563 | 4zel | B | 79 | ARG | p.Arg79Trp | P09172 |
| HOMO SAPIENS | 2 | 22563 | 4zel | B | 101 | VAL | p.Val101Met | P09172 |
| HOMO SAPIENS | 2 | 22563 | 4zel | B | 114 | ASP | p.Asp114Glu | P09172 |
| HOMO SAPIENS | 0 | 22563 | 4zel | B | 201 | ASN | p.Asn201Ser | P09172 |
| HOMO SAPIENS | 2 | 22563 | 4zel | B | 211 | ALA | p.Ala211Thr | P09172 |
| HOMO SAPIENS | -1 | 22563 | 4zel | B | 290 | ASP | p.Asp290Asn | P09172 |
| HOMO SAPIENS | 2 | 22563 | 4zel | B | 318 | ALA | p.Ala318Ser | P09172 |
| HOMO SAPIENS | 2 | 22563 | 4zel | B | 345 | ASP | p.Asp345Asn | P09172 |
| HOMO SAPIENS | 2 | 22563 | 4zel | B | 362 | ALA | p.Ala362Glu | P09172 |
| HOMO SAPIENS | 2 | 22563 | 4zel | B | 549 | ARG | p.Arg549Cys | P09172 |
| HOMO SAPIENS | 9 | 9780 | 1y6k | R | 36 | TYR | p.Tyr57Cys | Q13651 |
| HOMO SAPIENS | 1 | 9780 | 1y6k | R | 40 | LEU | p.Leu61Val | Q13651 |
| HOMO SAPIENS | 1 | 9780 | 1y6k | R | 63 | THR | p.Thr84Ile | Q13651 |
| HOMO SAPIENS | 9 | 9780 | 1y6k | R | 80 | ARG | p.Arg101Trp | Q13651 |
| HOMO SAPIENS | 0 | 9780 | 1y6k | R | 120 | GLY | p.Gly141Arg | Q13651 |
| HOMO SAPIENS | 0 | 9780 | 1y6k | R | 138 | SER | p.Ser159Gly | Q13651 |
| HOMO SAPIENS | 0 | 9780 | 1y6k | R | 203 | ILE | p.Ile224Val | Q13651 |
| HOMO SAPIENS | 0 | 16019 | 3rep | A | 211 | ARG | p.Arg211Cys | Q13418 |
| HOMO SAPIENS | 3 | 7879 | 5ec3 | A | 33 | ALA | p.Ala33Thr | P32754 |
| HOMO SAPIENS | 4 | 7879 | 5ec3 | A | 113 | ARG | p.Arg113Gln | P32754 |
| HOMO SAPIENS | 3 | 7879 | 5ec3 | A | 340 | VAL | p.Val340Leu | P32754 |
| HOMO SAPIENS | 0 | 8904 | 1qk9 | A | 6 | LYS | p.Lys82Arg | P51608 |
| HOMO SAPIENS | 4 | 8904 | 1qk9 | A | 24 | LEU | p.Leu100Val | P51608 |
| HOMO SAPIENS | 7 | 8904 | 1qk9 | A | 30 | ARG | p.Arg106Trp | P51608 |
| HOMO SAPIENS | 6 | 8904 | 1qk9 | A | 57 | ARG | p.Arg133Cys | P51608 |
| HOMO SAPIENS | 3 | 8904 | 1qk9 | A | 61 | GLU | p.Glu137Gly | P51608 |
| HOMO SAPIENS | 2 | 8904 | 1qk9 | A | 64 | ALA | p.Ala140Val | P51608 |
| HOMO SAPIENS | 5 | 8904 | 1qk9 | A | 79 | PHE | p.Phe155Ser | P51608 |
| HOMO SAPIENS | 3 | 8904 | 1qk9 | A | 82 | THR | p.Thr158Met | P51608 |
| HOMO SAPIENS | 1 | 23121 | 3e9k | A | 198 | THR | p.Thr198Ala | Q16719 |
| HOMO SAPIENS | -1 | 26090 | 3tje | F | 19 | LEU | p.Leu35Met | P25445 |
| HOMO SAPIENS | 8 | 26090 | 3tje | F | 106 | THR | p.Thr122Ile | P25445 |

| | | | | | | | | |
|---|---|---|---|---|---|---|---|---|
| HOMO SAPIENS | 2 | 36404 | 5cqr | A | 765 | GLY | p.Gly765Glu | O95163 |
| HOMO SAPIENS | 2 | 36404 | 5cqr | A | 816 | ILE | p.Ile816Leu | O95163 |
| HOMO SAPIENS | 2 | 36404 | 5cqr | A | 830 | ILE | p.Ile830Met | O95163 |
| HOMO SAPIENS | 5 | 36404 | 5cqr | A | 939 | TYR | p.Tyr939Ter | O95163 |
| HOMO SAPIENS | 2 | 36404 | 5cqr | A | 952 | LYS | p.Lys952Ile | O95163 |
| HOMO SAPIENS | 5 | 36404 | 5cqr | A | 1013 | GLY | p.Gly1013Ser | O95163 |
| HOMO SAPIENS | 5 | 36404 | 5cqr | A | 1072 | CYS | p.Cys1072Ser | O95163 |
| HOMO SAPIENS | -1 | 36404 | 5cqr | A | 1158 | PRO | p.Pro1158Leu | O95163 |
| HOMO SAPIENS | 1 | 34149 | 2k40 | A | 18 | ASN | p.Asn125Ser | Q9UBX0 |
| HOMO SAPIENS | 0 | 34149 | 2k40 | A | 42 | LEU | p.Glu149Lys | Q9UBX0 |
| HOMO SAPIENS | 9 | 34149 | 2k40 | A | 53 | ARG | p.Arg160Cys | Q9UBX0 |
| HOMO SAPIENS | 0 | 34149 | 2k40 | A | 63 | SER | p.Ser170Leu | Q9UBX0 |
| HOMO SAPIENS | 2 | 2468 | 3vp6 | A | 484 | LEU | p.Leu484Met | Q99259 |
| HOMO SAPIENS | 0 | 2468 | 3vp6 | A | 532 | ARG | p.Arg532Gln | Q99259 |
| HOMO SAPIENS | 5 | 1253 | 5xtd | S | 8 | GLY | p.Gly8Arg | O15239 |
| HOMO SAPIENS | 7 | 1253 | 5xtd | S | 32 | GLY | p.Gly32Arg | O15239 |
| HOMO SAPIENS | 7 | 1253 | 5xtd | S | 37 | ARG | p.Arg37Ser | O15239 |
| HOMO SAPIENS | -1 | 13292 | 3u1k | A | 121 | VAL | p.Ile121Val | Q8TCS8 |
| HOMO SAPIENS | 2 | 13292 | 3u1k | A | 165 | PRO | p.Pro165Ser | Q8TCS8 |
| HOMO SAPIENS | 2 | 13292 | 3u1k | A | 192 | ARG | p.Arg192Gln | Q8TCS8 |
| HOMO SAPIENS | 1 | 13292 | 3u1k | A | 315 | LYS | p.Lys315Arg | Q8TCS8 |
| HOMO SAPIENS | 4 | 13292 | 3u1k | A | 509 | VAL | p.Val509Ile | Q8TCS8 |
| HOMO SAPIENS | 7 | 13292 | 3u1k | A | 531 | THR | p.Thr531Arg | Q8TCS8 |
| HOMO SAPIENS | 0 | 13292 | 3u1k | A | 590 | ASN | p.Asn590Asp | Q8TCS8 |
| HOMO SAPIENS | 0 | 27048 | 4p5w | B | 184 | ARG | p.Arg184Ter | Q9UJS0 |
| HOMO SAPIENS | 0 | 27048 | 4p5w | B | 225 | SER | p.Ser225Ter | Q9UJS0 |
| HOMO SAPIENS | -1 | 27048 | 4p5w | B | 631 | ALA | p.Arg360Ter | Q9UJS0 |
| HOMO SAPIENS | 0 | 24240 | 4mdr | A | 318 | ARG | p.Arg318Ter | O00189 |
| HOMO SAPIENS | 0 | 24240 | 4mdr | A | 338 | ARG | p.Arg338Ter | O00189 |
| HOMO SAPIENS | 0 | 13429 | 4cg4 | A | 479 | PHE | p.Phe479Leu | O15553 |
| HOMO SAPIENS | 0 | 13429 | 4cg4 | A | 487 | VAL | p.Val487Leu | O15553 |
| HOMO SAPIENS | 3 | 13429 | 4cg4 | A | 653 | ARG | p.Arg653His | O15553 |
| HOMO SAPIENS | 1 | 13429 | 4cg4 | A | 680 | MET | p.Met680Ile | O15553 |
| HOMO SAPIENS | 1 | 13429 | 4cg4 | A | 688 | TYR | p.Tyr688Ter | O15553 |
| HOMO SAPIENS | 0 | 13429 | 4cg4 | A | 694 | MET | p.Met694Ile | O15553 |
| HOMO SAPIENS | 0 | 13429 | 4cg4 | A | 694 | MET | p.Met694Val | O15553 |
| HOMO SAPIENS | 1 | 13429 | 4cg4 | A | 726 | VAL | p.Val726Ala | O15553 |
| HOMO SAPIENS | 0 | 13429 | 4cg4 | A | 761 | ARG | p.Arg761His | O15553 |
| HOMO SAPIENS | 0 | 2079 | 4okh | A | 691 | GLY | p.Gly691Arg | P20807 |
| HOMO SAPIENS | 2 | 2079 | 4okh | A | 748 | ARG | p.Arg748Ter | P20807 |
| HOMO SAPIENS | 0 | 2079 | 4okh | A | 763 | TYR | p.Tyr763Cys | P20807 |
| HOMO SAPIENS | 1 | 2079 | 4okh | A | 769 | ARG | p.Arg769Gln | P20807 |
| HOMO SAPIENS | 9 | 2079 | 4okh | A | 777 | ILE | p.Ile777Val | P20807 |
| HOMO SAPIENS | 8 | 2079 | 4okh | A | 778 | ASP | p.Asp778His | P20807 |
| HOMO SAPIENS | 5 | 2079 | 4okh | A | 780 | ASP | p.Asp780His | P20807 |
| HOMO SAPIENS | 1 | 1200 | 2gi7 | A | 142 | ARG | p.Arg162Trp | Q9HCN6 |
| HOMO SAPIENS | 4 | 9199 | 3ezq | C | 260 | ASP | p.Asp260Val | P25445 |
| HOMO SAPIENS | 0 | 36522 | 5ms9 | A | 122 | ARG | p.Arg122Cys | P35555 |
| HOMO SAPIENS | 0 | 36522 | 5ms9 | A | 240 | ARG | p.Arg240Cys | P35555 |
| HOMO SAPIENS | 0 | 17448 | 3g2f | A | 225 | ARG | p.Arg225His | Q13873 |
| HOMO SAPIENS | 0 | 17448 | 3g2f | A | 347 | CYS | p.Cys347Tyr | Q13873 |
| HOMO SAPIENS | 3 | 17448 | 3g2f | A | 485 | ASP | p.Asp485Gly | Q13873 |
| HOMO SAPIENS | 7 | 17448 | 3g2f | A | 491 | ARG | p.Arg491Gln | Q13873 |
| HOMO SAPIENS | 7 | 17448 | 3g2f | A | 491 | ARG | p.Arg491Trp | Q13873 |

| | | | | | | | | |
|---|---|---|---|---|---|---|---|---|
| HOMO SAPIENS | 0 | 4861 | 4ux8 | A | 56 | LEU | p.Leu56Met | P07949 |
| HOMO SAPIENS | 1 | 4861 | 4ux8 | A | 88 | ILE | p.Ile88Leu | P07949 |
| HOMO SAPIENS | 0 | 4861 | 4ux8 | A | 114 | ARG | p.Arg114His | P07949 |
| HOMO SAPIENS | 5 | 4861 | 4ux8 | A | 180 | ARG | p.Arg180Gln | P07949 |
| HOMO SAPIENS | 0 | 4861 | 4ux8 | A | 208 | GLU | p.Glu208Asp | P07949 |
| HOMO SAPIENS | 0 | 4861 | 4ux8 | A | 373 | ALA | p.Ala373Val | P07949 |
| HOMO SAPIENS | 0 | 4861 | 4ux8 | A | 375 | LEU | p.Leu375Gln | P07949 |
| HOMO SAPIENS | -1 | 4861 | 4ux8 | A | 386 | ALA | p.Ala386Val | P07949 |
| HOMO SAPIENS | 0 | 4861 | 4ux8 | A | 446 | GLY | p.Gly446Arg | P07949 |
| HOMO SAPIENS | 0 | 4861 | 4ux8 | A | 480 | GLU | p.Glu480Lys | P07949 |
| HOMO SAPIENS | 0 | 4861 | 4ux8 | A | 489 | ASP | p.Asp489Asn | P07949 |
| HOMO SAPIENS | -1 | 4861 | 4ux8 | A | 510 | ALA | p.Ala510Val | P07949 |
| HOMO SAPIENS | -1 | 4861 | 4ux8 | A | 533 | GLY | p.Gly533Cys | P07949 |
| HOMO SAPIENS | -1 | 4861 | 4ux8 | A | 567 | ASP | p.Asp567Asn | P07949 |
| HOMO SAPIENS | -1 | 4861 | 4ux8 | A | 609 | CYS | p.Cys609Tyr | P07949 |
| HOMO SAPIENS | -1 | 4861 | 4ux8 | A | 609 | CYS | p.Cys609Gly | P07949 |
| HOMO SAPIENS | -1 | 4861 | 4ux8 | A | 611 | CYS | p.Cys611Trp | P07949 |
| HOMO SAPIENS | -1 | 4861 | 4ux8 | A | 618 | CYS | p.Cys618Gly | P07949 |
| HOMO SAPIENS | -1 | 4861 | 4ux8 | A | 618 | CYS | p.Cys618Phe | P07949 |
| HOMO SAPIENS | -1 | 4861 | 4ux8 | A | 620 | CYS | p.Cys620Gly | P07949 |
| HOMO SAPIENS | -1 | 4861 | 4ux8 | A | 620 | CYS | p.Cys620Ser | P07949 |
| HOMO SAPIENS | -1 | 4861 | 4ux8 | A | 620 | CYS | p.Cys620Trp | P07949 |
| HOMO SAPIENS | -1 | 4861 | 4ux8 | A | 623 | GLU | p.Glu623Lys | P07949 |
| HOMO SAPIENS | -1 | 4861 | 4ux8 | A | 631 | ASP | p.Asp631Gly | P07949 |
| HOMO SAPIENS | -1 | 4861 | 4ux8 | A | 631 | ASP | p.Asp631Glu | P07949 |
| HOMO SAPIENS | -1 | 4861 | 4ux8 | A | 634 | CYS | p.Cys634Gly | P07949 |
| HOMO SAPIENS | -1 | 4861 | 4ux8 | A | 634 | CYS | p.Cys634Phe | P07949 |
| HOMO SAPIENS | -1 | 4861 | 4ux8 | A | 634 | CYS | p.Cys634Trp | P07949 |
| HOMO SAPIENS | 0 | 36308 | 4nkq | B | 123 | PRO | p.Pro142Leu | P15509 |
| HOMO SAPIENS | 0 | 36308 | 4nkq | B | 145 | ARG | p.Arg164Gln | P15509 |
| HOMO SAPIENS | -1 | 36308 | 4nkq | B | 177 | GLY | p.Gly196Arg | P15509 |
| HOMO SAPIENS | 0 | 36006 | 1oqa | A | 107 | HIS | p.His1860Pro | P38398 |
| HOMO SAPIENS | 7 | 15394 | 2ozo | A | 465 | ARG | p.Arg465His | P43403 |
| HOMO SAPIENS | 7 | 15394 | 2ozo | A | 465 | ARG | p.Arg465Cys | P43403 |
| HOMO SAPIENS | 4 | 15394 | 2ozo | A | 518 | SER | p.Ser518Arg | P43403 |
| HOMO SAPIENS | 1 | 20813 | 3o5z | A | 19 | ARG | p.Arg19Trp | O00459 |
| HOMO SAPIENS | 2 | 4329 | 2bb5 | A | 5 | ILE | p.Ile23Val | P20062 |
| HOMO SAPIENS | 4 | 4329 | 2bb5 | A | 59 | LYS | p.Lys77Met | P20062 |
| HOMO SAPIENS | 4 | 4329 | 2bb5 | A | 197 | ARG | p.Arg215Trp | P20062 |
| HOMO SAPIENS | 1 | 4329 | 2bb5 | A | 241 | PRO | p.Arg259Pro | P20062 |
| HOMO SAPIENS | 2 | 4329 | 2bb5 | A | 315 | THR | p.Thr333Lys | P20062 |
| HOMO SAPIENS | 5 | 4329 | 2bb5 | A | 330 | SER | p.Ser348Phe | P20062 |
| HOMO SAPIENS | 4 | 4329 | 2bb5 | A | 358 | LEU | p.Leu376Ser | P20062 |
| HOMO SAPIENS | 5 | 4329 | 2bb5 | A | 381 | ARG | p.Arg399Gln | P20062 |
| HOMO SAPIENS | 4 | 18294 | 4r30 | A | 163 | ASN | p.Asn163Asp | O95278 |
| HOMO SAPIENS | 6 | 18294 | 4r30 | A | 227 | ALA | p.Ala227Val | O95278 |
| HOMO SAPIENS | 4 | 18294 | 4r30 | A | 241 | ARG | p.Arg241Ter | O95278 |
| HOMO SAPIENS | 4 | 18294 | 4r30 | A | 325 | ARG | p.Arg325His | O95278 |
| HOMO SAPIENS | 0 | 2632 | 4xem | A | 22 | GLU | p.Glu22Lys | P49588 |
| HOMO SAPIENS | 0 | 2632 | 4xem | A | 275 | GLY | p.Gly275Asp | P49588 |
| HOMO SAPIENS | 1 | 2632 | 4xem | A | 329 | ARG | p.Arg329His | P49588 |
| HOMO SAPIENS | 0 | 11431 | 5t4n | A | 284 | GLN | p.Gln305Glu | Q96QU1 |
| HOMO SAPIENS | 0 | 11431 | 5t4n | A | 315 | ARG | p.Arg336Ter | Q96QU1 |
| HOMO SAPIENS | 0 | 11431 | 5t4n | A | 322 | ARG | p.Arg343Lys | Q96QU1 |

| | | | | | | | | |
|---|---|---|---|---|---|---|---|---|
| HOMO SAPIENS | 1 | 11431 | 5t4n | A | 326 | LEU | p.Leu347Phe | Q96QU1 |
| HOMO SAPIENS | 0 | 11431 | 5t4n | A | 373 | TYR | p.Tyr394Cys | Q96QU1 |
| HOMO SAPIENS | 0 | 11431 | 5t4n | A | 398 | ASN | p.Asn419Ser | Q96QU1 |
| HOMO SAPIENS | 0 | 11431 | 5t4n | A | 414 | ALA | p.Asp435Ala | Q96QU1 |
| HOMO SAPIENS | 0 | 11431 | 5t4n | A | 433 | VAL | p.Val454Phe | Q96QU1 |
| HOMO SAPIENS | 1 | 11431 | 5t4n | A | 507 | VAL | p.Val528Asp | Q96QU1 |
| HOMO SAPIENS | 0 | 11431 | 5t4n | A | 547 | ALA | p.Ala568Ser | Q96QU1 |
| HOMO SAPIENS | 3 | 3194 | 5xtd | L | 84 | ARG | p.Arg84His | O43181 |
| HOMO SAPIENS | 1 | 3194 | 5xtd | L | 130 | THR | p.Thr130Ser | O43181 |
| HOMO SAPIENS | 8 | 3194 | 5xtd | L | 136 | SER | p.Ser136Ala | O43181 |
| HOMO SAPIENS | 0 | 40749 | 1nun | A | 106 | CYS | p.Cys106Phe | O15520 |
| HOMO SAPIENS | 9 | 40749 | 1nun | A | 156 | ILE | p.Ile156Arg | O15520 |
| HOMO SAPIENS | 1 | 40749 | 1nun | A | 193 | ARG | p.Arg193Ter | O15520 |
| HOMO SAPIENS | 0 | 40749 | 1nun | A | 207 | HIS | p.His207Pro | O15520 |
| HOMO SAPIENS | 0 | 7026 | 3n1f | D | 875 | SER | p.Ser875Gly | Q4KMG0 |
| HOMO SAPIENS | 6 | 23802 | 3rzv | A | 424 | ARG | p.Arg424Ter | O95630 |
| HOMO SAPIENS | 0 | 36720 | 1jm7 | A | 12 | GLN | p.Gln12Ter | P38398 |
| HOMO SAPIENS | 3 | 36720 | 1jm7 | A | 23 | GLU | p.Glu23Asp | P38398 |
| HOMO SAPIENS | 1 | 36720 | 1jm7 | A | 45 | LYS | p.Lys45Gln | P38398 |
| HOMO SAPIENS | 8 | 36720 | 1jm7 | A | 47 | CYS | p.Cys47Ter | P38398 |
| HOMO SAPIENS | 0 | 36720 | 1jm7 | A | 59 | SER | p.Ser59Ter | P38398 |
| HOMO SAPIENS | 8 | 36720 | 1jm7 | A | 61 | CYS | p.Cys61Gly | P38398 |
| HOMO SAPIENS | 8 | 36720 | 1jm7 | A | 63 | LEU | p.Leu63Ter | P38398 |
| HOMO SAPIENS | 7 | 36720 | 1jm7 | A | 64 | CYS | p.Cys64Gly | P38398 |
| HOMO SAPIENS | 0 | 36720 | 1jm7 | A | 101 | TYR | p.Tyr101Ter | P38398 |
| HOMO SAPIENS | 0 | 36720 | 1jm7 | A | null | TYR | p.Tyr105Cys | P38398 |
| HOMO SAPIENS | 0 | 28500 | 3kuz | A | 1217 | CYS | p.Cys1217Arg | O60486 |
| HOMO SAPIENS | 5 | 8097 | 3p56 | C | 69 | ARG | p.Arg69Trp | Q8TDP1 |
| HOMO SAPIENS | 5 | 8097 | 3p56 | C | 143 | LYS | p.Lys143Ile | Q8TDP1 |
| HOMO SAPIENS | 2 | 30073 | 3nmz | A | 380 | ARG | p.Arg380Gln | P25054 |
| HOMO SAPIENS | 1 | 22573 | 1cjy | A | 111 | SER | p.Ser111Pro | P47712 |
| HOMO SAPIENS | 0 | 22573 | 1cjy | A | 485 | ARG | p.Arg485His | P47712 |
| HOMO SAPIENS | 2 | 965 | 3E+077 | A | 99 | ALA | p.Ala99Val | Q9Y617 |
| HOMO SAPIENS | 2 | 965 | 3E+077 | A | 100 | ASP | p.Asp100Ala | Q9Y617 |
| HOMO SAPIENS | 4 | 1503 | 3kxc | A | 62 | ARG | p.Arg62Trp | O43617 |
| HOMO SAPIENS | 6 | 9418 | 1hdr | A | 22 | GLY | p.Gly23Asp | P09417 |
| HOMO SAPIENS | 2 | 9418 | 1hdr | A | 35 | TRP | p.Trp36Arg | P09417 |
| HOMO SAPIENS | 2 | 9418 | 1hdr | A | 107 | TRP | p.Trp108Gly | P09417 |
| HOMO SAPIENS | 6 | 9418 | 1hdr | A | 149 | TYR | p.Tyr150Cys | P09417 |
| HOMO SAPIENS | -1 | 12079 | 3tnu | B | 373 | GLY | p.Gly373Ala | P13647 |
| HOMO SAPIENS | 1 | 12079 | 3tnu | B | 463 | LEU | p.Leu463Pro | P13647 |
| HOMO SAPIENS | 1 | 12079 | 3tnu | B | 471 | ARG | p.Arg471Cys | P13647 |
| HOMO SAPIENS | 0 | 8135 | 3rbn | B | 89 | SER | p.Ser556Ile | P40692 |
| HOMO SAPIENS | 0 | 8135 | 3rbn | B | 111 | GLU | p.Glu578Gly | P40692 |
| HOMO SAPIENS | 2 | 8135 | 3rbn | B | 114 | PRO | p.Pro581Leu | P40692 |
| HOMO SAPIENS | 0 | 8135 | 3rbn | B | 133 | GLU | p.Glu600Gly | P40692 |
| HOMO SAPIENS | 2 | 8135 | 3rbn | B | 136 | PRO | p.Pro603Arg | P40692 |
| HOMO SAPIENS | 2 | 8135 | 3rbn | B | 136 | PRO | p.Pro603Leu | P40692 |
| HOMO SAPIENS | 7 | 8135 | 3rbn | B | 140 | LEU | p.Leu607His | P40692 |
| HOMO SAPIENS | 2 | 8135 | 3rbn | B | 151 | LYS | p.Lys618Ala | P40692 |
| HOMO SAPIENS | 2 | 8135 | 3rbn | B | 151 | LYS | p.Lys618Thr | P40692 |
| HOMO SAPIENS | 5 | 8135 | 3rbn | B | 181 | PRO | p.Pro648Ser | P40692 |
| HOMO SAPIENS | 2 | 8135 | 3rbn | B | 188 | ILE | p.Ile655Val | P40692 |
| HOMO SAPIENS | 2 | 8135 | 3rbn | B | 214 | ALA | p.Ala681Thr | P40692 |

| | | | | | | | | |
|---|---|---|---|---|---|---|---|---|
| HOMO SAPIENS | 2 | 8135 | 3rbn | B | 220 | ARG | p.Arg687Trp | P40692 |
| HOMO SAPIENS | 2 | 8135 | 3rbn | B | 222 | GLN | p.Gln689Arg | P40692 |
| HOMO SAPIENS | -1 | 8135 | 3rbn | B | 236 | GLN | p.Gln701Ter | P40692 |
| HOMO SAPIENS | 7 | 8135 | 3rbn | B | 251 | HIS | p.His718Tyr | P40692 |
| HOMO SAPIENS | 0 | 8135 | 3rbn | B | 284 | LYS | p.Lys751Arg | P40692 |
| HOMO SAPIENS | 1 | 21998 | 5d68 | A | 355 | LEU | p.Leu355Phe | O00522 |
| HOMO SAPIENS | 3 | 21998 | 5d68 | A | 380 | THR | p.Thr380Met | O00522 |
| HOMO SAPIENS | 0 | 21998 | 5d68 | A | 415 | ILE | p.Ile415Met | O00522 |
| HOMO SAPIENS | 0 | 33789 | 2m0c | A | 13 | ARG | p.Arg218Gln | Q9H161 |
| HOMO SAPIENS | 7 | 33789 | 2m0c | A | 52 | ARG | p.Arg257Thr | Q9H161 |
| HOMO SAPIENS | 0 | 33789 | 2m0c | A | 67 | ARG | p.Arg272Pro | Q9H161 |
| HOMO SAPIENS | 6 | 8263 | 2osg | A | 51 | GLY | p.Gly355Arg | Q9UDY2 |
| HOMO SAPIENS | 5 | 8263 | 2osg | A | 60 | THR | p.Thr364Met | Q9UDY2 |
| HOMO SAPIENS | 1 | 2475 | 5E+026 | B | 234 | THR | p.Thr234Ala | Q9BZ23 |
| HOMO SAPIENS | 2 | 2475 | 5E+026 | B | 264 | ARG | p.Arg264Trp | Q9BZ23 |
| HOMO SAPIENS | 4 | 2475 | 5E+026 | B | 286 | ARG | p.Arg286Cys | Q9BZ23 |
| HOMO SAPIENS | 0 | 2475 | 5E+026 | B | 357 | ARG | p.Arg357Trp | Q9BZ23 |
| HOMO SAPIENS | 2 | 2475 | 5E+026 | B | 481 | ARG | p.Arg481Ter | Q9BZ23 |
| HOMO SAPIENS | 8 | 2475 | 5E+026 | B | 521 | GLY | p.Gly521Arg | Q9BZ23 |
| HOMO SAPIENS | 2 | 2475 | 5E+026 | B | 528 | THR | p.Thr528Met | Q9BZ23 |
| HOMO SAPIENS | 1 | 3742 | 5xtd | E | 108 | HIS | p.His134Tyr | P56556 |
| HOMO SAPIENS | 1 | 16725 | 3g04 | C | 36 | ASP | p.Asp36His | P16473 |
| HOMO SAPIENS | 3 | 16725 | 3g04 | C | 52 | PRO | p.Pro52Thr | P16473 |
| HOMO SAPIENS | 1 | 16725 | 3g04 | C | 162 | PRO | p.Pro162Ala | P16473 |
| HOMO SAPIENS | -1 | 12366 | 5d2d | C | 750 | PRO | p.Pro750Leu | P13569 |
| HOMO SAPIENS | 0 | 12366 | 5d2d | C | 764 | ARG | p.Arg764Ter | P13569 |
| HOMO SAPIENS | 0 | 34845 | 2y23 | A | 1151 | ASP | p.Asp1151Glu | P52179 |
| HOMO SAPIENS | 0 | 34845 | 2y23 | A | 1180 | ASP | p.Asp1180Ala | P52179 |
| HOMO SAPIENS | 6 | 36740 | 3lbx | B | 2053 | ALA | p.Ala2053Pro | P11277 |
| HOMO SAPIENS | 0 | 19289 | 3ihj | A | 404 | ARG | p.Arg404Ter | Q8TD30 |
| HOMO SAPIENS | 5 | 1034 | 5xtd | Q | 228 | ARG | p.Arg228Gln | O75306 |
| HOMO SAPIENS | 5 | 1034 | 5xtd | Q | 229 | PRO | p.Pro229Gln | O75306 |
| HOMO SAPIENS | 4 | 1034 | 5xtd | Q | 323 | ARG | p.Arg323Gln | O75306 |
| HOMO SAPIENS | 4 | 1034 | 5xtd | Q | 352 | PRO | p.Pro352Ala | O75306 |
| HOMO SAPIENS | 5 | 1034 | 5xtd | Q | 413 | SER | p.Ser413Pro | O75306 |
| HOMO SAPIENS | 1 | 2772 | 2vaf | A | 1047 | THR | p.Thr66Ala | O14958 |
| HOMO SAPIENS | 5 | 2772 | 2vaf | A | 1057 | VAL | p.Val76Met | O14958 |
| HOMO SAPIENS | 0 | 2772 | 2vaf | A | 1148 | LEU | p.Leu167His | O14958 |
| HOMO SAPIENS | 1 | 2772 | 2vaf | A | 1149 | ILE | p.Ile168Thr | O14958 |
| HOMO SAPIENS | 8 | 2772 | 2vaf | A | 1225 | HIS | p.His244Arg | O14958 |
| HOMO SAPIENS | 8 | 2772 | 2vaf | A | 1225 | HIS | p.His244Tyr | O14958 |
| HOMO SAPIENS | 7 | 2772 | 2vaf | A | 1251 | ILE | p.Ile270Thr | O14958 |
| HOMO SAPIENS | 3 | 2772 | 2vaf | A | 1288 | ASP | p.Asp307His | O14958 |
| HOMO SAPIENS | 5 | 2772 | 2vaf | A | 1289 | PRO | p.Pro308Leu | O14958 |
| HOMO SAPIENS | 1 | 31885 | 2aex | A | 132 | ALA | p.Ala132Val | P36551 |
| HOMO SAPIENS | 1 | 31885 | 2aex | A | 272 | ASN | p.Asn272His | P36551 |
| HOMO SAPIENS | 8 | 31885 | 2aex | A | 279 | GLY | p.Gly279Arg | P36551 |
| HOMO SAPIENS | 5 | 31885 | 2aex | A | 294 | VAL | p.Val294Ile | P36551 |
| HOMO SAPIENS | 5 | 31885 | 2aex | A | 295 | HIS | p.His295Asp | P36551 |
| HOMO SAPIENS | 1 | 31885 | 2aex | A | 331 | ARG | p.Arg331Trp | P36551 |
| HOMO SAPIENS | 3 | 31885 | 2aex | A | 352 | ARG | p.Arg352Cys | P36551 |
| HOMO SAPIENS | 1 | 31885 | 2aex | A | 404 | LYS | p.Lys404Glu | P36551 |
| HOMO SAPIENS | 2 | 31885 | 2aex | A | 447 | ARG | p.Arg447Cys | P36551 |
| HOMO SAPIENS | 5 | 11721 | 2r3v | A | 20 | HIS | p.His20Pro | Q03426 |

| | | | | | | | | |
|---|---|---|---|---|---|---|---|---|
| HOMO SAPIENS | 1 | 11721 | 2r3v | A | 116 | TYR | p.Tyr116His | Q03426 |
| HOMO SAPIENS | 1 | 11721 | 2r3v | A | 148 | ALA | p.Ala148Thr | Q03426 |
| HOMO SAPIENS | 1 | 11721 | 2r3v | A | 167 | PRO | p.Pro167Leu | Q03426 |
| HOMO SAPIENS | 6 | 11721 | 2r3v | A | 203 | VAL | p.Val203Ala | Q03426 |
| HOMO SAPIENS | 2 | 11721 | 2r3v | A | 215 | ARG | p.Arg215Ter | Q03426 |
| HOMO SAPIENS | 2 | 11721 | 2r3v | A | 237 | THR | p.Thr237Ser | Q03426 |
| HOMO SAPIENS | 0 | 11721 | 2r3v | A | 268 | ILE | p.Ile268Thr | Q03426 |
| HOMO SAPIENS | 0 | 11721 | 2r3v | A | 268 | ILE | p.Ile268Val | Q03426 |
| HOMO SAPIENS | 0 | 11721 | 2r3v | A | 301 | ASN | p.Asn301Ser | Q03426 |
| HOMO SAPIENS | 6 | 11721 | 2r3v | A | 310 | VAL | p.Val310Met | Q03426 |
| HOMO SAPIENS | 5 | 11721 | 2r3v | A | 334 | ALA | p.Ala334Thr | Q03426 |
| HOMO SAPIENS | 2 | 11721 | 2r3v | A | 377 | VAL | p.Val377Ile | Q03426 |
| HOMO SAPIENS | 1 | 11721 | 2r3v | A | 386 | ASP | p.Asp386Asn | Q03426 |
| HOMO SAPIENS | 1 | 11721 | 2r3v | A | 388 | ARG | p.Arg388Ter | Q03426 |
| HOMO SAPIENS | 8 | 5010 | 2hgs | A | 125 | ARG | p.Arg125Cys | P48637 |
| HOMO SAPIENS | 1 | 5010 | 2hgs | A | 164 | ARG | p.Arg164Gln | P48637 |
| HOMO SAPIENS | 0 | 5010 | 2hgs | A | 219 | ASP | p.Asp219Gly | P48637 |
| HOMO SAPIENS | 7 | 5010 | 2hgs | A | 267 | ARG | p.Arg267Trp | P48637 |
| HOMO SAPIENS | 0 | 5010 | 2hgs | A | 283 | ARG | p.Arg283Cys | P48637 |
| HOMO SAPIENS | 0 | 6892 | 5erd | B | 7 | VAL | p.Val56Met | Q14126 |
| HOMO SAPIENS | 1 | 6892 | 5erd | B | 25 | HIS | p.His74Arg | Q14126 |
| HOMO SAPIENS | 0 | 6892 | 5erd | B | 40 | TYR | p.Tyr89Cys | Q14126 |
| HOMO SAPIENS | 0 | 6892 | 5erd | B | 97 | ARG | p.Arg146Leu | Q14126 |
| HOMO SAPIENS | 7 | 6892 | 5erd | B | 217 | ASN | p.Asn266Ser | Q14126 |
| HOMO SAPIENS | 0 | 6892 | 5erd | B | 243 | ARG | p.Arg292His | Q14126 |
| HOMO SAPIENS | 0 | 6892 | 5erd | B | 244 | ILE | p.Ile293Val | Q14126 |
| HOMO SAPIENS | 0 | 6892 | 5erd | B | 343 | VAL | p.Val392Ile | Q14126 |
| HOMO SAPIENS | 4 | 6892 | 5erd | B | 350 | GLU | p.Glu399Lys | Q14126 |
| HOMO SAPIENS | 0 | 6892 | 5erd | B | 410 | TYR | p.Tyr459Cys | Q14126 |
| HOMO SAPIENS | 0 | 6892 | 5erd | B | 466 | VAL | p.Val515Ile | Q14126 |
| HOMO SAPIENS | 0 | 6892 | 5erd | B | 482 | PHE | p.Phe531Cys | Q14126 |
| HOMO SAPIENS | 0 | 6892 | 5erd | B | 484 | VAL | p.Val533Ile | Q14126 |
| HOMO SAPIENS | 0 | 6892 | 5erd | B | 545 | LEU | p.Leu594Pro | Q14126 |
| HOMO SAPIENS | 0 | 1338 | 2ax4 | A | 58 | THR | p.Thr48Arg | O95340 |
| HOMO SAPIENS | 7 | 8675 | 2zkm | X | 385 | LYS | p.Lys385Arg | Q00722 |
| HOMO SAPIENS | 1 | 11769 | 2wya | B | 54 | VAL | p.Val54Met | P54868 |
| HOMO SAPIENS | 1 | 11769 | 2wya | B | 116 | ARG | p.Arg116Cys | P54868 |
| HOMO SAPIENS | 5 | 11769 | 2wya | B | 167 | TYR | p.Tyr167Cys | P54868 |
| HOMO SAPIENS | 1 | 11769 | 2wya | B | 174 | PHE | p.Phe174Leu | P54868 |
| HOMO SAPIENS | 0 | 11769 | 2wya | B | 212 | GLY | p.Gly212Arg | P54868 |
| HOMO SAPIENS | 1 | 11769 | 2wya | B | 288 | ARG | p.Arg288Ter | P54868 |
| HOMO SAPIENS | 0 | 11769 | 2wya | B | 375 | SER | p.Ser375Cys | P54868 |
| HOMO SAPIENS | 4 | 11769 | 2wya | B | 500 | ARG | p.Arg500His | P54868 |
| HOMO SAPIENS | 0 | 3102 | 2iwo | A | 1880 | ARG | p.Arg1908Lys | O75970 |
| HOMO SAPIENS | 6 | 14296 | 2m9w | A | 38 | GLY | p.Gly296Ser | P43694 |
| HOMO SAPIENS | 0 | 14296 | 2m9w | A | 58 | GLN | p.Gln316Glu | P43694 |
| HOMO SAPIENS | 2 | 704 | 5ekd | B | 239 | ARG | p.Arg239Ter | Q9UGM6 |
| HOMO SAPIENS | 2 | 704 | 5ekd | B | 264 | TYR | p.Tyr264Cys | Q9UGM6 |
| HOMO SAPIENS | 6 | 2228 | 1ey2 | A | 47 | SER | p.Ser47Ter | Q93099 |
| HOMO SAPIENS | 6 | 2228 | 1ey2 | A | 48 | ALA | p.Ala48Ser | Q93099 |
| HOMO SAPIENS | 6 | 2228 | 1ey2 | A | 53 | ARG | p.Arg53Gln | Q93099 |
| HOMO SAPIENS | 3 | 2228 | 1ey2 | A | 80 | HIS | p.Gln80His | Q93099 |
| HOMO SAPIENS | 0 | 2228 | 1ey2 | A | 120 | CYS | p.Cys120Trp | Q93099 |
| HOMO SAPIENS | 3 | 2228 | 1ey2 | A | 122 | ALA | p.Ala122Val | Q93099 |

| | | | | | | | | |
|---|---|---|---|---|---|---|---|---|
| HOMO SAPIENS | 3 | 2228 | 1ey2 | A | 161 | GLY | p.Gly161Arg | Q93099 |
| HOMO SAPIENS | 6 | 2228 | 1ey2 | A | 225 | ARG | p.Arg225His | Q93099 |
| HOMO SAPIENS | 0 | 2228 | 1ey2 | A | 230 | PRO | p.Pro230Ser | Q93099 |
| HOMO SAPIENS | 6 | 2228 | 1ey2 | A | 270 | GLY | p.Gly270Arg | Q93099 |
| HOMO SAPIENS | 6 | 2228 | 1ey2 | A | 330 | ARG | p.Arg330Ser | Q93099 |
| HOMO SAPIENS | -1 | 2228 | 1ey2 | A | 368 | MSE | p.Met368Val | Q93099 |
| HOMO SAPIENS | 6 | 2228 | 1ey2 | A | 371 | HIS | p.His371Arg | Q93099 |
| HOMO SAPIENS | 6 | 2228 | 1ey2 | A | 401 | GLU | p.Glu401Gln | Q93099 |
| HOMO SAPIENS | 5 | 34648 | 2k27 | A | 31 | ARG | p.Arg31His | Q06710 |
| HOMO SAPIENS | 7 | 34648 | 2k27 | A | 40 | GLN | p.Gln40Pro | Q06710 |
| HOMO SAPIENS | 5 | 34648 | 2k27 | A | 57 | CYS | p.Cys57Tyr | Q06710 |
| HOMO SAPIENS | 5 | 34648 | 2k27 | A | 62 | LEU | p.Leu62Arg | Q06710 |
| HOMO SAPIENS | 1 | 21399 | 3edu | A | 1819 | GLU | p.Glu1819Ter | P11277 |
| HOMO SAPIENS | 1 | 18153 | 1oey | J | 245 | TYR | p.Tyr245Ter | Q15080 |
| HOMO SAPIENS | 2 | 41567 | 3sw0 | X | 1050 | ASN | p.Asn1050Tyr | P08603 |
| HOMO SAPIENS | 5 | 41567 | 3sw0 | X | 1059 | ILE | p.Ile1059Thr | P08603 |
| HOMO SAPIENS | 0 | 36362 | 4wrm | A | 362 | HIS | p.His362Arg | P07333 |
| HOMO SAPIENS | -1 | 36362 | 4wrm | A | 413 | GLY | p.Gly413Ser | P07333 |
| HOMO SAPIENS | 0 | 31255 | 2w2j | A | 100 | SER | p.Ser100Pro | P35219 |
| HOMO SAPIENS | 0 | 31255 | 2w2j | A | 100 | SER | p.Ser100Leu | P35219 |
| HOMO SAPIENS | 5 | 31255 | 2w2j | A | 237 | ARG | p.Arg237Gln | P35219 |
| HOMO SAPIENS | -1 | 26450 | 1th1 | C | 1395 | SER | p.Ser1395Cys | P25054 |
| HOMO SAPIENS | -1 | 26450 | 1th1 | C | 1413 | MET | p.Met1413Val | P25054 |
| HOMO SAPIENS | -1 | 26450 | 1th1 | C | 1446 | ALA | p.Ala1446Thr | P25054 |
| HOMO SAPIENS | -1 | 26450 | 1th1 | C | 1454 | LYS | p.Lys1454Glu | P25054 |
| HOMO SAPIENS | 0 | 26450 | 1th1 | C | 1474 | ALA | p.Ala1474Thr | P25054 |
| HOMO SAPIENS | 1 | 981 | 5xtb | F | 14 | LEU | p.Leu14Val | O43678 |
| HOMO SAPIENS | 1 | 11069 | 3kq4 | A | 46 | SER | p.Ser46Leu | P27352 |
| HOMO SAPIENS | 1 | 11069 | 3kq4 | A | 97 | MET | p.Met97Thr | P27352 |
| HOMO SAPIENS | 1 | 11069 | 3kq4 | A | 255 | ASN | p.Asn255Ser | P27352 |
| HOMO SAPIENS | 1 | 11069 | 3kq4 | A | 330 | ASN | p.Asn330Lys | P27352 |
| HOMO SAPIENS | 0 | 41489 | 3erb | A | 53 | PRO | p.Pro73Leu | P06681 |
| HOMO SAPIENS | 6 | 8715 | 2bq8 | X | 23 | THR | p.Thr44Met | P13686 |
| HOMO SAPIENS | 0 | 8715 | 2bq8 | X | 88 | GLY | p.Gly109Arg | P13686 |
| HOMO SAPIENS | 6 | 8715 | 2bq8 | X | 127 | VAL | p.Val148Met | P13686 |
| HOMO SAPIENS | 2 | 8715 | 2bq8 | X | 179 | VAL | p.Val200Met | P13686 |
| HOMO SAPIENS | 6 | 8715 | 2bq8 | X | 251 | ARG | p.Arg272Cys | P13686 |
| HOMO SAPIENS | 7 | 8112 | 3p56 | A | 37 | GLY | p.Gly37Ser | O75792 |
| HOMO SAPIENS | -1 | 8112 | 3p56 | A | 202 | LEU | p.Leu202Ser | O75792 |
| HOMO SAPIENS | 1 | 8112 | 3p56 | A | 205 | ASP | p.Asp205Glu | O75792 |
| HOMO SAPIENS | 4 | 8112 | 3p56 | A | 212 | ASN | p.Asn212Ile | O75792 |
| HOMO SAPIENS | 0 | 8112 | 3p56 | A | 235 | ARG | p.Arg235Gln | O75792 |
| HOMO SAPIENS | 4 | 8112 | 3p56 | A | 240 | THR | p.Thr240Met | O75792 |
| HOMO SAPIENS | 0 | 8112 | 3p56 | A | 291 | ARG | p.Arg291His | O75792 |
| HOMO SAPIENS | 0 | 37603 | 2yqg | A | 10 | ARG | p.Arg49His | Q14126 |
| HOMO SAPIENS | 0 | 21178 | 3k1r | B | 458 | ASP | p.Asp458Val | Q495M9 |
| HOMO SAPIENS | -1 | 36828 | 4fq3 | B | 507 | GLY | p.Gly507Asp | P35637 |
| HOMO SAPIENS | 5 | 36828 | 4fq3 | B | 517 | HIS | p.His517Gln | P35637 |
| HOMO SAPIENS | 8 | 36828 | 4fq3 | B | 518 | ARG | p.Arg518Lys | P35637 |
| HOMO SAPIENS | 6 | 36828 | 4fq3 | B | 521 | ARG | p.Arg521Cys | P35637 |
| HOMO SAPIENS | 6 | 36828 | 4fq3 | B | 521 | ARG | p.Arg521His | P35637 |
| HOMO SAPIENS | 0 | 6363 | 3j7y | c | null | HIS | p.His13Arg | Q9H9J2 |
| HOMO SAPIENS | 2 | 6363 | 3j7y | c | 105 | ARG | p.Arg105Cys | Q9H9J2 |
| HOMO SAPIENS | 2 | 6363 | 3j7y | c | 138 | THR | p.Thr138Ile | Q9H9J2 |

| | | | | | | | | |
|---|---|---|---|---|---|---|---|---|
| HOMO SAPIENS | 1 | 6363 | 3j7y | c | 156 | LEU | p.Leu156Arg | Q9H9J2 |
| HOMO SAPIENS | 1 | 6782 | 5l46 | A | 109 | HIS | p.His109Arg | Q9UI17 |
| HOMO SAPIENS | 0 | 23951 | 5t76 | A | 175 | GLY | p.Gly931Ser | P49588 |
| HOMO SAPIENS | 8 | 12604 | 3p0l | A | 182 | ARG | p.Arg182Leu | P49675 |
| HOMO SAPIENS | 2 | 12604 | 3p0l | A | 187 | VAL | p.Val187Met | P49675 |
| HOMO SAPIENS | 2 | 12604 | 3p0l | A | 188 | ARG | p.Arg188Cys | P49675 |
| HOMO SAPIENS | 0 | 12604 | 3p0l | A | 193 | ARG | p.Arg193Ter | P49675 |
| HOMO SAPIENS | 9 | 12604 | 3p0l | A | 217 | ARG | p.Arg217Thr | P49675 |
| HOMO SAPIENS | 4 | 12604 | 3p0l | A | 218 | ALA | p.Ala218Val | P49675 |
| HOMO SAPIENS | 0 | 12604 | 3p0l | A | 250 | TRP | p.Trp250Ter | P49675 |
| HOMO SAPIENS | 1 | 12604 | 3p0l | A | 258 | GLN | p.Gln258Ter | P49675 |
| HOMO SAPIENS | -1 | 27765 | 4wkr | A | 4 | GLU | p.Glu4Lys | Q4G0J3 |
| HOMO SAPIENS | 2 | 3587 | 4kit | C | 2301 | PRO | p.Pro2301Thr | Q6P2Q9 |
| HOMO SAPIENS | 3 | 3587 | 4kit | C | 2304 | PHE | p.Phe2304Leu | Q6P2Q9 |
| HOMO SAPIENS | 3 | 3587 | 4kit | C | 2309 | HIS | p.His2309Pro | Q6P2Q9 |
| HOMO SAPIENS | 3 | 3587 | 4kit | C | 2310 | ARG | p.Arg2310Lys | Q6P2Q9 |
| HOMO SAPIENS | 0 | 9073 | 1n69 | B | 21 | ASN | p.Asn215His | P07602 |
| HOMO SAPIENS | 0 | 9073 | 1n69 | B | 23 | THR | p.Thr217Ile | P07602 |
| HOMO SAPIENS | 0 | 9073 | 1n69 | B | 47 | CYS | p.Cys241Ser | P07602 |
| HOMO SAPIENS | 0 | 475 | 2vd5 | A | 225 | THR | p.Thr225Met | Q09013 |
| HOMO SAPIENS | 1 | 9066 | 4mzv | A | 31 | ASN | p.Asn31Lys | P16422 |
| HOMO SAPIENS | 1 | 9066 | 4mzv | A | 47 | GLN | p.Gln47His | P16422 |
| HOMO SAPIENS | 3 | 9066 | 4mzv | A | 66 | CYS | p.Cys66Tyr | P16422 |
| HOMO SAPIENS | 1 | 9066 | 4mzv | A | 102 | SER | p.Ser102Gly | P16422 |
| HOMO SAPIENS | 5 | 9066 | 4mzv | A | 115 | THR | p.Met115Thr | P16422 |
| HOMO SAPIENS | 1 | 9066 | 4mzv | A | 138 | ARG | p.Arg138Ter | P16422 |
| HOMO SAPIENS | 1 | 9066 | 4mzv | A | 153 | ARG | p.Arg153Thr | P16422 |
| HOMO SAPIENS | 1 | 9066 | 4mzv | A | 165 | ALA | p.Ala165Thr | P16422 |
| HOMO SAPIENS | 8 | 1827 | 2nz2 | A | 14 | GLY | p.Gly14Ser | P00966 |
| HOMO SAPIENS | 5 | 1827 | 2nz2 | A | 18 | SER | p.Ser18Leu | P00966 |
| HOMO SAPIENS | 3 | 1827 | 2nz2 | A | 86 | ARG | p.Arg86Cys | P00966 |
| HOMO SAPIENS | 3 | 1827 | 2nz2 | A | 86 | ARG | p.Arg86His | P00966 |
| HOMO SAPIENS | 1 | 1827 | 2nz2 | A | 179 | TRP | p.Trp179Arg | P00966 |
| HOMO SAPIENS | 7 | 1827 | 2nz2 | A | 180 | SER | p.Ser180Asn | P00966 |
| HOMO SAPIENS | 7 | 1827 | 2nz2 | A | 191 | GLU | p.Glu191Lys | P00966 |
| HOMO SAPIENS | 4 | 1827 | 2nz2 | A | 256 | GLU | p.Glu256Lys | P00966 |
| HOMO SAPIENS | 4 | 1827 | 2nz2 | A | 263 | VAL | p.Val263Met | P00966 |
| HOMO SAPIENS | 5 | 1827 | 2nz2 | A | 265 | ARG | p.Arg265Cys | P00966 |
| HOMO SAPIENS | 5 | 1827 | 2nz2 | A | 268 | ILE | p.Ile268Met | P00966 |
| HOMO SAPIENS | 6 | 1827 | 2nz2 | A | 269 | VAL | p.Val269Met | P00966 |
| HOMO SAPIENS | 7 | 1827 | 2nz2 | A | 272 | ARG | p.Arg272Cys | P00966 |
| HOMO SAPIENS | 3 | 1827 | 2nz2 | A | 304 | ARG | p.Arg304Trp | P00966 |
| HOMO SAPIENS | 3 | 1827 | 2nz2 | A | 307 | ARG | p.Arg307Cys | P00966 |
| HOMO SAPIENS | 3 | 1827 | 2nz2 | A | 310 | LYS | p.Lys310Gln | P00966 |
| HOMO SAPIENS | 2 | 1827 | 2nz2 | A | 362 | GLY | p.Gly362Val | P00966 |
| HOMO SAPIENS | 2 | 1827 | 2nz2 | A | 363 | ARG | p.Arg363Trp | P00966 |
| HOMO SAPIENS | 2 | 1827 | 2nz2 | A | 363 | ARG | p.Arg363Gln | P00966 |
| HOMO SAPIENS | 1 | 1827 | 2nz2 | A | 390 | GLY | p.Gly390Arg | P00966 |
| HOMO SAPIENS | 0 | 29524 | 5vbn | B | 2165 | ARG | p.Arg2165His | Q07864 |
| HOMO SAPIENS | 2 | 29524 | 5vbn | B | 2255 | ILE | p.Ile2255Phe | Q07864 |
| HOMO SAPIENS | 2 | 29524 | 5vbn | B | 2274 | LEU | p.Leu2274Val | Q07864 |
| HOMO SAPIENS | 0 | 31897 | 3t0o | A | 184 | CYS | p.Cys184Arg | O00584 |
| HOMO SAPIENS | -1 | 31897 | 3t0o | A | 215 | GLU | p.Glu215Lys | O00584 |
| HOMO SAPIENS | 0 | 31897 | 3t0o | A | 236 | ARG | p.Arg236Trp | O00584 |

| | | | | | | | | |
|---|---|---|---|---|---|---|---|---|
| HOMO SAPIENS | 7 | 18490 | 4pcw | B | 31 | GLU | p.Glu32Ter | P20930 |
| HOMO SAPIENS | -1 | 39865 | 2h9e | L | 152 | GLY | p.Gly192Arg | P00742 |
| HOMO SAPIENS | 2 | 4309 | 2x0b | F | 174 | THR | p.Thr207Met | P01019 |
| HOMO SAPIENS | 3 | 4309 | 2x0b | F | 342 | ARG | p.Arg375Gln | P01019 |
| HOMO SAPIENS | 2 | 3094 | 1zll | B | 9 | ARG | p.Arg9Cys | P26678 |
| HOMO SAPIENS | 9 | 3094 | 1zll | B | 39 | LEU | p.Leu39Ter | P26678 |
| HOMO SAPIENS | 5 | 3094 | 1zll | B | 49 | VAL | p.Val49Met | P26678 |
| HOMO SAPIENS | -1 | 41498 | 3ffd | P | 8 | LEU | p.Leu44Pro | P12272 |
| HOMO SAPIENS | 1 | 20697 | 1igr | A | 407 | ARG | p.Arg437His | P08069 |
| HOMO SAPIENS | 0 | 20697 | 1igr | A | 416 | MET | p.Met446Val | P08069 |
| HOMO SAPIENS | -1 | 20697 | 1igr | A | 462 | VAL | p.Val492Ile | P08069 |
| HOMO SAPIENS | 0 | 3973 | 4cbz | A | 37 | LEU | p.Leu37Ser | P78504 |
| HOMO SAPIENS | 0 | 3973 | 4cbz | A | 45 | VAL | p.Val45Leu | P78504 |
| HOMO SAPIENS | 0 | 3973 | 4cbz | A | 134 | LEU | p.Leu134Met | P78504 |
| HOMO SAPIENS | -1 | 3973 | 4cbz | A | 146 | VAL | p.Val146Ile | P78504 |
| HOMO SAPIENS | 0 | 3973 | 4cbz | A | 176 | VAL | p.Val176Ile | P78504 |
| HOMO SAPIENS | 0 | 3973 | 4cbz | A | 184 | ARG | p.Arg184Cys | P78504 |
| HOMO SAPIENS | 6 | 1697 | 4hmy | S | 33 | ARG | p.Arg33Trp | Q96PC3 |
| HOMO SAPIENS | 2 | 23752 | 5j4o | A | 13 | ARG | p.Arg1611Ser | P02549 |
| HOMO SAPIENS | 0 | 23752 | 5j4o | A | 42 | ALA | p.Ala1640Gly | P02549 |
| HOMO SAPIENS | 0 | 23752 | 5j4o | A | 108 | HIS | p.His1706Gln | P02549 |
| HOMO SAPIENS | 6 | 23752 | 5j4o | A | 206 | LEU | p.Leu1804Phe | P02549 |
| HOMO SAPIENS | 8 | 531 | 2qz4 | A | 349 | GLY | p.Gly349Ser | Q9UQ90 |
| HOMO SAPIENS | 0 | 531 | 2qz4 | A | 486 | ARG | p.Arg486Gln | Q9UQ90 |
| HOMO SAPIENS | 1 | 531 | 2qz4 | A | 510 | ALA | p.Ala510Val | Q9UQ90 |
| HOMO SAPIENS | 0 | 531 | 2qz4 | A | 529 | ALA | p.Ala529Val | Q9UQ90 |
| HOMO SAPIENS | 0 | 531 | 2qz4 | A | 558 | LYS | p.Lys558Ter | Q9UQ90 |
| HOMO SAPIENS | 1 | 531 | 2qz4 | A | 559 | LYS | p.Lys559Ter | Q9UQ90 |
| HOMO SAPIENS | 2 | 7736 | 1fnh | A | 114 | TRP | p.Trp1834Arg | P02751 |
| HOMO SAPIENS | 0 | 7736 | 1fnh | A | 163 | LEU | p.Leu1883Arg | P02751 |
| HOMO SAPIENS | 3 | 20337 | 5wy5 | B | 264 | LEU | p.Leu264Phe | Q96MG7 |
| HOMO SAPIENS | 8 | 36096 | 2jku | A | 643 | GLY | p.Gly668Arg | P05165 |
| HOMO SAPIENS | 6 | 36096 | 2jku | A | 663 | CYS | p.Cys688Arg | P05165 |
| HOMO SAPIENS | 0 | 9850 | 3kbt | A | 1606 | TYR | p.Tyr1606Ter | P11277 |
| HOMO SAPIENS | 4 | 6876 | 4jkq | A | 217 | ILE | p.Ile217Val | Q92736 |
| HOMO SAPIENS | 5 | 6876 | 4jkq | A | 420 | ARG | p.Arg420Trp | Q92736 |
| HOMO SAPIENS | 2 | 27603 | 1itv | B | 62 | PRO | p.Arg574Pro | P14780 |
| HOMO SAPIENS | 4 | 27603 | 1itv | B | 156 | ARG | p.Arg668Gln | P14780 |
| HOMO SAPIENS | 2 | 22954 | 4f52 | E | 48 | ILE | p.Ile48Met | Q92990 |
| HOMO SAPIENS | 0 | 22954 | 4f52 | E | 91 | ASP | p.Asp91Asn | Q92990 |
| HOMO SAPIENS | 0 | 22954 | 4f52 | E | 146 | ALA | p.Ala146Thr | Q92990 |
| HOMO SAPIENS | 0 | 22954 | 4f52 | E | 181 | ILE | p.Ile181Lys | Q92990 |
| HOMO SAPIENS | 0 | 22954 | 4f52 | E | 194 | ASN | p.Asn194Ser | Q92990 |
| HOMO SAPIENS | 2 | 22954 | 4f52 | E | 254 | PRO | p.Pro254Arg | Q92990 |
| HOMO SAPIENS | 2 | 22954 | 4f52 | E | 300 | ILE | p.Ile300Thr | Q92990 |
| HOMO SAPIENS | 2 | 22954 | 4f52 | E | 336 | LEU | p.Leu336Ser | Q92990 |
| HOMO SAPIENS | 0 | 22954 | 4f52 | E | 353 | TYR | p.Tyr353His | Q92990 |
| HOMO SAPIENS | 0 | 22954 | 4f52 | E | 387 | MET | p.Met387Val | Q92990 |
| HOMO SAPIENS | 4 | 2946 | 5gl7 | A | 344 | TYR | p.Tyr344Cys | P56192 |
| HOMO SAPIENS | 0 | 2946 | 5gl7 | A | 370 | PHE | p.Phe370Leu | P56192 |
| HOMO SAPIENS | 0 | 2946 | 5gl7 | A | 393 | ALA | p.Ala393Thr | P56192 |
| HOMO SAPIENS | 0 | 2946 | 5gl7 | A | 523 | ILE | p.Ile523Thr | P56192 |
| HOMO SAPIENS | 0 | 2946 | 5gl7 | A | 567 | SER | p.Ser567Leu | P56192 |
| HOMO SAPIENS | 0 | 2946 | 5gl7 | A | 800 | PRO | p.Pro800Thr | P56192 |

| | | | | | | | | |
|---|---|---|---|---|---|---|---|---|
| HOMO SAPIENS | 3 | 34961 | 4wy4 | C | 44 | TYR | p.Tyr44His | O95721 |
| HOMO SAPIENS | 8 | 34961 | 4wy4 | C | 89 | GLU | p.Glu89Lys | O95721 |
| HOMO SAPIENS | 0 | 9427 | 4chl | B | 21 | ALA | p.Ala21Ser | O95571 |
| HOMO SAPIENS | 2 | 9427 | 4chl | B | 62 | ALA | p.Ala62Thr | O95571 |
| HOMO SAPIENS | 0 | 9427 | 4chl | B | 114 | ILE | p.Ile114Phe | O95571 |
| HOMO SAPIENS | 2 | 9427 | 4chl | B | 159 | ARG | p.Arg159His | O95571 |
| HOMO SAPIENS | 8 | 9427 | 4chl | B | 163 | ARG | p.Arg163Trp | O95571 |
| HOMO SAPIENS | 8 | 9427 | 4chl | B | 163 | ARG | p.Arg163Gln | O95571 |
| HOMO SAPIENS | 2 | 9427 | 4chl | B | 185 | LEU | p.Leu185Arg | O95571 |
| HOMO SAPIENS | 0 | 40917 | 2e8o | A | 11 | PRO | p.Pro26Leu | Q9Y3Z3 |
| HOMO SAPIENS | -1 | 39190 | 4gow | A | 556 | TYR | p.Tyr556Ter | P56696 |
| HOMO SAPIENS | 0 | 2399 | 2mkv | A | 8 | GLY | p.Gly10Ser | P54710 |
| HOMO SAPIENS | 1 | 2399 | 2mkv | A | 24 | VAL | p.Val26Ile | P54710 |
| HOMO SAPIENS | 4 | 2399 | 2mkv | A | 39 | GLY | p.Gly41Arg | P54710 |
| HOMO SAPIENS | 0 | 20801 | 2zej | B | 1441 | ARG | p.Arg1441Cys | Q5S007 |
| HOMO SAPIENS | 0 | 20801 | 2zej | B | 1441 | ARG | p.Arg1441His | Q5S007 |
| HOMO SAPIENS | 5 | 1019 | 5xtb | M | 62 | ARG | p.Arg62Gln | P28331 |
| HOMO SAPIENS | 1 | 1019 | 5xtb | M | 85 | ALA | p.Ala85Thr | P28331 |
| HOMO SAPIENS | 5 | 1019 | 5xtb | M | 231 | LEU | p.Leu231Val | P28331 |
| HOMO SAPIENS | 2 | 1019 | 5xtb | M | 241 | ARG | p.Arg241Trp | P28331 |
| HOMO SAPIENS | 1 | 1019 | 5xtb | M | 252 | ASP | p.Asp252Gly | P28331 |
| HOMO SAPIENS | 1 | 1019 | 5xtb | M | 408 | ARG | p.Arg408Cys | P28331 |
| HOMO SAPIENS | 2 | 1019 | 5xtb | M | 417 | ARG | p.Arg417Ser | P28331 |
| HOMO SAPIENS | 1 | 1019 | 5xtb | M | 557 | ARG | p.Arg557Ter | P28331 |
| HOMO SAPIENS | 0 | 1019 | 5xtb | M | 567 | ILE | p.Ile567Val | P28331 |
| HOMO SAPIENS | 1 | 1019 | 5xtb | M | 595 | THR | p.Thr595Ala | P28331 |
| HOMO SAPIENS | 1 | 1019 | 5xtb | M | 703 | ALA | p.Ala703Thr | P28331 |
| HOMO SAPIENS | -1 | 3437 | 5hyn | A | 185 | ASP | p.Asp185His | Q15910 |
| HOMO SAPIENS | -1 | 3437 | 5hyn | A | 482 | ALA | p.Ala482Thr | Q15910 |
| HOMO SAPIENS | 0 | 8201 | 2wfd | A | 373 | TYR | p.Tyr373Cys | Q9P2J5 |
| HOMO SAPIENS | 1 | 3590 | 3edi | A | 90 | VAL | p.Val238Ala | O43897 |
| HOMO SAPIENS | 6 | 4555 | 1ytq | A | 64 | ALA | p.Ala65Ser | P43320 |
| HOMO SAPIENS | 0 | 4555 | 1ytq | A | 194 | TRP | p.Trp195Gly | P43320 |
| HOMO SAPIENS | 0 | 1013 | 2nn6 | C | 272 | SER | p.Ser272Thr | Q96B26 |
| HOMO SAPIENS | -1 | 29026 | 5m73 | D | 530 | ILE | p.Ile530Asn | O76094 |
| HOMO SAPIENS | 8 | 2252 | 1d5v | A | 57 | SER | p.Ser125Leu | Q99958 |
| HOMO SAPIENS | 1 | 4337 | 3ecr | A | 29 | ARG | p.Arg26His | P08397 |
| HOMO SAPIENS | 2 | 4337 | 3ecr | A | 34 | ALA | p.Ala31Thr | P08397 |
| HOMO SAPIENS | 2 | 4337 | 3ecr | A | 37 | GLN | p.Gln34Lys | P08397 |
| HOMO SAPIENS | 1 | 4337 | 3ecr | A | 58 | ALA | p.Ala55Ser | P08397 |
| HOMO SAPIENS | 0 | 4337 | 3ecr | A | 84 | LEU | p.Leu81Pro | P08397 |
| HOMO SAPIENS | 0 | 4337 | 3ecr | A | 114 | GLY | p.Gly111Arg | P08397 |
| HOMO SAPIENS | 1 | 4337 | 3ecr | A | 119 | ARG | p.Arg116Trp | P08397 |
| HOMO SAPIENS | 1 | 4337 | 3ecr | A | 152 | ARG | p.Arg149Gln | P08397 |
| HOMO SAPIENS | 1 | 4337 | 3ecr | A | 170 | ARG | p.Arg167Trp | P08397 |
| HOMO SAPIENS | 1 | 4337 | 3ecr | A | 176 | ARG | p.Arg173Gln | P08397 |
| HOMO SAPIENS | 0 | 4337 | 3ecr | A | 180 | LEU | p.Leu177Arg | P08397 |
| HOMO SAPIENS | 0 | 4337 | 3ecr | A | 181 | ASP | p.Asp178Asn | P08397 |
| HOMO SAPIENS | 1 | 4337 | 3ecr | A | 198 | ARG | p.Arg195Cys | P08397 |
| HOMO SAPIENS | 0 | 4337 | 3ecr | A | 201 | TRP | p.Trp198Ter | P08397 |
| HOMO SAPIENS | 1 | 4337 | 3ecr | A | 204 | ARG | p.Arg201Trp | P08397 |
| HOMO SAPIENS | 1 | 4337 | 3ecr | A | 219 | GLY | p.Gly216Asp | P08397 |
| HOMO SAPIENS | 1 | 4337 | 3ecr | A | 226 | GLU | p.Glu223Lys | P08397 |
| HOMO SAPIENS | 0 | 4337 | 3ecr | A | 248 | LEU | p.Leu245Arg | P08397 |

| | | | | | | | | |
|---|---|---|---|---|---|---|---|---|
| HOMO SAPIENS | 1 | 4337 | 3ecr | A | 250 | CYS | p.Cys247Arg | P08397 |
| HOMO SAPIENS | 0 | 4337 | 3ecr | A | 253 | GLU | p.Glu250Lys | P08397 |
| HOMO SAPIENS | 0 | 4337 | 3ecr | A | 255 | ALA | p.Ala252Thr | P08397 |
| HOMO SAPIENS | 0 | 4337 | 3ecr | A | 255 | ALA | p.Ala252Val | P08397 |
| HOMO SAPIENS | 0 | 4337 | 3ecr | A | 259 | HIS | p.His256Asn | P08397 |
| HOMO SAPIENS | 0 | 4337 | 3ecr | A | 324 | ARG | p.Arg321His | P08397 |
| HOMO SAPIENS | -1 | 4337 | 3ecr | A | 362 | ASP | p.Asp359Asn | P08397 |
| HOMO SAPIENS | 1 | 1204 | 5lwv | A | 854 | ILE | p.Ala864Thr | O15294 |
| HOMO SAPIENS | 1 | 25016 | 1m4u | A | 35 | PRO | p.Pro35Arg | Q13253 |
| HOMO SAPIENS | 1 | 25016 | 1m4u | A | 35 | PRO | p.Pro35Ser | Q13253 |
| HOMO SAPIENS | 1 | 25016 | 1m4u | A | 167 | ARG | p.Arg167Gly | Q13253 |
| HOMO SAPIENS | 1 | 25016 | 1m4u | A | 184 | CYS | p.Cys184Tyr | Q13253 |
| HOMO SAPIENS | 0 | 25016 | 1m4u | A | 189 | GLY | p.Gly189Cys | Q13253 |
| HOMO SAPIENS | 1 | 25016 | 1m4u | A | 204 | ARG | p.Arg204Leu | Q13253 |
| HOMO SAPIENS | 1 | 25016 | 1m4u | A | 205 | TRP | p.Trp205Cys | Q13253 |
| HOMO SAPIENS | 1 | 25016 | 1m4u | A | 217 | TRP | p.Trp217Gly | Q13253 |
| HOMO SAPIENS | 0 | 25016 | 1m4u | A | 222 | TYR | p.Tyr222Cys | Q13253 |
| HOMO SAPIENS | 0 | 25016 | 1m4u | A | 222 | TYR | p.Tyr222Asp | Q13253 |
| HOMO SAPIENS | 1 | 25016 | 1m4u | A | 223 | PRO | p.Pro223Leu | Q13253 |
| HOMO SAPIENS | 0 | 1394 | 1um7 | A | 95 | ALA | p.Ala463Val | Q92796 |
| HOMO SAPIENS | 0 | 21294 | 4jpz | B | 1788 | GLU | p.Glu1788Asp | Q99250 |
| HOMO SAPIENS | 9 | 24068 | 2db6 | A | 47 | ARG | p.Arg119Cys | Q96MF2 |
| HOMO SAPIENS | 5 | 1869 | 4omt | A | 39 | ARG | p.Arg39Leu | P08237 |
| HOMO SAPIENS | 2 | 1869 | 4omt | A | 100 | ARG | p.Arg100Gln | P08237 |
| HOMO SAPIENS | 7 | 1869 | 4omt | A | 543 | ASP | p.Asp543Ala | P08237 |
| HOMO SAPIENS | 1 | 1869 | 4omt | A | 686 | TRP | p.Trp686Cys | P08237 |
| HOMO SAPIENS | 1 | 1869 | 4omt | A | 696 | ARG | p.Arg696His | P08237 |
| HOMO SAPIENS | 0 | 12163 | 4wxx | A | 495 | TYR | p.Tyr495Cys | P26358 |
| HOMO SAPIENS | 0 | 12163 | 4wxx | A | 528 | ILE | p.Ile528Met | P26358 |
| HOMO SAPIENS | 0 | 12391 | 1j1d | C | 36 | LYS | p.Lys36Gln | P19429 |
| HOMO SAPIENS | 0 | 12391 | 1j1d | C | 82 | PRO | p.Pro82Ser | P19429 |
| HOMO SAPIENS | 7 | 12391 | 1j1d | C | 102 | ALA | p.Ala102Thr | P19429 |
| HOMO SAPIENS | 1 | 5000 | 4cca | A | 17 | GLY | p.Gly17Arg | Q15833 |
| HOMO SAPIENS | 0 | 5000 | 4cca | A | 205 | VAL | p.Val205Ile | Q15833 |
| HOMO SAPIENS | 0 | 5000 | 4cca | A | 209 | LEU | p.Leu209Pro | Q15833 |
| HOMO SAPIENS | 1 | 5000 | 4cca | A | 459 | ARG | p.Arg459Trp | Q15833 |
| HOMO SAPIENS | 7 | 5000 | 4cca | A | 477 | PRO | p.Pro477Leu | Q15833 |
| HOMO SAPIENS | -1 | 5000 | 4cca | A | 526 | ILE | p.Ile526Val | Q15833 |
| HOMO SAPIENS | 0 | 5000 | 4cca | A | 541 | GLY | p.Gly541Ser | Q15833 |
| HOMO SAPIENS | 0 | 5000 | 4cca | A | 555 | ARG | p.Arg555Gly | Q15833 |
| HOMO SAPIENS | 5 | 28551 | 3vcm | P | 26 | ARG | p.Arg49Ter | P00797 |
| HOMO SAPIENS | 6 | 36453 | 5dzz | A | 2070 | ASP | p.Asp2070His | P15924 |
| HOMO SAPIENS | 0 | 36453 | 5dzz | A | 2166 | ARG | p.Arg2166Ter | P15924 |
| HOMO SAPIENS | 0 | 20268 | 2da6 | A | 96 | ASN | p.Asn321Ser | P35680 |
| HOMO SAPIENS | 1 | 1405 | 3iar | A | 8 | ASP | p.Asp8Asn | P00813 |
| HOMO SAPIENS | 8 | 1405 | 3iar | A | 15 | HIS | p.His15Asp | P00813 |
| HOMO SAPIENS | 6 | 1405 | 3iar | A | 74 | GLY | p.Gly74Val | P00813 |
| HOMO SAPIENS | 6 | 1405 | 3iar | A | 80 | LYS | p.Lys80Arg | P00813 |
| HOMO SAPIENS | 5 | 1405 | 3iar | A | 97 | TYR | p.Tyr97Cys | P00813 |
| HOMO SAPIENS | 5 | 1405 | 3iar | A | 101 | ARG | p.Arg101Leu | P00813 |
| HOMO SAPIENS | 5 | 1405 | 3iar | A | 101 | ARG | p.Arg101Trp | P00813 |
| HOMO SAPIENS | 5 | 1405 | 3iar | A | 106 | LEU | p.Leu106Val | P00813 |
| HOMO SAPIENS | 0 | 1405 | 3iar | A | 107 | LEU | p.Leu107Pro | P00813 |
| HOMO SAPIENS | 9 | 1405 | 3iar | A | 156 | ARG | p.Arg156Cys | P00813 |

| | | | | | | | | |
|---|---|---|---|---|---|---|---|---|
| HOMO SAPIENS | 6 | 1405 | 3iar | A | 211 | ARG | p.Arg211His | P00813 |
| HOMO SAPIENS | 0 | 1405 | 3iar | A | 216 | GLY | p.Gly216Arg | P00813 |
| HOMO SAPIENS | 0 | 1405 | 3iar | A | 291 | SER | p.Ser291Leu | P00813 |
| HOMO SAPIENS | 6 | 1405 | 3iar | A | 304 | LEU | p.Leu304Arg | P00813 |
| HOMO SAPIENS | 0 | 1405 | 3iar | A | 329 | ALA | p.Ala329Val | P00813 |
| HOMO SAPIENS | -1 | 31340 | 4atu | I | 186 | ARG | p.Arg186His | O43602 |
| HOMO SAPIENS | -1 | 31340 | 4atu | I | 192 | ARG | p.Arg192Trp | O43602 |
| HOMO SAPIENS | -1 | 31340 | 4atu | I | 196 | ARG | p.Arg196His | O43602 |
| HOMO SAPIENS | -1 | 31340 | 4atu | I | 203 | THR | p.Thr203Arg | O43602 |
| HOMO SAPIENS | -1 | 31340 | 4atu | I | 286 | ALA | p.Ala286Ser | O43602 |
| HOMO SAPIENS | 4 | 30734 | 1fsu | A | 95 | ARG | p.Arg95Gln | P15848 |
| HOMO SAPIENS | 0 | 30734 | 1fsu | A | 117 | CYS | p.Cys117Arg | P15848 |
| HOMO SAPIENS | 0 | 30734 | 1fsu | A | 137 | GLY | p.Gly137Val | P15848 |
| HOMO SAPIENS | 0 | 30734 | 1fsu | A | 210 | TYR | p.Tyr210Cys | P15848 |
| HOMO SAPIENS | 0 | 30734 | 1fsu | A | 236 | LEU | p.Leu236Pro | P15848 |
| HOMO SAPIENS | 1 | 30734 | 1fsu | A | 315 | ARG | p.Arg315Gln | P15848 |
| HOMO SAPIENS | 0 | 30734 | 1fsu | A | 358 | VAL | p.Val358Met | P15848 |
| HOMO SAPIENS | 2 | 30734 | 1fsu | A | 376 | VAL | p.Val376Met | P15848 |
| HOMO SAPIENS | 0 | 30734 | 1fsu | A | 384 | SER | p.Ser384Asn | P15848 |
| HOMO SAPIENS | 0 | 30734 | 1fsu | A | 393 | HIS | p.His393Pro | P15848 |
| HOMO SAPIENS | 0 | 30734 | 1fsu | A | 405 | CYS | p.Cys405Tyr | P15848 |
| HOMO SAPIENS | 0 | 30734 | 1fsu | A | 458 | ASN | p.Asn458Ser | P15848 |
| HOMO SAPIENS | 1 | 30734 | 1fsu | A | 484 | ARG | p.Arg484Gly | P15848 |
| HOMO SAPIENS | 0 | 38072 | 1wmv | A | 51 | PRO | p.Pro98Leu | Q9NZC7 |
| HOMO SAPIENS | -1 | 5674 | 1j1e | C | 185 | ASN | p.Asn185Lys | P19429 |
| HOMO SAPIENS | -1 | 5674 | 1j1e | C | 192 | ARG | p.Arg192His | P19429 |
| HOMO SAPIENS | -1 | 5674 | 1j1e | C | 196 | ASP | p.Asp196Asn | P19429 |
| HOMO SAPIENS | -1 | 5674 | 1j1e | C | 206 | LYS | p.Lys206Gln | P19429 |
| HOMO SAPIENS | 0 | 38067 | 1wlp | A | 13 | PRO | p.Pro156Gln | P13498 |
| HOMO SAPIENS | -1 | 20441 | 3vfd | A | 289 | HIS | p.His289Tyr | Q9UBP0 |
| HOMO SAPIENS | 2 | 20441 | 3vfd | A | 344 | ILE | p.Ile344Lys | Q9UBP0 |
| HOMO SAPIENS | 0 | 20441 | 3vfd | A | 362 | SER | p.Ser362Cys | Q9UBP0 |
| HOMO SAPIENS | 1 | 20441 | 3vfd | A | 386 | ASN | p.Asn386Ser | Q9UBP0 |
| HOMO SAPIENS | 9 | 20441 | 3vfd | A | 441 | ASP | p.Asp441Gly | Q9UBP0 |
| HOMO SAPIENS | -1 | 20441 | 3vfd | A | 448 | CYS | p.Cys448Tyr | Q9UBP0 |
| HOMO SAPIENS | 2 | 20441 | 3vfd | A | 500 | ARG | p.Arg499Cys | Q9UBP0 |
| HOMO SAPIENS | 1 | 20441 | 3vfd | A | 563 | ARG | p.Arg562Gly | Q9UBP0 |
| HOMO SAPIENS | 3 | 18471 | 4dx8 | J | 26 | ARG | p.Arg26Gln | O00522 |
| HOMO SAPIENS | -1 | 18471 | 4dx8 | J | 101 | GLY | p.Gly101Val | O00522 |
| HOMO SAPIENS | -1 | 34735 | 2ly4 | B | 83 | ALA | p.Ala83Glu | P04637 |
| HOMO SAPIENS | 0 | 31128 | 4cvh | A | 116 | ARG | p.Arg116Cys | A4D126 |
| HOMO SAPIENS | 8 | 31128 | 4cvh | A | 215 | GLN | p.Gln215Ter | A4D126 |
| HOMO SAPIENS | 0 | 31128 | 4cvh | A | 268 | ARG | p.Arg268Ter | A4D126 |
| HOMO SAPIENS | 0 | 31128 | 4cvh | A | 316 | THR | p.Thr316Lys | A4D126 |
| HOMO SAPIENS | 1 | 7164 | 3mt5 | A | 369 | ASP | p.Asp434Gly | Q12791 |
| HOMO SAPIENS | 2 | 26812 | 2wtk | F | 49 | TYR | p.Tyr49Asp | Q15831 |
| HOMO SAPIENS | 0 | 26812 | 2wtk | F | 67 | LEU | p.Leu67Pro | Q15831 |
| HOMO SAPIENS | 4 | 26812 | 2wtk | F | 104 | ARG | p.Arg104Ser | Q15831 |
| HOMO SAPIENS | 0 | 26812 | 2wtk | F | 135 | GLY | p.Gly135Arg | Q15831 |
| HOMO SAPIENS | 0 | 26812 | 2wtk | F | 163 | GLY | p.Gly163Asp | Q15831 |
| HOMO SAPIENS | 2 | 26812 | 2wtk | F | 194 | ALA | p.Asp194Asn | Q15831 |
| HOMO SAPIENS | 2 | 1357 | 5xtd | W | 57 | ARG | p.Arg57His | Q9P0J0 |
| HOMO SAPIENS | 0 | 28086 | 4nn2 | A | 229 | HIS | p.His229Arg | Q8IWS0 |
| HOMO SAPIENS | 0 | 28086 | 4nn2 | A | 234 | LYS | p.Lys234Glu | Q8IWS0 |

| | | | | | | | | |
|---|---|---|---|---|---|---|---|---|
| HOMO SAPIENS | 0 | 28086 | 4nn2 | A | 257 | ARG | p.Arg257Gly | Q8IWS0 |
| HOMO SAPIENS | 5 | 31182 | 2a1x | A | 176 | GLN | p.Gln176Lys | O14832 |
| HOMO SAPIENS | 1 | 31182 | 2a1x | A | 204 | GLY | p.Gly204Ser | O14832 |
| HOMO SAPIENS | 0 | 31182 | 2a1x | A | 269 | ASN | p.Asn269His | O14832 |
| HOMO SAPIENS | 3 | 31182 | 2a1x | A | 275 | ARG | p.Arg275Gln | O14832 |
| HOMO SAPIENS | 3 | 31182 | 2a1x | A | 275 | ARG | p.Arg275Trp | O14832 |
| HOMO SAPIENS | 4 | 2823 | 5xtb | P | 145 | THR | p.Thr145Ile | O75489 |
| HOMO SAPIENS | 5 | 2823 | 5xtb | P | 199 | ARG | p.Arg199Trp | O75489 |
| HOMO SAPIENS | 0 | 8645 | 2h5g | A | 372 | SER | p.Ser372Tyr | P54886 |
| HOMO SAPIENS | 3 | 8645 | 2h5g | A | 715 | ASP | p.Asp715His | P54886 |
| HOMO SAPIENS | 0 | 8645 | 2h5g | A | 765 | ARG | p.Arg765Gln | P54886 |
| HOMO SAPIENS | 0 | 8645 | 2h5g | A | 782 | TYR | p.Tyr782Cys | P54886 |
| HOMO SAPIENS | 0 | 8645 | 2h5g | A | 784 | HIS | p.His784Tyr | P54886 |
| HOMO SAPIENS | 1 | 13992 | 2cue | A | 39 | ARG | p.Arg242Thr | P26367 |
| HOMO SAPIENS | 9 | 13992 | 2cue | A | 55 | PHE | p.Phe258Ser | P26367 |
| HOMO SAPIENS | 0 | 30418 | 2xsw | A | 303 | VAL | p.Val303Met | Q9NRR6 |
| HOMO SAPIENS | 0 | 30418 | 2xsw | A | 315 | PRO | p.Pro315Leu | Q9NRR6 |
| HOMO SAPIENS | 0 | 30418 | 2xsw | A | 341 | GLY | p.Gly341Ser | Q9NRR6 |
| HOMO SAPIENS | 0 | 30418 | 2xsw | A | 378 | ARG | p.Arg378Cys | Q9NRR6 |
| HOMO SAPIENS | 0 | 30418 | 2xsw | A | 435 | ARG | p.Arg435Gln | Q9NRR6 |
| HOMO SAPIENS | 0 | 30418 | 2xsw | A | 515 | ARG | p.Arg515Trp | Q9NRR6 |
| HOMO SAPIENS | 1 | 30418 | 2xsw | A | 526 | PRO | p.Pro526Leu | Q9NRR6 |
| HOMO SAPIENS | 0 | 30418 | 2xsw | A | 563 | ARG | p.Arg563His | Q9NRR6 |
| HOMO SAPIENS | 2 | 30418 | 2xsw | A | 585 | ARG | p.Arg585His | Q9NRR6 |
| HOMO SAPIENS | 2 | 11167 | 4y99 | B | 205 | ARG | p.Arg215Leu | P45379 |
| HOMO SAPIENS | 5 | 11167 | 4y99 | B | 244 | GLU | p.Glu254Asp | P45379 |
| HOMO SAPIENS | 1 | 11167 | 4y99 | B | 253 | LYS | p.Lys263Arg | P45379 |
| HOMO SAPIENS | 8 | 11167 | 4y99 | B | 265 | ARG | p.Arg275Gln | P45379 |
| HOMO SAPIENS | 3 | 11167 | 4y99 | B | 270 | ASP | p.Asp280Asn | P45379 |
| HOMO SAPIENS | -1 | 11167 | 4y99 | B | 278 | ARG | p.Arg288Pro | P45379 |
| HOMO SAPIENS | -1 | 11167 | 4y99 | B | 286 | ARG | p.Arg296Cys | P45379 |
| HOMO SAPIENS | -1 | 11167 | 4y99 | B | 286 | ARG | p.Arg296His | P45379 |
| HOMO SAPIENS | 0 | 15993 | 4po6 | A | 197 | ARG | p.Arg197His | P29597 |
| HOMO SAPIENS | -1 | 15993 | 4po6 | A | 362 | VAL | p.Val362Phe | P29597 |
| HOMO SAPIENS | -1 | 15993 | 4po6 | A | 363 | GLY | p.Gly363Ser | P29597 |
| HOMO SAPIENS | 0 | 5646 | 5uak | A | 7 | GLU | p.Glu7Ter | P13569 |
| HOMO SAPIENS | 0 | 5646 | 5uak | A | 27 | GLY | p.Gly27Ter | P13569 |
| HOMO SAPIENS | 0 | 5646 | 5uak | A | 46 | ALA | p.Ala46Asp | P13569 |
| HOMO SAPIENS | 0 | 5646 | 5uak | A | 60 | GLU | p.Glu60Ter | P13569 |
| HOMO SAPIENS | 0 | 5646 | 5uak | A | 67 | PRO | p.Pro67Leu | P13569 |
| HOMO SAPIENS | 0 | 5646 | 5uak | A | 74 | ARG | p.Arg74Trp | P13569 |
| HOMO SAPIENS | 0 | 5646 | 5uak | A | 75 | ARG | p.Arg75Ter | P13569 |
| HOMO SAPIENS | 0 | 5646 | 5uak | A | 85 | GLY | p.Gly85Glu | P13569 |
| HOMO SAPIENS | 0 | 5646 | 5uak | A | 91 | GLY | p.Gly91Arg | P13569 |
| HOMO SAPIENS | 0 | 5646 | 5uak | A | 92 | GLU | p.Glu92Lys | P13569 |
| HOMO SAPIENS | 0 | 5646 | 5uak | A | 98 | GLN | p.Gln98Arg | P13569 |
| HOMO SAPIENS | 0 | 5646 | 5uak | A | 98 | GLN | p.Gln98Ter | P13569 |
| HOMO SAPIENS | 0 | 5646 | 5uak | A | 109 | TYR | p.Tyr109Cys | P13569 |
| HOMO SAPIENS | 0 | 5646 | 5uak | A | 110 | ASP | p.Asp110His | P13569 |
| HOMO SAPIENS | 0 | 5646 | 5uak | A | 117 | ARG | p.Arg117Cys | P13569 |
| HOMO SAPIENS | 0 | 5646 | 5uak | A | 117 | ARG | p.Arg117His | P13569 |
| HOMO SAPIENS | 0 | 5646 | 5uak | A | 125 | ILE | p.Ile125Thr | P13569 |
| HOMO SAPIENS | 0 | 5646 | 5uak | A | 148 | ILE | p.Ile148Thr | P13569 |
| HOMO SAPIENS | 0 | 5646 | 5uak | A | 170 | ARG | p.Arg170His | P13569 |

| | | | | | | | | |
|---|---|---|---|---|---|---|---|---|
| HOMO SAPIENS | 0 | 5646 | 5uak | A | 178 | GLY | p.Gly178Arg | P13569 |
| HOMO SAPIENS | 0 | 5646 | 5uak | A | 206 | LEU | p.Leu206Trp | P13569 |
| HOMO SAPIENS | 0 | 5646 | 5uak | A | 373 | ASP | p.Asp373Asn | P13569 |
| HOMO SAPIENS | -1 | 5646 | 5uak | A | 709 | ARG | p.Arg709Ter | P13569 |
| HOMO SAPIENS | -1 | 5646 | 5uak | A | 732 | LEU | p.Leu732Ter | P13569 |
| HOMO SAPIENS | -1 | 5646 | 5uak | A | 785 | ARG | p.Arg785Ter | P13569 |
| HOMO SAPIENS | -1 | 5646 | 5uak | A | 831 | GLU | p.Glu831Ter | P13569 |
| HOMO SAPIENS | -1 | 5646 | 5uak | A | 836 | ASP | p.Asp836Tyr | P13569 |
| HOMO SAPIENS | 0 | 5646 | 5uak | A | 851 | ARG | p.Arg851Ter | P13569 |
| HOMO SAPIENS | 0 | 5646 | 5uak | A | 913 | TYR | p.Tyr913Cys | P13569 |
| HOMO SAPIENS | 0 | 5646 | 5uak | A | 913 | TYR | p.Tyr913Ter | P13569 |
| HOMO SAPIENS | 0 | 5646 | 5uak | A | 945 | SER | p.Ser945Leu | P13569 |
| HOMO SAPIENS | 0 | 5646 | 5uak | A | 949 | HIS | p.His949Tyr | P13569 |
| HOMO SAPIENS | 0 | 5646 | 5uak | A | 970 | GLY | p.Gly970Asp | P13569 |
| HOMO SAPIENS | 0 | 5646 | 5uak | A | 1027 | ILE | p.Ile1027Thr | P13569 |
| HOMO SAPIENS | 0 | 5646 | 5uak | A | 1032 | TYR | p.Tyr1032Cys | P13569 |
| HOMO SAPIENS | 0 | 5646 | 5uak | A | 1061 | GLY | p.Gly1061Arg | P13569 |
| HOMO SAPIENS | 1 | 5646 | 5uak | A | 1065 | LEU | p.Leu1065Pro | P13569 |
| HOMO SAPIENS | 1 | 5646 | 5uak | A | 1066 | ARG | p.Arg1066Cys | P13569 |
| HOMO SAPIENS | 1 | 5646 | 5uak | A | 1066 | ARG | p.Arg1066His | P13569 |
| HOMO SAPIENS | 1 | 5646 | 5uak | A | 1067 | ALA | p.Ala1067Thr | P13569 |
| HOMO SAPIENS | 0 | 5646 | 5uak | A | 1069 | GLY | p.Gly1069Arg | P13569 |
| HOMO SAPIENS | 0 | 5646 | 5uak | A | 1070 | ARG | p.Arg1070Trp | P13569 |
| HOMO SAPIENS | 1 | 5646 | 5uak | A | 1071 | GLN | p.Gln1071Pro | P13569 |
| HOMO SAPIENS | 0 | 5646 | 5uak | A | 1074 | PHE | p.Phe1074Leu | P13569 |
| HOMO SAPIENS | 0 | 5646 | 5uak | A | 1077 | LEU | p.Leu1077Pro | P13569 |
| HOMO SAPIENS | 0 | 5646 | 5uak | A | 1089 | TRP | p.Trp1089Ter | P13569 |
| HOMO SAPIENS | 0 | 5646 | 5uak | A | 1092 | TYR | p.Tyr1092Ter | P13569 |
| HOMO SAPIENS | 0 | 5646 | 5uak | A | 1101 | MET | p.Met1101Lys | P13569 |
| HOMO SAPIENS | 0 | 5646 | 5uak | A | 1156 | LEU | p.Leu1156Phe | P13569 |
| HOMO SAPIENS | 0 | 5646 | 5uak | A | 1158 | ARG | p.Arg1158Ter | P13569 |
| HOMO SAPIENS | 0 | 5646 | 5uak | A | 1162 | ARG | p.Arg1162Leu | P13569 |
| HOMO SAPIENS | 0 | 5646 | 5uak | A | 1162 | ARG | p.Arg1162Ter | P13569 |
| HOMO SAPIENS | -1 | 5646 | 5uak | A | 1476 | GLN | p.Gln1476Ter | P13569 |
| HOMO SAPIENS | 0 | 22763 | 1imv | A | 52 | MET | p.Thr72Met | P36955 |
| HOMO SAPIENS | 0 | 22763 | 1imv | A | 301 | TYR | p.Tyr321Ter | P36955 |
| HOMO SAPIENS | 0 | 23179 | 5fql | A | 160 | PRO | p.Pro160Arg | P22304 |
| HOMO SAPIENS | 0 | 23179 | 5fql | A | 172 | ARG | p.Arg172Ter | P22304 |
| HOMO SAPIENS | 0 | 23179 | 5fql | A | 214 | THR | p.Thr214Met | P22304 |
| HOMO SAPIENS | 0 | 23179 | 5fql | A | 309 | THR | p.Thr309Ala | P22304 |
| HOMO SAPIENS | 0 | 23179 | 5fql | A | 333 | SER | p.Ser333Leu | P22304 |
| HOMO SAPIENS | 0 | 23179 | 5fql | A | 443 | ARG | p.Arg443Ter | P22304 |
| HOMO SAPIENS | 0 | 23179 | 5fql | A | 468 | ARG | p.Arg468Leu | P22304 |
| HOMO SAPIENS | 0 | 23179 | 5fql | A | 468 | ARG | p.Arg468Trp | P22304 |
| HOMO SAPIENS | 0 | 23179 | 5fql | A | 502 | TRP | p.Trp502Ser | P22304 |
| HOMO SAPIENS | 9 | 20989 | 1nfi | F | 185 | THR | p.Thr185Met | P25963 |
| HOMO SAPIENS | 4 | 20989 | 1nfi | F | 194 | GLY | p.Gly194Ala | P25963 |
| HOMO SAPIENS | 0 | 11538 | 3eab | A | 162 | VAL | p.Val162Ile | Q9UBP0 |
| HOMO SAPIENS | 0 | 33399 | 2p39 | A | 71 | SER | p.Ser71Gly | Q9GZV9 |
| HOMO SAPIENS | 3 | 33399 | 2p39 | A | 96 | MET | p.Met96Thr | Q9GZV9 |
| HOMO SAPIENS | 0 | 33399 | 2p39 | A | 129 | SER | p.Ser129Phe | Q9GZV9 |
| HOMO SAPIENS | -1 | 33399 | 2p39 | A | 172 | PRO | p.Pro172Leu | Q9GZV9 |
| HOMO SAPIENS | -1 | 33399 | 2p39 | A | 176 | ARG | p.Arg176Gln | Q9GZV9 |
| HOMO SAPIENS | -1 | 33399 | 2p39 | A | 179 | ARG | p.Arg179Trp | Q9GZV9 |

| | | | | | | | | |
|---|---|---|---|---|---|---|---|---|
| HOMO SAPIENS | -1 | 33399 | 2p39 | A | 179 | ARG | p.Arg179Gln | Q9GZV9 |
| HOMO SAPIENS | 0 | 16217 | 2dne | A | 79 | ILE | p.Ile163Leu | P10515 |
| HOMO SAPIENS | 8 | 7666 | 4kbq | A | 65 | ASN | p.Asn65Ser | Q9UNE7 |
| HOMO SAPIENS | 1 | 6407 | 1ikt | A | 71 | THR | p.Thr687Ile | P51659 |
| HOMO SAPIENS | 0 | 6407 | 1ikt | A | 112 | MET | p.Met728Val | P51659 |
| HOMO SAPIENS | 0 | 5849 | 4ihb | A | 67 | VAL | p.Val67Asp | O75923 |
| HOMO SAPIENS | 6 | 17703 | 5fcn | A | 20 | HIS | p.His99Gln | Q66GS9 |
| HOMO SAPIENS | 8 | 17703 | 5fcn | A | 33 | ARG | p.Arg112His | Q66GS9 |
| HOMO SAPIENS | 0 | 34095 | 2E+073 | A | 101 | HIS | p.His101Tyr | P05129 |
| HOMO SAPIENS | -1 | 41358 | 4liq | E | 507 | THR | p.Thr507Met | P07333 |
| HOMO SAPIENS | -1 | 21290 | 1fch | A | 235 | MET | p.Met272Arg | P50542 |
| HOMO SAPIENS | 5 | 23431 | 6pax | A | 23 | ARG | p.Arg26Gly | P26367 |
| HOMO SAPIENS | 1 | 23431 | 6pax | A | 61 | GLY | p.Gly64Val | P26367 |
| HOMO SAPIENS | 1 | 23431 | 6pax | A | 65 | PRO | p.Pro68Ser | P26367 |
| HOMO SAPIENS | 1 | 23431 | 6pax | A | 116 | SER | p.Ser119Arg | P26367 |
| HOMO SAPIENS | 9 | 23431 | 6pax | A | 123 | VAL | p.Val126Asp | P26367 |
| HOMO SAPIENS | 9 | 23431 | 6pax | A | 125 | ARG | p.Arg128Cys | P26367 |
| HOMO SAPIENS | 9 | 33003 | 4zrj | B | 535 | LEU | p.Leu535Pro | P35240 |
| HOMO SAPIENS | 7 | 33003 | 4zrj | B | 538 | GLN | p.Gln538Pro | P35240 |
| HOMO SAPIENS | 0 | 33003 | 4zrj | B | 585 | TRP | p.Ala585Thr | P35240 |
| HOMO SAPIENS | 0 | 3800 | 2nzj | A | 193 | VAL | p.Val193Met | O75628 |

| gene_name | max_cons | max_bs_id | max_pdb_id | max_chain_id |
|---|---|---|---|---|
| AKR1D1 | 0.306 | 22 | 3dop | A |
| AKR1D1 | 0.216 | 34 | 3cav | A |
| AKR1D1 | 0.276 | 21 | 3cot | A |
| AKR1D1 | 0.258 | 56 | 3cas | A |
| ALPP | None | None | None | None |
| BMPR1A | None | None | None | None |
| BMPR1A | None | None | None | None |
| FGFR2 | None | None | None | None |
| PROC | 0.587 | 3 | 1aut | C |
| PROC | 0.174 | 16 | 1aut | C |
| PROC | 0.007 | 73 | 1aut | C |
| PROC | 0.396 | 10 | 3f6u | H |
| PROC | 0.247 | 13 | 3f6u | H |
| PROC | 0.021 | 43 | 3f6u | H |
| PROC | 0.024 | 42 | 1aut | C |
| PROC | 0.191 | 15 | 3f6u | H |
| PROC | 0.09 | 42 | 3f6u | H |
| PROC | 0.587 | 3 | 1aut | C |
| PROC | 0.191 | 26 | 1aut | C |
| PROC | 0.003 | 107 | 3f6u | H |
| INS | 1 | 2 | 3zs2 | J |
| INS | 0.85 | 1 | 4ajx | D |
| INS | 1 | 1 | 3zs2 | J |
| INS | 1 | 1 | 3zs2 | J |
| FTL | 0.627 | 11 | 2fg8 | B |
| FTL | 0.313 | 62 | 2fg8 | G |
| FTL | 0.544 | 24 | 2fg8 | D |
| TNFSF11 | None | None | None | None |
| EHMT1 | None | None | None | None |
| EHMT1 | None | None | None | None |
| EHMT1 | None | None | None | None |
| CA2 | 0.939 | 1 | 4idr | X |
| CA2 | 0.939 | 1 | 4idr | X |
| CA2 | 0.902 | 5 | 4idr | X |
| CA2 | 0.902 | 5 | 4idr | X |
| SMAD4 | None | None | None | None |
| SMAD4 | None | None | None | None |
| TTR | 0.691 | 12 | 1ttb | B |
| TTR | 0.961 | 7 | 3grb | A |
| TTR | 0.891 | 4 | 3dk2 | B |
| TTR | 0.947 | 3 | 1tta | A |
| TTR | 0.923 | 1 | 3grb | B |
| TTR | 0.891 | 1 | 4y9b | B |
| TTR | 0.891 | 1 | 4y9b | B |
| TTR | 0.882 | 15 | 3grb | A |
| TTR | 0.909 | 5 | 4tl4 | A |
| TTR | 0.836 | 7 | 1z7j | B |
| TTR | 0.882 | 15 | 3grb | A |
| TTR | 0.891 | 5 | 3i9i | B |
| TTR | 0.902 | 9 | 3m1o | B |
| TTR | 0.902 | 9 | 3m1o | B |
| TTR | 0.902 | 9 | 3m1o | B |
| TTR | 0.636 | 16 | 1thc | A |

| | | | | |
|---|---|---|---|---|
| TTR | 0.902 | 9 | 3m1o | B |
| TTR | 0.836 | 7 | 1bze | A |
| TTR | 0.846 | 16 | 3gps | A |
| TTR | 0.846 | 16 | 3gps | A |
| TTR | 0.909 | 5 | 4tl4 | A |
| TTR | 0.909 | 5 | 4tl4 | A |
| TTR | 0.673 | 15 | 3dk2 | B |
| TTR | 0.945 | 4 | 4deu | A |
| TTR | 0.941 | 3 | 3did | B |
| TTR | 0.941 | 3 | 3did | B |
| TTR | 0.868 | 6 | 4l1t | A |
| TTR | 0.882 | 19 | 3gps | B |
| TTR | 0.891 | 4 | 3dk2 | B |
| TTR | 0.941 | 3 | 3did | B |
| TTR | 0.923 | 1 | 4hiq | A |
| TTR | 0.923 | 1 | 4hiq | A |
| TTR | 0.961 | 7 | 3grb | A |
| TTR | 0.839 | 7 | 1tsh | B |
| TTR | 0.891 | 6 | 1ttb | B |
| ALK | 0.682 | 1 | 4cd0 | A |
| ALK | 0.163 | 13 | 2xp2 | A |
| THRB | 0.155 | 15 | 2pin | B |
| THRB | 0.099 | 39 | 2pin | B |
| THRB | 0.155 | 15 | 2pin | B |
| THRB | 0.071 | 50 | 2pin | A |
| THRB | 0.155 | 15 | 2pin | B |
| THRB | 0.155 | 15 | 2pin | B |
| THRB | 0.063 | 32 | 4zo1 | X |
| THRB | None | None | None | None |
| THRB | None | None | None | None |
| THRB | None | None | None | None |
| THRB | None | None | None | None |
| THRB | None | None | None | None |
| THRB | None | None | None | None |
| THRB | 0.032 | 31 | 4zo1 | X |
| THRB | 0.022 | 47 | 3gws | X |
| THRB | None | None | None | None |
| THRB | 0.126 | 16 | 1q4x | A |
| THRB | 0.126 | 16 | 1q4x | A |
| THRB | 0.577 | 3 | 2j4a | A |
| IL2RG | None | None | None | None |
| IL2RG | None | None | None | None |
| IL2RG | None | None | None | None |
| IL2RG | 0.256 | 1 | 2b5i | C |
| IL2RG | None | None | None | None |
| ITGB3 | None | None | None | None |
| ITGB3 | None | None | None | None |
| ITGB3 | 0.294 | 1 | 1jv2 | B |
| ITGB3 | None | None | None | None |
| ITGB3 | None | None | None | None |
| ITGB3 | None | None | None | None |
| ITGB3 | None | None | None | None |
| ITGB3 | None | None | None | None |
| ITGB3 | None | None | None | None |

| | | | | |
|---|---|---|---|---|
| PKD2 | None | None | None | None |
| PKD2 | None | None | None | None |
| PKD2 | None | None | None | None |
| PKD2 | None | None | None | None |
| PKD2 | None | None | None | None |
| PKD2 | None | None | None | None |
| PKD2 | None | None | None | None |
| TP53 | 0.571 | 1 | 1saf | B |
| TP53 | 0.571 | 1 | 1saf | B |
| TP53 | None | None | None | None |
| TP53 | 0.585 | 15 | 4mzr | B |
| TP53 | 0.091 | 178 | 3q05 | C |
| TP53 | None | None | None | None |
| EGFR | None | None | None | None |
| EGFR | None | None | None | None |
| XRCC4 | None | None | None | None |
| XRCC4 | None | None | None | None |
| HBB | 0.438 | 2 | 2w6v | D |
| HBB | 0.467 | 2 | 3qje | D |
| HBB | 0.467 | 2 | 3qje | D |
| HBB | 0.303 | 15 | 2w72 | B |
| HBB | 0.303 | 21 | 2w72 | B |
| HBB | 0.332 | 7 | 1cbm | A |
| HBB | 0.421 | 11 | 3ic2 | B |
| HBB | 0.459 | 2 | 5ee4 | D |
| HBB | 0.459 | 2 | 5ee4 | D |
| HBB | 0.459 | 2 | 5ee4 | D |
| HBB | 0.535 | 3 | 3r5i | B |
| HBB | 0.383 | 3 | 1lfv | B |
| HBB | 0.383 | 3 | 1lfv | B |
| HBB | 0.402 | 11 | 1m9p | D |
| HBB | 0.256 | 24 | 4n7p | D |
| HBB | 0.459 | 2 | 5ee4 | D |
| HBB | 0.566 | 1 | 1cbm | B |
| HBB | 0.51 | 3 | 1abw | B |
| HBB | 0.392 | 2 | 1lfz | B |
| HBB | 0.474 | 1 | 2h35 | B |
| HBB | 0.474 | 1 | 2h35 | B |
| HBB | 0.38 | 2 | 1j7y | B |
| HBB | 0.508 | 7 | 3ic2 | B |
| HBB | 0.545 | 3 | 1bij | D |
| HBB | 0.6 | 1 | 2w72 | D |
| HBB | 0.332 | 4 | 1fn3 | D |
| HBB | 0.354 | 15 | 1bij | D |
| HBB | 0.308 | 15 | 4rol | D |
| HBB | 0.733 | 1 | 1yff | D |
| HBB | 0.692 | 2 | 1buw | B |
| RPS6KA3 | 0.016 | 83 | 4nw5 | A |
| RPS6KA3 | 0.555 | 1 | 4nw5 | A |
| RPS6KA3 | 0.226 | 6 | 4nw5 | A |
| RPS6KA3 | 0.016 | 63 | 5d9k | A |
| RPS6KA3 | 0.034 | 55 | 4nw5 | A |
| RPS6KA3 | 0.197 | 11 | 5d9k | A |
| GBA | 1 | 15 | 3gxd | B |

| | | | | |
|---|---|---|---|---|
| GBA | 1 | 55 | 3gxd | B |
| GBA | 1 | 15 | 3gxd | B |
| GBA | 0.75 | 45 | 3ril | A |
| GBA | 0.25 | 33 | 2nt0 | A |
| GBA | 0.25 | 33 | 2nt0 | A |
| GBA | 1 | 12 | 1ogs | A |
| GBA | 0.75 | 122 | 3gxi | A |
| GBA | 1 | 2 | 1ogs | A |
| GBA | 1 | 11 | 1ogs | A |
| GBA | 0.5 | 59 | 3gxf | B |
| GBA | 0.75 | 45 | 3ril | A |
| GBA | 1 | 18 | 1ogs | A |
| GBA | 1 | 3 | 1ogs | A |
| GBA | None | None | None | None |
| GBA | 1 | 2 | 1ogs | A |
| GBA | 0.25 | 35 | 2nt0 | A |
| GBA | 0.75 | 76 | 3gxm | C |
| GBA | 1 | 3 | 1ogs | A |
| GBA | 1 | 3 | 1ogs | A |
| GBA | 0.75 | 62 | 3ril | A |
| GBA | 1 | 28 | 1ogs | A |
| GBA | None | None | None | None |
| GBA | 0.5 | 78 | 3gxi | A |
| GBA | 0.5 | 92 | 3rik | B |
| GBA | 0.5 | 42 | 2xwe | A |
| GBA | 0.333 | 117 | 1ogs | A |
| GBA | 0.5 | 92 | 3rik | B |
| GBA | 0.75 | 49 | 3ril | A |
| GBA | 0.25 | 109 | 2f61 | A |
| GBA | 0.667 | 60 | 1ogs | A |
| GBA | None | None | None | None |
| GBA | 0.5 | 11 | 2nt1 | A |
| GBA | 0.75 | 21 | 3gxi | A |
| GBA | None | None | None | None |
| GBA | 0.5 | 35 | 2xwe | A |
| GBA | 0.75 | 25 | 3keh | B |
| GBA | 0.75 | 70 | 3ril | A |
| GBA | 0.75 | 70 | 3ril | A |
| GBA | 1 | 15 | 3gxd | B |
| GBA | 1 | 44 | 3gxd | B |
| GBA | 0.5 | 57 | 3gxf | D |
| GBA | 1 | 44 | 3gxd | B |
| GBA | 1 | 44 | 3gxd | B |
| GBA | None | None | None | None |
| GBA | 0.75 | 94 | 3gxi | A |
| GBA | 0.75 | 94 | 3gxi | A |
| IFIH1 | None | None | None | None |
| TGFBR2 | 0.008 | 82 | 5e8y | A |
| TGFBR2 | None | None | None | None |
| TGFBR2 | 0.143 | 9 | 5e8v | A |
| TGFBR2 | None | None | None | None |
| TGFBR2 | None | None | None | None |
| TGFBR2 | 0.017 | 51 | 5E+091 | A |
| TGFBR2 | 0.057 | 12 | 5e8v | A |

| | | | | | |
|---|---|---|---|---|---|
| TGFBR2 | None | None | None | | None |
| TGFBR2 | 0.169 | | 14 | 5E+092 | A |
| TGFBR2 | 0.742 | | 1 | 5E+091 | A |
| TGFBR2 | None | None | None | | None |
| TGFBR2 | None | None | None | | None |
| TGFBR2 | 0.206 | | 6 | 5E+092 | A |
| TGFBR2 | 0.676 | | 1 | 5e8v | A |
| TGFBR2 | 0.676 | | 1 | 5e8v | A |
| ACE | 0.684 | | 17 | 3nxq | A |
| ACE | 0.474 | | 25 | 4bxk | A |
| ACE | 0.263 | | 31 | 2c6f | A |
| ACE | None | None | None | | None |
| ACE | 0.211 | | 93 | 4ca6 | A |
| ACE | 0.211 | | 92 | 5ama | D |
| COL10A1 | None | None | None | | None |
| COL10A1 | None | None | None | | None |
| COL10A1 | None | None | None | | None |
| COL10A1 | 0.409 | | 1 | 1gr3 | A |
| COL10A1 | None | None | None | | None |
| COL10A1 | 0.318 | | 4 | 1gr3 | A |
| COL10A1 | None | None | None | | None |
| COL10A1 | 0.273 | | 6 | 1gr3 | A |
| COL10A1 | 0.318 | | 4 | 1gr3 | A |
| KLF1 | None | None | None | | None |
| C3 | None | None | None | | None |
| CTSC | 0.108 | | 49 | 3pdf | A |
| CTSC | 0.934 | | 1 | 2djg | C |
| CTSC | None | None | None | | None |
| LYZ | 0.877 | | 3 | 1bb5 | A |
| LYZ | 0.923 | | 1 | 1lz4 | A |
| LYZ | 0.852 | | 2 | 1ubz | A |
| DHODH | 0.111 | | 65 | 4zmg | A |
| DHODH | 0.207 | | 18 | 5mut | A |
| DHODH | 0.038 | | 63 | 4rr4 | A |
| DHODH | 0.077 | | 23 | 4rr4 | A |
| DHODH | 0.038 | | 76 | 4rr4 | A |
| DHODH | 0.107 | | 34 | 4ylw | A |
| DHODH | 0.154 | | 34 | 4rr4 | A |
| ADH1C | 0.527 | | 11 | 1ht0 | A |
| GDNF | None | None | None | | None |
| TNF | 0.5 | | 8 | 2e7a | B |
| ALDH4A1 | 0.636 | | 3 | 3v9h | A |
| ALDH4A1 | 0.105 | | 86 | 4oe5 | B |
| ALDH4A1 | 0.04 | | 42 | 3v9i | A |
| BRAF | 0.02 | | 44 | 2fb8 | A |
| BRAF | 0.02 | | 44 | 2fb8 | A |
| BRAF | 0.752 | | 2 | 3psb | A |
| BRAF | 0.752 | | 2 | 3psb | A |
| BRAF | 0.737 | | 1 | 3pri | A |
| BRAF | 0.737 | | 1 | 3pri | A |
| BRAF | 0.737 | | 1 | 3pri | A |
| BRAF | 0.737 | | 1 | 3prf | A |
| BRAF | 0.737 | | 1 | 3prf | A |
| BRAF | 0.752 | | 2 | 3psb | A |

| | | | | |
|---|---|---|---|---|
| BRAF | 0.752 | 2 | 3psb | A |
| BRAF | 0.752 | 2 | 3psb | A |
| BRAF | 0.752 | 2 | 3psb | A |
| BRAF | 0.555 | 3 | 5vam | A |
| BRAF | 0.663 | 1 | 4ehe | B |
| BRAF | 0.734 | 1 | 5csx | A |
| TBX5 | None | None | None | None |
| TBX5 | None | None | None | None |
| TBX5 | None | None | None | None |
| TBX5 | None | None | None | None |
| TBX5 | None | None | None | None |
| TBX5 | None | None | None | None |
| TBX5 | None | None | None | None |
| TBX5 | None | None | None | None |
| TBX5 | None | None | None | None |
| RB1 | None | None | None | None |
| RB1 | None | None | None | None |
| RB1 | None | None | None | None |
| RB1 | None | None | None | None |
| RB1 | None | None | None | None |
| RB1 | None | None | None | None |
| CLIC2 | 0.5 | 3 | 2per | A |
| CLIC2 | None | None | None | None |
| ALDH7A1 | 0.04 | 50 | 4zuk | C |
| ALDH7A1 | 0.026 | 58 | 4zvx | A |
| ALDH7A1 | 0.405 | 4 | 2j6l | D |
| ALDH7A1 | 0.301 | 14 | 4zuk | A |
| ALDH7A1 | 0.409 | 3 | 4zvw | C |
| ALDH7A1 | 0.082 | 43 | 4zvy | A |
| ALDH7A1 | 0.534 | 3 | 2j6l | A |
| ALDH7A1 | 0.191 | 13 | 2j6l | D |
| ALDH7A1 | 0.333 | 6 | 4x0t | A |
| ALDH7A1 | 0.024 | 98 | 4zuk | C |
| ALDH7A1 | 0.113 | 38 | 4zul | C |
| ALDH7A1 | 0.534 | 3 | 2j6l | A |
| ALDH7A1 | 0.024 | 68 | 4zvw | A |
| ALDH7A1 | None | None | None | None |
| ALDH7A1 | 0.632 | 1 | 4zuk | A |
| ALDH7A1 | 0.026 | 65 | 4zvy | B |
| KDR | 0.575 | 5 | 1y6a | A |
| MUT | None | None | None | None |
| MUT | None | None | None | None |
| MUT | None | None | None | None |
| MUT | None | None | None | None |
| MUT | None | None | None | None |
| MUT | None | None | None | None |
| MUT | None | None | None | None |
| MUT | None | None | None | None |
| MUT | None | None | None | None |
| MUT | None | None | None | None |
| MUT | None | None | None | None |
| MUT | None | None | None | None |
| MUT | None | None | None | None |
| MUT | None | None | None | None |

| | | | | | |
|---|---|---|---|---|---|
| MUT | None | None | None | | None |
| MUT | None | None | None | | None |
| MUT | | 0.6 | 1 | 3bic | A |
| MUT | | 0.6 | 2 | 2xiq | A |
| MUT | | 0.6 | 2 | 2xiq | A |
| MUT | None | None | None | | None |
| MUT | None | None | None | | None |
| MUT | | 0.4 | 1 | 2xiq | A |
| MUT | None | None | None | | None |
| MUT | None | None | None | | None |
| MUT | None | None | None | | None |
| MUT | None | None | None | | None |
| MUT | None | None | None | | None |
| MUT | None | None | None | | None |
| MUT | None | None | None | | None |
| MUT | None | None | None | | None |
| MUT | None | None | None | | None |
| MUT | None | None | None | | None |
| MUT | None | None | None | | None |
| MUT | None | None | None | | None |
| SOD1 | | 0.778 | 1 | 2gbu | A |
| SOD1 | | 0.727 | 4 | 4xcr | A |
| SOD1 | | 0.35 | 26 | 1n19 | A |
| SOD1 | | 0.621 | 12 | 1hl5 | H |
| NECTIN3 | None | None | None | | None |
| HFE | | 0.065 | 21 | 1de4 | D |
| HFE | | 0.065 | 21 | 1de4 | D |
| HFE | | 0.065 | 21 | 1de4 | D |
| HFE | | 0.5 | 1 | 1de4 | A |
| HFE | | 0.065 | 21 | 1de4 | D |
| HFE | | 0.065 | 21 | 1de4 | D |
| HFE | | 0.25 | 15 | 1a6z | A |
| EIF2AK3 | None | None | None | | None |
| EIF2AK3 | None | None | None | | None |
| TERT | None | None | None | | None |
| TERT | None | None | None | | None |
| SPR | None | None | None | | None |
| SPR | | 0.02 | 41 | 4hwk | A |
| SPR | | 0.036 | 47 | 4hwk | A |
| SPR | None | None | None | | None |
| SPR | None | None | None | | None |
| F7 | | 0.625 | 2 | 1klj | L |
| UNG | None | None | None | | None |
| MMP13 | | 0.356 | 22 | 2ozr | E |
| MMP13 | | 0.786 | 1 | 3tvc | A |
| MMP13 | | 0.766 | 1 | 3zxh | B |
| MMP13 | | 0.766 | 1 | 3zxh | B |
| CD207 | | 0.621 | 1 | 3kqg | B |
| F2 | | 1 | 2 | 1nrq | L |
| SPTBN2 | None | None | None | | None |
| PKLR | | 0.568 | 9 | 2vgb | A |
| PKLR | | 0.486 | 17 | 2vgg | D |
| PKLR | None | None | None | | None |
| PKLR | | 0.486 | 8 | 2vgg | D |

| | | | | |
|---|---|---|---|---|
| PKLR | 0.568 | 3 | 4ip7 | D |
| PKLR | 0.432 | 13 | 2vgi | A |
| PKLR | 0.378 | 45 | 4ip7 | B |
| PKLR | 0.324 | 25 | 2vgf | B |
| SACS | None | None | None | None |
| HRAS | 0.529 | 3 | 6q21 | A |
| HRAS | 0.529 | 3 | 6q21 | A |
| HRAS | 0.59 | 1 | 1gnq | A |
| HRAS | 0.59 | 1 | 1gnq | A |
| HRAS | 0.664 | 1 | 2lwi | A |
| HRAS | 0.742 | 1 | 1ctq | A |
| HRAS | 0.69 | 1 | 1xd2 | A |
| HRAS | 0.415 | 3 | 6q21 | D |
| HRAS | 0.529 | 3 | 6q21 | A |
| HRAS | 0.403 | 3 | 2n46 | A |
| HRAS | 0.521 | 4 | 4nyj | R |
| HRAS | 0.664 | 1 | 2lwi | A |
| HRAS | 0.664 | 1 | 2lwi | A |
| VHL | None | None | None | None |
| VHL | None | None | None | None |
| VHL | None | None | None | None |
| VHL | None | None | None | None |
| VHL | 0.5 | 22 | 4w9l | C |
| VHL | 0.5 | 22 | 5nvy | C |
| VHL | None | None | None | None |
| VHL | None | None | None | None |
| VHL | None | None | None | None |
| VHL | 0.5 | 2 | 5nvy | C |
| VHL | 0.5 | 1 | 5nvy | C |
| VHL | 0.5 | 1 | 5nvy | C |
| VHL | 0.5 | 4 | 5nvy | C |
| VHL | None | None | None | None |
| VHL | 0.5 | 6 | 5nvy | C |
| VHL | 0.5 | 2 | 5nvy | C |
| VHL | 0.5 | 8 | 3ztc | C |
| VHL | 0.5 | 2 | 5nvy | C |
| VHL | 0.5 | 2 | 5nvy | C |
| VHL | None | None | None | None |
| VHL | 0.5 | 7 | 4w9h | C |
| VHL | 0.5 | 2 | 5nvy | C |
| VHL | 0.5 | 2 | 5nvy | C |
| VHL | 0.5 | 19 | 4w9l | C |
| VHL | None | None | None | None |
| TTBK1 | None | None | None | None |
| RAD51 | 0.031 | 15 | 5h1c | B |
| ALB | None | None | None | None |
| ALB | 0.5 | 2 | 4n0u | D |
| ALB | None | None | None | None |
| ALB | None | None | None | None |
| MET | None | None | None | None |
| MET | None | None | None | None |
| MET | None | None | None | None |
| MET | None | None | None | None |
| MET | None | None | None | None |

| | | | | |
|---|---|---|---|---|
| MET | None | None | None | None |
| MET | None | None | None | None |
| MET | None | None | None | None |
| MET | None | None | None | None |
| CP | None | None | None | None |
| CP | None | None | None | None |
| PGM1 | None | None | None | None |
| PGM1 | 0.409 | 2 | 5f9c | A |
| PGM1 | 0.409 | 2 | 5f9c | A |
| PGM1 | None | None | None | None |
| PGM1 | None | None | None | None |
| PGM1 | None | None | None | None |
| PGM1 | None | None | None | None |
| NR5A1 | None | None | None | None |
| NR5A1 | 0.157 | 17 | 4qjr | A |
| NR5A1 | None | None | None | None |
| F9 | None | None | None | None |
| F9 | None | None | None | None |
| HPRT1 | 0.188 | 29 | 3ggj | B |
| HPRT1 | 0.125 | 29 | 1z7g | A |
| HPRT1 | 0.25 | 7 | 3gep | B |
| HPRT1 | 0.375 | 15 | 3gep | B |
| HPRT1 | 0.077 | 46 | 1bzy | D |
| HPRT1 | 0.871 | 3 | 4rac | B |
| HPRT1 | 0.806 | 2 | 4ran | D |
| HPRT1 | None | None | None | None |
| HPRT1 | 0.25 | 4 | 3gep | B |
| HPRT1 | 0.44 | 3 | 1d6n | B |
| HPRT1 | 0.154 | 31 | 1bzy | D |
| HPRT1 | 0.054 | 43 | 5brn | B |
| HPRT1 | 0.24 | 25 | 1d6n | B |
| HPRT1 | 0.703 | 4 | 5w8v | A |
| HPRT1 | 0.118 | 27 | 1z7g | B |
| HPRT1 | 0.321 | 8 | 4kn6 | A |
| HPRT1 | 0.312 | 27 | 3ggj | B |
| HPRT1 | 0.312 | 12 | 3gep | B |
| HPRT1 | 0.871 | 3 | 4rac | B |
| HPRT1 | 0.871 | 3 | 4rac | B |
| HPRT1 | 0.765 | 2 | 4rao | D |
| HPRT1 | 0.774 | 2 | 4rac | B |
| GPD1L | None | None | None | None |
| GPD1L | None | None | None | None |
| GPD1L | None | None | None | None |
| FGFR2 | None | None | None | None |
| FGFR2 | None | None | None | None |
| FGFR2 | None | None | None | None |
| FGFR2 | None | None | None | None |
| FGFR2 | None | None | None | None |
| FGFR2 | None | None | None | None |
| FGFR2 | None | None | None | None |
| FGFR2 | None | None | None | None |
| FGFR2 | None | None | None | None |
| FGFR2 | None | None | None | None |
| FGFR2 | None | None | None | None |

| | | | | |
|---|---|---|---|---|
| FGFR2 | None | None | None | None |
| FGFR2 | None | None | None | None |
| FGFR2 | None | None | None | None |
| FGFR2 | None | None | None | None |
| FGFR2 | None | None | None | None |
| FGFR2 | None | None | None | None |
| FGFR2 | None | None | None | None |
| FGFR2 | None | None | None | None |
| HSPD1 | None | None | None | None |
| HSPD1 | None | None | None | None |
| FGF8 | None | None | None | None |
| FGF8 | None | None | None | None |
| FGF8 | None | None | None | None |
| F9 | 0.438 | 1 | 5vyg | B |
| NCF4 | None | None | None | None |
| NCF4 | None | None | None | None |
| LDLR | None | None | None | None |
| LDLR | None | None | None | None |
| LDLR | 0.478 | 1 | 3gcx | E |
| LDLR | 0.478 | 1 | 3gcx | E |
| LDLR | 0.478 | 1 | 3gcx | E |
| LDLR | None | None | None | None |
| C3 | None | None | None | None |
| C3 | None | None | None | None |
| C3 | None | None | None | None |
| C3 | None | None | None | None |
| C3 | None | None | None | None |
| C3 | None | None | None | None |
| C3 | None | None | None | None |
| C3 | None | None | None | None |
| TULP1 | None | None | None | None |
| TULP1 | None | None | None | None |
| TULP1 | None | None | None | None |
| TULP1 | None | None | None | None |
| IKBKG | None | None | None | None |
| IKBKG | None | None | None | None |
| IKBKG | None | None | None | None |
| CD40 | None | None | None | None |
| CD40 | None | None | None | None |
| SLC9A3R1 | None | None | None | None |
| BCHE | 0.278 | 18 | 2xqg | A |
| BCHE | 0.439 | 5 | 2wil | B |
| BCHE | 0.544 | 11 | 1p0q | A |
| BCHE | 0.127 | 115 | 2pm8 | B |
| BCHE | 0.196 | 66 | 2wik | A |
| BCHE | 0.088 | 204 | 2pm8 | A |
| BCHE | 0.63 | 4 | 2xqg | A |
| BCHE | None | None | None | None |
| BCHE | 0.138 | 164 | 1xlu | A |
| BCHE | 0.255 | 18 | 2xqj | A |
| BCHE | 0.14 | 62 | 5dyt | A |
| BCKDHB | 0.833 | 1 | 1v11 | B |
| BCKDHB | 0.5 | 18 | 2bfc | B |
| BCKDHB | None | None | None | None |

| | | | | |
|---|---|---|---|---|
| BCKDHB | 0.667 | 11 | 2bfd | B |
| BCKDHB | 0.833 | 10 | 1olx | B |
| BCKDHB | 0.833 | 1 | 1v11 | B |
| BCKDHB | 0.833 | 1 | 1v11 | B |
| BCKDHB | 0.625 | 2 | 1x7z | B |
| BCKDHB | 0.429 | 16 | 1ols | B |
| BCKDHB | 0.333 | 99 | 1olx | B |
| BCKDHB | 0.333 | 64 | 1x7y | B |
| BCKDHB | 0.857 | 1 | 2bff | B |
| CDH1 | 0.278 | 8 | 3ff7 | B |
| CDH1 | 0.575 | 1 | 3ff8 | A |
| CDH1 | 0.543 | 1 | 3ff7 | A |
| CDH1 | None | None | None | None |
| CDH1 | 0.2 | 20 | 2o72 | A |
| CTNNB1 | None | None | None | None |
| AIRE | None | None | None | None |
| EDA | 1 | 1 | 1rj8 | A |
| EDA | None | None | None | None |
| EDA | None | None | None | None |
| EDA | None | None | None | None |
| EDA | None | None | None | None |
| SH2D1A | 0.444 | 5 | 1ka7 | A |
| SH2D1A | None | None | None | None |
| SH2D1A | None | None | None | None |
| GCK | 0.556 | 37 | 3idh | A |
| GCK | 0.333 | 32 | 4isf | A |
| GCK | None | None | None | None |
| GCK | None | None | None | None |
| GCK | 0.176 | 38 | 4isg | A |
| GCK | 0.611 | 13 | 3h1v | X |
| GCK | None | None | None | None |
| GCK | None | None | None | None |
| GCK | None | None | None | None |
| GCK | 0.778 | 5 | 3id8 | A |
| GCK | 0.833 | 1 | 4dhy | A |
| GCK | 0.389 | 6 | 4lc9 | B |
| GCK | 0.222 | 33 | 4mle | A |
| GCK | 0.611 | 6 | 3id8 | A |
| GCK | None | None | None | None |
| GCK | 0.167 | 33 | 4rch | A |
| GCK | None | None | None | None |
| GCK | 0.5 | 5 | 3vey | A |
| GCK | None | None | None | None |
| GCK | None | None | None | None |
| AQP2 | None | None | None | None |
| AQP2 | None | None | None | None |
| AQP2 | None | None | None | None |
| AQP2 | None | None | None | None |
| AQP2 | None | None | None | None |
| AQP2 | None | None | None | None |
| AQP2 | None | None | None | None |
| AQP2 | None | None | None | None |
| AQP2 | None | None | None | None |
| AQP2 | None | None | None | None |

| | | | | |
|---|---|---|---|---|
| AQP2 | None | None | None | None |
| AQP2 | None | None | None | None |
| AQP2 | None | None | None | None |
| SUMF1 | 0.75 | 6 | 1z70 | X |
| SUMF1 | 0.556 | 30 | 1y1i | X |
| SUMF1 | 0.5 | 38 | 2aij | X |
| SUMF1 | 0.625 | 18 | 2aij | X |
| SUMF1 | 0.875 | 35 | 2afy | X |
| SUMF1 | 0.667 | 19 | 1y1i | X |
| SUMF1 | 0.75 | 17 | 1y1j | X |
| SUMF1 | 0.75 | 17 | 1y1j | X |
| SUMF1 | 0.625 | 27 | 1y1g | X |
| FAM20A | None | None | None | None |
| FAM20A | None | None | None | None |
| FAM20A | None | None | None | None |
| FAM20A | None | None | None | None |
| NT5E | None | None | None | None |
| NT5E | None | None | None | None |
| NOS3 | 0.564 | 54 | 1m9k | B |
| ACTG1 | None | None | None | None |
| ACTG1 | None | None | None | None |
| ACTG1 | 0.229 | 48 | 5jlh | E |
| ACTG1 | 0.03 | 110 | 5jlh | A |
| ACTG1 | 0.571 | 8 | 5jlh | E |
| ACTG1 | None | None | None | None |
| FCGR1A | None | None | None | None |
| GPI | 0.304 | 64 | 1iri | A |
| GPI | 0.13 | 210 | 1jiq | A |
| GPI | 1 | 3 | 1jiq | A |
| GPI | None | None | None | None |
| GPI | 0.696 | 22 | 1iat | A |
| GPI | 0.522 | 22 | 1nuh | A |
| GPI | 0.696 | 27 | 1iat | A |
| GPI | 0.304 | 87 | 1jlh | B |
| GPI | 0.391 | 141 | 1jlh | B |
| CFH | None | None | None | None |
| OGG1 | 0.895 | 13 | 1m3h | A |
| TPK1 | None | None | None | None |
| TPK1 | None | None | None | None |
| TPK1 | None | None | None | None |
| TPK1 | None | None | None | None |
| SEC23A | None | None | None | None |
| SEC23A | None | None | None | None |
| TYRP1 | None | None | None | None |
| TYRP1 | None | None | None | None |
| TYRP1 | None | None | None | None |
| TYRP1 | 0.067 | 5 | 5m8p | A |
| TYRP1 | 0.3 | 5 | 5m8m | A |
| TYRP1 | 0.067 | 5 | 5m8p | A |
| CSTB | None | None | None | None |
| CSTB | None | None | None | None |
| CSTB | None | None | None | None |
| ACP1 | None | None | None | None |
| IL17F | None | None | None | None |

| | | | | |
|---|---|---|---|---|
| IL17F | None | None | None | None |
| IL17F | None | None | None | None |
| IL17F | None | None | None | None |
| IL17F | None | None | None | None |
| IDH2 | 0.5 | 2 | 5svn | B |
| IDH2 | 0.5 | 2 | 5svn | B |
| IDH2 | 0.48 | 13 | 5i96 | A |
| IDH2 | None | None | None | None |
| TP53 | 0.925 | 11 | 2ocj | A |
| TP53 | 0.8 | 16 | 2ahi | D |
| TP53 | 0.65 | 49 | 4ibq | A |
| TP53 | 0.725 | 27 | 5g4o | A |
| TP53 | 0.455 | 21 | 3q06 | A |
| TP53 | 0.9 | 1 | 4agp | A |
| TP53 | 0.9 | 1 | 4agp | A |
| TP53 | 0.925 | 10 | 4hje | A |
| TP53 | 0.7 | 29 | 3d05 | A |
| TP53 | 0.875 | 7 | 3kmd | B |
| TP53 | 0.875 | 7 | 3kmd | B |
| TP53 | 0.925 | 6 | 3kz8 | B |
| TP53 | 0.925 | 6 | 3kz8 | B |
| TP53 | 0.609 | 22 | 3q01 | B |
| TP53 | 0.925 | 5 | 3kmd | B |
| TP53 | 0.925 | 5 | 3kmd | B |
| TP53 | 0.925 | 8 | 3kz8 | B |
| TP53 | 0.925 | 8 | 3kz8 | B |
| TP53 | 0.925 | 1 | 3kmd | B |
| TP53 | 0.9 | 11 | 2pcx | A |
| TP53 | 0.9 | 1 | 5aol | A |
| TP53 | 0.85 | 5 | 5lgy | A |
| TP53 | 0.9 | 1 | 3q06 | B |
| TP53 | 0.725 | 19 | 5lgy | A |
| TP53 | 0.775 | 14 | 4mzi | A |
| TP53 | 0.825 | 8 | 1kzy | A |
| TP53 | 0.825 | 8 | 1kzy | A |
| TP53 | 0.8 | 11 | 4ibt | A |
| TP53 | 0.8 | 11 | 4ibt | A |
| TP53 | 0.725 | 15 | 5lgy | A |
| TP53 | 0.725 | 29 | 2h1l | O |
| TP53 | 0.875 | 7 | 3kmd | B |
| TP53 | 0.725 | 15 | 5lgy | A |
| TP53 | 0.8 | 10 | 4agp | A |
| TP53 | 0.8 | 10 | 4agp | A |
| TP53 | 0.8 | 10 | 4agp | A |
| TP53 | 0.8 | 10 | 4agp | A |
| TP53 | 0.675 | 35 | 4ibq | A |
| TP53 | 0.575 | 37 | 4agm | A |
| TP53 | 0.675 | 17 | 3kz8 | B |
| TP53 | 0.356 | 63 | 4mzr | C |
| TP53 | None | None | None | None |
| LMAN1 | None | None | None | None |
| CTSK | 0.75 | 2 | 4n79 | A |
| CTSK | 0.127 | 32 | 1ayw | A |
| CTSK | 0.413 | 12 | 4x6j | A |

| | | | | |
|---|---|---|---|---|
| FGB | None | None | None | None |
| FGB | None | None | None | None |
| FGB | 0.688 | 1 | 1fzf | E |
| FGB | 0.281 | 10 | 2ffd | B |
| ACADM | None | None | None | None |
| ACADM | None | None | None | None |
| ACADM | 0.218 | 11 | 1t9g | C |
| ACADM | 0.471 | 3 | 4p13 | D |
| ACADM | 0.127 | 10 | 1egc | C |
| ACADM | None | None | None | None |
| ACADM | 0.389 | 6 | 1t9g | D |
| ACADM | 0.074 | 38 | 1t9g | D |
| ACADM | 0.327 | 12 | 4p13 | C |
| ACADM | 0.389 | 6 | 1t9g | D |
| ACADM | 0.098 | 31 | 4p13 | D |
| ACADM | None | None | None | None |
| ACADM | None | None | None | None |
| ACADM | 0.291 | 14 | 1t9g | C |
| ACADM | 0.098 | 31 | 4p13 | D |
| ACADM | 0.481 | 2 | 4p13 | C |
| ACADM | 0.389 | 6 | 1t9g | D |
| ACADM | 0.109 | 24 | 2a1t | C |
| ACADM | None | None | None | None |
| ACADM | 0.104 | 32 | 4p13 | A |
| ACADM | 0.564 | 1 | 1t9g | C |
| ACADM | 0.255 | 10 | 1t9g | C |
| ACADM | 0.091 | 33 | 1t9g | C |
| ACADM | None | None | None | None |
| ACADM | 0.364 | 6 | 2a1t | A |
| ACADM | 0.312 | 12 | 4p13 | A |
| ACADM | 0.347 | 7 | 4p13 | B |
| PTS | None | None | None | None |
| PTS | 0.417 | 2 | 3i2b | E |
| PTS | None | None | None | None |
| PTS | 0.778 | 1 | 3i2b | D |
| PTS | None | None | None | None |
| PTS | None | None | None | None |
| PTS | None | None | None | None |
| PTS | None | None | None | None |
| PTS | None | None | None | None |
| INS | 0.75 | 2 | 3zqr | E |
| INS | 1 | 5 | 2l1z | A |
| INS | 1 | 5 | 2l1z | A |
| INS | 1 | 1 | 1his | A |
| FBP1 | 0.71 | 26 | 2y5l | C |
| FBP1 | 0.125 | 129 | 3a29 | C |
| FBP1 | 0.5 | 18 | 1fta | A |
| FBP1 | 0.7 | 10 | 2y5k | A |
| FBP1 | 0.516 | 38 | 3kc1 | B |
| FBP1 | 0.733 | 10 | 3kc1 | A |
| FBP1 | 0.633 | 43 | 2jjk | A |
| FBP1 | 0.8 | 11 | 2y5k | A |
| FBP1 | 0.548 | 41 | 2vt5 | D |
| FBP1 | 0.438 | 25 | 2fie | A |

| | | | | |
|---|---|---|---|---|
| FBP1 | 0.643 | 7 | 4mjo | B |
| KCNQ1 | None | None | None | None |
| KCNQ1 | None | None | None | None |
| KCNQ1 | None | None | None | None |
| UPP1 | 0.706 | 4 | 3euf | A |
| UMPS | 1 | 3 | 2qcl | A |
| UMPS | 0.556 | 59 | 2qcl | B |
| UMPS | 0.184 | 153 | 2v30 | B |
| DHFR | 0.167 | 16 | 3f91 | A |
| KHK | 0.667 | 2 | 2hw1 | A |
| TTPA | None | None | None | None |
| TTPA | None | None | None | None |
| TTPA | 0.5 | 1 | 1r5l | A |
| TTPA | None | None | None | None |
| MAP1LC3B | 0.387 | 8 | 3vtv | A |
| MAP1LC3B | None | None | None | None |
| ZAP70 | 0.317 | 1 | 2oq1 | A |
| PROC | 0.536 | 1 | 3f6u | L |
| SQSTM1 | None | None | None | None |
| AHCY | 0.19 | 113 | 4pfj | B |
| AHCY | 0.091 | 142 | 5w4b | E |
| PRNP | None | None | None | None |
| PRNP | None | None | None | None |
| PRNP | None | None | None | None |
| PRNP | None | None | None | None |
| PRNP | 0.277 | 1 | 1qlx | A |
| PRNP | None | None | None | None |
| PRNP | None | None | None | None |
| PRNP | 0.19 | 2 | 1e1p | A |
| PRNP | None | None | None | None |
| PRNP | 0.238 | 2 | 3hj5 | A |
| PRNP | 0.238 | 2 | 3hj5 | A |
| PRNP | None | None | None | None |
| PRNP | None | None | None | None |
| PRNP | 0.19 | 3 | 1qm3 | A |
| PRNP | None | None | None | None |
| PRNP | 0.277 | 1 | 1qlx | A |
| REN | 0.431 | 2 | 2v13 | A |
| REN | 0.431 | 2 | 2v13 | A |
| HINT1 | 0.556 | 2 | 5ed6 | B |
| HINT1 | 0.897 | 1 | 5km4 | A |
| HINT1 | 0.227 | 60 | 5ed6 | A |
| HINT1 | 0.667 | 11 | 5ed6 | B |
| HINT1 | 0.923 | 1 | 5ipc | A |
| OAT | 0.051 | 66 | 5vwo | B |
| OAT | 0.107 | 32 | 2can | B |
| OAT | 0.381 | 5 | 2can | B |
| OAT | 0.381 | 5 | 2can | B |
| OAT | 0.381 | 7 | 2byj | B |
| OAT | 0.071 | 20 | 2oat | C |
| OAT | 0.113 | 54 | 1oat | B |
| OAT | None | None | None | None |
| OAT | 0.278 | 12 | 2byj | A |
| OAT | None | None | None | None |

| | | | | |
|---|---|---|---|---|
| OAT | 0.675 | 2 | 2byl | B |
| OAT | 0.179 | 32 | 2oat | C |
| OAT | 0.179 | 32 | 2oat | C |
| OAT | 0.169 | 15 | 5vwo | A |
| OAT | 0.169 | 15 | 5vwo | A |
| OAT | 0.758 | 1 | 2byl | A |
| OAT | 0.375 | 5 | 2oat | C |
| OAT | 0.525 | 4 | 2byl | B |
| OAT | 0.429 | 6 | 2byj | B |
| OAT | None | None | None | None |
| OAT | 0.107 | 49 | 2can | B |
| OAT | 0.165 | 23 | 2byl | A |
| OAT | 0.087 | 70 | 1oat | B |
| OAT | 0.155 | 17 | 1gbn | C |
| OAT | 0.155 | 17 | 1gbn | C |
| OAT | 0.057 | 54 | 5vwo | C |
| OAT | 0.057 | 54 | 5vwo | C |
| OAT | None | None | None | None |
| CTSA | None | None | None | None |
| CTSA | None | None | None | None |
| PNP | 0.375 | 24 | 1ulb | A |
| PNP | 0.29 | 111 | 1yry | E |
| PNP | 0.406 | 43 | 3phb | E |
| PNP | 0.706 | 5 | 4ear | C |
| PNP | 0.367 | 28 | 3ggs | C |
| PNP | 0.5 | 17 | 1pf7 | E |
| PNP | 0.933 | 1 | 3ggs | C |
| PNP | 0.833 | 1 | 3d1v | A |
| MYL2 | None | None | None | None |
| MYL2 | None | None | None | None |
| ANG | 0.548 | 4 | 1k5b | A |
| ANG | 0.419 | 8 | 4ahi | A |
| ANG | 0.419 | 8 | 4ahi | A |
| ANG | 0.19 | 15 | 1k58 | A |
| ANG | 0.761 | 1 | 5eop | A |
| ANG | 0.576 | 4 | 5m9v | A |
| ANG | 0.579 | 4 | 5m9q | A |
| ANG | 0.14 | 34 | 4ahf | A |
| ANG | 0.112 | 34 | 5m9r | A |
| ANG | 0.52 | 5 | 5m9g | A |
| POLH | 0.241 | 101 | 4dl5 | A |
| POLH | 0.185 | 17 | 4j9l | A |
| POLH | 0.208 | 9 | 3si8 | A |
| IL2RA | None | None | None | None |
| TGFB1 | None | None | None | None |
| TP53 | None | None | None | None |
| TP53 | None | None | None | None |
| PAH | None | None | None | None |
| PAH | None | None | None | None |
| PAH | 0.44 | 11 | 2pah | A |
| PAH | 0.44 | 11 | 2pah | A |
| PAH | 0.32 | 19 | 6pah | A |
| PAH | 0.32 | 19 | 6pah | A |
| PAH | 0.48 | 2 | 1pah | A |

| | | | | |
|---|---|---|---|---|
| PAH | 0.48 | 2 | 1pah | A |
| PAH | 0.4 | 11 | 3pah | A |
| PAH | 0.2 | 51 | 1j8u | A |
| PAH | 0.2 | 51 | 1j8u | A |
| PAH | 0.36 | 7 | 2pah | B |
| PAH | 0.44 | 9 | 1dmw | A |
| PAH | 0.48 | 1 | 1dmw | A |
| PAH | 0.48 | 1 | 1dmw | A |
| PAH | 0.44 | 8 | 2pah | B |
| PAH | 0.44 | 8 | 2pah | B |
| PAH | 0.417 | 3 | 1j8t | A |
| PAH | 0.48 | 2 | 4anp | A |
| PAH | 0.48 | 1 | 1dmw | A |
| PAH | 0.48 | 4 | 2pah | B |
| PAH | 0.44 | 11 | 2pah | A |
| PAH | 0.48 | 4 | 1mmt | A |
| PAH | 0.68 | 6 | 2pah | A |
| PAH | 0.76 | 1 | 1tdw | A |
| PAH | 0.08 | 49 | 1j8u | A |
| PAH | 0.12 | 44 | 6pah | A |
| PAH | 0.4 | 14 | 1j8u | A |
| PAH | None | None | None | None |
| PAH | 0.08 | 49 | 1j8u | A |
| PAH | None | None | None | None |
| PAH | 0.2 | 43 | 3pah | A |
| PAH | 0.64 | 2 | 1lrm | A |
| PAH | 0.4 | 11 | 2pah | B |
| PAH | 0.28 | 31 | 4anp | A |
| PAH | 0.4 | 14 | 1j8u | A |
| PAH | 0.4 | 14 | 1j8u | A |
| PAH | 0.12 | 60 | 1j8u | A |
| PAH | 0.48 | 1 | 1dmw | A |
| PAH | 0.48 | 1 | 1dmw | A |
| PAH | 0.167 | 38 | 1j8t | A |
| HDAC8 | None | None | None | None |
| HDAC8 | 0.732 | 3 | 3rqd | A |
| VCP | 0.438 | 4 | 3tiw | A |
| VCP | 0.5 | 2 | 3qq8 | A |
| VCP | 0.5 | 2 | 3qq8 | A |
| VCP | None | None | None | None |
| VCP | None | None | None | None |
| VCP | None | None | None | None |
| VCP | None | None | None | None |
| B3GAT3 | None | None | None | None |
| B3GAT3 | None | None | None | None |
| CSF2RB | None | None | None | None |
| CSF2RB | None | None | None | None |
| CSF2RB | None | None | None | None |
| NF2 | 0.5 | 6 | 1h4r | B |
| NF2 | None | None | None | None |
| NF2 | None | None | None | None |
| PIK3CA | None | None | None | None |
| PHGDH | 0.019 | 40 | 5nzp | A |
| PHGDH | None | None | None | None |

| | | | | |
|---|---|---|---|---|
| PHGDH | 0.04 | 44 | 2g76 | A |
| PHGDH | 0.66 | 1 | 5n53 | A |
| POLH | None | None | None | None |
| XIAP | 0.5 | 7 | 5c7b | A |
| XIAP | None | None | None | None |
| PDHX | None | None | None | None |
| PDHX | None | None | None | None |
| PDHX | None | None | None | None |
| PDHX | None | None | None | None |
| PDHX | None | None | None | None |
| SAMHD1 | 0.5 | 12 | 5ao0 | A |
| SAMHD1 | 0.357 | 51 | 4zwe | D |
| SAMHD1 | 0.429 | 7 | 4tnr | D |
| SAMHD1 | None | None | None | None |
| SAMHD1 | 0.571 | 3 | 4qfy | D |
| SAMHD1 | None | None | None | None |
| SAMHD1 | 0.308 | 18 | 4qg2 | B |
| SAMHD1 | 0.286 | 57 | 5ao0 | B |
| CD36 | None | None | None | None |
| CD36 | None | None | None | None |
| CD36 | None | None | None | None |
| ASPH | None | None | None | None |
| CCT5 | None | None | None | None |
| CCT5 | None | None | None | None |
| CCT5 | None | None | None | None |
| SMN1 | None | None | None | None |
| SMN1 | None | None | None | None |
| SMN1 | None | None | None | None |
| SMN1 | None | None | None | None |
| ABCA1 | None | None | None | None |
| ABCA1 | None | None | None | None |
| ABCA1 | None | None | None | None |
| ABCA1 | None | None | None | None |
| ABCA1 | None | None | None | None |
| ABCA1 | None | None | None | None |
| ABCA1 | None | None | None | None |
| ABCA1 | None | None | None | None |
| ABCA1 | None | None | None | None |
| ABCA1 | None | None | None | None |
| ABCA1 | None | None | None | None |
| ABCA1 | None | None | None | None |
| ABCA1 | None | None | None | None |
| ABCA1 | None | None | None | None |
| ABCA1 | None | None | None | None |
| ABCA1 | None | None | None | None |
| ABCA1 | None | None | None | None |
| ABCA1 | None | None | None | None |
| ABCA1 | None | None | None | None |
| ABCA1 | None | None | None | None |
| ABCA1 | None | None | None | None |
| ABCA1 | None | None | None | None |
| ABCA1 | None | None | None | None |
| ABCA1 | None | None | None | None |
| ABCA1 | None | None | None | None |

| | | | | |
|---|---|---|---|---|
| ABCA1 | None | None | None | None |
| ABCA1 | None | None | None | None |
| ABCA1 | None | None | None | None |
| ABCA1 | None | None | None | None |
| ABCA1 | None | None | None | None |
| ABCA1 | None | None | None | None |
| ABCA1 | None | None | None | None |
| ABCA1 | None | None | None | None |
| ABCA1 | None | None | None | None |
| ABCA1 | None | None | None | None |
| ABCA1 | None | None | None | None |
| ERCC1 | None | None | None | None |
| FECH | 0.524 | 27 | 2po5 | A |
| FECH | 0.6 | 60 | 2hre | A |
| FECH | 0.636 | 20 | 2po7 | A |
| FECH | 0.476 | 61 | 4klr | A |
| FECH | 0.5 | 84 | 2hrc | A |
| FECH | 0.667 | 10 | 2po5 | A |
| FECH | 0.455 | 30 | 4kla | B |
| PTEN | None | None | None | None |
| PTEN | None | None | None | None |
| PTEN | None | None | None | None |
| PTEN | None | None | None | None |
| PTEN | None | None | None | None |
| PTEN | None | None | None | None |
| PTEN | None | None | None | None |
| PTEN | None | None | None | None |
| PTEN | None | None | None | None |
| PTEN | None | None | None | None |
| PTEN | None | None | None | None |
| PTEN | None | None | None | None |
| PTEN | None | None | None | None |
| PTEN | None | None | None | None |
| PTEN | None | None | None | None |
| PTEN | None | None | None | None |
| PTEN | None | None | None | None |
| PTEN | None | None | None | None |
| PTEN | None | None | None | None |
| PTEN | None | None | None | None |
| ACAT1 | 0.667 | 1 | 2ib9 | C |
| ACAT1 | 0.114 | 43 | 2ib8 | B |
| ACAT1 | None | None | None | None |
| ACAT1 | 0.667 | 1 | 2ib9 | C |
| ACAT1 | 0.667 | 1 | 2ib9 | C |
| ACAT1 | 0.472 | 11 | 2f2s | A |
| ACAT1 | 0.833 | 6 | 2ib7 | C |
| ACAT1 | 0.229 | 68 | 2ib8 | B |
| ACAT1 | 0.056 | 61 | 2ibw | D |
| ACAT1 | 0.528 | 9 | 2ibw | D |
| ACAT1 | 0.703 | 16 | 2ibu | A |
| CREBBP | 0.283 | 35 | 5mqg | A |
| ITGA2B | 0.083 | 83 | 3zdy | A |
| ITGA2B | 0.083 | 53 | 4cak | A |
| ITGA2B | 0.214 | 20 | 2vdn | A |

| | | | | |
|---|---|---|---|---|
| ITGA2B | 0.538 | 1 | 3nig | A |
| TSC1 | None | None | None | None |
| ADA2 | None | None | None | None |
| ADA2 | None | None | None | None |
| ADA2 | 0.75 | 1 | 3lgd | A |
| ADA2 | None | None | None | None |
| ADA2 | None | None | None | None |
| ADA2 | None | None | None | None |
| ADA2 | None | None | None | None |
| ADA2 | None | None | None | None |
| ADA2 | 1 | 1 | 3lgd | B |
| ADA2 | None | None | None | None |
| PGK1 | None | None | None | None |
| PGK1 | 0.536 | 8 | 2xe8 | A |
| PGK1 | 0.25 | 38 | 4axx | A |
| PGK1 | 0.214 | 33 | 2y3i | A |
| PGK1 | 0.571 | 3 | 2ybe | A |
| TUBA1A | 0.138 | 96 | 5jco | A |
| TUBA1A | 0.207 | 59 | 5jco | G |
| TUBA1A | 0.207 | 59 | 5jco | G |
| TUBA1A | 0.103 | 132 | 5jco | G |
| TUBA1A | 0.069 | 167 | 5jco | F |
| PTPN11 | None | None | None | None |
| PTPN11 | 0.237 | 25 | 4ohh | A |
| PTPN11 | 0.362 | 13 | 4ohl | A |
| PTPN11 | 0.327 | 15 | 4ohi | A |
| PTPN11 | 0.09 | 74 | 5ehp | A |
| PTPN11 | 0.22 | 4 | 4je4 | A |
| PTPN11 | 0.22 | 4 | 4je4 | A |
| PTPN11 | 0.22 | 4 | 4je4 | A |
| PTPN11 | 0.099 | 78 | 4ohh | A |
| PTPN11 | 0.404 | 5 | 3tl0 | A |
| PTPN11 | 0.404 | 5 | 3tl0 | A |
| PTPN11 | 0.404 | 5 | 3tl0 | A |
| PTPN11 | 0.11 | 7 | 4jeg | A |
| PTPN11 | 0.466 | 7 | 3zm1 | A |
| PTPN11 | 0.621 | 2 | 3b7o | A |
| PTPN11 | 0.776 | 1 | 3zm1 | A |
| PTPN11 | 0.453 | 5 | 4ohd | A |
| PTPN11 | 0.216 | 29 | 3b7o | A |
| PTPN11 | 0.216 | 29 | 3b7o | A |
| PTPN11 | 0.165 | 20 | 4ohd | A |
| PTPN11 | None | None | None | None |
| PTPN11 | 0.776 | 1 | 3zm1 | A |
| PTPN11 | 0.621 | 6 | 3zm1 | A |
| PTPN11 | 0.107 | 62 | 5ibs | A |
| PTPN11 | 0.216 | 29 | 3b7o | A |
| PTPN11 | 0.621 | 6 | 3zm1 | A |
| GLB1 | 0.333 | 34 | 3wez | A |
| GLB1 | 0.333 | 67 | 3wf4 | A |
| UQCRQ | None | None | None | None |
| VPS35 | None | None | None | None |
| NR3C1 | 0.107 | 39 | 4p6x | A |
| NR3C1 | 0.135 | 62 | 5g3j | A |

| | | | | |
|---|---|---|---|---|
| NR3C1 | None | None | None | None |
| NR3C1 | 0.359 | 21 | 1nhz | A |
| NR3C1 | 0.42 | 6 | 4mdd | A |
| NR3C1 | 0.177 | 20 | 3cld | B |
| NR3C1 | 0.041 | 74 | 3e7c | A |
| NR3C1 | 0.116 | 61 | 4udd | A |
| NR3C1 | 0.422 | 14 | 1nhz | A |
| APOA1 | None | None | None | None |
| APOA1 | None | None | None | None |
| APOA1 | None | None | None | None |
| APOA1 | None | None | None | None |
| APOA1 | None | None | None | None |
| APOA1 | None | None | None | None |
| APOA1 | None | None | None | None |
| BCKDHA | None | None | None | None |
| BCKDHA | 0.583 | 76 | 1x7x | A |
| BCKDHA | 0.963 | 1 | 1ols | A |
| BCKDHA | 0.696 | 3 | 1v1r | A |
| BCKDHA | 0.625 | 16 | 2beu | A |
| BCKDHA | None | None | None | None |
| BCKDHA | 0.652 | 11 | 1v1r | A |
| BCKDHA | 1 | 1 | 2beu | A |
| BCKDHA | 0.84 | 2 | 2j9f | A |
| BCKDHA | 0.556 | 42 | 1ols | A |
| BCKDHA | 0.609 | 90 | 1v1r | A |
| BCKDHA | 0.652 | 25 | 1v1r | A |
| BCKDHA | 0.346 | 5 | 1wci | A |
| BCKDHA | 0.625 | 60 | 2beu | A |
| BCKDHA | 0.625 | 28 | 2beu | A |
| TOP1 | None | None | None | None |
| ANTXR2 | None | None | None | None |
| ANTXR2 | None | None | None | None |
| ANTXR2 | None | None | None | None |
| TF | 0.625 | 35 | 5dyh | A |
| TF | 0.328 | 52 | 5dyh | A |
| FSHB | None | None | None | None |
| GALK1 | None | None | None | None |
| GALK1 | None | None | None | None |
| GALK1 | None | None | None | None |
| CYP17A1 | 0.119 | 7 | 5irq | A |
| CYP17A1 | 0.353 | 2 | 4nkx | A |
| CYP17A1 | 0.353 | 2 | 4nkx | A |
| CYP17A1 | 0.038 | 15 | 4nkv | A |
| CYP17A1 | 0.038 | 15 | 4nkv | A |
| CYP17A1 | 0.046 | 25 | 5irq | A |
| CYP17A1 | 0.244 | 5 | 4nkx | A |
| CYP17A1 | None | None | None | None |
| CYP17A1 | None | None | None | None |
| CYP17A1 | None | None | None | None |
| CYP17A1 | None | None | None | None |
| CYP17A1 | 0.071 | 13 | 4nkv | A |
| CYP17A1 | 0.071 | 13 | 4nkv | A |
| CYP17A1 | 0.194 | 8 | 4nkw | A |
| CYP17A1 | None | None | None | None |

| | | | | |
|---|---|---|---|---|
| CYP17A1 | None | None | None | None |
| CYP17A1 | None | None | None | None |
| CD44 | 0.667 | 24 | 1uuh | A |
| MDH2 | 0.426 | 5 | 4wlo | B |
| MDH2 | 0.088 | 44 | 4wlo | A |
| NDUFV1 | None | None | None | None |
| NDUFV1 | None | None | None | None |
| NDUFV1 | None | None | None | None |
| NDUFV1 | None | None | None | None |
| NDUFV1 | None | None | None | None |
| NDUFV1 | None | None | None | None |
| NDUFV1 | None | None | None | None |
| NDUFV1 | None | None | None | None |
| NDUFV1 | None | None | None | None |
| NDUFV1 | None | None | None | None |
| CTSC | None | None | None | None |
| CTSC | None | None | None | None |
| CTSC | 0.048 | 34 | 4oem | A |
| CTSC | 0.342 | 13 | 3pdf | A |
| CTSC | 0.643 | 1 | 4oel | A |
| CTSC | 0.136 | 24 | 4cdc | H |
| CTSC | None | None | None | None |
| VWF | None | None | None | None |
| VWF | None | None | None | None |
| VWF | None | None | None | None |
| VWF | None | None | None | None |
| VWF | None | None | None | None |
| VWF | None | None | None | None |
| VWF | None | None | None | None |
| ORC4 | None | None | None | None |
| ORC4 | None | None | None | None |
| CRYAB | None | None | None | None |
| CRYAB | None | None | None | None |
| CRYAB | None | None | None | None |
| MAP2K2 | None | None | None | None |
| MAP2K2 | None | None | None | None |
| MAP2K2 | 0.299 | 4 | 1s9i | A |
| MAP2K2 | 0.067 | 25 | 1s9i | A |
| MAP2K2 | 0.272 | 4 | 1s9i | B |
| MAP2K2 | None | None | None | None |
| MAP2K2 | None | None | None | None |
| MAP2K2 | None | None | None | None |
| MAP2K2 | None | None | None | None |
| CBL | None | None | None | None |
| CBL | None | None | None | None |
| CBL | None | None | None | None |
| CBL | None | None | None | None |
| CBL | None | None | None | None |
| CBL | None | None | None | None |
| CBL | None | None | None | None |
| PFN1 | None | None | None | None |
| PFN1 | 0.75 | 2 | 1fik | A |
| ACSM2A | 0.224 | 4 | 3eq6 | A |
| HSPG2 | None | None | None | None |

| | | | | |
|---|---|---|---|---|
| HSPG2 | None | None | None | None |
| HSPG2 | None | None | None | None |
| HSPG2 | None | None | None | None |
| PARK7 | 0.444 | 48 | 3cyf | A |
| PARK7 | 0.769 | 4 | 1pe0 | A |
| PARK7 | 0.808 | 2 | 1pe0 | A |
| PARK7 | 0.667 | 8 | 4rky | A |
| PARK7 | 0.889 | 1 | 3cz9 | A |
| PARK7 | 0.75 | 2 | 1pdw | B |
| INSR | None | None | None | None |
| INSR | None | None | None | None |
| INSR | None | None | None | None |
| INSR | None | None | None | None |
| INSR | None | None | None | None |
| DSC2 | None | None | None | None |
| DSC2 | None | None | None | None |
| DSC2 | None | None | None | None |
| DSC2 | None | None | None | None |
| DSC2 | None | None | None | None |
| DSC2 | None | None | None | None |
| DSC2 | None | None | None | None |
| DSC2 | None | None | None | None |
| DSC2 | None | None | None | None |
| DSC2 | None | None | None | None |
| MICU1 | None | None | None | None |
| MICU1 | None | None | None | None |
| ADAMTS13 | None | None | None | None |
| ADAMTS13 | None | None | None | None |
| ADAMTS13 | None | None | None | None |
| ADAMTS13 | None | None | None | None |
| ADAMTS13 | None | None | None | None |
| ADAMTS13 | None | None | None | None |
| ADAMTS13 | None | None | None | None |
| IRAK4 | 0.074 | 21 | 5uiu | A |
| IRAK4 | 0.074 | 21 | 5uiu | A |
| RNASEH2B | None | None | None | None |
| RNASEH2B | None | None | None | None |
| RNASEH2B | None | None | None | None |
| NTRK1 | None | None | None | None |
| MPO | 0.75 | 3 | 1d5l | C |
| MPO | 0.6 | 25 | 1myp | D |
| MPO | 0.25 | 86 | 1d5l | C |
| PEX3 | None | None | None | None |
| SERPINA1 | None | None | None | None |
| SERPINA1 | None | None | None | None |
| SERPINA1 | None | None | None | None |
| SERPINA1 | 0.113 | 3 | 1oo8 | A |
| SERPINA1 | None | None | None | None |
| SERPINA1 | None | None | None | None |
| SERPINA1 | 0.185 | 3 | 1qmb | A |
| SERPINA1 | 0.6 | 1 | 1ezx | A |
| SERPINA1 | 0.113 | 3 | 1oo8 | A |
| SERPINA1 | None | None | None | None |
| YARS2 | None | None | None | None |

| | | | | |
|---|---|---|---|---|
| YARS2 | None | None | None | None |
| YARS2 | None | None | None | None |
| YARS2 | None | None | None | None |
| YARS2 | None | None | None | None |
| YARS2 | 0.781 | 1 | 2pid | B |
| YARS2 | None | None | None | None |
| YARS2 | None | None | None | None |
| YARS2 | None | None | None | None |
| PDHA1 | None | None | None | None |
| PDHA1 | None | None | None | None |
| PDHA1 | None | None | None | None |
| PDHA1 | None | None | None | None |
| PDHA1 | None | None | None | None |
| PDHA1 | None | None | None | None |
| PDHA1 | None | None | None | None |
| PDHA1 | 0.96 | 1 | 3exh | C |
| PDHA1 | 0.3 | 5 | 1ni4 | A |
| PDHA1 | None | None | None | None |
| PDHA1 | None | None | None | None |
| CUL4B | None | None | None | None |
| AGXT | None | None | None | None |
| AGXT | None | None | None | None |
| AGXT | None | None | None | None |
| AGXT | 0.692 | 3 | 2yob | A |
| AGXT | 0.786 | 7 | 4kxk | C |
| AGXT | 0.538 | 21 | 3r9a | A |
| AGXT | 0.353 | 33 | 3r9a | C |
| AGXT | None | None | None | None |
| AGXT | None | None | None | None |
| AGXT | 0.611 | 4 | 5hhy | B |
| AGXT | 0.333 | 40 | 4cbs | A |
| AGXT | 0.333 | 59 | 5f9s | A |
| AGXT | 0.308 | 45 | 3r9a | A |
| AGXT | 0.294 | 55 | 3r9a | C |
| AGXT | 0.429 | 28 | 4kxk | C |
| AGXT | 0.684 | 1 | 1h0c | A |
| AGXT | 0.5 | 2 | 4cbr | A |
| AGXT | 0.538 | 19 | 4kyo | A |
| AGXT | 0.538 | 19 | 4kyo | A |
| AGXT | 0.75 | 2 | 5luc | B |
| AGXT | 0.722 | 3 | 5hhy | B |
| AGXT | 0.571 | 3 | 4kxk | C |
| AGXT | 0.571 | 5 | 4kxk | C |
| AGXT | None | None | None | None |
| AGXT | 0.778 | 1 | 5hhy | B |
| AGXT | 0.778 | 1 | 5hhy | B |
| AGXT | 0.421 | 12 | 1h0c | A |
| AGXT | 0.692 | 7 | 4kyo | A |
| AGXT | 0.5 | 18 | 4cbr | A |
| AGXT | 0.5 | 18 | 4cbr | A |
| AGXT | 0.615 | 13 | 3r9a | A |
| AGXT | 0.786 | 7 | 4kxk | C |
| AGXT | 0.846 | 1 | 4kxk | A |
| AGXT | 0.615 | 36 | 3r9a | A |

| | | | | | |
|---|---|---|---|---|---|
| AGXT | | 0.571 | 10 | 4kxk | C |
| SFTPD | | 0.509 | 2 | 3g83 | B |
| SFTPD | | 0.111 | 27 | 4E+052 | B |
| PIK3CA | None | None | | None | None |
| PIK3CA | None | None | | None | None |
| PIK3CA | None | None | | None | None |
| AUH | | 0.22 | 3 | 1hzd | E |
| AUH | None | None | | None | None |
| AUH | None | None | | None | None |
| AUH | | 0.293 | 16 | 2zqr | D |
| F11 | None | None | | None | None |
| F11 | None | None | | None | None |
| F11 | None | None | | None | None |
| F11 | None | None | | None | None |
| MEN1 | | 0.6 | 81 | 5db0 | A |
| MEN1 | | 0.8 | 54 | 4i80 | A |
| MEN1 | | 0.8 | 84 | 5db0 | A |
| MEN1 | | 0.8 | 30 | 5ddb | A |
| MEN1 | | 1 | 2 | 4i80 | A |
| MEN1 | | 0.8 | 17 | 5db0 | A |
| MEN1 | | 0.6 | 17 | 5ddb | A |
| MEN1 | | 0.6 | 31 | 3u88 | A |
| MEN1 | None | None | | None | None |
| MEN1 | | 0.6 | 56 | 5ddb | A |
| MYBPC1 | None | None | | None | None |
| MYBPC1 | None | None | | None | None |
| MYBPC1 | None | None | | None | None |
| PDCD10 | None | None | | None | None |
| SCARB2 | None | None | | None | None |
| SCARB2 | None | None | | None | None |
| SCARB2 | None | None | | None | None |
| SCARB2 | None | None | | None | None |
| SCARB2 | None | None | | None | None |
| TREX2 | None | None | | None | None |
| TBK1 | | 0.03 | 22 | 4im2 | A |
| TBK1 | None | None | | None | None |
| DLD | None | None | | None | None |
| DLD | None | None | | None | None |
| DLD | | 0.505 | 3 | 1zy8 | B |
| DLD | None | None | | None | None |
| DLD | | 0.24 | 11 | 1zy8 | A |
| DLD | | 0.078 | 48 | 3rnm | A |
| LDHA | | 0.802 | 1 | 4qt0 | A |
| TNNC1 | None | None | | None | None |
| TNNC1 | | 0.461 | 2 | 1wrl | A |
| TNNC1 | | 0.153 | 5 | 1mxl | C |
| SMAD3 | None | None | | None | None |
| HARS | None | None | | None | None |
| HARS | None | None | | None | None |
| TNNI3 | None | None | | None | None |
| TNNI3 | None | None | | None | None |
| TNNI3 | None | None | | None | None |
| TNNI3 | None | None | | None | None |
| TNNI3 | None | None | | None | None |

| | | | | |
|---|---|---|---|---|
| TNNI3 | None | None | None | None |
| ACKR1 | None | None | None | None |
| BUB1B | None | None | None | None |
| BUB1B | None | None | None | None |
| BUB1B | None | None | None | None |
| AR | 0.522 | 6 | 1xj7 | A |
| AR | 0.273 | 23 | 2ao6 | A |
| AR | 0.305 | 55 | 4olm | A |
| AR | 0.305 | 55 | 4olm | A |
| AR | 0.131 | 48 | 2ax6 | A |
| FTO | None | None | None | None |
| FTO | 0.333 | 1 | 4ie6 | A |
| TFAM | None | None | None | None |
| PYGL | None | None | None | None |
| PYGL | 0.778 | 3 | 1fa9 | A |
| PYGL | None | None | None | None |
| PYGL | 0.722 | 2 | 1fa9 | A |
| PYGL | 0.5 | 19 | 3dds | A |
| PYGL | 0.611 | 25 | 3ceh | B |
| PYGL | 0.611 | 32 | 1l7x | B |
| PYGL | 0.167 | 132 | 1em6 | B |
| PYGL | 0.722 | 2 | 1fc0 | B |
| PYGL | 0.778 | 3 | 1fa9 | A |
| PYGL | 0.278 | 167 | 1l7x | B |
| PYGL | 0.722 | 12 | 2ati | A |
| PYGL | 0.722 | 1 | 3ceh | B |
| PYGL | 0.611 | 11 | 3ceh | B |
| PYGL | 0.611 | 18 | 1fc0 | B |
| PYGL | 0.722 | 2 | 1fc0 | B |
| GALNS | None | None | None | None |
| GALNS | None | None | None | None |
| GALNS | None | None | None | None |
| GALNS | None | None | None | None |
| GALNS | None | None | None | None |
| GALNS | None | None | None | None |
| GALNS | None | None | None | None |
| GALNS | None | None | None | None |
| GALNS | None | None | None | None |
| GALNS | None | None | None | None |
| GALNS | None | None | None | None |
| GALNS | None | None | None | None |
| GALNS | None | None | None | None |
| GALNS | None | None | None | None |
| GALNS | None | None | None | None |
| GALNS | None | None | None | None |
| GALNS | None | None | None | None |
| GALNS | None | None | None | None |
| GALNS | None | None | None | None |
| GALNS | None | None | None | None |
| GALNS | None | None | None | None |
| GALNS | None | None | None | None |
| GALNS | None | None | None | None |
| GALNS | None | None | None | None |
| GALNS | None | None | None | None |

| | | | | |
|---|---|---|---|---|
| GALNS | None | None | None | None |
| GALNS | None | None | None | None |
| GALNS | None | None | None | None |
| GALNS | None | None | None | None |
| GALNS | None | None | None | None |
| GALNS | None | None | None | None |
| GALNS | None | None | None | None |
| GALNS | None | None | None | None |
| F10 | 0.304 | 21 | 1z6e | L |
| GLUD1 | None | None | None | None |
| GLUD1 | None | None | None | None |
| GLUD1 | None | None | None | None |
| GLUD1 | None | None | None | None |
| GLUD1 | None | None | None | None |
| GLUD1 | None | None | None | None |
| GLUD1 | None | None | None | None |
| GLUD1 | None | None | None | None |
| GLUD1 | None | None | None | None |
| GLUD1 | 0.094 | 5 | 1l1f | D |
| NOTCH1 | 0.528 | 3 | 3v79 | K |
| POGLUT1 | None | None | None | None |
| POGLUT1 | None | None | None | None |
| NR3C1 | 0.455 | 1 | 4hn5 | B |
| CASP8 | 0.535 | 6 | 1f9e | C |
| CASP8 | 0.592 | 8 | 1qdu | C |
| TRAP1 | None | None | None | None |
| FANCM | None | None | None | None |
| EPOR | None | None | None | None |
| EPOR | None | None | None | None |
| NDP | None | None | None | None |
| NDP | None | None | None | None |
| NDP | None | None | None | None |
| NDP | None | None | None | None |
| NDP | None | None | None | None |
| NDP | None | None | None | None |
| NDP | None | None | None | None |
| NDP | None | None | None | None |
| NDP | 0.238 | 1 | 5bqc | A |
| NDP | 0.238 | 1 | 5bqc | A |
| NDP | None | None | None | None |
| NDP | None | None | None | None |
| NDP | None | None | None | None |
| NDP | None | None | None | None |
| NDP | None | None | None | None |
| NDP | None | None | None | None |
| NDP | None | None | None | None |
| ROGDI | None | None | None | None |
| CYP19A1 | None | None | None | None |
| CYP19A1 | None | None | None | None |
| CYP19A1 | None | None | None | None |
| CYP19A1 | None | None | None | None |
| CYP19A1 | None | None | None | None |
| CYP19A1 | 0.487 | 1 | 3eqm | A |
| CYP19A1 | 0.487 | 1 | 3eqm | A |

| | | | | |
|---|---|---|---|---|
| CYP19A1 | 0.257 | 5 | 4kq8 | A |
| UBA5 | None | None | None | None |
| PRPS1 | None | None | None | None |
| PRPS1 | 0.316 | 13 | 4f8e | A |
| PRPS1 | None | None | None | None |
| PRPS1 | None | None | None | None |
| PRPS1 | None | None | None | None |
| PRPS1 | None | None | None | None |
| PRPS1 | None | None | None | None |
| PRPS1 | None | None | None | None |
| F2 | None | None | None | None |
| F2 | None | None | None | None |
| NTF4 | None | None | None | None |
| NTF4 | None | None | None | None |
| PSAP | None | None | None | None |
| PSAP | None | None | None | None |
| TAP1 | None | None | None | None |
| TAP1 | None | None | None | None |
| TAP1 | None | None | None | None |
| ARG1 | None | None | None | None |
| ARG1 | 0.714 | 1 | 2pha | A |
| ARG1 | None | None | None | None |
| ARG1 | 0.471 | 10 | 3th7 | A |
| ARG1 | 0.184 | 52 | 4gsz | A |
| ARG1 | 0.184 | 52 | 4gsz | A |
| CRYM | None | None | None | None |
| ADSL | None | None | None | None |
| ADSL | None | None | None | None |
| ADSL | None | None | None | None |
| ADSL | None | None | None | None |
| ADSL | None | None | None | None |
| ADSL | None | None | None | None |
| ADSL | 0.312 | 8 | 2vd6 | B |
| ADSL | 0.312 | 8 | 2vd6 | B |
| ADSL | None | None | None | None |
| ADSL | 0.353 | 5 | 2vd6 | D |
| ADSL | None | None | None | None |
| ADSL | 0.294 | 11 | 2vd6 | D |
| ADSL | 0.647 | 1 | 2vd6 | D |
| ADSL | 0.435 | 1 | 4ffx | A |
| ADSL | None | None | None | None |
| ADSL | None | None | None | None |
| ADSL | None | None | None | None |
| ADSL | None | None | None | None |
| ADSL | None | None | None | None |
| ADSL | None | None | None | None |
| VWF | None | None | None | None |
| PLA2G7 | None | None | None | None |
| MMP13 | None | None | None | None |
| MMP13 | None | None | None | None |
| MMP13 | None | None | None | None |
| MMP13 | None | None | None | None |
| POMGNT1 | None | None | None | None |
| POMGNT1 | None | None | None | None |

| | | | | |
|---|---|---|---|---|
| COL3A1 | None | None | None | None |
| COL3A1 | None | None | None | None |
| COL3A1 | None | None | None | None |
| COL3A1 | None | None | None | None |
| ACTN2 | None | None | None | None |
| ACTN2 | None | None | None | None |
| ACTN2 | None | None | None | None |
| ACTN2 | None | None | None | None |
| ACTN2 | None | None | None | None |
| ACTN2 | None | None | None | None |
| ACTN2 | None | None | None | None |
| UBA5 | None | None | None | None |
| ADAR | 0.75 | 1 | 1qbj | B |
| UMPS | None | None | None | None |
| UMPS | None | None | None | None |
| UMPS | None | None | None | None |
| TRNT1 | None | None | None | None |
| TRNT1 | None | None | None | None |
| NHEJ1 | None | None | None | None |
| NHEJ1 | None | None | None | None |
| NHEJ1 | None | None | None | None |
| C8A | None | None | None | None |
| GLUL | None | None | None | None |
| GLUL | 0.375 | 6 | 2qc8 | C |
| GLUL | 0.467 | 9 | 2qc8 | B |
| GYG1 | 0.615 | 45 | 3rmv | A |
| GYG1 | 0.923 | 1 | 3u2u | B |
| FLNA | None | None | None | None |
| FLNA | None | None | None | None |
| FLNA | None | None | None | None |
| FLNA | None | None | None | None |
| FLNA | None | None | None | None |
| FLNA | None | None | None | None |
| FLNA | None | None | None | None |
| TTN | None | None | None | None |
| TTN | None | None | None | None |
| TTN | None | None | None | None |
| TTN | None | None | None | None |
| SMS | None | None | None | None |
| SMS | None | None | None | None |
| SMS | None | None | None | None |
| C5 | None | None | None | None |
| SERPINC1 | None | None | None | None |
| SERPINC1 | None | None | None | None |
| SERPINC1 | None | None | None | None |
| SERPINC1 | None | None | None | None |
| SERPINC1 | None | None | None | None |
| SERPINC1 | None | None | None | None |
| SERPINC1 | None | None | None | None |
| SERPINC1 | None | None | None | None |
| SERPINC1 | None | None | None | None |
| SERPINC1 | 0.273 | 3 | 1ant | I |
| SERPINC1 | 0.262 | 2 | 2znh | B |
| SERPINC1 | None | None | None | None |

| | | | | |
|---|---|---|---|---|
| SERPINC1 | 0.136 | 8 | 2b5t | I |
| SERPINC1 | 0.157 | 7 | 3evj | I |
| SERPINC1 | None | None | None | None |
| SERPINC1 | None | None | None | None |
| SERPINC1 | None | None | None | None |
| SERPINC1 | None | None | None | None |
| SERPINC1 | None | None | None | None |
| SERPINC1 | None | None | None | None |
| SERPINC1 | None | None | None | None |
| ETFB | None | None | None | None |
| ETFB | None | None | None | None |
| ETFB | None | None | None | None |
| ETFB | None | None | None | None |
| ETFB | None | None | None | None |
| ETFB | None | None | None | None |
| DIABLO | None | None | None | None |
| DIABLO | None | None | None | None |
| DPYS | 0.333 | 24 | 2vr2 | A |
| DPYS | None | None | None | None |
| DPYS | None | None | None | None |
| DPYS | 0.533 | 7 | 2vr2 | A |
| DPYS | None | None | None | None |
| GNE | None | None | None | None |
| GNE | None | None | None | None |
| GNE | None | None | None | None |
| GNE | None | None | None | None |
| GNE | None | None | None | None |
| GNE | None | None | None | None |
| GNE | None | None | None | None |
| GNE | None | None | None | None |
| GNE | None | None | None | None |
| GNMT | None | None | None | None |
| GNMT | 0.2 | 12 | 2azt | B |
| CTH | 0.324 | 20 | 3cog | D |
| CTH | 0.214 | 15 | 2nmp | B |
| CTH | None | None | None | None |
| USH1C | None | None | None | None |
| PSEN1 | None | None | None | None |
| CHEK2 | None | None | None | None |
| CHEK2 | None | None | None | None |
| CHEK2 | None | None | None | None |
| CHEK2 | None | None | None | None |
| CHEK2 | None | None | None | None |
| CHEK2 | None | None | None | None |
| CHEK2 | None | None | None | None |
| CHEK2 | None | None | None | None |
| CHEK2 | None | None | None | None |
| SNCA | None | None | None | None |
| SNCA | None | None | None | None |
| GAN | None | None | None | None |
| GAN | None | None | None | None |
| GAN | None | None | None | None |
| SBDS | None | None | None | None |
| SBDS | None | None | None | None |

| | | | | | |
|---|---|---|---|---|---|
| SBDS | None | None | None | | None |
| SBDS | None | None | None | | None |
| SBDS | None | None | None | | None |
| SBDS | None | None | None | | None |
| MPO | | 1 | 3 | 3f9p | A |
| MPO | | 0.6 | 12 | 5mfa | A |
| KCNH2 | | 0.292 | 2 | 4hp9 | A |
| KCNH2 | None | None | None | | None |
| KCNH2 | | 0.75 | 1 | 4hp9 | A |
| KCNH2 | None | None | None | | None |
| TEK | None | None | None | | None |
| TEK | None | None | None | | None |
| TEK | None | None | None | | None |
| TEK | None | None | None | | None |
| NPC2 | None | None | None | | None |
| NPC2 | None | None | None | | None |
| NPC2 | None | None | None | | None |
| NPC2 | None | None | None | | None |
| NPC2 | None | None | None | | None |
| NPC2 | None | None | None | | None |
| NPC2 | None | None | None | | None |
| PLG | None | None | None | | None |
| PCSK9 | None | None | None | | None |
| SFTPC | None | None | None | | None |
| SFTPC | None | None | None | | None |
| SFTPC | None | None | None | | None |
| CD40LG | None | None | None | | None |
| CD40LG | None | None | None | | None |
| CD40LG | None | None | None | | None |
| CD40LG | None | None | None | | None |
| CD40LG | None | None | None | | None |
| TAT | None | None | None | | None |
| TAT | None | None | None | | None |
| TAT | None | None | None | | None |
| EP300 | None | None | None | | None |
| FARS2 | None | None | None | | None |
| FARS2 | None | None | None | | None |
| FARS2 | None | None | None | | None |
| FARS2 | None | None | None | | None |
| FARS2 | None | None | None | | None |
| FARS2 | None | None | None | | None |
| FARS2 | None | None | None | | None |
| FARS2 | None | None | None | | None |
| FARS2 | | 0.5 | 1 | 5mgw | A |
| FARS2 | None | None | None | | None |
| FARS2 | None | None | None | | None |
| FARS2 | None | None | None | | None |
| FARS2 | None | None | None | | None |
| MASP2 | None | None | None | | None |
| MASP2 | None | None | None | | None |
| MASP2 | None | None | None | | None |
| MASP2 | | 0.054 | 49 | 1q3x | A |
| STIM1 | None | None | None | | None |
| CHRM2 | None | None | None | | None |

| | | | | |
|---|---|---|---|---|
| MCFD2 | 0.206 | 9 | 3whu | B |
| MCFD2 | 0.574 | 1 | 4yge | B |
| MCFD2 | 0.623 | 1 | 3wnx | B |
| KITLG | None | None | None | None |
| F12 | 0.345 | 1 | 4bdx | A |
| F12 | 0.345 | 1 | 4bdx | A |
| ERAP1 | 0.381 | 5 | 3mdj | A |
| GTF2H5 | None | None | None | None |
| GTF2H5 | None | None | None | None |
| THBD | None | None | None | None |
| MRPL28 | None | None | None | None |
| MRPL28 | None | None | None | None |
| HTRA1 | None | None | None | None |
| HTRA1 | None | None | None | None |
| HTRA1 | None | None | None | None |
| HTRA1 | None | None | None | None |
| HTRA1 | 0.237 | 1 | 3nwu | B |
| HTRA1 | None | None | None | None |
| KARS | None | None | None | None |
| KARS | None | None | None | None |
| KARS | None | None | None | None |
| KARS | None | None | None | None |
| KARS | None | None | None | None |
| HK1 | 0.722 | 11 | 4fpb | B |
| GARS | None | None | None | None |
| GARS | None | None | None | None |
| GARS | None | None | None | None |
| GARS | None | None | None | None |
| GARS | None | None | None | None |
| GARS | None | None | None | None |
| GARS | None | None | None | None |
| IL12B | None | None | None | None |
| HADH | None | None | None | None |
| HADH | None | None | None | None |
| HADH | 0.52 | 4 | 1lsj | A |
| HADH | 0.458 | 3 | 1m75 | A |
| HADH | 0.52 | 4 | 1lsj | A |
| RRM2B | None | None | None | None |
| RRM2B | None | None | None | None |
| RRM2B | None | None | None | None |
| RRM2B | None | None | None | None |
| RRM2B | 0.28 | 4 | 4djn | B |
| RRM2B | None | None | None | None |
| RRM2B | 0.333 | 2 | 3hf1 | B |
| RRM2B | None | None | None | None |
| RRM2B | None | None | None | None |
| RRM2B | 0.286 | 8 | 2vux | A |
| RRM2B | None | None | None | None |
| RRM2B | None | None | None | None |
| RRM2B | None | None | None | None |
| RRM2B | None | None | None | None |
| RRM2B | None | None | None | None |
| MARVELD2 | None | None | None | None |
| TLR4 | None | None | None | None |

| | | | | |
|---|---|---|---|---|
| TTN | 0.146 | 1 | 3q5o | A |
| TTN | None | None | None | None |
| TNFRSF13B | None | None | None | None |
| POLG | None | None | None | None |
| POLG | None | None | None | None |
| POLG | None | None | None | None |
| POLG | None | None | None | None |
| POLG | None | None | None | None |
| POLG | None | None | None | None |
| POLG | None | None | None | None |
| POLG | None | None | None | None |
| POLG | None | None | None | None |
| POLG | None | None | None | None |
| POLG | None | None | None | None |
| POLG | None | None | None | None |
| POLG | None | None | None | None |
| POLG | None | None | None | None |
| POLG | None | None | None | None |
| POLG | None | None | None | None |
| POLG | None | None | None | None |
| POLG | None | None | None | None |
| POLG | None | None | None | None |
| POLG | None | None | None | None |
| POLG | None | None | None | None |
| POLG | None | None | None | None |
| POLG | None | None | None | None |
| POLG | None | None | None | None |
| POLG | None | None | None | None |
| POLG | None | None | None | None |
| POLG | None | None | None | None |
| POLG | 0.467 | 4 | 5c53 | A |
| POLG | 0.533 | 1 | 4ztu | A |
| POLG | None | None | None | None |
| POLG | None | None | None | None |
| POLG | None | None | None | None |
| POLG | None | None | None | None |
| POLG | None | None | None | None |
| POLG | 0.533 | 1 | 4ztu | A |
| POLG | 0.2 | 8 | 4ztu | A |
| POLG | 0.2 | 8 | 4ztu | A |
| POLG | None | None | None | None |
| POLG | None | None | None | None |
| POLG | None | None | None | None |
| POLG | None | None | None | None |
| POLG | None | None | None | None |
| POLG | None | None | None | None |
| POLG | None | None | None | None |
| POLG | None | None | None | None |
| POLG | None | None | None | None |
| POLG | 0.267 | 6 | 4ztu | A |
| POLG | None | None | None | None |
| POLG | None | None | None | None |
| UROD | 0.632 | 24 | 1jpk | A |

| | | | | |
|---|---|---|---|---|
| UROD | 0.579 | 33 | 1r3v | A |
| UROD | None | None | None | None |
| UROD | 0.526 | 49 | 1jph | A |
| UROD | None | None | None | None |
| UROD | 0.632 | 13 | 3gvw | A |
| UROD | 0.789 | 6 | 3gvw | A |
| UROD | 0.842 | 2 | 3gvv | A |
| UROD | 0.737 | 31 | 1r3q | A |
| UROD | 0.667 | 14 | 2q71 | A |
| PCK1 | 0.429 | 154 | 1khg | A |
| PCK1 | 0.053 | 121 | 1m51 | A |
| PCK1 | None | None | None | None |
| PCK1 | 0.167 | 187 | 1khf | A |
| PCK1 | 0.524 | 60 | 1nhx | A |
| HEXA | None | None | None | None |
| HEXA | None | None | None | None |
| HEXA | None | None | None | None |
| HEXA | None | None | None | None |
| HEXA | None | None | None | None |
| HEXA | 0.364 | 2 | 2gjx | A |
| HEXA | 0.364 | 2 | 2gjx | A |
| HEXA | None | None | None | None |
| HEXA | None | None | None | None |
| HEXA | None | None | None | None |
| HEXA | None | None | None | None |
| HEXA | 0.364 | 2 | 2gjx | A |
| HEXA | None | None | None | None |
| HEXA | None | None | None | None |
| HEXA | None | None | None | None |
| HEXA | 0.455 | 1 | 2gjx | A |
| HEXA | None | None | None | None |
| HEXA | None | None | None | None |
| HEXA | None | None | None | None |
| HEXA | None | None | None | None |
| HEXA | None | None | None | None |
| HEXA | None | None | None | None |
| HEXA | None | None | None | None |
| HEXA | None | None | None | None |
| HEXA | 0.273 | 6 | 2gjx | A |
| HEXA | None | None | None | None |
| HEXA | None | None | None | None |
| HEXA | 0.455 | 3 | 2gjx | A |
| HEXA | 0.455 | 3 | 2gjx | A |
| HEXA | None | None | None | None |
| HEXA | None | None | None | None |
| CFTR | None | None | None | None |
| CFTR | None | None | None | None |
| CFTR | None | None | None | None |
| CFTR | None | None | None | None |
| CFTR | None | None | None | None |
| CFTR | None | None | None | None |
| CFTR | None | None | None | None |
| CFTR | None | None | None | None |
| CFTR | None | None | None | None |

| | | | | |
|---|---|---|---|---|
| CFTR | None | None | None | None |
| CFTR | 0.462 | 1 | 2pzg | A |
| CFTR | 0.321 | 1 | 1xmj | A |
| CFTR | 0.462 | 2 | 2pzg | A |
| CFTR | None | None | None | None |
| CFTR | None | None | None | None |
| CFTR | None | None | None | None |
| CFTR | None | None | None | None |
| CFTR | None | None | None | None |
| CFTR | None | None | None | None |
| CFTR | None | None | None | None |
| CFTR | None | None | None | None |
| CFTR | None | None | None | None |
| CFTR | None | None | None | None |
| CFTR | None | None | None | None |
| CFTR | None | None | None | None |
| CFTR | 0.462 | 1 | 2pzg | A |
| CFTR | None | None | None | None |
| CFTR | 0.462 | 2 | 2pzg | A |
| CFTR | None | None | None | None |
| CFTR | None | None | None | None |
| CFTR | None | None | None | None |
| CFTR | 0.346 | 3 | 2pzg | A |
| CFTR | None | None | None | None |
| CFTR | None | None | None | None |
| FAN1 | None | None | None | None |
| CBS | None | None | None | None |
| CBS | None | None | None | None |
| CBS | None | None | None | None |
| CBS | 0.111 | 24 | 5mms | A |
| CBS | 0.406 | 6 | 4pcu | B |
| CBS | None | None | None | None |
| CBS | 0.469 | 6 | 4pcu | A |
| CBS | 0.597 | 3 | 5mms | C |
| CBS | 0.54 | 2 | 5mms | A |
| CBS | 0.54 | 2 | 5mms | A |
| CBS | 0.328 | 9 | 4pcu | A |
| CBS | None | None | None | None |
| CBS | 0.179 | 11 | 4l0d | A |
| CBS | 0.208 | 20 | 1m54 | B |
| CBS | 0.516 | 4 | 5mms | C |
| CBS | None | None | None | None |
| CBS | None | None | None | None |
| CBS | 0.103 | 21 | 4l28 | C |
| CBS | 0.698 | 2 | 1m54 | B |
| CBS | 0.122 | 11 | 4l0d | B |
| CBS | None | None | None | None |
| CBS | 0.19 | 12 | 4coo | A |
| CBS | None | None | None | None |
| CBS | None | None | None | None |
| CBS | None | None | None | None |
| CBS | None | None | None | None |
| CBS | 0.424 | 6 | 4coo | B |
| CBS | 0.143 | 14 | 1jbq | B |

| | | | | |
|---|---|---|---|---|
| CBS | 0.603 | 3 | 4coo | A |
| CBS | 0.603 | 3 | 4coo | A |
| CBS | None | None | None | None |
| CBS | None | None | None | None |
| CBS | None | None | None | None |
| CBS | None | None | None | None |
| CBS | None | None | None | None |
| CBS | None | None | None | None |
| TLR3 | None | None | None | None |
| LIG4 | None | None | None | None |
| LIG4 | None | None | None | None |
| ASPA | 0.769 | 1 | 2i3c | B |
| ASPA | 0.769 | 1 | 2i3c | B |
| ASPA | None | None | None | None |
| ASPA | None | None | None | None |
| ASPA | None | None | None | None |
| ASPA | None | None | None | None |
| ASPA | None | None | None | None |
| ASPA | None | None | None | None |
| ASPA | None | None | None | None |
| ASPA | None | None | None | None |
| ASPA | 0.583 | 2 | 4tnu | A |
| ASPA | 0.583 | 2 | 4tnu | A |
| ASPA | None | None | None | None |
| CFH | None | None | None | None |
| FCGR3A | None | None | None | None |
| FCGR3A | None | None | None | None |
| GRIN2A | 0.035 | 69 | 5kdt | A |
| GRIN2A | 0.658 | 1 | 5i2n | A |
| GRIN2A | None | None | None | None |
| GLA | 0.467 | 10 | 1r46 | B |
| GLA | 0.375 | 19 | 3hg3 | B |
| GLA | 0.375 | 14 | 3hg5 | B |
| GLA | 0.312 | 13 | 3tv8 | B |
| GLA | 0.375 | 14 | 3hg5 | B |
| GLA | 0.375 | 14 | 3hg5 | B |
| GLA | 0.267 | 22 | 3lxc | B |
| GLA | 0.647 | 2 | 3s5y | A |
| GLA | 0.562 | 20 | 3gxt | B |
| GLA | 0.353 | 21 | 3s5y | A |
| GLA | None | None | None | None |
| GLA | 0.75 | 3 | 3gxn | B |
| GLA | 0.75 | 3 | 3gxn | B |
| GLA | 0.438 | 7 | 1r47 | B |
| GLA | 0.375 | 3 | 3hg4 | B |
| GLA | 0.375 | 11 | 3hg2 | A |
| GLA | 0.188 | 40 | 3lxb | B |
| GLA | 0.25 | 42 | 3hg5 | A |
| GLA | 0.25 | 42 | 3hg5 | A |
| GLA | 0.312 | 13 | 3hg2 | B |
| GLA | 0.375 | 19 | 3hg3 | B |
| GLA | 0.4 | 5 | 3lxa | A |
| GLA | 0.214 | 50 | 3lxb | A |
| GLA | 0.214 | 50 | 3lxb | A |

| | | | | |
|---|---|---|---|---|
| GLA | 0.214 | 61 | 3lxb | A |
| GLA | 0.467 | 22 | 3gxn | A |
| GLA | 0.214 | 40 | 3lxb | A |
| GLA | 0.214 | 40 | 3lxb | A |
| GLA | 0.2 | 58 | 3lxc | B |
| GLA | 0.214 | 69 | 3lxb | A |
| GLA | 0.286 | 36 | 3lxb | A |
| SERPINA3 | None | None | None | None |
| MTAP | None | None | None | None |
| MTAP | 0.462 | 25 | 3oze | B |
| MTAP | None | None | None | None |
| BLMH | None | None | None | None |
| TUSC3 | None | None | None | None |
| ISCU | None | None | None | None |
| CYP11B2 | None | None | None | None |
| CYP11B2 | None | None | None | None |
| CYP11B2 | None | None | None | None |
| CYP11B2 | None | None | None | None |
| CYP11B2 | None | None | None | None |
| CYP11B2 | None | None | None | None |
| CYP11B2 | None | None | None | None |
| CYP11B2 | None | None | None | None |
| CYP11B2 | None | None | None | None |
| CYP11B2 | None | None | None | None |
| CYP11B2 | None | None | None | None |
| CYP11B2 | None | None | None | None |
| CYP11B2 | None | None | None | None |
| CYP11B2 | None | None | None | None |
| CYP11B2 | 0.049 | 19 | 4fdh | A |
| CYP11B2 | None | None | None | None |
| CYP11B2 | None | None | None | None |
| CYP11B2 | None | None | None | None |
| CYP11B2 | None | None | None | None |
| CYP11B2 | None | None | None | None |
| CRADD | None | None | None | None |
| CRADD | None | None | None | None |
| FGA | None | None | None | None |
| FGA | None | None | None | None |
| FGA | None | None | None | None |
| FGA | None | None | None | None |
| FGA | None | None | None | None |
| FGA | None | None | None | None |
| FGA | None | None | None | None |
| FGA | None | None | None | None |
| IYD | 1 | 1 | 4ttb | A |
| IYD | None | None | None | None |
| IYD | None | None | None | None |
| IYD | None | None | None | None |
| TCAP | None | None | None | None |
| TCAP | None | None | None | None |
| TCAP | None | None | None | None |
| TCAP | None | None | None | None |
| TCAP | None | None | None | None |
| CYP7A1 | None | None | None | None |

| | | | | | |
|---|---|---|---|---|---|
| TEK | None | None | None | | None |
| TEK | None | None | None | | None |
| TEK | None | None | None | | None |
| TEK | None | None | None | | None |
| TEK | None | None | None | | None |
| TEK | None | None | None | | None |
| TEK | None | None | None | | None |
| CD8A | None | None | None | | None |
| HERC2 | None | None | None | | None |
| NR1H3 | 0.537 | | 3 | 3ipq | A |
| GRHPR | None | None | None | | None |
| GRHPR | None | None | None | | None |
| GRHPR | 0.458 | | 2 | 2q50 | A |
| GRHPR | 0.458 | | 2 | 2q50 | A |
| GRHPR | 0.697 | | 1 | 2wwr | D |
| GRHPR | 0.697 | | 1 | 2wwr | D |
| GRHPR | None | None | None | | None |
| GBE1 | None | None | None | | None |
| GBE1 | None | None | None | | None |
| GBE1 | None | None | None | | None |
| GBE1 | None | None | None | | None |
| GBE1 | None | None | None | | None |
| GBE1 | None | None | None | | None |
| GBE1 | None | None | None | | None |
| GBE1 | None | None | None | | None |
| GBE1 | None | None | None | | None |
| GBE1 | None | None | None | | None |
| GBE1 | None | None | None | | None |
| GBE1 | None | None | None | | None |
| GBE1 | None | None | None | | None |
| GBE1 | None | None | None | | None |
| GBE1 | None | None | None | | None |
| GBE1 | None | None | None | | None |
| GBE1 | None | None | None | | None |
| GBE1 | None | None | None | | None |
| GBE1 | None | None | None | | None |
| GBE1 | None | None | None | | None |
| PC | None | None | None | | None |
| PC | 0.429 | | 1 | 3bg3 | A |
| PC | 0.05 | | 10 | 3bg9 | B |
| PC | 0.4 | | 2 | 3bg9 | B |
| PC | None | None | None | | None |
| PC | None | None | None | | None |
| PC | 0.4 | | 6 | 3bg3 | B |
| PC | None | None | None | | None |
| MSH6 | None | None | None | | None |
| MSH6 | None | None | None | | None |
| MSH6 | None | None | None | | None |
| MSH6 | None | None | None | | None |
| MSH6 | None | None | None | | None |
| MSH6 | None | None | None | | None |
| MSH6 | None | None | None | | None |
| MSH6 | None | None | None | | None |
| MSH6 | None | None | None | | None |

| | | | | | |
|---|---|---|---|---|---|
| MSH6 | None | None | None | | None |
| MSH6 | None | None | None | | None |
| MSH6 | None | None | None | | None |
| MSH6 | None | None | None | | None |
| MSH6 | None | None | None | | None |
| MSH6 | None | None | None | | None |
| MSH6 | None | None | None | | None |
| MSH6 | None | None | None | | None |
| MSH6 | None | None | None | | None |
| MSH6 | None | None | None | | None |
| MSH6 | None | None | None | | None |
| MSH6 | None | None | None | | None |
| MSH6 | None | None | None | | None |
| MSH6 | None | None | None | | None |
| MSH6 | None | None | None | | None |
| MSH6 | None | None | None | | None |
| MSH6 | None | None | None | | None |
| MSH6 | None | None | None | | None |
| MSH6 | None | None | None | | None |
| MSH6 | None | None | None | | None |
| MSH6 | None | None | None | | None |
| RNASEL | 0.32 | 6 | | 4g8l | A |
| ORC1 | None | None | None | | None |
| ORC1 | None | None | None | | None |
| ORC1 | None | None | None | | None |
| KCNH2 | None | None | None | | None |
| KCNH2 | None | None | None | | None |
| KCNH2 | None | None | None | | None |
| KCNH2 | None | None | None | | None |
| KCNH2 | None | None | None | | None |
| KCNH2 | None | None | None | | None |
| KCNH2 | None | None | None | | None |
| KCNH2 | None | None | None | | None |
| KCNH2 | None | None | None | | None |
| KCNH2 | None | None | None | | None |
| KCNH2 | None | None | None | | None |
| KCNH2 | None | None | None | | None |
| KCNH2 | None | None | None | | None |
| KCNH2 | None | None | None | | None |
| KCNH2 | None | None | None | | None |
| KCNH2 | None | None | None | | None |
| KCNH2 | None | None | None | | None |
| KCNH2 | None | None | None | | None |
| KCNH2 | None | None | None | | None |
| KCNH2 | None | None | None | | None |
| KCNH2 | None | None | None | | None |
| KCNH2 | None | None | None | | None |
| KCNH2 | None | None | None | | None |
| KCNH2 | None | None | None | | None |
| MAT1A | 0.429 | 3 | | 2obv | A |
| MAT1A | 0.429 | 14 | | 2obv | A |
| MAT1A | 0.571 | 1 | | 2obv | A |
| MAT1A | 0.571 | 1 | | 2obv | A |
| MAT1A | None | None | None | | None |

| | | | | |
|---|---|---|---|---|
| MAT1A | None | None | None | None |
| MAT1A | None | None | None | None |
| MAT1A | None | None | None | None |
| INSR | None | None | None | None |
| INSR | None | None | None | None |
| INSR | None | None | None | None |
| INSR | None | None | None | None |
| AMT | 0.545 | 5 | 1wsr | A |
| AMT | None | None | None | None |
| AMT | None | None | None | None |
| AMT | None | None | None | None |
| AMT | None | None | None | None |
| AMT | None | None | None | None |
| AMT | None | None | None | None |
| AMT | None | None | None | None |
| AMT | None | None | None | None |
| CSNK1D | None | None | None | None |
| SF3B1 | None | None | None | None |
| AGA | None | None | None | None |
| AGA | None | None | None | None |
| AGA | None | None | None | None |
| AGA | None | None | None | None |
| AGA | None | None | None | None |
| AGA | None | None | None | None |
| AGA | None | None | None | None |
| NECTIN4 | None | None | None | None |
| RS1 | None | None | None | None |
| RS1 | None | None | None | None |
| RS1 | None | None | None | None |
| RS1 | None | None | None | None |
| TBC1D7 | None | None | None | None |
| CTSA | None | None | None | None |
| CTSA | None | None | None | None |
| TKT | 0.273 | 20 | 4kxu | A |
| RP2 | None | None | None | None |
| RP2 | None | None | None | None |
| RP2 | None | None | None | None |
| RP2 | None | None | None | None |
| LYRM4 | None | None | None | None |
| SOS1 | None | None | None | None |
| SOS1 | None | None | None | None |
| SOS1 | None | None | None | None |
| SOS1 | None | None | None | None |
| SOS1 | None | None | None | None |
| SOS1 | None | None | None | None |
| HUWE1 | None | None | None | None |
| HUWE1 | None | None | None | None |
| TPI1 | 0.165 | 51 | 1wyi | B |
| TPI1 | 0.886 | 1 | 4unk | B |
| TPI1 | 0.146 | 43 | 1hti | A |
| MSH2 | None | None | None | None |
| MSH2 | None | None | None | None |
| MSH2 | None | None | None | None |
| MSH2 | None | None | None | None |

| | | | | |
|---|---|---|---|---|
| MSH2 | None | None | None | None |
| MSH2 | None | None | None | None |
| MSH2 | None | None | None | None |
| MSH2 | None | None | None | None |
| MSH2 | None | None | None | None |
| MSH2 | None | None | None | None |
| MSH2 | None | None | None | None |
| MSH2 | None | None | None | None |
| MSH2 | None | None | None | None |
| MSH2 | None | None | None | None |
| MSH2 | None | None | None | None |
| MSH2 | None | None | None | None |
| MSH2 | None | None | None | None |
| MSH2 | None | None | None | None |
| MSH2 | None | None | None | None |
| MSH2 | None | None | None | None |
| MSH2 | None | None | None | None |
| MSH2 | None | None | None | None |
| MSH2 | None | None | None | None |
| MSH2 | None | None | None | None |
| MSH2 | None | None | None | None |
| MSH2 | None | None | None | None |
| MSH2 | None | None | None | None |
| MSH2 | None | None | None | None |
| MSH2 | None | None | None | None |
| MSH2 | None | None | None | None |
| MSH2 | None | None | None | None |
| MSH2 | None | None | None | None |
| MSH2 | None | None | None | None |
| MSH2 | None | None | None | None |
| MSH2 | None | None | None | None |
| MSH2 | None | None | None | None |
| GALE | 0.172 | 6 | 1ek5 | A |
| GALE | 0.78 | 1 | 1i3n | A |
| GALE | 0.122 | 38 | 1ek6 | A |
| GALE | 0.106 | 41 | 1i3m | A |
| GALE | 0.106 | 36 | 1i3m | A |
| GALE | None | None | None | None |
| GALE | 0.176 | 29 | 1i3m | A |
| GALE | 0.259 | 5 | 1i3m | A |
| NSD2 | None | None | None | None |
| SRP72 | None | None | None | None |
| SRP72 | None | None | None | None |
| SRP72 | None | None | None | None |
| POLK | None | None | None | None |
| GALT | None | None | None | None |
| GALT | None | None | None | None |
| GALT | None | None | None | None |
| GALT | None | None | None | None |
| GALT | None | None | None | None |
| GALT | None | None | None | None |
| GALT | None | None | None | None |
| GALT | None | None | None | None |
| GALT | None | None | None | None |

| | | | | |
|---|---|---|---|---|
| GALT | None | None | None | None |
| GALT | None | None | None | None |
| GALT | None | None | None | None |
| GALT | None | None | None | None |
| GALT | None | None | None | None |
| GALT | None | None | None | None |
| GALT | None | None | None | None |
| GALT | None | None | None | None |
| GALT | None | None | None | None |
| GALT | None | None | None | None |
| GALT | None | None | None | None |
| GALT | None | None | None | None |
| GALT | None | None | None | None |
| GALT | None | None | None | None |
| GALT | None | None | None | None |
| GALT | None | None | None | None |
| GALT | None | None | None | None |
| GALT | None | None | None | None |
| GALT | None | None | None | None |
| GALT | None | None | None | None |
| GALT | None | None | None | None |
| GALT | None | None | None | None |
| GALT | None | None | None | None |
| GALT | None | None | None | None |
| GALT | None | None | None | None |
| GALT | None | None | None | None |
| GALT | None | None | None | None |
| GALT | None | None | None | None |
| PYCR1 | 0.75 | 11 | 5uav | A |
| PYCR1 | 0.75 | 11 | 5uav | A |
| PYCR1 | 0.5 | 2 | 2izz | A |
| PYCR1 | 0.556 | 2 | 5uax | B |
| F8 | None | None | None | None |
| F8 | None | None | None | None |
| F8 | None | None | None | None |
| F8 | None | None | None | None |
| F8 | None | None | None | None |
| F8 | None | None | None | None |
| F8 | None | None | None | None |
| F8 | None | None | None | None |
| F8 | None | None | None | None |
| F8 | None | None | None | None |
| F8 | None | None | None | None |
| F8 | None | None | None | None |
| F8 | None | None | None | None |
| F8 | None | None | None | None |
| F8 | None | None | None | None |
| F8 | None | None | None | None |
| ANK1 | None | None | None | None |
| CHAT | None | None | None | None |
| CHAT | None | None | None | None |
| CHAT | 0.615 | 8 | 2fy5 | A |
| CHAT | 0.538 | 2 | 2fy5 | A |
| CHAT | None | None | None | None |

| | | | | |
|---|---|---|---|---|
| CHAT | None | None | None | None |
| CHAT | None | None | None | None |
| CHAT | None | None | None | None |
| CHAT | None | None | None | None |
| CHAT | 0.538 | 6 | 2fy5 | A |
| CHAT | None | None | None | None |
| CHAT | None | None | None | None |
| CHAT | None | None | None | None |
| CHAT | None | None | None | None |
| CHAT | None | None | None | None |
| CHAT | None | None | None | None |
| CHAT | None | None | None | None |
| GHR | None | None | None | None |
| GHR | 0.312 | 2 | 3hhr | C |
| GHR | None | None | None | None |
| GHR | 0.312 | 1 | 3hhr | C |
| GHR | None | None | None | None |
| GHR | None | None | None | None |
| GHR | None | None | None | None |
| GHR | None | None | None | None |
| GHR | None | None | None | None |
| GHR | None | None | None | None |
| GHR | None | None | None | None |
| GHR | None | None | None | None |
| GHR | None | None | None | None |
| GHR | None | None | None | None |
| FZD4 | None | None | None | None |
| FZD4 | None | None | None | None |
| FZD4 | None | None | None | None |
| MTR | None | None | None | None |
| MTR | None | None | None | None |
| MTR | None | None | None | None |
| GM2A | None | None | None | None |
| GM2A | None | None | None | None |
| GM2A | None | None | None | None |
| GM2A | None | None | None | None |
| GM2A | None | None | None | None |
| CRYGD | 0.444 | 3 | 2g98 | A |
| CRYGD | None | None | None | None |
| CRYGD | 0.522 | 1 | 2kfb | A |
| CRYGD | 0.4 | 3 | 4gr7 | X |
| CRYGD | 0.235 | 9 | 1h4a | X |
| CRYGD | 0.333 | 9 | 1hk0 | X |
| CRYGD | None | None | None | None |
| CRYGD | None | None | None | None |
| CRYGD | 0.556 | 2 | 2g98 | A |
| TLR5 | None | None | None | None |
| HSD17B10 | 0.855 | 1 | 1u7t | B |
| HSD17B10 | 0.02 | 51 | 1u7t | C |
| HSD17B10 | 0.542 | 1 | 1so8 | A |
| HSD17B10 | 0.375 | 3 | 1so8 | A |
| TBC1D20 | None | None | None | None |
| FBN1 | None | None | None | None |
| FBN1 | None | None | None | None |

| | | | | |
|---|---|---|---|---|
| FBN1 | None | None | None | None |
| FBN1 | None | None | None | None |
| FBN1 | None | None | None | None |
| FBN1 | None | None | None | None |
| FBN1 | None | None | None | None |
| FXN | None | None | None | None |
| FXN | None | None | None | None |
| FXN | None | None | None | None |
| FXN | None | None | None | None |
| MYO5A | None | None | None | None |
| CTSK | 0.168 | 45 | 7pck | A |
| CTSK | None | None | None | None |
| SCN5A | None | None | None | None |
| SCN5A | None | None | None | None |
| SOS1 | None | None | None | None |
| SOS1 | None | None | None | None |
| SOS1 | None | None | None | None |
| SOS1 | None | None | None | None |
| SOS1 | None | None | None | None |
| SOS1 | None | None | None | None |
| SOS1 | None | None | None | None |
| SOS1 | None | None | None | None |
| SOS1 | None | None | None | None |
| SOS1 | None | None | None | None |
| SOS1 | None | None | None | None |
| SOS1 | None | None | None | None |
| GATM | None | None | None | None |
| GATM | None | None | None | None |
| GATM | 0.75 | 4 | 4jdw | A |
| GATM | None | None | None | None |
| GATM | None | None | None | None |
| GATM | 0.75 | 6 | 6jdw | A |
| GATM | 0.75 | 11 | 6jdw | A |
| MMP2 | 0.094 | 35 | 1ck7 | A |
| MMP2 | None | None | None | None |
| MMP2 | 0.407 | 7 | 1qib | A |
| MMP2 | 0.097 | 28 | 1gxd | A |
| NDUFB9 | None | None | None | None |
| NDUFB9 | None | None | None | None |
| GH1 | None | None | None | None |
| GH1 | None | None | None | None |
| GH1 | None | None | None | None |
| SLC6A4 | None | None | None | None |
| G6PD | None | None | None | None |
| G6PD | None | None | None | None |
| G6PD | None | None | None | None |
| G6PD | None | None | None | None |
| G6PD | None | None | None | None |
| G6PD | None | None | None | None |
| G6PD | None | None | None | None |
| G6PD | None | None | None | None |
| G6PD | 0.545 | 1 | 1qki | B |
| G6PD | None | None | None | None |
| G6PD | None | None | None | None |

| | | | | |
|---|---|---|---|---|
| G6PD | None | None | None | None |
| G6PD | None | None | None | None |
| G6PD | None | None | None | None |
| G6PD | None | None | None | None |
| G6PD | 0.455 | 1 | 1qki | D |
| G6PD | None | None | None | None |
| G6PD | None | None | None | None |
| G6PD | None | None | None | None |
| G6PD | None | None | None | None |
| CST3 | None | None | None | None |
| LMNA | None | None | None | None |
| LMNA | None | None | None | None |
| NDUFA9 | None | None | None | None |
| NDUFA9 | None | None | None | None |
| NDUFA9 | None | None | None | None |
| NDUFA9 | None | None | None | None |
| YARS | 0.581 | 1 | 5thl | A |
| YARS | None | None | None | None |
| YARS | None | None | None | None |
| YARS | None | None | None | None |
| EPHA2 | None | None | None | None |
| EPHA2 | None | None | None | None |
| EPHA2 | None | None | None | None |
| EPHA2 | None | None | None | None |
| DICER1 | None | None | None | None |
| DICER1 | None | None | None | None |
| SNX10 | None | None | None | None |
| SNX10 | None | None | None | None |
| TTN | None | None | None | None |
| TTN | None | None | None | None |
| GATA3 | None | None | None | None |
| C5 | None | None | None | None |
| THBD | None | None | None | None |
| BIN1 | None | None | None | None |
| BIN1 | None | None | None | None |
| SMARCAL1 | None | None | None | None |
| CFB | None | None | None | None |
| CYC1 | None | None | None | None |
| ANK2 | 0.006 | 30 | 4rlv | A |
| CHMP4B | None | None | None | None |
| CHMP4B | None | None | None | None |
| ERBB2 | None | None | None | None |
| ERBB2 | None | None | None | None |
| POLG2 | None | None | None | None |
| POLG2 | None | None | None | None |
| POLG2 | None | None | None | None |
| POLG2 | None | None | None | None |
| POLG2 | None | None | None | None |
| POLG2 | None | None | None | None |
| POLG2 | None | None | None | None |
| POLG2 | None | None | None | None |
| C3 | None | None | None | None |
| C3 | None | None | None | None |
| RPGR | None | None | None | None |

| | | | | |
|---|---|---|---|---|
| RPGR | None | None | None | None |
| RPGR | None | None | None | None |
| RPGR | None | None | None | None |
| RPGR | None | None | None | None |
| C1QA | None | None | None | None |
| PRKG1 | None | None | None | None |
| IGBP1 | None | None | None | None |
| ENO3 | 0.077 | 98 | 2xsx | B |
| ENO3 | None | None | None | None |
| ENO3 | 0.41 | 43 | 2xsx | A |
| ENO3 | 0.487 | 39 | 2xsx | A |
| ENO3 | 0.359 | 47 | 2xsx | B |
| ENO3 | 0.897 | 2 | 2xsx | B |
| MCPH1 | None | None | None | None |
| MCPH1 | None | None | None | None |
| MCPH1 | None | None | None | None |
| MCPH1 | None | None | None | None |
| MCPH1 | None | None | None | None |
| SNRNP200 | None | None | None | None |
| SNRNP200 | None | None | None | None |
| DNMT1 | None | None | None | None |
| DNMT1 | None | None | None | None |
| DNMT1 | None | None | None | None |
| AKT1 | 0.072 | 24 | 4ejn | A |
| NDUFS2 | None | None | None | None |
| ACVR2B | None | None | None | None |
| HMGCL | None | None | None | None |
| HMGCL | None | None | None | None |
| HMGCL | None | None | None | None |
| JUP | None | None | None | None |
| JUP | None | None | None | None |
| JUP | None | None | None | None |
| JUP | None | None | None | None |
| JUP | None | None | None | None |
| JUP | None | None | None | None |
| JUP | None | None | None | None |
| JUP | None | None | None | None |
| SPINK1 | None | None | None | None |
| SPINK1 | None | None | None | None |
| SPINK1 | None | None | None | None |
| VCL | None | None | None | None |
| VCL | None | None | None | None |
| NOTCH1 | None | None | None | None |
| NOTCH1 | None | None | None | None |
| NOTCH1 | None | None | None | None |
| NDUFB3 | None | None | None | None |
| IKBKG | None | None | None | None |
| GSN | 0.529 | 2 | 2fh1 | A |
| GSN | None | None | None | None |
| GSN | None | None | None | None |
| GSN | 0.5 | 3 | 1p8x | A |
| GSN | 0.263 | 13 | 2fh3 | A |
| GSN | None | None | None | None |
| GSN | None | None | None | None |

| | | | | |
|---|---|---|---|---|
| GSN | None | None | None | None |
| GSN | None | None | None | None |
| CLC | None | None | None | None |
| RAF1 | None | None | None | None |
| RAF1 | None | None | None | None |
| RAF1 | None | None | None | None |
| NDUFS8 | None | None | None | None |
| NDUFS8 | None | None | None | None |
| NDUFS8 | None | None | None | None |
| NDUFS8 | None | None | None | None |
| NDUFS8 | None | None | None | None |
| CASR | None | None | None | None |
| CASR | None | None | None | None |
| CASR | None | None | None | None |
| CASR | None | None | None | None |
| CASR | None | None | None | None |
| CASR | None | None | None | None |
| CASR | None | None | None | None |
| CASR | None | None | None | None |
| CASR | None | None | None | None |
| CASR | None | None | None | None |
| CASR | None | None | None | None |
| CASR | None | None | None | None |
| CASR | None | None | None | None |
| CASR | None | None | None | None |
| CASR | None | None | None | None |
| CASR | None | None | None | None |
| CASR | None | None | None | None |
| CASR | None | None | None | None |
| CASR | None | None | None | None |
| CASR | None | None | None | None |
| CASR | None | None | None | None |
| CASR | None | None | None | None |
| CASR | None | None | None | None |
| CASR | None | None | None | None |
| CASR | None | None | None | None |
| CASR | None | None | None | None |
| CASR | None | None | None | None |
| CASR | None | None | None | None |
| CASR | None | None | None | None |
| VCL | None | None | None | None |
| VCL | None | None | None | None |
| SMPD1 | None | None | None | None |
| SMPD1 | None | None | None | None |
| SMPD1 | None | None | None | None |
| KDM6A | None | None | None | None |
| KCNQ1 | None | None | None | None |
| KCNQ1 | None | None | None | None |
| KCNQ1 | None | None | None | None |
| KCNQ1 | None | None | None | None |
| KCNQ1 | None | None | None | None |
| KCNQ1 | None | None | None | None |
| NDUFA12 | None | None | None | None |
| GLRX5 | None | None | None | None |

| | | | | |
|---|---|---|---|---|
| PALB2 | None | None | None | None |
| PALB2 | None | None | None | None |
| PALB2 | None | None | None | None |
| PALB2 | None | None | None | None |
| PALB2 | None | None | None | None |
| PALB2 | None | None | None | None |
| PALB2 | None | None | None | None |
| PALB2 | None | None | None | None |
| PALB2 | None | None | None | None |
| PALB2 | None | None | None | None |
| ALDOB | 0.462 | 89 | 1qo5 | C |
| ALDOB | 0.731 | 2 | 1xdm | A |
| ALDOB | 0.731 | 1 | 1qo5 | B |
| ALDOB | 0.538 | 6 | 1xdm | A |
| ALDOB | 0.692 | 7 | 1qo5 | C |
| ALDOB | None | None | None | None |
| ALDOB | 0.577 | 25 | 1qo5 | B |
| GNE | None | None | None | None |
| GNE | None | None | None | None |
| GNE | None | None | None | None |
| GNE | None | None | None | None |
| GNE | None | None | None | None |
| GNE | None | None | None | None |
| GNE | None | None | None | None |
| GNE | None | None | None | None |
| GNE | None | None | None | None |
| GNE | None | None | None | None |
| GNE | None | None | None | None |
| GNE | None | None | None | None |
| CHRNB2 | None | None | None | None |
| CHRNB2 | None | None | None | None |
| ERCC3 | None | None | None | None |
| ERCC3 | None | None | None | None |
| ERCC3 | None | None | None | None |
| ERCC3 | None | None | None | None |
| ERCC3 | None | None | None | None |
| UNC119 | None | None | None | None |
| ATL1 | None | None | None | None |
| ATL1 | None | None | None | None |
| ATL1 | None | None | None | None |
| ATL1 | 0.8 | 1 | 3q5d | A |
| ATL1 | None | None | None | None |
| ATL1 | None | None | None | None |
| ATL1 | None | None | None | None |
| ATL1 | None | None | None | None |
| ATL1 | None | None | None | None |
| ATL1 | None | None | None | None |
| NR3C2 | 0.701 | 1 | 2oax | A |
| TBK1 | None | None | None | None |
| HYAL1 | None | None | None | None |
| HYAL1 | None | None | None | None |
| HYAL1 | None | None | None | None |
| HYAL1 | None | None | None | None |
| CTSD | 0.633 | 1 | 4obz | A |

| | | | | |
|---|---|---|---|---|
| PRKCG | None | None | None | None |
| TGFBR1 | None | None | None | None |
| XDH | 0.308 | 2 | 2e1q | A |
| XDH | None | None | None | None |
| XDH | 0.051 | 415 | 2e1q | A |
| XDH | 0.154 | 74 | 2e1q | A |
| XDH | 0.125 | 35 | 2e1q | B |
| XDH | None | None | None | None |
| NMNAT1 | 0.833 | 1 | 1kqn | E |
| NMNAT1 | None | None | None | None |
| NMNAT1 | 0.167 | 15 | 1kqo | B |
| NMNAT1 | 0.167 | 3 | 1kqo | B |
| NMNAT1 | 0.167 | 20 | 1kqo | B |
| NMNAT1 | 0.118 | 25 | 1kr2 | B |
| SERPING1 | None | None | None | None |
| SERPING1 | None | None | None | None |
| SERPING1 | None | None | None | None |
| SERPING1 | None | None | None | None |
| SERPING1 | None | None | None | None |
| PDZRN3 | None | None | None | None |
| FABP2 | None | None | None | None |
| IFNGR1 | 0.143 | 1 | 1jrh | I |
| IFNGR1 | 0.143 | 1 | 1jrh | I |
| TDP1 | None | None | None | None |
| TDP1 | 0.333 | 11 | 1rg1 | A |
| INS | 0.263 | 5 | 2lgb | B |
| INS | 0.769 | 1 | 2r34 | C |
| INS | 0.769 | 1 | 2r34 | C |
| PPOX | 0.857 | 1 | 4ivm | B |
| PPOX | None | None | None | None |
| PPOX | None | None | None | None |
| PPOX | None | None | None | None |
| PPOX | None | None | None | None |
| PPOX | None | None | None | None |
| PPOX | None | None | None | None |
| PPOX | None | None | None | None |
| PPOX | None | None | None | None |
| GFER | None | None | None | None |
| GFER | 0.769 | 4 | 3tk0 | A |
| GFER | None | None | None | None |
| GFER | None | None | None | None |
| LIFR | None | None | None | None |
| EGLN1 | 0.556 | 21 | 5a3u | C |
| EGLN1 | 0.889 | 6 | 5v18 | A |
| EGLN1 | 0.944 | 2 | 5a3u | C |
| GP1BA | None | None | None | None |
| GP1BA | None | None | None | None |
| GP1BA | None | None | None | None |
| GP1BA | None | None | None | None |
| GP1BA | None | None | None | None |
| MMADHC | None | None | None | None |
| MMADHC | None | None | None | None |
| MMADHC | None | None | None | None |
| MMADHC | None | None | None | None |

| | | | | |
|---|---|---|---|---|
| MMADHC | None | None | None | None |
| MMADHC | None | None | None | None |
| MMADHC | None | None | None | None |
| ACVRL1 | None | None | None | None |
| ACVRL1 | None | None | None | None |
| RFXANK | None | None | None | None |
| CHIT1 | None | None | None | None |
| UBR1 | None | None | None | None |
| BLM | None | None | None | None |
| BLM | None | None | None | None |
| BLM | None | None | None | None |
| BLM | None | None | None | None |
| BLM | None | None | None | None |
| BLM | None | None | None | None |
| BLM | None | None | None | None |
| BLM | None | None | None | None |
| BLM | None | None | None | None |
| BLM | None | None | None | None |
| BLM | None | None | None | None |
| BLM | None | None | None | None |
| BLM | None | None | None | None |
| VDR | 0.378 | 7 | 1kb4 | B |
| VDR | None | None | None | None |
| VDR | 0.378 | 4 | 1kb4 | A |
| VDR | 0.578 | 1 | 1kb2 | B |
| VDR | 0.578 | 1 | 1kb2 | B |
| VDR | 0.578 | 1 | 1kb2 | B |
| EPHA2 | None | None | None | None |
| EPHA2 | None | None | None | None |
| EPHA2 | None | None | None | None |
| EPHA2 | None | None | None | None |
| EPHA2 | None | None | None | None |
| EPHA2 | None | None | None | None |
| EPHA2 | None | None | None | None |
| EPHA2 | None | None | None | None |
| EPHA2 | None | None | None | None |
| EPHA2 | None | None | None | None |
| UBA1 | None | None | None | None |
| STAT1 | None | None | None | None |
| STAT1 | None | None | None | None |
| STAT1 | None | None | None | None |
| STAT1 | None | None | None | None |
| STAT1 | None | None | None | None |
| STAT1 | None | None | None | None |
| STAT1 | None | None | None | None |
| STAT1 | None | None | None | None |
| DNM1L | None | None | None | None |
| DNM1L | None | None | None | None |
| OTC | None | None | None | None |
| OTC | None | None | None | None |
| OTC | 0.071 | 7 | 1oth | A |
| OTC | None | None | None | None |
| OTC | None | None | None | None |
| OTC | None | None | None | None |

| | | | | | |
|---|---|---|---|---|---|
| OTC | | 0.021 | 14 | 1ep9 | A |
| OTC | None | None | | None | None |
| OTC | None | None | | None | None |
| OTC | | 0.281 | 3 | 1fvo | A |
| OTC | None | None | | None | None |
| OTC | None | None | | None | None |
| OTC | None | None | | None | None |
| OTC | None | None | | None | None |
| OTC | | 0.125 | 14 | 1c9y | A |
| OTC | | 0.125 | 14 | 1c9y | A |
| OTC | | 0.143 | 11 | 1oth | A |
| RARS | None | None | | None | None |
| RARS | None | None | | None | None |
| RARS | None | None | | None | None |
| SEPSECS | None | None | | None | None |
| SEPSECS | None | None | | None | None |
| SEPSECS | None | None | | None | None |
| SEPSECS | None | None | | None | None |
| SEPSECS | None | None | | None | None |
| BRCA1 | None | None | | None | None |
| BRCA1 | None | None | | None | None |
| BRCA1 | | 0.538 | 1 | 3pxc | X |
| BRCA1 | None | None | | None | None |
| BRCA1 | None | None | | None | None |
| BRCA1 | | 0.462 | 1 | 1t2u | A |
| BRCA1 | None | None | | None | None |
| BRCA1 | None | None | | None | None |
| BRCA1 | None | None | | None | None |
| BRCA1 | None | None | | None | None |
| BRCA1 | None | None | | None | None |
| BRCA1 | None | None | | None | None |
| BRCA1 | None | None | | None | None |
| BRCA1 | None | None | | None | None |
| BRCA1 | None | None | | None | None |
| BRCA1 | None | None | | None | None |
| BRCA1 | None | None | | None | None |
| BRCA1 | None | None | | None | None |
| BRCA1 | None | None | | None | None |
| BRCA1 | None | None | | None | None |
| BRCA1 | None | None | | None | None |
| BRCA1 | None | None | | None | None |
| CUBN | None | None | | None | None |
| ITGB3 | None | None | | None | None |
| MTRR | None | None | | None | None |
| MTRR | None | None | | None | None |
| MTRR | None | None | | None | None |
| MTRR | None | None | | None | None |
| MTRR | None | None | | None | None |
| MTRR | None | None | | None | None |
| MTRR | None | None | | None | None |
| MTRR | None | None | | None | None |
| IL7R | None | None | | None | None |
| IL7R | None | None | | None | None |
| RPS24 | None | None | | None | None |

| | | | | |
|---|---|---|---|---|
| RPS24 | None | None | None | None |
| HOGA1 | None | None | None | None |
| HOGA1 | 0.023 | 17 | 3s5o | A |
| HOGA1 | None | None | None | None |
| HOGA1 | None | None | None | None |
| HOGA1 | None | None | None | None |
| HOGA1 | None | None | None | None |
| HOGA1 | 0.791 | 1 | 3s5n | A |
| HOGA1 | None | None | None | None |
| HOGA1 | None | None | None | None |
| HOGA1 | None | None | None | None |
| HOGA1 | None | None | None | None |
| HOGA1 | 0.091 | 19 | 3s5o | A |
| HOGA1 | None | None | None | None |
| GJB2 | None | None | None | None |
| GJB2 | None | None | None | None |
| GJB2 | None | None | None | None |
| GJB2 | None | None | None | None |
| GJB2 | None | None | None | None |
| GJB2 | None | None | None | None |
| GJB2 | None | None | None | None |
| GJB2 | None | None | None | None |
| GJB2 | None | None | None | None |
| GJB2 | None | None | None | None |
| GJB2 | None | None | None | None |
| GJB2 | None | None | None | None |
| GJB2 | None | None | None | None |
| GJB2 | None | None | None | None |
| GJB2 | None | None | None | None |
| GJB2 | None | None | None | None |
| GJB2 | None | None | None | None |
| GJB2 | None | None | None | None |
| GJB2 | None | None | None | None |
| GJB2 | None | None | None | None |
| GJB2 | None | None | None | None |
| GJB2 | None | None | None | None |
| GJB2 | None | None | None | None |
| GJB2 | None | None | None | None |
| GJB2 | None | None | None | None |
| GJB2 | None | None | None | None |
| GJB2 | None | None | None | None |
| GJB2 | None | None | None | None |
| GJB2 | None | None | None | None |
| GJB2 | None | None | None | None |
| GJB2 | None | None | None | None |
| GJB2 | None | None | None | None |
| GJB2 | None | None | None | None |
| GJB2 | None | None | None | None |
| GJB2 | None | None | None | None |
| GJB2 | None | None | None | None |
| GJB2 | None | None | None | None |
| GJB2 | None | None | None | None |
| GJB2 | None | None | None | None |

| | | | | |
|---|---|---|---|---|
| GJB2 | None | None | None | None |
| GJB2 | None | None | None | None |
| GJB2 | None | None | None | None |
| GJB2 | None | None | None | None |
| GJB2 | None | None | None | None |
| GJB2 | None | None | None | None |
| GJB2 | None | None | None | None |
| GJB2 | None | None | None | None |
| GJB2 | None | None | None | None |
| GJB2 | None | None | None | None |
| GJB2 | None | None | None | None |
| GJB2 | None | None | None | None |
| GJB2 | None | None | None | None |
| GJB2 | None | None | None | None |
| CPS1 | None | None | None | None |
| CPS1 | 0.545 | 127 | 5dou | A |
| CPS1 | None | None | None | None |
| CPS1 | None | None | None | None |
| CPS1 | 0.364 | 19 | 5dou | A |
| CPS1 | 0.364 | 19 | 5dou | A |
| CPS1 | 0.182 | 108 | 5dou | A |
| CPS1 | None | None | None | None |
| CPS1 | None | None | None | None |
| CPS1 | None | None | None | None |
| PSPH | None | None | None | None |
| PSPH | None | None | None | None |
| PSPH | None | None | None | None |
| PSPH | None | None | None | None |
| PSPH | None | None | None | None |
| PSPH | None | None | None | None |
| PEPD | 0.512 | 2 | 5m4l | B |
| PEPD | 0.048 | 51 | 5m4j | A |
| PEPD | 0.045 | 58 | 5m4l | A |
| PEPD | 0.643 | 2 | 5m4j | B |
| PEPD | 0.643 | 2 | 5m4j | B |
| PEPD | 0.419 | 5 | 5m4l | B |
| PEPD | 0.106 | 31 | 2iw2 | A |
| PEPD | None | None | None | None |
| PEPD | 0.022 | 40 | 2okn | B |
| KDM5C | None | None | None | None |
| KDM5C | 0.222 | 117 | 5fwj | A |
| KDM5C | 0.333 | 55 | 5fwj | A |
| ITGB6 | 0.312 | 2 | 5ffo | B |
| ITGB6 | None | None | None | None |
| ITGB6 | None | None | None | None |
| DARS | None | None | None | None |
| DARS | None | None | None | None |
| DARS | None | None | None | None |
| DARS | None | None | None | None |
| BTK | None | None | None | None |
| BTK | None | None | None | None |
| BTK | None | None | None | None |
| HDAC4 | None | None | None | None |
| ITGA2B | None | None | None | None |

| | | | | |
|---|---|---|---|---|
| ITGA2B | None | None | None | None |
| ITGA2B | None | None | None | None |
| ITGA2B | None | None | None | None |
| PCSK9 | None | None | None | None |
| PCSK9 | None | None | None | None |
| PCSK9 | None | None | None | None |
| PCSK9 | 0.12 | 16 | 2pmw | B |
| MUTYH | None | None | None | None |
| MUTYH | None | None | None | None |
| MUTYH | None | None | None | None |
| MUTYH | None | None | None | None |
| MUTYH | None | None | None | None |
| MUTYH | 0.375 | 2 | 3n5n | X |
| MUTYH | None | None | None | None |
| MUTYH | None | None | None | None |
| MUTYH | None | None | None | None |
| MUTYH | 0.375 | 2 | 3n5n | X |
| MUTYH | 0.375 | 2 | 3n5n | X |
| MUTYH | None | None | None | None |
| MUTYH | 0.375 | 2 | 3n5n | X |
| MUTYH | None | None | None | None |
| MUTYH | None | None | None | None |
| MUTYH | None | None | None | None |
| MUTYH | None | None | None | None |
| MUTYH | None | None | None | None |
| MUTYH | None | None | None | None |
| MUTYH | None | None | None | None |
| MUTYH | None | None | None | None |
| MUTYH | None | None | None | None |
| MUTYH | None | None | None | None |
| MUTYH | None | None | None | None |
| SLC25A12 | None | None | None | None |
| GSN | None | None | None | None |
| IHH | 0.909 | 1 | 3n1p | B |
| IHH | None | None | None | None |
| IHH | None | None | None | None |
| IHH | 0.818 | 2 | 3n1p | B |
| IHH | 0.727 | 12 | 3n1p | B |
| IHH | 0.636 | 15 | 3n1p | B |
| IHH | 0.818 | 7 | 3n1p | B |
| CD247 | None | None | None | None |
| EP300 | 0.028 | 64 | 5lku | A |
| TTN | None | None | None | None |
| MRI1 | None | None | None | None |
| SGSH | None | None | None | None |
| SGSH | None | None | None | None |
| SGSH | None | None | None | None |
| SGSH | None | None | None | None |
| SGSH | None | None | None | None |
| SGSH | None | None | None | None |
| SGSH | None | None | None | None |
| SGSH | None | None | None | None |
| SGSH | None | None | None | None |
| SGSH | None | None | None | None |

| | | | | |
|---|---|---|---|---|
| SGSH | None | None | None | None |
| SGSH | None | None | None | None |
| SGSH | None | None | None | None |
| SGSH | None | None | None | None |
| SGSH | None | None | None | None |
| SGSH | None | None | None | None |
| SGSH | None | None | None | None |
| SGSH | None | None | None | None |
| SGSH | None | None | None | None |
| SGSH | None | None | None | None |
| SGSH | None | None | None | None |
| SGSH | None | None | None | None |
| CBL | None | None | None | None |
| CANT1 | None | None | None | None |
| CANT1 | None | None | None | None |
| CANT1 | None | None | None | None |
| CANT1 | None | None | None | None |
| CANT1 | None | None | None | None |
| CANT1 | None | None | None | None |
| CANT1 | None | None | None | None |
| SLC1A3 | None | None | None | None |
| SLC1A3 | None | None | None | None |
| SLC1A5 | None | None | None | None |
| SLC1A3 | None | None | None | None |
| SLC1A5 | None | None | None | None |
| SLC1A3 | None | None | None | None |
| SLC1A3 | None | None | None | None |
| CTSD | 0.423 | 15 | 4od9 | D |
| CTSD | 0.14 | 32 | 1lya | B |
| CTSD | None | None | None | None |
| CTSD | None | None | None | None |
| CTSD | None | None | None | None |
| MTOR | None | None | None | None |
| ILK | None | None | None | None |
| PCDH15 | None | None | None | None |
| PCDH15 | None | None | None | None |
| PCDH15 | None | None | None | None |
| PCDH15 | None | None | None | None |
| PCDH15 | None | None | None | None |
| PCDH15 | None | None | None | None |
| KLHL40 | None | None | None | None |
| KLHL40 | None | None | None | None |
| KLHL40 | None | None | None | None |
| KLHL40 | None | None | None | None |
| CSTA | None | None | None | None |
| CSTA | None | None | None | None |
| HNMT | None | None | None | None |
| HNMT | None | None | None | None |
| IFNGR2 | None | None | None | None |
| IFNGR2 | None | None | None | None |
| PEX5 | None | None | None | None |
| KIF7 | None | None | None | None |
| KIF7 | 0.185 | 18 | 4a14 | A |
| MYL3 | None | None | None | None |

| | | | | |
|---|---|---|---|---|
| MYL3 | None | None | None | None |
| MYL3 | None | None | None | None |
| MYL3 | None | None | None | None |
| MYL3 | None | None | None | None |
| MYL3 | None | None | None | None |
| MYL3 | None | None | None | None |
| NAGLU | None | None | None | None |
| NAGLU | None | None | None | None |
| NAGLU | None | None | None | None |
| NAGLU | None | None | None | None |
| NAGLU | None | None | None | None |
| NAGLU | None | None | None | None |
| NAGLU | None | None | None | None |
| NAGLU | None | None | None | None |
| NAGLU | None | None | None | None |
| NAGLU | None | None | None | None |
| NAGLU | None | None | None | None |
| NAGLU | None | None | None | None |
| NAGLU | None | None | None | None |
| TNFRSF13C | None | None | None | None |
| KIF21A | None | None | None | None |
| KIF21A | None | None | None | None |
| KIF21A | None | None | None | None |
| KIF21A | None | None | None | None |
| KIF21A | None | None | None | None |
| DOCK2 | None | None | None | None |
| DSP | None | None | None | None |
| CDKN2A | None | None | None | None |
| CDKN2A | None | None | None | None |
| CDKN2A | None | None | None | None |
| CDKN2A | None | None | None | None |
| CDKN2A | None | None | None | None |
| CDKN2A | None | None | None | None |
| CDKN2A | None | None | None | None |
| CDKN2A | None | None | None | None |
| CDKN2A | None | None | None | None |
| CDKN2A | None | None | None | None |
| CDKN2A | None | None | None | None |
| CDKN2A | None | None | None | None |
| CDKN2A | None | None | None | None |
| CDKN2A | None | None | None | None |
| VCL | None | None | None | None |
| TPP1 | None | None | None | None |
| TPP1 | None | None | None | None |
| TPP1 | None | None | None | None |
| TPP1 | None | None | None | None |
| TPP1 | None | None | None | None |
| TPP1 | None | None | None | None |
| TPP1 | None | None | None | None |
| TPP1 | None | None | None | None |
| TPP1 | None | None | None | None |
| TPP1 | None | None | None | None |
| TPP1 | None | None | None | None |
| TPP1 | None | None | None | None |

| | | | | |
|---|---|---|---|---|
| TPP1 | None | None | None | None |
| TPP1 | None | None | None | None |
| TPP1 | None | None | None | None |
| TPP1 | None | None | None | None |
| TPP1 | None | None | None | None |
| TPP1 | None | None | None | None |
| TPP1 | None | None | None | None |
| TPP1 | None | None | None | None |
| TPP1 | None | None | None | None |
| TPP1 | None | None | None | None |
| IL10RB | None | None | None | None |
| IL10RB | None | None | None | None |
| CCM2 | None | None | None | None |
| CCM2 | None | None | None | None |
| CCM2 | None | None | None | None |
| DCX | None | None | None | None |
| DCX | None | None | None | None |
| DCX | None | None | None | None |
| DCX | None | None | None | None |
| DCX | None | None | None | None |
| DCX | None | None | None | None |
| DCX | None | None | None | None |
| DCX | None | None | None | None |
| BRCA2 | None | None | None | None |
| NNT | None | None | None | None |
| C2 | None | None | None | None |
| C2 | None | None | None | None |
| C2 | None | None | None | None |
| C2 | None | None | None | None |
| WRN | None | None | None | None |
| WRN | None | None | None | None |
| F13A1 | None | None | None | None |
| F13A1 | None | None | None | None |
| F13A1 | None | None | None | None |
| F13A1 | None | None | None | None |
| TF | 0.188 | 24 | 3ve1 | B |
| TF | None | None | None | None |
| TF | 0.516 | 2 | 3ve1 | B |
| TF | 0.562 | 39 | 3skp | A |
| TF | 0.141 | 33 | 3v89 | B |
| ITPA | None | None | None | None |
| ITPA | None | None | None | None |
| SNCA | None | None | None | None |
| ITGB4 | None | None | None | None |
| ITGB4 | None | None | None | None |
| ITGB4 | None | None | None | None |
| LRP6 | 0.021 | 144 | 5air | B |
| LRP6 | None | None | None | None |
| LRP6 | 0.114 | 11 | 3s94 | A |
| PLG | None | None | None | None |
| PLG | 0.61 | 1 | 1i5k | A |
| TYROBP | None | None | None | None |
| UQCRC2 | None | None | None | None |
| GPSM2 | None | None | None | None |

| | | | | |
|---|---|---|---|---|
| GPSM2 | None | None | None | None |
| GPSM2 | None | None | None | None |
| SCN5A | None | None | None | None |
| SCN5A | None | None | None | None |
| SCN5A | None | None | None | None |
| SCN5A | None | None | None | None |
| SCN5A | None | None | None | None |
| ATIC | 0.6 | 6 | 1p4r | A |
| EZH2 | None | None | None | None |
| CPA1 | 0.387 | 33 | 4uez | A |
| SCN4B | None | None | None | None |
| SCN4B | None | None | None | None |
| MAN1B1 | None | None | None | None |
| MAN1B1 | 0.6 | 9 | 1fmi | A |
| MAN1B1 | None | None | None | None |
| MAN1B1 | None | None | None | None |
| CYCS | 0.74 | 1 | 2n9i | A |
| IDUA | None | None | None | None |
| IDUA | None | None | None | None |
| IDUA | None | None | None | None |
| IDUA | None | None | None | None |
| IDUA | None | None | None | None |
| IDUA | None | None | None | None |
| IDUA | None | None | None | None |
| IDUA | None | None | None | None |
| IDUA | None | None | None | None |
| IDUA | None | None | None | None |
| IDUA | None | None | None | None |
| IDUA | None | None | None | None |
| IDUA | None | None | None | None |
| IDUA | None | None | None | None |
| IDUA | None | None | None | None |
| IDUA | None | None | None | None |
| IDUA | None | None | None | None |
| IDUA | None | None | None | None |
| IDUA | None | None | None | None |
| IDUA | None | None | None | None |
| IDUA | None | None | None | None |
| IDUA | None | None | None | None |
| IDUA | None | None | None | None |
| IDUA | None | None | None | None |
| HIKESHI | None | None | None | None |
| APC | None | None | None | None |
| APC | None | None | None | None |
| APC | None | None | None | None |
| APP | None | None | None | None |
| CEP152 | None | None | None | None |
| GAA | 0.609 | 1 | 5kzx | A |
| GAA | 0.609 | 1 | 5kzx | A |
| GAA | None | None | None | None |
| GAA | 0.174 | 9 | 5kzx | A |
| GAA | 0.261 | 10 | 5kzx | A |
| GAA | 0.609 | 1 | 5kzx | A |

| | | | | |
|---|---|---|---|---|
| GAA | 0.609 | 1 | 5kzx | A |
| GAA | 0.609 | 1 | 5kzx | A |
| GAA | 0.609 | 1 | 5kzx | A |
| GAA | 0.609 | 1 | 5kzx | A |
| GAA | 0.609 | 1 | 5kzx | A |
| GAA | 0.609 | 1 | 5kzx | A |
| GAA | 0.609 | 1 | 5kzx | A |
| GAA | 0.609 | 1 | 5kzx | A |
| GAA | 0.609 | 1 | 5kzx | A |
| GAA | 0.609 | 1 | 5kzx | A |
| GAA | 0.609 | 1 | 5kzx | A |
| GAA | 0.609 | 1 | 5kzx | A |
| GAA | 0.609 | 1 | 5kzx | A |
| GAA | 0.609 | 1 | 5kzx | A |
| GAA | 0.609 | 1 | 5kzx | A |
| GAA | 0.609 | 1 | 5kzx | A |
| GAA | 0.609 | 1 | 5kzx | A |
| GAA | 0.609 | 1 | 5kzx | A |
| GAA | 0.609 | 1 | 5kzx | A |
| GAA | 0.609 | 1 | 5kzx | A |
| GAA | 0.609 | 1 | 5kzx | A |
| GAA | 0.609 | 1 | 5kzx | A |
| GAA | 0.609 | 1 | 5kzx | A |
| GAA | 0.609 | 1 | 5kzx | A |
| GAA | 0.609 | 1 | 5kzx | A |
| GAA | 0.609 | 1 | 5kzx | A |
| GAA | 0.609 | 1 | 5kzx | A |
| GAA | 0.609 | 1 | 5kzx | A |
| GAA | 0.609 | 1 | 5kzx | A |
| GAA | 0.609 | 1 | 5kzx | A |
| GAA | 0.609 | 1 | 5kzx | A |
| GAA | 0.609 | 1 | 5kzx | A |
| GAA | 0.609 | 1 | 5kzx | A |
| GAA | 0.609 | 1 | 5kzx | A |
| GAA | 0.609 | 1 | 5kzx | A |
| GAA | 0.609 | 1 | 5kzx | A |
| GAA | 0.609 | 1 | 5kzx | A |
| GAA | 0.609 | 1 | 5kzx | A |
| GAA | 0.609 | 1 | 5kzx | A |
| GAA | 0.609 | 1 | 5kzx | A |
| GAA | 0.609 | 1 | 5kzx | A |
| GAA | 0.609 | 1 | 5kzx | A |
| GAA | 0.609 | 1 | 5kzx | A |
| GAA | 0.261 | 10 | 5kzx | A |
| FOLR1 | None | None | None | None |
| DSP | None | None | None | None |
| DSP | None | None | None | None |
| DSP | None | None | None | None |
| DSP | None | None | None | None |
| DSP | None | None | None | None |
| CCM2 | None | None | None | None |
| FXN | None | None | None | None |
| FXN | None | None | None | None |
| COMP | None | None | None | None |
| COMP | None | None | None | None |

| | | | | | |
|---|---|---|---|---|---|
| COMP | None | None | | None | None |
| COMP | None | None | | None | None |
| COMP | None | None | | None | None |
| COMP | None | None | | None | None |
| COMP | None | None | | None | None |
| COMP | None | None | | None | None |
| COMP | None | None | | None | None |
| COMP | None | None | | None | None |
| COMP | None | None | | None | None |
| COMP | None | None | | None | None |
| AIFM1 | None | None | | None | None |
| AIFM1 | None | None | | None | None |
| AIFM1 | 0.229 | | 23 | 5fs8 | A |
| AIFM1 | None | None | | None | None |
| AIFM1 | 0.262 | | 40 | 5fs7 | A |
| AIFM1 | None | None | | None | None |
| AIFM1 | None | None | | None | None |
| MYH9 | None | None | | None | None |
| FHL2 | None | None | | None | None |
| FHL2 | None | None | | None | None |
| TNFRSF4 | None | None | | None | None |
| CFB | None | None | | None | None |
| CFB | None | None | | None | None |
| CFB | None | None | | None | None |
| CFB | None | None | | None | None |
| CFB | None | None | | None | None |
| CFB | None | None | | None | None |
| CFB | 0.258 | | 12 | 1dle | B |
| CFB | None | None | | None | None |
| CFB | None | None | | None | None |
| CFB | None | None | | None | None |
| MVD | None | None | | None | None |
| MVD | None | None | | None | None |
| HSD17B4 | None | None | | None | None |
| HSD17B4 | None | None | | None | None |
| HSD17B4 | None | None | | None | None |
| HSD17B4 | 0.429 | | 2 | 1s9c | A |
| HSD17B4 | None | None | | None | None |
| HSD17B4 | None | None | | None | None |
| F11 | None | None | | None | None |
| F11 | None | None | | None | None |
| F11 | None | None | | None | None |
| F11 | None | None | | None | None |
| F11 | None | None | | None | None |
| F11 | None | None | | None | None |
| F11 | None | None | | None | None |
| F11 | None | None | | None | None |
| F11 | None | None | | None | None |
| SPTA1 | None | None | | None | None |
| SPTA1 | None | None | | None | None |
| SPTA1 | None | None | | None | None |
| SPTA1 | None | None | | None | None |
| SPTA1 | None | None | | None | None |
| SPTA1 | None | None | | None | None |

| | | | | |
|---|---|---|---|---|
| GMPR | None | None | None | None |
| MITF | None | None | None | None |
| NPC1 | None | None | None | None |
| NPC1 | None | None | None | None |
| NPC1 | None | None | None | None |
| NPC1 | None | None | None | None |
| NPC1 | None | None | None | None |
| NPC1 | None | None | None | None |
| NPC1 | None | None | None | None |
| NPC1 | None | None | None | None |
| NPC1 | None | None | None | None |
| NPC1 | None | None | None | None |
| NPC1 | None | None | None | None |
| NPC1 | None | None | None | None |
| NPC1 | None | None | None | None |
| NPC1 | None | None | None | None |
| NPC1 | None | None | None | None |
| NPC1 | None | None | None | None |
| NPC1 | None | None | None | None |
| NPC1 | None | None | None | None |
| NPC1 | None | None | None | None |
| NPC1 | None | None | None | None |
| NPC1 | None | None | None | None |
| NPC1 | None | None | None | None |
| NPC1 | None | None | None | None |
| NPC1 | None | None | None | None |
| NPC1 | None | None | None | None |
| NPC1 | None | None | None | None |
| NPC1 | None | None | None | None |
| NPC1 | None | None | None | None |
| NPC1 | None | None | None | None |
| NPC1 | None | None | None | None |
| NPC1 | None | None | None | None |
| NPC1 | None | None | None | None |
| NPC1 | None | None | None | None |
| NPC1 | None | None | None | None |
| NPC1 | None | None | None | None |
| NPC1 | None | None | None | None |
| NPC1 | None | None | None | None |
| NPC1 | None | None | None | None |
| NPC1 | None | None | None | None |
| NPC1 | None | None | None | None |
| NPC1 | None | None | None | None |
| NPC1 | None | None | None | None |
| NPC1 | None | None | None | None |
| C3 | None | None | None | None |
| NLRX1 | None | None | None | None |
| IGLL1 | 0.244 | 2 | 2h3n | D |
| APRT | 0.769 | 4 | 1zn9 | B |
| APRT | 0.417 | 5 | 4x44 | A |
| APRT | None | None | None | None |
| APRT | 0.923 | 1 | 1zn8 | A |
| APRT | None | None | None | None |
| ZIC3 | None | None | None | None |

| | | | | |
|---|---|---|---|---|
| ZIC3 | None | None | None | None |
| KLHL3 | None | None | None | None |
| RPS6KA3 | None | None | None | None |
| RPS6KA3 | None | None | None | None |
| RPS6KA3 | None | None | None | None |
| F8 | None | None | None | None |
| F8 | None | None | None | None |
| F8 | None | None | None | None |
| F8 | None | None | None | None |
| F8 | None | None | None | None |
| F8 | None | None | None | None |
| F8 | None | None | None | None |
| F8 | None | None | None | None |
| F8 | None | None | None | None |
| F8 | None | None | None | None |
| F8 | None | None | None | None |
| F8 | None | None | None | None |
| F8 | None | None | None | None |
| F8 | None | None | None | None |
| F8 | None | None | None | None |
| F8 | None | None | None | None |
| F8 | None | None | None | None |
| ISG15 | None | None | None | None |
| EXOSC3 | None | None | None | None |
| EXOSC3 | None | None | None | None |
| EXOSC3 | None | None | None | None |
| NKX2-5 | 0.403 | 1 | 4s0h | B |
| NKX2-5 | None | None | None | None |
| NKX2-5 | 0.403 | 1 | 4s0h | B |
| NKX2-5 | 0.403 | 1 | 4s0h | B |
| FH | 0.207 | 44 | 30000 | A |
| FH | None | None | None | None |
| FH | None | None | None | None |
| FH | 0.304 | 30 | 30000 | D |
| FH | 0.621 | 4 | 30000 | A |
| FH | 0.462 | 6 | 30000 | C |
| FH | None | None | None | None |
| FH | None | None | None | None |
| FH | 0.64 | 6 | 5d6b | A |
| FH | 0.308 | 49 | 30000 | C |
| FH | 0.64 | 1 | 5d6b | A |
| FH | 0.64 | 1 | 5d6b | A |
| SNRNP200 | None | None | None | None |
| SNRNP200 | None | None | None | None |
| SNRNP200 | None | None | None | None |
| SNRNP200 | None | None | None | None |
| CHRNA1 | None | None | None | None |
| CHRNA1 | None | None | None | None |
| MYOM1 | None | None | None | None |
| MYOM1 | None | None | None | None |
| MYOM1 | None | None | None | None |
| MYOM1 | None | None | None | None |
| MYOM1 | None | None | None | None |
| MYOM1 | None | None | None | None |

| | | | | |
|---|---|---|---|---|
| LDLR | None | None | None | None |
| LDLR | None | None | None | None |
| LDLR | None | None | None | None |
| LDLR | None | None | None | None |
| LDLR | None | None | None | None |
| LDLR | None | None | None | None |
| EHMT1 | 0.286 | 8 | 3b7b | A |
| EHMT1 | 0.545 | 3 | 3b7b | A |
| EHMT1 | None | None | None | None |
| COL9A2 | None | None | None | None |
| HFE2 | None | None | None | None |
| HFE2 | None | None | None | None |
| CD3G | None | None | None | None |
| MMAB | None | None | None | None |
| MMAB | 0.5 | 3 | 2idx | C |
| MMAB | 0.45 | 7 | 2idx | C |
| MMAB | 0.65 | 3 | 2idx | A |
| MMAB | 0.65 | 3 | 2idx | A |
| MMAB | 0.7 | 2 | 2idx | B |
| MMAB | 0.7 | 2 | 2idx | B |
| MMAB | 0.35 | 4 | 2idx | A |
| MMAB | 0.4 | 11 | 2idx | C |
| MMAB | None | None | None | None |
| TAF1 | None | None | None | None |
| TAF1 | None | None | None | None |
| CD4 | None | None | None | None |
| QARS | None | None | None | None |
| QARS | None | None | None | None |
| QARS | None | None | None | None |
| QARS | None | None | None | None |
| QARS | None | None | None | None |
| FGFR3 | None | None | None | None |
| FGFR3 | None | None | None | None |
| FGFR3 | None | None | None | None |
| FGFR3 | None | None | None | None |
| FGFR3 | None | None | None | None |
| FGFR3 | None | None | None | None |
| FLNB | None | None | None | None |
| FLNB | None | None | None | None |
| FLNB | None | None | None | None |
| FLNB | None | None | None | None |
| FLNB | None | None | None | None |
| FLNB | None | None | None | None |
| FLNB | None | None | None | None |
| FLNB | None | None | None | None |
| IL21R | None | None | None | None |
| RB1 | None | None | None | None |
| RB1 | None | None | None | None |
| RB1 | None | None | None | None |
| NTRK1 | 0.533 | 1 | 1he7 | A |
| NOTCH1 | 0.045 | 11 | 4cuf | A |
| MYOM1 | None | None | None | None |
| MYOM1 | None | None | None | None |
| MYOM1 | None | None | None | None |

| | | | | |
|---|---|---|---|---|
| MYOM1 | None | None | None | None |
| MYOM1 | None | None | None | None |
| MYOM1 | None | None | None | None |
| COQ9 | None | None | None | None |
| COQ9 | None | None | None | None |
| CRBN | None | None | None | None |
| DCTN1 | None | None | None | None |
| DCTN1 | None | None | None | None |
| DCTN1 | None | None | None | None |
| DCTN1 | None | None | None | None |
| DCTN1 | None | None | None | None |
| DCTN1 | 0.333 | 1 | 2hkn | A |
| THAP1 | None | None | None | None |
| THAP1 | None | None | None | None |
| GNAS | None | None | None | None |
| GNAS | None | None | None | None |
| HNF4A | 0.018 | 27 | 4b7w | B |
| HNF4A | 0.016 | 63 | 4iqr | B |
| ALDH3A2 | None | None | None | None |
| ALDH3A2 | 0.039 | 38 | 4qgk | B |
| ALDH3A2 | 0.412 | 5 | 4qgk | B |
| ALDH3A2 | 0.557 | 2 | 4qgk | A |
| ALDH3A2 | None | None | None | None |
| ALDH3A2 | 0.569 | 3 | 4qgk | B |
| ALDH3A2 | None | None | None | None |
| ALDH3A2 | None | None | None | None |
| ALDH3A2 | None | None | None | None |
| HCN1 | None | None | None | None |
| PLEC | None | None | None | None |
| SCN5A | None | None | None | None |
| SCN5A | None | None | None | None |
| STAMBP | None | None | None | None |
| STAMBP | None | None | None | None |
| RNF125 | None | None | None | None |
| RNF125 | None | None | None | None |
| USH1C | None | None | None | None |
| OBSL1 | None | None | None | None |
| GRIN2B | None | None | None | None |
| GRIN2B | None | None | None | None |
| HCN4 | 0.186 | 6 | 4kl1 | C |
| HCN4 | 0.222 | 5 | 3u11 | A |
| TALDO1 | 0.192 | 51 | 1f05 | B |
| ITGB3 | None | None | None | None |
| CHKB | None | None | None | None |
| CHKB | None | None | None | None |
| CHKB | 0.333 | 7 | 3lq3 | A |
| CHKB | None | None | None | None |
| ACTN4 | None | None | None | None |
| ACTN4 | None | None | None | None |
| ACTN4 | None | None | None | None |
| ACTN4 | None | None | None | None |
| CDT1 | None | None | None | None |
| CDT1 | None | None | None | None |
| CDT1 | None | None | None | None |

| | | | | |
|---|---|---|---|---|
| CDT1 | None | None | None | None |
| CDT1 | None | None | None | None |
| CDT1 | None | None | None | None |
| MAPT | None | None | None | None |
| F5 | 0.517 | 4 | 1czs | A |
| F5 | None | None | None | None |
| F5 | None | None | None | None |
| ESRRB | 0.186 | 13 | 1lo1 | A |
| DGUOK | 0.188 | 23 | 2ocp | B |
| DGUOK | 0.471 | 8 | 2ocp | A |
| DGUOK | None | None | None | None |
| DGUOK | 0.375 | 6 | 2ocp | B |
| DGUOK | 0.765 | 1 | 2ocp | A |
| DGUOK | 0.438 | 11 | 2ocp | B |
| DGUOK | 0.294 | 19 | 2ocp | A |
| DGUOK | None | None | None | None |
| DGUOK | None | None | None | None |
| CENPJ | None | None | None | None |
| TTN | 0.041 | 5 | 2f8v | A |
| TTN | None | None | None | None |
| TTN | None | None | None | None |
| TTN | None | None | None | None |
| TNFRSF1A | None | None | None | None |
| TNFRSF1A | None | None | None | None |
| TNFRSF1A | None | None | None | None |
| TNFRSF1A | None | None | None | None |
| TNFRSF1A | None | None | None | None |
| TNFRSF1A | None | None | None | None |
| TNFRSF1A | None | None | None | None |
| TNFRSF1A | None | None | None | None |
| TNFRSF1A | None | None | None | None |
| TNFRSF1A | None | None | None | None |
| TNFRSF1A | None | None | None | None |
| TNFRSF1A | None | None | None | None |
| ASL | None | None | None | None |
| ASL | 0.625 | 25 | 1aos | A |
| ASL | None | None | None | None |
| ASL | 0.375 | 28 | 1k62 | A |
| ASL | 0.438 | 4 | 1k62 | B |
| ASL | 0.562 | 30 | 1k62 | A |
| ASL | 0.625 | 1 | 1aos | A |
| ASL | None | None | None | None |
| ASL | None | None | None | None |
| ASL | None | None | None | None |
| MYO7A | None | None | None | None |
| MYO7A | None | None | None | None |
| MYO7A | None | None | None | None |
| MYO7A | None | None | None | None |
| MYO7A | None | None | None | None |
| MYO7A | None | None | None | None |
| MYO7A | None | None | None | None |
| MYO7A | None | None | None | None |
| MYO7A | None | None | None | None |
| MYO7A | None | None | None | None |

| | | | | |
|---|---|---|---|---|
| MYO7A | None | None | None | None |
| MYO7A | None | None | None | None |
| MYO7A | None | None | None | None |
| MYO7A | None | None | None | None |
| MYO7A | None | None | None | None |
| MYO7A | None | None | None | None |
| MYO7A | None | None | None | None |
| MYO7A | None | None | None | None |
| MYO7A | None | None | None | None |
| MYO7A | None | None | None | None |
| APC | None | None | None | None |
| ALAD | None | None | None | None |
| ALAD | 0.143 | 21 | 1pv8 | A |
| ALAD | None | None | None | None |
| ALAD | 0.542 | 1 | 1E+051 | A |
| ALAD | None | None | None | None |
| ALAD | None | None | None | None |
| POR | None | None | None | None |
| CSF1R | None | None | None | None |
| CSF1R | None | None | None | None |
| CSF1R | None | None | None | None |
| CSF1R | None | None | None | None |
| CSF1R | None | None | None | None |
| CSF1R | None | None | None | None |
| CSF1R | None | None | None | None |
| CSF1R | None | None | None | None |
| CSF1R | None | None | None | None |
| CSF1R | None | None | None | None |
| CSF1R | None | None | None | None |
| VWF | None | None | None | None |
| VWF | None | None | None | None |
| PHF8 | 0.333 | 6 | 3k3n | A |
| ACY1 | None | None | None | None |
| SMN1 | None | None | None | None |
| SMN1 | None | None | None | None |
| DLAT | None | None | None | None |
| DLAT | None | None | None | None |
| B2M | None | None | None | None |
| SOX9 | None | None | None | None |
| SOX9 | None | None | None | None |
| SOX9 | None | None | None | None |
| SOX9 | None | None | None | None |
| SOX9 | None | None | None | None |
| CFH | None | None | None | None |
| FLNC | None | None | None | None |
| FLNC | None | None | None | None |
| FLNC | None | None | None | None |
| ITGB2 | None | None | None | None |
| ITGB2 | 0.261 | 2 | 5e6r | B |
| ITGB2 | None | None | None | None |
| ITGB2 | None | None | None | None |
| ITGB2 | None | None | None | None |
| ITGB2 | None | None | None | None |
| ITGB2 | 0.174 | 1 | 5e6u | B |

| | | | | |
|---|---|---|---|---|
| ITGB2 | None | None | None | None |
| ITGB2 | None | None | None | None |
| PSEN1 | None | None | None | None |
| PSEN1 | None | None | None | None |
| PSEN1 | None | None | None | None |
| PSEN1 | None | None | None | None |
| PSEN1 | None | None | None | None |
| PSEN1 | None | None | None | None |
| OXCT1 | None | None | None | None |
| OXCT1 | 0.385 | 24 | 3dlx | A |
| OXCT1 | None | None | None | None |
| OXCT1 | None | None | None | None |
| OXCT1 | None | None | None | None |
| OXCT1 | None | None | None | None |
| OXCT1 | None | None | None | None |
| DHH | None | None | None | None |
| PRKN | None | None | None | None |
| PRKN | None | None | None | None |
| PRKN | None | None | None | None |
| PRKN | None | None | None | None |
| PRKN | None | None | None | None |
| PRKN | None | None | None | None |
| PRKN | None | None | None | None |
| PRKN | None | None | None | None |
| PRKN | None | None | None | None |
| PRKN | None | None | None | None |
| PRKN | None | None | None | None |
| LCAT | None | None | None | None |
| LCAT | None | None | None | None |
| LCAT | None | None | None | None |
| LCAT | None | None | None | None |
| LCAT | None | None | None | None |
| LCAT | None | None | None | None |
| LCAT | None | None | None | None |
| LCAT | None | None | None | None |
| LCAT | None | None | None | None |
| LCAT | None | None | None | None |
| LCAT | None | None | None | None |
| LCAT | None | None | None | None |
| FOXP3 | None | None | None | None |
| FOXP3 | None | None | None | None |
| FOXP3 | None | None | None | None |
| FOXP3 | None | None | None | None |
| SUFU | None | None | None | None |
| HSPA9 | 0.571 | 14 | 4kbo | A |
| HSPA9 | None | None | None | None |
| SQSTM1 | None | None | None | None |
| DARS2 | None | None | None | None |
| DARS2 | None | None | None | None |
| DARS2 | None | None | None | None |
| DARS2 | None | None | None | None |
| DARS2 | 0.412 | 2 | 4ah6 | A |
| DARS2 | None | None | None | None |
| DARS2 | None | None | None | None |

| | | | | |
|---|---|---|---|---|
| DARS2 | None | None | None | None |
| DARS2 | None | None | None | None |
| DARS2 | None | None | None | None |
| DARS2 | None | None | None | None |
| DARS2 | None | None | None | None |
| NT5C3A | None | None | None | None |
| NT5C3A | 0.4 | 18 | 2vkq | A |
| NT5C3A | 0.4 | 7 | 2vkq | A |
| NT5C3A | 0.6 | 14 | 2vkq | A |
| ETFA | None | None | None | None |
| ETFA | None | None | None | None |
| ETFA | None | None | None | None |
| ETFA | None | None | None | None |
| ETFA | None | None | None | None |
| ETFA | None | None | None | None |
| ETFA | None | None | None | None |
| TTN | None | None | None | None |
| TTN | None | None | None | None |
| FLT4 | None | None | None | None |
| FGB | None | None | None | None |
| HPCA | None | None | None | None |
| TAP2 | None | None | None | None |
| TAP2 | None | None | None | None |
| PTPRC | None | None | None | None |
| SLC4A1 | None | None | None | None |
| SLC4A1 | None | None | None | None |
| SLC4A1 | None | None | None | None |
| SLC4A1 | None | None | None | None |
| SLC4A1 | None | None | None | None |
| SLC4A1 | None | None | None | None |
| MYH7 | None | None | None | None |
| MYH7 | None | None | None | None |
| MYH7 | None | None | None | None |
| MYH7 | None | None | None | None |
| MYH7 | None | None | None | None |
| MYH7 | None | None | None | None |
| MYH7 | None | None | None | None |
| MYH7 | None | None | None | None |
| MYH7 | None | None | None | None |
| PARN | None | None | None | None |
| PARN | None | None | None | None |
| NDUFS6 | None | None | None | None |
| VRK1 | None | None | None | None |
| VRK1 | None | None | None | None |
| VRK1 | 0.021 | 20 | 5ukf | A |
| VRK1 | 0.29 | 8 | 3op5 | A |
| VRK1 | None | None | None | None |
| VRK1 | None | None | None | None |
| COLEC11 | 0.229 | 23 | 4ymd | B |
| VHL | None | None | None | None |
| TTN | None | None | None | None |
| NCF2 | None | None | None | None |
| NCF2 | None | None | None | None |
| NCF2 | None | None | None | None |

| | | | | |
|---|---|---|---|---|
| NCF2 | None | None | None | None |
| NCF2 | None | None | None | None |
| RPS26 | None | None | None | None |
| CETP | None | None | None | None |
| CETP | None | None | None | None |
| CETP | None | None | None | None |
| CETP | None | None | None | None |
| CETP | None | None | None | None |
| CETP | None | None | None | None |
| CETP | None | None | None | None |
| CETP | None | None | None | None |
| CETP | None | None | None | None |
| CETP | None | None | None | None |
| CETP | None | None | None | None |
| CETP | None | None | None | None |
| CETP | None | None | None | None |
| FLNA | None | None | None | None |
| FLNA | None | None | None | None |
| FLNA | None | None | None | None |
| FLNA | None | None | None | None |
| FLNA | None | None | None | None |
| FLNA | None | None | None | None |
| FLNA | None | None | None | None |
| MYBPC3 | None | None | None | None |
| MYBPC3 | None | None | None | None |
| MYBPC3 | None | None | None | None |
| MYBPC3 | None | None | None | None |
| MYBPC3 | None | None | None | None |
| MYBPC3 | None | None | None | None |
| MYBPC3 | None | None | None | None |
| MYBPC3 | None | None | None | None |
| MYBPC3 | None | None | None | None |
| MYBPC3 | None | None | None | None |
| LSS | None | None | None | None |
| C6 | None | None | None | None |
| C6 | None | None | None | None |
| RLBP1 | None | None | None | None |
| RLBP1 | None | None | None | None |
| RLBP1 | None | None | None | None |
| RLBP1 | None | None | None | None |
| RLBP1 | None | None | None | None |
| RLBP1 | None | None | None | None |
| RLBP1 | None | None | None | None |
| RLBP1 | None | None | None | None |
| MMAA | None | None | None | None |
| MMAA | 0.3 | 1 | 2www | C |
| MMAA | 0.286 | 4 | 2www | D |
| MMAA | None | None | None | None |
| MMAA | None | None | None | None |
| MMAA | None | None | None | None |
| IL1RN | None | None | None | None |
| MASTL | None | None | None | None |
| FHL1 | None | None | None | None |
| FHL1 | None | None | None | None |

| | | | | |
|---|---|---|---|---|
| FHL1 | None | None | None | None |
| FHL1 | None | None | None | None |
| FHL1 | None | None | None | None |
| FHL1 | None | None | None | None |
| FHL1 | None | None | None | None |
| FHL1 | None | None | None | None |
| FHL1 | None | None | None | None |
| EIF2B1 | 0.636 | 1 | 3ecs | H |
| EIF2B1 | 0.636 | 1 | 3ecs | H |
| COQ8A | None | None | None | None |
| COQ8A | None | None | None | None |
| COQ8A | None | None | None | None |
| COQ8A | None | None | None | None |
| COQ8A | None | None | None | None |
| COQ8A | None | None | None | None |
| COQ8A | None | None | None | None |
| COQ8A | None | None | None | None |
| COQ8A | None | None | None | None |
| COQ8A | None | None | None | None |
| COQ8A | None | None | None | None |
| COQ8A | None | None | None | None |
| COQ8A | None | None | None | None |
| COQ8A | None | None | None | None |
| COQ8A | None | None | None | None |
| COQ8A | None | None | None | None |
| UNC119 | None | None | None | None |
| IGHMBP2 | None | None | None | None |
| JAG1 | None | None | None | None |
| JAG1 | None | None | None | None |
| JAG1 | None | None | None | None |
| GNB1 | None | None | None | None |
| GNB1 | 0.3 | 3 | 5ukl | B |
| FLNA | None | None | None | None |
| FLNA | None | None | None | None |
| FLNA | None | None | None | None |
| FLNA | None | None | None | None |
| FLNA | None | None | None | None |
| CREBBP | None | None | None | None |
| CSF3R | None | None | None | None |
| CSF3R | None | None | None | None |
| HIBCH | None | None | None | None |
| HIBCH | None | None | None | None |
| HIBCH | None | None | None | None |
| MMACHC | None | None | None | None |
| MMACHC | None | None | None | None |
| MMACHC | None | None | None | None |
| MMACHC | None | None | None | None |
| MMACHC | None | None | None | None |
| MMACHC | None | None | None | None |
| MMACHC | None | None | None | None |
| MMACHC | None | None | None | None |
| MMACHC | None | None | None | None |
| MMACHC | None | None | None | None |
| MMACHC | None | None | None | None |

| | | | | | |
|---|---|---|---|---|---|
| KLF1 | None | None | | None | None |
| NAGS | None | None | | None | None |
| NAGS | None | None | | None | None |
| NAGS | None | None | | None | None |
| PIK3R2 | None | None | | None | None |
| EDNRB | None | None | | None | None |
| EDNRB | None | None | | None | None |
| EDNRB | None | None | | None | None |
| EDNRB | None | None | | None | None |
| E | 0.156 | | 69 | 5glh | A |
| RUNX1 | None | None | | None | None |
| RUNX1 | None | None | | None | None |
| CTNNB1 | None | None | | None | None |
| CTNNB1 | None | None | | None | None |
| CTNNB1 | None | None | | None | None |
| CTNNB1 | None | None | | None | None |
| CTNNB1 | None | None | | None | None |
| CTNNB1 | None | None | | None | None |
| TTN | None | None | | None | None |
| TTN | None | None | | None | None |
| TYK2 | 0.035 | | 60 | 4oli | A |
| TYK2 | None | None | | None | None |
| TYK2 | 0.055 | | 20 | 5c03 | A |
| ERCC4 | None | None | | None | None |
| ERCC4 | None | None | | None | None |
| ERCC4 | None | None | | None | None |
| ERCC4 | None | None | | None | None |
| CFI | None | None | | None | None |
| CFI | None | None | | None | None |
| CFI | None | None | | None | None |
| CFI | None | None | | None | None |
| CFI | None | None | | None | None |
| CFI | None | None | | None | None |
| CFI | None | None | | None | None |
| CFI | None | None | | None | None |
| CFI | 0.094 | | 18 | 5o32 | J |
| CFI | 0.301 | | 5 | 2xrc | A |
| CFI | None | None | | None | None |
| CFI | 0.691 | | 2 | 5o32 | J |
| CFI | 0.691 | | 2 | 5o32 | J |
| CFI | 0.524 | | 5 | 5o32 | I |
| NOTCH3 | None | None | | None | None |
| NOTCH3 | None | None | | None | None |
| NOTCH3 | None | None | | None | None |
| NOTCH3 | None | None | | None | None |
| TBX1 | None | None | | None | None |
| TBX1 | None | None | | None | None |
| DDB2 | None | None | | None | None |
| DDB2 | None | None | | None | None |
| TP53 | None | None | | None | None |
| POMGNT1 | None | None | | None | None |
| POMGNT1 | None | None | | None | None |
| POMGNT1 | None | None | | None | None |
| POMGNT1 | None | None | | None | None |

| | | | | |
|---|---|---|---|---|
| POMGNT1 | None | None | None | None |
| POMGNT1 | None | None | None | None |
| POMGNT1 | None | None | None | None |
| POMGNT1 | None | None | None | None |
| POMGNT1 | None | None | None | None |
| POMGNT1 | None | None | None | None |
| POMGNT1 | None | None | None | None |
| POMGNT1 | None | None | None | None |
| POMGNT1 | None | None | None | None |
| FMR1 | None | None | None | None |
| FMR1 | None | None | None | None |
| SLC9A3R1 | None | None | None | None |
| HSPB1 | None | None | None | None |
| HSPB1 | None | None | None | None |
| HSPB1 | None | None | None | None |
| HSPB1 | None | None | None | None |
| HSPB1 | None | None | None | None |
| HSPB1 | None | None | None | None |
| HSPB1 | None | None | None | None |
| SOD3 | None | None | None | None |
| AGA | 1 | 2 | 1apz | D |
| AGA | None | None | None | None |
| AGA | None | None | None | None |
| NDUFS7 | None | None | None | None |
| NDUFS7 | None | None | None | None |
| SOX2 | None | None | None | None |
| CSRP3 | None | None | None | None |
| CSRP3 | None | None | None | None |
| PAH | None | None | None | None |
| PAH | None | None | None | None |
| PAH | None | None | None | None |
| PAH | None | None | None | None |
| CACNA1S | None | None | None | None |
| TRPV4 | None | None | None | None |
| TRPV4 | None | None | None | None |
| TRPV4 | None | None | None | None |
| TRPV4 | 0.58 | 3 | 4dx2 | A |
| TRPV4 | None | None | None | None |
| TRPV4 | None | None | None | None |
| TRPV4 | None | None | None | None |
| TRPV4 | None | None | None | None |
| TRPV4 | None | None | None | None |
| TRPV4 | None | None | None | None |
| TRPV4 | None | None | None | None |
| TRPV4 | None | None | None | None |
| TRPV4 | None | None | None | None |
| TNNI3 | None | None | None | None |
| MLYCD | None | None | None | None |
| MLYCD | None | None | None | None |
| MLYCD | None | None | None | None |
| MLYCD | None | None | None | None |
| AIP | None | None | None | None |
| AIP | None | None | None | None |
| ERCC2 | None | None | None | None |

| | | | | |
|---|---|---|---|---|
| ERCC2 | None | None | None | None |
| ERCC2 | None | None | None | None |
| ERCC2 | None | None | None | None |
| ERCC2 | None | None | None | None |
| ERCC2 | None | None | None | None |
| ERCC2 | None | None | None | None |
| ERCC2 | None | None | None | None |
| ERCC2 | None | None | None | None |
| ERCC2 | None | None | None | None |
| ERCC2 | None | None | None | None |
| ERCC2 | None | None | None | None |
| ERCC2 | None | None | None | None |
| ERCC2 | None | None | None | None |
| PUS1 | None | None | None | None |
| PUS1 | None | None | None | None |
| PUS1 | None | None | None | None |
| CASK | None | None | None | None |
| CASK | None | None | None | None |
| BLVRA | None | None | None | None |
| DICER1 | None | None | None | None |
| DICER1 | None | None | None | None |
| DICER1 | None | None | None | None |
| EIF2AK3 | None | None | None | None |
| EIF2AK3 | None | None | None | None |
| EIF2AK3 | None | None | None | None |
| KIT | None | None | None | None |
| KIT | None | None | None | None |
| KIT | None | None | None | None |
| KIT | None | None | None | None |
| KIT | None | None | None | None |
| KIT | None | None | None | None |
| DNMT3B | None | None | None | None |
| TGM1 | None | None | None | None |
| NDUFA10 | None | None | None | None |
| NDUFA10 | None | None | None | None |
| PRPF31 | None | None | None | None |
| PRPF31 | None | None | None | None |
| ECE1 | None | None | None | None |
| ECE1 | None | None | None | None |
| ECE1 | None | None | None | None |
| NDUFV2 | None | None | None | None |
| NDUFV2 | None | None | None | None |
| NDUFV2 | None | None | None | None |
| GHRHR | None | None | None | None |
| GHRHR | None | None | None | None |
| GHRHR | None | None | None | None |
| SLC4A1 | None | None | None | None |
| FGF9 | None | None | None | None |
| BARD1 | None | None | None | None |
| BARD1 | None | None | None | None |
| BARD1 | None | None | None | None |
| PTPN12 | 0.741 | 2 | 5j8r | D |
| MASP2 | None | None | None | None |
| MASP2 | None | None | None | None |

| | | | | |
|---|---|---|---|---|
| MASP2 | None | None | None | None |
| MASP2 | 0.118 | 3 | 1szb | A |
| HOXB1 | None | None | None | None |
| SLC4A1 | None | None | None | None |
| SLC4A1 | None | None | None | None |
| SLC4A1 | None | None | None | None |
| SLC4A1 | None | None | None | None |
| SLC4A1 | None | None | None | None |
| SLC4A1 | None | None | None | None |
| SLC4A1 | None | None | None | None |
| SLC4A1 | None | None | None | None |
| SLC4A1 | None | None | None | None |
| SLC4A1 | None | None | None | None |
| SLC4A1 | None | None | None | None |
| SLC4A1 | None | None | None | None |
| SLC4A1 | None | None | None | None |
| SLC4A1 | None | None | None | None |
| SLC4A1 | None | None | None | None |
| SLC4A1 | None | None | None | None |
| SLC4A1 | None | None | None | None |
| CBL | None | None | None | None |
| FAS | None | None | None | None |
| MTMR2 | None | None | None | None |
| MTMR2 | None | None | None | None |
| MTMR2 | None | None | None | None |
| MTMR2 | None | None | None | None |
| MTMR2 | None | None | None | None |
| MTMR2 | None | None | None | None |
| MTMR2 | None | None | None | None |
| MTMR2 | None | None | None | None |
| KLHL3 | None | None | None | None |
| KLHL3 | None | None | None | None |
| KLHL3 | 0.667 | 1 | 4ch9 | B |
| KLHL3 | 0.733 | 1 | 5nkp | A |
| KLHL3 | None | None | None | None |
| KLHL3 | None | None | None | None |
| RNASEH1 | None | None | None | None |
| ACTN1 | None | None | None | None |
| PHKG2 | 0.006 | 73 | 2y7j | D |
| PHKG2 | 0.006 | 143 | 2y7j | D |
| PHKG2 | 0.472 | 2 | 2y7j | A |
| NTRK1 | None | None | None | None |
| APOH | None | None | None | None |
| PMS2 | None | None | None | None |
| PMS2 | None | None | None | None |
| PMS2 | 0.797 | 1 | 1ea6 | A |
| PMS2 | None | None | None | None |
| PMS2 | None | None | None | None |
| PMS2 | None | None | None | None |
| PMS2 | 0.156 | 10 | 1ea6 | A |
| PMS2 | None | None | None | None |
| PMS2 | 0.188 | 8 | 1ea6 | A |
| PMS2 | None | None | None | None |
| PMS2 | None | None | None | None |

| | | | | |
|---|---|---|---|---|
| PMS2 | None | None | None | None |
| PMS2 | None | None | None | None |
| PMS2 | None | None | None | None |
| PMS2 | None | None | None | None |
| PMS2 | None | None | None | None |
| PMS2 | None | None | None | None |
| PMS2 | None | None | None | None |
| PMS2 | None | None | None | None |
| PMS2 | None | None | None | None |
| PMS2 | None | None | None | None |
| PMS2 | None | None | None | None |
| MOCS2 | None | None | None | None |
| MOCS2 | None | None | None | None |
| RPGRIP1 | None | None | None | None |
| RPGRIP1 | None | None | None | None |
| LIG4 | None | None | None | None |
| PROC | None | None | None | None |
| PROC | 0.111 | 34 | 1aut | C |
| NAT1 | None | None | None | None |
| NAT1 | None | None | None | None |
| PNPO | None | None | None | None |
| PNPO | 0.5 | 2 | 1nrg | A |
| PNPO | None | None | None | None |
| PNPO | None | None | None | None |
| PNPO | None | None | None | None |
| MRE11 | None | None | None | None |
| MRE11 | None | None | None | None |
| MRE11 | None | None | None | None |
| MRE11 | None | None | None | None |
| DDC | None | None | None | None |
| DDC | None | None | None | None |
| DDC | None | None | None | None |
| DDC | None | None | None | None |
| DDC | None | None | None | None |
| DDC | 0.909 | 7 | 3rch | B |
| DDC | 0.909 | 7 | 3rch | B |
| DDC | 0.643 | 4 | 3rbf | A |
| DDC | None | None | None | None |
| DDC | None | None | None | None |
| DDC | None | None | None | None |
| ANK2 | 0.2 | 7 | 4rlv | A |
| ANK2 | 0.242 | 5 | 4rlv | A |
| ANK2 | None | None | None | None |
| AK2 | None | None | None | None |
| AK2 | 0.241 | 12 | 2c9y | A |
| AK2 | 0.207 | 17 | 2c9y | A |
| AK2 | None | None | None | None |
| AK2 | 0.138 | 10 | 2c9y | A |
| AK2 | None | None | None | None |
| GPHN | None | None | None | None |
| HSD17B4 | 0.66 | 1 | 1zbq | A |
| HSD17B4 | 0.66 | 1 | 1zbq | A |
| HSD17B4 | None | None | None | None |
| HSD17B4 | 0.297 | 7 | 1zbq | B |

| | | | | |
|---|---|---|---|---|
| HSD17B4 | None | None | None | None |
| HSD17B4 | None | None | None | None |
| APTX | 0.6 | 5 | 4ndg | A |
| APTX | 0.6 | 5 | 4ndg | A |
| APTX | None | None | None | None |
| APTX | None | None | None | None |
| APTX | None | None | None | None |
| APTX | None | None | None | None |
| APTX | None | None | None | None |
| FANCL | None | None | None | None |
| FANCL | None | None | None | None |
| EMG1 | None | None | None | None |
| EMG1 | None | None | None | None |
| EZH2 | 0.205 | 1 | 4mi5 | A |
| ABCB6 | None | None | None | None |
| NCF2 | None | None | None | None |
| NCF2 | None | None | None | None |
| COL4A2 | None | None | None | None |
| COL4A2 | 1 | 1 | 1li1 | F |
| COL4A2 | None | None | None | None |
| COL4A2 | None | None | None | None |
| COL4A2 | 0.75 | 16 | 1li1 | C |
| SLX4 | None | None | None | None |
| SERPIND1 | None | None | None | None |
| FGFR3 | None | None | None | None |
| FGFR3 | None | None | None | None |
| FGFR3 | None | None | None | None |
| FGFR3 | None | None | None | None |
| FGFR3 | None | None | None | None |
| FGFR3 | None | None | None | None |
| TREM2 | None | None | None | None |
| TREM2 | None | None | None | None |
| TREM2 | None | None | None | None |
| TREM2 | None | None | None | None |
| TREM2 | None | None | None | None |
| TREM2 | None | None | None | None |
| TREM2 | None | None | None | None |
| LIG4 | None | None | None | None |
| LIG4 | None | None | None | None |
| LIG4 | None | None | None | None |
| LIG4 | None | None | None | None |
| LIG4 | None | None | None | None |
| LIG4 | None | None | None | None |
| ECHS1 | 0.069 | 25 | 2hw5 | C |
| ECHS1 | 0.279 | 4 | 2hw5 | A |
| ECHS1 | None | None | None | None |
| MTR | None | None | None | None |
| MTR | None | None | None | None |
| MTR | None | None | None | None |
| MTR | None | None | None | None |
| MTR | None | None | None | None |
| PMVK | None | None | None | None |
| PRNP | None | None | None | None |
| CHN1 | None | None | None | None |

| | | | | |
|---|---|---|---|---|
| CHN1 | None | None | None | None |
| CHN1 | None | None | None | None |
| CHN1 | 0.067 | 2 | 3cxl | A |
| CHN1 | None | None | None | None |
| CHN1 | None | None | None | None |
| CHN1 | 0.067 | 2 | 3cxl | A |
| CHN1 | None | None | None | None |
| SLC2A1 | None | None | None | None |
| SLC2A1 | None | None | None | None |
| SLC2A1 | None | None | None | None |
| SLC2A1 | None | None | None | None |
| SLC2A1 | None | None | None | None |
| SLC2A1 | None | None | None | None |
| SLC2A1 | None | None | None | None |
| SLC2A1 | None | None | None | None |
| SLC2A1 | None | None | None | None |
| SLC2A1 | None | None | None | None |
| SLC2A1 | None | None | None | None |
| SLC2A1 | None | None | None | None |
| SLC2A1 | None | None | None | None |
| SLC2A1 | None | None | None | None |
| SLC2A1 | None | None | None | None |
| SLC2A1 | None | None | None | None |
| SLC2A1 | None | None | None | None |
| PMM2 | None | None | None | None |
| PMM2 | None | None | None | None |
| PMM2 | None | None | None | None |
| PMM2 | None | None | None | None |
| PMM2 | None | None | None | None |
| PMM2 | None | None | None | None |
| PMM2 | None | None | None | None |
| PMM2 | None | None | None | None |
| PMM2 | None | None | None | None |
| PMM2 | None | None | None | None |
| PMM2 | None | None | None | None |
| PMM2 | None | None | None | None |
| PMM2 | None | None | None | None |
| PMM2 | None | None | None | None |
| PMM2 | None | None | None | None |
| PMM2 | None | None | None | None |
| PMM2 | None | None | None | None |
| PMM2 | None | None | None | None |
| PMM2 | None | None | None | None |
| PMM2 | None | None | None | None |
| PMM2 | None | None | None | None |
| PMM2 | None | None | None | None |
| PMM2 | None | None | None | None |
| PMM2 | None | None | None | None |
| PMM2 | None | None | None | None |
| PMM2 | None | None | None | None |
| PMM2 | None | None | None | None |
| PMM2 | None | None | None | None |
| CCKAR | None | None | None | None |
| MAN1B1 | None | None | None | None |

| | | | | | |
|---|---|---|---|---|---|
| MAN1B1 | None | None | None | | None |
| LAMA2 | None | None | None | | None |
| LAMA2 | None | None | None | | None |
| TBC1D4 | None | None | None | | None |
| TBC1D4 | None | None | None | | None |
| LDLR | None | None | None | | None |
| LDLR | None | None | None | | None |
| LDLR | None | None | None | | None |
| LDLR | None | None | None | | None |
| LDLR | None | None | None | | None |
| LDLR | None | None | None | | None |
| LDLR | None | None | None | | None |
| LDLR | None | None | None | | None |
| LDLR | None | None | None | | None |
| LDLR | None | None | None | | None |
| LDLR | None | None | None | | None |
| LDLR | None | None | None | | None |
| LDLR | None | None | None | | None |
| LDLR | None | None | None | | None |
| LDLR | None | None | None | | None |
| LDLR | None | None | None | | None |
| LDLR | None | None | None | | None |
| LDLR | None | None | None | | None |
| LDLR | None | None | None | | None |
| LDLR | None | None | None | | None |
| LDLR | None | None | None | | None |
| LDLR | None | None | None | | None |
| LDLR | None | None | None | | None |
| LDLR | None | None | None | | None |
| LDLR | None | None | None | | None |
| LDLR | None | None | None | | None |
| LDLR | None | None | None | | None |
| LDLR | None | None | None | | None |
| LDLR | None | None | None | | None |
| LDLR | None | None | None | | None |
| LDLR | None | None | None | | None |
| LDLR | None | None | None | | None |
| LDLR | None | None | None | | None |
| LDLR | None | None | None | | None |
| LDLR | None | None | None | | None |
| LDLR | None | None | None | | None |
| LDLR | None | None | None | | None |
| LDLR | None | None | None | | None |
| LDLR | 0.036 | | 6 | 1ijq | A |
| LDLR | 0.071 | | 7 | 3m0c | C |
| LDLR | 0.036 | | 6 | 1ijq | A |
| LDLR | 0.5 | | 1 | 1ijq | A |
| LDLR | None | None | None | | None |
| FANCM | None | None | None | | None |
| HAMP | None | None | None | | None |
| GUSB | 0.462 | | 8 | 3hn3 | A |
| GUSB | None | None | None | | None |
| GUSB | 0.45 | | 16 | 3hn3 | B |
| GUSB | None | None | None | | None |

| | | | | |
|---|---|---|---|---|
| GUSB | 0.105 | 30 | 1bhg | A |
| GUSB | 0.075 | 49 | 3hn3 | B |
| GUSB | None | None | None | None |
| GUSB | None | None | None | None |
| GUSB | 0.105 | 38 | 1bhg | A |
| GUSB | None | None | None | None |
| GUSB | 0.105 | 30 | 1bhg | A |
| GUSB | 0.088 | 53 | 1bhg | B |
| GUSB | None | None | None | None |
| LIG1 | None | None | None | None |
| LIG1 | None | None | None | None |
| MYH7 | None | None | None | None |
| MYH7 | None | None | None | None |
| MYH7 | None | None | None | None |
| MYH7 | None | None | None | None |
| MYH7 | None | None | None | None |
| MYH7 | None | None | None | None |
| MYH7 | None | None | None | None |
| MYH7 | None | None | None | None |
| MYH7 | None | None | None | None |
| MYH7 | None | None | None | None |
| MYH7 | None | None | None | None |
| MYH7 | None | None | None | None |
| MYH7 | None | None | None | None |
| MYH7 | None | None | None | None |
| MYH7 | None | None | None | None |
| MYH7 | None | None | None | None |
| MYH7 | None | None | None | None |
| MYH7 | None | None | None | None |
| MYH7 | None | None | None | None |
| MYH7 | None | None | None | None |
| MYH7 | None | None | None | None |
| MYH7 | None | None | None | None |
| MYH7 | None | None | None | None |
| MYH7 | None | None | None | None |
| MYH7 | None | None | None | None |
| MYH7 | None | None | None | None |
| MYH7 | None | None | None | None |
| MYH7 | None | None | None | None |
| MYH7 | None | None | None | None |
| MYH7 | None | None | None | None |
| MYH7 | None | None | None | None |
| MYH7 | None | None | None | None |
| MYH7 | None | None | None | None |
| MYH7 | None | None | None | None |
| HCN4 | None | None | None | None |
| RECQL4 | None | None | None | None |
| RECQL4 | None | None | None | None |
| RECQL4 | None | None | None | None |
| RECQL4 | None | None | None | None |
| RECQL4 | None | None | None | None |
| RECQL4 | None | None | None | None |
| RECQL4 | None | None | None | None |
| RECQL4 | None | None | None | None |

| | | | | |
|---|---|---|---|---|
| RECQL4 | None | None | None | None |
| RECQL4 | None | None | None | None |
| RECQL4 | None | None | None | None |
| RECQL4 | None | None | None | None |
| RECQL4 | None | None | None | None |
| CHEK2 | None | None | None | None |
| PPARG | None | None | None | None |
| ACADVL | None | None | None | None |
| ACADVL | 0.385 | 4 | 3b96 | A |
| ACADVL | None | None | None | None |
| ACADVL | None | None | None | None |
| ACADVL | None | None | None | None |
| ACADVL | None | None | None | None |
| ACADVL | None | None | None | None |
| ACADVL | 0.346 | 14 | 3b96 | A |
| ACADVL | 0.346 | 14 | 3b96 | A |
| ACADVL | None | None | None | None |
| ACADVL | None | None | None | None |
| ACADVL | 0.231 | 10 | 3b96 | A |
| ACADVL | None | None | None | None |
| ACADVL | 0.346 | 14 | 3b96 | A |
| ACADVL | 0.269 | 12 | 3b96 | A |
| ACADVL | 0.423 | 2 | 3b96 | A |
| ACADVL | None | None | None | None |
| ACADVL | 0.135 | 20 | 3b96 | A |
| ACADVL | 0.135 | 20 | 3b96 | A |
| ACADVL | None | None | None | None |
| ACADVL | None | None | None | None |
| ACADVL | None | None | None | None |
| APOA1 | None | None | None | None |
| APOA1 | None | None | None | None |
| APOA1 | None | None | None | None |
| APOA1 | None | None | None | None |
| APOA1 | None | None | None | None |
| DNMT3A | None | None | None | None |
| DNMT3A | None | None | None | None |
| APC | None | None | None | None |
| TMOD1 | None | None | None | None |
| USH1C | None | None | None | None |
| USH1C | None | None | None | None |
| USH1C | None | None | None | None |
| USH1C | None | None | None | None |
| ERCC6 | None | None | None | None |
| ERCC6 | None | None | None | None |
| ERCC6 | None | None | None | None |
| ERCC8 | None | None | None | None |
| ERCC8 | None | None | None | None |
| ERCC8 | None | None | None | None |
| DPAGT1 | None | None | None | None |
| DPAGT1 | None | None | None | None |
| DPAGT1 | None | None | None | None |
| DPAGT1 | None | None | None | None |
| DPAGT1 | None | None | None | None |
| DPAGT1 | None | None | None | None |

| | | | | |
|---|---|---|---|---|
| DPAGT1 | None | None | None | None |
| IL2RG | None | None | None | None |
| HNF1A | None | None | None | None |
| HNF4A | 0.016 | 63 | 4iqr | B |
| FLT4 | None | None | None | None |
| FLT4 | None | None | None | None |
| FLT4 | None | None | None | None |
| CD3D | None | None | None | None |
| CD3D | None | None | None | None |
| MCOLN1 | None | None | None | None |
| MCOLN1 | None | None | None | None |
| MOCS2 | None | None | None | None |
| MOCS2 | None | None | None | None |
| ABCG8 | None | None | None | None |
| ABCG8 | None | None | None | None |
| ABCG8 | None | None | None | None |
| ABCG8 | None | None | None | None |
| ABCG8 | None | None | None | None |
| ABCG8 | None | None | None | None |
| ABCG8 | None | None | None | None |
| ABCG8 | None | None | None | None |
| ABCG8 | None | None | None | None |
| ABCG8 | None | None | None | None |
| ABCG8 | None | None | None | None |
| ABCG8 | None | None | None | None |
| ABCG8 | None | None | None | None |
| ABCG8 | None | None | None | None |
| CTNNA1 | None | None | None | None |
| TYMP | None | None | None | None |
| TYMP | None | None | None | None |
| TYMP | None | None | None | None |
| TYMP | None | None | None | None |
| TYMP | None | None | None | None |
| TYMP | None | None | None | None |
| TYMP | None | None | None | None |
| TYMP | None | None | None | None |
| TYMP | None | None | None | None |
| TYMP | None | None | None | None |
| TYMP | None | None | None | None |
| TYMP | None | None | None | None |
| TYMP | None | None | None | None |
| GAMT | None | None | None | None |
| GAMT | None | None | None | None |
| GAMT | None | None | None | None |
| GAMT | None | None | None | None |
| GAMT | None | None | None | None |
| GAMT | None | None | None | None |
| GAMT | None | None | None | None |
| GAMT | None | None | None | None |
| GAMT | None | None | None | None |
| GAMT | None | None | None | None |
| GAMT | None | None | None | None |
| GLRA1 | None | None | None | None |
| GLRA1 | None | None | None | None |

| | | | | |
|---|---|---|---|---|
| GLRA1 | None | None | None | None |
| GLRA1 | None | None | None | None |
| GLRA1 | None | None | None | None |
| GLRA1 | None | None | None | None |
| GLRA1 | None | None | None | None |
| GLRA1 | None | None | None | None |
| GLRA1 | None | None | None | None |
| ADK | None | None | None | None |
| ACTN2 | None | None | None | None |
| UBE2T | 0.134 | 17 | 5ngz | A |
| TTN | None | None | None | None |
| ATP7B | None | None | None | None |
| ATP7B | None | None | None | None |
| ATP7B | None | None | None | None |
| ATP7B | None | None | None | None |
| ATP7B | None | None | None | None |
| ATP7B | None | None | None | None |
| ATP7B | None | None | None | None |
| ATP7B | None | None | None | None |
| ATP7B | None | None | None | None |
| ATP7B | None | None | None | None |
| ATP7B | None | None | None | None |
| ATP7B | None | None | None | None |
| ATP7B | None | None | None | None |
| CDKN1B | None | None | None | None |
| CDKN1B | None | None | None | None |
| CDKN1B | None | None | None | None |
| CDKN1B | None | None | None | None |
| CDKN1B | None | None | None | None |
| CDKN1B | None | None | None | None |
| MYOC | None | None | None | None |
| MYOC | None | None | None | None |
| MYOC | None | None | None | None |
| MYOC | None | None | None | None |
| MYOC | None | None | None | None |
| MYOC | None | None | None | None |
| MYOC | None | None | None | None |
| MYOC | None | None | None | None |
| MYOC | None | None | None | None |
| MYOC | None | None | None | None |
| MYOC | None | None | None | None |
| MYOC | None | None | None | None |
| MYOC | None | None | None | None |
| MYOC | None | None | None | None |
| MYOC | None | None | None | None |
| SPTAN1 | None | None | None | None |
| GC | None | None | None | None |
| GC | None | None | None | None |
| MAD1L1 | None | None | None | None |
| NDST1 | None | None | None | None |
| ARL2BP | None | None | None | None |
| TOR1A | None | None | None | None |
| C1S | 0.174 | 8 | 4j1y | A |
| ARL6 | 0.508 | 1 | 2h57 | A |

| | | | | |
|---|---|---|---|---|
| ARL6 | None | None | None | None |
| ARL6 | None | None | None | None |
| ARL6 | None | None | None | None |
| ARL6 | 0.056 | 20 | 2h57 | A |
| ARL6 | 0.056 | 20 | 2h57 | A |
| FGFR1 | None | None | None | None |
| KLHL7 | None | None | None | None |
| FLNC | None | None | None | None |
| PTPRC | None | None | None | None |
| GCH1 | None | None | None | None |
| GCH1 | None | None | None | None |
| GCH1 | 0.1 | 9 | 1fb1 | A |
| GCH1 | 0.1 | 5 | 1fb1 | A |
| GCH1 | None | None | None | None |
| GCH1 | 0.2 | 17 | 1fb1 | A |
| GCH1 | 0.1 | 8 | 1fb1 | A |
| GCH1 | None | None | None | None |
| GCH1 | None | None | None | None |
| GCH1 | None | None | None | None |
| GCH1 | 0.3 | 1 | 1fb1 | D |
| GCH1 | 0.2 | 3 | 1fb1 | D |
| GCH1 | 0.3 | 11 | 1fb1 | B |
| GCH1 | 0.1 | 10 | 1fb1 | A |
| CFH | None | None | None | None |
| COL2A1 | None | None | None | None |
| COL2A1 | None | None | None | None |
| OCRL | None | None | None | None |
| WRN | None | None | None | None |
| WRN | None | None | None | None |
| WRN | None | None | None | None |
| WRN | None | None | None | None |
| MASP1 | None | None | None | None |
| MASP1 | None | None | None | None |
| NSD1 | None | None | None | None |
| PITX2 | None | None | None | None |
| PITX2 | None | None | None | None |
| PITX2 | None | None | None | None |
| PITX2 | 0.362 | 2 | 2lkx | A |
| PITX2 | 0.362 | 2 | 2lkx | A |
| PITX2 | 0.362 | 2 | 2lkx | A |
| B4GALT7 | 0.917 | 1 | 4irq | A |
| B4GALT7 | None | None | None | None |
| B4GALT7 | None | None | None | None |
| B4GALT7 | None | None | None | None |
| B4GALT7 | None | None | None | None |
| FSHR | None | None | None | None |
| IGHMBP2 | None | None | None | None |
| IGHMBP2 | None | None | None | None |
| IGHMBP2 | None | None | None | None |
| IGHMBP2 | None | None | None | None |
| IGHMBP2 | None | None | None | None |
| IGHMBP2 | None | None | None | None |
| IGHMBP2 | None | None | None | None |
| IGHMBP2 | None | None | None | None |

| | | | | |
|---|---|---|---|---|
| IGHMBP2 | None | None | None | None |
| IGHMBP2 | None | None | None | None |
| IGHMBP2 | None | None | None | None |
| IGHMBP2 | None | None | None | None |
| IGHMBP2 | None | None | None | None |
| IGHMBP2 | None | None | None | None |
| IGHMBP2 | None | None | None | None |
| IGHMBP2 | None | None | None | None |
| TTN | None | None | None | None |
| TTN | None | None | None | None |
| IRAK4 | None | None | None | None |
| PROC | None | None | None | None |
| PROC | None | None | None | None |
| RAD50 | None | None | None | None |
| RAD50 | None | None | None | None |
| DYNC1H1 | None | None | None | None |
| DYNC1H1 | None | None | None | None |
| DYNC1H1 | None | None | None | None |
| DYNC1H1 | None | None | None | None |
| DYNC1H1 | None | None | None | None |
| DYNC1H1 | None | None | None | None |
| DYNC1H1 | None | None | None | None |
| DYNC1H1 | None | None | None | None |
| DYNC1H1 | None | None | None | None |
| DYNC1H1 | None | None | None | None |
| DYNC1H1 | None | None | None | None |
| CYB5R3 | None | None | None | None |
| CYB5R3 | 0.161 | 10 | 1umk | A |
| CYB5R3 | None | None | None | None |
| CYB5R3 | None | None | None | None |
| CYB5R3 | None | None | None | None |
| CYB5R3 | None | None | None | None |
| CYB5R3 | None | None | None | None |
| CYB5R3 | None | None | None | None |
| CYB5R3 | None | None | None | None |
| CYB5R3 | None | None | None | None |
| CYB5R3 | None | None | None | None |
| SPTB | None | None | None | None |
| FANCL | None | None | None | None |
| FAM126A | None | None | None | None |
| SEPT9 | None | None | None | None |
| TTN | None | None | None | None |
| TTN | None | None | None | None |
| TTN | 0.268 | 2 | 4jnw | A |
| CNTN1 | None | None | None | None |
| C1QTNF5 | None | None | None | None |
| VCL | None | None | None | None |
| VCL | None | None | None | None |
| VCL | None | None | None | None |
| VCL | None | None | None | None |
| VCL | None | None | None | None |
| VCL | None | None | None | None |
| PPT1 | None | None | None | None |
| PPT1 | None | None | None | None |

| | | | | |
|---|---|---|---|---|
| PPT1 | None | None | None | None |
| PPT1 | None | None | None | None |
| PPT1 | None | None | None | None |
| PPT1 | None | None | None | None |
| PPT1 | None | None | None | None |
| PPT1 | None | None | None | None |
| PPT1 | None | None | None | None |
| PPT1 | None | None | None | None |
| PPT1 | None | None | None | None |
| PPT1 | None | None | None | None |
| PPT1 | None | None | None | None |
| PPT1 | None | None | None | None |
| PPT1 | None | None | None | None |
| PPT1 | None | None | None | None |
| PPT1 | None | None | None | None |
| PPT1 | None | None | None | None |
| PPT1 | None | None | None | None |
| NOTCH2 | None | None | None | None |
| NOTCH2 | None | None | None | None |
| IFIH1 | None | None | None | None |
| IFIH1 | None | None | None | None |
| IKBKG | None | None | None | None |
| DBH | None | None | None | None |
| DBH | None | None | None | None |
| DBH | None | None | None | None |
| DBH | None | None | None | None |
| DBH | None | None | None | None |
| DBH | None | None | None | None |
| DBH | None | None | None | None |
| DBH | None | None | None | None |
| DBH | None | None | None | None |
| DBH | None | None | None | None |
| IL10RA | None | None | None | None |
| IL10RA | None | None | None | None |
| IL10RA | None | None | None | None |
| IL10RA | None | None | None | None |
| IL10RA | None | None | None | None |
| IL10RA | None | None | None | None |
| IL10RA | None | None | None | None |
| ILK | None | None | None | None |
| HPD | None | None | None | None |
| HPD | None | None | None | None |
| HPD | None | None | None | None |
| MECP2 | None | None | None | None |
| MECP2 | None | None | None | None |
| MECP2 | None | None | None | None |
| MECP2 | None | None | None | None |
| MECP2 | None | None | None | None |
| MECP2 | None | None | None | None |
| MECP2 | None | None | None | None |
| MECP2 | None | None | None | None |
| KYNU | None | None | None | None |
| FAS | None | None | None | None |
| FAS | None | None | None | None |

| | | | | |
|---|---|---|---|---|
| ELP1 | None | None | None | None |
| ELP1 | None | None | None | None |
| ELP1 | None | None | None | None |
| ELP1 | None | None | None | None |
| ELP1 | None | None | None | None |
| ELP1 | None | None | None | None |
| ELP1 | None | None | None | None |
| ELP1 | None | None | None | None |
| HESX1 | None | None | None | None |
| HESX1 | None | None | None | None |
| HESX1 | None | None | None | None |
| HESX1 | None | None | None | None |
| GAD1 | None | None | None | None |
| GAD1 | None | None | None | None |
| NDUFA1 | None | None | None | None |
| NDUFA1 | None | None | None | None |
| NDUFA1 | None | None | None | None |
| PNPT1 | None | None | None | None |
| PNPT1 | None | None | None | None |
| PNPT1 | None | None | None | None |
| PNPT1 | None | None | None | None |
| PNPT1 | 0.389 | 1 | 3u1k | A |
| PNPT1 | None | None | None | None |
| PNPT1 | None | None | None | None |
| SLC25A13 | None | None | None | None |
| SLC25A13 | None | None | None | None |
| SLC25A13 | None | None | None | None |
| AP4M1 | None | None | None | None |
| AP4M1 | None | None | None | None |
| MEFV | None | None | None | None |
| MEFV | None | None | None | None |
| MEFV | None | None | None | None |
| MEFV | None | None | None | None |
| MEFV | None | None | None | None |
| MEFV | None | None | None | None |
| MEFV | None | None | None | None |
| MEFV | None | None | None | None |
| MEFV | None | None | None | None |
| CAPN3 | None | None | None | None |
| CAPN3 | None | None | None | None |
| CAPN3 | None | None | None | None |
| CAPN3 | 0.158 | 14 | 4okh | A |
| CAPN3 | 0.474 | 2 | 4okh | A |
| CAPN3 | 0.474 | 2 | 4okh | A |
| CAPN3 | None | None | None | None |
| GP6 | None | None | None | None |
| FAS | None | None | None | None |
| FBN1 | None | None | None | None |
| FBN1 | None | None | None | None |
| BMPR2 | None | None | None | None |
| BMPR2 | None | None | None | None |
| BMPR2 | 0.031 | 38 | 3g2f | A |
| BMPR2 | 0.327 | 6 | 3g2f | A |
| BMPR2 | 0.327 | 6 | 3g2f | A |

| | | | | |
|---|---|---|---|---|
| RET | None | None | None | None |
| RET | None | None | None | None |
| RET | None | None | None | None |
| RET | None | None | None | None |
| RET | None | None | None | None |
| RET | None | None | None | None |
| RET | None | None | None | None |
| RET | None | None | None | None |
| RET | None | None | None | None |
| RET | None | None | None | None |
| RET | None | None | None | None |
| RET | None | None | None | None |
| RET | None | None | None | None |
| RET | None | None | None | None |
| RET | None | None | None | None |
| RET | None | None | None | None |
| RET | None | None | None | None |
| RET | None | None | None | None |
| RET | None | None | None | None |
| RET | None | None | None | None |
| RET | None | None | None | None |
| RET | None | None | None | None |
| RET | None | None | None | None |
| RET | None | None | None | None |
| RET | None | None | None | None |
| RET | None | None | None | None |
| RET | None | None | None | None |
| RET | None | None | None | None |
| CSF2RA | None | None | None | None |
| CSF2RA | None | None | None | None |
| CSF2RA | None | None | None | None |
| BRCA1 | None | None | None | None |
| ZAP70 | 0.636 | 1 | 2ozo | A |
| ZAP70 | 0.636 | 1 | 2ozo | A |
| ZAP70 | 0.26 | 9 | 2ozo | A |
| PIK3R2 | None | None | None | None |
| TCN2 | None | None | None | None |
| TCN2 | None | None | None | None |
| TCN2 | None | None | None | None |
| TCN2 | None | None | None | None |
| TCN2 | None | None | None | None |
| TCN2 | None | None | None | None |
| TCN2 | None | None | None | None |
| TCN2 | None | None | None | None |
| EPM2A | None | None | None | None |
| EPM2A | None | None | None | None |
| EPM2A | None | None | None | None |
| EPM2A | None | None | None | None |
| AARS | None | None | None | None |
| AARS | None | None | None | None |
| AARS | None | None | None | None |
| PCDH15 | None | None | None | None |
| PCDH15 | None | None | None | None |
| PCDH15 | None | None | None | None |

| | | | | |
|---|---|---|---|---|
| PCDH15 | None | None | None | None |
| PCDH15 | None | None | None | None |
| PCDH15 | None | None | None | None |
| PCDH15 | None | None | None | None |
| PCDH15 | None | None | None | None |
| PCDH15 | None | None | None | None |
| PCDH15 | None | None | None | None |
| NDUFS4 | None | None | None | None |
| NDUFS4 | None | None | None | None |
| NDUFS4 | None | None | None | None |
| FGF10 | None | None | None | None |
| FGF10 | 0.558 | 3 | 1nun | A |
| FGF10 | None | None | None | None |
| FGF10 | None | None | None | None |
| CDON | None | None | None | None |
| STAMBP | 0.909 | 2 | 3rzv | A |
| BRCA1 | None | None | None | None |
| BRCA1 | None | None | None | None |
| BRCA1 | None | None | None | None |
| BRCA1 | None | None | None | None |
| BRCA1 | None | None | None | None |
| BRCA1 | None | None | None | None |
| BRCA1 | None | None | None | None |
| BRCA1 | None | None | None | None |
| BRCA1 | None | None | None | None |
| BRCA1 | None | None | None | None |
| PLXNC1 | None | None | None | None |
| RNASEH2C | None | None | None | None |
| RNASEH2C | None | None | None | None |
| APC | None | None | None | None |
| PLA2G4A | None | None | None | None |
| PLA2G4A | None | None | None | None |
| PSAT1 | None | None | None | None |
| PSAT1 | None | None | None | None |
| TRAPPC3 | None | None | None | None |
| QDPR | None | None | None | None |
| QDPR | None | None | None | None |
| QDPR | None | None | None | None |
| QDPR | None | None | None | None |
| KRT5 | None | None | None | None |
| KRT5 | None | None | None | None |
| KRT5 | None | None | None | None |
| MLH1 | None | None | None | None |
| MLH1 | None | None | None | None |
| MLH1 | None | None | None | None |
| MLH1 | None | None | None | None |
| MLH1 | None | None | None | None |
| MLH1 | None | None | None | None |
| MLH1 | None | None | None | None |
| MLH1 | None | None | None | None |
| MLH1 | None | None | None | None |
| MLH1 | None | None | None | None |
| MLH1 | None | None | None | None |
| MLH1 | None | None | None | None |

| | | | | |
|---|---|---|---|---|
| MLH1 | None | None | None | None |
| MLH1 | None | None | None | None |
| MLH1 | None | None | None | None |
| MLH1 | None | None | None | None |
| MLH1 | None | None | None | None |
| KRIT1 | None | None | None | None |
| KRIT1 | None | None | None | None |
| KRIT1 | None | None | None | None |
| ALX4 | None | None | None | None |
| ALX4 | None | None | None | None |
| ALX4 | None | None | None | None |
| TJP2 | None | None | None | None |
| TJP2 | None | None | None | None |
| PANK2 | None | None | None | None |
| PANK2 | None | None | None | None |
| PANK2 | None | None | None | None |
| PANK2 | None | None | None | None |
| PANK2 | None | None | None | None |
| PANK2 | 0.333 | 2 | 5E+026 | B |
| PANK2 | None | None | None | None |
| NDUFA6 | None | None | None | None |
| TSHR | None | None | None | None |
| TSHR | None | None | None | None |
| TSHR | None | None | None | None |
| CFTR | None | None | None | None |
| CFTR | None | None | None | None |
| MYOM1 | None | None | None | None |
| MYOM1 | None | None | None | None |
| SPTB | None | None | None | None |
| GPT2 | None | None | None | None |
| NDUFS2 | None | None | None | None |
| NDUFS2 | None | None | None | None |
| NDUFS2 | None | None | None | None |
| NDUFS2 | None | None | None | None |
| NDUFS2 | None | None | None | None |
| CASQ2 | None | None | None | None |
| CASQ2 | None | None | None | None |
| CASQ2 | None | None | None | None |
| CASQ2 | None | None | None | None |
| CASQ2 | None | None | None | None |
| CASQ2 | None | None | None | None |
| CASQ2 | None | None | None | None |
| CASQ2 | None | None | None | None |
| CASQ2 | None | None | None | None |
| CPOX | None | None | None | None |
| CPOX | None | None | None | None |
| CPOX | 0.857 | 2 | 2aex | A |
| CPOX | None | None | None | None |
| CPOX | None | None | None | None |
| CPOX | None | None | None | None |
| CPOX | None | None | None | None |
| CPOX | None | None | None | None |
| CPOX | None | None | None | None |
| MVK | None | None | None | None |

| | | | | |
|---|---|---|---|---|
| MVK | None | None | None | None |
| MVK | None | None | None | None |
| MVK | None | None | None | None |
| MVK | None | None | None | None |
| MVK | None | None | None | None |
| MVK | None | None | None | None |
| MVK | None | None | None | None |
| MVK | None | None | None | None |
| MVK | None | None | None | None |
| MVK | None | None | None | None |
| MVK | None | None | None | None |
| MVK | None | None | None | None |
| MVK | None | None | None | None |
| MVK | None | None | None | None |
| GSS | None | None | None | None |
| GSS | None | None | None | None |
| GSS | None | None | None | None |
| GSS | None | None | None | None |
| GSS | None | None | None | None |
| DSG2 | None | None | None | None |
| DSG2 | None | None | None | None |
| DSG2 | None | None | None | None |
| DSG2 | None | None | None | None |
| DSG2 | None | None | None | None |
| DSG2 | None | None | None | None |
| DSG2 | None | None | None | None |
| DSG2 | None | None | None | None |
| DSG2 | None | None | None | None |
| DSG2 | None | None | None | None |
| DSG2 | None | None | None | None |
| DSG2 | None | None | None | None |
| DSG2 | None | None | None | None |
| DSG2 | None | None | None | None |
| PAPSS2 | None | None | None | None |
| PLCB2 | None | None | None | None |
| HMGCS2 | None | None | None | None |
| HMGCS2 | None | None | None | None |
| HMGCS2 | 0.938 | 1 | 2wya | C |
| HMGCS2 | None | None | None | None |
| HMGCS2 | None | None | None | None |
| HMGCS2 | None | None | None | None |
| HMGCS2 | 0.562 | 4 | 2wya | C |
| HMGCS2 | None | None | None | None |
| MPDZ | None | None | None | None |
| GATA4 | None | None | None | None |
| GATA4 | None | None | None | None |
| WARS2 | None | None | None | None |
| WARS2 | None | None | None | None |
| HGD | 1 | 47 | 1eyb | A |
| HGD | 1 | 47 | 1eyb | A |
| HGD | 1 | 70 | 1eyb | A |
| HGD | None | None | None | None |
| HGD | None | None | None | None |
| HGD | None | None | None | None |

| | | | | |
|---|---|---|---|---|
| HGD | None | None | None | None |
| HGD | 0.5 | 51 | 1ey2 | A |
| HGD | None | None | None | None |
| HGD | 1 | 1 | 1ey2 | A |
| HGD | 1 | 9 | 1ey2 | A |
| HGD | None | None | None | None |
| HGD | 1 | 10 | 1eyb | A |
| HGD | 1 | 7 | 1ey2 | A |
| PAX8 | None | None | None | None |
| PAX8 | None | None | None | None |
| PAX8 | None | None | None | None |
| PAX8 | None | None | None | None |
| SPTB | None | None | None | None |
| NCF4 | None | None | None | None |
| CFH | None | None | None | None |
| CFH | None | None | None | None |
| CSF1R | None | None | None | None |
| CSF1R | None | None | None | None |
| CA8 | None | None | None | None |
| CA8 | None | None | None | None |
| CA8 | 0.748 | 7 | 2w2j | A |
| APC | None | None | None | None |
| APC | None | None | None | None |
| APC | None | None | None | None |
| APC | None | None | None | None |
| APC | None | None | None | None |
| NDUFA2 | None | None | None | None |
| GIF | None | None | None | None |
| GIF | None | None | None | None |
| GIF | None | None | None | None |
| GIF | None | None | None | None |
| C2 | None | None | None | None |
| ACP5 | None | None | None | None |
| ACP5 | None | None | None | None |
| ACP5 | None | None | None | None |
| ACP5 | None | None | None | None |
| ACP5 | None | None | None | None |
| RNASEH2A | None | None | None | None |
| RNASEH2A | None | None | None | None |
| RNASEH2A | None | None | None | None |
| RNASEH2A | None | None | None | None |
| RNASEH2A | None | None | None | None |
| RNASEH2A | None | None | None | None |
| RNASEH2A | None | None | None | None |
| DSG2 | None | None | None | None |
| USH1G | None | None | None | None |
| FUS | None | None | None | None |
| FUS | None | None | None | None |
| FUS | None | None | None | None |
| FUS | None | None | None | None |
| FUS | None | None | None | None |
| MRPL44 | None | None | None | None |
| MRPL44 | None | None | None | None |
| MRPL44 | None | None | None | None |

| | | | | |
|---|---|---|---|---|
| MRPL44 | None | None | None | None |
| DMGDH | None | None | None | None |
| AARS | None | None | None | None |
| STAR | None | None | None | None |
| STAR | None | None | None | None |
| STAR | None | None | None | None |
| STAR | None | None | None | None |
| STAR | None | None | None | None |
| STAR | None | None | None | None |
| STAR | None | None | None | None |
| STAR | None | None | None | None |
| LARP7 | None | None | None | None |
| PRPF8 | None | None | None | None |
| PRPF8 | None | None | None | None |
| PRPF8 | None | None | None | None |
| PRPF8 | None | None | None | None |
| PSAP | None | None | None | None |
| PSAP | None | None | None | None |
| PSAP | None | None | None | None |
| DMPK | None | None | None | None |
| EPCAM | None | None | None | None |
| EPCAM | None | None | None | None |
| EPCAM | None | None | None | None |
| EPCAM | None | None | None | None |
| EPCAM | None | None | None | None |
| EPCAM | None | None | None | None |
| EPCAM | None | None | None | None |
| EPCAM | None | None | None | None |
| ASS1 | None | None | None | None |
| ASS1 | None | None | None | None |
| ASS1 | 0.571 | 3 | 2nz2 | A |
| ASS1 | 0.571 | 3 | 2nz2 | A |
| ASS1 | None | None | None | None |
| ASS1 | 0.857 | 20 | 2nz2 | A |
| ASS1 | 0.857 | 9 | 2nz2 | A |
| ASS1 | None | None | None | None |
| ASS1 | 0.429 | 13 | 2nz2 | A |
| ASS1 | 0.571 | 4 | 2nz2 | A |
| ASS1 | 0.857 | 9 | 2nz2 | A |
| ASS1 | 0.857 | 9 | 2nz2 | A |
| ASS1 | 0.857 | 9 | 2nz2 | A |
| ASS1 | None | None | None | None |
| ASS1 | None | None | None | None |
| ASS1 | None | None | None | None |
| ASS1 | None | None | None | None |
| ASS1 | None | None | None | None |
| ASS1 | None | None | None | None |
| ASS1 | None | None | None | None |
| POLE | None | None | None | None |
| POLE | None | None | None | None |
| POLE | None | None | None | None |
| RNASET2 | None | None | None | None |
| RNASET2 | None | None | None | None |
| RNASET2 | None | None | None | None |

| | | | | |
|---|---|---|---|---|
| FLG | 0.394 | | 2 4pcw | A |
| F10 | None | None | None | None |
| AGT | None | None | None | None |
| AGT | None | None | None | None |
| PLN | None | None | None | None |
| PLN | None | None | None | None |
| PLN | None | None | None | None |
| PTHLH | None | None | None | None |
| IGF1R | None | None | None | None |
| IGF1R | None | None | None | None |
| IGF1R | None | None | None | None |
| JAG1 | None | None | None | None |
| JAG1 | None | None | None | None |
| JAG1 | None | None | None | None |
| JAG1 | None | None | None | None |
| JAG1 | None | None | None | None |
| JAG1 | None | None | None | None |
| AP1S3 | None | None | None | None |
| SPTA1 | None | None | None | None |
| SPTA1 | None | None | None | None |
| SPTA1 | None | None | None | None |
| SPTA1 | None | None | None | None |
| SPG7 | None | None | None | None |
| SPG7 | None | None | None | None |
| SPG7 | None | None | None | None |
| SPG7 | None | None | None | None |
| SPG7 | None | None | None | None |
| SPG7 | None | None | None | None |
| FN1 | None | None | None | None |
| FN1 | None | None | None | None |
| NSMCE3 | None | None | None | None |
| PCCA | None | None | None | None |
| PCCA | None | None | None | None |
| SPTB | None | None | None | None |
| RYR2 | None | None | None | None |
| RYR2 | None | None | None | None |
| MMP9 | None | None | None | None |
| MMP9 | None | None | None | None |
| GLMN | None | None | None | None |
| GLMN | None | None | None | None |
| GLMN | None | None | None | None |
| GLMN | None | None | None | None |
| GLMN | None | None | None | None |
| GLMN | None | None | None | None |
| GLMN | None | None | None | None |
| GLMN | None | None | None | None |
| GLMN | None | None | None | None |
| GLMN | None | None | None | None |
| MARS | 0.333 | | 2 5goy | A |
| MARS | None | None | None | None |
| MARS | None | None | None | None |
| MARS | None | None | None | None |
| MARS | None | None | None | None |
| MARS | None | None | None | None |

| | | | | |
|---|---|---|---|---|
| SNAP29 | None | None | None | None |
| SNAP29 | None | None | None | None |
| ETHE1 | None | None | None | None |
| ETHE1 | None | None | None | None |
| ETHE1 | None | None | None | None |
| ETHE1 | None | None | None | None |
| ETHE1 | None | None | None | None |
| ETHE1 | None | None | None | None |
| ETHE1 | None | None | None | None |
| SAMHD1 | None | None | None | None |
| KCNQ4 | None | None | None | None |
| FXYD2 | None | None | None | None |
| FXYD2 | None | None | None | None |
| FXYD2 | None | None | None | None |
| LRRK2 | None | None | None | None |
| LRRK2 | None | None | None | None |
| NDUFS1 | None | None | None | None |
| NDUFS1 | None | None | None | None |
| NDUFS1 | None | None | None | None |
| NDUFS1 | None | None | None | None |
| NDUFS1 | None | None | None | None |
| NDUFS1 | None | None | None | None |
| NDUFS1 | None | None | None | None |
| NDUFS1 | None | None | None | None |
| NDUFS1 | None | None | None | None |
| NDUFS1 | None | None | None | None |
| NDUFS1 | None | None | None | None |
| EZH2 | None | None | None | None |
| EZH2 | None | None | None | None |
| LARS | None | None | None | None |
| TLL1 | None | None | None | None |
| CRYBB2 | None | None | None | None |
| CRYBB2 | None | None | None | None |
| EXOSC8 | None | None | None | None |
| SRP72 | None | None | None | None |
| FOXC2 | None | None | None | None |
| HMBS | None | None | None | None |
| HMBS | None | None | None | None |
| HMBS | None | None | None | None |
| HMBS | None | None | None | None |
| HMBS | None | None | None | None |
| HMBS | None | None | None | None |
| HMBS | None | None | None | None |
| HMBS | None | None | None | None |
| HMBS | None | None | None | None |
| HMBS | None | None | None | None |
| HMBS | None | None | None | None |
| HMBS | None | None | None | None |
| HMBS | None | None | None | None |
| HMBS | None | None | None | None |
| HMBS | None | None | None | None |
| HMBS | None | None | None | None |
| HMBS | None | None | None | None |
| HMBS | None | None | None | None |

| | | | | | |
|---|---|---|---|---|---|
| HMBS | None | None | | None | None |
| HMBS | None | None | | None | None |
| HMBS | None | None | | None | None |
| HMBS | None | None | | None | None |
| HMBS | None | None | | None | None |
| HMBS | None | None | | None | None |
| HMBS | None | None | | None | None |
| OGT | None | None | | None | None |
| NOG | None | None | | None | None |
| NOG | None | None | | None | None |
| NOG | None | None | | None | None |
| NOG | None | None | | None | None |
| NOG | None | None | | None | None |
| NOG | None | None | | None | None |
| NOG | None | None | | None | None |
| NOG | None | None | | None | None |
| NOG | None | None | | None | None |
| NOG | None | None | | None | None |
| NOG | None | None | | None | None |
| DLG3 | None | None | | None | None |
| SCN2A | None | None | | None | None |
| STAC3 | None | None | | None | None |
| PFKM | None | None | | None | None |
| PFKM | None | None | | None | None |
| PFKM | None | None | | None | None |
| PFKM | None | None | | None | None |
| PFKM | None | None | | None | None |
| DNMT1 | None | None | | None | None |
| DNMT1 | None | None | | None | None |
| TNNI3 | None | None | | None | None |
| TNNI3 | None | None | | None | None |
| TNNI3 | None | None | | None | None |
| STXBP2 | None | None | | None | None |
| STXBP2 | None | None | | None | None |
| STXBP2 | None | None | | None | None |
| STXBP2 | None | None | | None | None |
| STXBP2 | None | None | | None | None |
| STXBP2 | None | None | | None | None |
| STXBP2 | None | None | | None | None |
| STXBP2 | None | None | | None | None |
| REN | 0.083 | | 27 | 4amt | A |
| DSP | None | None | | None | None |
| DSP | None | None | | None | None |
| HNF1B | None | None | | None | None |
| ADA | None | None | | None | None |
| ADA | None | None | | None | None |
| ADA | 0.615 | | 7 | 3iar | A |
| ADA | None | None | | None | None |
| ADA | None | None | | None | None |
| ADA | 0.538 | | 5 | 3iar | A |
| ADA | 0.538 | | 5 | 3iar | A |
| ADA | None | None | | None | None |
| ADA | None | None | | None | None |
| ADA | 0.615 | | 1 | 3iar | A |

| | | | | | |
|---|---|---|---|---|---|
| ADA | | 0.462 | 6 | 3iar | A |
| ADA | | 0.615 | 1 | 3iar | A |
| ADA | None | None | | None | None |
| ADA | None | None | | None | None |
| ADA | None | None | | None | None |
| DCX | None | None | | None | None |
| DCX | None | None | | None | None |
| DCX | None | None | | None | None |
| DCX | None | None | | None | None |
| DCX | None | None | | None | None |
| ARSB | None | None | | None | None |
| ARSB | None | None | | None | None |
| ARSB | None | None | | None | None |
| ARSB | None | None | | None | None |
| ARSB | None | None | | None | None |
| ARSB | None | None | | None | None |
| ARSB | None | None | | None | None |
| ARSB | None | None | | None | None |
| ARSB | None | None | | None | None |
| ARSB | None | None | | None | None |
| ARSB | None | None | | None | None |
| ARSB | None | None | | None | None |
| ARSB | None | None | | None | None |
| WWOX | None | None | | None | None |
| TNNI3 | None | None | | None | None |
| TNNI3 | None | None | | None | None |
| TNNI3 | None | None | | None | None |
| TNNI3 | None | None | | None | None |
| CYBA | None | None | | None | None |
| SPAST | None | None | | None | None |
| SPAST | None | None | | None | None |
| SPAST | None | None | | None | None |
| SPAST | None | None | | None | None |
| SPAST | | 0.132 | 1 | 3vfd | A |
| SPAST | None | None | | None | None |
| SPAST | None | None | | None | None |
| SPAST | None | None | | None | None |
| KRIT1 | None | None | | None | None |
| KRIT1 | None | None | | None | None |
| TP53 | None | None | | None | None |
| ISPD | None | None | | None | None |
| ISPD | None | None | | None | None |
| ISPD | None | None | | None | None |
| ISPD | None | None | | None | None |
| KCNMA1 | None | None | | None | None |
| STK11 | | 0.002 | 58 | 2wtk | F |
| STK11 | None | None | | None | None |
| STK11 | | 0.034 | 32 | 2wtk | F |
| STK11 | None | None | | None | None |
| STK11 | None | None | | None | None |
| STK11 | | 0.412 | 2 | 2wtk | F |
| NDUFA13 | None | None | | None | None |
| PHF6 | None | None | | None | None |
| PHF6 | None | None | | None | None |

| | | | | |
|---|---|---|---|---|
| PHF6 | None | None | None | None |
| PHYH | None | None | None | None |
| PHYH | None | None | None | None |
| PHYH | None | None | None | None |
| PHYH | None | None | None | None |
| PHYH | None | None | None | None |
| NDUFS3 | None | None | None | None |
| NDUFS3 | None | None | None | None |
| ALDH18A1 | None | None | None | None |
| ALDH18A1 | None | None | None | None |
| ALDH18A1 | None | None | None | None |
| ALDH18A1 | None | None | None | None |
| ALDH18A1 | None | None | None | None |
| PAX6 | None | None | None | None |
| PAX6 | 0.339 | 1 | 2cue | A |
| INPP5E | None | None | None | None |
| INPP5E | None | None | None | None |
| INPP5E | None | None | None | None |
| INPP5E | None | None | None | None |
| INPP5E | None | None | None | None |
| INPP5E | None | None | None | None |
| INPP5E | None | None | None | None |
| INPP5E | None | None | None | None |
| INPP5E | None | None | None | None |
| TNNT2 | None | None | None | None |
| TNNT2 | None | None | None | None |
| TNNT2 | None | None | None | None |
| TNNT2 | None | None | None | None |
| TNNT2 | None | None | None | None |
| TNNT2 | None | None | None | None |
| TNNT2 | None | None | None | None |
| TNNT2 | None | None | None | None |
| TYK2 | None | None | None | None |
| TYK2 | None | None | None | None |
| TYK2 | None | None | None | None |
| CFTR | None | None | None | None |
| CFTR | None | None | None | None |
| CFTR | None | None | None | None |
| CFTR | None | None | None | None |
| CFTR | None | None | None | None |
| CFTR | None | None | None | None |
| CFTR | None | None | None | None |
| CFTR | None | None | None | None |
| CFTR | None | None | None | None |
| CFTR | None | None | None | None |
| CFTR | None | None | None | None |
| CFTR | None | None | None | None |
| CFTR | None | None | None | None |
| CFTR | None | None | None | None |
| CFTR | None | None | None | None |
| CFTR | None | None | None | None |
| CFTR | None | None | None | None |
| CFTR | None | None | None | None |
| CFTR | None | None | None | None |

| | | | | |
|---|---|---|---|---|
| CFTR | None | None | None | None |
| CFTR | None | None | None | None |
| CFTR | None | None | None | None |
| CFTR | None | None | None | None |
| CFTR | None | None | None | None |
| CFTR | None | None | None | None |
| CFTR | None | None | None | None |
| CFTR | None | None | None | None |
| CFTR | None | None | None | None |
| CFTR | None | None | None | None |
| CFTR | None | None | None | None |
| CFTR | None | None | None | None |
| CFTR | None | None | None | None |
| CFTR | None | None | None | None |
| CFTR | None | None | None | None |
| CFTR | None | None | None | None |
| CFTR | None | None | None | None |
| CFTR | None | None | None | None |
| CFTR | None | None | None | None |
| CFTR | None | None | None | None |
| CFTR | None | None | None | None |
| CFTR | None | None | None | None |
| CFTR | None | None | None | None |
| CFTR | None | None | None | None |
| CFTR | None | None | None | None |
| CFTR | None | None | None | None |
| CFTR | None | None | None | None |
| CFTR | None | None | None | None |
| CFTR | None | None | None | None |
| CFTR | None | None | None | None |
| CFTR | None | None | None | None |
| CFTR | None | None | None | None |
| CFTR | None | None | None | None |
| CFTR | None | None | None | None |
| SERPINF1 | None | None | None | None |
| SERPINF1 | None | None | None | None |
| IDS | None | None | None | None |
| IDS | None | None | None | None |
| IDS | None | None | None | None |
| IDS | None | None | None | None |
| IDS | None | None | None | None |
| IDS | None | None | None | None |
| IDS | None | None | None | None |
| IDS | None | None | None | None |
| IDS | None | None | None | None |
| NFKBIA | 0.7 | 1 | 1nfi | F |
| NFKBIA | None | None | None | None |
| SPAST | None | None | None | None |
| FGF23 | None | None | None | None |
| FGF23 | None | None | None | None |
| FGF23 | None | None | None | None |
| FGF23 | None | None | None | None |
| FGF23 | None | None | None | None |
| FGF23 | None | None | None | None |

| | | | | |
|---|---|---|---|---|
| FGF23 | None | None | None | None |
| DLAT | None | None | None | None |
| STUB1 | None | None | None | None |
| HSD17B4 | None | None | None | None |
| HSD17B4 | None | None | None | None |
| DYSF | None | None | None | None |
| CEP135 | None | None | None | None |
| CEP135 | None | None | None | None |
| PRKCG | None | None | None | None |
| CSF1R | None | None | None | None |
| PEX5 | None | None | None | None |
| PAX6 | None | None | None | None |
| PAX6 | None | None | None | None |
| PAX6 | None | None | None | None |
| PAX6 | None | None | None | None |
| PAX6 | None | None | None | None |
| PAX6 | None | None | None | None |
| NF2 | None | None | None | None |
| NF2 | None | None | None | None |
| NF2 | None | None | None | None |
| REM1 | None | None | None | None |

| source_id | classification | binding_site | binding_site_single_water | protein_binding_site |
|---|---|---|---|---|
| rs121918343 | pathogenic | 1 | 1 | 0 |
| rs267606649 | pathogenic | 1 | 1 | 1 |
| rs121918342 | pathogenic | 0 | 1 | 0 |
| rs267606650 | pathogenic | 1 | 1 | 1 |
| rs1130335 | benign | 1 | 1 | 1 |
| rs199476087 | pathogenic | 1 | 1 | 1 |
| rs772163112 | benign | 0 | 0 | 0 |
| rs755793 | benign | 0 | 0 | 0 |
| rs121918155 | pathogenic | 1 | 1 | 0 |
| rs121918156 | pathogenic | 1 | 1 | 0 |
| rs121918154 | pathogenic | 1 | 1 | 1 |
| rs121918151 | pathogenic | 1 | 1 | 1 |
| rs121918144 | pathogenic | 0 | 1 | 0 |
| rs121918146 | pathogenic | 1 | 1 | 1 |
| rs121918160 | pathogenic | 1 | 1 | 1 |
| rs121918150 | pathogenic | 1 | 1 | 1 |
| rs121918141 | pathogenic | 1 | 1 | 1 |
| rs142742242 | pathogenic | 1 | 1 | 1 |
| rs121918142 | pathogenic | 1 | 1 | 1 |
| rs121918157 | pathogenic | 1 | 1 | 1 |
| rs121918101 | pathogenic | 1 | 1 | 1 |
| rs121908260 | pathogenic | 1 | 1 | 1 |
| rs80356667 | pathogenic | 1 | 1 | 1 |
| rs80356668 | pathogenic | 1 | 1 | 1 |
| rs397514540 | pathogenic | 1 | 1 | 1 |
| rs104894685 | pathogenic | 0 | 1 | 0 |
| rs199869995 | pathogenic | 1 | 1 | 1 |
| rs121909072 | pathogenic | 0 | 0 | 0 |
| rs143669310 | benign | 1 | 1 | 1 |
| rs747347498 | benign | 0 | 0 | 0 |
| rs138283222 | benign | 0 | 0 | 0 |
| rs118203931 | pathogenic | 1 | 1 | 1 |
| rs118203933 | pathogenic | 1 | 1 | 0 |
| rs118203932 | pathogenic | 1 | 1 | 0 |
| rs2228063 | benign | 1 | 1 | 1 |
| rs377767360 | pathogenic | 0 | 0 | 0 |
| rs397518413 | pathogenic | 1 | 1 | 1 |
| rs1800458 | benign | 1 | 1 | 0 |
| rs121918083 | pathogenic | 1 | 1 | 1 |
| rs121918094 | pathogenic | 1 | 1 | 1 |
| rs121918098 | pathogenic | 1 | 1 | 1 |
| rs121918093 | pathogenic | 1 | 1 | 1 |
| rs79977247 | pathogenic | 1 | 1 | 1 |
| rs28933979 | pathogenic | 1 | 1 | 1 |
| rs121918068 | pathogenic | 1 | 1 | 1 |
| rs121918077 | pathogenic | 1 | 1 | 1 |
| rs11541796 | pathogenic | 1 | 1 | 1 |
| rs104894665 | pathogenic | 1 | 1 | 1 |
| rs121918090 | pathogenic | 1 | 1 | 1 |
| rs121918081 | pathogenic | 1 | 1 | 1 |
| rs121918080 | pathogenic | 1 | 1 | 1 |
| rs121918076 | pathogenic | 1 | 1 | 1 |
| rs121918097 | pathogenic | 1 | 1 | 1 |

| | | | | |
|---|---|---|---|---|
| rs121918079 | pathogenic | 1 | 1 | 1 |
| rs121918069 | pathogenic | 1 | 1 | 1 |
| rs121918086 | pathogenic | 1 | 1 | 1 |
| rs121918091 | pathogenic | 1 | 1 | 1 |
| rs121918085 | pathogenic | 1 | 1 | 1 |
| rs121918100 | pathogenic | 1 | 1 | 1 |
| rs121918084 | pathogenic | 1 | 1 | 1 |
| rs121918071 | pathogenic | 1 | 1 | 1 |
| rs121918072 | pathogenic | 1 | 1 | 1 |
| rs121918082 | pathogenic | 1 | 1 | 1 |
| rs121918087 | pathogenic | 1 | 1 | 1 |
| rs76410435 | benign | 1 | 1 | 1 |
| rs121918089 | pathogenic | 1 | 1 | 1 |
| rs121918073 | pathogenic | 1 | 1 | 1 |
| rs121918075 | pathogenic | 1 | 1 | 1 |
| rs121918088 | pathogenic | 1 | 1 | 1 |
| rs76992529 | pathogenic | 1 | 1 | 1 |
| rs536294863 | benign | 1 | 1 | 1 |
| rs730881162 | benign | 1 | 1 | 1 |
| rs113994087 | pathogenic | 1 | 1 | 1 |
| rs74716434 | benign | 1 | 1 | 1 |
| rs121918694 | pathogenic | 1 | 1 | 1 |
| rs121918707 | pathogenic | 1 | 1 | 1 |
| rs121918695 | pathogenic | 1 | 1 | 0 |
| rs121918690 | pathogenic | 1 | 1 | 0 |
| rs121918696 | pathogenic | 1 | 1 | 0 |
| rs121918693 | pathogenic | 1 | 1 | 0 |
| rs28999969 | pathogenic | 1 | 1 | 1 |
| rs121918697 | pathogenic | 1 | 1 | 1 |
| rs121918688 | pathogenic | 1 | 1 | 0 |
| rs28999970 | pathogenic | 1 | 1 | 0 |
| rs121918686 | pathogenic | 1 | 1 | 0 |
| rs28999971 | pathogenic | 1 | 1 | 1 |
| rs121918698 | pathogenic | 1 | 1 | 1 |
| rs121918691 | pathogenic | 1 | 1 | 0 |
| rs121918692 | pathogenic | 1 | 1 | 1 |
| rs121918703 | pathogenic | 1 | 1 | 1 |
| rs121918687 | pathogenic | 1 | 1 | 1 |
| rs28933408 | pathogenic | 1 | 1 | 1 |
| rs121918702 | pathogenic | 1 | 1 | 1 |
| rs17875899 | benign | 1 | 1 | 1 |
| rs111033620 | pathogenic | 1 | 1 | 1 |
| rs111033622 | pathogenic | 1 | 1 | 1 |
| rs111033621 | pathogenic | 1 | 1 | 1 |
| rs111033618 | pathogenic | 1 | 1 | 1 |
| rs5918 | benign | 0 | 0 | 0 |
| rs121918452 | pathogenic | 0 | 0 | 0 |
| rs121918445 | pathogenic | 1 | 1 | 1 |
| rs5917 | benign | 1 | 1 | 1 |
| rs121918444 | pathogenic | 1 | 1 | 1 |
| rs121918446 | pathogenic | 1 | 1 | 1 |
| rs121918449 | pathogenic | 0 | 0 | 0 |
| rs121918448 | benign | 0 | 0 | 0 |
| rs13306487 | benign | 1 | 1 | 1 |

| | | | | |
|---|---|---|---|---|
| rs1805044 | benign | 0 | 0 | 0 |
| rs117078377 | benign | 0 | 0 | 0 |
| rs138132026 | benign | 1 | 1 | 1 |
| rs75762896 | benign | 1 | 1 | 0 |
| rs121918043 | pathogenic | 1 | 1 | 0 |
| rs143581690 | benign | 1 | 1 | 0 |
| rs147654263 | benign | 0 | 0 | 0 |
| rs587782529 | pathogenic | 1 | 1 | 1 |
| rs121912664 | pathogenic | 1 | 1 | 1 |
| rs150293825 | benign | 1 | 1 | 1 |
| rs17882252 | benign | 1 | 1 | 1 |
| rs121912662 | pathogenic | 1 | 1 | 1 |
| rs35993958 | benign | 1 | 1 | 1 |
| rs606231253 | pathogenic | 1 | 1 | 1 |
| rs2227983 | benign | 1 | 1 | 1 |
| rs587779351 | pathogenic | 0 | 0 | 0 |
| rs779773463 | pathogenic | 1 | 1 | 1 |
| rs713040 | benign | 1 | 1 | 1 |
| rs334 | pathogenic | 1 | 1 | 1 |
| rs33930165 | pathogenic | 1 | 1 | 1 |
| rs33918131 | benign | 1 | 1 | 1 |
| rs63750783 | pathogenic | 1 | 1 | 1 |
| rs33986703 | pathogenic | 1 | 1 | 1 |
| rs33972047 | pathogenic | 1 | 1 | 1 |
| rs33936254 | pathogenic | 1 | 1 | 1 |
| rs33959855 | benign | 1 | 1 | 1 |
| rs33950507 | pathogenic | 1 | 1 | 1 |
| rs33916412 | pathogenic | 1 | 1 | 0 |
| rs1135071 | pathogenic | 1 | 1 | 1 |
| rs33960103 | pathogenic | 1 | 1 | 1 |
| rs11549407 | pathogenic | 1 | 1 | 1 |
| rs33922842 | pathogenic | 1 | 1 | 1 |
| rs35278874 | benign | 1 | 1 | 1 |
| rs34621955 | benign | 1 | 1 | 1 |
| rs33995148 | pathogenic | 1 | 1 | 1 |
| rs33945705 | benign | 1 | 1 | 1 |
| rs33993568 | benign | 1 | 1 | 1 |
| rs35553496 | benign | 1 | 1 | 1 |
| rs33913712 | pathogenic | 1 | 1 | 1 |
| rs33917785 | pathogenic | 1 | 1 | 1 |
| rs34515413 | benign | 1 | 1 | 1 |
| rs35256489 | pathogenic | 1 | 1 | 0 |
| rs34484056 | pathogenic | 1 | 1 | 1 |
| rs33947020 | benign | 1 | 1 | 1 |
| rs33946267 | pathogenic | 1 | 1 | 1 |
| rs33966761 | pathogenic | 1 | 1 | 1 |
| rs36020563 | benign | 1 | 1 | 1 |
| rs122454124 | pathogenic | 1 | 1 | 1 |
| rs122454126 | pathogenic | 1 | 1 | 0 |
| rs122454127 | pathogenic | 1 | 1 | 1 |
| rs122454130 | pathogenic | 1 | 1 | 1 |
| rs122454125 | pathogenic | 1 | 1 | 1 |
| rs122454131 | pathogenic | 1 | 1 | 0 |
| rs121908302 | pathogenic | 0 | 1 | 0 |

| | | | | |
|---|---|---|---|---|
| rs77829017 | pathogenic | 0 | 1 | 0 |
| rs1141814 | pathogenic | 0 | 1 | 0 |
| rs121908312 | pathogenic | 0 | 1 | 0 |
| rs79653797 | pathogenic | 1 | 1 | 0 |
| rs439898 | pathogenic | 1 | 1 | 0 |
| rs121908299 | pathogenic | 0 | 1 | 0 |
| rs80356763 | pathogenic | 1 | 1 | 1 |
| rs781152868 | pathogenic | 0 | 1 | 0 |
| rs121908297 | pathogenic | 0 | 1 | 0 |
| rs398123531 | benign | 0 | 1 | 0 |
| rs398123532 | pathogenic | 0 | 1 | 0 |
| rs364897 | pathogenic | 1 | 1 | 1 |
| rs1064644 | pathogenic | 0 | 1 | 0 |
| rs409652 | pathogenic | 1 | 1 | 1 |
| rs121908300 | pathogenic | 0 | 1 | 0 |
| rs381737 | pathogenic | 0 | 1 | 0 |
| rs74500255 | pathogenic | 0 | 1 | 0 |
| rs121908313 | pathogenic | 0 | 1 | 0 |
| rs367968666 | pathogenic | 0 | 1 | 0 |
| rs78973108 | pathogenic | 1 | 1 | 1 |
| rs121908298 | pathogenic | 1 | 1 | 1 |
| rs78396650 | pathogenic | 0 | 0 | 0 |
| rs78198234 | pathogenic | 1 | 1 | 0 |
| rs121908304 | pathogenic | 1 | 1 | 1 |
| rs76539814 | pathogenic | 1 | 1 | 1 |
| rs121908305 | pathogenic | 0 | 1 | 0 |
| rs121908306 | pathogenic | 1 | 1 | 0 |
| rs121908308 | pathogenic | 1 | 1 | 1 |
| rs121908307 | pathogenic | 0 | 1 | 0 |
| rs75548401 | benign | 0 | 1 | 0 |
| rs76763715 | pathogenic | 0 | 0 | 0 |
| rs121908314 | pathogenic | 0 | 1 | 0 |
| rs398123528 | pathogenic | 0 | 1 | 0 |
| rs121908311 | pathogenic | 0 | 0 | 0 |
| rs121908310 | pathogenic | 1 | 1 | 0 |
| rs74598136 | pathogenic | 0 | 1 | 0 |
| rs1064651 | pathogenic | 1 | 1 | 1 |
| rs77369218 | pathogenic | 1 | 1 | 1 |
| rs121908295 | pathogenic | 0 | 1 | 0 |
| rs421016 | pathogenic | 0 | 1 | 0 |
| rs368060 | pathogenic | 0 | 1 | 0 |
| rs80356771 | pathogenic | 1 | 1 | 1 |
| rs80356772 | pathogenic | 1 | 1 | 1 |
| rs121908301 | pathogenic | 0 | 0 | 0 |
| rs747506979 | pathogenic | 0 | 1 | 0 |
| rs75822236 | pathogenic | 0 | 1 | 0 |
| rs1990760 | benign | 1 | 1 | 0 |
| rs28934568 | pathogenic | 1 | 1 | 0 |
| rs34833812 | benign | 1 | 1 | 0 |
| rs193922666 | benign | 1 | 1 | 0 |
| rs104893812 | pathogenic | 1 | 1 | 0 |
| rs144701411 | benign | 0 | 0 | 0 |
| rs104893813 | pathogenic | 0 | 1 | 0 |
| rs199660234 | pathogenic | 1 | 1 | 0 |

| | | | | |
|---|---|---|---|---|
| rs104893814 | pathogenic | 1 | 1 | 0 |
| rs143095746 | benign | 1 | 1 | 1 |
| rs104893807 | pathogenic | 1 | 1 | 0 |
| rs104893811 | pathogenic | 0 | 0 | 0 |
| rs104893816 | pathogenic | 0 | 0 | 0 |
| rs121918714 | pathogenic | 0 | 1 | 0 |
| rs104893810 | pathogenic | 1 | 1 | 0 |
| rs104893815 | pathogenic | 1 | 1 | 0 |
| rs4303 | benign | 0 | 1 | 0 |
| rs121912704 | pathogenic | 0 | 1 | 0 |
| rs56394458 | benign | 1 | 1 | 0 |
| rs397514688 | pathogenic | 0 | 0 | 0 |
| rs367797185 | pathogenic | 1 | 1 | 0 |
| rs12709426 | benign | 0 | 1 | 0 |
| rs2228547 | benign | 0 | 0 | 0 |
| rs111033546 | pathogenic | 0 | 0 | 0 |
| rs111033553 | pathogenic | 0 | 0 | 0 |
| rs111033554 | pathogenic | 1 | 1 | 1 |
| rs111033544 | pathogenic | 1 | 1 | 1 |
| rs111033555 | pathogenic | 1 | 1 | 1 |
| rs111033545 | pathogenic | 1 | 1 | 1 |
| rs111033549 | pathogenic | 0 | 1 | 0 |
| rs111033552 | pathogenic | 1 | 1 | 1 |
| rs752204035 | benign | 0 | 0 | 0 |
| rs11569534 | benign | 0 | 0 | 0 |
| rs28937571 | pathogenic | 1 | 1 | 1 |
| rs104894215 | pathogenic | 1 | 1 | 1 |
| rs3888798 | benign | 1 | 1 | 1 |
| rs764263034 | benign | 1 | 1 | 1 |
| rs1800973 | benign | 1 | 1 | 1 |
| rs367623154 | benign | 1 | 1 | 1 |
| rs201230446 | pathogenic | 1 | 1 | 1 |
| rs267606766 | pathogenic | 0 | 1 | 0 |
| rs267606769 | pathogenic | 0 | 1 | 0 |
| rs267606767 | pathogenic | 1 | 1 | 0 |
| rs267606768 | pathogenic | 0 | 1 | 0 |
| rs61733129 | benign | 0 | 1 | 0 |
| rs201947120 | pathogenic | 1 | 1 | 1 |
| rs283413 | benign | 0 | 1 | 0 |
| rs36119840 | benign | 0 | 0 | 0 |
| rs281865419 | pathogenic | 1 | 1 | 1 |
| rs137852937 | pathogenic | 0 | 1 | 0 |
| rs2230709 | benign | 0 | 1 | 0 |
| rs6695033 | benign | 0 | 1 | 0 |
| rs180177032 | pathogenic | 1 | 1 | 1 |
| rs180177033 | pathogenic | 1 | 1 | 1 |
| rs121913351 | pathogenic | 1 | 1 | 1 |
| rs397507473 | pathogenic | 1 | 1 | 1 |
| rs121913355 | pathogenic | 1 | 1 | 1 |
| rs121913357 | pathogenic | 1 | 1 | 1 |
| rs180177036 | pathogenic | 1 | 1 | 1 |
| rs180177039 | pathogenic | 1 | 1 | 1 |
| rs180177038 | pathogenic | 1 | 1 | 1 |
| rs180177040 | pathogenic | 1 | 1 | 1 |

| | | | | |
|---|---|---|---|---|
| rs121913338 | pathogenic | 1 | 1 | 1 |
| rs121913366 | pathogenic | 1 | 1 | 1 |
| rs121913369 | pathogenic | 1 | 1 | 1 |
| rs113488022 | pathogenic | 1 | 1 | 1 |
| rs121913364 | pathogenic | 1 | 1 | 1 |
| rs180177042 | pathogenic | 1 | 1 | 1 |
| rs200461617 | benign | 0 | 0 | 0 |
| rs141609745 | benign | 0 | 0 | 0 |
| rs104894383 | pathogenic | 1 | 1 | 1 |
| rs104894384 | pathogenic | 1 | 1 | 1 |
| rs104894381 | pathogenic | 1 | 1 | 0 |
| rs147710408 | benign | 1 | 1 | 1 |
| rs77357563 | benign | 1 | 1 | 1 |
| rs104894378 | pathogenic | 0 | 0 | 0 |
| rs104894382 | pathogenic | 0 | 0 | 0 |
| rs137853292 | pathogenic | 1 | 1 | 1 |
| rs587778834 | pathogenic | 0 | 0 | 0 |
| rs137853294 | pathogenic | 1 | 1 | 1 |
| rs551747882 | benign | 0 | 0 | 0 |
| rs3092903 | benign | 1 | 1 | 1 |
| rs137853296 | pathogenic | 0 | 0 | 0 |
| rs398122917 | pathogenic | 0 | 1 | 0 |
| rs41304992 | benign | 0 | 0 | 0 |
| rs121912708 | pathogenic | 1 | 1 | 1 |
| rs140102105 | pathogenic | 1 | 1 | 1 |
| rs376917645 | pathogenic | 1 | 1 | 0 |
| rs372660425 | pathogenic | 1 | 1 | 0 |
| rs555896752 | pathogenic | 1 | 1 | 1 |
| rs777829351 | pathogenic | 1 | 1 | 0 |
| rs121912711 | pathogenic | 1 | 1 | 0 |
| rs754449549 | pathogenic | 0 | 1 | 0 |
| rs141701364 | benign | 0 | 1 | 0 |
| rs761011399 | benign | 1 | 1 | 1 |
| rs121912710 | benign | 1 | 1 | 0 |
| rs121912707 | pathogenic | 1 | 1 | 0 |
| rs747597620 | pathogenic | 0 | 1 | 0 |
| rs186558364 | pathogenic | 1 | 1 | 1 |
| rs556400964 | pathogenic | 1 | 1 | 1 |
| rs61757684 | benign | 1 | 1 | 1 |
| rs121917766 | pathogenic | 1 | 1 | 1 |
| rs121918248 | pathogenic | 0 | 0 | 0 |
| rs398123278 | pathogenic | 0 | 0 | 0 |
| rs121918251 | pathogenic | 1 | 1 | 1 |
| rs727504022 | pathogenic | 0 | 0 | 0 |
| rs190834116 | pathogenic | 1 | 1 | 1 |
| rs121918249 | pathogenic | 1 | 1 | 0 |
| rs121918257 | pathogenic | 1 | 1 | 0 |
| rs142068898 | benign | 0 | 0 | 0 |
| rs757000253 | pathogenic | 0 | 0 | 0 |
| rs200908035 | pathogenic | 1 | 1 | 0 |
| rs760782399 | pathogenic | 0 | 0 | 0 |
| rs778702777 | pathogenic | 0 | 0 | 0 |
| rs121918258 | pathogenic | 1 | 1 | 0 |
| rs121918256 | pathogenic | 0 | 0 | 0 |

| | | | | |
|---|---|---|---|---|
| rs200596762 | pathogenic | 1 | 1 | 0 |
| rs547709692 | pathogenic | 0 | 0 | 0 |
| rs758577372 | pathogenic | 0 | 1 | 0 |
| rs564069299 | pathogenic | 1 | 1 | 1 |
| rs772552898 | pathogenic | 1 | 1 | 1 |
| rs368790885 | pathogenic | 1 | 1 | 1 |
| rs121918250 | pathogenic | 0 | 0 | 0 |
| rs533755473 | pathogenic | 1 | 1 | 1 |
| rs753288303 | pathogenic | 1 | 1 | 1 |
| rs2229385 | benign | 0 | 0 | 0 |
| rs1141321 | benign | 0 | 0 | 0 |
| rs753564352 | pathogenic | 0 | 0 | 0 |
| rs775593146 | pathogenic | 0 | 0 | 0 |
| rs121918254 | pathogenic | 0 | 0 | 0 |
| rs8589 | benign | 1 | 1 | 0 |
| rs777758903 | pathogenic | 0 | 0 | 0 |
| rs140600746 | pathogenic | 1 | 1 | 0 |
| rs121918255 | pathogenic | 1 | 1 | 0 |
| rs121918252 | pathogenic | 0 | 0 | 0 |
| rs779990936 | pathogenic | 1 | 1 | 0 |
| rs121912455 | pathogenic | 1 | 1 | 1 |
| rs121912459 | pathogenic | 1 | 1 | 1 |
| rs74315452 | pathogenic | 1 | 1 | 1 |
| rs121912451 | pathogenic | 1 | 1 | 1 |
| rs79006549 | pathogenic | 0 | 0 | 0 |
| rs28934889 | benign | 0 | 1 | 0 |
| rs111033557 | benign | 1 | 1 | 1 |
| rs28934597 | pathogenic | 1 | 1 | 1 |
| rs28934596 | pathogenic | 1 | 1 | 1 |
| rs28934595 | pathogenic | 1 | 1 | 1 |
| rs146519482 | pathogenic | 1 | 1 | 1 |
| rs111033563 | pathogenic | 1 | 1 | 1 |
| rs75385605 | benign | 1 | 1 | 1 |
| rs121908569 | pathogenic | 1 | 1 | 1 |
| rs35719940 | benign | 0 | 0 | 0 |
| rs121918664 | pathogenic | 0 | 0 | 0 |
| rs104893665 | pathogenic | 1 | 1 | 1 |
| rs104893666 | pathogenic | 1 | 1 | 1 |
| rs200845457 | benign | 1 | 1 | 1 |
| rs779204655 | pathogenic | 1 | 1 | 1 |
| rs121917747 | pathogenic | 1 | 1 | 1 |
| rs6042 | benign | 1 | 1 | 1 |
| rs104894380 | pathogenic | 0 | 0 | 0 |
| rs369083541 | pathogenic | 1 | 1 | 1 |
| rs145243532 | benign | 1 | 1 | 1 |
| rs121909500 | pathogenic | 1 | 1 | 1 |
| rs140993310 | benign | 1 | 1 | 1 |
| rs200837270 | pathogenic | 1 | 1 | 1 |
| rs121918484 | pathogenic | 1 | 1 | 1 |
| rs121918306 | pathogenic | 0 | 0 | 0 |
| rs118204089 | pathogenic | 1 | 1 | 0 |
| rs118204083 | pathogenic | 1 | 1 | 0 |
| rs201953584 | pathogenic | 1 | 1 | 1 |
| rs74315362 | pathogenic | 1 | 1 | 1 |

| | | | | |
|---|---|---|---|---|
| rs118204084 | pathogenic | 1 | 1 | 0 |
| rs118204085 | pathogenic | 1 | 1 | 1 |
| rs113403872 | pathogenic | 1 | 1 | 0 |
| rs1052176 | benign | 1 | 1 | 1 |
| rs2031640 | benign | 0 | 0 | 0 |
| rs104894230 | pathogenic | 1 | 1 | 1 |
| rs104894229 | pathogenic | 1 | 1 | 1 |
| rs104894228 | pathogenic | 1 | 1 | 1 |
| rs104894226 | pathogenic | 1 | 1 | 1 |
| rs121917757 | pathogenic | 1 | 1 | 1 |
| rs121917758 | pathogenic | 1 | 1 | 1 |
| rs28933406 | pathogenic | 1 | 1 | 1 |
| rs121917756 | pathogenic | 1 | 1 | 1 |
| rs104894227 | pathogenic | 1 | 1 | 1 |
| rs369106578 | pathogenic | 1 | 1 | 1 |
| rs397517142 | benign | 1 | 1 | 1 |
| rs104894231 | pathogenic | 1 | 1 | 1 |
| rs121917759 | pathogenic | 1 | 1 | 1 |
| rs104893827 | pathogenic | 0 | 0 | 0 |
| rs104893826 | pathogenic | 1 | 1 | 1 |
| rs5030802 | pathogenic | 1 | 1 | 1 |
| rs5030827 | pathogenic | 1 | 1 | 0 |
| rs119103277 | pathogenic | 1 | 1 | 1 |
| rs5030808 | pathogenic | 0 | 1 | 0 |
| rs5030809 | pathogenic | 1 | 1 | 1 |
| rs765978945 | pathogenic | 1 | 1 | 1 |
| rs104893824 | pathogenic | 1 | 1 | 1 |
| rs104893831 | pathogenic | 0 | 1 | 0 |
| rs104893830 | pathogenic | 0 | 1 | 0 |
| rs119103278 | pathogenic | 0 | 1 | 0 |
| rs5030818 | pathogenic | 1 | 1 | 1 |
| rs397516444 | pathogenic | 1 | 1 | 1 |
| rs28940297 | pathogenic | 1 | 1 | 1 |
| rs267607170 | pathogenic | 1 | 1 | 1 |
| rs104893825 | pathogenic | 1 | 1 | 1 |
| rs5030820 | pathogenic | 0 | 1 | 0 |
| rs5030821 | pathogenic | 0 | 1 | 0 |
| rs5030823 | pathogenic | 1 | 1 | 1 |
| rs5030824 | pathogenic | 1 | 1 | 1 |
| rs28940301 | pathogenic | 0 | 1 | 0 |
| rs28940300 | pathogenic | 0 | 1 | 0 |
| rs28940298 | pathogenic | 0 | 1 | 0 |
| rs200019083 | benign | 0 | 0 | 0 |
| rs759748655 | pathogenic | 0 | 0 | 0 |
| rs121917739 | pathogenic | 1 | 1 | 1 |
| rs75353611 | pathogenic | 1 | 1 | 1 |
| rs79744198 | benign | 1 | 1 | 1 |
| rs79228041 | pathogenic | 1 | 1 | 1 |
| rs80106970 | benign | 1 | 1 | 1 |
| rs375353223 | benign | 0 | 0 | 0 |
| rs56311081 | benign | 0 | 0 | 0 |
| rs55985569 | benign | 1 | 1 | 1 |
| rs45483396 | benign | 0 | 0 | 0 |
| rs35284565 | benign | 1 | 1 | 1 |

| | | | | |
|---|---|---|---|---|
| rs34349517 | benign | 0 | 0 | 0 |
| rs35776110 | benign | 0 | 0 | 0 |
| rs33917957 | benign | 1 | 1 | 1 |
| rs587778445 | benign | 0 | 0 | 0 |
| rs34624984 | benign | 0 | 0 | 0 |
| rs701753 | benign | 0 | 0 | 0 |
| rs855314 | benign | 0 | 0 | 0 |
| rs121918371 | pathogenic | 1 | 1 | 0 |
| rs398122912 | pathogenic | 0 | 1 | 0 |
| rs1126728 | benign | 0 | 0 | 0 |
| rs587777404 | pathogenic | 1 | 1 | 0 |
| rs11208257 | benign | 0 | 0 | 0 |
| rs6676290 | benign | 0 | 0 | 0 |
| rs104894118 | pathogenic | 0 | 0 | 0 |
| rs121918655 | pathogenic | 1 | 1 | 1 |
| rs104894120 | pathogenic | 1 | 1 | 1 |
| rs137852238 | pathogenic | 0 | 0 | 0 |
| rs137852237 | pathogenic | 0 | 0 | 0 |
| rs137852499 | pathogenic | 1 | 1 | 1 |
| rs137852480 | pathogenic | 1 | 1 | 1 |
| rs137852494 | pathogenic | 1 | 1 | 1 |
| rs137852502 | pathogenic | 1 | 1 | 1 |
| rs137852500 | pathogenic | 1 | 1 | 1 |
| rs137852487 | pathogenic | 1 | 1 | 1 |
| rs137852481 | pathogenic | 1 | 1 | 1 |
| rs137852501 | pathogenic | 0 | 0 | 0 |
| rs137852478 | pathogenic | 1 | 1 | 1 |
| rs137852485 | pathogenic | 1 | 1 | 1 |
| rs137852482 | pathogenic | 1 | 1 | 0 |
| rs137852483 | pathogenic | 0 | 1 | 0 |
| rs137852477 | pathogenic | 1 | 1 | 0 |
| rs137852496 | pathogenic | 1 | 1 | 0 |
| rs137852484 | pathogenic | 0 | 1 | 0 |
| rs137852498 | pathogenic | 1 | 1 | 0 |
| rs137852497 | pathogenic | 1 | 1 | 0 |
| rs137852493 | pathogenic | 1 | 1 | 0 |
| rs137852504 | pathogenic | 1 | 1 | 1 |
| rs267606863 | pathogenic | 1 | 1 | 1 |
| rs137852486 | pathogenic | 1 | 1 | 1 |
| rs137852479 | pathogenic | 1 | 1 | 1 |
| rs112122950 | benign | 0 | 0 | 0 |
| rs72546648 | benign | 1 | 1 | 1 |
| rs72552291 | pathogenic | 0 | 0 | 0 |
| rs79184941 | pathogenic | 1 | 1 | 1 |
| rs77543610 | pathogenic | 1 | 1 | 1 |
| rs121918505 | pathogenic | 1 | 1 | 1 |
| rs776587763 | pathogenic | 1 | 1 | 1 |
| rs121918497 | pathogenic | 1 | 1 | 1 |
| rs121918499 | pathogenic | 1 | 1 | 1 |
| rs121918501 | pathogenic | 1 | 1 | 1 |
| rs121918500 | pathogenic | 0 | 0 | 0 |
| rs121918504 | pathogenic | 1 | 1 | 1 |
| rs121918510 | pathogenic | 1 | 1 | 1 |
| rs121918493 | pathogenic | 1 | 1 | 1 |

| | | | | |
|---|---|---|---|---|
| rs121918489 | pathogenic | 0 | 0 | 0 |
| rs121918495 | pathogenic | 1 | 1 | 1 |
| rs121918488 | pathogenic | 0 | 0 | 0 |
| rs121918496 | pathogenic | 0 | 0 | 0 |
| rs121918492 | pathogenic | 1 | 1 | 1 |
| rs121918494 | pathogenic | 1 | 1 | 1 |
| rs121918502 | pathogenic | 1 | 1 | 1 |
| rs121918490 | pathogenic | 1 | 1 | 1 |
| rs72466451 | pathogenic | 1 | 1 | 1 |
| rs66468541 | pathogenic | 0 | 0 | 0 |
| rs137852662 | pathogenic | 0 | 0 | 0 |
| rs137852663 | pathogenic | 0 | 0 | 0 |
| rs606231407 | pathogenic | 0 | 0 | 0 |
| rs137852233 | pathogenic | 1 | 1 | 1 |
| rs112306225 | benign | 0 | 0 | 0 |
| rs387906808 | pathogenic | 1 | 1 | 0 |
| rs730882094 | pathogenic | 1 | 1 | 1 |
| rs373869746 | pathogenic | 1 | 1 | 1 |
| rs777524402 | pathogenic | 1 | 1 | 1 |
| rs769737896 | pathogenic | 0 | 1 | 0 |
| rs13306515 | benign | 1 | 1 | 0 |
| rs746982741 | pathogenic | 0 | 0 | 0 |
| rs2230200 | benign | 1 | 1 | 1 |
| rs1047286 | benign | 0 | 0 | 0 |
| rs141737564 | benign | 0 | 0 | 0 |
| rs774826179 | benign | 0 | 0 | 0 |
| rs11569422 | benign | 1 | 1 | 1 |
| rs146613648 | benign | 0 | 0 | 0 |
| rs144432231 | benign | 0 | 0 | 0 |
| rs140655115 | benign | 0 | 0 | 0 |
| rs387906837 | pathogenic | 1 | 1 | 0 |
| rs121909076 | pathogenic | 1 | 1 | 0 |
| rs121909077 | pathogenic | 0 | 0 | 0 |
| rs121909074 | pathogenic | 0 | 0 | 0 |
| rs137853330 | pathogenic | 1 | 1 | 1 |
| rs137853331 | pathogenic | 1 | 1 | 1 |
| rs137853332 | pathogenic | 1 | 1 | 1 |
| rs28931586 | pathogenic | 1 | 1 | 0 |
| rs11569321 | benign | 1 | 1 | 1 |
| rs119486097 | benign | 0 | 0 | 0 |
| rs1799807 | pathogenic | 1 | 1 | 1 |
| rs201820739 | pathogenic | 1 | 1 | 0 |
| rs114706984 | pathogenic | 1 | 1 | 0 |
| rs140080572 | pathogenic | 1 | 1 | 0 |
| rs28933389 | pathogenic | 0 | 1 | 0 |
| rs16849700 | benign | 1 | 1 | 0 |
| rs115624085 | pathogenic | 1 | 1 | 1 |
| rs747196387 | pathogenic | 1 | 1 | 1 |
| rs104893684 | pathogenic | 1 | 1 | 0 |
| rs121918557 | pathogenic | 1 | 1 | 0 |
| rs121918556 | pathogenic | 1 | 1 | 1 |
| rs398124571 | pathogenic | 1 | 1 | 1 |
| rs776631396 | pathogenic | 0 | 1 | 0 |
| rs398124576 | pathogenic | 1 | 1 | 1 |

| | | | | |
|---|---|---|---|---|
| rs398124581 | pathogenic | 1 | 1 | 1 |
| rs398124582 | pathogenic | 1 | 1 | 1 |
| rs149766077 | pathogenic | 0 | 1 | 0 |
| rs79761867 | pathogenic | 0 | 1 | 0 |
| rs148905512 | pathogenic | 0 | 1 | 0 |
| rs121965004 | pathogenic | 1 | 1 | 1 |
| rs386834233 | pathogenic | 0 | 1 | 0 |
| rs398124598 | pathogenic | 0 | 1 | 0 |
| rs398124603 | pathogenic | 1 | 1 | 1 |
| rs546716073 | benign | 1 | 1 | 1 |
| rs200310662 | benign | 1 | 1 | 1 |
| rs115494727 | benign | 1 | 1 | 1 |
| rs781513008 | benign | 0 | 0 | 0 |
| rs587776398 | benign | 1 | 1 | 0 |
| rs35288908 | benign | 1 | 1 | 1 |
| rs150634562 | pathogenic | 1 | 1 | 0 |
| rs142948132 | pathogenic | 1 | 1 | 1 |
| rs132630321 | pathogenic | 1 | 1 | 1 |
| rs132630317 | pathogenic | 1 | 1 | 1 |
| rs132630320 | pathogenic | 0 | 0 | 0 |
| rs781394318 | pathogenic | 1 | 1 | 1 |
| rs111033624 | pathogenic | 1 | 1 | 1 |
| rs111033630 | pathogenic | 1 | 1 | 1 |
| rs111033627 | pathogenic | 1 | 1 | 1 |
| rs193922287 | pathogenic | 1 | 1 | 1 |
| rs759072800 | pathogenic | 0 | 1 | 0 |
| rs104894010 | pathogenic | 0 | 0 | 0 |
| rs193922296 | pathogenic | 0 | 0 | 0 |
| rs104894006 | pathogenic | 1 | 1 | 1 |
| rs193922311 | pathogenic | 1 | 1 | 0 |
| rs80356654 | pathogenic | 1 | 1 | 0 |
| rs144723656 | benign | 1 | 1 | 1 |
| rs148311934 | pathogenic | 1 | 1 | 0 |
| rs80356655 | pathogenic | 1 | 1 | 1 |
| rs769268803 | pathogenic | 1 | 1 | 0 |
| rs104894008 | pathogenic | 0 | 1 | 0 |
| rs104894005 | pathogenic | 1 | 1 | 1 |
| rs104894009 | pathogenic | 1 | 1 | 0 |
| rs193922340 | pathogenic | 0 | 0 | 0 |
| rs104894016 | pathogenic | 0 | 1 | 0 |
| rs193921338 | pathogenic | 0 | 0 | 0 |
| rs193922272 | pathogenic | 1 | 1 | 1 |
| rs104894012 | pathogenic | 1 | 1 | 0 |
| rs193922285 | pathogenic | 1 | 1 | 1 |
| rs104894336 | pathogenic | 1 | 1 | 1 |
| rs28931580 | pathogenic | 1 | 1 | 1 |
| rs104894326 | pathogenic | 1 | 1 | 1 |
| rs104894331 | pathogenic | 0 | 0 | 0 |
| rs104894338 | pathogenic | 0 | 0 | 0 |
| rs104894333 | pathogenic | 1 | 1 | 1 |
| rs104894330 | pathogenic | 1 | 1 | 1 |
| rs143886391 | benign | 1 | 1 | 1 |
| rs104894335 | pathogenic | 1 | 1 | 1 |
| rs104894337 | pathogenic | 0 | 0 | 0 |

| | | | | |
|---|---|---|---|---|
| rs104894328 | pathogenic | 0 | 0 | 0 |
| rs104894341 | pathogenic | 1 | 1 | 0 |
| rs104894329 | pathogenic | 0 | 0 | 0 |
| rs137852850 | pathogenic | 1 | 1 | 1 |
| rs137852854 | pathogenic | 0 | 1 | 0 |
| rs137852849 | pathogenic | 0 | 1 | 0 |
| rs137852848 | pathogenic | 1 | 1 | 1 |
| rs137852852 | pathogenic | 0 | 1 | 0 |
| rs137852853 | pathogenic | 0 | 1 | 0 |
| rs137852847 | pathogenic | 0 | 1 | 0 |
| rs137852846 | pathogenic | 0 | 1 | 0 |
| rs137852844 | pathogenic | 0 | 1 | 0 |
| rs144411158 | pathogenic | 0 | 0 | 0 |
| rs2302234 | benign | 0 | 0 | 0 |
| rs139620139 | pathogenic | 0 | 0 | 0 |
| rs146689929 | benign | 0 | 0 | 0 |
| rs373328681 | pathogenic | 1 | 1 | 0 |
| rs387906620 | pathogenic | 1 | 1 | 1 |
| rs1799983 | benign | 0 | 1 | 0 |
| rs28999111 | pathogenic | 1 | 1 | 1 |
| rs104894544 | pathogenic | 1 | 1 | 1 |
| rs104894546 | pathogenic | 1 | 1 | 1 |
| rs28999112 | pathogenic | 0 | 1 | 0 |
| rs104894545 | pathogenic | 1 | 1 | 1 |
| rs104894547 | pathogenic | 1 | 1 | 1 |
| rs74315310 | pathogenic | 0 | 0 | 0 |
| rs267606852 | pathogenic | 0 | 1 | 0 |
| rs137853586 | pathogenic | 0 | 1 | 0 |
| rs137853582 | pathogenic | 1 | 1 | 1 |
| rs61754634 | pathogenic | 1 | 1 | 1 |
| rs137853587 | pathogenic | 0 | 1 | 0 |
| rs267606851 | pathogenic | 1 | 1 | 1 |
| rs267606853 | pathogenic | 1 | 1 | 1 |
| rs137853584 | pathogenic | 1 | 1 | 1 |
| rs137853585 | pathogenic | 1 | 1 | 1 |
| rs460184 | pathogenic | 1 | 1 | 1 |
| rs104893751 | pathogenic | 0 | 1 | 0 |
| rs77358162 | benign | 0 | 0 | 0 |
| rs387906936 | pathogenic | 0 | 0 | 0 |
| rs769525399 | pathogenic | 0 | 0 | 0 |
| rs371271054 | pathogenic | 1 | 1 | 0 |
| rs118204000 | pathogenic | 0 | 0 | 0 |
| rs138568622 | pathogenic | 1 | 1 | 1 |
| rs34509359 | benign | 0 | 0 | 0 |
| rs104894130 | pathogenic | 1 | 1 | 0 |
| rs281865424 | pathogenic | 0 | 0 | 0 |
| rs121912778 | pathogenic | 1 | 1 | 0 |
| rs771055145 | pathogenic | 1 | 1 | 0 |
| rs776174514 | pathogenic | 1 | 1 | 0 |
| rs545986367 | pathogenic | 1 | 1 | 1 |
| rs570768038 | benign | 1 | 1 | 1 |
| rs74315442 | pathogenic | 1 | 1 | 1 |
| rs79716074 | benign | 1 | 1 | 1 |
| rs748486078 | pathogenic | 1 | 1 | 1 |

| | | | | |
|---|---|---|---|---|
| rs2397084 | benign | 1 | 1 | 1 |
| rs11465553 | benign | 1 | 1 | 1 |
| rs144576902 | benign | 1 | 1 | 1 |
| rs763780 | benign | 1 | 1 | 1 |
| rs267606870 | pathogenic | 1 | 1 | 0 |
| rs121913502 | pathogenic | 1 | 1 | 0 |
| rs118101777 | benign | 1 | 1 | 1 |
| rs118053940 | benign | 0 | 0 | 0 |
| rs368771578 | benign | 1 | 1 | 1 |
| rs11540654 | pathogenic | 1 | 1 | 1 |
| rs121912658 | pathogenic | 1 | 1 | 1 |
| rs28934873 | pathogenic | 0 | 1 | 0 |
| rs780442292 | pathogenic | 1 | 1 | 1 |
| rs758781593 | benign | 1 | 1 | 1 |
| rs28934875 | pathogenic | 1 | 1 | 1 |
| rs28934874 | pathogenic | 1 | 1 | 0 |
| rs587782705 | pathogenic | 1 | 1 | 1 |
| rs587780068 | pathogenic | 1 | 1 | 1 |
| rs587782144 | pathogenic | 1 | 1 | 1 |
| rs138729528 | pathogenic | 1 | 1 | 1 |
| rs28934578 | pathogenic | 1 | 1 | 1 |
| rs397514495 | pathogenic | 1 | 1 | 1 |
| rs121912665 | pathogenic | 1 | 1 | 1 |
| rs397516435 | pathogenic | 1 | 1 | 1 |
| rs587778720 | pathogenic | 1 | 1 | 1 |
| rs397516436 | pathogenic | 1 | 1 | 1 |
| rs587781386 | benign | 1 | 1 | 1 |
| rs121912666 | pathogenic | 1 | 1 | 0 |
| rs587780073 | pathogenic | 1 | 1 | 1 |
| rs587782664 | pathogenic | 1 | 1 | 1 |
| rs730882005 | pathogenic | 1 | 1 | 1 |
| rs28934573 | pathogenic | 1 | 1 | 1 |
| rs121912655 | pathogenic | 1 | 1 | 1 |
| rs28934575 | pathogenic | 1 | 1 | 1 |
| rs121912656 | pathogenic | 1 | 1 | 1 |
| rs11540652 | pathogenic | 1 | 1 | 1 |
| rs121912651 | pathogenic | 1 | 1 | 1 |
| rs730882027 | pathogenic | 1 | 1 | 1 |
| rs121912653 | pathogenic | 1 | 1 | 1 |
| rs121912652 | pathogenic | 1 | 1 | 1 |
| rs121912657 | pathogenic | 0 | 1 | 0 |
| rs121913343 | pathogenic | 1 | 1 | 1 |
| rs28934576 | pathogenic | 1 | 1 | 1 |
| rs763098116 | pathogenic | 1 | 1 | 1 |
| rs121912660 | pathogenic | 1 | 1 | 1 |
| rs28934574 | pathogenic | 1 | 1 | 1 |
| rs371409680 | pathogenic | 1 | 1 | 1 |
| rs121912667 | pathogenic | 1 | 1 | 1 |
| rs121912663 | pathogenic | 1 | 1 | 1 |
| rs201744589 | pathogenic | 0 | 0 | 0 |
| rs33926449 | benign | 0 | 0 | 0 |
| rs74315302 | pathogenic | 0 | 1 | 0 |
| rs202040269 | pathogenic | 1 | 1 | 1 |
| rs375958814 | pathogenic | 1 | 1 | 1 |

| | | | | |
|---|---|---|---|---|
| rs121909624 | pathogenic | 1 | 1 | 1 |
| rs121909621 | pathogenic | 1 | 1 | 1 |
| rs121909622 | pathogenic | 1 | 1 | 0 |
| rs4220 | benign | 1 | 1 | 1 |
| rs147559466 | pathogenic | 1 | 1 | 1 |
| rs757434857 | pathogenic | 1 | 1 | 1 |
| rs398123072 | pathogenic | 1 | 1 | 1 |
| rs121434280 | pathogenic | 0 | 1 | 0 |
| rs398123074 | pathogenic | 0 | 1 | 0 |
| rs762114560 | pathogenic | 1 | 1 | 1 |
| rs121434283 | pathogenic | 1 | 1 | 0 |
| rs121434277 | pathogenic | 0 | 1 | 0 |
| rs745844469 | pathogenic | 1 | 1 | 1 |
| rs121434279 | pathogenic | 1 | 1 | 0 |
| rs121434278 | pathogenic | 1 | 1 | 1 |
| rs373715782 | pathogenic | 1 | 1 | 1 |
| rs200724875 | pathogenic | 1 | 1 | 1 |
| rs149678400 | pathogenic | 0 | 1 | 0 |
| rs770273135 | pathogenic | 1 | 1 | 1 |
| rs121434276 | pathogenic | 0 | 1 | 0 |
| rs121434281 | pathogenic | 1 | 1 | 0 |
| rs768884003 | pathogenic | 1 | 1 | 1 |
| rs201375579 | pathogenic | 0 | 0 | 0 |
| rs121434274 | pathogenic | 1 | 1 | 0 |
| rs121434282 | pathogenic | 1 | 1 | 0 |
| rs779759347 | pathogenic | 0 | 1 | 0 |
| rs77931234 | pathogenic | 1 | 1 | 1 |
| rs148207467 | pathogenic | 1 | 1 | 1 |
| rs766140986 | pathogenic | 1 | 1 | 1 |
| rs121434275 | pathogenic | 1 | 1 | 1 |
| rs748110745 | pathogenic | 1 | 1 | 1 |
| rs104894274 | pathogenic | 1 | 1 | 1 |
| rs104894273 | pathogenic | 0 | 1 | 0 |
| rs104894278 | pathogenic | 1 | 1 | 1 |
| rs104894275 | pathogenic | 1 | 1 | 1 |
| rs104894277 | pathogenic | 1 | 1 | 1 |
| rs104894276 | pathogenic | 0 | 0 | 0 |
| rs104894280 | pathogenic | 1 | 1 | 1 |
| rs145882709 | pathogenic | 1 | 1 | 1 |
| rs104894279 | pathogenic | 1 | 1 | 1 |
| rs80356670 | pathogenic | 1 | 1 | 1 |
| rs121918102 | pathogenic | 1 | 1 | 1 |
| rs80356671 | pathogenic | 1 | 1 | 1 |
| rs80356672 | pathogenic | 1 | 1 | 1 |
| rs758609113 | pathogenic | 1 | 1 | 0 |
| rs121918188 | pathogenic | 0 | 1 | 0 |
| rs121918189 | pathogenic | 1 | 1 | 0 |
| rs121918191 | pathogenic | 1 | 1 | 1 |
| rs1769259 | benign | 1 | 1 | 1 |
| rs140222720 | benign | 1 | 1 | 1 |
| rs2297085 | benign | 1 | 1 | 1 |
| rs200679026 | pathogenic | 1 | 1 | 1 |
| rs121918192 | pathogenic | 0 | 1 | 0 |
| rs145771944 | benign | 0 | 1 | 0 |

| | | | | |
|---|---|---|---|---|
| rs201508880 | pathogenic | 0 | 1 | 0 |
| rs17221854 | pathogenic | 0 | 0 | 0 |
| rs120074189 | pathogenic | 1 | 1 | 1 |
| rs120074190 | pathogenic | 1 | 1 | 1 |
| rs758600283 | benign | 1 | 1 | 1 |
| rs200305064 | pathogenic | 1 | 1 | 1 |
| rs121917891 | pathogenic | 1 | 1 | 1 |
| rs3772809 | benign | 0 | 1 | 0 |
| rs387906619 | pathogenic | 1 | 1 | 0 |
| rs2304681 | benign | 0 | 1 | 0 |
| rs121917849 | pathogenic | 0 | 0 | 0 |
| rs143010236 | pathogenic | 0 | 0 | 0 |
| rs121917850 | pathogenic | 1 | 1 | 0 |
| rs35916840 | pathogenic | 1 | 1 | 0 |
| rs28939688 | pathogenic | 1 | 1 | 1 |
| rs11258194 | benign | 1 | 1 | 1 |
| rs199840952 | pathogenic | 1 | 1 | 1 |
| rs121918159 | pathogenic | 1 | 1 | 1 |
| rs200396166 | pathogenic | 0 | 0 | 0 |
| rs757966746 | pathogenic | 0 | 1 | 0 |
| rs121918608 | pathogenic | 0 | 1 | 0 |
| rs74315401 | pathogenic | 1 | 1 | 1 |
| rs11538758 | pathogenic | 0 | 0 | 0 |
| rs74315402 | pathogenic | 1 | 1 | 1 |
| rs1799990 | pathogenic | 1 | 1 | 1 |
| rs74315410 | pathogenic | 1 | 1 | 1 |
| rs150351644 | benign | 1 | 1 | 1 |
| rs16990018 | benign | 1 | 1 | 1 |
| rs74315403 | pathogenic | 1 | 1 | 1 |
| rs74315408 | pathogenic | 1 | 1 | 1 |
| rs74315411 | pathogenic | 1 | 1 | 1 |
| rs74315413 | pathogenic | 1 | 1 | 1 |
| rs74315405 | pathogenic | 1 | 1 | 1 |
| rs28933385 | pathogenic | 1 | 1 | 1 |
| rs74315412 | pathogenic | 1 | 1 | 1 |
| rs74315407 | pathogenic | 1 | 1 | 1 |
| rs74315406 | pathogenic | 1 | 1 | 1 |
| rs11571117 | benign | 1 | 1 | 1 |
| rs121917742 | pathogenic | 1 | 1 | 1 |
| rs149782619 | pathogenic | 1 | 1 | 1 |
| rs397514492 | pathogenic | 1 | 1 | 1 |
| rs397514489 | pathogenic | 0 | 1 | 0 |
| rs397514493 | pathogenic | 1 | 1 | 1 |
| rs373849532 | pathogenic | 1 | 1 | 1 |
| rs121965048 | pathogenic | 1 | 1 | 1 |
| rs121965037 | pathogenic | 1 | 1 | 1 |
| rs121965060 | pathogenic | 0 | 1 | 0 |
| rs386833603 | pathogenic | 1 | 1 | 1 |
| rs121965038 | pathogenic | 1 | 1 | 1 |
| rs121965039 | pathogenic | 1 | 1 | 1 |
| rs121965040 | pathogenic | 1 | 1 | 1 |
| rs267606925 | pathogenic | 1 | 1 | 1 |
| rs121965056 | pathogenic | 0 | 1 | 0 |
| rs121965059 | pathogenic | 0 | 0 | 0 |

| | | | | |
|---|---|---|---|---|
| rs386833615 | pathogenic | 1 | 1 | 0 |
| rs121965051 | pathogenic | 1 | 1 | 0 |
| rs121965046 | pathogenic | 0 | 1 | 0 |
| rs121965052 | pathogenic | 0 | 1 | 0 |
| rs386833617 | pathogenic | 0 | 1 | 0 |
| rs386833618 | pathogenic | 1 | 1 | 0 |
| rs121965041 | pathogenic | 0 | 1 | 0 |
| rs121965042 | pathogenic | 0 | 1 | 0 |
| rs121965057 | pathogenic | 1 | 1 | 1 |
| rs386833621 | pathogenic | 1 | 1 | 1 |
| rs121965049 | pathogenic | 1 | 1 | 1 |
| rs386833623 | pathogenic | 1 | 1 | 1 |
| rs121965047 | pathogenic | 1 | 1 | 1 |
| rs121965053 | pathogenic | 0 | 1 | 0 |
| rs121965045 | pathogenic | 0 | 1 | 0 |
| rs121965054 | pathogenic | 0 | 1 | 0 |
| rs121965043 | pathogenic | 0 | 1 | 0 |
| rs386833598 | pathogenic | 0 | 0 | 0 |
| rs137854545 | pathogenic | 1 | 1 | 1 |
| rs137854544 | pathogenic | 1 | 1 | 1 |
| rs104894455 | pathogenic | 0 | 1 | 0 |
| rs1049564 | benign | 0 | 1 | 0 |
| rs104894460 | pathogenic | 1 | 1 | 1 |
| rs104894453 | pathogenic | 1 | 1 | 1 |
| rs104894450 | pathogenic | 0 | 1 | 0 |
| rs104894454 | pathogenic | 0 | 1 | 0 |
| rs104894452 | pathogenic | 1 | 1 | 1 |
| rs104894451 | pathogenic | 0 | 1 | 0 |
| rs104894368 | pathogenic | 1 | 1 | 1 |
| rs730880950 | pathogenic | 1 | 1 | 1 |
| rs121909535 | pathogenic | 1 | 1 | 1 |
| rs121909537 | pathogenic | 1 | 1 | 1 |
| rs121909536 | pathogenic | 1 | 1 | 1 |
| rs121909542 | pathogenic | 1 | 1 | 1 |
| rs121909538 | pathogenic | 1 | 1 | 1 |
| rs121909539 | pathogenic | 1 | 1 | 1 |
| rs121909540 | pathogenic | 1 | 1 | 1 |
| rs17560 | benign | 1 | 1 | 1 |
| rs121909543 | pathogenic | 0 | 1 | 0 |
| rs121909544 | pathogenic | 1 | 1 | 1 |
| rs2307456 | benign | 0 | 1 | 0 |
| rs745778317 | pathogenic | 0 | 1 | 0 |
| rs759607901 | pathogenic | 0 | 1 | 0 |
| rs201188114 | benign | 0 | 0 | 0 |
| rs1800472 | benign | 0 | 0 | 0 |
| rs587780728 | benign | 1 | 1 | 1 |
| rs144386518 | benign | 1 | 1 | 1 |
| rs281865439 | pathogenic | 1 | 1 | 1 |
| rs140175796 | pathogenic | 1 | 1 | 1 |
| rs5030843 | pathogenic | 1 | 1 | 0 |
| rs75166491 | pathogenic | 1 | 1 | 0 |
| rs199475679 | pathogenic | 1 | 1 | 1 |
| rs281865440 | pathogenic | 1 | 1 | 1 |
| rs74486803 | pathogenic | 0 | 1 | 0 |

| | | | | |
|---|---|---|---|---|
| rs199475602 | pathogenic | 0 | 1 | 0 |
| rs5030844 | pathogenic | 1 | 1 | 0 |
| rs62514927 | pathogenic | 1 | 1 | 1 |
| rs62514928 | pathogenic | 1 | 1 | 1 |
| rs62516152 | pathogenic | 0 | 1 | 0 |
| rs62507348 | pathogenic | 1 | 1 | 1 |
| rs76687508 | pathogenic | 1 | 1 | 1 |
| rs62508730 | pathogenic | 1 | 1 | 1 |
| rs62508588 | pathogenic | 0 | 1 | 0 |
| rs5030846 | pathogenic | 0 | 1 | 0 |
| rs74503222 | pathogenic | 1 | 1 | 0 |
| rs5030847 | pathogenic | 1 | 1 | 0 |
| rs5030850 | pathogenic | 1 | 1 | 1 |
| rs778154939 | pathogenic | 0 | 1 | 0 |
| rs62514952 | pathogenic | 1 | 1 | 1 |
| rs78655458 | pathogenic | 1 | 1 | 1 |
| rs62508698 | pathogenic | 1 | 1 | 0 |
| rs5030851 | pathogenic | 1 | 1 | 0 |
| rs62642933 | pathogenic | 1 | 1 | 1 |
| rs5030853 | pathogenic | 1 | 1 | 1 |
| rs62642935 | pathogenic | 1 | 1 | 1 |
| rs199475615 | pathogenic | 0 | 0 | 0 |
| rs62516062 | pathogenic | 1 | 1 | 0 |
| rs62516092 | pathogenic | 0 | 0 | 0 |
| rs62516095 | pathogenic | 1 | 1 | 1 |
| rs62642937 | pathogenic | 1 | 1 | 1 |
| rs62516141 | pathogenic | 0 | 1 | 0 |
| rs5030856 | pathogenic | 1 | 1 | 1 |
| rs5030857 | pathogenic | 1 | 1 | 1 |
| rs5030858 | pathogenic | 1 | 1 | 1 |
| rs79931499 | pathogenic | 1 | 1 | 1 |
| rs5030860 | pathogenic | 1 | 1 | 1 |
| rs281865437 | pathogenic | 1 | 1 | 1 |
| rs62644499 | pathogenic | 1 | 1 | 1 |
| rs147689487 | benign | 0 | 0 | 0 |
| rs397515417 | pathogenic | 1 | 1 | 1 |
| rs121909332 | pathogenic | 1 | 1 | 1 |
| rs121909330 | pathogenic | 1 | 1 | 0 |
| rs121909329 | pathogenic | 1 | 1 | 0 |
| rs121909335 | pathogenic | 1 | 1 | 1 |
| rs387906789 | pathogenic | 1 | 1 | 1 |
| rs121909334 | pathogenic | 1 | 1 | 1 |
| rs121909331 | pathogenic | 1 | 1 | 1 |
| rs372487178 | pathogenic | 1 | 1 | 1 |
| rs387906937 | pathogenic | 1 | 1 | 0 |
| rs16845 | benign | 1 | 1 | 1 |
| rs672601313 | pathogenic | 1 | 1 | 1 |
| rs117805308 | benign | 0 | 0 | 0 |
| rs121434261 | pathogenic | 1 | 1 | 1 |
| rs141629512 | benign | 0 | 0 | 0 |
| rs199957176 | benign | 1 | 1 | 1 |
| rs2230461 | benign | 1 | 1 | 1 |
| rs764618040 | pathogenic | 1 | 1 | 1 |
| rs267606949 | pathogenic | 1 | 1 | 1 |

| | | | | |
|---|---|---|---|---|
| rs587777483 | pathogenic | 1 | 1 | 1 |
| rs267606947 | pathogenic | 1 | 1 | 1 |
| rs6941583 | benign | 0 | 0 | 0 |
| rs749597959 | benign | 1 | 1 | 1 |
| rs182340753 | benign | 1 | 1 | 1 |
| rs146678221 | benign | 0 | 0 | 0 |
| rs11539202 | benign | 0 | 0 | 0 |
| rs148723565 | benign | 0 | 0 | 0 |
| rs61752925 | benign | 1 | 1 | 1 |
| rs113309941 | pathogenic | 0 | 0 | 0 |
| rs121434520 | pathogenic | 1 | 1 | 1 |
| rs121434517 | pathogenic | 1 | 1 | 1 |
| rs267607027 | pathogenic | 1 | 1 | 1 |
| rs138603088 | pathogenic | 0 | 0 | 0 |
| rs121434516 | pathogenic | 1 | 1 | 0 |
| rs121434521 | pathogenic | 0 | 0 | 0 |
| rs145735112 | benign | 1 | 1 | 1 |
| rs121434519 | pathogenic | 1 | 1 | 1 |
| rs75326924 | pathogenic | 1 | 1 | 0 |
| rs142186404 | pathogenic | 0 | 0 | 0 |
| rs121918035 | pathogenic | 0 | 0 | 0 |
| rs374385878 | pathogenic | 1 | 1 | 0 |
| rs11557652 | benign | 0 | 0 | 0 |
| rs118203986 | pathogenic | 1 | 1 | 1 |
| rs141675330 | benign | 1 | 1 | 1 |
| rs104893927 | pathogenic | 0 | 0 | 0 |
| rs104893935 | pathogenic | 0 | 0 | 0 |
| rs104893933 | pathogenic | 1 | 1 | 0 |
| rs104893934 | pathogenic | 0 | 0 | 0 |
| rs145183203 | benign | 1 | 1 | 0 |
| rs2230806 | benign | 0 | 0 | 0 |
| rs9282541 | benign | 0 | 0 | 0 |
| rs559110055 | benign | 0 | 0 | 0 |
| rs200030513 | benign | 0 | 0 | 0 |
| rs9282543 | benign | 0 | 0 | 0 |
| rs147675550 | benign | 0 | 0 | 0 |
| rs137854496 | pathogenic | 0 | 0 | 0 |
| rs374190304 | benign | 0 | 0 | 0 |
| rs114620717 | benign | 0 | 0 | 0 |
| rs2066718 | benign | 0 | 0 | 0 |
| rs35819696 | benign | 0 | 0 | 0 |
| rs138880920 | benign | 0 | 0 | 0 |
| rs2066715 | benign | 0 | 0 | 0 |
| rs35207495 | benign | 0 | 0 | 0 |
| rs2066714 | benign | 0 | 0 | 0 |
| rs28937313 | pathogenic | 0 | 0 | 0 |
| rs28937314 | pathogenic | 0 | 0 | 0 |
| rs137854495 | pathogenic | 0 | 0 | 0 |
| rs140365800 | benign | 0 | 0 | 0 |
| rs13306073 | benign | 0 | 0 | 0 |
| rs28933692 | pathogenic | 0 | 0 | 0 |
| rs33918808 | benign | 0 | 0 | 0 |
| rs76881554 | benign | 0 | 0 | 0 |
| rs143180998 | benign | 0 | 0 | 0 |

| | | | | |
|---|---|---|---|---|
| rs137854500 | pathogenic | 0 | 0 | 0 |
| rs538614702 | benign | 0 | 0 | 0 |
| rs201879964 | benign | 0 | 0 | 0 |
| rs2230808 | benign | 0 | 0 | 0 |
| rs138422574 | benign | 0 | 0 | 0 |
| rs137854498 | pathogenic | 0 | 0 | 0 |
| rs150125857 | benign | 0 | 0 | 0 |
| rs146292819 | pathogenic | 0 | 0 | 0 |
| rs142688906 | benign | 0 | 0 | 0 |
| rs137854499 | pathogenic | 0 | 0 | 0 |
| rs34879708 | benign | 0 | 0 | 0 |
| rs121913028 | pathogenic | 1 | 1 | 1 |
| rs1041951 | benign | 1 | 1 | 0 |
| rs138714094 | benign | 1 | 1 | 0 |
| rs200650502 | benign | 0 | 1 | 0 |
| rs370708663 | pathogenic | 1 | 1 | 1 |
| rs118204040 | pathogenic | 0 | 1 | 0 |
| rs146899669 | pathogenic | 1 | 1 | 0 |
| rs118204039 | pathogenic | 0 | 1 | 0 |
| rs121909233 | pathogenic | 0 | 0 | 0 |
| rs121909225 | pathogenic | 1 | 1 | 0 |
| rs121909236 | pathogenic | 0 | 0 | 0 |
| rs398123317 | pathogenic | 0 | 0 | 0 |
| rs121909226 | pathogenic | 1 | 1 | 0 |
| rs121909238 | pathogenic | 1 | 1 | 0 |
| rs121909230 | pathogenic | 0 | 0 | 0 |
| rs121909237 | pathogenic | 0 | 0 | 0 |
| rs121909222 | pathogenic | 1 | 1 | 0 |
| rs121909223 | pathogenic | 1 | 1 | 0 |
| rs121909218 | pathogenic | 1 | 1 | 0 |
| rs121909229 | pathogenic | 1 | 1 | 0 |
| rs121909224 | pathogenic | 1 | 1 | 0 |
| rs121909241 | pathogenic | 0 | 0 | 0 |
| rs121909221 | pathogenic | 0 | 0 | 0 |
| rs121913294 | pathogenic | 0 | 0 | 0 |
| rs121909234 | pathogenic | 0 | 0 | 0 |
| rs121909240 | pathogenic | 0 | 0 | 0 |
| rs767623493 | benign | 0 | 0 | 0 |
| rs121909239 | pathogenic | 0 | 0 | 0 |
| rs120074145 | pathogenic | 1 | 1 | 1 |
| rs120074148 | pathogenic | 1 | 1 | 1 |
| rs762991875 | pathogenic | 0 | 0 | 0 |
| rs148639841 | pathogenic | 1 | 1 | 1 |
| rs199524907 | pathogenic | 1 | 1 | 1 |
| rs120074141 | pathogenic | 1 | 1 | 1 |
| rs120074144 | pathogenic | 0 | 1 | 0 |
| rs120074146 | pathogenic | 1 | 1 | 1 |
| rs120074147 | pathogenic | 0 | 1 | 0 |
| rs120074143 | pathogenic | 0 | 1 | 0 |
| rs120074140 | pathogenic | 1 | 1 | 0 |
| rs28937315 | pathogenic | 1 | 1 | 1 |
| rs76066357 | benign | 1 | 1 | 1 |
| rs137852911 | pathogenic | 0 | 1 | 0 |
| rs137852907 | pathogenic | 1 | 1 | 0 |

| | | | | |
|---|---|---|---|---|
| rs137852910 | pathogenic | 1 | 1 | 1 |
| rs537585211 | benign | 0 | 0 | 0 |
| rs202134424 | pathogenic | 0 | 0 | 0 |
| rs200930463 | pathogenic | 0 | 0 | 0 |
| rs587777241 | pathogenic | 1 | 1 | 0 |
| rs77563738 | pathogenic | 0 | 0 | 0 |
| rs750868279 | pathogenic | 1 | 1 | 0 |
| rs148936893 | pathogenic | 0 | 0 | 0 |
| rs146597836 | pathogenic | 0 | 0 | 0 |
| rs775440641 | pathogenic | 1 | 1 | 1 |
| rs770689762 | pathogenic | 0 | 1 | 0 |
| rs376785840 | pathogenic | 0 | 0 | 0 |
| rs137852531 | pathogenic | 0 | 0 | 0 |
| rs137852538 | pathogenic | 1 | 1 | 0 |
| rs137852529 | pathogenic | 0 | 1 | 0 |
| rs137852528 | pathogenic | 0 | 1 | 0 |
| rs137852535 | pathogenic | 1 | 1 | 0 |
| rs137853045 | pathogenic | 0 | 1 | 0 |
| rs137853046 | pathogenic | 1 | 1 | 1 |
| rs137853043 | pathogenic | 1 | 1 | 1 |
| rs137853044 | pathogenic | 1 | 1 | 1 |
| rs137853047 | pathogenic | 1 | 1 | 1 |
| rs267606990 | pathogenic | 1 | 1 | 1 |
| rs751437780 | pathogenic | 1 | 1 | 0 |
| rs121918461 | pathogenic | 1 | 1 | 1 |
| rs121918460 | pathogenic | 1 | 1 | 1 |
| rs121918459 | pathogenic | 1 | 1 | 0 |
| rs397507511 | pathogenic | 1 | 1 | 1 |
| rs121918454 | pathogenic | 1 | 1 | 1 |
| rs121918453 | pathogenic | 1 | 1 | 1 |
| rs121918462 | pathogenic | 1 | 1 | 0 |
| rs121918465 | pathogenic | 1 | 1 | 0 |
| rs121918464 | pathogenic | 1 | 1 | 0 |
| rs121918466 | pathogenic | 1 | 1 | 1 |
| rs397507521 | pathogenic | 1 | 1 | 1 |
| rs397507527 | pathogenic | 1 | 1 | 1 |
| rs777603059 | pathogenic | 0 | 1 | 0 |
| rs121918456 | pathogenic | 1 | 1 | 1 |
| rs121918463 | pathogenic | 1 | 1 | 1 |
| rs121918455 | pathogenic | 0 | 1 | 0 |
| rs28933386 | pathogenic | 0 | 1 | 0 |
| rs774356443 | benign | 1 | 1 | 1 |
| rs121918467 | pathogenic | 1 | 1 | 1 |
| rs121918468 | pathogenic | 1 | 1 | 1 |
| rs121918469 | pathogenic | 1 | 1 | 1 |
| rs397507539 | pathogenic | 1 | 1 | 0 |
| rs121918458 | pathogenic | 1 | 1 | 1 |
| rs121918470 | pathogenic | 1 | 1 | 1 |
| rs35289681 | benign | 0 | 1 | 0 |
| rs4302331 | benign | 0 | 1 | 0 |
| rs11544803 | pathogenic | 1 | 1 | 1 |
| rs116254156 | benign | 0 | 0 | 0 |
| rs104893909 | pathogenic | 1 | 1 | 1 |
| rs104893911 | pathogenic | 1 | 1 | 1 |

| | | | | |
|---|---|---|---|---|
| rs104893908 | pathogenic | 1 | 1 | 0 |
| rs258751 | benign | 1 | 1 | 1 |
| rs104893914 | pathogenic | 1 | 1 | 0 |
| rs121909727 | pathogenic | 1 | 1 | 1 |
| rs104893910 | pathogenic | 1 | 1 | 1 |
| rs121909726 | pathogenic | 1 | 1 | 1 |
| rs104893912 | pathogenic | 1 | 1 | 1 |
| rs138407155 | benign | 1 | 1 | 1 |
| rs574061789 | benign | 1 | 1 | 1 |
| rs121912718 | pathogenic | 1 | 1 | 1 |
| rs121912719 | pathogenic | 1 | 1 | 1 |
| rs140770089 | benign | 1 | 1 | 1 |
| rs28931573 | pathogenic | 1 | 1 | 1 |
| rs121912717 | pathogenic | 1 | 1 | 1 |
| rs376456598 | pathogenic | 0 | 0 | 0 |
| rs398123499 | pathogenic | 1 | 1 | 1 |
| rs773048903 | pathogenic | 1 | 1 | 0 |
| rs398123503 | pathogenic | 1 | 1 | 1 |
| rs375785084 | pathogenic | 0 | 1 | 0 |
| rs137852874 | pathogenic | 1 | 1 | 1 |
| rs373713279 | pathogenic | 1 | 1 | 1 |
| rs137852873 | pathogenic | 1 | 1 | 0 |
| rs367823977 | pathogenic | 1 | 1 | 1 |
| rs398123508 | pathogenic | 1 | 1 | 1 |
| rs137852871 | pathogenic | 1 | 1 | 1 |
| rs753698250 | pathogenic | 0 | 1 | 0 |
| rs182923857 | pathogenic | 1 | 1 | 1 |
| rs148090804 | benign | 1 | 1 | 1 |
| rs137852870 | pathogenic | 1 | 1 | 1 |
| rs267607131 | pathogenic | 1 | 1 | 0 |
| rs137852902 | pathogenic | 0 | 0 | 0 |
| rs61741646 | benign | 1 | 1 | 1 |
| rs137852905 | pathogenic | 0 | 0 | 0 |
| rs121918681 | pathogenic | 0 | 1 | 0 |
| rs8177238 | benign | 1 | 1 | 0 |
| rs5030776 | pathogenic | 1 | 1 | 1 |
| rs104894572 | pathogenic | 0 | 0 | 0 |
| rs104894576 | pathogenic | 0 | 0 | 0 |
| rs80084721 | pathogenic | 0 | 0 | 0 |
| rs104894146 | pathogenic | 0 | 1 | 0 |
| rs104894138 | pathogenic | 1 | 1 | 0 |
| rs104894153 | pathogenic | 1 | 1 | 0 |
| rs104894135 | pathogenic | 1 | 1 | 0 |
| rs104894147 | pathogenic | 1 | 1 | 0 |
| rs104894148 | pathogenic | 1 | 1 | 0 |
| rs104894154 | pathogenic | 1 | 1 | 0 |
| rs104894150 | pathogenic | 1 | 1 | 0 |
| rs104894136 | pathogenic | 1 | 1 | 0 |
| rs104894144 | pathogenic | 0 | 0 | 0 |
| rs104894137 | pathogenic | 0 | 0 | 0 |
| rs61754278 | pathogenic | 1 | 1 | 0 |
| rs104894149 | pathogenic | 1 | 1 | 0 |
| rs104894139 | pathogenic | 1 | 1 | 0 |
| rs104894142 | pathogenic | 0 | 0 | 0 |

| | | | | |
|---|---|---|---|---|
| rs104894143 | pathogenic | 0 | 0 | 0 |
| rs104894145 | pathogenic | 0 | 0 | 0 |
| rs121909545 | benign | 0 | 1 | 0 |
| rs782308462 | pathogenic | 1 | 1 | 1 |
| rs375002796 | pathogenic | 0 | 1 | 0 |
| rs201727685 | pathogenic | 0 | 0 | 0 |
| rs768050261 | pathogenic | 1 | 1 | 1 |
| rs780851340 | benign | 0 | 0 | 0 |
| rs139824241 | benign | 1 | 1 | 0 |
| rs757486575 | pathogenic | 1 | 1 | 0 |
| rs750831299 | pathogenic | 1 | 1 | 1 |
| rs767602961 | pathogenic | 1 | 1 | 1 |
| rs121913661 | pathogenic | 1 | 1 | 1 |
| rs777692271 | pathogenic | 1 | 1 | 1 |
| rs121913659 | pathogenic | 1 | 1 | 1 |
| rs587777533 | pathogenic | 1 | 1 | 1 |
| rs104894210 | pathogenic | 0 | 0 | 0 |
| rs104894216 | pathogenic | 1 | 1 | 1 |
| rs104894207 | pathogenic | 1 | 1 | 1 |
| rs104894208 | pathogenic | 0 | 1 | 0 |
| rs104894214 | pathogenic | 1 | 1 | 1 |
| rs104894211 | pathogenic | 1 | 1 | 1 |
| rs61749384 | pathogenic | 0 | 0 | 0 |
| rs61750071 | pathogenic | 1 | 1 | 1 |
| rs61750074 | pathogenic | 0 | 0 | 0 |
| rs11063988 | benign | 0 | 0 | 0 |
| rs216311 | benign | 0 | 0 | 0 |
| rs1800382 | benign | 1 | 1 | 1 |
| rs11063987 | benign | 1 | 1 | 1 |
| rs387906847 | pathogenic | 1 | 1 | 1 |
| rs73003466 | benign | 0 | 0 | 0 |
| rs149787233 | benign | 1 | 1 | 1 |
| rs144451841 | pathogenic | 1 | 1 | 1 |
| rs141638421 | pathogenic | 1 | 1 | 1 |
| rs121434497 | pathogenic | 0 | 0 | 0 |
| rs121434498 | pathogenic | 0 | 0 | 0 |
| rs267607230 | pathogenic | 1 | 1 | 0 |
| rs121434499 | pathogenic | 1 | 1 | 0 |
| rs546749336 | benign | 1 | 1 | 1 |
| rs138873805 | benign | 0 | 0 | 0 |
| rs730880522 | pathogenic | 0 | 0 | 0 |
| rs200371894 | benign | 0 | 0 | 0 |
| rs730880509 | benign | 0 | 0 | 0 |
| rs267606704 | pathogenic | 1 | 1 | 1 |
| rs267606706 | pathogenic | 0 | 0 | 0 |
| rs267606705 | pathogenic | 1 | 1 | 1 |
| rs267606707 | pathogenic | 0 | 0 | 0 |
| rs387906665 | pathogenic | 1 | 1 | 1 |
| rs267606708 | pathogenic | 1 | 1 | 1 |
| rs148368481 | benign | 0 | 0 | 0 |
| rs13204 | benign | 0 | 0 | 0 |
| rs140547520 | pathogenic | 1 | 1 | 1 |
| rs755234990 | benign | 1 | 1 | 0 |
| rs151184052 | benign | 0 | 0 | 0 |

| | | | | |
|---|---|---|---|---|
| rs3736358 | benign | 0 | 0 | 0 |
| rs3736360 | benign | 0 | 0 | 0 |
| rs1138469 | benign | 0 | 0 | 0 |
| rs74315351 | pathogenic | 0 | 1 | 0 |
| rs137853051 | pathogenic | 1 | 1 | 0 |
| rs74315353 | pathogenic | 0 | 1 | 0 |
| rs74315352 | pathogenic | 0 | 1 | 0 |
| rs74315354 | pathogenic | 1 | 1 | 1 |
| rs28938172 | pathogenic | 1 | 1 | 1 |
| rs52836744 | pathogenic | 0 | 0 | 0 |
| rs121913159 | pathogenic | 1 | 1 | 1 |
| rs121913145 | pathogenic | 1 | 1 | 1 |
| rs891087 | benign | 1 | 1 | 1 |
| rs9282757 | benign | 1 | 1 | 1 |
| rs397517404 | pathogenic | 1 | 1 | 1 |
| rs199918720 | benign | 1 | 1 | 0 |
| rs368299411 | benign | 1 | 1 | 0 |
| rs139399951 | benign | 1 | 1 | 0 |
| rs763981974 | benign | 1 | 1 | 0 |
| rs144242114 | benign | 0 | 0 | 0 |
| rs750965801 | benign | 0 | 0 | 0 |
| rs561310777 | benign | 0 | 0 | 0 |
| rs148185335 | benign | 0 | 0 | 0 |
| rs147742157 | benign | 0 | 0 | 0 |
| rs375664373 | pathogenic | 1 | 1 | 1 |
| rs777327250 | pathogenic | 0 | 0 | 0 |
| rs121908471 | pathogenic | 1 | 1 | 0 |
| rs2301612 | pathogenic | 1 | 1 | 0 |
| rs121908476 | pathogenic | 0 | 0 | 0 |
| rs36220239 | benign | 0 | 0 | 0 |
| rs121908473 | pathogenic | 0 | 0 | 0 |
| rs28647808 | benign | 0 | 0 | 0 |
| rs36090624 | benign | 0 | 0 | 0 |
| rs121908002 | pathogenic | 1 | 1 | 1 |
| rs4251545 | benign | 1 | 1 | 1 |
| rs78705382 | pathogenic | 0 | 0 | 0 |
| rs75184679 | pathogenic | 1 | 1 | 1 |
| rs74555752 | pathogenic | 0 | 0 | 0 |
| rs56320207 | benign | 1 | 1 | 1 |
| rs28730837 | pathogenic | 1 | 1 | 1 |
| rs119469014 | pathogenic | 1 | 1 | 1 |
| rs119468010 | pathogenic | 1 | 1 | 1 |
| rs35220041 | benign | 0 | 0 | 0 |
| rs762321137 | pathogenic | 1 | 1 | 1 |
| rs28931570 | pathogenic | 1 | 1 | 1 |
| rs709932 | benign | 1 | 1 | 1 |
| rs746531546 | benign | 1 | 1 | 1 |
| rs6647 | benign | 1 | 1 | 1 |
| rs141095970 | benign | 1 | 1 | 1 |
| rs143370956 | benign | 1 | 1 | 1 |
| rs28929474 | pathogenic | 1 | 1 | 1 |
| rs61761869 | pathogenic | 1 | 1 | 1 |
| rs1303 | benign | 1 | 1 | 1 |
| rs149447502 | benign | 0 | 0 | 0 |

| | | | | |
|---|---|---|---|---|
| rs267607180 | pathogenic | 1 | 1 | 1 |
| rs150304121 | benign | 0 | 0 | 0 |
| rs147630375 | benign | 1 | 1 | 1 |
| rs11539445 | benign | 1 | 1 | 1 |
| rs372098364 | pathogenic | 1 | 1 | 0 |
| rs760072286 | benign | 0 | 0 | 0 |
| rs745595833 | pathogenic | 1 | 1 | 1 |
| rs587777214 | pathogenic | 0 | 0 | 0 |
| rs150318528 | benign | 0 | 0 | 0 |
| rs137853254 | pathogenic | 1 | 1 | 1 |
| rs137853255 | pathogenic | 0 | 0 | 0 |
| rs137853253 | pathogenic | 0 | 0 | 0 |
| rs202166915 | benign | 0 | 0 | 0 |
| rs137853259 | pathogenic | 0 | 0 | 0 |
| rs2229137 | benign | 0 | 0 | 0 |
| rs137853258 | pathogenic | 1 | 1 | 0 |
| rs137853252 | pathogenic | 1 | 1 | 1 |
| rs137853256 | pathogenic | 0 | 0 | 0 |
| rs137853250 | pathogenic | 1 | 1 | 1 |
| rs121434615 | pathogenic | 0 | 0 | 0 |
| rs115014558 | benign | 1 | 1 | 1 |
| rs34116584 | benign | 1 | 1 | 1 |
| rs34885252 | benign | 1 | 1 | 1 |
| rs180177262 | pathogenic | 1 | 1 | 1 |
| rs180177162 | pathogenic | 1 | 1 | 1 |
| rs121908523 | pathogenic | 1 | 1 | 1 |
| rs180177170 | pathogenic | 1 | 1 | 1 |
| rs180177180 | pathogenic | 1 | 1 | 1 |
| rs767586362 | pathogenic | 1 | 1 | 1 |
| rs121908521 | pathogenic | 1 | 1 | 1 |
| rs180177203 | pathogenic | 1 | 1 | 1 |
| rs138025751 | pathogenic | 1 | 1 | 1 |
| rs180177210 | pathogenic | 0 | 1 | 0 |
| rs180177211 | pathogenic | 0 | 1 | 0 |
| rs121908524 | pathogenic | 1 | 1 | 0 |
| rs121908530 | pathogenic | 1 | 1 | 0 |
| rs180177227 | pathogenic | 0 | 1 | 0 |
| rs180177230 | pathogenic | 1 | 1 | 1 |
| rs121908529 | pathogenic | 0 | 1 | 0 |
| rs117195882 | benign | 1 | 1 | 0 |
| rs180177238 | pathogenic | 1 | 1 | 0 |
| rs34664134 | benign | 1 | 1 | 1 |
| rs180177246 | pathogenic | 0 | 1 | 0 |
| rs536352238 | pathogenic | 0 | 0 | 0 |
| rs121908520 | pathogenic | 1 | 1 | 0 |
| rs180177248 | pathogenic | 1 | 1 | 0 |
| rs180177252 | pathogenic | 1 | 1 | 1 |
| rs121908526 | pathogenic | 1 | 1 | 1 |
| rs121908525 | pathogenic | 1 | 1 | 1 |
| rs147106773 | benign | 1 | 1 | 1 |
| rs180177264 | pathogenic | 0 | 1 | 0 |
| rs146525143 | pathogenic | 1 | 1 | 1 |
| rs140992177 | benign | 1 | 1 | 1 |
| rs13408961 | benign | 0 | 1 | 0 |

| | | | | |
|---|---|---|---|---|
| rs4426527 | benign | 1 | 1 | 1 |
| rs3088308 | benign | 0 | 1 | 0 |
| rs4469829 | benign | 1 | 1 | 0 |
| rs104886003 | pathogenic | 1 | 1 | 1 |
| rs121913286 | pathogenic | 1 | 1 | 1 |
| rs121913279 | pathogenic | 1 | 1 | 1 |
| rs121434636 | pathogenic | 1 | 1 | 1 |
| rs755299132 | pathogenic | 1 | 1 | 1 |
| rs748318386 | pathogenic | 1 | 1 | 1 |
| rs387906757 | pathogenic | 1 | 1 | 1 |
| rs121965064 | pathogenic | 1 | 1 | 1 |
| rs28934608 | pathogenic | 0 | 0 | 0 |
| rs5967 | benign | 1 | 1 | 1 |
| rs773905328 | pathogenic | 0 | 0 | 0 |
| rs104894256 | pathogenic | 0 | 1 | 0 |
| rs28931612 | pathogenic | 0 | 1 | 0 |
| rs104894263 | pathogenic | 1 | 1 | 1 |
| rs607969 | benign | 1 | 1 | 0 |
| rs104894262 | pathogenic | 0 | 1 | 0 |
| rs104894268 | pathogenic | 1 | 1 | 1 |
| rs104894264 | pathogenic | 0 | 1 | 0 |
| rs104894259 | pathogenic | 1 | 1 | 0 |
| rs2959656 | benign | 0 | 0 | 0 |
| rs121913035 | pathogenic | 1 | 1 | 0 |
| rs139605474 | benign | 0 | 0 | 0 |
| rs3817552 | benign | 0 | 0 | 0 |
| rs150541115 | benign | 0 | 0 | 0 |
| rs562763010 | benign | 1 | 1 | 1 |
| rs200053119 | pathogenic | 0 | 0 | 0 |
| rs73826386 | benign | 0 | 0 | 0 |
| rs143655258 | benign | 0 | 0 | 0 |
| rs758857853 | pathogenic | 1 | 1 | 0 |
| rs2228380 | benign | 0 | 0 | 0 |
| rs141078733 | benign | 1 | 1 | 1 |
| rs55824172 | pathogenic | 1 | 1 | 1 |
| rs756751089 | pathogenic | 1 | 1 | 1 |
| rs121964990 | pathogenic | 0 | 0 | 0 |
| rs143555186 | benign | 0 | 0 | 0 |
| rs121964993 | pathogenic | 1 | 1 | 0 |
| rs121964992 | pathogenic | 1 | 1 | 1 |
| rs121964991 | pathogenic | 1 | 1 | 0 |
| rs121964988 | pathogenic | 1 | 1 | 1 |
| rs4687 | benign | 1 | 1 | 1 |
| rs267607125 | pathogenic | 1 | 1 | 1 |
| rs751682878 | benign | 1 | 1 | 1 |
| rs267607126 | pathogenic | 1 | 1 | 1 |
| rs768713596 | pathogenic | 0 | 0 | 0 |
| rs147288996 | benign | 0 | 0 | 0 |
| rs387906639 | pathogenic | 1 | 1 | 1 |
| rs730881069 | pathogenic | 1 | 1 | 1 |
| rs730881071 | pathogenic | 1 | 1 | 1 |
| rs397516348 | pathogenic | 0 | 0 | 0 |
| rs121917760 | pathogenic | 0 | 0 | 0 |
| rs104894724 | pathogenic | 1 | 1 | 1 |

| | | | | |
|---|---|---|---|---|
| rs397516349 | pathogenic | 1 | 1 | 1 |
| rs12075 | benign | 0 | 0 | 0 |
| rs147549987 | benign | 0 | 0 | 0 |
| rs56079734 | benign | 0 | 0 | 0 |
| rs1017842 | benign | 1 | 1 | 1 |
| rs137852593 | pathogenic | 1 | 1 | 1 |
| rs137852571 | pathogenic | 1 | 1 | 1 |
| rs137852591 | pathogenic | 1 | 1 | 1 |
| rs9332971 | pathogenic | 1 | 1 | 1 |
| rs755226547 | pathogenic | 1 | 1 | 1 |
| rs79206939 | benign | 0 | 0 | 0 |
| rs121918214 | pathogenic | 1 | 1 | 0 |
| rs757075712 | pathogenic | 1 | 1 | 0 |
| rs150483902 | benign | 1 | 1 | 1 |
| rs113993973 | pathogenic | 0 | 1 | 0 |
| rs946616 | benign | 0 | 0 | 0 |
| rs113993976 | pathogenic | 1 | 1 | 0 |
| rs151022264 | benign | 0 | 1 | 0 |
| rs113993977 | pathogenic | 1 | 1 | 0 |
| rs113993978 | pathogenic | 1 | 1 | 0 |
| rs2228499 | benign | 0 | 1 | 0 |
| rs113993979 | pathogenic | 1 | 1 | 0 |
| rs113993980 | pathogenic | 1 | 1 | 0 |
| rs113993983 | pathogenic | 0 | 1 | 0 |
| rs150547274 | pathogenic | 1 | 1 | 0 |
| rs113993984 | pathogenic | 1 | 1 | 0 |
| rs113993986 | pathogenic | 1 | 1 | 0 |
| rs34313873 | benign | 0 | 1 | 0 |
| rs113993988 | pathogenic | 1 | 1 | 0 |
| rs199638097 | pathogenic | 1 | 1 | 1 |
| rs118204447 | pathogenic | 1 | 1 | 1 |
| rs11862754 | benign | 0 | 0 | 0 |
| rs118204445 | pathogenic | 0 | 0 | 0 |
| rs118204441 | pathogenic | 0 | 0 | 0 |
| rs118204438 | pathogenic | 0 | 0 | 0 |
| rs118204436 | pathogenic | 0 | 0 | 0 |
| rs146093755 | pathogenic | 0 | 0 | 0 |
| rs781439830 | pathogenic | 0 | 0 | 0 |
| rs398123438 | pathogenic | 0 | 0 | 0 |
| rs398123439 | pathogenic | 0 | 0 | 0 |
| rs118204444 | pathogenic | 0 | 0 | 0 |
| rs7187889 | benign | 0 | 0 | 0 |
| rs118204435 | pathogenic | 1 | 1 | 0 |
| rs34745339 | benign | 0 | 0 | 0 |
| rs398123440 | pathogenic | 0 | 0 | 0 |
| rs118204442 | pathogenic | 0 | 0 | 0 |
| rs770053354 | pathogenic | 0 | 0 | 0 |
| rs118204448 | pathogenic | 0 | 0 | 0 |
| rs118204443 | pathogenic | 0 | 0 | 0 |
| rs118204446 | pathogenic | 0 | 0 | 0 |
| rs746756997 | pathogenic | 0 | 0 | 0 |
| rs267606838 | pathogenic | 0 | 0 | 0 |
| rs118204437 | pathogenic | 0 | 0 | 0 |
| rs398123429 | pathogenic | 0 | 0 | 0 |

| | | | | |
|---|---|---|---|---|
| rs398123430 | pathogenic | 0 | 0 | 0 |
| rs2303269 | benign | 0 | 0 | 0 |
| rs114703967 | benign | 0 | 0 | 0 |
| rs118204439 | pathogenic | 0 | 0 | 0 |
| rs151296605 | benign | 0 | 0 | 0 |
| rs118204440 | pathogenic | 0 | 0 | 0 |
| rs78127134 | benign | 0 | 0 | 0 |
| rs372893383 | pathogenic | 0 | 0 | 0 |
| rs61753266 | pathogenic | 1 | 1 | 1 |
| rs139579928 | benign | 1 | 1 | 1 |
| rs56275071 | pathogenic | 1 | 1 | 1 |
| rs121909736 | pathogenic | 1 | 1 | 0 |
| rs121909737 | pathogenic | 1 | 1 | 0 |
| rs121909735 | pathogenic | 1 | 1 | 0 |
| rs121909731 | pathogenic | 1 | 1 | 1 |
| rs121909734 | pathogenic | 1 | 1 | 1 |
| rs121909733 | pathogenic | 1 | 1 | 1 |
| rs121909732 | pathogenic | 1 | 1 | 1 |
| rs121909730 | pathogenic | 1 | 1 | 1 |
| rs202198360 | benign | 1 | 1 | 1 |
| rs587777294 | pathogenic | 1 | 1 | 0 |
| rs587777297 | pathogenic | 1 | 1 | 0 |
| rs104893913 | pathogenic | 1 | 1 | 0 |
| rs17860424 | pathogenic | 1 | 1 | 1 |
| rs1045485 | benign | 1 | 1 | 1 |
| rs13926 | benign | 0 | 0 | 0 |
| rs79343837 | benign | 0 | 0 | 0 |
| rs45516306 | benign | 1 | 1 | 1 |
| rs139849355 | benign | 1 | 1 | 1 |
| rs104894874 | pathogenic | 1 | 1 | 1 |
| rs104894870 | pathogenic | 1 | 1 | 1 |
| rs137852221 | pathogenic | 1 | 1 | 1 |
| rs104894869 | pathogenic | 1 | 1 | 1 |
| rs104894880 | pathogenic | 1 | 1 | 1 |
| rs104894872 | pathogenic | 1 | 1 | 1 |
| rs104894868 | pathogenic | 1 | 1 | 1 |
| rs104894867 | pathogenic | 1 | 1 | 1 |
| rs104894871 | pathogenic | 1 | 1 | 1 |
| rs104894877 | pathogenic | 1 | 1 | 1 |
| rs104894883 | pathogenic | 1 | 1 | 1 |
| rs104894875 | pathogenic | 1 | 1 | 1 |
| rs104894876 | pathogenic | 1 | 1 | 1 |
| rs104894878 | pathogenic | 1 | 1 | 1 |
| rs137852220 | pathogenic | 1 | 1 | 1 |
| rs28933684 | pathogenic | 1 | 1 | 1 |
| rs104894873 | pathogenic | 1 | 1 | 1 |
| rs138409264 | benign | 0 | 0 | 0 |
| rs2236722 | benign | 0 | 0 | 0 |
| rs28757184 | benign | 0 | 0 | 0 |
| rs121434538 | pathogenic | 0 | 0 | 0 |
| rs700519 | benign | 0 | 0 | 0 |
| rs80051519 | pathogenic | 0 | 0 | 0 |
| rs121434536 | pathogenic | 1 | 1 | 0 |
| rs121434534 | pathogenic | 1 | 1 | 0 |

| | | | | |
|---|---|---|---|---|
| rs78310315 | pathogenic | 1 | 1 | 0 |
| rs540839115 | pathogenic | 1 | 1 | 1 |
| rs80338731 | pathogenic | 1 | 1 | 1 |
| rs180177151 | pathogenic | 1 | 1 | 1 |
| rs180177152 | pathogenic | 0 | 0 | 0 |
| rs80338732 | pathogenic | 1 | 1 | 1 |
| rs80338675 | pathogenic | 1 | 1 | 1 |
| rs80338676 | pathogenic | 1 | 1 | 1 |
| rs180177153 | pathogenic | 0 | 0 | 0 |
| rs180177154 | pathogenic | 1 | 1 | 1 |
| rs5896 | benign | 1 | 1 | 1 |
| rs121918477 | pathogenic | 0 | 0 | 0 |
| rs121918428 | pathogenic | 1 | 1 | 1 |
| rs121918427 | pathogenic | 1 | 1 | 1 |
| rs544300820 | benign | 1 | 1 | 1 |
| rs121918110 | pathogenic | 1 | 1 | 1 |
| rs1057141 | benign | 0 | 0 | 0 |
| rs1135216 | benign | 1 | 1 | 1 |
| rs121917702 | pathogenic | 1 | 1 | 1 |
| rs28941474 | pathogenic | 0 | 0 | 0 |
| rs104893944 | pathogenic | 1 | 1 | 1 |
| rs104893947 | pathogenic | 1 | 1 | 0 |
| rs104893943 | pathogenic | 1 | 1 | 1 |
| rs104893942 | pathogenic | 1 | 1 | 1 |
| rs104893940 | pathogenic | 1 | 1 | 1 |
| rs34045013 | benign | 0 | 0 | 0 |
| rs143083947 | pathogenic | 0 | 0 | 0 |
| rs145786986 | benign | 0 | 0 | 0 |
| rs119450942 | pathogenic | 0 | 0 | 0 |
| rs374259530 | pathogenic | 0 | 0 | 0 |
| rs756210458 | pathogenic | 0 | 0 | 0 |
| rs11089991 | benign | 0 | 0 | 0 |
| rs28941471 | pathogenic | 0 | 1 | 0 |
| rs750614500 | pathogenic | 0 | 1 | 0 |
| rs199761158 | benign | 1 | 1 | 1 |
| rs119450944 | pathogenic | 1 | 1 | 1 |
| rs373458753 | pathogenic | 1 | 1 | 1 |
| rs202064195 | pathogenic | 1 | 1 | 1 |
| rs776496275 | pathogenic | 1 | 1 | 1 |
| rs761493155 | pathogenic | 1 | 1 | 1 |
| rs763542069 | pathogenic | 0 | 0 | 0 |
| rs119450941 | pathogenic | 0 | 0 | 0 |
| rs119450940 | pathogenic | 0 | 0 | 0 |
| rs777821034 | pathogenic | 1 | 1 | 1 |
| rs771121666 | pathogenic | 1 | 1 | 1 |
| rs372895468 | pathogenic | 1 | 1 | 1 |
| rs61750603 | pathogenic | 0 | 0 | 0 |
| rs76863441 | pathogenic | 0 | 0 | 0 |
| rs142601143 | benign | 1 | 1 | 1 |
| rs185832993 | benign | 1 | 1 | 0 |
| rs17860568 | benign | 0 | 0 | 0 |
| rs142064825 | benign | 0 | 0 | 0 |
| rs564076183 | benign | 0 | 0 | 0 |
| rs386834034 | pathogenic | 0 | 0 | 0 |

| | | | | |
|---|---|---|---|---|
| rs147663769 | benign | 0 | 0 | 0 |
| rs776478974 | benign | 0 | 0 | 0 |
| rs111840783 | benign | 0 | 0 | 0 |
| rs1516446 | benign | 1 | 1 | 1 |
| rs143749154 | benign | 1 | 1 | 1 |
| rs80257412 | benign | 0 | 0 | 0 |
| rs142943120 | benign | 0 | 0 | 0 |
| rs144553482 | benign | 0 | 0 | 0 |
| rs35997569 | benign | 0 | 0 | 0 |
| rs772909106 | pathogenic | 0 | 0 | 0 |
| rs149433837 | benign | 1 | 1 | 1 |
| rs532178791 | pathogenic | 0 | 0 | 0 |
| rs145588689 | pathogenic | 1 | 1 | 0 |
| rs17843776 | benign | 1 | 1 | 0 |
| rs121917890 | pathogenic | 1 | 1 | 1 |
| rs121917892 | pathogenic | 0 | 0 | 0 |
| rs606231289 | pathogenic | 0 | 0 | 0 |
| rs370011798 | pathogenic | 0 | 0 | 0 |
| rs34689457 | benign | 0 | 0 | 0 |
| rs118204451 | pathogenic | 0 | 0 | 0 |
| rs118204452 | pathogenic | 1 | 1 | 1 |
| rs652785 | benign | 0 | 0 | 0 |
| rs35378126 | benign | 0 | 0 | 0 |
| rs80358214 | pathogenic | 1 | 1 | 1 |
| rs80358215 | pathogenic | 0 | 1 | 0 |
| rs267606858 | pathogenic | 1 | 1 | 1 |
| rs143137713 | pathogenic | 1 | 1 | 1 |
| rs782168634 | benign | 0 | 0 | 0 |
| rs143873938 | benign | 0 | 0 | 0 |
| rs782776511 | benign | 0 | 0 | 0 |
| rs201093148 | benign | 0 | 0 | 0 |
| rs267606815 | pathogenic | 0 | 0 | 0 |
| rs137853311 | pathogenic | 0 | 0 | 0 |
| rs267606817 | pathogenic | 0 | 0 | 0 |
| rs72650006 | benign | 0 | 0 | 0 |
| rs727503638 | benign | 1 | 1 | 0 |
| rs189286381 | benign | 0 | 0 | 0 |
| rs376644553 | benign | 0 | 0 | 0 |
| rs121434610 | pathogenic | 1 | 1 | 1 |
| rs267607076 | pathogenic | 1 | 1 | 1 |
| rs201275730 | benign | 0 | 0 | 0 |
| rs121909587 | pathogenic | 0 | 0 | 0 |
| rs121909558 | pathogenic | 1 | 1 | 0 |
| rs28929469 | pathogenic | 1 | 1 | 0 |
| rs121909547 | pathogenic | 1 | 1 | 0 |
| rs121909552 | pathogenic | 1 | 1 | 0 |
| rs121909573 | pathogenic | 1 | 1 | 1 |
| rs121909567 | pathogenic | 1 | 1 | 0 |
| rs121909569 | pathogenic | 0 | 0 | 0 |
| rs121909563 | pathogenic | 1 | 1 | 0 |
| rs121909570 | pathogenic | 1 | 1 | 1 |
| rs121909571 | pathogenic | 1 | 1 | 0 |
| rs121909572 | pathogenic | 1 | 1 | 1 |
| rs121909565 | pathogenic | 1 | 1 | 1 |

| | | | | |
|---|---|---|---|---|
| rs121909557 | pathogenic | 1 | 1 | 1 |
| rs121909568 | pathogenic | 1 | 1 | 1 |
| rs121909566 | pathogenic | 1 | 1 | 1 |
| rs121909549 | pathogenic | 1 | 1 | 1 |
| rs121909554 | pathogenic | 1 | 1 | 1 |
| rs121909550 | pathogenic | 1 | 1 | 1 |
| rs121909546 | pathogenic | 1 | 1 | 0 |
| rs121909555 | pathogenic | 1 | 1 | 1 |
| rs121909564 | pathogenic | 1 | 1 | 1 |
| rs548046212 | pathogenic | 0 | 0 | 0 |
| rs147353781 | benign | 0 | 0 | 0 |
| rs104894678 | pathogenic | 1 | 1 | 1 |
| rs1130426 | benign | 0 | 0 | 0 |
| rs104894677 | pathogenic | 1 | 1 | 1 |
| rs141917423 | benign | 1 | 1 | 1 |
| rs574777883 | benign | 0 | 0 | 0 |
| rs150199226 | benign | 1 | 1 | 1 |
| rs199618701 | benign | 0 | 1 | 0 |
| rs121964923 | pathogenic | 0 | 0 | 0 |
| rs121964924 | pathogenic | 0 | 0 | 0 |
| rs201258823 | pathogenic | 0 | 1 | 0 |
| rs267606773 | pathogenic | 0 | 0 | 0 |
| rs35638832 | benign | 0 | 0 | 0 |
| rs121908632 | pathogenic | 1 | 1 | 1 |
| rs121908625 | pathogenic | 1 | 1 | 1 |
| rs748949603 | pathogenic | 0 | 0 | 0 |
| rs757523840 | pathogenic | 0 | 0 | 0 |
| rs62541771 | pathogenic | 0 | 0 | 0 |
| rs776384541 | pathogenic | 1 | 1 | 0 |
| rs121908627 | pathogenic | 1 | 1 | 0 |
| rs28937594 | pathogenic | 0 | 0 | 0 |
| rs121907888 | pathogenic | 0 | 0 | 0 |
| rs121907889 | pathogenic | 0 | 1 | 0 |
| rs28941785 | pathogenic | 1 | 1 | 1 |
| rs28941786 | pathogenic | 1 | 1 | 1 |
| rs1021737 | benign | 0 | 0 | 0 |
| rs121908370 | pathogenic | 1 | 1 | 1 |
| rs63750083 | pathogenic | 0 | 0 | 0 |
| rs17883862 | benign | 0 | 0 | 0 |
| rs587782070 | pathogenic | 1 | 1 | 1 |
| rs587781269 | pathogenic | 1 | 1 | 1 |
| rs28909982 | pathogenic | 1 | 1 | 1 |
| rs373648967 | benign | 1 | 1 | 1 |
| rs368570187 | benign | 1 | 1 | 1 |
| rs730881701 | pathogenic | 1 | 1 | 1 |
| rs137853007 | pathogenic | 0 | 0 | 0 |
| rs575910805 | benign | 0 | 0 | 0 |
| rs104893875 | pathogenic | 1 | 1 | 1 |
| rs104893877 | pathogenic | 1 | 1 | 1 |
| rs119485093 | pathogenic | 1 | 1 | 1 |
| rs119485092 | pathogenic | 0 | 0 | 0 |
| rs119485090 | pathogenic | 0 | 0 | 0 |
| rs28942099 | pathogenic | 1 | 1 | 1 |
| rs373730800 | pathogenic | 1 | 1 | 1 |

| | | | | |
|---|---|---|---|---|
| rs113993995 | pathogenic | 0 | 0 | 0 |
| rs201070132 | pathogenic | 0 | 0 | 0 |
| rs113993996 | pathogenic | 0 | 0 | 0 |
| rs113993998 | pathogenic | 0 | 0 | 0 |
| rs78950939 | pathogenic | 1 | 1 | 1 |
| rs56378716 | pathogenic | 1 | 1 | 1 |
| rs754921704 | benign | 0 | 1 | 0 |
| rs766379103 | pathogenic | 1 | 1 | 0 |
| rs121912515 | pathogenic | 0 | 1 | 0 |
| rs764831888 | benign | 1 | 1 | 0 |
| rs572527340 | benign | 0 | 0 | 0 |
| rs35969327 | benign | 0 | 0 | 0 |
| rs35814893 | benign | 0 | 0 | 0 |
| rs34032300 | benign | 0 | 0 | 0 |
| rs80358260 | pathogenic | 0 | 0 | 0 |
| rs151220873 | benign | 0 | 0 | 0 |
| rs80358261 | pathogenic | 1 | 1 | 0 |
| rs11694 | pathogenic | 0 | 0 | 0 |
| rs80358264 | pathogenic | 0 | 0 | 0 |
| rs80358266 | pathogenic | 1 | 1 | 0 |
| rs104894458 | pathogenic | 1 | 1 | 1 |
| rs121918028 | pathogenic | 0 | 0 | 0 |
| rs11583680 | pathogenic | 0 | 0 | 0 |
| rs4715 | benign | 1 | 1 | 1 |
| rs34957318 | benign | 0 | 0 | 0 |
| rs1124 | benign | 1 | 1 | 1 |
| rs104894778 | pathogenic | 1 | 1 | 1 |
| rs104894777 | pathogenic | 1 | 1 | 1 |
| rs104894769 | pathogenic | 1 | 1 | 1 |
| rs104894768 | pathogenic | 1 | 1 | 1 |
| rs104894771 | pathogenic | 1 | 1 | 1 |
| rs575372249 | benign | 1 | 1 | 1 |
| rs118203914 | pathogenic | 0 | 0 | 0 |
| rs118203916 | pathogenic | 1 | 1 | 0 |
| rs140154690 | benign | 0 | 0 | 0 |
| rs34382405 | benign | 0 | 0 | 0 |
| rs73718082 | benign | 0 | 0 | 0 |
| rs764932724 | benign | 0 | 0 | 0 |
| rs554931092 | pathogenic | 1 | 1 | 0 |
| rs202060864 | pathogenic | 0 | 0 | 0 |
| rs116567033 | benign | 0 | 0 | 0 |
| rs11243011 | benign | 0 | 0 | 0 |
| rs142073519 | pathogenic | 0 | 0 | 0 |
| rs397514611 | pathogenic | 1 | 1 | 0 |
| rs397514612 | pathogenic | 0 | 0 | 0 |
| rs372054960 | pathogenic | 0 | 0 | 0 |
| rs775690041 | pathogenic | 0 | 0 | 0 |
| rs148921184 | pathogenic | 0 | 0 | 0 |
| rs12711521 | benign | 0 | 0 | 0 |
| rs2273346 | benign | 0 | 0 | 0 |
| rs12085877 | benign | 1 | 1 | 1 |
| rs144471433 | benign | 1 | 1 | 1 |
| rs397514671 | pathogenic | 1 | 1 | 1 |
| rs76394680 | benign | 0 | 0 | 0 |

| | | | | |
|---|---|---|---|---|
| rs78289603 | pathogenic | 1 | 1 | 1 |
| rs137852913 | pathogenic | 1 | 1 | 1 |
| rs137852914 | pathogenic | 1 | 1 | 1 |
| rs121918653 | pathogenic | 1 | 1 | 1 |
| rs35515200 | benign | 1 | 1 | 0 |
| rs17876030 | benign | 1 | 1 | 1 |
| rs556623443 | benign | 1 | 1 | 1 |
| rs121434365 | pathogenic | 1 | 1 | 1 |
| rs121434364 | pathogenic | 0 | 0 | 0 |
| rs41400249 | benign | 0 | 0 | 0 |
| rs149440376 | benign | 1 | 1 | 1 |
| rs181590179 | benign | 0 | 0 | 0 |
| rs113993968 | pathogenic | 0 | 0 | 0 |
| rs587776445 | pathogenic | 1 | 1 | 1 |
| rs113993969 | pathogenic | 1 | 1 | 1 |
| rs113993970 | pathogenic | 1 | 1 | 1 |
| rs587776449 | pathogenic | 1 | 1 | 1 |
| rs113993971 | pathogenic | 1 | 1 | 1 |
| rs267607194 | pathogenic | 1 | 1 | 1 |
| rs397514745 | pathogenic | 0 | 0 | 0 |
| rs201650281 | pathogenic | 0 | 0 | 0 |
| rs150529876 | benign | 1 | 1 | 1 |
| rs77573084 | benign | 0 | 0 | 0 |
| rs398122379 | pathogenic | 1 | 1 | 0 |
| rs765478968 | benign | 0 | 0 | 0 |
| rs201399681 | benign | 0 | 0 | 0 |
| rs17159287 | benign | 0 | 0 | 0 |
| rs538571144 | benign | 0 | 0 | 0 |
| rs137852647 | pathogenic | 0 | 0 | 0 |
| rs201358272 | pathogenic | 1 | 1 | 1 |
| rs181251337 | benign | 0 | 0 | 0 |
| rs3213119 | benign | 0 | 0 | 0 |
| rs137853102 | pathogenic | 1 | 1 | 0 |
| rs4956145 | benign | 0 | 0 | 0 |
| rs146036912 | pathogenic | 1 | 1 | 1 |
| rs375717077 | pathogenic | 1 | 1 | 1 |
| rs137853103 | pathogenic | 1 | 1 | 1 |
| rs776184830 | pathogenic | 1 | 1 | 1 |
| rs200273673 | pathogenic | 1 | 1 | 1 |
| rs267607025 | pathogenic | 1 | 1 | 1 |
| rs267607024 | pathogenic | 1 | 1 | 1 |
| rs515726188 | pathogenic | 1 | 1 | 1 |
| rs147315735 | benign | 0 | 0 | 0 |
| rs121918308 | pathogenic | 1 | 1 | 0 |
| rs515726195 | pathogenic | 0 | 0 | 0 |
| rs515726196 | pathogenic | 1 | 1 | 0 |
| rs121918311 | pathogenic | 0 | 1 | 0 |
| rs121918309 | pathogenic | 0 | 0 | 0 |
| rs387906891 | pathogenic | 0 | 0 | 0 |
| rs182614164 | pathogenic | 1 | 1 | 0 |
| rs121918307 | pathogenic | 0 | 0 | 0 |
| rs201440849 | benign | 0 | 0 | 0 |
| rs118203957 | pathogenic | 0 | 0 | 0 |
| rs4986790 | benign | 0 | 0 | 0 |

| | | | | |
|---|---|---|---|---|
| rs281864928 | pathogenic | 1 | 1 | 0 |
| rs267607156 | pathogenic | 0 | 0 | 0 |
| rs55916807 | benign | 0 | 0 | 0 |
| rs749750052 | benign | 0 | 0 | 0 |
| rs74382477 | benign | 0 | 0 | 0 |
| rs201016638 | benign | 0 | 0 | 0 |
| rs376683989 | benign | 0 | 0 | 0 |
| rs144439703 | pathogenic | 0 | 0 | 0 |
| rs779122022 | benign | 0 | 0 | 0 |
| rs121918056 | pathogenic | 0 | 0 | 0 |
| rs113994093 | pathogenic | 1 | 1 | 1 |
| rs141367015 | pathogenic | 0 | 0 | 0 |
| rs775538075 | pathogenic | 0 | 0 | 0 |
| rs121918055 | pathogenic | 0 | 0 | 0 |
| rs754246294 | benign | 1 | 1 | 1 |
| rs2307447 | benign | 1 | 1 | 1 |
| rs113994096 | pathogenic | 1 | 1 | 1 |
| rs367610201 | pathogenic | 1 | 1 | 1 |
| rs758438414 | pathogenic | 1 | 1 | 0 |
| rs121918046 | pathogenic | 0 | 0 | 0 |
| rs201871736 | benign | 0 | 0 | 0 |
| rs773994204 | benign | 0 | 0 | 0 |
| rs2307450 | benign | 1 | 1 | 1 |
| rs200257554 | benign | 0 | 0 | 0 |
| rs185645212 | benign | 0 | 0 | 0 |
| rs121918054 | pathogenic | 0 | 0 | 0 |
| rs113994097 | pathogenic | 0 | 0 | 0 |
| rs202037973 | pathogenic | 0 | 0 | 0 |
| rs769827124 | pathogenic | 1 | 1 | 0 |
| rs41549716 | benign | 1 | 1 | 1 |
| rs113994098 | pathogenic | 1 | 1 | 0 |
| rs144500145 | pathogenic | 1 | 1 | 0 |
| rs121918053 | pathogenic | 1 | 1 | 0 |
| rs121918050 | pathogenic | 1 | 1 | 0 |
| rs368587966 | pathogenic | 0 | 0 | 0 |
| rs139590686 | pathogenic | 0 | 0 | 0 |
| rs121918048 | pathogenic | 1 | 1 | 0 |
| rs768653086 | pathogenic | 1 | 1 | 0 |
| rs113994099 | pathogenic | 1 | 1 | 0 |
| rs121918051 | pathogenic | 1 | 1 | 0 |
| rs753160398 | pathogenic | 1 | 1 | 0 |
| rs201477273 | pathogenic | 1 | 1 | 0 |
| rs142347031 | pathogenic | 0 | 0 | 0 |
| rs121918049 | pathogenic | 0 | 0 | 0 |
| rs267606959 | pathogenic | 1 | 1 | 0 |
| rs201732356 | pathogenic | 0 | 0 | 0 |
| rs368435864 | pathogenic | 0 | 0 | 0 |
| rs764036283 | pathogenic | 0 | 0 | 0 |
| rs2307442 | benign | 0 | 0 | 0 |
| rs2307440 | benign | 0 | 0 | 0 |
| rs548076633 | pathogenic | 0 | 1 | 0 |
| rs750792237 | benign | 0 | 0 | 0 |
| rs3087374 | benign | 0 | 0 | 0 |
| rs121918060 | pathogenic | 0 | 1 | 0 |

| | | | | |
|---|---|---|---|---|
| rs397514765 | pathogenic | 1 | 1 | 1 |
| rs121918063 | pathogenic | 0 | 0 | 0 |
| rs121918058 | pathogenic | 1 | 1 | 1 |
| rs121918064 | pathogenic | 0 | 0 | 0 |
| rs36033115 | benign | 0 | 1 | 0 |
| rs121918057 | pathogenic | 0 | 1 | 0 |
| rs121918065 | pathogenic | 1 | 1 | 1 |
| rs121918061 | pathogenic | 1 | 1 | 1 |
| rs121918066 | pathogenic | 0 | 1 | 0 |
| rs707555 | benign | 0 | 1 | 0 |
| rs8192708 | benign | 0 | 1 | 0 |
| rs11552145 | benign | 0 | 0 | 0 |
| rs1804160 | benign | 0 | 1 | 0 |
| rs34103538 | benign | 0 | 1 | 0 |
| rs751393950 | pathogenic | 1 | 1 | 1 |
| rs121907979 | pathogenic | 1 | 1 | 1 |
| rs121907975 | pathogenic | 1 | 1 | 1 |
| rs121907962 | pathogenic | 1 | 1 | 1 |
| rs121907957 | pathogenic | 0 | 0 | 0 |
| rs121907953 | pathogenic | 1 | 1 | 1 |
| rs28941770 | pathogenic | 1 | 1 | 1 |
| rs28941771 | pathogenic | 1 | 1 | 1 |
| rs121907969 | pathogenic | 1 | 1 | 1 |
| rs121907973 | pathogenic | 1 | 1 | 1 |
| rs1800429 | pathogenic | 0 | 0 | 0 |
| rs121907976 | pathogenic | 1 | 1 | 0 |
| rs121907961 | pathogenic | 0 | 0 | 0 |
| rs121907974 | pathogenic | 0 | 0 | 0 |
| rs121907959 | pathogenic | 0 | 0 | 0 |
| rs121907971 | pathogenic | 1 | 1 | 0 |
| rs121907954 | pathogenic | 0 | 0 | 0 |
| rs121907977 | pathogenic | 0 | 0 | 0 |
| rs121907963 | pathogenic | 1 | 1 | 0 |
| rs370266293 | pathogenic | 1 | 1 | 0 |
| rs1800430 | benign | 0 | 0 | 0 |
| rs121907958 | pathogenic | 0 | 0 | 0 |
| rs202173526 | pathogenic | 0 | 0 | 0 |
| rs121907978 | pathogenic | 0 | 0 | 0 |
| rs121907981 | pathogenic | 0 | 1 | 0 |
| rs145012038 | benign | 1 | 1 | 1 |
| rs121907968 | pathogenic | 1 | 1 | 1 |
| rs121907966 | pathogenic | 1 | 1 | 1 |
| rs121907956 | pathogenic | 1 | 1 | 1 |
| rs28942071 | pathogenic | 1 | 1 | 1 |
| rs121907955 | pathogenic | 1 | 1 | 1 |
| rs4727853 | benign | 0 | 0 | 0 |
| rs201880593 | benign | 0 | 0 | 0 |
| rs201434579 | benign | 1 | 1 | 1 |
| rs74551128 | pathogenic | 1 | 1 | 0 |
| rs193922500 | pathogenic | 0 | 0 | 0 |
| rs121909009 | pathogenic | 1 | 1 | 0 |
| rs121908805 | pathogenic | 1 | 1 | 0 |
| rs139573311 | pathogenic | 0 | 0 | 0 |
| rs213950 | benign | 0 | 0 | 0 |

| | | | | |
|---|---|---|---|---|
| rs79282516 | pathogenic | 0 | 0 | 0 |
| rs121909017 | pathogenic | 1 | 1 | 1 |
| rs77101217 | pathogenic | 1 | 1 | 1 |
| rs1800092 | pathogenic | 0 | 1 | 0 |
| rs77646904 | pathogenic | 0 | 0 | 0 |
| rs1800095 | benign | 1 | 1 | 1 |
| rs387906368 | pathogenic | 1 | 1 | 1 |
| rs113993959 | pathogenic | 1 | 1 | 1 |
| rs121908755 | pathogenic | 1 | 1 | 1 |
| rs121909005 | pathogenic | 1 | 1 | 1 |
| rs121908757 | pathogenic | 1 | 1 | 1 |
| rs75527207 | pathogenic | 1 | 1 | 0 |
| rs121909013 | pathogenic | 1 | 1 | 0 |
| rs121909044 | pathogenic | 1 | 1 | 0 |
| rs74597325 | pathogenic | 1 | 1 | 0 |
| rs75549581 | pathogenic | 0 | 0 | 0 |
| rs80055610 | pathogenic | 1 | 1 | 1 |
| rs1800097 | pathogenic | 0 | 0 | 0 |
| rs121909006 | pathogenic | 0 | 1 | 0 |
| rs397508276 | pathogenic | 0 | 0 | 0 |
| rs397508296 | pathogenic | 1 | 1 | 1 |
| rs139468767 | pathogenic | 0 | 0 | 0 |
| rs121908759 | pathogenic | 1 | 1 | 0 |
| rs121909033 | pathogenic | 1 | 1 | 0 |
| rs397508331 | pathogenic | 0 | 0 | 0 |
| rs387907279 | pathogenic | 1 | 1 | 1 |
| rs546530618 | benign | 0 | 0 | 0 |
| rs558259739 | benign | 0 | 0 | 0 |
| rs201827340 | benign | 0 | 0 | 0 |
| rs148865119 | pathogenic | 1 | 1 | 0 |
| rs2229413 | benign | 1 | 1 | 1 |
| rs34040148 | benign | 1 | 1 | 1 |
| rs121964964 | pathogenic | 1 | 1 | 1 |
| rs760214620 | pathogenic | 1 | 1 | 1 |
| rs775992753 | pathogenic | 1 | 1 | 1 |
| rs770095972 | pathogenic | 1 | 1 | 1 |
| rs781444670 | pathogenic | 1 | 1 | 1 |
| rs147474549 | pathogenic | 0 | 0 | 0 |
| rs121964965 | pathogenic | 0 | 1 | 0 |
| rs121964963 | pathogenic | 1 | 1 | 1 |
| rs755952006 | pathogenic | 1 | 1 | 1 |
| rs121964970 | pathogenic | 1 | 1 | 1 |
| rs121964973 | pathogenic | 1 | 1 | 1 |
| rs773734233 | pathogenic | 1 | 1 | 1 |
| rs758236584 | pathogenic | 1 | 1 | 1 |
| rs149119723 | pathogenic | 1 | 1 | 0 |
| rs5742905 | pathogenic | 0 | 0 | 0 |
| rs121964962 | pathogenic | 1 | 1 | 1 |
| rs781567152 | pathogenic | 1 | 1 | 0 |
| rs398123151 | pathogenic | 1 | 1 | 1 |
| rs760417941 | pathogenic | 1 | 1 | 1 |
| rs774926464 | pathogenic | 0 | 0 | 0 |
| rs121964972 | pathogenic | 1 | 1 | 0 |
| rs372010465 | pathogenic | 1 | 1 | 1 |

| | | | | |
|---|---|---|---|---|
| rs763036586 | pathogenic | 1 | 1 | 1 |
| rs769080151 | pathogenic | 1 | 1 | 1 |
| rs121964967 | pathogenic | 1 | 1 | 1 |
| rs28934892 | pathogenic | 1 | 1 | 0 |
| rs28934891 | pathogenic | 1 | 1 | 1 |
| rs121964971 | pathogenic | 1 | 1 | 1 |
| rs121964968 | pathogenic | 1 | 1 | 1 |
| rs150828989 | benign | 1 | 1 | 1 |
| rs3775291 | benign | 1 | 1 | 0 |
| rs104894419 | pathogenic | 1 | 1 | 1 |
| rs2232642 | benign | 0 | 0 | 0 |
| rs104894551 | pathogenic | 1 | 1 | 0 |
| rs766328537 | pathogenic | 0 | 1 | 0 |
| rs104894553 | pathogenic | 1 | 1 | 0 |
| rs80099330 | benign | 0 | 0 | 0 |
| rs199565861 | pathogenic | 0 | 0 | 0 |
| rs104894548 | pathogenic | 0 | 0 | 0 |
| rs104894549 | pathogenic | 0 | 0 | 0 |
| rs104894550 | pathogenic | 1 | 1 | 1 |
| rs12948217 | pathogenic | 1 | 1 | 1 |
| rs761064915 | pathogenic | 0 | 0 | 0 |
| rs28940279 | pathogenic | 1 | 1 | 1 |
| rs774323189 | pathogenic | 1 | 1 | 1 |
| rs28940574 | pathogenic | 0 | 0 | 0 |
| rs515299 | benign | 1 | 1 | 1 |
| rs10127939 | benign | 1 | 1 | 1 |
| rs396991 | benign | 1 | 1 | 1 |
| rs145956175 | benign | 0 | 1 | 0 |
| rs747838255 | pathogenic | 1 | 1 | 1 |
| rs762659685 | pathogenic | 1 | 1 | 0 |
| rs104894831 | pathogenic | 0 | 1 | 0 |
| rs104894848 | pathogenic | 1 | 1 | 0 |
| rs104894833 | pathogenic | 0 | 1 | 0 |
| rs781838005 | pathogenic | 0 | 1 | 0 |
| rs104894834 | pathogenic | 1 | 1 | 0 |
| rs372966991 | pathogenic | 1 | 1 | 0 |
| rs138886989 | benign | 1 | 1 | 1 |
| rs28935195 | pathogenic | 0 | 1 | 0 |
| rs28935196 | pathogenic | 0 | 1 | 0 |
| rs372416832 | pathogenic | 1 | 1 | 1 |
| rs28935197 | pathogenic | 1 | 1 | 0 |
| rs28935486 | pathogenic | 1 | 1 | 0 |
| rs28935487 | pathogenic | 1 | 1 | 0 |
| rs28935488 | pathogenic | 1 | 1 | 1 |
| rs28935495 | pathogenic | 1 | 1 | 1 |
| rs28935485 | pathogenic | 1 | 1 | 1 |
| rs140329381 | pathogenic | 0 | 1 | 0 |
| rs104894830 | pathogenic | 0 | 1 | 0 |
| rs104894846 | pathogenic | 0 | 1 | 0 |
| rs28935489 | pathogenic | 0 | 1 | 0 |
| rs104894828 | pathogenic | 1 | 1 | 0 |
| rs28935491 | pathogenic | 1 | 1 | 0 |
| rs104894832 | pathogenic | 1 | 1 | 0 |
| rs28935492 | pathogenic | 1 | 1 | 0 |

| | | | | |
|---|---|---|---|---|
| rs730880437 | benign | 1 | 1 | 0 |
| rs28935493 | pathogenic | 0 | 1 | 0 |
| rs28935494 | pathogenic | 1 | 1 | 1 |
| rs111422676 | pathogenic | 1 | 1 | 1 |
| rs397515869 | benign | 1 | 1 | 0 |
| rs782449839 | pathogenic | 0 | 1 | 0 |
| rs104894852 | pathogenic | 1 | 1 | 1 |
| rs116929575 | pathogenic | 1 | 1 | 1 |
| rs7023954 | benign | 0 | 0 | 0 |
| rs184520335 | benign | 1 | 1 | 1 |
| rs143080527 | benign | 0 | 0 | 0 |
| rs1050565 | benign | 0 | 0 | 0 |
| rs11545035 | benign | 0 | 0 | 0 |
| rs267607190 | pathogenic | 1 | 1 | 1 |
| rs200283987 | benign | 0 | 0 | 0 |
| rs372556807 | benign | 1 | 1 | 0 |
| rs563073392 | benign | 0 | 0 | 0 |
| rs4539 | benign | 0 | 0 | 0 |
| rs28931609 | pathogenic | 1 | 1 | 1 |
| rs121912978 | pathogenic | 1 | 1 | 1 |
| rs104894072 | pathogenic | 1 | 1 | 1 |
| rs551933154 | benign | 1 | 1 | 1 |
| rs752962897 | benign | 1 | 1 | 0 |
| rs121912977 | pathogenic | 1 | 1 | 0 |
| rs4537 | benign | 1 | 1 | 1 |
| rs201830462 | benign | 0 | 0 | 0 |
| rs4544 | benign | 0 | 0 | 0 |
| rs562670189 | benign | 0 | 0 | 0 |
| rs4538 | benign | 0 | 1 | 0 |
| rs61757294 | pathogenic | 0 | 0 | 0 |
| rs5313 | benign | 0 | 0 | 0 |
| rs4545 | benign | 0 | 0 | 0 |
| rs72554627 | pathogenic | 1 | 1 | 1 |
| rs72554626 | pathogenic | 1 | 1 | 1 |
| rs387906861 | pathogenic | 1 | 1 | 1 |
| rs749655461 | pathogenic | 1 | 1 | 1 |
| rs121909606 | pathogenic | 1 | 1 | 1 |
| rs121909607 | pathogenic | 1 | 1 | 1 |
| rs200203992 | benign | 0 | 0 | 0 |
| rs6050 | benign | 0 | 0 | 0 |
| rs184635235 | benign | 0 | 0 | 0 |
| rs139146037 | benign | 0 | 0 | 0 |
| rs121909612 | pathogenic | 0 | 0 | 0 |
| rs78506343 | pathogenic | 0 | 0 | 0 |
| rs121918138 | pathogenic | 1 | 1 | 1 |
| rs121918139 | pathogenic | 1 | 1 | 1 |
| rs121918140 | pathogenic | 0 | 0 | 0 |
| rs36063028 | benign | 1 | 1 | 1 |
| rs45495192 | benign | 1 | 1 | 1 |
| rs45458802 | benign | 1 | 1 | 1 |
| rs552865793 | pathogenic | 1 | 1 | 1 |
| rs753744791 | pathogenic | 1 | 1 | 1 |
| rs777518512 | benign | 1 | 1 | 1 |
| rs142708991 | benign | 1 | 1 | 0 |

| | | | | |
|---|---|---|---|---|
| rs746069957 | benign | 0 | 0 | 0 |
| rs1334811 | benign | 0 | 0 | 0 |
| rs138894008 | benign | 1 | 1 | 0 |
| rs35030851 | benign | 1 | 1 | 0 |
| rs35378598 | benign | 0 | 0 | 0 |
| rs35818764 | benign | 1 | 1 | 1 |
| rs4631561 | benign | 1 | 1 | 1 |
| rs121918660 | pathogenic | 0 | 0 | 0 |
| rs373175587 | benign | 0 | 0 | 0 |
| rs61731956 | pathogenic | 1 | 1 | 1 |
| rs180177305 | pathogenic | 0 | 0 | 0 |
| rs119490108 | pathogenic | 0 | 0 | 0 |
| rs180177312 | pathogenic | 1 | 1 | 0 |
| rs180177314 | pathogenic | 1 | 1 | 0 |
| rs180177322 | pathogenic | 1 | 1 | 1 |
| rs180177323 | pathogenic | 1 | 1 | 1 |
| rs180177325 | pathogenic | 0 | 0 | 0 |
| rs35196441 | benign | 0 | 0 | 0 |
| rs28763904 | benign | 0 | 0 | 0 |
| rs2229519 | benign | 0 | 0 | 0 |
| rs137852886 | pathogenic | 0 | 0 | 0 |
| rs137852892 | pathogenic | 0 | 0 | 0 |
| rs770427750 | pathogenic | 0 | 0 | 0 |
| rs137852887 | pathogenic | 0 | 0 | 0 |
| rs137852893 | pathogenic | 0 | 0 | 0 |
| rs80338671 | pathogenic | 1 | 1 | 0 |
| rs2228389 | benign | 0 | 0 | 0 |
| rs80338672 | pathogenic | 0 | 0 | 0 |
| rs201958741 | pathogenic | 0 | 0 | 0 |
| rs80338673 | pathogenic | 0 | 0 | 0 |
| rs137852888 | pathogenic | 0 | 0 | 0 |
| rs137852889 | pathogenic | 0 | 0 | 0 |
| rs137852894 | pathogenic | 0 | 0 | 0 |
| rs137852890 | pathogenic | 0 | 0 | 0 |
| rs137852891 | pathogenic | 0 | 0 | 0 |
| rs766935302 | pathogenic | 0 | 0 | 0 |
| rs193074572 | benign | 0 | 0 | 0 |
| rs555210692 | benign | 1 | 1 | 0 |
| rs119103242 | pathogenic | 0 | 1 | 0 |
| rs143725101 | benign | 0 | 1 | 0 |
| rs28940589 | pathogenic | 1 | 1 | 1 |
| rs113994145 | pathogenic | 0 | 0 | 0 |
| rs773872994 | benign | 0 | 0 | 0 |
| rs28940590 | pathogenic | 1 | 1 | 0 |
| rs145400751 | benign | 1 | 1 | 1 |
| rs587782331 | benign | 0 | 0 | 0 |
| rs730881787 | benign | 0 | 0 | 0 |
| rs587778531 | benign | 0 | 0 | 0 |
| rs781652274 | benign | 0 | 0 | 0 |
| rs55882234 | benign | 0 | 0 | 0 |
| rs2020908 | benign | 0 | 0 | 0 |
| rs63750741 | pathogenic | 1 | 1 | 1 |
| rs41295268 | benign | 1 | 1 | 1 |
| rs63750897 | benign | 1 | 1 | 0 |

| | | | | |
|---|---|---|---|---|
| rs63751005 | benign | 1 | 1 | 0 |
| rs587779223 | benign | 0 | 0 | 0 |
| rs202127474 | benign | 0 | 0 | 0 |
| rs374230313 | benign | 0 | 0 | 0 |
| rs2020912 | benign | 0 | 0 | 0 |
| rs63751017 | pathogenic | 0 | 0 | 0 |
| rs587779246 | pathogenic | 0 | 0 | 0 |
| rs63749919 | pathogenic | 0 | 0 | 0 |
| rs63750563 | pathogenic | 1 | 1 | 1 |
| rs63749999 | pathogenic | 0 | 0 | 0 |
| rs200492211 | pathogenic | 0 | 0 | 0 |
| rs576269342 | benign | 0 | 0 | 0 |
| rs267608075 | benign | 0 | 0 | 0 |
| rs149605979 | benign | 0 | 0 | 0 |
| rs63750252 | benign | 0 | 0 | 0 |
| rs754289472 | benign | 0 | 0 | 0 |
| rs63750119 | pathogenic | 0 | 0 | 0 |
| rs587782109 | benign | 0 | 0 | 0 |
| rs34625968 | benign | 1 | 1 | 1 |
| rs764786814 | benign | 1 | 1 | 1 |
| rs199594809 | benign | 1 | 1 | 1 |
| rs74315364 | pathogenic | 1 | 1 | 1 |
| rs201253919 | pathogenic | 0 | 0 | 0 |
| rs387906828 | pathogenic | 1 | 1 | 1 |
| rs34521609 | benign | 1 | 1 | 1 |
| rs199472894 | pathogenic | 0 | 0 | 0 |
| rs199472902 | pathogenic | 0 | 0 | 0 |
| rs121912505 | pathogenic | 0 | 0 | 0 |
| rs28928905 | pathogenic | 0 | 0 | 0 |
| rs374376640 | benign | 0 | 0 | 0 |
| rs202102276 | benign | 0 | 0 | 0 |
| rs199472918 | pathogenic | 0 | 0 | 0 |
| rs121912516 | pathogenic | 1 | 1 | 1 |
| rs121912504 | pathogenic | 0 | 0 | 0 |
| rs9333649 | pathogenic | 0 | 0 | 0 |
| rs121912508 | pathogenic | 0 | 0 | 0 |
| rs199473428 | pathogenic | 1 | 1 | 1 |
| rs104894021 | pathogenic | 0 | 0 | 0 |
| rs28928904 | pathogenic | 1 | 1 | 1 |
| rs199473039 | benign | 1 | 1 | 1 |
| rs199472958 | pathogenic | 1 | 1 | 1 |
| rs1137617 | pathogenic | 1 | 1 | 1 |
| rs199473532 | pathogenic | 1 | 1 | 1 |
| rs199472988 | pathogenic | 0 | 0 | 0 |
| rs189014161 | pathogenic | 0 | 0 | 0 |
| rs12720441 | pathogenic | 0 | 0 | 0 |
| rs199473001 | pathogenic | 1 | 1 | 0 |
| rs121912506 | pathogenic | 1 | 1 | 0 |
| rs773724817 | pathogenic | 1 | 1 | 1 |
| rs118204002 | pathogenic | 1 | 1 | 1 |
| rs138556525 | pathogenic | 1 | 1 | 1 |
| rs72558181 | pathogenic | 1 | 1 | 1 |
| rs118204005 | pathogenic | 1 | 1 | 1 |
| rs118204004 | pathogenic | 0 | 0 | 0 |

| | | | | |
|---|---|---|---|---|
| rs118204001 | pathogenic | 1 | 1 | 1 |
| rs118204006 | pathogenic | 1 | 1 | 1 |
| rs118204003 | pathogenic | 0 | 0 | 0 |
| rs121913147 | pathogenic | 1 | 1 | 0 |
| rs2229429 | benign | 1 | 1 | 0 |
| rs182552223 | benign | 0 | 0 | 0 |
| rs146588336 | benign | 0 | 0 | 0 |
| rs121964983 | pathogenic | 0 | 1 | 0 |
| rs121964982 | pathogenic | 0 | 0 | 0 |
| rs386833679 | pathogenic | 0 | 0 | 0 |
| rs386833680 | pathogenic | 0 | 0 | 0 |
| rs121964981 | pathogenic | 1 | 1 | 0 |
| rs121964984 | pathogenic | 0 | 0 | 0 |
| rs144971200 | benign | 0 | 0 | 0 |
| rs121964985 | pathogenic | 1 | 1 | 0 |
| rs141246107 | benign | 0 | 0 | 0 |
| rs104894561 | pathogenic | 1 | 1 | 0 |
| rs377023736 | pathogenic | 0 | 0 | 0 |
| rs121964907 | pathogenic | 1 | 1 | 1 |
| rs121964909 | pathogenic | 1 | 1 | 1 |
| rs765070743 | pathogenic | 0 | 0 | 0 |
| rs2228119 | benign | 0 | 0 | 0 |
| rs745976989 | pathogenic | 1 | 1 | 1 |
| rs121964904 | pathogenic | 0 | 0 | 0 |
| rs386833430 | pathogenic | 1 | 1 | 1 |
| rs267606992 | pathogenic | 0 | 0 | 0 |
| rs61752063 | pathogenic | 0 | 0 | 0 |
| rs61752067 | pathogenic | 1 | 1 | 1 |
| rs104894934 | pathogenic | 0 | 0 | 0 |
| rs104894929 | pathogenic | 1 | 1 | 1 |
| rs80189640 | benign | 0 | 0 | 0 |
| rs137854547 | pathogenic | 1 | 1 | 1 |
| rs137854540 | pathogenic | 1 | 1 | 1 |
| rs782092363 | pathogenic | 1 | 1 | 1 |
| rs147783915 | benign | 0 | 0 | 0 |
| rs28933687 | pathogenic | 1 | 1 | 1 |
| rs149503319 | benign | 0 | 0 | 0 |
| rs1805147 | benign | 0 | 0 | 0 |
| rs587777218 | pathogenic | 1 | 1 | 1 |
| rs200786705 | benign | 0 | 0 | 0 |
| rs56219475 | benign | 0 | 0 | 0 |
| rs587781172 | benign | 0 | 0 | 0 |
| rs757094189 | pathogenic | 0 | 0 | 0 |
| rs140811086 | benign | 1 | 1 | 1 |
| rs574088829 | pathogenic | 1 | 1 | 1 |
| rs121918525 | pathogenic | 0 | 0 | 0 |
| rs121918527 | pathogenic | 0 | 0 | 0 |
| rs121964848 | pathogenic | 0 | 1 | 0 |
| rs121964845 | pathogenic | 1 | 1 | 1 |
| rs121964846 | pathogenic | 0 | 1 | 0 |
| rs17217716 | benign | 1 | 1 | 1 |
| rs141711342 | benign | 0 | 0 | 0 |
| rs372619120 | benign | 0 | 0 | 0 |
| rs63750002 | benign | 1 | 1 | 1 |

| | | | | |
|---|---|---|---|---|
| rs193922373 | benign | 0 | 0 | 0 |
| rs41295286 | benign | 0 | 0 | 0 |
| rs17217772 | benign | 0 | 0 | 0 |
| rs145649774 | benign | 0 | 0 | 0 |
| rs63750214 | pathogenic | 0 | 0 | 0 |
| rs63750716 | benign | 0 | 0 | 0 |
| rs202026056 | benign | 0 | 0 | 0 |
| rs4987188 | benign | 1 | 1 | 0 |
| rs267607938 | benign | 0 | 0 | 0 |
| rs17224367 | benign | 0 | 0 | 0 |
| rs63750006 | benign | 0 | 0 | 0 |
| rs587781373 | benign | 0 | 0 | 0 |
| rs63751207 | pathogenic | 1 | 1 | 0 |
| rs730881759 | benign | 0 | 0 | 0 |
| rs372350768 | benign | 1 | 1 | 1 |
| rs55778204 | benign | 1 | 1 | 0 |
| rs63749910 | benign | 0 | 0 | 0 |
| rs63750844 | benign | 0 | 0 | 0 |
| rs28929483 | pathogenic | 0 | 0 | 0 |
| rs61756468 | benign | 0 | 0 | 0 |
| rs63750875 | pathogenic | 1 | 1 | 0 |
| rs776528054 | benign | 1 | 1 | 0 |
| rs751939698 | benign | 0 | 0 | 0 |
| rs587780685 | benign | 0 | 0 | 0 |
| rs730881763 | benign | 1 | 1 | 0 |
| rs373226409 | benign | 0 | 0 | 0 |
| rs202145681 | benign | 1 | 1 | 1 |
| rs63749841 | benign | 1 | 1 | 1 |
| rs63750757 | benign | 1 | 1 | 1 |
| rs587778527 | benign | 1 | 1 | 1 |
| rs63750849 | benign | 0 | 0 | 0 |
| rs587779969 | benign | 0 | 0 | 0 |
| rs121908046 | pathogenic | 1 | 1 | 1 |
| rs28940882 | pathogenic | 1 | 1 | 0 |
| rs28940883 | pathogenic | 1 | 1 | 1 |
| rs727503943 | pathogenic | 1 | 1 | 1 |
| rs137853859 | pathogenic | 1 | 1 | 1 |
| rs121908045 | pathogenic | 1 | 1 | 0 |
| rs137853861 | pathogenic | 1 | 1 | 0 |
| rs3180383 | pathogenic | 0 | 1 | 0 |
| rs144431814 | benign | 0 | 0 | 0 |
| rs139502866 | benign | 0 | 0 | 0 |
| rs17524437 | benign | 0 | 0 | 0 |
| rs760960779 | benign | 0 | 0 | 0 |
| rs148960463 | pathogenic | 0 | 0 | 0 |
| rs111033647 | pathogenic | 1 | 1 | 1 |
| rs111033648 | pathogenic | 0 | 0 | 0 |
| rs1800461 | benign | 0 | 0 | 0 |
| rs111033656 | pathogenic | 0 | 0 | 0 |
| rs111033663 | pathogenic | 1 | 1 | 0 |
| rs111033669 | pathogenic | 1 | 1 | 1 |
| rs111033690 | pathogenic | 0 | 0 | 0 |
| rs111033686 | pathogenic | 0 | 0 | 0 |
| rs111033687 | pathogenic | 0 | 0 | 0 |

| | | | | |
|---|---|---|---|---|
| rs111033695 | pathogenic | 0 | 0 | 0 |
| rs111033697 | pathogenic | 0 | 0 | 0 |
| rs111033693 | pathogenic | 0 | 0 | 0 |
| rs111033694 | pathogenic | 0 | 0 | 0 |
| rs111033721 | pathogenic | 1 | 1 | 1 |
| rs111033722 | pathogenic | 0 | 0 | 0 |
| rs111033725 | pathogenic | 1 | 1 | 0 |
| rs75391579 | pathogenic | 1 | 1 | 1 |
| rs111033728 | pathogenic | 1 | 1 | 1 |
| rs111033735 | pathogenic | 1 | 1 | 1 |
| rs111033739 | pathogenic | 1 | 1 | 1 |
| rs111033736 | pathogenic | 0 | 0 | 0 |
| rs111033744 | pathogenic | 1 | 1 | 1 |
| rs111033750 | pathogenic | 1 | 1 | 1 |
| rs111033752 | pathogenic | 0 | 0 | 0 |
| rs111033751 | pathogenic | 0 | 0 | 0 |
| rs111033846 | pathogenic | 0 | 0 | 0 |
| rs368166217 | pathogenic | 0 | 0 | 0 |
| rs111033764 | pathogenic | 0 | 0 | 0 |
| rs111033847 | benign | 0 | 0 | 0 |
| rs111033778 | pathogenic | 0 | 0 | 0 |
| rs111033773 | pathogenic | 0 | 0 | 0 |
| rs111033792 | pathogenic | 0 | 0 | 0 |
| rs367543266 | pathogenic | 1 | 1 | 1 |
| rs111033794 | pathogenic | 0 | 0 | 0 |
| rs111033802 | pathogenic | 0 | 0 | 0 |
| rs111033803 | pathogenic | 0 | 0 | 0 |
| rs111033809 | pathogenic | 1 | 1 | 0 |
| rs121918376 | pathogenic | 0 | 1 | 0 |
| rs121918377 | pathogenic | 0 | 1 | 0 |
| rs121918378 | pathogenic | 1 | 1 | 1 |
| rs121918374 | pathogenic | 1 | 1 | 1 |
| rs137852379 | pathogenic | 0 | 0 | 0 |
| rs137852388 | pathogenic | 1 | 1 | 1 |
| rs137852368 | pathogenic | 0 | 0 | 0 |
| rs28935499 | pathogenic | 0 | 0 | 0 |
| rs28933668 | pathogenic | 0 | 0 | 0 |
| rs28933669 | pathogenic | 0 | 0 | 0 |
| rs28933670 | pathogenic | 0 | 0 | 0 |
| rs28933671 | pathogenic | 0 | 0 | 0 |
| rs28933672 | pathogenic | 0 | 0 | 0 |
| rs28937272 | pathogenic | 0 | 0 | 0 |
| rs35383156 | benign | 0 | 0 | 0 |
| rs137852418 | pathogenic | 0 | 0 | 0 |
| rs137852419 | pathogenic | 0 | 0 | 0 |
| rs137852428 | pathogenic | 0 | 0 | 0 |
| rs137852436 | pathogenic | 0 | 0 | 0 |
| rs28937285 | pathogenic | 0 | 0 | 0 |
| rs35213384 | benign | 1 | 1 | 1 |
| rs3810950 | benign | 0 | 0 | 0 |
| rs201479289 | pathogenic | 0 | 0 | 0 |
| rs121912820 | pathogenic | 0 | 1 | 0 |
| rs121912815 | pathogenic | 0 | 1 | 0 |
| rs115510708 | benign | 0 | 0 | 0 |

| | | | | |
|---|---|---|---|---|
| rs868749 | benign | 0 | 0 | 0 |
| rs121912823 | pathogenic | 0 | 0 | 0 |
| rs115212829 | benign | 0 | 0 | 0 |
| rs121912822 | pathogenic | 0 | 0 | 0 |
| rs121912816 | pathogenic | 0 | 1 | 0 |
| rs4838544 | benign | 0 | 0 | 0 |
| rs121912818 | pathogenic | 0 | 0 | 0 |
| rs121912821 | pathogenic | 0 | 0 | 0 |
| rs121912817 | pathogenic | 0 | 0 | 0 |
| rs8178992 | benign | 0 | 0 | 0 |
| rs121912819 | pathogenic | 1 | 1 | 0 |
| rs80097077 | benign | 1 | 1 | 0 |
| rs121909359 | pathogenic | 1 | 1 | 0 |
| rs121909358 | pathogenic | 1 | 1 | 1 |
| rs121909361 | pathogenic | 1 | 1 | 1 |
| rs121909357 | pathogenic | 0 | 1 | 0 |
| rs121909365 | pathogenic | 1 | 1 | 1 |
| rs6413484 | pathogenic | 1 | 1 | 1 |
| rs121909373 | pathogenic | 1 | 1 | 1 |
| rs121909366 | pathogenic | 1 | 1 | 1 |
| rs121909367 | pathogenic | 1 | 1 | 1 |
| rs121909368 | pathogenic | 1 | 1 | 1 |
| rs121909369 | pathogenic | 0 | 0 | 0 |
| rs121909363 | pathogenic | 1 | 1 | 1 |
| rs45588036 | pathogenic | 1 | 1 | 1 |
| rs121909364 | pathogenic | 1 | 1 | 1 |
| rs80358282 | benign | 0 | 0 | 0 |
| rs80358284 | pathogenic | 1 | 1 | 1 |
| rs61735303 | benign | 0 | 0 | 0 |
| rs2229274 | benign | 0 | 0 | 0 |
| rs145006491 | benign | 0 | 0 | 0 |
| rs121913580 | pathogenic | 1 | 1 | 0 |
| rs730882196 | pathogenic | 0 | 0 | 0 |
| rs153477 | benign | 0 | 0 | 0 |
| rs153478 | benign | 1 | 1 | 0 |
| rs61740602 | benign | 1 | 1 | 0 |
| rs104893892 | pathogenic | 1 | 1 | 0 |
| rs121909595 | pathogenic | 1 | 1 | 1 |
| rs28931605 | pathogenic | 1 | 1 | 1 |
| rs121909597 | pathogenic | 0 | 1 | 0 |
| rs202233735 | pathogenic | 0 | 1 | 0 |
| rs121909596 | pathogenic | 1 | 1 | 1 |
| rs75156162 | benign | 0 | 1 | 0 |
| rs2305430 | benign | 1 | 1 | 1 |
| rs150318966 | benign | 0 | 0 | 0 |
| rs121909598 | pathogenic | 0 | 1 | 0 |
| rs5744175 | benign | 0 | 0 | 0 |
| rs28935476 | pathogenic | 1 | 1 | 1 |
| rs28935475 | pathogenic | 1 | 1 | 1 |
| rs122461163 | pathogenic | 1 | 1 | 1 |
| rs62626305 | pathogenic | 1 | 1 | 1 |
| rs587777157 | pathogenic | 0 | 0 | 0 |
| rs137854478 | pathogenic | 1 | 1 | 0 |
| rs137854465 | pathogenic | 0 | 0 | 0 |

| | | | | |
|---|---|---|---|---|
| rs137854470 | pathogenic | 0 | 0 | 0 |
| rs137854482 | pathogenic | 0 | 0 | 0 |
| rs137854473 | pathogenic | 0 | 0 | 0 |
| rs137854456 | pathogenic | 0 | 0 | 0 |
| rs140598 | benign | 0 | 0 | 0 |
| rs104894105 | pathogenic | 1 | 1 | 1 |
| rs104894107 | pathogenic | 1 | 1 | 1 |
| rs104894106 | pathogenic | 1 | 1 | 1 |
| rs74621026 | benign | 1 | 1 | 1 |
| rs2290332 | benign | 0 | 0 | 0 |
| rs780202604 | pathogenic | 1 | 1 | 1 |
| rs74315305 | pathogenic | 1 | 1 | 0 |
| rs766751878 | pathogenic | 1 | 1 | 1 |
| rs137854603 | pathogenic | 1 | 1 | 1 |
| rs139592595 | benign | 1 | 1 | 0 |
| rs150565592 | benign | 1 | 1 | 1 |
| rs763520126 | benign | 0 | 0 | 0 |
| rs55980502 | benign | 0 | 0 | 0 |
| rs137852812 | pathogenic | 0 | 0 | 0 |
| rs137852813 | pathogenic | 0 | 0 | 0 |
| rs190222208 | benign | 0 | 0 | 0 |
| rs138459502 | pathogenic | 0 | 0 | 0 |
| rs730881041 | pathogenic | 0 | 0 | 0 |
| rs267607080 | pathogenic | 0 | 0 | 0 |
| rs137852814 | pathogenic | 0 | 0 | 0 |
| rs267607079 | pathogenic | 0 | 0 | 0 |
| rs192378417 | benign | 0 | 0 | 0 |
| rs1288775 | benign | 0 | 0 | 0 |
| rs80338737 | pathogenic | 1 | 1 | 1 |
| rs397514708 | pathogenic | 0 | 0 | 0 |
| rs775933965 | pathogenic | 0 | 0 | 0 |
| rs747608698 | benign | 0 | 1 | 0 |
| rs1145086 | benign | 0 | 1 | 0 |
| rs121912953 | pathogenic | 1 | 1 | 0 |
| rs243849 | benign | 0 | 0 | 0 |
| rs121912955 | pathogenic | 1 | 1 | 1 |
| rs17859943 | benign | 0 | 1 | 0 |
| rs755557988 | benign | 1 | 1 | 1 |
| rs10195 | benign | 0 | 0 | 0 |
| rs61735357 | benign | 1 | 1 | 1 |
| rs137882374 | benign | 0 | 0 | 0 |
| rs5388 | benign | 0 | 0 | 0 |
| rs6352 | benign | 0 | 0 | 0 |
| rs76645461 | pathogenic | 1 | 1 | 0 |
| rs1050828 | pathogenic | 0 | 0 | 0 |
| rs1050829 | pathogenic | 0 | 0 | 0 |
| rs137852314 | pathogenic | 0 | 0 | 0 |
| rs137852331 | pathogenic | 0 | 0 | 0 |
| rs5030872 | pathogenic | 0 | 0 | 0 |
| rs267606836 | pathogenic | 0 | 0 | 0 |
| rs5030868 | pathogenic | 0 | 0 | 0 |
| rs137852330 | pathogenic | 0 | 1 | 0 |
| rs137852326 | pathogenic | 1 | 1 | 1 |
| rs137852318 | pathogenic | 1 | 1 | 1 |

| | | | | |
|---|---|---|---|---|
| rs74575103 | benign | 0 | 0 | 0 |
| rs137852327 | pathogenic | 1 | 1 | 1 |
| rs76723693 | pathogenic | 0 | 0 | 0 |
| rs5030869 | pathogenic | 0 | 0 | 0 |
| rs34193178 | pathogenic | 1 | 1 | 1 |
| rs137852316 | pathogenic | 1 | 1 | 0 |
| rs398123546 | pathogenic | 1 | 1 | 1 |
| rs72554665 | pathogenic | 0 | 0 | 0 |
| rs72554664 | pathogenic | 0 | 0 | 0 |
| rs28939068 | pathogenic | 1 | 1 | 1 |
| rs59301204 | pathogenic | 1 | 1 | 1 |
| rs143715750 | benign | 1 | 1 | 1 |
| rs35263902 | pathogenic | 1 | 1 | 1 |
| rs202214518 | benign | 1 | 1 | 1 |
| rs34076756 | benign | 0 | 0 | 0 |
| rs199592341 | pathogenic | 1 | 1 | 0 |
| rs121908833 | pathogenic | 1 | 1 | 1 |
| rs121908834 | pathogenic | 1 | 1 | 0 |
| rs138454151 | benign | 0 | 0 | 0 |
| rs141323599 | benign | 0 | 0 | 0 |
| rs137853200 | pathogenic | 0 | 0 | 0 |
| rs137853199 | pathogenic | 0 | 0 | 0 |
| rs139787163 | benign | 0 | 0 | 0 |
| rs61731097 | benign | 1 | 1 | 1 |
| rs61729795 | benign | 0 | 0 | 0 |
| rs149242330 | benign | 0 | 0 | 0 |
| rs587777490 | pathogenic | 1 | 1 | 1 |
| rs398123011 | pathogenic | 1 | 1 | 0 |
| rs775186117 | pathogenic | 0 | 0 | 0 |
| rs72677237 | benign | 0 | 0 | 0 |
| rs104894163 | pathogenic | 1 | 1 | 1 |
| rs121909588 | pathogenic | 0 | 0 | 0 |
| rs1042579 | benign | 0 | 0 | 0 |
| rs121909273 | pathogenic | 0 | 0 | 0 |
| rs121909274 | pathogenic | 0 | 0 | 0 |
| rs119473034 | pathogenic | 1 | 1 | 1 |
| rs398123065 | pathogenic | 0 | 0 | 0 |
| rs587777042 | pathogenic | 1 | 1 | 0 |
| rs761413864 | pathogenic | 1 | 1 | 1 |
| rs118203965 | pathogenic | 0 | 0 | 0 |
| rs118203966 | pathogenic | 0 | 0 | 0 |
| rs1801201 | benign | 1 | 1 | 0 |
| rs1136201 | benign | 1 | 1 | 1 |
| rs144148008 | benign | 0 | 0 | 0 |
| rs200118378 | benign | 1 | 1 | 1 |
| rs112984118 | benign | 1 | 1 | 1 |
| rs149446102 | benign | 1 | 1 | 1 |
| rs1427463 | benign | 0 | 0 | 0 |
| rs201936720 | benign | 1 | 1 | 1 |
| rs61751983 | benign | 1 | 1 | 1 |
| rs104894632 | pathogenic | 1 | 1 | 1 |
| rs149202905 | benign | 0 | 0 | 0 |
| rs2230210 | benign | 1 | 1 | 1 |
| rs111631988 | benign | 0 | 0 | 0 |

| | | | | |
|---|---|---|---|---|
| rs137852550 | pathogenic | 0 | 0 | 0 |
| rs5963403 | benign | 0 | 0 | 0 |
| rs138018739 | benign | 0 | 0 | 0 |
| rs62640590 | benign | 0 | 0 | 0 |
| rs121909581 | pathogenic | 1 | 1 | 1 |
| rs202017913 | benign | 1 | 1 | 1 |
| rs143163513 | benign | 0 | 0 | 0 |
| rs238238 | benign | 0 | 1 | 0 |
| rs143945974 | benign | 0 | 0 | 0 |
| rs238239 | benign | 1 | 1 | 1 |
| rs121918403 | pathogenic | 0 | 1 | 0 |
| rs61735456 | benign | 0 | 1 | 0 |
| rs121918404 | pathogenic | 1 | 1 | 0 |
| rs12674488 | benign | 0 | 0 | 0 |
| rs201599657 | pathogenic | 0 | 0 | 0 |
| rs1057090 | benign | 0 | 0 | 0 |
| rs1057091 | benign | 0 | 0 | 0 |
| rs587783740 | benign | 0 | 0 | 0 |
| rs75956769 | benign | 0 | 0 | 0 |
| rs201691299 | benign | 0 | 0 | 0 |
| rs62621087 | benign | 0 | 0 | 0 |
| rs201308454 | benign | 0 | 0 | 0 |
| rs145657360 | benign | 0 | 0 | 0 |
| rs121434592 | pathogenic | 1 | 1 | 0 |
| rs375650413 | benign | 0 | 0 | 0 |
| rs121434437 | pathogenic | 1 | 1 | 0 |
| rs763494292 | pathogenic | 0 | 0 | 0 |
| rs121964997 | pathogenic | 1 | 1 | 0 |
| rs121964996 | pathogenic | 0 | 0 | 0 |
| rs41283425 | benign | 1 | 1 | 1 |
| rs376289528 | benign | 1 | 1 | 1 |
| rs144171604 | benign | 1 | 1 | 1 |
| rs782440692 | pathogenic | 1 | 1 | 1 |
| rs782058451 | pathogenic | 1 | 1 | 1 |
| rs78437817 | benign | 0 | 0 | 0 |
| rs200327969 | benign | 1 | 1 | 1 |
| rs143043662 | benign | 0 | 0 | 0 |
| rs111966833 | benign | 1 | 1 | 1 |
| rs35523678 | benign | 0 | 0 | 0 |
| rs515726208 | pathogenic | 0 | 0 | 0 |
| rs397517239 | pathogenic | 1 | 1 | 1 |
| rs148966602 | benign | 1 | 1 | 1 |
| rs76371972 | benign | 1 | 1 | 1 |
| rs2229968 | benign | 1 | 1 | 1 |
| rs61751538 | benign | 1 | 1 | 1 |
| rs200800978 | pathogenic | 1 | 1 | 1 |
| rs148695964 | benign | 1 | 1 | 1 |
| rs140042418 | benign | 1 | 1 | 1 |
| rs116185403 | benign | 0 | 0 | 0 |
| rs142828669 | benign | 1 | 1 | 1 |
| rs139832048 | benign | 0 | 1 | 0 |
| rs58750568 | benign | 0 | 1 | 0 |
| rs77681311 | benign | 0 | 0 | 0 |
| rs151208452 | benign | 0 | 0 | 0 |

| | | | | |
|---|---|---|---|---|
| rs76463933 | benign | 0 | 0 | 0 |
| rs144434647 | benign | 0 | 0 | 0 |
| rs117361485 | benign | 0 | 0 | 0 |
| rs587777588 | pathogenic | 0 | 0 | 0 |
| rs80338796 | pathogenic | 0 | 0 | 0 |
| rs121434594 | pathogenic | 0 | 0 | 0 |
| rs146766138 | pathogenic | 1 | 1 | 1 |
| rs28939679 | pathogenic | 1 | 1 | 1 |
| rs748754134 | pathogenic | 1 | 1 | 0 |
| rs764276946 | pathogenic | 1 | 1 | 1 |
| rs111033588 | pathogenic | 1 | 1 | 1 |
| rs201633414 | pathogenic | 1 | 1 | 1 |
| rs121909262 | pathogenic | 1 | 1 | 1 |
| rs104893702 | pathogenic | 1 | 1 | 1 |
| rs121909265 | pathogenic | 1 | 1 | 1 |
| rs121909266 | pathogenic | 1 | 1 | 1 |
| rs193922439 | pathogenic | 1 | 1 | 0 |
| rs104893691 | pathogenic | 1 | 1 | 1 |
| rs104893695 | pathogenic | 0 | 0 | 0 |
| rs104893708 | pathogenic | 1 | 1 | 1 |
| rs121909260 | pathogenic | 0 | 0 | 0 |
| rs104893696 | pathogenic | 0 | 0 | 0 |
| rs121909267 | pathogenic | 0 | 0 | 0 |
| rs121909263 | pathogenic | 0 | 0 | 0 |
| rs121909264 | pathogenic | 0 | 0 | 0 |
| rs104893694 | pathogenic | 1 | 1 | 1 |
| rs121909268 | pathogenic | 1 | 1 | 1 |
| rs104893689 | pathogenic | 1 | 1 | 0 |
| rs104893707 | pathogenic | 1 | 1 | 0 |
| rs104893697 | pathogenic | 1 | 1 | 1 |
| rs28936684 | pathogenic | 1 | 1 | 1 |
| rs62269092 | benign | 1 | 1 | 0 |
| rs121909259 | pathogenic | 1 | 1 | 0 |
| rs548403340 | benign | 1 | 1 | 0 |
| rs142818334 | benign | 0 | 0 | 0 |
| rs12493789 | benign | 1 | 1 | 1 |
| rs193922423 | pathogenic | 1 | 1 | 0 |
| rs104893719 | pathogenic | 1 | 1 | 1 |
| rs117375173 | benign | 1 | 1 | 0 |
| rs104893712 | pathogenic | 0 | 0 | 0 |
| rs121917776 | pathogenic | 1 | 1 | 1 |
| rs767325003 | benign | 1 | 1 | 1 |
| rs35824453 | benign | 0 | 0 | 0 |
| rs141641266 | benign | 0 | 0 | 0 |
| rs1050239 | benign | 0 | 0 | 0 |
| rs141303384 | benign | 0 | 0 | 0 |
| rs12720458 | pathogenic | 0 | 0 | 0 |
| rs368507376 | pathogenic | 1 | 1 | 1 |
| rs17215500 | pathogenic | 1 | 1 | 1 |
| rs120074188 | pathogenic | 1 | 1 | 1 |
| rs397508097 | pathogenic | 1 | 1 | 1 |
| rs138551008 | pathogenic | 0 | 0 | 0 |
| rs387907139 | pathogenic | 1 | 1 | 1 |
| rs765487627 | pathogenic | 0 | 0 | 0 |

| | | | | |
|---|---|---|---|---|
| rs587778587 | benign | 0 | 0 | 0 |
| rs45568339 | benign | 0 | 0 | 0 |
| rs78179744 | benign | 0 | 0 | 0 |
| rs45624036 | benign | 0 | 0 | 0 |
| rs61755166 | benign | 0 | 0 | 0 |
| rs180177132 | pathogenic | 1 | 1 | 1 |
| rs587776527 | pathogenic | 0 | 0 | 0 |
| rs373783514 | benign | 0 | 0 | 0 |
| rs730881896 | benign | 0 | 0 | 0 |
| rs118203998 | pathogenic | 0 | 0 | 0 |
| rs41281039 | pathogenic | 1 | 1 | 1 |
| rs118204430 | pathogenic | 1 | 1 | 1 |
| rs1800546 | pathogenic | 1 | 1 | 0 |
| rs76917243 | pathogenic | 0 | 1 | 0 |
| rs78340951 | pathogenic | 0 | 1 | 0 |
| rs77718928 | pathogenic | 0 | 0 | 0 |
| rs369586696 | pathogenic | 0 | 1 | 0 |
| rs745517517 | pathogenic | 1 | 1 | 0 |
| rs372872777 | pathogenic | 0 | 0 | 0 |
| rs748704459 | pathogenic | 0 | 0 | 0 |
| rs769215411 | pathogenic | 0 | 0 | 0 |
| rs139425890 | pathogenic | 0 | 0 | 0 |
| rs779694939 | pathogenic | 1 | 1 | 0 |
| rs773729410 | pathogenic | 0 | 0 | 0 |
| rs121908623 | pathogenic | 1 | 1 | 1 |
| rs121908622 | pathogenic | 1 | 1 | 1 |
| rs121908621 | pathogenic | 1 | 1 | 1 |
| rs762106720 | pathogenic | 1 | 1 | 0 |
| rs199877522 | pathogenic | 0 | 0 | 0 |
| rs74315291 | pathogenic | 1 | 1 | 1 |
| rs79137415 | benign | 0 | 0 | 0 |
| rs121913045 | pathogenic | 0 | 0 | 0 |
| rs34295337 | pathogenic | 0 | 0 | 0 |
| rs121913046 | pathogenic | 0 | 0 | 0 |
| rs121913047 | pathogenic | 0 | 0 | 0 |
| rs121913048 | pathogenic | 0 | 0 | 0 |
| rs267607166 | pathogenic | 0 | 0 | 0 |
| rs17850684 | benign | 1 | 1 | 1 |
| rs119476051 | pathogenic | 1 | 1 | 1 |
| rs17850683 | benign | 1 | 1 | 1 |
| rs119476049 | pathogenic | 1 | 1 | 1 |
| rs119476046 | pathogenic | 1 | 1 | 1 |
| rs119476048 | pathogenic | 0 | 0 | 0 |
| rs119476047 | pathogenic | 1 | 1 | 1 |
| rs28939094 | pathogenic | 1 | 1 | 1 |
| rs119476050 | pathogenic | 0 | 0 | 0 |
| rs387906941 | pathogenic | 1 | 1 | 1 |
| rs41511344 | pathogenic | 1 | 1 | 0 |
| rs748112833 | pathogenic | 1 | 1 | 1 |
| rs74342080 | benign | 1 | 1 | 0 |
| rs116482870 | benign | 0 | 0 | 0 |
| rs104893743 | pathogenic | 0 | 0 | 0 |
| rs117179004 | benign | 0 | 0 | 0 |
| rs587779409 | pathogenic | 1 | 1 | 1 |

| | | | | |
|---|---|---|---|---|
| rs3745406 | benign | 1 | 1 | 0 |
| rs387906696 | pathogenic | 0 | 0 | 0 |
| rs72549369 | pathogenic | 1 | 1 | 0 |
| rs45523133 | benign | 0 | 0 | 0 |
| rs119460972 | pathogenic | 0 | 1 | 0 |
| rs34929837 | benign | 0 | 1 | 0 |
| rs17323225 | benign | 0 | 1 | 0 |
| rs17011368 | benign | 0 | 0 | 0 |
| rs748902766 | pathogenic | 1 | 1 | 1 |
| rs387907292 | pathogenic | 0 | 0 | 0 |
| rs387907293 | pathogenic | 1 | 1 | 0 |
| rs371526758 | pathogenic | 1 | 1 | 1 |
| rs142968179 | pathogenic | 1 | 1 | 1 |
| rs387907291 | pathogenic | 1 | 1 | 0 |
| rs121907949 | pathogenic | 0 | 0 | 0 |
| rs121907947 | pathogenic | 0 | 0 | 0 |
| rs121907950 | pathogenic | 1 | 1 | 1 |
| rs121907948 | pathogenic | 1 | 1 | 1 |
| rs4926 | benign | 0 | 0 | 0 |
| rs748809996 | pathogenic | 0 | 0 | 0 |
| rs1799883 | benign | 1 | 1 | 0 |
| rs104893974 | pathogenic | 0 | 1 | 0 |
| rs104893973 | pathogenic | 0 | 1 | 0 |
| rs34452707 | benign | 0 | 0 | 0 |
| rs119467003 | pathogenic | 1 | 1 | 1 |
| rs121908261 | pathogenic | 0 | 1 | 0 |
| rs28933985 | pathogenic | 1 | 1 | 1 |
| rs80356669 | pathogenic | 1 | 1 | 1 |
| rs28936677 | pathogenic | 1 | 1 | 0 |
| rs121918326 | pathogenic | 0 | 0 | 0 |
| rs121918324 | pathogenic | 1 | 1 | 0 |
| rs121918325 | pathogenic | 1 | 1 | 0 |
| rs41270025 | pathogenic | 1 | 1 | 0 |
| rs121918323 | pathogenic | 0 | 0 | 0 |
| rs12735723 | pathogenic | 1 | 1 | 0 |
| rs36013429 | benign | 0 | 0 | 0 |
| rs28936676 | pathogenic | 1 | 1 | 0 |
| rs771809901 | pathogenic | 1 | 1 | 1 |
| rs758350837 | benign | 1 | 1 | 0 |
| rs121908192 | pathogenic | 1 | 1 | 1 |
| rs147568500 | pathogenic | 1 | 1 | 1 |
| rs3729734 | benign | 0 | 0 | 0 |
| rs371536347 | benign | 1 | 1 | 1 |
| rs80358193 | pathogenic | 1 | 1 | 1 |
| rs119476044 | pathogenic | 1 | 1 | 1 |
| rs111292798 | benign | 1 | 1 | 1 |
| rs121908063 | pathogenic | 1 | 1 | 0 |
| rs121908065 | pathogenic | 1 | 1 | 1 |
| rs121908062 | pathogenic | 1 | 1 | 1 |
| rs121908064 | pathogenic | 1 | 1 | 1 |
| rs61746421 | benign | 1 | 1 | 1 |
| rs34886916 | benign | 1 | 1 | 1 |
| rs118204045 | pathogenic | 0 | 0 | 0 |
| rs138607412 | benign | 1 | 1 | 1 |

| | | | | |
|---|---|---|---|---|
| rs118204046 | pathogenic | 0 | 0 | 0 |
| rs118204048 | pathogenic | 0 | 0 | 0 |
| rs118204044 | pathogenic | 0 | 0 | 0 |
| rs149664056 | benign | 0 | 0 | 0 |
| rs121909285 | pathogenic | 0 | 0 | 0 |
| rs1802498 | benign | 0 | 0 | 0 |
| rs1065761 | benign | 0 | 0 | 0 |
| rs119477054 | pathogenic | 0 | 0 | 0 |
| rs12720097 | benign | 0 | 0 | 0 |
| rs747281324 | pathogenic | 1 | 1 | 0 |
| rs367543028 | pathogenic | 1 | 1 | 0 |
| rs146077918 | benign | 0 | 0 | 0 |
| rs142551229 | benign | 0 | 0 | 0 |
| rs367543039 | pathogenic | 0 | 0 | 0 |
| rs587779884 | pathogenic | 1 | 1 | 0 |
| rs137853153 | pathogenic | 0 | 0 | 0 |
| rs367543029 | pathogenic | 0 | 0 | 0 |
| rs587783037 | pathogenic | 1 | 1 | 0 |
| rs142928725 | benign | 0 | 0 | 0 |
| rs28385141 | benign | 0 | 0 | 0 |
| rs587779888 | pathogenic | 0 | 0 | 0 |
| rs121909801 | pathogenic | 1 | 1 | 1 |
| rs121909790 | pathogenic | 1 | 1 | 0 |
| rs121909797 | pathogenic | 1 | 1 | 0 |
| rs121909794 | pathogenic | 1 | 1 | 1 |
| rs121909791 | pathogenic | 1 | 1 | 1 |
| rs121909793 | pathogenic | 1 | 1 | 1 |
| rs147977279 | benign | 1 | 1 | 1 |
| rs147352564 | benign | 0 | 0 | 0 |
| rs368758045 | benign | 1 | 1 | 1 |
| rs114498261 | benign | 1 | 1 | 0 |
| rs35484156 | benign | 0 | 0 | 0 |
| rs139176878 | benign | 0 | 0 | 0 |
| rs11543934 | benign | 0 | 0 | 0 |
| rs34192549 | benign | 0 | 0 | 0 |
| rs568736295 | benign | 0 | 0 | 0 |
| rs55747232 | benign | 0 | 0 | 0 |
| rs781887272 | benign | 0 | 0 | 0 |
| rs146273341 | benign | 0 | 0 | 0 |
| rs41473544 | benign | 0 | 0 | 0 |
| rs137852680 | pathogenic | 0 | 0 | 0 |
| rs55891000 | benign | 0 | 0 | 0 |
| rs140351189 | benign | 0 | 0 | 0 |
| rs137852679 | pathogenic | 1 | 1 | 0 |
| rs137852678 | pathogenic | 0 | 0 | 0 |
| rs137852677 | pathogenic | 1 | 1 | 1 |
| rs138620818 | benign | 0 | 0 | 0 |
| rs121908531 | pathogenic | 0 | 0 | 0 |
| rs72554307 | pathogenic | 0 | 0 | 0 |
| rs72554308 | pathogenic | 0 | 0 | 0 |
| rs72554312 | pathogenic | 1 | 1 | 0 |
| rs67939655 | pathogenic | 0 | 0 | 0 |
| rs67486158 | pathogenic | 0 | 0 | 0 |
| rs72554338 | pathogenic | 1 | 1 | 1 |

| | | | | |
|---|---|---|---|---|
| rs1800324 | pathogenic | 1 | 1 | 1 |
| rs72554356 | pathogenic | 1 | 1 | 1 |
| rs66656800 | pathogenic | 1 | 1 | 1 |
| rs68026851 | pathogenic | 1 | 1 | 1 |
| rs72558412 | pathogenic | 0 | 0 | 0 |
| rs72558415 | pathogenic | 0 | 0 | 0 |
| rs72558423 | pathogenic | 0 | 0 | 0 |
| rs67120076 | pathogenic | 1 | 1 | 1 |
| rs1800328 | pathogenic | 1 | 1 | 1 |
| rs72558451 | pathogenic | 1 | 1 | 1 |
| rs72558454 | pathogenic | 1 | 1 | 1 |
| rs672601372 | pathogenic | 0 | 0 | 0 |
| rs139644798 | pathogenic | 0 | 0 | 0 |
| rs369398935 | pathogenic | 0 | 0 | 0 |
| rs267607035 | pathogenic | 0 | 0 | 0 |
| rs267607036 | pathogenic | 1 | 1 | 1 |
| rs2302566 | benign | 1 | 1 | 0 |
| rs115959591 | benign | 0 | 0 | 0 |
| rs773876739 | pathogenic | 0 | 0 | 0 |
| rs1799967 | benign | 0 | 0 | 0 |
| rs80356968 | benign | 0 | 0 | 0 |
| rs28897695 | benign | 1 | 1 | 1 |
| rs80357314 | benign | 0 | 0 | 0 |
| rs80357169 | benign | 0 | 0 | 0 |
| rs397507239 | benign | 1 | 1 | 1 |
| rs80187739 | pathogenic | 1 | 1 | 1 |
| rs41293459 | pathogenic | 1 | 1 | 1 |
| rs55770810 | pathogenic | 1 | 1 | 1 |
| rs80356860 | pathogenic | 0 | 0 | 0 |
| rs56195342 | benign | 0 | 0 | 0 |
| rs80357442 | benign | 0 | 0 | 0 |
| rs41293463 | pathogenic | 1 | 1 | 1 |
| rs80357069 | pathogenic | 0 | 0 | 0 |
| rs397509275 | benign | 0 | 0 | 0 |
| rs80356920 | benign | 0 | 0 | 0 |
| rs397509284 | pathogenic | 0 | 0 | 0 |
| rs80357286 | benign | 0 | 0 | 0 |
| rs80357274 | benign | 0 | 0 | 0 |
| rs765656957 | benign | 0 | 0 | 0 |
| rs755427809 | benign | 0 | 0 | 0 |
| rs80357322 | benign | 0 | 0 | 0 |
| rs121434430 | pathogenic | 0 | 0 | 0 |
| rs121918450 | pathogenic | 1 | 1 | 1 |
| rs1532268 | benign | 0 | 0 | 0 |
| rs2303080 | benign | 0 | 0 | 0 |
| rs10064631 | benign | 0 | 0 | 0 |
| rs162036 | benign | 0 | 0 | 0 |
| rs2287780 | benign | 0 | 0 | 0 |
| rs16879334 | benign | 0 | 0 | 0 |
| rs137853061 | pathogenic | 1 | 1 | 0 |
| rs10380 | benign | 0 | 0 | 0 |
| rs1494558 | benign | 0 | 0 | 0 |
| rs1494555 | benign | 0 | 0 | 0 |
| rs192994060 | benign | 0 | 0 | 0 |

| | | | | |
|---|---|---|---|---|
| rs147128920 | benign | 1 | 1 | 0 |
| rs201803986 | pathogenic | 0 | 0 | 0 |
| rs746419489 | pathogenic | 0 | 1 | 0 |
| rs764396564 | pathogenic | 0 | 0 | 0 |
| rs758304537 | pathogenic | 0 | 0 | 0 |
| rs267606763 | pathogenic | 0 | 0 | 0 |
| rs267606762 | pathogenic | 0 | 0 | 0 |
| rs150702945 | pathogenic | 1 | 1 | 1 |
| rs202047589 | pathogenic | 0 | 0 | 0 |
| rs755562733 | pathogenic | 0 | 0 | 0 |
| rs267606764 | pathogenic | 1 | 1 | 1 |
| rs138207257 | pathogenic | 1 | 1 | 0 |
| rs149150736 | pathogenic | 1 | 1 | 1 |
| rs777046879 | pathogenic | 0 | 0 | 0 |
| rs104894408 | pathogenic | 1 | 1 | 1 |
| rs1801002 | pathogenic | 1 | 1 | 1 |
| rs28929485 | pathogenic | 1 | 1 | 1 |
| rs104894396 | pathogenic | 1 | 1 | 1 |
| rs371024165 | pathogenic | 0 | 0 | 0 |
| rs35887622 | pathogenic | 1 | 1 | 1 |
| rs587783644 | pathogenic | 1 | 1 | 1 |
| rs72474224 | pathogenic | 1 | 1 | 1 |
| rs111033296 | pathogenic | 1 | 1 | 1 |
| rs104894407 | pathogenic | 1 | 1 | 1 |
| rs72561723 | pathogenic | 1 | 1 | 1 |
| rs104894398 | pathogenic | 1 | 1 | 1 |
| rs28931594 | pathogenic | 1 | 1 | 1 |
| rs104894412 | pathogenic | 1 | 1 | 1 |
| rs111033297 | pathogenic | 1 | 1 | 1 |
| rs104894404 | pathogenic | 1 | 1 | 1 |
| rs104894410 | pathogenic | 1 | 1 | 1 |
| rs104894403 | pathogenic | 1 | 1 | 1 |
| rs397516870 | benign | 0 | 0 | 0 |
| rs200023879 | pathogenic | 1 | 1 | 1 |
| rs121912968 | pathogenic | 0 | 0 | 0 |
| rs104894402 | pathogenic | 1 | 1 | 1 |
| rs28931593 | pathogenic | 1 | 1 | 1 |
| rs104894397 | pathogenic | 0 | 0 | 0 |
| rs80338944 | pathogenic | 0 | 0 | 0 |
| rs199883710 | pathogenic | 0 | 0 | 0 |
| rs727504302 | pathogenic | 0 | 0 | 0 |
| rs145216882 | benign | 1 | 1 | 1 |
| rs781534323 | pathogenic | 1 | 1 | 1 |
| rs111033218 | benign | 1 | 1 | 1 |
| rs104894409 | pathogenic | 0 | 0 | 0 |
| rs80338945 | pathogenic | 1 | 1 | 1 |
| rs111033299 | pathogenic | 0 | 0 | 0 |
| rs143343083 | pathogenic | 1 | 1 | 1 |
| rs80338946 | pathogenic | 0 | 0 | 0 |
| rs150529554 | pathogenic | 0 | 0 | 0 |
| rs111033295 | pathogenic | 0 | 0 | 0 |
| rs111033188 | benign | 0 | 0 | 0 |
| rs397516874 | pathogenic | 0 | 0 | 0 |
| rs76434661 | pathogenic | 0 | 0 | 0 |

| | | | | |
|---|---|---|---|---|
| rs80338948 | pathogenic | 0 | 0 | 0 |
| rs104894401 | pathogenic | 0 | 0 | 0 |
| rs767178508 | pathogenic | 0 | 0 | 0 |
| rs772264564 | pathogenic | 0 | 0 | 0 |
| rs28931592 | pathogenic | 0 | 0 | 0 |
| rs80338949 | pathogenic | 1 | 1 | 1 |
| rs376898963 | benign | 1 | 1 | 1 |
| rs111033360 | benign | 0 | 0 | 0 |
| rs774518779 | pathogenic | 1 | 1 | 1 |
| rs28931595 | pathogenic | 1 | 1 | 1 |
| rs771748289 | pathogenic | 0 | 0 | 0 |
| rs104894406 | pathogenic | 0 | 0 | 0 |
| rs76838169 | benign | 0 | 0 | 0 |
| rs111033294 | pathogenic | 0 | 0 | 0 |
| rs200214298 | pathogenic | 0 | 0 | 0 |
| rs28940283 | pathogenic | 0 | 1 | 0 |
| rs1047883 | benign | 0 | 0 | 0 |
| rs138254425 | benign | 0 | 0 | 0 |
| rs121912592 | pathogenic | 1 | 1 | 0 |
| rs369061090 | pathogenic | 1 | 1 | 0 |
| rs758724746 | pathogenic | 0 | 1 | 0 |
| rs121912595 | pathogenic | 0 | 0 | 0 |
| rs76340296 | benign | 0 | 0 | 0 |
| rs140578009 | benign | 1 | 1 | 1 |
| rs74445297 | benign | 0 | 0 | 0 |
| rs78599516 | benign | 1 | 1 | 0 |
| rs104894035 | pathogenic | 1 | 1 | 0 |
| rs79451216 | benign | 0 | 0 | 0 |
| rs104894036 | pathogenic | 0 | 0 | 0 |
| rs73343757 | benign | 0 | 0 | 0 |
| rs121917722 | pathogenic | 1 | 1 | 1 |
| rs747700126 | pathogenic | 0 | 1 | 0 |
| rs121917725 | pathogenic | 1 | 1 | 1 |
| rs121917721 | pathogenic | 1 | 1 | 1 |
| rs121917723 | pathogenic | 1 | 1 | 0 |
| rs17569 | benign | 1 | 1 | 1 |
| rs2230062 | benign | 1 | 1 | 1 |
| rs17570 | benign | 0 | 0 | 0 |
| rs121917724 | pathogenic | 0 | 1 | 0 |
| rs199422235 | pathogenic | 0 | 0 | 0 |
| rs199422237 | pathogenic | 1 | 1 | 1 |
| rs199422234 | pathogenic | 1 | 1 | 0 |
| rs140015315 | pathogenic | 1 | 1 | 1 |
| rs730880298 | pathogenic | 1 | 1 | 1 |
| rs779692470 | pathogenic | 0 | 0 | 0 |
| rs369152939 | pathogenic | 0 | 0 | 0 |
| rs527236040 | pathogenic | 0 | 0 | 0 |
| rs587776985 | pathogenic | 1 | 1 | 1 |
| rs147077598 | pathogenic | 1 | 1 | 1 |
| rs128620185 | pathogenic | 1 | 1 | 1 |
| rs128620189 | pathogenic | 0 | 0 | 0 |
| rs128621190 | pathogenic | 0 | 0 | 0 |
| rs61754648 | benign | 0 | 0 | 0 |
| rs137852906 | pathogenic | 0 | 0 | 0 |

| | | | | |
|---|---|---|---|---|
| rs76811038 | pathogenic | 0 | 0 | 0 |
| rs149468422 | benign | 0 | 0 | 0 |
| rs5911 | benign | 0 | 0 | 0 |
| rs28942112 | pathogenic | 1 | 1 | 1 |
| rs143275858 | pathogenic | 1 | 1 | 0 |
| rs28362261 | benign | 0 | 0 | 0 |
| rs28362263 | benign | 1 | 1 | 1 |
| rs558707786 | pathogenic | 0 | 0 | 0 |
| rs138775799 | pathogenic | 0 | 0 | 0 |
| rs748170941 | pathogenic | 0 | 0 | 0 |
| rs121908380 | pathogenic | 0 | 0 | 0 |
| rs765123255 | pathogenic | 1 | 1 | 1 |
| rs730881832 | pathogenic | 0 | 1 | 0 |
| rs587782165 | benign | 1 | 1 | 1 |
| rs762307622 | pathogenic | 0 | 0 | 0 |
| rs34612342 | pathogenic | 1 | 1 | 0 |
| rs143353451 | pathogenic | 0 | 1 | 0 |
| rs747993448 | pathogenic | 0 | 1 | 0 |
| rs376561094 | pathogenic | 0 | 0 | 0 |
| rs776487884 | pathogenic | 1 | 1 | 0 |
| rs34126013 | pathogenic | 1 | 1 | 0 |
| rs200495564 | pathogenic | 1 | 1 | 0 |
| rs140342925 | pathogenic | 1 | 1 | 0 |
| rs587780749 | pathogenic | 1 | 1 | 0 |
| rs773087549 | pathogenic | 1 | 1 | 0 |
| rs730881833 | pathogenic | 0 | 0 | 0 |
| rs374950566 | pathogenic | 1 | 1 | 0 |
| rs587780752 | benign | 0 | 0 | 0 |
| rs3219489 | benign | 0 | 0 | 0 |
| rs199742231 | benign | 0 | 0 | 0 |
| rs587780082 | pathogenic | 0 | 0 | 0 |
| rs35881803 | benign | 0 | 0 | 0 |
| rs41305623 | benign | 1 | 1 | 1 |
| rs121917856 | pathogenic | 1 | 1 | 1 |
| rs576284122 | benign | 0 | 0 | 0 |
| rs142036701 | benign | 0 | 0 | 0 |
| rs121917852 | pathogenic | 1 | 1 | 1 |
| rs121917854 | pathogenic | 1 | 1 | 1 |
| rs121917853 | pathogenic | 1 | 1 | 1 |
| rs121917857 | pathogenic | 0 | 1 | 0 |
| rs193922740 | pathogenic | 1 | 1 | 1 |
| rs565779970 | pathogenic | 1 | 1 | 1 |
| rs6723526 | benign | 0 | 0 | 0 |
| rs141094096 | pathogenic | 1 | 1 | 1 |
| rs104894637 | pathogenic | 0 | 0 | 0 |
| rs104894636 | pathogenic | 0 | 0 | 0 |
| rs104894642 | pathogenic | 0 | 0 | 0 |
| rs104894638 | pathogenic | 0 | 0 | 0 |
| rs753666460 | pathogenic | 0 | 0 | 0 |
| rs104894643 | pathogenic | 1 | 1 | 0 |
| rs34520362 | pathogenic | 0 | 0 | 0 |
| rs374621913 | pathogenic | 0 | 0 | 0 |
| rs113641837 | benign | 0 | 0 | 0 |
| rs104894635 | pathogenic | 0 | 0 | 0 |

| | | | | |
|---|---|---|---|---|
| rs143947056 | pathogenic | 0 | 0 | 0 |
| rs138504221 | pathogenic | 0 | 0 | 0 |
| rs9894254 | benign | 0 | 0 | 0 |
| rs104894640 | pathogenic | 1 | 1 | 1 |
| rs58786455 | benign | 0 | 0 | 0 |
| rs144143780 | pathogenic | 0 | 0 | 0 |
| rs62620232 | benign | 0 | 0 | 0 |
| rs34297805 | benign | 1 | 1 | 1 |
| rs104894641 | pathogenic | 0 | 0 | 0 |
| rs148876719 | benign | 0 | 0 | 0 |
| rs104894639 | pathogenic | 0 | 0 | 0 |
| rs7503034 | benign | 0 | 0 | 0 |
| rs201631570 | benign | 0 | 0 | 0 |
| rs150181226 | pathogenic | 0 | 0 | 0 |
| rs377546036 | pathogenic | 0 | 0 | 0 |
| rs267606700 | pathogenic | 0 | 0 | 0 |
| rs267606701 | pathogenic | 1 | 1 | 0 |
| rs267606699 | pathogenic | 1 | 1 | 0 |
| rs9903215 | benign | 0 | 0 | 0 |
| rs139486406 | benign | 0 | 0 | 0 |
| rs143791614 | benign | 0 | 0 | 0 |
| rs573483474 | benign | 0 | 0 | 0 |
| rs2032892 | benign | 0 | 0 | 0 |
| rs137852619 | pathogenic | 0 | 0 | 0 |
| rs200243548 | benign | 0 | 0 | 0 |
| rs115702388 | benign | 0 | 0 | 0 |
| rs138085358 | pathogenic | 0 | 0 | 0 |
| rs121912789 | pathogenic | 1 | 1 | 1 |
| rs763407972 | pathogenic | 1 | 1 | 1 |
| rs147278302 | benign | 0 | 0 | 0 |
| rs374540411 | benign | 0 | 0 | 0 |
| rs121912790 | pathogenic | 1 | 1 | 0 |
| rs752049917 | benign | 0 | 0 | 0 |
| rs139640491 | benign | 1 | 1 | 1 |
| rs138010738 | benign | 1 | 1 | 0 |
| rs201328768 | pathogenic | 1 | 1 | 0 |
| rs2135720 | benign | 1 | 1 | 0 |
| rs201816080 | benign | 1 | 1 | 0 |
| rs754391973 | pathogenic | 0 | 0 | 0 |
| rs139087859 | benign | 0 | 0 | 0 |
| rs6805421 | benign | 0 | 0 | 0 |
| rs146266900 | benign | 0 | 0 | 0 |
| rs397509419 | pathogenic | 0 | 0 | 0 |
| rs149980411 | benign | 0 | 0 | 0 |
| rs747711488 | pathogenic | 1 | 1 | 1 |
| rs149474339 | pathogenic | 1 | 1 | 1 |
| rs758252808 | pathogenic | 1 | 1 | 0 |
| rs745756308 | pathogenic | 0 | 0 | 0 |
| rs4986958 | benign | 0 | 0 | 0 |
| rs74315444 | pathogenic | 0 | 0 | 0 |
| rs61752137 | pathogenic | 0 | 0 | 0 |
| rs8179065 | benign | 1 | 1 | 1 |
| rs758361736 | pathogenic | 1 | 1 | 0 |
| rs139794067 | pathogenic | 1 | 1 | 1 |

| | | | | |
|---|---|---|---|---|
| rs730880954 | pathogenic | 1 | 1 | 1 |
| rs730880955 | pathogenic | 1 | 1 | 1 |
| rs150634297 | pathogenic | 0 | 0 | 0 |
| rs104893750 | pathogenic | 1 | 1 | 1 |
| rs104893748 | pathogenic | 1 | 1 | 1 |
| rs104893749 | pathogenic | 1 | 1 | 1 |
| rs753520553 | pathogenic | 1 | 1 | 0 |
| rs61737294 | benign | 0 | 0 | 0 |
| rs104894601 | pathogenic | 0 | 0 | 0 |
| rs376090795 | pathogenic | 0 | 0 | 0 |
| rs104894596 | pathogenic | 0 | 0 | 0 |
| rs104894595 | pathogenic | 0 | 0 | 0 |
| rs104894598 | pathogenic | 0 | 0 | 0 |
| rs104894597 | pathogenic | 0 | 0 | 0 |
| rs104894591 | pathogenic | 0 | 0 | 0 |
| rs104894593 | pathogenic | 0 | 0 | 0 |
| rs104894594 | pathogenic | 0 | 0 | 0 |
| rs104894590 | pathogenic | 0 | 0 | 0 |
| rs86312 | benign | 0 | 0 | 0 |
| rs77874543 | benign | 0 | 0 | 0 |
| rs121912590 | pathogenic | 1 | 1 | 1 |
| rs121912589 | pathogenic | 1 | 1 | 1 |
| rs121912586 | pathogenic | 1 | 1 | 1 |
| rs121912585 | pathogenic | 1 | 1 | 1 |
| rs121912587 | pathogenic | 1 | 1 | 1 |
| rs762909359 | pathogenic | 0 | 0 | 0 |
| rs28931610 | pathogenic | 0 | 0 | 0 |
| rs104894097 | pathogenic | 1 | 1 | 1 |
| rs775176191 | benign | 0 | 0 | 0 |
| rs199907548 | pathogenic | 0 | 0 | 0 |
| rs104894095 | pathogenic | 1 | 1 | 1 |
| rs104894109 | pathogenic | 1 | 1 | 1 |
| rs372266620 | benign | 1 | 1 | 1 |
| rs104894099 | pathogenic | 0 | 0 | 0 |
| rs137854599 | pathogenic | 1 | 1 | 1 |
| rs104894094 | pathogenic | 0 | 0 | 0 |
| rs104894104 | pathogenic | 0 | 0 | 0 |
| rs6413463 | benign | 1 | 1 | 1 |
| rs104894098 | pathogenic | 0 | 0 | 0 |
| rs6413464 | benign | 0 | 0 | 0 |
| rs3731249 | benign | 0 | 0 | 0 |
| rs16931179 | benign | 0 | 0 | 0 |
| rs2734715 | benign | 0 | 0 | 0 |
| rs759080581 | pathogenic | 0 | 0 | 0 |
| rs1800746 | benign | 0 | 0 | 0 |
| rs202189057 | pathogenic | 0 | 0 | 0 |
| rs121908204 | pathogenic | 0 | 0 | 0 |
| rs756564767 | pathogenic | 0 | 0 | 0 |
| rs34758634 | benign | 1 | 1 | 1 |
| rs28940573 | pathogenic | 0 | 0 | 0 |
| rs119455955 | pathogenic | 1 | 1 | 0 |
| rs752164603 | pathogenic | 1 | 1 | 0 |
| rs763162812 | pathogenic | 0 | 0 | 0 |
| rs119455957 | pathogenic | 0 | 0 | 0 |

| | | | | |
|---|---|---|---|---|
| rs119455958 | pathogenic | 1 | 1 | 0 |
| rs750428882 | pathogenic | 0 | 0 | 0 |
| rs141482368 | benign | 0 | 0 | 0 |
| rs119455953 | pathogenic | 0 | 0 | 0 |
| rs119455954 | pathogenic | 0 | 0 | 0 |
| rs112812685 | benign | 1 | 1 | 0 |
| rs138744051 | benign | 1 | 1 | 0 |
| rs121908200 | pathogenic | 0 | 0 | 0 |
| rs119455956 | pathogenic | 0 | 0 | 0 |
| rs78970155 | benign | 0 | 0 | 0 |
| rs2834167 | benign | 0 | 0 | 0 |
| rs387907326 | pathogenic | 1 | 1 | 1 |
| rs2107732 | benign | 0 | 0 | 0 |
| rs11552377 | benign | 0 | 0 | 0 |
| rs137852843 | pathogenic | 0 | 0 | 0 |
| rs104894783 | pathogenic | 0 | 0 | 0 |
| rs122457137 | pathogenic | 1 | 1 | 1 |
| rs104894779 | pathogenic | 1 | 1 | 1 |
| rs104894786 | pathogenic | 0 | 0 | 0 |
| rs104894784 | pathogenic | 0 | 0 | 0 |
| rs104894785 | pathogenic | 0 | 0 | 0 |
| rs201870761 | pathogenic | 0 | 0 | 0 |
| rs104894781 | pathogenic | 0 | 0 | 0 |
| rs28897728 | benign | 0 | 0 | 0 |
| rs370273690 | pathogenic | 1 | 1 | 0 |
| rs9332739 | benign | 0 | 0 | 0 |
| rs150878060 | benign | 0 | 0 | 0 |
| rs573509224 | benign | 0 | 0 | 0 |
| rs36221133 | benign | 0 | 0 | 0 |
| rs2230009 | benign | 0 | 0 | 0 |
| rs267607008 | pathogenic | 0 | 0 | 0 |
| rs5985 | benign | 1 | 1 | 1 |
| rs5982 | benign | 1 | 1 | 1 |
| rs5987 | benign | 0 | 0 | 0 |
| rs5988 | benign | 0 | 0 | 0 |
| rs121918680 | pathogenic | 1 | 1 | 0 |
| rs121918679 | pathogenic | 1 | 1 | 0 |
| rs7845 | benign | 1 | 1 | 0 |
| rs1049296 | benign | 1 | 1 | 1 |
| rs121918677 | pathogenic | 1 | 1 | 0 |
| rs200086262 | pathogenic | 1 | 1 | 0 |
| rs746930990 | pathogenic | 1 | 1 | 0 |
| rs104893878 | pathogenic | 1 | 1 | 1 |
| rs121912468 | pathogenic | 1 | 1 | 1 |
| rs121912467 | pathogenic | 1 | 1 | 1 |
| rs145351926 | benign | 1 | 1 | 1 |
| rs141212743 | pathogenic | 1 | 1 | 1 |
| rs397515473 | pathogenic | 1 | 1 | 0 |
| rs121918313 | pathogenic | 0 | 1 | 0 |
| rs73015965 | pathogenic | 0 | 0 | 0 |
| rs121918030 | pathogenic | 0 | 1 | 0 |
| rs77782321 | benign | 0 | 0 | 0 |
| rs374661051 | pathogenic | 1 | 1 | 1 |
| rs267606854 | pathogenic | 0 | 0 | 0 |

| | | | | |
|---|---|---|---|---|
| rs41279678 | benign | 0 | 0 | 0 |
| rs61754640 | benign | 0 | 0 | 0 |
| rs199473314 | pathogenic | 0 | 0 | 0 |
| rs137854601 | pathogenic | 0 | 0 | 0 |
| rs137854614 | pathogenic | 0 | 0 | 0 |
| rs137854615 | pathogenic | 0 | 0 | 0 |
| rs199473635 | pathogenic | 0 | 0 | 0 |
| rs121434478 | pathogenic | 1 | 1 | 1 |
| rs775407864 | pathogenic | 1 | 1 | 1 |
| rs12706927 | benign | 0 | 1 | 0 |
| rs775614261 | benign | 0 | 0 | 0 |
| rs768926261 | benign | 0 | 0 | 0 |
| rs114057640 | benign | 0 | 0 | 0 |
| rs387906886 | pathogenic | 1 | 1 | 0 |
| rs138658585 | benign | 0 | 0 | 0 |
| rs4880091 | benign | 0 | 0 | 0 |
| rs121918552 | pathogenic | 1 | 1 | 1 |
| rs10794537 | benign | 0 | 0 | 0 |
| rs121965022 | pathogenic | 1 | 1 | 0 |
| rs121965020 | pathogenic | 0 | 0 | 0 |
| rs758452450 | pathogenic | 0 | 0 | 0 |
| rs747981483 | benign | 0 | 0 | 0 |
| rs58037052 | benign | 0 | 0 | 0 |
| rs121965029 | pathogenic | 1 | 1 | 0 |
| rs147490060 | benign | 0 | 0 | 0 |
| rs3755955 | benign | 0 | 0 | 0 |
| rs148789453 | pathogenic | 0 | 0 | 0 |
| rs121965030 | pathogenic | 0 | 0 | 0 |
| rs76722191 | benign | 0 | 0 | 0 |
| rs199801029 | pathogenic | 0 | 0 | 0 |
| rs121965033 | pathogenic | 0 | 0 | 0 |
| rs368454909 | pathogenic | 1 | 1 | 0 |
| rs6831280 | benign | 0 | 0 | 0 |
| rs121965024 | pathogenic | 0 | 0 | 0 |
| rs121965019 | pathogenic | 0 | 0 | 0 |
| rs11934801 | benign | 0 | 0 | 0 |
| rs73066479 | benign | 0 | 0 | 0 |
| rs121965027 | pathogenic | 0 | 0 | 0 |
| rs121965026 | pathogenic | 1 | 1 | 0 |
| rs121965021 | pathogenic | 0 | 0 | 0 |
| rs398123257 | benign | 0 | 0 | 0 |
| rs121965031 | pathogenic | 0 | 0 | 0 |
| rs202003795 | pathogenic | 0 | 0 | 0 |
| rs559510809 | pathogenic | 0 | 0 | 0 |
| rs762936223 | benign | 0 | 0 | 0 |
| rs137854580 | pathogenic | 1 | 1 | 1 |
| rs112263157 | benign | 0 | 0 | 0 |
| rs2289181 | benign | 0 | 0 | 0 |
| rs138034915 | benign | 0 | 1 | 0 |
| rs528367092 | pathogenic | 0 | 1 | 0 |
| rs1042393 | benign | 0 | 0 | 0 |
| rs370950728 | pathogenic | 0 | 1 | 0 |
| rs530478036 | benign | 0 | 1 | 0 |
| rs374569672 | benign | 0 | 1 | 0 |

| | | | | |
|---|---|---|---|---|
| rs121907944 | pathogenic | 0 | 1 | 0 |
| rs200856561 | pathogenic | 0 | 1 | 0 |
| rs577915581 | pathogenic | 0 | 1 | 0 |
| rs201896815 | pathogenic | 0 | 1 | 0 |
| rs121907945 | pathogenic | 0 | 1 | 0 |
| rs121907940 | pathogenic | 0 | 1 | 0 |
| rs543300039 | pathogenic | 0 | 1 | 0 |
| rs121907936 | pathogenic | 0 | 1 | 0 |
| rs766074609 | pathogenic | 0 | 1 | 0 |
| rs755253527 | pathogenic | 0 | 1 | 0 |
| rs560575383 | pathogenic | 0 | 1 | 0 |
| rs528369909 | benign | 0 | 1 | 0 |
| rs770610356 | pathogenic | 0 | 1 | 0 |
| rs747610090 | pathogenic | 0 | 1 | 0 |
| rs145712232 | benign | 0 | 1 | 0 |
| rs7215458 | benign | 1 | 1 | 0 |
| rs535644999 | pathogenic | 0 | 1 | 0 |
| rs772883420 | pathogenic | 1 | 1 | 0 |
| rs398123169 | pathogenic | 0 | 1 | 0 |
| rs121907937 | pathogenic | 0 | 1 | 0 |
| rs121907941 | pathogenic | 1 | 1 | 0 |
| rs121907942 | pathogenic | 0 | 1 | 0 |
| rs779556619 | pathogenic | 0 | 1 | 0 |
| rs1800307 | benign | 0 | 1 | 0 |
| rs377544304 | pathogenic | 1 | 1 | 0 |
| rs549029029 | pathogenic | 0 | 1 | 0 |
| rs753269119 | pathogenic | 1 | 1 | 1 |
| rs757617999 | pathogenic | 0 | 1 | 0 |
| rs28937909 | pathogenic | 0 | 1 | 0 |
| rs28940868 | pathogenic | 0 | 1 | 0 |
| rs368438393 | pathogenic | 0 | 1 | 0 |
| rs536906561 | pathogenic | 0 | 1 | 0 |
| rs374143224 | pathogenic | 0 | 1 | 0 |
| rs778418246 | pathogenic | 0 | 1 | 0 |
| rs398123172 | pathogenic | 0 | 1 | 0 |
| rs121907938 | pathogenic | 0 | 1 | 0 |
| rs1800312 | pathogenic | 0 | 1 | 0 |
| rs1126690 | benign | 0 | 1 | 0 |
| rs1800314 | benign | 0 | 1 | 0 |
| rs369532274 | pathogenic | 0 | 1 | 0 |
| rs121907943 | pathogenic | 0 | 1 | 0 |
| rs780321415 | pathogenic | 0 | 1 | 0 |
| rs377286472 | benign | 0 | 1 | 0 |
| rs121918406 | pathogenic | 1 | 1 | 1 |
| rs147484870 | benign | 0 | 0 | 0 |
| rs193922671 | pathogenic | 0 | 0 | 0 |
| rs201397978 | benign | 1 | 1 | 1 |
| rs34884895 | benign | 1 | 1 | 1 |
| rs138329459 | benign | 0 | 0 | 0 |
| rs2289366 | benign | 1 | 1 | 1 |
| rs141858334 | benign | 1 | 1 | 1 |
| rs59907886 | benign | 1 | 1 | 1 |
| rs137852653 | pathogenic | 0 | 0 | 0 |
| rs137852652 | pathogenic | 0 | 0 | 0 |

| | | | | |
|---|---|---|---|---|
| rs137852656 | pathogenic | 0 | 0 | 0 |
| rs61739916 | benign | 0 | 0 | 0 |
| rs28936668 | pathogenic | 0 | 0 | 0 |
| rs137852651 | pathogenic | 0 | 0 | 0 |
| rs137852650 | pathogenic | 0 | 0 | 0 |
| rs28936669 | pathogenic | 0 | 0 | 0 |
| rs137852654 | pathogenic | 0 | 0 | 0 |
| rs28936368 | pathogenic | 1 | 1 | 0 |
| rs137852655 | pathogenic | 0 | 0 | 0 |
| rs61752496 | benign | 0 | 0 | 0 |
| rs750098055 | benign | 0 | 0 | 0 |
| rs764149793 | benign | 0 | 0 | 0 |
| rs724160017 | pathogenic | 0 | 1 | 0 |
| rs724160019 | pathogenic | 0 | 0 | 0 |
| rs202219398 | pathogenic | 1 | 1 | 0 |
| rs724160023 | pathogenic | 1 | 1 | 0 |
| rs761953453 | pathogenic | 0 | 0 | 0 |
| rs80338835 | pathogenic | 1 | 1 | 1 |
| rs56284940 | benign | 0 | 0 | 0 |
| rs74575749 | benign | 0 | 0 | 0 |
| rs587777075 | pathogenic | 1 | 1 | 1 |
| rs641153 | benign | 0 | 0 | 0 |
| rs12614 | benign | 0 | 0 | 0 |
| rs117314762 | benign | 0 | 0 | 0 |
| rs537478097 | benign | 1 | 1 | 1 |
| rs144812066 | benign | 0 | 0 | 0 |
| rs4151651 | benign | 0 | 0 | 0 |
| rs138207668 | benign | 1 | 1 | 1 |
| rs149101394 | benign | 1 | 1 | 1 |
| rs4151659 | benign | 1 | 1 | 1 |
| rs45484591 | benign | 1 | 1 | 1 |
| rs761991070 | pathogenic | 0 | 0 | 0 |
| rs755948940 | pathogenic | 0 | 0 | 0 |
| rs137853097 | pathogenic | 0 | 0 | 0 |
| rs775297352 | benign | 0 | 0 | 0 |
| rs28943591 | benign | 0 | 0 | 0 |
| rs191468413 | pathogenic | 1 | 1 | 0 |
| rs11539471 | benign | 1 | 1 | 1 |
| rs11205 | benign | 0 | 0 | 0 |
| rs768409400 | pathogenic | 0 | 0 | 0 |
| rs762013077 | pathogenic | 0 | 0 | 0 |
| rs768474112 | pathogenic | 0 | 0 | 0 |
| rs121965063 | pathogenic | 1 | 1 | 1 |
| rs143648758 | pathogenic | 0 | 0 | 0 |
| rs121965066 | pathogenic | 0 | 0 | 0 |
| rs752907087 | pathogenic | 0 | 0 | 0 |
| rs747702749 | pathogenic | 0 | 0 | 0 |
| rs754087775 | pathogenic | 0 | 0 | 0 |
| rs121918641 | pathogenic | 1 | 1 | 1 |
| rs121918642 | pathogenic | 1 | 1 | 1 |
| rs121918640 | pathogenic | 1 | 1 | 1 |
| rs121918637 | pathogenic | 1 | 1 | 1 |
| rs121918644 | pathogenic | 1 | 1 | 1 |
| rs16840544 | benign | 0 | 0 | 0 |

| | | | | |
|---|---|---|---|---|
| rs1042391 | benign | 1 | 1 | 1 |
| rs104893744 | pathogenic | 1 | 1 | 1 |
| rs120074136 | pathogenic | 0 | 0 | 0 |
| rs1805081 | benign | 1 | 1 | 0 |
| rs80358251 | benign | 0 | 0 | 0 |
| rs774588125 | benign | 0 | 0 | 0 |
| rs120074134 | pathogenic | 0 | 0 | 0 |
| rs139751448 | pathogenic | 0 | 0 | 0 |
| rs77080672 | benign | 0 | 0 | 0 |
| rs372445155 | pathogenic | 1 | 1 | 0 |
| rs13381670 | benign | 0 | 0 | 0 |
| rs1788799 | benign | 0 | 0 | 0 |
| rs376213990 | pathogenic | 0 | 0 | 0 |
| rs757475924 | pathogenic | 0 | 0 | 0 |
| rs80358253 | pathogenic | 0 | 0 | 0 |
| rs1805082 | benign | 0 | 0 | 0 |
| rs120074130 | pathogenic | 0 | 0 | 0 |
| rs768999208 | pathogenic | 0 | 0 | 0 |
| rs34302553 | benign | 0 | 0 | 0 |
| rs28940897 | pathogenic | 0 | 0 | 0 |
| rs1140458 | benign | 0 | 0 | 0 |
| rs143124972 | pathogenic | 0 | 0 | 0 |
| rs120074135 | pathogenic | 0 | 0 | 0 |
| rs543206298 | pathogenic | 0 | 0 | 0 |
| rs120074132 | pathogenic | 0 | 0 | 0 |
| rs759826138 | pathogenic | 0 | 0 | 0 |
| rs28942108 | pathogenic | 0 | 0 | 0 |
| rs80358254 | pathogenic | 0 | 0 | 0 |
| rs80358257 | pathogenic | 0 | 0 | 0 |
| rs781261962 | pathogenic | 0 | 0 | 0 |
| rs28942107 | pathogenic | 0 | 0 | 0 |
| rs80358258 | pathogenic | 0 | 0 | 0 |
| rs80358259 | pathogenic | 1 | 1 | 0 |
| rs141440861 | benign | 0 | 0 | 0 |
| rs28942106 | pathogenic | 0 | 0 | 0 |
| rs778878523 | pathogenic | 0 | 0 | 0 |
| rs28942105 | pathogenic | 0 | 0 | 0 |
| rs61731969 | benign | 0 | 0 | 0 |
| rs200444084 | pathogenic | 0 | 0 | 0 |
| rs113371321 | pathogenic | 0 | 0 | 0 |
| rs369098773 | pathogenic | 0 | 0 | 0 |
| rs35248744 | benign | 0 | 0 | 0 |
| rs758902805 | pathogenic | 0 | 0 | 0 |
| rs120074131 | pathogenic | 0 | 0 | 0 |
| rs1805084 | benign | 0 | 0 | 0 |
| rs117793540 | benign | 0 | 0 | 0 |
| rs530058063 | benign | 0 | 0 | 0 |
| rs1064422 | pathogenic | 1 | 1 | 1 |
| rs104894506 | pathogenic | 1 | 1 | 1 |
| rs104894507 | pathogenic | 1 | 1 | 1 |
| rs104894508 | pathogenic | 1 | 1 | 0 |
| rs28999113 | pathogenic | 1 | 1 | 0 |
| rs281860266 | pathogenic | 0 | 0 | 0 |
| rs104894961 | pathogenic | 1 | 1 | 0 |

| | | | | |
|---|---|---|---|---|
| rs122463168 | pathogenic | 0 | 0 | 0 |
| rs199469624 | pathogenic | 1 | 1 | 1 |
| rs150107747 | benign | 0 | 0 | 0 |
| rs144984628 | pathogenic | 0 | 0 | 0 |
| rs28935171 | pathogenic | 0 | 0 | 0 |
| rs137852443 | pathogenic | 1 | 1 | 0 |
| rs28933674 | pathogenic | 1 | 1 | 1 |
| rs28933675 | pathogenic | 1 | 1 | 1 |
| rs28933676 | pathogenic | 1 | 1 | 1 |
| rs28933677 | pathogenic | 1 | 1 | 1 |
| rs28933678 | pathogenic | 0 | 0 | 0 |
| rs28933679 | pathogenic | 1 | 1 | 1 |
| rs28933680 | pathogenic | 0 | 0 | 0 |
| rs28933681 | pathogenic | 0 | 0 | 0 |
| rs28933682 | pathogenic | 0 | 0 | 0 |
| rs28937294 | pathogenic | 1 | 1 | 0 |
| rs137852459 | pathogenic | 0 | 0 | 0 |
| rs137852461 | pathogenic | 0 | 0 | 0 |
| rs137852464 | pathogenic | 1 | 1 | 1 |
| rs137852465 | pathogenic | 1 | 1 | 1 |
| rs1800297 | benign | 1 | 1 | 0 |
| rs137852374 | pathogenic | 1 | 1 | 1 |
| rs672601312 | pathogenic | 1 | 1 | 1 |
| rs387907196 | pathogenic | 1 | 1 | 1 |
| rs148348866 | benign | 0 | 0 | 0 |
| rs141138948 | pathogenic | 1 | 1 | 1 |
| rs137852685 | pathogenic | 1 | 1 | 0 |
| rs104893900 | pathogenic | 1 | 1 | 0 |
| rs137852686 | pathogenic | 1 | 1 | 1 |
| rs104893906 | pathogenic | 1 | 1 | 0 |
| rs587782207 | pathogenic | 1 | 1 | 1 |
| rs75086406 | pathogenic | 0 | 0 | 0 |
| rs199822819 | pathogenic | 0 | 0 | 0 |
| rs398123166 | pathogenic | 1 | 1 | 1 |
| rs121913123 | pathogenic | 1 | 1 | 1 |
| rs772190176 | pathogenic | 0 | 1 | 0 |
| rs587782216 | pathogenic | 1 | 1 | 1 |
| rs398123159 | pathogenic | 1 | 1 | 1 |
| rs121913122 | pathogenic | 1 | 1 | 1 |
| rs755436052 | pathogenic | 1 | 1 | 1 |
| rs200796606 | pathogenic | 1 | 1 | 1 |
| rs2070080 | pathogenic | 0 | 1 | 0 |
| rs149616320 | benign | 0 | 0 | 0 |
| rs143529458 | benign | 0 | 0 | 0 |
| rs267607077 | pathogenic | 1 | 1 | 1 |
| rs143898031 | benign | 1 | 1 | 0 |
| rs148304857 | benign | 1 | 1 | 1 |
| rs137852809 | pathogenic | 0 | 0 | 0 |
| rs3765623 | benign | 0 | 0 | 0 |
| rs16944397 | benign | 0 | 0 | 0 |
| rs730880167 | benign | 0 | 0 | 0 |
| rs187108957 | benign | 0 | 0 | 0 |
| rs117342470 | benign | 0 | 0 | 0 |
| rs751200138 | pathogenic | 1 | 1 | 1 |

| | | | | |
|---|---|---|---|---|
| rs755799528 | pathogenic | 0 | 0 | 0 |
| rs537484504 | benign | 0 | 0 | 0 |
| rs748944640 | pathogenic | 0 | 0 | 0 |
| rs199774121 | pathogenic | 0 | 0 | 0 |
| rs766094434 | pathogenic | 1 | 1 | 0 |
| rs777321035 | pathogenic | 1 | 1 | 1 |
| rs146269774 | benign | 0 | 1 | 0 |
| rs144085805 | benign | 0 | 1 | 0 |
| rs367643028 | benign | 0 | 0 | 0 |
| rs42524 | benign | 0 | 0 | 0 |
| rs28940586 | pathogenic | 0 | 0 | 0 |
| rs74315327 | pathogenic | 1 | 1 | 1 |
| rs199676861 | pathogenic | 0 | 0 | 0 |
| rs115802744 | benign | 1 | 1 | 0 |
| rs752866643 | pathogenic | 1 | 1 | 1 |
| rs28941784 | pathogenic | 1 | 1 | 1 |
| rs398124434 | pathogenic | 1 | 1 | 1 |
| rs756414548 | pathogenic | 1 | 1 | 1 |
| rs376128990 | pathogenic | 1 | 1 | 1 |
| rs746219370 | pathogenic | 1 | 1 | 1 |
| rs756195708 | pathogenic | 1 | 1 | 1 |
| rs369296618 | pathogenic | 1 | 1 | 1 |
| rs9593 | benign | 1 | 1 | 1 |
| rs139510844 | benign | 0 | 0 | 0 |
| rs28382158 | benign | 0 | 0 | 0 |
| rs28919570 | pathogenic | 0 | 0 | 0 |
| rs587777331 | pathogenic | 0 | 0 | 0 |
| rs587777333 | pathogenic | 0 | 0 | 0 |
| rs587777332 | pathogenic | 0 | 0 | 0 |
| rs774980346 | pathogenic | 0 | 0 | 0 |
| rs370934093 | pathogenic | 0 | 0 | 0 |
| rs121913479 | pathogenic | 1 | 1 | 1 |
| rs121913484 | pathogenic | 0 | 0 | 0 |
| rs121913485 | pathogenic | 1 | 1 | 1 |
| rs28931614 | pathogenic | 1 | 1 | 1 |
| rs17881656 | benign | 1 | 1 | 1 |
| rs28931615 | pathogenic | 1 | 1 | 1 |
| rs147846832 | benign | 0 | 0 | 0 |
| rs80356506 | pathogenic | 0 | 0 | 0 |
| rs80356494 | pathogenic | 0 | 0 | 0 |
| rs121908894 | pathogenic | 0 | 0 | 0 |
| rs121908895 | pathogenic | 0 | 0 | 0 |
| rs80356508 | pathogenic | 0 | 0 | 0 |
| rs80356507 | pathogenic | 0 | 0 | 0 |
| rs121908896 | pathogenic | 0 | 0 | 0 |
| rs745372589 | pathogenic | 1 | 1 | 0 |
| rs183898408 | benign | 0 | 0 | 0 |
| rs200844292 | benign | 0 | 0 | 0 |
| rs146897002 | benign | 0 | 0 | 0 |
| rs121964869 | pathogenic | 1 | 1 | 1 |
| rs201768800 | benign | 1 | 1 | 1 |
| rs9807556 | benign | 1 | 1 | 1 |
| rs184721031 | benign | 0 | 0 | 0 |
| rs201104206 | benign | 0 | 0 | 0 |

| | | | | |
|---|---|---|---|---|
| rs188677538 | benign | 0 | 0 | 0 |
| rs149528866 | benign | 0 | 0 | 0 |
| rs115382168 | benign | 0 | 0 | 0 |
| rs78846023 | benign | 1 | 1 | 1 |
| rs61730662 | benign | 0 | 0 | 0 |
| rs121918368 | pathogenic | 0 | 0 | 0 |
| rs566433112 | pathogenic | 1 | 1 | 1 |
| rs121909342 | pathogenic | 0 | 0 | 0 |
| rs67586389 | pathogenic | 0 | 0 | 0 |
| rs72466485 | pathogenic | 0 | 0 | 0 |
| rs72466486 | pathogenic | 1 | 1 | 1 |
| rs72466487 | pathogenic | 1 | 1 | 1 |
| rs267607112 | pathogenic | 0 | 0 | 0 |
| rs118204013 | pathogenic | 0 | 0 | 0 |
| rs121913495 | pathogenic | 0 | 0 | 0 |
| rs11554273 | pathogenic | 0 | 0 | 0 |
| rs193922477 | pathogenic | 1 | 1 | 1 |
| rs193922479 | pathogenic | 1 | 1 | 1 |
| rs72547554 | pathogenic | 0 | 0 | 0 |
| rs72547561 | pathogenic | 1 | 1 | 1 |
| rs72547562 | pathogenic | 1 | 1 | 0 |
| rs115977487 | benign | 1 | 1 | 0 |
| rs72547564 | pathogenic | 0 | 0 | 0 |
| rs72547568 | pathogenic | 1 | 1 | 0 |
| rs72547569 | pathogenic | 1 | 1 | 1 |
| rs72547571 | pathogenic | 0 | 0 | 0 |
| rs72547575 | pathogenic | 1 | 1 | 1 |
| rs544994462 | benign | 0 | 0 | 0 |
| rs190470017 | benign | 0 | 0 | 0 |
| rs199473266 | pathogenic | 1 | 1 | 1 |
| rs137854602 | pathogenic | 0 | 0 | 0 |
| rs143739249 | pathogenic | 0 | 0 | 0 |
| rs397514697 | pathogenic | 1 | 1 | 1 |
| rs373764886 | pathogenic | 0 | 0 | 0 |
| rs370242930 | pathogenic | 0 | 0 | 0 |
| rs377145777 | pathogenic | 1 | 1 | 1 |
| rs191256186 | benign | 0 | 0 | 0 |
| rs150070901 | benign | 1 | 1 | 1 |
| rs138098032 | benign | 0 | 0 | 0 |
| rs117731813 | benign | 1 | 1 | 1 |
| rs104894488 | pathogenic | 1 | 1 | 1 |
| rs11302 | benign | 0 | 1 | 0 |
| rs121918447 | pathogenic | 1 | 1 | 1 |
| rs138205828 | benign | 1 | 1 | 0 |
| rs149858290 | benign | 0 | 0 | 0 |
| rs750764003 | pathogenic | 0 | 1 | 0 |
| rs141381896 | benign | 0 | 0 | 0 |
| rs149027682 | benign | 0 | 0 | 0 |
| rs121908415 | pathogenic | 0 | 0 | 0 |
| rs121908416 | pathogenic | 0 | 0 | 0 |
| rs121908417 | pathogenic | 0 | 0 | 0 |
| rs587780306 | benign | 0 | 0 | 0 |
| rs3218725 | benign | 0 | 0 | 0 |
| rs3218729 | benign | 0 | 0 | 0 |

| | | | | |
|---|---|---|---|---|
| rs387906917 | pathogenic | 0 | 0 | 0 |
| rs200652608 | pathogenic | 0 | 0 | 0 |
| rs3218721 | benign | 0 | 0 | 0 |
| rs63750869 | pathogenic | 1 | 1 | 1 |
| rs118203910 | pathogenic | 0 | 1 | 0 |
| rs9332701 | benign | 1 | 1 | 0 |
| rs6027 | benign | 0 | 0 | 0 |
| rs121909110 | pathogenic | 0 | 1 | 0 |
| rs762550967 | pathogenic | 1 | 1 | 0 |
| rs763615602 | pathogenic | 1 | 1 | 0 |
| rs144907403 | benign | 0 | 0 | 0 |
| rs104893631 | pathogenic | 1 | 1 | 0 |
| rs144181978 | pathogenic | 1 | 1 | 0 |
| rs74874677 | benign | 1 | 1 | 1 |
| rs104893632 | pathogenic | 0 | 1 | 0 |
| rs371342464 | benign | 1 | 1 | 1 |
| rs104893633 | pathogenic | 1 | 1 | 0 |
| rs201088712 | pathogenic | 0 | 0 | 0 |
| rs201857541 | benign | 1 | 1 | 1 |
| rs139517732 | pathogenic | 1 | 1 | 0 |
| rs191057824 | benign | 1 | 1 | 0 |
| rs35683768 | benign | 1 | 1 | 0 |
| rs104895271 | pathogenic | 0 | 0 | 0 |
| rs104895217 | pathogenic | 1 | 1 | 1 |
| rs104895223 | pathogenic | 1 | 1 | 1 |
| rs104895218 | pathogenic | 1 | 1 | 1 |
| rs104895225 | pathogenic | 1 | 1 | 1 |
| rs4149637 | benign | 1 | 1 | 1 |
| rs104895219 | pathogenic | 1 | 1 | 1 |
| rs104895220 | pathogenic | 1 | 1 | 1 |
| rs104895228 | pathogenic | 1 | 1 | 1 |
| rs104895221 | pathogenic | 1 | 1 | 1 |
| rs104895222 | pathogenic | 1 | 1 | 1 |
| rs104895247 | pathogenic | 0 | 0 | 0 |
| rs28940585 | pathogenic | 0 | 0 | 0 |
| rs561367199 | pathogenic | 1 | 1 | 1 |
| rs143793815 | benign | 0 | 0 | 0 |
| rs28941473 | pathogenic | 1 | 1 | 1 |
| rs751590073 | pathogenic | 1 | 1 | 1 |
| rs111407265 | benign | 1 | 1 | 1 |
| rs373697663 | pathogenic | 1 | 1 | 1 |
| rs369879957 | pathogenic | 0 | 0 | 0 |
| rs28941472 | pathogenic | 1 | 1 | 1 |
| rs28940287 | pathogenic | 1 | 1 | 1 |
| rs77625410 | benign | 0 | 0 | 0 |
| rs111033477 | benign | 0 | 0 | 0 |
| rs115123584 | benign | 0 | 0 | 0 |
| rs374576916 | benign | 0 | 0 | 0 |
| rs397516317 | pathogenic | 0 | 0 | 0 |
| rs368657015 | pathogenic | 0 | 0 | 0 |
| rs397516321 | pathogenic | 0 | 0 | 0 |
| rs397516322 | pathogenic | 0 | 0 | 0 |
| rs111033215 | pathogenic | 0 | 0 | 0 |
| rs199606180 | pathogenic | 0 | 0 | 0 |

| | | | | |
|---|---|---|---|---|
| rs111033192 | pathogenic | 0 | 0 | 0 |
| rs948962 | benign | 1 | 1 | 1 |
| rs142293185 | benign | 0 | 0 | 0 |
| rs376764423 | pathogenic | 0 | 0 | 0 |
| rs773844428 | pathogenic | 0 | 0 | 0 |
| rs186644871 | benign | 0 | 0 | 0 |
| rs200313391 | benign | 0 | 0 | 0 |
| rs377670513 | pathogenic | 0 | 0 | 0 |
| rs1132036 | benign | 0 | 0 | 0 |
| rs111033231 | benign | 0 | 0 | 0 |
| rs145945630 | pathogenic | 0 | 0 | 0 |
| rs121912984 | pathogenic | 1 | 1 | 1 |
| rs1800435 | benign | 0 | 1 | 0 |
| rs121912980 | pathogenic | 0 | 0 | 0 |
| rs121912982 | pathogenic | 1 | 1 | 1 |
| rs121912983 | pathogenic | 0 | 0 | 0 |
| rs121912981 | pathogenic | 0 | 0 | 0 |
| rs1057868 | benign | 0 | 0 | 0 |
| rs56048668 | benign | 0 | 0 | 0 |
| rs55865465 | benign | 0 | 0 | 0 |
| rs748096324 | benign | 1 | 1 | 0 |
| rs150475750 | benign | 0 | 0 | 0 |
| rs143025739 | benign | 1 | 1 | 0 |
| rs79702016 | benign | 1 | 1 | 0 |
| rs41338945 | benign | 0 | 0 | 0 |
| rs146406037 | benign | 1 | 1 | 1 |
| rs143274491 | benign | 0 | 0 | 0 |
| rs3829986 | benign | 0 | 0 | 0 |
| rs149168939 | benign | 0 | 0 | 0 |
| rs1800385 | benign | 0 | 0 | 0 |
| rs1800386 | pathogenic | 0 | 0 | 0 |
| rs121918524 | pathogenic | 1 | 1 | 1 |
| rs121912700 | pathogenic | 0 | 0 | 0 |
| rs104893930 | pathogenic | 1 | 1 | 1 |
| rs104893931 | pathogenic | 0 | 0 | 0 |
| rs116125936 | benign | 0 | 0 | 0 |
| rs627441 | benign | 0 | 0 | 0 |
| rs104894481 | pathogenic | 0 | 0 | 0 |
| rs137853129 | pathogenic | 0 | 0 | 0 |
| rs137853130 | pathogenic | 1 | 1 | 0 |
| rs28940282 | pathogenic | 0 | 0 | 0 |
| rs2229989 | benign | 0 | 0 | 0 |
| rs104894647 | pathogenic | 0 | 0 | 0 |
| rs1061170 | benign | 1 | 1 | 1 |
| rs576402053 | benign | 0 | 0 | 0 |
| rs34972246 | benign | 0 | 0 | 0 |
| rs192725607 | benign | 0 | 0 | 0 |
| rs137852615 | pathogenic | 1 | 1 | 0 |
| rs137852617 | pathogenic | 1 | 1 | 1 |
| rs137852611 | pathogenic | 0 | 0 | 0 |
| rs137852612 | pathogenic | 0 | 0 | 0 |
| rs137852614 | pathogenic | 1 | 1 | 1 |
| rs137852610 | pathogenic | 1 | 1 | 1 |
| rs137852618 | pathogenic | 1 | 1 | 1 |

| | | | | |
|---|---|---|---|---|
| rs137852613 | pathogenic | 1 | 1 | 0 |
| rs137852609 | pathogenic | 0 | 0 | 0 |
| rs63750306 | pathogenic | 1 | 1 | 1 |
| rs63750265 | pathogenic | 1 | 1 | 1 |
| rs63751287 | pathogenic | 1 | 1 | 1 |
| rs199723282 | benign | 0 | 0 | 0 |
| rs121917807 | pathogenic | 0 | 0 | 0 |
| rs121917809 | pathogenic | 0 | 0 | 0 |
| rs75134564 | pathogenic | 0 | 0 | 0 |
| rs267606930 | pathogenic | 1 | 1 | 0 |
| rs121909302 | pathogenic | 0 | 0 | 0 |
| rs121909303 | pathogenic | 0 | 0 | 0 |
| rs121909299 | pathogenic | 0 | 0 | 0 |
| rs121909301 | pathogenic | 0 | 0 | 0 |
| rs121909300 | pathogenic | 1 | 1 | 0 |
| rs111033589 | pathogenic | 0 | 0 | 0 |
| rs137853059 | pathogenic | 0 | 0 | 0 |
| rs137853057 | pathogenic | 1 | 1 | 1 |
| rs1801474 | benign | 0 | 0 | 0 |
| rs137853060 | pathogenic | 1 | 1 | 1 |
| rs137853058 | pathogenic | 1 | 1 | 1 |
| rs34424986 | pathogenic | 0 | 0 | 0 |
| rs199657839 | benign | 1 | 1 | 1 |
| rs56092260 | benign | 0 | 0 | 0 |
| rs1801582 | benign | 1 | 1 | 1 |
| rs1801334 | benign | 1 | 1 | 1 |
| rs191486604 | pathogenic | 0 | 0 | 0 |
| rs121908051 | pathogenic | 0 | 0 | 0 |
| rs121908055 | pathogenic | 1 | 1 | 0 |
| rs28940886 | pathogenic | 0 | 0 | 0 |
| rs121908050 | pathogenic | 0 | 0 | 0 |
| rs28940887 | pathogenic | 0 | 0 | 0 |
| rs387906300 | pathogenic | 0 | 0 | 0 |
| rs28942087 | pathogenic | 0 | 0 | 0 |
| rs121908049 | pathogenic | 1 | 1 | 0 |
| rs121908054 | pathogenic | 1 | 1 | 0 |
| rs121908048 | pathogenic | 0 | 0 | 0 |
| rs28940888 | pathogenic | 1 | 1 | 1 |
| rs121908053 | pathogenic | 1 | 1 | 0 |
| rs122467169 | pathogenic | 1 | 1 | 1 |
| rs122467172 | pathogenic | 1 | 1 | 1 |
| rs122467170 | pathogenic | 1 | 1 | 1 |
| rs28935477 | pathogenic | 1 | 1 | 0 |
| rs28942088 | pathogenic | 0 | 0 | 0 |
| rs751478142 | pathogenic | 1 | 1 | 0 |
| rs765368797 | pathogenic | 0 | 0 | 0 |
| rs147810437 | benign | 0 | 0 | 0 |
| rs121918209 | pathogenic | 1 | 1 | 1 |
| rs121918208 | pathogenic | 1 | 1 | 1 |
| rs121918210 | pathogenic | 1 | 1 | 1 |
| rs35515638 | benign | 1 | 1 | 1 |
| rs121918207 | pathogenic | 1 | 1 | 1 |
| rs141298312 | benign | 0 | 0 | 0 |
| rs121918211 | pathogenic | 0 | 0 | 0 |

| | | | | |
|---|---|---|---|---|
| rs149660059 | benign | 0 | 0 | 0 |
| rs200670286 | pathogenic | 1 | 1 | 1 |
| rs121918212 | pathogenic | 1 | 1 | 1 |
| rs121918213 | pathogenic | 1 | 1 | 1 |
| rs121918205 | pathogenic | 1 | 1 | 1 |
| rs104894025 | pathogenic | 1 | 1 | 1 |
| rs104894026 | pathogenic | 1 | 1 | 1 |
| rs104894028 | pathogenic | 0 | 1 | 0 |
| rs104894029 | pathogenic | 0 | 1 | 0 |
| rs119458971 | pathogenic | 0 | 0 | 0 |
| rs119458969 | pathogenic | 1 | 1 | 1 |
| rs1801591 | benign | 1 | 1 | 1 |
| rs140169311 | benign | 0 | 0 | 0 |
| rs753363278 | benign | 1 | 1 | 1 |
| rs199763682 | pathogenic | 0 | 0 | 0 |
| rs119458970 | pathogenic | 1 | 1 | 1 |
| rs763722868 | pathogenic | 0 | 0 | 0 |
| rs56404770 | benign | 0 | 0 | 0 |
| rs34221241 | benign | 0 | 0 | 0 |
| rs121909625 | pathogenic | 0 | 0 | 0 |
| rs550921485 | pathogenic | 0 | 0 | 0 |
| rs1800454 | benign | 0 | 0 | 0 |
| rs241447 | benign | 0 | 0 | 0 |
| rs2230606 | benign | 1 | 1 | 0 |
| rs5036 | pathogenic | 1 | 1 | 1 |
| rs13306788 | benign | 0 | 0 | 0 |
| rs28929480 | pathogenic | 0 | 0 | 0 |
| rs121912749 | pathogenic | 0 | 0 | 0 |
| rs56361140 | pathogenic | 0 | 0 | 0 |
| rs28931583 | pathogenic | 1 | 1 | 1 |
| rs36211715 | pathogenic | 1 | 1 | 1 |
| rs138049878 | pathogenic | 1 | 1 | 1 |
| rs727503253 | pathogenic | 1 | 1 | 1 |
| rs267606908 | pathogenic | 0 | 0 | 0 |
| rs121913631 | pathogenic | 1 | 1 | 1 |
| rs36211716 | benign | 1 | 1 | 1 |
| rs121913639 | pathogenic | 1 | 1 | 1 |
| rs730880761 | pathogenic | 1 | 1 | 1 |
| rs121913629 | pathogenic | 1 | 1 | 1 |
| rs754368658 | pathogenic | 0 | 0 | 0 |
| rs777090017 | pathogenic | 0 | 0 | 0 |
| rs267606913 | pathogenic | 1 | 1 | 1 |
| rs773138218 | pathogenic | 0 | 0 | 0 |
| rs371295780 | pathogenic | 0 | 0 | 0 |
| rs387906830 | pathogenic | 1 | 1 | 0 |
| rs771364038 | pathogenic | 0 | 1 | 0 |
| rs772731615 | pathogenic | 0 | 0 | 0 |
| rs137853063 | pathogenic | 0 | 0 | 0 |
| rs387907076 | pathogenic | 0 | 1 | 0 |
| rs35460768 | benign | 0 | 0 | 0 |
| rs2278196 | benign | 1 | 1 | 1 |
| rs119103275 | pathogenic | 0 | 0 | 0 |
| rs119103276 | pathogenic | 1 | 1 | 1 |
| rs374402066 | pathogenic | 1 | 1 | 1 |

| | | | | |
|---|---|---|---|---|
| rs119103274 | pathogenic | 0 | 0 | 0 |
| rs2274064 | benign | 0 | 0 | 0 |
| rs267607023 | pathogenic | 1 | 1 | 0 |
| rs142117489 | benign | 1 | 1 | 0 |
| rs371258270 | benign | 0 | 0 | 0 |
| rs34716057 | benign | 0 | 0 | 0 |
| rs201778606 | benign | 0 | 0 | 0 |
| rs201438792 | benign | 0 | 0 | 0 |
| rs140547417 | benign | 0 | 0 | 0 |
| rs5881 | benign | 0 | 0 | 0 |
| rs752298084 | benign | 0 | 0 | 0 |
| rs34855278 | benign | 0 | 0 | 0 |
| rs5880 | benign | 1 | 1 | 0 |
| rs5882 | benign | 0 | 0 | 0 |
| rs1800777 | benign | 0 | 0 | 0 |
| rs577301331 | benign | 0 | 0 | 0 |
| rs137853313 | pathogenic | 0 | 0 | 0 |
| rs28935169 | pathogenic | 0 | 0 | 0 |
| rs137853315 | pathogenic | 0 | 0 | 0 |
| rs137853314 | pathogenic | 0 | 0 | 0 |
| rs28935469 | pathogenic | 0 | 0 | 0 |
| rs137853318 | pathogenic | 0 | 0 | 0 |
| rs28935470 | pathogenic | 0 | 0 | 0 |
| rs3729986 | benign | 0 | 0 | 0 |
| rs730880619 | pathogenic | 0 | 0 | 0 |
| rs201012766 | benign | 0 | 0 | 0 |
| rs727503218 | pathogenic | 0 | 0 | 0 |
| rs11570052 | benign | 0 | 0 | 0 |
| rs730880623 | pathogenic | 0 | 0 | 0 |
| rs138753870 | benign | 0 | 0 | 0 |
| rs397516068 | pathogenic | 0 | 0 | 0 |
| rs3729989 | benign | 1 | 1 | 1 |
| rs397516074 | pathogenic | 0 | 0 | 0 |
| rs561449819 | pathogenic | 0 | 0 | 0 |
| rs121917779 | benign | 0 | 0 | 0 |
| rs142881576 | pathogenic | 0 | 0 | 0 |
| rs77384282 | benign | 0 | 0 | 0 |
| rs201865787 | benign | 0 | 0 | 0 |
| rs142244640 | benign | 0 | 0 | 0 |
| rs200488706 | benign | 0 | 0 | 0 |
| rs137853291 | pathogenic | 1 | 1 | 0 |
| rs28933990 | pathogenic | 1 | 1 | 0 |
| rs140570189 | benign | 0 | 0 | 0 |
| rs201588178 | benign | 0 | 0 | 0 |
| rs104893846 | pathogenic | 0 | 0 | 0 |
| rs754545360 | pathogenic | 0 | 1 | 0 |
| rs104893851 | pathogenic | 0 | 1 | 0 |
| rs104893849 | pathogenic | 0 | 0 | 0 |
| rs571038432 | pathogenic | 1 | 1 | 0 |
| rs2270655 | benign | 1 | 1 | 1 |
| rs315952 | benign | 0 | 0 | 0 |
| rs151176548 | benign | 0 | 0 | 0 |
| rs122459147 | pathogenic | 1 | 1 | 0 |
| rs122458140 | pathogenic | 0 | 0 | 0 |

| | | | | |
|---|---|---|---|---|
| rs122458142 | pathogenic | 1 | 1 | 0 |
| rs267606812 | pathogenic | 1 | 1 | 0 |
| rs267606813 | pathogenic | 1 | 1 | 0 |
| rs122458143 | pathogenic | 0 | 0 | 0 |
| rs122459146 | pathogenic | 0 | 0 | 0 |
| rs122458144 | pathogenic | 0 | 0 | 0 |
| rs122458145 | pathogenic | 0 | 0 | 0 |
| rs113994007 | pathogenic | 1 | 1 | 0 |
| rs758746181 | pathogenic | 1 | 1 | 1 |
| rs145034527 | pathogenic | 1 | 1 | 0 |
| rs119468006 | pathogenic | 1 | 1 | 0 |
| rs201908721 | pathogenic | 0 | 0 | 0 |
| rs755933881 | pathogenic | 1 | 1 | 0 |
| rs771578775 | pathogenic | 0 | 0 | 0 |
| rs17849927 | benign | 0 | 0 | 0 |
| rs139133094 | benign | 0 | 0 | 0 |
| rs12593 | benign | 0 | 0 | 0 |
| rs578189699 | pathogenic | 1 | 1 | 0 |
| rs119468008 | pathogenic | 0 | 0 | 0 |
| rs768524626 | benign | 0 | 0 | 0 |
| rs119468009 | pathogenic | 0 | 0 | 0 |
| rs119468004 | pathogenic | 0 | 0 | 0 |
| rs199874519 | pathogenic | 0 | 0 | 0 |
| rs376478331 | pathogenic | 0 | 0 | 0 |
| rs61995958 | pathogenic | 0 | 0 | 0 |
| rs199714731 | benign | 0 | 0 | 0 |
| rs149185954 | benign | 0 | 0 | 0 |
| rs121918353 | pathogenic | 0 | 0 | 0 |
| rs1131695 | benign | 0 | 0 | 0 |
| rs28939668 | pathogenic | 0 | 0 | 0 |
| rs758432471 | pathogenic | 1 | 1 | 1 |
| rs752746786 | pathogenic | 1 | 1 | 1 |
| rs28935471 | pathogenic | 0 | 0 | 0 |
| rs80338837 | pathogenic | 0 | 0 | 0 |
| rs137853312 | pathogenic | 0 | 0 | 0 |
| rs28935472 | pathogenic | 0 | 0 | 0 |
| rs28935473 | pathogenic | 0 | 0 | 0 |
| rs267606752 | pathogenic | 0 | 0 | 0 |
| rs3917973 | benign | 0 | 0 | 0 |
| rs3918018 | benign | 0 | 0 | 0 |
| rs757976755 | pathogenic | 0 | 0 | 0 |
| rs121918329 | pathogenic | 1 | 1 | 0 |
| rs770114459 | pathogenic | 0 | 0 | 0 |
| rs6662272 | benign | 0 | 0 | 0 |
| rs121918242 | pathogenic | 1 | 1 | 0 |
| rs121918240 | pathogenic | 1 | 1 | 0 |
| rs200094982 | pathogenic | 0 | 0 | 0 |
| rs121918241 | pathogenic | 0 | 0 | 0 |
| rs140522266 | pathogenic | 1 | 1 | 0 |
| rs121918243 | pathogenic | 1 | 1 | 0 |
| rs370596113 | pathogenic | 1 | 1 | 0 |
| rs398124295 | pathogenic | 1 | 1 | 0 |
| rs587776889 | pathogenic | 1 | 1 | 0 |
| rs747527726 | pathogenic | 1 | 1 | 0 |

| | | | | |
|---|---|---|---|---|
| rs112631212 | benign | 0 | 0 | 0 |
| rs104894605 | pathogenic | 0 | 0 | 0 |
| rs104894607 | pathogenic | 0 | 0 | 0 |
| rs104894606 | pathogenic | 0 | 0 | 0 |
| rs143873873 | benign | 0 | 0 | 0 |
| rs104894388 | pathogenic | 1 | 1 | 0 |
| rs104894391 | pathogenic | 1 | 1 | 0 |
| rs5350 | benign | 1 | 1 | 1 |
| rs104894390 | pathogenic | 1 | 1 | 1 |
| rs5346 | benign | 1 | 1 | 1 |
| rs121912500 | pathogenic | 0 | 0 | 0 |
| rs74315450 | pathogenic | 1 | 1 | 0 |
| rs121913396 | pathogenic | 1 | 1 | 1 |
| rs28931588 | pathogenic | 1 | 1 | 1 |
| rs121913400 | pathogenic | 0 | 0 | 0 |
| rs28931589 | pathogenic | 1 | 1 | 1 |
| rs121913403 | pathogenic | 0 | 0 | 0 |
| rs121913413 | pathogenic | 0 | 0 | 0 |
| rs760107623 | pathogenic | 1 | 1 | 0 |
| rs16866378 | benign | 1 | 1 | 0 |
| rs201025290 | pathogenic | 1 | 1 | 0 |
| rs12720356 | benign | 1 | 1 | 0 |
| rs55882956 | benign | 0 | 1 | 0 |
| rs4986933 | benign | 1 | 1 | 0 |
| rs2020957 | benign | 1 | 1 | 1 |
| rs1800124 | benign | 1 | 1 | 1 |
| rs150077735 | benign | 1 | 1 | 1 |
| rs201419000 | benign | 0 | 0 | 0 |
| rs772044176 | pathogenic | 0 | 0 | 0 |
| rs138346388 | benign | 0 | 0 | 0 |
| rs121964916 | pathogenic | 0 | 0 | 0 |
| rs112534524 | pathogenic | 0 | 0 | 0 |
| rs182078921 | benign | 1 | 1 | 1 |
| rs113273712 | benign | 0 | 0 | 0 |
| rs74817407 | benign | 1 | 1 | 1 |
| rs61733901 | benign | 1 | 1 | 1 |
| rs121964912 | pathogenic | 1 | 1 | 1 |
| rs41278047 | benign | 1 | 1 | 1 |
| rs7437875 | benign | 1 | 1 | 1 |
| rs113460688 | benign | 1 | 1 | 1 |
| rs754572081 | benign | 1 | 1 | 1 |
| rs201082692 | benign | 1 | 1 | 1 |
| rs202085803 | benign | 1 | 1 | 1 |
| rs141320511 | benign | 0 | 0 | 0 |
| rs78501403 | benign | 0 | 0 | 0 |
| rs28939675 | pathogenic | 0 | 0 | 0 |
| rs74315522 | pathogenic | 0 | 0 | 0 |
| rs121434639 | pathogenic | 1 | 1 | 0 |
| rs121434640 | pathogenic | 1 | 1 | 1 |
| rs757282628 | benign | 1 | 1 | 1 |
| rs386834010 | pathogenic | 0 | 0 | 0 |
| rs200863680 | pathogenic | 0 | 0 | 0 |
| rs193919336 | pathogenic | 1 | 1 | 0 |
| rs386834011 | pathogenic | 0 | 0 | 0 |

| | | | | |
|---|---|---|---|---|
| rs28940869 | pathogenic | 0 | 0 | 0 |
| rs386834014 | pathogenic | 1 | 1 | 0 |
| rs267606960 | pathogenic | 0 | 0 | 0 |
| rs28942068 | pathogenic | 0 | 0 | 0 |
| rs193919335 | pathogenic | 0 | 0 | 0 |
| rs74374973 | pathogenic | 0 | 0 | 0 |
| rs386834018 | pathogenic | 0 | 0 | 0 |
| rs267606962 | pathogenic | 0 | 0 | 0 |
| rs6659553 | benign | 0 | 0 | 0 |
| rs29281 | benign | 0 | 0 | 0 |
| rs782265090 | benign | 0 | 0 | 0 |
| rs35910969 | pathogenic | 0 | 0 | 0 |
| rs121909113 | pathogenic | 1 | 1 | 1 |
| rs29001571 | pathogenic | 1 | 1 | 1 |
| rs587781250 | pathogenic | 1 | 1 | 1 |
| rs28939680 | pathogenic | 1 | 1 | 1 |
| rs28939681 | pathogenic | 1 | 1 | 1 |
| rs121909112 | pathogenic | 1 | 1 | 1 |
| rs28937568 | pathogenic | 1 | 1 | 1 |
| rs1799895 | pathogenic | 0 | 0 | 0 |
| rs386833431 | pathogenic | 1 | 1 | 1 |
| rs121964905 | pathogenic | 1 | 1 | 1 |
| rs121964906 | pathogenic | 1 | 1 | 1 |
| rs11551664 | pathogenic | 1 | 1 | 1 |
| rs104894705 | pathogenic | 1 | 1 | 1 |
| rs104893805 | pathogenic | 0 | 0 | 0 |
| rs104894205 | pathogenic | 0 | 0 | 0 |
| rs104894204 | pathogenic | 1 | 1 | 0 |
| rs62642926 | pathogenic | 0 | 0 | 0 |
| rs5030841 | pathogenic | 1 | 1 | 1 |
| rs75193786 | pathogenic | 1 | 1 | 1 |
| rs62508682 | pathogenic | 0 | 0 | 0 |
| rs3850625 | benign | 1 | 1 | 1 |
| rs141908793 | benign | 0 | 0 | 0 |
| rs187864727 | benign | 0 | 0 | 0 |
| rs548909101 | benign | 0 | 0 | 0 |
| rs3825394 | benign | 0 | 1 | 0 |
| rs143548402 | benign | 0 | 0 | 0 |
| rs267607146 | pathogenic | 0 | 0 | 0 |
| rs267607144 | pathogenic | 0 | 0 | 0 |
| rs267607148 | pathogenic | 1 | 1 | 1 |
| rs267607143 | pathogenic | 0 | 0 | 0 |
| rs267607145 | pathogenic | 0 | 0 | 0 |
| rs121912636 | pathogenic | 1 | 1 | 1 |
| rs121912634 | pathogenic | 1 | 1 | 1 |
| rs570675468 | benign | 0 | 0 | 0 |
| rs368861241 | pathogenic | 1 | 1 | 1 |
| rs104894528 | pathogenic | 1 | 1 | 1 |
| rs77493891 | benign | 0 | 0 | 0 |
| rs62640903 | benign | 0 | 0 | 0 |
| rs149926091 | benign | 1 | 1 | 1 |
| rs641081 | benign | 0 | 0 | 0 |
| rs4930199 | benign | 0 | 0 | 0 |
| rs121913020 | pathogenic | 0 | 0 | 0 |

| | | | | |
|---|---|---|---|---|
| rs1799793 | benign | 0 | 0 | 0 |
| rs121913016 | pathogenic | 0 | 0 | 0 |
| rs121913025 | pathogenic | 0 | 0 | 0 |
| rs121913019 | pathogenic | 0 | 0 | 0 |
| rs376556895 | pathogenic | 1 | 1 | 1 |
| rs121913024 | pathogenic | 1 | 1 | 1 |
| rs121913021 | pathogenic | 0 | 0 | 0 |
| rs121913023 | pathogenic | 0 | 0 | 0 |
| rs41556519 | pathogenic | 0 | 0 | 0 |
| rs121913022 | pathogenic | 0 | 0 | 0 |
| rs144564120 | pathogenic | 0 | 0 | 0 |
| rs121913026 | pathogenic | 1 | 1 | 1 |
| rs121913018 | pathogenic | 1 | 1 | 1 |
| rs142954643 | benign | 0 | 0 | 0 |
| rs76655496 | benign | 0 | 0 | 0 |
| rs104894371 | pathogenic | 1 | 1 | 0 |
| rs137964936 | benign | 0 | 0 | 0 |
| rs759736132 | pathogenic | 0 | 0 | 0 |
| rs387906595 | pathogenic | 1 | 1 | 0 |
| rs768248216 | pathogenic | 0 | 0 | 0 |
| rs548231008 | benign | 0 | 0 | 0 |
| rs757094384 | benign | 0 | 0 | 0 |
| rs867529 | benign | 0 | 0 | 0 |
| rs13045 | benign | 1 | 1 | 1 |
| rs121908570 | pathogenic | 0 | 0 | 0 |
| rs201872586 | benign | 1 | 1 | 0 |
| rs149092990 | benign | 0 | 0 | 0 |
| rs115585711 | benign | 0 | 0 | 0 |
| rs137909416 | benign | 1 | 1 | 0 |
| rs143388949 | benign | 1 | 1 | 1 |
| rs73137716 | benign | 1 | 1 | 1 |
| rs121908947 | pathogenic | 1 | 1 | 1 |
| rs398122904 | pathogenic | 1 | 1 | 1 |
| rs35715497 | benign | 1 | 1 | 0 |
| rs35462421 | benign | 0 | 0 | 0 |
| rs119475043 | pathogenic | 0 | 0 | 0 |
| rs119475042 | pathogenic | 0 | 0 | 0 |
| rs1076669 | benign | 1 | 1 | 0 |
| rs141146885 | benign | 0 | 0 | 0 |
| rs3026906 | pathogenic | 0 | 0 | 0 |
| rs75362221 | benign | 1 | 1 | 1 |
| rs138472116 | benign | 1 | 1 | 1 |
| rs371040282 | pathogenic | 1 | 1 | 1 |
| rs4988496 | benign | 0 | 0 | 0 |
| rs121918117 | pathogenic | 1 | 1 | 1 |
| rs4988498 | benign | 0 | 0 | 0 |
| rs5035 | benign | 0 | 0 | 0 |
| rs121918322 | pathogenic | 1 | 1 | 1 |
| rs370359540 | benign | 0 | 0 | 0 |
| rs758972589 | pathogenic | 0 | 0 | 0 |
| rs144856889 | benign | 0 | 0 | 0 |
| rs121434623 | pathogenic | 1 | 1 | 1 |
| rs61735600 | benign | 0 | 0 | 0 |
| rs72550870 | benign | 1 | 1 | 1 |

| | | | | |
|---|---|---|---|---|
| rs56392418 | benign | 0 | 0 | 0 |
| rs41307788 | benign | 1 | 1 | 1 |
| rs387907239 | pathogenic | 1 | 1 | 0 |
| rs28931584 | pathogenic | 0 | 0 | 0 |
| rs121912745 | pathogenic | 0 | 0 | 0 |
| rs121912744 | pathogenic | 0 | 0 | 0 |
| rs121912754 | pathogenic | 1 | 1 | 1 |
| rs121912746 | pathogenic | 0 | 0 | 0 |
| rs121912757 | pathogenic | 0 | 0 | 0 |
| rs121912758 | pathogenic | 0 | 0 | 0 |
| rs121912748 | pathogenic | 0 | 0 | 0 |
| rs121912755 | pathogenic | 0 | 0 | 0 |
| rs121912741 | pathogenic | 0 | 0 | 0 |
| rs121912753 | pathogenic | 0 | 0 | 0 |
| rs121912750 | pathogenic | 0 | 0 | 0 |
| rs2285644 | pathogenic | 0 | 0 | 0 |
| rs121912751 | pathogenic | 0 | 0 | 0 |
| rs5026 | benign | 0 | 0 | 0 |
| rs121912759 | pathogenic | 0 | 0 | 0 |
| rs28931585 | pathogenic | 0 | 0 | 0 |
| rs17122769 | benign | 0 | 0 | 0 |
| rs28362322 | benign | 1 | 1 | 1 |
| rs3824874 | benign | 0 | 0 | 0 |
| rs146004831 | benign | 0 | 0 | 0 |
| rs142155860 | benign | 0 | 0 | 0 |
| rs121434404 | pathogenic | 0 | 0 | 0 |
| rs61735578 | benign | 1 | 1 | 1 |
| rs558018 | benign | 0 | 0 | 0 |
| rs61735577 | benign | 1 | 1 | 1 |
| rs76784113 | benign | 0 | 0 | 0 |
| rs199469640 | pathogenic | 0 | 0 | 0 |
| rs199469628 | pathogenic | 1 | 1 | 0 |
| rs199469629 | pathogenic | 0 | 1 | 0 |
| rs387907156 | pathogenic | 0 | 1 | 0 |
| rs199469643 | pathogenic | 0 | 0 | 0 |
| rs199469646 | pathogenic | 0 | 0 | 0 |
| rs766294940 | pathogenic | 0 | 0 | 0 |
| rs387907347 | pathogenic | 0 | 0 | 0 |
| rs137853590 | pathogenic | 1 | 1 | 1 |
| rs137853589 | pathogenic | 1 | 1 | 1 |
| rs137853588 | pathogenic | 1 | 1 | 1 |
| rs55909005 | benign | 0 | 0 | 0 |
| rs4581 | benign | 0 | 0 | 0 |
| rs63750123 | benign | 0 | 0 | 0 |
| rs10254120 | benign | 0 | 0 | 0 |
| rs121434629 | pathogenic | 1 | 1 | 0 |
| rs200313585 | benign | 0 | 0 | 0 |
| rs143162541 | benign | 1 | 1 | 0 |
| rs760615315 | benign | 0 | 0 | 0 |
| rs768488890 | benign | 1 | 1 | 1 |
| rs116373169 | benign | 1 | 1 | 0 |
| rs63750871 | pathogenic | 1 | 1 | 0 |
| rs116349687 | benign | 0 | 0 | 0 |
| rs375289386 | benign | 0 | 0 | 0 |

| | | | | |
|---|---|---|---|---|
| rs760228510 | pathogenic | 1 | 1 | 1 |
| rs587779343 | pathogenic | 1 | 1 | 1 |
| rs573125799 | benign | 0 | 0 | 0 |
| rs1805322 | benign | 0 | 0 | 0 |
| rs267608153 | pathogenic | 0 | 0 | 0 |
| rs139194813 | benign | 0 | 0 | 0 |
| rs530021751 | benign | 0 | 0 | 0 |
| rs200640585 | pathogenic | 0 | 0 | 0 |
| rs139438201 | benign | 0 | 0 | 0 |
| rs200513014 | benign | 0 | 0 | 0 |
| rs567102013 | benign | 0 | 0 | 0 |
| rs121908608 | pathogenic | 0 | 0 | 0 |
| rs121908606 | pathogenic | 0 | 0 | 0 |
| rs17103671 | benign | 0 | 0 | 0 |
| rs34725281 | benign | 0 | 0 | 0 |
| rs104894421 | pathogenic | 1 | 1 | 0 |
| rs121918145 | pathogenic | 1 | 1 | 1 |
| rs121918152 | pathogenic | 1 | 1 | 1 |
| rs56379106 | benign | 0 | 0 | 0 |
| rs4987076 | benign | 0 | 0 | 0 |
| rs370243877 | pathogenic | 0 | 0 | 0 |
| rs17679445 | benign | 1 | 1 | 1 |
| rs550423482 | pathogenic | 1 | 1 | 1 |
| rs104894629 | pathogenic | 1 | 1 | 1 |
| rs773450573 | pathogenic | 1 | 1 | 1 |
| rs116679717 | benign | 0 | 0 | 0 |
| rs372000848 | pathogenic | 0 | 0 | 0 |
| rs138595973 | benign | 0 | 0 | 0 |
| rs115244417 | benign | 0 | 0 | 0 |
| rs11575292 | benign | 0 | 0 | 0 |
| rs137853211 | pathogenic | 1 | 1 | 1 |
| rs137853207 | pathogenic | 0 | 0 | 0 |
| rs137853210 | pathogenic | 1 | 1 | 1 |
| rs6262 | benign | 0 | 0 | 0 |
| rs6263 | benign | 0 | 1 | 0 |
| rs137853208 | pathogenic | 0 | 1 | 0 |
| rs137853212 | pathogenic | 1 | 1 | 0 |
| rs137853209 | pathogenic | 1 | 1 | 1 |
| rs201951824 | pathogenic | 0 | 0 | 0 |
| rs11575542 | benign | 0 | 0 | 0 |
| rs199681788 | benign | 0 | 1 | 0 |
| rs748835902 | benign | 1 | 1 | 1 |
| rs753719949 | benign | 0 | 0 | 0 |
| rs267606647 | pathogenic | 0 | 0 | 0 |
| rs267606648 | pathogenic | 1 | 1 | 1 |
| rs267606643 | pathogenic | 0 | 1 | 0 |
| rs559947967 | pathogenic | 1 | 1 | 0 |
| rs267606645 | pathogenic | 1 | 1 | 1 |
| rs12116440 | benign | 0 | 0 | 0 |
| rs121908539 | pathogenic | 0 | 0 | 0 |
| rs137853096 | pathogenic | 1 | 1 | 0 |
| rs148363262 | benign | 1 | 1 | 0 |
| rs587777442 | pathogenic | 0 | 0 | 0 |
| rs25640 | benign | 1 | 1 | 1 |

| | | | | |
|---|---|---|---|---|
| rs28943589 | benign | 0 | 0 | 0 |
| rs1143650 | benign | 1 | 1 | 1 |
| rs150886026 | benign | 0 | 1 | 0 |
| rs121908133 | pathogenic | 1 | 1 | 0 |
| rs121908131 | pathogenic | 0 | 0 | 0 |
| rs754477154 | pathogenic | 0 | 0 | 0 |
| rs747334007 | benign | 0 | 0 | 0 |
| rs121908132 | pathogenic | 0 | 0 | 0 |
| rs104894103 | pathogenic | 1 | 1 | 0 |
| rs140088149 | benign | 0 | 0 | 0 |
| rs149803148 | benign | 0 | 0 | 0 |
| rs11064480 | benign | 0 | 0 | 0 |
| rs74435397 | pathogenic | 1 | 1 | 1 |
| rs587783626 | pathogenic | 1 | 1 | 1 |
| rs148211042 | pathogenic | 1 | 1 | 1 |
| rs17849502 | benign | 0 | 0 | 0 |
| rs13306575 | pathogenic | 1 | 1 | 1 |
| rs201426733 | benign | 0 | 0 | 0 |
| rs182278784 | benign | 1 | 1 | 1 |
| rs77786415 | benign | 1 | 1 | 1 |
| rs200192119 | benign | 1 | 1 | 1 |
| rs12877501 | benign | 1 | 1 | 1 |
| rs77985244 | benign | 1 | 1 | 1 |
| rs5907 | pathogenic | 1 | 1 | 0 |
| rs2305178 | benign | 0 | 0 | 0 |
| rs3135867 | benign | 0 | 0 | 0 |
| rs121913482 | pathogenic | 1 | 1 | 1 |
| rs121913483 | pathogenic | 1 | 1 | 1 |
| rs4647924 | pathogenic | 1 | 1 | 1 |
| rs121913111 | pathogenic | 1 | 1 | 1 |
| rs104894002 | pathogenic | 0 | 0 | 0 |
| rs104894001 | pathogenic | 0 | 0 | 0 |
| rs75932628 | benign | 0 | 0 | 0 |
| rs201258663 | pathogenic | 0 | 0 | 0 |
| rs104893998 | pathogenic | 0 | 0 | 0 |
| rs2234253 | benign | 1 | 1 | 0 |
| rs121908402 | pathogenic | 1 | 1 | 0 |
| rs1805389 | benign | 0 | 0 | 0 |
| rs1805388 | benign | 0 | 0 | 0 |
| rs748385144 | pathogenic | 0 | 0 | 0 |
| rs104894420 | pathogenic | 0 | 0 | 0 |
| rs104894418 | pathogenic | 0 | 0 | 0 |
| rs61731910 | benign | 0 | 0 | 0 |
| rs758723288 | pathogenic | 1 | 1 | 0 |
| rs201865375 | pathogenic | 1 | 1 | 0 |
| rs565090080 | pathogenic | 1 | 1 | 1 |
| rs113042166 | benign | 0 | 0 | 0 |
| rs61736326 | benign | 1 | 1 | 1 |
| rs12030699 | benign | 1 | 1 | 1 |
| rs121913578 | pathogenic | 1 | 1 | 1 |
| rs61739582 | benign | 0 | 0 | 0 |
| rs745983207 | pathogenic | 0 | 0 | 0 |
| rs763524380 | benign | 0 | 0 | 0 |
| rs121912792 | pathogenic | 0 | 0 | 0 |

| | | | | |
|---|---|---|---|---|
| rs121912793 | pathogenic | 0 | 0 | 0 |
| rs387906599 | pathogenic | 0 | 0 | 0 |
| rs121912794 | pathogenic | 0 | 1 | 0 |
| rs121912795 | pathogenic | 0 | 0 | 0 |
| rs121912796 | pathogenic | 1 | 1 | 0 |
| rs121912797 | pathogenic | 0 | 1 | 0 |
| rs121912798 | pathogenic | 1 | 1 | 1 |
| rs80359812 | pathogenic | 1 | 1 | 0 |
| rs776583130 | benign | 1 | 1 | 0 |
| rs80359814 | pathogenic | 1 | 1 | 0 |
| rs202060209 | pathogenic | 1 | 1 | 0 |
| rs267607061 | pathogenic | 1 | 1 | 0 |
| rs267607060 | pathogenic | 0 | 0 | 0 |
| rs80359816 | pathogenic | 1 | 1 | 0 |
| rs80359818 | pathogenic | 1 | 1 | 0 |
| rs387907313 | pathogenic | 0 | 0 | 0 |
| rs587784395 | benign | 0 | 0 | 0 |
| rs5811 | benign | 0 | 0 | 0 |
| rs121909740 | pathogenic | 1 | 1 | 0 |
| rs148518827 | benign | 1 | 1 | 0 |
| rs121909739 | pathogenic | 1 | 1 | 0 |
| rs780528770 | benign | 1 | 1 | 0 |
| rs13306758 | pathogenic | 0 | 0 | 0 |
| rs267607059 | pathogenic | 0 | 0 | 0 |
| rs61730638 | benign | 0 | 0 | 0 |
| rs104894533 | pathogenic | 0 | 0 | 0 |
| rs104894527 | pathogenic | 0 | 0 | 0 |
| rs200503569 | pathogenic | 1 | 1 | 1 |
| rs80338700 | pathogenic | 1 | 1 | 1 |
| rs765433263 | pathogenic | 1 | 1 | 1 |
| rs104894530 | pathogenic | 1 | 1 | 1 |
| rs80338701 | pathogenic | 1 | 1 | 1 |
| rs141498002 | pathogenic | 1 | 1 | 1 |
| rs104894525 | pathogenic | 0 | 0 | 0 |
| rs80338703 | pathogenic | 0 | 0 | 0 |
| rs28936415 | pathogenic | 1 | 1 | 0 |
| rs150719105 | pathogenic | 0 | 0 | 0 |
| rs148032587 | pathogenic | 0 | 0 | 0 |
| rs150577656 | pathogenic | 0 | 0 | 0 |
| rs190521996 | pathogenic | 0 | 0 | 0 |
| rs104894526 | pathogenic | 1 | 1 | 1 |
| rs80338704 | pathogenic | 0 | 0 | 0 |
| rs199562225 | pathogenic | 1 | 1 | 1 |
| rs532870929 | pathogenic | 0 | 0 | 0 |
| rs398123309 | pathogenic | 0 | 0 | 0 |
| rs78290141 | pathogenic | 0 | 0 | 0 |
| rs80338705 | pathogenic | 0 | 0 | 0 |
| rs104894531 | pathogenic | 1 | 1 | 1 |
| rs80338706 | pathogenic | 0 | 0 | 0 |
| rs80338707 | pathogenic | 0 | 0 | 0 |
| rs80338708 | pathogenic | 0 | 0 | 0 |
| rs80338709 | pathogenic | 0 | 0 | 0 |
| rs104893833 | benign | 0 | 0 | 0 |
| rs61744585 | benign | 0 | 0 | 0 |

| | | | | |
|---|---|---|---|---|
| rs73569515 | benign | 0 | 0 | 0 |
| rs398123373 | pathogenic | 0 | 0 | 0 |
| rs756854513 | pathogenic | 0 | 0 | 0 |
| rs58232698 | benign | 0 | 0 | 0 |
| rs9600455 | benign | 0 | 0 | 0 |
| rs201016593 | pathogenic | 0 | 0 | 0 |
| rs756039188 | pathogenic | 0 | 0 | 0 |
| rs121908024 | pathogenic | 0 | 0 | 0 |
| rs373144619 | benign | 0 | 0 | 0 |
| rs138078086 | benign | 0 | 0 | 0 |
| rs376207800 | pathogenic | 0 | 0 | 0 |
| rs749038326 | pathogenic | 1 | 1 | 0 |
| rs372845091 | benign | 0 | 0 | 0 |
| rs377271627 | pathogenic | 0 | 0 | 0 |
| rs563390335 | pathogenic | 0 | 0 | 0 |
| rs13306510 | benign | 0 | 0 | 0 |
| rs764042910 | pathogenic | 0 | 0 | 0 |
| rs201384282 | benign | 1 | 1 | 1 |
| rs730882086 | pathogenic | 1 | 1 | 1 |
| rs756613387 | pathogenic | 1 | 1 | 1 |
| rs121908028 | benign | 1 | 1 | 0 |
| rs773328511 | benign | 0 | 0 | 0 |
| rs749220643 | pathogenic | 1 | 1 | 0 |
| rs730882089 | pathogenic | 0 | 0 | 0 |
| rs146651743 | benign | 0 | 0 | 0 |
| rs148698650 | pathogenic | 0 | 0 | 0 |
| rs140241383 | benign | 0 | 0 | 0 |
| rs757252110 | pathogenic | 0 | 0 | 0 |
| rs13306512 | pathogenic | 1 | 1 | 1 |
| rs121908043 | pathogenic | 1 | 1 | 1 |
| rs773658037 | pathogenic | 0 | 0 | 0 |
| rs774439908 | pathogenic | 0 | 0 | 0 |
| rs779732323 | pathogenic | 1 | 1 | 0 |
| rs749780672 | pathogenic | 0 | 0 | 0 |
| rs375651668 | benign | 1 | 1 | 1 |
| rs193922568 | pathogenic | 1 | 1 | 0 |
| rs750363970 | benign | 1 | 1 | 0 |
| rs5932 | benign | 0 | 0 | 0 |
| rs755154048 | pathogenic | 1 | 1 | 0 |
| rs759876319 | pathogenic | 0 | 0 | 0 |
| rs764550980 | pathogenic | 1 | 1 | 0 |
| rs779940524 | pathogenic | 0 | 0 | 0 |
| rs563382937 | pathogenic | 1 | 1 | 1 |
| rs28942083 | pathogenic | 0 | 1 | 0 |
| rs770744861 | pathogenic | 0 | 1 | 0 |
| rs121908031 | pathogenic | 0 | 1 | 0 |
| rs776217028 | pathogenic | 0 | 1 | 0 |
| rs763625913 | pathogenic | 0 | 0 | 0 |
| rs3736772 | benign | 0 | 0 | 0 |
| rs104894696 | benign | 1 | 1 | 1 |
| rs121918177 | pathogenic | 0 | 1 | 0 |
| rs121918181 | pathogenic | 1 | 1 | 0 |
| rs121918174 | pathogenic | 0 | 1 | 0 |
| rs121918182 | pathogenic | 0 | 0 | 0 |

| | | | | |
|---|---|---|---|---|
| rs121918175 | pathogenic | 1 | 1 | 1 |
| rs121918185 | pathogenic | 1 | 1 | 1 |
| rs121918173 | pathogenic | 1 | 1 | 0 |
| rs121918178 | pathogenic | 1 | 1 | 1 |
| rs121918179 | pathogenic | 1 | 1 | 1 |
| rs121918183 | pathogenic | 0 | 0 | 0 |
| rs121918176 | pathogenic | 1 | 1 | 1 |
| rs121918172 | pathogenic | 0 | 1 | 0 |
| rs121918184 | pathogenic | 0 | 0 | 0 |
| rs121434560 | pathogenic | 1 | 1 | 0 |
| rs121434561 | pathogenic | 1 | 1 | 0 |
| rs3218716 | pathogenic | 1 | 1 | 1 |
| rs145532615 | benign | 0 | 0 | 0 |
| rs149840927 | pathogenic | 0 | 0 | 0 |
| rs730880762 | pathogenic | 0 | 0 | 0 |
| rs730880764 | pathogenic | 0 | 0 | 0 |
| rs730880765 | pathogenic | 0 | 0 | 0 |
| rs587782962 | pathogenic | 0 | 0 | 0 |
| rs141735183 | pathogenic | 0 | 0 | 0 |
| rs397516187 | pathogenic | 0 | 0 | 0 |
| rs727505026 | pathogenic | 0 | 0 | 0 |
| rs529700838 | pathogenic | 0 | 0 | 0 |
| rs540263945 | pathogenic | 0 | 0 | 0 |
| rs149103761 | benign | 0 | 0 | 0 |
| rs730880906 | pathogenic | 0 | 0 | 0 |
| rs730880785 | pathogenic | 0 | 0 | 0 |
| rs397516199 | pathogenic | 0 | 0 | 0 |
| rs730880725 | benign | 0 | 0 | 0 |
| rs730880796 | pathogenic | 0 | 0 | 0 |
| rs201307101 | benign | 0 | 0 | 0 |
| rs397516214 | pathogenic | 0 | 0 | 0 |
| rs3729823 | benign | 0 | 0 | 0 |
| rs397516218 | pathogenic | 0 | 0 | 0 |
| rs121913647 | pathogenic | 0 | 0 | 0 |
| rs373514686 | pathogenic | 0 | 0 | 0 |
| rs763538103 | pathogenic | 0 | 0 | 0 |
| rs121913650 | pathogenic | 0 | 0 | 0 |
| rs139222507 | pathogenic | 0 | 0 | 0 |
| rs369940645 | pathogenic | 0 | 0 | 0 |
| rs763073072 | pathogenic | 0 | 0 | 0 |
| rs145734640 | pathogenic | 0 | 0 | 0 |
| rs730880821 | pathogenic | 0 | 0 | 0 |
| rs28933098 | pathogenic | 0 | 0 | 0 |
| rs372381770 | pathogenic | 0 | 0 | 0 |
| rs146796870 | benign | 0 | 0 | 0 |
| rs104894485 | pathogenic | 1 | 1 | 1 |
| rs35407712 | benign | 0 | 0 | 0 |
| rs35842750 | benign | 0 | 0 | 0 |
| rs2721191 | benign | 0 | 0 | 0 |
| rs35098923 | benign | 0 | 0 | 0 |
| rs34293591 | benign | 1 | 1 | 0 |
| rs36023964 | benign | 0 | 0 | 0 |
| rs4251691 | benign | 0 | 0 | 0 |
| rs34666647 | benign | 0 | 0 | 0 |

| | | | | |
|---|---|---|---|---|
| rs4925828 | benign | 0 | 0 | 0 |
| rs35348691 | benign | 0 | 0 | 0 |
| rs36078464 | benign | 0 | 0 | 0 |
| rs34915097 | benign | 0 | 0 | 0 |
| rs35101495 | benign | 0 | 0 | 0 |
| rs141568342 | pathogenic | 0 | 0 | 0 |
| rs1800571 | pathogenic | 0 | 0 | 0 |
| rs140566084 | benign | 1 | 1 | 1 |
| rs727503791 | pathogenic | 1 | 1 | 0 |
| rs545215807 | pathogenic | 0 | 0 | 0 |
| rs113994168 | pathogenic | 0 | 0 | 0 |
| rs113994167 | pathogenic | 0 | 0 | 0 |
| rs200788251 | pathogenic | 0 | 0 | 0 |
| rs200573371 | pathogenic | 0 | 0 | 0 |
| rs771874163 | pathogenic | 1 | 1 | 1 |
| rs112406105 | pathogenic | 1 | 1 | 1 |
| rs118204015 | pathogenic | 1 | 1 | 1 |
| rs113994169 | pathogenic | 0 | 0 | 0 |
| rs118204018 | pathogenic | 0 | 1 | 0 |
| rs138834083 | pathogenic | 0 | 0 | 0 |
| rs533055438 | pathogenic | 1 | 1 | 1 |
| rs2309689 | pathogenic | 1 | 1 | 1 |
| rs138058572 | pathogenic | 1 | 1 | 1 |
| rs118204017 | pathogenic | 1 | 1 | 0 |
| rs766742117 | pathogenic | 1 | 1 | 1 |
| rs751995154 | pathogenic | 1 | 1 | 1 |
| rs200771970 | pathogenic | 0 | 0 | 0 |
| rs118204014 | pathogenic | 0 | 0 | 0 |
| rs148584617 | pathogenic | 1 | 1 | 1 |
| rs121912720 | pathogenic | 0 | 0 | 0 |
| rs121912721 | pathogenic | 0 | 0 | 0 |
| rs28929476 | pathogenic | 0 | 0 | 0 |
| rs28931574 | pathogenic | 1 | 1 | 1 |
| rs12718465 | benign | 0 | 0 | 0 |
| rs144689354 | pathogenic | 0 | 0 | 0 |
| rs149095705 | pathogenic | 0 | 0 | 0 |
| rs572839648 | benign | 1 | 1 | 1 |
| rs121909715 | pathogenic | 1 | 1 | 0 |
| rs34077456 | benign | 1 | 1 | 1 |
| rs397517876 | benign | 0 | 0 | 0 |
| rs1064074 | benign | 0 | 0 | 0 |
| rs142751309 | benign | 0 | 0 | 0 |
| rs116275562 | benign | 0 | 0 | 0 |
| rs148095899 | benign | 0 | 0 | 0 |
| rs751838040 | pathogenic | 0 | 0 | 0 |
| rs121434324 | pathogenic | 1 | 1 | 1 |
| rs143367518 | pathogenic | 0 | 0 | 0 |
| rs121434325 | pathogenic | 0 | 0 | 0 |
| rs397515328 | pathogenic | 0 | 0 | 0 |
| rs376039938 | pathogenic | 0 | 0 | 0 |
| rs387907243 | pathogenic | 0 | 0 | 0 |
| rs28934876 | pathogenic | 0 | 0 | 0 |
| rs387907245 | pathogenic | 0 | 0 | 0 |
| rs138544311 | benign | 0 | 0 | 0 |

| | | | | |
|---|---|---|---|---|
| rs643788 | benign | 0 | 0 | 0 |
| rs749977756 | benign | 0 | 0 | 0 |
| rs1800574 | benign | 0 | 0 | 0 |
| rs137853336 | pathogenic | 1 | 1 | 1 |
| rs111916034 | benign | 0 | 0 | 0 |
| rs307826 | benign | 0 | 0 | 0 |
| rs35874891 | benign | 0 | 0 | 0 |
| rs111033580 | pathogenic | 1 | 1 | 1 |
| rs111033581 | pathogenic | 1 | 1 | 1 |
| rs121908373 | pathogenic | 0 | 0 | 0 |
| rs767122713 | pathogenic | 0 | 0 | 0 |
| rs2233213 | benign | 0 | 0 | 0 |
| rs121908605 | pathogenic | 1 | 1 | 0 |
| rs11887534 | benign | 0 | 0 | 0 |
| rs4148211 | benign | 0 | 0 | 0 |
| rs80025980 | benign | 0 | 0 | 0 |
| rs137854891 | pathogenic | 1 | 1 | 0 |
| rs74358901 | benign | 0 | 0 | 0 |
| rs9282574 | benign | 0 | 0 | 0 |
| rs137852993 | pathogenic | 0 | 0 | 0 |
| rs137852990 | pathogenic | 0 | 0 | 0 |
| rs137852987 | pathogenic | 0 | 0 | 0 |
| rs4148217 | benign | 0 | 0 | 0 |
| rs140690030 | benign | 0 | 0 | 0 |
| rs137852992 | pathogenic | 0 | 0 | 0 |
| rs6544718 | benign | 0 | 0 | 0 |
| rs137852989 | pathogenic | 0 | 0 | 0 |
| rs150893072 | benign | 1 | 1 | 1 |
| rs28931613 | pathogenic | 1 | 1 | 1 |
| rs143789597 | benign | 1 | 1 | 1 |
| rs121913037 | pathogenic | 1 | 1 | 0 |
| rs188802138 | pathogenic | 1 | 1 | 0 |
| rs121913038 | pathogenic | 1 | 1 | 0 |
| rs146835206 | benign | 0 | 0 | 0 |
| rs121913041 | pathogenic | 1 | 1 | 1 |
| rs121913039 | pathogenic | 1 | 1 | 1 |
| rs149977726 | pathogenic | 1 | 1 | 0 |
| rs121913036 | pathogenic | 0 | 0 | 0 |
| rs121913040 | pathogenic | 0 | 0 | 0 |
| rs765007158 | benign | 0 | 0 | 0 |
| rs112723255 | benign | 0 | 0 | 0 |
| rs574164748 | benign | 0 | 0 | 0 |
| rs80338734 | pathogenic | 1 | 1 | 0 |
| rs104894694 | pathogenic | 1 | 1 | 0 |
| rs753198836 | pathogenic | 0 | 0 | 0 |
| rs760101382 | pathogenic | 0 | 0 | 0 |
| rs121909272 | pathogenic | 1 | 1 | 0 |
| rs370421531 | pathogenic | 0 | 0 | 0 |
| rs565109128 | benign | 0 | 0 | 0 |
| rs17851582 | benign | 0 | 0 | 0 |
| rs753228876 | benign | 0 | 0 | 0 |
| rs141471799 | benign | 0 | 0 | 0 |
| rs281864916 | pathogenic | 1 | 1 | 0 |
| rs121918409 | pathogenic | 1 | 1 | 1 |

| | | | | |
|---|---|---|---|---|
| rs121918413 | pathogenic | 1 | 1 | 1 |
| rs121918411 | pathogenic | 1 | 1 | 1 |
| rs121918408 | pathogenic | 1 | 1 | 1 |
| rs121918412 | pathogenic | 1 | 1 | 1 |
| rs121918410 | pathogenic | 1 | 1 | 1 |
| rs116474260 | benign | 1 | 1 | 1 |
| rs281864919 | pathogenic | 1 | 1 | 1 |
| rs397514452 | pathogenic | 1 | 1 | 0 |
| rs747843638 | benign | 0 | 0 | 0 |
| rs774357609 | pathogenic | 1 | 1 | 1 |
| rs201804005 | benign | 0 | 0 | 0 |
| rs746485916 | pathogenic | 0 | 0 | 0 |
| rs764131178 | pathogenic | 0 | 0 | 0 |
| rs374094065 | pathogenic | 0 | 0 | 0 |
| rs76151636 | pathogenic | 1 | 1 | 0 |
| rs753250853 | pathogenic | 0 | 0 | 0 |
| rs761632029 | pathogenic | 0 | 0 | 0 |
| rs775541743 | pathogenic | 0 | 0 | 0 |
| rs778749563 | pathogenic | 0 | 0 | 0 |
| rs60431989 | pathogenic | 1 | 1 | 0 |
| rs755554442 | pathogenic | 1 | 1 | 0 |
| rs749085322 | pathogenic | 0 | 0 | 0 |
| rs756029120 | pathogenic | 1 | 1 | 0 |
| rs137853279 | pathogenic | 1 | 1 | 0 |
| rs777354267 | pathogenic | 1 | 1 | 1 |
| rs142940486 | benign | 1 | 1 | 1 |
| rs140167393 | benign | 1 | 1 | 1 |
| rs2066827 | benign | 0 | 0 | 0 |
| rs765681672 | benign | 0 | 0 | 0 |
| rs137985549 | benign | 0 | 0 | 0 |
| rs74315340 | pathogenic | 1 | 1 | 1 |
| rs74315341 | pathogenic | 0 | 0 | 0 |
| rs74315335 | pathogenic | 0 | 0 | 0 |
| rs61730974 | benign | 0 | 0 | 0 |
| rs121909193 | pathogenic | 0 | 0 | 0 |
| rs74315334 | pathogenic | 0 | 0 | 0 |
| rs74315329 | pathogenic | 0 | 0 | 0 |
| rs74315330 | pathogenic | 0 | 0 | 0 |
| rs121909194 | pathogenic | 0 | 0 | 0 |
| rs28936694 | pathogenic | 0 | 0 | 0 |
| rs74315336 | pathogenic | 0 | 0 | 0 |
| rs74315338 | pathogenic | 1 | 1 | 1 |
| rs74315328 | pathogenic | 0 | 0 | 0 |
| rs74315331 | pathogenic | 0 | 0 | 0 |
| rs74315332 | pathogenic | 0 | 0 | 0 |
| rs141696651 | benign | 0 | 0 | 0 |
| rs7041 | benign | 0 | 0 | 0 |
| rs4588 | benign | 0 | 0 | 0 |
| rs121908981 | pathogenic | 1 | 1 | 1 |
| rs606231459 | pathogenic | 0 | 0 | 0 |
| rs398123053 | pathogenic | 0 | 0 | 0 |
| rs774259802 | benign | 0 | 0 | 0 |
| rs121909582 | pathogenic | 1 | 1 | 1 |
| rs104893680 | pathogenic | 1 | 1 | 1 |

| | | | | |
|---|---|---|---|---|
| rs137854907 | pathogenic | 0 | 0 | 0 |
| rs765715798 | pathogenic | 0 | 0 | 0 |
| rs104893678 | pathogenic | 0 | 0 | 0 |
| rs104893679 | pathogenic | 0 | 1 | 0 |
| rs104893681 | pathogenic | 1 | 1 | 0 |
| rs121909640 | pathogenic | 1 | 1 | 0 |
| rs780705654 | pathogenic | 0 | 0 | 0 |
| rs201672146 | benign | 1 | 1 | 0 |
| rs12136658 | benign | 0 | 0 | 0 |
| rs56127440 | benign | 0 | 0 | 0 |
| rs104894433 | pathogenic | 1 | 1 | 1 |
| rs104894435 | pathogenic | 0 | 1 | 0 |
| rs104894437 | pathogenic | 1 | 1 | 1 |
| rs104894441 | pathogenic | 1 | 1 | 1 |
| rs104894440 | pathogenic | 1 | 1 | 1 |
| rs104894445 | pathogenic | 1 | 1 | 1 |
| rs104894436 | pathogenic | 0 | 0 | 0 |
| rs104894438 | pathogenic | 0 | 0 | 0 |
| rs200891969 | pathogenic | 1 | 1 | 0 |
| rs104894443 | pathogenic | 1 | 1 | 1 |
| rs104894434 | pathogenic | 1 | 1 | 1 |
| rs41298442 | pathogenic | 1 | 1 | 1 |
| rs104894442 | pathogenic | 1 | 1 | 1 |
| rs35453854 | benign | 0 | 0 | 0 |
| rs121912898 | pathogenic | 0 | 0 | 0 |
| rs121912897 | pathogenic | 1 | 1 | 0 |
| rs138260625 | benign | 0 | 0 | 0 |
| rs147802438 | benign | 1 | 1 | 0 |
| rs146443310 | benign | 1 | 1 | 0 |
| rs747319628 | pathogenic | 1 | 1 | 0 |
| rs1801195 | benign | 0 | 0 | 0 |
| rs145045341 | benign | 1 | 1 | 0 |
| rs116001173 | benign | 1 | 1 | 0 |
| rs587784175 | pathogenic | 0 | 0 | 0 |
| rs104893857 | pathogenic | 0 | 0 | 0 |
| rs104893862 | pathogenic | 1 | 1 | 0 |
| rs104893858 | pathogenic | 1 | 1 | 1 |
| rs121909249 | pathogenic | 0 | 1 | 0 |
| rs121909248 | pathogenic | 1 | 1 | 1 |
| rs104893859 | pathogenic | 1 | 1 | 0 |
| rs187063864 | pathogenic | 1 | 1 | 0 |
| rs121917817 | pathogenic | 1 | 1 | 0 |
| rs121917818 | pathogenic | 0 | 0 | 0 |
| rs753594601 | pathogenic | 0 | 0 | 0 |
| rs28937869 | pathogenic | 1 | 1 | 0 |
| rs6165 | benign | 0 | 0 | 0 |
| rs200089714 | pathogenic | 1 | 1 | 0 |
| rs372000714 | pathogenic | 0 | 0 | 0 |
| rs2228206 | benign | 0 | 0 | 0 |
| rs560096 | benign | 0 | 0 | 0 |
| rs137852666 | pathogenic | 0 | 0 | 0 |
| rs10896380 | benign | 0 | 0 | 0 |
| rs773690764 | pathogenic | 0 | 0 | 0 |
| rs201060167 | pathogenic | 0 | 0 | 0 |

| | | | | |
|---|---|---|---|---|
| rs137852670 | pathogenic | 0 | 0 | 0 |
| rs780594709 | pathogenic | 0 | 0 | 0 |
| rs145226920 | pathogenic | 0 | 0 | 0 |
| rs137852665 | pathogenic | 0 | 0 | 0 |
| rs724159960 | pathogenic | 0 | 0 | 0 |
| rs756985703 | pathogenic | 0 | 0 | 0 |
| rs137852667 | pathogenic | 0 | 0 | 0 |
| rs151079750 | pathogenic | 1 | 1 | 0 |
| rs2306636 | benign | 0 | 0 | 0 |
| rs72647885 | benign | 1 | 1 | 0 |
| rs377584435 | pathogenic | 1 | 1 | 1 |
| rs757583846 | pathogenic | 1 | 1 | 0 |
| rs121918148 | pathogenic | 0 | 0 | 0 |
| rs149201802 | benign | 0 | 0 | 0 |
| rs587781904 | pathogenic | 1 | 1 | 1 |
| rs143421325 | benign | 0 | 0 | 0 |
| rs587781253 | pathogenic | 0 | 0 | 0 |
| rs587780331 | benign | 0 | 0 | 0 |
| rs151001016 | benign | 0 | 0 | 0 |
| rs150888094 | benign | 0 | 0 | 0 |
| rs149581331 | benign | 0 | 0 | 0 |
| rs774840535 | benign | 0 | 0 | 0 |
| rs138428684 | benign | 0 | 0 | 0 |
| rs149300055 | benign | 0 | 0 | 0 |
| rs571005578 | benign | 0 | 0 | 0 |
| rs200731839 | benign | 0 | 0 | 0 |
| rs121965007 | pathogenic | 0 | 0 | 0 |
| rs121965013 | pathogenic | 0 | 1 | 0 |
| rs121965009 | pathogenic | 0 | 0 | 0 |
| rs1800457 | benign | 1 | 1 | 0 |
| rs121965006 | pathogenic | 1 | 1 | 0 |
| rs121965008 | pathogenic | 0 | 0 | 0 |
| rs61732609 | pathogenic | 0 | 0 | 0 |
| rs121965011 | pathogenic | 0 | 0 | 0 |
| rs121965015 | pathogenic | 0 | 0 | 0 |
| rs121965018 | pathogenic | 1 | 1 | 0 |
| rs121965016 | pathogenic | 0 | 0 | 0 |
| rs77806 | benign | 0 | 0 | 0 |
| rs149731356 | benign | 0 | 0 | 0 |
| rs72549407 | pathogenic | 0 | 0 | 0 |
| rs200363929 | benign | 0 | 0 | 0 |
| rs376403708 | benign | 1 | 1 | 0 |
| rs56376197 | benign | 1 | 1 | 1 |
| rs200237973 | benign | 1 | 1 | 1 |
| rs548330075 | benign | 0 | 0 | 0 |
| rs148291350 | benign | 0 | 0 | 0 |
| rs71579353 | pathogenic | 0 | 0 | 0 |
| rs146278697 | benign | 0 | 0 | 0 |
| rs141033098 | benign | 0 | 0 | 0 |
| rs397517234 | pathogenic | 0 | 0 | 0 |
| rs71579374 | benign | 0 | 0 | 0 |
| rs140381835 | benign | 0 | 0 | 0 |
| rs137852702 | pathogenic | 0 | 0 | 0 |
| rs137852696 | pathogenic | 1 | 1 | 1 |

| | | | | |
|---|---|---|---|---|
| rs137852697 | pathogenic | 0 | 0 | 0 |
| rs386833639 | pathogenic | 0 | 0 | 0 |
| rs386833640 | pathogenic | 0 | 0 | 0 |
| rs386833641 | benign | 0 | 0 | 0 |
| rs137852701 | pathogenic | 0 | 0 | 0 |
| rs386833642 | pathogenic | 0 | 0 | 0 |
| rs137852695 | pathogenic | 0 | 0 | 0 |
| rs386833646 | pathogenic | 0 | 0 | 0 |
| rs137852700 | pathogenic | 1 | 1 | 0 |
| rs386833649 | pathogenic | 1 | 1 | 0 |
| rs386833650 | pathogenic | 0 | 0 | 0 |
| rs148412181 | pathogenic | 0 | 0 | 0 |
| rs386833655 | pathogenic | 0 | 0 | 0 |
| rs386833656 | pathogenic | 1 | 1 | 0 |
| rs137852698 | pathogenic | 0 | 0 | 0 |
| rs386833661 | pathogenic | 0 | 0 | 0 |
| rs386833670 | pathogenic | 0 | 0 | 0 |
| rs111033632 | pathogenic | 1 | 1 | 1 |
| rs34308573 | benign | 1 | 1 | 1 |
| rs376048533 | pathogenic | 1 | 1 | 0 |
| rs3747517 | benign | 1 | 1 | 0 |
| rs137853328 | pathogenic | 0 | 0 | 0 |
| rs77273740 | benign | 0 | 0 | 0 |
| rs267606760 | pathogenic | 0 | 0 | 0 |
| rs77576840 | pathogenic | 0 | 0 | 0 |
| rs45465204 | benign | 1 | 1 | 0 |
| rs5320 | benign | 0 | 0 | 0 |
| rs5324 | benign | 0 | 0 | 0 |
| rs4531 | benign | 0 | 0 | 0 |
| rs267606761 | pathogenic | 0 | 0 | 0 |
| rs75215331 | pathogenic | 0 | 0 | 0 |
| rs6271 | benign | 0 | 0 | 0 |
| rs201643277 | pathogenic | 0 | 0 | 0 |
| rs4252250 | benign | 0 | 0 | 0 |
| rs137853580 | pathogenic | 0 | 0 | 0 |
| rs368287711 | pathogenic | 0 | 0 | 0 |
| rs137853579 | pathogenic | 0 | 0 | 0 |
| rs3135932 | benign | 0 | 0 | 0 |
| rs2228055 | benign | 0 | 0 | 0 |
| rs140322345 | benign | 1 | 1 | 1 |
| rs1154510 | pathogenic | 0 | 0 | 0 |
| rs11833399 | benign | 0 | 0 | 0 |
| rs36023382 | benign | 1 | 1 | 1 |
| rs61754444 | benign | 0 | 0 | 0 |
| rs28935168 | pathogenic | 0 | 0 | 0 |
| rs28934907 | pathogenic | 0 | 0 | 0 |
| rs28934904 | pathogenic | 1 | 1 | 0 |
| rs61748392 | pathogenic | 1 | 1 | 0 |
| rs28934908 | pathogenic | 0 | 0 | 0 |
| rs28934905 | pathogenic | 0 | 0 | 0 |
| rs28934906 | pathogenic | 1 | 1 | 0 |
| rs606231307 | pathogenic | 0 | 0 | 0 |
| rs9333296 | benign | 0 | 0 | 0 |
| rs3218614 | benign | 0 | 0 | 0 |

| | | | | |
|---|---|---|---|---|
| rs2230792 | benign | 0 | 0 | 0 |
| rs2230793 | benign | 0 | 0 | 0 |
| rs2230794 | benign | 0 | 0 | 0 |
| rs749052963 | pathogenic | 0 | 0 | 0 |
| rs2230798 | benign | 0 | 0 | 0 |
| rs2230795 | benign | 0 | 0 | 0 |
| rs3204145 | benign | 0 | 0 | 0 |
| rs1538660 | benign | 0 | 0 | 0 |
| rs9878928 | benign | 0 | 0 | 0 |
| rs104893742 | pathogenic | 0 | 0 | 0 |
| rs28936702 | pathogenic | 0 | 0 | 0 |
| rs28936703 | pathogenic | 0 | 0 | 0 |
| rs769392 | benign | 0 | 0 | 0 |
| rs769402 | benign | 1 | 1 | 1 |
| rs104894884 | pathogenic | 1 | 1 | 1 |
| rs1801316 | benign | 1 | 1 | 1 |
| rs104894885 | pathogenic | 1 | 1 | 1 |
| rs782572 | benign | 0 | 0 | 0 |
| rs151166046 | pathogenic | 0 | 0 | 0 |
| rs772153760 | pathogenic | 1 | 1 | 0 |
| rs35405862 | benign | 0 | 0 | 0 |
| rs146571352 | pathogenic | 0 | 1 | 0 |
| rs374698153 | pathogenic | 0 | 0 | 0 |
| rs7594497 | benign | 0 | 0 | 0 |
| rs80338716 | pathogenic | 0 | 0 | 0 |
| rs80338719 | pathogenic | 1 | 1 | 1 |
| rs80338721 | pathogenic | 0 | 0 | 0 |
| rs730882249 | pathogenic | 0 | 0 | 0 |
| rs146262009 | pathogenic | 0 | 0 | 0 |
| rs104895083 | pathogenic | 1 | 1 | 1 |
| rs104895100 | benign | 1 | 1 | 1 |
| rs104895085 | pathogenic | 0 | 0 | 0 |
| rs28940580 | pathogenic | 0 | 0 | 0 |
| rs104895098 | pathogenic | 0 | 0 | 0 |
| rs28940578 | pathogenic | 0 | 0 | 0 |
| rs61752717 | pathogenic | 0 | 0 | 0 |
| rs28940579 | pathogenic | 0 | 0 | 0 |
| rs104895097 | pathogenic | 0 | 0 | 0 |
| rs140425651 | benign | 1 | 1 | 0 |
| rs768090444 | pathogenic | 0 | 0 | 0 |
| rs764459544 | pathogenic | 0 | 0 | 0 |
| rs80338802 | pathogenic | 1 | 1 | 1 |
| rs149969786 | pathogenic | 1 | 1 | 0 |
| rs115311625 | pathogenic | 1 | 1 | 1 |
| rs778768583 | pathogenic | 1 | 1 | 1 |
| rs892090 | benign | 0 | 0 | 0 |
| rs28929498 | pathogenic | 1 | 1 | 1 |
| rs137854467 | pathogenic | 0 | 0 | 0 |
| rs137854480 | pathogenic | 0 | 0 | 0 |
| rs148682262 | benign | 0 | 0 | 0 |
| rs137852744 | pathogenic | 0 | 0 | 0 |
| rs137852745 | pathogenic | 0 | 1 | 0 |
| rs137852749 | pathogenic | 0 | 1 | 0 |
| rs137852746 | pathogenic | 0 | 1 | 0 |

| | | | | |
|---|---|---|---|---|
| rs145633958 | benign | 0 | 0 | 0 |
| rs141679950 | benign | 0 | 0 | 0 |
| rs76397662 | benign | 1 | 1 | 1 |
| rs370736139 | benign | 0 | 0 | 0 |
| rs781750106 | benign | 0 | 0 | 0 |
| rs546866208 | benign | 0 | 0 | 0 |
| rs142338976 | benign | 0 | 0 | 0 |
| rs115272158 | benign | 0 | 0 | 0 |
| rs115423919 | benign | 0 | 0 | 0 |
| rs537874538 | benign | 0 | 0 | 0 |
| rs9282834 | benign | 0 | 0 | 0 |
| rs201745826 | benign | 0 | 0 | 0 |
| rs75873440 | pathogenic | 0 | 0 | 0 |
| rs147219360 | benign | 0 | 0 | 0 |
| rs77939446 | pathogenic | 0 | 0 | 0 |
| rs77558292 | pathogenic | 0 | 0 | 0 |
| rs80069458 | pathogenic | 0 | 0 | 0 |
| rs76262710 | pathogenic | 0 | 0 | 0 |
| rs79781594 | pathogenic | 0 | 0 | 0 |
| rs77316810 | pathogenic | 0 | 0 | 0 |
| rs77503355 | pathogenic | 0 | 0 | 0 |
| rs79890926 | pathogenic | 0 | 0 | 0 |
| rs377767402 | benign | 0 | 0 | 0 |
| rs121913308 | pathogenic | 0 | 0 | 0 |
| rs55846256 | benign | 0 | 0 | 0 |
| rs75076352 | pathogenic | 0 | 0 | 0 |
| rs75996173 | pathogenic | 0 | 0 | 0 |
| rs77709286 | pathogenic | 0 | 0 | 0 |
| rs151058706 | benign | 0 | 0 | 0 |
| rs138964358 | benign | 0 | 0 | 0 |
| rs137852353 | pathogenic | 1 | 1 | 0 |
| rs201196020 | benign | 0 | 0 | 0 |
| rs137853201 | pathogenic | 1 | 1 | 1 |
| rs113994174 | pathogenic | 1 | 1 | 1 |
| rs104893674 | pathogenic | 1 | 1 | 1 |
| rs779519249 | benign | 1 | 1 | 1 |
| rs9606756 | benign | 0 | 0 | 0 |
| rs75680863 | benign | 1 | 1 | 1 |
| rs35838082 | benign | 0 | 0 | 0 |
| rs1801198 | benign | 0 | 0 | 0 |
| rs117458738 | benign | 1 | 1 | 0 |
| rs9621049 | benign | 1 | 1 | 0 |
| rs1131603 | benign | 1 | 1 | 0 |
| rs4820889 | benign | 0 | 0 | 0 |
| rs777767978 | pathogenic | 1 | 1 | 0 |
| rs147399860 | benign | 1 | 1 | 1 |
| rs104893950 | pathogenic | 0 | 0 | 0 |
| rs777308612 | benign | 0 | 0 | 0 |
| rs151091410 | benign | 0 | 0 | 0 |
| rs11537667 | benign | 0 | 0 | 0 |
| rs267606621 | pathogenic | 1 | 1 | 0 |
| rs143058902 | benign | 0 | 0 | 0 |
| rs370261904 | pathogenic | 0 | 0 | 0 |
| rs141169746 | benign | 0 | 0 | 0 |

| | | | | |
|---|---|---|---|---|
| rs111033436 | benign | 0 | 0 | 0 |
| rs148533341 | benign | 0 | 0 | 0 |
| rs143827620 | benign | 0 | 0 | 0 |
| rs4935502 | benign | 0 | 0 | 0 |
| rs61735473 | benign | 0 | 0 | 0 |
| rs267606932 | pathogenic | 1 | 1 | 0 |
| rs61730754 | benign | 0 | 0 | 0 |
| rs374491359 | pathogenic | 1 | 1 | 1 |
| rs761800077 | benign | 1 | 1 | 1 |
| rs760391381 | benign | 1 | 1 | 1 |
| rs104893885 | pathogenic | 0 | 0 | 0 |
| rs104893886 | pathogenic | 1 | 1 | 1 |
| rs104893884 | pathogenic | 0 | 0 | 0 |
| rs147715509 | benign | 0 | 0 | 0 |
| rs115533243 | benign | 0 | 0 | 0 |
| rs397509390 | pathogenic | 0 | 1 | 0 |
| rs80357134 | pathogenic | 0 | 0 | 0 |
| rs766004110 | benign | 1 | 1 | 1 |
| rs769650474 | benign | 0 | 0 | 0 |
| rs398122635 | benign | 1 | 1 | 0 |
| rs199522616 | pathogenic | 0 | 0 | 0 |
| rs28897672 | pathogenic | 1 | 1 | 0 |
| rs80357086 | pathogenic | 1 | 1 | 1 |
| rs80357064 | pathogenic | 1 | 1 | 0 |
| rs80356936 | benign | 0 | 0 | 0 |
| rs28897673 | benign | 0 | 0 | 0 |
| rs146747324 | benign | 0 | 0 | 0 |
| rs78635798 | pathogenic | 0 | 0 | 0 |
| rs75146158 | pathogenic | 1 | 1 | 1 |
| rs587782886 | benign | 0 | 0 | 0 |
| rs121434634 | pathogenic | 0 | 0 | 0 |
| rs121434635 | pathogenic | 0 | 0 | 0 |
| rs587777778 | pathogenic | 0 | 0 | 0 |
| rs118203967 | pathogenic | 0 | 0 | 0 |
| rs751375244 | pathogenic | 1 | 1 | 1 |
| rs104893863 | pathogenic | 1 | 1 | 0 |
| rs104893865 | pathogenic | 0 | 0 | 0 |
| rs104893864 | pathogenic | 1 | 1 | 1 |
| rs104893866 | pathogenic | 1 | 1 | 0 |
| rs561738497 | benign | 0 | 0 | 0 |
| rs57599352 | pathogenic | 1 | 1 | 1 |
| rs267607448 | pathogenic | 1 | 1 | 1 |
| rs63751596 | pathogenic | 0 | 0 | 0 |
| rs63751612 | benign | 0 | 0 | 0 |
| rs63751684 | benign | 0 | 0 | 0 |
| rs267607861 | benign | 0 | 0 | 0 |
| rs35831931 | benign | 0 | 0 | 0 |
| rs63750876 | benign | 0 | 0 | 0 |
| rs41295284 | benign | 0 | 0 | 0 |
| rs35502531 | benign | 0 | 0 | 0 |
| rs63750449 | benign | 0 | 0 | 0 |
| rs63750899 | pathogenic | 0 | 0 | 0 |
| rs55907433 | benign | 0 | 0 | 0 |
| rs63750217 | pathogenic | 0 | 0 | 0 |

| | | | | |
|---|---|---|---|---|
| rs63751275 | pathogenic | 0 | 0 | 0 |
| rs63750702 | benign | 1 | 1 | 1 |
| rs63750114 | benign | 0 | 0 | 0 |
| rs2020873 | benign | 0 | 0 | 0 |
| rs140195825 | benign | 1 | 1 | 1 |
| rs182763805 | benign | 0 | 0 | 0 |
| rs533696377 | benign | 0 | 0 | 0 |
| rs41278788 | benign | 0 | 0 | 0 |
| rs104894193 | pathogenic | 1 | 1 | 0 |
| rs3824915 | benign | 1 | 1 | 1 |
| rs104894196 | pathogenic | 0 | 0 | 0 |
| rs78681604 | benign | 1 | 1 | 1 |
| rs77321498 | benign | 1 | 1 | 1 |
| rs137852965 | pathogenic | 0 | 0 | 0 |
| rs137852961 | pathogenic | 0 | 0 | 0 |
| rs137852962 | pathogenic | 1 | 1 | 0 |
| rs753376100 | pathogenic | 0 | 0 | 0 |
| rs137852968 | pathogenic | 1 | 1 | 1 |
| rs137852959 | pathogenic | 1 | 1 | 0 |
| rs137852967 | pathogenic | 0 | 0 | 0 |
| rs113437301 | benign | 1 | 1 | 1 |
| rs61747482 | benign | 1 | 1 | 1 |
| rs2234919 | benign | 1 | 1 | 0 |
| rs121908863 | pathogenic | 0 | 0 | 0 |
| rs140455771 | pathogenic | 0 | 0 | 0 |
| rs121908810 | pathogenic | 1 | 1 | 1 |
| rs143879853 | benign | 0 | 0 | 0 |
| rs188319622 | benign | 0 | 0 | 0 |
| rs121918645 | pathogenic | 0 | 0 | 0 |
| rs115352435 | pathogenic | 0 | 0 | 0 |
| rs121434427 | pathogenic | 1 | 1 | 1 |
| rs121434428 | pathogenic | 0 | 0 | 0 |
| rs35086265 | benign | 1 | 1 | 1 |
| rs11576415 | benign | 0 | 0 | 0 |
| rs121434429 | pathogenic | 1 | 1 | 1 |
| rs4074536 | benign | 1 | 1 | 1 |
| rs10801999 | benign | 0 | 0 | 0 |
| rs121434550 | pathogenic | 0 | 0 | 0 |
| rs148057999 | pathogenic | 0 | 0 | 0 |
| rs28730716 | benign | 0 | 0 | 0 |
| rs142036299 | benign | 0 | 0 | 0 |
| rs762381137 | pathogenic | 0 | 0 | 0 |
| rs121434549 | pathogenic | 0 | 0 | 0 |
| rs139228801 | pathogenic | 0 | 0 | 0 |
| rs147219463 | benign | 0 | 0 | 0 |
| rs1131857 | benign | 0 | 0 | 0 |
| rs121917874 | pathogenic | 0 | 1 | 0 |
| rs2228056 | benign | 0 | 0 | 0 |
| rs121917870 | pathogenic | 0 | 0 | 0 |
| rs121917866 | pathogenic | 0 | 0 | 0 |
| rs11921054 | benign | 0 | 0 | 0 |
| rs121917868 | pathogenic | 0 | 0 | 0 |
| rs28931603 | pathogenic | 0 | 0 | 0 |
| rs104895295 | pathogenic | 1 | 1 | 0 |

| | | | | |
|---|---|---|---|---|
| rs104895382 | pathogenic | 0 | 0 | 0 |
| rs104895298 | pathogenic | 0 | 0 | 0 |
| rs104895300 | pathogenic | 0 | 0 | 0 |
| rs104895332 | pathogenic | 0 | 0 | 0 |
| rs758026399 | pathogenic | 1 | 1 | 1 |
| rs104895366 | pathogenic | 0 | 0 | 0 |
| rs104895304 | pathogenic | 0 | 0 | 0 |
| rs759997079 | pathogenic | 0 | 0 | 0 |
| rs121917789 | pathogenic | 0 | 0 | 0 |
| rs104895319 | pathogenic | 0 | 0 | 0 |
| rs104895317 | pathogenic | 1 | 1 | 0 |
| rs28934897 | pathogenic | 0 | 0 | 0 |
| rs104895380 | benign | 0 | 0 | 0 |
| rs104895360 | pathogenic | 0 | 0 | 0 |
| rs28936396 | pathogenic | 1 | 1 | 0 |
| rs121909307 | pathogenic | 1 | 1 | 1 |
| rs28938472 | pathogenic | 1 | 1 | 0 |
| rs121909308 | pathogenic | 1 | 1 | 0 |
| rs121909309 | pathogenic | 0 | 0 | 0 |
| rs121913013 | benign | 1 | 1 | 1 |
| rs201855245 | benign | 1 | 1 | 1 |
| rs2230232 | benign | 1 | 1 | 0 |
| rs113451409 | benign | 0 | 0 | 0 |
| rs121913011 | pathogenic | 1 | 1 | 0 |
| rs185821167 | pathogenic | 1 | 1 | 0 |
| rs2230234 | benign | 0 | 0 | 0 |
| rs193922639 | benign | 1 | 1 | 0 |
| rs774863785 | pathogenic | 1 | 1 | 0 |
| rs576404380 | pathogenic | 1 | 1 | 0 |
| rs2230235 | benign | 1 | 1 | 0 |
| rs200484060 | pathogenic | 1 | 1 | 0 |
| rs199761749 | benign | 0 | 0 | 0 |
| rs199681901 | benign | 0 | 0 | 0 |
| rs121908951 | pathogenic | 0 | 0 | 0 |
| rs769251460 | benign | 0 | 0 | 0 |
| rs28937320 | pathogenic | 0 | 0 | 0 |
| rs200607527 | pathogenic | 0 | 0 | 0 |
| rs137852640 | pathogenic | 1 | 1 | 1 |
| rs137852636 | pathogenic | 1 | 1 | 1 |
| rs137852638 | pathogenic | 0 | 0 | 0 |
| rs142637231 | pathogenic | 0 | 0 | 0 |
| rs151328418 | benign | 0 | 1 | 0 |
| rs137852639 | pathogenic | 1 | 1 | 1 |
| rs34605667 | benign | 0 | 0 | 0 |
| rs104894073 | pathogenic | 1 | 1 | 0 |
| rs56298569 | pathogenic | 0 | 0 | 0 |
| rs757600616 | pathogenic | 0 | 0 | 0 |
| rs139194636 | pathogenic | 0 | 0 | 0 |
| rs369517993 | pathogenic | 1 | 1 | 1 |
| rs138846036 | benign | 1 | 1 | 1 |
| rs200808744 | pathogenic | 1 | 1 | 1 |
| rs2255543 | benign | 0 | 0 | 0 |
| rs149165166 | pathogenic | 0 | 0 | 0 |
| rs544956641 | pathogenic | 1 | 1 | 1 |

| | | | | |
|---|---|---|---|---|
| rs28941783 | pathogenic | 0 | 0 | 0 |
| rs562853291 | pathogenic | 1 | 1 | 1 |
| rs28942100 | pathogenic | 0 | 0 | 0 |
| rs120074174 | pathogenic | 1 | 1 | 1 |
| rs120074171 | pathogenic | 1 | 1 | 1 |
| rs120074173 | pathogenic | 0 | 0 | 0 |
| rs120074172 | pathogenic | 1 | 1 | 1 |
| rs767159114 | pathogenic | 1 | 1 | 1 |
| rs104893657 | pathogenic | 0 | 0 | 0 |
| rs104893656 | pathogenic | 0 | 0 | 0 |
| rs104893659 | pathogenic | 0 | 0 | 0 |
| rs104893658 | pathogenic | 0 | 0 | 0 |
| rs200386310 | pathogenic | 0 | 0 | 0 |
| rs2072712 | benign | 0 | 0 | 0 |
| rs35274867 | benign | 0 | 0 | 0 |
| rs35343172 | benign | 0 | 0 | 0 |
| rs10079250 | benign | 0 | 0 | 0 |
| rs34951517 | benign | 0 | 0 | 0 |
| rs267606695 | pathogenic | 0 | 0 | 0 |
| rs149391728 | benign | 0 | 0 | 0 |
| rs387906598 | pathogenic | 1 | 1 | 0 |
| rs137854578 | pathogenic | 0 | 0 | 0 |
| rs141519952 | benign | 0 | 0 | 0 |
| rs146572883 | benign | 0 | 0 | 0 |
| rs111866410 | benign | 0 | 0 | 0 |
| rs139387758 | benign | 1 | 1 | 1 |
| rs35672074 | benign | 1 | 1 | 1 |
| rs121434322 | pathogenic | 0 | 0 | 0 |
| rs150884181 | benign | 0 | 0 | 0 |
| rs35867471 | benign | 0 | 0 | 0 |
| rs2867802 | benign | 0 | 0 | 0 |
| rs137902889 | benign | 0 | 0 | 0 |
| rs369804864 | pathogenic | 0 | 0 | 0 |
| rs781050795 | pathogenic | 0 | 0 | 0 |
| rs2305799 | benign | 0 | 0 | 0 |
| rs2229531 | benign | 0 | 0 | 0 |
| rs147025508 | benign | 0 | 0 | 0 |
| rs76857106 | pathogenic | 0 | 0 | 0 |
| rs7247284 | benign | 0 | 0 | 0 |
| rs62619782 | benign | 0 | 0 | 0 |
| rs377244188 | pathogenic | 0 | 0 | 0 |
| rs75718910 | pathogenic | 0 | 0 | 0 |
| rs79843600 | pathogenic | 0 | 0 | 0 |
| rs75037667 | pathogenic | 1 | 1 | 1 |
| rs121913006 | pathogenic | 0 | 0 | 0 |
| rs397517925 | pathogenic | 1 | 1 | 1 |
| rs267606831 | pathogenic | 1 | 1 | 1 |
| rs121909667 | pathogenic | 1 | 1 | 1 |
| rs121909669 | pathogenic | 1 | 1 | 1 |
| rs121909668 | pathogenic | 1 | 1 | 1 |
| rs121909671 | pathogenic | 1 | 1 | 1 |
| rs534596036 | benign | 0 | 0 | 0 |
| rs144037629 | pathogenic | 0 | 0 | 0 |
| rs11546406 | benign | 0 | 0 | 0 |

| | | | | |
|---|---|---|---|---|
| rs143697995 | pathogenic | 0 | 0 | 0 |
| rs121908331 | pathogenic | 0 | 0 | 0 |
| rs149377346 | benign | 0 | 0 | 0 |
| rs104894086 | pathogenic | 1 | 1 | 0 |
| rs104894089 | pathogenic | 0 | 0 | 0 |
| rs104894090 | pathogenic | 1 | 1 | 0 |
| rs387907235 | pathogenic | 0 | 0 | 0 |
| rs137852689 | pathogenic | 0 | 0 | 0 |
| rs137852690 | pathogenic | 1 | 1 | 0 |
| rs104894087 | pathogenic | 0 | 0 | 0 |
| rs104894085 | pathogenic | 0 | 0 | 0 |
| rs79383654 | benign | 0 | 0 | 0 |
| rs121434239 | pathogenic | 0 | 0 | 0 |
| rs121434240 | pathogenic | 0 | 0 | 0 |
| rs121434236 | pathogenic | 0 | 0 | 0 |
| rs121434238 | pathogenic | 1 | 1 | 1 |
| rs121918107 | pathogenic | 1 | 1 | 1 |
| rs121918103 | pathogenic | 1 | 1 | 1 |
| rs121918104 | pathogenic | 1 | 1 | 1 |
| rs61747614 | benign | 1 | 1 | 1 |
| rs555329870 | benign | 0 | 0 | 0 |
| rs186094931 | benign | 0 | 0 | 0 |
| rs267606785 | pathogenic | 0 | 0 | 0 |
| rs34474955 | benign | 0 | 0 | 0 |
| rs1126497 | benign | 1 | 1 | 1 |
| rs397514661 | pathogenic | 0 | 0 | 0 |
| rs189732445 | benign | 0 | 0 | 0 |
| rs146685071 | benign | 0 | 0 | 0 |
| rs121908636 | pathogenic | 1 | 1 | 0 |
| rs121908643 | pathogenic | 0 | 0 | 0 |
| rs121908644 | pathogenic | 1 | 1 | 1 |
| rs575001023 | pathogenic | 1 | 1 | 1 |
| rs121908646 | pathogenic | 1 | 1 | 0 |
| rs121908638 | pathogenic | 1 | 1 | 1 |
| rs777828000 | pathogenic | 1 | 1 | 1 |
| rs74923032 | benign | 0 | 0 | 0 |
| rs192838388 | pathogenic | 1 | 1 | 1 |
| rs148918985 | pathogenic | 1 | 1 | 1 |
| rs773540764 | benign | 1 | 1 | 1 |
| rs370595480 | pathogenic | 1 | 1 | 1 |
| rs762387914 | pathogenic | 1 | 1 | 1 |
| rs121908642 | pathogenic | 1 | 1 | 1 |
| rs183276875 | pathogenic | 1 | 1 | 1 |
| rs121908648 | pathogenic | 1 | 1 | 1 |
| rs121908647 | pathogenic | 1 | 1 | 1 |
| rs121908640 | pathogenic | 1 | 1 | 1 |
| rs771937610 | pathogenic | 1 | 1 | 1 |
| rs121908641 | pathogenic | 1 | 1 | 1 |
| rs5745068 | benign | 0 | 0 | 0 |
| rs73155056 | benign | 1 | 1 | 1 |
| rs148788180 | benign | 0 | 0 | 0 |
| rs121918137 | pathogenic | 0 | 0 | 0 |
| rs41269593 | benign | 0 | 0 | 0 |
| rs11159 | benign | 1 | 1 | 0 |

| | | | | |
|---|---|---|---|---|
| rs114733570 | pathogenic | 1 | 1 | 0 |
| rs3211783 | benign | 0 | 0 | 0 |
| rs4762 | benign | 0 | 0 | 0 |
| rs74315283 | pathogenic | 0 | 0 | 0 |
| rs111033559 | pathogenic | 0 | 0 | 0 |
| rs111033560 | pathogenic | 1 | 1 | 1 |
| rs749962743 | benign | 0 | 0 | 0 |
| rs267606986 | pathogenic | 0 | 0 | 0 |
| rs34516635 | benign | 0 | 0 | 0 |
| rs117440569 | benign | 0 | 0 | 0 |
| rs61740877 | benign | 0 | 0 | 0 |
| rs121918352 | pathogenic | 0 | 0 | 0 |
| rs183974372 | benign | 0 | 0 | 0 |
| rs140511204 | benign | 1 | 1 | 0 |
| rs6040067 | benign | 0 | 0 | 0 |
| rs199674138 | benign | 0 | 0 | 0 |
| rs121918350 | pathogenic | 0 | 0 | 0 |
| rs138292988 | benign | 1 | 1 | 1 |
| rs145288947 | benign | 0 | 0 | 0 |
| rs369714529 | benign | 0 | 0 | 0 |
| rs200945419 | benign | 0 | 0 | 0 |
| rs116959874 | benign | 0 | 0 | 0 |
| rs141659620 | pathogenic | 1 | 1 | 0 |
| rs111475461 | benign | 0 | 0 | 0 |
| rs61755320 | pathogenic | 0 | 0 | 0 |
| rs748600162 | pathogenic | 1 | 1 | 1 |
| rs369227537 | pathogenic | 1 | 1 | 1 |
| rs372981030 | pathogenic | 1 | 1 | 1 |
| rs137854486 | pathogenic | 0 | 0 | 0 |
| rs137854487 | pathogenic | 0 | 0 | 0 |
| rs199905054 | pathogenic | 0 | 0 | 0 |
| rs771438170 | pathogenic | 0 | 0 | 0 |
| rs774949844 | pathogenic | 0 | 0 | 0 |
| rs4899145 | benign | 0 | 0 | 0 |
| rs200642525 | benign | 0 | 0 | 0 |
| rs190140598 | pathogenic | 0 | 0 | 0 |
| rs2250889 | benign | 0 | 0 | 0 |
| rs17577 | benign | 0 | 0 | 0 |
| rs142032681 | benign | 0 | 0 | 0 |
| rs144577963 | benign | 0 | 0 | 0 |
| rs61754623 | benign | 0 | 0 | 0 |
| rs139873200 | benign | 0 | 0 | 0 |
| rs532530906 | benign | 0 | 0 | 0 |
| rs145762716 | benign | 0 | 0 | 0 |
| rs34062020 | benign | 0 | 0 | 0 |
| rs35258161 | benign | 0 | 0 | 0 |
| rs149792649 | benign | 0 | 0 | 0 |
| rs78101774 | benign | 0 | 0 | 0 |
| rs766466297 | pathogenic | 0 | 1 | 0 |
| rs140467171 | pathogenic | 0 | 0 | 0 |
| rs141340466 | pathogenic | 0 | 0 | 0 |
| rs201555303 | pathogenic | 0 | 0 | 0 |
| rs143592405 | pathogenic | 0 | 0 | 0 |
| rs781249411 | pathogenic | 0 | 0 | 0 |

| | | | | |
|---|---|---|---|---|
| rs116644127 | benign | 0 | 0 | 0 |
| rs151146863 | benign | 1 | 1 | 1 |
| rs116440799 | benign | 0 | 0 | 0 |
| rs138958351 | benign | 0 | 0 | 0 |
| rs143319354 | benign | 0 | 0 | 0 |
| rs768669208 | pathogenic | 1 | 1 | 1 |
| rs28940289 | pathogenic | 1 | 1 | 0 |
| rs745656120 | pathogenic | 1 | 1 | 0 |
| rs387906987 | pathogenic | 0 | 0 | 0 |
| rs147240777 | benign | 0 | 0 | 0 |
| rs140945833 | benign | 0 | 0 | 0 |
| rs146614981 | benign | 0 | 0 | 0 |
| rs149878562 | benign | 0 | 0 | 0 |
| rs28938168 | pathogenic | 0 | 0 | 0 |
| rs33939927 | pathogenic | 1 | 1 | 1 |
| rs34995376 | pathogenic | 1 | 1 | 1 |
| rs201034481 | pathogenic | 1 | 1 | 1 |
| rs774069983 | benign | 0 | 0 | 0 |
| rs199422226 | pathogenic | 1 | 1 | 0 |
| rs199422225 | pathogenic | 1 | 1 | 1 |
| rs199422224 | pathogenic | 1 | 1 | 0 |
| rs149271416 | pathogenic | 1 | 1 | 0 |
| rs1801318 | benign | 1 | 1 | 1 |
| rs372691318 | pathogenic | 1 | 1 | 1 |
| rs147685849 | benign | 0 | 0 | 0 |
| rs387907199 | pathogenic | 0 | 0 | 0 |
| rs767122069 | pathogenic | 0 | 0 | 0 |
| rs2302427 | benign | 0 | 0 | 0 |
| rs201135441 | benign | 0 | 0 | 0 |
| rs201861847 | pathogenic | 0 | 0 | 0 |
| rs137852952 | pathogenic | 0 | 0 | 0 |
| rs16986560 | benign | 0 | 0 | 0 |
| rs147344332 | pathogenic | 1 | 1 | 1 |
| rs36027220 | pathogenic | 0 | 0 | 0 |
| rs192401229 | benign | 0 | 0 | 0 |
| rs121909106 | pathogenic | 1 | 1 | 0 |
| rs118204103 | pathogenic | 1 | 1 | 0 |
| rs118204104 | pathogenic | 1 | 1 | 0 |
| rs118204105 | pathogenic | 1 | 1 | 0 |
| rs118204106 | pathogenic | 0 | 0 | 0 |
| rs118204119 | pathogenic | 0 | 0 | 0 |
| rs118204107 | pathogenic | 0 | 0 | 0 |
| rs118204094 | pathogenic | 0 | 0 | 0 |
| rs118204098 | pathogenic | 1 | 1 | 0 |
| rs118204101 | pathogenic | 1 | 1 | 0 |
| rs118204096 | pathogenic | 1 | 1 | 0 |
| rs118204108 | pathogenic | 0 | 0 | 0 |
| rs536814318 | pathogenic | 0 | 0 | 0 |
| rs34413634 | pathogenic | 1 | 1 | 0 |
| rs118204100 | pathogenic | 0 | 0 | 0 |
| rs118204109 | pathogenic | 0 | 0 | 0 |
| rs118204116 | pathogenic | 1 | 1 | 0 |
| rs118204110 | pathogenic | 0 | 0 | 0 |
| rs118204099 | pathogenic | 0 | 0 | 0 |

| | | | | |
|---|---|---|---|---|
| rs118204111 | pathogenic | 0 | 0 | 0 |
| rs118204112 | pathogenic | 0 | 0 | 0 |
| rs118204113 | pathogenic | 0 | 0 | 0 |
| rs118204114 | pathogenic | 0 | 0 | 0 |
| rs118204115 | pathogenic | 0 | 0 | 0 |
| rs150428209 | benign | 0 | 0 | 0 |
| rs144949995 | benign | 0 | 0 | 0 |
| rs190023981 | benign | 0 | 0 | 0 |
| rs104894611 | pathogenic | 1 | 1 | 1 |
| rs28937580 | pathogenic | 1 | 1 | 1 |
| rs121908949 | pathogenic | 0 | 0 | 0 |
| rs104894612 | pathogenic | 1 | 1 | 1 |
| rs104894609 | pathogenic | 1 | 1 | 1 |
| rs104894610 | pathogenic | 1 | 1 | 1 |
| rs104894615 | pathogenic | 0 | 0 | 0 |
| rs104894603 | pathogenic | 0 | 0 | 0 |
| rs104894602 | pathogenic | 1 | 1 | 1 |
| rs121908948 | pathogenic | 1 | 1 | 1 |
| rs104894608 | pathogenic | 1 | 1 | 1 |
| rs371545673 | benign | 0 | 0 | 0 |
| rs199925238 | benign | 0 | 0 | 0 |
| rs146313451 | benign | 0 | 0 | 0 |
| rs121918193 | pathogenic | 1 | 1 | 0 |
| rs2228500 | benign | 0 | 0 | 0 |
| rs121918194 | pathogenic | 1 | 1 | 0 |
| rs121918196 | pathogenic | 0 | 0 | 0 |
| rs41291971 | benign | 0 | 0 | 0 |
| rs199473690 | pathogenic | 0 | 0 | 0 |
| rs2228613 | benign | 0 | 0 | 0 |
| rs267607130 | pathogenic | 1 | 1 | 1 |
| rs77615401 | benign | 1 | 1 | 1 |
| rs374618872 | benign | 1 | 1 | 1 |
| rs146165014 | benign | 0 | 0 | 0 |
| rs144586070 | benign | 0 | 0 | 0 |
| rs121918541 | pathogenic | 0 | 0 | 0 |
| rs142105943 | benign | 0 | 0 | 0 |
| rs121918540 | pathogenic | 0 | 0 | 0 |
| rs6791 | benign | 0 | 0 | 0 |
| rs61736587 | pathogenic | 0 | 0 | 0 |
| rs61736586 | benign | 0 | 0 | 0 |
| rs121917741 | pathogenic | 1 | 1 | 1 |
| rs41302885 | benign | 0 | 0 | 0 |
| rs141026028 | benign | 0 | 0 | 0 |
| rs193922493 | pathogenic | 0 | 0 | 0 |
| rs73598374 | benign | 0 | 0 | 0 |
| rs121908725 | pathogenic | 1 | 1 | 0 |
| rs199422328 | pathogenic | 0 | 1 | 0 |
| rs11555566 | benign | 1 | 1 | 1 |
| rs267606634 | pathogenic | 0 | 0 | 0 |
| rs121908714 | pathogenic | 1 | 1 | 0 |
| rs121908717 | pathogenic | 1 | 1 | 0 |
| rs267606635 | pathogenic | 1 | 1 | 0 |
| rs121908739 | pathogenic | 0 | 0 | 0 |
| rs121908735 | pathogenic | 0 | 1 | 0 |

| | | | | |
|---|---|---|---|---|
| rs121908716 | pathogenic | 0 | 1 | 0 |
| rs121908723 | pathogenic | 0 | 1 | 0 |
| rs121908721 | pathogenic | 0 | 0 | 0 |
| rs199422327 | pathogenic | 0 | 0 | 0 |
| rs121908715 | pathogenic | 0 | 0 | 0 |
| rs587783563 | pathogenic | 0 | 0 | 0 |
| rs104894780 | pathogenic | 0 | 0 | 0 |
| rs56030372 | pathogenic | 1 | 1 | 1 |
| rs104894782 | pathogenic | 1 | 1 | 1 |
| rs149495971 | benign | 0 | 0 | 0 |
| rs118203942 | pathogenic | 1 | 1 | 0 |
| rs118203939 | pathogenic | 0 | 0 | 0 |
| rs118203938 | pathogenic | 0 | 0 | 0 |
| rs118203943 | pathogenic | 0 | 0 | 0 |
| rs118203940 | pathogenic | 0 | 0 | 0 |
| rs727503809 | pathogenic | 0 | 0 | 0 |
| rs1065757 | benign | 0 | 0 | 0 |
| rs1071598 | benign | 0 | 0 | 0 |
| rs25414 | benign | 0 | 0 | 0 |
| rs118203944 | pathogenic | 0 | 0 | 0 |
| rs118203941 | pathogenic | 0 | 0 | 0 |
| rs144222167 | benign | 0 | 0 | 0 |
| rs201101343 | pathogenic | 0 | 0 | 0 |
| rs144601717 | benign | 0 | 0 | 0 |
| rs267607129 | pathogenic | 0 | 0 | 0 |
| rs104894729 | pathogenic | 0 | 0 | 0 |
| rs104894727 | pathogenic | 0 | 0 | 0 |
| rs104894725 | pathogenic | 0 | 0 | 0 |
| rs104894515 | pathogenic | 1 | 1 | 1 |
| rs74688377 | benign | 0 | 0 | 0 |
| rs121908513 | pathogenic | 1 | 1 | 0 |
| rs121908509 | pathogenic | 0 | 0 | 0 |
| rs121908514 | pathogenic | 1 | 1 | 0 |
| rs121908512 | pathogenic | 1 | 1 | 0 |
| rs121908510 | pathogenic | 0 | 0 | 0 |
| rs121908511 | pathogenic | 1 | 1 | 1 |
| rs121908518 | pathogenic | 1 | 1 | 1 |
| rs34358665 | benign | 0 | 0 | 0 |
| rs147568834 | benign | 0 | 0 | 0 |
| rs201717599 | benign | 0 | 0 | 0 |
| rs61744487 | benign | 0 | 0 | 0 |
| rs370627877 | pathogenic | 1 | 1 | 1 |
| rs368593151 | pathogenic | 1 | 1 | 1 |
| rs114363936 | benign | 0 | 0 | 0 |
| rs137853333 | pathogenic | 0 | 0 | 0 |
| rs137853080 | pathogenic | 1 | 1 | 1 |
| rs137853077 | pathogenic | 1 | 1 | 1 |
| rs780749732 | benign | 1 | 1 | 1 |
| rs137853081 | pathogenic | 1 | 1 | 0 |
| rs137853078 | pathogenic | 0 | 0 | 0 |
| rs121913315 | pathogenic | 1 | 1 | 1 |
| rs752513525 | pathogenic | 1 | 1 | 1 |
| rs104894918 | pathogenic | 0 | 0 | 0 |
| rs104894917 | pathogenic | 0 | 0 | 0 |

| | | | | |
|---|---|---|---|---|
| rs104894919 | pathogenic | 0 | 0 | 0 |
| rs28939672 | pathogenic | 0 | 0 | 0 |
| rs104894173 | pathogenic | 0 | 0 | 0 |
| rs104894179 | pathogenic | 0 | 0 | 0 |
| rs104894174 | pathogenic | 1 | 1 | 0 |
| rs104894178 | pathogenic | 1 | 1 | 0 |
| rs28939714 | pathogenic | 1 | 1 | 1 |
| rs104894270 | pathogenic | 1 | 1 | 1 |
| rs3765571 | benign | 0 | 0 | 0 |
| rs752669339 | pathogenic | 1 | 1 | 1 |
| rs537043237 | pathogenic | 1 | 1 | 1 |
| rs774047299 | pathogenic | 1 | 1 | 1 |
| rs121434583 | pathogenic | 1 | 1 | 1 |
| rs121907927 | pathogenic | 0 | 0 | 0 |
| rs121907925 | pathogenic | 0 | 1 | 0 |
| rs746212325 | pathogenic | 0 | 0 | 0 |
| rs754637179 | pathogenic | 0 | 0 | 0 |
| rs780882740 | pathogenic | 0 | 0 | 0 |
| rs121918130 | pathogenic | 0 | 0 | 0 |
| rs121918129 | pathogenic | 0 | 0 | 0 |
| rs13297509 | pathogenic | 0 | 0 | 0 |
| rs746867724 | pathogenic | 0 | 0 | 0 |
| rs121918128 | pathogenic | 0 | 0 | 0 |
| rs752300607 | pathogenic | 0 | 0 | 0 |
| rs121964860 | pathogenic | 0 | 0 | 0 |
| rs45466197 | pathogenic | 1 | 1 | 1 |
| rs3730238 | benign | 1 | 1 | 1 |
| rs730881125 | pathogenic | 1 | 1 | 1 |
| rs121964861 | pathogenic | 1 | 1 | 1 |
| rs397516484 | pathogenic | 0 | 0 | 0 |
| rs367785431 | pathogenic | 0 | 0 | 0 |
| rs141121678 | pathogenic | 0 | 0 | 0 |
| rs12720263 | benign | 0 | 0 | 0 |
| rs2304256 | benign | 0 | 0 | 0 |
| rs2304255 | benign | 0 | 0 | 0 |
| rs121909045 | pathogenic | 0 | 0 | 0 |
| rs397508796 | pathogenic | 0 | 0 | 0 |
| rs151020603 | pathogenic | 0 | 0 | 0 |
| rs77284892 | pathogenic | 0 | 0 | 0 |
| rs368505753 | pathogenic | 0 | 0 | 0 |
| rs115545701 | pathogenic | 0 | 0 | 0 |
| rs121908749 | pathogenic | 0 | 0 | 0 |
| rs75961395 | pathogenic | 0 | 0 | 0 |
| rs121908750 | pathogenic | 0 | 0 | 0 |
| rs121908751 | pathogenic | 0 | 0 | 0 |
| rs397508464 | pathogenic | 0 | 0 | 0 |
| rs397508461 | pathogenic | 0 | 0 | 0 |
| rs121909031 | pathogenic | 0 | 0 | 0 |
| rs113993958 | pathogenic | 0 | 0 | 0 |
| rs77834169 | pathogenic | 0 | 0 | 0 |
| rs78655421 | pathogenic | 0 | 0 | 0 |
| rs141723617 | benign | 0 | 0 | 0 |
| rs35516286 | benign | 0 | 0 | 0 |
| rs1800079 | pathogenic | 1 | 1 | 0 |

| | | | | |
|---|---|---|---|---|
| rs80282562 | pathogenic | 0 | 0 | 0 |
| rs121908752 | pathogenic | 0 | 0 | 0 |
| rs556880586 | pathogenic | 0 | 0 | 0 |
| rs121908760 | pathogenic | 0 | 0 | 0 |
| rs397508350 | pathogenic | 0 | 0 | 0 |
| rs374946172 | pathogenic | 0 | 0 | 0 |
| rs397508387 | pathogenic | 0 | 0 | 0 |
| rs201386642 | pathogenic | 0 | 0 | 0 |
| rs121909012 | pathogenic | 0 | 0 | 0 |
| rs121909008 | pathogenic | 0 | 0 | 0 |
| rs149790377 | pathogenic | 0 | 0 | 0 |
| rs397508442 | pathogenic | 0 | 0 | 0 |
| rs121909035 | pathogenic | 0 | 0 | 0 |
| rs386134230 | pathogenic | 0 | 0 | 0 |
| rs1800112 | benign | 0 | 0 | 0 |
| rs144055758 | pathogenic | 0 | 0 | 0 |
| rs142394380 | pathogenic | 0 | 0 | 0 |
| rs121909036 | pathogenic | 0 | 0 | 0 |
| rs78194216 | pathogenic | 0 | 0 | 0 |
| rs121909019 | pathogenic | 0 | 0 | 0 |
| rs121909020 | pathogenic | 0 | 0 | 0 |
| rs200321110 | pathogenic | 0 | 0 | 0 |
| rs202179988 | pathogenic | 0 | 0 | 0 |
| rs121909037 | pathogenic | 0 | 0 | 0 |
| rs186045772 | pathogenic | 0 | 0 | 0 |
| rs139304906 | pathogenic | 0 | 0 | 0 |
| rs78802634 | pathogenic | 0 | 0 | 0 |
| rs121908761 | pathogenic | 0 | 0 | 0 |
| rs36210737 | pathogenic | 0 | 0 | 0 |
| rs139729994 | pathogenic | 0 | 0 | 0 |
| rs79850223 | pathogenic | 0 | 0 | 0 |
| rs1800120 | benign | 0 | 0 | 0 |
| rs74767530 | pathogenic | 0 | 0 | 0 |
| rs374705585 | pathogenic | 0 | 0 | 0 |
| rs1136287 | benign | 0 | 0 | 0 |
| rs6828 | benign | 0 | 0 | 0 |
| rs104894856 | pathogenic | 0 | 0 | 0 |
| rs104894860 | pathogenic | 0 | 0 | 0 |
| rs61736892 | pathogenic | 0 | 0 | 0 |
| rs145807417 | benign | 0 | 0 | 0 |
| rs104894853 | pathogenic | 0 | 0 | 0 |
| rs199422227 | pathogenic | 0 | 0 | 0 |
| rs113993946 | pathogenic | 0 | 0 | 0 |
| rs199422231 | pathogenic | 0 | 0 | 0 |
| rs199422228 | pathogenic | 0 | 0 | 0 |
| rs142195196 | benign | 0 | 1 | 0 |
| rs148656104 | benign | 1 | 1 | 1 |
| rs141944844 | benign | 0 | 0 | 0 |
| rs104894342 | pathogenic | 0 | 0 | 0 |
| rs104894343 | pathogenic | 0 | 0 | 0 |
| rs104894344 | pathogenic | 0 | 0 | 0 |
| rs573322878 | benign | 0 | 0 | 0 |
| rs104894347 | pathogenic | 0 | 0 | 0 |
| rs28937882 | pathogenic | 0 | 0 | 0 |

| | | | | |
|---|---|---|---|---|
| rs193922702 | pathogenic | 0 | 0 | 0 |
| rs200786104 | benign | 0 | 0 | 0 |
| rs690016544 | pathogenic | 1 | 1 | 1 |
| rs28943592 | benign | 0 | 0 | 0 |
| rs28943594 | benign | 0 | 0 | 0 |
| rs121908957 | pathogenic | 0 | 0 | 0 |
| rs115408010 | benign | 1 | 1 | 1 |
| rs77591659 | benign | 0 | 0 | 0 |
| rs121918511 | pathogenic | 0 | 0 | 0 |
| rs538359948 | benign | 0 | 0 | 0 |
| rs76708142 | benign | 0 | 0 | 0 |
| rs121907913 | pathogenic | 1 | 1 | 0 |
| rs121907920 | pathogenic | 0 | 0 | 0 |
| rs121907923 | pathogenic | 1 | 1 | 0 |
| rs121907928 | pathogenic | 1 | 1 | 0 |
| rs121907919 | pathogenic | 1 | 1 | 0 |
| rs121907918 | pathogenic | 1 | 1 | 0 |
| rs74315493 | pathogenic | 1 | 1 | 1 |
| rs74315494 | pathogenic | 1 | 1 | 1 |
| rs145446060 | benign | 1 | 1 | 1 |
| rs550409143 | benign | 1 | 1 | 0 |

| compound_binding_site | cofactor_binding_site | nucleic_binding_site | ion_binding_site |
|---|---|---|---|
| 1 | 0 | 0 | 0 |
| 1 | 0 | 0 | 0 |
| 0 | 0 | 0 | 0 |
| 1 | 0 | 0 | 0 |
| 0 | 0 | 0 | 0 |
| 0 | 0 | 0 | 0 |
| 0 | 0 | 0 | 0 |
| 0 | 0 | 0 | 0 |
| 1 | 0 | 0 | 1 |
| 1 | 0 | 0 | 0 |
| 0 | 0 | 0 | 1 |
| 1 | 0 | 1 | 0 |
| 0 | 0 | 0 | 0 |
| 0 | 0 | 0 | 0 |
| 0 | 0 | 0 | 0 |
| 0 | 0 | 0 | 0 |
| 0 | 0 | 0 | 0 |
| 1 | 0 | 0 | 1 |
| 1 | 0 | 1 | 0 |
| 0 | 0 | 0 | 0 |
| 1 | 0 | 0 | 1 |
| 1 | 0 | 0 | 0 |
| 1 | 0 | 0 | 0 |
| 1 | 0 | 0 | 0 |
| 0 | 0 | 0 | 0 |
| 0 | 0 | 0 | 0 |
| 1 | 0 | 0 | 1 |
| 0 | 0 | 0 | 0 |
| 0 | 0 | 0 | 0 |
| 0 | 0 | 0 | 0 |
| 0 | 0 | 0 | 0 |
| 1 | 0 | 0 | 0 |
| 1 | 0 | 0 | 1 |
| 1 | 0 | 0 | 0 |
| 0 | 0 | 0 | 0 |
| 0 | 0 | 0 | 0 |
| 0 | 0 | 0 | 0 |
| 1 | 0 | 0 | 1 |
| 1 | 0 | 0 | 1 |
| 1 | 0 | 0 | 1 |
| 1 | 0 | 0 | 0 |
| 1 | 0 | 0 | 0 |
| 1 | 0 | 0 | 0 |
| 1 | 0 | 0 | 0 |
| 1 | 0 | 0 | 0 |
| 1 | 0 | 0 | 0 |
| 1 | 0 | 0 | 0 |
| 1 | 0 | 0 | 0 |
| 1 | 0 | 0 | 0 |
| 1 | 0 | 0 | 0 |
| 1 | 0 | 0 | 0 |
| 1 | 0 | 0 | 0 |
| 1 | 0 | 0 | 0 |

| | | | |
|---|---|---|---|
| 1 | 0 | 0 | 1 |
| 1 | 0 | 0 | 0 |
| 1 | 0 | 0 | 0 |
| 1 | 0 | 0 | 0 |
| 0 | 0 | 0 | 0 |
| 1 | 0 | 0 | 0 |
| 0 | 0 | 0 | 0 |
| 1 | 0 | 0 | 0 |
| 1 | 0 | 0 | 0 |
| 1 | 0 | 0 | 1 |
| 1 | 0 | 0 | 0 |
| 1 | 0 | 0 | 0 |
| 1 | 0 | 0 | 0 |
| 1 | 0 | 0 | 0 |
| 1 | 0 | 0 | 0 |
| 1 | 0 | 0 | 0 |
| 1 | 0 | 0 | 0 |
| 1 | 0 | 0 | 1 |
| 1 | 0 | 0 | 1 |
| 1 | 1 | 0 | 1 |
| 1 | 0 | 0 | 0 |
| 1 | 0 | 0 | 0 |
| 1 | 0 | 0 | 0 |
| 1 | 0 | 0 | 0 |
| 1 | 1 | 0 | 0 |
| 1 | 0 | 0 | 0 |
| 1 | 0 | 0 | 0 |
| 1 | 0 | 0 | 0 |
| 1 | 0 | 0 | 0 |
| 1 | 0 | 0 | 0 |
| 1 | 1 | 0 | 0 |
| 1 | 1 | 0 | 0 |
| 1 | 1 | 0 | 0 |
| 1 | 1 | 0 | 0 |
| 1 | 1 | 0 | 0 |
| 1 | 0 | 0 | 0 |
| 1 | 1 | 0 | 0 |
| 1 | 0 | 0 | 0 |
| 1 | 0 | 0 | 0 |
| 1 | 1 | 0 | 0 |
| 0 | 0 | 0 | 0 |
| 0 | 0 | 0 | 0 |
| 0 | 0 | 0 | 0 |
| 0 | 0 | 0 | 0 |
| 0 | 0 | 0 | 0 |
| 0 | 0 | 0 | 0 |
| 0 | 0 | 0 | 0 |
| 1 | 0 | 0 | 1 |
| 0 | 0 | 0 | 0 |
| 1 | 0 | 0 | 0 |
| 1 | 0 | 0 | 0 |
| 0 | 0 | 0 | 0 |
| 0 | 0 | 0 | 0 |
| 0 | 0 | 0 | 0 |

| | | | |
|---|---|---|---|
| 0 | 0 | 0 | 0 |
| 0 | 0 | 0 | 0 |
| 0 | 0 | 0 | 0 |
| 1 | 0 | 0 | 0 |
| 1 | 0 | 0 | 0 |
| 1 | 0 | 0 | 0 |
| 0 | 0 | 0 | 0 |
| 0 | 0 | 1 | 0 |
| 0 | 0 | 1 | 0 |
| 0 | 0 | 1 | 0 |
| 0 | 0 | 1 | 0 |
| 0 | 0 | 1 | 0 |
| 0 | 0 | 0 | 0 |
| 0 | 0 | 0 | 0 |
| 1 | 0 | 0 | 0 |
| 0 | 0 | 0 | 0 |
| 0 | 0 | 0 | 0 |
| 1 | 0 | 0 | 0 |
| 1 | 0 | 0 | 0 |
| 1 | 0 | 0 | 0 |
| 1 | 0 | 0 | 0 |
| 1 | 1 | 0 | 0 |
| 1 | 0 | 0 | 0 |
| 1 | 0 | 0 | 0 |
| 0 | 0 | 0 | 0 |
| 0 | 0 | 0 | 0 |
| 1 | 0 | 0 | 0 |
| 1 | 1 | 0 | 0 |
| 1 | 0 | 0 | 0 |
| 1 | 0 | 0 | 0 |
| 1 | 1 | 0 | 0 |
| 1 | 1 | 0 | 0 |
| 1 | 1 | 0 | 0 |
| 1 | 1 | 0 | 0 |
| 1 | 1 | 0 | 0 |
| 1 | 1 | 0 | 0 |
| 1 | 1 | 0 | 0 |
| 1 | 1 | 0 | 0 |
| 1 | 1 | 0 | 0 |
| 1 | 1 | 0 | 0 |
| 1 | 1 | 0 | 0 |
| 1 | 1 | 0 | 0 |
| 1 | 0 | 0 | 0 |
| 1 | 0 | 0 | 0 |
| 1 | 0 | 0 | 0 |
| 1 | 1 | 0 | 0 |
| 1 | 0 | 0 | 0 |
| 1 | 1 | 1 | 0 |
| 1 | 1 | 1 | 0 |
| 1 | 1 | 0 | 0 |
| 1 | 1 | 0 | 0 |
| 1 | 0 | 0 | 0 |
| 1 | 0 | 0 | 0 |
| 0 | 0 | 0 | 0 |

| | | | |
|---|---|---|---|
| 0 | 0 | 0 | 0 |
| 0 | 0 | 0 | 0 |
| 0 | 0 | 0 | 0 |
| 1 | 0 | 0 | 0 |
| 1 | 0 | 0 | 0 |
| 0 | 0 | 0 | 0 |
| 0 | 0 | 0 | 0 |
| 0 | 0 | 0 | 0 |
| 0 | 0 | 0 | 0 |
| 0 | 0 | 0 | 0 |
| 0 | 0 | 0 | 0 |
| 0 | 0 | 0 | 0 |
| 0 | 0 | 0 | 0 |
| 0 | 0 | 0 | 0 |
| 0 | 0 | 0 | 0 |
| 0 | 0 | 0 | 0 |
| 0 | 0 | 0 | 0 |
| 0 | 0 | 0 | 0 |
| 0 | 0 | 0 | 0 |
| 0 | 0 | 0 | 0 |
| 0 | 0 | 0 | 0 |
| 0 | 0 | 0 | 0 |
| 1 | 0 | 0 | 0 |
| 1 | 0 | 0 | 0 |
| 0 | 0 | 0 | 0 |
| 0 | 0 | 0 | 0 |
| 1 | 0 | 0 | 0 |
| 0 | 0 | 0 | 0 |
| 0 | 0 | 0 | 0 |
| 0 | 0 | 0 | 0 |
| 0 | 0 | 0 | 0 |
| 0 | 0 | 0 | 0 |
| 0 | 0 | 0 | 0 |
| 0 | 0 | 0 | 0 |
| 1 | 0 | 0 | 0 |
| 0 | 0 | 0 | 0 |
| 0 | 0 | 0 | 0 |
| 0 | 0 | 0 | 0 |
| 0 | 0 | 0 | 0 |
| 0 | 0 | 0 | 0 |
| 0 | 0 | 0 | 0 |
| 0 | 0 | 0 | 0 |
| 0 | 0 | 0 | 0 |
| 0 | 0 | 0 | 0 |
| 0 | 0 | 0 | 0 |
| 0 | 0 | 0 | 0 |
| 0 | 0 | 1 | 0 |
| 1 | 0 | 0 | 0 |
| 1 | 0 | 0 | 0 |
| 1 | 1 | 0 | 0 |
| 1 | 0 | 0 | 0 |
| 0 | 0 | 0 | 0 |
| 0 | 0 | 0 | 0 |
| 1 | 0 | 0 | 0 |

| | | | |
|---|---|---|---|
| 1 | 0 | 0 | 0 |
| 1 | 0 | 0 | 0 |
| 1 | 0 | 0 | 0 |
| 0 | 0 | 0 | 0 |
| 0 | 0 | 0 | 0 |
| 0 | 0 | 0 | 0 |
| 1 | 0 | 0 | 0 |
| 1 | 0 | 0 | 0 |
| 0 | 0 | 0 | 0 |
| 0 | 0 | 0 | 0 |
| 1 | 0 | 0 | 0 |
| 0 | 0 | 0 | 0 |
| 1 | 0 | 0 | 0 |
| 0 | 0 | 0 | 0 |
| 0 | 0 | 0 | 0 |
| 0 | 0 | 0 | 0 |
| 0 | 0 | 0 | 0 |
| 0 | 0 | 0 | 0 |
| 0 | 0 | 0 | 0 |
| 0 | 0 | 0 | 0 |
| 0 | 0 | 0 | 0 |
| 0 | 0 | 0 | 0 |
| 0 | 0 | 0 | 0 |
| 0 | 0 | 0 | 0 |
| 0 | 0 | 0 | 0 |
| 0 | 0 | 0 | 0 |
| 1 | 0 | 0 | 0 |
| 1 | 0 | 0 | 0 |
| 1 | 0 | 0 | 0 |
| 1 | 0 | 0 | 0 |
| 1 | 0 | 0 | 0 |
| 1 | 0 | 0 | 0 |
| 0 | 0 | 0 | 0 |
| 0 | 0 | 0 | 0 |
| 1 | 0 | 0 | 0 |
| 0 | 0 | 0 | 0 |
| 0 | 0 | 0 | 0 |
| 0 | 0 | 0 | 0 |
| 0 | 0 | 0 | 0 |
| 0 | 0 | 0 | 0 |
| 0 | 0 | 0 | 0 |
| 0 | 0 | 0 | 0 |
| 0 | 0 | 0 | 0 |
| 0 | 0 | 0 | 0 |
| 1 | 1 | 0 | 0 |
| 1 | 1 | 0 | 0 |
| 1 | 1 | 0 | 1 |
| 1 | 1 | 0 | 1 |
| 1 | 1 | 0 | 1 |
| 1 | 1 | 0 | 1 |
| 1 | 1 | 0 | 0 |
| 1 | 1 | 0 | 1 |
| 1 | 1 | 0 | 1 |
| 1 | 1 | 0 | 1 |

| | | | |
|---|---|---|---|
| 1 | 1 | 0 | 1 |
| 1 | 1 | 0 | 1 |
| 1 | 1 | 0 | 1 |
| 1 | 1 | 0 | 1 |
| 1 | 1 | 0 | 1 |
| 1 | 0 | 0 | 0 |
| 0 | 0 | 0 | 0 |
| 0 | 0 | 0 | 0 |
| 0 | 0 | 0 | 0 |
| 1 | 0 | 0 | 0 |
| 0 | 0 | 1 | 0 |
| 0 | 0 | 0 | 0 |
| 0 | 0 | 0 | 0 |
| 0 | 0 | 0 | 0 |
| 0 | 0 | 0 | 0 |
| 0 | 0 | 0 | 0 |
| 0 | 0 | 0 | 0 |
| 0 | 0 | 0 | 0 |
| 0 | 0 | 0 | 0 |
| 0 | 0 | 0 | 0 |
| 0 | 0 | 0 | 0 |
| 0 | 0 | 0 | 0 |
| 0 | 0 | 0 | 0 |
| 0 | 0 | 0 | 0 |
| 0 | 0 | 0 | 0 |
| 1 | 1 | 0 | 0 |
| 1 | 1 | 0 | 0 |
| 1 | 1 | 0 | 0 |
| 0 | 1 | 0 | 0 |
| 1 | 0 | 0 | 0 |
| 0 | 0 | 0 | 0 |
| 0 | 0 | 0 | 0 |
| 0 | 0 | 0 | 0 |
| 1 | 0 | 0 | 0 |
| 1 | 1 | 0 | 0 |
| 0 | 0 | 0 | 0 |
| 0 | 0 | 0 | 0 |
| 0 | 0 | 0 | 0 |
| 0 | 0 | 0 | 0 |
| 1 | 0 | 0 | 0 |
| 0 | 0 | 0 | 0 |
| 0 | 0 | 0 | 0 |
| 0 | 0 | 0 | 0 |
| 0 | 0 | 0 | 0 |
| 0 | 0 | 0 | 0 |
| 0 | 1 | 0 | 0 |
| 0 | 1 | 0 | 0 |
| 0 | 0 | 0 | 0 |
| 0 | 0 | 0 | 0 |
| 0 | 1 | 0 | 0 |
| 0 | 0 | 0 | 0 |
| 0 | 0 | 0 | 0 |
| 0 | 1 | 0 | 0 |
| 0 | 0 | 0 | 0 |

| | | | |
|---|---|---|---|
| 1 | 1 | 0 | 0 |
| 0 | 0 | 0 | 0 |
| 0 | 0 | 0 | 0 |
| 0 | 0 | 0 | 0 |
| 0 | 0 | 0 | 0 |
| 0 | 0 | 0 | 0 |
| 0 | 0 | 0 | 0 |
| 0 | 0 | 0 | 0 |
| 0 | 0 | 0 | 0 |
| 0 | 0 | 0 | 0 |
| 0 | 0 | 0 | 0 |
| 0 | 0 | 0 | 0 |
| 0 | 0 | 0 | 0 |
| 0 | 0 | 0 | 0 |
| 0 | 1 | 0 | 0 |
| 0 | 0 | 0 | 0 |
| 0 | 1 | 0 | 0 |
| 0 | 1 | 0 | 0 |
| 0 | 0 | 0 | 0 |
| 0 | 1 | 0 | 0 |
| 1 | 0 | 0 | 1 |
| 1 | 1 | 0 | 0 |
| 1 | 0 | 0 | 0 |
| 1 | 0 | 0 | 1 |
| 0 | 0 | 0 | 0 |
| 0 | 0 | 0 | 0 |
| 0 | 0 | 0 | 0 |
| 0 | 0 | 0 | 0 |
| 1 | 0 | 0 | 0 |
| 0 | 0 | 0 | 0 |
| 1 | 0 | 0 | 0 |
| 0 | 0 | 0 | 0 |
| 0 | 0 | 0 | 0 |
| 0 | 0 | 0 | 0 |
| 0 | 0 | 0 | 0 |
| 0 | 0 | 0 | 0 |
| 0 | 0 | 0 | 0 |
| 1 | 1 | 0 | 0 |
| 1 | 1 | 0 | 0 |
| 1 | 1 | 0 | 0 |
| 0 | 0 | 0 | 0 |
| 0 | 0 | 0 | 0 |
| 0 | 0 | 0 | 0 |
| 1 | 0 | 0 | 0 |
| 1 | 0 | 0 | 1 |
| 1 | 0 | 0 | 1 |
| 1 | 0 | 0 | 0 |
| 1 | 0 | 0 | 1 |
| 1 | 0 | 0 | 0 |
| 0 | 0 | 0 | 0 |
| 1 | 1 | 0 | 0 |
| 1 | 1 | 0 | 0 |
| 0 | 0 | 0 | 0 |
| 0 | 0 | 0 | 0 |

| | | | |
|---|---|---|---|
| 1 | 0 | 0 | 0 |
| 1 | 0 | 0 | 0 |
| 1 | 0 | 0 | 0 |
| 0 | 0 | 0 | 0 |
| 0 | 0 | 0 | 0 |
| 1 | 1 | 0 | 1 |
| 1 | 1 | 0 | 1 |
| 1 | 1 | 0 | 1 |
| 1 | 1 | 0 | 1 |
| 1 | 1 | 0 | 1 |
| 1 | 1 | 0 | 1 |
| 1 | 1 | 0 | 1 |
| 1 | 1 | 0 | 0 |
| 1 | 1 | 0 | 0 |
| 1 | 1 | 1 | 1 |
| 0 | 0 | 0 | 0 |
| 1 | 1 | 0 | 0 |
| 1 | 1 | 0 | 0 |
| 0 | 0 | 0 | 0 |
| 0 | 0 | 0 | 0 |
| 0 | 0 | 0 | 0 |
| 1 | 0 | 0 | 0 |
| 1 | 0 | 0 | 0 |
| 0 | 0 | 0 | 0 |
| 1 | 0 | 0 | 0 |
| 1 | 0 | 0 | 0 |
| 1 | 0 | 0 | 0 |
| 0 | 0 | 0 | 0 |
| 0 | 0 | 0 | 0 |
| 0 | 0 | 0 | 0 |
| 0 | 0 | 0 | 0 |
| 0 | 0 | 0 | 0 |
| 0 | 0 | 0 | 0 |
| 0 | 0 | 0 | 0 |
| 0 | 0 | 0 | 0 |
| 0 | 0 | 0 | 0 |
| 0 | 0 | 0 | 0 |
| 0 | 0 | 0 | 0 |
| 0 | 0 | 0 | 0 |
| 0 | 0 | 0 | 0 |
| 0 | 0 | 0 | 0 |
| 0 | 0 | 0 | 0 |
| 0 | 0 | 0 | 0 |
| 0 | 0 | 0 | 0 |
| 0 | 0 | 0 | 0 |
| 1 | 0 | 0 | 0 |
| 1 | 0 | 0 | 0 |
| 1 | 0 | 0 | 0 |
| 1 | 0 | 0 | 0 |
| 0 | 0 | 0 | 0 |
| 0 | 0 | 0 | 0 |
| 0 | 0 | 0 | 0 |
| 0 | 0 | 0 | 0 |
| 0 | 0 | 0 | 0 |

| | | | |
|---|---|---|---|
| 0 | 0 | 0 | 0 |
| 0 | 0 | 0 | 0 |
| 0 | 0 | 0 | 0 |
| 0 | 0 | 0 | 0 |
| 0 | 0 | 0 | 0 |
| 0 | 0 | 0 | 0 |
| 0 | 0 | 0 | 0 |
| 1 | 0 | 0 | 1 |
| 0 | 0 | 0 | 0 |
| 0 | 0 | 0 | 0 |
| 1 | 0 | 0 | 0 |
| 0 | 0 | 0 | 0 |
| 0 | 0 | 0 | 0 |
| 0 | 0 | 0 | 0 |
| 0 | 0 | 0 | 0 |
| 1 | 0 | 0 | 0 |
| 0 | 0 | 0 | 0 |
| 0 | 0 | 0 | 0 |
| 0 | 0 | 0 | 0 |
| 0 | 0 | 0 | 0 |
| 0 | 0 | 0 | 0 |
| 0 | 0 | 0 | 0 |
| 0 | 0 | 0 | 0 |
| 1 | 1 | 0 | 1 |
| 0 | 0 | 0 | 0 |
| 0 | 0 | 0 | 0 |
| 0 | 0 | 0 | 0 |
| 1 | 1 | 0 | 1 |
| 1 | 1 | 0 | 0 |
| 0 | 0 | 0 | 0 |
| 1 | 0 | 0 | 0 |
| 1 | 1 | 0 | 0 |
| 0 | 0 | 0 | 0 |
| 1 | 0 | 0 | 0 |
| 1 | 1 | 0 | 0 |
| 1 | 0 | 0 | 0 |
| 1 | 1 | 0 | 1 |
| 1 | 1 | 0 | 1 |
| 1 | 0 | 0 | 0 |
| 0 | 0 | 0 | 0 |
| 0 | 0 | 0 | 0 |
| 0 | 0 | 0 | 0 |
| 0 | 0 | 0 | 0 |
| 0 | 0 | 0 | 0 |
| 0 | 0 | 0 | 0 |
| 0 | 0 | 0 | 0 |
| 0 | 0 | 0 | 0 |
| 0 | 0 | 0 | 0 |
| 0 | 0 | 0 | 0 |
| 0 | 0 | 0 | 0 |
| 0 | 0 | 0 | 0 |
| 0 | 0 | 0 | 0 |
| 0 | 0 | 0 | 0 |
| 0 | 0 | 0 | 0 |

| | | | |
|---|---|---|---|
| 0 | 0 | 0 | 0 |
| 0 | 0 | 0 | 0 |
| 0 | 0 | 0 | 0 |
| 0 | 0 | 0 | 0 |
| 0 | 0 | 0 | 0 |
| 0 | 0 | 0 | 0 |
| 0 | 0 | 0 | 0 |
| 0 | 0 | 0 | 0 |
| 0 | 0 | 0 | 0 |
| 0 | 0 | 0 | 0 |
| 0 | 0 | 0 | 0 |
| 0 | 0 | 0 | 0 |
| 0 | 0 | 0 | 0 |
| 1 | 1 | 0 | 0 |
| 0 | 0 | 0 | 0 |
| 1 | 0 | 0 | 0 |
| 0 | 0 | 0 | 1 |
| 0 | 0 | 0 | 0 |
| 0 | 0 | 0 | 0 |
| 0 | 0 | 0 | 0 |
| 0 | 0 | 0 | 0 |
| 0 | 0 | 0 | 0 |
| 0 | 0 | 0 | 0 |
| 0 | 0 | 0 | 0 |
| 0 | 0 | 0 | 0 |
| 0 | 0 | 0 | 0 |
| 0 | 0 | 0 | 0 |
| 0 | 0 | 0 | 0 |
| 0 | 0 | 0 | 0 |
| 0 | 0 | 0 | 0 |
| 1 | 0 | 0 | 0 |
| 1 | 0 | 0 | 0 |
| 0 | 0 | 0 | 0 |
| 0 | 0 | 0 | 0 |
| 0 | 0 | 0 | 0 |
| 0 | 0 | 0 | 0 |
| 0 | 0 | 0 | 0 |
| 0 | 0 | 0 | 0 |
| 0 | 0 | 0 | 0 |
| 0 | 0 | 0 | 0 |
| 1 | 0 | 0 | 0 |
| 1 | 0 | 0 | 0 |
| 1 | 0 | 0 | 0 |
| 1 | 0 | 0 | 0 |
| 0 | 0 | 0 | 0 |
| 1 | 0 | 0 | 0 |
| 0 | 0 | 0 | 0 |
| 0 | 0 | 0 | 0 |
| 1 | 0 | 0 | 0 |
| 1 | 0 | 0 | 0 |
| 0 | 0 | 0 | 0 |
| 0 | 0 | 0 | 0 |
| 0 | 0 | 0 | 0 |
| 0 | 0 | 0 | 0 |

| | | | |
|---|---|---|---|
| 0 | 0 | 0 | 0 |
| 0 | 0 | 0 | 0 |
| 0 | 0 | 0 | 0 |
| 0 | 0 | 0 | 0 |
| 0 | 0 | 0 | 0 |
| 0 | 0 | 0 | 0 |
| 0 | 0 | 0 | 0 |
| 0 | 0 | 0 | 0 |
| 0 | 0 | 0 | 0 |
| 0 | 0 | 0 | 0 |
| 0 | 0 | 0 | 0 |
| 0 | 0 | 0 | 0 |
| 0 | 0 | 0 | 0 |
| 0 | 0 | 0 | 0 |
| 0 | 0 | 0 | 0 |
| 1 | 0 | 0 | 1 |
| 0 | 0 | 0 | 0 |
| 0 | 0 | 0 | 0 |
| 0 | 0 | 0 | 0 |
| 0 | 0 | 0 | 0 |
| 0 | 0 | 0 | 0 |
| 1 | 1 | 0 | 0 |
| 1 | 1 | 0 | 0 |
| 1 | 0 | 0 | 0 |
| 1 | 0 | 0 | 0 |
| 0 | 0 | 0 | 0 |
| 0 | 0 | 0 | 0 |
| 0 | 0 | 0 | 0 |
| 0 | 0 | 0 | 0 |
| 1 | 0 | 0 | 0 |
| 1 | 0 | 0 | 0 |
| 1 | 0 | 0 | 0 |
| 1 | 1 | 0 | 0 |
| 1 | 1 | 0 | 0 |
| 1 | 0 | 0 | 0 |
| 0 | 0 | 0 | 0 |
| 0 | 0 | 0 | 0 |
| 0 | 1 | 0 | 0 |
| 0 | 0 | 0 | 0 |
| 0 | 0 | 0 | 0 |
| 0 | 0 | 0 | 0 |
| 1 | 1 | 0 | 0 |
| 1 | 0 | 0 | 0 |
| 1 | 0 | 0 | 0 |
| 0 | 0 | 0 | 0 |
| 0 | 0 | 0 | 0 |
| 0 | 0 | 0 | 0 |
| 0 | 0 | 0 | 0 |
| 0 | 0 | 0 | 0 |
| 0 | 0 | 0 | 0 |
| 0 | 0 | 0 | 0 |
| 0 | 0 | 0 | 0 |
| 0 | 0 | 0 | 0 |
| 0 | 0 | 0 | 0 |

| | | | |
|---|---|---|---|
| 0 | 0 | 0 | 0 |
| 1 | 0 | 0 | 0 |
| 0 | 0 | 0 | 0 |
| 0 | 0 | 0 | 0 |
| 0 | 0 | 0 | 0 |
| 0 | 0 | 0 | 0 |
| 1 | 0 | 0 | 0 |
| 0 | 0 | 0 | 0 |
| 0 | 0 | 0 | 0 |
| 0 | 0 | 0 | 0 |
| 0 | 0 | 0 | 0 |
| 0 | 0 | 0 | 0 |
| 0 | 0 | 0 | 0 |
| 0 | 0 | 0 | 0 |
| 0 | 0 | 0 | 0 |
| 0 | 0 | 0 | 0 |
| 0 | 1 | 0 | 0 |
| 0 | 0 | 0 | 0 |
| 0 | 0 | 0 | 0 |
| 0 | 0 | 0 | 0 |
| 1 | 0 | 0 | 0 |
| 0 | 0 | 0 | 0 |
| 0 | 0 | 0 | 0 |
| 1 | 1 | 0 | 0 |
| 0 | 0 | 0 | 0 |
| 0 | 0 | 0 | 0 |
| 0 | 0 | 0 | 0 |
| 0 | 0 | 0 | 0 |
| 1 | 0 | 0 | 0 |
| 0 | 0 | 0 | 0 |
| 0 | 0 | 0 | 0 |
| 0 | 0 | 0 | 0 |
| 0 | 0 | 0 | 0 |
| 0 | 0 | 0 | 0 |
| 0 | 0 | 0 | 0 |
| 1 | 0 | 0 | 0 |
| 0 | 0 | 0 | 0 |
| 0 | 0 | 0 | 0 |
| 0 | 0 | 0 | 0 |
| 0 | 0 | 0 | 0 |
| 1 | 1 | 0 | 0 |
| 0 | 0 | 0 | 0 |
| 0 | 0 | 0 | 0 |
| 0 | 0 | 0 | 0 |
| 0 | 0 | 0 | 0 |
| 0 | 0 | 0 | 0 |
| 1 | 0 | 0 | 0 |
| 1 | 0 | 0 | 1 |
| 1 | 0 | 0 | 0 |
| 0 | 0 | 0 | 0 |
| 0 | 0 | 0 | 0 |
| 0 | 0 | 0 | 0 |
| 1 | 0 | 0 | 0 |
| 1 | 0 | 0 | 0 |

| | | | |
|---|---|---|---|
| 1 | 0 | 0 | 0 |
| 0 | 0 | 0 | 0 |
| 0 | 0 | 0 | 0 |
| 0 | 0 | 0 | 0 |
| 1 | 1 | 0 | 0 |
| 1 | 1 | 0 | 0 |
| 0 | 0 | 0 | 0 |
| 0 | 0 | 0 | 0 |
| 1 | 0 | 0 | 0 |
| 1 | 0 | 0 | 0 |
| 1 | 0 | 1 | 0 |
| 0 | 0 | 0 | 0 |
| 0 | 0 | 1 | 0 |
| 0 | 0 | 1 | 0 |
| 0 | 0 | 1 | 0 |
| 1 | 0 | 0 | 0 |
| 1 | 0 | 0 | 0 |
| 0 | 0 | 0 | 0 |
| 0 | 0 | 0 | 0 |
| 1 | 0 | 0 | 1 |
| 1 | 0 | 0 | 1 |
| 1 | 0 | 0 | 0 |
| 0 | 0 | 0 | 0 |
| 0 | 0 | 0 | 0 |
| 0 | 0 | 0 | 0 |
| 0 | 0 | 0 | 0 |
| 0 | 0 | 0 | 0 |
| 1 | 0 | 0 | 0 |
| 0 | 0 | 0 | 0 |
| 0 | 0 | 0 | 1 |
| 0 | 0 | 0 | 1 |
| 0 | 0 | 1 | 1 |
| 0 | 0 | 1 | 1 |
| 0 | 0 | 1 | 1 |
| 0 | 0 | 1 | 1 |
| 0 | 0 | 1 | 0 |
| 0 | 0 | 1 | 0 |
| 0 | 0 | 0 | 0 |
| 0 | 0 | 0 | 0 |
| 0 | 0 | 0 | 0 |
| 0 | 0 | 0 | 0 |
| 0 | 0 | 1 | 0 |
| 0 | 0 | 1 | 0 |
| 0 | 0 | 1 | 0 |
| 0 | 0 | 1 | 0 |
| 0 | 0 | 1 | 0 |
| 0 | 0 | 1 | 0 |
| 0 | 0 | 1 | 0 |
| 0 | 0 | 1 | 0 |
| 0 | 0 | 0 | 0 |
| 0 | 0 | 0 | 0 |
| 0 | 0 | 0 | 0 |
| 0 | 0 | 0 | 0 |
| 1 | 0 | 0 | 0 |

| | | | |
|---|---|---|---|
| 0 | 0 | 0 | 0 |
| 0 | 0 | 0 | 0 |
| 0 | 0 | 0 | 0 |
| 1 | 0 | 0 | 0 |
| 0 | 0 | 0 | 0 |
| 0 | 0 | 0 | 0 |
| 0 | 0 | 0 | 0 |
| 0 | 0 | 0 | 0 |
| 0 | 0 | 0 | 0 |
| 0 | 0 | 0 | 0 |
| 1 | 1 | 0 | 0 |
| 0 | 0 | 0 | 0 |
| 1 | 1 | 0 | 0 |
| 1 | 1 | 0 | 0 |
| 0 | 1 | 0 | 0 |
| 0 | 0 | 0 | 0 |
| 0 | 0 | 0 | 0 |
| 0 | 0 | 0 | 0 |
| 0 | 0 | 0 | 0 |
| 0 | 0 | 0 | 0 |
| 1 | 1 | 0 | 0 |
| 0 | 1 | 0 | 0 |
| 0 | 0 | 0 | 0 |
| 0 | 1 | 0 | 0 |
| 1 | 1 | 0 | 0 |
| 0 | 0 | 0 | 0 |
| 0 | 0 | 0 | 0 |
| 1 | 1 | 0 | 0 |
| 1 | 1 | 0 | 0 |
| 0 | 1 | 0 | 0 |
| 0 | 0 | 0 | 0 |
| 0 | 0 | 0 | 0 |
| 0 | 0 | 0 | 0 |
| 0 | 0 | 0 | 0 |
| 0 | 0 | 0 | 0 |
| 0 | 0 | 0 | 0 |
| 0 | 0 | 0 | 0 |
| 0 | 0 | 0 | 0 |
| 0 | 0 | 0 | 0 |
| 0 | 0 | 0 | 0 |
| 1 | 0 | 0 | 0 |
| 1 | 0 | 0 | 0 |
| 1 | 0 | 0 | 0 |
| 1 | 0 | 0 | 0 |
| 1 | 1 | 0 | 1 |
| 0 | 0 | 0 | 0 |
| 1 | 0 | 0 | 0 |
| 0 | 0 | 0 | 0 |
| 1 | 0 | 0 | 0 |
| 0 | 0 | 0 | 0 |
| 1 | 0 | 0 | 0 |
| 1 | 0 | 0 | 0 |
| 0 | 0 | 0 | 0 |
| 0 | 0 | 0 | 0 |

| | | | |
|---|---|---|---|
| 0 | 0 | 0 | 0 |
| 0 | 0 | 0 | 0 |
| 0 | 0 | 0 | 0 |
| 0 | 0 | 0 | 0 |
| 1 | 0 | 0 | 0 |
| 1 | 0 | 0 | 0 |
| 1 | 0 | 0 | 0 |
| 0 | 0 | 0 | 0 |
| 1 | 1 | 0 | 0 |
| 0 | 0 | 0 | 0 |
| 0 | 0 | 0 | 0 |
| 0 | 0 | 0 | 0 |
| 1 | 1 | 0 | 0 |
| 1 | 0 | 0 | 0 |
| 0 | 0 | 0 | 0 |
| 0 | 0 | 0 | 0 |
| 1 | 1 | 0 | 0 |
| 0 | 0 | 0 | 0 |
| 0 | 0 | 0 | 0 |
| 0 | 0 | 0 | 0 |
| 0 | 0 | 0 | 0 |
| 0 | 0 | 0 | 0 |
| 0 | 0 | 0 | 0 |
| 0 | 0 | 0 | 0 |
| 1 | 0 | 0 | 0 |
| 0 | 0 | 0 | 0 |
| 1 | 0 | 0 | 0 |
| 0 | 0 | 0 | 0 |
| 0 | 0 | 0 | 0 |
| 0 | 0 | 0 | 0 |
| 1 | 0 | 0 | 0 |
| 1 | 0 | 0 | 0 |
| 0 | 0 | 0 | 0 |
| 0 | 0 | 0 | 0 |
| 1 | 0 | 0 | 0 |
| 0 | 0 | 0 | 0 |
| 0 | 0 | 0 | 0 |
| 1 | 0 | 0 | 0 |
| 0 | 0 | 0 | 0 |
| 1 | 0 | 0 | 0 |
| 1 | 0 | 0 | 0 |
| 0 | 0 | 0 | 0 |
| 0 | 0 | 0 | 0 |
| 1 | 1 | 0 | 0 |
| 1 | 0 | 0 | 0 |
| 1 | 1 | 0 | 0 |
| 0 | 0 | 0 | 0 |
| 0 | 0 | 0 | 0 |
| 0 | 0 | 0 | 0 |
| 0 | 0 | 0 | 0 |
| 1 | 1 | 0 | 0 |
| 0 | 0 | 0 | 0 |
| 0 | 0 | 0 | 0 |
| 0 | 0 | 0 | 0 |

| | | | |
|---|---|---|---|
| 0 | 1 | 0 | 0 |
| 1 | 0 | 0 | 0 |
| 0 | 0 | 0 | 0 |
| 0 | 0 | 0 | 0 |
| 0 | 0 | 0 | 0 |
| 1 | 1 | 0 | 0 |
| 0 | 0 | 0 | 0 |
| 0 | 0 | 0 | 0 |
| 0 | 0 | 0 | 0 |
| 1 | 0 | 0 | 0 |
| 1 | 0 | 0 | 0 |
| 0 | 0 | 0 | 0 |
| 0 | 0 | 0 | 0 |
| 0 | 0 | 0 | 0 |
| 0 | 0 | 0 | 0 |
| 0 | 0 | 0 | 0 |
| 0 | 0 | 0 | 0 |
| 0 | 0 | 0 | 0 |
| 0 | 0 | 0 | 0 |
| 0 | 0 | 0 | 0 |
| 0 | 0 | 0 | 0 |
| 0 | 0 | 0 | 0 |
| 0 | 0 | 0 | 0 |
| 1 | 1 | 0 | 0 |
| 0 | 0 | 0 | 0 |
| 0 | 0 | 0 | 0 |
| 1 | 1 | 1 | 0 |
| 0 | 0 | 0 | 0 |
| 0 | 0 | 0 | 0 |
| 0 | 0 | 0 | 0 |
| 1 | 1 | 1 | 0 |
| 1 | 0 | 0 | 0 |
| 1 | 0 | 0 | 0 |
| 1 | 0 | 0 | 0 |
| 1 | 0 | 0 | 0 |
| 1 | 1 | 1 | 0 |
| 1 | 1 | 1 | 0 |
| 0 | 0 | 0 | 0 |
| 0 | 0 | 0 | 0 |
| 1 | 1 | 1 | 0 |
| 0 | 0 | 0 | 0 |
| 0 | 0 | 0 | 0 |
| 0 | 0 | 0 | 0 |
| 0 | 0 | 0 | 0 |
| 0 | 0 | 0 | 0 |
| 0 | 0 | 0 | 0 |
| 0 | 0 | 0 | 0 |
| 0 | 0 | 0 | 0 |
| 1 | 0 | 0 | 0 |
| 1 | 0 | 0 | 0 |
| 1 | 0 | 0 | 0 |
| 0 | 0 | 0 | 0 |
| 0 | 0 | 0 | 0 |
| 0 | 0 | 0 | 0 |

| | | | |
|---|---|---|---|
| 0 | 0 | 0 | 0 |
| 1 | 0 | 0 | 0 |
| 0 | 0 | 0 | 0 |
| 0 | 0 | 0 | 0 |
| 0 | 0 | 0 | 0 |
| 0 | 0 | 0 | 0 |
| 0 | 0 | 0 | 0 |
| 0 | 0 | 0 | 0 |
| 0 | 0 | 0 | 0 |
| 0 | 0 | 0 | 0 |
| 1 | 1 | 0 | 0 |
| 1 | 1 | 0 | 0 |
| 0 | 0 | 0 | 0 |
| 0 | 0 | 0 | 0 |
| 0 | 0 | 0 | 0 |
| 1 | 0 | 0 | 0 |
| 1 | 0 | 0 | 0 |
| 1 | 1 | 0 | 1 |
| 0 | 0 | 0 | 0 |
| 0 | 0 | 0 | 0 |
| 0 | 0 | 0 | 0 |
| 0 | 0 | 0 | 0 |
| 1 | 0 | 0 | 1 |
| 0 | 0 | 0 | 0 |
| 0 | 0 | 0 | 0 |
| 1 | 0 | 0 | 0 |
| 0 | 0 | 0 | 0 |
| 0 | 0 | 0 | 0 |
| 0 | 0 | 0 | 0 |
| 0 | 0 | 0 | 0 |
| 0 | 0 | 0 | 0 |
| 0 | 0 | 0 | 0 |
| 0 | 0 | 0 | 0 |
| 0 | 0 | 0 | 0 |
| 0 | 0 | 0 | 0 |
| 1 | 0 | 0 | 0 |
| 1 | 0 | 0 | 0 |
| 1 | 1 | 0 | 0 |
| 1 | 1 | 0 | 0 |
| 0 | 0 | 0 | 0 |
| 0 | 0 | 0 | 0 |
| 1 | 0 | 0 | 0 |
| 0 | 0 | 0 | 0 |
| 1 | 0 | 0 | 0 |
| 1 | 1 | 0 | 0 |
| 0 | 0 | 0 | 0 |
| 0 | 0 | 0 | 0 |
| 0 | 0 | 0 | 0 |
| 0 | 0 | 0 | 0 |
| 0 | 0 | 0 | 0 |
| 0 | 0 | 0 | 0 |
| 1 | 0 | 0 | 0 |
| 0 | 0 | 0 | 0 |
| 0 | 0 | 0 | 0 |

| | | | |
|---|---|---|---|
| 0 | 0 | 0 | 0 |
| 1 | 1 | 0 | 0 |
| 0 | 0 | 0 | 0 |
| 1 | 0 | 0 | 0 |
| 1 | 0 | 0 | 0 |
| 0 | 0 | 0 | 0 |
| 0 | 0 | 0 | 0 |
| 0 | 0 | 0 | 0 |
| 0 | 0 | 0 | 0 |
| 0 | 0 | 0 | 0 |
| 0 | 0 | 0 | 0 |
| 1 | 1 | 0 | 0 |
| 1 | 1 | 0 | 1 |
| 0 | 0 | 0 | 0 |
| 0 | 1 | 0 | 0 |
| 0 | 0 | 0 | 0 |
| 0 | 0 | 0 | 0 |
| 0 | 0 | 0 | 0 |
| 0 | 0 | 0 | 0 |
| 0 | 0 | 0 | 0 |
| 0 | 0 | 0 | 0 |
| 1 | 0 | 0 | 0 |
| 0 | 0 | 0 | 0 |
| 0 | 0 | 0 | 0 |
| 0 | 0 | 0 | 0 |
| 0 | 0 | 0 | 0 |
| 0 | 0 | 0 | 0 |
| 1 | 0 | 0 | 0 |
| 0 | 0 | 0 | 0 |
| 0 | 0 | 0 | 0 |
| 0 | 0 | 0 | 0 |
| 0 | 0 | 0 | 0 |
| 0 | 0 | 0 | 0 |
| 0 | 0 | 0 | 0 |
| 0 | 0 | 0 | 0 |
| 0 | 0 | 0 | 0 |
| 0 | 0 | 0 | 0 |
| 0 | 0 | 0 | 0 |
| 0 | 0 | 0 | 0 |
| 0 | 0 | 0 | 0 |
| 0 | 0 | 0 | 0 |
| 0 | 0 | 0 | 0 |
| 0 | 0 | 0 | 0 |
| 0 | 0 | 0 | 0 |
| 0 | 0 | 0 | 0 |
| 0 | 0 | 0 | 0 |
| 0 | 0 | 0 | 0 |
| 0 | 0 | 0 | 0 |
| 0 | 0 | 0 | 0 |
| 0 | 0 | 0 | 0 |
| 0 | 0 | 0 | 0 |
| 0 | 0 | 0 | 0 |
| 0 | 0 | 0 | 0 |
| 0 | 0 | 0 | 0 |

| | | | |
|---|---|---|---|
| 0 | 0 | 0 | 0 |
| 0 | 0 | 0 | 0 |
| 0 | 0 | 0 | 0 |
| 0 | 0 | 0 | 0 |
| 0 | 0 | 0 | 0 |
| 0 | 0 | 0 | 0 |
| 0 | 0 | 0 | 0 |
| 0 | 0 | 0 | 0 |
| 0 | 0 | 0 | 0 |
| 0 | 0 | 0 | 0 |
| 0 | 0 | 0 | 0 |
| 0 | 0 | 0 | 0 |
| 1 | 0 | 0 | 0 |
| 1 | 1 | 0 | 0 |
| 0 | 0 | 0 | 0 |
| 1 | 0 | 0 | 0 |
| 0 | 0 | 0 | 0 |
| 1 | 0 | 0 | 0 |
| 0 | 0 | 0 | 0 |
| 0 | 0 | 0 | 0 |
| 1 | 0 | 0 | 0 |
| 0 | 0 | 0 | 0 |
| 0 | 0 | 0 | 0 |
| 1 | 0 | 0 | 0 |
| 1 | 0 | 0 | 0 |
| 0 | 0 | 0 | 0 |
| 0 | 0 | 0 | 0 |
| 1 | 0 | 0 | 0 |
| 1 | 0 | 0 | 0 |
| 1 | 0 | 0 | 0 |
| 1 | 0 | 0 | 0 |
| 1 | 0 | 0 | 0 |
| 0 | 0 | 0 | 0 |
| 0 | 0 | 0 | 0 |
| 0 | 0 | 0 | 0 |
| 0 | 0 | 0 | 0 |
| 0 | 0 | 0 | 0 |
| 0 | 0 | 0 | 0 |
| 0 | 0 | 0 | 0 |
| 0 | 0 | 0 | 0 |
| 0 | 0 | 0 | 0 |
| 0 | 0 | 0 | 0 |
| 0 | 0 | 0 | 0 |
| 0 | 0 | 0 | 0 |
| 1 | 1 | 0 | 0 |
| 0 | 0 | 0 | 0 |
| 0 | 0 | 0 | 0 |
| 0 | 0 | 0 | 0 |
| 0 | 0 | 0 | 0 |
| 1 | 1 | 0 | 0 |
| 1 | 0 | 0 | 0 |
| 0 | 0 | 0 | 0 |
| 0 | 0 | 0 | 0 |
| 0 | 0 | 0 | 1 |

| | | | |
|---|---|---|---|
| 0 | 0 | 0 | 0 |
| 0 | 0 | 0 | 0 |
| 0 | 0 | 0 | 0 |
| 0 | 0 | 0 | 0 |
| 1 | 0 | 0 | 1 |
| 0 | 0 | 0 | 0 |
| 1 | 0 | 0 | 0 |
| 0 | 0 | 0 | 0 |
| 0 | 0 | 0 | 0 |
| 0 | 0 | 0 | 0 |
| 0 | 0 | 0 | 0 |
| 0 | 0 | 0 | 0 |
| 0 | 0 | 0 | 0 |
| 1 | 0 | 0 | 0 |
| 0 | 0 | 0 | 0 |
| 0 | 0 | 0 | 0 |
| 1 | 1 | 0 | 0 |
| 0 | 0 | 0 | 0 |
| 1 | 0 | 0 | 0 |
| 1 | 0 | 0 | 0 |
| 0 | 0 | 0 | 0 |
| 0 | 0 | 0 | 0 |
| 0 | 0 | 0 | 0 |
| 1 | 0 | 0 | 0 |
| 1 | 0 | 0 | 0 |
| 1 | 0 | 0 | 0 |
| 1 | 0 | 0 | 0 |
| 1 | 0 | 0 | 0 |
| 1 | 0 | 0 | 0 |
| 1 | 0 | 0 | 0 |
| 1 | 0 | 0 | 0 |
| 1 | 0 | 0 | 0 |
| 1 | 0 | 0 | 0 |
| 1 | 0 | 0 | 0 |
| 0 | 0 | 0 | 0 |
| 0 | 0 | 0 | 0 |
| 0 | 0 | 0 | 0 |
| 1 | 0 | 0 | 0 |
| 1 | 0 | 0 | 0 |
| 0 | 0 | 0 | 0 |
| 0 | 0 | 0 | 0 |
| 1 | 0 | 0 | 0 |
| 0 | 0 | 0 | 0 |
| 1 | 0 | 0 | 0 |
| 1 | 0 | 0 | 0 |
| 1 | 0 | 0 | 0 |
| 1 | 0 | 0 | 0 |
| 1 | 0 | 0 | 0 |
| 0 | 0 | 0 | 0 |
| 0 | 0 | 0 | 0 |
| 1 | 0 | 0 | 0 |
| 0 | 0 | 0 | 0 |
| 1 | 0 | 0 | 0 |
| 1 | 0 | 0 | 0 |

| | | | |
|---|---|---|---|
| 1 | 0 | 0 | 0 |
| 1 | 0 | 0 | 0 |
| 1 | 0 | 0 | 0 |
| 1 | 0 | 0 | 0 |
| 1 | 0 | 0 | 0 |
| 1 | 0 | 0 | 0 |
| 0 | 0 | 0 | 0 |
| 1 | 0 | 0 | 0 |
| 1 | 0 | 0 | 0 |
| 1 | 0 | 0 | 0 |
| 1 | 0 | 0 | 0 |
| 1 | 0 | 0 | 0 |
| 1 | 0 | 0 | 0 |
| 1 | 0 | 0 | 0 |
| 0 | 0 | 0 | 0 |
| 0 | 0 | 0 | 0 |
| 1 | 1 | 0 | 0 |
| 1 | 1 | 0 | 0 |
| 0 | 0 | 0 | 0 |
| 0 | 0 | 0 | 0 |
| 0 | 0 | 0 | 0 |
| 1 | 1 | 0 | 1 |
| 0 | 0 | 0 | 0 |
| 0 | 0 | 0 | 0 |
| 0 | 0 | 0 | 0 |
| 0 | 0 | 0 | 0 |
| 0 | 0 | 0 | 0 |
| 0 | 0 | 0 | 0 |
| 0 | 0 | 0 | 0 |
| 1 | 0 | 1 | 0 |
| 0 | 0 | 0 | 0 |
| 0 | 0 | 0 | 0 |
| 0 | 0 | 0 | 0 |
| 0 | 0 | 0 | 0 |
| 0 | 0 | 0 | 0 |
| 0 | 0 | 0 | 0 |
| 0 | 0 | 0 | 0 |
| 0 | 0 | 0 | 0 |
| 0 | 0 | 0 | 0 |
| 0 | 0 | 0 | 0 |
| 1 | 1 | 0 | 0 |
| 1 | 1 | 0 | 0 |
| 1 | 0 | 0 | 0 |
| 1 | 1 | 0 | 0 |
| 1 | 0 | 0 | 0 |
| 1 | 1 | 0 | 0 |
| 1 | 0 | 0 | 0 |
| 1 | 0 | 0 | 0 |
| 0 | 0 | 0 | 0 |
| 0 | 0 | 0 | 0 |
| 1 | 0 | 0 | 0 |
| 1 | 0 | 0 | 0 |
| 1 | 0 | 0 | 0 |
| 0 | 0 | 0 | 0 |

| | | | |
|---|---|---|---|
| 0 | 0 | 0 | 0 |
| 0 | 0 | 0 | 0 |
| 0 | 0 | 0 | 0 |
| 1 | 1 | 1 | 0 |
| 0 | 0 | 0 | 0 |
| 0 | 0 | 0 | 0 |
| 0 | 0 | 0 | 0 |
| 0 | 0 | 0 | 0 |
| 0 | 1 | 0 | 0 |
| 0 | 1 | 0 | 0 |
| 0 | 1 | 0 | 0 |
| 0 | 0 | 0 | 0 |
| 0 | 0 | 0 | 0 |
| 0 | 0 | 0 | 0 |
| 1 | 0 | 0 | 0 |
| 1 | 0 | 0 | 0 |
| 0 | 0 | 0 | 0 |
| 0 | 0 | 0 | 0 |
| 1 | 0 | 0 | 0 |
| 0 | 0 | 0 | 0 |
| 1 | 0 | 0 | 0 |
| 1 | 0 | 0 | 0 |
| 0 | 0 | 0 | 0 |
| 0 | 0 | 0 | 0 |
| 0 | 0 | 0 | 0 |
| 0 | 0 | 0 | 0 |
| 0 | 0 | 0 | 0 |
| 0 | 0 | 1 | 0 |
| 0 | 0 | 0 | 0 |
| 0 | 1 | 0 | 0 |
| 0 | 0 | 0 | 0 |
| 0 | 0 | 0 | 0 |
| 0 | 0 | 0 | 0 |
| 0 | 0 | 0 | 0 |
| 0 | 0 | 0 | 0 |
| 0 | 0 | 0 | 0 |
| 1 | 0 | 0 | 0 |
| 1 | 0 | 0 | 0 |
| 1 | 0 | 0 | 0 |
| 0 | 0 | 0 | 0 |
| 0 | 0 | 0 | 0 |
| 0 | 0 | 0 | 0 |
| 0 | 0 | 0 | 0 |
| 0 | 0 | 0 | 0 |
| 0 | 0 | 0 | 0 |
| 0 | 0 | 0 | 0 |
| 0 | 0 | 0 | 0 |
| 0 | 0 | 0 | 1 |
| 0 | 0 | 0 | 0 |
| 0 | 0 | 0 | 0 |
| 0 | 0 | 0 | 0 |
| 0 | 0 | 0 | 0 |
| 1 | 1 | 0 | 0 |
| 0 | 0 | 0 | 0 |

| | | | |
|---|---|---|---|
| 0 | 0 | 0 | 0 |
| 0 | 0 | 0 | 0 |
| 0 | 0 | 0 | 0 |
| 0 | 0 | 0 | 0 |
| 1 | 0 | 0 | 0 |
| 0 | 0 | 0 | 0 |
| 0 | 0 | 0 | 0 |
| 0 | 0 | 0 | 0 |
| 0 | 0 | 0 | 0 |
| 0 | 0 | 0 | 0 |
| 0 | 0 | 0 | 0 |
| 0 | 0 | 0 | 0 |
| 0 | 0 | 0 | 0 |
| 0 | 0 | 0 | 0 |
| 0 | 0 | 0 | 0 |
| 0 | 0 | 0 | 0 |
| 0 | 0 | 0 | 0 |
| 0 | 0 | 0 | 0 |
| 0 | 0 | 0 | 0 |
| 0 | 0 | 0 | 0 |
| 0 | 0 | 0 | 0 |
| 0 | 0 | 0 | 0 |
| 0 | 0 | 0 | 0 |
| 0 | 0 | 0 | 0 |
| 0 | 0 | 0 | 0 |
| 0 | 0 | 0 | 0 |
| 0 | 0 | 0 | 0 |
| 0 | 0 | 0 | 0 |
| 0 | 0 | 0 | 0 |
| 0 | 0 | 0 | 0 |
| 0 | 0 | 0 | 0 |
| 0 | 0 | 0 | 0 |
| 0 | 0 | 0 | 0 |
| 0 | 0 | 0 | 0 |
| 0 | 0 | 0 | 0 |
| 0 | 0 | 0 | 0 |
| 0 | 0 | 0 | 0 |
| 0 | 0 | 0 | 0 |
| 0 | 0 | 0 | 0 |
| 0 | 0 | 0 | 0 |
| 1 | 1 | 0 | 0 |
| 0 | 0 | 0 | 0 |
| 0 | 0 | 0 | 0 |
| 0 | 0 | 0 | 0 |
| 0 | 0 | 0 | 0 |
| 1 | 0 | 0 | 0 |
| 1 | 0 | 0 | 0 |
| 0 | 0 | 0 | 0 |
| 1 | 0 | 0 | 0 |
| 0 | 0 | 0 | 0 |
| 0 | 0 | 0 | 0 |
| 1 | 0 | 0 | 0 |
| 1 | 0 | 0 | 0 |
| 0 | 0 | 0 | 0 |

| | | | |
|---|---|---|---|
| 0 | 0 | 0 | 0 |
| 0 | 0 | 0 | 0 |
| 0 | 0 | 0 | 0 |
| 0 | 0 | 0 | 0 |
| 1 | 0 | 0 | 0 |
| 0 | 0 | 0 | 0 |
| 0 | 0 | 0 | 0 |
| 0 | 0 | 0 | 0 |
| 0 | 0 | 0 | 0 |
| 0 | 0 | 0 | 0 |
| 0 | 0 | 0 | 0 |
| 0 | 0 | 0 | 0 |
| 0 | 0 | 0 | 0 |
| 0 | 0 | 0 | 0 |
| 0 | 0 | 0 | 0 |
| 1 | 1 | 0 | 1 |
| 0 | 0 | 0 | 0 |
| 0 | 0 | 0 | 0 |
| 0 | 0 | 0 | 0 |
| 0 | 0 | 0 | 0 |
| 0 | 0 | 0 | 0 |
| 0 | 0 | 0 | 0 |
| 0 | 0 | 0 | 0 |
| 0 | 0 | 0 | 0 |
| 0 | 0 | 0 | 0 |
| 0 | 0 | 0 | 0 |
| 0 | 0 | 0 | 0 |
| 1 | 0 | 0 | 0 |
| 1 | 0 | 0 | 0 |
| 0 | 0 | 0 | 0 |
| 1 | 1 | 0 | 0 |
| 0 | 0 | 0 | 0 |
| 0 | 0 | 0 | 0 |
| 0 | 0 | 0 | 0 |
| 1 | 1 | 0 | 0 |
| 1 | 1 | 0 | 0 |
| 0 | 0 | 0 | 0 |
| 1 | 0 | 0 | 0 |
| 0 | 0 | 0 | 0 |
| 1 | 1 | 0 | 0 |
| 0 | 1 | 0 | 0 |
| 0 | 0 | 0 | 0 |
| 0 | 0 | 0 | 0 |
| 0 | 0 | 0 | 0 |
| 1 | 1 | 0 | 0 |
| 1 | 1 | 0 | 0 |
| 1 | 1 | 0 | 0 |
| 0 | 0 | 0 | 0 |
| 0 | 0 | 0 | 0 |
| 0 | 0 | 0 | 0 |
| 0 | 0 | 0 | 0 |
| 0 | 0 | 0 | 0 |
| 0 | 0 | 0 | 0 |
| 0 | 0 | 0 | 0 |

| | | | |
|---|---|---|---|
| 0 | 0 | 0 | 0 |
| 0 | 0 | 0 | 0 |
| 1 | 0 | 0 | 0 |
| 1 | 0 | 0 | 0 |
| 0 | 0 | 0 | 0 |
| 1 | 0 | 0 | 0 |
| 0 | 0 | 0 | 0 |
| 0 | 0 | 0 | 0 |
| 0 | 0 | 0 | 0 |
| 0 | 1 | 0 | 0 |
| 0 | 0 | 0 | 0 |
| 0 | 0 | 0 | 0 |
| 0 | 0 | 0 | 0 |
| 0 | 0 | 0 | 0 |
| 0 | 0 | 0 | 0 |
| 0 | 0 | 0 | 0 |
| 0 | 0 | 0 | 0 |
| 1 | 0 | 0 | 0 |
| 0 | 0 | 0 | 0 |
| 1 | 0 | 0 | 0 |
| 0 | 0 | 0 | 0 |
| 1 | 0 | 0 | 0 |
| 0 | 0 | 0 | 0 |
| 1 | 0 | 0 | 0 |
| 0 | 0 | 0 | 0 |
| 0 | 0 | 0 | 0 |
| 0 | 0 | 0 | 0 |
| 0 | 0 | 0 | 0 |
| 0 | 0 | 0 | 0 |
| 0 | 0 | 0 | 0 |
| 0 | 0 | 0 | 0 |
| 1 | 0 | 0 | 0 |
| 0 | 0 | 0 | 0 |
| 0 | 0 | 0 | 0 |
| 0 | 0 | 0 | 0 |
| 0 | 0 | 0 | 0 |
| 0 | 0 | 0 | 0 |
| 0 | 0 | 0 | 0 |
| 1 | 1 | 0 | 0 |
| 1 | 1 | 0 | 0 |
| 1 | 1 | 0 | 0 |
| 1 | 1 | 0 | 0 |
| 1 | 1 | 0 | 0 |
| 1 | 0 | 0 | 0 |
| 1 | 0 | 0 | 0 |
| 1 | 0 | 0 | 0 |
| 0 | 0 | 0 | 0 |
| 0 | 0 | 0 | 0 |
| 0 | 0 | 0 | 0 |
| 0 | 0 | 0 | 0 |
| 0 | 0 | 0 | 0 |
| 0 | 0 | 0 | 0 |
| 0 | 0 | 0 | 0 |
| 0 | 0 | 0 | 0 |

| | | | |
|---|---|---|---|
| 0 | 0 | 0 | 0 |
| 0 | 0 | 0 | 0 |
| 0 | 0 | 0 | 0 |
| 0 | 0 | 0 | 0 |
| 0 | 0 | 0 | 0 |
| 1 | 0 | 0 | 0 |
| 1 | 0 | 0 | 0 |
| 0 | 0 | 0 | 0 |
| 0 | 0 | 0 | 0 |
| 0 | 0 | 0 | 0 |
| 0 | 0 | 0 | 0 |
| 1 | 0 | 0 | 0 |
| 0 | 0 | 1 | 0 |
| 0 | 0 | 0 | 0 |
| 0 | 0 | 0 | 0 |
| 0 | 0 | 0 | 0 |
| 1 | 0 | 0 | 0 |
| 0 | 0 | 0 | 0 |
| 1 | 0 | 0 | 0 |
| 1 | 0 | 0 | 0 |
| 0 | 0 | 0 | 0 |
| 1 | 0 | 0 | 0 |
| 1 | 1 | 0 | 0 |
| 0 | 0 | 0 | 0 |
| 1 | 1 | 0 | 0 |
| 1 | 1 | 0 | 0 |
| 1 | 1 | 0 | 0 |
| 0 | 0 | 0 | 0 |
| 1 | 0 | 0 | 0 |
| 0 | 0 | 0 | 0 |
| 0 | 0 | 0 | 0 |
| 0 | 0 | 0 | 0 |
| 0 | 0 | 0 | 0 |
| 0 | 0 | 0 | 0 |
| 0 | 0 | 0 | 0 |
| 0 | 0 | 0 | 0 |
| 0 | 0 | 0 | 0 |
| 0 | 0 | 0 | 0 |
| 0 | 0 | 0 | 0 |
| 0 | 0 | 0 | 0 |
| 0 | 0 | 0 | 0 |
| 0 | 0 | 0 | 0 |
| 0 | 0 | 0 | 0 |
| 0 | 0 | 0 | 0 |
| 0 | 0 | 0 | 0 |
| 0 | 0 | 0 | 0 |
| 0 | 0 | 0 | 0 |
| 0 | 0 | 0 | 0 |
| 0 | 0 | 0 | 0 |
| 0 | 0 | 0 | 0 |
| 0 | 0 | 0 | 0 |
| 0 | 0 | 0 | 0 |
| 0 | 0 | 0 | 0 |
| 0 | 0 | 0 | 0 |

| | | | |
|---|---|---|---|
| 0 | 0 | 0 | 0 |
| 0 | 0 | 0 | 0 |
| 0 | 0 | 0 | 0 |
| 0 | 0 | 0 | 0 |
| 0 | 0 | 0 | 0 |
| 0 | 0 | 0 | 0 |
| 0 | 0 | 0 | 0 |
| 0 | 0 | 0 | 0 |
| 0 | 0 | 0 | 0 |
| 0 | 0 | 0 | 0 |
| 0 | 1 | 0 | 0 |
| 0 | 1 | 0 | 0 |
| 0 | 1 | 0 | 0 |
| 0 | 1 | 0 | 0 |
| 0 | 0 | 0 | 0 |
| 0 | 1 | 0 | 0 |
| 0 | 1 | 0 | 0 |
| 1 | 1 | 0 | 0 |
| 0 | 1 | 0 | 0 |
| 1 | 0 | 0 | 0 |
| 1 | 1 | 0 | 0 |
| 1 | 1 | 0 | 0 |
| 0 | 0 | 1 | 1 |
| 0 | 0 | 0 | 0 |
| 1 | 0 | 0 | 0 |
| 0 | 0 | 0 | 0 |
| 0 | 0 | 0 | 0 |
| 0 | 0 | 0 | 0 |
| 0 | 0 | 0 | 0 |
| 0 | 0 | 0 | 0 |
| 0 | 0 | 0 | 0 |
| 0 | 0 | 0 | 0 |
| 0 | 0 | 0 | 0 |
| 0 | 0 | 0 | 0 |
| 0 | 0 | 0 | 0 |
| 0 | 0 | 0 | 0 |
| 0 | 0 | 0 | 0 |
| 0 | 0 | 0 | 0 |
| 0 | 0 | 0 | 0 |
| 0 | 0 | 0 | 0 |
| 1 | 0 | 0 | 0 |
| 0 | 0 | 0 | 0 |
| 1 | 0 | 0 | 0 |
| 1 | 0 | 0 | 0 |
| 0 | 0 | 0 | 0 |
| 1 | 0 | 0 | 0 |
| 0 | 0 | 0 | 0 |
| 0 | 0 | 0 | 0 |
| 0 | 0 | 0 | 0 |
| 0 | 0 | 0 | 0 |
| 0 | 0 | 0 | 0 |
| 0 | 0 | 0 | 0 |
| 1 | 1 | 0 | 0 |
| 1 | 1 | 0 | 0 |

| | | | |
|---|---|---|---|
| 1 | 1 | 0 | 0 |
| 0 | 0 | 0 | 0 |
| 0 | 0 | 0 | 0 |
| 0 | 0 | 0 | 0 |
| 0 | 0 | 0 | 0 |
| 0 | 0 | 0 | 0 |
| 0 | 1 | 0 | 0 |
| 0 | 0 | 0 | 0 |
| 0 | 0 | 0 | 0 |
| 0 | 1 | 0 | 0 |
| 1 | 0 | 0 | 0 |
| 0 | 0 | 0 | 0 |
| 0 | 0 | 1 | 0 |
| 0 | 0 | 1 | 0 |
| 0 | 0 | 0 | 0 |
| 0 | 0 | 0 | 0 |
| 0 | 0 | 0 | 0 |
| 0 | 1 | 0 | 0 |
| 0 | 0 | 0 | 0 |
| 0 | 0 | 0 | 0 |
| 1 | 0 | 0 | 0 |
| 0 | 0 | 0 | 1 |
| 0 | 0 | 0 | 0 |
| 0 | 0 | 0 | 0 |
| 0 | 0 | 0 | 0 |
| 0 | 0 | 0 | 0 |
| 0 | 0 | 0 | 0 |
| 0 | 0 | 0 | 0 |
| 0 | 0 | 0 | 0 |
| 0 | 0 | 0 | 0 |
| 0 | 0 | 0 | 0 |
| 0 | 0 | 0 | 0 |
| 0 | 0 | 0 | 0 |
| 0 | 0 | 0 | 0 |
| 0 | 0 | 0 | 0 |
| 0 | 0 | 0 | 0 |
| 1 | 0 | 0 | 0 |
| 0 | 0 | 0 | 0 |
| 1 | 0 | 0 | 0 |
| 0 | 0 | 0 | 0 |
| 0 | 0 | 0 | 0 |
| 0 | 0 | 0 | 0 |
| 0 | 0 | 0 | 0 |
| 0 | 0 | 0 | 0 |
| 0 | 0 | 0 | 0 |
| 0 | 0 | 0 | 0 |
| 0 | 0 | 0 | 0 |
| 0 | 0 | 0 | 0 |
| 0 | 0 | 0 | 0 |
| 1 | 0 | 0 | 0 |
| 0 | 0 | 0 | 0 |
| 0 | 0 | 0 | 0 |
| 0 | 0 | 0 | 0 |
| 0 | 0 | 0 | 0 |

| | | | |
|---|---|---|---|
| 0 | 0 | 0 | 0 |
| 0 | 0 | 0 | 0 |
| 0 | 0 | 0 | 0 |
| 0 | 0 | 0 | 0 |
| 0 | 0 | 0 | 0 |
| 0 | 0 | 0 | 0 |
| 0 | 0 | 0 | 0 |
| 0 | 0 | 0 | 0 |
| 0 | 0 | 0 | 0 |
| 0 | 0 | 0 | 0 |
| 0 | 0 | 0 | 0 |
| 0 | 0 | 0 | 0 |
| 0 | 0 | 1 | 0 |
| 1 | 0 | 0 | 0 |
| 1 | 0 | 0 | 0 |
| 0 | 0 | 0 | 0 |
| 0 | 0 | 0 | 0 |
| 0 | 0 | 0 | 0 |
| 0 | 0 | 0 | 0 |
| 0 | 0 | 0 | 0 |
| 0 | 0 | 0 | 0 |
| 0 | 0 | 0 | 0 |
| 0 | 0 | 0 | 0 |
| 1 | 1 | 0 | 1 |
| 0 | 0 | 0 | 0 |
| 1 | 1 | 0 | 0 |
| 1 | 1 | 0 | 1 |
| 0 | 0 | 0 | 0 |
| 0 | 0 | 0 | 0 |
| 0 | 0 | 0 | 0 |
| 0 | 0 | 0 | 0 |
| 0 | 0 | 0 | 0 |
| 0 | 0 | 0 | 0 |
| 0 | 0 | 0 | 0 |
| 0 | 0 | 0 | 0 |
| 1 | 0 | 0 | 0 |
| 0 | 0 | 0 | 0 |
| 0 | 0 | 0 | 0 |
| 0 | 0 | 0 | 0 |
| 0 | 0 | 0 | 0 |
| 0 | 0 | 0 | 0 |
| 0 | 0 | 0 | 0 |
| 1 | 0 | 0 | 0 |
| 1 | 0 | 0 | 0 |
| 1 | 0 | 0 | 0 |
| 1 | 0 | 0 | 0 |
| 1 | 0 | 0 | 0 |
| 0 | 0 | 0 | 0 |
| 0 | 0 | 0 | 0 |
| 1 | 0 | 0 | 0 |
| 1 | 0 | 0 | 0 |
| 0 | 0 | 0 | 0 |
| 1 | 0 | 0 | 0 |
| 1 | 0 | 0 | 0 |

| | | | |
|---|---|---|---|
| 0 | 0 | 0 | 0 |
| 1 | 0 | 0 | 0 |
| 0 | 0 | 0 | 0 |
| 0 | 0 | 0 | 0 |
| 0 | 0 | 0 | 0 |
| 0 | 0 | 0 | 0 |
| 1 | 0 | 0 | 0 |
| 0 | 0 | 0 | 0 |
| 0 | 0 | 0 | 0 |
| 0 | 0 | 0 | 0 |
| 0 | 0 | 0 | 0 |
| 1 | 1 | 0 | 0 |
| 0 | 0 | 0 | 0 |
| 0 | 0 | 0 | 0 |
| 0 | 0 | 0 | 0 |
| 0 | 0 | 0 | 0 |
| 0 | 0 | 0 | 0 |
| 0 | 0 | 0 | 0 |
| 0 | 0 | 0 | 0 |
| 0 | 0 | 0 | 0 |
| 0 | 0 | 0 | 0 |
| 0 | 0 | 0 | 0 |
| 0 | 0 | 0 | 0 |
| 0 | 0 | 0 | 0 |
| 0 | 0 | 0 | 0 |
| 0 | 0 | 0 | 0 |
| 0 | 0 | 0 | 0 |
| 0 | 0 | 0 | 0 |
| 1 | 1 | 0 | 0 |
| 1 | 0 | 0 | 0 |
| 0 | 0 | 0 | 0 |
| 0 | 0 | 0 | 0 |
| 0 | 0 | 0 | 0 |
| 0 | 0 | 0 | 0 |
| 0 | 0 | 0 | 0 |
| 0 | 0 | 0 | 0 |
| 0 | 0 | 0 | 0 |
| 0 | 0 | 0 | 0 |
| 0 | 0 | 0 | 0 |
| 0 | 0 | 0 | 0 |
| 0 | 0 | 0 | 0 |
| 0 | 0 | 0 | 0 |
| 0 | 0 | 0 | 0 |
| 0 | 0 | 0 | 0 |
| 0 | 0 | 0 | 0 |
| 0 | 0 | 0 | 0 |
| 0 | 0 | 0 | 0 |
| 0 | 0 | 0 | 0 |
| 0 | 0 | 0 | 0 |
| 0 | 0 | 0 | 0 |
| 0 | 0 | 0 | 0 |
| 0 | 0 | 0 | 0 |
| 0 | 0 | 1 | 0 |
| 0 | 0 | 0 | 0 |

| | | | |
|---|---|---|---|
| 0 | 0 | 0 | 0 |
| 0 | 0 | 0 | 0 |
| 0 | 0 | 0 | 0 |
| 0 | 0 | 0 | 0 |
| 1 | 0 | 0 | 0 |
| 0 | 0 | 0 | 0 |
| 0 | 0 | 0 | 0 |
| 1 | 1 | 0 | 0 |
| 0 | 0 | 0 | 0 |
| 1 | 1 | 0 | 0 |
| 0 | 0 | 0 | 0 |
| 0 | 0 | 0 | 0 |
| 0 | 0 | 0 | 0 |
| 0 | 0 | 0 | 0 |
| 0 | 0 | 0 | 0 |
| 0 | 0 | 0 | 0 |
| 1 | 0 | 0 | 0 |
| 0 | 0 | 0 | 0 |
| 0 | 0 | 0 | 0 |
| 1 | 0 | 0 | 0 |
| 1 | 0 | 0 | 0 |
| 0 | 0 | 0 | 0 |
| 0 | 0 | 0 | 0 |
| 0 | 0 | 0 | 0 |
| 0 | 0 | 0 | 0 |
| 0 | 0 | 0 | 0 |
| 1 | 0 | 0 | 0 |
| 0 | 0 | 0 | 0 |
| 0 | 0 | 0 | 0 |
| 1 | 0 | 0 | 0 |
| 0 | 0 | 0 | 0 |
| 0 | 0 | 0 | 0 |
| 0 | 0 | 0 | 0 |
| 1 | 1 | 0 | 0 |
| 0 | 0 | 0 | 0 |
| 0 | 0 | 0 | 0 |
| 0 | 0 | 0 | 0 |
| 0 | 0 | 0 | 0 |
| 0 | 1 | 0 | 0 |
| 0 | 0 | 0 | 0 |
| 0 | 0 | 0 | 0 |
| 0 | 0 | 0 | 0 |
| 0 | 0 | 0 | 0 |
| 1 | 1 | 0 | 0 |
| 0 | 0 | 0 | 0 |
| 0 | 0 | 0 | 0 |
| 0 | 0 | 0 | 0 |
| 0 | 0 | 0 | 0 |
| 0 | 0 | 0 | 0 |
| 0 | 0 | 0 | 0 |
| 0 | 0 | 0 | 0 |
| 1 | 0 | 0 | 0 |
| 0 | 0 | 0 | 0 |
| 0 | 0 | 0 | 0 |

| | | | |
|---|---|---|---|
| 1 | 0 | 0 | 1 |
| 1 | 0 | 0 | 1 |
| 1 | 0 | 0 | 1 |
| 0 | 0 | 0 | 0 |
| 0 | 0 | 0 | 0 |
| 0 | 0 | 0 | 0 |
| 1 | 0 | 0 | 0 |
| 0 | 0 | 0 | 0 |
| 0 | 0 | 0 | 0 |
| 0 | 0 | 0 | 0 |
| 0 | 0 | 1 | 0 |
| 0 | 0 | 0 | 0 |
| 0 | 0 | 0 | 0 |
| 0 | 0 | 0 | 0 |
| 0 | 0 | 0 | 0 |
| 0 | 0 | 0 | 0 |
| 1 | 0 | 0 | 0 |
| 0 | 0 | 0 | 0 |
| 0 | 0 | 0 | 0 |
| 0 | 0 | 0 | 0 |
| 0 | 0 | 0 | 0 |
| 0 | 0 | 0 | 0 |
| 0 | 0 | 0 | 0 |
| 1 | 1 | 0 | 0 |
| 0 | 0 | 0 | 0 |
| 0 | 0 | 0 | 0 |
| 0 | 0 | 0 | 0 |
| 0 | 0 | 0 | 0 |
| 0 | 0 | 0 | 0 |
| 0 | 0 | 0 | 0 |
| 0 | 0 | 0 | 0 |
| 0 | 0 | 0 | 0 |
| 0 | 1 | 0 | 0 |
| 0 | 0 | 0 | 0 |
| 1 | 1 | 0 | 0 |
| 0 | 0 | 0 | 0 |
| 0 | 0 | 0 | 0 |
| 0 | 0 | 0 | 0 |
| 0 | 0 | 0 | 0 |
| 0 | 0 | 0 | 0 |
| 0 | 0 | 0 | 0 |
| 1 | 0 | 0 | 1 |
| 0 | 0 | 0 | 0 |
| 1 | 0 | 0 | 1 |
| 0 | 0 | 0 | 0 |
| 1 | 0 | 0 | 1 |
| 0 | 0 | 0 | 0 |
| 0 | 0 | 0 | 0 |
| 0 | 0 | 0 | 0 |
| 1 | 0 | 0 | 0 |
| 0 | 0 | 0 | 0 |
| 0 | 0 | 0 | 0 |
| 0 | 0 | 0 | 0 |
| 0 | 0 | 0 | 0 |

| | | | |
|---|---|---|---|
| 0 | 0 | 0 | 0 |
| 0 | 0 | 0 | 0 |
| 0 | 0 | 0 | 0 |
| 0 | 0 | 0 | 0 |
| 0 | 0 | 0 | 0 |
| 0 | 0 | 0 | 0 |
| 0 | 0 | 0 | 0 |
| 0 | 0 | 0 | 0 |
| 0 | 0 | 0 | 0 |
| 0 | 0 | 0 | 0 |
| 0 | 0 | 0 | 0 |
| 0 | 0 | 0 | 0 |
| 0 | 0 | 0 | 0 |
| 0 | 0 | 0 | 0 |
| 0 | 0 | 0 | 0 |
| 0 | 0 | 0 | 0 |
| 0 | 0 | 0 | 0 |
| 0 | 0 | 0 | 0 |
| 0 | 0 | 1 | 0 |
| 0 | 0 | 0 | 0 |
| 0 | 0 | 0 | 0 |
| 0 | 0 | 0 | 0 |
| 0 | 0 | 0 | 0 |
| 0 | 0 | 0 | 0 |
| 0 | 0 | 0 | 0 |
| 0 | 0 | 0 | 0 |
| 0 | 0 | 0 | 0 |
| 0 | 0 | 0 | 0 |
| 0 | 0 | 1 | 0 |
| 0 | 0 | 0 | 0 |
| 0 | 0 | 1 | 0 |
| 1 | 0 | 0 | 0 |
| 1 | 1 | 1 | 0 |
| 1 | 1 | 1 | 0 |
| 0 | 0 | 0 | 0 |
| 0 | 0 | 0 | 0 |
| 0 | 1 | 0 | 0 |
| 1 | 1 | 1 | 0 |
| 1 | 1 | 1 | 0 |
| 0 | 0 | 1 | 0 |
| 0 | 0 | 1 | 0 |
| 1 | 0 | 0 | 0 |
| 0 | 0 | 0 | 0 |
| 0 | 0 | 0 | 0 |
| 1 | 1 | 0 | 0 |
| 0 | 0 | 0 | 0 |
| 0 | 0 | 0 | 0 |
| 0 | 0 | 0 | 0 |
| 0 | 0 | 0 | 0 |
| 0 | 0 | 0 | 0 |
| 0 | 0 | 0 | 0 |
| 0 | 0 | 0 | 0 |
| 0 | 0 | 0 | 0 |
| 0 | 0 | 0 | 0 |

| | | | |
|---|---|---|---|
| 0 | 0 | 0 | 0 |
| 0 | 0 | 0 | 0 |
| 0 | 0 | 0 | 0 |
| 0 | 0 | 0 | 0 |
| 0 | 0 | 0 | 0 |
| 0 | 0 | 0 | 0 |
| 0 | 0 | 0 | 0 |
| 0 | 0 | 0 | 0 |
| 0 | 0 | 0 | 0 |
| 0 | 0 | 0 | 0 |
| 0 | 0 | 0 | 0 |
| 0 | 0 | 0 | 0 |
| 0 | 0 | 0 | 0 |
| 0 | 0 | 0 | 0 |
| 0 | 0 | 0 | 0 |
| 0 | 0 | 0 | 0 |
| 0 | 0 | 0 | 0 |
| 0 | 0 | 0 | 0 |
| 0 | 0 | 0 | 0 |
| 1 | 0 | 0 | 0 |
| 1 | 0 | 0 | 0 |
| 0 | 0 | 0 | 0 |
| 0 | 0 | 0 | 0 |
| 0 | 0 | 0 | 0 |
| 0 | 0 | 0 | 0 |
| 1 | 0 | 0 | 0 |
| 0 | 0 | 0 | 0 |
| 0 | 0 | 0 | 0 |
| 0 | 0 | 0 | 0 |
| 1 | 0 | 0 | 0 |
| 0 | 0 | 0 | 0 |
| 0 | 0 | 0 | 0 |
| 1 | 0 | 0 | 0 |
| 1 | 0 | 0 | 0 |
| 0 | 0 | 0 | 0 |
| 0 | 0 | 0 | 0 |
| 0 | 0 | 0 | 0 |
| 0 | 0 | 0 | 0 |
| 0 | 0 | 0 | 0 |
| 0 | 0 | 0 | 0 |
| 0 | 0 | 0 | 0 |
| 0 | 0 | 0 | 0 |
| 0 | 0 | 0 | 0 |
| 0 | 0 | 0 | 0 |
| 0 | 0 | 0 | 0 |
| 0 | 0 | 0 | 0 |
| 0 | 0 | 0 | 0 |
| 0 | 1 | 0 | 0 |
| 0 | 1 | 0 | 0 |
| 0 | 0 | 0 | 0 |
| 0 | 1 | 0 | 0 |
| 0 | 1 | 0 | 0 |
| 0 | 0 | 0 | 0 |
| 0 | 0 | 0 | 0 |

| | | | |
|---|---|---|---|
| 0 | 0 | 0 | 0 |
| 0 | 0 | 0 | 0 |
| 1 | 1 | 0 | 1 |
| 0 | 0 | 0 | 0 |
| 0 | 0 | 0 | 0 |
| 0 | 0 | 0 | 0 |
| 0 | 0 | 0 | 0 |
| 0 | 1 | 0 | 0 |
| 1 | 1 | 0 | 0 |
| 1 | 1 | 0 | 0 |
| 1 | 1 | 0 | 0 |
| 1 | 1 | 0 | 0 |
| 1 | 1 | 0 | 0 |
| 0 | 1 | 0 | 0 |
| 0 | 1 | 0 | 0 |
| 0 | 0 | 0 | 0 |
| 0 | 0 | 0 | 0 |
| 0 | 0 | 0 | 0 |
| 0 | 0 | 0 | 0 |
| 0 | 0 | 0 | 0 |
| 0 | 0 | 0 | 0 |
| 0 | 0 | 0 | 0 |
| 0 | 1 | 0 | 0 |
| 0 | 1 | 0 | 0 |
| 0 | 0 | 0 | 0 |
| 0 | 0 | 1 | 0 |
| 0 | 0 | 0 | 0 |
| 0 | 0 | 0 | 0 |
| 0 | 0 | 0 | 0 |
| 0 | 1 | 0 | 0 |
| 0 | 0 | 0 | 0 |
| 0 | 0 | 0 | 0 |
| 0 | 0 | 0 | 0 |
| 0 | 0 | 0 | 0 |
| 0 | 0 | 0 | 0 |
| 0 | 0 | 0 | 0 |
| 0 | 0 | 0 | 0 |
| 0 | 0 | 0 | 0 |
| 0 | 0 | 0 | 0 |
| 1 | 0 | 0 | 0 |
| 1 | 1 | 0 | 0 |
| 1 | 0 | 0 | 0 |
| 0 | 0 | 0 | 0 |
| 0 | 1 | 0 | 0 |
| 1 | 1 | 0 | 0 |
| 0 | 1 | 0 | 0 |
| 0 | 0 | 0 | 0 |
| 1 | 0 | 0 | 0 |
| 0 | 1 | 0 | 0 |
| 0 | 0 | 0 | 0 |
| 0 | 0 | 0 | 0 |
| 0 | 0 | 0 | 0 |
| 1 | 1 | 0 | 0 |
| 0 | 0 | 0 | 0 |

| | | | |
|---|---|---|---|
| 0 | 0 | 0 | 0 |
| 0 | 0 | 0 | 0 |
| 1 | 0 | 0 | 0 |
| 1 | 1 | 0 | 0 |
| 1 | 1 | 0 | 0 |
| 0 | 0 | 0 | 0 |
| 0 | 0 | 0 | 0 |
| 0 | 0 | 0 | 0 |
| 1 | 0 | 0 | 0 |
| 0 | 0 | 0 | 0 |
| 0 | 0 | 0 | 0 |
| 1 | 0 | 0 | 1 |
| 0 | 0 | 0 | 0 |
| 1 | 0 | 0 | 0 |
| 0 | 0 | 0 | 0 |
| 0 | 0 | 0 | 0 |
| 0 | 0 | 0 | 0 |
| 0 | 0 | 0 | 0 |
| 0 | 0 | 0 | 0 |
| 0 | 0 | 0 | 0 |
| 0 | 0 | 0 | 0 |
| 1 | 0 | 0 | 1 |
| 1 | 0 | 0 | 0 |
| 0 | 0 | 0 | 0 |
| 0 | 0 | 0 | 0 |
| 0 | 0 | 0 | 0 |
| 0 | 0 | 0 | 0 |
| 0 | 0 | 0 | 0 |
| 1 | 0 | 0 | 0 |
| 1 | 0 | 0 | 0 |
| 0 | 0 | 0 | 0 |
| 0 | 0 | 0 | 0 |
| 0 | 0 | 0 | 0 |
| 0 | 0 | 0 | 0 |
| 0 | 0 | 0 | 0 |
| 0 | 0 | 0 | 0 |
| 1 | 0 | 0 | 0 |
| 0 | 0 | 0 | 0 |
| 0 | 0 | 0 | 0 |
| 1 | 0 | 0 | 0 |
| 1 | 0 | 0 | 0 |
| 1 | 0 | 0 | 0 |
| 1 | 0 | 0 | 0 |
| 0 | 0 | 0 | 0 |
| 0 | 0 | 0 | 0 |
| 1 | 0 | 0 | 0 |
| 0 | 0 | 0 | 0 |
| 0 | 0 | 0 | 0 |
| 0 | 0 | 0 | 0 |
| 0 | 0 | 0 | 0 |
| 0 | 0 | 0 | 0 |
| 1 | 0 | 0 | 0 |
| 1 | 0 | 0 | 0 |
| 1 | 0 | 0 | 0 |

| | | | |
|---|---|---|---|
| 1 | 0 | 0 | 0 |
| 0 | 0 | 0 | 0 |
| 0 | 0 | 0 | 0 |
| 0 | 0 | 0 | 0 |
| 1 | 0 | 0 | 0 |
| 0 | 0 | 0 | 0 |
| 1 | 0 | 0 | 0 |
| 0 | 0 | 0 | 0 |
| 0 | 0 | 0 | 0 |
| 0 | 0 | 0 | 0 |
| 0 | 0 | 0 | 0 |
| 0 | 0 | 0 | 0 |
| 0 | 0 | 0 | 0 |
| 0 | 0 | 0 | 0 |
| 0 | 0 | 0 | 0 |
| 1 | 0 | 0 | 0 |
| 0 | 0 | 0 | 0 |
| 0 | 0 | 0 | 0 |
| 0 | 0 | 0 | 0 |
| 0 | 0 | 0 | 0 |
| 0 | 0 | 0 | 0 |
| 0 | 0 | 0 | 0 |
| 1 | 0 | 0 | 0 |
| 1 | 0 | 0 | 0 |
| 0 | 0 | 0 | 0 |
| 0 | 0 | 0 | 0 |
| 0 | 0 | 0 | 0 |
| 0 | 0 | 0 | 0 |
| 0 | 0 | 0 | 0 |
| 0 | 0 | 0 | 0 |
| 0 | 0 | 0 | 0 |
| 0 | 0 | 0 | 0 |
| 1 | 0 | 0 | 0 |
| 0 | 0 | 0 | 0 |
| 0 | 0 | 0 | 0 |
| 0 | 0 | 0 | 0 |
| 0 | 0 | 0 | 0 |
| 0 | 0 | 0 | 0 |
| 0 | 0 | 0 | 0 |
| 0 | 0 | 0 | 0 |
| 0 | 0 | 0 | 0 |
| 0 | 0 | 0 | 0 |
| 0 | 0 | 0 | 0 |
| 0 | 0 | 0 | 0 |
| 0 | 1 | 0 | 0 |
| 0 | 0 | 0 | 0 |
| 0 | 0 | 0 | 0 |
| 0 | 0 | 0 | 0 |
| 0 | 0 | 0 | 0 |
| 0 | 0 | 0 | 0 |
| 0 | 0 | 0 | 0 |
| 0 | 0 | 0 | 0 |
| 0 | 0 | 0 | 0 |
| 1 | 0 | 0 | 0 |

| | | | |
|---|---|---|---|
| 0 | 0 | 0 | 0 |
| 0 | 0 | 0 | 0 |
| 0 | 0 | 0 | 0 |
| 0 | 0 | 0 | 0 |
| 0 | 0 | 0 | 0 |
| 0 | 0 | 0 | 0 |
| 0 | 0 | 0 | 0 |
| 0 | 0 | 0 | 0 |
| 0 | 0 | 0 | 0 |
| 1 | 0 | 0 | 0 |
| 0 | 0 | 0 | 0 |
| 0 | 0 | 0 | 0 |
| 1 | 1 | 0 | 0 |
| 1 | 1 | 0 | 0 |
| 1 | 1 | 0 | 0 |
| 1 | 1 | 0 | 0 |
| 0 | 0 | 0 | 0 |
| 0 | 0 | 0 | 0 |
| 0 | 0 | 0 | 0 |
| 0 | 0 | 0 | 0 |
| 0 | 0 | 0 | 0 |
| 0 | 0 | 0 | 0 |
| 0 | 0 | 0 | 0 |
| 0 | 0 | 0 | 0 |
| 0 | 0 | 0 | 0 |
| 0 | 0 | 0 | 1 |
| 0 | 0 | 0 | 0 |
| 0 | 0 | 0 | 0 |
| 0 | 0 | 0 | 0 |
| 0 | 0 | 0 | 0 |
| 0 | 0 | 0 | 0 |
| 0 | 0 | 0 | 0 |
| 0 | 0 | 0 | 0 |
| 0 | 0 | 0 | 0 |
| 0 | 0 | 0 | 0 |
| 0 | 0 | 0 | 0 |
| 0 | 0 | 0 | 0 |
| 1 | 1 | 0 | 0 |
| 0 | 0 | 0 | 0 |
| 0 | 0 | 0 | 0 |
| 1 | 1 | 0 | 0 |
| 0 | 0 | 0 | 0 |
| 0 | 0 | 0 | 0 |
| 1 | 0 | 0 | 1 |
| 0 | 0 | 0 | 0 |
| 0 | 0 | 0 | 0 |
| 0 | 0 | 0 | 0 |
| 0 | 0 | 0 | 0 |
| 0 | 0 | 0 | 0 |
| 0 | 0 | 0 | 0 |
| 0 | 0 | 0 | 0 |
| 0 | 0 | 1 | 0 |
| 0 | 0 | 1 | 0 |
| 0 | 0 | 1 | 0 |

| | | | |
|---|---|---|---|
| 0 | 0 | 1 | 0 |
| 0 | 0 | 0 | 0 |
| 0 | 0 | 0 | 0 |
| 0 | 0 | 0 | 0 |
| 0 | 0 | 0 | 0 |
| 0 | 0 | 0 | 0 |
| 0 | 0 | 0 | 0 |
| 0 | 0 | 0 | 0 |
| 0 | 0 | 0 | 0 |
| 0 | 0 | 0 | 0 |
| 0 | 0 | 0 | 0 |
| 0 | 0 | 0 | 0 |
| 0 | 0 | 0 | 0 |
| 0 | 0 | 0 | 0 |
| 0 | 0 | 0 | 0 |
| 0 | 0 | 0 | 0 |
| 0 | 0 | 0 | 0 |
| 0 | 0 | 0 | 0 |
| 0 | 0 | 0 | 0 |
| 0 | 0 | 0 | 0 |
| 0 | 0 | 0 | 0 |
| 0 | 0 | 0 | 0 |
| 0 | 0 | 0 | 0 |
| 0 | 1 | 0 | 1 |
| 0 | 0 | 0 | 0 |
| 0 | 0 | 0 | 0 |
| 0 | 0 | 0 | 0 |
| 0 | 0 | 0 | 0 |
| 0 | 0 | 0 | 0 |
| 0 | 0 | 0 | 0 |
| 0 | 0 | 0 | 0 |
| 0 | 0 | 0 | 0 |
| 0 | 0 | 0 | 0 |
| 0 | 0 | 0 | 0 |
| 0 | 0 | 0 | 0 |
| 0 | 0 | 0 | 0 |
| 0 | 0 | 0 | 0 |
| 0 | 0 | 0 | 0 |
| 0 | 0 | 0 | 0 |
| 0 | 0 | 0 | 0 |
| 0 | 0 | 0 | 0 |
| 0 | 0 | 0 | 0 |
| 0 | 0 | 0 | 0 |
| 0 | 0 | 0 | 0 |
| 0 | 0 | 0 | 0 |
| 0 | 0 | 0 | 0 |
| 1 | 0 | 0 | 0 |
| 1 | 0 | 0 | 0 |
| 0 | 0 | 0 | 0 |
| 0 | 1 | 0 | 0 |
| 1 | 1 | 0 | 1 |
| 1 | 1 | 0 | 1 |
| 1 | 1 | 0 | 1 |
| 0 | 0 | 0 | 0 |

| | | | |
|---|---|---|---|
| 1 | 1 | 0 | 0 |
| 0 | 0 | 0 | 0 |
| 0 | 0 | 0 | 0 |
| 0 | 0 | 0 | 0 |
| 0 | 0 | 0 | 0 |
| 0 | 0 | 0 | 0 |
| 0 | 0 | 0 | 0 |
| 0 | 0 | 0 | 0 |
| 0 | 0 | 0 | 0 |
| 0 | 0 | 0 | 0 |
| 0 | 0 | 0 | 0 |
| 0 | 1 | 0 | 0 |
| 0 | 0 | 0 | 0 |
| 0 | 0 | 0 | 0 |
| 0 | 1 | 0 | 0 |
| 0 | 0 | 0 | 0 |
| 1 | 1 | 0 | 0 |
| 0 | 0 | 0 | 0 |
| 0 | 0 | 0 | 0 |
| 1 | 0 | 0 | 0 |
| 0 | 0 | 0 | 0 |
| 0 | 0 | 0 | 0 |
| 0 | 0 | 0 | 0 |
| 0 | 0 | 0 | 0 |
| 0 | 0 | 0 | 0 |
| 0 | 0 | 0 | 0 |
| 0 | 0 | 0 | 0 |
| 0 | 0 | 0 | 0 |
| 0 | 0 | 0 | 0 |
| 0 | 0 | 0 | 0 |
| 0 | 0 | 0 | 0 |
| 0 | 0 | 0 | 0 |
| 0 | 0 | 0 | 0 |
| 1 | 0 | 0 | 0 |
| 0 | 0 | 0 | 0 |
| 0 | 1 | 0 | 0 |
| 0 | 0 | 0 | 0 |
| 0 | 0 | 0 | 0 |
| 0 | 0 | 0 | 0 |
| 0 | 0 | 0 | 0 |
| 0 | 0 | 0 | 0 |
| 0 | 0 | 0 | 0 |
| 0 | 0 | 0 | 0 |
| 0 | 0 | 0 | 0 |
| 0 | 0 | 0 | 0 |
| 0 | 0 | 0 | 0 |
| 0 | 0 | 0 | 0 |
| 0 | 0 | 0 | 0 |
| 1 | 1 | 0 | 0 |
| 0 | 0 | 0 | 0 |
| 0 | 0 | 0 | 0 |
| 0 | 0 | 0 | 0 |
| 0 | 0 | 0 | 0 |
| 0 | 0 | 0 | 0 |

| | | | |
|---|---|---|---|
| 0 | 0 | 0 | 0 |
| 0 | 0 | 0 | 0 |
| 0 | 0 | 0 | 0 |
| 0 | 0 | 0 | 0 |
| 0 | 0 | 0 | 0 |
| 0 | 0 | 0 | 0 |
| 0 | 0 | 0 | 0 |
| 0 | 1 | 0 | 0 |
| 0 | 0 | 0 | 0 |
| 0 | 0 | 0 | 0 |
| 0 | 0 | 0 | 0 |
| 0 | 0 | 0 | 0 |
| 0 | 0 | 1 | 0 |
| 0 | 0 | 0 | 0 |
| 0 | 0 | 1 | 0 |
| 0 | 0 | 1 | 0 |
| 0 | 0 | 0 | 0 |
| 0 | 0 | 0 | 0 |
| 0 | 0 | 0 | 0 |
| 0 | 0 | 0 | 0 |
| 0 | 1 | 0 | 0 |
| 0 | 1 | 0 | 0 |
| 0 | 0 | 0 | 0 |
| 0 | 0 | 0 | 0 |
| 0 | 1 | 0 | 0 |
| 0 | 0 | 0 | 0 |
| 0 | 0 | 0 | 0 |
| 0 | 1 | 0 | 0 |
| 0 | 0 | 0 | 0 |
| 0 | 0 | 0 | 0 |
| 0 | 0 | 0 | 0 |
| 0 | 0 | 0 | 0 |
| 1 | 1 | 0 | 0 |
| 1 | 1 | 0 | 0 |
| 1 | 1 | 0 | 0 |
| 0 | 1 | 0 | 0 |
| 1 | 0 | 0 | 0 |
| 0 | 1 | 0 | 0 |
| 1 | 1 | 0 | 0 |
| 0 | 0 | 0 | 0 |
| 0 | 0 | 0 | 0 |
| 0 | 0 | 0 | 0 |
| 0 | 0 | 0 | 0 |
| 0 | 0 | 0 | 0 |
| 0 | 0 | 0 | 0 |
| 0 | 0 | 0 | 0 |
| 0 | 0 | 0 | 0 |
| 0 | 0 | 0 | 0 |
| 0 | 0 | 0 | 0 |
| 1 | 1 | 0 | 0 |
| 1 | 1 | 0 | 0 |
| 0 | 0 | 0 | 0 |
| 0 | 0 | 0 | 0 |
| 0 | 0 | 0 | 0 |

| | | | |
|---|---|---|---|
| 0 | 0 | 0 | 0 |
| 0 | 0 | 0 | 0 |
| 0 | 0 | 0 | 0 |
| 0 | 0 | 0 | 0 |
| 0 | 0 | 0 | 0 |
| 0 | 0 | 0 | 0 |
| 1 | 1 | 0 | 0 |
| 1 | 1 | 0 | 0 |
| 0 | 0 | 0 | 0 |
| 0 | 0 | 0 | 0 |
| 0 | 0 | 0 | 0 |
| 0 | 0 | 0 | 0 |
| 0 | 0 | 0 | 0 |
| 0 | 0 | 0 | 0 |
| 0 | 0 | 0 | 0 |
| 0 | 0 | 0 | 0 |
| 0 | 0 | 0 | 0 |
| 0 | 0 | 0 | 0 |
| 0 | 0 | 0 | 0 |
| 0 | 0 | 0 | 0 |
| 0 | 0 | 0 | 0 |
| 0 | 0 | 0 | 0 |
| 0 | 0 | 0 | 0 |
| 0 | 0 | 0 | 0 |
| 0 | 0 | 0 | 0 |
| 0 | 0 | 0 | 0 |
| 0 | 0 | 0 | 0 |
| 1 | 1 | 0 | 0 |
| 0 | 0 | 0 | 0 |
| 0 | 0 | 0 | 0 |
| 0 | 0 | 0 | 0 |
| 0 | 0 | 0 | 0 |
| 0 | 0 | 0 | 0 |
| 0 | 0 | 0 | 0 |
| 0 | 0 | 0 | 0 |
| 0 | 0 | 0 | 0 |
| 0 | 0 | 0 | 0 |
| 0 | 0 | 0 | 0 |
| 0 | 0 | 0 | 0 |
| 0 | 0 | 0 | 0 |
| 0 | 0 | 0 | 0 |
| 0 | 0 | 0 | 0 |
| 0 | 0 | 0 | 0 |
| 0 | 0 | 0 | 0 |
| 0 | 0 | 0 | 0 |
| 0 | 0 | 0 | 0 |
| 0 | 0 | 0 | 0 |
| 0 | 0 | 0 | 0 |
| 0 | 0 | 0 | 0 |
| 0 | 0 | 0 | 0 |
| 0 | 0 | 0 | 0 |
| 0 | 0 | 0 | 0 |
| 0 | 0 | 0 | 0 |
| 0 | 0 | 0 | 0 |

| | | | |
|---|---|---|---|
| 0 | 0 | 0 | 0 |
| 0 | 0 | 0 | 0 |
| 0 | 0 | 0 | 0 |
| 0 | 0 | 0 | 0 |
| 0 | 0 | 0 | 0 |
| 0 | 0 | 0 | 0 |
| 0 | 0 | 0 | 0 |
| 0 | 0 | 0 | 0 |
| 0 | 0 | 0 | 0 |
| 0 | 0 | 0 | 0 |
| 1 | 0 | 0 | 0 |
| 1 | 1 | 0 | 0 |
| 1 | 0 | 0 | 0 |
| 0 | 0 | 0 | 0 |
| 0 | 0 | 0 | 0 |
| 0 | 0 | 0 | 0 |
| 0 | 0 | 0 | 0 |
| 0 | 0 | 0 | 0 |
| 0 | 0 | 0 | 0 |
| 0 | 0 | 0 | 0 |
| 0 | 0 | 0 | 0 |
| 0 | 0 | 0 | 0 |
| 0 | 0 | 0 | 0 |
| 0 | 0 | 0 | 0 |
| 0 | 0 | 0 | 0 |
| 0 | 0 | 0 | 0 |
| 0 | 0 | 0 | 0 |
| 0 | 0 | 0 | 0 |
| 0 | 0 | 0 | 0 |
| 0 | 0 | 0 | 0 |
| 0 | 0 | 0 | 0 |
| 0 | 1 | 0 | 0 |
| 0 | 0 | 0 | 0 |
| 0 | 0 | 0 | 0 |
| 1 | 0 | 0 | 0 |
| 1 | 0 | 0 | 0 |
| 1 | 0 | 0 | 0 |
| 0 | 0 | 0 | 0 |
| 0 | 0 | 0 | 0 |
| 0 | 0 | 0 | 0 |
| 0 | 0 | 0 | 0 |
| 0 | 0 | 0 | 0 |
| 0 | 0 | 0 | 0 |
| 0 | 0 | 0 | 0 |
| 0 | 0 | 0 | 0 |
| 0 | 0 | 0 | 0 |
| 0 | 0 | 0 | 0 |
| 0 | 0 | 0 | 0 |
| 0 | 0 | 0 | 0 |
| 0 | 0 | 0 | 0 |
| 0 | 0 | 0 | 0 |
| 0 | 0 | 0 | 0 |
| 0 | 0 | 0 | 1 |
| 0 | 0 | 0 | 0 |

| | | | |
|---|---|---|---|
| 0 | 0 | 0 | 0 |
| 0 | 0 | 0 | 0 |
| 0 | 0 | 0 | 0 |
| 0 | 0 | 0 | 0 |
| 0 | 0 | 0 | 0 |
| 0 | 0 | 0 | 0 |
| 0 | 0 | 0 | 0 |
| 0 | 0 | 0 | 0 |
| 0 | 0 | 0 | 0 |
| 0 | 0 | 0 | 0 |
| 0 | 0 | 0 | 0 |
| 1 | 0 | 0 | 0 |
| 0 | 0 | 0 | 0 |
| 0 | 0 | 0 | 0 |
| 0 | 0 | 1 | 0 |
| 0 | 0 | 0 | 0 |
| 0 | 0 | 0 | 0 |
| 0 | 0 | 0 | 0 |
| 0 | 0 | 0 | 0 |
| 0 | 0 | 0 | 0 |
| 0 | 0 | 0 | 0 |
| 0 | 0 | 0 | 0 |
| 0 | 0 | 0 | 0 |
| 0 | 0 | 0 | 0 |
| 0 | 0 | 0 | 0 |
| 0 | 0 | 0 | 0 |
| 0 | 0 | 0 | 0 |
| 0 | 0 | 0 | 0 |
| 0 | 0 | 0 | 0 |
| 0 | 0 | 0 | 0 |
| 0 | 0 | 0 | 0 |
| 0 | 0 | 0 | 0 |
| 0 | 0 | 0 | 0 |
| 1 | 0 | 0 | 0 |
| 0 | 0 | 0 | 0 |
| 1 | 0 | 0 | 1 |
| 0 | 0 | 0 | 0 |
| 0 | 0 | 0 | 0 |
| 0 | 0 | 0 | 0 |
| 0 | 0 | 0 | 0 |
| 0 | 0 | 0 | 0 |
| 0 | 0 | 0 | 0 |
| 0 | 0 | 0 | 0 |
| 0 | 1 | 0 | 0 |
| 0 | 0 | 0 | 0 |
| 0 | 0 | 0 | 0 |
| 0 | 0 | 0 | 0 |
| 0 | 0 | 0 | 0 |
| 0 | 0 | 0 | 0 |
| 0 | 0 | 0 | 0 |
| 0 | 0 | 0 | 0 |
| 0 | 0 | 0 | 0 |
| 0 | 0 | 0 | 0 |
| 0 | 0 | 0 | 0 |

| | | | |
|---|---|---|---|
| 0 | 0 | 0 | 0 |
| 0 | 0 | 0 | 0 |
| 0 | 0 | 0 | 0 |
| 0 | 0 | 0 | 0 |
| 0 | 0 | 0 | 0 |
| 0 | 1 | 0 | 0 |
| 0 | 0 | 0 | 0 |
| 0 | 0 | 0 | 0 |
| 0 | 0 | 0 | 0 |
| 0 | 0 | 0 | 0 |
| 0 | 0 | 0 | 0 |
| 0 | 0 | 0 | 0 |
| 0 | 0 | 0 | 0 |
| 0 | 0 | 0 | 0 |
| 0 | 0 | 0 | 0 |
| 0 | 1 | 0 | 0 |
| 1 | 1 | 0 | 0 |
| 0 | 0 | 1 | 0 |
| 0 | 0 | 0 | 0 |
| 0 | 0 | 0 | 0 |
| 0 | 0 | 0 | 0 |
| 0 | 0 | 0 | 0 |
| 0 | 0 | 0 | 0 |
| 0 | 0 | 0 | 0 |
| 0 | 0 | 0 | 0 |
| 0 | 0 | 0 | 0 |
| 0 | 0 | 0 | 0 |
| 1 | 0 | 0 | 0 |
| 0 | 0 | 0 | 0 |
| 0 | 0 | 0 | 0 |
| 0 | 0 | 1 | 0 |
| 0 | 0 | 0 | 0 |
| 0 | 0 | 0 | 0 |
| 0 | 0 | 0 | 0 |
| 0 | 0 | 0 | 0 |
| 0 | 0 | 0 | 0 |
| 0 | 0 | 0 | 0 |
| 1 | 1 | 0 | 0 |
| 0 | 0 | 0 | 0 |
| 0 | 0 | 0 | 0 |
| 0 | 0 | 0 | 0 |
| 1 | 0 | 0 | 0 |
| 1 | 0 | 0 | 0 |
| 0 | 0 | 0 | 0 |
| 0 | 0 | 0 | 0 |
| 0 | 0 | 0 | 0 |
| 0 | 0 | 0 | 0 |
| 0 | 0 | 0 | 0 |
| 0 | 0 | 0 | 0 |
| 0 | 0 | 0 | 0 |
| 0 | 0 | 0 | 0 |
| 0 | 0 | 0 | 0 |
| 0 | 0 | 0 | 0 |
| 0 | 0 | 0 | 0 |

| | | | |
|---|---|---|---|
| 0 | 0 | 0 | 0 |
| 0 | 0 | 0 | 0 |
| 0 | 0 | 0 | 0 |
| 0 | 0 | 0 | 0 |
| 0 | 0 | 0 | 0 |
| 0 | 0 | 0 | 0 |
| 0 | 0 | 0 | 0 |
| 0 | 0 | 0 | 0 |
| 0 | 0 | 0 | 0 |
| 0 | 0 | 0 | 0 |
| 0 | 0 | 0 | 0 |
| 0 | 0 | 0 | 0 |
| 1 | 0 | 0 | 0 |
| 0 | 0 | 0 | 0 |
| 0 | 0 | 0 | 0 |
| 0 | 0 | 0 | 0 |
| 0 | 0 | 0 | 0 |
| 0 | 0 | 0 | 0 |
| 0 | 0 | 0 | 0 |
| 0 | 0 | 0 | 0 |
| 0 | 0 | 0 | 0 |
| 0 | 0 | 0 | 0 |
| 0 | 0 | 0 | 0 |
| 1 | 0 | 0 | 0 |
| 0 | 0 | 0 | 0 |
| 0 | 0 | 0 | 0 |
| 0 | 0 | 0 | 0 |
| 1 | 1 | 0 | 1 |
| 0 | 0 | 0 | 0 |
| 0 | 0 | 0 | 0 |
| 0 | 0 | 0 | 0 |
| 0 | 0 | 0 | 0 |
| 0 | 0 | 0 | 0 |
| 0 | 0 | 0 | 0 |
| 0 | 0 | 0 | 0 |
| 0 | 0 | 0 | 0 |
| 0 | 0 | 0 | 0 |
| 0 | 0 | 0 | 0 |
| 0 | 0 | 0 | 0 |
| 0 | 0 | 0 | 0 |
| 0 | 0 | 0 | 0 |
| 0 | 0 | 0 | 0 |
| 0 | 0 | 0 | 0 |
| 0 | 0 | 0 | 0 |
| 0 | 0 | 0 | 0 |
| 0 | 0 | 0 | 0 |
| 0 | 0 | 0 | 0 |
| 0 | 0 | 0 | 0 |
| 0 | 0 | 0 | 0 |
| 0 | 0 | 0 | 0 |
| 0 | 0 | 0 | 0 |
| 0 | 0 | 0 | 0 |
| 0 | 0 | 0 | 0 |
| 0 | 0 | 0 | 0 |

| | | | |
|---|---|---|---|
| 0 | 0 | 0 | 0 |
| 0 | 0 | 0 | 0 |
| 0 | 0 | 0 | 0 |
| 0 | 0 | 0 | 0 |
| 0 | 0 | 0 | 0 |
| 0 | 0 | 0 | 0 |
| 0 | 0 | 0 | 0 |
| 0 | 0 | 0 | 0 |
| 1 | 0 | 0 | 0 |
| 1 | 0 | 0 | 0 |
| 1 | 0 | 0 | 0 |
| 0 | 0 | 0 | 0 |
| 1 | 0 | 0 | 0 |
| 0 | 0 | 0 | 0 |
| 0 | 0 | 0 | 0 |
| 1 | 0 | 0 | 0 |
| 0 | 0 | 0 | 0 |
| 0 | 0 | 0 | 0 |
| 0 | 0 | 0 | 0 |
| 0 | 0 | 0 | 0 |
| 0 | 0 | 0 | 0 |
| 0 | 0 | 0 | 0 |
| 0 | 0 | 0 | 0 |
| 0 | 0 | 0 | 0 |
| 0 | 0 | 0 | 0 |
| 0 | 0 | 0 | 0 |
| 0 | 0 | 0 | 0 |
| 1 | 0 | 0 | 0 |
| 1 | 0 | 0 | 0 |
| 0 | 0 | 0 | 0 |
| 0 | 0 | 0 | 0 |
| 0 | 0 | 0 | 0 |
| 1 | 0 | 0 | 0 |
| 0 | 0 | 0 | 0 |
| 0 | 0 | 0 | 0 |
| 0 | 0 | 0 | 0 |
| 0 | 0 | 0 | 0 |
| 0 | 0 | 0 | 0 |
| 0 | 0 | 0 | 0 |
| 0 | 0 | 0 | 0 |
| 1 | 0 | 0 | 0 |
| 1 | 0 | 0 | 0 |
| 0 | 0 | 0 | 0 |
| 0 | 0 | 0 | 0 |
| 0 | 0 | 0 | 0 |
| 0 | 0 | 0 | 0 |
| 0 | 0 | 0 | 0 |
| 0 | 0 | 0 | 0 |
| 0 | 0 | 0 | 0 |
| 0 | 0 | 0 | 0 |
| 0 | 0 | 0 | 0 |
| 0 | 0 | 0 | 0 |
| 0 | 0 | 0 | 0 |
| 0 | 0 | 0 | 0 |

| | | | |
|---|---|---|---|
| 0 | 0 | 0 | 0 |
| 0 | 0 | 0 | 0 |
| 0 | 0 | 0 | 0 |
| 0 | 0 | 0 | 0 |
| 0 | 0 | 0 | 0 |
| 0 | 0 | 0 | 0 |
| 0 | 0 | 0 | 0 |
| 0 | 0 | 0 | 0 |
| 0 | 0 | 0 | 0 |
| 0 | 0 | 0 | 0 |
| 1 | 0 | 0 | 0 |
| 0 | 0 | 0 | 0 |
| 1 | 0 | 0 | 0 |
| 0 | 0 | 0 | 0 |
| 0 | 0 | 0 | 0 |
| 0 | 0 | 0 | 0 |
| 0 | 0 | 0 | 0 |
| 0 | 1 | 0 | 0 |
| 0 | 0 | 0 | 0 |
| 0 | 0 | 0 | 0 |
| 0 | 0 | 0 | 0 |
| 0 | 0 | 0 | 0 |
| 1 | 0 | 0 | 0 |
| 0 | 0 | 0 | 0 |
| 0 | 0 | 0 | 0 |
| 1 | 0 | 0 | 0 |
| 1 | 0 | 0 | 0 |
| 1 | 0 | 0 | 0 |
| 0 | 0 | 0 | 0 |
| 0 | 0 | 0 | 0 |
| 0 | 0 | 0 | 0 |
| 0 | 0 | 0 | 0 |
| 0 | 0 | 0 | 0 |
| 0 | 0 | 0 | 0 |
| 0 | 0 | 0 | 0 |
| 0 | 0 | 0 | 0 |
| 0 | 0 | 0 | 0 |
| 0 | 0 | 0 | 0 |
| 0 | 0 | 0 | 0 |
| 0 | 0 | 0 | 0 |
| 1 | 1 | 0 | 0 |
| 0 | 0 | 0 | 0 |
| 0 | 0 | 0 | 0 |
| 0 | 0 | 0 | 0 |
| 0 | 0 | 0 | 0 |
| 0 | 0 | 0 | 0 |
| 0 | 0 | 0 | 0 |
| 1 | 0 | 0 | 0 |
| 0 | 0 | 0 | 0 |
| 0 | 0 | 0 | 0 |
| 0 | 0 | 0 | 0 |
| 0 | 0 | 0 | 0 |
| 0 | 0 | 0 | 0 |
| 0 | 0 | 0 | 0 |

| | | | |
|---|---|---|---|
| 1 | 0 | 0 | 0 |
| 0 | 0 | 0 | 0 |
| 1 | 0 | 0 | 0 |
| 0 | 0 | 0 | 0 |
| 0 | 0 | 0 | 0 |
| 0 | 0 | 0 | 0 |
| 0 | 0 | 0 | 0 |
| 0 | 0 | 0 | 0 |
| 1 | 1 | 0 | 0 |
| 0 | 0 | 0 | 0 |
| 1 | 1 | 0 | 0 |
| 1 | 1 | 0 | 0 |
| 0 | 0 | 0 | 0 |
| 1 | 1 | 0 | 0 |
| 0 | 0 | 0 | 0 |
| 0 | 0 | 0 | 0 |
| 0 | 0 | 0 | 0 |
| 0 | 0 | 0 | 0 |
| 0 | 0 | 0 | 0 |
| 0 | 0 | 0 | 0 |
| 1 | 0 | 0 | 0 |
| 0 | 0 | 0 | 0 |
| 0 | 0 | 0 | 0 |
| 0 | 0 | 0 | 0 |
| 1 | 0 | 0 | 0 |
| 0 | 0 | 0 | 0 |
| 1 | 0 | 0 | 0 |
| 1 | 0 | 0 | 0 |
| 1 | 1 | 0 | 0 |
| 0 | 0 | 0 | 0 |
| 1 | 1 | 0 | 0 |
| 1 | 1 | 0 | 0 |
| 1 | 1 | 0 | 0 |
| 0 | 0 | 0 | 0 |
| 1 | 1 | 0 | 0 |
| 0 | 0 | 0 | 0 |
| 1 | 0 | 0 | 0 |
| 0 | 0 | 0 | 0 |
| 1 | 1 | 0 | 0 |
| 1 | 1 | 0 | 0 |
| 1 | 0 | 0 | 0 |
| 0 | 0 | 0 | 0 |
| 0 | 0 | 0 | 0 |
| 1 | 0 | 0 | 0 |
| 0 | 0 | 0 | 0 |
| 0 | 0 | 0 | 0 |
| 1 | 0 | 0 | 0 |
| 0 | 0 | 0 | 0 |
| 1 | 0 | 0 | 0 |
| 0 | 0 | 0 | 0 |
| 0 | 0 | 0 | 0 |
| 0 | 0 | 0 | 0 |
| 0 | 0 | 0 | 0 |
| 0 | 0 | 0 | 0 |

| | | | |
|---|---|---|---|
| 0 | 0 | 0 | 0 |
| 0 | 0 | 0 | 0 |
| 0 | 0 | 0 | 0 |
| 0 | 0 | 0 | 0 |
| 0 | 0 | 0 | 0 |
| 0 | 0 | 0 | 0 |
| 0 | 0 | 0 | 0 |
| 0 | 0 | 0 | 0 |
| 0 | 0 | 0 | 0 |
| 1 | 1 | 0 | 0 |
| 1 | 1 | 0 | 0 |
| 0 | 0 | 0 | 0 |
| 0 | 0 | 0 | 0 |
| 0 | 0 | 0 | 0 |
| 1 | 0 | 1 | 0 |
| 0 | 0 | 0 | 0 |
| 0 | 0 | 0 | 0 |
| 1 | 0 | 1 | 0 |
| 0 | 0 | 0 | 0 |
| 0 | 0 | 0 | 0 |
| 0 | 0 | 0 | 0 |
| 0 | 0 | 1 | 0 |
| 1 | 0 | 1 | 0 |
| 0 | 0 | 1 | 0 |
| 0 | 0 | 1 | 0 |
| 0 | 0 | 1 | 1 |
| 0 | 0 | 1 | 1 |
| 0 | 0 | 0 | 0 |
| 0 | 0 | 0 | 0 |
| 0 | 0 | 0 | 0 |
| 0 | 0 | 0 | 0 |
| 0 | 0 | 0 | 0 |
| 0 | 0 | 0 | 0 |
| 0 | 0 | 0 | 0 |
| 0 | 0 | 0 | 0 |
| 0 | 0 | 0 | 0 |
| 0 | 0 | 0 | 0 |
| 0 | 0 | 0 | 0 |
| 0 | 0 | 0 | 0 |
| 0 | 0 | 0 | 0 |
| 0 | 0 | 0 | 0 |
| 0 | 0 | 0 | 0 |
| 0 | 0 | 0 | 0 |
| 0 | 0 | 1 | 0 |
| 0 | 0 | 0 | 0 |
| 0 | 0 | 0 | 0 |
| 0 | 0 | 0 | 0 |
| 0 | 0 | 0 | 0 |
| 0 | 0 | 0 | 0 |
| 0 | 0 | 0 | 0 |
| 1 | 0 | 0 | 0 |
| 0 | 0 | 0 | 0 |
| 0 | 0 | 0 | 0 |
| 0 | 0 | 0 | 0 |

| | | | |
|---|---|---|---|
| 0 | 0 | 0 | 0 |
| 0 | 0 | 0 | 0 |
| 0 | 0 | 0 | 0 |
| 1 | 0 | 0 | 0 |
| 0 | 0 | 0 | 0 |
| 0 | 0 | 0 | 0 |
| 0 | 0 | 0 | 0 |
| 0 | 0 | 0 | 0 |
| 1 | 0 | 0 | 0 |
| 1 | 0 | 0 | 0 |
| 0 | 0 | 0 | 0 |
| 0 | 0 | 0 | 0 |
| 0 | 0 | 0 | 0 |
| 0 | 0 | 0 | 0 |
| 0 | 0 | 0 | 0 |
| 0 | 0 | 0 | 0 |
| 0 | 0 | 1 | 0 |
| 0 | 0 | 0 | 0 |
| 0 | 0 | 0 | 0 |
| 0 | 0 | 0 | 0 |
| 0 | 0 | 0 | 0 |
| 0 | 0 | 0 | 0 |
| 0 | 0 | 0 | 0 |
| 0 | 0 | 0 | 0 |
| 0 | 0 | 0 | 0 |
| 0 | 0 | 0 | 0 |
| 0 | 0 | 0 | 0 |
| 0 | 0 | 0 | 0 |
| 0 | 0 | 0 | 0 |
| 0 | 0 | 0 | 0 |
| 0 | 0 | 0 | 0 |
| 0 | 0 | 0 | 0 |
| 0 | 0 | 0 | 0 |
| 0 | 0 | 0 | 0 |
| 0 | 0 | 0 | 0 |
| 0 | 0 | 0 | 0 |
| 0 | 0 | 0 | 0 |
| 0 | 0 | 0 | 0 |
| 0 | 0 | 0 | 0 |
| 0 | 0 | 0 | 0 |
| 0 | 0 | 0 | 0 |
| 0 | 0 | 0 | 0 |
| 0 | 0 | 0 | 0 |
| 0 | 0 | 0 | 0 |
| 0 | 0 | 0 | 0 |
| 0 | 0 | 0 | 0 |
| 0 | 0 | 0 | 0 |
| 0 | 0 | 0 | 0 |
| 0 | 0 | 0 | 0 |
| 0 | 1 | 0 | 0 |
| 0 | 0 | 0 | 0 |
| 0 | 0 | 0 | 0 |
| 0 | 0 | 0 | 0 |
| 0 | 0 | 0 | 0 |

| | | | |
|---|---|---|---|
| 0 | 0 | 1 | 0 |
| 0 | 0 | 0 | 0 |
| 0 | 0 | 0 | 0 |
| 0 | 0 | 0 | 0 |
| 0 | 0 | 0 | 0 |
| 0 | 0 | 0 | 0 |
| 0 | 0 | 0 | 0 |
| 1 | 0 | 0 | 0 |
| 0 | 0 | 0 | 0 |
| 0 | 0 | 0 | 0 |
| 0 | 0 | 0 | 0 |
| 1 | 0 | 0 | 0 |
| 0 | 0 | 0 | 0 |
| 0 | 0 | 0 | 0 |
| 0 | 0 | 0 | 0 |
| 0 | 0 | 0 | 0 |
| 0 | 0 | 0 | 0 |
| 0 | 0 | 0 | 0 |
| 0 | 0 | 0 | 0 |
| 0 | 0 | 0 | 0 |
| 0 | 0 | 0 | 0 |
| 0 | 0 | 0 | 0 |
| 0 | 0 | 0 | 0 |
| 0 | 0 | 0 | 0 |
| 0 | 0 | 0 | 0 |
| 0 | 0 | 0 | 0 |
| 0 | 0 | 0 | 0 |
| 0 | 0 | 0 | 0 |
| 0 | 0 | 0 | 0 |
| 0 | 0 | 0 | 0 |
| 0 | 0 | 0 | 0 |
| 0 | 0 | 0 | 0 |
| 0 | 0 | 0 | 0 |
| 0 | 0 | 0 | 0 |
| 0 | 0 | 0 | 0 |
| 0 | 0 | 0 | 0 |
| 0 | 0 | 0 | 0 |
| 0 | 0 | 0 | 0 |
| 0 | 0 | 0 | 0 |
| 0 | 0 | 0 | 0 |
| 0 | 0 | 0 | 0 |
| 0 | 0 | 0 | 0 |
| 0 | 0 | 0 | 0 |
| 0 | 0 | 0 | 0 |
| 0 | 0 | 0 | 0 |
| 0 | 0 | 0 | 0 |
| 0 | 0 | 0 | 0 |
| 0 | 0 | 0 | 0 |
| 0 | 0 | 0 | 0 |
| 0 | 0 | 0 | 0 |
| 0 | 0 | 0 | 0 |
| 0 | 0 | 0 | 0 |
| 0 | 0 | 0 | 0 |
| 0 | 0 | 0 | 0 |

| | | | |
|---|---|---|---|
| 0 | 0 | 0 | 0 |
| 0 | 0 | 0 | 0 |
| 0 | 0 | 0 | 0 |
| 0 | 0 | 0 | 0 |
| 0 | 0 | 0 | 0 |
| 0 | 0 | 0 | 0 |
| 0 | 0 | 0 | 0 |
| 0 | 0 | 0 | 0 |
| 0 | 0 | 0 | 0 |
| 0 | 0 | 0 | 0 |
| 0 | 0 | 0 | 0 |
| 0 | 0 | 0 | 0 |
| 0 | 0 | 0 | 0 |
| 0 | 0 | 0 | 0 |
| 0 | 0 | 0 | 0 |
| 0 | 0 | 0 | 0 |
| 0 | 0 | 0 | 0 |
| 0 | 0 | 0 | 0 |
| 0 | 0 | 0 | 1 |
| 0 | 1 | 0 | 1 |
| 0 | 0 | 0 | 0 |
| 0 | 0 | 0 | 0 |
| 0 | 0 | 0 | 0 |
| 0 | 0 | 0 | 0 |
| 0 | 0 | 0 | 0 |
| 1 | 0 | 0 | 0 |
| 1 | 0 | 0 | 0 |
| 0 | 0 | 0 | 0 |
| 0 | 0 | 0 | 0 |
| 0 | 0 | 0 | 0 |
| 0 | 0 | 0 | 0 |
| 0 | 0 | 0 | 0 |
| 0 | 0 | 0 | 0 |
| 1 | 0 | 0 | 1 |
| 1 | 0 | 0 | 0 |
| 1 | 0 | 0 | 1 |
| 1 | 0 | 0 | 0 |
| 0 | 0 | 0 | 0 |
| 0 | 0 | 0 | 0 |
| 0 | 0 | 0 | 0 |
| 0 | 0 | 0 | 0 |
| 1 | 0 | 0 | 0 |
| 1 | 0 | 0 | 1 |
| 1 | 0 | 0 | 0 |
| 0 | 0 | 0 | 0 |
| 0 | 0 | 0 | 0 |
| 0 | 0 | 0 | 0 |
| 0 | 0 | 0 | 0 |
| 0 | 0 | 0 | 0 |
| 1 | 0 | 0 | 0 |
| 0 | 0 | 0 | 0 |
| 0 | 0 | 0 | 0 |
| 0 | 0 | 0 | 0 |
| 0 | 0 | 0 | 0 |

| | | | |
|---|---|---|---|
| 0 | 0 | 0 | 0 |
| 0 | 0 | 0 | 0 |
| 0 | 0 | 0 | 0 |
| 1 | 0 | 0 | 0 |
| 1 | 0 | 0 | 0 |
| 0 | 0 | 0 | 0 |
| 1 | 0 | 0 | 0 |
| 0 | 0 | 0 | 0 |
| 0 | 0 | 0 | 0 |
| 0 | 0 | 0 | 0 |
| 0 | 0 | 0 | 0 |
| 1 | 0 | 1 | 0 |
| 0 | 0 | 0 | 0 |
| 0 | 0 | 0 | 0 |
| 0 | 0 | 0 | 0 |
| 0 | 0 | 1 | 0 |
| 0 | 0 | 0 | 0 |
| 0 | 0 | 0 | 0 |
| 0 | 0 | 0 | 0 |
| 0 | 0 | 1 | 0 |
| 1 | 0 | 1 | 0 |
| 1 | 0 | 0 | 0 |
| 1 | 0 | 0 | 0 |
| 1 | 0 | 0 | 0 |
| 0 | 0 | 1 | 0 |
| 0 | 0 | 0 | 0 |
| 1 | 0 | 0 | 0 |
| 0 | 0 | 0 | 0 |
| 0 | 0 | 0 | 0 |
| 0 | 0 | 0 | 0 |
| 0 | 0 | 0 | 0 |
| 0 | 0 | 0 | 0 |
| 0 | 0 | 0 | 0 |
| 0 | 0 | 0 | 0 |
| 0 | 0 | 0 | 0 |
| 0 | 0 | 0 | 0 |
| 0 | 0 | 0 | 1 |
| 0 | 0 | 0 | 1 |
| 0 | 0 | 0 | 1 |
| 0 | 0 | 0 | 0 |
| 0 | 0 | 0 | 0 |
| 0 | 0 | 0 | 0 |
| 0 | 0 | 0 | 0 |
| 0 | 0 | 0 | 0 |
| 0 | 0 | 0 | 0 |
| 0 | 0 | 0 | 0 |
| 0 | 0 | 0 | 0 |
| 0 | 0 | 0 | 0 |
| 0 | 0 | 0 | 0 |
| 0 | 0 | 0 | 0 |
| 0 | 0 | 0 | 0 |
| 0 | 0 | 0 | 0 |
| 0 | 0 | 0 | 0 |
| 0 | 0 | 0 | 0 |

| | | | |
|---|---|---|---|
| 0 | 0 | 0 | 0 |
| 0 | 0 | 0 | 0 |
| 0 | 0 | 0 | 0 |
| 0 | 0 | 0 | 0 |
| 0 | 0 | 0 | 0 |
| 0 | 0 | 0 | 0 |
| 0 | 0 | 0 | 0 |
| 0 | 0 | 0 | 0 |
| 0 | 0 | 0 | 0 |
| 0 | 0 | 0 | 0 |
| 0 | 0 | 0 | 0 |
| 0 | 0 | 0 | 0 |
| 0 | 0 | 0 | 0 |
| 0 | 0 | 0 | 0 |
| 0 | 0 | 0 | 0 |
| 0 | 0 | 0 | 0 |
| 0 | 1 | 0 | 0 |
| 0 | 1 | 0 | 0 |
| 0 | 0 | 0 | 0 |
| 0 | 0 | 0 | 0 |
| 0 | 0 | 0 | 0 |
| 0 | 0 | 0 | 0 |
| 0 | 0 | 0 | 0 |
| 0 | 0 | 0 | 0 |
| 0 | 0 | 0 | 0 |
| 0 | 0 | 0 | 0 |
| 0 | 0 | 0 | 0 |
| 0 | 0 | 0 | 0 |
| 0 | 0 | 0 | 0 |
| 0 | 0 | 0 | 0 |
| 0 | 0 | 0 | 0 |
| 1 | 0 | 0 | 0 |
| 0 | 0 | 0 | 0 |
| 0 | 0 | 0 | 0 |
| 0 | 0 | 0 | 0 |
| 0 | 0 | 0 | 0 |
| 0 | 0 | 0 | 0 |
| 0 | 0 | 0 | 0 |
| 0 | 0 | 0 | 0 |
| 0 | 0 | 0 | 0 |
| 0 | 0 | 0 | 0 |
| 0 | 0 | 0 | 0 |
| 0 | 0 | 0 | 0 |
| 0 | 0 | 0 | 0 |
| 0 | 0 | 0 | 0 |
| 0 | 0 | 0 | 0 |
| 1 | 1 | 0 | 0 |
| 0 | 0 | 0 | 0 |
| 0 | 0 | 0 | 0 |
| 0 | 0 | 0 | 0 |
| 0 | 0 | 0 | 0 |
| 0 | 0 | 0 | 0 |
| 1 | 0 | 0 | 0 |
| 0 | 0 | 0 | 0 |

| | | | |
|---|---|---|---|
| 0 | 0 | 0 | 0 |
| 0 | 0 | 0 | 0 |
| 0 | 0 | 0 | 0 |
| 0 | 0 | 0 | 0 |
| 0 | 0 | 0 | 0 |
| 0 | 0 | 0 | 0 |
| 1 | 0 | 0 | 0 |
| 0 | 0 | 0 | 0 |
| 0 | 0 | 0 | 0 |
| 0 | 0 | 0 | 0 |
| 0 | 0 | 0 | 0 |
| 0 | 0 | 0 | 0 |
| 0 | 0 | 0 | 0 |
| 0 | 0 | 0 | 0 |
| 0 | 0 | 0 | 0 |
| 0 | 0 | 0 | 0 |
| 0 | 0 | 0 | 0 |
| 0 | 0 | 0 | 0 |
| 0 | 0 | 0 | 0 |
| 0 | 0 | 0 | 0 |
| 0 | 0 | 0 | 0 |
| 0 | 0 | 0 | 0 |
| 0 | 0 | 0 | 0 |
| 0 | 0 | 0 | 0 |
| 0 | 0 | 0 | 0 |
| 0 | 0 | 0 | 0 |
| 0 | 0 | 0 | 0 |
| 0 | 0 | 0 | 0 |
| 0 | 0 | 0 | 0 |
| 0 | 0 | 0 | 0 |
| 0 | 0 | 0 | 0 |
| 0 | 0 | 0 | 0 |
| 0 | 0 | 0 | 0 |
| 0 | 0 | 0 | 0 |
| 0 | 0 | 0 | 0 |
| 0 | 0 | 0 | 0 |
| 0 | 0 | 0 | 0 |
| 0 | 0 | 0 | 0 |
| 0 | 0 | 0 | 0 |
| 0 | 0 | 0 | 0 |
| 0 | 0 | 0 | 0 |
| 0 | 0 | 0 | 0 |
| 0 | 0 | 0 | 0 |
| 0 | 0 | 0 | 0 |
| 0 | 0 | 0 | 0 |
| 0 | 0 | 0 | 0 |
| 0 | 0 | 0 | 0 |
| 0 | 0 | 0 | 0 |
| 0 | 0 | 0 | 0 |
| 0 | 0 | 0 | 0 |
| 0 | 0 | 0 | 0 |
| 0 | 0 | 0 | 0 |
| 0 | 0 | 0 | 0 |
| 0 | 0 | 0 | 0 |

| | | | |
|---|---|---|---|
| 0 | 0 | 0 | 0 |
| 0 | 0 | 0 | 0 |
| 0 | 0 | 0 | 0 |
| 0 | 0 | 0 | 0 |
| 0 | 0 | 0 | 0 |
| 0 | 0 | 0 | 0 |
| 0 | 0 | 0 | 0 |
| 0 | 0 | 0 | 0 |
| 0 | 0 | 0 | 0 |
| 0 | 0 | 0 | 0 |
| 0 | 0 | 0 | 0 |
| 0 | 0 | 0 | 0 |
| 0 | 0 | 0 | 0 |
| 0 | 0 | 0 | 0 |
| 0 | 0 | 0 | 0 |
| 0 | 0 | 0 | 0 |
| 0 | 0 | 0 | 0 |
| 0 | 0 | 0 | 0 |
| 0 | 0 | 0 | 0 |
| 0 | 0 | 0 | 0 |
| 0 | 0 | 0 | 0 |
| 0 | 0 | 0 | 0 |
| 0 | 0 | 0 | 0 |
| 0 | 0 | 0 | 0 |
| 1 | 1 | 0 | 0 |
| 0 | 0 | 0 | 0 |
| 0 | 0 | 0 | 0 |
| 0 | 0 | 0 | 0 |
| 0 | 0 | 0 | 0 |
| 0 | 0 | 0 | 0 |
| 0 | 0 | 0 | 0 |
| 0 | 0 | 0 | 0 |
| 0 | 0 | 0 | 0 |
| 0 | 0 | 0 | 0 |
| 0 | 0 | 0 | 0 |
| 1 | 0 | 0 | 0 |
| 1 | 0 | 0 | 1 |
| 1 | 0 | 0 | 0 |
| 0 | 0 | 0 | 0 |
| 1 | 0 | 0 | 0 |
| 1 | 1 | 0 | 0 |
| 1 | 1 | 0 | 0 |
| 0 | 0 | 0 | 0 |
| 0 | 0 | 0 | 0 |
| 0 | 0 | 0 | 0 |
| 0 | 0 | 0 | 0 |
| 0 | 0 | 0 | 0 |
| 0 | 0 | 0 | 0 |
| 0 | 0 | 0 | 0 |
| 0 | 0 | 0 | 0 |
| 0 | 0 | 0 | 0 |
| 0 | 0 | 0 | 0 |
| 0 | 0 | 0 | 0 |
| 0 | 0 | 0 | 0 |

| | | | |
|---|---|---|---|
| 0 | 0 | 0 | 0 |
| 0 | 0 | 0 | 0 |
| 0 | 0 | 0 | 0 |
| 0 | 0 | 0 | 0 |
| 0 | 0 | 0 | 0 |
| 0 | 0 | 0 | 0 |
| 0 | 0 | 0 | 0 |
| 0 | 0 | 0 | 0 |
| 0 | 0 | 0 | 0 |
| 0 | 0 | 0 | 0 |
| 0 | 0 | 0 | 0 |
| 0 | 0 | 0 | 0 |
| 0 | 0 | 0 | 0 |
| 1 | 0 | 0 | 0 |
| 0 | 0 | 0 | 0 |
| 0 | 0 | 0 | 0 |
| 1 | 1 | 0 | 0 |
| 0 | 0 | 0 | 0 |
| 0 | 0 | 0 | 0 |
| 0 | 0 | 0 | 0 |
| 0 | 0 | 0 | 0 |
| 0 | 0 | 0 | 0 |
| 0 | 0 | 0 | 0 |
| 1 | 0 | 0 | 0 |
| 0 | 0 | 0 | 0 |
| 0 | 0 | 0 | 0 |
| 0 | 0 | 0 | 0 |
| 0 | 0 | 0 | 0 |
| 0 | 0 | 0 | 0 |
| 0 | 0 | 0 | 0 |
| 0 | 0 | 0 | 0 |
| 1 | 0 | 0 | 0 |
| 0 | 0 | 0 | 0 |
| 0 | 0 | 0 | 0 |
| 0 | 0 | 0 | 0 |
| 0 | 0 | 0 | 0 |
| 0 | 0 | 0 | 0 |
| 0 | 0 | 0 | 0 |
| 0 | 0 | 0 | 0 |
| 0 | 0 | 0 | 0 |
| 0 | 0 | 0 | 0 |
| 0 | 0 | 0 | 0 |
| 0 | 0 | 0 | 0 |
| 0 | 0 | 0 | 0 |
| 0 | 0 | 0 | 0 |
| 0 | 0 | 0 | 0 |
| 0 | 0 | 0 | 0 |
| 0 | 0 | 0 | 0 |
| 0 | 0 | 0 | 0 |
| 0 | 0 | 0 | 0 |
| 0 | 0 | 0 | 0 |
| 0 | 0 | 0 | 0 |
| 0 | 0 | 0 | 0 |
| 0 | 0 | 0 | 0 |

| | | | |
|---|---|---|---|
| 0 | 0 | 0 | 0 |
| 0 | 0 | 0 | 0 |
| 0 | 0 | 0 | 0 |
| 0 | 0 | 0 | 0 |
| 0 | 0 | 0 | 0 |
| 0 | 0 | 0 | 0 |
| 0 | 0 | 0 | 0 |
| 0 | 0 | 0 | 0 |
| 0 | 0 | 0 | 0 |
| 0 | 0 | 0 | 0 |
| 0 | 0 | 0 | 0 |
| 0 | 0 | 0 | 0 |
| 0 | 0 | 0 | 0 |
| 0 | 0 | 0 | 0 |
| 0 | 0 | 0 | 0 |
| 0 | 0 | 0 | 0 |
| 0 | 0 | 0 | 0 |
| 1 | 0 | 0 | 0 |
| 0 | 0 | 0 | 0 |
| 0 | 0 | 0 | 0 |
| 0 | 0 | 0 | 0 |
| 0 | 0 | 0 | 0 |
| 0 | 0 | 0 | 0 |
| 0 | 0 | 0 | 0 |
| 1 | 0 | 0 | 0 |
| 0 | 0 | 0 | 0 |
| 0 | 0 | 0 | 0 |
| 0 | 0 | 0 | 0 |
| 0 | 0 | 0 | 0 |
| 0 | 0 | 0 | 0 |
| 0 | 0 | 0 | 0 |
| 0 | 0 | 0 | 0 |
| 0 | 0 | 0 | 0 |
| 0 | 0 | 0 | 0 |
| 0 | 0 | 0 | 0 |
| 0 | 0 | 0 | 0 |
| 0 | 0 | 0 | 0 |
| 0 | 0 | 0 | 0 |
| 0 | 0 | 0 | 0 |
| 0 | 0 | 0 | 0 |
| 0 | 0 | 0 | 0 |
| 0 | 0 | 0 | 0 |
| 0 | 0 | 0 | 0 |
| 0 | 0 | 0 | 0 |
| 0 | 0 | 0 | 0 |
| 0 | 0 | 0 | 0 |
| 0 | 0 | 0 | 0 |
| 0 | 0 | 0 | 0 |
| 0 | 0 | 0 | 0 |
| 0 | 0 | 0 | 0 |
| 0 | 0 | 0 | 0 |
| 0 | 0 | 0 | 0 |
| 0 | 0 | 0 | 0 |
| 0 | 0 | 0 | 0 |

| | | | |
|---|---|---|---|
| 0 | 0 | 0 | 0 |
| 0 | 0 | 0 | 0 |
| 0 | 0 | 0 | 0 |
| 0 | 0 | 0 | 0 |
| 0 | 0 | 0 | 0 |
| 0 | 0 | 0 | 0 |
| 0 | 0 | 0 | 0 |
| 0 | 0 | 0 | 0 |
| 0 | 0 | 0 | 0 |
| 0 | 0 | 0 | 0 |
| 0 | 0 | 0 | 0 |
| 0 | 0 | 0 | 0 |
| 0 | 0 | 0 | 0 |
| 0 | 0 | 0 | 0 |
| 1 | 1 | 0 | 0 |
| 1 | 1 | 0 | 0 |
| 0 | 0 | 0 | 0 |
| 0 | 0 | 0 | 0 |
| 0 | 0 | 0 | 0 |
| 0 | 0 | 0 | 0 |
| 0 | 0 | 0 | 0 |
| 0 | 0 | 0 | 0 |
| 0 | 0 | 0 | 0 |
| 0 | 0 | 0 | 0 |
| 0 | 0 | 0 | 0 |
| 0 | 0 | 0 | 0 |
| 0 | 0 | 0 | 0 |
| 1 | 0 | 0 | 0 |
| 0 | 0 | 0 | 0 |
| 1 | 0 | 0 | 0 |
| 0 | 0 | 0 | 0 |
| 0 | 0 | 0 | 0 |
| 0 | 0 | 0 | 0 |
| 0 | 0 | 0 | 0 |
| 0 | 0 | 0 | 0 |
| 0 | 0 | 0 | 0 |
| 1 | 0 | 0 | 0 |
| 0 | 0 | 0 | 0 |
| 0 | 0 | 0 | 0 |
| 0 | 0 | 0 | 0 |
| 0 | 0 | 0 | 0 |
| 0 | 0 | 0 | 0 |
| 0 | 0 | 0 | 0 |
| 0 | 0 | 0 | 0 |
| 0 | 0 | 0 | 0 |
| 0 | 0 | 0 | 0 |
| 0 | 0 | 0 | 0 |
| 0 | 0 | 0 | 0 |
| 0 | 0 | 0 | 0 |
| 0 | 0 | 0 | 0 |
| 0 | 0 | 0 | 0 |
| 0 | 0 | 0 | 0 |
| 0 | 0 | 0 | 0 |
| 0 | 0 | 0 | 0 |

| | | | |
|---|---|---|---|
| 0 | 0 | 0 | 0 |
| 0 | 0 | 0 | 0 |
| 0 | 0 | 0 | 0 |
| 0 | 0 | 0 | 0 |
| 0 | 0 | 0 | 0 |
| 0 | 0 | 0 | 0 |
| 0 | 0 | 0 | 0 |
| 0 | 0 | 0 | 0 |
| 0 | 0 | 0 | 0 |
| 0 | 0 | 0 | 0 |
| 0 | 0 | 0 | 0 |
| 0 | 0 | 0 | 0 |
| 0 | 0 | 0 | 0 |
| 0 | 0 | 0 | 0 |
| 0 | 0 | 0 | 0 |
| 0 | 0 | 0 | 0 |
| 0 | 0 | 0 | 0 |
| 0 | 0 | 0 | 0 |
| 0 | 0 | 0 | 0 |
| 0 | 0 | 0 | 0 |
| 0 | 0 | 0 | 0 |
| 0 | 0 | 0 | 0 |
| 0 | 0 | 0 | 0 |
| 0 | 0 | 0 | 0 |
| 0 | 0 | 0 | 0 |
| 0 | 0 | 0 | 0 |
| 0 | 0 | 0 | 0 |
| 0 | 0 | 0 | 0 |
| 0 | 0 | 0 | 0 |
| 0 | 0 | 0 | 0 |
| 0 | 0 | 0 | 0 |
| 0 | 0 | 0 | 0 |
| 0 | 0 | 0 | 0 |
| 0 | 0 | 0 | 0 |
| 0 | 0 | 0 | 0 |
| 0 | 0 | 0 | 0 |
| 0 | 0 | 0 | 0 |
| 0 | 0 | 0 | 0 |
| 0 | 0 | 0 | 0 |
| 0 | 0 | 0 | 0 |
| 0 | 0 | 0 | 0 |
| 0 | 0 | 0 | 0 |
| 0 | 0 | 0 | 0 |
| 0 | 0 | 0 | 0 |
| 0 | 0 | 0 | 0 |
| 0 | 0 | 0 | 0 |
| 0 | 0 | 0 | 0 |
| 0 | 0 | 0 | 0 |
| 1 | 0 | 0 | 1 |
| 0 | 0 | 0 | 0 |
| 1 | 0 | 0 | 0 |
| 1 | 0 | 0 | 0 |
| 0 | 0 | 0 | 0 |
| 0 | 0 | 0 | 1 |

| | | | |
|---|---|---|---|
| 0 | 0 | 0 | 0 |
| 0 | 0 | 0 | 0 |
| 0 | 0 | 0 | 0 |
| 0 | 0 | 0 | 0 |
| 0 | 0 | 0 | 0 |
| 0 | 0 | 0 | 0 |
| 0 | 0 | 0 | 0 |
| 0 | 0 | 0 | 0 |
| 0 | 0 | 0 | 0 |
| 0 | 0 | 0 | 0 |
| 0 | 0 | 0 | 0 |
| 0 | 0 | 0 | 0 |
| 0 | 0 | 0 | 0 |
| 0 | 0 | 0 | 0 |
| 0 | 0 | 0 | 0 |
| 0 | 0 | 0 | 0 |
| 0 | 0 | 0 | 0 |
| 0 | 0 | 0 | 0 |
| 0 | 0 | 0 | 0 |
| 0 | 0 | 0 | 0 |
| 0 | 0 | 0 | 0 |
| 0 | 0 | 0 | 0 |
| 0 | 0 | 0 | 0 |
| 0 | 0 | 0 | 0 |
| 0 | 0 | 0 | 0 |
| 0 | 0 | 0 | 0 |
| 0 | 0 | 1 | 0 |
| 0 | 0 | 1 | 0 |
| 0 | 0 | 1 | 0 |
| 0 | 0 | 1 | 0 |
| 0 | 0 | 0 | 0 |
| 0 | 0 | 0 | 0 |
| 0 | 0 | 0 | 0 |
| 1 | 0 | 0 | 0 |
| 1 | 0 | 0 | 0 |
| 0 | 0 | 0 | 0 |
| 0 | 0 | 0 | 0 |
| 0 | 0 | 0 | 0 |
| 0 | 0 | 0 | 0 |
| 1 | 0 | 0 | 0 |
| 0 | 0 | 0 | 0 |
| 0 | 0 | 0 | 0 |
| 0 | 0 | 0 | 0 |
| 0 | 0 | 0 | 0 |
| 0 | 0 | 0 | 0 |
| 0 | 1 | 0 | 0 |
| 0 | 0 | 0 | 0 |
| 0 | 0 | 0 | 0 |
| 0 | 0 | 0 | 0 |
| 0 | 0 | 0 | 0 |
| 0 | 0 | 0 | 0 |
| 0 | 0 | 0 | 0 |
| 0 | 0 | 0 | 0 |
| 0 | 0 | 0 | 0 |

| | | | |
|---|---|---|---|
| 0 | 0 | 0 | 0 |
| 0 | 0 | 0 | 0 |
| 0 | 0 | 0 | 0 |
| 0 | 0 | 0 | 0 |
| 0 | 0 | 0 | 0 |
| 0 | 0 | 0 | 1 |
| 0 | 0 | 0 | 0 |
| 0 | 0 | 0 | 0 |
| 0 | 0 | 0 | 0 |
| 0 | 0 | 0 | 0 |
| 0 | 0 | 0 | 0 |
| 0 | 0 | 0 | 0 |
| 0 | 0 | 0 | 0 |
| 0 | 1 | 0 | 0 |
| 0 | 1 | 0 | 0 |
| 0 | 1 | 0 | 0 |
| 0 | 1 | 0 | 0 |
| 0 | 1 | 0 | 0 |
| 0 | 0 | 0 | 0 |
| 0 | 0 | 0 | 0 |
| 0 | 0 | 0 | 0 |
| 0 | 0 | 0 | 0 |
| 0 | 0 | 0 | 0 |
| 0 | 0 | 0 | 0 |
| 0 | 0 | 0 | 0 |
| 0 | 0 | 0 | 0 |
| 0 | 0 | 0 | 0 |
| 0 | 0 | 0 | 0 |
| 0 | 0 | 0 | 0 |
| 0 | 0 | 0 | 0 |
| 0 | 0 | 0 | 0 |
| 0 | 0 | 0 | 0 |
| 0 | 0 | 0 | 0 |
| 0 | 0 | 0 | 0 |
| 0 | 0 | 0 | 0 |
| 0 | 0 | 0 | 0 |
| 0 | 0 | 0 | 0 |
| 0 | 0 | 0 | 0 |
| 0 | 0 | 0 | 0 |
| 0 | 0 | 0 | 0 |
| 0 | 0 | 0 | 0 |
| 0 | 0 | 0 | 0 |
| 0 | 0 | 0 | 0 |
| 0 | 0 | 0 | 0 |
| 0 | 0 | 0 | 0 |
| 0 | 0 | 0 | 0 |
| 0 | 0 | 0 | 0 |
| 0 | 0 | 0 | 0 |
| 0 | 0 | 0 | 0 |
| 0 | 0 | 0 | 0 |
| 0 | 0 | 0 | 0 |
| 0 | 0 | 0 | 0 |
| 0 | 0 | 0 | 0 |
| 0 | 0 | 0 | 0 |

| | | | |
|---|---|---|---|
| 0 | 0 | 0 | 0 |
| 0 | 0 | 0 | 0 |
| 0 | 0 | 0 | 0 |
| 1 | 0 | 0 | 0 |
| 0 | 0 | 0 | 0 |
| 0 | 0 | 0 | 0 |
| 0 | 0 | 0 | 0 |
| 0 | 0 | 0 | 0 |
| 0 | 0 | 0 | 0 |
| 0 | 0 | 0 | 0 |
| 0 | 0 | 0 | 0 |
| 0 | 0 | 0 | 0 |
| 0 | 0 | 0 | 0 |
| 0 | 0 | 0 | 0 |
| 0 | 0 | 0 | 0 |
| 0 | 0 | 0 | 0 |
| 0 | 0 | 0 | 0 |
| 0 | 0 | 0 | 0 |
| 0 | 0 | 0 | 0 |
| 0 | 0 | 0 | 0 |
| 1 | 1 | 0 | 0 |
| 1 | 1 | 0 | 0 |
| 0 | 0 | 0 | 0 |
| 0 | 1 | 0 | 0 |
| 0 | 0 | 0 | 0 |
| 0 | 0 | 0 | 0 |
| 0 | 0 | 0 | 0 |
| 0 | 0 | 0 | 0 |
| 0 | 0 | 0 | 0 |
| 0 | 0 | 0 | 0 |
| 0 | 0 | 0 | 0 |
| 0 | 0 | 0 | 0 |
| 0 | 0 | 0 | 0 |
| 0 | 0 | 0 | 0 |
| 0 | 0 | 0 | 0 |
| 0 | 0 | 0 | 0 |
| 0 | 0 | 0 | 0 |
| 0 | 0 | 0 | 0 |
| 0 | 0 | 0 | 0 |
| 1 | 0 | 0 | 0 |
| 1 | 0 | 0 | 0 |
| 0 | 0 | 0 | 0 |
| 0 | 0 | 0 | 0 |
| 1 | 0 | 0 | 0 |
| 0 | 0 | 0 | 0 |
| 0 | 0 | 0 | 0 |
| 0 | 0 | 0 | 0 |
| 0 | 0 | 0 | 0 |
| 0 | 0 | 0 | 0 |
| 0 | 0 | 0 | 0 |
| 0 | 0 | 0 | 0 |
| 0 | 0 | 0 | 0 |
| 0 | 0 | 0 | 0 |
| 0 | 0 | 0 | 0 |

| | | | |
|---|---|---|---|
| 0 | 0 | 0 | 0 |
| 0 | 0 | 0 | 0 |
| 0 | 0 | 0 | 0 |
| 0 | 0 | 0 | 0 |
| 0 | 0 | 0 | 0 |
| 1 | 0 | 0 | 0 |
| 0 | 0 | 0 | 0 |
| 0 | 0 | 0 | 0 |
| 0 | 1 | 0 | 0 |
| 1 | 1 | 0 | 0 |
| 0 | 0 | 0 | 0 |
| 1 | 1 | 0 | 0 |
| 1 | 0 | 0 | 0 |
| 0 | 0 | 0 | 0 |
| 0 | 0 | 0 | 0 |
| 0 | 0 | 0 | 0 |
| 1 | 1 | 0 | 0 |
| 0 | 0 | 0 | 0 |
| 0 | 0 | 0 | 0 |
| 1 | 0 | 0 | 0 |
| 0 | 0 | 0 | 0 |
| 0 | 0 | 0 | 0 |
| 0 | 0 | 0 | 0 |
| 0 | 0 | 0 | 0 |
| 0 | 0 | 0 | 0 |
| 0 | 0 | 0 | 0 |
| 0 | 0 | 0 | 0 |
| 0 | 0 | 0 | 0 |
| 0 | 0 | 0 | 0 |
| 0 | 0 | 0 | 0 |
| 1 | 0 | 0 | 0 |
| 0 | 0 | 0 | 0 |
| 0 | 0 | 0 | 0 |
| 0 | 0 | 0 | 0 |
| 0 | 0 | 0 | 0 |
| 1 | 0 | 0 | 0 |
| 0 | 0 | 0 | 0 |
| 0 | 0 | 0 | 0 |
| 0 | 0 | 0 | 0 |
| 0 | 0 | 0 | 0 |
| 0 | 0 | 0 | 0 |
| 0 | 0 | 0 | 0 |
| 0 | 0 | 0 | 0 |
| 0 | 0 | 0 | 0 |
| 0 | 0 | 0 | 0 |
| 0 | 0 | 0 | 0 |
| 0 | 0 | 0 | 0 |
| 0 | 0 | 0 | 0 |
| 0 | 0 | 0 | 0 |
| 0 | 0 | 0 | 0 |
| 0 | 0 | 0 | 0 |
| 0 | 0 | 0 | 0 |
| 0 | 0 | 0 | 0 |
| 0 | 0 | 0 | 0 |

| | | | |
|---|---|---|---|
| 0 | 0 | 0 | 0 |
| 0 | 0 | 0 | 0 |
| 0 | 0 | 0 | 0 |
| 0 | 0 | 0 | 0 |
| 0 | 0 | 0 | 0 |
| 0 | 0 | 0 | 0 |
| 0 | 0 | 0 | 0 |
| 0 | 0 | 0 | 0 |
| 0 | 0 | 0 | 0 |
| 0 | 0 | 0 | 0 |
| 0 | 0 | 0 | 0 |
| 0 | 0 | 0 | 0 |
| 0 | 0 | 0 | 0 |
| 0 | 0 | 0 | 0 |
| 0 | 0 | 0 | 1 |
| 0 | 0 | 0 | 0 |
| 0 | 0 | 0 | 0 |
| 0 | 0 | 0 | 0 |
| 0 | 0 | 0 | 0 |
| 0 | 0 | 0 | 0 |
| 0 | 0 | 0 | 0 |
| 0 | 0 | 0 | 0 |
| 0 | 0 | 0 | 0 |
| 0 | 0 | 0 | 0 |
| 0 | 0 | 0 | 0 |
| 0 | 0 | 0 | 0 |
| 0 | 0 | 0 | 0 |
| 0 | 0 | 0 | 0 |
| 0 | 0 | 0 | 0 |
| 0 | 0 | 0 | 0 |
| 0 | 0 | 0 | 0 |
| 0 | 0 | 0 | 0 |
| 0 | 0 | 0 | 0 |
| 0 | 0 | 0 | 0 |
| 0 | 0 | 0 | 0 |
| 0 | 0 | 0 | 0 |
| 0 | 0 | 0 | 0 |
| 0 | 0 | 0 | 0 |
| 0 | 0 | 0 | 0 |
| 0 | 0 | 1 | 0 |
| 0 | 0 | 0 | 0 |
| 0 | 0 | 0 | 0 |
| 0 | 0 | 0 | 0 |
| 1 | 0 | 0 | 0 |
| 0 | 0 | 0 | 0 |
| 0 | 0 | 0 | 0 |
| 0 | 0 | 0 | 0 |
| 0 | 0 | 0 | 0 |
| 1 | 0 | 0 | 1 |
| 0 | 0 | 0 | 0 |
| 0 | 0 | 0 | 0 |
| 0 | 0 | 0 | 0 |
| 0 | 0 | 0 | 0 |
| 0 | 0 | 0 | 0 |

| | | | |
|---|---|---|---|
| 0 | 0 | 0 | 0 |
| 0 | 0 | 0 | 0 |
| 0 | 0 | 0 | 0 |
| 0 | 0 | 0 | 0 |
| 0 | 0 | 0 | 0 |
| 0 | 0 | 0 | 0 |
| 0 | 0 | 0 | 0 |
| 0 | 0 | 0 | 0 |
| 0 | 0 | 0 | 0 |
| 1 | 0 | 0 | 0 |
| 0 | 0 | 0 | 0 |
| 0 | 0 | 0 | 0 |
| 0 | 0 | 0 | 0 |
| 0 | 0 | 0 | 0 |
| 1 | 0 | 0 | 0 |
| 0 | 0 | 0 | 0 |
| 0 | 0 | 0 | 0 |
| 0 | 0 | 0 | 0 |
| 0 | 0 | 0 | 0 |
| 0 | 0 | 0 | 0 |
| 0 | 0 | 0 | 0 |
| 0 | 0 | 0 | 0 |
| 0 | 0 | 0 | 0 |
| 0 | 0 | 0 | 0 |
| 1 | 0 | 0 | 0 |
| 0 | 0 | 0 | 0 |
| 0 | 0 | 0 | 0 |
| 0 | 0 | 0 | 0 |
| 0 | 0 | 0 | 0 |
| 0 | 0 | 0 | 0 |
| 0 | 0 | 0 | 0 |
| 0 | 0 | 0 | 0 |
| 0 | 0 | 0 | 0 |
| 0 | 0 | 0 | 0 |
| 1 | 0 | 0 | 0 |
| 1 | 0 | 0 | 0 |
| 0 | 0 | 0 | 0 |
| 0 | 0 | 0 | 0 |
| 1 | 0 | 0 | 0 |
| 0 | 0 | 1 | 0 |
| 0 | 0 | 1 | 0 |
| 0 | 0 | 1 | 0 |
| 0 | 0 | 1 | 0 |
| 0 | 0 | 0 | 0 |
| 1 | 0 | 0 | 0 |
| 0 | 0 | 0 | 0 |
| 0 | 0 | 0 | 0 |
| 0 | 0 | 0 | 0 |
| 0 | 0 | 1 | 0 |
| 0 | 0 | 0 | 0 |
| 0 | 0 | 0 | 0 |
| 0 | 0 | 0 | 0 |
| 0 | 0 | 0 | 0 |
| 0 | 0 | 0 | 0 |

| | | | |
|---|---|---|---|
| 0 | 0 | 0 | 0 |
| 0 | 0 | 0 | 0 |
| 0 | 0 | 0 | 0 |
| 0 | 0 | 0 | 0 |
| 0 | 0 | 0 | 0 |
| 0 | 0 | 0 | 0 |
| 0 | 0 | 0 | 0 |
| 0 | 0 | 0 | 0 |
| 0 | 0 | 0 | 0 |
| 0 | 0 | 0 | 0 |
| 0 | 1 | 0 | 0 |
| 0 | 0 | 0 | 0 |
| 0 | 0 | 0 | 0 |
| 0 | 0 | 0 | 0 |
| 0 | 0 | 0 | 0 |
| 0 | 1 | 0 | 0 |
| 0 | 0 | 0 | 0 |
| 0 | 0 | 0 | 0 |
| 0 | 0 | 0 | 0 |
| 0 | 0 | 0 | 0 |
| 0 | 0 | 0 | 0 |
| 0 | 0 | 0 | 0 |
| 0 | 0 | 0 | 0 |
| 0 | 0 | 0 | 0 |
| 0 | 0 | 0 | 0 |
| 0 | 0 | 0 | 0 |
| 0 | 0 | 0 | 0 |
| 0 | 0 | 0 | 0 |
| 0 | 0 | 0 | 0 |
| 0 | 0 | 0 | 0 |
| 0 | 0 | 0 | 0 |
| 0 | 0 | 0 | 0 |
| 0 | 0 | 0 | 0 |
| 0 | 0 | 0 | 0 |
| 0 | 0 | 0 | 0 |
| 0 | 0 | 0 | 0 |
| 0 | 0 | 0 | 0 |
| 0 | 0 | 0 | 0 |
| 0 | 0 | 0 | 0 |
| 0 | 0 | 0 | 0 |
| 0 | 0 | 0 | 0 |
| 0 | 0 | 0 | 0 |
| 0 | 0 | 0 | 0 |
| 0 | 0 | 0 | 0 |
| 1 | 1 | 0 | 0 |
| 0 | 0 | 0 | 0 |
| 0 | 0 | 0 | 0 |
| 0 | 0 | 0 | 0 |
| 0 | 0 | 0 | 0 |
| 0 | 0 | 0 | 0 |
| 0 | 0 | 0 | 0 |
| 0 | 0 | 0 | 0 |
| 0 | 0 | 0 | 0 |
| 0 | 0 | 0 | 0 |

| | | | |
|---|---|---|---|
| 0 | 0 | 0 | 0 |
| 0 | 0 | 0 | 0 |
| 0 | 0 | 1 | 0 |
| 1 | 0 | 0 | 0 |
| 0 | 0 | 0 | 0 |
| 0 | 0 | 0 | 0 |
| 0 | 0 | 0 | 0 |
| 0 | 0 | 0 | 0 |
| 0 | 0 | 0 | 0 |
| 0 | 0 | 0 | 0 |
| 0 | 0 | 0 | 0 |
| 0 | 0 | 0 | 0 |
| 1 | 0 | 0 | 0 |
| 0 | 0 | 0 | 0 |
| 0 | 0 | 0 | 0 |
| 0 | 0 | 0 | 0 |
| 0 | 0 | 0 | 0 |
| 0 | 0 | 0 | 0 |
| 0 | 0 | 0 | 0 |
| 0 | 0 | 0 | 0 |
| 0 | 0 | 0 | 0 |
| 0 | 0 | 0 | 0 |
| 0 | 0 | 0 | 0 |
| 0 | 0 | 0 | 0 |
| 0 | 0 | 0 | 0 |
| 0 | 0 | 0 | 0 |
| 0 | 0 | 0 | 0 |
| 0 | 0 | 0 | 0 |
| 0 | 0 | 0 | 0 |
| 0 | 0 | 0 | 0 |
| 0 | 0 | 0 | 0 |
| 0 | 0 | 0 | 0 |
| 0 | 0 | 0 | 0 |
| 0 | 0 | 0 | 0 |
| 0 | 0 | 0 | 0 |
| 0 | 0 | 0 | 0 |
| 0 | 0 | 0 | 0 |
| 0 | 0 | 0 | 0 |
| 0 | 0 | 0 | 0 |
| 0 | 0 | 0 | 0 |
| 0 | 1 | 0 | 0 |
| 1 | 1 | 0 | 0 |
| 0 | 0 | 0 | 0 |
| 0 | 0 | 0 | 0 |
| 0 | 0 | 0 | 0 |
| 0 | 0 | 0 | 0 |
| 0 | 0 | 0 | 0 |
| 0 | 0 | 0 | 0 |
| 0 | 1 | 0 | 0 |
| 0 | 0 | 0 | 0 |
| 0 | 0 | 0 | 0 |
| 0 | 0 | 0 | 0 |
| 0 | 0 | 0 | 1 |
| 0 | 0 | 0 | 0 |

| | | | |
|---|---|---|---|
| 0 | 0 | 0 | 1 |
| 0 | 0 | 0 | 1 |
| 0 | 0 | 0 | 1 |
| 0 | 0 | 0 | 0 |
| 0 | 0 | 0 | 0 |
| 0 | 0 | 0 | 0 |
| 0 | 0 | 0 | 0 |
| 1 | 0 | 0 | 0 |
| 0 | 0 | 0 | 0 |
| 1 | 1 | 0 | 0 |
| 0 | 1 | 0 | 0 |
| 0 | 0 | 0 | 0 |
| 1 | 1 | 0 | 0 |
| 0 | 0 | 0 | 0 |
| 0 | 0 | 0 | 0 |
| 0 | 0 | 0 | 0 |
| 0 | 0 | 0 | 0 |
| 1 | 0 | 0 | 0 |
| 0 | 0 | 0 | 0 |
| 0 | 0 | 0 | 0 |
| 0 | 0 | 0 | 0 |
| 0 | 0 | 0 | 0 |
| 0 | 0 | 0 | 0 |
| 0 | 0 | 0 | 0 |
| 0 | 0 | 0 | 0 |
| 0 | 0 | 0 | 0 |
| 0 | 0 | 0 | 0 |
| 0 | 0 | 0 | 0 |
| 0 | 0 | 0 | 0 |
| 0 | 0 | 0 | 0 |
| 0 | 0 | 0 | 0 |
| 1 | 0 | 0 | 0 |
| 0 | 0 | 0 | 0 |
| 0 | 0 | 0 | 0 |
| 0 | 0 | 0 | 0 |
| 0 | 0 | 0 | 0 |
| 0 | 0 | 0 | 0 |
| 0 | 0 | 0 | 0 |
| 0 | 0 | 0 | 0 |
| 0 | 0 | 0 | 0 |
| 0 | 0 | 0 | 0 |
| 1 | 1 | 0 | 0 |
| 0 | 0 | 0 | 0 |
| 0 | 0 | 0 | 0 |
| 1 | 0 | 0 | 0 |
| 0 | 1 | 0 | 0 |
| 0 | 0 | 0 | 0 |
| 0 | 0 | 0 | 0 |
| 0 | 1 | 0 | 0 |
| 1 | 1 | 0 | 0 |
| 1 | 1 | 0 | 0 |
| 1 | 1 | 0 | 0 |
| 1 | 1 | 0 | 0 |
| 1 | 1 | 0 | 0 |

| | | | |
|---|---|---|---|
| 0 | 0 | 0 | 0 |
| 0 | 0 | 0 | 0 |
| 0 | 0 | 0 | 0 |
| 0 | 0 | 0 | 0 |
| 0 | 0 | 0 | 0 |
| 1 | 0 | 0 | 0 |
| 1 | 0 | 0 | 0 |
| 0 | 0 | 0 | 0 |
| 0 | 0 | 0 | 0 |
| 1 | 0 | 0 | 0 |
| 0 | 0 | 0 | 0 |
| 0 | 0 | 1 | 0 |
| 0 | 0 | 0 | 0 |
| 0 | 0 | 0 | 0 |
| 0 | 0 | 0 | 0 |
| 0 | 0 | 0 | 0 |
| 0 | 0 | 0 | 0 |
| 0 | 0 | 0 | 0 |
| 0 | 0 | 0 | 0 |
| 0 | 0 | 0 | 0 |
| 1 | 1 | 0 | 0 |
| 1 | 0 | 0 | 0 |
| 0 | 0 | 0 | 0 |
| 0 | 0 | 1 | 0 |
| 0 | 0 | 0 | 0 |
| 0 | 0 | 1 | 0 |
| 0 | 0 | 0 | 0 |
| 0 | 0 | 0 | 0 |
| 0 | 0 | 0 | 0 |
| 0 | 0 | 0 | 0 |
| 0 | 0 | 0 | 0 |
| 0 | 0 | 0 | 0 |
| 0 | 0 | 0 | 0 |
| 0 | 0 | 0 | 0 |
| 0 | 0 | 0 | 0 |
| 0 | 0 | 0 | 0 |
| 1 | 0 | 0 | 0 |
| 0 | 0 | 0 | 0 |
| 1 | 0 | 0 | 0 |
| 1 | 0 | 0 | 0 |
| 1 | 0 | 0 | 0 |
| 0 | 0 | 0 | 0 |
| 0 | 0 | 0 | 0 |
| 0 | 0 | 0 | 0 |
| 0 | 0 | 0 | 0 |
| 0 | 0 | 0 | 0 |
| 0 | 0 | 0 | 0 |
| 1 | 0 | 1 | 0 |
| 0 | 0 | 0 | 0 |
| 0 | 0 | 0 | 0 |
| 0 | 0 | 0 | 0 |
| 0 | 0 | 0 | 0 |
| 1 | 1 | 0 | 0 |
| 0 | 0 | 0 | 0 |

| | | | |
|---|---|---|---|
| 0 | 0 | 0 | 0 |
| 0 | 1 | 0 | 0 |
| 0 | 0 | 0 | 0 |
| 0 | 0 | 0 | 0 |
| 0 | 0 | 0 | 0 |
| 0 | 0 | 0 | 0 |
| 0 | 0 | 0 | 0 |
| 0 | 0 | 0 | 0 |
| 0 | 0 | 0 | 0 |
| 0 | 0 | 0 | 0 |
| 0 | 0 | 0 | 0 |
| 0 | 0 | 0 | 0 |
| 0 | 0 | 0 | 0 |
| 0 | 0 | 0 | 0 |
| 0 | 0 | 0 | 0 |
| 0 | 0 | 0 | 0 |
| 0 | 0 | 0 | 0 |
| 0 | 0 | 0 | 0 |
| 0 | 0 | 0 | 0 |
| 0 | 0 | 0 | 0 |
| 1 | 0 | 0 | 0 |
| 0 | 0 | 0 | 0 |
| 0 | 0 | 0 | 0 |
| 0 | 0 | 0 | 0 |
| 1 | 0 | 0 | 0 |
| 0 | 0 | 0 | 0 |
| 0 | 0 | 0 | 0 |
| 0 | 0 | 0 | 1 |
| 0 | 0 | 0 | 0 |
| 0 | 0 | 0 | 0 |
| 0 | 0 | 0 | 0 |
| 0 | 0 | 0 | 0 |
| 0 | 0 | 0 | 0 |
| 0 | 0 | 0 | 0 |
| 0 | 0 | 0 | 0 |
| 0 | 0 | 0 | 0 |
| 0 | 0 | 0 | 0 |
| 0 | 0 | 0 | 0 |
| 0 | 0 | 0 | 0 |
| 0 | 0 | 0 | 0 |
| 0 | 0 | 0 | 0 |
| 0 | 0 | 0 | 0 |
| 0 | 0 | 0 | 0 |
| 0 | 0 | 0 | 0 |
| 0 | 0 | 0 | 0 |
| 0 | 0 | 0 | 0 |
| 0 | 0 | 0 | 0 |
| 0 | 0 | 0 | 0 |
| 0 | 0 | 0 | 0 |
| 0 | 0 | 0 | 0 |
| 0 | 0 | 0 | 0 |
| 0 | 0 | 0 | 0 |
| 0 | 0 | 0 | 0 |
| 0 | 0 | 0 | 0 |

| | | | |
|---|---|---|---|
| 0 | 0 | 0 | 0 |
| 0 | 0 | 0 | 0 |
| 0 | 0 | 0 | 0 |
| 0 | 0 | 0 | 0 |
| 0 | 0 | 0 | 0 |
| 0 | 0 | 0 | 0 |
| 0 | 0 | 0 | 0 |
| 0 | 0 | 0 | 0 |
| 0 | 0 | 0 | 0 |
| 0 | 0 | 0 | 0 |
| 0 | 0 | 0 | 0 |
| 0 | 0 | 0 | 0 |
| 0 | 0 | 0 | 0 |
| 0 | 0 | 0 | 0 |
| 0 | 0 | 0 | 0 |
| 1 | 0 | 0 | 0 |
| 0 | 0 | 0 | 0 |
| 0 | 0 | 0 | 0 |
| 0 | 1 | 0 | 0 |
| 0 | 0 | 0 | 0 |
| 0 | 0 | 0 | 0 |
| 0 | 0 | 0 | 0 |
| 0 | 0 | 0 | 0 |
| 0 | 0 | 0 | 0 |
| 0 | 0 | 0 | 0 |
| 1 | 0 | 0 | 0 |
| 0 | 0 | 0 | 0 |
| 0 | 0 | 0 | 0 |
| 0 | 0 | 0 | 0 |
| 0 | 0 | 0 | 0 |
| 0 | 0 | 0 | 0 |
| 0 | 0 | 0 | 0 |
| 0 | 0 | 0 | 0 |
| 1 | 1 | 0 | 0 |
| 0 | 0 | 0 | 0 |
| 0 | 0 | 0 | 0 |
| 0 | 0 | 0 | 0 |
| 0 | 0 | 0 | 0 |
| 0 | 0 | 0 | 0 |
| 0 | 0 | 0 | 0 |
| 0 | 0 | 0 | 0 |
| 0 | 0 | 0 | 0 |
| 0 | 0 | 0 | 0 |
| 0 | 0 | 0 | 0 |
| 0 | 0 | 0 | 0 |
| 0 | 0 | 0 | 0 |
| 0 | 0 | 0 | 0 |
| 1 | 0 | 0 | 0 |
| 0 | 0 | 0 | 0 |
| 0 | 0 | 0 | 0 |
| 0 | 0 | 0 | 0 |
| 0 | 0 | 0 | 0 |
| 0 | 0 | 0 | 0 |
| 1 | 0 | 0 | 1 |

| | | | |
|---|---|---|---|
| 0 | 0 | 0 | 0 |
| 0 | 0 | 0 | 0 |
| 0 | 0 | 1 | 0 |
| 0 | 0 | 0 | 0 |
| 0 | 0 | 0 | 0 |
| 0 | 0 | 0 | 0 |
| 0 | 0 | 0 | 0 |
| 0 | 0 | 0 | 0 |
| 0 | 0 | 0 | 0 |
| 0 | 0 | 0 | 0 |
| 0 | 0 | 0 | 0 |
| 0 | 0 | 0 | 0 |
| 0 | 0 | 0 | 0 |
| 0 | 0 | 0 | 0 |
| 0 | 0 | 0 | 0 |
| 0 | 0 | 0 | 0 |
| 0 | 0 | 0 | 0 |
| 0 | 0 | 0 | 0 |
| 0 | 0 | 0 | 0 |
| 0 | 0 | 0 | 0 |
| 0 | 0 | 0 | 0 |
| 0 | 0 | 0 | 0 |
| 0 | 0 | 0 | 0 |
| 0 | 0 | 0 | 0 |
| 0 | 0 | 0 | 0 |
| 0 | 0 | 0 | 0 |
| 0 | 0 | 0 | 0 |
| 0 | 0 | 0 | 0 |
| 0 | 0 | 0 | 0 |
| 0 | 0 | 0 | 0 |
| 0 | 0 | 0 | 0 |
| 1 | 0 | 0 | 0 |
| 0 | 0 | 0 | 0 |
| 0 | 0 | 0 | 0 |
| 0 | 0 | 0 | 0 |
| 0 | 0 | 0 | 0 |
| 0 | 0 | 0 | 0 |
| 0 | 0 | 0 | 0 |
| 0 | 0 | 0 | 0 |
| 0 | 0 | 0 | 0 |
| 1 | 1 | 0 | 0 |
| 0 | 0 | 0 | 0 |
| 0 | 0 | 0 | 0 |
| 0 | 0 | 0 | 0 |
| 0 | 0 | 0 | 0 |
| 1 | 1 | 0 | 0 |
| 0 | 0 | 0 | 0 |
| 1 | 1 | 0 | 0 |
| 0 | 0 | 0 | 0 |
| 1 | 0 | 0 | 0 |
| 1 | 0 | 0 | 0 |
| 1 | 0 | 0 | 0 |
| 0 | 0 | 0 | 0 |
| 0 | 0 | 0 | 0 |

| | | | |
|---|---|---|---|
| 1 | 0 | 0 | 0 |
| 0 | 0 | 0 | 0 |
| 0 | 0 | 0 | 0 |
| 0 | 0 | 0 | 0 |
| 0 | 0 | 0 | 0 |
| 0 | 0 | 0 | 0 |
| 0 | 0 | 0 | 0 |
| 0 | 0 | 0 | 0 |
| 0 | 0 | 0 | 0 |
| 0 | 0 | 0 | 0 |
| 0 | 0 | 0 | 0 |
| 0 | 0 | 0 | 0 |
| 0 | 0 | 0 | 0 |
| 0 | 0 | 0 | 0 |
| 0 | 0 | 0 | 0 |
| 1 | 1 | 1 | 0 |
| 0 | 0 | 0 | 0 |
| 1 | 0 | 0 | 0 |
| 0 | 0 | 0 | 0 |
| 0 | 0 | 0 | 0 |
| 0 | 0 | 0 | 0 |
| 0 | 1 | 0 | 0 |
| 0 | 0 | 0 | 0 |
| 0 | 1 | 0 | 0 |
| 0 | 1 | 0 | 0 |
| 0 | 0 | 0 | 0 |
| 0 | 0 | 0 | 0 |
| 0 | 0 | 0 | 0 |
| 0 | 0 | 0 | 0 |
| 0 | 0 | 0 | 0 |
| 0 | 0 | 0 | 0 |
| 0 | 0 | 0 | 0 |
| 1 | 1 | 0 | 0 |
| 0 | 0 | 0 | 0 |
| 0 | 0 | 0 | 0 |
| 0 | 0 | 0 | 0 |
| 0 | 1 | 0 | 0 |
| 1 | 1 | 0 | 0 |
| 0 | 0 | 0 | 0 |
| 0 | 0 | 0 | 0 |
| 0 | 0 | 0 | 0 |
| 0 | 0 | 0 | 0 |
| 0 | 0 | 0 | 0 |
| 0 | 0 | 0 | 0 |
| 1 | 1 | 0 | 0 |
| 0 | 0 | 0 | 0 |
| 1 | 0 | 0 | 0 |
| 1 | 1 | 0 | 0 |
| 0 | 0 | 0 | 0 |
| 0 | 0 | 0 | 0 |
| 1 | 1 | 0 | 0 |
| 1 | 1 | 0 | 0 |
| 0 | 0 | 0 | 0 |
| 1 | 1 | 0 | 0 |

| | | | |
|---|---|---|---|
| 0 | 0 | 0 | 0 |
| 0 | 0 | 0 | 0 |
| 0 | 0 | 0 | 0 |
| 1 | 1 | 0 | 0 |
| 0 | 0 | 0 | 0 |
| 0 | 0 | 0 | 0 |
| 0 | 0 | 0 | 0 |
| 0 | 0 | 0 | 0 |
| 1 | 0 | 0 | 0 |
| 0 | 0 | 0 | 0 |
| 0 | 0 | 0 | 0 |
| 0 | 0 | 0 | 0 |
| 1 | 0 | 1 | 0 |
| 1 | 0 | 0 | 0 |
| 0 | 0 | 0 | 0 |
| 0 | 0 | 0 | 0 |
| 0 | 0 | 0 | 0 |
| 0 | 0 | 0 | 0 |
| 0 | 0 | 0 | 0 |
| 0 | 0 | 0 | 0 |
| 0 | 0 | 0 | 0 |
| 0 | 0 | 0 | 0 |
| 0 | 0 | 0 | 0 |
| 1 | 0 | 0 | 0 |
| 0 | 0 | 0 | 0 |
| 0 | 0 | 0 | 0 |
| 0 | 0 | 0 | 0 |
| 0 | 0 | 0 | 0 |
| 0 | 0 | 0 | 0 |
| 0 | 0 | 0 | 0 |
| 0 | 0 | 0 | 0 |
| 0 | 0 | 0 | 0 |
| 0 | 0 | 0 | 0 |
| 0 | 0 | 0 | 0 |
| 0 | 0 | 0 | 0 |
| 0 | 0 | 0 | 0 |
| 0 | 0 | 0 | 0 |
| 0 | 0 | 0 | 0 |
| 0 | 0 | 0 | 0 |
| 0 | 0 | 0 | 0 |
| 0 | 0 | 0 | 0 |
| 0 | 0 | 0 | 0 |
| 0 | 0 | 0 | 0 |
| 1 | 1 | 0 | 0 |
| 1 | 1 | 0 | 0 |
| 0 | 0 | 0 | 0 |
| 0 | 0 | 0 | 0 |
| 0 | 1 | 0 | 0 |
| 0 | 1 | 0 | 0 |
| 0 | 1 | 0 | 0 |
| 0 | 0 | 0 | 0 |
| 0 | 0 | 0 | 0 |
| 0 | 0 | 0 | 0 |
| 0 | 0 | 0 | 0 |

| | | | |
|---|---|---|---|
| 0 | 0 | 0 | 0 |
| 0 | 0 | 0 | 0 |
| 0 | 0 | 0 | 0 |
| 0 | 0 | 0 | 0 |
| 1 | 0 | 0 | 0 |
| 0 | 0 | 0 | 0 |
| 0 | 0 | 0 | 0 |
| 1 | 0 | 0 | 0 |
| 1 | 0 | 0 | 0 |
| 1 | 0 | 0 | 0 |
| 1 | 0 | 0 | 0 |
| 1 | 0 | 0 | 0 |
| 0 | 0 | 0 | 0 |
| 1 | 0 | 0 | 0 |
| 1 | 0 | 0 | 0 |
| 0 | 0 | 0 | 0 |
| 0 | 0 | 0 | 0 |
| 0 | 0 | 0 | 0 |
| 1 | 0 | 0 | 0 |
| 1 | 0 | 0 | 0 |
| 1 | 0 | 0 | 0 |
| 1 | 0 | 0 | 0 |
| 0 | 0 | 0 | 0 |
| 0 | 0 | 0 | 0 |
| 0 | 0 | 0 | 0 |
| 0 | 0 | 0 | 0 |
| 0 | 0 | 0 | 0 |
| 0 | 0 | 0 | 0 |
| 0 | 0 | 0 | 0 |
| 0 | 0 | 0 | 0 |
| 0 | 0 | 0 | 0 |
| 0 | 0 | 0 | 0 |
| 1 | 0 | 0 | 0 |
| 0 | 0 | 0 | 0 |
| 0 | 0 | 0 | 0 |
| 1 | 0 | 0 | 0 |
| 0 | 0 | 0 | 0 |
| 0 | 0 | 0 | 0 |
| 0 | 0 | 0 | 0 |
| 0 | 0 | 0 | 0 |
| 0 | 0 | 0 | 0 |
| 0 | 0 | 0 | 0 |
| 0 | 0 | 0 | 0 |
| 0 | 0 | 0 | 0 |
| 0 | 0 | 0 | 0 |
| 0 | 0 | 0 | 0 |
| 0 | 0 | 0 | 0 |
| 0 | 0 | 0 | 0 |
| 0 | 0 | 0 | 0 |
| 0 | 0 | 0 | 0 |
| 0 | 0 | 0 | 0 |
| 0 | 0 | 0 | 0 |
| 0 | 0 | 0 | 0 |
| 0 | 0 | 0 | 0 |

| | | | |
|---|---|---|---|
| 0 | 0 | 0 | 0 |
| 0 | 0 | 0 | 0 |
| 0 | 0 | 0 | 0 |
| 0 | 0 | 0 | 0 |
| 0 | 0 | 0 | 0 |
| 0 | 0 | 0 | 0 |
| 0 | 0 | 0 | 0 |
| 0 | 0 | 0 | 0 |
| 0 | 0 | 0 | 0 |
| 0 | 0 | 0 | 0 |
| 0 | 0 | 0 | 0 |
| 0 | 0 | 0 | 1 |
| 0 | 0 | 0 | 0 |
| 0 | 0 | 0 | 0 |
| 0 | 0 | 0 | 0 |
| 0 | 0 | 0 | 0 |
| 0 | 0 | 0 | 0 |
| 0 | 0 | 0 | 1 |
| 0 | 0 | 0 | 0 |
| 0 | 0 | 0 | 0 |
| 0 | 0 | 0 | 1 |
| 0 | 0 | 0 | 0 |
| 0 | 0 | 0 | 0 |
| 0 | 0 | 0 | 0 |
| 0 | 0 | 0 | 0 |
| 0 | 0 | 0 | 0 |
| 0 | 0 | 0 | 0 |
| 0 | 0 | 0 | 0 |
| 0 | 0 | 0 | 1 |
| 0 | 0 | 0 | 0 |
| 0 | 0 | 0 | 0 |
| 0 | 0 | 0 | 0 |
| 0 | 0 | 0 | 0 |
| 0 | 0 | 0 | 0 |
| 0 | 0 | 0 | 0 |
| 0 | 0 | 0 | 0 |
| 0 | 0 | 0 | 0 |
| 0 | 0 | 0 | 0 |
| 0 | 0 | 0 | 0 |
| 0 | 0 | 0 | 0 |
| 0 | 0 | 0 | 0 |
| 0 | 0 | 0 | 0 |
| 0 | 0 | 0 | 0 |
| 0 | 0 | 0 | 0 |
| 0 | 0 | 0 | 0 |
| 0 | 0 | 0 | 0 |
| 0 | 0 | 0 | 0 |
| 0 | 0 | 0 | 0 |
| 0 | 0 | 0 | 0 |
| 0 | 0 | 0 | 0 |
| 0 | 0 | 0 | 0 |
| 0 | 0 | 0 | 0 |
| 0 | 0 | 0 | 0 |
| 0 | 0 | 0 | 0 |

| | | | |
|---|---|---|---|
| 0 | 0 | 0 | 0 |
| 0 | 0 | 0 | 0 |
| 1 | 0 | 0 | 0 |
| 0 | 0 | 0 | 0 |
| 1 | 0 | 0 | 0 |
| 0 | 0 | 0 | 0 |
| 0 | 0 | 0 | 0 |
| 0 | 0 | 0 | 0 |
| 0 | 0 | 0 | 0 |
| 1 | 1 | 0 | 0 |
| 0 | 0 | 1 | 0 |
| 0 | 0 | 0 | 0 |
| 0 | 0 | 0 | 0 |
| 0 | 0 | 0 | 0 |
| 0 | 0 | 0 | 0 |
| 0 | 0 | 0 | 0 |
| 0 | 0 | 0 | 0 |
| 0 | 0 | 0 | 0 |
| 0 | 0 | 0 | 0 |
| 0 | 0 | 0 | 0 |
| 0 | 0 | 0 | 0 |
| 0 | 0 | 0 | 0 |
| 0 | 0 | 0 | 0 |
| 0 | 0 | 0 | 0 |
| 0 | 0 | 0 | 0 |
| 0 | 0 | 0 | 0 |
| 0 | 0 | 0 | 0 |
| 0 | 0 | 0 | 0 |
| 0 | 0 | 0 | 0 |
| 0 | 0 | 0 | 0 |
| 0 | 0 | 0 | 0 |
| 0 | 0 | 0 | 0 |
| 0 | 0 | 0 | 0 |
| 0 | 0 | 0 | 0 |
| 0 | 0 | 0 | 0 |
| 0 | 0 | 0 | 0 |
| 0 | 0 | 0 | 0 |
| 0 | 0 | 0 | 0 |
| 0 | 0 | 0 | 0 |
| 0 | 0 | 0 | 0 |
| 0 | 0 | 0 | 0 |
| 0 | 0 | 0 | 0 |
| 0 | 0 | 0 | 0 |
| 0 | 0 | 0 | 0 |
| 0 | 0 | 0 | 0 |
| 1 | 0 | 0 | 0 |
| 0 | 0 | 0 | 0 |
| 0 | 0 | 0 | 0 |
| 0 | 0 | 0 | 0 |
| 0 | 0 | 0 | 0 |
| 1 | 0 | 0 | 0 |
| 0 | 0 | 0 | 0 |
| 0 | 0 | 0 | 0 |
| 0 | 0 | 0 | 0 |

| | | | |
|---|---|---|---|
| 0 | 0 | 0 | 0 |
| 0 | 0 | 0 | 0 |
| 0 | 0 | 0 | 0 |
| 0 | 0 | 0 | 0 |
| 0 | 0 | 0 | 0 |
| 0 | 0 | 0 | 0 |
| 0 | 0 | 0 | 0 |
| 0 | 0 | 0 | 0 |
| 1 | 1 | 0 | 0 |
| 0 | 0 | 0 | 0 |
| 0 | 0 | 0 | 0 |
| 0 | 0 | 0 | 0 |
| 0 | 0 | 0 | 0 |
| 0 | 0 | 0 | 0 |
| 0 | 1 | 0 | 0 |
| 0 | 1 | 0 | 0 |
| 0 | 1 | 0 | 0 |
| 0 | 0 | 0 | 0 |
| 0 | 0 | 0 | 0 |
| 0 | 0 | 0 | 0 |
| 0 | 1 | 0 | 0 |
| 0 | 0 | 0 | 0 |
| 0 | 0 | 0 | 0 |
| 1 | 1 | 0 | 0 |
| 0 | 1 | 0 | 0 |
| 0 | 1 | 0 | 0 |
| 0 | 0 | 0 | 0 |
| 0 | 0 | 0 | 0 |
| 0 | 0 | 0 | 0 |
| 0 | 0 | 0 | 0 |
| 0 | 0 | 0 | 0 |
| 0 | 0 | 0 | 0 |
| 0 | 0 | 0 | 0 |
| 0 | 0 | 0 | 0 |
| 0 | 0 | 0 | 0 |
| 0 | 0 | 0 | 0 |
| 0 | 0 | 0 | 0 |
| 1 | 0 | 0 | 0 |
| 0 | 0 | 0 | 0 |
| 0 | 0 | 0 | 0 |
| 0 | 0 | 0 | 0 |
| 0 | 0 | 0 | 0 |
| 0 | 0 | 0 | 0 |
| 0 | 0 | 0 | 0 |
| 0 | 0 | 0 | 0 |
| 0 | 0 | 0 | 0 |
| 0 | 0 | 0 | 0 |
| 0 | 0 | 0 | 0 |
| 0 | 0 | 0 | 0 |
| 0 | 0 | 0 | 0 |
| 0 | 0 | 0 | 0 |
| 0 | 0 | 0 | 0 |
| 0 | 0 | 0 | 0 |
| 0 | 0 | 0 | 0 |

| | | | |
|---|---|---|---|
| 0 | 0 | 0 | 0 |
| 0 | 0 | 0 | 0 |
| 0 | 0 | 0 | 0 |
| 0 | 0 | 1 | 0 |
| 0 | 0 | 0 | 0 |
| 0 | 0 | 0 | 0 |
| 0 | 0 | 0 | 0 |
| 0 | 0 | 0 | 0 |
| 0 | 0 | 0 | 0 |
| 0 | 0 | 0 | 0 |
| 0 | 0 | 0 | 0 |
| 0 | 0 | 0 | 0 |
| 1 | 0 | 0 | 0 |
| 0 | 0 | 0 | 0 |
| 0 | 0 | 0 | 0 |
| 0 | 0 | 0 | 0 |
| 0 | 1 | 0 | 0 |
| 0 | 0 | 0 | 0 |
| 0 | 0 | 0 | 0 |
| 0 | 0 | 0 | 0 |
| 0 | 0 | 0 | 0 |
| 0 | 0 | 0 | 0 |
| 0 | 0 | 0 | 0 |
| 0 | 0 | 0 | 0 |
| 0 | 0 | 0 | 0 |
| 0 | 0 | 0 | 0 |
| 0 | 0 | 0 | 0 |
| 0 | 0 | 0 | 0 |
| 0 | 0 | 0 | 0 |
| 0 | 0 | 0 | 0 |
| 1 | 0 | 0 | 0 |
| 1 | 0 | 0 | 0 |
| 1 | 0 | 0 | 0 |
| 0 | 0 | 0 | 0 |
| 1 | 0 | 0 | 0 |
| 1 | 0 | 0 | 0 |
| 1 | 0 | 0 | 0 |
| 0 | 0 | 0 | 0 |
| 0 | 0 | 0 | 0 |
| 0 | 0 | 0 | 0 |
| 0 | 0 | 0 | 0 |
| 0 | 0 | 0 | 0 |
| 0 | 1 | 0 | 0 |
| 0 | 1 | 0 | 0 |
| 0 | 0 | 0 | 0 |
| 0 | 0 | 0 | 0 |
| 1 | 0 | 0 | 0 |
| 0 | 0 | 0 | 0 |
| 0 | 0 | 0 | 0 |
| 0 | 0 | 0 | 0 |
| 0 | 0 | 0 | 0 |
| 0 | 0 | 0 | 0 |
| 1 | 0 | 0 | 0 |
| 0 | 0 | 0 | 0 |

| | | | |
|---|---|---|---|
| 0 | 0 | 0 | 0 |
| 1 | 0 | 0 | 0 |
| 1 | 0 | 0 | 0 |
| 0 | 0 | 0 | 0 |
| 0 | 0 | 0 | 0 |
| 0 | 0 | 0 | 0 |
| 0 | 0 | 0 | 0 |
| 0 | 1 | 0 | 0 |
| 0 | 0 | 0 | 0 |
| 0 | 0 | 0 | 0 |
| 0 | 0 | 0 | 0 |
| 0 | 0 | 0 | 0 |
| 0 | 0 | 0 | 0 |
| 0 | 0 | 0 | 0 |
| 1 | 0 | 0 | 0 |
| 0 | 0 | 0 | 0 |
| 0 | 0 | 0 | 0 |
| 0 | 0 | 0 | 0 |
| 0 | 0 | 0 | 0 |
| 1 | 0 | 0 | 0 |
| 1 | 0 | 0 | 0 |
| 0 | 0 | 0 | 0 |
| 1 | 0 | 0 | 0 |
| 1 | 0 | 0 | 0 |
| 0 | 0 | 0 | 0 |
| 0 | 0 | 0 | 0 |
| 0 | 0 | 0 | 0 |
| 0 | 0 | 0 | 0 |
| 0 | 0 | 0 | 0 |
| 0 | 0 | 0 | 0 |
| 0 | 0 | 0 | 0 |
| 0 | 0 | 0 | 0 |
| 0 | 0 | 0 | 0 |
| 0 | 0 | 0 | 0 |
| 0 | 0 | 0 | 0 |
| 0 | 0 | 0 | 0 |
| 0 | 0 | 0 | 0 |
| 0 | 0 | 0 | 0 |
| 0 | 0 | 0 | 0 |
| 0 | 0 | 0 | 0 |
| 0 | 0 | 0 | 0 |
| 0 | 0 | 0 | 0 |
| 0 | 0 | 0 | 0 |
| 0 | 0 | 0 | 0 |
| 0 | 0 | 0 | 0 |
| 0 | 0 | 0 | 0 |
| 0 | 0 | 0 | 0 |
| 0 | 0 | 0 | 0 |
| 0 | 0 | 0 | 0 |
| 0 | 0 | 0 | 0 |
| 0 | 0 | 0 | 0 |
| 0 | 0 | 0 | 0 |
| 0 | 0 | 0 | 0 |
| 1 | 1 | 0 | 1 |

| | | | |
|---|---|---|---|
| 0 | 0 | 0 | 0 |
| 0 | 0 | 0 | 0 |
| 0 | 0 | 0 | 0 |
| 0 | 0 | 0 | 0 |
| 0 | 1 | 0 | 0 |
| 0 | 0 | 0 | 0 |
| 0 | 0 | 0 | 0 |
| 0 | 1 | 0 | 0 |
| 0 | 0 | 0 | 0 |
| 0 | 0 | 0 | 0 |
| 0 | 0 | 0 | 0 |
| 0 | 0 | 0 | 0 |
| 0 | 0 | 0 | 0 |
| 0 | 0 | 0 | 0 |
| 0 | 1 | 0 | 1 |
| 0 | 1 | 0 | 0 |
| 0 | 0 | 0 | 0 |
| 0 | 0 | 0 | 0 |
| 0 | 1 | 0 | 0 |
| 0 | 1 | 0 | 1 |
| 0 | 0 | 0 | 0 |
| 0 | 0 | 0 | 0 |
| 0 | 0 | 0 | 0 |
| 0 | 0 | 0 | 0 |
| 0 | 0 | 0 | 0 |
| 1 | 0 | 0 | 0 |
| 0 | 0 | 0 | 0 |
| 0 | 0 | 1 | 0 |
| 0 | 0 | 1 | 0 |
| 0 | 0 | 1 | 0 |
| 0 | 0 | 0 | 0 |
| 0 | 0 | 0 | 0 |
| 0 | 0 | 0 | 0 |
| 0 | 0 | 0 | 0 |
| 0 | 0 | 0 | 0 |
| 0 | 0 | 1 | 0 |
| 0 | 0 | 0 | 0 |
| 0 | 0 | 0 | 0 |
| 1 | 0 | 1 | 0 |
| 0 | 0 | 1 | 0 |
| 1 | 1 | 0 | 0 |
| 0 | 1 | 0 | 0 |
| 0 | 0 | 0 | 0 |
| 0 | 0 | 0 | 0 |
| 1 | 0 | 0 | 0 |
| 0 | 0 | 0 | 0 |
| 0 | 0 | 1 | 0 |
| 0 | 0 | 0 | 0 |
| 0 | 0 | 0 | 0 |
| 0 | 0 | 0 | 0 |
| 0 | 0 | 0 | 0 |
| 0 | 0 | 0 | 0 |
| 0 | 0 | 0 | 0 |
| 0 | 0 | 0 | 0 |

| | | | |
|---|---|---|---|
| 0 | 0 | 0 | 0 |
| 0 | 0 | 0 | 0 |
| 0 | 0 | 0 | 0 |
| 0 | 0 | 0 | 0 |
| 0 | 0 | 0 | 0 |
| 0 | 0 | 0 | 0 |
| 0 | 0 | 0 | 0 |
| 1 | 1 | 0 | 0 |
| 0 | 0 | 0 | 0 |
| 0 | 0 | 0 | 0 |
| 0 | 0 | 0 | 0 |
| 0 | 0 | 0 | 1 |
| 0 | 0 | 0 | 0 |
| 0 | 0 | 0 | 0 |
| 0 | 0 | 0 | 0 |
| 0 | 0 | 0 | 0 |
| 0 | 0 | 0 | 0 |
| 0 | 0 | 0 | 0 |
| 0 | 0 | 0 | 0 |
| 0 | 0 | 0 | 0 |
| 0 | 0 | 0 | 0 |
| 0 | 0 | 0 | 0 |
| 0 | 0 | 0 | 0 |
| 0 | 0 | 0 | 0 |
| 0 | 0 | 0 | 0 |
| 0 | 0 | 0 | 0 |
| 0 | 0 | 0 | 0 |
| 0 | 0 | 0 | 0 |
| 0 | 0 | 0 | 0 |
| 0 | 1 | 0 | 0 |
| 0 | 1 | 0 | 0 |
| 0 | 0 | 0 | 0 |
| 0 | 0 | 0 | 0 |
| 0 | 0 | 0 | 0 |
| 0 | 0 | 0 | 0 |
| 0 | 1 | 0 | 0 |
| 0 | 0 | 0 | 0 |
| 0 | 0 | 0 | 0 |
| 0 | 0 | 0 | 0 |
| 0 | 0 | 0 | 0 |
| 0 | 0 | 0 | 0 |
| 1 | 1 | 0 | 0 |
| 1 | 0 | 0 | 0 |
| 0 | 0 | 0 | 0 |
| 0 | 0 | 0 | 0 |
| 0 | 0 | 0 | 0 |
| 0 | 0 | 0 | 0 |
| 0 | 0 | 0 | 0 |
| 0 | 0 | 0 | 0 |
| 0 | 0 | 0 | 0 |
| 0 | 0 | 0 | 0 |
| 0 | 0 | 0 | 0 |
| 0 | 0 | 0 | 0 |
| 0 | 0 | 0 | 0 |

| | | | |
|---|---|---|---|
| 0 | 0 | 0 | 0 |
| 0 | 0 | 0 | 0 |
| 0 | 0 | 0 | 0 |
| 0 | 0 | 0 | 0 |
| 0 | 0 | 0 | 0 |
| 0 | 0 | 0 | 0 |
| 0 | 0 | 0 | 0 |
| 0 | 0 | 0 | 0 |
| 1 | 0 | 0 | 0 |
| 1 | 0 | 0 | 0 |
| 0 | 0 | 0 | 0 |
| 0 | 0 | 0 | 0 |
| 0 | 0 | 0 | 0 |
| 1 | 0 | 0 | 0 |
| 0 | 0 | 0 | 0 |
| 0 | 0 | 0 | 0 |
| 0 | 0 | 0 | 0 |
| 0 | 0 | 0 | 0 |
| 0 | 0 | 0 | 0 |
| 1 | 1 | 0 | 0 |
| 0 | 0 | 1 | 0 |
| 0 | 0 | 0 | 0 |
| 0 | 0 | 0 | 0 |
| 0 | 0 | 0 | 0 |
| 0 | 0 | 0 | 0 |
| 0 | 0 | 0 | 0 |
| 0 | 0 | 0 | 0 |
| 0 | 0 | 0 | 0 |
| 0 | 0 | 0 | 0 |
| 0 | 0 | 0 | 0 |
| 0 | 0 | 0 | 0 |
| 0 | 0 | 0 | 0 |
| 0 | 0 | 0 | 0 |
| 0 | 0 | 0 | 0 |
| 0 | 0 | 0 | 0 |
| 0 | 0 | 0 | 0 |
| 0 | 0 | 0 | 0 |
| 0 | 0 | 0 | 0 |
| 0 | 0 | 0 | 0 |
| 0 | 0 | 0 | 0 |
| 0 | 0 | 0 | 0 |
| 0 | 0 | 0 | 0 |
| 0 | 0 | 0 | 0 |
| 0 | 0 | 0 | 0 |
| 0 | 0 | 0 | 0 |
| 0 | 0 | 0 | 0 |
| 1 | 0 | 1 | 0 |
| 1 | 0 | 0 | 0 |
| 0 | 0 | 0 | 0 |
| 0 | 0 | 0 | 0 |
| 0 | 0 | 1 | 0 |
| 0 | 0 | 0 | 0 |
| 0 | 0 | 0 | 0 |
| 0 | 0 | 0 | 0 |

| | | | |
|---|---|---|---|
| 0 | 0 | 0 | 0 |
| 0 | 0 | 0 | 0 |
| 0 | 0 | 0 | 0 |
| 0 | 0 | 0 | 0 |
| 0 | 0 | 0 | 0 |
| 0 | 0 | 0 | 0 |
| 0 | 0 | 0 | 0 |
| 0 | 0 | 0 | 0 |
| 0 | 0 | 0 | 0 |
| 0 | 0 | 0 | 0 |
| 0 | 0 | 0 | 0 |
| 0 | 0 | 0 | 0 |
| 0 | 0 | 0 | 0 |
| 0 | 0 | 0 | 0 |
| 0 | 0 | 0 | 0 |
| 0 | 0 | 0 | 0 |
| 0 | 0 | 0 | 0 |
| 0 | 0 | 0 | 0 |
| 0 | 0 | 0 | 0 |
| 1 | 0 | 0 | 0 |
| 0 | 0 | 0 | 0 |
| 0 | 0 | 0 | 0 |
| 0 | 0 | 0 | 0 |
| 0 | 0 | 0 | 0 |
| 0 | 0 | 0 | 0 |
| 0 | 0 | 0 | 0 |
| 0 | 0 | 0 | 0 |
| 0 | 0 | 0 | 0 |
| 0 | 0 | 0 | 0 |
| 0 | 0 | 0 | 0 |
| 0 | 0 | 0 | 0 |
| 0 | 0 | 0 | 0 |
| 0 | 0 | 0 | 0 |
| 0 | 0 | 0 | 0 |
| 0 | 0 | 0 | 0 |
| 0 | 0 | 0 | 0 |
| 0 | 0 | 0 | 0 |
| 0 | 0 | 0 | 0 |
| 1 | 0 | 0 | 1 |
| 0 | 0 | 0 | 0 |
| 0 | 0 | 0 | 0 |
| 0 | 0 | 0 | 0 |
| 0 | 0 | 0 | 1 |
| 0 | 0 | 0 | 1 |
| 0 | 0 | 0 | 1 |
| 0 | 0 | 0 | 0 |
| 0 | 0 | 0 | 0 |
| 0 | 0 | 0 | 0 |
| 0 | 0 | 0 | 0 |
| 0 | 0 | 0 | 0 |
| 0 | 0 | 0 | 0 |
| 0 | 0 | 0 | 0 |
| 0 | 0 | 0 | 0 |
| 0 | 0 | 0 | 0 |

| | | | |
|---|---|---|---|
| 0 | 0 | 0 | 0 |
| 0 | 0 | 0 | 0 |
| 0 | 0 | 0 | 0 |
| 0 | 0 | 0 | 0 |
| 0 | 0 | 0 | 0 |
| 0 | 0 | 0 | 0 |
| 0 | 0 | 0 | 0 |
| 0 | 0 | 0 | 0 |
| 0 | 0 | 0 | 0 |
| 0 | 0 | 0 | 0 |
| 0 | 0 | 0 | 0 |
| 0 | 0 | 0 | 0 |
| 0 | 0 | 0 | 0 |
| 0 | 0 | 0 | 0 |
| 0 | 0 | 0 | 0 |
| 0 | 0 | 0 | 0 |
| 0 | 0 | 0 | 0 |
| 0 | 0 | 0 | 0 |
| 0 | 0 | 0 | 0 |
| 0 | 0 | 0 | 0 |
| 0 | 0 | 0 | 0 |
| 0 | 0 | 0 | 0 |
| 0 | 0 | 0 | 0 |
| 0 | 0 | 0 | 0 |
| 0 | 0 | 0 | 0 |
| 0 | 0 | 0 | 0 |
| 0 | 0 | 0 | 0 |
| 0 | 0 | 0 | 0 |
| 0 | 0 | 0 | 0 |
| 0 | 0 | 0 | 0 |
| 0 | 0 | 0 | 0 |
| 0 | 0 | 0 | 0 |
| 1 | 1 | 0 | 1 |
| 1 | 1 | 0 | 1 |
| 1 | 0 | 0 | 0 |
| 0 | 0 | 0 | 0 |
| 0 | 0 | 0 | 0 |
| 0 | 0 | 0 | 0 |
| 0 | 0 | 0 | 0 |
| 0 | 0 | 0 | 0 |
| 0 | 0 | 0 | 0 |
| 0 | 0 | 0 | 0 |
| 1 | 1 | 0 | 0 |
| 0 | 0 | 0 | 0 |
| 1 | 0 | 0 | 0 |
| 0 | 0 | 0 | 0 |
| 0 | 0 | 0 | 0 |
| 0 | 0 | 0 | 0 |
| 0 | 0 | 0 | 0 |
| 0 | 0 | 0 | 0 |
| 1 | 0 | 0 | 0 |
| 0 | 0 | 0 | 0 |
| 0 | 0 | 0 | 0 |
| 0 | 0 | 0 | 0 |

| | | | |
|---|---|---|---|
| 0 | 0 | 0 | 0 |
| 0 | 0 | 0 | 0 |
| 0 | 0 | 0 | 0 |
| 0 | 0 | 0 | 0 |
| 0 | 0 | 0 | 0 |
| 0 | 0 | 0 | 0 |
| 0 | 0 | 0 | 0 |
| 0 | 0 | 0 | 0 |
| 0 | 0 | 0 | 0 |
| 0 | 0 | 0 | 0 |
| 0 | 0 | 0 | 0 |
| 0 | 0 | 0 | 0 |
| 0 | 0 | 0 | 0 |
| 0 | 0 | 0 | 0 |
| 0 | 0 | 0 | 0 |
| 0 | 0 | 0 | 0 |
| 0 | 0 | 0 | 0 |
| 0 | 0 | 0 | 0 |
| 0 | 0 | 0 | 0 |
| 0 | 0 | 0 | 1 |
| 0 | 0 | 0 | 0 |
| 0 | 0 | 0 | 1 |
| 0 | 0 | 0 | 1 |
| 0 | 0 | 0 | 1 |
| 0 | 0 | 0 | 0 |
| 0 | 0 | 0 | 0 |
| 0 | 0 | 0 | 0 |
| 0 | 0 | 0 | 0 |
| 0 | 0 | 0 | 0 |
| 0 | 0 | 0 | 0 |
| 0 | 0 | 0 | 0 |
| 0 | 0 | 0 | 0 |
| 0 | 0 | 0 | 0 |
| 0 | 0 | 0 | 0 |
| 0 | 0 | 0 | 0 |
| 0 | 1 | 0 | 0 |
| 0 | 0 | 0 | 0 |
| 0 | 0 | 0 | 0 |
| 0 | 1 | 0 | 0 |
| 0 | 0 | 0 | 0 |
| 0 | 0 | 0 | 0 |
| 0 | 0 | 0 | 0 |
| 0 | 0 | 0 | 0 |
| 0 | 0 | 0 | 0 |
| 0 | 0 | 0 | 0 |
| 0 | 0 | 0 | 0 |
| 0 | 0 | 0 | 0 |
| 0 | 0 | 0 | 0 |
| 0 | 0 | 0 | 0 |
| 0 | 0 | 0 | 0 |
| 0 | 0 | 0 | 0 |
| 0 | 0 | 0 | 0 |
| 0 | 0 | 0 | 0 |
| 0 | 0 | 0 | 0 |

| | | | |
|---|---|---|---|
| 0 | 0 | 0 | 0 |
| 0 | 0 | 0 | 0 |
| 0 | 0 | 0 | 0 |
| 0 | 0 | 0 | 0 |
| 0 | 0 | 0 | 0 |
| 0 | 0 | 0 | 0 |
| 0 | 0 | 0 | 0 |
| 0 | 0 | 0 | 0 |
| 0 | 0 | 1 | 0 |
| 0 | 0 | 1 | 0 |
| 0 | 0 | 0 | 0 |
| 0 | 0 | 0 | 0 |
| 0 | 0 | 0 | 0 |
| 0 | 0 | 0 | 0 |
| 0 | 0 | 0 | 0 |
| 0 | 1 | 0 | 0 |
| 0 | 0 | 0 | 0 |
| 0 | 0 | 0 | 0 |
| 1 | 1 | 0 | 0 |
| 0 | 0 | 0 | 0 |
| 0 | 0 | 0 | 0 |
| 0 | 0 | 0 | 0 |
| 0 | 0 | 0 | 0 |
| 0 | 0 | 0 | 0 |
| 0 | 0 | 0 | 0 |
| 0 | 0 | 0 | 0 |
| 0 | 0 | 0 | 0 |
| 0 | 0 | 0 | 0 |
| 0 | 0 | 0 | 0 |
| 0 | 0 | 0 | 0 |
| 0 | 0 | 0 | 0 |
| 0 | 0 | 0 | 0 |
| 0 | 0 | 0 | 0 |
| 0 | 0 | 0 | 0 |
| 0 | 0 | 0 | 0 |
| 0 | 0 | 0 | 0 |
| 0 | 0 | 0 | 0 |
| 0 | 0 | 0 | 0 |
| 0 | 0 | 0 | 0 |
| 0 | 0 | 0 | 0 |
| 0 | 0 | 0 | 0 |
| 0 | 0 | 0 | 0 |
| 0 | 0 | 0 | 0 |
| 0 | 0 | 0 | 0 |
| 0 | 0 | 0 | 0 |
| 0 | 0 | 0 | 0 |
| 0 | 0 | 0 | 0 |
| 0 | 0 | 0 | 0 |
| 0 | 0 | 0 | 0 |
| 0 | 0 | 0 | 0 |
| 0 | 0 | 0 | 0 |
| 0 | 0 | 0 | 0 |
| 0 | 0 | 0 | 0 |
| 1 | 0 | 0 | 0 |

| | | | |
|---|---|---|---|
| 0 | 0 | 0 | 0 |
| 0 | 0 | 0 | 0 |
| 0 | 0 | 0 | 0 |
| 0 | 0 | 0 | 0 |
| 0 | 0 | 0 | 0 |
| 0 | 0 | 0 | 0 |
| 0 | 0 | 0 | 0 |
| 0 | 0 | 0 | 0 |
| 0 | 0 | 0 | 0 |
| 0 | 0 | 0 | 0 |
| 1 | 0 | 0 | 0 |
| 0 | 0 | 0 | 0 |
| 0 | 0 | 0 | 0 |
| 0 | 0 | 0 | 0 |
| 1 | 1 | 0 | 1 |
| 0 | 0 | 0 | 0 |
| 1 | 0 | 0 | 0 |
| 1 | 1 | 0 | 0 |
| 0 | 0 | 0 | 0 |
| 0 | 0 | 0 | 0 |
| 0 | 0 | 0 | 0 |
| 0 | 0 | 0 | 0 |
| 0 | 0 | 0 | 0 |
| 0 | 0 | 0 | 1 |
| 0 | 0 | 0 | 0 |
| 0 | 0 | 0 | 0 |
| 0 | 0 | 0 | 0 |
| 0 | 0 | 0 | 1 |
| 0 | 0 | 0 | 0 |
| 0 | 0 | 0 | 0 |
| 0 | 0 | 0 | 0 |
| 0 | 0 | 0 | 0 |
| 0 | 0 | 0 | 0 |
| 0 | 0 | 0 | 0 |
| 0 | 0 | 0 | 0 |
| 0 | 0 | 0 | 0 |
| 0 | 0 | 0 | 0 |
| 0 | 0 | 0 | 0 |
| 0 | 0 | 0 | 0 |
| 0 | 0 | 0 | 0 |
| 0 | 0 | 0 | 0 |
| 0 | 0 | 0 | 0 |
| 0 | 0 | 0 | 0 |
| 0 | 0 | 0 | 0 |
| 0 | 0 | 1 | 0 |
| 0 | 0 | 0 | 0 |
| 0 | 0 | 0 | 0 |
| 0 | 0 | 0 | 0 |
| 0 | 0 | 0 | 0 |
| 0 | 0 | 0 | 0 |
| 0 | 0 | 0 | 0 |
| 0 | 0 | 0 | 0 |
| 0 | 0 | 0 | 0 |
| 0 | 0 | 0 | 0 |

| | | | |
|---|---|---|---|
| 0 | 0 | 0 | 0 |
| 0 | 0 | 0 | 0 |
| 0 | 0 | 0 | 0 |
| 0 | 0 | 0 | 0 |
| 0 | 0 | 0 | 0 |
| 0 | 0 | 0 | 0 |
| 1 | 0 | 0 | 1 |
| 1 | 0 | 0 | 0 |
| 0 | 0 | 0 | 0 |
| 0 | 0 | 0 | 0 |
| 0 | 0 | 0 | 0 |
| 0 | 0 | 0 | 0 |
| 0 | 0 | 0 | 0 |
| 0 | 0 | 0 | 0 |
| 0 | 0 | 0 | 0 |
| 0 | 0 | 0 | 0 |
| 0 | 0 | 0 | 0 |
| 0 | 0 | 0 | 0 |
| 0 | 0 | 0 | 0 |
| 0 | 0 | 0 | 0 |
| 1 | 0 | 0 | 0 |
| 0 | 0 | 0 | 0 |
| 0 | 0 | 0 | 0 |
| 0 | 0 | 0 | 0 |
| 0 | 0 | 0 | 0 |
| 0 | 0 | 0 | 0 |
| 0 | 0 | 0 | 0 |
| 0 | 0 | 0 | 0 |
| 0 | 0 | 0 | 0 |
| 0 | 0 | 0 | 0 |
| 0 | 0 | 0 | 0 |
| 0 | 0 | 0 | 0 |
| 0 | 0 | 0 | 0 |
| 0 | 0 | 0 | 0 |
| 0 | 0 | 0 | 0 |
| 0 | 0 | 0 | 0 |
| 0 | 0 | 0 | 0 |
| 0 | 0 | 0 | 0 |
| 0 | 0 | 0 | 0 |
| 0 | 0 | 0 | 0 |
| 0 | 0 | 0 | 0 |
| 0 | 0 | 0 | 0 |
| 0 | 0 | 0 | 0 |
| 0 | 0 | 0 | 0 |
| 0 | 0 | 0 | 0 |
| 0 | 0 | 0 | 0 |
| 0 | 0 | 0 | 0 |
| 0 | 0 | 0 | 0 |
| 0 | 0 | 0 | 0 |
| 0 | 0 | 0 | 0 |
| 0 | 0 | 0 | 0 |
| 0 | 0 | 0 | 0 |
| 0 | 0 | 0 | 0 |
| 0 | 0 | 0 | 0 |

| | | | |
|---|---|---|---|
| 0 | 0 | 0 | 0 |
| 0 | 0 | 0 | 0 |
| 0 | 0 | 0 | 0 |
| 1 | 0 | 0 | 0 |
| 0 | 0 | 0 | 0 |
| 1 | 0 | 0 | 0 |
| 0 | 0 | 0 | 0 |
| 0 | 0 | 0 | 0 |
| 1 | 0 | 0 | 0 |
| 0 | 0 | 0 | 0 |
| 0 | 0 | 0 | 0 |
| 0 | 0 | 0 | 0 |
| 0 | 0 | 0 | 0 |
| 0 | 0 | 0 | 0 |
| 0 | 0 | 0 | 0 |
| 0 | 0 | 0 | 0 |
| 0 | 0 | 0 | 0 |
| 0 | 0 | 0 | 0 |
| 1 | 0 | 0 | 0 |
| 0 | 0 | 0 | 0 |
| 0 | 0 | 0 | 0 |
| 0 | 0 | 0 | 0 |
| 0 | 0 | 0 | 0 |
| 0 | 0 | 0 | 0 |
| 0 | 0 | 0 | 0 |
| 0 | 0 | 0 | 0 |
| 0 | 0 | 0 | 0 |
| 0 | 0 | 0 | 0 |
| 0 | 1 | 0 | 0 |
| 0 | 0 | 0 | 0 |
| 0 | 0 | 0 | 0 |
| 0 | 0 | 0 | 0 |
| 0 | 1 | 0 | 0 |
| 1 | 1 | 0 | 0 |
| 1 | 0 | 0 | 0 |
| 0 | 0 | 0 | 0 |
| 0 | 0 | 0 | 0 |
| 0 | 0 | 0 | 0 |
| 0 | 0 | 0 | 0 |
| 0 | 0 | 0 | 0 |
| 1 | 0 | 0 | 0 |
| 0 | 0 | 0 | 0 |
| 0 | 0 | 0 | 0 |
| 0 | 0 | 0 | 0 |
| 0 | 0 | 0 | 0 |
| 0 | 0 | 0 | 0 |
| 0 | 0 | 0 | 0 |
| 0 | 0 | 0 | 0 |
| 0 | 0 | 0 | 0 |
| 0 | 0 | 0 | 0 |
| 0 | 0 | 0 | 0 |
| 0 | 0 | 0 | 0 |
| 0 | 0 | 0 | 0 |
| 1 | 0 | 0 | 0 |

| | | | |
|---|---|---|---|
| 1 | 0 | 0 | 1 |
| 0 | 0 | 0 | 0 |
| 0 | 0 | 0 | 0 |
| 0 | 0 | 0 | 0 |
| 0 | 0 | 0 | 0 |
| 0 | 0 | 0 | 0 |
| 0 | 0 | 0 | 0 |
| 0 | 0 | 0 | 0 |
| 0 | 0 | 0 | 0 |
| 0 | 0 | 0 | 0 |
| 0 | 0 | 0 | 0 |
| 0 | 0 | 0 | 0 |
| 0 | 0 | 0 | 0 |
| 0 | 0 | 0 | 0 |
| 0 | 0 | 0 | 0 |
| 0 | 0 | 0 | 0 |
| 0 | 0 | 0 | 0 |
| 0 | 0 | 0 | 0 |
| 0 | 0 | 0 | 0 |
| 0 | 0 | 0 | 0 |
| 0 | 0 | 0 | 0 |
| 0 | 0 | 0 | 0 |
| 0 | 1 | 0 | 0 |
| 0 | 0 | 0 | 0 |
| 0 | 0 | 0 | 0 |
| 0 | 0 | 0 | 0 |
| 0 | 0 | 0 | 0 |
| 0 | 0 | 0 | 0 |
| 0 | 0 | 0 | 0 |
| 0 | 0 | 0 | 0 |
| 0 | 0 | 0 | 0 |
| 0 | 0 | 0 | 0 |
| 0 | 0 | 0 | 0 |
| 0 | 0 | 0 | 0 |
| 0 | 0 | 0 | 0 |
| 0 | 0 | 0 | 0 |
| 0 | 0 | 0 | 0 |
| 0 | 0 | 0 | 0 |
| 0 | 0 | 0 | 0 |
| 0 | 0 | 0 | 0 |
| 0 | 0 | 0 | 0 |
| 0 | 0 | 0 | 0 |
| 0 | 0 | 0 | 0 |
| 0 | 0 | 0 | 0 |
| 0 | 0 | 0 | 0 |
| 0 | 0 | 0 | 0 |
| 0 | 0 | 0 | 0 |
| 0 | 0 | 0 | 0 |
| 0 | 0 | 0 | 0 |
| 0 | 0 | 0 | 0 |
| 0 | 0 | 0 | 0 |
| 0 | 0 | 0 | 0 |
| 0 | 0 | 0 | 0 |
| 0 | 0 | 0 | 0 |

| | | | |
|---|---|---|---|
| 0 | 0 | 0 | 0 |
| 0 | 0 | 0 | 0 |
| 0 | 0 | 0 | 0 |
| 0 | 0 | 0 | 0 |
| 0 | 0 | 0 | 0 |
| 0 | 0 | 0 | 0 |
| 0 | 1 | 0 | 0 |
| 0 | 1 | 0 | 0 |
| 0 | 0 | 0 | 0 |
| 0 | 0 | 0 | 0 |
| 0 | 0 | 0 | 0 |
| 0 | 0 | 0 | 0 |
| 0 | 0 | 0 | 0 |
| 0 | 0 | 0 | 0 |
| 0 | 0 | 0 | 0 |
| 0 | 0 | 0 | 0 |
| 1 | 0 | 0 | 0 |
| 0 | 0 | 0 | 0 |
| 1 | 0 | 0 | 0 |
| 1 | 0 | 0 | 0 |
| 1 | 0 | 0 | 0 |
| 1 | 0 | 0 | 0 |
| 0 | 0 | 0 | 0 |
| 0 | 0 | 0 | 0 |
| 0 | 0 | 0 | 0 |
| 0 | 0 | 0 | 0 |
| 0 | 0 | 0 | 0 |
| 0 | 0 | 0 | 0 |
| 0 | 0 | 0 | 0 |
| 0 | 0 | 0 | 0 |
| 0 | 0 | 0 | 0 |
| 0 | 0 | 0 | 0 |
| 0 | 0 | 0 | 0 |
| 0 | 0 | 0 | 0 |
| 0 | 0 | 0 | 0 |
| 0 | 0 | 1 | 0 |
| 1 | 0 | 0 | 0 |
| 1 | 0 | 0 | 0 |
| 1 | 0 | 0 | 0 |
| 0 | 0 | 0 | 0 |
| 0 | 0 | 0 | 0 |
| 0 | 0 | 0 | 0 |
| 0 | 0 | 0 | 0 |
| 1 | 0 | 0 | 0 |
| 1 | 0 | 0 | 0 |
| 1 | 0 | 0 | 0 |
| 0 | 0 | 0 | 0 |
| 0 | 0 | 0 | 0 |
| 1 | 0 | 0 | 0 |
| 0 | 0 | 0 | 0 |
| 0 | 0 | 0 | 0 |
| 1 | 0 | 0 | 0 |
| 0 | 0 | 0 | 0 |
| 0 | 0 | 0 | 0 |

| | | | |
|---|---|---|---|
| 0 | 0 | 0 | 0 |
| 0 | 0 | 0 | 0 |
| 0 | 0 | 0 | 0 |
| 0 | 0 | 0 | 0 |
| 0 | 0 | 0 | 0 |
| 0 | 0 | 0 | 0 |
| 0 | 0 | 0 | 0 |
| 0 | 0 | 0 | 0 |
| 0 | 0 | 0 | 0 |
| 0 | 0 | 0 | 0 |
| 0 | 0 | 0 | 0 |
| 0 | 0 | 0 | 0 |
| 0 | 0 | 0 | 0 |
| 0 | 0 | 0 | 0 |
| 0 | 0 | 0 | 0 |
| 0 | 0 | 0 | 0 |
| 0 | 0 | 0 | 0 |
| 0 | 0 | 0 | 0 |
| 0 | 0 | 0 | 0 |
| 0 | 0 | 0 | 0 |
| 0 | 0 | 0 | 0 |
| 0 | 0 | 0 | 0 |
| 1 | 1 | 0 | 0 |
| 0 | 0 | 0 | 0 |
| 0 | 1 | 0 | 0 |
| 0 | 0 | 0 | 0 |
| 0 | 0 | 0 | 0 |
| 0 | 0 | 0 | 0 |
| 0 | 0 | 0 | 0 |
| 0 | 0 | 0 | 0 |
| 0 | 0 | 0 | 0 |
| 0 | 0 | 0 | 0 |
| 0 | 0 | 0 | 0 |
| 0 | 0 | 0 | 0 |
| 0 | 0 | 0 | 0 |
| 0 | 0 | 0 | 0 |
| 0 | 0 | 0 | 0 |
| 0 | 0 | 0 | 0 |
| 0 | 0 | 0 | 0 |
| 0 | 0 | 0 | 0 |
| 0 | 0 | 0 | 0 |
| 0 | 0 | 0 | 0 |
| 0 | 0 | 0 | 0 |
| 0 | 0 | 0 | 0 |
| 0 | 0 | 0 | 0 |
| 0 | 0 | 0 | 0 |
| 1 | 0 | 0 | 1 |
| 0 | 0 | 0 | 0 |
| 0 | 0 | 0 | 0 |
| 0 | 0 | 0 | 0 |
| 1 | 0 | 0 | 0 |
| 1 | 0 | 0 | 0 |
| 1 | 0 | 0 | 0 |
| 0 | 0 | 0 | 0 |
| 0 | 0 | 0 | 0 |

| | | | |
|---|---|---|---|
| 0 | 0 | 0 | 0 |
| 0 | 0 | 0 | 0 |
| 0 | 0 | 0 | 0 |
| 0 | 0 | 0 | 0 |
| 0 | 0 | 0 | 0 |
| 0 | 0 | 0 | 0 |
| 0 | 0 | 0 | 0 |
| 0 | 0 | 0 | 0 |
| 0 | 0 | 0 | 0 |
| 0 | 0 | 0 | 0 |
| 0 | 0 | 0 | 1 |
| 0 | 0 | 0 | 0 |
| 0 | 0 | 0 | 0 |
| 0 | 0 | 0 | 0 |
| 0 | 0 | 0 | 0 |
| 0 | 0 | 0 | 0 |
| 0 | 0 | 0 | 0 |
| 0 | 0 | 0 | 0 |
| 0 | 0 | 0 | 0 |
| 0 | 0 | 0 | 0 |
| 0 | 0 | 0 | 0 |
| 0 | 0 | 0 | 0 |
| 0 | 0 | 0 | 0 |
| 0 | 0 | 0 | 0 |
| 0 | 0 | 0 | 0 |
| 0 | 0 | 0 | 0 |
| 0 | 0 | 0 | 0 |
| 0 | 0 | 0 | 0 |
| 0 | 0 | 0 | 0 |
| 0 | 0 | 0 | 0 |
| 0 | 1 | 0 | 0 |
| 0 | 0 | 0 | 0 |
| 1 | 1 | 0 | 0 |
| 1 | 1 | 0 | 0 |
| 0 | 0 | 0 | 0 |
| 0 | 0 | 0 | 0 |
| 0 | 0 | 0 | 0 |
| 0 | 0 | 0 | 0 |
| 0 | 0 | 0 | 0 |
| 0 | 0 | 0 | 0 |
| 0 | 0 | 0 | 0 |
| 0 | 1 | 0 | 0 |
| 0 | 0 | 0 | 0 |
| 0 | 0 | 0 | 0 |
| 0 | 0 | 0 | 0 |
| 1 | 0 | 0 | 0 |
| 1 | 0 | 0 | 0 |
| 1 | 0 | 0 | 0 |
| 1 | 1 | 1 | 0 |
| 0 | 0 | 0 | 0 |
| 1 | 1 | 1 | 1 |
| 0 | 0 | 0 | 0 |
| 0 | 0 | 0 | 0 |
| 0 | 0 | 0 | 0 |

| | | | |
|---|---|---|---|
| 0 | 0 | 0 | 0 |
| 0 | 0 | 0 | 0 |
| 0 | 0 | 0 | 0 |
| 0 | 0 | 0 | 0 |
| 1 | 0 | 0 | 0 |
| 1 | 0 | 0 | 0 |
| 0 | 0 | 0 | 0 |
| 0 | 0 | 0 | 0 |
| 0 | 0 | 0 | 0 |
| 0 | 0 | 0 | 0 |
| 0 | 0 | 0 | 0 |
| 0 | 0 | 0 | 0 |
| 0 | 0 | 0 | 0 |
| 0 | 0 | 0 | 0 |
| 0 | 0 | 0 | 0 |
| 0 | 0 | 0 | 0 |
| 0 | 0 | 0 | 0 |
| 0 | 0 | 0 | 0 |
| 0 | 0 | 0 | 0 |
| 0 | 0 | 0 | 0 |
| 0 | 0 | 0 | 0 |
| 0 | 0 | 0 | 0 |
| 0 | 0 | 0 | 0 |
| 0 | 0 | 0 | 0 |
| 0 | 0 | 0 | 0 |
| 0 | 0 | 0 | 0 |
| 0 | 0 | 0 | 0 |
| 0 | 0 | 0 | 0 |
| 0 | 0 | 0 | 0 |
| 0 | 0 | 0 | 0 |
| 0 | 0 | 0 | 0 |
| 0 | 0 | 0 | 0 |
| 0 | 0 | 0 | 0 |
| 0 | 0 | 0 | 0 |
| 0 | 0 | 0 | 0 |
| 0 | 0 | 0 | 0 |
| 0 | 0 | 0 | 0 |
| 0 | 0 | 0 | 0 |
| 0 | 0 | 0 | 0 |
| 0 | 0 | 0 | 0 |
| 0 | 0 | 0 | 0 |
| 0 | 0 | 0 | 0 |
| 0 | 0 | 0 | 0 |
| 0 | 0 | 0 | 0 |
| 0 | 0 | 0 | 0 |
| 0 | 0 | 0 | 0 |
| 0 | 0 | 0 | 0 |
| 0 | 0 | 0 | 0 |
| 0 | 0 | 0 | 0 |
| 0 | 0 | 0 | 0 |
| 0 | 0 | 0 | 0 |
| 0 | 0 | 0 | 0 |
| 0 | 0 | 0 | 0 |
| 0 | 1 | 0 | 0 |

| | | | |
|---|---|---|---|
| 0 | 0 | 0 | 0 |
| 0 | 0 | 0 | 0 |
| 0 | 0 | 0 | 0 |
| 0 | 0 | 0 | 0 |
| 0 | 0 | 0 | 0 |
| 0 | 0 | 0 | 0 |
| 0 | 0 | 0 | 0 |
| 0 | 0 | 0 | 0 |
| 0 | 0 | 0 | 0 |
| 0 | 0 | 0 | 0 |
| 0 | 0 | 0 | 0 |
| 0 | 0 | 0 | 0 |
| 0 | 0 | 0 | 0 |
| 0 | 0 | 0 | 0 |
| 0 | 0 | 0 | 0 |
| 0 | 0 | 0 | 0 |
| 0 | 0 | 0 | 0 |
| 0 | 0 | 0 | 0 |
| 0 | 0 | 0 | 0 |
| 0 | 0 | 0 | 0 |
| 0 | 0 | 0 | 0 |
| 0 | 0 | 0 | 0 |
| 0 | 0 | 0 | 0 |
| 0 | 0 | 0 | 0 |
| 0 | 0 | 0 | 0 |
| 0 | 0 | 0 | 0 |
| 0 | 0 | 0 | 0 |
| 0 | 0 | 0 | 0 |
| 0 | 0 | 0 | 0 |
| 0 | 0 | 0 | 0 |
| 0 | 0 | 0 | 0 |
| 0 | 0 | 0 | 0 |
| 0 | 0 | 0 | 0 |
| 0 | 0 | 0 | 0 |
| 0 | 0 | 0 | 0 |
| 0 | 0 | 0 | 0 |
| 0 | 0 | 0 | 0 |
| 0 | 0 | 0 | 0 |
| 0 | 0 | 0 | 0 |
| 0 | 0 | 0 | 0 |
| 0 | 0 | 0 | 0 |
| 0 | 0 | 0 | 0 |
| 0 | 0 | 0 | 0 |
| 0 | 0 | 0 | 0 |
| 0 | 0 | 0 | 0 |
| 0 | 0 | 0 | 0 |
| 0 | 0 | 0 | 0 |
| 0 | 0 | 0 | 0 |
| 0 | 0 | 0 | 0 |
| 0 | 0 | 0 | 0 |
| 0 | 0 | 0 | 0 |
| 0 | 0 | 0 | 0 |
| 0 | 0 | 0 | 0 |
| 0 | 0 | 0 | 0 |

| | | | |
|---|---|---|---|
| 0 | 0 | 0 | 0 |
| 0 | 0 | 0 | 0 |
| 0 | 0 | 0 | 0 |
| 0 | 0 | 0 | 0 |
| 0 | 0 | 0 | 0 |
| 0 | 0 | 0 | 0 |
| 0 | 0 | 0 | 0 |
| 0 | 0 | 0 | 0 |
| 0 | 0 | 0 | 0 |
| 0 | 0 | 0 | 0 |
| 0 | 0 | 0 | 0 |
| 0 | 0 | 1 | 0 |
| 0 | 0 | 0 | 0 |
| 0 | 0 | 1 | 0 |
| 0 | 0 | 1 | 0 |
| 0 | 0 | 1 | 0 |
| 0 | 0 | 1 | 0 |
| 0 | 0 | 0 | 0 |
| 0 | 0 | 0 | 0 |
| 0 | 0 | 0 | 0 |
| 1 | 0 | 0 | 0 |

| glycan_binding_site | water_binding_site | water_only_binding_site | non_binding_site |
|---|---|---|---|
| 0 | 1 | 0 | 0 |
| 0 | 1 | 0 | 0 |
| 0 | 1 | 1 | 1 |
| 0 | 1 | 0 | 0 |
| 0 | 0 | 0 | 0 |
| 0 | 0 | 0 | 0 |
| 0 | 0 | 0 | 1 |
| 0 | 0 | 0 | 1 |
| 0 | 1 | 0 | 0 |
| 0 | 1 | 0 | 0 |
| 0 | 0 | 0 | 0 |
| 0 | 1 | 0 | 0 |
| 0 | 1 | 1 | 1 |
| 1 | 0 | 0 | 0 |
| 0 | 0 | 0 | 0 |
| 0 | 1 | 0 | 0 |
| 0 | 0 | 0 | 0 |
| 0 | 1 | 0 | 0 |
| 0 | 1 | 0 | 0 |
| 0 | 0 | 0 | 0 |
| 0 | 1 | 0 | 0 |
| 1 | 1 | 0 | 0 |
| 1 | 1 | 0 | 0 |
| 0 | 1 | 0 | 0 |
| 0 | 1 | 0 | 0 |
| 0 | 1 | 1 | 1 |
| 0 | 1 | 0 | 0 |
| 0 | 0 | 0 | 1 |
| 0 | 0 | 0 | 0 |
| 0 | 0 | 0 | 1 |
| 0 | 0 | 0 | 1 |
| 0 | 1 | 0 | 0 |
| 0 | 1 | 0 | 0 |
| 0 | 1 | 0 | 0 |
| 0 | 1 | 0 | 0 |
| 0 | 0 | 0 | 1 |
| 0 | 0 | 0 | 0 |
| 0 | 1 | 0 | 0 |
| 0 | 1 | 0 | 0 |
| 0 | 1 | 0 | 0 |
| 0 | 1 | 0 | 0 |
| 0 | 1 | 0 | 0 |
| 0 | 1 | 0 | 0 |
| 0 | 1 | 0 | 0 |
| 0 | 1 | 0 | 0 |
| 0 | 1 | 0 | 0 |
| 0 | 1 | 0 | 0 |
| 0 | 1 | 0 | 0 |
| 0 | 1 | 0 | 0 |
| 0 | 1 | 0 | 0 |
| 0 | 1 | 0 | 0 |
| 0 | 1 | 0 | 0 |
| 0 | 1 | 0 | 0 |

| | | | |
|---|---|---|---|
| 0 | 1 | 0 | 0 |
| 0 | 1 | 0 | 0 |
| 0 | 1 | 0 | 0 |
| 0 | 1 | 0 | 0 |
| 0 | 1 | 0 | 0 |
| 0 | 1 | 0 | 0 |
| 0 | 1 | 0 | 0 |
| 0 | 1 | 0 | 0 |
| 0 | 1 | 0 | 0 |
| 0 | 1 | 0 | 0 |
| 0 | 1 | 0 | 0 |
| 0 | 1 | 0 | 0 |
| 0 | 1 | 0 | 0 |
| 0 | 1 | 0 | 0 |
| 0 | 1 | 0 | 0 |
| 0 | 1 | 0 | 0 |
| 0 | 1 | 0 | 0 |
| 0 | 1 | 0 | 0 |
| 0 | 1 | 0 | 0 |
| 0 | 1 | 0 | 0 |
| 0 | 1 | 0 | 0 |
| 0 | 1 | 0 | 0 |
| 0 | 0 | 0 | 0 |
| 0 | 1 | 0 | 0 |
| 0 | 0 | 0 | 0 |
| 0 | 1 | 0 | 0 |
| 0 | 1 | 0 | 0 |
| 0 | 0 | 0 | 0 |
| 0 | 0 | 0 | 0 |
| 0 | 0 | 0 | 0 |
| 0 | 0 | 0 | 0 |
| 0 | 0 | 0 | 0 |
| 0 | 0 | 0 | 0 |
| 0 | 0 | 0 | 0 |
| 0 | 0 | 0 | 0 |
| 0 | 0 | 0 | 0 |
| 0 | 0 | 0 | 0 |
| 0 | 1 | 0 | 0 |
| 0 | 1 | 0 | 0 |
| 0 | 1 | 0 | 0 |
| 1 | 0 | 0 | 0 |
| 0 | 0 | 0 | 0 |
| 1 | 0 | 0 | 0 |
| 1 | 1 | 0 | 0 |
| 0 | 0 | 0 | 0 |
| 0 | 0 | 0 | 1 |
| 0 | 0 | 0 | 1 |
| 0 | 1 | 0 | 0 |
| 1 | 0 | 0 | 0 |
| 0 | 0 | 0 | 0 |
| 0 | 0 | 0 | 0 |
| 0 | 0 | 0 | 1 |
| 0 | 0 | 0 | 1 |
| 0 | 0 | 0 | 0 |

| | | | |
|---|---|---|---|
| 0 | 0 | 0 | 1 |
| 0 | 0 | 0 | 1 |
| 0 | 0 | 0 | 0 |
| 0 | 0 | 0 | 0 |
| 0 | 0 | 0 | 0 |
| 0 | 0 | 0 | 0 |
| 0 | 0 | 0 | 1 |
| 0 | 1 | 0 | 0 |
| 0 | 1 | 0 | 0 |
| 0 | 0 | 0 | 0 |
| 0 | 1 | 0 | 0 |
| 0 | 0 | 0 | 0 |
| 0 | 0 | 0 | 0 |
| 1 | 0 | 0 | 0 |
| 1 | 0 | 0 | 0 |
| 0 | 0 | 0 | 1 |
| 0 | 0 | 0 | 0 |
| 0 | 1 | 0 | 0 |
| 0 | 1 | 0 | 0 |
| 0 | 1 | 0 | 0 |
| 0 | 1 | 0 | 0 |
| 0 | 1 | 0 | 0 |
| 0 | 1 | 0 | 0 |
| 0 | 1 | 0 | 0 |
| 0 | 1 | 0 | 0 |
| 0 | 1 | 0 | 0 |
| 0 | 1 | 0 | 0 |
| 0 | 1 | 0 | 0 |
| 0 | 1 | 0 | 0 |
| 0 | 1 | 0 | 0 |
| 0 | 1 | 0 | 0 |
| 0 | 1 | 0 | 0 |
| 0 | 1 | 0 | 0 |
| 0 | 1 | 0 | 0 |
| 0 | 1 | 0 | 0 |
| 0 | 1 | 0 | 0 |
| 0 | 1 | 0 | 0 |
| 0 | 1 | 0 | 0 |
| 0 | 1 | 0 | 0 |
| 0 | 1 | 0 | 0 |
| 0 | 1 | 0 | 0 |
| 0 | 1 | 0 | 0 |
| 0 | 1 | 0 | 0 |
| 0 | 1 | 0 | 0 |
| 0 | 1 | 0 | 0 |
| 0 | 1 | 0 | 0 |
| 1 | 1 | 0 | 0 |
| 0 | 0 | 0 | 0 |
| 0 | 1 | 0 | 0 |
| 0 | 1 | 0 | 0 |
| 0 | 0 | 0 | 0 |
| 0 | 0 | 0 | 0 |
| 0 | 1 | 0 | 0 |
| 0 | 1 | 1 | 1 |

| | | | |
|---|---|---|---|
| 0 | 1 | 1 | 1 |
| 0 | 1 | 1 | 1 |
| 0 | 1 | 1 | 1 |
| 0 | 1 | 0 | 0 |
| 0 | 1 | 0 | 0 |
| 0 | 1 | 1 | 1 |
| 0 | 1 | 0 | 0 |
| 0 | 1 | 1 | 1 |
| 0 | 1 | 1 | 1 |
| 0 | 1 | 1 | 1 |
| 0 | 1 | 1 | 1 |
| 0 | 1 | 0 | 0 |
| 0 | 1 | 1 | 1 |
| 0 | 0 | 0 | 0 |
| 0 | 1 | 1 | 1 |
| 0 | 1 | 1 | 1 |
| 0 | 1 | 1 | 1 |
| 0 | 1 | 1 | 1 |
| 0 | 1 | 1 | 1 |
| 0 | 1 | 0 | 0 |
| 0 | 1 | 0 | 0 |
| 0 | 0 | 0 | 1 |
| 0 | 1 | 0 | 0 |
| 0 | 1 | 0 | 0 |
| 0 | 1 | 0 | 0 |
| 0 | 1 | 1 | 1 |
| 0 | 1 | 0 | 0 |
| 0 | 1 | 0 | 0 |
| 0 | 1 | 1 | 1 |
| 0 | 1 | 1 | 1 |
| 0 | 0 | 0 | 1 |
| 0 | 1 | 1 | 1 |
| 0 | 1 | 1 | 1 |
| 0 | 0 | 0 | 1 |
| 0 | 1 | 0 | 0 |
| 0 | 1 | 1 | 1 |
| 1 | 1 | 0 | 0 |
| 1 | 1 | 0 | 0 |
| 0 | 1 | 1 | 1 |
| 0 | 1 | 1 | 1 |
| 0 | 1 | 1 | 1 |
| 0 | 1 | 0 | 0 |
| 0 | 1 | 0 | 0 |
| 0 | 0 | 0 | 1 |
| 0 | 1 | 1 | 1 |
| 0 | 1 | 1 | 1 |
| 0 | 0 | 0 | 0 |
| 0 | 0 | 0 | 0 |
| 0 | 0 | 0 | 0 |
| 0 | 1 | 0 | 0 |
| 0 | 0 | 0 | 0 |
| 0 | 0 | 0 | 1 |
| 0 | 0 | 0 | 1 |
| 0 | 0 | 0 | 0 |

| | | | |
|---|---|---|---|
| 0 | 0 | 0 | 0 |
| 0 | 1 | 0 | 0 |
| 0 | 1 | 0 | 0 |
| 0 | 0 | 0 | 1 |
| 0 | 0 | 0 | 1 |
| 0 | 1 | 1 | 1 |
| 0 | 1 | 0 | 0 |
| 0 | 1 | 0 | 0 |
| 0 | 1 | 1 | 1 |
| 0 | 1 | 1 | 1 |
| 1 | 1 | 0 | 0 |
| 0 | 0 | 0 | 1 |
| 1 | 1 | 0 | 0 |
| 0 | 1 | 1 | 1 |
| 0 | 0 | 0 | 1 |
| 0 | 0 | 0 | 1 |
| 0 | 0 | 0 | 1 |
| 0 | 1 | 0 | 0 |
| 0 | 0 | 0 | 0 |
| 0 | 1 | 0 | 0 |
| 0 | 0 | 0 | 0 |
| 0 | 1 | 1 | 1 |
| 0 | 1 | 0 | 0 |
| 0 | 0 | 0 | 1 |
| 0 | 0 | 0 | 1 |
| 0 | 1 | 0 | 0 |
| 0 | 1 | 0 | 0 |
| 0 | 0 | 0 | 0 |
| 0 | 1 | 0 | 0 |
| 0 | 1 | 0 | 0 |
| 0 | 1 | 0 | 0 |
| 0 | 1 | 0 | 0 |
| 0 | 1 | 1 | 1 |
| 0 | 0 | 0 | 1 |
| 0 | 0 | 0 | 0 |
| 0 | 0 | 0 | 1 |
| 0 | 1 | 1 | 1 |
| 0 | 1 | 0 | 0 |
| 0 | 1 | 1 | 1 |
| 0 | 0 | 0 | 1 |
| 0 | 1 | 0 | 0 |
| 0 | 1 | 1 | 1 |
| 0 | 1 | 1 | 1 |
| 0 | 0 | 0 | 1 |
| 0 | 0 | 0 | 0 |
| 0 | 0 | 0 | 0 |
| 0 | 1 | 0 | 0 |
| 0 | 1 | 0 | 0 |
| 0 | 1 | 0 | 0 |
| 0 | 1 | 0 | 0 |
| 0 | 1 | 0 | 0 |
| 0 | 1 | 0 | 0 |
| 0 | 1 | 0 | 0 |
| 0 | 1 | 0 | 0 |

| | | | |
|---|---|---|---|
| 0 | 1 | 0 | 0 |
| 0 | 1 | 0 | 0 |
| 0 | 1 | 0 | 0 |
| 0 | 1 | 0 | 0 |
| 0 | 1 | 0 | 0 |
| 0 | 1 | 0 | 0 |
| 0 | 0 | 0 | 1 |
| 0 | 0 | 0 | 1 |
| 0 | 0 | 0 | 0 |
| 0 | 0 | 0 | 0 |
| 0 | 0 | 0 | 0 |
| 0 | 0 | 0 | 0 |
| 0 | 0 | 0 | 0 |
| 0 | 0 | 0 | 1 |
| 0 | 0 | 0 | 1 |
| 0 | 0 | 0 | 0 |
| 0 | 0 | 0 | 1 |
| 0 | 0 | 0 | 0 |
| 0 | 0 | 0 | 1 |
| 0 | 0 | 0 | 0 |
| 0 | 0 | 0 | 1 |
| 0 | 1 | 1 | 1 |
| 0 | 0 | 0 | 1 |
| 0 | 0 | 0 | 0 |
| 0 | 0 | 0 | 0 |
| 0 | 1 | 0 | 0 |
| 0 | 1 | 0 | 0 |
| 0 | 1 | 0 | 0 |
| 0 | 0 | 0 | 0 |
| 0 | 1 | 0 | 0 |
| 0 | 1 | 1 | 1 |
| 0 | 1 | 1 | 1 |
| 0 | 0 | 0 | 0 |
| 0 | 1 | 0 | 0 |
| 0 | 1 | 0 | 0 |
| 0 | 0 | 0 | 1 |
| 0 | 0 | 0 | 0 |
| 0 | 1 | 0 | 0 |
| 0 | 0 | 0 | 0 |
| 0 | 1 | 0 | 0 |
| 0 | 0 | 0 | 1 |
| 0 | 0 | 0 | 1 |
| 0 | 0 | 0 | 0 |
| 0 | 0 | 0 | 1 |
| 0 | 0 | 0 | 0 |
| 0 | 0 | 0 | 0 |
| 0 | 0 | 0 | 0 |
| 0 | 0 | 0 | 1 |
| 0 | 0 | 0 | 1 |
| 0 | 0 | 0 | 0 |
| 0 | 0 | 0 | 1 |
| 0 | 0 | 0 | 1 |
| 0 | 0 | 0 | 0 |
| 0 | 0 | 0 | 1 |

| | | | |
|---|---|---|---|
| 0 | 0 | 0 | 0 |
| 0 | 0 | 0 | 1 |
| 0 | 1 | 1 | 1 |
| 0 | 1 | 0 | 0 |
| 0 | 1 | 0 | 0 |
| 0 | 0 | 0 | 0 |
| 0 | 0 | 0 | 1 |
| 0 | 1 | 0 | 0 |
| 0 | 0 | 0 | 0 |
| 0 | 0 | 0 | 1 |
| 0 | 0 | 0 | 1 |
| 0 | 0 | 0 | 1 |
| 0 | 0 | 0 | 1 |
| 0 | 0 | 0 | 1 |
| 0 | 0 | 0 | 0 |
| 0 | 0 | 0 | 1 |
| 0 | 0 | 0 | 0 |
| 0 | 0 | 0 | 0 |
| 0 | 0 | 0 | 1 |
| 0 | 0 | 0 | 0 |
| 0 | 1 | 0 | 0 |
| 0 | 1 | 0 | 0 |
| 0 | 1 | 0 | 0 |
| 0 | 1 | 0 | 0 |
| 0 | 0 | 0 | 1 |
| 0 | 0 | 0 | 1 |
| 0 | 0 | 0 | 0 |
| 0 | 0 | 0 | 0 |
| 0 | 1 | 0 | 0 |
| 0 | 0 | 0 | 0 |
| 0 | 0 | 0 | 0 |
| 0 | 1 | 0 | 0 |
| 0 | 0 | 0 | 0 |
| 0 | 0 | 0 | 0 |
| 0 | 0 | 0 | 1 |
| 0 | 0 | 0 | 1 |
| 0 | 0 | 0 | 0 |
| 0 | 0 | 0 | 0 |
| 0 | 0 | 0 | 0 |
| 0 | 0 | 0 | 0 |
| 0 | 0 | 0 | 0 |
| 0 | 1 | 0 | 0 |
| 0 | 0 | 0 | 1 |
| 0 | 1 | 0 | 0 |
| 0 | 1 | 0 | 0 |
| 0 | 1 | 0 | 0 |
| 0 | 1 | 0 | 0 |
| 0 | 1 | 0 | 0 |
| 1 | 1 | 0 | 0 |
| 0 | 0 | 0 | 1 |
| 0 | 1 | 0 | 0 |
| 0 | 1 | 0 | 0 |
| 0 | 0 | 0 | 0 |
| 0 | 1 | 0 | 0 |

| | | | |
|---|---|---|---|
| 0 | 1 | 0 | 0 |
| 0 | 1 | 0 | 0 |
| 0 | 1 | 0 | 0 |
| 0 | 1 | 0 | 0 |
| 0 | 0 | 0 | 1 |
| 0 | 1 | 0 | 0 |
| 0 | 1 | 0 | 0 |
| 0 | 1 | 0 | 0 |
| 0 | 1 | 0 | 0 |
| 0 | 1 | 0 | 0 |
| 0 | 1 | 0 | 0 |
| 0 | 1 | 0 | 0 |
| 0 | 1 | 0 | 0 |
| 0 | 1 | 0 | 0 |
| 0 | 1 | 0 | 0 |
| 0 | 1 | 0 | 0 |
| 0 | 1 | 0 | 0 |
| 0 | 1 | 0 | 0 |
| 0 | 0 | 0 | 1 |
| 0 | 0 | 0 | 0 |
| 0 | 0 | 0 | 0 |
| 0 | 0 | 0 | 0 |
| 0 | 1 | 0 | 0 |
| 0 | 1 | 1 | 1 |
| 0 | 0 | 0 | 0 |
| 0 | 0 | 0 | 0 |
| 0 | 0 | 0 | 0 |
| 0 | 1 | 1 | 1 |
| 0 | 1 | 1 | 1 |
| 0 | 1 | 1 | 1 |
| 0 | 1 | 0 | 0 |
| 0 | 0 | 0 | 0 |
| 0 | 1 | 0 | 0 |
| 0 | 1 | 0 | 0 |
| 0 | 1 | 0 | 0 |
| 0 | 1 | 1 | 1 |
| 0 | 1 | 1 | 1 |
| 0 | 0 | 0 | 0 |
| 0 | 1 | 0 | 0 |
| 0 | 1 | 1 | 1 |
| 0 | 1 | 1 | 1 |
| 0 | 1 | 1 | 1 |
| 0 | 0 | 0 | 1 |
| 0 | 0 | 0 | 1 |
| 0 | 0 | 0 | 0 |
| 0 | 0 | 0 | 0 |
| 0 | 1 | 0 | 0 |
| 0 | 0 | 0 | 0 |
| 0 | 0 | 0 | 0 |
| 0 | 0 | 0 | 1 |
| 0 | 0 | 0 | 1 |
| 0 | 0 | 0 | 0 |
| 0 | 0 | 0 | 1 |
| 0 | 0 | 0 | 0 |

| | | | |
|---|---|---|---|
| 0 | 0 | 0 | 1 |
| 0 | 0 | 0 | 1 |
| 0 | 0 | 0 | 0 |
| 0 | 0 | 0 | 1 |
| 0 | 0 | 0 | 1 |
| 0 | 0 | 0 | 1 |
| 0 | 0 | 0 | 1 |
| 0 | 1 | 0 | 0 |
| 0 | 1 | 1 | 1 |
| 0 | 0 | 0 | 1 |
| 0 | 0 | 0 | 0 |
| 0 | 0 | 0 | 1 |
| 0 | 0 | 0 | 1 |
| 0 | 0 | 0 | 1 |
| 0 | 1 | 0 | 0 |
| 0 | 0 | 0 | 0 |
| 0 | 0 | 0 | 1 |
| 0 | 0 | 0 | 1 |
| 0 | 1 | 0 | 0 |
| 0 | 1 | 0 | 0 |
| 0 | 1 | 0 | 0 |
| 0 | 1 | 0 | 0 |
| 0 | 0 | 0 | 0 |
| 0 | 1 | 0 | 0 |
| 0 | 1 | 0 | 0 |
| 0 | 0 | 0 | 1 |
| 0 | 1 | 0 | 0 |
| 0 | 1 | 0 | 0 |
| 0 | 1 | 0 | 0 |
| 0 | 0 | 0 | 1 |
| 0 | 1 | 0 | 0 |
| 0 | 1 | 0 | 0 |
| 0 | 1 | 1 | 1 |
| 0 | 1 | 0 | 0 |
| 0 | 1 | 0 | 0 |
| 0 | 1 | 0 | 0 |
| 0 | 1 | 0 | 0 |
| 0 | 1 | 0 | 0 |
| 0 | 1 | 0 | 0 |
| 0 | 1 | 0 | 0 |
| 0 | 0 | 0 | 1 |
| 0 | 0 | 0 | 0 |
| 0 | 0 | 0 | 1 |
| 0 | 0 | 0 | 0 |
| 0 | 0 | 0 | 0 |
| 1 | 0 | 0 | 0 |
| 0 | 0 | 0 | 0 |
| 1 | 0 | 0 | 0 |
| 0 | 0 | 0 | 0 |
| 0 | 0 | 0 | 0 |
| 0 | 0 | 0 | 1 |
| 0 | 0 | 0 | 0 |
| 0 | 0 | 0 | 0 |
| 0 | 0 | 0 | 0 |

| | | | |
|---|---|---|---|
| 0 | 0 | 0 | 1 |
| 0 | 0 | 0 | 0 |
| 0 | 0 | 0 | 1 |
| 0 | 0 | 0 | 1 |
| 1 | 0 | 0 | 0 |
| 0 | 0 | 0 | 0 |
| 1 | 0 | 0 | 0 |
| 1 | 0 | 0 | 0 |
| 0 | 0 | 0 | 0 |
| 0 | 0 | 0 | 1 |
| 0 | 0 | 0 | 1 |
| 0 | 0 | 0 | 1 |
| 0 | 0 | 0 | 1 |
| 1 | 1 | 0 | 0 |
| 0 | 0 | 0 | 1 |
| 0 | 0 | 0 | 0 |
| 0 | 0 | 0 | 0 |
| 0 | 0 | 0 | 0 |
| 1 | 1 | 0 | 0 |
| 0 | 1 | 1 | 1 |
| 1 | 1 | 0 | 0 |
| 0 | 0 | 0 | 1 |
| 0 | 0 | 0 | 0 |
| 0 | 0 | 0 | 1 |
| 0 | 0 | 0 | 1 |
| 0 | 0 | 0 | 1 |
| 0 | 0 | 0 | 0 |
| 0 | 0 | 0 | 1 |
| 0 | 0 | 0 | 1 |
| 0 | 0 | 0 | 1 |
| 0 | 0 | 0 | 0 |
| 0 | 0 | 0 | 0 |
| 0 | 0 | 0 | 1 |
| 0 | 0 | 0 | 1 |
| 0 | 0 | 0 | 0 |
| 0 | 0 | 0 | 0 |
| 0 | 0 | 0 | 0 |
| 1 | 0 | 0 | 0 |
| 0 | 0 | 0 | 0 |
| 0 | 0 | 0 | 1 |
| 0 | 1 | 0 | 0 |
| 0 | 1 | 0 | 0 |
| 0 | 1 | 0 | 0 |
| 0 | 1 | 0 | 0 |
| 0 | 1 | 1 | 1 |
| 1 | 0 | 0 | 0 |
| 0 | 1 | 0 | 0 |
| 0 | 0 | 0 | 0 |
| 0 | 1 | 0 | 0 |
| 1 | 1 | 0 | 0 |
| 0 | 1 | 0 | 0 |
| 0 | 1 | 0 | 0 |
| 0 | 1 | 1 | 1 |
| 0 | 0 | 0 | 0 |

| | | | |
|---|---|---|---|
| 0 | 1 | 0 | 0 |
| 0 | 1 | 0 | 0 |
| 0 | 1 | 1 | 1 |
| 0 | 1 | 1 | 1 |
| 0 | 1 | 1 | 1 |
| 0 | 1 | 0 | 0 |
| 0 | 1 | 1 | 1 |
| 0 | 1 | 1 | 1 |
| 0 | 1 | 0 | 0 |
| 0 | 1 | 0 | 0 |
| 1 | 1 | 0 | 0 |
| 1 | 1 | 0 | 0 |
| 0 | 0 | 0 | 1 |
| 1 | 1 | 0 | 0 |
| 0 | 0 | 0 | 0 |
| 0 | 0 | 0 | 0 |
| 0 | 1 | 0 | 0 |
| 0 | 0 | 0 | 0 |
| 0 | 0 | 0 | 0 |
| 0 | 0 | 0 | 1 |
| 0 | 0 | 0 | 0 |
| 0 | 1 | 0 | 0 |
| 0 | 0 | 0 | 0 |
| 0 | 0 | 0 | 0 |
| 0 | 1 | 0 | 0 |
| 0 | 1 | 1 | 1 |
| 0 | 0 | 0 | 1 |
| 0 | 0 | 0 | 1 |
| 0 | 1 | 0 | 0 |
| 0 | 1 | 0 | 0 |
| 0 | 0 | 0 | 0 |
| 0 | 0 | 0 | 0 |
| 0 | 0 | 0 | 0 |
| 0 | 1 | 0 | 0 |
| 0 | 1 | 0 | 0 |
| 0 | 1 | 1 | 1 |
| 0 | 1 | 0 | 0 |
| 0 | 1 | 0 | 0 |
| 0 | 0 | 0 | 1 |
| 0 | 1 | 1 | 1 |
| 0 | 0 | 0 | 1 |
| 0 | 1 | 0 | 0 |
| 0 | 0 | 0 | 0 |
| 0 | 0 | 0 | 0 |
| 0 | 0 | 0 | 0 |
| 0 | 0 | 0 | 0 |
| 0 | 0 | 0 | 0 |
| 0 | 0 | 0 | 1 |
| 0 | 0 | 0 | 1 |
| 0 | 0 | 0 | 0 |
| 0 | 0 | 0 | 0 |
| 0 | 0 | 0 | 0 |
| 0 | 0 | 0 | 0 |
| 0 | 0 | 0 | 1 |

| | | | |
|---|---|---|---|
| 0 | 0 | 0 | 1 |
| 0 | 0 | 0 | 0 |
| 0 | 0 | 0 | 1 |
| 0 | 1 | 0 | 0 |
| 0 | 1 | 1 | 1 |
| 0 | 1 | 1 | 1 |
| 0 | 1 | 0 | 0 |
| 0 | 1 | 1 | 1 |
| 0 | 1 | 1 | 1 |
| 0 | 1 | 1 | 1 |
| 0 | 1 | 1 | 1 |
| 0 | 1 | 1 | 1 |
| 0 | 0 | 0 | 1 |
| 0 | 0 | 0 | 1 |
| 0 | 0 | 0 | 1 |
| 0 | 0 | 0 | 1 |
| 0 | 0 | 0 | 0 |
| 0 | 0 | 0 | 0 |
| 0 | 1 | 1 | 1 |
| 0 | 0 | 0 | 0 |
| 0 | 0 | 0 | 0 |
| 0 | 1 | 0 | 0 |
| 0 | 0 | 0 | 1 |
| 0 | 1 | 0 | 0 |
| 0 | 0 | 0 | 0 |
| 0 | 0 | 0 | 1 |
| 0 | 1 | 1 | 1 |
| 0 | 1 | 1 | 1 |
| 0 | 1 | 0 | 0 |
| 0 | 0 | 0 | 0 |
| 0 | 1 | 1 | 1 |
| 0 | 1 | 0 | 0 |
| 0 | 1 | 0 | 0 |
| 0 | 1 | 0 | 0 |
| 0 | 1 | 0 | 0 |
| 0 | 0 | 0 | 0 |
| 0 | 1 | 1 | 1 |
| 0 | 0 | 0 | 1 |
| 0 | 0 | 0 | 1 |
| 0 | 0 | 0 | 1 |
| 0 | 0 | 0 | 0 |
| 0 | 0 | 0 | 1 |
| 0 | 0 | 0 | 0 |
| 0 | 0 | 0 | 1 |
| 1 | 0 | 0 | 0 |
| 0 | 0 | 0 | 1 |
| 0 | 0 | 0 | 0 |
| 0 | 1 | 0 | 0 |
| 1 | 0 | 0 | 0 |
| 0 | 0 | 0 | 0 |
| 0 | 0 | 0 | 0 |
| 0 | 0 | 0 | 0 |
| 0 | 0 | 0 | 0 |
| 0 | 0 | 0 | 0 |

| | | | |
|---|---|---|---|
| 0 | 0 | 0 | 0 |
| 0 | 0 | 0 | 0 |
| 0 | 0 | 0 | 0 |
| 0 | 0 | 0 | 0 |
| 0 | 1 | 0 | 0 |
| 0 | 1 | 0 | 0 |
| 0 | 1 | 0 | 0 |
| 0 | 0 | 0 | 1 |
| 0 | 1 | 0 | 0 |
| 0 | 1 | 0 | 0 |
| 0 | 1 | 0 | 0 |
| 0 | 1 | 1 | 1 |
| 0 | 1 | 0 | 0 |
| 0 | 1 | 0 | 0 |
| 0 | 1 | 0 | 0 |
| 0 | 1 | 0 | 0 |
| 0 | 1 | 0 | 0 |
| 0 | 1 | 0 | 0 |
| 0 | 1 | 0 | 0 |
| 0 | 1 | 0 | 0 |
| 0 | 1 | 0 | 0 |
| 0 | 1 | 0 | 0 |
| 0 | 1 | 0 | 0 |
| 0 | 1 | 0 | 0 |
| 0 | 1 | 0 | 0 |
| 0 | 1 | 0 | 0 |
| 0 | 1 | 0 | 0 |
| 0 | 1 | 0 | 0 |
| 0 | 1 | 0 | 0 |
| 0 | 1 | 0 | 0 |
| 0 | 1 | 0 | 0 |
| 0 | 1 | 0 | 0 |
| 0 | 1 | 0 | 0 |
| 0 | 1 | 0 | 0 |
| 0 | 1 | 0 | 0 |
| 0 | 1 | 0 | 0 |
| 0 | 1 | 0 | 0 |
| 0 | 1 | 0 | 0 |
| 0 | 1 | 0 | 0 |
| 0 | 1 | 1 | 1 |
| 0 | 1 | 0 | 0 |
| 0 | 1 | 0 | 0 |
| 0 | 1 | 0 | 0 |
| 0 | 1 | 0 | 0 |
| 0 | 1 | 0 | 0 |
| 0 | 1 | 0 | 0 |
| 0 | 1 | 0 | 0 |
| 0 | 1 | 0 | 0 |
| 0 | 0 | 0 | 1 |
| 0 | 0 | 0 | 1 |
| 0 | 1 | 1 | 1 |
| 1 | 1 | 0 | 0 |
| 0 | 1 | 0 | 0 |

| | | | |
|---|---|---|---|
| 0 | 0 | 0 | 0 |
| 0 | 0 | 0 | 0 |
| 1 | 1 | 0 | 0 |
| 1 | 1 | 0 | 0 |
| 0 | 0 | 0 | 0 |
| 0 | 0 | 0 | 0 |
| 0 | 1 | 0 | 0 |
| 0 | 1 | 1 | 1 |
| 0 | 1 | 1 | 1 |
| 0 | 0 | 0 | 0 |
| 0 | 1 | 0 | 0 |
| 0 | 0 | 0 | 1 |
| 0 | 1 | 0 | 0 |
| 0 | 1 | 0 | 0 |
| 0 | 0 | 0 | 0 |
| 0 | 0 | 0 | 0 |
| 0 | 0 | 0 | 0 |
| 0 | 1 | 1 | 1 |
| 0 | 0 | 0 | 0 |
| 0 | 1 | 1 | 1 |
| 0 | 1 | 0 | 0 |
| 0 | 1 | 0 | 0 |
| 0 | 0 | 0 | 1 |
| 0 | 1 | 0 | 0 |
| 0 | 1 | 0 | 0 |
| 0 | 1 | 1 | 1 |
| 0 | 0 | 0 | 0 |
| 0 | 0 | 0 | 0 |
| 0 | 1 | 0 | 0 |
| 0 | 1 | 0 | 0 |
| 0 | 1 | 0 | 0 |
| 0 | 0 | 0 | 0 |
| 0 | 1 | 1 | 1 |
| 0 | 0 | 0 | 0 |
| 0 | 1 | 0 | 0 |
| 0 | 0 | 0 | 0 |
| 0 | 0 | 0 | 1 |
| 0 | 0 | 0 | 0 |
| 0 | 0 | 0 | 0 |
| 0 | 0 | 0 | 0 |
| 0 | 1 | 0 | 0 |
| 0 | 1 | 0 | 0 |
| 0 | 1 | 0 | 0 |
| 0 | 1 | 0 | 0 |
| 0 | 1 | 0 | 0 |
| 0 | 1 | 1 | 1 |
| 0 | 1 | 0 | 0 |
| 0 | 1 | 0 | 0 |
| 0 | 1 | 0 | 0 |
| 0 | 1 | 0 | 0 |
| 0 | 1 | 0 | 0 |
| 0 | 1 | 0 | 0 |
| 0 | 1 | 1 | 1 |
| 0 | 1 | 1 | 1 |

| | | | |
|---|---|---|---|
| 0 | 1 | 1 | 1 |
| 0 | 0 | 0 | 1 |
| 0 | 0 | 0 | 0 |
| 0 | 0 | 0 | 0 |
| 0 | 1 | 0 | 0 |
| 0 | 1 | 0 | 0 |
| 0 | 1 | 0 | 0 |
| 0 | 1 | 1 | 1 |
| 0 | 1 | 0 | 0 |
| 0 | 1 | 1 | 1 |
| 0 | 0 | 0 | 1 |
| 0 | 0 | 0 | 1 |
| 0 | 1 | 0 | 0 |
| 0 | 0 | 0 | 0 |
| 0 | 1 | 0 | 0 |
| 0 | 0 | 0 | 0 |
| 0 | 1 | 0 | 0 |
| 1 | 1 | 0 | 0 |
| 0 | 0 | 0 | 1 |
| 0 | 1 | 1 | 1 |
| 0 | 0 | 0 | 1 |
| 0 | 0 | 0 | 0 |
| 0 | 0 | 0 | 1 |
| 0 | 0 | 0 | 0 |
| 0 | 0 | 0 | 0 |
| 0 | 1 | 0 | 0 |
| 0 | 0 | 0 | 0 |
| 0 | 0 | 0 | 0 |
| 0 | 1 | 0 | 0 |
| 0 | 0 | 0 | 0 |
| 0 | 1 | 0 | 0 |
| 0 | 1 | 0 | 0 |
| 0 | 0 | 0 | 0 |
| 0 | 0 | 0 | 0 |
| 0 | 1 | 0 | 0 |
| 0 | 0 | 0 | 0 |
| 0 | 1 | 0 | 0 |
| 0 | 1 | 0 | 0 |
| 1 | 1 | 0 | 0 |
| 0 | 1 | 0 | 0 |
| 0 | 1 | 0 | 0 |
| 0 | 1 | 1 | 1 |
| 0 | 1 | 0 | 0 |
| 0 | 1 | 0 | 0 |
| 0 | 0 | 0 | 0 |
| 0 | 1 | 0 | 0 |
| 0 | 1 | 1 | 1 |
| 0 | 1 | 0 | 0 |
| 0 | 1 | 0 | 0 |
| 0 | 0 | 0 | 0 |
| 0 | 1 | 0 | 0 |
| 0 | 0 | 0 | 0 |
| 0 | 1 | 1 | 1 |
| 0 | 0 | 0 | 1 |

| | | | |
|---|---|---|---|
| 0 | 1 | 0 | 0 |
| 0 | 1 | 0 | 0 |
| 0 | 1 | 1 | 1 |
| 0 | 1 | 1 | 1 |
| 0 | 1 | 1 | 1 |
| 0 | 1 | 0 | 0 |
| 0 | 1 | 1 | 1 |
| 0 | 1 | 1 | 1 |
| 0 | 1 | 0 | 0 |
| 0 | 0 | 0 | 0 |
| 0 | 1 | 0 | 0 |
| 0 | 1 | 0 | 0 |
| 0 | 0 | 0 | 0 |
| 0 | 1 | 1 | 1 |
| 0 | 1 | 1 | 1 |
| 0 | 0 | 0 | 1 |
| 0 | 0 | 0 | 1 |
| 0 | 0 | 0 | 1 |
| 0 | 0 | 0 | 0 |
| 0 | 0 | 0 | 0 |
| 0 | 1 | 1 | 1 |
| 0 | 1 | 1 | 1 |
| 0 | 1 | 0 | 0 |
| 0 | 1 | 0 | 0 |
| 0 | 1 | 1 | 1 |
| 0 | 1 | 1 | 1 |
| 0 | 1 | 0 | 0 |
| 0 | 1 | 1 | 1 |
| 0 | 0 | 0 | 0 |
| 0 | 0 | 0 | 0 |
| 0 | 1 | 0 | 0 |
| 0 | 1 | 0 | 0 |
| 0 | 1 | 0 | 0 |
| 1 | 1 | 0 | 0 |
| 1 | 1 | 0 | 0 |
| 0 | 1 | 0 | 0 |
| 0 | 1 | 0 | 0 |
| 0 | 1 | 0 | 0 |
| 0 | 1 | 1 | 1 |
| 0 | 1 | 0 | 0 |
| 0 | 1 | 1 | 1 |
| 0 | 1 | 1 | 1 |
| 0 | 1 | 1 | 1 |
| 0 | 0 | 0 | 1 |
| 0 | 0 | 0 | 1 |
| 0 | 0 | 0 | 0 |
| 0 | 0 | 0 | 0 |
| 0 | 0 | 0 | 0 |
| 0 | 0 | 0 | 0 |
| 0 | 1 | 0 | 0 |
| 0 | 1 | 0 | 0 |
| 0 | 1 | 0 | 0 |
| 0 | 1 | 0 | 0 |
| 0 | 1 | 1 | 1 |

| | | | |
|---|---|---|---|
| 0 | 1 | 1 | 1 |
| 0 | 1 | 0 | 0 |
| 0 | 1 | 0 | 0 |
| 0 | 1 | 0 | 0 |
| 0 | 1 | 1 | 1 |
| 0 | 1 | 0 | 0 |
| 0 | 1 | 0 | 0 |
| 0 | 1 | 0 | 0 |
| 0 | 1 | 1 | 1 |
| 0 | 1 | 1 | 1 |
| 0 | 1 | 0 | 0 |
| 0 | 1 | 0 | 0 |
| 0 | 1 | 0 | 0 |
| 0 | 1 | 1 | 1 |
| 0 | 1 | 0 | 0 |
| 0 | 1 | 0 | 0 |
| 0 | 1 | 0 | 0 |
| 0 | 1 | 0 | 0 |
| 0 | 0 | 0 | 0 |
| 0 | 1 | 0 | 0 |
| 0 | 1 | 0 | 0 |
| 0 | 0 | 0 | 1 |
| 0 | 0 | 0 | 0 |
| 0 | 0 | 0 | 1 |
| 0 | 1 | 0 | 0 |
| 0 | 1 | 0 | 0 |
| 0 | 1 | 1 | 1 |
| 0 | 1 | 0 | 0 |
| 0 | 1 | 0 | 0 |
| 0 | 1 | 0 | 0 |
| 0 | 1 | 0 | 0 |
| 0 | 1 | 0 | 0 |
| 0 | 1 | 0 | 0 |
| 0 | 1 | 0 | 0 |
| 0 | 0 | 0 | 1 |
| 0 | 1 | 0 | 0 |
| 0 | 1 | 0 | 0 |
| 0 | 1 | 0 | 0 |
| 0 | 1 | 0 | 0 |
| 0 | 0 | 0 | 0 |
| 0 | 0 | 0 | 0 |
| 0 | 0 | 0 | 0 |
| 0 | 0 | 0 | 0 |
| 0 | 0 | 0 | 0 |
| 0 | 0 | 0 | 0 |
| 0 | 0 | 0 | 0 |
| 0 | 0 | 0 | 0 |
| 0 | 0 | 0 | 1 |
| 0 | 1 | 0 | 0 |
| 0 | 0 | 0 | 1 |
| 0 | 0 | 0 | 0 |
| 0 | 0 | 0 | 0 |
| 0 | 0 | 0 | 0 |
| 0 | 0 | 0 | 0 |

| | | | |
|---|---|---|---|
| 0 | 0 | 0 | 0 |
| 0 | 1 | 0 | 0 |
| 0 | 0 | 0 | 1 |
| 0 | 1 | 0 | 0 |
| 0 | 0 | 0 | 0 |
| 0 | 0 | 0 | 1 |
| 0 | 0 | 0 | 1 |
| 0 | 0 | 0 | 1 |
| 0 | 0 | 0 | 0 |
| 0 | 0 | 0 | 1 |
| 0 | 1 | 0 | 0 |
| 0 | 1 | 0 | 0 |
| 0 | 1 | 0 | 0 |
| 0 | 0 | 0 | 1 |
| 0 | 1 | 0 | 0 |
| 0 | 0 | 0 | 1 |
| 0 | 1 | 0 | 0 |
| 0 | 1 | 0 | 0 |
| 1 | 0 | 0 | 0 |
| 0 | 0 | 0 | 1 |
| 0 | 0 | 0 | 1 |
| 0 | 0 | 0 | 0 |
| 0 | 0 | 0 | 1 |
| 0 | 0 | 0 | 0 |
| 0 | 0 | 0 | 0 |
| 0 | 0 | 0 | 1 |
| 0 | 0 | 0 | 1 |
| 0 | 0 | 0 | 0 |
| 0 | 0 | 0 | 1 |
| 1 | 0 | 0 | 0 |
| 0 | 0 | 0 | 1 |
| 0 | 0 | 0 | 1 |
| 0 | 0 | 0 | 1 |
| 0 | 0 | 0 | 1 |
| 0 | 0 | 0 | 1 |
| 0 | 0 | 0 | 1 |
| 0 | 0 | 0 | 1 |
| 0 | 0 | 0 | 1 |
| 0 | 0 | 0 | 1 |
| 0 | 0 | 0 | 1 |
| 0 | 0 | 0 | 1 |
| 0 | 0 | 0 | 1 |
| 0 | 0 | 0 | 1 |
| 0 | 0 | 0 | 1 |
| 0 | 0 | 0 | 1 |
| 0 | 0 | 0 | 1 |
| 0 | 0 | 0 | 1 |
| 0 | 0 | 0 | 1 |
| 0 | 0 | 0 | 1 |
| 0 | 0 | 0 | 1 |
| 0 | 0 | 0 | 1 |
| 0 | 0 | 0 | 1 |
| 0 | 0 | 0 | 1 |
| 0 | 0 | 0 | 1 |

| | | | |
|---|---|---|---|
| 0 | 0 | 0 | 1 |
| 0 | 0 | 0 | 1 |
| 0 | 0 | 0 | 1 |
| 0 | 0 | 0 | 1 |
| 0 | 0 | 0 | 1 |
| 0 | 0 | 0 | 1 |
| 0 | 0 | 0 | 1 |
| 0 | 0 | 0 | 1 |
| 0 | 0 | 0 | 1 |
| 0 | 0 | 0 | 1 |
| 0 | 0 | 0 | 1 |
| 0 | 0 | 0 | 0 |
| 0 | 1 | 0 | 0 |
| 0 | 1 | 0 | 0 |
| 0 | 1 | 1 | 1 |
| 0 | 1 | 0 | 0 |
| 0 | 1 | 1 | 1 |
| 0 | 1 | 0 | 0 |
| 0 | 1 | 1 | 1 |
| 0 | 0 | 0 | 1 |
| 0 | 0 | 0 | 0 |
| 0 | 0 | 0 | 1 |
| 0 | 0 | 0 | 1 |
| 0 | 0 | 0 | 0 |
| 0 | 0 | 0 | 0 |
| 0 | 0 | 0 | 1 |
| 0 | 0 | 0 | 1 |
| 0 | 0 | 0 | 0 |
| 0 | 0 | 0 | 0 |
| 0 | 0 | 0 | 0 |
| 0 | 0 | 0 | 0 |
| 0 | 0 | 0 | 0 |
| 0 | 0 | 0 | 1 |
| 0 | 0 | 0 | 1 |
| 0 | 0 | 0 | 1 |
| 0 | 0 | 0 | 1 |
| 0 | 0 | 0 | 1 |
| 0 | 0 | 0 | 1 |
| 0 | 0 | 0 | 1 |
| 0 | 1 | 0 | 0 |
| 0 | 1 | 0 | 0 |
| 0 | 0 | 0 | 1 |
| 0 | 1 | 0 | 0 |
| 0 | 1 | 0 | 0 |
| 0 | 1 | 0 | 0 |
| 0 | 1 | 1 | 1 |
| 0 | 1 | 0 | 0 |
| 0 | 0 | 0 | 1 |
| 0 | 1 | 1 | 1 |
| 0 | 1 | 0 | 0 |
| 0 | 1 | 0 | 0 |
| 0 | 0 | 0 | 0 |
| 0 | 0 | 0 | 1 |
| 0 | 1 | 0 | 0 |

| | | | |
|---|---|---|---|
| 0 | 1 | 0 | 0 |
| 0 | 0 | 0 | 1 |
| 0 | 0 | 0 | 1 |
| 0 | 0 | 0 | 1 |
| 0 | 1 | 0 | 0 |
| 0 | 0 | 0 | 1 |
| 0 | 0 | 0 | 0 |
| 0 | 0 | 0 | 1 |
| 0 | 0 | 0 | 1 |
| 0 | 0 | 0 | 0 |
| 0 | 1 | 1 | 1 |
| 0 | 0 | 0 | 1 |
| 0 | 0 | 0 | 1 |
| 0 | 1 | 0 | 0 |
| 0 | 1 | 1 | 1 |
| 0 | 1 | 1 | 1 |
| 0 | 1 | 0 | 0 |
| 0 | 1 | 1 | 1 |
| 0 | 1 | 0 | 0 |
| 0 | 1 | 0 | 0 |
| 0 | 1 | 0 | 0 |
| 0 | 0 | 0 | 0 |
| 0 | 0 | 0 | 0 |
| 0 | 1 | 0 | 0 |
| 0 | 1 | 0 | 0 |
| 0 | 1 | 0 | 0 |
| 0 | 0 | 0 | 0 |
| 0 | 1 | 0 | 0 |
| 0 | 1 | 0 | 0 |
| 0 | 1 | 0 | 0 |
| 0 | 0 | 0 | 0 |
| 0 | 1 | 0 | 0 |
| 0 | 1 | 0 | 0 |
| 0 | 1 | 0 | 0 |
| 0 | 1 | 0 | 0 |
| 0 | 1 | 0 | 0 |
| 0 | 1 | 1 | 1 |
| 0 | 1 | 0 | 0 |
| 0 | 1 | 0 | 0 |
| 0 | 1 | 1 | 1 |
| 0 | 1 | 1 | 1 |
| 0 | 1 | 0 | 0 |
| 0 | 0 | 0 | 0 |
| 0 | 1 | 0 | 0 |
| 0 | 1 | 0 | 0 |
| 0 | 1 | 0 | 0 |
| 0 | 1 | 0 | 0 |
| 0 | 1 | 0 | 0 |
| 0 | 1 | 1 | 1 |
| 0 | 1 | 1 | 1 |
| 0 | 0 | 0 | 0 |
| 0 | 0 | 0 | 1 |
| 0 | 1 | 0 | 0 |
| 0 | 1 | 0 | 0 |

| | | | |
|---|---|---|---|
| 0 | 0 | 0 | 0 |
| 0 | 1 | 0 | 0 |
| 0 | 1 | 0 | 0 |
| 0 | 1 | 0 | 0 |
| 0 | 0 | 0 | 0 |
| 0 | 1 | 0 | 0 |
| 0 | 1 | 0 | 0 |
| 0 | 0 | 0 | 0 |
| 0 | 0 | 0 | 0 |
| 0 | 0 | 0 | 0 |
| 0 | 0 | 0 | 0 |
| 0 | 0 | 0 | 0 |
| 0 | 0 | 0 | 0 |
| 0 | 0 | 0 | 0 |
| 0 | 0 | 0 | 1 |
| 0 | 1 | 0 | 0 |
| 0 | 1 | 0 | 0 |
| 0 | 1 | 0 | 0 |
| 0 | 1 | 1 | 1 |
| 0 | 0 | 0 | 0 |
| 0 | 1 | 0 | 0 |
| 0 | 1 | 0 | 0 |
| 0 | 1 | 0 | 0 |
| 0 | 1 | 0 | 0 |
| 0 | 1 | 0 | 0 |
| 0 | 1 | 1 | 1 |
| 0 | 1 | 0 | 0 |
| 0 | 1 | 0 | 0 |
| 0 | 1 | 0 | 0 |
| 0 | 0 | 0 | 0 |
| 0 | 0 | 0 | 1 |
| 0 | 0 | 0 | 0 |
| 0 | 0 | 0 | 1 |
| 0 | 1 | 1 | 1 |
| 1 | 1 | 0 | 0 |
| 0 | 0 | 0 | 0 |
| 0 | 0 | 0 | 1 |
| 0 | 0 | 0 | 1 |
| 0 | 0 | 0 | 1 |
| 0 | 1 | 1 | 1 |
| 0 | 1 | 0 | 0 |
| 0 | 1 | 0 | 0 |
| 0 | 0 | 0 | 0 |
| 0 | 0 | 0 | 0 |
| 0 | 0 | 0 | 0 |
| 0 | 1 | 0 | 0 |
| 0 | 0 | 0 | 0 |
| 0 | 0 | 0 | 0 |
| 0 | 0 | 0 | 1 |
| 0 | 0 | 0 | 1 |
| 0 | 0 | 0 | 0 |
| 0 | 0 | 0 | 0 |
| 0 | 1 | 0 | 0 |
| 0 | 0 | 0 | 1 |

| | | | |
|---|---|---|---|
| 0 | 0 | 0 | 1 |
| 0 | 0 | 0 | 1 |
| 0 | 1 | 1 | 1 |
| 0 | 1 | 0 | 0 |
| 0 | 0 | 0 | 1 |
| 0 | 0 | 0 | 1 |
| 0 | 0 | 0 | 0 |
| 0 | 0 | 0 | 1 |
| 0 | 0 | 0 | 0 |
| 0 | 0 | 0 | 0 |
| 0 | 0 | 0 | 0 |
| 0 | 0 | 0 | 0 |
| 0 | 0 | 0 | 0 |
| 0 | 0 | 0 | 0 |
| 0 | 0 | 0 | 0 |
| 0 | 0 | 0 | 0 |
| 0 | 0 | 0 | 1 |
| 0 | 0 | 0 | 0 |
| 0 | 1 | 0 | 0 |
| 0 | 1 | 1 | 1 |
| 0 | 1 | 0 | 0 |
| 0 | 0 | 0 | 0 |
| 0 | 0 | 0 | 1 |
| 0 | 0 | 0 | 0 |
| 0 | 0 | 0 | 1 |
| 0 | 0 | 0 | 1 |
| 0 | 0 | 0 | 1 |
| 0 | 0 | 0 | 0 |
| 0 | 0 | 0 | 0 |
| 0 | 0 | 0 | 0 |
| 0 | 0 | 0 | 1 |
| 0 | 0 | 0 | 0 |
| 0 | 0 | 0 | 0 |
| 0 | 0 | 0 | 0 |
| 0 | 0 | 0 | 1 |
| 0 | 0 | 0 | 1 |
| 0 | 1 | 0 | 0 |
| 0 | 0 | 0 | 0 |
| 0 | 1 | 0 | 0 |
| 0 | 0 | 0 | 1 |
| 0 | 0 | 0 | 1 |
| 0 | 0 | 0 | 1 |
| 0 | 0 | 0 | 1 |
| 0 | 0 | 0 | 0 |
| 0 | 0 | 0 | 1 |
| 0 | 0 | 0 | 0 |
| 0 | 0 | 0 | 1 |
| 0 | 0 | 0 | 0 |
| 0 | 0 | 0 | 0 |
| 0 | 0 | 0 | 1 |
| 0 | 0 | 0 | 1 |
| 0 | 1 | 0 | 0 |
| 0 | 1 | 0 | 0 |
| 0 | 0 | 0 | 1 |

| | | | |
|---|---|---|---|
| 0 | 0 | 0 | 1 |
| 0 | 0 | 0 | 1 |
| 0 | 0 | 0 | 1 |
| 0 | 1 | 1 | 1 |
| 0 | 1 | 0 | 0 |
| 0 | 1 | 1 | 1 |
| 0 | 1 | 1 | 1 |
| 0 | 1 | 0 | 0 |
| 0 | 1 | 0 | 0 |
| 0 | 0 | 0 | 1 |
| 0 | 0 | 0 | 0 |
| 0 | 0 | 0 | 0 |
| 1 | 0 | 0 | 0 |
| 1 | 0 | 0 | 0 |
| 0 | 0 | 0 | 0 |
| 1 | 0 | 0 | 0 |
| 1 | 0 | 0 | 0 |
| 1 | 0 | 0 | 0 |
| 1 | 0 | 0 | 0 |
| 0 | 0 | 0 | 1 |
| 0 | 0 | 0 | 1 |
| 0 | 0 | 0 | 1 |
| 0 | 0 | 0 | 1 |
| 0 | 0 | 0 | 1 |
| 0 | 0 | 0 | 0 |
| 0 | 0 | 0 | 1 |
| 1 | 0 | 0 | 0 |
| 1 | 0 | 0 | 0 |
| 0 | 0 | 0 | 1 |
| 0 | 0 | 0 | 1 |
| 0 | 0 | 0 | 1 |
| 0 | 0 | 0 | 1 |
| 0 | 0 | 0 | 1 |
| 0 | 0 | 0 | 0 |
| 0 | 0 | 0 | 0 |
| 0 | 0 | 0 | 1 |
| 0 | 0 | 0 | 0 |
| 0 | 0 | 0 | 1 |
| 0 | 0 | 0 | 0 |
| 0 | 1 | 0 | 0 |
| 0 | 1 | 0 | 0 |
| 0 | 1 | 0 | 0 |
| 0 | 0 | 0 | 1 |
| 0 | 0 | 0 | 0 |
| 0 | 0 | 0 | 0 |
| 1 | 0 | 0 | 0 |
| 1 | 1 | 0 | 0 |
| 1 | 0 | 0 | 0 |
| 0 | 0 | 0 | 0 |
| 0 | 1 | 0 | 0 |
| 0 | 1 | 0 | 0 |
| 1 | 1 | 0 | 0 |
| 1 | 0 | 0 | 0 |
| 0 | 0 | 0 | 1 |

| | | | |
|---|---|---|---|
| 0 | 0 | 0 | 0 |
| 0 | 0 | 0 | 1 |
| 0 | 0 | 0 | 0 |
| 0 | 0 | 0 | 0 |
| 0 | 1 | 0 | 0 |
| 0 | 0 | 0 | 1 |
| 0 | 0 | 0 | 0 |
| 0 | 0 | 0 | 1 |
| 0 | 0 | 0 | 1 |
| 0 | 0 | 0 | 0 |
| 0 | 0 | 0 | 1 |
| 0 | 0 | 0 | 1 |
| 0 | 0 | 0 | 1 |
| 0 | 0 | 0 | 1 |
| 0 | 0 | 0 | 1 |
| 0 | 1 | 0 | 0 |
| 0 | 1 | 0 | 0 |
| 0 | 0 | 0 | 1 |
| 0 | 0 | 0 | 0 |
| 0 | 0 | 0 | 1 |
| 0 | 0 | 0 | 0 |
| 0 | 0 | 0 | 0 |
| 0 | 0 | 0 | 0 |
| 0 | 1 | 0 | 0 |
| 0 | 1 | 0 | 0 |
| 0 | 1 | 0 | 0 |
| 0 | 1 | 0 | 0 |
| 0 | 0 | 0 | 0 |
| 0 | 0 | 0 | 0 |
| 0 | 1 | 0 | 0 |
| 0 | 1 | 0 | 0 |
| 0 | 1 | 0 | 0 |
| 0 | 1 | 1 | 1 |
| 0 | 1 | 1 | 1 |
| 0 | 1 | 0 | 0 |
| 0 | 1 | 0 | 0 |
| 0 | 1 | 1 | 1 |
| 0 | 1 | 0 | 0 |
| 0 | 1 | 1 | 1 |
| 0 | 1 | 0 | 0 |
| 0 | 1 | 0 | 0 |
| 0 | 1 | 0 | 0 |
| 0 | 1 | 1 | 1 |
| 0 | 0 | 0 | 1 |
| 0 | 1 | 0 | 0 |
| 0 | 1 | 0 | 0 |
| 0 | 1 | 0 | 0 |
| 0 | 1 | 0 | 0 |
| 0 | 1 | 0 | 0 |
| 0 | 1 | 0 | 0 |
| 0 | 1 | 1 | 1 |
| 0 | 1 | 0 | 0 |
| 0 | 1 | 0 | 0 |
| 0 | 1 | 1 | 1 |

| | | | |
|---|---|---|---|
| 0 | 1 | 0 | 0 |
| 0 | 1 | 1 | 1 |
| 0 | 1 | 0 | 0 |
| 0 | 0 | 0 | 0 |
| 0 | 0 | 0 | 0 |
| 0 | 0 | 0 | 0 |
| 0 | 1 | 0 | 0 |
| 0 | 0 | 0 | 0 |
| 0 | 0 | 0 | 0 |
| 0 | 1 | 0 | 0 |
| 0 | 0 | 0 | 0 |
| 0 | 0 | 0 | 1 |
| 0 | 0 | 0 | 0 |
| 0 | 0 | 0 | 1 |
| 0 | 1 | 1 | 1 |
| 0 | 1 | 1 | 1 |
| 0 | 1 | 0 | 0 |
| 0 | 1 | 0 | 0 |
| 0 | 1 | 1 | 1 |
| 0 | 1 | 0 | 0 |
| 0 | 1 | 1 | 1 |
| 0 | 1 | 0 | 0 |
| 0 | 0 | 0 | 1 |
| 0 | 1 | 0 | 0 |
| 0 | 0 | 0 | 1 |
| 0 | 0 | 0 | 1 |
| 0 | 0 | 0 | 1 |
| 0 | 0 | 0 | 0 |
| 0 | 0 | 0 | 1 |
| 0 | 0 | 0 | 1 |
| 0 | 0 | 0 | 1 |
| 1 | 0 | 0 | 0 |
| 0 | 0 | 0 | 1 |
| 0 | 0 | 0 | 0 |
| 0 | 0 | 0 | 0 |
| 0 | 0 | 0 | 0 |
| 0 | 0 | 0 | 1 |
| 0 | 0 | 0 | 1 |
| 0 | 1 | 0 | 0 |
| 0 | 0 | 0 | 0 |
| 0 | 1 | 0 | 0 |
| 0 | 0 | 0 | 0 |
| 0 | 1 | 0 | 0 |
| 0 | 0 | 0 | 0 |
| 0 | 1 | 0 | 0 |
| 0 | 1 | 0 | 0 |
| 0 | 0 | 0 | 1 |
| 0 | 0 | 0 | 1 |
| 0 | 0 | 0 | 0 |
| 0 | 0 | 0 | 0 |
| 0 | 0 | 0 | 0 |
| 0 | 0 | 0 | 1 |
| 0 | 0 | 0 | 1 |
| 0 | 0 | 0 | 0 |

| | | | |
|---|---|---|---|
| 0 | 0 | 0 | 0 |
| 0 | 0 | 0 | 1 |
| 0 | 0 | 0 | 1 |
| 0 | 0 | 0 | 1 |
| 0 | 0 | 0 | 0 |
| 0 | 1 | 0 | 0 |
| 0 | 1 | 0 | 0 |
| 0 | 1 | 0 | 0 |
| 0 | 1 | 0 | 0 |
| 0 | 1 | 0 | 0 |
| 0 | 0 | 0 | 1 |
| 0 | 1 | 0 | 0 |
| 0 | 0 | 0 | 0 |
| 0 | 0 | 0 | 0 |
| 0 | 1 | 1 | 1 |
| 0 | 0 | 0 | 1 |
| 0 | 1 | 0 | 0 |
| 0 | 1 | 1 | 1 |
| 0 | 1 | 0 | 0 |
| 0 | 1 | 0 | 0 |
| 0 | 1 | 1 | 1 |
| 0 | 1 | 0 | 0 |
| 0 | 1 | 0 | 0 |
| 0 | 1 | 1 | 1 |
| 0 | 1 | 0 | 0 |
| 0 | 1 | 0 | 0 |
| 0 | 1 | 0 | 0 |
| 0 | 1 | 1 | 1 |
| 0 | 1 | 0 | 0 |
| 0 | 0 | 0 | 0 |
| 0 | 0 | 0 | 0 |
| 0 | 0 | 0 | 1 |
| 0 | 0 | 0 | 1 |
| 0 | 0 | 0 | 1 |
| 0 | 0 | 0 | 1 |
| 0 | 0 | 0 | 1 |
| 0 | 0 | 0 | 1 |
| 0 | 0 | 0 | 1 |
| 0 | 0 | 0 | 1 |
| 0 | 0 | 0 | 1 |
| 0 | 0 | 0 | 1 |
| 0 | 0 | 0 | 1 |
| 1 | 0 | 0 | 0 |
| 0 | 0 | 0 | 1 |
| 0 | 0 | 0 | 1 |
| 0 | 0 | 0 | 1 |
| 0 | 0 | 0 | 1 |
| 0 | 0 | 0 | 1 |
| 0 | 0 | 0 | 1 |
| 0 | 0 | 0 | 1 |
| 0 | 0 | 0 | 1 |
| 0 | 0 | 0 | 1 |
| 0 | 0 | 0 | 1 |
| 0 | 0 | 0 | 1 |

| | | | |
|---|---|---|---|
| 0 | 0 | 0 | 1 |
| 0 | 0 | 0 | 1 |
| 0 | 0 | 0 | 1 |
| 0 | 0 | 0 | 1 |
| 0 | 0 | 0 | 1 |
| 0 | 0 | 0 | 1 |
| 0 | 0 | 0 | 1 |
| 0 | 0 | 0 | 1 |
| 0 | 1 | 0 | 0 |
| 0 | 0 | 0 | 0 |
| 0 | 0 | 0 | 0 |
| 0 | 0 | 0 | 0 |
| 0 | 0 | 0 | 0 |
| 0 | 0 | 0 | 0 |
| 0 | 0 | 0 | 0 |
| 0 | 0 | 0 | 0 |
| 0 | 0 | 0 | 0 |
| 0 | 0 | 0 | 0 |
| 0 | 0 | 0 | 0 |
| 0 | 1 | 0 | 0 |
| 1 | 0 | 0 | 0 |
| 1 | 0 | 0 | 0 |
| 0 | 1 | 0 | 0 |
| 0 | 1 | 0 | 0 |
| 0 | 1 | 0 | 0 |
| 0 | 0 | 0 | 1 |
| 0 | 0 | 0 | 1 |
| 0 | 0 | 0 | 0 |
| 0 | 0 | 0 | 0 |
| 1 | 0 | 0 | 0 |
| 1 | 0 | 0 | 0 |
| 0 | 0 | 0 | 0 |
| 1 | 0 | 0 | 0 |
| 1 | 0 | 0 | 0 |
| 0 | 0 | 0 | 0 |
| 0 | 0 | 0 | 0 |
| 0 | 0 | 0 | 0 |
| 0 | 1 | 0 | 0 |
| 0 | 1 | 0 | 0 |
| 0 | 0 | 0 | 0 |
| 0 | 0 | 0 | 0 |
| 0 | 0 | 0 | 0 |
| 1 | 0 | 0 | 0 |
| 1 | 0 | 0 | 0 |
| 1 | 0 | 0 | 0 |
| 0 | 0 | 0 | 0 |
| 0 | 0 | 0 | 1 |
| 0 | 0 | 0 | 1 |
| 0 | 0 | 0 | 1 |
| 0 | 0 | 0 | 1 |
| 0 | 0 | 0 | 1 |
| 0 | 0 | 0 | 1 |
| 0 | 1 | 0 | 0 |
| 0 | 1 | 0 | 0 |

| | | | |
|---|---|---|---|
| 0 | 1 | 0 | 0 |
| 0 | 0 | 0 | 0 |
| 0 | 0 | 0 | 0 |
| 0 | 1 | 0 | 0 |
| 0 | 0 | 0 | 1 |
| 0 | 0 | 0 | 0 |
| 0 | 0 | 0 | 0 |
| 0 | 0 | 0 | 0 |
| 0 | 0 | 0 | 1 |
| 0 | 0 | 0 | 0 |
| 1 | 0 | 0 | 0 |
| 0 | 0 | 0 | 1 |
| 0 | 0 | 0 | 0 |
| 0 | 0 | 0 | 0 |
| 0 | 0 | 0 | 0 |
| 0 | 0 | 0 | 0 |
| 0 | 0 | 0 | 1 |
| 0 | 0 | 0 | 0 |
| 0 | 0 | 0 | 0 |
| 0 | 0 | 0 | 1 |
| 0 | 1 | 0 | 0 |
| 0 | 0 | 0 | 0 |
| 0 | 1 | 0 | 0 |
| 0 | 1 | 0 | 0 |
| 0 | 1 | 0 | 0 |
| 0 | 0 | 0 | 1 |
| 0 | 0 | 0 | 1 |
| 0 | 0 | 0 | 1 |
| 0 | 0 | 0 | 1 |
| 0 | 0 | 0 | 1 |
| 0 | 0 | 0 | 1 |
| 0 | 0 | 0 | 1 |
| 0 | 1 | 1 | 1 |
| 0 | 1 | 1 | 1 |
| 0 | 0 | 0 | 0 |
| 0 | 1 | 0 | 0 |
| 0 | 0 | 0 | 0 |
| 0 | 1 | 0 | 0 |
| 0 | 1 | 0 | 0 |
| 0 | 1 | 0 | 0 |
| 0 | 0 | 0 | 1 |
| 0 | 0 | 0 | 1 |
| 0 | 0 | 0 | 1 |
| 0 | 0 | 0 | 0 |
| 0 | 0 | 0 | 0 |
| 0 | 0 | 0 | 0 |
| 0 | 0 | 0 | 1 |
| 0 | 0 | 0 | 1 |
| 0 | 0 | 0 | 0 |
| 0 | 0 | 0 | 0 |
| 0 | 0 | 0 | 1 |
| 0 | 0 | 0 | 1 |
| 0 | 0 | 0 | 1 |
| 0 | 0 | 0 | 1 |

| | | | |
|---|---|---|---|
| 0 | 0 | 0 | 1 |
| 0 | 0 | 0 | 1 |
| 0 | 0 | 0 | 1 |
| 0 | 0 | 0 | 0 |
| 0 | 0 | 0 | 0 |
| 0 | 0 | 0 | 1 |
| 0 | 0 | 0 | 1 |
| 0 | 0 | 0 | 1 |
| 0 | 0 | 0 | 1 |
| 0 | 0 | 0 | 1 |
| 0 | 0 | 0 | 0 |
| 0 | 0 | 0 | 1 |
| 0 | 1 | 0 | 0 |
| 0 | 0 | 0 | 0 |
| 0 | 0 | 0 | 0 |
| 0 | 0 | 0 | 1 |
| 0 | 0 | 0 | 1 |
| 0 | 0 | 0 | 1 |
| 0 | 0 | 0 | 1 |
| 0 | 0 | 0 | 1 |
| 0 | 0 | 0 | 0 |
| 0 | 0 | 0 | 1 |
| 0 | 0 | 0 | 1 |
| 0 | 1 | 0 | 0 |
| 0 | 1 | 1 | 1 |
| 0 | 1 | 0 | 0 |
| 0 | 1 | 0 | 0 |
| 0 | 0 | 0 | 1 |
| 0 | 0 | 0 | 1 |
| 0 | 0 | 0 | 1 |
| 0 | 0 | 0 | 1 |
| 0 | 0 | 0 | 1 |
| 0 | 0 | 0 | 1 |
| 0 | 0 | 0 | 1 |
| 0 | 0 | 0 | 1 |
| 0 | 0 | 0 | 0 |
| 0 | 0 | 0 | 1 |
| 0 | 0 | 0 | 1 |
| 0 | 0 | 0 | 0 |
| 0 | 0 | 0 | 0 |
| 0 | 0 | 0 | 1 |
| 0 | 0 | 0 | 1 |
| 1 | 0 | 0 | 0 |
| 1 | 0 | 0 | 0 |
| 0 | 0 | 0 | 0 |
| 0 | 0 | 0 | 0 |
| 1 | 0 | 0 | 0 |
| 1 | 0 | 0 | 0 |
| 0 | 0 | 0 | 1 |
| 1 | 0 | 0 | 0 |
| 1 | 0 | 0 | 0 |
| 1 | 1 | 0 | 0 |
| 1 | 1 | 0 | 0 |
| 1 | 0 | 0 | 0 |

| | | | |
|---|---|---|---|
| 0 | 1 | 0 | 0 |
| 0 | 1 | 0 | 0 |
| 1 | 0 | 0 | 0 |
| 1 | 0 | 0 | 0 |
| 1 | 0 | 0 | 0 |
| 1 | 0 | 0 | 0 |
| 0 | 0 | 0 | 0 |
| 0 | 0 | 0 | 0 |
| 0 | 0 | 0 | 0 |
| 0 | 0 | 0 | 1 |
| 0 | 0 | 0 | 1 |
| 0 | 0 | 0 | 0 |
| 0 | 0 | 0 | 1 |
| 0 | 0 | 0 | 0 |
| 0 | 0 | 0 | 0 |
| 0 | 0 | 0 | 1 |
| 0 | 0 | 0 | 0 |
| 0 | 1 | 1 | 1 |
| 0 | 0 | 0 | 1 |
| 0 | 0 | 0 | 1 |
| 0 | 1 | 1 | 1 |
| 0 | 0 | 0 | 1 |
| 0 | 0 | 0 | 1 |
| 0 | 0 | 0 | 0 |
| 0 | 0 | 0 | 0 |
| 0 | 0 | 0 | 1 |
| 0 | 0 | 0 | 1 |
| 0 | 0 | 0 | 1 |
| 0 | 0 | 0 | 0 |
| 0 | 0 | 0 | 0 |
| 0 | 0 | 0 | 1 |
| 0 | 0 | 0 | 1 |
| 0 | 1 | 1 | 1 |
| 0 | 1 | 0 | 0 |
| 0 | 1 | 0 | 0 |
| 0 | 0 | 0 | 1 |
| 0 | 0 | 0 | 0 |
| 0 | 0 | 0 | 1 |
| 0 | 0 | 0 | 1 |
| 0 | 0 | 0 | 0 |
| 0 | 0 | 0 | 0 |
| 0 | 0 | 0 | 0 |
| 0 | 0 | 0 | 0 |
| 0 | 0 | 0 | 0 |
| 0 | 0 | 0 | 0 |
| 0 | 0 | 0 | 1 |
| 0 | 0 | 0 | 1 |
| 0 | 0 | 0 | 0 |
| 0 | 0 | 0 | 0 |
| 0 | 0 | 0 | 0 |
| 0 | 0 | 0 | 1 |
| 0 | 0 | 0 | 1 |
| 0 | 0 | 0 | 0 |
| 0 | 0 | 0 | 0 |

| | | | |
|---|---|---|---|
| 0 | 0 | 0 | 1 |
| 0 | 0 | 0 | 1 |
| 0 | 0 | 0 | 1 |
| 0 | 0 | 0 | 1 |
| 0 | 1 | 0 | 0 |
| 0 | 1 | 0 | 0 |
| 0 | 1 | 1 | 1 |
| 0 | 0 | 0 | 0 |
| 0 | 1 | 1 | 1 |
| 0 | 0 | 0 | 0 |
| 0 | 0 | 0 | 1 |
| 0 | 0 | 0 | 1 |
| 0 | 0 | 0 | 1 |
| 0 | 0 | 0 | 1 |
| 0 | 0 | 0 | 1 |
| 0 | 0 | 0 | 1 |
| 0 | 0 | 0 | 0 |
| 0 | 0 | 0 | 1 |
| 0 | 0 | 0 | 1 |
| 0 | 0 | 0 | 0 |
| 0 | 0 | 0 | 0 |
| 0 | 0 | 0 | 1 |
| 0 | 0 | 0 | 1 |
| 0 | 0 | 0 | 0 |
| 0 | 0 | 0 | 1 |
| 0 | 0 | 0 | 0 |
| 0 | 0 | 0 | 0 |
| 0 | 0 | 0 | 0 |
| 0 | 0 | 0 | 0 |
| 0 | 0 | 0 | 0 |
| 0 | 0 | 0 | 0 |
| 0 | 0 | 0 | 0 |
| 0 | 0 | 0 | 1 |
| 0 | 0 | 0 | 0 |
| 0 | 0 | 0 | 1 |
| 0 | 0 | 0 | 1 |
| 0 | 0 | 0 | 1 |
| 0 | 0 | 0 | 1 |
| 0 | 0 | 0 | 0 |
| 0 | 0 | 0 | 1 |
| 0 | 0 | 0 | 1 |
| 0 | 0 | 0 | 1 |
| 0 | 0 | 0 | 1 |
| 0 | 1 | 0 | 0 |
| 0 | 0 | 0 | 1 |
| 0 | 0 | 0 | 1 |
| 0 | 0 | 0 | 1 |
| 0 | 0 | 0 | 1 |
| 0 | 0 | 0 | 1 |
| 0 | 0 | 0 | 1 |
| 0 | 0 | 0 | 0 |
| 0 | 0 | 0 | 0 |
| 0 | 0 | 0 | 0 |
| 0 | 0 | 0 | 1 |

| | | | |
|---|---|---|---|
| 0 | 1 | 0 | 0 |
| 0 | 1 | 0 | 0 |
| 0 | 1 | 0 | 0 |
| 0 | 0 | 0 | 0 |
| 1 | 1 | 0 | 0 |
| 1 | 1 | 0 | 0 |
| 0 | 1 | 0 | 0 |
| 0 | 0 | 0 | 0 |
| 0 | 0 | 0 | 1 |
| 0 | 0 | 0 | 1 |
| 0 | 0 | 0 | 0 |
| 0 | 0 | 0 | 1 |
| 0 | 0 | 0 | 1 |
| 0 | 0 | 0 | 0 |
| 0 | 0 | 0 | 0 |
| 0 | 0 | 0 | 0 |
| 0 | 1 | 0 | 0 |
| 0 | 0 | 0 | 0 |
| 0 | 0 | 0 | 0 |
| 0 | 0 | 0 | 1 |
| 0 | 0 | 0 | 1 |
| 0 | 0 | 0 | 0 |
| 0 | 0 | 0 | 1 |
| 0 | 1 | 0 | 0 |
| 0 | 0 | 0 | 1 |
| 0 | 0 | 0 | 1 |
| 0 | 0 | 0 | 1 |
| 0 | 0 | 0 | 1 |
| 0 | 0 | 0 | 1 |
| 0 | 0 | 0 | 0 |
| 0 | 0 | 0 | 1 |
| 0 | 0 | 0 | 1 |
| 0 | 0 | 0 | 0 |
| 0 | 0 | 0 | 1 |
| 0 | 1 | 0 | 0 |
| 0 | 1 | 0 | 0 |
| 0 | 1 | 0 | 0 |
| 0 | 0 | 0 | 0 |
| 0 | 0 | 0 | 0 |
| 0 | 0 | 0 | 0 |
| 0 | 0 | 0 | 0 |
| 0 | 1 | 0 | 0 |
| 0 | 0 | 0 | 1 |
| 0 | 1 | 0 | 0 |
| 0 | 0 | 0 | 1 |
| 0 | 0 | 0 | 0 |
| 0 | 1 | 1 | 1 |
| 0 | 0 | 0 | 1 |
| 0 | 0 | 0 | 1 |
| 0 | 0 | 0 | 0 |
| 0 | 0 | 0 | 1 |
| 0 | 0 | 0 | 1 |
| 0 | 0 | 0 | 1 |
| 0 | 0 | 0 | 1 |

| | | | |
|---|---|---|---|
| 1 | 1 | 0 | 0 |
| 0 | 0 | 0 | 1 |
| 0 | 0 | 0 | 1 |
| 0 | 0 | 0 | 1 |
| 0 | 0 | 0 | 1 |
| 0 | 0 | 0 | 1 |
| 0 | 0 | 0 | 1 |
| 0 | 0 | 0 | 1 |
| 0 | 0 | 0 | 1 |
| 0 | 0 | 0 | 1 |
| 0 | 0 | 0 | 0 |
| 0 | 0 | 0 | 1 |
| 0 | 0 | 0 | 1 |
| 0 | 0 | 0 | 1 |
| 0 | 0 | 0 | 0 |
| 0 | 0 | 0 | 0 |
| 0 | 0 | 0 | 0 |
| 0 | 0 | 0 | 0 |
| 0 | 0 | 0 | 0 |
| 0 | 0 | 0 | 1 |
| 0 | 0 | 0 | 1 |
| 0 | 0 | 0 | 1 |
| 0 | 0 | 0 | 0 |
| 0 | 0 | 0 | 1 |
| 0 | 0 | 0 | 1 |
| 0 | 0 | 0 | 1 |
| 0 | 0 | 0 | 1 |
| 0 | 0 | 0 | 1 |
| 0 | 0 | 0 | 0 |
| 0 | 0 | 0 | 0 |
| 0 | 0 | 0 | 0 |
| 0 | 1 | 0 | 0 |
| 0 | 1 | 0 | 0 |
| 0 | 0 | 0 | 0 |
| 0 | 0 | 0 | 1 |
| 0 | 0 | 0 | 1 |
| 0 | 0 | 0 | 0 |
| 0 | 0 | 0 | 0 |
| 0 | 1 | 0 | 0 |
| 0 | 1 | 0 | 0 |
| 0 | 1 | 0 | 0 |
| 0 | 0 | 0 | 0 |
| 0 | 0 | 0 | 1 |
| 0 | 0 | 0 | 1 |
| 0 | 0 | 0 | 0 |
| 0 | 0 | 0 | 1 |
| 0 | 0 | 0 | 1 |
| 0 | 0 | 0 | 1 |
| 0 | 0 | 0 | 1 |
| 0 | 0 | 0 | 1 |
| 0 | 1 | 1 | 1 |
| 0 | 0 | 0 | 1 |
| 0 | 0 | 0 | 1 |
| 0 | 1 | 1 | 1 |

| | | | |
|---|---|---|---|
| 0 | 1 | 0 | 0 |
| 0 | 0 | 0 | 1 |
| 0 | 1 | 0 | 0 |
| 0 | 0 | 0 | 1 |
| 0 | 1 | 1 | 1 |
| 0 | 1 | 1 | 1 |
| 0 | 1 | 0 | 0 |
| 0 | 1 | 0 | 0 |
| 0 | 1 | 1 | 1 |
| 0 | 1 | 1 | 1 |
| 0 | 0 | 0 | 1 |
| 0 | 0 | 0 | 1 |
| 0 | 1 | 1 | 1 |
| 0 | 1 | 1 | 1 |
| 0 | 0 | 0 | 0 |
| 0 | 0 | 0 | 0 |
| 0 | 0 | 0 | 0 |
| 0 | 0 | 0 | 0 |
| 0 | 0 | 0 | 1 |
| 0 | 1 | 0 | 0 |
| 0 | 1 | 0 | 0 |
| 0 | 0 | 0 | 0 |
| 0 | 0 | 0 | 0 |
| 0 | 0 | 0 | 0 |
| 0 | 0 | 0 | 1 |
| 0 | 1 | 0 | 0 |
| 0 | 0 | 0 | 1 |
| 0 | 0 | 0 | 1 |
| 0 | 0 | 0 | 1 |
| 0 | 1 | 0 | 0 |
| 0 | 0 | 0 | 1 |
| 0 | 0 | 0 | 1 |
| 0 | 0 | 0 | 0 |
| 0 | 0 | 0 | 0 |
| 0 | 0 | 0 | 1 |
| 0 | 0 | 0 | 1 |
| 0 | 0 | 0 | 1 |
| 0 | 0 | 0 | 1 |
| 0 | 1 | 1 | 1 |
| 0 | 0 | 0 | 0 |
| 0 | 0 | 0 | 0 |
| 0 | 1 | 0 | 0 |
| 0 | 1 | 0 | 0 |
| 0 | 0 | 0 | 0 |
| 0 | 0 | 0 | 0 |
| 0 | 0 | 0 | 1 |
| 0 | 0 | 0 | 1 |
| 0 | 0 | 0 | 0 |
| 0 | 0 | 0 | 0 |
| 0 | 0 | 0 | 1 |
| 0 | 0 | 0 | 0 |
| 0 | 0 | 0 | 0 |
| 0 | 0 | 0 | 1 |
| 0 | 0 | 0 | 1 |

| | | | |
|---|---|---|---|
| 0 | 0 | 0 | 1 |
| 0 | 1 | 0 | 0 |
| 0 | 1 | 0 | 0 |
| 0 | 1 | 1 | 1 |
| 0 | 0 | 0 | 1 |
| 0 | 0 | 0 | 0 |
| 0 | 0 | 0 | 0 |
| 0 | 0 | 0 | 0 |
| 0 | 0 | 0 | 0 |
| 0 | 0 | 0 | 0 |
| 0 | 0 | 0 | 0 |
| 0 | 0 | 0 | 0 |
| 0 | 0 | 0 | 0 |
| 0 | 0 | 0 | 0 |
| 0 | 0 | 0 | 0 |
| 0 | 0 | 0 | 1 |
| 0 | 1 | 0 | 0 |
| 0 | 0 | 0 | 1 |
| 0 | 1 | 1 | 1 |
| 0 | 0 | 0 | 1 |
| 0 | 0 | 0 | 0 |
| 0 | 0 | 0 | 1 |
| 0 | 1 | 0 | 0 |
| 0 | 0 | 0 | 0 |
| 0 | 0 | 0 | 1 |
| 0 | 0 | 0 | 0 |
| 0 | 0 | 0 | 1 |
| 0 | 0 | 0 | 1 |
| 0 | 0 | 0 | 1 |
| 0 | 1 | 0 | 0 |
| 0 | 1 | 0 | 0 |
| 0 | 0 | 0 | 0 |
| 0 | 1 | 0 | 0 |
| 0 | 1 | 0 | 0 |
| 0 | 1 | 0 | 0 |
| 0 | 1 | 0 | 0 |
| 0 | 1 | 0 | 0 |
| 0 | 0 | 0 | 1 |
| 0 | 1 | 1 | 1 |
| 0 | 1 | 0 | 0 |
| 0 | 1 | 0 | 0 |
| 0 | 0 | 0 | 0 |
| 0 | 0 | 0 | 0 |
| 0 | 1 | 0 | 0 |
| 0 | 1 | 0 | 0 |
| 0 | 1 | 0 | 0 |
| 0 | 0 | 0 | 1 |
| 0 | 1 | 0 | 0 |
| 0 | 0 | 0 | 0 |
| 0 | 0 | 0 | 0 |
| 0 | 0 | 0 | 0 |
| 0 | 0 | 0 | 1 |
| 0 | 1 | 0 | 0 |
| 0 | 1 | 0 | 0 |

| | | | |
|---|---|---|---|
| 0 | 1 | 0 | 0 |
| 0 | 1 | 0 | 0 |
| 0 | 0 | 0 | 0 |
| 0 | 0 | 0 | 0 |
| 0 | 0 | 0 | 0 |
| 0 | 0 | 0 | 0 |
| 0 | 0 | 0 | 0 |
| 0 | 0 | 0 | 0 |
| 1 | 0 | 0 | 0 |
| 0 | 0 | 0 | 0 |
| 0 | 0 | 0 | 1 |
| 0 | 1 | 0 | 0 |
| 0 | 1 | 1 | 1 |
| 0 | 0 | 0 | 0 |
| 0 | 0 | 0 | 1 |
| 0 | 0 | 0 | 1 |
| 0 | 0 | 0 | 1 |
| 0 | 0 | 0 | 1 |
| 0 | 0 | 0 | 0 |
| 0 | 0 | 0 | 0 |
| 0 | 0 | 0 | 1 |
| 0 | 1 | 0 | 0 |
| 0 | 1 | 0 | 0 |
| 0 | 0 | 0 | 1 |
| 0 | 0 | 0 | 0 |
| 0 | 0 | 0 | 0 |
| 0 | 0 | 0 | 0 |
| 0 | 0 | 0 | 1 |
| 0 | 1 | 0 | 0 |
| 0 | 0 | 0 | 0 |
| 0 | 1 | 1 | 1 |
| 1 | 1 | 0 | 0 |
| 0 | 1 | 1 | 1 |
| 0 | 1 | 1 | 1 |
| 1 | 1 | 0 | 0 |
| 1 | 1 | 0 | 0 |
| 1 | 1 | 0 | 0 |
| 0 | 1 | 1 | 1 |
| 0 | 1 | 1 | 1 |
| 0 | 1 | 0 | 0 |
| 1 | 0 | 0 | 0 |
| 0 | 1 | 0 | 0 |
| 0 | 1 | 0 | 0 |
| 1 | 1 | 0 | 0 |
| 1 | 1 | 0 | 0 |
| 0 | 1 | 0 | 0 |
| 0 | 1 | 1 | 1 |
| 0 | 1 | 1 | 1 |
| 0 | 1 | 1 | 1 |
| 0 | 1 | 1 | 1 |
| 1 | 1 | 0 | 0 |
| 0 | 1 | 0 | 0 |
| 0 | 1 | 0 | 0 |
| 0 | 1 | 0 | 0 |

| | | | |
|---|---|---|---|
| 0 | 1 | 0 | 0 |
| 0 | 1 | 1 | 1 |
| 0 | 1 | 0 | 0 |
| 0 | 1 | 0 | 0 |
| 0 | 1 | 0 | 0 |
| 0 | 1 | 1 | 1 |
| 0 | 1 | 0 | 0 |
| 0 | 0 | 0 | 0 |
| 0 | 0 | 0 | 1 |
| 0 | 1 | 0 | 0 |
| 0 | 0 | 0 | 1 |
| 0 | 0 | 0 | 1 |
| 0 | 0 | 0 | 1 |
| 0 | 0 | 0 | 0 |
| 0 | 0 | 0 | 1 |
| 0 | 0 | 0 | 0 |
| 0 | 0 | 0 | 1 |
| 0 | 0 | 0 | 1 |
| 0 | 0 | 0 | 0 |
| 0 | 0 | 0 | 0 |
| 0 | 0 | 0 | 0 |
| 0 | 0 | 0 | 0 |
| 0 | 0 | 0 | 0 |
| 0 | 0 | 0 | 0 |
| 0 | 0 | 0 | 0 |
| 0 | 0 | 0 | 1 |
| 0 | 0 | 0 | 1 |
| 0 | 0 | 0 | 1 |
| 0 | 0 | 0 | 1 |
| 0 | 0 | 0 | 1 |
| 0 | 0 | 0 | 1 |
| 0 | 0 | 0 | 1 |
| 0 | 0 | 0 | 0 |
| 0 | 0 | 0 | 0 |
| 0 | 0 | 0 | 0 |
| 0 | 0 | 0 | 0 |
| 0 | 0 | 0 | 0 |
| 0 | 0 | 0 | 0 |
| 0 | 0 | 0 | 1 |
| 0 | 0 | 0 | 1 |
| 0 | 0 | 0 | 1 |
| 0 | 0 | 0 | 1 |
| 0 | 0 | 0 | 1 |
| 0 | 0 | 0 | 1 |
| 0 | 1 | 0 | 0 |
| 0 | 0 | 0 | 0 |
| 0 | 0 | 0 | 1 |
| 0 | 0 | 0 | 0 |
| 0 | 0 | 0 | 0 |
| 0 | 0 | 0 | 0 |
| 0 | 0 | 0 | 0 |
| 0 | 0 | 0 | 0 |
| 0 | 0 | 0 | 0 |
| 0 | 0 | 0 | 0 |

| | | | |
|---|---|---|---|
| 0 | 0 | 0 | 1 |
| 0 | 0 | 0 | 1 |
| 1 | 0 | 0 | 0 |
| 1 | 0 | 0 | 0 |
| 0 | 0 | 0 | 1 |
| 0 | 0 | 0 | 0 |
| 0 | 0 | 0 | 0 |
| 0 | 0 | 0 | 1 |
| 0 | 0 | 0 | 1 |
| 0 | 1 | 0 | 0 |
| 0 | 0 | 0 | 1 |
| 0 | 0 | 0 | 1 |
| 0 | 1 | 0 | 0 |
| 0 | 1 | 0 | 0 |
| 0 | 1 | 0 | 0 |
| 0 | 1 | 0 | 0 |
| 0 | 0 | 0 | 1 |
| 0 | 0 | 0 | 1 |
| 0 | 0 | 0 | 1 |
| 0 | 0 | 0 | 1 |
| 0 | 0 | 0 | 1 |
| 0 | 0 | 0 | 1 |
| 0 | 0 | 0 | 1 |
| 0 | 0 | 0 | 1 |
| 0 | 0 | 0 | 1 |
| 0 | 0 | 0 | 0 |
| 0 | 0 | 0 | 1 |
| 0 | 0 | 0 | 1 |
| 0 | 0 | 0 | 1 |
| 0 | 0 | 0 | 1 |
| 0 | 0 | 0 | 1 |
| 0 | 0 | 0 | 1 |
| 0 | 0 | 0 | 1 |
| 0 | 0 | 0 | 1 |
| 0 | 0 | 0 | 1 |
| 0 | 0 | 0 | 1 |
| 0 | 0 | 0 | 1 |
| 0 | 0 | 0 | 0 |
| 0 | 1 | 1 | 1 |
| 0 | 0 | 0 | 1 |
| 0 | 1 | 0 | 0 |
| 0 | 0 | 0 | 1 |
| 0 | 0 | 0 | 1 |
| 0 | 1 | 0 | 0 |
| 0 | 0 | 0 | 0 |
| 0 | 0 | 0 | 1 |
| 0 | 0 | 0 | 1 |
| 0 | 0 | 0 | 1 |
| 0 | 0 | 0 | 1 |
| 0 | 0 | 0 | 1 |
| 0 | 0 | 0 | 1 |
| 0 | 0 | 0 | 0 |
| 0 | 0 | 0 | 0 |
| 0 | 0 | 0 | 0 |

| | | | |
|---|---|---|---|
| 0 | 0 | 0 | 0 |
| 0 | 0 | 0 | 1 |
| 0 | 0 | 0 | 1 |
| 0 | 0 | 0 | 1 |
| 0 | 0 | 0 | 1 |
| 0 | 0 | 0 | 1 |
| 0 | 0 | 0 | 1 |
| 0 | 0 | 0 | 1 |
| 0 | 0 | 0 | 0 |
| 0 | 0 | 0 | 1 |
| 0 | 0 | 0 | 1 |
| 0 | 0 | 0 | 1 |
| 0 | 0 | 0 | 1 |
| 0 | 0 | 0 | 1 |
| 0 | 0 | 0 | 1 |
| 0 | 0 | 0 | 1 |
| 0 | 0 | 0 | 1 |
| 0 | 0 | 0 | 1 |
| 0 | 0 | 0 | 0 |
| 0 | 0 | 0 | 0 |
| 0 | 0 | 0 | 0 |
| 0 | 1 | 0 | 0 |
| 0 | 0 | 0 | 1 |
| 0 | 0 | 0 | 0 |
| 0 | 0 | 0 | 0 |
| 0 | 0 | 0 | 1 |
| 0 | 0 | 0 | 1 |
| 0 | 0 | 0 | 1 |
| 0 | 0 | 0 | 1 |
| 0 | 0 | 0 | 1 |
| 0 | 0 | 0 | 1 |
| 0 | 0 | 0 | 1 |
| 0 | 0 | 0 | 0 |
| 0 | 0 | 0 | 1 |
| 0 | 0 | 0 | 1 |
| 0 | 0 | 0 | 1 |
| 0 | 0 | 0 | 0 |
| 0 | 0 | 0 | 1 |
| 0 | 0 | 0 | 0 |
| 0 | 0 | 0 | 0 |
| 0 | 0 | 0 | 0 |
| 0 | 0 | 0 | 0 |
| 0 | 0 | 0 | 0 |
| 0 | 0 | 0 | 1 |
| 0 | 0 | 0 | 1 |
| 0 | 0 | 0 | 1 |
| 0 | 0 | 0 | 0 |
| 0 | 0 | 0 | 0 |
| 0 | 0 | 0 | 0 |
| 0 | 1 | 0 | 0 |
| 0 | 1 | 0 | 0 |
| 0 | 1 | 0 | 0 |
| 0 | 1 | 0 | 0 |
| 0 | 0 | 0 | 1 |

| | | | |
|---|---|---|---|
| 0 | 0 | 0 | 0 |
| 0 | 0 | 0 | 0 |
| 0 | 0 | 0 | 1 |
| 1 | 0 | 0 | 0 |
| 1 | 0 | 0 | 0 |
| 0 | 0 | 0 | 1 |
| 0 | 0 | 0 | 1 |
| 0 | 1 | 1 | 1 |
| 0 | 0 | 0 | 1 |
| 0 | 0 | 0 | 1 |
| 0 | 0 | 0 | 1 |
| 0 | 0 | 0 | 0 |
| 0 | 0 | 0 | 1 |
| 0 | 0 | 0 | 1 |
| 0 | 0 | 0 | 0 |
| 0 | 0 | 0 | 1 |
| 0 | 0 | 0 | 0 |
| 0 | 0 | 0 | 1 |
| 0 | 0 | 0 | 0 |
| 1 | 0 | 0 | 0 |
| 0 | 0 | 0 | 1 |
| 0 | 0 | 0 | 1 |
| 0 | 0 | 0 | 0 |
| 0 | 0 | 0 | 1 |
| 0 | 0 | 0 | 0 |
| 0 | 0 | 0 | 1 |
| 0 | 0 | 0 | 1 |
| 0 | 0 | 0 | 0 |
| 0 | 0 | 0 | 1 |
| 0 | 0 | 0 | 0 |
| 0 | 0 | 0 | 1 |
| 0 | 0 | 0 | 0 |
| 0 | 0 | 0 | 0 |
| 0 | 1 | 0 | 0 |
| 0 | 0 | 0 | 1 |
| 0 | 0 | 0 | 0 |
| 0 | 0 | 0 | 1 |
| 0 | 0 | 0 | 1 |
| 0 | 0 | 0 | 0 |
| 0 | 0 | 0 | 1 |
| 0 | 0 | 0 | 1 |
| 0 | 0 | 0 | 1 |
| 0 | 0 | 0 | 1 |
| 0 | 0 | 0 | 0 |
| 0 | 0 | 0 | 0 |
| 0 | 0 | 0 | 1 |
| 0 | 0 | 0 | 1 |
| 0 | 1 | 1 | 1 |
| 0 | 1 | 0 | 0 |
| 0 | 1 | 1 | 1 |
| 0 | 0 | 0 | 0 |
| 0 | 0 | 0 | 1 |
| 0 | 0 | 0 | 1 |
| 0 | 0 | 0 | 0 |

| | | | |
|---|---|---|---|
| 0 | 0 | 0 | 1 |
| 0 | 0 | 0 | 1 |
| 0 | 0 | 0 | 1 |
| 0 | 0 | 0 | 1 |
| 0 | 0 | 0 | 1 |
| 0 | 0 | 0 | 1 |
| 0 | 0 | 0 | 1 |
| 0 | 0 | 0 | 0 |
| 0 | 0 | 0 | 1 |
| 0 | 0 | 0 | 1 |
| 0 | 0 | 0 | 1 |
| 0 | 0 | 0 | 1 |
| 0 | 0 | 0 | 0 |
| 0 | 0 | 0 | 1 |
| 0 | 0 | 0 | 0 |
| 0 | 0 | 0 | 0 |
| 0 | 0 | 0 | 1 |
| 0 | 0 | 0 | 1 |
| 0 | 0 | 0 | 1 |
| 0 | 0 | 0 | 1 |
| 0 | 0 | 0 | 0 |
| 0 | 0 | 0 | 0 |
| 0 | 0 | 0 | 1 |
| 0 | 0 | 0 | 1 |
| 0 | 0 | 0 | 0 |
| 0 | 0 | 0 | 1 |
| 0 | 0 | 0 | 0 |
| 0 | 0 | 0 | 0 |
| 0 | 0 | 0 | 0 |
| 0 | 0 | 0 | 0 |
| 0 | 0 | 0 | 1 |
| 0 | 0 | 0 | 1 |
| 0 | 1 | 0 | 0 |
| 0 | 1 | 0 | 0 |
| 0 | 1 | 0 | 0 |
| 0 | 1 | 0 | 0 |
| 0 | 1 | 0 | 0 |
| 0 | 0 | 0 | 0 |
| 0 | 1 | 0 | 0 |
| 0 | 1 | 1 | 1 |
| 0 | 0 | 0 | 1 |
| 0 | 0 | 0 | 1 |
| 0 | 0 | 0 | 1 |
| 0 | 0 | 0 | 1 |
| 0 | 0 | 0 | 1 |
| 0 | 0 | 0 | 0 |
| 0 | 0 | 0 | 1 |
| 0 | 0 | 0 | 1 |
| 0 | 0 | 0 | 1 |
| 0 | 0 | 0 | 0 |
| 0 | 0 | 0 | 0 |
| 0 | 0 | 0 | 1 |
| 0 | 0 | 0 | 1 |
| 0 | 0 | 0 | 1 |

| | | | |
|---|---|---|---|
| 0 | 0 | 0 | 1 |
| 0 | 0 | 0 | 1 |
| 0 | 0 | 0 | 1 |
| 0 | 0 | 0 | 1 |
| 0 | 0 | 0 | 0 |
| 0 | 0 | 0 | 1 |
| 0 | 0 | 0 | 0 |
| 0 | 0 | 0 | 0 |
| 0 | 0 | 0 | 0 |
| 0 | 0 | 0 | 0 |
| 0 | 0 | 0 | 0 |
| 0 | 0 | 0 | 1 |
| 0 | 0 | 0 | 0 |
| 0 | 0 | 0 | 0 |
| 0 | 0 | 0 | 1 |
| 0 | 0 | 0 | 1 |
| 0 | 0 | 0 | 1 |
| 0 | 0 | 0 | 1 |
| 0 | 0 | 0 | 1 |
| 0 | 0 | 0 | 1 |
| 0 | 0 | 0 | 1 |
| 0 | 0 | 0 | 1 |
| 0 | 0 | 0 | 1 |
| 0 | 0 | 0 | 0 |
| 0 | 0 | 0 | 1 |
| 0 | 0 | 0 | 1 |
| 0 | 0 | 0 | 1 |
| 0 | 0 | 0 | 0 |
| 0 | 1 | 1 | 1 |
| 0 | 1 | 1 | 1 |
| 0 | 1 | 0 | 0 |
| 0 | 1 | 0 | 0 |
| 0 | 0 | 0 | 1 |
| 0 | 0 | 0 | 0 |
| 0 | 0 | 0 | 1 |
| 0 | 0 | 0 | 1 |
| 0 | 0 | 0 | 1 |
| 0 | 0 | 0 | 1 |
| 0 | 0 | 0 | 1 |
| 0 | 0 | 0 | 1 |
| 0 | 0 | 0 | 1 |
| 0 | 0 | 0 | 1 |
| 0 | 0 | 0 | 1 |
| 0 | 0 | 0 | 1 |
| 0 | 0 | 0 | 1 |
| 0 | 0 | 0 | 1 |
| 0 | 0 | 0 | 1 |
| 0 | 0 | 0 | 1 |
| 0 | 0 | 0 | 0 |
| 0 | 0 | 0 | 1 |
| 0 | 0 | 0 | 1 |
| 0 | 1 | 1 | 1 |
| 0 | 1 | 1 | 1 |
| 0 | 0 | 0 | 1 |

| | | | |
|---|---|---|---|
| 0 | 0 | 0 | 1 |
| 0 | 0 | 0 | 1 |
| 0 | 0 | 0 | 1 |
| 0 | 0 | 0 | 1 |
| 0 | 1 | 1 | 1 |
| 0 | 0 | 0 | 1 |
| 0 | 0 | 0 | 1 |
| 0 | 0 | 0 | 1 |
| 0 | 0 | 0 | 1 |
| 0 | 0 | 0 | 1 |
| 0 | 0 | 0 | 0 |
| 0 | 0 | 0 | 0 |
| 0 | 0 | 0 | 0 |
| 1 | 1 | 0 | 0 |
| 1 | 0 | 0 | 0 |
| 0 | 1 | 1 | 1 |
| 1 | 0 | 0 | 0 |
| 0 | 0 | 0 | 0 |
| 0 | 0 | 0 | 0 |
| 0 | 0 | 0 | 0 |
| 0 | 0 | 0 | 0 |
| 1 | 0 | 0 | 0 |
| 0 | 0 | 0 | 1 |
| 0 | 0 | 0 | 0 |
| 1 | 0 | 0 | 0 |
| 1 | 0 | 0 | 0 |
| 0 | 0 | 0 | 1 |
| 0 | 0 | 0 | 0 |
| 0 | 0 | 0 | 1 |
| 0 | 0 | 0 | 1 |
| 0 | 0 | 0 | 1 |
| 0 | 0 | 0 | 0 |
| 0 | 0 | 0 | 1 |
| 0 | 0 | 0 | 1 |
| 0 | 0 | 0 | 0 |
| 0 | 0 | 0 | 0 |
| 0 | 0 | 0 | 0 |
| 0 | 1 | 0 | 0 |
| 0 | 0 | 0 | 0 |
| 0 | 1 | 1 | 1 |
| 0 | 1 | 1 | 1 |
| 0 | 1 | 0 | 0 |
| 0 | 1 | 1 | 1 |
| 0 | 0 | 0 | 0 |
| 0 | 0 | 0 | 1 |
| 0 | 1 | 1 | 1 |
| 0 | 0 | 0 | 1 |
| 0 | 1 | 0 | 0 |
| 0 | 0 | 0 | 0 |
| 0 | 1 | 0 | 0 |
| 0 | 1 | 0 | 0 |
| 0 | 0 | 0 | 1 |
| 0 | 0 | 0 | 0 |
| 0 | 0 | 0 | 1 |

| | | | |
|---|---|---|---|
| 0 | 0 | 0 | 1 |
| 0 | 0 | 0 | 1 |
| 0 | 0 | 0 | 1 |
| 0 | 0 | 0 | 1 |
| 0 | 0 | 0 | 1 |
| 0 | 0 | 0 | 0 |
| 0 | 0 | 0 | 0 |
| 0 | 0 | 0 | 0 |
| 0 | 0 | 0 | 0 |
| 0 | 0 | 0 | 1 |
| 1 | 1 | 0 | 0 |
| 1 | 0 | 0 | 0 |
| 0 | 0 | 0 | 0 |
| 0 | 0 | 0 | 0 |
| 0 | 0 | 0 | 0 |
| 0 | 0 | 0 | 0 |
| 0 | 0 | 0 | 1 |
| 0 | 0 | 0 | 1 |
| 0 | 0 | 0 | 1 |
| 0 | 0 | 0 | 1 |
| 0 | 0 | 0 | 1 |
| 0 | 0 | 0 | 1 |
| 0 | 0 | 0 | 1 |
| 0 | 0 | 0 | 1 |
| 0 | 0 | 0 | 1 |
| 0 | 0 | 0 | 1 |
| 0 | 0 | 0 | 1 |
| 0 | 0 | 0 | 1 |
| 0 | 1 | 0 | 0 |
| 0 | 0 | 0 | 1 |
| 0 | 0 | 0 | 1 |
| 0 | 1 | 1 | 1 |
| 0 | 1 | 1 | 1 |
| 0 | 0 | 0 | 0 |
| 0 | 0 | 0 | 1 |
| 0 | 1 | 0 | 0 |
| 0 | 0 | 0 | 1 |
| 0 | 0 | 0 | 0 |
| 0 | 0 | 0 | 1 |
| 0 | 0 | 0 | 0 |
| 0 | 0 | 0 | 1 |
| 0 | 0 | 0 | 1 |
| 0 | 0 | 0 | 1 |
| 0 | 0 | 0 | 0 |
| 0 | 0 | 0 | 1 |
| 0 | 0 | 0 | 1 |
| 0 | 0 | 0 | 1 |
| 0 | 0 | 0 | 1 |
| 0 | 0 | 0 | 1 |
| 0 | 0 | 0 | 1 |
| 0 | 0 | 0 | 1 |
| 0 | 1 | 1 | 1 |
| 0 | 0 | 0 | 0 |
| 0 | 0 | 0 | 0 |

| | | | |
|---|---|---|---|
| 0 | 0 | 0 | 1 |
| 0 | 0 | 0 | 0 |
| 0 | 0 | 0 | 1 |
| 0 | 0 | 0 | 1 |
| 0 | 1 | 0 | 0 |
| 0 | 0 | 0 | 0 |
| 0 | 0 | 0 | 0 |
| 0 | 0 | 0 | 1 |
| 0 | 0 | 0 | 1 |
| 0 | 0 | 0 | 0 |
| 0 | 0 | 0 | 0 |
| 0 | 0 | 0 | 0 |
| 0 | 0 | 0 | 0 |
| 0 | 0 | 0 | 0 |
| 0 | 0 | 0 | 1 |
| 0 | 0 | 0 | 0 |
| 0 | 1 | 0 | 0 |
| 0 | 0 | 0 | 0 |
| 0 | 0 | 0 | 1 |
| 0 | 0 | 0 | 1 |
| 0 | 0 | 0 | 1 |
| 0 | 0 | 0 | 1 |
| 0 | 0 | 0 | 1 |
| 0 | 0 | 0 | 0 |
| 0 | 0 | 0 | 1 |
| 0 | 0 | 0 | 1 |
| 0 | 0 | 0 | 0 |
| 0 | 0 | 0 | 0 |
| 0 | 0 | 0 | 1 |
| 0 | 0 | 0 | 1 |
| 0 | 0 | 0 | 0 |
| 0 | 0 | 0 | 1 |
| 0 | 0 | 0 | 1 |
| 0 | 0 | 0 | 1 |
| 0 | 0 | 0 | 1 |
| 0 | 0 | 0 | 0 |
| 0 | 0 | 0 | 1 |
| 0 | 0 | 0 | 0 |
| 0 | 0 | 0 | 0 |
| 0 | 0 | 0 | 1 |
| 0 | 0 | 0 | 1 |
| 0 | 0 | 0 | 0 |
| 0 | 0 | 0 | 0 |
| 0 | 0 | 0 | 1 |
| 0 | 0 | 0 | 0 |
| 0 | 0 | 0 | 0 |
| 0 | 0 | 0 | 0 |
| 0 | 0 | 0 | 1 |
| 0 | 0 | 0 | 0 |
| 0 | 0 | 0 | 0 |
| 0 | 0 | 0 | 0 |
| 0 | 0 | 0 | 1 |
| 0 | 0 | 0 | 0 |
| 0 | 0 | 0 | 1 |

| | | | |
|---|---|---|---|
| 0 | 0 | 0 | 1 |
| 0 | 0 | 0 | 1 |
| 0 | 0 | 0 | 1 |
| 0 | 0 | 0 | 1 |
| 0 | 0 | 0 | 0 |
| 0 | 0 | 0 | 0 |
| 0 | 0 | 0 | 1 |
| 0 | 0 | 0 | 1 |
| 0 | 0 | 0 | 1 |
| 0 | 1 | 0 | 0 |
| 0 | 1 | 1 | 1 |
| 0 | 1 | 1 | 1 |
| 0 | 1 | 0 | 0 |
| 0 | 0 | 0 | 1 |
| 0 | 0 | 0 | 1 |
| 0 | 0 | 0 | 1 |
| 0 | 0 | 0 | 1 |
| 0 | 0 | 0 | 1 |
| 0 | 0 | 0 | 1 |
| 0 | 0 | 0 | 1 |
| 0 | 0 | 0 | 1 |
| 0 | 0 | 0 | 1 |
| 0 | 0 | 0 | 1 |
| 0 | 0 | 0 | 0 |
| 0 | 0 | 0 | 1 |
| 1 | 0 | 0 | 0 |
| 0 | 0 | 0 | 1 |
| 0 | 0 | 0 | 0 |
| 0 | 0 | 0 | 1 |
| 0 | 0 | 0 | 0 |
| 0 | 0 | 0 | 0 |
| 0 | 0 | 0 | 0 |
| 0 | 0 | 0 | 0 |
| 0 | 0 | 0 | 0 |
| 0 | 0 | 0 | 1 |
| 0 | 0 | 0 | 0 |
| 0 | 0 | 0 | 1 |
| 0 | 0 | 0 | 0 |
| 0 | 0 | 0 | 1 |
| 0 | 0 | 0 | 1 |
| 0 | 0 | 0 | 0 |
| 0 | 0 | 0 | 0 |
| 0 | 0 | 0 | 0 |
| 0 | 0 | 0 | 0 |
| 0 | 0 | 0 | 0 |
| 0 | 0 | 0 | 0 |
| 0 | 0 | 0 | 0 |
| 0 | 1 | 0 | 0 |
| 0 | 0 | 0 | 1 |
| 0 | 0 | 0 | 0 |
| 0 | 1 | 1 | 1 |
| 0 | 1 | 1 | 1 |
| 0 | 0 | 0 | 1 |
| 0 | 0 | 0 | 1 |

| | | | |
|---|---|---|---|
| 0 | 0 | 0 | 1 |
| 0 | 0 | 0 | 1 |
| 0 | 0 | 0 | 1 |
| 0 | 0 | 0 | 1 |
| 0 | 0 | 0 | 1 |
| 0 | 0 | 0 | 1 |
| 0 | 0 | 0 | 0 |
| 0 | 0 | 0 | 0 |
| 0 | 0 | 0 | 0 |
| 0 | 0 | 0 | 0 |
| 0 | 0 | 0 | 0 |
| 0 | 0 | 0 | 0 |
| 0 | 0 | 0 | 0 |
| 0 | 0 | 0 | 0 |
| 0 | 0 | 0 | 0 |
| 0 | 0 | 0 | 0 |
| 1 | 0 | 0 | 0 |
| 0 | 0 | 0 | 0 |
| 0 | 0 | 0 | 1 |
| 0 | 0 | 0 | 0 |
| 0 | 0 | 0 | 1 |
| 0 | 0 | 0 | 1 |
| 0 | 0 | 0 | 1 |
| 0 | 0 | 0 | 1 |
| 0 | 0 | 0 | 1 |
| 0 | 0 | 0 | 0 |
| 0 | 0 | 0 | 0 |
| 0 | 0 | 0 | 0 |
| 0 | 0 | 0 | 0 |
| 0 | 0 | 0 | 0 |
| 0 | 0 | 0 | 0 |
| 1 | 0 | 0 | 0 |
| 0 | 0 | 0 | 0 |
| 1 | 0 | 0 | 0 |
| 0 | 0 | 0 | 1 |
| 1 | 0 | 0 | 0 |
| 1 | 0 | 0 | 0 |
| 0 | 0 | 0 | 0 |
| 1 | 0 | 0 | 0 |
| 0 | 0 | 0 | 1 |
| 0 | 0 | 0 | 0 |
| 0 | 0 | 0 | 0 |
| 0 | 0 | 0 | 1 |
| 0 | 0 | 0 | 1 |
| 0 | 0 | 0 | 1 |
| 0 | 0 | 0 | 1 |
| 0 | 0 | 0 | 1 |
| 0 | 0 | 0 | 0 |
| 0 | 0 | 0 | 0 |
| 0 | 0 | 0 | 0 |
| 0 | 0 | 0 | 0 |
| 0 | 0 | 0 | 1 |
| 0 | 0 | 0 | 0 |
| 0 | 0 | 0 | 1 |

| | | | |
|---|---|---|---|
| 0 | 0 | 0 | 1 |
| 0 | 0 | 0 | 1 |
| 0 | 0 | 0 | 1 |
| 0 | 0 | 0 | 1 |
| 0 | 0 | 0 | 1 |
| 0 | 0 | 0 | 0 |
| 0 | 0 | 0 | 1 |
| 0 | 0 | 0 | 1 |
| 0 | 0 | 0 | 1 |
| 0 | 0 | 0 | 1 |
| 0 | 1 | 0 | 0 |
| 0 | 1 | 0 | 0 |
| 0 | 1 | 0 | 0 |
| 0 | 1 | 1 | 1 |
| 0 | 1 | 1 | 1 |
| 0 | 0 | 0 | 1 |
| 0 | 1 | 1 | 1 |
| 0 | 0 | 0 | 0 |
| 0 | 0 | 0 | 1 |
| 0 | 0 | 0 | 1 |
| 0 | 0 | 0 | 1 |
| 0 | 0 | 0 | 1 |
| 0 | 0 | 0 | 0 |
| 0 | 0 | 0 | 1 |
| 0 | 0 | 0 | 0 |
| 0 | 0 | 0 | 0 |
| 0 | 0 | 0 | 0 |
| 0 | 0 | 0 | 0 |
| 0 | 0 | 0 | 1 |
| 0 | 0 | 0 | 0 |
| 0 | 0 | 0 | 1 |
| 0 | 0 | 0 | 1 |
| 0 | 0 | 0 | 1 |
| 0 | 0 | 0 | 1 |
| 0 | 0 | 0 | 1 |
| 0 | 0 | 0 | 1 |
| 0 | 0 | 0 | 1 |
| 0 | 0 | 0 | 0 |
| 0 | 0 | 0 | 0 |
| 0 | 0 | 0 | 0 |
| 0 | 1 | 0 | 0 |
| 0 | 0 | 0 | 0 |
| 0 | 0 | 0 | 1 |
| 0 | 0 | 0 | 0 |
| 0 | 0 | 0 | 0 |
| 0 | 0 | 0 | 1 |
| 0 | 0 | 0 | 0 |
| 0 | 1 | 0 | 0 |
| 0 | 0 | 0 | 0 |
| 1 | 0 | 0 | 0 |
| 0 | 0 | 0 | 1 |
| 0 | 0 | 0 | 1 |
| 0 | 0 | 0 | 1 |
| 0 | 1 | 0 | 0 |

| | | | |
|---|---|---|---|
| 0 | 0 | 0 | 0 |
| 0 | 0 | 0 | 1 |
| 0 | 1 | 0 | 0 |
| 0 | 0 | 0 | 1 |
| 0 | 0 | 0 | 1 |
| 0 | 1 | 1 | 1 |
| 0 | 1 | 1 | 1 |
| 0 | 0 | 0 | 1 |
| 0 | 1 | 0 | 0 |
| 0 | 0 | 0 | 1 |
| 0 | 1 | 0 | 0 |
| 0 | 1 | 0 | 0 |
| 0 | 1 | 0 | 0 |
| 0 | 1 | 0 | 0 |
| 0 | 0 | 0 | 1 |
| 0 | 0 | 0 | 1 |
| 0 | 0 | 0 | 0 |
| 0 | 0 | 0 | 0 |
| 0 | 0 | 0 | 1 |
| 0 | 0 | 0 | 1 |
| 0 | 0 | 0 | 0 |
| 0 | 1 | 1 | 1 |
| 0 | 1 | 1 | 1 |
| 0 | 0 | 0 | 1 |
| 0 | 1 | 0 | 0 |
| 0 | 1 | 1 | 1 |
| 0 | 1 | 0 | 0 |
| 0 | 1 | 0 | 0 |
| 0 | 1 | 0 | 0 |
| 0 | 0 | 0 | 1 |
| 0 | 0 | 0 | 0 |
| 0 | 0 | 0 | 0 |
| 0 | 0 | 0 | 0 |
| 0 | 0 | 0 | 1 |
| 0 | 0 | 0 | 0 |
| 0 | 0 | 0 | 1 |
| 0 | 0 | 0 | 0 |
| 0 | 0 | 0 | 0 |
| 0 | 1 | 0 | 0 |
| 0 | 0 | 0 | 0 |
| 0 | 0 | 0 | 0 |
| 0 | 0 | 0 | 1 |
| 0 | 1 | 0 | 0 |
| 0 | 1 | 0 | 0 |
| 0 | 1 | 0 | 0 |
| 1 | 0 | 0 | 0 |
| 0 | 0 | 0 | 0 |
| 1 | 0 | 0 | 0 |
| 1 | 0 | 0 | 0 |
| 1 | 0 | 0 | 0 |
| 0 | 0 | 0 | 0 |
| 0 | 0 | 0 | 0 |
| 0 | 0 | 0 | 1 |
| 0 | 0 | 0 | 0 |

| | | | |
|---|---|---|---|
| 0 | 0 | 0 | 1 |
| 0 | 0 | 0 | 1 |
| 0 | 0 | 0 | 1 |
| 0 | 0 | 0 | 1 |
| 0 | 0 | 0 | 1 |
| 0 | 0 | 0 | 1 |
| 0 | 0 | 0 | 1 |
| 0 | 0 | 0 | 1 |
| 0 | 0 | 0 | 1 |
| 0 | 0 | 0 | 0 |
| 0 | 0 | 0 | 0 |
| 0 | 0 | 0 | 1 |
| 0 | 0 | 0 | 1 |
| 0 | 0 | 0 | 1 |
| 0 | 0 | 0 | 0 |
| 0 | 0 | 0 | 1 |
| 0 | 0 | 0 | 1 |
| 0 | 0 | 0 | 0 |
| 0 | 0 | 0 | 1 |
| 0 | 0 | 0 | 1 |
| 0 | 0 | 0 | 1 |
| 0 | 1 | 0 | 0 |
| 0 | 0 | 0 | 0 |
| 0 | 1 | 0 | 0 |
| 0 | 1 | 0 | 0 |
| 0 | 1 | 0 | 0 |
| 0 | 1 | 0 | 0 |
| 0 | 0 | 0 | 0 |
| 0 | 0 | 0 | 1 |
| 0 | 0 | 0 | 0 |
| 1 | 0 | 0 | 0 |
| 0 | 0 | 0 | 1 |
| 0 | 0 | 0 | 1 |
| 0 | 0 | 0 | 1 |
| 0 | 0 | 0 | 1 |
| 0 | 0 | 0 | 1 |
| 0 | 0 | 0 | 1 |
| 0 | 0 | 0 | 1 |
| 0 | 0 | 0 | 1 |
| 0 | 0 | 0 | 1 |
| 0 | 0 | 0 | 1 |
| 0 | 0 | 0 | 1 |
| 0 | 0 | 0 | 1 |
| 0 | 0 | 0 | 0 |
| 0 | 0 | 0 | 1 |
| 0 | 0 | 0 | 0 |
| 0 | 0 | 0 | 1 |
| 0 | 0 | 0 | 1 |
| 0 | 0 | 0 | 1 |
| 0 | 0 | 0 | 1 |
| 0 | 0 | 0 | 0 |
| 0 | 0 | 0 | 1 |
| 0 | 0 | 0 | 1 |
| 0 | 0 | 0 | 0 |

| | | | |
|---|---|---|---|
| 0 | 0 | 0 | 0 |
| 0 | 0 | 0 | 0 |
| 0 | 0 | 0 | 0 |
| 0 | 1 | 0 | 0 |
| 0 | 0 | 0 | 1 |
| 0 | 0 | 0 | 1 |
| 0 | 0 | 0 | 1 |
| 0 | 0 | 0 | 0 |
| 0 | 1 | 0 | 0 |
| 0 | 1 | 0 | 0 |
| 0 | 1 | 0 | 0 |
| 0 | 0 | 0 | 1 |
| 0 | 0 | 0 | 1 |
| 0 | 0 | 0 | 1 |
| 0 | 0 | 0 | 1 |
| 0 | 0 | 0 | 0 |
| 0 | 0 | 0 | 0 |
| 0 | 0 | 0 | 1 |
| 0 | 0 | 0 | 1 |
| 0 | 0 | 0 | 1 |
| 0 | 0 | 0 | 1 |
| 0 | 1 | 0 | 0 |
| 0 | 0 | 0 | 1 |
| 0 | 0 | 0 | 1 |
| 0 | 1 | 0 | 0 |
| 0 | 0 | 0 | 0 |
| 0 | 0 | 0 | 0 |
| 0 | 0 | 0 | 0 |
| 0 | 0 | 0 | 1 |
| 0 | 0 | 0 | 1 |
| 0 | 0 | 0 | 1 |
| 0 | 0 | 0 | 0 |
| 0 | 0 | 0 | 1 |
| 0 | 0 | 0 | 1 |
| 0 | 0 | 0 | 1 |
| 0 | 0 | 0 | 1 |
| 0 | 0 | 0 | 1 |
| 0 | 0 | 0 | 1 |
| 0 | 0 | 0 | 1 |
| 0 | 0 | 0 | 1 |
| 0 | 0 | 0 | 1 |
| 0 | 0 | 0 | 1 |
| 0 | 0 | 0 | 0 |
| 0 | 0 | 0 | 1 |
| 0 | 0 | 0 | 1 |
| 0 | 0 | 0 | 1 |
| 0 | 0 | 0 | 1 |
| 0 | 0 | 0 | 1 |
| 0 | 0 | 0 | 1 |
| 0 | 0 | 0 | 0 |
| 0 | 0 | 0 | 1 |
| 0 | 0 | 0 | 1 |
| 0 | 0 | 0 | 1 |
| 0 | 0 | 0 | 1 |

| | | | |
|---|---|---|---|
| 0 | 0 | 0 | 0 |
| 0 | 0 | 0 | 1 |
| 0 | 0 | 0 | 1 |
| 0 | 0 | 0 | 1 |
| 0 | 0 | 0 | 1 |
| 0 | 0 | 0 | 1 |
| 0 | 0 | 0 | 1 |
| 0 | 1 | 0 | 0 |
| 0 | 0 | 0 | 1 |
| 0 | 0 | 0 | 1 |
| 0 | 0 | 0 | 0 |
| 0 | 0 | 0 | 0 |
| 0 | 0 | 0 | 0 |
| 0 | 0 | 0 | 1 |
| 0 | 0 | 0 | 0 |
| 0 | 0 | 0 | 0 |
| 0 | 0 | 0 | 0 |
| 0 | 0 | 0 | 0 |
| 0 | 0 | 0 | 1 |
| 0 | 0 | 0 | 0 |
| 0 | 0 | 0 | 0 |
| 0 | 0 | 0 | 0 |
| 0 | 0 | 0 | 0 |
| 0 | 0 | 0 | 0 |
| 0 | 0 | 0 | 0 |
| 0 | 0 | 0 | 0 |
| 0 | 0 | 0 | 0 |
| 0 | 0 | 0 | 0 |
| 0 | 0 | 0 | 0 |
| 0 | 0 | 0 | 0 |
| 0 | 0 | 0 | 0 |
| 0 | 0 | 0 | 0 |
| 0 | 0 | 0 | 1 |
| 0 | 0 | 0 | 0 |
| 0 | 0 | 0 | 1 |
| 0 | 0 | 0 | 0 |
| 0 | 0 | 0 | 0 |
| 0 | 0 | 0 | 1 |
| 0 | 0 | 0 | 1 |
| 0 | 0 | 0 | 1 |
| 0 | 0 | 0 | 1 |
| 0 | 0 | 0 | 0 |
| 0 | 0 | 0 | 0 |
| 0 | 0 | 0 | 0 |
| 0 | 0 | 0 | 1 |
| 0 | 0 | 0 | 0 |
| 0 | 0 | 0 | 1 |
| 0 | 0 | 0 | 0 |
| 0 | 0 | 0 | 1 |
| 0 | 0 | 0 | 1 |
| 0 | 0 | 0 | 1 |
| 0 | 0 | 0 | 1 |
| 0 | 0 | 0 | 1 |
| 0 | 0 | 0 | 1 |

| | | | |
|---|---|---|---|
| 0 | 0 | 0 | 1 |
| 0 | 0 | 0 | 1 |
| 0 | 0 | 0 | 1 |
| 0 | 0 | 0 | 1 |
| 0 | 0 | 0 | 1 |
| 0 | 0 | 0 | 0 |
| 0 | 0 | 0 | 0 |
| 0 | 0 | 0 | 1 |
| 0 | 0 | 0 | 0 |
| 0 | 0 | 0 | 0 |
| 0 | 0 | 0 | 1 |
| 0 | 0 | 0 | 1 |
| 0 | 0 | 0 | 1 |
| 0 | 0 | 0 | 1 |
| 0 | 0 | 0 | 1 |
| 0 | 1 | 1 | 1 |
| 0 | 0 | 0 | 1 |
| 0 | 0 | 0 | 1 |
| 0 | 1 | 0 | 0 |
| 0 | 1 | 0 | 0 |
| 0 | 1 | 1 | 1 |
| 0 | 0 | 0 | 1 |
| 0 | 0 | 0 | 1 |
| 0 | 0 | 0 | 0 |
| 0 | 0 | 0 | 1 |
| 0 | 0 | 0 | 0 |
| 0 | 0 | 0 | 0 |
| 0 | 0 | 0 | 1 |
| 0 | 0 | 0 | 1 |
| 0 | 0 | 0 | 1 |
| 0 | 1 | 0 | 0 |
| 0 | 0 | 0 | 1 |
| 0 | 0 | 0 | 0 |
| 0 | 1 | 0 | 0 |
| 0 | 1 | 0 | 0 |
| 0 | 1 | 0 | 0 |
| 0 | 1 | 0 | 0 |
| 0 | 0 | 0 | 1 |
| 0 | 0 | 0 | 1 |
| 0 | 0 | 0 | 1 |
| 0 | 1 | 0 | 0 |
| 0 | 1 | 0 | 0 |
| 0 | 1 | 0 | 0 |
| 0 | 0 | 0 | 0 |
| 0 | 0 | 0 | 1 |
| 0 | 0 | 0 | 1 |
| 0 | 0 | 0 | 1 |
| 0 | 0 | 0 | 0 |
| 0 | 0 | 0 | 0 |
| 0 | 0 | 0 | 0 |
| 0 | 0 | 0 | 1 |
| 0 | 0 | 0 | 1 |
| 0 | 0 | 0 | 1 |
| 0 | 0 | 0 | 1 |

| | | | |
|---|---|---|---|
| 0 | 0 | 0 | 1 |
| 0 | 0 | 0 | 1 |
| 0 | 0 | 0 | 1 |
| 0 | 0 | 0 | 0 |
| 0 | 0 | 0 | 0 |
| 0 | 0 | 0 | 1 |
| 0 | 1 | 0 | 0 |
| 0 | 0 | 0 | 1 |
| 0 | 0 | 0 | 1 |
| 0 | 0 | 0 | 1 |
| 0 | 0 | 0 | 1 |
| 0 | 0 | 0 | 0 |
| 0 | 1 | 1 | 1 |
| 0 | 0 | 0 | 0 |
| 0 | 0 | 0 | 1 |
| 0 | 0 | 0 | 0 |
| 0 | 1 | 1 | 1 |
| 0 | 1 | 1 | 1 |
| 0 | 0 | 0 | 1 |
| 0 | 1 | 0 | 0 |
| 0 | 0 | 0 | 0 |
| 0 | 0 | 0 | 0 |
| 0 | 0 | 0 | 0 |
| 0 | 0 | 0 | 0 |
| 0 | 0 | 0 | 0 |
| 0 | 0 | 0 | 1 |
| 0 | 0 | 0 | 0 |
| 0 | 0 | 0 | 1 |
| 0 | 0 | 0 | 1 |
| 0 | 0 | 0 | 1 |
| 0 | 0 | 0 | 1 |
| 0 | 0 | 0 | 1 |
| 0 | 0 | 0 | 0 |
| 0 | 1 | 0 | 0 |
| 0 | 0 | 0 | 1 |
| 0 | 0 | 0 | 1 |
| 0 | 1 | 0 | 0 |
| 0 | 1 | 0 | 0 |
| 0 | 1 | 0 | 0 |
| 0 | 1 | 1 | 1 |
| 0 | 0 | 0 | 0 |
| 0 | 0 | 0 | 0 |
| 0 | 0 | 0 | 1 |
| 0 | 0 | 0 | 0 |
| 0 | 0 | 0 | 1 |
| 0 | 0 | 0 | 1 |
| 0 | 0 | 0 | 1 |
| 0 | 0 | 0 | 1 |
| 0 | 0 | 0 | 1 |
| 1 | 0 | 0 | 0 |
| 0 | 0 | 0 | 1 |
| 0 | 0 | 0 | 1 |
| 0 | 0 | 0 | 1 |
| 0 | 0 | 0 | 1 |

| | | | |
|---|---|---|---|
| 0 | 0 | 0 | 1 |
| 0 | 0 | 0 | 1 |
| 0 | 0 | 0 | 1 |
| 0 | 0 | 0 | 0 |
| 0 | 0 | 0 | 1 |
| 0 | 0 | 0 | 1 |
| 0 | 0 | 0 | 1 |
| 0 | 0 | 0 | 0 |
| 0 | 0 | 0 | 1 |
| 0 | 0 | 0 | 1 |
| 0 | 0 | 0 | 1 |
| 0 | 0 | 0 | 1 |
| 0 | 0 | 0 | 1 |
| 0 | 0 | 0 | 1 |
| 0 | 0 | 0 | 1 |
| 0 | 0 | 0 | 1 |
| 0 | 0 | 0 | 0 |
| 0 | 0 | 0 | 0 |
| 0 | 0 | 0 | 1 |
| 0 | 0 | 0 | 1 |
| 0 | 0 | 0 | 1 |
| 0 | 0 | 0 | 1 |
| 0 | 0 | 0 | 1 |
| 0 | 0 | 0 | 1 |
| 0 | 0 | 0 | 1 |
| 0 | 0 | 0 | 1 |
| 0 | 0 | 0 | 1 |
| 0 | 1 | 0 | 0 |
| 0 | 1 | 0 | 0 |
| 0 | 0 | 0 | 1 |
| 0 | 0 | 0 | 1 |
| 0 | 0 | 0 | 0 |
| 0 | 0 | 0 | 1 |
| 0 | 0 | 0 | 0 |
| 1 | 0 | 0 | 0 |
| 1 | 0 | 0 | 0 |
| 1 | 0 | 0 | 0 |
| 1 | 0 | 0 | 0 |
| 0 | 0 | 0 | 1 |
| 0 | 0 | 0 | 1 |
| 0 | 0 | 0 | 1 |
| 0 | 0 | 0 | 1 |
| 0 | 0 | 0 | 1 |
| 0 | 0 | 0 | 1 |
| 0 | 0 | 0 | 0 |
| 0 | 0 | 0 | 0 |
| 0 | 0 | 0 | 0 |
| 0 | 0 | 0 | 1 |
| 0 | 0 | 0 | 1 |
| 0 | 0 | 0 | 1 |
| 0 | 0 | 0 | 1 |
| 0 | 0 | 0 | 0 |
| 0 | 1 | 0 | 0 |
| 0 | 0 | 0 | 0 |

| | | | |
|---|---|---|---|
| 0 | 0 | 0 | 0 |
| 0 | 0 | 0 | 0 |
| 0 | 0 | 0 | 1 |
| 0 | 0 | 0 | 0 |
| 0 | 0 | 0 | 0 |
| 0 | 0 | 0 | 0 |
| 0 | 0 | 0 | 0 |
| 0 | 0 | 0 | 1 |
| 0 | 0 | 0 | 1 |
| 0 | 0 | 0 | 1 |
| 0 | 0 | 0 | 1 |
| 0 | 0 | 0 | 1 |
| 0 | 0 | 0 | 1 |
| 0 | 0 | 0 | 1 |
| 0 | 0 | 0 | 1 |
| 0 | 0 | 0 | 1 |
| 0 | 0 | 0 | 1 |
| 0 | 0 | 0 | 1 |
| 0 | 0 | 0 | 1 |
| 0 | 0 | 0 | 1 |
| 0 | 0 | 0 | 0 |
| 0 | 0 | 0 | 0 |
| 0 | 0 | 0 | 0 |
| 0 | 0 | 0 | 0 |
| 0 | 0 | 0 | 0 |
| 0 | 0 | 0 | 1 |
| 0 | 0 | 0 | 1 |
| 0 | 0 | 0 | 0 |
| 0 | 0 | 0 | 1 |
| 0 | 0 | 0 | 1 |
| 0 | 0 | 0 | 0 |
| 0 | 0 | 0 | 0 |
| 0 | 0 | 0 | 0 |
| 0 | 0 | 0 | 1 |
| 0 | 0 | 0 | 0 |
| 0 | 0 | 0 | 1 |
| 0 | 0 | 0 | 1 |
| 0 | 0 | 0 | 0 |
| 0 | 0 | 0 | 1 |
| 0 | 0 | 0 | 1 |
| 0 | 0 | 0 | 1 |
| 0 | 0 | 0 | 1 |
| 0 | 0 | 0 | 1 |
| 0 | 0 | 0 | 1 |
| 0 | 0 | 0 | 1 |
| 0 | 0 | 0 | 1 |
| 0 | 0 | 0 | 1 |
| 0 | 0 | 0 | 1 |
| 0 | 0 | 0 | 0 |
| 0 | 0 | 0 | 1 |
| 1 | 0 | 0 | 0 |
| 1 | 0 | 0 | 0 |
| 0 | 0 | 0 | 1 |
| 0 | 0 | 0 | 1 |

| | | | |
|---|---|---|---|
| 1 | 0 | 0 | 0 |
| 0 | 0 | 0 | 1 |
| 0 | 0 | 0 | 1 |
| 0 | 0 | 0 | 1 |
| 0 | 0 | 0 | 1 |
| 1 | 0 | 0 | 0 |
| 1 | 0 | 0 | 0 |
| 0 | 0 | 0 | 1 |
| 0 | 0 | 0 | 1 |
| 0 | 0 | 0 | 1 |
| 0 | 0 | 0 | 1 |
| 0 | 0 | 0 | 0 |
| 0 | 0 | 0 | 1 |
| 0 | 0 | 0 | 1 |
| 0 | 0 | 0 | 1 |
| 0 | 0 | 0 | 1 |
| 0 | 0 | 0 | 0 |
| 0 | 0 | 0 | 0 |
| 0 | 0 | 0 | 1 |
| 0 | 0 | 0 | 1 |
| 0 | 0 | 0 | 1 |
| 0 | 0 | 0 | 1 |
| 0 | 0 | 0 | 1 |
| 0 | 0 | 0 | 1 |
| 0 | 0 | 0 | 0 |
| 0 | 0 | 0 | 1 |
| 0 | 0 | 0 | 1 |
| 0 | 0 | 0 | 1 |
| 0 | 0 | 0 | 1 |
| 0 | 0 | 0 | 1 |
| 0 | 0 | 0 | 1 |
| 0 | 0 | 0 | 0 |
| 0 | 0 | 0 | 0 |
| 0 | 0 | 0 | 1 |
| 0 | 0 | 0 | 1 |
| 0 | 1 | 0 | 0 |
| 1 | 0 | 0 | 0 |
| 0 | 1 | 0 | 0 |
| 1 | 1 | 0 | 0 |
| 0 | 1 | 0 | 0 |
| 0 | 0 | 0 | 0 |
| 0 | 0 | 0 | 0 |
| 0 | 0 | 0 | 0 |
| 0 | 0 | 0 | 0 |
| 0 | 0 | 0 | 0 |
| 0 | 0 | 0 | 0 |
| 0 | 0 | 0 | 0 |
| 1 | 0 | 0 | 0 |
| 0 | 1 | 1 | 1 |
| 0 | 0 | 0 | 1 |
| 0 | 1 | 1 | 1 |
| 0 | 0 | 0 | 1 |
| 0 | 0 | 0 | 0 |
| 0 | 0 | 0 | 1 |

| | | | |
|---|---|---|---|
| 0 | 0 | 0 | 1 |
| 0 | 0 | 0 | 1 |
| 0 | 0 | 0 | 1 |
| 0 | 0 | 0 | 1 |
| 0 | 0 | 0 | 1 |
| 0 | 0 | 0 | 1 |
| 0 | 0 | 0 | 1 |
| 0 | 1 | 0 | 0 |
| 0 | 0 | 0 | 0 |
| 0 | 1 | 1 | 1 |
| 0 | 0 | 0 | 1 |
| 0 | 0 | 0 | 1 |
| 0 | 0 | 0 | 1 |
| 0 | 1 | 0 | 0 |
| 0 | 0 | 0 | 1 |
| 0 | 0 | 0 | 1 |
| 0 | 1 | 0 | 0 |
| 0 | 0 | 0 | 1 |
| 1 | 0 | 0 | 0 |
| 0 | 0 | 0 | 1 |
| 0 | 0 | 0 | 1 |
| 0 | 0 | 0 | 1 |
| 0 | 0 | 0 | 1 |
| 0 | 0 | 0 | 0 |
| 0 | 0 | 0 | 1 |
| 0 | 0 | 0 | 1 |
| 0 | 0 | 0 | 1 |
| 0 | 0 | 0 | 1 |
| 0 | 0 | 0 | 1 |
| 0 | 0 | 0 | 1 |
| 0 | 0 | 0 | 1 |
| 1 | 0 | 0 | 0 |
| 0 | 0 | 0 | 1 |
| 0 | 0 | 0 | 1 |
| 0 | 0 | 0 | 1 |
| 0 | 0 | 0 | 1 |
| 0 | 0 | 0 | 1 |
| 0 | 0 | 0 | 1 |
| 1 | 0 | 0 | 0 |
| 0 | 0 | 0 | 1 |
| 0 | 0 | 0 | 1 |
| 0 | 0 | 0 | 1 |
| 0 | 0 | 0 | 1 |
| 0 | 0 | 0 | 1 |
| 0 | 0 | 0 | 1 |
| 0 | 0 | 0 | 0 |
| 0 | 0 | 0 | 1 |
| 0 | 0 | 0 | 1 |
| 0 | 1 | 1 | 1 |
| 0 | 1 | 1 | 1 |
| 0 | 0 | 0 | 1 |
| 0 | 1 | 1 | 1 |
| 0 | 1 | 1 | 1 |
| 0 | 1 | 1 | 1 |

| | | | |
|---|---|---|---|
| 0 | 1 | 1 | 1 |
| 0 | 1 | 1 | 1 |
| 0 | 1 | 1 | 1 |
| 0 | 1 | 1 | 1 |
| 0 | 1 | 1 | 1 |
| 0 | 1 | 1 | 1 |
| 0 | 1 | 1 | 1 |
| 0 | 1 | 1 | 1 |
| 0 | 1 | 1 | 1 |
| 0 | 1 | 1 | 1 |
| 0 | 1 | 1 | 1 |
| 0 | 1 | 1 | 1 |
| 0 | 1 | 1 | 1 |
| 0 | 1 | 1 | 1 |
| 0 | 1 | 1 | 1 |
| 1 | 1 | 0 | 0 |
| 0 | 1 | 1 | 1 |
| 0 | 1 | 0 | 0 |
| 0 | 1 | 1 | 1 |
| 0 | 1 | 1 | 1 |
| 1 | 1 | 0 | 0 |
| 0 | 1 | 1 | 1 |
| 0 | 1 | 1 | 1 |
| 0 | 1 | 1 | 1 |
| 0 | 1 | 0 | 0 |
| 0 | 1 | 1 | 1 |
| 0 | 1 | 0 | 0 |
| 0 | 1 | 1 | 1 |
| 0 | 1 | 1 | 1 |
| 0 | 1 | 1 | 1 |
| 0 | 1 | 1 | 1 |
| 0 | 1 | 1 | 1 |
| 0 | 1 | 1 | 1 |
| 0 | 1 | 1 | 1 |
| 0 | 1 | 1 | 1 |
| 0 | 1 | 1 | 1 |
| 0 | 1 | 1 | 1 |
| 0 | 1 | 1 | 1 |
| 0 | 1 | 1 | 1 |
| 0 | 1 | 1 | 1 |
| 0 | 1 | 1 | 1 |
| 0 | 1 | 1 | 1 |
| 0 | 1 | 1 | 1 |
| 0 | 0 | 0 | 0 |
| 0 | 0 | 0 | 1 |
| 0 | 0 | 0 | 1 |
| 0 | 0 | 0 | 0 |
| 0 | 0 | 0 | 0 |
| 0 | 0 | 0 | 1 |
| 0 | 0 | 0 | 0 |
| 0 | 0 | 0 | 0 |
| 0 | 0 | 0 | 0 |
| 0 | 0 | 0 | 1 |
| 0 | 0 | 0 | 1 |

| | | | |
|---|---|---|---|
| 0 | 0 | 0 | 1 |
| 0 | 0 | 0 | 1 |
| 0 | 0 | 0 | 1 |
| 0 | 0 | 0 | 1 |
| 0 | 0 | 0 | 1 |
| 0 | 0 | 0 | 1 |
| 0 | 0 | 0 | 1 |
| 1 | 0 | 0 | 0 |
| 0 | 0 | 0 | 1 |
| 0 | 0 | 0 | 1 |
| 0 | 0 | 0 | 1 |
| 0 | 0 | 0 | 1 |
| 0 | 1 | 1 | 1 |
| 0 | 0 | 0 | 1 |
| 0 | 1 | 0 | 0 |
| 0 | 0 | 0 | 0 |
| 0 | 0 | 0 | 1 |
| 0 | 0 | 0 | 0 |
| 0 | 0 | 0 | 1 |
| 0 | 0 | 0 | 1 |
| 0 | 0 | 0 | 0 |
| 0 | 0 | 0 | 1 |
| 0 | 0 | 0 | 1 |
| 0 | 0 | 0 | 1 |
| 0 | 0 | 0 | 0 |
| 0 | 0 | 0 | 1 |
| 0 | 0 | 0 | 1 |
| 1 | 1 | 0 | 0 |
| 0 | 0 | 0 | 0 |
| 1 | 0 | 0 | 0 |
| 1 | 0 | 0 | 0 |
| 0 | 0 | 0 | 1 |
| 0 | 0 | 0 | 1 |
| 0 | 0 | 0 | 1 |
| 0 | 0 | 0 | 1 |
| 0 | 0 | 0 | 1 |
| 0 | 1 | 0 | 0 |
| 0 | 0 | 0 | 0 |
| 0 | 0 | 0 | 1 |
| 0 | 0 | 0 | 1 |
| 0 | 0 | 0 | 1 |
| 0 | 0 | 0 | 1 |
| 0 | 0 | 0 | 0 |
| 0 | 0 | 0 | 1 |
| 0 | 0 | 0 | 1 |
| 0 | 0 | 0 | 1 |
| 0 | 0 | 0 | 1 |
| 0 | 0 | 0 | 1 |
| 0 | 0 | 0 | 0 |
| 0 | 0 | 0 | 0 |
| 0 | 0 | 0 | 0 |
| 0 | 0 | 0 | 0 |
| 0 | 0 | 0 | 0 |
| 0 | 0 | 0 | 1 |

| | | | |
|---|---|---|---|
| 0 | 0 | 0 | 0 |
| 0 | 0 | 0 | 0 |
| 0 | 0 | 0 | 1 |
| 1 | 0 | 0 | 0 |
| 0 | 0 | 0 | 1 |
| 0 | 0 | 0 | 1 |
| 0 | 0 | 0 | 1 |
| 0 | 0 | 0 | 1 |
| 0 | 0 | 0 | 1 |
| 1 | 0 | 0 | 0 |
| 0 | 0 | 0 | 1 |
| 0 | 0 | 0 | 1 |
| 0 | 0 | 0 | 1 |
| 0 | 0 | 0 | 1 |
| 0 | 0 | 0 | 1 |
| 0 | 0 | 0 | 1 |
| 0 | 0 | 0 | 1 |
| 0 | 0 | 0 | 1 |
| 0 | 0 | 0 | 1 |
| 0 | 0 | 0 | 1 |
| 0 | 0 | 0 | 1 |
| 0 | 0 | 0 | 1 |
| 0 | 0 | 0 | 1 |
| 0 | 0 | 0 | 1 |
| 0 | 0 | 0 | 1 |
| 0 | 0 | 0 | 1 |
| 0 | 0 | 0 | 1 |
| 0 | 0 | 0 | 1 |
| 0 | 0 | 0 | 1 |
| 0 | 0 | 0 | 1 |
| 0 | 0 | 0 | 1 |
| 0 | 0 | 0 | 1 |
| 1 | 0 | 0 | 0 |
| 0 | 0 | 0 | 1 |
| 0 | 0 | 0 | 1 |
| 0 | 0 | 0 | 1 |
| 0 | 0 | 0 | 1 |
| 0 | 0 | 0 | 1 |
| 0 | 0 | 0 | 1 |
| 0 | 0 | 0 | 1 |
| 0 | 0 | 0 | 1 |
| 0 | 0 | 0 | 1 |
| 0 | 0 | 0 | 1 |
| 0 | 0 | 0 | 1 |
| 0 | 0 | 0 | 1 |
| 0 | 0 | 0 | 1 |
| 0 | 0 | 0 | 1 |
| 0 | 1 | 0 | 0 |
| 0 | 1 | 0 | 0 |
| 0 | 1 | 0 | 0 |
| 0 | 0 | 0 | 0 |
| 0 | 1 | 0 | 0 |
| 0 | 0 | 0 | 1 |
| 0 | 0 | 0 | 0 |

| | | | |
|---|---|---|---|
| 0 | 0 | 0 | 1 |
| 0 | 0 | 0 | 0 |
| 0 | 0 | 0 | 1 |
| 0 | 0 | 0 | 1 |
| 0 | 0 | 0 | 1 |
| 1 | 0 | 0 | 0 |
| 0 | 0 | 0 | 0 |
| 0 | 0 | 0 | 0 |
| 1 | 0 | 0 | 0 |
| 0 | 0 | 0 | 0 |
| 0 | 0 | 0 | 1 |
| 0 | 0 | 0 | 0 |
| 0 | 0 | 0 | 1 |
| 0 | 0 | 0 | 1 |
| 0 | 0 | 0 | 1 |
| 1 | 0 | 0 | 0 |
| 0 | 0 | 0 | 1 |
| 0 | 0 | 0 | 1 |
| 0 | 0 | 0 | 0 |
| 0 | 0 | 0 | 0 |
| 1 | 0 | 0 | 0 |
| 1 | 0 | 0 | 0 |
| 0 | 0 | 0 | 0 |
| 0 | 0 | 0 | 0 |
| 0 | 0 | 0 | 1 |
| 0 | 0 | 0 | 0 |
| 0 | 1 | 0 | 0 |
| 0 | 0 | 0 | 0 |
| 0 | 1 | 0 | 0 |
| 0 | 1 | 0 | 0 |
| 0 | 1 | 0 | 0 |
| 0 | 0 | 0 | 1 |
| 0 | 0 | 0 | 1 |
| 0 | 1 | 0 | 0 |
| 0 | 1 | 0 | 0 |
| 0 | 1 | 1 | 1 |
| 0 | 0 | 0 | 0 |
| 0 | 0 | 0 | 0 |
| 0 | 1 | 0 | 0 |
| 0 | 1 | 0 | 0 |
| 0 | 1 | 0 | 0 |
| 0 | 1 | 1 | 1 |
| 0 | 0 | 0 | 1 |
| 0 | 0 | 0 | 1 |
| 0 | 0 | 0 | 0 |
| 0 | 0 | 0 | 0 |
| 0 | 0 | 0 | 0 |
| 0 | 0 | 0 | 1 |
| 0 | 0 | 0 | 1 |
| 0 | 0 | 0 | 1 |
| 0 | 0 | 0 | 1 |
| 0 | 0 | 0 | 1 |
| 0 | 0 | 0 | 1 |
| 0 | 0 | 0 | 0 |

| | | | |
|---|---|---|---|
| 0 | 0 | 0 | 1 |
| 0 | 0 | 0 | 1 |
| 0 | 0 | 0 | 1 |
| 0 | 0 | 0 | 1 |
| 1 | 0 | 0 | 0 |
| 0 | 0 | 0 | 0 |
| 0 | 1 | 1 | 1 |
| 0 | 1 | 1 | 1 |
| 0 | 0 | 0 | 1 |
| 0 | 0 | 0 | 1 |
| 0 | 0 | 0 | 1 |
| 0 | 0 | 0 | 0 |
| 0 | 0 | 0 | 1 |
| 0 | 0 | 0 | 0 |
| 0 | 1 | 0 | 0 |
| 0 | 1 | 0 | 0 |
| 0 | 1 | 0 | 0 |
| 0 | 1 | 0 | 0 |
| 0 | 1 | 0 | 0 |
| 0 | 1 | 0 | 0 |
| 0 | 1 | 0 | 0 |
| 0 | 1 | 0 | 0 |
| 0 | 0 | 0 | 0 |
| 0 | 0 | 0 | 1 |
| 0 | 0 | 0 | 1 |
| 0 | 0 | 0 | 1 |
| 0 | 0 | 0 | 1 |
| 0 | 0 | 0 | 1 |
| 0 | 0 | 0 | 1 |
| 0 | 0 | 0 | 1 |
| 0 | 0 | 0 | 1 |
| 0 | 0 | 0 | 0 |
| 0 | 0 | 0 | 1 |
| 0 | 0 | 0 | 0 |
| 0 | 0 | 0 | 0 |
| 0 | 0 | 0 | 0 |
| 0 | 0 | 0 | 0 |
| 0 | 0 | 0 | 1 |
| 0 | 0 | 0 | 1 |
| 0 | 0 | 0 | 1 |
| 0 | 0 | 0 | 1 |
| 0 | 0 | 0 | 1 |
| 0 | 0 | 0 | 1 |
| 0 | 0 | 0 | 1 |
| 0 | 0 | 0 | 1 |
| 1 | 0 | 0 | 0 |
| 0 | 0 | 0 | 1 |
| 0 | 0 | 0 | 1 |
| 0 | 0 | 0 | 1 |
| 1 | 1 | 0 | 0 |
| 1 | 0 | 0 | 0 |
| 0 | 0 | 0 | 0 |
| 0 | 0 | 0 | 1 |
| 0 | 0 | 0 | 1 |

| | | | |
|---|---|---|---|
| 0 | 0 | 0 | 1 |
| 0 | 0 | 0 | 1 |
| 0 | 0 | 0 | 1 |
| 0 | 0 | 0 | 0 |
| 0 | 0 | 0 | 1 |
| 0 | 0 | 0 | 1 |
| 0 | 0 | 0 | 0 |
| 0 | 0 | 0 | 1 |
| 0 | 0 | 0 | 1 |
| 0 | 0 | 0 | 1 |
| 0 | 0 | 0 | 0 |
| 0 | 1 | 0 | 0 |
| 0 | 0 | 0 | 1 |
| 0 | 0 | 0 | 1 |
| 0 | 0 | 0 | 1 |
| 0 | 0 | 0 | 1 |
| 0 | 0 | 0 | 0 |
| 0 | 0 | 0 | 0 |
| 0 | 0 | 0 | 1 |
| 0 | 0 | 0 | 0 |
| 0 | 1 | 0 | 0 |
| 0 | 1 | 0 | 0 |
| 0 | 0 | 0 | 1 |
| 0 | 1 | 0 | 0 |
| 0 | 0 | 0 | 0 |
| 0 | 0 | 0 | 1 |
| 0 | 0 | 0 | 0 |
| 0 | 0 | 0 | 1 |
| 0 | 0 | 0 | 1 |
| 0 | 0 | 0 | 0 |
| 0 | 0 | 0 | 1 |
| 0 | 0 | 0 | 1 |
| 0 | 0 | 0 | 0 |
| 0 | 0 | 0 | 1 |
| 0 | 0 | 0 | 1 |
| 0 | 0 | 0 | 0 |
| 0 | 0 | 0 | 1 |
| 0 | 0 | 0 | 0 |
| 0 | 0 | 0 | 1 |
| 0 | 1 | 0 | 0 |
| 0 | 1 | 0 | 0 |
| 0 | 1 | 1 | 1 |
| 0 | 0 | 0 | 0 |
| 0 | 0 | 0 | 0 |
| 0 | 0 | 0 | 1 |
| 0 | 1 | 1 | 1 |
| 0 | 0 | 0 | 1 |
| 0 | 0 | 0 | 1 |
| 0 | 0 | 0 | 1 |
| 0 | 0 | 0 | 1 |
| 0 | 0 | 0 | 1 |
| 0 | 0 | 0 | 1 |
| 0 | 0 | 0 | 1 |
| 0 | 0 | 0 | 1 |

| | | | |
|---|---|---|---|
| 0 | 0 | 0 | 1 |
| 0 | 0 | 0 | 1 |
| 0 | 0 | 0 | 1 |
| 0 | 0 | 0 | 0 |
| 0 | 1 | 1 | 1 |
| 0 | 0 | 0 | 0 |
| 0 | 0 | 0 | 1 |
| 0 | 1 | 1 | 1 |
| 0 | 1 | 0 | 0 |
| 0 | 1 | 0 | 0 |
| 0 | 0 | 0 | 1 |
| 0 | 1 | 0 | 0 |
| 0 | 1 | 0 | 0 |
| 0 | 1 | 0 | 0 |
| 0 | 1 | 1 | 1 |
| 0 | 0 | 0 | 0 |
| 0 | 0 | 0 | 0 |
| 0 | 0 | 0 | 1 |
| 1 | 0 | 0 | 0 |
| 1 | 0 | 0 | 0 |
| 1 | 0 | 0 | 0 |
| 1 | 0 | 0 | 0 |
| 0 | 0 | 0 | 1 |
| 0 | 0 | 0 | 0 |
| 0 | 0 | 0 | 0 |
| 0 | 0 | 0 | 0 |
| 0 | 0 | 0 | 0 |
| 0 | 0 | 0 | 0 |
| 0 | 0 | 0 | 0 |
| 0 | 0 | 0 | 0 |
| 0 | 0 | 0 | 0 |
| 0 | 0 | 0 | 0 |
| 0 | 0 | 0 | 0 |
| 0 | 0 | 0 | 1 |
| 0 | 0 | 0 | 1 |
| 0 | 1 | 0 | 0 |
| 0 | 0 | 0 | 1 |
| 0 | 1 | 0 | 0 |
| 0 | 1 | 0 | 0 |
| 0 | 1 | 0 | 0 |
| 0 | 1 | 0 | 0 |
| 0 | 0 | 0 | 1 |
| 0 | 0 | 0 | 0 |
| 0 | 0 | 0 | 0 |
| 0 | 0 | 0 | 1 |
| 0 | 0 | 0 | 1 |
| 0 | 0 | 0 | 1 |
| 0 | 0 | 0 | 1 |
| 0 | 0 | 0 | 1 |
| 0 | 0 | 0 | 1 |
| 0 | 0 | 0 | 1 |
| 0 | 0 | 0 | 1 |
| 0 | 0 | 0 | 1 |
| 0 | 0 | 0 | 1 |

| | | | |
|---|---|---|---|
| 0 | 0 | 0 | 1 |
| 0 | 0 | 0 | 0 |
| 0 | 0 | 0 | 1 |
| 0 | 0 | 0 | 1 |
| 0 | 0 | 0 | 1 |
| 0 | 0 | 0 | 1 |
| 0 | 0 | 0 | 1 |
| 0 | 0 | 0 | 1 |
| 0 | 0 | 0 | 1 |
| 0 | 0 | 0 | 1 |
| 0 | 0 | 0 | 1 |
| 0 | 0 | 0 | 0 |
| 0 | 1 | 1 | 1 |
| 0 | 0 | 0 | 1 |
| 0 | 1 | 0 | 0 |
| 0 | 0 | 0 | 1 |
| 0 | 0 | 0 | 1 |
| 0 | 0 | 0 | 1 |
| 0 | 0 | 0 | 1 |
| 0 | 0 | 0 | 1 |
| 1 | 0 | 0 | 0 |
| 0 | 0 | 0 | 1 |
| 1 | 0 | 0 | 0 |
| 1 | 0 | 0 | 0 |
| 0 | 0 | 0 | 1 |
| 0 | 0 | 0 | 0 |
| 0 | 0 | 0 | 1 |
| 0 | 0 | 0 | 1 |
| 0 | 0 | 0 | 1 |
| 0 | 0 | 0 | 1 |
| 0 | 0 | 0 | 1 |
| 0 | 1 | 0 | 0 |
| 0 | 0 | 0 | 1 |
| 0 | 0 | 0 | 0 |
| 0 | 0 | 0 | 1 |
| 0 | 0 | 0 | 1 |
| 0 | 0 | 0 | 1 |
| 0 | 0 | 0 | 1 |
| 0 | 0 | 0 | 1 |
| 0 | 0 | 0 | 0 |
| 0 | 0 | 0 | 1 |
| 0 | 0 | 0 | 1 |
| 0 | 0 | 0 | 1 |
| 0 | 0 | 0 | 0 |
| 0 | 0 | 0 | 1 |
| 0 | 0 | 0 | 1 |
| 0 | 0 | 0 | 1 |
| 1 | 0 | 0 | 0 |
| 0 | 1 | 0 | 0 |
| 0 | 0 | 0 | 1 |
| 0 | 0 | 0 | 1 |
| 0 | 0 | 0 | 0 |
| 1 | 0 | 0 | 0 |
| 0 | 1 | 0 | 0 |

| | | | |
|---|---|---|---|
| 1 | 0 | 0 | 0 |
| 0 | 0 | 0 | 1 |
| 0 | 0 | 0 | 0 |
| 0 | 0 | 0 | 0 |
| 0 | 0 | 0 | 0 |
| 0 | 0 | 0 | 1 |
| 0 | 0 | 0 | 1 |
| 0 | 0 | 0 | 1 |
| 0 | 0 | 0 | 1 |
| 0 | 1 | 0 | 0 |
| 0 | 0 | 0 | 1 |
| 0 | 0 | 0 | 1 |
| 0 | 0 | 0 | 1 |
| 0 | 0 | 0 | 1 |
| 0 | 0 | 0 | 0 |
| 0 | 0 | 0 | 1 |
| 0 | 0 | 0 | 1 |
| 0 | 0 | 0 | 0 |
| 0 | 0 | 0 | 1 |
| 0 | 0 | 0 | 0 |
| 0 | 0 | 0 | 0 |
| 0 | 0 | 0 | 1 |
| 0 | 0 | 0 | 0 |
| 0 | 0 | 0 | 1 |
| 0 | 0 | 0 | 0 |
| 0 | 0 | 0 | 0 |
| 0 | 0 | 0 | 1 |
| 0 | 0 | 0 | 1 |
| 1 | 0 | 0 | 0 |
| 0 | 0 | 0 | 1 |
| 0 | 0 | 0 | 1 |
| 0 | 0 | 0 | 1 |
| 0 | 0 | 0 | 1 |
| 0 | 0 | 0 | 1 |
| 0 | 0 | 0 | 0 |
| 0 | 0 | 0 | 0 |
| 0 | 0 | 0 | 1 |
| 0 | 0 | 0 | 0 |
| 0 | 0 | 0 | 0 |
| 0 | 0 | 0 | 0 |
| 0 | 0 | 0 | 0 |
| 0 | 0 | 0 | 0 |
| 0 | 0 | 0 | 0 |
| 0 | 0 | 0 | 1 |
| 0 | 1 | 0 | 0 |
| 0 | 0 | 0 | 1 |
| 0 | 0 | 0 | 1 |
| 0 | 0 | 0 | 0 |
| 0 | 0 | 0 | 0 |
| 0 | 0 | 0 | 0 |
| 0 | 0 | 0 | 0 |
| 0 | 1 | 0 | 0 |
| 0 | 0 | 0 | 1 |
| 0 | 0 | 0 | 1 |

| | | | |
|---|---|---|---|
| 0 | 0 | 0 | 1 |
| 0 | 0 | 0 | 0 |
| 0 | 0 | 0 | 0 |
| 0 | 0 | 0 | 0 |
| 0 | 0 | 0 | 0 |
| 0 | 0 | 0 | 0 |
| 0 | 1 | 0 | 0 |
| 0 | 1 | 1 | 1 |
| 0 | 1 | 1 | 1 |
| 0 | 0 | 0 | 1 |
| 0 | 0 | 0 | 0 |
| 0 | 0 | 0 | 0 |
| 0 | 0 | 0 | 1 |
| 0 | 0 | 0 | 0 |
| 0 | 0 | 0 | 1 |
| 0 | 0 | 0 | 0 |
| 0 | 0 | 0 | 1 |
| 0 | 0 | 0 | 1 |
| 0 | 0 | 0 | 1 |
| 0 | 0 | 0 | 1 |
| 0 | 0 | 0 | 1 |
| 0 | 0 | 0 | 1 |
| 0 | 0 | 0 | 1 |
| 1 | 0 | 0 | 0 |
| 0 | 0 | 0 | 0 |
| 0 | 0 | 0 | 1 |
| 0 | 0 | 0 | 1 |
| 0 | 0 | 0 | 1 |
| 0 | 0 | 0 | 1 |
| 0 | 0 | 0 | 0 |
| 0 | 0 | 0 | 0 |
| 0 | 0 | 0 | 0 |
| 0 | 0 | 0 | 0 |
| 0 | 0 | 0 | 1 |
| 0 | 0 | 0 | 0 |
| 0 | 0 | 0 | 0 |
| 0 | 0 | 0 | 0 |
| 0 | 0 | 0 | 0 |
| 0 | 0 | 0 | 0 |
| 0 | 0 | 0 | 1 |
| 0 | 0 | 0 | 1 |
| 0 | 0 | 0 | 0 |
| 0 | 0 | 0 | 1 |
| 0 | 0 | 0 | 1 |
| 0 | 0 | 0 | 0 |
| 0 | 1 | 1 | 1 |
| 0 | 0 | 0 | 1 |
| 0 | 0 | 0 | 1 |
| 0 | 1 | 1 | 1 |
| 0 | 0 | 0 | 1 |
| 0 | 0 | 0 | 0 |
| 0 | 0 | 0 | 1 |
| 0 | 0 | 0 | 0 |
| 0 | 0 | 0 | 0 |

| | | | |
|---|---|---|---|
| 0 | 0 | 0 | 1 |
| 0 | 0 | 0 | 1 |
| 0 | 0 | 0 | 0 |
| 0 | 0 | 0 | 0 |
| 0 | 0 | 0 | 1 |
| 0 | 0 | 0 | 1 |
| 0 | 0 | 0 | 1 |
| 0 | 0 | 0 | 1 |
| 0 | 0 | 0 | 1 |
| 0 | 0 | 0 | 1 |
| 0 | 0 | 0 | 1 |
| 0 | 0 | 0 | 1 |
| 0 | 0 | 0 | 0 |
| 0 | 0 | 0 | 1 |
| 0 | 0 | 0 | 1 |
| 0 | 0 | 0 | 1 |
| 0 | 0 | 0 | 1 |
| 0 | 0 | 0 | 1 |
| 0 | 0 | 0 | 1 |
| 0 | 0 | 0 | 1 |
| 0 | 0 | 0 | 1 |
| 0 | 0 | 0 | 1 |
| 0 | 0 | 0 | 1 |
| 0 | 0 | 0 | 1 |
| 0 | 0 | 0 | 1 |
| 0 | 0 | 0 | 1 |
| 0 | 0 | 0 | 1 |
| 0 | 0 | 0 | 1 |
| 0 | 0 | 0 | 1 |
| 0 | 0 | 0 | 1 |
| 0 | 0 | 0 | 1 |
| 0 | 0 | 0 | 0 |
| 0 | 0 | 0 | 1 |
| 0 | 0 | 0 | 1 |
| 0 | 0 | 0 | 1 |
| 0 | 0 | 0 | 1 |
| 0 | 0 | 0 | 1 |
| 0 | 0 | 0 | 1 |
| 0 | 0 | 0 | 1 |
| 0 | 0 | 0 | 1 |
| 0 | 0 | 0 | 0 |
| 0 | 0 | 0 | 0 |
| 0 | 0 | 0 | 1 |
| 0 | 0 | 0 | 1 |
| 0 | 0 | 0 | 1 |
| 0 | 1 | 1 | 1 |
| 0 | 1 | 1 | 1 |
| 0 | 0 | 0 | 1 |
| 0 | 0 | 0 | 0 |
| 0 | 0 | 0 | 0 |
| 0 | 0 | 0 | 1 |
| 0 | 0 | 0 | 1 |
| 0 | 0 | 0 | 0 |
| 0 | 0 | 0 | 1 |

| | | | |
|---|---|---|---|
| 0 | 0 | 0 | 0 |
| 0 | 0 | 0 | 0 |
| 0 | 0 | 0 | 0 |
| 0 | 0 | 0 | 1 |
| 0 | 0 | 0 | 1 |
| 0 | 0 | 0 | 1 |
| 0 | 0 | 0 | 1 |
| 0 | 1 | 0 | 0 |
| 0 | 1 | 0 | 0 |
| 0 | 0 | 0 | 0 |
| 0 | 0 | 0 | 0 |
| 0 | 0 | 0 | 1 |
| 0 | 0 | 0 | 0 |
| 0 | 0 | 0 | 1 |
| 0 | 0 | 0 | 1 |
| 0 | 0 | 0 | 1 |
| 0 | 0 | 0 | 1 |
| 0 | 0 | 0 | 0 |
| 0 | 0 | 0 | 1 |
| 0 | 0 | 0 | 1 |
| 0 | 0 | 0 | 1 |
| 0 | 0 | 0 | 1 |
| 0 | 0 | 0 | 1 |
| 0 | 0 | 0 | 1 |
| 0 | 0 | 0 | 1 |
| 0 | 0 | 0 | 1 |
| 0 | 0 | 0 | 1 |
| 0 | 0 | 0 | 1 |
| 0 | 0 | 0 | 1 |
| 0 | 0 | 0 | 1 |
| 0 | 0 | 0 | 0 |
| 0 | 1 | 0 | 0 |
| 0 | 0 | 0 | 1 |
| 0 | 0 | 0 | 1 |
| 0 | 0 | 0 | 1 |
| 0 | 0 | 0 | 1 |
| 0 | 0 | 0 | 1 |
| 0 | 0 | 0 | 1 |
| 0 | 0 | 0 | 1 |
| 0 | 0 | 0 | 1 |
| 0 | 0 | 0 | 1 |
| 0 | 0 | 0 | 0 |
| 0 | 0 | 0 | 1 |
| 0 | 0 | 0 | 1 |
| 0 | 0 | 0 | 0 |
| 0 | 0 | 0 | 0 |
| 0 | 0 | 0 | 1 |
| 0 | 0 | 0 | 1 |
| 0 | 0 | 0 | 0 |
| 0 | 0 | 0 | 0 |
| 0 | 0 | 0 | 0 |
| 0 | 0 | 0 | 0 |
| 0 | 0 | 0 | 0 |
| 0 | 0 | 0 | 0 |

| | | | |
|---|---|---|---|
| 0 | 0 | 0 | 1 |
| 0 | 0 | 0 | 1 |
| 0 | 0 | 0 | 1 |
| 0 | 0 | 0 | 1 |
| 0 | 0 | 0 | 1 |
| 0 | 0 | 0 | 0 |
| 0 | 0 | 0 | 0 |
| 0 | 0 | 0 | 0 |
| 0 | 0 | 0 | 0 |
| 0 | 1 | 0 | 0 |
| 0 | 0 | 0 | 1 |
| 0 | 0 | 0 | 0 |
| 0 | 0 | 0 | 0 |
| 0 | 0 | 0 | 0 |
| 0 | 0 | 0 | 1 |
| 0 | 0 | 0 | 0 |
| 0 | 0 | 0 | 1 |
| 0 | 0 | 0 | 1 |
| 1 | 0 | 0 | 0 |
| 1 | 0 | 0 | 0 |
| 0 | 0 | 0 | 0 |
| 0 | 0 | 0 | 0 |
| 0 | 0 | 0 | 1 |
| 0 | 0 | 0 | 0 |
| 0 | 0 | 0 | 0 |
| 0 | 0 | 0 | 0 |
| 0 | 0 | 0 | 0 |
| 0 | 0 | 0 | 1 |
| 0 | 0 | 0 | 1 |
| 0 | 0 | 0 | 1 |
| 0 | 0 | 0 | 1 |
| 0 | 0 | 0 | 1 |
| 0 | 0 | 0 | 0 |
| 0 | 0 | 0 | 1 |
| 0 | 0 | 0 | 0 |
| 0 | 0 | 0 | 0 |
| 0 | 1 | 0 | 0 |
| 0 | 0 | 0 | 0 |
| 0 | 1 | 0 | 0 |
| 0 | 1 | 0 | 0 |
| 0 | 1 | 0 | 0 |
| 1 | 0 | 0 | 0 |
| 0 | 0 | 0 | 0 |
| 0 | 0 | 0 | 1 |
| 0 | 0 | 0 | 1 |
| 0 | 0 | 0 | 1 |
| 0 | 0 | 0 | 1 |
| 0 | 0 | 0 | 0 |
| 0 | 0 | 0 | 0 |
| 0 | 0 | 0 | 0 |
| 0 | 0 | 0 | 1 |
| 0 | 0 | 0 | 1 |
| 0 | 0 | 0 | 0 |
| 0 | 0 | 0 | 1 |

| | | | |
|---|---|---|---|
| 0 | 0 | 0 | 1 |
| 0 | 0 | 0 | 0 |
| 0 | 0 | 0 | 1 |
| 0 | 0 | 0 | 1 |
| 0 | 0 | 0 | 1 |
| 0 | 0 | 0 | 1 |
| 0 | 0 | 0 | 1 |
| 0 | 0 | 0 | 1 |
| 0 | 0 | 0 | 1 |
| 0 | 0 | 0 | 1 |
| 0 | 0 | 0 | 1 |
| 0 | 0 | 0 | 1 |
| 0 | 0 | 0 | 0 |
| 0 | 0 | 0 | 0 |
| 0 | 0 | 0 | 0 |
| 0 | 0 | 0 | 0 |
| 0 | 0 | 0 | 0 |
| 0 | 0 | 0 | 0 |
| 0 | 0 | 0 | 0 |
| 0 | 0 | 0 | 1 |
| 1 | 1 | 0 | 0 |
| 0 | 0 | 0 | 0 |
| 0 | 0 | 0 | 0 |
| 0 | 0 | 0 | 0 |
| 0 | 0 | 0 | 0 |
| 0 | 0 | 0 | 1 |
| 0 | 0 | 0 | 1 |
| 0 | 0 | 0 | 0 |
| 0 | 0 | 0 | 1 |
| 0 | 0 | 0 | 0 |
| 0 | 0 | 0 | 0 |
| 0 | 0 | 0 | 1 |
| 0 | 0 | 0 | 0 |
| 0 | 0 | 0 | 1 |
| 0 | 0 | 0 | 1 |
| 0 | 0 | 0 | 1 |
| 0 | 1 | 1 | 1 |
| 0 | 0 | 0 | 1 |
| 0 | 0 | 0 | 1 |
| 0 | 0 | 0 | 1 |
| 0 | 0 | 0 | 0 |
| 0 | 0 | 0 | 1 |
| 0 | 0 | 0 | 1 |
| 0 | 0 | 0 | 0 |
| 0 | 0 | 0 | 0 |
| 0 | 0 | 0 | 1 |
| 0 | 0 | 0 | 0 |
| 0 | 0 | 0 | 0 |
| 0 | 0 | 0 | 1 |
| 0 | 0 | 0 | 1 |
| 0 | 0 | 0 | 0 |
| 0 | 0 | 0 | 1 |
| 0 | 0 | 0 | 1 |
| 0 | 0 | 0 | 1 |

| | | | |
|---|---|---|---|
| 0 | 0 | 0 | 1 |
| 0 | 0 | 0 | 1 |
| 0 | 0 | 0 | 1 |
| 0 | 0 | 0 | 1 |
| 0 | 0 | 0 | 0 |
| 0 | 0 | 0 | 0 |
| 0 | 0 | 0 | 1 |
| 0 | 0 | 0 | 1 |
| 0 | 0 | 0 | 1 |
| 0 | 0 | 0 | 1 |
| 0 | 0 | 0 | 1 |
| 0 | 0 | 0 | 0 |
| 0 | 0 | 0 | 0 |
| 0 | 0 | 0 | 1 |
| 0 | 0 | 0 | 1 |
| 0 | 0 | 0 | 0 |
| 0 | 0 | 0 | 1 |
| 0 | 0 | 0 | 1 |
| 0 | 0 | 0 | 0 |
| 0 | 0 | 0 | 1 |
| 0 | 0 | 0 | 1 |
| 0 | 0 | 0 | 1 |
| 0 | 0 | 0 | 1 |
| 0 | 0 | 0 | 0 |
| 0 | 0 | 0 | 1 |
| 0 | 0 | 0 | 0 |
| 0 | 0 | 0 | 1 |
| 0 | 0 | 0 | 1 |
| 1 | 0 | 0 | 0 |
| 1 | 0 | 0 | 0 |
| 0 | 0 | 0 | 0 |
| 0 | 0 | 0 | 0 |
| 0 | 0 | 0 | 0 |
| 0 | 0 | 0 | 0 |
| 0 | 0 | 0 | 1 |
| 0 | 0 | 0 | 1 |
| 0 | 0 | 0 | 1 |
| 1 | 0 | 0 | 0 |
| 0 | 0 | 0 | 1 |
| 0 | 0 | 0 | 1 |
| 0 | 0 | 0 | 0 |
| 0 | 0 | 0 | 0 |
| 0 | 0 | 0 | 0 |
| 0 | 0 | 0 | 1 |
| 0 | 0 | 0 | 0 |
| 0 | 0 | 0 | 1 |
| 0 | 0 | 0 | 1 |
| 0 | 0 | 0 | 0 |
| 0 | 0 | 0 | 1 |
| 0 | 0 | 0 | 1 |
| 0 | 0 | 0 | 1 |
| 0 | 1 | 0 | 0 |
| 0 | 0 | 0 | 1 |
| 0 | 0 | 0 | 0 |

| | | | |
|---|---|---|---|
| 0 | 0 | 0 | 1 |
| 0 | 1 | 0 | 0 |
| 0 | 0 | 0 | 0 |
| 0 | 0 | 0 | 1 |
| 0 | 0 | 0 | 1 |
| 0 | 0 | 0 | 1 |
| 0 | 0 | 0 | 0 |
| 0 | 0 | 0 | 1 |
| 0 | 0 | 0 | 1 |
| 0 | 0 | 0 | 1 |
| 0 | 0 | 0 | 1 |
| 0 | 0 | 0 | 1 |
| 0 | 0 | 0 | 1 |
| 0 | 0 | 0 | 1 |
| 0 | 0 | 0 | 1 |
| 0 | 0 | 0 | 1 |
| 0 | 0 | 0 | 1 |
| 0 | 0 | 0 | 1 |
| 0 | 0 | 0 | 1 |
| 0 | 0 | 0 | 1 |
| 0 | 0 | 0 | 1 |
| 0 | 0 | 0 | 0 |
| 0 | 0 | 0 | 1 |
| 0 | 0 | 0 | 1 |
| 0 | 0 | 0 | 1 |
| 0 | 0 | 0 | 1 |
| 0 | 0 | 0 | 0 |
| 0 | 0 | 0 | 1 |
| 0 | 0 | 0 | 0 |
| 0 | 0 | 0 | 1 |
| 0 | 0 | 0 | 1 |
| 0 | 0 | 0 | 0 |
| 0 | 1 | 1 | 1 |
| 0 | 1 | 1 | 1 |
| 0 | 0 | 0 | 1 |
| 0 | 0 | 0 | 1 |
| 0 | 0 | 0 | 1 |
| 0 | 0 | 0 | 1 |
| 0 | 0 | 0 | 0 |
| 0 | 0 | 0 | 0 |
| 0 | 1 | 0 | 0 |
| 0 | 0 | 0 | 1 |
| 0 | 0 | 0 | 1 |
| 0 | 0 | 0 | 1 |
| 0 | 0 | 0 | 1 |
| 0 | 1 | 0 | 0 |
| 0 | 0 | 0 | 1 |
| 0 | 0 | 0 | 0 |
| 0 | 0 | 0 | 1 |
| 0 | 1 | 0 | 0 |
| 0 | 0 | 0 | 0 |
| 0 | 1 | 0 | 0 |
| 0 | 0 | 0 | 1 |
| 0 | 0 | 0 | 1 |

| | | | |
|---|---|---|---|
| 0 | 0 | 0 | 0 |
| 0 | 0 | 0 | 0 |
| 0 | 0 | 0 | 1 |
| 0 | 0 | 0 | 1 |
| 0 | 0 | 0 | 1 |
| 0 | 0 | 0 | 1 |
| 0 | 0 | 0 | 1 |
| 0 | 0 | 0 | 1 |
| 0 | 0 | 0 | 1 |
| 0 | 0 | 0 | 1 |
| 0 | 0 | 0 | 1 |
| 0 | 0 | 0 | 1 |
| 0 | 0 | 0 | 1 |
| 0 | 0 | 0 | 1 |
| 0 | 0 | 0 | 1 |
| 0 | 0 | 0 | 0 |
| 0 | 0 | 0 | 0 |
| 0 | 1 | 0 | 0 |
| 0 | 0 | 0 | 1 |
| 0 | 0 | 0 | 1 |
| 0 | 0 | 0 | 1 |
| 0 | 1 | 0 | 0 |
| 0 | 0 | 0 | 0 |
| 0 | 0 | 0 | 0 |
| 0 | 0 | 0 | 0 |
| 0 | 0 | 0 | 1 |
| 0 | 0 | 0 | 1 |
| 0 | 0 | 0 | 1 |
| 0 | 0 | 0 | 1 |
| 0 | 0 | 0 | 1 |
| 0 | 0 | 0 | 0 |
| 0 | 0 | 0 | 1 |
| 0 | 0 | 0 | 0 |
| 0 | 0 | 0 | 1 |
| 0 | 1 | 1 | 1 |
| 0 | 1 | 1 | 1 |
| 0 | 1 | 0 | 0 |
| 0 | 0 | 0 | 0 |
| 0 | 0 | 0 | 1 |
| 0 | 0 | 0 | 1 |
| 0 | 1 | 1 | 1 |
| 0 | 1 | 0 | 0 |
| 0 | 0 | 0 | 1 |
| 0 | 0 | 0 | 1 |
| 0 | 1 | 0 | 0 |
| 0 | 1 | 1 | 1 |
| 0 | 0 | 0 | 0 |
| 0 | 1 | 0 | 0 |
| 0 | 0 | 0 | 1 |
| 0 | 0 | 0 | 1 |
| 0 | 1 | 0 | 0 |
| 0 | 1 | 0 | 0 |
| 0 | 0 | 0 | 1 |
| 0 | 1 | 0 | 0 |

| | | | |
|---|---|---|---|
| 0 | 0 | 0 | 1 |
| 0 | 0 | 0 | 0 |
| 0 | 1 | 1 | 1 |
| 0 | 1 | 0 | 0 |
| 0 | 0 | 0 | 1 |
| 0 | 0 | 0 | 1 |
| 0 | 0 | 0 | 1 |
| 0 | 0 | 0 | 1 |
| 0 | 0 | 0 | 0 |
| 0 | 0 | 0 | 1 |
| 0 | 0 | 0 | 1 |
| 0 | 0 | 0 | 1 |
| 0 | 0 | 0 | 0 |
| 0 | 1 | 0 | 0 |
| 0 | 0 | 0 | 0 |
| 0 | 0 | 0 | 1 |
| 0 | 0 | 0 | 0 |
| 0 | 0 | 0 | 1 |
| 0 | 1 | 0 | 0 |
| 0 | 0 | 0 | 0 |
| 0 | 0 | 0 | 0 |
| 0 | 1 | 0 | 0 |
| 0 | 0 | 0 | 0 |
| 1 | 0 | 0 | 0 |
| 0 | 0 | 0 | 1 |
| 0 | 0 | 0 | 1 |
| 0 | 0 | 0 | 0 |
| 0 | 0 | 0 | 0 |
| 0 | 0 | 0 | 0 |
| 0 | 0 | 0 | 0 |
| 0 | 0 | 0 | 1 |
| 0 | 0 | 0 | 1 |
| 0 | 0 | 0 | 1 |
| 0 | 0 | 0 | 1 |
| 0 | 0 | 0 | 1 |
| 1 | 0 | 0 | 0 |
| 1 | 0 | 0 | 0 |
| 0 | 0 | 0 | 1 |
| 0 | 0 | 0 | 1 |
| 0 | 0 | 0 | 1 |
| 0 | 0 | 0 | 1 |
| 0 | 0 | 0 | 1 |
| 0 | 0 | 0 | 1 |
| 0 | 0 | 0 | 0 |
| 0 | 1 | 0 | 0 |
| 0 | 0 | 0 | 0 |
| 0 | 0 | 0 | 1 |
| 0 | 0 | 0 | 0 |
| 0 | 0 | 0 | 0 |
| 0 | 0 | 0 | 0 |
| 0 | 0 | 0 | 1 |
| 0 | 0 | 0 | 1 |
| 0 | 0 | 0 | 1 |
| 0 | 0 | 0 | 1 |

| | | | |
|---|---|---|---|
| 0 | 0 | 0 | 1 |
| 0 | 0 | 0 | 1 |
| 0 | 0 | 0 | 1 |
| 0 | 0 | 0 | 1 |
| 0 | 0 | 0 | 0 |
| 0 | 0 | 0 | 1 |
| 0 | 0 | 0 | 0 |
| 0 | 0 | 0 | 0 |
| 0 | 0 | 0 | 0 |
| 0 | 0 | 0 | 0 |
| 0 | 0 | 0 | 0 |
| 0 | 0 | 0 | 0 |
| 0 | 0 | 0 | 1 |
| 0 | 0 | 0 | 0 |
| 0 | 0 | 0 | 0 |
| 0 | 0 | 0 | 1 |
| 0 | 0 | 0 | 1 |
| 0 | 0 | 0 | 1 |
| 0 | 0 | 0 | 0 |
| 0 | 0 | 0 | 0 |
| 0 | 0 | 0 | 0 |
| 0 | 0 | 0 | 0 |
| 0 | 0 | 0 | 1 |
| 0 | 0 | 0 | 1 |
| 0 | 0 | 0 | 1 |
| 0 | 0 | 0 | 1 |
| 0 | 0 | 0 | 1 |
| 0 | 0 | 0 | 0 |
| 0 | 0 | 0 | 0 |
| 0 | 0 | 0 | 0 |
| 0 | 0 | 0 | 0 |
| 0 | 0 | 0 | 0 |
| 0 | 0 | 0 | 0 |
| 0 | 0 | 0 | 1 |
| 0 | 0 | 0 | 1 |
| 0 | 0 | 0 | 0 |
| 0 | 0 | 0 | 1 |
| 0 | 0 | 0 | 1 |
| 0 | 0 | 0 | 1 |
| 0 | 0 | 0 | 1 |
| 0 | 0 | 0 | 0 |
| 0 | 0 | 0 | 1 |
| 0 | 0 | 0 | 0 |
| 0 | 0 | 0 | 1 |
| 0 | 0 | 0 | 1 |
| 0 | 0 | 0 | 1 |
| 0 | 0 | 0 | 1 |
| 0 | 0 | 0 | 0 |
| 0 | 0 | 0 | 1 |
| 0 | 0 | 0 | 1 |
| 0 | 0 | 0 | 1 |
| 0 | 0 | 0 | 1 |
| 0 | 0 | 0 | 1 |
| 0 | 0 | 0 | 1 |

| | | | |
|---|---|---|---|
| 0 | 0 | 0 | 1 |
| 0 | 0 | 0 | 1 |
| 0 | 0 | 0 | 1 |
| 0 | 0 | 0 | 1 |
| 0 | 0 | 0 | 1 |
| 0 | 0 | 0 | 1 |
| 0 | 0 | 0 | 1 |
| 0 | 0 | 0 | 1 |
| 0 | 0 | 0 | 1 |
| 0 | 0 | 0 | 1 |
| 0 | 0 | 0 | 1 |
| 0 | 0 | 0 | 0 |
| 0 | 0 | 0 | 1 |
| 0 | 0 | 0 | 1 |
| 0 | 0 | 0 | 1 |
| 0 | 0 | 0 | 1 |
| 0 | 0 | 0 | 1 |
| 0 | 0 | 0 | 0 |
| 0 | 0 | 0 | 0 |
| 0 | 0 | 0 | 0 |
| 0 | 0 | 0 | 0 |
| 0 | 0 | 0 | 1 |
| 1 | 0 | 0 | 0 |
| 0 | 0 | 0 | 1 |
| 0 | 0 | 0 | 1 |
| 0 | 0 | 0 | 1 |
| 0 | 0 | 0 | 1 |
| 0 | 0 | 0 | 1 |
| 0 | 0 | 0 | 0 |
| 0 | 0 | 0 | 0 |
| 0 | 0 | 0 | 1 |
| 0 | 0 | 0 | 1 |
| 1 | 0 | 0 | 0 |
| 0 | 0 | 0 | 1 |
| 0 | 0 | 0 | 0 |
| 1 | 0 | 0 | 0 |
| 1 | 0 | 0 | 0 |
| 0 | 0 | 0 | 1 |
| 1 | 0 | 0 | 0 |
| 0 | 0 | 0 | 1 |
| 1 | 0 | 0 | 0 |
| 0 | 0 | 0 | 1 |
| 0 | 0 | 0 | 0 |
| 0 | 0 | 0 | 1 |
| 0 | 0 | 0 | 1 |
| 0 | 0 | 0 | 1 |
| 0 | 1 | 1 | 1 |
| 0 | 0 | 0 | 1 |
| 0 | 0 | 0 | 1 |
| 0 | 0 | 0 | 0 |
| 0 | 1 | 1 | 1 |
| 1 | 0 | 0 | 0 |
| 0 | 1 | 1 | 1 |
| 0 | 0 | 0 | 1 |

| | | | |
|---|---|---|---|
| 0 | 1 | 0 | 0 |
| 0 | 0 | 0 | 0 |
| 0 | 0 | 0 | 0 |
| 0 | 0 | 0 | 0 |
| 0 | 1 | 0 | 0 |
| 0 | 0 | 0 | 1 |
| 0 | 1 | 0 | 0 |
| 0 | 0 | 0 | 1 |
| 0 | 0 | 0 | 1 |
| 0 | 0 | 0 | 0 |
| 0 | 0 | 0 | 0 |
| 0 | 0 | 0 | 0 |
| 0 | 0 | 0 | 1 |
| 0 | 0 | 0 | 1 |
| 0 | 0 | 0 | 1 |
| 0 | 0 | 0 | 1 |
| 0 | 0 | 0 | 1 |
| 0 | 0 | 0 | 1 |
| 0 | 0 | 0 | 1 |
| 0 | 0 | 0 | 1 |
| 0 | 0 | 0 | 1 |
| 0 | 0 | 0 | 1 |
| 0 | 0 | 0 | 1 |
| 0 | 0 | 0 | 1 |
| 0 | 0 | 0 | 1 |
| 0 | 0 | 0 | 1 |
| 0 | 0 | 0 | 1 |
| 0 | 0 | 0 | 1 |
| 0 | 0 | 0 | 1 |
| 0 | 0 | 0 | 1 |
| 0 | 0 | 0 | 1 |
| 0 | 0 | 0 | 1 |
| 0 | 0 | 0 | 1 |
| 0 | 0 | 0 | 1 |
| 0 | 0 | 0 | 1 |
| 0 | 0 | 0 | 1 |
| 0 | 0 | 0 | 1 |
| 0 | 0 | 0 | 1 |
| 0 | 0 | 0 | 1 |
| 0 | 0 | 0 | 1 |
| 0 | 0 | 0 | 1 |
| 0 | 0 | 0 | 1 |
| 0 | 0 | 0 | 1 |
| 0 | 0 | 0 | 1 |
| 0 | 0 | 0 | 1 |
| 0 | 0 | 0 | 0 |
| 0 | 0 | 0 | 1 |
| 0 | 0 | 0 | 1 |
| 0 | 0 | 0 | 1 |
| 0 | 0 | 0 | 1 |
| 0 | 0 | 0 | 0 |
| 0 | 0 | 0 | 1 |
| 0 | 0 | 0 | 1 |
| 0 | 0 | 0 | 1 |

| | | | |
|---|---|---|---|
| 0 | 0 | 0 | 1 |
| 0 | 0 | 0 | 1 |
| 0 | 0 | 0 | 1 |
| 0 | 0 | 0 | 1 |
| 0 | 0 | 0 | 1 |
| 0 | 0 | 0 | 1 |
| 0 | 0 | 0 | 1 |
| 0 | 0 | 0 | 0 |
| 0 | 1 | 0 | 0 |
| 0 | 0 | 0 | 1 |
| 0 | 0 | 0 | 1 |
| 0 | 0 | 0 | 1 |
| 0 | 0 | 0 | 1 |
| 0 | 0 | 0 | 1 |
| 0 | 1 | 0 | 0 |
| 0 | 1 | 0 | 0 |
| 0 | 0 | 0 | 0 |
| 0 | 0 | 0 | 1 |
| 0 | 1 | 1 | 1 |
| 0 | 0 | 0 | 1 |
| 0 | 1 | 0 | 0 |
| 0 | 1 | 0 | 0 |
| 0 | 1 | 0 | 0 |
| 0 | 0 | 0 | 0 |
| 0 | 1 | 0 | 0 |
| 0 | 1 | 0 | 0 |
| 0 | 0 | 0 | 1 |
| 0 | 0 | 0 | 1 |
| 0 | 0 | 0 | 0 |
| 0 | 0 | 0 | 1 |
| 0 | 0 | 0 | 1 |
| 0 | 0 | 0 | 1 |
| 0 | 0 | 0 | 0 |
| 0 | 0 | 0 | 1 |
| 0 | 0 | 0 | 1 |
| 0 | 0 | 0 | 1 |
| 0 | 0 | 0 | 0 |
| 0 | 0 | 0 | 0 |
| 0 | 0 | 0 | 0 |
| 0 | 0 | 0 | 1 |
| 0 | 0 | 0 | 1 |
| 0 | 0 | 0 | 1 |
| 0 | 0 | 0 | 1 |
| 0 | 0 | 0 | 1 |
| 0 | 0 | 0 | 1 |
| 0 | 0 | 0 | 0 |
| 0 | 0 | 0 | 1 |
| 0 | 0 | 0 | 1 |
| 0 | 0 | 0 | 1 |
| 0 | 0 | 0 | 1 |
| 0 | 0 | 0 | 1 |
| 0 | 0 | 0 | 1 |
| 0 | 0 | 0 | 1 |
| 0 | 0 | 0 | 1 |

| | | | |
|---|---|---|---|
| 0 | 0 | 0 | 1 |
| 0 | 0 | 0 | 1 |
| 0 | 0 | 0 | 1 |
| 0 | 0 | 0 | 0 |
| 0 | 0 | 0 | 1 |
| 0 | 0 | 0 | 1 |
| 0 | 0 | 0 | 1 |
| 0 | 0 | 0 | 0 |
| 0 | 0 | 0 | 0 |
| 0 | 0 | 0 | 1 |
| 0 | 0 | 0 | 1 |
| 0 | 0 | 0 | 1 |
| 0 | 0 | 0 | 0 |
| 0 | 0 | 0 | 1 |
| 0 | 0 | 0 | 1 |
| 0 | 0 | 0 | 1 |
| 0 | 0 | 0 | 0 |
| 0 | 0 | 0 | 1 |
| 0 | 0 | 0 | 1 |
| 0 | 0 | 0 | 1 |
| 0 | 0 | 0 | 1 |
| 0 | 0 | 0 | 1 |
| 0 | 0 | 0 | 1 |
| 0 | 0 | 0 | 1 |
| 0 | 0 | 0 | 1 |
| 0 | 0 | 0 | 1 |
| 0 | 0 | 0 | 1 |
| 0 | 0 | 0 | 0 |
| 0 | 0 | 0 | 0 |
| 0 | 0 | 0 | 0 |
| 0 | 0 | 0 | 0 |
| 0 | 0 | 0 | 0 |
| 0 | 0 | 0 | 0 |
| 0 | 0 | 0 | 1 |
| 0 | 0 | 0 | 0 |
| 0 | 0 | 0 | 0 |
| 0 | 0 | 0 | 0 |
| 0 | 0 | 0 | 1 |
| 0 | 0 | 0 | 1 |
| 0 | 0 | 0 | 1 |
| 0 | 0 | 0 | 1 |
| 0 | 0 | 0 | 1 |
| 0 | 0 | 0 | 0 |
| 0 | 0 | 0 | 0 |
| 0 | 0 | 0 | 1 |
| 0 | 0 | 0 | 1 |
| 0 | 0 | 0 | 0 |
| 0 | 0 | 0 | 1 |
| 0 | 0 | 0 | 1 |
| 0 | 0 | 0 | 1 |
| 0 | 0 | 0 | 1 |
| 0 | 0 | 0 | 1 |
| 0 | 0 | 0 | 0 |
| 0 | 0 | 0 | 0 |

| | | | |
|---|---|---|---|
| 0 | 0 | 0 | 0 |
| 0 | 0 | 0 | 0 |
| 0 | 0 | 0 | 0 |
| 0 | 0 | 0 | 0 |
| 0 | 0 | 0 | 0 |
| 0 | 0 | 0 | 0 |
| 0 | 0 | 0 | 0 |
| 0 | 0 | 0 | 0 |
| 0 | 0 | 0 | 1 |
| 0 | 1 | 0 | 0 |
| 0 | 0 | 0 | 1 |
| 0 | 0 | 0 | 1 |
| 0 | 0 | 0 | 1 |
| 0 | 0 | 0 | 1 |
| 0 | 0 | 0 | 0 |
| 0 | 0 | 0 | 1 |
| 0 | 0 | 0 | 1 |
| 0 | 0 | 0 | 1 |
| 0 | 0 | 0 | 1 |
| 0 | 0 | 0 | 0 |
| 0 | 0 | 0 | 0 |
| 0 | 0 | 0 | 1 |
| 0 | 0 | 0 | 0 |
| 0 | 0 | 0 | 0 |
| 0 | 0 | 0 | 0 |
| 0 | 0 | 0 | 0 |
| 0 | 0 | 0 | 0 |
| 0 | 0 | 0 | 1 |
| 0 | 0 | 0 | 1 |
| 0 | 0 | 0 | 1 |
| 0 | 0 | 0 | 0 |
| 0 | 0 | 0 | 1 |
| 0 | 0 | 0 | 1 |
| 0 | 0 | 0 | 1 |
| 0 | 0 | 0 | 1 |
| 0 | 0 | 0 | 1 |
| 0 | 0 | 0 | 1 |
| 0 | 0 | 0 | 1 |
| 0 | 0 | 0 | 1 |
| 0 | 0 | 0 | 1 |
| 0 | 0 | 0 | 1 |
| 0 | 0 | 0 | 0 |
| 0 | 0 | 0 | 1 |
| 0 | 0 | 0 | 1 |
| 0 | 0 | 0 | 1 |
| 0 | 0 | 0 | 1 |
| 0 | 0 | 0 | 1 |
| 0 | 0 | 0 | 1 |
| 0 | 0 | 0 | 0 |
| 0 | 0 | 0 | 1 |
| 0 | 0 | 0 | 1 |
| 0 | 0 | 0 | 1 |
| 0 | 1 | 0 | 0 |
| 0 | 1 | 0 | 0 |

| | | | |
|---|---|---|---|
| 0 | 0 | 0 | 1 |
| 0 | 0 | 0 | 1 |
| 0 | 0 | 0 | 1 |
| 0 | 0 | 0 | 1 |
| 0 | 0 | 0 | 0 |
| 1 | 0 | 0 | 0 |
| 0 | 0 | 0 | 1 |
| 0 | 0 | 0 | 0 |
| 0 | 0 | 0 | 1 |
| 0 | 0 | 0 | 1 |
| 0 | 0 | 0 | 0 |
| 0 | 1 | 1 | 1 |
| 0 | 1 | 0 | 0 |
| 0 | 0 | 0 | 0 |
| 0 | 1 | 0 | 0 |
| 0 | 1 | 0 | 0 |
| 0 | 0 | 0 | 1 |
| 0 | 0 | 0 | 1 |
| 0 | 0 | 0 | 0 |
| 0 | 1 | 0 | 0 |
| 0 | 1 | 0 | 0 |
| 0 | 1 | 0 | 0 |
| 0 | 1 | 0 | 0 |
| 0 | 0 | 0 | 1 |
| 0 | 0 | 0 | 1 |
| 0 | 0 | 0 | 0 |
| 0 | 0 | 0 | 1 |
| 0 | 0 | 0 | 0 |
| 0 | 0 | 0 | 0 |
| 0 | 0 | 0 | 0 |
| 0 | 0 | 0 | 1 |
| 1 | 0 | 0 | 0 |
| 1 | 0 | 0 | 0 |
| 0 | 0 | 0 | 1 |
| 0 | 0 | 0 | 1 |
| 0 | 0 | 0 | 0 |
| 0 | 0 | 0 | 0 |
| 0 | 1 | 1 | 1 |
| 0 | 1 | 0 | 0 |
| 0 | 1 | 0 | 0 |
| 0 | 1 | 0 | 0 |
| 0 | 0 | 0 | 0 |
| 0 | 0 | 0 | 1 |
| 0 | 0 | 0 | 1 |
| 0 | 0 | 0 | 0 |
| 0 | 0 | 0 | 1 |
| 0 | 0 | 0 | 0 |
| 0 | 0 | 0 | 1 |
| 0 | 0 | 0 | 1 |
| 0 | 0 | 0 | 1 |
| 0 | 0 | 0 | 1 |
| 0 | 0 | 0 | 1 |
| 0 | 0 | 0 | 1 |
| 0 | 0 | 0 | 1 |

| | | | |
|---|---|---|---|
| 0 | 0 | 0 | 1 |
| 0 | 0 | 0 | 1 |
| 0 | 0 | 0 | 1 |
| 0 | 0 | 0 | 1 |
| 0 | 0 | 0 | 1 |
| 0 | 0 | 0 | 1 |
| 0 | 0 | 0 | 1 |
| 0 | 0 | 0 | 0 |
| 0 | 0 | 0 | 1 |
| 1 | 0 | 0 | 0 |
| 0 | 0 | 0 | 0 |
| 0 | 0 | 0 | 0 |
| 0 | 0 | 0 | 1 |
| 0 | 0 | 0 | 1 |
| 0 | 0 | 0 | 0 |
| 0 | 0 | 0 | 1 |
| 0 | 0 | 0 | 1 |
| 0 | 0 | 0 | 1 |
| 0 | 0 | 0 | 1 |
| 0 | 0 | 0 | 1 |
| 0 | 0 | 0 | 1 |
| 0 | 0 | 0 | 1 |
| 0 | 0 | 0 | 1 |
| 0 | 0 | 0 | 1 |
| 0 | 0 | 0 | 1 |
| 0 | 0 | 0 | 1 |
| 0 | 0 | 0 | 1 |
| 0 | 1 | 1 | 1 |
| 0 | 0 | 0 | 1 |
| 0 | 0 | 0 | 0 |
| 0 | 0 | 0 | 0 |
| 0 | 0 | 0 | 1 |
| 0 | 0 | 0 | 1 |
| 0 | 0 | 0 | 1 |
| 0 | 0 | 0 | 1 |
| 0 | 0 | 0 | 0 |
| 0 | 0 | 0 | 1 |
| 0 | 0 | 0 | 1 |
| 0 | 0 | 0 | 1 |
| 0 | 0 | 0 | 1 |
| 0 | 0 | 0 | 1 |
| 0 | 0 | 0 | 0 |
| 0 | 0 | 0 | 0 |
| 0 | 1 | 0 | 0 |
| 0 | 0 | 0 | 1 |
| 0 | 0 | 0 | 1 |
| 0 | 0 | 0 | 1 |
| 0 | 0 | 0 | 1 |
| 0 | 0 | 0 | 1 |
| 0 | 0 | 0 | 1 |
| 0 | 0 | 0 | 1 |
| 0 | 0 | 0 | 1 |
| 0 | 0 | 0 | 1 |
| 0 | 0 | 0 | 0 |

| | | | |
|---|---|---|---|
| 0 | 0 | 0 | 1 |
| 0 | 0 | 0 | 1 |
| 0 | 0 | 0 | 1 |
| 0 | 0 | 0 | 1 |
| 0 | 0 | 0 | 1 |
| 0 | 0 | 0 | 1 |
| 0 | 0 | 0 | 1 |
| 0 | 0 | 0 | 1 |
| 0 | 0 | 0 | 0 |
| 0 | 0 | 0 | 0 |
| 0 | 0 | 0 | 1 |
| 0 | 0 | 0 | 1 |
| 0 | 0 | 0 | 1 |
| 0 | 0 | 0 | 0 |
| 0 | 0 | 0 | 1 |
| 0 | 0 | 0 | 1 |
| 0 | 0 | 0 | 1 |
| 1 | 0 | 0 | 0 |
| 1 | 0 | 0 | 0 |
| 0 | 0 | 0 | 0 |
| 0 | 0 | 0 | 0 |
| 0 | 0 | 0 | 1 |
| 0 | 0 | 0 | 1 |
| 0 | 0 | 0 | 1 |
| 0 | 0 | 0 | 1 |
| 1 | 0 | 0 | 0 |
| 0 | 0 | 0 | 1 |
| 0 | 0 | 0 | 1 |
| 0 | 0 | 0 | 1 |
| 0 | 0 | 0 | 1 |
| 0 | 0 | 0 | 1 |
| 0 | 0 | 0 | 1 |
| 0 | 0 | 0 | 1 |
| 0 | 0 | 0 | 1 |
| 0 | 0 | 0 | 1 |
| 0 | 0 | 0 | 1 |
| 0 | 0 | 0 | 1 |
| 0 | 0 | 0 | 1 |
| 0 | 0 | 0 | 1 |
| 0 | 0 | 0 | 0 |
| 0 | 0 | 0 | 1 |
| 0 | 0 | 0 | 1 |
| 0 | 0 | 0 | 0 |
| 0 | 0 | 0 | 1 |
| 0 | 0 | 0 | 1 |
| 0 | 0 | 0 | 1 |
| 0 | 0 | 0 | 0 |
| 0 | 0 | 0 | 0 |
| 0 | 0 | 0 | 1 |
| 0 | 0 | 0 | 1 |
| 0 | 0 | 0 | 0 |
| 0 | 0 | 0 | 1 |
| 0 | 0 | 0 | 1 |
| 0 | 0 | 0 | 1 |

| | | | |
|---|---|---|---|
| 0 | 0 | 0 | 1 |
| 0 | 0 | 0 | 1 |
| 0 | 0 | 0 | 1 |
| 0 | 0 | 0 | 1 |
| 0 | 0 | 0 | 1 |
| 0 | 0 | 0 | 1 |
| 0 | 0 | 0 | 1 |
| 0 | 0 | 0 | 1 |
| 0 | 0 | 0 | 1 |
| 0 | 0 | 0 | 1 |
| 0 | 0 | 0 | 1 |
| 0 | 0 | 0 | 1 |
| 0 | 0 | 0 | 1 |
| 0 | 0 | 0 | 0 |
| 0 | 0 | 0 | 0 |
| 0 | 0 | 0 | 0 |
| 0 | 0 | 0 | 0 |
| 0 | 0 | 0 | 1 |
| 0 | 0 | 0 | 1 |
| 0 | 0 | 0 | 0 |
| 0 | 0 | 0 | 1 |
| 0 | 1 | 1 | 1 |
| 0 | 0 | 0 | 1 |
| 0 | 0 | 0 | 1 |
| 0 | 0 | 0 | 1 |
| 0 | 0 | 0 | 0 |
| 0 | 0 | 0 | 1 |
| 0 | 0 | 0 | 1 |
| 0 | 0 | 0 | 1 |
| 0 | 0 | 0 | 0 |
| 0 | 0 | 0 | 0 |
| 0 | 0 | 0 | 1 |
| 0 | 0 | 0 | 1 |
| 0 | 0 | 0 | 1 |
| 0 | 0 | 0 | 1 |
| 0 | 0 | 0 | 1 |
| 0 | 0 | 0 | 1 |
| 0 | 0 | 0 | 1 |
| 0 | 0 | 0 | 0 |
| 0 | 0 | 0 | 1 |
| 0 | 0 | 0 | 1 |
| 0 | 1 | 0 | 0 |
| 0 | 1 | 0 | 0 |
| 0 | 1 | 0 | 0 |
| 0 | 0 | 0 | 0 |
| 0 | 0 | 0 | 1 |
| 0 | 0 | 0 | 0 |
| 0 | 0 | 0 | 1 |
| 0 | 0 | 0 | 1 |
| 0 | 0 | 0 | 1 |
| 0 | 0 | 0 | 1 |
| 0 | 0 | 0 | 1 |
| 0 | 1 | 1 | 1 |
| 0 | 1 | 1 | 1 |

| | | | |
|---|---|---|---|
| 0 | 0 | 0 | 1 |
| 0 | 0 | 0 | 1 |
| 0 | 0 | 0 | 0 |
| 0 | 0 | 0 | 1 |
| 0 | 0 | 0 | 1 |
| 0 | 0 | 0 | 1 |
| 0 | 0 | 0 | 1 |
| 0 | 0 | 0 | 1 |
| 0 | 0 | 0 | 1 |
| 0 | 0 | 0 | 1 |
| 0 | 0 | 0 | 1 |
| 0 | 0 | 0 | 1 |
| 0 | 0 | 0 | 1 |
| 0 | 0 | 0 | 1 |
| 0 | 0 | 0 | 1 |
| 0 | 0 | 0 | 1 |
| 0 | 0 | 0 | 1 |
| 0 | 0 | 0 | 1 |
| 0 | 0 | 0 | 1 |
| 0 | 0 | 0 | 1 |
| 0 | 0 | 0 | 1 |
| 0 | 0 | 0 | 1 |
| 0 | 0 | 0 | 1 |
| 0 | 0 | 0 | 1 |
| 0 | 0 | 0 | 1 |
| 0 | 0 | 0 | 1 |
| 0 | 0 | 0 | 1 |
| 0 | 0 | 0 | 1 |
| 0 | 0 | 0 | 1 |
| 0 | 0 | 0 | 1 |
| 1 | 0 | 0 | 0 |
| 0 | 0 | 0 | 1 |
| 0 | 1 | 0 | 0 |
| 0 | 1 | 0 | 0 |
| 0 | 1 | 0 | 0 |
| 0 | 0 | 0 | 0 |
| 0 | 0 | 0 | 1 |
| 0 | 0 | 0 | 0 |
| 0 | 0 | 0 | 1 |
| 0 | 0 | 0 | 1 |
| 1 | 0 | 0 | 0 |
| 1 | 0 | 0 | 0 |
| 0 | 0 | 0 | 0 |
| 0 | 0 | 0 | 1 |
| 0 | 0 | 0 | 0 |
| 0 | 0 | 0 | 0 |
| 0 | 0 | 0 | 1 |
| 0 | 0 | 0 | 1 |
| 0 | 0 | 0 | 1 |
| 0 | 0 | 0 | 1 |
| 0 | 0 | 0 | 0 |
| 0 | 0 | 0 | 1 |
| 0 | 0 | 0 | 1 |
| 0 | 0 | 0 | 1 |

| | | | |
|---|---|---|---|
| 0 | 0 | 0 | 1 |
| 0 | 0 | 0 | 1 |
| 0 | 0 | 0 | 1 |
| 0 | 0 | 0 | 1 |
| 0 | 0 | 0 | 1 |
| 1 | 0 | 0 | 0 |
| 0 | 0 | 0 | 1 |
| 0 | 0 | 0 | 0 |
| 0 | 0 | 0 | 0 |
| 0 | 0 | 0 | 0 |
| 0 | 0 | 0 | 1 |
| 0 | 1 | 0 | 0 |
| 0 | 0 | 0 | 1 |
| 0 | 0 | 0 | 1 |
| 0 | 0 | 0 | 1 |
| 0 | 1 | 1 | 1 |
| 0 | 0 | 0 | 1 |
| 0 | 0 | 0 | 0 |
| 0 | 0 | 0 | 1 |
| 0 | 0 | 0 | 0 |
| 0 | 0 | 0 | 1 |
| 0 | 0 | 0 | 0 |
| 0 | 0 | 0 | 0 |
| 0 | 0 | 0 | 0 |
| 0 | 0 | 0 | 1 |
| 0 | 0 | 0 | 1 |
| 0 | 0 | 0 | 1 |
| 0 | 0 | 0 | 1 |
| 0 | 0 | 0 | 0 |
| 0 | 0 | 0 | 1 |
| 0 | 0 | 0 | 1 |
| 0 | 0 | 0 | 1 |
| 0 | 0 | 0 | 1 |
| 0 | 0 | 0 | 1 |
| 0 | 0 | 0 | 0 |
| 0 | 0 | 0 | 0 |
| 0 | 0 | 0 | 1 |
| 0 | 0 | 0 | 0 |
| 0 | 0 | 0 | 0 |
| 0 | 0 | 0 | 1 |
| 0 | 0 | 0 | 0 |
| 0 | 0 | 0 | 0 |
| 0 | 0 | 0 | 1 |
| 0 | 0 | 0 | 1 |
| 0 | 0 | 0 | 1 |
| 0 | 0 | 0 | 1 |
| 0 | 0 | 0 | 1 |
| 0 | 0 | 0 | 1 |
| 0 | 0 | 0 | 1 |
| 0 | 0 | 0 | 1 |
| 0 | 0 | 0 | 1 |
| 0 | 0 | 0 | 1 |
| 0 | 0 | 0 | 1 |
| 0 | 0 | 0 | 1 |

| | | | |
|---|---|---|---|
| 0 | 0 | 0 | 1 |
| 0 | 0 | 0 | 0 |
| 0 | 0 | 0 | 1 |
| 0 | 0 | 0 | 1 |
| 0 | 0 | 0 | 0 |
| 0 | 0 | 0 | 1 |
| 0 | 0 | 0 | 1 |
| 0 | 0 | 0 | 1 |
| 0 | 0 | 0 | 0 |
| 0 | 0 | 0 | 0 |
| 0 | 0 | 0 | 1 |
| 0 | 0 | 0 | 0 |
| 0 | 0 | 0 | 0 |
| 0 | 0 | 0 | 1 |
| 0 | 0 | 0 | 1 |
| 0 | 0 | 0 | 0 |
| 0 | 0 | 0 | 1 |
| 0 | 0 | 0 | 0 |
| 0 | 1 | 0 | 0 |
| 0 | 0 | 0 | 1 |
| 0 | 0 | 0 | 0 |
| 0 | 0 | 0 | 0 |
| 1 | 0 | 0 | 0 |
| 0 | 0 | 0 | 1 |
| 0 | 0 | 0 | 1 |
| 0 | 0 | 0 | 0 |
| 0 | 0 | 0 | 1 |
| 0 | 0 | 0 | 1 |
| 0 | 0 | 0 | 1 |
| 0 | 0 | 0 | 1 |
| 0 | 0 | 0 | 0 |
| 0 | 0 | 0 | 1 |
| 0 | 0 | 0 | 0 |
| 0 | 0 | 0 | 1 |
| 0 | 0 | 0 | 0 |
| 0 | 0 | 0 | 0 |
| 0 | 0 | 0 | 1 |
| 0 | 0 | 0 | 1 |
| 0 | 0 | 0 | 1 |
| 0 | 0 | 0 | 1 |
| 0 | 0 | 0 | 1 |
| 0 | 0 | 0 | 1 |
| 0 | 0 | 0 | 1 |
| 0 | 0 | 0 | 1 |
| 0 | 0 | 0 | 1 |
| 0 | 0 | 0 | 1 |
| 0 | 1 | 1 | 1 |
| 0 | 0 | 0 | 1 |
| 0 | 0 | 0 | 1 |
| 0 | 0 | 0 | 1 |
| 0 | 0 | 0 | 1 |
| 0 | 0 | 0 | 1 |
| 0 | 0 | 0 | 1 |
| 0 | 0 | 0 | 0 |

| | | | |
|---|---|---|---|
| 0 | 0 | 0 | 1 |
| 0 | 0 | 0 | 1 |
| 0 | 0 | 0 | 1 |
| 0 | 0 | 0 | 1 |
| 0 | 0 | 0 | 0 |
| 0 | 0 | 0 | 1 |
| 0 | 0 | 0 | 1 |
| 0 | 0 | 0 | 1 |
| 0 | 0 | 0 | 1 |
| 0 | 0 | 0 | 1 |
| 0 | 0 | 0 | 0 |
| 0 | 0 | 0 | 1 |
| 0 | 0 | 0 | 1 |
| 0 | 0 | 0 | 1 |
| 0 | 0 | 0 | 0 |
| 0 | 0 | 0 | 0 |
| 0 | 0 | 0 | 0 |
| 0 | 0 | 0 | 0 |
| 0 | 0 | 0 | 1 |
| 0 | 0 | 0 | 0 |
| 0 | 0 | 0 | 0 |
| 1 | 0 | 0 | 0 |
| 0 | 0 | 0 | 1 |
| 0 | 0 | 0 | 0 |
| 1 | 0 | 0 | 0 |
| 0 | 0 | 0 | 1 |
| 1 | 0 | 0 | 0 |
| 0 | 0 | 0 | 0 |
| 1 | 0 | 0 | 0 |
| 1 | 0 | 0 | 0 |
| 1 | 0 | 0 | 0 |
| 0 | 0 | 0 | 1 |
| 0 | 0 | 0 | 1 |
| 0 | 0 | 0 | 1 |
| 0 | 0 | 0 | 1 |
| 0 | 0 | 0 | 1 |
| 0 | 0 | 0 | 1 |
| 0 | 1 | 0 | 0 |
| 0 | 0 | 0 | 0 |
| 0 | 0 | 0 | 1 |
| 0 | 0 | 0 | 1 |
| 0 | 1 | 1 | 1 |
| 0 | 0 | 0 | 0 |
| 0 | 0 | 0 | 1 |
| 0 | 0 | 0 | 0 |
| 0 | 0 | 0 | 1 |
| 0 | 0 | 0 | 1 |
| 0 | 0 | 0 | 1 |
| 0 | 1 | 0 | 0 |
| 0 | 1 | 0 | 0 |
| 0 | 1 | 0 | 0 |
| 0 | 0 | 0 | 1 |
| 0 | 0 | 0 | 1 |
| 0 | 0 | 0 | 0 |

| | | | |
|---|---|---|---|
| 0 | 0 | 0 | 1 |
| 0 | 1 | 0 | 0 |
| 0 | 0 | 0 | 1 |
| 0 | 1 | 0 | 0 |
| 0 | 1 | 0 | 0 |
| 0 | 0 | 0 | 1 |
| 0 | 1 | 0 | 0 |
| 0 | 1 | 0 | 0 |
| 0 | 0 | 0 | 1 |
| 0 | 0 | 0 | 1 |
| 0 | 0 | 0 | 1 |
| 0 | 0 | 0 | 1 |
| 0 | 0 | 0 | 1 |
| 0 | 0 | 0 | 1 |
| 0 | 0 | 0 | 1 |
| 0 | 0 | 0 | 1 |
| 0 | 0 | 0 | 1 |
| 0 | 0 | 0 | 1 |
| 0 | 0 | 0 | 1 |
| 0 | 0 | 0 | 1 |
| 1 | 1 | 0 | 0 |
| 0 | 0 | 0 | 1 |
| 0 | 0 | 0 | 1 |
| 0 | 0 | 0 | 1 |
| 0 | 0 | 0 | 1 |
| 0 | 0 | 0 | 0 |
| 0 | 0 | 0 | 0 |
| 0 | 0 | 0 | 1 |
| 0 | 0 | 0 | 1 |
| 0 | 0 | 0 | 1 |
| 0 | 0 | 0 | 1 |
| 0 | 0 | 0 | 1 |
| 0 | 0 | 0 | 1 |
| 0 | 0 | 0 | 1 |
| 0 | 0 | 0 | 1 |
| 0 | 0 | 0 | 1 |
| 0 | 0 | 0 | 1 |
| 0 | 0 | 0 | 1 |
| 0 | 0 | 0 | 1 |
| 0 | 0 | 0 | 1 |
| 0 | 0 | 0 | 1 |
| 0 | 0 | 0 | 1 |
| 0 | 0 | 0 | 1 |
| 0 | 0 | 0 | 0 |
| 0 | 0 | 0 | 1 |
| 0 | 0 | 0 | 0 |
| 0 | 0 | 0 | 0 |
| 0 | 0 | 0 | 0 |
| 0 | 0 | 0 | 0 |
| 0 | 0 | 0 | 0 |
| 0 | 0 | 0 | 0 |
| 0 | 0 | 0 | 1 |
| 0 | 0 | 0 | 1 |
| 0 | 0 | 0 | 1 |

| | | | |
|---|---|---|---|
| 0 | 0 | 0 | 1 |
| 0 | 0 | 0 | 1 |
| 0 | 0 | 0 | 1 |
| 0 | 0 | 0 | 0 |
| 0 | 0 | 0 | 1 |
| 0 | 0 | 0 | 0 |
| 0 | 0 | 0 | 1 |
| 0 | 0 | 0 | 1 |
| 0 | 0 | 0 | 0 |
| 0 | 0 | 0 | 1 |
| 0 | 0 | 0 | 1 |
| 0 | 0 | 0 | 1 |
| 0 | 0 | 0 | 1 |
| 0 | 0 | 0 | 1 |
| 0 | 0 | 0 | 1 |
| 0 | 0 | 0 | 0 |
| 0 | 0 | 0 | 0 |
| 0 | 0 | 0 | 0 |
| 0 | 0 | 0 | 0 |
| 0 | 0 | 0 | 0 |
| 0 | 0 | 0 | 1 |
| 0 | 0 | 0 | 1 |
| 0 | 0 | 0 | 1 |
| 0 | 0 | 0 | 1 |
| 0 | 0 | 0 | 0 |
| 0 | 0 | 0 | 1 |
| 0 | 0 | 0 | 1 |
| 0 | 0 | 0 | 1 |
| 0 | 0 | 0 | 0 |
| 0 | 0 | 0 | 1 |
| 0 | 1 | 0 | 0 |
| 0 | 1 | 0 | 0 |
| 0 | 0 | 0 | 0 |
| 0 | 1 | 0 | 0 |
| 0 | 1 | 0 | 0 |
| 0 | 0 | 0 | 1 |
| 0 | 1 | 0 | 0 |
| 0 | 1 | 0 | 0 |
| 0 | 1 | 0 | 0 |
| 0 | 1 | 0 | 0 |
| 0 | 1 | 0 | 0 |
| 0 | 0 | 0 | 0 |
| 0 | 0 | 0 | 0 |
| 0 | 0 | 0 | 0 |
| 0 | 0 | 0 | 0 |
| 0 | 0 | 0 | 0 |
| 0 | 0 | 0 | 0 |
| 0 | 0 | 0 | 0 |
| 0 | 0 | 0 | 1 |
| 0 | 0 | 0 | 0 |
| 0 | 0 | 0 | 1 |
| 0 | 0 | 0 | 1 |
| 0 | 0 | 0 | 1 |
| 0 | 0 | 0 | 0 |

| | | | |
|---|---|---|---|
| 0 | 1 | 0 | 0 |
| 0 | 0 | 0 | 1 |
| 0 | 0 | 0 | 1 |
| 0 | 0 | 0 | 1 |
| 0 | 0 | 0 | 1 |
| 0 | 0 | 0 | 0 |
| 0 | 0 | 0 | 1 |
| 0 | 0 | 0 | 1 |
| 0 | 0 | 0 | 1 |
| 0 | 0 | 0 | 1 |
| 0 | 0 | 0 | 1 |
| 0 | 0 | 0 | 1 |
| 0 | 0 | 0 | 1 |
| 1 | 0 | 0 | 0 |
| 0 | 0 | 0 | 1 |
| 0 | 0 | 0 | 1 |
| 0 | 0 | 0 | 1 |
| 0 | 0 | 0 | 0 |
| 0 | 0 | 0 | 1 |
| 0 | 0 | 0 | 1 |
| 0 | 0 | 0 | 1 |
| 0 | 0 | 0 | 1 |
| 0 | 0 | 0 | 0 |
| 0 | 0 | 0 | 1 |
| 0 | 0 | 0 | 1 |
| 0 | 0 | 0 | 0 |
| 0 | 0 | 0 | 0 |
| 0 | 0 | 0 | 0 |
| 0 | 0 | 0 | 1 |
| 0 | 0 | 0 | 1 |
| 0 | 0 | 0 | 1 |
| 0 | 0 | 0 | 1 |
| 0 | 0 | 0 | 1 |
| 0 | 0 | 0 | 1 |
| 0 | 0 | 0 | 1 |
| 0 | 0 | 0 | 1 |
| 0 | 0 | 0 | 1 |
| 0 | 0 | 0 | 1 |
| 0 | 0 | 0 | 1 |
| 0 | 0 | 0 | 1 |
| 0 | 0 | 0 | 1 |
| 0 | 0 | 0 | 1 |
| 0 | 0 | 0 | 1 |
| 0 | 0 | 0 | 1 |
| 0 | 0 | 0 | 1 |
| 0 | 0 | 0 | 1 |
| 0 | 0 | 0 | 1 |
| 0 | 0 | 0 | 1 |
| 0 | 1 | 1 | 1 |
| 0 | 0 | 0 | 1 |
| 0 | 0 | 0 | 1 |
| 0 | 0 | 0 | 1 |
| 0 | 0 | 0 | 1 |
| 0 | 0 | 0 | 1 |

| | | | |
|---|---|---|---|
| 0 | 0 | 0 | 1 |
| 0 | 0 | 0 | 0 |
| 0 | 0 | 0 | 1 |
| 0 | 0 | 0 | 1 |
| 0 | 0 | 0 | 1 |
| 0 | 0 | 0 | 0 |
| 0 | 0 | 0 | 0 |
| 0 | 0 | 0 | 0 |
| 0 | 0 | 0 | 1 |
| 0 | 0 | 0 | 1 |
| 0 | 0 | 0 | 1 |
| 0 | 0 | 0 | 1 |
| 0 | 0 | 0 | 1 |
| 0 | 0 | 0 | 1 |
| 0 | 0 | 0 | 0 |
| 0 | 0 | 0 | 0 |
| 0 | 0 | 0 | 0 |
| 0 | 0 | 0 | 1 |
| 0 | 0 | 0 | 0 |
| 0 | 0 | 0 | 0 |
| 0 | 0 | 0 | 0 |
| 0 | 0 | 0 | 0 |
| 0 | 0 | 0 | 0 |
| 0 | 0 | 0 | 0 |
| 0 | 0 | 0 | 1 |
| 0 | 0 | 0 | 1 |
| 0 | 0 | 0 | 1 |
| 0 | 0 | 0 | 1 |
| 0 | 0 | 0 | 1 |
| 0 | 0 | 0 | 1 |
| 0 | 0 | 0 | 1 |
| 0 | 0 | 0 | 1 |
| 0 | 0 | 0 | 0 |
| 0 | 0 | 0 | 1 |
| 0 | 0 | 0 | 1 |
| 0 | 0 | 0 | 0 |
| 0 | 0 | 0 | 0 |
| 0 | 0 | 0 | 0 |
| 0 | 0 | 0 | 0 |
| 0 | 0 | 0 | 1 |
| 0 | 0 | 0 | 1 |
| 0 | 0 | 0 | 1 |
| 0 | 0 | 0 | 1 |
| 0 | 0 | 0 | 0 |
| 0 | 0 | 0 | 0 |
| 0 | 0 | 0 | 0 |
| 0 | 0 | 0 | 1 |
| 0 | 0 | 0 | 1 |
| 0 | 0 | 0 | 0 |
| 0 | 0 | 0 | 1 |
| 0 | 0 | 0 | 1 |
| 0 | 0 | 0 | 0 |
| 0 | 0 | 0 | 1 |
| 0 | 0 | 0 | 1 |

| | | | |
|---|---|---|---|
| 0 | 0 | 0 | 1 |
| 0 | 0 | 0 | 1 |
| 0 | 0 | 0 | 1 |
| 0 | 0 | 0 | 1 |
| 0 | 0 | 0 | 1 |
| 0 | 0 | 0 | 1 |
| 0 | 0 | 0 | 1 |
| 0 | 0 | 0 | 1 |
| 0 | 0 | 0 | 0 |
| 0 | 0 | 0 | 0 |
| 0 | 0 | 0 | 1 |
| 0 | 0 | 0 | 0 |
| 0 | 0 | 0 | 0 |
| 0 | 0 | 0 | 0 |
| 0 | 0 | 0 | 1 |
| 0 | 0 | 0 | 1 |
| 0 | 0 | 0 | 0 |
| 0 | 0 | 0 | 0 |
| 0 | 0 | 0 | 0 |
| 0 | 0 | 0 | 1 |
| 0 | 0 | 0 | 1 |
| 0 | 0 | 0 | 1 |
| 0 | 0 | 0 | 0 |
| 0 | 0 | 0 | 1 |
| 0 | 0 | 0 | 0 |
| 0 | 0 | 0 | 1 |
| 0 | 0 | 0 | 1 |
| 0 | 0 | 0 | 1 |
| 0 | 0 | 0 | 1 |
| 0 | 0 | 0 | 0 |
| 0 | 0 | 0 | 0 |
| 0 | 0 | 0 | 0 |
| 0 | 0 | 0 | 1 |
| 0 | 0 | 0 | 1 |
| 0 | 0 | 0 | 1 |
| 0 | 0 | 0 | 1 |
| 0 | 0 | 0 | 1 |
| 0 | 0 | 0 | 1 |
| 0 | 0 | 0 | 1 |
| 0 | 0 | 0 | 1 |
| 0 | 0 | 0 | 0 |
| 0 | 0 | 0 | 1 |
| 0 | 0 | 0 | 1 |
| 0 | 0 | 0 | 1 |
| 0 | 0 | 0 | 1 |
| 0 | 0 | 0 | 0 |
| 0 | 1 | 1 | 1 |
| 0 | 0 | 0 | 0 |
| 0 | 0 | 0 | 1 |
| 0 | 1 | 0 | 0 |
| 0 | 1 | 0 | 0 |
| 0 | 0 | 0 | 0 |
| 0 | 0 | 0 | 1 |
| 0 | 1 | 1 | 1 |

| | | | |
|---|---|---|---|
| 0 | 1 | 1 | 1 |
| 0 | 1 | 1 | 1 |
| 0 | 0 | 0 | 1 |
| 0 | 0 | 0 | 1 |
| 0 | 0 | 0 | 1 |
| 0 | 0 | 0 | 1 |
| 0 | 0 | 0 | 1 |
| 0 | 0 | 0 | 0 |
| 0 | 0 | 0 | 0 |
| 0 | 0 | 0 | 1 |
| 0 | 0 | 0 | 0 |
| 0 | 0 | 0 | 1 |
| 0 | 0 | 0 | 1 |
| 0 | 0 | 0 | 1 |
| 0 | 0 | 0 | 1 |
| 0 | 0 | 0 | 1 |
| 0 | 0 | 0 | 1 |
| 0 | 0 | 0 | 1 |
| 0 | 0 | 0 | 1 |
| 0 | 0 | 0 | 1 |
| 0 | 0 | 0 | 1 |
| 0 | 0 | 0 | 1 |
| 0 | 0 | 0 | 1 |
| 0 | 0 | 0 | 1 |
| 0 | 0 | 0 | 1 |
| 0 | 0 | 0 | 1 |
| 0 | 0 | 0 | 1 |
| 0 | 0 | 0 | 1 |
| 0 | 0 | 0 | 0 |
| 0 | 0 | 0 | 1 |
| 0 | 0 | 0 | 0 |
| 0 | 0 | 0 | 1 |
| 0 | 0 | 0 | 0 |
| 0 | 1 | 0 | 0 |
| 0 | 0 | 0 | 1 |
| 0 | 0 | 0 | 0 |
| 0 | 0 | 0 | 0 |
| 0 | 0 | 0 | 1 |
| 0 | 0 | 0 | 1 |
| 0 | 0 | 0 | 1 |
| 0 | 0 | 0 | 1 |
| 0 | 0 | 0 | 0 |
| 0 | 0 | 0 | 0 |
| 0 | 0 | 0 | 1 |
| 0 | 0 | 0 | 1 |
| 0 | 0 | 0 | 0 |
| 0 | 0 | 0 | 0 |
| 0 | 0 | 0 | 0 |
| 0 | 0 | 0 | 0 |
| 0 | 0 | 0 | 1 |
| 0 | 1 | 0 | 0 |
| 0 | 0 | 0 | 0 |
| 0 | 0 | 0 | 1 |
| 0 | 0 | 0 | 1 |

| | | | |
|---|---|---|---|
| 0 | 0 | 0 | 1 |
| 0 | 0 | 0 | 1 |
| 0 | 0 | 0 | 1 |
| 0 | 0 | 0 | 1 |
| 0 | 0 | 0 | 0 |
| 0 | 0 | 0 | 0 |
| 0 | 0 | 0 | 0 |
| 0 | 0 | 0 | 0 |
| 0 | 0 | 0 | 1 |
| 0 | 0 | 0 | 0 |
| 0 | 0 | 0 | 0 |
| 0 | 0 | 0 | 0 |
| 0 | 0 | 0 | 0 |
| 0 | 0 | 0 | 1 |
| 0 | 1 | 1 | 1 |
| 0 | 0 | 0 | 1 |
| 0 | 0 | 0 | 1 |
| 0 | 0 | 0 | 1 |
| 0 | 0 | 0 | 1 |
| 0 | 0 | 0 | 1 |
| 0 | 0 | 0 | 1 |
| 0 | 0 | 0 | 1 |
| 0 | 0 | 0 | 1 |
| 0 | 0 | 0 | 1 |
| 0 | 0 | 0 | 1 |
| 0 | 0 | 0 | 0 |
| 0 | 0 | 0 | 0 |
| 0 | 0 | 0 | 0 |
| 0 | 0 | 0 | 0 |
| 0 | 0 | 0 | 1 |
| 0 | 0 | 0 | 1 |
| 0 | 0 | 0 | 1 |
| 0 | 0 | 0 | 1 |
| 0 | 0 | 0 | 1 |
| 0 | 0 | 0 | 1 |
| 0 | 0 | 0 | 1 |
| 0 | 0 | 0 | 1 |
| 0 | 0 | 0 | 1 |
| 0 | 0 | 0 | 1 |
| 0 | 0 | 0 | 1 |
| 0 | 0 | 0 | 1 |
| 0 | 0 | 0 | 1 |
| 0 | 0 | 0 | 1 |
| 0 | 0 | 0 | 1 |
| 0 | 0 | 0 | 1 |
| 0 | 0 | 0 | 1 |
| 0 | 0 | 0 | 1 |
| 0 | 0 | 0 | 1 |
| 0 | 0 | 0 | 1 |
| 0 | 0 | 0 | 1 |
| 0 | 0 | 0 | 1 |
| 0 | 0 | 0 | 1 |
| 0 | 0 | 0 | 1 |
| 0 | 0 | 0 | 0 |

| | | | |
|---|---|---|---|
| 0 | 0 | 0 | 1 |
| 0 | 0 | 0 | 1 |
| 0 | 0 | 0 | 1 |
| 0 | 0 | 0 | 1 |
| 0 | 0 | 0 | 1 |
| 0 | 0 | 0 | 1 |
| 0 | 0 | 0 | 1 |
| 0 | 0 | 0 | 1 |
| 0 | 0 | 0 | 1 |
| 0 | 0 | 0 | 1 |
| 0 | 0 | 0 | 1 |
| 0 | 0 | 0 | 1 |
| 0 | 0 | 0 | 1 |
| 0 | 0 | 0 | 1 |
| 0 | 0 | 0 | 1 |
| 0 | 0 | 0 | 1 |
| 0 | 0 | 0 | 1 |
| 0 | 0 | 0 | 1 |
| 0 | 0 | 0 | 1 |
| 0 | 0 | 0 | 1 |
| 0 | 0 | 0 | 1 |
| 0 | 0 | 0 | 1 |
| 0 | 0 | 0 | 1 |
| 0 | 0 | 0 | 1 |
| 0 | 0 | 0 | 1 |
| 0 | 0 | 0 | 1 |
| 0 | 0 | 0 | 1 |
| 0 | 0 | 0 | 1 |
| 0 | 0 | 0 | 1 |
| 0 | 0 | 0 | 1 |
| 0 | 0 | 0 | 1 |
| 0 | 0 | 0 | 1 |
| 0 | 0 | 0 | 1 |
| 0 | 0 | 0 | 1 |
| 0 | 0 | 0 | 1 |
| 0 | 0 | 0 | 1 |
| 0 | 0 | 0 | 1 |
| 0 | 0 | 0 | 1 |
| 0 | 0 | 0 | 1 |
| 0 | 0 | 0 | 1 |
| 0 | 0 | 0 | 1 |
| 0 | 0 | 0 | 1 |
| 0 | 0 | 0 | 1 |
| 0 | 0 | 0 | 1 |
| 0 | 0 | 0 | 1 |
| 0 | 1 | 1 | 1 |
| 0 | 0 | 0 | 0 |
| 0 | 0 | 0 | 1 |
| 0 | 0 | 0 | 1 |
| 0 | 0 | 0 | 1 |
| 0 | 0 | 0 | 1 |
| 0 | 0 | 0 | 1 |
| 0 | 0 | 0 | 1 |
| 0 | 0 | 0 | 1 |

| | | | |
|---|---|---|---|
| 0 | 0 | 0 | 1 |
| 0 | 0 | 0 | 1 |
| 0 | 0 | 0 | 0 |
| 0 | 0 | 0 | 1 |
| 0 | 0 | 0 | 1 |
| 0 | 0 | 0 | 1 |
| 0 | 0 | 0 | 0 |
| 0 | 0 | 0 | 1 |
| 0 | 0 | 0 | 1 |
| 0 | 0 | 0 | 1 |
| 0 | 0 | 0 | 1 |
| 0 | 0 | 0 | 0 |
| 0 | 0 | 0 | 1 |
| 0 | 0 | 0 | 0 |
| 0 | 0 | 0 | 0 |
| 0 | 0 | 0 | 0 |
| 0 | 0 | 0 | 0 |
| 0 | 0 | 0 | 0 |
| 0 | 0 | 0 | 0 |
| 0 | 0 | 0 | 0 |
| 0 | 0 | 0 | 0 |

| surface_non_binding_site | non_binding_site_single_water | surface_non_binding_site_single_water |
|---|---|---|
| 0 | 0 | 0 |
| 0 | 0 | 0 |
| 0 | 0 | 0 |
| 0 | 0 | 0 |
| 0 | 0 | 0 |
| 0 | 0 | 0 |
| 1 | 1 | 1 |
| 1 | 1 | 1 |
| 0 | 0 | 0 |
| 0 | 0 | 0 |
| 0 | 0 | 0 |
| 0 | 0 | 0 |
| 0 | 0 | 0 |
| 0 | 0 | 0 |
| 0 | 0 | 0 |
| 0 | 0 | 0 |
| 0 | 0 | 0 |
| 0 | 0 | 0 |
| 0 | 0 | 0 |
| 0 | 0 | 0 |
| 0 | 0 | 0 |
| 0 | 0 | 0 |
| 0 | 0 | 0 |
| 0 | 0 | 0 |
| 0 | 0 | 0 |
| 1 | 0 | 0 |
| 0 | 0 | 0 |
| 1 | 1 | 1 |
| 0 | 0 | 0 |
| 0 | 1 | 0 |
| 0 | 1 | 0 |
| 0 | 0 | 0 |
| 0 | 0 | 0 |
| 0 | 0 | 0 |
| 0 | 0 | 0 |
| 1 | 1 | 1 |
| 0 | 0 | 0 |
| 0 | 0 | 0 |
| 0 | 0 | 0 |
| 0 | 0 | 0 |
| 0 | 0 | 0 |
| 0 | 0 | 0 |
| 0 | 0 | 0 |
| 0 | 0 | 0 |
| 0 | 0 | 0 |
| 0 | 0 | 0 |
| 0 | 0 | 0 |
| 0 | 0 | 0 |
| 0 | 0 | 0 |
| 0 | 0 | 0 |
| 0 | 0 | 0 |
| 0 | 0 | 0 |
| 0 | 0 | 0 |

| | | |
|---|---|---|
| 0 | 0 | 0 |
| 0 | 0 | 0 |
| 0 | 0 | 0 |
| 0 | 0 | 0 |
| 0 | 0 | 0 |
| 0 | 0 | 0 |
| 0 | 0 | 0 |
| 0 | 0 | 0 |
| 0 | 0 | 0 |
| 0 | 0 | 0 |
| 0 | 0 | 0 |
| 0 | 0 | 0 |
| 0 | 0 | 0 |
| 0 | 0 | 0 |
| 0 | 0 | 0 |
| 0 | 0 | 0 |
| 0 | 0 | 0 |
| 0 | 0 | 0 |
| 0 | 0 | 0 |
| 0 | 0 | 0 |
| 0 | 0 | 0 |
| 0 | 0 | 0 |
| 0 | 0 | 0 |
| 0 | 0 | 0 |
| 0 | 0 | 0 |
| 0 | 0 | 0 |
| 0 | 0 | 0 |
| 0 | 0 | 0 |
| 0 | 0 | 0 |
| 0 | 0 | 0 |
| 0 | 0 | 0 |
| 0 | 0 | 0 |
| 0 | 0 | 0 |
| 0 | 0 | 0 |
| 0 | 0 | 0 |
| 0 | 0 | 0 |
| 0 | 0 | 0 |
| 0 | 0 | 0 |
| 0 | 0 | 0 |
| 0 | 0 | 0 |
| 0 | 0 | 0 |
| 0 | 0 | 0 |
| 0 | 0 | 0 |
| 0 | 0 | 0 |
| 0 | 0 | 0 |
| 1 | 1 | 1 |
| 0 | 1 | 0 |
| 0 | 0 | 0 |
| 0 | 0 | 0 |
| 0 | 0 | 0 |
| 0 | 0 | 0 |
| 1 | 1 | 1 |
| 1 | 1 | 1 |
| 0 | 0 | 0 |

| | | |
|---|---|---|
| 0 | 1 | 0 |
| 0 | 1 | 0 |
| 0 | 0 | 0 |
| 0 | 0 | 0 |
| 0 | 0 | 0 |
| 0 | 0 | 0 |
| 0 | 1 | 0 |
| 0 | 0 | 0 |
| 0 | 0 | 0 |
| 0 | 0 | 0 |
| 0 | 0 | 0 |
| 0 | 0 | 0 |
| 0 | 0 | 0 |
| 0 | 0 | 0 |
| 0 | 0 | 0 |
| 1 | 1 | 1 |
| 0 | 0 | 0 |
| 0 | 0 | 0 |
| 0 | 0 | 0 |
| 0 | 0 | 0 |
| 0 | 0 | 0 |
| 0 | 0 | 0 |
| 0 | 0 | 0 |
| 0 | 0 | 0 |
| 0 | 0 | 0 |
| 0 | 0 | 0 |
| 0 | 0 | 0 |
| 0 | 0 | 0 |
| 0 | 0 | 0 |
| 0 | 0 | 0 |
| 0 | 0 | 0 |
| 0 | 0 | 0 |
| 0 | 0 | 0 |
| 0 | 0 | 0 |
| 0 | 0 | 0 |
| 0 | 0 | 0 |
| 0 | 0 | 0 |
| 0 | 0 | 0 |
| 0 | 0 | 0 |
| 0 | 0 | 0 |
| 0 | 0 | 0 |
| 0 | 0 | 0 |
| 0 | 0 | 0 |
| 0 | 0 | 0 |
| 0 | 0 | 0 |
| 0 | 0 | 0 |
| 0 | 0 | 0 |
| 0 | 0 | 0 |
| 0 | 0 | 0 |
| 0 | 0 | 0 |
| 0 | 0 | 0 |
| 0 | 0 | 0 |
| 0 | 0 | 0 |
| 0 | 0 | 0 |

| | | |
|---|---|---|
| 1 | 0 | 0 |
| 1 | 0 | 0 |
| 1 | 0 | 0 |
| 0 | 0 | 0 |
| 0 | 0 | 0 |
| 0 | 0 | 0 |
| 0 | 0 | 0 |
| 1 | 0 | 0 |
| 0 | 0 | 0 |
| 1 | 0 | 0 |
| 1 | 0 | 0 |
| 0 | 0 | 0 |
| 1 | 0 | 0 |
| 0 | 0 | 0 |
| 0 | 0 | 0 |
| 1 | 0 | 0 |
| 1 | 0 | 0 |
| 1 | 0 | 0 |
| 1 | 0 | 0 |
| 0 | 0 | 0 |
| 0 | 0 | 0 |
| 0 | 1 | 0 |
| 0 | 0 | 0 |
| 0 | 0 | 0 |
| 0 | 0 | 0 |
| 1 | 0 | 0 |
| 0 | 0 | 0 |
| 0 | 0 | 0 |
| 1 | 0 | 0 |
| 1 | 0 | 0 |
| 1 | 1 | 1 |
| 0 | 0 | 0 |
| 0 | 0 | 0 |
| 0 | 1 | 0 |
| 0 | 0 | 0 |
| 0 | 0 | 0 |
| 0 | 0 | 0 |
| 0 | 0 | 0 |
| 0 | 0 | 0 |
| 1 | 0 | 0 |
| 1 | 0 | 0 |
| 0 | 0 | 0 |
| 0 | 0 | 0 |
| 1 | 1 | 1 |
| 1 | 0 | 0 |
| 1 | 0 | 0 |
| 0 | 0 | 0 |
| 0 | 0 | 0 |
| 0 | 0 | 0 |
| 0 | 0 | 0 |
| 0 | 0 | 0 |
| 0 | 1 | 0 |
| 1 | 0 | 0 |
| 0 | 0 | 0 |

| | | |
|---|---|---|
| 0 | 0 | 0 |
| 0 | 0 | 0 |
| 0 | 0 | 0 |
| 1 | 1 | 1 |
| 1 | 1 | 1 |
| 1 | 0 | 0 |
| 0 | 0 | 0 |
| 0 | 0 | 0 |
| 1 | 0 | 0 |
| 1 | 0 | 0 |
| 0 | 0 | 0 |
| 0 | 1 | 0 |
| 0 | 0 | 0 |
| 1 | 0 | 0 |
| 0 | 1 | 0 |
| 1 | 1 | 1 |
| 1 | 1 | 1 |
| 0 | 0 | 0 |
| 0 | 0 | 0 |
| 0 | 0 | 0 |
| 0 | 0 | 0 |
| 1 | 0 | 0 |
| 0 | 0 | 0 |
| 0 | 1 | 0 |
| 1 | 1 | 1 |
| 0 | 0 | 0 |
| 0 | 0 | 0 |
| 0 | 0 | 0 |
| 0 | 0 | 0 |
| 0 | 0 | 0 |
| 0 | 0 | 0 |
| 0 | 0 | 0 |
| 1 | 0 | 0 |
| 1 | 0 | 0 |
| 0 | 0 | 0 |
| 1 | 0 | 0 |
| 0 | 0 | 0 |
| 0 | 0 | 0 |
| 1 | 0 | 0 |
| 0 | 1 | 0 |
| 0 | 0 | 0 |
| 1 | 0 | 0 |
| 1 | 0 | 0 |
| 1 | 0 | 0 |
| 0 | 0 | 0 |
| 0 | 0 | 0 |
| 0 | 0 | 0 |
| 0 | 0 | 0 |
| 0 | 0 | 0 |
| 0 | 0 | 0 |
| 0 | 0 | 0 |
| 0 | 0 | 0 |
| 0 | 0 | 0 |
| 0 | 0 | 0 |

| | | |
|---|---|---|
| 0 | 0 | 0 |
| 0 | 0 | 0 |
| 0 | 0 | 0 |
| 0 | 0 | 0 |
| 0 | 0 | 0 |
| 0 | 0 | 0 |
| 0 | 1 | 0 |
| 0 | 1 | 0 |
| 0 | 0 | 0 |
| 0 | 0 | 0 |
| 0 | 0 | 0 |
| 0 | 0 | 0 |
| 0 | 0 | 0 |
| 0 | 1 | 0 |
| 0 | 1 | 0 |
| 0 | 0 | 0 |
| 0 | 1 | 0 |
| 0 | 0 | 0 |
| 1 | 1 | 1 |
| 0 | 0 | 0 |
| 1 | 1 | 1 |
| 1 | 0 | 0 |
| 1 | 1 | 1 |
| 0 | 0 | 0 |
| 0 | 0 | 0 |
| 0 | 0 | 0 |
| 0 | 0 | 0 |
| 0 | 0 | 0 |
| 0 | 0 | 0 |
| 0 | 0 | 0 |
| 0 | 0 | 0 |
| 1 | 0 | 0 |
| 0 | 0 | 0 |
| 0 | 0 | 0 |
| 0 | 0 | 0 |
| 0 | 0 | 0 |
| 0 | 0 | 0 |
| 0 | 0 | 0 |
| 0 | 0 | 0 |
| 0 | 0 | 0 |
| 0 | 1 | 0 |
| 0 | 1 | 0 |
| 0 | 0 | 0 |
| 0 | 1 | 0 |
| 0 | 0 | 0 |
| 0 | 0 | 0 |
| 0 | 0 | 0 |
| 1 | 1 | 1 |
| 1 | 1 | 1 |
| 0 | 0 | 0 |
| 0 | 1 | 0 |
| 1 | 1 | 1 |
| 0 | 0 | 0 |
| 0 | 1 | 0 |

| | | |
|---|---|---|
| 0 | 0 | 0 |
| 1 | 1 | 1 |
| 0 | 0 | 0 |
| 0 | 0 | 0 |
| 0 | 0 | 0 |
| 0 | 0 | 0 |
| 0 | 1 | 0 |
| 0 | 0 | 0 |
| 0 | 0 | 0 |
| 1 | 1 | 1 |
| 1 | 1 | 1 |
| 1 | 1 | 1 |
| 1 | 1 | 1 |
| 1 | 1 | 1 |
| 0 | 0 | 0 |
| 1 | 1 | 1 |
| 0 | 0 | 0 |
| 0 | 0 | 0 |
| 1 | 1 | 1 |
| 0 | 0 | 0 |
| 0 | 0 | 0 |
| 0 | 0 | 0 |
| 0 | 0 | 0 |
| 0 | 0 | 0 |
| 1 | 1 | 1 |
| 1 | 0 | 0 |
| 0 | 0 | 0 |
| 0 | 0 | 0 |
| 0 | 0 | 0 |
| 0 | 0 | 0 |
| 0 | 0 | 0 |
| 0 | 0 | 0 |
| 0 | 0 | 0 |
| 0 | 0 | 0 |
| 1 | 1 | 1 |
| 1 | 1 | 1 |
| 0 | 0 | 0 |
| 0 | 0 | 0 |
| 0 | 0 | 0 |
| 0 | 0 | 0 |
| 0 | 0 | 0 |
| 0 | 0 | 0 |
| 0 | 1 | 0 |
| 0 | 0 | 0 |
| 0 | 0 | 0 |
| 0 | 0 | 0 |
| 0 | 0 | 0 |
| 0 | 0 | 0 |
| 0 | 0 | 0 |
| 1 | 1 | 1 |
| 0 | 0 | 0 |
| 0 | 0 | 0 |
| 0 | 0 | 0 |
| 0 | 0 | 0 |

| | | |
|---|---|---|
| 0 | 0 | 0 |
| 0 | 0 | 0 |
| 0 | 0 | 0 |
| 0 | 0 | 0 |
| 1 | 1 | 1 |
| 0 | 0 | 0 |
| 0 | 0 | 0 |
| 0 | 0 | 0 |
| 0 | 0 | 0 |
| 0 | 0 | 0 |
| 0 | 0 | 0 |
| 0 | 0 | 0 |
| 0 | 0 | 0 |
| 0 | 0 | 0 |
| 0 | 0 | 0 |
| 0 | 0 | 0 |
| 0 | 0 | 0 |
| 0 | 0 | 0 |
| 1 | 1 | 1 |
| 0 | 0 | 0 |
| 0 | 0 | 0 |
| 0 | 0 | 0 |
| 0 | 0 | 0 |
| 1 | 0 | 0 |
| 0 | 0 | 0 |
| 0 | 0 | 0 |
| 0 | 0 | 0 |
| 1 | 0 | 0 |
| 1 | 0 | 0 |
| 1 | 0 | 0 |
| 0 | 0 | 0 |
| 0 | 0 | 0 |
| 0 | 0 | 0 |
| 0 | 0 | 0 |
| 0 | 0 | 0 |
| 1 | 0 | 0 |
| 1 | 0 | 0 |
| 0 | 0 | 0 |
| 0 | 0 | 0 |
| 1 | 0 | 0 |
| 1 | 0 | 0 |
| 1 | 0 | 0 |
| 0 | 1 | 0 |
| 1 | 1 | 1 |
| 0 | 0 | 0 |
| 0 | 0 | 0 |
| 0 | 0 | 0 |
| 0 | 0 | 0 |
| 0 | 0 | 0 |
| 0 | 1 | 0 |
| 1 | 1 | 1 |
| 0 | 0 | 0 |
| 1 | 1 | 1 |
| 0 | 0 | 0 |

| | | |
|---|---|---|
| 1 | 1 | 1 |
| 1 | 1 | 1 |
| 0 | 0 | 0 |
| 1 | 1 | 1 |
| 1 | 1 | 1 |
| 1 | 1 | 1 |
| 1 | 1 | 1 |
| 0 | 0 | 0 |
| 1 | 0 | 0 |
| 1 | 1 | 1 |
| 0 | 0 | 0 |
| 1 | 1 | 1 |
| 1 | 1 | 1 |
| 0 | 1 | 0 |
| 0 | 0 | 0 |
| 0 | 0 | 0 |
| 1 | 1 | 1 |
| 1 | 1 | 1 |
| 0 | 0 | 0 |
| 0 | 0 | 0 |
| 0 | 0 | 0 |
| 0 | 0 | 0 |
| 0 | 0 | 0 |
| 0 | 0 | 0 |
| 0 | 0 | 0 |
| 1 | 1 | 1 |
| 0 | 0 | 0 |
| 0 | 0 | 0 |
| 0 | 0 | 0 |
| 1 | 0 | 0 |
| 0 | 0 | 0 |
| 0 | 0 | 0 |
| 0 | 0 | 0 |
| 0 | 0 | 0 |
| 0 | 0 | 0 |
| 0 | 0 | 0 |
| 0 | 0 | 0 |
| 0 | 0 | 0 |
| 0 | 0 | 0 |
| 0 | 0 | 0 |
| 0 | 1 | 0 |
| 0 | 0 | 0 |
| 1 | 1 | 1 |
| 0 | 0 | 0 |
| 0 | 0 | 0 |
| 0 | 0 | 0 |
| 0 | 0 | 0 |
| 0 | 0 | 0 |
| 0 | 0 | 0 |
| 0 | 0 | 0 |
| 0 | 1 | 0 |
| 0 | 0 | 0 |
| 0 | 0 | 0 |
| 0 | 0 | 0 |

| | | |
|---|---|---|
| 1 | 1 | 1 |
| 0 | 0 | 0 |
| 0 | 1 | 0 |
| 0 | 1 | 0 |
| 0 | 0 | 0 |
| 0 | 0 | 0 |
| 0 | 0 | 0 |
| 0 | 0 | 0 |
| 0 | 0 | 0 |
| 0 | 1 | 0 |
| 1 | 1 | 1 |
| 1 | 1 | 1 |
| 1 | 1 | 1 |
| 0 | 0 | 0 |
| 1 | 1 | 1 |
| 0 | 0 | 0 |
| 0 | 0 | 0 |
| 0 | 0 | 0 |
| 0 | 0 | 0 |
| 1 | 0 | 0 |
| 0 | 0 | 0 |
| 1 | 1 | 1 |
| 0 | 0 | 0 |
| 0 | 1 | 0 |
| 1 | 1 | 1 |
| 1 | 1 | 1 |
| 0 | 0 | 0 |
| 1 | 1 | 1 |
| 1 | 1 | 1 |
| 1 | 1 | 1 |
| 0 | 0 | 0 |
| 0 | 0 | 0 |
| 1 | 1 | 1 |
| 1 | 1 | 1 |
| 0 | 0 | 0 |
| 0 | 0 | 0 |
| 0 | 0 | 0 |
| 0 | 0 | 0 |
| 0 | 0 | 0 |
| 1 | 1 | 1 |
| 0 | 0 | 0 |
| 0 | 0 | 0 |
| 0 | 0 | 0 |
| 0 | 0 | 0 |
| 1 | 0 | 0 |
| 0 | 0 | 0 |
| 0 | 0 | 0 |
| 0 | 0 | 0 |
| 0 | 0 | 0 |
| 0 | 0 | 0 |
| 0 | 0 | 0 |
| 0 | 0 | 0 |
| 1 | 0 | 0 |
| 0 | 0 | 0 |

| | | |
|---|---|---|
| 0 | 0 | 0 |
| 0 | 0 | 0 |
| 0 | 0 | 0 |
| 0 | 0 | 0 |
| 0 | 0 | 0 |
| 0 | 0 | 0 |
| 0 | 0 | 0 |
| 1 | 0 | 0 |
| 0 | 0 | 0 |
| 0 | 0 | 0 |
| 0 | 0 | 0 |
| 0 | 0 | 0 |
| 1 | 1 | 1 |
| 0 | 0 | 0 |
| 0 | 0 | 0 |
| 0 | 0 | 0 |
| 0 | 0 | 0 |
| 0 | 0 | 0 |
| 0 | 0 | 0 |
| 1 | 1 | 1 |
| 0 | 0 | 0 |
| 0 | 0 | 0 |
| 0 | 0 | 0 |
| 0 | 0 | 0 |
| 0 | 0 | 0 |
| 1 | 0 | 0 |
| 1 | 1 | 1 |
| 0 | 1 | 0 |
| 0 | 0 | 0 |
| 0 | 0 | 0 |
| 0 | 0 | 0 |
| 0 | 0 | 0 |
| 0 | 0 | 0 |
| 0 | 0 | 0 |
| 0 | 0 | 0 |
| 1 | 0 | 0 |
| 0 | 0 | 0 |
| 0 | 0 | 0 |
| 1 | 1 | 1 |
| 1 | 0 | 0 |
| 0 | 1 | 0 |
| 0 | 0 | 0 |
| 0 | 0 | 0 |
| 0 | 0 | 0 |
| 0 | 0 | 0 |
| 0 | 0 | 0 |
| 0 | 0 | 0 |
| 1 | 1 | 1 |
| 1 | 1 | 1 |
| 0 | 0 | 0 |
| 0 | 0 | 0 |
| 0 | 0 | 0 |
| 0 | 0 | 0 |
| 1 | 1 | 1 |

| | | |
|---|---|---|
| 1 | 1 | 1 |
| 0 | 0 | 0 |
| 0 | 1 | 0 |
| 0 | 0 | 0 |
| 1 | 0 | 0 |
| 1 | 0 | 0 |
| 0 | 0 | 0 |
| 1 | 0 | 0 |
| 1 | 0 | 0 |
| 1 | 0 | 0 |
| 1 | 0 | 0 |
| 1 | 0 | 0 |
| 1 | 1 | 1 |
| 1 | 1 | 1 |
| 1 | 1 | 1 |
| 1 | 1 | 1 |
| 0 | 0 | 0 |
| 0 | 0 | 0 |
| 1 | 0 | 0 |
| 0 | 0 | 0 |
| 0 | 0 | 0 |
| 0 | 0 | 0 |
| 1 | 0 | 0 |
| 0 | 0 | 0 |
| 0 | 0 | 0 |
| 1 | 1 | 1 |
| 1 | 0 | 0 |
| 1 | 0 | 0 |
| 0 | 0 | 0 |
| 0 | 0 | 0 |
| 0 | 0 | 0 |
| 0 | 0 | 0 |
| 0 | 0 | 0 |
| 0 | 0 | 0 |
| 0 | 0 | 0 |
| 0 | 0 | 0 |
| 1 | 0 | 0 |
| 1 | 1 | 1 |
| 1 | 1 | 1 |
| 1 | 1 | 1 |
| 0 | 0 | 0 |
| 1 | 1 | 1 |
| 0 | 0 | 0 |
| 1 | 1 | 1 |
| 0 | 0 | 0 |
| 0 | 1 | 0 |
| 0 | 0 | 0 |
| 0 | 0 | 0 |
| 0 | 0 | 0 |
| 0 | 0 | 0 |
| 0 | 0 | 0 |
| 0 | 0 | 0 |
| 0 | 0 | 0 |
| 0 | 0 | 0 |

| | | |
|---|---|---|
| 0 | 0 | 0 |
| 0 | 0 | 0 |
| 0 | 0 | 0 |
| 0 | 0 | 0 |
| 0 | 0 | 0 |
| 0 | 0 | 0 |
| 0 | 0 | 0 |
| 1 | 1 | 1 |
| 0 | 0 | 0 |
| 0 | 0 | 0 |
| 0 | 0 | 0 |
| 1 | 0 | 0 |
| 0 | 0 | 0 |
| 0 | 0 | 0 |
| 0 | 0 | 0 |
| 0 | 0 | 0 |
| 0 | 0 | 0 |
| 0 | 0 | 0 |
| 0 | 0 | 0 |
| 0 | 0 | 0 |
| 0 | 0 | 0 |
| 0 | 0 | 0 |
| 0 | 0 | 0 |
| 0 | 0 | 0 |
| 0 | 0 | 0 |
| 0 | 0 | 0 |
| 0 | 0 | 0 |
| 0 | 0 | 0 |
| 0 | 0 | 0 |
| 0 | 0 | 0 |
| 0 | 0 | 0 |
| 0 | 0 | 0 |
| 0 | 0 | 0 |
| 0 | 0 | 0 |
| 0 | 0 | 0 |
| 0 | 0 | 0 |
| 0 | 0 | 0 |
| 0 | 0 | 0 |
| 0 | 0 | 0 |
| 0 | 0 | 0 |
| 1 | 0 | 0 |
| 0 | 0 | 0 |
| 0 | 0 | 0 |
| 0 | 0 | 0 |
| 0 | 0 | 0 |
| 0 | 0 | 0 |
| 0 | 0 | 0 |
| 0 | 0 | 0 |
| 0 | 0 | 0 |
| 0 | 1 | 0 |
| 0 | 1 | 0 |
| 1 | 0 | 0 |
| 0 | 0 | 0 |
| 0 | 0 | 0 |

| | | |
|---|---|---|
| 0 | 0 | 0 |
| 0 | 0 | 0 |
| 0 | 0 | 0 |
| 0 | 0 | 0 |
| 0 | 0 | 0 |
| 0 | 0 | 0 |
| 0 | 0 | 0 |
| 1 | 0 | 0 |
| 1 | 0 | 0 |
| 0 | 0 | 0 |
| 0 | 0 | 0 |
| 1 | 0 | 0 |
| 0 | 0 | 0 |
| 0 | 0 | 0 |
| 0 | 0 | 0 |
| 0 | 0 | 0 |
| 0 | 0 | 0 |
| 1 | 0 | 0 |
| 0 | 0 | 0 |
| 1 | 0 | 0 |
| 0 | 0 | 0 |
| 0 | 0 | 0 |
| 1 | 1 | 1 |
| 0 | 0 | 0 |
| 0 | 0 | 0 |
| 1 | 0 | 0 |
| 0 | 0 | 0 |
| 0 | 0 | 0 |
| 0 | 0 | 0 |
| 0 | 0 | 0 |
| 0 | 0 | 0 |
| 0 | 0 | 0 |
| 1 | 0 | 0 |
| 0 | 0 | 0 |
| 0 | 0 | 0 |
| 0 | 0 | 0 |
| 1 | 1 | 1 |
| 0 | 0 | 0 |
| 0 | 0 | 0 |
| 0 | 0 | 0 |
| 0 | 0 | 0 |
| 0 | 0 | 0 |
| 0 | 0 | 0 |
| 0 | 0 | 0 |
| 0 | 0 | 0 |
| 0 | 0 | 0 |
| 0 | 0 | 0 |
| 0 | 0 | 0 |
| 0 | 0 | 0 |
| 0 | 0 | 0 |
| 0 | 0 | 0 |
| 0 | 0 | 0 |
| 0 | 0 | 0 |
| 0 | 0 | 0 |

| | | |
|---|---|---|
| 1 | 0 | 0 |
| 0 | 1 | 0 |
| 0 | 0 | 0 |
| 0 | 0 | 0 |
| 0 | 0 | 0 |
| 0 | 0 | 0 |
| 0 | 0 | 0 |
| 0 | 0 | 0 |
| 0 | 0 | 0 |
| 1 | 0 | 0 |
| 0 | 1 | 0 |
| 1 | 1 | 1 |
| 0 | 0 | 0 |
| 0 | 0 | 0 |
| 0 | 0 | 0 |
| 0 | 0 | 0 |
| 0 | 0 | 0 |
| 0 | 0 | 0 |
| 1 | 1 | 1 |
| 1 | 0 | 0 |
| 1 | 0 | 0 |
| 0 | 0 | 0 |
| 0 | 1 | 0 |
| 0 | 0 | 0 |
| 0 | 0 | 0 |
| 0 | 0 | 0 |
| 0 | 0 | 0 |
| 0 | 0 | 0 |
| 0 | 0 | 0 |
| 0 | 0 | 0 |
| 0 | 0 | 0 |
| 0 | 0 | 0 |
| 0 | 0 | 0 |
| 0 | 0 | 0 |
| 0 | 0 | 0 |
| 0 | 0 | 0 |
| 0 | 0 | 0 |
| 0 | 0 | 0 |
| 0 | 0 | 0 |
| 0 | 0 | 0 |
| 0 | 0 | 0 |
| 1 | 0 | 0 |
| 0 | 0 | 0 |
| 0 | 0 | 0 |
| 0 | 0 | 0 |
| 0 | 0 | 0 |
| 1 | 0 | 0 |
| 0 | 0 | 0 |
| 0 | 0 | 0 |
| 0 | 0 | 0 |
| 0 | 0 | 0 |
| 0 | 0 | 0 |
| 1 | 0 | 0 |
| 0 | 1 | 0 |

| | | |
|---|---|---|
| 0 | 0 | 0 |
| 0 | 0 | 0 |
| 1 | 0 | 0 |
| 1 | 0 | 0 |
| 1 | 0 | 0 |
| 0 | 0 | 0 |
| 1 | 0 | 0 |
| 0 | 0 | 0 |
| 0 | 0 | 0 |
| 0 | 0 | 0 |
| 0 | 0 | 0 |
| 0 | 0 | 0 |
| 0 | 0 | 0 |
| 1 | 0 | 0 |
| 1 | 0 | 0 |
| 1 | 0 | 0 |
| 0 | 0 | 0 |
| 1 | 1 | 1 |
| 0 | 0 | 0 |
| 0 | 0 | 0 |
| 1 | 0 | 0 |
| 1 | 0 | 0 |
| 0 | 0 | 0 |
| 0 | 0 | 0 |
| 1 | 0 | 0 |
| 0 | 0 | 0 |
| 0 | 0 | 0 |
| 1 | 0 | 0 |
| 0 | 0 | 0 |
| 0 | 0 | 0 |
| 0 | 0 | 0 |
| 0 | 0 | 0 |
| 0 | 0 | 0 |
| 0 | 0 | 0 |
| 0 | 0 | 0 |
| 0 | 0 | 0 |
| 0 | 0 | 0 |
| 0 | 0 | 0 |
| 0 | 0 | 0 |
| 0 | 0 | 0 |
| 1 | 0 | 0 |
| 1 | 0 | 0 |
| 1 | 0 | 0 |
| 1 | 1 | 1 |
| 1 | 1 | 1 |
| 0 | 0 | 0 |
| 0 | 0 | 0 |
| 0 | 0 | 0 |
| 0 | 0 | 0 |
| 0 | 0 | 0 |
| 0 | 0 | 0 |
| 0 | 0 | 0 |
| 0 | 0 | 0 |
| 1 | 0 | 0 |

| | | |
|---|---|---|
| 1 | 0 | 0 |
| 0 | 0 | 0 |
| 0 | 0 | 0 |
| 0 | 0 | 0 |
| 1 | 0 | 0 |
| 0 | 0 | 0 |
| 0 | 0 | 0 |
| 0 | 0 | 0 |
| 1 | 0 | 0 |
| 1 | 0 | 0 |
| 0 | 0 | 0 |
| 0 | 0 | 0 |
| 0 | 0 | 0 |
| 1 | 0 | 0 |
| 0 | 0 | 0 |
| 0 | 0 | 0 |
| 0 | 0 | 0 |
| 0 | 0 | 0 |
| 0 | 0 | 0 |
| 0 | 0 | 0 |
| 0 | 0 | 0 |
| 1 | 1 | 1 |
| 0 | 0 | 0 |
| 0 | 1 | 0 |
| 0 | 0 | 0 |
| 0 | 0 | 0 |
| 0 | 0 | 0 |
| 0 | 0 | 0 |
| 0 | 0 | 0 |
| 0 | 0 | 0 |
| 0 | 0 | 0 |
| 0 | 0 | 0 |
| 0 | 0 | 0 |
| 0 | 0 | 0 |
| 0 | 1 | 0 |
| 0 | 0 | 0 |
| 0 | 0 | 0 |
| 0 | 0 | 0 |
| 0 | 0 | 0 |
| 0 | 0 | 0 |
| 0 | 0 | 0 |
| 0 | 0 | 0 |
| 0 | 0 | 0 |
| 0 | 0 | 0 |
| 0 | 0 | 0 |
| 0 | 0 | 0 |
| 0 | 0 | 0 |
| 1 | 1 | 1 |
| 0 | 0 | 0 |
| 1 | 1 | 1 |
| 0 | 0 | 0 |
| 0 | 0 | 0 |
| 0 | 0 | 0 |
| 0 | 0 | 0 |

| | | |
|---|---|---|
| 0 | 0 | 0 |
| 0 | 0 | 0 |
| 1 | 1 | 1 |
| 0 | 0 | 0 |
| 0 | 0 | 0 |
| 0 | 1 | 0 |
| 0 | 1 | 0 |
| 0 | 1 | 0 |
| 0 | 0 | 0 |
| 0 | 1 | 0 |
| 0 | 0 | 0 |
| 0 | 0 | 0 |
| 0 | 0 | 0 |
| 1 | 1 | 1 |
| 0 | 0 | 0 |
| 0 | 1 | 0 |
| 0 | 0 | 0 |
| 0 | 0 | 0 |
| 0 | 0 | 0 |
| 1 | 1 | 1 |
| 1 | 1 | 1 |
| 0 | 0 | 0 |
| 1 | 1 | 1 |
| 0 | 0 | 0 |
| 0 | 0 | 0 |
| 1 | 1 | 1 |
| 1 | 1 | 1 |
| 0 | 0 | 0 |
| 1 | 1 | 1 |
| 0 | 0 | 0 |
| 0 | 1 | 0 |
| 0 | 1 | 0 |
| 1 | 1 | 1 |
| 0 | 1 | 0 |
| 1 | 1 | 1 |
| 1 | 1 | 1 |
| 1 | 1 | 1 |
| 1 | 1 | 1 |
| 1 | 1 | 1 |
| 1 | 1 | 1 |
| 1 | 1 | 1 |
| 1 | 1 | 1 |
| 0 | 1 | 0 |
| 0 | 1 | 0 |
| 0 | 1 | 0 |
| 0 | 1 | 0 |
| 0 | 1 | 0 |
| 0 | 1 | 0 |
| 0 | 1 | 0 |
| 0 | 1 | 0 |
| 0 | 1 | 0 |
| 0 | 1 | 0 |
| 0 | 1 | 0 |
| 0 | 1 | 0 |

| | | |
|---|---|---|
| 0 | 1 | 0 |
| 1 | 1 | 1 |
| 1 | 1 | 1 |
| 1 | 1 | 1 |
| 1 | 1 | 1 |
| 1 | 1 | 1 |
| 1 | 1 | 1 |
| 0 | 1 | 0 |
| 0 | 1 | 0 |
| 0 | 1 | 0 |
| 0 | 1 | 0 |
| 0 | 0 | 0 |
| 0 | 0 | 0 |
| 0 | 0 | 0 |
| 1 | 0 | 0 |
| 0 | 0 | 0 |
| 1 | 0 | 0 |
| 0 | 0 | 0 |
| 1 | 0 | 0 |
| 1 | 1 | 1 |
| 0 | 0 | 0 |
| 1 | 1 | 1 |
| 1 | 1 | 1 |
| 0 | 0 | 0 |
| 0 | 0 | 0 |
| 1 | 1 | 1 |
| 1 | 1 | 1 |
| 0 | 0 | 0 |
| 0 | 0 | 0 |
| 0 | 0 | 0 |
| 0 | 0 | 0 |
| 0 | 0 | 0 |
| 0 | 1 | 0 |
| 0 | 1 | 0 |
| 1 | 1 | 1 |
| 1 | 1 | 1 |
| 1 | 1 | 1 |
| 1 | 1 | 1 |
| 1 | 1 | 1 |
| 0 | 0 | 0 |
| 0 | 0 | 0 |
| 0 | 1 | 0 |
| 0 | 0 | 0 |
| 0 | 0 | 0 |
| 0 | 0 | 0 |
| 1 | 0 | 0 |
| 0 | 0 | 0 |
| 1 | 0 | 0 |
| 1 | 0 | 0 |
| 0 | 0 | 0 |
| 0 | 0 | 0 |
| 0 | 0 | 0 |
| 0 | 0 | 0 |
| 0 | 0 | 0 |

| | | |
|---|---|---|
| 0 | 0 | 0 |
| 0 | 1 | 0 |
| 1 | 1 | 1 |
| 1 | 1 | 1 |
| 0 | 0 | 0 |
| 1 | 1 | 1 |
| 0 | 0 | 0 |
| 1 | 1 | 1 |
| 1 | 1 | 1 |
| 0 | 0 | 0 |
| 1 | 0 | 0 |
| 1 | 1 | 1 |
| 1 | 1 | 1 |
| 0 | 0 | 0 |
| 1 | 0 | 0 |
| 1 | 0 | 0 |
| 0 | 0 | 0 |
| 1 | 0 | 0 |
| 0 | 0 | 0 |
| 0 | 0 | 0 |
| 0 | 0 | 0 |
| 0 | 0 | 0 |
| 0 | 0 | 0 |
| 0 | 0 | 0 |
| 0 | 0 | 0 |
| 0 | 0 | 0 |
| 0 | 0 | 0 |
| 0 | 0 | 0 |
| 0 | 0 | 0 |
| 0 | 0 | 0 |
| 0 | 0 | 0 |
| 0 | 0 | 0 |
| 0 | 0 | 0 |
| 0 | 0 | 0 |
| 0 | 0 | 0 |
| 0 | 0 | 0 |
| 1 | 0 | 0 |
| 0 | 0 | 0 |
| 0 | 0 | 0 |
| 0 | 0 | 0 |
| 0 | 0 | 0 |
| 0 | 0 | 0 |
| 0 | 0 | 0 |
| 0 | 0 | 0 |
| 0 | 0 | 0 |
| 0 | 0 | 0 |
| 0 | 0 | 0 |
| 0 | 0 | 0 |
| 1 | 0 | 0 |
| 1 | 0 | 0 |
| 0 | 0 | 0 |
| 1 | 1 | 1 |
| 0 | 0 | 0 |
| 0 | 0 | 0 |

| | | |
|---|---|---|
| 0 | 0 | 0 |
| 0 | 0 | 0 |
| 0 | 0 | 0 |
| 0 | 0 | 0 |
| 0 | 0 | 0 |
| 0 | 0 | 0 |
| 0 | 0 | 0 |
| 0 | 0 | 0 |
| 0 | 0 | 0 |
| 0 | 0 | 0 |
| 0 | 0 | 0 |
| 0 | 0 | 0 |
| 0 | 0 | 0 |
| 0 | 0 | 0 |
| 0 | 1 | 0 |
| 0 | 0 | 0 |
| 0 | 0 | 0 |
| 0 | 0 | 0 |
| 1 | 0 | 0 |
| 0 | 0 | 0 |
| 0 | 0 | 0 |
| 0 | 0 | 0 |
| 0 | 0 | 0 |
| 0 | 0 | 0 |
| 0 | 0 | 0 |
| 1 | 0 | 0 |
| 0 | 0 | 0 |
| 0 | 0 | 0 |
| 0 | 0 | 0 |
| 0 | 0 | 0 |
| 1 | 1 | 1 |
| 0 | 0 | 0 |
| 1 | 1 | 1 |
| 1 | 0 | 0 |
| 0 | 0 | 0 |
| 0 | 0 | 0 |
| 1 | 1 | 1 |
| 1 | 1 | 1 |
| 1 | 1 | 1 |
| 1 | 0 | 0 |
| 0 | 0 | 0 |
| 0 | 0 | 0 |
| 0 | 0 | 0 |
| 0 | 0 | 0 |
| 0 | 0 | 0 |
| 0 | 0 | 0 |
| 0 | 0 | 0 |
| 0 | 0 | 0 |
| 1 | 1 | 1 |
| 1 | 1 | 1 |
| 0 | 0 | 0 |
| 0 | 0 | 0 |
| 0 | 0 | 0 |
| 0 | 1 | 0 |

| | | |
|---|---|---|
| 1 | 1 | 1 |
| 1 | 1 | 1 |
| 1 | 0 | 0 |
| 0 | 0 | 0 |
| 1 | 0 | 0 |
| 1 | 1 | 1 |
| 0 | 0 | 0 |
| 1 | 1 | 1 |
| 0 | 0 | 0 |
| 0 | 0 | 0 |
| 0 | 0 | 0 |
| 0 | 0 | 0 |
| 0 | 0 | 0 |
| 0 | 0 | 0 |
| 0 | 0 | 0 |
| 0 | 0 | 0 |
| 0 | 1 | 0 |
| 0 | 0 | 0 |
| 0 | 0 | 0 |
| 1 | 0 | 0 |
| 0 | 0 | 0 |
| 0 | 0 | 0 |
| 1 | 1 | 1 |
| 0 | 0 | 0 |
| 1 | 1 | 1 |
| 1 | 1 | 1 |
| 0 | 1 | 0 |
| 0 | 0 | 0 |
| 0 | 0 | 0 |
| 0 | 0 | 0 |
| 1 | 1 | 1 |
| 0 | 0 | 0 |
| 0 | 0 | 0 |
| 0 | 0 | 0 |
| 0 | 1 | 0 |
| 0 | 1 | 0 |
| 0 | 0 | 0 |
| 0 | 0 | 0 |
| 0 | 0 | 0 |
| 1 | 1 | 1 |
| 1 | 1 | 1 |
| 0 | 1 | 0 |
| 0 | 1 | 0 |
| 0 | 0 | 0 |
| 1 | 1 | 1 |
| 0 | 0 | 0 |
| 1 | 1 | 1 |
| 0 | 0 | 0 |
| 0 | 0 | 0 |
| 1 | 1 | 1 |
| 0 | 1 | 0 |
| 0 | 0 | 0 |
| 0 | 0 | 0 |
| 1 | 1 | 1 |

| | | |
|---|---|---|
| 1 | 1 | 1 |
| 1 | 1 | 1 |
| 0 | 1 | 0 |
| 0 | 0 | 0 |
| 0 | 0 | 0 |
| 1 | 0 | 0 |
| 1 | 0 | 0 |
| 0 | 0 | 0 |
| 0 | 0 | 0 |
| 1 | 1 | 1 |
| 0 | 0 | 0 |
| 0 | 0 | 0 |
| 0 | 0 | 0 |
| 0 | 0 | 0 |
| 0 | 0 | 0 |
| 0 | 0 | 0 |
| 0 | 0 | 0 |
| 0 | 0 | 0 |
| 0 | 0 | 0 |
| 0 | 1 | 0 |
| 1 | 1 | 1 |
| 1 | 1 | 1 |
| 1 | 1 | 1 |
| 1 | 1 | 1 |
| 0 | 0 | 0 |
| 0 | 1 | 0 |
| 0 | 0 | 0 |
| 0 | 0 | 0 |
| 1 | 1 | 1 |
| 1 | 1 | 1 |
| 1 | 1 | 1 |
| 1 | 1 | 1 |
| 1 | 1 | 1 |
| 0 | 0 | 0 |
| 0 | 0 | 0 |
| 1 | 1 | 1 |
| 0 | 0 | 0 |
| 0 | 1 | 0 |
| 0 | 0 | 0 |
| 0 | 0 | 0 |
| 0 | 0 | 0 |
| 0 | 0 | 0 |
| 0 | 1 | 0 |
| 0 | 0 | 0 |
| 0 | 0 | 0 |
| 0 | 0 | 0 |
| 0 | 0 | 0 |
| 0 | 0 | 0 |
| 0 | 0 | 0 |
| 0 | 0 | 0 |
| 0 | 0 | 0 |
| 0 | 0 | 0 |
| 0 | 0 | 0 |
| 0 | 1 | 0 |

| | | |
|---|---|---|
| 0 | 0 | 0 |
| 0 | 1 | 0 |
| 0 | 0 | 0 |
| 0 | 0 | 0 |
| 0 | 0 | 0 |
| 1 | 1 | 1 |
| 0 | 0 | 0 |
| 1 | 1 | 1 |
| 0 | 1 | 0 |
| 0 | 0 | 0 |
| 0 | 1 | 0 |
| 0 | 1 | 0 |
| 0 | 1 | 0 |
| 0 | 1 | 0 |
| 0 | 1 | 0 |
| 0 | 0 | 0 |
| 0 | 0 | 0 |
| 0 | 1 | 0 |
| 0 | 0 | 0 |
| 1 | 1 | 1 |
| 0 | 0 | 0 |
| 0 | 0 | 0 |
| 0 | 0 | 0 |
| 0 | 0 | 0 |
| 0 | 0 | 0 |
| 0 | 0 | 0 |
| 0 | 0 | 0 |
| 0 | 0 | 0 |
| 0 | 0 | 0 |
| 0 | 0 | 0 |
| 0 | 0 | 0 |
| 0 | 0 | 0 |
| 1 | 0 | 0 |
| 1 | 0 | 0 |
| 0 | 0 | 0 |
| 0 | 0 | 0 |
| 0 | 0 | 0 |
| 0 | 0 | 0 |
| 1 | 0 | 0 |
| 0 | 0 | 0 |
| 0 | 0 | 0 |
| 0 | 0 | 0 |
| 1 | 0 | 0 |
| 0 | 1 | 0 |
| 0 | 0 | 0 |
| 0 | 0 | 0 |
| 0 | 0 | 0 |
| 0 | 0 | 0 |
| 0 | 0 | 0 |
| 0 | 0 | 0 |
| 1 | 0 | 0 |
| 0 | 0 | 0 |
| 0 | 0 | 0 |
| 1 | 0 | 0 |

| | | |
|---|---|---|
| 0 | 0 | 0 |
| 1 | 0 | 0 |
| 0 | 0 | 0 |
| 0 | 0 | 0 |
| 0 | 0 | 0 |
| 0 | 0 | 0 |
| 0 | 0 | 0 |
| 0 | 0 | 0 |
| 0 | 0 | 0 |
| 0 | 0 | 0 |
| 0 | 0 | 0 |
| 1 | 1 | 1 |
| 0 | 0 | 0 |
| 1 | 1 | 1 |
| 1 | 0 | 0 |
| 1 | 0 | 0 |
| 0 | 0 | 0 |
| 0 | 0 | 0 |
| 0 | 0 | 0 |
| 0 | 0 | 0 |
| 1 | 0 | 0 |
| 0 | 0 | 0 |
| 0 | 1 | 0 |
| 0 | 0 | 0 |
| 1 | 1 | 1 |
| 1 | 1 | 1 |
| 1 | 1 | 1 |
| 0 | 0 | 0 |
| 1 | 1 | 1 |
| 1 | 1 | 1 |
| 1 | 1 | 1 |
| 0 | 0 | 0 |
| 1 | 1 | 1 |
| 0 | 0 | 0 |
| 0 | 0 | 0 |
| 0 | 0 | 0 |
| 0 | 1 | 0 |
| 1 | 1 | 1 |
| 0 | 0 | 0 |
| 0 | 0 | 0 |
| 0 | 0 | 0 |
| 0 | 0 | 0 |
| 0 | 0 | 0 |
| 0 | 0 | 0 |
| 0 | 0 | 0 |
| 0 | 0 | 0 |
| 1 | 1 | 1 |
| 1 | 1 | 1 |
| 0 | 0 | 0 |
| 0 | 0 | 0 |
| 0 | 0 | 0 |
| 1 | 1 | 1 |
| 1 | 1 | 1 |
| 0 | 0 | 0 |

| | | |
|---|---|---|
| 0 | 0 | 0 |
| 0 | 1 | 0 |
| 0 | 1 | 0 |
| 1 | 1 | 1 |
| 0 | 0 | 0 |
| 0 | 0 | 0 |
| 0 | 0 | 0 |
| 0 | 0 | 0 |
| 0 | 0 | 0 |
| 0 | 0 | 0 |
| 1 | 1 | 1 |
| 0 | 0 | 0 |
| 0 | 0 | 0 |
| 0 | 0 | 0 |
| 1 | 0 | 0 |
| 1 | 1 | 1 |
| 0 | 0 | 0 |
| 1 | 0 | 0 |
| 0 | 0 | 0 |
| 0 | 0 | 0 |
| 1 | 0 | 0 |
| 0 | 0 | 0 |
| 0 | 0 | 0 |
| 1 | 0 | 0 |
| 0 | 0 | 0 |
| 0 | 0 | 0 |
| 0 | 0 | 0 |
| 1 | 0 | 0 |
| 0 | 0 | 0 |
| 0 | 0 | 0 |
| 0 | 0 | 0 |
| 1 | 1 | 1 |
| 1 | 1 | 1 |
| 1 | 1 | 1 |
| 0 | 1 | 0 |
| 0 | 1 | 0 |
| 0 | 1 | 0 |
| 1 | 1 | 1 |
| 1 | 1 | 1 |
| 1 | 1 | 1 |
| 0 | 1 | 0 |
| 1 | 1 | 1 |
| 0 | 0 | 0 |
| 0 | 1 | 0 |
| 1 | 1 | 1 |
| 1 | 1 | 1 |
| 0 | 1 | 0 |
| 1 | 1 | 1 |
| 1 | 1 | 1 |
| 1 | 1 | 1 |
| 1 | 1 | 1 |
| 1 | 1 | 1 |
| 1 | 1 | 1 |
| 1 | 1 | 1 |

| | | |
|---|---|---|
| 0 | 1 | 0 |
| 0 | 1 | 0 |
| 1 | 1 | 1 |
| 1 | 1 | 1 |
| 1 | 1 | 1 |
| 1 | 1 | 1 |
| 1 | 1 | 1 |
| 1 | 1 | 1 |
| 0 | 0 | 0 |
| 0 | 0 | 0 |
| 0 | 0 | 0 |
| 0 | 0 | 0 |
| 0 | 0 | 0 |
| 0 | 0 | 0 |
| 0 | 0 | 0 |
| 0 | 0 | 0 |
| 0 | 0 | 0 |
| 0 | 0 | 0 |
| 0 | 0 | 0 |
| 0 | 0 | 0 |
| 0 | 0 | 0 |
| 0 | 0 | 0 |
| 0 | 0 | 0 |
| 0 | 0 | 0 |
| 0 | 0 | 0 |
| 1 | 1 | 1 |
| 1 | 1 | 1 |
| 0 | 0 | 0 |
| 0 | 0 | 0 |
| 0 | 0 | 0 |
| 0 | 0 | 0 |
| 0 | 0 | 0 |
| 0 | 0 | 0 |
| 0 | 0 | 0 |
| 0 | 0 | 0 |
| 0 | 0 | 0 |
| 0 | 0 | 0 |
| 0 | 0 | 0 |
| 0 | 0 | 0 |
| 0 | 0 | 0 |
| 0 | 0 | 0 |
| 0 | 0 | 0 |
| 0 | 0 | 0 |
| 0 | 0 | 0 |
| 0 | 0 | 0 |
| 0 | 0 | 0 |
| 1 | 1 | 1 |
| 0 | 1 | 0 |
| 1 | 1 | 1 |
| 1 | 1 | 1 |
| 1 | 1 | 1 |
| 1 | 1 | 1 |
| 0 | 0 | 0 |
| 0 | 0 | 0 |

| | | |
|---|---|---|
| 0 | 0 | 0 |
| 0 | 0 | 0 |
| 0 | 0 | 0 |
| 0 | 0 | 0 |
| 0 | 1 | 0 |
| 0 | 0 | 0 |
| 0 | 0 | 0 |
| 0 | 0 | 0 |
| 1 | 1 | 1 |
| 0 | 0 | 0 |
| 0 | 0 | 0 |
| 0 | 1 | 0 |
| 0 | 0 | 0 |
| 0 | 0 | 0 |
| 0 | 0 | 0 |
| 0 | 0 | 0 |
| 1 | 1 | 1 |
| 0 | 0 | 0 |
| 0 | 0 | 0 |
| 1 | 1 | 1 |
| 0 | 0 | 0 |
| 0 | 0 | 0 |
| 0 | 0 | 0 |
| 0 | 0 | 0 |
| 0 | 0 | 0 |
| 0 | 1 | 0 |
| 0 | 1 | 0 |
| 1 | 1 | 1 |
| 1 | 1 | 1 |
| 0 | 1 | 0 |
| 1 | 1 | 1 |
| 1 | 1 | 1 |
| 1 | 0 | 0 |
| 1 | 0 | 0 |
| 0 | 0 | 0 |
| 0 | 0 | 0 |
| 0 | 0 | 0 |
| 0 | 0 | 0 |
| 0 | 0 | 0 |
| 0 | 0 | 0 |
| 1 | 1 | 1 |
| 1 | 1 | 1 |
| 1 | 1 | 1 |
| 0 | 0 | 0 |
| 0 | 0 | 0 |
| 0 | 0 | 0 |
| 1 | 1 | 1 |
| 0 | 1 | 0 |
| 0 | 0 | 0 |
| 0 | 0 | 0 |
| 1 | 1 | 1 |
| 1 | 1 | 1 |
| 1 | 1 | 1 |
| 1 | 1 | 1 |

| | | |
|---|---|---|
| 0 | 1 | 0 |
| 1 | 1 | 1 |
| 0 | 1 | 0 |
| 0 | 0 | 0 |
| 0 | 0 | 0 |
| 1 | 1 | 1 |
| 1 | 1 | 1 |
| 1 | 1 | 1 |
| 1 | 1 | 1 |
| 1 | 1 | 1 |
| 0 | 0 | 0 |
| 0 | 1 | 0 |
| 0 | 0 | 0 |
| 0 | 0 | 0 |
| 0 | 0 | 0 |
| 1 | 1 | 1 |
| 1 | 1 | 1 |
| 0 | 1 | 0 |
| 1 | 1 | 1 |
| 1 | 1 | 1 |
| 0 | 0 | 0 |
| 1 | 1 | 1 |
| 1 | 1 | 1 |
| 0 | 0 | 0 |
| 1 | 0 | 0 |
| 0 | 0 | 0 |
| 0 | 0 | 0 |
| 1 | 1 | 1 |
| 1 | 1 | 1 |
| 1 | 1 | 1 |
| 1 | 1 | 1 |
| 1 | 1 | 1 |
| 1 | 1 | 1 |
| 1 | 1 | 1 |
| 1 | 1 | 1 |
| 0 | 0 | 0 |
| 1 | 1 | 1 |
| 1 | 1 | 1 |
| 0 | 0 | 0 |
| 0 | 0 | 0 |
| 1 | 1 | 1 |
| 0 | 1 | 0 |
| 0 | 0 | 0 |
| 0 | 0 | 0 |
| 0 | 0 | 0 |
| 0 | 0 | 0 |
| 0 | 0 | 0 |
| 0 | 0 | 0 |
| 1 | 1 | 1 |
| 0 | 0 | 0 |
| 0 | 0 | 0 |
| 0 | 0 | 0 |
| 0 | 0 | 0 |
| 0 | 0 | 0 |

| | | |
|---|---|---|
| 0 | 0 | 0 |
| 0 | 0 | 0 |
| 0 | 0 | 0 |
| 0 | 0 | 0 |
| 0 | 0 | 0 |
| 0 | 0 | 0 |
| 0 | 0 | 0 |
| 0 | 0 | 0 |
| 0 | 0 | 0 |
| 0 | 1 | 0 |
| 0 | 1 | 0 |
| 0 | 0 | 0 |
| 1 | 1 | 1 |
| 0 | 0 | 0 |
| 0 | 0 | 0 |
| 0 | 1 | 0 |
| 0 | 0 | 0 |
| 1 | 0 | 0 |
| 1 | 1 | 1 |
| 1 | 1 | 1 |
| 0 | 0 | 0 |
| 0 | 1 | 0 |
| 0 | 1 | 0 |
| 0 | 0 | 0 |
| 0 | 0 | 0 |
| 0 | 1 | 0 |
| 0 | 1 | 0 |
| 1 | 1 | 1 |
| 0 | 0 | 0 |
| 0 | 0 | 0 |
| 1 | 1 | 1 |
| 1 | 1 | 1 |
| 0 | 0 | 0 |
| 0 | 0 | 0 |
| 0 | 0 | 0 |
| 0 | 1 | 0 |
| 0 | 0 | 0 |
| 0 | 1 | 0 |
| 0 | 1 | 0 |
| 0 | 0 | 0 |
| 0 | 0 | 0 |
| 0 | 0 | 0 |
| 0 | 0 | 0 |
| 0 | 0 | 0 |
| 0 | 0 | 0 |
| 1 | 1 | 1 |
| 0 | 1 | 0 |
| 0 | 0 | 0 |
| 0 | 0 | 0 |
| 0 | 0 | 0 |
| 1 | 1 | 1 |
| 1 | 1 | 1 |
| 0 | 0 | 0 |
| 0 | 0 | 0 |

| | | |
|---|---|---|
| 1 | 1 | 1 |
| 1 | 1 | 1 |
| 1 | 1 | 1 |
| 1 | 1 | 1 |
| 0 | 0 | 0 |
| 0 | 0 | 0 |
| 1 | 0 | 0 |
| 0 | 0 | 0 |
| 1 | 0 | 0 |
| 0 | 0 | 0 |
| 1 | 1 | 1 |
| 1 | 1 | 1 |
| 1 | 1 | 1 |
| 1 | 1 | 1 |
| 1 | 1 | 1 |
| 1 | 1 | 1 |
| 0 | 0 | 0 |
| 1 | 1 | 1 |
| 1 | 1 | 1 |
| 0 | 0 | 0 |
| 0 | 0 | 0 |
| 0 | 1 | 0 |
| 0 | 1 | 0 |
| 0 | 0 | 0 |
| 0 | 1 | 0 |
| 0 | 0 | 0 |
| 0 | 0 | 0 |
| 0 | 0 | 0 |
| 0 | 0 | 0 |
| 0 | 0 | 0 |
| 0 | 0 | 0 |
| 0 | 0 | 0 |
| 0 | 1 | 0 |
| 0 | 0 | 0 |
| 1 | 1 | 1 |
| 1 | 1 | 1 |
| 1 | 1 | 1 |
| 1 | 1 | 1 |
| 0 | 0 | 0 |
| 1 | 1 | 1 |
| 1 | 1 | 1 |
| 1 | 1 | 1 |
| 1 | 1 | 1 |
| 0 | 0 | 0 |
| 1 | 1 | 1 |
| 1 | 1 | 1 |
| 1 | 1 | 1 |
| 1 | 1 | 1 |
| 1 | 1 | 1 |
| 1 | 1 | 1 |
| 0 | 0 | 0 |
| 0 | 0 | 0 |
| 0 | 0 | 0 |
| 0 | 1 | 0 |

| | | |
|---|---|---|
| 0 | 0 | 0 |
| 0 | 0 | 0 |
| 0 | 0 | 0 |
| 0 | 0 | 0 |
| 0 | 0 | 0 |
| 0 | 0 | 0 |
| 0 | 0 | 0 |
| 0 | 0 | 0 |
| 1 | 1 | 1 |
| 1 | 1 | 1 |
| 0 | 0 | 0 |
| 1 | 1 | 1 |
| 1 | 1 | 1 |
| 0 | 0 | 0 |
| 0 | 0 | 0 |
| 0 | 0 | 0 |
| 0 | 0 | 0 |
| 0 | 0 | 0 |
| 0 | 0 | 0 |
| 1 | 1 | 1 |
| 1 | 1 | 1 |
| 0 | 0 | 0 |
| 1 | 1 | 1 |
| 0 | 0 | 0 |
| 1 | 1 | 1 |
| 1 | 1 | 1 |
| 1 | 1 | 1 |
| 0 | 1 | 0 |
| 0 | 1 | 0 |
| 0 | 0 | 0 |
| 0 | 1 | 0 |
| 1 | 1 | 1 |
| 0 | 0 | 0 |
| 1 | 1 | 1 |
| 0 | 0 | 0 |
| 0 | 0 | 0 |
| 0 | 0 | 0 |
| 0 | 0 | 0 |
| 0 | 0 | 0 |
| 0 | 0 | 0 |
| 0 | 0 | 0 |
| 0 | 0 | 0 |
| 1 | 1 | 1 |
| 0 | 0 | 0 |
| 1 | 1 | 1 |
| 0 | 0 | 0 |
| 1 | 0 | 0 |
| 1 | 1 | 1 |
| 1 | 1 | 1 |
| 0 | 0 | 0 |
| 1 | 1 | 1 |
| 1 | 1 | 1 |
| 0 | 1 | 0 |
| 1 | 1 | 1 |

| | | |
|---|---|---|
| 0 | 0 | 0 |
| 0 | 1 | 0 |
| 1 | 1 | 1 |
| 0 | 1 | 0 |
| 0 | 1 | 0 |
| 0 | 1 | 0 |
| 0 | 1 | 0 |
| 1 | 1 | 1 |
| 1 | 1 | 1 |
| 1 | 1 | 1 |
| 0 | 0 | 0 |
| 1 | 1 | 1 |
| 1 | 1 | 1 |
| 0 | 1 | 0 |
| 0 | 0 | 0 |
| 0 | 0 | 0 |
| 0 | 0 | 0 |
| 0 | 0 | 0 |
| 0 | 0 | 0 |
| 0 | 1 | 0 |
| 0 | 1 | 0 |
| 1 | 1 | 1 |
| 0 | 0 | 0 |
| 0 | 1 | 0 |
| 0 | 1 | 0 |
| 0 | 1 | 0 |
| 1 | 1 | 1 |
| 1 | 1 | 1 |
| 0 | 0 | 0 |
| 0 | 0 | 0 |
| 0 | 0 | 0 |
| 0 | 0 | 0 |
| 0 | 0 | 0 |
| 0 | 0 | 0 |
| 1 | 1 | 1 |
| 1 | 1 | 1 |
| 0 | 0 | 0 |
| 0 | 0 | 0 |
| 0 | 0 | 0 |
| 0 | 0 | 0 |
| 0 | 0 | 0 |
| 0 | 0 | 0 |
| 1 | 1 | 1 |
| 1 | 1 | 1 |
| 0 | 0 | 0 |
| 1 | 1 | 1 |
| 1 | 1 | 1 |
| 1 | 1 | 1 |
| 1 | 1 | 1 |
| 1 | 1 | 1 |
| 1 | 0 | 0 |
| 0 | 1 | 0 |
| 0 | 1 | 0 |
| 1 | 0 | 0 |

| | | |
|---|---|---|
| 0 | 0 | 0 |
| 1 | 1 | 1 |
| 0 | 0 | 0 |
| 0 | 1 | 0 |
| 1 | 0 | 0 |
| 0 | 0 | 0 |
| 0 | 0 | 0 |
| 0 | 0 | 0 |
| 1 | 0 | 0 |
| 1 | 0 | 0 |
| 0 | 0 | 0 |
| 1 | 1 | 1 |
| 1 | 0 | 0 |
| 1 | 0 | 0 |
| 0 | 0 | 0 |
| 0 | 0 | 0 |
| 0 | 0 | 0 |
| 0 | 0 | 0 |
| 1 | 1 | 1 |
| 0 | 0 | 0 |
| 0 | 0 | 0 |
| 0 | 0 | 0 |
| 0 | 0 | 0 |
| 0 | 0 | 0 |
| 1 | 1 | 1 |
| 0 | 0 | 0 |
| 1 | 1 | 1 |
| 0 | 1 | 0 |
| 1 | 1 | 1 |
| 0 | 0 | 0 |
| 1 | 1 | 1 |
| 1 | 1 | 1 |
| 0 | 0 | 0 |
| 0 | 0 | 0 |
| 1 | 1 | 1 |
| 1 | 1 | 1 |
| 1 | 1 | 1 |
| 0 | 1 | 0 |
| 1 | 0 | 0 |
| 0 | 0 | 0 |
| 0 | 0 | 0 |
| 0 | 0 | 0 |
| 0 | 0 | 0 |
| 0 | 0 | 0 |
| 0 | 0 | 0 |
| 0 | 1 | 0 |
| 0 | 1 | 0 |
| 0 | 0 | 0 |
| 0 | 0 | 0 |
| 1 | 1 | 1 |
| 0 | 0 | 0 |
| 0 | 0 | 0 |
| 0 | 1 | 0 |
| 1 | 1 | 1 |

| | | |
|---|---|---|
| 1 | 1 | 1 |
| 0 | 0 | 0 |
| 0 | 0 | 0 |
| 1 | 0 | 0 |
| 1 | 1 | 1 |
| 0 | 0 | 0 |
| 0 | 0 | 0 |
| 0 | 0 | 0 |
| 0 | 0 | 0 |
| 0 | 0 | 0 |
| 0 | 0 | 0 |
| 0 | 0 | 0 |
| 0 | 0 | 0 |
| 0 | 0 | 0 |
| 0 | 0 | 0 |
| 0 | 1 | 0 |
| 0 | 0 | 0 |
| 1 | 1 | 1 |
| 1 | 0 | 0 |
| 1 | 1 | 1 |
| 0 | 0 | 0 |
| 1 | 1 | 1 |
| 0 | 0 | 0 |
| 0 | 0 | 0 |
| 1 | 1 | 1 |
| 0 | 0 | 0 |
| 0 | 1 | 0 |
| 0 | 1 | 0 |
| 0 | 1 | 0 |
| 0 | 0 | 0 |
| 0 | 0 | 0 |
| 0 | 0 | 0 |
| 0 | 0 | 0 |
| 0 | 0 | 0 |
| 0 | 0 | 0 |
| 0 | 0 | 0 |
| 0 | 0 | 0 |
| 1 | 1 | 1 |
| 1 | 0 | 0 |
| 0 | 0 | 0 |
| 0 | 0 | 0 |
| 0 | 0 | 0 |
| 0 | 0 | 0 |
| 0 | 0 | 0 |
| 0 | 0 | 0 |
| 0 | 0 | 0 |
| 1 | 1 | 1 |
| 0 | 0 | 0 |
| 0 | 0 | 0 |
| 0 | 0 | 0 |
| 0 | 0 | 0 |
| 0 | 1 | 0 |
| 0 | 0 | 0 |
| 0 | 0 | 0 |

| | | |
|---|---|---|
| 0 | 0 | 0 |
| 0 | 0 | 0 |
| 0 | 0 | 0 |
| 0 | 0 | 0 |
| 0 | 0 | 0 |
| 0 | 0 | 0 |
| 0 | 0 | 0 |
| 0 | 0 | 0 |
| 0 | 0 | 0 |
| 0 | 0 | 0 |
| 1 | 1 | 1 |
| 0 | 0 | 0 |
| 1 | 0 | 0 |
| 0 | 0 | 0 |
| 0 | 1 | 0 |
| 1 | 1 | 1 |
| 1 | 1 | 1 |
| 1 | 1 | 1 |
| 0 | 0 | 0 |
| 0 | 0 | 0 |
| 1 | 1 | 1 |
| 0 | 0 | 0 |
| 0 | 0 | 0 |
| 1 | 1 | 1 |
| 0 | 0 | 0 |
| 0 | 0 | 0 |
| 0 | 0 | 0 |
| 1 | 0 | 0 |
| 0 | 0 | 0 |
| 0 | 0 | 0 |
| 1 | 0 | 0 |
| 0 | 0 | 0 |
| 1 | 0 | 0 |
| 1 | 0 | 0 |
| 0 | 0 | 0 |
| 0 | 0 | 0 |
| 0 | 0 | 0 |
| 1 | 0 | 0 |
| 1 | 0 | 0 |
| 0 | 0 | 0 |
| 0 | 0 | 0 |
| 0 | 0 | 0 |
| 0 | 0 | 0 |
| 0 | 0 | 0 |
| 0 | 0 | 0 |
| 0 | 0 | 0 |
| 0 | 0 | 0 |
| 0 | 0 | 0 |
| 0 | 0 | 0 |
| 0 | 0 | 0 |
| 0 | 0 | 0 |
| 0 | 0 | 0 |
| 0 | 0 | 0 |
| 0 | 0 | 0 |

| | | |
|---|---|---|
| 0 | 0 | 0 |
| 1 | 0 | 0 |
| 0 | 0 | 0 |
| 0 | 0 | 0 |
| 0 | 0 | 0 |
| 1 | 0 | 0 |
| 0 | 0 | 0 |
| 0 | 0 | 0 |
| 1 | 1 | 1 |
| 0 | 0 | 0 |
| 0 | 1 | 0 |
| 1 | 1 | 1 |
| 1 | 1 | 1 |
| 0 | 0 | 0 |
| 1 | 1 | 1 |
| 0 | 0 | 0 |
| 1 | 1 | 1 |
| 1 | 1 | 1 |
| 0 | 0 | 0 |
| 0 | 0 | 0 |
| 0 | 0 | 0 |
| 0 | 0 | 0 |
| 0 | 0 | 0 |
| 0 | 0 | 0 |
| 0 | 0 | 0 |
| 0 | 1 | 0 |
| 1 | 1 | 1 |
| 1 | 1 | 1 |
| 1 | 0 | 0 |
| 1 | 1 | 1 |
| 1 | 1 | 1 |
| 0 | 1 | 0 |
| 0 | 0 | 0 |
| 0 | 0 | 0 |
| 0 | 0 | 0 |
| 0 | 0 | 0 |
| 0 | 0 | 0 |
| 0 | 0 | 0 |
| 0 | 1 | 0 |
| 0 | 1 | 0 |
| 0 | 1 | 0 |
| 0 | 1 | 0 |
| 0 | 1 | 0 |
| 0 | 1 | 0 |
| 0 | 0 | 0 |
| 0 | 0 | 0 |
| 0 | 1 | 0 |
| 0 | 0 | 0 |
| 0 | 0 | 0 |
| 0 | 0 | 0 |
| 0 | 0 | 0 |
| 0 | 0 | 0 |
| 0 | 0 | 0 |
| 0 | 0 | 0 |

| | | |
|---|---|---|
| 1 | 1 | 1 |
| 1 | 1 | 1 |
| 0 | 0 | 0 |
| 0 | 0 | 0 |
| 1 | 1 | 1 |
| 0 | 0 | 0 |
| 0 | 0 | 0 |
| 1 | 1 | 1 |
| 1 | 1 | 1 |
| 0 | 0 | 0 |
| 1 | 1 | 1 |
| 1 | 1 | 1 |
| 0 | 0 | 0 |
| 0 | 0 | 0 |
| 0 | 0 | 0 |
| 0 | 0 | 0 |
| 0 | 1 | 0 |
| 0 | 1 | 0 |
| 1 | 1 | 1 |
| 1 | 1 | 1 |
| 1 | 1 | 1 |
| 0 | 1 | 0 |
| 1 | 1 | 1 |
| 1 | 1 | 1 |
| 1 | 1 | 1 |
| 0 | 0 | 0 |
| 1 | 1 | 1 |
| 1 | 1 | 1 |
| 1 | 1 | 1 |
| 0 | 1 | 0 |
| 0 | 1 | 0 |
| 1 | 1 | 1 |
| 1 | 1 | 1 |
| 1 | 1 | 1 |
| 1 | 1 | 1 |
| 1 | 1 | 1 |
| 1 | 1 | 1 |
| 0 | 0 | 0 |
| 0 | 0 | 0 |
| 1 | 0 | 0 |
| 0 | 0 | 0 |
| 1 | 1 | 1 |
| 1 | 1 | 1 |
| 0 | 0 | 0 |
| 0 | 0 | 0 |
| 0 | 1 | 0 |
| 0 | 1 | 0 |
| 0 | 1 | 0 |
| 1 | 1 | 1 |
| 1 | 1 | 1 |
| 1 | 1 | 1 |
| 0 | 0 | 0 |
| 0 | 0 | 0 |
| 0 | 0 | 0 |

| | | |
|---|---|---|
| 0 | 0 | 0 |
| 1 | 1 | 1 |
| 1 | 1 | 1 |
| 1 | 1 | 1 |
| 1 | 1 | 1 |
| 1 | 1 | 1 |
| 1 | 1 | 1 |
| 1 | 1 | 1 |
| 0 | 0 | 0 |
| 1 | 1 | 1 |
| 1 | 1 | 1 |
| 1 | 1 | 1 |
| 0 | 1 | 0 |
| 0 | 1 | 0 |
| 1 | 1 | 1 |
| 1 | 1 | 1 |
| 0 | 1 | 0 |
| 0 | 1 | 0 |
| 0 | 0 | 0 |
| 0 | 0 | 0 |
| 0 | 0 | 0 |
| 0 | 0 | 0 |
| 0 | 1 | 0 |
| 0 | 0 | 0 |
| 0 | 0 | 0 |
| 1 | 1 | 1 |
| 1 | 1 | 1 |
| 1 | 1 | 1 |
| 1 | 1 | 1 |
| 1 | 1 | 1 |
| 1 | 1 | 1 |
| 1 | 1 | 1 |
| 0 | 0 | 0 |
| 1 | 1 | 1 |
| 1 | 1 | 1 |
| 0 | 1 | 0 |
| 0 | 0 | 0 |
| 1 | 1 | 1 |
| 0 | 0 | 0 |
| 0 | 0 | 0 |
| 0 | 0 | 0 |
| 0 | 0 | 0 |
| 0 | 0 | 0 |
| 1 | 1 | 1 |
| 1 | 1 | 1 |
| 1 | 1 | 1 |
| 0 | 0 | 0 |
| 0 | 0 | 0 |
| 0 | 0 | 0 |
| 0 | 0 | 0 |
| 0 | 0 | 0 |
| 0 | 0 | 0 |
| 0 | 0 | 0 |
| 0 | 1 | 0 |

| | | |
|---|---|---|
| 0 | 0 | 0 |
| 0 | 0 | 0 |
| 0 | 1 | 0 |
| 0 | 0 | 0 |
| 0 | 0 | 0 |
| 1 | 1 | 1 |
| 1 | 1 | 1 |
| 1 | 0 | 0 |
| 1 | 1 | 1 |
| 1 | 1 | 1 |
| 1 | 1 | 1 |
| 0 | 0 | 0 |
| 1 | 1 | 1 |
| 1 | 1 | 1 |
| 0 | 0 | 0 |
| 1 | 1 | 1 |
| 0 | 0 | 0 |
| 1 | 1 | 1 |
| 0 | 0 | 0 |
| 0 | 0 | 0 |
| 1 | 1 | 1 |
| 1 | 1 | 1 |
| 0 | 0 | 0 |
| 1 | 1 | 1 |
| 0 | 0 | 0 |
| 1 | 1 | 1 |
| 1 | 1 | 1 |
| 0 | 0 | 0 |
| 1 | 1 | 1 |
| 0 | 0 | 0 |
| 0 | 1 | 0 |
| 0 | 0 | 0 |
| 0 | 0 | 0 |
| 0 | 0 | 0 |
| 1 | 1 | 1 |
| 0 | 0 | 0 |
| 1 | 1 | 1 |
| 1 | 1 | 1 |
| 0 | 0 | 0 |
| 1 | 1 | 1 |
| 1 | 1 | 1 |
| 1 | 1 | 1 |
| 1 | 1 | 1 |
| 0 | 0 | 0 |
| 0 | 0 | 0 |
| 1 | 1 | 1 |
| 1 | 1 | 1 |
| 1 | 0 | 0 |
| 0 | 0 | 0 |
| 1 | 0 | 0 |
| 0 | 0 | 0 |
| 1 | 1 | 1 |
| 1 | 1 | 1 |
| 0 | 0 | 0 |

| | | |
|---|---|---|
| 1 | 1 | 1 |
| 1 | 1 | 1 |
| 1 | 1 | 1 |
| 1 | 1 | 1 |
| 0 | 1 | 0 |
| 1 | 1 | 1 |
| 1 | 1 | 1 |
| 0 | 0 | 0 |
| 1 | 1 | 1 |
| 1 | 1 | 1 |
| 1 | 1 | 1 |
| 1 | 1 | 1 |
| 0 | 0 | 0 |
| 1 | 1 | 1 |
| 0 | 0 | 0 |
| 0 | 0 | 0 |
| 1 | 1 | 1 |
| 1 | 1 | 1 |
| 1 | 1 | 1 |
| 1 | 1 | 1 |
| 0 | 0 | 0 |
| 0 | 0 | 0 |
| 1 | 1 | 1 |
| 1 | 1 | 1 |
| 0 | 0 | 0 |
| 1 | 1 | 1 |
| 0 | 0 | 0 |
| 0 | 0 | 0 |
| 0 | 0 | 0 |
| 0 | 0 | 0 |
| 0 | 1 | 0 |
| 0 | 1 | 0 |
| 0 | 0 | 0 |
| 0 | 0 | 0 |
| 0 | 0 | 0 |
| 0 | 0 | 0 |
| 0 | 0 | 0 |
| 0 | 0 | 0 |
| 0 | 0 | 0 |
| 1 | 0 | 0 |
| 1 | 1 | 1 |
| 0 | 1 | 0 |
| 0 | 1 | 0 |
| 0 | 1 | 0 |
| 0 | 1 | 0 |
| 0 | 0 | 0 |
| 0 | 1 | 0 |
| 0 | 1 | 0 |
| 1 | 1 | 1 |
| 0 | 0 | 0 |
| 0 | 0 | 0 |
| 1 | 1 | 1 |
| 1 | 1 | 1 |
| 1 | 1 | 1 |

| | | |
|---|---|---|
| 1 | 1 | 1 |
| 1 | 1 | 1 |
| 1 | 1 | 1 |
| 1 | 1 | 1 |
| 0 | 0 | 0 |
| 1 | 1 | 1 |
| 0 | 0 | 0 |
| 0 | 0 | 0 |
| 0 | 0 | 0 |
| 0 | 0 | 0 |
| 0 | 0 | 0 |
| 1 | 1 | 1 |
| 0 | 0 | 0 |
| 0 | 0 | 0 |
| 1 | 1 | 1 |
| 1 | 1 | 1 |
| 1 | 1 | 1 |
| 1 | 1 | 1 |
| 1 | 1 | 1 |
| 1 | 1 | 1 |
| 1 | 1 | 1 |
| 1 | 1 | 1 |
| 1 | 1 | 1 |
| 0 | 0 | 0 |
| 1 | 1 | 1 |
| 1 | 1 | 1 |
| 1 | 1 | 1 |
| 0 | 0 | 0 |
| 1 | 0 | 0 |
| 1 | 0 | 0 |
| 0 | 0 | 0 |
| 0 | 0 | 0 |
| 1 | 1 | 1 |
| 0 | 0 | 0 |
| 0 | 1 | 0 |
| 1 | 1 | 1 |
| 1 | 1 | 1 |
| 1 | 1 | 1 |
| 1 | 1 | 1 |
| 1 | 1 | 1 |
| 1 | 1 | 1 |
| 1 | 1 | 1 |
| 1 | 1 | 1 |
| 1 | 1 | 1 |
| 1 | 1 | 1 |
| 1 | 1 | 1 |
| 1 | 1 | 1 |
| 1 | 1 | 1 |
| 0 | 0 | 0 |
| 0 | 1 | 0 |
| 1 | 1 | 1 |
| 1 | 0 | 0 |
| 1 | 0 | 0 |
| 1 | 1 | 1 |

| | | |
|---|---|---|
| 1 | 1 | 1 |
| 0 | 1 | 0 |
| 1 | 1 | 1 |
| 0 | 1 | 0 |
| 1 | 0 | 0 |
| 1 | 1 | 1 |
| 1 | 1 | 1 |
| 1 | 1 | 1 |
| 1 | 1 | 1 |
| 1 | 1 | 1 |
| 0 | 0 | 0 |
| 0 | 0 | 0 |
| 0 | 0 | 0 |
| 0 | 0 | 0 |
| 0 | 0 | 0 |
| 1 | 0 | 0 |
| 0 | 0 | 0 |
| 0 | 0 | 0 |
| 0 | 0 | 0 |
| 0 | 0 | 0 |
| 0 | 0 | 0 |
| 0 | 0 | 0 |
| 0 | 1 | 0 |
| 0 | 0 | 0 |
| 0 | 0 | 0 |
| 0 | 0 | 0 |
| 1 | 1 | 1 |
| 0 | 0 | 0 |
| 0 | 1 | 0 |
| 0 | 1 | 0 |
| 0 | 1 | 0 |
| 0 | 0 | 0 |
| 1 | 1 | 1 |
| 1 | 1 | 1 |
| 0 | 0 | 0 |
| 0 | 0 | 0 |
| 0 | 0 | 0 |
| 0 | 0 | 0 |
| 0 | 0 | 0 |
| 1 | 0 | 0 |
| 1 | 0 | 0 |
| 0 | 0 | 0 |
| 1 | 0 | 0 |
| 0 | 0 | 0 |
| 1 | 1 | 1 |
| 1 | 0 | 0 |
| 1 | 1 | 1 |
| 0 | 0 | 0 |
| 0 | 0 | 0 |
| 0 | 0 | 0 |
| 0 | 0 | 0 |
| 1 | 1 | 1 |
| 0 | 0 | 0 |
| 1 | 1 | 1 |

| | | |
|---|---|---|
| 1 | 1 | 1 |
| 1 | 1 | 1 |
| 1 | 1 | 1 |
| 1 | 1 | 1 |
| 1 | 1 | 1 |
| 0 | 0 | 0 |
| 0 | 0 | 0 |
| 0 | 0 | 0 |
| 0 | 0 | 0 |
| 0 | 1 | 0 |
| 0 | 0 | 0 |
| 0 | 0 | 0 |
| 0 | 0 | 0 |
| 0 | 0 | 0 |
| 0 | 0 | 0 |
| 0 | 0 | 0 |
| 1 | 1 | 1 |
| 0 | 1 | 0 |
| 1 | 1 | 1 |
| 1 | 1 | 1 |
| 1 | 1 | 1 |
| 1 | 1 | 1 |
| 1 | 1 | 1 |
| 1 | 1 | 1 |
| 1 | 1 | 1 |
| 1 | 1 | 1 |
| 1 | 1 | 1 |
| 1 | 1 | 1 |
| 0 | 0 | 0 |
| 0 | 1 | 0 |
| 0 | 1 | 0 |
| 1 | 0 | 0 |
| 1 | 0 | 0 |
| 0 | 0 | 0 |
| 1 | 1 | 1 |
| 0 | 0 | 0 |
| 1 | 0 | 0 |
| 0 | 0 | 0 |
| 1 | 1 | 1 |
| 0 | 0 | 0 |
| 1 | 1 | 1 |
| 1 | 1 | 1 |
| 0 | 1 | 0 |
| 0 | 0 | 0 |
| 1 | 1 | 1 |
| 1 | 1 | 1 |
| 1 | 1 | 1 |
| 1 | 1 | 1 |
| 1 | 1 | 1 |
| 1 | 1 | 1 |
| 1 | 1 | 1 |
| 1 | 0 | 0 |
| 0 | 0 | 0 |
| 0 | 0 | 0 |

| | | |
|---|---|---|
| 1 | 1 | 1 |
| 0 | 0 | 0 |
| 1 | 1 | 1 |
| 1 | 1 | 1 |
| 0 | 0 | 0 |
| 0 | 0 | 0 |
| 0 | 0 | 0 |
| 1 | 1 | 1 |
| 1 | 1 | 1 |
| 0 | 0 | 0 |
| 0 | 0 | 0 |
| 0 | 0 | 0 |
| 0 | 0 | 0 |
| 0 | 0 | 0 |
| 1 | 1 | 1 |
| 0 | 0 | 0 |
| 0 | 0 | 0 |
| 0 | 0 | 0 |
| 1 | 1 | 1 |
| 1 | 1 | 1 |
| 1 | 1 | 1 |
| 1 | 1 | 1 |
| 1 | 1 | 1 |
| 0 | 0 | 0 |
| 0 | 1 | 0 |
| 0 | 1 | 0 |
| 0 | 0 | 0 |
| 0 | 0 | 0 |
| 1 | 1 | 1 |
| 1 | 1 | 1 |
| 0 | 0 | 0 |
| 1 | 1 | 1 |
| 1 | 1 | 1 |
| 0 | 1 | 0 |
| 1 | 1 | 1 |
| 0 | 0 | 0 |
| 0 | 1 | 0 |
| 0 | 0 | 0 |
| 0 | 0 | 0 |
| 0 | 1 | 0 |
| 0 | 1 | 0 |
| 0 | 0 | 0 |
| 0 | 0 | 0 |
| 0 | 1 | 0 |
| 0 | 0 | 0 |
| 0 | 0 | 0 |
| 0 | 0 | 0 |
| 0 | 1 | 0 |
| 0 | 0 | 0 |
| 0 | 0 | 0 |
| 0 | 0 | 0 |
| 1 | 1 | 1 |
| 0 | 0 | 0 |
| 1 | 1 | 1 |

| | | |
|---|---|---|
| 1 | 1 | 1 |
| 1 | 1 | 1 |
| 1 | 1 | 1 |
| 1 | 1 | 1 |
| 0 | 0 | 0 |
| 0 | 0 | 0 |
| 1 | 1 | 1 |
| 1 | 0 | 0 |
| 1 | 1 | 1 |
| 0 | 0 | 0 |
| 1 | 0 | 0 |
| 1 | 0 | 0 |
| 0 | 0 | 0 |
| 1 | 1 | 1 |
| 1 | 1 | 1 |
| 1 | 1 | 1 |
| 1 | 1 | 1 |
| 1 | 1 | 1 |
| 1 | 1 | 1 |
| 1 | 1 | 1 |
| 1 | 1 | 1 |
| 0 | 1 | 0 |
| 1 | 1 | 1 |
| 0 | 0 | 0 |
| 1 | 1 | 1 |
| 0 | 0 | 0 |
| 1 | 1 | 1 |
| 0 | 0 | 0 |
| 1 | 1 | 1 |
| 0 | 0 | 0 |
| 0 | 0 | 0 |
| 0 | 0 | 0 |
| 0 | 0 | 0 |
| 0 | 0 | 0 |
| 0 | 1 | 0 |
| 0 | 0 | 0 |
| 1 | 1 | 1 |
| 0 | 0 | 0 |
| 1 | 1 | 1 |
| 1 | 1 | 1 |
| 0 | 0 | 0 |
| 0 | 0 | 0 |
| 0 | 0 | 0 |
| 0 | 0 | 0 |
| 0 | 0 | 0 |
| 0 | 0 | 0 |
| 0 | 0 | 0 |
| 0 | 0 | 0 |
| 1 | 1 | 1 |
| 0 | 0 | 0 |
| 1 | 0 | 0 |
| 1 | 0 | 0 |
| 1 | 1 | 1 |
| 1 | 1 | 1 |

| | | |
|---|---|---|
| 1 | 1 | 1 |
| 1 | 1 | 1 |
| 1 | 1 | 1 |
| 0 | 1 | 0 |
| 0 | 1 | 0 |
| 0 | 1 | 0 |
| 0 | 0 | 0 |
| 0 | 0 | 0 |
| 0 | 0 | 0 |
| 0 | 0 | 0 |
| 0 | 0 | 0 |
| 0 | 0 | 0 |
| 0 | 0 | 0 |
| 0 | 0 | 0 |
| 0 | 0 | 0 |
| 0 | 0 | 0 |
| 0 | 0 | 0 |
| 0 | 0 | 0 |
| 1 | 1 | 1 |
| 0 | 0 | 0 |
| 0 | 1 | 0 |
| 0 | 1 | 0 |
| 0 | 1 | 0 |
| 1 | 1 | 1 |
| 0 | 1 | 0 |
| 0 | 0 | 0 |
| 0 | 0 | 0 |
| 0 | 0 | 0 |
| 0 | 0 | 0 |
| 0 | 0 | 0 |
| 0 | 0 | 0 |
| 0 | 0 | 0 |
| 0 | 0 | 0 |
| 0 | 0 | 0 |
| 1 | 1 | 1 |
| 0 | 0 | 0 |
| 0 | 0 | 0 |
| 0 | 0 | 0 |
| 0 | 0 | 0 |
| 0 | 1 | 0 |
| 0 | 0 | 0 |
| 0 | 0 | 0 |
| 1 | 1 | 1 |
| 0 | 1 | 0 |
| 1 | 1 | 1 |
| 0 | 1 | 0 |
| 0 | 1 | 0 |
| 0 | 0 | 0 |
| 0 | 0 | 0 |
| 0 | 0 | 0 |
| 0 | 0 | 0 |
| 0 | 1 | 0 |
| 0 | 0 | 0 |
| 1 | 1 | 1 |

| | | |
|---|---|---|
| 0 | 1 | 0 |
| 1 | 1 | 1 |
| 1 | 1 | 1 |
| 1 | 1 | 1 |
| 1 | 1 | 1 |
| 0 | 0 | 0 |
| 0 | 1 | 0 |
| 1 | 1 | 1 |
| 1 | 1 | 1 |
| 1 | 1 | 1 |
| 0 | 0 | 0 |
| 0 | 0 | 0 |
| 0 | 0 | 0 |
| 0 | 0 | 0 |
| 0 | 0 | 0 |
| 0 | 1 | 0 |
| 1 | 0 | 0 |
| 0 | 0 | 0 |
| 1 | 1 | 1 |
| 1 | 1 | 1 |
| 1 | 1 | 1 |
| 1 | 1 | 1 |
| 0 | 0 | 0 |
| 1 | 1 | 1 |
| 0 | 0 | 0 |
| 0 | 0 | 0 |
| 0 | 0 | 0 |
| 0 | 0 | 0 |
| 1 | 1 | 1 |
| 0 | 0 | 0 |
| 1 | 1 | 1 |
| 0 | 1 | 0 |
| 0 | 1 | 0 |
| 0 | 1 | 0 |
| 1 | 1 | 1 |
| 1 | 1 | 1 |
| 0 | 1 | 0 |
| 0 | 0 | 0 |
| 0 | 0 | 0 |
| 0 | 0 | 0 |
| 0 | 0 | 0 |
| 0 | 0 | 0 |
| 1 | 1 | 1 |
| 0 | 0 | 0 |
| 0 | 0 | 0 |
| 1 | 1 | 1 |
| 0 | 0 | 0 |
| 0 | 0 | 0 |
| 0 | 0 | 0 |
| 0 | 0 | 0 |
| 1 | 1 | 1 |
| 1 | 1 | 1 |
| 1 | 1 | 1 |
| 0 | 0 | 0 |

| | | |
|---|---|---|
| 0 | 0 | 0 |
| 1 | 1 | 1 |
| 0 | 0 | 0 |
| 0 | 1 | 0 |
| 1 | 0 | 0 |
| 1 | 0 | 0 |
| 1 | 0 | 0 |
| 1 | 1 | 1 |
| 0 | 0 | 0 |
| 1 | 1 | 1 |
| 0 | 0 | 0 |
| 0 | 0 | 0 |
| 0 | 0 | 0 |
| 0 | 0 | 0 |
| 1 | 1 | 1 |
| 1 | 1 | 1 |
| 0 | 0 | 0 |
| 0 | 0 | 0 |
| 0 | 1 | 0 |
| 1 | 1 | 1 |
| 0 | 0 | 0 |
| 1 | 0 | 0 |
| 1 | 0 | 0 |
| 1 | 1 | 1 |
| 0 | 0 | 0 |
| 1 | 0 | 0 |
| 0 | 0 | 0 |
| 0 | 0 | 0 |
| 0 | 0 | 0 |
| 1 | 1 | 1 |
| 0 | 0 | 0 |
| 0 | 0 | 0 |
| 0 | 0 | 0 |
| 1 | 1 | 1 |
| 0 | 0 | 0 |
| 1 | 1 | 1 |
| 0 | 0 | 0 |
| 0 | 0 | 0 |
| 0 | 0 | 0 |
| 0 | 0 | 0 |
| 0 | 0 | 0 |
| 1 | 1 | 1 |
| 0 | 0 | 0 |
| 0 | 0 | 0 |
| 0 | 0 | 0 |
| 0 | 0 | 0 |
| 0 | 0 | 0 |
| 0 | 0 | 0 |
| 0 | 0 | 0 |
| 0 | 0 | 0 |
| 0 | 0 | 0 |
| 0 | 0 | 0 |
| 1 | 1 | 1 |
| 0 | 0 | 0 |

| | | |
|---|---|---|
| 0 | 1 | 0 |
| 1 | 1 | 1 |
| 1 | 1 | 1 |
| 1 | 1 | 1 |
| 1 | 1 | 1 |
| 1 | 1 | 1 |
| 1 | 1 | 1 |
| 1 | 1 | 1 |
| 1 | 1 | 1 |
| 0 | 0 | 0 |
| 0 | 0 | 0 |
| 1 | 1 | 1 |
| 1 | 1 | 1 |
| 1 | 1 | 1 |
| 0 | 0 | 0 |
| 1 | 1 | 1 |
| 1 | 1 | 1 |
| 0 | 0 | 0 |
| 0 | 1 | 0 |
| 0 | 1 | 0 |
| 0 | 1 | 0 |
| 0 | 0 | 0 |
| 0 | 0 | 0 |
| 0 | 0 | 0 |
| 0 | 0 | 0 |
| 0 | 0 | 0 |
| 0 | 0 | 0 |
| 0 | 0 | 0 |
| 1 | 1 | 1 |
| 0 | 0 | 0 |
| 0 | 0 | 0 |
| 1 | 1 | 1 |
| 1 | 1 | 1 |
| 1 | 1 | 1 |
| 1 | 1 | 1 |
| 1 | 1 | 1 |
| 1 | 1 | 1 |
| 1 | 1 | 1 |
| 1 | 1 | 1 |
| 1 | 1 | 1 |
| 1 | 1 | 1 |
| 1 | 1 | 1 |
| 1 | 1 | 1 |
| 0 | 0 | 0 |
| 1 | 1 | 1 |
| 0 | 0 | 0 |
| 1 | 1 | 1 |
| 1 | 1 | 1 |
| 1 | 1 | 1 |
| 1 | 1 | 1 |
| 0 | 0 | 0 |
| 1 | 1 | 1 |
| 1 | 1 | 1 |
| 0 | 0 | 0 |

| | | |
|---|---|---|
| 0 | 0 | 0 |
| 0 | 0 | 0 |
| 0 | 0 | 0 |
| 0 | 0 | 0 |
| 1 | 1 | 1 |
| 0 | 1 | 0 |
| 1 | 1 | 1 |
| 0 | 0 | 0 |
| 0 | 0 | 0 |
| 0 | 0 | 0 |
| 0 | 0 | 0 |
| 1 | 1 | 1 |
| 1 | 1 | 1 |
| 1 | 1 | 1 |
| 1 | 1 | 1 |
| 0 | 0 | 0 |
| 0 | 0 | 0 |
| 0 | 1 | 0 |
| 0 | 1 | 0 |
| 1 | 1 | 1 |
| 1 | 1 | 1 |
| 0 | 0 | 0 |
| 1 | 1 | 1 |
| 0 | 1 | 0 |
| 0 | 0 | 0 |
| 0 | 0 | 0 |
| 0 | 0 | 0 |
| 0 | 0 | 0 |
| 1 | 1 | 1 |
| 1 | 1 | 1 |
| 1 | 1 | 1 |
| 0 | 0 | 0 |
| 1 | 1 | 1 |
| 1 | 1 | 1 |
| 1 | 1 | 1 |
| 1 | 1 | 1 |
| 1 | 1 | 1 |
| 1 | 1 | 1 |
| 1 | 1 | 1 |
| 1 | 1 | 1 |
| 1 | 1 | 1 |
| 1 | 1 | 1 |
| 0 | 0 | 0 |
| 0 | 1 | 0 |
| 0 | 1 | 0 |
| 1 | 1 | 1 |
| 0 | 1 | 0 |
| 1 | 1 | 1 |
| 1 | 1 | 1 |
| 0 | 0 | 0 |
| 1 | 1 | 1 |
| 1 | 1 | 1 |
| 1 | 1 | 1 |
| 1 | 1 | 1 |

| | | |
|---|---|---|
| 0 | 0 | 0 |
| 1 | 1 | 1 |
| 0 | 0 | 0 |
| 0 | 1 | 0 |
| 1 | 1 | 1 |
| 1 | 1 | 1 |
| 1 | 1 | 1 |
| 0 | 0 | 0 |
| 1 | 1 | 1 |
| 1 | 1 | 1 |
| 0 | 0 | 0 |
| 0 | 0 | 0 |
| 0 | 0 | 0 |
| 1 | 1 | 1 |
| 0 | 0 | 0 |
| 0 | 0 | 0 |
| 0 | 0 | 0 |
| 0 | 0 | 0 |
| 0 | 1 | 0 |
| 0 | 0 | 0 |
| 0 | 0 | 0 |
| 0 | 0 | 0 |
| 0 | 0 | 0 |
| 0 | 0 | 0 |
| 0 | 0 | 0 |
| 0 | 0 | 0 |
| 0 | 0 | 0 |
| 0 | 0 | 0 |
| 0 | 0 | 0 |
| 0 | 0 | 0 |
| 0 | 0 | 0 |
| 0 | 0 | 0 |
| 1 | 1 | 1 |
| 0 | 0 | 0 |
| 1 | 1 | 1 |
| 0 | 0 | 0 |
| 0 | 0 | 0 |
| 1 | 1 | 1 |
| 1 | 1 | 1 |
| 0 | 1 | 0 |
| 0 | 1 | 0 |
| 0 | 0 | 0 |
| 0 | 0 | 0 |
| 0 | 0 | 0 |
| 1 | 1 | 1 |
| 0 | 0 | 0 |
| 1 | 1 | 1 |
| 0 | 0 | 0 |
| 0 | 1 | 0 |
| 0 | 1 | 0 |
| 0 | 1 | 0 |
| 0 | 1 | 0 |
| 0 | 1 | 0 |
| 1 | 1 | 1 |

| | | |
|---|---|---|
| 1 | 1 | 1 |
| 1 | 1 | 1 |
| 0 | 1 | 0 |
| 1 | 1 | 1 |
| 1 | 1 | 1 |
| 0 | 0 | 0 |
| 0 | 0 | 0 |
| 1 | 1 | 1 |
| 0 | 0 | 0 |
| 0 | 0 | 0 |
| 1 | 1 | 1 |
| 0 | 1 | 0 |
| 1 | 1 | 1 |
| 1 | 1 | 1 |
| 1 | 1 | 1 |
| 0 | 0 | 0 |
| 1 | 1 | 1 |
| 1 | 1 | 1 |
| 0 | 0 | 0 |
| 0 | 0 | 0 |
| 0 | 0 | 0 |
| 1 | 1 | 1 |
| 1 | 1 | 1 |
| 0 | 0 | 0 |
| 1 | 1 | 1 |
| 0 | 0 | 0 |
| 0 | 0 | 0 |
| 1 | 1 | 1 |
| 1 | 1 | 1 |
| 1 | 1 | 1 |
| 0 | 0 | 0 |
| 1 | 0 | 0 |
| 0 | 0 | 0 |
| 0 | 0 | 0 |
| 0 | 0 | 0 |
| 0 | 0 | 0 |
| 0 | 0 | 0 |
| 1 | 1 | 1 |
| 0 | 0 | 0 |
| 0 | 1 | 0 |
| 0 | 0 | 0 |
| 0 | 0 | 0 |
| 0 | 0 | 0 |
| 0 | 0 | 0 |
| 1 | 1 | 1 |
| 0 | 1 | 0 |
| 0 | 1 | 0 |
| 0 | 0 | 0 |
| 0 | 0 | 0 |
| 0 | 0 | 0 |
| 1 | 1 | 1 |
| 0 | 1 | 0 |
| 1 | 1 | 1 |
| 1 | 1 | 1 |

| | | |
|---|---|---|
| 1 | 1 | 1 |
| 0 | 1 | 0 |
| 0 | 1 | 0 |
| 0 | 0 | 0 |
| 0 | 0 | 0 |
| 1 | 1 | 1 |
| 0 | 0 | 0 |
| 1 | 1 | 1 |
| 1 | 1 | 1 |
| 1 | 1 | 1 |
| 1 | 1 | 1 |
| 0 | 0 | 0 |
| 1 | 0 | 0 |
| 0 | 0 | 0 |
| 1 | 1 | 1 |
| 0 | 0 | 0 |
| 0 | 0 | 0 |
| 0 | 0 | 0 |
| 1 | 1 | 1 |
| 0 | 0 | 0 |
| 0 | 0 | 0 |
| 0 | 0 | 0 |
| 0 | 0 | 0 |
| 0 | 0 | 0 |
| 0 | 0 | 0 |
| 1 | 1 | 1 |
| 0 | 0 | 0 |
| 0 | 1 | 0 |
| 0 | 1 | 0 |
| 0 | 1 | 0 |
| 0 | 1 | 0 |
| 0 | 1 | 0 |
| 0 | 0 | 0 |
| 0 | 0 | 0 |
| 1 | 1 | 1 |
| 1 | 1 | 1 |
| 0 | 0 | 0 |
| 0 | 0 | 0 |
| 0 | 0 | 0 |
| 1 | 0 | 0 |
| 0 | 0 | 0 |
| 0 | 0 | 0 |
| 1 | 1 | 1 |
| 0 | 0 | 0 |
| 1 | 1 | 1 |
| 1 | 1 | 1 |
| 1 | 1 | 1 |
| 1 | 1 | 1 |
| 1 | 1 | 1 |
| 0 | 0 | 0 |
| 1 | 1 | 1 |
| 1 | 1 | 1 |
| 1 | 1 | 1 |
| 0 | 1 | 0 |

| | | |
|---|---|---|
| 1 | 1 | 1 |
| 1 | 1 | 1 |
| 1 | 1 | 1 |
| 0 | 0 | 0 |
| 1 | 1 | 1 |
| 1 | 1 | 1 |
| 0 | 1 | 0 |
| 0 | 0 | 0 |
| 1 | 1 | 1 |
| 1 | 1 | 1 |
| 1 | 1 | 1 |
| 1 | 1 | 1 |
| 1 | 1 | 1 |
| 0 | 1 | 0 |
| 0 | 1 | 0 |
| 0 | 1 | 0 |
| 0 | 0 | 0 |
| 0 | 0 | 0 |
| 0 | 1 | 0 |
| 1 | 1 | 1 |
| 0 | 1 | 0 |
| 1 | 1 | 1 |
| 0 | 1 | 0 |
| 1 | 1 | 1 |
| 1 | 1 | 1 |
| 1 | 1 | 1 |
| 1 | 1 | 1 |
| 0 | 0 | 0 |
| 0 | 0 | 0 |
| 1 | 1 | 1 |
| 1 | 1 | 1 |
| 0 | 0 | 0 |
| 0 | 1 | 0 |
| 0 | 0 | 0 |
| 0 | 0 | 0 |
| 0 | 0 | 0 |
| 0 | 0 | 0 |
| 0 | 0 | 0 |
| 1 | 1 | 1 |
| 1 | 1 | 1 |
| 1 | 1 | 1 |
| 0 | 1 | 0 |
| 1 | 1 | 1 |
| 1 | 1 | 1 |
| 0 | 0 | 0 |
| 0 | 0 | 0 |
| 0 | 0 | 0 |
| 1 | 1 | 1 |
| 1 | 1 | 1 |
| 1 | 1 | 1 |
| 1 | 1 | 1 |
| 0 | 0 | 0 |
| 0 | 0 | 0 |
| 0 | 0 | 0 |

| | | |
|---|---|---|
| 0 | 0 | 0 |
| 0 | 0 | 0 |
| 1 | 1 | 1 |
| 0 | 0 | 0 |
| 0 | 0 | 0 |
| 0 | 0 | 0 |
| 0 | 0 | 0 |
| 1 | 1 | 1 |
| 1 | 1 | 1 |
| 1 | 1 | 1 |
| 1 | 1 | 1 |
| 1 | 1 | 1 |
| 0 | 1 | 0 |
| 0 | 1 | 0 |
| 1 | 1 | 1 |
| 1 | 1 | 1 |
| 1 | 1 | 1 |
| 1 | 1 | 1 |
| 1 | 1 | 1 |
| 1 | 1 | 1 |
| 0 | 0 | 0 |
| 0 | 0 | 0 |
| 0 | 0 | 0 |
| 0 | 0 | 0 |
| 0 | 0 | 0 |
| 1 | 1 | 1 |
| 1 | 1 | 1 |
| 0 | 0 | 0 |
| 0 | 1 | 0 |
| 0 | 1 | 0 |
| 0 | 0 | 0 |
| 0 | 0 | 0 |
| 0 | 0 | 0 |
| 1 | 1 | 1 |
| 0 | 0 | 0 |
| 1 | 1 | 1 |
| 0 | 1 | 0 |
| 0 | 0 | 0 |
| 1 | 1 | 1 |
| 1 | 1 | 1 |
| 0 | 1 | 0 |
| 0 | 1 | 0 |
| 1 | 1 | 1 |
| 1 | 1 | 1 |
| 1 | 1 | 1 |
| 1 | 1 | 1 |
| 1 | 1 | 1 |
| 1 | 1 | 1 |
| 0 | 0 | 0 |
| 1 | 1 | 1 |
| 0 | 0 | 0 |
| 0 | 0 | 0 |
| 1 | 1 | 1 |
| 0 | 1 | 0 |

| | | |
|---|---|---|
| 0 | 0 | 0 |
| 1 | 1 | 1 |
| 1 | 1 | 1 |
| 1 | 1 | 1 |
| 1 | 1 | 1 |
| 0 | 0 | 0 |
| 0 | 0 | 0 |
| 1 | 1 | 1 |
| 0 | 1 | 0 |
| 1 | 1 | 1 |
| 1 | 1 | 1 |
| 0 | 0 | 0 |
| 0 | 1 | 0 |
| 1 | 1 | 1 |
| 0 | 1 | 0 |
| 1 | 1 | 1 |
| 0 | 0 | 0 |
| 0 | 0 | 0 |
| 1 | 1 | 1 |
| 1 | 1 | 1 |
| 1 | 1 | 1 |
| 1 | 1 | 1 |
| 1 | 1 | 1 |
| 1 | 1 | 1 |
| 0 | 0 | 0 |
| 1 | 1 | 1 |
| 1 | 1 | 1 |
| 1 | 1 | 1 |
| 1 | 1 | 1 |
| 1 | 1 | 1 |
| 0 | 1 | 0 |
| 0 | 0 | 0 |
| 0 | 0 | 0 |
| 0 | 1 | 0 |
| 1 | 1 | 1 |
| 0 | 0 | 0 |
| 0 | 0 | 0 |
| 0 | 0 | 0 |
| 0 | 0 | 0 |
| 0 | 0 | 0 |
| 0 | 0 | 0 |
| 0 | 0 | 0 |
| 0 | 0 | 0 |
| 0 | 0 | 0 |
| 0 | 0 | 0 |
| 0 | 0 | 0 |
| 0 | 0 | 0 |
| 0 | 0 | 0 |
| 1 | 0 | 0 |
| 1 | 1 | 1 |
| 1 | 0 | 0 |
| 1 | 1 | 1 |
| 0 | 0 | 0 |
| 1 | 1 | 1 |

| | | |
|---|---|---|
| 1 | 1 | 1 |
| 1 | 1 | 1 |
| 1 | 1 | 1 |
| 1 | 1 | 1 |
| 1 | 1 | 1 |
| 1 | 1 | 1 |
| 1 | 1 | 1 |
| 0 | 0 | 0 |
| 0 | 0 | 0 |
| 1 | 0 | 0 |
| 1 | 1 | 1 |
| 1 | 1 | 1 |
| 1 | 1 | 1 |
| 0 | 0 | 0 |
| 1 | 1 | 1 |
| 1 | 1 | 1 |
| 0 | 0 | 0 |
| 1 | 1 | 1 |
| 0 | 0 | 0 |
| 0 | 1 | 0 |
| 1 | 1 | 1 |
| 1 | 1 | 1 |
| 1 | 1 | 1 |
| 0 | 0 | 0 |
| 1 | 1 | 1 |
| 1 | 1 | 1 |
| 0 | 1 | 0 |
| 0 | 1 | 0 |
| 0 | 1 | 0 |
| 0 | 1 | 0 |
| 1 | 1 | 1 |
| 0 | 0 | 0 |
| 1 | 1 | 1 |
| 0 | 1 | 0 |
| 1 | 1 | 1 |
| 1 | 1 | 1 |
| 1 | 1 | 1 |
| 1 | 1 | 1 |
| 0 | 0 | 0 |
| 1 | 1 | 1 |
| 1 | 1 | 1 |
| 1 | 1 | 1 |
| 0 | 1 | 0 |
| 1 | 1 | 1 |
| 1 | 1 | 1 |
| 0 | 0 | 0 |
| 0 | 1 | 0 |
| 0 | 1 | 0 |
| 1 | 0 | 0 |
| 1 | 0 | 0 |
| 0 | 1 | 0 |
| 1 | 0 | 0 |
| 0 | 0 | 0 |
| 1 | 0 | 0 |

| | | |
|---|---|---|
| 1 | 0 | 0 |
| 1 | 0 | 0 |
| 1 | 0 | 0 |
| 1 | 0 | 0 |
| 0 | 0 | 0 |
| 0 | 0 | 0 |
| 0 | 0 | 0 |
| 0 | 0 | 0 |
| 1 | 0 | 0 |
| 1 | 0 | 0 |
| 1 | 0 | 0 |
| 1 | 0 | 0 |
| 1 | 0 | 0 |
| 0 | 0 | 0 |
| 1 | 0 | 0 |
| 0 | 0 | 0 |
| 1 | 0 | 0 |
| 0 | 0 | 0 |
| 1 | 0 | 0 |
| 0 | 0 | 0 |
| 0 | 0 | 0 |
| 1 | 0 | 0 |
| 1 | 0 | 0 |
| 0 | 0 | 0 |
| 0 | 0 | 0 |
| 1 | 0 | 0 |
| 0 | 0 | 0 |
| 0 | 0 | 0 |
| 0 | 0 | 0 |
| 1 | 0 | 0 |
| 1 | 0 | 0 |
| 0 | 0 | 0 |
| 1 | 0 | 0 |
| 1 | 0 | 0 |
| 1 | 0 | 0 |
| 1 | 0 | 0 |
| 1 | 0 | 0 |
| 1 | 0 | 0 |
| 1 | 0 | 0 |
| 1 | 0 | 0 |
| 1 | 0 | 0 |
| 1 | 0 | 0 |
| 1 | 0 | 0 |
| 0 | 0 | 0 |
| 1 | 1 | 1 |
| 1 | 1 | 1 |
| 0 | 0 | 0 |
| 0 | 0 | 0 |
| 0 | 1 | 0 |
| 0 | 0 | 0 |
| 0 | 0 | 0 |
| 0 | 0 | 0 |
| 1 | 1 | 1 |
| 1 | 1 | 1 |

| | | |
|---|---|---|
| 1 | 1 | 1 |
| 1 | 1 | 1 |
| 1 | 1 | 1 |
| 1 | 1 | 1 |
| 1 | 1 | 1 |
| 1 | 1 | 1 |
| 1 | 1 | 1 |
| 0 | 0 | 0 |
| 1 | 1 | 1 |
| 1 | 1 | 1 |
| 0 | 1 | 0 |
| 0 | 1 | 0 |
| 1 | 0 | 0 |
| 1 | 1 | 1 |
| 0 | 0 | 0 |
| 0 | 0 | 0 |
| 0 | 1 | 0 |
| 0 | 0 | 0 |
| 1 | 1 | 1 |
| 1 | 1 | 1 |
| 0 | 0 | 0 |
| 0 | 1 | 0 |
| 0 | 1 | 0 |
| 1 | 1 | 1 |
| 0 | 0 | 0 |
| 1 | 1 | 1 |
| 0 | 1 | 0 |
| 0 | 0 | 0 |
| 0 | 0 | 0 |
| 0 | 0 | 0 |
| 0 | 0 | 0 |
| 1 | 1 | 1 |
| 1 | 1 | 1 |
| 0 | 1 | 0 |
| 1 | 1 | 1 |
| 1 | 1 | 1 |
| 0 | 0 | 0 |
| 0 | 0 | 0 |
| 1 | 1 | 1 |
| 1 | 1 | 1 |
| 1 | 1 | 1 |
| 1 | 1 | 1 |
| 0 | 0 | 0 |
| 0 | 1 | 0 |
| 1 | 1 | 1 |
| 1 | 1 | 1 |
| 1 | 1 | 1 |
| 1 | 1 | 1 |
| 0 | 0 | 0 |
| 0 | 0 | 0 |
| 0 | 0 | 0 |
| 0 | 0 | 0 |
| 0 | 0 | 0 |
| 1 | 1 | 1 |

| | | |
|---|---|---|
| 0 | 0 | 0 |
| 0 | 0 | 0 |
| 0 | 1 | 0 |
| 0 | 0 | 0 |
| 1 | 1 | 1 |
| 0 | 1 | 0 |
| 0 | 1 | 0 |
| 1 | 1 | 1 |
| 1 | 1 | 1 |
| 0 | 0 | 0 |
| 1 | 1 | 1 |
| 0 | 1 | 0 |
| 1 | 1 | 1 |
| 0 | 1 | 0 |
| 0 | 1 | 0 |
| 0 | 1 | 0 |
| 0 | 1 | 0 |
| 1 | 1 | 1 |
| 1 | 1 | 1 |
| 0 | 1 | 0 |
| 0 | 1 | 0 |
| 0 | 1 | 0 |
| 0 | 1 | 0 |
| 0 | 1 | 0 |
| 0 | 1 | 0 |
| 0 | 1 | 0 |
| 0 | 1 | 0 |
| 0 | 1 | 0 |
| 0 | 1 | 0 |
| 0 | 1 | 0 |
| 1 | 1 | 1 |
| 1 | 1 | 1 |
| 0 | 0 | 0 |
| 1 | 1 | 1 |
| 1 | 1 | 1 |
| 1 | 1 | 1 |
| 1 | 1 | 1 |
| 0 | 1 | 0 |
| 1 | 1 | 1 |
| 1 | 1 | 1 |
| 1 | 1 | 1 |
| 1 | 1 | 1 |
| 1 | 1 | 1 |
| 0 | 1 | 0 |
| 0 | 1 | 0 |
| 1 | 1 | 1 |
| 1 | 1 | 1 |
| 0 | 0 | 0 |
| 0 | 0 | 0 |
| 0 | 0 | 0 |
| 0 | 0 | 0 |
| 0 | 0 | 0 |
| 1 | 1 | 1 |
| 0 | 0 | 0 |

| | | |
|---|---|---|
| 1 | 1 | 1 |
| 0 | 0 | 0 |
| 1 | 1 | 1 |
| 0 | 1 | 0 |
| 0 | 1 | 0 |
| 0 | 0 | 0 |
| 0 | 0 | 0 |
| 0 | 0 | 0 |
| 0 | 0 | 0 |
| 0 | 0 | 0 |
| 0 | 1 | 0 |
| 0 | 0 | 0 |
| 0 | 1 | 0 |
| 1 | 1 | 1 |
| 0 | 1 | 0 |
| 0 | 0 | 0 |
| 1 | 1 | 1 |
| 1 | 1 | 1 |
| 0 | 0 | 0 |
| 0 | 0 | 0 |
| 0 | 0 | 0 |
| 0 | 0 | 0 |
| 0 | 0 | 0 |
| 0 | 0 | 0 |
| 0 | 1 | 0 |
| 0 | 0 | 0 |
| 0 | 0 | 0 |
| 0 | 0 | 0 |
| 0 | 0 | 0 |
| 0 | 0 | 0 |
| 0 | 0 | 0 |
| 1 | 1 | 1 |
| 1 | 1 | 1 |
| 0 | 0 | 0 |
| 0 | 0 | 0 |
| 1 | 0 | 0 |
| 0 | 0 | 0 |
| 0 | 0 | 0 |
| 0 | 0 | 0 |
| 0 | 0 | 0 |
| 0 | 0 | 0 |
| 1 | 0 | 0 |
| 1 | 1 | 1 |
| 1 | 1 | 1 |
| 0 | 0 | 0 |
| 0 | 0 | 0 |
| 0 | 0 | 0 |
| 1 | 1 | 1 |
| 1 | 1 | 1 |
| 1 | 1 | 1 |
| 1 | 1 | 1 |
| 1 | 1 | 1 |
| 1 | 1 | 1 |
| 0 | 0 | 0 |

| | | |
|---|---|---|
| 1 | 1 | 1 |
| 1 | 1 | 1 |
| 1 | 1 | 1 |
| 1 | 1 | 1 |
| 0 | 0 | 0 |
| 0 | 0 | 0 |
| 1 | 0 | 0 |
| 1 | 0 | 0 |
| 0 | 1 | 0 |
| 1 | 1 | 1 |
| 1 | 1 | 1 |
| 0 | 0 | 0 |
| 1 | 1 | 1 |
| 0 | 0 | 0 |
| 0 | 0 | 0 |
| 0 | 0 | 0 |
| 0 | 0 | 0 |
| 0 | 0 | 0 |
| 0 | 0 | 0 |
| 0 | 0 | 0 |
| 0 | 0 | 0 |
| 0 | 0 | 0 |
| 0 | 0 | 0 |
| 0 | 1 | 0 |
| 0 | 1 | 0 |
| 0 | 1 | 0 |
| 1 | 1 | 1 |
| 1 | 1 | 1 |
| 1 | 1 | 1 |
| 1 | 1 | 1 |
| 1 | 1 | 1 |
| 0 | 0 | 0 |
| 1 | 1 | 1 |
| 0 | 0 | 0 |
| 0 | 0 | 0 |
| 0 | 0 | 0 |
| 0 | 0 | 0 |
| 1 | 1 | 1 |
| 1 | 1 | 1 |
| 0 | 1 | 0 |
| 1 | 1 | 1 |
| 1 | 1 | 1 |
| 1 | 1 | 1 |
| 0 | 1 | 0 |
| 1 | 1 | 1 |
| 0 | 0 | 0 |
| 1 | 1 | 1 |
| 1 | 1 | 1 |
| 1 | 1 | 1 |
| 0 | 0 | 0 |
| 0 | 0 | 0 |
| 0 | 0 | 0 |
| 0 | 1 | 0 |
| 0 | 1 | 0 |

| | | |
|---|---|---|
| 0 | 1 | 0 |
| 0 | 1 | 0 |
| 0 | 1 | 0 |
| 0 | 0 | 0 |
| 0 | 1 | 0 |
| 1 | 1 | 1 |
| 0 | 0 | 0 |
| 0 | 1 | 0 |
| 1 | 1 | 1 |
| 1 | 1 | 1 |
| 0 | 0 | 0 |
| 0 | 0 | 0 |
| 1 | 1 | 1 |
| 1 | 1 | 1 |
| 0 | 1 | 0 |
| 0 | 1 | 0 |
| 0 | 0 | 0 |
| 0 | 0 | 0 |
| 1 | 1 | 1 |
| 0 | 0 | 0 |
| 0 | 0 | 0 |
| 0 | 0 | 0 |
| 0 | 1 | 0 |
| 0 | 0 | 0 |
| 0 | 0 | 0 |
| 1 | 1 | 1 |
| 0 | 0 | 0 |
| 0 | 1 | 0 |
| 1 | 1 | 1 |
| 0 | 0 | 0 |
| 0 | 1 | 0 |
| 1 | 1 | 1 |
| 0 | 0 | 0 |
| 0 | 1 | 0 |
| 0 | 1 | 0 |
| 0 | 0 | 0 |
| 1 | 1 | 1 |
| 0 | 0 | 0 |
| 1 | 1 | 1 |
| 0 | 0 | 0 |
| 0 | 0 | 0 |
| 1 | 0 | 0 |
| 0 | 0 | 0 |
| 0 | 0 | 0 |
| 1 | 1 | 1 |
| 1 | 0 | 0 |
| 0 | 1 | 0 |
| 1 | 1 | 1 |
| 1 | 1 | 1 |
| 1 | 1 | 1 |
| 1 | 1 | 1 |
| 0 | 1 | 0 |
| 0 | 1 | 0 |
| 0 | 1 | 0 |

| | | |
|---|---|---|
| 0 | 1 | 0 |
| 0 | 1 | 0 |
| 0 | 1 | 0 |
| 0 | 0 | 0 |
| 1 | 0 | 0 |
| 0 | 0 | 0 |
| 1 | 1 | 1 |
| 1 | 0 | 0 |
| 0 | 0 | 0 |
| 0 | 0 | 0 |
| 1 | 1 | 1 |
| 0 | 0 | 0 |
| 0 | 0 | 0 |
| 0 | 0 | 0 |
| 1 | 0 | 0 |
| 0 | 0 | 0 |
| 0 | 0 | 0 |
| 0 | 1 | 0 |
| 0 | 0 | 0 |
| 0 | 0 | 0 |
| 0 | 0 | 0 |
| 0 | 0 | 0 |
| 1 | 1 | 1 |
| 0 | 0 | 0 |
| 0 | 0 | 0 |
| 0 | 0 | 0 |
| 0 | 0 | 0 |
| 0 | 0 | 0 |
| 0 | 0 | 0 |
| 0 | 0 | 0 |
| 0 | 0 | 0 |
| 0 | 0 | 0 |
| 0 | 0 | 0 |
| 0 | 1 | 0 |
| 0 | 1 | 0 |
| 0 | 0 | 0 |
| 1 | 1 | 1 |
| 0 | 0 | 0 |
| 0 | 0 | 0 |
| 0 | 0 | 0 |
| 0 | 0 | 0 |
| 1 | 1 | 1 |
| 0 | 0 | 0 |
| 0 | 0 | 0 |
| 1 | 1 | 1 |
| 0 | 1 | 0 |
| 1 | 1 | 1 |
| 1 | 1 | 1 |
| 1 | 1 | 1 |
| 0 | 1 | 0 |
| 1 | 1 | 1 |
| 1 | 1 | 1 |
| 1 | 1 | 1 |
| 1 | 1 | 1 |

| | | |
|---|---|---|
| 1 | 1 | 1 |
| 0 | 0 | 0 |
| 1 | 1 | 1 |
| 1 | 1 | 1 |
| 1 | 1 | 1 |
| 1 | 1 | 1 |
| 1 | 1 | 1 |
| 1 | 1 | 1 |
| 1 | 1 | 1 |
| 0 | 1 | 0 |
| 1 | 1 | 1 |
| 0 | 0 | 0 |
| 1 | 0 | 0 |
| 0 | 1 | 0 |
| 0 | 0 | 0 |
| 0 | 1 | 0 |
| 0 | 1 | 0 |
| 1 | 1 | 1 |
| 1 | 1 | 1 |
| 1 | 1 | 1 |
| 0 | 0 | 0 |
| 0 | 1 | 0 |
| 0 | 0 | 0 |
| 0 | 0 | 0 |
| 1 | 1 | 1 |
| 0 | 0 | 0 |
| 1 | 1 | 1 |
| 1 | 1 | 1 |
| 1 | 1 | 1 |
| 1 | 1 | 1 |
| 1 | 1 | 1 |
| 0 | 0 | 0 |
| 1 | 1 | 1 |
| 0 | 0 | 0 |
| 1 | 1 | 1 |
| 0 | 1 | 0 |
| 0 | 1 | 0 |
| 0 | 1 | 0 |
| 1 | 1 | 1 |
| 0 | 0 | 0 |
| 1 | 1 | 1 |
| 1 | 1 | 1 |
| 0 | 1 | 0 |
| 0 | 0 | 0 |
| 1 | 1 | 1 |
| 1 | 1 | 1 |
| 1 | 1 | 1 |
| 0 | 0 | 0 |
| 0 | 0 | 0 |
| 1 | 1 | 1 |
| 1 | 1 | 1 |
| 0 | 0 | 0 |
| 0 | 0 | 0 |
| 0 | 0 | 0 |

| | | |
|---|---|---|
| 0 | 0 | 0 |
| 1 | 1 | 1 |
| 0 | 0 | 0 |
| 0 | 0 | 0 |
| 0 | 0 | 0 |
| 1 | 1 | 1 |
| 1 | 1 | 1 |
| 0 | 1 | 0 |
| 1 | 1 | 1 |
| 0 | 0 | 0 |
| 1 | 1 | 1 |
| 0 | 1 | 0 |
| 1 | 1 | 1 |
| 1 | 1 | 1 |
| 0 | 0 | 0 |
| 0 | 1 | 0 |
| 1 | 1 | 1 |
| 0 | 0 | 0 |
| 1 | 1 | 1 |
| 0 | 0 | 0 |
| 0 | 0 | 0 |
| 1 | 1 | 1 |
| 0 | 0 | 0 |
| 1 | 1 | 1 |
| 0 | 0 | 0 |
| 0 | 0 | 0 |
| 1 | 1 | 1 |
| 0 | 1 | 0 |
| 0 | 0 | 0 |
| 1 | 1 | 1 |
| 1 | 1 | 1 |
| 1 | 1 | 1 |
| 1 | 1 | 1 |
| 0 | 1 | 0 |
| 0 | 0 | 0 |
| 0 | 0 | 0 |
| 1 | 1 | 1 |
| 0 | 0 | 0 |
| 0 | 0 | 0 |
| 0 | 0 | 0 |
| 0 | 0 | 0 |
| 0 | 0 | 0 |
| 0 | 0 | 0 |
| 0 | 1 | 0 |
| 0 | 0 | 0 |
| 1 | 1 | 1 |
| 0 | 1 | 0 |
| 0 | 0 | 0 |
| 0 | 0 | 0 |
| 0 | 0 | 0 |
| 0 | 0 | 0 |
| 0 | 0 | 0 |
| 1 | 1 | 1 |
| 1 | 1 | 1 |

| | | |
|---|---|---|
| 1 | 1 | 1 |
| 0 | 0 | 0 |
| 0 | 0 | 0 |
| 0 | 0 | 0 |
| 0 | 0 | 0 |
| 0 | 0 | 0 |
| 0 | 0 | 0 |
| 1 | 0 | 0 |
| 1 | 0 | 0 |
| 1 | 1 | 1 |
| 0 | 0 | 0 |
| 0 | 0 | 0 |
| 1 | 1 | 1 |
| 0 | 0 | 0 |
| 1 | 1 | 1 |
| 0 | 0 | 0 |
| 1 | 1 | 1 |
| 1 | 1 | 1 |
| 0 | 1 | 0 |
| 0 | 1 | 0 |
| 0 | 1 | 0 |
| 1 | 1 | 1 |
| 0 | 1 | 0 |
| 0 | 0 | 0 |
| 0 | 0 | 0 |
| 1 | 1 | 1 |
| 1 | 1 | 1 |
| 1 | 1 | 1 |
| 1 | 1 | 1 |
| 0 | 0 | 0 |
| 0 | 0 | 0 |
| 0 | 0 | 0 |
| 0 | 0 | 0 |
| 1 | 1 | 1 |
| 0 | 0 | 0 |
| 0 | 0 | 0 |
| 0 | 0 | 0 |
| 0 | 0 | 0 |
| 0 | 0 | 0 |
| 1 | 1 | 1 |
| 1 | 1 | 1 |
| 0 | 0 | 0 |
| 1 | 1 | 1 |
| 1 | 1 | 1 |
| 0 | 0 | 0 |
| 1 | 0 | 0 |
| 1 | 1 | 1 |
| 0 | 1 | 0 |
| 1 | 0 | 0 |
| 0 | 1 | 0 |
| 0 | 0 | 0 |
| 1 | 1 | 1 |
| 0 | 0 | 0 |
| 0 | 0 | 0 |

| | | |
|---|---|---|
| 0 | 1 | 0 |
| 1 | 1 | 1 |
| 0 | 0 | 0 |
| 0 | 0 | 0 |
| 1 | 1 | 1 |
| 1 | 1 | 1 |
| 1 | 1 | 1 |
| 1 | 1 | 1 |
| 1 | 1 | 1 |
| 1 | 1 | 1 |
| 1 | 1 | 1 |
| 1 | 1 | 1 |
| 0 | 0 | 0 |
| 1 | 1 | 1 |
| 1 | 1 | 1 |
| 1 | 1 | 1 |
| 1 | 1 | 1 |
| 1 | 1 | 1 |
| 1 | 1 | 1 |
| 1 | 1 | 1 |
| 1 | 1 | 1 |
| 1 | 1 | 1 |
| 1 | 1 | 1 |
| 1 | 1 | 1 |
| 1 | 1 | 1 |
| 1 | 1 | 1 |
| 0 | 1 | 0 |
| 1 | 1 | 1 |
| 1 | 1 | 1 |
| 1 | 1 | 1 |
| 1 | 1 | 1 |
| 0 | 0 | 0 |
| 1 | 1 | 1 |
| 0 | 1 | 0 |
| 1 | 1 | 1 |
| 1 | 1 | 1 |
| 0 | 1 | 0 |
| 1 | 1 | 1 |
| 1 | 1 | 1 |
| 1 | 1 | 1 |
| 0 | 0 | 0 |
| 0 | 0 | 0 |
| 1 | 1 | 1 |
| 1 | 1 | 1 |
| 1 | 1 | 1 |
| 1 | 0 | 0 |
| 1 | 0 | 0 |
| 1 | 1 | 1 |
| 0 | 0 | 0 |
| 0 | 0 | 0 |
| 1 | 1 | 1 |
| 0 | 1 | 0 |
| 0 | 0 | 0 |
| 1 | 1 | 1 |

| | | |
|---|---|---|
| 0 | 0 | 0 |
| 0 | 0 | 0 |
| 0 | 0 | 0 |
| 1 | 1 | 1 |
| 1 | 1 | 1 |
| 1 | 1 | 1 |
| 1 | 1 | 1 |
| 0 | 0 | 0 |
| 0 | 0 | 0 |
| 0 | 0 | 0 |
| 0 | 0 | 0 |
| 1 | 1 | 1 |
| 0 | 0 | 0 |
| 1 | 1 | 1 |
| 1 | 1 | 1 |
| 1 | 1 | 1 |
| 1 | 1 | 1 |
| 0 | 0 | 0 |
| 1 | 1 | 1 |
| 1 | 1 | 1 |
| 1 | 1 | 1 |
| 1 | 1 | 1 |
| 0 | 1 | 0 |
| 1 | 1 | 1 |
| 1 | 1 | 1 |
| 0 | 1 | 0 |
| 1 | 1 | 1 |
| 1 | 1 | 1 |
| 1 | 1 | 1 |
| 1 | 1 | 1 |
| 0 | 0 | 0 |
| 0 | 0 | 0 |
| 1 | 1 | 1 |
| 1 | 1 | 1 |
| 1 | 1 | 1 |
| 1 | 1 | 1 |
| 1 | 1 | 1 |
| 1 | 1 | 1 |
| 1 | 1 | 1 |
| 1 | 1 | 1 |
| 1 | 1 | 1 |
| 0 | 0 | 0 |
| 1 | 1 | 1 |
| 1 | 1 | 1 |
| 0 | 0 | 0 |
| 0 | 0 | 0 |
| 1 | 1 | 1 |
| 1 | 1 | 1 |
| 0 | 0 | 0 |
| 0 | 0 | 0 |
| 0 | 0 | 0 |
| 0 | 0 | 0 |
| 0 | 0 | 0 |
| 0 | 0 | 0 |

| | | |
|---|---|---|
| 1 | 1 | 1 |
| 1 | 1 | 1 |
| 1 | 1 | 1 |
| 1 | 1 | 1 |
| 1 | 1 | 1 |
| 0 | 0 | 0 |
| 0 | 0 | 0 |
| 0 | 0 | 0 |
| 0 | 0 | 0 |
| 0 | 0 | 0 |
| 0 | 1 | 0 |
| 0 | 0 | 0 |
| 0 | 0 | 0 |
| 0 | 0 | 0 |
| 0 | 1 | 0 |
| 0 | 0 | 0 |
| 0 | 1 | 0 |
| 0 | 1 | 0 |
| 0 | 0 | 0 |
| 0 | 0 | 0 |
| 0 | 0 | 0 |
| 0 | 0 | 0 |
| 1 | 0 | 0 |
| 0 | 0 | 0 |
| 0 | 0 | 0 |
| 0 | 0 | 0 |
| 0 | 0 | 0 |
| 1 | 1 | 1 |
| 1 | 1 | 1 |
| 1 | 1 | 1 |
| 1 | 1 | 1 |
| 1 | 1 | 1 |
| 0 | 0 | 0 |
| 0 | 1 | 0 |
| 0 | 0 | 0 |
| 0 | 0 | 0 |
| 0 | 0 | 0 |
| 0 | 0 | 0 |
| 0 | 0 | 0 |
| 0 | 0 | 0 |
| 0 | 0 | 0 |
| 0 | 0 | 0 |
| 0 | 0 | 0 |
| 1 | 1 | 1 |
| 1 | 1 | 1 |
| 1 | 1 | 1 |
| 1 | 1 | 1 |
| 0 | 0 | 0 |
| 0 | 0 | 0 |
| 0 | 0 | 0 |
| 1 | 1 | 1 |
| 1 | 1 | 1 |
| 0 | 0 | 0 |
| 1 | 1 | 1 |

| | | |
|---|---|---|
| 1 | 1 | 1 |
| 0 | 0 | 0 |
| 0 | 1 | 0 |
| 0 | 1 | 0 |
| 1 | 1 | 1 |
| 1 | 1 | 1 |
| 0 | 1 | 0 |
| 1 | 1 | 1 |
| 0 | 1 | 0 |
| 1 | 1 | 1 |
| 1 | 1 | 1 |
| 1 | 1 | 1 |
| 0 | 0 | 0 |
| 0 | 0 | 0 |
| 0 | 0 | 0 |
| 0 | 0 | 0 |
| 0 | 0 | 0 |
| 0 | 0 | 0 |
| 0 | 0 | 0 |
| 0 | 1 | 0 |
| 0 | 0 | 0 |
| 0 | 0 | 0 |
| 0 | 0 | 0 |
| 0 | 0 | 0 |
| 0 | 0 | 0 |
| 1 | 1 | 1 |
| 1 | 1 | 1 |
| 0 | 0 | 0 |
| 0 | 1 | 0 |
| 0 | 0 | 0 |
| 0 | 0 | 0 |
| 1 | 1 | 1 |
| 0 | 0 | 0 |
| 1 | 1 | 1 |
| 1 | 1 | 1 |
| 1 | 1 | 1 |
| 1 | 0 | 0 |
| 1 | 1 | 1 |
| 1 | 1 | 1 |
| 1 | 1 | 1 |
| 0 | 0 | 0 |
| 1 | 1 | 1 |
| 1 | 1 | 1 |
| 0 | 0 | 0 |
| 0 | 0 | 0 |
| 1 | 1 | 1 |
| 0 | 0 | 0 |
| 0 | 0 | 0 |
| 1 | 1 | 1 |
| 1 | 1 | 1 |
| 0 | 0 | 0 |
| 1 | 1 | 1 |
| 1 | 1 | 1 |
| 1 | 1 | 1 |

| | | |
|---|---|---|
| 1 | 1 | 1 |
| 1 | 1 | 1 |
| 1 | 1 | 1 |
| 1 | 1 | 1 |
| 0 | 0 | 0 |
| 0 | 0 | 0 |
| 1 | 1 | 1 |
| 1 | 1 | 1 |
| 1 | 1 | 1 |
| 0 | 1 | 0 |
| 1 | 1 | 1 |
| 0 | 0 | 0 |
| 0 | 0 | 0 |
| 1 | 1 | 1 |
| 1 | 1 | 1 |
| 0 | 0 | 0 |
| 0 | 1 | 0 |
| 1 | 1 | 1 |
| 0 | 0 | 0 |
| 1 | 1 | 1 |
| 1 | 1 | 1 |
| 1 | 1 | 1 |
| 1 | 1 | 1 |
| 0 | 0 | 0 |
| 1 | 1 | 1 |
| 0 | 0 | 0 |
| 1 | 1 | 1 |
| 1 | 1 | 1 |
| 0 | 0 | 0 |
| 0 | 0 | 0 |
| 0 | 0 | 0 |
| 0 | 0 | 0 |
| 0 | 0 | 0 |
| 0 | 0 | 0 |
| 1 | 1 | 1 |
| 1 | 1 | 1 |
| 1 | 1 | 1 |
| 0 | 0 | 0 |
| 1 | 1 | 1 |
| 1 | 1 | 1 |
| 0 | 0 | 0 |
| 0 | 0 | 0 |
| 0 | 0 | 0 |
| 1 | 1 | 1 |
| 0 | 0 | 0 |
| 0 | 1 | 0 |
| 0 | 1 | 0 |
| 0 | 0 | 0 |
| 1 | 1 | 1 |
| 1 | 1 | 1 |
| 1 | 1 | 1 |
| 0 | 0 | 0 |
| 1 | 1 | 1 |
| 0 | 0 | 0 |

| | | |
|---|---|---|
| 1 | 1 | 1 |
| 0 | 0 | 0 |
| 0 | 0 | 0 |
| 0 | 1 | 0 |
| 1 | 1 | 1 |
| 1 | 1 | 1 |
| 0 | 0 | 0 |
| 1 | 1 | 1 |
| 0 | 1 | 0 |
| 0 | 1 | 0 |
| 1 | 1 | 1 |
| 1 | 1 | 1 |
| 1 | 1 | 1 |
| 1 | 1 | 1 |
| 1 | 1 | 1 |
| 1 | 1 | 1 |
| 1 | 1 | 1 |
| 1 | 1 | 1 |
| 1 | 1 | 1 |
| 1 | 1 | 1 |
| 1 | 1 | 1 |
| 0 | 0 | 0 |
| 0 | 1 | 0 |
| 0 | 1 | 0 |
| 1 | 1 | 1 |
| 1 | 1 | 1 |
| 0 | 0 | 0 |
| 1 | 1 | 1 |
| 0 | 0 | 0 |
| 0 | 1 | 0 |
| 1 | 1 | 1 |
| 0 | 0 | 0 |
| 0 | 0 | 0 |
| 1 | 0 | 0 |
| 1 | 1 | 1 |
| 1 | 1 | 1 |
| 1 | 1 | 1 |
| 1 | 1 | 1 |
| 0 | 0 | 0 |
| 0 | 0 | 0 |
| 0 | 0 | 0 |
| 1 | 1 | 1 |
| 1 | 1 | 1 |
| 0 | 1 | 0 |
| 0 | 1 | 0 |
| 0 | 0 | 0 |
| 1 | 1 | 1 |
| 0 | 0 | 0 |
| 1 | 1 | 1 |
| 0 | 0 | 0 |
| 0 | 0 | 0 |
| 0 | 0 | 0 |
| 1 | 1 | 1 |
| 1 | 1 | 1 |

| | | |
|---|---|---|
| 0 | 0 | 0 |
| 0 | 0 | 0 |
| 1 | 1 | 1 |
| 1 | 1 | 1 |
| 1 | 1 | 1 |
| 1 | 1 | 1 |
| 1 | 1 | 1 |
| 1 | 1 | 1 |
| 1 | 1 | 1 |
| 1 | 1 | 1 |
| 1 | 1 | 1 |
| 1 | 1 | 1 |
| 1 | 1 | 1 |
| 1 | 1 | 1 |
| 1 | 1 | 1 |
| 0 | 0 | 0 |
| 0 | 0 | 0 |
| 0 | 0 | 0 |
| 1 | 1 | 1 |
| 1 | 1 | 1 |
| 0 | 1 | 0 |
| 0 | 0 | 0 |
| 0 | 0 | 0 |
| 0 | 0 | 0 |
| 0 | 0 | 0 |
| 1 | 1 | 1 |
| 1 | 1 | 1 |
| 1 | 1 | 1 |
| 1 | 1 | 1 |
| 1 | 1 | 1 |
| 0 | 0 | 0 |
| 0 | 1 | 0 |
| 0 | 0 | 0 |
| 1 | 1 | 1 |
| 1 | 0 | 0 |
| 1 | 0 | 0 |
| 0 | 0 | 0 |
| 0 | 0 | 0 |
| 0 | 1 | 0 |
| 1 | 1 | 1 |
| 1 | 0 | 0 |
| 0 | 0 | 0 |
| 1 | 1 | 1 |
| 0 | 1 | 0 |
| 0 | 0 | 0 |
| 1 | 0 | 0 |
| 0 | 0 | 0 |
| 0 | 0 | 0 |
| 1 | 1 | 1 |
| 0 | 1 | 0 |
| 0 | 0 | 0 |
| 0 | 0 | 0 |
| 1 | 1 | 1 |
| 0 | 0 | 0 |

| | | |
|---|---|---|
| 0 | 1 | 0 |
| 0 | 0 | 0 |
| 1 | 0 | 0 |
| 0 | 0 | 0 |
| 1 | 1 | 1 |
| 1 | 1 | 1 |
| 1 | 1 | 1 |
| 0 | 1 | 0 |
| 0 | 0 | 0 |
| 1 | 1 | 1 |
| 0 | 1 | 0 |
| 0 | 1 | 0 |
| 0 | 0 | 0 |
| 0 | 0 | 0 |
| 0 | 0 | 0 |
| 1 | 1 | 1 |
| 0 | 0 | 0 |
| 0 | 1 | 0 |
| 0 | 0 | 0 |
| 0 | 0 | 0 |
| 0 | 0 | 0 |
| 0 | 0 | 0 |
| 0 | 0 | 0 |
| 0 | 0 | 0 |
| 0 | 1 | 0 |
| 0 | 1 | 0 |
| 0 | 0 | 0 |
| 0 | 0 | 0 |
| 0 | 0 | 0 |
| 0 | 0 | 0 |
| 1 | 1 | 1 |
| 1 | 1 | 1 |
| 1 | 1 | 1 |
| 1 | 1 | 1 |
| 1 | 1 | 1 |
| 0 | 0 | 0 |
| 0 | 0 | 0 |
| 0 | 1 | 0 |
| 1 | 1 | 1 |
| 1 | 1 | 1 |
| 0 | 1 | 0 |
| 1 | 1 | 1 |
| 1 | 1 | 1 |
| 0 | 0 | 0 |
| 0 | 0 | 0 |
| 0 | 0 | 0 |
| 1 | 1 | 1 |
| 0 | 0 | 0 |
| 0 | 0 | 0 |
| 0 | 0 | 0 |
| 1 | 1 | 1 |
| 1 | 1 | 1 |
| 0 | 1 | 0 |
| 0 | 1 | 0 |

| | | |
|---|---|---|
| 0 | 1 | 0 |
| 1 | 1 | 1 |
| 1 | 0 | 0 |
| 1 | 1 | 1 |
| 0 | 0 | 0 |
| 1 | 0 | 0 |
| 0 | 0 | 0 |
| 0 | 0 | 0 |
| 0 | 0 | 0 |
| 0 | 0 | 0 |
| 0 | 0 | 0 |
| 0 | 0 | 0 |
| 0 | 1 | 0 |
| 0 | 0 | 0 |
| 0 | 0 | 0 |
| 1 | 1 | 1 |
| 1 | 1 | 1 |
| 1 | 1 | 1 |
| 0 | 0 | 0 |
| 0 | 0 | 0 |
| 0 | 0 | 0 |
| 0 | 0 | 0 |
| 0 | 1 | 0 |
| 0 | 1 | 0 |
| 1 | 1 | 1 |
| 1 | 1 | 1 |
| 1 | 1 | 1 |
| 0 | 0 | 0 |
| 0 | 0 | 0 |
| 0 | 0 | 0 |
| 0 | 0 | 0 |
| 0 | 0 | 0 |
| 0 | 0 | 0 |
| 0 | 1 | 0 |
| 1 | 1 | 1 |
| 0 | 0 | 0 |
| 1 | 1 | 1 |
| 1 | 1 | 1 |
| 1 | 1 | 1 |
| 1 | 1 | 1 |
| 0 | 0 | 0 |
| 1 | 1 | 1 |
| 0 | 0 | 0 |
| 0 | 1 | 0 |
| 1 | 1 | 1 |
| 1 | 1 | 1 |
| 1 | 1 | 1 |
| 0 | 0 | 0 |
| 1 | 1 | 1 |
| 1 | 1 | 1 |
| 1 | 1 | 1 |
| 1 | 1 | 1 |
| 1 | 1 | 1 |
| 0 | 1 | 0 |

| | | |
|---|---|---|
| 0 | 1 | 0 |
| 1 | 1 | 1 |
| 1 | 1 | 1 |
| 0 | 1 | 0 |
| 1 | 1 | 1 |
| 0 | 1 | 0 |
| 0 | 1 | 0 |
| 0 | 1 | 0 |
| 0 | 1 | 0 |
| 0 | 1 | 0 |
| 0 | 1 | 0 |
| 0 | 0 | 0 |
| 0 | 1 | 0 |
| 0 | 1 | 0 |
| 0 | 1 | 0 |
| 0 | 1 | 0 |
| 0 | 1 | 0 |
| 0 | 0 | 0 |
| 0 | 0 | 0 |
| 0 | 0 | 0 |
| 0 | 0 | 0 |
| 0 | 1 | 0 |
| 0 | 0 | 0 |
| 1 | 1 | 1 |
| 1 | 1 | 1 |
| 1 | 1 | 1 |
| 1 | 1 | 1 |
| 0 | 1 | 0 |
| 0 | 0 | 0 |
| 0 | 0 | 0 |
| 1 | 1 | 1 |
| 1 | 1 | 1 |
| 0 | 0 | 0 |
| 1 | 1 | 1 |
| 0 | 0 | 0 |
| 0 | 0 | 0 |
| 0 | 0 | 0 |
| 1 | 1 | 1 |
| 0 | 0 | 0 |
| 1 | 1 | 1 |
| 0 | 0 | 0 |
| 1 | 1 | 1 |
| 0 | 0 | 0 |
| 1 | 0 | 0 |
| 1 | 0 | 0 |
| 1 | 0 | 0 |
| 1 | 0 | 0 |
| 0 | 1 | 0 |
| 0 | 1 | 0 |
| 0 | 0 | 0 |
| 1 | 0 | 0 |
| 0 | 0 | 0 |
| 1 | 0 | 0 |
| 1 | 1 | 1 |

| | | |
|---|---|---|
| 0 | 0 | 0 |
| 0 | 0 | 0 |
| 0 | 0 | 0 |
| 0 | 0 | 0 |
| 0 | 0 | 0 |
| 1 | 1 | 1 |
| 0 | 0 | 0 |
| 0 | 0 | 0 |
| 1 | 1 | 1 |
| 0 | 0 | 0 |
| 0 | 0 | 0 |
| 0 | 0 | 0 |
| 0 | 1 | 0 |
| 0 | 1 | 0 |
| 0 | 1 | 0 |
| 0 | 1 | 0 |
| 0 | 1 | 0 |
| 0 | 1 | 0 |
| 0 | 1 | 0 |
| 0 | 1 | 0 |
| 0 | 1 | 0 |
| 0 | 1 | 0 |
| 0 | 1 | 0 |
| 0 | 1 | 0 |
| 0 | 1 | 0 |
| 0 | 1 | 0 |
| 0 | 1 | 0 |
| 0 | 1 | 0 |
| 0 | 1 | 0 |
| 0 | 1 | 0 |
| 0 | 1 | 0 |
| 0 | 1 | 0 |
| 0 | 1 | 0 |
| 0 | 1 | 0 |
| 0 | 1 | 0 |
| 0 | 1 | 0 |
| 0 | 1 | 0 |
| 0 | 1 | 0 |
| 0 | 1 | 0 |
| 0 | 1 | 0 |
| 0 | 1 | 0 |
| 0 | 1 | 0 |
| 0 | 1 | 0 |
| 0 | 1 | 0 |
| 0 | 1 | 0 |
| 0 | 0 | 0 |
| 1 | 1 | 1 |
| 1 | 1 | 1 |
| 1 | 1 | 1 |
| 1 | 1 | 1 |
| 0 | 0 | 0 |
| 1 | 1 | 1 |
| 1 | 1 | 1 |
| 0 | 1 | 0 |

| | | |
|---|---|---|
| 1 | 1 | 1 |
| 1 | 1 | 1 |
| 1 | 1 | 1 |
| 1 | 1 | 1 |
| 0 | 1 | 0 |
| 0 | 1 | 0 |
| 0 | 1 | 0 |
| 0 | 0 | 0 |
| 0 | 0 | 0 |
| 1 | 1 | 1 |
| 0 | 1 | 0 |
| 1 | 1 | 1 |
| 1 | 1 | 1 |
| 1 | 1 | 1 |
| 0 | 0 | 0 |
| 0 | 0 | 0 |
| 0 | 0 | 0 |
| 1 | 1 | 1 |
| 1 | 0 | 0 |
| 1 | 1 | 1 |
| 0 | 0 | 0 |
| 0 | 0 | 0 |
| 0 | 0 | 0 |
| 0 | 0 | 0 |
| 0 | 0 | 0 |
| 0 | 0 | 0 |
| 0 | 1 | 0 |
| 1 | 1 | 1 |
| 0 | 0 | 0 |
| 1 | 1 | 1 |
| 1 | 1 | 1 |
| 1 | 1 | 1 |
| 0 | 0 | 0 |
| 1 | 1 | 1 |
| 1 | 1 | 1 |
| 1 | 1 | 1 |
| 0 | 0 | 0 |
| 0 | 0 | 0 |
| 0 | 0 | 0 |
| 1 | 1 | 1 |
| 1 | 1 | 1 |
| 1 | 1 | 1 |
| 1 | 1 | 1 |
| 1 | 1 | 1 |
| 0 | 1 | 0 |
| 0 | 0 | 0 |
| 1 | 1 | 1 |
| 0 | 1 | 0 |
| 1 | 1 | 1 |
| 0 | 1 | 0 |
| 1 | 1 | 1 |
| 1 | 1 | 1 |
| 0 | 1 | 0 |
| 1 | 1 | 1 |

| | | |
|---|---|---|
| 1 | 1 | 1 |
| 0 | 1 | 0 |
| 1 | 1 | 1 |
| 0 | 0 | 0 |
| 1 | 1 | 1 |
| 1 | 1 | 1 |
| 1 | 1 | 1 |
| 0 | 0 | 0 |
| 0 | 0 | 0 |
| 1 | 1 | 1 |
| 1 | 1 | 1 |
| 1 | 1 | 1 |
| 0 | 0 | 0 |
| 0 | 1 | 0 |
| 0 | 1 | 0 |
| 0 | 1 | 0 |
| 0 | 0 | 0 |
| 1 | 1 | 1 |
| 1 | 1 | 1 |
| 1 | 1 | 1 |
| 1 | 1 | 1 |
| 1 | 1 | 1 |
| 1 | 1 | 1 |
| 1 | 1 | 1 |
| 0 | 1 | 0 |
| 1 | 1 | 1 |
| 0 | 1 | 0 |
| 0 | 0 | 0 |
| 0 | 0 | 0 |
| 0 | 0 | 0 |
| 0 | 0 | 0 |
| 0 | 0 | 0 |
| 0 | 0 | 0 |
| 0 | 1 | 0 |
| 0 | 0 | 0 |
| 0 | 0 | 0 |
| 0 | 0 | 0 |
| 1 | 1 | 1 |
| 1 | 1 | 1 |
| 1 | 1 | 1 |
| 1 | 1 | 1 |
| 0 | 1 | 0 |
| 0 | 0 | 0 |
| 0 | 0 | 0 |
| 1 | 1 | 1 |
| 1 | 1 | 1 |
| 0 | 0 | 0 |
| 1 | 1 | 1 |
| 1 | 1 | 1 |
| 1 | 1 | 1 |
| 1 | 1 | 1 |
| 1 | 1 | 1 |
| 0 | 0 | 0 |
| 0 | 0 | 0 |

| | | |
|---|---|---|
| 0 | 0 | 0 |
| 0 | 0 | 0 |
| 0 | 0 | 0 |
| 0 | 0 | 0 |
| 0 | 0 | 0 |
| 0 | 0 | 0 |
| 0 | 0 | 0 |
| 0 | 0 | 0 |
| 1 | 1 | 1 |
| 0 | 0 | 0 |
| 1 | 1 | 1 |
| 1 | 1 | 1 |
| 1 | 1 | 1 |
| 1 | 1 | 1 |
| 0 | 0 | 0 |
| 1 | 1 | 1 |
| 1 | 1 | 1 |
| 1 | 1 | 1 |
| 1 | 1 | 1 |
| 0 | 0 | 0 |
| 0 | 0 | 0 |
| 1 | 1 | 1 |
| 0 | 0 | 0 |
| 0 | 0 | 0 |
| 0 | 0 | 0 |
| 0 | 0 | 0 |
| 0 | 0 | 0 |
| 0 | 1 | 0 |
| 0 | 1 | 0 |
| 0 | 1 | 0 |
| 0 | 0 | 0 |
| 1 | 1 | 1 |
| 1 | 1 | 1 |
| 1 | 1 | 1 |
| 1 | 1 | 1 |
| 1 | 1 | 1 |
| 1 | 1 | 1 |
| 1 | 1 | 1 |
| 1 | 1 | 1 |
| 1 | 1 | 1 |
| 0 | 1 | 0 |
| 0 | 0 | 0 |
| 1 | 1 | 1 |
| 1 | 1 | 1 |
| 1 | 1 | 1 |
| 0 | 1 | 0 |
| 1 | 1 | 1 |
| 1 | 1 | 1 |
| 0 | 0 | 0 |
| 0 | 1 | 0 |
| 1 | 1 | 1 |
| 1 | 1 | 1 |
| 0 | 0 | 0 |
| 0 | 0 | 0 |

| | | |
|---|---|---|
| 1 | 1 | 1 |
| 1 | 1 | 1 |
| 0 | 1 | 0 |
| 1 | 0 | 0 |
| 0 | 0 | 0 |
| 0 | 0 | 0 |
| 0 | 1 | 0 |
| 0 | 0 | 0 |
| 1 | 1 | 1 |
| 1 | 1 | 1 |
| 0 | 0 | 0 |
| 1 | 0 | 0 |
| 0 | 0 | 0 |
| 0 | 0 | 0 |
| 0 | 0 | 0 |
| 0 | 0 | 0 |
| 1 | 1 | 1 |
| 1 | 1 | 1 |
| 0 | 0 | 0 |
| 0 | 0 | 0 |
| 0 | 0 | 0 |
| 0 | 0 | 0 |
| 0 | 0 | 0 |
| 1 | 1 | 1 |
| 1 | 1 | 1 |
| 0 | 0 | 0 |
| 1 | 1 | 1 |
| 0 | 0 | 0 |
| 0 | 0 | 0 |
| 0 | 0 | 0 |
| 0 | 1 | 0 |
| 0 | 0 | 0 |
| 0 | 0 | 0 |
| 1 | 1 | 1 |
| 1 | 1 | 1 |
| 0 | 0 | 0 |
| 0 | 0 | 0 |
| 1 | 0 | 0 |
| 0 | 0 | 0 |
| 0 | 0 | 0 |
| 0 | 0 | 0 |
| 0 | 0 | 0 |
| 0 | 1 | 0 |
| 1 | 1 | 1 |
| 0 | 0 | 0 |
| 0 | 1 | 0 |
| 0 | 0 | 0 |
| 1 | 1 | 1 |
| 1 | 1 | 1 |
| 1 | 1 | 1 |
| 1 | 1 | 1 |
| 1 | 1 | 1 |
| 0 | 1 | 0 |
| 1 | 1 | 1 |

| | | |
|---|---|---|
| 1 | 1 | 1 |
| 1 | 1 | 1 |
| 1 | 1 | 1 |
| 0 | 1 | 0 |
| 1 | 1 | 1 |
| 1 | 1 | 1 |
| 0 | 1 | 0 |
| 0 | 0 | 0 |
| 1 | 1 | 1 |
| 0 | 0 | 0 |
| 0 | 0 | 0 |
| 0 | 0 | 0 |
| 0 | 1 | 0 |
| 1 | 1 | 1 |
| 0 | 0 | 0 |
| 0 | 1 | 0 |
| 0 | 1 | 0 |
| 0 | 1 | 0 |
| 0 | 1 | 0 |
| 0 | 1 | 0 |
| 1 | 1 | 1 |
| 1 | 1 | 1 |
| 1 | 1 | 1 |
| 0 | 1 | 0 |
| 1 | 1 | 1 |
| 1 | 1 | 1 |
| 1 | 1 | 1 |
| 1 | 0 | 0 |
| 1 | 1 | 1 |
| 0 | 0 | 0 |
| 0 | 0 | 0 |
| 1 | 1 | 1 |
| 1 | 1 | 1 |
| 0 | 1 | 0 |
| 0 | 1 | 0 |
| 0 | 0 | 0 |
| 1 | 1 | 1 |
| 1 | 1 | 1 |
| 1 | 1 | 1 |
| 1 | 1 | 1 |
| 1 | 1 | 1 |
| 0 | 0 | 0 |
| 0 | 0 | 0 |
| 0 | 0 | 0 |
| 1 | 1 | 1 |
| 1 | 1 | 1 |
| 1 | 1 | 1 |
| 1 | 1 | 1 |
| 1 | 1 | 1 |
| 1 | 1 | 1 |
| 1 | 1 | 1 |
| 1 | 1 | 1 |
| 1 | 1 | 1 |
| 0 | 0 | 0 |

| | | |
|---|---|---|
| 1 | 1 | 1 |
| 1 | 1 | 1 |
| 1 | 1 | 1 |
| 1 | 1 | 1 |
| 1 | 1 | 1 |
| 1 | 1 | 1 |
| 1 | 1 | 1 |
| 0 | 1 | 0 |
| 0 | 0 | 0 |
| 0 | 0 | 0 |
| 1 | 1 | 1 |
| 0 | 1 | 0 |
| 0 | 1 | 0 |
| 0 | 0 | 0 |
| 1 | 1 | 1 |
| 1 | 1 | 1 |
| 1 | 1 | 1 |
| 0 | 0 | 0 |
| 0 | 0 | 0 |
| 0 | 0 | 0 |
| 0 | 0 | 0 |
| 0 | 1 | 0 |
| 1 | 1 | 1 |
| 1 | 1 | 1 |
| 1 | 1 | 1 |
| 0 | 0 | 0 |
| 1 | 1 | 1 |
| 0 | 1 | 0 |
| 1 | 1 | 1 |
| 1 | 1 | 1 |
| 1 | 1 | 1 |
| 1 | 1 | 1 |
| 1 | 1 | 1 |
| 1 | 1 | 1 |
| 1 | 1 | 1 |
| 1 | 1 | 1 |
| 1 | 1 | 1 |
| 1 | 1 | 1 |
| 1 | 1 | 1 |
| 0 | 0 | 0 |
| 1 | 1 | 1 |
| 1 | 1 | 1 |
| 0 | 0 | 0 |
| 1 | 1 | 1 |
| 1 | 1 | 1 |
| 1 | 1 | 1 |
| 0 | 0 | 0 |
| 0 | 0 | 0 |
| 1 | 1 | 1 |
| 1 | 1 | 1 |
| 0 | 0 | 0 |
| 1 | 1 | 1 |
| 0 | 1 | 0 |
| 1 | 1 | 1 |

| | | |
|---|---|---|
| 1 | 1 | 1 |
| 1 | 1 | 1 |
| 1 | 1 | 1 |
| 1 | 1 | 1 |
| 1 | 1 | 1 |
| 1 | 1 | 1 |
| 1 | 1 | 1 |
| 0 | 1 | 0 |
| 1 | 1 | 1 |
| 1 | 1 | 1 |
| 1 | 1 | 1 |
| 1 | 1 | 1 |
| 1 | 1 | 1 |
| 0 | 0 | 0 |
| 0 | 0 | 0 |
| 0 | 0 | 0 |
| 0 | 0 | 0 |
| 0 | 1 | 0 |
| 1 | 1 | 1 |
| 0 | 0 | 0 |
| 1 | 1 | 1 |
| 1 | 0 | 0 |
| 1 | 1 | 1 |
| 1 | 1 | 1 |
| 1 | 1 | 1 |
| 0 | 0 | 0 |
| 0 | 1 | 0 |
| 1 | 1 | 1 |
| 1 | 1 | 1 |
| 0 | 0 | 0 |
| 0 | 0 | 0 |
| 1 | 1 | 1 |
| 1 | 1 | 1 |
| 1 | 1 | 1 |
| 1 | 1 | 1 |
| 1 | 1 | 1 |
| 1 | 1 | 1 |
| 1 | 1 | 1 |
| 0 | 0 | 0 |
| 1 | 1 | 1 |
| 1 | 1 | 1 |
| 0 | 0 | 0 |
| 0 | 0 | 0 |
| 0 | 0 | 0 |
| 0 | 0 | 0 |
| 1 | 1 | 1 |
| 0 | 0 | 0 |
| 1 | 1 | 1 |
| 1 | 1 | 1 |
| 1 | 1 | 1 |
| 0 | 1 | 0 |
| 1 | 0 | 0 |
| 1 | 0 | 0 |
| 1 | 0 | 0 |

| | | |
|---|---|---|
| 1 | 1 | 1 |
| 1 | 1 | 1 |
| 0 | 0 | 0 |
| 1 | 1 | 1 |
| 1 | 1 | 1 |
| 1 | 1 | 1 |
| 1 | 1 | 1 |
| 0 | 1 | 0 |
| 1 | 1 | 1 |
| 1 | 1 | 1 |
| 1 | 1 | 1 |
| 0 | 1 | 0 |
| 0 | 1 | 0 |
| 0 | 1 | 0 |
| 0 | 1 | 0 |
| 0 | 1 | 0 |
| 0 | 1 | 0 |
| 0 | 1 | 0 |
| 0 | 1 | 0 |
| 0 | 1 | 0 |
| 0 | 1 | 0 |
| 0 | 1 | 0 |
| 0 | 1 | 0 |
| 0 | 1 | 0 |
| 0 | 1 | 0 |
| 0 | 1 | 0 |
| 0 | 1 | 0 |
| 0 | 1 | 0 |
| 1 | 1 | 1 |
| 1 | 1 | 1 |
| 0 | 0 | 0 |
| 1 | 1 | 1 |
| 0 | 0 | 0 |
| 0 | 0 | 0 |
| 0 | 0 | 0 |
| 0 | 0 | 0 |
| 1 | 1 | 1 |
| 0 | 0 | 0 |
| 1 | 1 | 1 |
| 1 | 1 | 1 |
| 0 | 0 | 0 |
| 0 | 0 | 0 |
| 0 | 0 | 0 |
| 1 | 1 | 1 |
| 0 | 0 | 0 |
| 0 | 0 | 0 |
| 1 | 1 | 1 |
| 1 | 1 | 1 |
| 1 | 1 | 1 |
| 1 | 1 | 1 |
| 0 | 0 | 0 |
| 1 | 1 | 1 |
| 1 | 1 | 1 |
| 1 | 1 | 1 |

| | | |
|---|---|---|
| 1 | 1 | 1 |
| 1 | 1 | 1 |
| 1 | 1 | 1 |
| 1 | 1 | 1 |
| 1 | 1 | 1 |
| 0 | 0 | 0 |
| 1 | 1 | 1 |
| 0 | 0 | 0 |
| 0 | 0 | 0 |
| 0 | 0 | 0 |
| 1 | 1 | 1 |
| 0 | 0 | 0 |
| 1 | 1 | 1 |
| 1 | 1 | 1 |
| 1 | 1 | 1 |
| 1 | 0 | 0 |
| 1 | 1 | 1 |
| 0 | 0 | 0 |
| 1 | 1 | 1 |
| 0 | 0 | 0 |
| 1 | 1 | 1 |
| 0 | 0 | 0 |
| 0 | 0 | 0 |
| 0 | 0 | 0 |
| 1 | 1 | 1 |
| 0 | 1 | 0 |
| 1 | 1 | 1 |
| 1 | 1 | 1 |
| 0 | 0 | 0 |
| 1 | 1 | 1 |
| 1 | 1 | 1 |
| 1 | 1 | 1 |
| 1 | 1 | 1 |
| 1 | 1 | 1 |
| 0 | 0 | 0 |
| 0 | 0 | 0 |
| 1 | 1 | 1 |
| 0 | 0 | 0 |
| 0 | 0 | 0 |
| 0 | 1 | 0 |
| 0 | 0 | 0 |
| 0 | 0 | 0 |
| 0 | 1 | 0 |
| 0 | 1 | 0 |
| 1 | 1 | 1 |
| 0 | 1 | 0 |
| 1 | 1 | 1 |
| 1 | 1 | 1 |
| 1 | 1 | 1 |
| 1 | 1 | 1 |
| 1 | 1 | 1 |
| 1 | 1 | 1 |
| 1 | 1 | 1 |
| 1 | 1 | 1 |

| | | |
|---|---|---|
| 1 | 1 | 1 |
| 0 | 0 | 0 |
| 0 | 1 | 0 |
| 1 | 1 | 1 |
| 0 | 0 | 0 |
| 1 | 1 | 1 |
| 1 | 1 | 1 |
| 1 | 1 | 1 |
| 0 | 0 | 0 |
| 0 | 0 | 0 |
| 1 | 1 | 1 |
| 0 | 0 | 0 |
| 0 | 0 | 0 |
| 1 | 1 | 1 |
| 1 | 1 | 1 |
| 0 | 0 | 0 |
| 0 | 1 | 0 |
| 0 | 0 | 0 |
| 0 | 0 | 0 |
| 1 | 1 | 1 |
| 0 | 0 | 0 |
| 0 | 0 | 0 |
| 0 | 0 | 0 |
| 1 | 1 | 1 |
| 0 | 1 | 0 |
| 0 | 0 | 0 |
| 1 | 1 | 1 |
| 1 | 1 | 1 |
| 1 | 1 | 1 |
| 1 | 1 | 1 |
| 0 | 0 | 0 |
| 1 | 1 | 1 |
| 0 | 0 | 0 |
| 1 | 1 | 1 |
| 0 | 0 | 0 |
| 0 | 0 | 0 |
| 1 | 1 | 1 |
| 1 | 1 | 1 |
| 1 | 1 | 1 |
| 1 | 1 | 1 |
| 1 | 1 | 1 |
| 1 | 1 | 1 |
| 1 | 1 | 1 |
| 1 | 1 | 1 |
| 1 | 1 | 1 |
| 1 | 1 | 1 |
| 1 | 0 | 0 |
| 1 | 1 | 1 |
| 1 | 1 | 1 |
| 1 | 1 | 1 |
| 1 | 1 | 1 |
| 1 | 1 | 1 |
| 1 | 1 | 1 |
| 0 | 0 | 0 |

| | | |
|---|---|---|
| 1 | 1 | 1 |
| 1 | 1 | 1 |
| 1 | 1 | 1 |
| 1 | 1 | 1 |
| 0 | 0 | 0 |
| 1 | 1 | 1 |
| 0 | 1 | 0 |
| 0 | 1 | 0 |
| 0 | 1 | 0 |
| 1 | 1 | 1 |
| 0 | 0 | 0 |
| 1 | 1 | 1 |
| 1 | 1 | 1 |
| 1 | 1 | 1 |
| 0 | 0 | 0 |
| 0 | 0 | 0 |
| 0 | 0 | 0 |
| 0 | 0 | 0 |
| 0 | 1 | 0 |
| 0 | 0 | 0 |
| 0 | 0 | 0 |
| 0 | 0 | 0 |
| 1 | 1 | 1 |
| 0 | 0 | 0 |
| 0 | 0 | 0 |
| 1 | 1 | 1 |
| 0 | 0 | 0 |
| 0 | 0 | 0 |
| 0 | 0 | 0 |
| 0 | 0 | 0 |
| 0 | 0 | 0 |
| 1 | 1 | 1 |
| 1 | 1 | 1 |
| 0 | 1 | 0 |
| 1 | 1 | 1 |
| 1 | 1 | 1 |
| 1 | 1 | 1 |
| 0 | 0 | 0 |
| 0 | 0 | 0 |
| 0 | 1 | 0 |
| 1 | 1 | 1 |
| 0 | 0 | 0 |
| 0 | 0 | 0 |
| 0 | 1 | 0 |
| 0 | 0 | 0 |
| 1 | 1 | 1 |
| 1 | 1 | 1 |
| 1 | 1 | 1 |
| 0 | 0 | 0 |
| 0 | 0 | 0 |
| 0 | 0 | 0 |
| 1 | 1 | 1 |
| 0 | 1 | 0 |
| 0 | 0 | 0 |

| | | |
|---|---|---|
| 1 | 1 | 1 |
| 0 | 0 | 0 |
| 1 | 1 | 1 |
| 0 | 0 | 0 |
| 0 | 0 | 0 |
| 0 | 1 | 0 |
| 0 | 0 | 0 |
| 0 | 0 | 0 |
| 1 | 1 | 1 |
| 1 | 1 | 1 |
| 1 | 1 | 1 |
| 1 | 1 | 1 |
| 1 | 1 | 1 |
| 1 | 1 | 1 |
| 1 | 1 | 1 |
| 1 | 1 | 1 |
| 1 | 1 | 1 |
| 0 | 1 | 0 |
| 1 | 1 | 1 |
| 1 | 1 | 1 |
| 0 | 0 | 0 |
| 0 | 1 | 0 |
| 0 | 1 | 0 |
| 0 | 1 | 0 |
| 0 | 1 | 0 |
| 0 | 0 | 0 |
| 0 | 0 | 0 |
| 1 | 1 | 1 |
| 1 | 1 | 1 |
| 1 | 1 | 1 |
| 1 | 1 | 1 |
| 1 | 1 | 1 |
| 1 | 1 | 1 |
| 0 | 1 | 0 |
| 1 | 1 | 1 |
| 1 | 1 | 1 |
| 1 | 1 | 1 |
| 1 | 1 | 1 |
| 0 | 1 | 0 |
| 1 | 1 | 1 |
| 1 | 1 | 1 |
| 0 | 1 | 0 |
| 1 | 1 | 1 |
| 0 | 0 | 0 |
| 1 | 1 | 1 |
| 0 | 0 | 0 |
| 0 | 0 | 0 |
| 0 | 0 | 0 |
| 0 | 0 | 0 |
| 0 | 0 | 0 |
| 0 | 0 | 0 |
| 0 | 1 | 0 |
| 1 | 1 | 1 |
| 1 | 1 | 1 |

| | | |
|---|---|---|
| 1 | 1 | 1 |
| 1 | 1 | 1 |
| 1 | 1 | 1 |
| 0 | 0 | 0 |
| 1 | 1 | 1 |
| 0 | 0 | 0 |
| 1 | 1 | 1 |
| 1 | 1 | 1 |
| 0 | 0 | 0 |
| 1 | 1 | 1 |
| 1 | 1 | 1 |
| 0 | 1 | 0 |
| 1 | 1 | 1 |
| 1 | 1 | 1 |
| 1 | 1 | 1 |
| 0 | 0 | 0 |
| 0 | 0 | 0 |
| 0 | 0 | 0 |
| 0 | 0 | 0 |
| 0 | 0 | 0 |
| 1 | 1 | 1 |
| 1 | 1 | 1 |
| 1 | 1 | 1 |
| 1 | 1 | 1 |
| 0 | 0 | 0 |
| 1 | 1 | 1 |
| 1 | 1 | 1 |
| 1 | 1 | 1 |
| 0 | 0 | 0 |
| 1 | 1 | 1 |
| 0 | 0 | 0 |
| 0 | 0 | 0 |
| 0 | 0 | 0 |
| 0 | 0 | 0 |
| 0 | 0 | 0 |
| 1 | 1 | 1 |
| 0 | 0 | 0 |
| 0 | 0 | 0 |
| 0 | 0 | 0 |
| 0 | 0 | 0 |
| 0 | 0 | 0 |
| 0 | 0 | 0 |
| 0 | 0 | 0 |
| 0 | 0 | 0 |
| 0 | 0 | 0 |
| 0 | 0 | 0 |
| 0 | 0 | 0 |
| 0 | 0 | 0 |
| 1 | 1 | 1 |
| 0 | 0 | 0 |
| 1 | 1 | 1 |
| 1 | 1 | 1 |
| 0 | 1 | 0 |
| 0 | 0 | 0 |

| | | |
|---|---|---|
| 0 | 0 | 0 |
| 0 | 1 | 0 |
| 1 | 1 | 1 |
| 1 | 1 | 1 |
| 1 | 1 | 1 |
| 0 | 0 | 0 |
| 1 | 1 | 1 |
| 0 | 1 | 0 |
| 1 | 1 | 1 |
| 0 | 1 | 0 |
| 0 | 1 | 0 |
| 0 | 1 | 0 |
| 1 | 1 | 1 |
| 0 | 0 | 0 |
| 0 | 1 | 0 |
| 1 | 1 | 1 |
| 1 | 1 | 1 |
| 0 | 0 | 0 |
| 1 | 1 | 1 |
| 1 | 1 | 1 |
| 1 | 1 | 1 |
| 1 | 1 | 1 |
| 0 | 0 | 0 |
| 1 | 1 | 1 |
| 1 | 1 | 1 |
| 0 | 0 | 0 |
| 0 | 0 | 0 |
| 0 | 0 | 0 |
| 1 | 1 | 1 |
| 0 | 1 | 0 |
| 1 | 1 | 1 |
| 1 | 1 | 1 |
| 1 | 1 | 1 |
| 1 | 1 | 1 |
| 1 | 1 | 1 |
| 1 | 1 | 1 |
| 1 | 1 | 1 |
| 1 | 1 | 1 |
| 1 | 1 | 1 |
| 1 | 1 | 1 |
| 1 | 1 | 1 |
| 1 | 1 | 1 |
| 1 | 1 | 1 |
| 1 | 1 | 1 |
| 1 | 1 | 1 |
| 1 | 1 | 1 |
| 1 | 1 | 1 |
| 1 | 1 | 1 |
| 1 | 0 | 0 |
| 1 | 1 | 1 |
| 1 | 1 | 1 |
| 1 | 1 | 1 |
| 1 | 1 | 1 |
| 1 | 1 | 1 |

| | | |
|---|---|---|
| 1 | 1 | 1 |
| 0 | 0 | 0 |
| 1 | 1 | 1 |
| 1 | 1 | 1 |
| 1 | 1 | 1 |
| 0 | 0 | 0 |
| 0 | 0 | 0 |
| 0 | 0 | 0 |
| 1 | 1 | 1 |
| 1 | 1 | 1 |
| 0 | 1 | 0 |
| 1 | 1 | 1 |
| 1 | 1 | 1 |
| 1 | 1 | 1 |
| 0 | 0 | 0 |
| 0 | 0 | 0 |
| 0 | 0 | 0 |
| 1 | 1 | 1 |
| 0 | 0 | 0 |
| 0 | 0 | 0 |
| 0 | 0 | 0 |
| 0 | 0 | 0 |
| 0 | 0 | 0 |
| 0 | 0 | 0 |
| 0 | 1 | 0 |
| 1 | 1 | 1 |
| 1 | 1 | 1 |
| 0 | 1 | 0 |
| 0 | 1 | 0 |
| 1 | 1 | 1 |
| 1 | 1 | 1 |
| 1 | 1 | 1 |
| 0 | 0 | 0 |
| 1 | 1 | 1 |
| 0 | 1 | 0 |
| 0 | 0 | 0 |
| 0 | 0 | 0 |
| 0 | 0 | 0 |
| 0 | 0 | 0 |
| 1 | 1 | 1 |
| 0 | 1 | 0 |
| 1 | 1 | 1 |
| 1 | 1 | 1 |
| 0 | 0 | 0 |
| 0 | 0 | 0 |
| 0 | 0 | 0 |
| 0 | 1 | 0 |
| 1 | 1 | 1 |
| 0 | 0 | 0 |
| 1 | 1 | 1 |
| 1 | 1 | 1 |
| 0 | 0 | 0 |
| 1 | 1 | 1 |
| 0 | 1 | 0 |

| | | |
|---|---|---|
| 1 | 1 | 1 |
| 0 | 1 | 0 |
| 1 | 1 | 1 |
| 1 | 1 | 1 |
| 1 | 1 | 1 |
| 1 | 1 | 1 |
| 0 | 1 | 0 |
| 1 | 1 | 1 |
| 0 | 0 | 0 |
| 0 | 0 | 0 |
| 1 | 1 | 1 |
| 0 | 0 | 0 |
| 0 | 0 | 0 |
| 0 | 0 | 0 |
| 1 | 1 | 1 |
| 1 | 1 | 1 |
| 0 | 0 | 0 |
| 0 | 0 | 0 |
| 0 | 0 | 0 |
| 1 | 1 | 1 |
| 1 | 1 | 1 |
| 1 | 1 | 1 |
| 0 | 0 | 0 |
| 1 | 1 | 1 |
| 0 | 0 | 0 |
| 1 | 1 | 1 |
| 1 | 1 | 1 |
| 1 | 1 | 1 |
| 1 | 1 | 1 |
| 0 | 0 | 0 |
| 0 | 0 | 0 |
| 0 | 0 | 0 |
| 1 | 1 | 1 |
| 0 | 1 | 0 |
| 1 | 1 | 1 |
| 1 | 1 | 1 |
| 1 | 1 | 1 |
| 0 | 1 | 0 |
| 0 | 1 | 0 |
| 0 | 1 | 0 |
| 0 | 0 | 0 |
| 1 | 1 | 1 |
| 1 | 1 | 1 |
| 1 | 1 | 1 |
| 1 | 1 | 1 |
| 0 | 0 | 0 |
| 1 | 0 | 0 |
| 0 | 0 | 0 |
| 1 | 1 | 1 |
| 0 | 0 | 0 |
| 0 | 0 | 0 |
| 0 | 0 | 0 |
| 0 | 1 | 0 |
| 1 | 0 | 0 |

| | | |
|---|---|---|
| 1 | 0 | 0 |
| 0 | 0 | 0 |
| 0 | 1 | 0 |
| 1 | 1 | 1 |
| 0 | 1 | 0 |
| 0 | 1 | 0 |
| 0 | 1 | 0 |
| 0 | 0 | 0 |
| 0 | 0 | 0 |
| 0 | 1 | 0 |
| 0 | 0 | 0 |
| 1 | 1 | 1 |
| 1 | 1 | 1 |
| 1 | 1 | 1 |
| 0 | 1 | 0 |
| 1 | 1 | 1 |
| 1 | 1 | 1 |
| 1 | 1 | 1 |
| 1 | 1 | 1 |
| 0 | 1 | 0 |
| 1 | 1 | 1 |
| 1 | 1 | 1 |
| 1 | 1 | 1 |
| 1 | 1 | 1 |
| 0 | 1 | 0 |
| 0 | 1 | 0 |
| 0 | 1 | 0 |
| 0 | 1 | 0 |
| 0 | 0 | 0 |
| 0 | 1 | 0 |
| 0 | 0 | 0 |
| 1 | 1 | 1 |
| 0 | 0 | 0 |
| 0 | 0 | 0 |
| 0 | 1 | 0 |
| 0 | 0 | 0 |
| 0 | 0 | 0 |
| 1 | 1 | 1 |
| 0 | 1 | 0 |
| 0 | 1 | 0 |
| 1 | 1 | 1 |
| 0 | 0 | 0 |
| 0 | 0 | 0 |
| 1 | 1 | 1 |
| 1 | 1 | 1 |
| 0 | 0 | 0 |
| 0 | 0 | 0 |
| 0 | 0 | 0 |
| 0 | 0 | 0 |
| 0 | 1 | 0 |
| 0 | 0 | 0 |
| 0 | 0 | 0 |
| 1 | 1 | 1 |
| 1 | 1 | 1 |

| | | |
|---|---|---|
| 1 | 1 | 1 |
| 1 | 1 | 1 |
| 1 | 1 | 1 |
| 1 | 1 | 1 |
| 0 | 0 | 0 |
| 0 | 0 | 0 |
| 0 | 0 | 0 |
| 0 | 0 | 0 |
| 1 | 1 | 1 |
| 0 | 0 | 0 |
| 0 | 0 | 0 |
| 0 | 0 | 0 |
| 0 | 0 | 0 |
| 1 | 1 | 1 |
| 1 | 0 | 0 |
| 0 | 1 | 0 |
| 1 | 1 | 1 |
| 1 | 1 | 1 |
| 1 | 1 | 1 |
| 0 | 1 | 0 |
| 1 | 1 | 1 |
| 1 | 1 | 1 |
| 1 | 1 | 1 |
| 1 | 1 | 1 |
| 1 | 1 | 1 |
| 0 | 0 | 0 |
| 0 | 0 | 0 |
| 0 | 0 | 0 |
| 0 | 0 | 0 |
| 0 | 1 | 0 |
| 0 | 1 | 0 |
| 0 | 1 | 0 |
| 1 | 1 | 1 |
| 0 | 1 | 0 |
| 0 | 1 | 0 |
| 1 | 1 | 1 |
| 1 | 1 | 1 |
| 1 | 1 | 1 |
| 1 | 1 | 1 |
| 1 | 1 | 1 |
| 1 | 1 | 1 |
| 1 | 1 | 1 |
| 1 | 1 | 1 |
| 1 | 1 | 1 |
| 1 | 1 | 1 |
| 1 | 1 | 1 |
| 1 | 1 | 1 |
| 1 | 1 | 1 |
| 1 | 1 | 1 |
| 1 | 1 | 1 |
| 1 | 1 | 1 |
| 1 | 1 | 1 |
| 1 | 1 | 1 |
| 0 | 0 | 0 |

| | | |
|---|---|---|
| 1 | 1 | 1 |
| 1 | 1 | 1 |
| 1 | 1 | 1 |
| 0 | 1 | 0 |
| 0 | 1 | 0 |
| 0 | 1 | 0 |
| 0 | 1 | 0 |
| 0 | 1 | 0 |
| 1 | 1 | 1 |
| 1 | 1 | 1 |
| 1 | 1 | 1 |
| 1 | 1 | 1 |
| 1 | 1 | 1 |
| 1 | 1 | 1 |
| 1 | 1 | 1 |
| 1 | 1 | 1 |
| 1 | 1 | 1 |
| 1 | 1 | 1 |
| 1 | 1 | 1 |
| 1 | 1 | 1 |
| 1 | 1 | 1 |
| 1 | 1 | 1 |
| 1 | 1 | 1 |
| 1 | 1 | 1 |
| 1 | 1 | 1 |
| 1 | 1 | 1 |
| 1 | 1 | 1 |
| 1 | 1 | 1 |
| 1 | 1 | 1 |
| 1 | 1 | 1 |
| 1 | 1 | 1 |
| 1 | 1 | 1 |
| 1 | 1 | 1 |
| 0 | 1 | 0 |
| 1 | 1 | 1 |
| 1 | 1 | 1 |
| 1 | 1 | 1 |
| 1 | 1 | 1 |
| 1 | 1 | 1 |
| 1 | 1 | 1 |
| 0 | 1 | 0 |
| 1 | 1 | 1 |
| 1 | 1 | 1 |
| 1 | 1 | 1 |
| 0 | 1 | 0 |
| 1 | 0 | 0 |
| 0 | 0 | 0 |
| 1 | 1 | 1 |
| 0 | 1 | 0 |
| 1 | 1 | 1 |
| 1 | 1 | 1 |
| 0 | 1 | 0 |
| 0 | 1 | 0 |
| 0 | 1 | 0 |

| | | |
|---|---|---|
| 0 | 1 | 0 |
| 1 | 1 | 1 |
| 0 | 0 | 0 |
| 1 | 1 | 1 |
| 1 | 1 | 1 |
| 1 | 1 | 1 |
| 0 | 0 | 0 |
| 1 | 1 | 1 |
| 1 | 1 | 1 |
| 0 | 1 | 0 |
| 0 | 1 | 0 |
| 0 | 0 | 0 |
| 1 | 1 | 1 |
| 0 | 0 | 0 |
| 0 | 0 | 0 |
| 0 | 0 | 0 |
| 0 | 0 | 0 |
| 0 | 0 | 0 |
| 0 | 0 | 0 |
| 0 | 0 | 0 |
| 0 | 0 | 0 |

| organism_scientific | evo_cons | cluster_id | pdb_id | chain_id | resi | resn | change | uniprot_id |
|---|---|---|---|---|---|---|---|---|
| HOMO SAPIENS | 5 | 87 | 3uzw | A | 106 | LEU | p.Leu106Phe | P51857 |
| HOMO SAPIENS | 3 | 87 | 3uzw | A | 133 | PRO | p.Pro133Arg | P51857 |
| HOMO SAPIENS | 0 | 87 | 3uzw | A | 198 | PRO | p.Pro198Leu | P51857 |
| HOMO SAPIENS | 7 | 87 | 3uzw | A | 261 | ARG | p.Arg261Cys | P51857 |
| HOMO SAPIENS | 0 | 4693 | 3mk1 | A | 3 | PRO | p.Pro25Leu | P05187 |
| HOMO SAPIENS | 8 | 4830 | 2h64 | B | 101 | CYS | p.Cys124Arg | P36894 |
| HOMO SAPIENS | 2 | 4830 | 2h64 | B | 116 | PRO | p.Pro139Ser | P36894 |
| HOMO SAPIENS | 0 | 6937 | 3dar | A | 186 | MET | p.Met186Thr | P21802 |
| HOMO SAPIENS | 3 | 5 | 1aut | C | 30 | GLN | p.Gln226His | P04070 |
| HOMO SAPIENS | 1 | 5 | 1aut | C | 68 | LEU | p.Leu265Phe | P04070 |
| HOMO SAPIENS | 0 | 5 | 1aut | C | 75 | ARG | p.Arg272Cys | P04070 |
| HOMO SAPIENS | 6 | 5 | 1aut | C | 92 | PRO | p.Pro289Leu | P04070 |
| HOMO SAPIENS | 0 | 5 | 1aut | C | 104 | ALA | p.Ala301Val | P04070 |
| HOMO SAPIENS | 1 | 5 | 1aut | C | 112 | ALA | p.Ala309Thr | P04070 |
| HOMO SAPIENS | 2 | 5 | 1aut | C | 115 | SER | p.Ser312Leu | P04070 |
| HOMO SAPIENS | 1 | 5 | 1aut | C | 133 | GLY | p.Gly334Ser | P04070 |
| HOMO SAPIENS | 0 | 5 | 1aut | C | 147 | ARG | p.Arg348Ter | P04070 |
| HOMO SAPIENS | 8 | 5 | 1aut | C | 194 | ASP | p.Asp401Asn | P04070 |
| HOMO SAPIENS | 7 | 5 | 1aut | C | 237 | TRP | p.Trp444Cys | P04070 |
| HOMO SAPIENS | 0 | 5 | 1aut | C | 238 | ILE | p.Ile445Met | P04070 |
| HOMO SAPIENS | 8 | 43 | 1aiy | F | 10 | HIS | p.His34Asp | P01308 |
| HOMO SAPIENS | 2 | 43 | 1aiy | F | 22 | ARG | p.Arg46Gln | P01308 |
| HOMO SAPIENS | 4 | 43 | 1aiy | F | 23 | GLY | p.Gly47Val | P01308 |
| HOMO SAPIENS | 7 | 43 | 1aiy | F | 24 | PHE | p.Phe48Ser | P01308 |
| HOMO SAPIENS | 5 | 24 | 2fg8 | A | 33 | THR | p.Thr30Ile | P02792 |
| HOMO SAPIENS | 4 | 24 | 2fg8 | A | 99 | ALA | p.Ala96Thr | P02792 |
| HOMO SAPIENS | 5 | 24 | 2fg8 | A | 107 | GLU | p.Glu104Ter | P02792 |
| HOMO SAPIENS | 3 | 5409 | 5bnq | A | 199 | MET | p.Met199Lys | O14788 |
| HOMO SAPIENS | -1 | 839 | 3swc | A | 959 | GLN | p.Gln990His | Q9H9B1 |
| HOMO SAPIENS | -1 | 839 | 3swc | A | 968 | ALA | p.Ala999Val | Q9H9B1 |
| HOMO SAPIENS | -1 | 839 | 3swc | A | 973 | SER | p.Ser1004Ile | Q9H9B1 |
| HOMO SAPIENS | 8 | 17 | 1eou | A | 18 | LYS | p.Lys18Glu | P00918 |
| HOMO SAPIENS | 2 | 17 | 1eou | A | 107 | HIS | p.His107Tyr | P00918 |
| HOMO SAPIENS | 1 | 17 | 1eou | A | 236 | PRO | p.Pro236His | P00918 |
| HOMO SAPIENS | 6 | 17 | 1eou | A | 252 | ASN | p.Asn252Asp | P00918 |
| HOMO SAPIENS | 2 | 9140 | 5c4v | A | 445 | ARG | p.Arg445Ter | Q13485 |
| HOMO SAPIENS | 7 | 9140 | 5c4v | A | 496 | ARG | p.Arg496Cys | Q13485 |
| HOMO SAPIENS | -1 | 42 | 4pwi | A | 6 | GLY | p.Gly26Ser | P02766 |
| HOMO SAPIENS | 6 | 42 | 4pwi | A | 10 | CYS | p.Cys30Arg | P02766 |
| HOMO SAPIENS | 8 | 42 | 4pwi | A | 12 | LEU | p.Leu32Pro | P02766 |
| HOMO SAPIENS | 9 | 42 | 4pwi | A | 18 | ASP | p.Asp38Gly | P02766 |
| HOMO SAPIENS | 8 | 42 | 4pwi | A | 20 | VAL | p.Val40Ile | P02766 |
| HOMO SAPIENS | 4 | 42 | 4pwi | A | 30 | MET | p.Val50Ala | P02766 |
| HOMO SAPIENS | 4 | 42 | 4pwi | A | 30 | MET | p.Val50Leu | P02766 |
| HOMO SAPIENS | 6 | 42 | 4pwi | A | 33 | PHE | p.Phe53Ile | P02766 |
| HOMO SAPIENS | 7 | 42 | 4pwi | A | 36 | ALA | p.Ala56Pro | P02766 |
| HOMO SAPIENS | 7 | 42 | 4pwi | A | 42 | GLU | p.Glu62Gly | P02766 |
| HOMO SAPIENS | 8 | 42 | 4pwi | A | 44 | PHE | p.Phe64Ser | P02766 |
| HOMO SAPIENS | 4 | 42 | 4pwi | A | 47 | GLY | p.Gly67Ala | P02766 |
| HOMO SAPIENS | 6 | 42 | 4pwi | A | 49 | THR | p.Thr69Ala | P02766 |
| HOMO SAPIENS | 6 | 42 | 4pwi | A | 50 | SER | p.Ser70Ile | P02766 |
| HOMO SAPIENS | 6 | 42 | 4pwi | A | 50 | SER | p.Ser70Arg | P02766 |
| HOMO SAPIENS | 0 | 42 | 4pwi | A | 53 | GLY | p.Gly73Glu | P02766 |

| | | | | | | | | |
|---|---|---|---|---|---|---|---|---|
| HOMO SAPIENS | 7 | 42 | 4pwi | A | 55 | LEU | p.Leu75Pro | P02766 |
| HOMO SAPIENS | 6 | 42 | 4pwi | A | 58 | LEU | p.Leu78His | P02766 |
| HOMO SAPIENS | 7 | 42 | 4pwi | A | 61 | GLU | p.Glu81Lys | P02766 |
| HOMO SAPIENS | 8 | 42 | 4pwi | A | 64 | PHE | p.Phe84Leu | P02766 |
| HOMO SAPIENS | 8 | 42 | 4pwi | A | 68 | ILE | p.Ile88Leu | P02766 |
| HOMO SAPIENS | 8 | 42 | 4pwi | A | 69 | TYR | p.Tyr89His | P02766 |
| HOMO SAPIENS | 4 | 42 | 4pwi | A | 71 | VAL | p.Val91Ala | P02766 |
| HOMO SAPIENS | 6 | 42 | 4pwi | A | 77 | SER | p.Ser97Tyr | P02766 |
| HOMO SAPIENS | 4 | 42 | 4pwi | A | 84 | ILE | p.Ile104Ser | P02766 |
| HOMO SAPIENS | 8 | 42 | 4pwi | A | 89 | GLU | p.Glu109Gln | P02766 |
| HOMO SAPIENS | 8 | 42 | 4pwi | A | 97 | ALA | p.Ala117Gly | P02766 |
| HOMO SAPIENS | 7 | 42 | 4pwi | A | 99 | ASP | p.Asp119Asn | P02766 |
| HOMO SAPIENS | 9 | 42 | 4pwi | A | 107 | ILE | p.Ile127Val | P02766 |
| HOMO SAPIENS | 9 | 42 | 4pwi | A | 111 | LEU | p.Leu131Met | P02766 |
| HOMO SAPIENS | 7 | 42 | 4pwi | A | 114 | TYR | p.Tyr134Cys | P02766 |
| HOMO SAPIENS | 7 | 42 | 4pwi | A | 114 | TYR | p.Tyr134His | P02766 |
| HOMO SAPIENS | 7 | 42 | 4pwi | A | 122 | VAL | p.Val142Ile | P02766 |
| HOMO SAPIENS | -1 | 42 | 4pwi | A | 126 | LYS | p.Lys146Arg | P02766 |
| HOMO SAPIENS | -1 | 42 | 4pwi | A | 127 | GLU | p.Glu147Ter | P02766 |
| HOMO SAPIENS | -1 | 1312 | 4dce | A | 1275 | ARG | p.Arg1275Gln | Q9UM73 |
| HOMO SAPIENS | -1 | 1312 | 4dce | A | 1280 | ALA | p.Ala1280Val | Q9UM73 |
| HOMO SAPIENS | 1 | 20 | 1y0x | X | 234 | ALA | p.Ala234Thr | P10828 |
| HOMO SAPIENS | 0 | 20 | 1y0x | X | 243 | ARG | p.Arg243Trp | P10828 |
| HOMO SAPIENS | 3 | 20 | 1y0x | X | 316 | ARG | p.Arg316His | P10828 |
| HOMO SAPIENS | 1 | 20 | 1y0x | X | 317 | ALA | p.Ala317Thr | P10828 |
| HOMO SAPIENS | 2 | 20 | 1y0x | X | 320 | ARG | p.Arg320Cys | P10828 |
| HOMO SAPIENS | 2 | 20 | 1y0x | X | 320 | ARG | p.Arg320His | P10828 |
| HOMO SAPIENS | 0 | 20 | 1y0x | X | 332 | GLY | p.Gly332Arg | P10828 |
| HOMO SAPIENS | 1 | 20 | 1y0x | X | 338 | ARG | p.Arg338Trp | P10828 |
| HOMO SAPIENS | 0 | 20 | 1y0x | X | 340 | GLN | p.Gln340His | P10828 |
| HOMO SAPIENS | 0 | 20 | 1y0x | X | 345 | GLY | p.Gly345Val | P10828 |
| HOMO SAPIENS | 0 | 20 | 1y0x | X | 345 | GLY | p.Gly345Ser | P10828 |
| HOMO SAPIENS | 0 | 20 | 1y0x | X | 347 | GLY | p.Gly347Glu | P10828 |
| HOMO SAPIENS | 1 | 20 | 1y0x | X | 438 | ARG | p.Arg438His | P10828 |
| HOMO SAPIENS | 0 | 20 | 1y0x | X | 442 | MET | p.Met442Val | P10828 |
| HOMO SAPIENS | 0 | 20 | 1y0x | X | 443 | LYS | p.Lys443Glu | P10828 |
| HOMO SAPIENS | 0 | 20 | 1y0x | X | 446 | CYS | p.Cys446Arg | P10828 |
| HOMO SAPIENS | 2 | 20 | 1y0x | X | 453 | PRO | p.Pro453His | P10828 |
| HOMO SAPIENS | 2 | 20 | 1y0x | X | 453 | PRO | p.Pro453Thr | P10828 |
| HOMO SAPIENS | 3 | 20 | 1y0x | X | 459 | PHE | p.Phe459Cys | P10828 |
| HOMO SAPIENS | 0 | 4409 | 3qb7 | D | 87 | GLU | p.Glu109Lys | P31785 |
| HOMO SAPIENS | 1 | 4409 | 3qb7 | D | 92 | GLY | p.Gly114Asp | P31785 |
| HOMO SAPIENS | 1 | 4409 | 3qb7 | D | 93 | CYS | p.Cys115Arg | P31785 |
| HOMO SAPIENS | 0 | 4409 | 3qb7 | D | 131 | ILE | p.Ile153Asn | P31785 |
| HOMO SAPIENS | 9 | 4409 | 3qb7 | D | 200 | ARG | p.Arg222Cys | P31785 |
| HOMO SAPIENS | 0 | 3891 | 3ije | B | 33 | LEU | p.Leu59Pro | P05106 |
| HOMO SAPIENS | 0 | 3891 | 3ije | B | 117 | LEU | p.Leu143Trp | P05106 |
| HOMO SAPIENS | 3 | 3891 | 3ije | B | 119 | ASP | p.Asp145Tyr | P05106 |
| HOMO SAPIENS | 2 | 3891 | 3ije | B | 143 | ARG | p.Arg169Gln | P05106 |
| HOMO SAPIENS | 2 | 3891 | 3ije | B | 214 | ARG | p.Arg240Gln | P05106 |
| HOMO SAPIENS | 2 | 3891 | 3ije | B | 214 | ARG | p.Arg240Trp | P05106 |
| HOMO SAPIENS | 2 | 3891 | 3ije | B | 374 | CYS | p.Cys400Tyr | P05106 |
| HOMO SAPIENS | 1 | 3891 | 3ije | B | 407 | PRO | p.Pro433Ala | P05106 |
| HOMO SAPIENS | 0 | 3891 | 3ije | B | 489 | ARG | p.Arg515Gln | P05106 |

| | | | | | | | | |
|---|---|---|---|---|---|---|---|---|
| HOMO SAPIENS | -1 | 9613 | 5mke | B | 28 | ARG | p.Arg28Pro | Q13563 |
| HOMO SAPIENS | -1 | 9613 | 5mke | B | 190 | ALA | p.Ala190Thr | Q13563 |
| HOMO SAPIENS | 4 | 9613 | 5mke | B | 262 | VAL | p.Val262Met | Q13563 |
| HOMO SAPIENS | 0 | 9613 | 5mke | B | 482 | PHE | p.Phe482Cys | Q13563 |
| HOMO SAPIENS | 4 | 9613 | 5mke | B | 511 | ASP | p.Asp511Val | Q13563 |
| HOMO SAPIENS | 5 | 9613 | 5mke | B | 516 | VAL | p.Val516Leu | Q13563 |
| HOMO SAPIENS | -1 | 9613 | 5mke | B | 807 | ARG | p.Arg807Gln | Q13563 |
| HOMO SAPIENS | 4 | 2428 | 1olh | C | 337 | ARG | p.Arg337Cys | P04637 |
| HOMO SAPIENS | 4 | 2428 | 1olh | C | 337 | ARG | p.Arg337His | P04637 |
| HOMO SAPIENS | 8 | 2428 | 1olh | C | 338 | PHE | p.Phe338Leu | P04637 |
| HOMO SAPIENS | 8 | 2428 | 1olh | C | 339 | GLU | p.Glu339Lys | P04637 |
| HOMO SAPIENS | 7 | 2428 | 1olh | C | 344 | LEU | p.Leu344Pro | P04637 |
| HOMO SAPIENS | 0 | 2428 | 1olh | C | 360 | GLY | p.Gly360Ala | P04637 |
| HOMO SAPIENS | 7 | 4867 | 1mox | B | 404 | GLY | p.Gly428Asp | P00533 |
| HOMO SAPIENS | 5 | 4867 | 1mox | B | 497 | ARG | p.Arg521Lys | P00533 |
| HOMO SAPIENS | 1 | 4630 | 1fu1 | A | 43 | TRP | p.Trp43Arg | Q13426 |
| HOMO SAPIENS | 2 | 4630 | 1fu1 | A | 161 | ARG | p.Arg161Ter | Q13426 |
| HOMO SAPIENS | 2 | 15 | 1lfy | B | 2 | HIS | p.His3Gln | P68871 |
| HOMO SAPIENS | 2 | 15 | 1lfy | B | 6 | GLU | p.Glu7Ala | P68871 |
| HOMO SAPIENS | 2 | 15 | 1lfy | B | 6 | GLU | p.Glu7Lys | P68871 |
| HOMO SAPIENS | 4 | 15 | 1lfy | B | 9 | SER | p.Ser10Cys | P68871 |
| HOMO SAPIENS | 6 | 15 | 1lfy | B | 15 | TRP | p.Trp16Ter | P68871 |
| HOMO SAPIENS | 3 | 15 | 1lfy | B | 17 | LYS | p.Lys18Glu | P68871 |
| HOMO SAPIENS | 6 | 15 | 1lfy | B | 19 | ASN | p.Asn20Ser | P68871 |
| HOMO SAPIENS | 6 | 15 | 1lfy | B | 22 | GLU | p.Glu23Ala | P68871 |
| HOMO SAPIENS | 6 | 15 | 1lfy | B | 22 | GLU | p.Glu23Lys | P68871 |
| HOMO SAPIENS | 8 | 15 | 1lfy | B | 26 | GLU | p.Glu27Lys | P68871 |
| HOMO SAPIENS | 8 | 15 | 1lfy | B | 28 | LEU | p.Leu29Pro | P68871 |
| HOMO SAPIENS | 8 | 15 | 1lfy | B | 30 | ARG | p.Arg31Ser | P68871 |
| HOMO SAPIENS | 8 | 15 | 1lfy | B | 30 | ARG | p.Arg31Lys | P68871 |
| HOMO SAPIENS | 6 | 15 | 1lfy | B | 39 | GLN | p.Gln40Ter | P68871 |
| HOMO SAPIENS | 2 | 15 | 1lfy | B | 43 | GLU | p.Glu44Gln | P68871 |
| HOMO SAPIENS | 6 | 15 | 1lfy | B | 57 | ASN | p.Asn58Lys | P68871 |
| HOMO SAPIENS | 6 | 15 | 1lfy | B | 59 | LYS | p.Lys60Asn | P68871 |
| HOMO SAPIENS | 8 | 15 | 1lfy | B | 61 | LYS | p.Lys62Glu | P68871 |
| HOMO SAPIENS | 4 | 15 | 1lfy | B | 73 | ASP | p.Asp74Tyr | P68871 |
| HOMO SAPIENS | 4 | 15 | 1lfy | B | 87 | THR | p.Thr88Ile | P68871 |
| HOMO SAPIENS | 4 | 15 | 1lfy | B | 87 | THR | p.Thr88Pro | P68871 |
| HOMO SAPIENS | 5 | 15 | 1lfy | B | 90 | GLU | p.Glu91Lys | P68871 |
| HOMO SAPIENS | 6 | 15 | 1lfy | B | 91 | LEU | p.Leu92Pro | P68871 |
| HOMO SAPIENS | 5 | 15 | 1lfy | B | 97 | HIS | p.His98Gln | P68871 |
| HOMO SAPIENS | 6 | 15 | 1lfy | B | 110 | LEU | p.Leu111Pro | P68871 |
| HOMO SAPIENS | 6 | 15 | 1lfy | B | 113 | VAL | p.Val114Glu | P68871 |
| HOMO SAPIENS | 2 | 15 | 1lfy | B | 119 | GLY | p.Gly120Ala | P68871 |
| HOMO SAPIENS | 4 | 15 | 1lfy | B | 121 | GLU | p.Glu122Gln | P68871 |
| HOMO SAPIENS | 5 | 15 | 1lfy | B | 134 | VAL | p.Val135Glu | P68871 |
| HOMO SAPIENS | 3 | 15 | 1lfy | B | 143 | HIS | p.His144Gln | P68871 |
| HOMO SAPIENS | 4 | 91 | 5d9k | A | 75 | GLY | p.Gly75Val | P51812 |
| HOMO SAPIENS | 5 | 91 | 5d9k | A | 82 | VAL | p.Val82Phe | P51812 |
| HOMO SAPIENS | 0 | 91 | 5d9k | A | 114 | ARG | p.Arg114Trp | P51812 |
| HOMO SAPIENS | 4 | 91 | 5d9k | A | 189 | ILE | p.Ile189Lys | P51812 |
| HOMO SAPIENS | -1 | 91 | 5d9k | A | 227 | SER | p.Ser227Ala | P51812 |
| HOMO SAPIENS | 1 | 91 | 5d9k | A | 268 | PHE | p.Phe268Ser | P51812 |
| HOMO SAPIENS | 0 | 954 | 2vt0 | A | 15 | VAL | p.Val54Leu | P04062 |

| | | | | | | | | |
|---|---|---|---|---|---|---|---|---|
| HOMO SAPIENS | 5 | 954 | 2vt0 | A | 46 | GLY | p.Gly85Glu | P04062 |
| HOMO SAPIENS | 5 | 954 | 2vt0 | A | 48 | ARG | p.Arg87Trp | P04062 |
| HOMO SAPIENS | 6 | 954 | 2vt0 | A | 79 | LYS | p.Lys118Asn | P04062 |
| HOMO SAPIENS | 0 | 954 | 2vt0 | A | 120 | ARG | p.Arg159Gln | P04062 |
| HOMO SAPIENS | 0 | 954 | 2vt0 | A | 120 | ARG | p.Arg159Trp | P04062 |
| HOMO SAPIENS | 0 | 954 | 2vt0 | A | 122 | PRO | p.Pro161Ser | P04062 |
| HOMO SAPIENS | 5 | 954 | 2vt0 | A | 131 | ARG | p.Arg170Leu | P04062 |
| HOMO SAPIENS | 5 | 954 | 2vt0 | A | 135 | TYR | p.Tyr174Cys | P04062 |
| HOMO SAPIENS | 0 | 954 | 2vt0 | A | 157 | LYS | p.Lys196Gln | P04062 |
| HOMO SAPIENS | 5 | 954 | 2vt0 | A | 163 | ARG | p.Arg202Gln | P04062 |
| HOMO SAPIENS | 5 | 954 | 2vt0 | A | 170 | ARG | p.Arg209Cys | P04062 |
| HOMO SAPIENS | 5 | 954 | 2vt0 | A | 188 | ASN | p.Asn227Ser | P04062 |
| HOMO SAPIENS | 3 | 954 | 2vt0 | A | 196 | SER | p.Ser235Pro | P04062 |
| HOMO SAPIENS | 5 | 954 | 2vt0 | A | 202 | GLY | p.Gly241Arg | P04062 |
| HOMO SAPIENS | 0 | 954 | 2vt0 | A | 212 | TYR | p.Tyr251His | P04062 |
| HOMO SAPIENS | 0 | 954 | 2vt0 | A | 213 | PHE | p.Phe252Ile | P04062 |
| HOMO SAPIENS | 5 | 954 | 2vt0 | A | 216 | PHE | p.Phe255Tyr | P04062 |
| HOMO SAPIENS | 6 | 954 | 2vt0 | A | 251 | PHE | p.Phe290Leu | P04062 |
| HOMO SAPIENS | 5 | 954 | 2vt0 | A | 255 | HIS | p.His294Gln | P04062 |
| HOMO SAPIENS | 6 | 954 | 2vt0 | A | 257 | ARG | p.Arg296Gln | P04062 |
| HOMO SAPIENS | 5 | 954 | 2vt0 | A | 289 | PRO | p.Pro328Leu | P04062 |
| HOMO SAPIENS | 0 | 954 | 2vt0 | A | 309 | ALA | p.Ala348Val | P04062 |
| HOMO SAPIENS | 0 | 954 | 2vt0 | A | 311 | HIS | p.His350Arg | P04062 |
| HOMO SAPIENS | 6 | 954 | 2vt0 | A | 312 | TRP | p.Trp351Cys | P04062 |
| HOMO SAPIENS | 5 | 954 | 2vt0 | A | 323 | THR | p.Thr362Ile | P04062 |
| HOMO SAPIENS | 5 | 954 | 2vt0 | A | 325 | GLY | p.Gly364Arg | P04062 |
| HOMO SAPIENS | 6 | 954 | 2vt0 | A | 342 | CYS | p.Cys381Gly | P04062 |
| HOMO SAPIENS | 5 | 954 | 2vt0 | A | 353 | ARG | p.Arg392Gly | P04062 |
| HOMO SAPIENS | 3 | 954 | 2vt0 | A | 364 | SER | p.Ser403Thr | P04062 |
| HOMO SAPIENS | 5 | 954 | 2vt0 | A | 369 | THR | p.Thr408Met | P04062 |
| HOMO SAPIENS | 3 | 954 | 2vt0 | A | 370 | ASN | p.Asn409Ser | P04062 |
| HOMO SAPIENS | 0 | 954 | 2vt0 | A | 371 | LEU | p.Leu410Val | P04062 |
| HOMO SAPIENS | 0 | 954 | 2vt0 | A | 375 | VAL | p.Val414Leu | P04062 |
| HOMO SAPIENS | 0 | 954 | 2vt0 | A | 377 | GLY | p.Gly416Ser | P04062 |
| HOMO SAPIENS | 8 | 954 | 2vt0 | A | 398 | VAL | p.Val437Phe | P04062 |
| HOMO SAPIENS | 0 | 954 | 2vt0 | A | 401 | PRO | p.Pro440Leu | P04062 |
| HOMO SAPIENS | 3 | 954 | 2vt0 | A | 409 | ASP | p.Asp448His | P04062 |
| HOMO SAPIENS | 3 | 954 | 2vt0 | A | 409 | ASP | p.Asp448Val | P04062 |
| HOMO SAPIENS | 0 | 954 | 2vt0 | A | 415 | PRO | p.Pro454Arg | P04062 |
| HOMO SAPIENS | 5 | 954 | 2vt0 | A | 444 | LEU | p.Leu483Arg | P04062 |
| HOMO SAPIENS | 5 | 954 | 2vt0 | A | 456 | ALA | p.Ala495Pro | P04062 |
| HOMO SAPIENS | 5 | 954 | 2vt0 | A | 463 | ARG | p.Arg502Cys | P04062 |
| HOMO SAPIENS | 5 | 954 | 2vt0 | A | 463 | ARG | p.Arg502His | P04062 |
| HOMO SAPIENS | 5 | 954 | 2vt0 | A | 478 | GLY | p.Gly517Ser | P04062 |
| HOMO SAPIENS | 5 | 954 | 2vt0 | A | 496 | ARG | p.Arg535Cys | P04062 |
| HOMO SAPIENS | 5 | 954 | 2vt0 | A | 496 | ARG | p.Arg535His | P04062 |
| HOMO SAPIENS | 1 | 9037 | 2rqb | A | 946 | ALA | p.Ala946Thr | Q9BYX4 |
| HOMO SAPIENS | 0 | 253 | 5E+092 | A | 308 | LEU | p.Leu308Pro | P37173 |
| HOMO SAPIENS | 0 | 253 | 5E+092 | A | 315 | THR | p.Thr315Met | P37173 |
| HOMO SAPIENS | 0 | 253 | 5E+092 | A | 328 | HIS | p.His328Gln | P37173 |
| HOMO SAPIENS | 0 | 253 | 5E+092 | A | 336 | TYR | p.Tyr336Asn | P37173 |
| HOMO SAPIENS | 0 | 253 | 5E+092 | A | 348 | ARG | p.Arg348Cys | P37173 |
| HOMO SAPIENS | 0 | 253 | 5E+092 | A | 355 | ALA | p.Ala355Pro | P37173 |
| HOMO SAPIENS | 1 | 253 | 5E+092 | A | 356 | ARG | p.Arg356Gly | P37173 |

| | | | | | | | | |
|---|---|---|---|---|---|---|---|---|
| HOMO SAPIENS | 0 | 253 | 5E+092 | A | 357 | GLY | p.Gly357Trp | P37173 |
| HOMO SAPIENS | 1 | 253 | 5E+092 | A | 403 | ARG | p.Arg403His | P37173 |
| HOMO SAPIENS | 5 | 253 | 5E+092 | A | 449 | SER | p.Ser449Phe | P37173 |
| HOMO SAPIENS | 0 | 253 | 5E+092 | A | 460 | ARG | p.Arg460Cys | P37173 |
| HOMO SAPIENS | 0 | 253 | 5E+092 | A | 460 | ARG | p.Arg460His | P37173 |
| HOMO SAPIENS | 2 | 253 | 5E+092 | A | 526 | GLU | p.Glu526Gln | P37173 |
| HOMO SAPIENS | 5 | 253 | 5E+092 | A | 528 | ARG | p.Arg528Cys | P37173 |
| HOMO SAPIENS | 5 | 253 | 5E+092 | A | 528 | ARG | p.Arg528His | P37173 |
| HOMO SAPIENS | 6 | 158 | 5amc | A | 231 | ARG | p.Arg260Leu | P12821 |
| HOMO SAPIENS | 7 | 158 | 5amc | A | 237 | TYR | p.Tyr266Ter | P12821 |
| HOMO SAPIENS | 6 | 158 | 5amc | A | 325 | GLY | p.Gly354Arg | P12821 |
| HOMO SAPIENS | 0 | 158 | 5amc | A | 467 | ARG | p.Arg496Ter | P12821 |
| HOMO SAPIENS | 4 | 158 | 5amc | A | 479 | ARG | p.Arg508Ter | P12821 |
| HOMO SAPIENS | 0 | 158 | 5amc | A | 563 | ASP | p.Asp592Gly | P12821 |
| HOMO SAPIENS | -1 | 33062 | 1gr3 | A | 545 | GLY | p.Gly545Arg | Q03692 |
| HOMO SAPIENS | 5 | 33062 | 1gr3 | A | 591 | CYS | p.Cys591Arg | Q03692 |
| HOMO SAPIENS | 2 | 33062 | 1gr3 | A | 595 | GLY | p.Gly595Glu | Q03692 |
| HOMO SAPIENS | 0 | 33062 | 1gr3 | A | 597 | TYR | p.Tyr597Cys | Q03692 |
| HOMO SAPIENS | 5 | 33062 | 1gr3 | A | 598 | TYR | p.Tyr598Asp | Q03692 |
| HOMO SAPIENS | 7 | 33062 | 1gr3 | A | 600 | SER | p.Ser600Pro | Q03692 |
| HOMO SAPIENS | 1 | 33062 | 1gr3 | A | 614 | LEU | p.Leu614Pro | Q03692 |
| HOMO SAPIENS | 6 | 33062 | 1gr3 | A | 651 | TRP | p.Trp651Arg | Q03692 |
| HOMO SAPIENS | 7 | 33062 | 1gr3 | A | 671 | SER | p.Ser671Pro | Q03692 |
| HOMO SAPIENS | -1 | 38257 | 2l2i | B | 87 | PRO | p.Pro87Ala | Q13351 |
| HOMO SAPIENS | 2 | 2036 | 1ghq | A | 231 | GLY | p.Gly1224Asp | P01024 |
| HOMO SAPIENS | 5 | 3594 | 4cde | F | 388 | TYR | p.Tyr412Cys | P53634 |
| HOMO SAPIENS | 9 | 3594 | 4cde | F | 405 | TRP | p.Trp429Cys | P53634 |
| HOMO SAPIENS | 0 | 3594 | 4cde | F | 429 | ILE | p.Ile453Val | P53634 |
| HOMO SAPIENS | 7 | 18 | 1jkb | A | 34 | TRP | p.Trp52Ter | P61626 |
| HOMO SAPIENS | 9 | 18 | 1jkb | A | 70 | THR | p.Thr88Asn | P61626 |
| HOMO SAPIENS | 8 | 18 | 1jkb | A | 121 | VAL | p.Val139Ile | P61626 |
| HOMO SAPIENS | 2 | 414 | 3fj6 | A | 136 | ARG | p.Arg135Cys | Q02127 |
| HOMO SAPIENS | 1 | 414 | 3fj6 | A | 153 | GLY | p.Gly152Arg | Q02127 |
| HOMO SAPIENS | 0 | 414 | 3fj6 | A | 200 | ARG | p.Arg199Cys | Q02127 |
| HOMO SAPIENS | 0 | 414 | 3fj6 | A | 203 | GLY | p.Gly202Ala | Q02127 |
| HOMO SAPIENS | 0 | 414 | 3fj6 | A | 245 | ARG | p.Arg244Trp | Q02127 |
| HOMO SAPIENS | 0 | 414 | 3fj6 | A | 342 | ALA | p.Ala341Val | Q02127 |
| HOMO SAPIENS | 1 | 414 | 3fj6 | A | 347 | ARG | p.Arg346Trp | Q02127 |
| HOMO SAPIENS | 6 | 103 | 1u3w | A | 77 | GLY | p.Gly78Ter | P00326 |
| HOMO SAPIENS | -1 | 8369 | 4ux8 | F | 16 | ARG | p.Arg93Trp | P39905 |
| HOMO SAPIENS | 0 | 366 | 2zjc | A | 32 | GLY | p.Arg108Trp | P01375 |
| HOMO SAPIENS | 0 | 1457 | 3v9i | A | 352 | LEU | p.Ser352Leu | P30038 |
| HOMO SAPIENS | 0 | 1457 | 3v9i | A | 470 | VAL | p.Val470Ile | P30038 |
| HOMO SAPIENS | 0 | 1457 | 3v9i | A | 473 | THR | p.Thr473Ala | P30038 |
| HOMO SAPIENS | 1 | 2 | 4wo5 | A | 462 | ARG | p.Arg462Ile | P15056 |
| HOMO SAPIENS | 3 | 2 | 4wo5 | A | 463 | ILE | p.Ile463Ser | P15056 |
| HOMO SAPIENS | 1 | 2 | 4wo5 | A | 466 | GLY | p.Gly466Val | P15056 |
| HOMO SAPIENS | 1 | 2 | 4wo5 | A | 468 | PHE | p.Phe468Ser | P15056 |
| HOMO SAPIENS | 2 | 2 | 4wo5 | A | 469 | GLY | p.Gly469Ala | P15056 |
| HOMO SAPIENS | 2 | 2 | 4wo5 | A | 469 | GLY | p.Gly469Arg | P15056 |
| HOMO SAPIENS | 2 | 2 | 4wo5 | A | 485 | LEU | p.Leu485Phe | P15056 |
| HOMO SAPIENS | 0 | 2 | 4wo5 | A | 501 | GLU | p.Glu501Gly | P15056 |
| HOMO SAPIENS | 0 | 2 | 4wo5 | A | 501 | GLU | p.Glu501Lys | P15056 |
| HOMO SAPIENS | 7 | 2 | 4wo5 | A | 581 | ASN | p.Asn581Asp | P15056 |

| | | | | | | | | |
|---|---|---|---|---|---|---|---|---|
| HOMO SAPIENS | 8 | 2 | 4wo5 | A | 594 | ASP | p.Asp594Gly | P15056 |
| HOMO SAPIENS | 0 | 2 | 4wo5 | A | 597 | LEU | p.Leu597Arg | P15056 |
| HOMO SAPIENS | 0 | 2 | 4wo5 | A | 597 | LEU | p.Leu597Val | P15056 |
| HOMO SAPIENS | 0 | 2 | 4wo5 | A | 600 | VAL | p.Val600Glu | P15056 |
| HOMO SAPIENS | 2 | 2 | 4wo5 | A | 601 | LYS | p.Lys601Glu | P15056 |
| HOMO SAPIENS | 8 | 2 | 4wo5 | A | 638 | ASP | p.Asp638Glu | P15056 |
| HOMO SAPIENS | -1 | 30366 | 5bqd | B | 19 | ALA | p.Ala19Ser | Q99593 |
| HOMO SAPIENS | -1 | 30366 | 5bqd | B | 23 | PRO | p.Pro23His | Q99593 |
| HOMO SAPIENS | 1 | 30366 | 5bqd | B | 49 | GLN | p.Gln49Lys | Q99593 |
| HOMO SAPIENS | 2 | 30366 | 5bqd | B | 54 | ILE | p.Ile54Thr | Q99593 |
| HOMO SAPIENS | 8 | 30366 | 5bqd | B | 80 | GLY | p.Gly80Arg | Q99593 |
| HOMO SAPIENS | 6 | 30366 | 5bqd | B | 106 | ILE | p.Ile106Val | Q99593 |
| HOMO SAPIENS | 7 | 30366 | 5bqd | B | 111 | ASP | p.Asp111Tyr | Q99593 |
| HOMO SAPIENS | -1 | 30366 | 5bqd | B | 237 | ARG | p.Arg237Gln | Q99593 |
| HOMO SAPIENS | -1 | 30366 | 5bqd | B | 237 | ARG | p.Arg237Trp | Q99593 |
| HOMO SAPIENS | 5 | 4789 | 1n4m | A | 567 | SER | p.Ser567Leu | P06400 |
| HOMO SAPIENS | 0 | 4789 | 1n4m | A | 658 | ALA | p.Ala658Asp | P06400 |
| HOMO SAPIENS | 7 | 4789 | 1n4m | A | 661 | ARG | p.Arg661Trp | P06400 |
| HOMO SAPIENS | 6 | 4789 | 1n4m | A | 668 | ARG | p.Arg668His | P06400 |
| HOMO SAPIENS | 8 | 4789 | 1n4m | A | 697 | ASP | p.Asp697Glu | P06400 |
| HOMO SAPIENS | 6 | 4789 | 1n4m | A | 712 | CYS | p.Cys712Arg | P06400 |
| HOMO SAPIENS | 1 | 1798 | 2r5g | A | 101 | HIS | p.His101Gln | O15247 |
| HOMO SAPIENS | 0 | 1798 | 2r5g | A | 160 | PRO | p.Pro160Ala | O15247 |
| HOMO SAPIENS | 2 | 22 | 4x0t | A | 82 | ARG | p.Arg110Ter | P49419 |
| HOMO SAPIENS | 0 | 22 | 4x0t | A | 149 | ALA | p.Ala177Pro | P49419 |
| HOMO SAPIENS | 4 | 22 | 4x0t | A | 164 | THR | p.Thr192Met | P49419 |
| HOMO SAPIENS | 9 | 22 | 4x0t | A | 167 | ASN | p.Asn195Ser | P49419 |
| HOMO SAPIENS | 0 | 22 | 4x0t | A | 175 | TRP | p.Trp203Gly | P49419 |
| HOMO SAPIENS | 0 | 22 | 4x0t | A | 194 | THR | p.Thr222Ala | P49419 |
| HOMO SAPIENS | 0 | 22 | 4x0t | A | 273 | ASN | p.Asn301Ile | P49419 |
| HOMO SAPIENS | 0 | 22 | 4x0t | A | 307 | ARG | p.Arg335Gln | P49419 |
| HOMO SAPIENS | 3 | 22 | 4x0t | A | 313 | SER | p.Ser341Asn | P49419 |
| HOMO SAPIENS | 0 | 22 | 4x0t | A | 329 | ILE | p.Ile357Val | P49419 |
| HOMO SAPIENS | 1 | 22 | 4x0t | A | 380 | TYR | p.Tyr408Ter | P49419 |
| HOMO SAPIENS | 9 | 22 | 4x0t | A | 399 | GLU | p.Glu427Gln | P49419 |
| HOMO SAPIENS | 0 | 22 | 4x0t | A | 406 | TYR | p.Tyr434Cys | P49419 |
| HOMO SAPIENS | 0 | 22 | 4x0t | A | 431 | ILE | p.Ile459Phe | P49419 |
| HOMO SAPIENS | 4 | 22 | 4x0t | A | 477 | GLY | p.Gly505Arg | P49419 |
| HOMO SAPIENS | 0 | 22 | 4x0t | A | 495 | THR | p.Thr523Ala | P49419 |
| HOMO SAPIENS | 8 | 11042 | 1y6b | A | 1145 | PRO | p.Pro1147Ser | P35968 |
| HOMO SAPIENS | -1 | 3867 | 2xij | A | 18 | GLN | p.Gln18Ter | P22033 |
| HOMO SAPIENS | -1 | 3867 | 2xij | A | 31 | ARG | p.Arg31Ter | P22033 |
| HOMO SAPIENS | 2 | 3867 | 2xij | A | 93 | ARG | p.Arg93His | P22033 |
| HOMO SAPIENS | 0 | 3867 | 2xij | A | 94 | GLY | p.Gly94Arg | P22033 |
| HOMO SAPIENS | 2 | 3867 | 2xij | A | 95 | PRO | p.Pro95Leu | P22033 |
| HOMO SAPIENS | 2 | 3867 | 2xij | A | 105 | TRP | p.Trp105Arg | P22033 |
| HOMO SAPIENS | 2 | 3867 | 2xij | A | 108 | ARG | p.Arg108Cys | P22033 |
| HOMO SAPIENS | 3 | 3867 | 2xij | A | 154 | ARG | p.Arg154His | P22033 |
| HOMO SAPIENS | 3 | 3867 | 2xij | A | 156 | ASP | p.Asp156Val | P22033 |
| HOMO SAPIENS | 0 | 3867 | 2xij | A | 189 | ASN | p.Asn189Ile | P22033 |
| HOMO SAPIENS | 0 | 3867 | 2xij | A | 191 | ALA | p.Ala191Glu | P22033 |
| HOMO SAPIENS | 0 | 3867 | 2xij | A | 203 | GLY | p.Gly203Arg | P22033 |
| HOMO SAPIENS | 2 | 3867 | 2xij | A | 215 | GLY | p.Gly215Ser | P22033 |
| HOMO SAPIENS | 0 | 3867 | 2xij | A | 219 | ASN | p.Asn219Tyr | P22033 |

| | | | | | | | | |
|---|---|---|---|---|---|---|---|---|
| HOMO SAPIENS | 3 | 3867 | 2xij | A | 228 | ARG | p.Arg228Ter | P22033 |
| HOMO SAPIENS | 1 | 3867 | 2xij | A | 297 | THR | p.Thr297Ile | P22033 |
| HOMO SAPIENS | 0 | 3867 | 2xij | A | 326 | ARG | p.Arg326Lys | P22033 |
| HOMO SAPIENS | 3 | 3867 | 2xij | A | 369 | ARG | p.Arg369His | P22033 |
| HOMO SAPIENS | 3 | 3867 | 2xij | A | 369 | ARG | p.Arg369Cys | P22033 |
| HOMO SAPIENS | 0 | 3867 | 2xij | A | 370 | THR | p.Thr370Pro | P22033 |
| HOMO SAPIENS | 0 | 3867 | 2xij | A | 377 | ALA | p.Ala377Glu | P22033 |
| HOMO SAPIENS | 2 | 3867 | 2xij | A | 426 | GLY | p.Gly426Ala | P22033 |
| HOMO SAPIENS | 2 | 3867 | 2xij | A | 427 | GLY | p.Gly427Asp | P22033 |
| HOMO SAPIENS | 1 | 3867 | 2xij | A | 499 | THR | p.Ala499Thr | P22033 |
| HOMO SAPIENS | 2 | 3867 | 2xij | A | 532 | ARG | p.Arg532His | P22033 |
| HOMO SAPIENS | 3 | 3867 | 2xij | A | 555 | ALA | p.Ala555Thr | P22033 |
| HOMO SAPIENS | 0 | 3867 | 2xij | A | 573 | PHE | p.Phe573Ser | P22033 |
| HOMO SAPIENS | 3 | 3867 | 2xij | A | 623 | GLY | p.Gly623Arg | P22033 |
| HOMO SAPIENS | 5 | 3867 | 2xij | A | 671 | VAL | p.Ile671Val | P22033 |
| HOMO SAPIENS | 4 | 3867 | 2xij | A | 694 | ARG | p.Arg694Trp | P22033 |
| HOMO SAPIENS | 1 | 3867 | 2xij | A | 700 | MET | p.Met700Lys | P22033 |
| HOMO SAPIENS | 8 | 3867 | 2xij | A | 703 | GLY | p.Gly703Arg | P22033 |
| HOMO SAPIENS | 0 | 3867 | 2xij | A | 717 | GLY | p.Gly717Val | P22033 |
| HOMO SAPIENS | 1 | 3867 | 2xij | A | 727 | ARG | p.Arg727Ter | P22033 |
| HOMO SAPIENS | -1 | 52 | 3k91 | B | 72 | GLY | p.Gly73Ser | P00441 |
| HOMO SAPIENS | 6 | 52 | 3k91 | B | 96 | ASP | p.Asp97Asn | P00441 |
| HOMO SAPIENS | 5 | 52 | 3k91 | B | 112 | ILE | p.Ile113Thr | P00441 |
| HOMO SAPIENS | -1 | 52 | 3k91 | B | 134 | SER | p.Ser135Asn | P00441 |
| HOMO SAPIENS | 0 | 3950 | 4fom | A | 296 | ASN | p.Asn296His | Q9NQS3 |
| HOMO SAPIENS | 1 | 10 | 1de4 | A | 31 | VAL | p.Val53Met | Q30201 |
| HOMO SAPIENS | 0 | 10 | 1de4 | A | 37 | VAL | p.Val59Met | Q30201 |
| HOMO SAPIENS | 0 | 10 | 1de4 | A | 71 | GLY | p.Gly93Arg | Q30201 |
| HOMO SAPIENS | 0 | 10 | 1de4 | A | 83 | ILE | p.Ile105Thr | Q30201 |
| HOMO SAPIENS | 0 | 10 | 1de4 | A | 105 | GLN | p.Gln127His | Q30201 |
| HOMO SAPIENS | 1 | 10 | 1de4 | A | 146 | GLU | p.Glu168Ter | Q30201 |
| HOMO SAPIENS | 2 | 10 | 1de4 | A | 261 | GLN | p.Gln283Pro | Q30201 |
| HOMO SAPIENS | 0 | 8038 | 4x7n | A | 586 | ILE | p.Ile586Leu | Q9NZJ5 |
| HOMO SAPIENS | 0 | 8038 | 4x7n | A | 588 | ARG | p.Arg588Gln | Q9NZJ5 |
| HOMO SAPIENS | 0 | 35175 | 5ugw | A | 1062 | ALA | p.Ala1062Thr | O14746 |
| HOMO SAPIENS | 2 | 35175 | 5ugw | A | 1090 | VAL | p.Val1090Met | O14746 |
| HOMO SAPIENS | 0 | 1722 | 4z3k | C | 147 | ARG | p.Arg150Gly | P35270 |
| HOMO SAPIENS | 0 | 1722 | 4z3k | C | 160 | PRO | p.Pro163Leu | P35270 |
| HOMO SAPIENS | 0 | 1722 | 4z3k | C | 207 | ARG | p.Arg210Trp | P35270 |
| HOMO SAPIENS | 0 | 1722 | 4z3k | C | 216 | ARG | p.Arg219Ter | P35270 |
| HOMO SAPIENS | 0 | 1722 | 4z3k | C | 248 | LYS | p.Lys251Ter | P35270 |
| HOMO SAPIENS | 1 | 2832 | 4x8v | L | 115 | HIS | p.His175Gln | P08709 |
| HOMO SAPIENS | 0 | 825 | 1emh | A | 242 | PHE | p.Phe251Ser | P13051 |
| HOMO SAPIENS | 0 | 122 | 5uwk | A | 109 | ARG | p.Arg109Ter | P45452 |
| HOMO SAPIENS | 1 | 122 | 5uwk | A | 170 | LYS | p.Lys170Thr | P45452 |
| HOMO SAPIENS | 9 | 122 | 5uwk | A | 232 | HIS | p.His232Asn | P45452 |
| HOMO SAPIENS | 5 | 122 | 5uwk | A | 257 | ASP | p.Asp257Val | P45452 |
| HOMO SAPIENS | 9 | 1074 | 3p5i | A | 264 | TRP | p.Trp264Arg | Q9UJ71 |
| HOMO SAPIENS | 0 | 146 | 3r3g | A | 14C | GLU | p.Glu352Lys | P00734 |
| HOMO SAPIENS | 3 | 9522 | 1wyq | A | 83 | LEU | p.Leu253Pro | O15020 |
| HOMO SAPIENS | 8 | 67 | 2vgb | A | 130 | SER | p.Ser130Tyr | P30613 |
| HOMO SAPIENS | 8 | 67 | 2vgb | A | 163 | ARG | p.Arg163Cys | P30613 |
| HOMO SAPIENS | 4 | 67 | 2vgb | A | 241 | GLU | p.Glu241Ter | P30613 |
| HOMO SAPIENS | 9 | 67 | 2vgb | A | 384 | THR | p.Thr384Met | P30613 |

| | | | | | | | | |
|---|---|---|---|---|---|---|---|---|
| HOMO SAPIENS | 0 | 67 | 2vgb | A | 421 | GLN | p.Gln421Lys | P30613 |
| HOMO SAPIENS | 6 | 67 | 2vgb | A | 479 | ARG | p.Arg479His | P30613 |
| HOMO SAPIENS | 8 | 67 | 2vgb | A | 510 | ARG | p.Arg510Gln | P30613 |
| HOMO SAPIENS | 4 | 67 | 2vgb | A | 569 | ARG | p.Arg569Trp | P30613 |
| HOMO SAPIENS | 0 | 9088 | 5v44 | A | 232 | ASN | p.Asn232Lys | Q9NZJ4 |
| HOMO SAPIENS | 4 | 16 | 3lbn | A | 12 | GLY | p.Gly12Val | P01112 |
| HOMO SAPIENS | 4 | 16 | 3lbn | A | 12 | GLY | p.Gly12Ser | P01112 |
| HOMO SAPIENS | 4 | 16 | 3lbn | A | 13 | GLY | p.Gly13Cys | P01112 |
| HOMO SAPIENS | 4 | 16 | 3lbn | A | 13 | GLY | p.Gly13Asp | P01112 |
| HOMO SAPIENS | 5 | 16 | 3lbn | A | 22 | GLN | p.Gln22Lys | P01112 |
| HOMO SAPIENS | 5 | 16 | 3lbn | A | 58 | THR | p.Thr58Ile | P01112 |
| HOMO SAPIENS | 3 | 16 | 3lbn | A | 61 | GLN | p.Gln61Lys | P01112 |
| HOMO SAPIENS | 1 | 16 | 3lbn | A | 63 | GLU | p.Glu63Lys | P01112 |
| HOMO SAPIENS | 9 | 16 | 3lbn | A | 117 | LYS | p.Lys117Arg | P01112 |
| HOMO SAPIENS | 6 | 16 | 3lbn | A | 123 | ARG | p.Arg123Cys | P01112 |
| HOMO SAPIENS | 1 | 16 | 3lbn | A | 138 | GLY | p.Gly138Ser | P01112 |
| HOMO SAPIENS | 9 | 16 | 3lbn | A | 146 | ALA | p.Ala146Thr | P01112 |
| HOMO SAPIENS | 9 | 16 | 3lbn | A | 146 | ALA | p.Ala146Val | P01112 |
| HOMO SAPIENS | 1 | 359 | 5nvy | C | 63 | LEU | p.Leu63Pro | P40337 |
| HOMO SAPIENS | 1 | 359 | 5nvy | C | 64 | ARG | p.Arg64Pro | P40337 |
| HOMO SAPIENS | 1 | 359 | 5nvy | C | 70 | GLU | p.Glu70Lys | P40337 |
| HOMO SAPIENS | 0 | 359 | 5nvy | C | 84 | VAL | p.Val84Leu | P40337 |
| HOMO SAPIENS | 1 | 359 | 5nvy | C | 88 | TRP | p.Trp88Ser | P40337 |
| HOMO SAPIENS | 1 | 359 | 5nvy | C | 93 | GLY | p.Gly93Cys | P40337 |
| HOMO SAPIENS | 1 | 359 | 5nvy | C | 98 | TYR | p.Tyr98His | P40337 |
| HOMO SAPIENS | 1 | 359 | 5nvy | C | 111 | SER | p.Ser111Arg | P40337 |
| HOMO SAPIENS | 1 | 359 | 5nvy | C | 112 | TYR | p.Tyr112His | P40337 |
| HOMO SAPIENS | 1 | 359 | 5nvy | C | 126 | ASP | p.Asp126Tyr | P40337 |
| HOMO SAPIENS | 1 | 359 | 5nvy | C | 130 | VAL | p.Val130Leu | P40337 |
| HOMO SAPIENS | 1 | 359 | 5nvy | C | 135 | LEU | p.Leu135Phe | P40337 |
| HOMO SAPIENS | 9 | 359 | 5nvy | C | 161 | ARG | p.Arg161Gly | P40337 |
| HOMO SAPIENS | 7 | 359 | 5nvy | C | 162 | CYS | p.Cys162Phe | P40337 |
| HOMO SAPIENS | 4 | 359 | 5nvy | C | 163 | LEU | p.Leu163Pro | P40337 |
| HOMO SAPIENS | 3 | 359 | 5nvy | C | 164 | GLN | p.Gln164Arg | P40337 |
| HOMO SAPIENS | 9 | 359 | 5nvy | C | 166 | VAL | p.Val166Phe | P40337 |
| HOMO SAPIENS | 7 | 359 | 5nvy | C | 167 | ARG | p.Arg167Gly | P40337 |
| HOMO SAPIENS | 7 | 359 | 5nvy | C | 167 | ARG | p.Arg167Gln | P40337 |
| HOMO SAPIENS | 1 | 359 | 5nvy | C | 183 | SER | p.Ser183Ter | P40337 |
| HOMO SAPIENS | 2 | 359 | 5nvy | C | 188 | LEU | p.Leu188Val | P40337 |
| HOMO SAPIENS | 1 | 359 | 5nvy | C | 191 | HIS | p.His191Asp | P40337 |
| HOMO SAPIENS | 1 | 359 | 5nvy | C | 192 | PRO | p.Pro192Ser | P40337 |
| HOMO SAPIENS | 1 | 359 | 5nvy | C | 200 | ARG | p.Arg200Trp | P40337 |
| HOMO SAPIENS | -1 | 359 | 5nvy | C | 211 | MET | p.Met211Leu | P40337 |
| HOMO SAPIENS | 0 | 14099 | 4btm | A | 282 | ARG | p.Arg258Gln | Q5TCY1 |
| HOMO SAPIENS | 0 | 696 | 5h1b | A | 150 | ARG | p.Arg150Gln | Q06609 |
| HOMO SAPIENS | -1 | 208 | 2bxl | A | 1 | ASP | p.Asp25Val | P02768 |
| HOMO SAPIENS | 0 | 208 | 2bxl | A | 269 | ASP | p.Asp293Gly | P02768 |
| HOMO SAPIENS | 0 | 208 | 2bxl | A | 570 | GLU | p.Glu594Lys | P02768 |
| HOMO SAPIENS | 0 | 208 | 2bxl | A | 573 | LYS | p.Lys597Glu | P02768 |
| HOMO SAPIENS | -1 | 12813 | 2uzx | B | 35 | MET | p.Met35Leu | P08581 |
| HOMO SAPIENS | 1 | 12813 | 2uzx | B | 156 | SER | p.Ser156Leu | P08581 |
| HOMO SAPIENS | 0 | 12813 | 2uzx | B | 168 | GLU | p.Glu168Asp | P08581 |
| HOMO SAPIENS | 2 | 12813 | 2uzx | B | 211 | LEU | p.Leu211Trp | P08581 |
| HOMO SAPIENS | 2 | 12813 | 2uzx | B | 218 | ARG | p.Arg218Ser | P08581 |

| | | | | | | | | |
|---|---|---|---|---|---|---|---|---|
| HOMO SAPIENS | 1 | 12813 | 2uzx | B | 238 | LEU | p.Leu238Ser | P08581 |
| HOMO SAPIENS | 4 | 12813 | 2uzx | B | 320 | ALA | p.Ala320Val | P08581 |
| HOMO SAPIENS | 1 | 12813 | 2uzx | B | 375 | ASN | p.Asn375Ser | P08581 |
| HOMO SAPIENS | 0 | 12813 | 2uzx | B | 575 | LEU | p.Leu575Ile | P08581 |
| HOMO SAPIENS | 1 | 10780 | 4ejx | A | 348 | ARG | p.Arg367Cys | P00450 |
| HOMO SAPIENS | 2 | 10780 | 4ejx | A | 525 | ASP | p.Asp544Glu | P00450 |
| HOMO SAPIENS | 1 | 1407 | 5jn5 | A | 88 | ILE | p.Ile88Val | P36871 |
| HOMO SAPIENS | 7 | 1407 | 5jn5 | A | 115 | THR | p.Thr115Ala | P36871 |
| HOMO SAPIENS | 3 | 1407 | 5jn5 | A | 121 | GLY | p.Gly121Arg | P36871 |
| HOMO SAPIENS | 2 | 1407 | 5jn5 | A | 221 | ARG | p.Arg221Cys | P36871 |
| HOMO SAPIENS | 6 | 1407 | 5jn5 | A | 263 | TYR | p.Asp263Tyr | P36871 |
| HOMO SAPIENS | 3 | 1407 | 5jn5 | A | 420 | TYR | p.Tyr420His | P36871 |
| HOMO SAPIENS | 0 | 1407 | 5jn5 | A | 501 | VAL | p.Val501Ile | P36871 |
| HOMO SAPIENS | -1 | 1878 | 1yow | A | 255 | ARG | p.Arg255Leu | Q13285 |
| HOMO SAPIENS | 6 | 1878 | 1yow | A | 293 | ASP | p.Asp293Asn | Q13285 |
| HOMO SAPIENS | 1 | 1878 | 1yow | A | 437 | LEU | p.Leu437Gln | Q13285 |
| HOMO SAPIENS | 0 | 3071 | 2wpm | E | 145 | ARG | p.Arg191His | P00740 |
| HOMO SAPIENS | 0 | 3071 | 2wpm | E | 145 | ARG | p.Arg191Cys | P00740 |
| HOMO SAPIENS | 3 | 589 | 3ggc | B | 15 | GLY | p.Gly16Ser | P00492 |
| HOMO SAPIENS | 4 | 589 | 3ggc | B | 40 | LEU | p.Leu41Pro | P00492 |
| HOMO SAPIENS | 2 | 589 | 3ggc | B | 50 | ARG | p.Arg51Gly | P00492 |
| HOMO SAPIENS | 5 | 589 | 3ggc | B | 51 | ASP | p.Asp52Gly | P00492 |
| HOMO SAPIENS | 0 | 589 | 3ggc | B | 57 | GLY | p.Gly58Arg | P00492 |
| HOMO SAPIENS | 6 | 589 | 3ggc | B | 69 | GLY | p.Gly70Glu | P00492 |
| HOMO SAPIENS | 1 | 589 | 3ggc | B | 73 | PHE | p.Phe74Leu | P00492 |
| HOMO SAPIENS | 0 | 589 | 3ggc | B | 77 | LEU | p.Leu78Val | P00492 |
| HOMO SAPIENS | 3 | 589 | 3ggc | B | 79 | ASP | p.Asp80Val | P00492 |
| HOMO SAPIENS | 0 | 589 | 3ggc | B | 103 | SER | p.Ser104Arg | P00492 |
| HOMO SAPIENS | 0 | 589 | 3ggc | B | 109 | SER | p.Ser110Leu | P00492 |
| HOMO SAPIENS | 1 | 589 | 3ggc | B | 129 | VAL | p.Val130Asp | P00492 |
| HOMO SAPIENS | 4 | 589 | 3ggc | B | 131 | ILE | p.Ile132Met | P00492 |
| HOMO SAPIENS | 7 | 589 | 3ggc | B | 142 | MET | p.Met143Lys | P00492 |
| HOMO SAPIENS | 0 | 589 | 3ggc | B | 160 | ALA | p.Ala161Ser | P00492 |
| HOMO SAPIENS | 6 | 589 | 3ggc | B | 167 | THR | p.Thr168Ile | P00492 |
| HOMO SAPIENS | 0 | 589 | 3ggc | B | 169 | ARG | p.Arg170Ter | P00492 |
| HOMO SAPIENS | 5 | 589 | 3ggc | B | 175 | PRO | p.Pro176Leu | P00492 |
| HOMO SAPIENS | 8 | 589 | 3ggc | B | 193 | ASP | p.Asp194Glu | P00492 |
| HOMO SAPIENS | 8 | 589 | 3ggc | B | 193 | ASP | p.Asp194Asn | P00492 |
| HOMO SAPIENS | 8 | 589 | 3ggc | B | 198 | PHE | p.Phe199Val | P00492 |
| HOMO SAPIENS | 8 | 589 | 3ggc | B | 200 | ASP | p.Asp201Gly | P00492 |
| HOMO SAPIENS | 0 | 4269 | 2pla | B | 174 | GLU | p.Glu174Lys | Q8N335 |
| HOMO SAPIENS | 1 | 4269 | 2pla | B | 220 | ARG | p.Arg220His | Q8N335 |
| HOMO SAPIENS | 4 | 4269 | 2pla | B | 280 | ALA | p.Ala280Val | Q8N335 |
| HOMO SAPIENS | 0 | 2712 | 1ii4 | E | 252 | TRP | p.Ser252Leu | P21802 |
| HOMO SAPIENS | 0 | 2712 | 1ii4 | E | 253 | PRO | p.Pro253Arg | P21802 |
| HOMO SAPIENS | 5 | 2712 | 1ii4 | E | 267 | SER | p.Ser267Pro | P21802 |
| HOMO SAPIENS | 0 | 2712 | 1ii4 | E | 278 | CYS | p.Cys278Phe | P21802 |
| HOMO SAPIENS | 0 | 2712 | 1ii4 | E | 289 | GLN | p.Gln289Pro | P21802 |
| HOMO SAPIENS | 0 | 2712 | 1ii4 | E | 290 | TRP | p.Trp290Cys | P21802 |
| HOMO SAPIENS | 0 | 2712 | 1ii4 | E | 290 | TRP | p.Trp290Arg | P21802 |
| HOMO SAPIENS | 0 | 2712 | 1ii4 | E | 292 | LYS | p.Lys292Glu | P21802 |
| HOMO SAPIENS | 0 | 2712 | 1ii4 | E | 315 | ALA | p.Ala315Ser | P21802 |
| HOMO SAPIENS | 0 | 2712 | 1ii4 | E | 321 | ASP | p.Asp321Ala | P21802 |
| HOMO SAPIENS | 1 | 2712 | 1ii4 | E | 328 | TYR | p.Tyr328Cys | P21802 |

| | | | | | | | | |
|---|---|---|---|---|---|---|---|---|
| HOMO SAPIENS | 9 | 2712 | 1ii4 | E | 340 | TYR | p.Tyr340His | P21802 |
| HOMO SAPIENS | 1 | 2712 | 1ii4 | E | 341 | THR | p.Thr341Pro | P21802 |
| HOMO SAPIENS | 0 | 2712 | 1ii4 | E | 342 | CYS | p.Cys342Arg | P21802 |
| HOMO SAPIENS | 0 | 2712 | 1ii4 | E | 342 | CYS | p.Cys342Trp | P21802 |
| HOMO SAPIENS | 0 | 2712 | 1ii4 | E | 344 | ALA | p.Ala344Gly | P21802 |
| HOMO SAPIENS | 0 | 2712 | 1ii4 | E | 347 | SER | p.Ser347Cys | P21802 |
| HOMO SAPIENS | 0 | 2712 | 1ii4 | E | 351 | SER | p.Ser351Cys | P21802 |
| HOMO SAPIENS | 1 | 2712 | 1ii4 | E | 354 | SER | p.Ser354Cys | P21802 |
| HOMO SAPIENS | 2 | 53 | 4pj1 | H | 5 | ASP | p.Asp29Gly | P10809 |
| HOMO SAPIENS | 0 | 53 | 4pj1 | H | 74 | VAL | p.Val98Ile | P10809 |
| HOMO SAPIENS | 0 | 29968 | 2fdb | M | 71 | LYS | p.Lys89Glu | P55075 |
| HOMO SAPIENS | 2 | 29968 | 2fdb | M | 98 | ARG | p.Arg116Gly | P55075 |
| HOMO SAPIENS | 2 | 29968 | 2fdb | M | 122 | GLY | p.Gly140Ser | P55075 |
| HOMO SAPIENS | 8 | 5061 | 5f85 | B | 60 | GLY | p.Gly106Ser | P00740 |
| HOMO SAPIENS | 0 | 35944 | 1h6h | A | 85 | THR | p.Thr85Asn | Q15080 |
| HOMO SAPIENS | 8 | 35944 | 1h6h | A | 105 | ARG | p.Arg105Gln | Q15080 |
| HOMO SAPIENS | 7 | 12016 | 1hj7 | A | 295 | ASN | p.Asn316Ser | P01130 |
| HOMO SAPIENS | 5 | 12016 | 1hj7 | A | 302 | GLY | p.Gly323Ser | P01130 |
| HOMO SAPIENS | 6 | 12016 | 1hj7 | A | 319 | CYS | p.Cys340Trp | P01130 |
| HOMO SAPIENS | 7 | 12016 | 1hj7 | A | 329 | ARG | p.Arg350Ter | P01130 |
| HOMO SAPIENS | 5 | 12016 | 1hj7 | A | 331 | CYS | p.Cys352Ter | P01130 |
| HOMO SAPIENS | 2 | 12016 | 1hj7 | A | 345 | GLN | p.Gln366Arg | P01130 |
| HOMO SAPIENS | 2 | 1481 | 2i07 | A | 239 | TYR | p.Tyr261Ter | P01024 |
| HOMO SAPIENS | -1 | 1481 | 2i07 | A | 292 | LEU | p.Pro314Leu | P01024 |
| HOMO SAPIENS | 2 | 1481 | 2i07 | A | 326 | ILE | p.Ile348Val | P01024 |
| HOMO SAPIENS | 2 | 1481 | 2i07 | A | 413 | GLU | p.Glu435Lys | P01024 |
| HOMO SAPIENS | 4 | 1481 | 2i07 | A | 447 | GLU | p.Glu469Asp | P01024 |
| HOMO SAPIENS | 2 | 1481 | 2i07 | A | 597 | VAL | p.Val619Met | P01024 |
| HOMO SAPIENS | 5 | 1481 | 2i07 | A | 603 | ILE | p.Ile625Phe | P01024 |
| HOMO SAPIENS | 0 | 1481 | 2i07 | A | 611 | LYS | p.Lys633Arg | P01024 |
| HOMO SAPIENS | 0 | 11123 | 3c5n | A | 315 | GLY | p.Gly368Trp | O00294 |
| HOMO SAPIENS | 8 | 11123 | 3c5n | A | 329 | PHE | p.Phe382Ser | O00294 |
| HOMO SAPIENS | 5 | 11123 | 3c5n | A | 429 | ARG | p.Arg482Trp | O00294 |
| HOMO SAPIENS | 8 | 11123 | 3c5n | A | 438 | PHE | p.Phe491Leu | O00294 |
| HOMO SAPIENS | 3 | 4398 | 3fx0 | B | 1288 | ALA | p.Ala288Gly | Q9Y6K9 |
| HOMO SAPIENS | 7 | 4398 | 3fx0 | B | 1315 | GLU | p.Glu315Ala | Q9Y6K9 |
| HOMO SAPIENS | 8 | 4398 | 3fx0 | B | 1319 | ARG | p.Arg319Gln | Q9Y6K9 |
| HOMO SAPIENS | 6 | 3989 | 3qd6 | S | 83 | CYS | p.Cys83Arg | P25942 |
| HOMO SAPIENS | 1 | 3989 | 3qd6 | S | 124 | SER | p.Ser124Leu | P25942 |
| HOMO SAPIENS | 2 | 24003 | 2m0v | A | 225 | GLU | p.Glu225Lys | O14745 |
| HOMO SAPIENS | 2 | 46 | 4aqd | A | 70 | ASP | p.Asp98Gly | P06276 |
| HOMO SAPIENS | 4 | 46 | 4aqd | A | 115 | GLY | p.Gly143Asp | P06276 |
| HOMO SAPIENS | 3 | 46 | 4aqd | A | 184 | ALA | p.Ala212Val | P06276 |
| HOMO SAPIENS | 0 | 46 | 4aqd | A | 225 | GLY | p.Gly253Ter | P06276 |
| HOMO SAPIENS | 2 | 46 | 4aqd | A | 243 | THR | p.Thr271Met | P06276 |
| HOMO SAPIENS | 2 | 46 | 4aqd | A | 255 | GLU | p.Glu283Asp | P06276 |
| HOMO SAPIENS | 5 | 46 | 4aqd | A | 267 | LYS | p.Lys295Arg | P06276 |
| HOMO SAPIENS | 3 | 46 | 4aqd | A | 271 | GLU | p.Glu299Ter | P06276 |
| HOMO SAPIENS | 2 | 46 | 4aqd | A | 307 | LEU | p.Leu335Pro | P06276 |
| HOMO SAPIENS | 1 | 46 | 4aqd | A | 330 | LEU | p.Leu358Ile | P06276 |
| HOMO SAPIENS | 0 | 46 | 4aqd | A | 497 | GLU | p.Glu525Val | P06276 |
| HOMO SAPIENS | 3 | 786 | 1ols | B | 51 | GLY | p.Gly101Asp | P21953 |
| HOMO SAPIENS | 2 | 786 | 1ols | B | 87 | ALA | p.Ala137Val | P21953 |
| HOMO SAPIENS | 0 | 786 | 1ols | B | 110 | ILE | p.Ile160Ser | P21953 |

| | | | | | | | | |
|---|---|---|---|---|---|---|---|---|
| HOMO SAPIENS | 4 | 786 | 1ols | B | 120 | ARG | p.Arg170Cys | P21953 |
| HOMO SAPIENS | 2 | 786 | 1ols | B | 126 | ASN | p.Asn176Tyr | P21953 |
| HOMO SAPIENS | 0 | 786 | 1ols | B | 133 | ARG | p.Arg183Gly | P21953 |
| HOMO SAPIENS | 0 | 786 | 1ols | B | 133 | ARG | p.Arg183Pro | P21953 |
| HOMO SAPIENS | 0 | 786 | 1ols | B | 135 | PRO | p.Pro185Leu | P21953 |
| HOMO SAPIENS | 3 | 786 | 1ols | B | 156 | HIS | p.His206Tyr | P21953 |
| HOMO SAPIENS | 0 | 786 | 1ols | B | 228 | GLY | p.Gly278Ser | P21953 |
| HOMO SAPIENS | 1 | 786 | 1ols | B | 235 | ARG | p.Arg285Ter | P21953 |
| HOMO SAPIENS | 6 | 786 | 1ols | B | 274 | ARG | p.Arg324Ter | P21953 |
| HOMO SAPIENS | 1 | 831 | 4zt1 | B | 48 | VAL | p.Val202Ile | P12830 |
| HOMO SAPIENS | 6 | 831 | 4zt1 | B | 70 | ARG | p.Arg224Cys | P12830 |
| HOMO SAPIENS | 5 | 831 | 4zt1 | B | 79 | HIS | p.His233Gln | P12830 |
| HOMO SAPIENS | 4 | 831 | 4zt1 | B | 120 | GLY | p.Gly274Ser | P12830 |
| HOMO SAPIENS | 7 | 831 | 4zt1 | B | 187 | TYR | p.Tyr341Ter | P12830 |
| HOMO SAPIENS | 0 | 8497 | 3sla | A | 287 | ASN | p.Asn287Ser | P35222 |
| HOMO SAPIENS | 7 | 23986 | 2ke1 | A | 301 | VAL | p.Val301Met | O43918 |
| HOMO SAPIENS | 0 | 4967 | 1rj7 | B | 334 | ARG | p.Arg334His | Q92838 |
| HOMO SAPIENS | 0 | 4967 | 1rj7 | B | 338 | THR | p.Thr338Met | Q92838 |
| HOMO SAPIENS | 3 | 4967 | 1rj7 | B | 349 | ALA | p.Ala349Thr | Q92838 |
| HOMO SAPIENS | 2 | 4967 | 1rj7 | B | 358 | GLN | p.Gln358Glu | Q92838 |
| HOMO SAPIENS | 1 | 4967 | 1rj7 | B | 375 | LYS | p.Lys375Glu | Q92838 |
| HOMO SAPIENS | 9 | 7623 | 1d4t | A | 32 | ARG | p.Arg32Thr | O60880 |
| HOMO SAPIENS | 6 | 7623 | 1d4t | A | 55 | ARG | p.Arg55Leu | O60880 |
| HOMO SAPIENS | 0 | 7623 | 1d4t | A | 68 | THR | p.Thr68Ile | O60880 |
| HOMO SAPIENS | 6 | 705 | 3idh | A | 59 | PRO | p.Pro59Ser | P35557 |
| HOMO SAPIENS | 5 | 705 | 3idh | A | 124 | ASP | p.Asp124Asn | P35557 |
| HOMO SAPIENS | 4 | 705 | 3idh | A | 131 | SER | p.Ser131Pro | P35557 |
| HOMO SAPIENS | 0 | 705 | 3idh | A | 147 | GLY | p.Gly147Asp | P35557 |
| HOMO SAPIENS | 5 | 705 | 3idh | A | 186 | ARG | p.Arg186Ter | P35557 |
| HOMO SAPIENS | 0 | 705 | 3idh | A | 202 | MET | p.Met202Thr | P35557 |
| HOMO SAPIENS | 2 | 705 | 3idh | A | 210 | MET | p.Met210Lys | P35557 |
| HOMO SAPIENS | 1 | 705 | 3idh | A | 215 | TYR | p.Tyr215Ter | P35557 |
| HOMO SAPIENS | 0 | 705 | 3idh | A | 226 | VAL | p.Val226Met | P35557 |
| HOMO SAPIENS | 9 | 705 | 3idh | A | 228 | THR | p.Thr228Met | P35557 |
| HOMO SAPIENS | 9 | 705 | 3idh | A | 256 | GLU | p.Glu256Lys | P35557 |
| HOMO SAPIENS | 2 | 705 | 3idh | A | 261 | GLY | p.Gly261Arg | P35557 |
| HOMO SAPIENS | 6 | 705 | 3idh | A | 279 | GLU | p.Glu279Gln | P35557 |
| HOMO SAPIENS | 7 | 705 | 3idh | A | 299 | GLY | p.Gly299Arg | P35557 |
| HOMO SAPIENS | 2 | 705 | 3idh | A | 318 | GLY | p.Gly318Trp | P35557 |
| HOMO SAPIENS | 4 | 705 | 3idh | A | 378 | ALA | p.Ala378Thr | P35557 |
| HOMO SAPIENS | 0 | 705 | 3idh | A | 387 | ALA | p.Ala387Glu | P35557 |
| HOMO SAPIENS | 5 | 705 | 3idh | A | 414 | LYS | p.Lys414Glu | P35557 |
| HOMO SAPIENS | 0 | 705 | 3idh | A | 455 | VAL | p.Val455Met | P35557 |
| HOMO SAPIENS | -1 | 705 | 3idh | A | 462 | MET | p.Met462Ile | P35557 |
| HOMO SAPIENS | 1 | 509 | 4nef | A | 22 | LEU | p.Leu22Val | P41181 |
| HOMO SAPIENS | 4 | 509 | 4nef | A | 57 | GLN | p.Gln57Pro | P41181 |
| HOMO SAPIENS | 8 | 509 | 4nef | A | 64 | GLY | p.Gly64Arg | P41181 |
| HOMO SAPIENS | 7 | 509 | 4nef | A | 68 | ASN | p.Asn68Ser | P41181 |
| HOMO SAPIENS | 1 | 509 | 4nef | A | 100 | GLY | p.Gly100Val | P41181 |
| HOMO SAPIENS | 0 | 509 | 4nef | A | 125 | THR | p.Thr125Met | P41181 |
| HOMO SAPIENS | 0 | 509 | 4nef | A | 126 | THR | p.Thr126Met | P41181 |
| HOMO SAPIENS | 4 | 509 | 4nef | A | 146 | PHE | p.Phe146Leu | P41181 |
| HOMO SAPIENS | 0 | 509 | 4nef | A | 175 | GLY | p.Gly175Arg | P41181 |
| HOMO SAPIENS | 5 | 509 | 4nef | A | 181 | CYS | p.Cys181Trp | P41181 |

| | | | | | | | | | |
|---|---|---|---|---|---|---|---|---|---|
| HOMO SAPIENS | 9 | 509 | 4nef | A | | 187 | ARG | p.Arg187Cys | P41181 |
| HOMO SAPIENS | 0 | 509 | 4nef | A | | 190 | ALA | p.Ala190Thr | P41181 |
| HOMO SAPIENS | 0 | 509 | 4nef | A | | 216 | SER | p.Ser216Pro | P41181 |
| HOMO SAPIENS | 4 | 2242 | 1z70 | X | | 1155 | SER | p.Ser155Pro | Q8NBK3 |
| HOMO SAPIENS | 3 | 2242 | 1z70 | X | | 1218 | CYS | p.Cys218Tyr | Q8NBK3 |
| HOMO SAPIENS | 0 | 2242 | 1z70 | X | | 1279 | ALA | p.Ala279Val | Q8NBK3 |
| HOMO SAPIENS | -1 | 2242 | 1z70 | X | | 1336 | OCS | p.Cys336Arg | Q8NBK3 |
| HOMO SAPIENS | 4 | 2242 | 1z70 | X | | 1345 | ARG | p.Arg345Cys | Q8NBK3 |
| HOMO SAPIENS | 2 | 2242 | 1z70 | X | | 1348 | ALA | p.Ala348Pro | Q8NBK3 |
| HOMO SAPIENS | 5 | 2242 | 1z70 | X | | 1349 | ARG | p.Arg349Gln | Q8NBK3 |
| HOMO SAPIENS | 5 | 2242 | 1z70 | X | | 1349 | ARG | p.Arg349Trp | Q8NBK3 |
| HOMO SAPIENS | 3 | 2242 | 1z70 | X | | 1359 | SER | p.Ser359Ter | Q8NBK3 |
| HOMO SAPIENS | 5 | 8269 | 5wrr | B | | 136 | ARG | p.Arg136Ter | Q96MK3 |
| HOMO SAPIENS | 1 | 8269 | 5wrr | B | | 332 | LYS | p.Asn332Lys | Q96MK3 |
| HOMO SAPIENS | 3 | 8269 | 5wrr | B | | 478 | ARG | p.Arg478Ter | Q96MK3 |
| HOMO SAPIENS | 5 | 8269 | 5wrr | B | | 504 | THR | p.Thr504Ile | Q96MK3 |
| HOMO SAPIENS | 5 | 1411 | 4h1s | B | | 221 | SER | p.Ser221Ter | P21589 |
| HOMO SAPIENS | 1 | 1411 | 4h1s | B | | 358 | CYS | p.Cys358Tyr | P21589 |
| HOMO SAPIENS | 2 | 23 | 5vvd | B | | 298 | GLU | p.Glu298Asp | P29474 |
| HOMO SAPIENS | 0 | 19 | 5jlh | E | | 88 | THR | p.Thr89Ile | P63261 |
| HOMO SAPIENS | 6 | 19 | 5jlh | E | | 117 | LYS | p.Lys118Met | P63261 |
| HOMO SAPIENS | 5 | 19 | 5jlh | E | | 263 | PRO | p.Pro264Leu | P63261 |
| HOMO SAPIENS | 1 | 19 | 5jlh | E | | 277 | THR | p.Thr278Ile | P63261 |
| HOMO SAPIENS | 6 | 19 | 5jlh | E | | 331 | PRO | p.Pro332Ala | P63261 |
| HOMO SAPIENS | 5 | 19 | 5jlh | E | | 369 | VAL | p.Val370Ala | P63261 |
| HOMO SAPIENS | 0 | 14306 | 4w4o | C | | 92 | ARG | p.Arg92Ter | P12314 |
| HOMO SAPIENS | 3 | 603 | 1nuh | A | | 4 | THR | p.Thr5Ile | P06744 |
| HOMO SAPIENS | 2 | 603 | 1nuh | A | | 19 | HIS | p.His20Pro | P06744 |
| HOMO SAPIENS | 8 | 603 | 1nuh | A | | 158 | GLY | p.Gly159Ser | P06744 |
| HOMO SAPIENS | 6 | 603 | 1nuh | A | | 223 | THR | p.Thr224Met | P06744 |
| HOMO SAPIENS | 0 | 603 | 1nuh | A | | 338 | LEU | p.Leu339Pro | P06744 |
| HOMO SAPIENS | 6 | 603 | 1nuh | A | | 342 | GLN | p.Gln343Arg | P06744 |
| HOMO SAPIENS | 6 | 603 | 1nuh | A | | 374 | THR | p.Thr375Arg | P06744 |
| HOMO SAPIENS | 4 | 603 | 1nuh | A | | 524 | ILE | p.Ile525Thr | P06744 |
| HOMO SAPIENS | 4 | 603 | 1nuh | A | | 538 | ASP | p.Asp539Asn | P06744 |
| HOMO SAPIENS | 1 | 3107 | 3kzj | A | | 1197 | VAL | p.Val1197Ala | P08603 |
| HOMO SAPIENS | 9 | 1748 | 2noh | A | | 46 | ARG | p.Arg46Gln | O15527 |
| HOMO SAPIENS | 2 | 11750 | 3s4y | A | | 33 | ARG | p.Arg33His | Q9H3S4 |
| HOMO SAPIENS | 2 | 11750 | 3s4y | A | | 40 | LEU | p.Leu40Pro | Q9H3S4 |
| HOMO SAPIENS | 4 | 11750 | 3s4y | A | | 142 | LEU | p.Leu142Phe | Q9H3S4 |
| HOMO SAPIENS | 5 | 11750 | 3s4y | A | | 219 | ASN | p.Asn219Ser | Q9H3S4 |
| HOMO SAPIENS | 1 | 2591 | 5vnj | A | | 382 | PHE | p.Phe382Leu | Q15436 |
| HOMO SAPIENS | 2 | 2591 | 5vnj | A | | 702 | MET | p.Met702Val | Q15436 |
| HOMO SAPIENS | 1 | 2426 | 5m8t | A | | 87 | ARG | p.Arg87Trp | P17643 |
| HOMO SAPIENS | 1 | 2426 | 5m8t | A | | 166 | SER | p.Ser166Ter | P17643 |
| HOMO SAPIENS | 0 | 2426 | 5m8t | A | | 356 | ARG | p.Arg356Gln | P17643 |
| HOMO SAPIENS | 2 | 2426 | 5m8t | A | | 374 | SER | p.Arg374Ter | P17643 |
| HOMO SAPIENS | 6 | 2426 | 5m8t | A | | 378 | ASN | p.Asn378Ser | P17643 |
| HOMO SAPIENS | 6 | 2426 | 5m8t | A | | 382 | LEU | p.Leu382Pro | P17643 |
| HOMO SAPIENS | 7 | 10097 | 4n6v | | 1 | 46 | GLN | p.Gln46Ter | P04080 |
| HOMO SAPIENS | 3 | 10097 | 4n6v | | 1 | 65 | VAL | p.Val65Ile | P04080 |
| HOMO SAPIENS | 8 | 10097 | 4n6v | | 1 | 68 | ARG | p.Arg68Ter | P04080 |
| HOMO SAPIENS | 2 | 859 | 4z9a | A | | 105 | GLN | p.Gln106Arg | P24666 |
| HOMO SAPIENS | 8 | 1420 | 5nan | F | | 95 | SER | p.Ser95Leu | Q96PD4 |

| | | | | | | | | |
|---|---|---|---|---|---|---|---|---|
| HOMO SAPIENS | 6 | 1420 | 5nan | F | 126 | GLU | p.Glu126Gly | Q96PD4 |
| HOMO SAPIENS | 8 | 1420 | 5nan | F | 155 | VAL | p.Val155Ile | Q96PD4 |
| HOMO SAPIENS | 6 | 1420 | 5nan | F | 157 | PRO | p.Pro157Ser | Q96PD4 |
| HOMO SAPIENS | -1 | 1420 | 5nan | F | 161 | HIS | p.His161Arg | Q96PD4 |
| HOMO SAPIENS | 3 | 288 | 5i96 | B | 140 | GLN | p.Arg140Gly | P48735 |
| HOMO SAPIENS | 3 | 288 | 5i96 | B | 140 | GLN | p.Arg140Gln | P48735 |
| HOMO SAPIENS | 2 | 288 | 5i96 | B | 261 | ARG | p.Arg261His | P48735 |
| HOMO SAPIENS | 2 | 288 | 5i96 | B | 435 | THR | p.Thr435Met | P48735 |
| HOMO SAPIENS | 9 | 120 | 1tup | B | 107 | TYR | p.Tyr107His | P04637 |
| HOMO SAPIENS | 8 | 120 | 1tup | B | 110 | ARG | p.Arg110Leu | P04637 |
| HOMO SAPIENS | 5 | 120 | 1tup | B | 120 | LYS | p.Lys120Glu | P04637 |
| HOMO SAPIENS | 4 | 120 | 1tup | B | 133 | MET | p.Met133Thr | P04637 |
| HOMO SAPIENS | 3 | 120 | 1tup | B | 134 | PHE | p.Phe134Cys | P04637 |
| HOMO SAPIENS | 8 | 120 | 1tup | B | 136 | GLN | p.Gln136His | P04637 |
| HOMO SAPIENS | 9 | 120 | 1tup | B | 138 | ALA | p.Ala138Pro | P04637 |
| HOMO SAPIENS | 9 | 120 | 1tup | B | 151 | PRO | p.Pro151Ser | P04637 |
| HOMO SAPIENS | 9 | 120 | 1tup | B | 152 | PRO | p.Pro152Leu | P04637 |
| HOMO SAPIENS | 9 | 120 | 1tup | B | 158 | ARG | p.Arg158Cys | P04637 |
| HOMO SAPIENS | 9 | 120 | 1tup | B | 158 | ARG | p.Arg158His | P04637 |
| HOMO SAPIENS | 9 | 120 | 1tup | B | 175 | ARG | p.Arg175Cys | P04637 |
| HOMO SAPIENS | 9 | 120 | 1tup | B | 175 | ARG | p.Arg175Leu | P04637 |
| HOMO SAPIENS | 8 | 120 | 1tup | B | 181 | ARG | p.Arg181Leu | P04637 |
| HOMO SAPIENS | 9 | 120 | 1tup | B | 189 | ALA | p.Ala189Val | P04637 |
| HOMO SAPIENS | 9 | 120 | 1tup | B | 196 | ARG | p.Arg196Gly | P04637 |
| HOMO SAPIENS | 9 | 120 | 1tup | B | 213 | ARG | p.Arg213Gln | P04637 |
| HOMO SAPIENS | 9 | 120 | 1tup | B | 213 | ARG | p.Arg213Ter | P04637 |
| HOMO SAPIENS | 9 | 120 | 1tup | B | 214 | HIS | p.His214Gln | P04637 |
| HOMO SAPIENS | 9 | 120 | 1tup | B | 220 | TYR | p.Tyr220Cys | P04637 |
| HOMO SAPIENS | 0 | 120 | 1tup | B | 234 | TYR | p.Tyr234Cys | P04637 |
| HOMO SAPIENS | 9 | 120 | 1tup | B | 237 | MET | p.Met237Ile | P04637 |
| HOMO SAPIENS | 0 | 120 | 1tup | B | 238 | CYS | p.Cys238Tyr | P04637 |
| HOMO SAPIENS | 9 | 120 | 1tup | B | 241 | SER | p.Ser241Phe | P04637 |
| HOMO SAPIENS | 9 | 120 | 1tup | B | 242 | CYS | p.Cys242Ser | P04637 |
| HOMO SAPIENS | 9 | 120 | 1tup | B | 245 | GLY | p.Gly245Cys | P04637 |
| HOMO SAPIENS | 9 | 120 | 1tup | B | 245 | GLY | p.Gly245Asp | P04637 |
| HOMO SAPIENS | 9 | 120 | 1tup | B | 248 | ARG | p.Arg248Leu | P04637 |
| HOMO SAPIENS | 9 | 120 | 1tup | B | 248 | ARG | p.Arg248Trp | P04637 |
| HOMO SAPIENS | 9 | 120 | 1tup | B | 251 | ILE | p.Ile251Ser | P04637 |
| HOMO SAPIENS | 9 | 120 | 1tup | B | 252 | LEU | p.Leu252Pro | P04637 |
| HOMO SAPIENS | 9 | 120 | 1tup | B | 258 | GLU | p.Glu258Lys | P04637 |
| HOMO SAPIENS | 8 | 120 | 1tup | B | 272 | VAL | p.Val272Leu | P04637 |
| HOMO SAPIENS | 9 | 120 | 1tup | B | 273 | ARG | p.Arg273Cys | P04637 |
| HOMO SAPIENS | 9 | 120 | 1tup | B | 273 | ARG | p.Arg273His | P04637 |
| HOMO SAPIENS | 9 | 120 | 1tup | B | 277 | CYS | p.Cys277Phe | P04637 |
| HOMO SAPIENS | 9 | 120 | 1tup | B | 280 | ARG | p.Arg280Ile | P04637 |
| HOMO SAPIENS | 9 | 120 | 1tup | B | 282 | ARG | p.Arg282Trp | P04637 |
| HOMO SAPIENS | 9 | 120 | 1tup | B | 283 | ARG | p.Arg283His | P04637 |
| HOMO SAPIENS | 9 | 120 | 1tup | B | 285 | GLU | p.Glu285Val | P04637 |
| HOMO SAPIENS | -1 | 120 | 1tup | B | 292 | LYS | p.Lys292Ile | P04637 |
| HOMO SAPIENS | -1 | 120 | 1tup | B | 298 | GLU | p.Glu298Lys | P04637 |
| HOMO SAPIENS | -1 | 2791 | 4ygb | A | 39 | VAL | p.Val39Ala | P49257 |
| HOMO SAPIENS | 2 | 106 | 4x6j | A | 32 | GLY | p.Gly146Arg | P43235 |
| HOMO SAPIENS | 0 | 106 | 4x6j | A | 76 | GLN | p.Gln190Ter | P43235 |
| HOMO SAPIENS | 7 | 106 | 4x6j | A | 198 | ARG | p.Arg312Ter | P43235 |

| | | | | | | | | |
|---|---|---|---|---|---|---|---|---|
| HOMO SAPIENS | 5 | 490 | 2hlo | B | 172 | LEU | p.Leu202Gln | P02675 |
| HOMO SAPIENS | 0 | 490 | 2hlo | B | 353 | LEU | p.Leu383Arg | P02675 |
| HOMO SAPIENS | 4 | 490 | 2hlo | B | 400 | GLY | p.Gly430Asp | P02675 |
| HOMO SAPIENS | 7 | 490 | 2hlo | B | 448 | ARG | p.Arg478Lys | P02675 |
| HOMO SAPIENS | 1 | 289 | 4p13 | B | 18 | GLU | p.Glu43Lys | P11310 |
| HOMO SAPIENS | 2 | 289 | 4p13 | B | 20 | GLN | p.Gln45Arg | P11310 |
| HOMO SAPIENS | 5 | 289 | 4p13 | B | 28 | ARG | p.Arg53Cys | P11310 |
| HOMO SAPIENS | 1 | 289 | 4p13 | B | 42 | TYR | p.Tyr67His | P11310 |
| HOMO SAPIENS | 1 | 289 | 4p13 | B | 53 | ILE | p.Ile78Thr | P11310 |
| HOMO SAPIENS | 1 | 289 | 4p13 | B | 59 | LEU | p.Leu84Phe | P11310 |
| HOMO SAPIENS | 2 | 289 | 4p13 | B | 96 | THR | p.Thr121Ile | P11310 |
| HOMO SAPIENS | 0 | 289 | 4p13 | B | 124 | MET | p.Met149Ile | P11310 |
| HOMO SAPIENS | 5 | 289 | 4p13 | B | 143 | ASP | p.Asp168Ala | P11310 |
| HOMO SAPIENS | 7 | 289 | 4p13 | B | 168 | THR | p.Thr193Ala | P11310 |
| HOMO SAPIENS | 1 | 289 | 4p13 | B | 170 | GLY | p.Gly195Arg | P11310 |
| HOMO SAPIENS | 2 | 289 | 4p13 | B | 181 | ARG | p.Arg206Cys | P11310 |
| HOMO SAPIENS | 2 | 289 | 4p13 | B | 181 | ARG | p.Arg206His | P11310 |
| HOMO SAPIENS | 8 | 289 | 4p13 | B | 203 | THR | p.Thr228Ile | P11310 |
| HOMO SAPIENS | 2 | 289 | 4p13 | B | 208 | ILE | p.Ile233Asn | P11310 |
| HOMO SAPIENS | 1 | 289 | 4p13 | B | 219 | CYS | p.Cys244Arg | P11310 |
| HOMO SAPIENS | 6 | 289 | 4p13 | B | 220 | SER | p.Ser245Leu | P11310 |
| HOMO SAPIENS | 5 | 289 | 4p13 | B | 228 | GLU | p.Glu253Lys | P11310 |
| HOMO SAPIENS | 2 | 289 | 4p13 | B | 241 | ASP | p.Asp266Gly | P11310 |
| HOMO SAPIENS | 1 | 289 | 4p13 | B | 242 | GLY | p.Gly267Arg | P11310 |
| HOMO SAPIENS | 7 | 289 | 4p13 | B | 256 | ARG | p.Arg281Thr | P11310 |
| HOMO SAPIENS | 4 | 289 | 4p13 | B | 269 | ARG | p.Arg294Thr | P11310 |
| HOMO SAPIENS | 4 | 289 | 4p13 | B | 304 | GLU | p.Lys329Glu | P11310 |
| HOMO SAPIENS | 1 | 289 | 4p13 | B | 324 | ARG | p.Arg349Ter | P11310 |
| HOMO SAPIENS | 3 | 289 | 4p13 | B | 326 | THR | p.Thr351Ile | P11310 |
| HOMO SAPIENS | 7 | 289 | 4p13 | B | 350 | ILE | p.Ile375Thr | P11310 |
| HOMO SAPIENS | 3 | 289 | 4p13 | B | 359 | GLU | p.Glu384Ter | P11310 |
| HOMO SAPIENS | 1 | 4563 | 3i2b | A | 16 | ARG | p.Arg16Cys | Q03393 |
| HOMO SAPIENS | 5 | 4563 | 3i2b | A | 25 | ARG | p.Arg25Gln | Q03393 |
| HOMO SAPIENS | 1 | 4563 | 3i2b | A | 47 | ASN | p.Asn47Asp | Q03393 |
| HOMO SAPIENS | 8 | 4563 | 3i2b | A | 52 | ASN | p.Asn52Ser | Q03393 |
| HOMO SAPIENS | 1 | 4563 | 3i2b | A | 56 | VAL | p.Val56Met | Q03393 |
| HOMO SAPIENS | 1 | 4563 | 3i2b | A | 87 | PRO | p.Pro87Ser | Q03393 |
| HOMO SAPIENS | 1 | 4563 | 3i2b | A | 96 | ASP | p.Asp96Asn | Q03393 |
| HOMO SAPIENS | 0 | 4563 | 3i2b | A | 99 | TYR | p.Tyr99Ter | Q03393 |
| HOMO SAPIENS | 2 | 4563 | 3i2b | A | 116 | ASP | p.Asp116Gly | Q03393 |
| HOMO SAPIENS | 2 | 41 | 3exx | A | 1 | GLY | p.Gly90Cys | P01308 |
| HOMO SAPIENS | 5 | 41 | 3exx | A | 3 | VAL | p.Val92Leu | P01308 |
| HOMO SAPIENS | 4 | 41 | 3exx | A | 7 | CYS | p.Cys96Ser | P01308 |
| HOMO SAPIENS | 9 | 41 | 3exx | A | 19 | TYR | p.Tyr108Cys | P01308 |
| HOMO SAPIENS | 9 | 131 | 4mjo | D | 118 | ASP | p.Asp119Asn | P09467 |
| HOMO SAPIENS | 0 | 131 | 4mjo | D | 163 | GLY | p.Gly164Ser | P09467 |
| HOMO SAPIENS | 0 | 131 | 4mjo | D | 176 | ALA | p.Ala177Asp | P09467 |
| HOMO SAPIENS | 5 | 131 | 4mjo | D | 193 | PHE | p.Phe194Ser | P09467 |
| HOMO SAPIENS | 4 | 131 | 4mjo | D | 217 | ARG | p.Arg218Lys | P09467 |
| HOMO SAPIENS | 4 | 131 | 4mjo | D | 223 | VAL | p.Val224Ile | P09467 |
| HOMO SAPIENS | 4 | 131 | 4mjo | D | 232 | PHE | p.Phe233Ile | P09467 |
| HOMO SAPIENS | 5 | 131 | 4mjo | D | 243 | ARG | p.Arg244Trp | P09467 |
| HOMO SAPIENS | 0 | 131 | 4mjo | D | 283 | PRO | p.Pro284Arg | P09467 |
| HOMO SAPIENS | 0 | 131 | 4mjo | D | 288 | MET | p.Met289Leu | P09467 |

| | | | | | | | | |
|---|---|---|---|---|---|---|---|---|
| HOMO SAPIENS | 0 | 131 | 4mjo | D | 323 | ASP | p.Asp324Asn | P09467 |
| HOMO SAPIENS | -1 | 13122 | 3hfc | A | 583 | ARG | p.Arg583Cys | P51787 |
| HOMO SAPIENS | 1 | 13122 | 3hfc | A | 587 | THR | p.Thr587Met | P51787 |
| HOMO SAPIENS | 0 | 13122 | 3hfc | A | 589 | GLY | p.Gly589Asp | P51787 |
| HOMO SAPIENS | 4 | 3002 | 3nbq | B | 94 | ARG | p.Arg94Cys | Q16831 |
| HOMO SAPIENS | 1 | 3616 | 3dbp | A | 97 | ILE | p.Ile286Asn | P11172 |
| HOMO SAPIENS | 0 | 3616 | 3dbp | A | 240 | GLY | p.Gly429Arg | P11172 |
| HOMO SAPIENS | 0 | 3616 | 3dbp | A | 257 | ILE | p.Ile446Val | P11172 |
| HOMO SAPIENS | 1 | 584 | 1drf | A | 79 | LEU | p.Leu80Phe | P00374 |
| HOMO SAPIENS | 3 | 1609 | 3qa2 | B | 49 | VAL | p.Val49Ile | P50053 |
| HOMO SAPIENS | 0 | 20560 | 5mug | A | 101 | HIS | p.His101Gln | P49638 |
| HOMO SAPIENS | 2 | 20560 | 5mug | A | 120 | ALA | p.Ala120Thr | P49638 |
| HOMO SAPIENS | 3 | 20560 | 5mug | A | 192 | ARG | p.Arg192His | P49638 |
| HOMO SAPIENS | 3 | 20560 | 5mug | A | 221 | ARG | p.Arg221Trp | P49638 |
| HOMO SAPIENS | 4 | 256 | 3vtv | A | 65 | THR | p.Glu50Lys | Q9GZQ8 |
| HOMO SAPIENS | 4 | 256 | 3vtv | A | 113 | VAL | p.Met98Lys | Q9GZQ8 |
| HOMO SAPIENS | 4 | 5397 | 1m61 | A | 192 | ARG | p.Arg192Trp | P43403 |
| HOMO SAPIENS | 5 | 39881 | 1aut | L | 107 | HIS | p.His149Pro | P04070 |
| HOMO SAPIENS | 5 | 4010 | 4uf8 | I | 33 | ALA | p.Ala33Val | Q13501 |
| HOMO SAPIENS | 3 | 313 | 5w49 | A | 98 | PRO | p.Pro98Leu | P23526 |
| HOMO SAPIENS | 2 | 313 | 5w49 | A | 143 | TYR | p.Tyr143Cys | P23526 |
| HOMO SAPIENS | -1 | 1643 | 2m8t | A | 102 | PRO | p.Pro102Leu | P04156 |
| HOMO SAPIENS | -1 | 1643 | 2m8t | A | 105 | PRO | p.Pro105Leu | P04156 |
| HOMO SAPIENS | -1 | 1643 | 2m8t | A | 117 | ALA | p.Ala117Val | P04156 |
| HOMO SAPIENS | 5 | 1643 | 2m8t | A | 129 | MET | p.Met129Val | P04156 |
| HOMO SAPIENS | 5 | 1643 | 2m8t | A | 131 | GLY | p.Gly131Val | P04156 |
| HOMO SAPIENS | 2 | 1643 | 2m8t | A | 142 | GLY | p.Gly142Ser | P04156 |
| HOMO SAPIENS | 3 | 1643 | 2m8t | A | 171 | ASN | p.Asn171Ser | P04156 |
| HOMO SAPIENS | 9 | 1643 | 2m8t | A | 178 | ASP | p.Asp178Asn | P04156 |
| HOMO SAPIENS | 9 | 1643 | 2m8t | A | 180 | VAL | p.Val180Ile | P04156 |
| HOMO SAPIENS | 9 | 1643 | 2m8t | A | 183 | THR | p.Thr183Ala | P04156 |
| HOMO SAPIENS | 9 | 1643 | 2m8t | A | 187 | HIS | p.His187Arg | P04156 |
| HOMO SAPIENS | 8 | 1643 | 2m8t | A | 198 | PHE | p.Phe198Ser | P04156 |
| HOMO SAPIENS | 9 | 1643 | 2m8t | A | 200 | GLU | p.Glu200Lys | P04156 |
| HOMO SAPIENS | 9 | 1643 | 2m8t | A | 208 | ARG | p.Arg208His | P04156 |
| HOMO SAPIENS | 3 | 1643 | 2m8t | A | 210 | VAL | p.Val210Ile | P04156 |
| HOMO SAPIENS | 6 | 1643 | 2m8t | A | 217 | GLN | p.Gln217Arg | P04156 |
| HOMO SAPIENS | 1 | 29 | 5kos | A | 151 | GLY | p.Gly217Arg | P00797 |
| HOMO SAPIENS | 1 | 29 | 5kos | A | 164 | ARG | p.Arg230Lys | P00797 |
| HOMO SAPIENS | 3 | 504 | 4zkv | B | 37 | ARG | p.Arg37Pro | P49773 |
| HOMO SAPIENS | 2 | 504 | 4zkv | B | 62 | GLN | p.Gln62Ter | P49773 |
| HOMO SAPIENS | 2 | 504 | 4zkv | B | 84 | CYS | p.Cys84Arg | P49773 |
| HOMO SAPIENS | 4 | 504 | 4zkv | B | 93 | GLY | p.Gly93Asp | P49773 |
| HOMO SAPIENS | 9 | 504 | 4zkv | B | 112 | HIS | p.His112Asn | P49773 |
| HOMO SAPIENS | 0 | 74 | 5vwo | C | 54 | ASN | p.Asn54Lys | P04181 |
| HOMO SAPIENS | 0 | 74 | 5vwo | C | 55 | TYR | p.Tyr55His | P04181 |
| HOMO SAPIENS | 2 | 74 | 5vwo | C | 90 | GLN | p.Gln90Glu | P04181 |
| HOMO SAPIENS | 6 | 74 | 5vwo | C | 91 | GLY | p.Gly91Glu | P04181 |
| HOMO SAPIENS | 2 | 74 | 5vwo | C | 93 | CYS | p.Cys93Phe | P04181 |
| HOMO SAPIENS | 1 | 74 | 5vwo | C | 154 | ARG | p.Arg154Leu | P04181 |
| HOMO SAPIENS | 0 | 74 | 5vwo | C | 180 | ARG | p.Arg180Thr | P04181 |
| HOMO SAPIENS | 0 | 74 | 5vwo | C | 199 | PRO | p.Pro199Gln | P04181 |
| HOMO SAPIENS | 4 | 74 | 5vwo | C | 209 | TYR | p.Tyr209Ter | P04181 |
| HOMO SAPIENS | 0 | 74 | 5vwo | C | 226 | ALA | p.Ala226Val | P04181 |

| | | | | | | | | |
|---|---|---|---|---|---|---|---|---|
| HOMO SAPIENS | 5 | 74 | 5vwo | C | 233 | GLN | p.Gln233Arg | P04181 |
| HOMO SAPIENS | 5 | 74 | 5vwo | C | 241 | PRO | p.Pro241Leu | P04181 |
| HOMO SAPIENS | 2 | 74 | 5vwo | C | 245 | TYR | p.Tyr245Cys | P04181 |
| HOMO SAPIENS | 6 | 74 | 5vwo | C | 250 | ARG | p.Arg250Pro | P04181 |
| HOMO SAPIENS | 6 | 74 | 5vwo | C | 250 | ARG | p.Arg250Ter | P04181 |
| HOMO SAPIENS | 8 | 74 | 5vwo | C | 267 | THR | p.Thr267Ile | P04181 |
| HOMO SAPIENS | 0 | 74 | 5vwo | C | 270 | ALA | p.Ala270Pro | P04181 |
| HOMO SAPIENS | 0 | 74 | 5vwo | C | 271 | ARG | p.Arg271Lys | P04181 |
| HOMO SAPIENS | 1 | 74 | 5vwo | C | 299 | TYR | p.Tyr299Ter | P04181 |
| HOMO SAPIENS | 0 | 74 | 5vwo | C | 318 | GLU | p.Glu318Lys | P04181 |
| HOMO SAPIENS | 1 | 74 | 5vwo | C | 319 | HIS | p.His319Tyr | P04181 |
| HOMO SAPIENS | 1 | 74 | 5vwo | C | 331 | ARG | p.Arg331Ter | P04181 |
| HOMO SAPIENS | 0 | 74 | 5vwo | C | 332 | VAL | p.Val332Met | P04181 |
| HOMO SAPIENS | 1 | 74 | 5vwo | C | 353 | GLY | p.Gly353Asp | P04181 |
| HOMO SAPIENS | 2 | 74 | 5vwo | C | 375 | GLY | p.Gly375Ala | P04181 |
| HOMO SAPIENS | 0 | 74 | 5vwo | C | 394 | CYS | p.Cys394Arg | P04181 |
| HOMO SAPIENS | 0 | 74 | 5vwo | C | 402 | LEU | p.Leu402Pro | P04181 |
| HOMO SAPIENS | 0 | 74 | 5vwo | C | 436 | ILE | p.Ile436Asn | P04181 |
| HOMO SAPIENS | 7 | 24356 | 4az0 | A | 104 | VAL | p.Val132Met | P10619 |
| HOMO SAPIENS | 4 | 24356 | 4az0 | A | 221 | TYR | p.Tyr249Asn | P10619 |
| HOMO SAPIENS | 3 | 245 | 1rsz | A | 24 | ARG | p.Arg24Ter | P00491 |
| HOMO SAPIENS | 3 | 245 | 1rsz | A | 51 | SER | p.Gly51Ser | P00491 |
| HOMO SAPIENS | 1 | 245 | 1rsz | A | 58 | ARG | p.Arg58Ter | P00491 |
| HOMO SAPIENS | 4 | 245 | 1rsz | A | 89 | GLU | p.Glu89Lys | P00491 |
| HOMO SAPIENS | 6 | 245 | 1rsz | A | 128 | ASP | p.Asp128Gly | P00491 |
| HOMO SAPIENS | 0 | 245 | 1rsz | A | 174 | ALA | p.Ala174Pro | P00491 |
| HOMO SAPIENS | 8 | 245 | 1rsz | A | 192 | TYR | p.Tyr192Cys | P00491 |
| HOMO SAPIENS | 8 | 245 | 1rsz | A | 234 | ARG | p.Arg234Pro | P00491 |
| HOMO SAPIENS | 0 | 37 | 5tby | E | 22 | GLU | p.Glu22Lys | P10916 |
| HOMO SAPIENS | 1 | 37 | 5tby | E | 86 | PHE | p.Phe86Ser | P10916 |
| HOMO SAPIENS | 9 | 71 | 5m9j | A | 12 | GLN | p.Gln36Leu | P03950 |
| HOMO SAPIENS | 3 | 71 | 5m9j | A | 17 | LYS | p.Lys41Glu | P03950 |
| HOMO SAPIENS | 3 | 71 | 5m9j | A | 17 | LYS | p.Lys41Ile | P03950 |
| HOMO SAPIENS | 4 | 71 | 5m9j | A | 28 | SER | p.Ser52Asn | P03950 |
| HOMO SAPIENS | 7 | 71 | 5m9j | A | 31 | ARG | p.Arg55Lys | P03950 |
| HOMO SAPIENS | 9 | 71 | 5m9j | A | 39 | CYS | p.Cys63Trp | P03950 |
| HOMO SAPIENS | 9 | 71 | 5m9j | A | 40 | LYS | p.Lys64Ile | P03950 |
| HOMO SAPIENS | 2 | 71 | 5m9j | A | 60 | GLU | p.Lys84Glu | P03950 |
| HOMO SAPIENS | 0 | 71 | 5m9j | A | 112 | PRO | p.Pro136Leu | P03950 |
| HOMO SAPIENS | 9 | 71 | 5m9j | A | 113 | VAL | p.Val137Ile | P03950 |
| HOMO SAPIENS | 3 | 326 | 5dqi | A | 209 | GLY | p.Gly209Val | Q9Y253 |
| HOMO SAPIENS | 1 | 326 | 5dqi | A | 242 | SER | p.Ser242Ter | Q9Y253 |
| HOMO SAPIENS | 1 | 326 | 5dqi | A | 303 | ARG | p.Arg303Ter | Q9Y253 |
| HOMO SAPIENS | 1 | 8203 | 3iu3 | K | 132 | VAL | p.Val153Ile | P01589 |
| HOMO SAPIENS | 0 | 4995 | 5ffo | H | 234 | THR | p.Thr263Ile | P01137 |
| HOMO SAPIENS | 5 | 38609 | 2ruk | A | 49 | ASP | p.Asp49His | P04637 |
| HOMO SAPIENS | 5 | 38609 | 2ruk | A | 58 | PRO | p.Pro58Arg | P04637 |
| HOMO SAPIENS | 0 | 1607 | 6pah | A | 117 | THR | p.Thr117Ile | P00439 |
| HOMO SAPIENS | 3 | 1607 | 6pah | A | 145 | ASP | p.Asp145Val | P00439 |
| HOMO SAPIENS | 5 | 1607 | 6pah | A | 158 | ARG | p.Arg158Gln | P00439 |
| HOMO SAPIENS | 5 | 1607 | 6pah | A | 158 | ARG | p.Arg158Trp | P00439 |
| HOMO SAPIENS | 5 | 1607 | 6pah | A | 169 | ARG | p.Arg169His | P00439 |
| HOMO SAPIENS | 5 | 1607 | 6pah | A | 169 | ARG | p.Arg169Cys | P00439 |
| HOMO SAPIENS | 3 | 1607 | 6pah | A | 176 | ARG | p.Arg176Leu | P00439 |

| | | | | | | | | |
|---|---|---|---|---|---|---|---|---|
| HOMO SAPIENS | 3 | 1607 | 6pah | A | 177 | VAL | p.Val177Leu | P00439 |
| HOMO SAPIENS | 1 | 1607 | 6pah | A | 194 | LEU | p.Leu194Pro | P00439 |
| HOMO SAPIENS | 1 | 1607 | 6pah | A | 204 | TYR | p.Tyr204Cys | P00439 |
| HOMO SAPIENS | 1 | 1607 | 6pah | A | 204 | TYR | p.Tyr204Ter | P00439 |
| HOMO SAPIENS | 2 | 1607 | 6pah | A | 230 | VAL | p.Val230Ile | P00439 |
| HOMO SAPIENS | 4 | 1607 | 6pah | A | 232 | GLN | p.Gln232Ter | P00439 |
| HOMO SAPIENS | 4 | 1607 | 6pah | A | 241 | ARG | p.Arg241Cys | P00439 |
| HOMO SAPIENS | 4 | 1607 | 6pah | A | 241 | ARG | p.Arg241His | P00439 |
| HOMO SAPIENS | 5 | 1607 | 6pah | A | 243 | ARG | p.Arg243Gln | P00439 |
| HOMO SAPIENS | 5 | 1607 | 6pah | A | 243 | ARG | p.Arg243Ter | P00439 |
| HOMO SAPIENS | 7 | 1607 | 6pah | A | 249 | LEU | p.Leu249Phe | P00439 |
| HOMO SAPIENS | 5 | 1607 | 6pah | A | 252 | ARG | p.Arg252Gly | P00439 |
| HOMO SAPIENS | 5 | 1607 | 6pah | A | 261 | ARG | p.Arg261Ter | P00439 |
| HOMO SAPIENS | 4 | 1607 | 6pah | A | 267 | GLN | p.Gln267Leu | P00439 |
| HOMO SAPIENS | 1 | 1607 | 6pah | A | 272 | GLY | p.Gly272Ter | P00439 |
| HOMO SAPIENS | 2 | 1607 | 6pah | A | 277 | TYR | p.Tyr277Asp | P00439 |
| HOMO SAPIENS | 7 | 1607 | 6pah | A | 280 | GLU | p.Glu280Lys | P00439 |
| HOMO SAPIENS | 8 | 1607 | 6pah | A | 281 | PRO | p.Pro281Leu | P00439 |
| HOMO SAPIENS | 0 | 1607 | 6pah | A | 299 | PHE | p.Phe299Cys | P00439 |
| HOMO SAPIENS | 5 | 1607 | 6pah | A | 300 | ALA | p.Ala300Ser | P00439 |
| HOMO SAPIENS | 5 | 1607 | 6pah | A | 309 | ALA | p.Ala309Asp | P00439 |
| HOMO SAPIENS | 4 | 1607 | 6pah | A | 320 | LYS | p.Lys320Asn | P00439 |
| HOMO SAPIENS | 0 | 1607 | 6pah | A | 345 | ALA | p.Ala345Ser | P00439 |
| HOMO SAPIENS | 0 | 1607 | 6pah | A | 348 | LEU | p.Leu348Val | P00439 |
| HOMO SAPIENS | 6 | 1607 | 6pah | A | 356 | TYR | p.Tyr356Ter | P00439 |
| HOMO SAPIENS | 2 | 1607 | 6pah | A | 380 | THR | p.Thr380Met | P00439 |
| HOMO SAPIENS | 0 | 1607 | 6pah | A | 386 | TYR | p.Tyr386Cys | P00439 |
| HOMO SAPIENS | 5 | 1607 | 6pah | A | 390 | GLU | p.Glu390Gly | P00439 |
| HOMO SAPIENS | 5 | 1607 | 6pah | A | 403 | ALA | p.Ala403Val | P00439 |
| HOMO SAPIENS | 4 | 1607 | 6pah | A | 408 | ARG | p.Arg408Trp | P00439 |
| HOMO SAPIENS | 3 | 1607 | 6pah | A | 413 | ARG | p.Arg413Pro | P00439 |
| HOMO SAPIENS | 4 | 1607 | 6pah | A | 414 | TYR | p.Tyr414Cys | P00439 |
| HOMO SAPIENS | 4 | 1607 | 6pah | A | 414 | TYR | p.Tyr414His | P00439 |
| HOMO SAPIENS | 4 | 1607 | 6pah | A | 415 | ASP | p.Asp415Asn | P00439 |
| HOMO SAPIENS | -1 | 676 | 5dc5 | A | 7 | PRO | p.Pro7Leu | Q9BY41 |
| HOMO SAPIENS | 0 | 676 | 5dc5 | A | 311 | THR | p.Thr311Met | Q9BY41 |
| HOMO SAPIENS | 1 | 124 | 5ftk | E | 95 | ARG | p.Arg95Gly | P55072 |
| HOMO SAPIENS | 1 | 124 | 5ftk | E | 155 | ARG | p.Arg155Cys | P55072 |
| HOMO SAPIENS | 1 | 124 | 5ftk | E | 155 | ARG | p.Arg155His | P55072 |
| HOMO SAPIENS | 1 | 124 | 5ftk | E | 159 | ARG | p.Arg159His | P55072 |
| HOMO SAPIENS | 1 | 124 | 5ftk | E | 159 | ARG | p.Arg159Gly | P55072 |
| HOMO SAPIENS | 0 | 124 | 5ftk | E | 191 | ARG | p.Arg191Gln | P55072 |
| HOMO SAPIENS | 0 | 124 | 5ftk | E | 232 | ALA | p.Ala232Glu | P55072 |
| HOMO SAPIENS | 5 | 5442 | 1fgg | B | 223 | GLY | p.Gly223Ser | O94766 |
| HOMO SAPIENS | 0 | 5442 | 1fgg | B | 277 | ARG | p.Arg277Gln | O94766 |
| HOMO SAPIENS | 0 | 9072 | 2gys | B | 225 | GLU | p.Glu249Gln | P32927 |
| HOMO SAPIENS | 0 | 9072 | 2gys | B | 247 | SER | p.Ser271Ter | P32927 |
| HOMO SAPIENS | 0 | 9072 | 2gys | B | 294 | ASP | p.Asp312Asn | P32927 |
| HOMO SAPIENS | 0 | 1484 | 4zrj | A | 62 | PHE | p.Phe62Ser | P35240 |
| HOMO SAPIENS | 3 | 1484 | 4zrj | A | 205 | MET | p.Met205Val | P35240 |
| HOMO SAPIENS | 3 | 1484 | 4zrj | A | 301 | ILE | p.Ile301Met | P35240 |
| HOMO SAPIENS | 4 | 40923 | 2enq | A | 68 | ILE | p.Ile391Met | P42336 |
| HOMO SAPIENS | 1 | 6179 | 5ofv | A | 125 | THR | p.Thr125Met | O43175 |
| HOMO SAPIENS | 0 | 6179 | 5ofv | A | 135 | ARG | p.Arg135Trp | O43175 |

| | | | | | | | | |
|---|---|---|---|---|---|---|---|---|
| HOMO SAPIENS | 1 | 6179 | 5ofv | A | 163 | ARG | p.Arg163Gln | O43175 |
| HOMO SAPIENS | 9 | 6179 | 5ofv | A | 261 | VAL | p.Val261Met | O43175 |
| HOMO SAPIENS | 7 | 34118 | 2i5o | A | 24 | MET | p.Met647Leu | Q9Y253 |
| HOMO SAPIENS | 8 | 3234 | 5c84 | A | 311 | LYS | p.Lys311Asn | P98170 |
| HOMO SAPIENS | 3 | 3234 | 5c84 | A | 321 | ALA | p.Ala321Gly | P98170 |
| HOMO SAPIENS | -1 | 15129 | 1zy8 | K | 1 | GLY | p.Gly54Ser | O00330 |
| HOMO SAPIENS | -1 | 15129 | 1zy8 | K | 48 | THR | p.Thr101Ala | O00330 |
| HOMO SAPIENS | -1 | 15129 | 1zy8 | K | 105 | GLU | p.Glu158Asp | O00330 |
| HOMO SAPIENS | 8 | 15129 | 1zy8 | K | 136 | ARG | p.Arg189His | O00330 |
| HOMO SAPIENS | -1 | 15129 | 1zy8 | K | 195 | GLN | p.Gln248Ter | O00330 |
| HOMO SAPIENS | 8 | 202 | 4bzb | A | 123 | HIS | p.His123Pro | Q9Y3Z3 |
| HOMO SAPIENS | 4 | 202 | 4bzb | A | 145 | ARG | p.Arg145Ter | Q9Y3Z3 |
| HOMO SAPIENS | 6 | 202 | 4bzb | A | 164 | ARG | p.Arg164Ter | Q9Y3Z3 |
| HOMO SAPIENS | 3 | 202 | 4bzb | A | 201 | ILE | p.Ile201Asn | Q9Y3Z3 |
| HOMO SAPIENS | 0 | 202 | 4bzb | A | 209 | GLY | p.Gly209Ser | Q9Y3Z3 |
| HOMO SAPIENS | 0 | 202 | 4bzb | A | 254 | MET | p.Met254Val | Q9Y3Z3 |
| HOMO SAPIENS | 5 | 202 | 4bzb | A | 531 | ARG | p.Arg531Ser | Q9Y3Z3 |
| HOMO SAPIENS | 6 | 202 | 4bzb | A | 548 | GLN | p.Gln548Ter | Q9Y3Z3 |
| HOMO SAPIENS | 3 | 30947 | 5lgd | A | 90 | PRO | p.Pro90Ser | P16671 |
| HOMO SAPIENS | 6 | 30947 | 5lgd | A | 254 | PHE | p.Phe254Leu | P16671 |
| HOMO SAPIENS | 0 | 30947 | 5lgd | A | 413 | ILE | p.Ile413Leu | P16671 |
| HOMO SAPIENS | 2 | 3906 | 5jtc | A | 735 | ARG | p.Arg735Trp | Q12797 |
| HOMO SAPIENS | 1 | 45 | 5uyx | C | 146 | GLU | p.Glu146Val | P48643 |
| HOMO SAPIENS | 1 | 45 | 5uyx | C | 147 | HIS | p.His147Arg | P48643 |
| HOMO SAPIENS | 0 | 45 | 5uyx | C | 362 | ILE | p.Ile362Met | P48643 |
| HOMO SAPIENS | 8 | 39596 | 1mhn | A | 95 | GLY | p.Gly95Arg | Q16637 |
| HOMO SAPIENS | 4 | 39596 | 1mhn | A | 111 | ALA | p.Ala111Gly | Q16637 |
| HOMO SAPIENS | 9 | 39596 | 1mhn | A | 116 | ILE | p.Ile116Phe | Q16637 |
| HOMO SAPIENS | 1 | 39596 | 1mhn | A | 136 | GLN | p.Gln136Glu | Q16637 |
| HOMO SAPIENS | 1 | 6408 | 5xjy | A | 85 | PRO | p.Pro85Leu | O95477 |
| HOMO SAPIENS | 0 | 6408 | 5xjy | A | 219 | ARG | p.Arg219Lys | O95477 |
| HOMO SAPIENS | 0 | 6408 | 5xjy | A | 230 | ARG | p.Arg230Cys | O95477 |
| HOMO SAPIENS | 1 | 6408 | 5xjy | A | 281 | MET | p.Met281Thr | O95477 |
| HOMO SAPIENS | -1 | 6408 | 5xjy | A | 343 | ALA | p.Ala343Gly | O95477 |
| HOMO SAPIENS | 1 | 6408 | 5xjy | A | 399 | VAL | p.Val399Ala | O95477 |
| HOMO SAPIENS | 1 | 6408 | 5xjy | A | 496 | ARG | p.Arg496Trp | O95477 |
| HOMO SAPIENS | 1 | 6408 | 5xjy | A | 590 | TRP | p.Trp590Ser | O95477 |
| HOMO SAPIENS | 1 | 6408 | 5xjy | A | 638 | ARG | p.Arg638Gln | O95477 |
| HOMO SAPIENS | 4 | 6408 | 5xjy | A | 697 | ALA | p.Ala697Thr | O95477 |
| HOMO SAPIENS | 1 | 6408 | 5xjy | A | 771 | VAL | p.Val771Met | O95477 |
| HOMO SAPIENS | 1 | 6408 | 5xjy | A | 774 | THR | p.Thr774Pro | O95477 |
| HOMO SAPIENS | 1 | 6408 | 5xjy | A | 776 | LYS | p.Lys776Asn | O95477 |
| HOMO SAPIENS | 0 | 6408 | 5xjy | A | 825 | VAL | p.Val825Ile | O95477 |
| HOMO SAPIENS | -1 | 6408 | 5xjy | A | 868 | GLU | p.Glu868Lys | O95477 |
| HOMO SAPIENS | -1 | 6408 | 5xjy | A | 883 | ILE | p.Ile883Met | O95477 |
| HOMO SAPIENS | 0 | 6408 | 5xjy | A | 935 | ASN | p.Asn935Ser | O95477 |
| HOMO SAPIENS | 0 | 6408 | 5xjy | A | 935 | ASN | p.Asn935His | O95477 |
| HOMO SAPIENS | 0 | 6408 | 5xjy | A | 937 | ALA | p.Ala937Val | O95477 |
| HOMO SAPIENS | 0 | 6408 | 5xjy | A | 1018 | ASP | p.Asp1018Gly | O95477 |
| HOMO SAPIENS | 0 | 6408 | 5xjy | A | 1096 | VAL | p.Val1096Ile | O95477 |
| HOMO SAPIENS | 0 | 6408 | 5xjy | A | 1099 | ASP | p.Asp1099Tyr | O95477 |
| HOMO SAPIENS | 0 | 6408 | 5xjy | A | 1172 | GLU | p.Glu1172Asp | O95477 |
| HOMO SAPIENS | 0 | 6408 | 5xjy | A | 1181 | SER | p.Ser1181Phe | O95477 |
| HOMO SAPIENS | 0 | 6408 | 5xjy | A | 1182 | ALA | p.Ala1182Thr | O95477 |

| | | | | | | | | |
|---|---|---|---|---|---|---|---|---|
| HOMO SAPIENS | -1 | 6408 | 5xjy | A | 1289 | ASP | p.Asp1289Asn | O95477 |
| HOMO SAPIENS | 0 | 6408 | 5xjy | A | 1399 | THR | p.Thr1399Ser | O95477 |
| HOMO SAPIENS | 1 | 6408 | 5xjy | A | 1408 | LEU | p.Leu1408Phe | O95477 |
| HOMO SAPIENS | 0 | 6408 | 5xjy | A | 1587 | LYS | p.Lys1587Arg | O95477 |
| HOMO SAPIENS | 1 | 6408 | 5xjy | A | 1674 | VAL | p.Val1674Ile | O95477 |
| HOMO SAPIENS | 1 | 6408 | 5xjy | A | 1680 | ARG | p.Arg1680Trp | O95477 |
| HOMO SAPIENS | 1 | 6408 | 5xjy | A | 1680 | ARG | p.Arg1680Gln | O95477 |
| HOMO SAPIENS | 0 | 6408 | 5xjy | A | 1800 | ASN | p.Asn1800His | O95477 |
| HOMO SAPIENS | -1 | 6408 | 5xjy | A | 1925 | ARG | p.Arg1925Gln | O95477 |
| HOMO SAPIENS | 0 | 6408 | 5xjy | A | 2009 | PHE | p.Phe2009Ser | O95477 |
| HOMO SAPIENS | -1 | 6408 | 5xjy | A | 2243 | ASP | p.Asp2243Glu | O95477 |
| HOMO SAPIENS | 4 | 23760 | 2mut | A | 231 | LEU | p.Phe231Leu | P07992 |
| HOMO SAPIENS | 6 | 1460 | 2hre | B | 96 | ARG | p.Arg96Gln | P22830 |
| HOMO SAPIENS | 8 | 1460 | 2hre | B | 129 | GLY | p.Gly129Arg | P22830 |
| HOMO SAPIENS | 9 | 1460 | 2hre | B | 172 | GLU | p.Glu172Lys | P22830 |
| HOMO SAPIENS | 4 | 1460 | 2hre | B | 285 | GLN | p.Gln285Arg | P22830 |
| HOMO SAPIENS | 5 | 1460 | 2hre | B | 362 | VAL | p.Val362Gly | P22830 |
| HOMO SAPIENS | 6 | 1460 | 2hre | B | 411 | CYS | p.Cys411Gly | P22830 |
| HOMO SAPIENS | 6 | 1460 | 2hre | B | 417 | PHE | p.Phe417Ser | P22830 |
| HOMO SAPIENS | 0 | 2210 | 5bug | A | 19 | ASP | p.Asp19Asn | P60484 |
| HOMO SAPIENS | 0 | 2210 | 5bug | A | 35 | MET | p.Met35Arg | P60484 |
| HOMO SAPIENS | 1 | 2210 | 5bug | A | 61 | HIS | p.His61Asp | P60484 |
| HOMO SAPIENS | 0 | 2210 | 5bug | A | 68 | TYR | p.Tyr68His | P60484 |
| HOMO SAPIENS | 8 | 2210 | 5bug | A | 70 | LEU | p.Leu70Pro | P60484 |
| HOMO SAPIENS | 6 | 2210 | 5bug | A | 93 | HIS | p.His93Arg | P60484 |
| HOMO SAPIENS | 1 | 2210 | 5bug | A | 112 | LEU | p.Leu112Pro | P60484 |
| HOMO SAPIENS | 0 | 2210 | 5bug | A | 121 | ALA | p.Ala121Gly | P60484 |
| HOMO SAPIENS | 3 | 2210 | 5bug | A | 123 | HIS | p.His123Arg | P60484 |
| HOMO SAPIENS | 7 | 2210 | 5bug | A | 124 | CYS | p.Cys124Arg | P60484 |
| HOMO SAPIENS | 9 | 2210 | 5bug | A | 129 | GLY | p.Gly129Glu | P60484 |
| HOMO SAPIENS | 9 | 2210 | 5bug | A | 130 | ARG | p.Arg130Gln | P60484 |
| HOMO SAPIENS | 9 | 2210 | 5bug | A | 130 | ARG | p.Arg130Gly | P60484 |
| HOMO SAPIENS | 0 | 2210 | 5bug | A | 132 | GLY | p.Gly132Val | P60484 |
| HOMO SAPIENS | 0 | 2210 | 5bug | A | 170 | SER | p.Ser170Arg | P60484 |
| HOMO SAPIENS | 1 | 2210 | 5bug | A | 173 | ARG | p.Arg173His | P60484 |
| HOMO SAPIENS | 0 | 2210 | 5bug | A | 217 | VAL | p.Val217Ile | P60484 |
| HOMO SAPIENS | 0 | 2210 | 5bug | A | 241 | PHE | p.Phe241Ser | P60484 |
| HOMO SAPIENS | 0 | 2210 | 5bug | A | 250 | CYS | p.Cys250Ter | P60484 |
| HOMO SAPIENS | 1 | 2210 | 5bug | A | 252 | ASP | p.Asp252Gly | P60484 |
| HOMO SAPIENS | 3 | 123 | 2ibu | A | 93 | ASN | p.Asn93Ser | P24752 |
| HOMO SAPIENS | 0 | 123 | 2ibu | A | 145 | GLN | p.Gln145Glu | P24752 |
| HOMO SAPIENS | 0 | 123 | 2ibu | A | 152 | GLY | p.Gly152Ala | P24752 |
| HOMO SAPIENS | 2 | 123 | 2ibu | A | 158 | ASN | p.Asn158Asp | P24752 |
| HOMO SAPIENS | 2 | 123 | 2ibu | A | 158 | ASN | p.Asn158Ser | P24752 |
| HOMO SAPIENS | 5 | 123 | 2ibu | A | 183 | GLY | p.Gly183Arg | P24752 |
| HOMO SAPIENS | 3 | 123 | 2ibu | A | 272 | GLN | p.Gln272Ter | P24752 |
| HOMO SAPIENS | 3 | 123 | 2ibu | A | 312 | ILE | p.Ile312Thr | P24752 |
| HOMO SAPIENS | 1 | 123 | 2ibu | A | 333 | ALA | p.Ala333Pro | P24752 |
| HOMO SAPIENS | 5 | 123 | 2ibu | A | 379 | GLY | p.Gly379Val | P24752 |
| HOMO SAPIENS | 0 | 123 | 2ibu | A | 380 | ALA | p.Ala380Thr | P24752 |
| HOMO SAPIENS | 2 | 369 | 5nrw | A | 1175 | TYR | p.Tyr1175Cys | Q92793 |
| HOMO SAPIENS | 2 | 830 | 2vc2 | A | 116 | LEU | p.Leu147Val | P08514 |
| HOMO SAPIENS | 0 | 830 | 2vc2 | A | 183 | LEU | p.Leu214Pro | P08514 |
| HOMO SAPIENS | 4 | 830 | 2vc2 | A | 242 | GLY | p.Gly273Asp | P08514 |

| | | | | | | | | |
|---|---|---|---|---|---|---|---|---|
| HOMO SAPIENS | 6 | 830 | 2vc2 | A | 324 | GLU | p.Glu355Lys | P08514 |
| HOMO SAPIENS | -1 | 10353 | 5ejc | E | 989 | ALA | p.Ala989Ser | Q92574 |
| HOMO SAPIENS | 0 | 15448 | 3lgg | A | 21 | GLY | p.Gly47Arg | Q9NZK5 |
| HOMO SAPIENS | 0 | 15448 | 3lgg | A | 21 | GLY | p.Gly47Val | Q9NZK5 |
| HOMO SAPIENS | 9 | 15448 | 3lgg | A | 86 | HIS | p.His112Gln | Q9NZK5 |
| HOMO SAPIENS | 0 | 15448 | 3lgg | A | 143 | ARG | p.Arg169Gln | Q9NZK5 |
| HOMO SAPIENS | 3 | 15448 | 3lgg | A | 221 | ALA | p.Ala247Val | Q9NZK5 |
| HOMO SAPIENS | 0 | 15448 | 3lgg | A | 225 | PRO | p.Pro251Leu | Q9NZK5 |
| HOMO SAPIENS | 0 | 15448 | 3lgg | A | 283 | MET | p.Met309Ile | Q9NZK5 |
| HOMO SAPIENS | 2 | 15448 | 3lgg | A | 334 | THR | p.Thr360Ala | Q9NZK5 |
| HOMO SAPIENS | 7 | 15448 | 3lgg | A | 357 | GLY | p.Gly383Ser | Q9NZK5 |
| HOMO SAPIENS | 0 | 15448 | 3lgg | A | 427 | TYR | p.Tyr453Cys | Q9NZK5 |
| HOMO SAPIENS | 0 | 872 | 2y3i | A | 87 | LEU | p.Leu88Pro | P00558 |
| HOMO SAPIENS | 1 | 872 | 2y3i | A | 163 | ASP | p.Asp164Val | P00558 |
| HOMO SAPIENS | 4 | 872 | 2y3i | A | 205 | ARG | p.Arg206Pro | P00558 |
| HOMO SAPIENS | 4 | 872 | 2y3i | A | 267 | ASP | p.Asp268Asn | P00558 |
| HOMO SAPIENS | 8 | 872 | 2y3i | A | 284 | ASP | p.Asp285Val | P00558 |
| HOMO SAPIENS | 4 | 8 | 5jco | B | 188 | ILE | p.Ile188Leu | Q71U36 |
| HOMO SAPIENS | 7 | 8 | 5jco | B | 263 | PRO | p.Pro263Thr | Q71U36 |
| HOMO SAPIENS | 6 | 8 | 5jco | B | 264 | ARG | p.Arg264Cys | Q71U36 |
| HOMO SAPIENS | 7 | 8 | 5jco | B | 402 | ARG | p.Arg402His | Q71U36 |
| HOMO SAPIENS | 7 | 8 | 5jco | B | 419 | SER | p.Ser419Leu | Q71U36 |
| HOMO SAPIENS | -1 | 880 | 4nwf | A | 2 | THR | p.Thr2Ile | Q06124 |
| HOMO SAPIENS | 2 | 880 | 4nwf | A | 58 | ASN | p.Asn58Ser | Q06124 |
| HOMO SAPIENS | 4 | 880 | 4nwf | A | 61 | ASP | p.Asp61Gly | Q06124 |
| HOMO SAPIENS | 2 | 880 | 4nwf | A | 62 | TYR | p.Tyr62Asp | Q06124 |
| HOMO SAPIENS | 3 | 880 | 4nwf | A | 63 | TYR | p.Tyr63Cys | Q06124 |
| HOMO SAPIENS | 2 | 880 | 4nwf | A | 69 | GLU | p.Glu69Gln | Q06124 |
| HOMO SAPIENS | 2 | 880 | 4nwf | A | 72 | ALA | p.Ala72Gly | Q06124 |
| HOMO SAPIENS | 2 | 880 | 4nwf | A | 72 | ALA | p.Ala72Ser | Q06124 |
| HOMO SAPIENS | 3 | 880 | 4nwf | A | 73 | THR | p.Thr73Ile | Q06124 |
| HOMO SAPIENS | 5 | 880 | 4nwf | A | 76 | GLU | p.Glu76Ala | Q06124 |
| HOMO SAPIENS | 5 | 880 | 4nwf | A | 76 | GLU | p.Glu76Lys | Q06124 |
| HOMO SAPIENS | 3 | 880 | 4nwf | A | 79 | GLN | p.Gln79Arg | Q06124 |
| HOMO SAPIENS | 5 | 880 | 4nwf | A | 152 | ARG | p.Arg152His | Q06124 |
| HOMO SAPIENS | 2 | 880 | 4nwf | A | 268 | GLY | p.Gly268Cys | Q06124 |
| HOMO SAPIENS | 9 | 880 | 4nwf | A | 276 | LYS | p.Lys276Arg | Q06124 |
| HOMO SAPIENS | 9 | 880 | 4nwf | A | 279 | TYR | p.Tyr279Cys | Q06124 |
| HOMO SAPIENS | 5 | 880 | 4nwf | A | 285 | PHE | p.Phe285Ser | Q06124 |
| HOMO SAPIENS | 0 | 880 | 4nwf | A | 308 | ASP | p.Asn308Ser | Q06124 |
| HOMO SAPIENS | 0 | 880 | 4nwf | A | 308 | ASP | p.Asn308Asp | Q06124 |
| HOMO SAPIENS | 5 | 880 | 4nwf | A | 392 | ALA | p.Ala392Thr | Q06124 |
| HOMO SAPIENS | 1 | 880 | 4nwf | A | 411 | THR | p.Thr415Met | Q06124 |
| HOMO SAPIENS | 0 | 880 | 4nwf | A | 461 | ALA | p.Ala465Thr | Q06124 |
| HOMO SAPIENS | 0 | 880 | 4nwf | A | 464 | GLY | p.Gly468Ala | Q06124 |
| HOMO SAPIENS | 1 | 880 | 4nwf | A | 491 | PRO | p.Pro495Ser | Q06124 |
| HOMO SAPIENS | 1 | 880 | 4nwf | A | 502 | SER | p.Ser506Thr | Q06124 |
| HOMO SAPIENS | 8 | 880 | 4nwf | A | 510 | GLN | p.Gln514Pro | Q06124 |
| HOMO SAPIENS | 4 | 1872 | 3wf1 | D | 109 | ARG | p.Arg109Trp | P16278 |
| HOMO SAPIENS | 0 | 1872 | 3wf1 | D | 521 | ARG | p.Arg521Cys | P16278 |
| HOMO SAPIENS | 5 | 910 | 5xte | A | 45 | SER | p.Ser45Phe | O14949 |
| HOMO SAPIENS | 2 | 9017 | 5f0k | A | 382 | LYS | p.Lys382Arg | Q96QK1 |
| HOMO SAPIENS | 1 | 710 | 4lsj | A | 559 | ILE | p.Ile559Asn | P04150 |
| HOMO SAPIENS | 5 | 710 | 4lsj | A | 571 | VAL | p.Val571Ala | P04150 |

| | | | | | | | | |
|---|---|---|---|---|---|---|---|---|
| HOMO SAPIENS | 0 | 710 | 4lsj | A | 641 | ASP | p.Asp641Val | P04150 |
| HOMO SAPIENS | 1 | 710 | 4lsj | A | 678 | ASP | p.Asp678Glu | P04150 |
| HOMO SAPIENS | 2 | 710 | 4lsj | A | 679 | GLY | p.Gly679Ser | P04150 |
| HOMO SAPIENS | 1 | 710 | 4lsj | A | 737 | PHE | p.Phe737Leu | P04150 |
| HOMO SAPIENS | 1 | 710 | 4lsj | A | 747 | ILE | p.Ile747Met | P04150 |
| HOMO SAPIENS | 1 | 710 | 4lsj | A | 753 | LEU | p.Leu753Phe | P04150 |
| HOMO SAPIENS | 1 | 710 | 4lsj | A | 773 | LEU | p.Leu773Pro | P04150 |
| HOMO SAPIENS | 1 | 2145 | 1av1 | B | 71 | PHE | p.Phe95Tyr | P02647 |
| HOMO SAPIENS | 1 | 2145 | 1av1 | B | 128 | GLU | p.Glu152Lys | P02647 |
| HOMO SAPIENS | 1 | 2145 | 1av1 | B | 136 | GLU | p.Glu160Lys | P02647 |
| HOMO SAPIENS | 2 | 2145 | 1av1 | B | 143 | PRO | p.Pro167Arg | P02647 |
| HOMO SAPIENS | 6 | 2145 | 1av1 | B | 164 | ALA | p.Ala188Ser | P02647 |
| HOMO SAPIENS | 7 | 2145 | 1av1 | B | 173 | ARG | p.Arg197Cys | P02647 |
| HOMO SAPIENS | 2 | 2145 | 1av1 | B | 198 | GLU | p.Glu222Lys | P02647 |
| HOMO SAPIENS | -1 | 3432 | 2beu | A | 1 | SER | p.Ser46Ter | P12694 |
| HOMO SAPIENS | 5 | 3432 | 2beu | A | 79 | ARG | p.Arg124Trp | P12694 |
| HOMO SAPIENS | 7 | 3432 | 2beu | A | 114 | ARG | p.Arg159Gln | P12694 |
| HOMO SAPIENS | 6 | 3432 | 2beu | A | 166 | THR | p.Thr211Met | P12694 |
| HOMO SAPIENS | 4 | 3432 | 2beu | A | 175 | ALA | p.Ala220Val | P12694 |
| HOMO SAPIENS | 5 | 3432 | 2beu | A | 204 | GLY | p.Gly249Ser | P12694 |
| HOMO SAPIENS | 4 | 3432 | 2beu | A | 209 | ALA | p.Ala254Asp | P12694 |
| HOMO SAPIENS | 6 | 3432 | 2beu | A | 220 | ARG | p.Arg265Trp | P12694 |
| HOMO SAPIENS | 6 | 3432 | 2beu | A | 234 | TYR | p.Tyr279Cys | P12694 |
| HOMO SAPIENS | 5 | 3432 | 2beu | A | 240 | ALA | p.Ala285Pro | P12694 |
| HOMO SAPIENS | 4 | 3432 | 2beu | A | 245 | GLY | p.Gly290Arg | P12694 |
| HOMO SAPIENS | 5 | 3432 | 2beu | A | 269 | ARG | p.Arg314Ter | P12694 |
| HOMO SAPIENS | 3 | 3432 | 2beu | A | 301 | ARG | p.Arg346Cys | P12694 |
| HOMO SAPIENS | 5 | 3432 | 2beu | A | 359 | ASN | p.Asn404Ser | P12694 |
| HOMO SAPIENS | 4 | 3432 | 2beu | A | 393 | TYR | p.Tyr438Asn | P12694 |
| HOMO SAPIENS | 8 | 4541 | 1rr8 | C | 533 | ASP | p.Asp533Gly | P11387 |
| HOMO SAPIENS | 2 | 3247 | 1tzn | d | 1105 | GLY | p.Gly105Asp | P58335 |
| HOMO SAPIENS | 3 | 3247 | 1tzn | d | 1111 | ARG | p.Arg111Cys | P58335 |
| HOMO SAPIENS | 2 | 3247 | 1tzn | d | 1189 | ILE | p.Ile189Thr | P58335 |
| HOMO SAPIENS | 9 | 467 | 1suv | C | 58 | ASP | p.Asp77Asn | P02787 |
| HOMO SAPIENS | 2 | 467 | 1suv | C | 277 | ASP | p.Asp296Gly | P02787 |
| HOMO SAPIENS | 9 | 6485 | 1xwd | B | 51 | CYS | p.Cys69Gly | P01225 |
| HOMO SAPIENS | 1 | 694 | 1wuu | A | 28 | PRO | p.Pro28Thr | P51570 |
| HOMO SAPIENS | 0 | 694 | 1wuu | A | 32 | VAL | p.Val32Met | P51570 |
| HOMO SAPIENS | 0 | 694 | 1wuu | A | 198 | ALA | p.Ala198Val | P51570 |
| HOMO SAPIENS | 0 | 143 | 4nkx | A | 93 | PHE | p.Phe93Cys | P05093 |
| HOMO SAPIENS | 3 | 143 | 4nkx | A | 96 | ARG | p.Arg96Trp | P05093 |
| HOMO SAPIENS | 3 | 143 | 4nkx | A | 96 | ARG | p.Arg96Gln | P05093 |
| HOMO SAPIENS | 0 | 143 | 4nkx | A | 106 | SER | p.Ser106Pro | P05093 |
| HOMO SAPIENS | 1 | 143 | 4nkx | A | 114 | PHE | p.Phe114Val | P05093 |
| HOMO SAPIENS | 1 | 143 | 4nkx | A | 116 | ASP | p.Asp116Val | P05093 |
| HOMO SAPIENS | 3 | 143 | 4nkx | A | 125 | ARG | p.Arg125Gln | P05093 |
| HOMO SAPIENS | 0 | 143 | 4nkx | A | 201 | TYR | p.Tyr201Asp | P05093 |
| HOMO SAPIENS | 0 | 143 | 4nkx | A | 239 | ARG | p.Arg239Ter | P05093 |
| HOMO SAPIENS | 0 | 143 | 4nkx | A | 329 | TYR | p.Tyr329Asp | P05093 |
| HOMO SAPIENS | 1 | 143 | 4nkx | A | 342 | PRO | p.Pro342Thr | P05093 |
| HOMO SAPIENS | 2 | 143 | 4nkx | A | 347 | ARG | p.Arg347His | P05093 |
| HOMO SAPIENS | 2 | 143 | 4nkx | A | 347 | ARG | p.Arg347Cys | P05093 |
| HOMO SAPIENS | 1 | 143 | 4nkx | A | 358 | ARG | p.Arg358Gln | P05093 |
| HOMO SAPIENS | 0 | 143 | 4nkx | A | 362 | ARG | p.Arg362Cys | P05093 |

| | | | | | | | | |
|---|---|---|---|---|---|---|---|---|
| HOMO SAPIENS | 1 | 143 | 4nkx | A | 406 | TRP | p.Trp406Arg | P05093 |
| HOMO SAPIENS | 0 | 143 | 4nkx | A | 428 | PRO | p.Pro428Leu | P05093 |
| HOMO SAPIENS | 6 | 1441 | 1uuh | A | 46 | ARG | p.Arg46Gly | P16070 |
| HOMO SAPIENS | 3 | 82 | 4wlv | D | 37 | GLY | p.Gly37Arg | P40926 |
| HOMO SAPIENS | 0 | 82 | 4wlv | D | 133 | PRO | p.Pro133Leu | P40926 |
| HOMO SAPIENS | 2 | 392 | 5xtd | A | 56 | SER | p.Ser56Pro | P49821 |
| HOMO SAPIENS | 2 | 392 | 5xtd | A | 59 | ARG | p.Arg59Ter | P49821 |
| HOMO SAPIENS | 2 | 392 | 5xtd | A | 74 | ASP | p.Asp74Asn | P49821 |
| HOMO SAPIENS | 4 | 392 | 5xtd | A | 116 | ASN | p.Asn116Lys | P49821 |
| HOMO SAPIENS | 4 | 392 | 5xtd | A | 117 | ALA | p.Ala117Thr | P49821 |
| HOMO SAPIENS | 5 | 392 | 5xtd | A | 122 | PRO | p.Pro122Leu | P49821 |
| HOMO SAPIENS | 3 | 392 | 5xtd | A | 160 | GLY | p.Gly160Glu | P49821 |
| HOMO SAPIENS | 4 | 392 | 5xtd | A | 214 | GLU | p.Glu214Lys | P49821 |
| HOMO SAPIENS | 2 | 392 | 5xtd | A | 357 | MET | p.Met357Thr | P49821 |
| HOMO SAPIENS | 4 | 392 | 5xtd | A | 423 | THR | p.Thr423Met | P49821 |
| HOMO SAPIENS | 0 | 22593 | 3pdf | A | 8 | TYR | p.Tyr32Ter | P53634 |
| HOMO SAPIENS | 0 | 22593 | 3pdf | A | 15 | TRP | p.Trp39Ser | P53634 |
| HOMO SAPIENS | 0 | 22593 | 3pdf | A | 103 | HIS | p.His127Pro | P53634 |
| HOMO SAPIENS | 9 | 22593 | 3pdf | A | 228 | GLN | p.Gln252Leu | P53634 |
| HOMO SAPIENS | 3 | 22593 | 3pdf | A | 262 | GLN | p.Gln286Arg | P53634 |
| HOMO SAPIENS | 4 | 22593 | 3pdf | A | 277 | GLY | p.Gly301Ser | P53634 |
| HOMO SAPIENS | 0 | 22593 | 3pdf | A | 323 | TYR | p.Tyr347Cys | P53634 |
| HOMO SAPIENS | 5 | 3461 | 4c2a | A | 1306 | GLN | p.Arg1306Trp | P04275 |
| HOMO SAPIENS | 5 | 3461 | 4c2a | A | 1374 | ARG | p.Arg1374Cys | P04275 |
| HOMO SAPIENS | 5 | 3461 | 4c2a | A | 1379 | ARG | p.Arg1379Cys | P04275 |
| HOMO SAPIENS | 4 | 3461 | 4c2a | A | 1380 | ILE | p.Ile1380Val | P04275 |
| HOMO SAPIENS | 0 | 3461 | 4c2a | A | 1381 | ALA | p.Thr1381Ala | P04275 |
| HOMO SAPIENS | 5 | 3461 | 4c2a | A | 1399 | ARG | p.Arg1399His | P04275 |
| HOMO SAPIENS | 5 | 3461 | 4c2a | A | 1435 | ASN | p.Asn1435Ser | P04275 |
| HOMO SAPIENS | 0 | 4533 | 5uj7 | C | 174 | TYR | p.Tyr174Cys | O43929 |
| HOMO SAPIENS | 0 | 4533 | 5uj7 | C | 202 | LEU | p.Leu202Val | O43929 |
| HOMO SAPIENS | 0 | 1374 | 3j07 | O | 39 | PRO | p.Pro39Leu | P02511 |
| HOMO SAPIENS | 0 | 1374 | 3j07 | O | 107 | ARG | p.Arg107His | P02511 |
| HOMO SAPIENS | 0 | 1374 | 3j07 | O | 157 | ARG | p.Arg157His | P02511 |
| HOMO SAPIENS | -1 | 942 | 1s9i | B | 57 | PHE | p.Phe57Cys | P36507 |
| HOMO SAPIENS | -1 | 942 | 1s9i | B | 57 | PHE | p.Phe57Val | P36507 |
| HOMO SAPIENS | 2 | 942 | 1s9i | B | 128 | PRO | p.Pro128Gln | P36507 |
| HOMO SAPIENS | 1 | 942 | 1s9i | B | 134 | TYR | p.Tyr134His | P36507 |
| HOMO SAPIENS | 1 | 942 | 1s9i | B | 188 | HIS | p.His188Gln | P36507 |
| HOMO SAPIENS | 0 | 942 | 1s9i | B | 262 | VAL | p.Val262Phe | P36507 |
| HOMO SAPIENS | 0 | 942 | 1s9i | B | 263 | GLY | p.Gly263Arg | P36507 |
| HOMO SAPIENS | -1 | 942 | 1s9i | B | 298 | PRO | p.Pro298Leu | P36507 |
| HOMO SAPIENS | -1 | 942 | 1s9i | B | 305 | VAL | p.Val305Ile | P36507 |
| HOMO SAPIENS | 0 | 3245 | 1fbv | A | 367 | GLN | p.Gln367Pro | P22681 |
| HOMO SAPIENS | 0 | 3245 | 1fbv | A | 371 | TYR | p.Tyr371His | P22681 |
| HOMO SAPIENS | 1 | 3245 | 1fbv | A | 382 | LYS | p.Lys382Glu | P22681 |
| HOMO SAPIENS | 2 | 3245 | 1fbv | A | 390 | ASP | p.Asp390Tyr | P22681 |
| HOMO SAPIENS | 9 | 3245 | 1fbv | A | 396 | CYS | p.Cys396Arg | P22681 |
| HOMO SAPIENS | 7 | 3245 | 1fbv | A | 420 | ARG | p.Arg420Gln | P22681 |
| HOMO SAPIENS | 1 | 3245 | 1fbv | A | 429 | ILE | p.Ile429Met | P22681 |
| HOMO SAPIENS | 0 | 2604 | 2pbd | P | 111 | LEU | p.Leu112Met | P07737 |
| HOMO SAPIENS | 3 | 2604 | 2pbd | P | 116 | GLU | p.Glu117Gly | P07737 |
| HOMO SAPIENS | 4 | 2013 | 3b7w | A | 472 | ARG | p.Arg472Gly | Q08AH3 |
| HOMO SAPIENS | 0 | 24346 | 3sh5 | A | 85 | ARG | p.Arg4281His | P98160 |

| | | | | | | | | |
|---|---|---|---|---|---|---|---|---|
| HOMO SAPIENS | 0 | 24346 | 3sh5 | A | 122 | ARG | p.Arg4318Trp | P98160 |
| HOMO SAPIENS | 0 | 24346 | 3sh5 | A | 135 | SER | p.Ser4331Asn | P98160 |
| HOMO SAPIENS | 0 | 24346 | 3sh5 | A | 136 | VAL | p.Val4332Ile | P98160 |
| HOMO SAPIENS | 0 | 707 | 4p2g | A | 26 | MET | p.Met26Ile | Q99497 |
| HOMO SAPIENS | 5 | 707 | 4p2g | A | 39 | ALA | p.Ala39Ser | Q99497 |
| HOMO SAPIENS | 5 | 707 | 4p2g | A | 64 | GLU | p.Glu64Asp | Q99497 |
| HOMO SAPIENS | 4 | 707 | 4p2g | A | 149 | ASP | p.Asp149Ala | Q99497 |
| HOMO SAPIENS | 6 | 707 | 4p2g | A | 163 | GLU | p.Glu163Lys | Q99497 |
| HOMO SAPIENS | 6 | 707 | 4p2g | A | 166 | LEU | p.Leu166Pro | Q99497 |
| HOMO SAPIENS | 3 | 5121 | 4xss | E | 31 | GLY | p.Gly58Arg | P06213 |
| HOMO SAPIENS | 2 | 5121 | 4xss | E | 119 | ILE | p.Ile146Met | P06213 |
| HOMO SAPIENS | 8 | 5121 | 4xss | E | 209 | HIS | p.His236Arg | P06213 |
| HOMO SAPIENS | 2 | 5121 | 4xss | E | 234 | ASP | p.Asp261Glu | P06213 |
| HOMO SAPIENS | 1 | 5121 | 4xss | E | 276 | GLN | p.Gln303His | P06213 |
| HOMO SAPIENS | 1 | 27790 | 5erp | B | 140 | THR | p.Thr275Met | Q02487 |
| HOMO SAPIENS | 0 | 27790 | 5erp | B | 150 | ILE | p.Ile285Thr | Q02487 |
| HOMO SAPIENS | 1 | 27790 | 5erp | B | 205 | THR | p.Thr340Ala | Q02487 |
| HOMO SAPIENS | 1 | 27790 | 5erp | B | 223 | THR | p.Thr358Ile | Q02487 |
| HOMO SAPIENS | 0 | 27790 | 5erp | B | 301 | GLY | p.Gly436Ala | Q02487 |
| HOMO SAPIENS | -1 | 27790 | 5erp | B | 315 | ARG | p.Arg450Ser | Q02487 |
| HOMO SAPIENS | 0 | 27790 | 5erp | B | 369 | SER | p.Ser504Gly | Q02487 |
| HOMO SAPIENS | 0 | 27790 | 5erp | B | 385 | ILE | p.Ile520Thr | Q02487 |
| HOMO SAPIENS | 0 | 27790 | 5erp | B | 461 | ALA | p.Ala596Val | Q02487 |
| HOMO SAPIENS | 0 | 27790 | 5erp | B | 503 | GLN | p.Gln638His | Q02487 |
| HOMO SAPIENS | 5 | 6599 | 4nsc | A | 129 | ARG | p.Arg129Leu | Q9BPX6 |
| HOMO SAPIENS | -1 | 6599 | 4nsc | A | 183 | GLN | p.Gln183Ter | Q9BPX6 |
| HOMO SAPIENS | 0 | 20420 | 3ghm | A | 398 | ARG | p.Arg398His | Q76LX8 |
| HOMO SAPIENS | 0 | 20420 | 3ghm | A | 448 | GLN | p.Gln448Glu | Q76LX8 |
| HOMO SAPIENS | 1 | 20420 | 3ghm | A | 449 | GLN | p.Gln449Ter | Q76LX8 |
| HOMO SAPIENS | 0 | 20420 | 3ghm | A | 456 | GLN | p.Gln456His | Q76LX8 |
| HOMO SAPIENS | 1 | 20420 | 3ghm | A | 528 | ARG | p.Arg528Gly | Q76LX8 |
| HOMO SAPIENS | 0 | 20420 | 3ghm | A | 618 | PRO | p.Pro618Ala | Q76LX8 |
| HOMO SAPIENS | 0 | 20420 | 3ghm | A | 625 | ARG | p.Arg625His | Q76LX8 |
| HOMO SAPIENS | 0 | 275 | 4rmz | A | 293 | GLN | p.Gln293Ter | Q9NWZ3 |
| HOMO SAPIENS | 0 | 275 | 4rmz | A | 428 | ALA | p.Ala428Thr | Q9NWZ3 |
| HOMO SAPIENS | 2 | 10458 | 3puf | B | 138 | LEU | p.Leu138Phe | Q5TBB1 |
| HOMO SAPIENS | 6 | 10458 | 3puf | B | 177 | ALA | p.Ala177Thr | Q5TBB1 |
| HOMO SAPIENS | -1 | 10458 | 3puf | B | 185 | VAL | p.Val185Gly | Q5TBB1 |
| HOMO SAPIENS | 0 | 26648 | 2n90 | A | 36 | ARG | p.Arg444Gln | P04629 |
| HOMO SAPIENS | 2 | 977 | 1dnw | D | 166 | ALA | p.Ala332Val | P05164 |
| HOMO SAPIENS | 4 | 977 | 1dnw | D | 333 | ARG | p.Arg499Cys | P05164 |
| HOMO SAPIENS | 1 | 977 | 1dnw | D | 403 | ARG | p.Arg569Trp | P05164 |
| HOMO SAPIENS | 0 | 25704 | 3ajb | A | 82 | GLN | p.Gln82Arg | P56589 |
| HOMO SAPIENS | 3 | 255 | 3t1p | A | 38 | TYR | p.Tyr62Ter | P01009 |
| HOMO SAPIENS | 3 | 255 | 3t1p | A | 39 | ARG | p.Arg63Cys | P01009 |
| HOMO SAPIENS | 0 | 255 | 3t1p | A | 101 | ARG | p.Arg125His | P01009 |
| HOMO SAPIENS | 0 | 255 | 3t1p | A | 212 | GLN | p.Gln236Arg | P01009 |
| HOMO SAPIENS | 0 | 255 | 3t1p | A | 213 | VAL | p.Val237Ala | P01009 |
| HOMO SAPIENS | 1 | 255 | 3t1p | A | 269 | HIS | p.His293Gln | P01009 |
| HOMO SAPIENS | 8 | 255 | 3t1p | A | 341 | ASP | p.Asp365Asn | P01009 |
| HOMO SAPIENS | 9 | 255 | 3t1p | A | 342 | GLU | p.Glu366Lys | P01009 |
| HOMO SAPIENS | 0 | 255 | 3t1p | A | 369 | PRO | p.Pro393Ser | P01009 |
| HOMO SAPIENS | 0 | 255 | 3t1p | A | 376 | GLU | p.Glu400Asp | P01009 |
| HOMO SAPIENS | -1 | 5175 | 3zxi | A | 35 | ALA | p.Ala35Asp | Q9Y2Z4 |

| | | | | | | | | | |
|---|---|---|---|---|---|---|---|---|---|
| HOMO SAPIENS | 0 | 5175 | 3zxi | A | | 52 | PHE | p.Phe52Leu | Q9Y2Z4 |
| HOMO SAPIENS | -1 | 5175 | 3zxi | A | | 68 | GLY | p.Gly68Ser | Q9Y2Z4 |
| HOMO SAPIENS | 0 | 5175 | 3zxi | A | | 179 | LYS | p.Lys179Gln | Q9Y2Z4 |
| HOMO SAPIENS | 0 | 5175 | 3zxi | A | | 191 | GLY | p.Gly191Val | Q9Y2Z4 |
| HOMO SAPIENS | 0 | 5175 | 3zxi | A | | 251 | ILE | p.Ile251Val | Q9Y2Z4 |
| HOMO SAPIENS | 2 | 5175 | 3zxi | A | | 295 | ASP | p.Asp295Asn | Q9Y2Z4 |
| HOMO SAPIENS | 0 | 5175 | 3zxi | A | | 311 | ASP | p.Asp311Glu | Q9Y2Z4 |
| HOMO SAPIENS | 0 | 5175 | 3zxi | A | | 360 | ARG | p.Arg360Ter | Q9Y2Z4 |
| HOMO SAPIENS | 1 | 3016 | 3exg | | 3 | 4 | ASP | p.Asp33His | P08559 |
| HOMO SAPIENS | 1 | 3016 | 3exg | | 3 | 176 | PHE | p.Phe205Leu | P08559 |
| HOMO SAPIENS | 1 | 3016 | 3exg | | 3 | 214 | TYR | p.Tyr243Asn | P08559 |
| HOMO SAPIENS | 1 | 3016 | 3exg | | 3 | 229 | ASP | p.Asp258Ala | P08559 |
| HOMO SAPIENS | 0 | 3016 | 3exg | | 3 | 233 | VAL | p.Val262Ile | P08559 |
| HOMO SAPIENS | 1 | 3016 | 3exg | | 3 | 234 | ARG | p.Arg263Gly | P08559 |
| HOMO SAPIENS | 0 | 3016 | 3exg | | 3 | 253 | MET | p.Met282Leu | P08559 |
| HOMO SAPIENS | 7 | 3016 | 3exg | | 3 | 259 | ARG | p.Arg288His | P08559 |
| HOMO SAPIENS | -1 | 3016 | 3exg | | 3 | 273 | ARG | p.Arg302Cys | P08559 |
| HOMO SAPIENS | 2 | 3016 | 3exg | | 3 | 286 | ASP | p.Asp315Asn | P08559 |
| HOMO SAPIENS | 1 | 3016 | 3exg | | 3 | 349 | ARG | p.Arg378His | P08559 |
| HOMO SAPIENS | 5 | 6729 | 4a0l | E | | 572 | ARG | p.Arg572Cys | Q13620 |
| HOMO SAPIENS | 0 | 1067 | 4kyo | A | | 9 | THR | p.Thr9Asn | P21549 |
| HOMO SAPIENS | 0 | 1067 | 4kyo | A | | 11 | PRO | p.Pro11Leu | P21549 |
| HOMO SAPIENS | 0 | 1067 | 4kyo | A | | 22 | ASN | p.Asn22Ser | P21549 |
| HOMO SAPIENS | 4 | 1067 | 4kyo | A | | 25 | LEU | p.Leu25Arg | P21549 |
| HOMO SAPIENS | 7 | 1067 | 4kyo | A | | 36 | ARG | p.Arg36His | P21549 |
| HOMO SAPIENS | 3 | 1067 | 4kyo | A | | 41 | GLY | p.Gly41Arg | P21549 |
| HOMO SAPIENS | 0 | 1067 | 4kyo | A | | 42 | GLY | p.Gly42Glu | P21549 |
| HOMO SAPIENS | 3 | 1067 | 4kyo | A | | 56 | ILE | p.Ile56Asn | P21549 |
| HOMO SAPIENS | 3 | 1067 | 4kyo | A | | 59 | GLU | p.Glu59Lys | P21549 |
| HOMO SAPIENS | 3 | 1067 | 4kyo | A | | 66 | TYR | p.Tyr66Ter | P21549 |
| HOMO SAPIENS | 3 | 1067 | 4kyo | A | | 111 | ARG | p.Arg111Leu | P21549 |
| HOMO SAPIENS | 7 | 1067 | 4kyo | A | | 118 | ARG | p.Arg118His | P21549 |
| HOMO SAPIENS | 3 | 1067 | 4kyo | A | | 122 | ARG | p.Arg122Ter | P21549 |
| HOMO SAPIENS | 3 | 1067 | 4kyo | A | | 124 | HIS | p.His124Pro | P21549 |
| HOMO SAPIENS | 0 | 1067 | 4kyo | A | | 152 | PHE | p.Phe152Ile | P21549 |
| HOMO SAPIENS | 1 | 1067 | 4kyo | A | | 156 | GLY | p.Gly156Arg | P21549 |
| HOMO SAPIENS | 0 | 1067 | 4kyo | A | | 161 | GLY | p.Gly161Arg | P21549 |
| HOMO SAPIENS | 3 | 1067 | 4kyo | A | | 166 | LEU | p.Leu166Pro | P21549 |
| HOMO SAPIENS | 5 | 1067 | 4kyo | A | | 170 | GLY | p.Gly170Arg | P21549 |
| HOMO SAPIENS | 5 | 1067 | 4kyo | A | | 186 | ALA | p.Ala186Val | P21549 |
| HOMO SAPIENS | 3 | 1067 | 4kyo | A | | 187 | SER | p.Ser187Phe | P21549 |
| HOMO SAPIENS | 3 | 1067 | 4kyo | A | | 197 | ARG | p.Arg197Pro | P21549 |
| HOMO SAPIENS | 5 | 1067 | 4kyo | A | | 201 | ASP | p.Asp201Glu | P21549 |
| HOMO SAPIENS | 0 | 1067 | 4kyo | A | | 202 | ILE | p.Ile202Asn | P21549 |
| HOMO SAPIENS | 1 | 1067 | 4kyo | A | | 205 | SER | p.Ser205Pro | P21549 |
| HOMO SAPIENS | 1 | 1067 | 4kyo | A | | 205 | SER | p.Ser205Ter | P21549 |
| HOMO SAPIENS | 2 | 1067 | 4kyo | A | | 216 | GLY | p.Gly216Arg | P21549 |
| HOMO SAPIENS | 5 | 1067 | 4kyo | A | | 233 | ARG | p.Arg233Cys | P21549 |
| HOMO SAPIENS | 5 | 1067 | 4kyo | A | | 244 | ILE | p.Ile244Thr | P21549 |
| HOMO SAPIENS | 5 | 1067 | 4kyo | A | | 244 | ILE | p.Ile244Met | P21549 |
| HOMO SAPIENS | 4 | 1067 | 4kyo | A | | 253 | CYS | p.Cys253Arg | P21549 |
| HOMO SAPIENS | 7 | 1067 | 4kyo | A | | 274 | GLU | p.Glu274Asp | P21549 |
| HOMO SAPIENS | 7 | 1067 | 4kyo | A | | 279 | ILE | p.Ile279Thr | P21549 |
| HOMO SAPIENS | 6 | 1067 | 4kyo | A | | 295 | ALA | p.Ala295Thr | P21549 |

| | | | | | | | | |
|---|---|---|---|---|---|---|---|---|
| HOMO SAPIENS | 5 | 1067 | 4kyo | A | 340 | ILE | p.Ile340Met | P21549 |
| HOMO SAPIENS | 1 | 790 | 2ric | C | 270 | SER | p.Ser290Thr | P35247 |
| HOMO SAPIENS | 0 | 790 | 2ric | C | 289 | GLU | p.Glu309Lys | P35247 |
| HOMO SAPIENS | 1 | 212 | 5sxe | A | 545 | GLU | p.Glu545Lys | P42336 |
| HOMO SAPIENS | 3 | 212 | 5sxe | A | 546 | GLN | p.Gln546Glu | P42336 |
| HOMO SAPIENS | 2 | 212 | 5sxe | A | 1047 | HIS | p.His1047Leu | P42336 |
| HOMO SAPIENS | 7 | 152 | 2zqq | F | 197 | ARG | p.Arg197Ter | Q13825 |
| HOMO SAPIENS | 0 | 152 | 2zqq | F | 248 | ALA | p.Ala248Thr | Q13825 |
| HOMO SAPIENS | 0 | 152 | 2zqq | F | 275 | ALA | p.Ala275Val | Q13825 |
| HOMO SAPIENS | 5 | 152 | 2zqq | F | 331 | LYS | p.Lys331Ter | Q13825 |
| HOMO SAPIENS | 6 | 17801 | 2j8l | A | 283 | PHE | p.Phe301Leu | P03951 |
| HOMO SAPIENS | 3 | 17801 | 2j8l | A | 308 | ARG | p.Arg326Cys | P03951 |
| HOMO SAPIENS | 0 | 17801 | 2j8l | A | 321 | CYS | p.Cys339Phe | P03951 |
| HOMO SAPIENS | 3 | 17801 | 2j8l | A | 351 | TYR | p.Tyr369Ter | P03951 |
| HOMO SAPIENS | 7 | 1802 | 5db0 | A | 22 | LEU | p.Leu22Arg | O00255 |
| HOMO SAPIENS | 7 | 1802 | 5db0 | A | 26 | GLU | p.Glu26Lys | O00255 |
| HOMO SAPIENS | 7 | 1802 | 5db0 | A | 139 | HIS | p.His139Asp | O00255 |
| HOMO SAPIENS | 0 | 1802 | 5db0 | A | 171 | ARG | p.Arg176Gln | O00255 |
| HOMO SAPIENS | 0 | 1802 | 5db0 | A | 184 | VAL | p.Val189Glu | O00255 |
| HOMO SAPIENS | 7 | 1802 | 5db0 | A | 255 | GLU | p.Glu260Lys | O00255 |
| HOMO SAPIENS | 4 | 1802 | 5db0 | A | 418 | ASP | p.Asp423Asn | O00255 |
| HOMO SAPIENS | 0 | 1802 | 5db0 | A | 436 | TRP | p.Trp441Arg | O00255 |
| HOMO SAPIENS | -1 | 1802 | 5db0 | A | 541 | ALA | p.Thr546Ala | O00255 |
| HOMO SAPIENS | 4 | 1802 | 5db0 | A | 552 | THR | p.Thr557Ser | O00255 |
| HOMO SAPIENS | 0 | 3450 | 2yuz | A | 36 | PRO | p.Pro461Gln | Q00872 |
| HOMO SAPIENS | 1 | 3450 | 2yuz | A | 56 | GLN | p.His481Gln | Q00872 |
| HOMO SAPIENS | 2 | 3450 | 2yuz | A | 68 | ALA | p.Ala493Val | Q00872 |
| HOMO SAPIENS | 0 | 3211 | 3rqg | A | 72 | VAL | p.Val72Leu | Q9BUL8 |
| HOMO SAPIENS | 2 | 4168 | 4q4f | A | 121 | ARG | p.Arg121Ter | Q14108 |
| HOMO SAPIENS | 2 | 4168 | 4q4f | A | 121 | ARG | p.Arg121Gln | Q14108 |
| HOMO SAPIENS | 2 | 4168 | 4q4f | A | 159 | MET | p.Met159Val | Q14108 |
| HOMO SAPIENS | 4 | 4168 | 4q4f | A | 363 | HIS | p.His363Asn | Q14108 |
| HOMO SAPIENS | 2 | 4168 | 4q4f | A | 396 | VAL | p.Val396Ile | Q14108 |
| HOMO SAPIENS | 0 | 1448 | 1y97 | A | 87 | ARG | p.Arg130Ter | Q9BQ50 |
| HOMO SAPIENS | 0 | 3444 | 4iwp | A | 151 | SER | p.Ser151Phe | Q9UHD2 |
| HOMO SAPIENS | 0 | 3444 | 4iwp | A | 401 | LYS | p.Lys401Glu | Q9UHD2 |
| HOMO SAPIENS | 0 | 116 | 1zy8 | B | 194 | GLY | p.Gly229Cys | P09622 |
| HOMO SAPIENS | 0 | 116 | 1zy8 | B | 255 | ASP | p.Asp290Gly | P09622 |
| HOMO SAPIENS | 2 | 116 | 1zy8 | B | 326 | MET | p.Met361Val | P09622 |
| HOMO SAPIENS | 0 | 116 | 1zy8 | B | 340 | GLU | p.Glu375Lys | P09622 |
| HOMO SAPIENS | 2 | 116 | 1zy8 | B | 358 | ILE | p.Ile393Thr | P09622 |
| HOMO SAPIENS | 1 | 116 | 1zy8 | B | 453 | PRO | p.Pro488Leu | P09622 |
| HOMO SAPIENS | 7 | 127 | 4qo7 | B | 160 | SER | p.Ser161Arg | P00338 |
| HOMO SAPIENS | 0 | 2025 | 1wrl | A | 8 | ALA | p.Ala8Val | P63316 |
| HOMO SAPIENS | 8 | 2025 | 1wrl | A | 24 | PHE | p.Phe24Leu | P63316 |
| HOMO SAPIENS | 0 | 2025 | 1wrl | A | 84 | SER | p.Cys84Tyr | P63316 |
| HOMO SAPIENS | 8 | 3791 | 1mhd | B | 93 | ARG | p.Arg93Ter | P84022 |
| HOMO SAPIENS | 0 | 1575 | 4phc | A | 205 | GLY | p.Gly205Asp | P12081 |
| HOMO SAPIENS | 0 | 1575 | 4phc | A | 454 | TYR | p.Tyr454Ser | P12081 |
| HOMO SAPIENS | 0 | 39620 | 1ozs | B | 135 | ARG | p.Arg136Gln | P19429 |
| HOMO SAPIENS | 5 | 39620 | 1ozs | B | 140 | ARG | p.Arg141Trp | P19429 |
| HOMO SAPIENS | 5 | 39620 | 1ozs | B | 142 | THR | p.Thr143Asn | P19429 |
| HOMO SAPIENS | 5 | 39620 | 1ozs | B | 143 | LEU | p.Leu144Gln | P19429 |
| HOMO SAPIENS | 5 | 39620 | 1ozs | B | 144 | ARG | p.Arg145Gly | P19429 |

| | | | | | | | | |
|---|---|---|---|---|---|---|---|---|
| HOMO SAPIENS | 5 | 39620 | 1ozs | B | 144 | ARG | p.Arg145Gln | P19429 |
| HOMO SAPIENS | -1 | 20934 | 4nuv | D | 42 | GLY | p.Gly42Asp | Q16570 |
| HOMO SAPIENS | -1 | 40565 | 5lcw | S | 4 | VAL | p.Val4Met | O60566 |
| HOMO SAPIENS | 0 | 40565 | 5lcw | S | 40 | THR | p.Thr40Met | O60566 |
| HOMO SAPIENS | 4 | 40565 | 5lcw | S | 198 | ALA | p.Glu390Asp | O60566 |
| HOMO SAPIENS | 4 | 758 | 4oiu | A | 726 | ARG | p.Arg727Leu | P10275 |
| HOMO SAPIENS | 4 | 758 | 4oiu | A | 730 | VAL | p.Val731Met | P10275 |
| HOMO SAPIENS | 2 | 758 | 4oiu | A | 798 | GLN | p.Gln799Glu | P10275 |
| HOMO SAPIENS | 3 | 758 | 4oiu | A | 855 | ARG | p.Arg856His | P10275 |
| HOMO SAPIENS | 1 | 758 | 4oiu | A | 886 | MET | p.Met887Val | P10275 |
| HOMO SAPIENS | 0 | 3303 | 4ie0 | A | 134 | ALA | p.Ala134Thr | Q9C0B1 |
| HOMO SAPIENS | 9 | 3303 | 4ie0 | A | 316 | ARG | p.Arg316Gln | Q9C0B1 |
| HOMO SAPIENS | 1 | 41502 | 3fgh | A | 136 | PRO | p.Pro178Leu | Q00059 |
| HOMO SAPIENS | 5 | 175 | 3ceh | B | 58 | THR | p.Thr59Lys | P06737 |
| HOMO SAPIENS | 8 | 175 | 3ceh | B | 93 | ARG | p.Arg94Ter | P06737 |
| HOMO SAPIENS | 5 | 175 | 3ceh | B | 221 | VAL | p.Val222Ile | P06737 |
| HOMO SAPIENS | 0 | 175 | 3ceh | B | 338 | ASN | p.Asn339Ser | P06737 |
| HOMO SAPIENS | 1 | 175 | 3ceh | B | 372 | PHE | p.Phe373Leu | P06737 |
| HOMO SAPIENS | 9 | 175 | 3ceh | B | 376 | ASN | p.Asn377Lys | P06737 |
| HOMO SAPIENS | 8 | 175 | 3ceh | B | 398 | ARG | p.Arg399Ter | P06737 |
| HOMO SAPIENS | 5 | 175 | 3ceh | B | 424 | ARG | p.Arg425Pro | P06737 |
| HOMO SAPIENS | 9 | 175 | 3ceh | B | 455 | VAL | p.Val456Met | P06737 |
| HOMO SAPIENS | 8 | 175 | 3ceh | B | 490 | ARG | p.Arg491Cys | P06737 |
| HOMO SAPIENS | 8 | 175 | 3ceh | B | 631 | ASN | p.Asn632Ile | P06737 |
| HOMO SAPIENS | 9 | 175 | 3ceh | B | 648 | TYR | p.Tyr649Ter | P06737 |
| HOMO SAPIENS | 9 | 175 | 3ceh | B | 672 | GLU | p.Glu673Lys | P06737 |
| HOMO SAPIENS | 9 | 175 | 3ceh | B | 674 | SER | p.Ser675Leu | P06737 |
| HOMO SAPIENS | 0 | 175 | 3ceh | B | 805 | ILE | p.Ile806Leu | P06737 |
| HOMO SAPIENS | 0 | 175 | 3ceh | B | 820 | TYR | p.Tyr821His | P06737 |
| HOMO SAPIENS | 1 | 6750 | 4fdj | A | 47 | GLY | p.Gly47Arg | P34059 |
| HOMO SAPIENS | 9 | 6750 | 4fdj | A | 60 | ASP | p.Asp60Asn | P34059 |
| HOMO SAPIENS | 5 | 6750 | 4fdj | A | 67 | LEU | p.Leu67Met | P34059 |
| HOMO SAPIENS | 0 | 6750 | 4fdj | A | 69 | PHE | p.Phe69Val | P34059 |
| HOMO SAPIENS | 3 | 6750 | 4fdj | A | 94 | ARG | p.Arg94Gly | P34059 |
| HOMO SAPIENS | 0 | 6750 | 4fdj | A | 113 | ILE | p.Ile113Phe | P34059 |
| HOMO SAPIENS | 0 | 6750 | 4fdj | A | 138 | VAL | p.Val138Ala | P34059 |
| HOMO SAPIENS | 0 | 6750 | 4fdj | A | 139 | GLY | p.Gly139Ser | P34059 |
| HOMO SAPIENS | 0 | 6750 | 4fdj | A | 151 | PRO | p.Pro151Thr | P34059 |
| HOMO SAPIENS | 0 | 6750 | 4fdj | A | 155 | GLY | p.Gly155Arg | P34059 |
| HOMO SAPIENS | 0 | 6750 | 4fdj | A | 159 | TRP | p.Trp159Ter | P34059 |
| HOMO SAPIENS | 0 | 6750 | 4fdj | A | 162 | SER | p.Ser162Phe | P34059 |
| HOMO SAPIENS | 0 | 6750 | 4fdj | A | 200 | THR | p.Thr200Met | P34059 |
| HOMO SAPIENS | 0 | 6750 | 4fdj | A | 204 | ASN | p.Asn204Lys | P34059 |
| HOMO SAPIENS | 0 | 6750 | 4fdj | A | 231 | ALA | p.Ala231Gly | P34059 |
| HOMO SAPIENS | 2 | 6750 | 4fdj | A | 235 | THR | p.Thr235Lys | P34059 |
| HOMO SAPIENS | 1 | 6750 | 4fdj | A | 259 | ARG | p.Arg259Gln | P34059 |
| HOMO SAPIENS | 0 | 6750 | 4fdj | A | 287 | SER | p.Ser287Leu | P34059 |
| HOMO SAPIENS | 0 | 6750 | 4fdj | A | 291 | ALA | p.Ala291Thr | P34059 |
| HOMO SAPIENS | 3 | 6750 | 4fdj | A | 301 | GLY | p.Gly301Cys | P34059 |
| HOMO SAPIENS | 0 | 6750 | 4fdj | A | 312 | THR | p.Thr312Ser | P34059 |
| HOMO SAPIENS | 0 | 6750 | 4fdj | A | 318 | MET | p.Met318Arg | P34059 |
| HOMO SAPIENS | 0 | 6750 | 4fdj | A | 340 | GLY | p.Gly340Asp | P34059 |
| HOMO SAPIENS | 0 | 6750 | 4fdj | A | 386 | ARG | p.Arg386Cys | P34059 |
| HOMO SAPIENS | 0 | 6750 | 4fdj | A | 391 | MET | p.Met391Val | P34059 |

| | | | | | | | | |
|---|---|---|---|---|---|---|---|---|
| HOMO SAPIENS | 0 | 6750 | 4fdj | A | 392 | ALA | p.Ala392Val | P34059 |
| HOMO SAPIENS | 0 | 6750 | 4fdj | A | 393 | ALA | p.Ala393Ser | P34059 |
| HOMO SAPIENS | 0 | 6750 | 4fdj | A | 459 | ALA | p.Ala459Val | P34059 |
| HOMO SAPIENS | 0 | 6750 | 4fdj | A | 473 | GLN | p.Gln473Ter | P34059 |
| HOMO SAPIENS | 2 | 6750 | 4fdj | A | 480 | VAL | p.Val480Phe | P34059 |
| HOMO SAPIENS | 0 | 6750 | 4fdj | A | 487 | ASN | p.Asn487Ser | P34059 |
| HOMO SAPIENS | 0 | 6750 | 4fdj | A | 488 | VAL | p.Val488Met | P34059 |
| HOMO SAPIENS | 0 | 6750 | 4fdj | A | 520 | TRP | p.Trp520Ter | P34059 |
| HOMO SAPIENS | 4 | 1088 | 2p3u | A | 102 | GLU | p.Glu142Lys | P00742 |
| HOMO SAPIENS | 6 | 370 | 1nr1 | D | 73 | ASP | p.Asp126His | P00367 |
| HOMO SAPIENS | 3 | 370 | 1nr1 | D | 221 | ARG | p.Arg274Cys | P00367 |
| HOMO SAPIENS | 2 | 370 | 1nr1 | D | 265 | ARG | p.Arg318Lys | P00367 |
| HOMO SAPIENS | 1 | 370 | 1nr1 | D | 269 | ARG | p.Arg322His | P00367 |
| HOMO SAPIENS | 1 | 370 | 1nr1 | D | 296 | GLU | p.Glu349Ala | P00367 |
| HOMO SAPIENS | 1 | 370 | 1nr1 | D | 445 | SER | p.Ser498Leu | P00367 |
| HOMO SAPIENS | 1 | 370 | 1nr1 | D | 446 | GLY | p.Gly499Asp | P00367 |
| HOMO SAPIENS | 1 | 370 | 1nr1 | D | 446 | GLY | p.Gly499Ser | P00367 |
| HOMO SAPIENS | 1 | 370 | 1nr1 | D | 448 | SER | p.Ser501Pro | P00367 |
| HOMO SAPIENS | 2 | 370 | 1nr1 | D | 454 | HIS | p.His507Tyr | P00367 |
| HOMO SAPIENS | 1 | 7720 | 2he0 | A | 147 | HIS | p.His2018Gln | P46531 |
| HOMO SAPIENS | 2 | 11844 | 5l0v | A | 218 | ARG | p.Arg218Ter | Q8NBL1 |
| HOMO SAPIENS | 2 | 11844 | 5l0v | A | 279 | ARG | p.Arg279Trp | Q8NBL1 |
| HOMO SAPIENS | 9 | 1485 | 5va0 | A | 477 | ARG | p.Arg477His | P04150 |
| HOMO SAPIENS | 0 | 235 | 2fun | B | 2248 | ARG | p.Arg248Trp | Q14790 |
| HOMO SAPIENS | 4 | 235 | 2fun | B | 2285 | HIS | p.Asp285His | Q14790 |
| HOMO SAPIENS | 0 | 4822 | 5y3n | A | 307 | ARG | p.Arg307Gly | Q12931 |
| HOMO SAPIENS | 8 | 32928 | 4m6w | A | 1886 | HIS | p.His1886Tyr | Q8IYD8 |
| HOMO SAPIENS | 0 | 2711 | 4y5v | F | 22 | GLY | p.Gly46Glu | P19235 |
| HOMO SAPIENS | -1 | 2711 | 4y5v | F | 166 | ALA | p.Ala190Ser | P19235 |
| HOMO SAPIENS | 4 | 5161 | 5bpu | A | 42 | HIS | p.His42Arg | Q00604 |
| HOMO SAPIENS | 0 | 5161 | 5bpu | A | 44 | TYR | p.Tyr44Cys | Q00604 |
| HOMO SAPIENS | 0 | 5161 | 5bpu | A | 45 | VAL | p.Val45Glu | Q00604 |
| HOMO SAPIENS | 0 | 5161 | 5bpu | A | 60 | VAL | p.Val60Glu | Q00604 |
| HOMO SAPIENS | 0 | 5161 | 5bpu | A | 61 | LEU | p.Leu61Phe | Q00604 |
| HOMO SAPIENS | 7 | 5161 | 5bpu | A | 69 | CYS | p.Cys69Ser | Q00604 |
| HOMO SAPIENS | 0 | 5161 | 5bpu | A | 75 | SER | p.Ser75Cys | Q00604 |
| HOMO SAPIENS | 0 | 5161 | 5bpu | A | 90 | ARG | p.Arg90Pro | Q00604 |
| HOMO SAPIENS | 9 | 5161 | 5bpu | A | 96 | CYS | p.Cys96Tyr | Q00604 |
| HOMO SAPIENS | 9 | 5161 | 5bpu | A | 96 | CYS | p.Cys96Trp | Q00604 |
| HOMO SAPIENS | 0 | 5161 | 5bpu | A | 101 | SER | p.Ser101Phe | Q00604 |
| HOMO SAPIENS | 0 | 5161 | 5bpu | A | 105 | ALA | p.Ala105Thr | Q00604 |
| HOMO SAPIENS | 0 | 5161 | 5bpu | A | 110 | CYS | p.Cys110Gly | Q00604 |
| HOMO SAPIENS | 0 | 5161 | 5bpu | A | 121 | ARG | p.Arg121Trp | Q00604 |
| HOMO SAPIENS | 0 | 5161 | 5bpu | A | 121 | ARG | p.Arg121Leu | Q00604 |
| HOMO SAPIENS | 0 | 5161 | 5bpu | A | 124 | LEU | p.Leu124Phe | Q00604 |
| HOMO SAPIENS | 8 | 5161 | 5bpu | A | 128 | CYS | p.Cys128Ter | Q00604 |
| HOMO SAPIENS | 1 | 13944 | 5xqh | A | 238 | ARG | p.Arg238His | Q9GZN7 |
| HOMO SAPIENS | -1 | 7508 | 5jl6 | A | 39 | TRP | p.Trp39Arg | P11511 |
| HOMO SAPIENS | 0 | 7508 | 5jl6 | A | 201 | THR | p.Thr201Met | P11511 |
| HOMO SAPIENS | 0 | 7508 | 5jl6 | A | 210 | GLU | p.Glu210Lys | P11511 |
| HOMO SAPIENS | 1 | 7508 | 5jl6 | A | 264 | ARG | p.Arg264Cys | P11511 |
| HOMO SAPIENS | 2 | 7508 | 5jl6 | A | 365 | ARG | p.Arg365Gln | P11511 |
| HOMO SAPIENS | 7 | 7508 | 5jl6 | A | 375 | ARG | p.Arg375Cys | P11511 |
| HOMO SAPIENS | 8 | 7508 | 5jl6 | A | 435 | ARG | p.Arg435Cys | P11511 |

| | | | | | | | | |
|---|---|---|---|---|---|---|---|---|
| HOMO SAPIENS | 9 | 7508 | 5jl6 | A | 437 | CYS | p.Cys437Tyr | P11511 |
| HOMO SAPIENS | -1 | 30206 | 5l95 | B | 246 | ARG | p.Arg246Ter | Q9GZZ9 |
| HOMO SAPIENS | 4 | 875 | 3efh | B | 43 | GLU | p.Glu43Asp | P60891 |
| HOMO SAPIENS | 5 | 875 | 3efh | B | 65 | ASP | p.Asp65Asn | P60891 |
| HOMO SAPIENS | 0 | 875 | 3efh | B | 87 | ALA | p.Ala87Thr | P60891 |
| HOMO SAPIENS | 2 | 875 | 3efh | B | 115 | MET | p.Met115Thr | P60891 |
| HOMO SAPIENS | 8 | 875 | 3efh | B | 133 | GLN | p.Gln133Pro | P60891 |
| HOMO SAPIENS | 8 | 875 | 3efh | B | 152 | LEU | p.Leu152Pro | P60891 |
| HOMO SAPIENS | 7 | 875 | 3efh | B | 290 | ILE | p.Ile290Thr | P60891 |
| HOMO SAPIENS | 6 | 875 | 3efh | B | 306 | GLY | p.Gly306Arg | P60891 |
| HOMO SAPIENS | 0 | 7978 | 5edm | A | 122 | MET | p.Thr165Met | P00734 |
| HOMO SAPIENS | -1 | 7978 | 5edm | A | 257 | ARG | p.Arg314Cys | P00734 |
| HOMO SAPIENS | 0 | 1716 | 1b98 | M | 126 | ARG | p.Arg206Gln | P34130 |
| HOMO SAPIENS | 0 | 1716 | 1b98 | M | 126 | ARG | p.Arg206Trp | P34130 |
| HOMO SAPIENS | 9 | 3960 | 2qyp | B | 28 | ILE | p.Ile338Val | P07602 |
| HOMO SAPIENS | 1 | 3960 | 2qyp | B | 39 | LEU | p.Leu349Pro | P07602 |
| HOMO SAPIENS | 0 | 31414 | 5u1d | A | 333 | ILE | p.Ile393Val | Q03518 |
| HOMO SAPIENS | 0 | 31414 | 5u1d | A | 637 | ASP | p.Asp697Gly | Q03518 |
| HOMO SAPIENS | 0 | 31414 | 5u1d | A | 659 | ARG | p.Arg719Gln | Q03518 |
| HOMO SAPIENS | 3 | 119 | 2pho | A | 11 | ILE | p.Ile11Thr | P05089 |
| HOMO SAPIENS | 8 | 119 | 2pho | A | 21 | ARG | p.Arg21Ter | P05089 |
| HOMO SAPIENS | 0 | 119 | 2pho | A | 122 | TRP | p.Trp122Ter | P05089 |
| HOMO SAPIENS | 6 | 119 | 2pho | A | 138 | GLY | p.Gly138Val | P05089 |
| HOMO SAPIENS | 5 | 119 | 2pho | A | 290 | THR | p.Thr290Ser | P05089 |
| HOMO SAPIENS | 5 | 119 | 2pho | A | 291 | ARG | p.Arg291Ter | P05089 |
| HOMO SAPIENS | 0 | 3732 | 2i99 | A | 247 | TYR | p.Tyr247Ter | Q14894 |
| HOMO SAPIENS | -1 | 1382 | 4ffx | A | 2 | ALA | p.Ala2Val | P30566 |
| HOMO SAPIENS | 1 | 1382 | 4ffx | A | 72 | ILE | p.Ile72Met | P30566 |
| HOMO SAPIENS | 1 | 1382 | 4ffx | A | 100 | PRO | p.Pro100Ala | P30566 |
| HOMO SAPIENS | 0 | 1382 | 4ffx | A | 114 | TYR | p.Tyr114His | P30566 |
| HOMO SAPIENS | 1 | 1382 | 4ffx | A | 141 | ARG | p.Arg141Trp | P30566 |
| HOMO SAPIENS | 0 | 1382 | 4ffx | A | 147 | LYS | p.Lys147Met | P30566 |
| HOMO SAPIENS | 3 | 1382 | 4ffx | A | 190 | ARG | p.Arg190Gln | P30566 |
| HOMO SAPIENS | 3 | 1382 | 4ffx | A | 190 | ARG | p.Arg190Ter | P30566 |
| HOMO SAPIENS | 0 | 1382 | 4ffx | A | 217 | HIS | p.His217Asp | P30566 |
| HOMO SAPIENS | 2 | 1382 | 4ffx | A | 246 | LYS | p.Lys246Glu | P30566 |
| HOMO SAPIENS | 5 | 1382 | 4ffx | A | 303 | ARG | p.Arg303Cys | P30566 |
| HOMO SAPIENS | 1 | 1382 | 4ffx | A | 318 | PRO | p.Pro318Leu | P30566 |
| HOMO SAPIENS | 6 | 1382 | 4ffx | A | 332 | ASP | p.Asp332His | P30566 |
| HOMO SAPIENS | 6 | 1382 | 4ffx | A | 337 | ARG | p.Arg337Ter | P30566 |
| HOMO SAPIENS | 1 | 1382 | 4ffx | A | 396 | ARG | p.Arg396His | P30566 |
| HOMO SAPIENS | 1 | 1382 | 4ffx | A | 426 | ARG | p.Arg426His | P30566 |
| HOMO SAPIENS | 0 | 1382 | 4ffx | A | 438 | SER | p.Ser438Pro | P30566 |
| HOMO SAPIENS | 1 | 1382 | 4ffx | A | 447 | SER | p.Ser447Pro | P30566 |
| HOMO SAPIENS | 2 | 1382 | 4ffx | A | 448 | SER | p.Ser448Pro | P30566 |
| HOMO SAPIENS | 2 | 1382 | 4ffx | A | 450 | THR | p.Thr450Ser | P30566 |
| HOMO SAPIENS | 0 | 6212 | 1ao3 | A | 46 | SER | p.Ser1731Thr | P04275 |
| HOMO SAPIENS | 0 | 1711 | 3f9c | A | 279 | VAL | p.Val279Phe | Q13093 |
| HOMO SAPIENS | 0 | 6112 | 4fvl | A | 312 | HIS | p.His312Arg | P45452 |
| HOMO SAPIENS | 0 | 6112 | 4fvl | A | 323 | THR | p.Thr323Met | P45452 |
| HOMO SAPIENS | 0 | 6112 | 4fvl | A | 390 | ASP | p.Asp390Gly | P45452 |
| HOMO SAPIENS | 0 | 6112 | 4fvl | A | 458 | ARG | p.Arg458Cys | P45452 |
| HOMO SAPIENS | 2 | 6180 | 5ggn | A | 97 | ARG | p.Arg97Trp | Q8WZA1 |
| HOMO SAPIENS | 2 | 6180 | 5ggn | A | 215 | ARG | p.Arg215Ter | Q8WZA1 |

| | | | | | | | | |
|---|---|---|---|---|---|---|---|---|
| HOMO SAPIENS | -1 | 5526 | 4ak3 | A | 11 | ASP | p.Asp1232Asn | P02461 |
| HOMO SAPIENS | 6 | 5526 | 4ak3 | A | 38 | ALA | p.Ala1259Thr | P02461 |
| HOMO SAPIENS | 0 | 5526 | 4ak3 | A | 92 | LYS | p.Lys1313Arg | P02461 |
| HOMO SAPIENS | 0 | 5526 | 4ak3 | A | 132 | GLN | p.His1353Gln | P02461 |
| HOMO SAPIENS | 4 | 16616 | 1hci | B | 433 | SER | p.Ser433Leu | P35609 |
| HOMO SAPIENS | 1 | 16616 | 1hci | B | 475 | ASP | p.Asp475Asn | P35609 |
| HOMO SAPIENS | 1 | 16616 | 1hci | B | 476 | ALA | p.Ala476Ser | P35609 |
| HOMO SAPIENS | 2 | 16616 | 1hci | B | 507 | MET | p.Met507Val | P35609 |
| HOMO SAPIENS | 1 | 16616 | 1hci | B | 604 | MET | p.Met604Val | P35609 |
| HOMO SAPIENS | 2 | 16616 | 1hci | B | 640 | ARG | p.Arg640His | P35609 |
| HOMO SAPIENS | 1 | 16616 | 1hci | B | 721 | ARG | p.Arg721Cys | P35609 |
| HOMO SAPIENS | -1 | 3402 | 5iaa | B | 57 | MET | p.Met57Val | Q9GZZ9 |
| HOMO SAPIENS | 7 | 3250 | 3irq | B | 193 | PRO | p.Pro193Ala | P55265 |
| HOMO SAPIENS | 2 | 15200 | 2wns | A | 30 | SER | p.Ser30Gly | P11172 |
| HOMO SAPIENS | 0 | 15200 | 2wns | A | 96 | ARG | p.Arg96Gly | P11172 |
| HOMO SAPIENS | 0 | 15200 | 2wns | A | 109 | VAL | p.Val109Gly | P11172 |
| HOMO SAPIENS | 2 | 15759 | 4x4w | B | 140 | LEU | p.Leu166Ser | Q96Q11 |
| HOMO SAPIENS | 0 | 15759 | 4x4w | B | 197 | ILE | p.Ile223Thr | Q96Q11 |
| HOMO SAPIENS | 0 | 3156 | 3w03 | B | 14 | ALA | p.Ala14Thr | Q9H9Q4 |
| HOMO SAPIENS | 0 | 3156 | 3w03 | B | 57 | ARG | p.Arg57Gly | Q9H9Q4 |
| HOMO SAPIENS | 0 | 3156 | 3w03 | B | 123 | CYS | p.Cys123Arg | Q9H9Q4 |
| HOMO SAPIENS | 0 | 11693 | 3ojy | A | 63 | GLN | p.Gln93Lys | P07357 |
| HOMO SAPIENS | 0 | 1421 | 2qc8 | E | 275 | GLU | p.Glu275Asp | P15104 |
| HOMO SAPIENS | 9 | 1421 | 2qc8 | E | 324 | ARG | p.Arg324Cys | P15104 |
| HOMO SAPIENS | 6 | 1421 | 2qc8 | E | 341 | ARG | p.Arg341Cys | P15104 |
| HOMO SAPIENS | 6 | 2850 | 3qvb | A | 83 | THR | p.Thr83Met | P46976 |
| HOMO SAPIENS | 8 | 2850 | 3qvb | A | 102 | ASP | p.Asp102His | P46976 |
| HOMO SAPIENS | 1 | 24314 | 4m9p | A | 480 | PRO | p.Pro480Leu | P21333 |
| HOMO SAPIENS | 0 | 24314 | 4m9p | A | 528 | VAL | p.Val528Met | P21333 |
| HOMO SAPIENS | 0 | 24314 | 4m9p | A | 590 | VAL | p.Val590Ile | P21333 |
| HOMO SAPIENS | 0 | 24314 | 4m9p | A | 605 | ASP | p.Asp605Asn | P21333 |
| HOMO SAPIENS | 1 | 24314 | 4m9p | A | 637 | PRO | p.Pro637Gln | P21333 |
| HOMO SAPIENS | 0 | 24314 | 4m9p | A | 656 | LEU | p.Leu656Phe | P21333 |
| HOMO SAPIENS | 2 | 24314 | 4m9p | A | 711 | VAL | p.Val711Asp | P21333 |
| HOMO SAPIENS | 0 | 4951 | 5joe | A | 40 | THR | p.Thr9621Ser | Q8WZ42 |
| HOMO SAPIENS | 1 | 4951 | 5joe | A | 41 | GLU | p.Glu9622Gln | Q8WZ42 |
| HOMO SAPIENS | 0 | 4951 | 5joe | A | 82 | ALA | p.Ala9663Ser | Q8WZ42 |
| HOMO SAPIENS | 4 | 4951 | 5joe | A | 89 | ILE | p.Ile9670Asn | Q8WZ42 |
| HOMO SAPIENS | 0 | 9664 | 3c6m | A | 56 | GLY | p.Gly56Ser | P52788 |
| HOMO SAPIENS | 0 | 9664 | 3c6m | A | 132 | VAL | p.Val132Gly | P52788 |
| HOMO SAPIENS | 1 | 9664 | 3c6m | A | 190 | THR | p.Thr190Ile | P52788 |
| HOMO SAPIENS | -1 | 12118 | 5hcd | B | 19 | GLN | p.Gln19Ter | P01031 |
| HOMO SAPIENS | 0 | 1583 | 2b4x | L | 7 | ILE | p.Ile39Asn | P01008 |
| HOMO SAPIENS | 0 | 1583 | 2b4x | L | 24 | ARG | p.Arg56Cys | P01008 |
| HOMO SAPIENS | 0 | 1583 | 2b4x | L | 47 | ARG | p.Arg79Cys | P01008 |
| HOMO SAPIENS | 0 | 1583 | 2b4x | L | 47 | ARG | p.Arg79His | P01008 |
| HOMO SAPIENS | 1 | 1583 | 2b4x | L | 95 | CYS | p.Cys127Arg | P01008 |
| HOMO SAPIENS | 1 | 1583 | 2b4x | L | 99 | LEU | p.Leu131Phe | P01008 |
| HOMO SAPIENS | 0 | 1583 | 2b4x | L | 116 | SER | p.Ser148Pro | P01008 |
| HOMO SAPIENS | 0 | 1583 | 2b4x | L | 129 | ARG | p.Arg161Gln | P01008 |
| HOMO SAPIENS | 0 | 1583 | 2b4x | L | 135 | ASN | p.Asn167Thr | P01008 |
| HOMO SAPIENS | 2 | 1583 | 2b4x | L | 187 | ASN | p.Asn219Asp | P01008 |
| HOMO SAPIENS | 0 | 1583 | 2b4x | L | 191 | SER | p.Ser223Pro | P01008 |
| HOMO SAPIENS | 2 | 1583 | 2b4x | L | 349 | SER | p.Ser381Pro | P01008 |

| | | | | | | | | |
|---|---|---|---|---|---|---|---|---|
| HOMO SAPIENS | 0 | 1583 | 2b4x | L | 382 | ALA | p.Ala414Thr | P01008 |
| HOMO SAPIENS | 0 | 1583 | 2b4x | L | 387 | ALA | p.Ala419Val | P01008 |
| HOMO SAPIENS | 0 | 1583 | 2b4x | L | 392 | GLY | p.Gly424Asp | P01008 |
| HOMO SAPIENS | 0 | 1583 | 2b4x | L | 393 | ARG | p.Arg425His | P01008 |
| HOMO SAPIENS | 0 | 1583 | 2b4x | L | 393 | ARG | p.Arg425Cys | P01008 |
| HOMO SAPIENS | 0 | 1583 | 2b4x | L | 394 | SER | p.Ser426Leu | P01008 |
| HOMO SAPIENS | 0 | 1583 | 2b4x | L | 404 | ALA | p.Ala436Thr | P01008 |
| HOMO SAPIENS | 2 | 1583 | 2b4x | L | 407 | PRO | p.Pro439Leu | P01008 |
| HOMO SAPIENS | 2 | 1583 | 2b4x | L | 429 | PRO | p.Pro461Leu | P01008 |
| HOMO SAPIENS | 0 | 11647 | 2a1t | S | 78 | ALA | p.Ala78Ser | P38117 |
| HOMO SAPIENS | 0 | 11647 | 2a1t | S | 98 | ARG | p.Arg98Cys | P38117 |
| HOMO SAPIENS | 8 | 11647 | 2a1t | S | 128 | ASP | p.Asp128Asn | P38117 |
| HOMO SAPIENS | 5 | 11647 | 2a1t | S | 154 | THR | p.Thr154Met | P38117 |
| HOMO SAPIENS | 6 | 11647 | 2a1t | S | 164 | ARG | p.Arg164Gln | P38117 |
| HOMO SAPIENS | 5 | 11647 | 2a1t | S | 174 | ARG | p.Arg174His | P38117 |
| HOMO SAPIENS | -1 | 11200 | 4tx5 | A | 4 | ILE | p.Ile59Val | Q9NR28 |
| HOMO SAPIENS | 0 | 11200 | 4tx5 | A | 169 | GLY | p.Gly224Arg | Q9NR28 |
| HOMO SAPIENS | 3 | 417 | 2vr2 | A | 6 | ARG | p.Arg6Gln | Q14117 |
| HOMO SAPIENS | 1 | 417 | 2vr2 | A | 334 | GLN | p.Gln334Arg | Q14117 |
| HOMO SAPIENS | 2 | 417 | 2vr2 | A | 360 | TRP | p.Trp360Arg | Q14117 |
| HOMO SAPIENS | 0 | 417 | 2vr2 | A | 379 | SER | p.Ser379Arg | Q14117 |
| HOMO SAPIENS | 0 | 417 | 2vr2 | A | 435 | GLY | p.Gly435Arg | Q14117 |
| HOMO SAPIENS | 0 | 12498 | 3eo3 | B | 423 | ILE | p.Ile423Val | Q9Y223 |
| HOMO SAPIENS | 5 | 12498 | 3eo3 | B | 572 | VAL | p.Val572Leu | Q9Y223 |
| HOMO SAPIENS | 0 | 12498 | 3eo3 | B | 576 | GLY | p.Gly576Glu | Q9Y223 |
| HOMO SAPIENS | 0 | 12498 | 3eo3 | B | 587 | ILE | p.Ile587Thr | Q9Y223 |
| HOMO SAPIENS | 0 | 12498 | 3eo3 | B | 615 | SER | p.Ser615Ter | Q9Y223 |
| HOMO SAPIENS | 1 | 12498 | 3eo3 | B | 631 | ALA | p.Ala631Val | Q9Y223 |
| HOMO SAPIENS | 1 | 12498 | 3eo3 | B | 669 | GLY | p.Gly669Arg | Q9Y223 |
| HOMO SAPIENS | 1 | 12498 | 3eo3 | B | 696 | VAL | p.Val696Met | Q9Y223 |
| HOMO SAPIENS | 2 | 12498 | 3eo3 | B | 712 | MET | p.Met712Thr | Q9Y223 |
| HOMO SAPIENS | 0 | 1568 | 1r74 | A | 49 | LEU | p.Leu50Pro | Q14749 |
| HOMO SAPIENS | 0 | 1568 | 1r74 | A | 176 | HIS | p.His177Asn | Q14749 |
| HOMO SAPIENS | 3 | 147 | 5eig | A | 67 | THR | p.Thr67Ile | P32929 |
| HOMO SAPIENS | 3 | 147 | 5eig | A | 240 | GLN | p.Gln240Glu | P32929 |
| HOMO SAPIENS | -1 | 147 | 5eig | A | 403 | SER | p.Ser403Ile | P32929 |
| HOMO SAPIENS | 8 | 20036 | 2lsr | A | 31 | ARG | p.Arg31Ter | Q9Y6N9 |
| HOMO SAPIENS | -1 | 41382 | 4uis | B | 431 | UNK | p.Ala431Glu | P49768 |
| HOMO SAPIENS | -1 | 8536 | 3i6w | B | 85 | PRO | p.Pro85Leu | O96017 |
| HOMO SAPIENS | 0 | 8536 | 3i6w | B | 93 | TRP | p.Trp93Ter | O96017 |
| HOMO SAPIENS | 0 | 8536 | 3i6w | B | 95 | ARG | p.Arg95Ter | O96017 |
| HOMO SAPIENS | 0 | 8536 | 3i6w | B | 117 | ARG | p.Arg117Gly | O96017 |
| HOMO SAPIENS | 0 | 8536 | 3i6w | B | 119 | LYS | p.Lys119Arg | O96017 |
| HOMO SAPIENS | 0 | 8536 | 3i6w | B | 137 | ARG | p.Arg137Gln | O96017 |
| HOMO SAPIENS | 0 | 8536 | 3i6w | B | 137 | ARG | p.Arg137Ter | O96017 |
| HOMO SAPIENS | 0 | 8536 | 3i6w | B | 145 | ARG | p.Arg145Trp | O96017 |
| HOMO SAPIENS | 0 | 8536 | 3i6w | B | 160 | ILE | p.Ile160Met | O96017 |
| HOMO SAPIENS | 0 | 41322 | 4bxl | C | 46 | GLU | p.Glu46Lys | P37840 |
| HOMO SAPIENS | 5 | 41322 | 4bxl | C | 53 | ALA | p.Ala53Thr | P37840 |
| HOMO SAPIENS | 2 | 24966 | 3hve | A | 15 | ARG | p.Arg15Ser | Q9H2C0 |
| HOMO SAPIENS | 1 | 24966 | 3hve | A | 138 | ARG | p.Arg138His | Q9H2C0 |
| HOMO SAPIENS | 0 | 24966 | 3hve | A | 201 | ARG | p.Arg201Ter | Q9H2C0 |
| HOMO SAPIENS | 0 | 11680 | 5an9 | J | 8 | ASN | p.Asn8Lys | Q9Y3A5 |
| HOMO SAPIENS | 1 | 11680 | 5an9 | J | 33 | LYS | p.Lys33Arg | Q9Y3A5 |

| | | | | | | | | |
|---|---|---|---|---|---|---|---|---|
| HOMO SAPIENS | 1 | 11680 | 5an9 | J | 126 | ARG | p.Arg126Thr | Q9Y3A5 |
| HOMO SAPIENS | 1 | 11680 | 5an9 | J | 130 | VAL | p.Val130Met | Q9Y3A5 |
| HOMO SAPIENS | 1 | 11680 | 5an9 | J | 169 | ARG | p.Arg169Cys | Q9Y3A5 |
| HOMO SAPIENS | 1 | 11680 | 5an9 | J | 218 | ARG | p.Arg218Ter | Q9Y3A5 |
| HOMO SAPIENS | 4 | 978 | 3zs0 | B | 7 | TYR | p.Tyr173Cys | P05164 |
| HOMO SAPIENS | 1 | 978 | 3zs0 | B | 85 | MET | p.Met251Thr | P05164 |
| HOMO SAPIENS | 4 | 5140 | 2l4r | A | 52 | CYS | p.Cys52Ter | Q12809 |
| HOMO SAPIENS | 4 | 5140 | 2l4r | A | 63 | PRO | p.Pro63His | Q12809 |
| HOMO SAPIENS | 0 | 5140 | 2l4r | A | 100 | ARG | p.Arg100Gly | Q12809 |
| HOMO SAPIENS | 4 | 5140 | 2l4r | A | 129 | PHE | p.Phe129Leu | Q12809 |
| HOMO SAPIENS | 0 | 25836 | 2gy7 | B | 103 | GLU | p.Glu103Asp | Q02763 |
| HOMO SAPIENS | 0 | 25836 | 2gy7 | B | 148 | ILE | p.Ile148Thr | Q02763 |
| HOMO SAPIENS | 0 | 25836 | 2gy7 | B | 226 | ALA | p.Ala226Val | Q02763 |
| HOMO SAPIENS | 0 | 25836 | 2gy7 | B | 391 | THR | p.Thr391Ile | Q02763 |
| HOMO SAPIENS | 4 | 12563 | 5kwy | D | 20 | GLU | p.Glu20Ter | P61916 |
| HOMO SAPIENS | 2 | 12563 | 5kwy | D | 30 | VAL | p.Val30Met | P61916 |
| HOMO SAPIENS | 4 | 12563 | 5kwy | D | 39 | VAL | p.Val39Met | P61916 |
| HOMO SAPIENS | 8 | 12563 | 5kwy | D | 67 | SER | p.Ser67Pro | P61916 |
| HOMO SAPIENS | 8 | 12563 | 5kwy | D | 99 | CYS | p.Cys99Arg | P61916 |
| HOMO SAPIENS | 2 | 12563 | 5kwy | D | 118 | GLU | p.Glu118Ter | P61916 |
| HOMO SAPIENS | 2 | 12563 | 5kwy | D | 120 | PRO | p.Pro120Ser | P61916 |
| HOMO SAPIENS | -1 | 1679 | 1pmk | A | -2 | VAL | p.Val374Phe | P00747 |
| HOMO SAPIENS | -1 | 5315 | 2w2n | P | 53 | ALA | p.Ala53Val | Q8NBP7 |
| HOMO SAPIENS | 8 | 13496 | 2yad | A | 138 | THR | p.Asn138Thr | P11686 |
| HOMO SAPIENS | -1 | 13496 | 2yad | A | 173 | ARG | p.Arg167Gln | P11686 |
| HOMO SAPIENS | 2 | 13496 | 2yad | A | 186 | SER | p.Asn186Ser | P11686 |
| HOMO SAPIENS | 0 | 4426 | 3qd6 | C | 123 | ALA | p.Ala123Glu | P29965 |
| HOMO SAPIENS | 1 | 4426 | 3qd6 | C | 140 | TRP | p.Trp140Gly | P29965 |
| HOMO SAPIENS | 0 | 4426 | 3qd6 | C | 155 | LEU | p.Leu155Pro | P29965 |
| HOMO SAPIENS | 6 | 4426 | 3qd6 | C | 227 | GLY | p.Gly227Val | P29965 |
| HOMO SAPIENS | 0 | 4426 | 3qd6 | C | 235 | ALA | p.Ala235Pro | P29965 |
| HOMO SAPIENS | 0 | 10734 | 3dyd | B | 45 | PRO | p.Pro45Ser | P17735 |
| HOMO SAPIENS | -1 | 10734 | 3dyd | B | 57 | ARG | p.Arg57Ter | P17735 |
| HOMO SAPIENS | 8 | 10734 | 3dyd | B | 417 | ARG | p.Arg417Ter | P17735 |
| HOMO SAPIENS | 0 | 6956 | 4pzs | A | 1600 | LEU | p.Leu1600Phe | Q09472 |
| HOMO SAPIENS | 0 | 9370 | 3hfv | A | 21 | SER | p.Ser57Cys | O95363 |
| HOMO SAPIENS | 1 | 9370 | 3hfv | A | 25 | ASP | p.Asp61Glu | O95363 |
| HOMO SAPIENS | 1 | 9370 | 3hfv | A | 130 | ALA | p.Ala166Thr | O95363 |
| HOMO SAPIENS | 5 | 9370 | 3hfv | A | 148 | ASP | p.Asp184Asn | O95363 |
| HOMO SAPIENS | 0 | 9370 | 3hfv | A | 187 | ARG | p.Arg223Cys | O95363 |
| HOMO SAPIENS | 0 | 9370 | 3hfv | A | 210 | THR | p.Thr246Met | O95363 |
| HOMO SAPIENS | 1 | 9370 | 3hfv | A | 244 | ASN | p.Asn280Ser | O95363 |
| HOMO SAPIENS | 2 | 9370 | 3hfv | A | 288 | TYR | p.Tyr324Cys | O95363 |
| HOMO SAPIENS | 5 | 9370 | 3hfv | A | 293 | ILE | p.Ile329Thr | O95363 |
| HOMO SAPIENS | 1 | 9370 | 3hfv | A | 355 | ASP | p.Asp391Val | O95363 |
| HOMO SAPIENS | 1 | 9370 | 3hfv | A | 371 | THR | p.Thr407Met | O95363 |
| HOMO SAPIENS | 1 | 9370 | 3hfv | A | 383 | ARG | p.Arg419Cys | O95363 |
| HOMO SAPIENS | 1 | 9370 | 3hfv | A | 387 | ARG | p.Arg423Gln | O95363 |
| HOMO SAPIENS | 0 | 14401 | 1q3x | A | 371 | ASP | p.Asp371Tyr | O00187 |
| HOMO SAPIENS | 0 | 14401 | 1q3x | A | 377 | VAL | p.Val377Ala | O00187 |
| HOMO SAPIENS | 0 | 14401 | 1q3x | A | 439 | ARG | p.Arg439His | O00187 |
| HOMO SAPIENS | 0 | 14401 | 1q3x | A | 577 | GLN | p.Gln577His | O00187 |
| HOMO SAPIENS | 2 | 18270 | 3teq | A | 429 | ARG | p.Arg429Cys | Q13586 |
| HOMO SAPIENS | -1 | 16081 | 4mqt | A | 231 | VAL | p.Val231Ile | P08172 |

| | | | | | | | | |
|---|---|---|---|---|---|---|---|---|
| HOMO SAPIENS | 9 | 6607 | 3whu | B | 81 | ASP | p.Asp81His | Q8NI22 |
| HOMO SAPIENS | 9 | 6607 | 3whu | B | 129 | ASP | p.Asp129Glu | Q8NI22 |
| HOMO SAPIENS | 9 | 6607 | 3whu | B | 136 | ILE | p.Ile136Thr | Q8NI22 |
| HOMO SAPIENS | 0 | 7208 | 2e9w | D | 11 | ASN | p.Asn36Ser | P21583 |
| HOMO SAPIENS | 0 | 24566 | 4bdx | A | 10 | LEU | p.Leu140Val | P00748 |
| HOMO SAPIENS | 5 | 24566 | 4bdx | A | 77 | PRO | p.Ala207Pro | P00748 |
| HOMO SAPIENS | 7 | 606 | 3qnf | A | 319 | MET | p.Met319Val | Q9NZ08 |
| HOMO SAPIENS | 8 | 4097 | 1ydl | A | 21 | LEU | p.Leu21Pro | Q6ZYL4 |
| HOMO SAPIENS | 6 | 4097 | 1ydl | A | 56 | ARG | p.Arg56Ter | Q6ZYL4 |
| HOMO SAPIENS | 8 | 36691 | 1dqb | A | 42 | ARG | p.Arg403Lys | P07204 |
| HOMO SAPIENS | 0 | 5642 | 3j7y | X | 59 | ARG | p.Arg59Pro | Q13084 |
| HOMO SAPIENS | 1 | 5642 | 3j7y | X | 204 | VAL | p.Val204Met | Q13084 |
| HOMO SAPIENS | 1 | 1119 | 3nwu | B | 252 | ALA | p.Ala252Thr | Q92743 |
| HOMO SAPIENS | 3 | 1119 | 3nwu | B | 274 | ARG | p.Arg274Gln | Q92743 |
| HOMO SAPIENS | 9 | 1119 | 3nwu | B | 297 | VAL | p.Val297Met | Q92743 |
| HOMO SAPIENS | -1 | 1119 | 3nwu | B | 302 | ARG | p.Arg302Ter | Q92743 |
| HOMO SAPIENS | 9 | 1119 | 3nwu | B | 321 | ALA | p.Ala321Thr | Q92743 |
| HOMO SAPIENS | -1 | 1119 | 3nwu | B | 370 | ARG | p.Arg370Ter | Q92743 |
| HOMO SAPIENS | 0 | 416 | 4ycu | B | 105 | LEU | p.Leu105His | Q15046 |
| HOMO SAPIENS | 0 | 416 | 4ycu | B | 145 | TYR | p.Tyr145His | Q15046 |
| HOMO SAPIENS | 0 | 416 | 4ycu | B | 200 | PRO | p.Pro200Leu | Q15046 |
| HOMO SAPIENS | 0 | 416 | 4ycu | B | 201 | TYR | p.Tyr201His | Q15046 |
| HOMO SAPIENS | 0 | 416 | 4ycu | B | 420 | ARG | p.Arg420Cys | Q15046 |
| HOMO SAPIENS | 9 | 3297 | 4fpb | A | 680 | THR | p.Thr680Ser | P19367 |
| HOMO SAPIENS | 0 | 6771 | 2zt6 | A | 201 | ALA | p.Ala255Val | P41250 |
| HOMO SAPIENS | 0 | 6771 | 2zt6 | A | 202 | ASP | p.Asp256His | P41250 |
| HOMO SAPIENS | 0 | 6771 | 2zt6 | A | 334 | ARG | p.Arg388Gln | P41250 |
| HOMO SAPIENS | -1 | 6771 | 2zt6 | A | 439 | ASN | p.Asn493Ser | P41250 |
| HOMO SAPIENS | -1 | 6771 | 2zt6 | A | 500 | ASP | p.Asp554Asn | P41250 |
| HOMO SAPIENS | 1 | 6771 | 2zt6 | A | 581 | SER | p.Ser635Leu | P41250 |
| HOMO SAPIENS | -1 | 6771 | 2zt6 | A | 684 | GLU | p.Glu738Lys | P41250 |
| HOMO SAPIENS | 0 | 2687 | 4grw | B | 276 | VAL | p.Val298Phe | P29460 |
| HOMO SAPIENS | 6 | 913 | 1m75 | B | 45 | ASP | p.Asp57Glu | Q16836 |
| HOMO SAPIENS | 2 | 913 | 1m75 | B | 74 | PRO | p.Leu86Pro | Q16836 |
| HOMO SAPIENS | 3 | 913 | 1m75 | B | 214 | TYR | p.Tyr226His | Q16836 |
| HOMO SAPIENS | 3 | 913 | 1m75 | B | 224 | ARG | p.Arg236Ter | Q16836 |
| HOMO SAPIENS | 4 | 913 | 1m75 | B | 246 | PRO | p.Pro258Leu | Q16836 |
| HOMO SAPIENS | 2 | 4713 | 4djn | B | 40 | ARG | p.Arg40Cys | Q7LG56 |
| HOMO SAPIENS | 1 | 4713 | 4djn | B | 41 | ARG | p.Arg41Pro | Q7LG56 |
| HOMO SAPIENS | 2 | 4713 | 4djn | B | 110 | ARG | p.Arg110His | Q7LG56 |
| HOMO SAPIENS | 2 | 4713 | 4djn | B | 121 | ARG | p.Arg121His | Q7LG56 |
| HOMO SAPIENS | 9 | 4713 | 4djn | B | 131 | GLU | p.Glu131Lys | Q7LG56 |
| HOMO SAPIENS | 2 | 4713 | 4djn | B | 182 | THR | p.Thr182Ala | Q7LG56 |
| HOMO SAPIENS | 6 | 4713 | 4djn | B | 194 | GLU | p.Glu194Lys | Q7LG56 |
| HOMO SAPIENS | 2 | 4713 | 4djn | B | 211 | ARG | p.Arg211Lys | Q7LG56 |
| HOMO SAPIENS | 0 | 4713 | 4djn | B | 224 | ILE | p.Ile224Ser | Q7LG56 |
| HOMO SAPIENS | 3 | 4713 | 4djn | B | 229 | GLY | p.Gly229Val | Q7LG56 |
| HOMO SAPIENS | 2 | 4713 | 4djn | B | 236 | CYS | p.Cys236Phe | Q7LG56 |
| HOMO SAPIENS | 0 | 4713 | 4djn | B | 273 | GLY | p.Gly273Ser | Q7LG56 |
| HOMO SAPIENS | -1 | 4713 | 4djn | B | 282 | MSE | p.Met282Ile | Q7LG56 |
| HOMO SAPIENS | 3 | 4713 | 4djn | B | 284 | GLN | p.Gln284Ter | Q7LG56 |
| HOMO SAPIENS | 3 | 4713 | 4djn | B | 305 | ALA | p.Ala305Ser | Q7LG56 |
| HOMO SAPIENS | -1 | 8974 | 5n7i | A | 500 | ARG | p.Arg500Ter | Q8N4S9 |
| HOMO SAPIENS | 0 | 17072 | 3fxi | B | 299 | ASP | p.Asp299Gly | O00206 |

| | | | | | | | | |
|---|---|---|---|---|---|---|---|---|
| HOMO SAPIENS | 1 | 14355 | 2wwk | T | 55 | ILE | p.Ile34306Asn | Q8WZ42 |
| HOMO SAPIENS | 0 | 14355 | 2wwk | T | 64 | LEU | p.Leu34315Pro | Q8WZ42 |
| HOMO SAPIENS | 2 | 17797 | 1xu1 | S | 72 | ARG | p.Arg72His | O14836 |
| HOMO SAPIENS | -1 | 10836 | 4ztu | A | 39 | GLN | p.Gln39Arg | P54098 |
| HOMO SAPIENS | -1 | 10836 | 4ztu | A | 42 | ARG | p.Arg42Gln | P54098 |
| HOMO SAPIENS | -1 | 10836 | 4ztu | A | 45 | GLN | p.Gln45Arg | P54098 |
| HOMO SAPIENS | -1 | 10836 | 4ztu | A | 52 | GLN | p.Gln52Glu | P54098 |
| HOMO SAPIENS | 0 | 10836 | 4ztu | A | 88 | PHE | p.Phe88Leu | P54098 |
| HOMO SAPIENS | 1 | 10836 | 4ztu | A | 194 | ALA | p.Ala194Val | P54098 |
| HOMO SAPIENS | 1 | 10836 | 4ztu | A | 227 | ARG | p.Arg227Trp | P54098 |
| HOMO SAPIENS | 0 | 10836 | 4ztu | A | 232 | ARG | p.Arg232His | P54098 |
| HOMO SAPIENS | 1 | 10836 | 4ztu | A | 285 | GLN | p.Gln285His | P54098 |
| HOMO SAPIENS | 1 | 10836 | 4ztu | A | 434 | TYR | p.Tyr434His | P54098 |
| HOMO SAPIENS | -1 | 10836 | 4ztu | A | 511 | SER | p.Ser511Asn | P54098 |
| HOMO SAPIENS | 0 | 10836 | 4ztu | A | 533 | CYS | p.Cys533Trp | P54098 |
| HOMO SAPIENS | 1 | 10836 | 4ztu | A | 546 | ARG | p.Arg546Cys | P54098 |
| HOMO SAPIENS | 1 | 10836 | 4ztu | A | 587 | PRO | p.Pro587Leu | P54098 |
| HOMO SAPIENS | 0 | 10836 | 4ztu | A | 603 | MET | p.Met603Thr | P54098 |
| HOMO SAPIENS | 0 | 10836 | 4ztu | A | 623 | LEU | p.Leu623Trp | P54098 |
| HOMO SAPIENS | -1 | 10836 | 4ztu | A | 627 | ARG | p.Arg627Trp | P54098 |
| HOMO SAPIENS | -1 | 10836 | 4ztu | A | 628 | ARG | p.Arg628Pro | P54098 |
| HOMO SAPIENS | 0 | 10836 | 4ztu | A | 635 | PRO | p.Pro635Leu | P54098 |
| HOMO SAPIENS | 0 | 10836 | 4ztu | A | 662 | GLU | p.Glu662Lys | P54098 |
| HOMO SAPIENS | -1 | 10836 | 4ztu | A | 674 | GLY | p.Gly674Asp | P54098 |
| HOMO SAPIENS | -1 | 10836 | 4ztu | A | 722 | ARG | p.Arg722His | P54098 |
| HOMO SAPIENS | -1 | 10836 | 4ztu | A | 737 | GLY | p.Gly737Arg | P54098 |
| HOMO SAPIENS | 0 | 10836 | 4ztu | A | 748 | TRP | p.Trp748Ser | P54098 |
| HOMO SAPIENS | 0 | 10836 | 4ztu | A | 749 | PHE | p.Phe749Ser | P54098 |
| HOMO SAPIENS | 1 | 10836 | 4ztu | A | 807 | ARG | p.Arg807Cys | P54098 |
| HOMO SAPIENS | 1 | 10836 | 4ztu | A | 831 | TYR | p.Tyr831Cys | P54098 |
| HOMO SAPIENS | 8 | 10836 | 4ztu | A | 848 | GLY | p.Gly848Ser | P54098 |
| HOMO SAPIENS | 1 | 10836 | 4ztu | A | 852 | ARG | p.Arg852Cys | P54098 |
| HOMO SAPIENS | 9 | 10836 | 4ztu | A | 853 | ARG | p.Arg853Trp | P54098 |
| HOMO SAPIENS | 1 | 10836 | 4ztu | A | 864 | ASN | p.Asn864Ser | P54098 |
| HOMO SAPIENS | 1 | 10836 | 4ztu | A | 879 | GLN | p.Gln879Arg | P54098 |
| HOMO SAPIENS | 1 | 10836 | 4ztu | A | 914 | THR | p.Thr914Pro | P54098 |
| HOMO SAPIENS | 6 | 10836 | 4ztu | A | 932 | HIS | p.His932Tyr | P54098 |
| HOMO SAPIENS | 2 | 10836 | 4ztu | A | 944 | GLU | p.Glu944Lys | P54098 |
| HOMO SAPIENS | 6 | 10836 | 4ztu | A | 955 | TYR | p.Tyr955Cys | P54098 |
| HOMO SAPIENS | 1 | 10836 | 4ztu | A | 957 | ALA | p.Ala957Ser | P54098 |
| HOMO SAPIENS | 1 | 10836 | 4ztu | A | 957 | ALA | p.Ala957Val | P54098 |
| HOMO SAPIENS | 1 | 10836 | 4ztu | A | 964 | ARG | p.Arg964Cys | P54098 |
| HOMO SAPIENS | 1 | 10836 | 4ztu | A | 966 | LEU | p.Leu966Arg | P54098 |
| HOMO SAPIENS | 0 | 10836 | 4ztu | A | 1051 | GLY | p.Gly1051Arg | P54098 |
| HOMO SAPIENS | 1 | 10836 | 4ztu | A | 1073 | PRO | p.Pro1073Leu | P54098 |
| HOMO SAPIENS | 1 | 10836 | 4ztu | A | 1096 | ARG | p.Arg1096Cys | P54098 |
| HOMO SAPIENS | 1 | 10836 | 4ztu | A | 1096 | ARG | p.Arg1096His | P54098 |
| HOMO SAPIENS | 0 | 10836 | 4ztu | A | 1116 | MET | p.Met1116Thr | P54098 |
| HOMO SAPIENS | 1 | 10836 | 4ztu | A | 1142 | ARG | p.Arg1142Trp | P54098 |
| HOMO SAPIENS | 1 | 10836 | 4ztu | A | 1146 | ARG | p.Arg1146Cys | P54098 |
| HOMO SAPIENS | 1 | 10836 | 4ztu | A | 1157 | ASN | p.Asn1157Ser | P54098 |
| HOMO SAPIENS | -1 | 10836 | 4ztu | A | 1234 | ARG | p.Arg1234Gln | P54098 |
| HOMO SAPIENS | -1 | 10836 | 4ztu | A | 1236 | GLN | p.Gln1236His | P54098 |
| HOMO SAPIENS | 5 | 2666 | 1jph | A | 62 | PRO | p.Pro62Leu | P06132 |

| | | | | | | | | |
|---|---|---|---|---|---|---|---|---|
| HOMO SAPIENS | 5 | 2666 | 1jph | A | 116 | GLN | p.Gln116Ter | P06132 |
| HOMO SAPIENS | 0 | 2666 | 1jph | A | 165 | MET | p.Met165Arg | P06132 |
| HOMO SAPIENS | 6 | 2666 | 1jph | A | 167 | GLU | p.Glu167Lys | P06132 |
| HOMO SAPIENS | 0 | 2666 | 1jph | A | 195 | LEU | p.Leu195Phe | P06132 |
| HOMO SAPIENS | 7 | 2666 | 1jph | A | 253 | LEU | p.Leu253Gln | P06132 |
| HOMO SAPIENS | 0 | 2666 | 1jph | A | 281 | GLY | p.Gly281Glu | P06132 |
| HOMO SAPIENS | 9 | 2666 | 1jph | A | 304 | ASN | p.Asn304Lys | P06132 |
| HOMO SAPIENS | 8 | 2666 | 1jph | A | 311 | TYR | p.Tyr311Cys | P06132 |
| HOMO SAPIENS | 5 | 2666 | 1jph | A | 332 | ARG | p.Arg332His | P06132 |
| HOMO SAPIENS | 1 | 889 | 1m51 | A | 184 | LEU | p.Val184Leu | P35558 |
| HOMO SAPIENS | 0 | 889 | 1m51 | A | 267 | VAL | p.Ile267Val | P35558 |
| HOMO SAPIENS | 2 | 889 | 1m51 | A | 276 | GLU | p.Glu276Lys | P35558 |
| HOMO SAPIENS | 6 | 889 | 1m51 | A | 368 | VAL | p.Val368Ile | P35558 |
| HOMO SAPIENS | 0 | 889 | 1m51 | A | 579 | SER | p.Ser579Arg | P35558 |
| HOMO SAPIENS | 1 | 4084 | 2gjx | A | 28 | GLN | p.Gln28Ter | P06865 |
| HOMO SAPIENS | 0 | 4084 | 2gjx | A | 39 | LEU | p.Leu39Arg | P06865 |
| HOMO SAPIENS | 0 | 4084 | 2gjx | A | 127 | LEU | p.Leu127Arg | P06865 |
| HOMO SAPIENS | 6 | 4084 | 2gjx | A | 137 | ARG | p.Arg137Ter | P06865 |
| HOMO SAPIENS | 6 | 4084 | 2gjx | A | 170 | ARG | p.Arg170Gln | P06865 |
| HOMO SAPIENS | 9 | 4084 | 2gjx | A | 178 | ARG | p.Arg178Cys | P06865 |
| HOMO SAPIENS | 9 | 4084 | 2gjx | A | 178 | ARG | p.Arg178His | P06865 |
| HOMO SAPIENS | 5 | 4084 | 2gjx | A | 180 | TYR | p.Tyr180His | P06865 |
| HOMO SAPIENS | 5 | 4084 | 2gjx | A | 180 | TYR | p.Tyr180Ter | P06865 |
| HOMO SAPIENS | 1 | 4084 | 2gjx | A | 197 | LYS | p.Lys197Thr | P06865 |
| HOMO SAPIENS | 0 | 4084 | 2gjx | A | 200 | VAL | p.Val200Met | P06865 |
| HOMO SAPIENS | 9 | 4084 | 2gjx | A | 204 | HIS | p.His204Arg | P06865 |
| HOMO SAPIENS | 5 | 4084 | 2gjx | A | 210 | SER | p.Ser210Phe | P06865 |
| HOMO SAPIENS | 0 | 4084 | 2gjx | A | 211 | PHE | p.Phe211Ser | P06865 |
| HOMO SAPIENS | 2 | 4084 | 2gjx | A | 250 | GLY | p.Gly250Asp | P06865 |
| HOMO SAPIENS | 9 | 4084 | 2gjx | A | 258 | ASP | p.Asp258His | P06865 |
| HOMO SAPIENS | 0 | 4084 | 2gjx | A | 269 | GLY | p.Gly269Ser | P06865 |
| HOMO SAPIENS | 0 | 4084 | 2gjx | A | 301 | MET | p.Met301Arg | P06865 |
| HOMO SAPIENS | 3 | 4084 | 2gjx | A | 393 | ARG | p.Arg393Ter | P06865 |
| HOMO SAPIENS | 3 | 4084 | 2gjx | A | 393 | ARG | p.Arg393Pro | P06865 |
| HOMO SAPIENS | 0 | 4084 | 2gjx | A | 399 | ASN | p.Asn399Asp | P06865 |
| HOMO SAPIENS | 2 | 4084 | 2gjx | A | 420 | TRP | p.Trp420Cys | P06865 |
| HOMO SAPIENS | 1 | 4084 | 2gjx | A | 434 | PHE | p.Phe434Leu | P06865 |
| HOMO SAPIENS | 0 | 4084 | 2gjx | A | 454 | GLY | p.Gly454Ser | P06865 |
| HOMO SAPIENS | 1 | 4084 | 2gjx | A | 474 | TRP | p.Trp474Cys | P06865 |
| HOMO SAPIENS | 0 | 4084 | 2gjx | A | 479 | ALA | p.Ala479Thr | P06865 |
| HOMO SAPIENS | 5 | 4084 | 2gjx | A | 485 | TRP | p.Trp485Arg | P06865 |
| HOMO SAPIENS | 4 | 4084 | 2gjx | A | 499 | ARG | p.Arg499Cys | P06865 |
| HOMO SAPIENS | 4 | 4084 | 2gjx | A | 499 | ARG | p.Arg499His | P06865 |
| HOMO SAPIENS | 2 | 4084 | 2gjx | A | 504 | ARG | p.Arg504Cys | P06865 |
| HOMO SAPIENS | 2 | 4084 | 2gjx | A | 504 | ARG | p.Arg504His | P06865 |
| HOMO SAPIENS | -1 | 1943 | 1xmj | A | 417 | ASN | p.Asn417Lys | P13569 |
| HOMO SAPIENS | -1 | 1943 | 1xmj | A | 422 | SER | p.Ser422Phe | P13569 |
| HOMO SAPIENS | 1 | 1943 | 1xmj | A | 438 | THR | p.Thr438Ala | P13569 |
| HOMO SAPIENS | 0 | 1943 | 1xmj | A | 455 | ALA | p.Ala455Glu | P13569 |
| HOMO SAPIENS | 4 | 1943 | 1xmj | A | 456 | VAL | p.Val456Ala | P13569 |
| HOMO SAPIENS | 9 | 1943 | 1xmj | A | 458 | GLY | p.Gly458Val | P13569 |
| HOMO SAPIENS | 8 | 1943 | 1xmj | A | 466 | SER | p.Ser466Ter | P13569 |
| HOMO SAPIENS | 0 | 1943 | 1xmj | A | 467 | LEU | p.Leu467Pro | P13569 |
| HOMO SAPIENS | 1 | 1943 | 1xmj | A | 470 | MET | p.Val470Met | P13569 |

| | | | | | | | | |
|---|---|---|---|---|---|---|---|---|
| HOMO SAPIENS | 5 | 1943 | 1xmj | A | 480 | GLY | p.Gly480Cys | P13569 |
| HOMO SAPIENS | 3 | 1943 | 1xmj | A | 492 | SER | p.Ser492Phe | P13569 |
| HOMO SAPIENS | 8 | 1943 | 1xmj | A | 493 | GLN | p.Gln493Ter | P13569 |
| HOMO SAPIENS | 5 | 1943 | 1xmj | A | 506 | ILE | p.Ile506Met | P13569 |
| HOMO SAPIENS | 3 | 1943 | 1xmj | A | 520 | VAL | p.Val520Phe | P13569 |
| HOMO SAPIENS | 1 | 1943 | 1xmj | A | 528 | GLU | p.Glu528Asp | P13569 |
| HOMO SAPIENS | 1 | 1943 | 1xmj | A | 534 | ALA | p.Ala534Glu | P13569 |
| HOMO SAPIENS | 4 | 1943 | 1xmj | A | 542 | GLY | p.Gly542Ter | P13569 |
| HOMO SAPIENS | 6 | 1943 | 1xmj | A | 549 | SER | p.Ser549Asn | P13569 |
| HOMO SAPIENS | 6 | 1943 | 1xmj | A | 549 | SER | p.Ser549Arg | P13569 |
| HOMO SAPIENS | 6 | 1943 | 1xmj | A | 549 | SER | p.Ser549Arg | P13569 |
| HOMO SAPIENS | 6 | 1943 | 1xmj | A | 551 | GLY | p.Gly551Asp | P13569 |
| HOMO SAPIENS | 6 | 1943 | 1xmj | A | 551 | GLY | p.Gly551Ser | P13569 |
| HOMO SAPIENS | 6 | 1943 | 1xmj | A | 553 | GLN | p.Arg553Gln | P13569 |
| HOMO SAPIENS | 6 | 1943 | 1xmj | A | 553 | GLN | p.Arg553Ter | P13569 |
| HOMO SAPIENS | 0 | 1943 | 1xmj | A | 559 | ALA | p.Ala559Thr | P13569 |
| HOMO SAPIENS | 7 | 1943 | 1xmj | A | 560 | ARG | p.Arg560Lys | P13569 |
| HOMO SAPIENS | 4 | 1943 | 1xmj | A | 562 | VAL | p.Val562Ile | P13569 |
| HOMO SAPIENS | 3 | 1943 | 1xmj | A | 563 | TYR | p.Tyr563Asn | P13569 |
| HOMO SAPIENS | 3 | 1943 | 1xmj | A | 569 | TYR | p.Tyr569Asp | P13569 |
| HOMO SAPIENS | 3 | 1943 | 1xmj | A | 585 | GLU | p.Glu585Ter | P13569 |
| HOMO SAPIENS | 1 | 1943 | 1xmj | A | 618 | ILE | p.Ile618Thr | P13569 |
| HOMO SAPIENS | 9 | 1943 | 1xmj | A | 622 | GLY | p.Gly622Asp | P13569 |
| HOMO SAPIENS | -1 | 1943 | 1xmj | A | 648 | ASP | p.Asp648Val | P13569 |
| HOMO SAPIENS | 0 | 1943 | 1xmj | A | 673 | GLY | p.Gly673Ter | P13569 |
| HOMO SAPIENS | 5 | 5111 | 4rec | A | 749 | ARG | p.Arg749Ter | Q9Y2M0 |
| HOMO SAPIENS | -1 | 4580 | 4l0d | B | 2 | GLY | p.Pro2Leu | P35520 |
| HOMO SAPIENS | -1 | 4580 | 4l0d | B | 12 | PRO | p.Pro12Ser | P35520 |
| HOMO SAPIENS | -1 | 4580 | 4l0d | B | 18 | ARG | p.Arg18Cys | P35520 |
| HOMO SAPIENS | 1 | 4580 | 4l0d | B | 49 | PRO | p.Pro49Leu | P35520 |
| HOMO SAPIENS | 2 | 4580 | 4l0d | B | 70 | PRO | p.Pro70Leu | P35520 |
| HOMO SAPIENS | 1 | 4580 | 4l0d | B | 102 | LYS | p.Lys102Gln | P35520 |
| HOMO SAPIENS | 7 | 4580 | 4l0d | B | 114 | ALA | p.Ala114Val | P35520 |
| HOMO SAPIENS | 5 | 4580 | 4l0d | B | 116 | GLY | p.Gly116Arg | P35520 |
| HOMO SAPIENS | 8 | 4580 | 4l0d | B | 121 | ARG | p.Arg121Cys | P35520 |
| HOMO SAPIENS | 8 | 4580 | 4l0d | B | 121 | ARG | p.Arg121His | P35520 |
| HOMO SAPIENS | 5 | 4580 | 4l0d | B | 125 | ARG | p.Arg125Gln | P35520 |
| HOMO SAPIENS | 4 | 4580 | 4l0d | B | 134 | GLY | p.Gly134Arg | P35520 |
| HOMO SAPIENS | 4 | 4580 | 4l0d | B | 139 | GLY | p.Gly139Arg | P35520 |
| HOMO SAPIENS | 0 | 4580 | 4l0d | B | 145 | PRO | p.Pro145Leu | P35520 |
| HOMO SAPIENS | 0 | 4580 | 4l0d | B | 148 | GLY | p.Gly148Arg | P35520 |
| HOMO SAPIENS | 0 | 4580 | 4l0d | B | 168 | VAL | p.Val168Met | P35520 |
| HOMO SAPIENS | 2 | 4580 | 4l0d | B | 191 | THR | p.Thr191Met | P35520 |
| HOMO SAPIENS | 5 | 4580 | 4l0d | B | 234 | ASP | p.Asp234Asn | P35520 |
| HOMO SAPIENS | 0 | 4580 | 4l0d | B | 257 | THR | p.Thr257Met | P35520 |
| HOMO SAPIENS | 4 | 4580 | 4l0d | B | 262 | THR | p.Thr262Met | P35520 |
| HOMO SAPIENS | 0 | 4580 | 4l0d | B | 278 | ILE | p.Ile278Thr | P35520 |
| HOMO SAPIENS | 2 | 4580 | 4l0d | B | 307 | GLY | p.Gly307Ser | P35520 |
| HOMO SAPIENS | 4 | 4580 | 4l0d | B | 320 | VAL | p.Val320Ala | P35520 |
| HOMO SAPIENS | 2 | 4580 | 4l0d | B | 336 | ARG | p.Arg336Cys | P35520 |
| HOMO SAPIENS | 2 | 4580 | 4l0d | B | 336 | ARG | p.Arg336His | P35520 |
| HOMO SAPIENS | 0 | 4580 | 4l0d | B | 351 | GLY | p.Gly351Arg | P35520 |
| HOMO SAPIENS | 0 | 4580 | 4l0d | B | 353 | THR | p.Thr353Met | P35520 |
| HOMO SAPIENS | 0 | 4580 | 4l0d | B | 371 | VAL | p.Val371Met | P35520 |

| | | | | | | | | |
|---|---|---|---|---|---|---|---|---|
| HOMO SAPIENS | 2 | 4580 | 4l0d | B | 379 | ARG | p.Arg379Gln | P35520 |
| HOMO SAPIENS | 2 | 4580 | 4l0d | B | 379 | ARG | p.Arg379Trp | P35520 |
| HOMO SAPIENS | 2 | 4580 | 4l0d | B | 384 | LYS | p.Lys384Glu | P35520 |
| HOMO SAPIENS | 0 | 4580 | 4l0d | B | 422 | PRO | p.Pro422Leu | P35520 |
| HOMO SAPIENS | 0 | 4580 | 4l0d | B | 444 | ASP | p.Asp444Asn | P35520 |
| HOMO SAPIENS | 0 | 4580 | 4l0d | B | 466 | SER | p.Ser466Leu | P35520 |
| HOMO SAPIENS | 0 | 4580 | 4l0d | B | 539 | LEU | p.Leu539Ser | P35520 |
| HOMO SAPIENS | 0 | 4580 | 4l0d | B | 548 | ARG | p.Arg548Gln | P35520 |
| HOMO SAPIENS | 1 | 6536 | 3ulu | A | 412 | LEU | p.Leu412Phe | O15455 |
| HOMO SAPIENS | 1 | 28493 | 3ii6 | X | 814 | ARG | p.Arg814Ter | P49917 |
| HOMO SAPIENS | 1 | 28493 | 3ii6 | X | 857 | ALA | p.Ala857Thr | P49917 |
| HOMO SAPIENS | 9 | 3543 | 2q51 | A | 24 | GLU | p.Glu24Gly | P45381 |
| HOMO SAPIENS | 1 | 3543 | 2q51 | A | 27 | GLY | p.Gly27Arg | P45381 |
| HOMO SAPIENS | 9 | 3543 | 2q51 | A | 71 | ARG | p.Arg71His | P45381 |
| HOMO SAPIENS | -1 | 3543 | 2q51 | A | 82 | MSE | p.Met82Thr | P45381 |
| HOMO SAPIENS | 0 | 3543 | 2q51 | A | 143 | ILE | p.Ile143Phe | P45381 |
| HOMO SAPIENS | 1 | 3543 | 2q51 | A | 152 | CYS | p.Cys152Arg | P45381 |
| HOMO SAPIENS | 2 | 3543 | 2q51 | A | 218 | CYS | p.Cys218Ter | P45381 |
| HOMO SAPIENS | 1 | 3543 | 2q51 | A | 231 | TYR | p.Tyr231Cys | P45381 |
| HOMO SAPIENS | 1 | 3543 | 2q51 | A | 231 | TYR | p.Tyr231Ter | P45381 |
| HOMO SAPIENS | 2 | 3543 | 2q51 | A | 274 | GLY | p.Gly274Arg | P45381 |
| HOMO SAPIENS | 6 | 3543 | 2q51 | A | 285 | GLU | p.Glu285Ala | P45381 |
| HOMO SAPIENS | 5 | 3543 | 2q51 | A | 287 | ALA | p.Ala287Thr | P45381 |
| HOMO SAPIENS | 2 | 3543 | 2q51 | A | 305 | ALA | p.Ala305Glu | P45381 |
| HOMO SAPIENS | 2 | 24687 | 1hfi | A | 25 | SER | p.Ser890Ile | P08603 |
| HOMO SAPIENS | 0 | 1508 | 3wn5 | C | 45 | LEU | p.Leu66Arg | P08637 |
| HOMO SAPIENS | 0 | 1508 | 3wn5 | C | 155 | VAL | p.Phe176Val | P08637 |
| HOMO SAPIENS | 0 | 750 | 5tpa | A | 54 | VAL | p.Val452Met | Q12879 |
| HOMO SAPIENS | 9 | 750 | 5tpa | A | 120 | ARG | p.Arg518Cys | Q12879 |
| HOMO SAPIENS | 0 | 750 | 5tpa | A | 199 | ALA | p.Ala716Thr | Q12879 |
| HOMO SAPIENS | 4 | 1363 | 3hg2 | B | 40 | PRO | p.Pro40Ser | P06280 |
| HOMO SAPIENS | 3 | 1363 | 3hg2 | B | 65 | SER | p.Ser65Thr | P06280 |
| HOMO SAPIENS | 4 | 1363 | 3hg2 | B | 66 | GLU | p.Glu66Gln | P06280 |
| HOMO SAPIENS | 2 | 1363 | 3hg2 | B | 80 | GLY | p.Gly80Asp | P06280 |
| HOMO SAPIENS | 3 | 1363 | 3hg2 | B | 112 | ARG | p.Arg112Cys | P06280 |
| HOMO SAPIENS | 3 | 1363 | 3hg2 | B | 112 | ARG | p.Arg112His | P06280 |
| HOMO SAPIENS | 0 | 1363 | 3hg2 | B | 139 | ASN | p.Asn139Ser | P06280 |
| HOMO SAPIENS | 7 | 1363 | 3hg2 | B | 156 | ALA | p.Ala156Thr | P06280 |
| HOMO SAPIENS | 5 | 1363 | 3hg2 | B | 162 | TRP | p.Trp162Arg | P06280 |
| HOMO SAPIENS | 5 | 1363 | 3hg2 | B | 207 | TYR | p.Tyr207His | P06280 |
| HOMO SAPIENS | 0 | 1363 | 3hg2 | B | 215 | ASN | p.Asn215Ser | P06280 |
| HOMO SAPIENS | 0 | 1363 | 3hg2 | B | 264 | ASP | p.Asp264Val | P06280 |
| HOMO SAPIENS | 5 | 1363 | 3hg2 | B | 266 | ASP | p.Asp266Val | P06280 |
| HOMO SAPIENS | 2 | 1363 | 3hg2 | B | 269 | VAL | p.Val269Ala | P06280 |
| HOMO SAPIENS | 3 | 1363 | 3hg2 | B | 272 | ASN | p.Asn272Ser | P06280 |
| HOMO SAPIENS | 3 | 1363 | 3hg2 | B | 279 | GLN | p.Gln279Glu | P06280 |
| HOMO SAPIENS | 0 | 1363 | 3hg2 | B | 289 | ILE | p.Ile289Val | P06280 |
| HOMO SAPIENS | 0 | 1363 | 3hg2 | B | 296 | MET | p.Met296Val | P06280 |
| HOMO SAPIENS | 0 | 1363 | 3hg2 | B | 296 | MET | p.Met296Ile | P06280 |
| HOMO SAPIENS | 0 | 1363 | 3hg2 | B | 297 | SER | p.Ser297Phe | P06280 |
| HOMO SAPIENS | 2 | 1363 | 3hg2 | B | 301 | ARG | p.Arg301Gln | P06280 |
| HOMO SAPIENS | 3 | 1363 | 3hg2 | B | 327 | GLN | p.Gln327Lys | P06280 |
| HOMO SAPIENS | 0 | 1363 | 3hg2 | B | 328 | GLY | p.Gly328Arg | P06280 |
| HOMO SAPIENS | 0 | 1363 | 3hg2 | B | 328 | GLY | p.Gly328Ala | P06280 |

| | | | | | | | | |
|---|---|---|---|---|---|---|---|---|
| HOMO SAPIENS | 2 | 1363 | 3hg2 | B | 331 | LEU | p.Leu331Phe | P06280 |
| HOMO SAPIENS | 0 | 1363 | 3hg2 | B | 342 | ARG | p.Arg342Gln | P06280 |
| HOMO SAPIENS | 1 | 1363 | 3hg2 | B | 361 | GLY | p.Gly361Arg | P06280 |
| HOMO SAPIENS | 1 | 1363 | 3hg2 | B | 363 | ARG | p.Arg363His | P06280 |
| HOMO SAPIENS | 1 | 1363 | 3hg2 | B | 385 | THR | p.Thr385Ala | P06280 |
| HOMO SAPIENS | 1 | 1363 | 3hg2 | B | 399 | TRP | p.Trp399Ser | P06280 |
| HOMO SAPIENS | 1 | 1363 | 3hg2 | B | 410 | THR | p.Thr410Ala | P06280 |
| HOMO SAPIENS | 2 | 3904 | 3caa | B | 385 | MET | p.Met414Val | P01011 |
| HOMO SAPIENS | 0 | 558 | 5eub | A | 56 | VAL | p.Val56Ile | Q13126 |
| HOMO SAPIENS | 0 | 558 | 5eub | A | 189 | TRP | p.Trp189Leu | Q13126 |
| HOMO SAPIENS | 0 | 558 | 5eub | A | 218 | ALA | p.Ala218Gly | Q13126 |
| HOMO SAPIENS | 1 | 3865 | 1cb5 | B | 443 | ILE | p.Ile443Val | Q13867 |
| HOMO SAPIENS | 2 | 14329 | 4m92 | A | 24 | ILE | p.Ile65Val | Q13454 |
| HOMO SAPIENS | 1 | 7381 | 5kz5 | k | 50 | GLY | p.Gly50Glu | Q9H1K1 |
| HOMO SAPIENS | 0 | 1102 | 4fdh | E | 86 | PRO | p.Pro86Ala | P19099 |
| HOMO SAPIENS | 0 | 1102 | 4fdh | E | 118 | ALA | p.Ala118Thr | P19099 |
| HOMO SAPIENS | 0 | 1102 | 4fdh | E | 159 | PRO | p.Pro159Leu | P19099 |
| HOMO SAPIENS | 0 | 1102 | 4fdh | E | 173 | LYS | p.Lys173Arg | P19099 |
| HOMO SAPIENS | 0 | 1102 | 4fdh | E | 181 | ARG | p.Arg181Trp | P19099 |
| HOMO SAPIENS | 0 | 1102 | 4fdh | E | 185 | THR | p.Thr185Ile | P19099 |
| HOMO SAPIENS | 0 | 1102 | 4fdh | E | 198 | GLU | p.Glu198Asp | P19099 |
| HOMO SAPIENS | 0 | 1102 | 4fdh | E | 214 | HIS | p.His214Asp | P19099 |
| HOMO SAPIENS | 0 | 1102 | 4fdh | E | 251 | LYS | p.Lys251Arg | P19099 |
| HOMO SAPIENS | 0 | 1102 | 4fdh | E | 255 | GLU | p.Glu255Ter | P19099 |
| HOMO SAPIENS | 0 | 1102 | 4fdh | E | 281 | ASN | p.Asn281Ser | P19099 |
| HOMO SAPIENS | 0 | 1102 | 4fdh | E | 320 | ALA | p.Ala320Val | P19099 |
| HOMO SAPIENS | 0 | 1102 | 4fdh | E | 339 | ILE | p.Ile339Thr | P19099 |
| HOMO SAPIENS | 0 | 1102 | 4fdh | E | 341 | ARG | p.Arg341Cys | P19099 |
| HOMO SAPIENS | 2 | 1102 | 4fdh | E | 374 | ARG | p.Arg374Trp | P19099 |
| HOMO SAPIENS | 0 | 1102 | 4fdh | E | 386 | VAL | p.Val386Ala | P19099 |
| HOMO SAPIENS | 0 | 1102 | 4fdh | E | 390 | LEU | p.Leu390Phe | P19099 |
| HOMO SAPIENS | -1 | 1102 | 4fdh | E | 435 | GLY | p.Gly435Ser | P19099 |
| HOMO SAPIENS | 0 | 1102 | 4fdh | E | 461 | LEU | p.Leu461Pro | P19099 |
| HOMO SAPIENS | 0 | 1102 | 4fdh | E | 498 | THR | p.Thr498Ala | P19099 |
| HOMO SAPIENS | 6 | 10292 | 2of5 | F | 128 | GLY | p.Gly128Arg | P78560 |
| HOMO SAPIENS | 6 | 10292 | 2of5 | F | 170 | ARG | p.Arg170Cys | P78560 |
| HOMO SAPIENS | -1 | 18965 | 3ghg | A | 16 | ARG | p.Arg35Cys | P02671 |
| HOMO SAPIENS | -1 | 18965 | 3ghg | A | 16 | ARG | p.Arg35His | P02671 |
| HOMO SAPIENS | -1 | 18965 | 3ghg | A | 283 | PRO | p.Pro302Ala | P02671 |
| HOMO SAPIENS | -1 | 18965 | 3ghg | A | 312 | THR | p.Thr331Ala | P02671 |
| HOMO SAPIENS | -1 | 18965 | 3ghg | A | 381 | SER | p.Ser400Phe | P02671 |
| HOMO SAPIENS | -1 | 18965 | 3ghg | A | 463 | VAL | p.Val482Met | P02671 |
| HOMO SAPIENS | -1 | 18965 | 3ghg | A | 526 | GLU | p.Glu545Val | P02671 |
| HOMO SAPIENS | -1 | 18965 | 3ghg | A | 554 | ARG | p.Arg573Leu | P02671 |
| HOMO SAPIENS | 8 | 3023 | 4ttb | A | 101 | ARG | p.Arg101Trp | Q6PHW0 |
| HOMO SAPIENS | 2 | 3023 | 4ttb | A | 116 | ILE | p.Ile116Thr | Q6PHW0 |
| HOMO SAPIENS | 0 | 3023 | 4ttb | A | 220 | ALA | p.Ala220Thr | Q6PHW0 |
| HOMO SAPIENS | 2 | 3023 | 4ttb | A | 271 | GLU | p.Glu271Lys | Q6PHW0 |
| HOMO SAPIENS | 3 | 33226 | 1ya5 | T | 11 | SER | p.Ser11Ter | O15273 |
| HOMO SAPIENS | 6 | 33226 | 1ya5 | T | 64 | SER | p.Ser64Leu | O15273 |
| HOMO SAPIENS | 6 | 33226 | 1ya5 | T | 70 | ARG | p.Arg70Gln | O15273 |
| HOMO SAPIENS | 3 | 33226 | 1ya5 | T | 75 | GLY | p.Gly75Ser | O15273 |
| HOMO SAPIENS | 0 | 33226 | 1ya5 | T | 87 | ARG | p.Arg87Trp | O15273 |
| HOMO SAPIENS | 2 | 5667 | 3dax | A | 398 | PRO | p.Pro398Ala | P22680 |

| | | | | | | | | |
|---|---|---|---|---|---|---|---|---|
| HOMO SAPIENS | 0 | 12861 | 5mya | A | 464 | ASN | p.Asn464His | Q02763 |
| HOMO SAPIENS | 2 | 12861 | 5mya | A | 486 | VAL | p.Val486Ile | Q02763 |
| HOMO SAPIENS | 0 | 12861 | 5mya | A | 522 | ARG | p.Arg522His | Q02763 |
| HOMO SAPIENS | 2 | 12861 | 5mya | A | 600 | VAL | p.Val600Leu | Q02763 |
| HOMO SAPIENS | 0 | 12861 | 5mya | A | 634 | LEU | p.Leu634Phe | Q02763 |
| HOMO SAPIENS | 0 | 12861 | 5mya | A | 684 | HIS | p.His684Gln | Q02763 |
| HOMO SAPIENS | 0 | 12861 | 5mya | A | 724 | ALA | p.Ala724Thr | Q02763 |
| HOMO SAPIENS | 0 | 7743 | 3qzw | G | 90 | GLY | p.Gly111Ser | P01732 |
| HOMO SAPIENS | 2 | 9498 | 3kci | A | 4267 | ASP | p.Asp4267Glu | O95714 |
| HOMO SAPIENS | 5 | 201 | 3ips | B | 415 | ARG | p.Arg415Gln | Q13133 |
| HOMO SAPIENS | 3 | 229 | 2q50 | A | 68 | LEU | p.Leu68Pro | Q9UBQ7 |
| HOMO SAPIENS | 2 | 229 | 2q50 | A | 99 | ARG | p.Arg99Ter | Q9UBQ7 |
| HOMO SAPIENS | 3 | 229 | 2q50 | A | 160 | GLY | p.Gly160Arg | Q9UBQ7 |
| HOMO SAPIENS | 5 | 229 | 2q50 | A | 165 | GLY | p.Gly165Asp | Q9UBQ7 |
| HOMO SAPIENS | 4 | 229 | 2q50 | A | 302 | ARG | p.Arg302Cys | Q9UBQ7 |
| HOMO SAPIENS | 4 | 229 | 2q50 | A | 302 | ARG | p.Arg302His | Q9UBQ7 |
| HOMO SAPIENS | -1 | 229 | 2q50 | A | 322 | MSE | p.Met322Arg | Q9UBQ7 |
| HOMO SAPIENS | -1 | 4519 | 5clw | A | 40 | PRO | p.Pro40Thr | Q04446 |
| HOMO SAPIENS | 1 | 4519 | 5clw | A | 59 | ILE | p.Ile59Thr | Q04446 |
| HOMO SAPIENS | 0 | 4519 | 5clw | A | 190 | ARG | p.Arg190Gly | Q04446 |
| HOMO SAPIENS | 1 | 4519 | 5clw | A | 224 | LEU | p.Leu224Pro | Q04446 |
| HOMO SAPIENS | 0 | 4519 | 5clw | A | 236 | GLN | p.Gln236His | Q04446 |
| HOMO SAPIENS | 1 | 4519 | 5clw | A | 254 | THR | p.Thr254Ala | Q04446 |
| HOMO SAPIENS | 0 | 4519 | 5clw | A | 257 | PHE | p.Phe257Leu | Q04446 |
| HOMO SAPIENS | 1 | 4519 | 5clw | A | 262 | ARG | p.Arg262Cys | Q04446 |
| HOMO SAPIENS | 3 | 4519 | 5clw | A | 329 | TYR | p.Tyr329Ser | Q04446 |
| HOMO SAPIENS | 0 | 4519 | 5clw | A | 507 | THR | p.Thr507Ala | Q04446 |
| HOMO SAPIENS | 0 | 4519 | 5clw | A | 515 | ARG | p.Arg515Cys | Q04446 |
| HOMO SAPIENS | 0 | 4519 | 5clw | A | 515 | ARG | p.Arg515His | Q04446 |
| HOMO SAPIENS | 0 | 4519 | 5clw | A | 524 | ARG | p.Arg524Gln | Q04446 |
| HOMO SAPIENS | 0 | 4519 | 5clw | A | 524 | ARG | p.Arg524Ter | Q04446 |
| HOMO SAPIENS | 0 | 4519 | 5clw | A | 545 | HIS | p.His545Arg | Q04446 |
| HOMO SAPIENS | 2 | 4519 | 5clw | A | 548 | TRP | p.Trp548Ter | Q04446 |
| HOMO SAPIENS | 0 | 4519 | 5clw | A | 592 | GLU | p.Glu592Ter | Q04446 |
| HOMO SAPIENS | 0 | 4519 | 5clw | A | 628 | HIS | p.His628Arg | Q04446 |
| HOMO SAPIENS | 0 | 4519 | 5clw | A | 637 | ARG | p.Arg637Ter | Q04446 |
| HOMO SAPIENS | 0 | 4519 | 5clw | A | 673 | ALA | p.Ala673Thr | Q04446 |
| HOMO SAPIENS | 1 | 10405 | 3bg3 | D | 507 | VAL | p.Val507Ile | P11498 |
| HOMO SAPIENS | 0 | 10405 | 3bg3 | D | 583 | ARG | p.Arg583Leu | P11498 |
| HOMO SAPIENS | 1 | 10405 | 3bg3 | D | 589 | LYS | p.Lys589Asn | P11498 |
| HOMO SAPIENS | 6 | 10405 | 3bg3 | D | 610 | ALA | p.Ala610Thr | P11498 |
| HOMO SAPIENS | 2 | 10405 | 3bg3 | D | 631 | ARG | p.Arg631Gln | P11498 |
| HOMO SAPIENS | 1 | 10405 | 3bg3 | D | 732 | ARG | p.Arg732Gln | P11498 |
| HOMO SAPIENS | 6 | 10405 | 3bg3 | D | 743 | MET | p.Met743Ile | P11498 |
| HOMO SAPIENS | 0 | 10405 | 3bg3 | D | 750 | THR | p.Thr750Met | P11498 |
| HOMO SAPIENS | -1 | 13751 | 2o8d | B | 350 | ALA | p.Ala350Val | P52701 |
| HOMO SAPIENS | -1 | 13751 | 2o8d | B | 352 | VAL | p.Val352Ile | P52701 |
| HOMO SAPIENS | -1 | 13751 | 2o8d | B | 355 | GLY | p.Gly355Ser | P52701 |
| HOMO SAPIENS | 0 | 13751 | 2o8d | B | 385 | ARG | p.Arg385Trp | P52701 |
| HOMO SAPIENS | 0 | 13751 | 2o8d | B | 390 | ASP | p.Asp390Glu | P52701 |
| HOMO SAPIENS | 0 | 13751 | 2o8d | B | 396 | LEU | p.Leu396Val | P52701 |
| HOMO SAPIENS | 0 | 13751 | 2o8d | B | 449 | LEU | p.Leu449Pro | P52701 |
| HOMO SAPIENS | 0 | 13751 | 2o8d | B | 468 | ARG | p.Arg468His | P52701 |
| HOMO SAPIENS | 0 | 13751 | 2o8d | B | 503 | SER | p.Ser503Cys | P52701 |

| | | | | | | | | |
|---|---|---|---|---|---|---|---|---|
| HOMO SAPIENS | 1 | 13751 | 2o8d | B | 509 | VAL | p.Val509Ala | P52701 |
| HOMO SAPIENS | 0 | 13751 | 2o8d | B | 675 | GLU | p.Glu675Asp | P52701 |
| HOMO SAPIENS | 0 | 13751 | 2o8d | B | 795 | ILE | p.Ile795Thr | P52701 |
| HOMO SAPIENS | 0 | 13751 | 2o8d | B | 850 | TYR | p.Tyr850Ter | P52701 |
| HOMO SAPIENS | 0 | 13751 | 2o8d | B | 878 | VAL | p.Val878Ala | P52701 |
| HOMO SAPIENS | 0 | 13751 | 2o8d | B | 911 | ARG | p.Arg911Ter | P52701 |
| HOMO SAPIENS | 0 | 13751 | 2o8d | B | 922 | ARG | p.Arg922Ter | P52701 |
| HOMO SAPIENS | 0 | 13751 | 2o8d | B | 969 | TYR | p.Tyr969Cys | P52701 |
| HOMO SAPIENS | 0 | 13751 | 2o8d | B | 1005 | ARG | p.Arg1005Ter | P52701 |
| HOMO SAPIENS | 0 | 13751 | 2o8d | B | 1035 | ARG | p.Arg1035Ter | P52701 |
| HOMO SAPIENS | 1 | 13751 | 2o8d | B | 1048 | GLN | p.Gln1048Ter | P52701 |
| HOMO SAPIENS | 1 | 13751 | 2o8d | B | 1051 | VAL | p.Val1051Ile | P52701 |
| HOMO SAPIENS | 0 | 13751 | 2o8d | B | 1054 | ILE | p.Ile1054Phe | P52701 |
| HOMO SAPIENS | 0 | 13751 | 2o8d | B | 1054 | ILE | p.Ile1054Met | P52701 |
| HOMO SAPIENS | 0 | 13751 | 2o8d | B | 1163 | GLU | p.Glu1163Val | P52701 |
| HOMO SAPIENS | 0 | 13751 | 2o8d | B | 1237 | GLU | p.Glu1237Asp | P52701 |
| HOMO SAPIENS | 0 | 13751 | 2o8d | B | 1242 | ARG | p.Arg1242His | P52701 |
| HOMO SAPIENS | -1 | 13751 | 2o8d | B | 1278 | PRO | p.Pro1278Thr | P52701 |
| HOMO SAPIENS | 0 | 13751 | 2o8d | B | 1304 | ARG | p.Arg1304Lys | P52701 |
| HOMO SAPIENS | 0 | 13751 | 2o8d | B | 1317 | HIS | p.His1317Gln | P52701 |
| HOMO SAPIENS | 0 | 13751 | 2o8d | B | 1329 | SER | p.Ser1329Leu | P52701 |
| HOMO SAPIENS | 3 | 13273 | 4g8k | A | 265 | GLU | p.Glu265Ter | Q05823 |
| HOMO SAPIENS | -1 | 11088 | 5uj7 | A | 666 | ARG | p.Arg666Trp | Q13415 |
| HOMO SAPIENS | 4 | 11088 | 5uj7 | A | 720 | ARG | p.Arg720Gln | Q13415 |
| HOMO SAPIENS | 0 | 11088 | 5uj7 | A | 816 | MET | p.Met816Thr | Q13415 |
| HOMO SAPIENS | 1 | 4352 | 5va1 | A | 421 | THR | p.Thr421Met | Q12809 |
| HOMO SAPIENS | 0 | 4352 | 5va1 | A | 451 | PRO | p.Pro451Leu | Q12809 |
| HOMO SAPIENS | 0 | 4352 | 5va1 | A | 470 | ASN | p.Asn470Asp | Q12809 |
| HOMO SAPIENS | 0 | 4352 | 5va1 | A | 490 | ALA | p.Ala490Thr | Q12809 |
| HOMO SAPIENS | 0 | 4352 | 5va1 | A | 491 | VAL | p.Val491Ile | Q12809 |
| HOMO SAPIENS | 1 | 4352 | 5va1 | A | 533 | VAL | p.Val533Gly | Q12809 |
| HOMO SAPIENS | 0 | 4352 | 5va1 | A | 552 | LEU | p.Leu552Ser | Q12809 |
| HOMO SAPIENS | 0 | 4352 | 5va1 | A | 558 | ALA | p.Ala558Pro | Q12809 |
| HOMO SAPIENS | 1 | 4352 | 5va1 | A | 561 | ALA | p.Ala561Val | Q12809 |
| HOMO SAPIENS | 0 | 4352 | 5va1 | A | 572 | GLY | p.Gly572Arg | Q12809 |
| HOMO SAPIENS | -1 | 4352 | 5va1 | A | 582 | ARG | p.Arg582Cys | Q12809 |
| HOMO SAPIENS | 0 | 4352 | 5va1 | A | 584 | GLY | p.Gly584Ser | Q12809 |
| HOMO SAPIENS | 0 | 4352 | 5va1 | A | 588 | ASN | p.Asn588Lys | Q12809 |
| HOMO SAPIENS | 0 | 4352 | 5va1 | A | 593 | ILE | p.Ile593Arg | Q12809 |
| HOMO SAPIENS | 1 | 4352 | 5va1 | A | 627 | PHE | p.Phe627Leu | Q12809 |
| HOMO SAPIENS | 0 | 4352 | 5va1 | A | 630 | VAL | p.Val630Leu | Q12809 |
| HOMO SAPIENS | 0 | 4352 | 5va1 | A | 652 | TYR | p.Tyr652Ter | Q12809 |
| HOMO SAPIENS | 0 | 4352 | 5va1 | A | 711 | ILE | p.Ile711Val | Q12809 |
| HOMO SAPIENS | 0 | 4352 | 5va1 | A | 735 | SER | p.Ser735Leu | Q12809 |
| HOMO SAPIENS | 0 | 4352 | 5va1 | A | 744 | ARG | p.Arg744Ter | Q12809 |
| HOMO SAPIENS | 0 | 4352 | 5va1 | A | 784 | ARG | p.Arg784Trp | Q12809 |
| HOMO SAPIENS | 1 | 4352 | 5va1 | A | 820 | GLY | p.Gly820Arg | Q12809 |
| HOMO SAPIENS | 5 | 4352 | 5va1 | A | 822 | VAL | p.Val822Met | Q12809 |
| HOMO SAPIENS | 1 | 4352 | 5va1 | A | 863 | ARG | p.Arg863Ter | Q12809 |
| HOMO SAPIENS | 5 | 303 | 2obv | A | 55 | ALA | p.Ala55Asp | Q00266 |
| HOMO SAPIENS | 7 | 303 | 2obv | A | 259 | ALA | p.Ala259Gly | Q00266 |
| HOMO SAPIENS | 9 | 303 | 2obv | A | 264 | ARG | p.Arg264His | Q00266 |
| HOMO SAPIENS | 9 | 303 | 2obv | A | 264 | ARG | p.Arg264Cys | Q00266 |
| HOMO SAPIENS | 0 | 303 | 2obv | A | 305 | LEU | p.Leu305Pro | Q00266 |

| | | | | | | | | |
|---|---|---|---|---|---|---|---|---|
| HOMO SAPIENS | 7 | 303 | 2obv | A | 322 | ILE | p.Ile322Met | Q00266 |
| HOMO SAPIENS | 0 | 303 | 2obv | A | 336 | GLY | p.Gly336Arg | Q00266 |
| HOMO SAPIENS | 0 | 303 | 2obv | A | 357 | PRO | p.Pro357Leu | Q00266 |
| HOMO SAPIENS | 0 | 2077 | 4zxb | E | 462 | ASN | p.Asn489Ser | P06213 |
| HOMO SAPIENS | -1 | 2077 | 4zxb | E | 519 | ASP | p.Asp546Glu | P06213 |
| HOMO SAPIENS | 2 | 2077 | 4zxb | E | 819 | THR | p.Thr858Ala | P06213 |
| HOMO SAPIENS | 0 | 2077 | 4zxb | E | 907 | ASP | p.Asp946Glu | P06213 |
| HOMO SAPIENS | 7 | 3274 | 1wsv | B | 14 | HIS | p.His42Arg | P48728 |
| HOMO SAPIENS | 0 | 3274 | 1wsv | B | 19 | GLY | p.Gly47Arg | P48728 |
| HOMO SAPIENS | 4 | 3274 | 1wsv | B | 45 | ARG | p.Arg73Cys | P48728 |
| HOMO SAPIENS | 1 | 3274 | 1wsv | B | 49 | SER | p.Ser77Leu | P48728 |
| HOMO SAPIENS | 3 | 3274 | 1wsv | B | 241 | GLY | p.Gly269Asp | P48728 |
| HOMO SAPIENS | 7 | 3274 | 1wsv | B | 248 | ASP | p.Asp276His | P48728 |
| HOMO SAPIENS | 0 | 3274 | 1wsv | B | 272 | MET | p.Met300Val | P48728 |
| HOMO SAPIENS | 4 | 3274 | 1wsv | B | 292 | ARG | p.Arg320His | P48728 |
| HOMO SAPIENS | 0 | 3274 | 1wsv | B | 354 | ARG | p.Arg382Gln | P48728 |
| HOMO SAPIENS | -1 | 435 | 3uyt | A | 44 | THR | p.Thr44Ala | P48730 |
| HOMO SAPIENS | 1 | 4698 | 5ife | C | 666 | LYS | p.Lys666Asn | O75533 |
| HOMO SAPIENS | 1 | 16917 | 1apy | C | 37 | GLY | p.Gly60Asp | P20933 |
| HOMO SAPIENS | 0 | 16917 | 1apy | C | 49 | SER | p.Ser72Pro | P20933 |
| HOMO SAPIENS | 0 | 16917 | 1apy | C | 84 | ARG | p.Arg107Ter | P20933 |
| HOMO SAPIENS | 0 | 16917 | 1apy | C | 126 | SER | p.Thr149Ser | P20933 |
| HOMO SAPIENS | 0 | 16917 | 1apy | C | 135 | TRP | p.Trp158Ter | P20933 |
| HOMO SAPIENS | 0 | 16917 | 1apy | C | 140 | CYS | p.Cys163Ser | P20933 |
| HOMO SAPIENS | 0 | 16917 | 1apy | C | 145 | TRP | p.Trp168Ter | P20933 |
| HOMO SAPIENS | 0 | 10413 | 4frw | A | 185 | THR | p.Thr185Met | Q96NY8 |
| HOMO SAPIENS | 0 | 5217 | 5n6w | G | 96 | TRP | p.Trp96Arg | O15537 |
| HOMO SAPIENS | 1 | 5217 | 5n6w | G | 102 | ARG | p.Arg102Trp | O15537 |
| HOMO SAPIENS | 0 | 5217 | 5n6w | G | 109 | GLY | p.Gly109Arg | O15537 |
| HOMO SAPIENS | -1 | 5217 | 5n6w | G | 223 | CYS | p.Cys223Arg | O15537 |
| HOMO SAPIENS | 0 | 9485 | 4z6y | E | 138 | ALA | p.Ala138Asp | Q9P0N9 |
| HOMO SAPIENS | 3 | 4504 | 4mwt | B | 411 | GLY | p.Gly439Ser | P10619 |
| HOMO SAPIENS | 3 | 4504 | 4mwt | B | 412 | PHE | p.Phe440Val | P10619 |
| HOMO SAPIENS | 6 | 6391 | 4kxy | A | 318 | ARG | p.Arg318Cys | P29401 |
| HOMO SAPIENS | 6 | 19546 | 2bx6 | A | 87 | THR | p.Thr87Ile | O75695 |
| HOMO SAPIENS | 1 | 19546 | 2bx6 | A | 118 | ARG | p.Arg118His | O75695 |
| HOMO SAPIENS | 1 | 19546 | 2bx6 | A | 201 | ILE | p.Ile201Val | O75695 |
| HOMO SAPIENS | 1 | 19546 | 2bx6 | A | 282 | ARG | p.Arg282Trp | O75695 |
| HOMO SAPIENS | 1 | 5265 | 5usr | H | 68 | ARG | p.Arg68Leu | Q9HD34 |
| HOMO SAPIENS | 2 | 1012 | 4nyi | S | 569 | LEU | p.Leu569Val | Q07889 |
| HOMO SAPIENS | 2 | 1012 | 4nyi | S | 655 | PRO | p.Pro655Leu | Q07889 |
| HOMO SAPIENS | 4 | 1012 | 4nyi | S | 663 | ILE | p.Ile663Met | Q07889 |
| HOMO SAPIENS | 2 | 1012 | 4nyi | S | 702 | TYR | p.Tyr702Cys | Q07889 |
| HOMO SAPIENS | 2 | 1012 | 4nyi | S | 708 | ALA | p.Ala708Thr | Q07889 |
| HOMO SAPIENS | 4 | 1012 | 4nyi | S | 733 | ILE | p.Ile733Phe | Q07889 |
| HOMO SAPIENS | 1 | 1848 | 3g1n | A | 4013 | ARG | p.Arg4013Trp | Q7Z6Z7 |
| HOMO SAPIENS | 1 | 1848 | 3g1n | A | 4187 | ARG | p.Arg4187Cys | Q7Z6Z7 |
| HOMO SAPIENS | 1 | 77 | 4pod | A | 41 | CYS | p.Cys79Tyr | P60174 |
| HOMO SAPIENS | 9 | 77 | 4pod | A | 104 | GLU | p.Glu142Asp | P60174 |
| HOMO SAPIENS | 1 | 77 | 4pod | A | 122 | GLY | p.Gly160Arg | P60174 |
| HOMO SAPIENS | -1 | 8672 | 3thz | A | 8 | THR | p.Thr8Met | P43246 |
| HOMO SAPIENS | 0 | 8672 | 3thz | A | 19 | PHE | p.Phe19Leu | P43246 |
| HOMO SAPIENS | 0 | 8672 | 3thz | A | 23 | PHE | p.Phe23Leu | P43246 |
| HOMO SAPIENS | 0 | 8672 | 3thz | A | 96 | ARG | p.Arg96His | P43246 |

| | | | | | | | | |
|---|---|---|---|---|---|---|---|---|
| HOMO SAPIENS | 0 | 8672 | 3thz | A | 102 | VAL | p.Val102Ile | P43246 |
| HOMO SAPIENS | 0 | 8672 | 3thz | A | 106 | ARG | p.Arg106Lys | P43246 |
| HOMO SAPIENS | 0 | 8672 | 3thz | A | 127 | ASN | p.Asn127Ser | P43246 |
| HOMO SAPIENS | 0 | 8672 | 3thz | A | 128 | LEU | p.Leu128Val | P43246 |
| HOMO SAPIENS | 0 | 8672 | 3thz | A | 163 | VAL | p.Val163Gly | P43246 |
| HOMO SAPIENS | 0 | 8672 | 3thz | A | 169 | ILE | p.Ile169Val | P43246 |
| HOMO SAPIENS | 0 | 8672 | 3thz | A | 315 | GLY | p.Gly315Asp | P43246 |
| HOMO SAPIENS | -1 | 8672 | 3thz | A | 322 | GLY | p.Gly322Asp | P43246 |
| HOMO SAPIENS | 0 | 8672 | 3thz | A | 331 | ASN | p.Asn331Asp | P43246 |
| HOMO SAPIENS | 1 | 8672 | 3thz | A | 390 | LEU | p.Leu390Phe | P43246 |
| HOMO SAPIENS | 0 | 8672 | 3thz | A | 419 | GLN | p.Gln419Lys | P43246 |
| HOMO SAPIENS | 0 | 8672 | 3thz | A | 447 | PHE | p.Phe447Leu | P43246 |
| HOMO SAPIENS | 0 | 8672 | 3thz | A | 524 | ARG | p.Arg524Pro | P43246 |
| HOMO SAPIENS | 0 | 8672 | 3thz | A | 539 | PHE | p.Phe539Leu | P43246 |
| HOMO SAPIENS | -1 | 8672 | 3thz | A | 546 | LYS | p.Lys546Asn | P43246 |
| HOMO SAPIENS | 0 | 8672 | 3thz | A | 564 | THR | p.Thr564Ala | P43246 |
| HOMO SAPIENS | 0 | 8672 | 3thz | A | 577 | ILE | p.Ile577Thr | P43246 |
| HOMO SAPIENS | 0 | 8672 | 3thz | A | 588 | TYR | p.Tyr588Ter | P43246 |
| HOMO SAPIENS | 0 | 8672 | 3thz | A | 622 | PRO | p.Pro622Leu | P43246 |
| HOMO SAPIENS | 0 | 8672 | 3thz | A | 629 | GLN | p.Gln629Arg | P43246 |
| HOMO SAPIENS | 0 | 8672 | 3thz | A | 636 | ALA | p.Ala636Pro | P43246 |
| HOMO SAPIENS | 2 | 8672 | 3thz | A | 642 | VAL | p.Val642Ile | P43246 |
| HOMO SAPIENS | 1 | 8672 | 3thz | A | 654 | ASP | p.Asp654Glu | P43246 |
| HOMO SAPIENS | 0 | 8672 | 3thz | A | 662 | GLN | p.Gln662His | P43246 |
| HOMO SAPIENS | 2 | 8672 | 3thz | A | 681 | GLN | p.Gln681His | P43246 |
| HOMO SAPIENS | 0 | 8672 | 3thz | A | 707 | CYS | p.Cys707Ser | P43246 |
| HOMO SAPIENS | 0 | 8672 | 3thz | A | 809 | GLU | p.Glu809Lys | P43246 |
| HOMO SAPIENS | 0 | 8672 | 3thz | A | 813 | MET | p.Met813Val | P43246 |
| HOMO SAPIENS | 1 | 8672 | 3thz | A | 834 | ALA | p.Ala834Thr | P43246 |
| HOMO SAPIENS | 0 | 8672 | 3thz | A | 849 | LEU | p.Leu849Val | P43246 |
| HOMO SAPIENS | -1 | 8672 | 3thz | A | 860 | SER | p.Ser860Leu | P43246 |
| HOMO SAPIENS | -1 | 8672 | 3thz | A | 934 | THR | p.Thr934Lys | P43246 |
| HOMO SAPIENS | 2 | 292 | 1i3k | A | 34 | ASN | p.Asn34Ser | Q14376 |
| HOMO SAPIENS | 7 | 292 | 1i3k | A | 90 | GLY | p.Gly90Glu | Q14376 |
| HOMO SAPIENS | 2 | 292 | 1i3k | A | 103 | ASP | p.Asp103Gly | Q14376 |
| HOMO SAPIENS | 2 | 292 | 1i3k | A | 136 | TYR | p.Tyr136Ter | Q14376 |
| HOMO SAPIENS | 0 | 292 | 1i3k | A | 169 | ARG | p.Arg169Trp | Q14376 |
| HOMO SAPIENS | 0 | 292 | 1i3k | A | 183 | LEU | p.Leu183Pro | Q14376 |
| HOMO SAPIENS | 0 | 292 | 1i3k | A | 302 | GLY | p.Gly302Asp | Q14376 |
| HOMO SAPIENS | 0 | 292 | 1i3k | A | 313 | LEU | p.Leu313Met | Q14376 |
| HOMO SAPIENS | 1 | 2084 | 5vc8 | A | 295 | GLN | p.Gln295Leu | O96028 |
| HOMO SAPIENS | -1 | 10157 | 5wrw | A | 7 | GLY | p.Gly7Ala | O76094 |
| HOMO SAPIENS | -1 | 10157 | 5wrw | A | 7 | GLY | p.Gly7Arg | O76094 |
| HOMO SAPIENS | -1 | 10157 | 5wrw | A | 8 | GLY | p.Gly8Ala | O76094 |
| HOMO SAPIENS | 0 | 3113 | 3in5 | A | 29 | GLU | p.Glu29Lys | Q9UBT6 |
| HOMO SAPIENS | 5 | 4546 | 5in3 | B | 44 | VAL | p.Val44Leu | P07902 |
| HOMO SAPIENS | -1 | 4546 | 5in3 | B | 51 | ARG | p.Arg51Leu | P07902 |
| HOMO SAPIENS | -1 | 4546 | 5in3 | B | 62 | LEU | p.Leu62Met | P07902 |
| HOMO SAPIENS | 4 | 4546 | 5in3 | B | 66 | PRO | p.Pro66His | P07902 |
| HOMO SAPIENS | 2 | 4546 | 5in3 | B | 74 | LEU | p.Leu74Pro | P07902 |
| HOMO SAPIENS | 5 | 4546 | 5in3 | B | 97 | ASN | p.Asn97Ser | P07902 |
| HOMO SAPIENS | 1 | 4546 | 5in3 | B | 135 | SER | p.Ser135Leu | P07902 |
| HOMO SAPIENS | 5 | 4546 | 5in3 | B | 138 | THR | p.Thr138Met | P07902 |
| HOMO SAPIENS | 7 | 4546 | 5in3 | B | 139 | LEU | p.Leu139Pro | P07902 |

| | | | | | | | | |
|---|---|---|---|---|---|---|---|---|
| HOMO SAPIENS | 2 | 4546 | 5in3 | B | 142 | MET | p.Met142Lys | P07902 |
| HOMO SAPIENS | 1 | 4546 | 5in3 | B | 143 | SER | p.Ser143Leu | P07902 |
| HOMO SAPIENS | 2 | 4546 | 5in3 | B | 148 | ARG | p.Arg148Gly | P07902 |
| HOMO SAPIENS | 2 | 4546 | 5in3 | B | 148 | ARG | p.Arg148Gln | P07902 |
| HOMO SAPIENS | 5 | 4546 | 5in3 | B | 183 | PRO | p.Pro183Thr | P07902 |
| HOMO SAPIENS | 5 | 4546 | 5in3 | B | 185 | PRO | p.Pro185His | P07902 |
| HOMO SAPIENS | 5 | 4546 | 5in3 | B | 186 | HIS | p.His186Tyr | P07902 |
| HOMO SAPIENS | 7 | 4546 | 5in3 | B | 188 | GLN | p.Gln188Arg | P07902 |
| HOMO SAPIENS | 5 | 4546 | 5in3 | B | 195 | LEU | p.Leu195Pro | P07902 |
| HOMO SAPIENS | 4 | 4546 | 5in3 | B | 201 | ARG | p.Arg201His | P07902 |
| HOMO SAPIENS | 4 | 4546 | 5in3 | B | 201 | ARG | p.Arg201Cys | P07902 |
| HOMO SAPIENS | 4 | 4546 | 5in3 | B | 203 | GLU | p.Glu203Lys | P07902 |
| HOMO SAPIENS | 2 | 4546 | 5in3 | B | 209 | TYR | p.Tyr209Cys | P07902 |
| HOMO SAPIENS | 5 | 4546 | 5in3 | B | 223 | ARG | p.Arg223Ser | P07902 |
| HOMO SAPIENS | 5 | 4546 | 5in3 | B | 226 | LEU | p.Leu226Pro | P07902 |
| HOMO SAPIENS | 5 | 4546 | 5in3 | B | 226 | LEU | p.Leu226Val | P07902 |
| HOMO SAPIENS | 2 | 4546 | 5in3 | B | 227 | LEU | p.Leu227Pro | P07902 |
| HOMO SAPIENS | 8 | 4546 | 5in3 | B | 258 | ARG | p.Arg258Cys | P07902 |
| HOMO SAPIENS | 1 | 4546 | 5in3 | B | 265 | PRO | p.Pro265Ala | P07902 |
| HOMO SAPIENS | 8 | 4546 | 5in3 | B | 268 | THR | p.Thr268Asn | P07902 |
| HOMO SAPIENS | 7 | 4546 | 5in3 | B | 278 | ILE | p.Ile278Asn | P07902 |
| HOMO SAPIENS | 5 | 4546 | 5in3 | B | 285 | LYS | p.Lys285Asn | P07902 |
| HOMO SAPIENS | 1 | 4546 | 5in3 | B | 319 | HIS | p.His319Gln | P07902 |
| HOMO SAPIENS | 4 | 4546 | 5in3 | B | 321 | HIS | p.His321Tyr | P07902 |
| HOMO SAPIENS | 4 | 4546 | 5in3 | B | 325 | PRO | p.Pro325Leu | P07902 |
| HOMO SAPIENS | 2 | 4546 | 5in3 | B | 328 | ARG | p.Arg328His | P07902 |
| HOMO SAPIENS | 2 | 4546 | 5in3 | B | 329 | SER | p.Ser329Phe | P07902 |
| HOMO SAPIENS | 5 | 4546 | 5in3 | B | 334 | LYS | p.Lys334Arg | P07902 |
| HOMO SAPIENS | 6 | 1290 | 5uav | C | 119 | ARG | p.Arg119Gly | P32322 |
| HOMO SAPIENS | 6 | 1290 | 5uav | C | 119 | ARG | p.Arg119His | P32322 |
| HOMO SAPIENS | 3 | 1290 | 5uav | C | 251 | ARG | p.Arg251His | P32322 |
| HOMO SAPIENS | 7 | 1290 | 5uav | C | 266 | ARG | p.Arg266Gln | P32322 |
| HOMO SAPIENS | 0 | 9618 | 2r7e | A | 22 | GLY | p.Gly41Cys | P00451 |
| HOMO SAPIENS | 2 | 9618 | 2r7e | A | 113 | GLU | p.Glu132Asp | P00451 |
| HOMO SAPIENS | -1 | 9618 | 2r7e | A | 336 | ARG | p.Arg355Ter | P00451 |
| HOMO SAPIENS | 0 | 9618 | 2r7e | A | 372 | ARG | p.Arg391His | P00451 |
| HOMO SAPIENS | 0 | 9618 | 2r7e | A | 373 | SER | p.Ser392Leu | P00451 |
| HOMO SAPIENS | 0 | 9618 | 2r7e | A | 373 | SER | p.Ser392Pro | P00451 |
| HOMO SAPIENS | 3 | 9618 | 2r7e | A | 386 | ILE | p.Ile405Ser | P00451 |
| HOMO SAPIENS | 6 | 9618 | 2r7e | A | 390 | GLU | p.Glu409Gly | P00451 |
| HOMO SAPIENS | 2 | 9618 | 2r7e | A | 412 | LEU | p.Leu431Phe | P00451 |
| HOMO SAPIENS | 5 | 9618 | 2r7e | A | 425 | LYS | p.Lys444Arg | P00451 |
| HOMO SAPIENS | 0 | 9618 | 2r7e | A | 484 | ARG | p.Arg503His | P00451 |
| HOMO SAPIENS | 2 | 9618 | 2r7e | A | 531 | ARG | p.Arg550His | P00451 |
| HOMO SAPIENS | 3 | 9618 | 2r7e | A | 535 | SER | p.Ser554Gly | P00451 |
| HOMO SAPIENS | 2 | 9618 | 2r7e | A | 593 | ARG | p.Arg612Cys | P00451 |
| HOMO SAPIENS | 0 | 9618 | 2r7e | A | 704 | ALA | p.Ala723Thr | P00451 |
| HOMO SAPIENS | 0 | 9618 | 2r7e | A | 720 | GLU | p.Glu739Lys | P00451 |
| HOMO SAPIENS | 1 | 13085 | 3ud1 | A | 1075 | ILE | p.Thr1075Ile | P16157 |
| HOMO SAPIENS | -1 | 3289 | 2fy4 | A | 2 | ALA | p.Ala120Thr | P28329 |
| HOMO SAPIENS | 7 | 3289 | 2fy4 | A | 18 | VAL | p.Val136Met | P28329 |
| HOMO SAPIENS | 5 | 3289 | 2fy4 | A | 92 | LEU | p.Leu210Pro | P28329 |
| HOMO SAPIENS | 2 | 3289 | 2fy4 | A | 93 | PRO | p.Pro211Ala | P28329 |
| HOMO SAPIENS | 4 | 3289 | 2fy4 | A | 131 | LEU | p.Leu249Val | P28329 |

| | | | | | | | | |
|---|---|---|---|---|---|---|---|---|
| HOMO SAPIENS | 2 | 3289 | 2fy4 | A | 181 | PRO | p.Pro299Leu | P28329 |
| HOMO SAPIENS | 0 | 3289 | 2fy4 | A | 218 | ILE | p.Ile336Thr | P28329 |
| HOMO SAPIENS | 2 | 3289 | 2fy4 | A | 261 | ASP | p.Asp379His | P28329 |
| HOMO SAPIENS | 0 | 3289 | 2fy4 | A | 302 | ARG | p.Arg420Cys | P28329 |
| HOMO SAPIENS | 5 | 3289 | 2fy4 | A | 323 | GLU | p.Glu441Lys | P28329 |
| HOMO SAPIENS | 0 | 3289 | 2fy4 | A | 343 | MET | p.Val461Met | P28329 |
| HOMO SAPIENS | 2 | 3289 | 2fy4 | A | 364 | ARG | p.Arg482Gly | P28329 |
| HOMO SAPIENS | 3 | 3289 | 2fy4 | A | 380 | SER | p.Ser498Leu | P28329 |
| HOMO SAPIENS | 6 | 3289 | 2fy4 | A | 388 | VAL | p.Val506Leu | P28329 |
| HOMO SAPIENS | 7 | 3289 | 2fy4 | A | 429 | HIS | p.His547Gln | P28329 |
| HOMO SAPIENS | 8 | 3289 | 2fy4 | A | 442 | ARG | p.Arg560His | P28329 |
| HOMO SAPIENS | 3 | 3289 | 2fy4 | A | 443 | ARG | p.Arg561Gln | P28329 |
| HOMO SAPIENS | 0 | 4137 | 3hhr | B | 38 | CYS | p.Cys56Ter | P10912 |
| HOMO SAPIENS | 0 | 4137 | 3hhr | B | 43 | ARG | p.Arg61Ter | P10912 |
| HOMO SAPIENS | 0 | 4137 | 3hhr | B | 44 | GLU | p.Glu62Lys | P10912 |
| HOMO SAPIENS | 0 | 4137 | 3hhr | B | 96 | PHE | p.Phe114Ser | P10912 |
| HOMO SAPIENS | 5 | 4137 | 3hhr | B | 131 | PRO | p.Pro149Gln | P10912 |
| HOMO SAPIENS | 0 | 4137 | 3hhr | B | 144 | VAL | p.Val162Ile | P10912 |
| HOMO SAPIENS | 0 | 4137 | 3hhr | B | 150 | HIS | p.His168Gln | P10912 |
| HOMO SAPIENS | 0 | 4137 | 3hhr | B | 152 | ASP | p.Asp170His | P10912 |
| HOMO SAPIENS | 0 | 4137 | 3hhr | B | 153 | ILE | p.Ile171Thr | P10912 |
| HOMO SAPIENS | 0 | 4137 | 3hhr | B | 154 | GLN | p.Gln172Pro | P10912 |
| HOMO SAPIENS | 0 | 4137 | 3hhr | B | 155 | VAL | p.Val173Gly | P10912 |
| HOMO SAPIENS | 0 | 4137 | 3hhr | B | 217 | ARG | p.Arg235Ter | P10912 |
| HOMO SAPIENS | 5 | 4137 | 3hhr | B | 224 | GLU | p.Glu242Asp | P10912 |
| HOMO SAPIENS | 5 | 4137 | 3hhr | B | 224 | GLU | p.Glu242Ter | P10912 |
| HOMO SAPIENS | 6 | 2493 | 5bpb | A | 69 | HIS | p.His69Tyr | Q9ULV1 |
| HOMO SAPIENS | 6 | 2493 | 5bpb | A | 105 | MET | p.Met105Val | Q9ULV1 |
| HOMO SAPIENS | -1 | 2493 | 5bpb | A | 168 | PRO | p.Pro168Ser | Q9ULV1 |
| HOMO SAPIENS | 0 | 32123 | 4ccz | A | 314 | ASP | p.Asp314Asn | Q99707 |
| HOMO SAPIENS | 0 | 32123 | 4ccz | A | 345 | VAL | p.Val345Phe | Q99707 |
| HOMO SAPIENS | 1 | 32123 | 4ccz | A | 585 | ARG | p.Arg585Ter | Q99707 |
| HOMO SAPIENS | 4 | 4618 | 2ag4 | A | 26 | PRO | p.Pro55Leu | P17900 |
| HOMO SAPIENS | 4 | 4618 | 2ag4 | A | 30 | VAL | p.Ile59Val | P17900 |
| HOMO SAPIENS | 2 | 4618 | 2ag4 | A | 40 | VAL | p.Met69Val | P17900 |
| HOMO SAPIENS | 2 | 4618 | 2ag4 | A | 124 | VAL | p.Val153Ala | P17900 |
| HOMO SAPIENS | 8 | 4618 | 2ag4 | A | 140 | ARG | p.Arg169Pro | P17900 |
| HOMO SAPIENS | 5 | 1082 | 1hk0 | X | 14 | ARG | p.Arg15Cys | P07320 |
| HOMO SAPIENS | 7 | 1082 | 1hk0 | X | 23 | PRO | p.Pro24Thr | P07320 |
| HOMO SAPIENS | 6 | 1082 | 1hk0 | X | 36 | ARG | p.Arg37Ser | P07320 |
| HOMO SAPIENS | 5 | 1082 | 1hk0 | X | 55 | TYR | p.Tyr56Ter | P07320 |
| HOMO SAPIENS | 3 | 1082 | 1hk0 | X | 58 | ARG | p.Arg59His | P07320 |
| HOMO SAPIENS | 7 | 1082 | 1hk0 | X | 88 | HIS | p.His88Gln | P07320 |
| HOMO SAPIENS | 6 | 1082 | 1hk0 | X | 95 | ARG | p.Arg95Ser | P07320 |
| HOMO SAPIENS | 5 | 1082 | 1hk0 | X | 126 | VAL | p.Val126Met | P07320 |
| HOMO SAPIENS | 8 | 1082 | 1hk0 | X | 157 | TRP | p.Trp157Ter | P07320 |
| HOMO SAPIENS | 0 | 22879 | 3j0a | B | 644 | ILE | p.Ile644Phe | O60602 |
| HOMO SAPIENS | 8 | 6 | 2o23 | B | 122 | LEU | p.Leu122Val | Q99714 |
| HOMO SAPIENS | 1 | 6 | 2o23 | B | 130 | ARG | p.Arg130Cys | Q99714 |
| HOMO SAPIENS | 1 | 6 | 2o23 | B | 247 | ASN | p.Asn247Ser | Q99714 |
| HOMO SAPIENS | 0 | 6 | 2o23 | B | 249 | GLU | p.Glu249Gln | Q99714 |
| HOMO SAPIENS | 1 | 12910 | 4hl4 | A | 67 | ARG | p.Arg67Ter | Q96BZ9 |
| HOMO SAPIENS | 8 | 16105 | 1lmj | A | 7 | GLU | p.Glu1073Lys | P35555 |
| HOMO SAPIENS | 8 | 16105 | 1lmj | A | 8 | CYS | p.Cys1074Arg | P35555 |

| | | | | | | | | |
|---|---|---|---|---|---|---|---|---|
| HOMO SAPIENS | 9 | 16105 | 1lmj | A | 51 | CYS | p.Cys1117Tyr | P35555 |
| HOMO SAPIENS | 9 | 16105 | 1lmj | A | 63 | CYS | p.Cys1129Tyr | P35555 |
| HOMO SAPIENS | 9 | 16105 | 1lmj | A | 65 | ASN | p.Asn1131Tyr | P35555 |
| HOMO SAPIENS | 4 | 16105 | 1lmj | A | 71 | ARG | p.Arg1137Pro | P35555 |
| HOMO SAPIENS | 1 | 16105 | 1lmj | A | 82 | PRO | p.Pro1148Ala | P35555 |
| HOMO SAPIENS | 0 | 4795 | 3s5f | A | 106 | LEU | p.Leu106Ser | Q16595 |
| HOMO SAPIENS | 0 | 4795 | 3s5f | A | 130 | GLY | p.Gly130Val | Q16595 |
| HOMO SAPIENS | 0 | 4795 | 3s5f | A | 154 | ILE | p.Ile154Phe | Q16595 |
| HOMO SAPIENS | -1 | 4795 | 3s5f | A | 209 | ASP | p.Asp209Gly | Q16595 |
| HOMO SAPIENS | -1 | 2884 | 4lli | A | 1655 | ASP | p.Asp1655Glu | Q9Y4I1 |
| HOMO SAPIENS | 0 | 1786 | 7pck | C | 56P | TYR | p.Tyr71Ter | P43235 |
| HOMO SAPIENS | 0 | 1786 | 7pck | C | 64P | GLY | p.Gly79Glu | P43235 |
| HOMO SAPIENS | 7 | 38286 | 2l53 | B | 1914 | ARG | p.Arg1914Gly | Q14524 |
| HOMO SAPIENS | 1 | 38286 | 2l53 | B | 1924 | ALA | p.Ala1924Thr | Q14524 |
| HOMO SAPIENS | 0 | 31747 | 3ksy | A | 25 | PRO | p.Pro25Ser | Q07889 |
| HOMO SAPIENS | 3 | 31747 | 3ksy | A | 37 | THR | p.Thr37Ala | Q07889 |
| HOMO SAPIENS | 0 | 31747 | 3ksy | A | 54 | GLN | p.Gln54His | Q07889 |
| HOMO SAPIENS | -1 | 31747 | 3ksy | A | 190 | ASP | p.Asp190Glu | Q07889 |
| HOMO SAPIENS | 2 | 31747 | 3ksy | A | 266 | THR | p.Thr266Lys | Q07889 |
| HOMO SAPIENS | 1 | 31747 | 3ksy | A | 269 | MET | p.Met269Arg | Q07889 |
| HOMO SAPIENS | 1 | 31747 | 3ksy | A | 340 | PRO | p.Pro340Ser | Q07889 |
| HOMO SAPIENS | 1 | 31747 | 3ksy | A | 423 | ASN | p.Asn423Lys | Q07889 |
| HOMO SAPIENS | 1 | 31747 | 3ksy | A | 424 | GLU | p.Glu424Lys | Q07889 |
| HOMO SAPIENS | 1 | 31747 | 3ksy | A | 432 | TRP | p.Trp432Arg | Q07889 |
| HOMO SAPIENS | 1 | 31747 | 3ksy | A | 552 | ARG | p.Arg552Gly | Q07889 |
| HOMO SAPIENS | 1 | 31747 | 3ksy | A | 552 | ARG | p.Arg552Ser | Q07889 |
| HOMO SAPIENS | 3 | 7529 | 6jdw | A | 93 | ILE | p.Ile93Met | P50440 |
| HOMO SAPIENS | 2 | 7529 | 6jdw | A | 110 | GLN | p.Gln110His | P50440 |
| HOMO SAPIENS | 5 | 7529 | 6jdw | A | 149 | TRP | p.Trp149Ter | P50440 |
| HOMO SAPIENS | 0 | 7529 | 6jdw | A | 169 | ARG | p.Arg169Ter | P50440 |
| HOMO SAPIENS | 0 | 7529 | 6jdw | A | 260 | ARG | p.Arg260Ter | P50440 |
| HOMO SAPIENS | 3 | 7529 | 6jdw | A | 344 | ILE | p.Ile344Val | P50440 |
| HOMO SAPIENS | 5 | 7529 | 6jdw | A | 418 | LEU | p.Leu418Ile | P50440 |
| HOMO SAPIENS | 1 | 14788 | 1eak | A | 101 | ARG | p.Arg101His | P08253 |
| HOMO SAPIENS | 0 | 14788 | 1eak | A | 383 | ASP | p.Asp383Glu | P08253 |
| HOMO SAPIENS | 7 | 14788 | 1eak | A | 404 | GLN | p.Glu404Lys | P08253 |
| HOMO SAPIENS | 2 | 14788 | 1eak | A | 447 | ALA | p.Ala447Val | P08253 |
| HOMO SAPIENS | 0 | 1255 | 5xtc | p | 8 | PRO | p.Pro8His | Q9Y6M9 |
| HOMO SAPIENS | 0 | 1255 | 5xtc | p | 146 | PRO | p.Pro146Ser | Q9Y6M9 |
| HOMO SAPIENS | 3 | 6721 | 1a22 | A | 25 | PHE | p.Phe51Tyr | P01241 |
| HOMO SAPIENS | 2 | 6721 | 1a22 | A | 95 | SER | p.Ser121Arg | P01241 |
| HOMO SAPIENS | 2 | 6721 | 1a22 | A | 110 | VAL | p.Val136Ile | P01241 |
| HOMO SAPIENS | 0 | 1380 | 5i6z | A | 605 | LYS | p.Lys605Asn | P31645 |
| HOMO SAPIENS | 0 | 941 | 1qki | D | 48 | ILE | p.Ile48Thr | P11413 |
| HOMO SAPIENS | 3 | 941 | 1qki | D | 68 | VAL | p.Val68Met | P11413 |
| HOMO SAPIENS | 2 | 941 | 1qki | D | 126 | ASN | p.Asn126Asp | P11413 |
| HOMO SAPIENS | 8 | 941 | 1qki | D | 163 | GLY | p.Gly163Ser | P11413 |
| HOMO SAPIENS | 6 | 941 | 1qki | D | 165 | ASN | p.Asn165Asp | P11413 |
| HOMO SAPIENS | 4 | 941 | 1qki | D | 181 | ASP | p.Asp181Val | P11413 |
| HOMO SAPIENS | 4 | 941 | 1qki | D | 182 | ARG | p.Arg182Trp | P11413 |
| HOMO SAPIENS | 5 | 941 | 1qki | D | 188 | SER | p.Ser188Phe | P11413 |
| HOMO SAPIENS | 9 | 941 | 1qki | D | 198 | ARG | p.Arg198Cys | P11413 |
| HOMO SAPIENS | 8 | 941 | 1qki | D | 213 | VAL | p.Val213Leu | P11413 |
| HOMO SAPIENS | 5 | 941 | 1qki | D | 282 | ASP | p.Asp282His | P11413 |

| | | | | | | | | |
|---|---|---|---|---|---|---|---|---|
| HOMO SAPIENS | 7 | 941 | 1qki | D | 285 | ARG | p.Arg285His | P11413 |
| HOMO SAPIENS | 5 | 941 | 1qki | D | 291 | VAL | p.Val291Met | P11413 |
| HOMO SAPIENS | 8 | 941 | 1qki | D | 323 | LEU | p.Leu323Pro | P11413 |
| HOMO SAPIENS | 3 | 941 | 1qki | D | 335 | ALA | p.Ala335Thr | P11413 |
| HOMO SAPIENS | 6 | 941 | 1qki | D | 350 | ASP | p.Asp350His | P11413 |
| HOMO SAPIENS | 5 | 941 | 1qki | D | 393 | ARG | p.Arg393His | P11413 |
| HOMO SAPIENS | 5 | 941 | 1qki | D | 454 | ARG | p.Arg454Cys | P11413 |
| HOMO SAPIENS | 2 | 941 | 1qki | D | 459 | LEU | p.Arg459Leu | P11413 |
| HOMO SAPIENS | 4 | 941 | 1qki | D | 463 | ARG | p.Arg463His | P11413 |
| HOMO SAPIENS | 4 | 2483 | 1tij | A | 68 | LEU | p.Leu94Gln | P01034 |
| HOMO SAPIENS | 2 | 20632 | 3v5b | A | 331 | ARG | p.Arg331Pro | P02545 |
| HOMO SAPIENS | 2 | 20632 | 3v5b | A | 374 | HIS | p.His374Gln | P02545 |
| HOMO SAPIENS | 4 | 13929 | 5xtd | J | 75 | ARG | p.Arg75His | Q16795 |
| HOMO SAPIENS | 3 | 13929 | 5xtd | J | 243 | VAL | p.Val243Ala | Q16795 |
| HOMO SAPIENS | 1 | 13929 | 5xtd | J | 294 | PRO | p.Pro294Leu | Q16795 |
| HOMO SAPIENS | 2 | 13929 | 5xtd | J | 321 | ARG | p.Arg321Pro | Q16795 |
| HOMO SAPIENS | 0 | 9411 | 5thl | A | 41 | ARG | p.Gly41Arg | P54577 |
| HOMO SAPIENS | 0 | 9411 | 5thl | A | 196 | GLU | p.Glu196Lys | P54577 |
| HOMO SAPIENS | 0 | 9411 | 5thl | A | 316 | ALA | p.Ala316Ser | P54577 |
| HOMO SAPIENS | 0 | 9411 | 5thl | A | 339 | ALA | p.Ala339Thr | P54577 |
| HOMO SAPIENS | 4 | 16273 | 2e8n | A | 46 | THR | p.Thr940Ile | P29317 |
| HOMO SAPIENS | 7 | 16273 | 2e8n | A | 54 | GLY | p.Gly948Trp | P29317 |
| HOMO SAPIENS | 6 | 16273 | 2e8n | A | 65 | ALA | p.Ala959Thr | P29317 |
| HOMO SAPIENS | 7 | 30387 | 2k9y | A | 541 | VAL | p.Val541Met | P29317 |
| HOMO SAPIENS | 0 | 6961 | 4nha | A | 843 | ILE | p.Ile853Val | Q9UPY3 |
| HOMO SAPIENS | -1 | 6961 | 4nha | A | 862 | ALA | p.Ala872Thr | Q9UPY3 |
| HOMO SAPIENS | 3 | 15708 | 4pzg | A | 16 | ARG | p.Arg16Ter | Q9Y5X0 |
| HOMO SAPIENS | 4 | 15708 | 4pzg | A | 51 | ARG | p.Arg51Gln | Q9Y5X0 |
| HOMO SAPIENS | 3 | 30166 | 4o00 | A | 44 | ARG | p.Arg14257Ter | Q8WZ42 |
| HOMO SAPIENS | 2 | 30166 | 4o00 | A | 54 | ARG | p.Arg14267His | Q8WZ42 |
| HOMO SAPIENS | 8 | 11158 | 4hca | A | 275 | TRP | p.Trp274Arg | P23771 |
| HOMO SAPIENS | 2 | 4328 | 4e0s | A | 1476 | ARG | p.Arg1476Ter | P01031 |
| HOMO SAPIENS | 0 | 11462 | 1dx5 | I | 455 | ALA | p.Ala473Val | P07204 |
| HOMO SAPIENS | -1 | 11150 | 2fic | A | 45 | LYS | p.Lys35Asn | O00499 |
| HOMO SAPIENS | 6 | 11150 | 2fic | A | 161 | ASP | p.Asp151Asn | O00499 |
| HOMO SAPIENS | 8 | 27382 | 4mqv | B | 17 | ARG | p.Arg17Ter | Q9NZC9 |
| HOMO SAPIENS | -1 | 37991 | 1q0p | A | 231 | GLN | p.Gln256Ter | P00751 |
| HOMO SAPIENS | 1 | 443 | 5xte | U | 215 | LEU | p.Leu215Phe | P08574 |
| HOMO SAPIENS | 8 | 21471 | 5y4e | A | 1461 | ASN | p.Thr692Ala | Q01484 |
| HOMO SAPIENS | -1 | 35443 | 3um3 | B | 129 | ASP | p.Asp129Val | Q9H444 |
| HOMO SAPIENS | -1 | 35443 | 3um3 | B | 161 | GLU | p.Glu161Lys | Q9H444 |
| HOMO SAPIENS | 6 | 28781 | 2n2a | B | 111 | ILE | p.Ile654Val | P04626 |
| HOMO SAPIENS | 7 | 28781 | 2n2a | B | 112 | ILE | p.Ile655Val | P04626 |
| HOMO SAPIENS | -1 | 3346 | 4ztz | C | 61 | GLY | p.Gly61Arg | Q9UHN1 |
| HOMO SAPIENS | 4 | 3346 | 4ztz | C | 103 | GLY | p.Gly103Ser | Q9UHN1 |
| HOMO SAPIENS | 4 | 3346 | 4ztz | C | 133 | HIS | p.His133Tyr | Q9UHN1 |
| HOMO SAPIENS | -1 | 3346 | 4ztz | C | 153 | LEU | p.Leu153Val | Q9UHN1 |
| HOMO SAPIENS | -1 | 3346 | 4ztz | C | 169 | ALA | p.Ala169Thr | Q9UHN1 |
| HOMO SAPIENS | 7 | 3346 | 4ztz | C | 369 | ARG | p.Arg369Gly | Q9UHN1 |
| HOMO SAPIENS | 4 | 3346 | 4ztz | C | 386 | ASP | p.Asp386Glu | Q9UHN1 |
| HOMO SAPIENS | 4 | 3346 | 4ztz | C | 451 | GLY | p.Gly451Glu | Q9UHN1 |
| HOMO SAPIENS | 4 | 2131 | 5fo9 | E | 1549 | LEU | p.Leu1549Met | P01024 |
| HOMO SAPIENS | 2 | 2131 | 5fo9 | E | 1619 | SER | p.Ser1619Arg | P01024 |
| HOMO SAPIENS | 3 | 12662 | 4jhp | C | 75 | ILE | p.Ile75Val | Q92834 |

| | | | | | | | | |
|---|---|---|---|---|---|---|---|---|
| HOMO SAPIENS | 1 | 12662 | 4jhp | C | 173 | GLY | p.Gly173Arg | Q92834 |
| HOMO SAPIENS | 0 | 12662 | 4jhp | C | 184 | GLN | p.Gln184His | Q92834 |
| HOMO SAPIENS | 3 | 12662 | 4jhp | C | 262 | ALA | p.Ala262Gly | Q92834 |
| HOMO SAPIENS | 3 | 12662 | 4jhp | C | 302 | CYS | p.Cys302Tyr | Q92834 |
| HOMO SAPIENS | 5 | 1093 | 2jg9 | A | 186 | GLN | p.Gln208Ter | P02745 |
| HOMO SAPIENS | 0 | 13181 | 4r4l | A | 12 | MET | p.Met13Ile | Q13976 |
| HOMO SAPIENS | 8 | 25939 | 4iyp | A | 106 | ARG | p.Arg106Gln | P78318 |
| HOMO SAPIENS | 1 | 217 | 2xsx | A | 71 | SER | p.Asn71Ser | P13929 |
| HOMO SAPIENS | 0 | 217 | 2xsx | A | 76 | ALA | p.Ala76Pro | P13929 |
| HOMO SAPIENS | 1 | 217 | 2xsx | A | 85 | ALA | p.Val85Ala | P13929 |
| HOMO SAPIENS | 8 | 217 | 2xsx | A | 156 | GLY | p.Gly156Asp | P13929 |
| HOMO SAPIENS | 5 | 217 | 2xsx | A | 332 | VAL | p.Val332Ile | P13929 |
| HOMO SAPIENS | 7 | 217 | 2xsx | A | 374 | GLY | p.Gly374Glu | P13929 |
| HOMO SAPIENS | 1 | 5508 | 3szm | A | 682 | THR | p.Thr682Asn | Q8NEM0 |
| HOMO SAPIENS | 0 | 5508 | 3szm | A | 715 | TRP | p.Trp715Ter | Q8NEM0 |
| HOMO SAPIENS | 1 | 5508 | 3szm | A | 761 | VAL | p.Ala761Val | Q8NEM0 |
| HOMO SAPIENS | 1 | 5508 | 3szm | A | 828 | PRO | p.Pro828Ser | Q8NEM0 |
| HOMO SAPIENS | 4 | 5508 | 3szm | A | 829 | GLU | p.Glu829Asp | Q8NEM0 |
| HOMO SAPIENS | 2 | 16353 | 2q0z | X | 171 | GLY | p.Gly1978Ala | O75643 |
| HOMO SAPIENS | 2 | 16353 | 2q0z | X | 188 | ALA | p.Ala1995Thr | O75643 |
| HOMO SAPIENS | 0 | 4127 | 3swr | A | 860 | GLY | p.Gly860Arg | P26358 |
| HOMO SAPIENS | -1 | 4127 | 3swr | A | 1122 | SER | p.Ser1122Cys | P26358 |
| HOMO SAPIENS | 0 | 4127 | 3swr | A | 1216 | ASP | p.Asp1216Glu | P26358 |
| HOMO SAPIENS | 0 | 6382 | 3o96 | A | 17 | GLU | p.Glu17Lys | P31749 |
| HOMO SAPIENS | 6 | 14314 | 5xtc | Q | 71 | PRO | p.Pro71Leu | O75306 |
| HOMO SAPIENS | 2 | 5886 | 4fao | E | 22 | ARG | p.Arg40His | Q13705 |
| HOMO SAPIENS | 1 | 2534 | 3mp4 | A | 37 | GLU | p.Glu37Ter | P35914 |
| HOMO SAPIENS | 3 | 2534 | 3mp4 | A | 41 | MET | p.Arg41Gln | P35914 |
| HOMO SAPIENS | 1 | 2534 | 3mp4 | A | 70 | VAL | p.Val70Leu | P35914 |
| HOMO SAPIENS | 2 | 2196 | 3ifq | A | 142 | ARG | p.Arg142His | P14923 |
| HOMO SAPIENS | 2 | 2196 | 3ifq | A | 153 | ASP | p.Asp153Glu | P14923 |
| HOMO SAPIENS | 3 | 2196 | 3ifq | A | 176 | ARG | p.Arg176Gln | P14923 |
| HOMO SAPIENS | 3 | 2196 | 3ifq | A | 265 | ARG | p.Arg265His | P14923 |
| HOMO SAPIENS | 2 | 2196 | 3ifq | A | 301 | GLU | p.Glu301Gly | P14923 |
| HOMO SAPIENS | 0 | 2196 | 3ifq | A | 456 | VAL | p.Val456Ile | P14923 |
| HOMO SAPIENS | 3 | 2196 | 3ifq | A | 603 | VAL | p.Val603Leu | P14923 |
| HOMO SAPIENS | 0 | 2196 | 3ifq | A | 648 | VAL | p.Val648Ile | P14923 |
| HOMO SAPIENS | 1 | 8746 | 1cgj | I | 32 | PRO | p.Pro55Ser | P00995 |
| HOMO SAPIENS | 1 | 8746 | 1cgj | I | 44 | ARG | p.Arg67His | P00995 |
| HOMO SAPIENS | 1 | 8746 | 1cgj | I | 44 | ARG | p.Arg67Cys | P00995 |
| HOMO SAPIENS | 3 | 3212 | 1syq | A | 105 | ARG | p.Arg105Ter | P18206 |
| HOMO SAPIENS | 3 | 3212 | 1syq | A | 126 | PHE | p.Phe126Leu | P18206 |
| HOMO SAPIENS | 4 | 10028 | 3eto | A | 1609 | ARG | p.Arg1608His | P46531 |
| HOMO SAPIENS | 1 | 10028 | 3eto | A | 1672 | VAL | p.Val1671Ile | P46531 |
| HOMO SAPIENS | 5 | 10028 | 3eto | A | 1692 | GLN | p.Gln1691His | P46531 |
| HOMO SAPIENS | 3 | 3749 | 5xtc | Z | 70 | GLY | p.Gly70Ter | O43676 |
| HOMO SAPIENS | 2 | 17039 | 3brv | B | 57 | GLU | p.Glu57Lys | Q9Y6K9 |
| HOMO SAPIENS | 8 | 3589 | 2fh3 | A | 433 | VAL | p.Val460Leu | P06396 |
| HOMO SAPIENS | 1 | 3589 | 2fh3 | A | 454 | ARG | p.Arg481Cys | P06396 |
| HOMO SAPIENS | 4 | 3589 | 2fh3 | A | 498 | ARG | p.Arg525His | P06396 |
| HOMO SAPIENS | 2 | 3589 | 2fh3 | A | 520 | TYR | p.Tyr547Cys | P06396 |
| HOMO SAPIENS | 1 | 3589 | 2fh3 | A | 529 | GLY | p.Gly556Arg | P06396 |
| HOMO SAPIENS | 2 | 3589 | 2fh3 | A | 536 | THR | p.Thr563Ser | P06396 |
| HOMO SAPIENS | 2 | 3589 | 2fh3 | A | 579 | VAL | p.Val606Met | P06396 |

| | | | | | | | | |
|---|---|---|---|---|---|---|---|---|
| HOMO SAPIENS | 1 | 3589 | 2fh3 | A | 589 | THR | p.Thr616Met | P06396 |
| HOMO SAPIENS | 5 | 3589 | 2fh3 | A | 668 | THR | p.Thr695Met | P06396 |
| HOMO SAPIENS | 0 | 148 | 1lcl | A | 130 | ILE | p.Ile130Ser | Q05315 |
| HOMO SAPIENS | -1 | 24727 | 4fj3 | P | 237 | ALA | p.Ala237Thr | P04049 |
| HOMO SAPIENS | 0 | 24727 | 4fj3 | P | 257 | SER | p.Ser257Leu | P04049 |
| HOMO SAPIENS | 0 | 24727 | 4fj3 | P | 261 | PRO | p.Pro261Ala | P04049 |
| HOMO SAPIENS | 0 | 2949 | 5xtb | B | 77 | ARG | p.Arg77Gly | O00217 |
| HOMO SAPIENS | 0 | 2949 | 5xtb | B | 79 | PRO | p.Pro79Leu | O00217 |
| HOMO SAPIENS | 0 | 2949 | 5xtb | B | 100 | ALA | p.Ala100Val | O00217 |
| HOMO SAPIENS | 1 | 2949 | 5xtb | B | 115 | LYS | p.Lys115Glu | O00217 |
| HOMO SAPIENS | 0 | 2949 | 5xtb | B | 138 | ARG | p.Arg138His | O00217 |
| HOMO SAPIENS | 1 | 3959 | 5k5s | B | 25 | ARG | p.Arg25Ter | P41180 |
| HOMO SAPIENS | 4 | 3959 | 5k5s | B | 39 | PRO | p.Pro39Ala | P41180 |
| HOMO SAPIENS | 0 | 3959 | 5k5s | B | 47 | LYS | p.Lys47Asn | P41180 |
| HOMO SAPIENS | 2 | 3959 | 5k5s | B | 62 | ARG | p.Arg62Met | P41180 |
| HOMO SAPIENS | 2 | 3959 | 5k5s | B | 66 | ARG | p.Arg66Cys | P41180 |
| HOMO SAPIENS | 4 | 3959 | 5k5s | B | 90 | ASN | p.Asn90Thr | P41180 |
| HOMO SAPIENS | 1 | 3959 | 5k5s | B | 116 | ALA | p.Ala116Thr | P41180 |
| HOMO SAPIENS | 1 | 3959 | 5k5s | B | 118 | ASN | p.Asn118Lys | P41180 |
| HOMO SAPIENS | -1 | 3959 | 5k5s | B | 125 | LEU | p.Leu125Pro | P41180 |
| HOMO SAPIENS | -1 | 3959 | 5k5s | B | 127 | GLU | p.Glu127Ala | P41180 |
| HOMO SAPIENS | -1 | 3959 | 5k5s | B | 128 | PHE | p.Phe128Leu | P41180 |
| HOMO SAPIENS | -1 | 3959 | 5k5s | B | 131 | CYS | p.Cys131Trp | P41180 |
| HOMO SAPIENS | 1 | 3959 | 5k5s | B | 138 | THR | p.Thr138Met | P41180 |
| HOMO SAPIENS | 0 | 3959 | 5k5s | B | 143 | GLY | p.Gly143Glu | P41180 |
| HOMO SAPIENS | 5 | 3959 | 5k5s | B | 151 | THR | p.Thr151Met | P41180 |
| HOMO SAPIENS | 4 | 3959 | 5k5s | B | 180 | PHE | p.Phe180Cys | P41180 |
| HOMO SAPIENS | 0 | 3959 | 5k5s | B | 185 | ARG | p.Arg185Gln | P41180 |
| HOMO SAPIENS | 0 | 3959 | 5k5s | B | 185 | ARG | p.Arg185Ter | P41180 |
| HOMO SAPIENS | 2 | 3959 | 5k5s | B | 191 | GLU | p.Glu191Lys | P41180 |
| HOMO SAPIENS | 2 | 3959 | 5k5s | B | 227 | ARG | p.Arg227Leu | P41180 |
| HOMO SAPIENS | 2 | 3959 | 5k5s | B | 250 | GLU | p.Glu250Lys | P41180 |
| HOMO SAPIENS | 6 | 3959 | 5k5s | B | 297 | GLU | p.Glu297Lys | P41180 |
| HOMO SAPIENS | 1 | 3959 | 5k5s | B | 336 | LYS | p.Lys336Asn | P41180 |
| HOMO SAPIENS | 6 | 3959 | 5k5s | B | 429 | HIS | p.His429Tyr | P41180 |
| HOMO SAPIENS | 0 | 3959 | 5k5s | B | 445 | THR | p.Thr445Ala | P41180 |
| HOMO SAPIENS | 3 | 3959 | 5k5s | B | 509 | GLY | p.Gly509Arg | P41180 |
| HOMO SAPIENS | 1 | 3959 | 5k5s | B | 553 | GLY | p.Gly553Arg | P41180 |
| HOMO SAPIENS | 3 | 3959 | 5k5s | B | 592 | ASN | p.Asn592Ser | P41180 |
| HOMO SAPIENS | -1 | 3959 | 5k5s | B | 604 | GLU | p.Glu604Lys | P41180 |
| HOMO SAPIENS | 4 | 2564 | 5l0c | C | 975 | ARG | p.Arg975Trp | P18206 |
| HOMO SAPIENS | 4 | 2564 | 5l0c | C | 975 | ARG | p.Arg975Leu | P18206 |
| HOMO SAPIENS | 1 | 5695 | 5i8r | A | 294 | ARG | p.Arg294Gln | P17405 |
| HOMO SAPIENS | 0 | 5695 | 5i8r | A | 485 | ALA | p.Ala485Val | P17405 |
| HOMO SAPIENS | 1 | 5695 | 5i8r | A | 506 | GLY | p.Gly506Arg | P17405 |
| HOMO SAPIENS | -1 | 1978 | 3avs | A | 1060 | HIS | p.His1060Leu | O15550 |
| HOMO SAPIENS | 0 | 11173 | 4v0c | B | 362 | LYS | p.Lys362Arg | P51787 |
| HOMO SAPIENS | 3 | 11173 | 4v0c | B | 381 | CYS | p.Cys381Tyr | P51787 |
| HOMO SAPIENS | 7 | 11173 | 4v0c | B | 518 | ARG | p.Arg518Gly | P51787 |
| HOMO SAPIENS | 2 | 11173 | 4v0c | B | 525 | ALA | p.Ala525Thr | P51787 |
| HOMO SAPIENS | 0 | 11173 | 4v0c | B | 530 | GLN | p.Gln530Ter | P51787 |
| HOMO SAPIENS | -1 | 11173 | 4v0c | B | 536 | TYR | p.Tyr536Ter | P51787 |
| HOMO SAPIENS | 2 | 1113 | 5xtd | N | 60 | ARG | p.Arg60Ter | Q9UI09 |
| HOMO SAPIENS | 3 | 6061 | 2mmz | A | 148 | LEU | p.Leu148Ser | Q86SX6 |

| | | | | | | | | |
|---|---|---|---|---|---|---|---|---|
| HOMO SAPIENS | -1 | 26081 | 3eu7 | A | 837 | GLU | p.Glu837Ter | Q86YC2 |
| HOMO SAPIENS | 0 | 26081 | 3eu7 | A | 864 | PRO | p.Pro864Ser | Q86YC2 |
| HOMO SAPIENS | 5 | 26081 | 3eu7 | A | 891 | CYS | p.Cys891Trp | Q86YC2 |
| HOMO SAPIENS | 0 | 26081 | 3eu7 | A | 932 | VAL | p.Val932Met | Q86YC2 |
| HOMO SAPIENS | 5 | 26081 | 3eu7 | A | 961 | LEU | p.Leu961Met | Q86YC2 |
| HOMO SAPIENS | 5 | 26081 | 3eu7 | A | 1038 | TRP | p.Trp1038Ter | Q86YC2 |
| HOMO SAPIENS | -1 | 26081 | 3eu7 | A | 1086 | ARG | p.Arg1086Ter | Q86YC2 |
| HOMO SAPIENS | 0 | 26081 | 3eu7 | A | 1122 | ASP | p.Asp1122Glu | Q86YC2 |
| HOMO SAPIENS | 5 | 26081 | 3eu7 | A | 1158 | HIS | p.His1158Gln | Q86YC2 |
| HOMO SAPIENS | 5 | 26081 | 3eu7 | A | 1183 | TYR | p.Tyr1183Ter | Q86YC2 |
| HOMO SAPIENS | 4 | 199 | 1qo5 | C | 45 | ARG | p.Arg46Trp | P05062 |
| HOMO SAPIENS | 5 | 199 | 1qo5 | C | 147 | TRP | p.Trp148Arg | P05062 |
| HOMO SAPIENS | 0 | 199 | 1qo5 | C | 149 | ALA | p.Ala150Pro | P05062 |
| HOMO SAPIENS | 0 | 199 | 1qo5 | C | 174 | ALA | p.Ala175Asp | P05062 |
| HOMO SAPIENS | 0 | 199 | 1qo5 | C | 334 | ASN | p.Asn335Lys | P05062 |
| HOMO SAPIENS | 0 | 199 | 1qo5 | C | 337 | ALA | p.Ala338Val | P05062 |
| HOMO SAPIENS | 5 | 199 | 1qo5 | C | 342 | TYR | p.Tyr343His | P05062 |
| HOMO SAPIENS | 0 | 6910 | 4zht | B | 59 | ARG | p.Arg59Ter | Q9Y223 |
| HOMO SAPIENS | 4 | 6910 | 4zht | B | 129 | ARG | p.Arg129Ter | Q9Y223 |
| HOMO SAPIENS | 4 | 6910 | 4zht | B | 129 | ARG | p.Arg129Gln | Q9Y223 |
| HOMO SAPIENS | 1 | 6910 | 4zht | B | 162 | ARG | p.Arg162Cys | Q9Y223 |
| HOMO SAPIENS | 0 | 6910 | 4zht | B | 176 | ASP | p.Asp176Val | Q9Y223 |
| HOMO SAPIENS | 0 | 6910 | 4zht | B | 216 | VAL | p.Val216Ala | Q9Y223 |
| HOMO SAPIENS | 0 | 6910 | 4zht | B | 246 | ARG | p.Arg246Trp | Q9Y223 |
| HOMO SAPIENS | 0 | 6910 | 4zht | B | 263 | ARG | p.Arg263Leu | Q9Y223 |
| HOMO SAPIENS | 0 | 6910 | 4zht | B | 266 | ARG | p.Arg266Gln | Q9Y223 |
| HOMO SAPIENS | 0 | 6910 | 4zht | B | 266 | ARG | p.Arg266Trp | Q9Y223 |
| HOMO SAPIENS | 0 | 6910 | 4zht | B | 277 | ARG | p.Arg277Cys | Q9Y223 |
| HOMO SAPIENS | 0 | 6910 | 4zht | B | 378 | ASP | p.Asp378Tyr | Q9Y223 |
| HOMO SAPIENS | 3 | 211 | 5kxi | C | 262 | VAL | p.Val287Leu | P17787 |
| HOMO SAPIENS | 0 | 211 | 5kxi | C | 369 | PHE | p.Phe478Leu | P17787 |
| HOMO SAPIENS | 0 | 2454 | 5iy9 | V | null | PHE | p.Phe99Ser | P19447 |
| HOMO SAPIENS | 0 | 2454 | 5iy9 | V | null | ARG | p.Arg109Ter | P19447 |
| HOMO SAPIENS | 0 | 2454 | 5iy9 | V | null | THR | p.Thr119Pro | P19447 |
| HOMO SAPIENS | 0 | 2454 | 5iy9 | V | 425 | ARG | p.Arg425Ter | P19447 |
| HOMO SAPIENS | 1 | 2454 | 5iy9 | V | 545 | GLN | p.Gln545Ter | P19447 |
| HOMO SAPIENS | -1 | 3805 | 3rbq | A | 57 | LYS | p.Lys57Ter | Q13432 |
| HOMO SAPIENS | 2 | 2649 | 4ido | A | 43 | ASP | p.Asp43Glu | Q8WXF7 |
| HOMO SAPIENS | 2 | 2649 | 4ido | A | 157 | LEU | p.Leu157Trp | Q8WXF7 |
| HOMO SAPIENS | 2 | 2649 | 4ido | A | 193 | PHE | p.Phe193Cys | Q8WXF7 |
| HOMO SAPIENS | 7 | 2649 | 4ido | A | 217 | ARG | p.Arg217Gln | Q8WXF7 |
| HOMO SAPIENS | 4 | 2649 | 4ido | A | 239 | ARG | p.Arg239Cys | Q8WXF7 |
| HOMO SAPIENS | 3 | 2649 | 4ido | A | 258 | HIS | p.His258Arg | Q8WXF7 |
| HOMO SAPIENS | 4 | 2649 | 4ido | A | 259 | SER | p.Ser259Tyr | Q8WXF7 |
| HOMO SAPIENS | 4 | 2649 | 4ido | A | 408 | MET | p.Met408Val | Q8WXF7 |
| HOMO SAPIENS | 2 | 2649 | 4ido | A | 415 | ARG | p.Arg415Trp | Q8WXF7 |
| HOMO SAPIENS | 3 | 2649 | 4ido | A | 416 | ARG | p.Arg416Cys | Q8WXF7 |
| HOMO SAPIENS | 5 | 2203 | 3wff | A | 810 | LEU | p.Ser810Leu | P08235 |
| HOMO SAPIENS | 4 | 12648 | 5ep6 | B | 696 | GLU | p.Glu696Lys | Q9UHD2 |
| HOMO SAPIENS | 0 | 30622 | 2pe4 | A | 90 | GLU | p.Glu90Asp | Q12794 |
| HOMO SAPIENS | 0 | 30622 | 2pe4 | A | 256 | GLY | p.Gly256Arg | Q12794 |
| HOMO SAPIENS | 1 | 30622 | 2pe4 | A | 268 | GLU | p.Glu268Lys | Q12794 |
| HOMO SAPIENS | 0 | 30622 | 2pe4 | A | 392 | THR | p.Thr392Met | Q12794 |
| HOMO SAPIENS | 5 | 5411 | 1lya | A | 93 | SER | p.Ser157Leu | P07339 |

| | | | | | | | | |
|---|---|---|---|---|---|---|---|---|
| HOMO SAPIENS | 3 | 5743 | 2uzp | A | 189 | ASN | p.Asn189Lys | P05129 |
| HOMO SAPIENS | 1 | 12529 | 3kfd | J | 21 | ASN | p.Asn45Ser | P36897 |
| HOMO SAPIENS | 1 | 1063 | 2e1q | B | 149 | ARG | p.Arg149Cys | P47989 |
| HOMO SAPIENS | -1 | 1063 | 2e1q | B | 172 | GLY | p.Gly172Arg | P47989 |
| HOMO SAPIENS | 4 | 1063 | 2e1q | B | 228 | ARG | p.Arg228Ter | P47989 |
| HOMO SAPIENS | 5 | 1063 | 2e1q | B | 395 | LYS | p.Lys395Met | P47989 |
| HOMO SAPIENS | 2 | 1063 | 2e1q | B | 646 | ILE | p.Ile646Val | P47989 |
| HOMO SAPIENS | 4 | 1063 | 2e1q | B | 703 | ILE | p.Ile703Val | P47989 |
| HOMO SAPIENS | 8 | 2525 | 1kqn | F | 18 | ASN | p.Asn18Ser | Q9HAN9 |
| HOMO SAPIENS | 0 | 2525 | 1kqn | F | 151 | VAL | p.Val151Phe | Q9HAN9 |
| HOMO SAPIENS | 0 | 2525 | 1kqn | F | 153 | LEU | p.Leu153Val | Q9HAN9 |
| HOMO SAPIENS | 0 | 2525 | 1kqn | F | 169 | TRP | p.Trp169Ter | Q9HAN9 |
| HOMO SAPIENS | 0 | 2525 | 1kqn | F | 207 | ARG | p.Arg207Trp | Q9HAN9 |
| HOMO SAPIENS | 0 | 2525 | 1kqn | F | 273 | ASN | p.Asn273Asp | Q9HAN9 |
| HOMO SAPIENS | 0 | 12299 | 5duq | A | 432 | VAL | p.Val454Glu | P05155 |
| HOMO SAPIENS | 0 | 12299 | 5duq | A | 436 | ALA | p.Ala458Thr | P05155 |
| HOMO SAPIENS | 0 | 12299 | 5duq | A | 443 | ALA | p.Ala465Val | P05155 |
| HOMO SAPIENS | 0 | 12299 | 5duq | A | 444 | ARG | p.Arg466His | P05155 |
| HOMO SAPIENS | 0 | 12299 | 5duq | A | 458 | MET | p.Val480Met | P05155 |
| HOMO SAPIENS | 0 | 327 | 1uhp | A | 36 | ASP | p.Asp274Asn | Q9UPQ7 |
| HOMO SAPIENS | 0 | 64 | 2mo5 | A | 54 | ALA | p.Ala55Thr | P12104 |
| HOMO SAPIENS | 2 | 35969 | 1jrh | I | 60 | CYS | p.Cys77Tyr | P15260 |
| HOMO SAPIENS | 4 | 35969 | 1jrh | I | 70 | ILE | p.Ile87Thr | P15260 |
| HOMO SAPIENS | 2 | 1068 | 1mu7 | A | 304 | ARG | p.Arg304Gln | Q9NUW8 |
| HOMO SAPIENS | 8 | 1068 | 1mu7 | A | 493 | HIS | p.His493Arg | Q9NUW8 |
| HOMO SAPIENS | 0 | 32360 | 2kqp | A | 31 | ARG | p.Arg55Cys | P01308 |
| HOMO SAPIENS | 0 | 32360 | 2kqp | A | 65 | ARG | p.Arg89His | P01308 |
| HOMO SAPIENS | 0 | 32360 | 2kqp | A | 65 | ARG | p.Arg89Cys | P01308 |
| HOMO SAPIENS | 6 | 10805 | 3nks | A | 12 | ILE | p.Ile12Thr | P50336 |
| HOMO SAPIENS | 0 | 10805 | 3nks | A | 20 | HIS | p.His20Pro | P50336 |
| HOMO SAPIENS | 0 | 10805 | 3nks | A | 59 | ARG | p.Arg59Trp | P50336 |
| HOMO SAPIENS | 0 | 10805 | 3nks | A | 168 | ARG | p.Arg168Cys | P50336 |
| HOMO SAPIENS | 0 | 10805 | 3nks | A | 168 | ARG | p.Arg168His | P50336 |
| HOMO SAPIENS | 2 | 10805 | 3nks | A | 232 | GLY | p.Gly232Arg | P50336 |
| HOMO SAPIENS | 3 | 10805 | 3nks | A | 256 | PRO | p.Pro256Arg | P50336 |
| HOMO SAPIENS | 0 | 10805 | 3nks | A | 304 | ARG | p.Arg304His | P50336 |
| HOMO SAPIENS | 0 | 10805 | 3nks | A | 349 | ASP | p.Asp349Ala | P50336 |
| HOMO SAPIENS | 9 | 7008 | 3u2l | A | 125 | GLN | p.Gln125Ter | P55789 |
| HOMO SAPIENS | 4 | 7008 | 3u2l | A | 172 | HIS | p.His172Asn | P55789 |
| HOMO SAPIENS | 4 | 7008 | 3u2l | A | 194 | ARG | p.Arg194His | P55789 |
| HOMO SAPIENS | 3 | 7008 | 3u2l | A | 198 | GLY | p.Gly198Ser | P55789 |
| HOMO SAPIENS | 0 | 20821 | 3e0g | A | 65 | HIS | p.His116Tyr | P42702 |
| HOMO SAPIENS | 3 | 1151 | 5l9v | A | 278 | ASP | p.Asp278Asn | Q9GZT9 |
| HOMO SAPIENS | 4 | 1151 | 5l9v | A | 317 | CYS | p.Pro317Arg | Q9GZT9 |
| HOMO SAPIENS | 5 | 1151 | 5l9v | A | 371 | ARG | p.Arg371His | Q9GZT9 |
| HOMO SAPIENS | 2 | 3935 | 1gwb | A | 36 | ARG | p.Arg36Gly | P07359 |
| HOMO SAPIENS | 1 | 3935 | 1gwb | A | 73 | LEU | p.Leu73Phe | P07359 |
| HOMO SAPIENS | 1 | 3935 | 1gwb | A | 172 | ALA | p.Ala172Val | P07359 |
| HOMO SAPIENS | 0 | 3935 | 1gwb | A | 249 | GLY | p.Gly249Val | P07359 |
| HOMO SAPIENS | 0 | 3935 | 1gwb | A | 255 | MET | p.Met255Val | P07359 |
| HOMO SAPIENS | 2 | 14541 | 5cuz | A | 138 | GLU | p.Glu138Lys | Q9H3L0 |
| HOMO SAPIENS | 2 | 14541 | 5cuz | A | 143 | SER | p.Ser143Ile | Q9H3L0 |
| HOMO SAPIENS | 4 | 14541 | 5cuz | A | 182 | THR | p.Thr182Asn | Q9H3L0 |
| HOMO SAPIENS | 6 | 14541 | 5cuz | A | 206 | ASN | p.Asn206Ile | Q9H3L0 |

| | | | | | | | | |
|---|---|---|---|---|---|---|---|---|
| HOMO SAPIENS | 0 | 14541 | 5cuz | A | 249 | TYR | p.Tyr249Cys | Q9H3L0 |
| HOMO SAPIENS | 4 | 14541 | 5cuz | A | 250 | ARG | p.Arg250Ter | Q9H3L0 |
| HOMO SAPIENS | 2 | 14541 | 5cuz | A | 259 | LEU | p.Leu259Pro | Q9H3L0 |
| HOMO SAPIENS | 0 | 4206 | 4fao | D | 30 | PRO | p.Pro30Ser | P37023 |
| HOMO SAPIENS | 1 | 4206 | 4fao | D | 50 | TRP | p.Trp50Cys | P37023 |
| HOMO SAPIENS | 0 | 16553 | 3v30 | A | 251 | GLN | p.Gln251Glu | O14593 |
| HOMO SAPIENS | 0 | 5012 | 5hbf | A | 442 | ALA | p.Ala442Gly | Q13231 |
| HOMO SAPIENS | 5 | 4844 | 5tdc | A | 136 | HIS | p.His136Arg | Q8IWV7 |
| HOMO SAPIENS | 0 | 15435 | 4cgz | A | 643 | ARG | p.Arg643His | P54132 |
| HOMO SAPIENS | 1 | 15435 | 4cgz | A | 672 | GLN | p.Gln672Arg | P54132 |
| HOMO SAPIENS | 1 | 15435 | 4cgz | A | 700 | GLN | p.Gln700Ter | P54132 |
| HOMO SAPIENS | 0 | 15435 | 4cgz | A | 707 | PRO | p.Pro707Ser | P54132 |
| HOMO SAPIENS | 0 | 15435 | 4cgz | A | 755 | LYS | p.Lys755Glu | P54132 |
| HOMO SAPIENS | 0 | 15435 | 4cgz | A | 881 | TRP | p.Trp881Ter | P54132 |
| HOMO SAPIENS | 2 | 15435 | 4cgz | A | 899 | ARG | p.Arg899Ter | P54132 |
| HOMO SAPIENS | 0 | 15435 | 4cgz | A | 1036 | CYS | p.Cys1036Phe | P54132 |
| HOMO SAPIENS | 0 | 15435 | 4cgz | A | 1055 | CYS | p.Cys1055Ser | P54132 |
| HOMO SAPIENS | 0 | 15435 | 4cgz | A | 1139 | ARG | p.Arg1139Ter | P54132 |
| HOMO SAPIENS | -1 | 15435 | 4cgz | A | 1198 | VAL | p.Val1198Met | P54132 |
| HOMO SAPIENS | -1 | 15435 | 4cgz | A | 1205 | VAL | p.Val1205Ile | P54132 |
| HOMO SAPIENS | 0 | 15435 | 4cgz | A | 1251 | GLU | p.Glu1251Gln | P54132 |
| HOMO SAPIENS | 0 | 11185 | 1kb4 | A | 30 | ARG | p.Arg30Ter | P11473 |
| HOMO SAPIENS | 5 | 11185 | 1kb4 | A | 33 | GLY | p.Gly33Asp | P11473 |
| HOMO SAPIENS | 5 | 11185 | 1kb4 | A | 46 | GLY | p.Gly46Asp | P11473 |
| HOMO SAPIENS | 6 | 11185 | 1kb4 | A | 50 | ARG | p.Arg50Gln | P11473 |
| HOMO SAPIENS | 9 | 11185 | 1kb4 | A | 73 | ARG | p.Arg73Gln | P11473 |
| HOMO SAPIENS | 9 | 11185 | 1kb4 | A | 80 | ARG | p.Arg80Gln | P11473 |
| HOMO SAPIENS | 0 | 5382 | 3fl7 | A | 41 | LEU | p.Leu41Val | P29317 |
| HOMO SAPIENS | 2 | 5382 | 3fl7 | A | 175 | ARG | p.Arg175Cys | P29317 |
| HOMO SAPIENS | 1 | 5382 | 3fl7 | A | 195 | ARG | p.Arg195Cys | P29317 |
| HOMO SAPIENS | 0 | 5382 | 3fl7 | A | 232 | ASP | p.Asp232Gly | P29317 |
| HOMO SAPIENS | 0 | 5382 | 3fl7 | A | 277 | SER | p.Ser277Leu | P29317 |
| HOMO SAPIENS | 0 | 5382 | 3fl7 | A | 315 | ARG | p.Arg315Gln | P29317 |
| HOMO SAPIENS | 7 | 5382 | 3fl7 | A | 350 | PRO | p.Pro350Thr | P29317 |
| HOMO SAPIENS | 0 | 5382 | 3fl7 | A | 391 | GLY | p.Gly391Arg | P29317 |
| HOMO SAPIENS | 0 | 5382 | 3fl7 | A | 418 | VAL | p.Val418Ile | P29317 |
| HOMO SAPIENS | 0 | 5382 | 3fl7 | A | 511 | THR | p.Thr511Met | P29317 |
| HOMO SAPIENS | 2 | 29168 | 4p22 | B | 144 | VAL | p.Val144Ile | P22314 |
| HOMO SAPIENS | 5 | 4348 | 1bf5 | A | 241 | ARG | p.Arg241Gln | P42224 |
| HOMO SAPIENS | 2 | 4348 | 1bf5 | A | 266 | VAL | p.Val266Ile | P42224 |
| HOMO SAPIENS | 4 | 4348 | 1bf5 | A | 320 | GLU | p.Glu320Gln | P42224 |
| HOMO SAPIENS | 1 | 4348 | 1bf5 | A | 372 | ASN | p.Asn372Lys | P42224 |
| HOMO SAPIENS | 1 | 4348 | 1bf5 | A | 447 | ASP | p.Asp447Glu | P42224 |
| HOMO SAPIENS | 2 | 4348 | 1bf5 | A | 463 | GLN | p.Gln463His | P42224 |
| HOMO SAPIENS | 4 | 4348 | 1bf5 | A | 600 | LEU | p.Leu600Pro | P42224 |
| HOMO SAPIENS | 0 | 4348 | 1bf5 | A | 706 | LEU | p.Leu706Ser | P42224 |
| HOMO SAPIENS | 4 | 4991 | 4bej | B | 141 | ASN | p.Asn141Ser | O00429 |
| HOMO SAPIENS | 0 | 4991 | 4bej | B | 395 | ALA | p.Ala395Asp | O00429 |
| HOMO SAPIENS | 2 | 11792 | 1fvo | A | 40 | ARG | p.Arg40Cys | P00480 |
| HOMO SAPIENS | 2 | 11792 | 1fvo | A | 40 | ARG | p.Arg40His | P00480 |
| HOMO SAPIENS | 0 | 11792 | 1fvo | A | 45 | LEU | p.Leu45Pro | P00480 |
| HOMO SAPIENS | 1 | 11792 | 1fvo | A | 47 | ASN | p.Asn47Ile | P00480 |
| HOMO SAPIENS | 0 | 11792 | 1fvo | A | 50 | GLY | p.Gly50Arg | P00480 |
| HOMO SAPIENS | 5 | 11792 | 1fvo | A | 87 | GLU | p.Glu87Lys | P00480 |

| | | | | | | | | |
|---|---|---|---|---|---|---|---|---|
| HOMO SAPIENS | 3 | 11792 | 1fvo | A | 111 | LEU | p.Leu111Pro | P00480 |
| HOMO SAPIENS | 1 | 11792 | 1fvo | A | 125 | THR | p.Thr125Met | P00480 |
| HOMO SAPIENS | 3 | 11792 | 1fvo | A | 129 | ARG | p.Arg129His | P00480 |
| HOMO SAPIENS | 9 | 11792 | 1fvo | A | 141 | ARG | p.Arg141Pro | P00480 |
| HOMO SAPIENS | 0 | 11792 | 1fvo | A | 206 | MET | p.Met206Arg | P00480 |
| HOMO SAPIENS | 0 | 11792 | 1fvo | A | 207 | SER | p.Ser207Arg | P00480 |
| HOMO SAPIENS | 0 | 11792 | 1fvo | A | 216 | GLN | p.Gln216Glu | P00480 |
| HOMO SAPIENS | 1 | 11792 | 1fvo | A | 225 | PRO | p.Pro225Leu | P00480 |
| HOMO SAPIENS | 0 | 11792 | 1fvo | A | 270 | ARG | p.Gln270Arg | P00480 |
| HOMO SAPIENS | 0 | 11792 | 1fvo | A | 270 | ARG | p.Gln270Ter | P00480 |
| HOMO SAPIENS | 2 | 11792 | 1fvo | A | 277 | ARG | p.Arg277Trp | P00480 |
| HOMO SAPIENS | 0 | 3721 | 4r3z | B | 2 | ASP | p.Asp2Gly | P54136 |
| HOMO SAPIENS | 0 | 3721 | 4r3z | B | 456 | SER | p.Ser456Leu | P54136 |
| HOMO SAPIENS | 5 | 3721 | 4r3z | B | 512 | ARG | p.Arg512Gln | P54136 |
| HOMO SAPIENS | 3 | 5749 | 3hl2 | A | 239 | ALA | p.Ala239Thr | Q9HD40 |
| HOMO SAPIENS | 1 | 5749 | 3hl2 | A | 334 | TYR | p.Tyr334Cys | Q9HD40 |
| HOMO SAPIENS | 1 | 5749 | 3hl2 | A | 452 | LYS | p.Lys452Asn | Q9HD40 |
| HOMO SAPIENS | -1 | 5749 | 3hl2 | A | 465 | ARG | p.Arg465Leu | Q9HD40 |
| HOMO SAPIENS | -1 | 5749 | 3hl2 | A | 489 | ASP | p.Asp489Gly | Q9HD40 |
| HOMO SAPIENS | 5 | 1721 | 3k15 | A | 1652 | MET | p.Met1652Ile | P38398 |
| HOMO SAPIENS | 5 | 1721 | 3k15 | A | 1652 | MET | p.Met1652Lys | P38398 |
| HOMO SAPIENS | 5 | 1721 | 3k15 | A | 1662 | PHE | p.Phe1662Cys | P38398 |
| HOMO SAPIENS | 5 | 1721 | 3k15 | A | 1664 | LEU | p.Leu1664Pro | P38398 |
| HOMO SAPIENS | 0 | 1721 | 3k15 | A | 1665 | VAL | p.Val1665Met | P38398 |
| HOMO SAPIENS | 5 | 1721 | 3k15 | A | 1690 | LYS | p.Lys1690Gln | P38398 |
| HOMO SAPIENS | 5 | 1721 | 3k15 | A | 1692 | ASP | p.Asp1692Asn | P38398 |
| HOMO SAPIENS | 5 | 1721 | 3k15 | A | 1699 | ARG | p.Arg1699Gln | P38398 |
| HOMO SAPIENS | 5 | 1721 | 3k15 | A | 1699 | ARG | p.Arg1699Trp | P38398 |
| HOMO SAPIENS | 3 | 1721 | 3k15 | A | 1706 | GLY | p.Gly1706Glu | P38398 |
| HOMO SAPIENS | 5 | 1721 | 3k15 | A | 1720 | THR | p.Thr1720Ala | P38398 |
| HOMO SAPIENS | 5 | 1721 | 3k15 | A | 1751 | ARG | p.Arg1751Pro | P38398 |
| HOMO SAPIENS | 4 | 1721 | 3k15 | A | 1775 | MET | p.Met1775Arg | P38398 |
| HOMO SAPIENS | 2 | 1721 | 3k15 | A | 1788 | GLY | p.Gly1788Val | P38398 |
| HOMO SAPIENS | 5 | 1721 | 3k15 | A | 1794 | GLU | p.Glu1794Asp | P38398 |
| HOMO SAPIENS | 5 | 1721 | 3k15 | A | 1804 | VAL | p.Val1804Asp | P38398 |
| HOMO SAPIENS | 3 | 1721 | 3k15 | A | 1815 | TRP | p.Trp1815Ter | P38398 |
| HOMO SAPIENS | 1 | 1721 | 3k15 | A | 1819 | ASN | p.Asn1819Ser | P38398 |
| HOMO SAPIENS | 5 | 1721 | 3k15 | A | 1856 | PRO | p.Pro1856Ser | P38398 |
| HOMO SAPIENS | 5 | 1721 | 3k15 | A | 1858 | ILE | p.Ile1858Leu | P38398 |
| HOMO SAPIENS | 5 | 1721 | 3k15 | A | 1858 | ILE | p.Ile1858Thr | P38398 |
| HOMO SAPIENS | 5 | 1721 | 3k15 | A | 1859 | PRO | p.Pro1859Arg | P38398 |
| HOMO SAPIENS | 2 | 22086 | 3kq4 | B | 1297 | PRO | p.Pro1297Leu | O60494 |
| HOMO SAPIENS | 1 | 12135 | 2k9j | B | 724 | ARG | p.Arg750Ter | P05106 |
| HOMO SAPIENS | -1 | 24027 | 2qtl | A | 175 | LEU | p.Ser202Leu | Q9UBK8 |
| HOMO SAPIENS | 0 | 24027 | 2qtl | A | 257 | SER | p.Ser284Thr | Q9UBK8 |
| HOMO SAPIENS | 0 | 24027 | 2qtl | A | 333 | LEU | p.Leu360Val | Q9UBK8 |
| HOMO SAPIENS | 0 | 24027 | 2qtl | A | 350 | LYS | p.Lys377Arg | Q9UBK8 |
| HOMO SAPIENS | 0 | 24027 | 2qtl | A | 415 | ARG | p.Arg442Cys | Q9UBK8 |
| HOMO SAPIENS | 2 | 24027 | 2qtl | A | 450 | PRO | p.Pro477Arg | Q9UBK8 |
| HOMO SAPIENS | 4 | 24027 | 2qtl | A | 487 | GLY | p.Gly514Arg | Q9UBK8 |
| HOMO SAPIENS | 1 | 24027 | 2qtl | A | 595 | HIS | p.His622Tyr | Q9UBK8 |
| HOMO SAPIENS | 0 | 5758 | 3di2 | B | 46 | THR | p.Thr66Ile | P16871 |
| HOMO SAPIENS | 1 | 5758 | 3di2 | B | 118 | VAL | p.Ile138Val | P16871 |
| HOMO SAPIENS | 4 | 848 | 5oa3 | Y | 79 | LEU | p.Leu79Val | P62847 |

| | | | | | | | | |
|---|---|---|---|---|---|---|---|---|
| HOMO SAPIENS | 2 | 848 | 5oa3 | Y | 124 | ASN | p.Asn124Ser | P62847 |
| HOMO SAPIENS | 0 | 81 | 3s5o | A | 36 | ALA | p.Ala36Glu | Q86XE5 |
| HOMO SAPIENS | 0 | 81 | 3s5o | A | 39 | TYR | p.Tyr39Ter | Q86XE5 |
| HOMO SAPIENS | 0 | 81 | 3s5o | A | 45 | PRO | p.Pro45Leu | Q86XE5 |
| HOMO SAPIENS | 0 | 81 | 3s5o | A | 70 | ARG | p.Arg70Ter | Q86XE5 |
| HOMO SAPIENS | 0 | 81 | 3s5o | A | 70 | ARG | p.Arg70Pro | Q86XE5 |
| HOMO SAPIENS | 1 | 81 | 3s5o | A | 97 | ARG | p.Arg97Cys | Q86XE5 |
| HOMO SAPIENS | 4 | 81 | 3s5o | A | 113 | GLU | p.Glu113Lys | Q86XE5 |
| HOMO SAPIENS | 1 | 81 | 3s5o | A | 190 | PRO | p.Pro190Leu | Q86XE5 |
| HOMO SAPIENS | 0 | 81 | 3s5o | A | 245 | VAL | p.Val245Phe | Q86XE5 |
| HOMO SAPIENS | 0 | 81 | 3s5o | A | 257 | CYS | p.Cys257Gly | Q86XE5 |
| HOMO SAPIENS | 0 | 81 | 3s5o | A | 287 | GLY | p.Gly287Val | Q86XE5 |
| HOMO SAPIENS | 7 | 81 | 3s5o | A | 303 | ARG | p.Arg303Cys | Q86XE5 |
| HOMO SAPIENS | 0 | 81 | 3s5o | A | 325 | GLY | p.Gly325Ser | Q86XE5 |
| HOMO SAPIENS | -1 | 622 | 5era | A | 12 | GLY | p.Gly12Arg | P29033 |
| HOMO SAPIENS | -1 | 622 | 5era | A | 12 | GLY | p.Gly12Asp | P29033 |
| HOMO SAPIENS | -1 | 622 | 5era | A | 17 | SER | p.Ser17Phe | P29033 |
| HOMO SAPIENS | 5 | 622 | 5era | A | 24 | TRP | p.Trp24Ter | P29033 |
| HOMO SAPIENS | 0 | 622 | 5era | A | 32 | ARG | p.Arg32Cys | P29033 |
| HOMO SAPIENS | 5 | 622 | 5era | A | 34 | MET | p.Met34Thr | P29033 |
| HOMO SAPIENS | 5 | 622 | 5era | A | 36 | LEU | p.Leu36Pro | P29033 |
| HOMO SAPIENS | 5 | 622 | 5era | A | 37 | VAL | p.Val37Ile | P29033 |
| HOMO SAPIENS | 5 | 622 | 5era | A | 40 | ALA | p.Ala40Glu | P29033 |
| HOMO SAPIENS | 5 | 622 | 5era | A | 44 | TRP | p.Trp44Cys | P29033 |
| HOMO SAPIENS | 5 | 622 | 5era | A | 45 | GLY | p.Gly45Glu | P29033 |
| HOMO SAPIENS | 5 | 622 | 5era | A | 47 | GLU | p.Glu47Ter | P29033 |
| HOMO SAPIENS | 5 | 622 | 5era | A | 50 | ASP | p.Asp50Asn | P29033 |
| HOMO SAPIENS | 5 | 622 | 5era | A | 54 | ASN | p.Asn54Lys | P29033 |
| HOMO SAPIENS | 5 | 622 | 5era | A | 57 | GLN | p.Gln57Ter | P29033 |
| HOMO SAPIENS | 5 | 622 | 5era | A | 59 | GLY | p.Gly59Ala | P29033 |
| HOMO SAPIENS | 5 | 622 | 5era | A | 59 | GLY | p.Gly59Ser | P29033 |
| HOMO SAPIENS | 5 | 622 | 5era | A | 66 | ASP | p.Asp66His | P29033 |
| HOMO SAPIENS | 5 | 622 | 5era | A | 68 | TYR | p.Tyr68Cys | P29033 |
| HOMO SAPIENS | 5 | 622 | 5era | A | 70 | PRO | p.Pro70Ala | P29033 |
| HOMO SAPIENS | 5 | 622 | 5era | A | 73 | HIS | p.His73Arg | P29033 |
| HOMO SAPIENS | 5 | 622 | 5era | A | 75 | ARG | p.Arg75Trp | P29033 |
| HOMO SAPIENS | 5 | 622 | 5era | A | 75 | ARG | p.Arg75Gln | P29033 |
| HOMO SAPIENS | 2 | 622 | 5era | A | 77 | TRP | p.Trp77Arg | P29033 |
| HOMO SAPIENS | 2 | 622 | 5era | A | 77 | TRP | p.Trp77Ter | P29033 |
| HOMO SAPIENS | 0 | 622 | 5era | A | 80 | GLN | p.Gln80Ter | P29033 |
| HOMO SAPIENS | 0 | 622 | 5era | A | 80 | GLN | p.Gln80Pro | P29033 |
| HOMO SAPIENS | 5 | 622 | 5era | A | 81 | LEU | p.Leu81Val | P29033 |
| HOMO SAPIENS | 5 | 622 | 5era | A | 82 | ILE | p.Ile82Met | P29033 |
| HOMO SAPIENS | 5 | 622 | 5era | A | 83 | PHE | p.Phe83Leu | P29033 |
| HOMO SAPIENS | 5 | 622 | 5era | A | 84 | VAL | p.Val84Leu | P29033 |
| HOMO SAPIENS | 5 | 622 | 5era | A | 90 | LEU | p.Leu90Pro | P29033 |
| HOMO SAPIENS | 5 | 622 | 5era | A | 95 | VAL | p.Val95Met | P29033 |
| HOMO SAPIENS | -1 | 622 | 5era | A | 100 | HIS | p.His100Tyr | P29033 |
| HOMO SAPIENS | -1 | 622 | 5era | A | 113 | SER | p.Ser113Arg | P29033 |
| HOMO SAPIENS | -1 | 622 | 5era | A | 119 | GLU | p.Glu119Lys | P29033 |
| HOMO SAPIENS | -1 | 622 | 5era | A | 122 | LYS | p.Lys122Ile | P29033 |
| HOMO SAPIENS | -1 | 622 | 5era | A | 123 | THR | p.Thr123Asn | P29033 |
| HOMO SAPIENS | -1 | 622 | 5era | A | 124 | GLN | p.Gln124Ter | P29033 |
| HOMO SAPIENS | 5 | 622 | 5era | A | 139 | SER | p.Ser139Asn | P29033 |

| | | | | | | | | |
|---|---|---|---|---|---|---|---|---|
| HOMO SAPIENS | 5 | 622 | 5era | A | 143 | ARG | p.Arg143Trp | P29033 |
| HOMO SAPIENS | 5 | 622 | 5era | A | 143 | ARG | p.Arg143Gln | P29033 |
| HOMO SAPIENS | 0 | 622 | 5era | A | 147 | GLU | p.Glu147Lys | P29033 |
| HOMO SAPIENS | 7 | 622 | 5era | A | 155 | TYR | p.Tyr155Ter | P29033 |
| HOMO SAPIENS | 2 | 622 | 5era | A | 159 | ASP | p.Asp159Val | P29033 |
| HOMO SAPIENS | 5 | 622 | 5era | A | 163 | MET | p.Met163Val | P29033 |
| HOMO SAPIENS | 5 | 622 | 5era | A | 165 | ARG | p.Arg165Trp | P29033 |
| HOMO SAPIENS | 5 | 622 | 5era | A | 167 | VAL | p.Val167Met | P29033 |
| HOMO SAPIENS | 5 | 622 | 5era | A | 169 | CYS | p.Cys169Tyr | P29033 |
| HOMO SAPIENS | 5 | 622 | 5era | A | 179 | ASP | p.Asp179Asn | P29033 |
| HOMO SAPIENS | 0 | 622 | 5era | A | 199 | SER | p.Ser199Phe | P29033 |
| HOMO SAPIENS | 0 | 622 | 5era | A | 202 | CYS | p.Cys202Phe | P29033 |
| HOMO SAPIENS | 5 | 622 | 5era | A | 203 | ILE | p.Ile203Thr | P29033 |
| HOMO SAPIENS | 5 | 622 | 5era | A | 206 | ASN | p.Asn206Ser | P29033 |
| HOMO SAPIENS | 0 | 8103 | 5dot | A | 163 | GLY | p.Gly163Ter | P31327 |
| HOMO SAPIENS | 0 | 8103 | 5dot | A | 337 | HIS | p.His337Arg | P31327 |
| HOMO SAPIENS | 3 | 8103 | 5dot | A | 344 | THR | p.Thr344Ala | P31327 |
| HOMO SAPIENS | 1 | 8103 | 5dot | A | 405 | THR | p.Thr405Asn | P31327 |
| HOMO SAPIENS | 5 | 8103 | 5dot | A | 544 | THR | p.Thr544Met | P31327 |
| HOMO SAPIENS | 5 | 8103 | 5dot | A | 716 | ASN | p.Asn716Lys | P31327 |
| HOMO SAPIENS | -1 | 8103 | 5dot | A | 780 | ARG | p.Arg780His | P31327 |
| HOMO SAPIENS | 0 | 8103 | 5dot | A | 982 | GLY | p.Gly982Asp | P31327 |
| HOMO SAPIENS | 3 | 8103 | 5dot | A | 1119 | ALA | p.Ala1119Ser | P31327 |
| HOMO SAPIENS | 1 | 8103 | 5dot | A | 1376 | GLY | p.Gly1376Ser | P31327 |
| HOMO SAPIENS | 4 | 6833 | 1l8l | B | 27 | ARG | p.Arg27Ser | P78330 |
| HOMO SAPIENS | 3 | 6833 | 1l8l | B | 32 | ASP | p.Asp32Gly | P78330 |
| HOMO SAPIENS | 3 | 6833 | 1l8l | B | 32 | ASP | p.Asp32Asn | P78330 |
| HOMO SAPIENS | 1 | 6833 | 1l8l | B | 49 | ARG | p.Arg49Trp | P78330 |
| HOMO SAPIENS | 0 | 6833 | 1l8l | B | 52 | MET | p.Met52Thr | P78330 |
| HOMO SAPIENS | 0 | 6833 | 1l8l | B | 83 | GLN | p.Gln83His | P78330 |
| HOMO SAPIENS | 6 | 810 | 2iw2 | A | 185 | ARG | p.Arg184Gln | P12955 |
| HOMO SAPIENS | 0 | 810 | 2iw2 | A | 213 | ALA | p.Ala212Pro | P12955 |
| HOMO SAPIENS | 1 | 810 | 2iw2 | A | 266 | ARG | p.Arg265Ter | P12955 |
| HOMO SAPIENS | 9 | 810 | 2iw2 | A | 277 | ASP | p.Asp276Asn | P12955 |
| HOMO SAPIENS | 0 | 810 | 2iw2 | A | 279 | GLY | p.Gly278Asp | P12955 |
| HOMO SAPIENS | 9 | 810 | 2iw2 | A | 378 | HIS | p.His377Gln | P12955 |
| HOMO SAPIENS | 0 | 810 | 2iw2 | A | 389 | ARG | p.Arg388His | P12955 |
| HOMO SAPIENS | 0 | 810 | 2iw2 | A | 436 | PHE | p.Leu435Phe | P12955 |
| HOMO SAPIENS | 0 | 810 | 2iw2 | A | 449 | GLY | p.Gly448Arg | P12955 |
| HOMO SAPIENS | -1 | 1530 | 5fwj | A | 388 | ALA | p.Ala388Pro | P41229 |
| HOMO SAPIENS | 4 | 1530 | 5fwj | A | 451 | SER | p.Ser451Arg | P41229 |
| HOMO SAPIENS | 3 | 1530 | 5fwj | A | 731 | LEU | p.Leu731Phe | P41229 |
| HOMO SAPIENS | 3 | 578 | 5net | B | 123 | ALA | p.Ala143Thr | P18564 |
| HOMO SAPIENS | 6 | 578 | 5net | B | 176 | PRO | p.Pro196Thr | P18564 |
| HOMO SAPIENS | 3 | 578 | 5net | B | 280 | GLU | p.Glu300Lys | P18564 |
| HOMO SAPIENS | -1 | 5247 | 4j15 | A | 274 | ALA | p.Ala274Val | P14868 |
| HOMO SAPIENS | -1 | 5247 | 4j15 | A | 280 | HIS | p.His280Leu | P14868 |
| HOMO SAPIENS | 2 | 5247 | 4j15 | A | 460 | ARG | p.Arg460His | P14868 |
| HOMO SAPIENS | 5 | 5247 | 4j15 | A | 494 | ARG | p.Arg494Cys | P14868 |
| HOMO SAPIENS | 7 | 2566 | 1b55 | A | 28 | ARG | p.Arg28His | Q06187 |
| HOMO SAPIENS | 5 | 2566 | 1b55 | A | 33 | THR | p.Thr33Pro | Q06187 |
| HOMO SAPIENS | 0 | 2566 | 1b55 | A | 113 | VAL | p.Val113Asp | Q06187 |
| HOMO SAPIENS | 0 | 356 | 5a2s | A | 786 | ALA | p.Ala786Thr | P56524 |
| HOMO SAPIENS | 5 | 2960 | 4cak | A | 553 | ARG | p.Arg584Ter | P08514 |

| | | | | | | | | |
|---|---|---|---|---|---|---|---|---|
| HOMO SAPIENS | 5 | 2960 | 4cak | A | 565 | ILE | p.Ile596Thr | P08514 |
| HOMO SAPIENS | -1 | 2960 | 4cak | A | 841 | LEU | p.Leu872Met | P08514 |
| HOMO SAPIENS | -1 | 2960 | 4cak | A | 843 | ILE | p.Ile874Ser | P08514 |
| HOMO SAPIENS | 0 | 5054 | 4k8r | B | 216 | PHE | p.Phe216Leu | Q8NBP7 |
| HOMO SAPIENS | 1 | 5054 | 4k8r | B | 417 | HIS | p.His417Gln | Q8NBP7 |
| HOMO SAPIENS | 1 | 5054 | 4k8r | B | 425 | ASN | p.Asn425Ser | Q8NBP7 |
| HOMO SAPIENS | 1 | 5054 | 4k8r | B | 443 | ALA | p.Ala443Thr | Q8NBP7 |
| HOMO SAPIENS | 0 | 27283 | 3n5n | X | 71 | HIS | p.His82Arg | Q9UIF7 |
| HOMO SAPIENS | 1 | 27283 | 3n5n | X | 83 | ARG | p.Arg94Ter | Q9UIF7 |
| HOMO SAPIENS | 8 | 27283 | 3n5n | X | 89 | TRP | p.Trp100Ter | Q9UIF7 |
| HOMO SAPIENS | 8 | 27283 | 3n5n | X | 90 | TYR | p.Tyr101Ter | Q9UIF7 |
| HOMO SAPIENS | 7 | 27283 | 3n5n | X | 95 | ARG | p.Arg106Trp | Q9UIF7 |
| HOMO SAPIENS | 7 | 27283 | 3n5n | X | 117 | TRP | p.Trp128Arg | Q9UIF7 |
| HOMO SAPIENS | 0 | 27283 | 3n5n | X | 137 | GLY | p.Gly148Arg | Q9UIF7 |
| HOMO SAPIENS | 5 | 27283 | 3n5n | X | 142 | TRP | p.Trp153Ter | Q9UIF7 |
| HOMO SAPIENS | 0 | 27283 | 3n5n | X | 165 | TYR | p.Tyr176Cys | Q9UIF7 |
| HOMO SAPIENS | 0 | 27283 | 3n5n | X | 168 | ARG | p.Arg179His | Q9UIF7 |
| HOMO SAPIENS | 0 | 27283 | 3n5n | X | 168 | ARG | p.Arg179Cys | Q9UIF7 |
| HOMO SAPIENS | 0 | 27283 | 3n5n | X | 196 | GLN | p.Gln207Ter | Q9UIF7 |
| HOMO SAPIENS | 8 | 27283 | 3n5n | X | 201 | VAL | p.Val212Met | Q9UIF7 |
| HOMO SAPIENS | 9 | 27283 | 3n5n | X | 227 | ARG | p.Arg238Trp | Q9UIF7 |
| HOMO SAPIENS | 9 | 27283 | 3n5n | X | 231 | ARG | p.Arg242Cys | Q9UIF7 |
| HOMO SAPIENS | 9 | 27283 | 3n5n | X | 231 | ARG | p.Arg242His | Q9UIF7 |
| HOMO SAPIENS | 3 | 27283 | 3n5n | X | 232 | VAL | p.Val243Phe | Q9UIF7 |
| HOMO SAPIENS | 3 | 27283 | 3n5n | X | 246 | GLN | p.Gln257Ter | Q9UIF7 |
| HOMO SAPIENS | 9 | 27283 | 3n5n | X | 272 | GLY | p.Gly283Glu | Q9UIF7 |
| HOMO SAPIENS | 9 | 27283 | 3n5n | X | 281 | PRO | p.Pro292Leu | Q9UIF7 |
| HOMO SAPIENS | -1 | 27283 | 3n5n | X | 314 | ASP | p.Asp325Glu | Q9UIF7 |
| HOMO SAPIENS | 0 | 27283 | 3n5n | X | 324 | GLN | p.Gln335His | Q9UIF7 |
| HOMO SAPIENS | 0 | 27283 | 3n5n | X | 324 | GLN | p.Gln335Arg | Q9UIF7 |
| HOMO SAPIENS | 0 | 27283 | 3n5n | X | 324 | GLN | p.Gln335Ter | Q9UIF7 |
| HOMO SAPIENS | -1 | 20852 | 4p60 | B | 243 | ARG | p.Arg243Lys | O75746 |
| HOMO SAPIENS | -1 | 1875 | 5dd2 | G | 152 | VAL | p.Val179Leu | P06396 |
| HOMO SAPIENS | 9 | 2126 | 3n1p | B | 46 | PRO | p.Pro46Leu | Q14623 |
| HOMO SAPIENS | 8 | 2126 | 3n1p | B | 76 | GLU | p.Glu76Ala | Q14623 |
| HOMO SAPIENS | 9 | 2126 | 3n1p | B | 77 | ARG | p.Arg77Ser | Q14623 |
| HOMO SAPIENS | 6 | 2126 | 3n1p | B | 95 | GLU | p.Glu95Lys | Q14623 |
| HOMO SAPIENS | 9 | 2126 | 3n1p | B | 100 | ASP | p.Asp100Glu | Q14623 |
| HOMO SAPIENS | 9 | 2126 | 3n1p | B | 131 | GLU | p.Glu131Lys | Q14623 |
| HOMO SAPIENS | 0 | 2126 | 3n1p | B | 190 | VAL | p.Val190Ala | Q14623 |
| HOMO SAPIENS | 5 | 40410 | 3ioz | B | 70 | GLN | p.Gln70Leu | P20963 |
| HOMO SAPIENS | 1 | 5754 | 5lkz | A | 1108 | THR | p.Tyr1191Ter | Q09472 |
| HOMO SAPIENS | 3 | 8159 | 1tiu | A | 3 | GLU | p.Lys12679Glu | Q8WZ42 |
| HOMO SAPIENS | 4 | 3275 | 4ldq | A | 210 | ARG | p.Arg210Gln | Q9BV20 |
| HOMO SAPIENS | 0 | 8143 | 4miv | B | 66 | SER | p.Ser66Trp | P51688 |
| HOMO SAPIENS | 1 | 8143 | 4miv | B | 74 | ARG | p.Arg74Cys | P51688 |
| HOMO SAPIENS | 0 | 8143 | 4miv | B | 128 | PRO | p.Pro128Leu | P51688 |
| HOMO SAPIENS | 1 | 8143 | 4miv | B | 150 | ARG | p.Arg150Gln | P51688 |
| HOMO SAPIENS | 0 | 8143 | 4miv | B | 191 | GLY | p.Gly191Arg | P51688 |
| HOMO SAPIENS | 0 | 8143 | 4miv | B | 206 | ARG | p.Arg206Pro | P51688 |
| HOMO SAPIENS | 0 | 8143 | 4miv | B | 225 | PHE | p.Phe225Leu | P51688 |
| HOMO SAPIENS | 0 | 8143 | 4miv | B | 233 | ARG | p.Arg233Ter | P51688 |
| HOMO SAPIENS | 0 | 8143 | 4miv | B | 234 | ALA | p.Ala234Gly | P51688 |
| HOMO SAPIENS | 0 | 8143 | 4miv | B | 245 | ARG | p.Arg245His | P51688 |

| | | | | | | | | |
|---|---|---|---|---|---|---|---|---|
| HOMO SAPIENS | 0 | 8143 | 4miv | B | 293 | PRO | p.Pro293Ser | P51688 |
| HOMO SAPIENS | 1 | 8143 | 4miv | B | 298 | SER | p.Ser298Pro | P51688 |
| HOMO SAPIENS | 0 | 8143 | 4miv | B | 361 | VAL | p.Val361Ile | P51688 |
| HOMO SAPIENS | 0 | 8143 | 4miv | B | 369 | GLU | p.Glu369Lys | P51688 |
| HOMO SAPIENS | 0 | 8143 | 4miv | B | 372 | MET | p.Met372Ile | P51688 |
| HOMO SAPIENS | 1 | 8143 | 4miv | B | 380 | GLN | p.Gln380Arg | P51688 |
| HOMO SAPIENS | 0 | 8143 | 4miv | B | 387 | VAL | p.Val387Met | P51688 |
| HOMO SAPIENS | 0 | 8143 | 4miv | B | 394 | MET | p.Met394Ile | P51688 |
| HOMO SAPIENS | 0 | 8143 | 4miv | B | 433 | ARG | p.Arg433Gln | P51688 |
| HOMO SAPIENS | 1 | 8143 | 4miv | B | 446 | HIS | p.His446Arg | P51688 |
| HOMO SAPIENS | 5 | 8143 | 4miv | B | 447 | GLU | p.Glu447Lys | P51688 |
| HOMO SAPIENS | 2 | 8143 | 4miv | B | 456 | ARG | p.Arg456His | P51688 |
| HOMO SAPIENS | 1 | 21880 | 2oo9 | B | 857 | LEU | p.Leu857Phe | P22681 |
| HOMO SAPIENS | 0 | 9467 | 1s1d | A | 154 | LEU | p.Leu224Pro | Q8WVQ1 |
| HOMO SAPIENS | 0 | 9467 | 1s1d | A | 156 | VAL | p.Val226Met | Q8WVQ1 |
| HOMO SAPIENS | 0 | 9467 | 1s1d | A | 229 | PRO | p.Pro299Leu | Q8WVQ1 |
| HOMO SAPIENS | 4 | 9467 | 1s1d | A | 230 | ARG | p.Arg300Cys | Q8WVQ1 |
| HOMO SAPIENS | 4 | 9467 | 1s1d | A | 230 | ARG | p.Arg300His | Q8WVQ1 |
| HOMO SAPIENS | 0 | 9467 | 1s1d | A | 253 | ALA | p.Ala323Thr | Q8WVQ1 |
| HOMO SAPIENS | 2 | 9467 | 1s1d | A | 305 | MET | p.Met375Leu | Q8WVQ1 |
| HOMO SAPIENS | -1 | 2361 | 5llm | A | 5 | ASN | p.Asn5Lys | P43003 |
| HOMO SAPIENS | 0 | 2361 | 5llm | A | 71 | TYR | p.Tyr71Cys | P43003 |
| HOMO SAPIENS | -1 | 2361 | 5llm | A | 211 | LYS | p.Glu219Asp | Q15758 |
| HOMO SAPIENS | 8 | 2361 | 5llm | A | 270 | PRO | p.Pro290Arg | P43003 |
| HOMO SAPIENS | 1 | 2361 | 5llm | A | 321 | THR | p.Ala329Thr | Q15758 |
| HOMO SAPIENS | 1 | 2361 | 5llm | A | 365 | ARG | p.Arg385His | P43003 |
| HOMO SAPIENS | 1 | 2361 | 5llm | A | 479 | ARG | p.Arg499Gln | P43003 |
| HOMO SAPIENS | 3 | 5412 | 4obz | D | 165 | PHE | p.Phe229Ile | P07339 |
| HOMO SAPIENS | 2 | 5412 | 4obz | D | 187 | ASP | p.Asp251Asn | P07339 |
| HOMO SAPIENS | 0 | 5412 | 4obz | D | 218 | GLY | p.Gly282Arg | P07339 |
| HOMO SAPIENS | 0 | 5412 | 4obz | D | 245 | ARG | p.Arg309His | P07339 |
| HOMO SAPIENS | 2 | 5412 | 4obz | D | 319 | TRP | p.Trp383Cys | P07339 |
| HOMO SAPIENS | 0 | 3544 | 5h64 | A | 277 | MET | p.Met277Ile | P42345 |
| HOMO SAPIENS | 0 | 14230 | 2kbx | A | 145 | ARG | p.Arg145Ser | Q13418 |
| HOMO SAPIENS | 2 | 34970 | 4xhz | A | 834 | ARG | p.Arg855Trp | Q96QU1 |
| HOMO SAPIENS | 4 | 34970 | 4xhz | A | 854 | SER | p.Ser875Ter | Q96QU1 |
| HOMO SAPIENS | 1 | 34970 | 4xhz | A | 908 | ARG | p.Arg929Gln | Q96QU1 |
| HOMO SAPIENS | 1 | 34970 | 4xhz | A | 941 | ARG | p.Arg962Cys | Q96QU1 |
| HOMO SAPIENS | 0 | 34970 | 4xhz | A | 970 | ARG | p.Arg991Ter | Q96QU1 |
| HOMO SAPIENS | 0 | 34970 | 4xhz | A | 976 | GLU | p.Glu997Gly | Q96QU1 |
| HOMO SAPIENS | 0 | 25734 | 4asc | A | 345 | SER | p.Asn345Ser | Q2TBA0 |
| HOMO SAPIENS | 0 | 25734 | 4asc | A | 514 | VAL | p.Val514Met | Q2TBA0 |
| HOMO SAPIENS | 6 | 25734 | 4asc | A | 528 | GLU | p.Glu528Lys | Q2TBA0 |
| HOMO SAPIENS | 0 | 25734 | 4asc | A | 580 | LEU | p.Leu580Ile | Q2TBA0 |
| HOMO SAPIENS | 8 | 2356 | 3kse | D | 22 | LYS | p.Lys22Ter | P01040 |
| HOMO SAPIENS | 1 | 2356 | 3kse | D | 58 | ARG | p.Arg58Ter | P01040 |
| HOMO SAPIENS | 2 | 6000 | 1jqd | A | 60 | GLY | p.Gly60Asp | P50135 |
| HOMO SAPIENS | 9 | 6000 | 1jqd | A | 208 | LEU | p.Leu208Pro | P50135 |
| HOMO SAPIENS | 0 | 14741 | 5eh1 | A | 58 | THR | p.Thr58Arg | P38484 |
| HOMO SAPIENS | 0 | 14741 | 5eh1 | A | 168 | THR | p.Thr168Asn | P38484 |
| HOMO SAPIENS | 1 | 4814 | 4kxk | D | 427 | ARG | p.Arg427Ter | P50542 |
| HOMO SAPIENS | 0 | 188 | 2xt3 | A | 49 | ASP | p.Asp52Asn | Q2M1P5 |
| HOMO SAPIENS | 3 | 188 | 2xt3 | A | 142 | TYR | p.Tyr145Ser | Q2M1P5 |
| HOMO SAPIENS | 9 | 8681 | 5tby | D | 57 | ALA | p.Ala57Asp | P08590 |

| | | | | | | | | |
|---|---|---|---|---|---|---|---|---|
| HOMO SAPIENS | 3 | 8681 | 5tby | D | 62 | ASP | p.Asp62Asn | P08590 |
| HOMO SAPIENS | 0 | 8681 | 5tby | D | 65 | PRO | p.Pro65Arg | P08590 |
| HOMO SAPIENS | 7 | 8681 | 5tby | D | 79 | VAL | p.Val79Ile | P08590 |
| HOMO SAPIENS | 0 | 8681 | 5tby | D | 143 | GLU | p.Glu143Lys | P08590 |
| HOMO SAPIENS | 0 | 8681 | 5tby | D | 149 | MET | p.Met149Val | P08590 |
| HOMO SAPIENS | 3 | 8681 | 5tby | D | 154 | ARG | p.Arg154His | P08590 |
| HOMO SAPIENS | 6 | 30984 | 4xwh | A | 140 | TYR | p.Tyr140Cys | P54802 |
| HOMO SAPIENS | 4 | 30984 | 4xwh | A | 141 | SER | p.Ser141Thr | P54802 |
| HOMO SAPIENS | 2 | 30984 | 4xwh | A | 234 | ARG | p.Arg234Cys | P54802 |
| HOMO SAPIENS | 2 | 30984 | 4xwh | A | 336 | GLU | p.Glu336Ter | P54802 |
| HOMO SAPIENS | 6 | 30984 | 4xwh | A | 482 | ARG | p.Arg482Trp | P54802 |
| HOMO SAPIENS | 2 | 30984 | 4xwh | A | 521 | PRO | p.Pro521Leu | P54802 |
| HOMO SAPIENS | 0 | 30984 | 4xwh | A | 565 | ARG | p.Arg565Gln | P54802 |
| HOMO SAPIENS | 0 | 30984 | 4xwh | A | 565 | ARG | p.Arg565Trp | P54802 |
| HOMO SAPIENS | 6 | 30984 | 4xwh | A | 626 | ARG | p.Arg626Ter | P54802 |
| HOMO SAPIENS | 6 | 30984 | 4xwh | A | 643 | ARG | p.Arg643His | P54802 |
| HOMO SAPIENS | 6 | 30984 | 4xwh | A | 643 | ARG | p.Arg643Cys | P54802 |
| HOMO SAPIENS | 2 | 30984 | 4xwh | A | 674 | ARG | p.Arg674His | P54802 |
| HOMO SAPIENS | 2 | 30984 | 4xwh | A | 737 | ARG | p.Arg737Gly | P54802 |
| HOMO SAPIENS | 5 | 10545 | 1oqe | N | 6 | PRO | p.Pro21Arg | Q96RJ3 |
| HOMO SAPIENS | 1 | 16995 | 5d3a | B | 12 | MET | p.Met947Arg | Q7Z4S6 |
| HOMO SAPIENS | 1 | 16995 | 5d3a | B | 12 | MET | p.Met947Val | Q7Z4S6 |
| HOMO SAPIENS | 2 | 16995 | 5d3a | B | 19 | ARG | p.Arg954Gln | Q7Z4S6 |
| HOMO SAPIENS | 2 | 16995 | 5d3a | B | 19 | ARG | p.Arg954Trp | Q7Z4S6 |
| HOMO SAPIENS | 0 | 16995 | 5d3a | B | 75 | ILE | p.Ile1010Thr | Q7Z4S6 |
| HOMO SAPIENS | 0 | 17091 | 2yin | B | 1324 | GLN | p.Gln1324Ter | Q92608 |
| HOMO SAPIENS | 5 | 27606 | 1lm7 | B | 2366 | ARG | p.Arg2366Cys | P15924 |
| HOMO SAPIENS | 2 | 15731 | 1bi7 | B | 24 | ARG | p.Arg24Pro | P42771 |
| HOMO SAPIENS | 0 | 15731 | 1bi7 | B | 28 | VAL | p.Val28Gly | P42771 |
| HOMO SAPIENS | 0 | 15731 | 1bi7 | B | 49 | ILE | p.Ile49Thr | P42771 |
| HOMO SAPIENS | 0 | 15731 | 1bi7 | B | 53 | MET | p.Met53Ile | P42771 |
| HOMO SAPIENS | 2 | 15731 | 1bi7 | B | 56 | SER | p.Ser56Ile | P42771 |
| HOMO SAPIENS | 1 | 15731 | 1bi7 | B | 57 | ALA | p.Ala57Val | P42771 |
| HOMO SAPIENS | 4 | 15731 | 1bi7 | B | 59 | VAL | p.Val59Gly | P42771 |
| HOMO SAPIENS | 9 | 15731 | 1bi7 | B | 89 | GLY | p.Gly89Asp | P42771 |
| HOMO SAPIENS | 0 | 15731 | 1bi7 | B | 101 | GLY | p.Gly101Trp | P42771 |
| HOMO SAPIENS | 0 | 15731 | 1bi7 | B | 114 | PRO | p.Pro114Ser | P42771 |
| HOMO SAPIENS | 9 | 15731 | 1bi7 | B | 123 | HIS | p.His123Gln | P42771 |
| HOMO SAPIENS | 8 | 15731 | 1bi7 | B | 126 | VAL | p.Val126Asp | P42771 |
| HOMO SAPIENS | 7 | 15731 | 1bi7 | B | 127 | ALA | p.Ala127Ser | P42771 |
| HOMO SAPIENS | -1 | 15731 | 1bi7 | B | 148 | ALA | p.Ala148Thr | P42771 |
| HOMO SAPIENS | -1 | 19858 | 3jbk | M | 866 | ALA | p.Ala934Val | P18206 |
| HOMO SAPIENS | 0 | 19484 | 3ee6 | B | 62 | SER | p.Ser62Leu | O14773 |
| HOMO SAPIENS | 1 | 19484 | 3ee6 | B | 66 | GLN | p.Gln66Ter | O14773 |
| HOMO SAPIENS | 0 | 19484 | 3ee6 | B | 100 | GLN | p.Gln100Arg | O14773 |
| HOMO SAPIENS | 0 | 19484 | 3ee6 | B | 104 | LEU | p.Leu104Ter | O14773 |
| HOMO SAPIENS | 0 | 19484 | 3ee6 | B | 127 | ARG | p.Arg127Gln | O14773 |
| HOMO SAPIENS | 0 | 19484 | 3ee6 | B | 127 | ARG | p.Arg127Ter | O14773 |
| HOMO SAPIENS | -1 | 19484 | 3ee6 | B | 185 | ARG | p.Arg185Cys | O14773 |
| HOMO SAPIENS | 0 | 19484 | 3ee6 | B | 206 | ARG | p.Arg206Cys | O14773 |
| HOMO SAPIENS | 0 | 19484 | 3ee6 | B | 208 | ARG | p.Arg208Ter | O14773 |
| HOMO SAPIENS | 0 | 19484 | 3ee6 | B | 214 | GLN | p.Gln214Ter | O14773 |
| HOMO SAPIENS | 1 | 19484 | 3ee6 | B | 276 | ASP | p.Asp276Val | O14773 |
| HOMO SAPIENS | 0 | 19484 | 3ee6 | B | 284 | GLY | p.Gly284Val | O14773 |

| | | | | | | | | |
|---|---|---|---|---|---|---|---|---|
| HOMO SAPIENS | 0 | 19484 | 3ee6 | B | 286 | ASN | p.Asn286Ser | O14773 |
| HOMO SAPIENS | 0 | 19484 | 3ee6 | B | 339 | ARG | p.Arg339Gly | O14773 |
| HOMO SAPIENS | 2 | 19484 | 3ee6 | B | 345 | MET | p.Met345Leu | O14773 |
| HOMO SAPIENS | 0 | 19484 | 3ee6 | B | 365 | CYS | p.Cys365Arg | O14773 |
| HOMO SAPIENS | 0 | 19484 | 3ee6 | B | 365 | CYS | p.Cys365Tyr | O14773 |
| HOMO SAPIENS | 0 | 19484 | 3ee6 | B | 373 | GLN | p.Gln373Glu | O14773 |
| HOMO SAPIENS | 6 | 19484 | 3ee6 | B | 418 | ARG | p.Arg418Gln | O14773 |
| HOMO SAPIENS | 7 | 19484 | 3ee6 | B | 422 | GLN | p.Gln422His | O14773 |
| HOMO SAPIENS | 0 | 19484 | 3ee6 | B | 447 | ARG | p.Arg447His | O14773 |
| HOMO SAPIENS | 0 | 19484 | 3ee6 | B | 466 | VAL | p.Val466Met | O14773 |
| HOMO SAPIENS | 0 | 4654 | 5t5w | A | 47 | LYS | p.Lys47Glu | Q08334 |
| HOMO SAPIENS | 0 | 4654 | 5t5w | A | 141 | GLU | p.Glu141Ter | Q08334 |
| HOMO SAPIENS | -1 | 18815 | 4wj7 | A | 53 | VAL | p.Val53Ile | Q9BSQ5 |
| HOMO SAPIENS | 0 | 18815 | 4wj7 | A | 120 | VAL | p.Val120Ile | Q9BSQ5 |
| HOMO SAPIENS | 0 | 18815 | 4wj7 | A | 198 | LEU | p.Leu198Arg | Q9BSQ5 |
| HOMO SAPIENS | 0 | 4602 | 1mjd | A | 47 | SER | p.Ser47Arg | O43602 |
| HOMO SAPIENS | 6 | 4602 | 1mjd | A | 59 | ARG | p.Arg59Leu | O43602 |
| HOMO SAPIENS | 7 | 4602 | 1mjd | A | 62 | ASP | p.Asp62Asn | O43602 |
| HOMO SAPIENS | 7 | 4602 | 1mjd | A | 71 | ALA | p.Ala71Ser | O43602 |
| HOMO SAPIENS | 7 | 4602 | 1mjd | A | 78 | ARG | p.Arg78Leu | O43602 |
| HOMO SAPIENS | 7 | 4602 | 1mjd | A | 89 | ARG | p.Arg89Gly | O43602 |
| HOMO SAPIENS | 8 | 4602 | 1mjd | A | 104 | ILE | p.Ile104Phe | O43602 |
| HOMO SAPIENS | 7 | 4602 | 1mjd | A | 125 | TYR | p.Tyr125His | O43602 |
| HOMO SAPIENS | 5 | 34467 | 1n0w | B | 1529 | GLY | p.Gly1529Arg | P51587 |
| HOMO SAPIENS | 8 | 5738 | 1u31 | A | 130 | ASN | p.Asn1009Lys | Q13423 |
| HOMO SAPIENS | 0 | 5760 | 2i6s | A | 298 | GLU | p.Glu318Asp | P06681 |
| HOMO SAPIENS | 0 | 5760 | 2i6s | A | 573 | ARG | p.Arg593Gln | P06681 |
| HOMO SAPIENS | 0 | 5760 | 2i6s | A | 592 | SER | p.Ser612Asn | P06681 |
| HOMO SAPIENS | 0 | 5760 | 2i6s | A | 621 | VAL | p.Val641Ala | P06681 |
| HOMO SAPIENS | 1 | 10977 | 2fby | A | 114 | VAL | p.Val114Ile | Q14191 |
| HOMO SAPIENS | 0 | 10977 | 2fby | A | 135 | LYS | p.Lys135Glu | Q14191 |
| HOMO SAPIENS | -1 | 1175 | 1qrk | A | 34 | VAL | p.Val35Leu | P00488 |
| HOMO SAPIENS | 4 | 1175 | 1qrk | A | 564 | PRO | p.Pro565Leu | P00488 |
| HOMO SAPIENS | 0 | 1175 | 1qrk | A | 650 | VAL | p.Val651Ile | P00488 |
| HOMO SAPIENS | 0 | 1175 | 1qrk | A | 651 | GLU | p.Gln652Glu | P00488 |
| HOMO SAPIENS | 3 | 909 | 4h0w | A | 375 | GLU | p.Glu394Lys | P02787 |
| HOMO SAPIENS | 0 | 909 | 4h0w | A | 458 | ALA | p.Ala477Pro | P02787 |
| HOMO SAPIENS | 6 | 909 | 4h0w | A | 515 | TYR | p.Tyr534Ter | P02787 |
| HOMO SAPIENS | 7 | 909 | 4h0w | A | 570 | PRO | p.Pro589Ser | P02787 |
| HOMO SAPIENS | 3 | 909 | 4h0w | A | 652 | GLY | p.Gly671Glu | P02787 |
| HOMO SAPIENS | 5 | 1964 | 2car | A | 151 | TRP | p.Trp151Ter | Q9BY32 |
| HOMO SAPIENS | 7 | 1964 | 2car | A | 178 | ARG | p.Arg178Cys | Q9BY32 |
| HOMO SAPIENS | 0 | 144 | 2n0a | J | 30 | ALA | p.Ala30Pro | P37840 |
| HOMO SAPIENS | 1 | 12288 | 4q58 | D | 1225 | ARG | p.Arg1225His | P16144 |
| HOMO SAPIENS | 2 | 12288 | 4q58 | D | 1281 | ARG | p.Arg1281Trp | P16144 |
| HOMO SAPIENS | 2 | 12288 | 4q58 | D | 1287 | ASN | p.Asn1287Lys | P16144 |
| HOMO SAPIENS | 0 | 5085 | 3s94 | A | 360 | ARG | p.Arg360His | O75581 |
| HOMO SAPIENS | 0 | 5085 | 3s94 | A | 433 | ASN | p.Asn433Ser | O75581 |
| HOMO SAPIENS | 2 | 5085 | 3s94 | A | 611 | ARG | p.Arg611Cys | O75581 |
| HOMO SAPIENS | 0 | 15452 | 4dur | A | 19 | LYS | p.Lys38Glu | P00747 |
| HOMO SAPIENS | 2 | 15452 | 4dur | A | 216 | ARG | p.Arg235His | P00747 |
| HOMO SAPIENS | 7 | 8406 | 4wo1 | D | 28 | VAL | p.Val55Leu | O43914 |
| HOMO SAPIENS | 2 | 895 | 5xte | K | 183 | ARG | p.Arg183Trp | P22695 |
| HOMO SAPIENS | 2 | 4191 | 4wnd | A | 120 | ARG | p.Arg127Ter | P81274 |

| | | | | | | | | |
|---|---|---|---|---|---|---|---|---|
| HOMO SAPIENS | 2 | 4191 | 4wnd | A | 120 | ARG | p.Arg127Gln | P81274 |
| HOMO SAPIENS | 1 | 4191 | 4wnd | A | 349 | GLY | p.Gly356Arg | P81274 |
| HOMO SAPIENS | 0 | 41734 | 2kbi | A | 1777 | VAL | p.Val1777Met | Q14524 |
| HOMO SAPIENS | 1 | 41734 | 2kbi | A | 1784 | GLU | p.Glu1784Lys | Q14524 |
| HOMO SAPIENS | 5 | 41734 | 2kbi | A | 1795 | TYR | p.Tyr1795Cys | Q14524 |
| HOMO SAPIENS | 5 | 41734 | 2kbi | A | 1795 | TYR | p.Tyr1795His | Q14524 |
| HOMO SAPIENS | 7 | 41734 | 2kbi | A | 1826 | ARG | p.Arg1826Cys | Q14524 |
| HOMO SAPIENS | 3 | 2119 | 1pl0 | A | 426 | LYS | p.Lys426Arg | P31939 |
| HOMO SAPIENS | 5 | 3475 | 5u62 | C | 50 | LEU | p.Leu50Ser | Q15910 |
| HOMO SAPIENS | 3 | 379 | 4uez | A | 200 | TYR | p.Tyr200Ter | P15085 |
| HOMO SAPIENS | 0 | 8195 | 5xaw | B | 67 | ARG | p.Arg67Trp | Q8IWT1 |
| HOMO SAPIENS | 0 | 8195 | 5xaw | B | 94 | THR | p.Thr94Met | Q8IWT1 |
| HOMO SAPIENS | 1 | 5711 | 1fo3 | A | 265 | ARG | p.Arg265His | Q9UKM7 |
| HOMO SAPIENS | 4 | 5711 | 1fo3 | A | 334 | ARG | p.Arg334Cys | Q9UKM7 |
| HOMO SAPIENS | 1 | 5711 | 1fo3 | A | 478 | LYS | p.Lys478Arg | Q9UKM7 |
| HOMO SAPIENS | 1 | 5711 | 1fo3 | A | 682 | ASP | p.Asp682Glu | Q9UKM7 |
| HOMO SAPIENS | 0 | 170 | 2n3y | A | 41 | GLY | p.Gly42Ser | P99999 |
| HOMO SAPIENS | 0 | 4122 | 3w81 | A | 33 | HIS | p.His33Gln | P35475 |
| HOMO SAPIENS | 0 | 4122 | 3w81 | A | 64 | TYR | p.Tyr64Ter | P35475 |
| HOMO SAPIENS | 0 | 4122 | 3w81 | A | 70 | GLN | p.Gln70Ter | P35475 |
| HOMO SAPIENS | 0 | 4122 | 3w81 | A | 75 | ALA | p.Ala75Thr | P35475 |
| HOMO SAPIENS | 0 | 4122 | 3w81 | A | 79 | ALA | p.Ala79Val | P35475 |
| HOMO SAPIENS | 0 | 4122 | 3w81 | A | 79 | ALA | p.Ala79Thr | P35475 |
| HOMO SAPIENS | 0 | 4122 | 3w81 | A | 89 | ARG | p.Arg89Gln | P35475 |
| HOMO SAPIENS | 0 | 4122 | 3w81 | A | 99 | THR | p.Thr99Ile | P35475 |
| HOMO SAPIENS | 0 | 4122 | 3w81 | A | 105 | ARG | p.Arg105Gln | P35475 |
| HOMO SAPIENS | 0 | 4122 | 3w81 | A | 238 | LEU | p.Leu238Gln | P35475 |
| HOMO SAPIENS | 0 | 4122 | 3w81 | A | 300 | ALA | p.Ala300Thr | P35475 |
| HOMO SAPIENS | 0 | 4122 | 3w81 | A | 322 | VAL | p.Val322Glu | P35475 |
| HOMO SAPIENS | 0 | 4122 | 3w81 | A | 327 | ALA | p.Ala327Pro | P35475 |
| HOMO SAPIENS | 0 | 4122 | 3w81 | A | 346 | LEU | p.Leu346Arg | P35475 |
| HOMO SAPIENS | 0 | 4122 | 3w81 | A | 349 | ASP | p.Asp349Asn | P35475 |
| HOMO SAPIENS | 0 | 4122 | 3w81 | A | 361 | ALA | p.Ala361Thr | P35475 |
| HOMO SAPIENS | 0 | 4122 | 3w81 | A | 366 | THR | p.Thr366Pro | P35475 |
| HOMO SAPIENS | 0 | 4122 | 3w81 | A | 402 | TRP | p.Trp402Ter | P35475 |
| HOMO SAPIENS | 0 | 4122 | 3w81 | A | 409 | GLY | p.Gly409Arg | P35475 |
| HOMO SAPIENS | 0 | 4122 | 3w81 | A | 454 | VAL | p.Val454Ile | P35475 |
| HOMO SAPIENS | 1 | 4122 | 3w81 | A | 490 | LEU | p.Leu490Pro | P35475 |
| HOMO SAPIENS | 1 | 4122 | 3w81 | A | 492 | ARG | p.Arg492Pro | P35475 |
| HOMO SAPIENS | 0 | 4122 | 3w81 | A | 533 | PRO | p.Pro533Arg | P35475 |
| HOMO SAPIENS | 0 | 4122 | 3w81 | A | 591 | ALA | p.Ala591Thr | P35475 |
| HOMO SAPIENS | 7 | 4122 | 3w81 | A | 619 | ARG | p.Arg619Gly | P35475 |
| HOMO SAPIENS | 0 | 15325 | 3wvz | B | 54 | VAL | p.Val54Leu | Q53FT3 |
| HOMO SAPIENS | 3 | 3228 | 5iz8 | A | 421 | TRP | p.Trp421Ter | P25054 |
| HOMO SAPIENS | 2 | 3228 | 5iz8 | A | 437 | PRO | p.Pro437Arg | P25054 |
| HOMO SAPIENS | 3 | 3228 | 5iz8 | A | 499 | ARG | p.Arg499Ter | P25054 |
| HOMO SAPIENS | -1 | 29229 | 5buo | A | 539 | SER | p.Ser614Gly | P05067 |
| HOMO SAPIENS | -1 | 34931 | 4n7v | C | 54 | SER | p.Ser54Leu | O94986 |
| HOMO SAPIENS | 0 | 2450 | 5kzx | A | 123 | GLY | p.Gly123Glu | P10253 |
| HOMO SAPIENS | 1 | 2450 | 5kzx | A | 190 | ARG | p.Arg190His | P10253 |
| HOMO SAPIENS | -1 | 2450 | 5kzx | A | 199 | HIS | p.His199Arg | P10253 |
| HOMO SAPIENS | 0 | 2450 | 5kzx | A | 219 | GLY | p.Gly219Arg | P10253 |
| HOMO SAPIENS | 0 | 2450 | 5kzx | A | 220 | VAL | p.Val220Leu | P10253 |
| HOMO SAPIENS | 0 | 2450 | 5kzx | A | 222 | VAL | p.Val222Met | P10253 |

| | | | | | | | | |
|---|---|---|---|---|---|---|---|---|
| HOMO SAPIENS | 0 | 2450 | 5kzx | A | 237 | ALA | p.Ala237Val | P10253 |
| HOMO SAPIENS | 0 | 2450 | 5kzx | A | 251 | SER | p.Ser251Leu | P10253 |
| HOMO SAPIENS | 0 | 2450 | 5kzx | A | 254 | SER | p.Ser254Leu | P10253 |
| HOMO SAPIENS | 0 | 2450 | 5kzx | A | 262 | GLU | p.Glu262Lys | P10253 |
| HOMO SAPIENS | 0 | 2450 | 5kzx | A | 293 | GLY | p.Gly293Arg | P10253 |
| HOMO SAPIENS | 0 | 2450 | 5kzx | A | 299 | LEU | p.Leu299Arg | P10253 |
| HOMO SAPIENS | 0 | 2450 | 5kzx | A | 309 | GLY | p.Gly309Arg | P10253 |
| HOMO SAPIENS | 0 | 2450 | 5kzx | A | 318 | MET | p.Met318Thr | P10253 |
| HOMO SAPIENS | 0 | 2450 | 5kzx | A | 355 | LEU | p.Leu355Pro | P10253 |
| HOMO SAPIENS | 1 | 2450 | 5kzx | A | 361 | PRO | p.Pro361Leu | P10253 |
| HOMO SAPIENS | 4 | 2450 | 5kzx | A | 408 | MET | p.Met408Val | P10253 |
| HOMO SAPIENS | 1 | 2450 | 5kzx | A | 429 | GLN | p.Gln429Glu | P10253 |
| HOMO SAPIENS | 4 | 2450 | 5kzx | A | 437 | ARG | p.Arg437Cys | P10253 |
| HOMO SAPIENS | 0 | 2450 | 5kzx | A | 439 | MET | p.Met439Lys | P10253 |
| HOMO SAPIENS | 1 | 2450 | 5kzx | A | 448 | SER | p.Ser448Thr | P10253 |
| HOMO SAPIENS | 0 | 2450 | 5kzx | A | 451 | PRO | p.Pro451Arg | P10253 |
| HOMO SAPIENS | 1 | 2450 | 5kzx | A | 459 | ASP | p.Asp459His | P10253 |
| HOMO SAPIENS | 8 | 2450 | 5kzx | A | 481 | TRP | p.Trp481Arg | P10253 |
| HOMO SAPIENS | 1 | 2450 | 5kzx | A | 489 | ASP | p.Asp489Asn | P10253 |
| HOMO SAPIENS | 0 | 2450 | 5kzx | A | 521 | GLU | p.Glu521Lys | P10253 |
| HOMO SAPIENS | 0 | 2450 | 5kzx | A | 529 | SER | p.Ser529Val | P10253 |
| HOMO SAPIENS | 0 | 2450 | 5kzx | A | 545 | PRO | p.Pro545Leu | P10253 |
| HOMO SAPIENS | 0 | 2450 | 5kzx | A | 552 | LEU | p.Leu552Pro | P10253 |
| HOMO SAPIENS | 0 | 2450 | 5kzx | A | 576 | GLY | p.Gly576Ser | P10253 |
| HOMO SAPIENS | 9 | 2450 | 5kzx | A | 600 | ARG | p.Arg600His | P10253 |
| HOMO SAPIENS | 8 | 2450 | 5kzx | A | 615 | GLY | p.Gly615Arg | P10253 |
| HOMO SAPIENS | 3 | 2450 | 5kzx | A | 619 | SER | p.Ser619Asn | P10253 |
| HOMO SAPIENS | 0 | 2450 | 5kzx | A | 638 | GLY | p.Gly638Trp | P10253 |
| HOMO SAPIENS | 0 | 2450 | 5kzx | A | 643 | GLY | p.Gly643Arg | P10253 |
| HOMO SAPIENS | 9 | 2450 | 5kzx | A | 645 | ASP | p.Asp645Glu | P10253 |
| HOMO SAPIENS | 9 | 2450 | 5kzx | A | 645 | ASP | p.Asp645His | P10253 |
| HOMO SAPIENS | 0 | 2450 | 5kzx | A | 648 | GLY | p.Gly648Ser | P10253 |
| HOMO SAPIENS | 5 | 2450 | 5kzx | A | 660 | ARG | p.Arg660His | P10253 |
| HOMO SAPIENS | 8 | 2450 | 5kzx | A | 672 | ARG | p.Arg672Gln | P10253 |
| HOMO SAPIENS | 0 | 2450 | 5kzx | A | 702 | ARG | p.Arg702Leu | P10253 |
| HOMO SAPIENS | 3 | 2450 | 5kzx | A | 725 | ARG | p.Arg725Trp | P10253 |
| HOMO SAPIENS | 0 | 2450 | 5kzx | A | 746 | TRP | p.Trp746Cys | P10253 |
| HOMO SAPIENS | 0 | 2450 | 5kzx | A | 780 | ILE | p.Val780Ile | P10253 |
| HOMO SAPIENS | 1 | 2450 | 5kzx | A | 816 | VAL | p.Val816Ile | P10253 |
| HOMO SAPIENS | 0 | 2450 | 5kzx | A | 838 | GLN | p.Gln838Ter | P10253 |
| HOMO SAPIENS | 2 | 2450 | 5kzx | A | 854 | ARG | p.Arg854Ter | P10253 |
| HOMO SAPIENS | 3 | 2450 | 5kzx | A | 870 | ARG | p.Arg870Ter | P10253 |
| HOMO SAPIENS | 0 | 2450 | 5kzx | A | 890 | VAL | p.Val890Leu | P10253 |
| HOMO SAPIENS | 6 | 1882 | 4km6 | A | 175 | CYS | p.Cys175Ter | P15328 |
| HOMO SAPIENS | 3 | 14767 | 1lm5 | A | 2628 | LEU | p.Leu2628Pro | P15924 |
| HOMO SAPIENS | 5 | 14767 | 1lm5 | A | 2655 | ALA | p.Ala2655Gly | P15924 |
| HOMO SAPIENS | 3 | 14767 | 1lm5 | A | 2731 | TYR | p.Tyr2731His | P15924 |
| HOMO SAPIENS | 1 | 14767 | 1lm5 | A | 2767 | THR | p.Thr2767Asn | P15924 |
| HOMO SAPIENS | -1 | 14767 | 1lm5 | A | 2819 | MET | p.Met2819Leu | P15924 |
| HOMO SAPIENS | -1 | 11246 | 4y5o | A | 289 | SER | p.Ser289Asn | Q9BSQ5 |
| HOMO SAPIENS | 0 | 7380 | 5kz5 | I | 60 | ARG | p.Arg60His | Q16595 |
| HOMO SAPIENS | 0 | 7380 | 5kz5 | I | 76 | MET | p.Met76Val | Q16595 |
| HOMO SAPIENS | 2 | 22187 | 3fby | A | 328 | CYS | p.Cys328Arg | P49747 |
| HOMO SAPIENS | 2 | 22187 | 3fby | A | 342 | ASP | p.Asp342Tyr | P49747 |

| | | | | | | | | |
|---|---|---|---|---|---|---|---|---|
| HOMO SAPIENS | 2 | 22187 | 3fby | A | 348 | CYS | p.Cys348Arg | P49747 |
| HOMO SAPIENS | 2 | 22187 | 3fby | A | 386 | ASN | p.Asn386Asp | P49747 |
| HOMO SAPIENS | 2 | 22187 | 3fby | A | 453 | ASN | p.Asn453Ser | P49747 |
| HOMO SAPIENS | 2 | 22187 | 3fby | A | 468 | CYS | p.Cys468Tyr | P49747 |
| HOMO SAPIENS | 3 | 22187 | 3fby | A | 472 | ASP | p.Asp472Tyr | P49747 |
| HOMO SAPIENS | 3 | 22187 | 3fby | A | 473 | ASP | p.Asp473Gly | P49747 |
| HOMO SAPIENS | 2 | 22187 | 3fby | A | 523 | ASN | p.Asn523Lys | P49747 |
| HOMO SAPIENS | 1 | 22187 | 3fby | A | 718 | ARG | p.Arg718Trp | P49747 |
| HOMO SAPIENS | 1 | 22187 | 3fby | A | 719 | GLY | p.Gly719Asp | P49747 |
| HOMO SAPIENS | 0 | 22187 | 3fby | A | 756 | GLN | p.Gln756Arg | P49747 |
| HOMO SAPIENS | -1 | 2784 | 5kvh | A | 88 | MET | p.Met88Val | O95831 |
| HOMO SAPIENS | -1 | 2784 | 5kvh | A | 96 | ASN | p.Asn96Ser | O95831 |
| HOMO SAPIENS | 0 | 2784 | 5kvh | A | 282 | THR | p.Thr282Met | O95831 |
| HOMO SAPIENS | 1 | 2784 | 5kvh | A | 366 | ASN | p.Asn366Ser | O95831 |
| HOMO SAPIENS | 3 | 2784 | 5kvh | A | 463 | ARG | p.Arg463Ile | O95831 |
| HOMO SAPIENS | 0 | 2784 | 5kvh | A | 498 | VAL | p.Val498Met | O95831 |
| HOMO SAPIENS | -1 | 2784 | 5kvh | A | 549 | ALA | p.Ala549Val | O95831 |
| HOMO SAPIENS | 0 | 38316 | 2lnk | C | 1933 | ARG | p.Arg1933Ter | P35579 |
| HOMO SAPIENS | 0 | 33808 | 2miu | A | 29 | VAL | p.Val29Met | Q14192 |
| HOMO SAPIENS | 1 | 33808 | 2miu | A | 37 | ALA | p.Ala37Ser | Q14192 |
| HOMO SAPIENS | 2 | 20994 | 2hey | R | 65 | ARG | p.Arg65Cys | P43489 |
| HOMO SAPIENS | -1 | 5687 | 2xwj | I | 7 | ARG | p.Arg32Gln | P00751 |
| HOMO SAPIENS | -1 | 5687 | 2xwj | I | 7 | ARG | p.Arg32Trp | P00751 |
| HOMO SAPIENS | 0 | 5687 | 2xwj | I | 49 | ARG | p.Arg74His | P00751 |
| HOMO SAPIENS | 0 | 5687 | 2xwj | I | 177 | ARG | p.Arg202Gly | P00751 |
| HOMO SAPIENS | 0 | 5687 | 2xwj | I | 217 | ILE | p.Ile242Leu | P00751 |
| HOMO SAPIENS | -1 | 5687 | 2xwj | I | 227 | GLY | p.Gly252Ser | P00751 |
| HOMO SAPIENS | 0 | 5687 | 2xwj | I | 483 | HIS | p.His508Gln | P00751 |
| HOMO SAPIENS | 0 | 5687 | 2xwj | I | 508 | LYS | p.Lys533Arg | P00751 |
| HOMO SAPIENS | 0 | 5687 | 2xwj | I | 540 | LYS | p.Lys565Glu | P00751 |
| HOMO SAPIENS | 0 | 5687 | 2xwj | I | 541 | GLU | p.Glu566Ala | P00751 |
| HOMO SAPIENS | 2 | 953 | 3d4j | A | 249 | PHE | p.Phe249Ser | P53602 |
| HOMO SAPIENS | 1 | 953 | 3d4j | A | 292 | ASN | p.Asn292Ser | P53602 |
| HOMO SAPIENS | 0 | 3300 | 1s9c | A | 140 | ASN | p.Asn457Tyr | P51659 |
| HOMO SAPIENS | 2 | 3300 | 1s9c | A | 167 | ILE | p.Ile484Val | P51659 |
| HOMO SAPIENS | 0 | 3300 | 1s9c | A | 174 | ALA | p.Ala491Thr | P51659 |
| HOMO SAPIENS | 8 | 3300 | 1s9c | A | 193 | ASP | p.Asp510Asn | P51659 |
| HOMO SAPIENS | 3 | 3300 | 1s9c | A | 194 | TRP | p.Trp511Arg | P51659 |
| HOMO SAPIENS | 0 | 3300 | 1s9c | A | 242 | VAL | p.Ile559Val | P51659 |
| HOMO SAPIENS | 0 | 10810 | 5i25 | A | 5 | GLN | p.Gln23Ter | P03951 |
| HOMO SAPIENS | 0 | 10810 | 5i25 | A | 55 | TRP | p.Trp73Ter | P03951 |
| HOMO SAPIENS | 0 | 10810 | 5i25 | A | 91 | ALA | p.Ala109Thr | P03951 |
| HOMO SAPIENS | 0 | 10810 | 5i25 | A | 117 | GLU | p.Glu135Ter | P03951 |
| HOMO SAPIENS | 0 | 10810 | 5i25 | A | 118 | CYS | p.Cys136Ter | P03951 |
| HOMO SAPIENS | 0 | 10810 | 5i25 | A | 128 | CYS | p.Cys146Ter | P03951 |
| HOMO SAPIENS | 0 | 10810 | 5i25 | A | 210 | ARG | p.Arg228Ter | P03951 |
| HOMO SAPIENS | 0 | 10810 | 5i25 | A | 226 | GLN | p.Gln244Ter | P03951 |
| HOMO SAPIENS | 0 | 10810 | 5i25 | A | 233 | GLN | p.Gln251Ter | P03951 |
| HOMO SAPIENS | 0 | 16668 | 3lbx | A | 28 | ARG | p.Arg28Leu | P02549 |
| HOMO SAPIENS | 0 | 16668 | 3lbx | A | 28 | ARG | p.Arg28Cys | P02549 |
| HOMO SAPIENS | 1 | 16668 | 3lbx | A | 41 | ARG | p.Arg41Trp | P02549 |
| HOMO SAPIENS | 1 | 16668 | 3lbx | A | 45 | ARG | p.Arg45Ser | P02549 |
| HOMO SAPIENS | 2 | 16668 | 3lbx | A | 48 | LYS | p.Lys48Arg | P02549 |
| HOMO SAPIENS | 1 | 16668 | 3lbx | A | 152 | ASP | p.Asp152Asn | P02549 |

| | | | | | | | | |
|---|---|---|---|---|---|---|---|---|
| HOMO SAPIENS | 0 | 226 | 2ble | A | 256 | ILE | p.Phe256Ile | P36959 |
| HOMO SAPIENS | 2 | 36808 | 4c7n | A | 3 | SER | p.Ser357Pro | O75030 |
| HOMO SAPIENS | 0 | 4092 | 5jnx | A | 113 | CYS | p.Cys113Arg | O15118 |
| HOMO SAPIENS | 4 | 4092 | 5jnx | A | 215 | HIS | p.His215Arg | O15118 |
| HOMO SAPIENS | 6 | 4092 | 5jnx | A | 237 | PRO | p.Pro237Ser | O15118 |
| HOMO SAPIENS | 0 | 4092 | 5jnx | A | 363 | ALA | p.Ala363Val | O15118 |
| HOMO SAPIENS | 0 | 4092 | 5jnx | A | 378 | VAL | p.Val378Ala | O15118 |
| HOMO SAPIENS | 8 | 4092 | 5jnx | A | 404 | ARG | p.Arg404Gln | O15118 |
| HOMO SAPIENS | 8 | 4092 | 5jnx | A | 411 | ARG | p.Arg411Gln | O15118 |
| HOMO SAPIENS | 2 | 4092 | 5jnx | A | 474 | PRO | p.Pro474Leu | O15118 |
| HOMO SAPIENS | 8 | 4092 | 5jnx | A | 511 | THR | p.Thr511Met | O15118 |
| HOMO SAPIENS | 0 | 4092 | 5jnx | A | 642 | MET | p.Met642Ile | O15118 |
| HOMO SAPIENS | 2 | 4092 | 5jnx | A | 664 | VAL | p.Val664Met | O15118 |
| HOMO SAPIENS | 0 | 4092 | 5jnx | A | 734 | SER | p.Ser734Ile | O15118 |
| HOMO SAPIENS | -1 | 4092 | 5jnx | A | 775 | GLN | p.Gln775Pro | O15118 |
| HOMO SAPIENS | 0 | 4092 | 5jnx | A | 858 | ILE | p.Ile858Val | O15118 |
| HOMO SAPIENS | 0 | 4092 | 5jnx | A | 889 | VAL | p.Val889Met | O15118 |
| HOMO SAPIENS | 0 | 4092 | 5jnx | A | 910 | GLY | p.Gly910Ser | O15118 |
| HOMO SAPIENS | 0 | 4092 | 5jnx | A | 911 | GLY | p.Gly911Ser | O15118 |
| HOMO SAPIENS | 0 | 4092 | 5jnx | A | 928 | GLN | p.Gln928Pro | O15118 |
| HOMO SAPIENS | 0 | 4092 | 5jnx | A | 931 | ASN | p.Asn931Lys | O15118 |
| HOMO SAPIENS | 0 | 4092 | 5jnx | A | 940 | SER | p.Ser940Leu | O15118 |
| HOMO SAPIENS | 0 | 4092 | 5jnx | A | 950 | VAL | p.Val950Met | O15118 |
| HOMO SAPIENS | 0 | 4092 | 5jnx | A | 954 | SER | p.Ser954Leu | O15118 |
| HOMO SAPIENS | 0 | 4092 | 5jnx | A | 958 | ARG | p.Arg958Gln | O15118 |
| HOMO SAPIENS | 0 | 4092 | 5jnx | A | 958 | ARG | p.Arg958Ter | O15118 |
| HOMO SAPIENS | -1 | 4092 | 5jnx | A | 978 | ARG | p.Arg978Cys | O15118 |
| HOMO SAPIENS | 0 | 4092 | 5jnx | A | 992 | GLY | p.Gly992Trp | O15118 |
| HOMO SAPIENS | 0 | 4092 | 5jnx | A | 1007 | PRO | p.Pro1007Ala | O15118 |
| HOMO SAPIENS | 0 | 4092 | 5jnx | A | 1019 | TYR | p.Tyr1019Cys | O15118 |
| HOMO SAPIENS | 2 | 4092 | 5jnx | A | 1035 | ALA | p.Ala1035Val | O15118 |
| HOMO SAPIENS | 2 | 4092 | 5jnx | A | 1054 | ALA | p.Ala1054Thr | O15118 |
| HOMO SAPIENS | 2 | 4092 | 5jnx | A | 1061 | ILE | p.Ile1061Thr | O15118 |
| HOMO SAPIENS | 0 | 4092 | 5jnx | A | 1073 | GLY | p.Gly1073Ser | O15118 |
| HOMO SAPIENS | 2 | 4092 | 5jnx | A | 1088 | TYR | p.Tyr1088Cys | O15118 |
| HOMO SAPIENS | 2 | 4092 | 5jnx | A | 1142 | MET | p.Met1142Thr | O15118 |
| HOMO SAPIENS | 2 | 4092 | 5jnx | A | 1156 | ASN | p.Asn1156Ser | O15118 |
| HOMO SAPIENS | 0 | 4092 | 5jnx | A | 1179 | MET | p.Met1179Val | O15118 |
| HOMO SAPIENS | 0 | 4092 | 5jnx | A | 1186 | ARG | p.Arg1186His | O15118 |
| HOMO SAPIENS | 0 | 4092 | 5jnx | A | 1187 | ALA | p.Ala1187Gly | O15118 |
| HOMO SAPIENS | 2 | 4092 | 5jnx | A | 1189 | GLU | p.Glu1189Gly | O15118 |
| HOMO SAPIENS | 2 | 4092 | 5jnx | A | 1200 | SER | p.Ser1200Gly | O15118 |
| HOMO SAPIENS | 2 | 4092 | 5jnx | A | 1205 | THR | p.Thr1205Lys | O15118 |
| HOMO SAPIENS | 0 | 4092 | 5jnx | A | 1213 | LEU | p.Leu1213Phe | O15118 |
| HOMO SAPIENS | -1 | 4092 | 5jnx | A | 1266 | ARG | p.Arg1266Gln | O15118 |
| HOMO SAPIENS | 9 | 16498 | 4i6o | A | 64 | ARG | p.Arg735Trp | P01024 |
| HOMO SAPIENS | 1 | 21902 | 3un9 | C | 751 | ARG | p.Arg751His | Q86UT6 |
| HOMO SAPIENS | 7 | 20391 | 2h32 | B | 105 | PRO | p.Pro142Leu | P15814 |
| HOMO SAPIENS | 6 | 1635 | 4x45 | A | 65 | ASP | p.Asp65Val | P07741 |
| HOMO SAPIENS | 1 | 1635 | 4x45 | A | 98 | TRP | p.Trp98Ter | P07741 |
| HOMO SAPIENS | 0 | 1635 | 4x45 | A | 110 | LEU | p.Leu110Pro | P07741 |
| HOMO SAPIENS | 7 | 1635 | 4x45 | A | 136 | MET | p.Met136Thr | P07741 |
| HOMO SAPIENS | 1 | 1635 | 4x45 | A | 150 | VAL | p.Val150Phe | P07741 |
| HOMO SAPIENS | 4 | 34810 | 2rpc | A | 16 | CYS | p.Cys253Ser | O60481 |

| | | | | | | | | |
|---|---|---|---|---|---|---|---|---|
| HOMO SAPIENS | 1 | 34810 | 2rpc | A | 18 | TRP | p.Trp255Gly | O60481 |
| HOMO SAPIENS | 1 | 14359 | 4hxi | A | 78 | MET | p.Met78Val | Q9UH77 |
| HOMO SAPIENS | 0 | 5342 | 4d9t | A | 628 | GLU | p.Glu628Asp | P51812 |
| HOMO SAPIENS | -1 | 5342 | 4d9t | A | 723 | ARG | p.Arg723His | P51812 |
| HOMO SAPIENS | -1 | 5342 | 4d9t | A | 729 | ARG | p.Arg729Gln | P51812 |
| HOMO SAPIENS | 1 | 9619 | 3j2s | B | 1781 | ARG | p.Arg1800Cys | P00451 |
| HOMO SAPIENS | 0 | 9619 | 3j2s | B | 1823 | MET | p.Met1842Ile | P00451 |
| HOMO SAPIENS | 1 | 9619 | 3j2s | B | 1825 | PRO | p.Pro1844Ser | P00451 |
| HOMO SAPIENS | 0 | 9619 | 3j2s | B | 1826 | THR | p.Thr1845Pro | P00451 |
| HOMO SAPIENS | 0 | 9619 | 3j2s | B | 1834 | ALA | p.Ala1853Val | P00451 |
| HOMO SAPIENS | 0 | 9619 | 3j2s | B | 1846 | ASP | p.Asp1865Asn | P00451 |
| HOMO SAPIENS | 1 | 9619 | 3j2s | B | 1848 | HIS | p.His1867Arg | P00451 |
| HOMO SAPIENS | 0 | 9619 | 3j2s | B | 1854 | PRO | p.Pro1873Arg | P00451 |
| HOMO SAPIENS | 1 | 9619 | 3j2s | B | 1885 | GLU | p.Glu1904Lys | P00451 |
| HOMO SAPIENS | 0 | 9619 | 3j2s | B | 1922 | ASN | p.Asn1941Ser | P00451 |
| HOMO SAPIENS | 1 | 9619 | 3j2s | B | 1941 | ARG | p.Arg1960Gln | P00451 |
| HOMO SAPIENS | 4 | 9619 | 3j2s | B | 2105 | TYR | p.Tyr2124Cys | P00451 |
| HOMO SAPIENS | 6 | 9619 | 3j2s | B | 2150 | ARG | p.Arg2169His | P00451 |
| HOMO SAPIENS | 0 | 9619 | 3j2s | B | 2159 | ARG | p.Arg2178Cys | P00451 |
| HOMO SAPIENS | 0 | 9619 | 3j2s | B | 2159 | ARG | p.Arg2178His | P00451 |
| HOMO SAPIENS | 2 | 9619 | 3j2s | B | 2238 | MET | p.Met2257Val | P00451 |
| HOMO SAPIENS | 2 | 9619 | 3j2s | B | 2300 | PRO | p.Pro2319Ser | P00451 |
| HOMO SAPIENS | 1 | 667 | 1z2m | A | 127 | GLU | p.Glu127Ter | P05161 |
| HOMO SAPIENS | 0 | 15717 | 2nn6 | G | 30 | GLY | p.Gly31Ala | Q9NQT5 |
| HOMO SAPIENS | -1 | 15717 | 2nn6 | G | 55 | ASN | p.Asn56His | Q9NQT5 |
| HOMO SAPIENS | 7 | 15717 | 2nn6 | G | 131 | ASP | p.Asp132Ala | Q9NQT5 |
| HOMO SAPIENS | 7 | 784 | 3rkq | A | 161 | ARG | p.Arg161Pro | P52952 |
| HOMO SAPIENS | 6 | 784 | 3rkq | A | 178 | THR | p.Thr178Met | P52952 |
| HOMO SAPIENS | 7 | 784 | 3rkq | A | 183 | LYS | p.Lys183Glu | P52952 |
| HOMO SAPIENS | 8 | 784 | 3rkq | A | 190 | ARG | p.Arg190Cys | P52952 |
| HOMO SAPIENS | 6 | 385 | 30000 | C | 70 | ALA | p.Ala70Thr | P07954 |
| HOMO SAPIENS | 2 | 385 | 30000 | C | 101 | ARG | p.Arg101Pro | P07954 |
| HOMO SAPIENS | 2 | 385 | 30000 | C | 174 | PRO | p.Pro174Arg | P07954 |
| HOMO SAPIENS | 6 | 385 | 30000 | C | 187 | SER | p.Ser187Ter | P07954 |
| HOMO SAPIENS | 9 | 385 | 30000 | C | 233 | ARG | p.Arg233His | P07954 |
| HOMO SAPIENS | 3 | 385 | 30000 | C | 273 | ALA | p.Ala273Thr | P07954 |
| HOMO SAPIENS | 4 | 385 | 30000 | C | 334 | SER | p.Ser334Arg | P07954 |
| HOMO SAPIENS | 6 | 385 | 30000 | C | 340 | ASN | p.Asn340Lys | P07954 |
| HOMO SAPIENS | 8 | 385 | 30000 | C | 343 | ARG | p.Arg343Ter | P07954 |
| HOMO SAPIENS | 3 | 385 | 30000 | C | 350 | ARG | p.Arg350Trp | P07954 |
| HOMO SAPIENS | 4 | 385 | 30000 | C | 376 | GLN | p.Gln376Pro | P07954 |
| HOMO SAPIENS | 6 | 385 | 30000 | C | 434 | CYS | p.Cys434Ter | P07954 |
| HOMO SAPIENS | 0 | 3316 | 4f91 | B | 934 | THR | p.Thr934Ala | O75643 |
| HOMO SAPIENS | 0 | 3316 | 4f91 | B | 1002 | ASN | p.Asn1002Ser | O75643 |
| HOMO SAPIENS | 0 | 3316 | 4f91 | B | 1087 | SER | p.Ser1087Leu | O75643 |
| HOMO SAPIENS | 4 | 3316 | 4f91 | B | 1389 | VAL | p.Val1389Ile | O75643 |
| HOMO SAPIENS | 0 | 619 | 5hbt | B | 195 | ASP | p.Asp240His | P02708 |
| HOMO SAPIENS | 1 | 619 | 5hbt | B | 209 | ARG | p.Arg254Leu | P02708 |
| HOMO SAPIENS | 0 | 18171 | 2y25 | B | 1408 | ASP | p.Asp1408Asn | P52179 |
| HOMO SAPIENS | 0 | 18171 | 2y25 | B | 1453 | MET | p.Met1453Thr | P52179 |
| HOMO SAPIENS | 0 | 18171 | 2y25 | B | 1459 | ILE | p.Ile1459Lys | P52179 |
| HOMO SAPIENS | 0 | 18171 | 2y25 | B | 1558 | PHE | p.Phe1558Leu | P52179 |
| HOMO SAPIENS | 0 | 18171 | 2y25 | B | 1573 | ARG | p.Arg1573Gln | P52179 |
| HOMO SAPIENS | 7 | 18171 | 2y25 | B | 1663 | VAL | p.Val1663Met | P52179 |

| | | | | | | | | |
|---|---|---|---|---|---|---|---|---|
| HOMO SAPIENS | 8 | 14262 | 2fcw | B | 117 | SER | p.Ser138Ter | P01130 |
| HOMO SAPIENS | 8 | 14262 | 2fcw | B | 118 | ASP | p.Asp139Glu | P01130 |
| HOMO SAPIENS | 9 | 14262 | 2fcw | B | 119 | GLU | p.Glu140Lys | P01130 |
| HOMO SAPIENS | 7 | 14262 | 2fcw | B | 122 | CYS | p.Cys143Ter | P01130 |
| HOMO SAPIENS | 6 | 14262 | 2fcw | B | 134 | CYS | p.Cys155Ter | P01130 |
| HOMO SAPIENS | 7 | 14262 | 2fcw | B | 147 | ASP | p.Asp168Glu | P01130 |
| HOMO SAPIENS | 4 | 2624 | 3b7b | A | 855 | VAL | p.Val886Met | Q9H9B1 |
| HOMO SAPIENS | 7 | 2624 | 3b7b | A | 868 | ILE | p.Ile899Val | Q9H9B1 |
| HOMO SAPIENS | 0 | 2624 | 3b7b | A | 892 | GLU | p.Glu923Lys | Q9H9B1 |
| HOMO SAPIENS | 0 | 4791 | 5ctd | B | 68 | LEU | p.Pro549Ala | Q14055 |
| HOMO SAPIENS | 5 | 460 | 4ui1 | D | 80 | CYS | p.Cys80Arg | Q6ZVN8 |
| HOMO SAPIENS | 3 | 460 | 4ui1 | D | 101 | LEU | p.Leu101Pro | Q6ZVN8 |
| HOMO SAPIENS | 0 | 40782 | 1sy6 | A | 47 | LYS | p.Lys69Ter | P09693 |
| HOMO SAPIENS | -1 | 1092 | 2idx | A | 62 | THR | p.Thr62Met | Q96EY8 |
| HOMO SAPIENS | 6 | 1092 | 2idx | A | 183 | HIS | p.His183Leu | Q96EY8 |
| HOMO SAPIENS | 7 | 1092 | 2idx | A | 186 | ARG | p.Arg186Trp | Q96EY8 |
| HOMO SAPIENS | 6 | 1092 | 2idx | A | 190 | ARG | p.Arg190Cys | Q96EY8 |
| HOMO SAPIENS | 6 | 1092 | 2idx | A | 190 | ARG | p.Arg190His | Q96EY8 |
| HOMO SAPIENS | 7 | 1092 | 2idx | A | 191 | ARG | p.Arg191Trp | Q96EY8 |
| HOMO SAPIENS | 7 | 1092 | 2idx | A | 191 | ARG | p.Arg191Gln | Q96EY8 |
| HOMO SAPIENS | 6 | 1092 | 2idx | A | 195 | ARG | p.Arg195His | Q96EY8 |
| HOMO SAPIENS | 1 | 1092 | 2idx | A | 234 | GLN | p.Gln234Ter | Q96EY8 |
| HOMO SAPIENS | 0 | 1092 | 2idx | A | 239 | MET | p.Met239Lys | Q96EY8 |
| HOMO SAPIENS | -1 | 6418 | 5fur | G | 11 | THR | p.Thr11Ile | P21675 |
| HOMO SAPIENS | -1 | 6418 | 5fur | G | 290 | LEU | p.Leu269Val | P21675 |
| HOMO SAPIENS | 0 | 6100 | 1wiq | A | 240 | ARG | p.Arg265Trp | P01730 |
| HOMO SAPIENS | 0 | 13757 | 4ye6 | A | 45 | GLY | p.Gly45Val | P47897 |
| HOMO SAPIENS | 1 | 13757 | 4ye6 | A | 57 | TYR | p.Tyr57His | P47897 |
| HOMO SAPIENS | 8 | 13757 | 4ye6 | A | 403 | ARG | p.Arg403Trp | P47897 |
| HOMO SAPIENS | 7 | 13757 | 4ye6 | A | 524 | ARG | p.Arg524Trp | P47897 |
| HOMO SAPIENS | 6 | 13757 | 4ye6 | A | 690 | ARG | p.Arg690Cys | P47897 |
| HOMO SAPIENS | 0 | 26958 | 2lzl | A | 370 | GLY | p.Gly370Cys | P22607 |
| HOMO SAPIENS | 0 | 26958 | 2lzl | A | 371 | SER | p.Ser371Cys | P22607 |
| HOMO SAPIENS | 0 | 26958 | 2lzl | A | 373 | TYR | p.Tyr373Cys | P22607 |
| HOMO SAPIENS | 0 | 26958 | 2lzl | A | 380 | GLY | p.Gly380Arg | P22607 |
| HOMO SAPIENS | 0 | 26958 | 2lzl | A | 384 | PHE | p.Phe384Leu | P22607 |
| HOMO SAPIENS | 5 | 26958 | 2lzl | A | 391 | ALA | p.Ala391Glu | P22607 |
| HOMO SAPIENS | 6 | 11325 | 2wa6 | A | 121 | TYR | p.Tyr121Phe | O75369 |
| HOMO SAPIENS | 2 | 11325 | 2wa6 | A | 161 | PHE | p.Phe161Cys | O75369 |
| HOMO SAPIENS | 0 | 11325 | 2wa6 | A | 171 | LEU | p.Leu171Arg | O75369 |
| HOMO SAPIENS | 2 | 11325 | 2wa6 | A | 173 | ALA | p.Ala173Val | O75369 |
| HOMO SAPIENS | 0 | 11325 | 2wa6 | A | 202 | MET | p.Met202Val | O75369 |
| HOMO SAPIENS | 3 | 11325 | 2wa6 | A | 227 | GLU | p.Glu227Lys | O75369 |
| HOMO SAPIENS | 0 | 11325 | 2wa6 | A | 234 | LEU | p.Leu234Val | O75369 |
| HOMO SAPIENS | 2 | 11325 | 2wa6 | A | 235 | SER | p.Ser235Pro | O75369 |
| HOMO SAPIENS | 9 | 7240 | 3tgx | A | 137 | TYR | p.Tyr156Cys | Q9HBE5 |
| HOMO SAPIENS | 0 | 35479 | 4elj | A | 307 | THR | p.Thr307Ile | P06400 |
| HOMO SAPIENS | 0 | 35479 | 4elj | A | 310 | GLY | p.Gly310Glu | P06400 |
| HOMO SAPIENS | 0 | 35479 | 4elj | A | 377 | THR | p.Thr377Ser | P06400 |
| HOMO SAPIENS | 9 | 36706 | 1he7 | A | 359 | TYR | p.Tyr359Cys | P04629 |
| HOMO SAPIENS | 4 | 6538 | 4d0e | A | 504 | ARG | p.Arg504His | P46531 |
| HOMO SAPIENS | 1 | 26518 | 5fm5 | M | 600 | GLU | p.Glu600Val | P52179 |
| HOMO SAPIENS | -1 | 26518 | 5fm5 | M | 651 | ARG | p.Arg651Gln | P52179 |
| HOMO SAPIENS | -1 | 26518 | 5fm5 | M | 660 | PRO | p.Pro660Ala | P52179 |

| | | | | | | | | |
|---|---|---|---|---|---|---|---|---|
| HOMO SAPIENS | -1 | 26518 | 5fm5 | M | 688 | THR | p.Thr688Ser | P52179 |
| HOMO SAPIENS | -1 | 26518 | 5fm5 | M | 704 | GLU | p.Glu704Lys | P52179 |
| HOMO SAPIENS | -1 | 26518 | 5fm5 | M | 727 | THR | p.Thr727Ala | P52179 |
| HOMO SAPIENS | 2 | 27054 | 4rhp | B | 209 | LEU | p.Leu209Val | O75208 |
| HOMO SAPIENS | -1 | 27054 | 4rhp | B | 288 | LYS | p.Lys288Asn | O75208 |
| HOMO SAPIENS | 5 | 2384 | 5fqd | B | 419 | ARG | p.Arg419Ter | Q96SW2 |
| HOMO SAPIENS | 4 | 5154 | 2hl3 | A | 56 | LYS | p.Lys56Arg | Q14203 |
| HOMO SAPIENS | 0 | 5154 | 2hl3 | A | 59 | GLY | p.Gly59Ser | Q14203 |
| HOMO SAPIENS | 0 | 5154 | 2hl3 | A | 71 | GLY | p.Gly71Ala | Q14203 |
| HOMO SAPIENS | 0 | 5154 | 2hl3 | A | 71 | GLY | p.Gly71Arg | Q14203 |
| HOMO SAPIENS | 7 | 5154 | 2hl3 | A | 72 | THR | p.Thr72Pro | Q14203 |
| HOMO SAPIENS | 2 | 5154 | 2hl3 | A | 74 | GLN | p.Gln74Pro | Q14203 |
| HOMO SAPIENS | 2 | 20039 | 2l1g | A | 9 | GLY | p.Gly9Cys | Q9NVV9 |
| HOMO SAPIENS | 5 | 20039 | 2l1g | A | 81 | PHE | p.Phe81Leu | Q9NVV9 |
| HOMO SAPIENS | -1 | 97 | 5uz7 | A | 201 | ARG | p.Arg201His | P63092 |
| HOMO SAPIENS | -1 | 97 | 5uz7 | A | 201 | ARG | p.Arg201Cys | P63092 |
| HOMO SAPIENS | 0 | 3257 | 4b7w | B | 278 | GLU | p.Glu278Asp | P41235 |
| HOMO SAPIENS | 1 | 3257 | 4b7w | B | 331 | ARG | p.Arg331Cys | P41235 |
| HOMO SAPIENS | 0 | 2814 | 4qgk | B | 10 | GLN | p.Gln10Ter | P51648 |
| HOMO SAPIENS | 0 | 2814 | 4qgk | B | 177 | ARG | p.Arg177Ter | P51648 |
| HOMO SAPIENS | 9 | 2814 | 4qgk | B | 184 | THR | p.Thr184Met | P51648 |
| HOMO SAPIENS | 1 | 2814 | 4qgk | B | 188 | ALA | p.Ala188Val | P51648 |
| HOMO SAPIENS | 0 | 2814 | 4qgk | B | 214 | CYS | p.Cys214Tyr | P51648 |
| HOMO SAPIENS | 0 | 2814 | 4qgk | B | 245 | ASP | p.Asp245Asn | P51648 |
| HOMO SAPIENS | 0 | 2814 | 4qgk | B | 266 | LYS | p.Lys266Asn | P51648 |
| HOMO SAPIENS | 0 | 2814 | 4qgk | B | 315 | PRO | p.Pro315Ser | P51648 |
| HOMO SAPIENS | 0 | 2814 | 4qgk | B | 386 | ASN | p.Asn386Ser | P51648 |
| HOMO SAPIENS | -1 | 5486 | 5u6o | B | 47 | GLY | p.Gly47Val | O60741 |
| HOMO SAPIENS | 1 | 21475 | 3pdy | A | 679 | THR | p.Thr789Ile | Q15149 |
| HOMO SAPIENS | 0 | 32890 | 4djc | B | 1501 | LEU | p.Leu1501Val | Q14524 |
| HOMO SAPIENS | -1 | 32890 | 4djc | B | 1512 | ARG | p.Arg1512Trp | Q14524 |
| HOMO SAPIENS | 0 | 26683 | 2xze | B | 38 | ARG | p.Arg38Cys | O95630 |
| HOMO SAPIENS | 0 | 26683 | 2xze | B | 100 | PHE | p.Phe100Tyr | O95630 |
| HOMO SAPIENS | -1 | 23175 | 5dka | A | 163 | SER | p.Ser163Leu | Q96EQ8 |
| HOMO SAPIENS | -1 | 23175 | 5dka | A | 174 | ARG | p.Arg174Cys | Q96EQ8 |
| HOMO SAPIENS | 6 | 21177 | 3k1r | A | 155 | ARG | p.Arg155Ter | Q9Y6N9 |
| HOMO SAPIENS | 0 | 1237 | 2wwm | O | 40 | PRO | p.Pro40Arg | O75147 |
| HOMO SAPIENS | 1 | 2543 | 5ewl | B | 64 | VAL | p.Val64Leu | Q13224 |
| HOMO SAPIENS | 2 | 2543 | 5ewl | B | 271 | ALA | p.Ala271Glu | Q13224 |
| HOMO SAPIENS | 3 | 34262 | 2mng | A | 613 | PHE | p.Phe613Leu | Q9Y3Q4 |
| HOMO SAPIENS | 7 | 34262 | 2mng | A | 672 | SER | p.Ser672Arg | Q9Y3Q4 |
| HOMO SAPIENS | 4 | 1667 | 1f05 | A | 321 | LYS | p.Lys321Met | P37837 |
| HOMO SAPIENS | 0 | 25179 | 2knc | B | 752 | SER | p.Ser778Pro | P05106 |
| HOMO SAPIENS | 0 | 2185 | 2ig7 | A | 50 | TYR | p.Tyr50Cys | Q9Y259 |
| HOMO SAPIENS | 0 | 2185 | 2ig7 | A | 224 | ASN | p.Asn224His | Q9Y259 |
| HOMO SAPIENS | 0 | 2185 | 2ig7 | A | 270 | TYR | p.Tyr270Ter | Q9Y259 |
| HOMO SAPIENS | 0 | 2185 | 2ig7 | A | 328 | GLN | p.Gln328Arg | Q9Y259 |
| HOMO SAPIENS | 3 | 2201 | 2r0o | A | 179 | PRO | p.Pro179Leu | O43707 |
| HOMO SAPIENS | 3 | 2201 | 2r0o | A | 255 | GLU | p.Lys255Glu | O43707 |
| HOMO SAPIENS | 3 | 2201 | 2r0o | A | 259 | THR | p.Thr259Ile | O43707 |
| HOMO SAPIENS | 4 | 2201 | 2r0o | A | 262 | SER | p.Ser262Pro | O43707 |
| HOMO SAPIENS | -1 | 31221 | 2wvr | C | 44 | THR | p.Thr44Pro | Q9H211 |
| HOMO SAPIENS | -1 | 31221 | 2wvr | C | 135 | ALA | p.Ala135Val | Q9H211 |
| HOMO SAPIENS | -1 | 31221 | 2wvr | C | 456 | GLU | p.Glu456Ala | Q9H211 |

| | | | | | | | | |
|---|---|---|---|---|---|---|---|---|
| HOMO SAPIENS | -1 | 31221 | 2wvr | C | 462 | ARG | p.Arg462Gln | Q9H211 |
| HOMO SAPIENS | -1 | 31221 | 2wvr | C | 468 | GLU | p.Glu468Lys | Q9H211 |
| HOMO SAPIENS | -1 | 31221 | 2wvr | C | 537 | ALA | p.Ala537Val | Q9H211 |
| HOMO SAPIENS | 5 | 2576 | 5o3l | C | 363 | VAL | p.Val680Ile | P10636 |
| HOMO SAPIENS | 5 | 1345 | 1czt | A | 37 | ARG | p.Arg2102Cys | P12259 |
| HOMO SAPIENS | 1 | 1345 | 1czt | A | 83 | MET | p.Met2148Thr | P12259 |
| HOMO SAPIENS | 1 | 1345 | 1czt | A | 157 | ASP | p.Asp2222Gly | P12259 |
| HOMO SAPIENS | 8 | 35985 | 1lo1 | A | 110 | ALA | p.Ala110Val | O95718 |
| HOMO SAPIENS | 1 | 617 | 2ocp | B | 44 | GLU | p.Glu44Lys | Q16854 |
| HOMO SAPIENS | 5 | 617 | 2ocp | B | 46 | ASN | p.Asn46Ser | Q16854 |
| HOMO SAPIENS | 0 | 617 | 2ocp | B | 59 | THR | p.Thr59Met | Q16854 |
| HOMO SAPIENS | 7 | 617 | 2ocp | B | 142 | ARG | p.Arg142Lys | Q16854 |
| HOMO SAPIENS | 3 | 617 | 2ocp | B | 154 | ASN | p.Asn154Lys | Q16854 |
| HOMO SAPIENS | 3 | 617 | 2ocp | B | 170 | GLN | p.Gln170Arg | Q16854 |
| HOMO SAPIENS | 3 | 617 | 2ocp | B | 227 | GLU | p.Glu227Lys | Q16854 |
| HOMO SAPIENS | 3 | 617 | 2ocp | B | 233 | LYS | p.Lys233Glu | Q16854 |
| HOMO SAPIENS | 3 | 617 | 2ocp | B | 255 | ASP | p.Asp255Tyr | Q16854 |
| HOMO SAPIENS | -1 | 38975 | 5itz | D | 421 | GLN | p.Gln421His | Q9HC77 |
| HOMO SAPIENS | 2 | 6235 | 2f8v | A | 15 | VAL | p.Val15Ile | Q8WZ42 |
| HOMO SAPIENS | 0 | 6235 | 2f8v | A | 54 | VAL | p.Val54Met | Q8WZ42 |
| HOMO SAPIENS | 0 | 6235 | 2f8v | A | 59 | SER | p.Ser59Arg | Q8WZ42 |
| HOMO SAPIENS | 0 | 6235 | 2f8v | A | 60 | ASP | p.Asp60Tyr | Q8WZ42 |
| HOMO SAPIENS | 0 | 12630 | 1ext | B | 12 | ASP | p.Asp41Glu | P19438 |
| HOMO SAPIENS | 3 | 12630 | 1ext | B | 30 | CYS | p.Cys59Arg | P19438 |
| HOMO SAPIENS | 3 | 12630 | 1ext | B | 30 | CYS | p.Cys59Ser | P19438 |
| HOMO SAPIENS | 5 | 12630 | 1ext | B | 33 | CYS | p.Cys62Tyr | P19438 |
| HOMO SAPIENS | 5 | 12630 | 1ext | B | 33 | CYS | p.Cys62Gly | P19438 |
| HOMO SAPIENS | 1 | 12630 | 1ext | B | 46 | PRO | p.Pro75Leu | P19438 |
| HOMO SAPIENS | 3 | 12630 | 1ext | B | 50 | THR | p.Thr79Met | P19438 |
| HOMO SAPIENS | 6 | 12630 | 1ext | B | 52 | CYS | p.Cys81Phe | P19438 |
| HOMO SAPIENS | 6 | 12630 | 1ext | B | 70 | CYS | p.Cys99Ser | P19438 |
| HOMO SAPIENS | 5 | 12630 | 1ext | B | 88 | CYS | p.Cys117Arg | P19438 |
| HOMO SAPIENS | 5 | 12630 | 1ext | B | 88 | CYS | p.Cys117Tyr | P19438 |
| HOMO SAPIENS | -1 | 12630 | 1ext | B | 170 | ILE | p.Ile199Asn | P19438 |
| HOMO SAPIENS | 0 | 994 | 1aos | B | 95 | ARG | p.Arg95Cys | P04424 |
| HOMO SAPIENS | 0 | 994 | 1aos | B | 111 | ARG | p.Arg111Gln | P04424 |
| HOMO SAPIENS | 0 | 994 | 1aos | B | 131 | THR | p.Thr131Met | P04424 |
| HOMO SAPIENS | 6 | 994 | 1aos | B | 178 | VAL | p.Val178Met | P04424 |
| HOMO SAPIENS | 9 | 994 | 1aos | B | 182 | ARG | p.Arg182Gln | P04424 |
| HOMO SAPIENS | 9 | 994 | 1aos | B | 186 | ARG | p.Arg186Trp | P04424 |
| HOMO SAPIENS | 8 | 994 | 1aos | B | 193 | ARG | p.Arg193Gln | P04424 |
| HOMO SAPIENS | 5 | 994 | 1aos | B | 217 | ARG | p.Arg217Ter | P04424 |
| HOMO SAPIENS | 0 | 994 | 1aos | B | 286 | GLN | p.Gln286Arg | P04424 |
| HOMO SAPIENS | 5 | 994 | 1aos | B | 379 | ARG | p.Arg379Cys | P04424 |
| HOMO SAPIENS | 4 | 16066 | 5mv9 | A | 1719 | TYR | p.Tyr1719Cys | Q13402 |
| HOMO SAPIENS | -1 | 16066 | 5mv9 | A | 1739 | ARG | p.Arg1739Ter | Q13402 |
| HOMO SAPIENS | 4 | 16066 | 5mv9 | A | 1775 | ILE | p.Ile1775Thr | Q13402 |
| HOMO SAPIENS | 2 | 16066 | 5mv9 | A | 1786 | SER | p.Ser1786Thr | Q13402 |
| HOMO SAPIENS | 4 | 16066 | 5mv9 | A | 1798 | GLN | p.Gln1798Ter | Q13402 |
| HOMO SAPIENS | 0 | 16066 | 5mv9 | A | 1858 | LEU | p.Leu1858Pro | Q13402 |
| HOMO SAPIENS | 6 | 16066 | 5mv9 | A | 1873 | ARG | p.Arg1873Trp | Q13402 |
| HOMO SAPIENS | 6 | 16066 | 5mv9 | A | 1873 | ARG | p.Arg1873Leu | Q13402 |
| HOMO SAPIENS | 6 | 16066 | 5mv9 | A | 1883 | ARG | p.Arg1883Gln | Q13402 |
| HOMO SAPIENS | 4 | 16066 | 5mv9 | A | 1887 | PRO | p.Pro1887Leu | Q13402 |

| | | | | | | | | |
|---|---|---|---|---|---|---|---|---|
| HOMO SAPIENS | 5 | 16066 | 5mv9 | A | 1942 | GLY | p.Gly1942Ter | Q13402 |
| HOMO SAPIENS | 5 | 16066 | 5mv9 | A | 1954 | LEU | p.Leu1954Ile | Q13402 |
| HOMO SAPIENS | 5 | 16066 | 5mv9 | A | 1956 | VAL | p.Val1956Ile | Q13402 |
| HOMO SAPIENS | 5 | 16066 | 5mv9 | A | 1967 | ARG | p.Arg1967Ter | Q13402 |
| HOMO SAPIENS | 5 | 16066 | 5mv9 | A | 1990 | GLN | p.Gln1990Ter | Q13402 |
| HOMO SAPIENS | 2 | 16066 | 5mv9 | A | 2005 | LYS | p.Lys2005Glu | Q13402 |
| HOMO SAPIENS | 1 | 16066 | 5mv9 | A | 2072 | VAL | p.Val2072Ile | Q13402 |
| HOMO SAPIENS | 4 | 16066 | 5mv9 | A | 2109 | THR | p.Thr2109Ile | Q13402 |
| HOMO SAPIENS | 4 | 16066 | 5mv9 | A | 2142 | ASP | p.Asp2142Asn | Q13402 |
| HOMO SAPIENS | -1 | 16066 | 5mv9 | A | 2214 | GLY | p.Gly2214Ser | Q13402 |
| HOMO SAPIENS | 7 | 27942 | 1deb | A | 24 | ARG | p.Arg24Ter | P25054 |
| HOMO SAPIENS | 0 | 876 | 5hnr | B | 12 | PHE | p.Phe12Leu | P13716 |
| HOMO SAPIENS | 0 | 876 | 5hnr | B | 59 | LYS | p.Lys59Asn | P13716 |
| HOMO SAPIENS | -1 | 876 | 5hnr | B | 133 | GLY | p.Gly133Arg | P13716 |
| HOMO SAPIENS | 2 | 876 | 5hnr | B | 240 | ARG | p.Arg240Trp | P13716 |
| HOMO SAPIENS | 0 | 876 | 5hnr | B | 274 | ALA | p.Ala274Thr | P13716 |
| HOMO SAPIENS | 0 | 876 | 5hnr | B | 275 | VAL | p.Val275Met | P13716 |
| HOMO SAPIENS | 0 | 1142 | 3qe2 | A | 503 | VAL | p.Ala500Val | P16435 |
| HOMO SAPIENS | 8 | 14358 | 4dkd | C | 32 | VAL | p.Val32Gly | P07333 |
| HOMO SAPIENS | 0 | 14358 | 4dkd | C | 60 | LEU | p.Leu60Val | P07333 |
| HOMO SAPIENS | 7 | 14358 | 4dkd | C | 75 | THR | p.Thr75Ile | P07333 |
| HOMO SAPIENS | -1 | 14358 | 4dkd | C | 90 | PRO | p.Pro90Thr | P07333 |
| HOMO SAPIENS | 0 | 14358 | 4dkd | C | 106 | ARG | p.Arg106Trp | P07333 |
| HOMO SAPIENS | 0 | 14358 | 4dkd | C | 241 | ASN | p.Asn241Asp | P07333 |
| HOMO SAPIENS | 0 | 14358 | 4dkd | C | 245 | ALA | p.Ala245Ser | P07333 |
| HOMO SAPIENS | 0 | 14358 | 4dkd | C | 255 | ASN | p.Asn255Ile | P07333 |
| HOMO SAPIENS | 0 | 14358 | 4dkd | C | 268 | VAL | p.Val268Ile | P07333 |
| HOMO SAPIENS | 0 | 14358 | 4dkd | C | 279 | VAL | p.Val279Met | P07333 |
| HOMO SAPIENS | 0 | 14358 | 4dkd | C | 294 | ARG | p.Arg294Gln | P07333 |
| HOMO SAPIENS | 5 | 8968 | 3ppy | A | 1565 | VAL | p.Val1565Leu | P04275 |
| HOMO SAPIENS | 4 | 8968 | 3ppy | A | 1584 | TYR | p.Tyr1584Cys | P04275 |
| HOMO SAPIENS | 5 | 5341 | 3k3n | A | 279 | PHE | p.Phe315Ser | Q9UPP1 |
| HOMO SAPIENS | 0 | 29463 | 1q7l | A | 197 | ARG | p.Arg197Trp | Q03154 |
| HOMO SAPIENS | 0 | 33761 | 2leh | B | 30 | ASP | p.Asp30Asn | Q16637 |
| HOMO SAPIENS | 6 | 33761 | 2leh | B | 44 | ASP | p.Asp44Val | Q16637 |
| HOMO SAPIENS | -1 | 12249 | 2pnr | C | 224 | GLU | p.Glu310Lys | P10515 |
| HOMO SAPIENS | -1 | 12249 | 2pnr | C | 232 | VAL | p.Val318Ala | P10515 |
| HOMO SAPIENS | -1 | 7 | 5whk | B | -9 | ALA | p.Ala11Pro | P61769 |
| HOMO SAPIENS | 8 | 20298 | 4euw | A | 154 | PHE | p.Phe154Leu | P48436 |
| HOMO SAPIENS | 9 | 20298 | 4euw | A | 158 | ALA | p.Ala158Thr | P48436 |
| HOMO SAPIENS | 5 | 20298 | 4euw | A | 165 | HIS | p.His165Tyr | P48436 |
| HOMO SAPIENS | 3 | 20298 | 4euw | A | 169 | HIS | p.His169Gln | P48436 |
| HOMO SAPIENS | 0 | 20298 | 4euw | A | 173 | LYS | p.Lys173Glu | P48436 |
| HOMO SAPIENS | 1 | 16465 | 2jgx | A | 402 | TYR | p.Tyr402His | P08603 |
| HOMO SAPIENS | 0 | 8993 | 3v8o | A | 659 | LEU | p.Leu659Arg | Q14315 |
| HOMO SAPIENS | 6 | 8993 | 3v8o | A | 693 | ASP | p.Asp693Ala | Q14315 |
| HOMO SAPIENS | 0 | 8993 | 3v8o | A | 709 | ALA | p.Ala709Thr | Q14315 |
| HOMO SAPIENS | 2 | 3852 | 5es4 | B | 106 | ASP | p.Asp128Asn | P05107 |
| HOMO SAPIENS | 2 | 3852 | 5es4 | B | 116 | SER | p.Ser138Pro | P05107 |
| HOMO SAPIENS | 1 | 3852 | 5es4 | B | 127 | LEU | p.Leu149Pro | P05107 |
| HOMO SAPIENS | 0 | 3852 | 5es4 | B | 147 | GLY | p.Gly169Arg | P05107 |
| HOMO SAPIENS | 3 | 3852 | 5es4 | B | 156 | PRO | p.Pro178Leu | P05107 |
| HOMO SAPIENS | 0 | 3852 | 5es4 | B | 174 | LYS | p.Lys196Thr | P05107 |
| HOMO SAPIENS | 2 | 3852 | 5es4 | B | 251 | GLY | p.Gly273Arg | P05107 |

| | | | | | | | | |
|---|---|---|---|---|---|---|---|---|
| HOMO SAPIENS | 2 | 3852 | 5es4 | B | 329 | ASN | p.Asn351Ser | P05107 |
| HOMO SAPIENS | 2 | 3852 | 5es4 | B | 571 | ARG | p.Arg593Cys | P05107 |
| HOMO SAPIENS | 7 | 5962 | 5fn2 | B | 146 | MET | p.Met146Leu | P49768 |
| HOMO SAPIENS | 3 | 5962 | 5fn2 | B | 166 | LEU | p.Leu166Pro | P49768 |
| HOMO SAPIENS | 1 | 5962 | 5fn2 | B | 233 | MET | p.Met233Leu | P49768 |
| HOMO SAPIENS | 1 | 5962 | 5fn2 | B | 261 | VAL | p.Val261Gly | P49768 |
| HOMO SAPIENS | 1 | 5962 | 5fn2 | B | 266 | GLY | p.Gly266Ser | P49768 |
| HOMO SAPIENS | -1 | 5962 | 5fn2 | B | 333 | ASP | p.Asp333Gly | P49768 |
| HOMO SAPIENS | 6 | 1918 | 3dlx | A | 58 | THR | p.Thr58Met | P55809 |
| HOMO SAPIENS | 5 | 1918 | 3dlx | A | 133 | VAL | p.Val133Glu | P55809 |
| HOMO SAPIENS | 5 | 1918 | 3dlx | A | 219 | GLY | p.Gly219Glu | P55809 |
| HOMO SAPIENS | 0 | 1918 | 3dlx | A | 221 | VAL | p.Val221Met | P55809 |
| HOMO SAPIENS | 5 | 1918 | 3dlx | A | 283 | SER | p.Ser283Ter | P55809 |
| HOMO SAPIENS | 4 | 1918 | 3dlx | A | 324 | GLY | p.Gly324Glu | P55809 |
| HOMO SAPIENS | 0 | 1918 | 3dlx | A | 456 | CYS | p.Cys456Phe | P55809 |
| HOMO SAPIENS | 0 | 9010 | 2wfr | A | 162 | LEU | p.Leu162Pro | O43323 |
| HOMO SAPIENS | 5 | 3770 | 5n38 | A | 56 | VAL | p.Val56Glu | O60260 |
| HOMO SAPIENS | 1 | 3770 | 5n38 | A | 161 | LYS | p.Lys161Asn | O60260 |
| HOMO SAPIENS | 1 | 3770 | 5n38 | A | 167 | SER | p.Ser167Asn | O60260 |
| HOMO SAPIENS | 1 | 3770 | 5n38 | A | 211 | LYS | p.Lys211Asn | O60260 |
| HOMO SAPIENS | 1 | 3770 | 5n38 | A | 212 | CYS | p.Cys212Tyr | O60260 |
| HOMO SAPIENS | 1 | 3770 | 5n38 | A | 275 | ARG | p.Arg275Trp | O60260 |
| HOMO SAPIENS | 2 | 3770 | 5n38 | A | 334 | ARG | p.Arg334Cys | O60260 |
| HOMO SAPIENS | 1 | 3770 | 5n38 | A | 366 | ARG | p.Arg366Trp | O60260 |
| HOMO SAPIENS | -1 | 3770 | 5n38 | A | 380 | VAL | p.Val380Leu | O60260 |
| HOMO SAPIENS | 1 | 3770 | 5n38 | A | 394 | ASP | p.Asp394Asn | O60260 |
| HOMO SAPIENS | 1 | 3770 | 5n38 | A | 430 | GLY | p.Gly430Asp | O60260 |
| HOMO SAPIENS | -1 | 2467 | 4xx1 | A | 10 | PRO | p.Pro34Leu | P04180 |
| HOMO SAPIENS | 4 | 2467 | 4xx1 | A | 83 | TYR | p.Tyr107Ter | P04180 |
| HOMO SAPIENS | 4 | 2467 | 4xx1 | A | 93 | ALA | p.Ala117Thr | P04180 |
| HOMO SAPIENS | 4 | 2467 | 4xx1 | A | 123 | THR | p.Thr147Ile | P04180 |
| HOMO SAPIENS | 4 | 2467 | 4xx1 | A | 135 | ARG | p.Arg159Trp | P04180 |
| HOMO SAPIENS | 3 | 2467 | 4xx1 | A | 158 | ARG | p.Arg182Cys | P04180 |
| HOMO SAPIENS | 0 | 2467 | 4xx1 | A | 209 | LEU | p.Leu233Pro | P04180 |
| HOMO SAPIENS | 1 | 2467 | 4xx1 | A | 228 | ASN | p.Asn252Lys | P04180 |
| HOMO SAPIENS | 4 | 2467 | 4xx1 | A | 252 | MET | p.Met276Lys | P04180 |
| HOMO SAPIENS | 3 | 2467 | 4xx1 | A | 293 | MET | p.Met317Ile | P04180 |
| HOMO SAPIENS | 5 | 2467 | 4xx1 | A | 321 | THR | p.Thr345Met | P04180 |
| HOMO SAPIENS | 9 | 2467 | 4xx1 | A | 347 | THR | p.Thr371Met | P04180 |
| HOMO SAPIENS | 1 | 5834 | 3qrf | G | 371 | PHE | p.Phe371Cys | Q9BZS1 |
| HOMO SAPIENS | 1 | 5834 | 3qrf | G | 373 | PHE | p.Phe373Ala | Q9BZS1 |
| HOMO SAPIENS | 9 | 5834 | 3qrf | G | 384 | ALA | p.Ala384Thr | Q9BZS1 |
| HOMO SAPIENS | 6 | 5834 | 3qrf | G | 397 | ARG | p.Arg397Trp | Q9BZS1 |
| HOMO SAPIENS | -1 | 11869 | 4km8 | A | 15 | PRO | p.Pro15Leu | Q9UMX1 |
| HOMO SAPIENS | 5 | 262 | 4kbo | A | 126 | ARG | p.Arg126Trp | P38646 |
| HOMO SAPIENS | 3 | 262 | 4kbo | A | 128 | TYR | p.Tyr128Cys | P38646 |
| HOMO SAPIENS | -1 | 11347 | 4uf9 | D | 117 | ALA | p.Ala117Val | Q13501 |
| HOMO SAPIENS | 0 | 2173 | 4ah6 | A | 45 | SER | p.Ser45Gly | Q6PI48 |
| HOMO SAPIENS | 2 | 2173 | 4ah6 | A | 152 | CYS | p.Cys152Phe | Q6PI48 |
| HOMO SAPIENS | 4 | 2173 | 4ah6 | A | 179 | ARG | p.Arg179His | Q6PI48 |
| HOMO SAPIENS | 0 | 2173 | 4ah6 | A | 196 | LYS | p.Lys196Arg | Q6PI48 |
| HOMO SAPIENS | 8 | 2173 | 4ah6 | A | 263 | ARG | p.Arg263Gln | Q6PI48 |
| HOMO SAPIENS | 1 | 2173 | 4ah6 | A | 338 | GLY | p.Gly338Glu | Q6PI48 |
| HOMO SAPIENS | 0 | 2173 | 4ah6 | A | 425 | GLU | p.Glu425Ter | Q6PI48 |

| | | | | | | | | |
|---|---|---|---|---|---|---|---|---|
| HOMO SAPIENS | 1 | 2173 | 4ah6 | A | 548 | LEU | p.Leu548Met | Q6PI48 |
| HOMO SAPIENS | 0 | 2173 | 4ah6 | A | 609 | ARG | p.Arg609Trp | Q6PI48 |
| HOMO SAPIENS | 2 | 2173 | 4ah6 | A | 613 | LEU | p.Leu613Phe | Q6PI48 |
| HOMO SAPIENS | 0 | 2173 | 4ah6 | A | 626 | LEU | p.Leu626Gln | Q6PI48 |
| HOMO SAPIENS | 0 | 2173 | 4ah6 | A | 626 | LEU | p.Leu626Val | Q6PI48 |
| HOMO SAPIENS | 7 | 3203 | 2cn1 | A | 87 | ASP | p.Asp137Val | Q9H0P0 |
| HOMO SAPIENS | 8 | 3203 | 2cn1 | A | 166 | GLN | p.Gln216Ter | Q9H0P0 |
| HOMO SAPIENS | 8 | 3203 | 2cn1 | A | 179 | ASN | p.Asn229Ser | Q9H0P0 |
| HOMO SAPIENS | 7 | 3203 | 2cn1 | A | 230 | GLY | p.Gly280Arg | Q9H0P0 |
| HOMO SAPIENS | 2 | 3424 | 2a1t | R | 116 | GLY | p.Gly116Arg | P13804 |
| HOMO SAPIENS | 2 | 3424 | 2a1t | R | 157 | VAL | p.Val157Gly | P13804 |
| HOMO SAPIENS | 2 | 3424 | 2a1t | R | 171 | THR | p.Thr171Ile | P13804 |
| HOMO SAPIENS | 0 | 3424 | 2a1t | R | 178 | THR | p.Thr178Arg | P13804 |
| HOMO SAPIENS | 0 | 3424 | 2a1t | R | 194 | VAL | p.Val194Met | P13804 |
| HOMO SAPIENS | 1 | 3424 | 2a1t | R | 209 | ARG | p.Arg209Ter | P13804 |
| HOMO SAPIENS | 6 | 3424 | 2a1t | R | 266 | THR | p.Thr266Met | P13804 |
| HOMO SAPIENS | 0 | 27598 | 1g1c | A | 26 | ARG | p.Arg2098Gly | Q8WZ42 |
| HOMO SAPIENS | 1 | 27598 | 1g1c | A | 46 | ILE | p.Ile2118Thr | Q8WZ42 |
| HOMO SAPIENS | 0 | 32885 | 4bsk | A | 149 | ASN | p.Asn149Asp | P35916 |
| HOMO SAPIENS | -1 | 8074 | 3ghg | B | 17 | ARG | p.Arg47Ter | P02675 |
| HOMO SAPIENS | -1 | 447 | 5g4p | A | 190 | ALA | p.Ala190Thr | P84074 |
| HOMO SAPIENS | 0 | 10761 | 5u1d | B | 379 | VAL | p.Val379Ile | Q03519 |
| HOMO SAPIENS | 0 | 10761 | 5u1d | B | 665 | THR | p.Thr665Ala | Q03519 |
| HOMO SAPIENS | 5 | 29251 | 5fn7 | A | 79 | ILE | p.Ile294Leu | P08575 |
| HOMO SAPIENS | -1 | 27727 | 4ky9 | A | 56 | LYS | p.Lys56Glu | P02730 |
| HOMO SAPIENS | 2 | 27727 | 4ky9 | A | 72 | GLU | p.Glu72Asp | P02730 |
| HOMO SAPIENS | 2 | 27727 | 4ky9 | A | 90 | GLU | p.Glu90Lys | P02730 |
| HOMO SAPIENS | 1 | 27727 | 4ky9 | A | 130 | GLY | p.Gly130Arg | P02730 |
| HOMO SAPIENS | 2 | 27727 | 4ky9 | A | 150 | ARG | p.Arg150Ter | P02730 |
| HOMO SAPIENS | 0 | 27727 | 4ky9 | A | 327 | PRO | p.Pro327Arg | P02730 |
| HOMO SAPIENS | 2 | 13202 | 2fxo | B | 870 | ARG | p.Arg870His | P12883 |
| HOMO SAPIENS | 2 | 13202 | 2fxo | B | 870 | ARG | p.Arg870Cys | P12883 |
| HOMO SAPIENS | 4 | 13202 | 2fxo | B | 904 | ARG | p.Arg904Cys | P12883 |
| HOMO SAPIENS | 4 | 13202 | 2fxo | B | 906 | ASP | p.Asp906Gly | P12883 |
| HOMO SAPIENS | 2 | 13202 | 2fxo | B | 908 | LEU | p.Leu908Val | P12883 |
| HOMO SAPIENS | 2 | 13202 | 2fxo | B | 923 | ASN | p.Asn923Lys | P12883 |
| HOMO SAPIENS | 7 | 13202 | 2fxo | B | 935 | GLU | p.Glu935Lys | P12883 |
| HOMO SAPIENS | 7 | 13202 | 2fxo | B | 935 | GLU | p.Glu935Val | P12883 |
| HOMO SAPIENS | 8 | 13202 | 2fxo | B | 949 | GLU | p.Glu949Lys | P12883 |
| HOMO SAPIENS | 0 | 13200 | 2a1s | A | 349 | ARG | p.Arg349Trp | O95453 |
| HOMO SAPIENS | 0 | 13200 | 2a1s | A | 421 | LYS | p.Lys421Arg | O95453 |
| HOMO SAPIENS | 6 | 3196 | 5xtb | T | 115 | CYS | p.Cys115Tyr | O75380 |
| HOMO SAPIENS | 0 | 1108 | 5ukf | A | 89 | ARG | p.Arg89Gln | Q99986 |
| HOMO SAPIENS | 0 | 1108 | 5ukf | A | 119 | HIS | p.His119Arg | Q99986 |
| HOMO SAPIENS | 0 | 1108 | 5ukf | A | 133 | ARG | p.Arg133Cys | Q99986 |
| HOMO SAPIENS | 0 | 1108 | 5ukf | A | 236 | VAL | p.Val236Leu | Q99986 |
| HOMO SAPIENS | 0 | 1108 | 5ukf | A | 321 | ARG | p.Arg321Cys | Q99986 |
| HOMO SAPIENS | -1 | 1108 | 5ukf | A | 358 | ARG | p.Arg358Ter | Q99986 |
| HOMO SAPIENS | 0 | 5842 | 4yli | C | 204 | GLY | p.Gly204Ser | Q9BWP8 |
| HOMO SAPIENS | -1 | 31362 | 4wqo | A | 25 | PRO | p.Pro25Leu | P40337 |
| HOMO SAPIENS | 2 | 10107 | 2j8h | A | 34 | VAL | p.Val31886Ile | Q8WZ42 |
| HOMO SAPIENS | 8 | 20190 | 1wm5 | A | 77 | ARG | p.Arg77Gln | P19878 |
| HOMO SAPIENS | 1 | 20190 | 1wm5 | A | 100 | GLN | p.Gln100Ter | P19878 |
| HOMO SAPIENS | 0 | 20190 | 1wm5 | A | 102 | ARG | p.Arg102Ter | P19878 |

| | | | | | | | | | |
|---|---|---|---|---|---|---|---|---|---|
| HOMO SAPIENS | 0 | 20190 | 1wm5 | A | | 128 | ALA | p.Ala128Val | P19878 |
| HOMO SAPIENS | 0 | 20190 | 1wm5 | A | | 181 | LYS | p.Lys181Arg | P19878 |
| HOMO SAPIENS | 7 | 1278 | 5a2q | a | | 33 | ASP | p.Asp33Asn | P62854 |
| HOMO SAPIENS | 3 | 19515 | 2obd | A | | 11 | ILE | p.Ile28Met | P11597 |
| HOMO SAPIENS | 3 | 19515 | 2obd | A | | 89 | VAL | p.Val106Met | P11597 |
| HOMO SAPIENS | 3 | 19515 | 2obd | A | | 137 | ARG | p.Arg154Trp | P11597 |
| HOMO SAPIENS | 3 | 19515 | 2obd | A | | 139 | ASP | p.Asp156Asn | P11597 |
| HOMO SAPIENS | 3 | 19515 | 2obd | A | | 204 | SER | p.Ser221Arg | P11597 |
| HOMO SAPIENS | 0 | 19515 | 2obd | A | | 297 | GLU | p.Glu314Lys | P11597 |
| HOMO SAPIENS | 0 | 19515 | 2obd | A | | 314 | GLY | p.Gly331Ser | P11597 |
| HOMO SAPIENS | 0 | 19515 | 2obd | A | | 332 | SER | p.Ser349Tyr | P11597 |
| HOMO SAPIENS | 0 | 19515 | 2obd | A | | 368 | VAL | p.Val385Met | P11597 |
| HOMO SAPIENS | 0 | 19515 | 2obd | A | | 373 | ALA | p.Ala390Pro | P11597 |
| HOMO SAPIENS | 3 | 19515 | 2obd | A | | 405 | ILE | p.Val422Ile | P11597 |
| HOMO SAPIENS | 0 | 19515 | 2obd | A | | 451 | ARG | p.Arg468Gln | P11597 |
| HOMO SAPIENS | 0 | 19515 | 2obd | A | | 451 | ARG | p.Arg468Ter | P11597 |
| HOMO SAPIENS | 0 | 6626 | 3hop | A | | 39 | ALA | p.Ala39Gly | P21333 |
| HOMO SAPIENS | 1 | 6626 | 3hop | A | | 82 | GLU | p.Glu82Val | P21333 |
| HOMO SAPIENS | 1 | 6626 | 3hop | A | | 128 | ALA | p.Ala128Val | P21333 |
| HOMO SAPIENS | 2 | 6626 | 3hop | A | | 203 | ASP | p.Asp203Tyr | P21333 |
| HOMO SAPIENS | 1 | 6626 | 3hop | A | | 207 | PRO | p.Pro207Leu | P21333 |
| HOMO SAPIENS | 2 | 6626 | 3hop | A | | 210 | CYS | p.Cys210Phe | P21333 |
| HOMO SAPIENS | 7 | 6626 | 3hop | A | | 254 | GLU | p.Glu254Lys | P21333 |
| HOMO SAPIENS | 4 | 4210 | 2avg | A | | 8 | VAL | p.Val158Met | Q14896 |
| HOMO SAPIENS | 6 | 4210 | 2avg | A | | 15 | GLU | p.Glu165Asp | Q14896 |
| HOMO SAPIENS | 4 | 4210 | 2avg | A | | 27 | ARG | p.Arg177His | Q14896 |
| HOMO SAPIENS | 0 | 4210 | 2avg | A | | 28 | VAL | p.Val178Met | Q14896 |
| HOMO SAPIENS | 6 | 4210 | 2avg | A | | 39 | VAL | p.Val189Ile | Q14896 |
| HOMO SAPIENS | 2 | 4210 | 2avg | A | | 52 | LYS | p.Lys202Gln | Q14896 |
| HOMO SAPIENS | 4 | 4210 | 2avg | A | | 67 | SER | p.Ser217Gly | Q14896 |
| HOMO SAPIENS | 6 | 4210 | 2avg | A | | 69 | VAL | p.Val219Leu | Q14896 |
| HOMO SAPIENS | 2 | 4210 | 2avg | A | | 86 | GLY | p.Ser236Gly | Q14896 |
| HOMO SAPIENS | 4 | 4210 | 2avg | A | | 108 | GLU | p.Glu258Lys | Q14896 |
| HOMO SAPIENS | 0 | 22237 | 1w6k | A | | 588 | GLY | p.Gly588Ser | P48449 |
| HOMO SAPIENS | 0 | 10765 | 4a5w | B | | 98 | GLU | p.Glu98Ala | P13671 |
| HOMO SAPIENS | 0 | 10765 | 4a5w | B | | 596 | ARG | p.Arg596Ter | P13671 |
| HOMO SAPIENS | -1 | 15515 | 3hx3 | A | | 11 | MSE | p.Met10Lys | P12271 |
| HOMO SAPIENS | 8 | 15515 | 3hx3 | A | | 64 | ARG | p.Arg64Gln | P12271 |
| HOMO SAPIENS | 0 | 15515 | 3hx3 | A | | 182 | PHE | p.Phe182Cys | P12271 |
| HOMO SAPIENS | 0 | 15515 | 3hx3 | A | | 216 | ARG | p.Arg216Gln | P12271 |
| HOMO SAPIENS | -1 | 15515 | 3hx3 | A | | 226 | MSE | p.Met226Lys | P12271 |
| HOMO SAPIENS | 0 | 15515 | 3hx3 | A | | 234 | TRP | p.Arg234Trp | P12271 |
| HOMO SAPIENS | 0 | 15515 | 3hx3 | A | | 273 | LEU | p.Leu273Phe | P12271 |
| HOMO SAPIENS | 0 | 15515 | 3hx3 | A | | 292 | THR | p.Thr292Met | P12271 |
| HOMO SAPIENS | 0 | 5152 | 2www | D | | 95 | GLN | p.Gln95Ter | Q8IVH4 |
| HOMO SAPIENS | 0 | 5152 | 2www | D | | 133 | GLN | p.Gln133Ter | Q8IVH4 |
| HOMO SAPIENS | 0 | 5152 | 2www | D | | 145 | ARG | p.Arg145Ter | Q8IVH4 |
| HOMO SAPIENS | 0 | 5152 | 2www | D | | 207 | TYR | p.Tyr207Cys | Q8IVH4 |
| HOMO SAPIENS | 8 | 5152 | 2www | D | | 330 | ARG | p.Arg330Ter | Q8IVH4 |
| HOMO SAPIENS | 0 | 5152 | 2www | D | | 363 | GLN | p.Gln363His | Q8IVH4 |
| HOMO SAPIENS | 1 | 1626 | 1ilr | | 1 | 105 | SER | p.Ser130Arg | P18510 |
| HOMO SAPIENS | -1 | 929 | 5loh | A | | 12 | GLY | p.Gly12Arg | Q96GX5 |
| HOMO SAPIENS | 9 | 3206 | 1x63 | A | | 21 | CYS | p.Cys104Arg | Q13642 |
| HOMO SAPIENS | 8 | 3206 | 1x63 | A | | 39 | TRP | p.Trp122Ser | Q13642 |

| | | | | | | | | |
|---|---|---|---|---|---|---|---|---|
| HOMO SAPIENS | 9 | 3206 | 1x63 | A | 40 | HIS | p.His123Tyr | Q13642 |
| HOMO SAPIENS | 9 | 3206 | 1x63 | A | 40 | HIS | p.His123Leu | Q13642 |
| HOMO SAPIENS | 9 | 3206 | 1x63 | A | 40 | HIS | p.His123Gln | Q13642 |
| HOMO SAPIENS | 3 | 3206 | 1x63 | A | 49 | CYS | p.Cys132Phe | Q13642 |
| HOMO SAPIENS | 3 | 3206 | 1x63 | A | 67 | CYS | p.Cys150Tyr | Q13642 |
| HOMO SAPIENS | 3 | 3206 | 1x63 | A | 70 | CYS | p.Cys153Arg | Q13642 |
| HOMO SAPIENS | 3 | 3206 | 1x63 | A | 70 | CYS | p.Cys153Tyr | Q13642 |
| HOMO SAPIENS | 5 | 1052 | 3ecs | E | 208 | ASN | p.Asn208Tyr | Q14232 |
| HOMO SAPIENS | 2 | 1052 | 3ecs | E | 275 | TYR | p.Tyr275Cys | Q14232 |
| HOMO SAPIENS | 0 | 24253 | 5i35 | A | 271 | ARG | p.Arg271Cys | Q8NI60 |
| HOMO SAPIENS | 0 | 24253 | 5i35 | A | 272 | GLY | p.Gly272Asp | Q8NI60 |
| HOMO SAPIENS | 0 | 24253 | 5i35 | A | 299 | ARG | p.Arg299Trp | Q8NI60 |
| HOMO SAPIENS | 1 | 24253 | 5i35 | A | 339 | ALA | p.Ala339Thr | Q8NI60 |
| HOMO SAPIENS | 0 | 24253 | 5i35 | A | 348 | ARG | p.Arg348Ter | Q8NI60 |
| HOMO SAPIENS | 0 | 24253 | 5i35 | A | 395 | ILE | p.Ile395Met | Q8NI60 |
| HOMO SAPIENS | 0 | 24253 | 5i35 | A | 396 | ASP | p.Asp396Glu | Q8NI60 |
| HOMO SAPIENS | 0 | 24253 | 5i35 | A | 480 | PHE | p.Phe480Leu | Q8NI60 |
| HOMO SAPIENS | 2 | 24253 | 5i35 | A | 511 | THR | p.Thr511Met | Q8NI60 |
| HOMO SAPIENS | 0 | 24253 | 5i35 | A | 514 | TYR | p.Tyr514Cys | Q8NI60 |
| HOMO SAPIENS | 0 | 24253 | 5i35 | A | 538 | ARG | p.Arg538Gln | Q8NI60 |
| HOMO SAPIENS | 0 | 24253 | 5i35 | A | 549 | GLY | p.Gly549Ser | Q8NI60 |
| HOMO SAPIENS | 0 | 24253 | 5i35 | A | 551 | GLU | p.Glu551Lys | Q8NI60 |
| HOMO SAPIENS | -1 | 24253 | 5i35 | A | 555 | MSE | p.Met555Ile | Q8NI60 |
| HOMO SAPIENS | 0 | 24253 | 5i35 | A | 568 | GLU | p.Glu568Ter | Q8NI60 |
| HOMO SAPIENS | 0 | 24253 | 5i35 | A | 602 | PRO | p.Pro602Arg | Q8NI60 |
| HOMO SAPIENS | -1 | 15505 | 4goj | C | 22 | GLY | p.Gly22Val | Q13432 |
| HOMO SAPIENS | 5 | 23902 | 2lrr | A | 765 | HIS | p.His765Gln | P38935 |
| HOMO SAPIENS | 1 | 29650 | 2vj2 | A | 234 | CYS | p.Cys234Tyr | P78504 |
| HOMO SAPIENS | 0 | 29650 | 2vj2 | A | 255 | TYR | p.Tyr255Ter | P78504 |
| HOMO SAPIENS | 3 | 29650 | 2vj2 | A | 274 | GLY | p.Gly274Asp | P78504 |
| HOMO SAPIENS | 1 | 105 | 5he0 | B | 77 | GLY | p.Gly77Ser | P62873 |
| HOMO SAPIENS | 2 | 105 | 5he0 | B | 80 | ILE | p.Ile80Thr | P62873 |
| HOMO SAPIENS | 3 | 4587 | 3rgh | A | 159 | ASP | p.Asp1159Ala | P21333 |
| HOMO SAPIENS | 3 | 4587 | 3rgh | A | 184 | ASP | p.Asp1184Glu | P21333 |
| HOMO SAPIENS | 2 | 4587 | 3rgh | A | 186 | SER | p.Ser1186Leu | P21333 |
| HOMO SAPIENS | 4 | 4587 | 3rgh | A | 188 | ALA | p.Ala1188Thr | P21333 |
| HOMO SAPIENS | 2 | 4587 | 3rgh | A | 199 | SER | p.Ser1199Leu | P21333 |
| HOMO SAPIENS | 0 | 20283 | 4n3w | A | 1278 | GLU | p.Glu1278Lys | Q92793 |
| HOMO SAPIENS | 1 | 14294 | 2d9q | B | 208 | MET | p.Met231Thr | Q99062 |
| HOMO SAPIENS | 6 | 14294 | 2d9q | B | 297 | ASP | p.Asp320Asn | Q99062 |
| HOMO SAPIENS | 0 | 11642 | 3bpt | A | 66 | ARG | p.Arg66Trp | Q6NVY1 |
| HOMO SAPIENS | 0 | 11642 | 3bpt | A | 122 | TYR | p.Tyr122Cys | Q6NVY1 |
| HOMO SAPIENS | 0 | 11642 | 3bpt | A | 345 | GLY | p.Gly345Ser | Q6NVY1 |
| HOMO SAPIENS | 1 | 3942 | 3sby | A | 60 | ASP | p.Asp60His | Q9Y4U1 |
| HOMO SAPIENS | 0 | 3942 | 3sby | A | 111 | ARG | p.Arg111Ter | Q9Y4U1 |
| HOMO SAPIENS | 1 | 3942 | 3sby | A | 116 | LEU | p.Leu116Pro | Q9Y4U1 |
| HOMO SAPIENS | 1 | 3942 | 3sby | A | 130 | TYR | p.Tyr130Cys | Q9Y4U1 |
| HOMO SAPIENS | 1 | 3942 | 3sby | A | 132 | ARG | p.Arg132Ter | Q9Y4U1 |
| HOMO SAPIENS | 1 | 3942 | 3sby | A | 147 | GLY | p.Gly147Ala | Q9Y4U1 |
| HOMO SAPIENS | 1 | 3942 | 3sby | A | 161 | ARG | p.Arg161Gln | Q9Y4U1 |
| HOMO SAPIENS | 1 | 3942 | 3sby | A | 161 | ARG | p.Arg161Ter | Q9Y4U1 |
| HOMO SAPIENS | 1 | 3942 | 3sby | A | 203 | TRP | p.Trp203Ter | Q9Y4U1 |
| HOMO SAPIENS | 1 | 3942 | 3sby | A | 203 | TRP | p.Trp203Ter | Q9Y4U1 |
| HOMO SAPIENS | 1 | 3942 | 3sby | A | 205 | TYR | p.Tyr205Ter | Q9Y4U1 |

| | | | | | | | | | |
|---|---|---|---|---|---|---|---|---|---|
| HOMO SAPIENS | 0 | 38340 | 2mbh | B | | 39 | MET | p.Met39Leu | Q13351 |
| HOMO SAPIENS | 2 | 7360 | 4k30 | A | | 430 | LEU | p.Leu430Pro | Q8N159 |
| HOMO SAPIENS | 2 | 7360 | 4k30 | A | | 433 | GLU | p.Glu433Asp | Q8N159 |
| HOMO SAPIENS | 7 | 7360 | 4k30 | A | | 484 | TRP | p.Trp484Arg | Q8N159 |
| HOMO SAPIENS | 1 | 41903 | 2xs6 | A | | 151 | PRO | p.Pro151Ser | O00459 |
| HOMO SAPIENS | 0 | 172 | 5glh | A | | 183 | ALA | p.Ala183Gly | P24530 |
| HOMO SAPIENS | 0 | 172 | 5glh | A | | 201 | ARG | p.Arg201Ter | P24530 |
| HOMO SAPIENS | 0 | 172 | 5glh | A | | 244 | THR | p.Thr244Met | P24530 |
| HOMO SAPIENS | 0 | 172 | 5glh | A | | 253 | ARG | p.Arg253Ter | P24530 |
| HOMO SAPIENS | 9 | 172 | 5glh | A | | 1017 | ILE | p.Leu17Phe | P00720 |
| HOMO SAPIENS | -1 | 3342 | 1ljm | A | | 1058 | HIS | p.His58Asn | Q01196 |
| HOMO SAPIENS | 4 | 3342 | 1ljm | A | | 1174 | ARG | p.Arg174Gln | Q01196 |
| HOMO SAPIENS | 5 | 25607 | 1p22 | C | | 32 | ASP | p.Asp32Ala | P35222 |
| HOMO SAPIENS | 5 | 25607 | 1p22 | C | | 32 | ASP | p.Asp32Tyr | P35222 |
| HOMO SAPIENS | -1 | 25607 | 1p22 | C | | 33 | SEP | p.Ser33Phe | P35222 |
| HOMO SAPIENS | 5 | 25607 | 1p22 | C | | 34 | GLY | p.Gly34Glu | P35222 |
| HOMO SAPIENS | -1 | 25607 | 1p22 | C | | 37 | SEP | p.Ser37Cys | P35222 |
| HOMO SAPIENS | -1 | 25607 | 1p22 | C | | 41 | THR | p.Thr41Ile | P35222 |
| HOMO SAPIENS | 1 | 48 | 4uow | | 9 | 60 | GLN | p.Arg34111His | Q8WZ42 |
| HOMO SAPIENS | 0 | 48 | 4uow | | 9 | 64 | LEU | p.Val34115Ala | Q8WZ42 |
| HOMO SAPIENS | 0 | 1127 | 5c01 | A | | 638 | ARG | p.Arg638Ter | P29597 |
| HOMO SAPIENS | 0 | 1127 | 5c01 | A | | 684 | ILE | p.Ile684Ser | P29597 |
| HOMO SAPIENS | 0 | 1127 | 5c01 | A | | 703 | ARG | p.Arg703Trp | P29597 |
| HOMO SAPIENS | 5 | 16389 | 2aq0 | B | | 128 | ALA | p.Ala860Asp | Q92889 |
| HOMO SAPIENS | 2 | 16389 | 2aq0 | B | | 141 | ILE | p.Ile873Val | Q92889 |
| HOMO SAPIENS | 5 | 16389 | 2aq0 | B | | 143 | GLU | p.Glu875Gly | Q92889 |
| HOMO SAPIENS | 0 | 16389 | 2aq0 | B | | 180 | GLY | p.Gly912Arg | Q92889 |
| HOMO SAPIENS | 0 | 15816 | 2xrc | A | | 89 | THR | p.Thr107Ala | P05156 |
| HOMO SAPIENS | 0 | 15816 | 2xrc | A | | 133 | ASN | p.Asn151Ser | P05156 |
| HOMO SAPIENS | 0 | 15816 | 2xrc | A | | 185 | THR | p.Thr203Ile | P05156 |
| HOMO SAPIENS | 0 | 15816 | 2xrc | A | | 225 | GLY | p.Gly243Asp | P05156 |
| HOMO SAPIENS | 0 | 15816 | 2xrc | A | | 243 | GLY | p.Gly261Ala | P05156 |
| HOMO SAPIENS | 0 | 15816 | 2xrc | A | | 269 | GLY | p.Gly287Arg | P05156 |
| HOMO SAPIENS | -1 | 15816 | 2xrc | A | | 288 | ILE | p.Ile306Val | P05156 |
| HOMO SAPIENS | -1 | 15816 | 2xrc | A | | 388 | ARG | p.Arg406His | P05156 |
| HOMO SAPIENS | 7 | 15816 | 2xrc | A | | 398 | ILE | p.Ile416Leu | P05156 |
| HOMO SAPIENS | 9 | 15816 | 2xrc | A | | 400 | HIS | p.His418Leu | P05156 |
| HOMO SAPIENS | 0 | 15816 | 2xrc | A | | 423 | LYS | p.Lys441Arg | P05156 |
| HOMO SAPIENS | -1 | 15816 | 2xrc | A | | 530 | GLU | p.Glu548Gln | P05156 |
| HOMO SAPIENS | 0 | 15816 | 2xrc | A | | 535 | PRO | p.Pro553Ala | P05156 |
| HOMO SAPIENS | 0 | 15816 | 2xrc | A | | 536 | GLU | p.Glu554Gly | P05156 |
| HOMO SAPIENS | 1 | 12385 | 4zlp | B | | 1450 | ALA | p.Ala1450Pro | Q9UM47 |
| HOMO SAPIENS | 7 | 12385 | 4zlp | B | | 1482 | GLY | p.Gly1482Arg | Q9UM47 |
| HOMO SAPIENS | 2 | 12385 | 4zlp | B | | 1518 | LEU | p.Leu1518Met | Q9UM47 |
| HOMO SAPIENS | 0 | 12385 | 4zlp | B | | 1560 | ARG | p.Arg1560Pro | Q9UM47 |
| HOMO SAPIENS | 0 | 1227 | 4a04 | A | | 148 | PHE | p.Phe148Tyr | O43435 |
| HOMO SAPIENS | 4 | 1227 | 4a04 | A | | 194 | HIS | p.His194Gln | O43435 |
| HOMO SAPIENS | 2 | 8017 | 3ei4 | B | | 244 | LYS | p.Lys244Glu | Q92466 |
| HOMO SAPIENS | 1 | 8017 | 3ei4 | B | | 273 | ARG | p.Arg273His | Q92466 |
| HOMO SAPIENS | 3 | 36112 | 2k8f | B | | 9 | SER | p.Ser9Arg | P04637 |
| HOMO SAPIENS | 2 | 12880 | 5ggf | A | | 265 | ARG | p.Arg265His | Q8WZA1 |
| HOMO SAPIENS | 2 | 12880 | 5ggf | A | | 287 | ILE | p.Ile287Ser | Q8WZA1 |
| HOMO SAPIENS | 2 | 12880 | 5ggf | A | | 311 | ARG | p.Arg311Gln | Q8WZA1 |
| HOMO SAPIENS | 2 | 12880 | 5ggf | A | | 425 | TRP | p.Trp425Ser | Q8WZA1 |

| | | | | | | | | |
|---|---|---|---|---|---|---|---|---|
| HOMO SAPIENS | 2 | 12880 | 5ggf | A | 442 | ARG | p.Arg442Cys | Q8WZA1 |
| HOMO SAPIENS | 2 | 12880 | 5ggf | A | 448 | GLY | p.Gly448Arg | Q8WZA1 |
| HOMO SAPIENS | 0 | 12880 | 5ggf | A | 490 | CYS | p.Cys490Tyr | Q8WZA1 |
| HOMO SAPIENS | 0 | 12880 | 5ggf | A | 493 | PRO | p.Pro493Arg | Q8WZA1 |
| HOMO SAPIENS | 3 | 12880 | 5ggf | A | 550 | SER | p.Ser550Asn | Q8WZA1 |
| HOMO SAPIENS | 1 | 12880 | 5ggf | A | 556 | ASP | p.Asp556Asn | Q8WZA1 |
| HOMO SAPIENS | 0 | 12880 | 5ggf | A | 580 | ARG | p.Arg580Ter | Q8WZA1 |
| HOMO SAPIENS | 2 | 12880 | 5ggf | A | 605 | ARG | p.Arg605Pro | Q8WZA1 |
| HOMO SAPIENS | 0 | 12880 | 5ggf | A | 623 | VAL | p.Met623Val | Q8WZA1 |
| HOMO SAPIENS | 1 | 10187 | 4qw2 | A | 145 | ALA | p.Ala145Ser | Q06787 |
| HOMO SAPIENS | 1 | 10187 | 4qw2 | A | 174 | ASN | p.Asn174Lys | Q06787 |
| HOMO SAPIENS | 0 | 24002 | 2m0t | A | 110 | LEU | p.Leu110Val | O14745 |
| HOMO SAPIENS | 9 | 3126 | 4mjh | A | 99 | LEU | p.Leu99Met | P04792 |
| HOMO SAPIENS | 1 | 3126 | 4mjh | A | 127 | ARG | p.Arg127Trp | P04792 |
| HOMO SAPIENS | 1 | 3126 | 4mjh | A | 127 | ARG | p.Arg127Leu | P04792 |
| HOMO SAPIENS | 1 | 3126 | 4mjh | A | 135 | SER | p.Ser135Phe | P04792 |
| HOMO SAPIENS | 2 | 3126 | 4mjh | A | 136 | ARG | p.Arg136Trp | P04792 |
| HOMO SAPIENS | 1 | 3126 | 4mjh | A | 140 | ARG | p.Arg140Gly | P04792 |
| HOMO SAPIENS | 0 | 3126 | 4mjh | A | 151 | THR | p.Thr151Ile | P04792 |
| HOMO SAPIENS | -1 | 15780 | 2jlp | A | 213 | ARG | p.Arg231Gly | P08294 |
| HOMO SAPIENS | 8 | 3578 | 1apz | D | 203 | GLY | p.Gly226Asp | P20933 |
| HOMO SAPIENS | 1 | 3578 | 1apz | D | 279 | GLY | p.Gly302Arg | P20933 |
| HOMO SAPIENS | 0 | 3578 | 1apz | D | 283 | CYS | p.Cys306Arg | P20933 |
| HOMO SAPIENS | 5 | 1335 | 5xtd | C | 115 | ARG | p.Arg115Cys | O75251 |
| HOMO SAPIENS | 0 | 1335 | 5xtd | C | 122 | VAL | p.Val122Met | O75251 |
| HOMO SAPIENS | 2 | 5063 | 2le4 | A | 37 | ARG | p.Arg74Pro | P48431 |
| HOMO SAPIENS | 1 | 25870 | 2o10 | A | 44 | LEU | p.Leu44Pro | P50461 |
| HOMO SAPIENS | 4 | 25870 | 2o10 | A | 58 | CYS | p.Cys58Gly | P50461 |
| HOMO SAPIENS | 0 | 18748 | 5fii | C | 39 | PHE | p.Phe39Leu | P00439 |
| HOMO SAPIENS | 6 | 18748 | 5fii | C | 48 | LEU | p.Leu48Ser | P00439 |
| HOMO SAPIENS | 2 | 18748 | 5fii | C | 65 | ILE | p.Ile65Asn | P00439 |
| HOMO SAPIENS | 3 | 18748 | 5fii | C | 95 | ILE | p.Ile95Phe | P00439 |
| HOMO SAPIENS | 1 | 14261 | 2vay | B | 1539 | ARG | p.Arg1539Cys | Q13698 |
| HOMO SAPIENS | 0 | 2494 | 4dx1 | A | 183 | GLU | p.Glu183Asp | Q9HBA0 |
| HOMO SAPIENS | 0 | 2494 | 4dx1 | A | 217 | ALA | p.Ala217Ser | Q9HBA0 |
| HOMO SAPIENS | 0 | 2494 | 4dx1 | A | 217 | ALA | p.Ala217Val | Q9HBA0 |
| HOMO SAPIENS | 4 | 2494 | 4dx1 | A | 224 | ARG | p.Arg224Trp | Q9HBA0 |
| HOMO SAPIENS | 4 | 2494 | 4dx1 | A | 254 | VAL | p.Val254Leu | Q9HBA0 |
| HOMO SAPIENS | 0 | 2494 | 4dx1 | A | 269 | ARG | p.Arg269Cys | Q9HBA0 |
| HOMO SAPIENS | 0 | 2494 | 4dx1 | A | 269 | ARG | p.Arg269His | Q9HBA0 |
| HOMO SAPIENS | 0 | 2494 | 4dx1 | A | 278 | GLU | p.Glu278Lys | Q9HBA0 |
| HOMO SAPIENS | 0 | 2494 | 4dx1 | A | 315 | ARG | p.Arg315Trp | Q9HBA0 |
| HOMO SAPIENS | 0 | 2494 | 4dx1 | A | 316 | ARG | p.Arg316Cys | Q9HBA0 |
| HOMO SAPIENS | 0 | 2494 | 4dx1 | A | 331 | ILE | p.Ile331Phe | Q9HBA0 |
| HOMO SAPIENS | 0 | 2494 | 4dx1 | A | 333 | ASP | p.Asp333Gly | Q9HBA0 |
| HOMO SAPIENS | 1 | 2494 | 4dx1 | A | 365 | VAL | p.Val365Met | Q9HBA0 |
| HOMO SAPIENS | 0 | 23998 | 2l1r | B | 161 | ARG | p.Arg162Trp | P19429 |
| HOMO SAPIENS | 0 | 8354 | 4f0x | D | 187 | SER | p.Ser187Ter | O95822 |
| HOMO SAPIENS | 2 | 8354 | 4f0x | D | 259 | GLY | p.Gly259Ala | O95822 |
| HOMO SAPIENS | 2 | 8354 | 4f0x | D | 284 | THR | p.Thr284Ala | O95822 |
| HOMO SAPIENS | 3 | 8354 | 4f0x | D | 309 | LYS | p.Lys309Gln | O95822 |
| HOMO SAPIENS | 0 | 21182 | 4apo | A | 228 | GLN | p.Gln228Lys | O00170 |
| HOMO SAPIENS | 1 | 21182 | 4apo | A | 307 | GLN | p.Gln307Arg | O00170 |
| HOMO SAPIENS | 0 | 2287 | 5of4 | B | 112 | ARG | p.Arg112His | P18074 |

| | | | | | | | | |
|---|---|---|---|---|---|---|---|---|
| HOMO SAPIENS | 0 | 2287 | 5of4 | B | 312 | ASP | p.Asp312Asn | P18074 |
| HOMO SAPIENS | 3 | 2287 | 5of4 | B | 461 | LEU | p.Leu461Val | P18074 |
| HOMO SAPIENS | 0 | 2287 | 5of4 | B | 485 | LEU | p.Leu485Pro | P18074 |
| HOMO SAPIENS | 1 | 2287 | 5of4 | B | 541 | SER | p.Ser541Arg | P18074 |
| HOMO SAPIENS | 0 | 2287 | 5of4 | B | 616 | ARG | p.Arg616Pro | P18074 |
| HOMO SAPIENS | 0 | 2287 | 5of4 | B | 616 | ARG | p.Arg616Trp | P18074 |
| HOMO SAPIENS | 1 | 2287 | 5of4 | B | 658 | ARG | p.Arg658Cys | P18074 |
| HOMO SAPIENS | 1 | 2287 | 5of4 | B | 681 | ASP | p.Asp681Asn | P18074 |
| HOMO SAPIENS | 1 | 2287 | 5of4 | B | 683 | ARG | p.Arg683Trp | P18074 |
| HOMO SAPIENS | 0 | 2287 | 5of4 | B | 713 | GLY | p.Gly713Arg | P18074 |
| HOMO SAPIENS | 0 | 2287 | 5of4 | B | 717 | ALA | p.Ala717Gly | P18074 |
| HOMO SAPIENS | 0 | 2287 | 5of4 | B | 722 | ARG | p.Arg722Trp | P18074 |
| HOMO SAPIENS | 0 | 2287 | 5of4 | B | 725 | ALA | p.Ala725Pro | P18074 |
| HOMO SAPIENS | 4 | 12188 | 4its | A | 122 | ARG | p.Arg122Gly | Q9Y606 |
| HOMO SAPIENS | 4 | 12188 | 4its | A | 133 | ASP | p.Asp133Asn | Q9Y606 |
| HOMO SAPIENS | 4 | 12188 | 4its | A | 144 | ARG | p.Arg144Trp | Q9Y606 |
| HOMO SAPIENS | 0 | 31529 | 1kgd | A | 761 | ARG | p.Arg766Gln | O14936 |
| HOMO SAPIENS | 1 | 31529 | 1kgd | A | 824 | GLN | p.Gln829Arg | O14936 |
| HOMO SAPIENS | 0 | 10906 | 2h63 | C | 18 | ARG | p.Arg18Ter | P53004 |
| HOMO SAPIENS | 9 | 27767 | 4wyq | A | 280 | LEU | p.Leu278Phe | Q9UPY3 |
| HOMO SAPIENS | 0 | 27767 | 4wyq | A | 297 | SER | p.Ser295Cys | Q9UPY3 |
| HOMO SAPIENS | 1 | 27767 | 4wyq | A | 334 | TYR | p.Tyr332Ter | Q9UPY3 |
| HOMO SAPIENS | 7 | 7775 | 5sv7 | C | 136 | SER | p.Ser136Cys | Q9NZJ5 |
| HOMO SAPIENS | 2 | 7775 | 5sv7 | C | 166 | GLN | p.Gln166Arg | Q9NZJ5 |
| HOMO SAPIENS | 2 | 7775 | 5sv7 | C | 332 | GLU | p.Glu332Ter | Q9NZJ5 |
| HOMO SAPIENS | 0 | 21449 | 2e9w | B | 84 | THR | p.Thr84Met | P10721 |
| HOMO SAPIENS | 0 | 21449 | 2e9w | B | 168 | ALA | p.Ala168Glu | P10721 |
| HOMO SAPIENS | 0 | 21449 | 2e9w | B | 178 | ALA | p.Ala178Thr | P10721 |
| HOMO SAPIENS | 0 | 21449 | 2e9w | B | 293 | ASN | p.Asn293Ser | P10721 |
| HOMO SAPIENS | -1 | 21449 | 2e9w | B | 318 | MET | p.Met318Leu | P10721 |
| HOMO SAPIENS | 0 | 21449 | 2e9w | B | 374 | VAL | p.Val374Ile | P10721 |
| HOMO SAPIENS | 6 | 3010 | 3flg | A | 66 | SER | p.Ser270Pro | Q9UBC3 |
| HOMO SAPIENS | 1 | 33918 | 2xzz | A | 760 | ARG | p.Arg760Ter | P22735 |
| HOMO SAPIENS | 2 | 1254 | 5xtc | w | 65 | ASN | p.Asn65Ser | O95299 |
| HOMO SAPIENS | 1 | 1254 | 5xtc | w | 238 | GLU | p.Glu238Lys | O95299 |
| HOMO SAPIENS | 3 | 10245 | 3siv | B | 194 | ALA | p.Ala194Glu | Q8WWY3 |
| HOMO SAPIENS | 4 | 10245 | 3siv | B | 216 | ALA | p.Ala216Pro | Q8WWY3 |
| HOMO SAPIENS | 0 | 2670 | 3dwb | A | 341 | THR | p.Thr341Ile | P42892 |
| HOMO SAPIENS | 1 | 2670 | 3dwb | A | 462 | SER | p.Ser462Asn | P42892 |
| HOMO SAPIENS | 1 | 2670 | 3dwb | A | 754 | ARG | p.Arg754Cys | P42892 |
| HOMO SAPIENS | 1 | 2961 | 5xtb | O | 40 | VAL | p.Val40Met | P19404 |
| HOMO SAPIENS | 5 | 2961 | 5xtb | O | 172 | ILE | p.Ile172Thr | P19404 |
| HOMO SAPIENS | 5 | 2961 | 5xtb | O | 190 | ASP | p.Asp190Gly | P19404 |
| HOMO SAPIENS | 5 | 29655 | 2xdg | A | 57 | ALA | p.Ala57Thr | Q02643 |
| HOMO SAPIENS | 4 | 29655 | 2xdg | A | 72 | GLU | p.Glu72Ter | Q02643 |
| HOMO SAPIENS | 0 | 29655 | 2xdg | A | 121 | GLU | p.Glu121Asp | Q02643 |
| HOMO SAPIENS | -1 | 17935 | 1hyn | P | 38 | ASP | p.Asp38Ala | P02730 |
| HOMO SAPIENS | 7 | 12825 | 1g82 | D | 99 | SER | p.Ser99Asn | P31371 |
| HOMO SAPIENS | 6 | 24927 | 1jm7 | B | 85 | VAL | p.Val85Leu | Q99728 |
| HOMO SAPIENS | 1 | 24927 | 1jm7 | B | 112 | ARG | p.Arg112Ter | Q99728 |
| HOMO SAPIENS | 0 | 24927 | 1jm7 | B | 116 | HIS | p.His116Tyr | Q99728 |
| HOMO SAPIENS | 7 | 92 | 5hde | A | 61 | LYS | p.Lys61Arg | Q05209 |
| HOMO SAPIENS | 0 | 17246 | 1szb | A | 84 | ARG | p.Arg99Gln | O00187 |
| HOMO SAPIENS | 1 | 17246 | 1szb | A | 105 | ASP | p.Asp120Gly | O00187 |

| | | | | | | | | |
|---|---|---|---|---|---|---|---|---|
| HOMO SAPIENS | 0 | 17246 | 1szb | A | 111 | PRO | p.Pro126Leu | O00187 |
| HOMO SAPIENS | 9 | 17246 | 1szb | A | 141 | CYS | p.Cys156Tyr | O00187 |
| HOMO SAPIENS | 0 | 32473 | 1b72 | A | 207 | ARG | p.Arg207Cys | P14653 |
| HOMO SAPIENS | 0 | 1920 | 4yzf | B | 488 | VAL | p.Val488Met | P02730 |
| HOMO SAPIENS | 0 | 1920 | 4yzf | B | 589 | ARG | p.Arg589Cys | P02730 |
| HOMO SAPIENS | 0 | 1920 | 4yzf | B | 589 | ARG | p.Arg589His | P02730 |
| HOMO SAPIENS | 0 | 1920 | 4yzf | B | 602 | ARG | p.Arg602Pro | P02730 |
| HOMO SAPIENS | 0 | 1920 | 4yzf | B | 613 | SER | p.Ser613Phe | P02730 |
| HOMO SAPIENS | -1 | 1920 | 4yzf | B | 646 | ARG | p.Arg646Gln | P02730 |
| HOMO SAPIENS | -1 | 1920 | 4yzf | B | 646 | ARG | p.Arg646Trp | P02730 |
| HOMO SAPIENS | 0 | 1920 | 4yzf | B | 701 | GLY | p.Gly701Asp | P02730 |
| HOMO SAPIENS | 0 | 1920 | 4yzf | B | 760 | ARG | p.Arg760Gln | P02730 |
| HOMO SAPIENS | 0 | 1920 | 4yzf | B | 771 | GLY | p.Gly771Asp | P02730 |
| HOMO SAPIENS | 0 | 1920 | 4yzf | B | 773 | SER | p.Ser773Pro | P02730 |
| HOMO SAPIENS | 0 | 1920 | 4yzf | B | 837 | THR | p.Thr837Ala | P02730 |
| HOMO SAPIENS | 0 | 1920 | 4yzf | B | 854 | PRO | p.Pro854Leu | P02730 |
| HOMO SAPIENS | 0 | 1920 | 4yzf | B | 858 | ALA | p.Ala858Asp | P02730 |
| HOMO SAPIENS | 0 | 1920 | 4yzf | B | 862 | VAL | p.Val862Ile | P02730 |
| HOMO SAPIENS | 0 | 1920 | 4yzf | B | 868 | PRO | p.Pro868Leu | P02730 |
| HOMO SAPIENS | 0 | 1920 | 4yzf | B | 870 | ARG | p.Arg870Trp | P02730 |
| HOMO SAPIENS | 0 | 9863 | 2juj | A | 54 | VAL | p.Val904Ile | P22681 |
| HOMO SAPIENS | 0 | 21934 | 2na7 | A | 184 | ILE | p.Ile184Val | P25445 |
| HOMO SAPIENS | -1 | 6746 | 1lw3 | A | 3 | THR | p.Lys3Thr | Q13614 |
| HOMO SAPIENS | -1 | 6746 | 1lw3 | A | 44 | VAL | p.Val44Ile | Q13614 |
| HOMO SAPIENS | 3 | 6746 | 1lw3 | A | 183 | ALA | p.Ala183Ser | Q13614 |
| HOMO SAPIENS | 4 | 6746 | 1lw3 | A | 482 | GLN | p.Gln482Ter | Q13614 |
| HOMO SAPIENS | 4 | 6746 | 1lw3 | A | 502 | GLU | p.Glu502Gln | Q13614 |
| HOMO SAPIENS | 3 | 6746 | 1lw3 | A | 545 | ASN | p.Asn545Ser | Q13614 |
| HOMO SAPIENS | 0 | 6746 | 1lw3 | A | 586 | ARG | p.Arg586Trp | Q13614 |
| HOMO SAPIENS | -1 | 6746 | 1lw3 | A | 602 | ALA | p.Ala602Gly | Q13614 |
| HOMO SAPIENS | 1 | 319 | 4ch9 | B | 336 | ARG | p.Arg336Ile | Q9UH77 |
| HOMO SAPIENS | 3 | 319 | 4ch9 | B | 340 | ALA | p.Ala340Val | Q9UH77 |
| HOMO SAPIENS | 0 | 319 | 4ch9 | B | 384 | ARG | p.Arg384Gln | Q9UH77 |
| HOMO SAPIENS | 5 | 319 | 4ch9 | B | 426 | PRO | p.Pro426Leu | Q9UH77 |
| HOMO SAPIENS | 3 | 319 | 4ch9 | B | 431 | ARG | p.Arg431Gln | Q9UH77 |
| HOMO SAPIENS | 2 | 319 | 4ch9 | B | 575 | ARG | p.Arg575Trp | Q9UH77 |
| HOMO SAPIENS | 0 | 574 | 2qkk | A | 142 | VAL | p.Val142Ile | O60930 |
| HOMO SAPIENS | 0 | 24021 | 2n8y | A | 752 | ARG | p.Arg752Gln | P12814 |
| HOMO SAPIENS | 0 | 18693 | 2y7j | B | 44 | ARG | p.Arg44Ter | P15735 |
| HOMO SAPIENS | 0 | 18693 | 2y7j | B | 106 | VAL | p.Val106Glu | P15735 |
| HOMO SAPIENS | 3 | 18693 | 2y7j | B | 189 | GLY | p.Gly189Glu | P15735 |
| HOMO SAPIENS | 0 | 17087 | 2ifg | A | 237 | THR | p.Thr237Met | P04629 |
| HOMO SAPIENS | 0 | 7980 | 1qub | A | 247 | VAL | p.Val266Leu | P02749 |
| HOMO SAPIENS | -1 | 2205 | 1h7u | A | 18 | ILE | p.Ile18Val | P54278 |
| HOMO SAPIENS | -1 | 2205 | 1h7u | A | 20 | ARG | p.Arg20Gln | P54278 |
| HOMO SAPIENS | 9 | 2205 | 1h7u | A | 46 | SER | p.Ser46Ile | P54278 |
| HOMO SAPIENS | 0 | 2205 | 1h7u | A | 60 | ASP | p.Asp60Glu | P54278 |
| HOMO SAPIENS | 0 | 2205 | 1h7u | A | 80 | PHE | p.Phe80Leu | P54278 |
| HOMO SAPIENS | 0 | 2205 | 1h7u | A | 118 | SER | p.Ser118Arg | P54278 |
| HOMO SAPIENS | 0 | 2205 | 1h7u | A | 126 | HIS | p.His126Gln | P54278 |
| HOMO SAPIENS | 0 | 2205 | 1h7u | A | 128 | SER | p.Ser128Leu | P54278 |
| HOMO SAPIENS | 1 | 2205 | 1h7u | A | 134 | ARG | p.Arg134Ter | P54278 |
| HOMO SAPIENS | 0 | 2205 | 1h7u | A | 166 | LEU | p.Leu166Pro | P54278 |
| HOMO SAPIENS | 1 | 2205 | 1h7u | A | 191 | TYR | p.Tyr191Cys | P54278 |

| | | | | | | | | |
|---|---|---|---|---|---|---|---|---|
| HOMO SAPIENS | 0 | 2205 | 1h7u | A | 211 | ARG | p.Arg211Ter | P54278 |
| HOMO SAPIENS | 0 | 2205 | 1h7u | A | 233 | GLN | p.Gln233Ter | P54278 |
| HOMO SAPIENS | 0 | 2205 | 1h7u | A | 255 | TYR | p.Tyr255Ter | P54278 |
| HOMO SAPIENS | 0 | 2205 | 1h7u | A | 277 | THR | p.Thr277Lys | P54278 |
| HOMO SAPIENS | 0 | 2205 | 1h7u | A | 301 | LYS | p.Lys301Asn | P54278 |
| HOMO SAPIENS | 0 | 2205 | 1h7u | A | 312 | MET | p.Met312Ile | P54278 |
| HOMO SAPIENS | 0 | 2205 | 1h7u | A | 312 | MET | p.Met312Thr | P54278 |
| HOMO SAPIENS | 0 | 2205 | 1h7u | A | 315 | ARG | p.Arg315Ter | P54278 |
| HOMO SAPIENS | 0 | 2205 | 1h7u | A | 318 | TYR | p.Tyr318Cys | P54278 |
| HOMO SAPIENS | 0 | 2205 | 1h7u | A | 335 | ASN | p.Asn335Ile | P54278 |
| HOMO SAPIENS | 0 | 2205 | 1h7u | A | 360 | ILE | p.Ile360Met | P54278 |
| HOMO SAPIENS | 6 | 20754 | 5mpo | B | 7 | VAL | p.Val7Phe | O96033 |
| HOMO SAPIENS | 4 | 20754 | 5mpo | B | 64 | VAL | p.Val64Ile | O96033 |
| HOMO SAPIENS | 5 | 34944 | 4qam | B | 1114 | ASP | p.Asp1114Gly | Q96KN7 |
| HOMO SAPIENS | 5 | 34944 | 4qam | B | 1240 | GLY | p.Gly1240Glu | Q96KN7 |
| HOMO SAPIENS | 9 | 40496 | 3vnn | A | 278 | ARG | p.Arg278His | P49917 |
| HOMO SAPIENS | 6 | 30131 | 4dt7 | E | 40 | PRO | p.Pro210Leu | P04070 |
| HOMO SAPIENS | -1 | 30131 | 4dt7 | E | 50 | ARG | p.Arg220Trp | P04070 |
| HOMO SAPIENS | 7 | 15311 | 2pqt | A | 64 | ARG | p.Arg64Trp | P18440 |
| HOMO SAPIENS | 2 | 15311 | 2pqt | A | 149 | VAL | p.Val149Ile | P18440 |
| HOMO SAPIENS | -1 | 19581 | 3hy8 | A | 33 | ASP | p.Asp33Val | Q9NVS9 |
| HOMO SAPIENS | 5 | 19581 | 3hy8 | A | 116 | ARG | p.Arg116Gln | Q9NVS9 |
| HOMO SAPIENS | 4 | 19581 | 3hy8 | A | 225 | ARG | p.Arg225His | Q9NVS9 |
| HOMO SAPIENS | 2 | 19581 | 3hy8 | A | 229 | TRP | p.Arg229Trp | Q9NVS9 |
| HOMO SAPIENS | 2 | 19581 | 3hy8 | A | 229 | TRP | p.Arg229Gln | Q9NVS9 |
| HOMO SAPIENS | 0 | 10266 | 3t1i | A | 41 | ASP | p.Asp41His | P49959 |
| HOMO SAPIENS | 1 | 10266 | 3t1i | A | 305 | ARG | p.Arg305Trp | P49959 |
| HOMO SAPIENS | 0 | 10266 | 3t1i | A | 308 | PHE | p.Phe308Leu | P49959 |
| HOMO SAPIENS | 0 | 10266 | 3t1i | A | 334 | SER | p.Ser334Arg | P49959 |
| HOMO SAPIENS | 1 | 1600 | 3rbf | A | 61 | GLU | p.Glu61Asp | P20711 |
| HOMO SAPIENS | 0 | 1600 | 3rbf | A | 91 | ALA | p.Ala91Val | P20711 |
| HOMO SAPIENS | -1 | 1600 | 3rbf | A | 102 | GLY | p.Gly102Ser | P20711 |
| HOMO SAPIENS | 3 | 1600 | 3rbf | A | 147 | SER | p.Ser147Arg | P20711 |
| HOMO SAPIENS | 1 | 1600 | 3rbf | A | 210 | PRO | p.Pro210Leu | P20711 |
| HOMO SAPIENS | 0 | 1600 | 3rbf | A | 217 | MET | p.Met217Val | P20711 |
| HOMO SAPIENS | 4 | 1600 | 3rbf | A | 250 | SER | p.Ser250Phe | P20711 |
| HOMO SAPIENS | 2 | 1600 | 3rbf | A | 275 | ALA | p.Ala275Thr | P20711 |
| HOMO SAPIENS | 1 | 1600 | 3rbf | A | 309 | PHE | p.Phe309Leu | P20711 |
| HOMO SAPIENS | -1 | 1600 | 3rbf | A | 347 | ARG | p.Arg347Gln | P20711 |
| HOMO SAPIENS | 0 | 1600 | 3rbf | A | 462 | ARG | p.Arg462Gln | P20711 |
| HOMO SAPIENS | 6 | 20830 | 5y4f | A | 714 | THR | p.Thr715Ile | Q01484 |
| HOMO SAPIENS | 7 | 20830 | 5y4f | A | 719 | ASP | p.Asp720Glu | Q01484 |
| HOMO SAPIENS | 0 | 20830 | 5y4f | A | 814 | VAL | p.Val815Ile | Q01484 |
| HOMO SAPIENS | -1 | 302 | 2c9y | A | 9 | GLU | p.Glu9Ter | P54819 |
| HOMO SAPIENS | 0 | 302 | 2c9y | A | 103 | ARG | p.Arg103Trp | P54819 |
| HOMO SAPIENS | 3 | 302 | 2c9y | A | 165 | ASP | p.Asp165Gly | P54819 |
| HOMO SAPIENS | 0 | 302 | 2c9y | A | 182 | ALA | p.Ala182Asp | P54819 |
| HOMO SAPIENS | 2 | 302 | 2c9y | A | 186 | ARG | p.Arg186Cys | P54819 |
| HOMO SAPIENS | 0 | 302 | 2c9y | A | 209 | ALA | p.Ala209Thr | P54819 |
| HOMO SAPIENS | -1 | 16920 | 1jlj | B | 10 | ASN | p.Asn10Tyr | Q9NQX3 |
| HOMO SAPIENS | 9 | 8462 | 1zbq | B | 16 | GLY | p.Gly16Ser | P51659 |
| HOMO SAPIENS | 5 | 8462 | 1zbq | B | 19 | ALA | p.Ala19Gly | P51659 |
| HOMO SAPIENS | 1 | 8462 | 1zbq | B | 34 | ALA | p.Ala34Val | P51659 |
| HOMO SAPIENS | 1 | 8462 | 1zbq | B | 106 | ARG | p.Arg106His | P51659 |

| | | | | | | | | |
|---|---|---|---|---|---|---|---|---|
| HOMO SAPIENS | 0 | 8462 | 1zbq | B | 140 | LYS | p.Lys140Asn | P51659 |
| HOMO SAPIENS | 0 | 8462 | 1zbq | B | 292 | THR | p.Thr292Ser | P51659 |
| HOMO SAPIENS | 0 | 4375 | 4ndi | A | 199 | ARG | p.Arg213His | Q7Z2E3 |
| HOMO SAPIENS | 3 | 4375 | 4ndi | A | 201 | HIS | p.His215Arg | Q7Z2E3 |
| HOMO SAPIENS | 0 | 4375 | 4ndi | A | 206 | PRO | p.Pro220Leu | Q7Z2E3 |
| HOMO SAPIENS | 2 | 4375 | 4ndi | A | 223 | LEU | p.Leu237Phe | Q7Z2E3 |
| HOMO SAPIENS | 0 | 4375 | 4ndi | A | 245 | ARG | p.Arg259His | Q7Z2E3 |
| HOMO SAPIENS | 0 | 4375 | 4ndi | A | 263 | VAL | p.Val277Gly | Q7Z2E3 |
| HOMO SAPIENS | 0 | 4375 | 4ndi | A | 279 | TRP | p.Trp293Ter | Q7Z2E3 |
| HOMO SAPIENS | 0 | 27004 | 4ccg | Y | 321 | ASP | p.Asp321Glu | Q9NW38 |
| HOMO SAPIENS | -1 | 27004 | 4ccg | Y | 372 | GLY | p.Gly372Ala | Q9NW38 |
| HOMO SAPIENS | -1 | 6383 | 5fai | A | 34 | ALA | p.Ala34Gly | Q92979 |
| HOMO SAPIENS | 8 | 6383 | 5fai | A | 86 | ASP | p.Asp86Gly | Q92979 |
| HOMO SAPIENS | 2 | 24585 | 4mi5 | A | 684 | ARG | p.Arg679Cys | Q15910 |
| HOMO SAPIENS | 0 | 15967 | 3nhb | A | 723 | ARG | p.Arg723Gln | Q9NP58 |
| HOMO SAPIENS | 5 | 18152 | 1oey | A | 389 | HIS | p.His389Gln | P19878 |
| HOMO SAPIENS | 5 | 18152 | 1oey | A | 395 | ARG | p.Arg395Trp | P19878 |
| HOMO SAPIENS | -1 | 1044 | 1li1 | F | 2 | VAL | p.Val1486Ile | P08572 |
| HOMO SAPIENS | 6 | 1044 | 1li1 | F | 37 | GLU | p.Glu1521Lys | P08572 |
| HOMO SAPIENS | 5 | 1044 | 1li1 | F | 157 | SER | p.Ser1639Leu | P08572 |
| HOMO SAPIENS | 5 | 1044 | 1li1 | F | 179 | ARG | p.Arg1662His | P08572 |
| HOMO SAPIENS | 5 | 1044 | 1li1 | F | 180 | GLY | p.Gly1663Arg | P08572 |
| HOMO SAPIENS | 6 | 20076 | 4uyi | A | 671 | LEU | p.Leu671Ser | Q8IY92 |
| HOMO SAPIENS | 0 | 19588 | 1jmj | A | 189 | ARG | p.Arg208His | P05546 |
| HOMO SAPIENS | -1 | 38023 | 1ry7 | B | 65 | GLY | p.Gly65Arg | P22607 |
| HOMO SAPIENS | -1 | 38023 | 1ry7 | B | 139 | ASP | p.Asp139Glu | P22607 |
| HOMO SAPIENS | 0 | 38023 | 1ry7 | B | 248 | ARG | p.Arg248Cys | P22607 |
| HOMO SAPIENS | 0 | 38023 | 1ry7 | B | 249 | SER | p.Ser249Cys | P22607 |
| HOMO SAPIENS | 0 | 38023 | 1ry7 | B | 250 | PRO | p.Pro250Arg | P22607 |
| HOMO SAPIENS | 0 | 38023 | 1ry7 | B | 322 | GLU | p.Glu322Lys | P22607 |
| HOMO SAPIENS | 5 | 16884 | 5eli | A | 33 | GLN | p.Gln33Ter | Q9NZC2 |
| HOMO SAPIENS | 0 | 16884 | 5eli | A | 44 | TRP | p.Trp44Ter | Q9NZC2 |
| HOMO SAPIENS | 0 | 16884 | 5eli | A | 47 | ARG | p.Arg47His | Q9NZC2 |
| HOMO SAPIENS | 0 | 16884 | 5eli | A | 66 | THR | p.Thr66Met | Q9NZC2 |
| HOMO SAPIENS | 0 | 16884 | 5eli | A | 78 | TRP | p.Trp78Ter | Q9NZC2 |
| HOMO SAPIENS | 1 | 16884 | 5eli | A | 96 | THR | p.Thr96Lys | Q9NZC2 |
| HOMO SAPIENS | 9 | 16884 | 5eli | A | 126 | VAL | p.Val126Gly | Q9NZC2 |
| HOMO SAPIENS | -1 | 12267 | 3w5o | A | 3 | ALA | p.Ala3Val | P49917 |
| HOMO SAPIENS | 1 | 12267 | 3w5o | A | 9 | THR | p.Thr9Ile | P49917 |
| HOMO SAPIENS | 2 | 12267 | 3w5o | A | 188 | ARG | p.Arg188Gln | P49917 |
| HOMO SAPIENS | 0 | 12267 | 3w5o | A | 469 | GLY | p.Gly469Glu | P49917 |
| HOMO SAPIENS | 1 | 12267 | 3w5o | A | 580 | ARG | p.Arg580Ter | P49917 |
| HOMO SAPIENS | 0 | 12267 | 3w5o | A | 600 | GLU | p.Glu600Lys | P49917 |
| HOMO SAPIENS | 9 | 2839 | 2hw5 | B | 54 | ARG | p.Arg54Cys | P30084 |
| HOMO SAPIENS | 9 | 2839 | 2hw5 | B | 59 | ASN | p.Asn59Ser | P30084 |
| HOMO SAPIENS | 0 | 2839 | 2hw5 | B | 273 | LYS | p.Lys273Glu | P30084 |
| HOMO SAPIENS | 1 | 12266 | 2o2k | B | 939 | GLY | p.Gly939Cys | Q99707 |
| HOMO SAPIENS | 5 | 12266 | 2o2k | B | 1164 | ARG | p.Arg1164His | Q99707 |
| HOMO SAPIENS | 1 | 12266 | 2o2k | B | 1166 | LEU | p.Leu1166Met | Q99707 |
| HOMO SAPIENS | 5 | 12266 | 2o2k | B | 1173 | PRO | p.Pro1173Leu | Q99707 |
| HOMO SAPIENS | 5 | 12266 | 2o2k | B | 1222 | ASN | p.Asn1222Ser | Q99707 |
| HOMO SAPIENS | 0 | 30891 | 3ch4 | B | 138 | ARG | p.Arg138Ter | Q15126 |
| HOMO SAPIENS | -1 | 4433 | 1qlz | A | 54 | GLY | p.Gly54Ser | P04156 |
| HOMO SAPIENS | 0 | 22568 | 3cxl | A | 20 | LEU | p.Leu20Phe | P15882 |

| | | | | | | | | |
|---|---|---|---|---|---|---|---|---|
| HOMO SAPIENS | 0 | 22568 | 3cxl | A | 126 | ILE | p.Ile126Met | P15882 |
| HOMO SAPIENS | 0 | 22568 | 3cxl | A | 141 | PRO | p.Pro141Leu | P15882 |
| HOMO SAPIENS | 0 | 22568 | 3cxl | A | 143 | TYR | p.Tyr143His | P15882 |
| HOMO SAPIENS | 0 | 22568 | 3cxl | A | 223 | ALA | p.Ala223Val | P15882 |
| HOMO SAPIENS | 0 | 22568 | 3cxl | A | 228 | GLY | p.Gly228Ser | P15882 |
| HOMO SAPIENS | 0 | 22568 | 3cxl | A | 252 | PRO | p.Pro252Gln | P15882 |
| HOMO SAPIENS | 0 | 22568 | 3cxl | A | 313 | GLU | p.Glu313Lys | P15882 |
| HOMO SAPIENS | 3 | 2668 | 5eqg | A | 34 | ASN | p.Asn34Ile | P11166 |
| HOMO SAPIENS | 3 | 2668 | 5eqg | A | 77 | MET | p.Met77Val | P11166 |
| HOMO SAPIENS | 6 | 2668 | 5eqg | A | 91 | GLY | p.Gly91Asp | P11166 |
| HOMO SAPIENS | 6 | 2668 | 5eqg | A | 92 | ARG | p.Arg92Trp | P11166 |
| HOMO SAPIENS | 6 | 2668 | 5eqg | A | 93 | ARG | p.Arg93Trp | P11166 |
| HOMO SAPIENS | 0 | 2668 | 5eqg | A | 95 | SER | p.Ser95Ile | P11166 |
| HOMO SAPIENS | 3 | 2668 | 5eqg | A | 126 | ARG | p.Arg126His | P11166 |
| HOMO SAPIENS | 3 | 2668 | 5eqg | A | 126 | ARG | p.Arg126Cys | P11166 |
| HOMO SAPIENS | 4 | 2668 | 5eqg | A | 232 | ARG | p.Arg232Cys | P11166 |
| HOMO SAPIENS | 3 | 2668 | 5eqg | A | 249 | ARG | p.Arg249Gln | P11166 |
| HOMO SAPIENS | 3 | 2668 | 5eqg | A | 255 | LYS | p.Lys255Arg | P11166 |
| HOMO SAPIENS | 0 | 2668 | 5eqg | A | 275 | ALA | p.Ala275Thr | P11166 |
| HOMO SAPIENS | 4 | 2668 | 5eqg | A | 299 | GLU | p.Glu299Lys | P11166 |
| HOMO SAPIENS | 6 | 2668 | 5eqg | A | 314 | GLY | p.Gly314Ser | P11166 |
| HOMO SAPIENS | 7 | 2668 | 5eqg | A | 315 | ILE | p.Ile315Met | P11166 |
| HOMO SAPIENS | -1 | 2668 | 5eqg | A | 458 | ARG | p.Arg458Trp | P11166 |
| HOMO SAPIENS | -1 | 2668 | 5eqg | A | 468 | ARG | p.Arg468Trp | P11166 |
| HOMO SAPIENS | 3 | 2291 | 2amy | A | 31 | PHE | p.Phe31Leu | O15305 |
| HOMO SAPIENS | 7 | 2291 | 2amy | A | 32 | LEU | p.Leu32Arg | O15305 |
| HOMO SAPIENS | 7 | 2291 | 2amy | A | 65 | ASP | p.Asp65Tyr | O15305 |
| HOMO SAPIENS | 1 | 2291 | 2amy | A | 108 | ALA | p.Ala108Val | O15305 |
| HOMO SAPIENS | 3 | 2291 | 2amy | A | 113 | PRO | p.Pro113Leu | O15305 |
| HOMO SAPIENS | 3 | 2291 | 2amy | A | 113 | PRO | p.Pro113Ser | O15305 |
| HOMO SAPIENS | 6 | 2291 | 2amy | A | 117 | GLY | p.Gly117Arg | O15305 |
| HOMO SAPIENS | 6 | 2291 | 2amy | A | 119 | PHE | p.Phe119Leu | O15305 |
| HOMO SAPIENS | 6 | 2291 | 2amy | A | 123 | ARG | p.Arg123Gln | O15305 |
| HOMO SAPIENS | 0 | 2291 | 2amy | A | 129 | VAL | p.Val129Met | O15305 |
| HOMO SAPIENS | 7 | 2291 | 2amy | A | 139 | GLU | p.Glu139Lys | O15305 |
| HOMO SAPIENS | 5 | 2291 | 2amy | A | 141 | ARG | p.Arg141His | O15305 |
| HOMO SAPIENS | 7 | 2291 | 2amy | A | 144 | PHE | p.Phe144Leu | O15305 |
| HOMO SAPIENS | 6 | 2291 | 2amy | A | 148 | ASP | p.Asp148Asn | O15305 |
| HOMO SAPIENS | 6 | 2291 | 2amy | A | 153 | ILE | p.Ile153Thr | O15305 |
| HOMO SAPIENS | 2 | 2291 | 2amy | A | 157 | PHE | p.Phe157Ser | O15305 |
| HOMO SAPIENS | 1 | 2291 | 2amy | A | 162 | ARG | p.Arg162Trp | O15305 |
| HOMO SAPIENS | 6 | 2291 | 2amy | A | 188 | ASP | p.Asp188Gly | O15305 |
| HOMO SAPIENS | 5 | 2291 | 2amy | A | 194 | ARG | p.Arg194Ter | O15305 |
| HOMO SAPIENS | 0 | 2291 | 2amy | A | 207 | PHE | p.Phe207Ser | O15305 |
| HOMO SAPIENS | 5 | 2291 | 2amy | A | 208 | GLY | p.Gly208Ala | O15305 |
| HOMO SAPIENS | 1 | 2291 | 2amy | A | 216 | ASN | p.Asn216Ile | O15305 |
| HOMO SAPIENS | 5 | 2291 | 2amy | A | 218 | HIS | p.His218Leu | O15305 |
| HOMO SAPIENS | 6 | 2291 | 2amy | A | 223 | ASP | p.Asp223Glu | O15305 |
| HOMO SAPIENS | 6 | 2291 | 2amy | A | 226 | THR | p.Thr226Ser | O15305 |
| HOMO SAPIENS | 5 | 2291 | 2amy | A | 231 | VAL | p.Val231Met | O15305 |
| HOMO SAPIENS | 1 | 2291 | 2amy | A | 237 | THR | p.Thr237Met | O15305 |
| HOMO SAPIENS | 1 | 2291 | 2amy | A | 241 | CYS | p.Cys241Ser | O15305 |
| HOMO SAPIENS | 5 | 35907 | 1d6g | A | 21 | GLY | p.Gly21Arg | P32238 |
| HOMO SAPIENS | -1 | 33209 | 1x9d | A | 197 | PRO | p.Pro197Leu | Q9UKM7 |

| | | | | | | | | |
|---|---|---|---|---|---|---|---|---|
| HOMO SAPIENS | -1 | 33209 | 1x9d | A | 201 | PRO | p.Pro201Leu | Q9UKM7 |
| HOMO SAPIENS | 6 | 21305 | 4yep | B | 1326 | ARG | p.Arg1326Ter | P24043 |
| HOMO SAPIENS | 6 | 21305 | 4yep | B | 1350 | ARG | p.Arg1350Ter | P24043 |
| HOMO SAPIENS | 0 | 21553 | 3qyb | A | 1120 | VAL | p.Val1119Ala | O60343 |
| HOMO SAPIENS | 2 | 21553 | 3qyb | A | 1148 | THR | p.Thr1147Met | O60343 |
| HOMO SAPIENS | -1 | 30645 | 3m0c | C | -17 | TRP | p.Trp4Ter | P01130 |
| HOMO SAPIENS | -1 | 30645 | 3m0c | C | -17 | TRP | p.Trp4Ter | P01130 |
| HOMO SAPIENS | -1 | 30645 | 3m0c | C | 12 | GLN | p.Gln33Ter | P01130 |
| HOMO SAPIENS | -1 | 30645 | 3m0c | C | 15 | ASP | p.Asp36Glu | P01130 |
| HOMO SAPIENS | -1 | 30645 | 3m0c | C | 28 | SER | p.Ser49Arg | P01130 |
| HOMO SAPIENS | -1 | 30645 | 3m0c | C | 41 | THR | p.Thr62Met | P01130 |
| HOMO SAPIENS | -1 | 30645 | 3m0c | C | 69 | ASP | p.Asp90Asn | P01130 |
| HOMO SAPIENS | -1 | 30645 | 3m0c | C | 76 | ASN | p.Asn97Lys | P01130 |
| HOMO SAPIENS | -1 | 30645 | 3m0c | C | 78 | SER | p.Ser99Ter | P01130 |
| HOMO SAPIENS | -1 | 30645 | 3m0c | C | 81 | GLN | p.Gln102Ter | P01130 |
| HOMO SAPIENS | -1 | 30645 | 3m0c | C | 84 | PRO | p.Pro105Ser | P01130 |
| HOMO SAPIENS | -1 | 30645 | 3m0c | C | 194 | ARG | p.Arg215Cys | P01130 |
| HOMO SAPIENS | -1 | 30645 | 3m0c | C | 198 | GLY | p.Gly219Asp | P01130 |
| HOMO SAPIENS | -1 | 30645 | 3m0c | C | 201 | CYS | p.Cys222Tyr | P01130 |
| HOMO SAPIENS | -1 | 30645 | 3m0c | C | 201 | CYS | p.Cys222Ter | P01130 |
| HOMO SAPIENS | -1 | 30645 | 3m0c | C | 206 | ASP | p.Asp227Glu | P01130 |
| HOMO SAPIENS | -1 | 30645 | 3m0c | C | 249 | CYS | p.Cys270Ter | P01130 |
| HOMO SAPIENS | -1 | 30645 | 3m0c | C | 250 | VAL | p.Val271Ile | P01130 |
| HOMO SAPIENS | 0 | 30645 | 3m0c | C | 255 | CYS | p.Cys276Tyr | P01130 |
| HOMO SAPIENS | 0 | 30645 | 3m0c | C | 255 | CYS | p.Cys276Ter | P01130 |
| HOMO SAPIENS | 0 | 30645 | 3m0c | C | 256 | GLU | p.Glu277Lys | P01130 |
| HOMO SAPIENS | 0 | 30645 | 3m0c | C | 265 | SER | p.Ser286Arg | P01130 |
| HOMO SAPIENS | 0 | 30645 | 3m0c | C | 284 | TRP | p.Trp305Ter | P01130 |
| HOMO SAPIENS | 0 | 30645 | 3m0c | C | 292 | CYS | p.Cys313Ter | P01130 |
| HOMO SAPIENS | 1 | 30645 | 3m0c | C | 385 | ARG | p.Arg406Trp | P01130 |
| HOMO SAPIENS | 0 | 30645 | 3m0c | C | 395 | ARG | p.Arg416Gln | P01130 |
| HOMO SAPIENS | 0 | 30645 | 3m0c | C | 398 | TYR | p.Tyr419Ter | P01130 |
| HOMO SAPIENS | 1 | 30645 | 3m0c | C | 415 | VAL | p.Val436Ala | P01130 |
| HOMO SAPIENS | 1 | 30645 | 3m0c | C | 424 | ASP | p.Asp445Glu | P01130 |
| HOMO SAPIENS | 1 | 30645 | 3m0c | C | 425 | LEU | p.Leu446Val | P01130 |
| HOMO SAPIENS | -1 | 30645 | 3m0c | C | 440 | GLY | p.Gly461Cys | P01130 |
| HOMO SAPIENS | -1 | 30645 | 3m0c | C | 441 | VAL | p.Val462Ile | P01130 |
| HOMO SAPIENS | 1 | 30645 | 3m0c | C | 447 | VAL | p.Val468Ile | P01130 |
| HOMO SAPIENS | 1 | 30645 | 3m0c | C | 489 | THR | p.Thr510Met | P01130 |
| HOMO SAPIENS | 0 | 30645 | 3m0c | C | 524 | GLY | p.Gly545Glu | P01130 |
| HOMO SAPIENS | 1 | 30645 | 3m0c | C | 615 | GLY | p.Gly636Ser | P01130 |
| HOMO SAPIENS | 1 | 30645 | 3m0c | C | 625 | LEU | p.Leu646Ile | P01130 |
| HOMO SAPIENS | 1 | 30645 | 3m0c | C | 630 | ASP | p.Asp651Gly | P01130 |
| HOMO SAPIENS | 3 | 30645 | 3m0c | C | 646 | CYS | p.Cys667Tyr | P01130 |
| HOMO SAPIENS | 4 | 30645 | 3m0c | C | 654 | GLY | p.Gly675Ser | P01130 |
| HOMO SAPIENS | 6 | 30645 | 3m0c | C | 660 | CYS | p.Cys681Ter | P01130 |
| HOMO SAPIENS | 5 | 30645 | 3m0c | C | 676 | ALA | p.Ala697Pro | P01130 |
| HOMO SAPIENS | -1 | 30645 | 3m0c | C | 749 | GLN | p.Gln770Ter | P01130 |
| HOMO SAPIENS | -1 | 41323 | 4bxo | A | 1812 | PRO | p.Pro1812Ala | Q8IYD8 |
| HOMO SAPIENS | 5 | 8335 | 3h0t | C | 12 | GLY | p.Gly71Asp | P81172 |
| HOMO SAPIENS | 6 | 609 | 1bhg | B | 148 | PRO | p.Pro148Ser | P08236 |
| HOMO SAPIENS | 1 | 609 | 1bhg | B | 176 | LEU | p.Leu176Phe | P08236 |
| HOMO SAPIENS | 6 | 609 | 1bhg | B | 216 | ARG | p.Arg216Trp | P08236 |
| HOMO SAPIENS | 0 | 609 | 1bhg | B | 350 | LYS | p.Lys350Asn | P08236 |

| | | | | | | | | |
|---|---|---|---|---|---|---|---|---|
| HOMO SAPIENS | 1 | 609 | 1bhg | B | 354 | ALA | p.Ala354Val | P08236 |
| HOMO SAPIENS | 1 | 609 | 1bhg | B | 357 | ARG | p.Arg357Ter | P08236 |
| HOMO SAPIENS | 0 | 609 | 1bhg | B | 382 | ARG | p.Arg382Cys | P08236 |
| HOMO SAPIENS | 0 | 609 | 1bhg | B | 495 | TYR | p.Tyr495Cys | P08236 |
| HOMO SAPIENS | 2 | 609 | 1bhg | B | 507 | TRP | p.Trp507Ter | P08236 |
| HOMO SAPIENS | 0 | 609 | 1bhg | B | 577 | ARG | p.Arg577Leu | P08236 |
| HOMO SAPIENS | 1 | 609 | 1bhg | B | 611 | ARG | p.Arg611Trp | P08236 |
| HOMO SAPIENS | 0 | 609 | 1bhg | B | 619 | ALA | p.Ala619Val | P08236 |
| HOMO SAPIENS | 0 | 609 | 1bhg | B | 627 | TRP | p.Trp627Cys | P08236 |
| HOMO SAPIENS | 5 | 3422 | 1x9n | A | 566 | GLU | p.Glu566Lys | P18858 |
| HOMO SAPIENS | 2 | 3422 | 1x9n | A | 771 | ARG | p.Arg771Trp | P18858 |
| HOMO SAPIENS | 0 | 3279 | 5tby | B | 797 | ALA | p.Ala797Thr | P12883 |
| HOMO SAPIENS | -1 | 3279 | 5tby | B | 982 | MET | p.Met982Thr | P12883 |
| HOMO SAPIENS | -1 | 3279 | 5tby | B | 992 | LEU | p.Leu992Met | P12883 |
| HOMO SAPIENS | -1 | 3279 | 5tby | B | 1004 | GLN | p.Gln1004Glu | P12883 |
| HOMO SAPIENS | -1 | 3279 | 5tby | B | 1013 | GLU | p.Glu1013Lys | P12883 |
| HOMO SAPIENS | -1 | 3279 | 5tby | B | 1032 | ASP | p.Asp1032Asn | P12883 |
| HOMO SAPIENS | -1 | 3279 | 5tby | B | 1053 | ARG | p.Arg1053Gln | P12883 |
| HOMO SAPIENS | -1 | 3279 | 5tby | B | 1117 | GLU | p.Glu1117Ter | P12883 |
| HOMO SAPIENS | -1 | 3279 | 5tby | B | 1193 | ARG | p.Arg1193His | P12883 |
| HOMO SAPIENS | -1 | 3279 | 5tby | B | 1205 | GLU | p.Glu1205Lys | P12883 |
| HOMO SAPIENS | -1 | 3279 | 5tby | B | 1207 | ILE | p.Ile1207Met | P12883 |
| HOMO SAPIENS | -1 | 3279 | 5tby | B | 1250 | ARG | p.Arg1250Pro | P12883 |
| HOMO SAPIENS | -1 | 3279 | 5tby | B | 1259 | HIS | p.His1259Gln | P12883 |
| HOMO SAPIENS | -1 | 3279 | 5tby | B | 1272 | ASP | p.Asp1272Asn | P12883 |
| HOMO SAPIENS | -1 | 3279 | 5tby | B | 1295 | GLU | p.Glu1295Lys | P12883 |
| HOMO SAPIENS | -1 | 3279 | 5tby | B | 1335 | SER | p.Ser1335Leu | P12883 |
| HOMO SAPIENS | -1 | 3279 | 5tby | B | 1399 | GLU | p.Glu1399Asp | P12883 |
| HOMO SAPIENS | -1 | 3279 | 5tby | B | 1413 | SER | p.Ser1413Ter | P12883 |
| HOMO SAPIENS | -1 | 3279 | 5tby | B | 1459 | LYS | p.Lys1459Asn | P12883 |
| HOMO SAPIENS | -1 | 3279 | 5tby | B | 1467 | LEU | p.Leu1467Val | P12883 |
| HOMO SAPIENS | -1 | 3279 | 5tby | B | 1491 | SER | p.Ser1491Cys | P12883 |
| HOMO SAPIENS | -1 | 3279 | 5tby | B | 1496 | GLU | p.Glu1496Ala | P12883 |
| HOMO SAPIENS | -1 | 3279 | 5tby | B | 1500 | ARG | p.Arg1500Pro | P12883 |
| HOMO SAPIENS | -1 | 3279 | 5tby | B | 1606 | ARG | p.Arg1606His | P12883 |
| HOMO SAPIENS | -1 | 3279 | 5tby | B | 1664 | ASN | p.Asn1664Lys | P12883 |
| HOMO SAPIENS | -1 | 3279 | 5tby | B | 1712 | ARG | p.Arg1712Trp | P12883 |
| HOMO SAPIENS | -1 | 3279 | 5tby | B | 1769 | LEU | p.Leu1769Met | P12883 |
| HOMO SAPIENS | -1 | 3279 | 5tby | B | 1808 | GLY | p.Gly1808Ser | P12883 |
| HOMO SAPIENS | -1 | 3279 | 5tby | B | 1818 | ARG | p.Arg1818Trp | P12883 |
| HOMO SAPIENS | -1 | 3279 | 5tby | B | 1820 | ARG | p.Arg1820Trp | P12883 |
| HOMO SAPIENS | -1 | 3279 | 5tby | B | 1844 | GLU | p.Glu1844Lys | P12883 |
| HOMO SAPIENS | -1 | 3279 | 5tby | B | 1845 | ARG | p.Arg1845Trp | P12883 |
| HOMO SAPIENS | -1 | 3279 | 5tby | B | 1854 | THR | p.Thr1854Met | P12883 |
| HOMO SAPIENS | -1 | 3279 | 5tby | B | 1909 | ARG | p.Arg1909Trp | P12883 |
| HOMO SAPIENS | 2 | 2260 | 4kl1 | A | 553 | ASP | p.Asp553Asn | Q9Y3Q4 |
| HOMO SAPIENS | 0 | 40570 | 5lst | A | 522 | ARG | p.Arg522Cys | O94761 |
| HOMO SAPIENS | 0 | 40570 | 5lst | A | 522 | ARG | p.Arg522His | O94761 |
| HOMO SAPIENS | 0 | 40570 | 5lst | A | 591 | PRO | p.Pro591Leu | O94761 |
| HOMO SAPIENS | 0 | 40570 | 5lst | A | 793 | PRO | p.Pro793Ser | O94761 |
| HOMO SAPIENS | 4 | 40570 | 5lst | A | 799 | VAL | p.Val799Met | O94761 |
| HOMO SAPIENS | 0 | 40570 | 5lst | A | 1004 | ARG | p.Arg1004Trp | O94761 |
| HOMO SAPIENS | 0 | 40570 | 5lst | A | 1005 | ARG | p.Arg1005Gln | O94761 |
| HOMO SAPIENS | 0 | 40570 | 5lst | A | 1021 | ARG | p.Arg1021Gln | O94761 |

| | | | | | | | | |
|---|---|---|---|---|---|---|---|---|
| HOMO SAPIENS | 0 | 40570 | 5lst | A | 1043 | LEU | p.Leu1043Pro | O94761 |
| HOMO SAPIENS | 0 | 40570 | 5lst | A | 1045 | ALA | p.Ala1045Thr | O94761 |
| HOMO SAPIENS | 0 | 40570 | 5lst | A | 1105 | GLY | p.Gly1105Asp | O94761 |
| HOMO SAPIENS | 0 | 40570 | 5lst | A | 1105 | GLY | p.Gly1105Ser | O94761 |
| HOMO SAPIENS | -1 | 40570 | 5lst | A | 1113 | GLY | p.Gly1113Arg | O94761 |
| HOMO SAPIENS | -1 | 18865 | 1gxc | A | 64 | GLU | p.Glu64Lys | O96017 |
| HOMO SAPIENS | -1 | 11673 | 3dzu | D | 85 | PRO | p.Pro113Gln | P37231 |
| HOMO SAPIENS | 0 | 19383 | 3b96 | A | 63 | LYS | p.Lys103Arg | P49748 |
| HOMO SAPIENS | 4 | 19383 | 3b96 | A | 140 | ALA | p.Ala180Thr | P49748 |
| HOMO SAPIENS | 1 | 19383 | 3b96 | A | 145 | GLY | p.Gly185Ser | P49748 |
| HOMO SAPIENS | 0 | 19383 | 3b96 | A | 220 | THR | p.Thr260Met | P49748 |
| HOMO SAPIENS | 5 | 19383 | 3b96 | A | 243 | VAL | p.Val283Ala | P49748 |
| HOMO SAPIENS | 0 | 19383 | 3b96 | A | 249 | GLY | p.Gly289Arg | P49748 |
| HOMO SAPIENS | 0 | 19383 | 3b96 | A | 254 | GLY | p.Gly294Glu | P49748 |
| HOMO SAPIENS | 7 | 19383 | 3b96 | A | 326 | ARG | p.Arg366Cys | P49748 |
| HOMO SAPIENS | 7 | 19383 | 3b96 | A | 326 | ARG | p.Arg366His | P49748 |
| HOMO SAPIENS | 5 | 19383 | 3b96 | A | 342 | LYS | p.Lys382Gln | P49748 |
| HOMO SAPIENS | 0 | 19383 | 3b96 | A | 369 | THR | p.Thr409Met | P49748 |
| HOMO SAPIENS | 5 | 19383 | 3b96 | A | 376 | ALA | p.Ala416Ser | P49748 |
| HOMO SAPIENS | 0 | 19383 | 3b96 | A | 385 | ALA | p.Ala425Thr | P49748 |
| HOMO SAPIENS | 5 | 19383 | 3b96 | A | 399 | GLY | p.Gly439Asp | P49748 |
| HOMO SAPIENS | 7 | 19383 | 3b96 | A | 401 | GLY | p.Gly441Asp | P49748 |
| HOMO SAPIENS | 9 | 19383 | 3b96 | A | 413 | ARG | p.Arg453Gln | P49748 |
| HOMO SAPIENS | 2 | 19383 | 3b96 | A | 418 | PHE | p.Phe458Leu | P49748 |
| HOMO SAPIENS | 6 | 19383 | 3b96 | A | 419 | ARG | p.Arg459Trp | P49748 |
| HOMO SAPIENS | 6 | 19383 | 3b96 | A | 419 | ARG | p.Arg459Gln | P49748 |
| HOMO SAPIENS | -1 | 19383 | 3b96 | A | 471 | ARG | p.Arg511Gln | P49748 |
| HOMO SAPIENS | 0 | 19383 | 3b96 | A | 573 | ARG | p.Arg613Trp | P49748 |
| HOMO SAPIENS | 0 | 19383 | 3b96 | A | 575 | ARG | p.Arg615Gln | P49748 |
| HOMO SAPIENS | 0 | 22286 | 3k2s | A | 3 | PRO | p.Pro27His | P02647 |
| HOMO SAPIENS | 0 | 22286 | 3k2s | A | 4 | PRO | p.Pro28Arg | P02647 |
| HOMO SAPIENS | 0 | 22286 | 3k2s | A | 10 | ARG | p.Arg34Leu | P02647 |
| HOMO SAPIENS | 0 | 22286 | 3k2s | A | 26 | GLY | p.Gly50Arg | P02647 |
| HOMO SAPIENS | 0 | 22286 | 3k2s | A | 37 | ALA | p.Ala61Thr | P02647 |
| HOMO SAPIENS | 1 | 1622 | 2qrv | D | 631 | ARG | p.Arg635Trp | Q9Y6K1 |
| HOMO SAPIENS | 1 | 1622 | 2qrv | D | 900 | PRO | p.Pro904Leu | Q9Y6K1 |
| HOMO SAPIENS | 6 | 40796 | 1v18 | B | 1509 | LEU | p.Leu1509Val | P25054 |
| HOMO SAPIENS | 1 | 22375 | 4pki | G | 1214 | LYS | p.Asp214Asn | P28289 |
| HOMO SAPIENS | 1 | 16307 | 5mv8 | B | 483 | ALA | p.Ala483Ser | Q9Y6N9 |
| HOMO SAPIENS | 6 | 16307 | 5mv8 | B | 506 | ASN | p.Asn506Lys | Q9Y6N9 |
| HOMO SAPIENS | 4 | 16307 | 5mv8 | B | 519 | ASP | p.Glu519Asp | Q9Y6N9 |
| HOMO SAPIENS | 0 | 16307 | 5mv8 | B | 530 | GLY | p.Gly530Arg | Q9Y6N9 |
| HOMO SAPIENS | 4 | 36221 | 4cvo | A | 127 | VAL | p.Val127Ile | Q03468 |
| HOMO SAPIENS | 7 | 36221 | 4cvo | A | 134 | ARG | p.Arg134Trp | Q03468 |
| HOMO SAPIENS | 0 | 36221 | 4cvo | A | 156 | GLN | p.Gln156Ter | Q03468 |
| HOMO SAPIENS | 0 | 10829 | 4a11 | B | 13 | GLU | p.Glu13Ter | Q13216 |
| HOMO SAPIENS | 1 | 10829 | 4a11 | B | 100 | TYR | p.Tyr100Ter | Q13216 |
| HOMO SAPIENS | 0 | 10829 | 4a11 | B | 160 | ALA | p.Ala160Val | Q13216 |
| HOMO SAPIENS | 1 | 6759 | 5lev | A | 29 | ILE | p.Ile29Phe | Q9H3H5 |
| HOMO SAPIENS | 0 | 6759 | 5lev | A | 108 | MET | p.Met108Ile | Q9H3H5 |
| HOMO SAPIENS | 2 | 6759 | 5lev | A | 117 | VAL | p.Val117Ile | Q9H3H5 |
| HOMO SAPIENS | 1 | 6759 | 5lev | A | 170 | TYR | p.Tyr170Cys | Q9H3H5 |
| HOMO SAPIENS | 0 | 6759 | 5lev | A | 264 | GLY | p.Val264Gly | Q9H3H5 |
| HOMO SAPIENS | 0 | 6759 | 5lev | A | 332 | PHE | p.Phe332Val | Q9H3H5 |

| | | | | | | | | |
|---|---|---|---|---|---|---|---|---|
| HOMO SAPIENS | 7 | 6759 | 5lev | A | 393 | ILE | p.Ile393Val | Q9H3H5 |
| HOMO SAPIENS | -1 | 33473 | 5m5e | C | 9 | ASN | p.Asn31Ser | P31785 |
| HOMO SAPIENS | 0 | 26155 | 1ic8 | B | 98 | ALA | p.Ala98Val | P20823 |
| HOMO SAPIENS | 0 | 15329 | 4iqr | B | 127 | ARG | p.Arg136Trp | P41235 |
| HOMO SAPIENS | 0 | 20725 | 4bsj | A | 350 | GLU | p.Glu350Val | P35916 |
| HOMO SAPIENS | 0 | 20725 | 4bsj | A | 494 | THR | p.Thr494Ala | P35916 |
| HOMO SAPIENS | 0 | 20725 | 4bsj | A | 527 | ASN | p.Asn527Ser | P35916 |
| HOMO SAPIENS | 6 | 29959 | 1xiw | B | 47 | ARG | p.Arg68Ter | P04234 |
| HOMO SAPIENS | 2 | 29959 | 1xiw | B | 72 | CYS | p.Cys93Ter | P04234 |
| HOMO SAPIENS | 3 | 20242 | 5tjc | A | 102 | ARG | p.Arg102Ter | Q9GZU1 |
| HOMO SAPIENS | 3 | 20242 | 5tjc | A | 232 | THR | p.Thr232Pro | Q9GZU1 |
| HOMO SAPIENS | 7 | 9590 | 4ap8 | A | 50 | THR | p.Thr50Ala | O96007 |
| HOMO SAPIENS | 4 | 9590 | 4ap8 | A | 168 | GLU | p.Glu168Lys | O96007 |
| HOMO SAPIENS | -1 | 4297 | 5do7 | B | 19 | ASP | p.Asp19His | Q9H221 |
| HOMO SAPIENS | -1 | 4297 | 5do7 | B | 54 | TYR | p.Tyr54Cys | Q9H221 |
| HOMO SAPIENS | -1 | 4297 | 5do7 | B | 80 | CYS | p.Cys80Tyr | Q9H221 |
| HOMO SAPIENS | 5 | 4297 | 5do7 | B | 107 | SER | p.Ser107Ter | Q9H221 |
| HOMO SAPIENS | 1 | 4297 | 5do7 | B | 164 | ARG | p.Arg164Pro | Q9H221 |
| HOMO SAPIENS | 0 | 4297 | 5do7 | B | 210 | VAL | p.Val210Met | Q9H221 |
| HOMO SAPIENS | 0 | 4297 | 5do7 | B | 231 | PRO | p.Pro231Thr | Q9H221 |
| HOMO SAPIENS | 0 | 4297 | 5do7 | B | 263 | ARG | p.Arg263Gln | Q9H221 |
| HOMO SAPIENS | 0 | 4297 | 5do7 | B | 361 | TRP | p.Trp361Ter | Q9H221 |
| HOMO SAPIENS | 0 | 4297 | 5do7 | B | 400 | THR | p.Thr400Lys | Q9H221 |
| HOMO SAPIENS | 0 | 4297 | 5do7 | B | 549 | ALA | p.Ala549Pro | Q9H221 |
| HOMO SAPIENS | 0 | 4297 | 5do7 | B | 596 | LEU | p.Leu596Arg | Q9H221 |
| HOMO SAPIENS | 0 | 4297 | 5do7 | B | 632 | VAL | p.Val632Ala | Q9H221 |
| HOMO SAPIENS | 0 | 4297 | 5do7 | B | 658 | TYR | p.Tyr658Ter | Q9H221 |
| HOMO SAPIENS | 2 | 9356 | 4igg | B | 206 | GLN | p.Gln206His | P35221 |
| HOMO SAPIENS | 6 | 2179 | 2wk6 | A | 44 | ARG | p.Arg44Gln | P19971 |
| HOMO SAPIENS | 6 | 2179 | 2wk6 | A | 81 | ARG | p.Arg81Gln | P19971 |
| HOMO SAPIENS | 6 | 2179 | 2wk6 | A | 145 | GLY | p.Gly145Arg | P19971 |
| HOMO SAPIENS | 7 | 2179 | 2wk6 | A | 146 | ARG | p.Arg146His | P19971 |
| HOMO SAPIENS | 7 | 2179 | 2wk6 | A | 153 | GLY | p.Gly153Ser | P19971 |
| HOMO SAPIENS | 0 | 2179 | 2wk6 | A | 173 | MET | p.Met173Ile | P19971 |
| HOMO SAPIENS | 6 | 2179 | 2wk6 | A | 202 | ARG | p.Arg202Lys | P19971 |
| HOMO SAPIENS | 6 | 2179 | 2wk6 | A | 208 | VAL | p.Val208Met | P19971 |
| HOMO SAPIENS | 5 | 2179 | 2wk6 | A | 222 | LYS | p.Lys222Arg | P19971 |
| HOMO SAPIENS | 5 | 2179 | 2wk6 | A | 289 | GLU | p.Glu289Ala | P19971 |
| HOMO SAPIENS | 2 | 2179 | 2wk6 | A | 311 | GLY | p.Gly311Arg | P19971 |
| HOMO SAPIENS | 2 | 2179 | 2wk6 | A | 376 | ARG | p.Arg376Pro | P19971 |
| HOMO SAPIENS | 2 | 2179 | 2wk6 | A | 465 | ALA | p.Ala465Thr | P19971 |
| HOMO SAPIENS | -1 | 11856 | 3orh | A | 3 | ALA | p.Ala3Ser | Q14353 |
| HOMO SAPIENS | 1 | 11856 | 3orh | A | 20 | TRP | p.Trp20Ser | Q14353 |
| HOMO SAPIENS | 1 | 11856 | 3orh | A | 50 | MET | p.Met50Leu | Q14353 |
| HOMO SAPIENS | 1 | 11856 | 3orh | A | 110 | VAL | p.Val110Phe | Q14353 |
| HOMO SAPIENS | 8 | 11856 | 3orh | A | 164 | GLY | p.Gly164Asp | Q14353 |
| HOMO SAPIENS | 1 | 11856 | 3orh | A | 169 | CYS | p.Cys169Tyr | Q14353 |
| HOMO SAPIENS | 1 | 11856 | 3orh | A | 174 | TRP | p.Trp174Ter | Q14353 |
| HOMO SAPIENS | 2 | 11856 | 3orh | A | 196 | ALA | p.Ala196Val | Q14353 |
| HOMO SAPIENS | 1 | 11856 | 3orh | A | 209 | THR | p.Thr209Met | Q14353 |
| HOMO SAPIENS | 1 | 11856 | 3orh | A | 219 | ASP | p.Asp219Asn | Q14353 |
| HOMO SAPIENS | 1 | 11856 | 3orh | A | 224 | ALA | p.Ala224Thr | Q14353 |
| HOMO SAPIENS | 1 | 13244 | 2m6i | D | 218 | ARG | p.Arg246Leu | P23415 |
| HOMO SAPIENS | 1 | 13244 | 2m6i | D | 244 | ILE | p.Ile272Asn | P23415 |

| | | | | | | | | |
|---|---|---|---|---|---|---|---|---|
| HOMO SAPIENS | 5 | 13244 | 2m6i | D | 250 | PRO | p.Pro278Thr | P23415 |
| HOMO SAPIENS | 6 | 13244 | 2m6i | D | 266 | GLN | p.Gln294His | P23415 |
| HOMO SAPIENS | 1 | 13244 | 2m6i | D | 271 | ARG | p.Arg299Leu | P23415 |
| HOMO SAPIENS | 1 | 13244 | 2m6i | D | 276 | LYS | p.Lys304Glu | P23415 |
| HOMO SAPIENS | 1 | 13244 | 2m6i | D | 279 | TYR | p.Tyr307Cys | P23415 |
| HOMO SAPIENS | 0 | 13244 | 2m6i | D | 316 | GLY | p.Gly378Ser | P23415 |
| HOMO SAPIENS | 1 | 13244 | 2m6i | D | 400 | ARG | p.Arg428His | P23415 |
| HOMO SAPIENS | 3 | 1967 | 2i6b | A | 301 | ALA | p.Ala318Glu | P55263 |
| HOMO SAPIENS | 4 | 11096 | 1h8b | A | 61 | ALA | p.Ala882Thr | P35609 |
| HOMO SAPIENS | 0 | 15423 | 5ngz | A | 2 | GLN | p.Gln2Glu | Q9NPD8 |
| HOMO SAPIENS | 3 | 30475 | 3lpw | A | 21 | SER | p.Ser22894Tyr | Q8WZ42 |
| HOMO SAPIENS | 6 | 32687 | 2arf | A | 1041 | ARG | p.Arg1041Trp | P35670 |
| HOMO SAPIENS | 8 | 32687 | 2arf | A | 1061 | GLY | p.Gly1061Glu | P35670 |
| HOMO SAPIENS | 6 | 32687 | 2arf | A | 1064 | GLU | p.Glu1064Ala | P35670 |
| HOMO SAPIENS | 2 | 32687 | 2arf | A | 1069 | HIS | p.His1069Gln | P35670 |
| HOMO SAPIENS | 2 | 32687 | 2arf | A | 1088 | LEU | p.Leu1088Ter | P35670 |
| HOMO SAPIENS | 0 | 32687 | 2arf | A | 1099 | GLY | p.Gly1099Ser | P35670 |
| HOMO SAPIENS | 4 | 32687 | 2arf | A | 1106 | VAL | p.Val1106Asp | P35670 |
| HOMO SAPIENS | 2 | 32687 | 2arf | A | 1142 | GLN | p.Gln1142His | P35670 |
| HOMO SAPIENS | 8 | 32687 | 2arf | A | 1148 | ILE | p.Ile1148Thr | P35670 |
| HOMO SAPIENS | 6 | 32687 | 2arf | A | 1151 | ARG | p.Arg1151Cys | P35670 |
| HOMO SAPIENS | 4 | 32687 | 2arf | A | 1169 | MET | p.Met1169Val | P35670 |
| HOMO SAPIENS | 4 | 32687 | 2arf | A | 1173 | GLU | p.Glu1173Lys | P35670 |
| HOMO SAPIENS | 2 | 32687 | 2arf | A | 1176 | GLY | p.Gly1176Arg | P35670 |
| HOMO SAPIENS | 3 | 15544 | 5uq3 | C | 575 | PRO | p.Pro69Leu | P46527 |
| HOMO SAPIENS | 6 | 15544 | 5uq3 | C | 592 | GLU | p.Glu86Asp | P46527 |
| HOMO SAPIENS | 6 | 15544 | 5uq3 | C | 599 | ARG | p.Arg93Trp | P46527 |
| HOMO SAPIENS | -1 | 15544 | 5uq3 | C | 615 | VAL | p.Val109Gly | P46527 |
| HOMO SAPIENS | -1 | 15544 | 5uq3 | C | 631 | SER | p.Ser125Pro | P46527 |
| HOMO SAPIENS | -1 | 15544 | 5uq3 | C | 639 | PRO | p.Pro133Ser | P46527 |
| HOMO SAPIENS | 2 | 2703 | 4wxq | A | 245 | CYS | p.Cys245Tyr | Q99972 |
| HOMO SAPIENS | 0 | 2703 | 4wxq | A | 252 | GLY | p.Gly252Arg | Q99972 |
| HOMO SAPIENS | 1 | 2703 | 4wxq | A | 337 | GLN | p.Gln337Arg | Q99972 |
| HOMO SAPIENS | 5 | 2703 | 4wxq | A | 347 | TYR | p.Tyr347Ter | Q99972 |
| HOMO SAPIENS | 3 | 2703 | 4wxq | A | 364 | GLY | p.Gly364Val | Q99972 |
| HOMO SAPIENS | 2 | 2703 | 4wxq | A | 367 | GLY | p.Gly367Arg | Q99972 |
| HOMO SAPIENS | 2 | 2703 | 4wxq | A | 368 | GLN | p.Gln368Ter | Q99972 |
| HOMO SAPIENS | 6 | 2703 | 4wxq | A | 370 | PRO | p.Pro370Leu | Q99972 |
| HOMO SAPIENS | 1 | 2703 | 4wxq | A | 380 | ASP | p.Asp380His | Q99972 |
| HOMO SAPIENS | 8 | 2703 | 4wxq | A | 399 | GLY | p.Gly399Val | Q99972 |
| HOMO SAPIENS | 0 | 2703 | 4wxq | A | 423 | LYS | p.Lys423Glu | Q99972 |
| HOMO SAPIENS | 3 | 2703 | 4wxq | A | 433 | CYS | p.Cys433Arg | Q99972 |
| HOMO SAPIENS | 3 | 2703 | 4wxq | A | 437 | TYR | p.Tyr437His | Q99972 |
| HOMO SAPIENS | 2 | 2703 | 4wxq | A | 477 | ILE | p.Ile477Ser | Q99972 |
| HOMO SAPIENS | 3 | 2703 | 4wxq | A | 480 | ASN | p.Asn480Lys | Q99972 |
| HOMO SAPIENS | -1 | 34100 | 2fot | C | 1171 | GLN | p.Gln1173Pro | Q13813 |
| HOMO SAPIENS | 0 | 8773 | 1kw2 | A | 432 | ASP | p.Asp432Glu | P02774 |
| HOMO SAPIENS | 0 | 8773 | 1kw2 | A | 436 | LYS | p.Lys436Thr | P02774 |
| HOMO SAPIENS | 0 | 8402 | 4dzo | B | 649 | TYR | p.Tyr649Ter | Q9Y6D9 |
| HOMO SAPIENS | 0 | 39962 | 1nst | A | 611 | GLY | p.Gly611Ser | P52848 |
| HOMO SAPIENS | 2 | 22578 | 3dof | B | 45 | MET | p.Met45Arg | Q9Y2Y0 |
| HOMO SAPIENS | 1 | 23290 | 5j1s | A | 292 | ILE | p.Ile292Thr | O14656 |
| HOMO SAPIENS | 4 | 9505 | 4j1y | A | 534 | ARG | p.Arg534Trp | P09871 |
| HOMO SAPIENS | 9 | 260 | 2h57 | A | 31 | THR | p.Thr31Met | Q9H0F7 |

| | | | | | | | | |
|---|---|---|---|---|---|---|---|---|
| HOMO SAPIENS | 0 | 260 | 2h57 | A | 91 | ILE | p.Ile91Thr | Q9H0F7 |
| HOMO SAPIENS | 0 | 260 | 2h57 | A | 121 | ARG | p.Arg121His | Q9H0F7 |
| HOMO SAPIENS | 0 | 260 | 2h57 | A | 122 | ARG | p.Arg122Ter | Q9H0F7 |
| HOMO SAPIENS | 2 | 260 | 2h57 | A | 169 | GLY | p.Gly169Ala | Q9H0F7 |
| HOMO SAPIENS | 4 | 260 | 2h57 | A | 170 | LEU | p.Leu170Trp | Q9H0F7 |
| HOMO SAPIENS | 4 | 34023 | 2cr3 | A | 18 | GLY | p.Gly48Ser | P11362 |
| HOMO SAPIENS | 0 | 37667 | 3ii7 | A | 420 | ARG | p.Arg420Cys | Q8IXQ5 |
| HOMO SAPIENS | 0 | 41720 | 2k9u | A | 68 | ARG | p.Arg2364His | Q14315 |
| HOMO SAPIENS | 1 | 29250 | 5fmv | A | 568 | HIS | p.His568Gln | P08575 |
| HOMO SAPIENS | 1 | 1280 | 1fb1 | A | 69 | PRO | p.Pro69Leu | P30793 |
| HOMO SAPIENS | 3 | 1280 | 1fb1 | A | 88 | ARG | p.Arg88Trp | P30793 |
| HOMO SAPIENS | 2 | 1280 | 1fb1 | A | 108 | GLY | p.Gly108Asp | P30793 |
| HOMO SAPIENS | 4 | 1280 | 1fb1 | A | 134 | ASP | p.Asp134Val | P30793 |
| HOMO SAPIENS | 6 | 1280 | 1fb1 | A | 135 | ILE | p.Ile135Lys | P30793 |
| HOMO SAPIENS | 9 | 1280 | 1fb1 | A | 144 | HIS | p.His144Pro | P30793 |
| HOMO SAPIENS | 9 | 1280 | 1fb1 | A | 184 | ARG | p.Arg184His | P30793 |
| HOMO SAPIENS | 1 | 1280 | 1fb1 | A | 196 | ALA | p.Ala196Ser | P30793 |
| HOMO SAPIENS | 3 | 1280 | 1fb1 | A | 201 | GLY | p.Gly201Glu | P30793 |
| HOMO SAPIENS | 2 | 1280 | 1fb1 | A | 204 | VAL | p.Val204Ile | P30793 |
| HOMO SAPIENS | 2 | 1280 | 1fb1 | A | 211 | MET | p.Met211Ile | P30793 |
| HOMO SAPIENS | 0 | 1280 | 1fb1 | A | 221 | MET | p.Met221Thr | P30793 |
| HOMO SAPIENS | 3 | 1280 | 1fb1 | A | 224 | LYS | p.Lys224Arg | P30793 |
| HOMO SAPIENS | 2 | 1280 | 1fb1 | A | 249 | ARG | p.Arg249Ser | P30793 |
| HOMO SAPIENS | 8 | 41352 | 4k12 | A | 47 | ILE | p.Ile551Thr | P08603 |
| HOMO SAPIENS | 9 | 20703 | 5nir | A | 57 | CYS | p.Cys57Tyr | P02458 |
| HOMO SAPIENS | 9 | 20703 | 5nir | A | 64 | CYS | p.Cys64Ter | P02458 |
| HOMO SAPIENS | 0 | 20087 | 4cmn | A | 299 | MET | p.Met299Ile | Q01968 |
| HOMO SAPIENS | 1 | 20819 | 3aaf | A | 979 | ILE | p.Ile979Met | Q14191 |
| HOMO SAPIENS | 1 | 20819 | 3aaf | A | 984 | LEU | p.Leu984Ile | Q14191 |
| HOMO SAPIENS | 3 | 20819 | 3aaf | A | 987 | ARG | p.Arg987Ter | Q14191 |
| HOMO SAPIENS | -1 | 20819 | 3aaf | A | 1074 | PHE | p.Leu1074Phe | Q14191 |
| HOMO SAPIENS | 0 | 3619 | 3dem | A | 206 | ASN | p.Asn225Ser | P48740 |
| HOMO SAPIENS | 1 | 3619 | 3dem | A | 269 | ARG | p.Arg288Leu | P48740 |
| HOMO SAPIENS | 0 | 3105 | 3ooi | A | 164 | TYR | p.Tyr2015His | Q96L73 |
| HOMO SAPIENS | 6 | 15828 | 2lkx | A | 16 | LEU | p.Leu100Gln | Q99697 |
| HOMO SAPIENS | 8 | 15828 | 2lkx | A | 24 | ARG | p.Arg108His | Q99697 |
| HOMO SAPIENS | 2 | 15828 | 2lkx | A | 30 | THR | p.Thr114Pro | Q99697 |
| HOMO SAPIENS | 8 | 15828 | 2lkx | A | 45 | VAL | p.Val129Leu | Q99697 |
| HOMO SAPIENS | 8 | 15828 | 2lkx | A | 46 | ARG | p.Arg130Trp | Q99697 |
| HOMO SAPIENS | 9 | 15828 | 2lkx | A | 53 | ARG | p.Arg137Pro | Q99697 |
| HOMO SAPIENS | 9 | 13017 | 4irq | A | 141 | ARG | p.Arg141Trp | Q9UBV7 |
| HOMO SAPIENS | 0 | 13017 | 4irq | A | 186 | ALA | p.Ala186Asp | Q9UBV7 |
| HOMO SAPIENS | 0 | 13017 | 4irq | A | 206 | LEU | p.Leu206Pro | Q9UBV7 |
| HOMO SAPIENS | 1 | 13017 | 4irq | A | 214 | CYS | p.Cys214Tyr | Q9UBV7 |
| HOMO SAPIENS | 0 | 13017 | 4irq | A | 270 | ARG | p.Arg270Cys | Q9UBV7 |
| HOMO SAPIENS | -1 | 13053 | 4mqw | X | 307 | ALA | p.Ala307Thr | P23945 |
| HOMO SAPIENS | 0 | 19406 | 4b3g | A | 43 | ARG | p.Arg43Ter | P38935 |
| HOMO SAPIENS | 0 | 19406 | 4b3g | A | 46 | CYS | p.Cys46Ter | P38935 |
| HOMO SAPIENS | 0 | 19406 | 4b3g | A | 75 | ALA | p.Ala75Thr | P38935 |
| HOMO SAPIENS | 0 | 19406 | 4b3g | A | 201 | LEU | p.Leu201Ser | P38935 |
| HOMO SAPIENS | 3 | 19406 | 4b3g | A | 213 | HIS | p.His213Arg | P38935 |
| HOMO SAPIENS | 0 | 19406 | 4b3g | A | 275 | ILE | p.Ile275Val | P38935 |
| HOMO SAPIENS | -1 | 19406 | 4b3g | A | 320 | ARG | p.Arg320Ter | P38935 |
| HOMO SAPIENS | 0 | 19406 | 4b3g | A | 361 | LEU | p.Leu361Pro | P38935 |

| | | | | | | | | |
|---|---|---|---|---|---|---|---|---|
| HOMO SAPIENS | 0 | 19406 | 4b3g | A | 369 | PHE | p.Phe369Leu | P38935 |
| HOMO SAPIENS | 0 | 19406 | 4b3g | A | 493 | THR | p.Thr493Ile | P38935 |
| HOMO SAPIENS | 0 | 19406 | 4b3g | A | 496 | CYS | p.Cys496Ter | P38935 |
| HOMO SAPIENS | 0 | 19406 | 4b3g | A | 514 | GLU | p.Glu514Lys | P38935 |
| HOMO SAPIENS | 0 | 19406 | 4b3g | A | 528 | ALA | p.Ala528Thr | P38935 |
| HOMO SAPIENS | 0 | 19406 | 4b3g | A | 531 | PRO | p.Pro531Thr | P38935 |
| HOMO SAPIENS | 0 | 19406 | 4b3g | A | 580 | VAL | p.Val580Ile | P38935 |
| HOMO SAPIENS | 6 | 19406 | 4b3g | A | 603 | ARG | p.Arg603His | P38935 |
| HOMO SAPIENS | 0 | 21501 | 5jde | A | 40 | SER | p.Ser2831Asn | Q8WZ42 |
| HOMO SAPIENS | 5 | 21501 | 5jde | A | 189 | ALA | p.Ala2980Thr | Q8WZ42 |
| HOMO SAPIENS | 6 | 15008 | 3mop | H | 12 | ARG | p.Arg12Cys | Q9NWZ3 |
| HOMO SAPIENS | 8 | 17455 | 3jtc | C | 15 | ARG | p.Arg57Trp | P04070 |
| HOMO SAPIENS | -1 | 17455 | 3jtc | C | 20 | CGU | p.Glu62Ala | P04070 |
| HOMO SAPIENS | 0 | 27795 | 5gox | B | 625 | TYR | p.Tyr625Ter | Q92878 |
| HOMO SAPIENS | 0 | 27795 | 5gox | B | 653 | SER | p.Ser653Ter | Q92878 |
| HOMO SAPIENS | -1 | 6558 | 5nug | A | 137 | VAL | p.Val137Ile | Q14204 |
| HOMO SAPIENS | -1 | 6558 | 5nug | A | 567 | ARG | p.Arg567His | Q14204 |
| HOMO SAPIENS | -1 | 6558 | 5nug | A | 1140 | THR | p.Thr1140Met | Q14204 |
| HOMO SAPIENS | -1 | 6558 | 5nug | A | 1991 | ASP | p.Asp1991Asn | Q14204 |
| HOMO SAPIENS | -1 | 6558 | 5nug | A | 2401 | LYS | p.Lys2401Asn | Q14204 |
| HOMO SAPIENS | 0 | 6558 | 5nug | A | 2488 | ARG | p.Arg2488His | Q14204 |
| HOMO SAPIENS | 0 | 6558 | 5nug | A | 3907 | HIS | p.His3907Gln | Q14204 |
| HOMO SAPIENS | 0 | 6558 | 5nug | A | 3981 | THR | p.Thr3981Arg | Q14204 |
| HOMO SAPIENS | 0 | 6558 | 5nug | A | 4394 | THR | p.Thr4394Met | Q14204 |
| HOMO SAPIENS | 0 | 6558 | 5nug | A | 4573 | ASN | p.Asn4573Ser | Q14204 |
| HOMO SAPIENS | 0 | 6558 | 5nug | A | 4603 | SER | p.Ser4603Ile | Q14204 |
| HOMO SAPIENS | 1 | 4244 | 1umk | A | 57 | ARG | p.Arg58Gln | P00387 |
| HOMO SAPIENS | 1 | 4244 | 1umk | A | 72 | LEU | p.Leu73Pro | P00387 |
| HOMO SAPIENS | 1 | 4244 | 1umk | A | 105 | VAL | p.Val106Met | P00387 |
| HOMO SAPIENS | 0 | 4244 | 1umk | A | 116 | THR | p.Thr117Ser | P00387 |
| HOMO SAPIENS | 8 | 4244 | 1umk | A | 127 | SER | p.Ser128Pro | P00387 |
| HOMO SAPIENS | 0 | 4244 | 1umk | A | 148 | LEU | p.Leu149Pro | P00387 |
| HOMO SAPIENS | 0 | 4244 | 1umk | A | 159 | ARG | p.Arg160Ter | P00387 |
| HOMO SAPIENS | 0 | 4244 | 1umk | A | 203 | CYS | p.Cys204Arg | P00387 |
| HOMO SAPIENS | 0 | 4244 | 1umk | A | 203 | CYS | p.Cys204Tyr | P00387 |
| HOMO SAPIENS | 2 | 4244 | 1umk | A | 239 | ASP | p.Asp240Gly | P00387 |
| HOMO SAPIENS | 0 | 4244 | 1umk | A | 291 | GLY | p.Gly292Asp | P00387 |
| HOMO SAPIENS | 4 | 34495 | 1s35 | A | 1151 | ASP | p.Asn1151Asp | P11277 |
| HOMO SAPIENS | 3 | 30110 | 3zqs | A | 224 | THR | p.Thr224Ala | Q9NW38 |
| HOMO SAPIENS | 3 | 17588 | 5dse | B | 53 | LEU | p.Leu53Pro | Q9BYI3 |
| HOMO SAPIENS | 0 | 1564 | 4yqf | B | 72 | HIS | p.His347Gln | Q9UHD8 |
| HOMO SAPIENS | 0 | 17673 | 4jnw | A | 32 | ARG | p.Arg32186His | Q8WZ42 |
| HOMO SAPIENS | 3 | 17673 | 4jnw | A | 140 | ILE | p.Ile32294Val | Q8WZ42 |
| HOMO SAPIENS | 5 | 17673 | 4jnw | A | 269 | ALA | p.Ala32423Thr | Q8WZ42 |
| HOMO SAPIENS | 3 | 12858 | 3s97 | C | 302 | LEU | p.Leu302Pro | Q12860 |
| HOMO SAPIENS | 0 | 6147 | 4nn0 | A | 114 | ARG | p.Arg114Gln | Q9BXJ0 |
| HOMO SAPIENS | 0 | 7549 | 1tr2 | A | 277 | LEU | p.Leu277Met | P18206 |
| HOMO SAPIENS | 1 | 7549 | 1tr2 | A | 413 | ALA | p.Ala413Thr | P18206 |
| HOMO SAPIENS | 1 | 7549 | 1tr2 | A | 519 | ILE | p.Ile519Leu | P18206 |
| HOMO SAPIENS | 5 | 7549 | 1tr2 | A | 547 | ARG | p.Arg547Ter | P18206 |
| HOMO SAPIENS | 2 | 7549 | 1tr2 | A | 636 | HIS | p.His636Arg | P18206 |
| HOMO SAPIENS | 2 | 7549 | 1tr2 | A | 824 | ILE | p.Ile824Met | P18206 |
| HOMO SAPIENS | 2 | 13900 | 3gro | A | 45 | CYS | p.Cys45Tyr | P50897 |
| HOMO SAPIENS | 2 | 13900 | 3gro | A | 75 | THR | p.Thr75Pro | P50897 |

| | | | | | | | | |
|---|---|---|---|---|---|---|---|---|
| HOMO SAPIENS | 2 | 13900 | 3gro | A | 79 | ASP | p.Asp79Gly | P50897 |
| HOMO SAPIENS | 5 | 13900 | 3gro | A | 91 | GLN | p.Gln91Pro | P50897 |
| HOMO SAPIENS | 5 | 13900 | 3gro | A | 96 | CYS | p.Cys96Tyr | P50897 |
| HOMO SAPIENS | 0 | 13900 | 3gro | A | 104 | LYS | p.Lys104Ter | P50897 |
| HOMO SAPIENS | 2 | 13900 | 3gro | A | 108 | GLY | p.Gly108Arg | P50897 |
| HOMO SAPIENS | 2 | 13900 | 3gro | A | 109 | TYR | p.Tyr109Asp | P50897 |
| HOMO SAPIENS | 0 | 13900 | 3gro | A | 122 | ARG | p.Arg122Trp | P50897 |
| HOMO SAPIENS | 0 | 13900 | 3gro | A | 138 | SER | p.Ser138Leu | P50897 |
| HOMO SAPIENS | 5 | 13900 | 3gro | A | 151 | ARG | p.Arg151Ter | P50897 |
| HOMO SAPIENS | 7 | 13900 | 3gro | A | 164 | ARG | p.Arg164Ter | P50897 |
| HOMO SAPIENS | 0 | 13900 | 3gro | A | 177 | GLN | p.Gln177Glu | P50897 |
| HOMO SAPIENS | 0 | 13900 | 3gro | A | 181 | VAL | p.Val181Leu | P50897 |
| HOMO SAPIENS | 0 | 13900 | 3gro | A | 184 | GLU | p.Glu184Lys | P50897 |
| HOMO SAPIENS | 5 | 13900 | 3gro | A | 186 | TRP | p.Trp186Ter | P50897 |
| HOMO SAPIENS | 0 | 13900 | 3gro | A | 219 | LEU | p.Leu219Gln | P50897 |
| HOMO SAPIENS | 0 | 13900 | 3gro | A | 222 | LEU | p.Leu222Pro | P50897 |
| HOMO SAPIENS | 5 | 13900 | 3gro | A | 296 | TRP | p.Trp296Ter | P50897 |
| HOMO SAPIENS | 6 | 35028 | 5mwb | A | 444 | CYS | p.Cys444Tyr | Q04721 |
| HOMO SAPIENS | 0 | 35028 | 5mwb | A | 514 | ASN | p.Asn514Ser | Q04721 |
| HOMO SAPIENS | 5 | 1483 | 4gl2 | A | 822 | ARG | p.Arg822Gln | Q9BYX4 |
| HOMO SAPIENS | 0 | 1483 | 4gl2 | A | 843 | ARG | p.His843Arg | Q9BYX4 |
| HOMO SAPIENS | -1 | 28838 | 3cl3 | D | 153 | LEU | p.Leu153Arg | Q9Y6K9 |
| HOMO SAPIENS | 2 | 22563 | 4zel | B | 79 | ARG | p.Arg79Trp | P09172 |
| HOMO SAPIENS | 2 | 22563 | 4zel | B | 101 | VAL | p.Val101Met | P09172 |
| HOMO SAPIENS | 2 | 22563 | 4zel | B | 114 | ASP | p.Asp114Glu | P09172 |
| HOMO SAPIENS | 0 | 22563 | 4zel | B | 201 | ASN | p.Asn201Ser | P09172 |
| HOMO SAPIENS | 2 | 22563 | 4zel | B | 211 | ALA | p.Ala211Thr | P09172 |
| HOMO SAPIENS | -1 | 22563 | 4zel | B | 290 | ASP | p.Asp290Asn | P09172 |
| HOMO SAPIENS | 2 | 22563 | 4zel | B | 318 | ALA | p.Ala318Ser | P09172 |
| HOMO SAPIENS | 2 | 22563 | 4zel | B | 345 | ASP | p.Asp345Asn | P09172 |
| HOMO SAPIENS | 2 | 22563 | 4zel | B | 362 | ALA | p.Ala362Glu | P09172 |
| HOMO SAPIENS | 2 | 22563 | 4zel | B | 549 | ARG | p.Arg549Cys | P09172 |
| HOMO SAPIENS | 9 | 9780 | 1y6k | R | 36 | TYR | p.Tyr57Cys | Q13651 |
| HOMO SAPIENS | 1 | 9780 | 1y6k | R | 40 | LEU | p.Leu61Val | Q13651 |
| HOMO SAPIENS | 1 | 9780 | 1y6k | R | 63 | THR | p.Thr84Ile | Q13651 |
| HOMO SAPIENS | 9 | 9780 | 1y6k | R | 80 | ARG | p.Arg101Trp | Q13651 |
| HOMO SAPIENS | 0 | 9780 | 1y6k | R | 120 | GLY | p.Gly141Arg | Q13651 |
| HOMO SAPIENS | 0 | 9780 | 1y6k | R | 138 | SER | p.Ser159Gly | Q13651 |
| HOMO SAPIENS | 0 | 9780 | 1y6k | R | 203 | ILE | p.Ile224Val | Q13651 |
| HOMO SAPIENS | 0 | 16019 | 3rep | A | 211 | ARG | p.Arg211Cys | Q13418 |
| HOMO SAPIENS | 3 | 7879 | 5ec3 | A | 33 | ALA | p.Ala33Thr | P32754 |
| HOMO SAPIENS | 4 | 7879 | 5ec3 | A | 113 | ARG | p.Arg113Gln | P32754 |
| HOMO SAPIENS | 3 | 7879 | 5ec3 | A | 340 | VAL | p.Val340Leu | P32754 |
| HOMO SAPIENS | 0 | 8904 | 1qk9 | A | 6 | LYS | p.Lys82Arg | P51608 |
| HOMO SAPIENS | 4 | 8904 | 1qk9 | A | 24 | LEU | p.Leu100Val | P51608 |
| HOMO SAPIENS | 7 | 8904 | 1qk9 | A | 30 | ARG | p.Arg106Trp | P51608 |
| HOMO SAPIENS | 6 | 8904 | 1qk9 | A | 57 | ARG | p.Arg133Cys | P51608 |
| HOMO SAPIENS | 3 | 8904 | 1qk9 | A | 61 | GLU | p.Glu137Gly | P51608 |
| HOMO SAPIENS | 2 | 8904 | 1qk9 | A | 64 | ALA | p.Ala140Val | P51608 |
| HOMO SAPIENS | 5 | 8904 | 1qk9 | A | 79 | PHE | p.Phe155Ser | P51608 |
| HOMO SAPIENS | 3 | 8904 | 1qk9 | A | 82 | THR | p.Thr158Met | P51608 |
| HOMO SAPIENS | 1 | 23121 | 3e9k | A | 198 | THR | p.Thr198Ala | Q16719 |
| HOMO SAPIENS | -1 | 26090 | 3tje | F | 19 | LEU | p.Leu35Met | P25445 |
| HOMO SAPIENS | 8 | 26090 | 3tje | F | 106 | THR | p.Thr122Ile | P25445 |

| | | | | | | | | |
|---|---|---|---|---|---|---|---|---|
| HOMO SAPIENS | 2 | 36404 | 5cqr | A | 765 | GLY | p.Gly765Glu | O95163 |
| HOMO SAPIENS | 2 | 36404 | 5cqr | A | 816 | ILE | p.Ile816Leu | O95163 |
| HOMO SAPIENS | 2 | 36404 | 5cqr | A | 830 | ILE | p.Ile830Met | O95163 |
| HOMO SAPIENS | 5 | 36404 | 5cqr | A | 939 | TYR | p.Tyr939Ter | O95163 |
| HOMO SAPIENS | 2 | 36404 | 5cqr | A | 952 | LYS | p.Lys952Ile | O95163 |
| HOMO SAPIENS | 5 | 36404 | 5cqr | A | 1013 | GLY | p.Gly1013Ser | O95163 |
| HOMO SAPIENS | 5 | 36404 | 5cqr | A | 1072 | CYS | p.Cys1072Ser | O95163 |
| HOMO SAPIENS | -1 | 36404 | 5cqr | A | 1158 | PRO | p.Pro1158Leu | O95163 |
| HOMO SAPIENS | 1 | 34149 | 2k40 | A | 18 | ASN | p.Asn125Ser | Q9UBX0 |
| HOMO SAPIENS | 0 | 34149 | 2k40 | A | 42 | LEU | p.Glu149Lys | Q9UBX0 |
| HOMO SAPIENS | 9 | 34149 | 2k40 | A | 53 | ARG | p.Arg160Cys | Q9UBX0 |
| HOMO SAPIENS | 0 | 34149 | 2k40 | A | 63 | SER | p.Ser170Leu | Q9UBX0 |
| HOMO SAPIENS | 2 | 2468 | 3vp6 | A | 484 | LEU | p.Leu484Met | Q99259 |
| HOMO SAPIENS | 0 | 2468 | 3vp6 | A | 532 | ARG | p.Arg532Gln | Q99259 |
| HOMO SAPIENS | 5 | 1253 | 5xtd | S | 8 | GLY | p.Gly8Arg | O15239 |
| HOMO SAPIENS | 7 | 1253 | 5xtd | S | 32 | GLY | p.Gly32Arg | O15239 |
| HOMO SAPIENS | 7 | 1253 | 5xtd | S | 37 | ARG | p.Arg37Ser | O15239 |
| HOMO SAPIENS | -1 | 13292 | 3u1k | A | 121 | VAL | p.Ile121Val | Q8TCS8 |
| HOMO SAPIENS | 2 | 13292 | 3u1k | A | 165 | PRO | p.Pro165Ser | Q8TCS8 |
| HOMO SAPIENS | 2 | 13292 | 3u1k | A | 192 | ARG | p.Arg192Gln | Q8TCS8 |
| HOMO SAPIENS | 1 | 13292 | 3u1k | A | 315 | LYS | p.Lys315Arg | Q8TCS8 |
| HOMO SAPIENS | 4 | 13292 | 3u1k | A | 509 | VAL | p.Val509Ile | Q8TCS8 |
| HOMO SAPIENS | 7 | 13292 | 3u1k | A | 531 | THR | p.Thr531Arg | Q8TCS8 |
| HOMO SAPIENS | 0 | 13292 | 3u1k | A | 590 | ASN | p.Asn590Asp | Q8TCS8 |
| HOMO SAPIENS | 0 | 27048 | 4p5w | B | 184 | ARG | p.Arg184Ter | Q9UJS0 |
| HOMO SAPIENS | 0 | 27048 | 4p5w | B | 225 | SER | p.Ser225Ter | Q9UJS0 |
| HOMO SAPIENS | -1 | 27048 | 4p5w | B | 631 | ALA | p.Arg360Ter | Q9UJS0 |
| HOMO SAPIENS | 0 | 24240 | 4mdr | A | 318 | ARG | p.Arg318Ter | O00189 |
| HOMO SAPIENS | 0 | 24240 | 4mdr | A | 338 | ARG | p.Arg338Ter | O00189 |
| HOMO SAPIENS | 0 | 13429 | 4cg4 | A | 479 | PHE | p.Phe479Leu | O15553 |
| HOMO SAPIENS | 0 | 13429 | 4cg4 | A | 487 | VAL | p.Val487Leu | O15553 |
| HOMO SAPIENS | 3 | 13429 | 4cg4 | A | 653 | ARG | p.Arg653His | O15553 |
| HOMO SAPIENS | 1 | 13429 | 4cg4 | A | 680 | MET | p.Met680Ile | O15553 |
| HOMO SAPIENS | 1 | 13429 | 4cg4 | A | 688 | TYR | p.Tyr688Ter | O15553 |
| HOMO SAPIENS | 0 | 13429 | 4cg4 | A | 694 | MET | p.Met694Ile | O15553 |
| HOMO SAPIENS | 0 | 13429 | 4cg4 | A | 694 | MET | p.Met694Val | O15553 |
| HOMO SAPIENS | 1 | 13429 | 4cg4 | A | 726 | VAL | p.Val726Ala | O15553 |
| HOMO SAPIENS | 0 | 13429 | 4cg4 | A | 761 | ARG | p.Arg761His | O15553 |
| HOMO SAPIENS | 0 | 2079 | 4okh | A | 691 | GLY | p.Gly691Arg | P20807 |
| HOMO SAPIENS | 2 | 2079 | 4okh | A | 748 | ARG | p.Arg748Ter | P20807 |
| HOMO SAPIENS | 0 | 2079 | 4okh | A | 763 | TYR | p.Tyr763Cys | P20807 |
| HOMO SAPIENS | 1 | 2079 | 4okh | A | 769 | ARG | p.Arg769Gln | P20807 |
| HOMO SAPIENS | 9 | 2079 | 4okh | A | 777 | ILE | p.Ile777Val | P20807 |
| HOMO SAPIENS | 8 | 2079 | 4okh | A | 778 | ASP | p.Asp778His | P20807 |
| HOMO SAPIENS | 5 | 2079 | 4okh | A | 780 | ASP | p.Asp780His | P20807 |
| HOMO SAPIENS | 1 | 1200 | 2gi7 | A | 142 | ARG | p.Arg162Trp | Q9HCN6 |
| HOMO SAPIENS | 4 | 9199 | 3ezq | C | 260 | ASP | p.Asp260Val | P25445 |
| HOMO SAPIENS | 0 | 36522 | 5ms9 | A | 122 | ARG | p.Arg122Cys | P35555 |
| HOMO SAPIENS | 0 | 36522 | 5ms9 | A | 240 | ARG | p.Arg240Cys | P35555 |
| HOMO SAPIENS | 0 | 17448 | 3g2f | A | 225 | ARG | p.Arg225His | Q13873 |
| HOMO SAPIENS | 0 | 17448 | 3g2f | A | 347 | CYS | p.Cys347Tyr | Q13873 |
| HOMO SAPIENS | 3 | 17448 | 3g2f | A | 485 | ASP | p.Asp485Gly | Q13873 |
| HOMO SAPIENS | 7 | 17448 | 3g2f | A | 491 | ARG | p.Arg491Gln | Q13873 |
| HOMO SAPIENS | 7 | 17448 | 3g2f | A | 491 | ARG | p.Arg491Trp | Q13873 |

| | | | | | | | | |
|---|---|---|---|---|---|---|---|---|
| HOMO SAPIENS | 0 | 4861 | 4ux8 | A | 56 | LEU | p.Leu56Met | P07949 |
| HOMO SAPIENS | 1 | 4861 | 4ux8 | A | 88 | ILE | p.Ile88Leu | P07949 |
| HOMO SAPIENS | 0 | 4861 | 4ux8 | A | 114 | ARG | p.Arg114His | P07949 |
| HOMO SAPIENS | 5 | 4861 | 4ux8 | A | 180 | ARG | p.Arg180Gln | P07949 |
| HOMO SAPIENS | 0 | 4861 | 4ux8 | A | 208 | GLU | p.Glu208Asp | P07949 |
| HOMO SAPIENS | 0 | 4861 | 4ux8 | A | 373 | ALA | p.Ala373Val | P07949 |
| HOMO SAPIENS | 0 | 4861 | 4ux8 | A | 375 | LEU | p.Leu375Gln | P07949 |
| HOMO SAPIENS | -1 | 4861 | 4ux8 | A | 386 | ALA | p.Ala386Val | P07949 |
| HOMO SAPIENS | 0 | 4861 | 4ux8 | A | 446 | GLY | p.Gly446Arg | P07949 |
| HOMO SAPIENS | 0 | 4861 | 4ux8 | A | 480 | GLU | p.Glu480Lys | P07949 |
| HOMO SAPIENS | 0 | 4861 | 4ux8 | A | 489 | ASP | p.Asp489Asn | P07949 |
| HOMO SAPIENS | -1 | 4861 | 4ux8 | A | 510 | ALA | p.Ala510Val | P07949 |
| HOMO SAPIENS | -1 | 4861 | 4ux8 | A | 533 | GLY | p.Gly533Cys | P07949 |
| HOMO SAPIENS | -1 | 4861 | 4ux8 | A | 567 | ASP | p.Asp567Asn | P07949 |
| HOMO SAPIENS | -1 | 4861 | 4ux8 | A | 609 | CYS | p.Cys609Tyr | P07949 |
| HOMO SAPIENS | -1 | 4861 | 4ux8 | A | 609 | CYS | p.Cys609Gly | P07949 |
| HOMO SAPIENS | -1 | 4861 | 4ux8 | A | 611 | CYS | p.Cys611Trp | P07949 |
| HOMO SAPIENS | -1 | 4861 | 4ux8 | A | 618 | CYS | p.Cys618Gly | P07949 |
| HOMO SAPIENS | -1 | 4861 | 4ux8 | A | 618 | CYS | p.Cys618Phe | P07949 |
| HOMO SAPIENS | -1 | 4861 | 4ux8 | A | 620 | CYS | p.Cys620Gly | P07949 |
| HOMO SAPIENS | -1 | 4861 | 4ux8 | A | 620 | CYS | p.Cys620Ser | P07949 |
| HOMO SAPIENS | -1 | 4861 | 4ux8 | A | 620 | CYS | p.Cys620Trp | P07949 |
| HOMO SAPIENS | -1 | 4861 | 4ux8 | A | 623 | GLU | p.Glu623Lys | P07949 |
| HOMO SAPIENS | -1 | 4861 | 4ux8 | A | 631 | ASP | p.Asp631Gly | P07949 |
| HOMO SAPIENS | -1 | 4861 | 4ux8 | A | 631 | ASP | p.Asp631Glu | P07949 |
| HOMO SAPIENS | -1 | 4861 | 4ux8 | A | 634 | CYS | p.Cys634Gly | P07949 |
| HOMO SAPIENS | -1 | 4861 | 4ux8 | A | 634 | CYS | p.Cys634Phe | P07949 |
| HOMO SAPIENS | -1 | 4861 | 4ux8 | A | 634 | CYS | p.Cys634Trp | P07949 |
| HOMO SAPIENS | 0 | 36308 | 4nkq | B | 123 | PRO | p.Pro142Leu | P15509 |
| HOMO SAPIENS | 0 | 36308 | 4nkq | B | 145 | ARG | p.Arg164Gln | P15509 |
| HOMO SAPIENS | -1 | 36308 | 4nkq | B | 177 | GLY | p.Gly196Arg | P15509 |
| HOMO SAPIENS | 0 | 36006 | 1oqa | A | 107 | HIS | p.His1860Pro | P38398 |
| HOMO SAPIENS | 7 | 15394 | 2ozo | A | 465 | ARG | p.Arg465His | P43403 |
| HOMO SAPIENS | 7 | 15394 | 2ozo | A | 465 | ARG | p.Arg465Cys | P43403 |
| HOMO SAPIENS | 4 | 15394 | 2ozo | A | 518 | SER | p.Ser518Arg | P43403 |
| HOMO SAPIENS | 1 | 20813 | 3o5z | A | 19 | ARG | p.Arg19Trp | O00459 |
| HOMO SAPIENS | 2 | 4329 | 2bb5 | A | 5 | ILE | p.Ile23Val | P20062 |
| HOMO SAPIENS | 4 | 4329 | 2bb5 | A | 59 | LYS | p.Lys77Met | P20062 |
| HOMO SAPIENS | 4 | 4329 | 2bb5 | A | 197 | ARG | p.Arg215Trp | P20062 |
| HOMO SAPIENS | 1 | 4329 | 2bb5 | A | 241 | PRO | p.Arg259Pro | P20062 |
| HOMO SAPIENS | 2 | 4329 | 2bb5 | A | 315 | THR | p.Thr333Lys | P20062 |
| HOMO SAPIENS | 5 | 4329 | 2bb5 | A | 330 | SER | p.Ser348Phe | P20062 |
| HOMO SAPIENS | 4 | 4329 | 2bb5 | A | 358 | LEU | p.Leu376Ser | P20062 |
| HOMO SAPIENS | 5 | 4329 | 2bb5 | A | 381 | ARG | p.Arg399Gln | P20062 |
| HOMO SAPIENS | 4 | 18294 | 4r30 | A | 163 | ASN | p.Asn163Asp | O95278 |
| HOMO SAPIENS | 6 | 18294 | 4r30 | A | 227 | ALA | p.Ala227Val | O95278 |
| HOMO SAPIENS | 4 | 18294 | 4r30 | A | 241 | ARG | p.Arg241Ter | O95278 |
| HOMO SAPIENS | 4 | 18294 | 4r30 | A | 325 | ARG | p.Arg325His | O95278 |
| HOMO SAPIENS | 0 | 2632 | 4xem | A | 22 | GLU | p.Glu22Lys | P49588 |
| HOMO SAPIENS | 0 | 2632 | 4xem | A | 275 | GLY | p.Gly275Asp | P49588 |
| HOMO SAPIENS | 1 | 2632 | 4xem | A | 329 | ARG | p.Arg329His | P49588 |
| HOMO SAPIENS | 0 | 11431 | 5t4n | A | 284 | GLN | p.Gln305Glu | Q96QU1 |
| HOMO SAPIENS | 0 | 11431 | 5t4n | A | 315 | ARG | p.Arg336Ter | Q96QU1 |
| HOMO SAPIENS | 0 | 11431 | 5t4n | A | 322 | ARG | p.Arg343Lys | Q96QU1 |

| | | | | | | | | |
|---|---|---|---|---|---|---|---|---|
| HOMO SAPIENS | 1 | 11431 | 5t4n | A | 326 | LEU | p.Leu347Phe | Q96QU1 |
| HOMO SAPIENS | 0 | 11431 | 5t4n | A | 373 | TYR | p.Tyr394Cys | Q96QU1 |
| HOMO SAPIENS | 0 | 11431 | 5t4n | A | 398 | ASN | p.Asn419Ser | Q96QU1 |
| HOMO SAPIENS | 0 | 11431 | 5t4n | A | 414 | ALA | p.Asp435Ala | Q96QU1 |
| HOMO SAPIENS | 0 | 11431 | 5t4n | A | 433 | VAL | p.Val454Phe | Q96QU1 |
| HOMO SAPIENS | 1 | 11431 | 5t4n | A | 507 | VAL | p.Val528Asp | Q96QU1 |
| HOMO SAPIENS | 0 | 11431 | 5t4n | A | 547 | ALA | p.Ala568Ser | Q96QU1 |
| HOMO SAPIENS | 3 | 3194 | 5xtd | L | 84 | ARG | p.Arg84His | O43181 |
| HOMO SAPIENS | 1 | 3194 | 5xtd | L | 130 | THR | p.Thr130Ser | O43181 |
| HOMO SAPIENS | 8 | 3194 | 5xtd | L | 136 | SER | p.Ser136Ala | O43181 |
| HOMO SAPIENS | 0 | 40749 | 1nun | A | 106 | CYS | p.Cys106Phe | O15520 |
| HOMO SAPIENS | 9 | 40749 | 1nun | A | 156 | ILE | p.Ile156Arg | O15520 |
| HOMO SAPIENS | 1 | 40749 | 1nun | A | 193 | ARG | p.Arg193Ter | O15520 |
| HOMO SAPIENS | 0 | 40749 | 1nun | A | 207 | HIS | p.His207Pro | O15520 |
| HOMO SAPIENS | 0 | 7026 | 3n1f | D | 875 | SER | p.Ser875Gly | Q4KMG0 |
| HOMO SAPIENS | 6 | 23802 | 3rzv | A | 424 | ARG | p.Arg424Ter | O95630 |
| HOMO SAPIENS | 0 | 36720 | 1jm7 | A | 12 | GLN | p.Gln12Ter | P38398 |
| HOMO SAPIENS | 3 | 36720 | 1jm7 | A | 23 | GLU | p.Glu23Asp | P38398 |
| HOMO SAPIENS | 1 | 36720 | 1jm7 | A | 45 | LYS | p.Lys45Gln | P38398 |
| HOMO SAPIENS | 8 | 36720 | 1jm7 | A | 47 | CYS | p.Cys47Ter | P38398 |
| HOMO SAPIENS | 0 | 36720 | 1jm7 | A | 59 | SER | p.Ser59Ter | P38398 |
| HOMO SAPIENS | 8 | 36720 | 1jm7 | A | 61 | CYS | p.Cys61Gly | P38398 |
| HOMO SAPIENS | 8 | 36720 | 1jm7 | A | 63 | LEU | p.Leu63Ter | P38398 |
| HOMO SAPIENS | 7 | 36720 | 1jm7 | A | 64 | CYS | p.Cys64Gly | P38398 |
| HOMO SAPIENS | 0 | 36720 | 1jm7 | A | 101 | TYR | p.Tyr101Ter | P38398 |
| HOMO SAPIENS | 0 | 36720 | 1jm7 | A | null | TYR | p.Tyr105Cys | P38398 |
| HOMO SAPIENS | 0 | 28500 | 3kuz | A | 1217 | CYS | p.Cys1217Arg | O60486 |
| HOMO SAPIENS | 5 | 8097 | 3p56 | C | 69 | ARG | p.Arg69Trp | Q8TDP1 |
| HOMO SAPIENS | 5 | 8097 | 3p56 | C | 143 | LYS | p.Lys143Ile | Q8TDP1 |
| HOMO SAPIENS | 2 | 30073 | 3nmz | A | 380 | ARG | p.Arg380Gln | P25054 |
| HOMO SAPIENS | 1 | 22573 | 1cjy | A | 111 | SER | p.Ser111Pro | P47712 |
| HOMO SAPIENS | 0 | 22573 | 1cjy | A | 485 | ARG | p.Arg485His | P47712 |
| HOMO SAPIENS | 2 | 965 | 3E+077 | A | 99 | ALA | p.Ala99Val | Q9Y617 |
| HOMO SAPIENS | 2 | 965 | 3E+077 | A | 100 | ASP | p.Asp100Ala | Q9Y617 |
| HOMO SAPIENS | 4 | 1503 | 3kxc | A | 62 | ARG | p.Arg62Trp | O43617 |
| HOMO SAPIENS | 6 | 9418 | 1hdr | A | 22 | GLY | p.Gly23Asp | P09417 |
| HOMO SAPIENS | 2 | 9418 | 1hdr | A | 35 | TRP | p.Trp36Arg | P09417 |
| HOMO SAPIENS | 2 | 9418 | 1hdr | A | 107 | TRP | p.Trp108Gly | P09417 |
| HOMO SAPIENS | 6 | 9418 | 1hdr | A | 149 | TYR | p.Tyr150Cys | P09417 |
| HOMO SAPIENS | -1 | 12079 | 3tnu | B | 373 | GLY | p.Gly373Ala | P13647 |
| HOMO SAPIENS | 1 | 12079 | 3tnu | B | 463 | LEU | p.Leu463Pro | P13647 |
| HOMO SAPIENS | 1 | 12079 | 3tnu | B | 471 | ARG | p.Arg471Cys | P13647 |
| HOMO SAPIENS | 0 | 8135 | 3rbn | B | 89 | SER | p.Ser556Ile | P40692 |
| HOMO SAPIENS | 0 | 8135 | 3rbn | B | 111 | GLU | p.Glu578Gly | P40692 |
| HOMO SAPIENS | 2 | 8135 | 3rbn | B | 114 | PRO | p.Pro581Leu | P40692 |
| HOMO SAPIENS | 0 | 8135 | 3rbn | B | 133 | GLU | p.Glu600Gly | P40692 |
| HOMO SAPIENS | 2 | 8135 | 3rbn | B | 136 | PRO | p.Pro603Arg | P40692 |
| HOMO SAPIENS | 2 | 8135 | 3rbn | B | 136 | PRO | p.Pro603Leu | P40692 |
| HOMO SAPIENS | 7 | 8135 | 3rbn | B | 140 | LEU | p.Leu607His | P40692 |
| HOMO SAPIENS | 2 | 8135 | 3rbn | B | 151 | LYS | p.Lys618Ala | P40692 |
| HOMO SAPIENS | 2 | 8135 | 3rbn | B | 151 | LYS | p.Lys618Thr | P40692 |
| HOMO SAPIENS | 5 | 8135 | 3rbn | B | 181 | PRO | p.Pro648Ser | P40692 |
| HOMO SAPIENS | 2 | 8135 | 3rbn | B | 188 | ILE | p.Ile655Val | P40692 |
| HOMO SAPIENS | 2 | 8135 | 3rbn | B | 214 | ALA | p.Ala681Thr | P40692 |

| | | | | | | | | |
|---|---|---|---|---|---|---|---|---|
| HOMO SAPIENS | 2 | 8135 | 3rbn | B | 220 | ARG | p.Arg687Trp | P40692 |
| HOMO SAPIENS | 2 | 8135 | 3rbn | B | 222 | GLN | p.Gln689Arg | P40692 |
| HOMO SAPIENS | -1 | 8135 | 3rbn | B | 236 | GLN | p.Gln701Ter | P40692 |
| HOMO SAPIENS | 7 | 8135 | 3rbn | B | 251 | HIS | p.His718Tyr | P40692 |
| HOMO SAPIENS | 0 | 8135 | 3rbn | B | 284 | LYS | p.Lys751Arg | P40692 |
| HOMO SAPIENS | 1 | 21998 | 5d68 | A | 355 | LEU | p.Leu355Phe | O00522 |
| HOMO SAPIENS | 3 | 21998 | 5d68 | A | 380 | THR | p.Thr380Met | O00522 |
| HOMO SAPIENS | 0 | 21998 | 5d68 | A | 415 | ILE | p.Ile415Met | O00522 |
| HOMO SAPIENS | 0 | 33789 | 2m0c | A | 13 | ARG | p.Arg218Gln | Q9H161 |
| HOMO SAPIENS | 7 | 33789 | 2m0c | A | 52 | ARG | p.Arg257Thr | Q9H161 |
| HOMO SAPIENS | 0 | 33789 | 2m0c | A | 67 | ARG | p.Arg272Pro | Q9H161 |
| HOMO SAPIENS | 6 | 8263 | 2osg | A | 51 | GLY | p.Gly355Arg | Q9UDY2 |
| HOMO SAPIENS | 5 | 8263 | 2osg | A | 60 | THR | p.Thr364Met | Q9UDY2 |
| HOMO SAPIENS | 1 | 2475 | 5E+026 | B | 234 | THR | p.Thr234Ala | Q9BZ23 |
| HOMO SAPIENS | 2 | 2475 | 5E+026 | B | 264 | ARG | p.Arg264Trp | Q9BZ23 |
| HOMO SAPIENS | 4 | 2475 | 5E+026 | B | 286 | ARG | p.Arg286Cys | Q9BZ23 |
| HOMO SAPIENS | 0 | 2475 | 5E+026 | B | 357 | ARG | p.Arg357Trp | Q9BZ23 |
| HOMO SAPIENS | 2 | 2475 | 5E+026 | B | 481 | ARG | p.Arg481Ter | Q9BZ23 |
| HOMO SAPIENS | 8 | 2475 | 5E+026 | B | 521 | GLY | p.Gly521Arg | Q9BZ23 |
| HOMO SAPIENS | 2 | 2475 | 5E+026 | B | 528 | THR | p.Thr528Met | Q9BZ23 |
| HOMO SAPIENS | 1 | 3742 | 5xtd | E | 108 | HIS | p.His134Tyr | P56556 |
| HOMO SAPIENS | 1 | 16725 | 3g04 | C | 36 | ASP | p.Asp36His | P16473 |
| HOMO SAPIENS | 3 | 16725 | 3g04 | C | 52 | PRO | p.Pro52Thr | P16473 |
| HOMO SAPIENS | 1 | 16725 | 3g04 | C | 162 | PRO | p.Pro162Ala | P16473 |
| HOMO SAPIENS | -1 | 12366 | 5d2d | C | 750 | PRO | p.Pro750Leu | P13569 |
| HOMO SAPIENS | 0 | 12366 | 5d2d | C | 764 | ARG | p.Arg764Ter | P13569 |
| HOMO SAPIENS | 0 | 34845 | 2y23 | A | 1151 | ASP | p.Asp1151Glu | P52179 |
| HOMO SAPIENS | 0 | 34845 | 2y23 | A | 1180 | ASP | p.Asp1180Ala | P52179 |
| HOMO SAPIENS | 6 | 36740 | 3lbx | B | 2053 | ALA | p.Ala2053Pro | P11277 |
| HOMO SAPIENS | 0 | 19289 | 3ihj | A | 404 | ARG | p.Arg404Ter | Q8TD30 |
| HOMO SAPIENS | 5 | 1034 | 5xtd | Q | 228 | ARG | p.Arg228Gln | O75306 |
| HOMO SAPIENS | 5 | 1034 | 5xtd | Q | 229 | PRO | p.Pro229Gln | O75306 |
| HOMO SAPIENS | 4 | 1034 | 5xtd | Q | 323 | ARG | p.Arg323Gln | O75306 |
| HOMO SAPIENS | 4 | 1034 | 5xtd | Q | 352 | PRO | p.Pro352Ala | O75306 |
| HOMO SAPIENS | 5 | 1034 | 5xtd | Q | 413 | SER | p.Ser413Pro | O75306 |
| HOMO SAPIENS | 1 | 2772 | 2vaf | A | 1047 | THR | p.Thr66Ala | O14958 |
| HOMO SAPIENS | 5 | 2772 | 2vaf | A | 1057 | VAL | p.Val76Met | O14958 |
| HOMO SAPIENS | 0 | 2772 | 2vaf | A | 1148 | LEU | p.Leu167His | O14958 |
| HOMO SAPIENS | 1 | 2772 | 2vaf | A | 1149 | ILE | p.Ile168Thr | O14958 |
| HOMO SAPIENS | 8 | 2772 | 2vaf | A | 1225 | HIS | p.His244Arg | O14958 |
| HOMO SAPIENS | 8 | 2772 | 2vaf | A | 1225 | HIS | p.His244Tyr | O14958 |
| HOMO SAPIENS | 7 | 2772 | 2vaf | A | 1251 | ILE | p.Ile270Thr | O14958 |
| HOMO SAPIENS | 3 | 2772 | 2vaf | A | 1288 | ASP | p.Asp307His | O14958 |
| HOMO SAPIENS | 5 | 2772 | 2vaf | A | 1289 | PRO | p.Pro308Leu | O14958 |
| HOMO SAPIENS | 1 | 31885 | 2aex | A | 132 | ALA | p.Ala132Val | P36551 |
| HOMO SAPIENS | 1 | 31885 | 2aex | A | 272 | ASN | p.Asn272His | P36551 |
| HOMO SAPIENS | 8 | 31885 | 2aex | A | 279 | GLY | p.Gly279Arg | P36551 |
| HOMO SAPIENS | 5 | 31885 | 2aex | A | 294 | VAL | p.Val294Ile | P36551 |
| HOMO SAPIENS | 5 | 31885 | 2aex | A | 295 | HIS | p.His295Asp | P36551 |
| HOMO SAPIENS | 1 | 31885 | 2aex | A | 331 | ARG | p.Arg331Trp | P36551 |
| HOMO SAPIENS | 3 | 31885 | 2aex | A | 352 | ARG | p.Arg352Cys | P36551 |
| HOMO SAPIENS | 1 | 31885 | 2aex | A | 404 | LYS | p.Lys404Glu | P36551 |
| HOMO SAPIENS | 2 | 31885 | 2aex | A | 447 | ARG | p.Arg447Cys | P36551 |
| HOMO SAPIENS | 5 | 11721 | 2r3v | A | 20 | HIS | p.His20Pro | Q03426 |

| | | | | | | | | |
|---|---|---|---|---|---|---|---|---|
| HOMO SAPIENS | 1 | 11721 | 2r3v | A | 116 | TYR | p.Tyr116His | Q03426 |
| HOMO SAPIENS | 1 | 11721 | 2r3v | A | 148 | ALA | p.Ala148Thr | Q03426 |
| HOMO SAPIENS | 1 | 11721 | 2r3v | A | 167 | PRO | p.Pro167Leu | Q03426 |
| HOMO SAPIENS | 6 | 11721 | 2r3v | A | 203 | VAL | p.Val203Ala | Q03426 |
| HOMO SAPIENS | 2 | 11721 | 2r3v | A | 215 | ARG | p.Arg215Ter | Q03426 |
| HOMO SAPIENS | 2 | 11721 | 2r3v | A | 237 | THR | p.Thr237Ser | Q03426 |
| HOMO SAPIENS | 0 | 11721 | 2r3v | A | 268 | ILE | p.Ile268Thr | Q03426 |
| HOMO SAPIENS | 0 | 11721 | 2r3v | A | 268 | ILE | p.Ile268Val | Q03426 |
| HOMO SAPIENS | 0 | 11721 | 2r3v | A | 301 | ASN | p.Asn301Ser | Q03426 |
| HOMO SAPIENS | 6 | 11721 | 2r3v | A | 310 | VAL | p.Val310Met | Q03426 |
| HOMO SAPIENS | 5 | 11721 | 2r3v | A | 334 | ALA | p.Ala334Thr | Q03426 |
| HOMO SAPIENS | 2 | 11721 | 2r3v | A | 377 | VAL | p.Val377Ile | Q03426 |
| HOMO SAPIENS | 1 | 11721 | 2r3v | A | 386 | ASP | p.Asp386Asn | Q03426 |
| HOMO SAPIENS | 1 | 11721 | 2r3v | A | 388 | ARG | p.Arg388Ter | Q03426 |
| HOMO SAPIENS | 8 | 5010 | 2hgs | A | 125 | ARG | p.Arg125Cys | P48637 |
| HOMO SAPIENS | 1 | 5010 | 2hgs | A | 164 | ARG | p.Arg164Gln | P48637 |
| HOMO SAPIENS | 0 | 5010 | 2hgs | A | 219 | ASP | p.Asp219Gly | P48637 |
| HOMO SAPIENS | 7 | 5010 | 2hgs | A | 267 | ARG | p.Arg267Trp | P48637 |
| HOMO SAPIENS | 0 | 5010 | 2hgs | A | 283 | ARG | p.Arg283Cys | P48637 |
| HOMO SAPIENS | 0 | 6892 | 5erd | B | 7 | VAL | p.Val56Met | Q14126 |
| HOMO SAPIENS | 1 | 6892 | 5erd | B | 25 | HIS | p.His74Arg | Q14126 |
| HOMO SAPIENS | 0 | 6892 | 5erd | B | 40 | TYR | p.Tyr89Cys | Q14126 |
| HOMO SAPIENS | 0 | 6892 | 5erd | B | 97 | ARG | p.Arg146Leu | Q14126 |
| HOMO SAPIENS | 7 | 6892 | 5erd | B | 217 | ASN | p.Asn266Ser | Q14126 |
| HOMO SAPIENS | 0 | 6892 | 5erd | B | 243 | ARG | p.Arg292His | Q14126 |
| HOMO SAPIENS | 0 | 6892 | 5erd | B | 244 | ILE | p.Ile293Val | Q14126 |
| HOMO SAPIENS | 0 | 6892 | 5erd | B | 343 | VAL | p.Val392Ile | Q14126 |
| HOMO SAPIENS | 4 | 6892 | 5erd | B | 350 | GLU | p.Glu399Lys | Q14126 |
| HOMO SAPIENS | 0 | 6892 | 5erd | B | 410 | TYR | p.Tyr459Cys | Q14126 |
| HOMO SAPIENS | 0 | 6892 | 5erd | B | 466 | VAL | p.Val515Ile | Q14126 |
| HOMO SAPIENS | 0 | 6892 | 5erd | B | 482 | PHE | p.Phe531Cys | Q14126 |
| HOMO SAPIENS | 0 | 6892 | 5erd | B | 484 | VAL | p.Val533Ile | Q14126 |
| HOMO SAPIENS | 0 | 6892 | 5erd | B | 545 | LEU | p.Leu594Pro | Q14126 |
| HOMO SAPIENS | 0 | 1338 | 2ax4 | A | 58 | THR | p.Thr48Arg | O95340 |
| HOMO SAPIENS | 7 | 8675 | 2zkm | X | 385 | LYS | p.Lys385Arg | Q00722 |
| HOMO SAPIENS | 1 | 11769 | 2wya | B | 54 | VAL | p.Val54Met | P54868 |
| HOMO SAPIENS | 1 | 11769 | 2wya | B | 116 | ARG | p.Arg116Cys | P54868 |
| HOMO SAPIENS | 5 | 11769 | 2wya | B | 167 | TYR | p.Tyr167Cys | P54868 |
| HOMO SAPIENS | 1 | 11769 | 2wya | B | 174 | PHE | p.Phe174Leu | P54868 |
| HOMO SAPIENS | 0 | 11769 | 2wya | B | 212 | GLY | p.Gly212Arg | P54868 |
| HOMO SAPIENS | 1 | 11769 | 2wya | B | 288 | ARG | p.Arg288Ter | P54868 |
| HOMO SAPIENS | 0 | 11769 | 2wya | B | 375 | SER | p.Ser375Cys | P54868 |
| HOMO SAPIENS | 4 | 11769 | 2wya | B | 500 | ARG | p.Arg500His | P54868 |
| HOMO SAPIENS | 0 | 3102 | 2iwo | A | 1880 | ARG | p.Arg1908Lys | O75970 |
| HOMO SAPIENS | 6 | 14296 | 2m9w | A | 38 | GLY | p.Gly296Ser | P43694 |
| HOMO SAPIENS | 0 | 14296 | 2m9w | A | 58 | GLN | p.Gln316Glu | P43694 |
| HOMO SAPIENS | 2 | 704 | 5ekd | B | 239 | ARG | p.Arg239Ter | Q9UGM6 |
| HOMO SAPIENS | 2 | 704 | 5ekd | B | 264 | TYR | p.Tyr264Cys | Q9UGM6 |
| HOMO SAPIENS | 6 | 2228 | 1ey2 | A | 47 | SER | p.Ser47Ter | Q93099 |
| HOMO SAPIENS | 6 | 2228 | 1ey2 | A | 48 | ALA | p.Ala48Ser | Q93099 |
| HOMO SAPIENS | 6 | 2228 | 1ey2 | A | 53 | ARG | p.Arg53Gln | Q93099 |
| HOMO SAPIENS | 3 | 2228 | 1ey2 | A | 80 | HIS | p.Gln80His | Q93099 |
| HOMO SAPIENS | 0 | 2228 | 1ey2 | A | 120 | CYS | p.Cys120Trp | Q93099 |
| HOMO SAPIENS | 3 | 2228 | 1ey2 | A | 122 | ALA | p.Ala122Val | Q93099 |

| | | | | | | | | |
|---|---|---|---|---|---|---|---|---|
| HOMO SAPIENS | 3 | 2228 | 1ey2 | A | 161 | GLY | p.Gly161Arg | Q93099 |
| HOMO SAPIENS | 6 | 2228 | 1ey2 | A | 225 | ARG | p.Arg225His | Q93099 |
| HOMO SAPIENS | 0 | 2228 | 1ey2 | A | 230 | PRO | p.Pro230Ser | Q93099 |
| HOMO SAPIENS | 6 | 2228 | 1ey2 | A | 270 | GLY | p.Gly270Arg | Q93099 |
| HOMO SAPIENS | 6 | 2228 | 1ey2 | A | 330 | ARG | p.Arg330Ser | Q93099 |
| HOMO SAPIENS | -1 | 2228 | 1ey2 | A | 368 | MSE | p.Met368Val | Q93099 |
| HOMO SAPIENS | 6 | 2228 | 1ey2 | A | 371 | HIS | p.His371Arg | Q93099 |
| HOMO SAPIENS | 6 | 2228 | 1ey2 | A | 401 | GLU | p.Glu401Gln | Q93099 |
| HOMO SAPIENS | 5 | 34648 | 2k27 | A | 31 | ARG | p.Arg31His | Q06710 |
| HOMO SAPIENS | 7 | 34648 | 2k27 | A | 40 | GLN | p.Gln40Pro | Q06710 |
| HOMO SAPIENS | 5 | 34648 | 2k27 | A | 57 | CYS | p.Cys57Tyr | Q06710 |
| HOMO SAPIENS | 5 | 34648 | 2k27 | A | 62 | LEU | p.Leu62Arg | Q06710 |
| HOMO SAPIENS | 1 | 21399 | 3edu | A | 1819 | GLU | p.Glu1819Ter | P11277 |
| HOMO SAPIENS | 1 | 18153 | 1oey | J | 245 | TYR | p.Tyr245Ter | Q15080 |
| HOMO SAPIENS | 2 | 41567 | 3sw0 | X | 1050 | ASN | p.Asn1050Tyr | P08603 |
| HOMO SAPIENS | 5 | 41567 | 3sw0 | X | 1059 | ILE | p.Ile1059Thr | P08603 |
| HOMO SAPIENS | 0 | 36362 | 4wrm | A | 362 | HIS | p.His362Arg | P07333 |
| HOMO SAPIENS | -1 | 36362 | 4wrm | A | 413 | GLY | p.Gly413Ser | P07333 |
| HOMO SAPIENS | 0 | 31255 | 2w2j | A | 100 | SER | p.Ser100Pro | P35219 |
| HOMO SAPIENS | 0 | 31255 | 2w2j | A | 100 | SER | p.Ser100Leu | P35219 |
| HOMO SAPIENS | 5 | 31255 | 2w2j | A | 237 | ARG | p.Arg237Gln | P35219 |
| HOMO SAPIENS | -1 | 26450 | 1th1 | C | 1395 | SER | p.Ser1395Cys | P25054 |
| HOMO SAPIENS | -1 | 26450 | 1th1 | C | 1413 | MET | p.Met1413Val | P25054 |
| HOMO SAPIENS | -1 | 26450 | 1th1 | C | 1446 | ALA | p.Ala1446Thr | P25054 |
| HOMO SAPIENS | -1 | 26450 | 1th1 | C | 1454 | LYS | p.Lys1454Glu | P25054 |
| HOMO SAPIENS | 0 | 26450 | 1th1 | C | 1474 | ALA | p.Ala1474Thr | P25054 |
| HOMO SAPIENS | 1 | 981 | 5xtb | F | 14 | LEU | p.Leu14Val | O43678 |
| HOMO SAPIENS | 1 | 11069 | 3kq4 | A | 46 | SER | p.Ser46Leu | P27352 |
| HOMO SAPIENS | 1 | 11069 | 3kq4 | A | 97 | MET | p.Met97Thr | P27352 |
| HOMO SAPIENS | 1 | 11069 | 3kq4 | A | 255 | ASN | p.Asn255Ser | P27352 |
| HOMO SAPIENS | 1 | 11069 | 3kq4 | A | 330 | ASN | p.Asn330Lys | P27352 |
| HOMO SAPIENS | 0 | 41489 | 3erb | A | 53 | PRO | p.Pro73Leu | P06681 |
| HOMO SAPIENS | 6 | 8715 | 2bq8 | X | 23 | THR | p.Thr44Met | P13686 |
| HOMO SAPIENS | 0 | 8715 | 2bq8 | X | 88 | GLY | p.Gly109Arg | P13686 |
| HOMO SAPIENS | 6 | 8715 | 2bq8 | X | 127 | VAL | p.Val148Met | P13686 |
| HOMO SAPIENS | 2 | 8715 | 2bq8 | X | 179 | VAL | p.Val200Met | P13686 |
| HOMO SAPIENS | 6 | 8715 | 2bq8 | X | 251 | ARG | p.Arg272Cys | P13686 |
| HOMO SAPIENS | 7 | 8112 | 3p56 | A | 37 | GLY | p.Gly37Ser | O75792 |
| HOMO SAPIENS | -1 | 8112 | 3p56 | A | 202 | LEU | p.Leu202Ser | O75792 |
| HOMO SAPIENS | 1 | 8112 | 3p56 | A | 205 | ASP | p.Asp205Glu | O75792 |
| HOMO SAPIENS | 4 | 8112 | 3p56 | A | 212 | ASN | p.Asn212Ile | O75792 |
| HOMO SAPIENS | 0 | 8112 | 3p56 | A | 235 | ARG | p.Arg235Gln | O75792 |
| HOMO SAPIENS | 4 | 8112 | 3p56 | A | 240 | THR | p.Thr240Met | O75792 |
| HOMO SAPIENS | 0 | 8112 | 3p56 | A | 291 | ARG | p.Arg291His | O75792 |
| HOMO SAPIENS | 0 | 37603 | 2yqg | A | 10 | ARG | p.Arg49His | Q14126 |
| HOMO SAPIENS | 0 | 21178 | 3k1r | B | 458 | ASP | p.Asp458Val | Q495M9 |
| HOMO SAPIENS | -1 | 36828 | 4fq3 | B | 507 | GLY | p.Gly507Asp | P35637 |
| HOMO SAPIENS | 5 | 36828 | 4fq3 | B | 517 | HIS | p.His517Gln | P35637 |
| HOMO SAPIENS | 8 | 36828 | 4fq3 | B | 518 | ARG | p.Arg518Lys | P35637 |
| HOMO SAPIENS | 6 | 36828 | 4fq3 | B | 521 | ARG | p.Arg521Cys | P35637 |
| HOMO SAPIENS | 6 | 36828 | 4fq3 | B | 521 | ARG | p.Arg521His | P35637 |
| HOMO SAPIENS | 0 | 6363 | 3j7y | c | null | HIS | p.His13Arg | Q9H9J2 |
| HOMO SAPIENS | 2 | 6363 | 3j7y | c | 105 | ARG | p.Arg105Cys | Q9H9J2 |
| HOMO SAPIENS | 2 | 6363 | 3j7y | c | 138 | THR | p.Thr138Ile | Q9H9J2 |

| | | | | | | | | |
|---|---|---|---|---|---|---|---|---|
| HOMO SAPIENS | 1 | 6363 | 3j7y | c | 156 | LEU | p.Leu156Arg | Q9H9J2 |
| HOMO SAPIENS | 1 | 6782 | 5l46 | A | 109 | HIS | p.His109Arg | Q9UI17 |
| HOMO SAPIENS | 0 | 23951 | 5t76 | A | 175 | GLY | p.Gly931Ser | P49588 |
| HOMO SAPIENS | 8 | 12604 | 3p0l | A | 182 | ARG | p.Arg182Leu | P49675 |
| HOMO SAPIENS | 2 | 12604 | 3p0l | A | 187 | VAL | p.Val187Met | P49675 |
| HOMO SAPIENS | 2 | 12604 | 3p0l | A | 188 | ARG | p.Arg188Cys | P49675 |
| HOMO SAPIENS | 0 | 12604 | 3p0l | A | 193 | ARG | p.Arg193Ter | P49675 |
| HOMO SAPIENS | 9 | 12604 | 3p0l | A | 217 | ARG | p.Arg217Thr | P49675 |
| HOMO SAPIENS | 4 | 12604 | 3p0l | A | 218 | ALA | p.Ala218Val | P49675 |
| HOMO SAPIENS | 0 | 12604 | 3p0l | A | 250 | TRP | p.Trp250Ter | P49675 |
| HOMO SAPIENS | 1 | 12604 | 3p0l | A | 258 | GLN | p.Gln258Ter | P49675 |
| HOMO SAPIENS | -1 | 27765 | 4wkr | A | 4 | GLU | p.Glu4Lys | Q4G0J3 |
| HOMO SAPIENS | 2 | 3587 | 4kit | C | 2301 | PRO | p.Pro2301Thr | Q6P2Q9 |
| HOMO SAPIENS | 3 | 3587 | 4kit | C | 2304 | PHE | p.Phe2304Leu | Q6P2Q9 |
| HOMO SAPIENS | 3 | 3587 | 4kit | C | 2309 | HIS | p.His2309Pro | Q6P2Q9 |
| HOMO SAPIENS | 3 | 3587 | 4kit | C | 2310 | ARG | p.Arg2310Lys | Q6P2Q9 |
| HOMO SAPIENS | 0 | 9073 | 1n69 | B | 21 | ASN | p.Asn215His | P07602 |
| HOMO SAPIENS | 0 | 9073 | 1n69 | B | 23 | THR | p.Thr217Ile | P07602 |
| HOMO SAPIENS | 0 | 9073 | 1n69 | B | 47 | CYS | p.Cys241Ser | P07602 |
| HOMO SAPIENS | 0 | 475 | 2vd5 | A | 225 | THR | p.Thr225Met | Q09013 |
| HOMO SAPIENS | 1 | 9066 | 4mzv | A | 31 | ASN | p.Asn31Lys | P16422 |
| HOMO SAPIENS | 1 | 9066 | 4mzv | A | 47 | GLN | p.Gln47His | P16422 |
| HOMO SAPIENS | 3 | 9066 | 4mzv | A | 66 | CYS | p.Cys66Tyr | P16422 |
| HOMO SAPIENS | 1 | 9066 | 4mzv | A | 102 | SER | p.Ser102Gly | P16422 |
| HOMO SAPIENS | 5 | 9066 | 4mzv | A | 115 | THR | p.Met115Thr | P16422 |
| HOMO SAPIENS | 1 | 9066 | 4mzv | A | 138 | ARG | p.Arg138Ter | P16422 |
| HOMO SAPIENS | 1 | 9066 | 4mzv | A | 153 | ARG | p.Arg153Thr | P16422 |
| HOMO SAPIENS | 1 | 9066 | 4mzv | A | 165 | ALA | p.Ala165Thr | P16422 |
| HOMO SAPIENS | 8 | 1827 | 2nz2 | A | 14 | GLY | p.Gly14Ser | P00966 |
| HOMO SAPIENS | 5 | 1827 | 2nz2 | A | 18 | SER | p.Ser18Leu | P00966 |
| HOMO SAPIENS | 3 | 1827 | 2nz2 | A | 86 | ARG | p.Arg86Cys | P00966 |
| HOMO SAPIENS | 3 | 1827 | 2nz2 | A | 86 | ARG | p.Arg86His | P00966 |
| HOMO SAPIENS | 1 | 1827 | 2nz2 | A | 179 | TRP | p.Trp179Arg | P00966 |
| HOMO SAPIENS | 7 | 1827 | 2nz2 | A | 180 | SER | p.Ser180Asn | P00966 |
| HOMO SAPIENS | 7 | 1827 | 2nz2 | A | 191 | GLU | p.Glu191Lys | P00966 |
| HOMO SAPIENS | 4 | 1827 | 2nz2 | A | 256 | GLU | p.Glu256Lys | P00966 |
| HOMO SAPIENS | 4 | 1827 | 2nz2 | A | 263 | VAL | p.Val263Met | P00966 |
| HOMO SAPIENS | 5 | 1827 | 2nz2 | A | 265 | ARG | p.Arg265Cys | P00966 |
| HOMO SAPIENS | 5 | 1827 | 2nz2 | A | 268 | ILE | p.Ile268Met | P00966 |
| HOMO SAPIENS | 6 | 1827 | 2nz2 | A | 269 | VAL | p.Val269Met | P00966 |
| HOMO SAPIENS | 7 | 1827 | 2nz2 | A | 272 | ARG | p.Arg272Cys | P00966 |
| HOMO SAPIENS | 3 | 1827 | 2nz2 | A | 304 | ARG | p.Arg304Trp | P00966 |
| HOMO SAPIENS | 3 | 1827 | 2nz2 | A | 307 | ARG | p.Arg307Cys | P00966 |
| HOMO SAPIENS | 3 | 1827 | 2nz2 | A | 310 | LYS | p.Lys310Gln | P00966 |
| HOMO SAPIENS | 2 | 1827 | 2nz2 | A | 362 | GLY | p.Gly362Val | P00966 |
| HOMO SAPIENS | 2 | 1827 | 2nz2 | A | 363 | ARG | p.Arg363Trp | P00966 |
| HOMO SAPIENS | 2 | 1827 | 2nz2 | A | 363 | ARG | p.Arg363Gln | P00966 |
| HOMO SAPIENS | 1 | 1827 | 2nz2 | A | 390 | GLY | p.Gly390Arg | P00966 |
| HOMO SAPIENS | 0 | 29524 | 5vbn | B | 2165 | ARG | p.Arg2165His | Q07864 |
| HOMO SAPIENS | 2 | 29524 | 5vbn | B | 2255 | ILE | p.Ile2255Phe | Q07864 |
| HOMO SAPIENS | 2 | 29524 | 5vbn | B | 2274 | LEU | p.Leu2274Val | Q07864 |
| HOMO SAPIENS | 0 | 31897 | 3t0o | A | 184 | CYS | p.Cys184Arg | O00584 |
| HOMO SAPIENS | -1 | 31897 | 3t0o | A | 215 | GLU | p.Glu215Lys | O00584 |
| HOMO SAPIENS | 0 | 31897 | 3t0o | A | 236 | ARG | p.Arg236Trp | O00584 |

| | | | | | | | | |
|---|---|---|---|---|---|---|---|---|
| HOMO SAPIENS | 7 | 18490 | 4pcw | B | 31 | GLU | p.Glu32Ter | P20930 |
| HOMO SAPIENS | -1 | 39865 | 2h9e | L | 152 | GLY | p.Gly192Arg | P00742 |
| HOMO SAPIENS | 2 | 4309 | 2x0b | F | 174 | THR | p.Thr207Met | P01019 |
| HOMO SAPIENS | 3 | 4309 | 2x0b | F | 342 | ARG | p.Arg375Gln | P01019 |
| HOMO SAPIENS | 2 | 3094 | 1zll | B | 9 | ARG | p.Arg9Cys | P26678 |
| HOMO SAPIENS | 9 | 3094 | 1zll | B | 39 | LEU | p.Leu39Ter | P26678 |
| HOMO SAPIENS | 5 | 3094 | 1zll | B | 49 | VAL | p.Val49Met | P26678 |
| HOMO SAPIENS | -1 | 41498 | 3ffd | P | 8 | LEU | p.Leu44Pro | P12272 |
| HOMO SAPIENS | 1 | 20697 | 1igr | A | 407 | ARG | p.Arg437His | P08069 |
| HOMO SAPIENS | 0 | 20697 | 1igr | A | 416 | MET | p.Met446Val | P08069 |
| HOMO SAPIENS | -1 | 20697 | 1igr | A | 462 | VAL | p.Val492Ile | P08069 |
| HOMO SAPIENS | 0 | 3973 | 4cbz | A | 37 | LEU | p.Leu37Ser | P78504 |
| HOMO SAPIENS | 0 | 3973 | 4cbz | A | 45 | VAL | p.Val45Leu | P78504 |
| HOMO SAPIENS | 0 | 3973 | 4cbz | A | 134 | LEU | p.Leu134Met | P78504 |
| HOMO SAPIENS | -1 | 3973 | 4cbz | A | 146 | VAL | p.Val146Ile | P78504 |
| HOMO SAPIENS | 0 | 3973 | 4cbz | A | 176 | VAL | p.Val176Ile | P78504 |
| HOMO SAPIENS | 0 | 3973 | 4cbz | A | 184 | ARG | p.Arg184Cys | P78504 |
| HOMO SAPIENS | 6 | 1697 | 4hmy | S | 33 | ARG | p.Arg33Trp | Q96PC3 |
| HOMO SAPIENS | 2 | 23752 | 5j4o | A | 13 | ARG | p.Arg1611Ser | P02549 |
| HOMO SAPIENS | 0 | 23752 | 5j4o | A | 42 | ALA | p.Ala1640Gly | P02549 |
| HOMO SAPIENS | 0 | 23752 | 5j4o | A | 108 | HIS | p.His1706Gln | P02549 |
| HOMO SAPIENS | 6 | 23752 | 5j4o | A | 206 | LEU | p.Leu1804Phe | P02549 |
| HOMO SAPIENS | 8 | 531 | 2qz4 | A | 349 | GLY | p.Gly349Ser | Q9UQ90 |
| HOMO SAPIENS | 0 | 531 | 2qz4 | A | 486 | ARG | p.Arg486Gln | Q9UQ90 |
| HOMO SAPIENS | 1 | 531 | 2qz4 | A | 510 | ALA | p.Ala510Val | Q9UQ90 |
| HOMO SAPIENS | 0 | 531 | 2qz4 | A | 529 | ALA | p.Ala529Val | Q9UQ90 |
| HOMO SAPIENS | 0 | 531 | 2qz4 | A | 558 | LYS | p.Lys558Ter | Q9UQ90 |
| HOMO SAPIENS | 1 | 531 | 2qz4 | A | 559 | LYS | p.Lys559Ter | Q9UQ90 |
| HOMO SAPIENS | 2 | 7736 | 1fnh | A | 114 | TRP | p.Trp1834Arg | P02751 |
| HOMO SAPIENS | 0 | 7736 | 1fnh | A | 163 | LEU | p.Leu1883Arg | P02751 |
| HOMO SAPIENS | 3 | 20337 | 5wy5 | B | 264 | LEU | p.Leu264Phe | Q96MG7 |
| HOMO SAPIENS | 8 | 36096 | 2jku | A | 643 | GLY | p.Gly668Arg | P05165 |
| HOMO SAPIENS | 6 | 36096 | 2jku | A | 663 | CYS | p.Cys688Arg | P05165 |
| HOMO SAPIENS | 0 | 9850 | 3kbt | A | 1606 | TYR | p.Tyr1606Ter | P11277 |
| HOMO SAPIENS | 4 | 6876 | 4jkq | A | 217 | ILE | p.Ile217Val | Q92736 |
| HOMO SAPIENS | 5 | 6876 | 4jkq | A | 420 | ARG | p.Arg420Trp | Q92736 |
| HOMO SAPIENS | 2 | 27603 | 1itv | B | 62 | PRO | p.Arg574Pro | P14780 |
| HOMO SAPIENS | 4 | 27603 | 1itv | B | 156 | ARG | p.Arg668Gln | P14780 |
| HOMO SAPIENS | 2 | 22954 | 4f52 | E | 48 | ILE | p.Ile48Met | Q92990 |
| HOMO SAPIENS | 0 | 22954 | 4f52 | E | 91 | ASP | p.Asp91Asn | Q92990 |
| HOMO SAPIENS | 0 | 22954 | 4f52 | E | 146 | ALA | p.Ala146Thr | Q92990 |
| HOMO SAPIENS | 0 | 22954 | 4f52 | E | 181 | ILE | p.Ile181Lys | Q92990 |
| HOMO SAPIENS | 0 | 22954 | 4f52 | E | 194 | ASN | p.Asn194Ser | Q92990 |
| HOMO SAPIENS | 2 | 22954 | 4f52 | E | 254 | PRO | p.Pro254Arg | Q92990 |
| HOMO SAPIENS | 2 | 22954 | 4f52 | E | 300 | ILE | p.Ile300Thr | Q92990 |
| HOMO SAPIENS | 2 | 22954 | 4f52 | E | 336 | LEU | p.Leu336Ser | Q92990 |
| HOMO SAPIENS | 0 | 22954 | 4f52 | E | 353 | TYR | p.Tyr353His | Q92990 |
| HOMO SAPIENS | 0 | 22954 | 4f52 | E | 387 | MET | p.Met387Val | Q92990 |
| HOMO SAPIENS | 4 | 2946 | 5gl7 | A | 344 | TYR | p.Tyr344Cys | P56192 |
| HOMO SAPIENS | 0 | 2946 | 5gl7 | A | 370 | PHE | p.Phe370Leu | P56192 |
| HOMO SAPIENS | 0 | 2946 | 5gl7 | A | 393 | ALA | p.Ala393Thr | P56192 |
| HOMO SAPIENS | 0 | 2946 | 5gl7 | A | 523 | ILE | p.Ile523Thr | P56192 |
| HOMO SAPIENS | 0 | 2946 | 5gl7 | A | 567 | SER | p.Ser567Leu | P56192 |
| HOMO SAPIENS | 0 | 2946 | 5gl7 | A | 800 | PRO | p.Pro800Thr | P56192 |

| | | | | | | | | |
|---|---|---|---|---|---|---|---|---|
| HOMO SAPIENS | 3 | 34961 | 4wy4 | C | 44 | TYR | p.Tyr44His | O95721 |
| HOMO SAPIENS | 8 | 34961 | 4wy4 | C | 89 | GLU | p.Glu89Lys | O95721 |
| HOMO SAPIENS | 0 | 9427 | 4chl | B | 21 | ALA | p.Ala21Ser | O95571 |
| HOMO SAPIENS | 2 | 9427 | 4chl | B | 62 | ALA | p.Ala62Thr | O95571 |
| HOMO SAPIENS | 0 | 9427 | 4chl | B | 114 | ILE | p.Ile114Phe | O95571 |
| HOMO SAPIENS | 2 | 9427 | 4chl | B | 159 | ARG | p.Arg159His | O95571 |
| HOMO SAPIENS | 8 | 9427 | 4chl | B | 163 | ARG | p.Arg163Trp | O95571 |
| HOMO SAPIENS | 8 | 9427 | 4chl | B | 163 | ARG | p.Arg163Gln | O95571 |
| HOMO SAPIENS | 2 | 9427 | 4chl | B | 185 | LEU | p.Leu185Arg | O95571 |
| HOMO SAPIENS | 0 | 40917 | 2e8o | A | 11 | PRO | p.Pro26Leu | Q9Y3Z3 |
| HOMO SAPIENS | -1 | 39190 | 4gow | A | 556 | TYR | p.Tyr556Ter | P56696 |
| HOMO SAPIENS | 0 | 2399 | 2mkv | A | 8 | GLY | p.Gly10Ser | P54710 |
| HOMO SAPIENS | 1 | 2399 | 2mkv | A | 24 | VAL | p.Val26Ile | P54710 |
| HOMO SAPIENS | 4 | 2399 | 2mkv | A | 39 | GLY | p.Gly41Arg | P54710 |
| HOMO SAPIENS | 0 | 20801 | 2zej | B | 1441 | ARG | p.Arg1441Cys | Q5S007 |
| HOMO SAPIENS | 0 | 20801 | 2zej | B | 1441 | ARG | p.Arg1441His | Q5S007 |
| HOMO SAPIENS | 5 | 1019 | 5xtb | M | 62 | ARG | p.Arg62Gln | P28331 |
| HOMO SAPIENS | 1 | 1019 | 5xtb | M | 85 | ALA | p.Ala85Thr | P28331 |
| HOMO SAPIENS | 5 | 1019 | 5xtb | M | 231 | LEU | p.Leu231Val | P28331 |
| HOMO SAPIENS | 2 | 1019 | 5xtb | M | 241 | ARG | p.Arg241Trp | P28331 |
| HOMO SAPIENS | 1 | 1019 | 5xtb | M | 252 | ASP | p.Asp252Gly | P28331 |
| HOMO SAPIENS | 1 | 1019 | 5xtb | M | 408 | ARG | p.Arg408Cys | P28331 |
| HOMO SAPIENS | 2 | 1019 | 5xtb | M | 417 | ARG | p.Arg417Ser | P28331 |
| HOMO SAPIENS | 1 | 1019 | 5xtb | M | 557 | ARG | p.Arg557Ter | P28331 |
| HOMO SAPIENS | 0 | 1019 | 5xtb | M | 567 | ILE | p.Ile567Val | P28331 |
| HOMO SAPIENS | 1 | 1019 | 5xtb | M | 595 | THR | p.Thr595Ala | P28331 |
| HOMO SAPIENS | 1 | 1019 | 5xtb | M | 703 | ALA | p.Ala703Thr | P28331 |
| HOMO SAPIENS | -1 | 3437 | 5hyn | A | 185 | ASP | p.Asp185His | Q15910 |
| HOMO SAPIENS | -1 | 3437 | 5hyn | A | 482 | ALA | p.Ala482Thr | Q15910 |
| HOMO SAPIENS | 0 | 8201 | 2wfd | A | 373 | TYR | p.Tyr373Cys | Q9P2J5 |
| HOMO SAPIENS | 1 | 3590 | 3edi | A | 90 | VAL | p.Val238Ala | O43897 |
| HOMO SAPIENS | 6 | 4555 | 1ytq | A | 64 | ALA | p.Ala65Ser | P43320 |
| HOMO SAPIENS | 0 | 4555 | 1ytq | A | 194 | TRP | p.Trp195Gly | P43320 |
| HOMO SAPIENS | 0 | 1013 | 2nn6 | C | 272 | SER | p.Ser272Thr | Q96B26 |
| HOMO SAPIENS | -1 | 29026 | 5m73 | D | 530 | ILE | p.Ile530Asn | O76094 |
| HOMO SAPIENS | 8 | 2252 | 1d5v | A | 57 | SER | p.Ser125Leu | Q99958 |
| HOMO SAPIENS | 1 | 4337 | 3ecr | A | 29 | ARG | p.Arg26His | P08397 |
| HOMO SAPIENS | 2 | 4337 | 3ecr | A | 34 | ALA | p.Ala31Thr | P08397 |
| HOMO SAPIENS | 2 | 4337 | 3ecr | A | 37 | GLN | p.Gln34Lys | P08397 |
| HOMO SAPIENS | 1 | 4337 | 3ecr | A | 58 | ALA | p.Ala55Ser | P08397 |
| HOMO SAPIENS | 0 | 4337 | 3ecr | A | 84 | LEU | p.Leu81Pro | P08397 |
| HOMO SAPIENS | 0 | 4337 | 3ecr | A | 114 | GLY | p.Gly111Arg | P08397 |
| HOMO SAPIENS | 1 | 4337 | 3ecr | A | 119 | ARG | p.Arg116Trp | P08397 |
| HOMO SAPIENS | 1 | 4337 | 3ecr | A | 152 | ARG | p.Arg149Gln | P08397 |
| HOMO SAPIENS | 1 | 4337 | 3ecr | A | 170 | ARG | p.Arg167Trp | P08397 |
| HOMO SAPIENS | 1 | 4337 | 3ecr | A | 176 | ARG | p.Arg173Gln | P08397 |
| HOMO SAPIENS | 0 | 4337 | 3ecr | A | 180 | LEU | p.Leu177Arg | P08397 |
| HOMO SAPIENS | 0 | 4337 | 3ecr | A | 181 | ASP | p.Asp178Asn | P08397 |
| HOMO SAPIENS | 1 | 4337 | 3ecr | A | 198 | ARG | p.Arg195Cys | P08397 |
| HOMO SAPIENS | 0 | 4337 | 3ecr | A | 201 | TRP | p.Trp198Ter | P08397 |
| HOMO SAPIENS | 1 | 4337 | 3ecr | A | 204 | ARG | p.Arg201Trp | P08397 |
| HOMO SAPIENS | 1 | 4337 | 3ecr | A | 219 | GLY | p.Gly216Asp | P08397 |
| HOMO SAPIENS | 1 | 4337 | 3ecr | A | 226 | GLU | p.Glu223Lys | P08397 |
| HOMO SAPIENS | 0 | 4337 | 3ecr | A | 248 | LEU | p.Leu245Arg | P08397 |

| | | | | | | | | |
|---|---|---|---|---|---|---|---|---|
| HOMO SAPIENS | 1 | 4337 | 3ecr | A | 250 | CYS | p.Cys247Arg | P08397 |
| HOMO SAPIENS | 0 | 4337 | 3ecr | A | 253 | GLU | p.Glu250Lys | P08397 |
| HOMO SAPIENS | 0 | 4337 | 3ecr | A | 255 | ALA | p.Ala252Thr | P08397 |
| HOMO SAPIENS | 0 | 4337 | 3ecr | A | 255 | ALA | p.Ala252Val | P08397 |
| HOMO SAPIENS | 0 | 4337 | 3ecr | A | 259 | HIS | p.His256Asn | P08397 |
| HOMO SAPIENS | 0 | 4337 | 3ecr | A | 324 | ARG | p.Arg321His | P08397 |
| HOMO SAPIENS | -1 | 4337 | 3ecr | A | 362 | ASP | p.Asp359Asn | P08397 |
| HOMO SAPIENS | 1 | 1204 | 5lwv | A | 854 | ILE | p.Ala864Thr | O15294 |
| HOMO SAPIENS | 1 | 25016 | 1m4u | A | 35 | PRO | p.Pro35Arg | Q13253 |
| HOMO SAPIENS | 1 | 25016 | 1m4u | A | 35 | PRO | p.Pro35Ser | Q13253 |
| HOMO SAPIENS | 1 | 25016 | 1m4u | A | 167 | ARG | p.Arg167Gly | Q13253 |
| HOMO SAPIENS | 1 | 25016 | 1m4u | A | 184 | CYS | p.Cys184Tyr | Q13253 |
| HOMO SAPIENS | 0 | 25016 | 1m4u | A | 189 | GLY | p.Gly189Cys | Q13253 |
| HOMO SAPIENS | 1 | 25016 | 1m4u | A | 204 | ARG | p.Arg204Leu | Q13253 |
| HOMO SAPIENS | 1 | 25016 | 1m4u | A | 205 | TRP | p.Trp205Cys | Q13253 |
| HOMO SAPIENS | 1 | 25016 | 1m4u | A | 217 | TRP | p.Trp217Gly | Q13253 |
| HOMO SAPIENS | 0 | 25016 | 1m4u | A | 222 | TYR | p.Tyr222Cys | Q13253 |
| HOMO SAPIENS | 0 | 25016 | 1m4u | A | 222 | TYR | p.Tyr222Asp | Q13253 |
| HOMO SAPIENS | 1 | 25016 | 1m4u | A | 223 | PRO | p.Pro223Leu | Q13253 |
| HOMO SAPIENS | 0 | 1394 | 1um7 | A | 95 | ALA | p.Ala463Val | Q92796 |
| HOMO SAPIENS | 0 | 21294 | 4jpz | B | 1788 | GLU | p.Glu1788Asp | Q99250 |
| HOMO SAPIENS | 9 | 24068 | 2db6 | A | 47 | ARG | p.Arg119Cys | Q96MF2 |
| HOMO SAPIENS | 5 | 1869 | 4omt | A | 39 | ARG | p.Arg39Leu | P08237 |
| HOMO SAPIENS | 2 | 1869 | 4omt | A | 100 | ARG | p.Arg100Gln | P08237 |
| HOMO SAPIENS | 7 | 1869 | 4omt | A | 543 | ASP | p.Asp543Ala | P08237 |
| HOMO SAPIENS | 1 | 1869 | 4omt | A | 686 | TRP | p.Trp686Cys | P08237 |
| HOMO SAPIENS | 1 | 1869 | 4omt | A | 696 | ARG | p.Arg696His | P08237 |
| HOMO SAPIENS | 0 | 12163 | 4wxx | A | 495 | TYR | p.Tyr495Cys | P26358 |
| HOMO SAPIENS | 0 | 12163 | 4wxx | A | 528 | ILE | p.Ile528Met | P26358 |
| HOMO SAPIENS | 0 | 12391 | 1j1d | C | 36 | LYS | p.Lys36Gln | P19429 |
| HOMO SAPIENS | 0 | 12391 | 1j1d | C | 82 | PRO | p.Pro82Ser | P19429 |
| HOMO SAPIENS | 7 | 12391 | 1j1d | C | 102 | ALA | p.Ala102Thr | P19429 |
| HOMO SAPIENS | 1 | 5000 | 4cca | A | 17 | GLY | p.Gly17Arg | Q15833 |
| HOMO SAPIENS | 0 | 5000 | 4cca | A | 205 | VAL | p.Val205Ile | Q15833 |
| HOMO SAPIENS | 0 | 5000 | 4cca | A | 209 | LEU | p.Leu209Pro | Q15833 |
| HOMO SAPIENS | 1 | 5000 | 4cca | A | 459 | ARG | p.Arg459Trp | Q15833 |
| HOMO SAPIENS | 7 | 5000 | 4cca | A | 477 | PRO | p.Pro477Leu | Q15833 |
| HOMO SAPIENS | -1 | 5000 | 4cca | A | 526 | ILE | p.Ile526Val | Q15833 |
| HOMO SAPIENS | 0 | 5000 | 4cca | A | 541 | GLY | p.Gly541Ser | Q15833 |
| HOMO SAPIENS | 0 | 5000 | 4cca | A | 555 | ARG | p.Arg555Gly | Q15833 |
| HOMO SAPIENS | 5 | 28551 | 3vcm | P | 26 | ARG | p.Arg49Ter | P00797 |
| HOMO SAPIENS | 6 | 36453 | 5dzz | A | 2070 | ASP | p.Asp2070His | P15924 |
| HOMO SAPIENS | 0 | 36453 | 5dzz | A | 2166 | ARG | p.Arg2166Ter | P15924 |
| HOMO SAPIENS | 0 | 20268 | 2da6 | A | 96 | ASN | p.Asn321Ser | P35680 |
| HOMO SAPIENS | 1 | 1405 | 3iar | A | 8 | ASP | p.Asp8Asn | P00813 |
| HOMO SAPIENS | 8 | 1405 | 3iar | A | 15 | HIS | p.His15Asp | P00813 |
| HOMO SAPIENS | 6 | 1405 | 3iar | A | 74 | GLY | p.Gly74Val | P00813 |
| HOMO SAPIENS | 6 | 1405 | 3iar | A | 80 | LYS | p.Lys80Arg | P00813 |
| HOMO SAPIENS | 5 | 1405 | 3iar | A | 97 | TYR | p.Tyr97Cys | P00813 |
| HOMO SAPIENS | 5 | 1405 | 3iar | A | 101 | ARG | p.Arg101Leu | P00813 |
| HOMO SAPIENS | 5 | 1405 | 3iar | A | 101 | ARG | p.Arg101Trp | P00813 |
| HOMO SAPIENS | 5 | 1405 | 3iar | A | 106 | LEU | p.Leu106Val | P00813 |
| HOMO SAPIENS | 0 | 1405 | 3iar | A | 107 | LEU | p.Leu107Pro | P00813 |
| HOMO SAPIENS | 9 | 1405 | 3iar | A | 156 | ARG | p.Arg156Cys | P00813 |

| | | | | | | | | |
|---|---|---|---|---|---|---|---|---|
| HOMO SAPIENS | 6 | 1405 | 3iar | A | 211 | ARG | p.Arg211His | P00813 |
| HOMO SAPIENS | 0 | 1405 | 3iar | A | 216 | GLY | p.Gly216Arg | P00813 |
| HOMO SAPIENS | 0 | 1405 | 3iar | A | 291 | SER | p.Ser291Leu | P00813 |
| HOMO SAPIENS | 6 | 1405 | 3iar | A | 304 | LEU | p.Leu304Arg | P00813 |
| HOMO SAPIENS | 0 | 1405 | 3iar | A | 329 | ALA | p.Ala329Val | P00813 |
| HOMO SAPIENS | -1 | 31340 | 4atu | I | 186 | ARG | p.Arg186His | O43602 |
| HOMO SAPIENS | -1 | 31340 | 4atu | I | 192 | ARG | p.Arg192Trp | O43602 |
| HOMO SAPIENS | -1 | 31340 | 4atu | I | 196 | ARG | p.Arg196His | O43602 |
| HOMO SAPIENS | -1 | 31340 | 4atu | I | 203 | THR | p.Thr203Arg | O43602 |
| HOMO SAPIENS | -1 | 31340 | 4atu | I | 286 | ALA | p.Ala286Ser | O43602 |
| HOMO SAPIENS | 4 | 30734 | 1fsu | A | 95 | ARG | p.Arg95Gln | P15848 |
| HOMO SAPIENS | 0 | 30734 | 1fsu | A | 117 | CYS | p.Cys117Arg | P15848 |
| HOMO SAPIENS | 0 | 30734 | 1fsu | A | 137 | GLY | p.Gly137Val | P15848 |
| HOMO SAPIENS | 0 | 30734 | 1fsu | A | 210 | TYR | p.Tyr210Cys | P15848 |
| HOMO SAPIENS | 0 | 30734 | 1fsu | A | 236 | LEU | p.Leu236Pro | P15848 |
| HOMO SAPIENS | 1 | 30734 | 1fsu | A | 315 | ARG | p.Arg315Gln | P15848 |
| HOMO SAPIENS | 0 | 30734 | 1fsu | A | 358 | VAL | p.Val358Met | P15848 |
| HOMO SAPIENS | 2 | 30734 | 1fsu | A | 376 | VAL | p.Val376Met | P15848 |
| HOMO SAPIENS | 0 | 30734 | 1fsu | A | 384 | SER | p.Ser384Asn | P15848 |
| HOMO SAPIENS | 0 | 30734 | 1fsu | A | 393 | HIS | p.His393Pro | P15848 |
| HOMO SAPIENS | 0 | 30734 | 1fsu | A | 405 | CYS | p.Cys405Tyr | P15848 |
| HOMO SAPIENS | 0 | 30734 | 1fsu | A | 458 | ASN | p.Asn458Ser | P15848 |
| HOMO SAPIENS | 1 | 30734 | 1fsu | A | 484 | ARG | p.Arg484Gly | P15848 |
| HOMO SAPIENS | 0 | 38072 | 1wmv | A | 51 | PRO | p.Pro98Leu | Q9NZC7 |
| HOMO SAPIENS | -1 | 5674 | 1j1e | C | 185 | ASN | p.Asn185Lys | P19429 |
| HOMO SAPIENS | -1 | 5674 | 1j1e | C | 192 | ARG | p.Arg192His | P19429 |
| HOMO SAPIENS | -1 | 5674 | 1j1e | C | 196 | ASP | p.Asp196Asn | P19429 |
| HOMO SAPIENS | -1 | 5674 | 1j1e | C | 206 | LYS | p.Lys206Gln | P19429 |
| HOMO SAPIENS | 0 | 38067 | 1wlp | A | 13 | PRO | p.Pro156Gln | P13498 |
| HOMO SAPIENS | -1 | 20441 | 3vfd | A | 289 | HIS | p.His289Tyr | Q9UBP0 |
| HOMO SAPIENS | 2 | 20441 | 3vfd | A | 344 | ILE | p.Ile344Lys | Q9UBP0 |
| HOMO SAPIENS | 0 | 20441 | 3vfd | A | 362 | SER | p.Ser362Cys | Q9UBP0 |
| HOMO SAPIENS | 1 | 20441 | 3vfd | A | 386 | ASN | p.Asn386Ser | Q9UBP0 |
| HOMO SAPIENS | 9 | 20441 | 3vfd | A | 441 | ASP | p.Asp441Gly | Q9UBP0 |
| HOMO SAPIENS | -1 | 20441 | 3vfd | A | 448 | CYS | p.Cys448Tyr | Q9UBP0 |
| HOMO SAPIENS | 2 | 20441 | 3vfd | A | 500 | ARG | p.Arg499Cys | Q9UBP0 |
| HOMO SAPIENS | 1 | 20441 | 3vfd | A | 563 | ARG | p.Arg562Gly | Q9UBP0 |
| HOMO SAPIENS | 3 | 18471 | 4dx8 | J | 26 | ARG | p.Arg26Gln | O00522 |
| HOMO SAPIENS | -1 | 18471 | 4dx8 | J | 101 | GLY | p.Gly101Val | O00522 |
| HOMO SAPIENS | -1 | 34735 | 2ly4 | B | 83 | ALA | p.Ala83Glu | P04637 |
| HOMO SAPIENS | 0 | 31128 | 4cvh | A | 116 | ARG | p.Arg116Cys | A4D126 |
| HOMO SAPIENS | 8 | 31128 | 4cvh | A | 215 | GLN | p.Gln215Ter | A4D126 |
| HOMO SAPIENS | 0 | 31128 | 4cvh | A | 268 | ARG | p.Arg268Ter | A4D126 |
| HOMO SAPIENS | 0 | 31128 | 4cvh | A | 316 | THR | p.Thr316Lys | A4D126 |
| HOMO SAPIENS | 1 | 7164 | 3mt5 | A | 369 | ASP | p.Asp434Gly | Q12791 |
| HOMO SAPIENS | 2 | 26812 | 2wtk | F | 49 | TYR | p.Tyr49Asp | Q15831 |
| HOMO SAPIENS | 0 | 26812 | 2wtk | F | 67 | LEU | p.Leu67Pro | Q15831 |
| HOMO SAPIENS | 4 | 26812 | 2wtk | F | 104 | ARG | p.Arg104Ser | Q15831 |
| HOMO SAPIENS | 0 | 26812 | 2wtk | F | 135 | GLY | p.Gly135Arg | Q15831 |
| HOMO SAPIENS | 0 | 26812 | 2wtk | F | 163 | GLY | p.Gly163Asp | Q15831 |
| HOMO SAPIENS | 2 | 26812 | 2wtk | F | 194 | ALA | p.Asp194Asn | Q15831 |
| HOMO SAPIENS | 2 | 1357 | 5xtd | W | 57 | ARG | p.Arg57His | Q9P0J0 |
| HOMO SAPIENS | 0 | 28086 | 4nn2 | A | 229 | HIS | p.His229Arg | Q8IWS0 |
| HOMO SAPIENS | 0 | 28086 | 4nn2 | A | 234 | LYS | p.Lys234Glu | Q8IWS0 |

| | | | | | | | | |
|---|---|---|---|---|---|---|---|---|
| HOMO SAPIENS | 0 | 28086 | 4nn2 | A | 257 | ARG | p.Arg257Gly | Q8IWS0 |
| HOMO SAPIENS | 5 | 31182 | 2a1x | A | 176 | GLN | p.Gln176Lys | O14832 |
| HOMO SAPIENS | 1 | 31182 | 2a1x | A | 204 | GLY | p.Gly204Ser | O14832 |
| HOMO SAPIENS | 0 | 31182 | 2a1x | A | 269 | ASN | p.Asn269His | O14832 |
| HOMO SAPIENS | 3 | 31182 | 2a1x | A | 275 | ARG | p.Arg275Gln | O14832 |
| HOMO SAPIENS | 3 | 31182 | 2a1x | A | 275 | ARG | p.Arg275Trp | O14832 |
| HOMO SAPIENS | 4 | 2823 | 5xtb | P | 145 | THR | p.Thr145Ile | O75489 |
| HOMO SAPIENS | 5 | 2823 | 5xtb | P | 199 | ARG | p.Arg199Trp | O75489 |
| HOMO SAPIENS | 0 | 8645 | 2h5g | A | 372 | SER | p.Ser372Tyr | P54886 |
| HOMO SAPIENS | 3 | 8645 | 2h5g | A | 715 | ASP | p.Asp715His | P54886 |
| HOMO SAPIENS | 0 | 8645 | 2h5g | A | 765 | ARG | p.Arg765Gln | P54886 |
| HOMO SAPIENS | 0 | 8645 | 2h5g | A | 782 | TYR | p.Tyr782Cys | P54886 |
| HOMO SAPIENS | 0 | 8645 | 2h5g | A | 784 | HIS | p.His784Tyr | P54886 |
| HOMO SAPIENS | 1 | 13992 | 2cue | A | 39 | ARG | p.Arg242Thr | P26367 |
| HOMO SAPIENS | 9 | 13992 | 2cue | A | 55 | PHE | p.Phe258Ser | P26367 |
| HOMO SAPIENS | 0 | 30418 | 2xsw | A | 303 | VAL | p.Val303Met | Q9NRR6 |
| HOMO SAPIENS | 0 | 30418 | 2xsw | A | 315 | PRO | p.Pro315Leu | Q9NRR6 |
| HOMO SAPIENS | 0 | 30418 | 2xsw | A | 341 | GLY | p.Gly341Ser | Q9NRR6 |
| HOMO SAPIENS | 0 | 30418 | 2xsw | A | 378 | ARG | p.Arg378Cys | Q9NRR6 |
| HOMO SAPIENS | 0 | 30418 | 2xsw | A | 435 | ARG | p.Arg435Gln | Q9NRR6 |
| HOMO SAPIENS | 0 | 30418 | 2xsw | A | 515 | ARG | p.Arg515Trp | Q9NRR6 |
| HOMO SAPIENS | 1 | 30418 | 2xsw | A | 526 | PRO | p.Pro526Leu | Q9NRR6 |
| HOMO SAPIENS | 0 | 30418 | 2xsw | A | 563 | ARG | p.Arg563His | Q9NRR6 |
| HOMO SAPIENS | 2 | 30418 | 2xsw | A | 585 | ARG | p.Arg585His | Q9NRR6 |
| HOMO SAPIENS | 2 | 11167 | 4y99 | B | 205 | ARG | p.Arg215Leu | P45379 |
| HOMO SAPIENS | 5 | 11167 | 4y99 | B | 244 | GLU | p.Glu254Asp | P45379 |
| HOMO SAPIENS | 1 | 11167 | 4y99 | B | 253 | LYS | p.Lys263Arg | P45379 |
| HOMO SAPIENS | 8 | 11167 | 4y99 | B | 265 | ARG | p.Arg275Gln | P45379 |
| HOMO SAPIENS | 3 | 11167 | 4y99 | B | 270 | ASP | p.Asp280Asn | P45379 |
| HOMO SAPIENS | -1 | 11167 | 4y99 | B | 278 | ARG | p.Arg288Pro | P45379 |
| HOMO SAPIENS | -1 | 11167 | 4y99 | B | 286 | ARG | p.Arg296Cys | P45379 |
| HOMO SAPIENS | -1 | 11167 | 4y99 | B | 286 | ARG | p.Arg296His | P45379 |
| HOMO SAPIENS | 0 | 15993 | 4po6 | A | 197 | ARG | p.Arg197His | P29597 |
| HOMO SAPIENS | -1 | 15993 | 4po6 | A | 362 | VAL | p.Val362Phe | P29597 |
| HOMO SAPIENS | -1 | 15993 | 4po6 | A | 363 | GLY | p.Gly363Ser | P29597 |
| HOMO SAPIENS | 0 | 5646 | 5uak | A | 7 | GLU | p.Glu7Ter | P13569 |
| HOMO SAPIENS | 0 | 5646 | 5uak | A | 27 | GLY | p.Gly27Ter | P13569 |
| HOMO SAPIENS | 0 | 5646 | 5uak | A | 46 | ALA | p.Ala46Asp | P13569 |
| HOMO SAPIENS | 0 | 5646 | 5uak | A | 60 | GLU | p.Glu60Ter | P13569 |
| HOMO SAPIENS | 0 | 5646 | 5uak | A | 67 | PRO | p.Pro67Leu | P13569 |
| HOMO SAPIENS | 0 | 5646 | 5uak | A | 74 | ARG | p.Arg74Trp | P13569 |
| HOMO SAPIENS | 0 | 5646 | 5uak | A | 75 | ARG | p.Arg75Ter | P13569 |
| HOMO SAPIENS | 0 | 5646 | 5uak | A | 85 | GLY | p.Gly85Glu | P13569 |
| HOMO SAPIENS | 0 | 5646 | 5uak | A | 91 | GLY | p.Gly91Arg | P13569 |
| HOMO SAPIENS | 0 | 5646 | 5uak | A | 92 | GLU | p.Glu92Lys | P13569 |
| HOMO SAPIENS | 0 | 5646 | 5uak | A | 98 | GLN | p.Gln98Arg | P13569 |
| HOMO SAPIENS | 0 | 5646 | 5uak | A | 98 | GLN | p.Gln98Ter | P13569 |
| HOMO SAPIENS | 0 | 5646 | 5uak | A | 109 | TYR | p.Tyr109Cys | P13569 |
| HOMO SAPIENS | 0 | 5646 | 5uak | A | 110 | ASP | p.Asp110His | P13569 |
| HOMO SAPIENS | 0 | 5646 | 5uak | A | 117 | ARG | p.Arg117Cys | P13569 |
| HOMO SAPIENS | 0 | 5646 | 5uak | A | 117 | ARG | p.Arg117His | P13569 |
| HOMO SAPIENS | 0 | 5646 | 5uak | A | 125 | ILE | p.Ile125Thr | P13569 |
| HOMO SAPIENS | 0 | 5646 | 5uak | A | 148 | ILE | p.Ile148Thr | P13569 |
| HOMO SAPIENS | 0 | 5646 | 5uak | A | 170 | ARG | p.Arg170His | P13569 |

| | | | | | | | | |
|---|---|---|---|---|---|---|---|---|
| HOMO SAPIENS | 0 | 5646 | 5uak | A | 178 | GLY | p.Gly178Arg | P13569 |
| HOMO SAPIENS | 0 | 5646 | 5uak | A | 206 | LEU | p.Leu206Trp | P13569 |
| HOMO SAPIENS | 0 | 5646 | 5uak | A | 373 | ASP | p.Asp373Asn | P13569 |
| HOMO SAPIENS | -1 | 5646 | 5uak | A | 709 | ARG | p.Arg709Ter | P13569 |
| HOMO SAPIENS | -1 | 5646 | 5uak | A | 732 | LEU | p.Leu732Ter | P13569 |
| HOMO SAPIENS | -1 | 5646 | 5uak | A | 785 | ARG | p.Arg785Ter | P13569 |
| HOMO SAPIENS | -1 | 5646 | 5uak | A | 831 | GLU | p.Glu831Ter | P13569 |
| HOMO SAPIENS | -1 | 5646 | 5uak | A | 836 | ASP | p.Asp836Tyr | P13569 |
| HOMO SAPIENS | 0 | 5646 | 5uak | A | 851 | ARG | p.Arg851Ter | P13569 |
| HOMO SAPIENS | 0 | 5646 | 5uak | A | 913 | TYR | p.Tyr913Cys | P13569 |
| HOMO SAPIENS | 0 | 5646 | 5uak | A | 913 | TYR | p.Tyr913Ter | P13569 |
| HOMO SAPIENS | 0 | 5646 | 5uak | A | 945 | SER | p.Ser945Leu | P13569 |
| HOMO SAPIENS | 0 | 5646 | 5uak | A | 949 | HIS | p.His949Tyr | P13569 |
| HOMO SAPIENS | 0 | 5646 | 5uak | A | 970 | GLY | p.Gly970Asp | P13569 |
| HOMO SAPIENS | 0 | 5646 | 5uak | A | 1027 | ILE | p.Ile1027Thr | P13569 |
| HOMO SAPIENS | 0 | 5646 | 5uak | A | 1032 | TYR | p.Tyr1032Cys | P13569 |
| HOMO SAPIENS | 0 | 5646 | 5uak | A | 1061 | GLY | p.Gly1061Arg | P13569 |
| HOMO SAPIENS | 1 | 5646 | 5uak | A | 1065 | LEU | p.Leu1065Pro | P13569 |
| HOMO SAPIENS | 1 | 5646 | 5uak | A | 1066 | ARG | p.Arg1066Cys | P13569 |
| HOMO SAPIENS | 1 | 5646 | 5uak | A | 1066 | ARG | p.Arg1066His | P13569 |
| HOMO SAPIENS | 1 | 5646 | 5uak | A | 1067 | ALA | p.Ala1067Thr | P13569 |
| HOMO SAPIENS | 0 | 5646 | 5uak | A | 1069 | GLY | p.Gly1069Arg | P13569 |
| HOMO SAPIENS | 0 | 5646 | 5uak | A | 1070 | ARG | p.Arg1070Trp | P13569 |
| HOMO SAPIENS | 1 | 5646 | 5uak | A | 1071 | GLN | p.Gln1071Pro | P13569 |
| HOMO SAPIENS | 0 | 5646 | 5uak | A | 1074 | PHE | p.Phe1074Leu | P13569 |
| HOMO SAPIENS | 0 | 5646 | 5uak | A | 1077 | LEU | p.Leu1077Pro | P13569 |
| HOMO SAPIENS | 0 | 5646 | 5uak | A | 1089 | TRP | p.Trp1089Ter | P13569 |
| HOMO SAPIENS | 0 | 5646 | 5uak | A | 1092 | TYR | p.Tyr1092Ter | P13569 |
| HOMO SAPIENS | 0 | 5646 | 5uak | A | 1101 | MET | p.Met1101Lys | P13569 |
| HOMO SAPIENS | 0 | 5646 | 5uak | A | 1156 | LEU | p.Leu1156Phe | P13569 |
| HOMO SAPIENS | 0 | 5646 | 5uak | A | 1158 | ARG | p.Arg1158Ter | P13569 |
| HOMO SAPIENS | 0 | 5646 | 5uak | A | 1162 | ARG | p.Arg1162Leu | P13569 |
| HOMO SAPIENS | 0 | 5646 | 5uak | A | 1162 | ARG | p.Arg1162Ter | P13569 |
| HOMO SAPIENS | -1 | 5646 | 5uak | A | 1476 | GLN | p.Gln1476Ter | P13569 |
| HOMO SAPIENS | 0 | 22763 | 1imv | A | 52 | MET | p.Thr72Met | P36955 |
| HOMO SAPIENS | 0 | 22763 | 1imv | A | 301 | TYR | p.Tyr321Ter | P36955 |
| HOMO SAPIENS | 0 | 23179 | 5fql | A | 160 | PRO | p.Pro160Arg | P22304 |
| HOMO SAPIENS | 0 | 23179 | 5fql | A | 172 | ARG | p.Arg172Ter | P22304 |
| HOMO SAPIENS | 0 | 23179 | 5fql | A | 214 | THR | p.Thr214Met | P22304 |
| HOMO SAPIENS | 0 | 23179 | 5fql | A | 309 | THR | p.Thr309Ala | P22304 |
| HOMO SAPIENS | 0 | 23179 | 5fql | A | 333 | SER | p.Ser333Leu | P22304 |
| HOMO SAPIENS | 0 | 23179 | 5fql | A | 443 | ARG | p.Arg443Ter | P22304 |
| HOMO SAPIENS | 0 | 23179 | 5fql | A | 468 | ARG | p.Arg468Leu | P22304 |
| HOMO SAPIENS | 0 | 23179 | 5fql | A | 468 | ARG | p.Arg468Trp | P22304 |
| HOMO SAPIENS | 0 | 23179 | 5fql | A | 502 | TRP | p.Trp502Ser | P22304 |
| HOMO SAPIENS | 9 | 20989 | 1nfi | F | 185 | THR | p.Thr185Met | P25963 |
| HOMO SAPIENS | 4 | 20989 | 1nfi | F | 194 | GLY | p.Gly194Ala | P25963 |
| HOMO SAPIENS | 0 | 11538 | 3eab | A | 162 | VAL | p.Val162Ile | Q9UBP0 |
| HOMO SAPIENS | 0 | 33399 | 2p39 | A | 71 | SER | p.Ser71Gly | Q9GZV9 |
| HOMO SAPIENS | 3 | 33399 | 2p39 | A | 96 | MET | p.Met96Thr | Q9GZV9 |
| HOMO SAPIENS | 0 | 33399 | 2p39 | A | 129 | SER | p.Ser129Phe | Q9GZV9 |
| HOMO SAPIENS | -1 | 33399 | 2p39 | A | 172 | PRO | p.Pro172Leu | Q9GZV9 |
| HOMO SAPIENS | -1 | 33399 | 2p39 | A | 176 | ARG | p.Arg176Gln | Q9GZV9 |
| HOMO SAPIENS | -1 | 33399 | 2p39 | A | 179 | ARG | p.Arg179Trp | Q9GZV9 |

| | | | | | | | | |
|---|---|---|---|---|---|---|---|---|
| HOMO SAPIENS | -1 | 33399 | 2p39 | A | 179 | ARG | p.Arg179Gln | Q9GZV9 |
| HOMO SAPIENS | 0 | 16217 | 2dne | A | 79 | ILE | p.Ile163Leu | P10515 |
| HOMO SAPIENS | 8 | 7666 | 4kbq | A | 65 | ASN | p.Asn65Ser | Q9UNE7 |
| HOMO SAPIENS | 1 | 6407 | 1ikt | A | 71 | THR | p.Thr687Ile | P51659 |
| HOMO SAPIENS | 0 | 6407 | 1ikt | A | 112 | MET | p.Met728Val | P51659 |
| HOMO SAPIENS | 0 | 5849 | 4ihb | A | 67 | VAL | p.Val67Asp | O75923 |
| HOMO SAPIENS | 6 | 17703 | 5fcn | A | 20 | HIS | p.His99Gln | Q66GS9 |
| HOMO SAPIENS | 8 | 17703 | 5fcn | A | 33 | ARG | p.Arg112His | Q66GS9 |
| HOMO SAPIENS | 0 | 34095 | 2E+073 | A | 101 | HIS | p.His101Tyr | P05129 |
| HOMO SAPIENS | -1 | 41358 | 4liq | E | 507 | THR | p.Thr507Met | P07333 |
| HOMO SAPIENS | -1 | 21290 | 1fch | A | 235 | MET | p.Met272Arg | P50542 |
| HOMO SAPIENS | 5 | 23431 | 6pax | A | 23 | ARG | p.Arg26Gly | P26367 |
| HOMO SAPIENS | 1 | 23431 | 6pax | A | 61 | GLY | p.Gly64Val | P26367 |
| HOMO SAPIENS | 1 | 23431 | 6pax | A | 65 | PRO | p.Pro68Ser | P26367 |
| HOMO SAPIENS | 1 | 23431 | 6pax | A | 116 | SER | p.Ser119Arg | P26367 |
| HOMO SAPIENS | 9 | 23431 | 6pax | A | 123 | VAL | p.Val126Asp | P26367 |
| HOMO SAPIENS | 9 | 23431 | 6pax | A | 125 | ARG | p.Arg128Cys | P26367 |
| HOMO SAPIENS | 9 | 33003 | 4zrj | B | 535 | LEU | p.Leu535Pro | P35240 |
| HOMO SAPIENS | 7 | 33003 | 4zrj | B | 538 | GLN | p.Gln538Pro | P35240 |
| HOMO SAPIENS | 0 | 33003 | 4zrj | B | 585 | TRP | p.Ala585Thr | P35240 |
| HOMO SAPIENS | 0 | 3800 | 2nzj | A | 193 | VAL | p.Val193Met | O75628 |

| gene_name | max_cons | max_bs_id | max_pdb_id | max_chain_id |
|---|---|---|---|---|
| AKR1D1 | 0.306 | 22 | 3dop | A |
| AKR1D1 | 0.216 | 34 | 3cav | A |
| AKR1D1 | 0.276 | 21 | 3cot | A |
| AKR1D1 | 0.258 | 56 | 3cas | A |
| ALPP | None | None | None | None |
| BMPR1A | None | None | None | None |
| BMPR1A | None | None | None | None |
| FGFR2 | None | None | None | None |
| PROC | 0.587 | 3 | 1aut | C |
| PROC | 0.174 | 16 | 1aut | C |
| PROC | None | None | None | None |
| PROC | 0.396 | 10 | 3f6u | H |
| PROC | 0.247 | 13 | 3f6u | H |
| PROC | None | None | None | None |
| PROC | None | None | None | None |
| PROC | 0.191 | 15 | 3f6u | H |
| PROC | None | None | None | None |
| PROC | 0.587 | 3 | 1aut | C |
| PROC | 0.191 | 26 | 1aut | C |
| PROC | None | None | None | None |
| INS | 1 | 2 | 3zs2 | J |
| INS | 0.85 | 1 | 4ajx | D |
| INS | 1 | 1 | 3zs2 | J |
| INS | 1 | 1 | 3zs2 | J |
| FTL | 0.627 | 11 | 2fg8 | B |
| FTL | 0.313 | 62 | 2fg8 | G |
| FTL | 0.544 | 24 | 2fg8 | D |
| TNFSF11 | None | None | None | None |
| EHMT1 | None | None | None | None |
| EHMT1 | None | None | None | None |
| EHMT1 | None | None | None | None |
| CA2 | 0.939 | 1 | 4idr | X |
| CA2 | 0.939 | 1 | 4idr | X |
| CA2 | 0.902 | 5 | 4idr | X |
| CA2 | 0.902 | 5 | 4idr | X |
| SMAD4 | None | None | None | None |
| SMAD4 | None | None | None | None |
| TTR | 0.691 | 12 | 1ttb | B |
| TTR | 0.961 | 7 | 3grb | A |
| TTR | 0.891 | 4 | 3dk2 | B |
| TTR | 0.947 | 3 | 1tta | A |
| TTR | 0.923 | 1 | 3grb | B |
| TTR | 0.891 | 1 | 4y9b | B |
| TTR | 0.891 | 1 | 4y9b | B |
| TTR | 0.882 | 15 | 3grb | A |
| TTR | 0.909 | 5 | 4tl4 | A |
| TTR | 0.836 | 7 | 1z7j | B |
| TTR | 0.882 | 15 | 3grb | A |
| TTR | 0.891 | 5 | 3i9i | B |
| TTR | 0.902 | 9 | 3m1o | B |
| TTR | 0.902 | 9 | 3m1o | B |
| TTR | 0.902 | 9 | 3m1o | B |
| TTR | 0.636 | 16 | 1thc | A |

| | | | | |
|---|---|---|---|---|
| TTR | 0.902 | 9 | 3m1o | B |
| TTR | 0.836 | 7 | 1bze | A |
| TTR | 0.846 | 16 | 3gps | A |
| TTR | 0.846 | 16 | 3gps | A |
| TTR | 0.909 | 5 | 4tl4 | A |
| TTR | 0.909 | 5 | 4tl4 | A |
| TTR | 0.673 | 15 | 3dk2 | B |
| TTR | 0.945 | 4 | 4deu | A |
| TTR | 0.941 | 3 | 3did | B |
| TTR | 0.941 | 3 | 3did | B |
| TTR | 0.868 | 6 | 4l1t | A |
| TTR | 0.882 | 19 | 3gps | B |
| TTR | 0.891 | 4 | 3dk2 | B |
| TTR | 0.941 | 3 | 3did | B |
| TTR | 0.923 | 1 | 4hiq | A |
| TTR | 0.923 | 1 | 4hiq | A |
| TTR | 0.961 | 7 | 3grb | A |
| TTR | 0.839 | 7 | 1tsh | B |
| TTR | 0.891 | 6 | 1ttb | B |
| ALK | 0.682 | 1 | 4cd0 | A |
| ALK | 0.163 | 13 | 2xp2 | A |
| THRB | 0.155 | 15 | 2pin | B |
| THRB | None | None | None | None |
| THRB | 0.155 | 15 | 2pin | B |
| THRB | None | None | None | None |
| THRB | 0.155 | 15 | 2pin | B |
| THRB | 0.155 | 15 | 2pin | B |
| THRB | None | None | None | None |
| THRB | None | None | None | None |
| THRB | None | None | None | None |
| THRB | None | None | None | None |
| THRB | None | None | None | None |
| THRB | None | None | None | None |
| THRB | None | None | None | None |
| THRB | None | None | None | None |
| THRB | None | None | None | None |
| THRB | None | None | None | None |
| THRB | 0.126 | 16 | 1q4x | A |
| THRB | 0.126 | 16 | 1q4x | A |
| THRB | 0.577 | 3 | 2j4a | A |
| IL2RG | None | None | None | None |
| IL2RG | None | None | None | None |
| IL2RG | None | None | None | None |
| IL2RG | 0.256 | 1 | 2b5i | C |
| IL2RG | None | None | None | None |
| ITGB3 | None | None | None | None |
| ITGB3 | None | None | None | None |
| ITGB3 | 0.294 | 1 | 1jv2 | B |
| ITGB3 | None | None | None | None |
| ITGB3 | None | None | None | None |
| ITGB3 | None | None | None | None |
| ITGB3 | None | None | None | None |
| ITGB3 | None | None | None | None |
| ITGB3 | None | None | None | None |

| | | | | |
|---|---|---|---|---|
| PKD2 | None | None | None | None |
| PKD2 | None | None | None | None |
| PKD2 | None | None | None | None |
| PKD2 | None | None | None | None |
| PKD2 | None | None | None | None |
| PKD2 | None | None | None | None |
| PKD2 | None | None | None | None |
| TP53 | 0.571 | 1 | 1saf | B |
| TP53 | 0.571 | 1 | 1saf | B |
| TP53 | None | None | None | None |
| TP53 | 0.585 | 15 | 4mzr | B |
| TP53 | None | None | None | None |
| TP53 | None | None | None | None |
| EGFR | None | None | None | None |
| EGFR | None | None | None | None |
| XRCC4 | None | None | None | None |
| XRCC4 | None | None | None | None |
| HBB | 0.438 | 2 | 2w6v | D |
| HBB | 0.467 | 2 | 3qje | D |
| HBB | 0.467 | 2 | 3qje | D |
| HBB | 0.303 | 15 | 2w72 | B |
| HBB | 0.303 | 21 | 2w72 | B |
| HBB | 0.332 | 7 | 1cbm | A |
| HBB | 0.421 | 11 | 3ic2 | B |
| HBB | 0.459 | 2 | 5ee4 | D |
| HBB | 0.459 | 2 | 5ee4 | D |
| HBB | 0.459 | 2 | 5ee4 | D |
| HBB | 0.535 | 3 | 3r5i | B |
| HBB | 0.383 | 3 | 1lfv | B |
| HBB | 0.383 | 3 | 1lfv | B |
| HBB | 0.402 | 11 | 1m9p | D |
| HBB | 0.256 | 24 | 4n7p | D |
| HBB | 0.459 | 2 | 5ee4 | D |
| HBB | 0.566 | 1 | 1cbm | B |
| HBB | 0.51 | 3 | 1abw | B |
| HBB | 0.392 | 2 | 1lfz | B |
| HBB | 0.474 | 1 | 2h35 | B |
| HBB | 0.474 | 1 | 2h35 | B |
| HBB | 0.38 | 2 | 1j7y | B |
| HBB | 0.508 | 7 | 3ic2 | B |
| HBB | 0.545 | 3 | 1bij | D |
| HBB | 0.6 | 1 | 2w72 | D |
| HBB | 0.332 | 4 | 1fn3 | D |
| HBB | 0.354 | 15 | 1bij | D |
| HBB | 0.308 | 15 | 4rol | D |
| HBB | 0.733 | 1 | 1yff | D |
| HBB | 0.692 | 2 | 1buw | B |
| RPS6KA3 | None | None | None | None |
| RPS6KA3 | 0.555 | 1 | 4nw5 | A |
| RPS6KA3 | 0.226 | 6 | 4nw5 | A |
| RPS6KA3 | None | None | None | None |
| RPS6KA3 | None | None | None | None |
| RPS6KA3 | 0.197 | 11 | 5d9k | A |
| GBA | 1 | 15 | 3gxd | B |

| | | | | |
|---|---|---|---|---|
| GBA | 1 | 55 | 3gxd | B |
| GBA | 1 | 15 | 3gxd | B |
| GBA | 0.75 | 45 | 3ril | A |
| GBA | 0.25 | 33 | 2nt0 | A |
| GBA | 0.25 | 33 | 2nt0 | A |
| GBA | 1 | 12 | 1ogs | A |
| GBA | 0.75 | 122 | 3gxi | A |
| GBA | 1 | 2 | 1ogs | A |
| GBA | 1 | 11 | 1ogs | A |
| GBA | 0.5 | 59 | 3gxf | B |
| GBA | 0.75 | 45 | 3ril | A |
| GBA | 1 | 18 | 1ogs | A |
| GBA | 1 | 3 | 1ogs | A |
| GBA | None | None | None | None |
| GBA | 1 | 2 | 1ogs | A |
| GBA | 0.25 | 35 | 2nt0 | A |
| GBA | 0.75 | 76 | 3gxm | C |
| GBA | 1 | 3 | 1ogs | A |
| GBA | 1 | 3 | 1ogs | A |
| GBA | 0.75 | 62 | 3ril | A |
| GBA | 1 | 28 | 1ogs | A |
| GBA | None | None | None | None |
| GBA | 0.5 | 78 | 3gxi | A |
| GBA | 0.5 | 92 | 3rik | B |
| GBA | 0.5 | 42 | 2xwe | A |
| GBA | 0.333 | 117 | 1ogs | A |
| GBA | 0.5 | 92 | 3rik | B |
| GBA | 0.75 | 49 | 3ril | A |
| GBA | 0.25 | 109 | 2f61 | A |
| GBA | 0.667 | 60 | 1ogs | A |
| GBA | None | None | None | None |
| GBA | 0.5 | 11 | 2nt1 | A |
| GBA | 0.75 | 21 | 3gxi | A |
| GBA | None | None | None | None |
| GBA | 0.5 | 35 | 2xwe | A |
| GBA | 0.75 | 25 | 3keh | B |
| GBA | 0.75 | 70 | 3ril | A |
| GBA | 0.75 | 70 | 3ril | A |
| GBA | 1 | 15 | 3gxd | B |
| GBA | 1 | 44 | 3gxd | B |
| GBA | 0.5 | 57 | 3gxf | D |
| GBA | 1 | 44 | 3gxd | B |
| GBA | 1 | 44 | 3gxd | B |
| GBA | None | None | None | None |
| GBA | 0.75 | 94 | 3gxi | A |
| GBA | 0.75 | 94 | 3gxi | A |
| IFIH1 | None | None | None | None |
| TGFBR2 | None | None | None | None |
| TGFBR2 | None | None | None | None |
| TGFBR2 | 0.143 | 9 | 5e8v | A |
| TGFBR2 | None | None | None | None |
| TGFBR2 | None | None | None | None |
| TGFBR2 | None | None | None | None |
| TGFBR2 | None | None | None | None |

| | | | | | |
|---|---|---|---|---|---|
| TGFBR2 | None | None | None | | None |
| TGFBR2 | 0.169 | 14 | | 5E+092 | A |
| TGFBR2 | 0.742 | 1 | | 5E+091 | A |
| TGFBR2 | None | None | None | | None |
| TGFBR2 | None | None | None | | None |
| TGFBR2 | 0.206 | 6 | | 5E+092 | A |
| TGFBR2 | 0.676 | 1 | 5e8v | | A |
| TGFBR2 | 0.676 | 1 | 5e8v | | A |
| ACE | 0.684 | 17 | 3nxq | | A |
| ACE | 0.474 | 25 | 4bxk | | A |
| ACE | 0.263 | 31 | 2c6f | | A |
| ACE | None | None | None | | None |
| ACE | 0.211 | 93 | 4ca6 | | A |
| ACE | 0.211 | 92 | 5ama | | D |
| COL10A1 | None | None | None | | None |
| COL10A1 | None | None | None | | None |
| COL10A1 | None | None | None | | None |
| COL10A1 | 0.409 | 1 | 1gr3 | | A |
| COL10A1 | None | None | None | | None |
| COL10A1 | 0.318 | 4 | 1gr3 | | A |
| COL10A1 | None | None | None | | None |
| COL10A1 | 0.273 | 6 | 1gr3 | | A |
| COL10A1 | 0.318 | 4 | 1gr3 | | A |
| KLF1 | None | None | None | | None |
| C3 | None | None | None | | None |
| CTSC | 0.108 | 49 | 3pdf | | A |
| CTSC | 0.934 | 1 | 2djg | | C |
| CTSC | None | None | None | | None |
| LYZ | 0.877 | 3 | 1bb5 | | A |
| LYZ | 0.923 | 1 | 1lz4 | | A |
| LYZ | 0.852 | 2 | 1ubz | | A |
| DHODH | 0.111 | 65 | 4zmg | | A |
| DHODH | 0.207 | 18 | 5mut | | A |
| DHODH | None | None | None | | None |
| DHODH | None | None | None | | None |
| DHODH | None | None | None | | None |
| DHODH | 0.107 | 34 | 4ylw | | A |
| DHODH | 0.154 | 34 | 4rr4 | | A |
| ADH1C | 0.527 | 11 | 1ht0 | | A |
| GDNF | None | None | None | | None |
| TNF | 0.5 | 8 | 2e7a | | B |
| ALDH4A1 | 0.636 | 3 | 3v9h | | A |
| ALDH4A1 | 0.105 | 86 | 4oe5 | | B |
| ALDH4A1 | None | None | None | | None |
| BRAF | None | None | None | | None |
| BRAF | None | None | None | | None |
| BRAF | 0.752 | 2 | 3psb | | A |
| BRAF | 0.752 | 2 | 3psb | | A |
| BRAF | 0.737 | 1 | 3pri | | A |
| BRAF | 0.737 | 1 | 3pri | | A |
| BRAF | 0.737 | 1 | 3pri | | A |
| BRAF | 0.737 | 1 | 3prf | | A |
| BRAF | 0.737 | 1 | 3prf | | A |
| BRAF | 0.752 | 2 | 3psb | | A |

| | | | | |
|---|---|---|---|---|
| BRAF | 0.752 | 2 | 3psb | A |
| BRAF | 0.752 | 2 | 3psb | A |
| BRAF | 0.752 | 2 | 3psb | A |
| BRAF | 0.555 | 3 | 5vam | A |
| BRAF | 0.663 | 1 | 4ehe | B |
| BRAF | 0.734 | 1 | 5csx | A |
| TBX5 | None | None | None | None |
| TBX5 | None | None | None | None |
| TBX5 | None | None | None | None |
| TBX5 | None | None | None | None |
| TBX5 | None | None | None | None |
| TBX5 | None | None | None | None |
| TBX5 | None | None | None | None |
| TBX5 | None | None | None | None |
| TBX5 | None | None | None | None |
| RB1 | None | None | None | None |
| RB1 | None | None | None | None |
| RB1 | None | None | None | None |
| RB1 | None | None | None | None |
| RB1 | None | None | None | None |
| RB1 | None | None | None | None |
| CLIC2 | 0.5 | 3 | 2per | A |
| CLIC2 | None | None | None | None |
| ALDH7A1 | None | None | None | None |
| ALDH7A1 | None | None | None | None |
| ALDH7A1 | 0.405 | 4 | 2j6l | D |
| ALDH7A1 | 0.301 | 14 | 4zuk | A |
| ALDH7A1 | 0.409 | 3 | 4zvw | C |
| ALDH7A1 | None | None | None | None |
| ALDH7A1 | 0.534 | 3 | 2j6l | A |
| ALDH7A1 | 0.191 | 13 | 2j6l | D |
| ALDH7A1 | 0.333 | 6 | 4x0t | A |
| ALDH7A1 | None | None | None | None |
| ALDH7A1 | 0.113 | 38 | 4zul | C |
| ALDH7A1 | 0.534 | 3 | 2j6l | A |
| ALDH7A1 | None | None | None | None |
| ALDH7A1 | None | None | None | None |
| ALDH7A1 | 0.632 | 1 | 4zuk | A |
| ALDH7A1 | None | None | None | None |
| KDR | 0.575 | 5 | 1y6a | A |
| MUT | None | None | None | None |
| MUT | None | None | None | None |
| MUT | None | None | None | None |
| MUT | None | None | None | None |
| MUT | None | None | None | None |
| MUT | None | None | None | None |
| MUT | None | None | None | None |
| MUT | None | None | None | None |
| MUT | None | None | None | None |
| MUT | None | None | None | None |
| MUT | None | None | None | None |
| MUT | None | None | None | None |
| MUT | None | None | None | None |
| MUT | None | None | None | None |

| | | | | |
|---|---|---|---|---|
| MUT | None | None | None | None |
| MUT | None | None | None | None |
| MUT | 0.6 | 1 | 3bic | A |
| MUT | 0.6 | 2 | 2xiq | A |
| MUT | 0.6 | 2 | 2xiq | A |
| MUT | None | None | None | None |
| MUT | None | None | None | None |
| MUT | 0.4 | 1 | 2xiq | A |
| MUT | None | None | None | None |
| MUT | None | None | None | None |
| MUT | None | None | None | None |
| MUT | None | None | None | None |
| MUT | None | None | None | None |
| MUT | None | None | None | None |
| MUT | None | None | None | None |
| MUT | None | None | None | None |
| MUT | None | None | None | None |
| MUT | None | None | None | None |
| MUT | None | None | None | None |
| MUT | None | None | None | None |
| SOD1 | 0.778 | 1 | 2gbu | A |
| SOD1 | 0.727 | 4 | 4xcr | A |
| SOD1 | 0.35 | 26 | 1n19 | A |
| SOD1 | 0.621 | 12 | 1hl5 | H |
| NECTIN3 | None | None | None | None |
| HFE | None | None | None | None |
| HFE | None | None | None | None |
| HFE | None | None | None | None |
| HFE | 0.5 | 1 | 1de4 | A |
| HFE | None | None | None | None |
| HFE | None | None | None | None |
| HFE | 0.25 | 15 | 1a6z | A |
| EIF2AK3 | None | None | None | None |
| EIF2AK3 | None | None | None | None |
| TERT | None | None | None | None |
| TERT | None | None | None | None |
| SPR | None | None | None | None |
| SPR | None | None | None | None |
| SPR | None | None | None | None |
| SPR | None | None | None | None |
| SPR | None | None | None | None |
| F7 | 0.625 | 2 | 1klj | L |
| UNG | None | None | None | None |
| MMP13 | 0.356 | 22 | 2ozr | E |
| MMP13 | 0.786 | 1 | 3tvc | A |
| MMP13 | 0.766 | 1 | 3zxh | B |
| MMP13 | 0.766 | 1 | 3zxh | B |
| CD207 | 0.621 | 1 | 3kqg | B |
| F2 | 1 | 2 | 1nrq | L |
| SPTBN2 | None | None | None | None |
| PKLR | 0.568 | 9 | 2vgb | A |
| PKLR | 0.486 | 17 | 2vgg | D |
| PKLR | None | None | None | None |
| PKLR | 0.486 | 8 | 2vgg | D |

| | | | | |
|---|---|---|---|---|
| PKLR | 0.568 | 3 | 4ip7 | D |
| PKLR | 0.432 | 13 | 2vgi | A |
| PKLR | 0.378 | 45 | 4ip7 | B |
| PKLR | 0.324 | 25 | 2vgf | B |
| SACS | None | None | None | None |
| HRAS | 0.529 | 3 | 6q21 | A |
| HRAS | 0.529 | 3 | 6q21 | A |
| HRAS | 0.59 | 1 | 1gnq | A |
| HRAS | 0.59 | 1 | 1gnq | A |
| HRAS | 0.664 | 1 | 2lwi | A |
| HRAS | 0.742 | 1 | 1ctq | A |
| HRAS | 0.69 | 1 | 1xd2 | A |
| HRAS | 0.415 | 3 | 6q21 | D |
| HRAS | 0.529 | 3 | 6q21 | A |
| HRAS | 0.403 | 3 | 2n46 | A |
| HRAS | 0.521 | 4 | 4nyj | R |
| HRAS | 0.664 | 1 | 2lwi | A |
| HRAS | 0.664 | 1 | 2lwi | A |
| VHL | None | None | None | None |
| VHL | None | None | None | None |
| VHL | None | None | None | None |
| VHL | None | None | None | None |
| VHL | 0.5 | 22 | 4w9l | C |
| VHL | 0.5 | 22 | 5nvy | C |
| VHL | None | None | None | None |
| VHL | None | None | None | None |
| VHL | None | None | None | None |
| VHL | 0.5 | 2 | 5nvy | C |
| VHL | 0.5 | 1 | 5nvy | C |
| VHL | 0.5 | 1 | 5nvy | C |
| VHL | 0.5 | 4 | 5nvy | C |
| VHL | None | None | None | None |
| VHL | 0.5 | 6 | 5nvy | C |
| VHL | 0.5 | 2 | 5nvy | C |
| VHL | 0.5 | 8 | 3ztc | C |
| VHL | 0.5 | 2 | 5nvy | C |
| VHL | 0.5 | 2 | 5nvy | C |
| VHL | None | None | None | None |
| VHL | 0.5 | 7 | 4w9h | C |
| VHL | 0.5 | 2 | 5nvy | C |
| VHL | 0.5 | 2 | 5nvy | C |
| VHL | 0.5 | 19 | 4w9l | C |
| VHL | None | None | None | None |
| TTBK1 | None | None | None | None |
| RAD51 | None | None | None | None |
| ALB | None | None | None | None |
| ALB | 0.5 | 2 | 4n0u | D |
| ALB | None | None | None | None |
| ALB | None | None | None | None |
| MET | None | None | None | None |
| MET | None | None | None | None |
| MET | None | None | None | None |
| MET | None | None | None | None |
| MET | None | None | None | None |

| | | | | |
|---|---|---|---|---|
| MET | None | None | None | None |
| MET | None | None | None | None |
| MET | None | None | None | None |
| MET | None | None | None | None |
| CP | None | None | None | None |
| CP | None | None | None | None |
| PGM1 | None | None | None | None |
| PGM1 | 0.409 | 2 | 5f9c | A |
| PGM1 | 0.409 | 2 | 5f9c | A |
| PGM1 | None | None | None | None |
| PGM1 | None | None | None | None |
| PGM1 | None | None | None | None |
| PGM1 | None | None | None | None |
| NR5A1 | None | None | None | None |
| NR5A1 | 0.157 | 17 | 4qjr | A |
| NR5A1 | None | None | None | None |
| F9 | None | None | None | None |
| F9 | None | None | None | None |
| HPRT1 | 0.188 | 29 | 3ggj | B |
| HPRT1 | 0.125 | 29 | 1z7g | A |
| HPRT1 | 0.25 | 7 | 3gep | B |
| HPRT1 | 0.375 | 15 | 3gep | B |
| HPRT1 | None | None | None | None |
| HPRT1 | 0.871 | 3 | 4rac | B |
| HPRT1 | 0.806 | 2 | 4ran | D |
| HPRT1 | None | None | None | None |
| HPRT1 | 0.25 | 4 | 3gep | B |
| HPRT1 | 0.44 | 3 | 1d6n | B |
| HPRT1 | 0.154 | 31 | 1bzy | D |
| HPRT1 | None | None | None | None |
| HPRT1 | 0.24 | 25 | 1d6n | B |
| HPRT1 | 0.703 | 4 | 5w8v | A |
| HPRT1 | 0.118 | 27 | 1z7g | B |
| HPRT1 | 0.321 | 8 | 4kn6 | A |
| HPRT1 | 0.312 | 27 | 3ggj | B |
| HPRT1 | 0.312 | 12 | 3gep | B |
| HPRT1 | 0.871 | 3 | 4rac | B |
| HPRT1 | 0.871 | 3 | 4rac | B |
| HPRT1 | 0.765 | 2 | 4rao | D |
| HPRT1 | 0.774 | 2 | 4rac | B |
| GPD1L | None | None | None | None |
| GPD1L | None | None | None | None |
| GPD1L | None | None | None | None |
| FGFR2 | None | None | None | None |
| FGFR2 | None | None | None | None |
| FGFR2 | None | None | None | None |
| FGFR2 | None | None | None | None |
| FGFR2 | None | None | None | None |
| FGFR2 | None | None | None | None |
| FGFR2 | None | None | None | None |
| FGFR2 | None | None | None | None |
| FGFR2 | None | None | None | None |
| FGFR2 | None | None | None | None |
| FGFR2 | None | None | None | None |

| | | | | |
|---|---|---|---|---|
| FGFR2 | None | None | None | None |
| FGFR2 | None | None | None | None |
| FGFR2 | None | None | None | None |
| FGFR2 | None | None | None | None |
| FGFR2 | None | None | None | None |
| FGFR2 | None | None | None | None |
| FGFR2 | None | None | None | None |
| FGFR2 | None | None | None | None |
| HSPD1 | None | None | None | None |
| HSPD1 | None | None | None | None |
| FGF8 | None | None | None | None |
| FGF8 | None | None | None | None |
| FGF8 | None | None | None | None |
| F9 | 0.438 | 1 | 5vyg | B |
| NCF4 | None | None | None | None |
| NCF4 | None | None | None | None |
| LDLR | None | None | None | None |
| LDLR | None | None | None | None |
| LDLR | 0.478 | 1 | 3gcx | E |
| LDLR | 0.478 | 1 | 3gcx | E |
| LDLR | 0.478 | 1 | 3gcx | E |
| LDLR | None | None | None | None |
| C3 | None | None | None | None |
| C3 | None | None | None | None |
| C3 | None | None | None | None |
| C3 | None | None | None | None |
| C3 | None | None | None | None |
| C3 | None | None | None | None |
| C3 | None | None | None | None |
| C3 | None | None | None | None |
| TULP1 | None | None | None | None |
| TULP1 | None | None | None | None |
| TULP1 | None | None | None | None |
| TULP1 | None | None | None | None |
| IKBKG | None | None | None | None |
| IKBKG | None | None | None | None |
| IKBKG | None | None | None | None |
| CD40 | None | None | None | None |
| CD40 | None | None | None | None |
| SLC9A3R1 | None | None | None | None |
| BCHE | 0.278 | 18 | 2xqg | A |
| BCHE | 0.439 | 5 | 2wil | B |
| BCHE | 0.544 | 11 | 1p0q | A |
| BCHE | 0.127 | 115 | 2pm8 | B |
| BCHE | 0.196 | 66 | 2wik | A |
| BCHE | None | None | None | None |
| BCHE | 0.63 | 4 | 2xqg | A |
| BCHE | None | None | None | None |
| BCHE | 0.138 | 164 | 1xlu | A |
| BCHE | 0.255 | 18 | 2xqj | A |
| BCHE | 0.14 | 62 | 5dyt | A |
| BCKDHB | 0.833 | 1 | 1v11 | B |
| BCKDHB | 0.5 | 18 | 2bfc | B |
| BCKDHB | None | None | None | None |

| | | | | |
|---|---|---|---|---|
| BCKDHB | 0.667 | 11 | 2bfd | B |
| BCKDHB | 0.833 | 10 | 1olx | B |
| BCKDHB | 0.833 | 1 | 1v11 | B |
| BCKDHB | 0.833 | 1 | 1v11 | B |
| BCKDHB | 0.625 | 2 | 1x7z | B |
| BCKDHB | 0.429 | 16 | 1ols | B |
| BCKDHB | 0.333 | 99 | 1olx | B |
| BCKDHB | 0.333 | 64 | 1x7y | B |
| BCKDHB | 0.857 | 1 | 2bff | B |
| CDH1 | 0.278 | 8 | 3ff7 | B |
| CDH1 | 0.575 | 1 | 3ff8 | A |
| CDH1 | 0.543 | 1 | 3ff7 | A |
| CDH1 | None | None | None | None |
| CDH1 | 0.2 | 20 | 2o72 | A |
| CTNNB1 | None | None | None | None |
| AIRE | None | None | None | None |
| EDA | 1 | 1 | 1rj8 | A |
| EDA | None | None | None | None |
| EDA | None | None | None | None |
| EDA | None | None | None | None |
| EDA | None | None | None | None |
| SH2D1A | 0.444 | 5 | 1ka7 | A |
| SH2D1A | None | None | None | None |
| SH2D1A | None | None | None | None |
| GCK | 0.556 | 37 | 3idh | A |
| GCK | 0.333 | 32 | 4isf | A |
| GCK | None | None | None | None |
| GCK | None | None | None | None |
| GCK | 0.176 | 38 | 4isg | A |
| GCK | 0.611 | 13 | 3h1v | X |
| GCK | None | None | None | None |
| GCK | None | None | None | None |
| GCK | None | None | None | None |
| GCK | 0.778 | 5 | 3id8 | A |
| GCK | 0.833 | 1 | 4dhy | A |
| GCK | 0.389 | 6 | 4lc9 | B |
| GCK | 0.222 | 33 | 4mle | A |
| GCK | 0.611 | 6 | 3id8 | A |
| GCK | None | None | None | None |
| GCK | 0.167 | 33 | 4rch | A |
| GCK | None | None | None | None |
| GCK | 0.5 | 5 | 3vey | A |
| GCK | None | None | None | None |
| GCK | None | None | None | None |
| AQP2 | None | None | None | None |
| AQP2 | None | None | None | None |
| AQP2 | None | None | None | None |
| AQP2 | None | None | None | None |
| AQP2 | None | None | None | None |
| AQP2 | None | None | None | None |
| AQP2 | None | None | None | None |
| AQP2 | None | None | None | None |
| AQP2 | None | None | None | None |
| AQP2 | None | None | None | None |

| | | | | |
|---|---|---|---|---|
| AQP2 | None | None | None | None |
| AQP2 | None | None | None | None |
| AQP2 | None | None | None | None |
| SUMF1 | 0.75 | 6 | 1z70 | X |
| SUMF1 | 0.556 | 30 | 1y1i | X |
| SUMF1 | 0.5 | 38 | 2aij | X |
| SUMF1 | 0.625 | 18 | 2aij | X |
| SUMF1 | 0.875 | 35 | 2afy | X |
| SUMF1 | 0.667 | 19 | 1y1i | X |
| SUMF1 | 0.75 | 17 | 1y1j | X |
| SUMF1 | 0.75 | 17 | 1y1j | X |
| SUMF1 | 0.625 | 27 | 1y1g | X |
| FAM20A | None | None | None | None |
| FAM20A | None | None | None | None |
| FAM20A | None | None | None | None |
| FAM20A | None | None | None | None |
| NT5E | None | None | None | None |
| NT5E | None | None | None | None |
| NOS3 | 0.564 | 54 | 1m9k | B |
| ACTG1 | None | None | None | None |
| ACTG1 | None | None | None | None |
| ACTG1 | 0.229 | 48 | 5jlh | E |
| ACTG1 | None | None | None | None |
| ACTG1 | 0.571 | 8 | 5jlh | E |
| ACTG1 | None | None | None | None |
| FCGR1A | None | None | None | None |
| GPI | 0.304 | 64 | 1iri | A |
| GPI | 0.13 | 210 | 1jiq | A |
| GPI | 1 | 3 | 1jiq | A |
| GPI | None | None | None | None |
| GPI | 0.696 | 22 | 1iat | A |
| GPI | 0.522 | 22 | 1nuh | A |
| GPI | 0.696 | 27 | 1iat | A |
| GPI | 0.304 | 87 | 1jlh | B |
| GPI | 0.391 | 141 | 1jlh | B |
| CFH | None | None | None | None |
| OGG1 | 0.895 | 13 | 1m3h | A |
| TPK1 | None | None | None | None |
| TPK1 | None | None | None | None |
| TPK1 | None | None | None | None |
| TPK1 | None | None | None | None |
| SEC23A | None | None | None | None |
| SEC23A | None | None | None | None |
| TYRP1 | None | None | None | None |
| TYRP1 | None | None | None | None |
| TYRP1 | None | None | None | None |
| TYRP1 | None | None | None | None |
| TYRP1 | 0.3 | 5 | 5m8m | A |
| TYRP1 | None | None | None | None |
| CSTB | None | None | None | None |
| CSTB | None | None | None | None |
| CSTB | None | None | None | None |
| ACP1 | None | None | None | None |
| IL17F | None | None | None | None |

| | | | | |
|---|---|---|---|---|
| IL17F | None | None | None | None |
| IL17F | None | None | None | None |
| IL17F | None | None | None | None |
| IL17F | None | None | None | None |
| IDH2 | 0.5 | 2 | 5svn | B |
| IDH2 | 0.5 | 2 | 5svn | B |
| IDH2 | 0.48 | 13 | 5i96 | A |
| IDH2 | None | None | None | None |
| TP53 | 0.925 | 11 | 2ocj | A |
| TP53 | 0.8 | 16 | 2ahi | D |
| TP53 | 0.65 | 49 | 4ibq | A |
| TP53 | 0.725 | 27 | 5g4o | A |
| TP53 | 0.455 | 21 | 3q06 | A |
| TP53 | 0.9 | 1 | 4agp | A |
| TP53 | 0.9 | 1 | 4agp | A |
| TP53 | 0.925 | 10 | 4hje | A |
| TP53 | 0.7 | 29 | 3d05 | A |
| TP53 | 0.875 | 7 | 3kmd | B |
| TP53 | 0.875 | 7 | 3kmd | B |
| TP53 | 0.925 | 6 | 3kz8 | B |
| TP53 | 0.925 | 6 | 3kz8 | B |
| TP53 | 0.609 | 22 | 3q01 | B |
| TP53 | 0.925 | 5 | 3kmd | B |
| TP53 | 0.925 | 5 | 3kmd | B |
| TP53 | 0.925 | 8 | 3kz8 | B |
| TP53 | 0.925 | 8 | 3kz8 | B |
| TP53 | 0.925 | 1 | 3kmd | B |
| TP53 | 0.9 | 11 | 2pcx | A |
| TP53 | 0.9 | 1 | 5aol | A |
| TP53 | 0.85 | 5 | 5lgy | A |
| TP53 | 0.9 | 1 | 3q06 | B |
| TP53 | 0.725 | 19 | 5lgy | A |
| TP53 | 0.775 | 14 | 4mzi | A |
| TP53 | 0.825 | 8 | 1kzy | A |
| TP53 | 0.825 | 8 | 1kzy | A |
| TP53 | 0.8 | 11 | 4ibt | A |
| TP53 | 0.8 | 11 | 4ibt | A |
| TP53 | 0.725 | 15 | 5lgy | A |
| TP53 | 0.725 | 29 | 2h1l | O |
| TP53 | 0.875 | 7 | 3kmd | B |
| TP53 | 0.725 | 15 | 5lgy | A |
| TP53 | 0.8 | 10 | 4agp | A |
| TP53 | 0.8 | 10 | 4agp | A |
| TP53 | 0.8 | 10 | 4agp | A |
| TP53 | 0.8 | 10 | 4agp | A |
| TP53 | 0.675 | 35 | 4ibq | A |
| TP53 | 0.575 | 37 | 4agm | A |
| TP53 | 0.675 | 17 | 3kz8 | B |
| TP53 | 0.356 | 63 | 4mzr | C |
| TP53 | None | None | None | None |
| LMAN1 | None | None | None | None |
| CTSK | 0.75 | 2 | 4n79 | A |
| CTSK | 0.127 | 32 | 1ayw | A |
| CTSK | 0.413 | 12 | 4x6j | A |

| | | | | |
|---|---|---|---|---|
| FGB | None | None | None | None |
| FGB | None | None | None | None |
| FGB | 0.688 | 1 | 1fzf | E |
| FGB | 0.281 | 10 | 2ffd | B |
| ACADM | None | None | None | None |
| ACADM | None | None | None | None |
| ACADM | 0.218 | 11 | 1t9g | C |
| ACADM | 0.471 | 3 | 4p13 | D |
| ACADM | 0.127 | 10 | 1egc | C |
| ACADM | None | None | None | None |
| ACADM | 0.389 | 6 | 1t9g | D |
| ACADM | None | None | None | None |
| ACADM | 0.327 | 12 | 4p13 | C |
| ACADM | 0.389 | 6 | 1t9g | D |
| ACADM | None | None | None | None |
| ACADM | None | None | None | None |
| ACADM | None | None | None | None |
| ACADM | 0.291 | 14 | 1t9g | C |
| ACADM | None | None | None | None |
| ACADM | 0.481 | 2 | 4p13 | C |
| ACADM | 0.389 | 6 | 1t9g | D |
| ACADM | 0.109 | 24 | 2a1t | C |
| ACADM | None | None | None | None |
| ACADM | 0.104 | 32 | 4p13 | A |
| ACADM | 0.564 | 1 | 1t9g | C |
| ACADM | 0.255 | 10 | 1t9g | C |
| ACADM | None | None | None | None |
| ACADM | None | None | None | None |
| ACADM | 0.364 | 6 | 2a1t | A |
| ACADM | 0.312 | 12 | 4p13 | A |
| ACADM | 0.347 | 7 | 4p13 | B |
| PTS | None | None | None | None |
| PTS | 0.417 | 2 | 3i2b | E |
| PTS | None | None | None | None |
| PTS | 0.778 | 1 | 3i2b | D |
| PTS | None | None | None | None |
| PTS | None | None | None | None |
| PTS | None | None | None | None |
| PTS | None | None | None | None |
| PTS | None | None | None | None |
| INS | 0.75 | 2 | 3zqr | E |
| INS | 1 | 5 | 2l1z | A |
| INS | 1 | 5 | 2l1z | A |
| INS | 1 | 1 | 1his | A |
| FBP1 | 0.71 | 26 | 2y5l | C |
| FBP1 | 0.125 | 129 | 3a29 | C |
| FBP1 | 0.5 | 18 | 1fta | A |
| FBP1 | 0.7 | 10 | 2y5k | A |
| FBP1 | 0.516 | 38 | 3kc1 | B |
| FBP1 | 0.733 | 10 | 3kc1 | A |
| FBP1 | 0.633 | 43 | 2jjk | A |
| FBP1 | 0.8 | 11 | 2y5k | A |
| FBP1 | 0.548 | 41 | 2vt5 | D |
| FBP1 | 0.438 | 25 | 2fie | A |

| | | | | |
|---|---|---|---|---|
| FBP1 | 0.643 | 7 | 4mjo | B |
| KCNQ1 | None | None | None | None |
| KCNQ1 | None | None | None | None |
| KCNQ1 | None | None | None | None |
| UPP1 | 0.706 | 4 | 3euf | A |
| UMPS | 1 | 3 | 2qcl | A |
| UMPS | 0.556 | 59 | 2qcl | B |
| UMPS | 0.184 | 153 | 2v30 | B |
| DHFR | 0.167 | 16 | 3f91 | A |
| KHK | 0.667 | 2 | 2hw1 | A |
| TTPA | None | None | None | None |
| TTPA | None | None | None | None |
| TTPA | 0.5 | 1 | 1r5l | A |
| TTPA | None | None | None | None |
| MAP1LC3B | 0.387 | 8 | 3vtv | A |
| MAP1LC3B | None | None | None | None |
| ZAP70 | 0.317 | 1 | 2oq1 | A |
| PROC | 0.536 | 1 | 3f6u | L |
| SQSTM1 | None | None | None | None |
| AHCY | 0.19 | 113 | 4pfj | B |
| AHCY | None | None | None | None |
| PRNP | None | None | None | None |
| PRNP | None | None | None | None |
| PRNP | None | None | None | None |
| PRNP | None | None | None | None |
| PRNP | 0.277 | 1 | 1qlx | A |
| PRNP | None | None | None | None |
| PRNP | None | None | None | None |
| PRNP | 0.19 | 2 | 1e1p | A |
| PRNP | None | None | None | None |
| PRNP | 0.238 | 2 | 3hj5 | A |
| PRNP | 0.238 | 2 | 3hj5 | A |
| PRNP | None | None | None | None |
| PRNP | None | None | None | None |
| PRNP | 0.19 | 3 | 1qm3 | A |
| PRNP | None | None | None | None |
| PRNP | 0.277 | 1 | 1qlx | A |
| REN | 0.431 | 2 | 2v13 | A |
| REN | 0.431 | 2 | 2v13 | A |
| HINT1 | 0.556 | 2 | 5ed6 | B |
| HINT1 | 0.897 | 1 | 5km4 | A |
| HINT1 | 0.227 | 60 | 5ed6 | A |
| HINT1 | 0.667 | 11 | 5ed6 | B |
| HINT1 | 0.923 | 1 | 5ipc | A |
| OAT | None | None | None | None |
| OAT | 0.107 | 32 | 2can | B |
| OAT | 0.381 | 5 | 2can | B |
| OAT | 0.381 | 5 | 2can | B |
| OAT | 0.381 | 7 | 2byj | B |
| OAT | None | None | None | None |
| OAT | 0.113 | 54 | 1oat | B |
| OAT | None | None | None | None |
| OAT | 0.278 | 12 | 2byj | A |
| OAT | None | None | None | None |

| | | | | |
|---|---|---|---|---|
| OAT | 0.675 | 2 | 2byl | B |
| OAT | 0.179 | 32 | 2oat | C |
| OAT | 0.179 | 32 | 2oat | C |
| OAT | 0.169 | 15 | 5vwo | A |
| OAT | 0.169 | 15 | 5vwo | A |
| OAT | 0.758 | 1 | 2byl | A |
| OAT | 0.375 | 5 | 2oat | C |
| OAT | 0.525 | 4 | 2byl | B |
| OAT | 0.429 | 6 | 2byj | B |
| OAT | None | None | None | None |
| OAT | 0.107 | 49 | 2can | B |
| OAT | 0.165 | 23 | 2byl | A |
| OAT | None | None | None | None |
| OAT | 0.155 | 17 | 1gbn | C |
| OAT | 0.155 | 17 | 1gbn | C |
| OAT | None | None | None | None |
| OAT | None | None | None | None |
| OAT | None | None | None | None |
| CTSA | None | None | None | None |
| CTSA | None | None | None | None |
| PNP | 0.375 | 24 | 1ulb | A |
| PNP | 0.29 | 111 | 1yry | E |
| PNP | 0.406 | 43 | 3phb | E |
| PNP | 0.706 | 5 | 4ear | C |
| PNP | 0.367 | 28 | 3ggs | C |
| PNP | 0.5 | 17 | 1pf7 | E |
| PNP | 0.933 | 1 | 3ggs | C |
| PNP | 0.833 | 1 | 3d1v | A |
| MYL2 | None | None | None | None |
| MYL2 | None | None | None | None |
| ANG | 0.548 | 4 | 1k5b | A |
| ANG | 0.419 | 8 | 4ahi | A |
| ANG | 0.419 | 8 | 4ahi | A |
| ANG | 0.19 | 15 | 1k58 | A |
| ANG | 0.761 | 1 | 5eop | A |
| ANG | 0.576 | 4 | 5m9v | A |
| ANG | 0.579 | 4 | 5m9q | A |
| ANG | 0.14 | 34 | 4ahf | A |
| ANG | 0.112 | 34 | 5m9r | A |
| ANG | 0.52 | 5 | 5m9g | A |
| POLH | 0.241 | 101 | 4dl5 | A |
| POLH | 0.185 | 17 | 4j9l | A |
| POLH | 0.208 | 9 | 3si8 | A |
| IL2RA | None | None | None | None |
| TGFB1 | None | None | None | None |
| TP53 | None | None | None | None |
| TP53 | None | None | None | None |
| PAH | None | None | None | None |
| PAH | None | None | None | None |
| PAH | 0.44 | 11 | 2pah | A |
| PAH | 0.44 | 11 | 2pah | A |
| PAH | 0.32 | 19 | 6pah | A |
| PAH | 0.32 | 19 | 6pah | A |
| PAH | 0.48 | 2 | 1pah | A |

| | | | | |
|---|---|---|---|---|
| PAH | 0.48 | 2 | 1pah | A |
| PAH | 0.4 | 11 | 3pah | A |
| PAH | 0.2 | 51 | 1j8u | A |
| PAH | 0.2 | 51 | 1j8u | A |
| PAH | 0.36 | 7 | 2pah | B |
| PAH | 0.44 | 9 | 1dmw | A |
| PAH | 0.48 | 1 | 1dmw | A |
| PAH | 0.48 | 1 | 1dmw | A |
| PAH | 0.44 | 8 | 2pah | B |
| PAH | 0.44 | 8 | 2pah | B |
| PAH | 0.417 | 3 | 1j8t | A |
| PAH | 0.48 | 2 | 4anp | A |
| PAH | 0.48 | 1 | 1dmw | A |
| PAH | 0.48 | 4 | 2pah | B |
| PAH | 0.44 | 11 | 2pah | A |
| PAH | 0.48 | 4 | 1mmt | A |
| PAH | 0.68 | 6 | 2pah | A |
| PAH | 0.76 | 1 | 1tdw | A |
| PAH | None | None | None | None |
| PAH | 0.12 | 44 | 6pah | A |
| PAH | 0.4 | 14 | 1j8u | A |
| PAH | None | None | None | None |
| PAH | None | None | None | None |
| PAH | None | None | None | None |
| PAH | 0.2 | 43 | 3pah | A |
| PAH | 0.64 | 2 | 1lrm | A |
| PAH | 0.4 | 11 | 2pah | B |
| PAH | 0.28 | 31 | 4anp | A |
| PAH | 0.4 | 14 | 1j8u | A |
| PAH | 0.4 | 14 | 1j8u | A |
| PAH | 0.12 | 60 | 1j8u | A |
| PAH | 0.48 | 1 | 1dmw | A |
| PAH | 0.48 | 1 | 1dmw | A |
| PAH | 0.167 | 38 | 1j8t | A |
| HDAC8 | None | None | None | None |
| HDAC8 | 0.732 | 3 | 3rqd | A |
| VCP | 0.438 | 4 | 3tiw | A |
| VCP | 0.5 | 2 | 3qq8 | A |
| VCP | 0.5 | 2 | 3qq8 | A |
| VCP | None | None | None | None |
| VCP | None | None | None | None |
| VCP | None | None | None | None |
| VCP | None | None | None | None |
| B3GAT3 | None | None | None | None |
| B3GAT3 | None | None | None | None |
| CSF2RB | None | None | None | None |
| CSF2RB | None | None | None | None |
| CSF2RB | None | None | None | None |
| NF2 | 0.5 | 6 | 1h4r | B |
| NF2 | None | None | None | None |
| NF2 | None | None | None | None |
| PIK3CA | None | None | None | None |
| PHGDH | None | None | None | None |
| PHGDH | None | None | None | None |

| | | | | |
|---|---|---|---|---|
| PHGDH | None | None | None | None |
| PHGDH | 0.66 | 1 | 5n53 | A |
| POLH | None | None | None | None |
| XIAP | 0.5 | 7 | 5c7b | A |
| XIAP | None | None | None | None |
| PDHX | None | None | None | None |
| PDHX | None | None | None | None |
| PDHX | None | None | None | None |
| PDHX | None | None | None | None |
| PDHX | None | None | None | None |
| SAMHD1 | 0.5 | 12 | 5ao0 | A |
| SAMHD1 | 0.357 | 51 | 4zwe | D |
| SAMHD1 | 0.429 | 7 | 4tnr | D |
| SAMHD1 | None | None | None | None |
| SAMHD1 | 0.571 | 3 | 4qfy | D |
| SAMHD1 | None | None | None | None |
| SAMHD1 | 0.308 | 18 | 4qg2 | B |
| SAMHD1 | 0.286 | 57 | 5ao0 | B |
| CD36 | None | None | None | None |
| CD36 | None | None | None | None |
| CD36 | None | None | None | None |
| ASPH | None | None | None | None |
| CCT5 | None | None | None | None |
| CCT5 | None | None | None | None |
| CCT5 | None | None | None | None |
| SMN1 | None | None | None | None |
| SMN1 | None | None | None | None |
| SMN1 | None | None | None | None |
| SMN1 | None | None | None | None |
| ABCA1 | None | None | None | None |
| ABCA1 | None | None | None | None |
| ABCA1 | None | None | None | None |
| ABCA1 | None | None | None | None |
| ABCA1 | None | None | None | None |
| ABCA1 | None | None | None | None |
| ABCA1 | None | None | None | None |
| ABCA1 | None | None | None | None |
| ABCA1 | None | None | None | None |
| ABCA1 | None | None | None | None |
| ABCA1 | None | None | None | None |
| ABCA1 | None | None | None | None |
| ABCA1 | None | None | None | None |
| ABCA1 | None | None | None | None |
| ABCA1 | None | None | None | None |
| ABCA1 | None | None | None | None |
| ABCA1 | None | None | None | None |
| ABCA1 | None | None | None | None |
| ABCA1 | None | None | None | None |
| ABCA1 | None | None | None | None |
| ABCA1 | None | None | None | None |
| ABCA1 | None | None | None | None |
| ABCA1 | None | None | None | None |
| ABCA1 | None | None | None | None |

| | | | | |
|---|---|---|---|---|
| ABCA1 | None | None | None | None |
| ABCA1 | None | None | None | None |
| ABCA1 | None | None | None | None |
| ABCA1 | None | None | None | None |
| ABCA1 | None | None | None | None |
| ABCA1 | None | None | None | None |
| ABCA1 | None | None | None | None |
| ABCA1 | None | None | None | None |
| ABCA1 | None | None | None | None |
| ABCA1 | None | None | None | None |
| ABCA1 | None | None | None | None |
| ERCC1 | None | None | None | None |
| FECH | 0.524 | 27 | 2po5 | A |
| FECH | 0.6 | 60 | 2hre | A |
| FECH | 0.636 | 20 | 2po7 | A |
| FECH | 0.476 | 61 | 4klr | A |
| FECH | 0.5 | 84 | 2hrc | A |
| FECH | 0.667 | 10 | 2po5 | A |
| FECH | 0.455 | 30 | 4kla | B |
| PTEN | None | None | None | None |
| PTEN | None | None | None | None |
| PTEN | None | None | None | None |
| PTEN | None | None | None | None |
| PTEN | None | None | None | None |
| PTEN | None | None | None | None |
| PTEN | None | None | None | None |
| PTEN | None | None | None | None |
| PTEN | None | None | None | None |
| PTEN | None | None | None | None |
| PTEN | None | None | None | None |
| PTEN | None | None | None | None |
| PTEN | None | None | None | None |
| PTEN | None | None | None | None |
| PTEN | None | None | None | None |
| PTEN | None | None | None | None |
| PTEN | None | None | None | None |
| PTEN | None | None | None | None |
| PTEN | None | None | None | None |
| PTEN | None | None | None | None |
| ACAT1 | 0.667 | 1 | 2ib9 | C |
| ACAT1 | 0.114 | 43 | 2ib8 | B |
| ACAT1 | None | None | None | None |
| ACAT1 | 0.667 | 1 | 2ib9 | C |
| ACAT1 | 0.667 | 1 | 2ib9 | C |
| ACAT1 | 0.472 | 11 | 2f2s | A |
| ACAT1 | 0.833 | 6 | 2ib7 | C |
| ACAT1 | 0.229 | 68 | 2ib8 | B |
| ACAT1 | None | None | None | None |
| ACAT1 | 0.528 | 9 | 2ibw | D |
| ACAT1 | 0.703 | 16 | 2ibu | A |
| CREBBP | 0.283 | 35 | 5mqg | A |
| ITGA2B | None | None | None | None |
| ITGA2B | None | None | None | None |
| ITGA2B | 0.214 | 20 | 2vdn | A |

| | | | | |
|---|---|---|---|---|
| ITGA2B | 0.538 | 1 | 3nig | A |
| TSC1 | None | None | None | None |
| ADA2 | None | None | None | None |
| ADA2 | None | None | None | None |
| ADA2 | 0.75 | 1 | 3lgd | A |
| ADA2 | None | None | None | None |
| ADA2 | None | None | None | None |
| ADA2 | None | None | None | None |
| ADA2 | None | None | None | None |
| ADA2 | None | None | None | None |
| ADA2 | 1 | 1 | 3lgd | B |
| ADA2 | None | None | None | None |
| PGK1 | None | None | None | None |
| PGK1 | 0.536 | 8 | 2xe8 | A |
| PGK1 | 0.25 | 38 | 4axx | A |
| PGK1 | 0.214 | 33 | 2y3i | A |
| PGK1 | 0.571 | 3 | 2ybe | A |
| TUBA1A | 0.138 | 96 | 5jco | A |
| TUBA1A | 0.207 | 59 | 5jco | G |
| TUBA1A | 0.207 | 59 | 5jco | G |
| TUBA1A | 0.103 | 132 | 5jco | G |
| TUBA1A | None | None | None | None |
| PTPN11 | None | None | None | None |
| PTPN11 | 0.237 | 25 | 4ohh | A |
| PTPN11 | 0.362 | 13 | 4ohl | A |
| PTPN11 | 0.327 | 15 | 4ohi | A |
| PTPN11 | None | None | None | None |
| PTPN11 | 0.22 | 4 | 4je4 | A |
| PTPN11 | 0.22 | 4 | 4je4 | A |
| PTPN11 | 0.22 | 4 | 4je4 | A |
| PTPN11 | None | None | None | None |
| PTPN11 | 0.404 | 5 | 3tl0 | A |
| PTPN11 | 0.404 | 5 | 3tl0 | A |
| PTPN11 | 0.404 | 5 | 3tl0 | A |
| PTPN11 | 0.11 | 7 | 4jeg | A |
| PTPN11 | 0.466 | 7 | 3zm1 | A |
| PTPN11 | 0.621 | 2 | 3b7o | A |
| PTPN11 | 0.776 | 1 | 3zm1 | A |
| PTPN11 | 0.453 | 5 | 4ohd | A |
| PTPN11 | 0.216 | 29 | 3b7o | A |
| PTPN11 | 0.216 | 29 | 3b7o | A |
| PTPN11 | 0.165 | 20 | 4ohd | A |
| PTPN11 | None | None | None | None |
| PTPN11 | 0.776 | 1 | 3zm1 | A |
| PTPN11 | 0.621 | 6 | 3zm1 | A |
| PTPN11 | 0.107 | 62 | 5ibs | A |
| PTPN11 | 0.216 | 29 | 3b7o | A |
| PTPN11 | 0.621 | 6 | 3zm1 | A |
| GLB1 | 0.333 | 34 | 3wez | A |
| GLB1 | 0.333 | 67 | 3wf4 | A |
| UQCRQ | None | None | None | None |
| VPS35 | None | None | None | None |
| NR3C1 | 0.107 | 39 | 4p6x | A |
| NR3C1 | 0.135 | 62 | 5g3j | A |

| | | | | |
|---|---|---|---|---|
| NR3C1 | None | None | None | None |
| NR3C1 | 0.359 | 21 | 1nhz | A |
| NR3C1 | 0.42 | 6 | 4mdd | A |
| NR3C1 | 0.177 | 20 | 3cld | B |
| NR3C1 | None | None | None | None |
| NR3C1 | 0.116 | 61 | 4udd | A |
| NR3C1 | 0.422 | 14 | 1nhz | A |
| APOA1 | None | None | None | None |
| APOA1 | None | None | None | None |
| APOA1 | None | None | None | None |
| APOA1 | None | None | None | None |
| APOA1 | None | None | None | None |
| APOA1 | None | None | None | None |
| APOA1 | None | None | None | None |
| BCKDHA | None | None | None | None |
| BCKDHA | 0.583 | 76 | 1x7x | A |
| BCKDHA | 0.963 | 1 | 1ols | A |
| BCKDHA | 0.696 | 3 | 1v1r | A |
| BCKDHA | 0.625 | 16 | 2beu | A |
| BCKDHA | None | None | None | None |
| BCKDHA | 0.652 | 11 | 1v1r | A |
| BCKDHA | 1 | 1 | 2beu | A |
| BCKDHA | 0.84 | 2 | 2j9f | A |
| BCKDHA | 0.556 | 42 | 1ols | A |
| BCKDHA | 0.609 | 90 | 1v1r | A |
| BCKDHA | 0.652 | 25 | 1v1r | A |
| BCKDHA | 0.346 | 5 | 1wci | A |
| BCKDHA | 0.625 | 60 | 2beu | A |
| BCKDHA | 0.625 | 28 | 2beu | A |
| TOP1 | None | None | None | None |
| ANTXR2 | None | None | None | None |
| ANTXR2 | None | None | None | None |
| ANTXR2 | None | None | None | None |
| TF | 0.625 | 35 | 5dyh | A |
| TF | 0.328 | 52 | 5dyh | A |
| FSHB | None | None | None | None |
| GALK1 | None | None | None | None |
| GALK1 | None | None | None | None |
| GALK1 | None | None | None | None |
| CYP17A1 | 0.119 | 7 | 5irq | A |
| CYP17A1 | 0.353 | 2 | 4nkx | A |
| CYP17A1 | 0.353 | 2 | 4nkx | A |
| CYP17A1 | None | None | None | None |
| CYP17A1 | None | None | None | None |
| CYP17A1 | None | None | None | None |
| CYP17A1 | 0.244 | 5 | 4nkx | A |
| CYP17A1 | None | None | None | None |
| CYP17A1 | None | None | None | None |
| CYP17A1 | None | None | None | None |
| CYP17A1 | None | None | None | None |
| CYP17A1 | None | None | None | None |
| CYP17A1 | None | None | None | None |
| CYP17A1 | 0.194 | 8 | 4nkw | A |
| CYP17A1 | None | None | None | None |

| | | | | |
|---|---|---|---|---|
| CYP17A1 | None | None | None | None |
| CYP17A1 | None | None | None | None |
| CD44 | 0.667 | 24 | 1uuh | A |
| MDH2 | 0.426 | 5 | 4wlo | B |
| MDH2 | None | None | None | None |
| NDUFV1 | None | None | None | None |
| NDUFV1 | None | None | None | None |
| NDUFV1 | None | None | None | None |
| NDUFV1 | None | None | None | None |
| NDUFV1 | None | None | None | None |
| NDUFV1 | None | None | None | None |
| NDUFV1 | None | None | None | None |
| NDUFV1 | None | None | None | None |
| NDUFV1 | None | None | None | None |
| NDUFV1 | None | None | None | None |
| CTSC | None | None | None | None |
| CTSC | None | None | None | None |
| CTSC | None | None | None | None |
| CTSC | 0.342 | 13 | 3pdf | A |
| CTSC | 0.643 | 1 | 4oel | A |
| CTSC | 0.136 | 24 | 4cdc | H |
| CTSC | None | None | None | None |
| VWF | None | None | None | None |
| VWF | None | None | None | None |
| VWF | None | None | None | None |
| VWF | None | None | None | None |
| VWF | None | None | None | None |
| VWF | None | None | None | None |
| VWF | None | None | None | None |
| ORC4 | None | None | None | None |
| ORC4 | None | None | None | None |
| CRYAB | None | None | None | None |
| CRYAB | None | None | None | None |
| CRYAB | None | None | None | None |
| MAP2K2 | None | None | None | None |
| MAP2K2 | None | None | None | None |
| MAP2K2 | 0.299 | 4 | 1s9i | A |
| MAP2K2 | None | None | None | None |
| MAP2K2 | 0.272 | 4 | 1s9i | B |
| MAP2K2 | None | None | None | None |
| MAP2K2 | None | None | None | None |
| MAP2K2 | None | None | None | None |
| MAP2K2 | None | None | None | None |
| CBL | None | None | None | None |
| CBL | None | None | None | None |
| CBL | None | None | None | None |
| CBL | None | None | None | None |
| CBL | None | None | None | None |
| CBL | None | None | None | None |
| CBL | None | None | None | None |
| PFN1 | None | None | None | None |
| PFN1 | 0.75 | 2 | 1fik | A |
| ACSM2A | 0.224 | 4 | 3eq6 | A |
| HSPG2 | None | None | None | None |

| | | | | |
|---|---|---|---|---|
| HSPG2 | None | None | None | None |
| HSPG2 | None | None | None | None |
| HSPG2 | None | None | None | None |
| PARK7 | 0.444 | 48 | 3cyf | A |
| PARK7 | 0.769 | 4 | 1pe0 | A |
| PARK7 | 0.808 | 2 | 1pe0 | A |
| PARK7 | 0.667 | 8 | 4rky | A |
| PARK7 | 0.889 | 1 | 3cz9 | A |
| PARK7 | 0.75 | 2 | 1pdw | B |
| INSR | None | None | None | None |
| INSR | None | None | None | None |
| INSR | None | None | None | None |
| INSR | None | None | None | None |
| INSR | None | None | None | None |
| DSC2 | None | None | None | None |
| DSC2 | None | None | None | None |
| DSC2 | None | None | None | None |
| DSC2 | None | None | None | None |
| DSC2 | None | None | None | None |
| DSC2 | None | None | None | None |
| DSC2 | None | None | None | None |
| DSC2 | None | None | None | None |
| DSC2 | None | None | None | None |
| DSC2 | None | None | None | None |
| MICU1 | None | None | None | None |
| MICU1 | None | None | None | None |
| ADAMTS13 | None | None | None | None |
| ADAMTS13 | None | None | None | None |
| ADAMTS13 | None | None | None | None |
| ADAMTS13 | None | None | None | None |
| ADAMTS13 | None | None | None | None |
| ADAMTS13 | None | None | None | None |
| ADAMTS13 | None | None | None | None |
| IRAK4 | None | None | None | None |
| IRAK4 | None | None | None | None |
| RNASEH2B | None | None | None | None |
| RNASEH2B | None | None | None | None |
| RNASEH2B | None | None | None | None |
| NTRK1 | None | None | None | None |
| MPO | 0.75 | 3 | 1d5l | C |
| MPO | 0.6 | 25 | 1myp | D |
| MPO | 0.25 | 86 | 1d5l | C |
| PEX3 | None | None | None | None |
| SERPINA1 | None | None | None | None |
| SERPINA1 | None | None | None | None |
| SERPINA1 | None | None | None | None |
| SERPINA1 | 0.113 | 3 | 1oo8 | A |
| SERPINA1 | None | None | None | None |
| SERPINA1 | None | None | None | None |
| SERPINA1 | 0.185 | 3 | 1qmb | A |
| SERPINA1 | 0.6 | 1 | 1ezx | A |
| SERPINA1 | 0.113 | 3 | 1oo8 | A |
| SERPINA1 | None | None | None | None |
| YARS2 | None | None | None | None |

| | | | | | |
|---|---|---|---|---|---|
| YARS2 | None | None | None | None | |
| YARS2 | None | None | None | None | |
| YARS2 | None | None | None | None | |
| YARS2 | None | None | None | None | |
| YARS2 | 0.781 | | 1 | 2pid | B |
| YARS2 | None | None | None | None | |
| YARS2 | None | None | None | None | |
| YARS2 | None | None | None | None | |
| PDHA1 | None | None | None | None | |
| PDHA1 | None | None | None | None | |
| PDHA1 | None | None | None | None | |
| PDHA1 | None | None | None | None | |
| PDHA1 | None | None | None | None | |
| PDHA1 | None | None | None | None | |
| PDHA1 | None | None | None | None | |
| PDHA1 | 0.96 | | 1 | 3exh | C |
| PDHA1 | 0.3 | | 5 | 1ni4 | A |
| PDHA1 | None | None | None | None | |
| PDHA1 | None | None | None | None | |
| CUL4B | None | None | None | None | |
| AGXT | None | None | None | None | |
| AGXT | None | None | None | None | |
| AGXT | None | None | None | None | |
| AGXT | 0.692 | | 3 | 2yob | A |
| AGXT | 0.786 | | 7 | 4kxk | C |
| AGXT | 0.538 | | 21 | 3r9a | A |
| AGXT | 0.353 | | 33 | 3r9a | C |
| AGXT | None | None | None | None | |
| AGXT | None | None | None | None | |
| AGXT | 0.611 | | 4 | 5hhy | B |
| AGXT | 0.333 | | 40 | 4cbs | A |
| AGXT | 0.333 | | 59 | 5f9s | A |
| AGXT | 0.308 | | 45 | 3r9a | A |
| AGXT | 0.294 | | 55 | 3r9a | C |
| AGXT | 0.429 | | 28 | 4kxk | C |
| AGXT | 0.684 | | 1 | 1h0c | A |
| AGXT | 0.5 | | 2 | 4cbr | A |
| AGXT | 0.538 | | 19 | 4kyo | A |
| AGXT | 0.538 | | 19 | 4kyo | A |
| AGXT | 0.75 | | 2 | 5luc | B |
| AGXT | 0.722 | | 3 | 5hhy | B |
| AGXT | 0.571 | | 3 | 4kxk | C |
| AGXT | 0.571 | | 5 | 4kxk | C |
| AGXT | None | None | None | None | |
| AGXT | 0.778 | | 1 | 5hhy | B |
| AGXT | 0.778 | | 1 | 5hhy | B |
| AGXT | 0.421 | | 12 | 1h0c | A |
| AGXT | 0.692 | | 7 | 4kyo | A |
| AGXT | 0.5 | | 18 | 4cbr | A |
| AGXT | 0.5 | | 18 | 4cbr | A |
| AGXT | 0.615 | | 13 | 3r9a | A |
| AGXT | 0.786 | | 7 | 4kxk | C |
| AGXT | 0.846 | | 1 | 4kxk | A |
| AGXT | 0.615 | | 36 | 3r9a | A |

| | | | | | | |
|---|---|---|---|---|---|---|
| AGXT | | 0.571 | 10 | 4kxk | | C |
| SFTPD | | 0.509 | 2 | 3g83 | | B |
| SFTPD | | 0.111 | 27 | | 4E+052 | B |
| PIK3CA | None | None | | None | | None |
| PIK3CA | None | None | | None | | None |
| PIK3CA | None | None | | None | | None |
| AUH | | 0.22 | 3 | 1hzd | | E |
| AUH | None | None | | None | | None |
| AUH | None | None | | None | | None |
| AUH | | 0.293 | 16 | 2zqr | | D |
| F11 | None | None | | None | | None |
| F11 | None | None | | None | | None |
| F11 | None | None | | None | | None |
| F11 | None | None | | None | | None |
| MEN1 | | 0.6 | 81 | 5db0 | | A |
| MEN1 | | 0.8 | 54 | 4i80 | | A |
| MEN1 | | 0.8 | 84 | 5db0 | | A |
| MEN1 | | 0.8 | 30 | 5ddb | | A |
| MEN1 | | 1 | 2 | 4i80 | | A |
| MEN1 | | 0.8 | 17 | 5db0 | | A |
| MEN1 | | 0.6 | 17 | 5ddb | | A |
| MEN1 | | 0.6 | 31 | 3u88 | | A |
| MEN1 | None | None | | None | | None |
| MEN1 | | 0.6 | 56 | 5ddb | | A |
| MYBPC1 | None | None | | None | | None |
| MYBPC1 | None | None | | None | | None |
| MYBPC1 | None | None | | None | | None |
| PDCD10 | None | None | | None | | None |
| SCARB2 | None | None | | None | | None |
| SCARB2 | None | None | | None | | None |
| SCARB2 | None | None | | None | | None |
| SCARB2 | None | None | | None | | None |
| SCARB2 | None | None | | None | | None |
| TREX2 | None | None | | None | | None |
| TBK1 | None | None | | None | | None |
| TBK1 | None | None | | None | | None |
| DLD | None | None | | None | | None |
| DLD | None | None | | None | | None |
| DLD | | 0.505 | 3 | 1zy8 | | B |
| DLD | None | None | | None | | None |
| DLD | | 0.24 | 11 | 1zy8 | | A |
| DLD | None | None | | None | | None |
| LDHA | | 0.802 | 1 | 4qt0 | | A |
| TNNC1 | None | None | | None | | None |
| TNNC1 | | 0.461 | 2 | 1wrl | | A |
| TNNC1 | | 0.153 | 5 | 1mxl | | C |
| SMAD3 | None | None | | None | | None |
| HARS | None | None | | None | | None |
| HARS | None | None | | None | | None |
| TNNI3 | None | None | | None | | None |
| TNNI3 | None | None | | None | | None |
| TNNI3 | None | None | | None | | None |
| TNNI3 | None | None | | None | | None |
| TNNI3 | None | None | | None | | None |

| | | | | |
|---|---|---|---|---|
| TNNI3 | None | None | None | None |
| ACKR1 | None | None | None | None |
| BUB1B | None | None | None | None |
| BUB1B | None | None | None | None |
| BUB1B | None | None | None | None |
| AR | 0.522 | 6 | 1xj7 | A |
| AR | 0.273 | 23 | 2ao6 | A |
| AR | 0.305 | 55 | 4olm | A |
| AR | 0.305 | 55 | 4olm | A |
| AR | 0.131 | 48 | 2ax6 | A |
| FTO | None | None | None | None |
| FTO | 0.333 | 1 | 4ie6 | A |
| TFAM | None | None | None | None |
| PYGL | None | None | None | None |
| PYGL | 0.778 | 3 | 1fa9 | A |
| PYGL | None | None | None | None |
| PYGL | 0.722 | 2 | 1fa9 | A |
| PYGL | 0.5 | 19 | 3dds | A |
| PYGL | 0.611 | 25 | 3ceh | B |
| PYGL | 0.611 | 32 | 1l7x | B |
| PYGL | 0.167 | 132 | 1em6 | B |
| PYGL | 0.722 | 2 | 1fc0 | B |
| PYGL | 0.778 | 3 | 1fa9 | A |
| PYGL | 0.278 | 167 | 1l7x | B |
| PYGL | 0.722 | 12 | 2ati | A |
| PYGL | 0.722 | 1 | 3ceh | B |
| PYGL | 0.611 | 11 | 3ceh | B |
| PYGL | 0.611 | 18 | 1fc0 | B |
| PYGL | 0.722 | 2 | 1fc0 | B |
| GALNS | None | None | None | None |
| GALNS | None | None | None | None |
| GALNS | None | None | None | None |
| GALNS | None | None | None | None |
| GALNS | None | None | None | None |
| GALNS | None | None | None | None |
| GALNS | None | None | None | None |
| GALNS | None | None | None | None |
| GALNS | None | None | None | None |
| GALNS | None | None | None | None |
| GALNS | None | None | None | None |
| GALNS | None | None | None | None |
| GALNS | None | None | None | None |
| GALNS | None | None | None | None |
| GALNS | None | None | None | None |
| GALNS | None | None | None | None |
| GALNS | None | None | None | None |
| GALNS | None | None | None | None |
| GALNS | None | None | None | None |
| GALNS | None | None | None | None |
| GALNS | None | None | None | None |
| GALNS | None | None | None | None |
| GALNS | None | None | None | None |
| GALNS | None | None | None | None |
| GALNS | None | None | None | None |

| | | | | | |
|---|---|---|---|---|---|
| GALNS | None | None | | None | None |
| GALNS | None | None | | None | None |
| GALNS | None | None | | None | None |
| GALNS | None | None | | None | None |
| GALNS | None | None | | None | None |
| GALNS | None | None | | None | None |
| GALNS | None | None | | None | None |
| GALNS | None | None | | None | None |
| F10 | 0.304 | | 21 | 1z6e | L |
| GLUD1 | None | None | | None | None |
| GLUD1 | None | None | | None | None |
| GLUD1 | None | None | | None | None |
| GLUD1 | None | None | | None | None |
| GLUD1 | None | None | | None | None |
| GLUD1 | None | None | | None | None |
| GLUD1 | None | None | | None | None |
| GLUD1 | None | None | | None | None |
| GLUD1 | None | None | | None | None |
| GLUD1 | None | None | | None | None |
| NOTCH1 | 0.528 | | 3 | 3v79 | K |
| POGLUT1 | None | None | | None | None |
| POGLUT1 | None | None | | None | None |
| NR3C1 | 0.455 | | 1 | 4hn5 | B |
| CASP8 | 0.535 | | 6 | 1f9e | C |
| CASP8 | 0.592 | | 8 | 1qdu | C |
| TRAP1 | None | None | | None | None |
| FANCM | None | None | | None | None |
| EPOR | None | None | | None | None |
| EPOR | None | None | | None | None |
| NDP | None | None | | None | None |
| NDP | None | None | | None | None |
| NDP | None | None | | None | None |
| NDP | None | None | | None | None |
| NDP | None | None | | None | None |
| NDP | None | None | | None | None |
| NDP | None | None | | None | None |
| NDP | None | None | | None | None |
| NDP | 0.238 | | 1 | 5bqc | A |
| NDP | 0.238 | | 1 | 5bqc | A |
| NDP | None | None | | None | None |
| NDP | None | None | | None | None |
| NDP | None | None | | None | None |
| NDP | None | None | | None | None |
| NDP | None | None | | None | None |
| NDP | None | None | | None | None |
| NDP | None | None | | None | None |
| ROGDI | None | None | | None | None |
| CYP19A1 | None | None | | None | None |
| CYP19A1 | None | None | | None | None |
| CYP19A1 | None | None | | None | None |
| CYP19A1 | None | None | | None | None |
| CYP19A1 | None | None | | None | None |
| CYP19A1 | 0.487 | | 1 | 3eqm | A |
| CYP19A1 | 0.487 | | 1 | 3eqm | A |

| | | | | | |
|---|---|---|---|---|---|
| CYP19A1 | | 0.257 | 5 | 4kq8 | A |
| UBA5 | None | None | | None | None |
| PRPS1 | None | None | | None | None |
| PRPS1 | | 0.316 | 13 | 4f8e | A |
| PRPS1 | None | None | | None | None |
| PRPS1 | None | None | | None | None |
| PRPS1 | None | None | | None | None |
| PRPS1 | None | None | | None | None |
| PRPS1 | None | None | | None | None |
| PRPS1 | None | None | | None | None |
| F2 | None | None | | None | None |
| F2 | None | None | | None | None |
| NTF4 | None | None | | None | None |
| NTF4 | None | None | | None | None |
| PSAP | None | None | | None | None |
| PSAP | None | None | | None | None |
| TAP1 | None | None | | None | None |
| TAP1 | None | None | | None | None |
| TAP1 | None | None | | None | None |
| ARG1 | None | None | | None | None |
| ARG1 | | 0.714 | 1 | 2pha | A |
| ARG1 | None | None | | None | None |
| ARG1 | | 0.471 | 10 | 3th7 | A |
| ARG1 | | 0.184 | 52 | 4gsz | A |
| ARG1 | | 0.184 | 52 | 4gsz | A |
| CRYM | None | None | | None | None |
| ADSL | None | None | | None | None |
| ADSL | None | None | | None | None |
| ADSL | None | None | | None | None |
| ADSL | None | None | | None | None |
| ADSL | None | None | | None | None |
| ADSL | None | None | | None | None |
| ADSL | | 0.312 | 8 | 2vd6 | B |
| ADSL | | 0.312 | 8 | 2vd6 | B |
| ADSL | None | None | | None | None |
| ADSL | | 0.353 | 5 | 2vd6 | D |
| ADSL | None | None | | None | None |
| ADSL | | 0.294 | 11 | 2vd6 | D |
| ADSL | | 0.647 | 1 | 2vd6 | D |
| ADSL | | 0.435 | 1 | 4ffx | A |
| ADSL | None | None | | None | None |
| ADSL | None | None | | None | None |
| ADSL | None | None | | None | None |
| ADSL | None | None | | None | None |
| ADSL | None | None | | None | None |
| ADSL | None | None | | None | None |
| VWF | None | None | | None | None |
| PLA2G7 | None | None | | None | None |
| MMP13 | None | None | | None | None |
| MMP13 | None | None | | None | None |
| MMP13 | None | None | | None | None |
| MMP13 | None | None | | None | None |
| POMGNT1 | None | None | | None | None |
| POMGNT1 | None | None | | None | None |

| | | | | |
|---|---|---|---|---|
| COL3A1 | None | None | None | None |
| COL3A1 | None | None | None | None |
| COL3A1 | None | None | None | None |
| COL3A1 | None | None | None | None |
| ACTN2 | None | None | None | None |
| ACTN2 | None | None | None | None |
| ACTN2 | None | None | None | None |
| ACTN2 | None | None | None | None |
| ACTN2 | None | None | None | None |
| ACTN2 | None | None | None | None |
| ACTN2 | None | None | None | None |
| UBA5 | None | None | None | None |
| ADAR | 0.75 | 1 | 1qbj | B |
| UMPS | None | None | None | None |
| UMPS | None | None | None | None |
| UMPS | None | None | None | None |
| TRNT1 | None | None | None | None |
| TRNT1 | None | None | None | None |
| NHEJ1 | None | None | None | None |
| NHEJ1 | None | None | None | None |
| NHEJ1 | None | None | None | None |
| C8A | None | None | None | None |
| GLUL | None | None | None | None |
| GLUL | 0.375 | 6 | 2qc8 | C |
| GLUL | 0.467 | 9 | 2qc8 | B |
| GYG1 | 0.615 | 45 | 3rmv | A |
| GYG1 | 0.923 | 1 | 3u2u | B |
| FLNA | None | None | None | None |
| FLNA | None | None | None | None |
| FLNA | None | None | None | None |
| FLNA | None | None | None | None |
| FLNA | None | None | None | None |
| FLNA | None | None | None | None |
| FLNA | None | None | None | None |
| TTN | None | None | None | None |
| TTN | None | None | None | None |
| TTN | None | None | None | None |
| TTN | None | None | None | None |
| SMS | None | None | None | None |
| SMS | None | None | None | None |
| SMS | None | None | None | None |
| C5 | None | None | None | None |
| SERPINC1 | None | None | None | None |
| SERPINC1 | None | None | None | None |
| SERPINC1 | None | None | None | None |
| SERPINC1 | None | None | None | None |
| SERPINC1 | None | None | None | None |
| SERPINC1 | None | None | None | None |
| SERPINC1 | None | None | None | None |
| SERPINC1 | None | None | None | None |
| SERPINC1 | None | None | None | None |
| SERPINC1 | 0.273 | 3 | 1ant | I |
| SERPINC1 | 0.262 | 2 | 2znh | B |
| SERPINC1 | None | None | None | None |

| | | | | |
|---|---|---|---|---|
| SERPINC1 | 0.136 | 8 | 2b5t | I |
| SERPINC1 | 0.157 | 7 | 3evj | I |
| SERPINC1 | None | None | None | None |
| SERPINC1 | None | None | None | None |
| SERPINC1 | None | None | None | None |
| SERPINC1 | None | None | None | None |
| SERPINC1 | None | None | None | None |
| SERPINC1 | None | None | None | None |
| SERPINC1 | None | None | None | None |
| ETFB | None | None | None | None |
| ETFB | None | None | None | None |
| ETFB | None | None | None | None |
| ETFB | None | None | None | None |
| ETFB | None | None | None | None |
| ETFB | None | None | None | None |
| DIABLO | None | None | None | None |
| DIABLO | None | None | None | None |
| DPYS | 0.333 | 24 | 2vr2 | A |
| DPYS | None | None | None | None |
| DPYS | None | None | None | None |
| DPYS | 0.533 | 7 | 2vr2 | A |
| DPYS | None | None | None | None |
| GNE | None | None | None | None |
| GNE | None | None | None | None |
| GNE | None | None | None | None |
| GNE | None | None | None | None |
| GNE | None | None | None | None |
| GNE | None | None | None | None |
| GNE | None | None | None | None |
| GNE | None | None | None | None |
| GNE | None | None | None | None |
| GNMT | None | None | None | None |
| GNMT | 0.2 | 12 | 2azt | B |
| CTH | 0.324 | 20 | 3cog | D |
| CTH | 0.214 | 15 | 2nmp | B |
| CTH | None | None | None | None |
| USH1C | None | None | None | None |
| PSEN1 | None | None | None | None |
| CHEK2 | None | None | None | None |
| CHEK2 | None | None | None | None |
| CHEK2 | None | None | None | None |
| CHEK2 | None | None | None | None |
| CHEK2 | None | None | None | None |
| CHEK2 | None | None | None | None |
| CHEK2 | None | None | None | None |
| CHEK2 | None | None | None | None |
| CHEK2 | None | None | None | None |
| SNCA | None | None | None | None |
| SNCA | None | None | None | None |
| GAN | None | None | None | None |
| GAN | None | None | None | None |
| GAN | None | None | None | None |
| SBDS | None | None | None | None |
| SBDS | None | None | None | None |

| | | | | |
|---|---|---|---|---|
| SBDS | None | None | None | None |
| SBDS | None | None | None | None |
| SBDS | None | None | None | None |
| SBDS | None | None | None | None |
| MPO | 1 | 3 | 3f9p | A |
| MPO | 0.6 | 12 | 5mfa | A |
| KCNH2 | 0.292 | 2 | 4hp9 | A |
| KCNH2 | None | None | None | None |
| KCNH2 | 0.75 | 1 | 4hp9 | A |
| KCNH2 | None | None | None | None |
| TEK | None | None | None | None |
| TEK | None | None | None | None |
| TEK | None | None | None | None |
| TEK | None | None | None | None |
| NPC2 | None | None | None | None |
| NPC2 | None | None | None | None |
| NPC2 | None | None | None | None |
| NPC2 | None | None | None | None |
| NPC2 | None | None | None | None |
| NPC2 | None | None | None | None |
| NPC2 | None | None | None | None |
| PLG | None | None | None | None |
| PCSK9 | None | None | None | None |
| SFTPC | None | None | None | None |
| SFTPC | None | None | None | None |
| SFTPC | None | None | None | None |
| CD40LG | None | None | None | None |
| CD40LG | None | None | None | None |
| CD40LG | None | None | None | None |
| CD40LG | None | None | None | None |
| CD40LG | None | None | None | None |
| TAT | None | None | None | None |
| TAT | None | None | None | None |
| TAT | None | None | None | None |
| EP300 | None | None | None | None |
| FARS2 | None | None | None | None |
| FARS2 | None | None | None | None |
| FARS2 | None | None | None | None |
| FARS2 | None | None | None | None |
| FARS2 | None | None | None | None |
| FARS2 | None | None | None | None |
| FARS2 | None | None | None | None |
| FARS2 | None | None | None | None |
| FARS2 | 0.5 | 1 | 5mgw | A |
| FARS2 | None | None | None | None |
| FARS2 | None | None | None | None |
| FARS2 | None | None | None | None |
| FARS2 | None | None | None | None |
| MASP2 | None | None | None | None |
| MASP2 | None | None | None | None |
| MASP2 | None | None | None | None |
| MASP2 | None | None | None | None |
| STIM1 | None | None | None | None |
| CHRM2 | None | None | None | None |

| | | | | |
|---|---|---|---|---|
| MCFD2 | 0.206 | 9 | 3whu | B |
| MCFD2 | 0.574 | 1 | 4yge | B |
| MCFD2 | 0.623 | 1 | 3wnx | B |
| KITLG | None | None | None | None |
| F12 | 0.345 | 1 | 4bdx | A |
| F12 | 0.345 | 1 | 4bdx | A |
| ERAP1 | 0.381 | 5 | 3mdj | A |
| GTF2H5 | None | None | None | None |
| GTF2H5 | None | None | None | None |
| THBD | None | None | None | None |
| MRPL28 | None | None | None | None |
| MRPL28 | None | None | None | None |
| HTRA1 | None | None | None | None |
| HTRA1 | None | None | None | None |
| HTRA1 | None | None | None | None |
| HTRA1 | None | None | None | None |
| HTRA1 | 0.237 | 1 | 3nwu | B |
| HTRA1 | None | None | None | None |
| KARS | None | None | None | None |
| KARS | None | None | None | None |
| KARS | None | None | None | None |
| KARS | None | None | None | None |
| KARS | None | None | None | None |
| HK1 | 0.722 | 11 | 4fpb | B |
| GARS | None | None | None | None |
| GARS | None | None | None | None |
| GARS | None | None | None | None |
| GARS | None | None | None | None |
| GARS | None | None | None | None |
| GARS | None | None | None | None |
| GARS | None | None | None | None |
| IL12B | None | None | None | None |
| HADH | None | None | None | None |
| HADH | None | None | None | None |
| HADH | 0.52 | 4 | 1lsj | A |
| HADH | 0.458 | 3 | 1m75 | A |
| HADH | 0.52 | 4 | 1lsj | A |
| RRM2B | None | None | None | None |
| RRM2B | None | None | None | None |
| RRM2B | None | None | None | None |
| RRM2B | None | None | None | None |
| RRM2B | 0.28 | 4 | 4djn | B |
| RRM2B | None | None | None | None |
| RRM2B | 0.333 | 2 | 3hf1 | B |
| RRM2B | None | None | None | None |
| RRM2B | None | None | None | None |
| RRM2B | 0.286 | 8 | 2vux | A |
| RRM2B | None | None | None | None |
| RRM2B | None | None | None | None |
| RRM2B | None | None | None | None |
| RRM2B | None | None | None | None |
| RRM2B | None | None | None | None |
| MARVELD2 | None | None | None | None |
| TLR4 | None | None | None | None |

| | | | | |
|---|---|---|---|---|
| TTN | 0.146 | 1 | 3q5o | A |
| TTN | None | None | None | None |
| TNFRSF13B | None | None | None | None |
| POLG | None | None | None | None |
| POLG | None | None | None | None |
| POLG | None | None | None | None |
| POLG | None | None | None | None |
| POLG | None | None | None | None |
| POLG | None | None | None | None |
| POLG | None | None | None | None |
| POLG | None | None | None | None |
| POLG | None | None | None | None |
| POLG | None | None | None | None |
| POLG | None | None | None | None |
| POLG | None | None | None | None |
| POLG | None | None | None | None |
| POLG | None | None | None | None |
| POLG | None | None | None | None |
| POLG | None | None | None | None |
| POLG | None | None | None | None |
| POLG | None | None | None | None |
| POLG | None | None | None | None |
| POLG | None | None | None | None |
| POLG | None | None | None | None |
| POLG | None | None | None | None |
| POLG | None | None | None | None |
| POLG | None | None | None | None |
| POLG | None | None | None | None |
| POLG | None | None | None | None |
| POLG | None | None | None | None |
| POLG | 0.467 | 4 | 5c53 | A |
| POLG | 0.533 | 1 | 4ztu | A |
| POLG | None | None | None | None |
| POLG | None | None | None | None |
| POLG | None | None | None | None |
| POLG | None | None | None | None |
| POLG | None | None | None | None |
| POLG | 0.533 | 1 | 4ztu | A |
| POLG | 0.2 | 8 | 4ztu | A |
| POLG | 0.2 | 8 | 4ztu | A |
| POLG | None | None | None | None |
| POLG | None | None | None | None |
| POLG | None | None | None | None |
| POLG | None | None | None | None |
| POLG | None | None | None | None |
| POLG | None | None | None | None |
| POLG | None | None | None | None |
| POLG | None | None | None | None |
| POLG | None | None | None | None |
| POLG | 0.267 | 6 | 4ztu | A |
| POLG | None | None | None | None |
| POLG | None | None | None | None |
| UROD | 0.632 | 24 | 1jpk | A |

| | | | | |
|---|---|---|---|---|
| UROD | 0.579 | 33 | 1r3v | A |
| UROD | None | None | None | None |
| UROD | 0.526 | 49 | 1jph | A |
| UROD | None | None | None | None |
| UROD | 0.632 | 13 | 3gvw | A |
| UROD | 0.789 | 6 | 3gvw | A |
| UROD | 0.842 | 2 | 3gvv | A |
| UROD | 0.737 | 31 | 1r3q | A |
| UROD | 0.667 | 14 | 2q71 | A |
| PCK1 | 0.429 | 154 | 1khg | A |
| PCK1 | None | None | None | None |
| PCK1 | None | None | None | None |
| PCK1 | 0.167 | 187 | 1khf | A |
| PCK1 | 0.524 | 60 | 1nhx | A |
| HEXA | None | None | None | None |
| HEXA | None | None | None | None |
| HEXA | None | None | None | None |
| HEXA | None | None | None | None |
| HEXA | None | None | None | None |
| HEXA | 0.364 | 2 | 2gjx | A |
| HEXA | 0.364 | 2 | 2gjx | A |
| HEXA | None | None | None | None |
| HEXA | None | None | None | None |
| HEXA | None | None | None | None |
| HEXA | None | None | None | None |
| HEXA | 0.364 | 2 | 2gjx | A |
| HEXA | None | None | None | None |
| HEXA | None | None | None | None |
| HEXA | None | None | None | None |
| HEXA | 0.455 | 1 | 2gjx | A |
| HEXA | None | None | None | None |
| HEXA | None | None | None | None |
| HEXA | None | None | None | None |
| HEXA | None | None | None | None |
| HEXA | None | None | None | None |
| HEXA | None | None | None | None |
| HEXA | None | None | None | None |
| HEXA | None | None | None | None |
| HEXA | 0.273 | 6 | 2gjx | A |
| HEXA | None | None | None | None |
| HEXA | None | None | None | None |
| HEXA | 0.455 | 3 | 2gjx | A |
| HEXA | 0.455 | 3 | 2gjx | A |
| HEXA | None | None | None | None |
| HEXA | None | None | None | None |
| CFTR | None | None | None | None |
| CFTR | None | None | None | None |
| CFTR | None | None | None | None |
| CFTR | None | None | None | None |
| CFTR | None | None | None | None |
| CFTR | None | None | None | None |
| CFTR | None | None | None | None |
| CFTR | None | None | None | None |
| CFTR | None | None | None | None |

| | | | | | |
|---|---|---|---|---|---|
| CFTR | None | None | None | None | |
| CFTR | 0.462 | | 1 | 2pzg | A |
| CFTR | 0.321 | | 1 | 1xmj | A |
| CFTR | 0.462 | | 2 | 2pzg | A |
| CFTR | None | None | None | None | |
| CFTR | None | None | None | None | |
| CFTR | None | None | None | None | |
| CFTR | None | None | None | None | |
| CFTR | None | None | None | None | |
| CFTR | None | None | None | None | |
| CFTR | None | None | None | None | |
| CFTR | None | None | None | None | |
| CFTR | None | None | None | None | |
| CFTR | None | None | None | None | |
| CFTR | None | None | None | None | |
| CFTR | None | None | None | None | |
| CFTR | 0.462 | | 1 | 2pzg | A |
| CFTR | None | None | None | None | |
| CFTR | 0.462 | | 2 | 2pzg | A |
| CFTR | None | None | None | None | |
| CFTR | None | None | None | None | |
| CFTR | None | None | None | None | |
| CFTR | 0.346 | | 3 | 2pzg | A |
| CFTR | None | None | None | None | |
| CFTR | None | None | None | None | |
| FAN1 | None | None | None | None | |
| CBS | None | None | None | None | |
| CBS | None | None | None | None | |
| CBS | None | None | None | None | |
| CBS | 0.111 | | 24 | 5mms | A |
| CBS | 0.406 | | 6 | 4pcu | B |
| CBS | None | None | None | None | |
| CBS | 0.469 | | 6 | 4pcu | A |
| CBS | 0.597 | | 3 | 5mms | C |
| CBS | 0.54 | | 2 | 5mms | A |
| CBS | 0.54 | | 2 | 5mms | A |
| CBS | 0.328 | | 9 | 4pcu | A |
| CBS | None | None | None | None | |
| CBS | 0.179 | | 11 | 4l0d | A |
| CBS | 0.208 | | 20 | 1m54 | B |
| CBS | 0.516 | | 4 | 5mms | C |
| CBS | None | None | None | None | |
| CBS | None | None | None | None | |
| CBS | 0.103 | | 21 | 4l28 | C |
| CBS | 0.698 | | 2 | 1m54 | B |
| CBS | 0.122 | | 11 | 4l0d | B |
| CBS | None | None | None | None | |
| CBS | 0.19 | | 12 | 4coo | A |
| CBS | None | None | None | None | |
| CBS | None | None | None | None | |
| CBS | None | None | None | None | |
| CBS | None | None | None | None | |
| CBS | 0.424 | | 6 | 4coo | B |
| CBS | 0.143 | | 14 | 1jbq | B |

| | | | | |
|---|---|---|---|---|
| CBS | 0.603 | 3 | 4coo | A |
| CBS | 0.603 | 3 | 4coo | A |
| CBS | None | None | None | None |
| CBS | None | None | None | None |
| CBS | None | None | None | None |
| CBS | None | None | None | None |
| CBS | None | None | None | None |
| CBS | None | None | None | None |
| TLR3 | None | None | None | None |
| LIG4 | None | None | None | None |
| LIG4 | None | None | None | None |
| ASPA | 0.769 | 1 | 2i3c | B |
| ASPA | 0.769 | 1 | 2i3c | B |
| ASPA | None | None | None | None |
| ASPA | None | None | None | None |
| ASPA | None | None | None | None |
| ASPA | None | None | None | None |
| ASPA | None | None | None | None |
| ASPA | None | None | None | None |
| ASPA | None | None | None | None |
| ASPA | None | None | None | None |
| ASPA | 0.583 | 2 | 4tnu | A |
| ASPA | 0.583 | 2 | 4tnu | A |
| ASPA | None | None | None | None |
| CFH | None | None | None | None |
| FCGR3A | None | None | None | None |
| FCGR3A | None | None | None | None |
| GRIN2A | None | None | None | None |
| GRIN2A | 0.658 | 1 | 5i2n | A |
| GRIN2A | None | None | None | None |
| GLA | 0.467 | 10 | 1r46 | B |
| GLA | 0.375 | 19 | 3hg3 | B |
| GLA | 0.375 | 14 | 3hg5 | B |
| GLA | 0.312 | 13 | 3tv8 | B |
| GLA | 0.375 | 14 | 3hg5 | B |
| GLA | 0.375 | 14 | 3hg5 | B |
| GLA | 0.267 | 22 | 3lxc | B |
| GLA | 0.647 | 2 | 3s5y | A |
| GLA | 0.562 | 20 | 3gxt | B |
| GLA | 0.353 | 21 | 3s5y | A |
| GLA | None | None | None | None |
| GLA | 0.75 | 3 | 3gxn | B |
| GLA | 0.75 | 3 | 3gxn | B |
| GLA | 0.438 | 7 | 1r47 | B |
| GLA | 0.375 | 3 | 3hg4 | B |
| GLA | 0.375 | 11 | 3hg2 | A |
| GLA | 0.188 | 40 | 3lxb | B |
| GLA | 0.25 | 42 | 3hg5 | A |
| GLA | 0.25 | 42 | 3hg5 | A |
| GLA | 0.312 | 13 | 3hg2 | B |
| GLA | 0.375 | 19 | 3hg3 | B |
| GLA | 0.4 | 5 | 3lxa | A |
| GLA | 0.214 | 50 | 3lxb | A |
| GLA | 0.214 | 50 | 3lxb | A |

| | | | | |
|---|---|---|---|---|
| GLA | 0.214 | 61 | 3lxb | A |
| GLA | 0.467 | 22 | 3gxn | A |
| GLA | 0.214 | 40 | 3lxb | A |
| GLA | 0.214 | 40 | 3lxb | A |
| GLA | 0.2 | 58 | 3lxc | B |
| GLA | 0.214 | 69 | 3lxb | A |
| GLA | 0.286 | 36 | 3lxb | A |
| SERPINA3 | None | None | None | None |
| MTAP | None | None | None | None |
| MTAP | 0.462 | 25 | 3oze | B |
| MTAP | None | None | None | None |
| BLMH | None | None | None | None |
| TUSC3 | None | None | None | None |
| ISCU | None | None | None | None |
| CYP11B2 | None | None | None | None |
| CYP11B2 | None | None | None | None |
| CYP11B2 | None | None | None | None |
| CYP11B2 | None | None | None | None |
| CYP11B2 | None | None | None | None |
| CYP11B2 | None | None | None | None |
| CYP11B2 | None | None | None | None |
| CYP11B2 | None | None | None | None |
| CYP11B2 | None | None | None | None |
| CYP11B2 | None | None | None | None |
| CYP11B2 | None | None | None | None |
| CYP11B2 | None | None | None | None |
| CYP11B2 | None | None | None | None |
| CYP11B2 | None | None | None | None |
| CYP11B2 | None | None | None | None |
| CYP11B2 | None | None | None | None |
| CYP11B2 | None | None | None | None |
| CYP11B2 | None | None | None | None |
| CYP11B2 | None | None | None | None |
| CRADD | None | None | None | None |
| CRADD | None | None | None | None |
| FGA | None | None | None | None |
| FGA | None | None | None | None |
| FGA | None | None | None | None |
| FGA | None | None | None | None |
| FGA | None | None | None | None |
| FGA | None | None | None | None |
| FGA | None | None | None | None |
| FGA | None | None | None | None |
| IYD | 1 | 1 | 4ttb | A |
| IYD | None | None | None | None |
| IYD | None | None | None | None |
| IYD | None | None | None | None |
| TCAP | None | None | None | None |
| TCAP | None | None | None | None |
| TCAP | None | None | None | None |
| TCAP | None | None | None | None |
| TCAP | None | None | None | None |
| CYP7A1 | None | None | None | None |

| | | | | |
|---|---|---|---|---|
| TEK | None | None | None | None |
| TEK | None | None | None | None |
| TEK | None | None | None | None |
| TEK | None | None | None | None |
| TEK | None | None | None | None |
| TEK | None | None | None | None |
| TEK | None | None | None | None |
| CD8A | None | None | None | None |
| HERC2 | None | None | None | None |
| NR1H3 | 0.537 | 3 | 3ipq | A |
| GRHPR | None | None | None | None |
| GRHPR | None | None | None | None |
| GRHPR | 0.458 | 2 | 2q50 | A |
| GRHPR | 0.458 | 2 | 2q50 | A |
| GRHPR | 0.697 | 1 | 2wwr | D |
| GRHPR | 0.697 | 1 | 2wwr | D |
| GRHPR | None | None | None | None |
| GBE1 | None | None | None | None |
| GBE1 | None | None | None | None |
| GBE1 | None | None | None | None |
| GBE1 | None | None | None | None |
| GBE1 | None | None | None | None |
| GBE1 | None | None | None | None |
| GBE1 | None | None | None | None |
| GBE1 | None | None | None | None |
| GBE1 | None | None | None | None |
| GBE1 | None | None | None | None |
| GBE1 | None | None | None | None |
| GBE1 | None | None | None | None |
| GBE1 | None | None | None | None |
| GBE1 | None | None | None | None |
| GBE1 | None | None | None | None |
| GBE1 | None | None | None | None |
| GBE1 | None | None | None | None |
| GBE1 | None | None | None | None |
| GBE1 | None | None | None | None |
| PC | None | None | None | None |
| PC | 0.429 | 1 | 3bg3 | A |
| PC | None | None | None | None |
| PC | 0.4 | 2 | 3bg9 | B |
| PC | None | None | None | None |
| PC | None | None | None | None |
| PC | 0.4 | 6 | 3bg3 | B |
| PC | None | None | None | None |
| MSH6 | None | None | None | None |
| MSH6 | None | None | None | None |
| MSH6 | None | None | None | None |
| MSH6 | None | None | None | None |
| MSH6 | None | None | None | None |
| MSH6 | None | None | None | None |
| MSH6 | None | None | None | None |
| MSH6 | None | None | None | None |
| MSH6 | None | None | None | None |

| | | | | | |
|---|---|---|---|---|---|
| MSH6 | None | None | None | | None |
| MSH6 | None | None | None | | None |
| MSH6 | None | None | None | | None |
| MSH6 | None | None | None | | None |
| MSH6 | None | None | None | | None |
| MSH6 | None | None | None | | None |
| MSH6 | None | None | None | | None |
| MSH6 | None | None | None | | None |
| MSH6 | None | None | None | | None |
| MSH6 | None | None | None | | None |
| MSH6 | None | None | None | | None |
| MSH6 | None | None | None | | None |
| MSH6 | None | None | None | | None |
| MSH6 | None | None | None | | None |
| MSH6 | None | None | None | | None |
| MSH6 | None | None | None | | None |
| MSH6 | None | None | None | | None |
| MSH6 | None | None | None | | None |
| MSH6 | None | None | None | | None |
| MSH6 | None | None | None | | None |
| MSH6 | None | None | None | | None |
| RNASEL | 0.32 | | 6 | 4g8l | A |
| ORC1 | None | None | None | | None |
| ORC1 | None | None | None | | None |
| ORC1 | None | None | None | | None |
| KCNH2 | None | None | None | | None |
| KCNH2 | None | None | None | | None |
| KCNH2 | None | None | None | | None |
| KCNH2 | None | None | None | | None |
| KCNH2 | None | None | None | | None |
| KCNH2 | None | None | None | | None |
| KCNH2 | None | None | None | | None |
| KCNH2 | None | None | None | | None |
| KCNH2 | None | None | None | | None |
| KCNH2 | None | None | None | | None |
| KCNH2 | None | None | None | | None |
| KCNH2 | None | None | None | | None |
| KCNH2 | None | None | None | | None |
| KCNH2 | None | None | None | | None |
| KCNH2 | None | None | None | | None |
| KCNH2 | None | None | None | | None |
| KCNH2 | None | None | None | | None |
| KCNH2 | None | None | None | | None |
| KCNH2 | None | None | None | | None |
| KCNH2 | None | None | None | | None |
| KCNH2 | None | None | None | | None |
| KCNH2 | None | None | None | | None |
| KCNH2 | None | None | None | | None |
| KCNH2 | None | None | None | | None |
| KCNH2 | None | None | None | | None |
| MAT1A | 0.429 | | 3 | 2obv | A |
| MAT1A | 0.429 | | 14 | 2obv | A |
| MAT1A | 0.571 | | 1 | 2obv | A |
| MAT1A | 0.571 | | 1 | 2obv | A |
| MAT1A | None | None | None | | None |

| | | | | |
|---|---|---|---|---|
| MAT1A | None | None | None | None |
| MAT1A | None | None | None | None |
| MAT1A | None | None | None | None |
| INSR | None | None | None | None |
| INSR | None | None | None | None |
| INSR | None | None | None | None |
| INSR | None | None | None | None |
| AMT | 0.545 | 5 | 1wsr | A |
| AMT | None | None | None | None |
| AMT | None | None | None | None |
| AMT | None | None | None | None |
| AMT | None | None | None | None |
| AMT | None | None | None | None |
| AMT | None | None | None | None |
| AMT | None | None | None | None |
| AMT | None | None | None | None |
| CSNK1D | None | None | None | None |
| SF3B1 | None | None | None | None |
| AGA | None | None | None | None |
| AGA | None | None | None | None |
| AGA | None | None | None | None |
| AGA | None | None | None | None |
| AGA | None | None | None | None |
| AGA | None | None | None | None |
| AGA | None | None | None | None |
| NECTIN4 | None | None | None | None |
| RS1 | None | None | None | None |
| RS1 | None | None | None | None |
| RS1 | None | None | None | None |
| RS1 | None | None | None | None |
| TBC1D7 | None | None | None | None |
| CTSA | None | None | None | None |
| CTSA | None | None | None | None |
| TKT | 0.273 | 20 | 4kxu | A |
| RP2 | None | None | None | None |
| RP2 | None | None | None | None |
| RP2 | None | None | None | None |
| RP2 | None | None | None | None |
| LYRM4 | None | None | None | None |
| SOS1 | None | None | None | None |
| SOS1 | None | None | None | None |
| SOS1 | None | None | None | None |
| SOS1 | None | None | None | None |
| SOS1 | None | None | None | None |
| SOS1 | None | None | None | None |
| HUWE1 | None | None | None | None |
| HUWE1 | None | None | None | None |
| TPI1 | 0.165 | 51 | 1wyi | B |
| TPI1 | 0.886 | 1 | 4unk | B |
| TPI1 | 0.146 | 43 | 1hti | A |
| MSH2 | None | None | None | None |
| MSH2 | None | None | None | None |
| MSH2 | None | None | None | None |
| MSH2 | None | None | None | None |

| | | | | |
|---|---|---|---|---|
| MSH2 | None | None | None | None |
| MSH2 | None | None | None | None |
| MSH2 | None | None | None | None |
| MSH2 | None | None | None | None |
| MSH2 | None | None | None | None |
| MSH2 | None | None | None | None |
| MSH2 | None | None | None | None |
| MSH2 | None | None | None | None |
| MSH2 | None | None | None | None |
| MSH2 | None | None | None | None |
| MSH2 | None | None | None | None |
| MSH2 | None | None | None | None |
| MSH2 | None | None | None | None |
| MSH2 | None | None | None | None |
| MSH2 | None | None | None | None |
| MSH2 | None | None | None | None |
| MSH2 | None | None | None | None |
| MSH2 | None | None | None | None |
| MSH2 | None | None | None | None |
| MSH2 | None | None | None | None |
| MSH2 | None | None | None | None |
| MSH2 | None | None | None | None |
| MSH2 | None | None | None | None |
| MSH2 | None | None | None | None |
| MSH2 | None | None | None | None |
| MSH2 | None | None | None | None |
| MSH2 | None | None | None | None |
| MSH2 | None | None | None | None |
| MSH2 | None | None | None | None |
| MSH2 | None | None | None | None |
| MSH2 | None | None | None | None |
| MSH2 | None | None | None | None |
| GALE | 0.172 | 6 | 1ek5 | A |
| GALE | 0.78 | 1 | 1i3n | A |
| GALE | 0.122 | 38 | 1ek6 | A |
| GALE | 0.106 | 41 | 1i3m | A |
| GALE | 0.106 | 36 | 1i3m | A |
| GALE | None | None | None | None |
| GALE | 0.176 | 29 | 1i3m | A |
| GALE | 0.259 | 5 | 1i3m | A |
| NSD2 | None | None | None | None |
| SRP72 | None | None | None | None |
| SRP72 | None | None | None | None |
| SRP72 | None | None | None | None |
| POLK | None | None | None | None |
| GALT | None | None | None | None |
| GALT | None | None | None | None |
| GALT | None | None | None | None |
| GALT | None | None | None | None |
| GALT | None | None | None | None |
| GALT | None | None | None | None |
| GALT | None | None | None | None |
| GALT | None | None | None | None |
| GALT | None | None | None | None |

| | | | | |
|---|---|---|---|---|
| GALT | None | None | None | None |
| GALT | None | None | None | None |
| GALT | None | None | None | None |
| GALT | None | None | None | None |
| GALT | None | None | None | None |
| GALT | None | None | None | None |
| GALT | None | None | None | None |
| GALT | None | None | None | None |
| GALT | None | None | None | None |
| GALT | None | None | None | None |
| GALT | None | None | None | None |
| GALT | None | None | None | None |
| GALT | None | None | None | None |
| GALT | None | None | None | None |
| GALT | None | None | None | None |
| GALT | None | None | None | None |
| GALT | None | None | None | None |
| GALT | None | None | None | None |
| GALT | None | None | None | None |
| GALT | None | None | None | None |
| GALT | None | None | None | None |
| GALT | None | None | None | None |
| GALT | None | None | None | None |
| GALT | None | None | None | None |
| GALT | None | None | None | None |
| GALT | None | None | None | None |
| GALT | None | None | None | None |
| GALT | None | None | None | None |
| PYCR1 | 0.75 | 11 | 5uav | A |
| PYCR1 | 0.75 | 11 | 5uav | A |
| PYCR1 | 0.5 | 2 | 2izz | A |
| PYCR1 | 0.556 | 2 | 5uax | B |
| F8 | None | None | None | None |
| F8 | None | None | None | None |
| F8 | None | None | None | None |
| F8 | None | None | None | None |
| F8 | None | None | None | None |
| F8 | None | None | None | None |
| F8 | None | None | None | None |
| F8 | None | None | None | None |
| F8 | None | None | None | None |
| F8 | None | None | None | None |
| F8 | None | None | None | None |
| F8 | None | None | None | None |
| F8 | None | None | None | None |
| F8 | None | None | None | None |
| F8 | None | None | None | None |
| F8 | None | None | None | None |
| ANK1 | None | None | None | None |
| CHAT | None | None | None | None |
| CHAT | None | None | None | None |
| CHAT | 0.615 | 8 | 2fy5 | A |
| CHAT | 0.538 | 2 | 2fy5 | A |
| CHAT | None | None | None | None |

| | | | | |
|---|---|---|---|---|
| CHAT | None | None | None | None |
| CHAT | None | None | None | None |
| CHAT | None | None | None | None |
| CHAT | None | None | None | None |
| CHAT | 0.538 | 6 | 2fy5 | A |
| CHAT | None | None | None | None |
| CHAT | None | None | None | None |
| CHAT | None | None | None | None |
| CHAT | None | None | None | None |
| CHAT | None | None | None | None |
| CHAT | None | None | None | None |
| CHAT | None | None | None | None |
| GHR | None | None | None | None |
| GHR | 0.312 | 2 | 3hhr | C |
| GHR | None | None | None | None |
| GHR | 0.312 | 1 | 3hhr | C |
| GHR | None | None | None | None |
| GHR | None | None | None | None |
| GHR | None | None | None | None |
| GHR | None | None | None | None |
| GHR | None | None | None | None |
| GHR | None | None | None | None |
| GHR | None | None | None | None |
| GHR | None | None | None | None |
| GHR | None | None | None | None |
| GHR | None | None | None | None |
| FZD4 | None | None | None | None |
| FZD4 | None | None | None | None |
| FZD4 | None | None | None | None |
| MTR | None | None | None | None |
| MTR | None | None | None | None |
| MTR | None | None | None | None |
| GM2A | None | None | None | None |
| GM2A | None | None | None | None |
| GM2A | None | None | None | None |
| GM2A | None | None | None | None |
| GM2A | None | None | None | None |
| CRYGD | 0.444 | 3 | 2g98 | A |
| CRYGD | None | None | None | None |
| CRYGD | 0.522 | 1 | 2kfb | A |
| CRYGD | 0.4 | 3 | 4gr7 | X |
| CRYGD | 0.235 | 9 | 1h4a | X |
| CRYGD | 0.333 | 9 | 1hk0 | X |
| CRYGD | None | None | None | None |
| CRYGD | None | None | None | None |
| CRYGD | 0.556 | 2 | 2g98 | A |
| TLR5 | None | None | None | None |
| HSD17B10 | 0.855 | 1 | 1u7t | B |
| HSD17B10 | None | None | None | None |
| HSD17B10 | 0.542 | 1 | 1so8 | A |
| HSD17B10 | 0.375 | 3 | 1so8 | A |
| TBC1D20 | None | None | None | None |
| FBN1 | None | None | None | None |
| FBN1 | None | None | None | None |

| | | | | |
|---|---|---|---|---|
| FBN1 | None | None | None | None |
| FBN1 | None | None | None | None |
| FBN1 | None | None | None | None |
| FBN1 | None | None | None | None |
| FBN1 | None | None | None | None |
| FXN | None | None | None | None |
| FXN | None | None | None | None |
| FXN | None | None | None | None |
| FXN | None | None | None | None |
| MYO5A | None | None | None | None |
| CTSK | 0.168 | 45 | 7pck | A |
| CTSK | None | None | None | None |
| SCN5A | None | None | None | None |
| SCN5A | None | None | None | None |
| SOS1 | None | None | None | None |
| SOS1 | None | None | None | None |
| SOS1 | None | None | None | None |
| SOS1 | None | None | None | None |
| SOS1 | None | None | None | None |
| SOS1 | None | None | None | None |
| SOS1 | None | None | None | None |
| SOS1 | None | None | None | None |
| SOS1 | None | None | None | None |
| SOS1 | None | None | None | None |
| SOS1 | None | None | None | None |
| SOS1 | None | None | None | None |
| GATM | None | None | None | None |
| GATM | None | None | None | None |
| GATM | 0.75 | 4 | 4jdw | A |
| GATM | None | None | None | None |
| GATM | None | None | None | None |
| GATM | 0.75 | 6 | 6jdw | A |
| GATM | 0.75 | 11 | 6jdw | A |
| MMP2 | None | None | None | None |
| MMP2 | None | None | None | None |
| MMP2 | 0.407 | 7 | 1qib | A |
| MMP2 | None | None | None | None |
| NDUFB9 | None | None | None | None |
| NDUFB9 | None | None | None | None |
| GH1 | None | None | None | None |
| GH1 | None | None | None | None |
| GH1 | None | None | None | None |
| SLC6A4 | None | None | None | None |
| G6PD | None | None | None | None |
| G6PD | None | None | None | None |
| G6PD | None | None | None | None |
| G6PD | None | None | None | None |
| G6PD | None | None | None | None |
| G6PD | None | None | None | None |
| G6PD | None | None | None | None |
| G6PD | None | None | None | None |
| G6PD | 0.545 | 1 | 1qki | B |
| G6PD | None | None | None | None |
| G6PD | None | None | None | None |

| | | | | |
|---|---|---|---|---|
| G6PD | None | None | None | None |
| G6PD | None | None | None | None |
| G6PD | None | None | None | None |
| G6PD | None | None | None | None |
| G6PD | 0.455 | 1 | 1qki | D |
| G6PD | None | None | None | None |
| G6PD | None | None | None | None |
| G6PD | None | None | None | None |
| G6PD | None | None | None | None |
| CST3 | None | None | None | None |
| LMNA | None | None | None | None |
| LMNA | None | None | None | None |
| NDUFA9 | None | None | None | None |
| NDUFA9 | None | None | None | None |
| NDUFA9 | None | None | None | None |
| NDUFA9 | None | None | None | None |
| YARS | 0.581 | 1 | 5thl | A |
| YARS | None | None | None | None |
| YARS | None | None | None | None |
| YARS | None | None | None | None |
| EPHA2 | None | None | None | None |
| EPHA2 | None | None | None | None |
| EPHA2 | None | None | None | None |
| EPHA2 | None | None | None | None |
| DICER1 | None | None | None | None |
| DICER1 | None | None | None | None |
| SNX10 | None | None | None | None |
| SNX10 | None | None | None | None |
| TTN | None | None | None | None |
| TTN | None | None | None | None |
| GATA3 | None | None | None | None |
| C5 | None | None | None | None |
| THBD | None | None | None | None |
| BIN1 | None | None | None | None |
| BIN1 | None | None | None | None |
| SMARCAL1 | None | None | None | None |
| CFB | None | None | None | None |
| CYC1 | None | None | None | None |
| ANK2 | None | None | None | None |
| CHMP4B | None | None | None | None |
| CHMP4B | None | None | None | None |
| ERBB2 | None | None | None | None |
| ERBB2 | None | None | None | None |
| POLG2 | None | None | None | None |
| POLG2 | None | None | None | None |
| POLG2 | None | None | None | None |
| POLG2 | None | None | None | None |
| POLG2 | None | None | None | None |
| POLG2 | None | None | None | None |
| POLG2 | None | None | None | None |
| POLG2 | None | None | None | None |
| C3 | None | None | None | None |
| C3 | None | None | None | None |
| RPGR | None | None | None | None |

| | | | | |
|---|---|---|---|---|
| RPGR | None | None | None | None |
| RPGR | None | None | None | None |
| RPGR | None | None | None | None |
| RPGR | None | None | None | None |
| C1QA | None | None | None | None |
| PRKG1 | None | None | None | None |
| IGBP1 | None | None | None | None |
| ENO3 | None | None | None | None |
| ENO3 | None | None | None | None |
| ENO3 | 0.41 | 43 | 2xsx | A |
| ENO3 | 0.487 | 39 | 2xsx | A |
| ENO3 | 0.359 | 47 | 2xsx | B |
| ENO3 | 0.897 | 2 | 2xsx | B |
| MCPH1 | None | None | None | None |
| MCPH1 | None | None | None | None |
| MCPH1 | None | None | None | None |
| MCPH1 | None | None | None | None |
| MCPH1 | None | None | None | None |
| SNRNP200 | None | None | None | None |
| SNRNP200 | None | None | None | None |
| DNMT1 | None | None | None | None |
| DNMT1 | None | None | None | None |
| DNMT1 | None | None | None | None |
| AKT1 | None | None | None | None |
| NDUFS2 | None | None | None | None |
| ACVR2B | None | None | None | None |
| HMGCL | None | None | None | None |
| HMGCL | None | None | None | None |
| HMGCL | None | None | None | None |
| JUP | None | None | None | None |
| JUP | None | None | None | None |
| JUP | None | None | None | None |
| JUP | None | None | None | None |
| JUP | None | None | None | None |
| JUP | None | None | None | None |
| JUP | None | None | None | None |
| JUP | None | None | None | None |
| SPINK1 | None | None | None | None |
| SPINK1 | None | None | None | None |
| SPINK1 | None | None | None | None |
| VCL | None | None | None | None |
| VCL | None | None | None | None |
| NOTCH1 | None | None | None | None |
| NOTCH1 | None | None | None | None |
| NOTCH1 | None | None | None | None |
| NDUFB3 | None | None | None | None |
| IKBKG | None | None | None | None |
| GSN | 0.529 | 2 | 2fh1 | A |
| GSN | None | None | None | None |
| GSN | None | None | None | None |
| GSN | 0.5 | 3 | 1p8x | A |
| GSN | 0.263 | 13 | 2fh3 | A |
| GSN | None | None | None | None |
| GSN | None | None | None | None |

| | | | | |
|---|---|---|---|---|
| GSN | None | None | None | None |
| GSN | None | None | None | None |
| CLC | None | None | None | None |
| RAF1 | None | None | None | None |
| RAF1 | None | None | None | None |
| RAF1 | None | None | None | None |
| NDUFS8 | None | None | None | None |
| NDUFS8 | None | None | None | None |
| NDUFS8 | None | None | None | None |
| NDUFS8 | None | None | None | None |
| NDUFS8 | None | None | None | None |
| CASR | None | None | None | None |
| CASR | None | None | None | None |
| CASR | None | None | None | None |
| CASR | None | None | None | None |
| CASR | None | None | None | None |
| CASR | None | None | None | None |
| CASR | None | None | None | None |
| CASR | None | None | None | None |
| CASR | None | None | None | None |
| CASR | None | None | None | None |
| CASR | None | None | None | None |
| CASR | None | None | None | None |
| CASR | None | None | None | None |
| CASR | None | None | None | None |
| CASR | None | None | None | None |
| CASR | None | None | None | None |
| CASR | None | None | None | None |
| CASR | None | None | None | None |
| CASR | None | None | None | None |
| CASR | None | None | None | None |
| CASR | None | None | None | None |
| CASR | None | None | None | None |
| CASR | None | None | None | None |
| CASR | None | None | None | None |
| CASR | None | None | None | None |
| CASR | None | None | None | None |
| CASR | None | None | None | None |
| CASR | None | None | None | None |
| CASR | None | None | None | None |
| VCL | None | None | None | None |
| VCL | None | None | None | None |
| SMPD1 | None | None | None | None |
| SMPD1 | None | None | None | None |
| SMPD1 | None | None | None | None |
| KDM6A | None | None | None | None |
| KCNQ1 | None | None | None | None |
| KCNQ1 | None | None | None | None |
| KCNQ1 | None | None | None | None |
| KCNQ1 | None | None | None | None |
| KCNQ1 | None | None | None | None |
| KCNQ1 | None | None | None | None |
| NDUFA12 | None | None | None | None |
| GLRX5 | None | None | None | None |

| | | | | |
|---|---|---|---|---|
| PALB2 | None | None | None | None |
| PALB2 | None | None | None | None |
| PALB2 | None | None | None | None |
| PALB2 | None | None | None | None |
| PALB2 | None | None | None | None |
| PALB2 | None | None | None | None |
| PALB2 | None | None | None | None |
| PALB2 | None | None | None | None |
| PALB2 | None | None | None | None |
| PALB2 | None | None | None | None |
| ALDOB | 0.462 | 89 | 1qo5 | C |
| ALDOB | 0.731 | 2 | 1xdm | A |
| ALDOB | 0.731 | 1 | 1qo5 | B |
| ALDOB | 0.538 | 6 | 1xdm | A |
| ALDOB | 0.692 | 7 | 1qo5 | C |
| ALDOB | None | None | None | None |
| ALDOB | 0.577 | 25 | 1qo5 | B |
| GNE | None | None | None | None |
| GNE | None | None | None | None |
| GNE | None | None | None | None |
| GNE | None | None | None | None |
| GNE | None | None | None | None |
| GNE | None | None | None | None |
| GNE | None | None | None | None |
| GNE | None | None | None | None |
| GNE | None | None | None | None |
| GNE | None | None | None | None |
| GNE | None | None | None | None |
| GNE | None | None | None | None |
| CHRNB2 | None | None | None | None |
| CHRNB2 | None | None | None | None |
| ERCC3 | None | None | None | None |
| ERCC3 | None | None | None | None |
| ERCC3 | None | None | None | None |
| ERCC3 | None | None | None | None |
| ERCC3 | None | None | None | None |
| UNC119 | None | None | None | None |
| ATL1 | None | None | None | None |
| ATL1 | None | None | None | None |
| ATL1 | None | None | None | None |
| ATL1 | 0.8 | 1 | 3q5d | A |
| ATL1 | None | None | None | None |
| ATL1 | None | None | None | None |
| ATL1 | None | None | None | None |
| ATL1 | None | None | None | None |
| ATL1 | None | None | None | None |
| ATL1 | None | None | None | None |
| NR3C2 | 0.701 | 1 | 2oax | A |
| TBK1 | None | None | None | None |
| HYAL1 | None | None | None | None |
| HYAL1 | None | None | None | None |
| HYAL1 | None | None | None | None |
| HYAL1 | None | None | None | None |
| CTSD | 0.633 | 1 | 4obz | A |

| | | | | |
|---|---|---|---|---|
| PRKCG | None | None | None | None |
| TGFBR1 | None | None | None | None |
| XDH | 0.308 | 2 | 2e1q | A |
| XDH | None | None | None | None |
| XDH | None | None | None | None |
| XDH | 0.154 | 74 | 2e1q | A |
| XDH | 0.125 | 35 | 2e1q | B |
| XDH | None | None | None | None |
| NMNAT1 | 0.833 | 1 | 1kqn | E |
| NMNAT1 | None | None | None | None |
| NMNAT1 | 0.167 | 15 | 1kqo | B |
| NMNAT1 | 0.167 | 3 | 1kqo | B |
| NMNAT1 | 0.167 | 20 | 1kqo | B |
| NMNAT1 | 0.118 | 25 | 1kr2 | B |
| SERPING1 | None | None | None | None |
| SERPING1 | None | None | None | None |
| SERPING1 | None | None | None | None |
| SERPING1 | None | None | None | None |
| SERPING1 | None | None | None | None |
| PDZRN3 | None | None | None | None |
| FABP2 | None | None | None | None |
| IFNGR1 | 0.143 | 1 | 1jrh | I |
| IFNGR1 | 0.143 | 1 | 1jrh | I |
| TDP1 | None | None | None | None |
| TDP1 | 0.333 | 11 | 1rg1 | A |
| INS | 0.263 | 5 | 2lgb | B |
| INS | 0.769 | 1 | 2r34 | C |
| INS | 0.769 | 1 | 2r34 | C |
| PPOX | 0.857 | 1 | 4ivm | B |
| PPOX | None | None | None | None |
| PPOX | None | None | None | None |
| PPOX | None | None | None | None |
| PPOX | None | None | None | None |
| PPOX | None | None | None | None |
| PPOX | None | None | None | None |
| PPOX | None | None | None | None |
| PPOX | None | None | None | None |
| GFER | None | None | None | None |
| GFER | 0.769 | 4 | 3tk0 | A |
| GFER | None | None | None | None |
| GFER | None | None | None | None |
| LIFR | None | None | None | None |
| EGLN1 | 0.556 | 21 | 5a3u | C |
| EGLN1 | 0.889 | 6 | 5v18 | A |
| EGLN1 | 0.944 | 2 | 5a3u | C |
| GP1BA | None | None | None | None |
| GP1BA | None | None | None | None |
| GP1BA | None | None | None | None |
| GP1BA | None | None | None | None |
| GP1BA | None | None | None | None |
| MMADHC | None | None | None | None |
| MMADHC | None | None | None | None |
| MMADHC | None | None | None | None |
| MMADHC | None | None | None | None |

| | | | | |
|---|---|---|---|---|
| MMADHC | None | None | None | None |
| MMADHC | None | None | None | None |
| MMADHC | None | None | None | None |
| ACVRL1 | None | None | None | None |
| ACVRL1 | None | None | None | None |
| RFXANK | None | None | None | None |
| CHIT1 | None | None | None | None |
| UBR1 | None | None | None | None |
| BLM | None | None | None | None |
| BLM | None | None | None | None |
| BLM | None | None | None | None |
| BLM | None | None | None | None |
| BLM | None | None | None | None |
| BLM | None | None | None | None |
| BLM | None | None | None | None |
| BLM | None | None | None | None |
| BLM | None | None | None | None |
| BLM | None | None | None | None |
| BLM | None | None | None | None |
| BLM | None | None | None | None |
| BLM | None | None | None | None |
| VDR | 0.378 | 7 | 1kb4 | B |
| VDR | None | None | None | None |
| VDR | 0.378 | 4 | 1kb4 | A |
| VDR | 0.578 | 1 | 1kb2 | B |
| VDR | 0.578 | 1 | 1kb2 | B |
| VDR | 0.578 | 1 | 1kb2 | B |
| EPHA2 | None | None | None | None |
| EPHA2 | None | None | None | None |
| EPHA2 | None | None | None | None |
| EPHA2 | None | None | None | None |
| EPHA2 | None | None | None | None |
| EPHA2 | None | None | None | None |
| EPHA2 | None | None | None | None |
| EPHA2 | None | None | None | None |
| EPHA2 | None | None | None | None |
| EPHA2 | None | None | None | None |
| UBA1 | None | None | None | None |
| STAT1 | None | None | None | None |
| STAT1 | None | None | None | None |
| STAT1 | None | None | None | None |
| STAT1 | None | None | None | None |
| STAT1 | None | None | None | None |
| STAT1 | None | None | None | None |
| STAT1 | None | None | None | None |
| STAT1 | None | None | None | None |
| DNM1L | None | None | None | None |
| DNM1L | None | None | None | None |
| OTC | None | None | None | None |
| OTC | None | None | None | None |
| OTC | None | None | None | None |
| OTC | None | None | None | None |
| OTC | None | None | None | None |
| OTC | None | None | None | None |

| | | | | |
|---|---|---|---|---|
| OTC | None | None | None | None |
| OTC | None | None | None | None |
| OTC | None | None | None | None |
| OTC | 0.281 | 3 | 1fvo | A |
| OTC | None | None | None | None |
| OTC | None | None | None | None |
| OTC | None | None | None | None |
| OTC | None | None | None | None |
| OTC | 0.125 | 14 | 1c9y | A |
| OTC | 0.125 | 14 | 1c9y | A |
| OTC | 0.143 | 11 | 1oth | A |
| RARS | None | None | None | None |
| RARS | None | None | None | None |
| RARS | None | None | None | None |
| SEPSECS | None | None | None | None |
| SEPSECS | None | None | None | None |
| SEPSECS | None | None | None | None |
| SEPSECS | None | None | None | None |
| SEPSECS | None | None | None | None |
| BRCA1 | None | None | None | None |
| BRCA1 | None | None | None | None |
| BRCA1 | 0.538 | 1 | 3pxc | X |
| BRCA1 | None | None | None | None |
| BRCA1 | None | None | None | None |
| BRCA1 | 0.462 | 1 | 1t2u | A |
| BRCA1 | None | None | None | None |
| BRCA1 | None | None | None | None |
| BRCA1 | None | None | None | None |
| BRCA1 | None | None | None | None |
| BRCA1 | None | None | None | None |
| BRCA1 | None | None | None | None |
| BRCA1 | None | None | None | None |
| BRCA1 | None | None | None | None |
| BRCA1 | None | None | None | None |
| BRCA1 | None | None | None | None |
| BRCA1 | None | None | None | None |
| BRCA1 | None | None | None | None |
| BRCA1 | None | None | None | None |
| BRCA1 | None | None | None | None |
| BRCA1 | None | None | None | None |
| BRCA1 | None | None | None | None |
| CUBN | None | None | None | None |
| ITGB3 | None | None | None | None |
| MTRR | None | None | None | None |
| MTRR | None | None | None | None |
| MTRR | None | None | None | None |
| MTRR | None | None | None | None |
| MTRR | None | None | None | None |
| MTRR | None | None | None | None |
| MTRR | None | None | None | None |
| MTRR | None | None | None | None |
| IL7R | None | None | None | None |
| IL7R | None | None | None | None |
| RPS24 | None | None | None | None |

| | | | | |
|---|---|---|---|---|
| RPS24 | None | None | None | None |
| HOGA1 | None | None | None | None |
| HOGA1 | None | None | None | None |
| HOGA1 | None | None | None | None |
| HOGA1 | None | None | None | None |
| HOGA1 | None | None | None | None |
| HOGA1 | None | None | None | None |
| HOGA1 | 0.791 | 1 | 3s5n | A |
| HOGA1 | None | None | None | None |
| HOGA1 | None | None | None | None |
| HOGA1 | None | None | None | None |
| HOGA1 | None | None | None | None |
| HOGA1 | None | None | None | None |
| HOGA1 | None | None | None | None |
| GJB2 | None | None | None | None |
| GJB2 | None | None | None | None |
| GJB2 | None | None | None | None |
| GJB2 | None | None | None | None |
| GJB2 | None | None | None | None |
| GJB2 | None | None | None | None |
| GJB2 | None | None | None | None |
| GJB2 | None | None | None | None |
| GJB2 | None | None | None | None |
| GJB2 | None | None | None | None |
| GJB2 | None | None | None | None |
| GJB2 | None | None | None | None |
| GJB2 | None | None | None | None |
| GJB2 | None | None | None | None |
| GJB2 | None | None | None | None |
| GJB2 | None | None | None | None |
| GJB2 | None | None | None | None |
| GJB2 | None | None | None | None |
| GJB2 | None | None | None | None |
| GJB2 | None | None | None | None |
| GJB2 | None | None | None | None |
| GJB2 | None | None | None | None |
| GJB2 | None | None | None | None |
| GJB2 | None | None | None | None |
| GJB2 | None | None | None | None |
| GJB2 | None | None | None | None |
| GJB2 | None | None | None | None |
| GJB2 | None | None | None | None |
| GJB2 | None | None | None | None |
| GJB2 | None | None | None | None |
| GJB2 | None | None | None | None |
| GJB2 | None | None | None | None |
| GJB2 | None | None | None | None |
| GJB2 | None | None | None | None |
| GJB2 | None | None | None | None |
| GJB2 | None | None | None | None |
| GJB2 | None | None | None | None |
| GJB2 | None | None | None | None |
| GJB2 | None | None | None | None |
| GJB2 | None | None | None | None |

| | | | | |
|---|---|---|---|---|
| GJB2 | None | None | None | None |
| GJB2 | None | None | None | None |
| GJB2 | None | None | None | None |
| GJB2 | None | None | None | None |
| GJB2 | None | None | None | None |
| GJB2 | None | None | None | None |
| GJB2 | None | None | None | None |
| GJB2 | None | None | None | None |
| GJB2 | None | None | None | None |
| GJB2 | None | None | None | None |
| GJB2 | None | None | None | None |
| GJB2 | None | None | None | None |
| GJB2 | None | None | None | None |
| GJB2 | None | None | None | None |
| CPS1 | None | None | None | None |
| CPS1 | 0.545 | 127 | 5dou | A |
| CPS1 | None | None | None | None |
| CPS1 | None | None | None | None |
| CPS1 | 0.364 | 19 | 5dou | A |
| CPS1 | 0.364 | 19 | 5dou | A |
| CPS1 | 0.182 | 108 | 5dou | A |
| CPS1 | None | None | None | None |
| CPS1 | None | None | None | None |
| CPS1 | None | None | None | None |
| PSPH | None | None | None | None |
| PSPH | None | None | None | None |
| PSPH | None | None | None | None |
| PSPH | None | None | None | None |
| PSPH | None | None | None | None |
| PSPH | None | None | None | None |
| PEPD | 0.512 | 2 | 5m4l | B |
| PEPD | None | None | None | None |
| PEPD | None | None | None | None |
| PEPD | 0.643 | 2 | 5m4j | B |
| PEPD | 0.643 | 2 | 5m4j | B |
| PEPD | 0.419 | 5 | 5m4l | B |
| PEPD | 0.106 | 31 | 2iw2 | A |
| PEPD | None | None | None | None |
| PEPD | None | None | None | None |
| KDM5C | None | None | None | None |
| KDM5C | 0.222 | 117 | 5fwj | A |
| KDM5C | 0.333 | 55 | 5fwj | A |
| ITGB6 | 0.312 | 2 | 5ffo | B |
| ITGB6 | None | None | None | None |
| ITGB6 | None | None | None | None |
| DARS | None | None | None | None |
| DARS | None | None | None | None |
| DARS | None | None | None | None |
| DARS | None | None | None | None |
| BTK | None | None | None | None |
| BTK | None | None | None | None |
| BTK | None | None | None | None |
| HDAC4 | None | None | None | None |
| ITGA2B | None | None | None | None |

| | | | | | |
|---|---|---|---|---|---|
| ITGA2B | None | None | | None | None |
| ITGA2B | None | None | | None | None |
| ITGA2B | None | None | | None | None |
| PCSK9 | None | None | | None | None |
| PCSK9 | None | None | | None | None |
| PCSK9 | None | None | | None | None |
| PCSK9 | 0.12 | | 16 | 2pmw | B |
| MUTYH | None | None | | None | None |
| MUTYH | None | None | | None | None |
| MUTYH | None | None | | None | None |
| MUTYH | None | None | | None | None |
| MUTYH | None | None | | None | None |
| MUTYH | 0.375 | | 2 | 3n5n | X |
| MUTYH | None | None | | None | None |
| MUTYH | None | None | | None | None |
| MUTYH | None | None | | None | None |
| MUTYH | 0.375 | | 2 | 3n5n | X |
| MUTYH | 0.375 | | 2 | 3n5n | X |
| MUTYH | None | None | | None | None |
| MUTYH | 0.375 | | 2 | 3n5n | X |
| MUTYH | None | None | | None | None |
| MUTYH | None | None | | None | None |
| MUTYH | None | None | | None | None |
| MUTYH | None | None | | None | None |
| MUTYH | None | None | | None | None |
| MUTYH | None | None | | None | None |
| MUTYH | None | None | | None | None |
| MUTYH | None | None | | None | None |
| MUTYH | None | None | | None | None |
| MUTYH | None | None | | None | None |
| MUTYH | None | None | | None | None |
| SLC25A12 | None | None | | None | None |
| GSN | None | None | | None | None |
| IHH | 0.909 | | 1 | 3n1p | B |
| IHH | None | None | | None | None |
| IHH | None | None | | None | None |
| IHH | 0.818 | | 2 | 3n1p | B |
| IHH | 0.727 | | 12 | 3n1p | B |
| IHH | 0.636 | | 15 | 3n1p | B |
| IHH | 0.818 | | 7 | 3n1p | B |
| CD247 | None | None | | None | None |
| EP300 | None | None | | None | None |
| TTN | None | None | | None | None |
| MRI1 | None | None | | None | None |
| SGSH | None | None | | None | None |
| SGSH | None | None | | None | None |
| SGSH | None | None | | None | None |
| SGSH | None | None | | None | None |
| SGSH | None | None | | None | None |
| SGSH | None | None | | None | None |
| SGSH | None | None | | None | None |
| SGSH | None | None | | None | None |
| SGSH | None | None | | None | None |
| SGSH | None | None | | None | None |

| | | | | |
|---|---|---|---|---|
| SGSH | None | None | None | None |
| SGSH | None | None | None | None |
| SGSH | None | None | None | None |
| SGSH | None | None | None | None |
| SGSH | None | None | None | None |
| SGSH | None | None | None | None |
| SGSH | None | None | None | None |
| SGSH | None | None | None | None |
| SGSH | None | None | None | None |
| SGSH | None | None | None | None |
| SGSH | None | None | None | None |
| SGSH | None | None | None | None |
| CBL | None | None | None | None |
| CANT1 | None | None | None | None |
| CANT1 | None | None | None | None |
| CANT1 | None | None | None | None |
| CANT1 | None | None | None | None |
| CANT1 | None | None | None | None |
| CANT1 | None | None | None | None |
| CANT1 | None | None | None | None |
| SLC1A3 | None | None | None | None |
| SLC1A3 | None | None | None | None |
| SLC1A5 | None | None | None | None |
| SLC1A3 | None | None | None | None |
| SLC1A5 | None | None | None | None |
| SLC1A3 | None | None | None | None |
| SLC1A3 | None | None | None | None |
| CTSD | 0.423 | 15 | 4od9 | D |
| CTSD | 0.14 | 32 | 1lya | B |
| CTSD | None | None | None | None |
| CTSD | None | None | None | None |
| CTSD | None | None | None | None |
| MTOR | None | None | None | None |
| ILK | None | None | None | None |
| PCDH15 | None | None | None | None |
| PCDH15 | None | None | None | None |
| PCDH15 | None | None | None | None |
| PCDH15 | None | None | None | None |
| PCDH15 | None | None | None | None |
| PCDH15 | None | None | None | None |
| KLHL40 | None | None | None | None |
| KLHL40 | None | None | None | None |
| KLHL40 | None | None | None | None |
| KLHL40 | None | None | None | None |
| CSTA | None | None | None | None |
| CSTA | None | None | None | None |
| HNMT | None | None | None | None |
| HNMT | None | None | None | None |
| IFNGR2 | None | None | None | None |
| IFNGR2 | None | None | None | None |
| PEX5 | None | None | None | None |
| KIF7 | None | None | None | None |
| KIF7 | 0.185 | 18 | 4a14 | A |
| MYL3 | None | None | None | None |

| | | | | |
|---|---|---|---|---|
| MYL3 | None | None | None | None |
| MYL3 | None | None | None | None |
| MYL3 | None | None | None | None |
| MYL3 | None | None | None | None |
| MYL3 | None | None | None | None |
| MYL3 | None | None | None | None |
| NAGLU | None | None | None | None |
| NAGLU | None | None | None | None |
| NAGLU | None | None | None | None |
| NAGLU | None | None | None | None |
| NAGLU | None | None | None | None |
| NAGLU | None | None | None | None |
| NAGLU | None | None | None | None |
| NAGLU | None | None | None | None |
| NAGLU | None | None | None | None |
| NAGLU | None | None | None | None |
| NAGLU | None | None | None | None |
| NAGLU | None | None | None | None |
| NAGLU | None | None | None | None |
| TNFRSF13C | None | None | None | None |
| KIF21A | None | None | None | None |
| KIF21A | None | None | None | None |
| KIF21A | None | None | None | None |
| KIF21A | None | None | None | None |
| KIF21A | None | None | None | None |
| DOCK2 | None | None | None | None |
| DSP | None | None | None | None |
| CDKN2A | None | None | None | None |
| CDKN2A | None | None | None | None |
| CDKN2A | None | None | None | None |
| CDKN2A | None | None | None | None |
| CDKN2A | None | None | None | None |
| CDKN2A | None | None | None | None |
| CDKN2A | None | None | None | None |
| CDKN2A | None | None | None | None |
| CDKN2A | None | None | None | None |
| CDKN2A | None | None | None | None |
| CDKN2A | None | None | None | None |
| CDKN2A | None | None | None | None |
| CDKN2A | None | None | None | None |
| CDKN2A | None | None | None | None |
| VCL | None | None | None | None |
| TPP1 | None | None | None | None |
| TPP1 | None | None | None | None |
| TPP1 | None | None | None | None |
| TPP1 | None | None | None | None |
| TPP1 | None | None | None | None |
| TPP1 | None | None | None | None |
| TPP1 | None | None | None | None |
| TPP1 | None | None | None | None |
| TPP1 | None | None | None | None |
| TPP1 | None | None | None | None |
| TPP1 | None | None | None | None |
| TPP1 | None | None | None | None |

| | | | | |
|---|---|---|---|---|
| TPP1 | None | None | None | None |
| TPP1 | None | None | None | None |
| TPP1 | None | None | None | None |
| TPP1 | None | None | None | None |
| TPP1 | None | None | None | None |
| TPP1 | None | None | None | None |
| TPP1 | None | None | None | None |
| TPP1 | None | None | None | None |
| TPP1 | None | None | None | None |
| TPP1 | None | None | None | None |
| IL10RB | None | None | None | None |
| IL10RB | None | None | None | None |
| CCM2 | None | None | None | None |
| CCM2 | None | None | None | None |
| CCM2 | None | None | None | None |
| DCX | None | None | None | None |
| DCX | None | None | None | None |
| DCX | None | None | None | None |
| DCX | None | None | None | None |
| DCX | None | None | None | None |
| DCX | None | None | None | None |
| DCX | None | None | None | None |
| DCX | None | None | None | None |
| BRCA2 | None | None | None | None |
| NNT | None | None | None | None |
| C2 | None | None | None | None |
| C2 | None | None | None | None |
| C2 | None | None | None | None |
| C2 | None | None | None | None |
| WRN | None | None | None | None |
| WRN | None | None | None | None |
| F13A1 | None | None | None | None |
| F13A1 | None | None | None | None |
| F13A1 | None | None | None | None |
| F13A1 | None | None | None | None |
| TF | 0.188 | 24 | 3ve1 | B |
| TF | None | None | None | None |
| TF | 0.516 | 2 | 3ve1 | B |
| TF | 0.562 | 39 | 3skp | A |
| TF | 0.141 | 33 | 3v89 | B |
| ITPA | None | None | None | None |
| ITPA | None | None | None | None |
| SNCA | None | None | None | None |
| ITGB4 | None | None | None | None |
| ITGB4 | None | None | None | None |
| ITGB4 | None | None | None | None |
| LRP6 | None | None | None | None |
| LRP6 | None | None | None | None |
| LRP6 | 0.114 | 11 | 3s94 | A |
| PLG | None | None | None | None |
| PLG | 0.61 | 1 | 1i5k | A |
| TYROBP | None | None | None | None |
| UQCRC2 | None | None | None | None |
| GPSM2 | None | None | None | None |

| | | | | |
|---|---|---|---|---|
| GPSM2 | None | None | None | None |
| GPSM2 | None | None | None | None |
| SCN5A | None | None | None | None |
| SCN5A | None | None | None | None |
| SCN5A | None | None | None | None |
| SCN5A | None | None | None | None |
| SCN5A | None | None | None | None |
| ATIC | 0.6 | 6 | 1p4r | A |
| EZH2 | None | None | None | None |
| CPA1 | 0.387 | 33 | 4uez | A |
| SCN4B | None | None | None | None |
| SCN4B | None | None | None | None |
| MAN1B1 | None | None | None | None |
| MAN1B1 | 0.6 | 9 | 1fmi | A |
| MAN1B1 | None | None | None | None |
| MAN1B1 | None | None | None | None |
| CYCS | 0.74 | 1 | 2n9i | A |
| IDUA | None | None | None | None |
| IDUA | None | None | None | None |
| IDUA | None | None | None | None |
| IDUA | None | None | None | None |
| IDUA | None | None | None | None |
| IDUA | None | None | None | None |
| IDUA | None | None | None | None |
| IDUA | None | None | None | None |
| IDUA | None | None | None | None |
| IDUA | None | None | None | None |
| IDUA | None | None | None | None |
| IDUA | None | None | None | None |
| IDUA | None | None | None | None |
| IDUA | None | None | None | None |
| IDUA | None | None | None | None |
| IDUA | None | None | None | None |
| IDUA | None | None | None | None |
| IDUA | None | None | None | None |
| IDUA | None | None | None | None |
| IDUA | None | None | None | None |
| IDUA | None | None | None | None |
| IDUA | None | None | None | None |
| IDUA | None | None | None | None |
| IDUA | None | None | None | None |
| IDUA | None | None | None | None |
| HIKESHI | None | None | None | None |
| APC | None | None | None | None |
| APC | None | None | None | None |
| APC | None | None | None | None |
| APP | None | None | None | None |
| CEP152 | None | None | None | None |
| GAA | 0.609 | 1 | 5kzx | A |
| GAA | 0.609 | 1 | 5kzx | A |
| GAA | None | None | None | None |
| GAA | 0.174 | 9 | 5kzx | A |
| GAA | 0.261 | 10 | 5kzx | A |
| GAA | 0.609 | 1 | 5kzx | A |

| | | | | |
|---|---|---|---|---|
| GAA | 0.609 | 1 | 5kzx | A |
| GAA | 0.609 | 1 | 5kzx | A |
| GAA | 0.609 | 1 | 5kzx | A |
| GAA | 0.609 | 1 | 5kzx | A |
| GAA | 0.609 | 1 | 5kzx | A |
| GAA | 0.609 | 1 | 5kzx | A |
| GAA | 0.609 | 1 | 5kzx | A |
| GAA | 0.609 | 1 | 5kzx | A |
| GAA | 0.609 | 1 | 5kzx | A |
| GAA | 0.609 | 1 | 5kzx | A |
| GAA | 0.609 | 1 | 5kzx | A |
| GAA | 0.609 | 1 | 5kzx | A |
| GAA | 0.609 | 1 | 5kzx | A |
| GAA | 0.609 | 1 | 5kzx | A |
| GAA | 0.609 | 1 | 5kzx | A |
| GAA | 0.609 | 1 | 5kzx | A |
| GAA | 0.609 | 1 | 5kzx | A |
| GAA | 0.609 | 1 | 5kzx | A |
| GAA | 0.609 | 1 | 5kzx | A |
| GAA | 0.609 | 1 | 5kzx | A |
| GAA | 0.609 | 1 | 5kzx | A |
| GAA | 0.609 | 1 | 5kzx | A |
| GAA | 0.609 | 1 | 5kzx | A |
| GAA | 0.609 | 1 | 5kzx | A |
| GAA | 0.609 | 1 | 5kzx | A |
| GAA | 0.609 | 1 | 5kzx | A |
| GAA | 0.609 | 1 | 5kzx | A |
| GAA | 0.609 | 1 | 5kzx | A |
| GAA | 0.609 | 1 | 5kzx | A |
| GAA | 0.609 | 1 | 5kzx | A |
| GAA | 0.609 | 1 | 5kzx | A |
| GAA | 0.609 | 1 | 5kzx | A |
| GAA | 0.609 | 1 | 5kzx | A |
| GAA | 0.609 | 1 | 5kzx | A |
| GAA | 0.609 | 1 | 5kzx | A |
| GAA | 0.609 | 1 | 5kzx | A |
| GAA | 0.609 | 1 | 5kzx | A |
| GAA | 0.609 | 1 | 5kzx | A |
| GAA | 0.609 | 1 | 5kzx | A |
| GAA | 0.609 | 1 | 5kzx | A |
| GAA | 0.609 | 1 | 5kzx | A |
| GAA | 0.609 | 1 | 5kzx | A |
| GAA | 0.261 | 10 | 5kzx | A |
| FOLR1 | None | None | None | None |
| DSP | None | None | None | None |
| DSP | None | None | None | None |
| DSP | None | None | None | None |
| DSP | None | None | None | None |
| DSP | None | None | None | None |
| CCM2 | None | None | None | None |
| FXN | None | None | None | None |
| FXN | None | None | None | None |
| COMP | None | None | None | None |
| COMP | None | None | None | None |

| | | | | |
|---|---|---|---|---|
| COMP | None | None | None | None |
| COMP | None | None | None | None |
| COMP | None | None | None | None |
| COMP | None | None | None | None |
| COMP | None | None | None | None |
| COMP | None | None | None | None |
| COMP | None | None | None | None |
| COMP | None | None | None | None |
| COMP | None | None | None | None |
| COMP | None | None | None | None |
| AIFM1 | None | None | None | None |
| AIFM1 | None | None | None | None |
| AIFM1 | 0.229 | 23 | 5fs8 | A |
| AIFM1 | None | None | None | None |
| AIFM1 | 0.262 | 40 | 5fs7 | A |
| AIFM1 | None | None | None | None |
| AIFM1 | None | None | None | None |
| MYH9 | None | None | None | None |
| FHL2 | None | None | None | None |
| FHL2 | None | None | None | None |
| TNFRSF4 | None | None | None | None |
| CFB | None | None | None | None |
| CFB | None | None | None | None |
| CFB | None | None | None | None |
| CFB | None | None | None | None |
| CFB | None | None | None | None |
| CFB | None | None | None | None |
| CFB | 0.258 | 12 | 1dle | B |
| CFB | None | None | None | None |
| CFB | None | None | None | None |
| CFB | None | None | None | None |
| MVD | None | None | None | None |
| MVD | None | None | None | None |
| HSD17B4 | None | None | None | None |
| HSD17B4 | None | None | None | None |
| HSD17B4 | None | None | None | None |
| HSD17B4 | 0.429 | 2 | 1s9c | A |
| HSD17B4 | None | None | None | None |
| HSD17B4 | None | None | None | None |
| F11 | None | None | None | None |
| F11 | None | None | None | None |
| F11 | None | None | None | None |
| F11 | None | None | None | None |
| F11 | None | None | None | None |
| F11 | None | None | None | None |
| F11 | None | None | None | None |
| F11 | None | None | None | None |
| F11 | None | None | None | None |
| SPTA1 | None | None | None | None |
| SPTA1 | None | None | None | None |
| SPTA1 | None | None | None | None |
| SPTA1 | None | None | None | None |
| SPTA1 | None | None | None | None |
| SPTA1 | None | None | None | None |

| | | | | |
|---|---|---|---|---|
| GMPR | None | None | None | None |
| MITF | None | None | None | None |
| NPC1 | None | None | None | None |
| NPC1 | None | None | None | None |
| NPC1 | None | None | None | None |
| NPC1 | None | None | None | None |
| NPC1 | None | None | None | None |
| NPC1 | None | None | None | None |
| NPC1 | None | None | None | None |
| NPC1 | None | None | None | None |
| NPC1 | None | None | None | None |
| NPC1 | None | None | None | None |
| NPC1 | None | None | None | None |
| NPC1 | None | None | None | None |
| NPC1 | None | None | None | None |
| NPC1 | None | None | None | None |
| NPC1 | None | None | None | None |
| NPC1 | None | None | None | None |
| NPC1 | None | None | None | None |
| NPC1 | None | None | None | None |
| NPC1 | None | None | None | None |
| NPC1 | None | None | None | None |
| NPC1 | None | None | None | None |
| NPC1 | None | None | None | None |
| NPC1 | None | None | None | None |
| NPC1 | None | None | None | None |
| NPC1 | None | None | None | None |
| NPC1 | None | None | None | None |
| NPC1 | None | None | None | None |
| NPC1 | None | None | None | None |
| NPC1 | None | None | None | None |
| NPC1 | None | None | None | None |
| NPC1 | None | None | None | None |
| NPC1 | None | None | None | None |
| NPC1 | None | None | None | None |
| NPC1 | None | None | None | None |
| NPC1 | None | None | None | None |
| NPC1 | None | None | None | None |
| NPC1 | None | None | None | None |
| NPC1 | None | None | None | None |
| NPC1 | None | None | None | None |
| NPC1 | None | None | None | None |
| NPC1 | None | None | None | None |
| NPC1 | None | None | None | None |
| NPC1 | None | None | None | None |
| C3 | None | None | None | None |
| NLRX1 | None | None | None | None |
| IGLL1 | 0.244 | 2 | 2h3n | D |
| APRT | 0.769 | 4 | 1zn9 | B |
| APRT | 0.417 | 5 | 4x44 | A |
| APRT | None | None | None | None |
| APRT | 0.923 | 1 | 1zn8 | A |
| APRT | None | None | None | None |
| ZIC3 | None | None | None | None |

| | | | | | |
|---|---|---|---|---|---|
| ZIC3 | None | None | None | | None |
| KLHL3 | None | None | None | | None |
| RPS6KA3 | None | None | None | | None |
| RPS6KA3 | None | None | None | | None |
| RPS6KA3 | None | None | None | | None |
| F8 | None | None | None | | None |
| F8 | None | None | None | | None |
| F8 | None | None | None | | None |
| F8 | None | None | None | | None |
| F8 | None | None | None | | None |
| F8 | None | None | None | | None |
| F8 | None | None | None | | None |
| F8 | None | None | None | | None |
| F8 | None | None | None | | None |
| F8 | None | None | None | | None |
| F8 | None | None | None | | None |
| F8 | None | None | None | | None |
| F8 | None | None | None | | None |
| F8 | None | None | None | | None |
| F8 | None | None | None | | None |
| F8 | None | None | None | | None |
| F8 | None | None | None | | None |
| ISG15 | None | None | None | | None |
| EXOSC3 | None | None | None | | None |
| EXOSC3 | None | None | None | | None |
| EXOSC3 | None | None | None | | None |
| NKX2-5 | 0.403 | 1 | 4s0h | | B |
| NKX2-5 | None | None | None | | None |
| NKX2-5 | 0.403 | 1 | 4s0h | | B |
| NKX2-5 | 0.403 | 1 | 4s0h | | B |
| FH | 0.207 | 44 | | 30000 | A |
| FH | None | None | None | | None |
| FH | None | None | None | | None |
| FH | 0.304 | 30 | | 30000 | D |
| FH | 0.621 | 4 | | 30000 | A |
| FH | 0.462 | 6 | | 30000 | C |
| FH | None | None | None | | None |
| FH | None | None | None | | None |
| FH | 0.64 | 6 | 5d6b | | A |
| FH | 0.308 | 49 | | 30000 | C |
| FH | 0.64 | 1 | 5d6b | | A |
| FH | 0.64 | 1 | 5d6b | | A |
| SNRNP200 | None | None | None | | None |
| SNRNP200 | None | None | None | | None |
| SNRNP200 | None | None | None | | None |
| SNRNP200 | None | None | None | | None |
| CHRNA1 | None | None | None | | None |
| CHRNA1 | None | None | None | | None |
| MYOM1 | None | None | None | | None |
| MYOM1 | None | None | None | | None |
| MYOM1 | None | None | None | | None |
| MYOM1 | None | None | None | | None |
| MYOM1 | None | None | None | | None |
| MYOM1 | None | None | None | | None |

| | | | | |
|---|---|---|---|---|
| LDLR | None | None | None | None |
| LDLR | None | None | None | None |
| LDLR | None | None | None | None |
| LDLR | None | None | None | None |
| LDLR | None | None | None | None |
| LDLR | None | None | None | None |
| EHMT1 | 0.286 | 8 | 3b7b | A |
| EHMT1 | 0.545 | 3 | 3b7b | A |
| EHMT1 | None | None | None | None |
| COL9A2 | None | None | None | None |
| HFE2 | None | None | None | None |
| HFE2 | None | None | None | None |
| CD3G | None | None | None | None |
| MMAB | None | None | None | None |
| MMAB | 0.5 | 3 | 2idx | C |
| MMAB | 0.45 | 7 | 2idx | C |
| MMAB | 0.65 | 3 | 2idx | A |
| MMAB | 0.65 | 3 | 2idx | A |
| MMAB | 0.7 | 2 | 2idx | B |
| MMAB | 0.7 | 2 | 2idx | B |
| MMAB | 0.35 | 4 | 2idx | A |
| MMAB | 0.4 | 11 | 2idx | C |
| MMAB | None | None | None | None |
| TAF1 | None | None | None | None |
| TAF1 | None | None | None | None |
| CD4 | None | None | None | None |
| QARS | None | None | None | None |
| QARS | None | None | None | None |
| QARS | None | None | None | None |
| QARS | None | None | None | None |
| QARS | None | None | None | None |
| FGFR3 | None | None | None | None |
| FGFR3 | None | None | None | None |
| FGFR3 | None | None | None | None |
| FGFR3 | None | None | None | None |
| FGFR3 | None | None | None | None |
| FGFR3 | None | None | None | None |
| FLNB | None | None | None | None |
| FLNB | None | None | None | None |
| FLNB | None | None | None | None |
| FLNB | None | None | None | None |
| FLNB | None | None | None | None |
| FLNB | None | None | None | None |
| FLNB | None | None | None | None |
| FLNB | None | None | None | None |
| IL21R | None | None | None | None |
| RB1 | None | None | None | None |
| RB1 | None | None | None | None |
| RB1 | None | None | None | None |
| NTRK1 | 0.533 | 1 | 1he7 | A |
| NOTCH1 | None | None | None | None |
| MYOM1 | None | None | None | None |
| MYOM1 | None | None | None | None |
| MYOM1 | None | None | None | None |

| | | | | |
|---|---|---|---|---|
| MYOM1 | None | None | None | None |
| MYOM1 | None | None | None | None |
| MYOM1 | None | None | None | None |
| COQ9 | None | None | None | None |
| COQ9 | None | None | None | None |
| CRBN | None | None | None | None |
| DCTN1 | None | None | None | None |
| DCTN1 | None | None | None | None |
| DCTN1 | None | None | None | None |
| DCTN1 | None | None | None | None |
| DCTN1 | None | None | None | None |
| DCTN1 | 0.333 | 1 | 2hkn | A |
| THAP1 | None | None | None | None |
| THAP1 | None | None | None | None |
| GNAS | None | None | None | None |
| GNAS | None | None | None | None |
| HNF4A | None | None | None | None |
| HNF4A | None | None | None | None |
| ALDH3A2 | None | None | None | None |
| ALDH3A2 | None | None | None | None |
| ALDH3A2 | 0.412 | 5 | 4qgk | B |
| ALDH3A2 | 0.557 | 2 | 4qgk | A |
| ALDH3A2 | None | None | None | None |
| ALDH3A2 | 0.569 | 3 | 4qgk | B |
| ALDH3A2 | None | None | None | None |
| ALDH3A2 | None | None | None | None |
| ALDH3A2 | None | None | None | None |
| HCN1 | None | None | None | None |
| PLEC | None | None | None | None |
| SCN5A | None | None | None | None |
| SCN5A | None | None | None | None |
| STAMBP | None | None | None | None |
| STAMBP | None | None | None | None |
| RNF125 | None | None | None | None |
| RNF125 | None | None | None | None |
| USH1C | None | None | None | None |
| OBSL1 | None | None | None | None |
| GRIN2B | None | None | None | None |
| GRIN2B | None | None | None | None |
| HCN4 | 0.186 | 6 | 4kl1 | C |
| HCN4 | 0.222 | 5 | 3u11 | A |
| TALDO1 | 0.192 | 51 | 1f05 | B |
| ITGB3 | None | None | None | None |
| CHKB | None | None | None | None |
| CHKB | None | None | None | None |
| CHKB | 0.333 | 7 | 3lq3 | A |
| CHKB | None | None | None | None |
| ACTN4 | None | None | None | None |
| ACTN4 | None | None | None | None |
| ACTN4 | None | None | None | None |
| ACTN4 | None | None | None | None |
| CDT1 | None | None | None | None |
| CDT1 | None | None | None | None |
| CDT1 | None | None | None | None |

| | | | | |
|---|---|---|---|---|
| CDT1 | None | None | None | None |
| CDT1 | None | None | None | None |
| CDT1 | None | None | None | None |
| MAPT | None | None | None | None |
| F5 | 0.517 | 4 | 1czs | A |
| F5 | None | None | None | None |
| F5 | None | None | None | None |
| ESRRB | 0.186 | 13 | 1lo1 | A |
| DGUOK | 0.188 | 23 | 2ocp | B |
| DGUOK | 0.471 | 8 | 2ocp | A |
| DGUOK | None | None | None | None |
| DGUOK | 0.375 | 6 | 2ocp | B |
| DGUOK | 0.765 | 1 | 2ocp | A |
| DGUOK | 0.438 | 11 | 2ocp | B |
| DGUOK | 0.294 | 19 | 2ocp | A |
| DGUOK | None | None | None | None |
| DGUOK | None | None | None | None |
| CENPJ | None | None | None | None |
| TTN | None | None | None | None |
| TTN | None | None | None | None |
| TTN | None | None | None | None |
| TTN | None | None | None | None |
| TNFRSF1A | None | None | None | None |
| TNFRSF1A | None | None | None | None |
| TNFRSF1A | None | None | None | None |
| TNFRSF1A | None | None | None | None |
| TNFRSF1A | None | None | None | None |
| TNFRSF1A | None | None | None | None |
| TNFRSF1A | None | None | None | None |
| TNFRSF1A | None | None | None | None |
| TNFRSF1A | None | None | None | None |
| TNFRSF1A | None | None | None | None |
| TNFRSF1A | None | None | None | None |
| TNFRSF1A | None | None | None | None |
| ASL | None | None | None | None |
| ASL | 0.625 | 25 | 1aos | A |
| ASL | None | None | None | None |
| ASL | 0.375 | 28 | 1k62 | A |
| ASL | 0.438 | 4 | 1k62 | B |
| ASL | 0.562 | 30 | 1k62 | A |
| ASL | 0.625 | 1 | 1aos | A |
| ASL | None | None | None | None |
| ASL | None | None | None | None |
| ASL | None | None | None | None |
| MYO7A | None | None | None | None |
| MYO7A | None | None | None | None |
| MYO7A | None | None | None | None |
| MYO7A | None | None | None | None |
| MYO7A | None | None | None | None |
| MYO7A | None | None | None | None |
| MYO7A | None | None | None | None |
| MYO7A | None | None | None | None |
| MYO7A | None | None | None | None |
| MYO7A | None | None | None | None |

| | | | | |
|---|---|---|---|---|
| MYO7A | None | None | None | None |
| MYO7A | None | None | None | None |
| MYO7A | None | None | None | None |
| MYO7A | None | None | None | None |
| MYO7A | None | None | None | None |
| MYO7A | None | None | None | None |
| MYO7A | None | None | None | None |
| MYO7A | None | None | None | None |
| MYO7A | None | None | None | None |
| MYO7A | None | None | None | None |
| APC | None | None | None | None |
| ALAD | None | None | None | None |
| ALAD | 0.143 | 21 | 1pv8 | A |
| ALAD | None | None | None | None |
| ALAD | 0.542 | 1 | 1E+051 | A |
| ALAD | None | None | None | None |
| ALAD | None | None | None | None |
| POR | None | None | None | None |
| CSF1R | None | None | None | None |
| CSF1R | None | None | None | None |
| CSF1R | None | None | None | None |
| CSF1R | None | None | None | None |
| CSF1R | None | None | None | None |
| CSF1R | None | None | None | None |
| CSF1R | None | None | None | None |
| CSF1R | None | None | None | None |
| CSF1R | None | None | None | None |
| CSF1R | None | None | None | None |
| CSF1R | None | None | None | None |
| VWF | None | None | None | None |
| VWF | None | None | None | None |
| PHF8 | 0.333 | 6 | 3k3n | A |
| ACY1 | None | None | None | None |
| SMN1 | None | None | None | None |
| SMN1 | None | None | None | None |
| DLAT | None | None | None | None |
| DLAT | None | None | None | None |
| B2M | None | None | None | None |
| SOX9 | None | None | None | None |
| SOX9 | None | None | None | None |
| SOX9 | None | None | None | None |
| SOX9 | None | None | None | None |
| SOX9 | None | None | None | None |
| CFH | None | None | None | None |
| FLNC | None | None | None | None |
| FLNC | None | None | None | None |
| FLNC | None | None | None | None |
| ITGB2 | None | None | None | None |
| ITGB2 | 0.261 | 2 | 5e6r | B |
| ITGB2 | None | None | None | None |
| ITGB2 | None | None | None | None |
| ITGB2 | None | None | None | None |
| ITGB2 | None | None | None | None |
| ITGB2 | 0.174 | 1 | 5e6u | B |

| | | | | |
|---|---|---|---|---|
| ITGB2 | None | None | None | None |
| ITGB2 | None | None | None | None |
| PSEN1 | None | None | None | None |
| PSEN1 | None | None | None | None |
| PSEN1 | None | None | None | None |
| PSEN1 | None | None | None | None |
| PSEN1 | None | None | None | None |
| PSEN1 | None | None | None | None |
| OXCT1 | None | None | None | None |
| OXCT1 | 0.385 | 24 | 3dlx | A |
| OXCT1 | None | None | None | None |
| OXCT1 | None | None | None | None |
| OXCT1 | None | None | None | None |
| OXCT1 | None | None | None | None |
| OXCT1 | None | None | None | None |
| DHH | None | None | None | None |
| PRKN | None | None | None | None |
| PRKN | None | None | None | None |
| PRKN | None | None | None | None |
| PRKN | None | None | None | None |
| PRKN | None | None | None | None |
| PRKN | None | None | None | None |
| PRKN | None | None | None | None |
| PRKN | None | None | None | None |
| PRKN | None | None | None | None |
| PRKN | None | None | None | None |
| PRKN | None | None | None | None |
| LCAT | None | None | None | None |
| LCAT | None | None | None | None |
| LCAT | None | None | None | None |
| LCAT | None | None | None | None |
| LCAT | None | None | None | None |
| LCAT | None | None | None | None |
| LCAT | None | None | None | None |
| LCAT | None | None | None | None |
| LCAT | None | None | None | None |
| LCAT | None | None | None | None |
| LCAT | None | None | None | None |
| LCAT | None | None | None | None |
| FOXP3 | None | None | None | None |
| FOXP3 | None | None | None | None |
| FOXP3 | None | None | None | None |
| FOXP3 | None | None | None | None |
| SUFU | None | None | None | None |
| HSPA9 | 0.571 | 14 | 4kbo | A |
| HSPA9 | None | None | None | None |
| SQSTM1 | None | None | None | None |
| DARS2 | None | None | None | None |
| DARS2 | None | None | None | None |
| DARS2 | None | None | None | None |
| DARS2 | None | None | None | None |
| DARS2 | 0.412 | 2 | 4ah6 | A |
| DARS2 | None | None | None | None |
| DARS2 | None | None | None | None |

| | | | | |
|---|---|---|---|---|
| DARS2 | None | None | None | None |
| DARS2 | None | None | None | None |
| DARS2 | None | None | None | None |
| DARS2 | None | None | None | None |
| DARS2 | None | None | None | None |
| NT5C3A | None | None | None | None |
| NT5C3A | 0.4 | 18 | 2vkq | A |
| NT5C3A | 0.4 | 7 | 2vkq | A |
| NT5C3A | 0.6 | 14 | 2vkq | A |
| ETFA | None | None | None | None |
| ETFA | None | None | None | None |
| ETFA | None | None | None | None |
| ETFA | None | None | None | None |
| ETFA | None | None | None | None |
| ETFA | None | None | None | None |
| ETFA | None | None | None | None |
| TTN | None | None | None | None |
| TTN | None | None | None | None |
| FLT4 | None | None | None | None |
| FGB | None | None | None | None |
| HPCA | None | None | None | None |
| TAP2 | None | None | None | None |
| TAP2 | None | None | None | None |
| PTPRC | None | None | None | None |
| SLC4A1 | None | None | None | None |
| SLC4A1 | None | None | None | None |
| SLC4A1 | None | None | None | None |
| SLC4A1 | None | None | None | None |
| SLC4A1 | None | None | None | None |
| SLC4A1 | None | None | None | None |
| MYH7 | None | None | None | None |
| MYH7 | None | None | None | None |
| MYH7 | None | None | None | None |
| MYH7 | None | None | None | None |
| MYH7 | None | None | None | None |
| MYH7 | None | None | None | None |
| MYH7 | None | None | None | None |
| MYH7 | None | None | None | None |
| MYH7 | None | None | None | None |
| PARN | None | None | None | None |
| PARN | None | None | None | None |
| NDUFS6 | None | None | None | None |
| VRK1 | None | None | None | None |
| VRK1 | None | None | None | None |
| VRK1 | None | None | None | None |
| VRK1 | 0.29 | 8 | 3op5 | A |
| VRK1 | None | None | None | None |
| VRK1 | None | None | None | None |
| COLEC11 | 0.229 | 23 | 4ymd | B |
| VHL | None | None | None | None |
| TTN | None | None | None | None |
| NCF2 | None | None | None | None |
| NCF2 | None | None | None | None |
| NCF2 | None | None | None | None |

| | | | | |
|---|---|---|---|---|
| NCF2 | None | None | None | None |
| NCF2 | None | None | None | None |
| RPS26 | None | None | None | None |
| CETP | None | None | None | None |
| CETP | None | None | None | None |
| CETP | None | None | None | None |
| CETP | None | None | None | None |
| CETP | None | None | None | None |
| CETP | None | None | None | None |
| CETP | None | None | None | None |
| CETP | None | None | None | None |
| CETP | None | None | None | None |
| CETP | None | None | None | None |
| CETP | None | None | None | None |
| CETP | None | None | None | None |
| CETP | None | None | None | None |
| FLNA | None | None | None | None |
| FLNA | None | None | None | None |
| FLNA | None | None | None | None |
| FLNA | None | None | None | None |
| FLNA | None | None | None | None |
| FLNA | None | None | None | None |
| FLNA | None | None | None | None |
| MYBPC3 | None | None | None | None |
| MYBPC3 | None | None | None | None |
| MYBPC3 | None | None | None | None |
| MYBPC3 | None | None | None | None |
| MYBPC3 | None | None | None | None |
| MYBPC3 | None | None | None | None |
| MYBPC3 | None | None | None | None |
| MYBPC3 | None | None | None | None |
| MYBPC3 | None | None | None | None |
| MYBPC3 | None | None | None | None |
| LSS | None | None | None | None |
| C6 | None | None | None | None |
| C6 | None | None | None | None |
| RLBP1 | None | None | None | None |
| RLBP1 | None | None | None | None |
| RLBP1 | None | None | None | None |
| RLBP1 | None | None | None | None |
| RLBP1 | None | None | None | None |
| RLBP1 | None | None | None | None |
| RLBP1 | None | None | None | None |
| RLBP1 | None | None | None | None |
| MMAA | None | None | None | None |
| MMAA | 0.3 | 1 | 2www | C |
| MMAA | 0.286 | 4 | 2www | D |
| MMAA | None | None | None | None |
| MMAA | None | None | None | None |
| MMAA | None | None | None | None |
| IL1RN | None | None | None | None |
| MASTL | None | None | None | None |
| FHL1 | None | None | None | None |
| FHL1 | None | None | None | None |

| | | | | |
|---|---|---|---|---|
| FHL1 | None | None | None | None |
| FHL1 | None | None | None | None |
| FHL1 | None | None | None | None |
| FHL1 | None | None | None | None |
| FHL1 | None | None | None | None |
| FHL1 | None | None | None | None |
| FHL1 | None | None | None | None |
| EIF2B1 | 0.636 | 1 | 3ecs | H |
| EIF2B1 | 0.636 | 1 | 3ecs | H |
| COQ8A | None | None | None | None |
| COQ8A | None | None | None | None |
| COQ8A | None | None | None | None |
| COQ8A | None | None | None | None |
| COQ8A | None | None | None | None |
| COQ8A | None | None | None | None |
| COQ8A | None | None | None | None |
| COQ8A | None | None | None | None |
| COQ8A | None | None | None | None |
| COQ8A | None | None | None | None |
| COQ8A | None | None | None | None |
| COQ8A | None | None | None | None |
| COQ8A | None | None | None | None |
| COQ8A | None | None | None | None |
| COQ8A | None | None | None | None |
| COQ8A | None | None | None | None |
| UNC119 | None | None | None | None |
| IGHMBP2 | None | None | None | None |
| JAG1 | None | None | None | None |
| JAG1 | None | None | None | None |
| JAG1 | None | None | None | None |
| GNB1 | None | None | None | None |
| GNB1 | 0.3 | 3 | 5ukl | B |
| FLNA | None | None | None | None |
| FLNA | None | None | None | None |
| FLNA | None | None | None | None |
| FLNA | None | None | None | None |
| FLNA | None | None | None | None |
| CREBBP | None | None | None | None |
| CSF3R | None | None | None | None |
| CSF3R | None | None | None | None |
| HIBCH | None | None | None | None |
| HIBCH | None | None | None | None |
| HIBCH | None | None | None | None |
| MMACHC | None | None | None | None |
| MMACHC | None | None | None | None |
| MMACHC | None | None | None | None |
| MMACHC | None | None | None | None |
| MMACHC | None | None | None | None |
| MMACHC | None | None | None | None |
| MMACHC | None | None | None | None |
| MMACHC | None | None | None | None |
| MMACHC | None | None | None | None |
| MMACHC | None | None | None | None |
| MMACHC | None | None | None | None |

| | | | | |
|---|---|---|---|---|
| KLF1 | None | None | None | None |
| NAGS | None | None | None | None |
| NAGS | None | None | None | None |
| NAGS | None | None | None | None |
| PIK3R2 | None | None | None | None |
| EDNRB | None | None | None | None |
| EDNRB | None | None | None | None |
| EDNRB | None | None | None | None |
| EDNRB | None | None | None | None |
| E | 0.156 | 69 | 5glh | A |
| RUNX1 | None | None | None | None |
| RUNX1 | None | None | None | None |
| CTNNB1 | None | None | None | None |
| CTNNB1 | None | None | None | None |
| CTNNB1 | None | None | None | None |
| CTNNB1 | None | None | None | None |
| CTNNB1 | None | None | None | None |
| CTNNB1 | None | None | None | None |
| TTN | None | None | None | None |
| TTN | None | None | None | None |
| TYK2 | None | None | None | None |
| TYK2 | None | None | None | None |
| TYK2 | None | None | None | None |
| ERCC4 | None | None | None | None |
| ERCC4 | None | None | None | None |
| ERCC4 | None | None | None | None |
| ERCC4 | None | None | None | None |
| CFI | None | None | None | None |
| CFI | None | None | None | None |
| CFI | None | None | None | None |
| CFI | None | None | None | None |
| CFI | None | None | None | None |
| CFI | None | None | None | None |
| CFI | None | None | None | None |
| CFI | None | None | None | None |
| CFI | None | None | None | None |
| CFI | 0.301 | 5 | 2xrc | A |
| CFI | None | None | None | None |
| CFI | 0.691 | 2 | 5o32 | J |
| CFI | 0.691 | 2 | 5o32 | J |
| CFI | 0.524 | 5 | 5o32 | I |
| NOTCH3 | None | None | None | None |
| NOTCH3 | None | None | None | None |
| NOTCH3 | None | None | None | None |
| NOTCH3 | None | None | None | None |
| TBX1 | None | None | None | None |
| TBX1 | None | None | None | None |
| DDB2 | None | None | None | None |
| DDB2 | None | None | None | None |
| TP53 | None | None | None | None |
| POMGNT1 | None | None | None | None |
| POMGNT1 | None | None | None | None |
| POMGNT1 | None | None | None | None |
| POMGNT1 | None | None | None | None |

| | | | | |
|---|---|---|---|---|
| POMGNT1 | None | None | None | None |
| POMGNT1 | None | None | None | None |
| POMGNT1 | None | None | None | None |
| POMGNT1 | None | None | None | None |
| POMGNT1 | None | None | None | None |
| POMGNT1 | None | None | None | None |
| POMGNT1 | None | None | None | None |
| POMGNT1 | None | None | None | None |
| POMGNT1 | None | None | None | None |
| FMR1 | None | None | None | None |
| FMR1 | None | None | None | None |
| SLC9A3R1 | None | None | None | None |
| HSPB1 | None | None | None | None |
| HSPB1 | None | None | None | None |
| HSPB1 | None | None | None | None |
| HSPB1 | None | None | None | None |
| HSPB1 | None | None | None | None |
| HSPB1 | None | None | None | None |
| HSPB1 | None | None | None | None |
| SOD3 | None | None | None | None |
| AGA | 1 | 2 | 1apz | D |
| AGA | None | None | None | None |
| AGA | None | None | None | None |
| NDUFS7 | None | None | None | None |
| NDUFS7 | None | None | None | None |
| SOX2 | None | None | None | None |
| CSRP3 | None | None | None | None |
| CSRP3 | None | None | None | None |
| PAH | None | None | None | None |
| PAH | None | None | None | None |
| PAH | None | None | None | None |
| PAH | None | None | None | None |
| CACNA1S | None | None | None | None |
| TRPV4 | None | None | None | None |
| TRPV4 | None | None | None | None |
| TRPV4 | None | None | None | None |
| TRPV4 | 0.58 | 3 | 4dx2 | A |
| TRPV4 | None | None | None | None |
| TRPV4 | None | None | None | None |
| TRPV4 | None | None | None | None |
| TRPV4 | None | None | None | None |
| TRPV4 | None | None | None | None |
| TRPV4 | None | None | None | None |
| TRPV4 | None | None | None | None |
| TRPV4 | None | None | None | None |
| TRPV4 | None | None | None | None |
| TNNI3 | None | None | None | None |
| MLYCD | None | None | None | None |
| MLYCD | None | None | None | None |
| MLYCD | None | None | None | None |
| MLYCD | None | None | None | None |
| AIP | None | None | None | None |
| AIP | None | None | None | None |
| ERCC2 | None | None | None | None |

| | | | | |
|---|---|---|---|---|
| ERCC2 | None | None | None | None |
| ERCC2 | None | None | None | None |
| ERCC2 | None | None | None | None |
| ERCC2 | None | None | None | None |
| ERCC2 | None | None | None | None |
| ERCC2 | None | None | None | None |
| ERCC2 | None | None | None | None |
| ERCC2 | None | None | None | None |
| ERCC2 | None | None | None | None |
| ERCC2 | None | None | None | None |
| ERCC2 | None | None | None | None |
| ERCC2 | None | None | None | None |
| ERCC2 | None | None | None | None |
| PUS1 | None | None | None | None |
| PUS1 | None | None | None | None |
| PUS1 | None | None | None | None |
| CASK | None | None | None | None |
| CASK | None | None | None | None |
| BLVRA | None | None | None | None |
| DICER1 | None | None | None | None |
| DICER1 | None | None | None | None |
| DICER1 | None | None | None | None |
| EIF2AK3 | None | None | None | None |
| EIF2AK3 | None | None | None | None |
| EIF2AK3 | None | None | None | None |
| KIT | None | None | None | None |
| KIT | None | None | None | None |
| KIT | None | None | None | None |
| KIT | None | None | None | None |
| KIT | None | None | None | None |
| KIT | None | None | None | None |
| DNMT3B | None | None | None | None |
| TGM1 | None | None | None | None |
| NDUFA10 | None | None | None | None |
| NDUFA10 | None | None | None | None |
| PRPF31 | None | None | None | None |
| PRPF31 | None | None | None | None |
| ECE1 | None | None | None | None |
| ECE1 | None | None | None | None |
| ECE1 | None | None | None | None |
| NDUFV2 | None | None | None | None |
| NDUFV2 | None | None | None | None |
| NDUFV2 | None | None | None | None |
| GHRHR | None | None | None | None |
| GHRHR | None | None | None | None |
| GHRHR | None | None | None | None |
| SLC4A1 | None | None | None | None |
| FGF9 | None | None | None | None |
| BARD1 | None | None | None | None |
| BARD1 | None | None | None | None |
| BARD1 | None | None | None | None |
| PTPN12 | 0.741 | 2 | 5j8r | D |
| MASP2 | None | None | None | None |
| MASP2 | None | None | None | None |

| | | | | |
|---|---|---|---|---|
| MASP2 | None | None | None | None |
| MASP2 | 0.118 | 3 | 1szb | A |
| HOXB1 | None | None | None | None |
| SLC4A1 | None | None | None | None |
| SLC4A1 | None | None | None | None |
| SLC4A1 | None | None | None | None |
| SLC4A1 | None | None | None | None |
| SLC4A1 | None | None | None | None |
| SLC4A1 | None | None | None | None |
| SLC4A1 | None | None | None | None |
| SLC4A1 | None | None | None | None |
| SLC4A1 | None | None | None | None |
| SLC4A1 | None | None | None | None |
| SLC4A1 | None | None | None | None |
| SLC4A1 | None | None | None | None |
| SLC4A1 | None | None | None | None |
| SLC4A1 | None | None | None | None |
| SLC4A1 | None | None | None | None |
| SLC4A1 | None | None | None | None |
| SLC4A1 | None | None | None | None |
| CBL | None | None | None | None |
| FAS | None | None | None | None |
| MTMR2 | None | None | None | None |
| MTMR2 | None | None | None | None |
| MTMR2 | None | None | None | None |
| MTMR2 | None | None | None | None |
| MTMR2 | None | None | None | None |
| MTMR2 | None | None | None | None |
| MTMR2 | None | None | None | None |
| MTMR2 | None | None | None | None |
| KLHL3 | None | None | None | None |
| KLHL3 | None | None | None | None |
| KLHL3 | 0.667 | 1 | 4ch9 | B |
| KLHL3 | 0.733 | 1 | 5nkp | A |
| KLHL3 | None | None | None | None |
| KLHL3 | None | None | None | None |
| RNASEH1 | None | None | None | None |
| ACTN1 | None | None | None | None |
| PHKG2 | None | None | None | None |
| PHKG2 | None | None | None | None |
| PHKG2 | 0.472 | 2 | 2y7j | A |
| NTRK1 | None | None | None | None |
| APOH | None | None | None | None |
| PMS2 | None | None | None | None |
| PMS2 | None | None | None | None |
| PMS2 | 0.797 | 1 | 1ea6 | A |
| PMS2 | None | None | None | None |
| PMS2 | None | None | None | None |
| PMS2 | None | None | None | None |
| PMS2 | 0.156 | 10 | 1ea6 | A |
| PMS2 | None | None | None | None |
| PMS2 | 0.188 | 8 | 1ea6 | A |
| PMS2 | None | None | None | None |
| PMS2 | None | None | None | None |

| | | | | |
|---|---|---|---|---|
| PMS2 | None | None | None | None |
| PMS2 | None | None | None | None |
| PMS2 | None | None | None | None |
| PMS2 | None | None | None | None |
| PMS2 | None | None | None | None |
| PMS2 | None | None | None | None |
| PMS2 | None | None | None | None |
| PMS2 | None | None | None | None |
| PMS2 | None | None | None | None |
| PMS2 | None | None | None | None |
| PMS2 | None | None | None | None |
| MOCS2 | None | None | None | None |
| MOCS2 | None | None | None | None |
| RPGRIP1 | None | None | None | None |
| RPGRIP1 | None | None | None | None |
| LIG4 | None | None | None | None |
| PROC | None | None | None | None |
| PROC | 0.111 | 34 | 1aut | C |
| NAT1 | None | None | None | None |
| NAT1 | None | None | None | None |
| PNPO | None | None | None | None |
| PNPO | 0.5 | 2 | 1nrg | A |
| PNPO | None | None | None | None |
| PNPO | None | None | None | None |
| PNPO | None | None | None | None |
| MRE11 | None | None | None | None |
| MRE11 | None | None | None | None |
| MRE11 | None | None | None | None |
| MRE11 | None | None | None | None |
| DDC | None | None | None | None |
| DDC | None | None | None | None |
| DDC | None | None | None | None |
| DDC | None | None | None | None |
| DDC | None | None | None | None |
| DDC | 0.909 | 7 | 3rch | B |
| DDC | 0.909 | 7 | 3rch | B |
| DDC | 0.643 | 4 | 3rbf | A |
| DDC | None | None | None | None |
| DDC | None | None | None | None |
| DDC | None | None | None | None |
| ANK2 | 0.2 | 7 | 4rlv | A |
| ANK2 | 0.242 | 5 | 4rlv | A |
| ANK2 | None | None | None | None |
| AK2 | None | None | None | None |
| AK2 | 0.241 | 12 | 2c9y | A |
| AK2 | 0.207 | 17 | 2c9y | A |
| AK2 | None | None | None | None |
| AK2 | 0.138 | 10 | 2c9y | A |
| AK2 | None | None | None | None |
| GPHN | None | None | None | None |
| HSD17B4 | 0.66 | 1 | 1zbq | A |
| HSD17B4 | 0.66 | 1 | 1zbq | A |
| HSD17B4 | None | None | None | None |
| HSD17B4 | 0.297 | 7 | 1zbq | B |

| | | | | |
|---|---|---|---|---|
| HSD17B4 | None | None | None | None |
| HSD17B4 | None | None | None | None |
| APTX | 0.6 | 5 | 4ndg | A |
| APTX | 0.6 | 5 | 4ndg | A |
| APTX | None | None | None | None |
| APTX | None | None | None | None |
| APTX | None | None | None | None |
| APTX | None | None | None | None |
| APTX | None | None | None | None |
| FANCL | None | None | None | None |
| FANCL | None | None | None | None |
| EMG1 | None | None | None | None |
| EMG1 | None | None | None | None |
| EZH2 | 0.205 | 1 | 4mi5 | A |
| ABCB6 | None | None | None | None |
| NCF2 | None | None | None | None |
| NCF2 | None | None | None | None |
| COL4A2 | None | None | None | None |
| COL4A2 | 1 | 1 | 1li1 | F |
| COL4A2 | None | None | None | None |
| COL4A2 | None | None | None | None |
| COL4A2 | 0.75 | 16 | 1li1 | C |
| SLX4 | None | None | None | None |
| SERPIND1 | None | None | None | None |
| FGFR3 | None | None | None | None |
| FGFR3 | None | None | None | None |
| FGFR3 | None | None | None | None |
| FGFR3 | None | None | None | None |
| FGFR3 | None | None | None | None |
| FGFR3 | None | None | None | None |
| TREM2 | None | None | None | None |
| TREM2 | None | None | None | None |
| TREM2 | None | None | None | None |
| TREM2 | None | None | None | None |
| TREM2 | None | None | None | None |
| TREM2 | None | None | None | None |
| TREM2 | None | None | None | None |
| LIG4 | None | None | None | None |
| LIG4 | None | None | None | None |
| LIG4 | None | None | None | None |
| LIG4 | None | None | None | None |
| LIG4 | None | None | None | None |
| LIG4 | None | None | None | None |
| ECHS1 | None | None | None | None |
| ECHS1 | 0.279 | 4 | 2hw5 | A |
| ECHS1 | None | None | None | None |
| MTR | None | None | None | None |
| MTR | None | None | None | None |
| MTR | None | None | None | None |
| MTR | None | None | None | None |
| MTR | None | None | None | None |
| PMVK | None | None | None | None |
| PRNP | None | None | None | None |
| CHN1 | None | None | None | None |

| | | | | |
|---|---|---|---|---|
| CHN1 | None | None | None | None |
| CHN1 | None | None | None | None |
| CHN1 | None | None | None | None |
| CHN1 | None | None | None | None |
| CHN1 | None | None | None | None |
| CHN1 | None | None | None | None |
| CHN1 | None | None | None | None |
| SLC2A1 | None | None | None | None |
| SLC2A1 | None | None | None | None |
| SLC2A1 | None | None | None | None |
| SLC2A1 | None | None | None | None |
| SLC2A1 | None | None | None | None |
| SLC2A1 | None | None | None | None |
| SLC2A1 | None | None | None | None |
| SLC2A1 | None | None | None | None |
| SLC2A1 | None | None | None | None |
| SLC2A1 | None | None | None | None |
| SLC2A1 | None | None | None | None |
| SLC2A1 | None | None | None | None |
| SLC2A1 | None | None | None | None |
| SLC2A1 | None | None | None | None |
| SLC2A1 | None | None | None | None |
| SLC2A1 | None | None | None | None |
| SLC2A1 | None | None | None | None |
| PMM2 | None | None | None | None |
| PMM2 | None | None | None | None |
| PMM2 | None | None | None | None |
| PMM2 | None | None | None | None |
| PMM2 | None | None | None | None |
| PMM2 | None | None | None | None |
| PMM2 | None | None | None | None |
| PMM2 | None | None | None | None |
| PMM2 | None | None | None | None |
| PMM2 | None | None | None | None |
| PMM2 | None | None | None | None |
| PMM2 | None | None | None | None |
| PMM2 | None | None | None | None |
| PMM2 | None | None | None | None |
| PMM2 | None | None | None | None |
| PMM2 | None | None | None | None |
| PMM2 | None | None | None | None |
| PMM2 | None | None | None | None |
| PMM2 | None | None | None | None |
| PMM2 | None | None | None | None |
| PMM2 | None | None | None | None |
| PMM2 | None | None | None | None |
| PMM2 | None | None | None | None |
| PMM2 | None | None | None | None |
| PMM2 | None | None | None | None |
| PMM2 | None | None | None | None |
| PMM2 | None | None | None | None |
| PMM2 | None | None | None | None |
| CCKAR | None | None | None | None |
| MAN1B1 | None | None | None | None |

| | | | | |
|---|---|---|---|---|
| MAN1B1 | None | None | None | None |
| LAMA2 | None | None | None | None |
| LAMA2 | None | None | None | None |
| TBC1D4 | None | None | None | None |
| TBC1D4 | None | None | None | None |
| LDLR | None | None | None | None |
| LDLR | None | None | None | None |
| LDLR | None | None | None | None |
| LDLR | None | None | None | None |
| LDLR | None | None | None | None |
| LDLR | None | None | None | None |
| LDLR | None | None | None | None |
| LDLR | None | None | None | None |
| LDLR | None | None | None | None |
| LDLR | None | None | None | None |
| LDLR | None | None | None | None |
| LDLR | None | None | None | None |
| LDLR | None | None | None | None |
| LDLR | None | None | None | None |
| LDLR | None | None | None | None |
| LDLR | None | None | None | None |
| LDLR | None | None | None | None |
| LDLR | None | None | None | None |
| LDLR | None | None | None | None |
| LDLR | None | None | None | None |
| LDLR | None | None | None | None |
| LDLR | None | None | None | None |
| LDLR | None | None | None | None |
| LDLR | None | None | None | None |
| LDLR | None | None | None | None |
| LDLR | None | None | None | None |
| LDLR | None | None | None | None |
| LDLR | None | None | None | None |
| LDLR | None | None | None | None |
| LDLR | None | None | None | None |
| LDLR | None | None | None | None |
| LDLR | None | None | None | None |
| LDLR | None | None | None | None |
| LDLR | None | None | None | None |
| LDLR | None | None | None | None |
| LDLR | None | None | None | None |
| LDLR | None | None | None | None |
| LDLR | None | None | None | None |
| LDLR | None | None | None | None |
| LDLR | None | None | None | None |
| LDLR | None | None | None | None |
| LDLR | None | None | None | None |
| LDLR | 0.5 | 1 | 1ijq | A |
| LDLR | None | None | None | None |
| FANCM | None | None | None | None |
| HAMP | None | None | None | None |
| GUSB | 0.462 | 8 | 3hn3 | A |
| GUSB | None | None | None | None |
| GUSB | 0.45 | 16 | 3hn3 | B |
| GUSB | None | None | None | None |

| | | | | |
|---|---|---|---|---|
| GUSB | 0.105 | 30 | 1bhg | A |
| GUSB | None | None | None | None |
| GUSB | None | None | None | None |
| GUSB | None | None | None | None |
| GUSB | 0.105 | 38 | 1bhg | A |
| GUSB | None | None | None | None |
| GUSB | 0.105 | 30 | 1bhg | A |
| GUSB | None | None | None | None |
| GUSB | None | None | None | None |
| LIG1 | None | None | None | None |
| LIG1 | None | None | None | None |
| MYH7 | None | None | None | None |
| MYH7 | None | None | None | None |
| MYH7 | None | None | None | None |
| MYH7 | None | None | None | None |
| MYH7 | None | None | None | None |
| MYH7 | None | None | None | None |
| MYH7 | None | None | None | None |
| MYH7 | None | None | None | None |
| MYH7 | None | None | None | None |
| MYH7 | None | None | None | None |
| MYH7 | None | None | None | None |
| MYH7 | None | None | None | None |
| MYH7 | None | None | None | None |
| MYH7 | None | None | None | None |
| MYH7 | None | None | None | None |
| MYH7 | None | None | None | None |
| MYH7 | None | None | None | None |
| MYH7 | None | None | None | None |
| MYH7 | None | None | None | None |
| MYH7 | None | None | None | None |
| MYH7 | None | None | None | None |
| MYH7 | None | None | None | None |
| MYH7 | None | None | None | None |
| MYH7 | None | None | None | None |
| MYH7 | None | None | None | None |
| MYH7 | None | None | None | None |
| MYH7 | None | None | None | None |
| MYH7 | None | None | None | None |
| MYH7 | None | None | None | None |
| MYH7 | None | None | None | None |
| MYH7 | None | None | None | None |
| MYH7 | None | None | None | None |
| MYH7 | None | None | None | None |
| MYH7 | None | None | None | None |
| MYH7 | None | None | None | None |
| HCN4 | None | None | None | None |
| RECQL4 | None | None | None | None |
| RECQL4 | None | None | None | None |
| RECQL4 | None | None | None | None |
| RECQL4 | None | None | None | None |
| RECQL4 | None | None | None | None |
| RECQL4 | None | None | None | None |
| RECQL4 | None | None | None | None |
| RECQL4 | None | None | None | None |

| | | | | |
|---|---|---|---|---|
| RECQL4 | None | None | None | None |
| RECQL4 | None | None | None | None |
| RECQL4 | None | None | None | None |
| RECQL4 | None | None | None | None |
| RECQL4 | None | None | None | None |
| CHEK2 | None | None | None | None |
| PPARG | None | None | None | None |
| ACADVL | None | None | None | None |
| ACADVL | 0.385 | 4 | 3b96 | A |
| ACADVL | None | None | None | None |
| ACADVL | None | None | None | None |
| ACADVL | None | None | None | None |
| ACADVL | None | None | None | None |
| ACADVL | None | None | None | None |
| ACADVL | 0.346 | 14 | 3b96 | A |
| ACADVL | 0.346 | 14 | 3b96 | A |
| ACADVL | None | None | None | None |
| ACADVL | None | None | None | None |
| ACADVL | 0.231 | 10 | 3b96 | A |
| ACADVL | None | None | None | None |
| ACADVL | 0.346 | 14 | 3b96 | A |
| ACADVL | 0.269 | 12 | 3b96 | A |
| ACADVL | 0.423 | 2 | 3b96 | A |
| ACADVL | None | None | None | None |
| ACADVL | 0.135 | 20 | 3b96 | A |
| ACADVL | 0.135 | 20 | 3b96 | A |
| ACADVL | None | None | None | None |
| ACADVL | None | None | None | None |
| ACADVL | None | None | None | None |
| APOA1 | None | None | None | None |
| APOA1 | None | None | None | None |
| APOA1 | None | None | None | None |
| APOA1 | None | None | None | None |
| APOA1 | None | None | None | None |
| DNMT3A | None | None | None | None |
| DNMT3A | None | None | None | None |
| APC | None | None | None | None |
| TMOD1 | None | None | None | None |
| USH1C | None | None | None | None |
| USH1C | None | None | None | None |
| USH1C | None | None | None | None |
| USH1C | None | None | None | None |
| ERCC6 | None | None | None | None |
| ERCC6 | None | None | None | None |
| ERCC6 | None | None | None | None |
| ERCC8 | None | None | None | None |
| ERCC8 | None | None | None | None |
| ERCC8 | None | None | None | None |
| DPAGT1 | None | None | None | None |
| DPAGT1 | None | None | None | None |
| DPAGT1 | None | None | None | None |
| DPAGT1 | None | None | None | None |
| DPAGT1 | None | None | None | None |
| DPAGT1 | None | None | None | None |

| | | | | |
|---|---|---|---|---|
| DPAGT1 | None | None | None | None |
| IL2RG | None | None | None | None |
| HNF1A | None | None | None | None |
| HNF4A | None | None | None | None |
| FLT4 | None | None | None | None |
| FLT4 | None | None | None | None |
| FLT4 | None | None | None | None |
| CD3D | None | None | None | None |
| CD3D | None | None | None | None |
| MCOLN1 | None | None | None | None |
| MCOLN1 | None | None | None | None |
| MOCS2 | None | None | None | None |
| MOCS2 | None | None | None | None |
| ABCG8 | None | None | None | None |
| ABCG8 | None | None | None | None |
| ABCG8 | None | None | None | None |
| ABCG8 | None | None | None | None |
| ABCG8 | None | None | None | None |
| ABCG8 | None | None | None | None |
| ABCG8 | None | None | None | None |
| ABCG8 | None | None | None | None |
| ABCG8 | None | None | None | None |
| ABCG8 | None | None | None | None |
| ABCG8 | None | None | None | None |
| ABCG8 | None | None | None | None |
| ABCG8 | None | None | None | None |
| ABCG8 | None | None | None | None |
| CTNNA1 | None | None | None | None |
| TYMP | None | None | None | None |
| TYMP | None | None | None | None |
| TYMP | None | None | None | None |
| TYMP | None | None | None | None |
| TYMP | None | None | None | None |
| TYMP | None | None | None | None |
| TYMP | None | None | None | None |
| TYMP | None | None | None | None |
| TYMP | None | None | None | None |
| TYMP | None | None | None | None |
| TYMP | None | None | None | None |
| TYMP | None | None | None | None |
| TYMP | None | None | None | None |
| GAMT | None | None | None | None |
| GAMT | None | None | None | None |
| GAMT | None | None | None | None |
| GAMT | None | None | None | None |
| GAMT | None | None | None | None |
| GAMT | None | None | None | None |
| GAMT | None | None | None | None |
| GAMT | None | None | None | None |
| GAMT | None | None | None | None |
| GAMT | None | None | None | None |
| GAMT | None | None | None | None |
| GLRA1 | None | None | None | None |
| GLRA1 | None | None | None | None |

| | | | | | |
|---|---|---|---|---|---|
| GLRA1 | None | None | | None | None |
| GLRA1 | None | None | | None | None |
| GLRA1 | None | None | | None | None |
| GLRA1 | None | None | | None | None |
| GLRA1 | None | None | | None | None |
| GLRA1 | None | None | | None | None |
| GLRA1 | None | None | | None | None |
| ADK | None | None | | None | None |
| ACTN2 | None | None | | None | None |
| UBE2T | 0.134 | | 17 | 5ngz | A |
| TTN | None | None | | None | None |
| ATP7B | None | None | | None | None |
| ATP7B | None | None | | None | None |
| ATP7B | None | None | | None | None |
| ATP7B | None | None | | None | None |
| ATP7B | None | None | | None | None |
| ATP7B | None | None | | None | None |
| ATP7B | None | None | | None | None |
| ATP7B | None | None | | None | None |
| ATP7B | None | None | | None | None |
| ATP7B | None | None | | None | None |
| ATP7B | None | None | | None | None |
| ATP7B | None | None | | None | None |
| ATP7B | None | None | | None | None |
| CDKN1B | None | None | | None | None |
| CDKN1B | None | None | | None | None |
| CDKN1B | None | None | | None | None |
| CDKN1B | None | None | | None | None |
| CDKN1B | None | None | | None | None |
| CDKN1B | None | None | | None | None |
| MYOC | None | None | | None | None |
| MYOC | None | None | | None | None |
| MYOC | None | None | | None | None |
| MYOC | None | None | | None | None |
| MYOC | None | None | | None | None |
| MYOC | None | None | | None | None |
| MYOC | None | None | | None | None |
| MYOC | None | None | | None | None |
| MYOC | None | None | | None | None |
| MYOC | None | None | | None | None |
| MYOC | None | None | | None | None |
| MYOC | None | None | | None | None |
| MYOC | None | None | | None | None |
| MYOC | None | None | | None | None |
| MYOC | None | None | | None | None |
| SPTAN1 | None | None | | None | None |
| GC | None | None | | None | None |
| GC | None | None | | None | None |
| MAD1L1 | None | None | | None | None |
| NDST1 | None | None | | None | None |
| ARL2BP | None | None | | None | None |
| TOR1A | None | None | | None | None |
| C1S | 0.174 | | 8 | 4j1y | A |
| ARL6 | 0.508 | | 1 | 2h57 | A |

| | | | | | |
|---|---|---|---|---|---|
| ARL6 | None | None | | None | None |
| ARL6 | None | None | | None | None |
| ARL6 | None | None | | None | None |
| ARL6 | None | None | | None | None |
| ARL6 | None | None | | None | None |
| FGFR1 | None | None | | None | None |
| KLHL7 | None | None | | None | None |
| FLNC | None | None | | None | None |
| PTPRC | None | None | | None | None |
| GCH1 | None | None | | None | None |
| GCH1 | None | None | | None | None |
| GCH1 | 0.1 | | 9 | 1fb1 | A |
| GCH1 | 0.1 | | 5 | 1fb1 | A |
| GCH1 | None | None | | None | None |
| GCH1 | 0.2 | | 17 | 1fb1 | A |
| GCH1 | 0.1 | | 8 | 1fb1 | A |
| GCH1 | None | None | | None | None |
| GCH1 | None | None | | None | None |
| GCH1 | None | None | | None | None |
| GCH1 | 0.3 | | 1 | 1fb1 | D |
| GCH1 | 0.2 | | 3 | 1fb1 | D |
| GCH1 | 0.3 | | 11 | 1fb1 | B |
| GCH1 | 0.1 | | 10 | 1fb1 | A |
| CFH | None | None | | None | None |
| COL2A1 | None | None | | None | None |
| COL2A1 | None | None | | None | None |
| OCRL | None | None | | None | None |
| WRN | None | None | | None | None |
| WRN | None | None | | None | None |
| WRN | None | None | | None | None |
| WRN | None | None | | None | None |
| MASP1 | None | None | | None | None |
| MASP1 | None | None | | None | None |
| NSD1 | None | None | | None | None |
| PITX2 | None | None | | None | None |
| PITX2 | None | None | | None | None |
| PITX2 | None | None | | None | None |
| PITX2 | 0.362 | | 2 | 2lkx | A |
| PITX2 | 0.362 | | 2 | 2lkx | A |
| PITX2 | 0.362 | | 2 | 2lkx | A |
| B4GALT7 | 0.917 | | 1 | 4irq | A |
| B4GALT7 | None | None | | None | None |
| B4GALT7 | None | None | | None | None |
| B4GALT7 | None | None | | None | None |
| B4GALT7 | None | None | | None | None |
| FSHR | None | None | | None | None |
| IGHMBP2 | None | None | | None | None |
| IGHMBP2 | None | None | | None | None |
| IGHMBP2 | None | None | | None | None |
| IGHMBP2 | None | None | | None | None |
| IGHMBP2 | None | None | | None | None |
| IGHMBP2 | None | None | | None | None |
| IGHMBP2 | None | None | | None | None |
| IGHMBP2 | None | None | | None | None |

| | | | | |
|---|---|---|---|---|
| IGHMBP2 | None | None | None | None |
| IGHMBP2 | None | None | None | None |
| IGHMBP2 | None | None | None | None |
| IGHMBP2 | None | None | None | None |
| IGHMBP2 | None | None | None | None |
| IGHMBP2 | None | None | None | None |
| IGHMBP2 | None | None | None | None |
| IGHMBP2 | None | None | None | None |
| TTN | None | None | None | None |
| TTN | None | None | None | None |
| IRAK4 | None | None | None | None |
| PROC | None | None | None | None |
| PROC | None | None | None | None |
| RAD50 | None | None | None | None |
| RAD50 | None | None | None | None |
| DYNC1H1 | None | None | None | None |
| DYNC1H1 | None | None | None | None |
| DYNC1H1 | None | None | None | None |
| DYNC1H1 | None | None | None | None |
| DYNC1H1 | None | None | None | None |
| DYNC1H1 | None | None | None | None |
| DYNC1H1 | None | None | None | None |
| DYNC1H1 | None | None | None | None |
| DYNC1H1 | None | None | None | None |
| DYNC1H1 | None | None | None | None |
| DYNC1H1 | None | None | None | None |
| CYB5R3 | None | None | None | None |
| CYB5R3 | 0.161 | 10 | 1umk | A |
| CYB5R3 | None | None | None | None |
| CYB5R3 | None | None | None | None |
| CYB5R3 | None | None | None | None |
| CYB5R3 | None | None | None | None |
| CYB5R3 | None | None | None | None |
| CYB5R3 | None | None | None | None |
| CYB5R3 | None | None | None | None |
| CYB5R3 | None | None | None | None |
| CYB5R3 | None | None | None | None |
| SPTB | None | None | None | None |
| FANCL | None | None | None | None |
| FAM126A | None | None | None | None |
| SEPT9 | None | None | None | None |
| TTN | None | None | None | None |
| TTN | None | None | None | None |
| TTN | 0.268 | 2 | 4jnw | A |
| CNTN1 | None | None | None | None |
| C1QTNF5 | None | None | None | None |
| VCL | None | None | None | None |
| VCL | None | None | None | None |
| VCL | None | None | None | None |
| VCL | None | None | None | None |
| VCL | None | None | None | None |
| VCL | None | None | None | None |
| PPT1 | None | None | None | None |
| PPT1 | None | None | None | None |

| | | | | |
|---|---|---|---|---|
| PPT1 | None | None | None | None |
| PPT1 | None | None | None | None |
| PPT1 | None | None | None | None |
| PPT1 | None | None | None | None |
| PPT1 | None | None | None | None |
| PPT1 | None | None | None | None |
| PPT1 | None | None | None | None |
| PPT1 | None | None | None | None |
| PPT1 | None | None | None | None |
| PPT1 | None | None | None | None |
| PPT1 | None | None | None | None |
| PPT1 | None | None | None | None |
| PPT1 | None | None | None | None |
| PPT1 | None | None | None | None |
| PPT1 | None | None | None | None |
| PPT1 | None | None | None | None |
| PPT1 | None | None | None | None |
| NOTCH2 | None | None | None | None |
| NOTCH2 | None | None | None | None |
| IFIH1 | None | None | None | None |
| IFIH1 | None | None | None | None |
| IKBKG | None | None | None | None |
| DBH | None | None | None | None |
| DBH | None | None | None | None |
| DBH | None | None | None | None |
| DBH | None | None | None | None |
| DBH | None | None | None | None |
| DBH | None | None | None | None |
| DBH | None | None | None | None |
| DBH | None | None | None | None |
| DBH | None | None | None | None |
| DBH | None | None | None | None |
| IL10RA | None | None | None | None |
| IL10RA | None | None | None | None |
| IL10RA | None | None | None | None |
| IL10RA | None | None | None | None |
| IL10RA | None | None | None | None |
| IL10RA | None | None | None | None |
| IL10RA | None | None | None | None |
| ILK | None | None | None | None |
| HPD | None | None | None | None |
| HPD | None | None | None | None |
| HPD | None | None | None | None |
| MECP2 | None | None | None | None |
| MECP2 | None | None | None | None |
| MECP2 | None | None | None | None |
| MECP2 | None | None | None | None |
| MECP2 | None | None | None | None |
| MECP2 | None | None | None | None |
| MECP2 | None | None | None | None |
| MECP2 | None | None | None | None |
| KYNU | None | None | None | None |
| FAS | None | None | None | None |
| FAS | None | None | None | None |

| | | | | |
|---|---|---|---|---|
| ELP1 | None | None | None | None |
| ELP1 | None | None | None | None |
| ELP1 | None | None | None | None |
| ELP1 | None | None | None | None |
| ELP1 | None | None | None | None |
| ELP1 | None | None | None | None |
| ELP1 | None | None | None | None |
| ELP1 | None | None | None | None |
| HESX1 | None | None | None | None |
| HESX1 | None | None | None | None |
| HESX1 | None | None | None | None |
| HESX1 | None | None | None | None |
| GAD1 | None | None | None | None |
| GAD1 | None | None | None | None |
| NDUFA1 | None | None | None | None |
| NDUFA1 | None | None | None | None |
| NDUFA1 | None | None | None | None |
| PNPT1 | None | None | None | None |
| PNPT1 | None | None | None | None |
| PNPT1 | None | None | None | None |
| PNPT1 | None | None | None | None |
| PNPT1 | 0.389 | 1 | 3u1k | A |
| PNPT1 | None | None | None | None |
| PNPT1 | None | None | None | None |
| SLC25A13 | None | None | None | None |
| SLC25A13 | None | None | None | None |
| SLC25A13 | None | None | None | None |
| AP4M1 | None | None | None | None |
| AP4M1 | None | None | None | None |
| MEFV | None | None | None | None |
| MEFV | None | None | None | None |
| MEFV | None | None | None | None |
| MEFV | None | None | None | None |
| MEFV | None | None | None | None |
| MEFV | None | None | None | None |
| MEFV | None | None | None | None |
| MEFV | None | None | None | None |
| MEFV | None | None | None | None |
| CAPN3 | None | None | None | None |
| CAPN3 | None | None | None | None |
| CAPN3 | None | None | None | None |
| CAPN3 | 0.158 | 14 | 4okh | A |
| CAPN3 | 0.474 | 2 | 4okh | A |
| CAPN3 | 0.474 | 2 | 4okh | A |
| CAPN3 | None | None | None | None |
| GP6 | None | None | None | None |
| FAS | None | None | None | None |
| FBN1 | None | None | None | None |
| FBN1 | None | None | None | None |
| BMPR2 | None | None | None | None |
| BMPR2 | None | None | None | None |
| BMPR2 | None | None | None | None |
| BMPR2 | 0.327 | 6 | 3g2f | A |
| BMPR2 | 0.327 | 6 | 3g2f | A |

| | | | | |
|---|---|---|---|---|
| RET | None | None | None | None |
| RET | None | None | None | None |
| RET | None | None | None | None |
| RET | None | None | None | None |
| RET | None | None | None | None |
| RET | None | None | None | None |
| RET | None | None | None | None |
| RET | None | None | None | None |
| RET | None | None | None | None |
| RET | None | None | None | None |
| RET | None | None | None | None |
| RET | None | None | None | None |
| RET | None | None | None | None |
| RET | None | None | None | None |
| RET | None | None | None | None |
| RET | None | None | None | None |
| RET | None | None | None | None |
| RET | None | None | None | None |
| RET | None | None | None | None |
| RET | None | None | None | None |
| RET | None | None | None | None |
| RET | None | None | None | None |
| RET | None | None | None | None |
| RET | None | None | None | None |
| RET | None | None | None | None |
| RET | None | None | None | None |
| RET | None | None | None | None |
| RET | None | None | None | None |
| CSF2RA | None | None | None | None |
| CSF2RA | None | None | None | None |
| CSF2RA | None | None | None | None |
| BRCA1 | None | None | None | None |
| ZAP70 | 0.636 | 1 | 2ozo | A |
| ZAP70 | 0.636 | 1 | 2ozo | A |
| ZAP70 | 0.26 | 9 | 2ozo | A |
| PIK3R2 | None | None | None | None |
| TCN2 | None | None | None | None |
| TCN2 | None | None | None | None |
| TCN2 | None | None | None | None |
| TCN2 | None | None | None | None |
| TCN2 | None | None | None | None |
| TCN2 | None | None | None | None |
| TCN2 | None | None | None | None |
| TCN2 | None | None | None | None |
| EPM2A | None | None | None | None |
| EPM2A | None | None | None | None |
| EPM2A | None | None | None | None |
| EPM2A | None | None | None | None |
| AARS | None | None | None | None |
| AARS | None | None | None | None |
| AARS | None | None | None | None |
| PCDH15 | None | None | None | None |
| PCDH15 | None | None | None | None |
| PCDH15 | None | None | None | None |

| | | | | |
|---|---|---|---|---|
| PCDH15 | None | None | None | None |
| PCDH15 | None | None | None | None |
| PCDH15 | None | None | None | None |
| PCDH15 | None | None | None | None |
| PCDH15 | None | None | None | None |
| PCDH15 | None | None | None | None |
| PCDH15 | None | None | None | None |
| NDUFS4 | None | None | None | None |
| NDUFS4 | None | None | None | None |
| NDUFS4 | None | None | None | None |
| FGF10 | None | None | None | None |
| FGF10 | 0.558 | 3 | 1nun | A |
| FGF10 | None | None | None | None |
| FGF10 | None | None | None | None |
| CDON | None | None | None | None |
| STAMBP | 0.909 | 2 | 3rzv | A |
| BRCA1 | None | None | None | None |
| BRCA1 | None | None | None | None |
| BRCA1 | None | None | None | None |
| BRCA1 | None | None | None | None |
| BRCA1 | None | None | None | None |
| BRCA1 | None | None | None | None |
| BRCA1 | None | None | None | None |
| BRCA1 | None | None | None | None |
| BRCA1 | None | None | None | None |
| BRCA1 | None | None | None | None |
| PLXNC1 | None | None | None | None |
| RNASEH2C | None | None | None | None |
| RNASEH2C | None | None | None | None |
| APC | None | None | None | None |
| PLA2G4A | None | None | None | None |
| PLA2G4A | None | None | None | None |
| PSAT1 | None | None | None | None |
| PSAT1 | None | None | None | None |
| TRAPPC3 | None | None | None | None |
| QDPR | None | None | None | None |
| QDPR | None | None | None | None |
| QDPR | None | None | None | None |
| QDPR | None | None | None | None |
| KRT5 | None | None | None | None |
| KRT5 | None | None | None | None |
| KRT5 | None | None | None | None |
| MLH1 | None | None | None | None |
| MLH1 | None | None | None | None |
| MLH1 | None | None | None | None |
| MLH1 | None | None | None | None |
| MLH1 | None | None | None | None |
| MLH1 | None | None | None | None |
| MLH1 | None | None | None | None |
| MLH1 | None | None | None | None |
| MLH1 | None | None | None | None |
| MLH1 | None | None | None | None |
| MLH1 | None | None | None | None |
| MLH1 | None | None | None | None |

| | | | | |
|---|---|---|---|---|
| MLH1 | None | None | None | None |
| MLH1 | None | None | None | None |
| MLH1 | None | None | None | None |
| MLH1 | None | None | None | None |
| MLH1 | None | None | None | None |
| KRIT1 | None | None | None | None |
| KRIT1 | None | None | None | None |
| KRIT1 | None | None | None | None |
| ALX4 | None | None | None | None |
| ALX4 | None | None | None | None |
| ALX4 | None | None | None | None |
| TJP2 | None | None | None | None |
| TJP2 | None | None | None | None |
| PANK2 | None | None | None | None |
| PANK2 | None | None | None | None |
| PANK2 | None | None | None | None |
| PANK2 | None | None | None | None |
| PANK2 | None | None | None | None |
| PANK2 | 0.333 | 2 | 5E+026 | B |
| PANK2 | None | None | None | None |
| NDUFA6 | None | None | None | None |
| TSHR | None | None | None | None |
| TSHR | None | None | None | None |
| TSHR | None | None | None | None |
| CFTR | None | None | None | None |
| CFTR | None | None | None | None |
| MYOM1 | None | None | None | None |
| MYOM1 | None | None | None | None |
| SPTB | None | None | None | None |
| GPT2 | None | None | None | None |
| NDUFS2 | None | None | None | None |
| NDUFS2 | None | None | None | None |
| NDUFS2 | None | None | None | None |
| NDUFS2 | None | None | None | None |
| NDUFS2 | None | None | None | None |
| CASQ2 | None | None | None | None |
| CASQ2 | None | None | None | None |
| CASQ2 | None | None | None | None |
| CASQ2 | None | None | None | None |
| CASQ2 | None | None | None | None |
| CASQ2 | None | None | None | None |
| CASQ2 | None | None | None | None |
| CASQ2 | None | None | None | None |
| CASQ2 | None | None | None | None |
| CPOX | None | None | None | None |
| CPOX | None | None | None | None |
| CPOX | 0.857 | 2 | 2aex | A |
| CPOX | None | None | None | None |
| CPOX | None | None | None | None |
| CPOX | None | None | None | None |
| CPOX | None | None | None | None |
| CPOX | None | None | None | None |
| CPOX | None | None | None | None |
| MVK | None | None | None | None |

| | | | | |
|---|---|---|---|---|
| MVK | None | None | None | None |
| MVK | None | None | None | None |
| MVK | None | None | None | None |
| MVK | None | None | None | None |
| MVK | None | None | None | None |
| MVK | None | None | None | None |
| MVK | None | None | None | None |
| MVK | None | None | None | None |
| MVK | None | None | None | None |
| MVK | None | None | None | None |
| MVK | None | None | None | None |
| MVK | None | None | None | None |
| MVK | None | None | None | None |
| MVK | None | None | None | None |
| GSS | None | None | None | None |
| GSS | None | None | None | None |
| GSS | None | None | None | None |
| GSS | None | None | None | None |
| GSS | None | None | None | None |
| DSG2 | None | None | None | None |
| DSG2 | None | None | None | None |
| DSG2 | None | None | None | None |
| DSG2 | None | None | None | None |
| DSG2 | None | None | None | None |
| DSG2 | None | None | None | None |
| DSG2 | None | None | None | None |
| DSG2 | None | None | None | None |
| DSG2 | None | None | None | None |
| DSG2 | None | None | None | None |
| DSG2 | None | None | None | None |
| DSG2 | None | None | None | None |
| DSG2 | None | None | None | None |
| DSG2 | None | None | None | None |
| PAPSS2 | None | None | None | None |
| PLCB2 | None | None | None | None |
| HMGCS2 | None | None | None | None |
| HMGCS2 | None | None | None | None |
| HMGCS2 | 0.938 | 1 | 2wya | C |
| HMGCS2 | None | None | None | None |
| HMGCS2 | None | None | None | None |
| HMGCS2 | None | None | None | None |
| HMGCS2 | 0.562 | 4 | 2wya | C |
| HMGCS2 | None | None | None | None |
| MPDZ | None | None | None | None |
| GATA4 | None | None | None | None |
| GATA4 | None | None | None | None |
| WARS2 | None | None | None | None |
| WARS2 | None | None | None | None |
| HGD | 1 | 47 | 1eyb | A |
| HGD | 1 | 47 | 1eyb | A |
| HGD | 1 | 70 | 1eyb | A |
| HGD | None | None | None | None |
| HGD | None | None | None | None |
| HGD | None | None | None | None |

| | | | | |
|---|---|---|---|---|
| HGD | None | None | None | None |
| HGD | 0.5 | 51 | 1ey2 | A |
| HGD | None | None | None | None |
| HGD | 1 | 1 | 1ey2 | A |
| HGD | 1 | 9 | 1ey2 | A |
| HGD | None | None | None | None |
| HGD | 1 | 10 | 1eyb | A |
| HGD | 1 | 7 | 1ey2 | A |
| PAX8 | None | None | None | None |
| PAX8 | None | None | None | None |
| PAX8 | None | None | None | None |
| PAX8 | None | None | None | None |
| SPTB | None | None | None | None |
| NCF4 | None | None | None | None |
| CFH | None | None | None | None |
| CFH | None | None | None | None |
| CSF1R | None | None | None | None |
| CSF1R | None | None | None | None |
| CA8 | None | None | None | None |
| CA8 | None | None | None | None |
| CA8 | 0.748 | 7 | 2w2j | A |
| APC | None | None | None | None |
| APC | None | None | None | None |
| APC | None | None | None | None |
| APC | None | None | None | None |
| APC | None | None | None | None |
| NDUFA2 | None | None | None | None |
| GIF | None | None | None | None |
| GIF | None | None | None | None |
| GIF | None | None | None | None |
| GIF | None | None | None | None |
| C2 | None | None | None | None |
| ACP5 | None | None | None | None |
| ACP5 | None | None | None | None |
| ACP5 | None | None | None | None |
| ACP5 | None | None | None | None |
| ACP5 | None | None | None | None |
| RNASEH2A | None | None | None | None |
| RNASEH2A | None | None | None | None |
| RNASEH2A | None | None | None | None |
| RNASEH2A | None | None | None | None |
| RNASEH2A | None | None | None | None |
| RNASEH2A | None | None | None | None |
| RNASEH2A | None | None | None | None |
| DSG2 | None | None | None | None |
| USH1G | None | None | None | None |
| FUS | None | None | None | None |
| FUS | None | None | None | None |
| FUS | None | None | None | None |
| FUS | None | None | None | None |
| FUS | None | None | None | None |
| MRPL44 | None | None | None | None |
| MRPL44 | None | None | None | None |
| MRPL44 | None | None | None | None |

| | | | | |
|---|---|---|---|---|
| MRPL44 | None | None | None | None |
| DMGDH | None | None | None | None |
| AARS | None | None | None | None |
| STAR | None | None | None | None |
| STAR | None | None | None | None |
| STAR | None | None | None | None |
| STAR | None | None | None | None |
| STAR | None | None | None | None |
| STAR | None | None | None | None |
| STAR | None | None | None | None |
| STAR | None | None | None | None |
| LARP7 | None | None | None | None |
| PRPF8 | None | None | None | None |
| PRPF8 | None | None | None | None |
| PRPF8 | None | None | None | None |
| PRPF8 | None | None | None | None |
| PSAP | None | None | None | None |
| PSAP | None | None | None | None |
| PSAP | None | None | None | None |
| DMPK | None | None | None | None |
| EPCAM | None | None | None | None |
| EPCAM | None | None | None | None |
| EPCAM | None | None | None | None |
| EPCAM | None | None | None | None |
| EPCAM | None | None | None | None |
| EPCAM | None | None | None | None |
| EPCAM | None | None | None | None |
| EPCAM | None | None | None | None |
| ASS1 | None | None | None | None |
| ASS1 | None | None | None | None |
| ASS1 | 0.571 | 3 | 2nz2 | A |
| ASS1 | 0.571 | 3 | 2nz2 | A |
| ASS1 | None | None | None | None |
| ASS1 | 0.857 | 20 | 2nz2 | A |
| ASS1 | 0.857 | 9 | 2nz2 | A |
| ASS1 | None | None | None | None |
| ASS1 | 0.429 | 13 | 2nz2 | A |
| ASS1 | 0.571 | 4 | 2nz2 | A |
| ASS1 | 0.857 | 9 | 2nz2 | A |
| ASS1 | 0.857 | 9 | 2nz2 | A |
| ASS1 | 0.857 | 9 | 2nz2 | A |
| ASS1 | None | None | None | None |
| ASS1 | None | None | None | None |
| ASS1 | None | None | None | None |
| ASS1 | None | None | None | None |
| ASS1 | None | None | None | None |
| ASS1 | None | None | None | None |
| ASS1 | None | None | None | None |
| POLE | None | None | None | None |
| POLE | None | None | None | None |
| POLE | None | None | None | None |
| RNASET2 | None | None | None | None |
| RNASET2 | None | None | None | None |
| RNASET2 | None | None | None | None |

| | | | | |
|---|---|---|---|---|
| FLG | 0.394 | 2 | 4pcw | A |
| F10 | None | None | None | None |
| AGT | None | None | None | None |
| AGT | None | None | None | None |
| PLN | None | None | None | None |
| PLN | None | None | None | None |
| PLN | None | None | None | None |
| PTHLH | None | None | None | None |
| IGF1R | None | None | None | None |
| IGF1R | None | None | None | None |
| IGF1R | None | None | None | None |
| JAG1 | None | None | None | None |
| JAG1 | None | None | None | None |
| JAG1 | None | None | None | None |
| JAG1 | None | None | None | None |
| JAG1 | None | None | None | None |
| JAG1 | None | None | None | None |
| AP1S3 | None | None | None | None |
| SPTA1 | None | None | None | None |
| SPTA1 | None | None | None | None |
| SPTA1 | None | None | None | None |
| SPTA1 | None | None | None | None |
| SPG7 | None | None | None | None |
| SPG7 | None | None | None | None |
| SPG7 | None | None | None | None |
| SPG7 | None | None | None | None |
| SPG7 | None | None | None | None |
| SPG7 | None | None | None | None |
| FN1 | None | None | None | None |
| FN1 | None | None | None | None |
| NSMCE3 | None | None | None | None |
| PCCA | None | None | None | None |
| PCCA | None | None | None | None |
| SPTB | None | None | None | None |
| RYR2 | None | None | None | None |
| RYR2 | None | None | None | None |
| MMP9 | None | None | None | None |
| MMP9 | None | None | None | None |
| GLMN | None | None | None | None |
| GLMN | None | None | None | None |
| GLMN | None | None | None | None |
| GLMN | None | None | None | None |
| GLMN | None | None | None | None |
| GLMN | None | None | None | None |
| GLMN | None | None | None | None |
| GLMN | None | None | None | None |
| GLMN | None | None | None | None |
| GLMN | None | None | None | None |
| MARS | 0.333 | 2 | 5goy | A |
| MARS | None | None | None | None |
| MARS | None | None | None | None |
| MARS | None | None | None | None |
| MARS | None | None | None | None |
| MARS | None | None | None | None |

| | | | | |
|---|---|---|---|---|
| SNAP29 | None | None | None | None |
| SNAP29 | None | None | None | None |
| ETHE1 | None | None | None | None |
| ETHE1 | None | None | None | None |
| ETHE1 | None | None | None | None |
| ETHE1 | None | None | None | None |
| ETHE1 | None | None | None | None |
| ETHE1 | None | None | None | None |
| ETHE1 | None | None | None | None |
| SAMHD1 | None | None | None | None |
| KCNQ4 | None | None | None | None |
| FXYD2 | None | None | None | None |
| FXYD2 | None | None | None | None |
| FXYD2 | None | None | None | None |
| LRRK2 | None | None | None | None |
| LRRK2 | None | None | None | None |
| NDUFS1 | None | None | None | None |
| NDUFS1 | None | None | None | None |
| NDUFS1 | None | None | None | None |
| NDUFS1 | None | None | None | None |
| NDUFS1 | None | None | None | None |
| NDUFS1 | None | None | None | None |
| NDUFS1 | None | None | None | None |
| NDUFS1 | None | None | None | None |
| NDUFS1 | None | None | None | None |
| NDUFS1 | None | None | None | None |
| NDUFS1 | None | None | None | None |
| EZH2 | None | None | None | None |
| EZH2 | None | None | None | None |
| LARS | None | None | None | None |
| TLL1 | None | None | None | None |
| CRYBB2 | None | None | None | None |
| CRYBB2 | None | None | None | None |
| EXOSC8 | None | None | None | None |
| SRP72 | None | None | None | None |
| FOXC2 | None | None | None | None |
| HMBS | None | None | None | None |
| HMBS | None | None | None | None |
| HMBS | None | None | None | None |
| HMBS | None | None | None | None |
| HMBS | None | None | None | None |
| HMBS | None | None | None | None |
| HMBS | None | None | None | None |
| HMBS | None | None | None | None |
| HMBS | None | None | None | None |
| HMBS | None | None | None | None |
| HMBS | None | None | None | None |
| HMBS | None | None | None | None |
| HMBS | None | None | None | None |
| HMBS | None | None | None | None |
| HMBS | None | None | None | None |
| HMBS | None | None | None | None |
| HMBS | None | None | None | None |
| HMBS | None | None | None | None |

| | | | | |
|---|---|---|---|---|
| HMBS | None | None | None | None |
| HMBS | None | None | None | None |
| HMBS | None | None | None | None |
| HMBS | None | None | None | None |
| HMBS | None | None | None | None |
| HMBS | None | None | None | None |
| HMBS | None | None | None | None |
| OGT | None | None | None | None |
| NOG | None | None | None | None |
| NOG | None | None | None | None |
| NOG | None | None | None | None |
| NOG | None | None | None | None |
| NOG | None | None | None | None |
| NOG | None | None | None | None |
| NOG | None | None | None | None |
| NOG | None | None | None | None |
| NOG | None | None | None | None |
| NOG | None | None | None | None |
| NOG | None | None | None | None |
| DLG3 | None | None | None | None |
| SCN2A | None | None | None | None |
| STAC3 | None | None | None | None |
| PFKM | None | None | None | None |
| PFKM | None | None | None | None |
| PFKM | None | None | None | None |
| PFKM | None | None | None | None |
| PFKM | None | None | None | None |
| DNMT1 | None | None | None | None |
| DNMT1 | None | None | None | None |
| TNNI3 | None | None | None | None |
| TNNI3 | None | None | None | None |
| TNNI3 | None | None | None | None |
| STXBP2 | None | None | None | None |
| STXBP2 | None | None | None | None |
| STXBP2 | None | None | None | None |
| STXBP2 | None | None | None | None |
| STXBP2 | None | None | None | None |
| STXBP2 | None | None | None | None |
| STXBP2 | None | None | None | None |
| STXBP2 | None | None | None | None |
| REN | None | None | None | None |
| DSP | None | None | None | None |
| DSP | None | None | None | None |
| HNF1B | None | None | None | None |
| ADA | None | None | None | None |
| ADA | None | None | None | None |
| ADA | 0.615 | 7 | 3iar | A |
| ADA | None | None | None | None |
| ADA | None | None | None | None |
| ADA | 0.538 | 5 | 3iar | A |
| ADA | 0.538 | 5 | 3iar | A |
| ADA | None | None | None | None |
| ADA | None | None | None | None |
| ADA | 0.615 | 1 | 3iar | A |

| | | | | |
|---|---|---|---|---|
| ADA | 0.462 | 6 | 3iar | A |
| ADA | 0.615 | 1 | 3iar | A |
| ADA | None | None | None | None |
| ADA | None | None | None | None |
| ADA | None | None | None | None |
| DCX | None | None | None | None |
| DCX | None | None | None | None |
| DCX | None | None | None | None |
| DCX | None | None | None | None |
| DCX | None | None | None | None |
| ARSB | None | None | None | None |
| ARSB | None | None | None | None |
| ARSB | None | None | None | None |
| ARSB | None | None | None | None |
| ARSB | None | None | None | None |
| ARSB | None | None | None | None |
| ARSB | None | None | None | None |
| ARSB | None | None | None | None |
| ARSB | None | None | None | None |
| ARSB | None | None | None | None |
| ARSB | None | None | None | None |
| ARSB | None | None | None | None |
| ARSB | None | None | None | None |
| WWOX | None | None | None | None |
| TNNI3 | None | None | None | None |
| TNNI3 | None | None | None | None |
| TNNI3 | None | None | None | None |
| TNNI3 | None | None | None | None |
| CYBA | None | None | None | None |
| SPAST | None | None | None | None |
| SPAST | None | None | None | None |
| SPAST | None | None | None | None |
| SPAST | None | None | None | None |
| SPAST | 0.132 | 1 | 3vfd | A |
| SPAST | None | None | None | None |
| SPAST | None | None | None | None |
| SPAST | None | None | None | None |
| KRIT1 | None | None | None | None |
| KRIT1 | None | None | None | None |
| TP53 | None | None | None | None |
| ISPD | None | None | None | None |
| ISPD | None | None | None | None |
| ISPD | None | None | None | None |
| ISPD | None | None | None | None |
| KCNMA1 | None | None | None | None |
| STK11 | None | None | None | None |
| STK11 | None | None | None | None |
| STK11 | None | None | None | None |
| STK11 | None | None | None | None |
| STK11 | None | None | None | None |
| STK11 | 0.412 | 2 | 2wtk | F |
| NDUFA13 | None | None | None | None |
| PHF6 | None | None | None | None |
| PHF6 | None | None | None | None |

| | | | | |
|---|---|---|---|---|
| PHF6 | None | None | None | None |
| PHYH | None | None | None | None |
| PHYH | None | None | None | None |
| PHYH | None | None | None | None |
| PHYH | None | None | None | None |
| PHYH | None | None | None | None |
| NDUFS3 | None | None | None | None |
| NDUFS3 | None | None | None | None |
| ALDH18A1 | None | None | None | None |
| ALDH18A1 | None | None | None | None |
| ALDH18A1 | None | None | None | None |
| ALDH18A1 | None | None | None | None |
| ALDH18A1 | None | None | None | None |
| PAX6 | None | None | None | None |
| PAX6 | 0.339 | 1 | 2cue | A |
| INPP5E | None | None | None | None |
| INPP5E | None | None | None | None |
| INPP5E | None | None | None | None |
| INPP5E | None | None | None | None |
| INPP5E | None | None | None | None |
| INPP5E | None | None | None | None |
| INPP5E | None | None | None | None |
| INPP5E | None | None | None | None |
| INPP5E | None | None | None | None |
| TNNT2 | None | None | None | None |
| TNNT2 | None | None | None | None |
| TNNT2 | None | None | None | None |
| TNNT2 | None | None | None | None |
| TNNT2 | None | None | None | None |
| TNNT2 | None | None | None | None |
| TNNT2 | None | None | None | None |
| TNNT2 | None | None | None | None |
| TYK2 | None | None | None | None |
| TYK2 | None | None | None | None |
| TYK2 | None | None | None | None |
| CFTR | None | None | None | None |
| CFTR | None | None | None | None |
| CFTR | None | None | None | None |
| CFTR | None | None | None | None |
| CFTR | None | None | None | None |
| CFTR | None | None | None | None |
| CFTR | None | None | None | None |
| CFTR | None | None | None | None |
| CFTR | None | None | None | None |
| CFTR | None | None | None | None |
| CFTR | None | None | None | None |
| CFTR | None | None | None | None |
| CFTR | None | None | None | None |
| CFTR | None | None | None | None |
| CFTR | None | None | None | None |
| CFTR | None | None | None | None |
| CFTR | None | None | None | None |
| CFTR | None | None | None | None |
| CFTR | None | None | None | None |

| | | | | |
|---|---|---|---|---|
| CFTR | None | None | None | None |
| CFTR | None | None | None | None |
| CFTR | None | None | None | None |
| CFTR | None | None | None | None |
| CFTR | None | None | None | None |
| CFTR | None | None | None | None |
| CFTR | None | None | None | None |
| CFTR | None | None | None | None |
| CFTR | None | None | None | None |
| CFTR | None | None | None | None |
| CFTR | None | None | None | None |
| CFTR | None | None | None | None |
| CFTR | None | None | None | None |
| CFTR | None | None | None | None |
| CFTR | None | None | None | None |
| CFTR | None | None | None | None |
| CFTR | None | None | None | None |
| CFTR | None | None | None | None |
| CFTR | None | None | None | None |
| CFTR | None | None | None | None |
| CFTR | None | None | None | None |
| CFTR | None | None | None | None |
| CFTR | None | None | None | None |
| CFTR | None | None | None | None |
| CFTR | None | None | None | None |
| CFTR | None | None | None | None |
| CFTR | None | None | None | None |
| CFTR | None | None | None | None |
| CFTR | None | None | None | None |
| CFTR | None | None | None | None |
| CFTR | None | None | None | None |
| CFTR | None | None | None | None |
| CFTR | None | None | None | None |
| CFTR | None | None | None | None |
| SERPINF1 | None | None | None | None |
| SERPINF1 | None | None | None | None |
| IDS | None | None | None | None |
| IDS | None | None | None | None |
| IDS | None | None | None | None |
| IDS | None | None | None | None |
| IDS | None | None | None | None |
| IDS | None | None | None | None |
| IDS | None | None | None | None |
| IDS | None | None | None | None |
| IDS | None | None | None | None |
| NFKBIA | 0.7 | 1 | 1nfi | F |
| NFKBIA | None | None | None | None |
| SPAST | None | None | None | None |
| FGF23 | None | None | None | None |
| FGF23 | None | None | None | None |
| FGF23 | None | None | None | None |
| FGF23 | None | None | None | None |
| FGF23 | None | None | None | None |
| FGF23 | None | None | None | None |

| | | | | |
|---|---|---|---|---|
| FGF23 | None | None | None | None |
| DLAT | None | None | None | None |
| STUB1 | None | None | None | None |
| HSD17B4 | None | None | None | None |
| HSD17B4 | None | None | None | None |
| DYSF | None | None | None | None |
| CEP135 | None | None | None | None |
| CEP135 | None | None | None | None |
| PRKCG | None | None | None | None |
| CSF1R | None | None | None | None |
| PEX5 | None | None | None | None |
| PAX6 | None | None | None | None |
| PAX6 | None | None | None | None |
| PAX6 | None | None | None | None |
| PAX6 | None | None | None | None |
| PAX6 | None | None | None | None |
| PAX6 | None | None | None | None |
| NF2 | None | None | None | None |
| NF2 | None | None | None | None |
| NF2 | None | None | None | None |
| REM1 | None | None | None | None |

| source_id | classification | binding_site | binding_site_single_water | protein_binding_site |
|---|---|---|---|---|
| rs121918343 | pathogenic | 1 | 1 | 0 |
| rs267606649 | pathogenic | 1 | 1 | 1 |
| rs121918342 | pathogenic | 0 | 1 | 0 |
| rs267606650 | pathogenic | 1 | 1 | 1 |
| rs1130335 | benign | 1 | 1 | 1 |
| rs199476087 | pathogenic | 1 | 1 | 1 |
| rs772163112 | benign | 0 | 0 | 0 |
| rs755793 | benign | 0 | 0 | 0 |
| rs121918155 | pathogenic | 1 | 1 | 0 |
| rs121918156 | pathogenic | 1 | 1 | 0 |
| rs121918154 | pathogenic | 1 | 1 | 1 |
| rs121918151 | pathogenic | 1 | 1 | 1 |
| rs121918144 | pathogenic | 0 | 1 | 0 |
| rs121918146 | pathogenic | 1 | 1 | 1 |
| rs121918160 | pathogenic | 1 | 1 | 1 |
| rs121918150 | pathogenic | 1 | 1 | 1 |
| rs121918141 | pathogenic | 1 | 1 | 1 |
| rs142742242 | pathogenic | 1 | 1 | 1 |
| rs121918142 | pathogenic | 1 | 1 | 1 |
| rs121918157 | pathogenic | 1 | 1 | 1 |
| rs121918101 | pathogenic | 1 | 1 | 1 |
| rs121908260 | pathogenic | 1 | 1 | 1 |
| rs80356667 | pathogenic | 1 | 1 | 1 |
| rs80356668 | pathogenic | 1 | 1 | 1 |
| rs397514540 | pathogenic | 1 | 1 | 1 |
| rs104894685 | pathogenic | 0 | 1 | 0 |
| rs199869995 | pathogenic | 1 | 1 | 1 |
| rs121909072 | pathogenic | 0 | 0 | 0 |
| rs143669310 | benign | 1 | 1 | 1 |
| rs747347498 | benign | 0 | 0 | 0 |
| rs138283222 | benign | 0 | 0 | 0 |
| rs118203931 | pathogenic | 1 | 1 | 1 |
| rs118203933 | pathogenic | 1 | 1 | 0 |
| rs118203932 | pathogenic | 1 | 1 | 0 |
| rs2228063 | benign | 1 | 1 | 1 |
| rs377767360 | pathogenic | 0 | 0 | 0 |
| rs397518413 | pathogenic | 1 | 1 | 1 |
| rs1800458 | benign | 1 | 1 | 0 |
| rs121918083 | pathogenic | 1 | 1 | 1 |
| rs121918094 | pathogenic | 1 | 1 | 1 |
| rs121918098 | pathogenic | 1 | 1 | 1 |
| rs121918093 | pathogenic | 1 | 1 | 1 |
| rs79977247 | pathogenic | 1 | 1 | 1 |
| rs28933979 | pathogenic | 1 | 1 | 1 |
| rs121918068 | pathogenic | 1 | 1 | 1 |
| rs121918077 | pathogenic | 1 | 1 | 1 |
| rs11541796 | pathogenic | 1 | 1 | 1 |
| rs104894665 | pathogenic | 1 | 1 | 1 |
| rs121918090 | pathogenic | 1 | 1 | 1 |
| rs121918081 | pathogenic | 1 | 1 | 1 |
| rs121918080 | pathogenic | 1 | 1 | 1 |
| rs121918076 | pathogenic | 1 | 1 | 1 |
| rs121918097 | pathogenic | 1 | 1 | 1 |

| | | | | |
|---|---|---|---|---|
| rs121918079 | pathogenic | 1 | 1 | 1 |
| rs121918069 | pathogenic | 1 | 1 | 1 |
| rs121918086 | pathogenic | 1 | 1 | 1 |
| rs121918091 | pathogenic | 1 | 1 | 1 |
| rs121918085 | pathogenic | 1 | 1 | 1 |
| rs121918100 | pathogenic | 1 | 1 | 1 |
| rs121918084 | pathogenic | 1 | 1 | 1 |
| rs121918071 | pathogenic | 1 | 1 | 1 |
| rs121918072 | pathogenic | 1 | 1 | 1 |
| rs121918082 | pathogenic | 1 | 1 | 1 |
| rs121918087 | pathogenic | 1 | 1 | 1 |
| rs76410435 | benign | 1 | 1 | 1 |
| rs121918089 | pathogenic | 1 | 1 | 1 |
| rs121918073 | pathogenic | 1 | 1 | 1 |
| rs121918075 | pathogenic | 1 | 1 | 1 |
| rs121918088 | pathogenic | 1 | 1 | 1 |
| rs76992529 | pathogenic | 1 | 1 | 1 |
| rs536294863 | benign | 1 | 1 | 1 |
| rs730881162 | benign | 1 | 1 | 1 |
| rs113994087 | pathogenic | 1 | 1 | 1 |
| rs74716434 | benign | 1 | 1 | 1 |
| rs121918694 | pathogenic | 1 | 1 | 1 |
| rs121918707 | pathogenic | 1 | 1 | 1 |
| rs121918695 | pathogenic | 1 | 1 | 0 |
| rs121918690 | pathogenic | 1 | 1 | 0 |
| rs121918696 | pathogenic | 1 | 1 | 0 |
| rs121918693 | pathogenic | 1 | 1 | 0 |
| rs28999969 | pathogenic | 1 | 1 | 1 |
| rs121918697 | pathogenic | 1 | 1 | 1 |
| rs121918688 | pathogenic | 1 | 1 | 0 |
| rs28999970 | pathogenic | 1 | 1 | 0 |
| rs121918686 | pathogenic | 1 | 1 | 0 |
| rs28999971 | pathogenic | 1 | 1 | 1 |
| rs121918698 | pathogenic | 1 | 1 | 1 |
| rs121918691 | pathogenic | 1 | 1 | 0 |
| rs121918692 | pathogenic | 1 | 1 | 1 |
| rs121918703 | pathogenic | 1 | 1 | 1 |
| rs121918687 | pathogenic | 1 | 1 | 1 |
| rs28933408 | pathogenic | 1 | 1 | 1 |
| rs121918702 | pathogenic | 1 | 1 | 1 |
| rs17875899 | benign | 1 | 1 | 1 |
| rs111033620 | pathogenic | 1 | 1 | 1 |
| rs111033622 | pathogenic | 1 | 1 | 1 |
| rs111033621 | pathogenic | 1 | 1 | 1 |
| rs111033618 | pathogenic | 1 | 1 | 1 |
| rs5918 | benign | 0 | 0 | 0 |
| rs121918452 | pathogenic | 0 | 0 | 0 |
| rs121918445 | pathogenic | 1 | 1 | 1 |
| rs5917 | benign | 1 | 1 | 1 |
| rs121918444 | pathogenic | 1 | 1 | 1 |
| rs121918446 | pathogenic | 1 | 1 | 1 |
| rs121918449 | pathogenic | 0 | 0 | 0 |
| rs121918448 | benign | 0 | 0 | 0 |
| rs13306487 | benign | 1 | 1 | 1 |

| | | | | |
|---|---|---|---|---|
| rs1805044 | benign | 0 | 0 | 0 |
| rs117078377 | benign | 0 | 0 | 0 |
| rs138132026 | benign | 1 | 1 | 1 |
| rs75762896 | benign | 1 | 1 | 0 |
| rs121918043 | pathogenic | 1 | 1 | 0 |
| rs143581690 | benign | 1 | 1 | 0 |
| rs147654263 | benign | 0 | 0 | 0 |
| rs587782529 | pathogenic | 1 | 1 | 1 |
| rs121912664 | pathogenic | 1 | 1 | 1 |
| rs150293825 | benign | 1 | 1 | 1 |
| rs17882252 | benign | 1 | 1 | 1 |
| rs121912662 | pathogenic | 1 | 1 | 1 |
| rs35993958 | benign | 1 | 1 | 1 |
| rs606231253 | pathogenic | 1 | 1 | 1 |
| rs2227983 | benign | 1 | 1 | 1 |
| rs587779351 | pathogenic | 0 | 0 | 0 |
| rs779773463 | pathogenic | 1 | 1 | 1 |
| rs713040 | benign | 1 | 1 | 1 |
| rs334 | pathogenic | 1 | 1 | 1 |
| rs33930165 | pathogenic | 1 | 1 | 1 |
| rs33918131 | benign | 1 | 1 | 1 |
| rs63750783 | pathogenic | 1 | 1 | 1 |
| rs33986703 | pathogenic | 1 | 1 | 1 |
| rs33972047 | pathogenic | 1 | 1 | 1 |
| rs33936254 | pathogenic | 1 | 1 | 1 |
| rs33959855 | benign | 1 | 1 | 1 |
| rs33950507 | pathogenic | 1 | 1 | 1 |
| rs33916412 | pathogenic | 1 | 1 | 0 |
| rs1135071 | pathogenic | 1 | 1 | 1 |
| rs33960103 | pathogenic | 1 | 1 | 1 |
| rs11549407 | pathogenic | 1 | 1 | 1 |
| rs33922842 | pathogenic | 1 | 1 | 1 |
| rs35278874 | benign | 1 | 1 | 1 |
| rs34621955 | benign | 1 | 1 | 1 |
| rs33995148 | pathogenic | 1 | 1 | 1 |
| rs33945705 | benign | 1 | 1 | 1 |
| rs33993568 | benign | 1 | 1 | 1 |
| rs35553496 | benign | 1 | 1 | 1 |
| rs33913712 | pathogenic | 1 | 1 | 1 |
| rs33917785 | pathogenic | 1 | 1 | 1 |
| rs34515413 | benign | 1 | 1 | 1 |
| rs35256489 | pathogenic | 1 | 1 | 0 |
| rs34484056 | pathogenic | 1 | 1 | 1 |
| rs33947020 | benign | 1 | 1 | 1 |
| rs33946267 | pathogenic | 1 | 1 | 1 |
| rs33966761 | pathogenic | 1 | 1 | 1 |
| rs36020563 | benign | 1 | 1 | 1 |
| rs122454124 | pathogenic | 1 | 1 | 1 |
| rs122454126 | pathogenic | 1 | 1 | 0 |
| rs122454127 | pathogenic | 1 | 1 | 1 |
| rs122454130 | pathogenic | 1 | 1 | 1 |
| rs122454125 | pathogenic | 1 | 1 | 1 |
| rs122454131 | pathogenic | 1 | 1 | 0 |
| rs121908302 | pathogenic | 0 | 1 | 0 |

| | | | | |
|---|---|---|---|---|
| rs77829017 | pathogenic | 0 | 1 | 0 |
| rs1141814 | pathogenic | 0 | 1 | 0 |
| rs121908312 | pathogenic | 0 | 1 | 0 |
| rs79653797 | pathogenic | 1 | 1 | 0 |
| rs439898 | pathogenic | 1 | 1 | 0 |
| rs121908299 | pathogenic | 0 | 1 | 0 |
| rs80356763 | pathogenic | 1 | 1 | 1 |
| rs781152868 | pathogenic | 0 | 1 | 0 |
| rs121908297 | pathogenic | 0 | 1 | 0 |
| rs398123531 | benign | 0 | 1 | 0 |
| rs398123532 | pathogenic | 0 | 1 | 0 |
| rs364897 | pathogenic | 1 | 1 | 1 |
| rs1064644 | pathogenic | 0 | 1 | 0 |
| rs409652 | pathogenic | 1 | 1 | 1 |
| rs121908300 | pathogenic | 0 | 1 | 0 |
| rs381737 | pathogenic | 0 | 1 | 0 |
| rs74500255 | pathogenic | 0 | 1 | 0 |
| rs121908313 | pathogenic | 0 | 1 | 0 |
| rs367968666 | pathogenic | 0 | 1 | 0 |
| rs78973108 | pathogenic | 1 | 1 | 1 |
| rs121908298 | pathogenic | 1 | 1 | 1 |
| rs78396650 | pathogenic | 0 | 0 | 0 |
| rs78198234 | pathogenic | 1 | 1 | 0 |
| rs121908304 | pathogenic | 1 | 1 | 1 |
| rs76539814 | pathogenic | 1 | 1 | 1 |
| rs121908305 | pathogenic | 0 | 1 | 0 |
| rs121908306 | pathogenic | 1 | 1 | 0 |
| rs121908308 | pathogenic | 1 | 1 | 1 |
| rs121908307 | pathogenic | 0 | 1 | 0 |
| rs75548401 | benign | 0 | 1 | 0 |
| rs76763715 | pathogenic | 0 | 0 | 0 |
| rs121908314 | pathogenic | 0 | 1 | 0 |
| rs398123528 | pathogenic | 0 | 1 | 0 |
| rs121908311 | pathogenic | 0 | 0 | 0 |
| rs121908310 | pathogenic | 1 | 1 | 0 |
| rs74598136 | pathogenic | 0 | 1 | 0 |
| rs1064651 | pathogenic | 1 | 1 | 1 |
| rs77369218 | pathogenic | 1 | 1 | 1 |
| rs121908295 | pathogenic | 0 | 1 | 0 |
| rs421016 | pathogenic | 0 | 1 | 0 |
| rs368060 | pathogenic | 0 | 1 | 0 |
| rs80356771 | pathogenic | 1 | 1 | 1 |
| rs80356772 | pathogenic | 1 | 1 | 1 |
| rs121908301 | pathogenic | 0 | 0 | 0 |
| rs747506979 | pathogenic | 0 | 1 | 0 |
| rs75822236 | pathogenic | 0 | 1 | 0 |
| rs1990760 | benign | 1 | 1 | 0 |
| rs28934568 | pathogenic | 1 | 1 | 0 |
| rs34833812 | benign | 1 | 1 | 0 |
| rs193922666 | benign | 1 | 1 | 0 |
| rs104893812 | pathogenic | 1 | 1 | 0 |
| rs144701411 | benign | 0 | 0 | 0 |
| rs104893813 | pathogenic | 0 | 1 | 0 |
| rs199660234 | pathogenic | 1 | 1 | 0 |

| | | | | |
|---|---|---|---|---|
| rs104893814 | pathogenic | 1 | 1 | 0 |
| rs143095746 | benign | 1 | 1 | 1 |
| rs104893807 | pathogenic | 1 | 1 | 0 |
| rs104893811 | pathogenic | 0 | 0 | 0 |
| rs104893816 | pathogenic | 0 | 0 | 0 |
| rs121918714 | pathogenic | 0 | 1 | 0 |
| rs104893810 | pathogenic | 1 | 1 | 0 |
| rs104893815 | pathogenic | 1 | 1 | 0 |
| rs4303 | benign | 0 | 1 | 0 |
| rs121912704 | pathogenic | 0 | 1 | 0 |
| rs56394458 | benign | 1 | 1 | 0 |
| rs397514688 | pathogenic | 0 | 0 | 0 |
| rs367797185 | pathogenic | 1 | 1 | 0 |
| rs12709426 | benign | 0 | 1 | 0 |
| rs2228547 | benign | 0 | 0 | 0 |
| rs111033546 | pathogenic | 0 | 0 | 0 |
| rs111033553 | pathogenic | 0 | 0 | 0 |
| rs111033554 | pathogenic | 1 | 1 | 1 |
| rs111033544 | pathogenic | 1 | 1 | 1 |
| rs111033555 | pathogenic | 1 | 1 | 1 |
| rs111033545 | pathogenic | 1 | 1 | 1 |
| rs111033549 | pathogenic | 0 | 1 | 0 |
| rs111033552 | pathogenic | 1 | 1 | 1 |
| rs752204035 | benign | 0 | 0 | 0 |
| rs11569534 | benign | 0 | 0 | 0 |
| rs28937571 | pathogenic | 1 | 1 | 1 |
| rs104894215 | pathogenic | 1 | 1 | 1 |
| rs3888798 | benign | 1 | 1 | 1 |
| rs764263034 | benign | 1 | 1 | 1 |
| rs1800973 | benign | 1 | 1 | 1 |
| rs367623154 | benign | 1 | 1 | 1 |
| rs201230446 | pathogenic | 1 | 1 | 1 |
| rs267606766 | pathogenic | 0 | 1 | 0 |
| rs267606769 | pathogenic | 0 | 1 | 0 |
| rs267606767 | pathogenic | 1 | 1 | 0 |
| rs267606768 | pathogenic | 0 | 1 | 0 |
| rs61733129 | benign | 0 | 1 | 0 |
| rs201947120 | pathogenic | 1 | 1 | 1 |
| rs283413 | benign | 0 | 1 | 0 |
| rs36119840 | benign | 0 | 0 | 0 |
| rs281865419 | pathogenic | 1 | 1 | 1 |
| rs137852937 | pathogenic | 0 | 1 | 0 |
| rs2230709 | benign | 0 | 1 | 0 |
| rs6695033 | benign | 0 | 1 | 0 |
| rs180177032 | pathogenic | 1 | 1 | 1 |
| rs180177033 | pathogenic | 1 | 1 | 1 |
| rs121913351 | pathogenic | 1 | 1 | 1 |
| rs397507473 | pathogenic | 1 | 1 | 1 |
| rs121913355 | pathogenic | 1 | 1 | 1 |
| rs121913357 | pathogenic | 1 | 1 | 1 |
| rs180177036 | pathogenic | 1 | 1 | 1 |
| rs180177039 | pathogenic | 1 | 1 | 1 |
| rs180177038 | pathogenic | 1 | 1 | 1 |
| rs180177040 | pathogenic | 1 | 1 | 1 |

| | | | | |
|---|---|---|---|---|
| rs121913338 | pathogenic | 1 | 1 | 1 |
| rs121913366 | pathogenic | 1 | 1 | 1 |
| rs121913369 | pathogenic | 1 | 1 | 1 |
| rs113488022 | pathogenic | 1 | 1 | 1 |
| rs121913364 | pathogenic | 1 | 1 | 1 |
| rs180177042 | pathogenic | 1 | 1 | 1 |
| rs200461617 | benign | 0 | 0 | 0 |
| rs141609745 | benign | 0 | 0 | 0 |
| rs104894383 | pathogenic | 1 | 1 | 1 |
| rs104894384 | pathogenic | 1 | 1 | 1 |
| rs104894381 | pathogenic | 1 | 1 | 0 |
| rs147710408 | benign | 1 | 1 | 1 |
| rs77357563 | benign | 1 | 1 | 1 |
| rs104894378 | pathogenic | 0 | 0 | 0 |
| rs104894382 | pathogenic | 0 | 0 | 0 |
| rs137853292 | pathogenic | 1 | 1 | 1 |
| rs587778834 | pathogenic | 0 | 0 | 0 |
| rs137853294 | pathogenic | 1 | 1 | 1 |
| rs551747882 | benign | 0 | 0 | 0 |
| rs3092903 | benign | 1 | 1 | 1 |
| rs137853296 | pathogenic | 0 | 0 | 0 |
| rs398122917 | pathogenic | 0 | 1 | 0 |
| rs41304992 | benign | 0 | 0 | 0 |
| rs121912708 | pathogenic | 1 | 1 | 1 |
| rs140102105 | pathogenic | 1 | 1 | 1 |
| rs376917645 | pathogenic | 1 | 1 | 0 |
| rs372660425 | pathogenic | 1 | 1 | 0 |
| rs555896752 | pathogenic | 1 | 1 | 1 |
| rs777829351 | pathogenic | 1 | 1 | 0 |
| rs121912711 | pathogenic | 1 | 1 | 0 |
| rs754449549 | pathogenic | 0 | 1 | 0 |
| rs141701364 | benign | 0 | 1 | 0 |
| rs761011399 | benign | 1 | 1 | 1 |
| rs121912710 | benign | 1 | 1 | 0 |
| rs121912707 | pathogenic | 1 | 1 | 0 |
| rs747597620 | pathogenic | 0 | 1 | 0 |
| rs186558364 | pathogenic | 1 | 1 | 1 |
| rs556400964 | pathogenic | 1 | 1 | 1 |
| rs61757684 | benign | 1 | 1 | 1 |
| rs121917766 | pathogenic | 1 | 1 | 1 |
| rs121918248 | pathogenic | 0 | 0 | 0 |
| rs398123278 | pathogenic | 0 | 0 | 0 |
| rs121918251 | pathogenic | 1 | 1 | 1 |
| rs727504022 | pathogenic | 0 | 0 | 0 |
| rs190834116 | pathogenic | 1 | 1 | 1 |
| rs121918249 | pathogenic | 1 | 1 | 0 |
| rs121918257 | pathogenic | 1 | 1 | 0 |
| rs142068898 | benign | 0 | 0 | 0 |
| rs757000253 | pathogenic | 0 | 0 | 0 |
| rs200908035 | pathogenic | 1 | 1 | 0 |
| rs760782399 | pathogenic | 0 | 0 | 0 |
| rs778702777 | pathogenic | 0 | 0 | 0 |
| rs121918258 | pathogenic | 1 | 1 | 0 |
| rs121918256 | pathogenic | 0 | 0 | 0 |

| | | | | |
|---|---|---|---|---|
| rs200596762 | pathogenic | 1 | 1 | 0 |
| rs547709692 | pathogenic | 0 | 0 | 0 |
| rs758577372 | pathogenic | 0 | 1 | 0 |
| rs564069299 | pathogenic | 1 | 1 | 1 |
| rs772552898 | pathogenic | 1 | 1 | 1 |
| rs368790885 | pathogenic | 1 | 1 | 1 |
| rs121918250 | pathogenic | 0 | 0 | 0 |
| rs533755473 | pathogenic | 1 | 1 | 1 |
| rs753288303 | pathogenic | 1 | 1 | 1 |
| rs2229385 | benign | 0 | 0 | 0 |
| rs1141321 | benign | 0 | 0 | 0 |
| rs753564352 | pathogenic | 0 | 0 | 0 |
| rs775593146 | pathogenic | 0 | 0 | 0 |
| rs121918254 | pathogenic | 0 | 0 | 0 |
| rs8589 | benign | 1 | 1 | 0 |
| rs777758903 | pathogenic | 0 | 0 | 0 |
| rs140600746 | pathogenic | 1 | 1 | 0 |
| rs121918255 | pathogenic | 1 | 1 | 0 |
| rs121918252 | pathogenic | 0 | 0 | 0 |
| rs779990936 | pathogenic | 1 | 1 | 0 |
| rs121912455 | pathogenic | 1 | 1 | 1 |
| rs121912459 | pathogenic | 1 | 1 | 1 |
| rs74315452 | pathogenic | 1 | 1 | 1 |
| rs121912451 | pathogenic | 1 | 1 | 1 |
| rs79006549 | pathogenic | 0 | 0 | 0 |
| rs28934889 | benign | 0 | 1 | 0 |
| rs111033557 | benign | 1 | 1 | 1 |
| rs28934597 | pathogenic | 1 | 1 | 1 |
| rs28934596 | pathogenic | 1 | 1 | 1 |
| rs28934595 | pathogenic | 1 | 1 | 1 |
| rs146519482 | pathogenic | 1 | 1 | 1 |
| rs111033563 | pathogenic | 1 | 1 | 1 |
| rs75385605 | benign | 1 | 1 | 1 |
| rs121908569 | pathogenic | 1 | 1 | 1 |
| rs35719940 | benign | 0 | 0 | 0 |
| rs121918664 | pathogenic | 0 | 0 | 0 |
| rs104893665 | pathogenic | 1 | 1 | 1 |
| rs104893666 | pathogenic | 1 | 1 | 1 |
| rs200845457 | benign | 1 | 1 | 1 |
| rs779204655 | pathogenic | 1 | 1 | 1 |
| rs121917747 | pathogenic | 1 | 1 | 1 |
| rs6042 | benign | 1 | 1 | 1 |
| rs104894380 | pathogenic | 0 | 0 | 0 |
| rs369083541 | pathogenic | 1 | 1 | 1 |
| rs145243532 | benign | 1 | 1 | 1 |
| rs121909500 | pathogenic | 1 | 1 | 1 |
| rs140993310 | benign | 1 | 1 | 1 |
| rs200837270 | pathogenic | 1 | 1 | 1 |
| rs121918484 | pathogenic | 1 | 1 | 1 |
| rs121918306 | pathogenic | 0 | 0 | 0 |
| rs118204089 | pathogenic | 1 | 1 | 0 |
| rs118204083 | pathogenic | 1 | 1 | 0 |
| rs201953584 | pathogenic | 1 | 1 | 1 |
| rs74315362 | pathogenic | 1 | 1 | 1 |

| | | | | |
|---|---|---|---|---|
| rs118204084 | pathogenic | 1 | 1 | 0 |
| rs118204085 | pathogenic | 1 | 1 | 1 |
| rs113403872 | pathogenic | 1 | 1 | 0 |
| rs1052176 | benign | 1 | 1 | 1 |
| rs2031640 | benign | 0 | 0 | 0 |
| rs104894230 | pathogenic | 1 | 1 | 1 |
| rs104894229 | pathogenic | 1 | 1 | 1 |
| rs104894228 | pathogenic | 1 | 1 | 1 |
| rs104894226 | pathogenic | 1 | 1 | 1 |
| rs121917757 | pathogenic | 1 | 1 | 1 |
| rs121917758 | pathogenic | 1 | 1 | 1 |
| rs28933406 | pathogenic | 1 | 1 | 1 |
| rs121917756 | pathogenic | 1 | 1 | 1 |
| rs104894227 | pathogenic | 1 | 1 | 1 |
| rs369106578 | pathogenic | 1 | 1 | 1 |
| rs397517142 | benign | 1 | 1 | 1 |
| rs104894231 | pathogenic | 1 | 1 | 1 |
| rs121917759 | pathogenic | 1 | 1 | 1 |
| rs104893827 | pathogenic | 0 | 0 | 0 |
| rs104893826 | pathogenic | 1 | 1 | 1 |
| rs5030802 | pathogenic | 1 | 1 | 1 |
| rs5030827 | pathogenic | 1 | 1 | 0 |
| rs119103277 | pathogenic | 1 | 1 | 1 |
| rs5030808 | pathogenic | 0 | 1 | 0 |
| rs5030809 | pathogenic | 1 | 1 | 1 |
| rs765978945 | pathogenic | 1 | 1 | 1 |
| rs104893824 | pathogenic | 1 | 1 | 1 |
| rs104893831 | pathogenic | 0 | 1 | 0 |
| rs104893830 | pathogenic | 0 | 1 | 0 |
| rs119103278 | pathogenic | 0 | 1 | 0 |
| rs5030818 | pathogenic | 1 | 1 | 1 |
| rs397516444 | pathogenic | 1 | 1 | 1 |
| rs28940297 | pathogenic | 1 | 1 | 1 |
| rs267607170 | pathogenic | 1 | 1 | 1 |
| rs104893825 | pathogenic | 1 | 1 | 1 |
| rs5030820 | pathogenic | 0 | 1 | 0 |
| rs5030821 | pathogenic | 0 | 1 | 0 |
| rs5030823 | pathogenic | 1 | 1 | 1 |
| rs5030824 | pathogenic | 1 | 1 | 1 |
| rs28940301 | pathogenic | 0 | 1 | 0 |
| rs28940300 | pathogenic | 0 | 1 | 0 |
| rs28940298 | pathogenic | 0 | 1 | 0 |
| rs200019083 | benign | 0 | 0 | 0 |
| rs759748655 | pathogenic | 0 | 0 | 0 |
| rs121917739 | pathogenic | 1 | 1 | 1 |
| rs75353611 | pathogenic | 1 | 1 | 1 |
| rs79744198 | benign | 1 | 1 | 1 |
| rs79228041 | pathogenic | 1 | 1 | 1 |
| rs80106970 | benign | 1 | 1 | 1 |
| rs375353223 | benign | 0 | 0 | 0 |
| rs56311081 | benign | 0 | 0 | 0 |
| rs55985569 | benign | 1 | 1 | 1 |
| rs45483396 | benign | 0 | 0 | 0 |
| rs35284565 | benign | 1 | 1 | 1 |

| | | | | |
|---|---|---|---|---|
| rs34349517 | benign | 0 | 0 | 0 |
| rs35776110 | benign | 0 | 0 | 0 |
| rs33917957 | benign | 1 | 1 | 1 |
| rs587778445 | benign | 0 | 0 | 0 |
| rs34624984 | benign | 0 | 0 | 0 |
| rs701753 | benign | 0 | 0 | 0 |
| rs855314 | benign | 0 | 0 | 0 |
| rs121918371 | pathogenic | 1 | 1 | 0 |
| rs398122912 | pathogenic | 0 | 1 | 0 |
| rs1126728 | benign | 0 | 0 | 0 |
| rs587777404 | pathogenic | 1 | 1 | 0 |
| rs11208257 | benign | 0 | 0 | 0 |
| rs6676290 | benign | 0 | 0 | 0 |
| rs104894118 | pathogenic | 0 | 0 | 0 |
| rs121918655 | pathogenic | 1 | 1 | 1 |
| rs104894120 | pathogenic | 1 | 1 | 1 |
| rs137852238 | pathogenic | 0 | 0 | 0 |
| rs137852237 | pathogenic | 0 | 0 | 0 |
| rs137852499 | pathogenic | 1 | 1 | 1 |
| rs137852480 | pathogenic | 1 | 1 | 1 |
| rs137852494 | pathogenic | 1 | 1 | 1 |
| rs137852502 | pathogenic | 1 | 1 | 1 |
| rs137852500 | pathogenic | 1 | 1 | 1 |
| rs137852487 | pathogenic | 1 | 1 | 1 |
| rs137852481 | pathogenic | 1 | 1 | 1 |
| rs137852501 | pathogenic | 0 | 0 | 0 |
| rs137852478 | pathogenic | 1 | 1 | 1 |
| rs137852485 | pathogenic | 1 | 1 | 1 |
| rs137852482 | pathogenic | 1 | 1 | 0 |
| rs137852483 | pathogenic | 0 | 1 | 0 |
| rs137852477 | pathogenic | 1 | 1 | 0 |
| rs137852496 | pathogenic | 1 | 1 | 0 |
| rs137852484 | pathogenic | 0 | 1 | 0 |
| rs137852498 | pathogenic | 1 | 1 | 0 |
| rs137852497 | pathogenic | 1 | 1 | 0 |
| rs137852493 | pathogenic | 1 | 1 | 0 |
| rs137852504 | pathogenic | 1 | 1 | 1 |
| rs267606863 | pathogenic | 1 | 1 | 1 |
| rs137852486 | pathogenic | 1 | 1 | 1 |
| rs137852479 | pathogenic | 1 | 1 | 1 |
| rs112122950 | benign | 0 | 0 | 0 |
| rs72546648 | benign | 1 | 1 | 1 |
| rs72552291 | pathogenic | 0 | 0 | 0 |
| rs79184941 | pathogenic | 1 | 1 | 1 |
| rs77543610 | pathogenic | 1 | 1 | 1 |
| rs121918505 | pathogenic | 1 | 1 | 1 |
| rs776587763 | pathogenic | 1 | 1 | 1 |
| rs121918497 | pathogenic | 1 | 1 | 1 |
| rs121918499 | pathogenic | 1 | 1 | 1 |
| rs121918501 | pathogenic | 1 | 1 | 1 |
| rs121918500 | pathogenic | 0 | 0 | 0 |
| rs121918504 | pathogenic | 1 | 1 | 1 |
| rs121918510 | pathogenic | 1 | 1 | 1 |
| rs121918493 | pathogenic | 1 | 1 | 1 |

| | | | | |
|---|---|---|---|---|
| rs121918489 | pathogenic | 0 | 0 | 0 |
| rs121918495 | pathogenic | 1 | 1 | 1 |
| rs121918488 | pathogenic | 0 | 0 | 0 |
| rs121918496 | pathogenic | 0 | 0 | 0 |
| rs121918492 | pathogenic | 1 | 1 | 1 |
| rs121918494 | pathogenic | 1 | 1 | 1 |
| rs121918502 | pathogenic | 1 | 1 | 1 |
| rs121918490 | pathogenic | 1 | 1 | 1 |
| rs72466451 | pathogenic | 1 | 1 | 1 |
| rs66468541 | pathogenic | 0 | 0 | 0 |
| rs137852662 | pathogenic | 0 | 0 | 0 |
| rs137852663 | pathogenic | 0 | 0 | 0 |
| rs606231407 | pathogenic | 0 | 0 | 0 |
| rs137852233 | pathogenic | 1 | 1 | 1 |
| rs112306225 | benign | 0 | 0 | 0 |
| rs387906808 | pathogenic | 1 | 1 | 0 |
| rs730882094 | pathogenic | 1 | 1 | 1 |
| rs373869746 | pathogenic | 1 | 1 | 1 |
| rs777524402 | pathogenic | 1 | 1 | 1 |
| rs769737896 | pathogenic | 0 | 1 | 0 |
| rs13306515 | benign | 1 | 1 | 0 |
| rs746982741 | pathogenic | 0 | 0 | 0 |
| rs2230200 | benign | 1 | 1 | 1 |
| rs1047286 | benign | 0 | 0 | 0 |
| rs141737564 | benign | 0 | 0 | 0 |
| rs774826179 | benign | 0 | 0 | 0 |
| rs11569422 | benign | 1 | 1 | 1 |
| rs146613648 | benign | 0 | 0 | 0 |
| rs144432231 | benign | 0 | 0 | 0 |
| rs140655115 | benign | 0 | 0 | 0 |
| rs387906837 | pathogenic | 1 | 1 | 0 |
| rs121909076 | pathogenic | 1 | 1 | 0 |
| rs121909077 | pathogenic | 0 | 0 | 0 |
| rs121909074 | pathogenic | 0 | 0 | 0 |
| rs137853330 | pathogenic | 1 | 1 | 1 |
| rs137853331 | pathogenic | 1 | 1 | 1 |
| rs137853332 | pathogenic | 1 | 1 | 1 |
| rs28931586 | pathogenic | 1 | 1 | 0 |
| rs11569321 | benign | 1 | 1 | 1 |
| rs119486097 | benign | 0 | 0 | 0 |
| rs1799807 | pathogenic | 1 | 1 | 1 |
| rs201820739 | pathogenic | 1 | 1 | 0 |
| rs114706984 | pathogenic | 1 | 1 | 0 |
| rs140080572 | pathogenic | 1 | 1 | 0 |
| rs28933389 | pathogenic | 0 | 1 | 0 |
| rs16849700 | benign | 1 | 1 | 0 |
| rs115624085 | pathogenic | 1 | 1 | 1 |
| rs747196387 | pathogenic | 1 | 1 | 1 |
| rs104893684 | pathogenic | 1 | 1 | 0 |
| rs121918557 | pathogenic | 1 | 1 | 0 |
| rs121918556 | pathogenic | 1 | 1 | 1 |
| rs398124571 | pathogenic | 1 | 1 | 1 |
| rs776631396 | pathogenic | 0 | 1 | 0 |
| rs398124576 | pathogenic | 1 | 1 | 1 |

| | | | | |
|---|---|---|---|---|
| rs398124581 | pathogenic | 1 | 1 | 1 |
| rs398124582 | pathogenic | 1 | 1 | 1 |
| rs149766077 | pathogenic | 0 | 1 | 0 |
| rs79761867 | pathogenic | 0 | 1 | 0 |
| rs148905512 | pathogenic | 0 | 1 | 0 |
| rs121965004 | pathogenic | 1 | 1 | 1 |
| rs386834233 | pathogenic | 0 | 1 | 0 |
| rs398124598 | pathogenic | 0 | 1 | 0 |
| rs398124603 | pathogenic | 1 | 1 | 1 |
| rs546716073 | benign | 1 | 1 | 1 |
| rs200310662 | benign | 1 | 1 | 1 |
| rs115494727 | benign | 1 | 1 | 1 |
| rs781513008 | benign | 0 | 0 | 0 |
| rs587776398 | benign | 1 | 1 | 0 |
| rs35288908 | benign | 1 | 1 | 1 |
| rs150634562 | pathogenic | 1 | 1 | 0 |
| rs142948132 | pathogenic | 1 | 1 | 1 |
| rs132630321 | pathogenic | 1 | 1 | 1 |
| rs132630317 | pathogenic | 1 | 1 | 1 |
| rs132630320 | pathogenic | 0 | 0 | 0 |
| rs781394318 | pathogenic | 1 | 1 | 1 |
| rs111033624 | pathogenic | 1 | 1 | 1 |
| rs111033630 | pathogenic | 1 | 1 | 1 |
| rs111033627 | pathogenic | 1 | 1 | 1 |
| rs193922287 | pathogenic | 1 | 1 | 1 |
| rs759072800 | pathogenic | 0 | 1 | 0 |
| rs104894010 | pathogenic | 0 | 0 | 0 |
| rs193922296 | pathogenic | 0 | 0 | 0 |
| rs104894006 | pathogenic | 1 | 1 | 1 |
| rs193922311 | pathogenic | 1 | 1 | 0 |
| rs80356654 | pathogenic | 1 | 1 | 0 |
| rs144723656 | benign | 1 | 1 | 1 |
| rs148311934 | pathogenic | 1 | 1 | 0 |
| rs80356655 | pathogenic | 1 | 1 | 1 |
| rs769268803 | pathogenic | 1 | 1 | 0 |
| rs104894008 | pathogenic | 0 | 1 | 0 |
| rs104894005 | pathogenic | 1 | 1 | 1 |
| rs104894009 | pathogenic | 1 | 1 | 0 |
| rs193922340 | pathogenic | 0 | 0 | 0 |
| rs104894016 | pathogenic | 0 | 1 | 0 |
| rs193921338 | pathogenic | 0 | 0 | 0 |
| rs193922272 | pathogenic | 1 | 1 | 1 |
| rs104894012 | pathogenic | 1 | 1 | 0 |
| rs193922285 | pathogenic | 1 | 1 | 1 |
| rs104894336 | pathogenic | 1 | 1 | 1 |
| rs28931580 | pathogenic | 1 | 1 | 1 |
| rs104894326 | pathogenic | 1 | 1 | 1 |
| rs104894331 | pathogenic | 0 | 0 | 0 |
| rs104894338 | pathogenic | 0 | 0 | 0 |
| rs104894333 | pathogenic | 1 | 1 | 1 |
| rs104894330 | pathogenic | 1 | 1 | 1 |
| rs143886391 | benign | 1 | 1 | 1 |
| rs104894335 | pathogenic | 1 | 1 | 1 |
| rs104894337 | pathogenic | 0 | 0 | 0 |

| | | | | |
|---|---|---|---|---|
| rs104894328 | pathogenic | 0 | 0 | 0 |
| rs104894341 | pathogenic | 1 | 1 | 0 |
| rs104894329 | pathogenic | 0 | 0 | 0 |
| rs137852850 | pathogenic | 1 | 1 | 1 |
| rs137852854 | pathogenic | 0 | 1 | 0 |
| rs137852849 | pathogenic | 0 | 1 | 0 |
| rs137852848 | pathogenic | 1 | 1 | 1 |
| rs137852852 | pathogenic | 0 | 1 | 0 |
| rs137852853 | pathogenic | 0 | 1 | 0 |
| rs137852847 | pathogenic | 0 | 1 | 0 |
| rs137852846 | pathogenic | 0 | 1 | 0 |
| rs137852844 | pathogenic | 0 | 1 | 0 |
| rs144411158 | pathogenic | 0 | 0 | 0 |
| rs2302234 | benign | 0 | 0 | 0 |
| rs139620139 | pathogenic | 0 | 0 | 0 |
| rs146689929 | benign | 0 | 0 | 0 |
| rs373328681 | pathogenic | 1 | 1 | 0 |
| rs387906620 | pathogenic | 1 | 1 | 1 |
| rs1799983 | benign | 0 | 1 | 0 |
| rs28999111 | pathogenic | 1 | 1 | 1 |
| rs104894544 | pathogenic | 1 | 1 | 1 |
| rs104894546 | pathogenic | 1 | 1 | 1 |
| rs28999112 | pathogenic | 0 | 1 | 0 |
| rs104894545 | pathogenic | 1 | 1 | 1 |
| rs104894547 | pathogenic | 1 | 1 | 1 |
| rs74315310 | pathogenic | 0 | 0 | 0 |
| rs267606852 | pathogenic | 0 | 1 | 0 |
| rs137853586 | pathogenic | 0 | 1 | 0 |
| rs137853582 | pathogenic | 1 | 1 | 1 |
| rs61754634 | pathogenic | 1 | 1 | 1 |
| rs137853587 | pathogenic | 0 | 1 | 0 |
| rs267606851 | pathogenic | 1 | 1 | 1 |
| rs267606853 | pathogenic | 1 | 1 | 1 |
| rs137853584 | pathogenic | 1 | 1 | 1 |
| rs137853585 | pathogenic | 1 | 1 | 1 |
| rs460184 | pathogenic | 1 | 1 | 1 |
| rs104893751 | pathogenic | 0 | 1 | 0 |
| rs77358162 | benign | 0 | 0 | 0 |
| rs387906936 | pathogenic | 0 | 0 | 0 |
| rs769525399 | pathogenic | 0 | 0 | 0 |
| rs371271054 | pathogenic | 1 | 1 | 0 |
| rs118204000 | pathogenic | 0 | 0 | 0 |
| rs138568622 | pathogenic | 1 | 1 | 1 |
| rs34509359 | benign | 0 | 0 | 0 |
| rs104894130 | pathogenic | 1 | 1 | 0 |
| rs281865424 | pathogenic | 0 | 0 | 0 |
| rs121912778 | pathogenic | 1 | 1 | 0 |
| rs771055145 | pathogenic | 1 | 1 | 0 |
| rs776174514 | pathogenic | 1 | 1 | 0 |
| rs545986367 | pathogenic | 1 | 1 | 1 |
| rs570768038 | benign | 1 | 1 | 1 |
| rs74315442 | pathogenic | 1 | 1 | 1 |
| rs79716074 | benign | 1 | 1 | 1 |
| rs748486078 | pathogenic | 1 | 1 | 1 |

| | | | | |
|---|---|---|---|---|
| rs2397084 | benign | 1 | 1 | 1 |
| rs11465553 | benign | 1 | 1 | 1 |
| rs144576902 | benign | 1 | 1 | 1 |
| rs763780 | benign | 1 | 1 | 1 |
| rs267606870 | pathogenic | 1 | 1 | 0 |
| rs121913502 | pathogenic | 1 | 1 | 0 |
| rs118101777 | benign | 1 | 1 | 1 |
| rs118053940 | benign | 0 | 0 | 0 |
| rs368771578 | benign | 1 | 1 | 1 |
| rs11540654 | pathogenic | 1 | 1 | 1 |
| rs121912658 | pathogenic | 1 | 1 | 1 |
| rs28934873 | pathogenic | 0 | 1 | 0 |
| rs780442292 | pathogenic | 1 | 1 | 1 |
| rs758781593 | benign | 1 | 1 | 1 |
| rs28934875 | pathogenic | 1 | 1 | 1 |
| rs28934874 | pathogenic | 1 | 1 | 0 |
| rs587782705 | pathogenic | 1 | 1 | 1 |
| rs587780068 | pathogenic | 1 | 1 | 1 |
| rs587782144 | pathogenic | 1 | 1 | 1 |
| rs138729528 | pathogenic | 1 | 1 | 1 |
| rs28934578 | pathogenic | 1 | 1 | 1 |
| rs397514495 | pathogenic | 1 | 1 | 1 |
| rs121912665 | pathogenic | 1 | 1 | 1 |
| rs397516435 | pathogenic | 1 | 1 | 1 |
| rs587778720 | pathogenic | 1 | 1 | 1 |
| rs397516436 | pathogenic | 1 | 1 | 1 |
| rs587781386 | benign | 1 | 1 | 1 |
| rs121912666 | pathogenic | 1 | 1 | 0 |
| rs587780073 | pathogenic | 1 | 1 | 1 |
| rs587782664 | pathogenic | 1 | 1 | 1 |
| rs730882005 | pathogenic | 1 | 1 | 1 |
| rs28934573 | pathogenic | 1 | 1 | 1 |
| rs121912655 | pathogenic | 1 | 1 | 1 |
| rs28934575 | pathogenic | 1 | 1 | 1 |
| rs121912656 | pathogenic | 1 | 1 | 1 |
| rs11540652 | pathogenic | 1 | 1 | 1 |
| rs121912651 | pathogenic | 1 | 1 | 1 |
| rs730882027 | pathogenic | 1 | 1 | 1 |
| rs121912653 | pathogenic | 1 | 1 | 1 |
| rs121912652 | pathogenic | 1 | 1 | 1 |
| rs121912657 | pathogenic | 0 | 1 | 0 |
| rs121913343 | pathogenic | 1 | 1 | 1 |
| rs28934576 | pathogenic | 1 | 1 | 1 |
| rs763098116 | pathogenic | 1 | 1 | 1 |
| rs121912660 | pathogenic | 1 | 1 | 1 |
| rs28934574 | pathogenic | 1 | 1 | 1 |
| rs371409680 | pathogenic | 1 | 1 | 1 |
| rs121912667 | pathogenic | 1 | 1 | 1 |
| rs121912663 | pathogenic | 1 | 1 | 1 |
| rs201744589 | pathogenic | 0 | 0 | 0 |
| rs33926449 | benign | 0 | 0 | 0 |
| rs74315302 | pathogenic | 0 | 1 | 0 |
| rs202040269 | pathogenic | 1 | 1 | 1 |
| rs375958814 | pathogenic | 1 | 1 | 1 |

| | | | | |
|---|---|---|---|---|
| rs121909624 | pathogenic | 1 | 1 | 1 |
| rs121909621 | pathogenic | 1 | 1 | 1 |
| rs121909622 | pathogenic | 1 | 1 | 0 |
| rs4220 | benign | 1 | 1 | 1 |
| rs147559466 | pathogenic | 1 | 1 | 1 |
| rs757434857 | pathogenic | 1 | 1 | 1 |
| rs398123072 | pathogenic | 1 | 1 | 1 |
| rs121434280 | pathogenic | 0 | 1 | 0 |
| rs398123074 | pathogenic | 0 | 1 | 0 |
| rs762114560 | pathogenic | 1 | 1 | 1 |
| rs121434283 | pathogenic | 1 | 1 | 0 |
| rs121434277 | pathogenic | 0 | 1 | 0 |
| rs745844469 | pathogenic | 1 | 1 | 1 |
| rs121434279 | pathogenic | 1 | 1 | 0 |
| rs121434278 | pathogenic | 1 | 1 | 1 |
| rs373715782 | pathogenic | 1 | 1 | 1 |
| rs200724875 | pathogenic | 1 | 1 | 1 |
| rs149678400 | pathogenic | 0 | 1 | 0 |
| rs770273135 | pathogenic | 1 | 1 | 1 |
| rs121434276 | pathogenic | 0 | 1 | 0 |
| rs121434281 | pathogenic | 1 | 1 | 0 |
| rs768884003 | pathogenic | 1 | 1 | 1 |
| rs201375579 | pathogenic | 0 | 0 | 0 |
| rs121434274 | pathogenic | 1 | 1 | 0 |
| rs121434282 | pathogenic | 1 | 1 | 0 |
| rs779759347 | pathogenic | 0 | 1 | 0 |
| rs77931234 | pathogenic | 1 | 1 | 1 |
| rs148207467 | pathogenic | 1 | 1 | 1 |
| rs766140986 | pathogenic | 1 | 1 | 1 |
| rs121434275 | pathogenic | 1 | 1 | 1 |
| rs748110745 | pathogenic | 1 | 1 | 1 |
| rs104894274 | pathogenic | 1 | 1 | 1 |
| rs104894273 | pathogenic | 0 | 1 | 0 |
| rs104894278 | pathogenic | 1 | 1 | 1 |
| rs104894275 | pathogenic | 1 | 1 | 1 |
| rs104894277 | pathogenic | 1 | 1 | 1 |
| rs104894276 | pathogenic | 0 | 0 | 0 |
| rs104894280 | pathogenic | 1 | 1 | 1 |
| rs145882709 | pathogenic | 1 | 1 | 1 |
| rs104894279 | pathogenic | 1 | 1 | 1 |
| rs80356670 | pathogenic | 1 | 1 | 1 |
| rs121918102 | pathogenic | 1 | 1 | 1 |
| rs80356671 | pathogenic | 1 | 1 | 1 |
| rs80356672 | pathogenic | 1 | 1 | 1 |
| rs758609113 | pathogenic | 1 | 1 | 0 |
| rs121918188 | pathogenic | 0 | 1 | 0 |
| rs121918189 | pathogenic | 1 | 1 | 0 |
| rs121918191 | pathogenic | 1 | 1 | 1 |
| rs1769259 | benign | 1 | 1 | 1 |
| rs140222720 | benign | 1 | 1 | 1 |
| rs2297085 | benign | 1 | 1 | 1 |
| rs200679026 | pathogenic | 1 | 1 | 1 |
| rs121918192 | pathogenic | 0 | 1 | 0 |
| rs145771944 | benign | 0 | 1 | 0 |

| | | | | |
|---|---|---|---|---|
| rs201508880 | pathogenic | 0 | 1 | 0 |
| rs17221854 | pathogenic | 0 | 0 | 0 |
| rs120074189 | pathogenic | 1 | 1 | 1 |
| rs120074190 | pathogenic | 1 | 1 | 1 |
| rs758600283 | benign | 1 | 1 | 1 |
| rs200305064 | pathogenic | 1 | 1 | 1 |
| rs121917891 | pathogenic | 1 | 1 | 1 |
| rs3772809 | benign | 0 | 1 | 0 |
| rs387906619 | pathogenic | 1 | 1 | 0 |
| rs2304681 | benign | 0 | 1 | 0 |
| rs121917849 | pathogenic | 0 | 0 | 0 |
| rs143010236 | pathogenic | 0 | 0 | 0 |
| rs121917850 | pathogenic | 1 | 1 | 0 |
| rs35916840 | pathogenic | 1 | 1 | 0 |
| rs28939688 | pathogenic | 1 | 1 | 1 |
| rs11258194 | benign | 1 | 1 | 1 |
| rs199840952 | pathogenic | 1 | 1 | 1 |
| rs121918159 | pathogenic | 1 | 1 | 1 |
| rs200396166 | pathogenic | 0 | 0 | 0 |
| rs757966746 | pathogenic | 0 | 1 | 0 |
| rs121918608 | pathogenic | 0 | 1 | 0 |
| rs74315401 | pathogenic | 1 | 1 | 1 |
| rs11538758 | pathogenic | 0 | 0 | 0 |
| rs74315402 | pathogenic | 1 | 1 | 1 |
| rs1799990 | pathogenic | 1 | 1 | 1 |
| rs74315410 | pathogenic | 1 | 1 | 1 |
| rs150351644 | benign | 1 | 1 | 1 |
| rs16990018 | benign | 1 | 1 | 1 |
| rs74315403 | pathogenic | 1 | 1 | 1 |
| rs74315408 | pathogenic | 1 | 1 | 1 |
| rs74315411 | pathogenic | 1 | 1 | 1 |
| rs74315413 | pathogenic | 1 | 1 | 1 |
| rs74315405 | pathogenic | 1 | 1 | 1 |
| rs28933385 | pathogenic | 1 | 1 | 1 |
| rs74315412 | pathogenic | 1 | 1 | 1 |
| rs74315407 | pathogenic | 1 | 1 | 1 |
| rs74315406 | pathogenic | 1 | 1 | 1 |
| rs11571117 | benign | 1 | 1 | 1 |
| rs121917742 | pathogenic | 1 | 1 | 1 |
| rs149782619 | pathogenic | 1 | 1 | 1 |
| rs397514492 | pathogenic | 1 | 1 | 1 |
| rs397514489 | pathogenic | 0 | 1 | 0 |
| rs397514493 | pathogenic | 1 | 1 | 1 |
| rs373849532 | pathogenic | 1 | 1 | 1 |
| rs121965048 | pathogenic | 1 | 1 | 1 |
| rs121965037 | pathogenic | 1 | 1 | 1 |
| rs121965060 | pathogenic | 0 | 1 | 0 |
| rs386833603 | pathogenic | 1 | 1 | 1 |
| rs121965038 | pathogenic | 1 | 1 | 1 |
| rs121965039 | pathogenic | 1 | 1 | 1 |
| rs121965040 | pathogenic | 1 | 1 | 1 |
| rs267606925 | pathogenic | 1 | 1 | 1 |
| rs121965056 | pathogenic | 0 | 1 | 0 |
| rs121965059 | pathogenic | 0 | 0 | 0 |

| | | | | |
|---|---|---|---|---|
| rs386833615 | pathogenic | 1 | 1 | 0 |
| rs121965051 | pathogenic | 1 | 1 | 0 |
| rs121965046 | pathogenic | 0 | 1 | 0 |
| rs121965052 | pathogenic | 0 | 1 | 0 |
| rs386833617 | pathogenic | 0 | 1 | 0 |
| rs386833618 | pathogenic | 1 | 1 | 0 |
| rs121965041 | pathogenic | 0 | 1 | 0 |
| rs121965042 | pathogenic | 0 | 1 | 0 |
| rs121965057 | pathogenic | 1 | 1 | 1 |
| rs386833621 | pathogenic | 1 | 1 | 1 |
| rs121965049 | pathogenic | 1 | 1 | 1 |
| rs386833623 | pathogenic | 1 | 1 | 1 |
| rs121965047 | pathogenic | 1 | 1 | 1 |
| rs121965053 | pathogenic | 0 | 1 | 0 |
| rs121965045 | pathogenic | 0 | 1 | 0 |
| rs121965054 | pathogenic | 0 | 1 | 0 |
| rs121965043 | pathogenic | 0 | 1 | 0 |
| rs386833598 | pathogenic | 0 | 0 | 0 |
| rs137854545 | pathogenic | 1 | 1 | 1 |
| rs137854544 | pathogenic | 1 | 1 | 1 |
| rs104894455 | pathogenic | 0 | 1 | 0 |
| rs1049564 | benign | 0 | 1 | 0 |
| rs104894460 | pathogenic | 1 | 1 | 1 |
| rs104894453 | pathogenic | 1 | 1 | 1 |
| rs104894450 | pathogenic | 0 | 1 | 0 |
| rs104894454 | pathogenic | 0 | 1 | 0 |
| rs104894452 | pathogenic | 1 | 1 | 1 |
| rs104894451 | pathogenic | 0 | 1 | 0 |
| rs104894368 | pathogenic | 1 | 1 | 1 |
| rs730880950 | pathogenic | 1 | 1 | 1 |
| rs121909535 | pathogenic | 1 | 1 | 1 |
| rs121909537 | pathogenic | 1 | 1 | 1 |
| rs121909536 | pathogenic | 1 | 1 | 1 |
| rs121909542 | pathogenic | 1 | 1 | 1 |
| rs121909538 | pathogenic | 1 | 1 | 1 |
| rs121909539 | pathogenic | 1 | 1 | 1 |
| rs121909540 | pathogenic | 1 | 1 | 1 |
| rs17560 | benign | 1 | 1 | 1 |
| rs121909543 | pathogenic | 0 | 1 | 0 |
| rs121909544 | pathogenic | 1 | 1 | 1 |
| rs2307456 | benign | 0 | 1 | 0 |
| rs745778317 | pathogenic | 0 | 1 | 0 |
| rs759607901 | pathogenic | 0 | 1 | 0 |
| rs201188114 | benign | 0 | 0 | 0 |
| rs1800472 | benign | 0 | 0 | 0 |
| rs587780728 | benign | 1 | 1 | 1 |
| rs144386518 | benign | 1 | 1 | 1 |
| rs281865439 | pathogenic | 1 | 1 | 1 |
| rs140175796 | pathogenic | 1 | 1 | 1 |
| rs5030843 | pathogenic | 1 | 1 | 0 |
| rs75166491 | pathogenic | 1 | 1 | 0 |
| rs199475679 | pathogenic | 1 | 1 | 1 |
| rs281865440 | pathogenic | 1 | 1 | 1 |
| rs74486803 | pathogenic | 0 | 1 | 0 |

| | | | | |
|---|---|---|---|---|
| rs199475602 | pathogenic | 0 | 1 | 0 |
| rs5030844 | pathogenic | 1 | 1 | 0 |
| rs62514927 | pathogenic | 1 | 1 | 1 |
| rs62514928 | pathogenic | 1 | 1 | 1 |
| rs62516152 | pathogenic | 0 | 1 | 0 |
| rs62507348 | pathogenic | 1 | 1 | 1 |
| rs76687508 | pathogenic | 1 | 1 | 1 |
| rs62508730 | pathogenic | 1 | 1 | 1 |
| rs62508588 | pathogenic | 0 | 1 | 0 |
| rs5030846 | pathogenic | 0 | 1 | 0 |
| rs74503222 | pathogenic | 1 | 1 | 0 |
| rs5030847 | pathogenic | 1 | 1 | 0 |
| rs5030850 | pathogenic | 1 | 1 | 1 |
| rs778154939 | pathogenic | 0 | 1 | 0 |
| rs62514952 | pathogenic | 1 | 1 | 1 |
| rs78655458 | pathogenic | 1 | 1 | 1 |
| rs62508698 | pathogenic | 1 | 1 | 0 |
| rs5030851 | pathogenic | 1 | 1 | 0 |
| rs62642933 | pathogenic | 1 | 1 | 1 |
| rs5030853 | pathogenic | 1 | 1 | 1 |
| rs62642935 | pathogenic | 1 | 1 | 1 |
| rs199475615 | pathogenic | 0 | 0 | 0 |
| rs62516062 | pathogenic | 1 | 1 | 0 |
| rs62516092 | pathogenic | 0 | 0 | 0 |
| rs62516095 | pathogenic | 1 | 1 | 1 |
| rs62642937 | pathogenic | 1 | 1 | 1 |
| rs62516141 | pathogenic | 0 | 1 | 0 |
| rs5030856 | pathogenic | 1 | 1 | 1 |
| rs5030857 | pathogenic | 1 | 1 | 1 |
| rs5030858 | pathogenic | 1 | 1 | 1 |
| rs79931499 | pathogenic | 1 | 1 | 1 |
| rs5030860 | pathogenic | 1 | 1 | 1 |
| rs281865437 | pathogenic | 1 | 1 | 1 |
| rs62644499 | pathogenic | 1 | 1 | 1 |
| rs147689487 | benign | 0 | 0 | 0 |
| rs397515417 | pathogenic | 1 | 1 | 1 |
| rs121909332 | pathogenic | 1 | 1 | 1 |
| rs121909330 | pathogenic | 1 | 1 | 0 |
| rs121909329 | pathogenic | 1 | 1 | 0 |
| rs121909335 | pathogenic | 1 | 1 | 1 |
| rs387906789 | pathogenic | 1 | 1 | 1 |
| rs121909334 | pathogenic | 1 | 1 | 1 |
| rs121909331 | pathogenic | 1 | 1 | 1 |
| rs372487178 | pathogenic | 1 | 1 | 1 |
| rs387906937 | pathogenic | 1 | 1 | 0 |
| rs16845 | benign | 1 | 1 | 1 |
| rs672601313 | pathogenic | 1 | 1 | 1 |
| rs117805308 | benign | 0 | 0 | 0 |
| rs121434261 | pathogenic | 1 | 1 | 1 |
| rs141629512 | benign | 0 | 0 | 0 |
| rs199957176 | benign | 1 | 1 | 1 |
| rs2230461 | benign | 1 | 1 | 1 |
| rs764618040 | pathogenic | 1 | 1 | 1 |
| rs267606949 | pathogenic | 1 | 1 | 1 |

| | | | | |
|---|---|---|---|---|
| rs587777483 | pathogenic | 1 | 1 | 1 |
| rs267606947 | pathogenic | 1 | 1 | 1 |
| rs6941583 | benign | 0 | 0 | 0 |
| rs749597959 | benign | 1 | 1 | 1 |
| rs182340753 | benign | 1 | 1 | 1 |
| rs146678221 | benign | 0 | 0 | 0 |
| rs11539202 | benign | 0 | 0 | 0 |
| rs148723565 | benign | 0 | 0 | 0 |
| rs61752925 | benign | 1 | 1 | 1 |
| rs113309941 | pathogenic | 0 | 0 | 0 |
| rs121434520 | pathogenic | 1 | 1 | 1 |
| rs121434517 | pathogenic | 1 | 1 | 1 |
| rs267607027 | pathogenic | 1 | 1 | 1 |
| rs138603088 | pathogenic | 0 | 0 | 0 |
| rs121434516 | pathogenic | 1 | 1 | 0 |
| rs121434521 | pathogenic | 0 | 0 | 0 |
| rs145735112 | benign | 1 | 1 | 1 |
| rs121434519 | pathogenic | 1 | 1 | 1 |
| rs75326924 | pathogenic | 1 | 1 | 0 |
| rs142186404 | pathogenic | 0 | 0 | 0 |
| rs121918035 | pathogenic | 0 | 0 | 0 |
| rs374385878 | pathogenic | 1 | 1 | 0 |
| rs11557652 | benign | 0 | 0 | 0 |
| rs118203986 | pathogenic | 1 | 1 | 1 |
| rs141675330 | benign | 1 | 1 | 1 |
| rs104893927 | pathogenic | 0 | 0 | 0 |
| rs104893935 | pathogenic | 0 | 0 | 0 |
| rs104893933 | pathogenic | 1 | 1 | 0 |
| rs104893934 | pathogenic | 0 | 0 | 0 |
| rs145183203 | benign | 1 | 1 | 0 |
| rs2230806 | benign | 0 | 0 | 0 |
| rs9282541 | benign | 0 | 0 | 0 |
| rs559110055 | benign | 0 | 0 | 0 |
| rs200030513 | benign | 0 | 0 | 0 |
| rs9282543 | benign | 0 | 0 | 0 |
| rs147675550 | benign | 0 | 0 | 0 |
| rs137854496 | pathogenic | 0 | 0 | 0 |
| rs374190304 | benign | 0 | 0 | 0 |
| rs114620717 | benign | 0 | 0 | 0 |
| rs2066718 | benign | 0 | 0 | 0 |
| rs35819696 | benign | 0 | 0 | 0 |
| rs138880920 | benign | 0 | 0 | 0 |
| rs2066715 | benign | 0 | 0 | 0 |
| rs35207495 | benign | 0 | 0 | 0 |
| rs2066714 | benign | 0 | 0 | 0 |
| rs28937313 | pathogenic | 0 | 0 | 0 |
| rs28937314 | pathogenic | 0 | 0 | 0 |
| rs137854495 | pathogenic | 0 | 0 | 0 |
| rs140365800 | benign | 0 | 0 | 0 |
| rs13306073 | benign | 0 | 0 | 0 |
| rs28933692 | pathogenic | 0 | 0 | 0 |
| rs33918808 | benign | 0 | 0 | 0 |
| rs76881554 | benign | 0 | 0 | 0 |
| rs143180998 | benign | 0 | 0 | 0 |

| | | | | |
|---|---|---|---|---|
| rs137854500 | pathogenic | 0 | 0 | 0 |
| rs538614702 | benign | 0 | 0 | 0 |
| rs201879964 | benign | 0 | 0 | 0 |
| rs2230808 | benign | 0 | 0 | 0 |
| rs138422574 | benign | 0 | 0 | 0 |
| rs137854498 | pathogenic | 0 | 0 | 0 |
| rs150125857 | benign | 0 | 0 | 0 |
| rs146292819 | pathogenic | 0 | 0 | 0 |
| rs142688906 | benign | 0 | 0 | 0 |
| rs137854499 | pathogenic | 0 | 0 | 0 |
| rs34879708 | benign | 0 | 0 | 0 |
| rs121913028 | pathogenic | 1 | 1 | 1 |
| rs1041951 | benign | 1 | 1 | 0 |
| rs138714094 | benign | 1 | 1 | 0 |
| rs200650502 | benign | 0 | 1 | 0 |
| rs370708663 | pathogenic | 1 | 1 | 1 |
| rs118204040 | pathogenic | 0 | 1 | 0 |
| rs146899669 | pathogenic | 1 | 1 | 0 |
| rs118204039 | pathogenic | 0 | 1 | 0 |
| rs121909233 | pathogenic | 0 | 0 | 0 |
| rs121909225 | pathogenic | 1 | 1 | 0 |
| rs121909236 | pathogenic | 0 | 0 | 0 |
| rs398123317 | pathogenic | 0 | 0 | 0 |
| rs121909226 | pathogenic | 1 | 1 | 0 |
| rs121909238 | pathogenic | 1 | 1 | 0 |
| rs121909230 | pathogenic | 0 | 0 | 0 |
| rs121909237 | pathogenic | 0 | 0 | 0 |
| rs121909222 | pathogenic | 1 | 1 | 0 |
| rs121909223 | pathogenic | 1 | 1 | 0 |
| rs121909218 | pathogenic | 1 | 1 | 0 |
| rs121909229 | pathogenic | 1 | 1 | 0 |
| rs121909224 | pathogenic | 1 | 1 | 0 |
| rs121909241 | pathogenic | 0 | 0 | 0 |
| rs121909221 | pathogenic | 0 | 0 | 0 |
| rs121913294 | pathogenic | 0 | 0 | 0 |
| rs121909234 | pathogenic | 0 | 0 | 0 |
| rs121909240 | pathogenic | 0 | 0 | 0 |
| rs767623493 | benign | 0 | 0 | 0 |
| rs121909239 | pathogenic | 0 | 0 | 0 |
| rs120074145 | pathogenic | 1 | 1 | 1 |
| rs120074148 | pathogenic | 1 | 1 | 1 |
| rs762991875 | pathogenic | 0 | 0 | 0 |
| rs148639841 | pathogenic | 1 | 1 | 1 |
| rs199524907 | pathogenic | 1 | 1 | 1 |
| rs120074141 | pathogenic | 1 | 1 | 1 |
| rs120074144 | pathogenic | 0 | 1 | 0 |
| rs120074146 | pathogenic | 1 | 1 | 1 |
| rs120074147 | pathogenic | 0 | 1 | 0 |
| rs120074143 | pathogenic | 0 | 1 | 0 |
| rs120074140 | pathogenic | 1 | 1 | 0 |
| rs28937315 | pathogenic | 1 | 1 | 1 |
| rs76066357 | benign | 1 | 1 | 1 |
| rs137852911 | pathogenic | 0 | 1 | 0 |
| rs137852907 | pathogenic | 1 | 1 | 0 |

| | | | | |
|---|---|---|---|---|
| rs137852910 | pathogenic | 1 | 1 | 1 |
| rs537585211 | benign | 0 | 0 | 0 |
| rs202134424 | pathogenic | 0 | 0 | 0 |
| rs200930463 | pathogenic | 0 | 0 | 0 |
| rs587777241 | pathogenic | 1 | 1 | 0 |
| rs77563738 | pathogenic | 0 | 0 | 0 |
| rs750868279 | pathogenic | 1 | 1 | 0 |
| rs148936893 | pathogenic | 0 | 0 | 0 |
| rs146597836 | pathogenic | 0 | 0 | 0 |
| rs775440641 | pathogenic | 1 | 1 | 1 |
| rs770689762 | pathogenic | 0 | 1 | 0 |
| rs376785840 | pathogenic | 0 | 0 | 0 |
| rs137852531 | pathogenic | 0 | 0 | 0 |
| rs137852538 | pathogenic | 1 | 1 | 0 |
| rs137852529 | pathogenic | 0 | 1 | 0 |
| rs137852528 | pathogenic | 0 | 1 | 0 |
| rs137852535 | pathogenic | 1 | 1 | 0 |
| rs137853045 | pathogenic | 0 | 1 | 0 |
| rs137853046 | pathogenic | 1 | 1 | 1 |
| rs137853043 | pathogenic | 1 | 1 | 1 |
| rs137853044 | pathogenic | 1 | 1 | 1 |
| rs137853047 | pathogenic | 1 | 1 | 1 |
| rs267606990 | pathogenic | 1 | 1 | 1 |
| rs751437780 | pathogenic | 1 | 1 | 0 |
| rs121918461 | pathogenic | 1 | 1 | 1 |
| rs121918460 | pathogenic | 1 | 1 | 1 |
| rs121918459 | pathogenic | 1 | 1 | 0 |
| rs397507511 | pathogenic | 1 | 1 | 1 |
| rs121918454 | pathogenic | 1 | 1 | 1 |
| rs121918453 | pathogenic | 1 | 1 | 1 |
| rs121918462 | pathogenic | 1 | 1 | 0 |
| rs121918465 | pathogenic | 1 | 1 | 0 |
| rs121918464 | pathogenic | 1 | 1 | 0 |
| rs121918466 | pathogenic | 1 | 1 | 1 |
| rs397507521 | pathogenic | 1 | 1 | 1 |
| rs397507527 | pathogenic | 1 | 1 | 1 |
| rs777603059 | pathogenic | 0 | 1 | 0 |
| rs121918456 | pathogenic | 1 | 1 | 1 |
| rs121918463 | pathogenic | 1 | 1 | 1 |
| rs121918455 | pathogenic | 0 | 1 | 0 |
| rs28933386 | pathogenic | 0 | 1 | 0 |
| rs774356443 | benign | 1 | 1 | 1 |
| rs121918467 | pathogenic | 1 | 1 | 1 |
| rs121918468 | pathogenic | 1 | 1 | 1 |
| rs121918469 | pathogenic | 1 | 1 | 1 |
| rs397507539 | pathogenic | 1 | 1 | 0 |
| rs121918458 | pathogenic | 1 | 1 | 1 |
| rs121918470 | pathogenic | 1 | 1 | 1 |
| rs35289681 | benign | 0 | 1 | 0 |
| rs4302331 | benign | 0 | 1 | 0 |
| rs11544803 | pathogenic | 1 | 1 | 1 |
| rs116254156 | benign | 0 | 0 | 0 |
| rs104893909 | pathogenic | 1 | 1 | 1 |
| rs104893911 | pathogenic | 1 | 1 | 1 |

| | | | | |
|---|---|---|---|---|
| rs104893908 | pathogenic | 1 | 1 | 0 |
| rs258751 | benign | 1 | 1 | 1 |
| rs104893914 | pathogenic | 1 | 1 | 0 |
| rs121909727 | pathogenic | 1 | 1 | 1 |
| rs104893910 | pathogenic | 1 | 1 | 1 |
| rs121909726 | pathogenic | 1 | 1 | 1 |
| rs104893912 | pathogenic | 1 | 1 | 1 |
| rs138407155 | benign | 1 | 1 | 1 |
| rs574061789 | benign | 1 | 1 | 1 |
| rs121912718 | pathogenic | 1 | 1 | 1 |
| rs121912719 | pathogenic | 1 | 1 | 1 |
| rs140770089 | benign | 1 | 1 | 1 |
| rs28931573 | pathogenic | 1 | 1 | 1 |
| rs121912717 | pathogenic | 1 | 1 | 1 |
| rs376456598 | pathogenic | 0 | 0 | 0 |
| rs398123499 | pathogenic | 1 | 1 | 1 |
| rs773048903 | pathogenic | 1 | 1 | 0 |
| rs398123503 | pathogenic | 1 | 1 | 1 |
| rs375785084 | pathogenic | 0 | 1 | 0 |
| rs137852874 | pathogenic | 1 | 1 | 1 |
| rs373713279 | pathogenic | 1 | 1 | 1 |
| rs137852873 | pathogenic | 1 | 1 | 0 |
| rs367823977 | pathogenic | 1 | 1 | 1 |
| rs398123508 | pathogenic | 1 | 1 | 1 |
| rs137852871 | pathogenic | 1 | 1 | 1 |
| rs753698250 | pathogenic | 0 | 1 | 0 |
| rs182923857 | pathogenic | 1 | 1 | 1 |
| rs148090804 | benign | 1 | 1 | 1 |
| rs137852870 | pathogenic | 1 | 1 | 1 |
| rs267607131 | pathogenic | 1 | 1 | 0 |
| rs137852902 | pathogenic | 0 | 0 | 0 |
| rs61741646 | benign | 1 | 1 | 1 |
| rs137852905 | pathogenic | 0 | 0 | 0 |
| rs121918681 | pathogenic | 0 | 1 | 0 |
| rs8177238 | benign | 1 | 1 | 0 |
| rs5030776 | pathogenic | 1 | 1 | 1 |
| rs104894572 | pathogenic | 0 | 0 | 0 |
| rs104894576 | pathogenic | 0 | 0 | 0 |
| rs80084721 | pathogenic | 0 | 0 | 0 |
| rs104894146 | pathogenic | 0 | 1 | 0 |
| rs104894138 | pathogenic | 1 | 1 | 0 |
| rs104894153 | pathogenic | 1 | 1 | 0 |
| rs104894135 | pathogenic | 1 | 1 | 0 |
| rs104894147 | pathogenic | 1 | 1 | 0 |
| rs104894148 | pathogenic | 1 | 1 | 0 |
| rs104894154 | pathogenic | 1 | 1 | 0 |
| rs104894150 | pathogenic | 1 | 1 | 0 |
| rs104894136 | pathogenic | 1 | 1 | 0 |
| rs104894144 | pathogenic | 0 | 0 | 0 |
| rs104894137 | pathogenic | 0 | 0 | 0 |
| rs61754278 | pathogenic | 1 | 1 | 0 |
| rs104894149 | pathogenic | 1 | 1 | 0 |
| rs104894139 | pathogenic | 1 | 1 | 0 |
| rs104894142 | pathogenic | 0 | 0 | 0 |

| | | | | |
|---|---|---|---|---|
| rs104894143 | pathogenic | 0 | 0 | 0 |
| rs104894145 | pathogenic | 0 | 0 | 0 |
| rs121909545 | benign | 0 | 1 | 0 |
| rs782308462 | pathogenic | 1 | 1 | 1 |
| rs375002796 | pathogenic | 0 | 1 | 0 |
| rs201727685 | pathogenic | 0 | 0 | 0 |
| rs768050261 | pathogenic | 1 | 1 | 1 |
| rs780851340 | benign | 0 | 0 | 0 |
| rs139824241 | benign | 1 | 1 | 0 |
| rs757486575 | pathogenic | 1 | 1 | 0 |
| rs750831299 | pathogenic | 1 | 1 | 1 |
| rs767602961 | pathogenic | 1 | 1 | 1 |
| rs121913661 | pathogenic | 1 | 1 | 1 |
| rs777692271 | pathogenic | 1 | 1 | 1 |
| rs121913659 | pathogenic | 1 | 1 | 1 |
| rs587777533 | pathogenic | 1 | 1 | 1 |
| rs104894210 | pathogenic | 0 | 0 | 0 |
| rs104894216 | pathogenic | 1 | 1 | 1 |
| rs104894207 | pathogenic | 1 | 1 | 1 |
| rs104894208 | pathogenic | 0 | 1 | 0 |
| rs104894214 | pathogenic | 1 | 1 | 1 |
| rs104894211 | pathogenic | 1 | 1 | 1 |
| rs61749384 | pathogenic | 0 | 0 | 0 |
| rs61750071 | pathogenic | 1 | 1 | 1 |
| rs61750074 | pathogenic | 0 | 0 | 0 |
| rs11063988 | benign | 0 | 0 | 0 |
| rs216311 | benign | 0 | 0 | 0 |
| rs1800382 | benign | 1 | 1 | 1 |
| rs11063987 | benign | 1 | 1 | 1 |
| rs387906847 | pathogenic | 1 | 1 | 1 |
| rs73003466 | benign | 0 | 0 | 0 |
| rs149787233 | benign | 1 | 1 | 1 |
| rs144451841 | pathogenic | 1 | 1 | 1 |
| rs141638421 | pathogenic | 1 | 1 | 1 |
| rs121434497 | pathogenic | 0 | 0 | 0 |
| rs121434498 | pathogenic | 0 | 0 | 0 |
| rs267607230 | pathogenic | 1 | 1 | 0 |
| rs121434499 | pathogenic | 1 | 1 | 0 |
| rs546749336 | benign | 1 | 1 | 1 |
| rs138873805 | benign | 0 | 0 | 0 |
| rs730880522 | pathogenic | 0 | 0 | 0 |
| rs200371894 | benign | 0 | 0 | 0 |
| rs730880509 | benign | 0 | 0 | 0 |
| rs267606704 | pathogenic | 1 | 1 | 1 |
| rs267606706 | pathogenic | 0 | 0 | 0 |
| rs267606705 | pathogenic | 1 | 1 | 1 |
| rs267606707 | pathogenic | 0 | 0 | 0 |
| rs387906665 | pathogenic | 1 | 1 | 1 |
| rs267606708 | pathogenic | 1 | 1 | 1 |
| rs148368481 | benign | 0 | 0 | 0 |
| rs13204 | benign | 0 | 0 | 0 |
| rs140547520 | pathogenic | 1 | 1 | 1 |
| rs755234990 | benign | 1 | 1 | 0 |
| rs151184052 | benign | 0 | 0 | 0 |

| | | | | |
|---|---|---|---|---|
| rs3736358 | benign | 0 | 0 | 0 |
| rs3736360 | benign | 0 | 0 | 0 |
| rs1138469 | benign | 0 | 0 | 0 |
| rs74315351 | pathogenic | 0 | 1 | 0 |
| rs137853051 | pathogenic | 1 | 1 | 0 |
| rs74315353 | pathogenic | 0 | 1 | 0 |
| rs74315352 | pathogenic | 0 | 1 | 0 |
| rs74315354 | pathogenic | 1 | 1 | 1 |
| rs28938172 | pathogenic | 1 | 1 | 1 |
| rs52836744 | pathogenic | 0 | 0 | 0 |
| rs121913159 | pathogenic | 1 | 1 | 1 |
| rs121913145 | pathogenic | 1 | 1 | 1 |
| rs891087 | benign | 1 | 1 | 1 |
| rs9282757 | benign | 1 | 1 | 1 |
| rs397517404 | pathogenic | 1 | 1 | 1 |
| rs199918720 | benign | 1 | 1 | 0 |
| rs368299411 | benign | 1 | 1 | 0 |
| rs139399951 | benign | 1 | 1 | 0 |
| rs763981974 | benign | 1 | 1 | 0 |
| rs144242114 | benign | 0 | 0 | 0 |
| rs750965801 | benign | 0 | 0 | 0 |
| rs561310777 | benign | 0 | 0 | 0 |
| rs148185335 | benign | 0 | 0 | 0 |
| rs147742157 | benign | 0 | 0 | 0 |
| rs375664373 | pathogenic | 1 | 1 | 1 |
| rs777327250 | pathogenic | 0 | 0 | 0 |
| rs121908471 | pathogenic | 1 | 1 | 0 |
| rs2301612 | pathogenic | 1 | 1 | 0 |
| rs121908476 | pathogenic | 0 | 0 | 0 |
| rs36220239 | benign | 0 | 0 | 0 |
| rs121908473 | pathogenic | 0 | 0 | 0 |
| rs28647808 | benign | 0 | 0 | 0 |
| rs36090624 | benign | 0 | 0 | 0 |
| rs121908002 | pathogenic | 1 | 1 | 1 |
| rs4251545 | benign | 1 | 1 | 1 |
| rs78705382 | pathogenic | 0 | 0 | 0 |
| rs75184679 | pathogenic | 1 | 1 | 1 |
| rs74555752 | pathogenic | 0 | 0 | 0 |
| rs56320207 | benign | 1 | 1 | 1 |
| rs28730837 | pathogenic | 1 | 1 | 1 |
| rs119469014 | pathogenic | 1 | 1 | 1 |
| rs119468010 | pathogenic | 1 | 1 | 1 |
| rs35220041 | benign | 0 | 0 | 0 |
| rs762321137 | pathogenic | 1 | 1 | 1 |
| rs28931570 | pathogenic | 1 | 1 | 1 |
| rs709932 | benign | 1 | 1 | 1 |
| rs746531546 | benign | 1 | 1 | 1 |
| rs6647 | benign | 1 | 1 | 1 |
| rs141095970 | benign | 1 | 1 | 1 |
| rs143370956 | benign | 1 | 1 | 1 |
| rs28929474 | pathogenic | 1 | 1 | 1 |
| rs61761869 | pathogenic | 1 | 1 | 1 |
| rs1303 | benign | 1 | 1 | 1 |
| rs149447502 | benign | 0 | 0 | 0 |

| | | | | |
|---|---|---|---|---|
| rs267607180 | pathogenic | 1 | 1 | 1 |
| rs150304121 | benign | 0 | 0 | 0 |
| rs147630375 | benign | 1 | 1 | 1 |
| rs11539445 | benign | 1 | 1 | 1 |
| rs372098364 | pathogenic | 1 | 1 | 0 |
| rs760072286 | benign | 0 | 0 | 0 |
| rs745595833 | pathogenic | 1 | 1 | 1 |
| rs587777214 | pathogenic | 0 | 0 | 0 |
| rs150318528 | benign | 0 | 0 | 0 |
| rs137853254 | pathogenic | 1 | 1 | 1 |
| rs137853255 | pathogenic | 0 | 0 | 0 |
| rs137853253 | pathogenic | 0 | 0 | 0 |
| rs202166915 | benign | 0 | 0 | 0 |
| rs137853259 | pathogenic | 0 | 0 | 0 |
| rs2229137 | benign | 0 | 0 | 0 |
| rs137853258 | pathogenic | 1 | 1 | 0 |
| rs137853252 | pathogenic | 1 | 1 | 1 |
| rs137853256 | pathogenic | 0 | 0 | 0 |
| rs137853250 | pathogenic | 1 | 1 | 1 |
| rs121434615 | pathogenic | 0 | 0 | 0 |
| rs115014558 | benign | 1 | 1 | 1 |
| rs34116584 | benign | 1 | 1 | 1 |
| rs34885252 | benign | 1 | 1 | 1 |
| rs180177262 | pathogenic | 1 | 1 | 1 |
| rs180177162 | pathogenic | 1 | 1 | 1 |
| rs121908523 | pathogenic | 1 | 1 | 1 |
| rs180177170 | pathogenic | 1 | 1 | 1 |
| rs180177180 | pathogenic | 1 | 1 | 1 |
| rs767586362 | pathogenic | 1 | 1 | 1 |
| rs121908521 | pathogenic | 1 | 1 | 1 |
| rs180177203 | pathogenic | 1 | 1 | 1 |
| rs138025751 | pathogenic | 1 | 1 | 1 |
| rs180177210 | pathogenic | 0 | 1 | 0 |
| rs180177211 | pathogenic | 0 | 1 | 0 |
| rs121908524 | pathogenic | 1 | 1 | 0 |
| rs121908530 | pathogenic | 1 | 1 | 0 |
| rs180177227 | pathogenic | 0 | 1 | 0 |
| rs180177230 | pathogenic | 1 | 1 | 1 |
| rs121908529 | pathogenic | 0 | 1 | 0 |
| rs117195882 | benign | 1 | 1 | 0 |
| rs180177238 | pathogenic | 1 | 1 | 0 |
| rs34664134 | benign | 1 | 1 | 1 |
| rs180177246 | pathogenic | 0 | 1 | 0 |
| rs536352238 | pathogenic | 0 | 0 | 0 |
| rs121908520 | pathogenic | 1 | 1 | 0 |
| rs180177248 | pathogenic | 1 | 1 | 0 |
| rs180177252 | pathogenic | 1 | 1 | 1 |
| rs121908526 | pathogenic | 1 | 1 | 1 |
| rs121908525 | pathogenic | 1 | 1 | 1 |
| rs147106773 | benign | 1 | 1 | 1 |
| rs180177264 | pathogenic | 0 | 1 | 0 |
| rs146525143 | pathogenic | 1 | 1 | 1 |
| rs140992177 | benign | 1 | 1 | 1 |
| rs13408961 | benign | 0 | 1 | 0 |

| | | | | |
|---|---|---|---|---|
| rs4426527 | benign | 1 | 1 | 1 |
| rs3088308 | benign | 0 | 1 | 0 |
| rs4469829 | benign | 1 | 1 | 0 |
| rs104886003 | pathogenic | 1 | 1 | 1 |
| rs121913286 | pathogenic | 1 | 1 | 1 |
| rs121913279 | pathogenic | 1 | 1 | 1 |
| rs121434636 | pathogenic | 1 | 1 | 1 |
| rs755299132 | pathogenic | 1 | 1 | 1 |
| rs748318386 | pathogenic | 1 | 1 | 1 |
| rs387906757 | pathogenic | 1 | 1 | 1 |
| rs121965064 | pathogenic | 1 | 1 | 1 |
| rs28934608 | pathogenic | 0 | 0 | 0 |
| rs5967 | benign | 1 | 1 | 1 |
| rs773905328 | pathogenic | 0 | 0 | 0 |
| rs104894256 | pathogenic | 0 | 1 | 0 |
| rs28931612 | pathogenic | 0 | 1 | 0 |
| rs104894263 | pathogenic | 1 | 1 | 1 |
| rs607969 | benign | 1 | 1 | 0 |
| rs104894262 | pathogenic | 0 | 1 | 0 |
| rs104894268 | pathogenic | 1 | 1 | 1 |
| rs104894264 | pathogenic | 0 | 1 | 0 |
| rs104894259 | pathogenic | 1 | 1 | 0 |
| rs2959656 | benign | 0 | 0 | 0 |
| rs121913035 | pathogenic | 1 | 1 | 0 |
| rs139605474 | benign | 0 | 0 | 0 |
| rs3817552 | benign | 0 | 0 | 0 |
| rs150541115 | benign | 0 | 0 | 0 |
| rs562763010 | benign | 1 | 1 | 1 |
| rs200053119 | pathogenic | 0 | 0 | 0 |
| rs73826386 | benign | 0 | 0 | 0 |
| rs143655258 | benign | 0 | 0 | 0 |
| rs758857853 | pathogenic | 1 | 1 | 0 |
| rs2228380 | benign | 0 | 0 | 0 |
| rs141078733 | benign | 1 | 1 | 1 |
| rs55824172 | pathogenic | 1 | 1 | 1 |
| rs756751089 | pathogenic | 1 | 1 | 1 |
| rs121964990 | pathogenic | 0 | 0 | 0 |
| rs143555186 | benign | 0 | 0 | 0 |
| rs121964993 | pathogenic | 1 | 1 | 0 |
| rs121964992 | pathogenic | 1 | 1 | 1 |
| rs121964991 | pathogenic | 1 | 1 | 0 |
| rs121964988 | pathogenic | 1 | 1 | 1 |
| rs4687 | benign | 1 | 1 | 1 |
| rs267607125 | pathogenic | 1 | 1 | 1 |
| rs751682878 | benign | 1 | 1 | 1 |
| rs267607126 | pathogenic | 1 | 1 | 1 |
| rs768713596 | pathogenic | 0 | 0 | 0 |
| rs147288996 | benign | 0 | 0 | 0 |
| rs387906639 | pathogenic | 1 | 1 | 1 |
| rs730881069 | pathogenic | 1 | 1 | 1 |
| rs730881071 | pathogenic | 1 | 1 | 1 |
| rs397516348 | pathogenic | 0 | 0 | 0 |
| rs121917760 | pathogenic | 0 | 0 | 0 |
| rs104894724 | pathogenic | 1 | 1 | 1 |

| | | | | |
|---|---|---|---|---|
| rs397516349 | pathogenic | 1 | 1 | 1 |
| rs12075 | benign | 0 | 0 | 0 |
| rs147549987 | benign | 0 | 0 | 0 |
| rs56079734 | benign | 0 | 0 | 0 |
| rs1017842 | benign | 1 | 1 | 1 |
| rs137852593 | pathogenic | 1 | 1 | 1 |
| rs137852571 | pathogenic | 1 | 1 | 1 |
| rs137852591 | pathogenic | 1 | 1 | 1 |
| rs9332971 | pathogenic | 1 | 1 | 1 |
| rs755226547 | pathogenic | 1 | 1 | 1 |
| rs79206939 | benign | 0 | 0 | 0 |
| rs121918214 | pathogenic | 1 | 1 | 0 |
| rs757075712 | pathogenic | 1 | 1 | 0 |
| rs150483902 | benign | 1 | 1 | 1 |
| rs113993973 | pathogenic | 0 | 1 | 0 |
| rs946616 | benign | 0 | 0 | 0 |
| rs113993976 | pathogenic | 1 | 1 | 0 |
| rs151022264 | benign | 0 | 1 | 0 |
| rs113993977 | pathogenic | 1 | 1 | 0 |
| rs113993978 | pathogenic | 1 | 1 | 0 |
| rs2228499 | benign | 0 | 1 | 0 |
| rs113993979 | pathogenic | 1 | 1 | 0 |
| rs113993980 | pathogenic | 1 | 1 | 0 |
| rs113993983 | pathogenic | 0 | 1 | 0 |
| rs150547274 | pathogenic | 1 | 1 | 0 |
| rs113993984 | pathogenic | 1 | 1 | 0 |
| rs113993986 | pathogenic | 1 | 1 | 0 |
| rs34313873 | benign | 0 | 1 | 0 |
| rs113993988 | pathogenic | 1 | 1 | 0 |
| rs199638097 | pathogenic | 1 | 1 | 1 |
| rs118204447 | pathogenic | 1 | 1 | 1 |
| rs11862754 | benign | 0 | 0 | 0 |
| rs118204445 | pathogenic | 0 | 0 | 0 |
| rs118204441 | pathogenic | 0 | 0 | 0 |
| rs118204438 | pathogenic | 0 | 0 | 0 |
| rs118204436 | pathogenic | 0 | 0 | 0 |
| rs146093755 | pathogenic | 0 | 0 | 0 |
| rs781439830 | pathogenic | 0 | 0 | 0 |
| rs398123438 | pathogenic | 0 | 0 | 0 |
| rs398123439 | pathogenic | 0 | 0 | 0 |
| rs118204444 | pathogenic | 0 | 0 | 0 |
| rs7187889 | benign | 0 | 0 | 0 |
| rs118204435 | pathogenic | 1 | 1 | 0 |
| rs34745339 | benign | 0 | 0 | 0 |
| rs398123440 | pathogenic | 0 | 0 | 0 |
| rs118204442 | pathogenic | 0 | 0 | 0 |
| rs770053354 | pathogenic | 0 | 0 | 0 |
| rs118204448 | pathogenic | 0 | 0 | 0 |
| rs118204443 | pathogenic | 0 | 0 | 0 |
| rs118204446 | pathogenic | 0 | 0 | 0 |
| rs746756997 | pathogenic | 0 | 0 | 0 |
| rs267606838 | pathogenic | 0 | 0 | 0 |
| rs118204437 | pathogenic | 0 | 0 | 0 |
| rs398123429 | pathogenic | 0 | 0 | 0 |

| | | | | |
|---|---|---|---|---|
| rs398123430 | pathogenic | 0 | 0 | 0 |
| rs2303269 | benign | 0 | 0 | 0 |
| rs114703967 | benign | 0 | 0 | 0 |
| rs118204439 | pathogenic | 0 | 0 | 0 |
| rs151296605 | benign | 0 | 0 | 0 |
| rs118204440 | pathogenic | 0 | 0 | 0 |
| rs78127134 | benign | 0 | 0 | 0 |
| rs372893383 | pathogenic | 0 | 0 | 0 |
| rs61753266 | pathogenic | 1 | 1 | 1 |
| rs139579928 | benign | 1 | 1 | 1 |
| rs56275071 | pathogenic | 1 | 1 | 1 |
| rs121909736 | pathogenic | 1 | 1 | 0 |
| rs121909737 | pathogenic | 1 | 1 | 0 |
| rs121909735 | pathogenic | 1 | 1 | 0 |
| rs121909731 | pathogenic | 1 | 1 | 1 |
| rs121909734 | pathogenic | 1 | 1 | 1 |
| rs121909733 | pathogenic | 1 | 1 | 1 |
| rs121909732 | pathogenic | 1 | 1 | 1 |
| rs121909730 | pathogenic | 1 | 1 | 1 |
| rs202198360 | benign | 1 | 1 | 1 |
| rs587777294 | pathogenic | 1 | 1 | 0 |
| rs587777297 | pathogenic | 1 | 1 | 0 |
| rs104893913 | pathogenic | 1 | 1 | 0 |
| rs17860424 | pathogenic | 1 | 1 | 1 |
| rs1045485 | benign | 1 | 1 | 1 |
| rs13926 | benign | 0 | 0 | 0 |
| rs79343837 | benign | 0 | 0 | 0 |
| rs45516306 | benign | 1 | 1 | 1 |
| rs139849355 | benign | 1 | 1 | 1 |
| rs104894874 | pathogenic | 1 | 1 | 1 |
| rs104894870 | pathogenic | 1 | 1 | 1 |
| rs137852221 | pathogenic | 1 | 1 | 1 |
| rs104894869 | pathogenic | 1 | 1 | 1 |
| rs104894880 | pathogenic | 1 | 1 | 1 |
| rs104894872 | pathogenic | 1 | 1 | 1 |
| rs104894868 | pathogenic | 1 | 1 | 1 |
| rs104894867 | pathogenic | 1 | 1 | 1 |
| rs104894871 | pathogenic | 1 | 1 | 1 |
| rs104894877 | pathogenic | 1 | 1 | 1 |
| rs104894883 | pathogenic | 1 | 1 | 1 |
| rs104894875 | pathogenic | 1 | 1 | 1 |
| rs104894876 | pathogenic | 1 | 1 | 1 |
| rs104894878 | pathogenic | 1 | 1 | 1 |
| rs137852220 | pathogenic | 1 | 1 | 1 |
| rs28933684 | pathogenic | 1 | 1 | 1 |
| rs104894873 | pathogenic | 1 | 1 | 1 |
| rs138409264 | benign | 0 | 0 | 0 |
| rs2236722 | benign | 0 | 0 | 0 |
| rs28757184 | benign | 0 | 0 | 0 |
| rs121434538 | pathogenic | 0 | 0 | 0 |
| rs700519 | benign | 0 | 0 | 0 |
| rs80051519 | pathogenic | 0 | 0 | 0 |
| rs121434536 | pathogenic | 1 | 1 | 0 |
| rs121434534 | pathogenic | 1 | 1 | 0 |

| | | | | |
|---|---|---|---|---|
| rs78310315 | pathogenic | 1 | 1 | 0 |
| rs540839115 | pathogenic | 1 | 1 | 1 |
| rs80338731 | pathogenic | 1 | 1 | 1 |
| rs180177151 | pathogenic | 1 | 1 | 1 |
| rs180177152 | pathogenic | 0 | 0 | 0 |
| rs80338732 | pathogenic | 1 | 1 | 1 |
| rs80338675 | pathogenic | 1 | 1 | 1 |
| rs80338676 | pathogenic | 1 | 1 | 1 |
| rs180177153 | pathogenic | 0 | 0 | 0 |
| rs180177154 | pathogenic | 1 | 1 | 1 |
| rs5896 | benign | 1 | 1 | 1 |
| rs121918477 | pathogenic | 0 | 0 | 0 |
| rs121918428 | pathogenic | 1 | 1 | 1 |
| rs121918427 | pathogenic | 1 | 1 | 1 |
| rs544300820 | benign | 1 | 1 | 1 |
| rs121918110 | pathogenic | 1 | 1 | 1 |
| rs1057141 | benign | 0 | 0 | 0 |
| rs1135216 | benign | 1 | 1 | 1 |
| rs121917702 | pathogenic | 1 | 1 | 1 |
| rs28941474 | pathogenic | 0 | 0 | 0 |
| rs104893944 | pathogenic | 1 | 1 | 1 |
| rs104893947 | pathogenic | 1 | 1 | 0 |
| rs104893943 | pathogenic | 1 | 1 | 1 |
| rs104893942 | pathogenic | 1 | 1 | 1 |
| rs104893940 | pathogenic | 1 | 1 | 1 |
| rs34045013 | benign | 0 | 0 | 0 |
| rs143083947 | pathogenic | 0 | 0 | 0 |
| rs145786986 | benign | 0 | 0 | 0 |
| rs119450942 | pathogenic | 0 | 0 | 0 |
| rs374259530 | pathogenic | 0 | 0 | 0 |
| rs756210458 | pathogenic | 0 | 0 | 0 |
| rs11089991 | benign | 0 | 0 | 0 |
| rs28941471 | pathogenic | 0 | 1 | 0 |
| rs750614500 | pathogenic | 0 | 1 | 0 |
| rs199761158 | benign | 1 | 1 | 1 |
| rs119450944 | pathogenic | 1 | 1 | 1 |
| rs373458753 | pathogenic | 1 | 1 | 1 |
| rs202064195 | pathogenic | 1 | 1 | 1 |
| rs776496275 | pathogenic | 1 | 1 | 1 |
| rs761493155 | pathogenic | 1 | 1 | 1 |
| rs763542069 | pathogenic | 0 | 0 | 0 |
| rs119450941 | pathogenic | 0 | 0 | 0 |
| rs119450940 | pathogenic | 0 | 0 | 0 |
| rs777821034 | pathogenic | 1 | 1 | 1 |
| rs771121666 | pathogenic | 1 | 1 | 1 |
| rs372895468 | pathogenic | 1 | 1 | 1 |
| rs61750603 | pathogenic | 0 | 0 | 0 |
| rs76863441 | pathogenic | 0 | 0 | 0 |
| rs142601143 | benign | 1 | 1 | 1 |
| rs185832993 | benign | 1 | 1 | 0 |
| rs17860568 | benign | 0 | 0 | 0 |
| rs142064825 | benign | 0 | 0 | 0 |
| rs564076183 | benign | 0 | 0 | 0 |
| rs386834034 | pathogenic | 0 | 0 | 0 |

| | | | | |
|---|---|---|---|---|
| rs147663769 | benign | 0 | 0 | 0 |
| rs776478974 | benign | 0 | 0 | 0 |
| rs111840783 | benign | 0 | 0 | 0 |
| rs1516446 | benign | 1 | 1 | 1 |
| rs143749154 | benign | 1 | 1 | 1 |
| rs80257412 | benign | 0 | 0 | 0 |
| rs142943120 | benign | 0 | 0 | 0 |
| rs144553482 | benign | 0 | 0 | 0 |
| rs35997569 | benign | 0 | 0 | 0 |
| rs772909106 | pathogenic | 0 | 0 | 0 |
| rs149433837 | benign | 1 | 1 | 1 |
| rs532178791 | pathogenic | 0 | 0 | 0 |
| rs145588689 | pathogenic | 1 | 1 | 0 |
| rs17843776 | benign | 1 | 1 | 0 |
| rs121917890 | pathogenic | 1 | 1 | 1 |
| rs121917892 | pathogenic | 0 | 0 | 0 |
| rs606231289 | pathogenic | 0 | 0 | 0 |
| rs370011798 | pathogenic | 0 | 0 | 0 |
| rs34689457 | benign | 0 | 0 | 0 |
| rs118204451 | pathogenic | 0 | 0 | 0 |
| rs118204452 | pathogenic | 1 | 1 | 1 |
| rs652785 | benign | 0 | 0 | 0 |
| rs35378126 | benign | 0 | 0 | 0 |
| rs80358214 | pathogenic | 1 | 1 | 1 |
| rs80358215 | pathogenic | 0 | 1 | 0 |
| rs267606858 | pathogenic | 1 | 1 | 1 |
| rs143137713 | pathogenic | 1 | 1 | 1 |
| rs782168634 | benign | 0 | 0 | 0 |
| rs143873938 | benign | 0 | 0 | 0 |
| rs782776511 | benign | 0 | 0 | 0 |
| rs201093148 | benign | 0 | 0 | 0 |
| rs267606815 | pathogenic | 0 | 0 | 0 |
| rs137853311 | pathogenic | 0 | 0 | 0 |
| rs267606817 | pathogenic | 0 | 0 | 0 |
| rs72650006 | benign | 0 | 0 | 0 |
| rs727503638 | benign | 1 | 1 | 0 |
| rs189286381 | benign | 0 | 0 | 0 |
| rs376644553 | benign | 0 | 0 | 0 |
| rs121434610 | pathogenic | 1 | 1 | 1 |
| rs267607076 | pathogenic | 1 | 1 | 1 |
| rs201275730 | benign | 0 | 0 | 0 |
| rs121909587 | pathogenic | 0 | 0 | 0 |
| rs121909558 | pathogenic | 1 | 1 | 0 |
| rs28929469 | pathogenic | 1 | 1 | 0 |
| rs121909547 | pathogenic | 1 | 1 | 0 |
| rs121909552 | pathogenic | 1 | 1 | 0 |
| rs121909573 | pathogenic | 1 | 1 | 1 |
| rs121909567 | pathogenic | 1 | 1 | 0 |
| rs121909569 | pathogenic | 0 | 0 | 0 |
| rs121909563 | pathogenic | 1 | 1 | 0 |
| rs121909570 | pathogenic | 1 | 1 | 1 |
| rs121909571 | pathogenic | 1 | 1 | 0 |
| rs121909572 | pathogenic | 1 | 1 | 1 |
| rs121909565 | pathogenic | 1 | 1 | 1 |

| | | | | |
|---|---|---|---|---|
| rs121909557 | pathogenic | 1 | 1 | 1 |
| rs121909568 | pathogenic | 1 | 1 | 1 |
| rs121909566 | pathogenic | 1 | 1 | 1 |
| rs121909549 | pathogenic | 1 | 1 | 1 |
| rs121909554 | pathogenic | 1 | 1 | 1 |
| rs121909550 | pathogenic | 1 | 1 | 1 |
| rs121909546 | pathogenic | 1 | 1 | 0 |
| rs121909555 | pathogenic | 1 | 1 | 1 |
| rs121909564 | pathogenic | 1 | 1 | 1 |
| rs548046212 | pathogenic | 0 | 0 | 0 |
| rs147353781 | benign | 0 | 0 | 0 |
| rs104894678 | pathogenic | 1 | 1 | 1 |
| rs1130426 | benign | 0 | 0 | 0 |
| rs104894677 | pathogenic | 1 | 1 | 1 |
| rs141917423 | benign | 1 | 1 | 1 |
| rs574777883 | benign | 0 | 0 | 0 |
| rs150199226 | benign | 1 | 1 | 1 |
| rs199618701 | benign | 0 | 1 | 0 |
| rs121964923 | pathogenic | 0 | 0 | 0 |
| rs121964924 | pathogenic | 0 | 0 | 0 |
| rs201258823 | pathogenic | 0 | 1 | 0 |
| rs267606773 | pathogenic | 0 | 0 | 0 |
| rs35638832 | benign | 0 | 0 | 0 |
| rs121908632 | pathogenic | 1 | 1 | 1 |
| rs121908625 | pathogenic | 1 | 1 | 1 |
| rs748949603 | pathogenic | 0 | 0 | 0 |
| rs757523840 | pathogenic | 0 | 0 | 0 |
| rs62541771 | pathogenic | 0 | 0 | 0 |
| rs776384541 | pathogenic | 1 | 1 | 0 |
| rs121908627 | pathogenic | 1 | 1 | 0 |
| rs28937594 | pathogenic | 0 | 0 | 0 |
| rs121907888 | pathogenic | 0 | 0 | 0 |
| rs121907889 | pathogenic | 0 | 1 | 0 |
| rs28941785 | pathogenic | 1 | 1 | 1 |
| rs28941786 | pathogenic | 1 | 1 | 1 |
| rs1021737 | benign | 0 | 0 | 0 |
| rs121908370 | pathogenic | 1 | 1 | 1 |
| rs63750083 | pathogenic | 0 | 0 | 0 |
| rs17883862 | benign | 0 | 0 | 0 |
| rs587782070 | pathogenic | 1 | 1 | 1 |
| rs587781269 | pathogenic | 1 | 1 | 1 |
| rs28909982 | pathogenic | 1 | 1 | 1 |
| rs373648967 | benign | 1 | 1 | 1 |
| rs368570187 | benign | 1 | 1 | 1 |
| rs730881701 | pathogenic | 1 | 1 | 1 |
| rs137853007 | pathogenic | 0 | 0 | 0 |
| rs575910805 | benign | 0 | 0 | 0 |
| rs104893875 | pathogenic | 1 | 1 | 1 |
| rs104893877 | pathogenic | 1 | 1 | 1 |
| rs119485093 | pathogenic | 1 | 1 | 1 |
| rs119485092 | pathogenic | 0 | 0 | 0 |
| rs119485090 | pathogenic | 0 | 0 | 0 |
| rs28942099 | pathogenic | 1 | 1 | 1 |
| rs373730800 | pathogenic | 1 | 1 | 1 |

| | | | | |
|---|---|---|---|---|
| rs113993995 | pathogenic | 0 | 0 | 0 |
| rs201070132 | pathogenic | 0 | 0 | 0 |
| rs113993996 | pathogenic | 0 | 0 | 0 |
| rs113993998 | pathogenic | 0 | 0 | 0 |
| rs78950939 | pathogenic | 1 | 1 | 1 |
| rs56378716 | pathogenic | 1 | 1 | 1 |
| rs754921704 | benign | 0 | 1 | 0 |
| rs766379103 | pathogenic | 1 | 1 | 0 |
| rs121912515 | pathogenic | 0 | 1 | 0 |
| rs764831888 | benign | 1 | 1 | 0 |
| rs572527340 | benign | 0 | 0 | 0 |
| rs35969327 | benign | 0 | 0 | 0 |
| rs35814893 | benign | 0 | 0 | 0 |
| rs34032300 | benign | 0 | 0 | 0 |
| rs80358260 | pathogenic | 0 | 0 | 0 |
| rs151220873 | benign | 0 | 0 | 0 |
| rs80358261 | pathogenic | 1 | 1 | 0 |
| rs11694 | pathogenic | 0 | 0 | 0 |
| rs80358264 | pathogenic | 0 | 0 | 0 |
| rs80358266 | pathogenic | 1 | 1 | 0 |
| rs104894458 | pathogenic | 1 | 1 | 1 |
| rs121918028 | pathogenic | 0 | 0 | 0 |
| rs11583680 | pathogenic | 0 | 0 | 0 |
| rs4715 | benign | 1 | 1 | 1 |
| rs34957318 | benign | 0 | 0 | 0 |
| rs1124 | benign | 1 | 1 | 1 |
| rs104894778 | pathogenic | 1 | 1 | 1 |
| rs104894777 | pathogenic | 1 | 1 | 1 |
| rs104894769 | pathogenic | 1 | 1 | 1 |
| rs104894768 | pathogenic | 1 | 1 | 1 |
| rs104894771 | pathogenic | 1 | 1 | 1 |
| rs575372249 | benign | 1 | 1 | 1 |
| rs118203914 | pathogenic | 0 | 0 | 0 |
| rs118203916 | pathogenic | 1 | 1 | 0 |
| rs140154690 | benign | 0 | 0 | 0 |
| rs34382405 | benign | 0 | 0 | 0 |
| rs73718082 | benign | 0 | 0 | 0 |
| rs764932724 | benign | 0 | 0 | 0 |
| rs554931092 | pathogenic | 1 | 1 | 0 |
| rs202060864 | pathogenic | 0 | 0 | 0 |
| rs116567033 | benign | 0 | 0 | 0 |
| rs11243011 | benign | 0 | 0 | 0 |
| rs142073519 | pathogenic | 0 | 0 | 0 |
| rs397514611 | pathogenic | 1 | 1 | 0 |
| rs397514612 | pathogenic | 0 | 0 | 0 |
| rs372054960 | pathogenic | 0 | 0 | 0 |
| rs775690041 | pathogenic | 0 | 0 | 0 |
| rs148921184 | pathogenic | 0 | 0 | 0 |
| rs12711521 | benign | 0 | 0 | 0 |
| rs2273346 | benign | 0 | 0 | 0 |
| rs12085877 | benign | 1 | 1 | 1 |
| rs144471433 | benign | 1 | 1 | 1 |
| rs397514671 | pathogenic | 1 | 1 | 1 |
| rs76394680 | benign | 0 | 0 | 0 |

| | | | | |
|---|---|---|---|---|
| rs78289603 | pathogenic | 1 | 1 | 1 |
| rs137852913 | pathogenic | 1 | 1 | 1 |
| rs137852914 | pathogenic | 1 | 1 | 1 |
| rs121918653 | pathogenic | 1 | 1 | 1 |
| rs35515200 | benign | 1 | 1 | 0 |
| rs17876030 | benign | 1 | 1 | 1 |
| rs556623443 | benign | 1 | 1 | 1 |
| rs121434365 | pathogenic | 1 | 1 | 1 |
| rs121434364 | pathogenic | 0 | 0 | 0 |
| rs41400249 | benign | 0 | 0 | 0 |
| rs149440376 | benign | 1 | 1 | 1 |
| rs181590179 | benign | 0 | 0 | 0 |
| rs113993968 | pathogenic | 0 | 0 | 0 |
| rs587776445 | pathogenic | 1 | 1 | 1 |
| rs113993969 | pathogenic | 1 | 1 | 1 |
| rs113993970 | pathogenic | 1 | 1 | 1 |
| rs587776449 | pathogenic | 1 | 1 | 1 |
| rs113993971 | pathogenic | 1 | 1 | 1 |
| rs267607194 | pathogenic | 1 | 1 | 1 |
| rs397514745 | pathogenic | 0 | 0 | 0 |
| rs201650281 | pathogenic | 0 | 0 | 0 |
| rs150529876 | benign | 1 | 1 | 1 |
| rs77573084 | benign | 0 | 0 | 0 |
| rs398122379 | pathogenic | 1 | 1 | 0 |
| rs765478968 | benign | 0 | 0 | 0 |
| rs201399681 | benign | 0 | 0 | 0 |
| rs17159287 | benign | 0 | 0 | 0 |
| rs538571144 | benign | 0 | 0 | 0 |
| rs137852647 | pathogenic | 0 | 0 | 0 |
| rs201358272 | pathogenic | 1 | 1 | 1 |
| rs181251337 | benign | 0 | 0 | 0 |
| rs3213119 | benign | 0 | 0 | 0 |
| rs137853102 | pathogenic | 1 | 1 | 0 |
| rs4956145 | benign | 0 | 0 | 0 |
| rs146036912 | pathogenic | 1 | 1 | 1 |
| rs375717077 | pathogenic | 1 | 1 | 1 |
| rs137853103 | pathogenic | 1 | 1 | 1 |
| rs776184830 | pathogenic | 1 | 1 | 1 |
| rs200273673 | pathogenic | 1 | 1 | 1 |
| rs267607025 | pathogenic | 1 | 1 | 1 |
| rs267607024 | pathogenic | 1 | 1 | 1 |
| rs515726188 | pathogenic | 1 | 1 | 1 |
| rs147315735 | benign | 0 | 0 | 0 |
| rs121918308 | pathogenic | 1 | 1 | 0 |
| rs515726195 | pathogenic | 0 | 0 | 0 |
| rs515726196 | pathogenic | 1 | 1 | 0 |
| rs121918311 | pathogenic | 0 | 1 | 0 |
| rs121918309 | pathogenic | 0 | 0 | 0 |
| rs387906891 | pathogenic | 0 | 0 | 0 |
| rs182614164 | pathogenic | 1 | 1 | 0 |
| rs121918307 | pathogenic | 0 | 0 | 0 |
| rs201440849 | benign | 0 | 0 | 0 |
| rs118203957 | pathogenic | 0 | 0 | 0 |
| rs4986790 | benign | 0 | 0 | 0 |

| | | | | |
|---|---|---|---|---|
| rs281864928 | pathogenic | 1 | 1 | 0 |
| rs267607156 | pathogenic | 0 | 0 | 0 |
| rs55916807 | benign | 0 | 0 | 0 |
| rs749750052 | benign | 0 | 0 | 0 |
| rs74382477 | benign | 0 | 0 | 0 |
| rs201016638 | benign | 0 | 0 | 0 |
| rs376683989 | benign | 0 | 0 | 0 |
| rs144439703 | pathogenic | 0 | 0 | 0 |
| rs779122022 | benign | 0 | 0 | 0 |
| rs121918056 | pathogenic | 0 | 0 | 0 |
| rs113994093 | pathogenic | 1 | 1 | 1 |
| rs141367015 | pathogenic | 0 | 0 | 0 |
| rs775538075 | pathogenic | 0 | 0 | 0 |
| rs121918055 | pathogenic | 0 | 0 | 0 |
| rs754246294 | benign | 1 | 1 | 1 |
| rs2307447 | benign | 1 | 1 | 1 |
| rs113994096 | pathogenic | 1 | 1 | 1 |
| rs367610201 | pathogenic | 1 | 1 | 1 |
| rs758438414 | pathogenic | 1 | 1 | 0 |
| rs121918046 | pathogenic | 0 | 0 | 0 |
| rs201871736 | benign | 0 | 0 | 0 |
| rs773994204 | benign | 0 | 0 | 0 |
| rs2307450 | benign | 1 | 1 | 1 |
| rs200257554 | benign | 0 | 0 | 0 |
| rs185645212 | benign | 0 | 0 | 0 |
| rs121918054 | pathogenic | 0 | 0 | 0 |
| rs113994097 | pathogenic | 0 | 0 | 0 |
| rs202037973 | pathogenic | 0 | 0 | 0 |
| rs769827124 | pathogenic | 1 | 1 | 0 |
| rs41549716 | benign | 1 | 1 | 1 |
| rs113994098 | pathogenic | 1 | 1 | 0 |
| rs144500145 | pathogenic | 1 | 1 | 0 |
| rs121918053 | pathogenic | 1 | 1 | 0 |
| rs121918050 | pathogenic | 1 | 1 | 0 |
| rs368587966 | pathogenic | 0 | 0 | 0 |
| rs139590686 | pathogenic | 0 | 0 | 0 |
| rs121918048 | pathogenic | 1 | 1 | 0 |
| rs768653086 | pathogenic | 1 | 1 | 0 |
| rs113994099 | pathogenic | 1 | 1 | 0 |
| rs121918051 | pathogenic | 1 | 1 | 0 |
| rs753160398 | pathogenic | 1 | 1 | 0 |
| rs201477273 | pathogenic | 1 | 1 | 0 |
| rs142347031 | pathogenic | 0 | 0 | 0 |
| rs121918049 | pathogenic | 0 | 0 | 0 |
| rs267606959 | pathogenic | 1 | 1 | 0 |
| rs201732356 | pathogenic | 0 | 0 | 0 |
| rs368435864 | pathogenic | 0 | 0 | 0 |
| rs764036283 | pathogenic | 0 | 0 | 0 |
| rs2307442 | benign | 0 | 0 | 0 |
| rs2307440 | benign | 0 | 0 | 0 |
| rs548076633 | pathogenic | 0 | 1 | 0 |
| rs750792237 | benign | 0 | 0 | 0 |
| rs3087374 | benign | 0 | 0 | 0 |
| rs121918060 | pathogenic | 0 | 1 | 0 |

| | | | | |
|---|---|---|---|---|
| rs397514765 | pathogenic | 1 | 1 | 1 |
| rs121918063 | pathogenic | 0 | 0 | 0 |
| rs121918058 | pathogenic | 1 | 1 | 1 |
| rs121918064 | pathogenic | 0 | 0 | 0 |
| rs36033115 | benign | 0 | 1 | 0 |
| rs121918057 | pathogenic | 0 | 1 | 0 |
| rs121918065 | pathogenic | 1 | 1 | 1 |
| rs121918061 | pathogenic | 1 | 1 | 1 |
| rs121918066 | pathogenic | 0 | 1 | 0 |
| rs707555 | benign | 0 | 1 | 0 |
| rs8192708 | benign | 0 | 1 | 0 |
| rs11552145 | benign | 0 | 0 | 0 |
| rs1804160 | benign | 0 | 1 | 0 |
| rs34103538 | benign | 0 | 1 | 0 |
| rs751393950 | pathogenic | 1 | 1 | 1 |
| rs121907979 | pathogenic | 1 | 1 | 1 |
| rs121907975 | pathogenic | 1 | 1 | 1 |
| rs121907962 | pathogenic | 1 | 1 | 1 |
| rs121907957 | pathogenic | 0 | 0 | 0 |
| rs121907953 | pathogenic | 1 | 1 | 1 |
| rs28941770 | pathogenic | 1 | 1 | 1 |
| rs28941771 | pathogenic | 1 | 1 | 1 |
| rs121907969 | pathogenic | 1 | 1 | 1 |
| rs121907973 | pathogenic | 1 | 1 | 1 |
| rs1800429 | pathogenic | 0 | 0 | 0 |
| rs121907976 | pathogenic | 1 | 1 | 0 |
| rs121907961 | pathogenic | 0 | 0 | 0 |
| rs121907974 | pathogenic | 0 | 0 | 0 |
| rs121907959 | pathogenic | 0 | 0 | 0 |
| rs121907971 | pathogenic | 1 | 1 | 0 |
| rs121907954 | pathogenic | 0 | 0 | 0 |
| rs121907977 | pathogenic | 0 | 0 | 0 |
| rs121907963 | pathogenic | 1 | 1 | 0 |
| rs370266293 | pathogenic | 1 | 1 | 0 |
| rs1800430 | benign | 0 | 0 | 0 |
| rs121907958 | pathogenic | 0 | 0 | 0 |
| rs202173526 | pathogenic | 0 | 0 | 0 |
| rs121907978 | pathogenic | 0 | 0 | 0 |
| rs121907981 | pathogenic | 0 | 1 | 0 |
| rs145012038 | benign | 1 | 1 | 1 |
| rs121907968 | pathogenic | 1 | 1 | 1 |
| rs121907966 | pathogenic | 1 | 1 | 1 |
| rs121907956 | pathogenic | 1 | 1 | 1 |
| rs28942071 | pathogenic | 1 | 1 | 1 |
| rs121907955 | pathogenic | 1 | 1 | 1 |
| rs4727853 | benign | 0 | 0 | 0 |
| rs201880593 | benign | 0 | 0 | 0 |
| rs201434579 | benign | 1 | 1 | 1 |
| rs74551128 | pathogenic | 1 | 1 | 0 |
| rs193922500 | pathogenic | 0 | 0 | 0 |
| rs121909009 | pathogenic | 1 | 1 | 0 |
| rs121908805 | pathogenic | 1 | 1 | 0 |
| rs139573311 | pathogenic | 0 | 0 | 0 |
| rs213950 | benign | 0 | 0 | 0 |

| | | | | |
|---|---|---|---|---|
| rs79282516 | pathogenic | 0 | 0 | 0 |
| rs121909017 | pathogenic | 1 | 1 | 1 |
| rs77101217 | pathogenic | 1 | 1 | 1 |
| rs1800092 | pathogenic | 0 | 1 | 0 |
| rs77646904 | pathogenic | 0 | 0 | 0 |
| rs1800095 | benign | 1 | 1 | 1 |
| rs387906368 | pathogenic | 1 | 1 | 1 |
| rs113993959 | pathogenic | 1 | 1 | 1 |
| rs121908755 | pathogenic | 1 | 1 | 1 |
| rs121909005 | pathogenic | 1 | 1 | 1 |
| rs121908757 | pathogenic | 1 | 1 | 1 |
| rs75527207 | pathogenic | 1 | 1 | 0 |
| rs121909013 | pathogenic | 1 | 1 | 0 |
| rs121909044 | pathogenic | 1 | 1 | 0 |
| rs74597325 | pathogenic | 1 | 1 | 0 |
| rs75549581 | pathogenic | 0 | 0 | 0 |
| rs80055610 | pathogenic | 1 | 1 | 1 |
| rs1800097 | pathogenic | 0 | 0 | 0 |
| rs121909006 | pathogenic | 0 | 1 | 0 |
| rs397508276 | pathogenic | 0 | 0 | 0 |
| rs397508296 | pathogenic | 1 | 1 | 1 |
| rs139468767 | pathogenic | 0 | 0 | 0 |
| rs121908759 | pathogenic | 1 | 1 | 0 |
| rs121909033 | pathogenic | 1 | 1 | 0 |
| rs397508331 | pathogenic | 0 | 0 | 0 |
| rs387907279 | pathogenic | 1 | 1 | 1 |
| rs546530618 | benign | 0 | 0 | 0 |
| rs558259739 | benign | 0 | 0 | 0 |
| rs201827340 | benign | 0 | 0 | 0 |
| rs148865119 | pathogenic | 1 | 1 | 0 |
| rs2229413 | benign | 1 | 1 | 1 |
| rs34040148 | benign | 1 | 1 | 1 |
| rs121964964 | pathogenic | 1 | 1 | 1 |
| rs760214620 | pathogenic | 1 | 1 | 1 |
| rs775992753 | pathogenic | 1 | 1 | 1 |
| rs770095972 | pathogenic | 1 | 1 | 1 |
| rs781444670 | pathogenic | 1 | 1 | 1 |
| rs147474549 | pathogenic | 0 | 0 | 0 |
| rs121964965 | pathogenic | 0 | 1 | 0 |
| rs121964963 | pathogenic | 1 | 1 | 1 |
| rs755952006 | pathogenic | 1 | 1 | 1 |
| rs121964970 | pathogenic | 1 | 1 | 1 |
| rs121964973 | pathogenic | 1 | 1 | 1 |
| rs773734233 | pathogenic | 1 | 1 | 1 |
| rs758236584 | pathogenic | 1 | 1 | 1 |
| rs149119723 | pathogenic | 1 | 1 | 0 |
| rs5742905 | pathogenic | 0 | 0 | 0 |
| rs121964962 | pathogenic | 1 | 1 | 1 |
| rs781567152 | pathogenic | 1 | 1 | 0 |
| rs398123151 | pathogenic | 1 | 1 | 1 |
| rs760417941 | pathogenic | 1 | 1 | 1 |
| rs774926464 | pathogenic | 0 | 0 | 0 |
| rs121964972 | pathogenic | 1 | 1 | 0 |
| rs372010465 | pathogenic | 1 | 1 | 1 |

| | | | | |
|---|---|---|---|---|
| rs763036586 | pathogenic | 1 | 1 | 1 |
| rs769080151 | pathogenic | 1 | 1 | 1 |
| rs121964967 | pathogenic | 1 | 1 | 1 |
| rs28934892 | pathogenic | 1 | 1 | 0 |
| rs28934891 | pathogenic | 1 | 1 | 1 |
| rs121964971 | pathogenic | 1 | 1 | 1 |
| rs121964968 | pathogenic | 1 | 1 | 1 |
| rs150828989 | benign | 1 | 1 | 1 |
| rs3775291 | benign | 1 | 1 | 0 |
| rs104894419 | pathogenic | 1 | 1 | 1 |
| rs2232642 | benign | 0 | 0 | 0 |
| rs104894551 | pathogenic | 1 | 1 | 0 |
| rs766328537 | pathogenic | 0 | 1 | 0 |
| rs104894553 | pathogenic | 1 | 1 | 0 |
| rs80099330 | benign | 0 | 0 | 0 |
| rs199565861 | pathogenic | 0 | 0 | 0 |
| rs104894548 | pathogenic | 0 | 0 | 0 |
| rs104894549 | pathogenic | 0 | 0 | 0 |
| rs104894550 | pathogenic | 1 | 1 | 1 |
| rs12948217 | pathogenic | 1 | 1 | 1 |
| rs761064915 | pathogenic | 0 | 0 | 0 |
| rs28940279 | pathogenic | 1 | 1 | 1 |
| rs774323189 | pathogenic | 1 | 1 | 1 |
| rs28940574 | pathogenic | 0 | 0 | 0 |
| rs515299 | benign | 1 | 1 | 1 |
| rs10127939 | benign | 1 | 1 | 1 |
| rs396991 | benign | 1 | 1 | 1 |
| rs145956175 | benign | 0 | 1 | 0 |
| rs747838255 | pathogenic | 1 | 1 | 1 |
| rs762659685 | pathogenic | 1 | 1 | 0 |
| rs104894831 | pathogenic | 0 | 1 | 0 |
| rs104894848 | pathogenic | 1 | 1 | 0 |
| rs104894833 | pathogenic | 0 | 1 | 0 |
| rs781838005 | pathogenic | 0 | 1 | 0 |
| rs104894834 | pathogenic | 1 | 1 | 0 |
| rs372966991 | pathogenic | 1 | 1 | 0 |
| rs138886989 | benign | 1 | 1 | 1 |
| rs28935195 | pathogenic | 0 | 1 | 0 |
| rs28935196 | pathogenic | 0 | 1 | 0 |
| rs372416832 | pathogenic | 1 | 1 | 1 |
| rs28935197 | pathogenic | 1 | 1 | 0 |
| rs28935486 | pathogenic | 1 | 1 | 0 |
| rs28935487 | pathogenic | 1 | 1 | 0 |
| rs28935488 | pathogenic | 1 | 1 | 1 |
| rs28935495 | pathogenic | 1 | 1 | 1 |
| rs28935485 | pathogenic | 1 | 1 | 1 |
| rs140329381 | pathogenic | 0 | 1 | 0 |
| rs104894830 | pathogenic | 0 | 1 | 0 |
| rs104894846 | pathogenic | 0 | 1 | 0 |
| rs28935489 | pathogenic | 0 | 1 | 0 |
| rs104894828 | pathogenic | 1 | 1 | 0 |
| rs28935491 | pathogenic | 1 | 1 | 0 |
| rs104894832 | pathogenic | 1 | 1 | 0 |
| rs28935492 | pathogenic | 1 | 1 | 0 |

| | | | | |
|---|---|---|---|---|
| rs730880437 | benign | 1 | 1 | 0 |
| rs28935493 | pathogenic | 0 | 1 | 0 |
| rs28935494 | pathogenic | 1 | 1 | 1 |
| rs111422676 | pathogenic | 1 | 1 | 1 |
| rs397515869 | benign | 1 | 1 | 0 |
| rs782449839 | pathogenic | 0 | 1 | 0 |
| rs104894852 | pathogenic | 1 | 1 | 1 |
| rs116929575 | pathogenic | 1 | 1 | 1 |
| rs7023954 | benign | 0 | 0 | 0 |
| rs184520335 | benign | 1 | 1 | 1 |
| rs143080527 | benign | 0 | 0 | 0 |
| rs1050565 | benign | 0 | 0 | 0 |
| rs11545035 | benign | 0 | 0 | 0 |
| rs267607190 | pathogenic | 1 | 1 | 1 |
| rs200283987 | benign | 0 | 0 | 0 |
| rs372556807 | benign | 1 | 1 | 0 |
| rs563073392 | benign | 0 | 0 | 0 |
| rs4539 | benign | 0 | 0 | 0 |
| rs28931609 | pathogenic | 1 | 1 | 1 |
| rs121912978 | pathogenic | 1 | 1 | 1 |
| rs104894072 | pathogenic | 1 | 1 | 1 |
| rs551933154 | benign | 1 | 1 | 1 |
| rs752962897 | benign | 1 | 1 | 0 |
| rs121912977 | pathogenic | 1 | 1 | 0 |
| rs4537 | benign | 1 | 1 | 1 |
| rs201830462 | benign | 0 | 0 | 0 |
| rs4544 | benign | 0 | 0 | 0 |
| rs562670189 | benign | 0 | 0 | 0 |
| rs4538 | benign | 0 | 1 | 0 |
| rs61757294 | pathogenic | 0 | 0 | 0 |
| rs5313 | benign | 0 | 0 | 0 |
| rs4545 | benign | 0 | 0 | 0 |
| rs72554627 | pathogenic | 1 | 1 | 1 |
| rs72554626 | pathogenic | 1 | 1 | 1 |
| rs387906861 | pathogenic | 1 | 1 | 1 |
| rs749655461 | pathogenic | 1 | 1 | 1 |
| rs121909606 | pathogenic | 1 | 1 | 1 |
| rs121909607 | pathogenic | 1 | 1 | 1 |
| rs200203992 | benign | 0 | 0 | 0 |
| rs6050 | benign | 0 | 0 | 0 |
| rs184635235 | benign | 0 | 0 | 0 |
| rs139146037 | benign | 0 | 0 | 0 |
| rs121909612 | pathogenic | 0 | 0 | 0 |
| rs78506343 | pathogenic | 0 | 0 | 0 |
| rs121918138 | pathogenic | 1 | 1 | 1 |
| rs121918139 | pathogenic | 1 | 1 | 1 |
| rs121918140 | pathogenic | 0 | 0 | 0 |
| rs36063028 | benign | 1 | 1 | 1 |
| rs45495192 | benign | 1 | 1 | 1 |
| rs45458802 | benign | 1 | 1 | 1 |
| rs552865793 | pathogenic | 1 | 1 | 1 |
| rs753744791 | pathogenic | 1 | 1 | 1 |
| rs777518512 | benign | 1 | 1 | 1 |
| rs142708991 | benign | 1 | 1 | 0 |

| | | | | |
|---|---|---|---|---|
| rs746069957 | benign | 0 | 0 | 0 |
| rs1334811 | benign | 0 | 0 | 0 |
| rs138894008 | benign | 1 | 1 | 0 |
| rs35030851 | benign | 1 | 1 | 0 |
| rs35378598 | benign | 0 | 0 | 0 |
| rs35818764 | benign | 1 | 1 | 1 |
| rs4631561 | benign | 1 | 1 | 1 |
| rs121918660 | pathogenic | 0 | 0 | 0 |
| rs373175587 | benign | 0 | 0 | 0 |
| rs61731956 | pathogenic | 1 | 1 | 1 |
| rs180177305 | pathogenic | 0 | 0 | 0 |
| rs119490108 | pathogenic | 0 | 0 | 0 |
| rs180177312 | pathogenic | 1 | 1 | 0 |
| rs180177314 | pathogenic | 1 | 1 | 0 |
| rs180177322 | pathogenic | 1 | 1 | 1 |
| rs180177323 | pathogenic | 1 | 1 | 1 |
| rs180177325 | pathogenic | 0 | 0 | 0 |
| rs35196441 | benign | 0 | 0 | 0 |
| rs28763904 | benign | 0 | 0 | 0 |
| rs2229519 | benign | 0 | 0 | 0 |
| rs137852886 | pathogenic | 0 | 0 | 0 |
| rs137852892 | pathogenic | 0 | 0 | 0 |
| rs770427750 | pathogenic | 0 | 0 | 0 |
| rs137852887 | pathogenic | 0 | 0 | 0 |
| rs137852893 | pathogenic | 0 | 0 | 0 |
| rs80338671 | pathogenic | 1 | 1 | 0 |
| rs2228389 | benign | 0 | 0 | 0 |
| rs80338672 | pathogenic | 0 | 0 | 0 |
| rs201958741 | pathogenic | 0 | 0 | 0 |
| rs80338673 | pathogenic | 0 | 0 | 0 |
| rs137852888 | pathogenic | 0 | 0 | 0 |
| rs137852889 | pathogenic | 0 | 0 | 0 |
| rs137852894 | pathogenic | 0 | 0 | 0 |
| rs137852890 | pathogenic | 0 | 0 | 0 |
| rs137852891 | pathogenic | 0 | 0 | 0 |
| rs766935302 | pathogenic | 0 | 0 | 0 |
| rs193074572 | benign | 0 | 0 | 0 |
| rs555210692 | benign | 1 | 1 | 0 |
| rs119103242 | pathogenic | 0 | 1 | 0 |
| rs143725101 | benign | 0 | 1 | 0 |
| rs28940589 | pathogenic | 1 | 1 | 1 |
| rs113994145 | pathogenic | 0 | 0 | 0 |
| rs773872994 | benign | 0 | 0 | 0 |
| rs28940590 | pathogenic | 1 | 1 | 0 |
| rs145400751 | benign | 1 | 1 | 1 |
| rs587782331 | benign | 0 | 0 | 0 |
| rs730881787 | benign | 0 | 0 | 0 |
| rs587778531 | benign | 0 | 0 | 0 |
| rs781652274 | benign | 0 | 0 | 0 |
| rs55882234 | benign | 0 | 0 | 0 |
| rs2020908 | benign | 0 | 0 | 0 |
| rs63750741 | pathogenic | 1 | 1 | 1 |
| rs41295268 | benign | 1 | 1 | 1 |
| rs63750897 | benign | 1 | 1 | 0 |

| | | | | |
|---|---|---|---|---|
| rs63751005 | benign | 1 | 1 | 0 |
| rs587779223 | benign | 0 | 0 | 0 |
| rs202127474 | benign | 0 | 0 | 0 |
| rs374230313 | benign | 0 | 0 | 0 |
| rs2020912 | benign | 0 | 0 | 0 |
| rs63751017 | pathogenic | 0 | 0 | 0 |
| rs587779246 | pathogenic | 0 | 0 | 0 |
| rs63749919 | pathogenic | 0 | 0 | 0 |
| rs63750563 | pathogenic | 1 | 1 | 1 |
| rs63749999 | pathogenic | 0 | 0 | 0 |
| rs200492211 | pathogenic | 0 | 0 | 0 |
| rs576269342 | benign | 0 | 0 | 0 |
| rs267608075 | benign | 0 | 0 | 0 |
| rs149605979 | benign | 0 | 0 | 0 |
| rs63750252 | benign | 0 | 0 | 0 |
| rs754289472 | benign | 0 | 0 | 0 |
| rs63750119 | pathogenic | 0 | 0 | 0 |
| rs587782109 | benign | 0 | 0 | 0 |
| rs34625968 | benign | 1 | 1 | 1 |
| rs764786814 | benign | 1 | 1 | 1 |
| rs199594809 | benign | 1 | 1 | 1 |
| rs74315364 | pathogenic | 1 | 1 | 1 |
| rs201253919 | pathogenic | 0 | 0 | 0 |
| rs387906828 | pathogenic | 1 | 1 | 1 |
| rs34521609 | benign | 1 | 1 | 1 |
| rs199472894 | pathogenic | 0 | 0 | 0 |
| rs199472902 | pathogenic | 0 | 0 | 0 |
| rs121912505 | pathogenic | 0 | 0 | 0 |
| rs28928905 | pathogenic | 0 | 0 | 0 |
| rs374376640 | benign | 0 | 0 | 0 |
| rs202102276 | benign | 0 | 0 | 0 |
| rs199472918 | pathogenic | 0 | 0 | 0 |
| rs121912516 | pathogenic | 1 | 1 | 1 |
| rs121912504 | pathogenic | 0 | 0 | 0 |
| rs9333649 | pathogenic | 0 | 0 | 0 |
| rs121912508 | pathogenic | 0 | 0 | 0 |
| rs199473428 | pathogenic | 1 | 1 | 1 |
| rs104894021 | pathogenic | 0 | 0 | 0 |
| rs28928904 | pathogenic | 1 | 1 | 1 |
| rs199473039 | benign | 1 | 1 | 1 |
| rs199472958 | pathogenic | 1 | 1 | 1 |
| rs1137617 | pathogenic | 1 | 1 | 1 |
| rs199473532 | pathogenic | 1 | 1 | 1 |
| rs199472988 | pathogenic | 0 | 0 | 0 |
| rs189014161 | pathogenic | 0 | 0 | 0 |
| rs12720441 | pathogenic | 0 | 0 | 0 |
| rs199473001 | pathogenic | 1 | 1 | 0 |
| rs121912506 | pathogenic | 1 | 1 | 0 |
| rs773724817 | pathogenic | 1 | 1 | 1 |
| rs118204002 | pathogenic | 1 | 1 | 1 |
| rs138556525 | pathogenic | 1 | 1 | 1 |
| rs72558181 | pathogenic | 1 | 1 | 1 |
| rs118204005 | pathogenic | 1 | 1 | 1 |
| rs118204004 | pathogenic | 0 | 0 | 0 |

| | | | | |
|---|---|---|---|---|
| rs118204001 | pathogenic | 1 | 1 | 1 |
| rs118204006 | pathogenic | 1 | 1 | 1 |
| rs118204003 | pathogenic | 0 | 0 | 0 |
| rs121913147 | pathogenic | 1 | 1 | 0 |
| rs2229429 | benign | 1 | 1 | 0 |
| rs182552223 | benign | 0 | 0 | 0 |
| rs146588336 | benign | 0 | 0 | 0 |
| rs121964983 | pathogenic | 0 | 1 | 0 |
| rs121964982 | pathogenic | 0 | 0 | 0 |
| rs386833679 | pathogenic | 0 | 0 | 0 |
| rs386833680 | pathogenic | 0 | 0 | 0 |
| rs121964981 | pathogenic | 1 | 1 | 0 |
| rs121964984 | pathogenic | 0 | 0 | 0 |
| rs144971200 | benign | 0 | 0 | 0 |
| rs121964985 | pathogenic | 1 | 1 | 0 |
| rs141246107 | benign | 0 | 0 | 0 |
| rs104894561 | pathogenic | 1 | 1 | 0 |
| rs377023736 | pathogenic | 0 | 0 | 0 |
| rs121964907 | pathogenic | 1 | 1 | 1 |
| rs121964909 | pathogenic | 1 | 1 | 1 |
| rs765070743 | pathogenic | 0 | 0 | 0 |
| rs2228119 | benign | 0 | 0 | 0 |
| rs745976989 | pathogenic | 1 | 1 | 1 |
| rs121964904 | pathogenic | 0 | 0 | 0 |
| rs386833430 | pathogenic | 1 | 1 | 1 |
| rs267606992 | pathogenic | 0 | 0 | 0 |
| rs61752063 | pathogenic | 0 | 0 | 0 |
| rs61752067 | pathogenic | 1 | 1 | 1 |
| rs104894934 | pathogenic | 0 | 0 | 0 |
| rs104894929 | pathogenic | 1 | 1 | 1 |
| rs80189640 | benign | 0 | 0 | 0 |
| rs137854547 | pathogenic | 1 | 1 | 1 |
| rs137854540 | pathogenic | 1 | 1 | 1 |
| rs782092363 | pathogenic | 1 | 1 | 1 |
| rs147783915 | benign | 0 | 0 | 0 |
| rs28933687 | pathogenic | 1 | 1 | 1 |
| rs149503319 | benign | 0 | 0 | 0 |
| rs1805147 | benign | 0 | 0 | 0 |
| rs587777218 | pathogenic | 1 | 1 | 1 |
| rs200786705 | benign | 0 | 0 | 0 |
| rs56219475 | benign | 0 | 0 | 0 |
| rs587781172 | benign | 0 | 0 | 0 |
| rs757094189 | pathogenic | 0 | 0 | 0 |
| rs140811086 | benign | 1 | 1 | 1 |
| rs574088829 | pathogenic | 1 | 1 | 1 |
| rs121918525 | pathogenic | 0 | 0 | 0 |
| rs121918527 | pathogenic | 0 | 0 | 0 |
| rs121964848 | pathogenic | 0 | 1 | 0 |
| rs121964845 | pathogenic | 1 | 1 | 1 |
| rs121964846 | pathogenic | 0 | 1 | 0 |
| rs17217716 | benign | 1 | 1 | 1 |
| rs141711342 | benign | 0 | 0 | 0 |
| rs372619120 | benign | 0 | 0 | 0 |
| rs63750002 | benign | 1 | 1 | 1 |

| | | | | |
|---|---|---|---|---|
| rs193922373 | benign | 0 | 0 | 0 |
| rs41295286 | benign | 0 | 0 | 0 |
| rs17217772 | benign | 0 | 0 | 0 |
| rs145649774 | benign | 0 | 0 | 0 |
| rs63750214 | pathogenic | 0 | 0 | 0 |
| rs63750716 | benign | 0 | 0 | 0 |
| rs202026056 | benign | 0 | 0 | 0 |
| rs4987188 | benign | 1 | 1 | 0 |
| rs267607938 | benign | 0 | 0 | 0 |
| rs17224367 | benign | 0 | 0 | 0 |
| rs63750006 | benign | 0 | 0 | 0 |
| rs587781373 | benign | 0 | 0 | 0 |
| rs63751207 | pathogenic | 1 | 1 | 0 |
| rs730881759 | benign | 0 | 0 | 0 |
| rs372350768 | benign | 1 | 1 | 1 |
| rs55778204 | benign | 1 | 1 | 0 |
| rs63749910 | benign | 0 | 0 | 0 |
| rs63750844 | benign | 0 | 0 | 0 |
| rs28929483 | pathogenic | 0 | 0 | 0 |
| rs61756468 | benign | 0 | 0 | 0 |
| rs63750875 | pathogenic | 1 | 1 | 0 |
| rs776528054 | benign | 1 | 1 | 0 |
| rs751939698 | benign | 0 | 0 | 0 |
| rs587780685 | benign | 0 | 0 | 0 |
| rs730881763 | benign | 1 | 1 | 0 |
| rs373226409 | benign | 0 | 0 | 0 |
| rs202145681 | benign | 1 | 1 | 1 |
| rs63749841 | benign | 1 | 1 | 1 |
| rs63750757 | benign | 1 | 1 | 1 |
| rs587778527 | benign | 1 | 1 | 1 |
| rs63750849 | benign | 0 | 0 | 0 |
| rs587779969 | benign | 0 | 0 | 0 |
| rs121908046 | pathogenic | 1 | 1 | 1 |
| rs28940882 | pathogenic | 1 | 1 | 0 |
| rs28940883 | pathogenic | 1 | 1 | 1 |
| rs727503943 | pathogenic | 1 | 1 | 1 |
| rs137853859 | pathogenic | 1 | 1 | 1 |
| rs121908045 | pathogenic | 1 | 1 | 0 |
| rs137853861 | pathogenic | 1 | 1 | 0 |
| rs3180383 | pathogenic | 0 | 1 | 0 |
| rs144431814 | benign | 0 | 0 | 0 |
| rs139502866 | benign | 0 | 0 | 0 |
| rs17524437 | benign | 0 | 0 | 0 |
| rs760960779 | benign | 0 | 0 | 0 |
| rs148960463 | pathogenic | 0 | 0 | 0 |
| rs111033647 | pathogenic | 1 | 1 | 1 |
| rs111033648 | pathogenic | 0 | 0 | 0 |
| rs1800461 | benign | 0 | 0 | 0 |
| rs111033656 | pathogenic | 0 | 0 | 0 |
| rs111033663 | pathogenic | 1 | 1 | 0 |
| rs111033669 | pathogenic | 1 | 1 | 1 |
| rs111033690 | pathogenic | 0 | 0 | 0 |
| rs111033686 | pathogenic | 0 | 0 | 0 |
| rs111033687 | pathogenic | 0 | 0 | 0 |

| | | | | |
|---|---|---|---|---|
| rs111033695 | pathogenic | 0 | 0 | 0 |
| rs111033697 | pathogenic | 0 | 0 | 0 |
| rs111033693 | pathogenic | 0 | 0 | 0 |
| rs111033694 | pathogenic | 0 | 0 | 0 |
| rs111033721 | pathogenic | 1 | 1 | 1 |
| rs111033722 | pathogenic | 0 | 0 | 0 |
| rs111033725 | pathogenic | 1 | 1 | 0 |
| rs75391579 | pathogenic | 1 | 1 | 1 |
| rs111033728 | pathogenic | 1 | 1 | 1 |
| rs111033735 | pathogenic | 1 | 1 | 1 |
| rs111033739 | pathogenic | 1 | 1 | 1 |
| rs111033736 | pathogenic | 0 | 0 | 0 |
| rs111033744 | pathogenic | 1 | 1 | 1 |
| rs111033750 | pathogenic | 1 | 1 | 1 |
| rs111033752 | pathogenic | 0 | 0 | 0 |
| rs111033751 | pathogenic | 0 | 0 | 0 |
| rs111033846 | pathogenic | 0 | 0 | 0 |
| rs368166217 | pathogenic | 0 | 0 | 0 |
| rs111033764 | pathogenic | 0 | 0 | 0 |
| rs111033847 | benign | 0 | 0 | 0 |
| rs111033778 | pathogenic | 0 | 0 | 0 |
| rs111033773 | pathogenic | 0 | 0 | 0 |
| rs111033792 | pathogenic | 0 | 0 | 0 |
| rs367543266 | pathogenic | 1 | 1 | 1 |
| rs111033794 | pathogenic | 0 | 0 | 0 |
| rs111033802 | pathogenic | 0 | 0 | 0 |
| rs111033803 | pathogenic | 0 | 0 | 0 |
| rs111033809 | pathogenic | 1 | 1 | 0 |
| rs121918376 | pathogenic | 0 | 1 | 0 |
| rs121918377 | pathogenic | 0 | 1 | 0 |
| rs121918378 | pathogenic | 1 | 1 | 1 |
| rs121918374 | pathogenic | 1 | 1 | 1 |
| rs137852379 | pathogenic | 0 | 0 | 0 |
| rs137852388 | pathogenic | 1 | 1 | 1 |
| rs137852368 | pathogenic | 0 | 0 | 0 |
| rs28935499 | pathogenic | 0 | 0 | 0 |
| rs28933668 | pathogenic | 0 | 0 | 0 |
| rs28933669 | pathogenic | 0 | 0 | 0 |
| rs28933670 | pathogenic | 0 | 0 | 0 |
| rs28933671 | pathogenic | 0 | 0 | 0 |
| rs28933672 | pathogenic | 0 | 0 | 0 |
| rs28937272 | pathogenic | 0 | 0 | 0 |
| rs35383156 | benign | 0 | 0 | 0 |
| rs137852418 | pathogenic | 0 | 0 | 0 |
| rs137852419 | pathogenic | 0 | 0 | 0 |
| rs137852428 | pathogenic | 0 | 0 | 0 |
| rs137852436 | pathogenic | 0 | 0 | 0 |
| rs28937285 | pathogenic | 0 | 0 | 0 |
| rs35213384 | benign | 1 | 1 | 1 |
| rs3810950 | benign | 0 | 0 | 0 |
| rs201479289 | pathogenic | 0 | 0 | 0 |
| rs121912820 | pathogenic | 0 | 1 | 0 |
| rs121912815 | pathogenic | 0 | 1 | 0 |
| rs115510708 | benign | 0 | 0 | 0 |

| | | | | |
|---|---|---|---|---|
| rs868749 | benign | 0 | 0 | 0 |
| rs121912823 | pathogenic | 0 | 0 | 0 |
| rs115212829 | benign | 0 | 0 | 0 |
| rs121912822 | pathogenic | 0 | 0 | 0 |
| rs121912816 | pathogenic | 0 | 1 | 0 |
| rs4838544 | benign | 0 | 0 | 0 |
| rs121912818 | pathogenic | 0 | 0 | 0 |
| rs121912821 | pathogenic | 0 | 0 | 0 |
| rs121912817 | pathogenic | 0 | 0 | 0 |
| rs8178992 | benign | 0 | 0 | 0 |
| rs121912819 | pathogenic | 1 | 1 | 0 |
| rs80097077 | benign | 1 | 1 | 0 |
| rs121909359 | pathogenic | 1 | 1 | 0 |
| rs121909358 | pathogenic | 1 | 1 | 1 |
| rs121909361 | pathogenic | 1 | 1 | 1 |
| rs121909357 | pathogenic | 0 | 1 | 0 |
| rs121909365 | pathogenic | 1 | 1 | 1 |
| rs6413484 | pathogenic | 1 | 1 | 1 |
| rs121909373 | pathogenic | 1 | 1 | 1 |
| rs121909366 | pathogenic | 1 | 1 | 1 |
| rs121909367 | pathogenic | 1 | 1 | 1 |
| rs121909368 | pathogenic | 1 | 1 | 1 |
| rs121909369 | pathogenic | 0 | 0 | 0 |
| rs121909363 | pathogenic | 1 | 1 | 1 |
| rs45588036 | pathogenic | 1 | 1 | 1 |
| rs121909364 | pathogenic | 1 | 1 | 1 |
| rs80358282 | benign | 0 | 0 | 0 |
| rs80358284 | pathogenic | 1 | 1 | 1 |
| rs61735303 | benign | 0 | 0 | 0 |
| rs2229274 | benign | 0 | 0 | 0 |
| rs145006491 | benign | 0 | 0 | 0 |
| rs121913580 | pathogenic | 1 | 1 | 0 |
| rs730882196 | pathogenic | 0 | 0 | 0 |
| rs153477 | benign | 0 | 0 | 0 |
| rs153478 | benign | 1 | 1 | 0 |
| rs61740602 | benign | 1 | 1 | 0 |
| rs104893892 | pathogenic | 1 | 1 | 0 |
| rs121909595 | pathogenic | 1 | 1 | 1 |
| rs28931605 | pathogenic | 1 | 1 | 1 |
| rs121909597 | pathogenic | 0 | 1 | 0 |
| rs202233735 | pathogenic | 0 | 1 | 0 |
| rs121909596 | pathogenic | 1 | 1 | 1 |
| rs75156162 | benign | 0 | 1 | 0 |
| rs2305430 | benign | 1 | 1 | 1 |
| rs150318966 | benign | 0 | 0 | 0 |
| rs121909598 | pathogenic | 0 | 1 | 0 |
| rs5744175 | benign | 0 | 0 | 0 |
| rs28935476 | pathogenic | 1 | 1 | 1 |
| rs28935475 | pathogenic | 1 | 1 | 1 |
| rs122461163 | pathogenic | 1 | 1 | 1 |
| rs62626305 | pathogenic | 1 | 1 | 1 |
| rs587777157 | pathogenic | 0 | 0 | 0 |
| rs137854478 | pathogenic | 1 | 1 | 0 |
| rs137854465 | pathogenic | 0 | 0 | 0 |

| | | | | |
|---|---|---|---|---|
| rs137854470 | pathogenic | 0 | 0 | 0 |
| rs137854482 | pathogenic | 0 | 0 | 0 |
| rs137854473 | pathogenic | 0 | 0 | 0 |
| rs137854456 | pathogenic | 0 | 0 | 0 |
| rs140598 | benign | 0 | 0 | 0 |
| rs104894105 | pathogenic | 1 | 1 | 1 |
| rs104894107 | pathogenic | 1 | 1 | 1 |
| rs104894106 | pathogenic | 1 | 1 | 1 |
| rs74621026 | benign | 1 | 1 | 1 |
| rs2290332 | benign | 0 | 0 | 0 |
| rs780202604 | pathogenic | 1 | 1 | 1 |
| rs74315305 | pathogenic | 1 | 1 | 0 |
| rs766751878 | pathogenic | 1 | 1 | 1 |
| rs137854603 | pathogenic | 1 | 1 | 1 |
| rs139592595 | benign | 1 | 1 | 0 |
| rs150565592 | benign | 1 | 1 | 1 |
| rs763520126 | benign | 0 | 0 | 0 |
| rs55980502 | benign | 0 | 0 | 0 |
| rs137852812 | pathogenic | 0 | 0 | 0 |
| rs137852813 | pathogenic | 0 | 0 | 0 |
| rs190222208 | benign | 0 | 0 | 0 |
| rs138459502 | pathogenic | 0 | 0 | 0 |
| rs730881041 | pathogenic | 0 | 0 | 0 |
| rs267607080 | pathogenic | 0 | 0 | 0 |
| rs137852814 | pathogenic | 0 | 0 | 0 |
| rs267607079 | pathogenic | 0 | 0 | 0 |
| rs192378417 | benign | 0 | 0 | 0 |
| rs1288775 | benign | 0 | 0 | 0 |
| rs80338737 | pathogenic | 1 | 1 | 1 |
| rs397514708 | pathogenic | 0 | 0 | 0 |
| rs775933965 | pathogenic | 0 | 0 | 0 |
| rs747608698 | benign | 0 | 1 | 0 |
| rs1145086 | benign | 0 | 1 | 0 |
| rs121912953 | pathogenic | 1 | 1 | 0 |
| rs243849 | benign | 0 | 0 | 0 |
| rs121912955 | pathogenic | 1 | 1 | 1 |
| rs17859943 | benign | 0 | 1 | 0 |
| rs755557988 | benign | 1 | 1 | 1 |
| rs10195 | benign | 0 | 0 | 0 |
| rs61735357 | benign | 1 | 1 | 1 |
| rs137882374 | benign | 0 | 0 | 0 |
| rs5388 | benign | 0 | 0 | 0 |
| rs6352 | benign | 0 | 0 | 0 |
| rs76645461 | pathogenic | 1 | 1 | 0 |
| rs1050828 | pathogenic | 0 | 0 | 0 |
| rs1050829 | pathogenic | 0 | 0 | 0 |
| rs137852314 | pathogenic | 0 | 0 | 0 |
| rs137852331 | pathogenic | 0 | 0 | 0 |
| rs5030872 | pathogenic | 0 | 0 | 0 |
| rs267606836 | pathogenic | 0 | 0 | 0 |
| rs5030868 | pathogenic | 0 | 0 | 0 |
| rs137852330 | pathogenic | 0 | 1 | 0 |
| rs137852326 | pathogenic | 1 | 1 | 1 |
| rs137852318 | pathogenic | 1 | 1 | 1 |

| | | | | |
|---|---|---|---|---|
| rs74575103 | benign | 0 | 0 | 0 |
| rs137852327 | pathogenic | 1 | 1 | 1 |
| rs76723693 | pathogenic | 0 | 0 | 0 |
| rs5030869 | pathogenic | 0 | 0 | 0 |
| rs34193178 | pathogenic | 1 | 1 | 1 |
| rs137852316 | pathogenic | 1 | 1 | 0 |
| rs398123546 | pathogenic | 1 | 1 | 1 |
| rs72554665 | pathogenic | 0 | 0 | 0 |
| rs72554664 | pathogenic | 0 | 0 | 0 |
| rs28939068 | pathogenic | 1 | 1 | 1 |
| rs59301204 | pathogenic | 1 | 1 | 1 |
| rs143715750 | benign | 1 | 1 | 1 |
| rs35263902 | pathogenic | 1 | 1 | 1 |
| rs202214518 | benign | 1 | 1 | 1 |
| rs34076756 | benign | 0 | 0 | 0 |
| rs199592341 | pathogenic | 1 | 1 | 0 |
| rs121908833 | pathogenic | 1 | 1 | 1 |
| rs121908834 | pathogenic | 1 | 1 | 0 |
| rs138454151 | benign | 0 | 0 | 0 |
| rs141323599 | benign | 0 | 0 | 0 |
| rs137853200 | pathogenic | 0 | 0 | 0 |
| rs137853199 | pathogenic | 0 | 0 | 0 |
| rs139787163 | benign | 0 | 0 | 0 |
| rs61731097 | benign | 1 | 1 | 1 |
| rs61729795 | benign | 0 | 0 | 0 |
| rs149242330 | benign | 0 | 0 | 0 |
| rs587777490 | pathogenic | 1 | 1 | 1 |
| rs398123011 | pathogenic | 1 | 1 | 0 |
| rs775186117 | pathogenic | 0 | 0 | 0 |
| rs72677237 | benign | 0 | 0 | 0 |
| rs104894163 | pathogenic | 1 | 1 | 1 |
| rs121909588 | pathogenic | 0 | 0 | 0 |
| rs1042579 | benign | 0 | 0 | 0 |
| rs121909273 | pathogenic | 0 | 0 | 0 |
| rs121909274 | pathogenic | 0 | 0 | 0 |
| rs119473034 | pathogenic | 1 | 1 | 1 |
| rs398123065 | pathogenic | 0 | 0 | 0 |
| rs587777042 | pathogenic | 1 | 1 | 0 |
| rs761413864 | pathogenic | 1 | 1 | 1 |
| rs118203965 | pathogenic | 0 | 0 | 0 |
| rs118203966 | pathogenic | 0 | 0 | 0 |
| rs1801201 | benign | 1 | 1 | 0 |
| rs1136201 | benign | 1 | 1 | 1 |
| rs144148008 | benign | 0 | 0 | 0 |
| rs200118378 | benign | 1 | 1 | 1 |
| rs112984118 | benign | 1 | 1 | 1 |
| rs149446102 | benign | 1 | 1 | 1 |
| rs1427463 | benign | 0 | 0 | 0 |
| rs201936720 | benign | 1 | 1 | 1 |
| rs61751983 | benign | 1 | 1 | 1 |
| rs104894632 | pathogenic | 1 | 1 | 1 |
| rs149202905 | benign | 0 | 0 | 0 |
| rs2230210 | benign | 1 | 1 | 1 |
| rs111631988 | benign | 0 | 0 | 0 |

| | | | | |
|---|---|---|---|---|
| rs137852550 | pathogenic | 0 | 0 | 0 |
| rs5963403 | benign | 0 | 0 | 0 |
| rs138018739 | benign | 0 | 0 | 0 |
| rs62640590 | benign | 0 | 0 | 0 |
| rs121909581 | pathogenic | 1 | 1 | 1 |
| rs202017913 | benign | 1 | 1 | 1 |
| rs143163513 | benign | 0 | 0 | 0 |
| rs238238 | benign | 0 | 1 | 0 |
| rs143945974 | benign | 0 | 0 | 0 |
| rs238239 | benign | 1 | 1 | 1 |
| rs121918403 | pathogenic | 0 | 1 | 0 |
| rs61735456 | benign | 0 | 1 | 0 |
| rs121918404 | pathogenic | 1 | 1 | 0 |
| rs12674488 | benign | 0 | 0 | 0 |
| rs201599657 | pathogenic | 0 | 0 | 0 |
| rs1057090 | benign | 0 | 0 | 0 |
| rs1057091 | benign | 0 | 0 | 0 |
| rs587783740 | benign | 0 | 0 | 0 |
| rs75956769 | benign | 0 | 0 | 0 |
| rs201691299 | benign | 0 | 0 | 0 |
| rs62621087 | benign | 0 | 0 | 0 |
| rs201308454 | benign | 0 | 0 | 0 |
| rs145657360 | benign | 0 | 0 | 0 |
| rs121434592 | pathogenic | 1 | 1 | 0 |
| rs375650413 | benign | 0 | 0 | 0 |
| rs121434437 | pathogenic | 1 | 1 | 0 |
| rs763494292 | pathogenic | 0 | 0 | 0 |
| rs121964997 | pathogenic | 1 | 1 | 0 |
| rs121964996 | pathogenic | 0 | 0 | 0 |
| rs41283425 | benign | 1 | 1 | 1 |
| rs376289528 | benign | 1 | 1 | 1 |
| rs144171604 | benign | 1 | 1 | 1 |
| rs782440692 | pathogenic | 1 | 1 | 1 |
| rs782058451 | pathogenic | 1 | 1 | 1 |
| rs78437817 | benign | 0 | 0 | 0 |
| rs200327969 | benign | 1 | 1 | 1 |
| rs143043662 | benign | 0 | 0 | 0 |
| rs111966833 | benign | 1 | 1 | 1 |
| rs35523678 | benign | 0 | 0 | 0 |
| rs515726208 | pathogenic | 0 | 0 | 0 |
| rs397517239 | pathogenic | 1 | 1 | 1 |
| rs148966602 | benign | 1 | 1 | 1 |
| rs76371972 | benign | 1 | 1 | 1 |
| rs2229968 | benign | 1 | 1 | 1 |
| rs61751538 | benign | 1 | 1 | 1 |
| rs200800978 | pathogenic | 1 | 1 | 1 |
| rs148695964 | benign | 1 | 1 | 1 |
| rs140042418 | benign | 1 | 1 | 1 |
| rs116185403 | benign | 0 | 0 | 0 |
| rs142828669 | benign | 1 | 1 | 1 |
| rs139832048 | benign | 0 | 1 | 0 |
| rs58750568 | benign | 0 | 1 | 0 |
| rs77681311 | benign | 0 | 0 | 0 |
| rs151208452 | benign | 0 | 0 | 0 |

| | | | | |
|---|---|---|---|---|
| rs76463933 | benign | 0 | 0 | 0 |
| rs144434647 | benign | 0 | 0 | 0 |
| rs117361485 | benign | 0 | 0 | 0 |
| rs587777588 | pathogenic | 0 | 0 | 0 |
| rs80338796 | pathogenic | 0 | 0 | 0 |
| rs121434594 | pathogenic | 0 | 0 | 0 |
| rs146766138 | pathogenic | 1 | 1 | 1 |
| rs28939679 | pathogenic | 1 | 1 | 1 |
| rs748754134 | pathogenic | 1 | 1 | 0 |
| rs764276946 | pathogenic | 1 | 1 | 1 |
| rs111033588 | pathogenic | 1 | 1 | 1 |
| rs201633414 | pathogenic | 1 | 1 | 1 |
| rs121909262 | pathogenic | 1 | 1 | 1 |
| rs104893702 | pathogenic | 1 | 1 | 1 |
| rs121909265 | pathogenic | 1 | 1 | 1 |
| rs121909266 | pathogenic | 1 | 1 | 1 |
| rs193922439 | pathogenic | 1 | 1 | 0 |
| rs104893691 | pathogenic | 1 | 1 | 1 |
| rs104893695 | pathogenic | 0 | 0 | 0 |
| rs104893708 | pathogenic | 1 | 1 | 1 |
| rs121909260 | pathogenic | 0 | 0 | 0 |
| rs104893696 | pathogenic | 0 | 0 | 0 |
| rs121909267 | pathogenic | 0 | 0 | 0 |
| rs121909263 | pathogenic | 0 | 0 | 0 |
| rs121909264 | pathogenic | 0 | 0 | 0 |
| rs104893694 | pathogenic | 1 | 1 | 1 |
| rs121909268 | pathogenic | 1 | 1 | 1 |
| rs104893689 | pathogenic | 1 | 1 | 0 |
| rs104893707 | pathogenic | 1 | 1 | 0 |
| rs104893697 | pathogenic | 1 | 1 | 1 |
| rs28936684 | pathogenic | 1 | 1 | 1 |
| rs62269092 | benign | 1 | 1 | 0 |
| rs121909259 | pathogenic | 1 | 1 | 0 |
| rs548403340 | benign | 1 | 1 | 0 |
| rs142818334 | benign | 0 | 0 | 0 |
| rs12493789 | benign | 1 | 1 | 1 |
| rs193922423 | pathogenic | 1 | 1 | 0 |
| rs104893719 | pathogenic | 1 | 1 | 1 |
| rs117375173 | benign | 1 | 1 | 0 |
| rs104893712 | pathogenic | 0 | 0 | 0 |
| rs121917776 | pathogenic | 1 | 1 | 1 |
| rs767325003 | benign | 1 | 1 | 1 |
| rs35824453 | benign | 0 | 0 | 0 |
| rs141641266 | benign | 0 | 0 | 0 |
| rs1050239 | benign | 0 | 0 | 0 |
| rs141303384 | benign | 0 | 0 | 0 |
| rs12720458 | pathogenic | 0 | 0 | 0 |
| rs368507376 | pathogenic | 1 | 1 | 1 |
| rs17215500 | pathogenic | 1 | 1 | 1 |
| rs120074188 | pathogenic | 1 | 1 | 1 |
| rs397508097 | pathogenic | 1 | 1 | 1 |
| rs138551008 | pathogenic | 0 | 0 | 0 |
| rs387907139 | pathogenic | 1 | 1 | 1 |
| rs765487627 | pathogenic | 0 | 0 | 0 |

| | | | | |
|---|---|---|---|---|
| rs587778587 | benign | 0 | 0 | 0 |
| rs45568339 | benign | 0 | 0 | 0 |
| rs78179744 | benign | 0 | 0 | 0 |
| rs45624036 | benign | 0 | 0 | 0 |
| rs61755166 | benign | 0 | 0 | 0 |
| rs180177132 | pathogenic | 1 | 1 | 1 |
| rs587776527 | pathogenic | 0 | 0 | 0 |
| rs373783514 | benign | 0 | 0 | 0 |
| rs730881896 | benign | 0 | 0 | 0 |
| rs118203998 | pathogenic | 0 | 0 | 0 |
| rs41281039 | pathogenic | 1 | 1 | 1 |
| rs118204430 | pathogenic | 1 | 1 | 1 |
| rs1800546 | pathogenic | 1 | 1 | 0 |
| rs76917243 | pathogenic | 0 | 1 | 0 |
| rs78340951 | pathogenic | 0 | 1 | 0 |
| rs77718928 | pathogenic | 0 | 0 | 0 |
| rs369586696 | pathogenic | 0 | 1 | 0 |
| rs745517517 | pathogenic | 1 | 1 | 0 |
| rs372872777 | pathogenic | 0 | 0 | 0 |
| rs748704459 | pathogenic | 0 | 0 | 0 |
| rs769215411 | pathogenic | 0 | 0 | 0 |
| rs139425890 | pathogenic | 0 | 0 | 0 |
| rs779694939 | pathogenic | 1 | 1 | 0 |
| rs773729410 | pathogenic | 0 | 0 | 0 |
| rs121908623 | pathogenic | 1 | 1 | 1 |
| rs121908622 | pathogenic | 1 | 1 | 1 |
| rs121908621 | pathogenic | 1 | 1 | 1 |
| rs762106720 | pathogenic | 1 | 1 | 0 |
| rs199877522 | pathogenic | 0 | 0 | 0 |
| rs74315291 | pathogenic | 1 | 1 | 1 |
| rs79137415 | benign | 0 | 0 | 0 |
| rs121913045 | pathogenic | 0 | 0 | 0 |
| rs34295337 | pathogenic | 0 | 0 | 0 |
| rs121913046 | pathogenic | 0 | 0 | 0 |
| rs121913047 | pathogenic | 0 | 0 | 0 |
| rs121913048 | pathogenic | 0 | 0 | 0 |
| rs267607166 | pathogenic | 0 | 0 | 0 |
| rs17850684 | benign | 1 | 1 | 1 |
| rs119476051 | pathogenic | 1 | 1 | 1 |
| rs17850683 | benign | 1 | 1 | 1 |
| rs119476049 | pathogenic | 1 | 1 | 1 |
| rs119476046 | pathogenic | 1 | 1 | 1 |
| rs119476048 | pathogenic | 0 | 0 | 0 |
| rs119476047 | pathogenic | 1 | 1 | 1 |
| rs28939094 | pathogenic | 1 | 1 | 1 |
| rs119476050 | pathogenic | 0 | 0 | 0 |
| rs387906941 | pathogenic | 1 | 1 | 1 |
| rs41511344 | pathogenic | 1 | 1 | 0 |
| rs748112833 | pathogenic | 1 | 1 | 1 |
| rs74342080 | benign | 1 | 1 | 0 |
| rs116482870 | benign | 0 | 0 | 0 |
| rs104893743 | pathogenic | 0 | 0 | 0 |
| rs117179004 | benign | 0 | 0 | 0 |
| rs587779409 | pathogenic | 1 | 1 | 1 |

| | | | | |
|---|---|---|---|---|
| rs3745406 | benign | 1 | 1 | 0 |
| rs387906696 | pathogenic | 0 | 0 | 0 |
| rs72549369 | pathogenic | 1 | 1 | 0 |
| rs45523133 | benign | 0 | 0 | 0 |
| rs119460972 | pathogenic | 0 | 1 | 0 |
| rs34929837 | benign | 0 | 1 | 0 |
| rs17323225 | benign | 0 | 1 | 0 |
| rs17011368 | benign | 0 | 0 | 0 |
| rs748902766 | pathogenic | 1 | 1 | 1 |
| rs387907292 | pathogenic | 0 | 0 | 0 |
| rs387907293 | pathogenic | 1 | 1 | 0 |
| rs371526758 | pathogenic | 1 | 1 | 1 |
| rs142968179 | pathogenic | 1 | 1 | 1 |
| rs387907291 | pathogenic | 1 | 1 | 0 |
| rs121907949 | pathogenic | 0 | 0 | 0 |
| rs121907947 | pathogenic | 0 | 0 | 0 |
| rs121907950 | pathogenic | 1 | 1 | 1 |
| rs121907948 | pathogenic | 1 | 1 | 1 |
| rs4926 | benign | 0 | 0 | 0 |
| rs748809996 | pathogenic | 0 | 0 | 0 |
| rs1799883 | benign | 1 | 1 | 0 |
| rs104893974 | pathogenic | 0 | 1 | 0 |
| rs104893973 | pathogenic | 0 | 1 | 0 |
| rs34452707 | benign | 0 | 0 | 0 |
| rs119467003 | pathogenic | 1 | 1 | 1 |
| rs121908261 | pathogenic | 0 | 1 | 0 |
| rs28933985 | pathogenic | 1 | 1 | 1 |
| rs80356669 | pathogenic | 1 | 1 | 1 |
| rs28936677 | pathogenic | 1 | 1 | 0 |
| rs121918326 | pathogenic | 0 | 0 | 0 |
| rs121918324 | pathogenic | 1 | 1 | 0 |
| rs121918325 | pathogenic | 1 | 1 | 0 |
| rs41270025 | pathogenic | 1 | 1 | 0 |
| rs121918323 | pathogenic | 0 | 0 | 0 |
| rs12735723 | pathogenic | 1 | 1 | 0 |
| rs36013429 | benign | 0 | 0 | 0 |
| rs28936676 | pathogenic | 1 | 1 | 0 |
| rs771809901 | pathogenic | 1 | 1 | 1 |
| rs758350837 | benign | 1 | 1 | 0 |
| rs121908192 | pathogenic | 1 | 1 | 1 |
| rs147568500 | pathogenic | 1 | 1 | 1 |
| rs3729734 | benign | 0 | 0 | 0 |
| rs371536347 | benign | 1 | 1 | 1 |
| rs80358193 | pathogenic | 1 | 1 | 1 |
| rs119476044 | pathogenic | 1 | 1 | 1 |
| rs111292798 | benign | 1 | 1 | 1 |
| rs121908063 | pathogenic | 1 | 1 | 0 |
| rs121908065 | pathogenic | 1 | 1 | 1 |
| rs121908062 | pathogenic | 1 | 1 | 1 |
| rs121908064 | pathogenic | 1 | 1 | 1 |
| rs61746421 | benign | 1 | 1 | 1 |
| rs34886916 | benign | 1 | 1 | 1 |
| rs118204045 | pathogenic | 0 | 0 | 0 |
| rs138607412 | benign | 1 | 1 | 1 |

| | | | | |
|---|---|---|---|---|
| rs118204046 | pathogenic | 0 | 0 | 0 |
| rs118204048 | pathogenic | 0 | 0 | 0 |
| rs118204044 | pathogenic | 0 | 0 | 0 |
| rs149664056 | benign | 0 | 0 | 0 |
| rs121909285 | pathogenic | 0 | 0 | 0 |
| rs1802498 | benign | 0 | 0 | 0 |
| rs1065761 | benign | 0 | 0 | 0 |
| rs119477054 | pathogenic | 0 | 0 | 0 |
| rs12720097 | benign | 0 | 0 | 0 |
| rs747281324 | pathogenic | 1 | 1 | 0 |
| rs367543028 | pathogenic | 1 | 1 | 0 |
| rs146077918 | benign | 0 | 0 | 0 |
| rs142551229 | benign | 0 | 0 | 0 |
| rs367543039 | pathogenic | 0 | 0 | 0 |
| rs587779884 | pathogenic | 1 | 1 | 0 |
| rs137853153 | pathogenic | 0 | 0 | 0 |
| rs367543029 | pathogenic | 0 | 0 | 0 |
| rs587783037 | pathogenic | 1 | 1 | 0 |
| rs142928725 | benign | 0 | 0 | 0 |
| rs28385141 | benign | 0 | 0 | 0 |
| rs587779888 | pathogenic | 0 | 0 | 0 |
| rs121909801 | pathogenic | 1 | 1 | 1 |
| rs121909790 | pathogenic | 1 | 1 | 0 |
| rs121909797 | pathogenic | 1 | 1 | 0 |
| rs121909794 | pathogenic | 1 | 1 | 1 |
| rs121909791 | pathogenic | 1 | 1 | 1 |
| rs121909793 | pathogenic | 1 | 1 | 1 |
| rs147977279 | benign | 1 | 1 | 1 |
| rs147352564 | benign | 0 | 0 | 0 |
| rs368758045 | benign | 1 | 1 | 1 |
| rs114498261 | benign | 1 | 1 | 0 |
| rs35484156 | benign | 0 | 0 | 0 |
| rs139176878 | benign | 0 | 0 | 0 |
| rs11543934 | benign | 0 | 0 | 0 |
| rs34192549 | benign | 0 | 0 | 0 |
| rs568736295 | benign | 0 | 0 | 0 |
| rs55747232 | benign | 0 | 0 | 0 |
| rs781887272 | benign | 0 | 0 | 0 |
| rs146273341 | benign | 0 | 0 | 0 |
| rs41473544 | benign | 0 | 0 | 0 |
| rs137852680 | pathogenic | 0 | 0 | 0 |
| rs55891000 | benign | 0 | 0 | 0 |
| rs140351189 | benign | 0 | 0 | 0 |
| rs137852679 | pathogenic | 1 | 1 | 0 |
| rs137852678 | pathogenic | 0 | 0 | 0 |
| rs137852677 | pathogenic | 1 | 1 | 1 |
| rs138620818 | benign | 0 | 0 | 0 |
| rs121908531 | pathogenic | 0 | 0 | 0 |
| rs72554307 | pathogenic | 0 | 0 | 0 |
| rs72554308 | pathogenic | 0 | 0 | 0 |
| rs72554312 | pathogenic | 1 | 1 | 0 |
| rs67939655 | pathogenic | 0 | 0 | 0 |
| rs67486158 | pathogenic | 0 | 0 | 0 |
| rs72554338 | pathogenic | 1 | 1 | 1 |

| | | | | |
|---|---|---|---|---|
| rs1800324 | pathogenic | 1 | 1 | 1 |
| rs72554356 | pathogenic | 1 | 1 | 1 |
| rs66656800 | pathogenic | 1 | 1 | 1 |
| rs68026851 | pathogenic | 1 | 1 | 1 |
| rs72558412 | pathogenic | 0 | 0 | 0 |
| rs72558415 | pathogenic | 0 | 0 | 0 |
| rs72558423 | pathogenic | 0 | 0 | 0 |
| rs67120076 | pathogenic | 1 | 1 | 1 |
| rs1800328 | pathogenic | 1 | 1 | 1 |
| rs72558451 | pathogenic | 1 | 1 | 1 |
| rs72558454 | pathogenic | 1 | 1 | 1 |
| rs672601372 | pathogenic | 0 | 0 | 0 |
| rs139644798 | pathogenic | 0 | 0 | 0 |
| rs369398935 | pathogenic | 0 | 0 | 0 |
| rs267607035 | pathogenic | 0 | 0 | 0 |
| rs267607036 | pathogenic | 1 | 1 | 1 |
| rs2302566 | benign | 1 | 1 | 0 |
| rs115959591 | benign | 0 | 0 | 0 |
| rs773876739 | pathogenic | 0 | 0 | 0 |
| rs1799967 | benign | 0 | 0 | 0 |
| rs80356968 | benign | 0 | 0 | 0 |
| rs28897695 | benign | 1 | 1 | 1 |
| rs80357314 | benign | 0 | 0 | 0 |
| rs80357169 | benign | 0 | 0 | 0 |
| rs397507239 | benign | 1 | 1 | 1 |
| rs80187739 | pathogenic | 1 | 1 | 1 |
| rs41293459 | pathogenic | 1 | 1 | 1 |
| rs55770810 | pathogenic | 1 | 1 | 1 |
| rs80356860 | pathogenic | 0 | 0 | 0 |
| rs56195342 | benign | 0 | 0 | 0 |
| rs80357442 | benign | 0 | 0 | 0 |
| rs41293463 | pathogenic | 1 | 1 | 1 |
| rs80357069 | pathogenic | 0 | 0 | 0 |
| rs397509275 | benign | 0 | 0 | 0 |
| rs80356920 | benign | 0 | 0 | 0 |
| rs397509284 | pathogenic | 0 | 0 | 0 |
| rs80357286 | benign | 0 | 0 | 0 |
| rs80357274 | benign | 0 | 0 | 0 |
| rs765656957 | benign | 0 | 0 | 0 |
| rs755427809 | benign | 0 | 0 | 0 |
| rs80357322 | benign | 0 | 0 | 0 |
| rs121434430 | pathogenic | 0 | 0 | 0 |
| rs121918450 | pathogenic | 1 | 1 | 1 |
| rs1532268 | benign | 0 | 0 | 0 |
| rs2303080 | benign | 0 | 0 | 0 |
| rs10064631 | benign | 0 | 0 | 0 |
| rs162036 | benign | 0 | 0 | 0 |
| rs2287780 | benign | 0 | 0 | 0 |
| rs16879334 | benign | 0 | 0 | 0 |
| rs137853061 | pathogenic | 1 | 1 | 0 |
| rs10380 | benign | 0 | 0 | 0 |
| rs1494558 | benign | 0 | 0 | 0 |
| rs1494555 | benign | 0 | 0 | 0 |
| rs192994060 | benign | 0 | 0 | 0 |

| | | | | |
|---|---|---|---|---|
| rs147128920 | benign | 1 | 1 | 0 |
| rs201803986 | pathogenic | 0 | 0 | 0 |
| rs746419489 | pathogenic | 0 | 1 | 0 |
| rs764396564 | pathogenic | 0 | 0 | 0 |
| rs758304537 | pathogenic | 0 | 0 | 0 |
| rs267606763 | pathogenic | 0 | 0 | 0 |
| rs267606762 | pathogenic | 0 | 0 | 0 |
| rs150702945 | pathogenic | 1 | 1 | 1 |
| rs202047589 | pathogenic | 0 | 0 | 0 |
| rs755562733 | pathogenic | 0 | 0 | 0 |
| rs267606764 | pathogenic | 1 | 1 | 1 |
| rs138207257 | pathogenic | 1 | 1 | 0 |
| rs149150736 | pathogenic | 1 | 1 | 1 |
| rs777046879 | pathogenic | 0 | 0 | 0 |
| rs104894408 | pathogenic | 1 | 1 | 1 |
| rs1801002 | pathogenic | 1 | 1 | 1 |
| rs28929485 | pathogenic | 1 | 1 | 1 |
| rs104894396 | pathogenic | 1 | 1 | 1 |
| rs371024165 | pathogenic | 0 | 0 | 0 |
| rs35887622 | pathogenic | 1 | 1 | 1 |
| rs587783644 | pathogenic | 1 | 1 | 1 |
| rs72474224 | pathogenic | 1 | 1 | 1 |
| rs111033296 | pathogenic | 1 | 1 | 1 |
| rs104894407 | pathogenic | 1 | 1 | 1 |
| rs72561723 | pathogenic | 1 | 1 | 1 |
| rs104894398 | pathogenic | 1 | 1 | 1 |
| rs28931594 | pathogenic | 1 | 1 | 1 |
| rs104894412 | pathogenic | 1 | 1 | 1 |
| rs111033297 | pathogenic | 1 | 1 | 1 |
| rs104894404 | pathogenic | 1 | 1 | 1 |
| rs104894410 | pathogenic | 1 | 1 | 1 |
| rs104894403 | pathogenic | 1 | 1 | 1 |
| rs397516870 | benign | 0 | 0 | 0 |
| rs200023879 | pathogenic | 1 | 1 | 1 |
| rs121912968 | pathogenic | 0 | 0 | 0 |
| rs104894402 | pathogenic | 1 | 1 | 1 |
| rs28931593 | pathogenic | 1 | 1 | 1 |
| rs104894397 | pathogenic | 0 | 0 | 0 |
| rs80338944 | pathogenic | 0 | 0 | 0 |
| rs199883710 | pathogenic | 0 | 0 | 0 |
| rs727504302 | pathogenic | 0 | 0 | 0 |
| rs145216882 | benign | 1 | 1 | 1 |
| rs781534323 | pathogenic | 1 | 1 | 1 |
| rs111033218 | benign | 1 | 1 | 1 |
| rs104894409 | pathogenic | 0 | 0 | 0 |
| rs80338945 | pathogenic | 1 | 1 | 1 |
| rs111033299 | pathogenic | 0 | 0 | 0 |
| rs143343083 | pathogenic | 1 | 1 | 1 |
| rs80338946 | pathogenic | 0 | 0 | 0 |
| rs150529554 | pathogenic | 0 | 0 | 0 |
| rs111033295 | pathogenic | 0 | 0 | 0 |
| rs111033188 | benign | 0 | 0 | 0 |
| rs397516874 | pathogenic | 0 | 0 | 0 |
| rs76434661 | pathogenic | 0 | 0 | 0 |

| | | | | |
|---|---|---|---|---|
| rs80338948 | pathogenic | 0 | 0 | 0 |
| rs104894401 | pathogenic | 0 | 0 | 0 |
| rs767178508 | pathogenic | 0 | 0 | 0 |
| rs772264564 | pathogenic | 0 | 0 | 0 |
| rs28931592 | pathogenic | 0 | 0 | 0 |
| rs80338949 | pathogenic | 1 | 1 | 1 |
| rs376898963 | benign | 1 | 1 | 1 |
| rs111033360 | benign | 0 | 0 | 0 |
| rs774518779 | pathogenic | 1 | 1 | 1 |
| rs28931595 | pathogenic | 1 | 1 | 1 |
| rs771748289 | pathogenic | 0 | 0 | 0 |
| rs104894406 | pathogenic | 0 | 0 | 0 |
| rs76838169 | benign | 0 | 0 | 0 |
| rs111033294 | pathogenic | 0 | 0 | 0 |
| rs200214298 | pathogenic | 0 | 0 | 0 |
| rs28940283 | pathogenic | 0 | 1 | 0 |
| rs1047883 | benign | 0 | 0 | 0 |
| rs138254425 | benign | 0 | 0 | 0 |
| rs121912592 | pathogenic | 1 | 1 | 0 |
| rs369061090 | pathogenic | 1 | 1 | 0 |
| rs758724746 | pathogenic | 0 | 1 | 0 |
| rs121912595 | pathogenic | 0 | 0 | 0 |
| rs76340296 | benign | 0 | 0 | 0 |
| rs140578009 | benign | 1 | 1 | 1 |
| rs74445297 | benign | 0 | 0 | 0 |
| rs78599516 | benign | 1 | 1 | 0 |
| rs104894035 | pathogenic | 1 | 1 | 0 |
| rs79451216 | benign | 0 | 0 | 0 |
| rs104894036 | pathogenic | 0 | 0 | 0 |
| rs73343757 | benign | 0 | 0 | 0 |
| rs121917722 | pathogenic | 1 | 1 | 1 |
| rs747700126 | pathogenic | 0 | 1 | 0 |
| rs121917725 | pathogenic | 1 | 1 | 1 |
| rs121917721 | pathogenic | 1 | 1 | 1 |
| rs121917723 | pathogenic | 1 | 1 | 0 |
| rs17569 | benign | 1 | 1 | 1 |
| rs2230062 | benign | 1 | 1 | 1 |
| rs17570 | benign | 0 | 0 | 0 |
| rs121917724 | pathogenic | 0 | 1 | 0 |
| rs199422235 | pathogenic | 0 | 0 | 0 |
| rs199422237 | pathogenic | 1 | 1 | 1 |
| rs199422234 | pathogenic | 1 | 1 | 0 |
| rs140015315 | pathogenic | 1 | 1 | 1 |
| rs730880298 | pathogenic | 1 | 1 | 1 |
| rs779692470 | pathogenic | 0 | 0 | 0 |
| rs369152939 | pathogenic | 0 | 0 | 0 |
| rs527236040 | pathogenic | 0 | 0 | 0 |
| rs587776985 | pathogenic | 1 | 1 | 1 |
| rs147077598 | pathogenic | 1 | 1 | 1 |
| rs128620185 | pathogenic | 1 | 1 | 1 |
| rs128620189 | pathogenic | 0 | 0 | 0 |
| rs128621190 | pathogenic | 0 | 0 | 0 |
| rs61754648 | benign | 0 | 0 | 0 |
| rs137852906 | pathogenic | 0 | 0 | 0 |

| | | | | |
|---|---|---|---|---|
| rs76811038 | pathogenic | 0 | 0 | 0 |
| rs149468422 | benign | 0 | 0 | 0 |
| rs5911 | benign | 0 | 0 | 0 |
| rs28942112 | pathogenic | 1 | 1 | 1 |
| rs143275858 | pathogenic | 1 | 1 | 0 |
| rs28362261 | benign | 0 | 0 | 0 |
| rs28362263 | benign | 1 | 1 | 1 |
| rs558707786 | pathogenic | 0 | 0 | 0 |
| rs138775799 | pathogenic | 0 | 0 | 0 |
| rs748170941 | pathogenic | 0 | 0 | 0 |
| rs121908380 | pathogenic | 0 | 0 | 0 |
| rs765123255 | pathogenic | 1 | 1 | 1 |
| rs730881832 | pathogenic | 0 | 1 | 0 |
| rs587782165 | benign | 1 | 1 | 1 |
| rs762307622 | pathogenic | 0 | 0 | 0 |
| rs34612342 | pathogenic | 1 | 1 | 0 |
| rs143353451 | pathogenic | 0 | 1 | 0 |
| rs747993448 | pathogenic | 0 | 1 | 0 |
| rs376561094 | pathogenic | 0 | 0 | 0 |
| rs776487884 | pathogenic | 1 | 1 | 0 |
| rs34126013 | pathogenic | 1 | 1 | 0 |
| rs200495564 | pathogenic | 1 | 1 | 0 |
| rs140342925 | pathogenic | 1 | 1 | 0 |
| rs587780749 | pathogenic | 1 | 1 | 0 |
| rs773087549 | pathogenic | 1 | 1 | 0 |
| rs730881833 | pathogenic | 0 | 0 | 0 |
| rs374950566 | pathogenic | 1 | 1 | 0 |
| rs587780752 | benign | 0 | 0 | 0 |
| rs3219489 | benign | 0 | 0 | 0 |
| rs199742231 | benign | 0 | 0 | 0 |
| rs587780082 | pathogenic | 0 | 0 | 0 |
| rs35881803 | benign | 0 | 0 | 0 |
| rs41305623 | benign | 1 | 1 | 1 |
| rs121917856 | pathogenic | 1 | 1 | 1 |
| rs576284122 | benign | 0 | 0 | 0 |
| rs142036701 | benign | 0 | 0 | 0 |
| rs121917852 | pathogenic | 1 | 1 | 1 |
| rs121917854 | pathogenic | 1 | 1 | 1 |
| rs121917853 | pathogenic | 1 | 1 | 1 |
| rs121917857 | pathogenic | 0 | 1 | 0 |
| rs193922740 | pathogenic | 1 | 1 | 1 |
| rs565779970 | pathogenic | 1 | 1 | 1 |
| rs6723526 | benign | 0 | 0 | 0 |
| rs141094096 | pathogenic | 1 | 1 | 1 |
| rs104894637 | pathogenic | 0 | 0 | 0 |
| rs104894636 | pathogenic | 0 | 0 | 0 |
| rs104894642 | pathogenic | 0 | 0 | 0 |
| rs104894638 | pathogenic | 0 | 0 | 0 |
| rs753666460 | pathogenic | 0 | 0 | 0 |
| rs104894643 | pathogenic | 1 | 1 | 0 |
| rs34520362 | pathogenic | 0 | 0 | 0 |
| rs374621913 | pathogenic | 0 | 0 | 0 |
| rs113641837 | benign | 0 | 0 | 0 |
| rs104894635 | pathogenic | 0 | 0 | 0 |

| | | | | |
|---|---|---|---|---|
| rs143947056 | pathogenic | 0 | 0 | 0 |
| rs138504221 | pathogenic | 0 | 0 | 0 |
| rs9894254 | benign | 0 | 0 | 0 |
| rs104894640 | pathogenic | 1 | 1 | 1 |
| rs58786455 | benign | 0 | 0 | 0 |
| rs144143780 | pathogenic | 0 | 0 | 0 |
| rs62620232 | benign | 0 | 0 | 0 |
| rs34297805 | benign | 1 | 1 | 1 |
| rs104894641 | pathogenic | 0 | 0 | 0 |
| rs148876719 | benign | 0 | 0 | 0 |
| rs104894639 | pathogenic | 0 | 0 | 0 |
| rs7503034 | benign | 0 | 0 | 0 |
| rs201631570 | benign | 0 | 0 | 0 |
| rs150181226 | pathogenic | 0 | 0 | 0 |
| rs377546036 | pathogenic | 0 | 0 | 0 |
| rs267606700 | pathogenic | 0 | 0 | 0 |
| rs267606701 | pathogenic | 1 | 1 | 0 |
| rs267606699 | pathogenic | 1 | 1 | 0 |
| rs9903215 | benign | 0 | 0 | 0 |
| rs139486406 | benign | 0 | 0 | 0 |
| rs143791614 | benign | 0 | 0 | 0 |
| rs573483474 | benign | 0 | 0 | 0 |
| rs2032892 | benign | 0 | 0 | 0 |
| rs137852619 | pathogenic | 0 | 0 | 0 |
| rs200243548 | benign | 0 | 0 | 0 |
| rs115702388 | benign | 0 | 0 | 0 |
| rs138085358 | pathogenic | 0 | 0 | 0 |
| rs121912789 | pathogenic | 1 | 1 | 1 |
| rs763407972 | pathogenic | 1 | 1 | 1 |
| rs147278302 | benign | 0 | 0 | 0 |
| rs374540411 | benign | 0 | 0 | 0 |
| rs121912790 | pathogenic | 1 | 1 | 0 |
| rs752049917 | benign | 0 | 0 | 0 |
| rs139640491 | benign | 1 | 1 | 1 |
| rs138010738 | benign | 1 | 1 | 0 |
| rs201328768 | pathogenic | 1 | 1 | 0 |
| rs2135720 | benign | 1 | 1 | 0 |
| rs201816080 | benign | 1 | 1 | 0 |
| rs754391973 | pathogenic | 0 | 0 | 0 |
| rs139087859 | benign | 0 | 0 | 0 |
| rs6805421 | benign | 0 | 0 | 0 |
| rs146266900 | benign | 0 | 0 | 0 |
| rs397509419 | pathogenic | 0 | 0 | 0 |
| rs149980411 | benign | 0 | 0 | 0 |
| rs747711488 | pathogenic | 1 | 1 | 1 |
| rs149474339 | pathogenic | 1 | 1 | 1 |
| rs758252808 | pathogenic | 1 | 1 | 0 |
| rs745756308 | pathogenic | 0 | 0 | 0 |
| rs4986958 | benign | 0 | 0 | 0 |
| rs74315444 | pathogenic | 0 | 0 | 0 |
| rs61752137 | pathogenic | 0 | 0 | 0 |
| rs8179065 | benign | 1 | 1 | 1 |
| rs758361736 | pathogenic | 1 | 1 | 0 |
| rs139794067 | pathogenic | 1 | 1 | 1 |

| | | | | |
|---|---|---|---|---|
| rs730880954 | pathogenic | 1 | 1 | 1 |
| rs730880955 | pathogenic | 1 | 1 | 1 |
| rs150634297 | pathogenic | 0 | 0 | 0 |
| rs104893750 | pathogenic | 1 | 1 | 1 |
| rs104893748 | pathogenic | 1 | 1 | 1 |
| rs104893749 | pathogenic | 1 | 1 | 1 |
| rs753520553 | pathogenic | 1 | 1 | 0 |
| rs61737294 | benign | 0 | 0 | 0 |
| rs104894601 | pathogenic | 0 | 0 | 0 |
| rs376090795 | pathogenic | 0 | 0 | 0 |
| rs104894596 | pathogenic | 0 | 0 | 0 |
| rs104894595 | pathogenic | 0 | 0 | 0 |
| rs104894598 | pathogenic | 0 | 0 | 0 |
| rs104894597 | pathogenic | 0 | 0 | 0 |
| rs104894591 | pathogenic | 0 | 0 | 0 |
| rs104894593 | pathogenic | 0 | 0 | 0 |
| rs104894594 | pathogenic | 0 | 0 | 0 |
| rs104894590 | pathogenic | 0 | 0 | 0 |
| rs86312 | benign | 0 | 0 | 0 |
| rs77874543 | benign | 0 | 0 | 0 |
| rs121912590 | pathogenic | 1 | 1 | 1 |
| rs121912589 | pathogenic | 1 | 1 | 1 |
| rs121912586 | pathogenic | 1 | 1 | 1 |
| rs121912585 | pathogenic | 1 | 1 | 1 |
| rs121912587 | pathogenic | 1 | 1 | 1 |
| rs762909359 | pathogenic | 0 | 0 | 0 |
| rs28931610 | pathogenic | 0 | 0 | 0 |
| rs104894097 | pathogenic | 1 | 1 | 1 |
| rs775176191 | benign | 0 | 0 | 0 |
| rs199907548 | pathogenic | 0 | 0 | 0 |
| rs104894095 | pathogenic | 1 | 1 | 1 |
| rs104894109 | pathogenic | 1 | 1 | 1 |
| rs372266620 | benign | 1 | 1 | 1 |
| rs104894099 | pathogenic | 0 | 0 | 0 |
| rs137854599 | pathogenic | 1 | 1 | 1 |
| rs104894094 | pathogenic | 0 | 0 | 0 |
| rs104894104 | pathogenic | 0 | 0 | 0 |
| rs6413463 | benign | 1 | 1 | 1 |
| rs104894098 | pathogenic | 0 | 0 | 0 |
| rs6413464 | benign | 0 | 0 | 0 |
| rs3731249 | benign | 0 | 0 | 0 |
| rs16931179 | benign | 0 | 0 | 0 |
| rs2734715 | benign | 0 | 0 | 0 |
| rs759080581 | pathogenic | 0 | 0 | 0 |
| rs1800746 | benign | 0 | 0 | 0 |
| rs202189057 | pathogenic | 0 | 0 | 0 |
| rs121908204 | pathogenic | 0 | 0 | 0 |
| rs756564767 | pathogenic | 0 | 0 | 0 |
| rs34758634 | benign | 1 | 1 | 1 |
| rs28940573 | pathogenic | 0 | 0 | 0 |
| rs119455955 | pathogenic | 1 | 1 | 0 |
| rs752164603 | pathogenic | 1 | 1 | 0 |
| rs763162812 | pathogenic | 0 | 0 | 0 |
| rs119455957 | pathogenic | 0 | 0 | 0 |

| | | | | |
|---|---|---|---|---|
| rs119455958 | pathogenic | 1 | 1 | 0 |
| rs750428882 | pathogenic | 0 | 0 | 0 |
| rs141482368 | benign | 0 | 0 | 0 |
| rs119455953 | pathogenic | 0 | 0 | 0 |
| rs119455954 | pathogenic | 0 | 0 | 0 |
| rs112812685 | benign | 1 | 1 | 0 |
| rs138744051 | benign | 1 | 1 | 0 |
| rs121908200 | pathogenic | 0 | 0 | 0 |
| rs119455956 | pathogenic | 0 | 0 | 0 |
| rs78970155 | benign | 0 | 0 | 0 |
| rs2834167 | benign | 0 | 0 | 0 |
| rs387907326 | pathogenic | 1 | 1 | 1 |
| rs2107732 | benign | 0 | 0 | 0 |
| rs11552377 | benign | 0 | 0 | 0 |
| rs137852843 | pathogenic | 0 | 0 | 0 |
| rs104894783 | pathogenic | 0 | 0 | 0 |
| rs122457137 | pathogenic | 1 | 1 | 1 |
| rs104894779 | pathogenic | 1 | 1 | 1 |
| rs104894786 | pathogenic | 0 | 0 | 0 |
| rs104894784 | pathogenic | 0 | 0 | 0 |
| rs104894785 | pathogenic | 0 | 0 | 0 |
| rs201870761 | pathogenic | 0 | 0 | 0 |
| rs104894781 | pathogenic | 0 | 0 | 0 |
| rs28897728 | benign | 0 | 0 | 0 |
| rs370273690 | pathogenic | 1 | 1 | 0 |
| rs9332739 | benign | 0 | 0 | 0 |
| rs150878060 | benign | 0 | 0 | 0 |
| rs573509224 | benign | 0 | 0 | 0 |
| rs36221133 | benign | 0 | 0 | 0 |
| rs2230009 | benign | 0 | 0 | 0 |
| rs267607008 | pathogenic | 0 | 0 | 0 |
| rs5985 | benign | 1 | 1 | 1 |
| rs5982 | benign | 1 | 1 | 1 |
| rs5987 | benign | 0 | 0 | 0 |
| rs5988 | benign | 0 | 0 | 0 |
| rs121918680 | pathogenic | 1 | 1 | 0 |
| rs121918679 | pathogenic | 1 | 1 | 0 |
| rs7845 | benign | 1 | 1 | 0 |
| rs1049296 | benign | 1 | 1 | 1 |
| rs121918677 | pathogenic | 1 | 1 | 0 |
| rs200086262 | pathogenic | 1 | 1 | 0 |
| rs746930990 | pathogenic | 1 | 1 | 0 |
| rs104893878 | pathogenic | 1 | 1 | 1 |
| rs121912468 | pathogenic | 1 | 1 | 1 |
| rs121912467 | pathogenic | 1 | 1 | 1 |
| rs145351926 | benign | 1 | 1 | 1 |
| rs141212743 | pathogenic | 1 | 1 | 1 |
| rs397515473 | pathogenic | 1 | 1 | 0 |
| rs121918313 | pathogenic | 0 | 1 | 0 |
| rs73015965 | pathogenic | 0 | 0 | 0 |
| rs121918030 | pathogenic | 0 | 1 | 0 |
| rs77782321 | benign | 0 | 0 | 0 |
| rs374661051 | pathogenic | 1 | 1 | 1 |
| rs267606854 | pathogenic | 0 | 0 | 0 |

| | | | | |
|---|---|---|---|---|
| rs41279678 | benign | 0 | 0 | 0 |
| rs61754640 | benign | 0 | 0 | 0 |
| rs199473314 | pathogenic | 0 | 0 | 0 |
| rs137854601 | pathogenic | 0 | 0 | 0 |
| rs137854614 | pathogenic | 0 | 0 | 0 |
| rs137854615 | pathogenic | 0 | 0 | 0 |
| rs199473635 | pathogenic | 0 | 0 | 0 |
| rs121434478 | pathogenic | 1 | 1 | 1 |
| rs775407864 | pathogenic | 1 | 1 | 1 |
| rs12706927 | benign | 0 | 1 | 0 |
| rs775614261 | benign | 0 | 0 | 0 |
| rs768926261 | benign | 0 | 0 | 0 |
| rs114057640 | benign | 0 | 0 | 0 |
| rs387906886 | pathogenic | 1 | 1 | 0 |
| rs138658585 | benign | 0 | 0 | 0 |
| rs4880091 | benign | 0 | 0 | 0 |
| rs121918552 | pathogenic | 1 | 1 | 1 |
| rs10794537 | benign | 0 | 0 | 0 |
| rs121965022 | pathogenic | 1 | 1 | 0 |
| rs121965020 | pathogenic | 0 | 0 | 0 |
| rs758452450 | pathogenic | 0 | 0 | 0 |
| rs747981483 | benign | 0 | 0 | 0 |
| rs58037052 | benign | 0 | 0 | 0 |
| rs121965029 | pathogenic | 1 | 1 | 0 |
| rs147490060 | benign | 0 | 0 | 0 |
| rs3755955 | benign | 0 | 0 | 0 |
| rs148789453 | pathogenic | 0 | 0 | 0 |
| rs121965030 | pathogenic | 0 | 0 | 0 |
| rs76722191 | benign | 0 | 0 | 0 |
| rs199801029 | pathogenic | 0 | 0 | 0 |
| rs121965033 | pathogenic | 0 | 0 | 0 |
| rs368454909 | pathogenic | 1 | 1 | 0 |
| rs6831280 | benign | 0 | 0 | 0 |
| rs121965024 | pathogenic | 0 | 0 | 0 |
| rs121965019 | pathogenic | 0 | 0 | 0 |
| rs11934801 | benign | 0 | 0 | 0 |
| rs73066479 | benign | 0 | 0 | 0 |
| rs121965027 | pathogenic | 0 | 0 | 0 |
| rs121965026 | pathogenic | 1 | 1 | 0 |
| rs121965021 | pathogenic | 0 | 0 | 0 |
| rs398123257 | benign | 0 | 0 | 0 |
| rs121965031 | pathogenic | 0 | 0 | 0 |
| rs202003795 | pathogenic | 0 | 0 | 0 |
| rs559510809 | pathogenic | 0 | 0 | 0 |
| rs762936223 | benign | 0 | 0 | 0 |
| rs137854580 | pathogenic | 1 | 1 | 1 |
| rs112263157 | benign | 0 | 0 | 0 |
| rs2289181 | benign | 0 | 0 | 0 |
| rs138034915 | benign | 0 | 1 | 0 |
| rs528367092 | pathogenic | 0 | 1 | 0 |
| rs1042393 | benign | 0 | 0 | 0 |
| rs370950728 | pathogenic | 0 | 1 | 0 |
| rs530478036 | benign | 0 | 1 | 0 |
| rs374569672 | benign | 0 | 1 | 0 |

| | | | | |
|---|---|---|---|---|
| rs121907944 | pathogenic | 0 | 1 | 0 |
| rs200856561 | pathogenic | 0 | 1 | 0 |
| rs577915581 | pathogenic | 0 | 1 | 0 |
| rs201896815 | pathogenic | 0 | 1 | 0 |
| rs121907945 | pathogenic | 0 | 1 | 0 |
| rs121907940 | pathogenic | 0 | 1 | 0 |
| rs543300039 | pathogenic | 0 | 1 | 0 |
| rs121907936 | pathogenic | 0 | 1 | 0 |
| rs766074609 | pathogenic | 0 | 1 | 0 |
| rs755253527 | pathogenic | 0 | 1 | 0 |
| rs560575383 | pathogenic | 0 | 1 | 0 |
| rs528369909 | benign | 0 | 1 | 0 |
| rs770610356 | pathogenic | 0 | 1 | 0 |
| rs747610090 | pathogenic | 0 | 1 | 0 |
| rs145712232 | benign | 0 | 1 | 0 |
| rs7215458 | benign | 1 | 1 | 0 |
| rs535644999 | pathogenic | 0 | 1 | 0 |
| rs772883420 | pathogenic | 1 | 1 | 0 |
| rs398123169 | pathogenic | 0 | 1 | 0 |
| rs121907937 | pathogenic | 0 | 1 | 0 |
| rs121907941 | pathogenic | 1 | 1 | 0 |
| rs121907942 | pathogenic | 0 | 1 | 0 |
| rs779556619 | pathogenic | 0 | 1 | 0 |
| rs1800307 | benign | 0 | 1 | 0 |
| rs377544304 | pathogenic | 1 | 1 | 0 |
| rs549029029 | pathogenic | 0 | 1 | 0 |
| rs753269119 | pathogenic | 1 | 1 | 1 |
| rs757617999 | pathogenic | 0 | 1 | 0 |
| rs28937909 | pathogenic | 0 | 1 | 0 |
| rs28940868 | pathogenic | 0 | 1 | 0 |
| rs368438393 | pathogenic | 0 | 1 | 0 |
| rs536906561 | pathogenic | 0 | 1 | 0 |
| rs374143224 | pathogenic | 0 | 1 | 0 |
| rs778418246 | pathogenic | 0 | 1 | 0 |
| rs398123172 | pathogenic | 0 | 1 | 0 |
| rs121907938 | pathogenic | 0 | 1 | 0 |
| rs1800312 | pathogenic | 0 | 1 | 0 |
| rs1126690 | benign | 0 | 1 | 0 |
| rs1800314 | benign | 0 | 1 | 0 |
| rs369532274 | pathogenic | 0 | 1 | 0 |
| rs121907943 | pathogenic | 0 | 1 | 0 |
| rs780321415 | pathogenic | 0 | 1 | 0 |
| rs377286472 | benign | 0 | 1 | 0 |
| rs121918406 | pathogenic | 1 | 1 | 1 |
| rs147484870 | benign | 0 | 0 | 0 |
| rs193922671 | pathogenic | 0 | 0 | 0 |
| rs201397978 | benign | 1 | 1 | 1 |
| rs34884895 | benign | 1 | 1 | 1 |
| rs138329459 | benign | 0 | 0 | 0 |
| rs2289366 | benign | 1 | 1 | 1 |
| rs141858334 | benign | 1 | 1 | 1 |
| rs59907886 | benign | 1 | 1 | 1 |
| rs137852653 | pathogenic | 0 | 0 | 0 |
| rs137852652 | pathogenic | 0 | 0 | 0 |

| | | | | |
|---|---|---|---|---|
| rs137852656 | pathogenic | 0 | 0 | 0 |
| rs61739916 | benign | 0 | 0 | 0 |
| rs28936668 | pathogenic | 0 | 0 | 0 |
| rs137852651 | pathogenic | 0 | 0 | 0 |
| rs137852650 | pathogenic | 0 | 0 | 0 |
| rs28936669 | pathogenic | 0 | 0 | 0 |
| rs137852654 | pathogenic | 0 | 0 | 0 |
| rs28936368 | pathogenic | 1 | 1 | 0 |
| rs137852655 | pathogenic | 0 | 0 | 0 |
| rs61752496 | benign | 0 | 0 | 0 |
| rs750098055 | benign | 0 | 0 | 0 |
| rs764149793 | benign | 0 | 0 | 0 |
| rs724160017 | pathogenic | 0 | 1 | 0 |
| rs724160019 | pathogenic | 0 | 0 | 0 |
| rs202219398 | pathogenic | 1 | 1 | 0 |
| rs724160023 | pathogenic | 1 | 1 | 0 |
| rs761953453 | pathogenic | 0 | 0 | 0 |
| rs80338835 | pathogenic | 1 | 1 | 1 |
| rs56284940 | benign | 0 | 0 | 0 |
| rs74575749 | benign | 0 | 0 | 0 |
| rs587777075 | pathogenic | 1 | 1 | 1 |
| rs641153 | benign | 0 | 0 | 0 |
| rs12614 | benign | 0 | 0 | 0 |
| rs117314762 | benign | 0 | 0 | 0 |
| rs537478097 | benign | 1 | 1 | 1 |
| rs144812066 | benign | 0 | 0 | 0 |
| rs4151651 | benign | 0 | 0 | 0 |
| rs138207668 | benign | 1 | 1 | 1 |
| rs149101394 | benign | 1 | 1 | 1 |
| rs4151659 | benign | 1 | 1 | 1 |
| rs45484591 | benign | 1 | 1 | 1 |
| rs761991070 | pathogenic | 0 | 0 | 0 |
| rs755948940 | pathogenic | 0 | 0 | 0 |
| rs137853097 | pathogenic | 0 | 0 | 0 |
| rs775297352 | benign | 0 | 0 | 0 |
| rs28943591 | benign | 0 | 0 | 0 |
| rs191468413 | pathogenic | 1 | 1 | 0 |
| rs11539471 | benign | 1 | 1 | 1 |
| rs11205 | benign | 0 | 0 | 0 |
| rs768409400 | pathogenic | 0 | 0 | 0 |
| rs762013077 | pathogenic | 0 | 0 | 0 |
| rs768474112 | pathogenic | 0 | 0 | 0 |
| rs121965063 | pathogenic | 1 | 1 | 1 |
| rs143648758 | pathogenic | 0 | 0 | 0 |
| rs121965066 | pathogenic | 0 | 0 | 0 |
| rs752907087 | pathogenic | 0 | 0 | 0 |
| rs747702749 | pathogenic | 0 | 0 | 0 |
| rs754087775 | pathogenic | 0 | 0 | 0 |
| rs121918641 | pathogenic | 1 | 1 | 1 |
| rs121918642 | pathogenic | 1 | 1 | 1 |
| rs121918640 | pathogenic | 1 | 1 | 1 |
| rs121918637 | pathogenic | 1 | 1 | 1 |
| rs121918644 | pathogenic | 1 | 1 | 1 |
| rs16840544 | benign | 0 | 0 | 0 |

| | | | | |
|---|---|---|---|---|
| rs1042391 | benign | 1 | 1 | 1 |
| rs104893744 | pathogenic | 1 | 1 | 1 |
| rs120074136 | pathogenic | 0 | 0 | 0 |
| rs1805081 | benign | 1 | 1 | 0 |
| rs80358251 | benign | 0 | 0 | 0 |
| rs774588125 | benign | 0 | 0 | 0 |
| rs120074134 | pathogenic | 0 | 0 | 0 |
| rs139751448 | pathogenic | 0 | 0 | 0 |
| rs77080672 | benign | 0 | 0 | 0 |
| rs372445155 | pathogenic | 1 | 1 | 0 |
| rs13381670 | benign | 0 | 0 | 0 |
| rs1788799 | benign | 0 | 0 | 0 |
| rs376213990 | pathogenic | 0 | 0 | 0 |
| rs757475924 | pathogenic | 0 | 0 | 0 |
| rs80358253 | pathogenic | 0 | 0 | 0 |
| rs1805082 | benign | 0 | 0 | 0 |
| rs120074130 | pathogenic | 0 | 0 | 0 |
| rs768999208 | pathogenic | 0 | 0 | 0 |
| rs34302553 | benign | 0 | 0 | 0 |
| rs28940897 | pathogenic | 0 | 0 | 0 |
| rs1140458 | benign | 0 | 0 | 0 |
| rs143124972 | pathogenic | 0 | 0 | 0 |
| rs120074135 | pathogenic | 0 | 0 | 0 |
| rs543206298 | pathogenic | 0 | 0 | 0 |
| rs120074132 | pathogenic | 0 | 0 | 0 |
| rs759826138 | pathogenic | 0 | 0 | 0 |
| rs28942108 | pathogenic | 0 | 0 | 0 |
| rs80358254 | pathogenic | 0 | 0 | 0 |
| rs80358257 | pathogenic | 0 | 0 | 0 |
| rs781261962 | pathogenic | 0 | 0 | 0 |
| rs28942107 | pathogenic | 0 | 0 | 0 |
| rs80358258 | pathogenic | 0 | 0 | 0 |
| rs80358259 | pathogenic | 1 | 1 | 0 |
| rs141440861 | benign | 0 | 0 | 0 |
| rs28942106 | pathogenic | 0 | 0 | 0 |
| rs778878523 | pathogenic | 0 | 0 | 0 |
| rs28942105 | pathogenic | 0 | 0 | 0 |
| rs61731969 | benign | 0 | 0 | 0 |
| rs200444084 | pathogenic | 0 | 0 | 0 |
| rs113371321 | pathogenic | 0 | 0 | 0 |
| rs369098773 | pathogenic | 0 | 0 | 0 |
| rs35248744 | benign | 0 | 0 | 0 |
| rs758902805 | pathogenic | 0 | 0 | 0 |
| rs120074131 | pathogenic | 0 | 0 | 0 |
| rs1805084 | benign | 0 | 0 | 0 |
| rs117793540 | benign | 0 | 0 | 0 |
| rs530058063 | benign | 0 | 0 | 0 |
| rs1064422 | pathogenic | 1 | 1 | 1 |
| rs104894506 | pathogenic | 1 | 1 | 1 |
| rs104894507 | pathogenic | 1 | 1 | 1 |
| rs104894508 | pathogenic | 1 | 1 | 0 |
| rs28999113 | pathogenic | 1 | 1 | 0 |
| rs281860266 | pathogenic | 0 | 0 | 0 |
| rs104894961 | pathogenic | 1 | 1 | 0 |

| | | | | |
|---|---|---|---|---|
| rs122463168 | pathogenic | 0 | 0 | 0 |
| rs199469624 | pathogenic | 1 | 1 | 1 |
| rs150107747 | benign | 0 | 0 | 0 |
| rs144984628 | pathogenic | 0 | 0 | 0 |
| rs28935171 | pathogenic | 0 | 0 | 0 |
| rs137852443 | pathogenic | 1 | 1 | 0 |
| rs28933674 | pathogenic | 1 | 1 | 1 |
| rs28933675 | pathogenic | 1 | 1 | 1 |
| rs28933676 | pathogenic | 1 | 1 | 1 |
| rs28933677 | pathogenic | 1 | 1 | 1 |
| rs28933678 | pathogenic | 0 | 0 | 0 |
| rs28933679 | pathogenic | 1 | 1 | 1 |
| rs28933680 | pathogenic | 0 | 0 | 0 |
| rs28933681 | pathogenic | 0 | 0 | 0 |
| rs28933682 | pathogenic | 0 | 0 | 0 |
| rs28937294 | pathogenic | 1 | 1 | 0 |
| rs137852459 | pathogenic | 0 | 0 | 0 |
| rs137852461 | pathogenic | 0 | 0 | 0 |
| rs137852464 | pathogenic | 1 | 1 | 1 |
| rs137852465 | pathogenic | 1 | 1 | 1 |
| rs1800297 | benign | 1 | 1 | 0 |
| rs137852374 | pathogenic | 1 | 1 | 1 |
| rs672601312 | pathogenic | 1 | 1 | 1 |
| rs387907196 | pathogenic | 1 | 1 | 1 |
| rs148348866 | benign | 0 | 0 | 0 |
| rs141138948 | pathogenic | 1 | 1 | 1 |
| rs137852685 | pathogenic | 1 | 1 | 0 |
| rs104893900 | pathogenic | 1 | 1 | 0 |
| rs137852686 | pathogenic | 1 | 1 | 1 |
| rs104893906 | pathogenic | 1 | 1 | 0 |
| rs587782207 | pathogenic | 1 | 1 | 1 |
| rs75086406 | pathogenic | 0 | 0 | 0 |
| rs199822819 | pathogenic | 0 | 0 | 0 |
| rs398123166 | pathogenic | 1 | 1 | 1 |
| rs121913123 | pathogenic | 1 | 1 | 1 |
| rs772190176 | pathogenic | 0 | 1 | 0 |
| rs587782216 | pathogenic | 1 | 1 | 1 |
| rs398123159 | pathogenic | 1 | 1 | 1 |
| rs121913122 | pathogenic | 1 | 1 | 1 |
| rs755436052 | pathogenic | 1 | 1 | 1 |
| rs200796606 | pathogenic | 1 | 1 | 1 |
| rs2070080 | pathogenic | 0 | 1 | 0 |
| rs149616320 | benign | 0 | 0 | 0 |
| rs143529458 | benign | 0 | 0 | 0 |
| rs267607077 | pathogenic | 1 | 1 | 1 |
| rs143898031 | benign | 1 | 1 | 0 |
| rs148304857 | benign | 1 | 1 | 1 |
| rs137852809 | pathogenic | 0 | 0 | 0 |
| rs3765623 | benign | 0 | 0 | 0 |
| rs16944397 | benign | 0 | 0 | 0 |
| rs730880167 | benign | 0 | 0 | 0 |
| rs187108957 | benign | 0 | 0 | 0 |
| rs117342470 | benign | 0 | 0 | 0 |
| rs751200138 | pathogenic | 1 | 1 | 1 |

| | | | | |
|---|---|---|---|---|
| rs755799528 | pathogenic | 0 | 0 | 0 |
| rs537484504 | benign | 0 | 0 | 0 |
| rs748944640 | pathogenic | 0 | 0 | 0 |
| rs199774121 | pathogenic | 0 | 0 | 0 |
| rs766094434 | pathogenic | 1 | 1 | 0 |
| rs777321035 | pathogenic | 1 | 1 | 1 |
| rs146269774 | benign | 0 | 1 | 0 |
| rs144085805 | benign | 0 | 1 | 0 |
| rs367643028 | benign | 0 | 0 | 0 |
| rs42524 | benign | 0 | 0 | 0 |
| rs28940586 | pathogenic | 0 | 0 | 0 |
| rs74315327 | pathogenic | 1 | 1 | 1 |
| rs199676861 | pathogenic | 0 | 0 | 0 |
| rs115802744 | benign | 1 | 1 | 0 |
| rs752866643 | pathogenic | 1 | 1 | 1 |
| rs28941784 | pathogenic | 1 | 1 | 1 |
| rs398124434 | pathogenic | 1 | 1 | 1 |
| rs756414548 | pathogenic | 1 | 1 | 1 |
| rs376128990 | pathogenic | 1 | 1 | 1 |
| rs746219370 | pathogenic | 1 | 1 | 1 |
| rs756195708 | pathogenic | 1 | 1 | 1 |
| rs369296618 | pathogenic | 1 | 1 | 1 |
| rs9593 | benign | 1 | 1 | 1 |
| rs139510844 | benign | 0 | 0 | 0 |
| rs28382158 | benign | 0 | 0 | 0 |
| rs28919570 | pathogenic | 0 | 0 | 0 |
| rs587777331 | pathogenic | 0 | 0 | 0 |
| rs587777333 | pathogenic | 0 | 0 | 0 |
| rs587777332 | pathogenic | 0 | 0 | 0 |
| rs774980346 | pathogenic | 0 | 0 | 0 |
| rs370934093 | pathogenic | 0 | 0 | 0 |
| rs121913479 | pathogenic | 1 | 1 | 1 |
| rs121913484 | pathogenic | 0 | 0 | 0 |
| rs121913485 | pathogenic | 1 | 1 | 1 |
| rs28931614 | pathogenic | 1 | 1 | 1 |
| rs17881656 | benign | 1 | 1 | 1 |
| rs28931615 | pathogenic | 1 | 1 | 1 |
| rs147846832 | benign | 0 | 0 | 0 |
| rs80356506 | pathogenic | 0 | 0 | 0 |
| rs80356494 | pathogenic | 0 | 0 | 0 |
| rs121908894 | pathogenic | 0 | 0 | 0 |
| rs121908895 | pathogenic | 0 | 0 | 0 |
| rs80356508 | pathogenic | 0 | 0 | 0 |
| rs80356507 | pathogenic | 0 | 0 | 0 |
| rs121908896 | pathogenic | 0 | 0 | 0 |
| rs745372589 | pathogenic | 1 | 1 | 0 |
| rs183898408 | benign | 0 | 0 | 0 |
| rs200844292 | benign | 0 | 0 | 0 |
| rs146897002 | benign | 0 | 0 | 0 |
| rs121964869 | pathogenic | 1 | 1 | 1 |
| rs201768800 | benign | 1 | 1 | 1 |
| rs9807556 | benign | 1 | 1 | 1 |
| rs184721031 | benign | 0 | 0 | 0 |
| rs201104206 | benign | 0 | 0 | 0 |

| | | | | |
|---|---|---|---|---|
| rs188677538 | benign | 0 | 0 | 0 |
| rs149528866 | benign | 0 | 0 | 0 |
| rs115382168 | benign | 0 | 0 | 0 |
| rs78846023 | benign | 1 | 1 | 1 |
| rs61730662 | benign | 0 | 0 | 0 |
| rs121918368 | pathogenic | 0 | 0 | 0 |
| rs566433112 | pathogenic | 1 | 1 | 1 |
| rs121909342 | pathogenic | 0 | 0 | 0 |
| rs67586389 | pathogenic | 0 | 0 | 0 |
| rs72466485 | pathogenic | 0 | 0 | 0 |
| rs72466486 | pathogenic | 1 | 1 | 1 |
| rs72466487 | pathogenic | 1 | 1 | 1 |
| rs267607112 | pathogenic | 0 | 0 | 0 |
| rs118204013 | pathogenic | 0 | 0 | 0 |
| rs121913495 | pathogenic | 0 | 0 | 0 |
| rs11554273 | pathogenic | 0 | 0 | 0 |
| rs193922477 | pathogenic | 1 | 1 | 1 |
| rs193922479 | pathogenic | 1 | 1 | 1 |
| rs72547554 | pathogenic | 0 | 0 | 0 |
| rs72547561 | pathogenic | 1 | 1 | 1 |
| rs72547562 | pathogenic | 1 | 1 | 0 |
| rs115977487 | benign | 1 | 1 | 0 |
| rs72547564 | pathogenic | 0 | 0 | 0 |
| rs72547568 | pathogenic | 1 | 1 | 0 |
| rs72547569 | pathogenic | 1 | 1 | 1 |
| rs72547571 | pathogenic | 0 | 0 | 0 |
| rs72547575 | pathogenic | 1 | 1 | 1 |
| rs544994462 | benign | 0 | 0 | 0 |
| rs190470017 | benign | 0 | 0 | 0 |
| rs199473266 | pathogenic | 1 | 1 | 1 |
| rs137854602 | pathogenic | 0 | 0 | 0 |
| rs143739249 | pathogenic | 0 | 0 | 0 |
| rs397514697 | pathogenic | 1 | 1 | 1 |
| rs373764886 | pathogenic | 0 | 0 | 0 |
| rs370242930 | pathogenic | 0 | 0 | 0 |
| rs377145777 | pathogenic | 1 | 1 | 1 |
| rs191256186 | benign | 0 | 0 | 0 |
| rs150070901 | benign | 1 | 1 | 1 |
| rs138098032 | benign | 0 | 0 | 0 |
| rs117731813 | benign | 1 | 1 | 1 |
| rs104894488 | pathogenic | 1 | 1 | 1 |
| rs11302 | benign | 0 | 1 | 0 |
| rs121918447 | pathogenic | 1 | 1 | 1 |
| rs138205828 | benign | 1 | 1 | 0 |
| rs149858290 | benign | 0 | 0 | 0 |
| rs750764003 | pathogenic | 0 | 1 | 0 |
| rs141381896 | benign | 0 | 0 | 0 |
| rs149027682 | benign | 0 | 0 | 0 |
| rs121908415 | pathogenic | 0 | 0 | 0 |
| rs121908416 | pathogenic | 0 | 0 | 0 |
| rs121908417 | pathogenic | 0 | 0 | 0 |
| rs587780306 | benign | 0 | 0 | 0 |
| rs3218725 | benign | 0 | 0 | 0 |
| rs3218729 | benign | 0 | 0 | 0 |

| | | | | |
|---|---|---|---|---|
| rs387906917 | pathogenic | 0 | 0 | 0 |
| rs200652608 | pathogenic | 0 | 0 | 0 |
| rs3218721 | benign | 0 | 0 | 0 |
| rs63750869 | pathogenic | 1 | 1 | 1 |
| rs118203910 | pathogenic | 0 | 1 | 0 |
| rs9332701 | benign | 1 | 1 | 0 |
| rs6027 | benign | 0 | 0 | 0 |
| rs121909110 | pathogenic | 0 | 1 | 0 |
| rs762550967 | pathogenic | 1 | 1 | 0 |
| rs763615602 | pathogenic | 1 | 1 | 0 |
| rs144907403 | benign | 0 | 0 | 0 |
| rs104893631 | pathogenic | 1 | 1 | 0 |
| rs144181978 | pathogenic | 1 | 1 | 0 |
| rs74874677 | benign | 1 | 1 | 1 |
| rs104893632 | pathogenic | 0 | 1 | 0 |
| rs371342464 | benign | 1 | 1 | 1 |
| rs104893633 | pathogenic | 1 | 1 | 0 |
| rs201088712 | pathogenic | 0 | 0 | 0 |
| rs201857541 | benign | 1 | 1 | 1 |
| rs139517732 | pathogenic | 1 | 1 | 0 |
| rs191057824 | benign | 1 | 1 | 0 |
| rs35683768 | benign | 1 | 1 | 0 |
| rs104895271 | pathogenic | 0 | 0 | 0 |
| rs104895217 | pathogenic | 1 | 1 | 1 |
| rs104895223 | pathogenic | 1 | 1 | 1 |
| rs104895218 | pathogenic | 1 | 1 | 1 |
| rs104895225 | pathogenic | 1 | 1 | 1 |
| rs4149637 | benign | 1 | 1 | 1 |
| rs104895219 | pathogenic | 1 | 1 | 1 |
| rs104895220 | pathogenic | 1 | 1 | 1 |
| rs104895228 | pathogenic | 1 | 1 | 1 |
| rs104895221 | pathogenic | 1 | 1 | 1 |
| rs104895222 | pathogenic | 1 | 1 | 1 |
| rs104895247 | pathogenic | 0 | 0 | 0 |
| rs28940585 | pathogenic | 0 | 0 | 0 |
| rs561367199 | pathogenic | 1 | 1 | 1 |
| rs143793815 | benign | 0 | 0 | 0 |
| rs28941473 | pathogenic | 1 | 1 | 1 |
| rs751590073 | pathogenic | 1 | 1 | 1 |
| rs111407265 | benign | 1 | 1 | 1 |
| rs373697663 | pathogenic | 1 | 1 | 1 |
| rs369879957 | pathogenic | 0 | 0 | 0 |
| rs28941472 | pathogenic | 1 | 1 | 1 |
| rs28940287 | pathogenic | 1 | 1 | 1 |
| rs77625410 | benign | 0 | 0 | 0 |
| rs111033477 | benign | 0 | 0 | 0 |
| rs115123584 | benign | 0 | 0 | 0 |
| rs374576916 | benign | 0 | 0 | 0 |
| rs397516317 | pathogenic | 0 | 0 | 0 |
| rs368657015 | pathogenic | 0 | 0 | 0 |
| rs397516321 | pathogenic | 0 | 0 | 0 |
| rs397516322 | pathogenic | 0 | 0 | 0 |
| rs111033215 | pathogenic | 0 | 0 | 0 |
| rs199606180 | pathogenic | 0 | 0 | 0 |

| | | | | |
|---|---|---|---|---|
| rs111033192 | pathogenic | 0 | 0 | 0 |
| rs948962 | benign | 1 | 1 | 1 |
| rs142293185 | benign | 0 | 0 | 0 |
| rs376764423 | pathogenic | 0 | 0 | 0 |
| rs773844428 | pathogenic | 0 | 0 | 0 |
| rs186644871 | benign | 0 | 0 | 0 |
| rs200313391 | benign | 0 | 0 | 0 |
| rs377670513 | pathogenic | 0 | 0 | 0 |
| rs1132036 | benign | 0 | 0 | 0 |
| rs111033231 | benign | 0 | 0 | 0 |
| rs145945630 | pathogenic | 0 | 0 | 0 |
| rs121912984 | pathogenic | 1 | 1 | 1 |
| rs1800435 | benign | 0 | 1 | 0 |
| rs121912980 | pathogenic | 0 | 0 | 0 |
| rs121912982 | pathogenic | 1 | 1 | 1 |
| rs121912983 | pathogenic | 0 | 0 | 0 |
| rs121912981 | pathogenic | 0 | 0 | 0 |
| rs1057868 | benign | 0 | 0 | 0 |
| rs56048668 | benign | 0 | 0 | 0 |
| rs55865465 | benign | 0 | 0 | 0 |
| rs748096324 | benign | 1 | 1 | 0 |
| rs150475750 | benign | 0 | 0 | 0 |
| rs143025739 | benign | 1 | 1 | 0 |
| rs79702016 | benign | 1 | 1 | 0 |
| rs41338945 | benign | 0 | 0 | 0 |
| rs146406037 | benign | 1 | 1 | 1 |
| rs143274491 | benign | 0 | 0 | 0 |
| rs3829986 | benign | 0 | 0 | 0 |
| rs149168939 | benign | 0 | 0 | 0 |
| rs1800385 | benign | 0 | 0 | 0 |
| rs1800386 | pathogenic | 0 | 0 | 0 |
| rs121918524 | pathogenic | 1 | 1 | 1 |
| rs121912700 | pathogenic | 0 | 0 | 0 |
| rs104893930 | pathogenic | 1 | 1 | 1 |
| rs104893931 | pathogenic | 0 | 0 | 0 |
| rs116125936 | benign | 0 | 0 | 0 |
| rs627441 | benign | 0 | 0 | 0 |
| rs104894481 | pathogenic | 0 | 0 | 0 |
| rs137853129 | pathogenic | 0 | 0 | 0 |
| rs137853130 | pathogenic | 1 | 1 | 0 |
| rs28940282 | pathogenic | 0 | 0 | 0 |
| rs2229989 | benign | 0 | 0 | 0 |
| rs104894647 | pathogenic | 0 | 0 | 0 |
| rs1061170 | benign | 1 | 1 | 1 |
| rs576402053 | benign | 0 | 0 | 0 |
| rs34972246 | benign | 0 | 0 | 0 |
| rs192725607 | benign | 0 | 0 | 0 |
| rs137852615 | pathogenic | 1 | 1 | 0 |
| rs137852617 | pathogenic | 1 | 1 | 1 |
| rs137852611 | pathogenic | 0 | 0 | 0 |
| rs137852612 | pathogenic | 0 | 0 | 0 |
| rs137852614 | pathogenic | 1 | 1 | 1 |
| rs137852610 | pathogenic | 1 | 1 | 1 |
| rs137852618 | pathogenic | 1 | 1 | 1 |

| | | | | |
|---|---|---|---|---|
| rs137852613 | pathogenic | 1 | 1 | 0 |
| rs137852609 | pathogenic | 0 | 0 | 0 |
| rs63750306 | pathogenic | 1 | 1 | 1 |
| rs63750265 | pathogenic | 1 | 1 | 1 |
| rs63751287 | pathogenic | 1 | 1 | 1 |
| rs199723282 | benign | 0 | 0 | 0 |
| rs121917807 | pathogenic | 0 | 0 | 0 |
| rs121917809 | pathogenic | 0 | 0 | 0 |
| rs75134564 | pathogenic | 0 | 0 | 0 |
| rs267606930 | pathogenic | 1 | 1 | 0 |
| rs121909302 | pathogenic | 0 | 0 | 0 |
| rs121909303 | pathogenic | 0 | 0 | 0 |
| rs121909299 | pathogenic | 0 | 0 | 0 |
| rs121909301 | pathogenic | 0 | 0 | 0 |
| rs121909300 | pathogenic | 1 | 1 | 0 |
| rs111033589 | pathogenic | 0 | 0 | 0 |
| rs137853059 | pathogenic | 0 | 0 | 0 |
| rs137853057 | pathogenic | 1 | 1 | 1 |
| rs1801474 | benign | 0 | 0 | 0 |
| rs137853060 | pathogenic | 1 | 1 | 1 |
| rs137853058 | pathogenic | 1 | 1 | 1 |
| rs34424986 | pathogenic | 0 | 0 | 0 |
| rs199657839 | benign | 1 | 1 | 1 |
| rs56092260 | benign | 0 | 0 | 0 |
| rs1801582 | benign | 1 | 1 | 1 |
| rs1801334 | benign | 1 | 1 | 1 |
| rs191486604 | pathogenic | 0 | 0 | 0 |
| rs121908051 | pathogenic | 0 | 0 | 0 |
| rs121908055 | pathogenic | 1 | 1 | 0 |
| rs28940886 | pathogenic | 0 | 0 | 0 |
| rs121908050 | pathogenic | 0 | 0 | 0 |
| rs28940887 | pathogenic | 0 | 0 | 0 |
| rs387906300 | pathogenic | 0 | 0 | 0 |
| rs28942087 | pathogenic | 0 | 0 | 0 |
| rs121908049 | pathogenic | 1 | 1 | 0 |
| rs121908054 | pathogenic | 1 | 1 | 0 |
| rs121908048 | pathogenic | 0 | 0 | 0 |
| rs28940888 | pathogenic | 1 | 1 | 1 |
| rs121908053 | pathogenic | 1 | 1 | 0 |
| rs122467169 | pathogenic | 1 | 1 | 1 |
| rs122467172 | pathogenic | 1 | 1 | 1 |
| rs122467170 | pathogenic | 1 | 1 | 1 |
| rs28935477 | pathogenic | 1 | 1 | 0 |
| rs28942088 | pathogenic | 0 | 0 | 0 |
| rs751478142 | pathogenic | 1 | 1 | 0 |
| rs765368797 | pathogenic | 0 | 0 | 0 |
| rs147810437 | benign | 0 | 0 | 0 |
| rs121918209 | pathogenic | 1 | 1 | 1 |
| rs121918208 | pathogenic | 1 | 1 | 1 |
| rs121918210 | pathogenic | 1 | 1 | 1 |
| rs35515638 | benign | 1 | 1 | 1 |
| rs121918207 | pathogenic | 1 | 1 | 1 |
| rs141298312 | benign | 0 | 0 | 0 |
| rs121918211 | pathogenic | 0 | 0 | 0 |

| | | | | |
|---|---|---|---|---|
| rs149660059 | benign | 0 | 0 | 0 |
| rs200670286 | pathogenic | 1 | 1 | 1 |
| rs121918212 | pathogenic | 1 | 1 | 1 |
| rs121918213 | pathogenic | 1 | 1 | 1 |
| rs121918205 | pathogenic | 1 | 1 | 1 |
| rs104894025 | pathogenic | 1 | 1 | 1 |
| rs104894026 | pathogenic | 1 | 1 | 1 |
| rs104894028 | pathogenic | 0 | 1 | 0 |
| rs104894029 | pathogenic | 0 | 1 | 0 |
| rs119458971 | pathogenic | 0 | 0 | 0 |
| rs119458969 | pathogenic | 1 | 1 | 1 |
| rs1801591 | benign | 1 | 1 | 1 |
| rs140169311 | benign | 0 | 0 | 0 |
| rs753363278 | benign | 1 | 1 | 1 |
| rs199763682 | pathogenic | 0 | 0 | 0 |
| rs119458970 | pathogenic | 1 | 1 | 1 |
| rs763722868 | pathogenic | 0 | 0 | 0 |
| rs56404770 | benign | 0 | 0 | 0 |
| rs34221241 | benign | 0 | 0 | 0 |
| rs121909625 | pathogenic | 0 | 0 | 0 |
| rs550921485 | pathogenic | 0 | 0 | 0 |
| rs1800454 | benign | 0 | 0 | 0 |
| rs241447 | benign | 0 | 0 | 0 |
| rs2230606 | benign | 1 | 1 | 0 |
| rs5036 | pathogenic | 1 | 1 | 1 |
| rs13306788 | benign | 0 | 0 | 0 |
| rs28929480 | pathogenic | 0 | 0 | 0 |
| rs121912749 | pathogenic | 0 | 0 | 0 |
| rs56361140 | pathogenic | 0 | 0 | 0 |
| rs28931583 | pathogenic | 1 | 1 | 1 |
| rs36211715 | pathogenic | 1 | 1 | 1 |
| rs138049878 | pathogenic | 1 | 1 | 1 |
| rs727503253 | pathogenic | 1 | 1 | 1 |
| rs267606908 | pathogenic | 0 | 0 | 0 |
| rs121913631 | pathogenic | 1 | 1 | 1 |
| rs36211716 | benign | 1 | 1 | 1 |
| rs121913639 | pathogenic | 1 | 1 | 1 |
| rs730880761 | pathogenic | 1 | 1 | 1 |
| rs121913629 | pathogenic | 1 | 1 | 1 |
| rs754368658 | pathogenic | 0 | 0 | 0 |
| rs777090017 | pathogenic | 0 | 0 | 0 |
| rs267606913 | pathogenic | 1 | 1 | 1 |
| rs773138218 | pathogenic | 0 | 0 | 0 |
| rs371295780 | pathogenic | 0 | 0 | 0 |
| rs387906830 | pathogenic | 1 | 1 | 0 |
| rs771364038 | pathogenic | 0 | 1 | 0 |
| rs772731615 | pathogenic | 0 | 0 | 0 |
| rs137853063 | pathogenic | 0 | 0 | 0 |
| rs387907076 | pathogenic | 0 | 1 | 0 |
| rs35460768 | benign | 0 | 0 | 0 |
| rs2278196 | benign | 1 | 1 | 1 |
| rs119103275 | pathogenic | 0 | 0 | 0 |
| rs119103276 | pathogenic | 1 | 1 | 1 |
| rs374402066 | pathogenic | 1 | 1 | 1 |

| | | | | |
|---|---|---|---|---|
| rs119103274 | pathogenic | 0 | 0 | 0 |
| rs2274064 | benign | 0 | 0 | 0 |
| rs267607023 | pathogenic | 1 | 1 | 0 |
| rs142117489 | benign | 1 | 1 | 0 |
| rs371258270 | benign | 0 | 0 | 0 |
| rs34716057 | benign | 0 | 0 | 0 |
| rs201778606 | benign | 0 | 0 | 0 |
| rs201438792 | benign | 0 | 0 | 0 |
| rs140547417 | benign | 0 | 0 | 0 |
| rs5881 | benign | 0 | 0 | 0 |
| rs752298084 | benign | 0 | 0 | 0 |
| rs34855278 | benign | 0 | 0 | 0 |
| rs5880 | benign | 1 | 1 | 0 |
| rs5882 | benign | 0 | 0 | 0 |
| rs1800777 | benign | 0 | 0 | 0 |
| rs577301331 | benign | 0 | 0 | 0 |
| rs137853313 | pathogenic | 0 | 0 | 0 |
| rs28935169 | pathogenic | 0 | 0 | 0 |
| rs137853315 | pathogenic | 0 | 0 | 0 |
| rs137853314 | pathogenic | 0 | 0 | 0 |
| rs28935469 | pathogenic | 0 | 0 | 0 |
| rs137853318 | pathogenic | 0 | 0 | 0 |
| rs28935470 | pathogenic | 0 | 0 | 0 |
| rs3729986 | benign | 0 | 0 | 0 |
| rs730880619 | pathogenic | 0 | 0 | 0 |
| rs201012766 | benign | 0 | 0 | 0 |
| rs727503218 | pathogenic | 0 | 0 | 0 |
| rs11570052 | benign | 0 | 0 | 0 |
| rs730880623 | pathogenic | 0 | 0 | 0 |
| rs138753870 | benign | 0 | 0 | 0 |
| rs397516068 | pathogenic | 0 | 0 | 0 |
| rs3729989 | benign | 1 | 1 | 1 |
| rs397516074 | pathogenic | 0 | 0 | 0 |
| rs561449819 | pathogenic | 0 | 0 | 0 |
| rs121917779 | benign | 0 | 0 | 0 |
| rs142881576 | pathogenic | 0 | 0 | 0 |
| rs77384282 | benign | 0 | 0 | 0 |
| rs201865787 | benign | 0 | 0 | 0 |
| rs142244640 | benign | 0 | 0 | 0 |
| rs200488706 | benign | 0 | 0 | 0 |
| rs137853291 | pathogenic | 1 | 1 | 0 |
| rs28933990 | pathogenic | 1 | 1 | 0 |
| rs140570189 | benign | 0 | 0 | 0 |
| rs201588178 | benign | 0 | 0 | 0 |
| rs104893846 | pathogenic | 0 | 0 | 0 |
| rs754545360 | pathogenic | 0 | 1 | 0 |
| rs104893851 | pathogenic | 0 | 1 | 0 |
| rs104893849 | pathogenic | 0 | 0 | 0 |
| rs571038432 | pathogenic | 1 | 1 | 0 |
| rs2270655 | benign | 1 | 1 | 1 |
| rs315952 | benign | 0 | 0 | 0 |
| rs151176548 | benign | 0 | 0 | 0 |
| rs122459147 | pathogenic | 1 | 1 | 0 |
| rs122458140 | pathogenic | 0 | 0 | 0 |

| | | | | |
|---|---|---|---|---|
| rs122458142 | pathogenic | 1 | 1 | 0 |
| rs267606812 | pathogenic | 1 | 1 | 0 |
| rs267606813 | pathogenic | 1 | 1 | 0 |
| rs122458143 | pathogenic | 0 | 0 | 0 |
| rs122459146 | pathogenic | 0 | 0 | 0 |
| rs122458144 | pathogenic | 0 | 0 | 0 |
| rs122458145 | pathogenic | 0 | 0 | 0 |
| rs113994007 | pathogenic | 1 | 1 | 0 |
| rs758746181 | pathogenic | 1 | 1 | 1 |
| rs145034527 | pathogenic | 1 | 1 | 0 |
| rs119468006 | pathogenic | 1 | 1 | 0 |
| rs201908721 | pathogenic | 0 | 0 | 0 |
| rs755933881 | pathogenic | 1 | 1 | 0 |
| rs771578775 | pathogenic | 0 | 0 | 0 |
| rs17849927 | benign | 0 | 0 | 0 |
| rs139133094 | benign | 0 | 0 | 0 |
| rs12593 | benign | 0 | 0 | 0 |
| rs578189699 | pathogenic | 1 | 1 | 0 |
| rs119468008 | pathogenic | 0 | 0 | 0 |
| rs768524626 | benign | 0 | 0 | 0 |
| rs119468009 | pathogenic | 0 | 0 | 0 |
| rs119468004 | pathogenic | 0 | 0 | 0 |
| rs199874519 | pathogenic | 0 | 0 | 0 |
| rs376478331 | pathogenic | 0 | 0 | 0 |
| rs61995958 | pathogenic | 0 | 0 | 0 |
| rs199714731 | benign | 0 | 0 | 0 |
| rs149185954 | benign | 0 | 0 | 0 |
| rs121918353 | pathogenic | 0 | 0 | 0 |
| rs1131695 | benign | 0 | 0 | 0 |
| rs28939668 | pathogenic | 0 | 0 | 0 |
| rs758432471 | pathogenic | 1 | 1 | 1 |
| rs752746786 | pathogenic | 1 | 1 | 1 |
| rs28935471 | pathogenic | 0 | 0 | 0 |
| rs80338837 | pathogenic | 0 | 0 | 0 |
| rs137853312 | pathogenic | 0 | 0 | 0 |
| rs28935472 | pathogenic | 0 | 0 | 0 |
| rs28935473 | pathogenic | 0 | 0 | 0 |
| rs267606752 | pathogenic | 0 | 0 | 0 |
| rs3917973 | benign | 0 | 0 | 0 |
| rs3918018 | benign | 0 | 0 | 0 |
| rs757976755 | pathogenic | 0 | 0 | 0 |
| rs121918329 | pathogenic | 1 | 1 | 0 |
| rs770114459 | pathogenic | 0 | 0 | 0 |
| rs6662272 | benign | 0 | 0 | 0 |
| rs121918242 | pathogenic | 1 | 1 | 0 |
| rs121918240 | pathogenic | 1 | 1 | 0 |
| rs200094982 | pathogenic | 0 | 0 | 0 |
| rs121918241 | pathogenic | 0 | 0 | 0 |
| rs140522266 | pathogenic | 1 | 1 | 0 |
| rs121918243 | pathogenic | 1 | 1 | 0 |
| rs370596113 | pathogenic | 1 | 1 | 0 |
| rs398124295 | pathogenic | 1 | 1 | 0 |
| rs587776889 | pathogenic | 1 | 1 | 0 |
| rs747527726 | pathogenic | 1 | 1 | 0 |

| | | | | |
|---|---|---|---|---|
| rs112631212 | benign | 0 | 0 | 0 |
| rs104894605 | pathogenic | 0 | 0 | 0 |
| rs104894607 | pathogenic | 0 | 0 | 0 |
| rs104894606 | pathogenic | 0 | 0 | 0 |
| rs143873873 | benign | 0 | 0 | 0 |
| rs104894388 | pathogenic | 1 | 1 | 0 |
| rs104894391 | pathogenic | 1 | 1 | 0 |
| rs5350 | benign | 1 | 1 | 1 |
| rs104894390 | pathogenic | 1 | 1 | 1 |
| rs5346 | benign | 1 | 1 | 1 |
| rs121912500 | pathogenic | 0 | 0 | 0 |
| rs74315450 | pathogenic | 1 | 1 | 0 |
| rs121913396 | pathogenic | 1 | 1 | 1 |
| rs28931588 | pathogenic | 1 | 1 | 1 |
| rs121913400 | pathogenic | 0 | 0 | 0 |
| rs28931589 | pathogenic | 1 | 1 | 1 |
| rs121913403 | pathogenic | 0 | 0 | 0 |
| rs121913413 | pathogenic | 0 | 0 | 0 |
| rs760107623 | pathogenic | 1 | 1 | 0 |
| rs16866378 | benign | 1 | 1 | 0 |
| rs201025290 | pathogenic | 1 | 1 | 0 |
| rs12720356 | benign | 1 | 1 | 0 |
| rs55882956 | benign | 0 | 1 | 0 |
| rs4986933 | benign | 1 | 1 | 0 |
| rs2020957 | benign | 1 | 1 | 1 |
| rs1800124 | benign | 1 | 1 | 1 |
| rs150077735 | benign | 1 | 1 | 1 |
| rs201419000 | benign | 0 | 0 | 0 |
| rs772044176 | pathogenic | 0 | 0 | 0 |
| rs138346388 | benign | 0 | 0 | 0 |
| rs121964916 | pathogenic | 0 | 0 | 0 |
| rs112534524 | pathogenic | 0 | 0 | 0 |
| rs182078921 | benign | 1 | 1 | 1 |
| rs113273712 | benign | 0 | 0 | 0 |
| rs74817407 | benign | 1 | 1 | 1 |
| rs61733901 | benign | 1 | 1 | 1 |
| rs121964912 | pathogenic | 1 | 1 | 1 |
| rs41278047 | benign | 1 | 1 | 1 |
| rs7437875 | benign | 1 | 1 | 1 |
| rs113460688 | benign | 1 | 1 | 1 |
| rs754572081 | benign | 1 | 1 | 1 |
| rs201082692 | benign | 1 | 1 | 1 |
| rs202085803 | benign | 1 | 1 | 1 |
| rs141320511 | benign | 0 | 0 | 0 |
| rs78501403 | benign | 0 | 0 | 0 |
| rs28939675 | pathogenic | 0 | 0 | 0 |
| rs74315522 | pathogenic | 0 | 0 | 0 |
| rs121434639 | pathogenic | 1 | 1 | 0 |
| rs121434640 | pathogenic | 1 | 1 | 1 |
| rs757282628 | benign | 1 | 1 | 1 |
| rs386834010 | pathogenic | 0 | 0 | 0 |
| rs200863680 | pathogenic | 0 | 0 | 0 |
| rs193919336 | pathogenic | 1 | 1 | 0 |
| rs386834011 | pathogenic | 0 | 0 | 0 |

| | | | | |
|---|---|---|---|---|
| rs28940869 | pathogenic | 0 | 0 | 0 |
| rs386834014 | pathogenic | 1 | 1 | 0 |
| rs267606960 | pathogenic | 0 | 0 | 0 |
| rs28942068 | pathogenic | 0 | 0 | 0 |
| rs193919335 | pathogenic | 0 | 0 | 0 |
| rs74374973 | pathogenic | 0 | 0 | 0 |
| rs386834018 | pathogenic | 0 | 0 | 0 |
| rs267606962 | pathogenic | 0 | 0 | 0 |
| rs6659553 | benign | 0 | 0 | 0 |
| rs29281 | benign | 0 | 0 | 0 |
| rs782265090 | benign | 0 | 0 | 0 |
| rs35910969 | pathogenic | 0 | 0 | 0 |
| rs121909113 | pathogenic | 1 | 1 | 1 |
| rs29001571 | pathogenic | 1 | 1 | 1 |
| rs587781250 | pathogenic | 1 | 1 | 1 |
| rs28939680 | pathogenic | 1 | 1 | 1 |
| rs28939681 | pathogenic | 1 | 1 | 1 |
| rs121909112 | pathogenic | 1 | 1 | 1 |
| rs28937568 | pathogenic | 1 | 1 | 1 |
| rs1799895 | pathogenic | 0 | 0 | 0 |
| rs386833431 | pathogenic | 1 | 1 | 1 |
| rs121964905 | pathogenic | 1 | 1 | 1 |
| rs121964906 | pathogenic | 1 | 1 | 1 |
| rs11551664 | pathogenic | 1 | 1 | 1 |
| rs104894705 | pathogenic | 1 | 1 | 1 |
| rs104893805 | pathogenic | 0 | 0 | 0 |
| rs104894205 | pathogenic | 0 | 0 | 0 |
| rs104894204 | pathogenic | 1 | 1 | 0 |
| rs62642926 | pathogenic | 0 | 0 | 0 |
| rs5030841 | pathogenic | 1 | 1 | 1 |
| rs75193786 | pathogenic | 1 | 1 | 1 |
| rs62508682 | pathogenic | 0 | 0 | 0 |
| rs3850625 | benign | 1 | 1 | 1 |
| rs141908793 | benign | 0 | 0 | 0 |
| rs187864727 | benign | 0 | 0 | 0 |
| rs548909101 | benign | 0 | 0 | 0 |
| rs3825394 | benign | 0 | 1 | 0 |
| rs143548402 | benign | 0 | 0 | 0 |
| rs267607146 | pathogenic | 0 | 0 | 0 |
| rs267607144 | pathogenic | 0 | 0 | 0 |
| rs267607148 | pathogenic | 1 | 1 | 1 |
| rs267607143 | pathogenic | 0 | 0 | 0 |
| rs267607145 | pathogenic | 0 | 0 | 0 |
| rs121912636 | pathogenic | 1 | 1 | 1 |
| rs121912634 | pathogenic | 1 | 1 | 1 |
| rs570675468 | benign | 0 | 0 | 0 |
| rs368861241 | pathogenic | 1 | 1 | 1 |
| rs104894528 | pathogenic | 1 | 1 | 1 |
| rs77493891 | benign | 0 | 0 | 0 |
| rs62640903 | benign | 0 | 0 | 0 |
| rs149926091 | benign | 1 | 1 | 1 |
| rs641081 | benign | 0 | 0 | 0 |
| rs4930199 | benign | 0 | 0 | 0 |
| rs121913020 | pathogenic | 0 | 0 | 0 |

| | | | | |
|---|---|---|---|---|
| rs1799793 | benign | 0 | 0 | 0 |
| rs121913016 | pathogenic | 0 | 0 | 0 |
| rs121913025 | pathogenic | 0 | 0 | 0 |
| rs121913019 | pathogenic | 0 | 0 | 0 |
| rs376556895 | pathogenic | 1 | 1 | 1 |
| rs121913024 | pathogenic | 1 | 1 | 1 |
| rs121913021 | pathogenic | 0 | 0 | 0 |
| rs121913023 | pathogenic | 0 | 0 | 0 |
| rs41556519 | pathogenic | 0 | 0 | 0 |
| rs121913022 | pathogenic | 0 | 0 | 0 |
| rs144564120 | pathogenic | 0 | 0 | 0 |
| rs121913026 | pathogenic | 1 | 1 | 1 |
| rs121913018 | pathogenic | 1 | 1 | 1 |
| rs142954643 | benign | 0 | 0 | 0 |
| rs76655496 | benign | 0 | 0 | 0 |
| rs104894371 | pathogenic | 1 | 1 | 0 |
| rs137964936 | benign | 0 | 0 | 0 |
| rs759736132 | pathogenic | 0 | 0 | 0 |
| rs387906595 | pathogenic | 1 | 1 | 0 |
| rs768248216 | pathogenic | 0 | 0 | 0 |
| rs548231008 | benign | 0 | 0 | 0 |
| rs757094384 | benign | 0 | 0 | 0 |
| rs867529 | benign | 0 | 0 | 0 |
| rs13045 | benign | 1 | 1 | 1 |
| rs121908570 | pathogenic | 0 | 0 | 0 |
| rs201872586 | benign | 1 | 1 | 0 |
| rs149092990 | benign | 0 | 0 | 0 |
| rs115585711 | benign | 0 | 0 | 0 |
| rs137909416 | benign | 1 | 1 | 0 |
| rs143388949 | benign | 1 | 1 | 1 |
| rs73137716 | benign | 1 | 1 | 1 |
| rs121908947 | pathogenic | 1 | 1 | 1 |
| rs398122904 | pathogenic | 1 | 1 | 1 |
| rs35715497 | benign | 1 | 1 | 0 |
| rs35462421 | benign | 0 | 0 | 0 |
| rs119475043 | pathogenic | 0 | 0 | 0 |
| rs119475042 | pathogenic | 0 | 0 | 0 |
| rs1076669 | benign | 1 | 1 | 0 |
| rs141146885 | benign | 0 | 0 | 0 |
| rs3026906 | pathogenic | 0 | 0 | 0 |
| rs75362221 | benign | 1 | 1 | 1 |
| rs138472116 | benign | 1 | 1 | 1 |
| rs371040282 | pathogenic | 1 | 1 | 1 |
| rs4988496 | benign | 0 | 0 | 0 |
| rs121918117 | pathogenic | 1 | 1 | 1 |
| rs4988498 | benign | 0 | 0 | 0 |
| rs5035 | benign | 0 | 0 | 0 |
| rs121918322 | pathogenic | 1 | 1 | 1 |
| rs370359540 | benign | 0 | 0 | 0 |
| rs758972589 | pathogenic | 0 | 0 | 0 |
| rs144856889 | benign | 0 | 0 | 0 |
| rs121434623 | pathogenic | 1 | 1 | 1 |
| rs61735600 | benign | 0 | 0 | 0 |
| rs72550870 | benign | 1 | 1 | 1 |

| | | | | |
|---|---|---|---|---|
| rs56392418 | benign | 0 | 0 | 0 |
| rs41307788 | benign | 1 | 1 | 1 |
| rs387907239 | pathogenic | 1 | 1 | 0 |
| rs28931584 | pathogenic | 0 | 0 | 0 |
| rs121912745 | pathogenic | 0 | 0 | 0 |
| rs121912744 | pathogenic | 0 | 0 | 0 |
| rs121912754 | pathogenic | 1 | 1 | 1 |
| rs121912746 | pathogenic | 0 | 0 | 0 |
| rs121912757 | pathogenic | 0 | 0 | 0 |
| rs121912758 | pathogenic | 0 | 0 | 0 |
| rs121912748 | pathogenic | 0 | 0 | 0 |
| rs121912755 | pathogenic | 0 | 0 | 0 |
| rs121912741 | pathogenic | 0 | 0 | 0 |
| rs121912753 | pathogenic | 0 | 0 | 0 |
| rs121912750 | pathogenic | 0 | 0 | 0 |
| rs2285644 | pathogenic | 0 | 0 | 0 |
| rs121912751 | pathogenic | 0 | 0 | 0 |
| rs5026 | benign | 0 | 0 | 0 |
| rs121912759 | pathogenic | 0 | 0 | 0 |
| rs28931585 | pathogenic | 0 | 0 | 0 |
| rs17122769 | benign | 0 | 0 | 0 |
| rs28362322 | benign | 1 | 1 | 1 |
| rs3824874 | benign | 0 | 0 | 0 |
| rs146004831 | benign | 0 | 0 | 0 |
| rs142155860 | benign | 0 | 0 | 0 |
| rs121434404 | pathogenic | 0 | 0 | 0 |
| rs61735578 | benign | 1 | 1 | 1 |
| rs558018 | benign | 0 | 0 | 0 |
| rs61735577 | benign | 1 | 1 | 1 |
| rs76784113 | benign | 0 | 0 | 0 |
| rs199469640 | pathogenic | 0 | 0 | 0 |
| rs199469628 | pathogenic | 1 | 1 | 0 |
| rs199469629 | pathogenic | 0 | 1 | 0 |
| rs387907156 | pathogenic | 0 | 1 | 0 |
| rs199469643 | pathogenic | 0 | 0 | 0 |
| rs199469646 | pathogenic | 0 | 0 | 0 |
| rs766294940 | pathogenic | 0 | 0 | 0 |
| rs387907347 | pathogenic | 0 | 0 | 0 |
| rs137853590 | pathogenic | 1 | 1 | 1 |
| rs137853589 | pathogenic | 1 | 1 | 1 |
| rs137853588 | pathogenic | 1 | 1 | 1 |
| rs55909005 | benign | 0 | 0 | 0 |
| rs4581 | benign | 0 | 0 | 0 |
| rs63750123 | benign | 0 | 0 | 0 |
| rs10254120 | benign | 0 | 0 | 0 |
| rs121434629 | pathogenic | 1 | 1 | 0 |
| rs200313585 | benign | 0 | 0 | 0 |
| rs143162541 | benign | 1 | 1 | 0 |
| rs760615315 | benign | 0 | 0 | 0 |
| rs768488890 | benign | 1 | 1 | 1 |
| rs116373169 | benign | 1 | 1 | 0 |
| rs63750871 | pathogenic | 1 | 1 | 0 |
| rs116349687 | benign | 0 | 0 | 0 |
| rs375289386 | benign | 0 | 0 | 0 |

| | | | | |
|---|---|---|---|---|
| rs760228510 | pathogenic | 1 | 1 | 1 |
| rs587779343 | pathogenic | 1 | 1 | 1 |
| rs573125799 | benign | 0 | 0 | 0 |
| rs1805322 | benign | 0 | 0 | 0 |
| rs267608153 | pathogenic | 0 | 0 | 0 |
| rs139194813 | benign | 0 | 0 | 0 |
| rs530021751 | benign | 0 | 0 | 0 |
| rs200640585 | pathogenic | 0 | 0 | 0 |
| rs139438201 | benign | 0 | 0 | 0 |
| rs200513014 | benign | 0 | 0 | 0 |
| rs567102013 | benign | 0 | 0 | 0 |
| rs121908608 | pathogenic | 0 | 0 | 0 |
| rs121908606 | pathogenic | 0 | 0 | 0 |
| rs17103671 | benign | 0 | 0 | 0 |
| rs34725281 | benign | 0 | 0 | 0 |
| rs104894421 | pathogenic | 1 | 1 | 0 |
| rs121918145 | pathogenic | 1 | 1 | 1 |
| rs121918152 | pathogenic | 1 | 1 | 1 |
| rs56379106 | benign | 0 | 0 | 0 |
| rs4987076 | benign | 0 | 0 | 0 |
| rs370243877 | pathogenic | 0 | 0 | 0 |
| rs17679445 | benign | 1 | 1 | 1 |
| rs550423482 | pathogenic | 1 | 1 | 1 |
| rs104894629 | pathogenic | 1 | 1 | 1 |
| rs773450573 | pathogenic | 1 | 1 | 1 |
| rs116679717 | benign | 0 | 0 | 0 |
| rs372000848 | pathogenic | 0 | 0 | 0 |
| rs138595973 | benign | 0 | 0 | 0 |
| rs115244417 | benign | 0 | 0 | 0 |
| rs11575292 | benign | 0 | 0 | 0 |
| rs137853211 | pathogenic | 1 | 1 | 1 |
| rs137853207 | pathogenic | 0 | 0 | 0 |
| rs137853210 | pathogenic | 1 | 1 | 1 |
| rs6262 | benign | 0 | 0 | 0 |
| rs6263 | benign | 0 | 1 | 0 |
| rs137853208 | pathogenic | 0 | 1 | 0 |
| rs137853212 | pathogenic | 1 | 1 | 0 |
| rs137853209 | pathogenic | 1 | 1 | 1 |
| rs201951824 | pathogenic | 0 | 0 | 0 |
| rs11575542 | benign | 0 | 0 | 0 |
| rs199681788 | benign | 0 | 1 | 0 |
| rs748835902 | benign | 1 | 1 | 1 |
| rs753719949 | benign | 0 | 0 | 0 |
| rs267606647 | pathogenic | 0 | 0 | 0 |
| rs267606648 | pathogenic | 1 | 1 | 1 |
| rs267606643 | pathogenic | 0 | 1 | 0 |
| rs559947967 | pathogenic | 1 | 1 | 0 |
| rs267606645 | pathogenic | 1 | 1 | 1 |
| rs12116440 | benign | 0 | 0 | 0 |
| rs121908539 | pathogenic | 0 | 0 | 0 |
| rs137853096 | pathogenic | 1 | 1 | 0 |
| rs148363262 | benign | 1 | 1 | 0 |
| rs587777442 | pathogenic | 0 | 0 | 0 |
| rs25640 | benign | 1 | 1 | 1 |

| | | | | |
|---|---|---|---|---|
| rs28943589 | benign | 0 | 0 | 0 |
| rs1143650 | benign | 1 | 1 | 1 |
| rs150886026 | benign | 0 | 1 | 0 |
| rs121908133 | pathogenic | 1 | 1 | 0 |
| rs121908131 | pathogenic | 0 | 0 | 0 |
| rs754477154 | pathogenic | 0 | 0 | 0 |
| rs747334007 | benign | 0 | 0 | 0 |
| rs121908132 | pathogenic | 0 | 0 | 0 |
| rs104894103 | pathogenic | 1 | 1 | 0 |
| rs140088149 | benign | 0 | 0 | 0 |
| rs149803148 | benign | 0 | 0 | 0 |
| rs11064480 | benign | 0 | 0 | 0 |
| rs74435397 | pathogenic | 1 | 1 | 1 |
| rs587783626 | pathogenic | 1 | 1 | 1 |
| rs148211042 | pathogenic | 1 | 1 | 1 |
| rs17849502 | benign | 0 | 0 | 0 |
| rs13306575 | pathogenic | 1 | 1 | 1 |
| rs201426733 | benign | 0 | 0 | 0 |
| rs182278784 | benign | 1 | 1 | 1 |
| rs77786415 | benign | 1 | 1 | 1 |
| rs200192119 | benign | 1 | 1 | 1 |
| rs12877501 | benign | 1 | 1 | 1 |
| rs77985244 | benign | 1 | 1 | 1 |
| rs5907 | pathogenic | 1 | 1 | 0 |
| rs2305178 | benign | 0 | 0 | 0 |
| rs3135867 | benign | 0 | 0 | 0 |
| rs121913482 | pathogenic | 1 | 1 | 1 |
| rs121913483 | pathogenic | 1 | 1 | 1 |
| rs4647924 | pathogenic | 1 | 1 | 1 |
| rs121913111 | pathogenic | 1 | 1 | 1 |
| rs104894002 | pathogenic | 0 | 0 | 0 |
| rs104894001 | pathogenic | 0 | 0 | 0 |
| rs75932628 | benign | 0 | 0 | 0 |
| rs201258663 | pathogenic | 0 | 0 | 0 |
| rs104893998 | pathogenic | 0 | 0 | 0 |
| rs2234253 | benign | 1 | 1 | 0 |
| rs121908402 | pathogenic | 1 | 1 | 0 |
| rs1805389 | benign | 0 | 0 | 0 |
| rs1805388 | benign | 0 | 0 | 0 |
| rs748385144 | pathogenic | 0 | 0 | 0 |
| rs104894420 | pathogenic | 0 | 0 | 0 |
| rs104894418 | pathogenic | 0 | 0 | 0 |
| rs61731910 | benign | 0 | 0 | 0 |
| rs758723288 | pathogenic | 1 | 1 | 0 |
| rs201865375 | pathogenic | 1 | 1 | 0 |
| rs565090080 | pathogenic | 1 | 1 | 1 |
| rs113042166 | benign | 0 | 0 | 0 |
| rs61736326 | benign | 1 | 1 | 1 |
| rs12030699 | benign | 1 | 1 | 1 |
| rs121913578 | pathogenic | 1 | 1 | 1 |
| rs61739582 | benign | 0 | 0 | 0 |
| rs745983207 | pathogenic | 0 | 0 | 0 |
| rs763524380 | benign | 0 | 0 | 0 |
| rs121912792 | pathogenic | 0 | 0 | 0 |

| | | | | |
|---|---|---|---|---|
| rs121912793 | pathogenic | 0 | 0 | 0 |
| rs387906599 | pathogenic | 0 | 0 | 0 |
| rs121912794 | pathogenic | 0 | 1 | 0 |
| rs121912795 | pathogenic | 0 | 0 | 0 |
| rs121912796 | pathogenic | 1 | 1 | 0 |
| rs121912797 | pathogenic | 0 | 1 | 0 |
| rs121912798 | pathogenic | 1 | 1 | 1 |
| rs80359812 | pathogenic | 1 | 1 | 0 |
| rs776583130 | benign | 1 | 1 | 0 |
| rs80359814 | pathogenic | 1 | 1 | 0 |
| rs202060209 | pathogenic | 1 | 1 | 0 |
| rs267607061 | pathogenic | 1 | 1 | 0 |
| rs267607060 | pathogenic | 0 | 0 | 0 |
| rs80359816 | pathogenic | 1 | 1 | 0 |
| rs80359818 | pathogenic | 1 | 1 | 0 |
| rs387907313 | pathogenic | 0 | 0 | 0 |
| rs587784395 | benign | 0 | 0 | 0 |
| rs5811 | benign | 0 | 0 | 0 |
| rs121909740 | pathogenic | 1 | 1 | 0 |
| rs148518827 | benign | 1 | 1 | 0 |
| rs121909739 | pathogenic | 1 | 1 | 0 |
| rs780528770 | benign | 1 | 1 | 0 |
| rs13306758 | pathogenic | 0 | 0 | 0 |
| rs267607059 | pathogenic | 0 | 0 | 0 |
| rs61730638 | benign | 0 | 0 | 0 |
| rs104894533 | pathogenic | 0 | 0 | 0 |
| rs104894527 | pathogenic | 0 | 0 | 0 |
| rs200503569 | pathogenic | 1 | 1 | 1 |
| rs80338700 | pathogenic | 1 | 1 | 1 |
| rs765433263 | pathogenic | 1 | 1 | 1 |
| rs104894530 | pathogenic | 1 | 1 | 1 |
| rs80338701 | pathogenic | 1 | 1 | 1 |
| rs141498002 | pathogenic | 1 | 1 | 1 |
| rs104894525 | pathogenic | 0 | 0 | 0 |
| rs80338703 | pathogenic | 0 | 0 | 0 |
| rs28936415 | pathogenic | 1 | 1 | 0 |
| rs150719105 | pathogenic | 0 | 0 | 0 |
| rs148032587 | pathogenic | 0 | 0 | 0 |
| rs150577656 | pathogenic | 0 | 0 | 0 |
| rs190521996 | pathogenic | 0 | 0 | 0 |
| rs104894526 | pathogenic | 1 | 1 | 1 |
| rs80338704 | pathogenic | 0 | 0 | 0 |
| rs199562225 | pathogenic | 1 | 1 | 1 |
| rs532870929 | pathogenic | 0 | 0 | 0 |
| rs398123309 | pathogenic | 0 | 0 | 0 |
| rs78290141 | pathogenic | 0 | 0 | 0 |
| rs80338705 | pathogenic | 0 | 0 | 0 |
| rs104894531 | pathogenic | 1 | 1 | 1 |
| rs80338706 | pathogenic | 0 | 0 | 0 |
| rs80338707 | pathogenic | 0 | 0 | 0 |
| rs80338708 | pathogenic | 0 | 0 | 0 |
| rs80338709 | pathogenic | 0 | 0 | 0 |
| rs104893833 | benign | 0 | 0 | 0 |
| rs61744585 | benign | 0 | 0 | 0 |

| | | | | |
|---|---|---|---|---|
| rs73569515 | benign | 0 | 0 | 0 |
| rs398123373 | pathogenic | 0 | 0 | 0 |
| rs756854513 | pathogenic | 0 | 0 | 0 |
| rs58232698 | benign | 0 | 0 | 0 |
| rs9600455 | benign | 0 | 0 | 0 |
| rs201016593 | pathogenic | 0 | 0 | 0 |
| rs756039188 | pathogenic | 0 | 0 | 0 |
| rs121908024 | pathogenic | 0 | 0 | 0 |
| rs373144619 | benign | 0 | 0 | 0 |
| rs138078086 | benign | 0 | 0 | 0 |
| rs376207800 | pathogenic | 0 | 0 | 0 |
| rs749038326 | pathogenic | 1 | 1 | 0 |
| rs372845091 | benign | 0 | 0 | 0 |
| rs377271627 | pathogenic | 0 | 0 | 0 |
| rs563390335 | pathogenic | 0 | 0 | 0 |
| rs13306510 | benign | 0 | 0 | 0 |
| rs764042910 | pathogenic | 0 | 0 | 0 |
| rs201384282 | benign | 1 | 1 | 1 |
| rs730882086 | pathogenic | 1 | 1 | 1 |
| rs756613387 | pathogenic | 1 | 1 | 1 |
| rs121908028 | benign | 1 | 1 | 0 |
| rs773328511 | benign | 0 | 0 | 0 |
| rs749220643 | pathogenic | 1 | 1 | 0 |
| rs730882089 | pathogenic | 0 | 0 | 0 |
| rs146651743 | benign | 0 | 0 | 0 |
| rs148698650 | pathogenic | 0 | 0 | 0 |
| rs140241383 | benign | 0 | 0 | 0 |
| rs757252110 | pathogenic | 0 | 0 | 0 |
| rs13306512 | pathogenic | 1 | 1 | 1 |
| rs121908043 | pathogenic | 1 | 1 | 1 |
| rs773658037 | pathogenic | 0 | 0 | 0 |
| rs774439908 | pathogenic | 0 | 0 | 0 |
| rs779732323 | pathogenic | 1 | 1 | 0 |
| rs749780672 | pathogenic | 0 | 0 | 0 |
| rs375651668 | benign | 1 | 1 | 1 |
| rs193922568 | pathogenic | 1 | 1 | 0 |
| rs750363970 | benign | 1 | 1 | 0 |
| rs5932 | benign | 0 | 0 | 0 |
| rs755154048 | pathogenic | 1 | 1 | 0 |
| rs759876319 | pathogenic | 0 | 0 | 0 |
| rs764550980 | pathogenic | 1 | 1 | 0 |
| rs779940524 | pathogenic | 0 | 0 | 0 |
| rs563382937 | pathogenic | 1 | 1 | 1 |
| rs28942083 | pathogenic | 0 | 1 | 0 |
| rs770744861 | pathogenic | 0 | 1 | 0 |
| rs121908031 | pathogenic | 0 | 1 | 0 |
| rs776217028 | pathogenic | 0 | 1 | 0 |
| rs763625913 | pathogenic | 0 | 0 | 0 |
| rs3736772 | benign | 0 | 0 | 0 |
| rs104894696 | benign | 1 | 1 | 1 |
| rs121918177 | pathogenic | 0 | 1 | 0 |
| rs121918181 | pathogenic | 1 | 1 | 0 |
| rs121918174 | pathogenic | 0 | 1 | 0 |
| rs121918182 | pathogenic | 0 | 0 | 0 |

| | | | | |
|---|---|---|---|---|
| rs121918175 | pathogenic | 1 | 1 | 1 |
| rs121918185 | pathogenic | 1 | 1 | 1 |
| rs121918173 | pathogenic | 1 | 1 | 0 |
| rs121918178 | pathogenic | 1 | 1 | 1 |
| rs121918179 | pathogenic | 1 | 1 | 1 |
| rs121918183 | pathogenic | 0 | 0 | 0 |
| rs121918176 | pathogenic | 1 | 1 | 1 |
| rs121918172 | pathogenic | 0 | 1 | 0 |
| rs121918184 | pathogenic | 0 | 0 | 0 |
| rs121434560 | pathogenic | 1 | 1 | 0 |
| rs121434561 | pathogenic | 1 | 1 | 0 |
| rs3218716 | pathogenic | 1 | 1 | 1 |
| rs145532615 | benign | 0 | 0 | 0 |
| rs149840927 | pathogenic | 0 | 0 | 0 |
| rs730880762 | pathogenic | 0 | 0 | 0 |
| rs730880764 | pathogenic | 0 | 0 | 0 |
| rs730880765 | pathogenic | 0 | 0 | 0 |
| rs587782962 | pathogenic | 0 | 0 | 0 |
| rs141735183 | pathogenic | 0 | 0 | 0 |
| rs397516187 | pathogenic | 0 | 0 | 0 |
| rs727505026 | pathogenic | 0 | 0 | 0 |
| rs529700838 | pathogenic | 0 | 0 | 0 |
| rs540263945 | pathogenic | 0 | 0 | 0 |
| rs149103761 | benign | 0 | 0 | 0 |
| rs730880906 | pathogenic | 0 | 0 | 0 |
| rs730880785 | pathogenic | 0 | 0 | 0 |
| rs397516199 | pathogenic | 0 | 0 | 0 |
| rs730880725 | benign | 0 | 0 | 0 |
| rs730880796 | pathogenic | 0 | 0 | 0 |
| rs201307101 | benign | 0 | 0 | 0 |
| rs397516214 | pathogenic | 0 | 0 | 0 |
| rs3729823 | benign | 0 | 0 | 0 |
| rs397516218 | pathogenic | 0 | 0 | 0 |
| rs121913647 | pathogenic | 0 | 0 | 0 |
| rs373514686 | pathogenic | 0 | 0 | 0 |
| rs763538103 | pathogenic | 0 | 0 | 0 |
| rs121913650 | pathogenic | 0 | 0 | 0 |
| rs139222507 | pathogenic | 0 | 0 | 0 |
| rs369940645 | pathogenic | 0 | 0 | 0 |
| rs763073072 | pathogenic | 0 | 0 | 0 |
| rs145734640 | pathogenic | 0 | 0 | 0 |
| rs730880821 | pathogenic | 0 | 0 | 0 |
| rs28933098 | pathogenic | 0 | 0 | 0 |
| rs372381770 | pathogenic | 0 | 0 | 0 |
| rs146796870 | benign | 0 | 0 | 0 |
| rs104894485 | pathogenic | 1 | 1 | 1 |
| rs35407712 | benign | 0 | 0 | 0 |
| rs35842750 | benign | 0 | 0 | 0 |
| rs2721191 | benign | 0 | 0 | 0 |
| rs35098923 | benign | 0 | 0 | 0 |
| rs34293591 | benign | 1 | 1 | 0 |
| rs36023964 | benign | 0 | 0 | 0 |
| rs4251691 | benign | 0 | 0 | 0 |
| rs34666647 | benign | 0 | 0 | 0 |

| | | | | |
|---|---|---|---|---|
| rs4925828 | benign | 0 | 0 | 0 |
| rs35348691 | benign | 0 | 0 | 0 |
| rs36078464 | benign | 0 | 0 | 0 |
| rs34915097 | benign | 0 | 0 | 0 |
| rs35101495 | benign | 0 | 0 | 0 |
| rs141568342 | pathogenic | 0 | 0 | 0 |
| rs1800571 | pathogenic | 0 | 0 | 0 |
| rs140566084 | benign | 1 | 1 | 1 |
| rs727503791 | pathogenic | 1 | 1 | 0 |
| rs545215807 | pathogenic | 0 | 0 | 0 |
| rs113994168 | pathogenic | 0 | 0 | 0 |
| rs113994167 | pathogenic | 0 | 0 | 0 |
| rs200788251 | pathogenic | 0 | 0 | 0 |
| rs200573371 | pathogenic | 0 | 0 | 0 |
| rs771874163 | pathogenic | 1 | 1 | 1 |
| rs112406105 | pathogenic | 1 | 1 | 1 |
| rs118204015 | pathogenic | 1 | 1 | 1 |
| rs113994169 | pathogenic | 0 | 0 | 0 |
| rs118204018 | pathogenic | 0 | 1 | 0 |
| rs138834083 | pathogenic | 0 | 0 | 0 |
| rs533055438 | pathogenic | 1 | 1 | 1 |
| rs2309689 | pathogenic | 1 | 1 | 1 |
| rs138058572 | pathogenic | 1 | 1 | 1 |
| rs118204017 | pathogenic | 1 | 1 | 0 |
| rs766742117 | pathogenic | 1 | 1 | 1 |
| rs751995154 | pathogenic | 1 | 1 | 1 |
| rs200771970 | pathogenic | 0 | 0 | 0 |
| rs118204014 | pathogenic | 0 | 0 | 0 |
| rs148584617 | pathogenic | 1 | 1 | 1 |
| rs121912720 | pathogenic | 0 | 0 | 0 |
| rs121912721 | pathogenic | 0 | 0 | 0 |
| rs28929476 | pathogenic | 0 | 0 | 0 |
| rs28931574 | pathogenic | 1 | 1 | 1 |
| rs12718465 | benign | 0 | 0 | 0 |
| rs144689354 | pathogenic | 0 | 0 | 0 |
| rs149095705 | pathogenic | 0 | 0 | 0 |
| rs572839648 | benign | 1 | 1 | 1 |
| rs121909715 | pathogenic | 1 | 1 | 0 |
| rs34077456 | benign | 1 | 1 | 1 |
| rs397517876 | benign | 0 | 0 | 0 |
| rs1064074 | benign | 0 | 0 | 0 |
| rs142751309 | benign | 0 | 0 | 0 |
| rs116275562 | benign | 0 | 0 | 0 |
| rs148095899 | benign | 0 | 0 | 0 |
| rs751838040 | pathogenic | 0 | 0 | 0 |
| rs121434324 | pathogenic | 1 | 1 | 1 |
| rs143367518 | pathogenic | 0 | 0 | 0 |
| rs121434325 | pathogenic | 0 | 0 | 0 |
| rs397515328 | pathogenic | 0 | 0 | 0 |
| rs376039938 | pathogenic | 0 | 0 | 0 |
| rs387907243 | pathogenic | 0 | 0 | 0 |
| rs28934876 | pathogenic | 0 | 0 | 0 |
| rs387907245 | pathogenic | 0 | 0 | 0 |
| rs138544311 | benign | 0 | 0 | 0 |

| | | | | |
|---|---|---|---|---|
| rs643788 | benign | 0 | 0 | 0 |
| rs749977756 | benign | 0 | 0 | 0 |
| rs1800574 | benign | 0 | 0 | 0 |
| rs137853336 | pathogenic | 1 | 1 | 1 |
| rs111916034 | benign | 0 | 0 | 0 |
| rs307826 | benign | 0 | 0 | 0 |
| rs35874891 | benign | 0 | 0 | 0 |
| rs111033580 | pathogenic | 1 | 1 | 1 |
| rs111033581 | pathogenic | 1 | 1 | 1 |
| rs121908373 | pathogenic | 0 | 0 | 0 |
| rs767122713 | pathogenic | 0 | 0 | 0 |
| rs2233213 | benign | 0 | 0 | 0 |
| rs121908605 | pathogenic | 1 | 1 | 0 |
| rs11887534 | benign | 0 | 0 | 0 |
| rs4148211 | benign | 0 | 0 | 0 |
| rs80025980 | benign | 0 | 0 | 0 |
| rs137854891 | pathogenic | 1 | 1 | 0 |
| rs74358901 | benign | 0 | 0 | 0 |
| rs9282574 | benign | 0 | 0 | 0 |
| rs137852993 | pathogenic | 0 | 0 | 0 |
| rs137852990 | pathogenic | 0 | 0 | 0 |
| rs137852987 | pathogenic | 0 | 0 | 0 |
| rs4148217 | benign | 0 | 0 | 0 |
| rs140690030 | benign | 0 | 0 | 0 |
| rs137852992 | pathogenic | 0 | 0 | 0 |
| rs6544718 | benign | 0 | 0 | 0 |
| rs137852989 | pathogenic | 0 | 0 | 0 |
| rs150893072 | benign | 1 | 1 | 1 |
| rs28931613 | pathogenic | 1 | 1 | 1 |
| rs143789597 | benign | 1 | 1 | 1 |
| rs121913037 | pathogenic | 1 | 1 | 0 |
| rs188802138 | pathogenic | 1 | 1 | 0 |
| rs121913038 | pathogenic | 1 | 1 | 0 |
| rs146835206 | benign | 0 | 0 | 0 |
| rs121913041 | pathogenic | 1 | 1 | 1 |
| rs121913039 | pathogenic | 1 | 1 | 1 |
| rs149977726 | pathogenic | 1 | 1 | 0 |
| rs121913036 | pathogenic | 0 | 0 | 0 |
| rs121913040 | pathogenic | 0 | 0 | 0 |
| rs765007158 | benign | 0 | 0 | 0 |
| rs112723255 | benign | 0 | 0 | 0 |
| rs574164748 | benign | 0 | 0 | 0 |
| rs80338734 | pathogenic | 1 | 1 | 0 |
| rs104894694 | pathogenic | 1 | 1 | 0 |
| rs753198836 | pathogenic | 0 | 0 | 0 |
| rs760101382 | pathogenic | 0 | 0 | 0 |
| rs121909272 | pathogenic | 1 | 1 | 0 |
| rs370421531 | pathogenic | 0 | 0 | 0 |
| rs565109128 | benign | 0 | 0 | 0 |
| rs17851582 | benign | 0 | 0 | 0 |
| rs753228876 | benign | 0 | 0 | 0 |
| rs141471799 | benign | 0 | 0 | 0 |
| rs281864916 | pathogenic | 1 | 1 | 0 |
| rs121918409 | pathogenic | 1 | 1 | 1 |

| | | | | |
|---|---|---|---|---|
| rs121918413 | pathogenic | 1 | 1 | 1 |
| rs121918411 | pathogenic | 1 | 1 | 1 |
| rs121918408 | pathogenic | 1 | 1 | 1 |
| rs121918412 | pathogenic | 1 | 1 | 1 |
| rs121918410 | pathogenic | 1 | 1 | 1 |
| rs116474260 | benign | 1 | 1 | 1 |
| rs281864919 | pathogenic | 1 | 1 | 1 |
| rs397514452 | pathogenic | 1 | 1 | 0 |
| rs747843638 | benign | 0 | 0 | 0 |
| rs774357609 | pathogenic | 1 | 1 | 1 |
| rs201804005 | benign | 0 | 0 | 0 |
| rs746485916 | pathogenic | 0 | 0 | 0 |
| rs764131178 | pathogenic | 0 | 0 | 0 |
| rs374094065 | pathogenic | 0 | 0 | 0 |
| rs76151636 | pathogenic | 1 | 1 | 0 |
| rs753250853 | pathogenic | 0 | 0 | 0 |
| rs761632029 | pathogenic | 0 | 0 | 0 |
| rs775541743 | pathogenic | 0 | 0 | 0 |
| rs778749563 | pathogenic | 0 | 0 | 0 |
| rs60431989 | pathogenic | 1 | 1 | 0 |
| rs755554442 | pathogenic | 1 | 1 | 0 |
| rs749085322 | pathogenic | 0 | 0 | 0 |
| rs756029120 | pathogenic | 1 | 1 | 0 |
| rs137853279 | pathogenic | 1 | 1 | 0 |
| rs777354267 | pathogenic | 1 | 1 | 1 |
| rs142940486 | benign | 1 | 1 | 1 |
| rs140167393 | benign | 1 | 1 | 1 |
| rs2066827 | benign | 0 | 0 | 0 |
| rs765681672 | benign | 0 | 0 | 0 |
| rs137985549 | benign | 0 | 0 | 0 |
| rs74315340 | pathogenic | 1 | 1 | 1 |
| rs74315341 | pathogenic | 0 | 0 | 0 |
| rs74315335 | pathogenic | 0 | 0 | 0 |
| rs61730974 | benign | 0 | 0 | 0 |
| rs121909193 | pathogenic | 0 | 0 | 0 |
| rs74315334 | pathogenic | 0 | 0 | 0 |
| rs74315329 | pathogenic | 0 | 0 | 0 |
| rs74315330 | pathogenic | 0 | 0 | 0 |
| rs121909194 | pathogenic | 0 | 0 | 0 |
| rs28936694 | pathogenic | 0 | 0 | 0 |
| rs74315336 | pathogenic | 0 | 0 | 0 |
| rs74315338 | pathogenic | 1 | 1 | 1 |
| rs74315328 | pathogenic | 0 | 0 | 0 |
| rs74315331 | pathogenic | 0 | 0 | 0 |
| rs74315332 | pathogenic | 0 | 0 | 0 |
| rs141696651 | benign | 0 | 0 | 0 |
| rs7041 | benign | 0 | 0 | 0 |
| rs4588 | benign | 0 | 0 | 0 |
| rs121908981 | pathogenic | 1 | 1 | 1 |
| rs606231459 | pathogenic | 0 | 0 | 0 |
| rs398123053 | pathogenic | 0 | 0 | 0 |
| rs774259802 | benign | 0 | 0 | 0 |
| rs121909582 | pathogenic | 1 | 1 | 1 |
| rs104893680 | pathogenic | 1 | 1 | 1 |

| | | | | |
|---|---|---|---|---|
| rs137854907 | pathogenic | 0 | 0 | 0 |
| rs765715798 | pathogenic | 0 | 0 | 0 |
| rs104893678 | pathogenic | 0 | 0 | 0 |
| rs104893679 | pathogenic | 0 | 1 | 0 |
| rs104893681 | pathogenic | 1 | 1 | 0 |
| rs121909640 | pathogenic | 1 | 1 | 0 |
| rs780705654 | pathogenic | 0 | 0 | 0 |
| rs201672146 | benign | 1 | 1 | 0 |
| rs12136658 | benign | 0 | 0 | 0 |
| rs56127440 | benign | 0 | 0 | 0 |
| rs104894433 | pathogenic | 1 | 1 | 1 |
| rs104894435 | pathogenic | 0 | 1 | 0 |
| rs104894437 | pathogenic | 1 | 1 | 1 |
| rs104894441 | pathogenic | 1 | 1 | 1 |
| rs104894440 | pathogenic | 1 | 1 | 1 |
| rs104894445 | pathogenic | 1 | 1 | 1 |
| rs104894436 | pathogenic | 0 | 0 | 0 |
| rs104894438 | pathogenic | 0 | 0 | 0 |
| rs200891969 | pathogenic | 1 | 1 | 0 |
| rs104894443 | pathogenic | 1 | 1 | 1 |
| rs104894434 | pathogenic | 1 | 1 | 1 |
| rs41298442 | pathogenic | 1 | 1 | 1 |
| rs104894442 | pathogenic | 1 | 1 | 1 |
| rs35453854 | benign | 0 | 0 | 0 |
| rs121912898 | pathogenic | 0 | 0 | 0 |
| rs121912897 | pathogenic | 1 | 1 | 0 |
| rs138260625 | benign | 0 | 0 | 0 |
| rs147802438 | benign | 1 | 1 | 0 |
| rs146443310 | benign | 1 | 1 | 0 |
| rs747319628 | pathogenic | 1 | 1 | 0 |
| rs1801195 | benign | 0 | 0 | 0 |
| rs145045341 | benign | 1 | 1 | 0 |
| rs116001173 | benign | 1 | 1 | 0 |
| rs587784175 | pathogenic | 0 | 0 | 0 |
| rs104893857 | pathogenic | 0 | 0 | 0 |
| rs104893862 | pathogenic | 1 | 1 | 0 |
| rs104893858 | pathogenic | 1 | 1 | 1 |
| rs121909249 | pathogenic | 0 | 1 | 0 |
| rs121909248 | pathogenic | 1 | 1 | 1 |
| rs104893859 | pathogenic | 1 | 1 | 0 |
| rs187063864 | pathogenic | 1 | 1 | 0 |
| rs121917817 | pathogenic | 1 | 1 | 0 |
| rs121917818 | pathogenic | 0 | 0 | 0 |
| rs753594601 | pathogenic | 0 | 0 | 0 |
| rs28937869 | pathogenic | 1 | 1 | 0 |
| rs6165 | benign | 0 | 0 | 0 |
| rs200089714 | pathogenic | 1 | 1 | 0 |
| rs372000714 | pathogenic | 0 | 0 | 0 |
| rs2228206 | benign | 0 | 0 | 0 |
| rs560096 | benign | 0 | 0 | 0 |
| rs137852666 | pathogenic | 0 | 0 | 0 |
| rs10896380 | benign | 0 | 0 | 0 |
| rs773690764 | pathogenic | 0 | 0 | 0 |
| rs201060167 | pathogenic | 0 | 0 | 0 |

| | | | | |
|---|---|---|---|---|
| rs137852670 | pathogenic | 0 | 0 | 0 |
| rs780594709 | pathogenic | 0 | 0 | 0 |
| rs145226920 | pathogenic | 0 | 0 | 0 |
| rs137852665 | pathogenic | 0 | 0 | 0 |
| rs724159960 | pathogenic | 0 | 0 | 0 |
| rs756985703 | pathogenic | 0 | 0 | 0 |
| rs137852667 | pathogenic | 0 | 0 | 0 |
| rs151079750 | pathogenic | 1 | 1 | 0 |
| rs2306636 | benign | 0 | 0 | 0 |
| rs72647885 | benign | 1 | 1 | 0 |
| rs377584435 | pathogenic | 1 | 1 | 1 |
| rs757583846 | pathogenic | 1 | 1 | 0 |
| rs121918148 | pathogenic | 0 | 0 | 0 |
| rs149201802 | benign | 0 | 0 | 0 |
| rs587781904 | pathogenic | 1 | 1 | 1 |
| rs143421325 | benign | 0 | 0 | 0 |
| rs587781253 | pathogenic | 0 | 0 | 0 |
| rs587780331 | benign | 0 | 0 | 0 |
| rs151001016 | benign | 0 | 0 | 0 |
| rs150888094 | benign | 0 | 0 | 0 |
| rs149581331 | benign | 0 | 0 | 0 |
| rs774840535 | benign | 0 | 0 | 0 |
| rs138428684 | benign | 0 | 0 | 0 |
| rs149300055 | benign | 0 | 0 | 0 |
| rs571005578 | benign | 0 | 0 | 0 |
| rs200731839 | benign | 0 | 0 | 0 |
| rs121965007 | pathogenic | 0 | 0 | 0 |
| rs121965013 | pathogenic | 0 | 1 | 0 |
| rs121965009 | pathogenic | 0 | 0 | 0 |
| rs1800457 | benign | 1 | 1 | 0 |
| rs121965006 | pathogenic | 1 | 1 | 0 |
| rs121965008 | pathogenic | 0 | 0 | 0 |
| rs61732609 | pathogenic | 0 | 0 | 0 |
| rs121965011 | pathogenic | 0 | 0 | 0 |
| rs121965015 | pathogenic | 0 | 0 | 0 |
| rs121965018 | pathogenic | 1 | 1 | 0 |
| rs121965016 | pathogenic | 0 | 0 | 0 |
| rs77806 | benign | 0 | 0 | 0 |
| rs149731356 | benign | 0 | 0 | 0 |
| rs72549407 | pathogenic | 0 | 0 | 0 |
| rs200363929 | benign | 0 | 0 | 0 |
| rs376403708 | benign | 1 | 1 | 0 |
| rs56376197 | benign | 1 | 1 | 1 |
| rs200237973 | benign | 1 | 1 | 1 |
| rs548330075 | benign | 0 | 0 | 0 |
| rs148291350 | benign | 0 | 0 | 0 |
| rs71579353 | pathogenic | 0 | 0 | 0 |
| rs146278697 | benign | 0 | 0 | 0 |
| rs141033098 | benign | 0 | 0 | 0 |
| rs397517234 | pathogenic | 0 | 0 | 0 |
| rs71579374 | benign | 0 | 0 | 0 |
| rs140381835 | benign | 0 | 0 | 0 |
| rs137852702 | pathogenic | 0 | 0 | 0 |
| rs137852696 | pathogenic | 1 | 1 | 1 |

| | | | | |
|---|---|---|---|---|
| rs137852697 | pathogenic | 0 | 0 | 0 |
| rs386833639 | pathogenic | 0 | 0 | 0 |
| rs386833640 | pathogenic | 0 | 0 | 0 |
| rs386833641 | benign | 0 | 0 | 0 |
| rs137852701 | pathogenic | 0 | 0 | 0 |
| rs386833642 | pathogenic | 0 | 0 | 0 |
| rs137852695 | pathogenic | 0 | 0 | 0 |
| rs386833646 | pathogenic | 0 | 0 | 0 |
| rs137852700 | pathogenic | 1 | 1 | 0 |
| rs386833649 | pathogenic | 1 | 1 | 0 |
| rs386833650 | pathogenic | 0 | 0 | 0 |
| rs148412181 | pathogenic | 0 | 0 | 0 |
| rs386833655 | pathogenic | 0 | 0 | 0 |
| rs386833656 | pathogenic | 1 | 1 | 0 |
| rs137852698 | pathogenic | 0 | 0 | 0 |
| rs386833661 | pathogenic | 0 | 0 | 0 |
| rs386833670 | pathogenic | 0 | 0 | 0 |
| rs111033632 | pathogenic | 1 | 1 | 1 |
| rs34308573 | benign | 1 | 1 | 1 |
| rs376048533 | pathogenic | 1 | 1 | 0 |
| rs3747517 | benign | 1 | 1 | 0 |
| rs137853328 | pathogenic | 0 | 0 | 0 |
| rs77273740 | benign | 0 | 0 | 0 |
| rs267606760 | pathogenic | 0 | 0 | 0 |
| rs77576840 | pathogenic | 0 | 0 | 0 |
| rs45465204 | benign | 1 | 1 | 0 |
| rs5320 | benign | 0 | 0 | 0 |
| rs5324 | benign | 0 | 0 | 0 |
| rs4531 | benign | 0 | 0 | 0 |
| rs267606761 | pathogenic | 0 | 0 | 0 |
| rs75215331 | pathogenic | 0 | 0 | 0 |
| rs6271 | benign | 0 | 0 | 0 |
| rs201643277 | pathogenic | 0 | 0 | 0 |
| rs4252250 | benign | 0 | 0 | 0 |
| rs137853580 | pathogenic | 0 | 0 | 0 |
| rs368287711 | pathogenic | 0 | 0 | 0 |
| rs137853579 | pathogenic | 0 | 0 | 0 |
| rs3135932 | benign | 0 | 0 | 0 |
| rs2228055 | benign | 0 | 0 | 0 |
| rs140322345 | benign | 1 | 1 | 1 |
| rs1154510 | pathogenic | 0 | 0 | 0 |
| rs11833399 | benign | 0 | 0 | 0 |
| rs36023382 | benign | 1 | 1 | 1 |
| rs61754444 | benign | 0 | 0 | 0 |
| rs28935168 | pathogenic | 0 | 0 | 0 |
| rs28934907 | pathogenic | 0 | 0 | 0 |
| rs28934904 | pathogenic | 1 | 1 | 0 |
| rs61748392 | pathogenic | 1 | 1 | 0 |
| rs28934908 | pathogenic | 0 | 0 | 0 |
| rs28934905 | pathogenic | 0 | 0 | 0 |
| rs28934906 | pathogenic | 1 | 1 | 0 |
| rs606231307 | pathogenic | 0 | 0 | 0 |
| rs9333296 | benign | 0 | 0 | 0 |
| rs3218614 | benign | 0 | 0 | 0 |

| | | | | |
|---|---|---|---|---|
| rs2230792 | benign | 0 | 0 | 0 |
| rs2230793 | benign | 0 | 0 | 0 |
| rs2230794 | benign | 0 | 0 | 0 |
| rs749052963 | pathogenic | 0 | 0 | 0 |
| rs2230798 | benign | 0 | 0 | 0 |
| rs2230795 | benign | 0 | 0 | 0 |
| rs3204145 | benign | 0 | 0 | 0 |
| rs1538660 | benign | 0 | 0 | 0 |
| rs9878928 | benign | 0 | 0 | 0 |
| rs104893742 | pathogenic | 0 | 0 | 0 |
| rs28936702 | pathogenic | 0 | 0 | 0 |
| rs28936703 | pathogenic | 0 | 0 | 0 |
| rs769392 | benign | 0 | 0 | 0 |
| rs769402 | benign | 1 | 1 | 1 |
| rs104894884 | pathogenic | 1 | 1 | 1 |
| rs1801316 | benign | 1 | 1 | 1 |
| rs104894885 | pathogenic | 1 | 1 | 1 |
| rs782572 | benign | 0 | 0 | 0 |
| rs151166046 | pathogenic | 0 | 0 | 0 |
| rs772153760 | pathogenic | 1 | 1 | 0 |
| rs35405862 | benign | 0 | 0 | 0 |
| rs146571352 | pathogenic | 0 | 1 | 0 |
| rs374698153 | pathogenic | 0 | 0 | 0 |
| rs7594497 | benign | 0 | 0 | 0 |
| rs80338716 | pathogenic | 0 | 0 | 0 |
| rs80338719 | pathogenic | 1 | 1 | 1 |
| rs80338721 | pathogenic | 0 | 0 | 0 |
| rs730882249 | pathogenic | 0 | 0 | 0 |
| rs146262009 | pathogenic | 0 | 0 | 0 |
| rs104895083 | pathogenic | 1 | 1 | 1 |
| rs104895100 | benign | 1 | 1 | 1 |
| rs104895085 | pathogenic | 0 | 0 | 0 |
| rs28940580 | pathogenic | 0 | 0 | 0 |
| rs104895098 | pathogenic | 0 | 0 | 0 |
| rs28940578 | pathogenic | 0 | 0 | 0 |
| rs61752717 | pathogenic | 0 | 0 | 0 |
| rs28940579 | pathogenic | 0 | 0 | 0 |
| rs104895097 | pathogenic | 0 | 0 | 0 |
| rs140425651 | benign | 1 | 1 | 0 |
| rs768090444 | pathogenic | 0 | 0 | 0 |
| rs764459544 | pathogenic | 0 | 0 | 0 |
| rs80338802 | pathogenic | 1 | 1 | 1 |
| rs149969786 | pathogenic | 1 | 1 | 0 |
| rs115311625 | pathogenic | 1 | 1 | 1 |
| rs778768583 | pathogenic | 1 | 1 | 1 |
| rs892090 | benign | 0 | 0 | 0 |
| rs28929498 | pathogenic | 1 | 1 | 1 |
| rs137854467 | pathogenic | 0 | 0 | 0 |
| rs137854480 | pathogenic | 0 | 0 | 0 |
| rs148682262 | benign | 0 | 0 | 0 |
| rs137852744 | pathogenic | 0 | 0 | 0 |
| rs137852745 | pathogenic | 0 | 1 | 0 |
| rs137852749 | pathogenic | 0 | 1 | 0 |
| rs137852746 | pathogenic | 0 | 1 | 0 |

| | | | | |
|---|---|---|---|---|
| rs145633958 | benign | 0 | 0 | 0 |
| rs141679950 | benign | 0 | 0 | 0 |
| rs76397662 | benign | 1 | 1 | 1 |
| rs370736139 | benign | 0 | 0 | 0 |
| rs781750106 | benign | 0 | 0 | 0 |
| rs546866208 | benign | 0 | 0 | 0 |
| rs142338976 | benign | 0 | 0 | 0 |
| rs115272158 | benign | 0 | 0 | 0 |
| rs115423919 | benign | 0 | 0 | 0 |
| rs537874538 | benign | 0 | 0 | 0 |
| rs9282834 | benign | 0 | 0 | 0 |
| rs201745826 | benign | 0 | 0 | 0 |
| rs75873440 | pathogenic | 0 | 0 | 0 |
| rs147219360 | benign | 0 | 0 | 0 |
| rs77939446 | pathogenic | 0 | 0 | 0 |
| rs77558292 | pathogenic | 0 | 0 | 0 |
| rs80069458 | pathogenic | 0 | 0 | 0 |
| rs76262710 | pathogenic | 0 | 0 | 0 |
| rs79781594 | pathogenic | 0 | 0 | 0 |
| rs77316810 | pathogenic | 0 | 0 | 0 |
| rs77503355 | pathogenic | 0 | 0 | 0 |
| rs79890926 | pathogenic | 0 | 0 | 0 |
| rs377767402 | benign | 0 | 0 | 0 |
| rs121913308 | pathogenic | 0 | 0 | 0 |
| rs55846256 | benign | 0 | 0 | 0 |
| rs75076352 | pathogenic | 0 | 0 | 0 |
| rs75996173 | pathogenic | 0 | 0 | 0 |
| rs77709286 | pathogenic | 0 | 0 | 0 |
| rs151058706 | benign | 0 | 0 | 0 |
| rs138964358 | benign | 0 | 0 | 0 |
| rs137852353 | pathogenic | 1 | 1 | 0 |
| rs201196020 | benign | 0 | 0 | 0 |
| rs137853201 | pathogenic | 1 | 1 | 1 |
| rs113994174 | pathogenic | 1 | 1 | 1 |
| rs104893674 | pathogenic | 1 | 1 | 1 |
| rs779519249 | benign | 1 | 1 | 1 |
| rs9606756 | benign | 0 | 0 | 0 |
| rs75680863 | benign | 1 | 1 | 1 |
| rs35838082 | benign | 0 | 0 | 0 |
| rs1801198 | benign | 0 | 0 | 0 |
| rs117458738 | benign | 1 | 1 | 0 |
| rs9621049 | benign | 1 | 1 | 0 |
| rs1131603 | benign | 1 | 1 | 0 |
| rs4820889 | benign | 0 | 0 | 0 |
| rs777767978 | pathogenic | 1 | 1 | 0 |
| rs147399860 | benign | 1 | 1 | 1 |
| rs104893950 | pathogenic | 0 | 0 | 0 |
| rs777308612 | benign | 0 | 0 | 0 |
| rs151091410 | benign | 0 | 0 | 0 |
| rs11537667 | benign | 0 | 0 | 0 |
| rs267606621 | pathogenic | 1 | 1 | 0 |
| rs143058902 | benign | 0 | 0 | 0 |
| rs370261904 | pathogenic | 0 | 0 | 0 |
| rs141169746 | benign | 0 | 0 | 0 |

| | | | | |
|---|---|---|---|---|
| rs111033436 | benign | 0 | 0 | 0 |
| rs148533341 | benign | 0 | 0 | 0 |
| rs143827620 | benign | 0 | 0 | 0 |
| rs4935502 | benign | 0 | 0 | 0 |
| rs61735473 | benign | 0 | 0 | 0 |
| rs267606932 | pathogenic | 1 | 1 | 0 |
| rs61730754 | benign | 0 | 0 | 0 |
| rs374491359 | pathogenic | 1 | 1 | 1 |
| rs761800077 | benign | 1 | 1 | 1 |
| rs760391381 | benign | 1 | 1 | 1 |
| rs104893885 | pathogenic | 0 | 0 | 0 |
| rs104893886 | pathogenic | 1 | 1 | 1 |
| rs104893884 | pathogenic | 0 | 0 | 0 |
| rs147715509 | benign | 0 | 0 | 0 |
| rs115533243 | benign | 0 | 0 | 0 |
| rs397509390 | pathogenic | 0 | 1 | 0 |
| rs80357134 | pathogenic | 0 | 0 | 0 |
| rs766004110 | benign | 1 | 1 | 1 |
| rs769650474 | benign | 0 | 0 | 0 |
| rs398122635 | benign | 1 | 1 | 0 |
| rs199522616 | pathogenic | 0 | 0 | 0 |
| rs28897672 | pathogenic | 1 | 1 | 0 |
| rs80357086 | pathogenic | 1 | 1 | 1 |
| rs80357064 | pathogenic | 1 | 1 | 0 |
| rs80356936 | benign | 0 | 0 | 0 |
| rs28897673 | benign | 0 | 0 | 0 |
| rs146747324 | benign | 0 | 0 | 0 |
| rs78635798 | pathogenic | 0 | 0 | 0 |
| rs75146158 | pathogenic | 1 | 1 | 1 |
| rs587782886 | benign | 0 | 0 | 0 |
| rs121434634 | pathogenic | 0 | 0 | 0 |
| rs121434635 | pathogenic | 0 | 0 | 0 |
| rs587777778 | pathogenic | 0 | 0 | 0 |
| rs118203967 | pathogenic | 0 | 0 | 0 |
| rs751375244 | pathogenic | 1 | 1 | 1 |
| rs104893863 | pathogenic | 1 | 1 | 0 |
| rs104893865 | pathogenic | 0 | 0 | 0 |
| rs104893864 | pathogenic | 1 | 1 | 1 |
| rs104893866 | pathogenic | 1 | 1 | 0 |
| rs561738497 | benign | 0 | 0 | 0 |
| rs57599352 | pathogenic | 1 | 1 | 1 |
| rs267607448 | pathogenic | 1 | 1 | 1 |
| rs63751596 | pathogenic | 0 | 0 | 0 |
| rs63751612 | benign | 0 | 0 | 0 |
| rs63751684 | benign | 0 | 0 | 0 |
| rs267607861 | benign | 0 | 0 | 0 |
| rs35831931 | benign | 0 | 0 | 0 |
| rs63750876 | benign | 0 | 0 | 0 |
| rs41295284 | benign | 0 | 0 | 0 |
| rs35502531 | benign | 0 | 0 | 0 |
| rs63750449 | benign | 0 | 0 | 0 |
| rs63750899 | pathogenic | 0 | 0 | 0 |
| rs55907433 | benign | 0 | 0 | 0 |
| rs63750217 | pathogenic | 0 | 0 | 0 |

| | | | | |
|---|---|---|---|---|
| rs63751275 | pathogenic | 0 | 0 | 0 |
| rs63750702 | benign | 1 | 1 | 1 |
| rs63750114 | benign | 0 | 0 | 0 |
| rs2020873 | benign | 0 | 0 | 0 |
| rs140195825 | benign | 1 | 1 | 1 |
| rs182763805 | benign | 0 | 0 | 0 |
| rs533696377 | benign | 0 | 0 | 0 |
| rs41278788 | benign | 0 | 0 | 0 |
| rs104894193 | pathogenic | 1 | 1 | 0 |
| rs3824915 | benign | 1 | 1 | 1 |
| rs104894196 | pathogenic | 0 | 0 | 0 |
| rs78681604 | benign | 1 | 1 | 1 |
| rs77321498 | benign | 1 | 1 | 1 |
| rs137852965 | pathogenic | 0 | 0 | 0 |
| rs137852961 | pathogenic | 0 | 0 | 0 |
| rs137852962 | pathogenic | 1 | 1 | 0 |
| rs753376100 | pathogenic | 0 | 0 | 0 |
| rs137852968 | pathogenic | 1 | 1 | 1 |
| rs137852959 | pathogenic | 1 | 1 | 0 |
| rs137852967 | pathogenic | 0 | 0 | 0 |
| rs113437301 | benign | 1 | 1 | 1 |
| rs61747482 | benign | 1 | 1 | 1 |
| rs2234919 | benign | 1 | 1 | 0 |
| rs121908863 | pathogenic | 0 | 0 | 0 |
| rs140455771 | pathogenic | 0 | 0 | 0 |
| rs121908810 | pathogenic | 1 | 1 | 1 |
| rs143879853 | benign | 0 | 0 | 0 |
| rs188319622 | benign | 0 | 0 | 0 |
| rs121918645 | pathogenic | 0 | 0 | 0 |
| rs115352435 | pathogenic | 0 | 0 | 0 |
| rs121434427 | pathogenic | 1 | 1 | 1 |
| rs121434428 | pathogenic | 0 | 0 | 0 |
| rs35086265 | benign | 1 | 1 | 1 |
| rs11576415 | benign | 0 | 0 | 0 |
| rs121434429 | pathogenic | 1 | 1 | 1 |
| rs4074536 | benign | 1 | 1 | 1 |
| rs10801999 | benign | 0 | 0 | 0 |
| rs121434550 | pathogenic | 0 | 0 | 0 |
| rs148057999 | pathogenic | 0 | 0 | 0 |
| rs28730716 | benign | 0 | 0 | 0 |
| rs142036299 | benign | 0 | 0 | 0 |
| rs762381137 | pathogenic | 0 | 0 | 0 |
| rs121434549 | pathogenic | 0 | 0 | 0 |
| rs139228801 | pathogenic | 0 | 0 | 0 |
| rs147219463 | benign | 0 | 0 | 0 |
| rs1131857 | benign | 0 | 0 | 0 |
| rs121917874 | pathogenic | 0 | 1 | 0 |
| rs2228056 | benign | 0 | 0 | 0 |
| rs121917870 | pathogenic | 0 | 0 | 0 |
| rs121917866 | pathogenic | 0 | 0 | 0 |
| rs11921054 | benign | 0 | 0 | 0 |
| rs121917868 | pathogenic | 0 | 0 | 0 |
| rs28931603 | pathogenic | 0 | 0 | 0 |
| rs104895295 | pathogenic | 1 | 1 | 0 |

| | | | | |
|---|---|---|---|---|
| rs104895382 | pathogenic | 0 | 0 | 0 |
| rs104895298 | pathogenic | 0 | 0 | 0 |
| rs104895300 | pathogenic | 0 | 0 | 0 |
| rs104895332 | pathogenic | 0 | 0 | 0 |
| rs758026399 | pathogenic | 1 | 1 | 1 |
| rs104895366 | pathogenic | 0 | 0 | 0 |
| rs104895304 | pathogenic | 0 | 0 | 0 |
| rs759997079 | pathogenic | 0 | 0 | 0 |
| rs121917789 | pathogenic | 0 | 0 | 0 |
| rs104895319 | pathogenic | 0 | 0 | 0 |
| rs104895317 | pathogenic | 1 | 1 | 0 |
| rs28934897 | pathogenic | 0 | 0 | 0 |
| rs104895380 | benign | 0 | 0 | 0 |
| rs104895360 | pathogenic | 0 | 0 | 0 |
| rs28936396 | pathogenic | 1 | 1 | 0 |
| rs121909307 | pathogenic | 1 | 1 | 1 |
| rs28938472 | pathogenic | 1 | 1 | 0 |
| rs121909308 | pathogenic | 1 | 1 | 0 |
| rs121909309 | pathogenic | 0 | 0 | 0 |
| rs121913013 | benign | 1 | 1 | 1 |
| rs201855245 | benign | 1 | 1 | 1 |
| rs2230232 | benign | 1 | 1 | 0 |
| rs113451409 | benign | 0 | 0 | 0 |
| rs121913011 | pathogenic | 1 | 1 | 0 |
| rs185821167 | pathogenic | 1 | 1 | 0 |
| rs2230234 | benign | 0 | 0 | 0 |
| rs193922639 | benign | 1 | 1 | 0 |
| rs774863785 | pathogenic | 1 | 1 | 0 |
| rs576404380 | pathogenic | 1 | 1 | 0 |
| rs2230235 | benign | 1 | 1 | 0 |
| rs200484060 | pathogenic | 1 | 1 | 0 |
| rs199761749 | benign | 0 | 0 | 0 |
| rs199681901 | benign | 0 | 0 | 0 |
| rs121908951 | pathogenic | 0 | 0 | 0 |
| rs769251460 | benign | 0 | 0 | 0 |
| rs28937320 | pathogenic | 0 | 0 | 0 |
| rs200607527 | pathogenic | 0 | 0 | 0 |
| rs137852640 | pathogenic | 1 | 1 | 1 |
| rs137852636 | pathogenic | 1 | 1 | 1 |
| rs137852638 | pathogenic | 0 | 0 | 0 |
| rs142637231 | pathogenic | 0 | 0 | 0 |
| rs151328418 | benign | 0 | 1 | 0 |
| rs137852639 | pathogenic | 1 | 1 | 1 |
| rs34605667 | benign | 0 | 0 | 0 |
| rs104894073 | pathogenic | 1 | 1 | 0 |
| rs56298569 | pathogenic | 0 | 0 | 0 |
| rs757600616 | pathogenic | 0 | 0 | 0 |
| rs139194636 | pathogenic | 0 | 0 | 0 |
| rs369517993 | pathogenic | 1 | 1 | 1 |
| rs138846036 | benign | 1 | 1 | 1 |
| rs200808744 | pathogenic | 1 | 1 | 1 |
| rs2255543 | benign | 0 | 0 | 0 |
| rs149165166 | pathogenic | 0 | 0 | 0 |
| rs544956641 | pathogenic | 1 | 1 | 1 |

| | | | | |
|---|---|---|---|---|
| rs28941783 | pathogenic | 0 | 0 | 0 |
| rs562853291 | pathogenic | 1 | 1 | 1 |
| rs28942100 | pathogenic | 0 | 0 | 0 |
| rs120074174 | pathogenic | 1 | 1 | 1 |
| rs120074171 | pathogenic | 1 | 1 | 1 |
| rs120074173 | pathogenic | 0 | 0 | 0 |
| rs120074172 | pathogenic | 1 | 1 | 1 |
| rs767159114 | pathogenic | 1 | 1 | 1 |
| rs104893657 | pathogenic | 0 | 0 | 0 |
| rs104893656 | pathogenic | 0 | 0 | 0 |
| rs104893659 | pathogenic | 0 | 0 | 0 |
| rs104893658 | pathogenic | 0 | 0 | 0 |
| rs200386310 | pathogenic | 0 | 0 | 0 |
| rs2072712 | benign | 0 | 0 | 0 |
| rs35274867 | benign | 0 | 0 | 0 |
| rs35343172 | benign | 0 | 0 | 0 |
| rs10079250 | benign | 0 | 0 | 0 |
| rs34951517 | benign | 0 | 0 | 0 |
| rs267606695 | pathogenic | 0 | 0 | 0 |
| rs149391728 | benign | 0 | 0 | 0 |
| rs387906598 | pathogenic | 1 | 1 | 0 |
| rs137854578 | pathogenic | 0 | 0 | 0 |
| rs141519952 | benign | 0 | 0 | 0 |
| rs146572883 | benign | 0 | 0 | 0 |
| rs111866410 | benign | 0 | 0 | 0 |
| rs139387758 | benign | 1 | 1 | 1 |
| rs35672074 | benign | 1 | 1 | 1 |
| rs121434322 | pathogenic | 0 | 0 | 0 |
| rs150884181 | benign | 0 | 0 | 0 |
| rs35867471 | benign | 0 | 0 | 0 |
| rs2867802 | benign | 0 | 0 | 0 |
| rs137902889 | benign | 0 | 0 | 0 |
| rs369804864 | pathogenic | 0 | 0 | 0 |
| rs781050795 | pathogenic | 0 | 0 | 0 |
| rs2305799 | benign | 0 | 0 | 0 |
| rs2229531 | benign | 0 | 0 | 0 |
| rs147025508 | benign | 0 | 0 | 0 |
| rs76857106 | pathogenic | 0 | 0 | 0 |
| rs7247284 | benign | 0 | 0 | 0 |
| rs62619782 | benign | 0 | 0 | 0 |
| rs377244188 | pathogenic | 0 | 0 | 0 |
| rs75718910 | pathogenic | 0 | 0 | 0 |
| rs79843600 | pathogenic | 0 | 0 | 0 |
| rs75037667 | pathogenic | 1 | 1 | 1 |
| rs121913006 | pathogenic | 0 | 0 | 0 |
| rs397517925 | pathogenic | 1 | 1 | 1 |
| rs267606831 | pathogenic | 1 | 1 | 1 |
| rs121909667 | pathogenic | 1 | 1 | 1 |
| rs121909669 | pathogenic | 1 | 1 | 1 |
| rs121909668 | pathogenic | 1 | 1 | 1 |
| rs121909671 | pathogenic | 1 | 1 | 1 |
| rs534596036 | benign | 0 | 0 | 0 |
| rs144037629 | pathogenic | 0 | 0 | 0 |
| rs11546406 | benign | 0 | 0 | 0 |

| | | | | |
|---|---|---|---|---|
| rs143697995 | pathogenic | 0 | 0 | 0 |
| rs121908331 | pathogenic | 0 | 0 | 0 |
| rs149377346 | benign | 0 | 0 | 0 |
| rs104894086 | pathogenic | 1 | 1 | 0 |
| rs104894089 | pathogenic | 0 | 0 | 0 |
| rs104894090 | pathogenic | 1 | 1 | 0 |
| rs387907235 | pathogenic | 0 | 0 | 0 |
| rs137852689 | pathogenic | 0 | 0 | 0 |
| rs137852690 | pathogenic | 1 | 1 | 0 |
| rs104894087 | pathogenic | 0 | 0 | 0 |
| rs104894085 | pathogenic | 0 | 0 | 0 |
| rs79383654 | benign | 0 | 0 | 0 |
| rs121434239 | pathogenic | 0 | 0 | 0 |
| rs121434240 | pathogenic | 0 | 0 | 0 |
| rs121434236 | pathogenic | 0 | 0 | 0 |
| rs121434238 | pathogenic | 1 | 1 | 1 |
| rs121918107 | pathogenic | 1 | 1 | 1 |
| rs121918103 | pathogenic | 1 | 1 | 1 |
| rs121918104 | pathogenic | 1 | 1 | 1 |
| rs61747614 | benign | 1 | 1 | 1 |
| rs555329870 | benign | 0 | 0 | 0 |
| rs186094931 | benign | 0 | 0 | 0 |
| rs267606785 | pathogenic | 0 | 0 | 0 |
| rs34474955 | benign | 0 | 0 | 0 |
| rs1126497 | benign | 1 | 1 | 1 |
| rs397514661 | pathogenic | 0 | 0 | 0 |
| rs189732445 | benign | 0 | 0 | 0 |
| rs146685071 | benign | 0 | 0 | 0 |
| rs121908636 | pathogenic | 1 | 1 | 0 |
| rs121908643 | pathogenic | 0 | 0 | 0 |
| rs121908644 | pathogenic | 1 | 1 | 1 |
| rs575001023 | pathogenic | 1 | 1 | 1 |
| rs121908646 | pathogenic | 1 | 1 | 0 |
| rs121908638 | pathogenic | 1 | 1 | 1 |
| rs777828000 | pathogenic | 1 | 1 | 1 |
| rs74923032 | benign | 0 | 0 | 0 |
| rs192838388 | pathogenic | 1 | 1 | 1 |
| rs148918985 | pathogenic | 1 | 1 | 1 |
| rs773540764 | benign | 1 | 1 | 1 |
| rs370595480 | pathogenic | 1 | 1 | 1 |
| rs762387914 | pathogenic | 1 | 1 | 1 |
| rs121908642 | pathogenic | 1 | 1 | 1 |
| rs183276875 | pathogenic | 1 | 1 | 1 |
| rs121908648 | pathogenic | 1 | 1 | 1 |
| rs121908647 | pathogenic | 1 | 1 | 1 |
| rs121908640 | pathogenic | 1 | 1 | 1 |
| rs771937610 | pathogenic | 1 | 1 | 1 |
| rs121908641 | pathogenic | 1 | 1 | 1 |
| rs5745068 | benign | 0 | 0 | 0 |
| rs73155056 | benign | 1 | 1 | 1 |
| rs148788180 | benign | 0 | 0 | 0 |
| rs121918137 | pathogenic | 0 | 0 | 0 |
| rs41269593 | benign | 0 | 0 | 0 |
| rs11159 | benign | 1 | 1 | 0 |

| | | | | |
|---|---|---|---|---|
| rs114733570 | pathogenic | 1 | 1 | 0 |
| rs3211783 | benign | 0 | 0 | 0 |
| rs4762 | benign | 0 | 0 | 0 |
| rs74315283 | pathogenic | 0 | 0 | 0 |
| rs111033559 | pathogenic | 0 | 0 | 0 |
| rs111033560 | pathogenic | 1 | 1 | 1 |
| rs749962743 | benign | 0 | 0 | 0 |
| rs267606986 | pathogenic | 0 | 0 | 0 |
| rs34516635 | benign | 0 | 0 | 0 |
| rs117440569 | benign | 0 | 0 | 0 |
| rs61740877 | benign | 0 | 0 | 0 |
| rs121918352 | pathogenic | 0 | 0 | 0 |
| rs183974372 | benign | 0 | 0 | 0 |
| rs140511204 | benign | 1 | 1 | 0 |
| rs6040067 | benign | 0 | 0 | 0 |
| rs199674138 | benign | 0 | 0 | 0 |
| rs121918350 | pathogenic | 0 | 0 | 0 |
| rs138292988 | benign | 1 | 1 | 1 |
| rs145288947 | benign | 0 | 0 | 0 |
| rs369714529 | benign | 0 | 0 | 0 |
| rs200945419 | benign | 0 | 0 | 0 |
| rs116959874 | benign | 0 | 0 | 0 |
| rs141659620 | pathogenic | 1 | 1 | 0 |
| rs111475461 | benign | 0 | 0 | 0 |
| rs61755320 | pathogenic | 0 | 0 | 0 |
| rs748600162 | pathogenic | 1 | 1 | 1 |
| rs369227537 | pathogenic | 1 | 1 | 1 |
| rs372981030 | pathogenic | 1 | 1 | 1 |
| rs137854486 | pathogenic | 0 | 0 | 0 |
| rs137854487 | pathogenic | 0 | 0 | 0 |
| rs199905054 | pathogenic | 0 | 0 | 0 |
| rs771438170 | pathogenic | 0 | 0 | 0 |
| rs774949844 | pathogenic | 0 | 0 | 0 |
| rs4899145 | benign | 0 | 0 | 0 |
| rs200642525 | benign | 0 | 0 | 0 |
| rs190140598 | pathogenic | 0 | 0 | 0 |
| rs2250889 | benign | 0 | 0 | 0 |
| rs17577 | benign | 0 | 0 | 0 |
| rs142032681 | benign | 0 | 0 | 0 |
| rs144577963 | benign | 0 | 0 | 0 |
| rs61754623 | benign | 0 | 0 | 0 |
| rs139873200 | benign | 0 | 0 | 0 |
| rs532530906 | benign | 0 | 0 | 0 |
| rs145762716 | benign | 0 | 0 | 0 |
| rs34062020 | benign | 0 | 0 | 0 |
| rs35258161 | benign | 0 | 0 | 0 |
| rs149792649 | benign | 0 | 0 | 0 |
| rs78101774 | benign | 0 | 0 | 0 |
| rs766466297 | pathogenic | 0 | 1 | 0 |
| rs140467171 | pathogenic | 0 | 0 | 0 |
| rs141340466 | pathogenic | 0 | 0 | 0 |
| rs201555303 | pathogenic | 0 | 0 | 0 |
| rs143592405 | pathogenic | 0 | 0 | 0 |
| rs781249411 | pathogenic | 0 | 0 | 0 |

| | | | | |
|---|---|---|---|---|
| rs116644127 | benign | 0 | 0 | 0 |
| rs151146863 | benign | 1 | 1 | 1 |
| rs116440799 | benign | 0 | 0 | 0 |
| rs138958351 | benign | 0 | 0 | 0 |
| rs143319354 | benign | 0 | 0 | 0 |
| rs768669208 | pathogenic | 1 | 1 | 1 |
| rs28940289 | pathogenic | 1 | 1 | 0 |
| rs745656120 | pathogenic | 1 | 1 | 0 |
| rs387906987 | pathogenic | 0 | 0 | 0 |
| rs147240777 | benign | 0 | 0 | 0 |
| rs140945833 | benign | 0 | 0 | 0 |
| rs146614981 | benign | 0 | 0 | 0 |
| rs149878562 | benign | 0 | 0 | 0 |
| rs28938168 | pathogenic | 0 | 0 | 0 |
| rs33939927 | pathogenic | 1 | 1 | 1 |
| rs34995376 | pathogenic | 1 | 1 | 1 |
| rs201034481 | pathogenic | 1 | 1 | 1 |
| rs774069983 | benign | 0 | 0 | 0 |
| rs199422226 | pathogenic | 1 | 1 | 0 |
| rs199422225 | pathogenic | 1 | 1 | 1 |
| rs199422224 | pathogenic | 1 | 1 | 0 |
| rs149271416 | pathogenic | 1 | 1 | 0 |
| rs1801318 | benign | 1 | 1 | 1 |
| rs372691318 | pathogenic | 1 | 1 | 1 |
| rs147685849 | benign | 0 | 0 | 0 |
| rs387907199 | pathogenic | 0 | 0 | 0 |
| rs767122069 | pathogenic | 0 | 0 | 0 |
| rs2302427 | benign | 0 | 0 | 0 |
| rs201135441 | benign | 0 | 0 | 0 |
| rs201861847 | pathogenic | 0 | 0 | 0 |
| rs137852952 | pathogenic | 0 | 0 | 0 |
| rs16986560 | benign | 0 | 0 | 0 |
| rs147344332 | pathogenic | 1 | 1 | 1 |
| rs36027220 | pathogenic | 0 | 0 | 0 |
| rs192401229 | benign | 0 | 0 | 0 |
| rs121909106 | pathogenic | 1 | 1 | 0 |
| rs118204103 | pathogenic | 1 | 1 | 0 |
| rs118204104 | pathogenic | 1 | 1 | 0 |
| rs118204105 | pathogenic | 1 | 1 | 0 |
| rs118204106 | pathogenic | 0 | 0 | 0 |
| rs118204119 | pathogenic | 0 | 0 | 0 |
| rs118204107 | pathogenic | 0 | 0 | 0 |
| rs118204094 | pathogenic | 0 | 0 | 0 |
| rs118204098 | pathogenic | 1 | 1 | 0 |
| rs118204101 | pathogenic | 1 | 1 | 0 |
| rs118204096 | pathogenic | 1 | 1 | 0 |
| rs118204108 | pathogenic | 0 | 0 | 0 |
| rs536814318 | pathogenic | 0 | 0 | 0 |
| rs34413634 | pathogenic | 1 | 1 | 0 |
| rs118204100 | pathogenic | 0 | 0 | 0 |
| rs118204109 | pathogenic | 0 | 0 | 0 |
| rs118204116 | pathogenic | 1 | 1 | 0 |
| rs118204110 | pathogenic | 0 | 0 | 0 |
| rs118204099 | pathogenic | 0 | 0 | 0 |

| | | | | |
|---|---|---|---|---|
| rs118204111 | pathogenic | 0 | 0 | 0 |
| rs118204112 | pathogenic | 0 | 0 | 0 |
| rs118204113 | pathogenic | 0 | 0 | 0 |
| rs118204114 | pathogenic | 0 | 0 | 0 |
| rs118204115 | pathogenic | 0 | 0 | 0 |
| rs150428209 | benign | 0 | 0 | 0 |
| rs144949995 | benign | 0 | 0 | 0 |
| rs190023981 | benign | 0 | 0 | 0 |
| rs104894611 | pathogenic | 1 | 1 | 1 |
| rs28937580 | pathogenic | 1 | 1 | 1 |
| rs121908949 | pathogenic | 0 | 0 | 0 |
| rs104894612 | pathogenic | 1 | 1 | 1 |
| rs104894609 | pathogenic | 1 | 1 | 1 |
| rs104894610 | pathogenic | 1 | 1 | 1 |
| rs104894615 | pathogenic | 0 | 0 | 0 |
| rs104894603 | pathogenic | 0 | 0 | 0 |
| rs104894602 | pathogenic | 1 | 1 | 1 |
| rs121908948 | pathogenic | 1 | 1 | 1 |
| rs104894608 | pathogenic | 1 | 1 | 1 |
| rs371545673 | benign | 0 | 0 | 0 |
| rs199925238 | benign | 0 | 0 | 0 |
| rs146313451 | benign | 0 | 0 | 0 |
| rs121918193 | pathogenic | 1 | 1 | 0 |
| rs2228500 | benign | 0 | 0 | 0 |
| rs121918194 | pathogenic | 1 | 1 | 0 |
| rs121918196 | pathogenic | 0 | 0 | 0 |
| rs41291971 | benign | 0 | 0 | 0 |
| rs199473690 | pathogenic | 0 | 0 | 0 |
| rs2228613 | benign | 0 | 0 | 0 |
| rs267607130 | pathogenic | 1 | 1 | 1 |
| rs77615401 | benign | 1 | 1 | 1 |
| rs374618872 | benign | 1 | 1 | 1 |
| rs146165014 | benign | 0 | 0 | 0 |
| rs144586070 | benign | 0 | 0 | 0 |
| rs121918541 | pathogenic | 0 | 0 | 0 |
| rs142105943 | benign | 0 | 0 | 0 |
| rs121918540 | pathogenic | 0 | 0 | 0 |
| rs6791 | benign | 0 | 0 | 0 |
| rs61736587 | pathogenic | 0 | 0 | 0 |
| rs61736586 | benign | 0 | 0 | 0 |
| rs121917741 | pathogenic | 1 | 1 | 1 |
| rs41302885 | benign | 0 | 0 | 0 |
| rs141026028 | benign | 0 | 0 | 0 |
| rs193922493 | pathogenic | 0 | 0 | 0 |
| rs73598374 | benign | 0 | 0 | 0 |
| rs121908725 | pathogenic | 1 | 1 | 0 |
| rs199422328 | pathogenic | 0 | 1 | 0 |
| rs11555566 | benign | 1 | 1 | 1 |
| rs267606634 | pathogenic | 0 | 0 | 0 |
| rs121908714 | pathogenic | 1 | 1 | 0 |
| rs121908717 | pathogenic | 1 | 1 | 0 |
| rs267606635 | pathogenic | 1 | 1 | 0 |
| rs121908739 | pathogenic | 0 | 0 | 0 |
| rs121908735 | pathogenic | 0 | 1 | 0 |

| | | | | |
|---|---|---|---|---|
| rs121908716 | pathogenic | 0 | 1 | 0 |
| rs121908723 | pathogenic | 0 | 1 | 0 |
| rs121908721 | pathogenic | 0 | 0 | 0 |
| rs199422327 | pathogenic | 0 | 0 | 0 |
| rs121908715 | pathogenic | 0 | 0 | 0 |
| rs587783563 | pathogenic | 0 | 0 | 0 |
| rs104894780 | pathogenic | 0 | 0 | 0 |
| rs56030372 | pathogenic | 1 | 1 | 1 |
| rs104894782 | pathogenic | 1 | 1 | 1 |
| rs149495971 | benign | 0 | 0 | 0 |
| rs118203942 | pathogenic | 1 | 1 | 0 |
| rs118203939 | pathogenic | 0 | 0 | 0 |
| rs118203938 | pathogenic | 0 | 0 | 0 |
| rs118203943 | pathogenic | 0 | 0 | 0 |
| rs118203940 | pathogenic | 0 | 0 | 0 |
| rs727503809 | pathogenic | 0 | 0 | 0 |
| rs1065757 | benign | 0 | 0 | 0 |
| rs1071598 | benign | 0 | 0 | 0 |
| rs25414 | benign | 0 | 0 | 0 |
| rs118203944 | pathogenic | 0 | 0 | 0 |
| rs118203941 | pathogenic | 0 | 0 | 0 |
| rs144222167 | benign | 0 | 0 | 0 |
| rs201101343 | pathogenic | 0 | 0 | 0 |
| rs144601717 | benign | 0 | 0 | 0 |
| rs267607129 | pathogenic | 0 | 0 | 0 |
| rs104894729 | pathogenic | 0 | 0 | 0 |
| rs104894727 | pathogenic | 0 | 0 | 0 |
| rs104894725 | pathogenic | 0 | 0 | 0 |
| rs104894515 | pathogenic | 1 | 1 | 1 |
| rs74688377 | benign | 0 | 0 | 0 |
| rs121908513 | pathogenic | 1 | 1 | 0 |
| rs121908509 | pathogenic | 0 | 0 | 0 |
| rs121908514 | pathogenic | 1 | 1 | 0 |
| rs121908512 | pathogenic | 1 | 1 | 0 |
| rs121908510 | pathogenic | 0 | 0 | 0 |
| rs121908511 | pathogenic | 1 | 1 | 1 |
| rs121908518 | pathogenic | 1 | 1 | 1 |
| rs34358665 | benign | 0 | 0 | 0 |
| rs147568834 | benign | 0 | 0 | 0 |
| rs201717599 | benign | 0 | 0 | 0 |
| rs61744487 | benign | 0 | 0 | 0 |
| rs370627877 | pathogenic | 1 | 1 | 1 |
| rs368593151 | pathogenic | 1 | 1 | 1 |
| rs114363936 | benign | 0 | 0 | 0 |
| rs137853333 | pathogenic | 0 | 0 | 0 |
| rs137853080 | pathogenic | 1 | 1 | 1 |
| rs137853077 | pathogenic | 1 | 1 | 1 |
| rs780749732 | benign | 1 | 1 | 1 |
| rs137853081 | pathogenic | 1 | 1 | 0 |
| rs137853078 | pathogenic | 0 | 0 | 0 |
| rs121913315 | pathogenic | 1 | 1 | 1 |
| rs752513525 | pathogenic | 1 | 1 | 1 |
| rs104894918 | pathogenic | 0 | 0 | 0 |
| rs104894917 | pathogenic | 0 | 0 | 0 |

| | | | | |
|---|---|---|---|---|
| rs104894919 | pathogenic | 0 | 0 | 0 |
| rs28939672 | pathogenic | 0 | 0 | 0 |
| rs104894173 | pathogenic | 0 | 0 | 0 |
| rs104894179 | pathogenic | 0 | 0 | 0 |
| rs104894174 | pathogenic | 1 | 1 | 0 |
| rs104894178 | pathogenic | 1 | 1 | 0 |
| rs28939714 | pathogenic | 1 | 1 | 1 |
| rs104894270 | pathogenic | 1 | 1 | 1 |
| rs3765571 | benign | 0 | 0 | 0 |
| rs752669339 | pathogenic | 1 | 1 | 1 |
| rs537043237 | pathogenic | 1 | 1 | 1 |
| rs774047299 | pathogenic | 1 | 1 | 1 |
| rs121434583 | pathogenic | 1 | 1 | 1 |
| rs121907927 | pathogenic | 0 | 0 | 0 |
| rs121907925 | pathogenic | 0 | 1 | 0 |
| rs746212325 | pathogenic | 0 | 0 | 0 |
| rs754637179 | pathogenic | 0 | 0 | 0 |
| rs780882740 | pathogenic | 0 | 0 | 0 |
| rs121918130 | pathogenic | 0 | 0 | 0 |
| rs121918129 | pathogenic | 0 | 0 | 0 |
| rs13297509 | pathogenic | 0 | 0 | 0 |
| rs746867724 | pathogenic | 0 | 0 | 0 |
| rs121918128 | pathogenic | 0 | 0 | 0 |
| rs752300607 | pathogenic | 0 | 0 | 0 |
| rs121964860 | pathogenic | 0 | 0 | 0 |
| rs45466197 | pathogenic | 1 | 1 | 1 |
| rs3730238 | benign | 1 | 1 | 1 |
| rs730881125 | pathogenic | 1 | 1 | 1 |
| rs121964861 | pathogenic | 1 | 1 | 1 |
| rs397516484 | pathogenic | 0 | 0 | 0 |
| rs367785431 | pathogenic | 0 | 0 | 0 |
| rs141121678 | pathogenic | 0 | 0 | 0 |
| rs12720263 | benign | 0 | 0 | 0 |
| rs2304256 | benign | 0 | 0 | 0 |
| rs2304255 | benign | 0 | 0 | 0 |
| rs121909045 | pathogenic | 0 | 0 | 0 |
| rs397508796 | pathogenic | 0 | 0 | 0 |
| rs151020603 | pathogenic | 0 | 0 | 0 |
| rs77284892 | pathogenic | 0 | 0 | 0 |
| rs368505753 | pathogenic | 0 | 0 | 0 |
| rs115545701 | pathogenic | 0 | 0 | 0 |
| rs121908749 | pathogenic | 0 | 0 | 0 |
| rs75961395 | pathogenic | 0 | 0 | 0 |
| rs121908750 | pathogenic | 0 | 0 | 0 |
| rs121908751 | pathogenic | 0 | 0 | 0 |
| rs397508464 | pathogenic | 0 | 0 | 0 |
| rs397508461 | pathogenic | 0 | 0 | 0 |
| rs121909031 | pathogenic | 0 | 0 | 0 |
| rs113993958 | pathogenic | 0 | 0 | 0 |
| rs77834169 | pathogenic | 0 | 0 | 0 |
| rs78655421 | pathogenic | 0 | 0 | 0 |
| rs141723617 | benign | 0 | 0 | 0 |
| rs35516286 | benign | 0 | 0 | 0 |
| rs1800079 | pathogenic | 1 | 1 | 0 |

| | | | | |
|---|---|---|---|---|
| rs80282562 | pathogenic | 0 | 0 | 0 |
| rs121908752 | pathogenic | 0 | 0 | 0 |
| rs556880586 | pathogenic | 0 | 0 | 0 |
| rs121908760 | pathogenic | 0 | 0 | 0 |
| rs397508350 | pathogenic | 0 | 0 | 0 |
| rs374946172 | pathogenic | 0 | 0 | 0 |
| rs397508387 | pathogenic | 0 | 0 | 0 |
| rs201386642 | pathogenic | 0 | 0 | 0 |
| rs121909012 | pathogenic | 0 | 0 | 0 |
| rs121909008 | pathogenic | 0 | 0 | 0 |
| rs149790377 | pathogenic | 0 | 0 | 0 |
| rs397508442 | pathogenic | 0 | 0 | 0 |
| rs121909035 | pathogenic | 0 | 0 | 0 |
| rs386134230 | pathogenic | 0 | 0 | 0 |
| rs1800112 | benign | 0 | 0 | 0 |
| rs144055758 | pathogenic | 0 | 0 | 0 |
| rs142394380 | pathogenic | 0 | 0 | 0 |
| rs121909036 | pathogenic | 0 | 0 | 0 |
| rs78194216 | pathogenic | 0 | 0 | 0 |
| rs121909019 | pathogenic | 0 | 0 | 0 |
| rs121909020 | pathogenic | 0 | 0 | 0 |
| rs200321110 | pathogenic | 0 | 0 | 0 |
| rs202179988 | pathogenic | 0 | 0 | 0 |
| rs121909037 | pathogenic | 0 | 0 | 0 |
| rs186045772 | pathogenic | 0 | 0 | 0 |
| rs139304906 | pathogenic | 0 | 0 | 0 |
| rs78802634 | pathogenic | 0 | 0 | 0 |
| rs121908761 | pathogenic | 0 | 0 | 0 |
| rs36210737 | pathogenic | 0 | 0 | 0 |
| rs139729994 | pathogenic | 0 | 0 | 0 |
| rs79850223 | pathogenic | 0 | 0 | 0 |
| rs1800120 | benign | 0 | 0 | 0 |
| rs74767530 | pathogenic | 0 | 0 | 0 |
| rs374705585 | pathogenic | 0 | 0 | 0 |
| rs1136287 | benign | 0 | 0 | 0 |
| rs6828 | benign | 0 | 0 | 0 |
| rs104894856 | pathogenic | 0 | 0 | 0 |
| rs104894860 | pathogenic | 0 | 0 | 0 |
| rs61736892 | pathogenic | 0 | 0 | 0 |
| rs145807417 | benign | 0 | 0 | 0 |
| rs104894853 | pathogenic | 0 | 0 | 0 |
| rs199422227 | pathogenic | 0 | 0 | 0 |
| rs113993946 | pathogenic | 0 | 0 | 0 |
| rs199422231 | pathogenic | 0 | 0 | 0 |
| rs199422228 | pathogenic | 0 | 0 | 0 |
| rs142195196 | benign | 0 | 1 | 0 |
| rs148656104 | benign | 1 | 1 | 1 |
| rs141944844 | benign | 0 | 0 | 0 |
| rs104894342 | pathogenic | 0 | 0 | 0 |
| rs104894343 | pathogenic | 0 | 0 | 0 |
| rs104894344 | pathogenic | 0 | 0 | 0 |
| rs573322878 | benign | 0 | 0 | 0 |
| rs104894347 | pathogenic | 0 | 0 | 0 |
| rs28937882 | pathogenic | 0 | 0 | 0 |

| | | | | |
|---|---|---|---|---|
| rs193922702 | pathogenic | 0 | 0 | 0 |
| rs200786104 | benign | 0 | 0 | 0 |
| rs690016544 | pathogenic | 1 | 1 | 1 |
| rs28943592 | benign | 0 | 0 | 0 |
| rs28943594 | benign | 0 | 0 | 0 |
| rs121908957 | pathogenic | 0 | 0 | 0 |
| rs115408010 | benign | 1 | 1 | 1 |
| rs77591659 | benign | 0 | 0 | 0 |
| rs121918511 | pathogenic | 0 | 0 | 0 |
| rs538359948 | benign | 0 | 0 | 0 |
| rs76708142 | benign | 0 | 0 | 0 |
| rs121907913 | pathogenic | 1 | 1 | 0 |
| rs121907920 | pathogenic | 0 | 0 | 0 |
| rs121907923 | pathogenic | 1 | 1 | 0 |
| rs121907928 | pathogenic | 1 | 1 | 0 |
| rs121907919 | pathogenic | 1 | 1 | 0 |
| rs121907918 | pathogenic | 1 | 1 | 0 |
| rs74315493 | pathogenic | 1 | 1 | 1 |
| rs74315494 | pathogenic | 1 | 1 | 1 |
| rs145446060 | benign | 1 | 1 | 1 |
| rs550409143 | benign | 1 | 1 | 0 |

| compound_binding_site | cofactor_binding_site | nucleic_binding_site | ion_binding_site |
|---|---|---|---|
| 1 | 0 | 0 | 0 |
| 1 | 0 | 0 | 0 |
| 0 | 0 | 0 | 0 |
| 1 | 0 | 0 | 0 |
| 0 | 0 | 0 | 0 |
| 0 | 0 | 0 | 0 |
| 0 | 0 | 0 | 0 |
| 0 | 0 | 0 | 0 |
| 1 | 0 | 0 | 1 |
| 1 | 0 | 0 | 0 |
| 0 | 0 | 0 | 1 |
| 1 | 0 | 1 | 0 |
| 0 | 0 | 0 | 0 |
| 0 | 0 | 0 | 0 |
| 0 | 0 | 0 | 0 |
| 0 | 0 | 0 | 0 |
| 0 | 0 | 0 | 0 |
| 1 | 0 | 0 | 1 |
| 1 | 0 | 1 | 0 |
| 0 | 0 | 0 | 0 |
| 1 | 0 | 0 | 1 |
| 1 | 0 | 0 | 0 |
| 1 | 0 | 0 | 0 |
| 1 | 0 | 0 | 0 |
| 0 | 0 | 0 | 0 |
| 0 | 0 | 0 | 0 |
| 1 | 0 | 0 | 1 |
| 0 | 0 | 0 | 0 |
| 0 | 0 | 0 | 0 |
| 0 | 0 | 0 | 0 |
| 0 | 0 | 0 | 0 |
| 1 | 0 | 0 | 0 |
| 1 | 0 | 0 | 1 |
| 1 | 0 | 0 | 0 |
| 0 | 0 | 0 | 0 |
| 0 | 0 | 0 | 0 |
| 0 | 0 | 0 | 0 |
| 1 | 0 | 0 | 1 |
| 1 | 0 | 0 | 1 |
| 1 | 0 | 0 | 1 |
| 1 | 0 | 0 | 0 |
| 1 | 0 | 0 | 0 |
| 1 | 0 | 0 | 0 |
| 1 | 0 | 0 | 0 |
| 1 | 0 | 0 | 0 |
| 1 | 0 | 0 | 0 |
| 1 | 0 | 0 | 0 |
| 1 | 0 | 0 | 0 |
| 1 | 0 | 0 | 0 |
| 1 | 0 | 0 | 0 |
| 1 | 0 | 0 | 0 |
| 1 | 0 | 0 | 0 |
| 1 | 0 | 0 | 0 |

| | | | |
|---|---|---|---|
| 1 | 0 | 0 | 1 |
| 1 | 0 | 0 | 0 |
| 1 | 0 | 0 | 0 |
| 1 | 0 | 0 | 0 |
| 0 | 0 | 0 | 0 |
| 1 | 0 | 0 | 0 |
| 0 | 0 | 0 | 0 |
| 1 | 0 | 0 | 0 |
| 1 | 0 | 0 | 0 |
| 1 | 0 | 0 | 1 |
| 1 | 0 | 0 | 0 |
| 1 | 0 | 0 | 0 |
| 1 | 0 | 0 | 0 |
| 1 | 0 | 0 | 0 |
| 1 | 0 | 0 | 0 |
| 1 | 0 | 0 | 0 |
| 1 | 0 | 0 | 0 |
| 1 | 0 | 0 | 1 |
| 1 | 0 | 0 | 1 |
| 1 | 1 | 0 | 1 |
| 1 | 0 | 0 | 0 |
| 1 | 0 | 0 | 0 |
| 1 | 0 | 0 | 0 |
| 1 | 0 | 0 | 0 |
| 1 | 1 | 0 | 0 |
| 1 | 0 | 0 | 0 |
| 1 | 0 | 0 | 0 |
| 1 | 0 | 0 | 0 |
| 1 | 0 | 0 | 0 |
| 1 | 0 | 0 | 0 |
| 1 | 1 | 0 | 0 |
| 1 | 1 | 0 | 0 |
| 1 | 1 | 0 | 0 |
| 1 | 1 | 0 | 0 |
| 1 | 1 | 0 | 0 |
| 1 | 0 | 0 | 0 |
| 1 | 1 | 0 | 0 |
| 1 | 0 | 0 | 0 |
| 1 | 0 | 0 | 0 |
| 1 | 1 | 0 | 0 |
| 0 | 0 | 0 | 0 |
| 0 | 0 | 0 | 0 |
| 0 | 0 | 0 | 0 |
| 0 | 0 | 0 | 0 |
| 0 | 0 | 0 | 0 |
| 0 | 0 | 0 | 0 |
| 0 | 0 | 0 | 0 |
| 1 | 0 | 0 | 1 |
| 0 | 0 | 0 | 0 |
| 1 | 0 | 0 | 0 |
| 1 | 0 | 0 | 0 |
| 0 | 0 | 0 | 0 |
| 0 | 0 | 0 | 0 |
| 0 | 0 | 0 | 0 |

| | | | |
|---|---|---|---|
| 0 | 0 | 0 | 0 |
| 0 | 0 | 0 | 0 |
| 0 | 0 | 0 | 0 |
| 1 | 0 | 0 | 0 |
| 1 | 0 | 0 | 0 |
| 1 | 0 | 0 | 0 |
| 0 | 0 | 0 | 0 |
| 0 | 0 | 1 | 0 |
| 0 | 0 | 1 | 0 |
| 0 | 0 | 1 | 0 |
| 0 | 0 | 1 | 0 |
| 0 | 0 | 1 | 0 |
| 0 | 0 | 0 | 0 |
| 0 | 0 | 0 | 0 |
| 1 | 0 | 0 | 0 |
| 0 | 0 | 0 | 0 |
| 0 | 0 | 0 | 0 |
| 1 | 0 | 0 | 0 |
| 1 | 0 | 0 | 0 |
| 1 | 0 | 0 | 0 |
| 1 | 0 | 0 | 0 |
| 1 | 1 | 0 | 0 |
| 1 | 0 | 0 | 0 |
| 1 | 0 | 0 | 0 |
| 0 | 0 | 0 | 0 |
| 0 | 0 | 0 | 0 |
| 1 | 0 | 0 | 0 |
| 1 | 1 | 0 | 0 |
| 1 | 0 | 0 | 0 |
| 1 | 0 | 0 | 0 |
| 1 | 1 | 0 | 0 |
| 1 | 1 | 0 | 0 |
| 1 | 1 | 0 | 0 |
| 1 | 1 | 0 | 0 |
| 1 | 1 | 0 | 0 |
| 1 | 1 | 0 | 0 |
| 1 | 1 | 0 | 0 |
| 1 | 1 | 0 | 0 |
| 1 | 1 | 0 | 0 |
| 1 | 1 | 0 | 0 |
| 1 | 1 | 0 | 0 |
| 1 | 1 | 0 | 0 |
| 1 | 0 | 0 | 0 |
| 1 | 0 | 0 | 0 |
| 1 | 0 | 0 | 0 |
| 1 | 1 | 0 | 0 |
| 1 | 0 | 0 | 0 |
| 1 | 1 | 1 | 0 |
| 1 | 1 | 1 | 0 |
| 1 | 1 | 0 | 0 |
| 1 | 1 | 0 | 0 |
| 1 | 0 | 0 | 0 |
| 1 | 0 | 0 | 0 |
| 0 | 0 | 0 | 0 |

| | | | |
|---|---|---|---|
| 0 | 0 | 0 | 0 |
| 0 | 0 | 0 | 0 |
| 0 | 0 | 0 | 0 |
| 1 | 0 | 0 | 0 |
| 1 | 0 | 0 | 0 |
| 0 | 0 | 0 | 0 |
| 0 | 0 | 0 | 0 |
| 0 | 0 | 0 | 0 |
| 0 | 0 | 0 | 0 |
| 0 | 0 | 0 | 0 |
| 0 | 0 | 0 | 0 |
| 0 | 0 | 0 | 0 |
| 0 | 0 | 0 | 0 |
| 0 | 0 | 0 | 0 |
| 0 | 0 | 0 | 0 |
| 0 | 0 | 0 | 0 |
| 0 | 0 | 0 | 0 |
| 0 | 0 | 0 | 0 |
| 0 | 0 | 0 | 0 |
| 0 | 0 | 0 | 0 |
| 0 | 0 | 0 | 0 |
| 0 | 0 | 0 | 0 |
| 1 | 0 | 0 | 0 |
| 1 | 0 | 0 | 0 |
| 0 | 0 | 0 | 0 |
| 0 | 0 | 0 | 0 |
| 1 | 0 | 0 | 0 |
| 0 | 0 | 0 | 0 |
| 0 | 0 | 0 | 0 |
| 0 | 0 | 0 | 0 |
| 0 | 0 | 0 | 0 |
| 0 | 0 | 0 | 0 |
| 0 | 0 | 0 | 0 |
| 0 | 0 | 0 | 0 |
| 1 | 0 | 0 | 0 |
| 0 | 0 | 0 | 0 |
| 0 | 0 | 0 | 0 |
| 0 | 0 | 0 | 0 |
| 0 | 0 | 0 | 0 |
| 0 | 0 | 0 | 0 |
| 0 | 0 | 0 | 0 |
| 0 | 0 | 0 | 0 |
| 0 | 0 | 0 | 0 |
| 0 | 0 | 0 | 0 |
| 0 | 0 | 0 | 0 |
| 0 | 0 | 0 | 0 |
| 0 | 0 | 1 | 0 |
| 1 | 0 | 0 | 0 |
| 1 | 0 | 0 | 0 |
| 1 | 1 | 0 | 0 |
| 1 | 0 | 0 | 0 |
| 0 | 0 | 0 | 0 |
| 0 | 0 | 0 | 0 |
| 1 | 0 | 0 | 0 |

| | | | |
|---|---|---|---|
| 1 | 0 | 0 | 0 |
| 1 | 0 | 0 | 0 |
| 1 | 0 | 0 | 0 |
| 0 | 0 | 0 | 0 |
| 0 | 0 | 0 | 0 |
| 0 | 0 | 0 | 0 |
| 1 | 0 | 0 | 0 |
| 1 | 0 | 0 | 0 |
| 0 | 0 | 0 | 0 |
| 0 | 0 | 0 | 0 |
| 1 | 0 | 0 | 0 |
| 0 | 0 | 0 | 0 |
| 1 | 0 | 0 | 0 |
| 0 | 0 | 0 | 0 |
| 0 | 0 | 0 | 0 |
| 0 | 0 | 0 | 0 |
| 0 | 0 | 0 | 0 |
| 0 | 0 | 0 | 0 |
| 0 | 0 | 0 | 0 |
| 0 | 0 | 0 | 0 |
| 0 | 0 | 0 | 0 |
| 0 | 0 | 0 | 0 |
| 0 | 0 | 0 | 0 |
| 0 | 0 | 0 | 0 |
| 0 | 0 | 0 | 0 |
| 0 | 0 | 0 | 0 |
| 1 | 0 | 0 | 0 |
| 1 | 0 | 0 | 0 |
| 1 | 0 | 0 | 0 |
| 1 | 0 | 0 | 0 |
| 1 | 0 | 0 | 0 |
| 1 | 0 | 0 | 0 |
| 0 | 0 | 0 | 0 |
| 0 | 0 | 0 | 0 |
| 1 | 0 | 0 | 0 |
| 0 | 0 | 0 | 0 |
| 0 | 0 | 0 | 0 |
| 0 | 0 | 0 | 0 |
| 0 | 0 | 0 | 0 |
| 0 | 0 | 0 | 0 |
| 0 | 0 | 0 | 0 |
| 0 | 0 | 0 | 0 |
| 0 | 0 | 0 | 0 |
| 0 | 0 | 0 | 0 |
| 1 | 1 | 0 | 0 |
| 1 | 1 | 0 | 0 |
| 1 | 1 | 0 | 1 |
| 1 | 1 | 0 | 1 |
| 1 | 1 | 0 | 1 |
| 1 | 1 | 0 | 1 |
| 1 | 1 | 0 | 0 |
| 1 | 1 | 0 | 1 |
| 1 | 1 | 0 | 1 |
| 1 | 1 | 0 | 1 |

| | | | |
|---|---|---|---|
| 1 | 1 | 0 | 1 |
| 1 | 1 | 0 | 1 |
| 1 | 1 | 0 | 1 |
| 1 | 1 | 0 | 1 |
| 1 | 1 | 0 | 1 |
| 1 | 0 | 0 | 0 |
| 0 | 0 | 0 | 0 |
| 0 | 0 | 0 | 0 |
| 0 | 0 | 0 | 0 |
| 1 | 0 | 0 | 0 |
| 0 | 0 | 1 | 0 |
| 0 | 0 | 0 | 0 |
| 0 | 0 | 0 | 0 |
| 0 | 0 | 0 | 0 |
| 0 | 0 | 0 | 0 |
| 0 | 0 | 0 | 0 |
| 0 | 0 | 0 | 0 |
| 0 | 0 | 0 | 0 |
| 0 | 0 | 0 | 0 |
| 0 | 0 | 0 | 0 |
| 0 | 0 | 0 | 0 |
| 0 | 0 | 0 | 0 |
| 0 | 0 | 0 | 0 |
| 0 | 0 | 0 | 0 |
| 0 | 0 | 0 | 0 |
| 1 | 1 | 0 | 0 |
| 1 | 1 | 0 | 0 |
| 1 | 1 | 0 | 0 |
| 0 | 1 | 0 | 0 |
| 1 | 0 | 0 | 0 |
| 0 | 0 | 0 | 0 |
| 0 | 0 | 0 | 0 |
| 0 | 0 | 0 | 0 |
| 1 | 0 | 0 | 0 |
| 1 | 1 | 0 | 0 |
| 0 | 0 | 0 | 0 |
| 0 | 0 | 0 | 0 |
| 0 | 0 | 0 | 0 |
| 0 | 0 | 0 | 0 |
| 1 | 0 | 0 | 0 |
| 0 | 0 | 0 | 0 |
| 0 | 0 | 0 | 0 |
| 0 | 0 | 0 | 0 |
| 0 | 0 | 0 | 0 |
| 0 | 0 | 0 | 0 |
| 0 | 1 | 0 | 0 |
| 0 | 1 | 0 | 0 |
| 0 | 0 | 0 | 0 |
| 0 | 0 | 0 | 0 |
| 0 | 1 | 0 | 0 |
| 0 | 0 | 0 | 0 |
| 0 | 0 | 0 | 0 |
| 0 | 1 | 0 | 0 |
| 0 | 0 | 0 | 0 |

| | | | |
|---|---|---|---|
| 1 | 1 | 0 | 0 |
| 0 | 0 | 0 | 0 |
| 0 | 0 | 0 | 0 |
| 0 | 0 | 0 | 0 |
| 0 | 0 | 0 | 0 |
| 0 | 0 | 0 | 0 |
| 0 | 0 | 0 | 0 |
| 0 | 0 | 0 | 0 |
| 0 | 0 | 0 | 0 |
| 0 | 0 | 0 | 0 |
| 0 | 0 | 0 | 0 |
| 0 | 0 | 0 | 0 |
| 0 | 0 | 0 | 0 |
| 0 | 0 | 0 | 0 |
| 0 | 1 | 0 | 0 |
| 0 | 0 | 0 | 0 |
| 0 | 1 | 0 | 0 |
| 0 | 1 | 0 | 0 |
| 0 | 0 | 0 | 0 |
| 0 | 1 | 0 | 0 |
| 1 | 0 | 0 | 1 |
| 1 | 1 | 0 | 0 |
| 1 | 0 | 0 | 0 |
| 1 | 0 | 0 | 1 |
| 0 | 0 | 0 | 0 |
| 0 | 0 | 0 | 0 |
| 0 | 0 | 0 | 0 |
| 0 | 0 | 0 | 0 |
| 1 | 0 | 0 | 0 |
| 0 | 0 | 0 | 0 |
| 1 | 0 | 0 | 0 |
| 0 | 0 | 0 | 0 |
| 0 | 0 | 0 | 0 |
| 0 | 0 | 0 | 0 |
| 0 | 0 | 0 | 0 |
| 0 | 0 | 0 | 0 |
| 0 | 0 | 0 | 0 |
| 1 | 1 | 0 | 0 |
| 1 | 1 | 0 | 0 |
| 1 | 1 | 0 | 0 |
| 0 | 0 | 0 | 0 |
| 0 | 0 | 0 | 0 |
| 0 | 0 | 0 | 0 |
| 1 | 0 | 0 | 0 |
| 1 | 0 | 0 | 1 |
| 1 | 0 | 0 | 1 |
| 1 | 0 | 0 | 0 |
| 1 | 0 | 0 | 1 |
| 1 | 0 | 0 | 0 |
| 0 | 0 | 0 | 0 |
| 1 | 1 | 0 | 0 |
| 1 | 1 | 0 | 0 |
| 0 | 0 | 0 | 0 |
| 0 | 0 | 0 | 0 |

| | | | |
|---|---|---|---|
| 1 | 0 | 0 | 0 |
| 1 | 0 | 0 | 0 |
| 1 | 0 | 0 | 0 |
| 0 | 0 | 0 | 0 |
| 0 | 0 | 0 | 0 |
| 1 | 1 | 0 | 1 |
| 1 | 1 | 0 | 1 |
| 1 | 1 | 0 | 1 |
| 1 | 1 | 0 | 1 |
| 1 | 1 | 0 | 1 |
| 1 | 1 | 0 | 1 |
| 1 | 1 | 0 | 1 |
| 1 | 1 | 0 | 0 |
| 1 | 1 | 0 | 0 |
| 1 | 1 | 1 | 1 |
| 0 | 0 | 0 | 0 |
| 1 | 1 | 0 | 0 |
| 1 | 1 | 0 | 0 |
| 0 | 0 | 0 | 0 |
| 0 | 0 | 0 | 0 |
| 0 | 0 | 0 | 0 |
| 1 | 0 | 0 | 0 |
| 1 | 0 | 0 | 0 |
| 0 | 0 | 0 | 0 |
| 1 | 0 | 0 | 0 |
| 1 | 0 | 0 | 0 |
| 1 | 0 | 0 | 0 |
| 0 | 0 | 0 | 0 |
| 0 | 0 | 0 | 0 |
| 0 | 0 | 0 | 0 |
| 0 | 0 | 0 | 0 |
| 0 | 0 | 0 | 0 |
| 0 | 0 | 0 | 0 |
| 0 | 0 | 0 | 0 |
| 0 | 0 | 0 | 0 |
| 0 | 0 | 0 | 0 |
| 0 | 0 | 0 | 0 |
| 0 | 0 | 0 | 0 |
| 0 | 0 | 0 | 0 |
| 0 | 0 | 0 | 0 |
| 0 | 0 | 0 | 0 |
| 0 | 0 | 0 | 0 |
| 0 | 0 | 0 | 0 |
| 0 | 0 | 0 | 0 |
| 0 | 0 | 0 | 0 |
| 1 | 0 | 0 | 0 |
| 1 | 0 | 0 | 0 |
| 1 | 0 | 0 | 0 |
| 1 | 0 | 0 | 0 |
| 0 | 0 | 0 | 0 |
| 0 | 0 | 0 | 0 |
| 0 | 0 | 0 | 0 |
| 0 | 0 | 0 | 0 |
| 0 | 0 | 0 | 0 |

| | | | |
|---|---|---|---|
| 0 | 0 | 0 | 0 |
| 0 | 0 | 0 | 0 |
| 0 | 0 | 0 | 0 |
| 0 | 0 | 0 | 0 |
| 0 | 0 | 0 | 0 |
| 0 | 0 | 0 | 0 |
| 0 | 0 | 0 | 0 |
| 1 | 0 | 0 | 1 |
| 0 | 0 | 0 | 0 |
| 0 | 0 | 0 | 0 |
| 1 | 0 | 0 | 0 |
| 0 | 0 | 0 | 0 |
| 0 | 0 | 0 | 0 |
| 0 | 0 | 0 | 0 |
| 0 | 0 | 0 | 0 |
| 1 | 0 | 0 | 0 |
| 0 | 0 | 0 | 0 |
| 0 | 0 | 0 | 0 |
| 0 | 0 | 0 | 0 |
| 0 | 0 | 0 | 0 |
| 0 | 0 | 0 | 0 |
| 0 | 0 | 0 | 0 |
| 0 | 0 | 0 | 0 |
| 1 | 1 | 0 | 1 |
| 0 | 0 | 0 | 0 |
| 0 | 0 | 0 | 0 |
| 0 | 0 | 0 | 0 |
| 1 | 1 | 0 | 1 |
| 1 | 1 | 0 | 0 |
| 0 | 0 | 0 | 0 |
| 1 | 0 | 0 | 0 |
| 1 | 1 | 0 | 0 |
| 0 | 0 | 0 | 0 |
| 1 | 0 | 0 | 0 |
| 1 | 1 | 0 | 0 |
| 1 | 0 | 0 | 0 |
| 1 | 1 | 0 | 1 |
| 1 | 1 | 0 | 1 |
| 1 | 0 | 0 | 0 |
| 0 | 0 | 0 | 0 |
| 0 | 0 | 0 | 0 |
| 0 | 0 | 0 | 0 |
| 0 | 0 | 0 | 0 |
| 0 | 0 | 0 | 0 |
| 0 | 0 | 0 | 0 |
| 0 | 0 | 0 | 0 |
| 0 | 0 | 0 | 0 |
| 0 | 0 | 0 | 0 |
| 0 | 0 | 0 | 0 |
| 0 | 0 | 0 | 0 |
| 0 | 0 | 0 | 0 |
| 0 | 0 | 0 | 0 |
| 0 | 0 | 0 | 0 |
| 0 | 0 | 0 | 0 |

| | | | |
|---|---|---|---|
| 0 | 0 | 0 | 0 |
| 0 | 0 | 0 | 0 |
| 0 | 0 | 0 | 0 |
| 0 | 0 | 0 | 0 |
| 0 | 0 | 0 | 0 |
| 0 | 0 | 0 | 0 |
| 0 | 0 | 0 | 0 |
| 0 | 0 | 0 | 0 |
| 0 | 0 | 0 | 0 |
| 0 | 0 | 0 | 0 |
| 0 | 0 | 0 | 0 |
| 0 | 0 | 0 | 0 |
| 0 | 0 | 0 | 0 |
| 1 | 1 | 0 | 0 |
| 0 | 0 | 0 | 0 |
| 1 | 0 | 0 | 0 |
| 0 | 0 | 0 | 1 |
| 0 | 0 | 0 | 0 |
| 0 | 0 | 0 | 0 |
| 0 | 0 | 0 | 0 |
| 0 | 0 | 0 | 0 |
| 0 | 0 | 0 | 0 |
| 0 | 0 | 0 | 0 |
| 0 | 0 | 0 | 0 |
| 0 | 0 | 0 | 0 |
| 0 | 0 | 0 | 0 |
| 0 | 0 | 0 | 0 |
| 0 | 0 | 0 | 0 |
| 0 | 0 | 0 | 0 |
| 0 | 0 | 0 | 0 |
| 1 | 0 | 0 | 0 |
| 1 | 0 | 0 | 0 |
| 0 | 0 | 0 | 0 |
| 0 | 0 | 0 | 0 |
| 0 | 0 | 0 | 0 |
| 0 | 0 | 0 | 0 |
| 0 | 0 | 0 | 0 |
| 0 | 0 | 0 | 0 |
| 0 | 0 | 0 | 0 |
| 0 | 0 | 0 | 0 |
| 1 | 0 | 0 | 0 |
| 1 | 0 | 0 | 0 |
| 1 | 0 | 0 | 0 |
| 1 | 0 | 0 | 0 |
| 0 | 0 | 0 | 0 |
| 1 | 0 | 0 | 0 |
| 0 | 0 | 0 | 0 |
| 0 | 0 | 0 | 0 |
| 1 | 0 | 0 | 0 |
| 1 | 0 | 0 | 0 |
| 0 | 0 | 0 | 0 |
| 0 | 0 | 0 | 0 |
| 0 | 0 | 0 | 0 |
| 0 | 0 | 0 | 0 |

| | | | |
|---|---|---|---|
| 0 | 0 | 0 | 0 |
| 0 | 0 | 0 | 0 |
| 0 | 0 | 0 | 0 |
| 0 | 0 | 0 | 0 |
| 0 | 0 | 0 | 0 |
| 0 | 0 | 0 | 0 |
| 0 | 0 | 0 | 0 |
| 0 | 0 | 0 | 0 |
| 0 | 0 | 0 | 0 |
| 0 | 0 | 0 | 0 |
| 0 | 0 | 0 | 0 |
| 0 | 0 | 0 | 0 |
| 0 | 0 | 0 | 0 |
| 0 | 0 | 0 | 0 |
| 0 | 0 | 0 | 0 |
| 1 | 0 | 0 | 1 |
| 0 | 0 | 0 | 0 |
| 0 | 0 | 0 | 0 |
| 0 | 0 | 0 | 0 |
| 0 | 0 | 0 | 0 |
| 0 | 0 | 0 | 0 |
| 1 | 1 | 0 | 0 |
| 1 | 1 | 0 | 0 |
| 1 | 0 | 0 | 0 |
| 1 | 0 | 0 | 0 |
| 0 | 0 | 0 | 0 |
| 0 | 0 | 0 | 0 |
| 0 | 0 | 0 | 0 |
| 0 | 0 | 0 | 0 |
| 1 | 0 | 0 | 0 |
| 1 | 0 | 0 | 0 |
| 1 | 0 | 0 | 0 |
| 1 | 1 | 0 | 0 |
| 1 | 1 | 0 | 0 |
| 1 | 0 | 0 | 0 |
| 0 | 0 | 0 | 0 |
| 0 | 0 | 0 | 0 |
| 0 | 1 | 0 | 0 |
| 0 | 0 | 0 | 0 |
| 0 | 0 | 0 | 0 |
| 0 | 0 | 0 | 0 |
| 1 | 1 | 0 | 0 |
| 1 | 0 | 0 | 0 |
| 1 | 0 | 0 | 0 |
| 0 | 0 | 0 | 0 |
| 0 | 0 | 0 | 0 |
| 0 | 0 | 0 | 0 |
| 0 | 0 | 0 | 0 |
| 0 | 0 | 0 | 0 |
| 0 | 0 | 0 | 0 |
| 0 | 0 | 0 | 0 |
| 0 | 0 | 0 | 0 |
| 0 | 0 | 0 | 0 |
| 0 | 0 | 0 | 0 |

| | | | |
|---|---|---|---|
| 0 | 0 | 0 | 0 |
| 1 | 0 | 0 | 0 |
| 0 | 0 | 0 | 0 |
| 0 | 0 | 0 | 0 |
| 0 | 0 | 0 | 0 |
| 0 | 0 | 0 | 0 |
| 1 | 0 | 0 | 0 |
| 0 | 0 | 0 | 0 |
| 0 | 0 | 0 | 0 |
| 0 | 0 | 0 | 0 |
| 0 | 0 | 0 | 0 |
| 0 | 0 | 0 | 0 |
| 0 | 0 | 0 | 0 |
| 0 | 0 | 0 | 0 |
| 0 | 0 | 0 | 0 |
| 0 | 0 | 0 | 0 |
| 0 | 1 | 0 | 0 |
| 0 | 0 | 0 | 0 |
| 0 | 0 | 0 | 0 |
| 0 | 0 | 0 | 0 |
| 1 | 0 | 0 | 0 |
| 0 | 0 | 0 | 0 |
| 0 | 0 | 0 | 0 |
| 1 | 1 | 0 | 0 |
| 0 | 0 | 0 | 0 |
| 0 | 0 | 0 | 0 |
| 0 | 0 | 0 | 0 |
| 0 | 0 | 0 | 0 |
| 1 | 0 | 0 | 0 |
| 0 | 0 | 0 | 0 |
| 0 | 0 | 0 | 0 |
| 0 | 0 | 0 | 0 |
| 0 | 0 | 0 | 0 |
| 0 | 0 | 0 | 0 |
| 0 | 0 | 0 | 0 |
| 1 | 0 | 0 | 0 |
| 0 | 0 | 0 | 0 |
| 0 | 0 | 0 | 0 |
| 0 | 0 | 0 | 0 |
| 0 | 0 | 0 | 0 |
| 1 | 1 | 0 | 0 |
| 0 | 0 | 0 | 0 |
| 0 | 0 | 0 | 0 |
| 0 | 0 | 0 | 0 |
| 0 | 0 | 0 | 0 |
| 0 | 0 | 0 | 0 |
| 1 | 0 | 0 | 0 |
| 1 | 0 | 0 | 1 |
| 1 | 0 | 0 | 0 |
| 0 | 0 | 0 | 0 |
| 0 | 0 | 0 | 0 |
| 0 | 0 | 0 | 0 |
| 1 | 0 | 0 | 0 |
| 1 | 0 | 0 | 0 |

| | | | |
|---|---|---|---|
| 1 | 0 | 0 | 0 |
| 0 | 0 | 0 | 0 |
| 0 | 0 | 0 | 0 |
| 0 | 0 | 0 | 0 |
| 1 | 1 | 0 | 0 |
| 1 | 1 | 0 | 0 |
| 0 | 0 | 0 | 0 |
| 0 | 0 | 0 | 0 |
| 1 | 0 | 0 | 0 |
| 1 | 0 | 0 | 0 |
| 1 | 0 | 1 | 0 |
| 0 | 0 | 0 | 0 |
| 0 | 0 | 1 | 0 |
| 0 | 0 | 1 | 0 |
| 0 | 0 | 1 | 0 |
| 1 | 0 | 0 | 0 |
| 1 | 0 | 0 | 0 |
| 0 | 0 | 0 | 0 |
| 0 | 0 | 0 | 0 |
| 1 | 0 | 0 | 1 |
| 1 | 0 | 0 | 1 |
| 1 | 0 | 0 | 0 |
| 0 | 0 | 0 | 0 |
| 0 | 0 | 0 | 0 |
| 0 | 0 | 0 | 0 |
| 0 | 0 | 0 | 0 |
| 0 | 0 | 0 | 0 |
| 1 | 0 | 0 | 0 |
| 0 | 0 | 0 | 0 |
| 0 | 0 | 0 | 1 |
| 0 | 0 | 0 | 1 |
| 0 | 0 | 1 | 1 |
| 0 | 0 | 1 | 1 |
| 0 | 0 | 1 | 1 |
| 0 | 0 | 1 | 1 |
| 0 | 0 | 1 | 0 |
| 0 | 0 | 1 | 0 |
| 0 | 0 | 0 | 0 |
| 0 | 0 | 0 | 0 |
| 0 | 0 | 0 | 0 |
| 0 | 0 | 0 | 0 |
| 0 | 0 | 1 | 0 |
| 0 | 0 | 1 | 0 |
| 0 | 0 | 1 | 0 |
| 0 | 0 | 1 | 0 |
| 0 | 0 | 1 | 0 |
| 0 | 0 | 1 | 0 |
| 0 | 0 | 1 | 0 |
| 0 | 0 | 1 | 0 |
| 0 | 0 | 0 | 0 |
| 0 | 0 | 0 | 0 |
| 0 | 0 | 0 | 0 |
| 0 | 0 | 0 | 0 |
| 1 | 0 | 0 | 0 |

| | | | |
|---|---|---|---|
| 0 | 0 | 0 | 0 |
| 0 | 0 | 0 | 0 |
| 0 | 0 | 0 | 0 |
| 1 | 0 | 0 | 0 |
| 0 | 0 | 0 | 0 |
| 0 | 0 | 0 | 0 |
| 0 | 0 | 0 | 0 |
| 0 | 0 | 0 | 0 |
| 0 | 0 | 0 | 0 |
| 0 | 0 | 0 | 0 |
| 1 | 1 | 0 | 0 |
| 0 | 0 | 0 | 0 |
| 1 | 1 | 0 | 0 |
| 1 | 1 | 0 | 0 |
| 0 | 1 | 0 | 0 |
| 0 | 0 | 0 | 0 |
| 0 | 0 | 0 | 0 |
| 0 | 0 | 0 | 0 |
| 0 | 0 | 0 | 0 |
| 0 | 0 | 0 | 0 |
| 1 | 1 | 0 | 0 |
| 0 | 1 | 0 | 0 |
| 0 | 0 | 0 | 0 |
| 0 | 1 | 0 | 0 |
| 1 | 1 | 0 | 0 |
| 0 | 0 | 0 | 0 |
| 0 | 0 | 0 | 0 |
| 1 | 1 | 0 | 0 |
| 1 | 1 | 0 | 0 |
| 0 | 1 | 0 | 0 |
| 0 | 0 | 0 | 0 |
| 0 | 0 | 0 | 0 |
| 0 | 0 | 0 | 0 |
| 0 | 0 | 0 | 0 |
| 0 | 0 | 0 | 0 |
| 0 | 0 | 0 | 0 |
| 0 | 0 | 0 | 0 |
| 0 | 0 | 0 | 0 |
| 0 | 0 | 0 | 0 |
| 0 | 0 | 0 | 0 |
| 1 | 0 | 0 | 0 |
| 1 | 0 | 0 | 0 |
| 1 | 0 | 0 | 0 |
| 1 | 0 | 0 | 0 |
| 1 | 1 | 0 | 1 |
| 0 | 0 | 0 | 0 |
| 1 | 0 | 0 | 0 |
| 0 | 0 | 0 | 0 |
| 1 | 0 | 0 | 0 |
| 0 | 0 | 0 | 0 |
| 1 | 0 | 0 | 0 |
| 1 | 0 | 0 | 0 |
| 0 | 0 | 0 | 0 |
| 0 | 0 | 0 | 0 |

| | | | |
|---|---|---|---|
| 0 | 0 | 0 | 0 |
| 0 | 0 | 0 | 0 |
| 0 | 0 | 0 | 0 |
| 0 | 0 | 0 | 0 |
| 1 | 0 | 0 | 0 |
| 1 | 0 | 0 | 0 |
| 1 | 0 | 0 | 0 |
| 0 | 0 | 0 | 0 |
| 1 | 1 | 0 | 0 |
| 0 | 0 | 0 | 0 |
| 0 | 0 | 0 | 0 |
| 0 | 0 | 0 | 0 |
| 1 | 1 | 0 | 0 |
| 1 | 0 | 0 | 0 |
| 0 | 0 | 0 | 0 |
| 0 | 0 | 0 | 0 |
| 1 | 1 | 0 | 0 |
| 0 | 0 | 0 | 0 |
| 0 | 0 | 0 | 0 |
| 0 | 0 | 0 | 0 |
| 0 | 0 | 0 | 0 |
| 0 | 0 | 0 | 0 |
| 0 | 0 | 0 | 0 |
| 0 | 0 | 0 | 0 |
| 1 | 0 | 0 | 0 |
| 0 | 0 | 0 | 0 |
| 1 | 0 | 0 | 0 |
| 0 | 0 | 0 | 0 |
| 0 | 0 | 0 | 0 |
| 0 | 0 | 0 | 0 |
| 1 | 0 | 0 | 0 |
| 1 | 0 | 0 | 0 |
| 0 | 0 | 0 | 0 |
| 0 | 0 | 0 | 0 |
| 1 | 0 | 0 | 0 |
| 0 | 0 | 0 | 0 |
| 0 | 0 | 0 | 0 |
| 1 | 0 | 0 | 0 |
| 0 | 0 | 0 | 0 |
| 1 | 0 | 0 | 0 |
| 1 | 0 | 0 | 0 |
| 0 | 0 | 0 | 0 |
| 0 | 0 | 0 | 0 |
| 1 | 1 | 0 | 0 |
| 1 | 0 | 0 | 0 |
| 1 | 1 | 0 | 0 |
| 0 | 0 | 0 | 0 |
| 0 | 0 | 0 | 0 |
| 0 | 0 | 0 | 0 |
| 0 | 0 | 0 | 0 |
| 1 | 1 | 0 | 0 |
| 0 | 0 | 0 | 0 |
| 0 | 0 | 0 | 0 |
| 0 | 0 | 0 | 0 |

| | | | |
|---|---|---|---|
| 0 | 1 | 0 | 0 |
| 1 | 0 | 0 | 0 |
| 0 | 0 | 0 | 0 |
| 0 | 0 | 0 | 0 |
| 0 | 0 | 0 | 0 |
| 1 | 1 | 0 | 0 |
| 0 | 0 | 0 | 0 |
| 0 | 0 | 0 | 0 |
| 0 | 0 | 0 | 0 |
| 1 | 0 | 0 | 0 |
| 1 | 0 | 0 | 0 |
| 0 | 0 | 0 | 0 |
| 0 | 0 | 0 | 0 |
| 0 | 0 | 0 | 0 |
| 0 | 0 | 0 | 0 |
| 0 | 0 | 0 | 0 |
| 0 | 0 | 0 | 0 |
| 0 | 0 | 0 | 0 |
| 0 | 0 | 0 | 0 |
| 0 | 0 | 0 | 0 |
| 0 | 0 | 0 | 0 |
| 0 | 0 | 0 | 0 |
| 0 | 0 | 0 | 0 |
| 1 | 1 | 0 | 0 |
| 0 | 0 | 0 | 0 |
| 0 | 0 | 0 | 0 |
| 1 | 1 | 1 | 0 |
| 0 | 0 | 0 | 0 |
| 0 | 0 | 0 | 0 |
| 0 | 0 | 0 | 0 |
| 1 | 1 | 1 | 0 |
| 1 | 0 | 0 | 0 |
| 1 | 0 | 0 | 0 |
| 1 | 0 | 0 | 0 |
| 1 | 0 | 0 | 0 |
| 1 | 1 | 1 | 0 |
| 1 | 1 | 1 | 0 |
| 0 | 0 | 0 | 0 |
| 0 | 0 | 0 | 0 |
| 1 | 1 | 1 | 0 |
| 0 | 0 | 0 | 0 |
| 0 | 0 | 0 | 0 |
| 0 | 0 | 0 | 0 |
| 0 | 0 | 0 | 0 |
| 0 | 0 | 0 | 0 |
| 0 | 0 | 0 | 0 |
| 0 | 0 | 0 | 0 |
| 0 | 0 | 0 | 0 |
| 1 | 0 | 0 | 0 |
| 1 | 0 | 0 | 0 |
| 1 | 0 | 0 | 0 |
| 0 | 0 | 0 | 0 |
| 0 | 0 | 0 | 0 |
| 0 | 0 | 0 | 0 |

| | | | |
|---|---|---|---|
| 0 | 0 | 0 | 0 |
| 1 | 0 | 0 | 0 |
| 0 | 0 | 0 | 0 |
| 0 | 0 | 0 | 0 |
| 0 | 0 | 0 | 0 |
| 0 | 0 | 0 | 0 |
| 0 | 0 | 0 | 0 |
| 0 | 0 | 0 | 0 |
| 0 | 0 | 0 | 0 |
| 0 | 0 | 0 | 0 |
| 1 | 1 | 0 | 0 |
| 1 | 1 | 0 | 0 |
| 0 | 0 | 0 | 0 |
| 0 | 0 | 0 | 0 |
| 0 | 0 | 0 | 0 |
| 1 | 0 | 0 | 0 |
| 1 | 0 | 0 | 0 |
| 1 | 1 | 0 | 1 |
| 0 | 0 | 0 | 0 |
| 0 | 0 | 0 | 0 |
| 0 | 0 | 0 | 0 |
| 0 | 0 | 0 | 0 |
| 1 | 0 | 0 | 1 |
| 0 | 0 | 0 | 0 |
| 0 | 0 | 0 | 0 |
| 1 | 0 | 0 | 0 |
| 0 | 0 | 0 | 0 |
| 0 | 0 | 0 | 0 |
| 0 | 0 | 0 | 0 |
| 0 | 0 | 0 | 0 |
| 0 | 0 | 0 | 0 |
| 0 | 0 | 0 | 0 |
| 0 | 0 | 0 | 0 |
| 0 | 0 | 0 | 0 |
| 0 | 0 | 0 | 0 |
| 1 | 0 | 0 | 0 |
| 1 | 0 | 0 | 0 |
| 1 | 1 | 0 | 0 |
| 1 | 1 | 0 | 0 |
| 0 | 0 | 0 | 0 |
| 0 | 0 | 0 | 0 |
| 1 | 0 | 0 | 0 |
| 0 | 0 | 0 | 0 |
| 1 | 0 | 0 | 0 |
| 1 | 1 | 0 | 0 |
| 0 | 0 | 0 | 0 |
| 0 | 0 | 0 | 0 |
| 0 | 0 | 0 | 0 |
| 0 | 0 | 0 | 0 |
| 0 | 0 | 0 | 0 |
| 0 | 0 | 0 | 0 |
| 1 | 0 | 0 | 0 |
| 0 | 0 | 0 | 0 |
| 0 | 0 | 0 | 0 |

| | | | |
|---|---|---|---|
| 0 | 0 | 0 | 0 |
| 1 | 1 | 0 | 0 |
| 0 | 0 | 0 | 0 |
| 1 | 0 | 0 | 0 |
| 1 | 0 | 0 | 0 |
| 0 | 0 | 0 | 0 |
| 0 | 0 | 0 | 0 |
| 0 | 0 | 0 | 0 |
| 0 | 0 | 0 | 0 |
| 0 | 0 | 0 | 0 |
| 0 | 0 | 0 | 0 |
| 1 | 1 | 0 | 0 |
| 1 | 1 | 0 | 1 |
| 0 | 0 | 0 | 0 |
| 0 | 1 | 0 | 0 |
| 0 | 0 | 0 | 0 |
| 0 | 0 | 0 | 0 |
| 0 | 0 | 0 | 0 |
| 0 | 0 | 0 | 0 |
| 0 | 0 | 0 | 0 |
| 0 | 0 | 0 | 0 |
| 1 | 0 | 0 | 0 |
| 0 | 0 | 0 | 0 |
| 0 | 0 | 0 | 0 |
| 0 | 0 | 0 | 0 |
| 0 | 0 | 0 | 0 |
| 0 | 0 | 0 | 0 |
| 1 | 0 | 0 | 0 |
| 0 | 0 | 0 | 0 |
| 0 | 0 | 0 | 0 |
| 0 | 0 | 0 | 0 |
| 0 | 0 | 0 | 0 |
| 0 | 0 | 0 | 0 |
| 0 | 0 | 0 | 0 |
| 0 | 0 | 0 | 0 |
| 0 | 0 | 0 | 0 |
| 0 | 0 | 0 | 0 |
| 0 | 0 | 0 | 0 |
| 0 | 0 | 0 | 0 |
| 0 | 0 | 0 | 0 |
| 0 | 0 | 0 | 0 |
| 0 | 0 | 0 | 0 |
| 0 | 0 | 0 | 0 |
| 0 | 0 | 0 | 0 |
| 0 | 0 | 0 | 0 |
| 0 | 0 | 0 | 0 |
| 0 | 0 | 0 | 0 |
| 0 | 0 | 0 | 0 |
| 0 | 0 | 0 | 0 |
| 0 | 0 | 0 | 0 |
| 0 | 0 | 0 | 0 |
| 0 | 0 | 0 | 0 |
| 0 | 0 | 0 | 0 |
| 0 | 0 | 0 | 0 |

| | | | |
|---|---|---|---|
| 0 | 0 | 0 | 0 |
| 0 | 0 | 0 | 0 |
| 0 | 0 | 0 | 0 |
| 0 | 0 | 0 | 0 |
| 0 | 0 | 0 | 0 |
| 0 | 0 | 0 | 0 |
| 0 | 0 | 0 | 0 |
| 0 | 0 | 0 | 0 |
| 0 | 0 | 0 | 0 |
| 0 | 0 | 0 | 0 |
| 0 | 0 | 0 | 0 |
| 0 | 0 | 0 | 0 |
| 1 | 0 | 0 | 0 |
| 1 | 1 | 0 | 0 |
| 0 | 0 | 0 | 0 |
| 1 | 0 | 0 | 0 |
| 0 | 0 | 0 | 0 |
| 1 | 0 | 0 | 0 |
| 0 | 0 | 0 | 0 |
| 0 | 0 | 0 | 0 |
| 1 | 0 | 0 | 0 |
| 0 | 0 | 0 | 0 |
| 0 | 0 | 0 | 0 |
| 1 | 0 | 0 | 0 |
| 1 | 0 | 0 | 0 |
| 0 | 0 | 0 | 0 |
| 0 | 0 | 0 | 0 |
| 1 | 0 | 0 | 0 |
| 1 | 0 | 0 | 0 |
| 1 | 0 | 0 | 0 |
| 1 | 0 | 0 | 0 |
| 1 | 0 | 0 | 0 |
| 0 | 0 | 0 | 0 |
| 0 | 0 | 0 | 0 |
| 0 | 0 | 0 | 0 |
| 0 | 0 | 0 | 0 |
| 0 | 0 | 0 | 0 |
| 0 | 0 | 0 | 0 |
| 0 | 0 | 0 | 0 |
| 0 | 0 | 0 | 0 |
| 0 | 0 | 0 | 0 |
| 0 | 0 | 0 | 0 |
| 0 | 0 | 0 | 0 |
| 0 | 0 | 0 | 0 |
| 1 | 1 | 0 | 0 |
| 0 | 0 | 0 | 0 |
| 0 | 0 | 0 | 0 |
| 0 | 0 | 0 | 0 |
| 0 | 0 | 0 | 0 |
| 1 | 1 | 0 | 0 |
| 1 | 0 | 0 | 0 |
| 0 | 0 | 0 | 0 |
| 0 | 0 | 0 | 0 |
| 0 | 0 | 0 | 1 |

| | | | |
|---|---|---|---|
| 0 | 0 | 0 | 0 |
| 0 | 0 | 0 | 0 |
| 0 | 0 | 0 | 0 |
| 0 | 0 | 0 | 0 |
| 1 | 0 | 0 | 1 |
| 0 | 0 | 0 | 0 |
| 1 | 0 | 0 | 0 |
| 0 | 0 | 0 | 0 |
| 0 | 0 | 0 | 0 |
| 0 | 0 | 0 | 0 |
| 0 | 0 | 0 | 0 |
| 0 | 0 | 0 | 0 |
| 0 | 0 | 0 | 0 |
| 1 | 0 | 0 | 0 |
| 0 | 0 | 0 | 0 |
| 0 | 0 | 0 | 0 |
| 1 | 1 | 0 | 0 |
| 0 | 0 | 0 | 0 |
| 1 | 0 | 0 | 0 |
| 1 | 0 | 0 | 0 |
| 0 | 0 | 0 | 0 |
| 0 | 0 | 0 | 0 |
| 0 | 0 | 0 | 0 |
| 1 | 0 | 0 | 0 |
| 1 | 0 | 0 | 0 |
| 1 | 0 | 0 | 0 |
| 1 | 0 | 0 | 0 |
| 1 | 0 | 0 | 0 |
| 1 | 0 | 0 | 0 |
| 1 | 0 | 0 | 0 |
| 1 | 0 | 0 | 0 |
| 1 | 0 | 0 | 0 |
| 1 | 0 | 0 | 0 |
| 1 | 0 | 0 | 0 |
| 0 | 0 | 0 | 0 |
| 0 | 0 | 0 | 0 |
| 0 | 0 | 0 | 0 |
| 1 | 0 | 0 | 0 |
| 1 | 0 | 0 | 0 |
| 0 | 0 | 0 | 0 |
| 0 | 0 | 0 | 0 |
| 1 | 0 | 0 | 0 |
| 0 | 0 | 0 | 0 |
| 1 | 0 | 0 | 0 |
| 1 | 0 | 0 | 0 |
| 1 | 0 | 0 | 0 |
| 1 | 0 | 0 | 0 |
| 1 | 0 | 0 | 0 |
| 0 | 0 | 0 | 0 |
| 0 | 0 | 0 | 0 |
| 1 | 0 | 0 | 0 |
| 0 | 0 | 0 | 0 |
| 1 | 0 | 0 | 0 |
| 1 | 0 | 0 | 0 |

| | | | |
|---|---|---|---|
| 1 | 0 | 0 | 0 |
| 1 | 0 | 0 | 0 |
| 1 | 0 | 0 | 0 |
| 1 | 0 | 0 | 0 |
| 1 | 0 | 0 | 0 |
| 1 | 0 | 0 | 0 |
| 0 | 0 | 0 | 0 |
| 1 | 0 | 0 | 0 |
| 1 | 0 | 0 | 0 |
| 1 | 0 | 0 | 0 |
| 1 | 0 | 0 | 0 |
| 1 | 0 | 0 | 0 |
| 1 | 0 | 0 | 0 |
| 1 | 0 | 0 | 0 |
| 0 | 0 | 0 | 0 |
| 0 | 0 | 0 | 0 |
| 1 | 1 | 0 | 0 |
| 1 | 1 | 0 | 0 |
| 0 | 0 | 0 | 0 |
| 0 | 0 | 0 | 0 |
| 0 | 0 | 0 | 0 |
| 1 | 1 | 0 | 1 |
| 0 | 0 | 0 | 0 |
| 0 | 0 | 0 | 0 |
| 0 | 0 | 0 | 0 |
| 0 | 0 | 0 | 0 |
| 0 | 0 | 0 | 0 |
| 0 | 0 | 0 | 0 |
| 0 | 0 | 0 | 0 |
| 1 | 0 | 1 | 0 |
| 0 | 0 | 0 | 0 |
| 0 | 0 | 0 | 0 |
| 0 | 0 | 0 | 0 |
| 0 | 0 | 0 | 0 |
| 0 | 0 | 0 | 0 |
| 0 | 0 | 0 | 0 |
| 0 | 0 | 0 | 0 |
| 0 | 0 | 0 | 0 |
| 0 | 0 | 0 | 0 |
| 0 | 0 | 0 | 0 |
| 1 | 1 | 0 | 0 |
| 1 | 1 | 0 | 0 |
| 1 | 0 | 0 | 0 |
| 1 | 1 | 0 | 0 |
| 1 | 0 | 0 | 0 |
| 1 | 1 | 0 | 0 |
| 1 | 0 | 0 | 0 |
| 1 | 0 | 0 | 0 |
| 0 | 0 | 0 | 0 |
| 0 | 0 | 0 | 0 |
| 1 | 0 | 0 | 0 |
| 1 | 0 | 0 | 0 |
| 1 | 0 | 0 | 0 |
| 0 | 0 | 0 | 0 |

| | | | |
|---|---|---|---|
| 0 | 0 | 0 | 0 |
| 0 | 0 | 0 | 0 |
| 0 | 0 | 0 | 0 |
| 1 | 1 | 1 | 0 |
| 0 | 0 | 0 | 0 |
| 0 | 0 | 0 | 0 |
| 0 | 0 | 0 | 0 |
| 0 | 0 | 0 | 0 |
| 0 | 1 | 0 | 0 |
| 0 | 1 | 0 | 0 |
| 0 | 1 | 0 | 0 |
| 0 | 0 | 0 | 0 |
| 0 | 0 | 0 | 0 |
| 0 | 0 | 0 | 0 |
| 1 | 0 | 0 | 0 |
| 1 | 0 | 0 | 0 |
| 0 | 0 | 0 | 0 |
| 0 | 0 | 0 | 0 |
| 1 | 0 | 0 | 0 |
| 0 | 0 | 0 | 0 |
| 1 | 0 | 0 | 0 |
| 1 | 0 | 0 | 0 |
| 0 | 0 | 0 | 0 |
| 0 | 0 | 0 | 0 |
| 0 | 0 | 0 | 0 |
| 0 | 0 | 0 | 0 |
| 0 | 0 | 0 | 0 |
| 0 | 0 | 1 | 0 |
| 0 | 0 | 0 | 0 |
| 0 | 1 | 0 | 0 |
| 0 | 0 | 0 | 0 |
| 0 | 0 | 0 | 0 |
| 0 | 0 | 0 | 0 |
| 0 | 0 | 0 | 0 |
| 0 | 0 | 0 | 0 |
| 0 | 0 | 0 | 0 |
| 1 | 0 | 0 | 0 |
| 1 | 0 | 0 | 0 |
| 1 | 0 | 0 | 0 |
| 0 | 0 | 0 | 0 |
| 0 | 0 | 0 | 0 |
| 0 | 0 | 0 | 0 |
| 0 | 0 | 0 | 0 |
| 0 | 0 | 0 | 0 |
| 0 | 0 | 0 | 0 |
| 0 | 0 | 0 | 0 |
| 0 | 0 | 0 | 0 |
| 0 | 0 | 0 | 1 |
| 0 | 0 | 0 | 0 |
| 0 | 0 | 0 | 0 |
| 0 | 0 | 0 | 0 |
| 0 | 0 | 0 | 0 |
| 1 | 1 | 0 | 0 |
| 0 | 0 | 0 | 0 |

| | | | |
|---|---|---|---|
| 0 | 0 | 0 | 0 |
| 0 | 0 | 0 | 0 |
| 0 | 0 | 0 | 0 |
| 0 | 0 | 0 | 0 |
| 1 | 0 | 0 | 0 |
| 0 | 0 | 0 | 0 |
| 0 | 0 | 0 | 0 |
| 0 | 0 | 0 | 0 |
| 0 | 0 | 0 | 0 |
| 0 | 0 | 0 | 0 |
| 0 | 0 | 0 | 0 |
| 0 | 0 | 0 | 0 |
| 0 | 0 | 0 | 0 |
| 0 | 0 | 0 | 0 |
| 0 | 0 | 0 | 0 |
| 0 | 0 | 0 | 0 |
| 0 | 0 | 0 | 0 |
| 0 | 0 | 0 | 0 |
| 0 | 0 | 0 | 0 |
| 0 | 0 | 0 | 0 |
| 0 | 0 | 0 | 0 |
| 0 | 0 | 0 | 0 |
| 0 | 0 | 0 | 0 |
| 0 | 0 | 0 | 0 |
| 0 | 0 | 0 | 0 |
| 0 | 0 | 0 | 0 |
| 0 | 0 | 0 | 0 |
| 0 | 0 | 0 | 0 |
| 0 | 0 | 0 | 0 |
| 0 | 0 | 0 | 0 |
| 0 | 0 | 0 | 0 |
| 0 | 0 | 0 | 0 |
| 0 | 0 | 0 | 0 |
| 0 | 0 | 0 | 0 |
| 0 | 0 | 0 | 0 |
| 0 | 0 | 0 | 0 |
| 0 | 0 | 0 | 0 |
| 0 | 0 | 0 | 0 |
| 0 | 0 | 0 | 0 |
| 0 | 0 | 0 | 0 |
| 1 | 1 | 0 | 0 |
| 0 | 0 | 0 | 0 |
| 0 | 0 | 0 | 0 |
| 0 | 0 | 0 | 0 |
| 0 | 0 | 0 | 0 |
| 1 | 0 | 0 | 0 |
| 1 | 0 | 0 | 0 |
| 0 | 0 | 0 | 0 |
| 1 | 0 | 0 | 0 |
| 0 | 0 | 0 | 0 |
| 0 | 0 | 0 | 0 |
| 1 | 0 | 0 | 0 |
| 1 | 0 | 0 | 0 |
| 0 | 0 | 0 | 0 |

| | | | |
|---|---|---|---|
| 0 | 0 | 0 | 0 |
| 0 | 0 | 0 | 0 |
| 0 | 0 | 0 | 0 |
| 0 | 0 | 0 | 0 |
| 1 | 0 | 0 | 0 |
| 0 | 0 | 0 | 0 |
| 0 | 0 | 0 | 0 |
| 0 | 0 | 0 | 0 |
| 0 | 0 | 0 | 0 |
| 0 | 0 | 0 | 0 |
| 0 | 0 | 0 | 0 |
| 0 | 0 | 0 | 0 |
| 0 | 0 | 0 | 0 |
| 0 | 0 | 0 | 0 |
| 0 | 0 | 0 | 0 |
| 1 | 1 | 0 | 1 |
| 0 | 0 | 0 | 0 |
| 0 | 0 | 0 | 0 |
| 0 | 0 | 0 | 0 |
| 0 | 0 | 0 | 0 |
| 0 | 0 | 0 | 0 |
| 0 | 0 | 0 | 0 |
| 0 | 0 | 0 | 0 |
| 0 | 0 | 0 | 0 |
| 0 | 0 | 0 | 0 |
| 0 | 0 | 0 | 0 |
| 0 | 0 | 0 | 0 |
| 1 | 0 | 0 | 0 |
| 1 | 0 | 0 | 0 |
| 0 | 0 | 0 | 0 |
| 1 | 1 | 0 | 0 |
| 0 | 0 | 0 | 0 |
| 0 | 0 | 0 | 0 |
| 0 | 0 | 0 | 0 |
| 1 | 1 | 0 | 0 |
| 1 | 1 | 0 | 0 |
| 0 | 0 | 0 | 0 |
| 1 | 0 | 0 | 0 |
| 0 | 0 | 0 | 0 |
| 1 | 1 | 0 | 0 |
| 0 | 1 | 0 | 0 |
| 0 | 0 | 0 | 0 |
| 0 | 0 | 0 | 0 |
| 0 | 0 | 0 | 0 |
| 1 | 1 | 0 | 0 |
| 1 | 1 | 0 | 0 |
| 1 | 1 | 0 | 0 |
| 0 | 0 | 0 | 0 |
| 0 | 0 | 0 | 0 |
| 0 | 0 | 0 | 0 |
| 0 | 0 | 0 | 0 |
| 0 | 0 | 0 | 0 |
| 0 | 0 | 0 | 0 |
| 0 | 0 | 0 | 0 |

| | | | |
|---|---|---|---|
| 0 | 0 | 0 | 0 |
| 0 | 0 | 0 | 0 |
| 1 | 0 | 0 | 0 |
| 1 | 0 | 0 | 0 |
| 0 | 0 | 0 | 0 |
| 1 | 0 | 0 | 0 |
| 0 | 0 | 0 | 0 |
| 0 | 0 | 0 | 0 |
| 0 | 0 | 0 | 0 |
| 0 | 1 | 0 | 0 |
| 0 | 0 | 0 | 0 |
| 0 | 0 | 0 | 0 |
| 0 | 0 | 0 | 0 |
| 0 | 0 | 0 | 0 |
| 0 | 0 | 0 | 0 |
| 0 | 0 | 0 | 0 |
| 0 | 0 | 0 | 0 |
| 1 | 0 | 0 | 0 |
| 0 | 0 | 0 | 0 |
| 1 | 0 | 0 | 0 |
| 0 | 0 | 0 | 0 |
| 1 | 0 | 0 | 0 |
| 0 | 0 | 0 | 0 |
| 1 | 0 | 0 | 0 |
| 0 | 0 | 0 | 0 |
| 0 | 0 | 0 | 0 |
| 0 | 0 | 0 | 0 |
| 0 | 0 | 0 | 0 |
| 0 | 0 | 0 | 0 |
| 0 | 0 | 0 | 0 |
| 0 | 0 | 0 | 0 |
| 1 | 0 | 0 | 0 |
| 0 | 0 | 0 | 0 |
| 0 | 0 | 0 | 0 |
| 0 | 0 | 0 | 0 |
| 0 | 0 | 0 | 0 |
| 0 | 0 | 0 | 0 |
| 0 | 0 | 0 | 0 |
| 1 | 1 | 0 | 0 |
| 1 | 1 | 0 | 0 |
| 1 | 1 | 0 | 0 |
| 1 | 1 | 0 | 0 |
| 1 | 1 | 0 | 0 |
| 1 | 0 | 0 | 0 |
| 1 | 0 | 0 | 0 |
| 1 | 0 | 0 | 0 |
| 0 | 0 | 0 | 0 |
| 0 | 0 | 0 | 0 |
| 0 | 0 | 0 | 0 |
| 0 | 0 | 0 | 0 |
| 0 | 0 | 0 | 0 |
| 0 | 0 | 0 | 0 |
| 0 | 0 | 0 | 0 |
| 0 | 0 | 0 | 0 |

| | | | |
|---|---|---|---|
| 0 | 0 | 0 | 0 |
| 0 | 0 | 0 | 0 |
| 0 | 0 | 0 | 0 |
| 0 | 0 | 0 | 0 |
| 0 | 0 | 0 | 0 |
| 1 | 0 | 0 | 0 |
| 1 | 0 | 0 | 0 |
| 0 | 0 | 0 | 0 |
| 0 | 0 | 0 | 0 |
| 0 | 0 | 0 | 0 |
| 0 | 0 | 0 | 0 |
| 1 | 0 | 0 | 0 |
| 0 | 0 | 1 | 0 |
| 0 | 0 | 0 | 0 |
| 0 | 0 | 0 | 0 |
| 0 | 0 | 0 | 0 |
| 1 | 0 | 0 | 0 |
| 0 | 0 | 0 | 0 |
| 1 | 0 | 0 | 0 |
| 1 | 0 | 0 | 0 |
| 0 | 0 | 0 | 0 |
| 1 | 0 | 0 | 0 |
| 1 | 1 | 0 | 0 |
| 0 | 0 | 0 | 0 |
| 1 | 1 | 0 | 0 |
| 1 | 1 | 0 | 0 |
| 1 | 1 | 0 | 0 |
| 0 | 0 | 0 | 0 |
| 1 | 0 | 0 | 0 |
| 0 | 0 | 0 | 0 |
| 0 | 0 | 0 | 0 |
| 0 | 0 | 0 | 0 |
| 0 | 0 | 0 | 0 |
| 0 | 0 | 0 | 0 |
| 0 | 0 | 0 | 0 |
| 0 | 0 | 0 | 0 |
| 0 | 0 | 0 | 0 |
| 0 | 0 | 0 | 0 |
| 0 | 0 | 0 | 0 |
| 0 | 0 | 0 | 0 |
| 0 | 0 | 0 | 0 |
| 0 | 0 | 0 | 0 |
| 0 | 0 | 0 | 0 |
| 0 | 0 | 0 | 0 |
| 0 | 0 | 0 | 0 |
| 0 | 0 | 0 | 0 |
| 0 | 0 | 0 | 0 |
| 0 | 0 | 0 | 0 |
| 0 | 0 | 0 | 0 |
| 0 | 0 | 0 | 0 |
| 0 | 0 | 0 | 0 |
| 0 | 0 | 0 | 0 |
| 0 | 0 | 0 | 0 |
| 0 | 0 | 0 | 0 |

| | | | |
|---|---|---|---|
| 0 | 0 | 0 | 0 |
| 0 | 0 | 0 | 0 |
| 0 | 0 | 0 | 0 |
| 0 | 0 | 0 | 0 |
| 0 | 0 | 0 | 0 |
| 0 | 0 | 0 | 0 |
| 0 | 0 | 0 | 0 |
| 0 | 0 | 0 | 0 |
| 0 | 0 | 0 | 0 |
| 0 | 0 | 0 | 0 |
| 0 | 1 | 0 | 0 |
| 0 | 1 | 0 | 0 |
| 0 | 1 | 0 | 0 |
| 0 | 1 | 0 | 0 |
| 0 | 0 | 0 | 0 |
| 0 | 1 | 0 | 0 |
| 0 | 1 | 0 | 0 |
| 1 | 1 | 0 | 0 |
| 0 | 1 | 0 | 0 |
| 1 | 0 | 0 | 0 |
| 1 | 1 | 0 | 0 |
| 1 | 1 | 0 | 0 |
| 0 | 0 | 1 | 1 |
| 0 | 0 | 0 | 0 |
| 1 | 0 | 0 | 0 |
| 0 | 0 | 0 | 0 |
| 0 | 0 | 0 | 0 |
| 0 | 0 | 0 | 0 |
| 0 | 0 | 0 | 0 |
| 0 | 0 | 0 | 0 |
| 0 | 0 | 0 | 0 |
| 0 | 0 | 0 | 0 |
| 0 | 0 | 0 | 0 |
| 0 | 0 | 0 | 0 |
| 0 | 0 | 0 | 0 |
| 0 | 0 | 0 | 0 |
| 0 | 0 | 0 | 0 |
| 0 | 0 | 0 | 0 |
| 0 | 0 | 0 | 0 |
| 0 | 0 | 0 | 0 |
| 1 | 0 | 0 | 0 |
| 0 | 0 | 0 | 0 |
| 1 | 0 | 0 | 0 |
| 1 | 0 | 0 | 0 |
| 0 | 0 | 0 | 0 |
| 1 | 0 | 0 | 0 |
| 0 | 0 | 0 | 0 |
| 0 | 0 | 0 | 0 |
| 0 | 0 | 0 | 0 |
| 0 | 0 | 0 | 0 |
| 0 | 0 | 0 | 0 |
| 0 | 0 | 0 | 0 |
| 1 | 1 | 0 | 0 |
| 1 | 1 | 0 | 0 |

| | | | |
|---|---|---|---|
| 1 | 1 | 0 | 0 |
| 0 | 0 | 0 | 0 |
| 0 | 0 | 0 | 0 |
| 0 | 0 | 0 | 0 |
| 0 | 0 | 0 | 0 |
| 0 | 0 | 0 | 0 |
| 0 | 1 | 0 | 0 |
| 0 | 0 | 0 | 0 |
| 0 | 0 | 0 | 0 |
| 0 | 1 | 0 | 0 |
| 1 | 0 | 0 | 0 |
| 0 | 0 | 0 | 0 |
| 0 | 0 | 1 | 0 |
| 0 | 0 | 1 | 0 |
| 0 | 0 | 0 | 0 |
| 0 | 0 | 0 | 0 |
| 0 | 0 | 0 | 0 |
| 0 | 1 | 0 | 0 |
| 0 | 0 | 0 | 0 |
| 0 | 0 | 0 | 0 |
| 1 | 0 | 0 | 0 |
| 0 | 0 | 0 | 1 |
| 0 | 0 | 0 | 0 |
| 0 | 0 | 0 | 0 |
| 0 | 0 | 0 | 0 |
| 0 | 0 | 0 | 0 |
| 0 | 0 | 0 | 0 |
| 0 | 0 | 0 | 0 |
| 0 | 0 | 0 | 0 |
| 0 | 0 | 0 | 0 |
| 0 | 0 | 0 | 0 |
| 0 | 0 | 0 | 0 |
| 0 | 0 | 0 | 0 |
| 0 | 0 | 0 | 0 |
| 0 | 0 | 0 | 0 |
| 0 | 0 | 0 | 0 |
| 1 | 0 | 0 | 0 |
| 0 | 0 | 0 | 0 |
| 1 | 0 | 0 | 0 |
| 0 | 0 | 0 | 0 |
| 0 | 0 | 0 | 0 |
| 0 | 0 | 0 | 0 |
| 0 | 0 | 0 | 0 |
| 0 | 0 | 0 | 0 |
| 0 | 0 | 0 | 0 |
| 0 | 0 | 0 | 0 |
| 0 | 0 | 0 | 0 |
| 0 | 0 | 0 | 0 |
| 0 | 0 | 0 | 0 |
| 1 | 0 | 0 | 0 |
| 0 | 0 | 0 | 0 |
| 0 | 0 | 0 | 0 |
| 0 | 0 | 0 | 0 |
| 0 | 0 | 0 | 0 |

| | | | |
|---|---|---|---|
| 0 | 0 | 0 | 0 |
| 0 | 0 | 0 | 0 |
| 0 | 0 | 0 | 0 |
| 0 | 0 | 0 | 0 |
| 0 | 0 | 0 | 0 |
| 0 | 0 | 0 | 0 |
| 0 | 0 | 0 | 0 |
| 0 | 0 | 0 | 0 |
| 0 | 0 | 0 | 0 |
| 0 | 0 | 0 | 0 |
| 0 | 0 | 0 | 0 |
| 0 | 0 | 0 | 0 |
| 0 | 0 | 1 | 0 |
| 1 | 0 | 0 | 0 |
| 1 | 0 | 0 | 0 |
| 0 | 0 | 0 | 0 |
| 0 | 0 | 0 | 0 |
| 0 | 0 | 0 | 0 |
| 0 | 0 | 0 | 0 |
| 0 | 0 | 0 | 0 |
| 0 | 0 | 0 | 0 |
| 0 | 0 | 0 | 0 |
| 0 | 0 | 0 | 0 |
| 1 | 1 | 0 | 1 |
| 0 | 0 | 0 | 0 |
| 1 | 1 | 0 | 0 |
| 1 | 1 | 0 | 1 |
| 0 | 0 | 0 | 0 |
| 0 | 0 | 0 | 0 |
| 0 | 0 | 0 | 0 |
| 0 | 0 | 0 | 0 |
| 0 | 0 | 0 | 0 |
| 0 | 0 | 0 | 0 |
| 0 | 0 | 0 | 0 |
| 0 | 0 | 0 | 0 |
| 1 | 0 | 0 | 0 |
| 0 | 0 | 0 | 0 |
| 0 | 0 | 0 | 0 |
| 0 | 0 | 0 | 0 |
| 0 | 0 | 0 | 0 |
| 0 | 0 | 0 | 0 |
| 0 | 0 | 0 | 0 |
| 1 | 0 | 0 | 0 |
| 1 | 0 | 0 | 0 |
| 1 | 0 | 0 | 0 |
| 1 | 0 | 0 | 0 |
| 1 | 0 | 0 | 0 |
| 0 | 0 | 0 | 0 |
| 0 | 0 | 0 | 0 |
| 1 | 0 | 0 | 0 |
| 1 | 0 | 0 | 0 |
| 0 | 0 | 0 | 0 |
| 1 | 0 | 0 | 0 |
| 1 | 0 | 0 | 0 |

| | | | |
|---|---|---|---|
| 0 | 0 | 0 | 0 |
| 1 | 0 | 0 | 0 |
| 0 | 0 | 0 | 0 |
| 0 | 0 | 0 | 0 |
| 0 | 0 | 0 | 0 |
| 0 | 0 | 0 | 0 |
| 1 | 0 | 0 | 0 |
| 0 | 0 | 0 | 0 |
| 0 | 0 | 0 | 0 |
| 0 | 0 | 0 | 0 |
| 0 | 0 | 0 | 0 |
| 1 | 1 | 0 | 0 |
| 0 | 0 | 0 | 0 |
| 0 | 0 | 0 | 0 |
| 0 | 0 | 0 | 0 |
| 0 | 0 | 0 | 0 |
| 0 | 0 | 0 | 0 |
| 0 | 0 | 0 | 0 |
| 0 | 0 | 0 | 0 |
| 0 | 0 | 0 | 0 |
| 0 | 0 | 0 | 0 |
| 0 | 0 | 0 | 0 |
| 0 | 0 | 0 | 0 |
| 0 | 0 | 0 | 0 |
| 0 | 0 | 0 | 0 |
| 0 | 0 | 0 | 0 |
| 0 | 0 | 0 | 0 |
| 0 | 0 | 0 | 0 |
| 1 | 1 | 0 | 0 |
| 1 | 0 | 0 | 0 |
| 0 | 0 | 0 | 0 |
| 0 | 0 | 0 | 0 |
| 0 | 0 | 0 | 0 |
| 0 | 0 | 0 | 0 |
| 0 | 0 | 0 | 0 |
| 0 | 0 | 0 | 0 |
| 0 | 0 | 0 | 0 |
| 0 | 0 | 0 | 0 |
| 0 | 0 | 0 | 0 |
| 0 | 0 | 0 | 0 |
| 0 | 0 | 0 | 0 |
| 0 | 0 | 0 | 0 |
| 0 | 0 | 0 | 0 |
| 0 | 0 | 0 | 0 |
| 0 | 0 | 0 | 0 |
| 0 | 0 | 0 | 0 |
| 0 | 0 | 0 | 0 |
| 0 | 0 | 0 | 0 |
| 0 | 0 | 0 | 0 |
| 0 | 0 | 0 | 0 |
| 0 | 0 | 0 | 0 |
| 0 | 0 | 0 | 0 |
| 0 | 0 | 1 | 0 |
| 0 | 0 | 0 | 0 |

| | | | |
|---|---|---|---|
| 0 | 0 | 0 | 0 |
| 0 | 0 | 0 | 0 |
| 0 | 0 | 0 | 0 |
| 0 | 0 | 0 | 0 |
| 1 | 0 | 0 | 0 |
| 0 | 0 | 0 | 0 |
| 0 | 0 | 0 | 0 |
| 1 | 1 | 0 | 0 |
| 0 | 0 | 0 | 0 |
| 1 | 1 | 0 | 0 |
| 0 | 0 | 0 | 0 |
| 0 | 0 | 0 | 0 |
| 0 | 0 | 0 | 0 |
| 0 | 0 | 0 | 0 |
| 0 | 0 | 0 | 0 |
| 0 | 0 | 0 | 0 |
| 1 | 0 | 0 | 0 |
| 0 | 0 | 0 | 0 |
| 0 | 0 | 0 | 0 |
| 1 | 0 | 0 | 0 |
| 1 | 0 | 0 | 0 |
| 0 | 0 | 0 | 0 |
| 0 | 0 | 0 | 0 |
| 0 | 0 | 0 | 0 |
| 0 | 0 | 0 | 0 |
| 0 | 0 | 0 | 0 |
| 1 | 0 | 0 | 0 |
| 0 | 0 | 0 | 0 |
| 0 | 0 | 0 | 0 |
| 1 | 0 | 0 | 0 |
| 0 | 0 | 0 | 0 |
| 0 | 0 | 0 | 0 |
| 0 | 0 | 0 | 0 |
| 1 | 1 | 0 | 0 |
| 0 | 0 | 0 | 0 |
| 0 | 0 | 0 | 0 |
| 0 | 0 | 0 | 0 |
| 0 | 0 | 0 | 0 |
| 0 | 1 | 0 | 0 |
| 0 | 0 | 0 | 0 |
| 0 | 0 | 0 | 0 |
| 0 | 0 | 0 | 0 |
| 0 | 0 | 0 | 0 |
| 1 | 1 | 0 | 0 |
| 0 | 0 | 0 | 0 |
| 0 | 0 | 0 | 0 |
| 0 | 0 | 0 | 0 |
| 0 | 0 | 0 | 0 |
| 0 | 0 | 0 | 0 |
| 0 | 0 | 0 | 0 |
| 0 | 0 | 0 | 0 |
| 1 | 0 | 0 | 0 |
| 0 | 0 | 0 | 0 |
| 0 | 0 | 0 | 0 |

| | | | |
|---|---|---|---|
| 1 | 0 | 0 | 1 |
| 1 | 0 | 0 | 1 |
| 1 | 0 | 0 | 1 |
| 0 | 0 | 0 | 0 |
| 0 | 0 | 0 | 0 |
| 0 | 0 | 0 | 0 |
| 1 | 0 | 0 | 0 |
| 0 | 0 | 0 | 0 |
| 0 | 0 | 0 | 0 |
| 0 | 0 | 0 | 0 |
| 0 | 0 | 1 | 0 |
| 0 | 0 | 0 | 0 |
| 0 | 0 | 0 | 0 |
| 0 | 0 | 0 | 0 |
| 0 | 0 | 0 | 0 |
| 0 | 0 | 0 | 0 |
| 1 | 0 | 0 | 0 |
| 0 | 0 | 0 | 0 |
| 0 | 0 | 0 | 0 |
| 0 | 0 | 0 | 0 |
| 0 | 0 | 0 | 0 |
| 0 | 0 | 0 | 0 |
| 0 | 0 | 0 | 0 |
| 1 | 1 | 0 | 0 |
| 0 | 0 | 0 | 0 |
| 0 | 0 | 0 | 0 |
| 0 | 0 | 0 | 0 |
| 0 | 0 | 0 | 0 |
| 0 | 0 | 0 | 0 |
| 0 | 0 | 0 | 0 |
| 0 | 0 | 0 | 0 |
| 0 | 0 | 0 | 0 |
| 0 | 1 | 0 | 0 |
| 0 | 0 | 0 | 0 |
| 1 | 1 | 0 | 0 |
| 0 | 0 | 0 | 0 |
| 0 | 0 | 0 | 0 |
| 0 | 0 | 0 | 0 |
| 0 | 0 | 0 | 0 |
| 0 | 0 | 0 | 0 |
| 0 | 0 | 0 | 0 |
| 1 | 0 | 0 | 1 |
| 0 | 0 | 0 | 0 |
| 1 | 0 | 0 | 1 |
| 0 | 0 | 0 | 0 |
| 1 | 0 | 0 | 1 |
| 0 | 0 | 0 | 0 |
| 0 | 0 | 0 | 0 |
| 0 | 0 | 0 | 0 |
| 1 | 0 | 0 | 0 |
| 0 | 0 | 0 | 0 |
| 0 | 0 | 0 | 0 |
| 0 | 0 | 0 | 0 |
| 0 | 0 | 0 | 0 |

| | | | |
|---|---|---|---|
| 0 | 0 | 0 | 0 |
| 0 | 0 | 0 | 0 |
| 0 | 0 | 0 | 0 |
| 0 | 0 | 0 | 0 |
| 0 | 0 | 0 | 0 |
| 0 | 0 | 0 | 0 |
| 0 | 0 | 0 | 0 |
| 0 | 0 | 0 | 0 |
| 0 | 0 | 0 | 0 |
| 0 | 0 | 0 | 0 |
| 0 | 0 | 0 | 0 |
| 0 | 0 | 0 | 0 |
| 0 | 0 | 0 | 0 |
| 0 | 0 | 0 | 0 |
| 0 | 0 | 0 | 0 |
| 0 | 0 | 0 | 0 |
| 0 | 0 | 0 | 0 |
| 0 | 0 | 0 | 0 |
| 0 | 0 | 1 | 0 |
| 0 | 0 | 0 | 0 |
| 0 | 0 | 0 | 0 |
| 0 | 0 | 0 | 0 |
| 0 | 0 | 0 | 0 |
| 0 | 0 | 0 | 0 |
| 0 | 0 | 0 | 0 |
| 0 | 0 | 0 | 0 |
| 0 | 0 | 0 | 0 |
| 0 | 0 | 0 | 0 |
| 0 | 0 | 1 | 0 |
| 0 | 0 | 0 | 0 |
| 0 | 0 | 1 | 0 |
| 1 | 0 | 0 | 0 |
| 1 | 1 | 1 | 0 |
| 1 | 1 | 1 | 0 |
| 0 | 0 | 0 | 0 |
| 0 | 0 | 0 | 0 |
| 0 | 1 | 0 | 0 |
| 1 | 1 | 1 | 0 |
| 1 | 1 | 1 | 0 |
| 0 | 0 | 1 | 0 |
| 0 | 0 | 1 | 0 |
| 1 | 0 | 0 | 0 |
| 0 | 0 | 0 | 0 |
| 0 | 0 | 0 | 0 |
| 1 | 1 | 0 | 0 |
| 0 | 0 | 0 | 0 |
| 0 | 0 | 0 | 0 |
| 0 | 0 | 0 | 0 |
| 0 | 0 | 0 | 0 |
| 0 | 0 | 0 | 0 |
| 0 | 0 | 0 | 0 |
| 0 | 0 | 0 | 0 |
| 0 | 0 | 0 | 0 |
| 0 | 0 | 0 | 0 |

| | | | |
|---|---|---|---|
| 0 | 0 | 0 | 0 |
| 0 | 0 | 0 | 0 |
| 0 | 0 | 0 | 0 |
| 0 | 0 | 0 | 0 |
| 0 | 0 | 0 | 0 |
| 0 | 0 | 0 | 0 |
| 0 | 0 | 0 | 0 |
| 0 | 0 | 0 | 0 |
| 0 | 0 | 0 | 0 |
| 0 | 0 | 0 | 0 |
| 0 | 0 | 0 | 0 |
| 0 | 0 | 0 | 0 |
| 0 | 0 | 0 | 0 |
| 0 | 0 | 0 | 0 |
| 0 | 0 | 0 | 0 |
| 0 | 0 | 0 | 0 |
| 0 | 0 | 0 | 0 |
| 0 | 0 | 0 | 0 |
| 0 | 0 | 0 | 0 |
| 1 | 0 | 0 | 0 |
| 1 | 0 | 0 | 0 |
| 0 | 0 | 0 | 0 |
| 0 | 0 | 0 | 0 |
| 0 | 0 | 0 | 0 |
| 0 | 0 | 0 | 0 |
| 1 | 0 | 0 | 0 |
| 0 | 0 | 0 | 0 |
| 0 | 0 | 0 | 0 |
| 0 | 0 | 0 | 0 |
| 1 | 0 | 0 | 0 |
| 0 | 0 | 0 | 0 |
| 0 | 0 | 0 | 0 |
| 1 | 0 | 0 | 0 |
| 1 | 0 | 0 | 0 |
| 0 | 0 | 0 | 0 |
| 0 | 0 | 0 | 0 |
| 0 | 0 | 0 | 0 |
| 0 | 0 | 0 | 0 |
| 0 | 0 | 0 | 0 |
| 0 | 0 | 0 | 0 |
| 0 | 0 | 0 | 0 |
| 0 | 0 | 0 | 0 |
| 0 | 0 | 0 | 0 |
| 0 | 0 | 0 | 0 |
| 0 | 0 | 0 | 0 |
| 0 | 0 | 0 | 0 |
| 0 | 0 | 0 | 0 |
| 0 | 1 | 0 | 0 |
| 0 | 1 | 0 | 0 |
| 0 | 0 | 0 | 0 |
| 0 | 1 | 0 | 0 |
| 0 | 1 | 0 | 0 |
| 0 | 0 | 0 | 0 |
| 0 | 0 | 0 | 0 |

| | | | |
|---|---|---|---|
| 0 | 0 | 0 | 0 |
| 0 | 0 | 0 | 0 |
| 1 | 1 | 0 | 1 |
| 0 | 0 | 0 | 0 |
| 0 | 0 | 0 | 0 |
| 0 | 0 | 0 | 0 |
| 0 | 0 | 0 | 0 |
| 0 | 1 | 0 | 0 |
| 1 | 1 | 0 | 0 |
| 1 | 1 | 0 | 0 |
| 1 | 1 | 0 | 0 |
| 1 | 1 | 0 | 0 |
| 1 | 1 | 0 | 0 |
| 0 | 1 | 0 | 0 |
| 0 | 1 | 0 | 0 |
| 0 | 0 | 0 | 0 |
| 0 | 0 | 0 | 0 |
| 0 | 0 | 0 | 0 |
| 0 | 0 | 0 | 0 |
| 0 | 0 | 0 | 0 |
| 0 | 0 | 0 | 0 |
| 0 | 0 | 0 | 0 |
| 0 | 1 | 0 | 0 |
| 0 | 1 | 0 | 0 |
| 0 | 0 | 0 | 0 |
| 0 | 0 | 1 | 0 |
| 0 | 0 | 0 | 0 |
| 0 | 0 | 0 | 0 |
| 0 | 0 | 0 | 0 |
| 0 | 1 | 0 | 0 |
| 0 | 0 | 0 | 0 |
| 0 | 0 | 0 | 0 |
| 0 | 0 | 0 | 0 |
| 0 | 0 | 0 | 0 |
| 0 | 0 | 0 | 0 |
| 0 | 0 | 0 | 0 |
| 0 | 0 | 0 | 0 |
| 0 | 0 | 0 | 0 |
| 0 | 0 | 0 | 0 |
| 1 | 0 | 0 | 0 |
| 1 | 1 | 0 | 0 |
| 1 | 0 | 0 | 0 |
| 0 | 0 | 0 | 0 |
| 0 | 1 | 0 | 0 |
| 1 | 1 | 0 | 0 |
| 0 | 1 | 0 | 0 |
| 0 | 0 | 0 | 0 |
| 1 | 0 | 0 | 0 |
| 0 | 1 | 0 | 0 |
| 0 | 0 | 0 | 0 |
| 0 | 0 | 0 | 0 |
| 0 | 0 | 0 | 0 |
| 1 | 1 | 0 | 0 |
| 0 | 0 | 0 | 0 |

| | | | |
|---|---|---|---|
| 0 | 0 | 0 | 0 |
| 0 | 0 | 0 | 0 |
| 1 | 0 | 0 | 0 |
| 1 | 1 | 0 | 0 |
| 1 | 1 | 0 | 0 |
| 0 | 0 | 0 | 0 |
| 0 | 0 | 0 | 0 |
| 0 | 0 | 0 | 0 |
| 1 | 0 | 0 | 0 |
| 0 | 0 | 0 | 0 |
| 0 | 0 | 0 | 0 |
| 1 | 0 | 0 | 1 |
| 0 | 0 | 0 | 0 |
| 1 | 0 | 0 | 0 |
| 0 | 0 | 0 | 0 |
| 0 | 0 | 0 | 0 |
| 0 | 0 | 0 | 0 |
| 0 | 0 | 0 | 0 |
| 0 | 0 | 0 | 0 |
| 0 | 0 | 0 | 0 |
| 0 | 0 | 0 | 0 |
| 1 | 0 | 0 | 1 |
| 1 | 0 | 0 | 0 |
| 0 | 0 | 0 | 0 |
| 0 | 0 | 0 | 0 |
| 0 | 0 | 0 | 0 |
| 0 | 0 | 0 | 0 |
| 0 | 0 | 0 | 0 |
| 1 | 0 | 0 | 0 |
| 1 | 0 | 0 | 0 |
| 0 | 0 | 0 | 0 |
| 0 | 0 | 0 | 0 |
| 0 | 0 | 0 | 0 |
| 0 | 0 | 0 | 0 |
| 0 | 0 | 0 | 0 |
| 0 | 0 | 0 | 0 |
| 1 | 0 | 0 | 0 |
| 0 | 0 | 0 | 0 |
| 0 | 0 | 0 | 0 |
| 1 | 0 | 0 | 0 |
| 1 | 0 | 0 | 0 |
| 1 | 0 | 0 | 0 |
| 1 | 0 | 0 | 0 |
| 0 | 0 | 0 | 0 |
| 0 | 0 | 0 | 0 |
| 1 | 0 | 0 | 0 |
| 0 | 0 | 0 | 0 |
| 0 | 0 | 0 | 0 |
| 0 | 0 | 0 | 0 |
| 0 | 0 | 0 | 0 |
| 0 | 0 | 0 | 0 |
| 1 | 0 | 0 | 0 |
| 1 | 0 | 0 | 0 |
| 1 | 0 | 0 | 0 |

| | | | |
|---|---|---|---|
| 1 | 0 | 0 | 0 |
| 0 | 0 | 0 | 0 |
| 0 | 0 | 0 | 0 |
| 0 | 0 | 0 | 0 |
| 1 | 0 | 0 | 0 |
| 0 | 0 | 0 | 0 |
| 1 | 0 | 0 | 0 |
| 0 | 0 | 0 | 0 |
| 0 | 0 | 0 | 0 |
| 0 | 0 | 0 | 0 |
| 0 | 0 | 0 | 0 |
| 0 | 0 | 0 | 0 |
| 0 | 0 | 0 | 0 |
| 0 | 0 | 0 | 0 |
| 0 | 0 | 0 | 0 |
| 1 | 0 | 0 | 0 |
| 0 | 0 | 0 | 0 |
| 0 | 0 | 0 | 0 |
| 0 | 0 | 0 | 0 |
| 0 | 0 | 0 | 0 |
| 0 | 0 | 0 | 0 |
| 0 | 0 | 0 | 0 |
| 1 | 0 | 0 | 0 |
| 1 | 0 | 0 | 0 |
| 0 | 0 | 0 | 0 |
| 0 | 0 | 0 | 0 |
| 0 | 0 | 0 | 0 |
| 0 | 0 | 0 | 0 |
| 0 | 0 | 0 | 0 |
| 0 | 0 | 0 | 0 |
| 0 | 0 | 0 | 0 |
| 0 | 0 | 0 | 0 |
| 1 | 0 | 0 | 0 |
| 0 | 0 | 0 | 0 |
| 0 | 0 | 0 | 0 |
| 0 | 0 | 0 | 0 |
| 0 | 0 | 0 | 0 |
| 0 | 0 | 0 | 0 |
| 0 | 0 | 0 | 0 |
| 0 | 0 | 0 | 0 |
| 0 | 0 | 0 | 0 |
| 0 | 0 | 0 | 0 |
| 0 | 0 | 0 | 0 |
| 0 | 0 | 0 | 0 |
| 0 | 1 | 0 | 0 |
| 0 | 0 | 0 | 0 |
| 0 | 0 | 0 | 0 |
| 0 | 0 | 0 | 0 |
| 0 | 0 | 0 | 0 |
| 0 | 0 | 0 | 0 |
| 0 | 0 | 0 | 0 |
| 0 | 0 | 0 | 0 |
| 0 | 0 | 0 | 0 |
| 1 | 0 | 0 | 0 |

| | | | |
|---|---|---|---|
| 0 | 0 | 0 | 0 |
| 0 | 0 | 0 | 0 |
| 0 | 0 | 0 | 0 |
| 0 | 0 | 0 | 0 |
| 0 | 0 | 0 | 0 |
| 0 | 0 | 0 | 0 |
| 0 | 0 | 0 | 0 |
| 0 | 0 | 0 | 0 |
| 0 | 0 | 0 | 0 |
| 1 | 0 | 0 | 0 |
| 0 | 0 | 0 | 0 |
| 0 | 0 | 0 | 0 |
| 1 | 1 | 0 | 0 |
| 1 | 1 | 0 | 0 |
| 1 | 1 | 0 | 0 |
| 1 | 1 | 0 | 0 |
| 0 | 0 | 0 | 0 |
| 0 | 0 | 0 | 0 |
| 0 | 0 | 0 | 0 |
| 0 | 0 | 0 | 0 |
| 0 | 0 | 0 | 0 |
| 0 | 0 | 0 | 0 |
| 0 | 0 | 0 | 0 |
| 0 | 0 | 0 | 0 |
| 0 | 0 | 0 | 0 |
| 0 | 0 | 0 | 1 |
| 0 | 0 | 0 | 0 |
| 0 | 0 | 0 | 0 |
| 0 | 0 | 0 | 0 |
| 0 | 0 | 0 | 0 |
| 0 | 0 | 0 | 0 |
| 0 | 0 | 0 | 0 |
| 0 | 0 | 0 | 0 |
| 0 | 0 | 0 | 0 |
| 0 | 0 | 0 | 0 |
| 0 | 0 | 0 | 0 |
| 0 | 0 | 0 | 0 |
| 1 | 1 | 0 | 0 |
| 0 | 0 | 0 | 0 |
| 0 | 0 | 0 | 0 |
| 1 | 1 | 0 | 0 |
| 0 | 0 | 0 | 0 |
| 0 | 0 | 0 | 0 |
| 1 | 0 | 0 | 1 |
| 0 | 0 | 0 | 0 |
| 0 | 0 | 0 | 0 |
| 0 | 0 | 0 | 0 |
| 0 | 0 | 0 | 0 |
| 0 | 0 | 0 | 0 |
| 0 | 0 | 0 | 0 |
| 0 | 0 | 0 | 0 |
| 0 | 0 | 1 | 0 |
| 0 | 0 | 1 | 0 |
| 0 | 0 | 1 | 0 |

| | | | |
|---|---|---|---|
| 0 | 0 | 1 | 0 |
| 0 | 0 | 0 | 0 |
| 0 | 0 | 0 | 0 |
| 0 | 0 | 0 | 0 |
| 0 | 0 | 0 | 0 |
| 0 | 0 | 0 | 0 |
| 0 | 0 | 0 | 0 |
| 0 | 0 | 0 | 0 |
| 0 | 0 | 0 | 0 |
| 0 | 0 | 0 | 0 |
| 0 | 0 | 0 | 0 |
| 0 | 0 | 0 | 0 |
| 0 | 0 | 0 | 0 |
| 0 | 0 | 0 | 0 |
| 0 | 0 | 0 | 0 |
| 0 | 0 | 0 | 0 |
| 0 | 0 | 0 | 0 |
| 0 | 0 | 0 | 0 |
| 0 | 0 | 0 | 0 |
| 0 | 0 | 0 | 0 |
| 0 | 0 | 0 | 0 |
| 0 | 0 | 0 | 0 |
| 0 | 0 | 0 | 0 |
| 0 | 1 | 0 | 1 |
| 0 | 0 | 0 | 0 |
| 0 | 0 | 0 | 0 |
| 0 | 0 | 0 | 0 |
| 0 | 0 | 0 | 0 |
| 0 | 0 | 0 | 0 |
| 0 | 0 | 0 | 0 |
| 0 | 0 | 0 | 0 |
| 0 | 0 | 0 | 0 |
| 0 | 0 | 0 | 0 |
| 0 | 0 | 0 | 0 |
| 0 | 0 | 0 | 0 |
| 0 | 0 | 0 | 0 |
| 0 | 0 | 0 | 0 |
| 0 | 0 | 0 | 0 |
| 0 | 0 | 0 | 0 |
| 0 | 0 | 0 | 0 |
| 0 | 0 | 0 | 0 |
| 0 | 0 | 0 | 0 |
| 0 | 0 | 0 | 0 |
| 0 | 0 | 0 | 0 |
| 0 | 0 | 0 | 0 |
| 0 | 0 | 0 | 0 |
| 1 | 0 | 0 | 0 |
| 1 | 0 | 0 | 0 |
| 0 | 0 | 0 | 0 |
| 0 | 1 | 0 | 0 |
| 1 | 1 | 0 | 1 |
| 1 | 1 | 0 | 1 |
| 1 | 1 | 0 | 1 |
| 0 | 0 | 0 | 0 |

| | | | |
|---|---|---|---|
| 1 | 1 | 0 | 0 |
| 0 | 0 | 0 | 0 |
| 0 | 0 | 0 | 0 |
| 0 | 0 | 0 | 0 |
| 0 | 0 | 0 | 0 |
| 0 | 0 | 0 | 0 |
| 0 | 0 | 0 | 0 |
| 0 | 0 | 0 | 0 |
| 0 | 0 | 0 | 0 |
| 0 | 0 | 0 | 0 |
| 0 | 0 | 0 | 0 |
| 0 | 1 | 0 | 0 |
| 0 | 0 | 0 | 0 |
| 0 | 0 | 0 | 0 |
| 0 | 1 | 0 | 0 |
| 0 | 0 | 0 | 0 |
| 1 | 1 | 0 | 0 |
| 0 | 0 | 0 | 0 |
| 0 | 0 | 0 | 0 |
| 1 | 0 | 0 | 0 |
| 0 | 0 | 0 | 0 |
| 0 | 0 | 0 | 0 |
| 0 | 0 | 0 | 0 |
| 0 | 0 | 0 | 0 |
| 0 | 0 | 0 | 0 |
| 0 | 0 | 0 | 0 |
| 0 | 0 | 0 | 0 |
| 0 | 0 | 0 | 0 |
| 0 | 0 | 0 | 0 |
| 0 | 0 | 0 | 0 |
| 0 | 0 | 0 | 0 |
| 0 | 0 | 0 | 0 |
| 0 | 0 | 0 | 0 |
| 1 | 0 | 0 | 0 |
| 0 | 0 | 0 | 0 |
| 0 | 1 | 0 | 0 |
| 0 | 0 | 0 | 0 |
| 0 | 0 | 0 | 0 |
| 0 | 0 | 0 | 0 |
| 0 | 0 | 0 | 0 |
| 0 | 0 | 0 | 0 |
| 0 | 0 | 0 | 0 |
| 0 | 0 | 0 | 0 |
| 0 | 0 | 0 | 0 |
| 0 | 0 | 0 | 0 |
| 0 | 0 | 0 | 0 |
| 0 | 0 | 0 | 0 |
| 0 | 0 | 0 | 0 |
| 1 | 1 | 0 | 0 |
| 0 | 0 | 0 | 0 |
| 0 | 0 | 0 | 0 |
| 0 | 0 | 0 | 0 |
| 0 | 0 | 0 | 0 |
| 0 | 0 | 0 | 0 |

| | | | |
|---|---|---|---|
| 0 | 0 | 0 | 0 |
| 0 | 0 | 0 | 0 |
| 0 | 0 | 0 | 0 |
| 0 | 0 | 0 | 0 |
| 0 | 0 | 0 | 0 |
| 0 | 0 | 0 | 0 |
| 0 | 0 | 0 | 0 |
| 0 | 1 | 0 | 0 |
| 0 | 0 | 0 | 0 |
| 0 | 0 | 0 | 0 |
| 0 | 0 | 0 | 0 |
| 0 | 0 | 0 | 0 |
| 0 | 0 | 1 | 0 |
| 0 | 0 | 0 | 0 |
| 0 | 0 | 1 | 0 |
| 0 | 0 | 1 | 0 |
| 0 | 0 | 0 | 0 |
| 0 | 0 | 0 | 0 |
| 0 | 0 | 0 | 0 |
| 0 | 0 | 0 | 0 |
| 0 | 1 | 0 | 0 |
| 0 | 1 | 0 | 0 |
| 0 | 0 | 0 | 0 |
| 0 | 0 | 0 | 0 |
| 0 | 1 | 0 | 0 |
| 0 | 0 | 0 | 0 |
| 0 | 0 | 0 | 0 |
| 0 | 1 | 0 | 0 |
| 0 | 0 | 0 | 0 |
| 0 | 0 | 0 | 0 |
| 0 | 0 | 0 | 0 |
| 0 | 0 | 0 | 0 |
| 1 | 1 | 0 | 0 |
| 1 | 1 | 0 | 0 |
| 1 | 1 | 0 | 0 |
| 0 | 1 | 0 | 0 |
| 1 | 0 | 0 | 0 |
| 0 | 1 | 0 | 0 |
| 1 | 1 | 0 | 0 |
| 0 | 0 | 0 | 0 |
| 0 | 0 | 0 | 0 |
| 0 | 0 | 0 | 0 |
| 0 | 0 | 0 | 0 |
| 0 | 0 | 0 | 0 |
| 0 | 0 | 0 | 0 |
| 0 | 0 | 0 | 0 |
| 0 | 0 | 0 | 0 |
| 0 | 0 | 0 | 0 |
| 0 | 0 | 0 | 0 |
| 1 | 1 | 0 | 0 |
| 1 | 1 | 0 | 0 |
| 0 | 0 | 0 | 0 |
| 0 | 0 | 0 | 0 |
| 0 | 0 | 0 | 0 |

| | | | |
|---|---|---|---|
| 0 | 0 | 0 | 0 |
| 0 | 0 | 0 | 0 |
| 0 | 0 | 0 | 0 |
| 0 | 0 | 0 | 0 |
| 0 | 0 | 0 | 0 |
| 0 | 0 | 0 | 0 |
| 1 | 1 | 0 | 0 |
| 1 | 1 | 0 | 0 |
| 0 | 0 | 0 | 0 |
| 0 | 0 | 0 | 0 |
| 0 | 0 | 0 | 0 |
| 0 | 0 | 0 | 0 |
| 0 | 0 | 0 | 0 |
| 0 | 0 | 0 | 0 |
| 0 | 0 | 0 | 0 |
| 0 | 0 | 0 | 0 |
| 0 | 0 | 0 | 0 |
| 0 | 0 | 0 | 0 |
| 0 | 0 | 0 | 0 |
| 0 | 0 | 0 | 0 |
| 0 | 0 | 0 | 0 |
| 0 | 0 | 0 | 0 |
| 0 | 0 | 0 | 0 |
| 0 | 0 | 0 | 0 |
| 0 | 0 | 0 | 0 |
| 0 | 0 | 0 | 0 |
| 0 | 0 | 0 | 0 |
| 1 | 1 | 0 | 0 |
| 0 | 0 | 0 | 0 |
| 0 | 0 | 0 | 0 |
| 0 | 0 | 0 | 0 |
| 0 | 0 | 0 | 0 |
| 0 | 0 | 0 | 0 |
| 0 | 0 | 0 | 0 |
| 0 | 0 | 0 | 0 |
| 0 | 0 | 0 | 0 |
| 0 | 0 | 0 | 0 |
| 0 | 0 | 0 | 0 |
| 0 | 0 | 0 | 0 |
| 0 | 0 | 0 | 0 |
| 0 | 0 | 0 | 0 |
| 0 | 0 | 0 | 0 |
| 0 | 0 | 0 | 0 |
| 0 | 0 | 0 | 0 |
| 0 | 0 | 0 | 0 |
| 0 | 0 | 0 | 0 |
| 0 | 0 | 0 | 0 |
| 0 | 0 | 0 | 0 |
| 0 | 0 | 0 | 0 |
| 0 | 0 | 0 | 0 |
| 0 | 0 | 0 | 0 |
| 0 | 0 | 0 | 0 |
| 0 | 0 | 0 | 0 |
| 0 | 0 | 0 | 0 |

| | | | |
|---|---|---|---|
| 0 | 0 | 0 | 0 |
| 0 | 0 | 0 | 0 |
| 0 | 0 | 0 | 0 |
| 0 | 0 | 0 | 0 |
| 0 | 0 | 0 | 0 |
| 0 | 0 | 0 | 0 |
| 0 | 0 | 0 | 0 |
| 0 | 0 | 0 | 0 |
| 0 | 0 | 0 | 0 |
| 0 | 0 | 0 | 0 |
| 1 | 0 | 0 | 0 |
| 1 | 1 | 0 | 0 |
| 1 | 0 | 0 | 0 |
| 0 | 0 | 0 | 0 |
| 0 | 0 | 0 | 0 |
| 0 | 0 | 0 | 0 |
| 0 | 0 | 0 | 0 |
| 0 | 0 | 0 | 0 |
| 0 | 0 | 0 | 0 |
| 0 | 0 | 0 | 0 |
| 0 | 0 | 0 | 0 |
| 0 | 0 | 0 | 0 |
| 0 | 0 | 0 | 0 |
| 0 | 0 | 0 | 0 |
| 0 | 0 | 0 | 0 |
| 0 | 0 | 0 | 0 |
| 0 | 0 | 0 | 0 |
| 0 | 0 | 0 | 0 |
| 0 | 0 | 0 | 0 |
| 0 | 0 | 0 | 0 |
| 0 | 1 | 0 | 0 |
| 0 | 0 | 0 | 0 |
| 0 | 0 | 0 | 0 |
| 1 | 0 | 0 | 0 |
| 1 | 0 | 0 | 0 |
| 1 | 0 | 0 | 0 |
| 0 | 0 | 0 | 0 |
| 0 | 0 | 0 | 0 |
| 0 | 0 | 0 | 0 |
| 0 | 0 | 0 | 0 |
| 0 | 0 | 0 | 0 |
| 0 | 0 | 0 | 0 |
| 0 | 0 | 0 | 0 |
| 0 | 0 | 0 | 0 |
| 0 | 0 | 0 | 0 |
| 0 | 0 | 0 | 0 |
| 0 | 0 | 0 | 0 |
| 0 | 0 | 0 | 0 |
| 0 | 0 | 0 | 0 |
| 0 | 0 | 0 | 0 |
| 0 | 0 | 0 | 0 |
| 0 | 0 | 0 | 0 |
| 0 | 0 | 0 | 1 |
| 0 | 0 | 0 | 0 |

| | | | |
|---|---|---|---|
| 0 | 0 | 0 | 0 |
| 0 | 0 | 0 | 0 |
| 0 | 0 | 0 | 0 |
| 0 | 0 | 0 | 0 |
| 0 | 0 | 0 | 0 |
| 0 | 0 | 0 | 0 |
| 0 | 0 | 0 | 0 |
| 0 | 0 | 0 | 0 |
| 0 | 0 | 0 | 0 |
| 0 | 0 | 0 | 0 |
| 0 | 0 | 0 | 0 |
| 1 | 0 | 0 | 0 |
| 0 | 0 | 0 | 0 |
| 0 | 0 | 0 | 0 |
| 0 | 0 | 1 | 0 |
| 0 | 0 | 0 | 0 |
| 0 | 0 | 0 | 0 |
| 0 | 0 | 0 | 0 |
| 0 | 0 | 0 | 0 |
| 0 | 0 | 0 | 0 |
| 0 | 0 | 0 | 0 |
| 0 | 0 | 0 | 0 |
| 0 | 0 | 0 | 0 |
| 0 | 0 | 0 | 0 |
| 0 | 0 | 0 | 0 |
| 0 | 0 | 0 | 0 |
| 0 | 0 | 0 | 0 |
| 0 | 0 | 0 | 0 |
| 0 | 0 | 0 | 0 |
| 0 | 0 | 0 | 0 |
| 0 | 0 | 0 | 0 |
| 0 | 0 | 0 | 0 |
| 0 | 0 | 0 | 0 |
| 1 | 0 | 0 | 0 |
| 0 | 0 | 0 | 0 |
| 1 | 0 | 0 | 1 |
| 0 | 0 | 0 | 0 |
| 0 | 0 | 0 | 0 |
| 0 | 0 | 0 | 0 |
| 0 | 0 | 0 | 0 |
| 0 | 0 | 0 | 0 |
| 0 | 0 | 0 | 0 |
| 0 | 0 | 0 | 0 |
| 0 | 1 | 0 | 0 |
| 0 | 0 | 0 | 0 |
| 0 | 0 | 0 | 0 |
| 0 | 0 | 0 | 0 |
| 0 | 0 | 0 | 0 |
| 0 | 0 | 0 | 0 |
| 0 | 0 | 0 | 0 |
| 0 | 0 | 0 | 0 |
| 0 | 0 | 0 | 0 |
| 0 | 0 | 0 | 0 |
| 0 | 0 | 0 | 0 |

| | | | |
|---|---|---|---|
| 0 | 0 | 0 | 0 |
| 0 | 0 | 0 | 0 |
| 0 | 0 | 0 | 0 |
| 0 | 0 | 0 | 0 |
| 0 | 0 | 0 | 0 |
| 0 | 1 | 0 | 0 |
| 0 | 0 | 0 | 0 |
| 0 | 0 | 0 | 0 |
| 0 | 0 | 0 | 0 |
| 0 | 0 | 0 | 0 |
| 0 | 0 | 0 | 0 |
| 0 | 0 | 0 | 0 |
| 0 | 0 | 0 | 0 |
| 0 | 0 | 0 | 0 |
| 0 | 0 | 0 | 0 |
| 0 | 1 | 0 | 0 |
| 1 | 1 | 0 | 0 |
| 0 | 0 | 1 | 0 |
| 0 | 0 | 0 | 0 |
| 0 | 0 | 0 | 0 |
| 0 | 0 | 0 | 0 |
| 0 | 0 | 0 | 0 |
| 0 | 0 | 0 | 0 |
| 0 | 0 | 0 | 0 |
| 0 | 0 | 0 | 0 |
| 0 | 0 | 0 | 0 |
| 0 | 0 | 0 | 0 |
| 1 | 0 | 0 | 0 |
| 0 | 0 | 0 | 0 |
| 0 | 0 | 0 | 0 |
| 0 | 0 | 1 | 0 |
| 0 | 0 | 0 | 0 |
| 0 | 0 | 0 | 0 |
| 0 | 0 | 0 | 0 |
| 0 | 0 | 0 | 0 |
| 0 | 0 | 0 | 0 |
| 0 | 0 | 0 | 0 |
| 1 | 1 | 0 | 0 |
| 0 | 0 | 0 | 0 |
| 0 | 0 | 0 | 0 |
| 0 | 0 | 0 | 0 |
| 1 | 0 | 0 | 0 |
| 1 | 0 | 0 | 0 |
| 0 | 0 | 0 | 0 |
| 0 | 0 | 0 | 0 |
| 0 | 0 | 0 | 0 |
| 0 | 0 | 0 | 0 |
| 0 | 0 | 0 | 0 |
| 0 | 0 | 0 | 0 |
| 0 | 0 | 0 | 0 |
| 0 | 0 | 0 | 0 |
| 0 | 0 | 0 | 0 |
| 0 | 0 | 0 | 0 |
| 0 | 0 | 0 | 0 |

| | | | |
|---|---|---|---|
| 0 | 0 | 0 | 0 |
| 0 | 0 | 0 | 0 |
| 0 | 0 | 0 | 0 |
| 0 | 0 | 0 | 0 |
| 0 | 0 | 0 | 0 |
| 0 | 0 | 0 | 0 |
| 0 | 0 | 0 | 0 |
| 0 | 0 | 0 | 0 |
| 0 | 0 | 0 | 0 |
| 0 | 0 | 0 | 0 |
| 0 | 0 | 0 | 0 |
| 0 | 0 | 0 | 0 |
| 1 | 0 | 0 | 0 |
| 0 | 0 | 0 | 0 |
| 0 | 0 | 0 | 0 |
| 0 | 0 | 0 | 0 |
| 0 | 0 | 0 | 0 |
| 0 | 0 | 0 | 0 |
| 0 | 0 | 0 | 0 |
| 0 | 0 | 0 | 0 |
| 0 | 0 | 0 | 0 |
| 0 | 0 | 0 | 0 |
| 0 | 0 | 0 | 0 |
| 1 | 0 | 0 | 0 |
| 0 | 0 | 0 | 0 |
| 0 | 0 | 0 | 0 |
| 0 | 0 | 0 | 0 |
| 1 | 1 | 0 | 1 |
| 0 | 0 | 0 | 0 |
| 0 | 0 | 0 | 0 |
| 0 | 0 | 0 | 0 |
| 0 | 0 | 0 | 0 |
| 0 | 0 | 0 | 0 |
| 0 | 0 | 0 | 0 |
| 0 | 0 | 0 | 0 |
| 0 | 0 | 0 | 0 |
| 0 | 0 | 0 | 0 |
| 0 | 0 | 0 | 0 |
| 0 | 0 | 0 | 0 |
| 0 | 0 | 0 | 0 |
| 0 | 0 | 0 | 0 |
| 0 | 0 | 0 | 0 |
| 0 | 0 | 0 | 0 |
| 0 | 0 | 0 | 0 |
| 0 | 0 | 0 | 0 |
| 0 | 0 | 0 | 0 |
| 0 | 0 | 0 | 0 |
| 0 | 0 | 0 | 0 |
| 0 | 0 | 0 | 0 |
| 0 | 0 | 0 | 0 |
| 0 | 0 | 0 | 0 |
| 0 | 0 | 0 | 0 |
| 0 | 0 | 0 | 0 |
| 0 | 0 | 0 | 0 |

| | | | |
|---|---|---|---|
| 0 | 0 | 0 | 0 |
| 0 | 0 | 0 | 0 |
| 0 | 0 | 0 | 0 |
| 0 | 0 | 0 | 0 |
| 0 | 0 | 0 | 0 |
| 0 | 0 | 0 | 0 |
| 0 | 0 | 0 | 0 |
| 0 | 0 | 0 | 0 |
| 1 | 0 | 0 | 0 |
| 1 | 0 | 0 | 0 |
| 1 | 0 | 0 | 0 |
| 0 | 0 | 0 | 0 |
| 1 | 0 | 0 | 0 |
| 0 | 0 | 0 | 0 |
| 0 | 0 | 0 | 0 |
| 1 | 0 | 0 | 0 |
| 0 | 0 | 0 | 0 |
| 0 | 0 | 0 | 0 |
| 0 | 0 | 0 | 0 |
| 0 | 0 | 0 | 0 |
| 0 | 0 | 0 | 0 |
| 0 | 0 | 0 | 0 |
| 0 | 0 | 0 | 0 |
| 0 | 0 | 0 | 0 |
| 0 | 0 | 0 | 0 |
| 0 | 0 | 0 | 0 |
| 0 | 0 | 0 | 0 |
| 1 | 0 | 0 | 0 |
| 1 | 0 | 0 | 0 |
| 0 | 0 | 0 | 0 |
| 0 | 0 | 0 | 0 |
| 0 | 0 | 0 | 0 |
| 1 | 0 | 0 | 0 |
| 0 | 0 | 0 | 0 |
| 0 | 0 | 0 | 0 |
| 0 | 0 | 0 | 0 |
| 0 | 0 | 0 | 0 |
| 0 | 0 | 0 | 0 |
| 0 | 0 | 0 | 0 |
| 0 | 0 | 0 | 0 |
| 1 | 0 | 0 | 0 |
| 1 | 0 | 0 | 0 |
| 0 | 0 | 0 | 0 |
| 0 | 0 | 0 | 0 |
| 0 | 0 | 0 | 0 |
| 0 | 0 | 0 | 0 |
| 0 | 0 | 0 | 0 |
| 0 | 0 | 0 | 0 |
| 0 | 0 | 0 | 0 |
| 0 | 0 | 0 | 0 |
| 0 | 0 | 0 | 0 |
| 0 | 0 | 0 | 0 |
| 0 | 0 | 0 | 0 |
| 0 | 0 | 0 | 0 |

| | | | |
|---|---|---|---|
| 0 | 0 | 0 | 0 |
| 0 | 0 | 0 | 0 |
| 0 | 0 | 0 | 0 |
| 0 | 0 | 0 | 0 |
| 0 | 0 | 0 | 0 |
| 0 | 0 | 0 | 0 |
| 0 | 0 | 0 | 0 |
| 0 | 0 | 0 | 0 |
| 0 | 0 | 0 | 0 |
| 0 | 0 | 0 | 0 |
| 1 | 0 | 0 | 0 |
| 0 | 0 | 0 | 0 |
| 1 | 0 | 0 | 0 |
| 0 | 0 | 0 | 0 |
| 0 | 0 | 0 | 0 |
| 0 | 0 | 0 | 0 |
| 0 | 0 | 0 | 0 |
| 0 | 1 | 0 | 0 |
| 0 | 0 | 0 | 0 |
| 0 | 0 | 0 | 0 |
| 0 | 0 | 0 | 0 |
| 0 | 0 | 0 | 0 |
| 1 | 0 | 0 | 0 |
| 0 | 0 | 0 | 0 |
| 0 | 0 | 0 | 0 |
| 1 | 0 | 0 | 0 |
| 1 | 0 | 0 | 0 |
| 1 | 0 | 0 | 0 |
| 0 | 0 | 0 | 0 |
| 0 | 0 | 0 | 0 |
| 0 | 0 | 0 | 0 |
| 0 | 0 | 0 | 0 |
| 0 | 0 | 0 | 0 |
| 0 | 0 | 0 | 0 |
| 0 | 0 | 0 | 0 |
| 0 | 0 | 0 | 0 |
| 0 | 0 | 0 | 0 |
| 0 | 0 | 0 | 0 |
| 0 | 0 | 0 | 0 |
| 0 | 0 | 0 | 0 |
| 1 | 1 | 0 | 0 |
| 0 | 0 | 0 | 0 |
| 0 | 0 | 0 | 0 |
| 0 | 0 | 0 | 0 |
| 0 | 0 | 0 | 0 |
| 0 | 0 | 0 | 0 |
| 0 | 0 | 0 | 0 |
| 1 | 0 | 0 | 0 |
| 0 | 0 | 0 | 0 |
| 0 | 0 | 0 | 0 |
| 0 | 0 | 0 | 0 |
| 0 | 0 | 0 | 0 |
| 0 | 0 | 0 | 0 |
| 0 | 0 | 0 | 0 |

| | | | |
|---|---|---|---|
| 1 | 0 | 0 | 0 |
| 0 | 0 | 0 | 0 |
| 1 | 0 | 0 | 0 |
| 0 | 0 | 0 | 0 |
| 0 | 0 | 0 | 0 |
| 0 | 0 | 0 | 0 |
| 0 | 0 | 0 | 0 |
| 0 | 0 | 0 | 0 |
| 1 | 1 | 0 | 0 |
| 0 | 0 | 0 | 0 |
| 1 | 1 | 0 | 0 |
| 1 | 1 | 0 | 0 |
| 0 | 0 | 0 | 0 |
| 1 | 1 | 0 | 0 |
| 0 | 0 | 0 | 0 |
| 0 | 0 | 0 | 0 |
| 0 | 0 | 0 | 0 |
| 0 | 0 | 0 | 0 |
| 0 | 0 | 0 | 0 |
| 0 | 0 | 0 | 0 |
| 1 | 0 | 0 | 0 |
| 0 | 0 | 0 | 0 |
| 0 | 0 | 0 | 0 |
| 0 | 0 | 0 | 0 |
| 1 | 0 | 0 | 0 |
| 0 | 0 | 0 | 0 |
| 1 | 0 | 0 | 0 |
| 1 | 0 | 0 | 0 |
| 1 | 1 | 0 | 0 |
| 0 | 0 | 0 | 0 |
| 1 | 1 | 0 | 0 |
| 1 | 1 | 0 | 0 |
| 1 | 1 | 0 | 0 |
| 0 | 0 | 0 | 0 |
| 1 | 1 | 0 | 0 |
| 0 | 0 | 0 | 0 |
| 1 | 0 | 0 | 0 |
| 0 | 0 | 0 | 0 |
| 1 | 1 | 0 | 0 |
| 1 | 1 | 0 | 0 |
| 1 | 0 | 0 | 0 |
| 0 | 0 | 0 | 0 |
| 0 | 0 | 0 | 0 |
| 1 | 0 | 0 | 0 |
| 0 | 0 | 0 | 0 |
| 0 | 0 | 0 | 0 |
| 1 | 0 | 0 | 0 |
| 0 | 0 | 0 | 0 |
| 1 | 0 | 0 | 0 |
| 0 | 0 | 0 | 0 |
| 0 | 0 | 0 | 0 |
| 0 | 0 | 0 | 0 |
| 0 | 0 | 0 | 0 |
| 0 | 0 | 0 | 0 |

| | | | |
|---|---|---|---|
| 0 | 0 | 0 | 0 |
| 0 | 0 | 0 | 0 |
| 0 | 0 | 0 | 0 |
| 0 | 0 | 0 | 0 |
| 0 | 0 | 0 | 0 |
| 0 | 0 | 0 | 0 |
| 0 | 0 | 0 | 0 |
| 0 | 0 | 0 | 0 |
| 0 | 0 | 0 | 0 |
| 1 | 1 | 0 | 0 |
| 1 | 1 | 0 | 0 |
| 0 | 0 | 0 | 0 |
| 0 | 0 | 0 | 0 |
| 0 | 0 | 0 | 0 |
| 1 | 0 | 1 | 0 |
| 0 | 0 | 0 | 0 |
| 0 | 0 | 0 | 0 |
| 1 | 0 | 1 | 0 |
| 0 | 0 | 0 | 0 |
| 0 | 0 | 0 | 0 |
| 0 | 0 | 0 | 0 |
| 0 | 0 | 1 | 0 |
| 1 | 0 | 1 | 0 |
| 0 | 0 | 1 | 0 |
| 0 | 0 | 1 | 0 |
| 0 | 0 | 1 | 1 |
| 0 | 0 | 1 | 1 |
| 0 | 0 | 0 | 0 |
| 0 | 0 | 0 | 0 |
| 0 | 0 | 0 | 0 |
| 0 | 0 | 0 | 0 |
| 0 | 0 | 0 | 0 |
| 0 | 0 | 0 | 0 |
| 0 | 0 | 0 | 0 |
| 0 | 0 | 0 | 0 |
| 0 | 0 | 0 | 0 |
| 0 | 0 | 0 | 0 |
| 0 | 0 | 0 | 0 |
| 0 | 0 | 0 | 0 |
| 0 | 0 | 0 | 0 |
| 0 | 0 | 0 | 0 |
| 0 | 0 | 0 | 0 |
| 0 | 0 | 0 | 0 |
| 0 | 0 | 1 | 0 |
| 0 | 0 | 0 | 0 |
| 0 | 0 | 0 | 0 |
| 0 | 0 | 0 | 0 |
| 0 | 0 | 0 | 0 |
| 0 | 0 | 0 | 0 |
| 0 | 0 | 0 | 0 |
| 1 | 0 | 0 | 0 |
| 0 | 0 | 0 | 0 |
| 0 | 0 | 0 | 0 |
| 0 | 0 | 0 | 0 |

| | | | |
|---|---|---|---|
| 0 | 0 | 0 | 0 |
| 0 | 0 | 0 | 0 |
| 0 | 0 | 0 | 0 |
| 1 | 0 | 0 | 0 |
| 0 | 0 | 0 | 0 |
| 0 | 0 | 0 | 0 |
| 0 | 0 | 0 | 0 |
| 0 | 0 | 0 | 0 |
| 1 | 0 | 0 | 0 |
| 1 | 0 | 0 | 0 |
| 0 | 0 | 0 | 0 |
| 0 | 0 | 0 | 0 |
| 0 | 0 | 0 | 0 |
| 0 | 0 | 0 | 0 |
| 0 | 0 | 0 | 0 |
| 0 | 0 | 0 | 0 |
| 0 | 0 | 1 | 0 |
| 0 | 0 | 0 | 0 |
| 0 | 0 | 0 | 0 |
| 0 | 0 | 0 | 0 |
| 0 | 0 | 0 | 0 |
| 0 | 0 | 0 | 0 |
| 0 | 0 | 0 | 0 |
| 0 | 0 | 0 | 0 |
| 0 | 0 | 0 | 0 |
| 0 | 0 | 0 | 0 |
| 0 | 0 | 0 | 0 |
| 0 | 0 | 0 | 0 |
| 0 | 0 | 0 | 0 |
| 0 | 0 | 0 | 0 |
| 0 | 0 | 0 | 0 |
| 0 | 0 | 0 | 0 |
| 0 | 0 | 0 | 0 |
| 0 | 0 | 0 | 0 |
| 0 | 0 | 0 | 0 |
| 0 | 0 | 0 | 0 |
| 0 | 0 | 0 | 0 |
| 0 | 0 | 0 | 0 |
| 0 | 0 | 0 | 0 |
| 0 | 0 | 0 | 0 |
| 0 | 0 | 0 | 0 |
| 0 | 0 | 0 | 0 |
| 0 | 0 | 0 | 0 |
| 0 | 0 | 0 | 0 |
| 0 | 0 | 0 | 0 |
| 0 | 0 | 0 | 0 |
| 0 | 0 | 0 | 0 |
| 0 | 0 | 0 | 0 |
| 0 | 0 | 0 | 0 |
| 0 | 1 | 0 | 0 |
| 0 | 0 | 0 | 0 |
| 0 | 0 | 0 | 0 |
| 0 | 0 | 0 | 0 |
| 0 | 0 | 0 | 0 |

| | | | |
|---|---|---|---|
| 0 | 0 | 1 | 0 |
| 0 | 0 | 0 | 0 |
| 0 | 0 | 0 | 0 |
| 0 | 0 | 0 | 0 |
| 0 | 0 | 0 | 0 |
| 0 | 0 | 0 | 0 |
| 0 | 0 | 0 | 0 |
| 1 | 0 | 0 | 0 |
| 0 | 0 | 0 | 0 |
| 0 | 0 | 0 | 0 |
| 0 | 0 | 0 | 0 |
| 1 | 0 | 0 | 0 |
| 0 | 0 | 0 | 0 |
| 0 | 0 | 0 | 0 |
| 0 | 0 | 0 | 0 |
| 0 | 0 | 0 | 0 |
| 0 | 0 | 0 | 0 |
| 0 | 0 | 0 | 0 |
| 0 | 0 | 0 | 0 |
| 0 | 0 | 0 | 0 |
| 0 | 0 | 0 | 0 |
| 0 | 0 | 0 | 0 |
| 0 | 0 | 0 | 0 |
| 0 | 0 | 0 | 0 |
| 0 | 0 | 0 | 0 |
| 0 | 0 | 0 | 0 |
| 0 | 0 | 0 | 0 |
| 0 | 0 | 0 | 0 |
| 0 | 0 | 0 | 0 |
| 0 | 0 | 0 | 0 |
| 0 | 0 | 0 | 0 |
| 0 | 0 | 0 | 0 |
| 0 | 0 | 0 | 0 |
| 0 | 0 | 0 | 0 |
| 0 | 0 | 0 | 0 |
| 0 | 0 | 0 | 0 |
| 0 | 0 | 0 | 0 |
| 0 | 0 | 0 | 0 |
| 0 | 0 | 0 | 0 |
| 0 | 0 | 0 | 0 |
| 0 | 0 | 0 | 0 |
| 0 | 0 | 0 | 0 |
| 0 | 0 | 0 | 0 |
| 0 | 0 | 0 | 0 |
| 0 | 0 | 0 | 0 |
| 0 | 0 | 0 | 0 |
| 0 | 0 | 0 | 0 |
| 0 | 0 | 0 | 0 |
| 0 | 0 | 0 | 0 |
| 0 | 0 | 0 | 0 |
| 0 | 0 | 0 | 0 |
| 0 | 0 | 0 | 0 |
| 0 | 0 | 0 | 0 |
| 0 | 0 | 0 | 0 |

| | | | |
|---|---|---|---|
| 0 | 0 | 0 | 0 |
| 0 | 0 | 0 | 0 |
| 0 | 0 | 0 | 0 |
| 0 | 0 | 0 | 0 |
| 0 | 0 | 0 | 0 |
| 0 | 0 | 0 | 0 |
| 0 | 0 | 0 | 0 |
| 0 | 0 | 0 | 0 |
| 0 | 0 | 0 | 0 |
| 0 | 0 | 0 | 0 |
| 0 | 0 | 0 | 0 |
| 0 | 0 | 0 | 0 |
| 0 | 0 | 0 | 0 |
| 0 | 0 | 0 | 0 |
| 0 | 0 | 0 | 0 |
| 0 | 0 | 0 | 0 |
| 0 | 0 | 0 | 0 |
| 0 | 0 | 0 | 0 |
| 0 | 0 | 0 | 1 |
| 0 | 1 | 0 | 1 |
| 0 | 0 | 0 | 0 |
| 0 | 0 | 0 | 0 |
| 0 | 0 | 0 | 0 |
| 0 | 0 | 0 | 0 |
| 0 | 0 | 0 | 0 |
| 1 | 0 | 0 | 0 |
| 1 | 0 | 0 | 0 |
| 0 | 0 | 0 | 0 |
| 0 | 0 | 0 | 0 |
| 0 | 0 | 0 | 0 |
| 0 | 0 | 0 | 0 |
| 0 | 0 | 0 | 0 |
| 0 | 0 | 0 | 0 |
| 1 | 0 | 0 | 1 |
| 1 | 0 | 0 | 0 |
| 1 | 0 | 0 | 1 |
| 1 | 0 | 0 | 0 |
| 0 | 0 | 0 | 0 |
| 0 | 0 | 0 | 0 |
| 0 | 0 | 0 | 0 |
| 0 | 0 | 0 | 0 |
| 1 | 0 | 0 | 0 |
| 1 | 0 | 0 | 1 |
| 1 | 0 | 0 | 0 |
| 0 | 0 | 0 | 0 |
| 0 | 0 | 0 | 0 |
| 0 | 0 | 0 | 0 |
| 0 | 0 | 0 | 0 |
| 0 | 0 | 0 | 0 |
| 1 | 0 | 0 | 0 |
| 0 | 0 | 0 | 0 |
| 0 | 0 | 0 | 0 |
| 0 | 0 | 0 | 0 |
| 0 | 0 | 0 | 0 |

| | | | |
|---|---|---|---|
| 0 | 0 | 0 | 0 |
| 0 | 0 | 0 | 0 |
| 0 | 0 | 0 | 0 |
| 1 | 0 | 0 | 0 |
| 1 | 0 | 0 | 0 |
| 0 | 0 | 0 | 0 |
| 1 | 0 | 0 | 0 |
| 0 | 0 | 0 | 0 |
| 0 | 0 | 0 | 0 |
| 0 | 0 | 0 | 0 |
| 0 | 0 | 0 | 0 |
| 1 | 0 | 1 | 0 |
| 0 | 0 | 0 | 0 |
| 0 | 0 | 0 | 0 |
| 0 | 0 | 0 | 0 |
| 0 | 0 | 1 | 0 |
| 0 | 0 | 0 | 0 |
| 0 | 0 | 0 | 0 |
| 0 | 0 | 0 | 0 |
| 0 | 0 | 1 | 0 |
| 1 | 0 | 1 | 0 |
| 1 | 0 | 0 | 0 |
| 1 | 0 | 0 | 0 |
| 1 | 0 | 0 | 0 |
| 0 | 0 | 1 | 0 |
| 0 | 0 | 0 | 0 |
| 1 | 0 | 0 | 0 |
| 0 | 0 | 0 | 0 |
| 0 | 0 | 0 | 0 |
| 0 | 0 | 0 | 0 |
| 0 | 0 | 0 | 0 |
| 0 | 0 | 0 | 0 |
| 0 | 0 | 0 | 0 |
| 0 | 0 | 0 | 0 |
| 0 | 0 | 0 | 0 |
| 0 | 0 | 0 | 0 |
| 0 | 0 | 0 | 1 |
| 0 | 0 | 0 | 1 |
| 0 | 0 | 0 | 1 |
| 0 | 0 | 0 | 0 |
| 0 | 0 | 0 | 0 |
| 0 | 0 | 0 | 0 |
| 0 | 0 | 0 | 0 |
| 0 | 0 | 0 | 0 |
| 0 | 0 | 0 | 0 |
| 0 | 0 | 0 | 0 |
| 0 | 0 | 0 | 0 |
| 0 | 0 | 0 | 0 |
| 0 | 0 | 0 | 0 |
| 0 | 0 | 0 | 0 |
| 0 | 0 | 0 | 0 |
| 0 | 0 | 0 | 0 |
| 0 | 0 | 0 | 0 |
| 0 | 0 | 0 | 0 |

| | | | |
|---|---|---|---|
| 0 | 0 | 0 | 0 |
| 0 | 0 | 0 | 0 |
| 0 | 0 | 0 | 0 |
| 0 | 0 | 0 | 0 |
| 0 | 0 | 0 | 0 |
| 0 | 0 | 0 | 0 |
| 0 | 0 | 0 | 0 |
| 0 | 0 | 0 | 0 |
| 0 | 0 | 0 | 0 |
| 0 | 0 | 0 | 0 |
| 0 | 0 | 0 | 0 |
| 0 | 0 | 0 | 0 |
| 0 | 0 | 0 | 0 |
| 0 | 0 | 0 | 0 |
| 0 | 0 | 0 | 0 |
| 0 | 0 | 0 | 0 |
| 0 | 1 | 0 | 0 |
| 0 | 1 | 0 | 0 |
| 0 | 0 | 0 | 0 |
| 0 | 0 | 0 | 0 |
| 0 | 0 | 0 | 0 |
| 0 | 0 | 0 | 0 |
| 0 | 0 | 0 | 0 |
| 0 | 0 | 0 | 0 |
| 0 | 0 | 0 | 0 |
| 0 | 0 | 0 | 0 |
| 0 | 0 | 0 | 0 |
| 0 | 0 | 0 | 0 |
| 0 | 0 | 0 | 0 |
| 0 | 0 | 0 | 0 |
| 0 | 0 | 0 | 0 |
| 1 | 0 | 0 | 0 |
| 0 | 0 | 0 | 0 |
| 0 | 0 | 0 | 0 |
| 0 | 0 | 0 | 0 |
| 0 | 0 | 0 | 0 |
| 0 | 0 | 0 | 0 |
| 0 | 0 | 0 | 0 |
| 0 | 0 | 0 | 0 |
| 0 | 0 | 0 | 0 |
| 0 | 0 | 0 | 0 |
| 0 | 0 | 0 | 0 |
| 0 | 0 | 0 | 0 |
| 0 | 0 | 0 | 0 |
| 0 | 0 | 0 | 0 |
| 0 | 0 | 0 | 0 |
| 1 | 1 | 0 | 0 |
| 0 | 0 | 0 | 0 |
| 0 | 0 | 0 | 0 |
| 0 | 0 | 0 | 0 |
| 0 | 0 | 0 | 0 |
| 0 | 0 | 0 | 0 |
| 1 | 0 | 0 | 0 |
| 0 | 0 | 0 | 0 |

| | | | |
|---|---|---|---|
| 0 | 0 | 0 | 0 |
| 0 | 0 | 0 | 0 |
| 0 | 0 | 0 | 0 |
| 0 | 0 | 0 | 0 |
| 0 | 0 | 0 | 0 |
| 0 | 0 | 0 | 0 |
| 1 | 0 | 0 | 0 |
| 0 | 0 | 0 | 0 |
| 0 | 0 | 0 | 0 |
| 0 | 0 | 0 | 0 |
| 0 | 0 | 0 | 0 |
| 0 | 0 | 0 | 0 |
| 0 | 0 | 0 | 0 |
| 0 | 0 | 0 | 0 |
| 0 | 0 | 0 | 0 |
| 0 | 0 | 0 | 0 |
| 0 | 0 | 0 | 0 |
| 0 | 0 | 0 | 0 |
| 0 | 0 | 0 | 0 |
| 0 | 0 | 0 | 0 |
| 0 | 0 | 0 | 0 |
| 0 | 0 | 0 | 0 |
| 0 | 0 | 0 | 0 |
| 0 | 0 | 0 | 0 |
| 0 | 0 | 0 | 0 |
| 0 | 0 | 0 | 0 |
| 0 | 0 | 0 | 0 |
| 0 | 0 | 0 | 0 |
| 0 | 0 | 0 | 0 |
| 0 | 0 | 0 | 0 |
| 0 | 0 | 0 | 0 |
| 0 | 0 | 0 | 0 |
| 0 | 0 | 0 | 0 |
| 0 | 0 | 0 | 0 |
| 0 | 0 | 0 | 0 |
| 0 | 0 | 0 | 0 |
| 0 | 0 | 0 | 0 |
| 0 | 0 | 0 | 0 |
| 0 | 0 | 0 | 0 |
| 0 | 0 | 0 | 0 |
| 0 | 0 | 0 | 0 |
| 0 | 0 | 0 | 0 |
| 0 | 0 | 0 | 0 |
| 0 | 0 | 0 | 0 |
| 0 | 0 | 0 | 0 |
| 0 | 0 | 0 | 0 |
| 0 | 0 | 0 | 0 |
| 0 | 0 | 0 | 0 |
| 0 | 0 | 0 | 0 |
| 0 | 0 | 0 | 0 |
| 0 | 0 | 0 | 0 |
| 0 | 0 | 0 | 0 |
| 0 | 0 | 0 | 0 |
| 0 | 0 | 0 | 0 |

| | | | |
|---|---|---|---|
| 0 | 0 | 0 | 0 |
| 0 | 0 | 0 | 0 |
| 0 | 0 | 0 | 0 |
| 0 | 0 | 0 | 0 |
| 0 | 0 | 0 | 0 |
| 0 | 0 | 0 | 0 |
| 0 | 0 | 0 | 0 |
| 0 | 0 | 0 | 0 |
| 0 | 0 | 0 | 0 |
| 0 | 0 | 0 | 0 |
| 0 | 0 | 0 | 0 |
| 0 | 0 | 0 | 0 |
| 0 | 0 | 0 | 0 |
| 0 | 0 | 0 | 0 |
| 0 | 0 | 0 | 0 |
| 0 | 0 | 0 | 0 |
| 0 | 0 | 0 | 0 |
| 0 | 0 | 0 | 0 |
| 0 | 0 | 0 | 0 |
| 0 | 0 | 0 | 0 |
| 0 | 0 | 0 | 0 |
| 0 | 0 | 0 | 0 |
| 0 | 0 | 0 | 0 |
| 0 | 0 | 0 | 0 |
| 1 | 1 | 0 | 0 |
| 0 | 0 | 0 | 0 |
| 0 | 0 | 0 | 0 |
| 0 | 0 | 0 | 0 |
| 0 | 0 | 0 | 0 |
| 0 | 0 | 0 | 0 |
| 0 | 0 | 0 | 0 |
| 0 | 0 | 0 | 0 |
| 0 | 0 | 0 | 0 |
| 0 | 0 | 0 | 0 |
| 0 | 0 | 0 | 0 |
| 1 | 0 | 0 | 0 |
| 1 | 0 | 0 | 1 |
| 1 | 0 | 0 | 0 |
| 0 | 0 | 0 | 0 |
| 1 | 0 | 0 | 0 |
| 1 | 1 | 0 | 0 |
| 1 | 1 | 0 | 0 |
| 0 | 0 | 0 | 0 |
| 0 | 0 | 0 | 0 |
| 0 | 0 | 0 | 0 |
| 0 | 0 | 0 | 0 |
| 0 | 0 | 0 | 0 |
| 0 | 0 | 0 | 0 |
| 0 | 0 | 0 | 0 |
| 0 | 0 | 0 | 0 |
| 0 | 0 | 0 | 0 |
| 0 | 0 | 0 | 0 |
| 0 | 0 | 0 | 0 |
| 0 | 0 | 0 | 0 |

| | | | |
|---|---|---|---|
| 0 | 0 | 0 | 0 |
| 0 | 0 | 0 | 0 |
| 0 | 0 | 0 | 0 |
| 0 | 0 | 0 | 0 |
| 0 | 0 | 0 | 0 |
| 0 | 0 | 0 | 0 |
| 0 | 0 | 0 | 0 |
| 0 | 0 | 0 | 0 |
| 0 | 0 | 0 | 0 |
| 0 | 0 | 0 | 0 |
| 0 | 0 | 0 | 0 |
| 0 | 0 | 0 | 0 |
| 0 | 0 | 0 | 0 |
| 1 | 0 | 0 | 0 |
| 0 | 0 | 0 | 0 |
| 0 | 0 | 0 | 0 |
| 1 | 1 | 0 | 0 |
| 0 | 0 | 0 | 0 |
| 0 | 0 | 0 | 0 |
| 0 | 0 | 0 | 0 |
| 0 | 0 | 0 | 0 |
| 0 | 0 | 0 | 0 |
| 0 | 0 | 0 | 0 |
| 1 | 0 | 0 | 0 |
| 0 | 0 | 0 | 0 |
| 0 | 0 | 0 | 0 |
| 0 | 0 | 0 | 0 |
| 0 | 0 | 0 | 0 |
| 0 | 0 | 0 | 0 |
| 0 | 0 | 0 | 0 |
| 0 | 0 | 0 | 0 |
| 1 | 0 | 0 | 0 |
| 0 | 0 | 0 | 0 |
| 0 | 0 | 0 | 0 |
| 0 | 0 | 0 | 0 |
| 0 | 0 | 0 | 0 |
| 0 | 0 | 0 | 0 |
| 0 | 0 | 0 | 0 |
| 0 | 0 | 0 | 0 |
| 0 | 0 | 0 | 0 |
| 0 | 0 | 0 | 0 |
| 0 | 0 | 0 | 0 |
| 0 | 0 | 0 | 0 |
| 0 | 0 | 0 | 0 |
| 0 | 0 | 0 | 0 |
| 0 | 0 | 0 | 0 |
| 0 | 0 | 0 | 0 |
| 0 | 0 | 0 | 0 |
| 0 | 0 | 0 | 0 |
| 0 | 0 | 0 | 0 |
| 0 | 0 | 0 | 0 |
| 0 | 0 | 0 | 0 |
| 0 | 0 | 0 | 0 |
| 0 | 0 | 0 | 0 |

| | | | |
|---|---|---|---|
| 0 | 0 | 0 | 0 |
| 0 | 0 | 0 | 0 |
| 0 | 0 | 0 | 0 |
| 0 | 0 | 0 | 0 |
| 0 | 0 | 0 | 0 |
| 0 | 0 | 0 | 0 |
| 0 | 0 | 0 | 0 |
| 0 | 0 | 0 | 0 |
| 0 | 0 | 0 | 0 |
| 0 | 0 | 0 | 0 |
| 0 | 0 | 0 | 0 |
| 0 | 0 | 0 | 0 |
| 0 | 0 | 0 | 0 |
| 0 | 0 | 0 | 0 |
| 0 | 0 | 0 | 0 |
| 0 | 0 | 0 | 0 |
| 0 | 0 | 0 | 0 |
| 1 | 0 | 0 | 0 |
| 0 | 0 | 0 | 0 |
| 0 | 0 | 0 | 0 |
| 0 | 0 | 0 | 0 |
| 0 | 0 | 0 | 0 |
| 0 | 0 | 0 | 0 |
| 0 | 0 | 0 | 0 |
| 1 | 0 | 0 | 0 |
| 0 | 0 | 0 | 0 |
| 0 | 0 | 0 | 0 |
| 0 | 0 | 0 | 0 |
| 0 | 0 | 0 | 0 |
| 0 | 0 | 0 | 0 |
| 0 | 0 | 0 | 0 |
| 0 | 0 | 0 | 0 |
| 0 | 0 | 0 | 0 |
| 0 | 0 | 0 | 0 |
| 0 | 0 | 0 | 0 |
| 0 | 0 | 0 | 0 |
| 0 | 0 | 0 | 0 |
| 0 | 0 | 0 | 0 |
| 0 | 0 | 0 | 0 |
| 0 | 0 | 0 | 0 |
| 0 | 0 | 0 | 0 |
| 0 | 0 | 0 | 0 |
| 0 | 0 | 0 | 0 |
| 0 | 0 | 0 | 0 |
| 0 | 0 | 0 | 0 |
| 0 | 0 | 0 | 0 |
| 0 | 0 | 0 | 0 |
| 0 | 0 | 0 | 0 |
| 0 | 0 | 0 | 0 |
| 0 | 0 | 0 | 0 |
| 0 | 0 | 0 | 0 |
| 0 | 0 | 0 | 0 |
| 0 | 0 | 0 | 0 |
| 0 | 0 | 0 | 0 |

| | | | |
|---|---|---|---|
| 0 | 0 | 0 | 0 |
| 0 | 0 | 0 | 0 |
| 0 | 0 | 0 | 0 |
| 0 | 0 | 0 | 0 |
| 0 | 0 | 0 | 0 |
| 0 | 0 | 0 | 0 |
| 0 | 0 | 0 | 0 |
| 0 | 0 | 0 | 0 |
| 0 | 0 | 0 | 0 |
| 0 | 0 | 0 | 0 |
| 0 | 0 | 0 | 0 |
| 0 | 0 | 0 | 0 |
| 0 | 0 | 0 | 0 |
| 0 | 0 | 0 | 0 |
| 1 | 1 | 0 | 0 |
| 1 | 1 | 0 | 0 |
| 0 | 0 | 0 | 0 |
| 0 | 0 | 0 | 0 |
| 0 | 0 | 0 | 0 |
| 0 | 0 | 0 | 0 |
| 0 | 0 | 0 | 0 |
| 0 | 0 | 0 | 0 |
| 0 | 0 | 0 | 0 |
| 0 | 0 | 0 | 0 |
| 0 | 0 | 0 | 0 |
| 0 | 0 | 0 | 0 |
| 0 | 0 | 0 | 0 |
| 1 | 0 | 0 | 0 |
| 0 | 0 | 0 | 0 |
| 1 | 0 | 0 | 0 |
| 0 | 0 | 0 | 0 |
| 0 | 0 | 0 | 0 |
| 0 | 0 | 0 | 0 |
| 0 | 0 | 0 | 0 |
| 0 | 0 | 0 | 0 |
| 0 | 0 | 0 | 0 |
| 1 | 0 | 0 | 0 |
| 0 | 0 | 0 | 0 |
| 0 | 0 | 0 | 0 |
| 0 | 0 | 0 | 0 |
| 0 | 0 | 0 | 0 |
| 0 | 0 | 0 | 0 |
| 0 | 0 | 0 | 0 |
| 0 | 0 | 0 | 0 |
| 0 | 0 | 0 | 0 |
| 0 | 0 | 0 | 0 |
| 0 | 0 | 0 | 0 |
| 0 | 0 | 0 | 0 |
| 0 | 0 | 0 | 0 |
| 0 | 0 | 0 | 0 |
| 0 | 0 | 0 | 0 |
| 0 | 0 | 0 | 0 |
| 0 | 0 | 0 | 0 |
| 0 | 0 | 0 | 0 |

| | | | |
|---|---|---|---|
| 0 | 0 | 0 | 0 |
| 0 | 0 | 0 | 0 |
| 0 | 0 | 0 | 0 |
| 0 | 0 | 0 | 0 |
| 0 | 0 | 0 | 0 |
| 0 | 0 | 0 | 0 |
| 0 | 0 | 0 | 0 |
| 0 | 0 | 0 | 0 |
| 0 | 0 | 0 | 0 |
| 0 | 0 | 0 | 0 |
| 0 | 0 | 0 | 0 |
| 0 | 0 | 0 | 0 |
| 0 | 0 | 0 | 0 |
| 0 | 0 | 0 | 0 |
| 0 | 0 | 0 | 0 |
| 0 | 0 | 0 | 0 |
| 0 | 0 | 0 | 0 |
| 0 | 0 | 0 | 0 |
| 0 | 0 | 0 | 0 |
| 0 | 0 | 0 | 0 |
| 0 | 0 | 0 | 0 |
| 0 | 0 | 0 | 0 |
| 0 | 0 | 0 | 0 |
| 0 | 0 | 0 | 0 |
| 0 | 0 | 0 | 0 |
| 0 | 0 | 0 | 0 |
| 0 | 0 | 0 | 0 |
| 0 | 0 | 0 | 0 |
| 0 | 0 | 0 | 0 |
| 0 | 0 | 0 | 0 |
| 0 | 0 | 0 | 0 |
| 0 | 0 | 0 | 0 |
| 0 | 0 | 0 | 0 |
| 0 | 0 | 0 | 0 |
| 0 | 0 | 0 | 0 |
| 0 | 0 | 0 | 0 |
| 0 | 0 | 0 | 0 |
| 0 | 0 | 0 | 0 |
| 0 | 0 | 0 | 0 |
| 0 | 0 | 0 | 0 |
| 0 | 0 | 0 | 0 |
| 0 | 0 | 0 | 0 |
| 0 | 0 | 0 | 0 |
| 0 | 0 | 0 | 0 |
| 0 | 0 | 0 | 0 |
| 0 | 0 | 0 | 0 |
| 0 | 0 | 0 | 0 |
| 0 | 0 | 0 | 0 |
| 1 | 0 | 0 | 1 |
| 0 | 0 | 0 | 0 |
| 1 | 0 | 0 | 0 |
| 1 | 0 | 0 | 0 |
| 0 | 0 | 0 | 0 |
| 0 | 0 | 0 | 1 |

| | | | |
|---|---|---|---|
| 0 | 0 | 0 | 0 |
| 0 | 0 | 0 | 0 |
| 0 | 0 | 0 | 0 |
| 0 | 0 | 0 | 0 |
| 0 | 0 | 0 | 0 |
| 0 | 0 | 0 | 0 |
| 0 | 0 | 0 | 0 |
| 0 | 0 | 0 | 0 |
| 0 | 0 | 0 | 0 |
| 0 | 0 | 0 | 0 |
| 0 | 0 | 0 | 0 |
| 0 | 0 | 0 | 0 |
| 0 | 0 | 0 | 0 |
| 0 | 0 | 0 | 0 |
| 0 | 0 | 0 | 0 |
| 0 | 0 | 0 | 0 |
| 0 | 0 | 0 | 0 |
| 0 | 0 | 0 | 0 |
| 0 | 0 | 0 | 0 |
| 0 | 0 | 0 | 0 |
| 0 | 0 | 0 | 0 |
| 0 | 0 | 0 | 0 |
| 0 | 0 | 0 | 0 |
| 0 | 0 | 0 | 0 |
| 0 | 0 | 0 | 0 |
| 0 | 0 | 0 | 0 |
| 0 | 0 | 1 | 0 |
| 0 | 0 | 1 | 0 |
| 0 | 0 | 1 | 0 |
| 0 | 0 | 1 | 0 |
| 0 | 0 | 0 | 0 |
| 0 | 0 | 0 | 0 |
| 0 | 0 | 0 | 0 |
| 1 | 0 | 0 | 0 |
| 1 | 0 | 0 | 0 |
| 0 | 0 | 0 | 0 |
| 0 | 0 | 0 | 0 |
| 0 | 0 | 0 | 0 |
| 0 | 0 | 0 | 0 |
| 1 | 0 | 0 | 0 |
| 0 | 0 | 0 | 0 |
| 0 | 0 | 0 | 0 |
| 0 | 0 | 0 | 0 |
| 0 | 0 | 0 | 0 |
| 0 | 0 | 0 | 0 |
| 0 | 1 | 0 | 0 |
| 0 | 0 | 0 | 0 |
| 0 | 0 | 0 | 0 |
| 0 | 0 | 0 | 0 |
| 0 | 0 | 0 | 0 |
| 0 | 0 | 0 | 0 |
| 0 | 0 | 0 | 0 |
| 0 | 0 | 0 | 0 |
| 0 | 0 | 0 | 0 |

| | | | |
|---|---|---|---|
| 0 | 0 | 0 | 0 |
| 0 | 0 | 0 | 0 |
| 0 | 0 | 0 | 0 |
| 0 | 0 | 0 | 0 |
| 0 | 0 | 0 | 0 |
| 0 | 0 | 0 | 1 |
| 0 | 0 | 0 | 0 |
| 0 | 0 | 0 | 0 |
| 0 | 0 | 0 | 0 |
| 0 | 0 | 0 | 0 |
| 0 | 0 | 0 | 0 |
| 0 | 0 | 0 | 0 |
| 0 | 0 | 0 | 0 |
| 0 | 1 | 0 | 0 |
| 0 | 1 | 0 | 0 |
| 0 | 1 | 0 | 0 |
| 0 | 1 | 0 | 0 |
| 0 | 1 | 0 | 0 |
| 0 | 0 | 0 | 0 |
| 0 | 0 | 0 | 0 |
| 0 | 0 | 0 | 0 |
| 0 | 0 | 0 | 0 |
| 0 | 0 | 0 | 0 |
| 0 | 0 | 0 | 0 |
| 0 | 0 | 0 | 0 |
| 0 | 0 | 0 | 0 |
| 0 | 0 | 0 | 0 |
| 0 | 0 | 0 | 0 |
| 0 | 0 | 0 | 0 |
| 0 | 0 | 0 | 0 |
| 0 | 0 | 0 | 0 |
| 0 | 0 | 0 | 0 |
| 0 | 0 | 0 | 0 |
| 0 | 0 | 0 | 0 |
| 0 | 0 | 0 | 0 |
| 0 | 0 | 0 | 0 |
| 0 | 0 | 0 | 0 |
| 0 | 0 | 0 | 0 |
| 0 | 0 | 0 | 0 |
| 0 | 0 | 0 | 0 |
| 0 | 0 | 0 | 0 |
| 0 | 0 | 0 | 0 |
| 0 | 0 | 0 | 0 |
| 0 | 0 | 0 | 0 |
| 0 | 0 | 0 | 0 |
| 0 | 0 | 0 | 0 |
| 0 | 0 | 0 | 0 |
| 0 | 0 | 0 | 0 |
| 0 | 0 | 0 | 0 |
| 0 | 0 | 0 | 0 |
| 0 | 0 | 0 | 0 |
| 0 | 0 | 0 | 0 |
| 0 | 0 | 0 | 0 |
| 0 | 0 | 0 | 0 |

| | | | |
|---|---|---|---|
| 0 | 0 | 0 | 0 |
| 0 | 0 | 0 | 0 |
| 0 | 0 | 0 | 0 |
| 1 | 0 | 0 | 0 |
| 0 | 0 | 0 | 0 |
| 0 | 0 | 0 | 0 |
| 0 | 0 | 0 | 0 |
| 0 | 0 | 0 | 0 |
| 0 | 0 | 0 | 0 |
| 0 | 0 | 0 | 0 |
| 0 | 0 | 0 | 0 |
| 0 | 0 | 0 | 0 |
| 0 | 0 | 0 | 0 |
| 0 | 0 | 0 | 0 |
| 0 | 0 | 0 | 0 |
| 0 | 0 | 0 | 0 |
| 0 | 0 | 0 | 0 |
| 0 | 0 | 0 | 0 |
| 0 | 0 | 0 | 0 |
| 0 | 0 | 0 | 0 |
| 1 | 1 | 0 | 0 |
| 1 | 1 | 0 | 0 |
| 0 | 0 | 0 | 0 |
| 0 | 1 | 0 | 0 |
| 0 | 0 | 0 | 0 |
| 0 | 0 | 0 | 0 |
| 0 | 0 | 0 | 0 |
| 0 | 0 | 0 | 0 |
| 0 | 0 | 0 | 0 |
| 0 | 0 | 0 | 0 |
| 0 | 0 | 0 | 0 |
| 0 | 0 | 0 | 0 |
| 0 | 0 | 0 | 0 |
| 0 | 0 | 0 | 0 |
| 0 | 0 | 0 | 0 |
| 0 | 0 | 0 | 0 |
| 0 | 0 | 0 | 0 |
| 0 | 0 | 0 | 0 |
| 0 | 0 | 0 | 0 |
| 1 | 0 | 0 | 0 |
| 1 | 0 | 0 | 0 |
| 0 | 0 | 0 | 0 |
| 0 | 0 | 0 | 0 |
| 1 | 0 | 0 | 0 |
| 0 | 0 | 0 | 0 |
| 0 | 0 | 0 | 0 |
| 0 | 0 | 0 | 0 |
| 0 | 0 | 0 | 0 |
| 0 | 0 | 0 | 0 |
| 0 | 0 | 0 | 0 |
| 0 | 0 | 0 | 0 |
| 0 | 0 | 0 | 0 |
| 0 | 0 | 0 | 0 |
| 0 | 0 | 0 | 0 |

| | | | |
|---|---|---|---|
| 0 | 0 | 0 | 0 |
| 0 | 0 | 0 | 0 |
| 0 | 0 | 0 | 0 |
| 0 | 0 | 0 | 0 |
| 0 | 0 | 0 | 0 |
| 1 | 0 | 0 | 0 |
| 0 | 0 | 0 | 0 |
| 0 | 0 | 0 | 0 |
| 0 | 1 | 0 | 0 |
| 1 | 1 | 0 | 0 |
| 0 | 0 | 0 | 0 |
| 1 | 1 | 0 | 0 |
| 1 | 0 | 0 | 0 |
| 0 | 0 | 0 | 0 |
| 0 | 0 | 0 | 0 |
| 0 | 0 | 0 | 0 |
| 1 | 1 | 0 | 0 |
| 0 | 0 | 0 | 0 |
| 0 | 0 | 0 | 0 |
| 1 | 0 | 0 | 0 |
| 0 | 0 | 0 | 0 |
| 0 | 0 | 0 | 0 |
| 0 | 0 | 0 | 0 |
| 0 | 0 | 0 | 0 |
| 0 | 0 | 0 | 0 |
| 0 | 0 | 0 | 0 |
| 0 | 0 | 0 | 0 |
| 0 | 0 | 0 | 0 |
| 0 | 0 | 0 | 0 |
| 0 | 0 | 0 | 0 |
| 1 | 0 | 0 | 0 |
| 0 | 0 | 0 | 0 |
| 0 | 0 | 0 | 0 |
| 0 | 0 | 0 | 0 |
| 0 | 0 | 0 | 0 |
| 1 | 0 | 0 | 0 |
| 0 | 0 | 0 | 0 |
| 0 | 0 | 0 | 0 |
| 0 | 0 | 0 | 0 |
| 0 | 0 | 0 | 0 |
| 0 | 0 | 0 | 0 |
| 0 | 0 | 0 | 0 |
| 0 | 0 | 0 | 0 |
| 0 | 0 | 0 | 0 |
| 0 | 0 | 0 | 0 |
| 0 | 0 | 0 | 0 |
| 0 | 0 | 0 | 0 |
| 0 | 0 | 0 | 0 |
| 0 | 0 | 0 | 0 |
| 0 | 0 | 0 | 0 |
| 0 | 0 | 0 | 0 |
| 0 | 0 | 0 | 0 |
| 0 | 0 | 0 | 0 |
| 0 | 0 | 0 | 0 |

| | | | |
|---|---|---|---|
| 0 | 0 | 0 | 0 |
| 0 | 0 | 0 | 0 |
| 0 | 0 | 0 | 0 |
| 0 | 0 | 0 | 0 |
| 0 | 0 | 0 | 0 |
| 0 | 0 | 0 | 0 |
| 0 | 0 | 0 | 0 |
| 0 | 0 | 0 | 0 |
| 0 | 0 | 0 | 0 |
| 0 | 0 | 0 | 0 |
| 0 | 0 | 0 | 0 |
| 0 | 0 | 0 | 0 |
| 0 | 0 | 0 | 0 |
| 0 | 0 | 0 | 0 |
| 0 | 0 | 0 | 1 |
| 0 | 0 | 0 | 0 |
| 0 | 0 | 0 | 0 |
| 0 | 0 | 0 | 0 |
| 0 | 0 | 0 | 0 |
| 0 | 0 | 0 | 0 |
| 0 | 0 | 0 | 0 |
| 0 | 0 | 0 | 0 |
| 0 | 0 | 0 | 0 |
| 0 | 0 | 0 | 0 |
| 0 | 0 | 0 | 0 |
| 0 | 0 | 0 | 0 |
| 0 | 0 | 0 | 0 |
| 0 | 0 | 0 | 0 |
| 0 | 0 | 0 | 0 |
| 0 | 0 | 0 | 0 |
| 0 | 0 | 0 | 0 |
| 0 | 0 | 0 | 0 |
| 0 | 0 | 0 | 0 |
| 0 | 0 | 0 | 0 |
| 0 | 0 | 0 | 0 |
| 0 | 0 | 0 | 0 |
| 0 | 0 | 0 | 0 |
| 0 | 0 | 0 | 0 |
| 0 | 0 | 0 | 0 |
| 0 | 0 | 1 | 0 |
| 0 | 0 | 0 | 0 |
| 0 | 0 | 0 | 0 |
| 0 | 0 | 0 | 0 |
| 1 | 0 | 0 | 0 |
| 0 | 0 | 0 | 0 |
| 0 | 0 | 0 | 0 |
| 0 | 0 | 0 | 0 |
| 0 | 0 | 0 | 0 |
| 1 | 0 | 0 | 1 |
| 0 | 0 | 0 | 0 |
| 0 | 0 | 0 | 0 |
| 0 | 0 | 0 | 0 |
| 0 | 0 | 0 | 0 |
| 0 | 0 | 0 | 0 |

| | | | |
|---|---|---|---|
| 0 | 0 | 0 | 0 |
| 0 | 0 | 0 | 0 |
| 0 | 0 | 0 | 0 |
| 0 | 0 | 0 | 0 |
| 0 | 0 | 0 | 0 |
| 0 | 0 | 0 | 0 |
| 0 | 0 | 0 | 0 |
| 0 | 0 | 0 | 0 |
| 0 | 0 | 0 | 0 |
| 1 | 0 | 0 | 0 |
| 0 | 0 | 0 | 0 |
| 0 | 0 | 0 | 0 |
| 0 | 0 | 0 | 0 |
| 0 | 0 | 0 | 0 |
| 1 | 0 | 0 | 0 |
| 0 | 0 | 0 | 0 |
| 0 | 0 | 0 | 0 |
| 0 | 0 | 0 | 0 |
| 0 | 0 | 0 | 0 |
| 0 | 0 | 0 | 0 |
| 0 | 0 | 0 | 0 |
| 0 | 0 | 0 | 0 |
| 0 | 0 | 0 | 0 |
| 0 | 0 | 0 | 0 |
| 1 | 0 | 0 | 0 |
| 0 | 0 | 0 | 0 |
| 0 | 0 | 0 | 0 |
| 0 | 0 | 0 | 0 |
| 0 | 0 | 0 | 0 |
| 0 | 0 | 0 | 0 |
| 0 | 0 | 0 | 0 |
| 0 | 0 | 0 | 0 |
| 0 | 0 | 0 | 0 |
| 0 | 0 | 0 | 0 |
| 1 | 0 | 0 | 0 |
| 1 | 0 | 0 | 0 |
| 0 | 0 | 0 | 0 |
| 0 | 0 | 0 | 0 |
| 1 | 0 | 0 | 0 |
| 0 | 0 | 1 | 0 |
| 0 | 0 | 1 | 0 |
| 0 | 0 | 1 | 0 |
| 0 | 0 | 1 | 0 |
| 0 | 0 | 0 | 0 |
| 1 | 0 | 0 | 0 |
| 0 | 0 | 0 | 0 |
| 0 | 0 | 0 | 0 |
| 0 | 0 | 0 | 0 |
| 0 | 0 | 1 | 0 |
| 0 | 0 | 0 | 0 |
| 0 | 0 | 0 | 0 |
| 0 | 0 | 0 | 0 |
| 0 | 0 | 0 | 0 |
| 0 | 0 | 0 | 0 |

| | | | |
|---|---|---|---|
| 0 | 0 | 0 | 0 |
| 0 | 0 | 0 | 0 |
| 0 | 0 | 0 | 0 |
| 0 | 0 | 0 | 0 |
| 0 | 0 | 0 | 0 |
| 0 | 0 | 0 | 0 |
| 0 | 0 | 0 | 0 |
| 0 | 0 | 0 | 0 |
| 0 | 0 | 0 | 0 |
| 0 | 0 | 0 | 0 |
| 0 | 1 | 0 | 0 |
| 0 | 0 | 0 | 0 |
| 0 | 0 | 0 | 0 |
| 0 | 0 | 0 | 0 |
| 0 | 0 | 0 | 0 |
| 0 | 1 | 0 | 0 |
| 0 | 0 | 0 | 0 |
| 0 | 0 | 0 | 0 |
| 0 | 0 | 0 | 0 |
| 0 | 0 | 0 | 0 |
| 0 | 0 | 0 | 0 |
| 0 | 0 | 0 | 0 |
| 0 | 0 | 0 | 0 |
| 0 | 0 | 0 | 0 |
| 0 | 0 | 0 | 0 |
| 0 | 0 | 0 | 0 |
| 0 | 0 | 0 | 0 |
| 0 | 0 | 0 | 0 |
| 0 | 0 | 0 | 0 |
| 0 | 0 | 0 | 0 |
| 0 | 0 | 0 | 0 |
| 0 | 0 | 0 | 0 |
| 0 | 0 | 0 | 0 |
| 0 | 0 | 0 | 0 |
| 0 | 0 | 0 | 0 |
| 0 | 0 | 0 | 0 |
| 0 | 0 | 0 | 0 |
| 0 | 0 | 0 | 0 |
| 0 | 0 | 0 | 0 |
| 0 | 0 | 0 | 0 |
| 0 | 0 | 0 | 0 |
| 0 | 0 | 0 | 0 |
| 0 | 0 | 0 | 0 |
| 0 | 0 | 0 | 0 |
| 1 | 1 | 0 | 0 |
| 0 | 0 | 0 | 0 |
| 0 | 0 | 0 | 0 |
| 0 | 0 | 0 | 0 |
| 0 | 0 | 0 | 0 |
| 0 | 0 | 0 | 0 |
| 0 | 0 | 0 | 0 |
| 0 | 0 | 0 | 0 |
| 0 | 0 | 0 | 0 |
| 0 | 0 | 0 | 0 |

| | | | |
|---|---|---|---|
| 0 | 0 | 0 | 0 |
| 0 | 0 | 0 | 0 |
| 0 | 0 | 1 | 0 |
| 1 | 0 | 0 | 0 |
| 0 | 0 | 0 | 0 |
| 0 | 0 | 0 | 0 |
| 0 | 0 | 0 | 0 |
| 0 | 0 | 0 | 0 |
| 0 | 0 | 0 | 0 |
| 0 | 0 | 0 | 0 |
| 0 | 0 | 0 | 0 |
| 0 | 0 | 0 | 0 |
| 1 | 0 | 0 | 0 |
| 0 | 0 | 0 | 0 |
| 0 | 0 | 0 | 0 |
| 0 | 0 | 0 | 0 |
| 0 | 0 | 0 | 0 |
| 0 | 0 | 0 | 0 |
| 0 | 0 | 0 | 0 |
| 0 | 0 | 0 | 0 |
| 0 | 0 | 0 | 0 |
| 0 | 0 | 0 | 0 |
| 0 | 0 | 0 | 0 |
| 0 | 0 | 0 | 0 |
| 0 | 0 | 0 | 0 |
| 0 | 0 | 0 | 0 |
| 0 | 0 | 0 | 0 |
| 0 | 0 | 0 | 0 |
| 0 | 0 | 0 | 0 |
| 0 | 0 | 0 | 0 |
| 0 | 0 | 0 | 0 |
| 0 | 0 | 0 | 0 |
| 0 | 0 | 0 | 0 |
| 0 | 0 | 0 | 0 |
| 0 | 0 | 0 | 0 |
| 0 | 0 | 0 | 0 |
| 0 | 0 | 0 | 0 |
| 0 | 0 | 0 | 0 |
| 0 | 0 | 0 | 0 |
| 0 | 0 | 0 | 0 |
| 0 | 0 | 0 | 0 |
| 0 | 1 | 0 | 0 |
| 1 | 1 | 0 | 0 |
| 0 | 0 | 0 | 0 |
| 0 | 0 | 0 | 0 |
| 0 | 0 | 0 | 0 |
| 0 | 0 | 0 | 0 |
| 0 | 0 | 0 | 0 |
| 0 | 0 | 0 | 0 |
| 0 | 1 | 0 | 0 |
| 0 | 0 | 0 | 0 |
| 0 | 0 | 0 | 0 |
| 0 | 0 | 0 | 0 |
| 0 | 0 | 0 | 1 |
| 0 | 0 | 0 | 0 |

| | | | |
|---|---|---|---|
| 0 | 0 | 0 | 1 |
| 0 | 0 | 0 | 1 |
| 0 | 0 | 0 | 1 |
| 0 | 0 | 0 | 0 |
| 0 | 0 | 0 | 0 |
| 0 | 0 | 0 | 0 |
| 0 | 0 | 0 | 0 |
| 1 | 0 | 0 | 0 |
| 0 | 0 | 0 | 0 |
| 1 | 1 | 0 | 0 |
| 0 | 1 | 0 | 0 |
| 0 | 0 | 0 | 0 |
| 1 | 1 | 0 | 0 |
| 0 | 0 | 0 | 0 |
| 0 | 0 | 0 | 0 |
| 0 | 0 | 0 | 0 |
| 0 | 0 | 0 | 0 |
| 1 | 0 | 0 | 0 |
| 0 | 0 | 0 | 0 |
| 0 | 0 | 0 | 0 |
| 0 | 0 | 0 | 0 |
| 0 | 0 | 0 | 0 |
| 0 | 0 | 0 | 0 |
| 0 | 0 | 0 | 0 |
| 0 | 0 | 0 | 0 |
| 0 | 0 | 0 | 0 |
| 0 | 0 | 0 | 0 |
| 0 | 0 | 0 | 0 |
| 0 | 0 | 0 | 0 |
| 0 | 0 | 0 | 0 |
| 0 | 0 | 0 | 0 |
| 1 | 0 | 0 | 0 |
| 0 | 0 | 0 | 0 |
| 0 | 0 | 0 | 0 |
| 0 | 0 | 0 | 0 |
| 0 | 0 | 0 | 0 |
| 0 | 0 | 0 | 0 |
| 0 | 0 | 0 | 0 |
| 0 | 0 | 0 | 0 |
| 0 | 0 | 0 | 0 |
| 0 | 0 | 0 | 0 |
| 1 | 1 | 0 | 0 |
| 0 | 0 | 0 | 0 |
| 0 | 0 | 0 | 0 |
| 1 | 0 | 0 | 0 |
| 0 | 1 | 0 | 0 |
| 0 | 0 | 0 | 0 |
| 0 | 0 | 0 | 0 |
| 0 | 1 | 0 | 0 |
| 1 | 1 | 0 | 0 |
| 1 | 1 | 0 | 0 |
| 1 | 1 | 0 | 0 |
| 1 | 1 | 0 | 0 |
| 1 | 1 | 0 | 0 |

| | | | |
|---|---|---|---|
| 0 | 0 | 0 | 0 |
| 0 | 0 | 0 | 0 |
| 0 | 0 | 0 | 0 |
| 0 | 0 | 0 | 0 |
| 0 | 0 | 0 | 0 |
| 1 | 0 | 0 | 0 |
| 1 | 0 | 0 | 0 |
| 0 | 0 | 0 | 0 |
| 0 | 0 | 0 | 0 |
| 1 | 0 | 0 | 0 |
| 0 | 0 | 0 | 0 |
| 0 | 0 | 1 | 0 |
| 0 | 0 | 0 | 0 |
| 0 | 0 | 0 | 0 |
| 0 | 0 | 0 | 0 |
| 0 | 0 | 0 | 0 |
| 0 | 0 | 0 | 0 |
| 0 | 0 | 0 | 0 |
| 0 | 0 | 0 | 0 |
| 0 | 0 | 0 | 0 |
| 1 | 1 | 0 | 0 |
| 1 | 0 | 0 | 0 |
| 0 | 0 | 0 | 0 |
| 0 | 0 | 1 | 0 |
| 0 | 0 | 0 | 0 |
| 0 | 0 | 1 | 0 |
| 0 | 0 | 0 | 0 |
| 0 | 0 | 0 | 0 |
| 0 | 0 | 0 | 0 |
| 0 | 0 | 0 | 0 |
| 0 | 0 | 0 | 0 |
| 0 | 0 | 0 | 0 |
| 0 | 0 | 0 | 0 |
| 0 | 0 | 0 | 0 |
| 0 | 0 | 0 | 0 |
| 0 | 0 | 0 | 0 |
| 1 | 0 | 0 | 0 |
| 0 | 0 | 0 | 0 |
| 1 | 0 | 0 | 0 |
| 1 | 0 | 0 | 0 |
| 1 | 0 | 0 | 0 |
| 0 | 0 | 0 | 0 |
| 0 | 0 | 0 | 0 |
| 0 | 0 | 0 | 0 |
| 0 | 0 | 0 | 0 |
| 0 | 0 | 0 | 0 |
| 0 | 0 | 0 | 0 |
| 1 | 0 | 1 | 0 |
| 0 | 0 | 0 | 0 |
| 0 | 0 | 0 | 0 |
| 0 | 0 | 0 | 0 |
| 0 | 0 | 0 | 0 |
| 1 | 1 | 0 | 0 |
| 0 | 0 | 0 | 0 |

| | | | |
|---|---|---|---|
| 0 | 0 | 0 | 0 |
| 0 | 1 | 0 | 0 |
| 0 | 0 | 0 | 0 |
| 0 | 0 | 0 | 0 |
| 0 | 0 | 0 | 0 |
| 0 | 0 | 0 | 0 |
| 0 | 0 | 0 | 0 |
| 0 | 0 | 0 | 0 |
| 0 | 0 | 0 | 0 |
| 0 | 0 | 0 | 0 |
| 0 | 0 | 0 | 0 |
| 0 | 0 | 0 | 0 |
| 0 | 0 | 0 | 0 |
| 0 | 0 | 0 | 0 |
| 0 | 0 | 0 | 0 |
| 0 | 0 | 0 | 0 |
| 0 | 0 | 0 | 0 |
| 0 | 0 | 0 | 0 |
| 0 | 0 | 0 | 0 |
| 0 | 0 | 0 | 0 |
| 1 | 0 | 0 | 0 |
| 0 | 0 | 0 | 0 |
| 0 | 0 | 0 | 0 |
| 0 | 0 | 0 | 0 |
| 1 | 0 | 0 | 0 |
| 0 | 0 | 0 | 0 |
| 0 | 0 | 0 | 0 |
| 0 | 0 | 0 | 1 |
| 0 | 0 | 0 | 0 |
| 0 | 0 | 0 | 0 |
| 0 | 0 | 0 | 0 |
| 0 | 0 | 0 | 0 |
| 0 | 0 | 0 | 0 |
| 0 | 0 | 0 | 0 |
| 0 | 0 | 0 | 0 |
| 0 | 0 | 0 | 0 |
| 0 | 0 | 0 | 0 |
| 0 | 0 | 0 | 0 |
| 0 | 0 | 0 | 0 |
| 0 | 0 | 0 | 0 |
| 0 | 0 | 0 | 0 |
| 0 | 0 | 0 | 0 |
| 0 | 0 | 0 | 0 |
| 0 | 0 | 0 | 0 |
| 0 | 0 | 0 | 0 |
| 0 | 0 | 0 | 0 |
| 0 | 0 | 0 | 0 |
| 0 | 0 | 0 | 0 |
| 0 | 0 | 0 | 0 |
| 0 | 0 | 0 | 0 |
| 0 | 0 | 0 | 0 |
| 0 | 0 | 0 | 0 |
| 0 | 0 | 0 | 0 |
| 0 | 0 | 0 | 0 |

| | | | |
|---|---|---|---|
| 0 | 0 | 0 | 0 |
| 0 | 0 | 0 | 0 |
| 0 | 0 | 0 | 0 |
| 0 | 0 | 0 | 0 |
| 0 | 0 | 0 | 0 |
| 0 | 0 | 0 | 0 |
| 0 | 0 | 0 | 0 |
| 0 | 0 | 0 | 0 |
| 0 | 0 | 0 | 0 |
| 0 | 0 | 0 | 0 |
| 0 | 0 | 0 | 0 |
| 0 | 0 | 0 | 0 |
| 0 | 0 | 0 | 0 |
| 0 | 0 | 0 | 0 |
| 0 | 0 | 0 | 0 |
| 1 | 0 | 0 | 0 |
| 0 | 0 | 0 | 0 |
| 0 | 0 | 0 | 0 |
| 0 | 1 | 0 | 0 |
| 0 | 0 | 0 | 0 |
| 0 | 0 | 0 | 0 |
| 0 | 0 | 0 | 0 |
| 0 | 0 | 0 | 0 |
| 0 | 0 | 0 | 0 |
| 0 | 0 | 0 | 0 |
| 1 | 0 | 0 | 0 |
| 0 | 0 | 0 | 0 |
| 0 | 0 | 0 | 0 |
| 0 | 0 | 0 | 0 |
| 0 | 0 | 0 | 0 |
| 0 | 0 | 0 | 0 |
| 0 | 0 | 0 | 0 |
| 0 | 0 | 0 | 0 |
| 1 | 1 | 0 | 0 |
| 0 | 0 | 0 | 0 |
| 0 | 0 | 0 | 0 |
| 0 | 0 | 0 | 0 |
| 0 | 0 | 0 | 0 |
| 0 | 0 | 0 | 0 |
| 0 | 0 | 0 | 0 |
| 0 | 0 | 0 | 0 |
| 0 | 0 | 0 | 0 |
| 0 | 0 | 0 | 0 |
| 0 | 0 | 0 | 0 |
| 0 | 0 | 0 | 0 |
| 0 | 0 | 0 | 0 |
| 0 | 0 | 0 | 0 |
| 1 | 0 | 0 | 0 |
| 0 | 0 | 0 | 0 |
| 0 | 0 | 0 | 0 |
| 0 | 0 | 0 | 0 |
| 0 | 0 | 0 | 0 |
| 0 | 0 | 0 | 0 |
| 1 | 0 | 0 | 1 |

| | | | |
|---|---|---|---|
| 0 | 0 | 0 | 0 |
| 0 | 0 | 0 | 0 |
| 0 | 0 | 1 | 0 |
| 0 | 0 | 0 | 0 |
| 0 | 0 | 0 | 0 |
| 0 | 0 | 0 | 0 |
| 0 | 0 | 0 | 0 |
| 0 | 0 | 0 | 0 |
| 0 | 0 | 0 | 0 |
| 0 | 0 | 0 | 0 |
| 0 | 0 | 0 | 0 |
| 0 | 0 | 0 | 0 |
| 0 | 0 | 0 | 0 |
| 0 | 0 | 0 | 0 |
| 0 | 0 | 0 | 0 |
| 0 | 0 | 0 | 0 |
| 0 | 0 | 0 | 0 |
| 0 | 0 | 0 | 0 |
| 0 | 0 | 0 | 0 |
| 0 | 0 | 0 | 0 |
| 0 | 0 | 0 | 0 |
| 0 | 0 | 0 | 0 |
| 0 | 0 | 0 | 0 |
| 0 | 0 | 0 | 0 |
| 0 | 0 | 0 | 0 |
| 0 | 0 | 0 | 0 |
| 0 | 0 | 0 | 0 |
| 0 | 0 | 0 | 0 |
| 0 | 0 | 0 | 0 |
| 0 | 0 | 0 | 0 |
| 0 | 0 | 0 | 0 |
| 1 | 0 | 0 | 0 |
| 0 | 0 | 0 | 0 |
| 0 | 0 | 0 | 0 |
| 0 | 0 | 0 | 0 |
| 0 | 0 | 0 | 0 |
| 0 | 0 | 0 | 0 |
| 0 | 0 | 0 | 0 |
| 0 | 0 | 0 | 0 |
| 0 | 0 | 0 | 0 |
| 1 | 1 | 0 | 0 |
| 0 | 0 | 0 | 0 |
| 0 | 0 | 0 | 0 |
| 0 | 0 | 0 | 0 |
| 0 | 0 | 0 | 0 |
| 1 | 1 | 0 | 0 |
| 0 | 0 | 0 | 0 |
| 1 | 1 | 0 | 0 |
| 0 | 0 | 0 | 0 |
| 1 | 0 | 0 | 0 |
| 1 | 0 | 0 | 0 |
| 1 | 0 | 0 | 0 |
| 0 | 0 | 0 | 0 |
| 0 | 0 | 0 | 0 |

| | | | |
|---|---|---|---|
| 1 | 0 | 0 | 0 |
| 0 | 0 | 0 | 0 |
| 0 | 0 | 0 | 0 |
| 0 | 0 | 0 | 0 |
| 0 | 0 | 0 | 0 |
| 0 | 0 | 0 | 0 |
| 0 | 0 | 0 | 0 |
| 0 | 0 | 0 | 0 |
| 0 | 0 | 0 | 0 |
| 0 | 0 | 0 | 0 |
| 0 | 0 | 0 | 0 |
| 0 | 0 | 0 | 0 |
| 0 | 0 | 0 | 0 |
| 0 | 0 | 0 | 0 |
| 0 | 0 | 0 | 0 |
| 1 | 1 | 1 | 0 |
| 0 | 0 | 0 | 0 |
| 1 | 0 | 0 | 0 |
| 0 | 0 | 0 | 0 |
| 0 | 0 | 0 | 0 |
| 0 | 0 | 0 | 0 |
| 0 | 1 | 0 | 0 |
| 0 | 0 | 0 | 0 |
| 0 | 1 | 0 | 0 |
| 0 | 1 | 0 | 0 |
| 0 | 0 | 0 | 0 |
| 0 | 0 | 0 | 0 |
| 0 | 0 | 0 | 0 |
| 0 | 0 | 0 | 0 |
| 0 | 0 | 0 | 0 |
| 0 | 0 | 0 | 0 |
| 0 | 0 | 0 | 0 |
| 1 | 1 | 0 | 0 |
| 0 | 0 | 0 | 0 |
| 0 | 0 | 0 | 0 |
| 0 | 0 | 0 | 0 |
| 0 | 1 | 0 | 0 |
| 1 | 1 | 0 | 0 |
| 0 | 0 | 0 | 0 |
| 0 | 0 | 0 | 0 |
| 0 | 0 | 0 | 0 |
| 0 | 0 | 0 | 0 |
| 0 | 0 | 0 | 0 |
| 0 | 0 | 0 | 0 |
| 1 | 1 | 0 | 0 |
| 0 | 0 | 0 | 0 |
| 1 | 0 | 0 | 0 |
| 1 | 1 | 0 | 0 |
| 0 | 0 | 0 | 0 |
| 0 | 0 | 0 | 0 |
| 1 | 1 | 0 | 0 |
| 1 | 1 | 0 | 0 |
| 0 | 0 | 0 | 0 |
| 1 | 1 | 0 | 0 |

| | | | |
|---|---|---|---|
| 0 | 0 | 0 | 0 |
| 0 | 0 | 0 | 0 |
| 0 | 0 | 0 | 0 |
| 1 | 1 | 0 | 0 |
| 0 | 0 | 0 | 0 |
| 0 | 0 | 0 | 0 |
| 0 | 0 | 0 | 0 |
| 0 | 0 | 0 | 0 |
| 1 | 0 | 0 | 0 |
| 0 | 0 | 0 | 0 |
| 0 | 0 | 0 | 0 |
| 0 | 0 | 0 | 0 |
| 1 | 0 | 1 | 0 |
| 1 | 0 | 0 | 0 |
| 0 | 0 | 0 | 0 |
| 0 | 0 | 0 | 0 |
| 0 | 0 | 0 | 0 |
| 0 | 0 | 0 | 0 |
| 0 | 0 | 0 | 0 |
| 0 | 0 | 0 | 0 |
| 0 | 0 | 0 | 0 |
| 0 | 0 | 0 | 0 |
| 0 | 0 | 0 | 0 |
| 1 | 0 | 0 | 0 |
| 0 | 0 | 0 | 0 |
| 0 | 0 | 0 | 0 |
| 0 | 0 | 0 | 0 |
| 0 | 0 | 0 | 0 |
| 0 | 0 | 0 | 0 |
| 0 | 0 | 0 | 0 |
| 0 | 0 | 0 | 0 |
| 0 | 0 | 0 | 0 |
| 0 | 0 | 0 | 0 |
| 0 | 0 | 0 | 0 |
| 0 | 0 | 0 | 0 |
| 0 | 0 | 0 | 0 |
| 0 | 0 | 0 | 0 |
| 0 | 0 | 0 | 0 |
| 0 | 0 | 0 | 0 |
| 0 | 0 | 0 | 0 |
| 0 | 0 | 0 | 0 |
| 0 | 0 | 0 | 0 |
| 0 | 0 | 0 | 0 |
| 1 | 1 | 0 | 0 |
| 1 | 1 | 0 | 0 |
| 0 | 0 | 0 | 0 |
| 0 | 0 | 0 | 0 |
| 0 | 1 | 0 | 0 |
| 0 | 1 | 0 | 0 |
| 0 | 1 | 0 | 0 |
| 0 | 0 | 0 | 0 |
| 0 | 0 | 0 | 0 |
| 0 | 0 | 0 | 0 |
| 0 | 0 | 0 | 0 |

| | | | |
|---|---|---|---|
| 0 | 0 | 0 | 0 |
| 0 | 0 | 0 | 0 |
| 0 | 0 | 0 | 0 |
| 0 | 0 | 0 | 0 |
| 1 | 0 | 0 | 0 |
| 0 | 0 | 0 | 0 |
| 0 | 0 | 0 | 0 |
| 1 | 0 | 0 | 0 |
| 1 | 0 | 0 | 0 |
| 1 | 0 | 0 | 0 |
| 1 | 0 | 0 | 0 |
| 1 | 0 | 0 | 0 |
| 0 | 0 | 0 | 0 |
| 1 | 0 | 0 | 0 |
| 1 | 0 | 0 | 0 |
| 0 | 0 | 0 | 0 |
| 0 | 0 | 0 | 0 |
| 0 | 0 | 0 | 0 |
| 1 | 0 | 0 | 0 |
| 1 | 0 | 0 | 0 |
| 1 | 0 | 0 | 0 |
| 1 | 0 | 0 | 0 |
| 0 | 0 | 0 | 0 |
| 0 | 0 | 0 | 0 |
| 0 | 0 | 0 | 0 |
| 0 | 0 | 0 | 0 |
| 0 | 0 | 0 | 0 |
| 0 | 0 | 0 | 0 |
| 0 | 0 | 0 | 0 |
| 0 | 0 | 0 | 0 |
| 0 | 0 | 0 | 0 |
| 0 | 0 | 0 | 0 |
| 1 | 0 | 0 | 0 |
| 0 | 0 | 0 | 0 |
| 0 | 0 | 0 | 0 |
| 1 | 0 | 0 | 0 |
| 0 | 0 | 0 | 0 |
| 0 | 0 | 0 | 0 |
| 0 | 0 | 0 | 0 |
| 0 | 0 | 0 | 0 |
| 0 | 0 | 0 | 0 |
| 0 | 0 | 0 | 0 |
| 0 | 0 | 0 | 0 |
| 0 | 0 | 0 | 0 |
| 0 | 0 | 0 | 0 |
| 0 | 0 | 0 | 0 |
| 0 | 0 | 0 | 0 |
| 0 | 0 | 0 | 0 |
| 0 | 0 | 0 | 0 |
| 0 | 0 | 0 | 0 |
| 0 | 0 | 0 | 0 |
| 0 | 0 | 0 | 0 |
| 0 | 0 | 0 | 0 |
| 0 | 0 | 0 | 0 |

| | | | |
|---|---|---|---|
| 0 | 0 | 0 | 0 |
| 0 | 0 | 0 | 0 |
| 0 | 0 | 0 | 0 |
| 0 | 0 | 0 | 0 |
| 0 | 0 | 0 | 0 |
| 0 | 0 | 0 | 0 |
| 0 | 0 | 0 | 0 |
| 0 | 0 | 0 | 0 |
| 0 | 0 | 0 | 0 |
| 0 | 0 | 0 | 0 |
| 0 | 0 | 0 | 0 |
| 0 | 0 | 0 | 1 |
| 0 | 0 | 0 | 0 |
| 0 | 0 | 0 | 0 |
| 0 | 0 | 0 | 0 |
| 0 | 0 | 0 | 0 |
| 0 | 0 | 0 | 0 |
| 0 | 0 | 0 | 1 |
| 0 | 0 | 0 | 0 |
| 0 | 0 | 0 | 0 |
| 0 | 0 | 0 | 1 |
| 0 | 0 | 0 | 0 |
| 0 | 0 | 0 | 0 |
| 0 | 0 | 0 | 0 |
| 0 | 0 | 0 | 0 |
| 0 | 0 | 0 | 0 |
| 0 | 0 | 0 | 0 |
| 0 | 0 | 0 | 0 |
| 0 | 0 | 0 | 1 |
| 0 | 0 | 0 | 0 |
| 0 | 0 | 0 | 0 |
| 0 | 0 | 0 | 0 |
| 0 | 0 | 0 | 0 |
| 0 | 0 | 0 | 0 |
| 0 | 0 | 0 | 0 |
| 0 | 0 | 0 | 0 |
| 0 | 0 | 0 | 0 |
| 0 | 0 | 0 | 0 |
| 0 | 0 | 0 | 0 |
| 0 | 0 | 0 | 0 |
| 0 | 0 | 0 | 0 |
| 0 | 0 | 0 | 0 |
| 0 | 0 | 0 | 0 |
| 0 | 0 | 0 | 0 |
| 0 | 0 | 0 | 0 |
| 0 | 0 | 0 | 0 |
| 0 | 0 | 0 | 0 |
| 0 | 0 | 0 | 0 |
| 0 | 0 | 0 | 0 |
| 0 | 0 | 0 | 0 |
| 0 | 0 | 0 | 0 |
| 0 | 0 | 0 | 0 |
| 0 | 0 | 0 | 0 |
| 0 | 0 | 0 | 0 |

| | | | |
|---|---|---|---|
| 0 | 0 | 0 | 0 |
| 0 | 0 | 0 | 0 |
| 1 | 0 | 0 | 0 |
| 0 | 0 | 0 | 0 |
| 1 | 0 | 0 | 0 |
| 0 | 0 | 0 | 0 |
| 0 | 0 | 0 | 0 |
| 0 | 0 | 0 | 0 |
| 0 | 0 | 0 | 0 |
| 1 | 1 | 0 | 0 |
| 0 | 0 | 1 | 0 |
| 0 | 0 | 0 | 0 |
| 0 | 0 | 0 | 0 |
| 0 | 0 | 0 | 0 |
| 0 | 0 | 0 | 0 |
| 0 | 0 | 0 | 0 |
| 0 | 0 | 0 | 0 |
| 0 | 0 | 0 | 0 |
| 0 | 0 | 0 | 0 |
| 0 | 0 | 0 | 0 |
| 0 | 0 | 0 | 0 |
| 0 | 0 | 0 | 0 |
| 0 | 0 | 0 | 0 |
| 0 | 0 | 0 | 0 |
| 0 | 0 | 0 | 0 |
| 0 | 0 | 0 | 0 |
| 0 | 0 | 0 | 0 |
| 0 | 0 | 0 | 0 |
| 0 | 0 | 0 | 0 |
| 0 | 0 | 0 | 0 |
| 0 | 0 | 0 | 0 |
| 0 | 0 | 0 | 0 |
| 0 | 0 | 0 | 0 |
| 0 | 0 | 0 | 0 |
| 0 | 0 | 0 | 0 |
| 0 | 0 | 0 | 0 |
| 0 | 0 | 0 | 0 |
| 0 | 0 | 0 | 0 |
| 0 | 0 | 0 | 0 |
| 0 | 0 | 0 | 0 |
| 0 | 0 | 0 | 0 |
| 0 | 0 | 0 | 0 |
| 0 | 0 | 0 | 0 |
| 0 | 0 | 0 | 0 |
| 0 | 0 | 0 | 0 |
| 1 | 0 | 0 | 0 |
| 0 | 0 | 0 | 0 |
| 0 | 0 | 0 | 0 |
| 0 | 0 | 0 | 0 |
| 0 | 0 | 0 | 0 |
| 1 | 0 | 0 | 0 |
| 0 | 0 | 0 | 0 |
| 0 | 0 | 0 | 0 |
| 0 | 0 | 0 | 0 |

| | | | |
|---|---|---|---|
| 0 | 0 | 0 | 0 |
| 0 | 0 | 0 | 0 |
| 0 | 0 | 0 | 0 |
| 0 | 0 | 0 | 0 |
| 0 | 0 | 0 | 0 |
| 0 | 0 | 0 | 0 |
| 0 | 0 | 0 | 0 |
| 0 | 0 | 0 | 0 |
| 1 | 1 | 0 | 0 |
| 0 | 0 | 0 | 0 |
| 0 | 0 | 0 | 0 |
| 0 | 0 | 0 | 0 |
| 0 | 0 | 0 | 0 |
| 0 | 0 | 0 | 0 |
| 0 | 1 | 0 | 0 |
| 0 | 1 | 0 | 0 |
| 0 | 1 | 0 | 0 |
| 0 | 0 | 0 | 0 |
| 0 | 0 | 0 | 0 |
| 0 | 0 | 0 | 0 |
| 0 | 1 | 0 | 0 |
| 0 | 0 | 0 | 0 |
| 0 | 0 | 0 | 0 |
| 1 | 1 | 0 | 0 |
| 0 | 1 | 0 | 0 |
| 0 | 1 | 0 | 0 |
| 0 | 0 | 0 | 0 |
| 0 | 0 | 0 | 0 |
| 0 | 0 | 0 | 0 |
| 0 | 0 | 0 | 0 |
| 0 | 0 | 0 | 0 |
| 0 | 0 | 0 | 0 |
| 0 | 0 | 0 | 0 |
| 0 | 0 | 0 | 0 |
| 0 | 0 | 0 | 0 |
| 0 | 0 | 0 | 0 |
| 0 | 0 | 0 | 0 |
| 1 | 0 | 0 | 0 |
| 0 | 0 | 0 | 0 |
| 0 | 0 | 0 | 0 |
| 0 | 0 | 0 | 0 |
| 0 | 0 | 0 | 0 |
| 0 | 0 | 0 | 0 |
| 0 | 0 | 0 | 0 |
| 0 | 0 | 0 | 0 |
| 0 | 0 | 0 | 0 |
| 0 | 0 | 0 | 0 |
| 0 | 0 | 0 | 0 |
| 0 | 0 | 0 | 0 |
| 0 | 0 | 0 | 0 |
| 0 | 0 | 0 | 0 |
| 0 | 0 | 0 | 0 |
| 0 | 0 | 0 | 0 |
| 0 | 0 | 0 | 0 |

| | | | |
|---|---|---|---|
| 0 | 0 | 0 | 0 |
| 0 | 0 | 0 | 0 |
| 0 | 0 | 0 | 0 |
| 0 | 0 | 1 | 0 |
| 0 | 0 | 0 | 0 |
| 0 | 0 | 0 | 0 |
| 0 | 0 | 0 | 0 |
| 0 | 0 | 0 | 0 |
| 0 | 0 | 0 | 0 |
| 0 | 0 | 0 | 0 |
| 0 | 0 | 0 | 0 |
| 0 | 0 | 0 | 0 |
| 1 | 0 | 0 | 0 |
| 0 | 0 | 0 | 0 |
| 0 | 0 | 0 | 0 |
| 0 | 0 | 0 | 0 |
| 0 | 1 | 0 | 0 |
| 0 | 0 | 0 | 0 |
| 0 | 0 | 0 | 0 |
| 0 | 0 | 0 | 0 |
| 0 | 0 | 0 | 0 |
| 0 | 0 | 0 | 0 |
| 0 | 0 | 0 | 0 |
| 0 | 0 | 0 | 0 |
| 0 | 0 | 0 | 0 |
| 0 | 0 | 0 | 0 |
| 0 | 0 | 0 | 0 |
| 0 | 0 | 0 | 0 |
| 0 | 0 | 0 | 0 |
| 0 | 0 | 0 | 0 |
| 1 | 0 | 0 | 0 |
| 1 | 0 | 0 | 0 |
| 1 | 0 | 0 | 0 |
| 0 | 0 | 0 | 0 |
| 1 | 0 | 0 | 0 |
| 1 | 0 | 0 | 0 |
| 1 | 0 | 0 | 0 |
| 0 | 0 | 0 | 0 |
| 0 | 0 | 0 | 0 |
| 0 | 0 | 0 | 0 |
| 0 | 0 | 0 | 0 |
| 0 | 0 | 0 | 0 |
| 0 | 1 | 0 | 0 |
| 0 | 1 | 0 | 0 |
| 0 | 0 | 0 | 0 |
| 0 | 0 | 0 | 0 |
| 1 | 0 | 0 | 0 |
| 0 | 0 | 0 | 0 |
| 0 | 0 | 0 | 0 |
| 0 | 0 | 0 | 0 |
| 0 | 0 | 0 | 0 |
| 0 | 0 | 0 | 0 |
| 1 | 0 | 0 | 0 |
| 0 | 0 | 0 | 0 |

| | | | |
|---|---|---|---|
| 0 | 0 | 0 | 0 |
| 1 | 0 | 0 | 0 |
| 1 | 0 | 0 | 0 |
| 0 | 0 | 0 | 0 |
| 0 | 0 | 0 | 0 |
| 0 | 0 | 0 | 0 |
| 0 | 0 | 0 | 0 |
| 0 | 1 | 0 | 0 |
| 0 | 0 | 0 | 0 |
| 0 | 0 | 0 | 0 |
| 0 | 0 | 0 | 0 |
| 0 | 0 | 0 | 0 |
| 0 | 0 | 0 | 0 |
| 0 | 0 | 0 | 0 |
| 1 | 0 | 0 | 0 |
| 0 | 0 | 0 | 0 |
| 0 | 0 | 0 | 0 |
| 0 | 0 | 0 | 0 |
| 0 | 0 | 0 | 0 |
| 1 | 0 | 0 | 0 |
| 1 | 0 | 0 | 0 |
| 0 | 0 | 0 | 0 |
| 1 | 0 | 0 | 0 |
| 1 | 0 | 0 | 0 |
| 0 | 0 | 0 | 0 |
| 0 | 0 | 0 | 0 |
| 0 | 0 | 0 | 0 |
| 0 | 0 | 0 | 0 |
| 0 | 0 | 0 | 0 |
| 0 | 0 | 0 | 0 |
| 0 | 0 | 0 | 0 |
| 0 | 0 | 0 | 0 |
| 0 | 0 | 0 | 0 |
| 0 | 0 | 0 | 0 |
| 0 | 0 | 0 | 0 |
| 0 | 0 | 0 | 0 |
| 0 | 0 | 0 | 0 |
| 0 | 0 | 0 | 0 |
| 0 | 0 | 0 | 0 |
| 0 | 0 | 0 | 0 |
| 0 | 0 | 0 | 0 |
| 0 | 0 | 0 | 0 |
| 0 | 0 | 0 | 0 |
| 0 | 0 | 0 | 0 |
| 0 | 0 | 0 | 0 |
| 0 | 0 | 0 | 0 |
| 0 | 0 | 0 | 0 |
| 0 | 0 | 0 | 0 |
| 0 | 0 | 0 | 0 |
| 0 | 0 | 0 | 0 |
| 0 | 0 | 0 | 0 |
| 0 | 0 | 0 | 0 |
| 0 | 0 | 0 | 0 |
| 1 | 1 | 0 | 1 |

| | | | |
|---|---|---|---|
| 0 | 0 | 0 | 0 |
| 0 | 0 | 0 | 0 |
| 0 | 0 | 0 | 0 |
| 0 | 0 | 0 | 0 |
| 0 | 1 | 0 | 0 |
| 0 | 0 | 0 | 0 |
| 0 | 0 | 0 | 0 |
| 0 | 1 | 0 | 0 |
| 0 | 0 | 0 | 0 |
| 0 | 0 | 0 | 0 |
| 0 | 0 | 0 | 0 |
| 0 | 0 | 0 | 0 |
| 0 | 0 | 0 | 0 |
| 0 | 0 | 0 | 0 |
| 0 | 1 | 0 | 1 |
| 0 | 1 | 0 | 0 |
| 0 | 0 | 0 | 0 |
| 0 | 0 | 0 | 0 |
| 0 | 1 | 0 | 0 |
| 0 | 1 | 0 | 1 |
| 0 | 0 | 0 | 0 |
| 0 | 0 | 0 | 0 |
| 0 | 0 | 0 | 0 |
| 0 | 0 | 0 | 0 |
| 0 | 0 | 0 | 0 |
| 1 | 0 | 0 | 0 |
| 0 | 0 | 0 | 0 |
| 0 | 0 | 1 | 0 |
| 0 | 0 | 1 | 0 |
| 0 | 0 | 1 | 0 |
| 0 | 0 | 0 | 0 |
| 0 | 0 | 0 | 0 |
| 0 | 0 | 0 | 0 |
| 0 | 0 | 0 | 0 |
| 0 | 0 | 0 | 0 |
| 0 | 0 | 1 | 0 |
| 0 | 0 | 0 | 0 |
| 0 | 0 | 0 | 0 |
| 1 | 0 | 1 | 0 |
| 0 | 0 | 1 | 0 |
| 1 | 1 | 0 | 0 |
| 0 | 1 | 0 | 0 |
| 0 | 0 | 0 | 0 |
| 0 | 0 | 0 | 0 |
| 1 | 0 | 0 | 0 |
| 0 | 0 | 0 | 0 |
| 0 | 0 | 1 | 0 |
| 0 | 0 | 0 | 0 |
| 0 | 0 | 0 | 0 |
| 0 | 0 | 0 | 0 |
| 0 | 0 | 0 | 0 |
| 0 | 0 | 0 | 0 |
| 0 | 0 | 0 | 0 |
| 0 | 0 | 0 | 0 |

| | | | |
|---|---|---|---|
| 0 | 0 | 0 | 0 |
| 0 | 0 | 0 | 0 |
| 0 | 0 | 0 | 0 |
| 0 | 0 | 0 | 0 |
| 0 | 0 | 0 | 0 |
| 0 | 0 | 0 | 0 |
| 0 | 0 | 0 | 0 |
| 1 | 1 | 0 | 0 |
| 0 | 0 | 0 | 0 |
| 0 | 0 | 0 | 0 |
| 0 | 0 | 0 | 0 |
| 0 | 0 | 0 | 1 |
| 0 | 0 | 0 | 0 |
| 0 | 0 | 0 | 0 |
| 0 | 0 | 0 | 0 |
| 0 | 0 | 0 | 0 |
| 0 | 0 | 0 | 0 |
| 0 | 0 | 0 | 0 |
| 0 | 0 | 0 | 0 |
| 0 | 0 | 0 | 0 |
| 0 | 0 | 0 | 0 |
| 0 | 0 | 0 | 0 |
| 0 | 0 | 0 | 0 |
| 0 | 0 | 0 | 0 |
| 0 | 0 | 0 | 0 |
| 0 | 0 | 0 | 0 |
| 0 | 0 | 0 | 0 |
| 0 | 0 | 0 | 0 |
| 0 | 0 | 0 | 0 |
| 0 | 1 | 0 | 0 |
| 0 | 1 | 0 | 0 |
| 0 | 0 | 0 | 0 |
| 0 | 0 | 0 | 0 |
| 0 | 0 | 0 | 0 |
| 0 | 0 | 0 | 0 |
| 0 | 1 | 0 | 0 |
| 0 | 0 | 0 | 0 |
| 0 | 0 | 0 | 0 |
| 0 | 0 | 0 | 0 |
| 0 | 0 | 0 | 0 |
| 0 | 0 | 0 | 0 |
| 1 | 1 | 0 | 0 |
| 1 | 0 | 0 | 0 |
| 0 | 0 | 0 | 0 |
| 0 | 0 | 0 | 0 |
| 0 | 0 | 0 | 0 |
| 0 | 0 | 0 | 0 |
| 0 | 0 | 0 | 0 |
| 0 | 0 | 0 | 0 |
| 0 | 0 | 0 | 0 |
| 0 | 0 | 0 | 0 |
| 0 | 0 | 0 | 0 |
| 0 | 0 | 0 | 0 |
| 0 | 0 | 0 | 0 |

| | | | |
|---|---|---|---|
| 0 | 0 | 0 | 0 |
| 0 | 0 | 0 | 0 |
| 0 | 0 | 0 | 0 |
| 0 | 0 | 0 | 0 |
| 0 | 0 | 0 | 0 |
| 0 | 0 | 0 | 0 |
| 0 | 0 | 0 | 0 |
| 0 | 0 | 0 | 0 |
| 1 | 0 | 0 | 0 |
| 1 | 0 | 0 | 0 |
| 0 | 0 | 0 | 0 |
| 0 | 0 | 0 | 0 |
| 0 | 0 | 0 | 0 |
| 1 | 0 | 0 | 0 |
| 0 | 0 | 0 | 0 |
| 0 | 0 | 0 | 0 |
| 0 | 0 | 0 | 0 |
| 0 | 0 | 0 | 0 |
| 0 | 0 | 0 | 0 |
| 1 | 1 | 0 | 0 |
| 0 | 0 | 1 | 0 |
| 0 | 0 | 0 | 0 |
| 0 | 0 | 0 | 0 |
| 0 | 0 | 0 | 0 |
| 0 | 0 | 0 | 0 |
| 0 | 0 | 0 | 0 |
| 0 | 0 | 0 | 0 |
| 0 | 0 | 0 | 0 |
| 0 | 0 | 0 | 0 |
| 0 | 0 | 0 | 0 |
| 0 | 0 | 0 | 0 |
| 0 | 0 | 0 | 0 |
| 0 | 0 | 0 | 0 |
| 0 | 0 | 0 | 0 |
| 0 | 0 | 0 | 0 |
| 0 | 0 | 0 | 0 |
| 0 | 0 | 0 | 0 |
| 0 | 0 | 0 | 0 |
| 0 | 0 | 0 | 0 |
| 0 | 0 | 0 | 0 |
| 0 | 0 | 0 | 0 |
| 0 | 0 | 0 | 0 |
| 0 | 0 | 0 | 0 |
| 0 | 0 | 0 | 0 |
| 0 | 0 | 0 | 0 |
| 0 | 0 | 0 | 0 |
| 1 | 0 | 1 | 0 |
| 1 | 0 | 0 | 0 |
| 0 | 0 | 0 | 0 |
| 0 | 0 | 0 | 0 |
| 0 | 0 | 1 | 0 |
| 0 | 0 | 0 | 0 |
| 0 | 0 | 0 | 0 |
| 0 | 0 | 0 | 0 |

| | | | |
|---|---|---|---|
| 0 | 0 | 0 | 0 |
| 0 | 0 | 0 | 0 |
| 0 | 0 | 0 | 0 |
| 0 | 0 | 0 | 0 |
| 0 | 0 | 0 | 0 |
| 0 | 0 | 0 | 0 |
| 0 | 0 | 0 | 0 |
| 0 | 0 | 0 | 0 |
| 0 | 0 | 0 | 0 |
| 0 | 0 | 0 | 0 |
| 0 | 0 | 0 | 0 |
| 0 | 0 | 0 | 0 |
| 0 | 0 | 0 | 0 |
| 0 | 0 | 0 | 0 |
| 0 | 0 | 0 | 0 |
| 0 | 0 | 0 | 0 |
| 0 | 0 | 0 | 0 |
| 0 | 0 | 0 | 0 |
| 0 | 0 | 0 | 0 |
| 1 | 0 | 0 | 0 |
| 0 | 0 | 0 | 0 |
| 0 | 0 | 0 | 0 |
| 0 | 0 | 0 | 0 |
| 0 | 0 | 0 | 0 |
| 0 | 0 | 0 | 0 |
| 0 | 0 | 0 | 0 |
| 0 | 0 | 0 | 0 |
| 0 | 0 | 0 | 0 |
| 0 | 0 | 0 | 0 |
| 0 | 0 | 0 | 0 |
| 0 | 0 | 0 | 0 |
| 0 | 0 | 0 | 0 |
| 0 | 0 | 0 | 0 |
| 0 | 0 | 0 | 0 |
| 0 | 0 | 0 | 0 |
| 0 | 0 | 0 | 0 |
| 0 | 0 | 0 | 0 |
| 0 | 0 | 0 | 0 |
| 1 | 0 | 0 | 1 |
| 0 | 0 | 0 | 0 |
| 0 | 0 | 0 | 0 |
| 0 | 0 | 0 | 0 |
| 0 | 0 | 0 | 1 |
| 0 | 0 | 0 | 1 |
| 0 | 0 | 0 | 1 |
| 0 | 0 | 0 | 0 |
| 0 | 0 | 0 | 0 |
| 0 | 0 | 0 | 0 |
| 0 | 0 | 0 | 0 |
| 0 | 0 | 0 | 0 |
| 0 | 0 | 0 | 0 |
| 0 | 0 | 0 | 0 |
| 0 | 0 | 0 | 0 |
| 0 | 0 | 0 | 0 |

| | | | |
|---|---|---|---|
| 0 | 0 | 0 | 0 |
| 0 | 0 | 0 | 0 |
| 0 | 0 | 0 | 0 |
| 0 | 0 | 0 | 0 |
| 0 | 0 | 0 | 0 |
| 0 | 0 | 0 | 0 |
| 0 | 0 | 0 | 0 |
| 0 | 0 | 0 | 0 |
| 0 | 0 | 0 | 0 |
| 0 | 0 | 0 | 0 |
| 0 | 0 | 0 | 0 |
| 0 | 0 | 0 | 0 |
| 0 | 0 | 0 | 0 |
| 0 | 0 | 0 | 0 |
| 0 | 0 | 0 | 0 |
| 0 | 0 | 0 | 0 |
| 0 | 0 | 0 | 0 |
| 0 | 0 | 0 | 0 |
| 0 | 0 | 0 | 0 |
| 0 | 0 | 0 | 0 |
| 0 | 0 | 0 | 0 |
| 0 | 0 | 0 | 0 |
| 0 | 0 | 0 | 0 |
| 0 | 0 | 0 | 0 |
| 0 | 0 | 0 | 0 |
| 0 | 0 | 0 | 0 |
| 0 | 0 | 0 | 0 |
| 0 | 0 | 0 | 0 |
| 0 | 0 | 0 | 0 |
| 0 | 0 | 0 | 0 |
| 0 | 0 | 0 | 0 |
| 0 | 0 | 0 | 0 |
| 1 | 1 | 0 | 1 |
| 1 | 1 | 0 | 1 |
| 1 | 0 | 0 | 0 |
| 0 | 0 | 0 | 0 |
| 0 | 0 | 0 | 0 |
| 0 | 0 | 0 | 0 |
| 0 | 0 | 0 | 0 |
| 0 | 0 | 0 | 0 |
| 0 | 0 | 0 | 0 |
| 0 | 0 | 0 | 0 |
| 1 | 1 | 0 | 0 |
| 0 | 0 | 0 | 0 |
| 1 | 0 | 0 | 0 |
| 0 | 0 | 0 | 0 |
| 0 | 0 | 0 | 0 |
| 0 | 0 | 0 | 0 |
| 0 | 0 | 0 | 0 |
| 0 | 0 | 0 | 0 |
| 1 | 0 | 0 | 0 |
| 0 | 0 | 0 | 0 |
| 0 | 0 | 0 | 0 |
| 0 | 0 | 0 | 0 |

| | | | |
|---|---|---|---|
| 0 | 0 | 0 | 0 |
| 0 | 0 | 0 | 0 |
| 0 | 0 | 0 | 0 |
| 0 | 0 | 0 | 0 |
| 0 | 0 | 0 | 0 |
| 0 | 0 | 0 | 0 |
| 0 | 0 | 0 | 0 |
| 0 | 0 | 0 | 0 |
| 0 | 0 | 0 | 0 |
| 0 | 0 | 0 | 0 |
| 0 | 0 | 0 | 0 |
| 0 | 0 | 0 | 0 |
| 0 | 0 | 0 | 0 |
| 0 | 0 | 0 | 0 |
| 0 | 0 | 0 | 0 |
| 0 | 0 | 0 | 0 |
| 0 | 0 | 0 | 0 |
| 0 | 0 | 0 | 0 |
| 0 | 0 | 0 | 0 |
| 0 | 0 | 0 | 1 |
| 0 | 0 | 0 | 0 |
| 0 | 0 | 0 | 1 |
| 0 | 0 | 0 | 1 |
| 0 | 0 | 0 | 1 |
| 0 | 0 | 0 | 0 |
| 0 | 0 | 0 | 0 |
| 0 | 0 | 0 | 0 |
| 0 | 0 | 0 | 0 |
| 0 | 0 | 0 | 0 |
| 0 | 0 | 0 | 0 |
| 0 | 0 | 0 | 0 |
| 0 | 0 | 0 | 0 |
| 0 | 0 | 0 | 0 |
| 0 | 0 | 0 | 0 |
| 0 | 0 | 0 | 0 |
| 0 | 1 | 0 | 0 |
| 0 | 0 | 0 | 0 |
| 0 | 0 | 0 | 0 |
| 0 | 1 | 0 | 0 |
| 0 | 0 | 0 | 0 |
| 0 | 0 | 0 | 0 |
| 0 | 0 | 0 | 0 |
| 0 | 0 | 0 | 0 |
| 0 | 0 | 0 | 0 |
| 0 | 0 | 0 | 0 |
| 0 | 0 | 0 | 0 |
| 0 | 0 | 0 | 0 |
| 0 | 0 | 0 | 0 |
| 0 | 0 | 0 | 0 |
| 0 | 0 | 0 | 0 |
| 0 | 0 | 0 | 0 |
| 0 | 0 | 0 | 0 |
| 0 | 0 | 0 | 0 |
| 0 | 0 | 0 | 0 |

| | | | |
|---|---|---|---|
| 0 | 0 | 0 | 0 |
| 0 | 0 | 0 | 0 |
| 0 | 0 | 0 | 0 |
| 0 | 0 | 0 | 0 |
| 0 | 0 | 0 | 0 |
| 0 | 0 | 0 | 0 |
| 0 | 0 | 0 | 0 |
| 0 | 0 | 0 | 0 |
| 0 | 0 | 1 | 0 |
| 0 | 0 | 1 | 0 |
| 0 | 0 | 0 | 0 |
| 0 | 0 | 0 | 0 |
| 0 | 0 | 0 | 0 |
| 0 | 0 | 0 | 0 |
| 0 | 0 | 0 | 0 |
| 0 | 1 | 0 | 0 |
| 0 | 0 | 0 | 0 |
| 0 | 0 | 0 | 0 |
| 1 | 1 | 0 | 0 |
| 0 | 0 | 0 | 0 |
| 0 | 0 | 0 | 0 |
| 0 | 0 | 0 | 0 |
| 0 | 0 | 0 | 0 |
| 0 | 0 | 0 | 0 |
| 0 | 0 | 0 | 0 |
| 0 | 0 | 0 | 0 |
| 0 | 0 | 0 | 0 |
| 0 | 0 | 0 | 0 |
| 0 | 0 | 0 | 0 |
| 0 | 0 | 0 | 0 |
| 0 | 0 | 0 | 0 |
| 0 | 0 | 0 | 0 |
| 0 | 0 | 0 | 0 |
| 0 | 0 | 0 | 0 |
| 0 | 0 | 0 | 0 |
| 0 | 0 | 0 | 0 |
| 0 | 0 | 0 | 0 |
| 0 | 0 | 0 | 0 |
| 0 | 0 | 0 | 0 |
| 0 | 0 | 0 | 0 |
| 0 | 0 | 0 | 0 |
| 0 | 0 | 0 | 0 |
| 0 | 0 | 0 | 0 |
| 0 | 0 | 0 | 0 |
| 0 | 0 | 0 | 0 |
| 0 | 0 | 0 | 0 |
| 0 | 0 | 0 | 0 |
| 0 | 0 | 0 | 0 |
| 0 | 0 | 0 | 0 |
| 0 | 0 | 0 | 0 |
| 0 | 0 | 0 | 0 |
| 0 | 0 | 0 | 0 |
| 0 | 0 | 0 | 0 |
| 1 | 0 | 0 | 0 |

| | | | |
|---|---|---|---|
| 0 | 0 | 0 | 0 |
| 0 | 0 | 0 | 0 |
| 0 | 0 | 0 | 0 |
| 0 | 0 | 0 | 0 |
| 0 | 0 | 0 | 0 |
| 0 | 0 | 0 | 0 |
| 0 | 0 | 0 | 0 |
| 0 | 0 | 0 | 0 |
| 0 | 0 | 0 | 0 |
| 0 | 0 | 0 | 0 |
| 1 | 0 | 0 | 0 |
| 0 | 0 | 0 | 0 |
| 0 | 0 | 0 | 0 |
| 0 | 0 | 0 | 0 |
| 1 | 1 | 0 | 1 |
| 0 | 0 | 0 | 0 |
| 1 | 0 | 0 | 0 |
| 1 | 1 | 0 | 0 |
| 0 | 0 | 0 | 0 |
| 0 | 0 | 0 | 0 |
| 0 | 0 | 0 | 0 |
| 0 | 0 | 0 | 0 |
| 0 | 0 | 0 | 0 |
| 0 | 0 | 0 | 1 |
| 0 | 0 | 0 | 0 |
| 0 | 0 | 0 | 0 |
| 0 | 0 | 0 | 0 |
| 0 | 0 | 0 | 1 |
| 0 | 0 | 0 | 0 |
| 0 | 0 | 0 | 0 |
| 0 | 0 | 0 | 0 |
| 0 | 0 | 0 | 0 |
| 0 | 0 | 0 | 0 |
| 0 | 0 | 0 | 0 |
| 0 | 0 | 0 | 0 |
| 0 | 0 | 0 | 0 |
| 0 | 0 | 0 | 0 |
| 0 | 0 | 0 | 0 |
| 0 | 0 | 0 | 0 |
| 0 | 0 | 0 | 0 |
| 0 | 0 | 0 | 0 |
| 0 | 0 | 0 | 0 |
| 0 | 0 | 0 | 0 |
| 0 | 0 | 0 | 0 |
| 0 | 0 | 1 | 0 |
| 0 | 0 | 0 | 0 |
| 0 | 0 | 0 | 0 |
| 0 | 0 | 0 | 0 |
| 0 | 0 | 0 | 0 |
| 0 | 0 | 0 | 0 |
| 0 | 0 | 0 | 0 |
| 0 | 0 | 0 | 0 |
| 0 | 0 | 0 | 0 |
| 0 | 0 | 0 | 0 |

| | | | |
|---|---|---|---|
| 0 | 0 | 0 | 0 |
| 0 | 0 | 0 | 0 |
| 0 | 0 | 0 | 0 |
| 0 | 0 | 0 | 0 |
| 0 | 0 | 0 | 0 |
| 0 | 0 | 0 | 0 |
| 1 | 0 | 0 | 1 |
| 1 | 0 | 0 | 0 |
| 0 | 0 | 0 | 0 |
| 0 | 0 | 0 | 0 |
| 0 | 0 | 0 | 0 |
| 0 | 0 | 0 | 0 |
| 0 | 0 | 0 | 0 |
| 0 | 0 | 0 | 0 |
| 0 | 0 | 0 | 0 |
| 0 | 0 | 0 | 0 |
| 0 | 0 | 0 | 0 |
| 0 | 0 | 0 | 0 |
| 0 | 0 | 0 | 0 |
| 0 | 0 | 0 | 0 |
| 1 | 0 | 0 | 0 |
| 0 | 0 | 0 | 0 |
| 0 | 0 | 0 | 0 |
| 0 | 0 | 0 | 0 |
| 0 | 0 | 0 | 0 |
| 0 | 0 | 0 | 0 |
| 0 | 0 | 0 | 0 |
| 0 | 0 | 0 | 0 |
| 0 | 0 | 0 | 0 |
| 0 | 0 | 0 | 0 |
| 0 | 0 | 0 | 0 |
| 0 | 0 | 0 | 0 |
| 0 | 0 | 0 | 0 |
| 0 | 0 | 0 | 0 |
| 0 | 0 | 0 | 0 |
| 0 | 0 | 0 | 0 |
| 0 | 0 | 0 | 0 |
| 0 | 0 | 0 | 0 |
| 0 | 0 | 0 | 0 |
| 0 | 0 | 0 | 0 |
| 0 | 0 | 0 | 0 |
| 0 | 0 | 0 | 0 |
| 0 | 0 | 0 | 0 |
| 0 | 0 | 0 | 0 |
| 0 | 0 | 0 | 0 |
| 0 | 0 | 0 | 0 |
| 0 | 0 | 0 | 0 |
| 0 | 0 | 0 | 0 |
| 0 | 0 | 0 | 0 |
| 0 | 0 | 0 | 0 |
| 0 | 0 | 0 | 0 |
| 0 | 0 | 0 | 0 |
| 0 | 0 | 0 | 0 |
| 0 | 0 | 0 | 0 |

| | | | |
|---|---|---|---|
| 0 | 0 | 0 | 0 |
| 0 | 0 | 0 | 0 |
| 0 | 0 | 0 | 0 |
| 1 | 0 | 0 | 0 |
| 0 | 0 | 0 | 0 |
| 1 | 0 | 0 | 0 |
| 0 | 0 | 0 | 0 |
| 0 | 0 | 0 | 0 |
| 1 | 0 | 0 | 0 |
| 0 | 0 | 0 | 0 |
| 0 | 0 | 0 | 0 |
| 0 | 0 | 0 | 0 |
| 0 | 0 | 0 | 0 |
| 0 | 0 | 0 | 0 |
| 0 | 0 | 0 | 0 |
| 0 | 0 | 0 | 0 |
| 0 | 0 | 0 | 0 |
| 0 | 0 | 0 | 0 |
| 1 | 0 | 0 | 0 |
| 0 | 0 | 0 | 0 |
| 0 | 0 | 0 | 0 |
| 0 | 0 | 0 | 0 |
| 0 | 0 | 0 | 0 |
| 0 | 0 | 0 | 0 |
| 0 | 0 | 0 | 0 |
| 0 | 0 | 0 | 0 |
| 0 | 0 | 0 | 0 |
| 0 | 0 | 0 | 0 |
| 0 | 1 | 0 | 0 |
| 0 | 0 | 0 | 0 |
| 0 | 0 | 0 | 0 |
| 0 | 0 | 0 | 0 |
| 0 | 1 | 0 | 0 |
| 1 | 1 | 0 | 0 |
| 1 | 0 | 0 | 0 |
| 0 | 0 | 0 | 0 |
| 0 | 0 | 0 | 0 |
| 0 | 0 | 0 | 0 |
| 0 | 0 | 0 | 0 |
| 0 | 0 | 0 | 0 |
| 1 | 0 | 0 | 0 |
| 0 | 0 | 0 | 0 |
| 0 | 0 | 0 | 0 |
| 0 | 0 | 0 | 0 |
| 0 | 0 | 0 | 0 |
| 0 | 0 | 0 | 0 |
| 0 | 0 | 0 | 0 |
| 0 | 0 | 0 | 0 |
| 0 | 0 | 0 | 0 |
| 0 | 0 | 0 | 0 |
| 0 | 0 | 0 | 0 |
| 0 | 0 | 0 | 0 |
| 0 | 0 | 0 | 0 |
| 1 | 0 | 0 | 0 |

| | | | |
|---|---|---|---|
| 1 | 0 | 0 | 1 |
| 0 | 0 | 0 | 0 |
| 0 | 0 | 0 | 0 |
| 0 | 0 | 0 | 0 |
| 0 | 0 | 0 | 0 |
| 0 | 0 | 0 | 0 |
| 0 | 0 | 0 | 0 |
| 0 | 0 | 0 | 0 |
| 0 | 0 | 0 | 0 |
| 0 | 0 | 0 | 0 |
| 0 | 0 | 0 | 0 |
| 0 | 0 | 0 | 0 |
| 0 | 0 | 0 | 0 |
| 0 | 0 | 0 | 0 |
| 0 | 0 | 0 | 0 |
| 0 | 0 | 0 | 0 |
| 0 | 0 | 0 | 0 |
| 0 | 0 | 0 | 0 |
| 0 | 0 | 0 | 0 |
| 0 | 0 | 0 | 0 |
| 0 | 0 | 0 | 0 |
| 0 | 0 | 0 | 0 |
| 0 | 1 | 0 | 0 |
| 0 | 0 | 0 | 0 |
| 0 | 0 | 0 | 0 |
| 0 | 0 | 0 | 0 |
| 0 | 0 | 0 | 0 |
| 0 | 0 | 0 | 0 |
| 0 | 0 | 0 | 0 |
| 0 | 0 | 0 | 0 |
| 0 | 0 | 0 | 0 |
| 0 | 0 | 0 | 0 |
| 0 | 0 | 0 | 0 |
| 0 | 0 | 0 | 0 |
| 0 | 0 | 0 | 0 |
| 0 | 0 | 0 | 0 |
| 0 | 0 | 0 | 0 |
| 0 | 0 | 0 | 0 |
| 0 | 0 | 0 | 0 |
| 0 | 0 | 0 | 0 |
| 0 | 0 | 0 | 0 |
| 0 | 0 | 0 | 0 |
| 0 | 0 | 0 | 0 |
| 0 | 0 | 0 | 0 |
| 0 | 0 | 0 | 0 |
| 0 | 0 | 0 | 0 |
| 0 | 0 | 0 | 0 |
| 0 | 0 | 0 | 0 |
| 0 | 0 | 0 | 0 |
| 0 | 0 | 0 | 0 |
| 0 | 0 | 0 | 0 |
| 0 | 0 | 0 | 0 |
| 0 | 0 | 0 | 0 |
| 0 | 0 | 0 | 0 |

| | | | |
|---|---|---|---|
| 0 | 0 | 0 | 0 |
| 0 | 0 | 0 | 0 |
| 0 | 0 | 0 | 0 |
| 0 | 0 | 0 | 0 |
| 0 | 0 | 0 | 0 |
| 0 | 0 | 0 | 0 |
| 0 | 1 | 0 | 0 |
| 0 | 1 | 0 | 0 |
| 0 | 0 | 0 | 0 |
| 0 | 0 | 0 | 0 |
| 0 | 0 | 0 | 0 |
| 0 | 0 | 0 | 0 |
| 0 | 0 | 0 | 0 |
| 0 | 0 | 0 | 0 |
| 0 | 0 | 0 | 0 |
| 0 | 0 | 0 | 0 |
| 1 | 0 | 0 | 0 |
| 0 | 0 | 0 | 0 |
| 1 | 0 | 0 | 0 |
| 1 | 0 | 0 | 0 |
| 1 | 0 | 0 | 0 |
| 1 | 0 | 0 | 0 |
| 0 | 0 | 0 | 0 |
| 0 | 0 | 0 | 0 |
| 0 | 0 | 0 | 0 |
| 0 | 0 | 0 | 0 |
| 0 | 0 | 0 | 0 |
| 0 | 0 | 0 | 0 |
| 0 | 0 | 0 | 0 |
| 0 | 0 | 0 | 0 |
| 0 | 0 | 0 | 0 |
| 0 | 0 | 0 | 0 |
| 0 | 0 | 0 | 0 |
| 0 | 0 | 0 | 0 |
| 0 | 0 | 0 | 0 |
| 0 | 0 | 1 | 0 |
| 1 | 0 | 0 | 0 |
| 1 | 0 | 0 | 0 |
| 1 | 0 | 0 | 0 |
| 0 | 0 | 0 | 0 |
| 0 | 0 | 0 | 0 |
| 0 | 0 | 0 | 0 |
| 0 | 0 | 0 | 0 |
| 1 | 0 | 0 | 0 |
| 1 | 0 | 0 | 0 |
| 1 | 0 | 0 | 0 |
| 0 | 0 | 0 | 0 |
| 0 | 0 | 0 | 0 |
| 1 | 0 | 0 | 0 |
| 0 | 0 | 0 | 0 |
| 0 | 0 | 0 | 0 |
| 1 | 0 | 0 | 0 |
| 0 | 0 | 0 | 0 |
| 0 | 0 | 0 | 0 |

| | | | |
|---|---|---|---|
| 0 | 0 | 0 | 0 |
| 0 | 0 | 0 | 0 |
| 0 | 0 | 0 | 0 |
| 0 | 0 | 0 | 0 |
| 0 | 0 | 0 | 0 |
| 0 | 0 | 0 | 0 |
| 0 | 0 | 0 | 0 |
| 0 | 0 | 0 | 0 |
| 0 | 0 | 0 | 0 |
| 0 | 0 | 0 | 0 |
| 0 | 0 | 0 | 0 |
| 0 | 0 | 0 | 0 |
| 0 | 0 | 0 | 0 |
| 0 | 0 | 0 | 0 |
| 0 | 0 | 0 | 0 |
| 0 | 0 | 0 | 0 |
| 0 | 0 | 0 | 0 |
| 0 | 0 | 0 | 0 |
| 0 | 0 | 0 | 0 |
| 0 | 0 | 0 | 0 |
| 0 | 0 | 0 | 0 |
| 0 | 0 | 0 | 0 |
| 1 | 1 | 0 | 0 |
| 0 | 0 | 0 | 0 |
| 0 | 1 | 0 | 0 |
| 0 | 0 | 0 | 0 |
| 0 | 0 | 0 | 0 |
| 0 | 0 | 0 | 0 |
| 0 | 0 | 0 | 0 |
| 0 | 0 | 0 | 0 |
| 0 | 0 | 0 | 0 |
| 0 | 0 | 0 | 0 |
| 0 | 0 | 0 | 0 |
| 0 | 0 | 0 | 0 |
| 0 | 0 | 0 | 0 |
| 0 | 0 | 0 | 0 |
| 0 | 0 | 0 | 0 |
| 0 | 0 | 0 | 0 |
| 0 | 0 | 0 | 0 |
| 0 | 0 | 0 | 0 |
| 0 | 0 | 0 | 0 |
| 0 | 0 | 0 | 0 |
| 0 | 0 | 0 | 0 |
| 0 | 0 | 0 | 0 |
| 0 | 0 | 0 | 0 |
| 1 | 0 | 0 | 1 |
| 0 | 0 | 0 | 0 |
| 0 | 0 | 0 | 0 |
| 0 | 0 | 0 | 0 |
| 1 | 0 | 0 | 0 |
| 1 | 0 | 0 | 0 |
| 1 | 0 | 0 | 0 |
| 0 | 0 | 0 | 0 |
| 0 | 0 | 0 | 0 |

| | | | |
|---|---|---|---|
| 0 | 0 | 0 | 0 |
| 0 | 0 | 0 | 0 |
| 0 | 0 | 0 | 0 |
| 0 | 0 | 0 | 0 |
| 0 | 0 | 0 | 0 |
| 0 | 0 | 0 | 0 |
| 0 | 0 | 0 | 0 |
| 0 | 0 | 0 | 0 |
| 0 | 0 | 0 | 0 |
| 0 | 0 | 0 | 0 |
| 0 | 0 | 0 | 1 |
| 0 | 0 | 0 | 0 |
| 0 | 0 | 0 | 0 |
| 0 | 0 | 0 | 0 |
| 0 | 0 | 0 | 0 |
| 0 | 0 | 0 | 0 |
| 0 | 0 | 0 | 0 |
| 0 | 0 | 0 | 0 |
| 0 | 0 | 0 | 0 |
| 0 | 0 | 0 | 0 |
| 0 | 0 | 0 | 0 |
| 0 | 0 | 0 | 0 |
| 0 | 0 | 0 | 0 |
| 0 | 0 | 0 | 0 |
| 0 | 0 | 0 | 0 |
| 0 | 0 | 0 | 0 |
| 0 | 0 | 0 | 0 |
| 0 | 0 | 0 | 0 |
| 0 | 0 | 0 | 0 |
| 0 | 0 | 0 | 0 |
| 0 | 1 | 0 | 0 |
| 0 | 0 | 0 | 0 |
| 1 | 1 | 0 | 0 |
| 1 | 1 | 0 | 0 |
| 0 | 0 | 0 | 0 |
| 0 | 0 | 0 | 0 |
| 0 | 0 | 0 | 0 |
| 0 | 0 | 0 | 0 |
| 0 | 0 | 0 | 0 |
| 0 | 0 | 0 | 0 |
| 0 | 0 | 0 | 0 |
| 0 | 1 | 0 | 0 |
| 0 | 0 | 0 | 0 |
| 0 | 0 | 0 | 0 |
| 0 | 0 | 0 | 0 |
| 1 | 0 | 0 | 0 |
| 1 | 0 | 0 | 0 |
| 1 | 0 | 0 | 0 |
| 1 | 1 | 1 | 0 |
| 0 | 0 | 0 | 0 |
| 1 | 1 | 1 | 1 |
| 0 | 0 | 0 | 0 |
| 0 | 0 | 0 | 0 |
| 0 | 0 | 0 | 0 |

| | | | |
|---|---|---|---|
| 0 | 0 | 0 | 0 |
| 0 | 0 | 0 | 0 |
| 0 | 0 | 0 | 0 |
| 0 | 0 | 0 | 0 |
| 1 | 0 | 0 | 0 |
| 1 | 0 | 0 | 0 |
| 0 | 0 | 0 | 0 |
| 0 | 0 | 0 | 0 |
| 0 | 0 | 0 | 0 |
| 0 | 0 | 0 | 0 |
| 0 | 0 | 0 | 0 |
| 0 | 0 | 0 | 0 |
| 0 | 0 | 0 | 0 |
| 0 | 0 | 0 | 0 |
| 0 | 0 | 0 | 0 |
| 0 | 0 | 0 | 0 |
| 0 | 0 | 0 | 0 |
| 0 | 0 | 0 | 0 |
| 0 | 0 | 0 | 0 |
| 0 | 0 | 0 | 0 |
| 0 | 0 | 0 | 0 |
| 0 | 0 | 0 | 0 |
| 0 | 0 | 0 | 0 |
| 0 | 0 | 0 | 0 |
| 0 | 0 | 0 | 0 |
| 0 | 0 | 0 | 0 |
| 0 | 0 | 0 | 0 |
| 0 | 0 | 0 | 0 |
| 0 | 0 | 0 | 0 |
| 0 | 0 | 0 | 0 |
| 0 | 0 | 0 | 0 |
| 0 | 0 | 0 | 0 |
| 0 | 0 | 0 | 0 |
| 0 | 0 | 0 | 0 |
| 0 | 0 | 0 | 0 |
| 0 | 0 | 0 | 0 |
| 0 | 0 | 0 | 0 |
| 0 | 0 | 0 | 0 |
| 0 | 0 | 0 | 0 |
| 0 | 0 | 0 | 0 |
| 0 | 0 | 0 | 0 |
| 0 | 0 | 0 | 0 |
| 0 | 0 | 0 | 0 |
| 0 | 0 | 0 | 0 |
| 0 | 0 | 0 | 0 |
| 0 | 0 | 0 | 0 |
| 0 | 0 | 0 | 0 |
| 0 | 0 | 0 | 0 |
| 0 | 0 | 0 | 0 |
| 0 | 0 | 0 | 0 |
| 0 | 0 | 0 | 0 |
| 0 | 0 | 0 | 0 |
| 0 | 0 | 0 | 0 |
| 0 | 1 | 0 | 0 |

| | | | |
|---|---|---|---|
| 0 | 0 | 0 | 0 |
| 0 | 0 | 0 | 0 |
| 0 | 0 | 0 | 0 |
| 0 | 0 | 0 | 0 |
| 0 | 0 | 0 | 0 |
| 0 | 0 | 0 | 0 |
| 0 | 0 | 0 | 0 |
| 0 | 0 | 0 | 0 |
| 0 | 0 | 0 | 0 |
| 0 | 0 | 0 | 0 |
| 0 | 0 | 0 | 0 |
| 0 | 0 | 0 | 0 |
| 0 | 0 | 0 | 0 |
| 0 | 0 | 0 | 0 |
| 0 | 0 | 0 | 0 |
| 0 | 0 | 0 | 0 |
| 0 | 0 | 0 | 0 |
| 0 | 0 | 0 | 0 |
| 0 | 0 | 0 | 0 |
| 0 | 0 | 0 | 0 |
| 0 | 0 | 0 | 0 |
| 0 | 0 | 0 | 0 |
| 0 | 0 | 0 | 0 |
| 0 | 0 | 0 | 0 |
| 0 | 0 | 0 | 0 |
| 0 | 0 | 0 | 0 |
| 0 | 0 | 0 | 0 |
| 0 | 0 | 0 | 0 |
| 0 | 0 | 0 | 0 |
| 0 | 0 | 0 | 0 |
| 0 | 0 | 0 | 0 |
| 0 | 0 | 0 | 0 |
| 0 | 0 | 0 | 0 |
| 0 | 0 | 0 | 0 |
| 0 | 0 | 0 | 0 |
| 0 | 0 | 0 | 0 |
| 0 | 0 | 0 | 0 |
| 0 | 0 | 0 | 0 |
| 0 | 0 | 0 | 0 |
| 0 | 0 | 0 | 0 |
| 0 | 0 | 0 | 0 |
| 0 | 0 | 0 | 0 |
| 0 | 0 | 0 | 0 |
| 0 | 0 | 0 | 0 |
| 0 | 0 | 0 | 0 |
| 0 | 0 | 0 | 0 |
| 0 | 0 | 0 | 0 |
| 0 | 0 | 0 | 0 |
| 0 | 0 | 0 | 0 |
| 0 | 0 | 0 | 0 |
| 0 | 0 | 0 | 0 |
| 0 | 0 | 0 | 0 |
| 0 | 0 | 0 | 0 |
| 0 | 0 | 0 | 0 |

| | | | |
|---|---|---|---|
| 0 | 0 | 0 | 0 |
| 0 | 0 | 0 | 0 |
| 0 | 0 | 0 | 0 |
| 0 | 0 | 0 | 0 |
| 0 | 0 | 0 | 0 |
| 0 | 0 | 0 | 0 |
| 0 | 0 | 0 | 0 |
| 0 | 0 | 0 | 0 |
| 0 | 0 | 0 | 0 |
| 0 | 0 | 0 | 0 |
| 0 | 0 | 0 | 0 |
| 0 | 0 | 1 | 0 |
| 0 | 0 | 0 | 0 |
| 0 | 0 | 1 | 0 |
| 0 | 0 | 1 | 0 |
| 0 | 0 | 1 | 0 |
| 0 | 0 | 1 | 0 |
| 0 | 0 | 0 | 0 |
| 0 | 0 | 0 | 0 |
| 0 | 0 | 0 | 0 |
| 1 | 0 | 0 | 0 |

| glycan_binding_site | water_binding_site | water_only_binding_site | non_binding_site |
|---|---|---|---|
| 0 | 1 | 0 | 0 |
| 0 | 1 | 0 | 0 |
| 0 | 1 | 1 | 1 |
| 0 | 1 | 0 | 0 |
| 0 | 0 | 0 | 0 |
| 0 | 0 | 0 | 0 |
| 0 | 0 | 0 | 1 |
| 0 | 0 | 0 | 1 |
| 0 | 1 | 0 | 0 |
| 0 | 0 | 0 | 0 |
| 0 | 0 | 0 | 0 |
| 0 | 1 | 0 | 0 |
| 0 | 1 | 1 | 1 |
| 1 | 0 | 0 | 0 |
| 0 | 0 | 0 | 0 |
| 0 | 0 | 0 | 0 |
| 0 | 0 | 0 | 0 |
| 0 | 1 | 0 | 0 |
| 0 | 0 | 0 | 0 |
| 0 | 0 | 0 | 0 |
| 0 | 1 | 0 | 0 |
| 1 | 1 | 0 | 0 |
| 1 | 1 | 0 | 0 |
| 0 | 1 | 0 | 0 |
| 0 | 1 | 0 | 0 |
| 0 | 1 | 1 | 1 |
| 0 | 1 | 0 | 0 |
| 0 | 0 | 0 | 1 |
| 0 | 0 | 0 | 0 |
| 0 | 0 | 0 | 1 |
| 0 | 0 | 0 | 1 |
| 0 | 1 | 0 | 0 |
| 0 | 1 | 0 | 0 |
| 0 | 1 | 0 | 0 |
| 0 | 1 | 0 | 0 |
| 0 | 0 | 0 | 1 |
| 0 | 0 | 0 | 0 |
| 0 | 1 | 0 | 0 |
| 0 | 1 | 0 | 0 |
| 0 | 1 | 0 | 0 |
| 0 | 1 | 0 | 0 |
| 0 | 1 | 0 | 0 |
| 0 | 1 | 0 | 0 |
| 0 | 1 | 0 | 0 |
| 0 | 1 | 0 | 0 |
| 0 | 1 | 0 | 0 |
| 0 | 1 | 0 | 0 |
| 0 | 1 | 0 | 0 |
| 0 | 1 | 0 | 0 |
| 0 | 1 | 0 | 0 |
| 0 | 1 | 0 | 0 |
| 0 | 1 | 0 | 0 |
| 0 | 1 | 0 | 0 |

| | | | |
|---|---|---|---|
| 0 | 1 | 0 | 0 |
| 0 | 1 | 0 | 0 |
| 0 | 1 | 0 | 0 |
| 0 | 1 | 0 | 0 |
| 0 | 1 | 0 | 0 |
| 0 | 1 | 0 | 0 |
| 0 | 1 | 0 | 0 |
| 0 | 1 | 0 | 0 |
| 0 | 1 | 0 | 0 |
| 0 | 1 | 0 | 0 |
| 0 | 1 | 0 | 0 |
| 0 | 1 | 0 | 0 |
| 0 | 1 | 0 | 0 |
| 0 | 1 | 0 | 0 |
| 0 | 1 | 0 | 0 |
| 0 | 1 | 0 | 0 |
| 0 | 1 | 0 | 0 |
| 0 | 1 | 0 | 0 |
| 0 | 1 | 0 | 0 |
| 0 | 1 | 0 | 0 |
| 0 | 0 | 0 | 0 |
| 0 | 0 | 0 | 0 |
| 0 | 0 | 0 | 0 |
| 0 | 0 | 0 | 0 |
| 0 | 0 | 0 | 0 |
| 0 | 0 | 0 | 0 |
| 0 | 0 | 0 | 0 |
| 0 | 0 | 0 | 0 |
| 0 | 0 | 0 | 0 |
| 0 | 0 | 0 | 0 |
| 0 | 0 | 0 | 0 |
| 0 | 0 | 0 | 0 |
| 0 | 0 | 0 | 0 |
| 0 | 0 | 0 | 0 |
| 0 | 0 | 0 | 0 |
| 0 | 0 | 0 | 0 |
| 0 | 0 | 0 | 0 |
| 0 | 0 | 0 | 0 |
| 0 | 0 | 0 | 0 |
| 0 | 1 | 0 | 0 |
| 1 | 0 | 0 | 0 |
| 0 | 0 | 0 | 0 |
| 1 | 0 | 0 | 0 |
| 1 | 1 | 0 | 0 |
| 0 | 0 | 0 | 0 |
| 0 | 0 | 0 | 1 |
| 0 | 0 | 0 | 1 |
| 0 | 1 | 0 | 0 |
| 1 | 0 | 0 | 0 |
| 0 | 0 | 0 | 0 |
| 0 | 0 | 0 | 0 |
| 0 | 0 | 0 | 1 |
| 0 | 0 | 0 | 1 |
| 0 | 0 | 0 | 0 |

| | | | |
|---|---|---|---|
| 0 | 0 | 0 | 1 |
| 0 | 0 | 0 | 1 |
| 0 | 0 | 0 | 0 |
| 0 | 0 | 0 | 0 |
| 0 | 0 | 0 | 0 |
| 0 | 0 | 0 | 0 |
| 0 | 0 | 0 | 1 |
| 0 | 1 | 0 | 0 |
| 0 | 1 | 0 | 0 |
| 0 | 0 | 0 | 0 |
| 0 | 1 | 0 | 0 |
| 0 | 0 | 0 | 0 |
| 0 | 0 | 0 | 0 |
| 1 | 0 | 0 | 0 |
| 1 | 0 | 0 | 0 |
| 0 | 0 | 0 | 1 |
| 0 | 0 | 0 | 0 |
| 0 | 1 | 0 | 0 |
| 0 | 1 | 0 | 0 |
| 0 | 1 | 0 | 0 |
| 0 | 1 | 0 | 0 |
| 0 | 1 | 0 | 0 |
| 0 | 1 | 0 | 0 |
| 0 | 1 | 0 | 0 |
| 0 | 1 | 0 | 0 |
| 0 | 1 | 0 | 0 |
| 0 | 1 | 0 | 0 |
| 0 | 1 | 0 | 0 |
| 0 | 1 | 0 | 0 |
| 0 | 1 | 0 | 0 |
| 0 | 1 | 0 | 0 |
| 0 | 1 | 0 | 0 |
| 0 | 1 | 0 | 0 |
| 0 | 1 | 0 | 0 |
| 0 | 1 | 0 | 0 |
| 0 | 1 | 0 | 0 |
| 0 | 1 | 0 | 0 |
| 0 | 1 | 0 | 0 |
| 0 | 1 | 0 | 0 |
| 0 | 1 | 0 | 0 |
| 0 | 1 | 0 | 0 |
| 0 | 1 | 0 | 0 |
| 0 | 1 | 0 | 0 |
| 0 | 1 | 0 | 0 |
| 0 | 1 | 0 | 0 |
| 0 | 1 | 0 | 0 |
| 1 | 1 | 0 | 0 |
| 0 | 0 | 0 | 0 |
| 0 | 1 | 0 | 0 |
| 0 | 1 | 0 | 0 |
| 0 | 0 | 0 | 0 |
| 0 | 0 | 0 | 0 |
| 0 | 0 | 0 | 0 |
| 0 | 1 | 1 | 1 |

| | | | |
|---|---|---|---|
| 0 | 1 | 1 | 1 |
| 0 | 1 | 1 | 1 |
| 0 | 1 | 1 | 1 |
| 0 | 1 | 0 | 0 |
| 0 | 1 | 0 | 0 |
| 0 | 1 | 1 | 1 |
| 0 | 1 | 0 | 0 |
| 0 | 1 | 1 | 1 |
| 0 | 1 | 1 | 1 |
| 0 | 1 | 1 | 1 |
| 0 | 1 | 1 | 1 |
| 0 | 1 | 0 | 0 |
| 0 | 1 | 1 | 1 |
| 0 | 0 | 0 | 0 |
| 0 | 1 | 1 | 1 |
| 0 | 1 | 1 | 1 |
| 0 | 1 | 1 | 1 |
| 0 | 1 | 1 | 1 |
| 0 | 1 | 1 | 1 |
| 0 | 1 | 0 | 0 |
| 0 | 1 | 0 | 0 |
| 0 | 0 | 0 | 1 |
| 0 | 1 | 0 | 0 |
| 0 | 1 | 0 | 0 |
| 0 | 1 | 0 | 0 |
| 0 | 1 | 1 | 1 |
| 0 | 1 | 0 | 0 |
| 0 | 1 | 0 | 0 |
| 0 | 1 | 1 | 1 |
| 0 | 1 | 1 | 1 |
| 0 | 0 | 0 | 1 |
| 0 | 1 | 1 | 1 |
| 0 | 1 | 1 | 1 |
| 0 | 0 | 0 | 1 |
| 0 | 1 | 0 | 0 |
| 0 | 1 | 1 | 1 |
| 1 | 1 | 0 | 0 |
| 1 | 1 | 0 | 0 |
| 0 | 1 | 1 | 1 |
| 0 | 1 | 1 | 1 |
| 0 | 1 | 1 | 1 |
| 0 | 1 | 0 | 0 |
| 0 | 1 | 0 | 0 |
| 0 | 0 | 0 | 1 |
| 0 | 1 | 1 | 1 |
| 0 | 1 | 1 | 1 |
| 0 | 0 | 0 | 0 |
| 0 | 0 | 0 | 0 |
| 0 | 0 | 0 | 0 |
| 0 | 0 | 0 | 0 |
| 0 | 0 | 0 | 0 |
| 0 | 0 | 0 | 1 |
| 0 | 0 | 0 | 1 |
| 0 | 0 | 0 | 0 |

| | | | |
|---|---|---|---|
| 0 | 0 | 0 | 0 |
| 0 | 0 | 0 | 0 |
| 0 | 1 | 0 | 0 |
| 0 | 0 | 0 | 1 |
| 0 | 0 | 0 | 1 |
| 0 | 1 | 1 | 1 |
| 0 | 1 | 0 | 0 |
| 0 | 1 | 0 | 0 |
| 0 | 1 | 1 | 1 |
| 0 | 1 | 1 | 1 |
| 1 | 1 | 0 | 0 |
| 0 | 0 | 0 | 1 |
| 1 | 1 | 0 | 0 |
| 0 | 1 | 1 | 1 |
| 0 | 0 | 0 | 1 |
| 0 | 0 | 0 | 1 |
| 0 | 0 | 0 | 1 |
| 0 | 1 | 0 | 0 |
| 0 | 0 | 0 | 0 |
| 0 | 1 | 0 | 0 |
| 0 | 0 | 0 | 0 |
| 0 | 1 | 1 | 1 |
| 0 | 1 | 0 | 0 |
| 0 | 0 | 0 | 1 |
| 0 | 0 | 0 | 1 |
| 0 | 0 | 0 | 0 |
| 0 | 1 | 0 | 0 |
| 0 | 0 | 0 | 0 |
| 0 | 1 | 0 | 0 |
| 0 | 1 | 0 | 0 |
| 0 | 1 | 0 | 0 |
| 0 | 0 | 0 | 0 |
| 0 | 1 | 1 | 1 |
| 0 | 0 | 0 | 1 |
| 0 | 0 | 0 | 0 |
| 0 | 0 | 0 | 1 |
| 0 | 0 | 0 | 1 |
| 0 | 0 | 0 | 0 |
| 0 | 1 | 1 | 1 |
| 0 | 0 | 0 | 1 |
| 0 | 1 | 0 | 0 |
| 0 | 1 | 1 | 1 |
| 0 | 0 | 0 | 1 |
| 0 | 0 | 0 | 1 |
| 0 | 0 | 0 | 0 |
| 0 | 0 | 0 | 0 |
| 0 | 1 | 0 | 0 |
| 0 | 1 | 0 | 0 |
| 0 | 1 | 0 | 0 |
| 0 | 1 | 0 | 0 |
| 0 | 1 | 0 | 0 |
| 0 | 1 | 0 | 0 |
| 0 | 1 | 0 | 0 |
| 0 | 1 | 0 | 0 |

| | | | |
|---|---|---|---|
| 0 | 1 | 0 | 0 |
| 0 | 1 | 0 | 0 |
| 0 | 1 | 0 | 0 |
| 0 | 1 | 0 | 0 |
| 0 | 1 | 0 | 0 |
| 0 | 1 | 0 | 0 |
| 0 | 0 | 0 | 1 |
| 0 | 0 | 0 | 1 |
| 0 | 0 | 0 | 0 |
| 0 | 0 | 0 | 0 |
| 0 | 0 | 0 | 0 |
| 0 | 0 | 0 | 0 |
| 0 | 0 | 0 | 0 |
| 0 | 0 | 0 | 1 |
| 0 | 0 | 0 | 1 |
| 0 | 0 | 0 | 0 |
| 0 | 0 | 0 | 1 |
| 0 | 0 | 0 | 0 |
| 0 | 0 | 0 | 1 |
| 0 | 0 | 0 | 0 |
| 0 | 0 | 0 | 1 |
| 0 | 1 | 1 | 1 |
| 0 | 0 | 0 | 1 |
| 0 | 0 | 0 | 0 |
| 0 | 0 | 0 | 0 |
| 0 | 1 | 0 | 0 |
| 0 | 1 | 0 | 0 |
| 0 | 1 | 0 | 0 |
| 0 | 0 | 0 | 0 |
| 0 | 1 | 0 | 0 |
| 0 | 0 | 0 | 1 |
| 0 | 1 | 1 | 1 |
| 0 | 0 | 0 | 0 |
| 0 | 0 | 0 | 0 |
| 0 | 1 | 0 | 0 |
| 0 | 0 | 0 | 1 |
| 0 | 0 | 0 | 0 |
| 0 | 1 | 0 | 0 |
| 0 | 0 | 0 | 0 |
| 0 | 1 | 0 | 0 |
| 0 | 0 | 0 | 1 |
| 0 | 0 | 0 | 1 |
| 0 | 0 | 0 | 0 |
| 0 | 0 | 0 | 1 |
| 0 | 0 | 0 | 0 |
| 0 | 0 | 0 | 0 |
| 0 | 0 | 0 | 0 |
| 0 | 0 | 0 | 1 |
| 0 | 0 | 0 | 1 |
| 0 | 0 | 0 | 0 |
| 0 | 0 | 0 | 1 |
| 0 | 0 | 0 | 1 |
| 0 | 0 | 0 | 0 |
| 0 | 0 | 0 | 1 |

| | | | |
|---|---|---|---|
| 0 | 0 | 0 | 0 |
| 0 | 0 | 0 | 1 |
| 0 | 1 | 1 | 1 |
| 0 | 1 | 0 | 0 |
| 0 | 1 | 0 | 0 |
| 0 | 0 | 0 | 0 |
| 0 | 0 | 0 | 1 |
| 0 | 1 | 0 | 0 |
| 0 | 0 | 0 | 0 |
| 0 | 0 | 0 | 1 |
| 0 | 0 | 0 | 1 |
| 0 | 0 | 0 | 1 |
| 0 | 0 | 0 | 1 |
| 0 | 0 | 0 | 1 |
| 0 | 0 | 0 | 0 |
| 0 | 0 | 0 | 1 |
| 0 | 0 | 0 | 0 |
| 0 | 0 | 0 | 0 |
| 0 | 0 | 0 | 1 |
| 0 | 0 | 0 | 0 |
| 0 | 1 | 0 | 0 |
| 0 | 1 | 0 | 0 |
| 0 | 1 | 0 | 0 |
| 0 | 1 | 0 | 0 |
| 0 | 0 | 0 | 1 |
| 0 | 0 | 0 | 1 |
| 0 | 0 | 0 | 0 |
| 0 | 0 | 0 | 0 |
| 0 | 1 | 0 | 0 |
| 0 | 0 | 0 | 0 |
| 0 | 0 | 0 | 0 |
| 0 | 1 | 0 | 0 |
| 0 | 0 | 0 | 0 |
| 0 | 0 | 0 | 0 |
| 0 | 0 | 0 | 1 |
| 0 | 0 | 0 | 1 |
| 0 | 0 | 0 | 0 |
| 0 | 0 | 0 | 0 |
| 0 | 0 | 0 | 0 |
| 0 | 0 | 0 | 0 |
| 0 | 0 | 0 | 0 |
| 0 | 1 | 0 | 0 |
| 0 | 0 | 0 | 1 |
| 0 | 1 | 0 | 0 |
| 0 | 1 | 0 | 0 |
| 0 | 1 | 0 | 0 |
| 0 | 1 | 0 | 0 |
| 0 | 1 | 0 | 0 |
| 1 | 1 | 0 | 0 |
| 0 | 0 | 0 | 1 |
| 0 | 1 | 0 | 0 |
| 0 | 1 | 0 | 0 |
| 0 | 0 | 0 | 0 |
| 0 | 1 | 0 | 0 |

| | | | |
|---|---|---|---|
| 0 | 1 | 0 | 0 |
| 0 | 1 | 0 | 0 |
| 0 | 1 | 0 | 0 |
| 0 | 1 | 0 | 0 |
| 0 | 0 | 0 | 1 |
| 0 | 1 | 0 | 0 |
| 0 | 1 | 0 | 0 |
| 0 | 1 | 0 | 0 |
| 0 | 1 | 0 | 0 |
| 0 | 1 | 0 | 0 |
| 0 | 1 | 0 | 0 |
| 0 | 1 | 0 | 0 |
| 0 | 1 | 0 | 0 |
| 0 | 1 | 0 | 0 |
| 0 | 1 | 0 | 0 |
| 0 | 1 | 0 | 0 |
| 0 | 1 | 0 | 0 |
| 0 | 1 | 0 | 0 |
| 0 | 0 | 0 | 1 |
| 0 | 0 | 0 | 0 |
| 0 | 0 | 0 | 0 |
| 0 | 0 | 0 | 0 |
| 0 | 1 | 0 | 0 |
| 0 | 1 | 1 | 1 |
| 0 | 0 | 0 | 0 |
| 0 | 0 | 0 | 0 |
| 0 | 0 | 0 | 0 |
| 0 | 1 | 1 | 1 |
| 0 | 1 | 1 | 1 |
| 0 | 1 | 1 | 1 |
| 0 | 1 | 0 | 0 |
| 0 | 0 | 0 | 0 |
| 0 | 1 | 0 | 0 |
| 0 | 1 | 0 | 0 |
| 0 | 1 | 0 | 0 |
| 0 | 1 | 1 | 1 |
| 0 | 1 | 1 | 1 |
| 0 | 0 | 0 | 0 |
| 0 | 1 | 0 | 0 |
| 0 | 1 | 1 | 1 |
| 0 | 1 | 1 | 1 |
| 0 | 1 | 1 | 1 |
| 0 | 0 | 0 | 1 |
| 0 | 0 | 0 | 1 |
| 0 | 0 | 0 | 0 |
| 0 | 0 | 0 | 0 |
| 0 | 1 | 0 | 0 |
| 0 | 0 | 0 | 0 |
| 0 | 0 | 0 | 0 |
| 0 | 0 | 0 | 1 |
| 0 | 0 | 0 | 1 |
| 0 | 0 | 0 | 0 |
| 0 | 0 | 0 | 1 |
| 0 | 0 | 0 | 0 |

| | | | |
|---|---|---|---|
| 0 | 0 | 0 | 1 |
| 0 | 0 | 0 | 1 |
| 0 | 0 | 0 | 0 |
| 0 | 0 | 0 | 1 |
| 0 | 0 | 0 | 1 |
| 0 | 0 | 0 | 1 |
| 0 | 0 | 0 | 1 |
| 0 | 1 | 0 | 0 |
| 0 | 1 | 1 | 1 |
| 0 | 0 | 0 | 1 |
| 0 | 0 | 0 | 0 |
| 0 | 0 | 0 | 1 |
| 0 | 0 | 0 | 1 |
| 0 | 0 | 0 | 1 |
| 0 | 0 | 0 | 0 |
| 0 | 0 | 0 | 0 |
| 0 | 0 | 0 | 1 |
| 0 | 0 | 0 | 1 |
| 0 | 0 | 0 | 0 |
| 0 | 0 | 0 | 0 |
| 0 | 1 | 0 | 0 |
| 0 | 1 | 0 | 0 |
| 0 | 0 | 0 | 0 |
| 0 | 1 | 0 | 0 |
| 0 | 1 | 0 | 0 |
| 0 | 0 | 0 | 1 |
| 0 | 1 | 0 | 0 |
| 0 | 1 | 0 | 0 |
| 0 | 0 | 0 | 0 |
| 0 | 0 | 0 | 1 |
| 0 | 1 | 0 | 0 |
| 0 | 1 | 0 | 0 |
| 0 | 0 | 0 | 1 |
| 0 | 1 | 0 | 0 |
| 0 | 1 | 0 | 0 |
| 0 | 1 | 0 | 0 |
| 0 | 1 | 0 | 0 |
| 0 | 1 | 0 | 0 |
| 0 | 1 | 0 | 0 |
| 0 | 1 | 0 | 0 |
| 0 | 0 | 0 | 1 |
| 0 | 0 | 0 | 0 |
| 0 | 0 | 0 | 1 |
| 0 | 0 | 0 | 0 |
| 0 | 0 | 0 | 0 |
| 1 | 0 | 0 | 0 |
| 0 | 0 | 0 | 0 |
| 1 | 0 | 0 | 0 |
| 0 | 0 | 0 | 0 |
| 0 | 0 | 0 | 0 |
| 0 | 0 | 0 | 1 |
| 0 | 0 | 0 | 0 |
| 0 | 0 | 0 | 0 |
| 0 | 0 | 0 | 0 |

| | | | |
|---|---|---|---|
| 0 | 0 | 0 | 1 |
| 0 | 0 | 0 | 0 |
| 0 | 0 | 0 | 1 |
| 0 | 0 | 0 | 1 |
| 1 | 0 | 0 | 0 |
| 0 | 0 | 0 | 0 |
| 1 | 0 | 0 | 0 |
| 1 | 0 | 0 | 0 |
| 0 | 0 | 0 | 0 |
| 0 | 0 | 0 | 1 |
| 0 | 0 | 0 | 1 |
| 0 | 0 | 0 | 1 |
| 0 | 0 | 0 | 1 |
| 1 | 1 | 0 | 0 |
| 0 | 0 | 0 | 1 |
| 0 | 0 | 0 | 0 |
| 0 | 0 | 0 | 0 |
| 0 | 0 | 0 | 0 |
| 1 | 1 | 0 | 0 |
| 0 | 1 | 1 | 1 |
| 1 | 1 | 0 | 0 |
| 0 | 0 | 0 | 1 |
| 0 | 0 | 0 | 0 |
| 0 | 0 | 0 | 1 |
| 0 | 0 | 0 | 1 |
| 0 | 0 | 0 | 1 |
| 0 | 0 | 0 | 0 |
| 0 | 0 | 0 | 1 |
| 0 | 0 | 0 | 1 |
| 0 | 0 | 0 | 1 |
| 0 | 0 | 0 | 0 |
| 0 | 0 | 0 | 0 |
| 0 | 0 | 0 | 1 |
| 0 | 0 | 0 | 1 |
| 0 | 0 | 0 | 0 |
| 0 | 0 | 0 | 0 |
| 0 | 0 | 0 | 0 |
| 1 | 0 | 0 | 0 |
| 0 | 0 | 0 | 0 |
| 0 | 0 | 0 | 1 |
| 0 | 1 | 0 | 0 |
| 0 | 1 | 0 | 0 |
| 0 | 1 | 0 | 0 |
| 0 | 0 | 0 | 0 |
| 0 | 0 | 0 | 1 |
| 1 | 0 | 0 | 0 |
| 0 | 1 | 0 | 0 |
| 0 | 0 | 0 | 0 |
| 0 | 0 | 0 | 0 |
| 1 | 1 | 0 | 0 |
| 0 | 0 | 0 | 0 |
| 0 | 1 | 0 | 0 |
| 0 | 1 | 1 | 1 |
| 0 | 0 | 0 | 0 |

| | | | |
|---|---|---|---|
| 0 | 1 | 0 | 0 |
| 0 | 1 | 0 | 0 |
| 0 | 1 | 1 | 1 |
| 0 | 1 | 1 | 1 |
| 0 | 1 | 1 | 1 |
| 0 | 1 | 0 | 0 |
| 0 | 1 | 1 | 1 |
| 0 | 1 | 1 | 1 |
| 0 | 1 | 0 | 0 |
| 0 | 1 | 0 | 0 |
| 1 | 1 | 0 | 0 |
| 1 | 1 | 0 | 0 |
| 0 | 0 | 0 | 1 |
| 1 | 1 | 0 | 0 |
| 0 | 0 | 0 | 0 |
| 0 | 0 | 0 | 0 |
| 0 | 1 | 0 | 0 |
| 0 | 0 | 0 | 0 |
| 0 | 0 | 0 | 0 |
| 0 | 0 | 0 | 1 |
| 0 | 0 | 0 | 0 |
| 0 | 1 | 0 | 0 |
| 0 | 0 | 0 | 0 |
| 0 | 0 | 0 | 0 |
| 0 | 1 | 0 | 0 |
| 0 | 1 | 1 | 1 |
| 0 | 0 | 0 | 1 |
| 0 | 0 | 0 | 1 |
| 0 | 0 | 0 | 0 |
| 0 | 1 | 0 | 0 |
| 0 | 0 | 0 | 0 |
| 0 | 0 | 0 | 0 |
| 0 | 0 | 0 | 0 |
| 0 | 1 | 0 | 0 |
| 0 | 1 | 0 | 0 |
| 0 | 1 | 1 | 1 |
| 0 | 1 | 0 | 0 |
| 0 | 1 | 0 | 0 |
| 0 | 0 | 0 | 1 |
| 0 | 0 | 0 | 1 |
| 0 | 0 | 0 | 1 |
| 0 | 1 | 0 | 0 |
| 0 | 0 | 0 | 0 |
| 0 | 0 | 0 | 0 |
| 0 | 0 | 0 | 0 |
| 0 | 0 | 0 | 0 |
| 0 | 0 | 0 | 0 |
| 0 | 0 | 0 | 1 |
| 0 | 0 | 0 | 1 |
| 0 | 0 | 0 | 0 |
| 0 | 0 | 0 | 0 |
| 0 | 0 | 0 | 0 |
| 0 | 0 | 0 | 0 |
| 0 | 0 | 0 | 1 |

| | | | |
|---|---|---|---|
| 0 | 0 | 0 | 1 |
| 0 | 0 | 0 | 0 |
| 0 | 0 | 0 | 1 |
| 0 | 1 | 0 | 0 |
| 0 | 1 | 1 | 1 |
| 0 | 1 | 1 | 1 |
| 0 | 1 | 0 | 0 |
| 0 | 1 | 1 | 1 |
| 0 | 1 | 1 | 1 |
| 0 | 1 | 1 | 1 |
| 0 | 1 | 1 | 1 |
| 0 | 1 | 1 | 1 |
| 0 | 0 | 0 | 1 |
| 0 | 0 | 0 | 1 |
| 0 | 0 | 0 | 1 |
| 0 | 0 | 0 | 1 |
| 0 | 0 | 0 | 0 |
| 0 | 0 | 0 | 0 |
| 0 | 1 | 1 | 1 |
| 0 | 0 | 0 | 0 |
| 0 | 0 | 0 | 0 |
| 0 | 1 | 0 | 0 |
| 0 | 0 | 0 | 1 |
| 0 | 1 | 0 | 0 |
| 0 | 0 | 0 | 0 |
| 0 | 0 | 0 | 1 |
| 0 | 1 | 1 | 1 |
| 0 | 0 | 0 | 1 |
| 0 | 1 | 0 | 0 |
| 0 | 0 | 0 | 0 |
| 0 | 1 | 1 | 1 |
| 0 | 1 | 0 | 0 |
| 0 | 1 | 0 | 0 |
| 0 | 1 | 0 | 0 |
| 0 | 1 | 0 | 0 |
| 0 | 0 | 0 | 0 |
| 0 | 1 | 1 | 1 |
| 0 | 0 | 0 | 1 |
| 0 | 0 | 0 | 1 |
| 0 | 0 | 0 | 1 |
| 0 | 0 | 0 | 0 |
| 0 | 0 | 0 | 1 |
| 0 | 0 | 0 | 0 |
| 0 | 0 | 0 | 1 |
| 1 | 0 | 0 | 0 |
| 0 | 0 | 0 | 1 |
| 0 | 0 | 0 | 0 |
| 0 | 1 | 0 | 0 |
| 1 | 0 | 0 | 0 |
| 0 | 0 | 0 | 0 |
| 0 | 0 | 0 | 0 |
| 0 | 0 | 0 | 0 |
| 0 | 0 | 0 | 0 |
| 0 | 0 | 0 | 0 |

| | | | |
|---|---|---|---|
| 0 | 0 | 0 | 0 |
| 0 | 0 | 0 | 0 |
| 0 | 0 | 0 | 0 |
| 0 | 0 | 0 | 0 |
| 0 | 1 | 0 | 0 |
| 0 | 1 | 0 | 0 |
| 0 | 1 | 0 | 0 |
| 0 | 0 | 0 | 1 |
| 0 | 1 | 0 | 0 |
| 0 | 1 | 0 | 0 |
| 0 | 1 | 0 | 0 |
| 0 | 1 | 1 | 1 |
| 0 | 1 | 0 | 0 |
| 0 | 1 | 0 | 0 |
| 0 | 1 | 0 | 0 |
| 0 | 1 | 0 | 0 |
| 0 | 1 | 0 | 0 |
| 0 | 1 | 0 | 0 |
| 0 | 1 | 0 | 0 |
| 0 | 1 | 0 | 0 |
| 0 | 1 | 0 | 0 |
| 0 | 1 | 0 | 0 |
| 0 | 1 | 0 | 0 |
| 0 | 1 | 0 | 0 |
| 0 | 1 | 0 | 0 |
| 0 | 1 | 0 | 0 |
| 0 | 1 | 0 | 0 |
| 0 | 1 | 0 | 0 |
| 0 | 1 | 0 | 0 |
| 0 | 1 | 0 | 0 |
| 0 | 1 | 0 | 0 |
| 0 | 1 | 0 | 0 |
| 0 | 1 | 0 | 0 |
| 0 | 1 | 0 | 0 |
| 0 | 1 | 0 | 0 |
| 0 | 1 | 0 | 0 |
| 0 | 1 | 0 | 0 |
| 0 | 1 | 0 | 0 |
| 0 | 1 | 0 | 0 |
| 0 | 1 | 1 | 1 |
| 0 | 1 | 0 | 0 |
| 0 | 1 | 0 | 0 |
| 0 | 1 | 0 | 0 |
| 0 | 1 | 0 | 0 |
| 0 | 1 | 0 | 0 |
| 0 | 1 | 0 | 0 |
| 0 | 1 | 0 | 0 |
| 0 | 1 | 0 | 0 |
| 0 | 0 | 0 | 1 |
| 0 | 0 | 0 | 1 |
| 0 | 1 | 1 | 1 |
| 1 | 0 | 0 | 0 |
| 0 | 1 | 0 | 0 |

| | | | |
|---|---|---|---|
| 0 | 0 | 0 | 0 |
| 0 | 0 | 0 | 0 |
| 1 | 1 | 0 | 0 |
| 1 | 1 | 0 | 0 |
| 0 | 0 | 0 | 0 |
| 0 | 0 | 0 | 0 |
| 0 | 1 | 0 | 0 |
| 0 | 1 | 1 | 1 |
| 0 | 0 | 0 | 1 |
| 0 | 0 | 0 | 0 |
| 0 | 1 | 0 | 0 |
| 0 | 0 | 0 | 1 |
| 0 | 1 | 0 | 0 |
| 0 | 1 | 0 | 0 |
| 0 | 0 | 0 | 0 |
| 0 | 0 | 0 | 0 |
| 0 | 0 | 0 | 0 |
| 0 | 1 | 1 | 1 |
| 0 | 0 | 0 | 0 |
| 0 | 1 | 1 | 1 |
| 0 | 1 | 0 | 0 |
| 0 | 0 | 0 | 0 |
| 0 | 0 | 0 | 1 |
| 0 | 0 | 0 | 0 |
| 0 | 1 | 0 | 0 |
| 0 | 1 | 1 | 1 |
| 0 | 0 | 0 | 0 |
| 0 | 0 | 0 | 0 |
| 0 | 1 | 0 | 0 |
| 0 | 1 | 0 | 0 |
| 0 | 1 | 0 | 0 |
| 0 | 0 | 0 | 0 |
| 0 | 1 | 1 | 1 |
| 0 | 0 | 0 | 0 |
| 0 | 1 | 0 | 0 |
| 0 | 0 | 0 | 0 |
| 0 | 0 | 0 | 1 |
| 0 | 0 | 0 | 0 |
| 0 | 0 | 0 | 0 |
| 0 | 0 | 0 | 0 |
| 0 | 1 | 0 | 0 |
| 0 | 1 | 0 | 0 |
| 0 | 1 | 0 | 0 |
| 0 | 1 | 0 | 0 |
| 0 | 1 | 0 | 0 |
| 0 | 0 | 0 | 1 |
| 0 | 1 | 0 | 0 |
| 0 | 1 | 0 | 0 |
| 0 | 1 | 0 | 0 |
| 0 | 1 | 0 | 0 |
| 0 | 1 | 0 | 0 |
| 0 | 1 | 0 | 0 |
| 0 | 1 | 1 | 1 |
| 0 | 1 | 1 | 1 |

| | | | |
|---|---|---|---|
| 0 | 1 | 1 | 1 |
| 0 | 0 | 0 | 1 |
| 0 | 0 | 0 | 0 |
| 0 | 0 | 0 | 0 |
| 0 | 1 | 0 | 0 |
| 0 | 1 | 0 | 0 |
| 0 | 1 | 0 | 0 |
| 0 | 0 | 0 | 1 |
| 0 | 0 | 0 | 0 |
| 0 | 1 | 1 | 1 |
| 0 | 0 | 0 | 1 |
| 0 | 0 | 0 | 1 |
| 0 | 1 | 0 | 0 |
| 0 | 0 | 0 | 0 |
| 0 | 1 | 0 | 0 |
| 0 | 0 | 0 | 0 |
| 0 | 1 | 0 | 0 |
| 1 | 1 | 0 | 0 |
| 0 | 0 | 0 | 1 |
| 0 | 0 | 0 | 1 |
| 0 | 0 | 0 | 1 |
| 0 | 0 | 0 | 0 |
| 0 | 0 | 0 | 1 |
| 0 | 0 | 0 | 0 |
| 0 | 0 | 0 | 0 |
| 0 | 1 | 0 | 0 |
| 0 | 0 | 0 | 0 |
| 0 | 0 | 0 | 0 |
| 0 | 0 | 0 | 0 |
| 0 | 0 | 0 | 0 |
| 0 | 1 | 0 | 0 |
| 0 | 1 | 0 | 0 |
| 0 | 0 | 0 | 0 |
| 0 | 0 | 0 | 0 |
| 0 | 0 | 0 | 0 |
| 0 | 0 | 0 | 0 |
| 0 | 1 | 0 | 0 |
| 0 | 1 | 0 | 0 |
| 1 | 1 | 0 | 0 |
| 0 | 1 | 0 | 0 |
| 0 | 1 | 0 | 0 |
| 0 | 1 | 1 | 1 |
| 0 | 1 | 0 | 0 |
| 0 | 1 | 0 | 0 |
| 0 | 0 | 0 | 0 |
| 0 | 0 | 0 | 0 |
| 0 | 1 | 1 | 1 |
| 0 | 1 | 0 | 0 |
| 0 | 1 | 0 | 0 |
| 0 | 0 | 0 | 0 |
| 0 | 0 | 0 | 0 |
| 0 | 0 | 0 | 0 |
| 0 | 1 | 1 | 1 |
| 0 | 0 | 0 | 1 |

| | | | |
|---|---|---|---|
| 0 | 1 | 0 | 0 |
| 0 | 0 | 0 | 0 |
| 0 | 0 | 0 | 1 |
| 0 | 0 | 0 | 1 |
| 0 | 0 | 0 | 1 |
| 0 | 1 | 0 | 0 |
| 0 | 1 | 1 | 1 |
| 0 | 1 | 1 | 1 |
| 0 | 1 | 0 | 0 |
| 0 | 0 | 0 | 0 |
| 0 | 0 | 0 | 0 |
| 0 | 0 | 0 | 0 |
| 0 | 0 | 0 | 0 |
| 0 | 0 | 0 | 1 |
| 0 | 0 | 0 | 1 |
| 0 | 0 | 0 | 1 |
| 0 | 0 | 0 | 1 |
| 0 | 0 | 0 | 1 |
| 0 | 0 | 0 | 0 |
| 0 | 0 | 0 | 0 |
| 0 | 1 | 1 | 1 |
| 0 | 1 | 1 | 1 |
| 0 | 1 | 0 | 0 |
| 0 | 1 | 0 | 0 |
| 0 | 1 | 1 | 1 |
| 0 | 1 | 1 | 1 |
| 0 | 1 | 0 | 0 |
| 0 | 1 | 1 | 1 |
| 0 | 0 | 0 | 0 |
| 0 | 0 | 0 | 0 |
| 0 | 1 | 0 | 0 |
| 0 | 1 | 0 | 0 |
| 0 | 1 | 0 | 0 |
| 1 | 0 | 0 | 0 |
| 1 | 1 | 0 | 0 |
| 0 | 1 | 0 | 0 |
| 0 | 1 | 0 | 0 |
| 0 | 0 | 0 | 0 |
| 0 | 0 | 0 | 1 |
| 0 | 1 | 0 | 0 |
| 0 | 1 | 1 | 1 |
| 0 | 0 | 0 | 1 |
| 0 | 1 | 1 | 1 |
| 0 | 0 | 0 | 1 |
| 0 | 0 | 0 | 1 |
| 0 | 0 | 0 | 0 |
| 0 | 0 | 0 | 0 |
| 0 | 0 | 0 | 0 |
| 0 | 0 | 0 | 0 |
| 0 | 1 | 0 | 0 |
| 0 | 1 | 0 | 0 |
| 0 | 1 | 0 | 0 |
| 0 | 1 | 0 | 0 |
| 0 | 1 | 1 | 1 |

| | | | |
|---|---|---|---|
| 0 | 1 | 1 | 1 |
| 0 | 1 | 0 | 0 |
| 0 | 1 | 0 | 0 |
| 0 | 1 | 0 | 0 |
| 0 | 1 | 1 | 1 |
| 0 | 1 | 0 | 0 |
| 0 | 1 | 0 | 0 |
| 0 | 1 | 0 | 0 |
| 0 | 1 | 1 | 1 |
| 0 | 1 | 1 | 1 |
| 0 | 1 | 0 | 0 |
| 0 | 1 | 0 | 0 |
| 0 | 1 | 0 | 0 |
| 0 | 1 | 1 | 1 |
| 0 | 1 | 0 | 0 |
| 0 | 1 | 0 | 0 |
| 0 | 1 | 0 | 0 |
| 0 | 1 | 0 | 0 |
| 0 | 0 | 0 | 0 |
| 0 | 0 | 0 | 0 |
| 0 | 1 | 0 | 0 |
| 0 | 0 | 0 | 1 |
| 0 | 0 | 0 | 0 |
| 0 | 0 | 0 | 1 |
| 0 | 1 | 0 | 0 |
| 0 | 1 | 0 | 0 |
| 0 | 1 | 1 | 1 |
| 0 | 1 | 0 | 0 |
| 0 | 1 | 0 | 0 |
| 0 | 1 | 0 | 0 |
| 0 | 0 | 0 | 0 |
| 0 | 1 | 0 | 0 |
| 0 | 1 | 0 | 0 |
| 0 | 0 | 0 | 0 |
| 0 | 0 | 0 | 1 |
| 0 | 1 | 0 | 0 |
| 0 | 1 | 0 | 0 |
| 0 | 1 | 0 | 0 |
| 0 | 1 | 0 | 0 |
| 0 | 0 | 0 | 0 |
| 0 | 0 | 0 | 0 |
| 0 | 0 | 0 | 0 |
| 0 | 0 | 0 | 0 |
| 0 | 0 | 0 | 0 |
| 0 | 0 | 0 | 0 |
| 0 | 0 | 0 | 0 |
| 0 | 0 | 0 | 0 |
| 0 | 0 | 0 | 1 |
| 0 | 1 | 0 | 0 |
| 0 | 0 | 0 | 1 |
| 0 | 0 | 0 | 0 |
| 0 | 0 | 0 | 0 |
| 0 | 0 | 0 | 0 |
| 0 | 0 | 0 | 0 |

| | | | |
|---|---|---|---|
| 0 | 0 | 0 | 0 |
| 0 | 1 | 0 | 0 |
| 0 | 0 | 0 | 1 |
| 0 | 1 | 0 | 0 |
| 0 | 0 | 0 | 0 |
| 0 | 0 | 0 | 1 |
| 0 | 0 | 0 | 1 |
| 0 | 0 | 0 | 1 |
| 0 | 0 | 0 | 0 |
| 0 | 0 | 0 | 1 |
| 0 | 1 | 0 | 0 |
| 0 | 1 | 0 | 0 |
| 0 | 1 | 0 | 0 |
| 0 | 0 | 0 | 1 |
| 0 | 1 | 0 | 0 |
| 0 | 0 | 0 | 1 |
| 0 | 1 | 0 | 0 |
| 0 | 1 | 0 | 0 |
| 1 | 0 | 0 | 0 |
| 0 | 0 | 0 | 1 |
| 0 | 0 | 0 | 1 |
| 0 | 0 | 0 | 0 |
| 0 | 0 | 0 | 1 |
| 0 | 0 | 0 | 0 |
| 0 | 0 | 0 | 0 |
| 0 | 0 | 0 | 1 |
| 0 | 0 | 0 | 1 |
| 0 | 0 | 0 | 0 |
| 0 | 0 | 0 | 1 |
| 1 | 0 | 0 | 0 |
| 0 | 0 | 0 | 1 |
| 0 | 0 | 0 | 1 |
| 0 | 0 | 0 | 1 |
| 0 | 0 | 0 | 1 |
| 0 | 0 | 0 | 1 |
| 0 | 0 | 0 | 1 |
| 0 | 0 | 0 | 1 |
| 0 | 0 | 0 | 1 |
| 0 | 0 | 0 | 1 |
| 0 | 0 | 0 | 1 |
| 0 | 0 | 0 | 1 |
| 0 | 0 | 0 | 1 |
| 0 | 0 | 0 | 1 |
| 0 | 0 | 0 | 1 |
| 0 | 0 | 0 | 1 |
| 0 | 0 | 0 | 1 |
| 0 | 0 | 0 | 1 |
| 0 | 0 | 0 | 1 |
| 0 | 0 | 0 | 1 |
| 0 | 0 | 0 | 1 |
| 0 | 0 | 0 | 1 |
| 0 | 0 | 0 | 1 |
| 0 | 0 | 0 | 1 |
| 0 | 0 | 0 | 1 |

| | | | |
|---|---|---|---|
| 0 | 0 | 0 | 1 |
| 0 | 0 | 0 | 1 |
| 0 | 0 | 0 | 1 |
| 0 | 0 | 0 | 1 |
| 0 | 0 | 0 | 1 |
| 0 | 0 | 0 | 1 |
| 0 | 0 | 0 | 1 |
| 0 | 0 | 0 | 1 |
| 0 | 0 | 0 | 1 |
| 0 | 0 | 0 | 1 |
| 0 | 0 | 0 | 1 |
| 0 | 0 | 0 | 0 |
| 0 | 1 | 0 | 0 |
| 0 | 1 | 0 | 0 |
| 0 | 1 | 1 | 1 |
| 0 | 1 | 0 | 0 |
| 0 | 1 | 1 | 1 |
| 0 | 1 | 0 | 0 |
| 0 | 1 | 1 | 1 |
| 0 | 0 | 0 | 1 |
| 0 | 0 | 0 | 0 |
| 0 | 0 | 0 | 1 |
| 0 | 0 | 0 | 1 |
| 0 | 0 | 0 | 0 |
| 0 | 0 | 0 | 0 |
| 0 | 0 | 0 | 1 |
| 0 | 0 | 0 | 1 |
| 0 | 0 | 0 | 0 |
| 0 | 0 | 0 | 0 |
| 0 | 0 | 0 | 0 |
| 0 | 0 | 0 | 0 |
| 0 | 0 | 0 | 0 |
| 0 | 0 | 0 | 1 |
| 0 | 0 | 0 | 1 |
| 0 | 0 | 0 | 1 |
| 0 | 0 | 0 | 1 |
| 0 | 0 | 0 | 1 |
| 0 | 0 | 0 | 1 |
| 0 | 0 | 0 | 1 |
| 0 | 1 | 0 | 0 |
| 0 | 0 | 0 | 0 |
| 0 | 0 | 0 | 1 |
| 0 | 1 | 0 | 0 |
| 0 | 1 | 0 | 0 |
| 0 | 1 | 0 | 0 |
| 0 | 1 | 1 | 1 |
| 0 | 1 | 0 | 0 |
| 0 | 0 | 0 | 1 |
| 0 | 1 | 1 | 1 |
| 0 | 1 | 0 | 0 |
| 0 | 1 | 0 | 0 |
| 0 | 0 | 0 | 0 |
| 0 | 0 | 0 | 1 |
| 0 | 1 | 0 | 0 |

| | | | |
|---|---|---|---|
| 0 | 1 | 0 | 0 |
| 0 | 0 | 0 | 1 |
| 0 | 0 | 0 | 1 |
| 0 | 0 | 0 | 1 |
| 0 | 1 | 0 | 0 |
| 0 | 0 | 0 | 1 |
| 0 | 0 | 0 | 0 |
| 0 | 0 | 0 | 1 |
| 0 | 0 | 0 | 1 |
| 0 | 0 | 0 | 0 |
| 0 | 1 | 1 | 1 |
| 0 | 0 | 0 | 1 |
| 0 | 0 | 0 | 1 |
| 0 | 1 | 0 | 0 |
| 0 | 1 | 1 | 1 |
| 0 | 1 | 1 | 1 |
| 0 | 1 | 0 | 0 |
| 0 | 0 | 0 | 1 |
| 0 | 1 | 0 | 0 |
| 0 | 1 | 0 | 0 |
| 0 | 0 | 0 | 0 |
| 0 | 0 | 0 | 0 |
| 0 | 0 | 0 | 0 |
| 0 | 1 | 0 | 0 |
| 0 | 1 | 0 | 0 |
| 0 | 1 | 0 | 0 |
| 0 | 0 | 0 | 0 |
| 0 | 1 | 0 | 0 |
| 0 | 1 | 0 | 0 |
| 0 | 1 | 0 | 0 |
| 0 | 0 | 0 | 0 |
| 0 | 1 | 0 | 0 |
| 0 | 1 | 0 | 0 |
| 0 | 1 | 0 | 0 |
| 0 | 0 | 0 | 0 |
| 0 | 1 | 0 | 0 |
| 0 | 1 | 1 | 1 |
| 0 | 1 | 0 | 0 |
| 0 | 1 | 0 | 0 |
| 0 | 1 | 1 | 1 |
| 0 | 1 | 1 | 1 |
| 0 | 0 | 0 | 0 |
| 0 | 0 | 0 | 0 |
| 0 | 1 | 0 | 0 |
| 0 | 1 | 0 | 0 |
| 0 | 0 | 0 | 0 |
| 0 | 1 | 0 | 0 |
| 0 | 1 | 0 | 0 |
| 0 | 1 | 1 | 1 |
| 0 | 1 | 1 | 1 |
| 0 | 0 | 0 | 0 |
| 0 | 0 | 0 | 1 |
| 0 | 0 | 0 | 0 |
| 0 | 0 | 0 | 0 |

| | | | |
|---|---|---|---|
| 0 | 0 | 0 | 0 |
| 0 | 1 | 0 | 0 |
| 0 | 1 | 0 | 0 |
| 0 | 0 | 0 | 0 |
| 0 | 0 | 0 | 0 |
| 0 | 0 | 0 | 0 |
| 0 | 1 | 0 | 0 |
| 0 | 0 | 0 | 0 |
| 0 | 0 | 0 | 0 |
| 0 | 0 | 0 | 0 |
| 0 | 0 | 0 | 0 |
| 0 | 0 | 0 | 0 |
| 0 | 0 | 0 | 0 |
| 0 | 0 | 0 | 0 |
| 0 | 0 | 0 | 1 |
| 0 | 1 | 0 | 0 |
| 0 | 1 | 0 | 0 |
| 0 | 1 | 0 | 0 |
| 0 | 1 | 1 | 1 |
| 0 | 0 | 0 | 0 |
| 0 | 1 | 0 | 0 |
| 0 | 1 | 0 | 0 |
| 0 | 1 | 0 | 0 |
| 0 | 1 | 0 | 0 |
| 0 | 1 | 0 | 0 |
| 0 | 1 | 1 | 1 |
| 0 | 1 | 0 | 0 |
| 0 | 1 | 0 | 0 |
| 0 | 1 | 0 | 0 |
| 0 | 0 | 0 | 0 |
| 0 | 0 | 0 | 1 |
| 0 | 0 | 0 | 0 |
| 0 | 0 | 0 | 1 |
| 0 | 1 | 1 | 1 |
| 1 | 1 | 0 | 0 |
| 0 | 0 | 0 | 0 |
| 0 | 0 | 0 | 1 |
| 0 | 0 | 0 | 1 |
| 0 | 0 | 0 | 1 |
| 0 | 0 | 0 | 1 |
| 0 | 1 | 0 | 0 |
| 0 | 1 | 0 | 0 |
| 0 | 0 | 0 | 0 |
| 0 | 0 | 0 | 0 |
| 0 | 0 | 0 | 0 |
| 0 | 1 | 0 | 0 |
| 0 | 0 | 0 | 0 |
| 0 | 0 | 0 | 0 |
| 0 | 0 | 0 | 1 |
| 0 | 0 | 0 | 1 |
| 0 | 0 | 0 | 0 |
| 0 | 0 | 0 | 0 |
| 0 | 0 | 0 | 0 |
| 0 | 0 | 0 | 1 |

| | | | |
|---|---|---|---|
| 0 | 0 | 0 | 1 |
| 0 | 0 | 0 | 1 |
| 0 | 1 | 1 | 1 |
| 0 | 1 | 0 | 0 |
| 0 | 0 | 0 | 1 |
| 0 | 0 | 0 | 1 |
| 0 | 0 | 0 | 0 |
| 0 | 0 | 0 | 1 |
| 0 | 0 | 0 | 0 |
| 0 | 0 | 0 | 0 |
| 0 | 0 | 0 | 0 |
| 0 | 0 | 0 | 0 |
| 0 | 0 | 0 | 0 |
| 0 | 0 | 0 | 0 |
| 0 | 0 | 0 | 0 |
| 0 | 0 | 0 | 0 |
| 0 | 0 | 0 | 1 |
| 0 | 0 | 0 | 0 |
| 0 | 1 | 0 | 0 |
| 0 | 1 | 1 | 1 |
| 0 | 0 | 0 | 0 |
| 0 | 0 | 0 | 0 |
| 0 | 0 | 0 | 1 |
| 0 | 0 | 0 | 0 |
| 0 | 0 | 0 | 1 |
| 0 | 0 | 0 | 1 |
| 0 | 0 | 0 | 1 |
| 0 | 0 | 0 | 0 |
| 0 | 0 | 0 | 0 |
| 0 | 0 | 0 | 0 |
| 0 | 0 | 0 | 1 |
| 0 | 0 | 0 | 0 |
| 0 | 0 | 0 | 0 |
| 0 | 0 | 0 | 0 |
| 0 | 0 | 0 | 1 |
| 0 | 0 | 0 | 1 |
| 0 | 1 | 0 | 0 |
| 0 | 0 | 0 | 0 |
| 0 | 1 | 0 | 0 |
| 0 | 0 | 0 | 1 |
| 0 | 0 | 0 | 1 |
| 0 | 0 | 0 | 1 |
| 0 | 0 | 0 | 1 |
| 0 | 0 | 0 | 0 |
| 0 | 0 | 0 | 1 |
| 0 | 0 | 0 | 0 |
| 0 | 0 | 0 | 1 |
| 0 | 0 | 0 | 0 |
| 0 | 0 | 0 | 0 |
| 0 | 0 | 0 | 1 |
| 0 | 0 | 0 | 1 |
| 0 | 1 | 0 | 0 |
| 0 | 1 | 0 | 0 |
| 0 | 0 | 0 | 1 |

| | | | |
|---|---|---|---|
| 0 | 0 | 0 | 1 |
| 0 | 0 | 0 | 1 |
| 0 | 0 | 0 | 1 |
| 0 | 1 | 1 | 1 |
| 0 | 1 | 0 | 0 |
| 0 | 1 | 1 | 1 |
| 0 | 1 | 1 | 1 |
| 0 | 1 | 0 | 0 |
| 0 | 1 | 0 | 0 |
| 0 | 0 | 0 | 1 |
| 0 | 0 | 0 | 0 |
| 0 | 0 | 0 | 0 |
| 1 | 0 | 0 | 0 |
| 1 | 0 | 0 | 0 |
| 0 | 0 | 0 | 0 |
| 1 | 0 | 0 | 0 |
| 1 | 0 | 0 | 0 |
| 1 | 0 | 0 | 0 |
| 1 | 0 | 0 | 0 |
| 0 | 0 | 0 | 1 |
| 0 | 0 | 0 | 1 |
| 0 | 0 | 0 | 1 |
| 0 | 0 | 0 | 1 |
| 0 | 0 | 0 | 1 |
| 0 | 0 | 0 | 0 |
| 0 | 0 | 0 | 1 |
| 1 | 0 | 0 | 0 |
| 1 | 0 | 0 | 0 |
| 0 | 0 | 0 | 1 |
| 0 | 0 | 0 | 1 |
| 0 | 0 | 0 | 1 |
| 0 | 0 | 0 | 1 |
| 0 | 0 | 0 | 1 |
| 0 | 0 | 0 | 0 |
| 0 | 0 | 0 | 0 |
| 0 | 0 | 0 | 1 |
| 0 | 0 | 0 | 0 |
| 0 | 0 | 0 | 1 |
| 0 | 0 | 0 | 0 |
| 0 | 1 | 0 | 0 |
| 0 | 1 | 0 | 0 |
| 0 | 1 | 0 | 0 |
| 0 | 0 | 0 | 1 |
| 0 | 0 | 0 | 0 |
| 0 | 0 | 0 | 0 |
| 1 | 0 | 0 | 0 |
| 1 | 0 | 0 | 0 |
| 1 | 0 | 0 | 0 |
| 0 | 0 | 0 | 0 |
| 0 | 0 | 0 | 0 |
| 0 | 1 | 0 | 0 |
| 1 | 0 | 0 | 0 |
| 1 | 0 | 0 | 0 |
| 0 | 0 | 0 | 1 |

| | | | |
|---|---|---|---|
| 0 | 0 | 0 | 0 |
| 0 | 0 | 0 | 1 |
| 0 | 0 | 0 | 0 |
| 0 | 0 | 0 | 0 |
| 0 | 1 | 0 | 0 |
| 0 | 0 | 0 | 1 |
| 0 | 0 | 0 | 0 |
| 0 | 0 | 0 | 1 |
| 0 | 0 | 0 | 1 |
| 0 | 0 | 0 | 0 |
| 0 | 0 | 0 | 1 |
| 0 | 0 | 0 | 1 |
| 0 | 0 | 0 | 1 |
| 0 | 0 | 0 | 1 |
| 0 | 0 | 0 | 1 |
| 0 | 1 | 0 | 0 |
| 0 | 1 | 0 | 0 |
| 0 | 0 | 0 | 1 |
| 0 | 0 | 0 | 0 |
| 0 | 0 | 0 | 1 |
| 0 | 0 | 0 | 0 |
| 0 | 0 | 0 | 0 |
| 0 | 0 | 0 | 0 |
| 0 | 1 | 0 | 0 |
| 0 | 1 | 0 | 0 |
| 0 | 1 | 0 | 0 |
| 0 | 1 | 0 | 0 |
| 0 | 0 | 0 | 0 |
| 0 | 0 | 0 | 0 |
| 0 | 1 | 0 | 0 |
| 0 | 1 | 0 | 0 |
| 0 | 1 | 0 | 0 |
| 0 | 1 | 1 | 1 |
| 0 | 1 | 1 | 1 |
| 0 | 1 | 0 | 0 |
| 0 | 1 | 0 | 0 |
| 0 | 1 | 1 | 1 |
| 0 | 1 | 0 | 0 |
| 0 | 1 | 1 | 1 |
| 0 | 1 | 0 | 0 |
| 0 | 1 | 0 | 0 |
| 0 | 1 | 0 | 0 |
| 0 | 1 | 1 | 1 |
| 0 | 0 | 0 | 1 |
| 0 | 1 | 0 | 0 |
| 0 | 1 | 0 | 0 |
| 0 | 1 | 0 | 0 |
| 0 | 1 | 0 | 0 |
| 0 | 1 | 0 | 0 |
| 0 | 1 | 0 | 0 |
| 0 | 1 | 1 | 1 |
| 0 | 1 | 0 | 0 |
| 0 | 1 | 0 | 0 |
| 0 | 1 | 1 | 1 |

| | | | |
|---|---|---|---|
| 0 | 1 | 0 | 0 |
| 0 | 1 | 1 | 1 |
| 0 | 0 | 0 | 0 |
| 0 | 0 | 0 | 0 |
| 0 | 0 | 0 | 0 |
| 0 | 0 | 0 | 0 |
| 0 | 1 | 0 | 0 |
| 0 | 0 | 0 | 0 |
| 0 | 0 | 0 | 0 |
| 0 | 1 | 0 | 0 |
| 0 | 0 | 0 | 0 |
| 0 | 0 | 0 | 1 |
| 0 | 0 | 0 | 0 |
| 0 | 0 | 0 | 1 |
| 0 | 1 | 1 | 1 |
| 0 | 1 | 1 | 1 |
| 0 | 1 | 0 | 0 |
| 0 | 1 | 0 | 0 |
| 0 | 1 | 1 | 1 |
| 0 | 1 | 0 | 0 |
| 0 | 1 | 1 | 1 |
| 0 | 1 | 0 | 0 |
| 0 | 0 | 0 | 1 |
| 0 | 1 | 0 | 0 |
| 0 | 0 | 0 | 1 |
| 0 | 0 | 0 | 1 |
| 0 | 0 | 0 | 1 |
| 0 | 0 | 0 | 0 |
| 0 | 0 | 0 | 1 |
| 0 | 0 | 0 | 1 |
| 0 | 0 | 0 | 1 |
| 1 | 0 | 0 | 0 |
| 0 | 0 | 0 | 1 |
| 0 | 0 | 0 | 0 |
| 0 | 0 | 0 | 0 |
| 0 | 0 | 0 | 0 |
| 0 | 0 | 0 | 1 |
| 0 | 0 | 0 | 1 |
| 0 | 1 | 0 | 0 |
| 0 | 0 | 0 | 0 |
| 0 | 1 | 0 | 0 |
| 0 | 0 | 0 | 0 |
| 0 | 1 | 0 | 0 |
| 0 | 0 | 0 | 0 |
| 0 | 1 | 0 | 0 |
| 0 | 0 | 0 | 0 |
| 0 | 0 | 0 | 1 |
| 0 | 0 | 0 | 1 |
| 0 | 0 | 0 | 0 |
| 0 | 0 | 0 | 0 |
| 0 | 0 | 0 | 0 |
| 0 | 0 | 0 | 1 |
| 0 | 0 | 0 | 1 |
| 0 | 0 | 0 | 0 |

| | | | |
|---|---|---|---|
| 0 | 0 | 0 | 0 |
| 0 | 0 | 0 | 1 |
| 0 | 0 | 0 | 1 |
| 0 | 0 | 0 | 1 |
| 0 | 0 | 0 | 0 |
| 0 | 1 | 0 | 0 |
| 0 | 1 | 0 | 0 |
| 0 | 1 | 0 | 0 |
| 0 | 1 | 0 | 0 |
| 0 | 0 | 0 | 0 |
| 0 | 0 | 0 | 1 |
| 0 | 1 | 0 | 0 |
| 0 | 0 | 0 | 0 |
| 0 | 0 | 0 | 0 |
| 0 | 1 | 1 | 1 |
| 0 | 0 | 0 | 1 |
| 0 | 1 | 0 | 0 |
| 0 | 1 | 1 | 1 |
| 0 | 1 | 0 | 0 |
| 0 | 1 | 0 | 0 |
| 0 | 0 | 0 | 1 |
| 0 | 1 | 0 | 0 |
| 0 | 1 | 0 | 0 |
| 0 | 1 | 1 | 1 |
| 0 | 1 | 0 | 0 |
| 0 | 1 | 0 | 0 |
| 0 | 1 | 0 | 0 |
| 0 | 1 | 1 | 1 |
| 0 | 1 | 0 | 0 |
| 0 | 0 | 0 | 0 |
| 0 | 0 | 0 | 0 |
| 0 | 0 | 0 | 1 |
| 0 | 0 | 0 | 1 |
| 0 | 0 | 0 | 1 |
| 0 | 0 | 0 | 1 |
| 0 | 0 | 0 | 1 |
| 0 | 0 | 0 | 1 |
| 0 | 0 | 0 | 1 |
| 0 | 0 | 0 | 1 |
| 0 | 0 | 0 | 1 |
| 0 | 0 | 0 | 1 |
| 0 | 0 | 0 | 1 |
| 1 | 0 | 0 | 0 |
| 0 | 0 | 0 | 1 |
| 0 | 0 | 0 | 1 |
| 0 | 0 | 0 | 1 |
| 0 | 0 | 0 | 1 |
| 0 | 0 | 0 | 1 |
| 0 | 0 | 0 | 1 |
| 0 | 0 | 0 | 1 |
| 0 | 0 | 0 | 1 |
| 0 | 0 | 0 | 1 |
| 0 | 0 | 0 | 1 |
| 0 | 0 | 0 | 1 |

| | | | |
|---|---|---|---|
| 0 | 0 | 0 | 1 |
| 0 | 0 | 0 | 1 |
| 0 | 0 | 0 | 1 |
| 0 | 0 | 0 | 1 |
| 0 | 0 | 0 | 1 |
| 0 | 0 | 0 | 1 |
| 0 | 0 | 0 | 1 |
| 0 | 0 | 0 | 1 |
| 0 | 1 | 0 | 0 |
| 0 | 0 | 0 | 0 |
| 0 | 0 | 0 | 0 |
| 0 | 0 | 0 | 0 |
| 0 | 0 | 0 | 0 |
| 0 | 0 | 0 | 0 |
| 0 | 0 | 0 | 0 |
| 0 | 0 | 0 | 0 |
| 0 | 0 | 0 | 0 |
| 0 | 0 | 0 | 0 |
| 0 | 0 | 0 | 0 |
| 0 | 1 | 0 | 0 |
| 1 | 0 | 0 | 0 |
| 1 | 0 | 0 | 0 |
| 0 | 1 | 0 | 0 |
| 0 | 1 | 0 | 0 |
| 0 | 1 | 0 | 0 |
| 0 | 0 | 0 | 1 |
| 0 | 0 | 0 | 1 |
| 0 | 0 | 0 | 0 |
| 0 | 0 | 0 | 0 |
| 1 | 0 | 0 | 0 |
| 1 | 0 | 0 | 0 |
| 0 | 0 | 0 | 0 |
| 1 | 0 | 0 | 0 |
| 1 | 0 | 0 | 0 |
| 0 | 0 | 0 | 0 |
| 0 | 0 | 0 | 0 |
| 0 | 0 | 0 | 0 |
| 0 | 1 | 0 | 0 |
| 0 | 1 | 0 | 0 |
| 0 | 0 | 0 | 0 |
| 0 | 0 | 0 | 0 |
| 0 | 0 | 0 | 0 |
| 1 | 0 | 0 | 0 |
| 1 | 0 | 0 | 0 |
| 1 | 0 | 0 | 0 |
| 0 | 0 | 0 | 0 |
| 0 | 0 | 0 | 1 |
| 0 | 0 | 0 | 1 |
| 0 | 0 | 0 | 1 |
| 0 | 0 | 0 | 1 |
| 0 | 0 | 0 | 1 |
| 0 | 0 | 0 | 1 |
| 0 | 1 | 0 | 0 |
| 0 | 1 | 0 | 0 |

| | | | |
|---|---|---|---|
| 0 | 1 | 0 | 0 |
| 0 | 0 | 0 | 0 |
| 0 | 0 | 0 | 0 |
| 0 | 1 | 0 | 0 |
| 0 | 0 | 0 | 1 |
| 0 | 0 | 0 | 0 |
| 0 | 0 | 0 | 0 |
| 0 | 0 | 0 | 0 |
| 0 | 0 | 0 | 1 |
| 0 | 0 | 0 | 0 |
| 1 | 0 | 0 | 0 |
| 0 | 0 | 0 | 1 |
| 0 | 0 | 0 | 0 |
| 0 | 0 | 0 | 0 |
| 0 | 0 | 0 | 0 |
| 0 | 0 | 0 | 0 |
| 0 | 0 | 0 | 1 |
| 0 | 0 | 0 | 0 |
| 0 | 0 | 0 | 0 |
| 0 | 0 | 0 | 1 |
| 0 | 1 | 0 | 0 |
| 0 | 0 | 0 | 0 |
| 0 | 1 | 0 | 0 |
| 0 | 0 | 0 | 0 |
| 0 | 0 | 0 | 0 |
| 0 | 0 | 0 | 1 |
| 0 | 0 | 0 | 1 |
| 0 | 0 | 0 | 1 |
| 0 | 0 | 0 | 1 |
| 0 | 0 | 0 | 1 |
| 0 | 0 | 0 | 1 |
| 0 | 0 | 0 | 1 |
| 0 | 1 | 1 | 1 |
| 0 | 1 | 1 | 1 |
| 0 | 0 | 0 | 0 |
| 0 | 1 | 0 | 0 |
| 0 | 0 | 0 | 0 |
| 0 | 1 | 0 | 0 |
| 0 | 1 | 0 | 0 |
| 0 | 1 | 0 | 0 |
| 0 | 0 | 0 | 1 |
| 0 | 0 | 0 | 1 |
| 0 | 0 | 0 | 1 |
| 0 | 0 | 0 | 0 |
| 0 | 0 | 0 | 0 |
| 0 | 0 | 0 | 0 |
| 0 | 0 | 0 | 1 |
| 0 | 0 | 0 | 1 |
| 0 | 0 | 0 | 0 |
| 0 | 0 | 0 | 0 |
| 0 | 0 | 0 | 1 |
| 0 | 0 | 0 | 1 |
| 0 | 0 | 0 | 1 |
| 0 | 0 | 0 | 1 |

| | | | |
|---|---|---|---|
| 0 | 0 | 0 | 1 |
| 0 | 0 | 0 | 1 |
| 0 | 0 | 0 | 1 |
| 0 | 0 | 0 | 0 |
| 0 | 0 | 0 | 0 |
| 0 | 0 | 0 | 1 |
| 0 | 0 | 0 | 1 |
| 0 | 0 | 0 | 1 |
| 0 | 0 | 0 | 1 |
| 0 | 0 | 0 | 1 |
| 0 | 0 | 0 | 0 |
| 0 | 0 | 0 | 1 |
| 0 | 1 | 0 | 0 |
| 0 | 0 | 0 | 0 |
| 0 | 0 | 0 | 0 |
| 0 | 0 | 0 | 1 |
| 0 | 0 | 0 | 1 |
| 0 | 0 | 0 | 1 |
| 0 | 0 | 0 | 1 |
| 0 | 0 | 0 | 1 |
| 0 | 0 | 0 | 0 |
| 0 | 0 | 0 | 1 |
| 0 | 0 | 0 | 1 |
| 0 | 1 | 0 | 0 |
| 0 | 1 | 1 | 1 |
| 0 | 1 | 0 | 0 |
| 0 | 1 | 0 | 0 |
| 0 | 0 | 0 | 1 |
| 0 | 0 | 0 | 1 |
| 0 | 0 | 0 | 1 |
| 0 | 0 | 0 | 1 |
| 0 | 0 | 0 | 1 |
| 0 | 0 | 0 | 1 |
| 0 | 0 | 0 | 1 |
| 0 | 0 | 0 | 1 |
| 0 | 0 | 0 | 0 |
| 0 | 0 | 0 | 1 |
| 0 | 0 | 0 | 1 |
| 0 | 0 | 0 | 0 |
| 0 | 0 | 0 | 0 |
| 0 | 0 | 0 | 1 |
| 0 | 0 | 0 | 1 |
| 1 | 0 | 0 | 0 |
| 1 | 0 | 0 | 0 |
| 0 | 0 | 0 | 0 |
| 0 | 0 | 0 | 0 |
| 1 | 0 | 0 | 0 |
| 1 | 0 | 0 | 0 |
| 0 | 0 | 0 | 1 |
| 1 | 0 | 0 | 0 |
| 1 | 0 | 0 | 0 |
| 1 | 1 | 0 | 0 |
| 1 | 1 | 0 | 0 |
| 1 | 0 | 0 | 0 |

| | | | |
|---|---|---|---|
| 0 | 0 | 0 | 0 |
| 0 | 0 | 0 | 0 |
| 1 | 0 | 0 | 0 |
| 1 | 0 | 0 | 0 |
| 1 | 0 | 0 | 0 |
| 1 | 0 | 0 | 0 |
| 0 | 0 | 0 | 0 |
| 0 | 0 | 0 | 0 |
| 0 | 0 | 0 | 0 |
| 0 | 0 | 0 | 1 |
| 0 | 0 | 0 | 1 |
| 0 | 0 | 0 | 0 |
| 0 | 0 | 0 | 1 |
| 0 | 0 | 0 | 0 |
| 0 | 0 | 0 | 0 |
| 0 | 0 | 0 | 1 |
| 0 | 0 | 0 | 0 |
| 0 | 1 | 1 | 1 |
| 0 | 0 | 0 | 1 |
| 0 | 0 | 0 | 1 |
| 0 | 1 | 1 | 1 |
| 0 | 0 | 0 | 1 |
| 0 | 0 | 0 | 1 |
| 0 | 0 | 0 | 0 |
| 0 | 0 | 0 | 0 |
| 0 | 0 | 0 | 1 |
| 0 | 0 | 0 | 1 |
| 0 | 0 | 0 | 1 |
| 0 | 0 | 0 | 0 |
| 0 | 0 | 0 | 0 |
| 0 | 0 | 0 | 1 |
| 0 | 0 | 0 | 1 |
| 0 | 1 | 1 | 1 |
| 0 | 1 | 0 | 0 |
| 0 | 1 | 0 | 0 |
| 0 | 0 | 0 | 1 |
| 0 | 0 | 0 | 0 |
| 0 | 0 | 0 | 1 |
| 0 | 0 | 0 | 1 |
| 0 | 0 | 0 | 0 |
| 0 | 0 | 0 | 0 |
| 0 | 0 | 0 | 0 |
| 0 | 0 | 0 | 0 |
| 0 | 0 | 0 | 0 |
| 0 | 0 | 0 | 0 |
| 0 | 0 | 0 | 1 |
| 0 | 0 | 0 | 1 |
| 0 | 0 | 0 | 0 |
| 0 | 0 | 0 | 0 |
| 0 | 0 | 0 | 0 |
| 0 | 0 | 0 | 1 |
| 0 | 0 | 0 | 1 |
| 0 | 0 | 0 | 0 |
| 0 | 0 | 0 | 0 |

| | | | |
|---|---|---|---|
| 0 | 0 | 0 | 1 |
| 0 | 0 | 0 | 1 |
| 0 | 0 | 0 | 1 |
| 0 | 0 | 0 | 1 |
| 0 | 1 | 0 | 0 |
| 0 | 1 | 0 | 0 |
| 0 | 1 | 1 | 1 |
| 0 | 0 | 0 | 0 |
| 0 | 1 | 1 | 1 |
| 0 | 0 | 0 | 0 |
| 0 | 0 | 0 | 1 |
| 0 | 0 | 0 | 1 |
| 0 | 0 | 0 | 1 |
| 0 | 0 | 0 | 1 |
| 0 | 0 | 0 | 1 |
| 0 | 0 | 0 | 1 |
| 0 | 0 | 0 | 0 |
| 0 | 0 | 0 | 1 |
| 0 | 0 | 0 | 1 |
| 0 | 0 | 0 | 0 |
| 0 | 0 | 0 | 0 |
| 0 | 0 | 0 | 1 |
| 0 | 0 | 0 | 1 |
| 0 | 0 | 0 | 0 |
| 0 | 0 | 0 | 1 |
| 0 | 0 | 0 | 0 |
| 0 | 0 | 0 | 0 |
| 0 | 0 | 0 | 0 |
| 0 | 0 | 0 | 0 |
| 0 | 0 | 0 | 0 |
| 0 | 0 | 0 | 0 |
| 0 | 0 | 0 | 0 |
| 0 | 0 | 0 | 1 |
| 0 | 0 | 0 | 0 |
| 0 | 0 | 0 | 1 |
| 0 | 0 | 0 | 1 |
| 0 | 0 | 0 | 1 |
| 0 | 0 | 0 | 1 |
| 0 | 0 | 0 | 0 |
| 0 | 0 | 0 | 1 |
| 0 | 0 | 0 | 1 |
| 0 | 0 | 0 | 1 |
| 0 | 0 | 0 | 1 |
| 0 | 1 | 0 | 0 |
| 0 | 0 | 0 | 1 |
| 0 | 0 | 0 | 1 |
| 0 | 0 | 0 | 1 |
| 0 | 0 | 0 | 1 |
| 0 | 0 | 0 | 1 |
| 0 | 0 | 0 | 1 |
| 0 | 0 | 0 | 0 |
| 0 | 0 | 0 | 0 |
| 0 | 0 | 0 | 0 |
| 0 | 0 | 0 | 1 |

| | | | |
|---|---|---|---|
| 0 | 1 | 0 | 0 |
| 0 | 1 | 0 | 0 |
| 0 | 1 | 0 | 0 |
| 0 | 0 | 0 | 0 |
| 1 | 1 | 0 | 0 |
| 1 | 1 | 0 | 0 |
| 0 | 1 | 0 | 0 |
| 0 | 0 | 0 | 0 |
| 0 | 0 | 0 | 1 |
| 0 | 0 | 0 | 1 |
| 0 | 0 | 0 | 0 |
| 0 | 0 | 0 | 1 |
| 0 | 0 | 0 | 1 |
| 0 | 0 | 0 | 0 |
| 0 | 0 | 0 | 0 |
| 0 | 0 | 0 | 0 |
| 0 | 1 | 0 | 0 |
| 0 | 0 | 0 | 0 |
| 0 | 0 | 0 | 0 |
| 0 | 0 | 0 | 1 |
| 0 | 0 | 0 | 1 |
| 0 | 0 | 0 | 0 |
| 0 | 0 | 0 | 1 |
| 0 | 1 | 0 | 0 |
| 0 | 0 | 0 | 1 |
| 0 | 0 | 0 | 1 |
| 0 | 0 | 0 | 1 |
| 0 | 0 | 0 | 1 |
| 0 | 0 | 0 | 1 |
| 0 | 0 | 0 | 0 |
| 0 | 0 | 0 | 1 |
| 0 | 0 | 0 | 1 |
| 0 | 0 | 0 | 0 |
| 0 | 0 | 0 | 1 |
| 0 | 1 | 0 | 0 |
| 0 | 1 | 0 | 0 |
| 0 | 1 | 0 | 0 |
| 0 | 0 | 0 | 0 |
| 0 | 0 | 0 | 0 |
| 0 | 0 | 0 | 0 |
| 0 | 0 | 0 | 0 |
| 0 | 1 | 0 | 0 |
| 0 | 0 | 0 | 1 |
| 0 | 1 | 0 | 0 |
| 0 | 0 | 0 | 1 |
| 0 | 0 | 0 | 0 |
| 0 | 1 | 1 | 1 |
| 0 | 0 | 0 | 1 |
| 0 | 0 | 0 | 1 |
| 0 | 0 | 0 | 0 |
| 0 | 0 | 0 | 1 |
| 0 | 0 | 0 | 1 |
| 0 | 0 | 0 | 1 |
| 0 | 0 | 0 | 1 |

| | | | |
|---|---|---|---|
| 1 | 0 | 0 | 0 |
| 0 | 0 | 0 | 1 |
| 0 | 0 | 0 | 1 |
| 0 | 0 | 0 | 1 |
| 0 | 0 | 0 | 1 |
| 0 | 0 | 0 | 1 |
| 0 | 0 | 0 | 1 |
| 0 | 0 | 0 | 1 |
| 0 | 0 | 0 | 1 |
| 0 | 0 | 0 | 1 |
| 0 | 0 | 0 | 0 |
| 0 | 0 | 0 | 1 |
| 0 | 0 | 0 | 1 |
| 0 | 0 | 0 | 1 |
| 0 | 0 | 0 | 0 |
| 0 | 0 | 0 | 0 |
| 0 | 0 | 0 | 0 |
| 0 | 0 | 0 | 0 |
| 0 | 0 | 0 | 0 |
| 0 | 0 | 0 | 1 |
| 0 | 0 | 0 | 1 |
| 0 | 0 | 0 | 1 |
| 0 | 0 | 0 | 0 |
| 0 | 0 | 0 | 1 |
| 0 | 0 | 0 | 1 |
| 0 | 0 | 0 | 1 |
| 0 | 0 | 0 | 1 |
| 0 | 0 | 0 | 1 |
| 0 | 0 | 0 | 0 |
| 0 | 0 | 0 | 0 |
| 0 | 0 | 0 | 0 |
| 0 | 1 | 0 | 0 |
| 0 | 1 | 0 | 0 |
| 0 | 0 | 0 | 0 |
| 0 | 0 | 0 | 1 |
| 0 | 0 | 0 | 1 |
| 0 | 0 | 0 | 0 |
| 0 | 0 | 0 | 0 |
| 0 | 1 | 0 | 0 |
| 0 | 1 | 0 | 0 |
| 0 | 1 | 0 | 0 |
| 0 | 0 | 0 | 0 |
| 0 | 0 | 0 | 1 |
| 0 | 0 | 0 | 1 |
| 0 | 0 | 0 | 0 |
| 0 | 0 | 0 | 1 |
| 0 | 0 | 0 | 1 |
| 0 | 0 | 0 | 1 |
| 0 | 0 | 0 | 1 |
| 0 | 0 | 0 | 1 |
| 0 | 1 | 1 | 1 |
| 0 | 0 | 0 | 1 |
| 0 | 0 | 0 | 1 |
| 0 | 1 | 1 | 1 |

| | | | |
|---|---|---|---|
| 0 | 1 | 0 | 0 |
| 0 | 0 | 0 | 1 |
| 0 | 1 | 0 | 0 |
| 0 | 0 | 0 | 1 |
| 0 | 1 | 1 | 1 |
| 0 | 1 | 1 | 1 |
| 0 | 1 | 0 | 0 |
| 0 | 1 | 0 | 0 |
| 0 | 1 | 1 | 1 |
| 0 | 1 | 1 | 1 |
| 0 | 0 | 0 | 1 |
| 0 | 0 | 0 | 1 |
| 0 | 0 | 0 | 1 |
| 0 | 1 | 1 | 1 |
| 0 | 0 | 0 | 0 |
| 0 | 0 | 0 | 0 |
| 0 | 0 | 0 | 0 |
| 0 | 0 | 0 | 0 |
| 0 | 0 | 0 | 1 |
| 0 | 1 | 0 | 0 |
| 0 | 1 | 0 | 0 |
| 0 | 0 | 0 | 0 |
| 0 | 0 | 0 | 0 |
| 0 | 0 | 0 | 0 |
| 0 | 0 | 0 | 1 |
| 0 | 1 | 0 | 0 |
| 0 | 0 | 0 | 1 |
| 0 | 0 | 0 | 1 |
| 0 | 0 | 0 | 1 |
| 0 | 1 | 0 | 0 |
| 0 | 0 | 0 | 1 |
| 0 | 0 | 0 | 1 |
| 0 | 0 | 0 | 0 |
| 0 | 0 | 0 | 0 |
| 0 | 0 | 0 | 1 |
| 0 | 0 | 0 | 1 |
| 0 | 0 | 0 | 1 |
| 0 | 0 | 0 | 1 |
| 0 | 1 | 1 | 1 |
| 0 | 0 | 0 | 0 |
| 0 | 0 | 0 | 0 |
| 0 | 1 | 0 | 0 |
| 0 | 1 | 0 | 0 |
| 0 | 0 | 0 | 0 |
| 0 | 0 | 0 | 0 |
| 0 | 0 | 0 | 1 |
| 0 | 0 | 0 | 1 |
| 0 | 0 | 0 | 0 |
| 0 | 0 | 0 | 0 |
| 0 | 0 | 0 | 1 |
| 0 | 0 | 0 | 0 |
| 0 | 0 | 0 | 0 |
| 0 | 0 | 0 | 1 |
| 0 | 0 | 0 | 1 |

| | | | |
|---|---|---|---|
| 0 | 0 | 0 | 1 |
| 0 | 1 | 0 | 0 |
| 0 | 1 | 0 | 0 |
| 0 | 1 | 1 | 1 |
| 0 | 0 | 0 | 1 |
| 0 | 0 | 0 | 0 |
| 0 | 0 | 0 | 0 |
| 0 | 0 | 0 | 0 |
| 0 | 0 | 0 | 0 |
| 0 | 0 | 0 | 0 |
| 0 | 0 | 0 | 0 |
| 0 | 0 | 0 | 0 |
| 0 | 0 | 0 | 0 |
| 0 | 0 | 0 | 0 |
| 0 | 0 | 0 | 0 |
| 0 | 0 | 0 | 1 |
| 0 | 1 | 0 | 0 |
| 0 | 0 | 0 | 1 |
| 0 | 1 | 1 | 1 |
| 0 | 0 | 0 | 1 |
| 0 | 0 | 0 | 0 |
| 0 | 0 | 0 | 1 |
| 0 | 1 | 0 | 0 |
| 0 | 0 | 0 | 0 |
| 0 | 0 | 0 | 1 |
| 0 | 0 | 0 | 0 |
| 0 | 0 | 0 | 1 |
| 0 | 0 | 0 | 1 |
| 0 | 0 | 0 | 1 |
| 0 | 0 | 0 | 0 |
| 0 | 1 | 0 | 0 |
| 0 | 0 | 0 | 0 |
| 0 | 1 | 0 | 0 |
| 0 | 1 | 0 | 0 |
| 0 | 1 | 0 | 0 |
| 0 | 1 | 0 | 0 |
| 0 | 1 | 0 | 0 |
| 0 | 0 | 0 | 1 |
| 0 | 0 | 0 | 1 |
| 0 | 1 | 0 | 0 |
| 0 | 1 | 0 | 0 |
| 0 | 0 | 0 | 0 |
| 0 | 0 | 0 | 0 |
| 0 | 0 | 0 | 0 |
| 0 | 1 | 0 | 0 |
| 0 | 0 | 0 | 0 |
| 0 | 0 | 0 | 1 |
| 0 | 0 | 0 | 0 |
| 0 | 0 | 0 | 0 |
| 0 | 0 | 0 | 0 |
| 0 | 0 | 0 | 0 |
| 0 | 0 | 0 | 1 |
| 0 | 1 | 0 | 0 |
| 0 | 0 | 0 | 0 |

| | | | |
|---|---|---|---|
| 0 | 1 | 0 | 0 |
| 0 | 1 | 0 | 0 |
| 0 | 0 | 0 | 0 |
| 0 | 0 | 0 | 0 |
| 0 | 0 | 0 | 0 |
| 0 | 0 | 0 | 0 |
| 0 | 0 | 0 | 0 |
| 0 | 0 | 0 | 0 |
| 1 | 0 | 0 | 0 |
| 0 | 0 | 0 | 0 |
| 0 | 0 | 0 | 1 |
| 0 | 1 | 0 | 0 |
| 0 | 1 | 1 | 1 |
| 0 | 0 | 0 | 0 |
| 0 | 0 | 0 | 1 |
| 0 | 0 | 0 | 1 |
| 0 | 0 | 0 | 1 |
| 0 | 0 | 0 | 1 |
| 0 | 0 | 0 | 0 |
| 0 | 0 | 0 | 0 |
| 0 | 0 | 0 | 1 |
| 0 | 1 | 0 | 0 |
| 0 | 1 | 0 | 0 |
| 0 | 0 | 0 | 1 |
| 0 | 0 | 0 | 0 |
| 0 | 0 | 0 | 0 |
| 0 | 0 | 0 | 0 |
| 0 | 0 | 0 | 1 |
| 0 | 1 | 0 | 0 |
| 0 | 0 | 0 | 0 |
| 0 | 1 | 1 | 1 |
| 1 | 1 | 0 | 0 |
| 0 | 1 | 1 | 1 |
| 0 | 1 | 1 | 1 |
| 1 | 1 | 0 | 0 |
| 1 | 1 | 0 | 0 |
| 1 | 1 | 0 | 0 |
| 0 | 1 | 1 | 1 |
| 0 | 1 | 1 | 1 |
| 0 | 1 | 0 | 0 |
| 1 | 0 | 0 | 0 |
| 0 | 1 | 0 | 0 |
| 0 | 1 | 0 | 0 |
| 1 | 1 | 0 | 0 |
| 1 | 1 | 0 | 0 |
| 0 | 1 | 0 | 0 |
| 0 | 0 | 0 | 1 |
| 0 | 1 | 1 | 1 |
| 0 | 1 | 1 | 1 |
| 0 | 1 | 1 | 1 |
| 1 | 1 | 0 | 0 |
| 0 | 1 | 0 | 0 |
| 0 | 1 | 0 | 0 |
| 0 | 1 | 0 | 0 |

| | | | |
|---|---|---|---|
| 0 | 1 | 0 | 0 |
| 0 | 1 | 1 | 1 |
| 0 | 1 | 0 | 0 |
| 0 | 1 | 0 | 0 |
| 0 | 1 | 0 | 0 |
| 0 | 1 | 1 | 1 |
| 0 | 1 | 0 | 0 |
| 0 | 0 | 0 | 0 |
| 0 | 0 | 0 | 1 |
| 0 | 1 | 0 | 0 |
| 0 | 0 | 0 | 1 |
| 0 | 0 | 0 | 1 |
| 0 | 0 | 0 | 1 |
| 0 | 0 | 0 | 0 |
| 0 | 0 | 0 | 1 |
| 0 | 0 | 0 | 0 |
| 0 | 0 | 0 | 1 |
| 0 | 0 | 0 | 1 |
| 0 | 0 | 0 | 0 |
| 0 | 0 | 0 | 0 |
| 0 | 0 | 0 | 0 |
| 0 | 0 | 0 | 0 |
| 0 | 0 | 0 | 0 |
| 0 | 0 | 0 | 0 |
| 0 | 0 | 0 | 0 |
| 0 | 0 | 0 | 1 |
| 0 | 0 | 0 | 1 |
| 0 | 0 | 0 | 1 |
| 0 | 0 | 0 | 1 |
| 0 | 0 | 0 | 1 |
| 0 | 0 | 0 | 1 |
| 0 | 0 | 0 | 1 |
| 0 | 0 | 0 | 0 |
| 0 | 0 | 0 | 0 |
| 0 | 0 | 0 | 0 |
| 0 | 0 | 0 | 0 |
| 0 | 0 | 0 | 0 |
| 0 | 0 | 0 | 0 |
| 0 | 0 | 0 | 1 |
| 0 | 0 | 0 | 1 |
| 0 | 0 | 0 | 1 |
| 0 | 0 | 0 | 1 |
| 0 | 0 | 0 | 1 |
| 0 | 0 | 0 | 1 |
| 0 | 1 | 0 | 0 |
| 0 | 0 | 0 | 0 |
| 0 | 0 | 0 | 1 |
| 0 | 0 | 0 | 0 |
| 0 | 0 | 0 | 0 |
| 0 | 0 | 0 | 0 |
| 0 | 0 | 0 | 0 |
| 0 | 0 | 0 | 0 |
| 0 | 0 | 0 | 0 |
| 0 | 0 | 0 | 0 |

| | | | |
|---|---|---|---|
| 0 | 0 | 0 | 1 |
| 0 | 0 | 0 | 1 |
| 1 | 0 | 0 | 0 |
| 1 | 0 | 0 | 0 |
| 0 | 0 | 0 | 1 |
| 0 | 0 | 0 | 0 |
| 0 | 0 | 0 | 0 |
| 0 | 0 | 0 | 1 |
| 0 | 0 | 0 | 1 |
| 0 | 1 | 0 | 0 |
| 0 | 0 | 0 | 1 |
| 0 | 0 | 0 | 1 |
| 0 | 1 | 0 | 0 |
| 0 | 1 | 0 | 0 |
| 0 | 1 | 0 | 0 |
| 0 | 1 | 0 | 0 |
| 0 | 0 | 0 | 1 |
| 0 | 0 | 0 | 1 |
| 0 | 0 | 0 | 1 |
| 0 | 0 | 0 | 1 |
| 0 | 0 | 0 | 1 |
| 0 | 0 | 0 | 1 |
| 0 | 0 | 0 | 1 |
| 0 | 0 | 0 | 1 |
| 0 | 0 | 0 | 1 |
| 0 | 0 | 0 | 0 |
| 0 | 0 | 0 | 1 |
| 0 | 0 | 0 | 1 |
| 0 | 0 | 0 | 1 |
| 0 | 0 | 0 | 1 |
| 0 | 0 | 0 | 1 |
| 0 | 0 | 0 | 1 |
| 0 | 0 | 0 | 1 |
| 0 | 0 | 0 | 1 |
| 0 | 0 | 0 | 1 |
| 0 | 0 | 0 | 1 |
| 0 | 0 | 0 | 1 |
| 0 | 0 | 0 | 0 |
| 0 | 1 | 1 | 1 |
| 0 | 0 | 0 | 1 |
| 0 | 1 | 0 | 0 |
| 0 | 0 | 0 | 1 |
| 0 | 0 | 0 | 1 |
| 0 | 1 | 0 | 0 |
| 0 | 0 | 0 | 0 |
| 0 | 0 | 0 | 1 |
| 0 | 0 | 0 | 1 |
| 0 | 0 | 0 | 1 |
| 0 | 0 | 0 | 1 |
| 0 | 0 | 0 | 1 |
| 0 | 0 | 0 | 1 |
| 0 | 0 | 0 | 0 |
| 0 | 0 | 0 | 0 |
| 0 | 0 | 0 | 0 |

| | | | |
|---|---|---|---|
| 0 | 0 | 0 | 0 |
| 0 | 0 | 0 | 1 |
| 0 | 0 | 0 | 1 |
| 0 | 0 | 0 | 1 |
| 0 | 0 | 0 | 1 |
| 0 | 0 | 0 | 1 |
| 0 | 0 | 0 | 1 |
| 0 | 0 | 0 | 1 |
| 0 | 0 | 0 | 0 |
| 0 | 0 | 0 | 1 |
| 0 | 0 | 0 | 1 |
| 0 | 0 | 0 | 1 |
| 0 | 0 | 0 | 1 |
| 0 | 0 | 0 | 1 |
| 0 | 0 | 0 | 1 |
| 0 | 0 | 0 | 1 |
| 0 | 0 | 0 | 1 |
| 0 | 0 | 0 | 1 |
| 0 | 0 | 0 | 0 |
| 0 | 0 | 0 | 0 |
| 0 | 0 | 0 | 0 |
| 0 | 1 | 0 | 0 |
| 0 | 0 | 0 | 1 |
| 0 | 0 | 0 | 0 |
| 0 | 0 | 0 | 0 |
| 0 | 0 | 0 | 1 |
| 0 | 0 | 0 | 1 |
| 0 | 0 | 0 | 1 |
| 0 | 0 | 0 | 1 |
| 0 | 0 | 0 | 1 |
| 0 | 0 | 0 | 1 |
| 0 | 0 | 0 | 1 |
| 0 | 0 | 0 | 0 |
| 0 | 0 | 0 | 1 |
| 0 | 0 | 0 | 1 |
| 0 | 0 | 0 | 1 |
| 0 | 0 | 0 | 0 |
| 0 | 0 | 0 | 1 |
| 0 | 0 | 0 | 0 |
| 0 | 0 | 0 | 0 |
| 0 | 0 | 0 | 0 |
| 0 | 0 | 0 | 0 |
| 0 | 0 | 0 | 0 |
| 0 | 0 | 0 | 1 |
| 0 | 0 | 0 | 1 |
| 0 | 0 | 0 | 1 |
| 0 | 0 | 0 | 0 |
| 0 | 0 | 0 | 0 |
| 0 | 0 | 0 | 0 |
| 0 | 1 | 0 | 0 |
| 0 | 1 | 0 | 0 |
| 0 | 1 | 0 | 0 |
| 0 | 1 | 0 | 0 |
| 0 | 0 | 0 | 1 |

| | | | |
|---|---|---|---|
| 0 | 0 | 0 | 0 |
| 0 | 0 | 0 | 0 |
| 0 | 0 | 0 | 1 |
| 1 | 0 | 0 | 0 |
| 1 | 0 | 0 | 0 |
| 0 | 0 | 0 | 1 |
| 0 | 0 | 0 | 1 |
| 0 | 1 | 1 | 1 |
| 0 | 0 | 0 | 1 |
| 0 | 0 | 0 | 1 |
| 0 | 0 | 0 | 1 |
| 0 | 0 | 0 | 0 |
| 0 | 0 | 0 | 1 |
| 0 | 0 | 0 | 1 |
| 0 | 0 | 0 | 0 |
| 0 | 0 | 0 | 1 |
| 0 | 0 | 0 | 0 |
| 0 | 0 | 0 | 1 |
| 0 | 0 | 0 | 0 |
| 1 | 0 | 0 | 0 |
| 0 | 0 | 0 | 1 |
| 0 | 0 | 0 | 1 |
| 0 | 0 | 0 | 0 |
| 0 | 0 | 0 | 1 |
| 0 | 0 | 0 | 0 |
| 0 | 0 | 0 | 1 |
| 0 | 0 | 0 | 1 |
| 0 | 0 | 0 | 0 |
| 0 | 0 | 0 | 1 |
| 0 | 0 | 0 | 0 |
| 0 | 0 | 0 | 1 |
| 0 | 0 | 0 | 0 |
| 0 | 0 | 0 | 0 |
| 0 | 1 | 0 | 0 |
| 0 | 0 | 0 | 1 |
| 0 | 0 | 0 | 0 |
| 0 | 0 | 0 | 1 |
| 0 | 0 | 0 | 1 |
| 0 | 0 | 0 | 0 |
| 0 | 0 | 0 | 1 |
| 0 | 0 | 0 | 1 |
| 0 | 0 | 0 | 1 |
| 0 | 0 | 0 | 1 |
| 0 | 0 | 0 | 0 |
| 0 | 0 | 0 | 0 |
| 0 | 0 | 0 | 1 |
| 0 | 0 | 0 | 1 |
| 0 | 0 | 0 | 1 |
| 0 | 1 | 0 | 0 |
| 0 | 0 | 0 | 1 |
| 0 | 0 | 0 | 0 |
| 0 | 0 | 0 | 1 |
| 0 | 0 | 0 | 1 |
| 0 | 0 | 0 | 0 |

| | | | |
|---|---|---|---|
| 0 | 0 | 0 | 1 |
| 0 | 0 | 0 | 1 |
| 0 | 0 | 0 | 1 |
| 0 | 0 | 0 | 1 |
| 0 | 0 | 0 | 1 |
| 0 | 0 | 0 | 1 |
| 0 | 0 | 0 | 1 |
| 0 | 0 | 0 | 0 |
| 0 | 0 | 0 | 1 |
| 0 | 0 | 0 | 1 |
| 0 | 0 | 0 | 1 |
| 0 | 0 | 0 | 1 |
| 0 | 0 | 0 | 0 |
| 0 | 0 | 0 | 1 |
| 0 | 0 | 0 | 0 |
| 0 | 0 | 0 | 0 |
| 0 | 0 | 0 | 1 |
| 0 | 0 | 0 | 1 |
| 0 | 0 | 0 | 1 |
| 0 | 0 | 0 | 1 |
| 0 | 0 | 0 | 0 |
| 0 | 0 | 0 | 0 |
| 0 | 0 | 0 | 1 |
| 0 | 0 | 0 | 1 |
| 0 | 0 | 0 | 0 |
| 0 | 0 | 0 | 1 |
| 0 | 0 | 0 | 0 |
| 0 | 0 | 0 | 0 |
| 0 | 0 | 0 | 0 |
| 0 | 0 | 0 | 0 |
| 0 | 0 | 0 | 1 |
| 0 | 0 | 0 | 1 |
| 0 | 0 | 0 | 0 |
| 0 | 1 | 0 | 0 |
| 0 | 0 | 0 | 0 |
| 0 | 0 | 0 | 0 |
| 0 | 0 | 0 | 0 |
| 0 | 0 | 0 | 0 |
| 0 | 0 | 0 | 0 |
| 0 | 1 | 1 | 1 |
| 0 | 0 | 0 | 1 |
| 0 | 0 | 0 | 1 |
| 0 | 0 | 0 | 1 |
| 0 | 0 | 0 | 1 |
| 0 | 0 | 0 | 1 |
| 0 | 0 | 0 | 0 |
| 0 | 0 | 0 | 1 |
| 0 | 0 | 0 | 1 |
| 0 | 0 | 0 | 1 |
| 0 | 0 | 0 | 0 |
| 0 | 0 | 0 | 0 |
| 0 | 0 | 0 | 1 |
| 0 | 0 | 0 | 1 |
| 0 | 0 | 0 | 1 |

| | | | |
|---|---|---|---|
| 0 | 0 | 0 | 1 |
| 0 | 0 | 0 | 1 |
| 0 | 0 | 0 | 1 |
| 0 | 0 | 0 | 1 |
| 0 | 0 | 0 | 0 |
| 0 | 0 | 0 | 1 |
| 0 | 0 | 0 | 0 |
| 0 | 0 | 0 | 0 |
| 0 | 0 | 0 | 0 |
| 0 | 0 | 0 | 0 |
| 0 | 0 | 0 | 0 |
| 0 | 0 | 0 | 1 |
| 0 | 0 | 0 | 0 |
| 0 | 0 | 0 | 0 |
| 0 | 0 | 0 | 1 |
| 0 | 0 | 0 | 1 |
| 0 | 0 | 0 | 1 |
| 0 | 0 | 0 | 1 |
| 0 | 0 | 0 | 1 |
| 0 | 0 | 0 | 1 |
| 0 | 0 | 0 | 1 |
| 0 | 0 | 0 | 1 |
| 0 | 0 | 0 | 1 |
| 0 | 0 | 0 | 0 |
| 0 | 0 | 0 | 1 |
| 0 | 0 | 0 | 1 |
| 0 | 0 | 0 | 1 |
| 0 | 0 | 0 | 0 |
| 0 | 1 | 1 | 1 |
| 0 | 1 | 1 | 1 |
| 0 | 1 | 0 | 0 |
| 0 | 1 | 0 | 0 |
| 0 | 0 | 0 | 1 |
| 0 | 0 | 0 | 0 |
| 0 | 0 | 0 | 1 |
| 0 | 0 | 0 | 1 |
| 0 | 0 | 0 | 1 |
| 0 | 0 | 0 | 1 |
| 0 | 0 | 0 | 1 |
| 0 | 0 | 0 | 1 |
| 0 | 0 | 0 | 1 |
| 0 | 0 | 0 | 1 |
| 0 | 0 | 0 | 1 |
| 0 | 0 | 0 | 1 |
| 0 | 0 | 0 | 1 |
| 0 | 0 | 0 | 1 |
| 0 | 0 | 0 | 1 |
| 0 | 0 | 0 | 1 |
| 0 | 0 | 0 | 0 |
| 0 | 0 | 0 | 1 |
| 0 | 0 | 0 | 1 |
| 0 | 1 | 1 | 1 |
| 0 | 1 | 1 | 1 |
| 0 | 0 | 0 | 1 |

| | | | |
|---|---|---|---|
| 0 | 0 | 0 | 1 |
| 0 | 0 | 0 | 1 |
| 0 | 0 | 0 | 1 |
| 0 | 0 | 0 | 1 |
| 0 | 1 | 1 | 1 |
| 0 | 0 | 0 | 1 |
| 0 | 0 | 0 | 1 |
| 0 | 0 | 0 | 1 |
| 0 | 0 | 0 | 1 |
| 0 | 0 | 0 | 1 |
| 0 | 0 | 0 | 0 |
| 0 | 0 | 0 | 0 |
| 0 | 0 | 0 | 0 |
| 1 | 1 | 0 | 0 |
| 1 | 0 | 0 | 0 |
| 0 | 1 | 1 | 1 |
| 1 | 0 | 0 | 0 |
| 0 | 0 | 0 | 0 |
| 0 | 0 | 0 | 0 |
| 0 | 0 | 0 | 0 |
| 0 | 0 | 0 | 0 |
| 1 | 0 | 0 | 0 |
| 0 | 0 | 0 | 1 |
| 0 | 0 | 0 | 0 |
| 1 | 0 | 0 | 0 |
| 1 | 0 | 0 | 0 |
| 0 | 0 | 0 | 1 |
| 0 | 0 | 0 | 0 |
| 0 | 0 | 0 | 1 |
| 0 | 0 | 0 | 1 |
| 0 | 0 | 0 | 1 |
| 0 | 0 | 0 | 0 |
| 0 | 0 | 0 | 1 |
| 0 | 0 | 0 | 1 |
| 0 | 0 | 0 | 0 |
| 0 | 0 | 0 | 0 |
| 0 | 0 | 0 | 0 |
| 0 | 1 | 0 | 0 |
| 0 | 0 | 0 | 0 |
| 0 | 1 | 1 | 1 |
| 0 | 1 | 1 | 1 |
| 0 | 1 | 0 | 0 |
| 0 | 1 | 1 | 1 |
| 0 | 0 | 0 | 0 |
| 0 | 0 | 0 | 1 |
| 0 | 1 | 1 | 1 |
| 0 | 0 | 0 | 1 |
| 0 | 1 | 0 | 0 |
| 0 | 0 | 0 | 0 |
| 0 | 1 | 0 | 0 |
| 0 | 1 | 0 | 0 |
| 0 | 0 | 0 | 1 |
| 0 | 0 | 0 | 0 |
| 0 | 0 | 0 | 1 |

| | | | |
|---|---|---|---|
| 0 | 0 | 0 | 1 |
| 0 | 0 | 0 | 1 |
| 0 | 0 | 0 | 1 |
| 0 | 0 | 0 | 1 |
| 0 | 0 | 0 | 1 |
| 0 | 0 | 0 | 0 |
| 0 | 0 | 0 | 0 |
| 0 | 0 | 0 | 0 |
| 0 | 0 | 0 | 0 |
| 0 | 0 | 0 | 1 |
| 1 | 0 | 0 | 0 |
| 1 | 0 | 0 | 0 |
| 0 | 0 | 0 | 0 |
| 0 | 0 | 0 | 0 |
| 0 | 0 | 0 | 0 |
| 0 | 0 | 0 | 0 |
| 0 | 0 | 0 | 1 |
| 0 | 0 | 0 | 1 |
| 0 | 0 | 0 | 1 |
| 0 | 0 | 0 | 1 |
| 0 | 0 | 0 | 1 |
| 0 | 0 | 0 | 1 |
| 0 | 0 | 0 | 1 |
| 0 | 0 | 0 | 1 |
| 0 | 0 | 0 | 1 |
| 0 | 0 | 0 | 1 |
| 0 | 0 | 0 | 1 |
| 0 | 0 | 0 | 1 |
| 0 | 1 | 0 | 0 |
| 0 | 0 | 0 | 1 |
| 0 | 0 | 0 | 1 |
| 0 | 1 | 1 | 1 |
| 0 | 1 | 1 | 1 |
| 0 | 0 | 0 | 0 |
| 0 | 0 | 0 | 1 |
| 0 | 1 | 0 | 0 |
| 0 | 0 | 0 | 1 |
| 0 | 0 | 0 | 0 |
| 0 | 0 | 0 | 1 |
| 0 | 0 | 0 | 0 |
| 0 | 0 | 0 | 1 |
| 0 | 0 | 0 | 1 |
| 0 | 0 | 0 | 1 |
| 0 | 0 | 0 | 0 |
| 0 | 0 | 0 | 1 |
| 0 | 0 | 0 | 1 |
| 0 | 0 | 0 | 1 |
| 0 | 0 | 0 | 1 |
| 0 | 0 | 0 | 1 |
| 0 | 0 | 0 | 1 |
| 0 | 0 | 0 | 1 |
| 0 | 1 | 1 | 1 |
| 0 | 0 | 0 | 0 |
| 0 | 0 | 0 | 0 |

| | | | |
|---|---|---|---|
| 0 | 0 | 0 | 1 |
| 0 | 0 | 0 | 0 |
| 0 | 0 | 0 | 1 |
| 0 | 0 | 0 | 1 |
| 0 | 1 | 0 | 0 |
| 0 | 0 | 0 | 0 |
| 0 | 0 | 0 | 0 |
| 0 | 0 | 0 | 1 |
| 0 | 0 | 0 | 1 |
| 0 | 0 | 0 | 0 |
| 0 | 0 | 0 | 0 |
| 0 | 0 | 0 | 0 |
| 0 | 0 | 0 | 0 |
| 0 | 0 | 0 | 0 |
| 0 | 0 | 0 | 1 |
| 0 | 0 | 0 | 0 |
| 0 | 1 | 0 | 0 |
| 0 | 0 | 0 | 0 |
| 0 | 0 | 0 | 1 |
| 0 | 0 | 0 | 1 |
| 0 | 0 | 0 | 1 |
| 0 | 0 | 0 | 1 |
| 0 | 0 | 0 | 1 |
| 0 | 0 | 0 | 0 |
| 0 | 0 | 0 | 1 |
| 0 | 0 | 0 | 1 |
| 0 | 0 | 0 | 0 |
| 0 | 0 | 0 | 0 |
| 0 | 0 | 0 | 1 |
| 0 | 0 | 0 | 1 |
| 0 | 0 | 0 | 0 |
| 0 | 0 | 0 | 1 |
| 0 | 0 | 0 | 1 |
| 0 | 0 | 0 | 1 |
| 0 | 0 | 0 | 1 |
| 0 | 0 | 0 | 0 |
| 0 | 0 | 0 | 1 |
| 0 | 0 | 0 | 0 |
| 0 | 0 | 0 | 0 |
| 0 | 0 | 0 | 1 |
| 0 | 0 | 0 | 1 |
| 0 | 0 | 0 | 0 |
| 0 | 0 | 0 | 0 |
| 0 | 0 | 0 | 1 |
| 0 | 0 | 0 | 0 |
| 0 | 0 | 0 | 0 |
| 0 | 0 | 0 | 0 |
| 0 | 0 | 0 | 1 |
| 0 | 0 | 0 | 0 |
| 0 | 0 | 0 | 0 |
| 0 | 0 | 0 | 0 |
| 0 | 0 | 0 | 1 |
| 0 | 0 | 0 | 0 |
| 0 | 0 | 0 | 1 |

| | | | |
|---|---|---|---|
| 0 | 0 | 0 | 1 |
| 0 | 0 | 0 | 1 |
| 0 | 0 | 0 | 1 |
| 0 | 0 | 0 | 1 |
| 0 | 0 | 0 | 0 |
| 0 | 0 | 0 | 0 |
| 0 | 0 | 0 | 1 |
| 0 | 0 | 0 | 1 |
| 0 | 0 | 0 | 1 |
| 0 | 1 | 0 | 0 |
| 0 | 1 | 1 | 1 |
| 0 | 1 | 1 | 1 |
| 0 | 1 | 0 | 0 |
| 0 | 0 | 0 | 1 |
| 0 | 0 | 0 | 1 |
| 0 | 0 | 0 | 1 |
| 0 | 0 | 0 | 1 |
| 0 | 0 | 0 | 1 |
| 0 | 0 | 0 | 1 |
| 0 | 0 | 0 | 1 |
| 0 | 0 | 0 | 1 |
| 0 | 0 | 0 | 1 |
| 0 | 0 | 0 | 1 |
| 0 | 0 | 0 | 0 |
| 0 | 0 | 0 | 1 |
| 1 | 0 | 0 | 0 |
| 0 | 0 | 0 | 1 |
| 0 | 0 | 0 | 0 |
| 0 | 0 | 0 | 1 |
| 0 | 0 | 0 | 0 |
| 0 | 0 | 0 | 0 |
| 0 | 0 | 0 | 0 |
| 0 | 0 | 0 | 0 |
| 0 | 0 | 0 | 0 |
| 0 | 0 | 0 | 1 |
| 0 | 0 | 0 | 0 |
| 0 | 0 | 0 | 1 |
| 0 | 0 | 0 | 0 |
| 0 | 0 | 0 | 1 |
| 0 | 0 | 0 | 1 |
| 0 | 0 | 0 | 0 |
| 0 | 0 | 0 | 0 |
| 0 | 0 | 0 | 0 |
| 0 | 0 | 0 | 0 |
| 0 | 0 | 0 | 0 |
| 0 | 0 | 0 | 0 |
| 0 | 0 | 0 | 0 |
| 0 | 1 | 0 | 0 |
| 0 | 0 | 0 | 1 |
| 0 | 0 | 0 | 0 |
| 0 | 1 | 1 | 1 |
| 0 | 1 | 1 | 1 |
| 0 | 0 | 0 | 1 |
| 0 | 0 | 0 | 1 |

| | | | |
|---|---|---|---|
| 0 | 0 | 0 | 1 |
| 0 | 0 | 0 | 1 |
| 0 | 0 | 0 | 1 |
| 0 | 0 | 0 | 1 |
| 0 | 0 | 0 | 1 |
| 0 | 0 | 0 | 1 |
| 0 | 0 | 0 | 0 |
| 0 | 0 | 0 | 0 |
| 0 | 0 | 0 | 0 |
| 0 | 0 | 0 | 0 |
| 0 | 0 | 0 | 0 |
| 0 | 0 | 0 | 0 |
| 0 | 0 | 0 | 0 |
| 0 | 0 | 0 | 0 |
| 0 | 0 | 0 | 0 |
| 0 | 0 | 0 | 0 |
| 1 | 0 | 0 | 0 |
| 0 | 0 | 0 | 0 |
| 0 | 0 | 0 | 1 |
| 0 | 0 | 0 | 0 |
| 0 | 0 | 0 | 1 |
| 0 | 0 | 0 | 1 |
| 0 | 0 | 0 | 1 |
| 0 | 0 | 0 | 1 |
| 0 | 0 | 0 | 1 |
| 0 | 0 | 0 | 0 |
| 0 | 0 | 0 | 0 |
| 0 | 0 | 0 | 0 |
| 0 | 0 | 0 | 0 |
| 0 | 0 | 0 | 0 |
| 0 | 0 | 0 | 0 |
| 1 | 0 | 0 | 0 |
| 0 | 0 | 0 | 0 |
| 1 | 0 | 0 | 0 |
| 0 | 0 | 0 | 1 |
| 1 | 0 | 0 | 0 |
| 1 | 0 | 0 | 0 |
| 0 | 0 | 0 | 0 |
| 1 | 0 | 0 | 0 |
| 0 | 0 | 0 | 1 |
| 0 | 0 | 0 | 0 |
| 0 | 0 | 0 | 0 |
| 0 | 0 | 0 | 1 |
| 0 | 0 | 0 | 1 |
| 0 | 0 | 0 | 1 |
| 0 | 0 | 0 | 1 |
| 0 | 0 | 0 | 1 |
| 0 | 0 | 0 | 0 |
| 0 | 0 | 0 | 0 |
| 0 | 0 | 0 | 0 |
| 0 | 0 | 0 | 0 |
| 0 | 0 | 0 | 1 |
| 0 | 0 | 0 | 0 |
| 0 | 0 | 0 | 1 |

| | | | |
|---|---|---|---|
| 0 | 0 | 0 | 1 |
| 0 | 0 | 0 | 1 |
| 0 | 0 | 0 | 1 |
| 0 | 0 | 0 | 1 |
| 0 | 0 | 0 | 1 |
| 0 | 0 | 0 | 0 |
| 0 | 0 | 0 | 1 |
| 0 | 0 | 0 | 1 |
| 0 | 0 | 0 | 1 |
| 0 | 0 | 0 | 1 |
| 0 | 1 | 0 | 0 |
| 0 | 1 | 0 | 0 |
| 0 | 1 | 0 | 0 |
| 0 | 1 | 1 | 1 |
| 0 | 1 | 1 | 1 |
| 0 | 0 | 0 | 1 |
| 0 | 1 | 1 | 1 |
| 0 | 0 | 0 | 0 |
| 0 | 0 | 0 | 1 |
| 0 | 0 | 0 | 1 |
| 0 | 0 | 0 | 1 |
| 0 | 0 | 0 | 1 |
| 0 | 0 | 0 | 0 |
| 0 | 0 | 0 | 1 |
| 0 | 0 | 0 | 0 |
| 0 | 0 | 0 | 0 |
| 0 | 0 | 0 | 0 |
| 0 | 0 | 0 | 0 |
| 0 | 0 | 0 | 1 |
| 0 | 0 | 0 | 0 |
| 0 | 0 | 0 | 1 |
| 0 | 0 | 0 | 1 |
| 0 | 0 | 0 | 1 |
| 0 | 0 | 0 | 1 |
| 0 | 0 | 0 | 1 |
| 0 | 0 | 0 | 1 |
| 0 | 0 | 0 | 1 |
| 0 | 0 | 0 | 0 |
| 0 | 0 | 0 | 0 |
| 0 | 0 | 0 | 0 |
| 0 | 1 | 0 | 0 |
| 0 | 0 | 0 | 0 |
| 0 | 0 | 0 | 1 |
| 0 | 0 | 0 | 0 |
| 0 | 0 | 0 | 0 |
| 0 | 0 | 0 | 1 |
| 0 | 0 | 0 | 0 |
| 0 | 1 | 0 | 0 |
| 0 | 0 | 0 | 0 |
| 1 | 0 | 0 | 0 |
| 0 | 0 | 0 | 1 |
| 0 | 0 | 0 | 1 |
| 0 | 0 | 0 | 1 |
| 0 | 1 | 0 | 0 |

| | | | |
|---|---|---|---|
| 0 | 0 | 0 | 0 |
| 0 | 0 | 0 | 1 |
| 0 | 1 | 0 | 0 |
| 0 | 0 | 0 | 1 |
| 0 | 0 | 0 | 1 |
| 0 | 0 | 0 | 1 |
| 0 | 0 | 0 | 1 |
| 0 | 0 | 0 | 1 |
| 0 | 1 | 0 | 0 |
| 0 | 0 | 0 | 1 |
| 0 | 0 | 0 | 0 |
| 0 | 0 | 0 | 0 |
| 0 | 0 | 0 | 0 |
| 0 | 0 | 0 | 0 |
| 0 | 0 | 0 | 1 |
| 0 | 0 | 0 | 1 |
| 0 | 0 | 0 | 0 |
| 0 | 0 | 0 | 0 |
| 0 | 0 | 0 | 1 |
| 0 | 0 | 0 | 1 |
| 0 | 0 | 0 | 0 |
| 0 | 0 | 0 | 1 |
| 0 | 0 | 0 | 1 |
| 0 | 0 | 0 | 1 |
| 0 | 1 | 0 | 0 |
| 0 | 1 | 1 | 1 |
| 0 | 1 | 0 | 0 |
| 0 | 1 | 0 | 0 |
| 0 | 1 | 0 | 0 |
| 0 | 0 | 0 | 1 |
| 0 | 0 | 0 | 0 |
| 0 | 0 | 0 | 0 |
| 0 | 0 | 0 | 0 |
| 0 | 0 | 0 | 1 |
| 0 | 0 | 0 | 0 |
| 0 | 0 | 0 | 1 |
| 0 | 0 | 0 | 0 |
| 0 | 0 | 0 | 0 |
| 0 | 1 | 0 | 0 |
| 0 | 0 | 0 | 0 |
| 0 | 0 | 0 | 0 |
| 0 | 0 | 0 | 1 |
| 0 | 1 | 0 | 0 |
| 0 | 1 | 0 | 0 |
| 0 | 1 | 0 | 0 |
| 1 | 0 | 0 | 0 |
| 0 | 0 | 0 | 0 |
| 1 | 0 | 0 | 0 |
| 1 | 0 | 0 | 0 |
| 1 | 0 | 0 | 0 |
| 0 | 0 | 0 | 0 |
| 0 | 0 | 0 | 0 |
| 0 | 0 | 0 | 1 |
| 0 | 0 | 0 | 0 |

| | | | |
|---|---|---|---|
| 0 | 0 | 0 | 1 |
| 0 | 0 | 0 | 1 |
| 0 | 0 | 0 | 1 |
| 0 | 0 | 0 | 1 |
| 0 | 0 | 0 | 1 |
| 0 | 0 | 0 | 1 |
| 0 | 0 | 0 | 1 |
| 0 | 0 | 0 | 1 |
| 0 | 0 | 0 | 1 |
| 0 | 0 | 0 | 0 |
| 0 | 0 | 0 | 0 |
| 0 | 0 | 0 | 1 |
| 0 | 0 | 0 | 1 |
| 0 | 0 | 0 | 1 |
| 0 | 0 | 0 | 0 |
| 0 | 0 | 0 | 1 |
| 0 | 0 | 0 | 1 |
| 0 | 0 | 0 | 0 |
| 0 | 0 | 0 | 1 |
| 0 | 0 | 0 | 1 |
| 0 | 0 | 0 | 1 |
| 0 | 1 | 0 | 0 |
| 0 | 0 | 0 | 0 |
| 0 | 1 | 0 | 0 |
| 0 | 1 | 0 | 0 |
| 0 | 1 | 0 | 0 |
| 0 | 1 | 0 | 0 |
| 0 | 0 | 0 | 0 |
| 0 | 0 | 0 | 1 |
| 0 | 0 | 0 | 0 |
| 1 | 0 | 0 | 0 |
| 0 | 0 | 0 | 1 |
| 0 | 0 | 0 | 1 |
| 0 | 0 | 0 | 1 |
| 0 | 0 | 0 | 1 |
| 0 | 0 | 0 | 1 |
| 0 | 0 | 0 | 1 |
| 0 | 0 | 0 | 1 |
| 0 | 0 | 0 | 1 |
| 0 | 0 | 0 | 1 |
| 0 | 0 | 0 | 1 |
| 0 | 0 | 0 | 1 |
| 0 | 0 | 0 | 1 |
| 0 | 0 | 0 | 0 |
| 0 | 0 | 0 | 1 |
| 0 | 0 | 0 | 0 |
| 0 | 0 | 0 | 1 |
| 0 | 0 | 0 | 1 |
| 0 | 0 | 0 | 1 |
| 0 | 0 | 0 | 1 |
| 0 | 0 | 0 | 0 |
| 0 | 0 | 0 | 1 |
| 0 | 0 | 0 | 1 |
| 0 | 0 | 0 | 0 |

| | | | |
|---|---|---|---|
| 0 | 0 | 0 | 0 |
| 0 | 0 | 0 | 0 |
| 0 | 0 | 0 | 0 |
| 0 | 1 | 0 | 0 |
| 0 | 0 | 0 | 1 |
| 0 | 0 | 0 | 1 |
| 0 | 0 | 0 | 1 |
| 0 | 0 | 0 | 0 |
| 0 | 0 | 0 | 0 |
| 0 | 0 | 0 | 0 |
| 0 | 0 | 0 | 0 |
| 0 | 0 | 0 | 1 |
| 0 | 0 | 0 | 1 |
| 0 | 0 | 0 | 1 |
| 0 | 0 | 0 | 1 |
| 0 | 0 | 0 | 0 |
| 0 | 0 | 0 | 0 |
| 0 | 0 | 0 | 1 |
| 0 | 0 | 0 | 1 |
| 0 | 0 | 0 | 1 |
| 0 | 0 | 0 | 1 |
| 0 | 1 | 0 | 0 |
| 0 | 0 | 0 | 1 |
| 0 | 0 | 0 | 1 |
| 0 | 1 | 0 | 0 |
| 0 | 0 | 0 | 0 |
| 0 | 0 | 0 | 0 |
| 0 | 0 | 0 | 0 |
| 0 | 0 | 0 | 1 |
| 0 | 0 | 0 | 1 |
| 0 | 0 | 0 | 1 |
| 0 | 0 | 0 | 0 |
| 0 | 0 | 0 | 1 |
| 0 | 0 | 0 | 1 |
| 0 | 0 | 0 | 1 |
| 0 | 0 | 0 | 1 |
| 0 | 0 | 0 | 1 |
| 0 | 0 | 0 | 1 |
| 0 | 0 | 0 | 1 |
| 0 | 0 | 0 | 1 |
| 0 | 0 | 0 | 1 |
| 0 | 0 | 0 | 1 |
| 0 | 0 | 0 | 0 |
| 0 | 0 | 0 | 1 |
| 0 | 0 | 0 | 1 |
| 0 | 0 | 0 | 1 |
| 0 | 0 | 0 | 1 |
| 0 | 0 | 0 | 1 |
| 0 | 0 | 0 | 1 |
| 0 | 0 | 0 | 0 |
| 0 | 0 | 0 | 1 |
| 0 | 0 | 0 | 1 |
| 0 | 0 | 0 | 1 |
| 0 | 0 | 0 | 1 |

| | | | |
|---|---|---|---|
| 0 | 0 | 0 | 0 |
| 0 | 0 | 0 | 1 |
| 0 | 0 | 0 | 1 |
| 0 | 0 | 0 | 1 |
| 0 | 0 | 0 | 1 |
| 0 | 0 | 0 | 1 |
| 0 | 0 | 0 | 1 |
| 0 | 1 | 0 | 0 |
| 0 | 0 | 0 | 1 |
| 0 | 0 | 0 | 1 |
| 0 | 0 | 0 | 0 |
| 0 | 0 | 0 | 0 |
| 0 | 0 | 0 | 0 |
| 0 | 0 | 0 | 1 |
| 0 | 0 | 0 | 0 |
| 0 | 0 | 0 | 0 |
| 0 | 0 | 0 | 0 |
| 0 | 0 | 0 | 0 |
| 0 | 0 | 0 | 1 |
| 0 | 0 | 0 | 0 |
| 0 | 0 | 0 | 0 |
| 0 | 0 | 0 | 0 |
| 0 | 0 | 0 | 0 |
| 0 | 0 | 0 | 0 |
| 0 | 0 | 0 | 0 |
| 0 | 0 | 0 | 0 |
| 0 | 0 | 0 | 0 |
| 0 | 0 | 0 | 0 |
| 0 | 0 | 0 | 0 |
| 0 | 0 | 0 | 0 |
| 0 | 0 | 0 | 0 |
| 0 | 0 | 0 | 0 |
| 0 | 0 | 0 | 1 |
| 0 | 0 | 0 | 0 |
| 0 | 0 | 0 | 1 |
| 0 | 0 | 0 | 0 |
| 0 | 0 | 0 | 0 |
| 0 | 0 | 0 | 1 |
| 0 | 0 | 0 | 1 |
| 0 | 0 | 0 | 1 |
| 0 | 0 | 0 | 1 |
| 0 | 0 | 0 | 0 |
| 0 | 0 | 0 | 0 |
| 0 | 0 | 0 | 0 |
| 0 | 0 | 0 | 1 |
| 0 | 0 | 0 | 0 |
| 0 | 0 | 0 | 1 |
| 0 | 0 | 0 | 0 |
| 0 | 0 | 0 | 1 |
| 0 | 0 | 0 | 1 |
| 0 | 0 | 0 | 1 |
| 0 | 0 | 0 | 1 |
| 0 | 0 | 0 | 1 |
| 0 | 0 | 0 | 1 |

| | | | |
|---|---|---|---|
| 0 | 0 | 0 | 1 |
| 0 | 0 | 0 | 1 |
| 0 | 0 | 0 | 1 |
| 0 | 0 | 0 | 1 |
| 0 | 0 | 0 | 1 |
| 0 | 0 | 0 | 0 |
| 0 | 0 | 0 | 0 |
| 0 | 0 | 0 | 1 |
| 0 | 0 | 0 | 0 |
| 0 | 0 | 0 | 0 |
| 0 | 0 | 0 | 1 |
| 0 | 0 | 0 | 1 |
| 0 | 0 | 0 | 1 |
| 0 | 0 | 0 | 1 |
| 0 | 0 | 0 | 1 |
| 0 | 1 | 1 | 1 |
| 0 | 0 | 0 | 1 |
| 0 | 0 | 0 | 1 |
| 0 | 1 | 0 | 0 |
| 0 | 1 | 0 | 0 |
| 0 | 0 | 0 | 1 |
| 0 | 0 | 0 | 1 |
| 0 | 0 | 0 | 1 |
| 0 | 0 | 0 | 0 |
| 0 | 0 | 0 | 1 |
| 0 | 0 | 0 | 0 |
| 0 | 0 | 0 | 0 |
| 0 | 0 | 0 | 1 |
| 0 | 0 | 0 | 1 |
| 0 | 0 | 0 | 1 |
| 0 | 1 | 0 | 0 |
| 0 | 0 | 0 | 1 |
| 0 | 0 | 0 | 0 |
| 0 | 1 | 0 | 0 |
| 0 | 1 | 0 | 0 |
| 0 | 1 | 0 | 0 |
| 0 | 0 | 0 | 0 |
| 0 | 0 | 0 | 1 |
| 0 | 0 | 0 | 1 |
| 0 | 0 | 0 | 1 |
| 0 | 1 | 0 | 0 |
| 0 | 1 | 0 | 0 |
| 0 | 1 | 0 | 0 |
| 0 | 0 | 0 | 0 |
| 0 | 0 | 0 | 1 |
| 0 | 0 | 0 | 1 |
| 0 | 0 | 0 | 1 |
| 0 | 0 | 0 | 0 |
| 0 | 0 | 0 | 0 |
| 0 | 0 | 0 | 0 |
| 0 | 0 | 0 | 1 |
| 0 | 0 | 0 | 1 |
| 0 | 0 | 0 | 1 |
| 0 | 0 | 0 | 1 |

| | | | |
|---|---|---|---|
| 0 | 0 | 0 | 1 |
| 0 | 0 | 0 | 1 |
| 0 | 0 | 0 | 1 |
| 0 | 0 | 0 | 0 |
| 0 | 0 | 0 | 0 |
| 0 | 0 | 0 | 1 |
| 0 | 0 | 0 | 0 |
| 0 | 0 | 0 | 1 |
| 0 | 0 | 0 | 1 |
| 0 | 0 | 0 | 1 |
| 0 | 0 | 0 | 1 |
| 0 | 0 | 0 | 0 |
| 0 | 1 | 1 | 1 |
| 0 | 0 | 0 | 0 |
| 0 | 0 | 0 | 1 |
| 0 | 0 | 0 | 0 |
| 0 | 1 | 1 | 1 |
| 0 | 1 | 1 | 1 |
| 0 | 0 | 0 | 1 |
| 0 | 1 | 0 | 0 |
| 0 | 0 | 0 | 0 |
| 0 | 0 | 0 | 0 |
| 0 | 0 | 0 | 0 |
| 0 | 0 | 0 | 0 |
| 0 | 0 | 0 | 0 |
| 0 | 0 | 0 | 1 |
| 0 | 0 | 0 | 0 |
| 0 | 0 | 0 | 1 |
| 0 | 0 | 0 | 1 |
| 0 | 0 | 0 | 1 |
| 0 | 0 | 0 | 1 |
| 0 | 0 | 0 | 1 |
| 0 | 0 | 0 | 0 |
| 0 | 1 | 0 | 0 |
| 0 | 0 | 0 | 1 |
| 0 | 0 | 0 | 1 |
| 0 | 1 | 0 | 0 |
| 0 | 1 | 0 | 0 |
| 0 | 1 | 0 | 0 |
| 0 | 1 | 1 | 1 |
| 0 | 0 | 0 | 0 |
| 0 | 0 | 0 | 0 |
| 0 | 0 | 0 | 1 |
| 0 | 0 | 0 | 0 |
| 0 | 0 | 0 | 1 |
| 0 | 0 | 0 | 1 |
| 0 | 0 | 0 | 1 |
| 0 | 0 | 0 | 1 |
| 0 | 0 | 0 | 1 |
| 1 | 0 | 0 | 0 |
| 0 | 0 | 0 | 1 |
| 0 | 0 | 0 | 1 |
| 0 | 0 | 0 | 1 |
| 0 | 0 | 0 | 1 |

| | | | |
|---|---|---|---|
| 0 | 0 | 0 | 1 |
| 0 | 0 | 0 | 1 |
| 0 | 0 | 0 | 1 |
| 0 | 0 | 0 | 0 |
| 0 | 0 | 0 | 1 |
| 0 | 0 | 0 | 1 |
| 0 | 0 | 0 | 1 |
| 0 | 0 | 0 | 0 |
| 0 | 0 | 0 | 1 |
| 0 | 0 | 0 | 1 |
| 0 | 0 | 0 | 1 |
| 0 | 0 | 0 | 1 |
| 0 | 0 | 0 | 1 |
| 0 | 0 | 0 | 1 |
| 0 | 0 | 0 | 1 |
| 0 | 0 | 0 | 1 |
| 0 | 0 | 0 | 0 |
| 0 | 0 | 0 | 0 |
| 0 | 0 | 0 | 1 |
| 0 | 0 | 0 | 1 |
| 0 | 0 | 0 | 1 |
| 0 | 0 | 0 | 1 |
| 0 | 0 | 0 | 1 |
| 0 | 0 | 0 | 1 |
| 0 | 0 | 0 | 1 |
| 0 | 0 | 0 | 1 |
| 0 | 0 | 0 | 1 |
| 0 | 1 | 0 | 0 |
| 0 | 0 | 0 | 0 |
| 0 | 0 | 0 | 1 |
| 0 | 0 | 0 | 1 |
| 0 | 0 | 0 | 0 |
| 0 | 0 | 0 | 1 |
| 0 | 0 | 0 | 0 |
| 1 | 0 | 0 | 0 |
| 1 | 0 | 0 | 0 |
| 1 | 0 | 0 | 0 |
| 1 | 0 | 0 | 0 |
| 0 | 0 | 0 | 1 |
| 0 | 0 | 0 | 1 |
| 0 | 0 | 0 | 1 |
| 0 | 0 | 0 | 1 |
| 0 | 0 | 0 | 1 |
| 0 | 0 | 0 | 1 |
| 0 | 0 | 0 | 0 |
| 0 | 0 | 0 | 0 |
| 0 | 0 | 0 | 0 |
| 0 | 0 | 0 | 1 |
| 0 | 0 | 0 | 1 |
| 0 | 0 | 0 | 1 |
| 0 | 0 | 0 | 1 |
| 0 | 0 | 0 | 0 |
| 0 | 0 | 0 | 0 |
| 0 | 0 | 0 | 0 |

| | | | |
|---|---|---|---|
| 0 | 0 | 0 | 0 |
| 0 | 0 | 0 | 0 |
| 0 | 0 | 0 | 1 |
| 0 | 0 | 0 | 0 |
| 0 | 0 | 0 | 0 |
| 0 | 0 | 0 | 0 |
| 0 | 0 | 0 | 0 |
| 0 | 0 | 0 | 1 |
| 0 | 0 | 0 | 1 |
| 0 | 0 | 0 | 1 |
| 0 | 0 | 0 | 1 |
| 0 | 0 | 0 | 1 |
| 0 | 0 | 0 | 1 |
| 0 | 0 | 0 | 1 |
| 0 | 0 | 0 | 1 |
| 0 | 0 | 0 | 1 |
| 0 | 0 | 0 | 1 |
| 0 | 0 | 0 | 1 |
| 0 | 0 | 0 | 1 |
| 0 | 0 | 0 | 1 |
| 0 | 0 | 0 | 0 |
| 0 | 0 | 0 | 0 |
| 0 | 0 | 0 | 0 |
| 0 | 0 | 0 | 0 |
| 0 | 0 | 0 | 0 |
| 0 | 0 | 0 | 1 |
| 0 | 0 | 0 | 1 |
| 0 | 0 | 0 | 0 |
| 0 | 0 | 0 | 1 |
| 0 | 0 | 0 | 1 |
| 0 | 0 | 0 | 0 |
| 0 | 0 | 0 | 0 |
| 0 | 0 | 0 | 0 |
| 0 | 0 | 0 | 1 |
| 0 | 0 | 0 | 0 |
| 0 | 0 | 0 | 1 |
| 0 | 0 | 0 | 1 |
| 0 | 0 | 0 | 0 |
| 0 | 0 | 0 | 1 |
| 0 | 0 | 0 | 1 |
| 0 | 0 | 0 | 1 |
| 0 | 0 | 0 | 1 |
| 0 | 0 | 0 | 1 |
| 0 | 0 | 0 | 1 |
| 0 | 0 | 0 | 1 |
| 0 | 0 | 0 | 1 |
| 0 | 0 | 0 | 1 |
| 0 | 0 | 0 | 1 |
| 0 | 0 | 0 | 0 |
| 0 | 0 | 0 | 1 |
| 1 | 0 | 0 | 0 |
| 1 | 0 | 0 | 0 |
| 0 | 0 | 0 | 1 |
| 0 | 0 | 0 | 1 |

| | | | |
|---|---|---|---|
| 1 | 0 | 0 | 0 |
| 0 | 0 | 0 | 1 |
| 0 | 0 | 0 | 1 |
| 0 | 0 | 0 | 1 |
| 0 | 0 | 0 | 1 |
| 1 | 0 | 0 | 0 |
| 1 | 0 | 0 | 0 |
| 0 | 0 | 0 | 1 |
| 0 | 0 | 0 | 1 |
| 0 | 0 | 0 | 1 |
| 0 | 0 | 0 | 1 |
| 0 | 0 | 0 | 0 |
| 0 | 0 | 0 | 1 |
| 0 | 0 | 0 | 1 |
| 0 | 0 | 0 | 1 |
| 0 | 0 | 0 | 1 |
| 0 | 0 | 0 | 0 |
| 0 | 0 | 0 | 0 |
| 0 | 0 | 0 | 1 |
| 0 | 0 | 0 | 1 |
| 0 | 0 | 0 | 1 |
| 0 | 0 | 0 | 1 |
| 0 | 0 | 0 | 1 |
| 0 | 0 | 0 | 1 |
| 0 | 0 | 0 | 0 |
| 0 | 0 | 0 | 1 |
| 0 | 0 | 0 | 1 |
| 0 | 0 | 0 | 1 |
| 0 | 0 | 0 | 1 |
| 0 | 0 | 0 | 1 |
| 0 | 0 | 0 | 1 |
| 0 | 0 | 0 | 0 |
| 0 | 0 | 0 | 0 |
| 0 | 0 | 0 | 1 |
| 0 | 0 | 0 | 1 |
| 0 | 0 | 0 | 0 |
| 1 | 0 | 0 | 0 |
| 0 | 1 | 0 | 0 |
| 1 | 1 | 0 | 0 |
| 0 | 0 | 0 | 0 |
| 0 | 0 | 0 | 0 |
| 0 | 0 | 0 | 0 |
| 0 | 0 | 0 | 0 |
| 0 | 0 | 0 | 0 |
| 0 | 0 | 0 | 0 |
| 0 | 0 | 0 | 0 |
| 0 | 0 | 0 | 0 |
| 1 | 0 | 0 | 0 |
| 0 | 0 | 0 | 1 |
| 0 | 0 | 0 | 1 |
| 0 | 1 | 1 | 1 |
| 0 | 0 | 0 | 1 |
| 0 | 0 | 0 | 0 |
| 0 | 0 | 0 | 1 |

| | | | |
|---|---|---|---|
| 0 | 0 | 0 | 1 |
| 0 | 0 | 0 | 1 |
| 0 | 0 | 0 | 1 |
| 0 | 0 | 0 | 1 |
| 0 | 0 | 0 | 1 |
| 0 | 0 | 0 | 1 |
| 0 | 0 | 0 | 1 |
| 0 | 1 | 0 | 0 |
| 0 | 0 | 0 | 0 |
| 0 | 1 | 1 | 1 |
| 0 | 0 | 0 | 1 |
| 0 | 0 | 0 | 1 |
| 0 | 0 | 0 | 1 |
| 0 | 1 | 0 | 0 |
| 0 | 0 | 0 | 1 |
| 0 | 0 | 0 | 1 |
| 0 | 1 | 0 | 0 |
| 0 | 0 | 0 | 1 |
| 1 | 0 | 0 | 0 |
| 0 | 0 | 0 | 1 |
| 0 | 0 | 0 | 1 |
| 0 | 0 | 0 | 1 |
| 0 | 0 | 0 | 1 |
| 0 | 0 | 0 | 0 |
| 0 | 0 | 0 | 1 |
| 0 | 0 | 0 | 1 |
| 0 | 0 | 0 | 1 |
| 0 | 0 | 0 | 1 |
| 0 | 0 | 0 | 1 |
| 0 | 0 | 0 | 1 |
| 0 | 0 | 0 | 1 |
| 1 | 0 | 0 | 0 |
| 0 | 0 | 0 | 1 |
| 0 | 0 | 0 | 1 |
| 0 | 0 | 0 | 1 |
| 0 | 0 | 0 | 1 |
| 0 | 0 | 0 | 1 |
| 0 | 0 | 0 | 1 |
| 1 | 0 | 0 | 0 |
| 0 | 0 | 0 | 1 |
| 0 | 0 | 0 | 1 |
| 0 | 0 | 0 | 1 |
| 0 | 0 | 0 | 1 |
| 0 | 0 | 0 | 1 |
| 0 | 0 | 0 | 1 |
| 0 | 0 | 0 | 0 |
| 0 | 0 | 0 | 1 |
| 0 | 0 | 0 | 1 |
| 0 | 1 | 1 | 1 |
| 0 | 1 | 1 | 1 |
| 0 | 0 | 0 | 1 |
| 0 | 0 | 0 | 1 |
| 0 | 1 | 1 | 1 |
| 0 | 1 | 1 | 1 |

| | | | |
|---|---|---|---|
| 0 | 1 | 1 | 1 |
| 0 | 1 | 1 | 1 |
| 0 | 1 | 1 | 1 |
| 0 | 1 | 1 | 1 |
| 0 | 1 | 1 | 1 |
| 0 | 1 | 1 | 1 |
| 0 | 1 | 1 | 1 |
| 0 | 1 | 1 | 1 |
| 0 | 1 | 1 | 1 |
| 0 | 1 | 1 | 1 |
| 0 | 1 | 1 | 1 |
| 0 | 1 | 1 | 1 |
| 0 | 1 | 1 | 1 |
| 0 | 1 | 1 | 1 |
| 0 | 1 | 1 | 1 |
| 1 | 1 | 0 | 0 |
| 0 | 1 | 1 | 1 |
| 0 | 1 | 0 | 0 |
| 0 | 1 | 1 | 1 |
| 0 | 1 | 1 | 1 |
| 1 | 1 | 0 | 0 |
| 0 | 1 | 1 | 1 |
| 0 | 1 | 1 | 1 |
| 0 | 1 | 1 | 1 |
| 0 | 1 | 0 | 0 |
| 0 | 1 | 1 | 1 |
| 0 | 1 | 0 | 0 |
| 0 | 1 | 1 | 1 |
| 0 | 1 | 1 | 1 |
| 0 | 1 | 1 | 1 |
| 0 | 1 | 1 | 1 |
| 0 | 1 | 1 | 1 |
| 0 | 1 | 1 | 1 |
| 0 | 1 | 1 | 1 |
| 0 | 1 | 1 | 1 |
| 0 | 1 | 1 | 1 |
| 0 | 1 | 1 | 1 |
| 0 | 1 | 1 | 1 |
| 0 | 1 | 1 | 1 |
| 0 | 1 | 1 | 1 |
| 0 | 1 | 1 | 1 |
| 0 | 1 | 1 | 1 |
| 0 | 1 | 1 | 1 |
| 0 | 0 | 0 | 0 |
| 0 | 0 | 0 | 1 |
| 0 | 0 | 0 | 1 |
| 0 | 0 | 0 | 0 |
| 0 | 0 | 0 | 0 |
| 0 | 0 | 0 | 1 |
| 0 | 0 | 0 | 0 |
| 0 | 0 | 0 | 0 |
| 0 | 0 | 0 | 0 |
| 0 | 0 | 0 | 1 |
| 0 | 0 | 0 | 1 |

| | | | |
|---|---|---|---|
| 0 | 0 | 0 | 1 |
| 0 | 0 | 0 | 1 |
| 0 | 0 | 0 | 1 |
| 0 | 0 | 0 | 1 |
| 0 | 0 | 0 | 1 |
| 0 | 0 | 0 | 1 |
| 0 | 0 | 0 | 1 |
| 1 | 0 | 0 | 0 |
| 0 | 0 | 0 | 1 |
| 0 | 0 | 0 | 1 |
| 0 | 0 | 0 | 1 |
| 0 | 0 | 0 | 1 |
| 0 | 1 | 1 | 1 |
| 0 | 0 | 0 | 1 |
| 0 | 1 | 0 | 0 |
| 0 | 0 | 0 | 0 |
| 0 | 0 | 0 | 1 |
| 0 | 0 | 0 | 0 |
| 0 | 0 | 0 | 1 |
| 0 | 0 | 0 | 1 |
| 0 | 0 | 0 | 0 |
| 0 | 0 | 0 | 1 |
| 0 | 0 | 0 | 1 |
| 0 | 0 | 0 | 1 |
| 0 | 0 | 0 | 0 |
| 0 | 0 | 0 | 1 |
| 0 | 0 | 0 | 1 |
| 1 | 1 | 0 | 0 |
| 0 | 0 | 0 | 0 |
| 1 | 0 | 0 | 0 |
| 1 | 0 | 0 | 0 |
| 0 | 0 | 0 | 1 |
| 0 | 0 | 0 | 1 |
| 0 | 0 | 0 | 1 |
| 0 | 0 | 0 | 1 |
| 0 | 0 | 0 | 1 |
| 0 | 1 | 0 | 0 |
| 0 | 0 | 0 | 0 |
| 0 | 0 | 0 | 1 |
| 0 | 0 | 0 | 1 |
| 0 | 0 | 0 | 1 |
| 0 | 0 | 0 | 1 |
| 0 | 0 | 0 | 0 |
| 0 | 0 | 0 | 1 |
| 0 | 0 | 0 | 1 |
| 0 | 0 | 0 | 1 |
| 0 | 0 | 0 | 1 |
| 0 | 0 | 0 | 1 |
| 0 | 0 | 0 | 0 |
| 0 | 0 | 0 | 0 |
| 0 | 0 | 0 | 0 |
| 0 | 0 | 0 | 0 |
| 0 | 0 | 0 | 0 |
| 0 | 0 | 0 | 1 |

| | | | |
|---|---|---|---|
| 0 | 0 | 0 | 0 |
| 0 | 0 | 0 | 0 |
| 0 | 0 | 0 | 1 |
| 1 | 0 | 0 | 0 |
| 0 | 0 | 0 | 1 |
| 0 | 0 | 0 | 1 |
| 0 | 0 | 0 | 1 |
| 0 | 0 | 0 | 1 |
| 0 | 0 | 0 | 1 |
| 1 | 0 | 0 | 0 |
| 0 | 0 | 0 | 1 |
| 0 | 0 | 0 | 1 |
| 0 | 0 | 0 | 1 |
| 0 | 0 | 0 | 1 |
| 0 | 0 | 0 | 1 |
| 0 | 0 | 0 | 1 |
| 0 | 0 | 0 | 1 |
| 0 | 0 | 0 | 1 |
| 0 | 0 | 0 | 1 |
| 0 | 0 | 0 | 1 |
| 0 | 0 | 0 | 1 |
| 0 | 0 | 0 | 1 |
| 0 | 0 | 0 | 1 |
| 0 | 0 | 0 | 1 |
| 0 | 0 | 0 | 1 |
| 0 | 0 | 0 | 1 |
| 0 | 0 | 0 | 1 |
| 0 | 0 | 0 | 1 |
| 0 | 0 | 0 | 1 |
| 0 | 0 | 0 | 1 |
| 0 | 0 | 0 | 1 |
| 0 | 0 | 0 | 1 |
| 1 | 0 | 0 | 0 |
| 0 | 0 | 0 | 1 |
| 0 | 0 | 0 | 1 |
| 0 | 0 | 0 | 1 |
| 0 | 0 | 0 | 1 |
| 0 | 0 | 0 | 1 |
| 0 | 0 | 0 | 1 |
| 0 | 0 | 0 | 1 |
| 0 | 0 | 0 | 1 |
| 0 | 0 | 0 | 1 |
| 0 | 0 | 0 | 1 |
| 0 | 0 | 0 | 1 |
| 0 | 0 | 0 | 1 |
| 0 | 0 | 0 | 1 |
| 0 | 0 | 0 | 1 |
| 0 | 1 | 0 | 0 |
| 0 | 1 | 0 | 0 |
| 0 | 1 | 0 | 0 |
| 0 | 0 | 0 | 0 |
| 0 | 1 | 0 | 0 |
| 0 | 0 | 0 | 1 |
| 0 | 0 | 0 | 0 |

| | | | |
|---|---|---|---|
| 0 | 0 | 0 | 1 |
| 0 | 0 | 0 | 0 |
| 0 | 0 | 0 | 1 |
| 0 | 0 | 0 | 1 |
| 0 | 0 | 0 | 1 |
| 1 | 0 | 0 | 0 |
| 0 | 0 | 0 | 0 |
| 0 | 0 | 0 | 0 |
| 1 | 0 | 0 | 0 |
| 0 | 0 | 0 | 0 |
| 0 | 0 | 0 | 1 |
| 0 | 0 | 0 | 0 |
| 0 | 0 | 0 | 1 |
| 0 | 0 | 0 | 1 |
| 0 | 0 | 0 | 1 |
| 1 | 0 | 0 | 0 |
| 0 | 0 | 0 | 1 |
| 0 | 0 | 0 | 1 |
| 0 | 0 | 0 | 0 |
| 0 | 0 | 0 | 0 |
| 1 | 0 | 0 | 0 |
| 1 | 0 | 0 | 0 |
| 0 | 0 | 0 | 0 |
| 0 | 0 | 0 | 0 |
| 0 | 0 | 0 | 1 |
| 0 | 0 | 0 | 0 |
| 0 | 1 | 0 | 0 |
| 0 | 0 | 0 | 0 |
| 0 | 1 | 0 | 0 |
| 0 | 1 | 0 | 0 |
| 0 | 1 | 0 | 0 |
| 0 | 0 | 0 | 1 |
| 0 | 0 | 0 | 1 |
| 0 | 1 | 0 | 0 |
| 0 | 1 | 0 | 0 |
| 0 | 1 | 1 | 1 |
| 0 | 0 | 0 | 0 |
| 0 | 0 | 0 | 0 |
| 0 | 1 | 0 | 0 |
| 0 | 1 | 0 | 0 |
| 0 | 1 | 0 | 0 |
| 0 | 1 | 1 | 1 |
| 0 | 0 | 0 | 1 |
| 0 | 0 | 0 | 1 |
| 0 | 0 | 0 | 0 |
| 0 | 0 | 0 | 0 |
| 0 | 0 | 0 | 0 |
| 0 | 0 | 0 | 1 |
| 0 | 0 | 0 | 1 |
| 0 | 0 | 0 | 1 |
| 0 | 0 | 0 | 1 |
| 0 | 0 | 0 | 1 |
| 0 | 0 | 0 | 1 |
| 0 | 0 | 0 | 0 |

| | | | |
|---|---|---|---|
| 0 | 0 | 0 | 1 |
| 0 | 0 | 0 | 1 |
| 0 | 0 | 0 | 1 |
| 0 | 0 | 0 | 1 |
| 1 | 0 | 0 | 0 |
| 0 | 0 | 0 | 0 |
| 0 | 1 | 1 | 1 |
| 0 | 1 | 1 | 1 |
| 0 | 0 | 0 | 1 |
| 0 | 0 | 0 | 1 |
| 0 | 0 | 0 | 1 |
| 0 | 0 | 0 | 0 |
| 0 | 0 | 0 | 1 |
| 0 | 0 | 0 | 0 |
| 0 | 1 | 0 | 0 |
| 0 | 1 | 0 | 0 |
| 0 | 1 | 0 | 0 |
| 0 | 1 | 0 | 0 |
| 0 | 1 | 0 | 0 |
| 0 | 1 | 0 | 0 |
| 0 | 1 | 0 | 0 |
| 0 | 1 | 0 | 0 |
| 0 | 0 | 0 | 0 |
| 0 | 0 | 0 | 1 |
| 0 | 0 | 0 | 1 |
| 0 | 0 | 0 | 1 |
| 0 | 0 | 0 | 1 |
| 0 | 0 | 0 | 1 |
| 0 | 0 | 0 | 1 |
| 0 | 0 | 0 | 1 |
| 0 | 0 | 0 | 1 |
| 0 | 0 | 0 | 0 |
| 0 | 0 | 0 | 1 |
| 0 | 0 | 0 | 0 |
| 0 | 0 | 0 | 0 |
| 0 | 0 | 0 | 0 |
| 0 | 0 | 0 | 0 |
| 0 | 0 | 0 | 1 |
| 0 | 0 | 0 | 1 |
| 0 | 0 | 0 | 1 |
| 0 | 0 | 0 | 1 |
| 0 | 0 | 0 | 1 |
| 0 | 0 | 0 | 1 |
| 0 | 0 | 0 | 1 |
| 0 | 0 | 0 | 1 |
| 1 | 0 | 0 | 0 |
| 0 | 0 | 0 | 1 |
| 0 | 0 | 0 | 1 |
| 0 | 0 | 0 | 1 |
| 1 | 1 | 0 | 0 |
| 1 | 0 | 0 | 0 |
| 0 | 0 | 0 | 0 |
| 0 | 0 | 0 | 1 |
| 0 | 0 | 0 | 1 |

| | | | |
|---|---|---|---|
| 0 | 0 | 0 | 1 |
| 0 | 0 | 0 | 1 |
| 0 | 0 | 0 | 1 |
| 0 | 0 | 0 | 0 |
| 0 | 0 | 0 | 1 |
| 0 | 0 | 0 | 1 |
| 0 | 0 | 0 | 0 |
| 0 | 0 | 0 | 1 |
| 0 | 0 | 0 | 1 |
| 0 | 0 | 0 | 1 |
| 0 | 0 | 0 | 0 |
| 0 | 1 | 0 | 0 |
| 0 | 0 | 0 | 1 |
| 0 | 0 | 0 | 1 |
| 0 | 0 | 0 | 1 |
| 0 | 0 | 0 | 1 |
| 0 | 0 | 0 | 0 |
| 0 | 0 | 0 | 0 |
| 0 | 0 | 0 | 1 |
| 0 | 0 | 0 | 0 |
| 0 | 1 | 0 | 0 |
| 0 | 1 | 0 | 0 |
| 0 | 0 | 0 | 1 |
| 0 | 1 | 0 | 0 |
| 0 | 0 | 0 | 0 |
| 0 | 0 | 0 | 1 |
| 0 | 0 | 0 | 0 |
| 0 | 0 | 0 | 1 |
| 0 | 0 | 0 | 1 |
| 0 | 0 | 0 | 0 |
| 0 | 0 | 0 | 1 |
| 0 | 0 | 0 | 1 |
| 0 | 0 | 0 | 0 |
| 0 | 0 | 0 | 1 |
| 0 | 0 | 0 | 1 |
| 0 | 0 | 0 | 0 |
| 0 | 0 | 0 | 1 |
| 0 | 0 | 0 | 0 |
| 0 | 0 | 0 | 1 |
| 0 | 0 | 0 | 0 |
| 0 | 1 | 0 | 0 |
| 0 | 0 | 0 | 1 |
| 0 | 0 | 0 | 0 |
| 0 | 0 | 0 | 0 |
| 0 | 0 | 0 | 1 |
| 0 | 1 | 1 | 1 |
| 0 | 0 | 0 | 1 |
| 0 | 0 | 0 | 1 |
| 0 | 0 | 0 | 1 |
| 0 | 0 | 0 | 1 |
| 0 | 0 | 0 | 1 |
| 0 | 0 | 0 | 1 |
| 0 | 0 | 0 | 1 |
| 0 | 0 | 0 | 1 |

| | | | |
|---|---|---|---|
| 0 | 0 | 0 | 1 |
| 0 | 0 | 0 | 1 |
| 0 | 0 | 0 | 1 |
| 0 | 0 | 0 | 0 |
| 0 | 1 | 1 | 1 |
| 0 | 0 | 0 | 0 |
| 0 | 0 | 0 | 1 |
| 0 | 0 | 0 | 1 |
| 0 | 0 | 0 | 0 |
| 0 | 1 | 0 | 0 |
| 0 | 0 | 0 | 1 |
| 0 | 1 | 0 | 0 |
| 0 | 1 | 0 | 0 |
| 0 | 1 | 0 | 0 |
| 0 | 1 | 1 | 1 |
| 0 | 0 | 0 | 0 |
| 0 | 0 | 0 | 0 |
| 0 | 0 | 0 | 1 |
| 1 | 0 | 0 | 0 |
| 1 | 0 | 0 | 0 |
| 1 | 0 | 0 | 0 |
| 1 | 0 | 0 | 0 |
| 0 | 0 | 0 | 1 |
| 0 | 0 | 0 | 0 |
| 0 | 0 | 0 | 0 |
| 0 | 0 | 0 | 0 |
| 0 | 0 | 0 | 0 |
| 0 | 0 | 0 | 0 |
| 0 | 0 | 0 | 0 |
| 0 | 0 | 0 | 0 |
| 0 | 0 | 0 | 0 |
| 0 | 0 | 0 | 0 |
| 0 | 0 | 0 | 0 |
| 0 | 0 | 0 | 1 |
| 0 | 0 | 0 | 1 |
| 0 | 1 | 0 | 0 |
| 0 | 0 | 0 | 1 |
| 0 | 1 | 0 | 0 |
| 0 | 1 | 0 | 0 |
| 0 | 1 | 0 | 0 |
| 0 | 1 | 0 | 0 |
| 0 | 0 | 0 | 1 |
| 0 | 0 | 0 | 0 |
| 0 | 0 | 0 | 0 |
| 0 | 0 | 0 | 1 |
| 0 | 0 | 0 | 1 |
| 0 | 0 | 0 | 1 |
| 0 | 0 | 0 | 1 |
| 0 | 0 | 0 | 1 |
| 0 | 0 | 0 | 1 |
| 0 | 0 | 0 | 1 |
| 0 | 0 | 0 | 1 |
| 0 | 0 | 0 | 1 |
| 0 | 0 | 0 | 1 |

| | | | |
|---|---|---|---|
| 0 | 0 | 0 | 1 |
| 0 | 0 | 0 | 0 |
| 0 | 0 | 0 | 1 |
| 0 | 0 | 0 | 1 |
| 0 | 0 | 0 | 1 |
| 0 | 0 | 0 | 1 |
| 0 | 0 | 0 | 1 |
| 0 | 0 | 0 | 1 |
| 0 | 0 | 0 | 1 |
| 0 | 0 | 0 | 1 |
| 0 | 0 | 0 | 1 |
| 0 | 0 | 0 | 0 |
| 0 | 0 | 0 | 1 |
| 0 | 0 | 0 | 1 |
| 0 | 1 | 0 | 0 |
| 0 | 0 | 0 | 1 |
| 0 | 0 | 0 | 1 |
| 0 | 0 | 0 | 1 |
| 0 | 0 | 0 | 1 |
| 0 | 0 | 0 | 1 |
| 1 | 0 | 0 | 0 |
| 0 | 0 | 0 | 1 |
| 1 | 0 | 0 | 0 |
| 1 | 0 | 0 | 0 |
| 0 | 0 | 0 | 1 |
| 0 | 0 | 0 | 0 |
| 0 | 0 | 0 | 1 |
| 0 | 0 | 0 | 1 |
| 0 | 0 | 0 | 1 |
| 0 | 0 | 0 | 1 |
| 0 | 0 | 0 | 1 |
| 0 | 1 | 0 | 0 |
| 0 | 0 | 0 | 1 |
| 0 | 0 | 0 | 0 |
| 0 | 0 | 0 | 1 |
| 0 | 0 | 0 | 1 |
| 0 | 0 | 0 | 1 |
| 0 | 0 | 0 | 1 |
| 0 | 0 | 0 | 1 |
| 0 | 0 | 0 | 0 |
| 0 | 0 | 0 | 1 |
| 0 | 0 | 0 | 1 |
| 0 | 0 | 0 | 1 |
| 0 | 0 | 0 | 0 |
| 0 | 0 | 0 | 1 |
| 0 | 0 | 0 | 1 |
| 0 | 0 | 0 | 1 |
| 1 | 0 | 0 | 0 |
| 0 | 1 | 0 | 0 |
| 0 | 0 | 0 | 1 |
| 0 | 0 | 0 | 1 |
| 0 | 0 | 0 | 0 |
| 1 | 0 | 0 | 0 |
| 0 | 0 | 0 | 0 |

| | | | |
|---|---|---|---|
| 1 | 0 | 0 | 0 |
| 0 | 0 | 0 | 1 |
| 0 | 0 | 0 | 0 |
| 0 | 0 | 0 | 0 |
| 0 | 0 | 0 | 0 |
| 0 | 0 | 0 | 1 |
| 0 | 0 | 0 | 1 |
| 0 | 0 | 0 | 1 |
| 0 | 0 | 0 | 1 |
| 0 | 1 | 0 | 0 |
| 0 | 0 | 0 | 1 |
| 0 | 0 | 0 | 1 |
| 0 | 0 | 0 | 1 |
| 0 | 0 | 0 | 1 |
| 0 | 0 | 0 | 0 |
| 0 | 0 | 0 | 1 |
| 0 | 0 | 0 | 1 |
| 0 | 0 | 0 | 0 |
| 0 | 0 | 0 | 1 |
| 0 | 0 | 0 | 0 |
| 0 | 0 | 0 | 0 |
| 0 | 0 | 0 | 1 |
| 0 | 0 | 0 | 0 |
| 0 | 0 | 0 | 1 |
| 0 | 0 | 0 | 0 |
| 0 | 0 | 0 | 0 |
| 0 | 0 | 0 | 1 |
| 0 | 0 | 0 | 1 |
| 1 | 0 | 0 | 0 |
| 0 | 0 | 0 | 1 |
| 0 | 0 | 0 | 1 |
| 0 | 0 | 0 | 1 |
| 0 | 0 | 0 | 1 |
| 0 | 0 | 0 | 1 |
| 0 | 0 | 0 | 0 |
| 0 | 0 | 0 | 0 |
| 0 | 0 | 0 | 1 |
| 0 | 0 | 0 | 0 |
| 0 | 0 | 0 | 0 |
| 0 | 0 | 0 | 0 |
| 0 | 0 | 0 | 0 |
| 0 | 0 | 0 | 0 |
| 0 | 0 | 0 | 0 |
| 0 | 0 | 0 | 1 |
| 0 | 1 | 0 | 0 |
| 0 | 0 | 0 | 1 |
| 0 | 0 | 0 | 1 |
| 0 | 0 | 0 | 0 |
| 0 | 0 | 0 | 0 |
| 0 | 0 | 0 | 0 |
| 0 | 0 | 0 | 0 |
| 0 | 1 | 0 | 0 |
| 0 | 0 | 0 | 1 |
| 0 | 0 | 0 | 1 |

| | | | |
|---|---|---|---|
| 0 | 0 | 0 | 1 |
| 0 | 0 | 0 | 0 |
| 0 | 0 | 0 | 0 |
| 0 | 0 | 0 | 0 |
| 0 | 0 | 0 | 0 |
| 0 | 0 | 0 | 0 |
| 0 | 1 | 0 | 0 |
| 0 | 1 | 1 | 1 |
| 0 | 1 | 1 | 1 |
| 0 | 0 | 0 | 1 |
| 0 | 0 | 0 | 0 |
| 0 | 0 | 0 | 0 |
| 0 | 0 | 0 | 1 |
| 0 | 0 | 0 | 0 |
| 0 | 0 | 0 | 1 |
| 0 | 0 | 0 | 0 |
| 0 | 0 | 0 | 1 |
| 0 | 0 | 0 | 1 |
| 0 | 0 | 0 | 1 |
| 0 | 0 | 0 | 1 |
| 0 | 0 | 0 | 1 |
| 0 | 0 | 0 | 1 |
| 0 | 0 | 0 | 1 |
| 1 | 0 | 0 | 0 |
| 0 | 0 | 0 | 0 |
| 0 | 0 | 0 | 1 |
| 0 | 0 | 0 | 1 |
| 0 | 0 | 0 | 1 |
| 0 | 0 | 0 | 1 |
| 0 | 0 | 0 | 0 |
| 0 | 0 | 0 | 0 |
| 0 | 0 | 0 | 0 |
| 0 | 0 | 0 | 0 |
| 0 | 0 | 0 | 1 |
| 0 | 0 | 0 | 0 |
| 0 | 0 | 0 | 0 |
| 0 | 0 | 0 | 0 |
| 0 | 0 | 0 | 0 |
| 0 | 0 | 0 | 0 |
| 0 | 0 | 0 | 1 |
| 0 | 0 | 0 | 1 |
| 0 | 0 | 0 | 0 |
| 0 | 0 | 0 | 1 |
| 0 | 0 | 0 | 1 |
| 0 | 0 | 0 | 0 |
| 0 | 1 | 1 | 1 |
| 0 | 0 | 0 | 1 |
| 0 | 0 | 0 | 1 |
| 0 | 1 | 1 | 1 |
| 0 | 0 | 0 | 1 |
| 0 | 0 | 0 | 0 |
| 0 | 0 | 0 | 1 |
| 0 | 0 | 0 | 0 |
| 0 | 0 | 0 | 0 |

| | | | |
|---|---|---|---|
| 0 | 0 | 0 | 1 |
| 0 | 0 | 0 | 1 |
| 0 | 0 | 0 | 0 |
| 0 | 0 | 0 | 0 |
| 0 | 0 | 0 | 1 |
| 0 | 0 | 0 | 1 |
| 0 | 0 | 0 | 1 |
| 0 | 0 | 0 | 1 |
| 0 | 0 | 0 | 1 |
| 0 | 0 | 0 | 1 |
| 0 | 0 | 0 | 1 |
| 0 | 0 | 0 | 1 |
| 0 | 0 | 0 | 0 |
| 0 | 0 | 0 | 1 |
| 0 | 0 | 0 | 1 |
| 0 | 0 | 0 | 1 |
| 0 | 0 | 0 | 1 |
| 0 | 0 | 0 | 1 |
| 0 | 0 | 0 | 1 |
| 0 | 0 | 0 | 1 |
| 0 | 0 | 0 | 1 |
| 0 | 0 | 0 | 1 |
| 0 | 0 | 0 | 1 |
| 0 | 0 | 0 | 1 |
| 0 | 0 | 0 | 1 |
| 0 | 0 | 0 | 1 |
| 0 | 0 | 0 | 1 |
| 0 | 0 | 0 | 1 |
| 0 | 0 | 0 | 1 |
| 0 | 0 | 0 | 1 |
| 0 | 0 | 0 | 1 |
| 0 | 0 | 0 | 0 |
| 0 | 0 | 0 | 1 |
| 0 | 0 | 0 | 1 |
| 0 | 0 | 0 | 1 |
| 0 | 0 | 0 | 1 |
| 0 | 0 | 0 | 1 |
| 0 | 0 | 0 | 1 |
| 0 | 0 | 0 | 1 |
| 0 | 0 | 0 | 1 |
| 0 | 0 | 0 | 0 |
| 0 | 0 | 0 | 0 |
| 0 | 0 | 0 | 1 |
| 0 | 0 | 0 | 1 |
| 0 | 0 | 0 | 1 |
| 0 | 1 | 1 | 1 |
| 0 | 1 | 1 | 1 |
| 0 | 0 | 0 | 1 |
| 0 | 0 | 0 | 0 |
| 0 | 0 | 0 | 0 |
| 0 | 0 | 0 | 1 |
| 0 | 0 | 0 | 1 |
| 0 | 0 | 0 | 0 |
| 0 | 0 | 0 | 1 |

| | | | |
|---|---|---|---|
| 0 | 0 | 0 | 0 |
| 0 | 0 | 0 | 0 |
| 0 | 0 | 0 | 0 |
| 0 | 0 | 0 | 1 |
| 0 | 0 | 0 | 1 |
| 0 | 0 | 0 | 1 |
| 0 | 0 | 0 | 1 |
| 0 | 1 | 0 | 0 |
| 0 | 1 | 0 | 0 |
| 0 | 0 | 0 | 0 |
| 0 | 0 | 0 | 0 |
| 0 | 0 | 0 | 1 |
| 0 | 0 | 0 | 0 |
| 0 | 0 | 0 | 1 |
| 0 | 0 | 0 | 1 |
| 0 | 0 | 0 | 1 |
| 0 | 0 | 0 | 1 |
| 0 | 0 | 0 | 0 |
| 0 | 0 | 0 | 1 |
| 0 | 0 | 0 | 1 |
| 0 | 0 | 0 | 1 |
| 0 | 0 | 0 | 1 |
| 0 | 0 | 0 | 1 |
| 0 | 0 | 0 | 1 |
| 0 | 0 | 0 | 1 |
| 0 | 0 | 0 | 1 |
| 0 | 0 | 0 | 1 |
| 0 | 0 | 0 | 1 |
| 0 | 0 | 0 | 1 |
| 0 | 0 | 0 | 1 |
| 0 | 0 | 0 | 0 |
| 0 | 1 | 0 | 0 |
| 0 | 0 | 0 | 1 |
| 0 | 0 | 0 | 1 |
| 0 | 0 | 0 | 1 |
| 0 | 0 | 0 | 1 |
| 0 | 0 | 0 | 1 |
| 0 | 0 | 0 | 1 |
| 0 | 0 | 0 | 1 |
| 0 | 0 | 0 | 1 |
| 0 | 0 | 0 | 1 |
| 0 | 0 | 0 | 0 |
| 0 | 0 | 0 | 1 |
| 0 | 0 | 0 | 1 |
| 0 | 0 | 0 | 0 |
| 0 | 0 | 0 | 0 |
| 0 | 0 | 0 | 1 |
| 0 | 0 | 0 | 1 |
| 0 | 0 | 0 | 0 |
| 0 | 0 | 0 | 0 |
| 0 | 0 | 0 | 0 |
| 0 | 0 | 0 | 0 |
| 0 | 0 | 0 | 0 |
| 0 | 0 | 0 | 0 |

| | | | |
|---|---|---|---|
| 0 | 0 | 0 | 1 |
| 0 | 0 | 0 | 1 |
| 0 | 0 | 0 | 1 |
| 0 | 0 | 0 | 1 |
| 0 | 0 | 0 | 1 |
| 0 | 0 | 0 | 0 |
| 0 | 0 | 0 | 0 |
| 0 | 0 | 0 | 0 |
| 0 | 0 | 0 | 0 |
| 0 | 0 | 0 | 0 |
| 0 | 0 | 0 | 1 |
| 0 | 0 | 0 | 0 |
| 0 | 0 | 0 | 0 |
| 0 | 0 | 0 | 0 |
| 0 | 0 | 0 | 1 |
| 0 | 0 | 0 | 0 |
| 0 | 0 | 0 | 1 |
| 0 | 0 | 0 | 1 |
| 1 | 0 | 0 | 0 |
| 1 | 0 | 0 | 0 |
| 0 | 0 | 0 | 0 |
| 0 | 0 | 0 | 0 |
| 0 | 0 | 0 | 1 |
| 0 | 0 | 0 | 0 |
| 0 | 0 | 0 | 0 |
| 0 | 0 | 0 | 0 |
| 0 | 0 | 0 | 0 |
| 0 | 0 | 0 | 1 |
| 0 | 0 | 0 | 1 |
| 0 | 0 | 0 | 1 |
| 0 | 0 | 0 | 1 |
| 0 | 0 | 0 | 1 |
| 0 | 0 | 0 | 0 |
| 0 | 0 | 0 | 1 |
| 0 | 0 | 0 | 0 |
| 0 | 0 | 0 | 0 |
| 0 | 1 | 0 | 0 |
| 0 | 0 | 0 | 0 |
| 0 | 1 | 0 | 0 |
| 0 | 1 | 0 | 0 |
| 0 | 1 | 0 | 0 |
| 1 | 0 | 0 | 0 |
| 0 | 0 | 0 | 0 |
| 0 | 0 | 0 | 1 |
| 0 | 0 | 0 | 1 |
| 0 | 0 | 0 | 1 |
| 0 | 0 | 0 | 1 |
| 0 | 0 | 0 | 0 |
| 0 | 0 | 0 | 0 |
| 0 | 0 | 0 | 0 |
| 0 | 0 | 0 | 1 |
| 0 | 0 | 0 | 1 |
| 0 | 0 | 0 | 0 |
| 0 | 0 | 0 | 1 |

| | | | |
|---|---|---|---|
| 0 | 0 | 0 | 1 |
| 0 | 0 | 0 | 0 |
| 0 | 0 | 0 | 1 |
| 0 | 0 | 0 | 1 |
| 0 | 0 | 0 | 1 |
| 0 | 0 | 0 | 1 |
| 0 | 0 | 0 | 1 |
| 0 | 0 | 0 | 1 |
| 0 | 0 | 0 | 1 |
| 0 | 0 | 0 | 1 |
| 0 | 0 | 0 | 1 |
| 0 | 0 | 0 | 1 |
| 0 | 0 | 0 | 0 |
| 0 | 0 | 0 | 0 |
| 0 | 0 | 0 | 0 |
| 0 | 0 | 0 | 0 |
| 0 | 0 | 0 | 0 |
| 0 | 0 | 0 | 0 |
| 0 | 0 | 0 | 0 |
| 0 | 0 | 0 | 1 |
| 1 | 1 | 0 | 0 |
| 0 | 0 | 0 | 0 |
| 0 | 0 | 0 | 0 |
| 0 | 0 | 0 | 0 |
| 0 | 0 | 0 | 0 |
| 0 | 0 | 0 | 1 |
| 0 | 0 | 0 | 1 |
| 0 | 0 | 0 | 0 |
| 0 | 0 | 0 | 1 |
| 0 | 0 | 0 | 0 |
| 0 | 0 | 0 | 0 |
| 0 | 0 | 0 | 1 |
| 0 | 0 | 0 | 0 |
| 0 | 0 | 0 | 1 |
| 0 | 0 | 0 | 1 |
| 0 | 0 | 0 | 1 |
| 0 | 1 | 1 | 1 |
| 0 | 0 | 0 | 1 |
| 0 | 0 | 0 | 1 |
| 0 | 0 | 0 | 1 |
| 0 | 0 | 0 | 0 |
| 0 | 0 | 0 | 1 |
| 0 | 0 | 0 | 1 |
| 0 | 0 | 0 | 0 |
| 0 | 0 | 0 | 0 |
| 0 | 0 | 0 | 1 |
| 0 | 0 | 0 | 0 |
| 0 | 0 | 0 | 0 |
| 0 | 0 | 0 | 1 |
| 0 | 0 | 0 | 1 |
| 0 | 0 | 0 | 0 |
| 0 | 0 | 0 | 1 |
| 0 | 0 | 0 | 1 |
| 0 | 0 | 0 | 1 |

| | | | |
|---|---|---|---|
| 0 | 0 | 0 | 1 |
| 0 | 0 | 0 | 1 |
| 0 | 0 | 0 | 1 |
| 0 | 0 | 0 | 1 |
| 0 | 0 | 0 | 0 |
| 0 | 0 | 0 | 0 |
| 0 | 0 | 0 | 1 |
| 0 | 0 | 0 | 1 |
| 0 | 0 | 0 | 1 |
| 0 | 0 | 0 | 1 |
| 0 | 0 | 0 | 1 |
| 0 | 0 | 0 | 0 |
| 0 | 0 | 0 | 0 |
| 0 | 0 | 0 | 1 |
| 0 | 0 | 0 | 1 |
| 0 | 0 | 0 | 0 |
| 0 | 0 | 0 | 1 |
| 0 | 0 | 0 | 1 |
| 0 | 0 | 0 | 0 |
| 0 | 0 | 0 | 1 |
| 0 | 0 | 0 | 1 |
| 0 | 0 | 0 | 1 |
| 0 | 0 | 0 | 1 |
| 0 | 0 | 0 | 0 |
| 0 | 0 | 0 | 1 |
| 0 | 0 | 0 | 0 |
| 0 | 0 | 0 | 1 |
| 0 | 0 | 0 | 1 |
| 1 | 0 | 0 | 0 |
| 1 | 0 | 0 | 0 |
| 0 | 0 | 0 | 0 |
| 0 | 0 | 0 | 0 |
| 0 | 0 | 0 | 0 |
| 0 | 0 | 0 | 0 |
| 0 | 0 | 0 | 1 |
| 0 | 0 | 0 | 1 |
| 0 | 0 | 0 | 1 |
| 1 | 0 | 0 | 0 |
| 0 | 0 | 0 | 1 |
| 0 | 0 | 0 | 1 |
| 0 | 0 | 0 | 0 |
| 0 | 0 | 0 | 0 |
| 0 | 0 | 0 | 0 |
| 0 | 0 | 0 | 1 |
| 0 | 0 | 0 | 0 |
| 0 | 0 | 0 | 1 |
| 0 | 0 | 0 | 1 |
| 0 | 0 | 0 | 0 |
| 0 | 0 | 0 | 1 |
| 0 | 0 | 0 | 1 |
| 0 | 0 | 0 | 1 |
| 0 | 1 | 0 | 0 |
| 0 | 0 | 0 | 1 |
| 0 | 0 | 0 | 0 |

| | | | |
|---|---|---|---|
| 0 | 0 | 0 | 1 |
| 0 | 0 | 0 | 0 |
| 0 | 0 | 0 | 0 |
| 0 | 0 | 0 | 1 |
| 0 | 0 | 0 | 1 |
| 0 | 0 | 0 | 1 |
| 0 | 0 | 0 | 0 |
| 0 | 0 | 0 | 1 |
| 0 | 0 | 0 | 1 |
| 0 | 0 | 0 | 1 |
| 0 | 0 | 0 | 1 |
| 0 | 0 | 0 | 1 |
| 0 | 0 | 0 | 1 |
| 0 | 0 | 0 | 1 |
| 0 | 0 | 0 | 1 |
| 0 | 0 | 0 | 1 |
| 0 | 0 | 0 | 1 |
| 0 | 0 | 0 | 1 |
| 0 | 0 | 0 | 1 |
| 0 | 0 | 0 | 1 |
| 0 | 0 | 0 | 1 |
| 0 | 0 | 0 | 0 |
| 0 | 0 | 0 | 1 |
| 0 | 0 | 0 | 1 |
| 0 | 0 | 0 | 1 |
| 0 | 0 | 0 | 1 |
| 0 | 0 | 0 | 0 |
| 0 | 0 | 0 | 1 |
| 0 | 0 | 0 | 0 |
| 0 | 0 | 0 | 1 |
| 0 | 0 | 0 | 1 |
| 0 | 0 | 0 | 0 |
| 0 | 1 | 1 | 1 |
| 0 | 1 | 1 | 1 |
| 0 | 0 | 0 | 1 |
| 0 | 0 | 0 | 1 |
| 0 | 0 | 0 | 1 |
| 0 | 0 | 0 | 1 |
| 0 | 0 | 0 | 0 |
| 0 | 0 | 0 | 0 |
| 0 | 1 | 0 | 0 |
| 0 | 0 | 0 | 1 |
| 0 | 0 | 0 | 1 |
| 0 | 0 | 0 | 1 |
| 0 | 0 | 0 | 1 |
| 0 | 1 | 0 | 0 |
| 0 | 0 | 0 | 1 |
| 0 | 0 | 0 | 0 |
| 0 | 0 | 0 | 1 |
| 0 | 0 | 0 | 0 |
| 0 | 0 | 0 | 0 |
| 0 | 0 | 0 | 0 |
| 0 | 0 | 0 | 1 |
| 0 | 0 | 0 | 1 |

| | | | |
|---|---|---|---|
| 0 | 0 | 0 | 0 |
| 0 | 0 | 0 | 0 |
| 0 | 0 | 0 | 1 |
| 0 | 0 | 0 | 1 |
| 0 | 0 | 0 | 1 |
| 0 | 0 | 0 | 1 |
| 0 | 0 | 0 | 1 |
| 0 | 0 | 0 | 1 |
| 0 | 0 | 0 | 1 |
| 0 | 0 | 0 | 1 |
| 0 | 0 | 0 | 1 |
| 0 | 0 | 0 | 1 |
| 0 | 0 | 0 | 1 |
| 0 | 0 | 0 | 1 |
| 0 | 0 | 0 | 1 |
| 0 | 0 | 0 | 0 |
| 0 | 0 | 0 | 0 |
| 0 | 0 | 0 | 0 |
| 0 | 0 | 0 | 1 |
| 0 | 0 | 0 | 1 |
| 0 | 0 | 0 | 1 |
| 0 | 1 | 0 | 0 |
| 0 | 0 | 0 | 0 |
| 0 | 0 | 0 | 0 |
| 0 | 0 | 0 | 0 |
| 0 | 0 | 0 | 1 |
| 0 | 0 | 0 | 1 |
| 0 | 0 | 0 | 1 |
| 0 | 0 | 0 | 1 |
| 0 | 0 | 0 | 1 |
| 0 | 0 | 0 | 0 |
| 0 | 0 | 0 | 1 |
| 0 | 0 | 0 | 0 |
| 0 | 0 | 0 | 1 |
| 0 | 1 | 1 | 1 |
| 0 | 1 | 1 | 1 |
| 0 | 1 | 0 | 0 |
| 0 | 0 | 0 | 0 |
| 0 | 0 | 0 | 1 |
| 0 | 0 | 0 | 1 |
| 0 | 1 | 1 | 1 |
| 0 | 1 | 0 | 0 |
| 0 | 0 | 0 | 1 |
| 0 | 0 | 0 | 1 |
| 0 | 1 | 0 | 0 |
| 0 | 1 | 1 | 1 |
| 0 | 0 | 0 | 0 |
| 0 | 0 | 0 | 0 |
| 0 | 0 | 0 | 1 |
| 0 | 0 | 0 | 1 |
| 0 | 1 | 0 | 0 |
| 0 | 1 | 0 | 0 |
| 0 | 0 | 0 | 1 |
| 0 | 1 | 0 | 0 |

| | | | |
|---|---|---|---|
| 0 | 0 | 0 | 1 |
| 0 | 0 | 0 | 0 |
| 0 | 1 | 1 | 1 |
| 0 | 1 | 0 | 0 |
| 0 | 0 | 0 | 1 |
| 0 | 0 | 0 | 1 |
| 0 | 0 | 0 | 1 |
| 0 | 0 | 0 | 1 |
| 0 | 0 | 0 | 0 |
| 0 | 0 | 0 | 1 |
| 0 | 0 | 0 | 1 |
| 0 | 0 | 0 | 1 |
| 0 | 0 | 0 | 0 |
| 0 | 1 | 0 | 0 |
| 0 | 0 | 0 | 0 |
| 0 | 0 | 0 | 1 |
| 0 | 0 | 0 | 0 |
| 0 | 0 | 0 | 1 |
| 0 | 1 | 0 | 0 |
| 0 | 0 | 0 | 0 |
| 0 | 0 | 0 | 0 |
| 0 | 1 | 0 | 0 |
| 0 | 0 | 0 | 0 |
| 1 | 0 | 0 | 0 |
| 0 | 0 | 0 | 1 |
| 0 | 0 | 0 | 1 |
| 0 | 0 | 0 | 0 |
| 0 | 0 | 0 | 0 |
| 0 | 0 | 0 | 0 |
| 0 | 0 | 0 | 0 |
| 0 | 0 | 0 | 1 |
| 0 | 0 | 0 | 1 |
| 0 | 0 | 0 | 1 |
| 0 | 0 | 0 | 1 |
| 0 | 0 | 0 | 1 |
| 1 | 0 | 0 | 0 |
| 1 | 0 | 0 | 0 |
| 0 | 0 | 0 | 1 |
| 0 | 0 | 0 | 1 |
| 0 | 0 | 0 | 1 |
| 0 | 0 | 0 | 1 |
| 0 | 0 | 0 | 1 |
| 0 | 0 | 0 | 1 |
| 0 | 0 | 0 | 0 |
| 0 | 1 | 0 | 0 |
| 0 | 0 | 0 | 0 |
| 0 | 0 | 0 | 1 |
| 0 | 0 | 0 | 0 |
| 0 | 0 | 0 | 0 |
| 0 | 0 | 0 | 0 |
| 0 | 0 | 0 | 1 |
| 0 | 0 | 0 | 1 |
| 0 | 0 | 0 | 1 |
| 0 | 0 | 0 | 1 |

| | | | |
|---|---|---|---|
| 0 | 0 | 0 | 1 |
| 0 | 0 | 0 | 1 |
| 0 | 0 | 0 | 1 |
| 0 | 0 | 0 | 1 |
| 0 | 0 | 0 | 0 |
| 0 | 0 | 0 | 1 |
| 0 | 0 | 0 | 0 |
| 0 | 0 | 0 | 0 |
| 0 | 0 | 0 | 0 |
| 0 | 0 | 0 | 0 |
| 0 | 0 | 0 | 0 |
| 0 | 0 | 0 | 0 |
| 0 | 0 | 0 | 1 |
| 0 | 0 | 0 | 0 |
| 0 | 0 | 0 | 0 |
| 0 | 0 | 0 | 1 |
| 0 | 0 | 0 | 1 |
| 0 | 0 | 0 | 1 |
| 0 | 0 | 0 | 0 |
| 0 | 0 | 0 | 0 |
| 0 | 0 | 0 | 0 |
| 0 | 0 | 0 | 0 |
| 0 | 0 | 0 | 1 |
| 0 | 0 | 0 | 1 |
| 0 | 0 | 0 | 1 |
| 0 | 0 | 0 | 1 |
| 0 | 0 | 0 | 1 |
| 0 | 0 | 0 | 0 |
| 0 | 0 | 0 | 0 |
| 0 | 0 | 0 | 0 |
| 0 | 0 | 0 | 0 |
| 0 | 0 | 0 | 0 |
| 0 | 0 | 0 | 0 |
| 0 | 0 | 0 | 1 |
| 0 | 0 | 0 | 1 |
| 0 | 0 | 0 | 0 |
| 0 | 0 | 0 | 1 |
| 0 | 0 | 0 | 1 |
| 0 | 0 | 0 | 1 |
| 0 | 0 | 0 | 1 |
| 0 | 0 | 0 | 0 |
| 0 | 0 | 0 | 1 |
| 0 | 0 | 0 | 0 |
| 0 | 0 | 0 | 1 |
| 0 | 0 | 0 | 1 |
| 0 | 0 | 0 | 1 |
| 0 | 0 | 0 | 1 |
| 0 | 0 | 0 | 0 |
| 0 | 0 | 0 | 1 |
| 0 | 0 | 0 | 1 |
| 0 | 0 | 0 | 1 |
| 0 | 0 | 0 | 1 |
| 0 | 0 | 0 | 1 |
| 0 | 0 | 0 | 1 |

| | | | |
|---|---|---|---|
| 0 | 0 | 0 | 1 |
| 0 | 0 | 0 | 1 |
| 0 | 0 | 0 | 1 |
| 0 | 0 | 0 | 1 |
| 0 | 0 | 0 | 1 |
| 0 | 0 | 0 | 1 |
| 0 | 0 | 0 | 1 |
| 0 | 0 | 0 | 1 |
| 0 | 0 | 0 | 1 |
| 0 | 0 | 0 | 1 |
| 0 | 0 | 0 | 1 |
| 0 | 0 | 0 | 0 |
| 0 | 0 | 0 | 1 |
| 0 | 0 | 0 | 1 |
| 0 | 0 | 0 | 1 |
| 0 | 0 | 0 | 1 |
| 0 | 0 | 0 | 1 |
| 0 | 0 | 0 | 0 |
| 0 | 0 | 0 | 0 |
| 0 | 0 | 0 | 0 |
| 0 | 0 | 0 | 0 |
| 0 | 0 | 0 | 1 |
| 1 | 0 | 0 | 0 |
| 0 | 0 | 0 | 1 |
| 0 | 0 | 0 | 1 |
| 0 | 0 | 0 | 1 |
| 0 | 0 | 0 | 1 |
| 0 | 0 | 0 | 1 |
| 0 | 0 | 0 | 0 |
| 0 | 0 | 0 | 0 |
| 0 | 0 | 0 | 1 |
| 0 | 0 | 0 | 1 |
| 1 | 0 | 0 | 0 |
| 0 | 0 | 0 | 1 |
| 0 | 0 | 0 | 0 |
| 1 | 0 | 0 | 0 |
| 1 | 0 | 0 | 0 |
| 0 | 0 | 0 | 1 |
| 1 | 0 | 0 | 0 |
| 0 | 0 | 0 | 1 |
| 1 | 0 | 0 | 0 |
| 0 | 0 | 0 | 1 |
| 0 | 0 | 0 | 0 |
| 0 | 0 | 0 | 1 |
| 0 | 0 | 0 | 1 |
| 0 | 0 | 0 | 1 |
| 0 | 1 | 1 | 1 |
| 0 | 0 | 0 | 1 |
| 0 | 0 | 0 | 1 |
| 0 | 0 | 0 | 0 |
| 0 | 1 | 1 | 1 |
| 1 | 0 | 0 | 0 |
| 0 | 1 | 1 | 1 |
| 0 | 0 | 0 | 1 |

| | | | |
|---|---|---|---|
| 0 | 0 | 0 | 0 |
| 0 | 0 | 0 | 0 |
| 0 | 0 | 0 | 0 |
| 0 | 0 | 0 | 0 |
| 0 | 0 | 0 | 0 |
| 0 | 0 | 0 | 1 |
| 0 | 0 | 0 | 0 |
| 0 | 0 | 0 | 1 |
| 0 | 0 | 0 | 1 |
| 0 | 0 | 0 | 0 |
| 0 | 0 | 0 | 0 |
| 0 | 0 | 0 | 0 |
| 0 | 0 | 0 | 1 |
| 0 | 0 | 0 | 1 |
| 0 | 0 | 0 | 1 |
| 0 | 0 | 0 | 1 |
| 0 | 0 | 0 | 1 |
| 0 | 0 | 0 | 1 |
| 0 | 0 | 0 | 1 |
| 0 | 0 | 0 | 1 |
| 0 | 0 | 0 | 1 |
| 0 | 0 | 0 | 1 |
| 0 | 0 | 0 | 1 |
| 0 | 0 | 0 | 1 |
| 0 | 0 | 0 | 1 |
| 0 | 0 | 0 | 1 |
| 0 | 0 | 0 | 1 |
| 0 | 0 | 0 | 1 |
| 0 | 0 | 0 | 1 |
| 0 | 0 | 0 | 1 |
| 0 | 0 | 0 | 1 |
| 0 | 0 | 0 | 1 |
| 0 | 0 | 0 | 1 |
| 0 | 0 | 0 | 1 |
| 0 | 0 | 0 | 1 |
| 0 | 0 | 0 | 1 |
| 0 | 0 | 0 | 1 |
| 0 | 0 | 0 | 1 |
| 0 | 0 | 0 | 1 |
| 0 | 0 | 0 | 1 |
| 0 | 0 | 0 | 1 |
| 0 | 0 | 0 | 1 |
| 0 | 0 | 0 | 1 |
| 0 | 0 | 0 | 1 |
| 0 | 0 | 0 | 1 |
| 0 | 0 | 0 | 0 |
| 0 | 0 | 0 | 1 |
| 0 | 0 | 0 | 1 |
| 0 | 0 | 0 | 1 |
| 0 | 0 | 0 | 1 |
| 0 | 0 | 0 | 0 |
| 0 | 0 | 0 | 1 |
| 0 | 0 | 0 | 1 |
| 0 | 0 | 0 | 1 |

| | | | |
|---|---|---|---|
| 0 | 0 | 0 | 1 |
| 0 | 0 | 0 | 1 |
| 0 | 0 | 0 | 1 |
| 0 | 0 | 0 | 1 |
| 0 | 0 | 0 | 1 |
| 0 | 0 | 0 | 1 |
| 0 | 0 | 0 | 1 |
| 0 | 0 | 0 | 0 |
| 0 | 1 | 0 | 0 |
| 0 | 0 | 0 | 1 |
| 0 | 0 | 0 | 1 |
| 0 | 0 | 0 | 1 |
| 0 | 0 | 0 | 1 |
| 0 | 0 | 0 | 1 |
| 0 | 1 | 0 | 0 |
| 0 | 1 | 0 | 0 |
| 0 | 0 | 0 | 0 |
| 0 | 0 | 0 | 1 |
| 0 | 1 | 1 | 1 |
| 0 | 0 | 0 | 1 |
| 0 | 1 | 0 | 0 |
| 0 | 1 | 0 | 0 |
| 0 | 1 | 0 | 0 |
| 0 | 0 | 0 | 0 |
| 0 | 0 | 0 | 0 |
| 0 | 0 | 0 | 0 |
| 0 | 0 | 0 | 1 |
| 0 | 0 | 0 | 1 |
| 0 | 0 | 0 | 0 |
| 0 | 0 | 0 | 1 |
| 0 | 0 | 0 | 1 |
| 0 | 0 | 0 | 1 |
| 0 | 0 | 0 | 0 |
| 0 | 0 | 0 | 1 |
| 0 | 0 | 0 | 1 |
| 0 | 0 | 0 | 1 |
| 0 | 0 | 0 | 0 |
| 0 | 0 | 0 | 0 |
| 0 | 0 | 0 | 0 |
| 0 | 0 | 0 | 1 |
| 0 | 0 | 0 | 1 |
| 0 | 0 | 0 | 1 |
| 0 | 0 | 0 | 1 |
| 0 | 0 | 0 | 1 |
| 0 | 0 | 0 | 1 |
| 0 | 0 | 0 | 0 |
| 0 | 0 | 0 | 1 |
| 0 | 0 | 0 | 1 |
| 0 | 0 | 0 | 1 |
| 0 | 0 | 0 | 1 |
| 0 | 0 | 0 | 1 |
| 0 | 0 | 0 | 1 |
| 0 | 0 | 0 | 1 |
| 0 | 0 | 0 | 1 |

| | | | |
|---|---|---|---|
| 0 | 0 | 0 | 1 |
| 0 | 0 | 0 | 1 |
| 0 | 0 | 0 | 1 |
| 0 | 0 | 0 | 0 |
| 0 | 0 | 0 | 1 |
| 0 | 0 | 0 | 1 |
| 0 | 0 | 0 | 1 |
| 0 | 0 | 0 | 0 |
| 0 | 0 | 0 | 0 |
| 0 | 0 | 0 | 1 |
| 0 | 0 | 0 | 1 |
| 0 | 0 | 0 | 1 |
| 0 | 0 | 0 | 0 |
| 0 | 0 | 0 | 1 |
| 0 | 0 | 0 | 1 |
| 0 | 0 | 0 | 1 |
| 0 | 0 | 0 | 0 |
| 0 | 0 | 0 | 1 |
| 0 | 0 | 0 | 1 |
| 0 | 0 | 0 | 1 |
| 0 | 0 | 0 | 1 |
| 0 | 0 | 0 | 1 |
| 0 | 0 | 0 | 1 |
| 0 | 0 | 0 | 1 |
| 0 | 0 | 0 | 1 |
| 0 | 0 | 0 | 1 |
| 0 | 0 | 0 | 1 |
| 0 | 0 | 0 | 0 |
| 0 | 0 | 0 | 0 |
| 0 | 0 | 0 | 0 |
| 0 | 0 | 0 | 0 |
| 0 | 0 | 0 | 0 |
| 0 | 0 | 0 | 0 |
| 0 | 0 | 0 | 1 |
| 0 | 0 | 0 | 0 |
| 0 | 0 | 0 | 0 |
| 0 | 0 | 0 | 0 |
| 0 | 0 | 0 | 1 |
| 0 | 0 | 0 | 1 |
| 0 | 0 | 0 | 1 |
| 0 | 0 | 0 | 1 |
| 0 | 0 | 0 | 1 |
| 0 | 0 | 0 | 0 |
| 0 | 0 | 0 | 0 |
| 0 | 0 | 0 | 1 |
| 0 | 0 | 0 | 1 |
| 0 | 0 | 0 | 0 |
| 0 | 0 | 0 | 1 |
| 0 | 0 | 0 | 1 |
| 0 | 0 | 0 | 1 |
| 0 | 0 | 0 | 1 |
| 0 | 0 | 0 | 1 |
| 0 | 0 | 0 | 0 |
| 0 | 0 | 0 | 0 |

| | | | |
|---|---|---|---|
| 0 | 0 | 0 | 0 |
| 0 | 0 | 0 | 0 |
| 0 | 0 | 0 | 0 |
| 0 | 0 | 0 | 0 |
| 0 | 0 | 0 | 0 |
| 0 | 0 | 0 | 0 |
| 0 | 0 | 0 | 0 |
| 0 | 0 | 0 | 0 |
| 0 | 0 | 0 | 1 |
| 0 | 0 | 0 | 0 |
| 0 | 0 | 0 | 1 |
| 0 | 0 | 0 | 1 |
| 0 | 0 | 0 | 1 |
| 0 | 0 | 0 | 1 |
| 0 | 0 | 0 | 0 |
| 0 | 0 | 0 | 1 |
| 0 | 0 | 0 | 1 |
| 0 | 0 | 0 | 1 |
| 0 | 0 | 0 | 1 |
| 0 | 0 | 0 | 0 |
| 0 | 0 | 0 | 0 |
| 0 | 0 | 0 | 1 |
| 0 | 0 | 0 | 0 |
| 0 | 0 | 0 | 0 |
| 0 | 0 | 0 | 0 |
| 0 | 0 | 0 | 0 |
| 0 | 0 | 0 | 0 |
| 0 | 0 | 0 | 1 |
| 0 | 0 | 0 | 1 |
| 0 | 0 | 0 | 1 |
| 0 | 0 | 0 | 0 |
| 0 | 0 | 0 | 1 |
| 0 | 0 | 0 | 1 |
| 0 | 0 | 0 | 1 |
| 0 | 0 | 0 | 1 |
| 0 | 0 | 0 | 1 |
| 0 | 0 | 0 | 1 |
| 0 | 0 | 0 | 1 |
| 0 | 0 | 0 | 1 |
| 0 | 0 | 0 | 1 |
| 0 | 0 | 0 | 1 |
| 0 | 0 | 0 | 0 |
| 0 | 0 | 0 | 1 |
| 0 | 0 | 0 | 1 |
| 0 | 0 | 0 | 1 |
| 0 | 0 | 0 | 1 |
| 0 | 0 | 0 | 1 |
| 0 | 0 | 0 | 1 |
| 0 | 0 | 0 | 0 |
| 0 | 0 | 0 | 1 |
| 0 | 0 | 0 | 1 |
| 0 | 0 | 0 | 1 |
| 0 | 0 | 0 | 0 |
| 0 | 1 | 0 | 0 |

| | | | |
|---|---|---|---|
| 0 | 0 | 0 | 1 |
| 0 | 0 | 0 | 1 |
| 0 | 0 | 0 | 1 |
| 0 | 0 | 0 | 1 |
| 0 | 0 | 0 | 0 |
| 1 | 0 | 0 | 0 |
| 0 | 0 | 0 | 1 |
| 0 | 0 | 0 | 0 |
| 0 | 0 | 0 | 1 |
| 0 | 0 | 0 | 1 |
| 0 | 0 | 0 | 0 |
| 0 | 0 | 0 | 1 |
| 0 | 0 | 0 | 0 |
| 0 | 0 | 0 | 0 |
| 0 | 1 | 0 | 0 |
| 0 | 0 | 0 | 0 |
| 0 | 0 | 0 | 1 |
| 0 | 0 | 0 | 1 |
| 0 | 0 | 0 | 0 |
| 0 | 1 | 0 | 0 |
| 0 | 1 | 0 | 0 |
| 0 | 1 | 0 | 0 |
| 0 | 0 | 0 | 0 |
| 0 | 0 | 0 | 1 |
| 0 | 0 | 0 | 1 |
| 0 | 0 | 0 | 0 |
| 0 | 0 | 0 | 1 |
| 0 | 0 | 0 | 0 |
| 0 | 0 | 0 | 0 |
| 0 | 0 | 0 | 0 |
| 0 | 0 | 0 | 1 |
| 1 | 0 | 0 | 0 |
| 1 | 0 | 0 | 0 |
| 0 | 0 | 0 | 1 |
| 0 | 0 | 0 | 1 |
| 0 | 0 | 0 | 0 |
| 0 | 0 | 0 | 0 |
| 0 | 1 | 1 | 1 |
| 0 | 1 | 0 | 0 |
| 0 | 1 | 0 | 0 |
| 0 | 1 | 0 | 0 |
| 0 | 0 | 0 | 0 |
| 0 | 0 | 0 | 1 |
| 0 | 0 | 0 | 1 |
| 0 | 0 | 0 | 0 |
| 0 | 0 | 0 | 1 |
| 0 | 0 | 0 | 0 |
| 0 | 0 | 0 | 1 |
| 0 | 0 | 0 | 1 |
| 0 | 0 | 0 | 1 |
| 0 | 0 | 0 | 1 |
| 0 | 0 | 0 | 1 |
| 0 | 0 | 0 | 1 |
| 0 | 0 | 0 | 1 |

| | | | |
|---|---|---|---|
| 0 | 0 | 0 | 1 |
| 0 | 0 | 0 | 1 |
| 0 | 0 | 0 | 1 |
| 0 | 0 | 0 | 1 |
| 0 | 0 | 0 | 1 |
| 0 | 0 | 0 | 1 |
| 0 | 0 | 0 | 1 |
| 0 | 0 | 0 | 0 |
| 0 | 0 | 0 | 1 |
| 1 | 0 | 0 | 0 |
| 0 | 0 | 0 | 0 |
| 0 | 0 | 0 | 0 |
| 0 | 0 | 0 | 1 |
| 0 | 0 | 0 | 1 |
| 0 | 0 | 0 | 0 |
| 0 | 0 | 0 | 1 |
| 0 | 0 | 0 | 1 |
| 0 | 0 | 0 | 1 |
| 0 | 0 | 0 | 1 |
| 0 | 0 | 0 | 1 |
| 0 | 0 | 0 | 1 |
| 0 | 0 | 0 | 1 |
| 0 | 0 | 0 | 1 |
| 0 | 0 | 0 | 1 |
| 0 | 0 | 0 | 1 |
| 0 | 0 | 0 | 1 |
| 0 | 0 | 0 | 1 |
| 0 | 0 | 0 | 1 |
| 0 | 0 | 0 | 1 |
| 0 | 0 | 0 | 0 |
| 0 | 0 | 0 | 0 |
| 0 | 0 | 0 | 1 |
| 0 | 0 | 0 | 1 |
| 0 | 0 | 0 | 1 |
| 0 | 0 | 0 | 1 |
| 0 | 0 | 0 | 0 |
| 0 | 0 | 0 | 1 |
| 0 | 0 | 0 | 1 |
| 0 | 0 | 0 | 1 |
| 0 | 0 | 0 | 1 |
| 0 | 0 | 0 | 1 |
| 0 | 0 | 0 | 0 |
| 0 | 0 | 0 | 0 |
| 0 | 1 | 0 | 0 |
| 0 | 0 | 0 | 1 |
| 0 | 0 | 0 | 1 |
| 0 | 0 | 0 | 1 |
| 0 | 0 | 0 | 1 |
| 0 | 0 | 0 | 1 |
| 0 | 0 | 0 | 1 |
| 0 | 0 | 0 | 1 |
| 0 | 0 | 0 | 1 |
| 0 | 0 | 0 | 1 |
| 0 | 0 | 0 | 0 |

| | | | |
|---|---|---|---|
| 0 | 0 | 0 | 1 |
| 0 | 0 | 0 | 1 |
| 0 | 0 | 0 | 1 |
| 0 | 0 | 0 | 1 |
| 0 | 0 | 0 | 1 |
| 0 | 0 | 0 | 1 |
| 0 | 0 | 0 | 1 |
| 0 | 0 | 0 | 1 |
| 0 | 0 | 0 | 0 |
| 0 | 0 | 0 | 0 |
| 0 | 0 | 0 | 1 |
| 0 | 0 | 0 | 1 |
| 0 | 0 | 0 | 1 |
| 0 | 0 | 0 | 0 |
| 0 | 0 | 0 | 1 |
| 0 | 0 | 0 | 1 |
| 0 | 0 | 0 | 1 |
| 1 | 0 | 0 | 0 |
| 1 | 0 | 0 | 0 |
| 0 | 0 | 0 | 0 |
| 0 | 0 | 0 | 0 |
| 0 | 0 | 0 | 1 |
| 0 | 0 | 0 | 1 |
| 0 | 0 | 0 | 1 |
| 0 | 0 | 0 | 1 |
| 1 | 0 | 0 | 0 |
| 0 | 0 | 0 | 1 |
| 0 | 0 | 0 | 1 |
| 0 | 0 | 0 | 1 |
| 0 | 0 | 0 | 1 |
| 0 | 0 | 0 | 1 |
| 0 | 0 | 0 | 1 |
| 0 | 0 | 0 | 1 |
| 0 | 0 | 0 | 1 |
| 0 | 0 | 0 | 1 |
| 0 | 0 | 0 | 1 |
| 0 | 0 | 0 | 1 |
| 0 | 0 | 0 | 1 |
| 0 | 0 | 0 | 1 |
| 0 | 0 | 0 | 0 |
| 0 | 0 | 0 | 1 |
| 0 | 0 | 0 | 1 |
| 0 | 0 | 0 | 0 |
| 0 | 0 | 0 | 1 |
| 0 | 0 | 0 | 1 |
| 0 | 0 | 0 | 1 |
| 0 | 0 | 0 | 0 |
| 0 | 0 | 0 | 0 |
| 0 | 0 | 0 | 1 |
| 0 | 0 | 0 | 1 |
| 0 | 0 | 0 | 0 |
| 0 | 0 | 0 | 1 |
| 0 | 0 | 0 | 1 |
| 0 | 0 | 0 | 1 |

| | | | |
|---|---|---|---|
| 0 | 0 | 0 | 1 |
| 0 | 0 | 0 | 1 |
| 0 | 0 | 0 | 1 |
| 0 | 0 | 0 | 1 |
| 0 | 0 | 0 | 1 |
| 0 | 0 | 0 | 1 |
| 0 | 0 | 0 | 1 |
| 0 | 0 | 0 | 1 |
| 0 | 0 | 0 | 1 |
| 0 | 0 | 0 | 1 |
| 0 | 0 | 0 | 1 |
| 0 | 0 | 0 | 1 |
| 0 | 0 | 0 | 1 |
| 0 | 0 | 0 | 0 |
| 0 | 0 | 0 | 0 |
| 0 | 0 | 0 | 0 |
| 0 | 0 | 0 | 0 |
| 0 | 0 | 0 | 1 |
| 0 | 0 | 0 | 1 |
| 0 | 0 | 0 | 0 |
| 0 | 0 | 0 | 1 |
| 0 | 1 | 1 | 1 |
| 0 | 0 | 0 | 1 |
| 0 | 0 | 0 | 1 |
| 0 | 0 | 0 | 1 |
| 0 | 0 | 0 | 0 |
| 0 | 0 | 0 | 1 |
| 0 | 0 | 0 | 1 |
| 0 | 0 | 0 | 1 |
| 0 | 0 | 0 | 0 |
| 0 | 0 | 0 | 0 |
| 0 | 0 | 0 | 1 |
| 0 | 0 | 0 | 1 |
| 0 | 0 | 0 | 1 |
| 0 | 0 | 0 | 1 |
| 0 | 0 | 0 | 1 |
| 0 | 0 | 0 | 1 |
| 0 | 0 | 0 | 1 |
| 0 | 0 | 0 | 0 |
| 0 | 0 | 0 | 1 |
| 0 | 0 | 0 | 1 |
| 0 | 0 | 0 | 0 |
| 0 | 1 | 0 | 0 |
| 0 | 1 | 0 | 0 |
| 0 | 0 | 0 | 0 |
| 0 | 0 | 0 | 1 |
| 0 | 0 | 0 | 0 |
| 0 | 0 | 0 | 1 |
| 0 | 0 | 0 | 1 |
| 0 | 0 | 0 | 1 |
| 0 | 0 | 0 | 1 |
| 0 | 0 | 0 | 1 |
| 0 | 1 | 1 | 1 |
| 0 | 1 | 1 | 1 |

| | | | |
|---|---|---|---|
| 0 | 0 | 0 | 1 |
| 0 | 0 | 0 | 1 |
| 0 | 0 | 0 | 0 |
| 0 | 0 | 0 | 1 |
| 0 | 0 | 0 | 1 |
| 0 | 0 | 0 | 1 |
| 0 | 0 | 0 | 1 |
| 0 | 0 | 0 | 1 |
| 0 | 0 | 0 | 1 |
| 0 | 0 | 0 | 1 |
| 0 | 0 | 0 | 1 |
| 0 | 0 | 0 | 1 |
| 0 | 0 | 0 | 1 |
| 0 | 0 | 0 | 1 |
| 0 | 0 | 0 | 1 |
| 0 | 0 | 0 | 1 |
| 0 | 0 | 0 | 1 |
| 0 | 0 | 0 | 1 |
| 0 | 0 | 0 | 1 |
| 0 | 0 | 0 | 1 |
| 0 | 0 | 0 | 1 |
| 0 | 0 | 0 | 1 |
| 0 | 0 | 0 | 1 |
| 0 | 0 | 0 | 1 |
| 0 | 0 | 0 | 1 |
| 0 | 0 | 0 | 1 |
| 0 | 0 | 0 | 1 |
| 0 | 0 | 0 | 1 |
| 0 | 0 | 0 | 1 |
| 0 | 0 | 0 | 1 |
| 1 | 0 | 0 | 0 |
| 0 | 0 | 0 | 1 |
| 0 | 1 | 0 | 0 |
| 0 | 1 | 0 | 0 |
| 0 | 1 | 0 | 0 |
| 0 | 0 | 0 | 0 |
| 0 | 0 | 0 | 1 |
| 0 | 0 | 0 | 0 |
| 0 | 0 | 0 | 1 |
| 0 | 0 | 0 | 1 |
| 1 | 0 | 0 | 0 |
| 1 | 0 | 0 | 0 |
| 0 | 0 | 0 | 0 |
| 0 | 0 | 0 | 1 |
| 0 | 0 | 0 | 0 |
| 0 | 0 | 0 | 0 |
| 0 | 0 | 0 | 1 |
| 0 | 0 | 0 | 1 |
| 0 | 0 | 0 | 1 |
| 0 | 0 | 0 | 1 |
| 0 | 0 | 0 | 0 |
| 0 | 0 | 0 | 1 |
| 0 | 0 | 0 | 1 |
| 0 | 0 | 0 | 1 |

| | | | |
|---|---|---|---|
| 0 | 0 | 0 | 1 |
| 0 | 0 | 0 | 1 |
| 0 | 0 | 0 | 1 |
| 0 | 0 | 0 | 1 |
| 0 | 0 | 0 | 1 |
| 1 | 0 | 0 | 0 |
| 0 | 0 | 0 | 1 |
| 0 | 0 | 0 | 0 |
| 0 | 0 | 0 | 0 |
| 0 | 0 | 0 | 0 |
| 0 | 0 | 0 | 1 |
| 0 | 1 | 0 | 0 |
| 0 | 0 | 0 | 1 |
| 0 | 0 | 0 | 1 |
| 0 | 0 | 0 | 1 |
| 0 | 1 | 1 | 1 |
| 0 | 0 | 0 | 1 |
| 0 | 0 | 0 | 0 |
| 0 | 0 | 0 | 1 |
| 0 | 0 | 0 | 0 |
| 0 | 0 | 0 | 1 |
| 0 | 0 | 0 | 0 |
| 0 | 0 | 0 | 0 |
| 0 | 0 | 0 | 0 |
| 0 | 0 | 0 | 1 |
| 0 | 0 | 0 | 1 |
| 0 | 0 | 0 | 1 |
| 0 | 0 | 0 | 1 |
| 0 | 0 | 0 | 0 |
| 0 | 0 | 0 | 1 |
| 0 | 0 | 0 | 1 |
| 0 | 0 | 0 | 1 |
| 0 | 0 | 0 | 1 |
| 0 | 0 | 0 | 1 |
| 0 | 0 | 0 | 0 |
| 0 | 0 | 0 | 0 |
| 0 | 0 | 0 | 1 |
| 0 | 0 | 0 | 0 |
| 0 | 0 | 0 | 0 |
| 0 | 0 | 0 | 1 |
| 0 | 0 | 0 | 0 |
| 0 | 0 | 0 | 0 |
| 0 | 0 | 0 | 1 |
| 0 | 0 | 0 | 1 |
| 0 | 0 | 0 | 1 |
| 0 | 0 | 0 | 1 |
| 0 | 0 | 0 | 1 |
| 0 | 0 | 0 | 1 |
| 0 | 0 | 0 | 1 |
| 0 | 0 | 0 | 1 |
| 0 | 0 | 0 | 1 |
| 0 | 0 | 0 | 1 |
| 0 | 0 | 0 | 1 |
| 0 | 0 | 0 | 1 |

| | | | |
|---|---|---|---|
| 0 | 0 | 0 | 1 |
| 0 | 0 | 0 | 0 |
| 0 | 0 | 0 | 1 |
| 0 | 0 | 0 | 1 |
| 0 | 0 | 0 | 0 |
| 0 | 0 | 0 | 1 |
| 0 | 0 | 0 | 1 |
| 0 | 0 | 0 | 1 |
| 0 | 0 | 0 | 0 |
| 0 | 0 | 0 | 0 |
| 0 | 0 | 0 | 1 |
| 0 | 0 | 0 | 0 |
| 0 | 0 | 0 | 0 |
| 0 | 0 | 0 | 1 |
| 0 | 0 | 0 | 1 |
| 0 | 0 | 0 | 0 |
| 0 | 0 | 0 | 1 |
| 0 | 0 | 0 | 0 |
| 0 | 1 | 0 | 0 |
| 0 | 0 | 0 | 1 |
| 0 | 0 | 0 | 0 |
| 0 | 0 | 0 | 0 |
| 1 | 0 | 0 | 0 |
| 0 | 0 | 0 | 1 |
| 0 | 0 | 0 | 1 |
| 0 | 0 | 0 | 0 |
| 0 | 0 | 0 | 1 |
| 0 | 0 | 0 | 1 |
| 0 | 0 | 0 | 1 |
| 0 | 0 | 0 | 1 |
| 0 | 0 | 0 | 0 |
| 0 | 0 | 0 | 1 |
| 0 | 0 | 0 | 0 |
| 0 | 0 | 0 | 1 |
| 0 | 0 | 0 | 0 |
| 0 | 0 | 0 | 0 |
| 0 | 0 | 0 | 1 |
| 0 | 0 | 0 | 1 |
| 0 | 0 | 0 | 1 |
| 0 | 0 | 0 | 1 |
| 0 | 0 | 0 | 1 |
| 0 | 0 | 0 | 1 |
| 0 | 0 | 0 | 1 |
| 0 | 0 | 0 | 1 |
| 0 | 0 | 0 | 1 |
| 0 | 0 | 0 | 1 |
| 0 | 1 | 1 | 1 |
| 0 | 0 | 0 | 1 |
| 0 | 0 | 0 | 1 |
| 0 | 0 | 0 | 1 |
| 0 | 0 | 0 | 1 |
| 0 | 0 | 0 | 1 |
| 0 | 0 | 0 | 1 |
| 0 | 0 | 0 | 0 |

| | | | |
|---|---|---|---|
| 0 | 0 | 0 | 1 |
| 0 | 0 | 0 | 1 |
| 0 | 0 | 0 | 1 |
| 0 | 0 | 0 | 1 |
| 0 | 0 | 0 | 0 |
| 0 | 0 | 0 | 1 |
| 0 | 0 | 0 | 1 |
| 0 | 0 | 0 | 1 |
| 0 | 0 | 0 | 1 |
| 0 | 0 | 0 | 1 |
| 0 | 0 | 0 | 0 |
| 0 | 0 | 0 | 1 |
| 0 | 0 | 0 | 1 |
| 0 | 0 | 0 | 1 |
| 0 | 0 | 0 | 0 |
| 0 | 0 | 0 | 0 |
| 0 | 0 | 0 | 0 |
| 0 | 0 | 0 | 0 |
| 0 | 0 | 0 | 1 |
| 0 | 0 | 0 | 0 |
| 0 | 0 | 0 | 0 |
| 1 | 0 | 0 | 0 |
| 0 | 0 | 0 | 1 |
| 0 | 0 | 0 | 0 |
| 1 | 0 | 0 | 0 |
| 0 | 0 | 0 | 1 |
| 1 | 0 | 0 | 0 |
| 0 | 0 | 0 | 0 |
| 1 | 0 | 0 | 0 |
| 1 | 0 | 0 | 0 |
| 1 | 0 | 0 | 0 |
| 0 | 0 | 0 | 1 |
| 0 | 0 | 0 | 1 |
| 0 | 0 | 0 | 1 |
| 0 | 0 | 0 | 1 |
| 0 | 0 | 0 | 1 |
| 0 | 0 | 0 | 1 |
| 0 | 1 | 0 | 0 |
| 0 | 0 | 0 | 0 |
| 0 | 0 | 0 | 1 |
| 0 | 0 | 0 | 1 |
| 0 | 1 | 1 | 1 |
| 0 | 0 | 0 | 0 |
| 0 | 0 | 0 | 1 |
| 0 | 0 | 0 | 0 |
| 0 | 0 | 0 | 1 |
| 0 | 0 | 0 | 1 |
| 0 | 0 | 0 | 1 |
| 0 | 1 | 0 | 0 |
| 0 | 1 | 0 | 0 |
| 0 | 1 | 0 | 0 |
| 0 | 0 | 0 | 1 |
| 0 | 0 | 0 | 1 |
| 0 | 0 | 0 | 0 |

| | | | |
|---|---|---|---|
| 0 | 0 | 0 | 1 |
| 0 | 1 | 0 | 0 |
| 0 | 0 | 0 | 1 |
| 0 | 1 | 0 | 0 |
| 0 | 1 | 0 | 0 |
| 0 | 0 | 0 | 1 |
| 0 | 1 | 0 | 0 |
| 0 | 1 | 0 | 0 |
| 0 | 0 | 0 | 1 |
| 0 | 0 | 0 | 1 |
| 0 | 0 | 0 | 1 |
| 0 | 0 | 0 | 1 |
| 0 | 0 | 0 | 1 |
| 0 | 0 | 0 | 1 |
| 0 | 0 | 0 | 1 |
| 0 | 0 | 0 | 1 |
| 0 | 0 | 0 | 1 |
| 0 | 0 | 0 | 1 |
| 0 | 0 | 0 | 1 |
| 0 | 0 | 0 | 1 |
| 1 | 1 | 0 | 0 |
| 0 | 0 | 0 | 1 |
| 0 | 0 | 0 | 1 |
| 0 | 0 | 0 | 1 |
| 0 | 0 | 0 | 1 |
| 0 | 0 | 0 | 0 |
| 0 | 0 | 0 | 0 |
| 0 | 0 | 0 | 1 |
| 0 | 0 | 0 | 1 |
| 0 | 0 | 0 | 1 |
| 0 | 0 | 0 | 1 |
| 0 | 0 | 0 | 1 |
| 0 | 0 | 0 | 1 |
| 0 | 0 | 0 | 1 |
| 0 | 0 | 0 | 1 |
| 0 | 0 | 0 | 1 |
| 0 | 0 | 0 | 1 |
| 0 | 0 | 0 | 1 |
| 0 | 0 | 0 | 1 |
| 0 | 0 | 0 | 1 |
| 0 | 0 | 0 | 1 |
| 0 | 0 | 0 | 1 |
| 0 | 0 | 0 | 1 |
| 0 | 0 | 0 | 0 |
| 0 | 0 | 0 | 1 |
| 0 | 0 | 0 | 0 |
| 0 | 0 | 0 | 0 |
| 0 | 0 | 0 | 0 |
| 0 | 0 | 0 | 0 |
| 0 | 0 | 0 | 0 |
| 0 | 0 | 0 | 0 |
| 0 | 0 | 0 | 1 |
| 0 | 0 | 0 | 1 |
| 0 | 0 | 0 | 1 |

| | | | |
|---|---|---|---|
| 0 | 0 | 0 | 1 |
| 0 | 0 | 0 | 1 |
| 0 | 0 | 0 | 1 |
| 0 | 0 | 0 | 0 |
| 0 | 0 | 0 | 1 |
| 0 | 0 | 0 | 0 |
| 0 | 0 | 0 | 1 |
| 0 | 0 | 0 | 1 |
| 0 | 0 | 0 | 0 |
| 0 | 0 | 0 | 1 |
| 0 | 0 | 0 | 1 |
| 0 | 0 | 0 | 1 |
| 0 | 0 | 0 | 1 |
| 0 | 0 | 0 | 1 |
| 0 | 0 | 0 | 1 |
| 0 | 0 | 0 | 0 |
| 0 | 0 | 0 | 0 |
| 0 | 0 | 0 | 0 |
| 0 | 0 | 0 | 0 |
| 0 | 0 | 0 | 0 |
| 0 | 0 | 0 | 1 |
| 0 | 0 | 0 | 1 |
| 0 | 0 | 0 | 1 |
| 0 | 0 | 0 | 1 |
| 0 | 0 | 0 | 0 |
| 0 | 0 | 0 | 1 |
| 0 | 0 | 0 | 1 |
| 0 | 0 | 0 | 1 |
| 0 | 0 | 0 | 0 |
| 0 | 0 | 0 | 1 |
| 0 | 1 | 0 | 0 |
| 0 | 1 | 0 | 0 |
| 0 | 0 | 0 | 0 |
| 0 | 1 | 0 | 0 |
| 0 | 1 | 0 | 0 |
| 0 | 0 | 0 | 1 |
| 0 | 1 | 0 | 0 |
| 0 | 1 | 0 | 0 |
| 0 | 1 | 0 | 0 |
| 0 | 1 | 0 | 0 |
| 0 | 1 | 0 | 0 |
| 0 | 0 | 0 | 0 |
| 0 | 0 | 0 | 0 |
| 0 | 0 | 0 | 0 |
| 0 | 0 | 0 | 0 |
| 0 | 0 | 0 | 0 |
| 0 | 0 | 0 | 0 |
| 0 | 0 | 0 | 0 |
| 0 | 0 | 0 | 1 |
| 0 | 0 | 0 | 0 |
| 0 | 0 | 0 | 1 |
| 0 | 0 | 0 | 1 |
| 0 | 0 | 0 | 1 |
| 0 | 0 | 0 | 0 |

| | | | |
|---|---|---|---|
| 0 | 1 | 0 | 0 |
| 0 | 0 | 0 | 1 |
| 0 | 0 | 0 | 1 |
| 0 | 0 | 0 | 1 |
| 0 | 0 | 0 | 1 |
| 0 | 0 | 0 | 0 |
| 0 | 0 | 0 | 1 |
| 0 | 0 | 0 | 1 |
| 0 | 0 | 0 | 1 |
| 0 | 0 | 0 | 1 |
| 0 | 0 | 0 | 1 |
| 0 | 0 | 0 | 1 |
| 0 | 0 | 0 | 1 |
| 1 | 0 | 0 | 0 |
| 0 | 0 | 0 | 1 |
| 0 | 0 | 0 | 1 |
| 0 | 0 | 0 | 1 |
| 0 | 0 | 0 | 0 |
| 0 | 0 | 0 | 1 |
| 0 | 0 | 0 | 1 |
| 0 | 0 | 0 | 1 |
| 0 | 0 | 0 | 1 |
| 0 | 0 | 0 | 0 |
| 0 | 0 | 0 | 1 |
| 0 | 0 | 0 | 1 |
| 0 | 0 | 0 | 0 |
| 0 | 0 | 0 | 0 |
| 0 | 0 | 0 | 0 |
| 0 | 0 | 0 | 1 |
| 0 | 0 | 0 | 1 |
| 0 | 0 | 0 | 1 |
| 0 | 0 | 0 | 1 |
| 0 | 0 | 0 | 1 |
| 0 | 0 | 0 | 1 |
| 0 | 0 | 0 | 1 |
| 0 | 0 | 0 | 1 |
| 0 | 0 | 0 | 1 |
| 0 | 0 | 0 | 1 |
| 0 | 0 | 0 | 1 |
| 0 | 0 | 0 | 1 |
| 0 | 0 | 0 | 1 |
| 0 | 0 | 0 | 1 |
| 0 | 0 | 0 | 1 |
| 0 | 0 | 0 | 1 |
| 0 | 0 | 0 | 1 |
| 0 | 0 | 0 | 1 |
| 0 | 0 | 0 | 1 |
| 0 | 0 | 0 | 1 |
| 0 | 1 | 1 | 1 |
| 0 | 0 | 0 | 1 |
| 0 | 0 | 0 | 1 |
| 0 | 0 | 0 | 1 |
| 0 | 0 | 0 | 1 |
| 0 | 0 | 0 | 1 |

| | | | |
|---|---|---|---|
| 0 | 0 | 0 | 1 |
| 0 | 0 | 0 | 0 |
| 0 | 0 | 0 | 1 |
| 0 | 0 | 0 | 1 |
| 0 | 0 | 0 | 1 |
| 0 | 0 | 0 | 0 |
| 0 | 0 | 0 | 0 |
| 0 | 0 | 0 | 0 |
| 0 | 0 | 0 | 1 |
| 0 | 0 | 0 | 1 |
| 0 | 0 | 0 | 1 |
| 0 | 0 | 0 | 1 |
| 0 | 0 | 0 | 1 |
| 0 | 0 | 0 | 1 |
| 0 | 0 | 0 | 0 |
| 0 | 0 | 0 | 0 |
| 0 | 0 | 0 | 0 |
| 0 | 0 | 0 | 1 |
| 0 | 0 | 0 | 0 |
| 0 | 0 | 0 | 0 |
| 0 | 0 | 0 | 0 |
| 0 | 0 | 0 | 0 |
| 0 | 0 | 0 | 0 |
| 0 | 0 | 0 | 0 |
| 0 | 0 | 0 | 1 |
| 0 | 0 | 0 | 1 |
| 0 | 0 | 0 | 1 |
| 0 | 0 | 0 | 1 |
| 0 | 0 | 0 | 1 |
| 0 | 0 | 0 | 1 |
| 0 | 0 | 0 | 1 |
| 0 | 0 | 0 | 1 |
| 0 | 0 | 0 | 0 |
| 0 | 0 | 0 | 1 |
| 0 | 0 | 0 | 1 |
| 0 | 0 | 0 | 0 |
| 0 | 0 | 0 | 0 |
| 0 | 0 | 0 | 0 |
| 0 | 0 | 0 | 0 |
| 0 | 0 | 0 | 1 |
| 0 | 0 | 0 | 1 |
| 0 | 0 | 0 | 1 |
| 0 | 0 | 0 | 1 |
| 0 | 0 | 0 | 0 |
| 0 | 0 | 0 | 0 |
| 0 | 0 | 0 | 0 |
| 0 | 0 | 0 | 1 |
| 0 | 0 | 0 | 1 |
| 0 | 0 | 0 | 0 |
| 0 | 0 | 0 | 1 |
| 0 | 0 | 0 | 1 |
| 0 | 0 | 0 | 0 |
| 0 | 0 | 0 | 1 |
| 0 | 0 | 0 | 1 |

| | | | |
|---|---|---|---|
| 0 | 0 | 0 | 1 |
| 0 | 0 | 0 | 1 |
| 0 | 0 | 0 | 1 |
| 0 | 0 | 0 | 1 |
| 0 | 0 | 0 | 1 |
| 0 | 0 | 0 | 1 |
| 0 | 0 | 0 | 1 |
| 0 | 0 | 0 | 1 |
| 0 | 0 | 0 | 0 |
| 0 | 0 | 0 | 0 |
| 0 | 0 | 0 | 1 |
| 0 | 0 | 0 | 0 |
| 0 | 0 | 0 | 0 |
| 0 | 0 | 0 | 0 |
| 0 | 0 | 0 | 1 |
| 0 | 0 | 0 | 1 |
| 0 | 0 | 0 | 0 |
| 0 | 0 | 0 | 0 |
| 0 | 0 | 0 | 0 |
| 0 | 0 | 0 | 1 |
| 0 | 0 | 0 | 1 |
| 0 | 0 | 0 | 1 |
| 0 | 0 | 0 | 0 |
| 0 | 0 | 0 | 1 |
| 0 | 0 | 0 | 0 |
| 0 | 0 | 0 | 1 |
| 0 | 0 | 0 | 1 |
| 0 | 0 | 0 | 1 |
| 0 | 0 | 0 | 1 |
| 0 | 0 | 0 | 0 |
| 0 | 0 | 0 | 0 |
| 0 | 0 | 0 | 0 |
| 0 | 0 | 0 | 1 |
| 0 | 0 | 0 | 1 |
| 0 | 0 | 0 | 1 |
| 0 | 0 | 0 | 1 |
| 0 | 0 | 0 | 1 |
| 0 | 0 | 0 | 1 |
| 0 | 0 | 0 | 1 |
| 0 | 0 | 0 | 1 |
| 0 | 0 | 0 | 0 |
| 0 | 0 | 0 | 1 |
| 0 | 0 | 0 | 1 |
| 0 | 0 | 0 | 1 |
| 0 | 0 | 0 | 1 |
| 0 | 0 | 0 | 0 |
| 0 | 1 | 1 | 1 |
| 0 | 0 | 0 | 0 |
| 0 | 0 | 0 | 1 |
| 0 | 1 | 0 | 0 |
| 0 | 1 | 0 | 0 |
| 0 | 0 | 0 | 0 |
| 0 | 0 | 0 | 1 |
| 0 | 1 | 1 | 1 |

| | | | |
|---|---|---|---|
| 0 | 1 | 1 | 1 |
| 0 | 1 | 1 | 1 |
| 0 | 0 | 0 | 1 |
| 0 | 0 | 0 | 1 |
| 0 | 0 | 0 | 1 |
| 0 | 0 | 0 | 1 |
| 0 | 0 | 0 | 1 |
| 0 | 0 | 0 | 0 |
| 0 | 0 | 0 | 0 |
| 0 | 0 | 0 | 1 |
| 0 | 0 | 0 | 0 |
| 0 | 0 | 0 | 1 |
| 0 | 0 | 0 | 1 |
| 0 | 0 | 0 | 1 |
| 0 | 0 | 0 | 1 |
| 0 | 0 | 0 | 1 |
| 0 | 0 | 0 | 1 |
| 0 | 0 | 0 | 1 |
| 0 | 0 | 0 | 1 |
| 0 | 0 | 0 | 1 |
| 0 | 0 | 0 | 1 |
| 0 | 0 | 0 | 1 |
| 0 | 0 | 0 | 1 |
| 0 | 0 | 0 | 1 |
| 0 | 0 | 0 | 1 |
| 0 | 0 | 0 | 1 |
| 0 | 0 | 0 | 1 |
| 0 | 0 | 0 | 1 |
| 0 | 0 | 0 | 0 |
| 0 | 0 | 0 | 1 |
| 0 | 0 | 0 | 0 |
| 0 | 0 | 0 | 1 |
| 0 | 0 | 0 | 0 |
| 0 | 0 | 0 | 0 |
| 0 | 0 | 0 | 1 |
| 0 | 0 | 0 | 0 |
| 0 | 0 | 0 | 0 |
| 0 | 0 | 0 | 1 |
| 0 | 0 | 0 | 1 |
| 0 | 0 | 0 | 1 |
| 0 | 0 | 0 | 1 |
| 0 | 0 | 0 | 0 |
| 0 | 0 | 0 | 0 |
| 0 | 0 | 0 | 1 |
| 0 | 0 | 0 | 1 |
| 0 | 0 | 0 | 0 |
| 0 | 0 | 0 | 0 |
| 0 | 0 | 0 | 0 |
| 0 | 0 | 0 | 0 |
| 0 | 0 | 0 | 1 |
| 0 | 1 | 0 | 0 |
| 0 | 0 | 0 | 0 |
| 0 | 0 | 0 | 1 |
| 0 | 0 | 0 | 1 |

| | | | |
|---|---|---|---|
| 0 | 0 | 0 | 1 |
| 0 | 0 | 0 | 1 |
| 0 | 0 | 0 | 1 |
| 0 | 0 | 0 | 1 |
| 0 | 0 | 0 | 0 |
| 0 | 0 | 0 | 0 |
| 0 | 0 | 0 | 0 |
| 0 | 0 | 0 | 0 |
| 0 | 0 | 0 | 1 |
| 0 | 0 | 0 | 0 |
| 0 | 0 | 0 | 0 |
| 0 | 0 | 0 | 0 |
| 0 | 0 | 0 | 0 |
| 0 | 0 | 0 | 1 |
| 0 | 1 | 1 | 1 |
| 0 | 0 | 0 | 1 |
| 0 | 0 | 0 | 1 |
| 0 | 0 | 0 | 1 |
| 0 | 0 | 0 | 1 |
| 0 | 0 | 0 | 1 |
| 0 | 0 | 0 | 1 |
| 0 | 0 | 0 | 1 |
| 0 | 0 | 0 | 1 |
| 0 | 0 | 0 | 1 |
| 0 | 0 | 0 | 1 |
| 0 | 0 | 0 | 0 |
| 0 | 0 | 0 | 0 |
| 0 | 0 | 0 | 0 |
| 0 | 0 | 0 | 0 |
| 0 | 0 | 0 | 1 |
| 0 | 0 | 0 | 1 |
| 0 | 0 | 0 | 1 |
| 0 | 0 | 0 | 1 |
| 0 | 0 | 0 | 1 |
| 0 | 0 | 0 | 1 |
| 0 | 0 | 0 | 1 |
| 0 | 0 | 0 | 1 |
| 0 | 0 | 0 | 1 |
| 0 | 0 | 0 | 1 |
| 0 | 0 | 0 | 1 |
| 0 | 0 | 0 | 1 |
| 0 | 0 | 0 | 1 |
| 0 | 0 | 0 | 1 |
| 0 | 0 | 0 | 1 |
| 0 | 0 | 0 | 1 |
| 0 | 0 | 0 | 1 |
| 0 | 0 | 0 | 1 |
| 0 | 0 | 0 | 1 |
| 0 | 0 | 0 | 1 |
| 0 | 0 | 0 | 1 |
| 0 | 0 | 0 | 1 |
| 0 | 0 | 0 | 1 |
| 0 | 0 | 0 | 1 |
| 0 | 0 | 0 | 0 |

| | | | |
|---|---|---|---|
| 0 | 0 | 0 | 1 |
| 0 | 0 | 0 | 1 |
| 0 | 0 | 0 | 1 |
| 0 | 0 | 0 | 1 |
| 0 | 0 | 0 | 1 |
| 0 | 0 | 0 | 1 |
| 0 | 0 | 0 | 1 |
| 0 | 0 | 0 | 1 |
| 0 | 0 | 0 | 1 |
| 0 | 0 | 0 | 1 |
| 0 | 0 | 0 | 1 |
| 0 | 0 | 0 | 1 |
| 0 | 0 | 0 | 1 |
| 0 | 0 | 0 | 1 |
| 0 | 0 | 0 | 1 |
| 0 | 0 | 0 | 1 |
| 0 | 0 | 0 | 1 |
| 0 | 0 | 0 | 1 |
| 0 | 0 | 0 | 1 |
| 0 | 0 | 0 | 1 |
| 0 | 0 | 0 | 1 |
| 0 | 0 | 0 | 1 |
| 0 | 0 | 0 | 1 |
| 0 | 0 | 0 | 1 |
| 0 | 0 | 0 | 1 |
| 0 | 0 | 0 | 1 |
| 0 | 0 | 0 | 1 |
| 0 | 0 | 0 | 1 |
| 0 | 0 | 0 | 1 |
| 0 | 0 | 0 | 1 |
| 0 | 0 | 0 | 1 |
| 0 | 0 | 0 | 1 |
| 0 | 0 | 0 | 1 |
| 0 | 0 | 0 | 1 |
| 0 | 0 | 0 | 1 |
| 0 | 0 | 0 | 1 |
| 0 | 0 | 0 | 1 |
| 0 | 0 | 0 | 1 |
| 0 | 0 | 0 | 1 |
| 0 | 0 | 0 | 1 |
| 0 | 0 | 0 | 1 |
| 0 | 0 | 0 | 1 |
| 0 | 0 | 0 | 1 |
| 0 | 0 | 0 | 1 |
| 0 | 0 | 0 | 1 |
| 0 | 1 | 1 | 1 |
| 0 | 0 | 0 | 0 |
| 0 | 0 | 0 | 1 |
| 0 | 0 | 0 | 1 |
| 0 | 0 | 0 | 1 |
| 0 | 0 | 0 | 1 |
| 0 | 0 | 0 | 1 |
| 0 | 0 | 0 | 1 |
| 0 | 0 | 0 | 1 |

| | | | |
|---|---|---|---|
| 0 | 0 | 0 | 1 |
| 0 | 0 | 0 | 1 |
| 0 | 0 | 0 | 0 |
| 0 | 0 | 0 | 1 |
| 0 | 0 | 0 | 1 |
| 0 | 0 | 0 | 1 |
| 0 | 0 | 0 | 0 |
| 0 | 0 | 0 | 1 |
| 0 | 0 | 0 | 1 |
| 0 | 0 | 0 | 1 |
| 0 | 0 | 0 | 1 |
| 0 | 0 | 0 | 0 |
| 0 | 0 | 0 | 1 |
| 0 | 0 | 0 | 0 |
| 0 | 0 | 0 | 0 |
| 0 | 0 | 0 | 0 |
| 0 | 0 | 0 | 0 |
| 0 | 0 | 0 | 0 |
| 0 | 0 | 0 | 0 |
| 0 | 0 | 0 | 0 |
| 0 | 0 | 0 | 0 |

| surface_non_binding_site | non_binding_site_single_water | surface_non_binding_site_single_water |
|---|---|---|
| 0 | 0 | 0 |
| 0 | 0 | 0 |
| 0 | 0 | 0 |
| 0 | 0 | 0 |
| 0 | 0 | 0 |
| 0 | 0 | 0 |
| 1 | 1 | 1 |
| 1 | 1 | 1 |
| 0 | 0 | 0 |
| 0 | 0 | 0 |
| 0 | 0 | 0 |
| 0 | 0 | 0 |
| 0 | 0 | 0 |
| 0 | 0 | 0 |
| 0 | 0 | 0 |
| 0 | 0 | 0 |
| 0 | 0 | 0 |
| 0 | 0 | 0 |
| 0 | 0 | 0 |
| 0 | 0 | 0 |
| 0 | 0 | 0 |
| 0 | 0 | 0 |
| 0 | 0 | 0 |
| 0 | 0 | 0 |
| 0 | 0 | 0 |
| 1 | 0 | 0 |
| 0 | 0 | 0 |
| 1 | 1 | 1 |
| 0 | 0 | 0 |
| 0 | 1 | 0 |
| 0 | 1 | 0 |
| 0 | 0 | 0 |
| 0 | 0 | 0 |
| 0 | 0 | 0 |
| 0 | 0 | 0 |
| 1 | 1 | 1 |
| 0 | 0 | 0 |
| 0 | 0 | 0 |
| 0 | 0 | 0 |
| 0 | 0 | 0 |
| 0 | 0 | 0 |
| 0 | 0 | 0 |
| 0 | 0 | 0 |
| 0 | 0 | 0 |
| 0 | 0 | 0 |
| 0 | 0 | 0 |
| 0 | 0 | 0 |
| 0 | 0 | 0 |
| 0 | 0 | 0 |
| 0 | 0 | 0 |
| 0 | 0 | 0 |
| 0 | 0 | 0 |
| 0 | 0 | 0 |

| | | |
|---|---|---|
| 0 | 0 | 0 |
| 0 | 0 | 0 |
| 0 | 0 | 0 |
| 0 | 0 | 0 |
| 0 | 0 | 0 |
| 0 | 0 | 0 |
| 0 | 0 | 0 |
| 0 | 0 | 0 |
| 0 | 0 | 0 |
| 0 | 0 | 0 |
| 0 | 0 | 0 |
| 0 | 0 | 0 |
| 0 | 0 | 0 |
| 0 | 0 | 0 |
| 0 | 0 | 0 |
| 0 | 0 | 0 |
| 0 | 0 | 0 |
| 0 | 0 | 0 |
| 0 | 0 | 0 |
| 0 | 0 | 0 |
| 0 | 0 | 0 |
| 0 | 0 | 0 |
| 0 | 0 | 0 |
| 0 | 0 | 0 |
| 0 | 0 | 0 |
| 0 | 0 | 0 |
| 0 | 0 | 0 |
| 0 | 0 | 0 |
| 0 | 0 | 0 |
| 0 | 0 | 0 |
| 0 | 0 | 0 |
| 0 | 0 | 0 |
| 0 | 0 | 0 |
| 0 | 0 | 0 |
| 0 | 0 | 0 |
| 0 | 0 | 0 |
| 0 | 0 | 0 |
| 0 | 0 | 0 |
| 0 | 0 | 0 |
| 0 | 0 | 0 |
| 0 | 0 | 0 |
| 0 | 0 | 0 |
| 0 | 0 | 0 |
| 0 | 0 | 0 |
| 0 | 0 | 0 |
| 1 | 1 | 1 |
| 0 | 1 | 0 |
| 0 | 0 | 0 |
| 0 | 0 | 0 |
| 0 | 0 | 0 |
| 0 | 0 | 0 |
| 1 | 1 | 1 |
| 1 | 1 | 1 |
| 0 | 0 | 0 |

| | | |
|---|---|---|
| 0 | 1 | 0 |
| 0 | 1 | 0 |
| 0 | 0 | 0 |
| 0 | 0 | 0 |
| 0 | 0 | 0 |
| 0 | 0 | 0 |
| 0 | 1 | 0 |
| 0 | 0 | 0 |
| 0 | 0 | 0 |
| 0 | 0 | 0 |
| 0 | 0 | 0 |
| 0 | 0 | 0 |
| 0 | 0 | 0 |
| 0 | 0 | 0 |
| 0 | 0 | 0 |
| 1 | 1 | 1 |
| 0 | 0 | 0 |
| 0 | 0 | 0 |
| 0 | 0 | 0 |
| 0 | 0 | 0 |
| 0 | 0 | 0 |
| 0 | 0 | 0 |
| 0 | 0 | 0 |
| 0 | 0 | 0 |
| 0 | 0 | 0 |
| 0 | 0 | 0 |
| 0 | 0 | 0 |
| 0 | 0 | 0 |
| 0 | 0 | 0 |
| 0 | 0 | 0 |
| 0 | 0 | 0 |
| 0 | 0 | 0 |
| 0 | 0 | 0 |
| 0 | 0 | 0 |
| 0 | 0 | 0 |
| 0 | 0 | 0 |
| 0 | 0 | 0 |
| 0 | 0 | 0 |
| 0 | 0 | 0 |
| 0 | 0 | 0 |
| 0 | 0 | 0 |
| 0 | 0 | 0 |
| 0 | 0 | 0 |
| 0 | 0 | 0 |
| 0 | 0 | 0 |
| 0 | 0 | 0 |
| 0 | 0 | 0 |
| 0 | 0 | 0 |
| 0 | 0 | 0 |
| 0 | 0 | 0 |
| 0 | 0 | 0 |
| 0 | 0 | 0 |
| 0 | 0 | 0 |
| 0 | 0 | 0 |

| | | |
|---|---|---|
| 1 | 0 | 0 |
| 1 | 0 | 0 |
| 1 | 0 | 0 |
| 0 | 0 | 0 |
| 0 | 0 | 0 |
| 0 | 0 | 0 |
| 0 | 0 | 0 |
| 1 | 0 | 0 |
| 0 | 0 | 0 |
| 1 | 0 | 0 |
| 1 | 0 | 0 |
| 0 | 0 | 0 |
| 1 | 0 | 0 |
| 0 | 0 | 0 |
| 0 | 0 | 0 |
| 1 | 0 | 0 |
| 1 | 0 | 0 |
| 1 | 0 | 0 |
| 1 | 0 | 0 |
| 0 | 0 | 0 |
| 0 | 0 | 0 |
| 0 | 1 | 0 |
| 0 | 0 | 0 |
| 0 | 0 | 0 |
| 0 | 0 | 0 |
| 1 | 0 | 0 |
| 0 | 0 | 0 |
| 0 | 0 | 0 |
| 1 | 0 | 0 |
| 1 | 0 | 0 |
| 1 | 1 | 1 |
| 0 | 0 | 0 |
| 0 | 0 | 0 |
| 0 | 1 | 0 |
| 0 | 0 | 0 |
| 0 | 0 | 0 |
| 0 | 0 | 0 |
| 0 | 0 | 0 |
| 0 | 0 | 0 |
| 1 | 0 | 0 |
| 1 | 0 | 0 |
| 0 | 0 | 0 |
| 0 | 0 | 0 |
| 1 | 1 | 1 |
| 1 | 0 | 0 |
| 1 | 0 | 0 |
| 0 | 0 | 0 |
| 0 | 0 | 0 |
| 0 | 0 | 0 |
| 0 | 0 | 0 |
| 0 | 0 | 0 |
| 0 | 1 | 0 |
| 1 | 0 | 0 |
| 0 | 0 | 0 |

| | | |
|---|---|---|
| 0 | 0 | 0 |
| 0 | 0 | 0 |
| 0 | 0 | 0 |
| 1 | 1 | 1 |
| 1 | 1 | 1 |
| 1 | 0 | 0 |
| 0 | 0 | 0 |
| 0 | 0 | 0 |
| 1 | 0 | 0 |
| 1 | 0 | 0 |
| 0 | 0 | 0 |
| 0 | 1 | 0 |
| 0 | 0 | 0 |
| 1 | 0 | 0 |
| 0 | 1 | 0 |
| 1 | 1 | 1 |
| 1 | 1 | 1 |
| 0 | 0 | 0 |
| 0 | 0 | 0 |
| 0 | 0 | 0 |
| 0 | 0 | 0 |
| 1 | 0 | 0 |
| 0 | 0 | 0 |
| 0 | 1 | 0 |
| 1 | 1 | 1 |
| 0 | 0 | 0 |
| 0 | 0 | 0 |
| 0 | 0 | 0 |
| 0 | 0 | 0 |
| 0 | 0 | 0 |
| 0 | 0 | 0 |
| 0 | 0 | 0 |
| 1 | 0 | 0 |
| 1 | 0 | 0 |
| 0 | 0 | 0 |
| 1 | 0 | 0 |
| 0 | 0 | 0 |
| 0 | 0 | 0 |
| 1 | 0 | 0 |
| 0 | 1 | 0 |
| 0 | 0 | 0 |
| 1 | 0 | 0 |
| 1 | 0 | 0 |
| 1 | 0 | 0 |
| 0 | 0 | 0 |
| 0 | 0 | 0 |
| 0 | 0 | 0 |
| 0 | 0 | 0 |
| 0 | 0 | 0 |
| 0 | 0 | 0 |
| 0 | 0 | 0 |
| 0 | 0 | 0 |
| 0 | 0 | 0 |
| 0 | 0 | 0 |

| | | |
|---|---|---|
| 0 | 0 | 0 |
| 0 | 0 | 0 |
| 0 | 0 | 0 |
| 0 | 0 | 0 |
| 0 | 0 | 0 |
| 0 | 0 | 0 |
| 0 | 1 | 0 |
| 0 | 1 | 0 |
| 0 | 0 | 0 |
| 0 | 0 | 0 |
| 0 | 0 | 0 |
| 0 | 0 | 0 |
| 0 | 0 | 0 |
| 0 | 1 | 0 |
| 0 | 1 | 0 |
| 0 | 0 | 0 |
| 0 | 1 | 0 |
| 0 | 0 | 0 |
| 1 | 1 | 1 |
| 0 | 0 | 0 |
| 1 | 1 | 1 |
| 1 | 0 | 0 |
| 1 | 1 | 1 |
| 0 | 0 | 0 |
| 0 | 0 | 0 |
| 0 | 0 | 0 |
| 0 | 0 | 0 |
| 0 | 0 | 0 |
| 0 | 0 | 0 |
| 0 | 0 | 0 |
| 0 | 0 | 0 |
| 1 | 0 | 0 |
| 0 | 0 | 0 |
| 0 | 0 | 0 |
| 0 | 0 | 0 |
| 0 | 0 | 0 |
| 0 | 0 | 0 |
| 0 | 0 | 0 |
| 0 | 0 | 0 |
| 0 | 0 | 0 |
| 0 | 1 | 0 |
| 0 | 1 | 0 |
| 0 | 0 | 0 |
| 0 | 1 | 0 |
| 0 | 0 | 0 |
| 0 | 0 | 0 |
| 0 | 0 | 0 |
| 1 | 1 | 1 |
| 1 | 1 | 1 |
| 0 | 0 | 0 |
| 0 | 1 | 0 |
| 1 | 1 | 1 |
| 0 | 0 | 0 |
| 0 | 1 | 0 |

| | | |
|---|---|---|
| 0 | 0 | 0 |
| 1 | 1 | 1 |
| 0 | 0 | 0 |
| 0 | 0 | 0 |
| 0 | 0 | 0 |
| 0 | 0 | 0 |
| 0 | 1 | 0 |
| 0 | 0 | 0 |
| 0 | 0 | 0 |
| 1 | 1 | 1 |
| 1 | 1 | 1 |
| 1 | 1 | 1 |
| 1 | 1 | 1 |
| 1 | 1 | 1 |
| 0 | 0 | 0 |
| 1 | 1 | 1 |
| 0 | 0 | 0 |
| 0 | 0 | 0 |
| 1 | 1 | 1 |
| 0 | 0 | 0 |
| 0 | 0 | 0 |
| 0 | 0 | 0 |
| 0 | 0 | 0 |
| 0 | 0 | 0 |
| 1 | 1 | 1 |
| 1 | 0 | 0 |
| 0 | 0 | 0 |
| 0 | 0 | 0 |
| 0 | 0 | 0 |
| 0 | 0 | 0 |
| 0 | 0 | 0 |
| 0 | 0 | 0 |
| 0 | 0 | 0 |
| 0 | 0 | 0 |
| 1 | 1 | 1 |
| 1 | 1 | 1 |
| 0 | 0 | 0 |
| 0 | 0 | 0 |
| 0 | 0 | 0 |
| 0 | 0 | 0 |
| 0 | 0 | 0 |
| 0 | 0 | 0 |
| 0 | 1 | 0 |
| 0 | 0 | 0 |
| 0 | 0 | 0 |
| 0 | 0 | 0 |
| 0 | 0 | 0 |
| 0 | 0 | 0 |
| 0 | 0 | 0 |
| 1 | 1 | 1 |
| 0 | 0 | 0 |
| 0 | 0 | 0 |
| 0 | 0 | 0 |
| 0 | 0 | 0 |

| | | |
|---|---|---|
| 0 | 0 | 0 |
| 0 | 0 | 0 |
| 0 | 0 | 0 |
| 0 | 0 | 0 |
| 1 | 1 | 1 |
| 0 | 0 | 0 |
| 0 | 0 | 0 |
| 0 | 0 | 0 |
| 0 | 0 | 0 |
| 0 | 0 | 0 |
| 0 | 0 | 0 |
| 0 | 0 | 0 |
| 0 | 0 | 0 |
| 0 | 0 | 0 |
| 0 | 0 | 0 |
| 0 | 0 | 0 |
| 0 | 0 | 0 |
| 0 | 0 | 0 |
| 1 | 1 | 1 |
| 0 | 0 | 0 |
| 0 | 0 | 0 |
| 0 | 0 | 0 |
| 0 | 0 | 0 |
| 1 | 0 | 0 |
| 0 | 0 | 0 |
| 0 | 0 | 0 |
| 0 | 0 | 0 |
| 1 | 0 | 0 |
| 1 | 0 | 0 |
| 1 | 0 | 0 |
| 0 | 0 | 0 |
| 0 | 0 | 0 |
| 0 | 0 | 0 |
| 0 | 0 | 0 |
| 0 | 0 | 0 |
| 1 | 0 | 0 |
| 1 | 0 | 0 |
| 0 | 0 | 0 |
| 0 | 0 | 0 |
| 1 | 0 | 0 |
| 1 | 0 | 0 |
| 1 | 0 | 0 |
| 0 | 1 | 0 |
| 1 | 1 | 1 |
| 0 | 0 | 0 |
| 0 | 0 | 0 |
| 0 | 0 | 0 |
| 0 | 0 | 0 |
| 0 | 0 | 0 |
| 0 | 1 | 0 |
| 1 | 1 | 1 |
| 0 | 0 | 0 |
| 1 | 1 | 1 |
| 0 | 0 | 0 |

| | | |
|---|---|---|
| 1 | 1 | 1 |
| 1 | 1 | 1 |
| 0 | 0 | 0 |
| 1 | 1 | 1 |
| 1 | 1 | 1 |
| 1 | 1 | 1 |
| 1 | 1 | 1 |
| 0 | 0 | 0 |
| 1 | 0 | 0 |
| 1 | 1 | 1 |
| 0 | 0 | 0 |
| 1 | 1 | 1 |
| 1 | 1 | 1 |
| 0 | 1 | 0 |
| 0 | 0 | 0 |
| 0 | 0 | 0 |
| 1 | 1 | 1 |
| 1 | 1 | 1 |
| 0 | 0 | 0 |
| 0 | 0 | 0 |
| 0 | 0 | 0 |
| 0 | 0 | 0 |
| 0 | 0 | 0 |
| 0 | 0 | 0 |
| 0 | 0 | 0 |
| 1 | 1 | 1 |
| 0 | 0 | 0 |
| 0 | 0 | 0 |
| 0 | 0 | 0 |
| 1 | 0 | 0 |
| 0 | 0 | 0 |
| 0 | 0 | 0 |
| 0 | 0 | 0 |
| 0 | 0 | 0 |
| 0 | 0 | 0 |
| 0 | 0 | 0 |
| 0 | 0 | 0 |
| 0 | 0 | 0 |
| 0 | 0 | 0 |
| 0 | 0 | 0 |
| 0 | 1 | 0 |
| 0 | 0 | 0 |
| 1 | 1 | 1 |
| 0 | 0 | 0 |
| 0 | 0 | 0 |
| 0 | 0 | 0 |
| 0 | 0 | 0 |
| 0 | 0 | 0 |
| 0 | 0 | 0 |
| 0 | 0 | 0 |
| 0 | 1 | 0 |
| 0 | 0 | 0 |
| 0 | 0 | 0 |
| 0 | 0 | 0 |

| | | |
|---|---|---|
| 1 | 1 | 1 |
| 0 | 0 | 0 |
| 0 | 1 | 0 |
| 0 | 1 | 0 |
| 0 | 0 | 0 |
| 0 | 0 | 0 |
| 0 | 0 | 0 |
| 0 | 0 | 0 |
| 0 | 0 | 0 |
| 0 | 1 | 0 |
| 1 | 1 | 1 |
| 1 | 1 | 1 |
| 1 | 1 | 1 |
| 0 | 0 | 0 |
| 1 | 1 | 1 |
| 0 | 0 | 0 |
| 0 | 0 | 0 |
| 0 | 0 | 0 |
| 0 | 0 | 0 |
| 1 | 0 | 0 |
| 0 | 0 | 0 |
| 1 | 1 | 1 |
| 0 | 0 | 0 |
| 0 | 1 | 0 |
| 1 | 1 | 1 |
| 1 | 1 | 1 |
| 0 | 0 | 0 |
| 1 | 1 | 1 |
| 1 | 1 | 1 |
| 1 | 1 | 1 |
| 0 | 0 | 0 |
| 0 | 0 | 0 |
| 1 | 1 | 1 |
| 1 | 1 | 1 |
| 0 | 0 | 0 |
| 0 | 0 | 0 |
| 0 | 0 | 0 |
| 0 | 0 | 0 |
| 0 | 0 | 0 |
| 1 | 1 | 1 |
| 0 | 0 | 0 |
| 0 | 0 | 0 |
| 0 | 0 | 0 |
| 0 | 0 | 0 |
| 1 | 0 | 0 |
| 0 | 0 | 0 |
| 0 | 0 | 0 |
| 0 | 0 | 0 |
| 0 | 0 | 0 |
| 0 | 0 | 0 |
| 0 | 0 | 0 |
| 0 | 0 | 0 |
| 1 | 0 | 0 |
| 0 | 0 | 0 |

| | | |
|---|---|---|
| 0 | 0 | 0 |
| 0 | 0 | 0 |
| 0 | 0 | 0 |
| 0 | 0 | 0 |
| 0 | 0 | 0 |
| 0 | 0 | 0 |
| 0 | 0 | 0 |
| 1 | 0 | 0 |
| 0 | 0 | 0 |
| 0 | 0 | 0 |
| 0 | 0 | 0 |
| 0 | 0 | 0 |
| 1 | 1 | 1 |
| 0 | 0 | 0 |
| 0 | 0 | 0 |
| 0 | 0 | 0 |
| 0 | 0 | 0 |
| 0 | 0 | 0 |
| 0 | 0 | 0 |
| 1 | 1 | 1 |
| 0 | 0 | 0 |
| 0 | 0 | 0 |
| 0 | 0 | 0 |
| 0 | 0 | 0 |
| 0 | 0 | 0 |
| 1 | 0 | 0 |
| 1 | 1 | 1 |
| 0 | 1 | 0 |
| 0 | 0 | 0 |
| 0 | 0 | 0 |
| 0 | 0 | 0 |
| 0 | 0 | 0 |
| 0 | 0 | 0 |
| 0 | 0 | 0 |
| 0 | 0 | 0 |
| 1 | 0 | 0 |
| 0 | 0 | 0 |
| 0 | 0 | 0 |
| 1 | 1 | 1 |
| 1 | 0 | 0 |
| 0 | 1 | 0 |
| 0 | 0 | 0 |
| 0 | 0 | 0 |
| 0 | 0 | 0 |
| 0 | 0 | 0 |
| 0 | 0 | 0 |
| 0 | 0 | 0 |
| 1 | 1 | 1 |
| 1 | 1 | 1 |
| 0 | 0 | 0 |
| 0 | 0 | 0 |
| 0 | 0 | 0 |
| 0 | 0 | 0 |
| 1 | 1 | 1 |

| | | |
|---|---|---|
| 1 | 1 | 1 |
| 0 | 0 | 0 |
| 0 | 1 | 0 |
| 0 | 0 | 0 |
| 1 | 0 | 0 |
| 1 | 0 | 0 |
| 0 | 0 | 0 |
| 1 | 0 | 0 |
| 1 | 0 | 0 |
| 1 | 0 | 0 |
| 1 | 0 | 0 |
| 1 | 0 | 0 |
| 1 | 1 | 1 |
| 1 | 1 | 1 |
| 1 | 1 | 1 |
| 1 | 1 | 1 |
| 0 | 0 | 0 |
| 0 | 0 | 0 |
| 1 | 0 | 0 |
| 0 | 0 | 0 |
| 0 | 0 | 0 |
| 0 | 0 | 0 |
| 1 | 0 | 0 |
| 0 | 0 | 0 |
| 0 | 0 | 0 |
| 1 | 1 | 1 |
| 1 | 0 | 0 |
| 1 | 0 | 0 |
| 0 | 0 | 0 |
| 0 | 0 | 0 |
| 0 | 0 | 0 |
| 0 | 0 | 0 |
| 0 | 0 | 0 |
| 0 | 0 | 0 |
| 0 | 0 | 0 |
| 0 | 0 | 0 |
| 1 | 0 | 0 |
| 1 | 1 | 1 |
| 1 | 1 | 1 |
| 1 | 1 | 1 |
| 0 | 0 | 0 |
| 1 | 1 | 1 |
| 0 | 0 | 0 |
| 1 | 1 | 1 |
| 0 | 0 | 0 |
| 0 | 1 | 0 |
| 0 | 0 | 0 |
| 0 | 0 | 0 |
| 0 | 0 | 0 |
| 0 | 0 | 0 |
| 0 | 0 | 0 |
| 0 | 0 | 0 |
| 0 | 0 | 0 |
| 0 | 0 | 0 |

| | | |
|---|---|---|
| 0 | 0 | 0 |
| 0 | 0 | 0 |
| 0 | 0 | 0 |
| 0 | 0 | 0 |
| 0 | 0 | 0 |
| 0 | 0 | 0 |
| 0 | 0 | 0 |
| 1 | 1 | 1 |
| 0 | 0 | 0 |
| 0 | 0 | 0 |
| 0 | 0 | 0 |
| 1 | 0 | 0 |
| 0 | 0 | 0 |
| 0 | 0 | 0 |
| 0 | 0 | 0 |
| 0 | 0 | 0 |
| 0 | 0 | 0 |
| 0 | 0 | 0 |
| 0 | 0 | 0 |
| 0 | 0 | 0 |
| 0 | 0 | 0 |
| 0 | 0 | 0 |
| 0 | 0 | 0 |
| 0 | 0 | 0 |
| 0 | 0 | 0 |
| 0 | 0 | 0 |
| 0 | 0 | 0 |
| 0 | 0 | 0 |
| 0 | 0 | 0 |
| 0 | 0 | 0 |
| 0 | 0 | 0 |
| 0 | 0 | 0 |
| 0 | 0 | 0 |
| 0 | 0 | 0 |
| 0 | 0 | 0 |
| 0 | 0 | 0 |
| 0 | 0 | 0 |
| 0 | 0 | 0 |
| 0 | 0 | 0 |
| 1 | 0 | 0 |
| 0 | 0 | 0 |
| 0 | 0 | 0 |
| 0 | 0 | 0 |
| 0 | 0 | 0 |
| 0 | 0 | 0 |
| 0 | 0 | 0 |
| 0 | 0 | 0 |
| 0 | 0 | 0 |
| 0 | 1 | 0 |
| 0 | 1 | 0 |
| 1 | 0 | 0 |
| 0 | 0 | 0 |
| 0 | 0 | 0 |

| | | |
|---|---|---|
| 0 | 0 | 0 |
| 0 | 0 | 0 |
| 0 | 0 | 0 |
| 0 | 0 | 0 |
| 0 | 0 | 0 |
| 0 | 0 | 0 |
| 0 | 0 | 0 |
| 1 | 0 | 0 |
| 1 | 0 | 0 |
| 0 | 0 | 0 |
| 0 | 0 | 0 |
| 1 | 0 | 0 |
| 0 | 0 | 0 |
| 0 | 0 | 0 |
| 0 | 0 | 0 |
| 0 | 0 | 0 |
| 0 | 0 | 0 |
| 1 | 0 | 0 |
| 0 | 0 | 0 |
| 1 | 0 | 0 |
| 0 | 0 | 0 |
| 0 | 0 | 0 |
| 1 | 1 | 1 |
| 0 | 0 | 0 |
| 0 | 0 | 0 |
| 1 | 0 | 0 |
| 0 | 0 | 0 |
| 0 | 0 | 0 |
| 0 | 0 | 0 |
| 0 | 0 | 0 |
| 0 | 0 | 0 |
| 0 | 0 | 0 |
| 1 | 0 | 0 |
| 0 | 0 | 0 |
| 0 | 0 | 0 |
| 0 | 0 | 0 |
| 1 | 1 | 1 |
| 0 | 0 | 0 |
| 0 | 0 | 0 |
| 0 | 0 | 0 |
| 0 | 0 | 0 |
| 0 | 0 | 0 |
| 0 | 0 | 0 |
| 0 | 0 | 0 |
| 0 | 0 | 0 |
| 0 | 0 | 0 |
| 0 | 0 | 0 |
| 0 | 0 | 0 |
| 0 | 0 | 0 |
| 0 | 0 | 0 |
| 0 | 0 | 0 |
| 0 | 0 | 0 |
| 0 | 0 | 0 |
| 0 | 0 | 0 |

| | | |
|---|---|---|
| 1 | 0 | 0 |
| 0 | 1 | 0 |
| 0 | 0 | 0 |
| 0 | 0 | 0 |
| 0 | 0 | 0 |
| 0 | 0 | 0 |
| 0 | 0 | 0 |
| 0 | 0 | 0 |
| 0 | 0 | 0 |
| 1 | 0 | 0 |
| 0 | 1 | 0 |
| 1 | 1 | 1 |
| 0 | 0 | 0 |
| 0 | 0 | 0 |
| 0 | 0 | 0 |
| 0 | 0 | 0 |
| 0 | 0 | 0 |
| 0 | 0 | 0 |
| 1 | 1 | 1 |
| 1 | 0 | 0 |
| 1 | 0 | 0 |
| 0 | 0 | 0 |
| 0 | 1 | 0 |
| 0 | 0 | 0 |
| 0 | 0 | 0 |
| 0 | 0 | 0 |
| 0 | 0 | 0 |
| 0 | 0 | 0 |
| 0 | 0 | 0 |
| 0 | 0 | 0 |
| 0 | 0 | 0 |
| 0 | 0 | 0 |
| 0 | 0 | 0 |
| 0 | 0 | 0 |
| 0 | 0 | 0 |
| 0 | 0 | 0 |
| 0 | 0 | 0 |
| 0 | 0 | 0 |
| 0 | 0 | 0 |
| 0 | 0 | 0 |
| 0 | 0 | 0 |
| 1 | 0 | 0 |
| 0 | 0 | 0 |
| 0 | 0 | 0 |
| 0 | 0 | 0 |
| 0 | 0 | 0 |
| 1 | 0 | 0 |
| 0 | 0 | 0 |
| 0 | 0 | 0 |
| 0 | 0 | 0 |
| 0 | 0 | 0 |
| 0 | 0 | 0 |
| 1 | 0 | 0 |
| 0 | 1 | 0 |

| | | |
|---|---|---|
| 0 | 0 | 0 |
| 0 | 0 | 0 |
| 1 | 0 | 0 |
| 1 | 0 | 0 |
| 1 | 0 | 0 |
| 0 | 0 | 0 |
| 1 | 0 | 0 |
| 0 | 0 | 0 |
| 0 | 0 | 0 |
| 0 | 0 | 0 |
| 0 | 0 | 0 |
| 0 | 0 | 0 |
| 0 | 0 | 0 |
| 1 | 0 | 0 |
| 1 | 0 | 0 |
| 1 | 0 | 0 |
| 0 | 0 | 0 |
| 1 | 1 | 1 |
| 0 | 0 | 0 |
| 0 | 0 | 0 |
| 1 | 0 | 0 |
| 1 | 0 | 0 |
| 0 | 0 | 0 |
| 0 | 0 | 0 |
| 1 | 0 | 0 |
| 0 | 0 | 0 |
| 0 | 0 | 0 |
| 1 | 0 | 0 |
| 0 | 0 | 0 |
| 0 | 0 | 0 |
| 0 | 0 | 0 |
| 0 | 0 | 0 |
| 0 | 0 | 0 |
| 0 | 0 | 0 |
| 0 | 0 | 0 |
| 0 | 0 | 0 |
| 0 | 0 | 0 |
| 0 | 0 | 0 |
| 0 | 0 | 0 |
| 0 | 0 | 0 |
| 1 | 0 | 0 |
| 1 | 0 | 0 |
| 1 | 0 | 0 |
| 1 | 1 | 1 |
| 1 | 1 | 1 |
| 0 | 0 | 0 |
| 0 | 0 | 0 |
| 0 | 0 | 0 |
| 0 | 0 | 0 |
| 0 | 0 | 0 |
| 0 | 0 | 0 |
| 0 | 0 | 0 |
| 0 | 0 | 0 |
| 1 | 0 | 0 |

| | | |
|---|---|---|
| 1 | 0 | 0 |
| 0 | 0 | 0 |
| 0 | 0 | 0 |
| 0 | 0 | 0 |
| 1 | 0 | 0 |
| 0 | 0 | 0 |
| 0 | 0 | 0 |
| 0 | 0 | 0 |
| 1 | 0 | 0 |
| 1 | 0 | 0 |
| 0 | 0 | 0 |
| 0 | 0 | 0 |
| 0 | 0 | 0 |
| 1 | 0 | 0 |
| 0 | 0 | 0 |
| 0 | 0 | 0 |
| 0 | 0 | 0 |
| 0 | 0 | 0 |
| 0 | 0 | 0 |
| 0 | 0 | 0 |
| 0 | 0 | 0 |
| 1 | 1 | 1 |
| 0 | 0 | 0 |
| 0 | 1 | 0 |
| 0 | 0 | 0 |
| 0 | 0 | 0 |
| 0 | 0 | 0 |
| 0 | 0 | 0 |
| 0 | 0 | 0 |
| 0 | 0 | 0 |
| 0 | 0 | 0 |
| 0 | 0 | 0 |
| 0 | 0 | 0 |
| 0 | 0 | 0 |
| 0 | 1 | 0 |
| 0 | 0 | 0 |
| 0 | 0 | 0 |
| 0 | 0 | 0 |
| 0 | 0 | 0 |
| 0 | 0 | 0 |
| 0 | 0 | 0 |
| 0 | 0 | 0 |
| 0 | 0 | 0 |
| 0 | 0 | 0 |
| 0 | 0 | 0 |
| 0 | 0 | 0 |
| 0 | 0 | 0 |
| 1 | 1 | 1 |
| 0 | 0 | 0 |
| 1 | 1 | 1 |
| 0 | 0 | 0 |
| 0 | 0 | 0 |
| 0 | 0 | 0 |
| 0 | 0 | 0 |

| | | |
|---|---|---|
| 0 | 0 | 0 |
| 0 | 0 | 0 |
| 1 | 1 | 1 |
| 0 | 0 | 0 |
| 0 | 0 | 0 |
| 0 | 1 | 0 |
| 0 | 1 | 0 |
| 0 | 1 | 0 |
| 0 | 0 | 0 |
| 0 | 1 | 0 |
| 0 | 0 | 0 |
| 0 | 0 | 0 |
| 0 | 0 | 0 |
| 1 | 1 | 1 |
| 0 | 0 | 0 |
| 0 | 1 | 0 |
| 0 | 0 | 0 |
| 0 | 0 | 0 |
| 0 | 0 | 0 |
| 1 | 1 | 1 |
| 1 | 1 | 1 |
| 0 | 0 | 0 |
| 1 | 1 | 1 |
| 0 | 0 | 0 |
| 0 | 0 | 0 |
| 1 | 1 | 1 |
| 1 | 1 | 1 |
| 0 | 0 | 0 |
| 1 | 1 | 1 |
| 0 | 0 | 0 |
| 0 | 1 | 0 |
| 0 | 1 | 0 |
| 1 | 1 | 1 |
| 0 | 1 | 0 |
| 1 | 1 | 1 |
| 1 | 1 | 1 |
| 1 | 1 | 1 |
| 1 | 1 | 1 |
| 1 | 1 | 1 |
| 1 | 1 | 1 |
| 1 | 1 | 1 |
| 1 | 1 | 1 |
| 0 | 1 | 0 |
| 0 | 1 | 0 |
| 0 | 1 | 0 |
| 0 | 1 | 0 |
| 0 | 1 | 0 |
| 0 | 1 | 0 |
| 0 | 1 | 0 |
| 0 | 1 | 0 |
| 0 | 1 | 0 |
| 0 | 1 | 0 |
| 0 | 1 | 0 |
| 0 | 1 | 0 |

| | | |
|---|---|---|
| 0 | 1 | 0 |
| 1 | 1 | 1 |
| 1 | 1 | 1 |
| 1 | 1 | 1 |
| 1 | 1 | 1 |
| 1 | 1 | 1 |
| 1 | 1 | 1 |
| 0 | 1 | 0 |
| 0 | 1 | 0 |
| 0 | 1 | 0 |
| 0 | 1 | 0 |
| 0 | 0 | 0 |
| 0 | 0 | 0 |
| 0 | 0 | 0 |
| 1 | 0 | 0 |
| 0 | 0 | 0 |
| 1 | 0 | 0 |
| 0 | 0 | 0 |
| 1 | 0 | 0 |
| 1 | 1 | 1 |
| 0 | 0 | 0 |
| 1 | 1 | 1 |
| 1 | 1 | 1 |
| 0 | 0 | 0 |
| 0 | 0 | 0 |
| 1 | 1 | 1 |
| 1 | 1 | 1 |
| 0 | 0 | 0 |
| 0 | 0 | 0 |
| 0 | 0 | 0 |
| 0 | 0 | 0 |
| 0 | 0 | 0 |
| 0 | 1 | 0 |
| 0 | 1 | 0 |
| 1 | 1 | 1 |
| 1 | 1 | 1 |
| 1 | 1 | 1 |
| 1 | 1 | 1 |
| 1 | 1 | 1 |
| 0 | 0 | 0 |
| 0 | 0 | 0 |
| 0 | 1 | 0 |
| 0 | 0 | 0 |
| 0 | 0 | 0 |
| 0 | 0 | 0 |
| 1 | 0 | 0 |
| 0 | 0 | 0 |
| 1 | 0 | 0 |
| 1 | 0 | 0 |
| 0 | 0 | 0 |
| 0 | 0 | 0 |
| 0 | 0 | 0 |
| 0 | 0 | 0 |
| 0 | 0 | 0 |

| | | |
|---|---|---|
| 0 | 0 | 0 |
| 0 | 1 | 0 |
| 1 | 1 | 1 |
| 1 | 1 | 1 |
| 0 | 0 | 0 |
| 1 | 1 | 1 |
| 0 | 0 | 0 |
| 1 | 1 | 1 |
| 1 | 1 | 1 |
| 0 | 0 | 0 |
| 1 | 0 | 0 |
| 1 | 1 | 1 |
| 1 | 1 | 1 |
| 0 | 0 | 0 |
| 1 | 0 | 0 |
| 1 | 0 | 0 |
| 0 | 0 | 0 |
| 1 | 0 | 0 |
| 0 | 0 | 0 |
| 0 | 0 | 0 |
| 0 | 0 | 0 |
| 0 | 0 | 0 |
| 0 | 0 | 0 |
| 0 | 0 | 0 |
| 0 | 0 | 0 |
| 0 | 0 | 0 |
| 0 | 0 | 0 |
| 0 | 0 | 0 |
| 0 | 0 | 0 |
| 0 | 0 | 0 |
| 0 | 0 | 0 |
| 0 | 0 | 0 |
| 0 | 0 | 0 |
| 0 | 0 | 0 |
| 0 | 0 | 0 |
| 0 | 0 | 0 |
| 1 | 0 | 0 |
| 0 | 0 | 0 |
| 0 | 0 | 0 |
| 0 | 0 | 0 |
| 0 | 0 | 0 |
| 0 | 0 | 0 |
| 0 | 0 | 0 |
| 0 | 0 | 0 |
| 0 | 0 | 0 |
| 0 | 0 | 0 |
| 0 | 0 | 0 |
| 0 | 0 | 0 |
| 1 | 0 | 0 |
| 1 | 0 | 0 |
| 0 | 0 | 0 |
| 1 | 1 | 1 |
| 0 | 0 | 0 |
| 0 | 0 | 0 |

| | | |
|---|---|---|
| 0 | 0 | 0 |
| 0 | 0 | 0 |
| 0 | 0 | 0 |
| 0 | 0 | 0 |
| 0 | 0 | 0 |
| 0 | 0 | 0 |
| 0 | 0 | 0 |
| 0 | 0 | 0 |
| 0 | 0 | 0 |
| 0 | 0 | 0 |
| 0 | 0 | 0 |
| 0 | 0 | 0 |
| 0 | 0 | 0 |
| 0 | 0 | 0 |
| 0 | 1 | 0 |
| 0 | 0 | 0 |
| 0 | 0 | 0 |
| 0 | 0 | 0 |
| 1 | 0 | 0 |
| 0 | 0 | 0 |
| 0 | 0 | 0 |
| 0 | 0 | 0 |
| 0 | 0 | 0 |
| 0 | 0 | 0 |
| 0 | 0 | 0 |
| 1 | 0 | 0 |
| 0 | 0 | 0 |
| 0 | 0 | 0 |
| 0 | 0 | 0 |
| 0 | 0 | 0 |
| 1 | 1 | 1 |
| 0 | 0 | 0 |
| 1 | 1 | 1 |
| 1 | 0 | 0 |
| 0 | 0 | 0 |
| 0 | 0 | 0 |
| 1 | 1 | 1 |
| 1 | 1 | 1 |
| 1 | 1 | 1 |
| 1 | 0 | 0 |
| 0 | 0 | 0 |
| 0 | 0 | 0 |
| 0 | 0 | 0 |
| 0 | 0 | 0 |
| 0 | 0 | 0 |
| 0 | 0 | 0 |
| 0 | 0 | 0 |
| 0 | 0 | 0 |
| 1 | 1 | 1 |
| 1 | 1 | 1 |
| 0 | 0 | 0 |
| 0 | 0 | 0 |
| 0 | 0 | 0 |
| 0 | 1 | 0 |

| | | |
|---|---|---|
| 1 | 1 | 1 |
| 1 | 1 | 1 |
| 1 | 0 | 0 |
| 0 | 0 | 0 |
| 1 | 0 | 0 |
| 1 | 1 | 1 |
| 0 | 0 | 0 |
| 1 | 1 | 1 |
| 0 | 0 | 0 |
| 0 | 0 | 0 |
| 0 | 0 | 0 |
| 0 | 0 | 0 |
| 0 | 0 | 0 |
| 0 | 0 | 0 |
| 0 | 0 | 0 |
| 0 | 0 | 0 |
| 0 | 1 | 0 |
| 0 | 0 | 0 |
| 0 | 0 | 0 |
| 1 | 0 | 0 |
| 0 | 0 | 0 |
| 0 | 0 | 0 |
| 1 | 1 | 1 |
| 0 | 0 | 0 |
| 1 | 1 | 1 |
| 1 | 1 | 1 |
| 0 | 1 | 0 |
| 0 | 0 | 0 |
| 0 | 0 | 0 |
| 0 | 0 | 0 |
| 1 | 1 | 1 |
| 0 | 0 | 0 |
| 0 | 0 | 0 |
| 0 | 0 | 0 |
| 0 | 1 | 0 |
| 0 | 1 | 0 |
| 0 | 0 | 0 |
| 0 | 0 | 0 |
| 0 | 0 | 0 |
| 1 | 1 | 1 |
| 1 | 1 | 1 |
| 0 | 1 | 0 |
| 0 | 1 | 0 |
| 0 | 0 | 0 |
| 1 | 1 | 1 |
| 0 | 0 | 0 |
| 1 | 1 | 1 |
| 0 | 0 | 0 |
| 0 | 0 | 0 |
| 1 | 1 | 1 |
| 0 | 1 | 0 |
| 0 | 0 | 0 |
| 0 | 0 | 0 |
| 1 | 1 | 1 |

| | | |
|---|---|---|
| 1 | 1 | 1 |
| 1 | 1 | 1 |
| 0 | 1 | 0 |
| 0 | 0 | 0 |
| 0 | 0 | 0 |
| 1 | 0 | 0 |
| 1 | 0 | 0 |
| 0 | 0 | 0 |
| 0 | 0 | 0 |
| 1 | 1 | 1 |
| 0 | 0 | 0 |
| 0 | 0 | 0 |
| 0 | 0 | 0 |
| 0 | 0 | 0 |
| 0 | 0 | 0 |
| 0 | 0 | 0 |
| 0 | 0 | 0 |
| 0 | 0 | 0 |
| 0 | 0 | 0 |
| 0 | 1 | 0 |
| 1 | 1 | 1 |
| 1 | 1 | 1 |
| 1 | 1 | 1 |
| 1 | 1 | 1 |
| 0 | 0 | 0 |
| 0 | 1 | 0 |
| 0 | 0 | 0 |
| 0 | 0 | 0 |
| 1 | 1 | 1 |
| 1 | 1 | 1 |
| 1 | 1 | 1 |
| 1 | 1 | 1 |
| 1 | 1 | 1 |
| 0 | 0 | 0 |
| 0 | 0 | 0 |
| 1 | 1 | 1 |
| 0 | 0 | 0 |
| 0 | 1 | 0 |
| 0 | 0 | 0 |
| 0 | 0 | 0 |
| 0 | 0 | 0 |
| 0 | 0 | 0 |
| 0 | 1 | 0 |
| 0 | 0 | 0 |
| 0 | 0 | 0 |
| 0 | 0 | 0 |
| 0 | 0 | 0 |
| 0 | 0 | 0 |
| 0 | 0 | 0 |
| 0 | 0 | 0 |
| 0 | 0 | 0 |
| 0 | 0 | 0 |
| 0 | 0 | 0 |
| 0 | 1 | 0 |

| | | |
|---|---|---|
| 0 | 0 | 0 |
| 0 | 1 | 0 |
| 0 | 0 | 0 |
| 0 | 0 | 0 |
| 0 | 0 | 0 |
| 1 | 1 | 1 |
| 0 | 0 | 0 |
| 1 | 1 | 1 |
| 0 | 1 | 0 |
| 0 | 0 | 0 |
| 0 | 1 | 0 |
| 0 | 1 | 0 |
| 0 | 1 | 0 |
| 0 | 1 | 0 |
| 0 | 1 | 0 |
| 0 | 0 | 0 |
| 0 | 0 | 0 |
| 0 | 1 | 0 |
| 0 | 0 | 0 |
| 1 | 1 | 1 |
| 0 | 0 | 0 |
| 0 | 0 | 0 |
| 0 | 0 | 0 |
| 0 | 0 | 0 |
| 0 | 0 | 0 |
| 0 | 0 | 0 |
| 0 | 0 | 0 |
| 0 | 0 | 0 |
| 0 | 0 | 0 |
| 0 | 0 | 0 |
| 0 | 0 | 0 |
| 0 | 0 | 0 |
| 1 | 0 | 0 |
| 1 | 0 | 0 |
| 0 | 0 | 0 |
| 0 | 0 | 0 |
| 0 | 0 | 0 |
| 0 | 0 | 0 |
| 1 | 0 | 0 |
| 0 | 0 | 0 |
| 0 | 0 | 0 |
| 0 | 0 | 0 |
| 1 | 0 | 0 |
| 0 | 1 | 0 |
| 0 | 0 | 0 |
| 0 | 0 | 0 |
| 0 | 0 | 0 |
| 0 | 0 | 0 |
| 0 | 0 | 0 |
| 0 | 0 | 0 |
| 1 | 0 | 0 |
| 0 | 0 | 0 |
| 0 | 0 | 0 |
| 1 | 0 | 0 |

| | | |
|---|---|---|
| 0 | 0 | 0 |
| 1 | 0 | 0 |
| 0 | 0 | 0 |
| 0 | 0 | 0 |
| 0 | 0 | 0 |
| 0 | 0 | 0 |
| 0 | 0 | 0 |
| 0 | 0 | 0 |
| 0 | 0 | 0 |
| 0 | 0 | 0 |
| 0 | 0 | 0 |
| 1 | 1 | 1 |
| 0 | 0 | 0 |
| 1 | 1 | 1 |
| 1 | 0 | 0 |
| 1 | 0 | 0 |
| 0 | 0 | 0 |
| 0 | 0 | 0 |
| 0 | 0 | 0 |
| 0 | 0 | 0 |
| 1 | 0 | 0 |
| 0 | 0 | 0 |
| 0 | 1 | 0 |
| 0 | 0 | 0 |
| 1 | 1 | 1 |
| 1 | 1 | 1 |
| 1 | 1 | 1 |
| 0 | 0 | 0 |
| 1 | 1 | 1 |
| 1 | 1 | 1 |
| 1 | 1 | 1 |
| 0 | 0 | 0 |
| 1 | 1 | 1 |
| 0 | 0 | 0 |
| 0 | 0 | 0 |
| 0 | 0 | 0 |
| 0 | 1 | 0 |
| 1 | 1 | 1 |
| 0 | 0 | 0 |
| 0 | 0 | 0 |
| 0 | 0 | 0 |
| 0 | 0 | 0 |
| 0 | 0 | 0 |
| 0 | 0 | 0 |
| 0 | 0 | 0 |
| 0 | 0 | 0 |
| 1 | 1 | 1 |
| 1 | 1 | 1 |
| 0 | 0 | 0 |
| 0 | 0 | 0 |
| 0 | 0 | 0 |
| 1 | 1 | 1 |
| 1 | 1 | 1 |
| 0 | 0 | 0 |

| | | |
|---|---|---|
| 0 | 0 | 0 |
| 0 | 1 | 0 |
| 0 | 1 | 0 |
| 1 | 1 | 1 |
| 0 | 0 | 0 |
| 0 | 0 | 0 |
| 0 | 0 | 0 |
| 0 | 0 | 0 |
| 0 | 0 | 0 |
| 0 | 0 | 0 |
| 1 | 1 | 1 |
| 0 | 0 | 0 |
| 0 | 0 | 0 |
| 0 | 0 | 0 |
| 1 | 0 | 0 |
| 1 | 1 | 1 |
| 0 | 0 | 0 |
| 1 | 0 | 0 |
| 0 | 0 | 0 |
| 0 | 0 | 0 |
| 1 | 0 | 0 |
| 0 | 0 | 0 |
| 0 | 0 | 0 |
| 1 | 0 | 0 |
| 0 | 0 | 0 |
| 0 | 0 | 0 |
| 0 | 0 | 0 |
| 1 | 0 | 0 |
| 0 | 0 | 0 |
| 0 | 0 | 0 |
| 0 | 0 | 0 |
| 1 | 1 | 1 |
| 1 | 1 | 1 |
| 1 | 1 | 1 |
| 0 | 1 | 0 |
| 0 | 1 | 0 |
| 0 | 1 | 0 |
| 1 | 1 | 1 |
| 1 | 1 | 1 |
| 1 | 1 | 1 |
| 0 | 1 | 0 |
| 1 | 1 | 1 |
| 0 | 0 | 0 |
| 0 | 1 | 0 |
| 1 | 1 | 1 |
| 1 | 1 | 1 |
| 0 | 1 | 0 |
| 1 | 1 | 1 |
| 1 | 1 | 1 |
| 1 | 1 | 1 |
| 1 | 1 | 1 |
| 1 | 1 | 1 |
| 1 | 1 | 1 |
| 1 | 1 | 1 |

| | | |
|---|---|---|
| 0 | 1 | 0 |
| 0 | 1 | 0 |
| 1 | 1 | 1 |
| 1 | 1 | 1 |
| 1 | 1 | 1 |
| 1 | 1 | 1 |
| 1 | 1 | 1 |
| 1 | 1 | 1 |
| 0 | 0 | 0 |
| 0 | 0 | 0 |
| 0 | 0 | 0 |
| 0 | 0 | 0 |
| 0 | 0 | 0 |
| 0 | 0 | 0 |
| 0 | 0 | 0 |
| 0 | 0 | 0 |
| 0 | 0 | 0 |
| 0 | 0 | 0 |
| 0 | 0 | 0 |
| 0 | 0 | 0 |
| 0 | 0 | 0 |
| 0 | 0 | 0 |
| 0 | 0 | 0 |
| 0 | 0 | 0 |
| 0 | 0 | 0 |
| 1 | 1 | 1 |
| 1 | 1 | 1 |
| 0 | 0 | 0 |
| 0 | 0 | 0 |
| 0 | 0 | 0 |
| 0 | 0 | 0 |
| 0 | 0 | 0 |
| 0 | 0 | 0 |
| 0 | 0 | 0 |
| 0 | 0 | 0 |
| 0 | 0 | 0 |
| 0 | 0 | 0 |
| 0 | 0 | 0 |
| 0 | 0 | 0 |
| 0 | 0 | 0 |
| 0 | 0 | 0 |
| 0 | 0 | 0 |
| 0 | 0 | 0 |
| 0 | 0 | 0 |
| 0 | 0 | 0 |
| 0 | 0 | 0 |
| 1 | 1 | 1 |
| 0 | 1 | 0 |
| 1 | 1 | 1 |
| 1 | 1 | 1 |
| 1 | 1 | 1 |
| 1 | 1 | 1 |
| 0 | 0 | 0 |
| 0 | 0 | 0 |

| | | |
|---|---|---|
| 0 | 0 | 0 |
| 0 | 0 | 0 |
| 0 | 0 | 0 |
| 0 | 0 | 0 |
| 0 | 1 | 0 |
| 0 | 0 | 0 |
| 0 | 0 | 0 |
| 0 | 0 | 0 |
| 1 | 1 | 1 |
| 0 | 0 | 0 |
| 0 | 0 | 0 |
| 0 | 1 | 0 |
| 0 | 0 | 0 |
| 0 | 0 | 0 |
| 0 | 0 | 0 |
| 0 | 0 | 0 |
| 1 | 1 | 1 |
| 0 | 0 | 0 |
| 0 | 0 | 0 |
| 1 | 1 | 1 |
| 0 | 0 | 0 |
| 0 | 0 | 0 |
| 0 | 0 | 0 |
| 0 | 0 | 0 |
| 0 | 0 | 0 |
| 0 | 1 | 0 |
| 0 | 1 | 0 |
| 1 | 1 | 1 |
| 1 | 1 | 1 |
| 0 | 1 | 0 |
| 1 | 1 | 1 |
| 1 | 1 | 1 |
| 1 | 0 | 0 |
| 1 | 0 | 0 |
| 0 | 0 | 0 |
| 0 | 0 | 0 |
| 0 | 0 | 0 |
| 0 | 0 | 0 |
| 0 | 0 | 0 |
| 0 | 0 | 0 |
| 1 | 1 | 1 |
| 1 | 1 | 1 |
| 1 | 1 | 1 |
| 0 | 0 | 0 |
| 0 | 0 | 0 |
| 0 | 0 | 0 |
| 1 | 1 | 1 |
| 0 | 1 | 0 |
| 0 | 0 | 0 |
| 0 | 0 | 0 |
| 1 | 1 | 1 |
| 1 | 1 | 1 |
| 1 | 1 | 1 |
| 1 | 1 | 1 |

| | | |
|---|---|---|
| 0 | 1 | 0 |
| 1 | 1 | 1 |
| 0 | 1 | 0 |
| 0 | 0 | 0 |
| 0 | 0 | 0 |
| 1 | 1 | 1 |
| 1 | 1 | 1 |
| 1 | 1 | 1 |
| 1 | 1 | 1 |
| 1 | 1 | 1 |
| 0 | 0 | 0 |
| 0 | 1 | 0 |
| 0 | 0 | 0 |
| 0 | 0 | 0 |
| 0 | 0 | 0 |
| 1 | 1 | 1 |
| 1 | 1 | 1 |
| 0 | 1 | 0 |
| 1 | 1 | 1 |
| 1 | 1 | 1 |
| 0 | 0 | 0 |
| 1 | 1 | 1 |
| 1 | 1 | 1 |
| 0 | 0 | 0 |
| 1 | 0 | 0 |
| 0 | 0 | 0 |
| 0 | 0 | 0 |
| 1 | 1 | 1 |
| 1 | 1 | 1 |
| 1 | 1 | 1 |
| 1 | 1 | 1 |
| 1 | 1 | 1 |
| 1 | 1 | 1 |
| 1 | 1 | 1 |
| 1 | 1 | 1 |
| 0 | 0 | 0 |
| 1 | 1 | 1 |
| 1 | 1 | 1 |
| 0 | 0 | 0 |
| 0 | 0 | 0 |
| 1 | 1 | 1 |
| 0 | 1 | 0 |
| 0 | 0 | 0 |
| 0 | 0 | 0 |
| 0 | 0 | 0 |
| 0 | 0 | 0 |
| 0 | 0 | 0 |
| 0 | 0 | 0 |
| 1 | 1 | 1 |
| 0 | 0 | 0 |
| 0 | 0 | 0 |
| 0 | 0 | 0 |
| 0 | 0 | 0 |
| 0 | 0 | 0 |

| | | |
|---|---|---|
| 0 | 0 | 0 |
| 0 | 0 | 0 |
| 0 | 0 | 0 |
| 0 | 0 | 0 |
| 0 | 0 | 0 |
| 0 | 0 | 0 |
| 0 | 0 | 0 |
| 0 | 0 | 0 |
| 0 | 0 | 0 |
| 0 | 1 | 0 |
| 0 | 1 | 0 |
| 0 | 0 | 0 |
| 1 | 1 | 1 |
| 0 | 0 | 0 |
| 0 | 0 | 0 |
| 0 | 1 | 0 |
| 0 | 0 | 0 |
| 1 | 0 | 0 |
| 1 | 1 | 1 |
| 1 | 1 | 1 |
| 0 | 0 | 0 |
| 0 | 1 | 0 |
| 0 | 1 | 0 |
| 0 | 0 | 0 |
| 0 | 0 | 0 |
| 0 | 1 | 0 |
| 0 | 1 | 0 |
| 1 | 1 | 1 |
| 0 | 0 | 0 |
| 0 | 0 | 0 |
| 1 | 1 | 1 |
| 1 | 1 | 1 |
| 0 | 0 | 0 |
| 0 | 0 | 0 |
| 0 | 0 | 0 |
| 0 | 1 | 0 |
| 0 | 0 | 0 |
| 0 | 1 | 0 |
| 0 | 1 | 0 |
| 0 | 0 | 0 |
| 0 | 0 | 0 |
| 0 | 0 | 0 |
| 0 | 0 | 0 |
| 0 | 0 | 0 |
| 0 | 0 | 0 |
| 1 | 1 | 1 |
| 0 | 1 | 0 |
| 0 | 0 | 0 |
| 0 | 0 | 0 |
| 0 | 0 | 0 |
| 1 | 1 | 1 |
| 1 | 1 | 1 |
| 0 | 0 | 0 |
| 0 | 0 | 0 |

| | | |
|---|---|---|
| 1 | 1 | 1 |
| 1 | 1 | 1 |
| 1 | 1 | 1 |
| 1 | 1 | 1 |
| 0 | 0 | 0 |
| 0 | 0 | 0 |
| 1 | 0 | 0 |
| 0 | 0 | 0 |
| 1 | 0 | 0 |
| 0 | 0 | 0 |
| 1 | 1 | 1 |
| 1 | 1 | 1 |
| 1 | 1 | 1 |
| 1 | 1 | 1 |
| 1 | 1 | 1 |
| 1 | 1 | 1 |
| 0 | 0 | 0 |
| 1 | 1 | 1 |
| 1 | 1 | 1 |
| 0 | 0 | 0 |
| 0 | 0 | 0 |
| 0 | 1 | 0 |
| 0 | 1 | 0 |
| 0 | 0 | 0 |
| 0 | 1 | 0 |
| 0 | 0 | 0 |
| 0 | 0 | 0 |
| 0 | 0 | 0 |
| 0 | 0 | 0 |
| 0 | 0 | 0 |
| 0 | 0 | 0 |
| 0 | 0 | 0 |
| 0 | 1 | 0 |
| 0 | 0 | 0 |
| 1 | 1 | 1 |
| 1 | 1 | 1 |
| 1 | 1 | 1 |
| 1 | 1 | 1 |
| 0 | 0 | 0 |
| 1 | 1 | 1 |
| 1 | 1 | 1 |
| 1 | 1 | 1 |
| 1 | 1 | 1 |
| 0 | 0 | 0 |
| 1 | 1 | 1 |
| 1 | 1 | 1 |
| 1 | 1 | 1 |
| 1 | 1 | 1 |
| 1 | 1 | 1 |
| 1 | 1 | 1 |
| 0 | 0 | 0 |
| 0 | 0 | 0 |
| 0 | 0 | 0 |
| 0 | 1 | 0 |

| | | |
|---|---|---|
| 0 | 0 | 0 |
| 0 | 0 | 0 |
| 0 | 0 | 0 |
| 0 | 0 | 0 |
| 0 | 0 | 0 |
| 0 | 0 | 0 |
| 0 | 0 | 0 |
| 0 | 0 | 0 |
| 1 | 1 | 1 |
| 1 | 1 | 1 |
| 0 | 0 | 0 |
| 1 | 1 | 1 |
| 1 | 1 | 1 |
| 0 | 0 | 0 |
| 0 | 0 | 0 |
| 0 | 0 | 0 |
| 0 | 0 | 0 |
| 0 | 0 | 0 |
| 0 | 0 | 0 |
| 1 | 1 | 1 |
| 1 | 1 | 1 |
| 0 | 0 | 0 |
| 1 | 1 | 1 |
| 0 | 0 | 0 |
| 1 | 1 | 1 |
| 1 | 1 | 1 |
| 1 | 1 | 1 |
| 0 | 1 | 0 |
| 0 | 1 | 0 |
| 0 | 0 | 0 |
| 0 | 1 | 0 |
| 1 | 1 | 1 |
| 0 | 0 | 0 |
| 1 | 1 | 1 |
| 0 | 0 | 0 |
| 0 | 0 | 0 |
| 0 | 0 | 0 |
| 0 | 0 | 0 |
| 0 | 0 | 0 |
| 0 | 0 | 0 |
| 0 | 0 | 0 |
| 0 | 0 | 0 |
| 1 | 1 | 1 |
| 0 | 0 | 0 |
| 1 | 1 | 1 |
| 0 | 0 | 0 |
| 1 | 0 | 0 |
| 1 | 1 | 1 |
| 1 | 1 | 1 |
| 0 | 0 | 0 |
| 1 | 1 | 1 |
| 1 | 1 | 1 |
| 0 | 1 | 0 |
| 1 | 1 | 1 |

| | | |
|---|---|---|
| 0 | 0 | 0 |
| 0 | 1 | 0 |
| 1 | 1 | 1 |
| 0 | 1 | 0 |
| 0 | 1 | 0 |
| 0 | 1 | 0 |
| 0 | 1 | 0 |
| 1 | 1 | 1 |
| 1 | 1 | 1 |
| 1 | 1 | 1 |
| 0 | 0 | 0 |
| 1 | 1 | 1 |
| 1 | 1 | 1 |
| 0 | 1 | 0 |
| 0 | 0 | 0 |
| 0 | 0 | 0 |
| 0 | 0 | 0 |
| 0 | 0 | 0 |
| 0 | 0 | 0 |
| 0 | 1 | 0 |
| 0 | 1 | 0 |
| 1 | 1 | 1 |
| 0 | 0 | 0 |
| 0 | 1 | 0 |
| 0 | 1 | 0 |
| 0 | 1 | 0 |
| 1 | 1 | 1 |
| 1 | 1 | 1 |
| 0 | 0 | 0 |
| 0 | 0 | 0 |
| 0 | 0 | 0 |
| 0 | 0 | 0 |
| 0 | 0 | 0 |
| 0 | 0 | 0 |
| 1 | 1 | 1 |
| 1 | 1 | 1 |
| 0 | 0 | 0 |
| 0 | 0 | 0 |
| 0 | 0 | 0 |
| 0 | 0 | 0 |
| 0 | 0 | 0 |
| 0 | 0 | 0 |
| 1 | 1 | 1 |
| 1 | 1 | 1 |
| 0 | 0 | 0 |
| 1 | 1 | 1 |
| 1 | 1 | 1 |
| 1 | 1 | 1 |
| 1 | 1 | 1 |
| 1 | 1 | 1 |
| 1 | 0 | 0 |
| 0 | 1 | 0 |
| 0 | 1 | 0 |
| 1 | 0 | 0 |

| | | |
|---|---|---|
| 0 | 0 | 0 |
| 1 | 1 | 1 |
| 0 | 0 | 0 |
| 0 | 1 | 0 |
| 1 | 0 | 0 |
| 0 | 0 | 0 |
| 0 | 0 | 0 |
| 0 | 0 | 0 |
| 1 | 0 | 0 |
| 1 | 0 | 0 |
| 0 | 0 | 0 |
| 1 | 1 | 1 |
| 1 | 0 | 0 |
| 1 | 0 | 0 |
| 0 | 0 | 0 |
| 0 | 0 | 0 |
| 0 | 0 | 0 |
| 0 | 0 | 0 |
| 1 | 1 | 1 |
| 0 | 0 | 0 |
| 0 | 0 | 0 |
| 0 | 0 | 0 |
| 0 | 0 | 0 |
| 0 | 0 | 0 |
| 1 | 1 | 1 |
| 0 | 0 | 0 |
| 1 | 1 | 1 |
| 0 | 1 | 0 |
| 1 | 1 | 1 |
| 0 | 0 | 0 |
| 1 | 1 | 1 |
| 1 | 1 | 1 |
| 0 | 0 | 0 |
| 0 | 0 | 0 |
| 1 | 1 | 1 |
| 1 | 1 | 1 |
| 1 | 1 | 1 |
| 0 | 1 | 0 |
| 1 | 0 | 0 |
| 0 | 0 | 0 |
| 0 | 0 | 0 |
| 0 | 0 | 0 |
| 0 | 0 | 0 |
| 0 | 0 | 0 |
| 0 | 0 | 0 |
| 0 | 1 | 0 |
| 0 | 1 | 0 |
| 0 | 0 | 0 |
| 0 | 0 | 0 |
| 1 | 1 | 1 |
| 0 | 0 | 0 |
| 0 | 0 | 0 |
| 0 | 1 | 0 |
| 1 | 1 | 1 |

| | | |
|---|---|---|
| 1 | 1 | 1 |
| 0 | 0 | 0 |
| 0 | 0 | 0 |
| 1 | 0 | 0 |
| 1 | 1 | 1 |
| 0 | 0 | 0 |
| 0 | 0 | 0 |
| 0 | 0 | 0 |
| 0 | 0 | 0 |
| 0 | 0 | 0 |
| 0 | 0 | 0 |
| 0 | 0 | 0 |
| 0 | 0 | 0 |
| 0 | 0 | 0 |
| 0 | 0 | 0 |
| 0 | 1 | 0 |
| 0 | 0 | 0 |
| 1 | 1 | 1 |
| 1 | 0 | 0 |
| 1 | 1 | 1 |
| 0 | 0 | 0 |
| 1 | 1 | 1 |
| 0 | 0 | 0 |
| 0 | 0 | 0 |
| 1 | 1 | 1 |
| 0 | 0 | 0 |
| 0 | 1 | 0 |
| 0 | 1 | 0 |
| 0 | 1 | 0 |
| 0 | 0 | 0 |
| 0 | 0 | 0 |
| 0 | 0 | 0 |
| 0 | 0 | 0 |
| 0 | 0 | 0 |
| 0 | 0 | 0 |
| 0 | 0 | 0 |
| 0 | 0 | 0 |
| 1 | 1 | 1 |
| 1 | 0 | 0 |
| 0 | 0 | 0 |
| 0 | 0 | 0 |
| 0 | 0 | 0 |
| 0 | 0 | 0 |
| 0 | 0 | 0 |
| 0 | 0 | 0 |
| 0 | 0 | 0 |
| 1 | 1 | 1 |
| 0 | 0 | 0 |
| 0 | 0 | 0 |
| 0 | 0 | 0 |
| 0 | 0 | 0 |
| 0 | 1 | 0 |
| 0 | 0 | 0 |
| 0 | 0 | 0 |

| | | |
|---|---|---|
| 0 | 0 | 0 |
| 0 | 0 | 0 |
| 0 | 0 | 0 |
| 0 | 0 | 0 |
| 0 | 0 | 0 |
| 0 | 0 | 0 |
| 0 | 0 | 0 |
| 0 | 0 | 0 |
| 0 | 0 | 0 |
| 0 | 0 | 0 |
| 1 | 1 | 1 |
| 0 | 0 | 0 |
| 1 | 0 | 0 |
| 0 | 0 | 0 |
| 0 | 1 | 0 |
| 1 | 1 | 1 |
| 1 | 1 | 1 |
| 1 | 1 | 1 |
| 0 | 0 | 0 |
| 0 | 0 | 0 |
| 1 | 1 | 1 |
| 0 | 0 | 0 |
| 0 | 0 | 0 |
| 1 | 1 | 1 |
| 0 | 0 | 0 |
| 0 | 0 | 0 |
| 0 | 0 | 0 |
| 1 | 0 | 0 |
| 0 | 0 | 0 |
| 0 | 0 | 0 |
| 1 | 0 | 0 |
| 0 | 0 | 0 |
| 1 | 0 | 0 |
| 1 | 0 | 0 |
| 0 | 0 | 0 |
| 0 | 0 | 0 |
| 0 | 0 | 0 |
| 1 | 0 | 0 |
| 1 | 0 | 0 |
| 0 | 0 | 0 |
| 0 | 0 | 0 |
| 0 | 0 | 0 |
| 0 | 0 | 0 |
| 0 | 0 | 0 |
| 0 | 0 | 0 |
| 0 | 0 | 0 |
| 0 | 0 | 0 |
| 0 | 0 | 0 |
| 0 | 0 | 0 |
| 0 | 0 | 0 |
| 0 | 0 | 0 |
| 0 | 0 | 0 |
| 0 | 0 | 0 |
| 0 | 0 | 0 |

| | | |
|---|---|---|
| 0 | 0 | 0 |
| 1 | 0 | 0 |
| 0 | 0 | 0 |
| 0 | 0 | 0 |
| 0 | 0 | 0 |
| 1 | 0 | 0 |
| 0 | 0 | 0 |
| 0 | 0 | 0 |
| 1 | 1 | 1 |
| 0 | 0 | 0 |
| 0 | 1 | 0 |
| 1 | 1 | 1 |
| 1 | 1 | 1 |
| 0 | 0 | 0 |
| 1 | 1 | 1 |
| 0 | 0 | 0 |
| 1 | 1 | 1 |
| 1 | 1 | 1 |
| 0 | 0 | 0 |
| 0 | 0 | 0 |
| 0 | 0 | 0 |
| 0 | 0 | 0 |
| 0 | 0 | 0 |
| 0 | 0 | 0 |
| 0 | 0 | 0 |
| 0 | 1 | 0 |
| 1 | 1 | 1 |
| 1 | 1 | 1 |
| 1 | 0 | 0 |
| 1 | 1 | 1 |
| 1 | 1 | 1 |
| 0 | 1 | 0 |
| 0 | 0 | 0 |
| 0 | 0 | 0 |
| 0 | 0 | 0 |
| 0 | 0 | 0 |
| 0 | 0 | 0 |
| 0 | 0 | 0 |
| 0 | 1 | 0 |
| 0 | 1 | 0 |
| 0 | 1 | 0 |
| 0 | 1 | 0 |
| 0 | 1 | 0 |
| 0 | 1 | 0 |
| 0 | 0 | 0 |
| 0 | 0 | 0 |
| 0 | 1 | 0 |
| 0 | 0 | 0 |
| 0 | 0 | 0 |
| 0 | 0 | 0 |
| 0 | 0 | 0 |
| 0 | 0 | 0 |
| 0 | 0 | 0 |
| 0 | 0 | 0 |

| | | |
|---|---|---|
| 1 | 1 | 1 |
| 1 | 1 | 1 |
| 0 | 0 | 0 |
| 0 | 0 | 0 |
| 1 | 1 | 1 |
| 0 | 0 | 0 |
| 0 | 0 | 0 |
| 1 | 1 | 1 |
| 1 | 1 | 1 |
| 0 | 0 | 0 |
| 1 | 1 | 1 |
| 1 | 1 | 1 |
| 0 | 0 | 0 |
| 0 | 0 | 0 |
| 0 | 0 | 0 |
| 0 | 0 | 0 |
| 0 | 1 | 0 |
| 0 | 1 | 0 |
| 1 | 1 | 1 |
| 1 | 1 | 1 |
| 1 | 1 | 1 |
| 0 | 1 | 0 |
| 1 | 1 | 1 |
| 1 | 1 | 1 |
| 1 | 1 | 1 |
| 0 | 0 | 0 |
| 1 | 1 | 1 |
| 1 | 1 | 1 |
| 1 | 1 | 1 |
| 0 | 1 | 0 |
| 0 | 1 | 0 |
| 1 | 1 | 1 |
| 1 | 1 | 1 |
| 1 | 1 | 1 |
| 1 | 1 | 1 |
| 1 | 1 | 1 |
| 1 | 1 | 1 |
| 0 | 0 | 0 |
| 0 | 0 | 0 |
| 1 | 0 | 0 |
| 0 | 0 | 0 |
| 1 | 1 | 1 |
| 1 | 1 | 1 |
| 0 | 0 | 0 |
| 0 | 0 | 0 |
| 0 | 1 | 0 |
| 0 | 1 | 0 |
| 0 | 1 | 0 |
| 1 | 1 | 1 |
| 1 | 1 | 1 |
| 1 | 1 | 1 |
| 0 | 0 | 0 |
| 0 | 0 | 0 |
| 0 | 0 | 0 |

| | | |
|---|---|---|
| 0 | 0 | 0 |
| 1 | 1 | 1 |
| 1 | 1 | 1 |
| 1 | 1 | 1 |
| 1 | 1 | 1 |
| 1 | 1 | 1 |
| 1 | 1 | 1 |
| 1 | 1 | 1 |
| 0 | 0 | 0 |
| 1 | 1 | 1 |
| 1 | 1 | 1 |
| 1 | 1 | 1 |
| 0 | 1 | 0 |
| 0 | 1 | 0 |
| 1 | 1 | 1 |
| 1 | 1 | 1 |
| 0 | 1 | 0 |
| 0 | 1 | 0 |
| 0 | 0 | 0 |
| 0 | 0 | 0 |
| 0 | 0 | 0 |
| 0 | 0 | 0 |
| 0 | 1 | 0 |
| 0 | 0 | 0 |
| 0 | 0 | 0 |
| 1 | 1 | 1 |
| 1 | 1 | 1 |
| 1 | 1 | 1 |
| 1 | 1 | 1 |
| 1 | 1 | 1 |
| 1 | 1 | 1 |
| 1 | 1 | 1 |
| 0 | 0 | 0 |
| 1 | 1 | 1 |
| 1 | 1 | 1 |
| 0 | 1 | 0 |
| 0 | 0 | 0 |
| 1 | 1 | 1 |
| 0 | 0 | 0 |
| 0 | 0 | 0 |
| 0 | 0 | 0 |
| 0 | 0 | 0 |
| 0 | 0 | 0 |
| 1 | 1 | 1 |
| 1 | 1 | 1 |
| 1 | 1 | 1 |
| 0 | 0 | 0 |
| 0 | 0 | 0 |
| 0 | 0 | 0 |
| 0 | 0 | 0 |
| 0 | 0 | 0 |
| 0 | 0 | 0 |
| 0 | 0 | 0 |
| 0 | 1 | 0 |

| | | |
|---|---|---|
| 0 | 0 | 0 |
| 0 | 0 | 0 |
| 0 | 1 | 0 |
| 0 | 0 | 0 |
| 0 | 0 | 0 |
| 1 | 1 | 1 |
| 1 | 1 | 1 |
| 1 | 0 | 0 |
| 1 | 1 | 1 |
| 1 | 1 | 1 |
| 1 | 1 | 1 |
| 0 | 0 | 0 |
| 1 | 1 | 1 |
| 1 | 1 | 1 |
| 0 | 0 | 0 |
| 1 | 1 | 1 |
| 0 | 0 | 0 |
| 1 | 1 | 1 |
| 0 | 0 | 0 |
| 0 | 0 | 0 |
| 1 | 1 | 1 |
| 1 | 1 | 1 |
| 0 | 0 | 0 |
| 1 | 1 | 1 |
| 0 | 0 | 0 |
| 1 | 1 | 1 |
| 1 | 1 | 1 |
| 0 | 0 | 0 |
| 1 | 1 | 1 |
| 0 | 0 | 0 |
| 0 | 1 | 0 |
| 0 | 0 | 0 |
| 0 | 0 | 0 |
| 0 | 0 | 0 |
| 1 | 1 | 1 |
| 0 | 0 | 0 |
| 1 | 1 | 1 |
| 1 | 1 | 1 |
| 0 | 0 | 0 |
| 1 | 1 | 1 |
| 1 | 1 | 1 |
| 1 | 1 | 1 |
| 1 | 1 | 1 |
| 0 | 0 | 0 |
| 0 | 0 | 0 |
| 1 | 1 | 1 |
| 1 | 1 | 1 |
| 1 | 0 | 0 |
| 0 | 0 | 0 |
| 1 | 0 | 0 |
| 0 | 0 | 0 |
| 1 | 1 | 1 |
| 1 | 1 | 1 |
| 0 | 0 | 0 |

| | | |
|---|---|---|
| 1 | 1 | 1 |
| 1 | 1 | 1 |
| 1 | 1 | 1 |
| 1 | 1 | 1 |
| 0 | 1 | 0 |
| 1 | 1 | 1 |
| 1 | 1 | 1 |
| 0 | 0 | 0 |
| 1 | 1 | 1 |
| 1 | 1 | 1 |
| 1 | 1 | 1 |
| 1 | 1 | 1 |
| 0 | 0 | 0 |
| 1 | 1 | 1 |
| 0 | 0 | 0 |
| 0 | 0 | 0 |
| 1 | 1 | 1 |
| 1 | 1 | 1 |
| 1 | 1 | 1 |
| 1 | 1 | 1 |
| 0 | 0 | 0 |
| 0 | 0 | 0 |
| 1 | 1 | 1 |
| 1 | 1 | 1 |
| 0 | 0 | 0 |
| 1 | 1 | 1 |
| 0 | 0 | 0 |
| 0 | 0 | 0 |
| 0 | 0 | 0 |
| 0 | 0 | 0 |
| 0 | 1 | 0 |
| 0 | 1 | 0 |
| 0 | 0 | 0 |
| 0 | 0 | 0 |
| 0 | 0 | 0 |
| 0 | 0 | 0 |
| 0 | 0 | 0 |
| 0 | 0 | 0 |
| 0 | 0 | 0 |
| 1 | 0 | 0 |
| 1 | 1 | 1 |
| 0 | 1 | 0 |
| 0 | 1 | 0 |
| 0 | 1 | 0 |
| 0 | 1 | 0 |
| 0 | 0 | 0 |
| 0 | 1 | 0 |
| 0 | 1 | 0 |
| 1 | 1 | 1 |
| 0 | 0 | 0 |
| 0 | 0 | 0 |
| 1 | 1 | 1 |
| 1 | 1 | 1 |
| 1 | 1 | 1 |

| | | |
|---|---|---|
| 1 | 1 | 1 |
| 1 | 1 | 1 |
| 1 | 1 | 1 |
| 1 | 1 | 1 |
| 0 | 0 | 0 |
| 1 | 1 | 1 |
| 0 | 0 | 0 |
| 0 | 0 | 0 |
| 0 | 0 | 0 |
| 0 | 0 | 0 |
| 0 | 0 | 0 |
| 1 | 1 | 1 |
| 0 | 0 | 0 |
| 0 | 0 | 0 |
| 1 | 1 | 1 |
| 1 | 1 | 1 |
| 1 | 1 | 1 |
| 1 | 1 | 1 |
| 1 | 1 | 1 |
| 1 | 1 | 1 |
| 1 | 1 | 1 |
| 1 | 1 | 1 |
| 1 | 1 | 1 |
| 0 | 0 | 0 |
| 1 | 1 | 1 |
| 1 | 1 | 1 |
| 1 | 1 | 1 |
| 0 | 0 | 0 |
| 1 | 0 | 0 |
| 1 | 0 | 0 |
| 0 | 0 | 0 |
| 0 | 0 | 0 |
| 1 | 1 | 1 |
| 0 | 0 | 0 |
| 0 | 1 | 0 |
| 1 | 1 | 1 |
| 1 | 1 | 1 |
| 1 | 1 | 1 |
| 1 | 1 | 1 |
| 1 | 1 | 1 |
| 1 | 1 | 1 |
| 1 | 1 | 1 |
| 1 | 1 | 1 |
| 1 | 1 | 1 |
| 1 | 1 | 1 |
| 1 | 1 | 1 |
| 1 | 1 | 1 |
| 1 | 1 | 1 |
| 0 | 0 | 0 |
| 0 | 1 | 0 |
| 1 | 1 | 1 |
| 1 | 0 | 0 |
| 1 | 0 | 0 |
| 1 | 1 | 1 |

| | | |
|---|---|---|
| 1 | 1 | 1 |
| 0 | 1 | 0 |
| 1 | 1 | 1 |
| 0 | 1 | 0 |
| 1 | 0 | 0 |
| 1 | 1 | 1 |
| 1 | 1 | 1 |
| 1 | 1 | 1 |
| 1 | 1 | 1 |
| 1 | 1 | 1 |
| 0 | 0 | 0 |
| 0 | 0 | 0 |
| 0 | 0 | 0 |
| 0 | 0 | 0 |
| 0 | 0 | 0 |
| 1 | 0 | 0 |
| 0 | 0 | 0 |
| 0 | 0 | 0 |
| 0 | 0 | 0 |
| 0 | 0 | 0 |
| 0 | 0 | 0 |
| 0 | 0 | 0 |
| 0 | 1 | 0 |
| 0 | 0 | 0 |
| 0 | 0 | 0 |
| 0 | 0 | 0 |
| 1 | 1 | 1 |
| 0 | 0 | 0 |
| 0 | 1 | 0 |
| 0 | 1 | 0 |
| 0 | 1 | 0 |
| 0 | 0 | 0 |
| 1 | 1 | 1 |
| 1 | 1 | 1 |
| 0 | 0 | 0 |
| 0 | 0 | 0 |
| 0 | 0 | 0 |
| 0 | 0 | 0 |
| 0 | 0 | 0 |
| 1 | 0 | 0 |
| 1 | 0 | 0 |
| 0 | 0 | 0 |
| 1 | 0 | 0 |
| 0 | 0 | 0 |
| 1 | 1 | 1 |
| 1 | 0 | 0 |
| 1 | 1 | 1 |
| 0 | 0 | 0 |
| 0 | 0 | 0 |
| 0 | 0 | 0 |
| 0 | 0 | 0 |
| 1 | 1 | 1 |
| 0 | 0 | 0 |
| 1 | 1 | 1 |

| | | |
|---|---|---|
| 1 | 1 | 1 |
| 1 | 1 | 1 |
| 1 | 1 | 1 |
| 1 | 1 | 1 |
| 1 | 1 | 1 |
| 0 | 0 | 0 |
| 0 | 0 | 0 |
| 0 | 0 | 0 |
| 0 | 0 | 0 |
| 0 | 1 | 0 |
| 0 | 0 | 0 |
| 0 | 0 | 0 |
| 0 | 0 | 0 |
| 0 | 0 | 0 |
| 0 | 0 | 0 |
| 0 | 0 | 0 |
| 1 | 1 | 1 |
| 0 | 1 | 0 |
| 1 | 1 | 1 |
| 1 | 1 | 1 |
| 1 | 1 | 1 |
| 1 | 1 | 1 |
| 1 | 1 | 1 |
| 1 | 1 | 1 |
| 1 | 1 | 1 |
| 1 | 1 | 1 |
| 1 | 1 | 1 |
| 1 | 1 | 1 |
| 0 | 0 | 0 |
| 0 | 1 | 0 |
| 0 | 1 | 0 |
| 1 | 0 | 0 |
| 1 | 0 | 0 |
| 0 | 0 | 0 |
| 1 | 1 | 1 |
| 0 | 0 | 0 |
| 1 | 0 | 0 |
| 0 | 0 | 0 |
| 1 | 1 | 1 |
| 0 | 0 | 0 |
| 1 | 1 | 1 |
| 1 | 1 | 1 |
| 0 | 1 | 0 |
| 0 | 0 | 0 |
| 1 | 1 | 1 |
| 1 | 1 | 1 |
| 1 | 1 | 1 |
| 1 | 1 | 1 |
| 1 | 1 | 1 |
| 1 | 1 | 1 |
| 1 | 1 | 1 |
| 1 | 0 | 0 |
| 0 | 0 | 0 |
| 0 | 0 | 0 |

| | | |
|---|---|---|
| 1 | 1 | 1 |
| 0 | 0 | 0 |
| 1 | 1 | 1 |
| 1 | 1 | 1 |
| 0 | 0 | 0 |
| 0 | 0 | 0 |
| 0 | 0 | 0 |
| 1 | 1 | 1 |
| 1 | 1 | 1 |
| 0 | 0 | 0 |
| 0 | 0 | 0 |
| 0 | 0 | 0 |
| 0 | 0 | 0 |
| 0 | 0 | 0 |
| 1 | 1 | 1 |
| 0 | 0 | 0 |
| 0 | 0 | 0 |
| 0 | 0 | 0 |
| 1 | 1 | 1 |
| 1 | 1 | 1 |
| 1 | 1 | 1 |
| 1 | 1 | 1 |
| 1 | 1 | 1 |
| 0 | 0 | 0 |
| 0 | 1 | 0 |
| 0 | 1 | 0 |
| 0 | 0 | 0 |
| 0 | 0 | 0 |
| 1 | 1 | 1 |
| 1 | 1 | 1 |
| 0 | 0 | 0 |
| 1 | 1 | 1 |
| 1 | 1 | 1 |
| 0 | 1 | 0 |
| 1 | 1 | 1 |
| 0 | 0 | 0 |
| 0 | 1 | 0 |
| 0 | 0 | 0 |
| 0 | 0 | 0 |
| 0 | 1 | 0 |
| 0 | 1 | 0 |
| 0 | 0 | 0 |
| 0 | 0 | 0 |
| 0 | 1 | 0 |
| 0 | 0 | 0 |
| 0 | 0 | 0 |
| 0 | 0 | 0 |
| 0 | 1 | 0 |
| 0 | 0 | 0 |
| 0 | 0 | 0 |
| 0 | 0 | 0 |
| 1 | 1 | 1 |
| 0 | 0 | 0 |
| 1 | 1 | 1 |

| | | |
|---|---|---|
| 1 | 1 | 1 |
| 1 | 1 | 1 |
| 1 | 1 | 1 |
| 1 | 1 | 1 |
| 0 | 0 | 0 |
| 0 | 0 | 0 |
| 1 | 1 | 1 |
| 1 | 0 | 0 |
| 1 | 1 | 1 |
| 0 | 0 | 0 |
| 1 | 0 | 0 |
| 1 | 0 | 0 |
| 0 | 0 | 0 |
| 1 | 1 | 1 |
| 1 | 1 | 1 |
| 1 | 1 | 1 |
| 1 | 1 | 1 |
| 1 | 1 | 1 |
| 1 | 1 | 1 |
| 1 | 1 | 1 |
| 1 | 1 | 1 |
| 0 | 1 | 0 |
| 1 | 1 | 1 |
| 0 | 0 | 0 |
| 1 | 1 | 1 |
| 0 | 0 | 0 |
| 1 | 1 | 1 |
| 0 | 0 | 0 |
| 1 | 1 | 1 |
| 0 | 0 | 0 |
| 0 | 0 | 0 |
| 0 | 0 | 0 |
| 0 | 0 | 0 |
| 0 | 0 | 0 |
| 0 | 1 | 0 |
| 0 | 0 | 0 |
| 1 | 1 | 1 |
| 0 | 0 | 0 |
| 1 | 1 | 1 |
| 1 | 1 | 1 |
| 0 | 0 | 0 |
| 0 | 0 | 0 |
| 0 | 0 | 0 |
| 0 | 0 | 0 |
| 0 | 0 | 0 |
| 0 | 0 | 0 |
| 0 | 0 | 0 |
| 0 | 0 | 0 |
| 1 | 1 | 1 |
| 0 | 0 | 0 |
| 1 | 0 | 0 |
| 1 | 0 | 0 |
| 1 | 1 | 1 |
| 1 | 1 | 1 |

| | | |
|---|---|---|
| 1 | 1 | 1 |
| 1 | 1 | 1 |
| 1 | 1 | 1 |
| 0 | 1 | 0 |
| 0 | 1 | 0 |
| 0 | 1 | 0 |
| 0 | 0 | 0 |
| 0 | 0 | 0 |
| 0 | 0 | 0 |
| 0 | 0 | 0 |
| 0 | 0 | 0 |
| 0 | 0 | 0 |
| 0 | 0 | 0 |
| 0 | 0 | 0 |
| 0 | 0 | 0 |
| 0 | 0 | 0 |
| 0 | 0 | 0 |
| 0 | 0 | 0 |
| 1 | 1 | 1 |
| 0 | 0 | 0 |
| 0 | 1 | 0 |
| 0 | 1 | 0 |
| 0 | 1 | 0 |
| 1 | 1 | 1 |
| 0 | 1 | 0 |
| 0 | 0 | 0 |
| 0 | 0 | 0 |
| 0 | 0 | 0 |
| 0 | 0 | 0 |
| 0 | 0 | 0 |
| 0 | 0 | 0 |
| 0 | 0 | 0 |
| 0 | 0 | 0 |
| 0 | 0 | 0 |
| 1 | 1 | 1 |
| 0 | 0 | 0 |
| 0 | 0 | 0 |
| 0 | 0 | 0 |
| 0 | 0 | 0 |
| 0 | 1 | 0 |
| 0 | 0 | 0 |
| 0 | 0 | 0 |
| 1 | 1 | 1 |
| 0 | 1 | 0 |
| 1 | 1 | 1 |
| 0 | 1 | 0 |
| 0 | 1 | 0 |
| 0 | 0 | 0 |
| 0 | 0 | 0 |
| 0 | 0 | 0 |
| 0 | 0 | 0 |
| 0 | 1 | 0 |
| 0 | 0 | 0 |
| 1 | 1 | 1 |

| | | |
|---|---|---|
| 0 | 1 | 0 |
| 1 | 1 | 1 |
| 1 | 1 | 1 |
| 1 | 1 | 1 |
| 1 | 1 | 1 |
| 0 | 0 | 0 |
| 0 | 1 | 0 |
| 1 | 1 | 1 |
| 1 | 1 | 1 |
| 1 | 1 | 1 |
| 0 | 0 | 0 |
| 0 | 0 | 0 |
| 0 | 0 | 0 |
| 0 | 0 | 0 |
| 0 | 0 | 0 |
| 0 | 1 | 0 |
| 1 | 0 | 0 |
| 0 | 0 | 0 |
| 1 | 1 | 1 |
| 1 | 1 | 1 |
| 1 | 1 | 1 |
| 1 | 1 | 1 |
| 0 | 0 | 0 |
| 1 | 1 | 1 |
| 0 | 0 | 0 |
| 0 | 0 | 0 |
| 0 | 0 | 0 |
| 0 | 0 | 0 |
| 1 | 1 | 1 |
| 0 | 0 | 0 |
| 1 | 1 | 1 |
| 0 | 1 | 0 |
| 0 | 1 | 0 |
| 0 | 1 | 0 |
| 1 | 1 | 1 |
| 1 | 1 | 1 |
| 0 | 1 | 0 |
| 0 | 0 | 0 |
| 0 | 0 | 0 |
| 0 | 0 | 0 |
| 0 | 0 | 0 |
| 0 | 0 | 0 |
| 1 | 1 | 1 |
| 0 | 0 | 0 |
| 0 | 0 | 0 |
| 1 | 1 | 1 |
| 0 | 0 | 0 |
| 0 | 0 | 0 |
| 0 | 0 | 0 |
| 0 | 0 | 0 |
| 1 | 1 | 1 |
| 1 | 1 | 1 |
| 1 | 1 | 1 |
| 0 | 0 | 0 |

| | | |
|---|---|---|
| 0 | 0 | 0 |
| 1 | 1 | 1 |
| 0 | 0 | 0 |
| 0 | 1 | 0 |
| 1 | 0 | 0 |
| 1 | 0 | 0 |
| 1 | 0 | 0 |
| 1 | 1 | 1 |
| 0 | 0 | 0 |
| 1 | 1 | 1 |
| 0 | 0 | 0 |
| 0 | 0 | 0 |
| 0 | 0 | 0 |
| 0 | 0 | 0 |
| 1 | 1 | 1 |
| 1 | 1 | 1 |
| 0 | 0 | 0 |
| 0 | 0 | 0 |
| 0 | 1 | 0 |
| 1 | 1 | 1 |
| 0 | 0 | 0 |
| 1 | 0 | 0 |
| 1 | 0 | 0 |
| 1 | 1 | 1 |
| 0 | 0 | 0 |
| 1 | 0 | 0 |
| 0 | 0 | 0 |
| 0 | 0 | 0 |
| 0 | 0 | 0 |
| 1 | 1 | 1 |
| 0 | 0 | 0 |
| 0 | 0 | 0 |
| 0 | 0 | 0 |
| 1 | 1 | 1 |
| 0 | 0 | 0 |
| 1 | 1 | 1 |
| 0 | 0 | 0 |
| 0 | 0 | 0 |
| 0 | 0 | 0 |
| 0 | 0 | 0 |
| 0 | 0 | 0 |
| 1 | 1 | 1 |
| 0 | 0 | 0 |
| 0 | 0 | 0 |
| 0 | 0 | 0 |
| 0 | 0 | 0 |
| 0 | 0 | 0 |
| 0 | 0 | 0 |
| 0 | 0 | 0 |
| 0 | 0 | 0 |
| 0 | 0 | 0 |
| 0 | 0 | 0 |
| 1 | 1 | 1 |
| 0 | 0 | 0 |

| | | |
|---|---|---|
| 0 | 1 | 0 |
| 1 | 1 | 1 |
| 1 | 1 | 1 |
| 1 | 1 | 1 |
| 1 | 1 | 1 |
| 1 | 1 | 1 |
| 1 | 1 | 1 |
| 1 | 1 | 1 |
| 1 | 1 | 1 |
| 0 | 0 | 0 |
| 0 | 0 | 0 |
| 1 | 1 | 1 |
| 1 | 1 | 1 |
| 1 | 1 | 1 |
| 0 | 0 | 0 |
| 1 | 1 | 1 |
| 1 | 1 | 1 |
| 0 | 0 | 0 |
| 0 | 1 | 0 |
| 0 | 1 | 0 |
| 0 | 1 | 0 |
| 0 | 0 | 0 |
| 0 | 0 | 0 |
| 0 | 0 | 0 |
| 0 | 0 | 0 |
| 0 | 0 | 0 |
| 0 | 0 | 0 |
| 0 | 0 | 0 |
| 1 | 1 | 1 |
| 0 | 0 | 0 |
| 0 | 0 | 0 |
| 1 | 1 | 1 |
| 1 | 1 | 1 |
| 1 | 1 | 1 |
| 1 | 1 | 1 |
| 1 | 1 | 1 |
| 1 | 1 | 1 |
| 1 | 1 | 1 |
| 1 | 1 | 1 |
| 1 | 1 | 1 |
| 1 | 1 | 1 |
| 1 | 1 | 1 |
| 1 | 1 | 1 |
| 0 | 0 | 0 |
| 1 | 1 | 1 |
| 0 | 0 | 0 |
| 1 | 1 | 1 |
| 1 | 1 | 1 |
| 1 | 1 | 1 |
| 1 | 1 | 1 |
| 0 | 0 | 0 |
| 1 | 1 | 1 |
| 1 | 1 | 1 |
| 0 | 0 | 0 |

| | | |
|---|---|---|
| 0 | 0 | 0 |
| 0 | 0 | 0 |
| 0 | 0 | 0 |
| 0 | 0 | 0 |
| 1 | 1 | 1 |
| 0 | 1 | 0 |
| 1 | 1 | 1 |
| 0 | 0 | 0 |
| 0 | 0 | 0 |
| 0 | 0 | 0 |
| 0 | 0 | 0 |
| 1 | 1 | 1 |
| 1 | 1 | 1 |
| 1 | 1 | 1 |
| 1 | 1 | 1 |
| 0 | 0 | 0 |
| 0 | 0 | 0 |
| 0 | 1 | 0 |
| 0 | 1 | 0 |
| 1 | 1 | 1 |
| 1 | 1 | 1 |
| 0 | 0 | 0 |
| 1 | 1 | 1 |
| 0 | 1 | 0 |
| 0 | 0 | 0 |
| 0 | 0 | 0 |
| 0 | 0 | 0 |
| 0 | 0 | 0 |
| 1 | 1 | 1 |
| 1 | 1 | 1 |
| 1 | 1 | 1 |
| 0 | 0 | 0 |
| 1 | 1 | 1 |
| 1 | 1 | 1 |
| 1 | 1 | 1 |
| 1 | 1 | 1 |
| 1 | 1 | 1 |
| 1 | 1 | 1 |
| 1 | 1 | 1 |
| 1 | 1 | 1 |
| 1 | 1 | 1 |
| 1 | 1 | 1 |
| 0 | 0 | 0 |
| 0 | 1 | 0 |
| 0 | 1 | 0 |
| 1 | 1 | 1 |
| 0 | 1 | 0 |
| 1 | 1 | 1 |
| 1 | 1 | 1 |
| 0 | 0 | 0 |
| 1 | 1 | 1 |
| 1 | 1 | 1 |
| 1 | 1 | 1 |
| 1 | 1 | 1 |

| | | |
|---|---|---|
| 0 | 0 | 0 |
| 1 | 1 | 1 |
| 0 | 0 | 0 |
| 0 | 1 | 0 |
| 1 | 1 | 1 |
| 1 | 1 | 1 |
| 1 | 1 | 1 |
| 0 | 0 | 0 |
| 1 | 1 | 1 |
| 1 | 1 | 1 |
| 0 | 0 | 0 |
| 0 | 0 | 0 |
| 0 | 0 | 0 |
| 1 | 1 | 1 |
| 0 | 0 | 0 |
| 0 | 0 | 0 |
| 0 | 0 | 0 |
| 0 | 0 | 0 |
| 0 | 1 | 0 |
| 0 | 0 | 0 |
| 0 | 0 | 0 |
| 0 | 0 | 0 |
| 0 | 0 | 0 |
| 0 | 0 | 0 |
| 0 | 0 | 0 |
| 0 | 0 | 0 |
| 0 | 0 | 0 |
| 0 | 0 | 0 |
| 0 | 0 | 0 |
| 0 | 0 | 0 |
| 0 | 0 | 0 |
| 0 | 0 | 0 |
| 1 | 1 | 1 |
| 0 | 0 | 0 |
| 1 | 1 | 1 |
| 0 | 0 | 0 |
| 0 | 0 | 0 |
| 1 | 1 | 1 |
| 1 | 1 | 1 |
| 0 | 1 | 0 |
| 0 | 1 | 0 |
| 0 | 0 | 0 |
| 0 | 0 | 0 |
| 0 | 0 | 0 |
| 1 | 1 | 1 |
| 0 | 0 | 0 |
| 1 | 1 | 1 |
| 0 | 0 | 0 |
| 0 | 1 | 0 |
| 0 | 1 | 0 |
| 0 | 1 | 0 |
| 0 | 1 | 0 |
| 0 | 1 | 0 |
| 1 | 1 | 1 |

| | | |
|---|---|---|
| 1 | 1 | 1 |
| 1 | 1 | 1 |
| 0 | 1 | 0 |
| 1 | 1 | 1 |
| 1 | 1 | 1 |
| 0 | 0 | 0 |
| 0 | 0 | 0 |
| 1 | 1 | 1 |
| 0 | 0 | 0 |
| 0 | 0 | 0 |
| 1 | 1 | 1 |
| 0 | 1 | 0 |
| 1 | 1 | 1 |
| 1 | 1 | 1 |
| 1 | 1 | 1 |
| 0 | 0 | 0 |
| 1 | 1 | 1 |
| 1 | 1 | 1 |
| 0 | 0 | 0 |
| 0 | 0 | 0 |
| 0 | 0 | 0 |
| 1 | 1 | 1 |
| 1 | 1 | 1 |
| 0 | 0 | 0 |
| 1 | 1 | 1 |
| 0 | 0 | 0 |
| 0 | 0 | 0 |
| 1 | 1 | 1 |
| 1 | 1 | 1 |
| 1 | 1 | 1 |
| 0 | 0 | 0 |
| 1 | 0 | 0 |
| 0 | 0 | 0 |
| 0 | 0 | 0 |
| 0 | 0 | 0 |
| 0 | 0 | 0 |
| 0 | 0 | 0 |
| 1 | 1 | 1 |
| 0 | 0 | 0 |
| 0 | 1 | 0 |
| 0 | 0 | 0 |
| 0 | 0 | 0 |
| 0 | 0 | 0 |
| 0 | 0 | 0 |
| 1 | 1 | 1 |
| 0 | 1 | 0 |
| 0 | 1 | 0 |
| 0 | 0 | 0 |
| 0 | 0 | 0 |
| 0 | 0 | 0 |
| 1 | 1 | 1 |
| 0 | 1 | 0 |
| 1 | 1 | 1 |
| 1 | 1 | 1 |

| | | |
|---|---|---|
| 1 | 1 | 1 |
| 0 | 1 | 0 |
| 0 | 1 | 0 |
| 0 | 0 | 0 |
| 0 | 0 | 0 |
| 1 | 1 | 1 |
| 0 | 0 | 0 |
| 1 | 1 | 1 |
| 1 | 1 | 1 |
| 1 | 1 | 1 |
| 1 | 1 | 1 |
| 0 | 0 | 0 |
| 1 | 0 | 0 |
| 0 | 0 | 0 |
| 1 | 1 | 1 |
| 0 | 0 | 0 |
| 0 | 0 | 0 |
| 0 | 0 | 0 |
| 1 | 1 | 1 |
| 0 | 0 | 0 |
| 0 | 0 | 0 |
| 0 | 0 | 0 |
| 0 | 0 | 0 |
| 0 | 0 | 0 |
| 0 | 0 | 0 |
| 1 | 1 | 1 |
| 0 | 0 | 0 |
| 0 | 1 | 0 |
| 0 | 1 | 0 |
| 0 | 1 | 0 |
| 0 | 1 | 0 |
| 0 | 1 | 0 |
| 0 | 0 | 0 |
| 0 | 0 | 0 |
| 1 | 1 | 1 |
| 1 | 1 | 1 |
| 0 | 0 | 0 |
| 0 | 0 | 0 |
| 0 | 0 | 0 |
| 1 | 0 | 0 |
| 0 | 0 | 0 |
| 0 | 0 | 0 |
| 1 | 1 | 1 |
| 0 | 0 | 0 |
| 1 | 1 | 1 |
| 1 | 1 | 1 |
| 1 | 1 | 1 |
| 1 | 1 | 1 |
| 1 | 1 | 1 |
| 0 | 0 | 0 |
| 1 | 1 | 1 |
| 1 | 1 | 1 |
| 1 | 1 | 1 |
| 0 | 1 | 0 |

| | | |
|---|---|---|
| 1 | 1 | 1 |
| 1 | 1 | 1 |
| 1 | 1 | 1 |
| 0 | 0 | 0 |
| 1 | 1 | 1 |
| 1 | 1 | 1 |
| 0 | 1 | 0 |
| 0 | 0 | 0 |
| 1 | 1 | 1 |
| 1 | 1 | 1 |
| 1 | 1 | 1 |
| 1 | 1 | 1 |
| 1 | 1 | 1 |
| 0 | 1 | 0 |
| 0 | 1 | 0 |
| 0 | 1 | 0 |
| 0 | 0 | 0 |
| 0 | 0 | 0 |
| 0 | 1 | 0 |
| 1 | 1 | 1 |
| 0 | 1 | 0 |
| 1 | 1 | 1 |
| 0 | 1 | 0 |
| 1 | 1 | 1 |
| 1 | 1 | 1 |
| 1 | 1 | 1 |
| 1 | 1 | 1 |
| 0 | 0 | 0 |
| 0 | 0 | 0 |
| 1 | 1 | 1 |
| 1 | 1 | 1 |
| 0 | 0 | 0 |
| 0 | 1 | 0 |
| 0 | 0 | 0 |
| 0 | 0 | 0 |
| 0 | 0 | 0 |
| 0 | 0 | 0 |
| 0 | 0 | 0 |
| 1 | 1 | 1 |
| 1 | 1 | 1 |
| 1 | 1 | 1 |
| 0 | 1 | 0 |
| 1 | 1 | 1 |
| 1 | 1 | 1 |
| 0 | 0 | 0 |
| 0 | 0 | 0 |
| 0 | 0 | 0 |
| 1 | 1 | 1 |
| 1 | 1 | 1 |
| 1 | 1 | 1 |
| 1 | 1 | 1 |
| 0 | 0 | 0 |
| 0 | 0 | 0 |
| 0 | 0 | 0 |

| | | |
|---|---|---|
| 0 | 0 | 0 |
| 0 | 0 | 0 |
| 1 | 1 | 1 |
| 0 | 0 | 0 |
| 0 | 0 | 0 |
| 0 | 0 | 0 |
| 0 | 0 | 0 |
| 1 | 1 | 1 |
| 1 | 1 | 1 |
| 1 | 1 | 1 |
| 1 | 1 | 1 |
| 1 | 1 | 1 |
| 0 | 1 | 0 |
| 0 | 1 | 0 |
| 1 | 1 | 1 |
| 1 | 1 | 1 |
| 1 | 1 | 1 |
| 1 | 1 | 1 |
| 1 | 1 | 1 |
| 1 | 1 | 1 |
| 0 | 0 | 0 |
| 0 | 0 | 0 |
| 0 | 0 | 0 |
| 0 | 0 | 0 |
| 0 | 0 | 0 |
| 1 | 1 | 1 |
| 1 | 1 | 1 |
| 0 | 0 | 0 |
| 0 | 1 | 0 |
| 0 | 1 | 0 |
| 0 | 0 | 0 |
| 0 | 0 | 0 |
| 0 | 0 | 0 |
| 1 | 1 | 1 |
| 0 | 0 | 0 |
| 1 | 1 | 1 |
| 0 | 1 | 0 |
| 0 | 0 | 0 |
| 1 | 1 | 1 |
| 1 | 1 | 1 |
| 0 | 1 | 0 |
| 0 | 1 | 0 |
| 1 | 1 | 1 |
| 1 | 1 | 1 |
| 1 | 1 | 1 |
| 1 | 1 | 1 |
| 1 | 1 | 1 |
| 1 | 1 | 1 |
| 0 | 0 | 0 |
| 1 | 1 | 1 |
| 0 | 0 | 0 |
| 0 | 0 | 0 |
| 1 | 1 | 1 |
| 0 | 1 | 0 |

| | | |
|---|---|---|
| 0 | 0 | 0 |
| 1 | 1 | 1 |
| 1 | 1 | 1 |
| 1 | 1 | 1 |
| 1 | 1 | 1 |
| 0 | 0 | 0 |
| 0 | 0 | 0 |
| 1 | 1 | 1 |
| 0 | 1 | 0 |
| 1 | 1 | 1 |
| 1 | 1 | 1 |
| 0 | 0 | 0 |
| 0 | 1 | 0 |
| 1 | 1 | 1 |
| 0 | 1 | 0 |
| 1 | 1 | 1 |
| 0 | 0 | 0 |
| 0 | 0 | 0 |
| 1 | 1 | 1 |
| 1 | 1 | 1 |
| 1 | 1 | 1 |
| 1 | 1 | 1 |
| 1 | 1 | 1 |
| 1 | 1 | 1 |
| 0 | 0 | 0 |
| 1 | 1 | 1 |
| 1 | 1 | 1 |
| 1 | 1 | 1 |
| 1 | 1 | 1 |
| 1 | 1 | 1 |
| 0 | 1 | 0 |
| 0 | 0 | 0 |
| 0 | 0 | 0 |
| 0 | 1 | 0 |
| 1 | 1 | 1 |
| 0 | 0 | 0 |
| 0 | 0 | 0 |
| 0 | 0 | 0 |
| 0 | 0 | 0 |
| 0 | 0 | 0 |
| 0 | 0 | 0 |
| 0 | 0 | 0 |
| 0 | 0 | 0 |
| 0 | 0 | 0 |
| 0 | 0 | 0 |
| 0 | 0 | 0 |
| 0 | 0 | 0 |
| 0 | 0 | 0 |
| 1 | 0 | 0 |
| 1 | 1 | 1 |
| 1 | 0 | 0 |
| 1 | 1 | 1 |
| 0 | 0 | 0 |
| 1 | 1 | 1 |

| | | |
|---|---|---|
| 1 | 1 | 1 |
| 1 | 1 | 1 |
| 1 | 1 | 1 |
| 1 | 1 | 1 |
| 1 | 1 | 1 |
| 1 | 1 | 1 |
| 1 | 1 | 1 |
| 0 | 0 | 0 |
| 0 | 0 | 0 |
| 1 | 0 | 0 |
| 1 | 1 | 1 |
| 1 | 1 | 1 |
| 1 | 1 | 1 |
| 0 | 0 | 0 |
| 1 | 1 | 1 |
| 1 | 1 | 1 |
| 0 | 0 | 0 |
| 1 | 1 | 1 |
| 0 | 0 | 0 |
| 0 | 1 | 0 |
| 1 | 1 | 1 |
| 1 | 1 | 1 |
| 1 | 1 | 1 |
| 0 | 0 | 0 |
| 1 | 1 | 1 |
| 1 | 1 | 1 |
| 0 | 1 | 0 |
| 0 | 1 | 0 |
| 0 | 1 | 0 |
| 0 | 1 | 0 |
| 1 | 1 | 1 |
| 0 | 0 | 0 |
| 1 | 1 | 1 |
| 0 | 1 | 0 |
| 1 | 1 | 1 |
| 1 | 1 | 1 |
| 1 | 1 | 1 |
| 1 | 1 | 1 |
| 0 | 0 | 0 |
| 1 | 1 | 1 |
| 1 | 1 | 1 |
| 1 | 1 | 1 |
| 0 | 1 | 0 |
| 1 | 1 | 1 |
| 1 | 1 | 1 |
| 0 | 0 | 0 |
| 0 | 1 | 0 |
| 0 | 1 | 0 |
| 1 | 0 | 0 |
| 1 | 0 | 0 |
| 0 | 1 | 0 |
| 1 | 0 | 0 |
| 0 | 0 | 0 |
| 1 | 0 | 0 |

| | | |
|---|---|---|
| 1 | 0 | 0 |
| 1 | 0 | 0 |
| 1 | 0 | 0 |
| 1 | 0 | 0 |
| 0 | 0 | 0 |
| 0 | 0 | 0 |
| 0 | 0 | 0 |
| 0 | 0 | 0 |
| 1 | 0 | 0 |
| 1 | 0 | 0 |
| 1 | 0 | 0 |
| 1 | 0 | 0 |
| 1 | 0 | 0 |
| 0 | 0 | 0 |
| 1 | 0 | 0 |
| 0 | 0 | 0 |
| 1 | 0 | 0 |
| 0 | 0 | 0 |
| 1 | 0 | 0 |
| 0 | 0 | 0 |
| 0 | 0 | 0 |
| 1 | 0 | 0 |
| 1 | 0 | 0 |
| 0 | 0 | 0 |
| 0 | 0 | 0 |
| 1 | 0 | 0 |
| 0 | 0 | 0 |
| 0 | 0 | 0 |
| 0 | 0 | 0 |
| 1 | 0 | 0 |
| 1 | 0 | 0 |
| 0 | 0 | 0 |
| 1 | 0 | 0 |
| 1 | 0 | 0 |
| 1 | 0 | 0 |
| 1 | 0 | 0 |
| 1 | 0 | 0 |
| 1 | 0 | 0 |
| 1 | 0 | 0 |
| 1 | 0 | 0 |
| 1 | 0 | 0 |
| 1 | 0 | 0 |
| 1 | 0 | 0 |
| 0 | 0 | 0 |
| 1 | 1 | 1 |
| 1 | 1 | 1 |
| 0 | 0 | 0 |
| 0 | 0 | 0 |
| 0 | 1 | 0 |
| 0 | 0 | 0 |
| 0 | 0 | 0 |
| 0 | 0 | 0 |
| 1 | 1 | 1 |
| 1 | 1 | 1 |

| | | |
|---|---|---|
| 1 | 1 | 1 |
| 1 | 1 | 1 |
| 1 | 1 | 1 |
| 1 | 1 | 1 |
| 1 | 1 | 1 |
| 1 | 1 | 1 |
| 1 | 1 | 1 |
| 0 | 0 | 0 |
| 1 | 1 | 1 |
| 1 | 1 | 1 |
| 0 | 1 | 0 |
| 0 | 1 | 0 |
| 1 | 0 | 0 |
| 1 | 1 | 1 |
| 0 | 0 | 0 |
| 0 | 0 | 0 |
| 0 | 1 | 0 |
| 0 | 0 | 0 |
| 1 | 1 | 1 |
| 1 | 1 | 1 |
| 0 | 0 | 0 |
| 0 | 1 | 0 |
| 0 | 1 | 0 |
| 1 | 1 | 1 |
| 0 | 0 | 0 |
| 1 | 1 | 1 |
| 0 | 1 | 0 |
| 0 | 0 | 0 |
| 0 | 0 | 0 |
| 0 | 0 | 0 |
| 0 | 0 | 0 |
| 1 | 1 | 1 |
| 1 | 1 | 1 |
| 0 | 1 | 0 |
| 1 | 1 | 1 |
| 1 | 1 | 1 |
| 0 | 0 | 0 |
| 0 | 0 | 0 |
| 1 | 1 | 1 |
| 1 | 1 | 1 |
| 1 | 1 | 1 |
| 1 | 1 | 1 |
| 0 | 0 | 0 |
| 0 | 1 | 0 |
| 1 | 1 | 1 |
| 1 | 1 | 1 |
| 1 | 1 | 1 |
| 1 | 1 | 1 |
| 0 | 0 | 0 |
| 0 | 0 | 0 |
| 0 | 0 | 0 |
| 0 | 0 | 0 |
| 0 | 0 | 0 |
| 1 | 1 | 1 |

| | | |
|---|---|---|
| 0 | 0 | 0 |
| 0 | 0 | 0 |
| 0 | 1 | 0 |
| 0 | 0 | 0 |
| 1 | 1 | 1 |
| 0 | 1 | 0 |
| 0 | 1 | 0 |
| 1 | 1 | 1 |
| 1 | 1 | 1 |
| 0 | 0 | 0 |
| 1 | 1 | 1 |
| 0 | 1 | 0 |
| 1 | 1 | 1 |
| 0 | 1 | 0 |
| 0 | 1 | 0 |
| 0 | 1 | 0 |
| 0 | 1 | 0 |
| 1 | 1 | 1 |
| 1 | 1 | 1 |
| 0 | 1 | 0 |
| 0 | 1 | 0 |
| 0 | 1 | 0 |
| 0 | 1 | 0 |
| 0 | 1 | 0 |
| 0 | 1 | 0 |
| 0 | 1 | 0 |
| 0 | 1 | 0 |
| 0 | 1 | 0 |
| 0 | 1 | 0 |
| 0 | 1 | 0 |
| 1 | 1 | 1 |
| 1 | 1 | 1 |
| 0 | 0 | 0 |
| 1 | 1 | 1 |
| 1 | 1 | 1 |
| 1 | 1 | 1 |
| 1 | 1 | 1 |
| 0 | 1 | 0 |
| 1 | 1 | 1 |
| 1 | 1 | 1 |
| 1 | 1 | 1 |
| 1 | 1 | 1 |
| 1 | 1 | 1 |
| 0 | 1 | 0 |
| 0 | 1 | 0 |
| 1 | 1 | 1 |
| 1 | 1 | 1 |
| 0 | 0 | 0 |
| 0 | 0 | 0 |
| 0 | 0 | 0 |
| 0 | 0 | 0 |
| 0 | 0 | 0 |
| 1 | 1 | 1 |
| 0 | 0 | 0 |

| | | |
|---|---|---|
| 1 | 1 | 1 |
| 0 | 0 | 0 |
| 1 | 1 | 1 |
| 0 | 1 | 0 |
| 0 | 1 | 0 |
| 0 | 0 | 0 |
| 0 | 0 | 0 |
| 0 | 0 | 0 |
| 0 | 0 | 0 |
| 0 | 0 | 0 |
| 0 | 1 | 0 |
| 0 | 0 | 0 |
| 0 | 1 | 0 |
| 1 | 1 | 1 |
| 0 | 1 | 0 |
| 0 | 0 | 0 |
| 1 | 1 | 1 |
| 1 | 1 | 1 |
| 0 | 0 | 0 |
| 0 | 0 | 0 |
| 0 | 0 | 0 |
| 0 | 0 | 0 |
| 0 | 0 | 0 |
| 0 | 0 | 0 |
| 0 | 1 | 0 |
| 0 | 0 | 0 |
| 0 | 0 | 0 |
| 0 | 0 | 0 |
| 0 | 0 | 0 |
| 0 | 0 | 0 |
| 0 | 0 | 0 |
| 1 | 1 | 1 |
| 1 | 1 | 1 |
| 0 | 0 | 0 |
| 0 | 0 | 0 |
| 1 | 0 | 0 |
| 0 | 0 | 0 |
| 0 | 0 | 0 |
| 0 | 0 | 0 |
| 0 | 0 | 0 |
| 0 | 0 | 0 |
| 1 | 0 | 0 |
| 1 | 1 | 1 |
| 1 | 1 | 1 |
| 0 | 0 | 0 |
| 0 | 0 | 0 |
| 0 | 0 | 0 |
| 1 | 1 | 1 |
| 1 | 1 | 1 |
| 1 | 1 | 1 |
| 1 | 1 | 1 |
| 1 | 1 | 1 |
| 1 | 1 | 1 |
| 0 | 0 | 0 |

| | | |
|---|---|---|
| 1 | 1 | 1 |
| 1 | 1 | 1 |
| 1 | 1 | 1 |
| 1 | 1 | 1 |
| 0 | 0 | 0 |
| 0 | 0 | 0 |
| 1 | 0 | 0 |
| 1 | 0 | 0 |
| 0 | 1 | 0 |
| 1 | 1 | 1 |
| 1 | 1 | 1 |
| 0 | 0 | 0 |
| 1 | 1 | 1 |
| 0 | 0 | 0 |
| 0 | 0 | 0 |
| 0 | 0 | 0 |
| 0 | 0 | 0 |
| 0 | 0 | 0 |
| 0 | 0 | 0 |
| 0 | 0 | 0 |
| 0 | 0 | 0 |
| 0 | 0 | 0 |
| 0 | 0 | 0 |
| 0 | 1 | 0 |
| 0 | 1 | 0 |
| 0 | 1 | 0 |
| 1 | 1 | 1 |
| 1 | 1 | 1 |
| 1 | 1 | 1 |
| 1 | 1 | 1 |
| 1 | 1 | 1 |
| 0 | 0 | 0 |
| 1 | 1 | 1 |
| 0 | 0 | 0 |
| 0 | 0 | 0 |
| 0 | 0 | 0 |
| 0 | 0 | 0 |
| 1 | 1 | 1 |
| 1 | 1 | 1 |
| 0 | 1 | 0 |
| 1 | 1 | 1 |
| 1 | 1 | 1 |
| 1 | 1 | 1 |
| 0 | 1 | 0 |
| 1 | 1 | 1 |
| 0 | 0 | 0 |
| 1 | 1 | 1 |
| 1 | 1 | 1 |
| 1 | 1 | 1 |
| 0 | 0 | 0 |
| 0 | 0 | 0 |
| 0 | 0 | 0 |
| 0 | 1 | 0 |
| 0 | 1 | 0 |

| | | |
|---|---|---|
| 0 | 1 | 0 |
| 0 | 1 | 0 |
| 0 | 1 | 0 |
| 0 | 0 | 0 |
| 0 | 1 | 0 |
| 1 | 1 | 1 |
| 0 | 0 | 0 |
| 0 | 1 | 0 |
| 1 | 1 | 1 |
| 1 | 1 | 1 |
| 0 | 0 | 0 |
| 0 | 0 | 0 |
| 1 | 1 | 1 |
| 1 | 1 | 1 |
| 0 | 1 | 0 |
| 0 | 1 | 0 |
| 0 | 0 | 0 |
| 0 | 0 | 0 |
| 1 | 1 | 1 |
| 0 | 0 | 0 |
| 0 | 0 | 0 |
| 0 | 0 | 0 |
| 0 | 1 | 0 |
| 0 | 0 | 0 |
| 0 | 0 | 0 |
| 1 | 1 | 1 |
| 0 | 0 | 0 |
| 0 | 1 | 0 |
| 1 | 1 | 1 |
| 0 | 0 | 0 |
| 0 | 1 | 0 |
| 1 | 1 | 1 |
| 0 | 0 | 0 |
| 0 | 1 | 0 |
| 0 | 1 | 0 |
| 0 | 0 | 0 |
| 1 | 1 | 1 |
| 0 | 0 | 0 |
| 1 | 1 | 1 |
| 0 | 0 | 0 |
| 0 | 0 | 0 |
| 1 | 0 | 0 |
| 0 | 0 | 0 |
| 0 | 0 | 0 |
| 1 | 1 | 1 |
| 1 | 0 | 0 |
| 0 | 1 | 0 |
| 1 | 1 | 1 |
| 1 | 1 | 1 |
| 1 | 1 | 1 |
| 1 | 1 | 1 |
| 0 | 1 | 0 |
| 0 | 1 | 0 |
| 0 | 1 | 0 |

| | | |
|---|---|---|
| 0 | 1 | 0 |
| 0 | 1 | 0 |
| 0 | 1 | 0 |
| 0 | 0 | 0 |
| 1 | 0 | 0 |
| 0 | 0 | 0 |
| 1 | 1 | 1 |
| 1 | 0 | 0 |
| 0 | 0 | 0 |
| 0 | 0 | 0 |
| 1 | 1 | 1 |
| 0 | 0 | 0 |
| 0 | 0 | 0 |
| 0 | 0 | 0 |
| 1 | 0 | 0 |
| 0 | 0 | 0 |
| 0 | 0 | 0 |
| 0 | 1 | 0 |
| 0 | 0 | 0 |
| 0 | 0 | 0 |
| 0 | 0 | 0 |
| 0 | 0 | 0 |
| 1 | 1 | 1 |
| 0 | 0 | 0 |
| 0 | 0 | 0 |
| 0 | 0 | 0 |
| 0 | 0 | 0 |
| 0 | 0 | 0 |
| 0 | 0 | 0 |
| 0 | 0 | 0 |
| 0 | 0 | 0 |
| 0 | 0 | 0 |
| 0 | 0 | 0 |
| 0 | 1 | 0 |
| 0 | 1 | 0 |
| 0 | 0 | 0 |
| 1 | 1 | 1 |
| 0 | 0 | 0 |
| 0 | 0 | 0 |
| 0 | 0 | 0 |
| 0 | 0 | 0 |
| 1 | 1 | 1 |
| 0 | 0 | 0 |
| 0 | 0 | 0 |
| 1 | 1 | 1 |
| 0 | 1 | 0 |
| 1 | 1 | 1 |
| 1 | 1 | 1 |
| 1 | 1 | 1 |
| 0 | 1 | 0 |
| 1 | 1 | 1 |
| 1 | 1 | 1 |
| 1 | 1 | 1 |
| 1 | 1 | 1 |

| | | |
|---|---|---|
| 1 | 1 | 1 |
| 0 | 0 | 0 |
| 1 | 1 | 1 |
| 1 | 1 | 1 |
| 1 | 1 | 1 |
| 1 | 1 | 1 |
| 1 | 1 | 1 |
| 1 | 1 | 1 |
| 1 | 1 | 1 |
| 0 | 1 | 0 |
| 1 | 1 | 1 |
| 0 | 0 | 0 |
| 1 | 0 | 0 |
| 0 | 1 | 0 |
| 0 | 0 | 0 |
| 0 | 1 | 0 |
| 0 | 1 | 0 |
| 1 | 1 | 1 |
| 1 | 1 | 1 |
| 1 | 1 | 1 |
| 0 | 0 | 0 |
| 0 | 1 | 0 |
| 0 | 0 | 0 |
| 0 | 0 | 0 |
| 1 | 1 | 1 |
| 0 | 0 | 0 |
| 1 | 1 | 1 |
| 1 | 1 | 1 |
| 1 | 1 | 1 |
| 1 | 1 | 1 |
| 1 | 1 | 1 |
| 0 | 0 | 0 |
| 1 | 1 | 1 |
| 0 | 0 | 0 |
| 1 | 1 | 1 |
| 0 | 1 | 0 |
| 0 | 1 | 0 |
| 0 | 1 | 0 |
| 1 | 1 | 1 |
| 0 | 0 | 0 |
| 1 | 1 | 1 |
| 1 | 1 | 1 |
| 0 | 1 | 0 |
| 0 | 0 | 0 |
| 1 | 1 | 1 |
| 1 | 1 | 1 |
| 1 | 1 | 1 |
| 0 | 0 | 0 |
| 0 | 0 | 0 |
| 1 | 1 | 1 |
| 1 | 1 | 1 |
| 0 | 0 | 0 |
| 0 | 0 | 0 |
| 0 | 0 | 0 |

| | | |
|---|---|---|
| 0 | 0 | 0 |
| 1 | 1 | 1 |
| 0 | 0 | 0 |
| 0 | 0 | 0 |
| 0 | 0 | 0 |
| 1 | 1 | 1 |
| 1 | 1 | 1 |
| 0 | 1 | 0 |
| 1 | 1 | 1 |
| 0 | 0 | 0 |
| 1 | 1 | 1 |
| 0 | 1 | 0 |
| 1 | 1 | 1 |
| 1 | 1 | 1 |
| 0 | 0 | 0 |
| 0 | 1 | 0 |
| 1 | 1 | 1 |
| 0 | 0 | 0 |
| 1 | 1 | 1 |
| 0 | 0 | 0 |
| 0 | 0 | 0 |
| 1 | 1 | 1 |
| 0 | 0 | 0 |
| 1 | 1 | 1 |
| 0 | 0 | 0 |
| 0 | 0 | 0 |
| 1 | 1 | 1 |
| 0 | 1 | 0 |
| 0 | 0 | 0 |
| 1 | 1 | 1 |
| 1 | 1 | 1 |
| 1 | 1 | 1 |
| 1 | 1 | 1 |
| 0 | 1 | 0 |
| 0 | 0 | 0 |
| 0 | 0 | 0 |
| 1 | 1 | 1 |
| 0 | 0 | 0 |
| 0 | 0 | 0 |
| 0 | 0 | 0 |
| 0 | 0 | 0 |
| 0 | 0 | 0 |
| 0 | 0 | 0 |
| 0 | 1 | 0 |
| 0 | 0 | 0 |
| 1 | 1 | 1 |
| 0 | 1 | 0 |
| 0 | 0 | 0 |
| 0 | 0 | 0 |
| 0 | 0 | 0 |
| 0 | 0 | 0 |
| 0 | 0 | 0 |
| 1 | 1 | 1 |
| 1 | 1 | 1 |

| | | |
|---|---|---|
| 1 | 1 | 1 |
| 0 | 0 | 0 |
| 0 | 0 | 0 |
| 0 | 0 | 0 |
| 0 | 0 | 0 |
| 0 | 0 | 0 |
| 0 | 0 | 0 |
| 1 | 0 | 0 |
| 1 | 0 | 0 |
| 1 | 1 | 1 |
| 0 | 0 | 0 |
| 0 | 0 | 0 |
| 1 | 1 | 1 |
| 0 | 0 | 0 |
| 1 | 1 | 1 |
| 0 | 0 | 0 |
| 1 | 1 | 1 |
| 1 | 1 | 1 |
| 0 | 1 | 0 |
| 0 | 1 | 0 |
| 0 | 1 | 0 |
| 1 | 1 | 1 |
| 0 | 1 | 0 |
| 0 | 0 | 0 |
| 0 | 0 | 0 |
| 1 | 1 | 1 |
| 1 | 1 | 1 |
| 1 | 1 | 1 |
| 1 | 1 | 1 |
| 0 | 0 | 0 |
| 0 | 0 | 0 |
| 0 | 0 | 0 |
| 0 | 0 | 0 |
| 1 | 1 | 1 |
| 0 | 0 | 0 |
| 0 | 0 | 0 |
| 0 | 0 | 0 |
| 0 | 0 | 0 |
| 0 | 0 | 0 |
| 1 | 1 | 1 |
| 1 | 1 | 1 |
| 0 | 0 | 0 |
| 1 | 1 | 1 |
| 1 | 1 | 1 |
| 0 | 0 | 0 |
| 1 | 0 | 0 |
| 1 | 1 | 1 |
| 0 | 1 | 0 |
| 1 | 0 | 0 |
| 0 | 1 | 0 |
| 0 | 0 | 0 |
| 1 | 1 | 1 |
| 0 | 0 | 0 |
| 0 | 0 | 0 |

| | | |
|---|---|---|
| 0 | 1 | 0 |
| 1 | 1 | 1 |
| 0 | 0 | 0 |
| 0 | 0 | 0 |
| 1 | 1 | 1 |
| 1 | 1 | 1 |
| 1 | 1 | 1 |
| 1 | 1 | 1 |
| 1 | 1 | 1 |
| 1 | 1 | 1 |
| 1 | 1 | 1 |
| 1 | 1 | 1 |
| 0 | 0 | 0 |
| 1 | 1 | 1 |
| 1 | 1 | 1 |
| 1 | 1 | 1 |
| 1 | 1 | 1 |
| 1 | 1 | 1 |
| 1 | 1 | 1 |
| 1 | 1 | 1 |
| 1 | 1 | 1 |
| 1 | 1 | 1 |
| 1 | 1 | 1 |
| 1 | 1 | 1 |
| 1 | 1 | 1 |
| 1 | 1 | 1 |
| 0 | 1 | 0 |
| 1 | 1 | 1 |
| 1 | 1 | 1 |
| 1 | 1 | 1 |
| 1 | 1 | 1 |
| 0 | 0 | 0 |
| 1 | 1 | 1 |
| 0 | 1 | 0 |
| 1 | 1 | 1 |
| 1 | 1 | 1 |
| 0 | 1 | 0 |
| 1 | 1 | 1 |
| 1 | 1 | 1 |
| 1 | 1 | 1 |
| 0 | 0 | 0 |
| 0 | 0 | 0 |
| 1 | 1 | 1 |
| 1 | 1 | 1 |
| 1 | 1 | 1 |
| 1 | 0 | 0 |
| 1 | 0 | 0 |
| 1 | 1 | 1 |
| 0 | 0 | 0 |
| 0 | 0 | 0 |
| 1 | 1 | 1 |
| 0 | 1 | 0 |
| 0 | 0 | 0 |
| 1 | 1 | 1 |

| | | |
|---|---|---|
| 0 | 0 | 0 |
| 0 | 0 | 0 |
| 0 | 0 | 0 |
| 1 | 1 | 1 |
| 1 | 1 | 1 |
| 1 | 1 | 1 |
| 1 | 1 | 1 |
| 0 | 0 | 0 |
| 0 | 0 | 0 |
| 0 | 0 | 0 |
| 0 | 0 | 0 |
| 1 | 1 | 1 |
| 0 | 0 | 0 |
| 1 | 1 | 1 |
| 1 | 1 | 1 |
| 1 | 1 | 1 |
| 1 | 1 | 1 |
| 0 | 0 | 0 |
| 1 | 1 | 1 |
| 1 | 1 | 1 |
| 1 | 1 | 1 |
| 1 | 1 | 1 |
| 0 | 1 | 0 |
| 1 | 1 | 1 |
| 1 | 1 | 1 |
| 0 | 1 | 0 |
| 1 | 1 | 1 |
| 1 | 1 | 1 |
| 1 | 1 | 1 |
| 1 | 1 | 1 |
| 0 | 0 | 0 |
| 0 | 0 | 0 |
| 1 | 1 | 1 |
| 1 | 1 | 1 |
| 1 | 1 | 1 |
| 1 | 1 | 1 |
| 1 | 1 | 1 |
| 1 | 1 | 1 |
| 1 | 1 | 1 |
| 1 | 1 | 1 |
| 1 | 1 | 1 |
| 0 | 0 | 0 |
| 1 | 1 | 1 |
| 1 | 1 | 1 |
| 0 | 0 | 0 |
| 0 | 0 | 0 |
| 1 | 1 | 1 |
| 1 | 1 | 1 |
| 0 | 0 | 0 |
| 0 | 0 | 0 |
| 0 | 0 | 0 |
| 0 | 0 | 0 |
| 0 | 0 | 0 |
| 0 | 0 | 0 |

| | | |
|---|---|---|
| 1 | 1 | 1 |
| 1 | 1 | 1 |
| 1 | 1 | 1 |
| 1 | 1 | 1 |
| 1 | 1 | 1 |
| 0 | 0 | 0 |
| 0 | 0 | 0 |
| 0 | 0 | 0 |
| 0 | 0 | 0 |
| 0 | 0 | 0 |
| 0 | 1 | 0 |
| 0 | 0 | 0 |
| 0 | 0 | 0 |
| 0 | 0 | 0 |
| 0 | 1 | 0 |
| 0 | 0 | 0 |
| 0 | 1 | 0 |
| 0 | 1 | 0 |
| 0 | 0 | 0 |
| 0 | 0 | 0 |
| 0 | 0 | 0 |
| 0 | 0 | 0 |
| 1 | 0 | 0 |
| 0 | 0 | 0 |
| 0 | 0 | 0 |
| 0 | 0 | 0 |
| 0 | 0 | 0 |
| 1 | 1 | 1 |
| 1 | 1 | 1 |
| 1 | 1 | 1 |
| 1 | 1 | 1 |
| 1 | 1 | 1 |
| 0 | 0 | 0 |
| 0 | 1 | 0 |
| 0 | 0 | 0 |
| 0 | 0 | 0 |
| 0 | 0 | 0 |
| 0 | 0 | 0 |
| 0 | 0 | 0 |
| 0 | 0 | 0 |
| 0 | 0 | 0 |
| 0 | 0 | 0 |
| 0 | 0 | 0 |
| 1 | 1 | 1 |
| 1 | 1 | 1 |
| 1 | 1 | 1 |
| 1 | 1 | 1 |
| 0 | 0 | 0 |
| 0 | 0 | 0 |
| 0 | 0 | 0 |
| 1 | 1 | 1 |
| 1 | 1 | 1 |
| 0 | 0 | 0 |
| 1 | 1 | 1 |

| | | |
|---|---|---|
| 1 | 1 | 1 |
| 0 | 0 | 0 |
| 0 | 1 | 0 |
| 0 | 1 | 0 |
| 1 | 1 | 1 |
| 1 | 1 | 1 |
| 0 | 1 | 0 |
| 1 | 1 | 1 |
| 0 | 1 | 0 |
| 1 | 1 | 1 |
| 1 | 1 | 1 |
| 1 | 1 | 1 |
| 0 | 0 | 0 |
| 0 | 0 | 0 |
| 0 | 0 | 0 |
| 0 | 0 | 0 |
| 0 | 0 | 0 |
| 0 | 0 | 0 |
| 0 | 0 | 0 |
| 0 | 1 | 0 |
| 0 | 0 | 0 |
| 0 | 0 | 0 |
| 0 | 0 | 0 |
| 0 | 0 | 0 |
| 0 | 0 | 0 |
| 1 | 1 | 1 |
| 1 | 1 | 1 |
| 0 | 0 | 0 |
| 0 | 1 | 0 |
| 0 | 0 | 0 |
| 0 | 0 | 0 |
| 1 | 1 | 1 |
| 0 | 0 | 0 |
| 1 | 1 | 1 |
| 1 | 1 | 1 |
| 1 | 1 | 1 |
| 1 | 0 | 0 |
| 1 | 1 | 1 |
| 1 | 1 | 1 |
| 1 | 1 | 1 |
| 0 | 0 | 0 |
| 1 | 1 | 1 |
| 1 | 1 | 1 |
| 0 | 0 | 0 |
| 0 | 0 | 0 |
| 1 | 1 | 1 |
| 0 | 0 | 0 |
| 0 | 0 | 0 |
| 1 | 1 | 1 |
| 1 | 1 | 1 |
| 0 | 0 | 0 |
| 1 | 1 | 1 |
| 1 | 1 | 1 |
| 1 | 1 | 1 |

| | | |
|---|---|---|
| 1 | 1 | 1 |
| 1 | 1 | 1 |
| 1 | 1 | 1 |
| 1 | 1 | 1 |
| 0 | 0 | 0 |
| 0 | 0 | 0 |
| 1 | 1 | 1 |
| 1 | 1 | 1 |
| 1 | 1 | 1 |
| 0 | 1 | 0 |
| 1 | 1 | 1 |
| 0 | 0 | 0 |
| 0 | 0 | 0 |
| 1 | 1 | 1 |
| 1 | 1 | 1 |
| 0 | 0 | 0 |
| 0 | 1 | 0 |
| 1 | 1 | 1 |
| 0 | 0 | 0 |
| 1 | 1 | 1 |
| 1 | 1 | 1 |
| 1 | 1 | 1 |
| 1 | 1 | 1 |
| 0 | 0 | 0 |
| 1 | 1 | 1 |
| 0 | 0 | 0 |
| 1 | 1 | 1 |
| 1 | 1 | 1 |
| 0 | 0 | 0 |
| 0 | 0 | 0 |
| 0 | 0 | 0 |
| 0 | 0 | 0 |
| 0 | 0 | 0 |
| 0 | 0 | 0 |
| 1 | 1 | 1 |
| 1 | 1 | 1 |
| 1 | 1 | 1 |
| 0 | 0 | 0 |
| 1 | 1 | 1 |
| 1 | 1 | 1 |
| 0 | 0 | 0 |
| 0 | 0 | 0 |
| 0 | 0 | 0 |
| 1 | 1 | 1 |
| 0 | 0 | 0 |
| 0 | 1 | 0 |
| 0 | 1 | 0 |
| 0 | 0 | 0 |
| 1 | 1 | 1 |
| 1 | 1 | 1 |
| 1 | 1 | 1 |
| 0 | 0 | 0 |
| 1 | 1 | 1 |
| 0 | 0 | 0 |

| | | |
|---|---|---|
| 1 | 1 | 1 |
| 0 | 0 | 0 |
| 0 | 0 | 0 |
| 0 | 1 | 0 |
| 1 | 1 | 1 |
| 1 | 1 | 1 |
| 0 | 0 | 0 |
| 1 | 1 | 1 |
| 0 | 1 | 0 |
| 0 | 1 | 0 |
| 1 | 1 | 1 |
| 1 | 1 | 1 |
| 1 | 1 | 1 |
| 1 | 1 | 1 |
| 1 | 1 | 1 |
| 1 | 1 | 1 |
| 1 | 1 | 1 |
| 1 | 1 | 1 |
| 1 | 1 | 1 |
| 1 | 1 | 1 |
| 1 | 1 | 1 |
| 0 | 0 | 0 |
| 0 | 1 | 0 |
| 0 | 1 | 0 |
| 1 | 1 | 1 |
| 1 | 1 | 1 |
| 0 | 0 | 0 |
| 1 | 1 | 1 |
| 0 | 0 | 0 |
| 0 | 1 | 0 |
| 1 | 1 | 1 |
| 0 | 0 | 0 |
| 0 | 0 | 0 |
| 1 | 0 | 0 |
| 1 | 1 | 1 |
| 1 | 1 | 1 |
| 1 | 1 | 1 |
| 1 | 1 | 1 |
| 0 | 0 | 0 |
| 0 | 0 | 0 |
| 0 | 0 | 0 |
| 1 | 1 | 1 |
| 1 | 1 | 1 |
| 0 | 1 | 0 |
| 0 | 1 | 0 |
| 0 | 0 | 0 |
| 1 | 1 | 1 |
| 0 | 0 | 0 |
| 1 | 1 | 1 |
| 0 | 0 | 0 |
| 0 | 0 | 0 |
| 0 | 0 | 0 |
| 1 | 1 | 1 |
| 1 | 1 | 1 |

| | | |
|---|---|---|
| 0 | 0 | 0 |
| 0 | 0 | 0 |
| 1 | 1 | 1 |
| 1 | 1 | 1 |
| 1 | 1 | 1 |
| 1 | 1 | 1 |
| 1 | 1 | 1 |
| 1 | 1 | 1 |
| 1 | 1 | 1 |
| 1 | 1 | 1 |
| 1 | 1 | 1 |
| 1 | 1 | 1 |
| 1 | 1 | 1 |
| 1 | 1 | 1 |
| 1 | 1 | 1 |
| 0 | 0 | 0 |
| 0 | 0 | 0 |
| 0 | 0 | 0 |
| 1 | 1 | 1 |
| 1 | 1 | 1 |
| 0 | 1 | 0 |
| 0 | 0 | 0 |
| 0 | 0 | 0 |
| 0 | 0 | 0 |
| 0 | 0 | 0 |
| 1 | 1 | 1 |
| 1 | 1 | 1 |
| 1 | 1 | 1 |
| 1 | 1 | 1 |
| 1 | 1 | 1 |
| 0 | 0 | 0 |
| 0 | 1 | 0 |
| 0 | 0 | 0 |
| 1 | 1 | 1 |
| 1 | 0 | 0 |
| 1 | 0 | 0 |
| 0 | 0 | 0 |
| 0 | 0 | 0 |
| 0 | 1 | 0 |
| 1 | 1 | 1 |
| 1 | 0 | 0 |
| 0 | 0 | 0 |
| 1 | 1 | 1 |
| 0 | 1 | 0 |
| 0 | 0 | 0 |
| 1 | 0 | 0 |
| 0 | 0 | 0 |
| 0 | 0 | 0 |
| 1 | 1 | 1 |
| 0 | 1 | 0 |
| 0 | 0 | 0 |
| 0 | 0 | 0 |
| 1 | 1 | 1 |
| 0 | 0 | 0 |

| | | |
|---|---|---|
| 0 | 1 | 0 |
| 0 | 0 | 0 |
| 1 | 0 | 0 |
| 0 | 0 | 0 |
| 1 | 1 | 1 |
| 1 | 1 | 1 |
| 1 | 1 | 1 |
| 0 | 1 | 0 |
| 0 | 0 | 0 |
| 1 | 1 | 1 |
| 0 | 1 | 0 |
| 0 | 1 | 0 |
| 0 | 0 | 0 |
| 0 | 0 | 0 |
| 0 | 0 | 0 |
| 1 | 1 | 1 |
| 0 | 0 | 0 |
| 0 | 1 | 0 |
| 0 | 0 | 0 |
| 0 | 0 | 0 |
| 0 | 0 | 0 |
| 0 | 0 | 0 |
| 0 | 0 | 0 |
| 0 | 0 | 0 |
| 0 | 1 | 0 |
| 0 | 1 | 0 |
| 0 | 0 | 0 |
| 0 | 0 | 0 |
| 0 | 0 | 0 |
| 0 | 0 | 0 |
| 1 | 1 | 1 |
| 1 | 1 | 1 |
| 1 | 1 | 1 |
| 1 | 1 | 1 |
| 1 | 1 | 1 |
| 0 | 0 | 0 |
| 0 | 0 | 0 |
| 0 | 1 | 0 |
| 1 | 1 | 1 |
| 1 | 1 | 1 |
| 0 | 1 | 0 |
| 1 | 1 | 1 |
| 1 | 1 | 1 |
| 0 | 0 | 0 |
| 0 | 0 | 0 |
| 0 | 0 | 0 |
| 1 | 1 | 1 |
| 0 | 0 | 0 |
| 0 | 0 | 0 |
| 0 | 0 | 0 |
| 1 | 1 | 1 |
| 1 | 1 | 1 |
| 0 | 1 | 0 |
| 0 | 1 | 0 |

| | | |
|---|---|---|
| 0 | 1 | 0 |
| 1 | 1 | 1 |
| 1 | 0 | 0 |
| 1 | 1 | 1 |
| 0 | 0 | 0 |
| 1 | 0 | 0 |
| 0 | 0 | 0 |
| 0 | 0 | 0 |
| 0 | 0 | 0 |
| 0 | 0 | 0 |
| 0 | 0 | 0 |
| 0 | 0 | 0 |
| 0 | 1 | 0 |
| 0 | 0 | 0 |
| 0 | 0 | 0 |
| 1 | 1 | 1 |
| 1 | 1 | 1 |
| 1 | 1 | 1 |
| 0 | 0 | 0 |
| 0 | 0 | 0 |
| 0 | 0 | 0 |
| 0 | 0 | 0 |
| 0 | 1 | 0 |
| 0 | 1 | 0 |
| 1 | 1 | 1 |
| 1 | 1 | 1 |
| 1 | 1 | 1 |
| 0 | 0 | 0 |
| 0 | 0 | 0 |
| 0 | 0 | 0 |
| 0 | 0 | 0 |
| 0 | 0 | 0 |
| 0 | 0 | 0 |
| 0 | 1 | 0 |
| 1 | 1 | 1 |
| 0 | 0 | 0 |
| 1 | 1 | 1 |
| 1 | 1 | 1 |
| 1 | 1 | 1 |
| 1 | 1 | 1 |
| 0 | 0 | 0 |
| 1 | 1 | 1 |
| 0 | 0 | 0 |
| 0 | 1 | 0 |
| 1 | 1 | 1 |
| 1 | 1 | 1 |
| 1 | 1 | 1 |
| 0 | 0 | 0 |
| 1 | 1 | 1 |
| 1 | 1 | 1 |
| 1 | 1 | 1 |
| 1 | 1 | 1 |
| 1 | 1 | 1 |
| 0 | 1 | 0 |

| | | |
|---|---|---|
| 0 | 1 | 0 |
| 1 | 1 | 1 |
| 1 | 1 | 1 |
| 0 | 1 | 0 |
| 1 | 1 | 1 |
| 0 | 1 | 0 |
| 0 | 1 | 0 |
| 0 | 1 | 0 |
| 0 | 1 | 0 |
| 0 | 1 | 0 |
| 0 | 1 | 0 |
| 0 | 0 | 0 |
| 0 | 1 | 0 |
| 0 | 1 | 0 |
| 0 | 1 | 0 |
| 0 | 1 | 0 |
| 0 | 1 | 0 |
| 0 | 0 | 0 |
| 0 | 0 | 0 |
| 0 | 0 | 0 |
| 0 | 0 | 0 |
| 0 | 1 | 0 |
| 0 | 0 | 0 |
| 1 | 1 | 1 |
| 1 | 1 | 1 |
| 1 | 1 | 1 |
| 1 | 1 | 1 |
| 0 | 1 | 0 |
| 0 | 0 | 0 |
| 0 | 0 | 0 |
| 1 | 1 | 1 |
| 1 | 1 | 1 |
| 0 | 0 | 0 |
| 1 | 1 | 1 |
| 0 | 0 | 0 |
| 0 | 0 | 0 |
| 0 | 0 | 0 |
| 1 | 1 | 1 |
| 0 | 0 | 0 |
| 1 | 1 | 1 |
| 0 | 0 | 0 |
| 1 | 1 | 1 |
| 0 | 0 | 0 |
| 1 | 0 | 0 |
| 1 | 0 | 0 |
| 1 | 0 | 0 |
| 1 | 0 | 0 |
| 0 | 1 | 0 |
| 0 | 1 | 0 |
| 0 | 0 | 0 |
| 1 | 0 | 0 |
| 0 | 0 | 0 |
| 1 | 0 | 0 |
| 1 | 1 | 1 |

| | | |
|---|---|---|
| 0 | 0 | 0 |
| 0 | 0 | 0 |
| 0 | 0 | 0 |
| 0 | 0 | 0 |
| 0 | 0 | 0 |
| 1 | 1 | 1 |
| 0 | 0 | 0 |
| 0 | 0 | 0 |
| 1 | 1 | 1 |
| 0 | 0 | 0 |
| 0 | 0 | 0 |
| 0 | 0 | 0 |
| 0 | 1 | 0 |
| 0 | 1 | 0 |
| 0 | 1 | 0 |
| 0 | 1 | 0 |
| 0 | 1 | 0 |
| 0 | 1 | 0 |
| 0 | 1 | 0 |
| 0 | 1 | 0 |
| 0 | 1 | 0 |
| 0 | 1 | 0 |
| 0 | 1 | 0 |
| 0 | 1 | 0 |
| 0 | 1 | 0 |
| 0 | 1 | 0 |
| 0 | 1 | 0 |
| 0 | 1 | 0 |
| 0 | 1 | 0 |
| 0 | 1 | 0 |
| 0 | 1 | 0 |
| 0 | 1 | 0 |
| 0 | 1 | 0 |
| 0 | 1 | 0 |
| 0 | 1 | 0 |
| 0 | 1 | 0 |
| 0 | 1 | 0 |
| 0 | 1 | 0 |
| 0 | 1 | 0 |
| 0 | 1 | 0 |
| 0 | 1 | 0 |
| 0 | 1 | 0 |
| 0 | 1 | 0 |
| 0 | 1 | 0 |
| 0 | 1 | 0 |
| 0 | 0 | 0 |
| 1 | 1 | 1 |
| 1 | 1 | 1 |
| 1 | 1 | 1 |
| 1 | 1 | 1 |
| 0 | 0 | 0 |
| 1 | 1 | 1 |
| 1 | 1 | 1 |
| 0 | 1 | 0 |

| | | |
|---|---|---|
| 1 | 1 | 1 |
| 1 | 1 | 1 |
| 1 | 1 | 1 |
| 1 | 1 | 1 |
| 0 | 1 | 0 |
| 0 | 1 | 0 |
| 0 | 1 | 0 |
| 0 | 0 | 0 |
| 0 | 0 | 0 |
| 1 | 1 | 1 |
| 0 | 1 | 0 |
| 1 | 1 | 1 |
| 1 | 1 | 1 |
| 1 | 1 | 1 |
| 0 | 0 | 0 |
| 0 | 0 | 0 |
| 0 | 0 | 0 |
| 1 | 1 | 1 |
| 1 | 0 | 0 |
| 1 | 1 | 1 |
| 0 | 0 | 0 |
| 0 | 0 | 0 |
| 0 | 0 | 0 |
| 0 | 0 | 0 |
| 0 | 0 | 0 |
| 0 | 0 | 0 |
| 0 | 1 | 0 |
| 1 | 1 | 1 |
| 0 | 0 | 0 |
| 1 | 1 | 1 |
| 1 | 1 | 1 |
| 1 | 1 | 1 |
| 0 | 0 | 0 |
| 1 | 1 | 1 |
| 1 | 1 | 1 |
| 1 | 1 | 1 |
| 0 | 0 | 0 |
| 0 | 0 | 0 |
| 0 | 0 | 0 |
| 1 | 1 | 1 |
| 1 | 1 | 1 |
| 1 | 1 | 1 |
| 1 | 1 | 1 |
| 1 | 1 | 1 |
| 0 | 1 | 0 |
| 0 | 0 | 0 |
| 1 | 1 | 1 |
| 0 | 1 | 0 |
| 1 | 1 | 1 |
| 0 | 1 | 0 |
| 1 | 1 | 1 |
| 1 | 1 | 1 |
| 0 | 1 | 0 |
| 1 | 1 | 1 |

| | | |
|---|---|---|
| 1 | 1 | 1 |
| 0 | 1 | 0 |
| 1 | 1 | 1 |
| 0 | 0 | 0 |
| 1 | 1 | 1 |
| 1 | 1 | 1 |
| 1 | 1 | 1 |
| 0 | 0 | 0 |
| 0 | 0 | 0 |
| 1 | 1 | 1 |
| 1 | 1 | 1 |
| 1 | 1 | 1 |
| 0 | 0 | 0 |
| 0 | 1 | 0 |
| 0 | 1 | 0 |
| 0 | 1 | 0 |
| 0 | 0 | 0 |
| 1 | 1 | 1 |
| 1 | 1 | 1 |
| 1 | 1 | 1 |
| 1 | 1 | 1 |
| 1 | 1 | 1 |
| 1 | 1 | 1 |
| 1 | 1 | 1 |
| 0 | 1 | 0 |
| 1 | 1 | 1 |
| 0 | 1 | 0 |
| 0 | 0 | 0 |
| 0 | 0 | 0 |
| 0 | 0 | 0 |
| 0 | 0 | 0 |
| 0 | 0 | 0 |
| 0 | 0 | 0 |
| 0 | 1 | 0 |
| 0 | 0 | 0 |
| 0 | 0 | 0 |
| 0 | 0 | 0 |
| 1 | 1 | 1 |
| 1 | 1 | 1 |
| 1 | 1 | 1 |
| 1 | 1 | 1 |
| 0 | 1 | 0 |
| 0 | 0 | 0 |
| 0 | 0 | 0 |
| 1 | 1 | 1 |
| 1 | 1 | 1 |
| 0 | 0 | 0 |
| 1 | 1 | 1 |
| 1 | 1 | 1 |
| 1 | 1 | 1 |
| 1 | 1 | 1 |
| 1 | 1 | 1 |
| 0 | 0 | 0 |
| 0 | 0 | 0 |

| | | |
|---|---|---|
| 0 | 0 | 0 |
| 0 | 0 | 0 |
| 0 | 0 | 0 |
| 0 | 0 | 0 |
| 0 | 0 | 0 |
| 0 | 0 | 0 |
| 0 | 0 | 0 |
| 0 | 0 | 0 |
| 1 | 1 | 1 |
| 0 | 0 | 0 |
| 1 | 1 | 1 |
| 1 | 1 | 1 |
| 1 | 1 | 1 |
| 1 | 1 | 1 |
| 0 | 0 | 0 |
| 1 | 1 | 1 |
| 1 | 1 | 1 |
| 1 | 1 | 1 |
| 1 | 1 | 1 |
| 0 | 0 | 0 |
| 0 | 0 | 0 |
| 1 | 1 | 1 |
| 0 | 0 | 0 |
| 0 | 0 | 0 |
| 0 | 0 | 0 |
| 0 | 0 | 0 |
| 0 | 0 | 0 |
| 0 | 1 | 0 |
| 0 | 1 | 0 |
| 0 | 1 | 0 |
| 0 | 0 | 0 |
| 1 | 1 | 1 |
| 1 | 1 | 1 |
| 1 | 1 | 1 |
| 1 | 1 | 1 |
| 1 | 1 | 1 |
| 1 | 1 | 1 |
| 1 | 1 | 1 |
| 1 | 1 | 1 |
| 1 | 1 | 1 |
| 0 | 1 | 0 |
| 0 | 0 | 0 |
| 1 | 1 | 1 |
| 1 | 1 | 1 |
| 1 | 1 | 1 |
| 0 | 1 | 0 |
| 1 | 1 | 1 |
| 1 | 1 | 1 |
| 0 | 0 | 0 |
| 0 | 1 | 0 |
| 1 | 1 | 1 |
| 1 | 1 | 1 |
| 0 | 0 | 0 |
| 0 | 0 | 0 |

| | | |
|---|---|---|
| 1 | 1 | 1 |
| 1 | 1 | 1 |
| 0 | 1 | 0 |
| 1 | 0 | 0 |
| 0 | 0 | 0 |
| 0 | 0 | 0 |
| 0 | 1 | 0 |
| 0 | 0 | 0 |
| 1 | 1 | 1 |
| 1 | 1 | 1 |
| 0 | 0 | 0 |
| 1 | 0 | 0 |
| 0 | 0 | 0 |
| 0 | 0 | 0 |
| 0 | 0 | 0 |
| 0 | 0 | 0 |
| 1 | 1 | 1 |
| 1 | 1 | 1 |
| 0 | 0 | 0 |
| 0 | 0 | 0 |
| 0 | 0 | 0 |
| 0 | 0 | 0 |
| 0 | 0 | 0 |
| 1 | 1 | 1 |
| 1 | 1 | 1 |
| 0 | 0 | 0 |
| 1 | 1 | 1 |
| 0 | 0 | 0 |
| 0 | 0 | 0 |
| 0 | 0 | 0 |
| 0 | 1 | 0 |
| 0 | 0 | 0 |
| 0 | 0 | 0 |
| 1 | 1 | 1 |
| 1 | 1 | 1 |
| 0 | 0 | 0 |
| 0 | 0 | 0 |
| 1 | 0 | 0 |
| 0 | 0 | 0 |
| 0 | 0 | 0 |
| 0 | 0 | 0 |
| 0 | 0 | 0 |
| 0 | 1 | 0 |
| 1 | 1 | 1 |
| 0 | 0 | 0 |
| 0 | 1 | 0 |
| 0 | 0 | 0 |
| 1 | 1 | 1 |
| 1 | 1 | 1 |
| 1 | 1 | 1 |
| 1 | 1 | 1 |
| 1 | 1 | 1 |
| 0 | 1 | 0 |
| 1 | 1 | 1 |

| | | |
|---|---|---|
| 1 | 1 | 1 |
| 1 | 1 | 1 |
| 1 | 1 | 1 |
| 0 | 1 | 0 |
| 1 | 1 | 1 |
| 1 | 1 | 1 |
| 0 | 1 | 0 |
| 0 | 0 | 0 |
| 1 | 1 | 1 |
| 0 | 0 | 0 |
| 0 | 0 | 0 |
| 0 | 0 | 0 |
| 0 | 1 | 0 |
| 1 | 1 | 1 |
| 0 | 0 | 0 |
| 0 | 1 | 0 |
| 0 | 1 | 0 |
| 0 | 1 | 0 |
| 0 | 1 | 0 |
| 0 | 1 | 0 |
| 1 | 1 | 1 |
| 1 | 1 | 1 |
| 1 | 1 | 1 |
| 0 | 1 | 0 |
| 1 | 1 | 1 |
| 1 | 1 | 1 |
| 1 | 1 | 1 |
| 1 | 0 | 0 |
| 1 | 1 | 1 |
| 0 | 0 | 0 |
| 0 | 0 | 0 |
| 1 | 1 | 1 |
| 1 | 1 | 1 |
| 0 | 1 | 0 |
| 0 | 1 | 0 |
| 0 | 0 | 0 |
| 1 | 1 | 1 |
| 1 | 1 | 1 |
| 1 | 1 | 1 |
| 1 | 1 | 1 |
| 1 | 1 | 1 |
| 0 | 0 | 0 |
| 0 | 0 | 0 |
| 0 | 0 | 0 |
| 1 | 1 | 1 |
| 1 | 1 | 1 |
| 1 | 1 | 1 |
| 1 | 1 | 1 |
| 1 | 1 | 1 |
| 1 | 1 | 1 |
| 1 | 1 | 1 |
| 1 | 1 | 1 |
| 1 | 1 | 1 |
| 0 | 0 | 0 |

| | | |
|---|---|---|
| 1 | 1 | 1 |
| 1 | 1 | 1 |
| 1 | 1 | 1 |
| 1 | 1 | 1 |
| 1 | 1 | 1 |
| 1 | 1 | 1 |
| 1 | 1 | 1 |
| 0 | 1 | 0 |
| 0 | 0 | 0 |
| 0 | 0 | 0 |
| 1 | 1 | 1 |
| 0 | 1 | 0 |
| 0 | 1 | 0 |
| 0 | 0 | 0 |
| 1 | 1 | 1 |
| 1 | 1 | 1 |
| 1 | 1 | 1 |
| 0 | 0 | 0 |
| 0 | 0 | 0 |
| 0 | 0 | 0 |
| 0 | 0 | 0 |
| 0 | 1 | 0 |
| 1 | 1 | 1 |
| 1 | 1 | 1 |
| 1 | 1 | 1 |
| 0 | 0 | 0 |
| 1 | 1 | 1 |
| 0 | 1 | 0 |
| 1 | 1 | 1 |
| 1 | 1 | 1 |
| 1 | 1 | 1 |
| 1 | 1 | 1 |
| 1 | 1 | 1 |
| 1 | 1 | 1 |
| 1 | 1 | 1 |
| 1 | 1 | 1 |
| 1 | 1 | 1 |
| 1 | 1 | 1 |
| 1 | 1 | 1 |
| 0 | 0 | 0 |
| 1 | 1 | 1 |
| 1 | 1 | 1 |
| 0 | 0 | 0 |
| 1 | 1 | 1 |
| 1 | 1 | 1 |
| 1 | 1 | 1 |
| 0 | 0 | 0 |
| 0 | 0 | 0 |
| 1 | 1 | 1 |
| 1 | 1 | 1 |
| 0 | 0 | 0 |
| 1 | 1 | 1 |
| 0 | 1 | 0 |
| 1 | 1 | 1 |

| | | |
|---|---|---|
| 1 | 1 | 1 |
| 1 | 1 | 1 |
| 1 | 1 | 1 |
| 1 | 1 | 1 |
| 1 | 1 | 1 |
| 1 | 1 | 1 |
| 1 | 1 | 1 |
| 0 | 1 | 0 |
| 1 | 1 | 1 |
| 1 | 1 | 1 |
| 1 | 1 | 1 |
| 1 | 1 | 1 |
| 1 | 1 | 1 |
| 0 | 0 | 0 |
| 0 | 0 | 0 |
| 0 | 0 | 0 |
| 0 | 0 | 0 |
| 0 | 1 | 0 |
| 1 | 1 | 1 |
| 0 | 0 | 0 |
| 1 | 1 | 1 |
| 1 | 0 | 0 |
| 1 | 1 | 1 |
| 1 | 1 | 1 |
| 1 | 1 | 1 |
| 0 | 0 | 0 |
| 0 | 1 | 0 |
| 1 | 1 | 1 |
| 1 | 1 | 1 |
| 0 | 0 | 0 |
| 0 | 0 | 0 |
| 1 | 1 | 1 |
| 1 | 1 | 1 |
| 1 | 1 | 1 |
| 1 | 1 | 1 |
| 1 | 1 | 1 |
| 1 | 1 | 1 |
| 1 | 1 | 1 |
| 0 | 0 | 0 |
| 1 | 1 | 1 |
| 1 | 1 | 1 |
| 0 | 0 | 0 |
| 0 | 0 | 0 |
| 0 | 0 | 0 |
| 0 | 0 | 0 |
| 1 | 1 | 1 |
| 0 | 0 | 0 |
| 1 | 1 | 1 |
| 1 | 1 | 1 |
| 1 | 1 | 1 |
| 0 | 1 | 0 |
| 1 | 0 | 0 |
| 1 | 0 | 0 |
| 1 | 0 | 0 |

| | | |
|---|---|---|
| 1 | 1 | 1 |
| 1 | 1 | 1 |
| 0 | 0 | 0 |
| 1 | 1 | 1 |
| 1 | 1 | 1 |
| 1 | 1 | 1 |
| 1 | 1 | 1 |
| 0 | 1 | 0 |
| 1 | 1 | 1 |
| 1 | 1 | 1 |
| 1 | 1 | 1 |
| 0 | 1 | 0 |
| 0 | 1 | 0 |
| 0 | 1 | 0 |
| 0 | 1 | 0 |
| 0 | 1 | 0 |
| 0 | 1 | 0 |
| 0 | 1 | 0 |
| 0 | 1 | 0 |
| 0 | 1 | 0 |
| 0 | 1 | 0 |
| 0 | 1 | 0 |
| 0 | 1 | 0 |
| 0 | 1 | 0 |
| 0 | 1 | 0 |
| 0 | 1 | 0 |
| 0 | 1 | 0 |
| 0 | 1 | 0 |
| 1 | 1 | 1 |
| 1 | 1 | 1 |
| 0 | 0 | 0 |
| 1 | 1 | 1 |
| 0 | 0 | 0 |
| 0 | 0 | 0 |
| 0 | 0 | 0 |
| 0 | 0 | 0 |
| 1 | 1 | 1 |
| 0 | 0 | 0 |
| 1 | 1 | 1 |
| 1 | 1 | 1 |
| 0 | 0 | 0 |
| 0 | 0 | 0 |
| 0 | 0 | 0 |
| 1 | 1 | 1 |
| 0 | 0 | 0 |
| 0 | 0 | 0 |
| 1 | 1 | 1 |
| 1 | 1 | 1 |
| 1 | 1 | 1 |
| 1 | 1 | 1 |
| 0 | 0 | 0 |
| 1 | 1 | 1 |
| 1 | 1 | 1 |
| 1 | 1 | 1 |

| | | |
|---|---|---|
| 1 | 1 | 1 |
| 1 | 1 | 1 |
| 1 | 1 | 1 |
| 1 | 1 | 1 |
| 1 | 1 | 1 |
| 0 | 0 | 0 |
| 1 | 1 | 1 |
| 0 | 0 | 0 |
| 0 | 0 | 0 |
| 0 | 0 | 0 |
| 1 | 1 | 1 |
| 0 | 0 | 0 |
| 1 | 1 | 1 |
| 1 | 1 | 1 |
| 1 | 1 | 1 |
| 1 | 0 | 0 |
| 1 | 1 | 1 |
| 0 | 0 | 0 |
| 1 | 1 | 1 |
| 0 | 0 | 0 |
| 1 | 1 | 1 |
| 0 | 0 | 0 |
| 0 | 0 | 0 |
| 0 | 0 | 0 |
| 1 | 1 | 1 |
| 0 | 1 | 0 |
| 1 | 1 | 1 |
| 1 | 1 | 1 |
| 0 | 0 | 0 |
| 1 | 1 | 1 |
| 1 | 1 | 1 |
| 1 | 1 | 1 |
| 1 | 1 | 1 |
| 1 | 1 | 1 |
| 0 | 0 | 0 |
| 0 | 0 | 0 |
| 1 | 1 | 1 |
| 0 | 0 | 0 |
| 0 | 0 | 0 |
| 0 | 1 | 0 |
| 0 | 0 | 0 |
| 0 | 0 | 0 |
| 0 | 1 | 0 |
| 0 | 1 | 0 |
| 1 | 1 | 1 |
| 0 | 1 | 0 |
| 1 | 1 | 1 |
| 1 | 1 | 1 |
| 1 | 1 | 1 |
| 1 | 1 | 1 |
| 1 | 1 | 1 |
| 1 | 1 | 1 |
| 1 | 1 | 1 |
| 1 | 1 | 1 |

| | | |
|---|---|---|
| 1 | 1 | 1 |
| 0 | 0 | 0 |
| 0 | 1 | 0 |
| 1 | 1 | 1 |
| 0 | 0 | 0 |
| 1 | 1 | 1 |
| 1 | 1 | 1 |
| 1 | 1 | 1 |
| 0 | 0 | 0 |
| 0 | 0 | 0 |
| 1 | 1 | 1 |
| 0 | 0 | 0 |
| 0 | 0 | 0 |
| 1 | 1 | 1 |
| 1 | 1 | 1 |
| 0 | 0 | 0 |
| 0 | 1 | 0 |
| 0 | 0 | 0 |
| 0 | 0 | 0 |
| 1 | 1 | 1 |
| 0 | 0 | 0 |
| 0 | 0 | 0 |
| 0 | 0 | 0 |
| 1 | 1 | 1 |
| 0 | 1 | 0 |
| 0 | 0 | 0 |
| 1 | 1 | 1 |
| 1 | 1 | 1 |
| 1 | 1 | 1 |
| 1 | 1 | 1 |
| 0 | 0 | 0 |
| 1 | 1 | 1 |
| 0 | 0 | 0 |
| 1 | 1 | 1 |
| 0 | 0 | 0 |
| 0 | 0 | 0 |
| 1 | 1 | 1 |
| 1 | 1 | 1 |
| 1 | 1 | 1 |
| 1 | 1 | 1 |
| 1 | 1 | 1 |
| 1 | 1 | 1 |
| 1 | 1 | 1 |
| 1 | 1 | 1 |
| 1 | 1 | 1 |
| 1 | 1 | 1 |
| 1 | 0 | 0 |
| 1 | 1 | 1 |
| 1 | 1 | 1 |
| 1 | 1 | 1 |
| 1 | 1 | 1 |
| 1 | 1 | 1 |
| 1 | 1 | 1 |
| 0 | 0 | 0 |

| | | |
|---|---|---|
| 1 | 1 | 1 |
| 1 | 1 | 1 |
| 1 | 1 | 1 |
| 1 | 1 | 1 |
| 0 | 0 | 0 |
| 1 | 1 | 1 |
| 0 | 1 | 0 |
| 0 | 1 | 0 |
| 0 | 1 | 0 |
| 1 | 1 | 1 |
| 0 | 0 | 0 |
| 1 | 1 | 1 |
| 1 | 1 | 1 |
| 1 | 1 | 1 |
| 0 | 0 | 0 |
| 0 | 0 | 0 |
| 0 | 0 | 0 |
| 0 | 0 | 0 |
| 0 | 1 | 0 |
| 0 | 0 | 0 |
| 0 | 0 | 0 |
| 0 | 0 | 0 |
| 1 | 1 | 1 |
| 0 | 0 | 0 |
| 0 | 0 | 0 |
| 1 | 1 | 1 |
| 0 | 0 | 0 |
| 0 | 0 | 0 |
| 0 | 0 | 0 |
| 0 | 0 | 0 |
| 0 | 0 | 0 |
| 1 | 1 | 1 |
| 1 | 1 | 1 |
| 0 | 1 | 0 |
| 1 | 1 | 1 |
| 1 | 1 | 1 |
| 1 | 1 | 1 |
| 0 | 0 | 0 |
| 0 | 0 | 0 |
| 0 | 1 | 0 |
| 1 | 1 | 1 |
| 0 | 0 | 0 |
| 0 | 0 | 0 |
| 0 | 1 | 0 |
| 0 | 0 | 0 |
| 1 | 1 | 1 |
| 1 | 1 | 1 |
| 1 | 1 | 1 |
| 0 | 0 | 0 |
| 0 | 0 | 0 |
| 0 | 0 | 0 |
| 1 | 1 | 1 |
| 0 | 1 | 0 |
| 0 | 0 | 0 |

| | | |
|---|---|---|
| 1 | 1 | 1 |
| 0 | 0 | 0 |
| 1 | 1 | 1 |
| 0 | 0 | 0 |
| 0 | 0 | 0 |
| 0 | 1 | 0 |
| 0 | 0 | 0 |
| 0 | 0 | 0 |
| 1 | 1 | 1 |
| 1 | 1 | 1 |
| 1 | 1 | 1 |
| 1 | 1 | 1 |
| 1 | 1 | 1 |
| 1 | 1 | 1 |
| 1 | 1 | 1 |
| 1 | 1 | 1 |
| 1 | 1 | 1 |
| 0 | 1 | 0 |
| 1 | 1 | 1 |
| 1 | 1 | 1 |
| 0 | 0 | 0 |
| 0 | 1 | 0 |
| 0 | 1 | 0 |
| 0 | 1 | 0 |
| 0 | 1 | 0 |
| 0 | 0 | 0 |
| 0 | 0 | 0 |
| 1 | 1 | 1 |
| 1 | 1 | 1 |
| 1 | 1 | 1 |
| 1 | 1 | 1 |
| 1 | 1 | 1 |
| 1 | 1 | 1 |
| 0 | 1 | 0 |
| 1 | 1 | 1 |
| 1 | 1 | 1 |
| 1 | 1 | 1 |
| 1 | 1 | 1 |
| 0 | 1 | 0 |
| 1 | 1 | 1 |
| 1 | 1 | 1 |
| 0 | 1 | 0 |
| 1 | 1 | 1 |
| 0 | 0 | 0 |
| 1 | 1 | 1 |
| 0 | 0 | 0 |
| 0 | 0 | 0 |
| 0 | 0 | 0 |
| 0 | 0 | 0 |
| 0 | 0 | 0 |
| 0 | 0 | 0 |
| 0 | 1 | 0 |
| 1 | 1 | 1 |
| 1 | 1 | 1 |

| | | |
|---|---|---|
| 1 | 1 | 1 |
| 1 | 1 | 1 |
| 1 | 1 | 1 |
| 0 | 0 | 0 |
| 1 | 1 | 1 |
| 0 | 0 | 0 |
| 1 | 1 | 1 |
| 1 | 1 | 1 |
| 0 | 0 | 0 |
| 1 | 1 | 1 |
| 1 | 1 | 1 |
| 0 | 1 | 0 |
| 1 | 1 | 1 |
| 1 | 1 | 1 |
| 1 | 1 | 1 |
| 0 | 0 | 0 |
| 0 | 0 | 0 |
| 0 | 0 | 0 |
| 0 | 0 | 0 |
| 0 | 0 | 0 |
| 1 | 1 | 1 |
| 1 | 1 | 1 |
| 1 | 1 | 1 |
| 1 | 1 | 1 |
| 0 | 0 | 0 |
| 1 | 1 | 1 |
| 1 | 1 | 1 |
| 1 | 1 | 1 |
| 0 | 0 | 0 |
| 1 | 1 | 1 |
| 0 | 0 | 0 |
| 0 | 0 | 0 |
| 0 | 0 | 0 |
| 0 | 0 | 0 |
| 0 | 0 | 0 |
| 1 | 1 | 1 |
| 0 | 0 | 0 |
| 0 | 0 | 0 |
| 0 | 0 | 0 |
| 0 | 0 | 0 |
| 0 | 0 | 0 |
| 0 | 0 | 0 |
| 0 | 0 | 0 |
| 0 | 0 | 0 |
| 0 | 0 | 0 |
| 0 | 0 | 0 |
| 0 | 0 | 0 |
| 0 | 0 | 0 |
| 1 | 1 | 1 |
| 0 | 0 | 0 |
| 1 | 1 | 1 |
| 1 | 1 | 1 |
| 0 | 1 | 0 |
| 0 | 0 | 0 |

| | | |
|---|---|---|
| 0 | 0 | 0 |
| 0 | 1 | 0 |
| 1 | 1 | 1 |
| 1 | 1 | 1 |
| 1 | 1 | 1 |
| 0 | 0 | 0 |
| 1 | 1 | 1 |
| 0 | 1 | 0 |
| 1 | 1 | 1 |
| 0 | 1 | 0 |
| 0 | 1 | 0 |
| 0 | 1 | 0 |
| 1 | 1 | 1 |
| 0 | 0 | 0 |
| 0 | 1 | 0 |
| 1 | 1 | 1 |
| 1 | 1 | 1 |
| 0 | 0 | 0 |
| 1 | 1 | 1 |
| 1 | 1 | 1 |
| 1 | 1 | 1 |
| 1 | 1 | 1 |
| 0 | 0 | 0 |
| 1 | 1 | 1 |
| 1 | 1 | 1 |
| 0 | 0 | 0 |
| 0 | 0 | 0 |
| 0 | 0 | 0 |
| 1 | 1 | 1 |
| 0 | 1 | 0 |
| 1 | 1 | 1 |
| 1 | 1 | 1 |
| 1 | 1 | 1 |
| 1 | 1 | 1 |
| 1 | 1 | 1 |
| 1 | 1 | 1 |
| 1 | 1 | 1 |
| 1 | 1 | 1 |
| 1 | 1 | 1 |
| 1 | 1 | 1 |
| 1 | 1 | 1 |
| 1 | 1 | 1 |
| 1 | 1 | 1 |
| 1 | 1 | 1 |
| 1 | 1 | 1 |
| 1 | 1 | 1 |
| 1 | 1 | 1 |
| 1 | 1 | 1 |
| 1 | 0 | 0 |
| 1 | 1 | 1 |
| 1 | 1 | 1 |
| 1 | 1 | 1 |
| 1 | 1 | 1 |
| 1 | 1 | 1 |

| | | |
|---|---|---|
| 1 | 1 | 1 |
| 0 | 0 | 0 |
| 1 | 1 | 1 |
| 1 | 1 | 1 |
| 1 | 1 | 1 |
| 0 | 0 | 0 |
| 0 | 0 | 0 |
| 0 | 0 | 0 |
| 1 | 1 | 1 |
| 1 | 1 | 1 |
| 0 | 1 | 0 |
| 1 | 1 | 1 |
| 1 | 1 | 1 |
| 1 | 1 | 1 |
| 0 | 0 | 0 |
| 0 | 0 | 0 |
| 0 | 0 | 0 |
| 1 | 1 | 1 |
| 0 | 0 | 0 |
| 0 | 0 | 0 |
| 0 | 0 | 0 |
| 0 | 0 | 0 |
| 0 | 0 | 0 |
| 0 | 0 | 0 |
| 0 | 1 | 0 |
| 1 | 1 | 1 |
| 1 | 1 | 1 |
| 0 | 1 | 0 |
| 0 | 1 | 0 |
| 1 | 1 | 1 |
| 1 | 1 | 1 |
| 1 | 1 | 1 |
| 0 | 0 | 0 |
| 1 | 1 | 1 |
| 0 | 1 | 0 |
| 0 | 0 | 0 |
| 0 | 0 | 0 |
| 0 | 0 | 0 |
| 0 | 0 | 0 |
| 1 | 1 | 1 |
| 0 | 1 | 0 |
| 1 | 1 | 1 |
| 1 | 1 | 1 |
| 0 | 0 | 0 |
| 0 | 0 | 0 |
| 0 | 0 | 0 |
| 0 | 1 | 0 |
| 1 | 1 | 1 |
| 0 | 0 | 0 |
| 1 | 1 | 1 |
| 1 | 1 | 1 |
| 0 | 0 | 0 |
| 1 | 1 | 1 |
| 0 | 1 | 0 |

| | | |
|---|---|---|
| 1 | 1 | 1 |
| 0 | 1 | 0 |
| 1 | 1 | 1 |
| 1 | 1 | 1 |
| 1 | 1 | 1 |
| 1 | 1 | 1 |
| 0 | 1 | 0 |
| 1 | 1 | 1 |
| 0 | 0 | 0 |
| 0 | 0 | 0 |
| 1 | 1 | 1 |
| 0 | 0 | 0 |
| 0 | 0 | 0 |
| 0 | 0 | 0 |
| 1 | 1 | 1 |
| 1 | 1 | 1 |
| 0 | 0 | 0 |
| 0 | 0 | 0 |
| 0 | 0 | 0 |
| 1 | 1 | 1 |
| 1 | 1 | 1 |
| 1 | 1 | 1 |
| 0 | 0 | 0 |
| 1 | 1 | 1 |
| 0 | 0 | 0 |
| 1 | 1 | 1 |
| 1 | 1 | 1 |
| 1 | 1 | 1 |
| 1 | 1 | 1 |
| 0 | 0 | 0 |
| 0 | 0 | 0 |
| 0 | 0 | 0 |
| 1 | 1 | 1 |
| 0 | 1 | 0 |
| 1 | 1 | 1 |
| 1 | 1 | 1 |
| 1 | 1 | 1 |
| 0 | 1 | 0 |
| 0 | 1 | 0 |
| 0 | 1 | 0 |
| 0 | 0 | 0 |
| 1 | 1 | 1 |
| 1 | 1 | 1 |
| 1 | 1 | 1 |
| 1 | 1 | 1 |
| 0 | 0 | 0 |
| 1 | 0 | 0 |
| 0 | 0 | 0 |
| 1 | 1 | 1 |
| 0 | 0 | 0 |
| 0 | 0 | 0 |
| 0 | 0 | 0 |
| 0 | 1 | 0 |
| 1 | 0 | 0 |

| | | |
|---|---|---|
| 1 | 0 | 0 |
| 0 | 0 | 0 |
| 0 | 1 | 0 |
| 1 | 1 | 1 |
| 0 | 1 | 0 |
| 0 | 1 | 0 |
| 0 | 1 | 0 |
| 0 | 0 | 0 |
| 0 | 0 | 0 |
| 0 | 1 | 0 |
| 0 | 0 | 0 |
| 1 | 1 | 1 |
| 1 | 1 | 1 |
| 1 | 1 | 1 |
| 0 | 1 | 0 |
| 1 | 1 | 1 |
| 1 | 1 | 1 |
| 1 | 1 | 1 |
| 1 | 1 | 1 |
| 0 | 1 | 0 |
| 1 | 1 | 1 |
| 1 | 1 | 1 |
| 1 | 1 | 1 |
| 1 | 1 | 1 |
| 0 | 1 | 0 |
| 0 | 1 | 0 |
| 0 | 1 | 0 |
| 0 | 1 | 0 |
| 0 | 0 | 0 |
| 0 | 1 | 0 |
| 0 | 0 | 0 |
| 1 | 1 | 1 |
| 0 | 0 | 0 |
| 0 | 0 | 0 |
| 0 | 1 | 0 |
| 0 | 0 | 0 |
| 0 | 0 | 0 |
| 1 | 1 | 1 |
| 0 | 1 | 0 |
| 0 | 1 | 0 |
| 1 | 1 | 1 |
| 0 | 0 | 0 |
| 0 | 0 | 0 |
| 1 | 1 | 1 |
| 1 | 1 | 1 |
| 0 | 0 | 0 |
| 0 | 0 | 0 |
| 0 | 0 | 0 |
| 0 | 0 | 0 |
| 0 | 1 | 0 |
| 0 | 0 | 0 |
| 0 | 0 | 0 |
| 1 | 1 | 1 |
| 1 | 1 | 1 |

| | | |
|---|---|---|
| 1 | 1 | 1 |
| 1 | 1 | 1 |
| 1 | 1 | 1 |
| 1 | 1 | 1 |
| 0 | 0 | 0 |
| 0 | 0 | 0 |
| 0 | 0 | 0 |
| 0 | 0 | 0 |
| 1 | 1 | 1 |
| 0 | 0 | 0 |
| 0 | 0 | 0 |
| 0 | 0 | 0 |
| 0 | 0 | 0 |
| 1 | 1 | 1 |
| 1 | 0 | 0 |
| 0 | 1 | 0 |
| 1 | 1 | 1 |
| 1 | 1 | 1 |
| 1 | 1 | 1 |
| 0 | 1 | 0 |
| 1 | 1 | 1 |
| 1 | 1 | 1 |
| 1 | 1 | 1 |
| 1 | 1 | 1 |
| 1 | 1 | 1 |
| 0 | 0 | 0 |
| 0 | 0 | 0 |
| 0 | 0 | 0 |
| 0 | 0 | 0 |
| 0 | 1 | 0 |
| 0 | 1 | 0 |
| 0 | 1 | 0 |
| 1 | 1 | 1 |
| 0 | 1 | 0 |
| 0 | 1 | 0 |
| 1 | 1 | 1 |
| 1 | 1 | 1 |
| 1 | 1 | 1 |
| 1 | 1 | 1 |
| 1 | 1 | 1 |
| 1 | 1 | 1 |
| 1 | 1 | 1 |
| 1 | 1 | 1 |
| 1 | 1 | 1 |
| 1 | 1 | 1 |
| 1 | 1 | 1 |
| 1 | 1 | 1 |
| 1 | 1 | 1 |
| 1 | 1 | 1 |
| 1 | 1 | 1 |
| 1 | 1 | 1 |
| 1 | 1 | 1 |
| 1 | 1 | 1 |
| 1 | 1 | 1 |
| 0 | 0 | 0 |

| | | |
|---|---|---|
| 1 | 1 | 1 |
| 1 | 1 | 1 |
| 1 | 1 | 1 |
| 0 | 1 | 0 |
| 0 | 1 | 0 |
| 0 | 1 | 0 |
| 0 | 1 | 0 |
| 0 | 1 | 0 |
| 1 | 1 | 1 |
| 1 | 1 | 1 |
| 1 | 1 | 1 |
| 1 | 1 | 1 |
| 1 | 1 | 1 |
| 1 | 1 | 1 |
| 1 | 1 | 1 |
| 1 | 1 | 1 |
| 1 | 1 | 1 |
| 1 | 1 | 1 |
| 1 | 1 | 1 |
| 1 | 1 | 1 |
| 1 | 1 | 1 |
| 1 | 1 | 1 |
| 1 | 1 | 1 |
| 1 | 1 | 1 |
| 1 | 1 | 1 |
| 1 | 1 | 1 |
| 1 | 1 | 1 |
| 1 | 1 | 1 |
| 1 | 1 | 1 |
| 1 | 1 | 1 |
| 1 | 1 | 1 |
| 1 | 1 | 1 |
| 1 | 1 | 1 |
| 0 | 1 | 0 |
| 1 | 1 | 1 |
| 1 | 1 | 1 |
| 1 | 1 | 1 |
| 1 | 1 | 1 |
| 1 | 1 | 1 |
| 1 | 1 | 1 |
| 0 | 1 | 0 |
| 1 | 1 | 1 |
| 1 | 1 | 1 |
| 1 | 1 | 1 |
| 0 | 1 | 0 |
| 1 | 0 | 0 |
| 0 | 0 | 0 |
| 1 | 1 | 1 |
| 0 | 1 | 0 |
| 1 | 1 | 1 |
| 1 | 1 | 1 |
| 0 | 1 | 0 |
| 0 | 1 | 0 |
| 0 | 1 | 0 |

| | | |
|---|---|---|
| 0 | 1 | 0 |
| 1 | 1 | 1 |
| 0 | 0 | 0 |
| 1 | 1 | 1 |
| 1 | 1 | 1 |
| 1 | 1 | 1 |
| 0 | 0 | 0 |
| 1 | 1 | 1 |
| 1 | 1 | 1 |
| 0 | 1 | 0 |
| 0 | 1 | 0 |
| 0 | 0 | 0 |
| 1 | 1 | 1 |
| 0 | 0 | 0 |
| 0 | 0 | 0 |
| 0 | 0 | 0 |
| 0 | 0 | 0 |
| 0 | 0 | 0 |
| 0 | 0 | 0 |
| 0 | 0 | 0 |
| 0 | 0 | 0 |

| organism_scientific | evo_cons | cluster_id | pdb_id | chain_id | resi | resn | change | uniprot_id |
|---|---|---|---|---|---|---|---|---|
| HOMO SAPIENS | 5 | 87 | 3uzw | A | 106 | LEU | p.Leu106Phe | P51857 |
| HOMO SAPIENS | 3 | 87 | 3uzw | A | 133 | PRO | p.Pro133Arg | P51857 |
| HOMO SAPIENS | 0 | 87 | 3uzw | A | 198 | PRO | p.Pro198Leu | P51857 |
| HOMO SAPIENS | 7 | 87 | 3uzw | A | 261 | ARG | p.Arg261Cys | P51857 |
| HOMO SAPIENS | 0 | 4693 | 3mk1 | A | 3 | PRO | p.Pro25Leu | P05187 |
| HOMO SAPIENS | 8 | 4830 | 2h64 | B | 101 | CYS | p.Cys124Arg | P36894 |
| HOMO SAPIENS | 2 | 4830 | 2h64 | B | 116 | PRO | p.Pro139Ser | P36894 |
| HOMO SAPIENS | 0 | 6937 | 3dar | A | 186 | MET | p.Met186Thr | P21802 |
| HOMO SAPIENS | 3 | 5 | 1aut | C | 30 | GLN | p.Gln226His | P04070 |
| HOMO SAPIENS | 1 | 5 | 1aut | C | 68 | LEU | p.Leu265Phe | P04070 |
| HOMO SAPIENS | 0 | 5 | 1aut | C | 75 | ARG | p.Arg272Cys | P04070 |
| HOMO SAPIENS | 6 | 5 | 1aut | C | 92 | PRO | p.Pro289Leu | P04070 |
| HOMO SAPIENS | 0 | 5 | 1aut | C | 104 | ALA | p.Ala301Val | P04070 |
| HOMO SAPIENS | 1 | 5 | 1aut | C | 112 | ALA | p.Ala309Thr | P04070 |
| HOMO SAPIENS | 2 | 5 | 1aut | C | 115 | SER | p.Ser312Leu | P04070 |
| HOMO SAPIENS | 1 | 5 | 1aut | C | 133 | GLY | p.Gly334Ser | P04070 |
| HOMO SAPIENS | 0 | 5 | 1aut | C | 147 | ARG | p.Arg348Ter | P04070 |
| HOMO SAPIENS | 8 | 5 | 1aut | C | 194 | ASP | p.Asp401Asn | P04070 |
| HOMO SAPIENS | 7 | 5 | 1aut | C | 237 | TRP | p.Trp444Cys | P04070 |
| HOMO SAPIENS | 0 | 5 | 1aut | C | 238 | ILE | p.Ile445Met | P04070 |
| HOMO SAPIENS | 8 | 43 | 1aiy | F | 10 | HIS | p.His34Asp | P01308 |
| HOMO SAPIENS | 2 | 43 | 1aiy | F | 22 | ARG | p.Arg46Gln | P01308 |
| HOMO SAPIENS | 4 | 43 | 1aiy | F | 23 | GLY | p.Gly47Val | P01308 |
| HOMO SAPIENS | 7 | 43 | 1aiy | F | 24 | PHE | p.Phe48Ser | P01308 |
| HOMO SAPIENS | 5 | 24 | 2fg8 | A | 33 | THR | p.Thr30Ile | P02792 |
| HOMO SAPIENS | 4 | 24 | 2fg8 | A | 99 | ALA | p.Ala96Thr | P02792 |
| HOMO SAPIENS | 5 | 24 | 2fg8 | A | 107 | GLU | p.Glu104Ter | P02792 |
| HOMO SAPIENS | 3 | 5409 | 5bnq | A | 199 | MET | p.Met199Lys | O14788 |
| HOMO SAPIENS | -1 | 839 | 3swc | A | 959 | GLN | p.Gln990His | Q9H9B1 |
| HOMO SAPIENS | -1 | 839 | 3swc | A | 968 | ALA | p.Ala999Val | Q9H9B1 |
| HOMO SAPIENS | -1 | 839 | 3swc | A | 973 | SER | p.Ser1004Ile | Q9H9B1 |
| HOMO SAPIENS | 8 | 17 | 1eou | A | 18 | LYS | p.Lys18Glu | P00918 |
| HOMO SAPIENS | 2 | 17 | 1eou | A | 107 | HIS | p.His107Tyr | P00918 |
| HOMO SAPIENS | 1 | 17 | 1eou | A | 236 | PRO | p.Pro236His | P00918 |
| HOMO SAPIENS | 6 | 17 | 1eou | A | 252 | ASN | p.Asn252Asp | P00918 |
| HOMO SAPIENS | 2 | 9140 | 5c4v | A | 445 | ARG | p.Arg445Ter | Q13485 |
| HOMO SAPIENS | 7 | 9140 | 5c4v | A | 496 | ARG | p.Arg496Cys | Q13485 |
| HOMO SAPIENS | -1 | 42 | 4pwi | A | 6 | GLY | p.Gly26Ser | P02766 |
| HOMO SAPIENS | 6 | 42 | 4pwi | A | 10 | CYS | p.Cys30Arg | P02766 |
| HOMO SAPIENS | 8 | 42 | 4pwi | A | 12 | LEU | p.Leu32Pro | P02766 |
| HOMO SAPIENS | 9 | 42 | 4pwi | A | 18 | ASP | p.Asp38Gly | P02766 |
| HOMO SAPIENS | 8 | 42 | 4pwi | A | 20 | VAL | p.Val40Ile | P02766 |
| HOMO SAPIENS | 4 | 42 | 4pwi | A | 30 | MET | p.Val50Ala | P02766 |
| HOMO SAPIENS | 4 | 42 | 4pwi | A | 30 | MET | p.Val50Leu | P02766 |
| HOMO SAPIENS | 6 | 42 | 4pwi | A | 33 | PHE | p.Phe53Ile | P02766 |
| HOMO SAPIENS | 7 | 42 | 4pwi | A | 36 | ALA | p.Ala56Pro | P02766 |
| HOMO SAPIENS | 7 | 42 | 4pwi | A | 42 | GLU | p.Glu62Gly | P02766 |
| HOMO SAPIENS | 8 | 42 | 4pwi | A | 44 | PHE | p.Phe64Ser | P02766 |
| HOMO SAPIENS | 4 | 42 | 4pwi | A | 47 | GLY | p.Gly67Ala | P02766 |
| HOMO SAPIENS | 6 | 42 | 4pwi | A | 49 | THR | p.Thr69Ala | P02766 |
| HOMO SAPIENS | 6 | 42 | 4pwi | A | 50 | SER | p.Ser70Ile | P02766 |
| HOMO SAPIENS | 6 | 42 | 4pwi | A | 50 | SER | p.Ser70Arg | P02766 |
| HOMO SAPIENS | 0 | 42 | 4pwi | A | 53 | GLY | p.Gly73Glu | P02766 |

| | | | | | | | | |
|---|---|---|---|---|---|---|---|---|
| HOMO SAPIENS | 7 | 42 | 4pwi | A | 55 | LEU | p.Leu75Pro | P02766 |
| HOMO SAPIENS | 6 | 42 | 4pwi | A | 58 | LEU | p.Leu78His | P02766 |
| HOMO SAPIENS | 7 | 42 | 4pwi | A | 61 | GLU | p.Glu81Lys | P02766 |
| HOMO SAPIENS | 8 | 42 | 4pwi | A | 64 | PHE | p.Phe84Leu | P02766 |
| HOMO SAPIENS | 8 | 42 | 4pwi | A | 68 | ILE | p.Ile88Leu | P02766 |
| HOMO SAPIENS | 8 | 42 | 4pwi | A | 69 | TYR | p.Tyr89His | P02766 |
| HOMO SAPIENS | 4 | 42 | 4pwi | A | 71 | VAL | p.Val91Ala | P02766 |
| HOMO SAPIENS | 6 | 42 | 4pwi | A | 77 | SER | p.Ser97Tyr | P02766 |
| HOMO SAPIENS | 4 | 42 | 4pwi | A | 84 | ILE | p.Ile104Ser | P02766 |
| HOMO SAPIENS | 8 | 42 | 4pwi | A | 89 | GLU | p.Glu109Gln | P02766 |
| HOMO SAPIENS | 8 | 42 | 4pwi | A | 97 | ALA | p.Ala117Gly | P02766 |
| HOMO SAPIENS | 7 | 42 | 4pwi | A | 99 | ASP | p.Asp119Asn | P02766 |
| HOMO SAPIENS | 9 | 42 | 4pwi | A | 107 | ILE | p.Ile127Val | P02766 |
| HOMO SAPIENS | 9 | 42 | 4pwi | A | 111 | LEU | p.Leu131Met | P02766 |
| HOMO SAPIENS | 7 | 42 | 4pwi | A | 114 | TYR | p.Tyr134Cys | P02766 |
| HOMO SAPIENS | 7 | 42 | 4pwi | A | 114 | TYR | p.Tyr134His | P02766 |
| HOMO SAPIENS | 7 | 42 | 4pwi | A | 122 | VAL | p.Val142Ile | P02766 |
| HOMO SAPIENS | -1 | 42 | 4pwi | A | 126 | LYS | p.Lys146Arg | P02766 |
| HOMO SAPIENS | -1 | 42 | 4pwi | A | 127 | GLU | p.Glu147Ter | P02766 |
| HOMO SAPIENS | -1 | 1312 | 4dce | A | 1275 | ARG | p.Arg1275Gln | Q9UM73 |
| HOMO SAPIENS | -1 | 1312 | 4dce | A | 1280 | ALA | p.Ala1280Val | Q9UM73 |
| HOMO SAPIENS | 1 | 20 | 1y0x | X | 234 | ALA | p.Ala234Thr | P10828 |
| HOMO SAPIENS | 0 | 20 | 1y0x | X | 243 | ARG | p.Arg243Trp | P10828 |
| HOMO SAPIENS | 3 | 20 | 1y0x | X | 316 | ARG | p.Arg316His | P10828 |
| HOMO SAPIENS | 1 | 20 | 1y0x | X | 317 | ALA | p.Ala317Thr | P10828 |
| HOMO SAPIENS | 2 | 20 | 1y0x | X | 320 | ARG | p.Arg320Cys | P10828 |
| HOMO SAPIENS | 2 | 20 | 1y0x | X | 320 | ARG | p.Arg320His | P10828 |
| HOMO SAPIENS | 0 | 20 | 1y0x | X | 332 | GLY | p.Gly332Arg | P10828 |
| HOMO SAPIENS | 1 | 20 | 1y0x | X | 338 | ARG | p.Arg338Trp | P10828 |
| HOMO SAPIENS | 0 | 20 | 1y0x | X | 340 | GLN | p.Gln340His | P10828 |
| HOMO SAPIENS | 0 | 20 | 1y0x | X | 345 | GLY | p.Gly345Val | P10828 |
| HOMO SAPIENS | 0 | 20 | 1y0x | X | 345 | GLY | p.Gly345Ser | P10828 |
| HOMO SAPIENS | 0 | 20 | 1y0x | X | 347 | GLY | p.Gly347Glu | P10828 |
| HOMO SAPIENS | 1 | 20 | 1y0x | X | 438 | ARG | p.Arg438His | P10828 |
| HOMO SAPIENS | 0 | 20 | 1y0x | X | 442 | MET | p.Met442Val | P10828 |
| HOMO SAPIENS | 0 | 20 | 1y0x | X | 443 | LYS | p.Lys443Glu | P10828 |
| HOMO SAPIENS | 0 | 20 | 1y0x | X | 446 | CYS | p.Cys446Arg | P10828 |
| HOMO SAPIENS | 2 | 20 | 1y0x | X | 453 | PRO | p.Pro453His | P10828 |
| HOMO SAPIENS | 2 | 20 | 1y0x | X | 453 | PRO | p.Pro453Thr | P10828 |
| HOMO SAPIENS | 3 | 20 | 1y0x | X | 459 | PHE | p.Phe459Cys | P10828 |
| HOMO SAPIENS | 0 | 4409 | 3qb7 | D | 87 | GLU | p.Glu109Lys | P31785 |
| HOMO SAPIENS | 1 | 4409 | 3qb7 | D | 92 | GLY | p.Gly114Asp | P31785 |
| HOMO SAPIENS | 1 | 4409 | 3qb7 | D | 93 | CYS | p.Cys115Arg | P31785 |
| HOMO SAPIENS | 0 | 4409 | 3qb7 | D | 131 | ILE | p.Ile153Asn | P31785 |
| HOMO SAPIENS | 9 | 4409 | 3qb7 | D | 200 | ARG | p.Arg222Cys | P31785 |
| HOMO SAPIENS | 0 | 3891 | 3ije | B | 33 | LEU | p.Leu59Pro | P05106 |
| HOMO SAPIENS | 0 | 3891 | 3ije | B | 117 | LEU | p.Leu143Trp | P05106 |
| HOMO SAPIENS | 3 | 3891 | 3ije | B | 119 | ASP | p.Asp145Tyr | P05106 |
| HOMO SAPIENS | 2 | 3891 | 3ije | B | 143 | ARG | p.Arg169Gln | P05106 |
| HOMO SAPIENS | 2 | 3891 | 3ije | B | 214 | ARG | p.Arg240Gln | P05106 |
| HOMO SAPIENS | 2 | 3891 | 3ije | B | 214 | ARG | p.Arg240Trp | P05106 |
| HOMO SAPIENS | 2 | 3891 | 3ije | B | 374 | CYS | p.Cys400Tyr | P05106 |
| HOMO SAPIENS | 1 | 3891 | 3ije | B | 407 | PRO | p.Pro433Ala | P05106 |
| HOMO SAPIENS | 0 | 3891 | 3ije | B | 489 | ARG | p.Arg515Gln | P05106 |

| | | | | | | | | |
|---|---|---|---|---|---|---|---|---|
| HOMO SAPIENS | -1 | 9613 | 5mke | B | 28 | ARG | p.Arg28Pro | Q13563 |
| HOMO SAPIENS | -1 | 9613 | 5mke | B | 190 | ALA | p.Ala190Thr | Q13563 |
| HOMO SAPIENS | 4 | 9613 | 5mke | B | 262 | VAL | p.Val262Met | Q13563 |
| HOMO SAPIENS | 0 | 9613 | 5mke | B | 482 | PHE | p.Phe482Cys | Q13563 |
| HOMO SAPIENS | 4 | 9613 | 5mke | B | 511 | ASP | p.Asp511Val | Q13563 |
| HOMO SAPIENS | 5 | 9613 | 5mke | B | 516 | VAL | p.Val516Leu | Q13563 |
| HOMO SAPIENS | -1 | 9613 | 5mke | B | 807 | ARG | p.Arg807Gln | Q13563 |
| HOMO SAPIENS | 4 | 2428 | 1olh | C | 337 | ARG | p.Arg337Cys | P04637 |
| HOMO SAPIENS | 4 | 2428 | 1olh | C | 337 | ARG | p.Arg337His | P04637 |
| HOMO SAPIENS | 8 | 2428 | 1olh | C | 338 | PHE | p.Phe338Leu | P04637 |
| HOMO SAPIENS | 8 | 2428 | 1olh | C | 339 | GLU | p.Glu339Lys | P04637 |
| HOMO SAPIENS | 7 | 2428 | 1olh | C | 344 | LEU | p.Leu344Pro | P04637 |
| HOMO SAPIENS | 0 | 2428 | 1olh | C | 360 | GLY | p.Gly360Ala | P04637 |
| HOMO SAPIENS | 7 | 4867 | 1mox | B | 404 | GLY | p.Gly428Asp | P00533 |
| HOMO SAPIENS | 5 | 4867 | 1mox | B | 497 | ARG | p.Arg521Lys | P00533 |
| HOMO SAPIENS | 1 | 4630 | 1fu1 | A | 43 | TRP | p.Trp43Arg | Q13426 |
| HOMO SAPIENS | 2 | 4630 | 1fu1 | A | 161 | ARG | p.Arg161Ter | Q13426 |
| HOMO SAPIENS | 2 | 15 | 1lfy | B | 2 | HIS | p.His3Gln | P68871 |
| HOMO SAPIENS | 2 | 15 | 1lfy | B | 6 | GLU | p.Glu7Ala | P68871 |
| HOMO SAPIENS | 2 | 15 | 1lfy | B | 6 | GLU | p.Glu7Lys | P68871 |
| HOMO SAPIENS | 4 | 15 | 1lfy | B | 9 | SER | p.Ser10Cys | P68871 |
| HOMO SAPIENS | 6 | 15 | 1lfy | B | 15 | TRP | p.Trp16Ter | P68871 |
| HOMO SAPIENS | 3 | 15 | 1lfy | B | 17 | LYS | p.Lys18Glu | P68871 |
| HOMO SAPIENS | 6 | 15 | 1lfy | B | 19 | ASN | p.Asn20Ser | P68871 |
| HOMO SAPIENS | 6 | 15 | 1lfy | B | 22 | GLU | p.Glu23Ala | P68871 |
| HOMO SAPIENS | 6 | 15 | 1lfy | B | 22 | GLU | p.Glu23Lys | P68871 |
| HOMO SAPIENS | 8 | 15 | 1lfy | B | 26 | GLU | p.Glu27Lys | P68871 |
| HOMO SAPIENS | 8 | 15 | 1lfy | B | 28 | LEU | p.Leu29Pro | P68871 |
| HOMO SAPIENS | 8 | 15 | 1lfy | B | 30 | ARG | p.Arg31Ser | P68871 |
| HOMO SAPIENS | 8 | 15 | 1lfy | B | 30 | ARG | p.Arg31Lys | P68871 |
| HOMO SAPIENS | 6 | 15 | 1lfy | B | 39 | GLN | p.Gln40Ter | P68871 |
| HOMO SAPIENS | 2 | 15 | 1lfy | B | 43 | GLU | p.Glu44Gln | P68871 |
| HOMO SAPIENS | 6 | 15 | 1lfy | B | 57 | ASN | p.Asn58Lys | P68871 |
| HOMO SAPIENS | 6 | 15 | 1lfy | B | 59 | LYS | p.Lys60Asn | P68871 |
| HOMO SAPIENS | 8 | 15 | 1lfy | B | 61 | LYS | p.Lys62Glu | P68871 |
| HOMO SAPIENS | 4 | 15 | 1lfy | B | 73 | ASP | p.Asp74Tyr | P68871 |
| HOMO SAPIENS | 4 | 15 | 1lfy | B | 87 | THR | p.Thr88Ile | P68871 |
| HOMO SAPIENS | 4 | 15 | 1lfy | B | 87 | THR | p.Thr88Pro | P68871 |
| HOMO SAPIENS | 5 | 15 | 1lfy | B | 90 | GLU | p.Glu91Lys | P68871 |
| HOMO SAPIENS | 6 | 15 | 1lfy | B | 91 | LEU | p.Leu92Pro | P68871 |
| HOMO SAPIENS | 5 | 15 | 1lfy | B | 97 | HIS | p.His98Gln | P68871 |
| HOMO SAPIENS | 6 | 15 | 1lfy | B | 110 | LEU | p.Leu111Pro | P68871 |
| HOMO SAPIENS | 6 | 15 | 1lfy | B | 113 | VAL | p.Val114Glu | P68871 |
| HOMO SAPIENS | 2 | 15 | 1lfy | B | 119 | GLY | p.Gly120Ala | P68871 |
| HOMO SAPIENS | 4 | 15 | 1lfy | B | 121 | GLU | p.Glu122Gln | P68871 |
| HOMO SAPIENS | 5 | 15 | 1lfy | B | 134 | VAL | p.Val135Glu | P68871 |
| HOMO SAPIENS | 3 | 15 | 1lfy | B | 143 | HIS | p.His144Gln | P68871 |
| HOMO SAPIENS | 4 | 91 | 5d9k | A | 75 | GLY | p.Gly75Val | P51812 |
| HOMO SAPIENS | 5 | 91 | 5d9k | A | 82 | VAL | p.Val82Phe | P51812 |
| HOMO SAPIENS | 0 | 91 | 5d9k | A | 114 | ARG | p.Arg114Trp | P51812 |
| HOMO SAPIENS | 4 | 91 | 5d9k | A | 189 | ILE | p.Ile189Lys | P51812 |
| HOMO SAPIENS | -1 | 91 | 5d9k | A | 227 | SER | p.Ser227Ala | P51812 |
| HOMO SAPIENS | 1 | 91 | 5d9k | A | 268 | PHE | p.Phe268Ser | P51812 |
| HOMO SAPIENS | 0 | 954 | 2vt0 | A | 15 | VAL | p.Val54Leu | P04062 |

| | | | | | | | | |
|---|---|---|---|---|---|---|---|---|
| HOMO SAPIENS | 5 | 954 | 2vt0 | A | 46 | GLY | p.Gly85Glu | P04062 |
| HOMO SAPIENS | 5 | 954 | 2vt0 | A | 48 | ARG | p.Arg87Trp | P04062 |
| HOMO SAPIENS | 6 | 954 | 2vt0 | A | 79 | LYS | p.Lys118Asn | P04062 |
| HOMO SAPIENS | 0 | 954 | 2vt0 | A | 120 | ARG | p.Arg159Gln | P04062 |
| HOMO SAPIENS | 0 | 954 | 2vt0 | A | 120 | ARG | p.Arg159Trp | P04062 |
| HOMO SAPIENS | 0 | 954 | 2vt0 | A | 122 | PRO | p.Pro161Ser | P04062 |
| HOMO SAPIENS | 5 | 954 | 2vt0 | A | 131 | ARG | p.Arg170Leu | P04062 |
| HOMO SAPIENS | 5 | 954 | 2vt0 | A | 135 | TYR | p.Tyr174Cys | P04062 |
| HOMO SAPIENS | 0 | 954 | 2vt0 | A | 157 | LYS | p.Lys196Gln | P04062 |
| HOMO SAPIENS | 5 | 954 | 2vt0 | A | 163 | ARG | p.Arg202Gln | P04062 |
| HOMO SAPIENS | 5 | 954 | 2vt0 | A | 170 | ARG | p.Arg209Cys | P04062 |
| HOMO SAPIENS | 5 | 954 | 2vt0 | A | 188 | ASN | p.Asn227Ser | P04062 |
| HOMO SAPIENS | 3 | 954 | 2vt0 | A | 196 | SER | p.Ser235Pro | P04062 |
| HOMO SAPIENS | 5 | 954 | 2vt0 | A | 202 | GLY | p.Gly241Arg | P04062 |
| HOMO SAPIENS | 0 | 954 | 2vt0 | A | 212 | TYR | p.Tyr251His | P04062 |
| HOMO SAPIENS | 0 | 954 | 2vt0 | A | 213 | PHE | p.Phe252Ile | P04062 |
| HOMO SAPIENS | 5 | 954 | 2vt0 | A | 216 | PHE | p.Phe255Tyr | P04062 |
| HOMO SAPIENS | 6 | 954 | 2vt0 | A | 251 | PHE | p.Phe290Leu | P04062 |
| HOMO SAPIENS | 5 | 954 | 2vt0 | A | 255 | HIS | p.His294Gln | P04062 |
| HOMO SAPIENS | 6 | 954 | 2vt0 | A | 257 | ARG | p.Arg296Gln | P04062 |
| HOMO SAPIENS | 5 | 954 | 2vt0 | A | 289 | PRO | p.Pro328Leu | P04062 |
| HOMO SAPIENS | 0 | 954 | 2vt0 | A | 309 | ALA | p.Ala348Val | P04062 |
| HOMO SAPIENS | 0 | 954 | 2vt0 | A | 311 | HIS | p.His350Arg | P04062 |
| HOMO SAPIENS | 6 | 954 | 2vt0 | A | 312 | TRP | p.Trp351Cys | P04062 |
| HOMO SAPIENS | 5 | 954 | 2vt0 | A | 323 | THR | p.Thr362Ile | P04062 |
| HOMO SAPIENS | 5 | 954 | 2vt0 | A | 325 | GLY | p.Gly364Arg | P04062 |
| HOMO SAPIENS | 6 | 954 | 2vt0 | A | 342 | CYS | p.Cys381Gly | P04062 |
| HOMO SAPIENS | 5 | 954 | 2vt0 | A | 353 | ARG | p.Arg392Gly | P04062 |
| HOMO SAPIENS | 3 | 954 | 2vt0 | A | 364 | SER | p.Ser403Thr | P04062 |
| HOMO SAPIENS | 5 | 954 | 2vt0 | A | 369 | THR | p.Thr408Met | P04062 |
| HOMO SAPIENS | 3 | 954 | 2vt0 | A | 370 | ASN | p.Asn409Ser | P04062 |
| HOMO SAPIENS | 0 | 954 | 2vt0 | A | 371 | LEU | p.Leu410Val | P04062 |
| HOMO SAPIENS | 0 | 954 | 2vt0 | A | 375 | VAL | p.Val414Leu | P04062 |
| HOMO SAPIENS | 0 | 954 | 2vt0 | A | 377 | GLY | p.Gly416Ser | P04062 |
| HOMO SAPIENS | 8 | 954 | 2vt0 | A | 398 | VAL | p.Val437Phe | P04062 |
| HOMO SAPIENS | 0 | 954 | 2vt0 | A | 401 | PRO | p.Pro440Leu | P04062 |
| HOMO SAPIENS | 3 | 954 | 2vt0 | A | 409 | ASP | p.Asp448His | P04062 |
| HOMO SAPIENS | 3 | 954 | 2vt0 | A | 409 | ASP | p.Asp448Val | P04062 |
| HOMO SAPIENS | 0 | 954 | 2vt0 | A | 415 | PRO | p.Pro454Arg | P04062 |
| HOMO SAPIENS | 5 | 954 | 2vt0 | A | 444 | LEU | p.Leu483Arg | P04062 |
| HOMO SAPIENS | 5 | 954 | 2vt0 | A | 456 | ALA | p.Ala495Pro | P04062 |
| HOMO SAPIENS | 5 | 954 | 2vt0 | A | 463 | ARG | p.Arg502Cys | P04062 |
| HOMO SAPIENS | 5 | 954 | 2vt0 | A | 463 | ARG | p.Arg502His | P04062 |
| HOMO SAPIENS | 5 | 954 | 2vt0 | A | 478 | GLY | p.Gly517Ser | P04062 |
| HOMO SAPIENS | 5 | 954 | 2vt0 | A | 496 | ARG | p.Arg535Cys | P04062 |
| HOMO SAPIENS | 5 | 954 | 2vt0 | A | 496 | ARG | p.Arg535His | P04062 |
| HOMO SAPIENS | 1 | 9037 | 2rqb | A | 946 | ALA | p.Ala946Thr | Q9BYX4 |
| HOMO SAPIENS | 0 | 253 | 5E+092 | A | 308 | LEU | p.Leu308Pro | P37173 |
| HOMO SAPIENS | 0 | 253 | 5E+092 | A | 315 | THR | p.Thr315Met | P37173 |
| HOMO SAPIENS | 0 | 253 | 5E+092 | A | 328 | HIS | p.His328Gln | P37173 |
| HOMO SAPIENS | 0 | 253 | 5E+092 | A | 336 | TYR | p.Tyr336Asn | P37173 |
| HOMO SAPIENS | 0 | 253 | 5E+092 | A | 348 | ARG | p.Arg348Cys | P37173 |
| HOMO SAPIENS | 0 | 253 | 5E+092 | A | 355 | ALA | p.Ala355Pro | P37173 |
| HOMO SAPIENS | 1 | 253 | 5E+092 | A | 356 | ARG | p.Arg356Gly | P37173 |

| | | | | | | | | |
|---|---|---|---|---|---|---|---|---|
| HOMO SAPIENS | 0 | 253 | 5E+092 | A | 357 | GLY | p.Gly357Trp | P37173 |
| HOMO SAPIENS | 1 | 253 | 5E+092 | A | 403 | ARG | p.Arg403His | P37173 |
| HOMO SAPIENS | 5 | 253 | 5E+092 | A | 449 | SER | p.Ser449Phe | P37173 |
| HOMO SAPIENS | 0 | 253 | 5E+092 | A | 460 | ARG | p.Arg460Cys | P37173 |
| HOMO SAPIENS | 0 | 253 | 5E+092 | A | 460 | ARG | p.Arg460His | P37173 |
| HOMO SAPIENS | 2 | 253 | 5E+092 | A | 526 | GLU | p.Glu526Gln | P37173 |
| HOMO SAPIENS | 5 | 253 | 5E+092 | A | 528 | ARG | p.Arg528Cys | P37173 |
| HOMO SAPIENS | 5 | 253 | 5E+092 | A | 528 | ARG | p.Arg528His | P37173 |
| HOMO SAPIENS | 6 | 158 | 5amc | A | 231 | ARG | p.Arg260Leu | P12821 |
| HOMO SAPIENS | 7 | 158 | 5amc | A | 237 | TYR | p.Tyr266Ter | P12821 |
| HOMO SAPIENS | 6 | 158 | 5amc | A | 325 | GLY | p.Gly354Arg | P12821 |
| HOMO SAPIENS | 0 | 158 | 5amc | A | 467 | ARG | p.Arg496Ter | P12821 |
| HOMO SAPIENS | 4 | 158 | 5amc | A | 479 | ARG | p.Arg508Ter | P12821 |
| HOMO SAPIENS | 0 | 158 | 5amc | A | 563 | ASP | p.Asp592Gly | P12821 |
| HOMO SAPIENS | -1 | 33062 | 1gr3 | A | 545 | GLY | p.Gly545Arg | Q03692 |
| HOMO SAPIENS | 5 | 33062 | 1gr3 | A | 591 | CYS | p.Cys591Arg | Q03692 |
| HOMO SAPIENS | 2 | 33062 | 1gr3 | A | 595 | GLY | p.Gly595Glu | Q03692 |
| HOMO SAPIENS | 0 | 33062 | 1gr3 | A | 597 | TYR | p.Tyr597Cys | Q03692 |
| HOMO SAPIENS | 5 | 33062 | 1gr3 | A | 598 | TYR | p.Tyr598Asp | Q03692 |
| HOMO SAPIENS | 7 | 33062 | 1gr3 | A | 600 | SER | p.Ser600Pro | Q03692 |
| HOMO SAPIENS | 1 | 33062 | 1gr3 | A | 614 | LEU | p.Leu614Pro | Q03692 |
| HOMO SAPIENS | 6 | 33062 | 1gr3 | A | 651 | TRP | p.Trp651Arg | Q03692 |
| HOMO SAPIENS | 7 | 33062 | 1gr3 | A | 671 | SER | p.Ser671Pro | Q03692 |
| HOMO SAPIENS | -1 | 38257 | 2l2i | B | 87 | PRO | p.Pro87Ala | Q13351 |
| HOMO SAPIENS | 2 | 2036 | 1ghq | A | 231 | GLY | p.Gly1224Asp | P01024 |
| HOMO SAPIENS | 5 | 3594 | 4cde | F | 388 | TYR | p.Tyr412Cys | P53634 |
| HOMO SAPIENS | 9 | 3594 | 4cde | F | 405 | TRP | p.Trp429Cys | P53634 |
| HOMO SAPIENS | 0 | 3594 | 4cde | F | 429 | ILE | p.Ile453Val | P53634 |
| HOMO SAPIENS | 7 | 18 | 1jkb | A | 34 | TRP | p.Trp52Ter | P61626 |
| HOMO SAPIENS | 9 | 18 | 1jkb | A | 70 | THR | p.Thr88Asn | P61626 |
| HOMO SAPIENS | 8 | 18 | 1jkb | A | 121 | VAL | p.Val139Ile | P61626 |
| HOMO SAPIENS | 2 | 414 | 3fj6 | A | 136 | ARG | p.Arg135Cys | Q02127 |
| HOMO SAPIENS | 1 | 414 | 3fj6 | A | 153 | GLY | p.Gly152Arg | Q02127 |
| HOMO SAPIENS | 0 | 414 | 3fj6 | A | 200 | ARG | p.Arg199Cys | Q02127 |
| HOMO SAPIENS | 0 | 414 | 3fj6 | A | 203 | GLY | p.Gly202Ala | Q02127 |
| HOMO SAPIENS | 0 | 414 | 3fj6 | A | 245 | ARG | p.Arg244Trp | Q02127 |
| HOMO SAPIENS | 0 | 414 | 3fj6 | A | 342 | ALA | p.Ala341Val | Q02127 |
| HOMO SAPIENS | 1 | 414 | 3fj6 | A | 347 | ARG | p.Arg346Trp | Q02127 |
| HOMO SAPIENS | 6 | 103 | 1u3w | A | 77 | GLY | p.Gly78Ter | P00326 |
| HOMO SAPIENS | -1 | 8369 | 4ux8 | F | 16 | ARG | p.Arg93Trp | P39905 |
| HOMO SAPIENS | 0 | 366 | 2zjc | A | 32 | GLY | p.Arg108Trp | P01375 |
| HOMO SAPIENS | 0 | 1457 | 3v9i | A | 352 | LEU | p.Ser352Leu | P30038 |
| HOMO SAPIENS | 0 | 1457 | 3v9i | A | 470 | VAL | p.Val470Ile | P30038 |
| HOMO SAPIENS | 0 | 1457 | 3v9i | A | 473 | THR | p.Thr473Ala | P30038 |
| HOMO SAPIENS | 1 | 2 | 4wo5 | A | 462 | ARG | p.Arg462Ile | P15056 |
| HOMO SAPIENS | 3 | 2 | 4wo5 | A | 463 | ILE | p.Ile463Ser | P15056 |
| HOMO SAPIENS | 1 | 2 | 4wo5 | A | 466 | GLY | p.Gly466Val | P15056 |
| HOMO SAPIENS | 1 | 2 | 4wo5 | A | 468 | PHE | p.Phe468Ser | P15056 |
| HOMO SAPIENS | 2 | 2 | 4wo5 | A | 469 | GLY | p.Gly469Ala | P15056 |
| HOMO SAPIENS | 2 | 2 | 4wo5 | A | 469 | GLY | p.Gly469Arg | P15056 |
| HOMO SAPIENS | 2 | 2 | 4wo5 | A | 485 | LEU | p.Leu485Phe | P15056 |
| HOMO SAPIENS | 0 | 2 | 4wo5 | A | 501 | GLU | p.Glu501Gly | P15056 |
| HOMO SAPIENS | 0 | 2 | 4wo5 | A | 501 | GLU | p.Glu501Lys | P15056 |
| HOMO SAPIENS | 7 | 2 | 4wo5 | A | 581 | ASN | p.Asn581Asp | P15056 |

| | | | | | | | | |
|---|---|---|---|---|---|---|---|---|
| HOMO SAPIENS | 8 | 2 | 4wo5 | A | 594 | ASP | p.Asp594Gly | P15056 |
| HOMO SAPIENS | 0 | 2 | 4wo5 | A | 597 | LEU | p.Leu597Arg | P15056 |
| HOMO SAPIENS | 0 | 2 | 4wo5 | A | 597 | LEU | p.Leu597Val | P15056 |
| HOMO SAPIENS | 0 | 2 | 4wo5 | A | 600 | VAL | p.Val600Glu | P15056 |
| HOMO SAPIENS | 2 | 2 | 4wo5 | A | 601 | LYS | p.Lys601Glu | P15056 |
| HOMO SAPIENS | 8 | 2 | 4wo5 | A | 638 | ASP | p.Asp638Glu | P15056 |
| HOMO SAPIENS | -1 | 30366 | 5bqd | B | 19 | ALA | p.Ala19Ser | Q99593 |
| HOMO SAPIENS | -1 | 30366 | 5bqd | B | 23 | PRO | p.Pro23His | Q99593 |
| HOMO SAPIENS | 1 | 30366 | 5bqd | B | 49 | GLN | p.Gln49Lys | Q99593 |
| HOMO SAPIENS | 2 | 30366 | 5bqd | B | 54 | ILE | p.Ile54Thr | Q99593 |
| HOMO SAPIENS | 8 | 30366 | 5bqd | B | 80 | GLY | p.Gly80Arg | Q99593 |
| HOMO SAPIENS | 6 | 30366 | 5bqd | B | 106 | ILE | p.Ile106Val | Q99593 |
| HOMO SAPIENS | 7 | 30366 | 5bqd | B | 111 | ASP | p.Asp111Tyr | Q99593 |
| HOMO SAPIENS | -1 | 30366 | 5bqd | B | 237 | ARG | p.Arg237Gln | Q99593 |
| HOMO SAPIENS | -1 | 30366 | 5bqd | B | 237 | ARG | p.Arg237Trp | Q99593 |
| HOMO SAPIENS | 5 | 4789 | 1n4m | A | 567 | SER | p.Ser567Leu | P06400 |
| HOMO SAPIENS | 0 | 4789 | 1n4m | A | 658 | ALA | p.Ala658Asp | P06400 |
| HOMO SAPIENS | 7 | 4789 | 1n4m | A | 661 | ARG | p.Arg661Trp | P06400 |
| HOMO SAPIENS | 6 | 4789 | 1n4m | A | 668 | ARG | p.Arg668His | P06400 |
| HOMO SAPIENS | 8 | 4789 | 1n4m | A | 697 | ASP | p.Asp697Glu | P06400 |
| HOMO SAPIENS | 6 | 4789 | 1n4m | A | 712 | CYS | p.Cys712Arg | P06400 |
| HOMO SAPIENS | 1 | 1798 | 2r5g | A | 101 | HIS | p.His101Gln | O15247 |
| HOMO SAPIENS | 0 | 1798 | 2r5g | A | 160 | PRO | p.Pro160Ala | O15247 |
| HOMO SAPIENS | 2 | 22 | 4x0t | A | 82 | ARG | p.Arg110Ter | P49419 |
| HOMO SAPIENS | 0 | 22 | 4x0t | A | 149 | ALA | p.Ala177Pro | P49419 |
| HOMO SAPIENS | 4 | 22 | 4x0t | A | 164 | THR | p.Thr192Met | P49419 |
| HOMO SAPIENS | 9 | 22 | 4x0t | A | 167 | ASN | p.Asn195Ser | P49419 |
| HOMO SAPIENS | 0 | 22 | 4x0t | A | 175 | TRP | p.Trp203Gly | P49419 |
| HOMO SAPIENS | 0 | 22 | 4x0t | A | 194 | THR | p.Thr222Ala | P49419 |
| HOMO SAPIENS | 0 | 22 | 4x0t | A | 273 | ASN | p.Asn301Ile | P49419 |
| HOMO SAPIENS | 0 | 22 | 4x0t | A | 307 | ARG | p.Arg335Gln | P49419 |
| HOMO SAPIENS | 3 | 22 | 4x0t | A | 313 | SER | p.Ser341Asn | P49419 |
| HOMO SAPIENS | 0 | 22 | 4x0t | A | 329 | ILE | p.Ile357Val | P49419 |
| HOMO SAPIENS | 1 | 22 | 4x0t | A | 380 | TYR | p.Tyr408Ter | P49419 |
| HOMO SAPIENS | 9 | 22 | 4x0t | A | 399 | GLU | p.Glu427Gln | P49419 |
| HOMO SAPIENS | 0 | 22 | 4x0t | A | 406 | TYR | p.Tyr434Cys | P49419 |
| HOMO SAPIENS | 0 | 22 | 4x0t | A | 431 | ILE | p.Ile459Phe | P49419 |
| HOMO SAPIENS | 4 | 22 | 4x0t | A | 477 | GLY | p.Gly505Arg | P49419 |
| HOMO SAPIENS | 0 | 22 | 4x0t | A | 495 | THR | p.Thr523Ala | P49419 |
| HOMO SAPIENS | 8 | 11042 | 1y6b | A | 1145 | PRO | p.Pro1147Ser | P35968 |
| HOMO SAPIENS | -1 | 3867 | 2xij | A | 18 | GLN | p.Gln18Ter | P22033 |
| HOMO SAPIENS | -1 | 3867 | 2xij | A | 31 | ARG | p.Arg31Ter | P22033 |
| HOMO SAPIENS | 2 | 3867 | 2xij | A | 93 | ARG | p.Arg93His | P22033 |
| HOMO SAPIENS | 0 | 3867 | 2xij | A | 94 | GLY | p.Gly94Arg | P22033 |
| HOMO SAPIENS | 2 | 3867 | 2xij | A | 95 | PRO | p.Pro95Leu | P22033 |
| HOMO SAPIENS | 2 | 3867 | 2xij | A | 105 | TRP | p.Trp105Arg | P22033 |
| HOMO SAPIENS | 2 | 3867 | 2xij | A | 108 | ARG | p.Arg108Cys | P22033 |
| HOMO SAPIENS | 3 | 3867 | 2xij | A | 154 | ARG | p.Arg154His | P22033 |
| HOMO SAPIENS | 3 | 3867 | 2xij | A | 156 | ASP | p.Asp156Val | P22033 |
| HOMO SAPIENS | 0 | 3867 | 2xij | A | 189 | ASN | p.Asn189Ile | P22033 |
| HOMO SAPIENS | 0 | 3867 | 2xij | A | 191 | ALA | p.Ala191Glu | P22033 |
| HOMO SAPIENS | 0 | 3867 | 2xij | A | 203 | GLY | p.Gly203Arg | P22033 |
| HOMO SAPIENS | 2 | 3867 | 2xij | A | 215 | GLY | p.Gly215Ser | P22033 |
| HOMO SAPIENS | 0 | 3867 | 2xij | A | 219 | ASN | p.Asn219Tyr | P22033 |

| | | | | | | | | |
|---|---|---|---|---|---|---|---|---|
| HOMO SAPIENS | 3 | 3867 | 2xij | A | 228 | ARG | p.Arg228Ter | P22033 |
| HOMO SAPIENS | 1 | 3867 | 2xij | A | 297 | THR | p.Thr297Ile | P22033 |
| HOMO SAPIENS | 0 | 3867 | 2xij | A | 326 | ARG | p.Arg326Lys | P22033 |
| HOMO SAPIENS | 3 | 3867 | 2xij | A | 369 | ARG | p.Arg369His | P22033 |
| HOMO SAPIENS | 3 | 3867 | 2xij | A | 369 | ARG | p.Arg369Cys | P22033 |
| HOMO SAPIENS | 0 | 3867 | 2xij | A | 370 | THR | p.Thr370Pro | P22033 |
| HOMO SAPIENS | 0 | 3867 | 2xij | A | 377 | ALA | p.Ala377Glu | P22033 |
| HOMO SAPIENS | 2 | 3867 | 2xij | A | 426 | GLY | p.Gly426Ala | P22033 |
| HOMO SAPIENS | 2 | 3867 | 2xij | A | 427 | GLY | p.Gly427Asp | P22033 |
| HOMO SAPIENS | 1 | 3867 | 2xij | A | 499 | THR | p.Ala499Thr | P22033 |
| HOMO SAPIENS | 2 | 3867 | 2xij | A | 532 | ARG | p.Arg532His | P22033 |
| HOMO SAPIENS | 3 | 3867 | 2xij | A | 555 | ALA | p.Ala555Thr | P22033 |
| HOMO SAPIENS | 0 | 3867 | 2xij | A | 573 | PHE | p.Phe573Ser | P22033 |
| HOMO SAPIENS | 3 | 3867 | 2xij | A | 623 | GLY | p.Gly623Arg | P22033 |
| HOMO SAPIENS | 5 | 3867 | 2xij | A | 671 | VAL | p.Ile671Val | P22033 |
| HOMO SAPIENS | 4 | 3867 | 2xij | A | 694 | ARG | p.Arg694Trp | P22033 |
| HOMO SAPIENS | 1 | 3867 | 2xij | A | 700 | MET | p.Met700Lys | P22033 |
| HOMO SAPIENS | 8 | 3867 | 2xij | A | 703 | GLY | p.Gly703Arg | P22033 |
| HOMO SAPIENS | 0 | 3867 | 2xij | A | 717 | GLY | p.Gly717Val | P22033 |
| HOMO SAPIENS | 1 | 3867 | 2xij | A | 727 | ARG | p.Arg727Ter | P22033 |
| HOMO SAPIENS | -1 | 52 | 3k91 | B | 72 | GLY | p.Gly73Ser | P00441 |
| HOMO SAPIENS | 6 | 52 | 3k91 | B | 96 | ASP | p.Asp97Asn | P00441 |
| HOMO SAPIENS | 5 | 52 | 3k91 | B | 112 | ILE | p.Ile113Thr | P00441 |
| HOMO SAPIENS | -1 | 52 | 3k91 | B | 134 | SER | p.Ser135Asn | P00441 |
| HOMO SAPIENS | 0 | 3950 | 4fom | A | 296 | ASN | p.Asn296His | Q9NQS3 |
| HOMO SAPIENS | 1 | 10 | 1de4 | A | 31 | VAL | p.Val53Met | Q30201 |
| HOMO SAPIENS | 0 | 10 | 1de4 | A | 37 | VAL | p.Val59Met | Q30201 |
| HOMO SAPIENS | 0 | 10 | 1de4 | A | 71 | GLY | p.Gly93Arg | Q30201 |
| HOMO SAPIENS | 0 | 10 | 1de4 | A | 83 | ILE | p.Ile105Thr | Q30201 |
| HOMO SAPIENS | 0 | 10 | 1de4 | A | 105 | GLN | p.Gln127His | Q30201 |
| HOMO SAPIENS | 1 | 10 | 1de4 | A | 146 | GLU | p.Glu168Ter | Q30201 |
| HOMO SAPIENS | 2 | 10 | 1de4 | A | 261 | GLN | p.Gln283Pro | Q30201 |
| HOMO SAPIENS | 0 | 8038 | 4x7n | A | 586 | ILE | p.Ile586Leu | Q9NZJ5 |
| HOMO SAPIENS | 0 | 8038 | 4x7n | A | 588 | ARG | p.Arg588Gln | Q9NZJ5 |
| HOMO SAPIENS | 0 | 35175 | 5ugw | A | 1062 | ALA | p.Ala1062Thr | O14746 |
| HOMO SAPIENS | 2 | 35175 | 5ugw | A | 1090 | VAL | p.Val1090Met | O14746 |
| HOMO SAPIENS | 0 | 1722 | 4z3k | C | 147 | ARG | p.Arg150Gly | P35270 |
| HOMO SAPIENS | 0 | 1722 | 4z3k | C | 160 | PRO | p.Pro163Leu | P35270 |
| HOMO SAPIENS | 0 | 1722 | 4z3k | C | 207 | ARG | p.Arg210Trp | P35270 |
| HOMO SAPIENS | 0 | 1722 | 4z3k | C | 216 | ARG | p.Arg219Ter | P35270 |
| HOMO SAPIENS | 0 | 1722 | 4z3k | C | 248 | LYS | p.Lys251Ter | P35270 |
| HOMO SAPIENS | 1 | 2832 | 4x8v | L | 115 | HIS | p.His175Gln | P08709 |
| HOMO SAPIENS | 0 | 825 | 1emh | A | 242 | PHE | p.Phe251Ser | P13051 |
| HOMO SAPIENS | 0 | 122 | 5uwk | A | 109 | ARG | p.Arg109Ter | P45452 |
| HOMO SAPIENS | 1 | 122 | 5uwk | A | 170 | LYS | p.Lys170Thr | P45452 |
| HOMO SAPIENS | 9 | 122 | 5uwk | A | 232 | HIS | p.His232Asn | P45452 |
| HOMO SAPIENS | 5 | 122 | 5uwk | A | 257 | ASP | p.Asp257Val | P45452 |
| HOMO SAPIENS | 9 | 1074 | 3p5i | A | 264 | TRP | p.Trp264Arg | Q9UJ71 |
| HOMO SAPIENS | 0 | 146 | 3r3g | A | 14C | GLU | p.Glu352Lys | P00734 |
| HOMO SAPIENS | 3 | 9522 | 1wyq | A | 83 | LEU | p.Leu253Pro | O15020 |
| HOMO SAPIENS | 8 | 67 | 2vgb | A | 130 | SER | p.Ser130Tyr | P30613 |
| HOMO SAPIENS | 8 | 67 | 2vgb | A | 163 | ARG | p.Arg163Cys | P30613 |
| HOMO SAPIENS | 4 | 67 | 2vgb | A | 241 | GLU | p.Glu241Ter | P30613 |
| HOMO SAPIENS | 9 | 67 | 2vgb | A | 384 | THR | p.Thr384Met | P30613 |

| | | | | | | | | |
|---|---|---|---|---|---|---|---|---|
| HOMO SAPIENS | 0 | 67 | 2vgb | A | 421 | GLN | p.Gln421Lys | P30613 |
| HOMO SAPIENS | 6 | 67 | 2vgb | A | 479 | ARG | p.Arg479His | P30613 |
| HOMO SAPIENS | 8 | 67 | 2vgb | A | 510 | ARG | p.Arg510Gln | P30613 |
| HOMO SAPIENS | 4 | 67 | 2vgb | A | 569 | ARG | p.Arg569Trp | P30613 |
| HOMO SAPIENS | 0 | 9088 | 5v44 | A | 232 | ASN | p.Asn232Lys | Q9NZJ4 |
| HOMO SAPIENS | 4 | 16 | 3lbn | A | 12 | GLY | p.Gly12Val | P01112 |
| HOMO SAPIENS | 4 | 16 | 3lbn | A | 12 | GLY | p.Gly12Ser | P01112 |
| HOMO SAPIENS | 4 | 16 | 3lbn | A | 13 | GLY | p.Gly13Cys | P01112 |
| HOMO SAPIENS | 4 | 16 | 3lbn | A | 13 | GLY | p.Gly13Asp | P01112 |
| HOMO SAPIENS | 5 | 16 | 3lbn | A | 22 | GLN | p.Gln22Lys | P01112 |
| HOMO SAPIENS | 5 | 16 | 3lbn | A | 58 | THR | p.Thr58Ile | P01112 |
| HOMO SAPIENS | 3 | 16 | 3lbn | A | 61 | GLN | p.Gln61Lys | P01112 |
| HOMO SAPIENS | 1 | 16 | 3lbn | A | 63 | GLU | p.Glu63Lys | P01112 |
| HOMO SAPIENS | 9 | 16 | 3lbn | A | 117 | LYS | p.Lys117Arg | P01112 |
| HOMO SAPIENS | 6 | 16 | 3lbn | A | 123 | ARG | p.Arg123Cys | P01112 |
| HOMO SAPIENS | 1 | 16 | 3lbn | A | 138 | GLY | p.Gly138Ser | P01112 |
| HOMO SAPIENS | 9 | 16 | 3lbn | A | 146 | ALA | p.Ala146Thr | P01112 |
| HOMO SAPIENS | 9 | 16 | 3lbn | A | 146 | ALA | p.Ala146Val | P01112 |
| HOMO SAPIENS | 1 | 359 | 5nvy | C | 63 | LEU | p.Leu63Pro | P40337 |
| HOMO SAPIENS | 1 | 359 | 5nvy | C | 64 | ARG | p.Arg64Pro | P40337 |
| HOMO SAPIENS | 1 | 359 | 5nvy | C | 70 | GLU | p.Glu70Lys | P40337 |
| HOMO SAPIENS | 0 | 359 | 5nvy | C | 84 | VAL | p.Val84Leu | P40337 |
| HOMO SAPIENS | 1 | 359 | 5nvy | C | 88 | TRP | p.Trp88Ser | P40337 |
| HOMO SAPIENS | 1 | 359 | 5nvy | C | 93 | GLY | p.Gly93Cys | P40337 |
| HOMO SAPIENS | 1 | 359 | 5nvy | C | 98 | TYR | p.Tyr98His | P40337 |
| HOMO SAPIENS | 1 | 359 | 5nvy | C | 111 | SER | p.Ser111Arg | P40337 |
| HOMO SAPIENS | 1 | 359 | 5nvy | C | 112 | TYR | p.Tyr112His | P40337 |
| HOMO SAPIENS | 1 | 359 | 5nvy | C | 126 | ASP | p.Asp126Tyr | P40337 |
| HOMO SAPIENS | 1 | 359 | 5nvy | C | 130 | VAL | p.Val130Leu | P40337 |
| HOMO SAPIENS | 1 | 359 | 5nvy | C | 135 | LEU | p.Leu135Phe | P40337 |
| HOMO SAPIENS | 9 | 359 | 5nvy | C | 161 | ARG | p.Arg161Gly | P40337 |
| HOMO SAPIENS | 7 | 359 | 5nvy | C | 162 | CYS | p.Cys162Phe | P40337 |
| HOMO SAPIENS | 4 | 359 | 5nvy | C | 163 | LEU | p.Leu163Pro | P40337 |
| HOMO SAPIENS | 3 | 359 | 5nvy | C | 164 | GLN | p.Gln164Arg | P40337 |
| HOMO SAPIENS | 9 | 359 | 5nvy | C | 166 | VAL | p.Val166Phe | P40337 |
| HOMO SAPIENS | 7 | 359 | 5nvy | C | 167 | ARG | p.Arg167Gly | P40337 |
| HOMO SAPIENS | 7 | 359 | 5nvy | C | 167 | ARG | p.Arg167Gln | P40337 |
| HOMO SAPIENS | 1 | 359 | 5nvy | C | 183 | SER | p.Ser183Ter | P40337 |
| HOMO SAPIENS | 2 | 359 | 5nvy | C | 188 | LEU | p.Leu188Val | P40337 |
| HOMO SAPIENS | 1 | 359 | 5nvy | C | 191 | HIS | p.His191Asp | P40337 |
| HOMO SAPIENS | 1 | 359 | 5nvy | C | 192 | PRO | p.Pro192Ser | P40337 |
| HOMO SAPIENS | 1 | 359 | 5nvy | C | 200 | ARG | p.Arg200Trp | P40337 |
| HOMO SAPIENS | -1 | 359 | 5nvy | C | 211 | MET | p.Met211Leu | P40337 |
| HOMO SAPIENS | 0 | 14099 | 4btm | A | 282 | ARG | p.Arg258Gln | Q5TCY1 |
| HOMO SAPIENS | 0 | 696 | 5h1b | A | 150 | ARG | p.Arg150Gln | Q06609 |
| HOMO SAPIENS | -1 | 208 | 2bxl | A | 1 | ASP | p.Asp25Val | P02768 |
| HOMO SAPIENS | 0 | 208 | 2bxl | A | 269 | ASP | p.Asp293Gly | P02768 |
| HOMO SAPIENS | 0 | 208 | 2bxl | A | 570 | GLU | p.Glu594Lys | P02768 |
| HOMO SAPIENS | 0 | 208 | 2bxl | A | 573 | LYS | p.Lys597Glu | P02768 |
| HOMO SAPIENS | -1 | 12813 | 2uzx | B | 35 | MET | p.Met35Leu | P08581 |
| HOMO SAPIENS | 1 | 12813 | 2uzx | B | 156 | SER | p.Ser156Leu | P08581 |
| HOMO SAPIENS | 0 | 12813 | 2uzx | B | 168 | GLU | p.Glu168Asp | P08581 |
| HOMO SAPIENS | 2 | 12813 | 2uzx | B | 211 | LEU | p.Leu211Trp | P08581 |
| HOMO SAPIENS | 2 | 12813 | 2uzx | B | 218 | ARG | p.Arg218Ser | P08581 |

| | | | | | | | | |
|---|---|---|---|---|---|---|---|---|
| HOMO SAPIENS | 1 | 12813 | 2uzx | B | 238 | LEU | p.Leu238Ser | P08581 |
| HOMO SAPIENS | 4 | 12813 | 2uzx | B | 320 | ALA | p.Ala320Val | P08581 |
| HOMO SAPIENS | 1 | 12813 | 2uzx | B | 375 | ASN | p.Asn375Ser | P08581 |
| HOMO SAPIENS | 0 | 12813 | 2uzx | B | 575 | LEU | p.Leu575Ile | P08581 |
| HOMO SAPIENS | 1 | 10780 | 4ejx | A | 348 | ARG | p.Arg367Cys | P00450 |
| HOMO SAPIENS | 2 | 10780 | 4ejx | A | 525 | ASP | p.Asp544Glu | P00450 |
| HOMO SAPIENS | 1 | 1407 | 5jn5 | A | 88 | ILE | p.Ile88Val | P36871 |
| HOMO SAPIENS | 7 | 1407 | 5jn5 | A | 115 | THR | p.Thr115Ala | P36871 |
| HOMO SAPIENS | 3 | 1407 | 5jn5 | A | 121 | GLY | p.Gly121Arg | P36871 |
| HOMO SAPIENS | 2 | 1407 | 5jn5 | A | 221 | ARG | p.Arg221Cys | P36871 |
| HOMO SAPIENS | 6 | 1407 | 5jn5 | A | 263 | TYR | p.Asp263Tyr | P36871 |
| HOMO SAPIENS | 3 | 1407 | 5jn5 | A | 420 | TYR | p.Tyr420His | P36871 |
| HOMO SAPIENS | 0 | 1407 | 5jn5 | A | 501 | VAL | p.Val501Ile | P36871 |
| HOMO SAPIENS | -1 | 1878 | 1yow | A | 255 | ARG | p.Arg255Leu | Q13285 |
| HOMO SAPIENS | 6 | 1878 | 1yow | A | 293 | ASP | p.Asp293Asn | Q13285 |
| HOMO SAPIENS | 1 | 1878 | 1yow | A | 437 | LEU | p.Leu437Gln | Q13285 |
| HOMO SAPIENS | 0 | 3071 | 2wpm | E | 145 | ARG | p.Arg191His | P00740 |
| HOMO SAPIENS | 0 | 3071 | 2wpm | E | 145 | ARG | p.Arg191Cys | P00740 |
| HOMO SAPIENS | 3 | 589 | 3ggc | B | 15 | GLY | p.Gly16Ser | P00492 |
| HOMO SAPIENS | 4 | 589 | 3ggc | B | 40 | LEU | p.Leu41Pro | P00492 |
| HOMO SAPIENS | 2 | 589 | 3ggc | B | 50 | ARG | p.Arg51Gly | P00492 |
| HOMO SAPIENS | 5 | 589 | 3ggc | B | 51 | ASP | p.Asp52Gly | P00492 |
| HOMO SAPIENS | 0 | 589 | 3ggc | B | 57 | GLY | p.Gly58Arg | P00492 |
| HOMO SAPIENS | 6 | 589 | 3ggc | B | 69 | GLY | p.Gly70Glu | P00492 |
| HOMO SAPIENS | 1 | 589 | 3ggc | B | 73 | PHE | p.Phe74Leu | P00492 |
| HOMO SAPIENS | 0 | 589 | 3ggc | B | 77 | LEU | p.Leu78Val | P00492 |
| HOMO SAPIENS | 3 | 589 | 3ggc | B | 79 | ASP | p.Asp80Val | P00492 |
| HOMO SAPIENS | 0 | 589 | 3ggc | B | 103 | SER | p.Ser104Arg | P00492 |
| HOMO SAPIENS | 0 | 589 | 3ggc | B | 109 | SER | p.Ser110Leu | P00492 |
| HOMO SAPIENS | 1 | 589 | 3ggc | B | 129 | VAL | p.Val130Asp | P00492 |
| HOMO SAPIENS | 4 | 589 | 3ggc | B | 131 | ILE | p.Ile132Met | P00492 |
| HOMO SAPIENS | 7 | 589 | 3ggc | B | 142 | MET | p.Met143Lys | P00492 |
| HOMO SAPIENS | 0 | 589 | 3ggc | B | 160 | ALA | p.Ala161Ser | P00492 |
| HOMO SAPIENS | 6 | 589 | 3ggc | B | 167 | THR | p.Thr168Ile | P00492 |
| HOMO SAPIENS | 0 | 589 | 3ggc | B | 169 | ARG | p.Arg170Ter | P00492 |
| HOMO SAPIENS | 5 | 589 | 3ggc | B | 175 | PRO | p.Pro176Leu | P00492 |
| HOMO SAPIENS | 8 | 589 | 3ggc | B | 193 | ASP | p.Asp194Glu | P00492 |
| HOMO SAPIENS | 8 | 589 | 3ggc | B | 193 | ASP | p.Asp194Asn | P00492 |
| HOMO SAPIENS | 8 | 589 | 3ggc | B | 198 | PHE | p.Phe199Val | P00492 |
| HOMO SAPIENS | 8 | 589 | 3ggc | B | 200 | ASP | p.Asp201Gly | P00492 |
| HOMO SAPIENS | 0 | 4269 | 2pla | B | 174 | GLU | p.Glu174Lys | Q8N335 |
| HOMO SAPIENS | 1 | 4269 | 2pla | B | 220 | ARG | p.Arg220His | Q8N335 |
| HOMO SAPIENS | 4 | 4269 | 2pla | B | 280 | ALA | p.Ala280Val | Q8N335 |
| HOMO SAPIENS | 0 | 2712 | 1ii4 | E | 252 | TRP | p.Ser252Leu | P21802 |
| HOMO SAPIENS | 0 | 2712 | 1ii4 | E | 253 | PRO | p.Pro253Arg | P21802 |
| HOMO SAPIENS | 5 | 2712 | 1ii4 | E | 267 | SER | p.Ser267Pro | P21802 |
| HOMO SAPIENS | 0 | 2712 | 1ii4 | E | 278 | CYS | p.Cys278Phe | P21802 |
| HOMO SAPIENS | 0 | 2712 | 1ii4 | E | 289 | GLN | p.Gln289Pro | P21802 |
| HOMO SAPIENS | 0 | 2712 | 1ii4 | E | 290 | TRP | p.Trp290Cys | P21802 |
| HOMO SAPIENS | 0 | 2712 | 1ii4 | E | 290 | TRP | p.Trp290Arg | P21802 |
| HOMO SAPIENS | 0 | 2712 | 1ii4 | E | 292 | LYS | p.Lys292Glu | P21802 |
| HOMO SAPIENS | 0 | 2712 | 1ii4 | E | 315 | ALA | p.Ala315Ser | P21802 |
| HOMO SAPIENS | 0 | 2712 | 1ii4 | E | 321 | ASP | p.Asp321Ala | P21802 |
| HOMO SAPIENS | 1 | 2712 | 1ii4 | E | 328 | TYR | p.Tyr328Cys | P21802 |

| | | | | | | | | |
|---|---|---|---|---|---|---|---|---|
| HOMO SAPIENS | 9 | 2712 | 1ii4 | E | 340 | TYR | p.Tyr340His | P21802 |
| HOMO SAPIENS | 1 | 2712 | 1ii4 | E | 341 | THR | p.Thr341Pro | P21802 |
| HOMO SAPIENS | 0 | 2712 | 1ii4 | E | 342 | CYS | p.Cys342Arg | P21802 |
| HOMO SAPIENS | 0 | 2712 | 1ii4 | E | 342 | CYS | p.Cys342Trp | P21802 |
| HOMO SAPIENS | 0 | 2712 | 1ii4 | E | 344 | ALA | p.Ala344Gly | P21802 |
| HOMO SAPIENS | 0 | 2712 | 1ii4 | E | 347 | SER | p.Ser347Cys | P21802 |
| HOMO SAPIENS | 0 | 2712 | 1ii4 | E | 351 | SER | p.Ser351Cys | P21802 |
| HOMO SAPIENS | 1 | 2712 | 1ii4 | E | 354 | SER | p.Ser354Cys | P21802 |
| HOMO SAPIENS | 2 | 53 | 4pj1 | H | 5 | ASP | p.Asp29Gly | P10809 |
| HOMO SAPIENS | 0 | 53 | 4pj1 | H | 74 | VAL | p.Val98Ile | P10809 |
| HOMO SAPIENS | 0 | 29968 | 2fdb | M | 71 | LYS | p.Lys89Glu | P55075 |
| HOMO SAPIENS | 2 | 29968 | 2fdb | M | 98 | ARG | p.Arg116Gly | P55075 |
| HOMO SAPIENS | 2 | 29968 | 2fdb | M | 122 | GLY | p.Gly140Ser | P55075 |
| HOMO SAPIENS | 8 | 5061 | 5f85 | B | 60 | GLY | p.Gly106Ser | P00740 |
| HOMO SAPIENS | 0 | 35944 | 1h6h | A | 85 | THR | p.Thr85Asn | Q15080 |
| HOMO SAPIENS | 8 | 35944 | 1h6h | A | 105 | ARG | p.Arg105Gln | Q15080 |
| HOMO SAPIENS | 7 | 12016 | 1hj7 | A | 295 | ASN | p.Asn316Ser | P01130 |
| HOMO SAPIENS | 5 | 12016 | 1hj7 | A | 302 | GLY | p.Gly323Ser | P01130 |
| HOMO SAPIENS | 6 | 12016 | 1hj7 | A | 319 | CYS | p.Cys340Trp | P01130 |
| HOMO SAPIENS | 7 | 12016 | 1hj7 | A | 329 | ARG | p.Arg350Ter | P01130 |
| HOMO SAPIENS | 5 | 12016 | 1hj7 | A | 331 | CYS | p.Cys352Ter | P01130 |
| HOMO SAPIENS | 2 | 12016 | 1hj7 | A | 345 | GLN | p.Gln366Arg | P01130 |
| HOMO SAPIENS | 2 | 1481 | 2i07 | A | 239 | TYR | p.Tyr261Ter | P01024 |
| HOMO SAPIENS | -1 | 1481 | 2i07 | A | 292 | LEU | p.Pro314Leu | P01024 |
| HOMO SAPIENS | 2 | 1481 | 2i07 | A | 326 | ILE | p.Ile348Val | P01024 |
| HOMO SAPIENS | 2 | 1481 | 2i07 | A | 413 | GLU | p.Glu435Lys | P01024 |
| HOMO SAPIENS | 4 | 1481 | 2i07 | A | 447 | GLU | p.Glu469Asp | P01024 |
| HOMO SAPIENS | 2 | 1481 | 2i07 | A | 597 | VAL | p.Val619Met | P01024 |
| HOMO SAPIENS | 5 | 1481 | 2i07 | A | 603 | ILE | p.Ile625Phe | P01024 |
| HOMO SAPIENS | 0 | 1481 | 2i07 | A | 611 | LYS | p.Lys633Arg | P01024 |
| HOMO SAPIENS | 0 | 11123 | 3c5n | A | 315 | GLY | p.Gly368Trp | O00294 |
| HOMO SAPIENS | 8 | 11123 | 3c5n | A | 329 | PHE | p.Phe382Ser | O00294 |
| HOMO SAPIENS | 5 | 11123 | 3c5n | A | 429 | ARG | p.Arg482Trp | O00294 |
| HOMO SAPIENS | 8 | 11123 | 3c5n | A | 438 | PHE | p.Phe491Leu | O00294 |
| HOMO SAPIENS | 3 | 4398 | 3fx0 | B | 1288 | ALA | p.Ala288Gly | Q9Y6K9 |
| HOMO SAPIENS | 7 | 4398 | 3fx0 | B | 1315 | GLU | p.Glu315Ala | Q9Y6K9 |
| HOMO SAPIENS | 8 | 4398 | 3fx0 | B | 1319 | ARG | p.Arg319Gln | Q9Y6K9 |
| HOMO SAPIENS | 6 | 3989 | 3qd6 | S | 83 | CYS | p.Cys83Arg | P25942 |
| HOMO SAPIENS | 1 | 3989 | 3qd6 | S | 124 | SER | p.Ser124Leu | P25942 |
| HOMO SAPIENS | 2 | 24003 | 2m0v | A | 225 | GLU | p.Glu225Lys | O14745 |
| HOMO SAPIENS | 2 | 46 | 4aqd | A | 70 | ASP | p.Asp98Gly | P06276 |
| HOMO SAPIENS | 4 | 46 | 4aqd | A | 115 | GLY | p.Gly143Asp | P06276 |
| HOMO SAPIENS | 3 | 46 | 4aqd | A | 184 | ALA | p.Ala212Val | P06276 |
| HOMO SAPIENS | 0 | 46 | 4aqd | A | 225 | GLY | p.Gly253Ter | P06276 |
| HOMO SAPIENS | 2 | 46 | 4aqd | A | 243 | THR | p.Thr271Met | P06276 |
| HOMO SAPIENS | 2 | 46 | 4aqd | A | 255 | GLU | p.Glu283Asp | P06276 |
| HOMO SAPIENS | 5 | 46 | 4aqd | A | 267 | LYS | p.Lys295Arg | P06276 |
| HOMO SAPIENS | 3 | 46 | 4aqd | A | 271 | GLU | p.Glu299Ter | P06276 |
| HOMO SAPIENS | 2 | 46 | 4aqd | A | 307 | LEU | p.Leu335Pro | P06276 |
| HOMO SAPIENS | 1 | 46 | 4aqd | A | 330 | LEU | p.Leu358Ile | P06276 |
| HOMO SAPIENS | 0 | 46 | 4aqd | A | 497 | GLU | p.Glu525Val | P06276 |
| HOMO SAPIENS | 3 | 786 | 1ols | B | 51 | GLY | p.Gly101Asp | P21953 |
| HOMO SAPIENS | 2 | 786 | 1ols | B | 87 | ALA | p.Ala137Val | P21953 |
| HOMO SAPIENS | 0 | 786 | 1ols | B | 110 | ILE | p.Ile160Ser | P21953 |

| | | | | | | | | |
|---|---|---|---|---|---|---|---|---|
| HOMO SAPIENS | 4 | 786 | 1ols | B | 120 | ARG | p.Arg170Cys | P21953 |
| HOMO SAPIENS | 2 | 786 | 1ols | B | 126 | ASN | p.Asn176Tyr | P21953 |
| HOMO SAPIENS | 0 | 786 | 1ols | B | 133 | ARG | p.Arg183Gly | P21953 |
| HOMO SAPIENS | 0 | 786 | 1ols | B | 133 | ARG | p.Arg183Pro | P21953 |
| HOMO SAPIENS | 0 | 786 | 1ols | B | 135 | PRO | p.Pro185Leu | P21953 |
| HOMO SAPIENS | 3 | 786 | 1ols | B | 156 | HIS | p.His206Tyr | P21953 |
| HOMO SAPIENS | 0 | 786 | 1ols | B | 228 | GLY | p.Gly278Ser | P21953 |
| HOMO SAPIENS | 1 | 786 | 1ols | B | 235 | ARG | p.Arg285Ter | P21953 |
| HOMO SAPIENS | 6 | 786 | 1ols | B | 274 | ARG | p.Arg324Ter | P21953 |
| HOMO SAPIENS | 1 | 831 | 4zt1 | B | 48 | VAL | p.Val202Ile | P12830 |
| HOMO SAPIENS | 6 | 831 | 4zt1 | B | 70 | ARG | p.Arg224Cys | P12830 |
| HOMO SAPIENS | 5 | 831 | 4zt1 | B | 79 | HIS | p.His233Gln | P12830 |
| HOMO SAPIENS | 4 | 831 | 4zt1 | B | 120 | GLY | p.Gly274Ser | P12830 |
| HOMO SAPIENS | 7 | 831 | 4zt1 | B | 187 | TYR | p.Tyr341Ter | P12830 |
| HOMO SAPIENS | 0 | 8497 | 3sla | A | 287 | ASN | p.Asn287Ser | P35222 |
| HOMO SAPIENS | 7 | 23986 | 2ke1 | A | 301 | VAL | p.Val301Met | O43918 |
| HOMO SAPIENS | 0 | 4967 | 1rj7 | B | 334 | ARG | p.Arg334His | Q92838 |
| HOMO SAPIENS | 0 | 4967 | 1rj7 | B | 338 | THR | p.Thr338Met | Q92838 |
| HOMO SAPIENS | 3 | 4967 | 1rj7 | B | 349 | ALA | p.Ala349Thr | Q92838 |
| HOMO SAPIENS | 2 | 4967 | 1rj7 | B | 358 | GLN | p.Gln358Glu | Q92838 |
| HOMO SAPIENS | 1 | 4967 | 1rj7 | B | 375 | LYS | p.Lys375Glu | Q92838 |
| HOMO SAPIENS | 9 | 7623 | 1d4t | A | 32 | ARG | p.Arg32Thr | O60880 |
| HOMO SAPIENS | 6 | 7623 | 1d4t | A | 55 | ARG | p.Arg55Leu | O60880 |
| HOMO SAPIENS | 0 | 7623 | 1d4t | A | 68 | THR | p.Thr68Ile | O60880 |
| HOMO SAPIENS | 6 | 705 | 3idh | A | 59 | PRO | p.Pro59Ser | P35557 |
| HOMO SAPIENS | 5 | 705 | 3idh | A | 124 | ASP | p.Asp124Asn | P35557 |
| HOMO SAPIENS | 4 | 705 | 3idh | A | 131 | SER | p.Ser131Pro | P35557 |
| HOMO SAPIENS | 0 | 705 | 3idh | A | 147 | GLY | p.Gly147Asp | P35557 |
| HOMO SAPIENS | 5 | 705 | 3idh | A | 186 | ARG | p.Arg186Ter | P35557 |
| HOMO SAPIENS | 0 | 705 | 3idh | A | 202 | MET | p.Met202Thr | P35557 |
| HOMO SAPIENS | 2 | 705 | 3idh | A | 210 | MET | p.Met210Lys | P35557 |
| HOMO SAPIENS | 1 | 705 | 3idh | A | 215 | TYR | p.Tyr215Ter | P35557 |
| HOMO SAPIENS | 0 | 705 | 3idh | A | 226 | VAL | p.Val226Met | P35557 |
| HOMO SAPIENS | 9 | 705 | 3idh | A | 228 | THR | p.Thr228Met | P35557 |
| HOMO SAPIENS | 9 | 705 | 3idh | A | 256 | GLU | p.Glu256Lys | P35557 |
| HOMO SAPIENS | 2 | 705 | 3idh | A | 261 | GLY | p.Gly261Arg | P35557 |
| HOMO SAPIENS | 6 | 705 | 3idh | A | 279 | GLU | p.Glu279Gln | P35557 |
| HOMO SAPIENS | 7 | 705 | 3idh | A | 299 | GLY | p.Gly299Arg | P35557 |
| HOMO SAPIENS | 2 | 705 | 3idh | A | 318 | GLY | p.Gly318Trp | P35557 |
| HOMO SAPIENS | 4 | 705 | 3idh | A | 378 | ALA | p.Ala378Thr | P35557 |
| HOMO SAPIENS | 0 | 705 | 3idh | A | 387 | ALA | p.Ala387Glu | P35557 |
| HOMO SAPIENS | 5 | 705 | 3idh | A | 414 | LYS | p.Lys414Glu | P35557 |
| HOMO SAPIENS | 0 | 705 | 3idh | A | 455 | VAL | p.Val455Met | P35557 |
| HOMO SAPIENS | -1 | 705 | 3idh | A | 462 | MET | p.Met462Ile | P35557 |
| HOMO SAPIENS | 1 | 509 | 4nef | A | 22 | LEU | p.Leu22Val | P41181 |
| HOMO SAPIENS | 4 | 509 | 4nef | A | 57 | GLN | p.Gln57Pro | P41181 |
| HOMO SAPIENS | 8 | 509 | 4nef | A | 64 | GLY | p.Gly64Arg | P41181 |
| HOMO SAPIENS | 7 | 509 | 4nef | A | 68 | ASN | p.Asn68Ser | P41181 |
| HOMO SAPIENS | 1 | 509 | 4nef | A | 100 | GLY | p.Gly100Val | P41181 |
| HOMO SAPIENS | 0 | 509 | 4nef | A | 125 | THR | p.Thr125Met | P41181 |
| HOMO SAPIENS | 0 | 509 | 4nef | A | 126 | THR | p.Thr126Met | P41181 |
| HOMO SAPIENS | 4 | 509 | 4nef | A | 146 | PHE | p.Phe146Leu | P41181 |
| HOMO SAPIENS | 0 | 509 | 4nef | A | 175 | GLY | p.Gly175Arg | P41181 |
| HOMO SAPIENS | 5 | 509 | 4nef | A | 181 | CYS | p.Cys181Trp | P41181 |

| | | | | | | | | | |
|---|---|---|---|---|---|---|---|---|---|
| HOMO SAPIENS | 9 | 509 | 4nef | A | | 187 | ARG | p.Arg187Cys | P41181 |
| HOMO SAPIENS | 0 | 509 | 4nef | A | | 190 | ALA | p.Ala190Thr | P41181 |
| HOMO SAPIENS | 0 | 509 | 4nef | A | | 216 | SER | p.Ser216Pro | P41181 |
| HOMO SAPIENS | 4 | 2242 | 1z70 | X | | 1155 | SER | p.Ser155Pro | Q8NBK3 |
| HOMO SAPIENS | 3 | 2242 | 1z70 | X | | 1218 | CYS | p.Cys218Tyr | Q8NBK3 |
| HOMO SAPIENS | 0 | 2242 | 1z70 | X | | 1279 | ALA | p.Ala279Val | Q8NBK3 |
| HOMO SAPIENS | -1 | 2242 | 1z70 | X | | 1336 | OCS | p.Cys336Arg | Q8NBK3 |
| HOMO SAPIENS | 4 | 2242 | 1z70 | X | | 1345 | ARG | p.Arg345Cys | Q8NBK3 |
| HOMO SAPIENS | 2 | 2242 | 1z70 | X | | 1348 | ALA | p.Ala348Pro | Q8NBK3 |
| HOMO SAPIENS | 5 | 2242 | 1z70 | X | | 1349 | ARG | p.Arg349Gln | Q8NBK3 |
| HOMO SAPIENS | 5 | 2242 | 1z70 | X | | 1349 | ARG | p.Arg349Trp | Q8NBK3 |
| HOMO SAPIENS | 3 | 2242 | 1z70 | X | | 1359 | SER | p.Ser359Ter | Q8NBK3 |
| HOMO SAPIENS | 5 | 8269 | 5wrr | B | | 136 | ARG | p.Arg136Ter | Q96MK3 |
| HOMO SAPIENS | 1 | 8269 | 5wrr | B | | 332 | LYS | p.Asn332Lys | Q96MK3 |
| HOMO SAPIENS | 3 | 8269 | 5wrr | B | | 478 | ARG | p.Arg478Ter | Q96MK3 |
| HOMO SAPIENS | 5 | 8269 | 5wrr | B | | 504 | THR | p.Thr504Ile | Q96MK3 |
| HOMO SAPIENS | 5 | 1411 | 4h1s | B | | 221 | SER | p.Ser221Ter | P21589 |
| HOMO SAPIENS | 1 | 1411 | 4h1s | B | | 358 | CYS | p.Cys358Tyr | P21589 |
| HOMO SAPIENS | 2 | 23 | 5vvd | B | | 298 | GLU | p.Glu298Asp | P29474 |
| HOMO SAPIENS | 0 | 19 | 5jlh | E | | 88 | THR | p.Thr89Ile | P63261 |
| HOMO SAPIENS | 6 | 19 | 5jlh | E | | 117 | LYS | p.Lys118Met | P63261 |
| HOMO SAPIENS | 5 | 19 | 5jlh | E | | 263 | PRO | p.Pro264Leu | P63261 |
| HOMO SAPIENS | 1 | 19 | 5jlh | E | | 277 | THR | p.Thr278Ile | P63261 |
| HOMO SAPIENS | 6 | 19 | 5jlh | E | | 331 | PRO | p.Pro332Ala | P63261 |
| HOMO SAPIENS | 5 | 19 | 5jlh | E | | 369 | VAL | p.Val370Ala | P63261 |
| HOMO SAPIENS | 0 | 14306 | 4w4o | C | | 92 | ARG | p.Arg92Ter | P12314 |
| HOMO SAPIENS | 3 | 603 | 1nuh | A | | 4 | THR | p.Thr5Ile | P06744 |
| HOMO SAPIENS | 2 | 603 | 1nuh | A | | 19 | HIS | p.His20Pro | P06744 |
| HOMO SAPIENS | 8 | 603 | 1nuh | A | | 158 | GLY | p.Gly159Ser | P06744 |
| HOMO SAPIENS | 6 | 603 | 1nuh | A | | 223 | THR | p.Thr224Met | P06744 |
| HOMO SAPIENS | 0 | 603 | 1nuh | A | | 338 | LEU | p.Leu339Pro | P06744 |
| HOMO SAPIENS | 6 | 603 | 1nuh | A | | 342 | GLN | p.Gln343Arg | P06744 |
| HOMO SAPIENS | 6 | 603 | 1nuh | A | | 374 | THR | p.Thr375Arg | P06744 |
| HOMO SAPIENS | 4 | 603 | 1nuh | A | | 524 | ILE | p.Ile525Thr | P06744 |
| HOMO SAPIENS | 4 | 603 | 1nuh | A | | 538 | ASP | p.Asp539Asn | P06744 |
| HOMO SAPIENS | 1 | 3107 | 3kzj | A | | 1197 | VAL | p.Val1197Ala | P08603 |
| HOMO SAPIENS | 9 | 1748 | 2noh | A | | 46 | ARG | p.Arg46Gln | O15527 |
| HOMO SAPIENS | 2 | 11750 | 3s4y | A | | 33 | ARG | p.Arg33His | Q9H3S4 |
| HOMO SAPIENS | 2 | 11750 | 3s4y | A | | 40 | LEU | p.Leu40Pro | Q9H3S4 |
| HOMO SAPIENS | 4 | 11750 | 3s4y | A | | 142 | LEU | p.Leu142Phe | Q9H3S4 |
| HOMO SAPIENS | 5 | 11750 | 3s4y | A | | 219 | ASN | p.Asn219Ser | Q9H3S4 |
| HOMO SAPIENS | 1 | 2591 | 5vnj | A | | 382 | PHE | p.Phe382Leu | Q15436 |
| HOMO SAPIENS | 2 | 2591 | 5vnj | A | | 702 | MET | p.Met702Val | Q15436 |
| HOMO SAPIENS | 1 | 2426 | 5m8t | A | | 87 | ARG | p.Arg87Trp | P17643 |
| HOMO SAPIENS | 1 | 2426 | 5m8t | A | | 166 | SER | p.Ser166Ter | P17643 |
| HOMO SAPIENS | 0 | 2426 | 5m8t | A | | 356 | ARG | p.Arg356Gln | P17643 |
| HOMO SAPIENS | 2 | 2426 | 5m8t | A | | 374 | SER | p.Arg374Ter | P17643 |
| HOMO SAPIENS | 6 | 2426 | 5m8t | A | | 378 | ASN | p.Asn378Ser | P17643 |
| HOMO SAPIENS | 6 | 2426 | 5m8t | A | | 382 | LEU | p.Leu382Pro | P17643 |
| HOMO SAPIENS | 7 | 10097 | 4n6v | | 1 | 46 | GLN | p.Gln46Ter | P04080 |
| HOMO SAPIENS | 3 | 10097 | 4n6v | | 1 | 65 | VAL | p.Val65Ile | P04080 |
| HOMO SAPIENS | 8 | 10097 | 4n6v | | 1 | 68 | ARG | p.Arg68Ter | P04080 |
| HOMO SAPIENS | 2 | 859 | 4z9a | A | | 105 | GLN | p.Gln106Arg | P24666 |
| HOMO SAPIENS | 8 | 1420 | 5nan | F | | 95 | SER | p.Ser95Leu | Q96PD4 |

| | | | | | | | | |
|---|---|---|---|---|---|---|---|---|
| HOMO SAPIENS | 6 | 1420 | 5nan | F | 126 | GLU | p.Glu126Gly | Q96PD4 |
| HOMO SAPIENS | 8 | 1420 | 5nan | F | 155 | VAL | p.Val155Ile | Q96PD4 |
| HOMO SAPIENS | 6 | 1420 | 5nan | F | 157 | PRO | p.Pro157Ser | Q96PD4 |
| HOMO SAPIENS | -1 | 1420 | 5nan | F | 161 | HIS | p.His161Arg | Q96PD4 |
| HOMO SAPIENS | 3 | 288 | 5i96 | B | 140 | GLN | p.Arg140Gly | P48735 |
| HOMO SAPIENS | 3 | 288 | 5i96 | B | 140 | GLN | p.Arg140Gln | P48735 |
| HOMO SAPIENS | 2 | 288 | 5i96 | B | 261 | ARG | p.Arg261His | P48735 |
| HOMO SAPIENS | 2 | 288 | 5i96 | B | 435 | THR | p.Thr435Met | P48735 |
| HOMO SAPIENS | 9 | 120 | 1tup | B | 107 | TYR | p.Tyr107His | P04637 |
| HOMO SAPIENS | 8 | 120 | 1tup | B | 110 | ARG | p.Arg110Leu | P04637 |
| HOMO SAPIENS | 5 | 120 | 1tup | B | 120 | LYS | p.Lys120Glu | P04637 |
| HOMO SAPIENS | 4 | 120 | 1tup | B | 133 | MET | p.Met133Thr | P04637 |
| HOMO SAPIENS | 3 | 120 | 1tup | B | 134 | PHE | p.Phe134Cys | P04637 |
| HOMO SAPIENS | 8 | 120 | 1tup | B | 136 | GLN | p.Gln136His | P04637 |
| HOMO SAPIENS | 9 | 120 | 1tup | B | 138 | ALA | p.Ala138Pro | P04637 |
| HOMO SAPIENS | 9 | 120 | 1tup | B | 151 | PRO | p.Pro151Ser | P04637 |
| HOMO SAPIENS | 9 | 120 | 1tup | B | 152 | PRO | p.Pro152Leu | P04637 |
| HOMO SAPIENS | 9 | 120 | 1tup | B | 158 | ARG | p.Arg158Cys | P04637 |
| HOMO SAPIENS | 9 | 120 | 1tup | B | 158 | ARG | p.Arg158His | P04637 |
| HOMO SAPIENS | 9 | 120 | 1tup | B | 175 | ARG | p.Arg175Cys | P04637 |
| HOMO SAPIENS | 9 | 120 | 1tup | B | 175 | ARG | p.Arg175Leu | P04637 |
| HOMO SAPIENS | 8 | 120 | 1tup | B | 181 | ARG | p.Arg181Leu | P04637 |
| HOMO SAPIENS | 9 | 120 | 1tup | B | 189 | ALA | p.Ala189Val | P04637 |
| HOMO SAPIENS | 9 | 120 | 1tup | B | 196 | ARG | p.Arg196Gly | P04637 |
| HOMO SAPIENS | 9 | 120 | 1tup | B | 213 | ARG | p.Arg213Gln | P04637 |
| HOMO SAPIENS | 9 | 120 | 1tup | B | 213 | ARG | p.Arg213Ter | P04637 |
| HOMO SAPIENS | 9 | 120 | 1tup | B | 214 | HIS | p.His214Gln | P04637 |
| HOMO SAPIENS | 9 | 120 | 1tup | B | 220 | TYR | p.Tyr220Cys | P04637 |
| HOMO SAPIENS | 0 | 120 | 1tup | B | 234 | TYR | p.Tyr234Cys | P04637 |
| HOMO SAPIENS | 9 | 120 | 1tup | B | 237 | MET | p.Met237Ile | P04637 |
| HOMO SAPIENS | 0 | 120 | 1tup | B | 238 | CYS | p.Cys238Tyr | P04637 |
| HOMO SAPIENS | 9 | 120 | 1tup | B | 241 | SER | p.Ser241Phe | P04637 |
| HOMO SAPIENS | 9 | 120 | 1tup | B | 242 | CYS | p.Cys242Ser | P04637 |
| HOMO SAPIENS | 9 | 120 | 1tup | B | 245 | GLY | p.Gly245Cys | P04637 |
| HOMO SAPIENS | 9 | 120 | 1tup | B | 245 | GLY | p.Gly245Asp | P04637 |
| HOMO SAPIENS | 9 | 120 | 1tup | B | 248 | ARG | p.Arg248Leu | P04637 |
| HOMO SAPIENS | 9 | 120 | 1tup | B | 248 | ARG | p.Arg248Trp | P04637 |
| HOMO SAPIENS | 9 | 120 | 1tup | B | 251 | ILE | p.Ile251Ser | P04637 |
| HOMO SAPIENS | 9 | 120 | 1tup | B | 252 | LEU | p.Leu252Pro | P04637 |
| HOMO SAPIENS | 9 | 120 | 1tup | B | 258 | GLU | p.Glu258Lys | P04637 |
| HOMO SAPIENS | 8 | 120 | 1tup | B | 272 | VAL | p.Val272Leu | P04637 |
| HOMO SAPIENS | 9 | 120 | 1tup | B | 273 | ARG | p.Arg273Cys | P04637 |
| HOMO SAPIENS | 9 | 120 | 1tup | B | 273 | ARG | p.Arg273His | P04637 |
| HOMO SAPIENS | 9 | 120 | 1tup | B | 277 | CYS | p.Cys277Phe | P04637 |
| HOMO SAPIENS | 9 | 120 | 1tup | B | 280 | ARG | p.Arg280Ile | P04637 |
| HOMO SAPIENS | 9 | 120 | 1tup | B | 282 | ARG | p.Arg282Trp | P04637 |
| HOMO SAPIENS | 9 | 120 | 1tup | B | 283 | ARG | p.Arg283His | P04637 |
| HOMO SAPIENS | 9 | 120 | 1tup | B | 285 | GLU | p.Glu285Val | P04637 |
| HOMO SAPIENS | -1 | 120 | 1tup | B | 292 | LYS | p.Lys292Ile | P04637 |
| HOMO SAPIENS | -1 | 120 | 1tup | B | 298 | GLU | p.Glu298Lys | P04637 |
| HOMO SAPIENS | -1 | 2791 | 4ygb | A | 39 | VAL | p.Val39Ala | P49257 |
| HOMO SAPIENS | 2 | 106 | 4x6j | A | 32 | GLY | p.Gly146Arg | P43235 |
| HOMO SAPIENS | 0 | 106 | 4x6j | A | 76 | GLN | p.Gln190Ter | P43235 |
| HOMO SAPIENS | 7 | 106 | 4x6j | A | 198 | ARG | p.Arg312Ter | P43235 |

| | | | | | | | | |
|---|---|---|---|---|---|---|---|---|
| HOMO SAPIENS | 5 | 490 | 2hlo | B | 172 | LEU | p.Leu202Gln | P02675 |
| HOMO SAPIENS | 0 | 490 | 2hlo | B | 353 | LEU | p.Leu383Arg | P02675 |
| HOMO SAPIENS | 4 | 490 | 2hlo | B | 400 | GLY | p.Gly430Asp | P02675 |
| HOMO SAPIENS | 7 | 490 | 2hlo | B | 448 | ARG | p.Arg478Lys | P02675 |
| HOMO SAPIENS | 1 | 289 | 4p13 | B | 18 | GLU | p.Glu43Lys | P11310 |
| HOMO SAPIENS | 2 | 289 | 4p13 | B | 20 | GLN | p.Gln45Arg | P11310 |
| HOMO SAPIENS | 5 | 289 | 4p13 | B | 28 | ARG | p.Arg53Cys | P11310 |
| HOMO SAPIENS | 1 | 289 | 4p13 | B | 42 | TYR | p.Tyr67His | P11310 |
| HOMO SAPIENS | 1 | 289 | 4p13 | B | 53 | ILE | p.Ile78Thr | P11310 |
| HOMO SAPIENS | 1 | 289 | 4p13 | B | 59 | LEU | p.Leu84Phe | P11310 |
| HOMO SAPIENS | 2 | 289 | 4p13 | B | 96 | THR | p.Thr121Ile | P11310 |
| HOMO SAPIENS | 0 | 289 | 4p13 | B | 124 | MET | p.Met149Ile | P11310 |
| HOMO SAPIENS | 5 | 289 | 4p13 | B | 143 | ASP | p.Asp168Ala | P11310 |
| HOMO SAPIENS | 7 | 289 | 4p13 | B | 168 | THR | p.Thr193Ala | P11310 |
| HOMO SAPIENS | 1 | 289 | 4p13 | B | 170 | GLY | p.Gly195Arg | P11310 |
| HOMO SAPIENS | 2 | 289 | 4p13 | B | 181 | ARG | p.Arg206Cys | P11310 |
| HOMO SAPIENS | 2 | 289 | 4p13 | B | 181 | ARG | p.Arg206His | P11310 |
| HOMO SAPIENS | 8 | 289 | 4p13 | B | 203 | THR | p.Thr228Ile | P11310 |
| HOMO SAPIENS | 2 | 289 | 4p13 | B | 208 | ILE | p.Ile233Asn | P11310 |
| HOMO SAPIENS | 1 | 289 | 4p13 | B | 219 | CYS | p.Cys244Arg | P11310 |
| HOMO SAPIENS | 6 | 289 | 4p13 | B | 220 | SER | p.Ser245Leu | P11310 |
| HOMO SAPIENS | 5 | 289 | 4p13 | B | 228 | GLU | p.Glu253Lys | P11310 |
| HOMO SAPIENS | 2 | 289 | 4p13 | B | 241 | ASP | p.Asp266Gly | P11310 |
| HOMO SAPIENS | 1 | 289 | 4p13 | B | 242 | GLY | p.Gly267Arg | P11310 |
| HOMO SAPIENS | 7 | 289 | 4p13 | B | 256 | ARG | p.Arg281Thr | P11310 |
| HOMO SAPIENS | 4 | 289 | 4p13 | B | 269 | ARG | p.Arg294Thr | P11310 |
| HOMO SAPIENS | 4 | 289 | 4p13 | B | 304 | GLU | p.Lys329Glu | P11310 |
| HOMO SAPIENS | 1 | 289 | 4p13 | B | 324 | ARG | p.Arg349Ter | P11310 |
| HOMO SAPIENS | 3 | 289 | 4p13 | B | 326 | THR | p.Thr351Ile | P11310 |
| HOMO SAPIENS | 7 | 289 | 4p13 | B | 350 | ILE | p.Ile375Thr | P11310 |
| HOMO SAPIENS | 3 | 289 | 4p13 | B | 359 | GLU | p.Glu384Ter | P11310 |
| HOMO SAPIENS | 1 | 4563 | 3i2b | A | 16 | ARG | p.Arg16Cys | Q03393 |
| HOMO SAPIENS | 5 | 4563 | 3i2b | A | 25 | ARG | p.Arg25Gln | Q03393 |
| HOMO SAPIENS | 1 | 4563 | 3i2b | A | 47 | ASN | p.Asn47Asp | Q03393 |
| HOMO SAPIENS | 8 | 4563 | 3i2b | A | 52 | ASN | p.Asn52Ser | Q03393 |
| HOMO SAPIENS | 1 | 4563 | 3i2b | A | 56 | VAL | p.Val56Met | Q03393 |
| HOMO SAPIENS | 1 | 4563 | 3i2b | A | 87 | PRO | p.Pro87Ser | Q03393 |
| HOMO SAPIENS | 1 | 4563 | 3i2b | A | 96 | ASP | p.Asp96Asn | Q03393 |
| HOMO SAPIENS | 0 | 4563 | 3i2b | A | 99 | TYR | p.Tyr99Ter | Q03393 |
| HOMO SAPIENS | 2 | 4563 | 3i2b | A | 116 | ASP | p.Asp116Gly | Q03393 |
| HOMO SAPIENS | 2 | 41 | 3exx | A | 1 | GLY | p.Gly90Cys | P01308 |
| HOMO SAPIENS | 5 | 41 | 3exx | A | 3 | VAL | p.Val92Leu | P01308 |
| HOMO SAPIENS | 4 | 41 | 3exx | A | 7 | CYS | p.Cys96Ser | P01308 |
| HOMO SAPIENS | 9 | 41 | 3exx | A | 19 | TYR | p.Tyr108Cys | P01308 |
| HOMO SAPIENS | 9 | 131 | 4mjo | D | 118 | ASP | p.Asp119Asn | P09467 |
| HOMO SAPIENS | 0 | 131 | 4mjo | D | 163 | GLY | p.Gly164Ser | P09467 |
| HOMO SAPIENS | 0 | 131 | 4mjo | D | 176 | ALA | p.Ala177Asp | P09467 |
| HOMO SAPIENS | 5 | 131 | 4mjo | D | 193 | PHE | p.Phe194Ser | P09467 |
| HOMO SAPIENS | 4 | 131 | 4mjo | D | 217 | ARG | p.Arg218Lys | P09467 |
| HOMO SAPIENS | 4 | 131 | 4mjo | D | 223 | VAL | p.Val224Ile | P09467 |
| HOMO SAPIENS | 4 | 131 | 4mjo | D | 232 | PHE | p.Phe233Ile | P09467 |
| HOMO SAPIENS | 5 | 131 | 4mjo | D | 243 | ARG | p.Arg244Trp | P09467 |
| HOMO SAPIENS | 0 | 131 | 4mjo | D | 283 | PRO | p.Pro284Arg | P09467 |
| HOMO SAPIENS | 0 | 131 | 4mjo | D | 288 | MET | p.Met289Leu | P09467 |

| | | | | | | | | |
|---|---|---|---|---|---|---|---|---|
| HOMO SAPIENS | 0 | 131 | 4mjo | D | 323 | ASP | p.Asp324Asn | P09467 |
| HOMO SAPIENS | -1 | 13122 | 3hfc | A | 583 | ARG | p.Arg583Cys | P51787 |
| HOMO SAPIENS | 1 | 13122 | 3hfc | A | 587 | THR | p.Thr587Met | P51787 |
| HOMO SAPIENS | 0 | 13122 | 3hfc | A | 589 | GLY | p.Gly589Asp | P51787 |
| HOMO SAPIENS | 4 | 3002 | 3nbq | B | 94 | ARG | p.Arg94Cys | Q16831 |
| HOMO SAPIENS | 1 | 3616 | 3dbp | A | 97 | ILE | p.Ile286Asn | P11172 |
| HOMO SAPIENS | 0 | 3616 | 3dbp | A | 240 | GLY | p.Gly429Arg | P11172 |
| HOMO SAPIENS | 0 | 3616 | 3dbp | A | 257 | ILE | p.Ile446Val | P11172 |
| HOMO SAPIENS | 1 | 584 | 1drf | A | 79 | LEU | p.Leu80Phe | P00374 |
| HOMO SAPIENS | 3 | 1609 | 3qa2 | B | 49 | VAL | p.Val49Ile | P50053 |
| HOMO SAPIENS | 0 | 20560 | 5mug | A | 101 | HIS | p.His101Gln | P49638 |
| HOMO SAPIENS | 2 | 20560 | 5mug | A | 120 | ALA | p.Ala120Thr | P49638 |
| HOMO SAPIENS | 3 | 20560 | 5mug | A | 192 | ARG | p.Arg192His | P49638 |
| HOMO SAPIENS | 3 | 20560 | 5mug | A | 221 | ARG | p.Arg221Trp | P49638 |
| HOMO SAPIENS | 4 | 256 | 3vtv | A | 65 | THR | p.Glu50Lys | Q9GZQ8 |
| HOMO SAPIENS | 4 | 256 | 3vtv | A | 113 | VAL | p.Met98Lys | Q9GZQ8 |
| HOMO SAPIENS | 4 | 5397 | 1m61 | A | 192 | ARG | p.Arg192Trp | P43403 |
| HOMO SAPIENS | 5 | 39881 | 1aut | L | 107 | HIS | p.His149Pro | P04070 |
| HOMO SAPIENS | 5 | 4010 | 4uf8 | I | 33 | ALA | p.Ala33Val | Q13501 |
| HOMO SAPIENS | 3 | 313 | 5w49 | A | 98 | PRO | p.Pro98Leu | P23526 |
| HOMO SAPIENS | 2 | 313 | 5w49 | A | 143 | TYR | p.Tyr143Cys | P23526 |
| HOMO SAPIENS | -1 | 1643 | 2m8t | A | 102 | PRO | p.Pro102Leu | P04156 |
| HOMO SAPIENS | -1 | 1643 | 2m8t | A | 105 | PRO | p.Pro105Leu | P04156 |
| HOMO SAPIENS | -1 | 1643 | 2m8t | A | 117 | ALA | p.Ala117Val | P04156 |
| HOMO SAPIENS | 5 | 1643 | 2m8t | A | 129 | MET | p.Met129Val | P04156 |
| HOMO SAPIENS | 5 | 1643 | 2m8t | A | 131 | GLY | p.Gly131Val | P04156 |
| HOMO SAPIENS | 2 | 1643 | 2m8t | A | 142 | GLY | p.Gly142Ser | P04156 |
| HOMO SAPIENS | 3 | 1643 | 2m8t | A | 171 | ASN | p.Asn171Ser | P04156 |
| HOMO SAPIENS | 9 | 1643 | 2m8t | A | 178 | ASP | p.Asp178Asn | P04156 |
| HOMO SAPIENS | 9 | 1643 | 2m8t | A | 180 | VAL | p.Val180Ile | P04156 |
| HOMO SAPIENS | 9 | 1643 | 2m8t | A | 183 | THR | p.Thr183Ala | P04156 |
| HOMO SAPIENS | 9 | 1643 | 2m8t | A | 187 | HIS | p.His187Arg | P04156 |
| HOMO SAPIENS | 8 | 1643 | 2m8t | A | 198 | PHE | p.Phe198Ser | P04156 |
| HOMO SAPIENS | 9 | 1643 | 2m8t | A | 200 | GLU | p.Glu200Lys | P04156 |
| HOMO SAPIENS | 9 | 1643 | 2m8t | A | 208 | ARG | p.Arg208His | P04156 |
| HOMO SAPIENS | 3 | 1643 | 2m8t | A | 210 | VAL | p.Val210Ile | P04156 |
| HOMO SAPIENS | 6 | 1643 | 2m8t | A | 217 | GLN | p.Gln217Arg | P04156 |
| HOMO SAPIENS | 1 | 29 | 5kos | A | 151 | GLY | p.Gly217Arg | P00797 |
| HOMO SAPIENS | 1 | 29 | 5kos | A | 164 | ARG | p.Arg230Lys | P00797 |
| HOMO SAPIENS | 3 | 504 | 4zkv | B | 37 | ARG | p.Arg37Pro | P49773 |
| HOMO SAPIENS | 2 | 504 | 4zkv | B | 62 | GLN | p.Gln62Ter | P49773 |
| HOMO SAPIENS | 2 | 504 | 4zkv | B | 84 | CYS | p.Cys84Arg | P49773 |
| HOMO SAPIENS | 4 | 504 | 4zkv | B | 93 | GLY | p.Gly93Asp | P49773 |
| HOMO SAPIENS | 9 | 504 | 4zkv | B | 112 | HIS | p.His112Asn | P49773 |
| HOMO SAPIENS | 0 | 74 | 5vwo | C | 54 | ASN | p.Asn54Lys | P04181 |
| HOMO SAPIENS | 0 | 74 | 5vwo | C | 55 | TYR | p.Tyr55His | P04181 |
| HOMO SAPIENS | 2 | 74 | 5vwo | C | 90 | GLN | p.Gln90Glu | P04181 |
| HOMO SAPIENS | 6 | 74 | 5vwo | C | 91 | GLY | p.Gly91Glu | P04181 |
| HOMO SAPIENS | 2 | 74 | 5vwo | C | 93 | CYS | p.Cys93Phe | P04181 |
| HOMO SAPIENS | 1 | 74 | 5vwo | C | 154 | ARG | p.Arg154Leu | P04181 |
| HOMO SAPIENS | 0 | 74 | 5vwo | C | 180 | ARG | p.Arg180Thr | P04181 |
| HOMO SAPIENS | 0 | 74 | 5vwo | C | 199 | PRO | p.Pro199Gln | P04181 |
| HOMO SAPIENS | 4 | 74 | 5vwo | C | 209 | TYR | p.Tyr209Ter | P04181 |
| HOMO SAPIENS | 0 | 74 | 5vwo | C | 226 | ALA | p.Ala226Val | P04181 |

| | | | | | | | | |
|---|---|---|---|---|---|---|---|---|
| HOMO SAPIENS | 5 | 74 | 5vwo | C | 233 | GLN | p.Gln233Arg | P04181 |
| HOMO SAPIENS | 5 | 74 | 5vwo | C | 241 | PRO | p.Pro241Leu | P04181 |
| HOMO SAPIENS | 2 | 74 | 5vwo | C | 245 | TYR | p.Tyr245Cys | P04181 |
| HOMO SAPIENS | 6 | 74 | 5vwo | C | 250 | ARG | p.Arg250Pro | P04181 |
| HOMO SAPIENS | 6 | 74 | 5vwo | C | 250 | ARG | p.Arg250Ter | P04181 |
| HOMO SAPIENS | 8 | 74 | 5vwo | C | 267 | THR | p.Thr267Ile | P04181 |
| HOMO SAPIENS | 0 | 74 | 5vwo | C | 270 | ALA | p.Ala270Pro | P04181 |
| HOMO SAPIENS | 0 | 74 | 5vwo | C | 271 | ARG | p.Arg271Lys | P04181 |
| HOMO SAPIENS | 1 | 74 | 5vwo | C | 299 | TYR | p.Tyr299Ter | P04181 |
| HOMO SAPIENS | 0 | 74 | 5vwo | C | 318 | GLU | p.Glu318Lys | P04181 |
| HOMO SAPIENS | 1 | 74 | 5vwo | C | 319 | HIS | p.His319Tyr | P04181 |
| HOMO SAPIENS | 1 | 74 | 5vwo | C | 331 | ARG | p.Arg331Ter | P04181 |
| HOMO SAPIENS | 0 | 74 | 5vwo | C | 332 | VAL | p.Val332Met | P04181 |
| HOMO SAPIENS | 1 | 74 | 5vwo | C | 353 | GLY | p.Gly353Asp | P04181 |
| HOMO SAPIENS | 2 | 74 | 5vwo | C | 375 | GLY | p.Gly375Ala | P04181 |
| HOMO SAPIENS | 0 | 74 | 5vwo | C | 394 | CYS | p.Cys394Arg | P04181 |
| HOMO SAPIENS | 0 | 74 | 5vwo | C | 402 | LEU | p.Leu402Pro | P04181 |
| HOMO SAPIENS | 0 | 74 | 5vwo | C | 436 | ILE | p.Ile436Asn | P04181 |
| HOMO SAPIENS | 7 | 24356 | 4az0 | A | 104 | VAL | p.Val132Met | P10619 |
| HOMO SAPIENS | 4 | 24356 | 4az0 | A | 221 | TYR | p.Tyr249Asn | P10619 |
| HOMO SAPIENS | 3 | 245 | 1rsz | A | 24 | ARG | p.Arg24Ter | P00491 |
| HOMO SAPIENS | 3 | 245 | 1rsz | A | 51 | SER | p.Gly51Ser | P00491 |
| HOMO SAPIENS | 1 | 245 | 1rsz | A | 58 | ARG | p.Arg58Ter | P00491 |
| HOMO SAPIENS | 4 | 245 | 1rsz | A | 89 | GLU | p.Glu89Lys | P00491 |
| HOMO SAPIENS | 6 | 245 | 1rsz | A | 128 | ASP | p.Asp128Gly | P00491 |
| HOMO SAPIENS | 0 | 245 | 1rsz | A | 174 | ALA | p.Ala174Pro | P00491 |
| HOMO SAPIENS | 8 | 245 | 1rsz | A | 192 | TYR | p.Tyr192Cys | P00491 |
| HOMO SAPIENS | 8 | 245 | 1rsz | A | 234 | ARG | p.Arg234Pro | P00491 |
| HOMO SAPIENS | 0 | 37 | 5tby | E | 22 | GLU | p.Glu22Lys | P10916 |
| HOMO SAPIENS | 1 | 37 | 5tby | E | 86 | PHE | p.Phe86Ser | P10916 |
| HOMO SAPIENS | 9 | 71 | 5m9j | A | 12 | GLN | p.Gln36Leu | P03950 |
| HOMO SAPIENS | 3 | 71 | 5m9j | A | 17 | LYS | p.Lys41Glu | P03950 |
| HOMO SAPIENS | 3 | 71 | 5m9j | A | 17 | LYS | p.Lys41Ile | P03950 |
| HOMO SAPIENS | 4 | 71 | 5m9j | A | 28 | SER | p.Ser52Asn | P03950 |
| HOMO SAPIENS | 7 | 71 | 5m9j | A | 31 | ARG | p.Arg55Lys | P03950 |
| HOMO SAPIENS | 9 | 71 | 5m9j | A | 39 | CYS | p.Cys63Trp | P03950 |
| HOMO SAPIENS | 9 | 71 | 5m9j | A | 40 | LYS | p.Lys64Ile | P03950 |
| HOMO SAPIENS | 2 | 71 | 5m9j | A | 60 | GLU | p.Lys84Glu | P03950 |
| HOMO SAPIENS | 0 | 71 | 5m9j | A | 112 | PRO | p.Pro136Leu | P03950 |
| HOMO SAPIENS | 9 | 71 | 5m9j | A | 113 | VAL | p.Val137Ile | P03950 |
| HOMO SAPIENS | 3 | 326 | 5dqi | A | 209 | GLY | p.Gly209Val | Q9Y253 |
| HOMO SAPIENS | 1 | 326 | 5dqi | A | 242 | SER | p.Ser242Ter | Q9Y253 |
| HOMO SAPIENS | 1 | 326 | 5dqi | A | 303 | ARG | p.Arg303Ter | Q9Y253 |
| HOMO SAPIENS | 1 | 8203 | 3iu3 | K | 132 | VAL | p.Val153Ile | P01589 |
| HOMO SAPIENS | 0 | 4995 | 5ffo | H | 234 | THR | p.Thr263Ile | P01137 |
| HOMO SAPIENS | 5 | 38609 | 2ruk | A | 49 | ASP | p.Asp49His | P04637 |
| HOMO SAPIENS | 5 | 38609 | 2ruk | A | 58 | PRO | p.Pro58Arg | P04637 |
| HOMO SAPIENS | 0 | 1607 | 6pah | A | 117 | THR | p.Thr117Ile | P00439 |
| HOMO SAPIENS | 3 | 1607 | 6pah | A | 145 | ASP | p.Asp145Val | P00439 |
| HOMO SAPIENS | 5 | 1607 | 6pah | A | 158 | ARG | p.Arg158Gln | P00439 |
| HOMO SAPIENS | 5 | 1607 | 6pah | A | 158 | ARG | p.Arg158Trp | P00439 |
| HOMO SAPIENS | 5 | 1607 | 6pah | A | 169 | ARG | p.Arg169His | P00439 |
| HOMO SAPIENS | 5 | 1607 | 6pah | A | 169 | ARG | p.Arg169Cys | P00439 |
| HOMO SAPIENS | 3 | 1607 | 6pah | A | 176 | ARG | p.Arg176Leu | P00439 |

| | | | | | | | | |
|---|---|---|---|---|---|---|---|---|
| HOMO SAPIENS | 3 | 1607 | 6pah | A | 177 | VAL | p.Val177Leu | P00439 |
| HOMO SAPIENS | 1 | 1607 | 6pah | A | 194 | LEU | p.Leu194Pro | P00439 |
| HOMO SAPIENS | 1 | 1607 | 6pah | A | 204 | TYR | p.Tyr204Cys | P00439 |
| HOMO SAPIENS | 1 | 1607 | 6pah | A | 204 | TYR | p.Tyr204Ter | P00439 |
| HOMO SAPIENS | 2 | 1607 | 6pah | A | 230 | VAL | p.Val230Ile | P00439 |
| HOMO SAPIENS | 4 | 1607 | 6pah | A | 232 | GLN | p.Gln232Ter | P00439 |
| HOMO SAPIENS | 4 | 1607 | 6pah | A | 241 | ARG | p.Arg241Cys | P00439 |
| HOMO SAPIENS | 4 | 1607 | 6pah | A | 241 | ARG | p.Arg241His | P00439 |
| HOMO SAPIENS | 5 | 1607 | 6pah | A | 243 | ARG | p.Arg243Gln | P00439 |
| HOMO SAPIENS | 5 | 1607 | 6pah | A | 243 | ARG | p.Arg243Ter | P00439 |
| HOMO SAPIENS | 7 | 1607 | 6pah | A | 249 | LEU | p.Leu249Phe | P00439 |
| HOMO SAPIENS | 5 | 1607 | 6pah | A | 252 | ARG | p.Arg252Gly | P00439 |
| HOMO SAPIENS | 5 | 1607 | 6pah | A | 261 | ARG | p.Arg261Ter | P00439 |
| HOMO SAPIENS | 4 | 1607 | 6pah | A | 267 | GLN | p.Gln267Leu | P00439 |
| HOMO SAPIENS | 1 | 1607 | 6pah | A | 272 | GLY | p.Gly272Ter | P00439 |
| HOMO SAPIENS | 2 | 1607 | 6pah | A | 277 | TYR | p.Tyr277Asp | P00439 |
| HOMO SAPIENS | 7 | 1607 | 6pah | A | 280 | GLU | p.Glu280Lys | P00439 |
| HOMO SAPIENS | 8 | 1607 | 6pah | A | 281 | PRO | p.Pro281Leu | P00439 |
| HOMO SAPIENS | 0 | 1607 | 6pah | A | 299 | PHE | p.Phe299Cys | P00439 |
| HOMO SAPIENS | 5 | 1607 | 6pah | A | 300 | ALA | p.Ala300Ser | P00439 |
| HOMO SAPIENS | 5 | 1607 | 6pah | A | 309 | ALA | p.Ala309Asp | P00439 |
| HOMO SAPIENS | 4 | 1607 | 6pah | A | 320 | LYS | p.Lys320Asn | P00439 |
| HOMO SAPIENS | 0 | 1607 | 6pah | A | 345 | ALA | p.Ala345Ser | P00439 |
| HOMO SAPIENS | 0 | 1607 | 6pah | A | 348 | LEU | p.Leu348Val | P00439 |
| HOMO SAPIENS | 6 | 1607 | 6pah | A | 356 | TYR | p.Tyr356Ter | P00439 |
| HOMO SAPIENS | 2 | 1607 | 6pah | A | 380 | THR | p.Thr380Met | P00439 |
| HOMO SAPIENS | 0 | 1607 | 6pah | A | 386 | TYR | p.Tyr386Cys | P00439 |
| HOMO SAPIENS | 5 | 1607 | 6pah | A | 390 | GLU | p.Glu390Gly | P00439 |
| HOMO SAPIENS | 5 | 1607 | 6pah | A | 403 | ALA | p.Ala403Val | P00439 |
| HOMO SAPIENS | 4 | 1607 | 6pah | A | 408 | ARG | p.Arg408Trp | P00439 |
| HOMO SAPIENS | 3 | 1607 | 6pah | A | 413 | ARG | p.Arg413Pro | P00439 |
| HOMO SAPIENS | 4 | 1607 | 6pah | A | 414 | TYR | p.Tyr414Cys | P00439 |
| HOMO SAPIENS | 4 | 1607 | 6pah | A | 414 | TYR | p.Tyr414His | P00439 |
| HOMO SAPIENS | 4 | 1607 | 6pah | A | 415 | ASP | p.Asp415Asn | P00439 |
| HOMO SAPIENS | -1 | 676 | 5dc5 | A | 7 | PRO | p.Pro7Leu | Q9BY41 |
| HOMO SAPIENS | 0 | 676 | 5dc5 | A | 311 | THR | p.Thr311Met | Q9BY41 |
| HOMO SAPIENS | 1 | 124 | 5ftk | E | 95 | ARG | p.Arg95Gly | P55072 |
| HOMO SAPIENS | 1 | 124 | 5ftk | E | 155 | ARG | p.Arg155Cys | P55072 |
| HOMO SAPIENS | 1 | 124 | 5ftk | E | 155 | ARG | p.Arg155His | P55072 |
| HOMO SAPIENS | 1 | 124 | 5ftk | E | 159 | ARG | p.Arg159His | P55072 |
| HOMO SAPIENS | 1 | 124 | 5ftk | E | 159 | ARG | p.Arg159Gly | P55072 |
| HOMO SAPIENS | 0 | 124 | 5ftk | E | 191 | ARG | p.Arg191Gln | P55072 |
| HOMO SAPIENS | 0 | 124 | 5ftk | E | 232 | ALA | p.Ala232Glu | P55072 |
| HOMO SAPIENS | 5 | 5442 | 1fgg | B | 223 | GLY | p.Gly223Ser | O94766 |
| HOMO SAPIENS | 0 | 5442 | 1fgg | B | 277 | ARG | p.Arg277Gln | O94766 |
| HOMO SAPIENS | 0 | 9072 | 2gys | B | 225 | GLU | p.Glu249Gln | P32927 |
| HOMO SAPIENS | 0 | 9072 | 2gys | B | 247 | SER | p.Ser271Ter | P32927 |
| HOMO SAPIENS | 0 | 9072 | 2gys | B | 294 | ASP | p.Asp312Asn | P32927 |
| HOMO SAPIENS | 0 | 1484 | 4zrj | A | 62 | PHE | p.Phe62Ser | P35240 |
| HOMO SAPIENS | 3 | 1484 | 4zrj | A | 205 | MET | p.Met205Val | P35240 |
| HOMO SAPIENS | 3 | 1484 | 4zrj | A | 301 | ILE | p.Ile301Met | P35240 |
| HOMO SAPIENS | 4 | 40923 | 2enq | A | 68 | ILE | p.Ile391Met | P42336 |
| HOMO SAPIENS | 1 | 6179 | 5ofv | A | 125 | THR | p.Thr125Met | O43175 |
| HOMO SAPIENS | 0 | 6179 | 5ofv | A | 135 | ARG | p.Arg135Trp | O43175 |

| | | | | | | | | |
|---|---|---|---|---|---|---|---|---|
| HOMO SAPIENS | 1 | 6179 | 5ofv | A | 163 | ARG | p.Arg163Gln | O43175 |
| HOMO SAPIENS | 9 | 6179 | 5ofv | A | 261 | VAL | p.Val261Met | O43175 |
| HOMO SAPIENS | 7 | 34118 | 2i5o | A | 24 | MET | p.Met647Leu | Q9Y253 |
| HOMO SAPIENS | 8 | 3234 | 5c84 | A | 311 | LYS | p.Lys311Asn | P98170 |
| HOMO SAPIENS | 3 | 3234 | 5c84 | A | 321 | ALA | p.Ala321Gly | P98170 |
| HOMO SAPIENS | -1 | 15129 | 1zy8 | K | 1 | GLY | p.Gly54Ser | O00330 |
| HOMO SAPIENS | -1 | 15129 | 1zy8 | K | 48 | THR | p.Thr101Ala | O00330 |
| HOMO SAPIENS | -1 | 15129 | 1zy8 | K | 105 | GLU | p.Glu158Asp | O00330 |
| HOMO SAPIENS | 8 | 15129 | 1zy8 | K | 136 | ARG | p.Arg189His | O00330 |
| HOMO SAPIENS | -1 | 15129 | 1zy8 | K | 195 | GLN | p.Gln248Ter | O00330 |
| HOMO SAPIENS | 8 | 202 | 4bzb | A | 123 | HIS | p.His123Pro | Q9Y3Z3 |
| HOMO SAPIENS | 4 | 202 | 4bzb | A | 145 | ARG | p.Arg145Ter | Q9Y3Z3 |
| HOMO SAPIENS | 6 | 202 | 4bzb | A | 164 | ARG | p.Arg164Ter | Q9Y3Z3 |
| HOMO SAPIENS | 3 | 202 | 4bzb | A | 201 | ILE | p.Ile201Asn | Q9Y3Z3 |
| HOMO SAPIENS | 0 | 202 | 4bzb | A | 209 | GLY | p.Gly209Ser | Q9Y3Z3 |
| HOMO SAPIENS | 0 | 202 | 4bzb | A | 254 | MET | p.Met254Val | Q9Y3Z3 |
| HOMO SAPIENS | 5 | 202 | 4bzb | A | 531 | ARG | p.Arg531Ser | Q9Y3Z3 |
| HOMO SAPIENS | 6 | 202 | 4bzb | A | 548 | GLN | p.Gln548Ter | Q9Y3Z3 |
| HOMO SAPIENS | 3 | 30947 | 5lgd | A | 90 | PRO | p.Pro90Ser | P16671 |
| HOMO SAPIENS | 6 | 30947 | 5lgd | A | 254 | PHE | p.Phe254Leu | P16671 |
| HOMO SAPIENS | 0 | 30947 | 5lgd | A | 413 | ILE | p.Ile413Leu | P16671 |
| HOMO SAPIENS | 2 | 3906 | 5jtc | A | 735 | ARG | p.Arg735Trp | Q12797 |
| HOMO SAPIENS | 1 | 45 | 5uyx | C | 146 | GLU | p.Glu146Val | P48643 |
| HOMO SAPIENS | 1 | 45 | 5uyx | C | 147 | HIS | p.His147Arg | P48643 |
| HOMO SAPIENS | 0 | 45 | 5uyx | C | 362 | ILE | p.Ile362Met | P48643 |
| HOMO SAPIENS | 8 | 39596 | 1mhn | A | 95 | GLY | p.Gly95Arg | Q16637 |
| HOMO SAPIENS | 4 | 39596 | 1mhn | A | 111 | ALA | p.Ala111Gly | Q16637 |
| HOMO SAPIENS | 9 | 39596 | 1mhn | A | 116 | ILE | p.Ile116Phe | Q16637 |
| HOMO SAPIENS | 1 | 39596 | 1mhn | A | 136 | GLN | p.Gln136Glu | Q16637 |
| HOMO SAPIENS | 1 | 6408 | 5xjy | A | 85 | PRO | p.Pro85Leu | O95477 |
| HOMO SAPIENS | 0 | 6408 | 5xjy | A | 219 | ARG | p.Arg219Lys | O95477 |
| HOMO SAPIENS | 0 | 6408 | 5xjy | A | 230 | ARG | p.Arg230Cys | O95477 |
| HOMO SAPIENS | 1 | 6408 | 5xjy | A | 281 | MET | p.Met281Thr | O95477 |
| HOMO SAPIENS | -1 | 6408 | 5xjy | A | 343 | ALA | p.Ala343Gly | O95477 |
| HOMO SAPIENS | 1 | 6408 | 5xjy | A | 399 | VAL | p.Val399Ala | O95477 |
| HOMO SAPIENS | 1 | 6408 | 5xjy | A | 496 | ARG | p.Arg496Trp | O95477 |
| HOMO SAPIENS | 1 | 6408 | 5xjy | A | 590 | TRP | p.Trp590Ser | O95477 |
| HOMO SAPIENS | 1 | 6408 | 5xjy | A | 638 | ARG | p.Arg638Gln | O95477 |
| HOMO SAPIENS | 4 | 6408 | 5xjy | A | 697 | ALA | p.Ala697Thr | O95477 |
| HOMO SAPIENS | 1 | 6408 | 5xjy | A | 771 | VAL | p.Val771Met | O95477 |
| HOMO SAPIENS | 1 | 6408 | 5xjy | A | 774 | THR | p.Thr774Pro | O95477 |
| HOMO SAPIENS | 1 | 6408 | 5xjy | A | 776 | LYS | p.Lys776Asn | O95477 |
| HOMO SAPIENS | 0 | 6408 | 5xjy | A | 825 | VAL | p.Val825Ile | O95477 |
| HOMO SAPIENS | -1 | 6408 | 5xjy | A | 868 | GLU | p.Glu868Lys | O95477 |
| HOMO SAPIENS | -1 | 6408 | 5xjy | A | 883 | ILE | p.Ile883Met | O95477 |
| HOMO SAPIENS | 0 | 6408 | 5xjy | A | 935 | ASN | p.Asn935Ser | O95477 |
| HOMO SAPIENS | 0 | 6408 | 5xjy | A | 935 | ASN | p.Asn935His | O95477 |
| HOMO SAPIENS | 0 | 6408 | 5xjy | A | 937 | ALA | p.Ala937Val | O95477 |
| HOMO SAPIENS | 0 | 6408 | 5xjy | A | 1018 | ASP | p.Asp1018Gly | O95477 |
| HOMO SAPIENS | 0 | 6408 | 5xjy | A | 1096 | VAL | p.Val1096Ile | O95477 |
| HOMO SAPIENS | 0 | 6408 | 5xjy | A | 1099 | ASP | p.Asp1099Tyr | O95477 |
| HOMO SAPIENS | 0 | 6408 | 5xjy | A | 1172 | GLU | p.Glu1172Asp | O95477 |
| HOMO SAPIENS | 0 | 6408 | 5xjy | A | 1181 | SER | p.Ser1181Phe | O95477 |
| HOMO SAPIENS | 0 | 6408 | 5xjy | A | 1182 | ALA | p.Ala1182Thr | O95477 |

| | | | | | | | | |
|---|---|---|---|---|---|---|---|---|
| HOMO SAPIENS | -1 | 6408 | 5xjy | A | 1289 | ASP | p.Asp1289Asn | O95477 |
| HOMO SAPIENS | 0 | 6408 | 5xjy | A | 1399 | THR | p.Thr1399Ser | O95477 |
| HOMO SAPIENS | 1 | 6408 | 5xjy | A | 1408 | LEU | p.Leu1408Phe | O95477 |
| HOMO SAPIENS | 0 | 6408 | 5xjy | A | 1587 | LYS | p.Lys1587Arg | O95477 |
| HOMO SAPIENS | 1 | 6408 | 5xjy | A | 1674 | VAL | p.Val1674Ile | O95477 |
| HOMO SAPIENS | 1 | 6408 | 5xjy | A | 1680 | ARG | p.Arg1680Trp | O95477 |
| HOMO SAPIENS | 1 | 6408 | 5xjy | A | 1680 | ARG | p.Arg1680Gln | O95477 |
| HOMO SAPIENS | 0 | 6408 | 5xjy | A | 1800 | ASN | p.Asn1800His | O95477 |
| HOMO SAPIENS | -1 | 6408 | 5xjy | A | 1925 | ARG | p.Arg1925Gln | O95477 |
| HOMO SAPIENS | 0 | 6408 | 5xjy | A | 2009 | PHE | p.Phe2009Ser | O95477 |
| HOMO SAPIENS | -1 | 6408 | 5xjy | A | 2243 | ASP | p.Asp2243Glu | O95477 |
| HOMO SAPIENS | 4 | 23760 | 2mut | A | 231 | LEU | p.Phe231Leu | P07992 |
| HOMO SAPIENS | 6 | 1460 | 2hre | B | 96 | ARG | p.Arg96Gln | P22830 |
| HOMO SAPIENS | 8 | 1460 | 2hre | B | 129 | GLY | p.Gly129Arg | P22830 |
| HOMO SAPIENS | 9 | 1460 | 2hre | B | 172 | GLU | p.Glu172Lys | P22830 |
| HOMO SAPIENS | 4 | 1460 | 2hre | B | 285 | GLN | p.Gln285Arg | P22830 |
| HOMO SAPIENS | 5 | 1460 | 2hre | B | 362 | VAL | p.Val362Gly | P22830 |
| HOMO SAPIENS | 6 | 1460 | 2hre | B | 411 | CYS | p.Cys411Gly | P22830 |
| HOMO SAPIENS | 6 | 1460 | 2hre | B | 417 | PHE | p.Phe417Ser | P22830 |
| HOMO SAPIENS | 0 | 2210 | 5bug | A | 19 | ASP | p.Asp19Asn | P60484 |
| HOMO SAPIENS | 0 | 2210 | 5bug | A | 35 | MET | p.Met35Arg | P60484 |
| HOMO SAPIENS | 1 | 2210 | 5bug | A | 61 | HIS | p.His61Asp | P60484 |
| HOMO SAPIENS | 0 | 2210 | 5bug | A | 68 | TYR | p.Tyr68His | P60484 |
| HOMO SAPIENS | 8 | 2210 | 5bug | A | 70 | LEU | p.Leu70Pro | P60484 |
| HOMO SAPIENS | 6 | 2210 | 5bug | A | 93 | HIS | p.His93Arg | P60484 |
| HOMO SAPIENS | 1 | 2210 | 5bug | A | 112 | LEU | p.Leu112Pro | P60484 |
| HOMO SAPIENS | 0 | 2210 | 5bug | A | 121 | ALA | p.Ala121Gly | P60484 |
| HOMO SAPIENS | 3 | 2210 | 5bug | A | 123 | HIS | p.His123Arg | P60484 |
| HOMO SAPIENS | 7 | 2210 | 5bug | A | 124 | CYS | p.Cys124Arg | P60484 |
| HOMO SAPIENS | 9 | 2210 | 5bug | A | 129 | GLY | p.Gly129Glu | P60484 |
| HOMO SAPIENS | 9 | 2210 | 5bug | A | 130 | ARG | p.Arg130Gln | P60484 |
| HOMO SAPIENS | 9 | 2210 | 5bug | A | 130 | ARG | p.Arg130Gly | P60484 |
| HOMO SAPIENS | 0 | 2210 | 5bug | A | 132 | GLY | p.Gly132Val | P60484 |
| HOMO SAPIENS | 0 | 2210 | 5bug | A | 170 | SER | p.Ser170Arg | P60484 |
| HOMO SAPIENS | 1 | 2210 | 5bug | A | 173 | ARG | p.Arg173His | P60484 |
| HOMO SAPIENS | 0 | 2210 | 5bug | A | 217 | VAL | p.Val217Ile | P60484 |
| HOMO SAPIENS | 0 | 2210 | 5bug | A | 241 | PHE | p.Phe241Ser | P60484 |
| HOMO SAPIENS | 0 | 2210 | 5bug | A | 250 | CYS | p.Cys250Ter | P60484 |
| HOMO SAPIENS | 1 | 2210 | 5bug | A | 252 | ASP | p.Asp252Gly | P60484 |
| HOMO SAPIENS | 3 | 123 | 2ibu | A | 93 | ASN | p.Asn93Ser | P24752 |
| HOMO SAPIENS | 0 | 123 | 2ibu | A | 145 | GLN | p.Gln145Glu | P24752 |
| HOMO SAPIENS | 0 | 123 | 2ibu | A | 152 | GLY | p.Gly152Ala | P24752 |
| HOMO SAPIENS | 2 | 123 | 2ibu | A | 158 | ASN | p.Asn158Asp | P24752 |
| HOMO SAPIENS | 2 | 123 | 2ibu | A | 158 | ASN | p.Asn158Ser | P24752 |
| HOMO SAPIENS | 5 | 123 | 2ibu | A | 183 | GLY | p.Gly183Arg | P24752 |
| HOMO SAPIENS | 3 | 123 | 2ibu | A | 272 | GLN | p.Gln272Ter | P24752 |
| HOMO SAPIENS | 3 | 123 | 2ibu | A | 312 | ILE | p.Ile312Thr | P24752 |
| HOMO SAPIENS | 1 | 123 | 2ibu | A | 333 | ALA | p.Ala333Pro | P24752 |
| HOMO SAPIENS | 5 | 123 | 2ibu | A | 379 | GLY | p.Gly379Val | P24752 |
| HOMO SAPIENS | 0 | 123 | 2ibu | A | 380 | ALA | p.Ala380Thr | P24752 |
| HOMO SAPIENS | 2 | 369 | 5nrw | A | 1175 | TYR | p.Tyr1175Cys | Q92793 |
| HOMO SAPIENS | 2 | 830 | 2vc2 | A | 116 | LEU | p.Leu147Val | P08514 |
| HOMO SAPIENS | 0 | 830 | 2vc2 | A | 183 | LEU | p.Leu214Pro | P08514 |
| HOMO SAPIENS | 4 | 830 | 2vc2 | A | 242 | GLY | p.Gly273Asp | P08514 |

| | | | | | | | | |
|---|---|---|---|---|---|---|---|---|
| HOMO SAPIENS | 6 | 830 | 2vc2 | A | 324 | GLU | p.Glu355Lys | P08514 |
| HOMO SAPIENS | -1 | 10353 | 5ejc | E | 989 | ALA | p.Ala989Ser | Q92574 |
| HOMO SAPIENS | 0 | 15448 | 3lgg | A | 21 | GLY | p.Gly47Arg | Q9NZK5 |
| HOMO SAPIENS | 0 | 15448 | 3lgg | A | 21 | GLY | p.Gly47Val | Q9NZK5 |
| HOMO SAPIENS | 9 | 15448 | 3lgg | A | 86 | HIS | p.His112Gln | Q9NZK5 |
| HOMO SAPIENS | 0 | 15448 | 3lgg | A | 143 | ARG | p.Arg169Gln | Q9NZK5 |
| HOMO SAPIENS | 3 | 15448 | 3lgg | A | 221 | ALA | p.Ala247Val | Q9NZK5 |
| HOMO SAPIENS | 0 | 15448 | 3lgg | A | 225 | PRO | p.Pro251Leu | Q9NZK5 |
| HOMO SAPIENS | 0 | 15448 | 3lgg | A | 283 | MET | p.Met309Ile | Q9NZK5 |
| HOMO SAPIENS | 2 | 15448 | 3lgg | A | 334 | THR | p.Thr360Ala | Q9NZK5 |
| HOMO SAPIENS | 7 | 15448 | 3lgg | A | 357 | GLY | p.Gly383Ser | Q9NZK5 |
| HOMO SAPIENS | 0 | 15448 | 3lgg | A | 427 | TYR | p.Tyr453Cys | Q9NZK5 |
| HOMO SAPIENS | 0 | 872 | 2y3i | A | 87 | LEU | p.Leu88Pro | P00558 |
| HOMO SAPIENS | 1 | 872 | 2y3i | A | 163 | ASP | p.Asp164Val | P00558 |
| HOMO SAPIENS | 4 | 872 | 2y3i | A | 205 | ARG | p.Arg206Pro | P00558 |
| HOMO SAPIENS | 4 | 872 | 2y3i | A | 267 | ASP | p.Asp268Asn | P00558 |
| HOMO SAPIENS | 8 | 872 | 2y3i | A | 284 | ASP | p.Asp285Val | P00558 |
| HOMO SAPIENS | 4 | 8 | 5jco | B | 188 | ILE | p.Ile188Leu | Q71U36 |
| HOMO SAPIENS | 7 | 8 | 5jco | B | 263 | PRO | p.Pro263Thr | Q71U36 |
| HOMO SAPIENS | 6 | 8 | 5jco | B | 264 | ARG | p.Arg264Cys | Q71U36 |
| HOMO SAPIENS | 7 | 8 | 5jco | B | 402 | ARG | p.Arg402His | Q71U36 |
| HOMO SAPIENS | 7 | 8 | 5jco | B | 419 | SER | p.Ser419Leu | Q71U36 |
| HOMO SAPIENS | -1 | 880 | 4nwf | A | 2 | THR | p.Thr2Ile | Q06124 |
| HOMO SAPIENS | 2 | 880 | 4nwf | A | 58 | ASN | p.Asn58Ser | Q06124 |
| HOMO SAPIENS | 4 | 880 | 4nwf | A | 61 | ASP | p.Asp61Gly | Q06124 |
| HOMO SAPIENS | 2 | 880 | 4nwf | A | 62 | TYR | p.Tyr62Asp | Q06124 |
| HOMO SAPIENS | 3 | 880 | 4nwf | A | 63 | TYR | p.Tyr63Cys | Q06124 |
| HOMO SAPIENS | 2 | 880 | 4nwf | A | 69 | GLU | p.Glu69Gln | Q06124 |
| HOMO SAPIENS | 2 | 880 | 4nwf | A | 72 | ALA | p.Ala72Gly | Q06124 |
| HOMO SAPIENS | 2 | 880 | 4nwf | A | 72 | ALA | p.Ala72Ser | Q06124 |
| HOMO SAPIENS | 3 | 880 | 4nwf | A | 73 | THR | p.Thr73Ile | Q06124 |
| HOMO SAPIENS | 5 | 880 | 4nwf | A | 76 | GLU | p.Glu76Ala | Q06124 |
| HOMO SAPIENS | 5 | 880 | 4nwf | A | 76 | GLU | p.Glu76Lys | Q06124 |
| HOMO SAPIENS | 3 | 880 | 4nwf | A | 79 | GLN | p.Gln79Arg | Q06124 |
| HOMO SAPIENS | 5 | 880 | 4nwf | A | 152 | ARG | p.Arg152His | Q06124 |
| HOMO SAPIENS | 2 | 880 | 4nwf | A | 268 | GLY | p.Gly268Cys | Q06124 |
| HOMO SAPIENS | 9 | 880 | 4nwf | A | 276 | LYS | p.Lys276Arg | Q06124 |
| HOMO SAPIENS | 9 | 880 | 4nwf | A | 279 | TYR | p.Tyr279Cys | Q06124 |
| HOMO SAPIENS | 5 | 880 | 4nwf | A | 285 | PHE | p.Phe285Ser | Q06124 |
| HOMO SAPIENS | 0 | 880 | 4nwf | A | 308 | ASP | p.Asn308Ser | Q06124 |
| HOMO SAPIENS | 0 | 880 | 4nwf | A | 308 | ASP | p.Asn308Asp | Q06124 |
| HOMO SAPIENS | 5 | 880 | 4nwf | A | 392 | ALA | p.Ala392Thr | Q06124 |
| HOMO SAPIENS | 1 | 880 | 4nwf | A | 411 | THR | p.Thr415Met | Q06124 |
| HOMO SAPIENS | 0 | 880 | 4nwf | A | 461 | ALA | p.Ala465Thr | Q06124 |
| HOMO SAPIENS | 0 | 880 | 4nwf | A | 464 | GLY | p.Gly468Ala | Q06124 |
| HOMO SAPIENS | 1 | 880 | 4nwf | A | 491 | PRO | p.Pro495Ser | Q06124 |
| HOMO SAPIENS | 1 | 880 | 4nwf | A | 502 | SER | p.Ser506Thr | Q06124 |
| HOMO SAPIENS | 8 | 880 | 4nwf | A | 510 | GLN | p.Gln514Pro | Q06124 |
| HOMO SAPIENS | 4 | 1872 | 3wf1 | D | 109 | ARG | p.Arg109Trp | P16278 |
| HOMO SAPIENS | 0 | 1872 | 3wf1 | D | 521 | ARG | p.Arg521Cys | P16278 |
| HOMO SAPIENS | 5 | 910 | 5xte | A | 45 | SER | p.Ser45Phe | O14949 |
| HOMO SAPIENS | 2 | 9017 | 5f0k | A | 382 | LYS | p.Lys382Arg | Q96QK1 |
| HOMO SAPIENS | 1 | 710 | 4lsj | A | 559 | ILE | p.Ile559Asn | P04150 |
| HOMO SAPIENS | 5 | 710 | 4lsj | A | 571 | VAL | p.Val571Ala | P04150 |

| | | | | | | | | |
|---|---|---|---|---|---|---|---|---|
| HOMO SAPIENS | 0 | 710 | 4lsj | A | 641 | ASP | p.Asp641Val | P04150 |
| HOMO SAPIENS | 1 | 710 | 4lsj | A | 678 | ASP | p.Asp678Glu | P04150 |
| HOMO SAPIENS | 2 | 710 | 4lsj | A | 679 | GLY | p.Gly679Ser | P04150 |
| HOMO SAPIENS | 1 | 710 | 4lsj | A | 737 | PHE | p.Phe737Leu | P04150 |
| HOMO SAPIENS | 1 | 710 | 4lsj | A | 747 | ILE | p.Ile747Met | P04150 |
| HOMO SAPIENS | 1 | 710 | 4lsj | A | 753 | LEU | p.Leu753Phe | P04150 |
| HOMO SAPIENS | 1 | 710 | 4lsj | A | 773 | LEU | p.Leu773Pro | P04150 |
| HOMO SAPIENS | 1 | 2145 | 1av1 | B | 71 | PHE | p.Phe95Tyr | P02647 |
| HOMO SAPIENS | 1 | 2145 | 1av1 | B | 128 | GLU | p.Glu152Lys | P02647 |
| HOMO SAPIENS | 1 | 2145 | 1av1 | B | 136 | GLU | p.Glu160Lys | P02647 |
| HOMO SAPIENS | 2 | 2145 | 1av1 | B | 143 | PRO | p.Pro167Arg | P02647 |
| HOMO SAPIENS | 6 | 2145 | 1av1 | B | 164 | ALA | p.Ala188Ser | P02647 |
| HOMO SAPIENS | 7 | 2145 | 1av1 | B | 173 | ARG | p.Arg197Cys | P02647 |
| HOMO SAPIENS | 2 | 2145 | 1av1 | B | 198 | GLU | p.Glu222Lys | P02647 |
| HOMO SAPIENS | -1 | 3432 | 2beu | A | 1 | SER | p.Ser46Ter | P12694 |
| HOMO SAPIENS | 5 | 3432 | 2beu | A | 79 | ARG | p.Arg124Trp | P12694 |
| HOMO SAPIENS | 7 | 3432 | 2beu | A | 114 | ARG | p.Arg159Gln | P12694 |
| HOMO SAPIENS | 6 | 3432 | 2beu | A | 166 | THR | p.Thr211Met | P12694 |
| HOMO SAPIENS | 4 | 3432 | 2beu | A | 175 | ALA | p.Ala220Val | P12694 |
| HOMO SAPIENS | 5 | 3432 | 2beu | A | 204 | GLY | p.Gly249Ser | P12694 |
| HOMO SAPIENS | 4 | 3432 | 2beu | A | 209 | ALA | p.Ala254Asp | P12694 |
| HOMO SAPIENS | 6 | 3432 | 2beu | A | 220 | ARG | p.Arg265Trp | P12694 |
| HOMO SAPIENS | 6 | 3432 | 2beu | A | 234 | TYR | p.Tyr279Cys | P12694 |
| HOMO SAPIENS | 5 | 3432 | 2beu | A | 240 | ALA | p.Ala285Pro | P12694 |
| HOMO SAPIENS | 4 | 3432 | 2beu | A | 245 | GLY | p.Gly290Arg | P12694 |
| HOMO SAPIENS | 5 | 3432 | 2beu | A | 269 | ARG | p.Arg314Ter | P12694 |
| HOMO SAPIENS | 3 | 3432 | 2beu | A | 301 | ARG | p.Arg346Cys | P12694 |
| HOMO SAPIENS | 5 | 3432 | 2beu | A | 359 | ASN | p.Asn404Ser | P12694 |
| HOMO SAPIENS | 4 | 3432 | 2beu | A | 393 | TYR | p.Tyr438Asn | P12694 |
| HOMO SAPIENS | 8 | 4541 | 1rr8 | C | 533 | ASP | p.Asp533Gly | P11387 |
| HOMO SAPIENS | 2 | 3247 | 1tzn | d | 1105 | GLY | p.Gly105Asp | P58335 |
| HOMO SAPIENS | 3 | 3247 | 1tzn | d | 1111 | ARG | p.Arg111Cys | P58335 |
| HOMO SAPIENS | 2 | 3247 | 1tzn | d | 1189 | ILE | p.Ile189Thr | P58335 |
| HOMO SAPIENS | 9 | 467 | 1suv | C | 58 | ASP | p.Asp77Asn | P02787 |
| HOMO SAPIENS | 2 | 467 | 1suv | C | 277 | ASP | p.Asp296Gly | P02787 |
| HOMO SAPIENS | 9 | 6485 | 1xwd | B | 51 | CYS | p.Cys69Gly | P01225 |
| HOMO SAPIENS | 1 | 694 | 1wuu | A | 28 | PRO | p.Pro28Thr | P51570 |
| HOMO SAPIENS | 0 | 694 | 1wuu | A | 32 | VAL | p.Val32Met | P51570 |
| HOMO SAPIENS | 0 | 694 | 1wuu | A | 198 | ALA | p.Ala198Val | P51570 |
| HOMO SAPIENS | 0 | 143 | 4nkx | A | 93 | PHE | p.Phe93Cys | P05093 |
| HOMO SAPIENS | 3 | 143 | 4nkx | A | 96 | ARG | p.Arg96Trp | P05093 |
| HOMO SAPIENS | 3 | 143 | 4nkx | A | 96 | ARG | p.Arg96Gln | P05093 |
| HOMO SAPIENS | 0 | 143 | 4nkx | A | 106 | SER | p.Ser106Pro | P05093 |
| HOMO SAPIENS | 1 | 143 | 4nkx | A | 114 | PHE | p.Phe114Val | P05093 |
| HOMO SAPIENS | 1 | 143 | 4nkx | A | 116 | ASP | p.Asp116Val | P05093 |
| HOMO SAPIENS | 3 | 143 | 4nkx | A | 125 | ARG | p.Arg125Gln | P05093 |
| HOMO SAPIENS | 0 | 143 | 4nkx | A | 201 | TYR | p.Tyr201Asp | P05093 |
| HOMO SAPIENS | 0 | 143 | 4nkx | A | 239 | ARG | p.Arg239Ter | P05093 |
| HOMO SAPIENS | 0 | 143 | 4nkx | A | 329 | TYR | p.Tyr329Asp | P05093 |
| HOMO SAPIENS | 1 | 143 | 4nkx | A | 342 | PRO | p.Pro342Thr | P05093 |
| HOMO SAPIENS | 2 | 143 | 4nkx | A | 347 | ARG | p.Arg347His | P05093 |
| HOMO SAPIENS | 2 | 143 | 4nkx | A | 347 | ARG | p.Arg347Cys | P05093 |
| HOMO SAPIENS | 1 | 143 | 4nkx | A | 358 | ARG | p.Arg358Gln | P05093 |
| HOMO SAPIENS | 0 | 143 | 4nkx | A | 362 | ARG | p.Arg362Cys | P05093 |

| | | | | | | | | |
|---|---|---|---|---|---|---|---|---|
| HOMO SAPIENS | 1 | 143 | 4nkx | A | 406 | TRP | p.Trp406Arg | P05093 |
| HOMO SAPIENS | 0 | 143 | 4nkx | A | 428 | PRO | p.Pro428Leu | P05093 |
| HOMO SAPIENS | 6 | 1441 | 1uuh | A | 46 | ARG | p.Arg46Gly | P16070 |
| HOMO SAPIENS | 3 | 82 | 4wlv | D | 37 | GLY | p.Gly37Arg | P40926 |
| HOMO SAPIENS | 0 | 82 | 4wlv | D | 133 | PRO | p.Pro133Leu | P40926 |
| HOMO SAPIENS | 2 | 392 | 5xtd | A | 56 | SER | p.Ser56Pro | P49821 |
| HOMO SAPIENS | 2 | 392 | 5xtd | A | 59 | ARG | p.Arg59Ter | P49821 |
| HOMO SAPIENS | 2 | 392 | 5xtd | A | 74 | ASP | p.Asp74Asn | P49821 |
| HOMO SAPIENS | 4 | 392 | 5xtd | A | 116 | ASN | p.Asn116Lys | P49821 |
| HOMO SAPIENS | 4 | 392 | 5xtd | A | 117 | ALA | p.Ala117Thr | P49821 |
| HOMO SAPIENS | 5 | 392 | 5xtd | A | 122 | PRO | p.Pro122Leu | P49821 |
| HOMO SAPIENS | 3 | 392 | 5xtd | A | 160 | GLY | p.Gly160Glu | P49821 |
| HOMO SAPIENS | 4 | 392 | 5xtd | A | 214 | GLU | p.Glu214Lys | P49821 |
| HOMO SAPIENS | 2 | 392 | 5xtd | A | 357 | MET | p.Met357Thr | P49821 |
| HOMO SAPIENS | 4 | 392 | 5xtd | A | 423 | THR | p.Thr423Met | P49821 |
| HOMO SAPIENS | 0 | 22593 | 3pdf | A | 8 | TYR | p.Tyr32Ter | P53634 |
| HOMO SAPIENS | 0 | 22593 | 3pdf | A | 15 | TRP | p.Trp39Ser | P53634 |
| HOMO SAPIENS | 0 | 22593 | 3pdf | A | 103 | HIS | p.His127Pro | P53634 |
| HOMO SAPIENS | 9 | 22593 | 3pdf | A | 228 | GLN | p.Gln252Leu | P53634 |
| HOMO SAPIENS | 3 | 22593 | 3pdf | A | 262 | GLN | p.Gln286Arg | P53634 |
| HOMO SAPIENS | 4 | 22593 | 3pdf | A | 277 | GLY | p.Gly301Ser | P53634 |
| HOMO SAPIENS | 0 | 22593 | 3pdf | A | 323 | TYR | p.Tyr347Cys | P53634 |
| HOMO SAPIENS | 5 | 3461 | 4c2a | A | 1306 | GLN | p.Arg1306Trp | P04275 |
| HOMO SAPIENS | 5 | 3461 | 4c2a | A | 1374 | ARG | p.Arg1374Cys | P04275 |
| HOMO SAPIENS | 5 | 3461 | 4c2a | A | 1379 | ARG | p.Arg1379Cys | P04275 |
| HOMO SAPIENS | 4 | 3461 | 4c2a | A | 1380 | ILE | p.Ile1380Val | P04275 |
| HOMO SAPIENS | 0 | 3461 | 4c2a | A | 1381 | ALA | p.Thr1381Ala | P04275 |
| HOMO SAPIENS | 5 | 3461 | 4c2a | A | 1399 | ARG | p.Arg1399His | P04275 |
| HOMO SAPIENS | 5 | 3461 | 4c2a | A | 1435 | ASN | p.Asn1435Ser | P04275 |
| HOMO SAPIENS | 0 | 4533 | 5uj7 | C | 174 | TYR | p.Tyr174Cys | O43929 |
| HOMO SAPIENS | 0 | 4533 | 5uj7 | C | 202 | LEU | p.Leu202Val | O43929 |
| HOMO SAPIENS | 0 | 1374 | 3j07 | O | 39 | PRO | p.Pro39Leu | P02511 |
| HOMO SAPIENS | 0 | 1374 | 3j07 | O | 107 | ARG | p.Arg107His | P02511 |
| HOMO SAPIENS | 0 | 1374 | 3j07 | O | 157 | ARG | p.Arg157His | P02511 |
| HOMO SAPIENS | -1 | 942 | 1s9i | B | 57 | PHE | p.Phe57Cys | P36507 |
| HOMO SAPIENS | -1 | 942 | 1s9i | B | 57 | PHE | p.Phe57Val | P36507 |
| HOMO SAPIENS | 2 | 942 | 1s9i | B | 128 | PRO | p.Pro128Gln | P36507 |
| HOMO SAPIENS | 1 | 942 | 1s9i | B | 134 | TYR | p.Tyr134His | P36507 |
| HOMO SAPIENS | 1 | 942 | 1s9i | B | 188 | HIS | p.His188Gln | P36507 |
| HOMO SAPIENS | 0 | 942 | 1s9i | B | 262 | VAL | p.Val262Phe | P36507 |
| HOMO SAPIENS | 0 | 942 | 1s9i | B | 263 | GLY | p.Gly263Arg | P36507 |
| HOMO SAPIENS | -1 | 942 | 1s9i | B | 298 | PRO | p.Pro298Leu | P36507 |
| HOMO SAPIENS | -1 | 942 | 1s9i | B | 305 | VAL | p.Val305Ile | P36507 |
| HOMO SAPIENS | 0 | 3245 | 1fbv | A | 367 | GLN | p.Gln367Pro | P22681 |
| HOMO SAPIENS | 0 | 3245 | 1fbv | A | 371 | TYR | p.Tyr371His | P22681 |
| HOMO SAPIENS | 1 | 3245 | 1fbv | A | 382 | LYS | p.Lys382Glu | P22681 |
| HOMO SAPIENS | 2 | 3245 | 1fbv | A | 390 | ASP | p.Asp390Tyr | P22681 |
| HOMO SAPIENS | 9 | 3245 | 1fbv | A | 396 | CYS | p.Cys396Arg | P22681 |
| HOMO SAPIENS | 7 | 3245 | 1fbv | A | 420 | ARG | p.Arg420Gln | P22681 |
| HOMO SAPIENS | 1 | 3245 | 1fbv | A | 429 | ILE | p.Ile429Met | P22681 |
| HOMO SAPIENS | 0 | 2604 | 2pbd | P | 111 | LEU | p.Leu112Met | P07737 |
| HOMO SAPIENS | 3 | 2604 | 2pbd | P | 116 | GLU | p.Glu117Gly | P07737 |
| HOMO SAPIENS | 4 | 2013 | 3b7w | A | 472 | ARG | p.Arg472Gly | Q08AH3 |
| HOMO SAPIENS | 0 | 24346 | 3sh5 | A | 85 | ARG | p.Arg4281His | P98160 |

| | | | | | | | | |
|---|---|---|---|---|---|---|---|---|
| HOMO SAPIENS | 0 | 24346 | 3sh5 | A | 122 | ARG | p.Arg4318Trp | P98160 |
| HOMO SAPIENS | 0 | 24346 | 3sh5 | A | 135 | SER | p.Ser4331Asn | P98160 |
| HOMO SAPIENS | 0 | 24346 | 3sh5 | A | 136 | VAL | p.Val4332Ile | P98160 |
| HOMO SAPIENS | 0 | 707 | 4p2g | A | 26 | MET | p.Met26Ile | Q99497 |
| HOMO SAPIENS | 5 | 707 | 4p2g | A | 39 | ALA | p.Ala39Ser | Q99497 |
| HOMO SAPIENS | 5 | 707 | 4p2g | A | 64 | GLU | p.Glu64Asp | Q99497 |
| HOMO SAPIENS | 4 | 707 | 4p2g | A | 149 | ASP | p.Asp149Ala | Q99497 |
| HOMO SAPIENS | 6 | 707 | 4p2g | A | 163 | GLU | p.Glu163Lys | Q99497 |
| HOMO SAPIENS | 6 | 707 | 4p2g | A | 166 | LEU | p.Leu166Pro | Q99497 |
| HOMO SAPIENS | 3 | 5121 | 4xss | E | 31 | GLY | p.Gly58Arg | P06213 |
| HOMO SAPIENS | 2 | 5121 | 4xss | E | 119 | ILE | p.Ile146Met | P06213 |
| HOMO SAPIENS | 8 | 5121 | 4xss | E | 209 | HIS | p.His236Arg | P06213 |
| HOMO SAPIENS | 2 | 5121 | 4xss | E | 234 | ASP | p.Asp261Glu | P06213 |
| HOMO SAPIENS | 1 | 5121 | 4xss | E | 276 | GLN | p.Gln303His | P06213 |
| HOMO SAPIENS | 1 | 27790 | 5erp | B | 140 | THR | p.Thr275Met | Q02487 |
| HOMO SAPIENS | 0 | 27790 | 5erp | B | 150 | ILE | p.Ile285Thr | Q02487 |
| HOMO SAPIENS | 1 | 27790 | 5erp | B | 205 | THR | p.Thr340Ala | Q02487 |
| HOMO SAPIENS | 1 | 27790 | 5erp | B | 223 | THR | p.Thr358Ile | Q02487 |
| HOMO SAPIENS | 0 | 27790 | 5erp | B | 301 | GLY | p.Gly436Ala | Q02487 |
| HOMO SAPIENS | -1 | 27790 | 5erp | B | 315 | ARG | p.Arg450Ser | Q02487 |
| HOMO SAPIENS | 0 | 27790 | 5erp | B | 369 | SER | p.Ser504Gly | Q02487 |
| HOMO SAPIENS | 0 | 27790 | 5erp | B | 385 | ILE | p.Ile520Thr | Q02487 |
| HOMO SAPIENS | 0 | 27790 | 5erp | B | 461 | ALA | p.Ala596Val | Q02487 |
| HOMO SAPIENS | 0 | 27790 | 5erp | B | 503 | GLN | p.Gln638His | Q02487 |
| HOMO SAPIENS | 5 | 6599 | 4nsc | A | 129 | ARG | p.Arg129Leu | Q9BPX6 |
| HOMO SAPIENS | -1 | 6599 | 4nsc | A | 183 | GLN | p.Gln183Ter | Q9BPX6 |
| HOMO SAPIENS | 0 | 20420 | 3ghm | A | 398 | ARG | p.Arg398His | Q76LX8 |
| HOMO SAPIENS | 0 | 20420 | 3ghm | A | 448 | GLN | p.Gln448Glu | Q76LX8 |
| HOMO SAPIENS | 1 | 20420 | 3ghm | A | 449 | GLN | p.Gln449Ter | Q76LX8 |
| HOMO SAPIENS | 0 | 20420 | 3ghm | A | 456 | GLN | p.Gln456His | Q76LX8 |
| HOMO SAPIENS | 1 | 20420 | 3ghm | A | 528 | ARG | p.Arg528Gly | Q76LX8 |
| HOMO SAPIENS | 0 | 20420 | 3ghm | A | 618 | PRO | p.Pro618Ala | Q76LX8 |
| HOMO SAPIENS | 0 | 20420 | 3ghm | A | 625 | ARG | p.Arg625His | Q76LX8 |
| HOMO SAPIENS | 0 | 275 | 4rmz | A | 293 | GLN | p.Gln293Ter | Q9NWZ3 |
| HOMO SAPIENS | 0 | 275 | 4rmz | A | 428 | ALA | p.Ala428Thr | Q9NWZ3 |
| HOMO SAPIENS | 2 | 10458 | 3puf | B | 138 | LEU | p.Leu138Phe | Q5TBB1 |
| HOMO SAPIENS | 6 | 10458 | 3puf | B | 177 | ALA | p.Ala177Thr | Q5TBB1 |
| HOMO SAPIENS | -1 | 10458 | 3puf | B | 185 | VAL | p.Val185Gly | Q5TBB1 |
| HOMO SAPIENS | 0 | 26648 | 2n90 | A | 36 | ARG | p.Arg444Gln | P04629 |
| HOMO SAPIENS | 2 | 977 | 1dnw | D | 166 | ALA | p.Ala332Val | P05164 |
| HOMO SAPIENS | 4 | 977 | 1dnw | D | 333 | ARG | p.Arg499Cys | P05164 |
| HOMO SAPIENS | 1 | 977 | 1dnw | D | 403 | ARG | p.Arg569Trp | P05164 |
| HOMO SAPIENS | 0 | 25704 | 3ajb | A | 82 | GLN | p.Gln82Arg | P56589 |
| HOMO SAPIENS | 3 | 255 | 3t1p | A | 38 | TYR | p.Tyr62Ter | P01009 |
| HOMO SAPIENS | 3 | 255 | 3t1p | A | 39 | ARG | p.Arg63Cys | P01009 |
| HOMO SAPIENS | 0 | 255 | 3t1p | A | 101 | ARG | p.Arg125His | P01009 |
| HOMO SAPIENS | 0 | 255 | 3t1p | A | 212 | GLN | p.Gln236Arg | P01009 |
| HOMO SAPIENS | 0 | 255 | 3t1p | A | 213 | VAL | p.Val237Ala | P01009 |
| HOMO SAPIENS | 1 | 255 | 3t1p | A | 269 | HIS | p.His293Gln | P01009 |
| HOMO SAPIENS | 8 | 255 | 3t1p | A | 341 | ASP | p.Asp365Asn | P01009 |
| HOMO SAPIENS | 9 | 255 | 3t1p | A | 342 | GLU | p.Glu366Lys | P01009 |
| HOMO SAPIENS | 0 | 255 | 3t1p | A | 369 | PRO | p.Pro393Ser | P01009 |
| HOMO SAPIENS | 0 | 255 | 3t1p | A | 376 | GLU | p.Glu400Asp | P01009 |
| HOMO SAPIENS | -1 | 5175 | 3zxi | A | 35 | ALA | p.Ala35Asp | Q9Y2Z4 |

| | | | | | | | | | |
|---|---|---|---|---|---|---|---|---|---|
| HOMO SAPIENS | 0 | 5175 | 3zxi | A | | 52 | PHE | p.Phe52Leu | Q9Y2Z4 |
| HOMO SAPIENS | -1 | 5175 | 3zxi | A | | 68 | GLY | p.Gly68Ser | Q9Y2Z4 |
| HOMO SAPIENS | 0 | 5175 | 3zxi | A | | 179 | LYS | p.Lys179Gln | Q9Y2Z4 |
| HOMO SAPIENS | 0 | 5175 | 3zxi | A | | 191 | GLY | p.Gly191Val | Q9Y2Z4 |
| HOMO SAPIENS | 0 | 5175 | 3zxi | A | | 251 | ILE | p.Ile251Val | Q9Y2Z4 |
| HOMO SAPIENS | 2 | 5175 | 3zxi | A | | 295 | ASP | p.Asp295Asn | Q9Y2Z4 |
| HOMO SAPIENS | 0 | 5175 | 3zxi | A | | 311 | ASP | p.Asp311Glu | Q9Y2Z4 |
| HOMO SAPIENS | 0 | 5175 | 3zxi | A | | 360 | ARG | p.Arg360Ter | Q9Y2Z4 |
| HOMO SAPIENS | 1 | 3016 | 3exg | | 3 | 4 | ASP | p.Asp33His | P08559 |
| HOMO SAPIENS | 1 | 3016 | 3exg | | 3 | 176 | PHE | p.Phe205Leu | P08559 |
| HOMO SAPIENS | 1 | 3016 | 3exg | | 3 | 214 | TYR | p.Tyr243Asn | P08559 |
| HOMO SAPIENS | 1 | 3016 | 3exg | | 3 | 229 | ASP | p.Asp258Ala | P08559 |
| HOMO SAPIENS | 0 | 3016 | 3exg | | 3 | 233 | VAL | p.Val262Ile | P08559 |
| HOMO SAPIENS | 1 | 3016 | 3exg | | 3 | 234 | ARG | p.Arg263Gly | P08559 |
| HOMO SAPIENS | 0 | 3016 | 3exg | | 3 | 253 | MET | p.Met282Leu | P08559 |
| HOMO SAPIENS | 7 | 3016 | 3exg | | 3 | 259 | ARG | p.Arg288His | P08559 |
| HOMO SAPIENS | -1 | 3016 | 3exg | | 3 | 273 | ARG | p.Arg302Cys | P08559 |
| HOMO SAPIENS | 2 | 3016 | 3exg | | 3 | 286 | ASP | p.Asp315Asn | P08559 |
| HOMO SAPIENS | 1 | 3016 | 3exg | | 3 | 349 | ARG | p.Arg378His | P08559 |
| HOMO SAPIENS | 5 | 6729 | 4a0l | E | | 572 | ARG | p.Arg572Cys | Q13620 |
| HOMO SAPIENS | 0 | 1067 | 4kyo | A | | 9 | THR | p.Thr9Asn | P21549 |
| HOMO SAPIENS | 0 | 1067 | 4kyo | A | | 11 | PRO | p.Pro11Leu | P21549 |
| HOMO SAPIENS | 0 | 1067 | 4kyo | A | | 22 | ASN | p.Asn22Ser | P21549 |
| HOMO SAPIENS | 4 | 1067 | 4kyo | A | | 25 | LEU | p.Leu25Arg | P21549 |
| HOMO SAPIENS | 7 | 1067 | 4kyo | A | | 36 | ARG | p.Arg36His | P21549 |
| HOMO SAPIENS | 3 | 1067 | 4kyo | A | | 41 | GLY | p.Gly41Arg | P21549 |
| HOMO SAPIENS | 0 | 1067 | 4kyo | A | | 42 | GLY | p.Gly42Glu | P21549 |
| HOMO SAPIENS | 3 | 1067 | 4kyo | A | | 56 | ILE | p.Ile56Asn | P21549 |
| HOMO SAPIENS | 3 | 1067 | 4kyo | A | | 59 | GLU | p.Glu59Lys | P21549 |
| HOMO SAPIENS | 3 | 1067 | 4kyo | A | | 66 | TYR | p.Tyr66Ter | P21549 |
| HOMO SAPIENS | 3 | 1067 | 4kyo | A | | 111 | ARG | p.Arg111Leu | P21549 |
| HOMO SAPIENS | 7 | 1067 | 4kyo | A | | 118 | ARG | p.Arg118His | P21549 |
| HOMO SAPIENS | 3 | 1067 | 4kyo | A | | 122 | ARG | p.Arg122Ter | P21549 |
| HOMO SAPIENS | 3 | 1067 | 4kyo | A | | 124 | HIS | p.His124Pro | P21549 |
| HOMO SAPIENS | 0 | 1067 | 4kyo | A | | 152 | PHE | p.Phe152Ile | P21549 |
| HOMO SAPIENS | 1 | 1067 | 4kyo | A | | 156 | GLY | p.Gly156Arg | P21549 |
| HOMO SAPIENS | 0 | 1067 | 4kyo | A | | 161 | GLY | p.Gly161Arg | P21549 |
| HOMO SAPIENS | 3 | 1067 | 4kyo | A | | 166 | LEU | p.Leu166Pro | P21549 |
| HOMO SAPIENS | 5 | 1067 | 4kyo | A | | 170 | GLY | p.Gly170Arg | P21549 |
| HOMO SAPIENS | 5 | 1067 | 4kyo | A | | 186 | ALA | p.Ala186Val | P21549 |
| HOMO SAPIENS | 3 | 1067 | 4kyo | A | | 187 | SER | p.Ser187Phe | P21549 |
| HOMO SAPIENS | 3 | 1067 | 4kyo | A | | 197 | ARG | p.Arg197Pro | P21549 |
| HOMO SAPIENS | 5 | 1067 | 4kyo | A | | 201 | ASP | p.Asp201Glu | P21549 |
| HOMO SAPIENS | 0 | 1067 | 4kyo | A | | 202 | ILE | p.Ile202Asn | P21549 |
| HOMO SAPIENS | 1 | 1067 | 4kyo | A | | 205 | SER | p.Ser205Pro | P21549 |
| HOMO SAPIENS | 1 | 1067 | 4kyo | A | | 205 | SER | p.Ser205Ter | P21549 |
| HOMO SAPIENS | 2 | 1067 | 4kyo | A | | 216 | GLY | p.Gly216Arg | P21549 |
| HOMO SAPIENS | 5 | 1067 | 4kyo | A | | 233 | ARG | p.Arg233Cys | P21549 |
| HOMO SAPIENS | 5 | 1067 | 4kyo | A | | 244 | ILE | p.Ile244Thr | P21549 |
| HOMO SAPIENS | 5 | 1067 | 4kyo | A | | 244 | ILE | p.Ile244Met | P21549 |
| HOMO SAPIENS | 4 | 1067 | 4kyo | A | | 253 | CYS | p.Cys253Arg | P21549 |
| HOMO SAPIENS | 7 | 1067 | 4kyo | A | | 274 | GLU | p.Glu274Asp | P21549 |
| HOMO SAPIENS | 7 | 1067 | 4kyo | A | | 279 | ILE | p.Ile279Thr | P21549 |
| HOMO SAPIENS | 6 | 1067 | 4kyo | A | | 295 | ALA | p.Ala295Thr | P21549 |

| | | | | | | | | |
|---|---|---|---|---|---|---|---|---|
| HOMO SAPIENS | 5 | 1067 | 4kyo | A | 340 | ILE | p.Ile340Met | P21549 |
| HOMO SAPIENS | 1 | 790 | 2ric | C | 270 | SER | p.Ser290Thr | P35247 |
| HOMO SAPIENS | 0 | 790 | 2ric | C | 289 | GLU | p.Glu309Lys | P35247 |
| HOMO SAPIENS | 1 | 212 | 5sxe | A | 545 | GLU | p.Glu545Lys | P42336 |
| HOMO SAPIENS | 3 | 212 | 5sxe | A | 546 | GLN | p.Gln546Glu | P42336 |
| HOMO SAPIENS | 2 | 212 | 5sxe | A | 1047 | HIS | p.His1047Leu | P42336 |
| HOMO SAPIENS | 7 | 152 | 2zqq | F | 197 | ARG | p.Arg197Ter | Q13825 |
| HOMO SAPIENS | 0 | 152 | 2zqq | F | 248 | ALA | p.Ala248Thr | Q13825 |
| HOMO SAPIENS | 0 | 152 | 2zqq | F | 275 | ALA | p.Ala275Val | Q13825 |
| HOMO SAPIENS | 5 | 152 | 2zqq | F | 331 | LYS | p.Lys331Ter | Q13825 |
| HOMO SAPIENS | 6 | 17801 | 2j8l | A | 283 | PHE | p.Phe301Leu | P03951 |
| HOMO SAPIENS | 3 | 17801 | 2j8l | A | 308 | ARG | p.Arg326Cys | P03951 |
| HOMO SAPIENS | 0 | 17801 | 2j8l | A | 321 | CYS | p.Cys339Phe | P03951 |
| HOMO SAPIENS | 3 | 17801 | 2j8l | A | 351 | TYR | p.Tyr369Ter | P03951 |
| HOMO SAPIENS | 7 | 1802 | 5db0 | A | 22 | LEU | p.Leu22Arg | O00255 |
| HOMO SAPIENS | 7 | 1802 | 5db0 | A | 26 | GLU | p.Glu26Lys | O00255 |
| HOMO SAPIENS | 7 | 1802 | 5db0 | A | 139 | HIS | p.His139Asp | O00255 |
| HOMO SAPIENS | 0 | 1802 | 5db0 | A | 171 | ARG | p.Arg176Gln | O00255 |
| HOMO SAPIENS | 0 | 1802 | 5db0 | A | 184 | VAL | p.Val189Glu | O00255 |
| HOMO SAPIENS | 7 | 1802 | 5db0 | A | 255 | GLU | p.Glu260Lys | O00255 |
| HOMO SAPIENS | 4 | 1802 | 5db0 | A | 418 | ASP | p.Asp423Asn | O00255 |
| HOMO SAPIENS | 0 | 1802 | 5db0 | A | 436 | TRP | p.Trp441Arg | O00255 |
| HOMO SAPIENS | -1 | 1802 | 5db0 | A | 541 | ALA | p.Thr546Ala | O00255 |
| HOMO SAPIENS | 4 | 1802 | 5db0 | A | 552 | THR | p.Thr557Ser | O00255 |
| HOMO SAPIENS | 0 | 3450 | 2yuz | A | 36 | PRO | p.Pro461Gln | Q00872 |
| HOMO SAPIENS | 1 | 3450 | 2yuz | A | 56 | GLN | p.His481Gln | Q00872 |
| HOMO SAPIENS | 2 | 3450 | 2yuz | A | 68 | ALA | p.Ala493Val | Q00872 |
| HOMO SAPIENS | 0 | 3211 | 3rqg | A | 72 | VAL | p.Val72Leu | Q9BUL8 |
| HOMO SAPIENS | 2 | 4168 | 4q4f | A | 121 | ARG | p.Arg121Ter | Q14108 |
| HOMO SAPIENS | 2 | 4168 | 4q4f | A | 121 | ARG | p.Arg121Gln | Q14108 |
| HOMO SAPIENS | 2 | 4168 | 4q4f | A | 159 | MET | p.Met159Val | Q14108 |
| HOMO SAPIENS | 4 | 4168 | 4q4f | A | 363 | HIS | p.His363Asn | Q14108 |
| HOMO SAPIENS | 2 | 4168 | 4q4f | A | 396 | VAL | p.Val396Ile | Q14108 |
| HOMO SAPIENS | 0 | 1448 | 1y97 | A | 87 | ARG | p.Arg130Ter | Q9BQ50 |
| HOMO SAPIENS | 0 | 3444 | 4iwp | A | 151 | SER | p.Ser151Phe | Q9UHD2 |
| HOMO SAPIENS | 0 | 3444 | 4iwp | A | 401 | LYS | p.Lys401Glu | Q9UHD2 |
| HOMO SAPIENS | 0 | 116 | 1zy8 | B | 194 | GLY | p.Gly229Cys | P09622 |
| HOMO SAPIENS | 0 | 116 | 1zy8 | B | 255 | ASP | p.Asp290Gly | P09622 |
| HOMO SAPIENS | 2 | 116 | 1zy8 | B | 326 | MET | p.Met361Val | P09622 |
| HOMO SAPIENS | 0 | 116 | 1zy8 | B | 340 | GLU | p.Glu375Lys | P09622 |
| HOMO SAPIENS | 2 | 116 | 1zy8 | B | 358 | ILE | p.Ile393Thr | P09622 |
| HOMO SAPIENS | 1 | 116 | 1zy8 | B | 453 | PRO | p.Pro488Leu | P09622 |
| HOMO SAPIENS | 7 | 127 | 4qo7 | B | 160 | SER | p.Ser161Arg | P00338 |
| HOMO SAPIENS | 0 | 2025 | 1wrl | A | 8 | ALA | p.Ala8Val | P63316 |
| HOMO SAPIENS | 8 | 2025 | 1wrl | A | 24 | PHE | p.Phe24Leu | P63316 |
| HOMO SAPIENS | 0 | 2025 | 1wrl | A | 84 | SER | p.Cys84Tyr | P63316 |
| HOMO SAPIENS | 8 | 3791 | 1mhd | B | 93 | ARG | p.Arg93Ter | P84022 |
| HOMO SAPIENS | 0 | 1575 | 4phc | A | 205 | GLY | p.Gly205Asp | P12081 |
| HOMO SAPIENS | 0 | 1575 | 4phc | A | 454 | TYR | p.Tyr454Ser | P12081 |
| HOMO SAPIENS | 0 | 39620 | 1ozs | B | 135 | ARG | p.Arg136Gln | P19429 |
| HOMO SAPIENS | 5 | 39620 | 1ozs | B | 140 | ARG | p.Arg141Trp | P19429 |
| HOMO SAPIENS | 5 | 39620 | 1ozs | B | 142 | THR | p.Thr143Asn | P19429 |
| HOMO SAPIENS | 5 | 39620 | 1ozs | B | 143 | LEU | p.Leu144Gln | P19429 |
| HOMO SAPIENS | 5 | 39620 | 1ozs | B | 144 | ARG | p.Arg145Gly | P19429 |

| | | | | | | | | |
|---|---|---|---|---|---|---|---|---|
| HOMO SAPIENS | 5 | 39620 | 1ozs | B | 144 | ARG | p.Arg145Gln | P19429 |
| HOMO SAPIENS | -1 | 20934 | 4nuv | D | 42 | GLY | p.Gly42Asp | Q16570 |
| HOMO SAPIENS | -1 | 40565 | 5lcw | S | 4 | VAL | p.Val4Met | O60566 |
| HOMO SAPIENS | 0 | 40565 | 5lcw | S | 40 | THR | p.Thr40Met | O60566 |
| HOMO SAPIENS | 4 | 40565 | 5lcw | S | 198 | ALA | p.Glu390Asp | O60566 |
| HOMO SAPIENS | 4 | 758 | 4oiu | A | 726 | ARG | p.Arg727Leu | P10275 |
| HOMO SAPIENS | 4 | 758 | 4oiu | A | 730 | VAL | p.Val731Met | P10275 |
| HOMO SAPIENS | 2 | 758 | 4oiu | A | 798 | GLN | p.Gln799Glu | P10275 |
| HOMO SAPIENS | 3 | 758 | 4oiu | A | 855 | ARG | p.Arg856His | P10275 |
| HOMO SAPIENS | 1 | 758 | 4oiu | A | 886 | MET | p.Met887Val | P10275 |
| HOMO SAPIENS | 0 | 3303 | 4ie0 | A | 134 | ALA | p.Ala134Thr | Q9C0B1 |
| HOMO SAPIENS | 9 | 3303 | 4ie0 | A | 316 | ARG | p.Arg316Gln | Q9C0B1 |
| HOMO SAPIENS | 1 | 41502 | 3fgh | A | 136 | PRO | p.Pro178Leu | Q00059 |
| HOMO SAPIENS | 5 | 175 | 3ceh | B | 58 | THR | p.Thr59Lys | P06737 |
| HOMO SAPIENS | 8 | 175 | 3ceh | B | 93 | ARG | p.Arg94Ter | P06737 |
| HOMO SAPIENS | 5 | 175 | 3ceh | B | 221 | VAL | p.Val222Ile | P06737 |
| HOMO SAPIENS | 0 | 175 | 3ceh | B | 338 | ASN | p.Asn339Ser | P06737 |
| HOMO SAPIENS | 1 | 175 | 3ceh | B | 372 | PHE | p.Phe373Leu | P06737 |
| HOMO SAPIENS | 9 | 175 | 3ceh | B | 376 | ASN | p.Asn377Lys | P06737 |
| HOMO SAPIENS | 8 | 175 | 3ceh | B | 398 | ARG | p.Arg399Ter | P06737 |
| HOMO SAPIENS | 5 | 175 | 3ceh | B | 424 | ARG | p.Arg425Pro | P06737 |
| HOMO SAPIENS | 9 | 175 | 3ceh | B | 455 | VAL | p.Val456Met | P06737 |
| HOMO SAPIENS | 8 | 175 | 3ceh | B | 490 | ARG | p.Arg491Cys | P06737 |
| HOMO SAPIENS | 8 | 175 | 3ceh | B | 631 | ASN | p.Asn632Ile | P06737 |
| HOMO SAPIENS | 9 | 175 | 3ceh | B | 648 | TYR | p.Tyr649Ter | P06737 |
| HOMO SAPIENS | 9 | 175 | 3ceh | B | 672 | GLU | p.Glu673Lys | P06737 |
| HOMO SAPIENS | 9 | 175 | 3ceh | B | 674 | SER | p.Ser675Leu | P06737 |
| HOMO SAPIENS | 0 | 175 | 3ceh | B | 805 | ILE | p.Ile806Leu | P06737 |
| HOMO SAPIENS | 0 | 175 | 3ceh | B | 820 | TYR | p.Tyr821His | P06737 |
| HOMO SAPIENS | 1 | 6750 | 4fdj | A | 47 | GLY | p.Gly47Arg | P34059 |
| HOMO SAPIENS | 9 | 6750 | 4fdj | A | 60 | ASP | p.Asp60Asn | P34059 |
| HOMO SAPIENS | 5 | 6750 | 4fdj | A | 67 | LEU | p.Leu67Met | P34059 |
| HOMO SAPIENS | 0 | 6750 | 4fdj | A | 69 | PHE | p.Phe69Val | P34059 |
| HOMO SAPIENS | 3 | 6750 | 4fdj | A | 94 | ARG | p.Arg94Gly | P34059 |
| HOMO SAPIENS | 0 | 6750 | 4fdj | A | 113 | ILE | p.Ile113Phe | P34059 |
| HOMO SAPIENS | 0 | 6750 | 4fdj | A | 138 | VAL | p.Val138Ala | P34059 |
| HOMO SAPIENS | 0 | 6750 | 4fdj | A | 139 | GLY | p.Gly139Ser | P34059 |
| HOMO SAPIENS | 0 | 6750 | 4fdj | A | 151 | PRO | p.Pro151Thr | P34059 |
| HOMO SAPIENS | 0 | 6750 | 4fdj | A | 155 | GLY | p.Gly155Arg | P34059 |
| HOMO SAPIENS | 0 | 6750 | 4fdj | A | 159 | TRP | p.Trp159Ter | P34059 |
| HOMO SAPIENS | 0 | 6750 | 4fdj | A | 162 | SER | p.Ser162Phe | P34059 |
| HOMO SAPIENS | 0 | 6750 | 4fdj | A | 200 | THR | p.Thr200Met | P34059 |
| HOMO SAPIENS | 0 | 6750 | 4fdj | A | 204 | ASN | p.Asn204Lys | P34059 |
| HOMO SAPIENS | 0 | 6750 | 4fdj | A | 231 | ALA | p.Ala231Gly | P34059 |
| HOMO SAPIENS | 2 | 6750 | 4fdj | A | 235 | THR | p.Thr235Lys | P34059 |
| HOMO SAPIENS | 1 | 6750 | 4fdj | A | 259 | ARG | p.Arg259Gln | P34059 |
| HOMO SAPIENS | 0 | 6750 | 4fdj | A | 287 | SER | p.Ser287Leu | P34059 |
| HOMO SAPIENS | 0 | 6750 | 4fdj | A | 291 | ALA | p.Ala291Thr | P34059 |
| HOMO SAPIENS | 3 | 6750 | 4fdj | A | 301 | GLY | p.Gly301Cys | P34059 |
| HOMO SAPIENS | 0 | 6750 | 4fdj | A | 312 | THR | p.Thr312Ser | P34059 |
| HOMO SAPIENS | 0 | 6750 | 4fdj | A | 318 | MET | p.Met318Arg | P34059 |
| HOMO SAPIENS | 0 | 6750 | 4fdj | A | 340 | GLY | p.Gly340Asp | P34059 |
| HOMO SAPIENS | 0 | 6750 | 4fdj | A | 386 | ARG | p.Arg386Cys | P34059 |
| HOMO SAPIENS | 0 | 6750 | 4fdj | A | 391 | MET | p.Met391Val | P34059 |

| | | | | | | | | |
|---|---|---|---|---|---|---|---|---|
| HOMO SAPIENS | 0 | 6750 | 4fdj | A | 392 | ALA | p.Ala392Val | P34059 |
| HOMO SAPIENS | 0 | 6750 | 4fdj | A | 393 | ALA | p.Ala393Ser | P34059 |
| HOMO SAPIENS | 0 | 6750 | 4fdj | A | 459 | ALA | p.Ala459Val | P34059 |
| HOMO SAPIENS | 0 | 6750 | 4fdj | A | 473 | GLN | p.Gln473Ter | P34059 |
| HOMO SAPIENS | 2 | 6750 | 4fdj | A | 480 | VAL | p.Val480Phe | P34059 |
| HOMO SAPIENS | 0 | 6750 | 4fdj | A | 487 | ASN | p.Asn487Ser | P34059 |
| HOMO SAPIENS | 0 | 6750 | 4fdj | A | 488 | VAL | p.Val488Met | P34059 |
| HOMO SAPIENS | 0 | 6750 | 4fdj | A | 520 | TRP | p.Trp520Ter | P34059 |
| HOMO SAPIENS | 4 | 1088 | 2p3u | A | 102 | GLU | p.Glu142Lys | P00742 |
| HOMO SAPIENS | 6 | 370 | 1nr1 | D | 73 | ASP | p.Asp126His | P00367 |
| HOMO SAPIENS | 3 | 370 | 1nr1 | D | 221 | ARG | p.Arg274Cys | P00367 |
| HOMO SAPIENS | 2 | 370 | 1nr1 | D | 265 | ARG | p.Arg318Lys | P00367 |
| HOMO SAPIENS | 1 | 370 | 1nr1 | D | 269 | ARG | p.Arg322His | P00367 |
| HOMO SAPIENS | 1 | 370 | 1nr1 | D | 296 | GLU | p.Glu349Ala | P00367 |
| HOMO SAPIENS | 1 | 370 | 1nr1 | D | 445 | SER | p.Ser498Leu | P00367 |
| HOMO SAPIENS | 1 | 370 | 1nr1 | D | 446 | GLY | p.Gly499Asp | P00367 |
| HOMO SAPIENS | 1 | 370 | 1nr1 | D | 446 | GLY | p.Gly499Ser | P00367 |
| HOMO SAPIENS | 1 | 370 | 1nr1 | D | 448 | SER | p.Ser501Pro | P00367 |
| HOMO SAPIENS | 2 | 370 | 1nr1 | D | 454 | HIS | p.His507Tyr | P00367 |
| HOMO SAPIENS | 1 | 7720 | 2he0 | A | 147 | HIS | p.His2018Gln | P46531 |
| HOMO SAPIENS | 2 | 11844 | 5l0v | A | 218 | ARG | p.Arg218Ter | Q8NBL1 |
| HOMO SAPIENS | 2 | 11844 | 5l0v | A | 279 | ARG | p.Arg279Trp | Q8NBL1 |
| HOMO SAPIENS | 9 | 1485 | 5va0 | A | 477 | ARG | p.Arg477His | P04150 |
| HOMO SAPIENS | 0 | 235 | 2fun | B | 2248 | ARG | p.Arg248Trp | Q14790 |
| HOMO SAPIENS | 4 | 235 | 2fun | B | 2285 | HIS | p.Asp285His | Q14790 |
| HOMO SAPIENS | 0 | 4822 | 5y3n | A | 307 | ARG | p.Arg307Gly | Q12931 |
| HOMO SAPIENS | 8 | 32928 | 4m6w | A | 1886 | HIS | p.His1886Tyr | Q8IYD8 |
| HOMO SAPIENS | 0 | 2711 | 4y5v | F | 22 | GLY | p.Gly46Glu | P19235 |
| HOMO SAPIENS | -1 | 2711 | 4y5v | F | 166 | ALA | p.Ala190Ser | P19235 |
| HOMO SAPIENS | 4 | 5161 | 5bpu | A | 42 | HIS | p.His42Arg | Q00604 |
| HOMO SAPIENS | 0 | 5161 | 5bpu | A | 44 | TYR | p.Tyr44Cys | Q00604 |
| HOMO SAPIENS | 0 | 5161 | 5bpu | A | 45 | VAL | p.Val45Glu | Q00604 |
| HOMO SAPIENS | 0 | 5161 | 5bpu | A | 60 | VAL | p.Val60Glu | Q00604 |
| HOMO SAPIENS | 0 | 5161 | 5bpu | A | 61 | LEU | p.Leu61Phe | Q00604 |
| HOMO SAPIENS | 7 | 5161 | 5bpu | A | 69 | CYS | p.Cys69Ser | Q00604 |
| HOMO SAPIENS | 0 | 5161 | 5bpu | A | 75 | SER | p.Ser75Cys | Q00604 |
| HOMO SAPIENS | 0 | 5161 | 5bpu | A | 90 | ARG | p.Arg90Pro | Q00604 |
| HOMO SAPIENS | 9 | 5161 | 5bpu | A | 96 | CYS | p.Cys96Tyr | Q00604 |
| HOMO SAPIENS | 9 | 5161 | 5bpu | A | 96 | CYS | p.Cys96Trp | Q00604 |
| HOMO SAPIENS | 0 | 5161 | 5bpu | A | 101 | SER | p.Ser101Phe | Q00604 |
| HOMO SAPIENS | 0 | 5161 | 5bpu | A | 105 | ALA | p.Ala105Thr | Q00604 |
| HOMO SAPIENS | 0 | 5161 | 5bpu | A | 110 | CYS | p.Cys110Gly | Q00604 |
| HOMO SAPIENS | 0 | 5161 | 5bpu | A | 121 | ARG | p.Arg121Trp | Q00604 |
| HOMO SAPIENS | 0 | 5161 | 5bpu | A | 121 | ARG | p.Arg121Leu | Q00604 |
| HOMO SAPIENS | 0 | 5161 | 5bpu | A | 124 | LEU | p.Leu124Phe | Q00604 |
| HOMO SAPIENS | 8 | 5161 | 5bpu | A | 128 | CYS | p.Cys128Ter | Q00604 |
| HOMO SAPIENS | 1 | 13944 | 5xqh | A | 238 | ARG | p.Arg238His | Q9GZN7 |
| HOMO SAPIENS | -1 | 7508 | 5jl6 | A | 39 | TRP | p.Trp39Arg | P11511 |
| HOMO SAPIENS | 0 | 7508 | 5jl6 | A | 201 | THR | p.Thr201Met | P11511 |
| HOMO SAPIENS | 0 | 7508 | 5jl6 | A | 210 | GLU | p.Glu210Lys | P11511 |
| HOMO SAPIENS | 1 | 7508 | 5jl6 | A | 264 | ARG | p.Arg264Cys | P11511 |
| HOMO SAPIENS | 2 | 7508 | 5jl6 | A | 365 | ARG | p.Arg365Gln | P11511 |
| HOMO SAPIENS | 7 | 7508 | 5jl6 | A | 375 | ARG | p.Arg375Cys | P11511 |
| HOMO SAPIENS | 8 | 7508 | 5jl6 | A | 435 | ARG | p.Arg435Cys | P11511 |

| | | | | | | | | |
|---|---|---|---|---|---|---|---|---|
| HOMO SAPIENS | 9 | 7508 | 5jl6 | A | 437 | CYS | p.Cys437Tyr | P11511 |
| HOMO SAPIENS | -1 | 30206 | 5l95 | B | 246 | ARG | p.Arg246Ter | Q9GZZ9 |
| HOMO SAPIENS | 4 | 875 | 3efh | B | 43 | GLU | p.Glu43Asp | P60891 |
| HOMO SAPIENS | 5 | 875 | 3efh | B | 65 | ASP | p.Asp65Asn | P60891 |
| HOMO SAPIENS | 0 | 875 | 3efh | B | 87 | ALA | p.Ala87Thr | P60891 |
| HOMO SAPIENS | 2 | 875 | 3efh | B | 115 | MET | p.Met115Thr | P60891 |
| HOMO SAPIENS | 8 | 875 | 3efh | B | 133 | GLN | p.Gln133Pro | P60891 |
| HOMO SAPIENS | 8 | 875 | 3efh | B | 152 | LEU | p.Leu152Pro | P60891 |
| HOMO SAPIENS | 7 | 875 | 3efh | B | 290 | ILE | p.Ile290Thr | P60891 |
| HOMO SAPIENS | 6 | 875 | 3efh | B | 306 | GLY | p.Gly306Arg | P60891 |
| HOMO SAPIENS | 0 | 7978 | 5edm | A | 122 | MET | p.Thr165Met | P00734 |
| HOMO SAPIENS | -1 | 7978 | 5edm | A | 257 | ARG | p.Arg314Cys | P00734 |
| HOMO SAPIENS | 0 | 1716 | 1b98 | M | 126 | ARG | p.Arg206Gln | P34130 |
| HOMO SAPIENS | 0 | 1716 | 1b98 | M | 126 | ARG | p.Arg206Trp | P34130 |
| HOMO SAPIENS | 9 | 3960 | 2qyp | B | 28 | ILE | p.Ile338Val | P07602 |
| HOMO SAPIENS | 1 | 3960 | 2qyp | B | 39 | LEU | p.Leu349Pro | P07602 |
| HOMO SAPIENS | 0 | 31414 | 5u1d | A | 333 | ILE | p.Ile393Val | Q03518 |
| HOMO SAPIENS | 0 | 31414 | 5u1d | A | 637 | ASP | p.Asp697Gly | Q03518 |
| HOMO SAPIENS | 0 | 31414 | 5u1d | A | 659 | ARG | p.Arg719Gln | Q03518 |
| HOMO SAPIENS | 3 | 119 | 2pho | A | 11 | ILE | p.Ile11Thr | P05089 |
| HOMO SAPIENS | 8 | 119 | 2pho | A | 21 | ARG | p.Arg21Ter | P05089 |
| HOMO SAPIENS | 0 | 119 | 2pho | A | 122 | TRP | p.Trp122Ter | P05089 |
| HOMO SAPIENS | 6 | 119 | 2pho | A | 138 | GLY | p.Gly138Val | P05089 |
| HOMO SAPIENS | 5 | 119 | 2pho | A | 290 | THR | p.Thr290Ser | P05089 |
| HOMO SAPIENS | 5 | 119 | 2pho | A | 291 | ARG | p.Arg291Ter | P05089 |
| HOMO SAPIENS | 0 | 3732 | 2i99 | A | 247 | TYR | p.Tyr247Ter | Q14894 |
| HOMO SAPIENS | -1 | 1382 | 4ffx | A | 2 | ALA | p.Ala2Val | P30566 |
| HOMO SAPIENS | 1 | 1382 | 4ffx | A | 72 | ILE | p.Ile72Met | P30566 |
| HOMO SAPIENS | 1 | 1382 | 4ffx | A | 100 | PRO | p.Pro100Ala | P30566 |
| HOMO SAPIENS | 0 | 1382 | 4ffx | A | 114 | TYR | p.Tyr114His | P30566 |
| HOMO SAPIENS | 1 | 1382 | 4ffx | A | 141 | ARG | p.Arg141Trp | P30566 |
| HOMO SAPIENS | 0 | 1382 | 4ffx | A | 147 | LYS | p.Lys147Met | P30566 |
| HOMO SAPIENS | 3 | 1382 | 4ffx | A | 190 | ARG | p.Arg190Gln | P30566 |
| HOMO SAPIENS | 3 | 1382 | 4ffx | A | 190 | ARG | p.Arg190Ter | P30566 |
| HOMO SAPIENS | 0 | 1382 | 4ffx | A | 217 | HIS | p.His217Asp | P30566 |
| HOMO SAPIENS | 2 | 1382 | 4ffx | A | 246 | LYS | p.Lys246Glu | P30566 |
| HOMO SAPIENS | 5 | 1382 | 4ffx | A | 303 | ARG | p.Arg303Cys | P30566 |
| HOMO SAPIENS | 1 | 1382 | 4ffx | A | 318 | PRO | p.Pro318Leu | P30566 |
| HOMO SAPIENS | 6 | 1382 | 4ffx | A | 332 | ASP | p.Asp332His | P30566 |
| HOMO SAPIENS | 6 | 1382 | 4ffx | A | 337 | ARG | p.Arg337Ter | P30566 |
| HOMO SAPIENS | 1 | 1382 | 4ffx | A | 396 | ARG | p.Arg396His | P30566 |
| HOMO SAPIENS | 1 | 1382 | 4ffx | A | 426 | ARG | p.Arg426His | P30566 |
| HOMO SAPIENS | 0 | 1382 | 4ffx | A | 438 | SER | p.Ser438Pro | P30566 |
| HOMO SAPIENS | 1 | 1382 | 4ffx | A | 447 | SER | p.Ser447Pro | P30566 |
| HOMO SAPIENS | 2 | 1382 | 4ffx | A | 448 | SER | p.Ser448Pro | P30566 |
| HOMO SAPIENS | 2 | 1382 | 4ffx | A | 450 | THR | p.Thr450Ser | P30566 |
| HOMO SAPIENS | 0 | 6212 | 1ao3 | A | 46 | SER | p.Ser1731Thr | P04275 |
| HOMO SAPIENS | 0 | 1711 | 3f9c | A | 279 | VAL | p.Val279Phe | Q13093 |
| HOMO SAPIENS | 0 | 6112 | 4fvl | A | 312 | HIS | p.His312Arg | P45452 |
| HOMO SAPIENS | 0 | 6112 | 4fvl | A | 323 | THR | p.Thr323Met | P45452 |
| HOMO SAPIENS | 0 | 6112 | 4fvl | A | 390 | ASP | p.Asp390Gly | P45452 |
| HOMO SAPIENS | 0 | 6112 | 4fvl | A | 458 | ARG | p.Arg458Cys | P45452 |
| HOMO SAPIENS | 2 | 6180 | 5ggn | A | 97 | ARG | p.Arg97Trp | Q8WZA1 |
| HOMO SAPIENS | 2 | 6180 | 5ggn | A | 215 | ARG | p.Arg215Ter | Q8WZA1 |

| | | | | | | | | |
|---|---|---|---|---|---|---|---|---|
| HOMO SAPIENS | -1 | 5526 | 4ak3 | A | 11 | ASP | p.Asp1232Asn | P02461 |
| HOMO SAPIENS | 6 | 5526 | 4ak3 | A | 38 | ALA | p.Ala1259Thr | P02461 |
| HOMO SAPIENS | 0 | 5526 | 4ak3 | A | 92 | LYS | p.Lys1313Arg | P02461 |
| HOMO SAPIENS | 0 | 5526 | 4ak3 | A | 132 | GLN | p.His1353Gln | P02461 |
| HOMO SAPIENS | 4 | 16616 | 1hci | B | 433 | SER | p.Ser433Leu | P35609 |
| HOMO SAPIENS | 1 | 16616 | 1hci | B | 475 | ASP | p.Asp475Asn | P35609 |
| HOMO SAPIENS | 1 | 16616 | 1hci | B | 476 | ALA | p.Ala476Ser | P35609 |
| HOMO SAPIENS | 2 | 16616 | 1hci | B | 507 | MET | p.Met507Val | P35609 |
| HOMO SAPIENS | 1 | 16616 | 1hci | B | 604 | MET | p.Met604Val | P35609 |
| HOMO SAPIENS | 2 | 16616 | 1hci | B | 640 | ARG | p.Arg640His | P35609 |
| HOMO SAPIENS | 1 | 16616 | 1hci | B | 721 | ARG | p.Arg721Cys | P35609 |
| HOMO SAPIENS | -1 | 3402 | 5iaa | B | 57 | MET | p.Met57Val | Q9GZZ9 |
| HOMO SAPIENS | 7 | 3250 | 3irq | B | 193 | PRO | p.Pro193Ala | P55265 |
| HOMO SAPIENS | 2 | 15200 | 2wns | A | 30 | SER | p.Ser30Gly | P11172 |
| HOMO SAPIENS | 0 | 15200 | 2wns | A | 96 | ARG | p.Arg96Gly | P11172 |
| HOMO SAPIENS | 0 | 15200 | 2wns | A | 109 | VAL | p.Val109Gly | P11172 |
| HOMO SAPIENS | 2 | 15759 | 4x4w | B | 140 | LEU | p.Leu166Ser | Q96Q11 |
| HOMO SAPIENS | 0 | 15759 | 4x4w | B | 197 | ILE | p.Ile223Thr | Q96Q11 |
| HOMO SAPIENS | 0 | 3156 | 3w03 | B | 14 | ALA | p.Ala14Thr | Q9H9Q4 |
| HOMO SAPIENS | 0 | 3156 | 3w03 | B | 57 | ARG | p.Arg57Gly | Q9H9Q4 |
| HOMO SAPIENS | 0 | 3156 | 3w03 | B | 123 | CYS | p.Cys123Arg | Q9H9Q4 |
| HOMO SAPIENS | 0 | 11693 | 3ojy | A | 63 | GLN | p.Gln93Lys | P07357 |
| HOMO SAPIENS | 0 | 1421 | 2qc8 | E | 275 | GLU | p.Glu275Asp | P15104 |
| HOMO SAPIENS | 9 | 1421 | 2qc8 | E | 324 | ARG | p.Arg324Cys | P15104 |
| HOMO SAPIENS | 6 | 1421 | 2qc8 | E | 341 | ARG | p.Arg341Cys | P15104 |
| HOMO SAPIENS | 6 | 2850 | 3qvb | A | 83 | THR | p.Thr83Met | P46976 |
| HOMO SAPIENS | 8 | 2850 | 3qvb | A | 102 | ASP | p.Asp102His | P46976 |
| HOMO SAPIENS | 1 | 24314 | 4m9p | A | 480 | PRO | p.Pro480Leu | P21333 |
| HOMO SAPIENS | 0 | 24314 | 4m9p | A | 528 | VAL | p.Val528Met | P21333 |
| HOMO SAPIENS | 0 | 24314 | 4m9p | A | 590 | VAL | p.Val590Ile | P21333 |
| HOMO SAPIENS | 0 | 24314 | 4m9p | A | 605 | ASP | p.Asp605Asn | P21333 |
| HOMO SAPIENS | 1 | 24314 | 4m9p | A | 637 | PRO | p.Pro637Gln | P21333 |
| HOMO SAPIENS | 0 | 24314 | 4m9p | A | 656 | LEU | p.Leu656Phe | P21333 |
| HOMO SAPIENS | 2 | 24314 | 4m9p | A | 711 | VAL | p.Val711Asp | P21333 |
| HOMO SAPIENS | 0 | 4951 | 5joe | A | 40 | THR | p.Thr9621Ser | Q8WZ42 |
| HOMO SAPIENS | 1 | 4951 | 5joe | A | 41 | GLU | p.Glu9622Gln | Q8WZ42 |
| HOMO SAPIENS | 0 | 4951 | 5joe | A | 82 | ALA | p.Ala9663Ser | Q8WZ42 |
| HOMO SAPIENS | 4 | 4951 | 5joe | A | 89 | ILE | p.Ile9670Asn | Q8WZ42 |
| HOMO SAPIENS | 0 | 9664 | 3c6m | A | 56 | GLY | p.Gly56Ser | P52788 |
| HOMO SAPIENS | 0 | 9664 | 3c6m | A | 132 | VAL | p.Val132Gly | P52788 |
| HOMO SAPIENS | 1 | 9664 | 3c6m | A | 190 | THR | p.Thr190Ile | P52788 |
| HOMO SAPIENS | -1 | 12118 | 5hcd | B | 19 | GLN | p.Gln19Ter | P01031 |
| HOMO SAPIENS | 0 | 1583 | 2b4x | L | 7 | ILE | p.Ile39Asn | P01008 |
| HOMO SAPIENS | 0 | 1583 | 2b4x | L | 24 | ARG | p.Arg56Cys | P01008 |
| HOMO SAPIENS | 0 | 1583 | 2b4x | L | 47 | ARG | p.Arg79Cys | P01008 |
| HOMO SAPIENS | 0 | 1583 | 2b4x | L | 47 | ARG | p.Arg79His | P01008 |
| HOMO SAPIENS | 1 | 1583 | 2b4x | L | 95 | CYS | p.Cys127Arg | P01008 |
| HOMO SAPIENS | 1 | 1583 | 2b4x | L | 99 | LEU | p.Leu131Phe | P01008 |
| HOMO SAPIENS | 0 | 1583 | 2b4x | L | 116 | SER | p.Ser148Pro | P01008 |
| HOMO SAPIENS | 0 | 1583 | 2b4x | L | 129 | ARG | p.Arg161Gln | P01008 |
| HOMO SAPIENS | 0 | 1583 | 2b4x | L | 135 | ASN | p.Asn167Thr | P01008 |
| HOMO SAPIENS | 2 | 1583 | 2b4x | L | 187 | ASN | p.Asn219Asp | P01008 |
| HOMO SAPIENS | 0 | 1583 | 2b4x | L | 191 | SER | p.Ser223Pro | P01008 |
| HOMO SAPIENS | 2 | 1583 | 2b4x | L | 349 | SER | p.Ser381Pro | P01008 |

| | | | | | | | | |
|---|---|---|---|---|---|---|---|---|
| HOMO SAPIENS | 0 | 1583 | 2b4x | L | 382 | ALA | p.Ala414Thr | P01008 |
| HOMO SAPIENS | 0 | 1583 | 2b4x | L | 387 | ALA | p.Ala419Val | P01008 |
| HOMO SAPIENS | 0 | 1583 | 2b4x | L | 392 | GLY | p.Gly424Asp | P01008 |
| HOMO SAPIENS | 0 | 1583 | 2b4x | L | 393 | ARG | p.Arg425His | P01008 |
| HOMO SAPIENS | 0 | 1583 | 2b4x | L | 393 | ARG | p.Arg425Cys | P01008 |
| HOMO SAPIENS | 0 | 1583 | 2b4x | L | 394 | SER | p.Ser426Leu | P01008 |
| HOMO SAPIENS | 0 | 1583 | 2b4x | L | 404 | ALA | p.Ala436Thr | P01008 |
| HOMO SAPIENS | 2 | 1583 | 2b4x | L | 407 | PRO | p.Pro439Leu | P01008 |
| HOMO SAPIENS | 2 | 1583 | 2b4x | L | 429 | PRO | p.Pro461Leu | P01008 |
| HOMO SAPIENS | 0 | 11647 | 2a1t | S | 78 | ALA | p.Ala78Ser | P38117 |
| HOMO SAPIENS | 0 | 11647 | 2a1t | S | 98 | ARG | p.Arg98Cys | P38117 |
| HOMO SAPIENS | 8 | 11647 | 2a1t | S | 128 | ASP | p.Asp128Asn | P38117 |
| HOMO SAPIENS | 5 | 11647 | 2a1t | S | 154 | THR | p.Thr154Met | P38117 |
| HOMO SAPIENS | 6 | 11647 | 2a1t | S | 164 | ARG | p.Arg164Gln | P38117 |
| HOMO SAPIENS | 5 | 11647 | 2a1t | S | 174 | ARG | p.Arg174His | P38117 |
| HOMO SAPIENS | -1 | 11200 | 4tx5 | A | 4 | ILE | p.Ile59Val | Q9NR28 |
| HOMO SAPIENS | 0 | 11200 | 4tx5 | A | 169 | GLY | p.Gly224Arg | Q9NR28 |
| HOMO SAPIENS | 3 | 417 | 2vr2 | A | 6 | ARG | p.Arg6Gln | Q14117 |
| HOMO SAPIENS | 1 | 417 | 2vr2 | A | 334 | GLN | p.Gln334Arg | Q14117 |
| HOMO SAPIENS | 2 | 417 | 2vr2 | A | 360 | TRP | p.Trp360Arg | Q14117 |
| HOMO SAPIENS | 0 | 417 | 2vr2 | A | 379 | SER | p.Ser379Arg | Q14117 |
| HOMO SAPIENS | 0 | 417 | 2vr2 | A | 435 | GLY | p.Gly435Arg | Q14117 |
| HOMO SAPIENS | 0 | 12498 | 3eo3 | B | 423 | ILE | p.Ile423Val | Q9Y223 |
| HOMO SAPIENS | 5 | 12498 | 3eo3 | B | 572 | VAL | p.Val572Leu | Q9Y223 |
| HOMO SAPIENS | 0 | 12498 | 3eo3 | B | 576 | GLY | p.Gly576Glu | Q9Y223 |
| HOMO SAPIENS | 0 | 12498 | 3eo3 | B | 587 | ILE | p.Ile587Thr | Q9Y223 |
| HOMO SAPIENS | 0 | 12498 | 3eo3 | B | 615 | SER | p.Ser615Ter | Q9Y223 |
| HOMO SAPIENS | 1 | 12498 | 3eo3 | B | 631 | ALA | p.Ala631Val | Q9Y223 |
| HOMO SAPIENS | 1 | 12498 | 3eo3 | B | 669 | GLY | p.Gly669Arg | Q9Y223 |
| HOMO SAPIENS | 1 | 12498 | 3eo3 | B | 696 | VAL | p.Val696Met | Q9Y223 |
| HOMO SAPIENS | 2 | 12498 | 3eo3 | B | 712 | MET | p.Met712Thr | Q9Y223 |
| HOMO SAPIENS | 0 | 1568 | 1r74 | A | 49 | LEU | p.Leu50Pro | Q14749 |
| HOMO SAPIENS | 0 | 1568 | 1r74 | A | 176 | HIS | p.His177Asn | Q14749 |
| HOMO SAPIENS | 3 | 147 | 5eig | A | 67 | THR | p.Thr67Ile | P32929 |
| HOMO SAPIENS | 3 | 147 | 5eig | A | 240 | GLN | p.Gln240Glu | P32929 |
| HOMO SAPIENS | -1 | 147 | 5eig | A | 403 | SER | p.Ser403Ile | P32929 |
| HOMO SAPIENS | 8 | 20036 | 2lsr | A | 31 | ARG | p.Arg31Ter | Q9Y6N9 |
| HOMO SAPIENS | -1 | 41382 | 4uis | B | 431 | UNK | p.Ala431Glu | P49768 |
| HOMO SAPIENS | -1 | 8536 | 3i6w | B | 85 | PRO | p.Pro85Leu | O96017 |
| HOMO SAPIENS | 0 | 8536 | 3i6w | B | 93 | TRP | p.Trp93Ter | O96017 |
| HOMO SAPIENS | 0 | 8536 | 3i6w | B | 95 | ARG | p.Arg95Ter | O96017 |
| HOMO SAPIENS | 0 | 8536 | 3i6w | B | 117 | ARG | p.Arg117Gly | O96017 |
| HOMO SAPIENS | 0 | 8536 | 3i6w | B | 119 | LYS | p.Lys119Arg | O96017 |
| HOMO SAPIENS | 0 | 8536 | 3i6w | B | 137 | ARG | p.Arg137Gln | O96017 |
| HOMO SAPIENS | 0 | 8536 | 3i6w | B | 137 | ARG | p.Arg137Ter | O96017 |
| HOMO SAPIENS | 0 | 8536 | 3i6w | B | 145 | ARG | p.Arg145Trp | O96017 |
| HOMO SAPIENS | 0 | 8536 | 3i6w | B | 160 | ILE | p.Ile160Met | O96017 |
| HOMO SAPIENS | 0 | 41322 | 4bxl | C | 46 | GLU | p.Glu46Lys | P37840 |
| HOMO SAPIENS | 5 | 41322 | 4bxl | C | 53 | ALA | p.Ala53Thr | P37840 |
| HOMO SAPIENS | 2 | 24966 | 3hve | A | 15 | ARG | p.Arg15Ser | Q9H2C0 |
| HOMO SAPIENS | 1 | 24966 | 3hve | A | 138 | ARG | p.Arg138His | Q9H2C0 |
| HOMO SAPIENS | 0 | 24966 | 3hve | A | 201 | ARG | p.Arg201Ter | Q9H2C0 |
| HOMO SAPIENS | 0 | 11680 | 5an9 | J | 8 | ASN | p.Asn8Lys | Q9Y3A5 |
| HOMO SAPIENS | 1 | 11680 | 5an9 | J | 33 | LYS | p.Lys33Arg | Q9Y3A5 |

| | | | | | | | | |
|---|---|---|---|---|---|---|---|---|
| HOMO SAPIENS | 1 | 11680 | 5an9 | J | 126 | ARG | p.Arg126Thr | Q9Y3A5 |
| HOMO SAPIENS | 1 | 11680 | 5an9 | J | 130 | VAL | p.Val130Met | Q9Y3A5 |
| HOMO SAPIENS | 1 | 11680 | 5an9 | J | 169 | ARG | p.Arg169Cys | Q9Y3A5 |
| HOMO SAPIENS | 1 | 11680 | 5an9 | J | 218 | ARG | p.Arg218Ter | Q9Y3A5 |
| HOMO SAPIENS | 4 | 978 | 3zs0 | B | 7 | TYR | p.Tyr173Cys | P05164 |
| HOMO SAPIENS | 1 | 978 | 3zs0 | B | 85 | MET | p.Met251Thr | P05164 |
| HOMO SAPIENS | 4 | 5140 | 2l4r | A | 52 | CYS | p.Cys52Ter | Q12809 |
| HOMO SAPIENS | 4 | 5140 | 2l4r | A | 63 | PRO | p.Pro63His | Q12809 |
| HOMO SAPIENS | 0 | 5140 | 2l4r | A | 100 | ARG | p.Arg100Gly | Q12809 |
| HOMO SAPIENS | 4 | 5140 | 2l4r | A | 129 | PHE | p.Phe129Leu | Q12809 |
| HOMO SAPIENS | 0 | 25836 | 2gy7 | B | 103 | GLU | p.Glu103Asp | Q02763 |
| HOMO SAPIENS | 0 | 25836 | 2gy7 | B | 148 | ILE | p.Ile148Thr | Q02763 |
| HOMO SAPIENS | 0 | 25836 | 2gy7 | B | 226 | ALA | p.Ala226Val | Q02763 |
| HOMO SAPIENS | 0 | 25836 | 2gy7 | B | 391 | THR | p.Thr391Ile | Q02763 |
| HOMO SAPIENS | 4 | 12563 | 5kwy | D | 20 | GLU | p.Glu20Ter | P61916 |
| HOMO SAPIENS | 2 | 12563 | 5kwy | D | 30 | VAL | p.Val30Met | P61916 |
| HOMO SAPIENS | 4 | 12563 | 5kwy | D | 39 | VAL | p.Val39Met | P61916 |
| HOMO SAPIENS | 8 | 12563 | 5kwy | D | 67 | SER | p.Ser67Pro | P61916 |
| HOMO SAPIENS | 8 | 12563 | 5kwy | D | 99 | CYS | p.Cys99Arg | P61916 |
| HOMO SAPIENS | 2 | 12563 | 5kwy | D | 118 | GLU | p.Glu118Ter | P61916 |
| HOMO SAPIENS | 2 | 12563 | 5kwy | D | 120 | PRO | p.Pro120Ser | P61916 |
| HOMO SAPIENS | -1 | 1679 | 1pmk | A | -2 | VAL | p.Val374Phe | P00747 |
| HOMO SAPIENS | -1 | 5315 | 2w2n | P | 53 | ALA | p.Ala53Val | Q8NBP7 |
| HOMO SAPIENS | 8 | 13496 | 2yad | A | 138 | THR | p.Asn138Thr | P11686 |
| HOMO SAPIENS | -1 | 13496 | 2yad | A | 173 | ARG | p.Arg167Gln | P11686 |
| HOMO SAPIENS | 2 | 13496 | 2yad | A | 186 | SER | p.Asn186Ser | P11686 |
| HOMO SAPIENS | 0 | 4426 | 3qd6 | C | 123 | ALA | p.Ala123Glu | P29965 |
| HOMO SAPIENS | 1 | 4426 | 3qd6 | C | 140 | TRP | p.Trp140Gly | P29965 |
| HOMO SAPIENS | 0 | 4426 | 3qd6 | C | 155 | LEU | p.Leu155Pro | P29965 |
| HOMO SAPIENS | 6 | 4426 | 3qd6 | C | 227 | GLY | p.Gly227Val | P29965 |
| HOMO SAPIENS | 0 | 4426 | 3qd6 | C | 235 | ALA | p.Ala235Pro | P29965 |
| HOMO SAPIENS | 0 | 10734 | 3dyd | B | 45 | PRO | p.Pro45Ser | P17735 |
| HOMO SAPIENS | -1 | 10734 | 3dyd | B | 57 | ARG | p.Arg57Ter | P17735 |
| HOMO SAPIENS | 8 | 10734 | 3dyd | B | 417 | ARG | p.Arg417Ter | P17735 |
| HOMO SAPIENS | 0 | 6956 | 4pzs | A | 1600 | LEU | p.Leu1600Phe | Q09472 |
| HOMO SAPIENS | 0 | 9370 | 3hfv | A | 21 | SER | p.Ser57Cys | O95363 |
| HOMO SAPIENS | 1 | 9370 | 3hfv | A | 25 | ASP | p.Asp61Glu | O95363 |
| HOMO SAPIENS | 1 | 9370 | 3hfv | A | 130 | ALA | p.Ala166Thr | O95363 |
| HOMO SAPIENS | 5 | 9370 | 3hfv | A | 148 | ASP | p.Asp184Asn | O95363 |
| HOMO SAPIENS | 0 | 9370 | 3hfv | A | 187 | ARG | p.Arg223Cys | O95363 |
| HOMO SAPIENS | 0 | 9370 | 3hfv | A | 210 | THR | p.Thr246Met | O95363 |
| HOMO SAPIENS | 1 | 9370 | 3hfv | A | 244 | ASN | p.Asn280Ser | O95363 |
| HOMO SAPIENS | 2 | 9370 | 3hfv | A | 288 | TYR | p.Tyr324Cys | O95363 |
| HOMO SAPIENS | 5 | 9370 | 3hfv | A | 293 | ILE | p.Ile329Thr | O95363 |
| HOMO SAPIENS | 1 | 9370 | 3hfv | A | 355 | ASP | p.Asp391Val | O95363 |
| HOMO SAPIENS | 1 | 9370 | 3hfv | A | 371 | THR | p.Thr407Met | O95363 |
| HOMO SAPIENS | 1 | 9370 | 3hfv | A | 383 | ARG | p.Arg419Cys | O95363 |
| HOMO SAPIENS | 1 | 9370 | 3hfv | A | 387 | ARG | p.Arg423Gln | O95363 |
| HOMO SAPIENS | 0 | 14401 | 1q3x | A | 371 | ASP | p.Asp371Tyr | O00187 |
| HOMO SAPIENS | 0 | 14401 | 1q3x | A | 377 | VAL | p.Val377Ala | O00187 |
| HOMO SAPIENS | 0 | 14401 | 1q3x | A | 439 | ARG | p.Arg439His | O00187 |
| HOMO SAPIENS | 0 | 14401 | 1q3x | A | 577 | GLN | p.Gln577His | O00187 |
| HOMO SAPIENS | 2 | 18270 | 3teq | A | 429 | ARG | p.Arg429Cys | Q13586 |
| HOMO SAPIENS | -1 | 16081 | 4mqt | A | 231 | VAL | p.Val231Ile | P08172 |

| | | | | | | | | |
|---|---|---|---|---|---|---|---|---|
| HOMO SAPIENS | 9 | 6607 | 3whu | B | 81 | ASP | p.Asp81His | Q8NI22 |
| HOMO SAPIENS | 9 | 6607 | 3whu | B | 129 | ASP | p.Asp129Glu | Q8NI22 |
| HOMO SAPIENS | 9 | 6607 | 3whu | B | 136 | ILE | p.Ile136Thr | Q8NI22 |
| HOMO SAPIENS | 0 | 7208 | 2e9w | D | 11 | ASN | p.Asn36Ser | P21583 |
| HOMO SAPIENS | 0 | 24566 | 4bdx | A | 10 | LEU | p.Leu140Val | P00748 |
| HOMO SAPIENS | 5 | 24566 | 4bdx | A | 77 | PRO | p.Ala207Pro | P00748 |
| HOMO SAPIENS | 7 | 606 | 3qnf | A | 319 | MET | p.Met319Val | Q9NZ08 |
| HOMO SAPIENS | 8 | 4097 | 1ydl | A | 21 | LEU | p.Leu21Pro | Q6ZYL4 |
| HOMO SAPIENS | 6 | 4097 | 1ydl | A | 56 | ARG | p.Arg56Ter | Q6ZYL4 |
| HOMO SAPIENS | 8 | 36691 | 1dqb | A | 42 | ARG | p.Arg403Lys | P07204 |
| HOMO SAPIENS | 0 | 5642 | 3j7y | X | 59 | ARG | p.Arg59Pro | Q13084 |
| HOMO SAPIENS | 1 | 5642 | 3j7y | X | 204 | VAL | p.Val204Met | Q13084 |
| HOMO SAPIENS | 1 | 1119 | 3nwu | B | 252 | ALA | p.Ala252Thr | Q92743 |
| HOMO SAPIENS | 3 | 1119 | 3nwu | B | 274 | ARG | p.Arg274Gln | Q92743 |
| HOMO SAPIENS | 9 | 1119 | 3nwu | B | 297 | VAL | p.Val297Met | Q92743 |
| HOMO SAPIENS | -1 | 1119 | 3nwu | B | 302 | ARG | p.Arg302Ter | Q92743 |
| HOMO SAPIENS | 9 | 1119 | 3nwu | B | 321 | ALA | p.Ala321Thr | Q92743 |
| HOMO SAPIENS | -1 | 1119 | 3nwu | B | 370 | ARG | p.Arg370Ter | Q92743 |
| HOMO SAPIENS | 0 | 416 | 4ycu | B | 105 | LEU | p.Leu105His | Q15046 |
| HOMO SAPIENS | 0 | 416 | 4ycu | B | 145 | TYR | p.Tyr145His | Q15046 |
| HOMO SAPIENS | 0 | 416 | 4ycu | B | 200 | PRO | p.Pro200Leu | Q15046 |
| HOMO SAPIENS | 0 | 416 | 4ycu | B | 201 | TYR | p.Tyr201His | Q15046 |
| HOMO SAPIENS | 0 | 416 | 4ycu | B | 420 | ARG | p.Arg420Cys | Q15046 |
| HOMO SAPIENS | 9 | 3297 | 4fpb | A | 680 | THR | p.Thr680Ser | P19367 |
| HOMO SAPIENS | 0 | 6771 | 2zt6 | A | 201 | ALA | p.Ala255Val | P41250 |
| HOMO SAPIENS | 0 | 6771 | 2zt6 | A | 202 | ASP | p.Asp256His | P41250 |
| HOMO SAPIENS | 0 | 6771 | 2zt6 | A | 334 | ARG | p.Arg388Gln | P41250 |
| HOMO SAPIENS | -1 | 6771 | 2zt6 | A | 439 | ASN | p.Asn493Ser | P41250 |
| HOMO SAPIENS | -1 | 6771 | 2zt6 | A | 500 | ASP | p.Asp554Asn | P41250 |
| HOMO SAPIENS | 1 | 6771 | 2zt6 | A | 581 | SER | p.Ser635Leu | P41250 |
| HOMO SAPIENS | -1 | 6771 | 2zt6 | A | 684 | GLU | p.Glu738Lys | P41250 |
| HOMO SAPIENS | 0 | 2687 | 4grw | B | 276 | VAL | p.Val298Phe | P29460 |
| HOMO SAPIENS | 6 | 913 | 1m75 | B | 45 | ASP | p.Asp57Glu | Q16836 |
| HOMO SAPIENS | 2 | 913 | 1m75 | B | 74 | PRO | p.Leu86Pro | Q16836 |
| HOMO SAPIENS | 3 | 913 | 1m75 | B | 214 | TYR | p.Tyr226His | Q16836 |
| HOMO SAPIENS | 3 | 913 | 1m75 | B | 224 | ARG | p.Arg236Ter | Q16836 |
| HOMO SAPIENS | 4 | 913 | 1m75 | B | 246 | PRO | p.Pro258Leu | Q16836 |
| HOMO SAPIENS | 2 | 4713 | 4djn | B | 40 | ARG | p.Arg40Cys | Q7LG56 |
| HOMO SAPIENS | 1 | 4713 | 4djn | B | 41 | ARG | p.Arg41Pro | Q7LG56 |
| HOMO SAPIENS | 2 | 4713 | 4djn | B | 110 | ARG | p.Arg110His | Q7LG56 |
| HOMO SAPIENS | 2 | 4713 | 4djn | B | 121 | ARG | p.Arg121His | Q7LG56 |
| HOMO SAPIENS | 9 | 4713 | 4djn | B | 131 | GLU | p.Glu131Lys | Q7LG56 |
| HOMO SAPIENS | 2 | 4713 | 4djn | B | 182 | THR | p.Thr182Ala | Q7LG56 |
| HOMO SAPIENS | 6 | 4713 | 4djn | B | 194 | GLU | p.Glu194Lys | Q7LG56 |
| HOMO SAPIENS | 2 | 4713 | 4djn | B | 211 | ARG | p.Arg211Lys | Q7LG56 |
| HOMO SAPIENS | 0 | 4713 | 4djn | B | 224 | ILE | p.Ile224Ser | Q7LG56 |
| HOMO SAPIENS | 3 | 4713 | 4djn | B | 229 | GLY | p.Gly229Val | Q7LG56 |
| HOMO SAPIENS | 2 | 4713 | 4djn | B | 236 | CYS | p.Cys236Phe | Q7LG56 |
| HOMO SAPIENS | 0 | 4713 | 4djn | B | 273 | GLY | p.Gly273Ser | Q7LG56 |
| HOMO SAPIENS | -1 | 4713 | 4djn | B | 282 | MSE | p.Met282Ile | Q7LG56 |
| HOMO SAPIENS | 3 | 4713 | 4djn | B | 284 | GLN | p.Gln284Ter | Q7LG56 |
| HOMO SAPIENS | 3 | 4713 | 4djn | B | 305 | ALA | p.Ala305Ser | Q7LG56 |
| HOMO SAPIENS | -1 | 8974 | 5n7i | A | 500 | ARG | p.Arg500Ter | Q8N4S9 |
| HOMO SAPIENS | 0 | 17072 | 3fxi | B | 299 | ASP | p.Asp299Gly | O00206 |

| | | | | | | | | |
|---|---|---|---|---|---|---|---|---|
| HOMO SAPIENS | 1 | 14355 | 2wwk | T | 55 | ILE | p.Ile34306Asn | Q8WZ42 |
| HOMO SAPIENS | 0 | 14355 | 2wwk | T | 64 | LEU | p.Leu34315Pro | Q8WZ42 |
| HOMO SAPIENS | 2 | 17797 | 1xu1 | S | 72 | ARG | p.Arg72His | O14836 |
| HOMO SAPIENS | -1 | 10836 | 4ztu | A | 39 | GLN | p.Gln39Arg | P54098 |
| HOMO SAPIENS | -1 | 10836 | 4ztu | A | 42 | ARG | p.Arg42Gln | P54098 |
| HOMO SAPIENS | -1 | 10836 | 4ztu | A | 45 | GLN | p.Gln45Arg | P54098 |
| HOMO SAPIENS | -1 | 10836 | 4ztu | A | 52 | GLN | p.Gln52Glu | P54098 |
| HOMO SAPIENS | 0 | 10836 | 4ztu | A | 88 | PHE | p.Phe88Leu | P54098 |
| HOMO SAPIENS | 1 | 10836 | 4ztu | A | 194 | ALA | p.Ala194Val | P54098 |
| HOMO SAPIENS | 1 | 10836 | 4ztu | A | 227 | ARG | p.Arg227Trp | P54098 |
| HOMO SAPIENS | 0 | 10836 | 4ztu | A | 232 | ARG | p.Arg232His | P54098 |
| HOMO SAPIENS | 1 | 10836 | 4ztu | A | 285 | GLN | p.Gln285His | P54098 |
| HOMO SAPIENS | 1 | 10836 | 4ztu | A | 434 | TYR | p.Tyr434His | P54098 |
| HOMO SAPIENS | -1 | 10836 | 4ztu | A | 511 | SER | p.Ser511Asn | P54098 |
| HOMO SAPIENS | 0 | 10836 | 4ztu | A | 533 | CYS | p.Cys533Trp | P54098 |
| HOMO SAPIENS | 1 | 10836 | 4ztu | A | 546 | ARG | p.Arg546Cys | P54098 |
| HOMO SAPIENS | 1 | 10836 | 4ztu | A | 587 | PRO | p.Pro587Leu | P54098 |
| HOMO SAPIENS | 0 | 10836 | 4ztu | A | 603 | MET | p.Met603Thr | P54098 |
| HOMO SAPIENS | 0 | 10836 | 4ztu | A | 623 | LEU | p.Leu623Trp | P54098 |
| HOMO SAPIENS | -1 | 10836 | 4ztu | A | 627 | ARG | p.Arg627Trp | P54098 |
| HOMO SAPIENS | -1 | 10836 | 4ztu | A | 628 | ARG | p.Arg628Pro | P54098 |
| HOMO SAPIENS | 0 | 10836 | 4ztu | A | 635 | PRO | p.Pro635Leu | P54098 |
| HOMO SAPIENS | 0 | 10836 | 4ztu | A | 662 | GLU | p.Glu662Lys | P54098 |
| HOMO SAPIENS | -1 | 10836 | 4ztu | A | 674 | GLY | p.Gly674Asp | P54098 |
| HOMO SAPIENS | -1 | 10836 | 4ztu | A | 722 | ARG | p.Arg722His | P54098 |
| HOMO SAPIENS | -1 | 10836 | 4ztu | A | 737 | GLY | p.Gly737Arg | P54098 |
| HOMO SAPIENS | 0 | 10836 | 4ztu | A | 748 | TRP | p.Trp748Ser | P54098 |
| HOMO SAPIENS | 0 | 10836 | 4ztu | A | 749 | PHE | p.Phe749Ser | P54098 |
| HOMO SAPIENS | 1 | 10836 | 4ztu | A | 807 | ARG | p.Arg807Cys | P54098 |
| HOMO SAPIENS | 1 | 10836 | 4ztu | A | 831 | TYR | p.Tyr831Cys | P54098 |
| HOMO SAPIENS | 8 | 10836 | 4ztu | A | 848 | GLY | p.Gly848Ser | P54098 |
| HOMO SAPIENS | 1 | 10836 | 4ztu | A | 852 | ARG | p.Arg852Cys | P54098 |
| HOMO SAPIENS | 9 | 10836 | 4ztu | A | 853 | ARG | p.Arg853Trp | P54098 |
| HOMO SAPIENS | 1 | 10836 | 4ztu | A | 864 | ASN | p.Asn864Ser | P54098 |
| HOMO SAPIENS | 1 | 10836 | 4ztu | A | 879 | GLN | p.Gln879Arg | P54098 |
| HOMO SAPIENS | 1 | 10836 | 4ztu | A | 914 | THR | p.Thr914Pro | P54098 |
| HOMO SAPIENS | 6 | 10836 | 4ztu | A | 932 | HIS | p.His932Tyr | P54098 |
| HOMO SAPIENS | 2 | 10836 | 4ztu | A | 944 | GLU | p.Glu944Lys | P54098 |
| HOMO SAPIENS | 6 | 10836 | 4ztu | A | 955 | TYR | p.Tyr955Cys | P54098 |
| HOMO SAPIENS | 1 | 10836 | 4ztu | A | 957 | ALA | p.Ala957Ser | P54098 |
| HOMO SAPIENS | 1 | 10836 | 4ztu | A | 957 | ALA | p.Ala957Val | P54098 |
| HOMO SAPIENS | 1 | 10836 | 4ztu | A | 964 | ARG | p.Arg964Cys | P54098 |
| HOMO SAPIENS | 1 | 10836 | 4ztu | A | 966 | LEU | p.Leu966Arg | P54098 |
| HOMO SAPIENS | 0 | 10836 | 4ztu | A | 1051 | GLY | p.Gly1051Arg | P54098 |
| HOMO SAPIENS | 1 | 10836 | 4ztu | A | 1073 | PRO | p.Pro1073Leu | P54098 |
| HOMO SAPIENS | 1 | 10836 | 4ztu | A | 1096 | ARG | p.Arg1096Cys | P54098 |
| HOMO SAPIENS | 1 | 10836 | 4ztu | A | 1096 | ARG | p.Arg1096His | P54098 |
| HOMO SAPIENS | 0 | 10836 | 4ztu | A | 1116 | MET | p.Met1116Thr | P54098 |
| HOMO SAPIENS | 1 | 10836 | 4ztu | A | 1142 | ARG | p.Arg1142Trp | P54098 |
| HOMO SAPIENS | 1 | 10836 | 4ztu | A | 1146 | ARG | p.Arg1146Cys | P54098 |
| HOMO SAPIENS | 1 | 10836 | 4ztu | A | 1157 | ASN | p.Asn1157Ser | P54098 |
| HOMO SAPIENS | -1 | 10836 | 4ztu | A | 1234 | ARG | p.Arg1234Gln | P54098 |
| HOMO SAPIENS | -1 | 10836 | 4ztu | A | 1236 | GLN | p.Gln1236His | P54098 |
| HOMO SAPIENS | 5 | 2666 | 1jph | A | 62 | PRO | p.Pro62Leu | P06132 |

| | | | | | | | | |
|---|---|---|---|---|---|---|---|---|
| HOMO SAPIENS | 5 | 2666 | 1jph | A | 116 | GLN | p.Gln116Ter | P06132 |
| HOMO SAPIENS | 0 | 2666 | 1jph | A | 165 | MET | p.Met165Arg | P06132 |
| HOMO SAPIENS | 6 | 2666 | 1jph | A | 167 | GLU | p.Glu167Lys | P06132 |
| HOMO SAPIENS | 0 | 2666 | 1jph | A | 195 | LEU | p.Leu195Phe | P06132 |
| HOMO SAPIENS | 7 | 2666 | 1jph | A | 253 | LEU | p.Leu253Gln | P06132 |
| HOMO SAPIENS | 0 | 2666 | 1jph | A | 281 | GLY | p.Gly281Glu | P06132 |
| HOMO SAPIENS | 9 | 2666 | 1jph | A | 304 | ASN | p.Asn304Lys | P06132 |
| HOMO SAPIENS | 8 | 2666 | 1jph | A | 311 | TYR | p.Tyr311Cys | P06132 |
| HOMO SAPIENS | 5 | 2666 | 1jph | A | 332 | ARG | p.Arg332His | P06132 |
| HOMO SAPIENS | 1 | 889 | 1m51 | A | 184 | LEU | p.Val184Leu | P35558 |
| HOMO SAPIENS | 0 | 889 | 1m51 | A | 267 | VAL | p.Ile267Val | P35558 |
| HOMO SAPIENS | 2 | 889 | 1m51 | A | 276 | GLU | p.Glu276Lys | P35558 |
| HOMO SAPIENS | 6 | 889 | 1m51 | A | 368 | VAL | p.Val368Ile | P35558 |
| HOMO SAPIENS | 0 | 889 | 1m51 | A | 579 | SER | p.Ser579Arg | P35558 |
| HOMO SAPIENS | 1 | 4084 | 2gjx | A | 28 | GLN | p.Gln28Ter | P06865 |
| HOMO SAPIENS | 0 | 4084 | 2gjx | A | 39 | LEU | p.Leu39Arg | P06865 |
| HOMO SAPIENS | 0 | 4084 | 2gjx | A | 127 | LEU | p.Leu127Arg | P06865 |
| HOMO SAPIENS | 6 | 4084 | 2gjx | A | 137 | ARG | p.Arg137Ter | P06865 |
| HOMO SAPIENS | 6 | 4084 | 2gjx | A | 170 | ARG | p.Arg170Gln | P06865 |
| HOMO SAPIENS | 9 | 4084 | 2gjx | A | 178 | ARG | p.Arg178Cys | P06865 |
| HOMO SAPIENS | 9 | 4084 | 2gjx | A | 178 | ARG | p.Arg178His | P06865 |
| HOMO SAPIENS | 5 | 4084 | 2gjx | A | 180 | TYR | p.Tyr180His | P06865 |
| HOMO SAPIENS | 5 | 4084 | 2gjx | A | 180 | TYR | p.Tyr180Ter | P06865 |
| HOMO SAPIENS | 1 | 4084 | 2gjx | A | 197 | LYS | p.Lys197Thr | P06865 |
| HOMO SAPIENS | 0 | 4084 | 2gjx | A | 200 | VAL | p.Val200Met | P06865 |
| HOMO SAPIENS | 9 | 4084 | 2gjx | A | 204 | HIS | p.His204Arg | P06865 |
| HOMO SAPIENS | 5 | 4084 | 2gjx | A | 210 | SER | p.Ser210Phe | P06865 |
| HOMO SAPIENS | 0 | 4084 | 2gjx | A | 211 | PHE | p.Phe211Ser | P06865 |
| HOMO SAPIENS | 2 | 4084 | 2gjx | A | 250 | GLY | p.Gly250Asp | P06865 |
| HOMO SAPIENS | 9 | 4084 | 2gjx | A | 258 | ASP | p.Asp258His | P06865 |
| HOMO SAPIENS | 0 | 4084 | 2gjx | A | 269 | GLY | p.Gly269Ser | P06865 |
| HOMO SAPIENS | 0 | 4084 | 2gjx | A | 301 | MET | p.Met301Arg | P06865 |
| HOMO SAPIENS | 3 | 4084 | 2gjx | A | 393 | ARG | p.Arg393Ter | P06865 |
| HOMO SAPIENS | 3 | 4084 | 2gjx | A | 393 | ARG | p.Arg393Pro | P06865 |
| HOMO SAPIENS | 0 | 4084 | 2gjx | A | 399 | ASN | p.Asn399Asp | P06865 |
| HOMO SAPIENS | 2 | 4084 | 2gjx | A | 420 | TRP | p.Trp420Cys | P06865 |
| HOMO SAPIENS | 1 | 4084 | 2gjx | A | 434 | PHE | p.Phe434Leu | P06865 |
| HOMO SAPIENS | 0 | 4084 | 2gjx | A | 454 | GLY | p.Gly454Ser | P06865 |
| HOMO SAPIENS | 1 | 4084 | 2gjx | A | 474 | TRP | p.Trp474Cys | P06865 |
| HOMO SAPIENS | 0 | 4084 | 2gjx | A | 479 | ALA | p.Ala479Thr | P06865 |
| HOMO SAPIENS | 5 | 4084 | 2gjx | A | 485 | TRP | p.Trp485Arg | P06865 |
| HOMO SAPIENS | 4 | 4084 | 2gjx | A | 499 | ARG | p.Arg499Cys | P06865 |
| HOMO SAPIENS | 4 | 4084 | 2gjx | A | 499 | ARG | p.Arg499His | P06865 |
| HOMO SAPIENS | 2 | 4084 | 2gjx | A | 504 | ARG | p.Arg504Cys | P06865 |
| HOMO SAPIENS | 2 | 4084 | 2gjx | A | 504 | ARG | p.Arg504His | P06865 |
| HOMO SAPIENS | -1 | 1943 | 1xmj | A | 417 | ASN | p.Asn417Lys | P13569 |
| HOMO SAPIENS | -1 | 1943 | 1xmj | A | 422 | SER | p.Ser422Phe | P13569 |
| HOMO SAPIENS | 1 | 1943 | 1xmj | A | 438 | THR | p.Thr438Ala | P13569 |
| HOMO SAPIENS | 0 | 1943 | 1xmj | A | 455 | ALA | p.Ala455Glu | P13569 |
| HOMO SAPIENS | 4 | 1943 | 1xmj | A | 456 | VAL | p.Val456Ala | P13569 |
| HOMO SAPIENS | 9 | 1943 | 1xmj | A | 458 | GLY | p.Gly458Val | P13569 |
| HOMO SAPIENS | 8 | 1943 | 1xmj | A | 466 | SER | p.Ser466Ter | P13569 |
| HOMO SAPIENS | 0 | 1943 | 1xmj | A | 467 | LEU | p.Leu467Pro | P13569 |
| HOMO SAPIENS | 1 | 1943 | 1xmj | A | 470 | MET | p.Val470Met | P13569 |

| | | | | | | | | |
|---|---|---|---|---|---|---|---|---|
| HOMO SAPIENS | 5 | 1943 | 1xmj | A | 480 | GLY | p.Gly480Cys | P13569 |
| HOMO SAPIENS | 3 | 1943 | 1xmj | A | 492 | SER | p.Ser492Phe | P13569 |
| HOMO SAPIENS | 8 | 1943 | 1xmj | A | 493 | GLN | p.Gln493Ter | P13569 |
| HOMO SAPIENS | 5 | 1943 | 1xmj | A | 506 | ILE | p.Ile506Met | P13569 |
| HOMO SAPIENS | 3 | 1943 | 1xmj | A | 520 | VAL | p.Val520Phe | P13569 |
| HOMO SAPIENS | 1 | 1943 | 1xmj | A | 528 | GLU | p.Glu528Asp | P13569 |
| HOMO SAPIENS | 1 | 1943 | 1xmj | A | 534 | ALA | p.Ala534Glu | P13569 |
| HOMO SAPIENS | 4 | 1943 | 1xmj | A | 542 | GLY | p.Gly542Ter | P13569 |
| HOMO SAPIENS | 6 | 1943 | 1xmj | A | 549 | SER | p.Ser549Asn | P13569 |
| HOMO SAPIENS | 6 | 1943 | 1xmj | A | 549 | SER | p.Ser549Arg | P13569 |
| HOMO SAPIENS | 6 | 1943 | 1xmj | A | 549 | SER | p.Ser549Arg | P13569 |
| HOMO SAPIENS | 6 | 1943 | 1xmj | A | 551 | GLY | p.Gly551Asp | P13569 |
| HOMO SAPIENS | 6 | 1943 | 1xmj | A | 551 | GLY | p.Gly551Ser | P13569 |
| HOMO SAPIENS | 6 | 1943 | 1xmj | A | 553 | GLN | p.Arg553Gln | P13569 |
| HOMO SAPIENS | 6 | 1943 | 1xmj | A | 553 | GLN | p.Arg553Ter | P13569 |
| HOMO SAPIENS | 0 | 1943 | 1xmj | A | 559 | ALA | p.Ala559Thr | P13569 |
| HOMO SAPIENS | 7 | 1943 | 1xmj | A | 560 | ARG | p.Arg560Lys | P13569 |
| HOMO SAPIENS | 4 | 1943 | 1xmj | A | 562 | VAL | p.Val562Ile | P13569 |
| HOMO SAPIENS | 3 | 1943 | 1xmj | A | 563 | TYR | p.Tyr563Asn | P13569 |
| HOMO SAPIENS | 3 | 1943 | 1xmj | A | 569 | TYR | p.Tyr569Asp | P13569 |
| HOMO SAPIENS | 3 | 1943 | 1xmj | A | 585 | GLU | p.Glu585Ter | P13569 |
| HOMO SAPIENS | 1 | 1943 | 1xmj | A | 618 | ILE | p.Ile618Thr | P13569 |
| HOMO SAPIENS | 9 | 1943 | 1xmj | A | 622 | GLY | p.Gly622Asp | P13569 |
| HOMO SAPIENS | -1 | 1943 | 1xmj | A | 648 | ASP | p.Asp648Val | P13569 |
| HOMO SAPIENS | 0 | 1943 | 1xmj | A | 673 | GLY | p.Gly673Ter | P13569 |
| HOMO SAPIENS | 5 | 5111 | 4rec | A | 749 | ARG | p.Arg749Ter | Q9Y2M0 |
| HOMO SAPIENS | -1 | 4580 | 4l0d | B | 2 | GLY | p.Pro2Leu | P35520 |
| HOMO SAPIENS | -1 | 4580 | 4l0d | B | 12 | PRO | p.Pro12Ser | P35520 |
| HOMO SAPIENS | -1 | 4580 | 4l0d | B | 18 | ARG | p.Arg18Cys | P35520 |
| HOMO SAPIENS | 1 | 4580 | 4l0d | B | 49 | PRO | p.Pro49Leu | P35520 |
| HOMO SAPIENS | 2 | 4580 | 4l0d | B | 70 | PRO | p.Pro70Leu | P35520 |
| HOMO SAPIENS | 1 | 4580 | 4l0d | B | 102 | LYS | p.Lys102Gln | P35520 |
| HOMO SAPIENS | 7 | 4580 | 4l0d | B | 114 | ALA | p.Ala114Val | P35520 |
| HOMO SAPIENS | 5 | 4580 | 4l0d | B | 116 | GLY | p.Gly116Arg | P35520 |
| HOMO SAPIENS | 8 | 4580 | 4l0d | B | 121 | ARG | p.Arg121Cys | P35520 |
| HOMO SAPIENS | 8 | 4580 | 4l0d | B | 121 | ARG | p.Arg121His | P35520 |
| HOMO SAPIENS | 5 | 4580 | 4l0d | B | 125 | ARG | p.Arg125Gln | P35520 |
| HOMO SAPIENS | 4 | 4580 | 4l0d | B | 134 | GLY | p.Gly134Arg | P35520 |
| HOMO SAPIENS | 4 | 4580 | 4l0d | B | 139 | GLY | p.Gly139Arg | P35520 |
| HOMO SAPIENS | 0 | 4580 | 4l0d | B | 145 | PRO | p.Pro145Leu | P35520 |
| HOMO SAPIENS | 0 | 4580 | 4l0d | B | 148 | GLY | p.Gly148Arg | P35520 |
| HOMO SAPIENS | 0 | 4580 | 4l0d | B | 168 | VAL | p.Val168Met | P35520 |
| HOMO SAPIENS | 2 | 4580 | 4l0d | B | 191 | THR | p.Thr191Met | P35520 |
| HOMO SAPIENS | 5 | 4580 | 4l0d | B | 234 | ASP | p.Asp234Asn | P35520 |
| HOMO SAPIENS | 0 | 4580 | 4l0d | B | 257 | THR | p.Thr257Met | P35520 |
| HOMO SAPIENS | 4 | 4580 | 4l0d | B | 262 | THR | p.Thr262Met | P35520 |
| HOMO SAPIENS | 0 | 4580 | 4l0d | B | 278 | ILE | p.Ile278Thr | P35520 |
| HOMO SAPIENS | 2 | 4580 | 4l0d | B | 307 | GLY | p.Gly307Ser | P35520 |
| HOMO SAPIENS | 4 | 4580 | 4l0d | B | 320 | VAL | p.Val320Ala | P35520 |
| HOMO SAPIENS | 2 | 4580 | 4l0d | B | 336 | ARG | p.Arg336Cys | P35520 |
| HOMO SAPIENS | 2 | 4580 | 4l0d | B | 336 | ARG | p.Arg336His | P35520 |
| HOMO SAPIENS | 0 | 4580 | 4l0d | B | 351 | GLY | p.Gly351Arg | P35520 |
| HOMO SAPIENS | 0 | 4580 | 4l0d | B | 353 | THR | p.Thr353Met | P35520 |
| HOMO SAPIENS | 0 | 4580 | 4l0d | B | 371 | VAL | p.Val371Met | P35520 |

| | | | | | | | | |
|---|---|---|---|---|---|---|---|---|
| HOMO SAPIENS | 2 | 4580 | 4l0d | B | 379 | ARG | p.Arg379Gln | P35520 |
| HOMO SAPIENS | 2 | 4580 | 4l0d | B | 379 | ARG | p.Arg379Trp | P35520 |
| HOMO SAPIENS | 2 | 4580 | 4l0d | B | 384 | LYS | p.Lys384Glu | P35520 |
| HOMO SAPIENS | 0 | 4580 | 4l0d | B | 422 | PRO | p.Pro422Leu | P35520 |
| HOMO SAPIENS | 0 | 4580 | 4l0d | B | 444 | ASP | p.Asp444Asn | P35520 |
| HOMO SAPIENS | 0 | 4580 | 4l0d | B | 466 | SER | p.Ser466Leu | P35520 |
| HOMO SAPIENS | 0 | 4580 | 4l0d | B | 539 | LEU | p.Leu539Ser | P35520 |
| HOMO SAPIENS | 0 | 4580 | 4l0d | B | 548 | ARG | p.Arg548Gln | P35520 |
| HOMO SAPIENS | 1 | 6536 | 3ulu | A | 412 | LEU | p.Leu412Phe | O15455 |
| HOMO SAPIENS | 1 | 28493 | 3ii6 | X | 814 | ARG | p.Arg814Ter | P49917 |
| HOMO SAPIENS | 1 | 28493 | 3ii6 | X | 857 | ALA | p.Ala857Thr | P49917 |
| HOMO SAPIENS | 9 | 3543 | 2q51 | A | 24 | GLU | p.Glu24Gly | P45381 |
| HOMO SAPIENS | 1 | 3543 | 2q51 | A | 27 | GLY | p.Gly27Arg | P45381 |
| HOMO SAPIENS | 9 | 3543 | 2q51 | A | 71 | ARG | p.Arg71His | P45381 |
| HOMO SAPIENS | -1 | 3543 | 2q51 | A | 82 | MSE | p.Met82Thr | P45381 |
| HOMO SAPIENS | 0 | 3543 | 2q51 | A | 143 | ILE | p.Ile143Phe | P45381 |
| HOMO SAPIENS | 1 | 3543 | 2q51 | A | 152 | CYS | p.Cys152Arg | P45381 |
| HOMO SAPIENS | 2 | 3543 | 2q51 | A | 218 | CYS | p.Cys218Ter | P45381 |
| HOMO SAPIENS | 1 | 3543 | 2q51 | A | 231 | TYR | p.Tyr231Cys | P45381 |
| HOMO SAPIENS | 1 | 3543 | 2q51 | A | 231 | TYR | p.Tyr231Ter | P45381 |
| HOMO SAPIENS | 2 | 3543 | 2q51 | A | 274 | GLY | p.Gly274Arg | P45381 |
| HOMO SAPIENS | 6 | 3543 | 2q51 | A | 285 | GLU | p.Glu285Ala | P45381 |
| HOMO SAPIENS | 5 | 3543 | 2q51 | A | 287 | ALA | p.Ala287Thr | P45381 |
| HOMO SAPIENS | 2 | 3543 | 2q51 | A | 305 | ALA | p.Ala305Glu | P45381 |
| HOMO SAPIENS | 2 | 24687 | 1hfi | A | 25 | SER | p.Ser890Ile | P08603 |
| HOMO SAPIENS | 0 | 1508 | 3wn5 | C | 45 | LEU | p.Leu66Arg | P08637 |
| HOMO SAPIENS | 0 | 1508 | 3wn5 | C | 155 | VAL | p.Phe176Val | P08637 |
| HOMO SAPIENS | 0 | 750 | 5tpa | A | 54 | VAL | p.Val452Met | Q12879 |
| HOMO SAPIENS | 9 | 750 | 5tpa | A | 120 | ARG | p.Arg518Cys | Q12879 |
| HOMO SAPIENS | 0 | 750 | 5tpa | A | 199 | ALA | p.Ala716Thr | Q12879 |
| HOMO SAPIENS | 4 | 1363 | 3hg2 | B | 40 | PRO | p.Pro40Ser | P06280 |
| HOMO SAPIENS | 3 | 1363 | 3hg2 | B | 65 | SER | p.Ser65Thr | P06280 |
| HOMO SAPIENS | 4 | 1363 | 3hg2 | B | 66 | GLU | p.Glu66Gln | P06280 |
| HOMO SAPIENS | 2 | 1363 | 3hg2 | B | 80 | GLY | p.Gly80Asp | P06280 |
| HOMO SAPIENS | 3 | 1363 | 3hg2 | B | 112 | ARG | p.Arg112Cys | P06280 |
| HOMO SAPIENS | 3 | 1363 | 3hg2 | B | 112 | ARG | p.Arg112His | P06280 |
| HOMO SAPIENS | 0 | 1363 | 3hg2 | B | 139 | ASN | p.Asn139Ser | P06280 |
| HOMO SAPIENS | 7 | 1363 | 3hg2 | B | 156 | ALA | p.Ala156Thr | P06280 |
| HOMO SAPIENS | 5 | 1363 | 3hg2 | B | 162 | TRP | p.Trp162Arg | P06280 |
| HOMO SAPIENS | 5 | 1363 | 3hg2 | B | 207 | TYR | p.Tyr207His | P06280 |
| HOMO SAPIENS | 0 | 1363 | 3hg2 | B | 215 | ASN | p.Asn215Ser | P06280 |
| HOMO SAPIENS | 0 | 1363 | 3hg2 | B | 264 | ASP | p.Asp264Val | P06280 |
| HOMO SAPIENS | 5 | 1363 | 3hg2 | B | 266 | ASP | p.Asp266Val | P06280 |
| HOMO SAPIENS | 2 | 1363 | 3hg2 | B | 269 | VAL | p.Val269Ala | P06280 |
| HOMO SAPIENS | 3 | 1363 | 3hg2 | B | 272 | ASN | p.Asn272Ser | P06280 |
| HOMO SAPIENS | 3 | 1363 | 3hg2 | B | 279 | GLN | p.Gln279Glu | P06280 |
| HOMO SAPIENS | 0 | 1363 | 3hg2 | B | 289 | ILE | p.Ile289Val | P06280 |
| HOMO SAPIENS | 0 | 1363 | 3hg2 | B | 296 | MET | p.Met296Val | P06280 |
| HOMO SAPIENS | 0 | 1363 | 3hg2 | B | 296 | MET | p.Met296Ile | P06280 |
| HOMO SAPIENS | 0 | 1363 | 3hg2 | B | 297 | SER | p.Ser297Phe | P06280 |
| HOMO SAPIENS | 2 | 1363 | 3hg2 | B | 301 | ARG | p.Arg301Gln | P06280 |
| HOMO SAPIENS | 3 | 1363 | 3hg2 | B | 327 | GLN | p.Gln327Lys | P06280 |
| HOMO SAPIENS | 0 | 1363 | 3hg2 | B | 328 | GLY | p.Gly328Arg | P06280 |
| HOMO SAPIENS | 0 | 1363 | 3hg2 | B | 328 | GLY | p.Gly328Ala | P06280 |

| | | | | | | | | |
|---|---|---|---|---|---|---|---|---|
| HOMO SAPIENS | 2 | 1363 | 3hg2 | B | 331 | LEU | p.Leu331Phe | P06280 |
| HOMO SAPIENS | 0 | 1363 | 3hg2 | B | 342 | ARG | p.Arg342Gln | P06280 |
| HOMO SAPIENS | 1 | 1363 | 3hg2 | B | 361 | GLY | p.Gly361Arg | P06280 |
| HOMO SAPIENS | 1 | 1363 | 3hg2 | B | 363 | ARG | p.Arg363His | P06280 |
| HOMO SAPIENS | 1 | 1363 | 3hg2 | B | 385 | THR | p.Thr385Ala | P06280 |
| HOMO SAPIENS | 1 | 1363 | 3hg2 | B | 399 | TRP | p.Trp399Ser | P06280 |
| HOMO SAPIENS | 1 | 1363 | 3hg2 | B | 410 | THR | p.Thr410Ala | P06280 |
| HOMO SAPIENS | 2 | 3904 | 3caa | B | 385 | MET | p.Met414Val | P01011 |
| HOMO SAPIENS | 0 | 558 | 5eub | A | 56 | VAL | p.Val56Ile | Q13126 |
| HOMO SAPIENS | 0 | 558 | 5eub | A | 189 | TRP | p.Trp189Leu | Q13126 |
| HOMO SAPIENS | 0 | 558 | 5eub | A | 218 | ALA | p.Ala218Gly | Q13126 |
| HOMO SAPIENS | 1 | 3865 | 1cb5 | B | 443 | ILE | p.Ile443Val | Q13867 |
| HOMO SAPIENS | 2 | 14329 | 4m92 | A | 24 | ILE | p.Ile65Val | Q13454 |
| HOMO SAPIENS | 1 | 7381 | 5kz5 | k | 50 | GLY | p.Gly50Glu | Q9H1K1 |
| HOMO SAPIENS | 0 | 1102 | 4fdh | E | 86 | PRO | p.Pro86Ala | P19099 |
| HOMO SAPIENS | 0 | 1102 | 4fdh | E | 118 | ALA | p.Ala118Thr | P19099 |
| HOMO SAPIENS | 0 | 1102 | 4fdh | E | 159 | PRO | p.Pro159Leu | P19099 |
| HOMO SAPIENS | 0 | 1102 | 4fdh | E | 173 | LYS | p.Lys173Arg | P19099 |
| HOMO SAPIENS | 0 | 1102 | 4fdh | E | 181 | ARG | p.Arg181Trp | P19099 |
| HOMO SAPIENS | 0 | 1102 | 4fdh | E | 185 | THR | p.Thr185Ile | P19099 |
| HOMO SAPIENS | 0 | 1102 | 4fdh | E | 198 | GLU | p.Glu198Asp | P19099 |
| HOMO SAPIENS | 0 | 1102 | 4fdh | E | 214 | HIS | p.His214Asp | P19099 |
| HOMO SAPIENS | 0 | 1102 | 4fdh | E | 251 | LYS | p.Lys251Arg | P19099 |
| HOMO SAPIENS | 0 | 1102 | 4fdh | E | 255 | GLU | p.Glu255Ter | P19099 |
| HOMO SAPIENS | 0 | 1102 | 4fdh | E | 281 | ASN | p.Asn281Ser | P19099 |
| HOMO SAPIENS | 0 | 1102 | 4fdh | E | 320 | ALA | p.Ala320Val | P19099 |
| HOMO SAPIENS | 0 | 1102 | 4fdh | E | 339 | ILE | p.Ile339Thr | P19099 |
| HOMO SAPIENS | 0 | 1102 | 4fdh | E | 341 | ARG | p.Arg341Cys | P19099 |
| HOMO SAPIENS | 2 | 1102 | 4fdh | E | 374 | ARG | p.Arg374Trp | P19099 |
| HOMO SAPIENS | 0 | 1102 | 4fdh | E | 386 | VAL | p.Val386Ala | P19099 |
| HOMO SAPIENS | 0 | 1102 | 4fdh | E | 390 | LEU | p.Leu390Phe | P19099 |
| HOMO SAPIENS | -1 | 1102 | 4fdh | E | 435 | GLY | p.Gly435Ser | P19099 |
| HOMO SAPIENS | 0 | 1102 | 4fdh | E | 461 | LEU | p.Leu461Pro | P19099 |
| HOMO SAPIENS | 0 | 1102 | 4fdh | E | 498 | THR | p.Thr498Ala | P19099 |
| HOMO SAPIENS | 6 | 10292 | 2of5 | F | 128 | GLY | p.Gly128Arg | P78560 |
| HOMO SAPIENS | 6 | 10292 | 2of5 | F | 170 | ARG | p.Arg170Cys | P78560 |
| HOMO SAPIENS | -1 | 18965 | 3ghg | A | 16 | ARG | p.Arg35Cys | P02671 |
| HOMO SAPIENS | -1 | 18965 | 3ghg | A | 16 | ARG | p.Arg35His | P02671 |
| HOMO SAPIENS | -1 | 18965 | 3ghg | A | 283 | PRO | p.Pro302Ala | P02671 |
| HOMO SAPIENS | -1 | 18965 | 3ghg | A | 312 | THR | p.Thr331Ala | P02671 |
| HOMO SAPIENS | -1 | 18965 | 3ghg | A | 381 | SER | p.Ser400Phe | P02671 |
| HOMO SAPIENS | -1 | 18965 | 3ghg | A | 463 | VAL | p.Val482Met | P02671 |
| HOMO SAPIENS | -1 | 18965 | 3ghg | A | 526 | GLU | p.Glu545Val | P02671 |
| HOMO SAPIENS | -1 | 18965 | 3ghg | A | 554 | ARG | p.Arg573Leu | P02671 |
| HOMO SAPIENS | 8 | 3023 | 4ttb | A | 101 | ARG | p.Arg101Trp | Q6PHW0 |
| HOMO SAPIENS | 2 | 3023 | 4ttb | A | 116 | ILE | p.Ile116Thr | Q6PHW0 |
| HOMO SAPIENS | 0 | 3023 | 4ttb | A | 220 | ALA | p.Ala220Thr | Q6PHW0 |
| HOMO SAPIENS | 2 | 3023 | 4ttb | A | 271 | GLU | p.Glu271Lys | Q6PHW0 |
| HOMO SAPIENS | 3 | 33226 | 1ya5 | T | 11 | SER | p.Ser11Ter | O15273 |
| HOMO SAPIENS | 6 | 33226 | 1ya5 | T | 64 | SER | p.Ser64Leu | O15273 |
| HOMO SAPIENS | 6 | 33226 | 1ya5 | T | 70 | ARG | p.Arg70Gln | O15273 |
| HOMO SAPIENS | 3 | 33226 | 1ya5 | T | 75 | GLY | p.Gly75Ser | O15273 |
| HOMO SAPIENS | 0 | 33226 | 1ya5 | T | 87 | ARG | p.Arg87Trp | O15273 |
| HOMO SAPIENS | 2 | 5667 | 3dax | A | 398 | PRO | p.Pro398Ala | P22680 |

| | | | | | | | | |
|---|---|---|---|---|---|---|---|---|
| HOMO SAPIENS | 0 | 12861 | 5mya | A | 464 | ASN | p.Asn464His | Q02763 |
| HOMO SAPIENS | 2 | 12861 | 5mya | A | 486 | VAL | p.Val486Ile | Q02763 |
| HOMO SAPIENS | 0 | 12861 | 5mya | A | 522 | ARG | p.Arg522His | Q02763 |
| HOMO SAPIENS | 2 | 12861 | 5mya | A | 600 | VAL | p.Val600Leu | Q02763 |
| HOMO SAPIENS | 0 | 12861 | 5mya | A | 634 | LEU | p.Leu634Phe | Q02763 |
| HOMO SAPIENS | 0 | 12861 | 5mya | A | 684 | HIS | p.His684Gln | Q02763 |
| HOMO SAPIENS | 0 | 12861 | 5mya | A | 724 | ALA | p.Ala724Thr | Q02763 |
| HOMO SAPIENS | 0 | 7743 | 3qzw | G | 90 | GLY | p.Gly111Ser | P01732 |
| HOMO SAPIENS | 2 | 9498 | 3kci | A | 4267 | ASP | p.Asp4267Glu | O95714 |
| HOMO SAPIENS | 5 | 201 | 3ips | B | 415 | ARG | p.Arg415Gln | Q13133 |
| HOMO SAPIENS | 3 | 229 | 2q50 | A | 68 | LEU | p.Leu68Pro | Q9UBQ7 |
| HOMO SAPIENS | 2 | 229 | 2q50 | A | 99 | ARG | p.Arg99Ter | Q9UBQ7 |
| HOMO SAPIENS | 3 | 229 | 2q50 | A | 160 | GLY | p.Gly160Arg | Q9UBQ7 |
| HOMO SAPIENS | 5 | 229 | 2q50 | A | 165 | GLY | p.Gly165Asp | Q9UBQ7 |
| HOMO SAPIENS | 4 | 229 | 2q50 | A | 302 | ARG | p.Arg302Cys | Q9UBQ7 |
| HOMO SAPIENS | 4 | 229 | 2q50 | A | 302 | ARG | p.Arg302His | Q9UBQ7 |
| HOMO SAPIENS | -1 | 229 | 2q50 | A | 322 | MSE | p.Met322Arg | Q9UBQ7 |
| HOMO SAPIENS | -1 | 4519 | 5clw | A | 40 | PRO | p.Pro40Thr | Q04446 |
| HOMO SAPIENS | 1 | 4519 | 5clw | A | 59 | ILE | p.Ile59Thr | Q04446 |
| HOMO SAPIENS | 0 | 4519 | 5clw | A | 190 | ARG | p.Arg190Gly | Q04446 |
| HOMO SAPIENS | 1 | 4519 | 5clw | A | 224 | LEU | p.Leu224Pro | Q04446 |
| HOMO SAPIENS | 0 | 4519 | 5clw | A | 236 | GLN | p.Gln236His | Q04446 |
| HOMO SAPIENS | 1 | 4519 | 5clw | A | 254 | THR | p.Thr254Ala | Q04446 |
| HOMO SAPIENS | 0 | 4519 | 5clw | A | 257 | PHE | p.Phe257Leu | Q04446 |
| HOMO SAPIENS | 1 | 4519 | 5clw | A | 262 | ARG | p.Arg262Cys | Q04446 |
| HOMO SAPIENS | 3 | 4519 | 5clw | A | 329 | TYR | p.Tyr329Ser | Q04446 |
| HOMO SAPIENS | 0 | 4519 | 5clw | A | 507 | THR | p.Thr507Ala | Q04446 |
| HOMO SAPIENS | 0 | 4519 | 5clw | A | 515 | ARG | p.Arg515Cys | Q04446 |
| HOMO SAPIENS | 0 | 4519 | 5clw | A | 515 | ARG | p.Arg515His | Q04446 |
| HOMO SAPIENS | 0 | 4519 | 5clw | A | 524 | ARG | p.Arg524Gln | Q04446 |
| HOMO SAPIENS | 0 | 4519 | 5clw | A | 524 | ARG | p.Arg524Ter | Q04446 |
| HOMO SAPIENS | 0 | 4519 | 5clw | A | 545 | HIS | p.His545Arg | Q04446 |
| HOMO SAPIENS | 2 | 4519 | 5clw | A | 548 | TRP | p.Trp548Ter | Q04446 |
| HOMO SAPIENS | 0 | 4519 | 5clw | A | 592 | GLU | p.Glu592Ter | Q04446 |
| HOMO SAPIENS | 0 | 4519 | 5clw | A | 628 | HIS | p.His628Arg | Q04446 |
| HOMO SAPIENS | 0 | 4519 | 5clw | A | 637 | ARG | p.Arg637Ter | Q04446 |
| HOMO SAPIENS | 0 | 4519 | 5clw | A | 673 | ALA | p.Ala673Thr | Q04446 |
| HOMO SAPIENS | 1 | 10405 | 3bg3 | D | 507 | VAL | p.Val507Ile | P11498 |
| HOMO SAPIENS | 0 | 10405 | 3bg3 | D | 583 | ARG | p.Arg583Leu | P11498 |
| HOMO SAPIENS | 1 | 10405 | 3bg3 | D | 589 | LYS | p.Lys589Asn | P11498 |
| HOMO SAPIENS | 6 | 10405 | 3bg3 | D | 610 | ALA | p.Ala610Thr | P11498 |
| HOMO SAPIENS | 2 | 10405 | 3bg3 | D | 631 | ARG | p.Arg631Gln | P11498 |
| HOMO SAPIENS | 1 | 10405 | 3bg3 | D | 732 | ARG | p.Arg732Gln | P11498 |
| HOMO SAPIENS | 6 | 10405 | 3bg3 | D | 743 | MET | p.Met743Ile | P11498 |
| HOMO SAPIENS | 0 | 10405 | 3bg3 | D | 750 | THR | p.Thr750Met | P11498 |
| HOMO SAPIENS | -1 | 13751 | 2o8d | B | 350 | ALA | p.Ala350Val | P52701 |
| HOMO SAPIENS | -1 | 13751 | 2o8d | B | 352 | VAL | p.Val352Ile | P52701 |
| HOMO SAPIENS | -1 | 13751 | 2o8d | B | 355 | GLY | p.Gly355Ser | P52701 |
| HOMO SAPIENS | 0 | 13751 | 2o8d | B | 385 | ARG | p.Arg385Trp | P52701 |
| HOMO SAPIENS | 0 | 13751 | 2o8d | B | 390 | ASP | p.Asp390Glu | P52701 |
| HOMO SAPIENS | 0 | 13751 | 2o8d | B | 396 | LEU | p.Leu396Val | P52701 |
| HOMO SAPIENS | 0 | 13751 | 2o8d | B | 449 | LEU | p.Leu449Pro | P52701 |
| HOMO SAPIENS | 0 | 13751 | 2o8d | B | 468 | ARG | p.Arg468His | P52701 |
| HOMO SAPIENS | 0 | 13751 | 2o8d | B | 503 | SER | p.Ser503Cys | P52701 |

| | | | | | | | | |
|---|---|---|---|---|---|---|---|---|
| HOMO SAPIENS | 1 | 13751 | 2o8d | B | 509 | VAL | p.Val509Ala | P52701 |
| HOMO SAPIENS | 0 | 13751 | 2o8d | B | 675 | GLU | p.Glu675Asp | P52701 |
| HOMO SAPIENS | 0 | 13751 | 2o8d | B | 795 | ILE | p.Ile795Thr | P52701 |
| HOMO SAPIENS | 0 | 13751 | 2o8d | B | 850 | TYR | p.Tyr850Ter | P52701 |
| HOMO SAPIENS | 0 | 13751 | 2o8d | B | 878 | VAL | p.Val878Ala | P52701 |
| HOMO SAPIENS | 0 | 13751 | 2o8d | B | 911 | ARG | p.Arg911Ter | P52701 |
| HOMO SAPIENS | 0 | 13751 | 2o8d | B | 922 | ARG | p.Arg922Ter | P52701 |
| HOMO SAPIENS | 0 | 13751 | 2o8d | B | 969 | TYR | p.Tyr969Cys | P52701 |
| HOMO SAPIENS | 0 | 13751 | 2o8d | B | 1005 | ARG | p.Arg1005Ter | P52701 |
| HOMO SAPIENS | 0 | 13751 | 2o8d | B | 1035 | ARG | p.Arg1035Ter | P52701 |
| HOMO SAPIENS | 1 | 13751 | 2o8d | B | 1048 | GLN | p.Gln1048Ter | P52701 |
| HOMO SAPIENS | 1 | 13751 | 2o8d | B | 1051 | VAL | p.Val1051Ile | P52701 |
| HOMO SAPIENS | 0 | 13751 | 2o8d | B | 1054 | ILE | p.Ile1054Phe | P52701 |
| HOMO SAPIENS | 0 | 13751 | 2o8d | B | 1054 | ILE | p.Ile1054Met | P52701 |
| HOMO SAPIENS | 0 | 13751 | 2o8d | B | 1163 | GLU | p.Glu1163Val | P52701 |
| HOMO SAPIENS | 0 | 13751 | 2o8d | B | 1237 | GLU | p.Glu1237Asp | P52701 |
| HOMO SAPIENS | 0 | 13751 | 2o8d | B | 1242 | ARG | p.Arg1242His | P52701 |
| HOMO SAPIENS | -1 | 13751 | 2o8d | B | 1278 | PRO | p.Pro1278Thr | P52701 |
| HOMO SAPIENS | 0 | 13751 | 2o8d | B | 1304 | ARG | p.Arg1304Lys | P52701 |
| HOMO SAPIENS | 0 | 13751 | 2o8d | B | 1317 | HIS | p.His1317Gln | P52701 |
| HOMO SAPIENS | 0 | 13751 | 2o8d | B | 1329 | SER | p.Ser1329Leu | P52701 |
| HOMO SAPIENS | 3 | 13273 | 4g8k | A | 265 | GLU | p.Glu265Ter | Q05823 |
| HOMO SAPIENS | -1 | 11088 | 5uj7 | A | 666 | ARG | p.Arg666Trp | Q13415 |
| HOMO SAPIENS | 4 | 11088 | 5uj7 | A | 720 | ARG | p.Arg720Gln | Q13415 |
| HOMO SAPIENS | 0 | 11088 | 5uj7 | A | 816 | MET | p.Met816Thr | Q13415 |
| HOMO SAPIENS | 1 | 4352 | 5va1 | A | 421 | THR | p.Thr421Met | Q12809 |
| HOMO SAPIENS | 0 | 4352 | 5va1 | A | 451 | PRO | p.Pro451Leu | Q12809 |
| HOMO SAPIENS | 0 | 4352 | 5va1 | A | 470 | ASN | p.Asn470Asp | Q12809 |
| HOMO SAPIENS | 0 | 4352 | 5va1 | A | 490 | ALA | p.Ala490Thr | Q12809 |
| HOMO SAPIENS | 0 | 4352 | 5va1 | A | 491 | VAL | p.Val491Ile | Q12809 |
| HOMO SAPIENS | 1 | 4352 | 5va1 | A | 533 | VAL | p.Val533Gly | Q12809 |
| HOMO SAPIENS | 0 | 4352 | 5va1 | A | 552 | LEU | p.Leu552Ser | Q12809 |
| HOMO SAPIENS | 0 | 4352 | 5va1 | A | 558 | ALA | p.Ala558Pro | Q12809 |
| HOMO SAPIENS | 1 | 4352 | 5va1 | A | 561 | ALA | p.Ala561Val | Q12809 |
| HOMO SAPIENS | 0 | 4352 | 5va1 | A | 572 | GLY | p.Gly572Arg | Q12809 |
| HOMO SAPIENS | -1 | 4352 | 5va1 | A | 582 | ARG | p.Arg582Cys | Q12809 |
| HOMO SAPIENS | 0 | 4352 | 5va1 | A | 584 | GLY | p.Gly584Ser | Q12809 |
| HOMO SAPIENS | 0 | 4352 | 5va1 | A | 588 | ASN | p.Asn588Lys | Q12809 |
| HOMO SAPIENS | 0 | 4352 | 5va1 | A | 593 | ILE | p.Ile593Arg | Q12809 |
| HOMO SAPIENS | 1 | 4352 | 5va1 | A | 627 | PHE | p.Phe627Leu | Q12809 |
| HOMO SAPIENS | 0 | 4352 | 5va1 | A | 630 | VAL | p.Val630Leu | Q12809 |
| HOMO SAPIENS | 0 | 4352 | 5va1 | A | 652 | TYR | p.Tyr652Ter | Q12809 |
| HOMO SAPIENS | 0 | 4352 | 5va1 | A | 711 | ILE | p.Ile711Val | Q12809 |
| HOMO SAPIENS | 0 | 4352 | 5va1 | A | 735 | SER | p.Ser735Leu | Q12809 |
| HOMO SAPIENS | 0 | 4352 | 5va1 | A | 744 | ARG | p.Arg744Ter | Q12809 |
| HOMO SAPIENS | 0 | 4352 | 5va1 | A | 784 | ARG | p.Arg784Trp | Q12809 |
| HOMO SAPIENS | 1 | 4352 | 5va1 | A | 820 | GLY | p.Gly820Arg | Q12809 |
| HOMO SAPIENS | 5 | 4352 | 5va1 | A | 822 | VAL | p.Val822Met | Q12809 |
| HOMO SAPIENS | 1 | 4352 | 5va1 | A | 863 | ARG | p.Arg863Ter | Q12809 |
| HOMO SAPIENS | 5 | 303 | 2obv | A | 55 | ALA | p.Ala55Asp | Q00266 |
| HOMO SAPIENS | 7 | 303 | 2obv | A | 259 | ALA | p.Ala259Gly | Q00266 |
| HOMO SAPIENS | 9 | 303 | 2obv | A | 264 | ARG | p.Arg264His | Q00266 |
| HOMO SAPIENS | 9 | 303 | 2obv | A | 264 | ARG | p.Arg264Cys | Q00266 |
| HOMO SAPIENS | 0 | 303 | 2obv | A | 305 | LEU | p.Leu305Pro | Q00266 |

| | | | | | | | | |
|---|---|---|---|---|---|---|---|---|
| HOMO SAPIENS | 7 | 303 | 2obv | A | 322 | ILE | p.Ile322Met | Q00266 |
| HOMO SAPIENS | 0 | 303 | 2obv | A | 336 | GLY | p.Gly336Arg | Q00266 |
| HOMO SAPIENS | 0 | 303 | 2obv | A | 357 | PRO | p.Pro357Leu | Q00266 |
| HOMO SAPIENS | 0 | 2077 | 4zxb | E | 462 | ASN | p.Asn489Ser | P06213 |
| HOMO SAPIENS | -1 | 2077 | 4zxb | E | 519 | ASP | p.Asp546Glu | P06213 |
| HOMO SAPIENS | 2 | 2077 | 4zxb | E | 819 | THR | p.Thr858Ala | P06213 |
| HOMO SAPIENS | 0 | 2077 | 4zxb | E | 907 | ASP | p.Asp946Glu | P06213 |
| HOMO SAPIENS | 7 | 3274 | 1wsv | B | 14 | HIS | p.His42Arg | P48728 |
| HOMO SAPIENS | 0 | 3274 | 1wsv | B | 19 | GLY | p.Gly47Arg | P48728 |
| HOMO SAPIENS | 4 | 3274 | 1wsv | B | 45 | ARG | p.Arg73Cys | P48728 |
| HOMO SAPIENS | 1 | 3274 | 1wsv | B | 49 | SER | p.Ser77Leu | P48728 |
| HOMO SAPIENS | 3 | 3274 | 1wsv | B | 241 | GLY | p.Gly269Asp | P48728 |
| HOMO SAPIENS | 7 | 3274 | 1wsv | B | 248 | ASP | p.Asp276His | P48728 |
| HOMO SAPIENS | 0 | 3274 | 1wsv | B | 272 | MET | p.Met300Val | P48728 |
| HOMO SAPIENS | 4 | 3274 | 1wsv | B | 292 | ARG | p.Arg320His | P48728 |
| HOMO SAPIENS | 0 | 3274 | 1wsv | B | 354 | ARG | p.Arg382Gln | P48728 |
| HOMO SAPIENS | -1 | 435 | 3uyt | A | 44 | THR | p.Thr44Ala | P48730 |
| HOMO SAPIENS | 1 | 4698 | 5ife | C | 666 | LYS | p.Lys666Asn | O75533 |
| HOMO SAPIENS | 1 | 16917 | 1apy | C | 37 | GLY | p.Gly60Asp | P20933 |
| HOMO SAPIENS | 0 | 16917 | 1apy | C | 49 | SER | p.Ser72Pro | P20933 |
| HOMO SAPIENS | 0 | 16917 | 1apy | C | 84 | ARG | p.Arg107Ter | P20933 |
| HOMO SAPIENS | 0 | 16917 | 1apy | C | 126 | SER | p.Thr149Ser | P20933 |
| HOMO SAPIENS | 0 | 16917 | 1apy | C | 135 | TRP | p.Trp158Ter | P20933 |
| HOMO SAPIENS | 0 | 16917 | 1apy | C | 140 | CYS | p.Cys163Ser | P20933 |
| HOMO SAPIENS | 0 | 16917 | 1apy | C | 145 | TRP | p.Trp168Ter | P20933 |
| HOMO SAPIENS | 0 | 10413 | 4frw | A | 185 | THR | p.Thr185Met | Q96NY8 |
| HOMO SAPIENS | 0 | 5217 | 5n6w | G | 96 | TRP | p.Trp96Arg | O15537 |
| HOMO SAPIENS | 1 | 5217 | 5n6w | G | 102 | ARG | p.Arg102Trp | O15537 |
| HOMO SAPIENS | 0 | 5217 | 5n6w | G | 109 | GLY | p.Gly109Arg | O15537 |
| HOMO SAPIENS | -1 | 5217 | 5n6w | G | 223 | CYS | p.Cys223Arg | O15537 |
| HOMO SAPIENS | 0 | 9485 | 4z6y | E | 138 | ALA | p.Ala138Asp | Q9P0N9 |
| HOMO SAPIENS | 3 | 4504 | 4mwt | B | 411 | GLY | p.Gly439Ser | P10619 |
| HOMO SAPIENS | 3 | 4504 | 4mwt | B | 412 | PHE | p.Phe440Val | P10619 |
| HOMO SAPIENS | 6 | 6391 | 4kxy | A | 318 | ARG | p.Arg318Cys | P29401 |
| HOMO SAPIENS | 6 | 19546 | 2bx6 | A | 87 | THR | p.Thr87Ile | O75695 |
| HOMO SAPIENS | 1 | 19546 | 2bx6 | A | 118 | ARG | p.Arg118His | O75695 |
| HOMO SAPIENS | 1 | 19546 | 2bx6 | A | 201 | ILE | p.Ile201Val | O75695 |
| HOMO SAPIENS | 1 | 19546 | 2bx6 | A | 282 | ARG | p.Arg282Trp | O75695 |
| HOMO SAPIENS | 1 | 5265 | 5usr | H | 68 | ARG | p.Arg68Leu | Q9HD34 |
| HOMO SAPIENS | 2 | 1012 | 4nyi | S | 569 | LEU | p.Leu569Val | Q07889 |
| HOMO SAPIENS | 2 | 1012 | 4nyi | S | 655 | PRO | p.Pro655Leu | Q07889 |
| HOMO SAPIENS | 4 | 1012 | 4nyi | S | 663 | ILE | p.Ile663Met | Q07889 |
| HOMO SAPIENS | 2 | 1012 | 4nyi | S | 702 | TYR | p.Tyr702Cys | Q07889 |
| HOMO SAPIENS | 2 | 1012 | 4nyi | S | 708 | ALA | p.Ala708Thr | Q07889 |
| HOMO SAPIENS | 4 | 1012 | 4nyi | S | 733 | ILE | p.Ile733Phe | Q07889 |
| HOMO SAPIENS | 1 | 1848 | 3g1n | A | 4013 | ARG | p.Arg4013Trp | Q7Z6Z7 |
| HOMO SAPIENS | 1 | 1848 | 3g1n | A | 4187 | ARG | p.Arg4187Cys | Q7Z6Z7 |
| HOMO SAPIENS | 1 | 77 | 4pod | A | 41 | CYS | p.Cys79Tyr | P60174 |
| HOMO SAPIENS | 9 | 77 | 4pod | A | 104 | GLU | p.Glu142Asp | P60174 |
| HOMO SAPIENS | 1 | 77 | 4pod | A | 122 | GLY | p.Gly160Arg | P60174 |
| HOMO SAPIENS | -1 | 8672 | 3thz | A | 8 | THR | p.Thr8Met | P43246 |
| HOMO SAPIENS | 0 | 8672 | 3thz | A | 19 | PHE | p.Phe19Leu | P43246 |
| HOMO SAPIENS | 0 | 8672 | 3thz | A | 23 | PHE | p.Phe23Leu | P43246 |
| HOMO SAPIENS | 0 | 8672 | 3thz | A | 96 | ARG | p.Arg96His | P43246 |

| | | | | | | | | |
|---|---|---|---|---|---|---|---|---|
| HOMO SAPIENS | 0 | 8672 | 3thz | A | 102 | VAL | p.Val102Ile | P43246 |
| HOMO SAPIENS | 0 | 8672 | 3thz | A | 106 | ARG | p.Arg106Lys | P43246 |
| HOMO SAPIENS | 0 | 8672 | 3thz | A | 127 | ASN | p.Asn127Ser | P43246 |
| HOMO SAPIENS | 0 | 8672 | 3thz | A | 128 | LEU | p.Leu128Val | P43246 |
| HOMO SAPIENS | 0 | 8672 | 3thz | A | 163 | VAL | p.Val163Gly | P43246 |
| HOMO SAPIENS | 0 | 8672 | 3thz | A | 169 | ILE | p.Ile169Val | P43246 |
| HOMO SAPIENS | 0 | 8672 | 3thz | A | 315 | GLY | p.Gly315Asp | P43246 |
| HOMO SAPIENS | -1 | 8672 | 3thz | A | 322 | GLY | p.Gly322Asp | P43246 |
| HOMO SAPIENS | 0 | 8672 | 3thz | A | 331 | ASN | p.Asn331Asp | P43246 |
| HOMO SAPIENS | 1 | 8672 | 3thz | A | 390 | LEU | p.Leu390Phe | P43246 |
| HOMO SAPIENS | 0 | 8672 | 3thz | A | 419 | GLN | p.Gln419Lys | P43246 |
| HOMO SAPIENS | 0 | 8672 | 3thz | A | 447 | PHE | p.Phe447Leu | P43246 |
| HOMO SAPIENS | 0 | 8672 | 3thz | A | 524 | ARG | p.Arg524Pro | P43246 |
| HOMO SAPIENS | 0 | 8672 | 3thz | A | 539 | PHE | p.Phe539Leu | P43246 |
| HOMO SAPIENS | -1 | 8672 | 3thz | A | 546 | LYS | p.Lys546Asn | P43246 |
| HOMO SAPIENS | 0 | 8672 | 3thz | A | 564 | THR | p.Thr564Ala | P43246 |
| HOMO SAPIENS | 0 | 8672 | 3thz | A | 577 | ILE | p.Ile577Thr | P43246 |
| HOMO SAPIENS | 0 | 8672 | 3thz | A | 588 | TYR | p.Tyr588Ter | P43246 |
| HOMO SAPIENS | 0 | 8672 | 3thz | A | 622 | PRO | p.Pro622Leu | P43246 |
| HOMO SAPIENS | 0 | 8672 | 3thz | A | 629 | GLN | p.Gln629Arg | P43246 |
| HOMO SAPIENS | 0 | 8672 | 3thz | A | 636 | ALA | p.Ala636Pro | P43246 |
| HOMO SAPIENS | 2 | 8672 | 3thz | A | 642 | VAL | p.Val642Ile | P43246 |
| HOMO SAPIENS | 1 | 8672 | 3thz | A | 654 | ASP | p.Asp654Glu | P43246 |
| HOMO SAPIENS | 0 | 8672 | 3thz | A | 662 | GLN | p.Gln662His | P43246 |
| HOMO SAPIENS | 2 | 8672 | 3thz | A | 681 | GLN | p.Gln681His | P43246 |
| HOMO SAPIENS | 0 | 8672 | 3thz | A | 707 | CYS | p.Cys707Ser | P43246 |
| HOMO SAPIENS | 0 | 8672 | 3thz | A | 809 | GLU | p.Glu809Lys | P43246 |
| HOMO SAPIENS | 0 | 8672 | 3thz | A | 813 | MET | p.Met813Val | P43246 |
| HOMO SAPIENS | 1 | 8672 | 3thz | A | 834 | ALA | p.Ala834Thr | P43246 |
| HOMO SAPIENS | 0 | 8672 | 3thz | A | 849 | LEU | p.Leu849Val | P43246 |
| HOMO SAPIENS | -1 | 8672 | 3thz | A | 860 | SER | p.Ser860Leu | P43246 |
| HOMO SAPIENS | -1 | 8672 | 3thz | A | 934 | THR | p.Thr934Lys | P43246 |
| HOMO SAPIENS | 2 | 292 | 1i3k | A | 34 | ASN | p.Asn34Ser | Q14376 |
| HOMO SAPIENS | 7 | 292 | 1i3k | A | 90 | GLY | p.Gly90Glu | Q14376 |
| HOMO SAPIENS | 2 | 292 | 1i3k | A | 103 | ASP | p.Asp103Gly | Q14376 |
| HOMO SAPIENS | 2 | 292 | 1i3k | A | 136 | TYR | p.Tyr136Ter | Q14376 |
| HOMO SAPIENS | 0 | 292 | 1i3k | A | 169 | ARG | p.Arg169Trp | Q14376 |
| HOMO SAPIENS | 0 | 292 | 1i3k | A | 183 | LEU | p.Leu183Pro | Q14376 |
| HOMO SAPIENS | 0 | 292 | 1i3k | A | 302 | GLY | p.Gly302Asp | Q14376 |
| HOMO SAPIENS | 0 | 292 | 1i3k | A | 313 | LEU | p.Leu313Met | Q14376 |
| HOMO SAPIENS | 1 | 2084 | 5vc8 | A | 295 | GLN | p.Gln295Leu | O96028 |
| HOMO SAPIENS | -1 | 10157 | 5wrw | A | 7 | GLY | p.Gly7Ala | O76094 |
| HOMO SAPIENS | -1 | 10157 | 5wrw | A | 7 | GLY | p.Gly7Arg | O76094 |
| HOMO SAPIENS | -1 | 10157 | 5wrw | A | 8 | GLY | p.Gly8Ala | O76094 |
| HOMO SAPIENS | 0 | 3113 | 3in5 | A | 29 | GLU | p.Glu29Lys | Q9UBT6 |
| HOMO SAPIENS | 5 | 4546 | 5in3 | B | 44 | VAL | p.Val44Leu | P07902 |
| HOMO SAPIENS | -1 | 4546 | 5in3 | B | 51 | ARG | p.Arg51Leu | P07902 |
| HOMO SAPIENS | -1 | 4546 | 5in3 | B | 62 | LEU | p.Leu62Met | P07902 |
| HOMO SAPIENS | 4 | 4546 | 5in3 | B | 66 | PRO | p.Pro66His | P07902 |
| HOMO SAPIENS | 2 | 4546 | 5in3 | B | 74 | LEU | p.Leu74Pro | P07902 |
| HOMO SAPIENS | 5 | 4546 | 5in3 | B | 97 | ASN | p.Asn97Ser | P07902 |
| HOMO SAPIENS | 1 | 4546 | 5in3 | B | 135 | SER | p.Ser135Leu | P07902 |
| HOMO SAPIENS | 5 | 4546 | 5in3 | B | 138 | THR | p.Thr138Met | P07902 |
| HOMO SAPIENS | 7 | 4546 | 5in3 | B | 139 | LEU | p.Leu139Pro | P07902 |

| | | | | | | | | |
|---|---|---|---|---|---|---|---|---|
| HOMO SAPIENS | 2 | 4546 | 5in3 | B | 142 | MET | p.Met142Lys | P07902 |
| HOMO SAPIENS | 1 | 4546 | 5in3 | B | 143 | SER | p.Ser143Leu | P07902 |
| HOMO SAPIENS | 2 | 4546 | 5in3 | B | 148 | ARG | p.Arg148Gly | P07902 |
| HOMO SAPIENS | 2 | 4546 | 5in3 | B | 148 | ARG | p.Arg148Gln | P07902 |
| HOMO SAPIENS | 5 | 4546 | 5in3 | B | 183 | PRO | p.Pro183Thr | P07902 |
| HOMO SAPIENS | 5 | 4546 | 5in3 | B | 185 | PRO | p.Pro185His | P07902 |
| HOMO SAPIENS | 5 | 4546 | 5in3 | B | 186 | HIS | p.His186Tyr | P07902 |
| HOMO SAPIENS | 7 | 4546 | 5in3 | B | 188 | GLN | p.Gln188Arg | P07902 |
| HOMO SAPIENS | 5 | 4546 | 5in3 | B | 195 | LEU | p.Leu195Pro | P07902 |
| HOMO SAPIENS | 4 | 4546 | 5in3 | B | 201 | ARG | p.Arg201His | P07902 |
| HOMO SAPIENS | 4 | 4546 | 5in3 | B | 201 | ARG | p.Arg201Cys | P07902 |
| HOMO SAPIENS | 4 | 4546 | 5in3 | B | 203 | GLU | p.Glu203Lys | P07902 |
| HOMO SAPIENS | 2 | 4546 | 5in3 | B | 209 | TYR | p.Tyr209Cys | P07902 |
| HOMO SAPIENS | 5 | 4546 | 5in3 | B | 223 | ARG | p.Arg223Ser | P07902 |
| HOMO SAPIENS | 5 | 4546 | 5in3 | B | 226 | LEU | p.Leu226Pro | P07902 |
| HOMO SAPIENS | 5 | 4546 | 5in3 | B | 226 | LEU | p.Leu226Val | P07902 |
| HOMO SAPIENS | 2 | 4546 | 5in3 | B | 227 | LEU | p.Leu227Pro | P07902 |
| HOMO SAPIENS | 8 | 4546 | 5in3 | B | 258 | ARG | p.Arg258Cys | P07902 |
| HOMO SAPIENS | 1 | 4546 | 5in3 | B | 265 | PRO | p.Pro265Ala | P07902 |
| HOMO SAPIENS | 8 | 4546 | 5in3 | B | 268 | THR | p.Thr268Asn | P07902 |
| HOMO SAPIENS | 7 | 4546 | 5in3 | B | 278 | ILE | p.Ile278Asn | P07902 |
| HOMO SAPIENS | 5 | 4546 | 5in3 | B | 285 | LYS | p.Lys285Asn | P07902 |
| HOMO SAPIENS | 1 | 4546 | 5in3 | B | 319 | HIS | p.His319Gln | P07902 |
| HOMO SAPIENS | 4 | 4546 | 5in3 | B | 321 | HIS | p.His321Tyr | P07902 |
| HOMO SAPIENS | 4 | 4546 | 5in3 | B | 325 | PRO | p.Pro325Leu | P07902 |
| HOMO SAPIENS | 2 | 4546 | 5in3 | B | 328 | ARG | p.Arg328His | P07902 |
| HOMO SAPIENS | 2 | 4546 | 5in3 | B | 329 | SER | p.Ser329Phe | P07902 |
| HOMO SAPIENS | 5 | 4546 | 5in3 | B | 334 | LYS | p.Lys334Arg | P07902 |
| HOMO SAPIENS | 6 | 1290 | 5uav | C | 119 | ARG | p.Arg119Gly | P32322 |
| HOMO SAPIENS | 6 | 1290 | 5uav | C | 119 | ARG | p.Arg119His | P32322 |
| HOMO SAPIENS | 3 | 1290 | 5uav | C | 251 | ARG | p.Arg251His | P32322 |
| HOMO SAPIENS | 7 | 1290 | 5uav | C | 266 | ARG | p.Arg266Gln | P32322 |
| HOMO SAPIENS | 0 | 9618 | 2r7e | A | 22 | GLY | p.Gly41Cys | P00451 |
| HOMO SAPIENS | 2 | 9618 | 2r7e | A | 113 | GLU | p.Glu132Asp | P00451 |
| HOMO SAPIENS | -1 | 9618 | 2r7e | A | 336 | ARG | p.Arg355Ter | P00451 |
| HOMO SAPIENS | 0 | 9618 | 2r7e | A | 372 | ARG | p.Arg391His | P00451 |
| HOMO SAPIENS | 0 | 9618 | 2r7e | A | 373 | SER | p.Ser392Leu | P00451 |
| HOMO SAPIENS | 0 | 9618 | 2r7e | A | 373 | SER | p.Ser392Pro | P00451 |
| HOMO SAPIENS | 3 | 9618 | 2r7e | A | 386 | ILE | p.Ile405Ser | P00451 |
| HOMO SAPIENS | 6 | 9618 | 2r7e | A | 390 | GLU | p.Glu409Gly | P00451 |
| HOMO SAPIENS | 2 | 9618 | 2r7e | A | 412 | LEU | p.Leu431Phe | P00451 |
| HOMO SAPIENS | 5 | 9618 | 2r7e | A | 425 | LYS | p.Lys444Arg | P00451 |
| HOMO SAPIENS | 0 | 9618 | 2r7e | A | 484 | ARG | p.Arg503His | P00451 |
| HOMO SAPIENS | 2 | 9618 | 2r7e | A | 531 | ARG | p.Arg550His | P00451 |
| HOMO SAPIENS | 3 | 9618 | 2r7e | A | 535 | SER | p.Ser554Gly | P00451 |
| HOMO SAPIENS | 2 | 9618 | 2r7e | A | 593 | ARG | p.Arg612Cys | P00451 |
| HOMO SAPIENS | 0 | 9618 | 2r7e | A | 704 | ALA | p.Ala723Thr | P00451 |
| HOMO SAPIENS | 0 | 9618 | 2r7e | A | 720 | GLU | p.Glu739Lys | P00451 |
| HOMO SAPIENS | 1 | 13085 | 3ud1 | A | 1075 | ILE | p.Thr1075Ile | P16157 |
| HOMO SAPIENS | -1 | 3289 | 2fy4 | A | 2 | ALA | p.Ala120Thr | P28329 |
| HOMO SAPIENS | 7 | 3289 | 2fy4 | A | 18 | VAL | p.Val136Met | P28329 |
| HOMO SAPIENS | 5 | 3289 | 2fy4 | A | 92 | LEU | p.Leu210Pro | P28329 |
| HOMO SAPIENS | 2 | 3289 | 2fy4 | A | 93 | PRO | p.Pro211Ala | P28329 |
| HOMO SAPIENS | 4 | 3289 | 2fy4 | A | 131 | LEU | p.Leu249Val | P28329 |

| | | | | | | | | |
|---|---|---|---|---|---|---|---|---|
| HOMO SAPIENS | 2 | 3289 | 2fy4 | A | 181 | PRO | p.Pro299Leu | P28329 |
| HOMO SAPIENS | 0 | 3289 | 2fy4 | A | 218 | ILE | p.Ile336Thr | P28329 |
| HOMO SAPIENS | 2 | 3289 | 2fy4 | A | 261 | ASP | p.Asp379His | P28329 |
| HOMO SAPIENS | 0 | 3289 | 2fy4 | A | 302 | ARG | p.Arg420Cys | P28329 |
| HOMO SAPIENS | 5 | 3289 | 2fy4 | A | 323 | GLU | p.Glu441Lys | P28329 |
| HOMO SAPIENS | 0 | 3289 | 2fy4 | A | 343 | MET | p.Val461Met | P28329 |
| HOMO SAPIENS | 2 | 3289 | 2fy4 | A | 364 | ARG | p.Arg482Gly | P28329 |
| HOMO SAPIENS | 3 | 3289 | 2fy4 | A | 380 | SER | p.Ser498Leu | P28329 |
| HOMO SAPIENS | 6 | 3289 | 2fy4 | A | 388 | VAL | p.Val506Leu | P28329 |
| HOMO SAPIENS | 7 | 3289 | 2fy4 | A | 429 | HIS | p.His547Gln | P28329 |
| HOMO SAPIENS | 8 | 3289 | 2fy4 | A | 442 | ARG | p.Arg560His | P28329 |
| HOMO SAPIENS | 3 | 3289 | 2fy4 | A | 443 | ARG | p.Arg561Gln | P28329 |
| HOMO SAPIENS | 0 | 4137 | 3hhr | B | 38 | CYS | p.Cys56Ter | P10912 |
| HOMO SAPIENS | 0 | 4137 | 3hhr | B | 43 | ARG | p.Arg61Ter | P10912 |
| HOMO SAPIENS | 0 | 4137 | 3hhr | B | 44 | GLU | p.Glu62Lys | P10912 |
| HOMO SAPIENS | 0 | 4137 | 3hhr | B | 96 | PHE | p.Phe114Ser | P10912 |
| HOMO SAPIENS | 5 | 4137 | 3hhr | B | 131 | PRO | p.Pro149Gln | P10912 |
| HOMO SAPIENS | 0 | 4137 | 3hhr | B | 144 | VAL | p.Val162Ile | P10912 |
| HOMO SAPIENS | 0 | 4137 | 3hhr | B | 150 | HIS | p.His168Gln | P10912 |
| HOMO SAPIENS | 0 | 4137 | 3hhr | B | 152 | ASP | p.Asp170His | P10912 |
| HOMO SAPIENS | 0 | 4137 | 3hhr | B | 153 | ILE | p.Ile171Thr | P10912 |
| HOMO SAPIENS | 0 | 4137 | 3hhr | B | 154 | GLN | p.Gln172Pro | P10912 |
| HOMO SAPIENS | 0 | 4137 | 3hhr | B | 155 | VAL | p.Val173Gly | P10912 |
| HOMO SAPIENS | 0 | 4137 | 3hhr | B | 217 | ARG | p.Arg235Ter | P10912 |
| HOMO SAPIENS | 5 | 4137 | 3hhr | B | 224 | GLU | p.Glu242Asp | P10912 |
| HOMO SAPIENS | 5 | 4137 | 3hhr | B | 224 | GLU | p.Glu242Ter | P10912 |
| HOMO SAPIENS | 6 | 2493 | 5bpb | A | 69 | HIS | p.His69Tyr | Q9ULV1 |
| HOMO SAPIENS | 6 | 2493 | 5bpb | A | 105 | MET | p.Met105Val | Q9ULV1 |
| HOMO SAPIENS | -1 | 2493 | 5bpb | A | 168 | PRO | p.Pro168Ser | Q9ULV1 |
| HOMO SAPIENS | 0 | 32123 | 4ccz | A | 314 | ASP | p.Asp314Asn | Q99707 |
| HOMO SAPIENS | 0 | 32123 | 4ccz | A | 345 | VAL | p.Val345Phe | Q99707 |
| HOMO SAPIENS | 1 | 32123 | 4ccz | A | 585 | ARG | p.Arg585Ter | Q99707 |
| HOMO SAPIENS | 4 | 4618 | 2ag4 | A | 26 | PRO | p.Pro55Leu | P17900 |
| HOMO SAPIENS | 4 | 4618 | 2ag4 | A | 30 | VAL | p.Ile59Val | P17900 |
| HOMO SAPIENS | 2 | 4618 | 2ag4 | A | 40 | VAL | p.Met69Val | P17900 |
| HOMO SAPIENS | 2 | 4618 | 2ag4 | A | 124 | VAL | p.Val153Ala | P17900 |
| HOMO SAPIENS | 8 | 4618 | 2ag4 | A | 140 | ARG | p.Arg169Pro | P17900 |
| HOMO SAPIENS | 5 | 1082 | 1hk0 | X | 14 | ARG | p.Arg15Cys | P07320 |
| HOMO SAPIENS | 7 | 1082 | 1hk0 | X | 23 | PRO | p.Pro24Thr | P07320 |
| HOMO SAPIENS | 6 | 1082 | 1hk0 | X | 36 | ARG | p.Arg37Ser | P07320 |
| HOMO SAPIENS | 5 | 1082 | 1hk0 | X | 55 | TYR | p.Tyr56Ter | P07320 |
| HOMO SAPIENS | 3 | 1082 | 1hk0 | X | 58 | ARG | p.Arg59His | P07320 |
| HOMO SAPIENS | 7 | 1082 | 1hk0 | X | 88 | HIS | p.His88Gln | P07320 |
| HOMO SAPIENS | 6 | 1082 | 1hk0 | X | 95 | ARG | p.Arg95Ser | P07320 |
| HOMO SAPIENS | 5 | 1082 | 1hk0 | X | 126 | VAL | p.Val126Met | P07320 |
| HOMO SAPIENS | 8 | 1082 | 1hk0 | X | 157 | TRP | p.Trp157Ter | P07320 |
| HOMO SAPIENS | 0 | 22879 | 3j0a | B | 644 | ILE | p.Ile644Phe | O60602 |
| HOMO SAPIENS | 8 | 6 | 2o23 | B | 122 | LEU | p.Leu122Val | Q99714 |
| HOMO SAPIENS | 1 | 6 | 2o23 | B | 130 | ARG | p.Arg130Cys | Q99714 |
| HOMO SAPIENS | 1 | 6 | 2o23 | B | 247 | ASN | p.Asn247Ser | Q99714 |
| HOMO SAPIENS | 0 | 6 | 2o23 | B | 249 | GLU | p.Glu249Gln | Q99714 |
| HOMO SAPIENS | 1 | 12910 | 4hl4 | A | 67 | ARG | p.Arg67Ter | Q96BZ9 |
| HOMO SAPIENS | 8 | 16105 | 1lmj | A | 7 | GLU | p.Glu1073Lys | P35555 |
| HOMO SAPIENS | 8 | 16105 | 1lmj | A | 8 | CYS | p.Cys1074Arg | P35555 |

| | | | | | | | | |
|---|---|---|---|---|---|---|---|---|
| HOMO SAPIENS | 9 | 16105 | 1lmj | A | 51 | CYS | p.Cys1117Tyr | P35555 |
| HOMO SAPIENS | 9 | 16105 | 1lmj | A | 63 | CYS | p.Cys1129Tyr | P35555 |
| HOMO SAPIENS | 9 | 16105 | 1lmj | A | 65 | ASN | p.Asn1131Tyr | P35555 |
| HOMO SAPIENS | 4 | 16105 | 1lmj | A | 71 | ARG | p.Arg1137Pro | P35555 |
| HOMO SAPIENS | 1 | 16105 | 1lmj | A | 82 | PRO | p.Pro1148Ala | P35555 |
| HOMO SAPIENS | 0 | 4795 | 3s5f | A | 106 | LEU | p.Leu106Ser | Q16595 |
| HOMO SAPIENS | 0 | 4795 | 3s5f | A | 130 | GLY | p.Gly130Val | Q16595 |
| HOMO SAPIENS | 0 | 4795 | 3s5f | A | 154 | ILE | p.Ile154Phe | Q16595 |
| HOMO SAPIENS | -1 | 4795 | 3s5f | A | 209 | ASP | p.Asp209Gly | Q16595 |
| HOMO SAPIENS | -1 | 2884 | 4lli | A | 1655 | ASP | p.Asp1655Glu | Q9Y4I1 |
| HOMO SAPIENS | 0 | 1786 | 7pck | C | 56P | TYR | p.Tyr71Ter | P43235 |
| HOMO SAPIENS | 0 | 1786 | 7pck | C | 64P | GLY | p.Gly79Glu | P43235 |
| HOMO SAPIENS | 7 | 38286 | 2l53 | B | 1914 | ARG | p.Arg1914Gly | Q14524 |
| HOMO SAPIENS | 1 | 38286 | 2l53 | B | 1924 | ALA | p.Ala1924Thr | Q14524 |
| HOMO SAPIENS | 0 | 31747 | 3ksy | A | 25 | PRO | p.Pro25Ser | Q07889 |
| HOMO SAPIENS | 3 | 31747 | 3ksy | A | 37 | THR | p.Thr37Ala | Q07889 |
| HOMO SAPIENS | 0 | 31747 | 3ksy | A | 54 | GLN | p.Gln54His | Q07889 |
| HOMO SAPIENS | -1 | 31747 | 3ksy | A | 190 | ASP | p.Asp190Glu | Q07889 |
| HOMO SAPIENS | 2 | 31747 | 3ksy | A | 266 | THR | p.Thr266Lys | Q07889 |
| HOMO SAPIENS | 1 | 31747 | 3ksy | A | 269 | MET | p.Met269Arg | Q07889 |
| HOMO SAPIENS | 1 | 31747 | 3ksy | A | 340 | PRO | p.Pro340Ser | Q07889 |
| HOMO SAPIENS | 1 | 31747 | 3ksy | A | 423 | ASN | p.Asn423Lys | Q07889 |
| HOMO SAPIENS | 1 | 31747 | 3ksy | A | 424 | GLU | p.Glu424Lys | Q07889 |
| HOMO SAPIENS | 1 | 31747 | 3ksy | A | 432 | TRP | p.Trp432Arg | Q07889 |
| HOMO SAPIENS | 1 | 31747 | 3ksy | A | 552 | ARG | p.Arg552Gly | Q07889 |
| HOMO SAPIENS | 1 | 31747 | 3ksy | A | 552 | ARG | p.Arg552Ser | Q07889 |
| HOMO SAPIENS | 3 | 7529 | 6jdw | A | 93 | ILE | p.Ile93Met | P50440 |
| HOMO SAPIENS | 2 | 7529 | 6jdw | A | 110 | GLN | p.Gln110His | P50440 |
| HOMO SAPIENS | 5 | 7529 | 6jdw | A | 149 | TRP | p.Trp149Ter | P50440 |
| HOMO SAPIENS | 0 | 7529 | 6jdw | A | 169 | ARG | p.Arg169Ter | P50440 |
| HOMO SAPIENS | 0 | 7529 | 6jdw | A | 260 | ARG | p.Arg260Ter | P50440 |
| HOMO SAPIENS | 3 | 7529 | 6jdw | A | 344 | ILE | p.Ile344Val | P50440 |
| HOMO SAPIENS | 5 | 7529 | 6jdw | A | 418 | LEU | p.Leu418Ile | P50440 |
| HOMO SAPIENS | 1 | 14788 | 1eak | A | 101 | ARG | p.Arg101His | P08253 |
| HOMO SAPIENS | 0 | 14788 | 1eak | A | 383 | ASP | p.Asp383Glu | P08253 |
| HOMO SAPIENS | 7 | 14788 | 1eak | A | 404 | GLN | p.Glu404Lys | P08253 |
| HOMO SAPIENS | 2 | 14788 | 1eak | A | 447 | ALA | p.Ala447Val | P08253 |
| HOMO SAPIENS | 0 | 1255 | 5xtc | p | 8 | PRO | p.Pro8His | Q9Y6M9 |
| HOMO SAPIENS | 0 | 1255 | 5xtc | p | 146 | PRO | p.Pro146Ser | Q9Y6M9 |
| HOMO SAPIENS | 3 | 6721 | 1a22 | A | 25 | PHE | p.Phe51Tyr | P01241 |
| HOMO SAPIENS | 2 | 6721 | 1a22 | A | 95 | SER | p.Ser121Arg | P01241 |
| HOMO SAPIENS | 2 | 6721 | 1a22 | A | 110 | VAL | p.Val136Ile | P01241 |
| HOMO SAPIENS | 0 | 1380 | 5i6z | A | 605 | LYS | p.Lys605Asn | P31645 |
| HOMO SAPIENS | 0 | 941 | 1qki | D | 48 | ILE | p.Ile48Thr | P11413 |
| HOMO SAPIENS | 3 | 941 | 1qki | D | 68 | VAL | p.Val68Met | P11413 |
| HOMO SAPIENS | 2 | 941 | 1qki | D | 126 | ASN | p.Asn126Asp | P11413 |
| HOMO SAPIENS | 8 | 941 | 1qki | D | 163 | GLY | p.Gly163Ser | P11413 |
| HOMO SAPIENS | 6 | 941 | 1qki | D | 165 | ASN | p.Asn165Asp | P11413 |
| HOMO SAPIENS | 4 | 941 | 1qki | D | 181 | ASP | p.Asp181Val | P11413 |
| HOMO SAPIENS | 4 | 941 | 1qki | D | 182 | ARG | p.Arg182Trp | P11413 |
| HOMO SAPIENS | 5 | 941 | 1qki | D | 188 | SER | p.Ser188Phe | P11413 |
| HOMO SAPIENS | 9 | 941 | 1qki | D | 198 | ARG | p.Arg198Cys | P11413 |
| HOMO SAPIENS | 8 | 941 | 1qki | D | 213 | VAL | p.Val213Leu | P11413 |
| HOMO SAPIENS | 5 | 941 | 1qki | D | 282 | ASP | p.Asp282His | P11413 |

| | | | | | | | | |
|---|---|---|---|---|---|---|---|---|
| HOMO SAPIENS | 7 | 941 | 1qki | D | 285 | ARG | p.Arg285His | P11413 |
| HOMO SAPIENS | 5 | 941 | 1qki | D | 291 | VAL | p.Val291Met | P11413 |
| HOMO SAPIENS | 8 | 941 | 1qki | D | 323 | LEU | p.Leu323Pro | P11413 |
| HOMO SAPIENS | 3 | 941 | 1qki | D | 335 | ALA | p.Ala335Thr | P11413 |
| HOMO SAPIENS | 6 | 941 | 1qki | D | 350 | ASP | p.Asp350His | P11413 |
| HOMO SAPIENS | 5 | 941 | 1qki | D | 393 | ARG | p.Arg393His | P11413 |
| HOMO SAPIENS | 5 | 941 | 1qki | D | 454 | ARG | p.Arg454Cys | P11413 |
| HOMO SAPIENS | 2 | 941 | 1qki | D | 459 | LEU | p.Arg459Leu | P11413 |
| HOMO SAPIENS | 4 | 941 | 1qki | D | 463 | ARG | p.Arg463His | P11413 |
| HOMO SAPIENS | 4 | 2483 | 1tij | A | 68 | LEU | p.Leu94Gln | P01034 |
| HOMO SAPIENS | 2 | 20632 | 3v5b | A | 331 | ARG | p.Arg331Pro | P02545 |
| HOMO SAPIENS | 2 | 20632 | 3v5b | A | 374 | HIS | p.His374Gln | P02545 |
| HOMO SAPIENS | 4 | 13929 | 5xtd | J | 75 | ARG | p.Arg75His | Q16795 |
| HOMO SAPIENS | 3 | 13929 | 5xtd | J | 243 | VAL | p.Val243Ala | Q16795 |
| HOMO SAPIENS | 1 | 13929 | 5xtd | J | 294 | PRO | p.Pro294Leu | Q16795 |
| HOMO SAPIENS | 2 | 13929 | 5xtd | J | 321 | ARG | p.Arg321Pro | Q16795 |
| HOMO SAPIENS | 0 | 9411 | 5thl | A | 41 | ARG | p.Gly41Arg | P54577 |
| HOMO SAPIENS | 0 | 9411 | 5thl | A | 196 | GLU | p.Glu196Lys | P54577 |
| HOMO SAPIENS | 0 | 9411 | 5thl | A | 316 | ALA | p.Ala316Ser | P54577 |
| HOMO SAPIENS | 0 | 9411 | 5thl | A | 339 | ALA | p.Ala339Thr | P54577 |
| HOMO SAPIENS | 4 | 16273 | 2e8n | A | 46 | THR | p.Thr940Ile | P29317 |
| HOMO SAPIENS | 7 | 16273 | 2e8n | A | 54 | GLY | p.Gly948Trp | P29317 |
| HOMO SAPIENS | 6 | 16273 | 2e8n | A | 65 | ALA | p.Ala959Thr | P29317 |
| HOMO SAPIENS | 7 | 30387 | 2k9y | A | 541 | VAL | p.Val541Met | P29317 |
| HOMO SAPIENS | 0 | 6961 | 4nha | A | 843 | ILE | p.Ile853Val | Q9UPY3 |
| HOMO SAPIENS | -1 | 6961 | 4nha | A | 862 | ALA | p.Ala872Thr | Q9UPY3 |
| HOMO SAPIENS | 3 | 15708 | 4pzg | A | 16 | ARG | p.Arg16Ter | Q9Y5X0 |
| HOMO SAPIENS | 4 | 15708 | 4pzg | A | 51 | ARG | p.Arg51Gln | Q9Y5X0 |
| HOMO SAPIENS | 3 | 30166 | 4o00 | A | 44 | ARG | p.Arg14257Ter | Q8WZ42 |
| HOMO SAPIENS | 2 | 30166 | 4o00 | A | 54 | ARG | p.Arg14267His | Q8WZ42 |
| HOMO SAPIENS | 8 | 11158 | 4hca | A | 275 | TRP | p.Trp274Arg | P23771 |
| HOMO SAPIENS | 2 | 4328 | 4e0s | A | 1476 | ARG | p.Arg1476Ter | P01031 |
| HOMO SAPIENS | 0 | 11462 | 1dx5 | I | 455 | ALA | p.Ala473Val | P07204 |
| HOMO SAPIENS | -1 | 11150 | 2fic | A | 45 | LYS | p.Lys35Asn | O00499 |
| HOMO SAPIENS | 6 | 11150 | 2fic | A | 161 | ASP | p.Asp151Asn | O00499 |
| HOMO SAPIENS | 8 | 27382 | 4mqv | B | 17 | ARG | p.Arg17Ter | Q9NZC9 |
| HOMO SAPIENS | -1 | 37991 | 1q0p | A | 231 | GLN | p.Gln256Ter | P00751 |
| HOMO SAPIENS | 1 | 443 | 5xte | U | 215 | LEU | p.Leu215Phe | P08574 |
| HOMO SAPIENS | 8 | 21471 | 5y4e | A | 1461 | ASN | p.Thr692Ala | Q01484 |
| HOMO SAPIENS | -1 | 35443 | 3um3 | B | 129 | ASP | p.Asp129Val | Q9H444 |
| HOMO SAPIENS | -1 | 35443 | 3um3 | B | 161 | GLU | p.Glu161Lys | Q9H444 |
| HOMO SAPIENS | 6 | 28781 | 2n2a | B | 111 | ILE | p.Ile654Val | P04626 |
| HOMO SAPIENS | 7 | 28781 | 2n2a | B | 112 | ILE | p.Ile655Val | P04626 |
| HOMO SAPIENS | -1 | 3346 | 4ztz | C | 61 | GLY | p.Gly61Arg | Q9UHN1 |
| HOMO SAPIENS | 4 | 3346 | 4ztz | C | 103 | GLY | p.Gly103Ser | Q9UHN1 |
| HOMO SAPIENS | 4 | 3346 | 4ztz | C | 133 | HIS | p.His133Tyr | Q9UHN1 |
| HOMO SAPIENS | -1 | 3346 | 4ztz | C | 153 | LEU | p.Leu153Val | Q9UHN1 |
| HOMO SAPIENS | -1 | 3346 | 4ztz | C | 169 | ALA | p.Ala169Thr | Q9UHN1 |
| HOMO SAPIENS | 7 | 3346 | 4ztz | C | 369 | ARG | p.Arg369Gly | Q9UHN1 |
| HOMO SAPIENS | 4 | 3346 | 4ztz | C | 386 | ASP | p.Asp386Glu | Q9UHN1 |
| HOMO SAPIENS | 4 | 3346 | 4ztz | C | 451 | GLY | p.Gly451Glu | Q9UHN1 |
| HOMO SAPIENS | 4 | 2131 | 5fo9 | E | 1549 | LEU | p.Leu1549Met | P01024 |
| HOMO SAPIENS | 2 | 2131 | 5fo9 | E | 1619 | SER | p.Ser1619Arg | P01024 |
| HOMO SAPIENS | 3 | 12662 | 4jhp | C | 75 | ILE | p.Ile75Val | Q92834 |

| | | | | | | | | |
|---|---|---|---|---|---|---|---|---|
| HOMO SAPIENS | 1 | 12662 | 4jhp | C | 173 | GLY | p.Gly173Arg | Q92834 |
| HOMO SAPIENS | 0 | 12662 | 4jhp | C | 184 | GLN | p.Gln184His | Q92834 |
| HOMO SAPIENS | 3 | 12662 | 4jhp | C | 262 | ALA | p.Ala262Gly | Q92834 |
| HOMO SAPIENS | 3 | 12662 | 4jhp | C | 302 | CYS | p.Cys302Tyr | Q92834 |
| HOMO SAPIENS | 5 | 1093 | 2jg9 | A | 186 | GLN | p.Gln208Ter | P02745 |
| HOMO SAPIENS | 0 | 13181 | 4r4l | A | 12 | MET | p.Met13Ile | Q13976 |
| HOMO SAPIENS | 8 | 25939 | 4iyp | A | 106 | ARG | p.Arg106Gln | P78318 |
| HOMO SAPIENS | 1 | 217 | 2xsx | A | 71 | SER | p.Asn71Ser | P13929 |
| HOMO SAPIENS | 0 | 217 | 2xsx | A | 76 | ALA | p.Ala76Pro | P13929 |
| HOMO SAPIENS | 1 | 217 | 2xsx | A | 85 | ALA | p.Val85Ala | P13929 |
| HOMO SAPIENS | 8 | 217 | 2xsx | A | 156 | GLY | p.Gly156Asp | P13929 |
| HOMO SAPIENS | 5 | 217 | 2xsx | A | 332 | VAL | p.Val332Ile | P13929 |
| HOMO SAPIENS | 7 | 217 | 2xsx | A | 374 | GLY | p.Gly374Glu | P13929 |
| HOMO SAPIENS | 1 | 5508 | 3szm | A | 682 | THR | p.Thr682Asn | Q8NEM0 |
| HOMO SAPIENS | 0 | 5508 | 3szm | A | 715 | TRP | p.Trp715Ter | Q8NEM0 |
| HOMO SAPIENS | 1 | 5508 | 3szm | A | 761 | VAL | p.Ala761Val | Q8NEM0 |
| HOMO SAPIENS | 1 | 5508 | 3szm | A | 828 | PRO | p.Pro828Ser | Q8NEM0 |
| HOMO SAPIENS | 4 | 5508 | 3szm | A | 829 | GLU | p.Glu829Asp | Q8NEM0 |
| HOMO SAPIENS | 2 | 16353 | 2q0z | X | 171 | GLY | p.Gly1978Ala | O75643 |
| HOMO SAPIENS | 2 | 16353 | 2q0z | X | 188 | ALA | p.Ala1995Thr | O75643 |
| HOMO SAPIENS | 0 | 4127 | 3swr | A | 860 | GLY | p.Gly860Arg | P26358 |
| HOMO SAPIENS | -1 | 4127 | 3swr | A | 1122 | SER | p.Ser1122Cys | P26358 |
| HOMO SAPIENS | 0 | 4127 | 3swr | A | 1216 | ASP | p.Asp1216Glu | P26358 |
| HOMO SAPIENS | 0 | 6382 | 3o96 | A | 17 | GLU | p.Glu17Lys | P31749 |
| HOMO SAPIENS | 6 | 14314 | 5xtc | Q | 71 | PRO | p.Pro71Leu | O75306 |
| HOMO SAPIENS | 2 | 5886 | 4fao | E | 22 | ARG | p.Arg40His | Q13705 |
| HOMO SAPIENS | 1 | 2534 | 3mp4 | A | 37 | GLU | p.Glu37Ter | P35914 |
| HOMO SAPIENS | 3 | 2534 | 3mp4 | A | 41 | MET | p.Arg41Gln | P35914 |
| HOMO SAPIENS | 1 | 2534 | 3mp4 | A | 70 | VAL | p.Val70Leu | P35914 |
| HOMO SAPIENS | 2 | 2196 | 3ifq | A | 142 | ARG | p.Arg142His | P14923 |
| HOMO SAPIENS | 2 | 2196 | 3ifq | A | 153 | ASP | p.Asp153Glu | P14923 |
| HOMO SAPIENS | 3 | 2196 | 3ifq | A | 176 | ARG | p.Arg176Gln | P14923 |
| HOMO SAPIENS | 3 | 2196 | 3ifq | A | 265 | ARG | p.Arg265His | P14923 |
| HOMO SAPIENS | 2 | 2196 | 3ifq | A | 301 | GLU | p.Glu301Gly | P14923 |
| HOMO SAPIENS | 0 | 2196 | 3ifq | A | 456 | VAL | p.Val456Ile | P14923 |
| HOMO SAPIENS | 3 | 2196 | 3ifq | A | 603 | VAL | p.Val603Leu | P14923 |
| HOMO SAPIENS | 0 | 2196 | 3ifq | A | 648 | VAL | p.Val648Ile | P14923 |
| HOMO SAPIENS | 1 | 8746 | 1cgj | I | 32 | PRO | p.Pro55Ser | P00995 |
| HOMO SAPIENS | 1 | 8746 | 1cgj | I | 44 | ARG | p.Arg67His | P00995 |
| HOMO SAPIENS | 1 | 8746 | 1cgj | I | 44 | ARG | p.Arg67Cys | P00995 |
| HOMO SAPIENS | 3 | 3212 | 1syq | A | 105 | ARG | p.Arg105Ter | P18206 |
| HOMO SAPIENS | 3 | 3212 | 1syq | A | 126 | PHE | p.Phe126Leu | P18206 |
| HOMO SAPIENS | 4 | 10028 | 3eto | A | 1609 | ARG | p.Arg1608His | P46531 |
| HOMO SAPIENS | 1 | 10028 | 3eto | A | 1672 | VAL | p.Val1671Ile | P46531 |
| HOMO SAPIENS | 5 | 10028 | 3eto | A | 1692 | GLN | p.Gln1691His | P46531 |
| HOMO SAPIENS | 3 | 3749 | 5xtc | Z | 70 | GLY | p.Gly70Ter | O43676 |
| HOMO SAPIENS | 2 | 17039 | 3brv | B | 57 | GLU | p.Glu57Lys | Q9Y6K9 |
| HOMO SAPIENS | 8 | 3589 | 2fh3 | A | 433 | VAL | p.Val460Leu | P06396 |
| HOMO SAPIENS | 1 | 3589 | 2fh3 | A | 454 | ARG | p.Arg481Cys | P06396 |
| HOMO SAPIENS | 4 | 3589 | 2fh3 | A | 498 | ARG | p.Arg525His | P06396 |
| HOMO SAPIENS | 2 | 3589 | 2fh3 | A | 520 | TYR | p.Tyr547Cys | P06396 |
| HOMO SAPIENS | 1 | 3589 | 2fh3 | A | 529 | GLY | p.Gly556Arg | P06396 |
| HOMO SAPIENS | 2 | 3589 | 2fh3 | A | 536 | THR | p.Thr563Ser | P06396 |
| HOMO SAPIENS | 2 | 3589 | 2fh3 | A | 579 | VAL | p.Val606Met | P06396 |

| | | | | | | | | |
|---|---|---|---|---|---|---|---|---|
| HOMO SAPIENS | 1 | 3589 | 2fh3 | A | 589 | THR | p.Thr616Met | P06396 |
| HOMO SAPIENS | 5 | 3589 | 2fh3 | A | 668 | THR | p.Thr695Met | P06396 |
| HOMO SAPIENS | 0 | 148 | 1lcl | A | 130 | ILE | p.Ile130Ser | Q05315 |
| HOMO SAPIENS | -1 | 24727 | 4fj3 | P | 237 | ALA | p.Ala237Thr | P04049 |
| HOMO SAPIENS | 0 | 24727 | 4fj3 | P | 257 | SER | p.Ser257Leu | P04049 |
| HOMO SAPIENS | 0 | 24727 | 4fj3 | P | 261 | PRO | p.Pro261Ala | P04049 |
| HOMO SAPIENS | 0 | 2949 | 5xtb | B | 77 | ARG | p.Arg77Gly | O00217 |
| HOMO SAPIENS | 0 | 2949 | 5xtb | B | 79 | PRO | p.Pro79Leu | O00217 |
| HOMO SAPIENS | 0 | 2949 | 5xtb | B | 100 | ALA | p.Ala100Val | O00217 |
| HOMO SAPIENS | 1 | 2949 | 5xtb | B | 115 | LYS | p.Lys115Glu | O00217 |
| HOMO SAPIENS | 0 | 2949 | 5xtb | B | 138 | ARG | p.Arg138His | O00217 |
| HOMO SAPIENS | 1 | 3959 | 5k5s | B | 25 | ARG | p.Arg25Ter | P41180 |
| HOMO SAPIENS | 4 | 3959 | 5k5s | B | 39 | PRO | p.Pro39Ala | P41180 |
| HOMO SAPIENS | 0 | 3959 | 5k5s | B | 47 | LYS | p.Lys47Asn | P41180 |
| HOMO SAPIENS | 2 | 3959 | 5k5s | B | 62 | ARG | p.Arg62Met | P41180 |
| HOMO SAPIENS | 2 | 3959 | 5k5s | B | 66 | ARG | p.Arg66Cys | P41180 |
| HOMO SAPIENS | 4 | 3959 | 5k5s | B | 90 | ASN | p.Asn90Thr | P41180 |
| HOMO SAPIENS | 1 | 3959 | 5k5s | B | 116 | ALA | p.Ala116Thr | P41180 |
| HOMO SAPIENS | 1 | 3959 | 5k5s | B | 118 | ASN | p.Asn118Lys | P41180 |
| HOMO SAPIENS | -1 | 3959 | 5k5s | B | 125 | LEU | p.Leu125Pro | P41180 |
| HOMO SAPIENS | -1 | 3959 | 5k5s | B | 127 | GLU | p.Glu127Ala | P41180 |
| HOMO SAPIENS | -1 | 3959 | 5k5s | B | 128 | PHE | p.Phe128Leu | P41180 |
| HOMO SAPIENS | -1 | 3959 | 5k5s | B | 131 | CYS | p.Cys131Trp | P41180 |
| HOMO SAPIENS | 1 | 3959 | 5k5s | B | 138 | THR | p.Thr138Met | P41180 |
| HOMO SAPIENS | 0 | 3959 | 5k5s | B | 143 | GLY | p.Gly143Glu | P41180 |
| HOMO SAPIENS | 5 | 3959 | 5k5s | B | 151 | THR | p.Thr151Met | P41180 |
| HOMO SAPIENS | 4 | 3959 | 5k5s | B | 180 | PHE | p.Phe180Cys | P41180 |
| HOMO SAPIENS | 0 | 3959 | 5k5s | B | 185 | ARG | p.Arg185Gln | P41180 |
| HOMO SAPIENS | 0 | 3959 | 5k5s | B | 185 | ARG | p.Arg185Ter | P41180 |
| HOMO SAPIENS | 2 | 3959 | 5k5s | B | 191 | GLU | p.Glu191Lys | P41180 |
| HOMO SAPIENS | 2 | 3959 | 5k5s | B | 227 | ARG | p.Arg227Leu | P41180 |
| HOMO SAPIENS | 2 | 3959 | 5k5s | B | 250 | GLU | p.Glu250Lys | P41180 |
| HOMO SAPIENS | 6 | 3959 | 5k5s | B | 297 | GLU | p.Glu297Lys | P41180 |
| HOMO SAPIENS | 1 | 3959 | 5k5s | B | 336 | LYS | p.Lys336Asn | P41180 |
| HOMO SAPIENS | 6 | 3959 | 5k5s | B | 429 | HIS | p.His429Tyr | P41180 |
| HOMO SAPIENS | 0 | 3959 | 5k5s | B | 445 | THR | p.Thr445Ala | P41180 |
| HOMO SAPIENS | 3 | 3959 | 5k5s | B | 509 | GLY | p.Gly509Arg | P41180 |
| HOMO SAPIENS | 1 | 3959 | 5k5s | B | 553 | GLY | p.Gly553Arg | P41180 |
| HOMO SAPIENS | 3 | 3959 | 5k5s | B | 592 | ASN | p.Asn592Ser | P41180 |
| HOMO SAPIENS | -1 | 3959 | 5k5s | B | 604 | GLU | p.Glu604Lys | P41180 |
| HOMO SAPIENS | 4 | 2564 | 5l0c | C | 975 | ARG | p.Arg975Trp | P18206 |
| HOMO SAPIENS | 4 | 2564 | 5l0c | C | 975 | ARG | p.Arg975Leu | P18206 |
| HOMO SAPIENS | 1 | 5695 | 5i8r | A | 294 | ARG | p.Arg294Gln | P17405 |
| HOMO SAPIENS | 0 | 5695 | 5i8r | A | 485 | ALA | p.Ala485Val | P17405 |
| HOMO SAPIENS | 1 | 5695 | 5i8r | A | 506 | GLY | p.Gly506Arg | P17405 |
| HOMO SAPIENS | -1 | 1978 | 3avs | A | 1060 | HIS | p.His1060Leu | O15550 |
| HOMO SAPIENS | 0 | 11173 | 4v0c | B | 362 | LYS | p.Lys362Arg | P51787 |
| HOMO SAPIENS | 3 | 11173 | 4v0c | B | 381 | CYS | p.Cys381Tyr | P51787 |
| HOMO SAPIENS | 7 | 11173 | 4v0c | B | 518 | ARG | p.Arg518Gly | P51787 |
| HOMO SAPIENS | 2 | 11173 | 4v0c | B | 525 | ALA | p.Ala525Thr | P51787 |
| HOMO SAPIENS | 0 | 11173 | 4v0c | B | 530 | GLN | p.Gln530Ter | P51787 |
| HOMO SAPIENS | -1 | 11173 | 4v0c | B | 536 | TYR | p.Tyr536Ter | P51787 |
| HOMO SAPIENS | 2 | 1113 | 5xtd | N | 60 | ARG | p.Arg60Ter | Q9UI09 |
| HOMO SAPIENS | 3 | 6061 | 2mmz | A | 148 | LEU | p.Leu148Ser | Q86SX6 |

| | | | | | | | | |
|---|---|---|---|---|---|---|---|---|
| HOMO SAPIENS | -1 | 26081 | 3eu7 | A | 837 | GLU | p.Glu837Ter | Q86YC2 |
| HOMO SAPIENS | 0 | 26081 | 3eu7 | A | 864 | PRO | p.Pro864Ser | Q86YC2 |
| HOMO SAPIENS | 5 | 26081 | 3eu7 | A | 891 | CYS | p.Cys891Trp | Q86YC2 |
| HOMO SAPIENS | 0 | 26081 | 3eu7 | A | 932 | VAL | p.Val932Met | Q86YC2 |
| HOMO SAPIENS | 5 | 26081 | 3eu7 | A | 961 | LEU | p.Leu961Met | Q86YC2 |
| HOMO SAPIENS | 5 | 26081 | 3eu7 | A | 1038 | TRP | p.Trp1038Ter | Q86YC2 |
| HOMO SAPIENS | -1 | 26081 | 3eu7 | A | 1086 | ARG | p.Arg1086Ter | Q86YC2 |
| HOMO SAPIENS | 0 | 26081 | 3eu7 | A | 1122 | ASP | p.Asp1122Glu | Q86YC2 |
| HOMO SAPIENS | 5 | 26081 | 3eu7 | A | 1158 | HIS | p.His1158Gln | Q86YC2 |
| HOMO SAPIENS | 5 | 26081 | 3eu7 | A | 1183 | TYR | p.Tyr1183Ter | Q86YC2 |
| HOMO SAPIENS | 4 | 199 | 1qo5 | C | 45 | ARG | p.Arg46Trp | P05062 |
| HOMO SAPIENS | 5 | 199 | 1qo5 | C | 147 | TRP | p.Trp148Arg | P05062 |
| HOMO SAPIENS | 0 | 199 | 1qo5 | C | 149 | ALA | p.Ala150Pro | P05062 |
| HOMO SAPIENS | 0 | 199 | 1qo5 | C | 174 | ALA | p.Ala175Asp | P05062 |
| HOMO SAPIENS | 0 | 199 | 1qo5 | C | 334 | ASN | p.Asn335Lys | P05062 |
| HOMO SAPIENS | 0 | 199 | 1qo5 | C | 337 | ALA | p.Ala338Val | P05062 |
| HOMO SAPIENS | 5 | 199 | 1qo5 | C | 342 | TYR | p.Tyr343His | P05062 |
| HOMO SAPIENS | 0 | 6910 | 4zht | B | 59 | ARG | p.Arg59Ter | Q9Y223 |
| HOMO SAPIENS | 4 | 6910 | 4zht | B | 129 | ARG | p.Arg129Ter | Q9Y223 |
| HOMO SAPIENS | 4 | 6910 | 4zht | B | 129 | ARG | p.Arg129Gln | Q9Y223 |
| HOMO SAPIENS | 1 | 6910 | 4zht | B | 162 | ARG | p.Arg162Cys | Q9Y223 |
| HOMO SAPIENS | 0 | 6910 | 4zht | B | 176 | ASP | p.Asp176Val | Q9Y223 |
| HOMO SAPIENS | 0 | 6910 | 4zht | B | 216 | VAL | p.Val216Ala | Q9Y223 |
| HOMO SAPIENS | 0 | 6910 | 4zht | B | 246 | ARG | p.Arg246Trp | Q9Y223 |
| HOMO SAPIENS | 0 | 6910 | 4zht | B | 263 | ARG | p.Arg263Leu | Q9Y223 |
| HOMO SAPIENS | 0 | 6910 | 4zht | B | 266 | ARG | p.Arg266Gln | Q9Y223 |
| HOMO SAPIENS | 0 | 6910 | 4zht | B | 266 | ARG | p.Arg266Trp | Q9Y223 |
| HOMO SAPIENS | 0 | 6910 | 4zht | B | 277 | ARG | p.Arg277Cys | Q9Y223 |
| HOMO SAPIENS | 0 | 6910 | 4zht | B | 378 | ASP | p.Asp378Tyr | Q9Y223 |
| HOMO SAPIENS | 3 | 211 | 5kxi | C | 262 | VAL | p.Val287Leu | P17787 |
| HOMO SAPIENS | 0 | 211 | 5kxi | C | 369 | PHE | p.Phe478Leu | P17787 |
| HOMO SAPIENS | 0 | 2454 | 5iy9 | V | null | PHE | p.Phe99Ser | P19447 |
| HOMO SAPIENS | 0 | 2454 | 5iy9 | V | null | ARG | p.Arg109Ter | P19447 |
| HOMO SAPIENS | 0 | 2454 | 5iy9 | V | null | THR | p.Thr119Pro | P19447 |
| HOMO SAPIENS | 0 | 2454 | 5iy9 | V | 425 | ARG | p.Arg425Ter | P19447 |
| HOMO SAPIENS | 1 | 2454 | 5iy9 | V | 545 | GLN | p.Gln545Ter | P19447 |
| HOMO SAPIENS | -1 | 3805 | 3rbq | A | 57 | LYS | p.Lys57Ter | Q13432 |
| HOMO SAPIENS | 2 | 2649 | 4ido | A | 43 | ASP | p.Asp43Glu | Q8WXF7 |
| HOMO SAPIENS | 2 | 2649 | 4ido | A | 157 | LEU | p.Leu157Trp | Q8WXF7 |
| HOMO SAPIENS | 2 | 2649 | 4ido | A | 193 | PHE | p.Phe193Cys | Q8WXF7 |
| HOMO SAPIENS | 7 | 2649 | 4ido | A | 217 | ARG | p.Arg217Gln | Q8WXF7 |
| HOMO SAPIENS | 4 | 2649 | 4ido | A | 239 | ARG | p.Arg239Cys | Q8WXF7 |
| HOMO SAPIENS | 3 | 2649 | 4ido | A | 258 | HIS | p.His258Arg | Q8WXF7 |
| HOMO SAPIENS | 4 | 2649 | 4ido | A | 259 | SER | p.Ser259Tyr | Q8WXF7 |
| HOMO SAPIENS | 4 | 2649 | 4ido | A | 408 | MET | p.Met408Val | Q8WXF7 |
| HOMO SAPIENS | 2 | 2649 | 4ido | A | 415 | ARG | p.Arg415Trp | Q8WXF7 |
| HOMO SAPIENS | 3 | 2649 | 4ido | A | 416 | ARG | p.Arg416Cys | Q8WXF7 |
| HOMO SAPIENS | 5 | 2203 | 3wff | A | 810 | LEU | p.Ser810Leu | P08235 |
| HOMO SAPIENS | 4 | 12648 | 5ep6 | B | 696 | GLU | p.Glu696Lys | Q9UHD2 |
| HOMO SAPIENS | 0 | 30622 | 2pe4 | A | 90 | GLU | p.Glu90Asp | Q12794 |
| HOMO SAPIENS | 0 | 30622 | 2pe4 | A | 256 | GLY | p.Gly256Arg | Q12794 |
| HOMO SAPIENS | 1 | 30622 | 2pe4 | A | 268 | GLU | p.Glu268Lys | Q12794 |
| HOMO SAPIENS | 0 | 30622 | 2pe4 | A | 392 | THR | p.Thr392Met | Q12794 |
| HOMO SAPIENS | 5 | 5411 | 1lya | A | 93 | SER | p.Ser157Leu | P07339 |

| | | | | | | | | |
|---|---|---|---|---|---|---|---|---|
| HOMO SAPIENS | 3 | 5743 | 2uzp | A | 189 | ASN | p.Asn189Lys | P05129 |
| HOMO SAPIENS | 1 | 12529 | 3kfd | J | 21 | ASN | p.Asn45Ser | P36897 |
| HOMO SAPIENS | 1 | 1063 | 2e1q | B | 149 | ARG | p.Arg149Cys | P47989 |
| HOMO SAPIENS | -1 | 1063 | 2e1q | B | 172 | GLY | p.Gly172Arg | P47989 |
| HOMO SAPIENS | 4 | 1063 | 2e1q | B | 228 | ARG | p.Arg228Ter | P47989 |
| HOMO SAPIENS | 5 | 1063 | 2e1q | B | 395 | LYS | p.Lys395Met | P47989 |
| HOMO SAPIENS | 2 | 1063 | 2e1q | B | 646 | ILE | p.Ile646Val | P47989 |
| HOMO SAPIENS | 4 | 1063 | 2e1q | B | 703 | ILE | p.Ile703Val | P47989 |
| HOMO SAPIENS | 8 | 2525 | 1kqn | F | 18 | ASN | p.Asn18Ser | Q9HAN9 |
| HOMO SAPIENS | 0 | 2525 | 1kqn | F | 151 | VAL | p.Val151Phe | Q9HAN9 |
| HOMO SAPIENS | 0 | 2525 | 1kqn | F | 153 | LEU | p.Leu153Val | Q9HAN9 |
| HOMO SAPIENS | 0 | 2525 | 1kqn | F | 169 | TRP | p.Trp169Ter | Q9HAN9 |
| HOMO SAPIENS | 0 | 2525 | 1kqn | F | 207 | ARG | p.Arg207Trp | Q9HAN9 |
| HOMO SAPIENS | 0 | 2525 | 1kqn | F | 273 | ASN | p.Asn273Asp | Q9HAN9 |
| HOMO SAPIENS | 0 | 12299 | 5duq | A | 432 | VAL | p.Val454Glu | P05155 |
| HOMO SAPIENS | 0 | 12299 | 5duq | A | 436 | ALA | p.Ala458Thr | P05155 |
| HOMO SAPIENS | 0 | 12299 | 5duq | A | 443 | ALA | p.Ala465Val | P05155 |
| HOMO SAPIENS | 0 | 12299 | 5duq | A | 444 | ARG | p.Arg466His | P05155 |
| HOMO SAPIENS | 0 | 12299 | 5duq | A | 458 | MET | p.Val480Met | P05155 |
| HOMO SAPIENS | 0 | 327 | 1uhp | A | 36 | ASP | p.Asp274Asn | Q9UPQ7 |
| HOMO SAPIENS | 0 | 64 | 2mo5 | A | 54 | ALA | p.Ala55Thr | P12104 |
| HOMO SAPIENS | 2 | 35969 | 1jrh | I | 60 | CYS | p.Cys77Tyr | P15260 |
| HOMO SAPIENS | 4 | 35969 | 1jrh | I | 70 | ILE | p.Ile87Thr | P15260 |
| HOMO SAPIENS | 2 | 1068 | 1mu7 | A | 304 | ARG | p.Arg304Gln | Q9NUW8 |
| HOMO SAPIENS | 8 | 1068 | 1mu7 | A | 493 | HIS | p.His493Arg | Q9NUW8 |
| HOMO SAPIENS | 0 | 32360 | 2kqp | A | 31 | ARG | p.Arg55Cys | P01308 |
| HOMO SAPIENS | 0 | 32360 | 2kqp | A | 65 | ARG | p.Arg89His | P01308 |
| HOMO SAPIENS | 0 | 32360 | 2kqp | A | 65 | ARG | p.Arg89Cys | P01308 |
| HOMO SAPIENS | 6 | 10805 | 3nks | A | 12 | ILE | p.Ile12Thr | P50336 |
| HOMO SAPIENS | 0 | 10805 | 3nks | A | 20 | HIS | p.His20Pro | P50336 |
| HOMO SAPIENS | 0 | 10805 | 3nks | A | 59 | ARG | p.Arg59Trp | P50336 |
| HOMO SAPIENS | 0 | 10805 | 3nks | A | 168 | ARG | p.Arg168Cys | P50336 |
| HOMO SAPIENS | 0 | 10805 | 3nks | A | 168 | ARG | p.Arg168His | P50336 |
| HOMO SAPIENS | 2 | 10805 | 3nks | A | 232 | GLY | p.Gly232Arg | P50336 |
| HOMO SAPIENS | 3 | 10805 | 3nks | A | 256 | PRO | p.Pro256Arg | P50336 |
| HOMO SAPIENS | 0 | 10805 | 3nks | A | 304 | ARG | p.Arg304His | P50336 |
| HOMO SAPIENS | 0 | 10805 | 3nks | A | 349 | ASP | p.Asp349Ala | P50336 |
| HOMO SAPIENS | 9 | 7008 | 3u2l | A | 125 | GLN | p.Gln125Ter | P55789 |
| HOMO SAPIENS | 4 | 7008 | 3u2l | A | 172 | HIS | p.His172Asn | P55789 |
| HOMO SAPIENS | 4 | 7008 | 3u2l | A | 194 | ARG | p.Arg194His | P55789 |
| HOMO SAPIENS | 3 | 7008 | 3u2l | A | 198 | GLY | p.Gly198Ser | P55789 |
| HOMO SAPIENS | 0 | 20821 | 3e0g | A | 65 | HIS | p.His116Tyr | P42702 |
| HOMO SAPIENS | 3 | 1151 | 5l9v | A | 278 | ASP | p.Asp278Asn | Q9GZT9 |
| HOMO SAPIENS | 4 | 1151 | 5l9v | A | 317 | CYS | p.Pro317Arg | Q9GZT9 |
| HOMO SAPIENS | 5 | 1151 | 5l9v | A | 371 | ARG | p.Arg371His | Q9GZT9 |
| HOMO SAPIENS | 2 | 3935 | 1gwb | A | 36 | ARG | p.Arg36Gly | P07359 |
| HOMO SAPIENS | 1 | 3935 | 1gwb | A | 73 | LEU | p.Leu73Phe | P07359 |
| HOMO SAPIENS | 1 | 3935 | 1gwb | A | 172 | ALA | p.Ala172Val | P07359 |
| HOMO SAPIENS | 0 | 3935 | 1gwb | A | 249 | GLY | p.Gly249Val | P07359 |
| HOMO SAPIENS | 0 | 3935 | 1gwb | A | 255 | MET | p.Met255Val | P07359 |
| HOMO SAPIENS | 2 | 14541 | 5cuz | A | 138 | GLU | p.Glu138Lys | Q9H3L0 |
| HOMO SAPIENS | 2 | 14541 | 5cuz | A | 143 | SER | p.Ser143Ile | Q9H3L0 |
| HOMO SAPIENS | 4 | 14541 | 5cuz | A | 182 | THR | p.Thr182Asn | Q9H3L0 |
| HOMO SAPIENS | 6 | 14541 | 5cuz | A | 206 | ASN | p.Asn206Ile | Q9H3L0 |

| | | | | | | | | |
|---|---|---|---|---|---|---|---|---|
| HOMO SAPIENS | 0 | 14541 | 5cuz | A | 249 | TYR | p.Tyr249Cys | Q9H3L0 |
| HOMO SAPIENS | 4 | 14541 | 5cuz | A | 250 | ARG | p.Arg250Ter | Q9H3L0 |
| HOMO SAPIENS | 2 | 14541 | 5cuz | A | 259 | LEU | p.Leu259Pro | Q9H3L0 |
| HOMO SAPIENS | 0 | 4206 | 4fao | D | 30 | PRO | p.Pro30Ser | P37023 |
| HOMO SAPIENS | 1 | 4206 | 4fao | D | 50 | TRP | p.Trp50Cys | P37023 |
| HOMO SAPIENS | 0 | 16553 | 3v30 | A | 251 | GLN | p.Gln251Glu | O14593 |
| HOMO SAPIENS | 0 | 5012 | 5hbf | A | 442 | ALA | p.Ala442Gly | Q13231 |
| HOMO SAPIENS | 5 | 4844 | 5tdc | A | 136 | HIS | p.His136Arg | Q8IWV7 |
| HOMO SAPIENS | 0 | 15435 | 4cgz | A | 643 | ARG | p.Arg643His | P54132 |
| HOMO SAPIENS | 1 | 15435 | 4cgz | A | 672 | GLN | p.Gln672Arg | P54132 |
| HOMO SAPIENS | 1 | 15435 | 4cgz | A | 700 | GLN | p.Gln700Ter | P54132 |
| HOMO SAPIENS | 0 | 15435 | 4cgz | A | 707 | PRO | p.Pro707Ser | P54132 |
| HOMO SAPIENS | 0 | 15435 | 4cgz | A | 755 | LYS | p.Lys755Glu | P54132 |
| HOMO SAPIENS | 0 | 15435 | 4cgz | A | 881 | TRP | p.Trp881Ter | P54132 |
| HOMO SAPIENS | 2 | 15435 | 4cgz | A | 899 | ARG | p.Arg899Ter | P54132 |
| HOMO SAPIENS | 0 | 15435 | 4cgz | A | 1036 | CYS | p.Cys1036Phe | P54132 |
| HOMO SAPIENS | 0 | 15435 | 4cgz | A | 1055 | CYS | p.Cys1055Ser | P54132 |
| HOMO SAPIENS | 0 | 15435 | 4cgz | A | 1139 | ARG | p.Arg1139Ter | P54132 |
| HOMO SAPIENS | -1 | 15435 | 4cgz | A | 1198 | VAL | p.Val1198Met | P54132 |
| HOMO SAPIENS | -1 | 15435 | 4cgz | A | 1205 | VAL | p.Val1205Ile | P54132 |
| HOMO SAPIENS | 0 | 15435 | 4cgz | A | 1251 | GLU | p.Glu1251Gln | P54132 |
| HOMO SAPIENS | 0 | 11185 | 1kb4 | A | 30 | ARG | p.Arg30Ter | P11473 |
| HOMO SAPIENS | 5 | 11185 | 1kb4 | A | 33 | GLY | p.Gly33Asp | P11473 |
| HOMO SAPIENS | 5 | 11185 | 1kb4 | A | 46 | GLY | p.Gly46Asp | P11473 |
| HOMO SAPIENS | 6 | 11185 | 1kb4 | A | 50 | ARG | p.Arg50Gln | P11473 |
| HOMO SAPIENS | 9 | 11185 | 1kb4 | A | 73 | ARG | p.Arg73Gln | P11473 |
| HOMO SAPIENS | 9 | 11185 | 1kb4 | A | 80 | ARG | p.Arg80Gln | P11473 |
| HOMO SAPIENS | 0 | 5382 | 3fl7 | A | 41 | LEU | p.Leu41Val | P29317 |
| HOMO SAPIENS | 2 | 5382 | 3fl7 | A | 175 | ARG | p.Arg175Cys | P29317 |
| HOMO SAPIENS | 1 | 5382 | 3fl7 | A | 195 | ARG | p.Arg195Cys | P29317 |
| HOMO SAPIENS | 0 | 5382 | 3fl7 | A | 232 | ASP | p.Asp232Gly | P29317 |
| HOMO SAPIENS | 0 | 5382 | 3fl7 | A | 277 | SER | p.Ser277Leu | P29317 |
| HOMO SAPIENS | 0 | 5382 | 3fl7 | A | 315 | ARG | p.Arg315Gln | P29317 |
| HOMO SAPIENS | 7 | 5382 | 3fl7 | A | 350 | PRO | p.Pro350Thr | P29317 |
| HOMO SAPIENS | 0 | 5382 | 3fl7 | A | 391 | GLY | p.Gly391Arg | P29317 |
| HOMO SAPIENS | 0 | 5382 | 3fl7 | A | 418 | VAL | p.Val418Ile | P29317 |
| HOMO SAPIENS | 0 | 5382 | 3fl7 | A | 511 | THR | p.Thr511Met | P29317 |
| HOMO SAPIENS | 2 | 29168 | 4p22 | B | 144 | VAL | p.Val144Ile | P22314 |
| HOMO SAPIENS | 5 | 4348 | 1bf5 | A | 241 | ARG | p.Arg241Gln | P42224 |
| HOMO SAPIENS | 2 | 4348 | 1bf5 | A | 266 | VAL | p.Val266Ile | P42224 |
| HOMO SAPIENS | 4 | 4348 | 1bf5 | A | 320 | GLU | p.Glu320Gln | P42224 |
| HOMO SAPIENS | 1 | 4348 | 1bf5 | A | 372 | ASN | p.Asn372Lys | P42224 |
| HOMO SAPIENS | 1 | 4348 | 1bf5 | A | 447 | ASP | p.Asp447Glu | P42224 |
| HOMO SAPIENS | 2 | 4348 | 1bf5 | A | 463 | GLN | p.Gln463His | P42224 |
| HOMO SAPIENS | 4 | 4348 | 1bf5 | A | 600 | LEU | p.Leu600Pro | P42224 |
| HOMO SAPIENS | 0 | 4348 | 1bf5 | A | 706 | LEU | p.Leu706Ser | P42224 |
| HOMO SAPIENS | 4 | 4991 | 4bej | B | 141 | ASN | p.Asn141Ser | O00429 |
| HOMO SAPIENS | 0 | 4991 | 4bej | B | 395 | ALA | p.Ala395Asp | O00429 |
| HOMO SAPIENS | 2 | 11792 | 1fvo | A | 40 | ARG | p.Arg40Cys | P00480 |
| HOMO SAPIENS | 2 | 11792 | 1fvo | A | 40 | ARG | p.Arg40His | P00480 |
| HOMO SAPIENS | 0 | 11792 | 1fvo | A | 45 | LEU | p.Leu45Pro | P00480 |
| HOMO SAPIENS | 1 | 11792 | 1fvo | A | 47 | ASN | p.Asn47Ile | P00480 |
| HOMO SAPIENS | 0 | 11792 | 1fvo | A | 50 | GLY | p.Gly50Arg | P00480 |
| HOMO SAPIENS | 5 | 11792 | 1fvo | A | 87 | GLU | p.Glu87Lys | P00480 |

| | | | | | | | | |
|---|---|---|---|---|---|---|---|---|
| HOMO SAPIENS | 3 | 11792 | 1fvo | A | 111 | LEU | p.Leu111Pro | P00480 |
| HOMO SAPIENS | 1 | 11792 | 1fvo | A | 125 | THR | p.Thr125Met | P00480 |
| HOMO SAPIENS | 3 | 11792 | 1fvo | A | 129 | ARG | p.Arg129His | P00480 |
| HOMO SAPIENS | 9 | 11792 | 1fvo | A | 141 | ARG | p.Arg141Pro | P00480 |
| HOMO SAPIENS | 0 | 11792 | 1fvo | A | 206 | MET | p.Met206Arg | P00480 |
| HOMO SAPIENS | 0 | 11792 | 1fvo | A | 207 | SER | p.Ser207Arg | P00480 |
| HOMO SAPIENS | 0 | 11792 | 1fvo | A | 216 | GLN | p.Gln216Glu | P00480 |
| HOMO SAPIENS | 1 | 11792 | 1fvo | A | 225 | PRO | p.Pro225Leu | P00480 |
| HOMO SAPIENS | 0 | 11792 | 1fvo | A | 270 | ARG | p.Gln270Arg | P00480 |
| HOMO SAPIENS | 0 | 11792 | 1fvo | A | 270 | ARG | p.Gln270Ter | P00480 |
| HOMO SAPIENS | 2 | 11792 | 1fvo | A | 277 | ARG | p.Arg277Trp | P00480 |
| HOMO SAPIENS | 0 | 3721 | 4r3z | B | 2 | ASP | p.Asp2Gly | P54136 |
| HOMO SAPIENS | 0 | 3721 | 4r3z | B | 456 | SER | p.Ser456Leu | P54136 |
| HOMO SAPIENS | 5 | 3721 | 4r3z | B | 512 | ARG | p.Arg512Gln | P54136 |
| HOMO SAPIENS | 3 | 5749 | 3hl2 | A | 239 | ALA | p.Ala239Thr | Q9HD40 |
| HOMO SAPIENS | 1 | 5749 | 3hl2 | A | 334 | TYR | p.Tyr334Cys | Q9HD40 |
| HOMO SAPIENS | 1 | 5749 | 3hl2 | A | 452 | LYS | p.Lys452Asn | Q9HD40 |
| HOMO SAPIENS | -1 | 5749 | 3hl2 | A | 465 | ARG | p.Arg465Leu | Q9HD40 |
| HOMO SAPIENS | -1 | 5749 | 3hl2 | A | 489 | ASP | p.Asp489Gly | Q9HD40 |
| HOMO SAPIENS | 5 | 1721 | 3k15 | A | 1652 | MET | p.Met1652Ile | P38398 |
| HOMO SAPIENS | 5 | 1721 | 3k15 | A | 1652 | MET | p.Met1652Lys | P38398 |
| HOMO SAPIENS | 5 | 1721 | 3k15 | A | 1662 | PHE | p.Phe1662Cys | P38398 |
| HOMO SAPIENS | 5 | 1721 | 3k15 | A | 1664 | LEU | p.Leu1664Pro | P38398 |
| HOMO SAPIENS | 0 | 1721 | 3k15 | A | 1665 | VAL | p.Val1665Met | P38398 |
| HOMO SAPIENS | 5 | 1721 | 3k15 | A | 1690 | LYS | p.Lys1690Gln | P38398 |
| HOMO SAPIENS | 5 | 1721 | 3k15 | A | 1692 | ASP | p.Asp1692Asn | P38398 |
| HOMO SAPIENS | 5 | 1721 | 3k15 | A | 1699 | ARG | p.Arg1699Gln | P38398 |
| HOMO SAPIENS | 5 | 1721 | 3k15 | A | 1699 | ARG | p.Arg1699Trp | P38398 |
| HOMO SAPIENS | 3 | 1721 | 3k15 | A | 1706 | GLY | p.Gly1706Glu | P38398 |
| HOMO SAPIENS | 5 | 1721 | 3k15 | A | 1720 | THR | p.Thr1720Ala | P38398 |
| HOMO SAPIENS | 5 | 1721 | 3k15 | A | 1751 | ARG | p.Arg1751Pro | P38398 |
| HOMO SAPIENS | 4 | 1721 | 3k15 | A | 1775 | MET | p.Met1775Arg | P38398 |
| HOMO SAPIENS | 2 | 1721 | 3k15 | A | 1788 | GLY | p.Gly1788Val | P38398 |
| HOMO SAPIENS | 5 | 1721 | 3k15 | A | 1794 | GLU | p.Glu1794Asp | P38398 |
| HOMO SAPIENS | 5 | 1721 | 3k15 | A | 1804 | VAL | p.Val1804Asp | P38398 |
| HOMO SAPIENS | 3 | 1721 | 3k15 | A | 1815 | TRP | p.Trp1815Ter | P38398 |
| HOMO SAPIENS | 1 | 1721 | 3k15 | A | 1819 | ASN | p.Asn1819Ser | P38398 |
| HOMO SAPIENS | 5 | 1721 | 3k15 | A | 1856 | PRO | p.Pro1856Ser | P38398 |
| HOMO SAPIENS | 5 | 1721 | 3k15 | A | 1858 | ILE | p.Ile1858Leu | P38398 |
| HOMO SAPIENS | 5 | 1721 | 3k15 | A | 1858 | ILE | p.Ile1858Thr | P38398 |
| HOMO SAPIENS | 5 | 1721 | 3k15 | A | 1859 | PRO | p.Pro1859Arg | P38398 |
| HOMO SAPIENS | 2 | 22086 | 3kq4 | B | 1297 | PRO | p.Pro1297Leu | O60494 |
| HOMO SAPIENS | 1 | 12135 | 2k9j | B | 724 | ARG | p.Arg750Ter | P05106 |
| HOMO SAPIENS | -1 | 24027 | 2qtl | A | 175 | LEU | p.Ser202Leu | Q9UBK8 |
| HOMO SAPIENS | 0 | 24027 | 2qtl | A | 257 | SER | p.Ser284Thr | Q9UBK8 |
| HOMO SAPIENS | 0 | 24027 | 2qtl | A | 333 | LEU | p.Leu360Val | Q9UBK8 |
| HOMO SAPIENS | 0 | 24027 | 2qtl | A | 350 | LYS | p.Lys377Arg | Q9UBK8 |
| HOMO SAPIENS | 0 | 24027 | 2qtl | A | 415 | ARG | p.Arg442Cys | Q9UBK8 |
| HOMO SAPIENS | 2 | 24027 | 2qtl | A | 450 | PRO | p.Pro477Arg | Q9UBK8 |
| HOMO SAPIENS | 4 | 24027 | 2qtl | A | 487 | GLY | p.Gly514Arg | Q9UBK8 |
| HOMO SAPIENS | 1 | 24027 | 2qtl | A | 595 | HIS | p.His622Tyr | Q9UBK8 |
| HOMO SAPIENS | 0 | 5758 | 3di2 | B | 46 | THR | p.Thr66Ile | P16871 |
| HOMO SAPIENS | 1 | 5758 | 3di2 | B | 118 | VAL | p.Ile138Val | P16871 |
| HOMO SAPIENS | 4 | 848 | 5oa3 | Y | 79 | LEU | p.Leu79Val | P62847 |

| | | | | | | | | |
|---|---|---|---|---|---|---|---|---|
| HOMO SAPIENS | 2 | 848 | 5oa3 | Y | 124 | ASN | p.Asn124Ser | P62847 |
| HOMO SAPIENS | 0 | 81 | 3s5o | A | 36 | ALA | p.Ala36Glu | Q86XE5 |
| HOMO SAPIENS | 0 | 81 | 3s5o | A | 39 | TYR | p.Tyr39Ter | Q86XE5 |
| HOMO SAPIENS | 0 | 81 | 3s5o | A | 45 | PRO | p.Pro45Leu | Q86XE5 |
| HOMO SAPIENS | 0 | 81 | 3s5o | A | 70 | ARG | p.Arg70Ter | Q86XE5 |
| HOMO SAPIENS | 0 | 81 | 3s5o | A | 70 | ARG | p.Arg70Pro | Q86XE5 |
| HOMO SAPIENS | 1 | 81 | 3s5o | A | 97 | ARG | p.Arg97Cys | Q86XE5 |
| HOMO SAPIENS | 4 | 81 | 3s5o | A | 113 | GLU | p.Glu113Lys | Q86XE5 |
| HOMO SAPIENS | 1 | 81 | 3s5o | A | 190 | PRO | p.Pro190Leu | Q86XE5 |
| HOMO SAPIENS | 0 | 81 | 3s5o | A | 245 | VAL | p.Val245Phe | Q86XE5 |
| HOMO SAPIENS | 0 | 81 | 3s5o | A | 257 | CYS | p.Cys257Gly | Q86XE5 |
| HOMO SAPIENS | 0 | 81 | 3s5o | A | 287 | GLY | p.Gly287Val | Q86XE5 |
| HOMO SAPIENS | 7 | 81 | 3s5o | A | 303 | ARG | p.Arg303Cys | Q86XE5 |
| HOMO SAPIENS | 0 | 81 | 3s5o | A | 325 | GLY | p.Gly325Ser | Q86XE5 |
| HOMO SAPIENS | -1 | 622 | 5era | A | 12 | GLY | p.Gly12Arg | P29033 |
| HOMO SAPIENS | -1 | 622 | 5era | A | 12 | GLY | p.Gly12Asp | P29033 |
| HOMO SAPIENS | -1 | 622 | 5era | A | 17 | SER | p.Ser17Phe | P29033 |
| HOMO SAPIENS | 5 | 622 | 5era | A | 24 | TRP | p.Trp24Ter | P29033 |
| HOMO SAPIENS | 0 | 622 | 5era | A | 32 | ARG | p.Arg32Cys | P29033 |
| HOMO SAPIENS | 5 | 622 | 5era | A | 34 | MET | p.Met34Thr | P29033 |
| HOMO SAPIENS | 5 | 622 | 5era | A | 36 | LEU | p.Leu36Pro | P29033 |
| HOMO SAPIENS | 5 | 622 | 5era | A | 37 | VAL | p.Val37Ile | P29033 |
| HOMO SAPIENS | 5 | 622 | 5era | A | 40 | ALA | p.Ala40Glu | P29033 |
| HOMO SAPIENS | 5 | 622 | 5era | A | 44 | TRP | p.Trp44Cys | P29033 |
| HOMO SAPIENS | 5 | 622 | 5era | A | 45 | GLY | p.Gly45Glu | P29033 |
| HOMO SAPIENS | 5 | 622 | 5era | A | 47 | GLU | p.Glu47Ter | P29033 |
| HOMO SAPIENS | 5 | 622 | 5era | A | 50 | ASP | p.Asp50Asn | P29033 |
| HOMO SAPIENS | 5 | 622 | 5era | A | 54 | ASN | p.Asn54Lys | P29033 |
| HOMO SAPIENS | 5 | 622 | 5era | A | 57 | GLN | p.Gln57Ter | P29033 |
| HOMO SAPIENS | 5 | 622 | 5era | A | 59 | GLY | p.Gly59Ala | P29033 |
| HOMO SAPIENS | 5 | 622 | 5era | A | 59 | GLY | p.Gly59Ser | P29033 |
| HOMO SAPIENS | 5 | 622 | 5era | A | 66 | ASP | p.Asp66His | P29033 |
| HOMO SAPIENS | 5 | 622 | 5era | A | 68 | TYR | p.Tyr68Cys | P29033 |
| HOMO SAPIENS | 5 | 622 | 5era | A | 70 | PRO | p.Pro70Ala | P29033 |
| HOMO SAPIENS | 5 | 622 | 5era | A | 73 | HIS | p.His73Arg | P29033 |
| HOMO SAPIENS | 5 | 622 | 5era | A | 75 | ARG | p.Arg75Trp | P29033 |
| HOMO SAPIENS | 5 | 622 | 5era | A | 75 | ARG | p.Arg75Gln | P29033 |
| HOMO SAPIENS | 2 | 622 | 5era | A | 77 | TRP | p.Trp77Arg | P29033 |
| HOMO SAPIENS | 2 | 622 | 5era | A | 77 | TRP | p.Trp77Ter | P29033 |
| HOMO SAPIENS | 0 | 622 | 5era | A | 80 | GLN | p.Gln80Ter | P29033 |
| HOMO SAPIENS | 0 | 622 | 5era | A | 80 | GLN | p.Gln80Pro | P29033 |
| HOMO SAPIENS | 5 | 622 | 5era | A | 81 | LEU | p.Leu81Val | P29033 |
| HOMO SAPIENS | 5 | 622 | 5era | A | 82 | ILE | p.Ile82Met | P29033 |
| HOMO SAPIENS | 5 | 622 | 5era | A | 83 | PHE | p.Phe83Leu | P29033 |
| HOMO SAPIENS | 5 | 622 | 5era | A | 84 | VAL | p.Val84Leu | P29033 |
| HOMO SAPIENS | 5 | 622 | 5era | A | 90 | LEU | p.Leu90Pro | P29033 |
| HOMO SAPIENS | 5 | 622 | 5era | A | 95 | VAL | p.Val95Met | P29033 |
| HOMO SAPIENS | -1 | 622 | 5era | A | 100 | HIS | p.His100Tyr | P29033 |
| HOMO SAPIENS | -1 | 622 | 5era | A | 113 | SER | p.Ser113Arg | P29033 |
| HOMO SAPIENS | -1 | 622 | 5era | A | 119 | GLU | p.Glu119Lys | P29033 |
| HOMO SAPIENS | -1 | 622 | 5era | A | 122 | LYS | p.Lys122Ile | P29033 |
| HOMO SAPIENS | -1 | 622 | 5era | A | 123 | THR | p.Thr123Asn | P29033 |
| HOMO SAPIENS | -1 | 622 | 5era | A | 124 | GLN | p.Gln124Ter | P29033 |
| HOMO SAPIENS | 5 | 622 | 5era | A | 139 | SER | p.Ser139Asn | P29033 |

| | | | | | | | | |
|---|---|---|---|---|---|---|---|---|
| HOMO SAPIENS | 5 | 622 | 5era | A | 143 | ARG | p.Arg143Trp | P29033 |
| HOMO SAPIENS | 5 | 622 | 5era | A | 143 | ARG | p.Arg143Gln | P29033 |
| HOMO SAPIENS | 0 | 622 | 5era | A | 147 | GLU | p.Glu147Lys | P29033 |
| HOMO SAPIENS | 7 | 622 | 5era | A | 155 | TYR | p.Tyr155Ter | P29033 |
| HOMO SAPIENS | 2 | 622 | 5era | A | 159 | ASP | p.Asp159Val | P29033 |
| HOMO SAPIENS | 5 | 622 | 5era | A | 163 | MET | p.Met163Val | P29033 |
| HOMO SAPIENS | 5 | 622 | 5era | A | 165 | ARG | p.Arg165Trp | P29033 |
| HOMO SAPIENS | 5 | 622 | 5era | A | 167 | VAL | p.Val167Met | P29033 |
| HOMO SAPIENS | 5 | 622 | 5era | A | 169 | CYS | p.Cys169Tyr | P29033 |
| HOMO SAPIENS | 5 | 622 | 5era | A | 179 | ASP | p.Asp179Asn | P29033 |
| HOMO SAPIENS | 0 | 622 | 5era | A | 199 | SER | p.Ser199Phe | P29033 |
| HOMO SAPIENS | 0 | 622 | 5era | A | 202 | CYS | p.Cys202Phe | P29033 |
| HOMO SAPIENS | 5 | 622 | 5era | A | 203 | ILE | p.Ile203Thr | P29033 |
| HOMO SAPIENS | 5 | 622 | 5era | A | 206 | ASN | p.Asn206Ser | P29033 |
| HOMO SAPIENS | 0 | 8103 | 5dot | A | 163 | GLY | p.Gly163Ter | P31327 |
| HOMO SAPIENS | 0 | 8103 | 5dot | A | 337 | HIS | p.His337Arg | P31327 |
| HOMO SAPIENS | 3 | 8103 | 5dot | A | 344 | THR | p.Thr344Ala | P31327 |
| HOMO SAPIENS | 1 | 8103 | 5dot | A | 405 | THR | p.Thr405Asn | P31327 |
| HOMO SAPIENS | 5 | 8103 | 5dot | A | 544 | THR | p.Thr544Met | P31327 |
| HOMO SAPIENS | 5 | 8103 | 5dot | A | 716 | ASN | p.Asn716Lys | P31327 |
| HOMO SAPIENS | -1 | 8103 | 5dot | A | 780 | ARG | p.Arg780His | P31327 |
| HOMO SAPIENS | 0 | 8103 | 5dot | A | 982 | GLY | p.Gly982Asp | P31327 |
| HOMO SAPIENS | 3 | 8103 | 5dot | A | 1119 | ALA | p.Ala1119Ser | P31327 |
| HOMO SAPIENS | 1 | 8103 | 5dot | A | 1376 | GLY | p.Gly1376Ser | P31327 |
| HOMO SAPIENS | 4 | 6833 | 1l8l | B | 27 | ARG | p.Arg27Ser | P78330 |
| HOMO SAPIENS | 3 | 6833 | 1l8l | B | 32 | ASP | p.Asp32Gly | P78330 |
| HOMO SAPIENS | 3 | 6833 | 1l8l | B | 32 | ASP | p.Asp32Asn | P78330 |
| HOMO SAPIENS | 1 | 6833 | 1l8l | B | 49 | ARG | p.Arg49Trp | P78330 |
| HOMO SAPIENS | 0 | 6833 | 1l8l | B | 52 | MET | p.Met52Thr | P78330 |
| HOMO SAPIENS | 0 | 6833 | 1l8l | B | 83 | GLN | p.Gln83His | P78330 |
| HOMO SAPIENS | 6 | 810 | 2iw2 | A | 185 | ARG | p.Arg184Gln | P12955 |
| HOMO SAPIENS | 0 | 810 | 2iw2 | A | 213 | ALA | p.Ala212Pro | P12955 |
| HOMO SAPIENS | 1 | 810 | 2iw2 | A | 266 | ARG | p.Arg265Ter | P12955 |
| HOMO SAPIENS | 9 | 810 | 2iw2 | A | 277 | ASP | p.Asp276Asn | P12955 |
| HOMO SAPIENS | 0 | 810 | 2iw2 | A | 279 | GLY | p.Gly278Asp | P12955 |
| HOMO SAPIENS | 9 | 810 | 2iw2 | A | 378 | HIS | p.His377Gln | P12955 |
| HOMO SAPIENS | 0 | 810 | 2iw2 | A | 389 | ARG | p.Arg388His | P12955 |
| HOMO SAPIENS | 0 | 810 | 2iw2 | A | 436 | PHE | p.Leu435Phe | P12955 |
| HOMO SAPIENS | 0 | 810 | 2iw2 | A | 449 | GLY | p.Gly448Arg | P12955 |
| HOMO SAPIENS | -1 | 1530 | 5fwj | A | 388 | ALA | p.Ala388Pro | P41229 |
| HOMO SAPIENS | 4 | 1530 | 5fwj | A | 451 | SER | p.Ser451Arg | P41229 |
| HOMO SAPIENS | 3 | 1530 | 5fwj | A | 731 | LEU | p.Leu731Phe | P41229 |
| HOMO SAPIENS | 3 | 578 | 5net | B | 123 | ALA | p.Ala143Thr | P18564 |
| HOMO SAPIENS | 6 | 578 | 5net | B | 176 | PRO | p.Pro196Thr | P18564 |
| HOMO SAPIENS | 3 | 578 | 5net | B | 280 | GLU | p.Glu300Lys | P18564 |
| HOMO SAPIENS | -1 | 5247 | 4j15 | A | 274 | ALA | p.Ala274Val | P14868 |
| HOMO SAPIENS | -1 | 5247 | 4j15 | A | 280 | HIS | p.His280Leu | P14868 |
| HOMO SAPIENS | 2 | 5247 | 4j15 | A | 460 | ARG | p.Arg460His | P14868 |
| HOMO SAPIENS | 5 | 5247 | 4j15 | A | 494 | ARG | p.Arg494Cys | P14868 |
| HOMO SAPIENS | 7 | 2566 | 1b55 | A | 28 | ARG | p.Arg28His | Q06187 |
| HOMO SAPIENS | 5 | 2566 | 1b55 | A | 33 | THR | p.Thr33Pro | Q06187 |
| HOMO SAPIENS | 0 | 2566 | 1b55 | A | 113 | VAL | p.Val113Asp | Q06187 |
| HOMO SAPIENS | 0 | 356 | 5a2s | A | 786 | ALA | p.Ala786Thr | P56524 |
| HOMO SAPIENS | 5 | 2960 | 4cak | A | 553 | ARG | p.Arg584Ter | P08514 |

| | | | | | | | | |
|---|---|---|---|---|---|---|---|---|
| HOMO SAPIENS | 5 | 2960 | 4cak | A | 565 | ILE | p.Ile596Thr | P08514 |
| HOMO SAPIENS | -1 | 2960 | 4cak | A | 841 | LEU | p.Leu872Met | P08514 |
| HOMO SAPIENS | -1 | 2960 | 4cak | A | 843 | ILE | p.Ile874Ser | P08514 |
| HOMO SAPIENS | 0 | 5054 | 4k8r | B | 216 | PHE | p.Phe216Leu | Q8NBP7 |
| HOMO SAPIENS | 1 | 5054 | 4k8r | B | 417 | HIS | p.His417Gln | Q8NBP7 |
| HOMO SAPIENS | 1 | 5054 | 4k8r | B | 425 | ASN | p.Asn425Ser | Q8NBP7 |
| HOMO SAPIENS | 1 | 5054 | 4k8r | B | 443 | ALA | p.Ala443Thr | Q8NBP7 |
| HOMO SAPIENS | 0 | 27283 | 3n5n | X | 71 | HIS | p.His82Arg | Q9UIF7 |
| HOMO SAPIENS | 1 | 27283 | 3n5n | X | 83 | ARG | p.Arg94Ter | Q9UIF7 |
| HOMO SAPIENS | 8 | 27283 | 3n5n | X | 89 | TRP | p.Trp100Ter | Q9UIF7 |
| HOMO SAPIENS | 8 | 27283 | 3n5n | X | 90 | TYR | p.Tyr101Ter | Q9UIF7 |
| HOMO SAPIENS | 7 | 27283 | 3n5n | X | 95 | ARG | p.Arg106Trp | Q9UIF7 |
| HOMO SAPIENS | 7 | 27283 | 3n5n | X | 117 | TRP | p.Trp128Arg | Q9UIF7 |
| HOMO SAPIENS | 0 | 27283 | 3n5n | X | 137 | GLY | p.Gly148Arg | Q9UIF7 |
| HOMO SAPIENS | 5 | 27283 | 3n5n | X | 142 | TRP | p.Trp153Ter | Q9UIF7 |
| HOMO SAPIENS | 0 | 27283 | 3n5n | X | 165 | TYR | p.Tyr176Cys | Q9UIF7 |
| HOMO SAPIENS | 0 | 27283 | 3n5n | X | 168 | ARG | p.Arg179His | Q9UIF7 |
| HOMO SAPIENS | 0 | 27283 | 3n5n | X | 168 | ARG | p.Arg179Cys | Q9UIF7 |
| HOMO SAPIENS | 0 | 27283 | 3n5n | X | 196 | GLN | p.Gln207Ter | Q9UIF7 |
| HOMO SAPIENS | 8 | 27283 | 3n5n | X | 201 | VAL | p.Val212Met | Q9UIF7 |
| HOMO SAPIENS | 9 | 27283 | 3n5n | X | 227 | ARG | p.Arg238Trp | Q9UIF7 |
| HOMO SAPIENS | 9 | 27283 | 3n5n | X | 231 | ARG | p.Arg242Cys | Q9UIF7 |
| HOMO SAPIENS | 9 | 27283 | 3n5n | X | 231 | ARG | p.Arg242His | Q9UIF7 |
| HOMO SAPIENS | 3 | 27283 | 3n5n | X | 232 | VAL | p.Val243Phe | Q9UIF7 |
| HOMO SAPIENS | 3 | 27283 | 3n5n | X | 246 | GLN | p.Gln257Ter | Q9UIF7 |
| HOMO SAPIENS | 9 | 27283 | 3n5n | X | 272 | GLY | p.Gly283Glu | Q9UIF7 |
| HOMO SAPIENS | 9 | 27283 | 3n5n | X | 281 | PRO | p.Pro292Leu | Q9UIF7 |
| HOMO SAPIENS | -1 | 27283 | 3n5n | X | 314 | ASP | p.Asp325Glu | Q9UIF7 |
| HOMO SAPIENS | 0 | 27283 | 3n5n | X | 324 | GLN | p.Gln335His | Q9UIF7 |
| HOMO SAPIENS | 0 | 27283 | 3n5n | X | 324 | GLN | p.Gln335Arg | Q9UIF7 |
| HOMO SAPIENS | 0 | 27283 | 3n5n | X | 324 | GLN | p.Gln335Ter | Q9UIF7 |
| HOMO SAPIENS | -1 | 20852 | 4p60 | B | 243 | ARG | p.Arg243Lys | O75746 |
| HOMO SAPIENS | -1 | 1875 | 5dd2 | G | 152 | VAL | p.Val179Leu | P06396 |
| HOMO SAPIENS | 9 | 2126 | 3n1p | B | 46 | PRO | p.Pro46Leu | Q14623 |
| HOMO SAPIENS | 8 | 2126 | 3n1p | B | 76 | GLU | p.Glu76Ala | Q14623 |
| HOMO SAPIENS | 9 | 2126 | 3n1p | B | 77 | ARG | p.Arg77Ser | Q14623 |
| HOMO SAPIENS | 6 | 2126 | 3n1p | B | 95 | GLU | p.Glu95Lys | Q14623 |
| HOMO SAPIENS | 9 | 2126 | 3n1p | B | 100 | ASP | p.Asp100Glu | Q14623 |
| HOMO SAPIENS | 9 | 2126 | 3n1p | B | 131 | GLU | p.Glu131Lys | Q14623 |
| HOMO SAPIENS | 0 | 2126 | 3n1p | B | 190 | VAL | p.Val190Ala | Q14623 |
| HOMO SAPIENS | 5 | 40410 | 3ioz | B | 70 | GLN | p.Gln70Leu | P20963 |
| HOMO SAPIENS | 1 | 5754 | 5lkz | A | 1108 | THR | p.Tyr1191Ter | Q09472 |
| HOMO SAPIENS | 3 | 8159 | 1tiu | A | 3 | GLU | p.Lys12679Glu | Q8WZ42 |
| HOMO SAPIENS | 4 | 3275 | 4ldq | A | 210 | ARG | p.Arg210Gln | Q9BV20 |
| HOMO SAPIENS | 0 | 8143 | 4miv | B | 66 | SER | p.Ser66Trp | P51688 |
| HOMO SAPIENS | 1 | 8143 | 4miv | B | 74 | ARG | p.Arg74Cys | P51688 |
| HOMO SAPIENS | 0 | 8143 | 4miv | B | 128 | PRO | p.Pro128Leu | P51688 |
| HOMO SAPIENS | 1 | 8143 | 4miv | B | 150 | ARG | p.Arg150Gln | P51688 |
| HOMO SAPIENS | 0 | 8143 | 4miv | B | 191 | GLY | p.Gly191Arg | P51688 |
| HOMO SAPIENS | 0 | 8143 | 4miv | B | 206 | ARG | p.Arg206Pro | P51688 |
| HOMO SAPIENS | 0 | 8143 | 4miv | B | 225 | PHE | p.Phe225Leu | P51688 |
| HOMO SAPIENS | 0 | 8143 | 4miv | B | 233 | ARG | p.Arg233Ter | P51688 |
| HOMO SAPIENS | 0 | 8143 | 4miv | B | 234 | ALA | p.Ala234Gly | P51688 |
| HOMO SAPIENS | 0 | 8143 | 4miv | B | 245 | ARG | p.Arg245His | P51688 |

| | | | | | | | | |
|---|---|---|---|---|---|---|---|---|
| HOMO SAPIENS | 0 | 8143 | 4miv | B | 293 | PRO | p.Pro293Ser | P51688 |
| HOMO SAPIENS | 1 | 8143 | 4miv | B | 298 | SER | p.Ser298Pro | P51688 |
| HOMO SAPIENS | 0 | 8143 | 4miv | B | 361 | VAL | p.Val361Ile | P51688 |
| HOMO SAPIENS | 0 | 8143 | 4miv | B | 369 | GLU | p.Glu369Lys | P51688 |
| HOMO SAPIENS | 0 | 8143 | 4miv | B | 372 | MET | p.Met372Ile | P51688 |
| HOMO SAPIENS | 1 | 8143 | 4miv | B | 380 | GLN | p.Gln380Arg | P51688 |
| HOMO SAPIENS | 0 | 8143 | 4miv | B | 387 | VAL | p.Val387Met | P51688 |
| HOMO SAPIENS | 0 | 8143 | 4miv | B | 394 | MET | p.Met394Ile | P51688 |
| HOMO SAPIENS | 0 | 8143 | 4miv | B | 433 | ARG | p.Arg433Gln | P51688 |
| HOMO SAPIENS | 1 | 8143 | 4miv | B | 446 | HIS | p.His446Arg | P51688 |
| HOMO SAPIENS | 5 | 8143 | 4miv | B | 447 | GLU | p.Glu447Lys | P51688 |
| HOMO SAPIENS | 2 | 8143 | 4miv | B | 456 | ARG | p.Arg456His | P51688 |
| HOMO SAPIENS | 1 | 21880 | 2oo9 | B | 857 | LEU | p.Leu857Phe | P22681 |
| HOMO SAPIENS | 0 | 9467 | 1s1d | A | 154 | LEU | p.Leu224Pro | Q8WVQ1 |
| HOMO SAPIENS | 0 | 9467 | 1s1d | A | 156 | VAL | p.Val226Met | Q8WVQ1 |
| HOMO SAPIENS | 0 | 9467 | 1s1d | A | 229 | PRO | p.Pro299Leu | Q8WVQ1 |
| HOMO SAPIENS | 4 | 9467 | 1s1d | A | 230 | ARG | p.Arg300Cys | Q8WVQ1 |
| HOMO SAPIENS | 4 | 9467 | 1s1d | A | 230 | ARG | p.Arg300His | Q8WVQ1 |
| HOMO SAPIENS | 0 | 9467 | 1s1d | A | 253 | ALA | p.Ala323Thr | Q8WVQ1 |
| HOMO SAPIENS | 2 | 9467 | 1s1d | A | 305 | MET | p.Met375Leu | Q8WVQ1 |
| HOMO SAPIENS | -1 | 2361 | 5llm | A | 5 | ASN | p.Asn5Lys | P43003 |
| HOMO SAPIENS | 0 | 2361 | 5llm | A | 71 | TYR | p.Tyr71Cys | P43003 |
| HOMO SAPIENS | -1 | 2361 | 5llm | A | 211 | LYS | p.Glu219Asp | Q15758 |
| HOMO SAPIENS | 8 | 2361 | 5llm | A | 270 | PRO | p.Pro290Arg | P43003 |
| HOMO SAPIENS | 1 | 2361 | 5llm | A | 321 | THR | p.Ala329Thr | Q15758 |
| HOMO SAPIENS | 1 | 2361 | 5llm | A | 365 | ARG | p.Arg385His | P43003 |
| HOMO SAPIENS | 1 | 2361 | 5llm | A | 479 | ARG | p.Arg499Gln | P43003 |
| HOMO SAPIENS | 3 | 5412 | 4obz | D | 165 | PHE | p.Phe229Ile | P07339 |
| HOMO SAPIENS | 2 | 5412 | 4obz | D | 187 | ASP | p.Asp251Asn | P07339 |
| HOMO SAPIENS | 0 | 5412 | 4obz | D | 218 | GLY | p.Gly282Arg | P07339 |
| HOMO SAPIENS | 0 | 5412 | 4obz | D | 245 | ARG | p.Arg309His | P07339 |
| HOMO SAPIENS | 2 | 5412 | 4obz | D | 319 | TRP | p.Trp383Cys | P07339 |
| HOMO SAPIENS | 0 | 3544 | 5h64 | A | 277 | MET | p.Met277Ile | P42345 |
| HOMO SAPIENS | 0 | 14230 | 2kbx | A | 145 | ARG | p.Arg145Ser | Q13418 |
| HOMO SAPIENS | 2 | 34970 | 4xhz | A | 834 | ARG | p.Arg855Trp | Q96QU1 |
| HOMO SAPIENS | 4 | 34970 | 4xhz | A | 854 | SER | p.Ser875Ter | Q96QU1 |
| HOMO SAPIENS | 1 | 34970 | 4xhz | A | 908 | ARG | p.Arg929Gln | Q96QU1 |
| HOMO SAPIENS | 1 | 34970 | 4xhz | A | 941 | ARG | p.Arg962Cys | Q96QU1 |
| HOMO SAPIENS | 0 | 34970 | 4xhz | A | 970 | ARG | p.Arg991Ter | Q96QU1 |
| HOMO SAPIENS | 0 | 34970 | 4xhz | A | 976 | GLU | p.Glu997Gly | Q96QU1 |
| HOMO SAPIENS | 0 | 25734 | 4asc | A | 345 | SER | p.Asn345Ser | Q2TBA0 |
| HOMO SAPIENS | 0 | 25734 | 4asc | A | 514 | VAL | p.Val514Met | Q2TBA0 |
| HOMO SAPIENS | 6 | 25734 | 4asc | A | 528 | GLU | p.Glu528Lys | Q2TBA0 |
| HOMO SAPIENS | 0 | 25734 | 4asc | A | 580 | LEU | p.Leu580Ile | Q2TBA0 |
| HOMO SAPIENS | 8 | 2356 | 3kse | D | 22 | LYS | p.Lys22Ter | P01040 |
| HOMO SAPIENS | 1 | 2356 | 3kse | D | 58 | ARG | p.Arg58Ter | P01040 |
| HOMO SAPIENS | 2 | 6000 | 1jqd | A | 60 | GLY | p.Gly60Asp | P50135 |
| HOMO SAPIENS | 9 | 6000 | 1jqd | A | 208 | LEU | p.Leu208Pro | P50135 |
| HOMO SAPIENS | 0 | 14741 | 5eh1 | A | 58 | THR | p.Thr58Arg | P38484 |
| HOMO SAPIENS | 0 | 14741 | 5eh1 | A | 168 | THR | p.Thr168Asn | P38484 |
| HOMO SAPIENS | 1 | 4814 | 4kxk | D | 427 | ARG | p.Arg427Ter | P50542 |
| HOMO SAPIENS | 0 | 188 | 2xt3 | A | 49 | ASP | p.Asp52Asn | Q2M1P5 |
| HOMO SAPIENS | 3 | 188 | 2xt3 | A | 142 | TYR | p.Tyr145Ser | Q2M1P5 |
| HOMO SAPIENS | 9 | 8681 | 5tby | D | 57 | ALA | p.Ala57Asp | P08590 |

| | | | | | | | | |
|---|---|---|---|---|---|---|---|---|
| HOMO SAPIENS | 3 | 8681 | 5tby | D | 62 | ASP | p.Asp62Asn | P08590 |
| HOMO SAPIENS | 0 | 8681 | 5tby | D | 65 | PRO | p.Pro65Arg | P08590 |
| HOMO SAPIENS | 7 | 8681 | 5tby | D | 79 | VAL | p.Val79Ile | P08590 |
| HOMO SAPIENS | 0 | 8681 | 5tby | D | 143 | GLU | p.Glu143Lys | P08590 |
| HOMO SAPIENS | 0 | 8681 | 5tby | D | 149 | MET | p.Met149Val | P08590 |
| HOMO SAPIENS | 3 | 8681 | 5tby | D | 154 | ARG | p.Arg154His | P08590 |
| HOMO SAPIENS | 6 | 30984 | 4xwh | A | 140 | TYR | p.Tyr140Cys | P54802 |
| HOMO SAPIENS | 4 | 30984 | 4xwh | A | 141 | SER | p.Ser141Thr | P54802 |
| HOMO SAPIENS | 2 | 30984 | 4xwh | A | 234 | ARG | p.Arg234Cys | P54802 |
| HOMO SAPIENS | 2 | 30984 | 4xwh | A | 336 | GLU | p.Glu336Ter | P54802 |
| HOMO SAPIENS | 6 | 30984 | 4xwh | A | 482 | ARG | p.Arg482Trp | P54802 |
| HOMO SAPIENS | 2 | 30984 | 4xwh | A | 521 | PRO | p.Pro521Leu | P54802 |
| HOMO SAPIENS | 0 | 30984 | 4xwh | A | 565 | ARG | p.Arg565Gln | P54802 |
| HOMO SAPIENS | 0 | 30984 | 4xwh | A | 565 | ARG | p.Arg565Trp | P54802 |
| HOMO SAPIENS | 6 | 30984 | 4xwh | A | 626 | ARG | p.Arg626Ter | P54802 |
| HOMO SAPIENS | 6 | 30984 | 4xwh | A | 643 | ARG | p.Arg643His | P54802 |
| HOMO SAPIENS | 6 | 30984 | 4xwh | A | 643 | ARG | p.Arg643Cys | P54802 |
| HOMO SAPIENS | 2 | 30984 | 4xwh | A | 674 | ARG | p.Arg674His | P54802 |
| HOMO SAPIENS | 2 | 30984 | 4xwh | A | 737 | ARG | p.Arg737Gly | P54802 |
| HOMO SAPIENS | 5 | 10545 | 1oqe | N | 6 | PRO | p.Pro21Arg | Q96RJ3 |
| HOMO SAPIENS | 1 | 16995 | 5d3a | B | 12 | MET | p.Met947Arg | Q7Z4S6 |
| HOMO SAPIENS | 1 | 16995 | 5d3a | B | 12 | MET | p.Met947Val | Q7Z4S6 |
| HOMO SAPIENS | 2 | 16995 | 5d3a | B | 19 | ARG | p.Arg954Gln | Q7Z4S6 |
| HOMO SAPIENS | 2 | 16995 | 5d3a | B | 19 | ARG | p.Arg954Trp | Q7Z4S6 |
| HOMO SAPIENS | 0 | 16995 | 5d3a | B | 75 | ILE | p.Ile1010Thr | Q7Z4S6 |
| HOMO SAPIENS | 0 | 17091 | 2yin | B | 1324 | GLN | p.Gln1324Ter | Q92608 |
| HOMO SAPIENS | 5 | 27606 | 1lm7 | B | 2366 | ARG | p.Arg2366Cys | P15924 |
| HOMO SAPIENS | 2 | 15731 | 1bi7 | B | 24 | ARG | p.Arg24Pro | P42771 |
| HOMO SAPIENS | 0 | 15731 | 1bi7 | B | 28 | VAL | p.Val28Gly | P42771 |
| HOMO SAPIENS | 0 | 15731 | 1bi7 | B | 49 | ILE | p.Ile49Thr | P42771 |
| HOMO SAPIENS | 0 | 15731 | 1bi7 | B | 53 | MET | p.Met53Ile | P42771 |
| HOMO SAPIENS | 2 | 15731 | 1bi7 | B | 56 | SER | p.Ser56Ile | P42771 |
| HOMO SAPIENS | 1 | 15731 | 1bi7 | B | 57 | ALA | p.Ala57Val | P42771 |
| HOMO SAPIENS | 4 | 15731 | 1bi7 | B | 59 | VAL | p.Val59Gly | P42771 |
| HOMO SAPIENS | 9 | 15731 | 1bi7 | B | 89 | GLY | p.Gly89Asp | P42771 |
| HOMO SAPIENS | 0 | 15731 | 1bi7 | B | 101 | GLY | p.Gly101Trp | P42771 |
| HOMO SAPIENS | 0 | 15731 | 1bi7 | B | 114 | PRO | p.Pro114Ser | P42771 |
| HOMO SAPIENS | 9 | 15731 | 1bi7 | B | 123 | HIS | p.His123Gln | P42771 |
| HOMO SAPIENS | 8 | 15731 | 1bi7 | B | 126 | VAL | p.Val126Asp | P42771 |
| HOMO SAPIENS | 7 | 15731 | 1bi7 | B | 127 | ALA | p.Ala127Ser | P42771 |
| HOMO SAPIENS | -1 | 15731 | 1bi7 | B | 148 | ALA | p.Ala148Thr | P42771 |
| HOMO SAPIENS | -1 | 19858 | 3jbk | M | 866 | ALA | p.Ala934Val | P18206 |
| HOMO SAPIENS | 0 | 19484 | 3ee6 | B | 62 | SER | p.Ser62Leu | O14773 |
| HOMO SAPIENS | 1 | 19484 | 3ee6 | B | 66 | GLN | p.Gln66Ter | O14773 |
| HOMO SAPIENS | 0 | 19484 | 3ee6 | B | 100 | GLN | p.Gln100Arg | O14773 |
| HOMO SAPIENS | 0 | 19484 | 3ee6 | B | 104 | LEU | p.Leu104Ter | O14773 |
| HOMO SAPIENS | 0 | 19484 | 3ee6 | B | 127 | ARG | p.Arg127Gln | O14773 |
| HOMO SAPIENS | 0 | 19484 | 3ee6 | B | 127 | ARG | p.Arg127Ter | O14773 |
| HOMO SAPIENS | -1 | 19484 | 3ee6 | B | 185 | ARG | p.Arg185Cys | O14773 |
| HOMO SAPIENS | 0 | 19484 | 3ee6 | B | 206 | ARG | p.Arg206Cys | O14773 |
| HOMO SAPIENS | 0 | 19484 | 3ee6 | B | 208 | ARG | p.Arg208Ter | O14773 |
| HOMO SAPIENS | 0 | 19484 | 3ee6 | B | 214 | GLN | p.Gln214Ter | O14773 |
| HOMO SAPIENS | 1 | 19484 | 3ee6 | B | 276 | ASP | p.Asp276Val | O14773 |
| HOMO SAPIENS | 0 | 19484 | 3ee6 | B | 284 | GLY | p.Gly284Val | O14773 |

| | | | | | | | | |
|---|---|---|---|---|---|---|---|---|
| HOMO SAPIENS | 0 | 19484 | 3ee6 | B | 286 | ASN | p.Asn286Ser | O14773 |
| HOMO SAPIENS | 0 | 19484 | 3ee6 | B | 339 | ARG | p.Arg339Gly | O14773 |
| HOMO SAPIENS | 2 | 19484 | 3ee6 | B | 345 | MET | p.Met345Leu | O14773 |
| HOMO SAPIENS | 0 | 19484 | 3ee6 | B | 365 | CYS | p.Cys365Arg | O14773 |
| HOMO SAPIENS | 0 | 19484 | 3ee6 | B | 365 | CYS | p.Cys365Tyr | O14773 |
| HOMO SAPIENS | 0 | 19484 | 3ee6 | B | 373 | GLN | p.Gln373Glu | O14773 |
| HOMO SAPIENS | 6 | 19484 | 3ee6 | B | 418 | ARG | p.Arg418Gln | O14773 |
| HOMO SAPIENS | 7 | 19484 | 3ee6 | B | 422 | GLN | p.Gln422His | O14773 |
| HOMO SAPIENS | 0 | 19484 | 3ee6 | B | 447 | ARG | p.Arg447His | O14773 |
| HOMO SAPIENS | 0 | 19484 | 3ee6 | B | 466 | VAL | p.Val466Met | O14773 |
| HOMO SAPIENS | 0 | 4654 | 5t5w | A | 47 | LYS | p.Lys47Glu | Q08334 |
| HOMO SAPIENS | 0 | 4654 | 5t5w | A | 141 | GLU | p.Glu141Ter | Q08334 |
| HOMO SAPIENS | -1 | 18815 | 4wj7 | A | 53 | VAL | p.Val53Ile | Q9BSQ5 |
| HOMO SAPIENS | 0 | 18815 | 4wj7 | A | 120 | VAL | p.Val120Ile | Q9BSQ5 |
| HOMO SAPIENS | 0 | 18815 | 4wj7 | A | 198 | LEU | p.Leu198Arg | Q9BSQ5 |
| HOMO SAPIENS | 0 | 4602 | 1mjd | A | 47 | SER | p.Ser47Arg | O43602 |
| HOMO SAPIENS | 6 | 4602 | 1mjd | A | 59 | ARG | p.Arg59Leu | O43602 |
| HOMO SAPIENS | 7 | 4602 | 1mjd | A | 62 | ASP | p.Asp62Asn | O43602 |
| HOMO SAPIENS | 7 | 4602 | 1mjd | A | 71 | ALA | p.Ala71Ser | O43602 |
| HOMO SAPIENS | 7 | 4602 | 1mjd | A | 78 | ARG | p.Arg78Leu | O43602 |
| HOMO SAPIENS | 7 | 4602 | 1mjd | A | 89 | ARG | p.Arg89Gly | O43602 |
| HOMO SAPIENS | 8 | 4602 | 1mjd | A | 104 | ILE | p.Ile104Phe | O43602 |
| HOMO SAPIENS | 7 | 4602 | 1mjd | A | 125 | TYR | p.Tyr125His | O43602 |
| HOMO SAPIENS | 5 | 34467 | 1n0w | B | 1529 | GLY | p.Gly1529Arg | P51587 |
| HOMO SAPIENS | 8 | 5738 | 1u31 | A | 130 | ASN | p.Asn1009Lys | Q13423 |
| HOMO SAPIENS | 0 | 5760 | 2i6s | A | 298 | GLU | p.Glu318Asp | P06681 |
| HOMO SAPIENS | 0 | 5760 | 2i6s | A | 573 | ARG | p.Arg593Gln | P06681 |
| HOMO SAPIENS | 0 | 5760 | 2i6s | A | 592 | SER | p.Ser612Asn | P06681 |
| HOMO SAPIENS | 0 | 5760 | 2i6s | A | 621 | VAL | p.Val641Ala | P06681 |
| HOMO SAPIENS | 1 | 10977 | 2fby | A | 114 | VAL | p.Val114Ile | Q14191 |
| HOMO SAPIENS | 0 | 10977 | 2fby | A | 135 | LYS | p.Lys135Glu | Q14191 |
| HOMO SAPIENS | -1 | 1175 | 1qrk | A | 34 | VAL | p.Val35Leu | P00488 |
| HOMO SAPIENS | 4 | 1175 | 1qrk | A | 564 | PRO | p.Pro565Leu | P00488 |
| HOMO SAPIENS | 0 | 1175 | 1qrk | A | 650 | VAL | p.Val651Ile | P00488 |
| HOMO SAPIENS | 0 | 1175 | 1qrk | A | 651 | GLU | p.Gln652Glu | P00488 |
| HOMO SAPIENS | 3 | 909 | 4h0w | A | 375 | GLU | p.Glu394Lys | P02787 |
| HOMO SAPIENS | 0 | 909 | 4h0w | A | 458 | ALA | p.Ala477Pro | P02787 |
| HOMO SAPIENS | 6 | 909 | 4h0w | A | 515 | TYR | p.Tyr534Ter | P02787 |
| HOMO SAPIENS | 7 | 909 | 4h0w | A | 570 | PRO | p.Pro589Ser | P02787 |
| HOMO SAPIENS | 3 | 909 | 4h0w | A | 652 | GLY | p.Gly671Glu | P02787 |
| HOMO SAPIENS | 5 | 1964 | 2car | A | 151 | TRP | p.Trp151Ter | Q9BY32 |
| HOMO SAPIENS | 7 | 1964 | 2car | A | 178 | ARG | p.Arg178Cys | Q9BY32 |
| HOMO SAPIENS | 0 | 144 | 2n0a | J | 30 | ALA | p.Ala30Pro | P37840 |
| HOMO SAPIENS | 1 | 12288 | 4q58 | D | 1225 | ARG | p.Arg1225His | P16144 |
| HOMO SAPIENS | 2 | 12288 | 4q58 | D | 1281 | ARG | p.Arg1281Trp | P16144 |
| HOMO SAPIENS | 2 | 12288 | 4q58 | D | 1287 | ASN | p.Asn1287Lys | P16144 |
| HOMO SAPIENS | 0 | 5085 | 3s94 | A | 360 | ARG | p.Arg360His | O75581 |
| HOMO SAPIENS | 0 | 5085 | 3s94 | A | 433 | ASN | p.Asn433Ser | O75581 |
| HOMO SAPIENS | 2 | 5085 | 3s94 | A | 611 | ARG | p.Arg611Cys | O75581 |
| HOMO SAPIENS | 0 | 15452 | 4dur | A | 19 | LYS | p.Lys38Glu | P00747 |
| HOMO SAPIENS | 2 | 15452 | 4dur | A | 216 | ARG | p.Arg235His | P00747 |
| HOMO SAPIENS | 7 | 8406 | 4wo1 | D | 28 | VAL | p.Val55Leu | O43914 |
| HOMO SAPIENS | 2 | 895 | 5xte | K | 183 | ARG | p.Arg183Trp | P22695 |
| HOMO SAPIENS | 2 | 4191 | 4wnd | A | 120 | ARG | p.Arg127Ter | P81274 |

| | | | | | | | | |
|---|---|---|---|---|---|---|---|---|
| HOMO SAPIENS | 2 | 4191 | 4wnd | A | 120 | ARG | p.Arg127Gln | P81274 |
| HOMO SAPIENS | 1 | 4191 | 4wnd | A | 349 | GLY | p.Gly356Arg | P81274 |
| HOMO SAPIENS | 0 | 41734 | 2kbi | A | 1777 | VAL | p.Val1777Met | Q14524 |
| HOMO SAPIENS | 1 | 41734 | 2kbi | A | 1784 | GLU | p.Glu1784Lys | Q14524 |
| HOMO SAPIENS | 5 | 41734 | 2kbi | A | 1795 | TYR | p.Tyr1795Cys | Q14524 |
| HOMO SAPIENS | 5 | 41734 | 2kbi | A | 1795 | TYR | p.Tyr1795His | Q14524 |
| HOMO SAPIENS | 7 | 41734 | 2kbi | A | 1826 | ARG | p.Arg1826Cys | Q14524 |
| HOMO SAPIENS | 3 | 2119 | 1pl0 | A | 426 | LYS | p.Lys426Arg | P31939 |
| HOMO SAPIENS | 5 | 3475 | 5u62 | C | 50 | LEU | p.Leu50Ser | Q15910 |
| HOMO SAPIENS | 3 | 379 | 4uez | A | 200 | TYR | p.Tyr200Ter | P15085 |
| HOMO SAPIENS | 0 | 8195 | 5xaw | B | 67 | ARG | p.Arg67Trp | Q8IWT1 |
| HOMO SAPIENS | 0 | 8195 | 5xaw | B | 94 | THR | p.Thr94Met | Q8IWT1 |
| HOMO SAPIENS | 1 | 5711 | 1fo3 | A | 265 | ARG | p.Arg265His | Q9UKM7 |
| HOMO SAPIENS | 4 | 5711 | 1fo3 | A | 334 | ARG | p.Arg334Cys | Q9UKM7 |
| HOMO SAPIENS | 1 | 5711 | 1fo3 | A | 478 | LYS | p.Lys478Arg | Q9UKM7 |
| HOMO SAPIENS | 1 | 5711 | 1fo3 | A | 682 | ASP | p.Asp682Glu | Q9UKM7 |
| HOMO SAPIENS | 0 | 170 | 2n3y | A | 41 | GLY | p.Gly42Ser | P99999 |
| HOMO SAPIENS | 0 | 4122 | 3w81 | A | 33 | HIS | p.His33Gln | P35475 |
| HOMO SAPIENS | 0 | 4122 | 3w81 | A | 64 | TYR | p.Tyr64Ter | P35475 |
| HOMO SAPIENS | 0 | 4122 | 3w81 | A | 70 | GLN | p.Gln70Ter | P35475 |
| HOMO SAPIENS | 0 | 4122 | 3w81 | A | 75 | ALA | p.Ala75Thr | P35475 |
| HOMO SAPIENS | 0 | 4122 | 3w81 | A | 79 | ALA | p.Ala79Val | P35475 |
| HOMO SAPIENS | 0 | 4122 | 3w81 | A | 79 | ALA | p.Ala79Thr | P35475 |
| HOMO SAPIENS | 0 | 4122 | 3w81 | A | 89 | ARG | p.Arg89Gln | P35475 |
| HOMO SAPIENS | 0 | 4122 | 3w81 | A | 99 | THR | p.Thr99Ile | P35475 |
| HOMO SAPIENS | 0 | 4122 | 3w81 | A | 105 | ARG | p.Arg105Gln | P35475 |
| HOMO SAPIENS | 0 | 4122 | 3w81 | A | 238 | LEU | p.Leu238Gln | P35475 |
| HOMO SAPIENS | 0 | 4122 | 3w81 | A | 300 | ALA | p.Ala300Thr | P35475 |
| HOMO SAPIENS | 0 | 4122 | 3w81 | A | 322 | VAL | p.Val322Glu | P35475 |
| HOMO SAPIENS | 0 | 4122 | 3w81 | A | 327 | ALA | p.Ala327Pro | P35475 |
| HOMO SAPIENS | 0 | 4122 | 3w81 | A | 346 | LEU | p.Leu346Arg | P35475 |
| HOMO SAPIENS | 0 | 4122 | 3w81 | A | 349 | ASP | p.Asp349Asn | P35475 |
| HOMO SAPIENS | 0 | 4122 | 3w81 | A | 361 | ALA | p.Ala361Thr | P35475 |
| HOMO SAPIENS | 0 | 4122 | 3w81 | A | 366 | THR | p.Thr366Pro | P35475 |
| HOMO SAPIENS | 0 | 4122 | 3w81 | A | 402 | TRP | p.Trp402Ter | P35475 |
| HOMO SAPIENS | 0 | 4122 | 3w81 | A | 409 | GLY | p.Gly409Arg | P35475 |
| HOMO SAPIENS | 0 | 4122 | 3w81 | A | 454 | VAL | p.Val454Ile | P35475 |
| HOMO SAPIENS | 1 | 4122 | 3w81 | A | 490 | LEU | p.Leu490Pro | P35475 |
| HOMO SAPIENS | 1 | 4122 | 3w81 | A | 492 | ARG | p.Arg492Pro | P35475 |
| HOMO SAPIENS | 0 | 4122 | 3w81 | A | 533 | PRO | p.Pro533Arg | P35475 |
| HOMO SAPIENS | 0 | 4122 | 3w81 | A | 591 | ALA | p.Ala591Thr | P35475 |
| HOMO SAPIENS | 7 | 4122 | 3w81 | A | 619 | ARG | p.Arg619Gly | P35475 |
| HOMO SAPIENS | 0 | 15325 | 3wvz | B | 54 | VAL | p.Val54Leu | Q53FT3 |
| HOMO SAPIENS | 3 | 3228 | 5iz8 | A | 421 | TRP | p.Trp421Ter | P25054 |
| HOMO SAPIENS | 2 | 3228 | 5iz8 | A | 437 | PRO | p.Pro437Arg | P25054 |
| HOMO SAPIENS | 3 | 3228 | 5iz8 | A | 499 | ARG | p.Arg499Ter | P25054 |
| HOMO SAPIENS | -1 | 29229 | 5buo | A | 539 | SER | p.Ser614Gly | P05067 |
| HOMO SAPIENS | -1 | 34931 | 4n7v | C | 54 | SER | p.Ser54Leu | O94986 |
| HOMO SAPIENS | 0 | 2450 | 5kzx | A | 123 | GLY | p.Gly123Glu | P10253 |
| HOMO SAPIENS | 1 | 2450 | 5kzx | A | 190 | ARG | p.Arg190His | P10253 |
| HOMO SAPIENS | -1 | 2450 | 5kzx | A | 199 | HIS | p.His199Arg | P10253 |
| HOMO SAPIENS | 0 | 2450 | 5kzx | A | 219 | GLY | p.Gly219Arg | P10253 |
| HOMO SAPIENS | 0 | 2450 | 5kzx | A | 220 | VAL | p.Val220Leu | P10253 |
| HOMO SAPIENS | 0 | 2450 | 5kzx | A | 222 | VAL | p.Val222Met | P10253 |

| | | | | | | | | |
|---|---|---|---|---|---|---|---|---|
| HOMO SAPIENS | 0 | 2450 | 5kzx | A | 237 | ALA | p.Ala237Val | P10253 |
| HOMO SAPIENS | 0 | 2450 | 5kzx | A | 251 | SER | p.Ser251Leu | P10253 |
| HOMO SAPIENS | 0 | 2450 | 5kzx | A | 254 | SER | p.Ser254Leu | P10253 |
| HOMO SAPIENS | 0 | 2450 | 5kzx | A | 262 | GLU | p.Glu262Lys | P10253 |
| HOMO SAPIENS | 0 | 2450 | 5kzx | A | 293 | GLY | p.Gly293Arg | P10253 |
| HOMO SAPIENS | 0 | 2450 | 5kzx | A | 299 | LEU | p.Leu299Arg | P10253 |
| HOMO SAPIENS | 0 | 2450 | 5kzx | A | 309 | GLY | p.Gly309Arg | P10253 |
| HOMO SAPIENS | 0 | 2450 | 5kzx | A | 318 | MET | p.Met318Thr | P10253 |
| HOMO SAPIENS | 0 | 2450 | 5kzx | A | 355 | LEU | p.Leu355Pro | P10253 |
| HOMO SAPIENS | 1 | 2450 | 5kzx | A | 361 | PRO | p.Pro361Leu | P10253 |
| HOMO SAPIENS | 4 | 2450 | 5kzx | A | 408 | MET | p.Met408Val | P10253 |
| HOMO SAPIENS | 1 | 2450 | 5kzx | A | 429 | GLN | p.Gln429Glu | P10253 |
| HOMO SAPIENS | 4 | 2450 | 5kzx | A | 437 | ARG | p.Arg437Cys | P10253 |
| HOMO SAPIENS | 0 | 2450 | 5kzx | A | 439 | MET | p.Met439Lys | P10253 |
| HOMO SAPIENS | 1 | 2450 | 5kzx | A | 448 | SER | p.Ser448Thr | P10253 |
| HOMO SAPIENS | 0 | 2450 | 5kzx | A | 451 | PRO | p.Pro451Arg | P10253 |
| HOMO SAPIENS | 1 | 2450 | 5kzx | A | 459 | ASP | p.Asp459His | P10253 |
| HOMO SAPIENS | 8 | 2450 | 5kzx | A | 481 | TRP | p.Trp481Arg | P10253 |
| HOMO SAPIENS | 1 | 2450 | 5kzx | A | 489 | ASP | p.Asp489Asn | P10253 |
| HOMO SAPIENS | 0 | 2450 | 5kzx | A | 521 | GLU | p.Glu521Lys | P10253 |
| HOMO SAPIENS | 0 | 2450 | 5kzx | A | 529 | SER | p.Ser529Val | P10253 |
| HOMO SAPIENS | 0 | 2450 | 5kzx | A | 545 | PRO | p.Pro545Leu | P10253 |
| HOMO SAPIENS | 0 | 2450 | 5kzx | A | 552 | LEU | p.Leu552Pro | P10253 |
| HOMO SAPIENS | 0 | 2450 | 5kzx | A | 576 | GLY | p.Gly576Ser | P10253 |
| HOMO SAPIENS | 9 | 2450 | 5kzx | A | 600 | ARG | p.Arg600His | P10253 |
| HOMO SAPIENS | 8 | 2450 | 5kzx | A | 615 | GLY | p.Gly615Arg | P10253 |
| HOMO SAPIENS | 3 | 2450 | 5kzx | A | 619 | SER | p.Ser619Asn | P10253 |
| HOMO SAPIENS | 0 | 2450 | 5kzx | A | 638 | GLY | p.Gly638Trp | P10253 |
| HOMO SAPIENS | 0 | 2450 | 5kzx | A | 643 | GLY | p.Gly643Arg | P10253 |
| HOMO SAPIENS | 9 | 2450 | 5kzx | A | 645 | ASP | p.Asp645Glu | P10253 |
| HOMO SAPIENS | 9 | 2450 | 5kzx | A | 645 | ASP | p.Asp645His | P10253 |
| HOMO SAPIENS | 0 | 2450 | 5kzx | A | 648 | GLY | p.Gly648Ser | P10253 |
| HOMO SAPIENS | 5 | 2450 | 5kzx | A | 660 | ARG | p.Arg660His | P10253 |
| HOMO SAPIENS | 8 | 2450 | 5kzx | A | 672 | ARG | p.Arg672Gln | P10253 |
| HOMO SAPIENS | 0 | 2450 | 5kzx | A | 702 | ARG | p.Arg702Leu | P10253 |
| HOMO SAPIENS | 3 | 2450 | 5kzx | A | 725 | ARG | p.Arg725Trp | P10253 |
| HOMO SAPIENS | 0 | 2450 | 5kzx | A | 746 | TRP | p.Trp746Cys | P10253 |
| HOMO SAPIENS | 0 | 2450 | 5kzx | A | 780 | ILE | p.Val780Ile | P10253 |
| HOMO SAPIENS | 1 | 2450 | 5kzx | A | 816 | VAL | p.Val816Ile | P10253 |
| HOMO SAPIENS | 0 | 2450 | 5kzx | A | 838 | GLN | p.Gln838Ter | P10253 |
| HOMO SAPIENS | 2 | 2450 | 5kzx | A | 854 | ARG | p.Arg854Ter | P10253 |
| HOMO SAPIENS | 3 | 2450 | 5kzx | A | 870 | ARG | p.Arg870Ter | P10253 |
| HOMO SAPIENS | 0 | 2450 | 5kzx | A | 890 | VAL | p.Val890Leu | P10253 |
| HOMO SAPIENS | 6 | 1882 | 4km6 | A | 175 | CYS | p.Cys175Ter | P15328 |
| HOMO SAPIENS | 3 | 14767 | 1lm5 | A | 2628 | LEU | p.Leu2628Pro | P15924 |
| HOMO SAPIENS | 5 | 14767 | 1lm5 | A | 2655 | ALA | p.Ala2655Gly | P15924 |
| HOMO SAPIENS | 3 | 14767 | 1lm5 | A | 2731 | TYR | p.Tyr2731His | P15924 |
| HOMO SAPIENS | 1 | 14767 | 1lm5 | A | 2767 | THR | p.Thr2767Asn | P15924 |
| HOMO SAPIENS | -1 | 14767 | 1lm5 | A | 2819 | MET | p.Met2819Leu | P15924 |
| HOMO SAPIENS | -1 | 11246 | 4y5o | A | 289 | SER | p.Ser289Asn | Q9BSQ5 |
| HOMO SAPIENS | 0 | 7380 | 5kz5 | I | 60 | ARG | p.Arg60His | Q16595 |
| HOMO SAPIENS | 0 | 7380 | 5kz5 | I | 76 | MET | p.Met76Val | Q16595 |
| HOMO SAPIENS | 2 | 22187 | 3fby | A | 328 | CYS | p.Cys328Arg | P49747 |
| HOMO SAPIENS | 2 | 22187 | 3fby | A | 342 | ASP | p.Asp342Tyr | P49747 |

| | | | | | | | | |
|---|---|---|---|---|---|---|---|---|
| HOMO SAPIENS | 2 | 22187 | 3fby | A | 348 | CYS | p.Cys348Arg | P49747 |
| HOMO SAPIENS | 2 | 22187 | 3fby | A | 386 | ASN | p.Asn386Asp | P49747 |
| HOMO SAPIENS | 2 | 22187 | 3fby | A | 453 | ASN | p.Asn453Ser | P49747 |
| HOMO SAPIENS | 2 | 22187 | 3fby | A | 468 | CYS | p.Cys468Tyr | P49747 |
| HOMO SAPIENS | 3 | 22187 | 3fby | A | 472 | ASP | p.Asp472Tyr | P49747 |
| HOMO SAPIENS | 3 | 22187 | 3fby | A | 473 | ASP | p.Asp473Gly | P49747 |
| HOMO SAPIENS | 2 | 22187 | 3fby | A | 523 | ASN | p.Asn523Lys | P49747 |
| HOMO SAPIENS | 1 | 22187 | 3fby | A | 718 | ARG | p.Arg718Trp | P49747 |
| HOMO SAPIENS | 1 | 22187 | 3fby | A | 719 | GLY | p.Gly719Asp | P49747 |
| HOMO SAPIENS | 0 | 22187 | 3fby | A | 756 | GLN | p.Gln756Arg | P49747 |
| HOMO SAPIENS | -1 | 2784 | 5kvh | A | 88 | MET | p.Met88Val | O95831 |
| HOMO SAPIENS | -1 | 2784 | 5kvh | A | 96 | ASN | p.Asn96Ser | O95831 |
| HOMO SAPIENS | 0 | 2784 | 5kvh | A | 282 | THR | p.Thr282Met | O95831 |
| HOMO SAPIENS | 1 | 2784 | 5kvh | A | 366 | ASN | p.Asn366Ser | O95831 |
| HOMO SAPIENS | 3 | 2784 | 5kvh | A | 463 | ARG | p.Arg463Ile | O95831 |
| HOMO SAPIENS | 0 | 2784 | 5kvh | A | 498 | VAL | p.Val498Met | O95831 |
| HOMO SAPIENS | -1 | 2784 | 5kvh | A | 549 | ALA | p.Ala549Val | O95831 |
| HOMO SAPIENS | 0 | 38316 | 2lnk | C | 1933 | ARG | p.Arg1933Ter | P35579 |
| HOMO SAPIENS | 0 | 33808 | 2miu | A | 29 | VAL | p.Val29Met | Q14192 |
| HOMO SAPIENS | 1 | 33808 | 2miu | A | 37 | ALA | p.Ala37Ser | Q14192 |
| HOMO SAPIENS | 2 | 20994 | 2hey | R | 65 | ARG | p.Arg65Cys | P43489 |
| HOMO SAPIENS | -1 | 5687 | 2xwj | I | 7 | ARG | p.Arg32Gln | P00751 |
| HOMO SAPIENS | -1 | 5687 | 2xwj | I | 7 | ARG | p.Arg32Trp | P00751 |
| HOMO SAPIENS | 0 | 5687 | 2xwj | I | 49 | ARG | p.Arg74His | P00751 |
| HOMO SAPIENS | 0 | 5687 | 2xwj | I | 177 | ARG | p.Arg202Gly | P00751 |
| HOMO SAPIENS | 0 | 5687 | 2xwj | I | 217 | ILE | p.Ile242Leu | P00751 |
| HOMO SAPIENS | -1 | 5687 | 2xwj | I | 227 | GLY | p.Gly252Ser | P00751 |
| HOMO SAPIENS | 0 | 5687 | 2xwj | I | 483 | HIS | p.His508Gln | P00751 |
| HOMO SAPIENS | 0 | 5687 | 2xwj | I | 508 | LYS | p.Lys533Arg | P00751 |
| HOMO SAPIENS | 0 | 5687 | 2xwj | I | 540 | LYS | p.Lys565Glu | P00751 |
| HOMO SAPIENS | 0 | 5687 | 2xwj | I | 541 | GLU | p.Glu566Ala | P00751 |
| HOMO SAPIENS | 2 | 953 | 3d4j | A | 249 | PHE | p.Phe249Ser | P53602 |
| HOMO SAPIENS | 1 | 953 | 3d4j | A | 292 | ASN | p.Asn292Ser | P53602 |
| HOMO SAPIENS | 0 | 3300 | 1s9c | A | 140 | ASN | p.Asn457Tyr | P51659 |
| HOMO SAPIENS | 2 | 3300 | 1s9c | A | 167 | ILE | p.Ile484Val | P51659 |
| HOMO SAPIENS | 0 | 3300 | 1s9c | A | 174 | ALA | p.Ala491Thr | P51659 |
| HOMO SAPIENS | 8 | 3300 | 1s9c | A | 193 | ASP | p.Asp510Asn | P51659 |
| HOMO SAPIENS | 3 | 3300 | 1s9c | A | 194 | TRP | p.Trp511Arg | P51659 |
| HOMO SAPIENS | 0 | 3300 | 1s9c | A | 242 | VAL | p.Ile559Val | P51659 |
| HOMO SAPIENS | 0 | 10810 | 5i25 | A | 5 | GLN | p.Gln23Ter | P03951 |
| HOMO SAPIENS | 0 | 10810 | 5i25 | A | 55 | TRP | p.Trp73Ter | P03951 |
| HOMO SAPIENS | 0 | 10810 | 5i25 | A | 91 | ALA | p.Ala109Thr | P03951 |
| HOMO SAPIENS | 0 | 10810 | 5i25 | A | 117 | GLU | p.Glu135Ter | P03951 |
| HOMO SAPIENS | 0 | 10810 | 5i25 | A | 118 | CYS | p.Cys136Ter | P03951 |
| HOMO SAPIENS | 0 | 10810 | 5i25 | A | 128 | CYS | p.Cys146Ter | P03951 |
| HOMO SAPIENS | 0 | 10810 | 5i25 | A | 210 | ARG | p.Arg228Ter | P03951 |
| HOMO SAPIENS | 0 | 10810 | 5i25 | A | 226 | GLN | p.Gln244Ter | P03951 |
| HOMO SAPIENS | 0 | 10810 | 5i25 | A | 233 | GLN | p.Gln251Ter | P03951 |
| HOMO SAPIENS | 0 | 16668 | 3lbx | A | 28 | ARG | p.Arg28Leu | P02549 |
| HOMO SAPIENS | 0 | 16668 | 3lbx | A | 28 | ARG | p.Arg28Cys | P02549 |
| HOMO SAPIENS | 1 | 16668 | 3lbx | A | 41 | ARG | p.Arg41Trp | P02549 |
| HOMO SAPIENS | 1 | 16668 | 3lbx | A | 45 | ARG | p.Arg45Ser | P02549 |
| HOMO SAPIENS | 2 | 16668 | 3lbx | A | 48 | LYS | p.Lys48Arg | P02549 |
| HOMO SAPIENS | 1 | 16668 | 3lbx | A | 152 | ASP | p.Asp152Asn | P02549 |

| | | | | | | | | |
|---|---|---|---|---|---|---|---|---|
| HOMO SAPIENS | 0 | 226 | 2ble | A | 256 | ILE | p.Phe256Ile | P36959 |
| HOMO SAPIENS | 2 | 36808 | 4c7n | A | 3 | SER | p.Ser357Pro | O75030 |
| HOMO SAPIENS | 0 | 4092 | 5jnx | A | 113 | CYS | p.Cys113Arg | O15118 |
| HOMO SAPIENS | 4 | 4092 | 5jnx | A | 215 | HIS | p.His215Arg | O15118 |
| HOMO SAPIENS | 6 | 4092 | 5jnx | A | 237 | PRO | p.Pro237Ser | O15118 |
| HOMO SAPIENS | 0 | 4092 | 5jnx | A | 363 | ALA | p.Ala363Val | O15118 |
| HOMO SAPIENS | 0 | 4092 | 5jnx | A | 378 | VAL | p.Val378Ala | O15118 |
| HOMO SAPIENS | 8 | 4092 | 5jnx | A | 404 | ARG | p.Arg404Gln | O15118 |
| HOMO SAPIENS | 8 | 4092 | 5jnx | A | 411 | ARG | p.Arg411Gln | O15118 |
| HOMO SAPIENS | 2 | 4092 | 5jnx | A | 474 | PRO | p.Pro474Leu | O15118 |
| HOMO SAPIENS | 8 | 4092 | 5jnx | A | 511 | THR | p.Thr511Met | O15118 |
| HOMO SAPIENS | 0 | 4092 | 5jnx | A | 642 | MET | p.Met642Ile | O15118 |
| HOMO SAPIENS | 2 | 4092 | 5jnx | A | 664 | VAL | p.Val664Met | O15118 |
| HOMO SAPIENS | 0 | 4092 | 5jnx | A | 734 | SER | p.Ser734Ile | O15118 |
| HOMO SAPIENS | -1 | 4092 | 5jnx | A | 775 | GLN | p.Gln775Pro | O15118 |
| HOMO SAPIENS | 0 | 4092 | 5jnx | A | 858 | ILE | p.Ile858Val | O15118 |
| HOMO SAPIENS | 0 | 4092 | 5jnx | A | 889 | VAL | p.Val889Met | O15118 |
| HOMO SAPIENS | 0 | 4092 | 5jnx | A | 910 | GLY | p.Gly910Ser | O15118 |
| HOMO SAPIENS | 0 | 4092 | 5jnx | A | 911 | GLY | p.Gly911Ser | O15118 |
| HOMO SAPIENS | 0 | 4092 | 5jnx | A | 928 | GLN | p.Gln928Pro | O15118 |
| HOMO SAPIENS | 0 | 4092 | 5jnx | A | 931 | ASN | p.Asn931Lys | O15118 |
| HOMO SAPIENS | 0 | 4092 | 5jnx | A | 940 | SER | p.Ser940Leu | O15118 |
| HOMO SAPIENS | 0 | 4092 | 5jnx | A | 950 | VAL | p.Val950Met | O15118 |
| HOMO SAPIENS | 0 | 4092 | 5jnx | A | 954 | SER | p.Ser954Leu | O15118 |
| HOMO SAPIENS | 0 | 4092 | 5jnx | A | 958 | ARG | p.Arg958Gln | O15118 |
| HOMO SAPIENS | 0 | 4092 | 5jnx | A | 958 | ARG | p.Arg958Ter | O15118 |
| HOMO SAPIENS | -1 | 4092 | 5jnx | A | 978 | ARG | p.Arg978Cys | O15118 |
| HOMO SAPIENS | 0 | 4092 | 5jnx | A | 992 | GLY | p.Gly992Trp | O15118 |
| HOMO SAPIENS | 0 | 4092 | 5jnx | A | 1007 | PRO | p.Pro1007Ala | O15118 |
| HOMO SAPIENS | 0 | 4092 | 5jnx | A | 1019 | TYR | p.Tyr1019Cys | O15118 |
| HOMO SAPIENS | 2 | 4092 | 5jnx | A | 1035 | ALA | p.Ala1035Val | O15118 |
| HOMO SAPIENS | 2 | 4092 | 5jnx | A | 1054 | ALA | p.Ala1054Thr | O15118 |
| HOMO SAPIENS | 2 | 4092 | 5jnx | A | 1061 | ILE | p.Ile1061Thr | O15118 |
| HOMO SAPIENS | 0 | 4092 | 5jnx | A | 1073 | GLY | p.Gly1073Ser | O15118 |
| HOMO SAPIENS | 2 | 4092 | 5jnx | A | 1088 | TYR | p.Tyr1088Cys | O15118 |
| HOMO SAPIENS | 2 | 4092 | 5jnx | A | 1142 | MET | p.Met1142Thr | O15118 |
| HOMO SAPIENS | 2 | 4092 | 5jnx | A | 1156 | ASN | p.Asn1156Ser | O15118 |
| HOMO SAPIENS | 0 | 4092 | 5jnx | A | 1179 | MET | p.Met1179Val | O15118 |
| HOMO SAPIENS | 0 | 4092 | 5jnx | A | 1186 | ARG | p.Arg1186His | O15118 |
| HOMO SAPIENS | 0 | 4092 | 5jnx | A | 1187 | ALA | p.Ala1187Gly | O15118 |
| HOMO SAPIENS | 2 | 4092 | 5jnx | A | 1189 | GLU | p.Glu1189Gly | O15118 |
| HOMO SAPIENS | 2 | 4092 | 5jnx | A | 1200 | SER | p.Ser1200Gly | O15118 |
| HOMO SAPIENS | 2 | 4092 | 5jnx | A | 1205 | THR | p.Thr1205Lys | O15118 |
| HOMO SAPIENS | 0 | 4092 | 5jnx | A | 1213 | LEU | p.Leu1213Phe | O15118 |
| HOMO SAPIENS | -1 | 4092 | 5jnx | A | 1266 | ARG | p.Arg1266Gln | O15118 |
| HOMO SAPIENS | 9 | 16498 | 4i6o | A | 64 | ARG | p.Arg735Trp | P01024 |
| HOMO SAPIENS | 1 | 21902 | 3un9 | C | 751 | ARG | p.Arg751His | Q86UT6 |
| HOMO SAPIENS | 7 | 20391 | 2h32 | B | 105 | PRO | p.Pro142Leu | P15814 |
| HOMO SAPIENS | 6 | 1635 | 4x45 | A | 65 | ASP | p.Asp65Val | P07741 |
| HOMO SAPIENS | 1 | 1635 | 4x45 | A | 98 | TRP | p.Trp98Ter | P07741 |
| HOMO SAPIENS | 0 | 1635 | 4x45 | A | 110 | LEU | p.Leu110Pro | P07741 |
| HOMO SAPIENS | 7 | 1635 | 4x45 | A | 136 | MET | p.Met136Thr | P07741 |
| HOMO SAPIENS | 1 | 1635 | 4x45 | A | 150 | VAL | p.Val150Phe | P07741 |
| HOMO SAPIENS | 4 | 34810 | 2rpc | A | 16 | CYS | p.Cys253Ser | O60481 |

| | | | | | | | | |
|---|---|---|---|---|---|---|---|---|
| HOMO SAPIENS | 1 | 34810 | 2rpc | A | 18 | TRP | p.Trp255Gly | O60481 |
| HOMO SAPIENS | 1 | 14359 | 4hxi | A | 78 | MET | p.Met78Val | Q9UH77 |
| HOMO SAPIENS | 0 | 5342 | 4d9t | A | 628 | GLU | p.Glu628Asp | P51812 |
| HOMO SAPIENS | -1 | 5342 | 4d9t | A | 723 | ARG | p.Arg723His | P51812 |
| HOMO SAPIENS | -1 | 5342 | 4d9t | A | 729 | ARG | p.Arg729Gln | P51812 |
| HOMO SAPIENS | 1 | 9619 | 3j2s | B | 1781 | ARG | p.Arg1800Cys | P00451 |
| HOMO SAPIENS | 0 | 9619 | 3j2s | B | 1823 | MET | p.Met1842Ile | P00451 |
| HOMO SAPIENS | 1 | 9619 | 3j2s | B | 1825 | PRO | p.Pro1844Ser | P00451 |
| HOMO SAPIENS | 0 | 9619 | 3j2s | B | 1826 | THR | p.Thr1845Pro | P00451 |
| HOMO SAPIENS | 0 | 9619 | 3j2s | B | 1834 | ALA | p.Ala1853Val | P00451 |
| HOMO SAPIENS | 0 | 9619 | 3j2s | B | 1846 | ASP | p.Asp1865Asn | P00451 |
| HOMO SAPIENS | 1 | 9619 | 3j2s | B | 1848 | HIS | p.His1867Arg | P00451 |
| HOMO SAPIENS | 0 | 9619 | 3j2s | B | 1854 | PRO | p.Pro1873Arg | P00451 |
| HOMO SAPIENS | 1 | 9619 | 3j2s | B | 1885 | GLU | p.Glu1904Lys | P00451 |
| HOMO SAPIENS | 0 | 9619 | 3j2s | B | 1922 | ASN | p.Asn1941Ser | P00451 |
| HOMO SAPIENS | 1 | 9619 | 3j2s | B | 1941 | ARG | p.Arg1960Gln | P00451 |
| HOMO SAPIENS | 4 | 9619 | 3j2s | B | 2105 | TYR | p.Tyr2124Cys | P00451 |
| HOMO SAPIENS | 6 | 9619 | 3j2s | B | 2150 | ARG | p.Arg2169His | P00451 |
| HOMO SAPIENS | 0 | 9619 | 3j2s | B | 2159 | ARG | p.Arg2178Cys | P00451 |
| HOMO SAPIENS | 0 | 9619 | 3j2s | B | 2159 | ARG | p.Arg2178His | P00451 |
| HOMO SAPIENS | 2 | 9619 | 3j2s | B | 2238 | MET | p.Met2257Val | P00451 |
| HOMO SAPIENS | 2 | 9619 | 3j2s | B | 2300 | PRO | p.Pro2319Ser | P00451 |
| HOMO SAPIENS | 1 | 667 | 1z2m | A | 127 | GLU | p.Glu127Ter | P05161 |
| HOMO SAPIENS | 0 | 15717 | 2nn6 | G | 30 | GLY | p.Gly31Ala | Q9NQT5 |
| HOMO SAPIENS | -1 | 15717 | 2nn6 | G | 55 | ASN | p.Asn56His | Q9NQT5 |
| HOMO SAPIENS | 7 | 15717 | 2nn6 | G | 131 | ASP | p.Asp132Ala | Q9NQT5 |
| HOMO SAPIENS | 7 | 784 | 3rkq | A | 161 | ARG | p.Arg161Pro | P52952 |
| HOMO SAPIENS | 6 | 784 | 3rkq | A | 178 | THR | p.Thr178Met | P52952 |
| HOMO SAPIENS | 7 | 784 | 3rkq | A | 183 | LYS | p.Lys183Glu | P52952 |
| HOMO SAPIENS | 8 | 784 | 3rkq | A | 190 | ARG | p.Arg190Cys | P52952 |
| HOMO SAPIENS | 6 | 385 | 30000 | C | 70 | ALA | p.Ala70Thr | P07954 |
| HOMO SAPIENS | 2 | 385 | 30000 | C | 101 | ARG | p.Arg101Pro | P07954 |
| HOMO SAPIENS | 2 | 385 | 30000 | C | 174 | PRO | p.Pro174Arg | P07954 |
| HOMO SAPIENS | 6 | 385 | 30000 | C | 187 | SER | p.Ser187Ter | P07954 |
| HOMO SAPIENS | 9 | 385 | 30000 | C | 233 | ARG | p.Arg233His | P07954 |
| HOMO SAPIENS | 3 | 385 | 30000 | C | 273 | ALA | p.Ala273Thr | P07954 |
| HOMO SAPIENS | 4 | 385 | 30000 | C | 334 | SER | p.Ser334Arg | P07954 |
| HOMO SAPIENS | 6 | 385 | 30000 | C | 340 | ASN | p.Asn340Lys | P07954 |
| HOMO SAPIENS | 8 | 385 | 30000 | C | 343 | ARG | p.Arg343Ter | P07954 |
| HOMO SAPIENS | 3 | 385 | 30000 | C | 350 | ARG | p.Arg350Trp | P07954 |
| HOMO SAPIENS | 4 | 385 | 30000 | C | 376 | GLN | p.Gln376Pro | P07954 |
| HOMO SAPIENS | 6 | 385 | 30000 | C | 434 | CYS | p.Cys434Ter | P07954 |
| HOMO SAPIENS | 0 | 3316 | 4f91 | B | 934 | THR | p.Thr934Ala | O75643 |
| HOMO SAPIENS | 0 | 3316 | 4f91 | B | 1002 | ASN | p.Asn1002Ser | O75643 |
| HOMO SAPIENS | 0 | 3316 | 4f91 | B | 1087 | SER | p.Ser1087Leu | O75643 |
| HOMO SAPIENS | 4 | 3316 | 4f91 | B | 1389 | VAL | p.Val1389Ile | O75643 |
| HOMO SAPIENS | 0 | 619 | 5hbt | B | 195 | ASP | p.Asp240His | P02708 |
| HOMO SAPIENS | 1 | 619 | 5hbt | B | 209 | ARG | p.Arg254Leu | P02708 |
| HOMO SAPIENS | 0 | 18171 | 2y25 | B | 1408 | ASP | p.Asp1408Asn | P52179 |
| HOMO SAPIENS | 0 | 18171 | 2y25 | B | 1453 | MET | p.Met1453Thr | P52179 |
| HOMO SAPIENS | 0 | 18171 | 2y25 | B | 1459 | ILE | p.Ile1459Lys | P52179 |
| HOMO SAPIENS | 0 | 18171 | 2y25 | B | 1558 | PHE | p.Phe1558Leu | P52179 |
| HOMO SAPIENS | 0 | 18171 | 2y25 | B | 1573 | ARG | p.Arg1573Gln | P52179 |
| HOMO SAPIENS | 7 | 18171 | 2y25 | B | 1663 | VAL | p.Val1663Met | P52179 |

| | | | | | | | | |
|---|---|---|---|---|---|---|---|---|
| HOMO SAPIENS | 8 | 14262 | 2fcw | B | 117 | SER | p.Ser138Ter | P01130 |
| HOMO SAPIENS | 8 | 14262 | 2fcw | B | 118 | ASP | p.Asp139Glu | P01130 |
| HOMO SAPIENS | 9 | 14262 | 2fcw | B | 119 | GLU | p.Glu140Lys | P01130 |
| HOMO SAPIENS | 7 | 14262 | 2fcw | B | 122 | CYS | p.Cys143Ter | P01130 |
| HOMO SAPIENS | 6 | 14262 | 2fcw | B | 134 | CYS | p.Cys155Ter | P01130 |
| HOMO SAPIENS | 7 | 14262 | 2fcw | B | 147 | ASP | p.Asp168Glu | P01130 |
| HOMO SAPIENS | 4 | 2624 | 3b7b | A | 855 | VAL | p.Val886Met | Q9H9B1 |
| HOMO SAPIENS | 7 | 2624 | 3b7b | A | 868 | ILE | p.Ile899Val | Q9H9B1 |
| HOMO SAPIENS | 0 | 2624 | 3b7b | A | 892 | GLU | p.Glu923Lys | Q9H9B1 |
| HOMO SAPIENS | 0 | 4791 | 5ctd | B | 68 | LEU | p.Pro549Ala | Q14055 |
| HOMO SAPIENS | 5 | 460 | 4ui1 | D | 80 | CYS | p.Cys80Arg | Q6ZVN8 |
| HOMO SAPIENS | 3 | 460 | 4ui1 | D | 101 | LEU | p.Leu101Pro | Q6ZVN8 |
| HOMO SAPIENS | 0 | 40782 | 1sy6 | A | 47 | LYS | p.Lys69Ter | P09693 |
| HOMO SAPIENS | -1 | 1092 | 2idx | A | 62 | THR | p.Thr62Met | Q96EY8 |
| HOMO SAPIENS | 6 | 1092 | 2idx | A | 183 | HIS | p.His183Leu | Q96EY8 |
| HOMO SAPIENS | 7 | 1092 | 2idx | A | 186 | ARG | p.Arg186Trp | Q96EY8 |
| HOMO SAPIENS | 6 | 1092 | 2idx | A | 190 | ARG | p.Arg190Cys | Q96EY8 |
| HOMO SAPIENS | 6 | 1092 | 2idx | A | 190 | ARG | p.Arg190His | Q96EY8 |
| HOMO SAPIENS | 7 | 1092 | 2idx | A | 191 | ARG | p.Arg191Trp | Q96EY8 |
| HOMO SAPIENS | 7 | 1092 | 2idx | A | 191 | ARG | p.Arg191Gln | Q96EY8 |
| HOMO SAPIENS | 6 | 1092 | 2idx | A | 195 | ARG | p.Arg195His | Q96EY8 |
| HOMO SAPIENS | 1 | 1092 | 2idx | A | 234 | GLN | p.Gln234Ter | Q96EY8 |
| HOMO SAPIENS | 0 | 1092 | 2idx | A | 239 | MET | p.Met239Lys | Q96EY8 |
| HOMO SAPIENS | -1 | 6418 | 5fur | G | 11 | THR | p.Thr11Ile | P21675 |
| HOMO SAPIENS | -1 | 6418 | 5fur | G | 290 | LEU | p.Leu269Val | P21675 |
| HOMO SAPIENS | 0 | 6100 | 1wiq | A | 240 | ARG | p.Arg265Trp | P01730 |
| HOMO SAPIENS | 0 | 13757 | 4ye6 | A | 45 | GLY | p.Gly45Val | P47897 |
| HOMO SAPIENS | 1 | 13757 | 4ye6 | A | 57 | TYR | p.Tyr57His | P47897 |
| HOMO SAPIENS | 8 | 13757 | 4ye6 | A | 403 | ARG | p.Arg403Trp | P47897 |
| HOMO SAPIENS | 7 | 13757 | 4ye6 | A | 524 | ARG | p.Arg524Trp | P47897 |
| HOMO SAPIENS | 6 | 13757 | 4ye6 | A | 690 | ARG | p.Arg690Cys | P47897 |
| HOMO SAPIENS | 0 | 26958 | 2lzl | A | 370 | GLY | p.Gly370Cys | P22607 |
| HOMO SAPIENS | 0 | 26958 | 2lzl | A | 371 | SER | p.Ser371Cys | P22607 |
| HOMO SAPIENS | 0 | 26958 | 2lzl | A | 373 | TYR | p.Tyr373Cys | P22607 |
| HOMO SAPIENS | 0 | 26958 | 2lzl | A | 380 | GLY | p.Gly380Arg | P22607 |
| HOMO SAPIENS | 0 | 26958 | 2lzl | A | 384 | PHE | p.Phe384Leu | P22607 |
| HOMO SAPIENS | 5 | 26958 | 2lzl | A | 391 | ALA | p.Ala391Glu | P22607 |
| HOMO SAPIENS | 6 | 11325 | 2wa6 | A | 121 | TYR | p.Tyr121Phe | O75369 |
| HOMO SAPIENS | 2 | 11325 | 2wa6 | A | 161 | PHE | p.Phe161Cys | O75369 |
| HOMO SAPIENS | 0 | 11325 | 2wa6 | A | 171 | LEU | p.Leu171Arg | O75369 |
| HOMO SAPIENS | 2 | 11325 | 2wa6 | A | 173 | ALA | p.Ala173Val | O75369 |
| HOMO SAPIENS | 0 | 11325 | 2wa6 | A | 202 | MET | p.Met202Val | O75369 |
| HOMO SAPIENS | 3 | 11325 | 2wa6 | A | 227 | GLU | p.Glu227Lys | O75369 |
| HOMO SAPIENS | 0 | 11325 | 2wa6 | A | 234 | LEU | p.Leu234Val | O75369 |
| HOMO SAPIENS | 2 | 11325 | 2wa6 | A | 235 | SER | p.Ser235Pro | O75369 |
| HOMO SAPIENS | 9 | 7240 | 3tgx | A | 137 | TYR | p.Tyr156Cys | Q9HBE5 |
| HOMO SAPIENS | 0 | 35479 | 4elj | A | 307 | THR | p.Thr307Ile | P06400 |
| HOMO SAPIENS | 0 | 35479 | 4elj | A | 310 | GLY | p.Gly310Glu | P06400 |
| HOMO SAPIENS | 0 | 35479 | 4elj | A | 377 | THR | p.Thr377Ser | P06400 |
| HOMO SAPIENS | 9 | 36706 | 1he7 | A | 359 | TYR | p.Tyr359Cys | P04629 |
| HOMO SAPIENS | 4 | 6538 | 4d0e | A | 504 | ARG | p.Arg504His | P46531 |
| HOMO SAPIENS | 1 | 26518 | 5fm5 | M | 600 | GLU | p.Glu600Val | P52179 |
| HOMO SAPIENS | -1 | 26518 | 5fm5 | M | 651 | ARG | p.Arg651Gln | P52179 |
| HOMO SAPIENS | -1 | 26518 | 5fm5 | M | 660 | PRO | p.Pro660Ala | P52179 |

| | | | | | | | | |
|---|---|---|---|---|---|---|---|---|
| HOMO SAPIENS | -1 | 26518 | 5fm5 | M | 688 | THR | p.Thr688Ser | P52179 |
| HOMO SAPIENS | -1 | 26518 | 5fm5 | M | 704 | GLU | p.Glu704Lys | P52179 |
| HOMO SAPIENS | -1 | 26518 | 5fm5 | M | 727 | THR | p.Thr727Ala | P52179 |
| HOMO SAPIENS | 2 | 27054 | 4rhp | B | 209 | LEU | p.Leu209Val | O75208 |
| HOMO SAPIENS | -1 | 27054 | 4rhp | B | 288 | LYS | p.Lys288Asn | O75208 |
| HOMO SAPIENS | 5 | 2384 | 5fqd | B | 419 | ARG | p.Arg419Ter | Q96SW2 |
| HOMO SAPIENS | 4 | 5154 | 2hl3 | A | 56 | LYS | p.Lys56Arg | Q14203 |
| HOMO SAPIENS | 0 | 5154 | 2hl3 | A | 59 | GLY | p.Gly59Ser | Q14203 |
| HOMO SAPIENS | 0 | 5154 | 2hl3 | A | 71 | GLY | p.Gly71Ala | Q14203 |
| HOMO SAPIENS | 0 | 5154 | 2hl3 | A | 71 | GLY | p.Gly71Arg | Q14203 |
| HOMO SAPIENS | 7 | 5154 | 2hl3 | A | 72 | THR | p.Thr72Pro | Q14203 |
| HOMO SAPIENS | 2 | 5154 | 2hl3 | A | 74 | GLN | p.Gln74Pro | Q14203 |
| HOMO SAPIENS | 2 | 20039 | 2l1g | A | 9 | GLY | p.Gly9Cys | Q9NVV9 |
| HOMO SAPIENS | 5 | 20039 | 2l1g | A | 81 | PHE | p.Phe81Leu | Q9NVV9 |
| HOMO SAPIENS | -1 | 97 | 5uz7 | A | 201 | ARG | p.Arg201His | P63092 |
| HOMO SAPIENS | -1 | 97 | 5uz7 | A | 201 | ARG | p.Arg201Cys | P63092 |
| HOMO SAPIENS | 0 | 3257 | 4b7w | B | 278 | GLU | p.Glu278Asp | P41235 |
| HOMO SAPIENS | 1 | 3257 | 4b7w | B | 331 | ARG | p.Arg331Cys | P41235 |
| HOMO SAPIENS | 0 | 2814 | 4qgk | B | 10 | GLN | p.Gln10Ter | P51648 |
| HOMO SAPIENS | 0 | 2814 | 4qgk | B | 177 | ARG | p.Arg177Ter | P51648 |
| HOMO SAPIENS | 9 | 2814 | 4qgk | B | 184 | THR | p.Thr184Met | P51648 |
| HOMO SAPIENS | 1 | 2814 | 4qgk | B | 188 | ALA | p.Ala188Val | P51648 |
| HOMO SAPIENS | 0 | 2814 | 4qgk | B | 214 | CYS | p.Cys214Tyr | P51648 |
| HOMO SAPIENS | 0 | 2814 | 4qgk | B | 245 | ASP | p.Asp245Asn | P51648 |
| HOMO SAPIENS | 0 | 2814 | 4qgk | B | 266 | LYS | p.Lys266Asn | P51648 |
| HOMO SAPIENS | 0 | 2814 | 4qgk | B | 315 | PRO | p.Pro315Ser | P51648 |
| HOMO SAPIENS | 0 | 2814 | 4qgk | B | 386 | ASN | p.Asn386Ser | P51648 |
| HOMO SAPIENS | -1 | 5486 | 5u6o | B | 47 | GLY | p.Gly47Val | O60741 |
| HOMO SAPIENS | 1 | 21475 | 3pdy | A | 679 | THR | p.Thr789Ile | Q15149 |
| HOMO SAPIENS | 0 | 32890 | 4djc | B | 1501 | LEU | p.Leu1501Val | Q14524 |
| HOMO SAPIENS | -1 | 32890 | 4djc | B | 1512 | ARG | p.Arg1512Trp | Q14524 |
| HOMO SAPIENS | 0 | 26683 | 2xze | B | 38 | ARG | p.Arg38Cys | O95630 |
| HOMO SAPIENS | 0 | 26683 | 2xze | B | 100 | PHE | p.Phe100Tyr | O95630 |
| HOMO SAPIENS | -1 | 23175 | 5dka | A | 163 | SER | p.Ser163Leu | Q96EQ8 |
| HOMO SAPIENS | -1 | 23175 | 5dka | A | 174 | ARG | p.Arg174Cys | Q96EQ8 |
| HOMO SAPIENS | 6 | 21177 | 3k1r | A | 155 | ARG | p.Arg155Ter | Q9Y6N9 |
| HOMO SAPIENS | 0 | 1237 | 2wwm | O | 40 | PRO | p.Pro40Arg | O75147 |
| HOMO SAPIENS | 1 | 2543 | 5ewl | B | 64 | VAL | p.Val64Leu | Q13224 |
| HOMO SAPIENS | 2 | 2543 | 5ewl | B | 271 | ALA | p.Ala271Glu | Q13224 |
| HOMO SAPIENS | 3 | 34262 | 2mng | A | 613 | PHE | p.Phe613Leu | Q9Y3Q4 |
| HOMO SAPIENS | 7 | 34262 | 2mng | A | 672 | SER | p.Ser672Arg | Q9Y3Q4 |
| HOMO SAPIENS | 4 | 1667 | 1f05 | A | 321 | LYS | p.Lys321Met | P37837 |
| HOMO SAPIENS | 0 | 25179 | 2knc | B | 752 | SER | p.Ser778Pro | P05106 |
| HOMO SAPIENS | 0 | 2185 | 2ig7 | A | 50 | TYR | p.Tyr50Cys | Q9Y259 |
| HOMO SAPIENS | 0 | 2185 | 2ig7 | A | 224 | ASN | p.Asn224His | Q9Y259 |
| HOMO SAPIENS | 0 | 2185 | 2ig7 | A | 270 | TYR | p.Tyr270Ter | Q9Y259 |
| HOMO SAPIENS | 0 | 2185 | 2ig7 | A | 328 | GLN | p.Gln328Arg | Q9Y259 |
| HOMO SAPIENS | 3 | 2201 | 2r0o | A | 179 | PRO | p.Pro179Leu | O43707 |
| HOMO SAPIENS | 3 | 2201 | 2r0o | A | 255 | GLU | p.Lys255Glu | O43707 |
| HOMO SAPIENS | 3 | 2201 | 2r0o | A | 259 | THR | p.Thr259Ile | O43707 |
| HOMO SAPIENS | 4 | 2201 | 2r0o | A | 262 | SER | p.Ser262Pro | O43707 |
| HOMO SAPIENS | -1 | 31221 | 2wvr | C | 44 | THR | p.Thr44Pro | Q9H211 |
| HOMO SAPIENS | -1 | 31221 | 2wvr | C | 135 | ALA | p.Ala135Val | Q9H211 |
| HOMO SAPIENS | -1 | 31221 | 2wvr | C | 456 | GLU | p.Glu456Ala | Q9H211 |

| | | | | | | | | |
|---|---|---|---|---|---|---|---|---|
| HOMO SAPIENS | -1 | 31221 | 2wvr | C | 462 | ARG | p.Arg462Gln | Q9H211 |
| HOMO SAPIENS | -1 | 31221 | 2wvr | C | 468 | GLU | p.Glu468Lys | Q9H211 |
| HOMO SAPIENS | -1 | 31221 | 2wvr | C | 537 | ALA | p.Ala537Val | Q9H211 |
| HOMO SAPIENS | 5 | 2576 | 5o3l | C | 363 | VAL | p.Val680Ile | P10636 |
| HOMO SAPIENS | 5 | 1345 | 1czt | A | 37 | ARG | p.Arg2102Cys | P12259 |
| HOMO SAPIENS | 1 | 1345 | 1czt | A | 83 | MET | p.Met2148Thr | P12259 |
| HOMO SAPIENS | 1 | 1345 | 1czt | A | 157 | ASP | p.Asp2222Gly | P12259 |
| HOMO SAPIENS | 8 | 35985 | 1lo1 | A | 110 | ALA | p.Ala110Val | O95718 |
| HOMO SAPIENS | 1 | 617 | 2ocp | B | 44 | GLU | p.Glu44Lys | Q16854 |
| HOMO SAPIENS | 5 | 617 | 2ocp | B | 46 | ASN | p.Asn46Ser | Q16854 |
| HOMO SAPIENS | 0 | 617 | 2ocp | B | 59 | THR | p.Thr59Met | Q16854 |
| HOMO SAPIENS | 7 | 617 | 2ocp | B | 142 | ARG | p.Arg142Lys | Q16854 |
| HOMO SAPIENS | 3 | 617 | 2ocp | B | 154 | ASN | p.Asn154Lys | Q16854 |
| HOMO SAPIENS | 3 | 617 | 2ocp | B | 170 | GLN | p.Gln170Arg | Q16854 |
| HOMO SAPIENS | 3 | 617 | 2ocp | B | 227 | GLU | p.Glu227Lys | Q16854 |
| HOMO SAPIENS | 3 | 617 | 2ocp | B | 233 | LYS | p.Lys233Glu | Q16854 |
| HOMO SAPIENS | 3 | 617 | 2ocp | B | 255 | ASP | p.Asp255Tyr | Q16854 |
| HOMO SAPIENS | -1 | 38975 | 5itz | D | 421 | GLN | p.Gln421His | Q9HC77 |
| HOMO SAPIENS | 2 | 6235 | 2f8v | A | 15 | VAL | p.Val15Ile | Q8WZ42 |
| HOMO SAPIENS | 0 | 6235 | 2f8v | A | 54 | VAL | p.Val54Met | Q8WZ42 |
| HOMO SAPIENS | 0 | 6235 | 2f8v | A | 59 | SER | p.Ser59Arg | Q8WZ42 |
| HOMO SAPIENS | 0 | 6235 | 2f8v | A | 60 | ASP | p.Asp60Tyr | Q8WZ42 |
| HOMO SAPIENS | 0 | 12630 | 1ext | B | 12 | ASP | p.Asp41Glu | P19438 |
| HOMO SAPIENS | 3 | 12630 | 1ext | B | 30 | CYS | p.Cys59Arg | P19438 |
| HOMO SAPIENS | 3 | 12630 | 1ext | B | 30 | CYS | p.Cys59Ser | P19438 |
| HOMO SAPIENS | 5 | 12630 | 1ext | B | 33 | CYS | p.Cys62Tyr | P19438 |
| HOMO SAPIENS | 5 | 12630 | 1ext | B | 33 | CYS | p.Cys62Gly | P19438 |
| HOMO SAPIENS | 1 | 12630 | 1ext | B | 46 | PRO | p.Pro75Leu | P19438 |
| HOMO SAPIENS | 3 | 12630 | 1ext | B | 50 | THR | p.Thr79Met | P19438 |
| HOMO SAPIENS | 6 | 12630 | 1ext | B | 52 | CYS | p.Cys81Phe | P19438 |
| HOMO SAPIENS | 6 | 12630 | 1ext | B | 70 | CYS | p.Cys99Ser | P19438 |
| HOMO SAPIENS | 5 | 12630 | 1ext | B | 88 | CYS | p.Cys117Arg | P19438 |
| HOMO SAPIENS | 5 | 12630 | 1ext | B | 88 | CYS | p.Cys117Tyr | P19438 |
| HOMO SAPIENS | -1 | 12630 | 1ext | B | 170 | ILE | p.Ile199Asn | P19438 |
| HOMO SAPIENS | 0 | 994 | 1aos | B | 95 | ARG | p.Arg95Cys | P04424 |
| HOMO SAPIENS | 0 | 994 | 1aos | B | 111 | ARG | p.Arg111Gln | P04424 |
| HOMO SAPIENS | 0 | 994 | 1aos | B | 131 | THR | p.Thr131Met | P04424 |
| HOMO SAPIENS | 6 | 994 | 1aos | B | 178 | VAL | p.Val178Met | P04424 |
| HOMO SAPIENS | 9 | 994 | 1aos | B | 182 | ARG | p.Arg182Gln | P04424 |
| HOMO SAPIENS | 9 | 994 | 1aos | B | 186 | ARG | p.Arg186Trp | P04424 |
| HOMO SAPIENS | 8 | 994 | 1aos | B | 193 | ARG | p.Arg193Gln | P04424 |
| HOMO SAPIENS | 5 | 994 | 1aos | B | 217 | ARG | p.Arg217Ter | P04424 |
| HOMO SAPIENS | 0 | 994 | 1aos | B | 286 | GLN | p.Gln286Arg | P04424 |
| HOMO SAPIENS | 5 | 994 | 1aos | B | 379 | ARG | p.Arg379Cys | P04424 |
| HOMO SAPIENS | 4 | 16066 | 5mv9 | A | 1719 | TYR | p.Tyr1719Cys | Q13402 |
| HOMO SAPIENS | -1 | 16066 | 5mv9 | A | 1739 | ARG | p.Arg1739Ter | Q13402 |
| HOMO SAPIENS | 4 | 16066 | 5mv9 | A | 1775 | ILE | p.Ile1775Thr | Q13402 |
| HOMO SAPIENS | 2 | 16066 | 5mv9 | A | 1786 | SER | p.Ser1786Thr | Q13402 |
| HOMO SAPIENS | 4 | 16066 | 5mv9 | A | 1798 | GLN | p.Gln1798Ter | Q13402 |
| HOMO SAPIENS | 0 | 16066 | 5mv9 | A | 1858 | LEU | p.Leu1858Pro | Q13402 |
| HOMO SAPIENS | 6 | 16066 | 5mv9 | A | 1873 | ARG | p.Arg1873Trp | Q13402 |
| HOMO SAPIENS | 6 | 16066 | 5mv9 | A | 1873 | ARG | p.Arg1873Leu | Q13402 |
| HOMO SAPIENS | 6 | 16066 | 5mv9 | A | 1883 | ARG | p.Arg1883Gln | Q13402 |
| HOMO SAPIENS | 4 | 16066 | 5mv9 | A | 1887 | PRO | p.Pro1887Leu | Q13402 |

| | | | | | | | | |
|---|---|---|---|---|---|---|---|---|
| HOMO SAPIENS | 5 | 16066 | 5mv9 | A | 1942 | GLY | p.Gly1942Ter | Q13402 |
| HOMO SAPIENS | 5 | 16066 | 5mv9 | A | 1954 | LEU | p.Leu1954Ile | Q13402 |
| HOMO SAPIENS | 5 | 16066 | 5mv9 | A | 1956 | VAL | p.Val1956Ile | Q13402 |
| HOMO SAPIENS | 5 | 16066 | 5mv9 | A | 1967 | ARG | p.Arg1967Ter | Q13402 |
| HOMO SAPIENS | 5 | 16066 | 5mv9 | A | 1990 | GLN | p.Gln1990Ter | Q13402 |
| HOMO SAPIENS | 2 | 16066 | 5mv9 | A | 2005 | LYS | p.Lys2005Glu | Q13402 |
| HOMO SAPIENS | 1 | 16066 | 5mv9 | A | 2072 | VAL | p.Val2072Ile | Q13402 |
| HOMO SAPIENS | 4 | 16066 | 5mv9 | A | 2109 | THR | p.Thr2109Ile | Q13402 |
| HOMO SAPIENS | 4 | 16066 | 5mv9 | A | 2142 | ASP | p.Asp2142Asn | Q13402 |
| HOMO SAPIENS | -1 | 16066 | 5mv9 | A | 2214 | GLY | p.Gly2214Ser | Q13402 |
| HOMO SAPIENS | 7 | 27942 | 1deb | A | 24 | ARG | p.Arg24Ter | P25054 |
| HOMO SAPIENS | 0 | 876 | 5hnr | B | 12 | PHE | p.Phe12Leu | P13716 |
| HOMO SAPIENS | 0 | 876 | 5hnr | B | 59 | LYS | p.Lys59Asn | P13716 |
| HOMO SAPIENS | -1 | 876 | 5hnr | B | 133 | GLY | p.Gly133Arg | P13716 |
| HOMO SAPIENS | 2 | 876 | 5hnr | B | 240 | ARG | p.Arg240Trp | P13716 |
| HOMO SAPIENS | 0 | 876 | 5hnr | B | 274 | ALA | p.Ala274Thr | P13716 |
| HOMO SAPIENS | 0 | 876 | 5hnr | B | 275 | VAL | p.Val275Met | P13716 |
| HOMO SAPIENS | 0 | 1142 | 3qe2 | A | 503 | VAL | p.Ala500Val | P16435 |
| HOMO SAPIENS | 8 | 14358 | 4dkd | C | 32 | VAL | p.Val32Gly | P07333 |
| HOMO SAPIENS | 0 | 14358 | 4dkd | C | 60 | LEU | p.Leu60Val | P07333 |
| HOMO SAPIENS | 7 | 14358 | 4dkd | C | 75 | THR | p.Thr75Ile | P07333 |
| HOMO SAPIENS | -1 | 14358 | 4dkd | C | 90 | PRO | p.Pro90Thr | P07333 |
| HOMO SAPIENS | 0 | 14358 | 4dkd | C | 106 | ARG | p.Arg106Trp | P07333 |
| HOMO SAPIENS | 0 | 14358 | 4dkd | C | 241 | ASN | p.Asn241Asp | P07333 |
| HOMO SAPIENS | 0 | 14358 | 4dkd | C | 245 | ALA | p.Ala245Ser | P07333 |
| HOMO SAPIENS | 0 | 14358 | 4dkd | C | 255 | ASN | p.Asn255Ile | P07333 |
| HOMO SAPIENS | 0 | 14358 | 4dkd | C | 268 | VAL | p.Val268Ile | P07333 |
| HOMO SAPIENS | 0 | 14358 | 4dkd | C | 279 | VAL | p.Val279Met | P07333 |
| HOMO SAPIENS | 0 | 14358 | 4dkd | C | 294 | ARG | p.Arg294Gln | P07333 |
| HOMO SAPIENS | 5 | 8968 | 3ppy | A | 1565 | VAL | p.Val1565Leu | P04275 |
| HOMO SAPIENS | 4 | 8968 | 3ppy | A | 1584 | TYR | p.Tyr1584Cys | P04275 |
| HOMO SAPIENS | 5 | 5341 | 3k3n | A | 279 | PHE | p.Phe315Ser | Q9UPP1 |
| HOMO SAPIENS | 0 | 29463 | 1q7l | A | 197 | ARG | p.Arg197Trp | Q03154 |
| HOMO SAPIENS | 0 | 33761 | 2leh | B | 30 | ASP | p.Asp30Asn | Q16637 |
| HOMO SAPIENS | 6 | 33761 | 2leh | B | 44 | ASP | p.Asp44Val | Q16637 |
| HOMO SAPIENS | -1 | 12249 | 2pnr | C | 224 | GLU | p.Glu310Lys | P10515 |
| HOMO SAPIENS | -1 | 12249 | 2pnr | C | 232 | VAL | p.Val318Ala | P10515 |
| HOMO SAPIENS | -1 | 7 | 5whk | B | -9 | ALA | p.Ala11Pro | P61769 |
| HOMO SAPIENS | 8 | 20298 | 4euw | A | 154 | PHE | p.Phe154Leu | P48436 |
| HOMO SAPIENS | 9 | 20298 | 4euw | A | 158 | ALA | p.Ala158Thr | P48436 |
| HOMO SAPIENS | 5 | 20298 | 4euw | A | 165 | HIS | p.His165Tyr | P48436 |
| HOMO SAPIENS | 3 | 20298 | 4euw | A | 169 | HIS | p.His169Gln | P48436 |
| HOMO SAPIENS | 0 | 20298 | 4euw | A | 173 | LYS | p.Lys173Glu | P48436 |
| HOMO SAPIENS | 1 | 16465 | 2jgx | A | 402 | TYR | p.Tyr402His | P08603 |
| HOMO SAPIENS | 0 | 8993 | 3v8o | A | 659 | LEU | p.Leu659Arg | Q14315 |
| HOMO SAPIENS | 6 | 8993 | 3v8o | A | 693 | ASP | p.Asp693Ala | Q14315 |
| HOMO SAPIENS | 0 | 8993 | 3v8o | A | 709 | ALA | p.Ala709Thr | Q14315 |
| HOMO SAPIENS | 2 | 3852 | 5es4 | B | 106 | ASP | p.Asp128Asn | P05107 |
| HOMO SAPIENS | 2 | 3852 | 5es4 | B | 116 | SER | p.Ser138Pro | P05107 |
| HOMO SAPIENS | 1 | 3852 | 5es4 | B | 127 | LEU | p.Leu149Pro | P05107 |
| HOMO SAPIENS | 0 | 3852 | 5es4 | B | 147 | GLY | p.Gly169Arg | P05107 |
| HOMO SAPIENS | 3 | 3852 | 5es4 | B | 156 | PRO | p.Pro178Leu | P05107 |
| HOMO SAPIENS | 0 | 3852 | 5es4 | B | 174 | LYS | p.Lys196Thr | P05107 |
| HOMO SAPIENS | 2 | 3852 | 5es4 | B | 251 | GLY | p.Gly273Arg | P05107 |

| | | | | | | | | |
|---|---|---|---|---|---|---|---|---|
| HOMO SAPIENS | 2 | 3852 | 5es4 | B | 329 | ASN | p.Asn351Ser | P05107 |
| HOMO SAPIENS | 2 | 3852 | 5es4 | B | 571 | ARG | p.Arg593Cys | P05107 |
| HOMO SAPIENS | 7 | 5962 | 5fn2 | B | 146 | MET | p.Met146Leu | P49768 |
| HOMO SAPIENS | 3 | 5962 | 5fn2 | B | 166 | LEU | p.Leu166Pro | P49768 |
| HOMO SAPIENS | 1 | 5962 | 5fn2 | B | 233 | MET | p.Met233Leu | P49768 |
| HOMO SAPIENS | 1 | 5962 | 5fn2 | B | 261 | VAL | p.Val261Gly | P49768 |
| HOMO SAPIENS | 1 | 5962 | 5fn2 | B | 266 | GLY | p.Gly266Ser | P49768 |
| HOMO SAPIENS | -1 | 5962 | 5fn2 | B | 333 | ASP | p.Asp333Gly | P49768 |
| HOMO SAPIENS | 6 | 1918 | 3dlx | A | 58 | THR | p.Thr58Met | P55809 |
| HOMO SAPIENS | 5 | 1918 | 3dlx | A | 133 | VAL | p.Val133Glu | P55809 |
| HOMO SAPIENS | 5 | 1918 | 3dlx | A | 219 | GLY | p.Gly219Glu | P55809 |
| HOMO SAPIENS | 0 | 1918 | 3dlx | A | 221 | VAL | p.Val221Met | P55809 |
| HOMO SAPIENS | 5 | 1918 | 3dlx | A | 283 | SER | p.Ser283Ter | P55809 |
| HOMO SAPIENS | 4 | 1918 | 3dlx | A | 324 | GLY | p.Gly324Glu | P55809 |
| HOMO SAPIENS | 0 | 1918 | 3dlx | A | 456 | CYS | p.Cys456Phe | P55809 |
| HOMO SAPIENS | 0 | 9010 | 2wfr | A | 162 | LEU | p.Leu162Pro | O43323 |
| HOMO SAPIENS | 5 | 3770 | 5n38 | A | 56 | VAL | p.Val56Glu | O60260 |
| HOMO SAPIENS | 1 | 3770 | 5n38 | A | 161 | LYS | p.Lys161Asn | O60260 |
| HOMO SAPIENS | 1 | 3770 | 5n38 | A | 167 | SER | p.Ser167Asn | O60260 |
| HOMO SAPIENS | 1 | 3770 | 5n38 | A | 211 | LYS | p.Lys211Asn | O60260 |
| HOMO SAPIENS | 1 | 3770 | 5n38 | A | 212 | CYS | p.Cys212Tyr | O60260 |
| HOMO SAPIENS | 1 | 3770 | 5n38 | A | 275 | ARG | p.Arg275Trp | O60260 |
| HOMO SAPIENS | 2 | 3770 | 5n38 | A | 334 | ARG | p.Arg334Cys | O60260 |
| HOMO SAPIENS | 1 | 3770 | 5n38 | A | 366 | ARG | p.Arg366Trp | O60260 |
| HOMO SAPIENS | -1 | 3770 | 5n38 | A | 380 | VAL | p.Val380Leu | O60260 |
| HOMO SAPIENS | 1 | 3770 | 5n38 | A | 394 | ASP | p.Asp394Asn | O60260 |
| HOMO SAPIENS | 1 | 3770 | 5n38 | A | 430 | GLY | p.Gly430Asp | O60260 |
| HOMO SAPIENS | -1 | 2467 | 4xx1 | A | 10 | PRO | p.Pro34Leu | P04180 |
| HOMO SAPIENS | 4 | 2467 | 4xx1 | A | 83 | TYR | p.Tyr107Ter | P04180 |
| HOMO SAPIENS | 4 | 2467 | 4xx1 | A | 93 | ALA | p.Ala117Thr | P04180 |
| HOMO SAPIENS | 4 | 2467 | 4xx1 | A | 123 | THR | p.Thr147Ile | P04180 |
| HOMO SAPIENS | 4 | 2467 | 4xx1 | A | 135 | ARG | p.Arg159Trp | P04180 |
| HOMO SAPIENS | 3 | 2467 | 4xx1 | A | 158 | ARG | p.Arg182Cys | P04180 |
| HOMO SAPIENS | 0 | 2467 | 4xx1 | A | 209 | LEU | p.Leu233Pro | P04180 |
| HOMO SAPIENS | 1 | 2467 | 4xx1 | A | 228 | ASN | p.Asn252Lys | P04180 |
| HOMO SAPIENS | 4 | 2467 | 4xx1 | A | 252 | MET | p.Met276Lys | P04180 |
| HOMO SAPIENS | 3 | 2467 | 4xx1 | A | 293 | MET | p.Met317Ile | P04180 |
| HOMO SAPIENS | 5 | 2467 | 4xx1 | A | 321 | THR | p.Thr345Met | P04180 |
| HOMO SAPIENS | 9 | 2467 | 4xx1 | A | 347 | THR | p.Thr371Met | P04180 |
| HOMO SAPIENS | 1 | 5834 | 3qrf | G | 371 | PHE | p.Phe371Cys | Q9BZS1 |
| HOMO SAPIENS | 1 | 5834 | 3qrf | G | 373 | PHE | p.Phe373Ala | Q9BZS1 |
| HOMO SAPIENS | 9 | 5834 | 3qrf | G | 384 | ALA | p.Ala384Thr | Q9BZS1 |
| HOMO SAPIENS | 6 | 5834 | 3qrf | G | 397 | ARG | p.Arg397Trp | Q9BZS1 |
| HOMO SAPIENS | -1 | 11869 | 4km8 | A | 15 | PRO | p.Pro15Leu | Q9UMX1 |
| HOMO SAPIENS | 5 | 262 | 4kbo | A | 126 | ARG | p.Arg126Trp | P38646 |
| HOMO SAPIENS | 3 | 262 | 4kbo | A | 128 | TYR | p.Tyr128Cys | P38646 |
| HOMO SAPIENS | -1 | 11347 | 4uf9 | D | 117 | ALA | p.Ala117Val | Q13501 |
| HOMO SAPIENS | 0 | 2173 | 4ah6 | A | 45 | SER | p.Ser45Gly | Q6PI48 |
| HOMO SAPIENS | 2 | 2173 | 4ah6 | A | 152 | CYS | p.Cys152Phe | Q6PI48 |
| HOMO SAPIENS | 4 | 2173 | 4ah6 | A | 179 | ARG | p.Arg179His | Q6PI48 |
| HOMO SAPIENS | 0 | 2173 | 4ah6 | A | 196 | LYS | p.Lys196Arg | Q6PI48 |
| HOMO SAPIENS | 8 | 2173 | 4ah6 | A | 263 | ARG | p.Arg263Gln | Q6PI48 |
| HOMO SAPIENS | 1 | 2173 | 4ah6 | A | 338 | GLY | p.Gly338Glu | Q6PI48 |
| HOMO SAPIENS | 0 | 2173 | 4ah6 | A | 425 | GLU | p.Glu425Ter | Q6PI48 |

| | | | | | | | | |
|---|---|---|---|---|---|---|---|---|
| HOMO SAPIENS | 1 | 2173 | 4ah6 | A | 548 | LEU | p.Leu548Met | Q6PI48 |
| HOMO SAPIENS | 0 | 2173 | 4ah6 | A | 609 | ARG | p.Arg609Trp | Q6PI48 |
| HOMO SAPIENS | 2 | 2173 | 4ah6 | A | 613 | LEU | p.Leu613Phe | Q6PI48 |
| HOMO SAPIENS | 0 | 2173 | 4ah6 | A | 626 | LEU | p.Leu626Gln | Q6PI48 |
| HOMO SAPIENS | 0 | 2173 | 4ah6 | A | 626 | LEU | p.Leu626Val | Q6PI48 |
| HOMO SAPIENS | 7 | 3203 | 2cn1 | A | 87 | ASP | p.Asp137Val | Q9H0P0 |
| HOMO SAPIENS | 8 | 3203 | 2cn1 | A | 166 | GLN | p.Gln216Ter | Q9H0P0 |
| HOMO SAPIENS | 8 | 3203 | 2cn1 | A | 179 | ASN | p.Asn229Ser | Q9H0P0 |
| HOMO SAPIENS | 7 | 3203 | 2cn1 | A | 230 | GLY | p.Gly280Arg | Q9H0P0 |
| HOMO SAPIENS | 2 | 3424 | 2a1t | R | 116 | GLY | p.Gly116Arg | P13804 |
| HOMO SAPIENS | 2 | 3424 | 2a1t | R | 157 | VAL | p.Val157Gly | P13804 |
| HOMO SAPIENS | 2 | 3424 | 2a1t | R | 171 | THR | p.Thr171Ile | P13804 |
| HOMO SAPIENS | 0 | 3424 | 2a1t | R | 178 | THR | p.Thr178Arg | P13804 |
| HOMO SAPIENS | 0 | 3424 | 2a1t | R | 194 | VAL | p.Val194Met | P13804 |
| HOMO SAPIENS | 1 | 3424 | 2a1t | R | 209 | ARG | p.Arg209Ter | P13804 |
| HOMO SAPIENS | 6 | 3424 | 2a1t | R | 266 | THR | p.Thr266Met | P13804 |
| HOMO SAPIENS | 0 | 27598 | 1g1c | A | 26 | ARG | p.Arg2098Gly | Q8WZ42 |
| HOMO SAPIENS | 1 | 27598 | 1g1c | A | 46 | ILE | p.Ile2118Thr | Q8WZ42 |
| HOMO SAPIENS | 0 | 32885 | 4bsk | A | 149 | ASN | p.Asn149Asp | P35916 |
| HOMO SAPIENS | -1 | 8074 | 3ghg | B | 17 | ARG | p.Arg47Ter | P02675 |
| HOMO SAPIENS | -1 | 447 | 5g4p | A | 190 | ALA | p.Ala190Thr | P84074 |
| HOMO SAPIENS | 0 | 10761 | 5u1d | B | 379 | VAL | p.Val379Ile | Q03519 |
| HOMO SAPIENS | 0 | 10761 | 5u1d | B | 665 | THR | p.Thr665Ala | Q03519 |
| HOMO SAPIENS | 5 | 29251 | 5fn7 | A | 79 | ILE | p.Ile294Leu | P08575 |
| HOMO SAPIENS | -1 | 27727 | 4ky9 | A | 56 | LYS | p.Lys56Glu | P02730 |
| HOMO SAPIENS | 2 | 27727 | 4ky9 | A | 72 | GLU | p.Glu72Asp | P02730 |
| HOMO SAPIENS | 2 | 27727 | 4ky9 | A | 90 | GLU | p.Glu90Lys | P02730 |
| HOMO SAPIENS | 1 | 27727 | 4ky9 | A | 130 | GLY | p.Gly130Arg | P02730 |
| HOMO SAPIENS | 2 | 27727 | 4ky9 | A | 150 | ARG | p.Arg150Ter | P02730 |
| HOMO SAPIENS | 0 | 27727 | 4ky9 | A | 327 | PRO | p.Pro327Arg | P02730 |
| HOMO SAPIENS | 2 | 13202 | 2fxo | B | 870 | ARG | p.Arg870His | P12883 |
| HOMO SAPIENS | 2 | 13202 | 2fxo | B | 870 | ARG | p.Arg870Cys | P12883 |
| HOMO SAPIENS | 4 | 13202 | 2fxo | B | 904 | ARG | p.Arg904Cys | P12883 |
| HOMO SAPIENS | 4 | 13202 | 2fxo | B | 906 | ASP | p.Asp906Gly | P12883 |
| HOMO SAPIENS | 2 | 13202 | 2fxo | B | 908 | LEU | p.Leu908Val | P12883 |
| HOMO SAPIENS | 2 | 13202 | 2fxo | B | 923 | ASN | p.Asn923Lys | P12883 |
| HOMO SAPIENS | 7 | 13202 | 2fxo | B | 935 | GLU | p.Glu935Lys | P12883 |
| HOMO SAPIENS | 7 | 13202 | 2fxo | B | 935 | GLU | p.Glu935Val | P12883 |
| HOMO SAPIENS | 8 | 13202 | 2fxo | B | 949 | GLU | p.Glu949Lys | P12883 |
| HOMO SAPIENS | 0 | 13200 | 2a1s | A | 349 | ARG | p.Arg349Trp | O95453 |
| HOMO SAPIENS | 0 | 13200 | 2a1s | A | 421 | LYS | p.Lys421Arg | O95453 |
| HOMO SAPIENS | 6 | 3196 | 5xtb | T | 115 | CYS | p.Cys115Tyr | O75380 |
| HOMO SAPIENS | 0 | 1108 | 5ukf | A | 89 | ARG | p.Arg89Gln | Q99986 |
| HOMO SAPIENS | 0 | 1108 | 5ukf | A | 119 | HIS | p.His119Arg | Q99986 |
| HOMO SAPIENS | 0 | 1108 | 5ukf | A | 133 | ARG | p.Arg133Cys | Q99986 |
| HOMO SAPIENS | 0 | 1108 | 5ukf | A | 236 | VAL | p.Val236Leu | Q99986 |
| HOMO SAPIENS | 0 | 1108 | 5ukf | A | 321 | ARG | p.Arg321Cys | Q99986 |
| HOMO SAPIENS | -1 | 1108 | 5ukf | A | 358 | ARG | p.Arg358Ter | Q99986 |
| HOMO SAPIENS | 0 | 5842 | 4yli | C | 204 | GLY | p.Gly204Ser | Q9BWP8 |
| HOMO SAPIENS | -1 | 31362 | 4wqo | A | 25 | PRO | p.Pro25Leu | P40337 |
| HOMO SAPIENS | 2 | 10107 | 2j8h | A | 34 | VAL | p.Val31886Ile | Q8WZ42 |
| HOMO SAPIENS | 8 | 20190 | 1wm5 | A | 77 | ARG | p.Arg77Gln | P19878 |
| HOMO SAPIENS | 1 | 20190 | 1wm5 | A | 100 | GLN | p.Gln100Ter | P19878 |
| HOMO SAPIENS | 0 | 20190 | 1wm5 | A | 102 | ARG | p.Arg102Ter | P19878 |

| | | | | | | | | | |
|---|---|---|---|---|---|---|---|---|---|
| HOMO SAPIENS | 0 | 20190 | 1wm5 | A | | 128 | ALA | p.Ala128Val | P19878 |
| HOMO SAPIENS | 0 | 20190 | 1wm5 | A | | 181 | LYS | p.Lys181Arg | P19878 |
| HOMO SAPIENS | 7 | 1278 | 5a2q | a | | 33 | ASP | p.Asp33Asn | P62854 |
| HOMO SAPIENS | 3 | 19515 | 2obd | A | | 11 | ILE | p.Ile28Met | P11597 |
| HOMO SAPIENS | 3 | 19515 | 2obd | A | | 89 | VAL | p.Val106Met | P11597 |
| HOMO SAPIENS | 3 | 19515 | 2obd | A | | 137 | ARG | p.Arg154Trp | P11597 |
| HOMO SAPIENS | 3 | 19515 | 2obd | A | | 139 | ASP | p.Asp156Asn | P11597 |
| HOMO SAPIENS | 3 | 19515 | 2obd | A | | 204 | SER | p.Ser221Arg | P11597 |
| HOMO SAPIENS | 0 | 19515 | 2obd | A | | 297 | GLU | p.Glu314Lys | P11597 |
| HOMO SAPIENS | 0 | 19515 | 2obd | A | | 314 | GLY | p.Gly331Ser | P11597 |
| HOMO SAPIENS | 0 | 19515 | 2obd | A | | 332 | SER | p.Ser349Tyr | P11597 |
| HOMO SAPIENS | 0 | 19515 | 2obd | A | | 368 | VAL | p.Val385Met | P11597 |
| HOMO SAPIENS | 0 | 19515 | 2obd | A | | 373 | ALA | p.Ala390Pro | P11597 |
| HOMO SAPIENS | 3 | 19515 | 2obd | A | | 405 | ILE | p.Val422Ile | P11597 |
| HOMO SAPIENS | 0 | 19515 | 2obd | A | | 451 | ARG | p.Arg468Gln | P11597 |
| HOMO SAPIENS | 0 | 19515 | 2obd | A | | 451 | ARG | p.Arg468Ter | P11597 |
| HOMO SAPIENS | 0 | 6626 | 3hop | A | | 39 | ALA | p.Ala39Gly | P21333 |
| HOMO SAPIENS | 1 | 6626 | 3hop | A | | 82 | GLU | p.Glu82Val | P21333 |
| HOMO SAPIENS | 1 | 6626 | 3hop | A | | 128 | ALA | p.Ala128Val | P21333 |
| HOMO SAPIENS | 2 | 6626 | 3hop | A | | 203 | ASP | p.Asp203Tyr | P21333 |
| HOMO SAPIENS | 1 | 6626 | 3hop | A | | 207 | PRO | p.Pro207Leu | P21333 |
| HOMO SAPIENS | 2 | 6626 | 3hop | A | | 210 | CYS | p.Cys210Phe | P21333 |
| HOMO SAPIENS | 7 | 6626 | 3hop | A | | 254 | GLU | p.Glu254Lys | P21333 |
| HOMO SAPIENS | 4 | 4210 | 2avg | A | | 8 | VAL | p.Val158Met | Q14896 |
| HOMO SAPIENS | 6 | 4210 | 2avg | A | | 15 | GLU | p.Glu165Asp | Q14896 |
| HOMO SAPIENS | 4 | 4210 | 2avg | A | | 27 | ARG | p.Arg177His | Q14896 |
| HOMO SAPIENS | 0 | 4210 | 2avg | A | | 28 | VAL | p.Val178Met | Q14896 |
| HOMO SAPIENS | 6 | 4210 | 2avg | A | | 39 | VAL | p.Val189Ile | Q14896 |
| HOMO SAPIENS | 2 | 4210 | 2avg | A | | 52 | LYS | p.Lys202Gln | Q14896 |
| HOMO SAPIENS | 4 | 4210 | 2avg | A | | 67 | SER | p.Ser217Gly | Q14896 |
| HOMO SAPIENS | 6 | 4210 | 2avg | A | | 69 | VAL | p.Val219Leu | Q14896 |
| HOMO SAPIENS | 2 | 4210 | 2avg | A | | 86 | GLY | p.Ser236Gly | Q14896 |
| HOMO SAPIENS | 4 | 4210 | 2avg | A | | 108 | GLU | p.Glu258Lys | Q14896 |
| HOMO SAPIENS | 0 | 22237 | 1w6k | A | | 588 | GLY | p.Gly588Ser | P48449 |
| HOMO SAPIENS | 0 | 10765 | 4a5w | B | | 98 | GLU | p.Glu98Ala | P13671 |
| HOMO SAPIENS | 0 | 10765 | 4a5w | B | | 596 | ARG | p.Arg596Ter | P13671 |
| HOMO SAPIENS | -1 | 15515 | 3hx3 | A | | 11 | MSE | p.Met10Lys | P12271 |
| HOMO SAPIENS | 8 | 15515 | 3hx3 | A | | 64 | ARG | p.Arg64Gln | P12271 |
| HOMO SAPIENS | 0 | 15515 | 3hx3 | A | | 182 | PHE | p.Phe182Cys | P12271 |
| HOMO SAPIENS | 0 | 15515 | 3hx3 | A | | 216 | ARG | p.Arg216Gln | P12271 |
| HOMO SAPIENS | -1 | 15515 | 3hx3 | A | | 226 | MSE | p.Met226Lys | P12271 |
| HOMO SAPIENS | 0 | 15515 | 3hx3 | A | | 234 | TRP | p.Arg234Trp | P12271 |
| HOMO SAPIENS | 0 | 15515 | 3hx3 | A | | 273 | LEU | p.Leu273Phe | P12271 |
| HOMO SAPIENS | 0 | 15515 | 3hx3 | A | | 292 | THR | p.Thr292Met | P12271 |
| HOMO SAPIENS | 0 | 5152 | 2www | D | | 95 | GLN | p.Gln95Ter | Q8IVH4 |
| HOMO SAPIENS | 0 | 5152 | 2www | D | | 133 | GLN | p.Gln133Ter | Q8IVH4 |
| HOMO SAPIENS | 0 | 5152 | 2www | D | | 145 | ARG | p.Arg145Ter | Q8IVH4 |
| HOMO SAPIENS | 0 | 5152 | 2www | D | | 207 | TYR | p.Tyr207Cys | Q8IVH4 |
| HOMO SAPIENS | 8 | 5152 | 2www | D | | 330 | ARG | p.Arg330Ter | Q8IVH4 |
| HOMO SAPIENS | 0 | 5152 | 2www | D | | 363 | GLN | p.Gln363His | Q8IVH4 |
| HOMO SAPIENS | 1 | 1626 | 1ilr | | 1 | 105 | SER | p.Ser130Arg | P18510 |
| HOMO SAPIENS | -1 | 929 | 5loh | A | | 12 | GLY | p.Gly12Arg | Q96GX5 |
| HOMO SAPIENS | 9 | 3206 | 1x63 | A | | 21 | CYS | p.Cys104Arg | Q13642 |
| HOMO SAPIENS | 8 | 3206 | 1x63 | A | | 39 | TRP | p.Trp122Ser | Q13642 |

| | | | | | | | | |
|---|---|---|---|---|---|---|---|---|
| HOMO SAPIENS | 9 | 3206 | 1x63 | A | 40 | HIS | p.His123Tyr | Q13642 |
| HOMO SAPIENS | 9 | 3206 | 1x63 | A | 40 | HIS | p.His123Leu | Q13642 |
| HOMO SAPIENS | 9 | 3206 | 1x63 | A | 40 | HIS | p.His123Gln | Q13642 |
| HOMO SAPIENS | 3 | 3206 | 1x63 | A | 49 | CYS | p.Cys132Phe | Q13642 |
| HOMO SAPIENS | 3 | 3206 | 1x63 | A | 67 | CYS | p.Cys150Tyr | Q13642 |
| HOMO SAPIENS | 3 | 3206 | 1x63 | A | 70 | CYS | p.Cys153Arg | Q13642 |
| HOMO SAPIENS | 3 | 3206 | 1x63 | A | 70 | CYS | p.Cys153Tyr | Q13642 |
| HOMO SAPIENS | 5 | 1052 | 3ecs | E | 208 | ASN | p.Asn208Tyr | Q14232 |
| HOMO SAPIENS | 2 | 1052 | 3ecs | E | 275 | TYR | p.Tyr275Cys | Q14232 |
| HOMO SAPIENS | 0 | 24253 | 5i35 | A | 271 | ARG | p.Arg271Cys | Q8NI60 |
| HOMO SAPIENS | 0 | 24253 | 5i35 | A | 272 | GLY | p.Gly272Asp | Q8NI60 |
| HOMO SAPIENS | 0 | 24253 | 5i35 | A | 299 | ARG | p.Arg299Trp | Q8NI60 |
| HOMO SAPIENS | 1 | 24253 | 5i35 | A | 339 | ALA | p.Ala339Thr | Q8NI60 |
| HOMO SAPIENS | 0 | 24253 | 5i35 | A | 348 | ARG | p.Arg348Ter | Q8NI60 |
| HOMO SAPIENS | 0 | 24253 | 5i35 | A | 395 | ILE | p.Ile395Met | Q8NI60 |
| HOMO SAPIENS | 0 | 24253 | 5i35 | A | 396 | ASP | p.Asp396Glu | Q8NI60 |
| HOMO SAPIENS | 0 | 24253 | 5i35 | A | 480 | PHE | p.Phe480Leu | Q8NI60 |
| HOMO SAPIENS | 2 | 24253 | 5i35 | A | 511 | THR | p.Thr511Met | Q8NI60 |
| HOMO SAPIENS | 0 | 24253 | 5i35 | A | 514 | TYR | p.Tyr514Cys | Q8NI60 |
| HOMO SAPIENS | 0 | 24253 | 5i35 | A | 538 | ARG | p.Arg538Gln | Q8NI60 |
| HOMO SAPIENS | 0 | 24253 | 5i35 | A | 549 | GLY | p.Gly549Ser | Q8NI60 |
| HOMO SAPIENS | 0 | 24253 | 5i35 | A | 551 | GLU | p.Glu551Lys | Q8NI60 |
| HOMO SAPIENS | -1 | 24253 | 5i35 | A | 555 | MSE | p.Met555Ile | Q8NI60 |
| HOMO SAPIENS | 0 | 24253 | 5i35 | A | 568 | GLU | p.Glu568Ter | Q8NI60 |
| HOMO SAPIENS | 0 | 24253 | 5i35 | A | 602 | PRO | p.Pro602Arg | Q8NI60 |
| HOMO SAPIENS | -1 | 15505 | 4goj | C | 22 | GLY | p.Gly22Val | Q13432 |
| HOMO SAPIENS | 5 | 23902 | 2lrr | A | 765 | HIS | p.His765Gln | P38935 |
| HOMO SAPIENS | 1 | 29650 | 2vj2 | A | 234 | CYS | p.Cys234Tyr | P78504 |
| HOMO SAPIENS | 0 | 29650 | 2vj2 | A | 255 | TYR | p.Tyr255Ter | P78504 |
| HOMO SAPIENS | 3 | 29650 | 2vj2 | A | 274 | GLY | p.Gly274Asp | P78504 |
| HOMO SAPIENS | 1 | 105 | 5he0 | B | 77 | GLY | p.Gly77Ser | P62873 |
| HOMO SAPIENS | 2 | 105 | 5he0 | B | 80 | ILE | p.Ile80Thr | P62873 |
| HOMO SAPIENS | 3 | 4587 | 3rgh | A | 159 | ASP | p.Asp1159Ala | P21333 |
| HOMO SAPIENS | 3 | 4587 | 3rgh | A | 184 | ASP | p.Asp1184Glu | P21333 |
| HOMO SAPIENS | 2 | 4587 | 3rgh | A | 186 | SER | p.Ser1186Leu | P21333 |
| HOMO SAPIENS | 4 | 4587 | 3rgh | A | 188 | ALA | p.Ala1188Thr | P21333 |
| HOMO SAPIENS | 2 | 4587 | 3rgh | A | 199 | SER | p.Ser1199Leu | P21333 |
| HOMO SAPIENS | 0 | 20283 | 4n3w | A | 1278 | GLU | p.Glu1278Lys | Q92793 |
| HOMO SAPIENS | 1 | 14294 | 2d9q | B | 208 | MET | p.Met231Thr | Q99062 |
| HOMO SAPIENS | 6 | 14294 | 2d9q | B | 297 | ASP | p.Asp320Asn | Q99062 |
| HOMO SAPIENS | 0 | 11642 | 3bpt | A | 66 | ARG | p.Arg66Trp | Q6NVY1 |
| HOMO SAPIENS | 0 | 11642 | 3bpt | A | 122 | TYR | p.Tyr122Cys | Q6NVY1 |
| HOMO SAPIENS | 0 | 11642 | 3bpt | A | 345 | GLY | p.Gly345Ser | Q6NVY1 |
| HOMO SAPIENS | 1 | 3942 | 3sby | A | 60 | ASP | p.Asp60His | Q9Y4U1 |
| HOMO SAPIENS | 0 | 3942 | 3sby | A | 111 | ARG | p.Arg111Ter | Q9Y4U1 |
| HOMO SAPIENS | 1 | 3942 | 3sby | A | 116 | LEU | p.Leu116Pro | Q9Y4U1 |
| HOMO SAPIENS | 1 | 3942 | 3sby | A | 130 | TYR | p.Tyr130Cys | Q9Y4U1 |
| HOMO SAPIENS | 1 | 3942 | 3sby | A | 132 | ARG | p.Arg132Ter | Q9Y4U1 |
| HOMO SAPIENS | 1 | 3942 | 3sby | A | 147 | GLY | p.Gly147Ala | Q9Y4U1 |
| HOMO SAPIENS | 1 | 3942 | 3sby | A | 161 | ARG | p.Arg161Gln | Q9Y4U1 |
| HOMO SAPIENS | 1 | 3942 | 3sby | A | 161 | ARG | p.Arg161Ter | Q9Y4U1 |
| HOMO SAPIENS | 1 | 3942 | 3sby | A | 203 | TRP | p.Trp203Ter | Q9Y4U1 |
| HOMO SAPIENS | 1 | 3942 | 3sby | A | 203 | TRP | p.Trp203Ter | Q9Y4U1 |
| HOMO SAPIENS | 1 | 3942 | 3sby | A | 205 | TYR | p.Tyr205Ter | Q9Y4U1 |

| | | | | | | | | | |
|---|---|---|---|---|---|---|---|---|---|
| HOMO SAPIENS | 0 | 38340 | 2mbh | B | | 39 | MET | p.Met39Leu | Q13351 |
| HOMO SAPIENS | 2 | 7360 | 4k30 | A | | 430 | LEU | p.Leu430Pro | Q8N159 |
| HOMO SAPIENS | 2 | 7360 | 4k30 | A | | 433 | GLU | p.Glu433Asp | Q8N159 |
| HOMO SAPIENS | 7 | 7360 | 4k30 | A | | 484 | TRP | p.Trp484Arg | Q8N159 |
| HOMO SAPIENS | 1 | 41903 | 2xs6 | A | | 151 | PRO | p.Pro151Ser | O00459 |
| HOMO SAPIENS | 0 | 172 | 5glh | A | | 183 | ALA | p.Ala183Gly | P24530 |
| HOMO SAPIENS | 0 | 172 | 5glh | A | | 201 | ARG | p.Arg201Ter | P24530 |
| HOMO SAPIENS | 0 | 172 | 5glh | A | | 244 | THR | p.Thr244Met | P24530 |
| HOMO SAPIENS | 0 | 172 | 5glh | A | | 253 | ARG | p.Arg253Ter | P24530 |
| HOMO SAPIENS | 9 | 172 | 5glh | A | | 1017 | ILE | p.Leu17Phe | P00720 |
| HOMO SAPIENS | -1 | 3342 | 1ljm | A | | 1058 | HIS | p.His58Asn | Q01196 |
| HOMO SAPIENS | 4 | 3342 | 1ljm | A | | 1174 | ARG | p.Arg174Gln | Q01196 |
| HOMO SAPIENS | 5 | 25607 | 1p22 | C | | 32 | ASP | p.Asp32Ala | P35222 |
| HOMO SAPIENS | 5 | 25607 | 1p22 | C | | 32 | ASP | p.Asp32Tyr | P35222 |
| HOMO SAPIENS | -1 | 25607 | 1p22 | C | | 33 | SEP | p.Ser33Phe | P35222 |
| HOMO SAPIENS | 5 | 25607 | 1p22 | C | | 34 | GLY | p.Gly34Glu | P35222 |
| HOMO SAPIENS | -1 | 25607 | 1p22 | C | | 37 | SEP | p.Ser37Cys | P35222 |
| HOMO SAPIENS | -1 | 25607 | 1p22 | C | | 41 | THR | p.Thr41Ile | P35222 |
| HOMO SAPIENS | 1 | 48 | 4uow | | 9 | 60 | GLN | p.Arg34111His | Q8WZ42 |
| HOMO SAPIENS | 0 | 48 | 4uow | | 9 | 64 | LEU | p.Val34115Ala | Q8WZ42 |
| HOMO SAPIENS | 0 | 1127 | 5c01 | A | | 638 | ARG | p.Arg638Ter | P29597 |
| HOMO SAPIENS | 0 | 1127 | 5c01 | A | | 684 | ILE | p.Ile684Ser | P29597 |
| HOMO SAPIENS | 0 | 1127 | 5c01 | A | | 703 | ARG | p.Arg703Trp | P29597 |
| HOMO SAPIENS | 5 | 16389 | 2aq0 | B | | 128 | ALA | p.Ala860Asp | Q92889 |
| HOMO SAPIENS | 2 | 16389 | 2aq0 | B | | 141 | ILE | p.Ile873Val | Q92889 |
| HOMO SAPIENS | 5 | 16389 | 2aq0 | B | | 143 | GLU | p.Glu875Gly | Q92889 |
| HOMO SAPIENS | 0 | 16389 | 2aq0 | B | | 180 | GLY | p.Gly912Arg | Q92889 |
| HOMO SAPIENS | 0 | 15816 | 2xrc | A | | 89 | THR | p.Thr107Ala | P05156 |
| HOMO SAPIENS | 0 | 15816 | 2xrc | A | | 133 | ASN | p.Asn151Ser | P05156 |
| HOMO SAPIENS | 0 | 15816 | 2xrc | A | | 185 | THR | p.Thr203Ile | P05156 |
| HOMO SAPIENS | 0 | 15816 | 2xrc | A | | 225 | GLY | p.Gly243Asp | P05156 |
| HOMO SAPIENS | 0 | 15816 | 2xrc | A | | 243 | GLY | p.Gly261Ala | P05156 |
| HOMO SAPIENS | 0 | 15816 | 2xrc | A | | 269 | GLY | p.Gly287Arg | P05156 |
| HOMO SAPIENS | -1 | 15816 | 2xrc | A | | 288 | ILE | p.Ile306Val | P05156 |
| HOMO SAPIENS | -1 | 15816 | 2xrc | A | | 388 | ARG | p.Arg406His | P05156 |
| HOMO SAPIENS | 7 | 15816 | 2xrc | A | | 398 | ILE | p.Ile416Leu | P05156 |
| HOMO SAPIENS | 9 | 15816 | 2xrc | A | | 400 | HIS | p.His418Leu | P05156 |
| HOMO SAPIENS | 0 | 15816 | 2xrc | A | | 423 | LYS | p.Lys441Arg | P05156 |
| HOMO SAPIENS | -1 | 15816 | 2xrc | A | | 530 | GLU | p.Glu548Gln | P05156 |
| HOMO SAPIENS | 0 | 15816 | 2xrc | A | | 535 | PRO | p.Pro553Ala | P05156 |
| HOMO SAPIENS | 0 | 15816 | 2xrc | A | | 536 | GLU | p.Glu554Gly | P05156 |
| HOMO SAPIENS | 1 | 12385 | 4zlp | B | | 1450 | ALA | p.Ala1450Pro | Q9UM47 |
| HOMO SAPIENS | 7 | 12385 | 4zlp | B | | 1482 | GLY | p.Gly1482Arg | Q9UM47 |
| HOMO SAPIENS | 2 | 12385 | 4zlp | B | | 1518 | LEU | p.Leu1518Met | Q9UM47 |
| HOMO SAPIENS | 0 | 12385 | 4zlp | B | | 1560 | ARG | p.Arg1560Pro | Q9UM47 |
| HOMO SAPIENS | 0 | 1227 | 4a04 | A | | 148 | PHE | p.Phe148Tyr | O43435 |
| HOMO SAPIENS | 4 | 1227 | 4a04 | A | | 194 | HIS | p.His194Gln | O43435 |
| HOMO SAPIENS | 2 | 8017 | 3ei4 | B | | 244 | LYS | p.Lys244Glu | Q92466 |
| HOMO SAPIENS | 1 | 8017 | 3ei4 | B | | 273 | ARG | p.Arg273His | Q92466 |
| HOMO SAPIENS | 3 | 36112 | 2k8f | B | | 9 | SER | p.Ser9Arg | P04637 |
| HOMO SAPIENS | 2 | 12880 | 5ggf | A | | 265 | ARG | p.Arg265His | Q8WZA1 |
| HOMO SAPIENS | 2 | 12880 | 5ggf | A | | 287 | ILE | p.Ile287Ser | Q8WZA1 |
| HOMO SAPIENS | 2 | 12880 | 5ggf | A | | 311 | ARG | p.Arg311Gln | Q8WZA1 |
| HOMO SAPIENS | 2 | 12880 | 5ggf | A | | 425 | TRP | p.Trp425Ser | Q8WZA1 |

| | | | | | | | | |
|---|---|---|---|---|---|---|---|---|
| HOMO SAPIENS | 2 | 12880 | 5ggf | A | 442 | ARG | p.Arg442Cys | Q8WZA1 |
| HOMO SAPIENS | 2 | 12880 | 5ggf | A | 448 | GLY | p.Gly448Arg | Q8WZA1 |
| HOMO SAPIENS | 0 | 12880 | 5ggf | A | 490 | CYS | p.Cys490Tyr | Q8WZA1 |
| HOMO SAPIENS | 0 | 12880 | 5ggf | A | 493 | PRO | p.Pro493Arg | Q8WZA1 |
| HOMO SAPIENS | 3 | 12880 | 5ggf | A | 550 | SER | p.Ser550Asn | Q8WZA1 |
| HOMO SAPIENS | 1 | 12880 | 5ggf | A | 556 | ASP | p.Asp556Asn | Q8WZA1 |
| HOMO SAPIENS | 0 | 12880 | 5ggf | A | 580 | ARG | p.Arg580Ter | Q8WZA1 |
| HOMO SAPIENS | 2 | 12880 | 5ggf | A | 605 | ARG | p.Arg605Pro | Q8WZA1 |
| HOMO SAPIENS | 0 | 12880 | 5ggf | A | 623 | VAL | p.Met623Val | Q8WZA1 |
| HOMO SAPIENS | 1 | 10187 | 4qw2 | A | 145 | ALA | p.Ala145Ser | Q06787 |
| HOMO SAPIENS | 1 | 10187 | 4qw2 | A | 174 | ASN | p.Asn174Lys | Q06787 |
| HOMO SAPIENS | 0 | 24002 | 2m0t | A | 110 | LEU | p.Leu110Val | O14745 |
| HOMO SAPIENS | 9 | 3126 | 4mjh | A | 99 | LEU | p.Leu99Met | P04792 |
| HOMO SAPIENS | 1 | 3126 | 4mjh | A | 127 | ARG | p.Arg127Trp | P04792 |
| HOMO SAPIENS | 1 | 3126 | 4mjh | A | 127 | ARG | p.Arg127Leu | P04792 |
| HOMO SAPIENS | 1 | 3126 | 4mjh | A | 135 | SER | p.Ser135Phe | P04792 |
| HOMO SAPIENS | 2 | 3126 | 4mjh | A | 136 | ARG | p.Arg136Trp | P04792 |
| HOMO SAPIENS | 1 | 3126 | 4mjh | A | 140 | ARG | p.Arg140Gly | P04792 |
| HOMO SAPIENS | 0 | 3126 | 4mjh | A | 151 | THR | p.Thr151Ile | P04792 |
| HOMO SAPIENS | -1 | 15780 | 2jlp | A | 213 | ARG | p.Arg231Gly | P08294 |
| HOMO SAPIENS | 8 | 3578 | 1apz | D | 203 | GLY | p.Gly226Asp | P20933 |
| HOMO SAPIENS | 1 | 3578 | 1apz | D | 279 | GLY | p.Gly302Arg | P20933 |
| HOMO SAPIENS | 0 | 3578 | 1apz | D | 283 | CYS | p.Cys306Arg | P20933 |
| HOMO SAPIENS | 5 | 1335 | 5xtd | C | 115 | ARG | p.Arg115Cys | O75251 |
| HOMO SAPIENS | 0 | 1335 | 5xtd | C | 122 | VAL | p.Val122Met | O75251 |
| HOMO SAPIENS | 2 | 5063 | 2le4 | A | 37 | ARG | p.Arg74Pro | P48431 |
| HOMO SAPIENS | 1 | 25870 | 2o10 | A | 44 | LEU | p.Leu44Pro | P50461 |
| HOMO SAPIENS | 4 | 25870 | 2o10 | A | 58 | CYS | p.Cys58Gly | P50461 |
| HOMO SAPIENS | 0 | 18748 | 5fii | C | 39 | PHE | p.Phe39Leu | P00439 |
| HOMO SAPIENS | 6 | 18748 | 5fii | C | 48 | LEU | p.Leu48Ser | P00439 |
| HOMO SAPIENS | 2 | 18748 | 5fii | C | 65 | ILE | p.Ile65Asn | P00439 |
| HOMO SAPIENS | 3 | 18748 | 5fii | C | 95 | ILE | p.Ile95Phe | P00439 |
| HOMO SAPIENS | 1 | 14261 | 2vay | B | 1539 | ARG | p.Arg1539Cys | Q13698 |
| HOMO SAPIENS | 0 | 2494 | 4dx1 | A | 183 | GLU | p.Glu183Asp | Q9HBA0 |
| HOMO SAPIENS | 0 | 2494 | 4dx1 | A | 217 | ALA | p.Ala217Ser | Q9HBA0 |
| HOMO SAPIENS | 0 | 2494 | 4dx1 | A | 217 | ALA | p.Ala217Val | Q9HBA0 |
| HOMO SAPIENS | 4 | 2494 | 4dx1 | A | 224 | ARG | p.Arg224Trp | Q9HBA0 |
| HOMO SAPIENS | 4 | 2494 | 4dx1 | A | 254 | VAL | p.Val254Leu | Q9HBA0 |
| HOMO SAPIENS | 0 | 2494 | 4dx1 | A | 269 | ARG | p.Arg269Cys | Q9HBA0 |
| HOMO SAPIENS | 0 | 2494 | 4dx1 | A | 269 | ARG | p.Arg269His | Q9HBA0 |
| HOMO SAPIENS | 0 | 2494 | 4dx1 | A | 278 | GLU | p.Glu278Lys | Q9HBA0 |
| HOMO SAPIENS | 0 | 2494 | 4dx1 | A | 315 | ARG | p.Arg315Trp | Q9HBA0 |
| HOMO SAPIENS | 0 | 2494 | 4dx1 | A | 316 | ARG | p.Arg316Cys | Q9HBA0 |
| HOMO SAPIENS | 0 | 2494 | 4dx1 | A | 331 | ILE | p.Ile331Phe | Q9HBA0 |
| HOMO SAPIENS | 0 | 2494 | 4dx1 | A | 333 | ASP | p.Asp333Gly | Q9HBA0 |
| HOMO SAPIENS | 1 | 2494 | 4dx1 | A | 365 | VAL | p.Val365Met | Q9HBA0 |
| HOMO SAPIENS | 0 | 23998 | 2l1r | B | 161 | ARG | p.Arg162Trp | P19429 |
| HOMO SAPIENS | 0 | 8354 | 4f0x | D | 187 | SER | p.Ser187Ter | O95822 |
| HOMO SAPIENS | 2 | 8354 | 4f0x | D | 259 | GLY | p.Gly259Ala | O95822 |
| HOMO SAPIENS | 2 | 8354 | 4f0x | D | 284 | THR | p.Thr284Ala | O95822 |
| HOMO SAPIENS | 3 | 8354 | 4f0x | D | 309 | LYS | p.Lys309Gln | O95822 |
| HOMO SAPIENS | 0 | 21182 | 4apo | A | 228 | GLN | p.Gln228Lys | O00170 |
| HOMO SAPIENS | 1 | 21182 | 4apo | A | 307 | GLN | p.Gln307Arg | O00170 |
| HOMO SAPIENS | 0 | 2287 | 5of4 | B | 112 | ARG | p.Arg112His | P18074 |

| | | | | | | | | |
|---|---|---|---|---|---|---|---|---|
| HOMO SAPIENS | 0 | 2287 | 5of4 | B | 312 | ASP | p.Asp312Asn | P18074 |
| HOMO SAPIENS | 3 | 2287 | 5of4 | B | 461 | LEU | p.Leu461Val | P18074 |
| HOMO SAPIENS | 0 | 2287 | 5of4 | B | 485 | LEU | p.Leu485Pro | P18074 |
| HOMO SAPIENS | 1 | 2287 | 5of4 | B | 541 | SER | p.Ser541Arg | P18074 |
| HOMO SAPIENS | 0 | 2287 | 5of4 | B | 616 | ARG | p.Arg616Pro | P18074 |
| HOMO SAPIENS | 0 | 2287 | 5of4 | B | 616 | ARG | p.Arg616Trp | P18074 |
| HOMO SAPIENS | 1 | 2287 | 5of4 | B | 658 | ARG | p.Arg658Cys | P18074 |
| HOMO SAPIENS | 1 | 2287 | 5of4 | B | 681 | ASP | p.Asp681Asn | P18074 |
| HOMO SAPIENS | 1 | 2287 | 5of4 | B | 683 | ARG | p.Arg683Trp | P18074 |
| HOMO SAPIENS | 0 | 2287 | 5of4 | B | 713 | GLY | p.Gly713Arg | P18074 |
| HOMO SAPIENS | 0 | 2287 | 5of4 | B | 717 | ALA | p.Ala717Gly | P18074 |
| HOMO SAPIENS | 0 | 2287 | 5of4 | B | 722 | ARG | p.Arg722Trp | P18074 |
| HOMO SAPIENS | 0 | 2287 | 5of4 | B | 725 | ALA | p.Ala725Pro | P18074 |
| HOMO SAPIENS | 4 | 12188 | 4its | A | 122 | ARG | p.Arg122Gly | Q9Y606 |
| HOMO SAPIENS | 4 | 12188 | 4its | A | 133 | ASP | p.Asp133Asn | Q9Y606 |
| HOMO SAPIENS | 4 | 12188 | 4its | A | 144 | ARG | p.Arg144Trp | Q9Y606 |
| HOMO SAPIENS | 0 | 31529 | 1kgd | A | 761 | ARG | p.Arg766Gln | O14936 |
| HOMO SAPIENS | 1 | 31529 | 1kgd | A | 824 | GLN | p.Gln829Arg | O14936 |
| HOMO SAPIENS | 0 | 10906 | 2h63 | C | 18 | ARG | p.Arg18Ter | P53004 |
| HOMO SAPIENS | 9 | 27767 | 4wyq | A | 280 | LEU | p.Leu278Phe | Q9UPY3 |
| HOMO SAPIENS | 0 | 27767 | 4wyq | A | 297 | SER | p.Ser295Cys | Q9UPY3 |
| HOMO SAPIENS | 1 | 27767 | 4wyq | A | 334 | TYR | p.Tyr332Ter | Q9UPY3 |
| HOMO SAPIENS | 7 | 7775 | 5sv7 | C | 136 | SER | p.Ser136Cys | Q9NZJ5 |
| HOMO SAPIENS | 2 | 7775 | 5sv7 | C | 166 | GLN | p.Gln166Arg | Q9NZJ5 |
| HOMO SAPIENS | 2 | 7775 | 5sv7 | C | 332 | GLU | p.Glu332Ter | Q9NZJ5 |
| HOMO SAPIENS | 0 | 21449 | 2e9w | B | 84 | THR | p.Thr84Met | P10721 |
| HOMO SAPIENS | 0 | 21449 | 2e9w | B | 168 | ALA | p.Ala168Glu | P10721 |
| HOMO SAPIENS | 0 | 21449 | 2e9w | B | 178 | ALA | p.Ala178Thr | P10721 |
| HOMO SAPIENS | 0 | 21449 | 2e9w | B | 293 | ASN | p.Asn293Ser | P10721 |
| HOMO SAPIENS | -1 | 21449 | 2e9w | B | 318 | MET | p.Met318Leu | P10721 |
| HOMO SAPIENS | 0 | 21449 | 2e9w | B | 374 | VAL | p.Val374Ile | P10721 |
| HOMO SAPIENS | 6 | 3010 | 3flg | A | 66 | SER | p.Ser270Pro | Q9UBC3 |
| HOMO SAPIENS | 1 | 33918 | 2xzz | A | 760 | ARG | p.Arg760Ter | P22735 |
| HOMO SAPIENS | 2 | 1254 | 5xtc | w | 65 | ASN | p.Asn65Ser | O95299 |
| HOMO SAPIENS | 1 | 1254 | 5xtc | w | 238 | GLU | p.Glu238Lys | O95299 |
| HOMO SAPIENS | 3 | 10245 | 3siv | B | 194 | ALA | p.Ala194Glu | Q8WWY3 |
| HOMO SAPIENS | 4 | 10245 | 3siv | B | 216 | ALA | p.Ala216Pro | Q8WWY3 |
| HOMO SAPIENS | 0 | 2670 | 3dwb | A | 341 | THR | p.Thr341Ile | P42892 |
| HOMO SAPIENS | 1 | 2670 | 3dwb | A | 462 | SER | p.Ser462Asn | P42892 |
| HOMO SAPIENS | 1 | 2670 | 3dwb | A | 754 | ARG | p.Arg754Cys | P42892 |
| HOMO SAPIENS | 1 | 2961 | 5xtb | O | 40 | VAL | p.Val40Met | P19404 |
| HOMO SAPIENS | 5 | 2961 | 5xtb | O | 172 | ILE | p.Ile172Thr | P19404 |
| HOMO SAPIENS | 5 | 2961 | 5xtb | O | 190 | ASP | p.Asp190Gly | P19404 |
| HOMO SAPIENS | 5 | 29655 | 2xdg | A | 57 | ALA | p.Ala57Thr | Q02643 |
| HOMO SAPIENS | 4 | 29655 | 2xdg | A | 72 | GLU | p.Glu72Ter | Q02643 |
| HOMO SAPIENS | 0 | 29655 | 2xdg | A | 121 | GLU | p.Glu121Asp | Q02643 |
| HOMO SAPIENS | -1 | 17935 | 1hyn | P | 38 | ASP | p.Asp38Ala | P02730 |
| HOMO SAPIENS | 7 | 12825 | 1g82 | D | 99 | SER | p.Ser99Asn | P31371 |
| HOMO SAPIENS | 6 | 24927 | 1jm7 | B | 85 | VAL | p.Val85Leu | Q99728 |
| HOMO SAPIENS | 1 | 24927 | 1jm7 | B | 112 | ARG | p.Arg112Ter | Q99728 |
| HOMO SAPIENS | 0 | 24927 | 1jm7 | B | 116 | HIS | p.His116Tyr | Q99728 |
| HOMO SAPIENS | 7 | 92 | 5hde | A | 61 | LYS | p.Lys61Arg | Q05209 |
| HOMO SAPIENS | 0 | 17246 | 1szb | A | 84 | ARG | p.Arg99Gln | O00187 |
| HOMO SAPIENS | 1 | 17246 | 1szb | A | 105 | ASP | p.Asp120Gly | O00187 |

| | | | | | | | | |
|---|---|---|---|---|---|---|---|---|
| HOMO SAPIENS | 0 | 17246 | 1szb | A | 111 | PRO | p.Pro126Leu | O00187 |
| HOMO SAPIENS | 9 | 17246 | 1szb | A | 141 | CYS | p.Cys156Tyr | O00187 |
| HOMO SAPIENS | 0 | 32473 | 1b72 | A | 207 | ARG | p.Arg207Cys | P14653 |
| HOMO SAPIENS | 0 | 1920 | 4yzf | B | 488 | VAL | p.Val488Met | P02730 |
| HOMO SAPIENS | 0 | 1920 | 4yzf | B | 589 | ARG | p.Arg589Cys | P02730 |
| HOMO SAPIENS | 0 | 1920 | 4yzf | B | 589 | ARG | p.Arg589His | P02730 |
| HOMO SAPIENS | 0 | 1920 | 4yzf | B | 602 | ARG | p.Arg602Pro | P02730 |
| HOMO SAPIENS | 0 | 1920 | 4yzf | B | 613 | SER | p.Ser613Phe | P02730 |
| HOMO SAPIENS | -1 | 1920 | 4yzf | B | 646 | ARG | p.Arg646Gln | P02730 |
| HOMO SAPIENS | -1 | 1920 | 4yzf | B | 646 | ARG | p.Arg646Trp | P02730 |
| HOMO SAPIENS | 0 | 1920 | 4yzf | B | 701 | GLY | p.Gly701Asp | P02730 |
| HOMO SAPIENS | 0 | 1920 | 4yzf | B | 760 | ARG | p.Arg760Gln | P02730 |
| HOMO SAPIENS | 0 | 1920 | 4yzf | B | 771 | GLY | p.Gly771Asp | P02730 |
| HOMO SAPIENS | 0 | 1920 | 4yzf | B | 773 | SER | p.Ser773Pro | P02730 |
| HOMO SAPIENS | 0 | 1920 | 4yzf | B | 837 | THR | p.Thr837Ala | P02730 |
| HOMO SAPIENS | 0 | 1920 | 4yzf | B | 854 | PRO | p.Pro854Leu | P02730 |
| HOMO SAPIENS | 0 | 1920 | 4yzf | B | 858 | ALA | p.Ala858Asp | P02730 |
| HOMO SAPIENS | 0 | 1920 | 4yzf | B | 862 | VAL | p.Val862Ile | P02730 |
| HOMO SAPIENS | 0 | 1920 | 4yzf | B | 868 | PRO | p.Pro868Leu | P02730 |
| HOMO SAPIENS | 0 | 1920 | 4yzf | B | 870 | ARG | p.Arg870Trp | P02730 |
| HOMO SAPIENS | 0 | 9863 | 2juj | A | 54 | VAL | p.Val904Ile | P22681 |
| HOMO SAPIENS | 0 | 21934 | 2na7 | A | 184 | ILE | p.Ile184Val | P25445 |
| HOMO SAPIENS | -1 | 6746 | 1lw3 | A | 3 | THR | p.Lys3Thr | Q13614 |
| HOMO SAPIENS | -1 | 6746 | 1lw3 | A | 44 | VAL | p.Val44Ile | Q13614 |
| HOMO SAPIENS | 3 | 6746 | 1lw3 | A | 183 | ALA | p.Ala183Ser | Q13614 |
| HOMO SAPIENS | 4 | 6746 | 1lw3 | A | 482 | GLN | p.Gln482Ter | Q13614 |
| HOMO SAPIENS | 4 | 6746 | 1lw3 | A | 502 | GLU | p.Glu502Gln | Q13614 |
| HOMO SAPIENS | 3 | 6746 | 1lw3 | A | 545 | ASN | p.Asn545Ser | Q13614 |
| HOMO SAPIENS | 0 | 6746 | 1lw3 | A | 586 | ARG | p.Arg586Trp | Q13614 |
| HOMO SAPIENS | -1 | 6746 | 1lw3 | A | 602 | ALA | p.Ala602Gly | Q13614 |
| HOMO SAPIENS | 1 | 319 | 4ch9 | B | 336 | ARG | p.Arg336Ile | Q9UH77 |
| HOMO SAPIENS | 3 | 319 | 4ch9 | B | 340 | ALA | p.Ala340Val | Q9UH77 |
| HOMO SAPIENS | 0 | 319 | 4ch9 | B | 384 | ARG | p.Arg384Gln | Q9UH77 |
| HOMO SAPIENS | 5 | 319 | 4ch9 | B | 426 | PRO | p.Pro426Leu | Q9UH77 |
| HOMO SAPIENS | 3 | 319 | 4ch9 | B | 431 | ARG | p.Arg431Gln | Q9UH77 |
| HOMO SAPIENS | 2 | 319 | 4ch9 | B | 575 | ARG | p.Arg575Trp | Q9UH77 |
| HOMO SAPIENS | 0 | 574 | 2qkk | A | 142 | VAL | p.Val142Ile | O60930 |
| HOMO SAPIENS | 0 | 24021 | 2n8y | A | 752 | ARG | p.Arg752Gln | P12814 |
| HOMO SAPIENS | 0 | 18693 | 2y7j | B | 44 | ARG | p.Arg44Ter | P15735 |
| HOMO SAPIENS | 0 | 18693 | 2y7j | B | 106 | VAL | p.Val106Glu | P15735 |
| HOMO SAPIENS | 3 | 18693 | 2y7j | B | 189 | GLY | p.Gly189Glu | P15735 |
| HOMO SAPIENS | 0 | 17087 | 2ifg | A | 237 | THR | p.Thr237Met | P04629 |
| HOMO SAPIENS | 0 | 7980 | 1qub | A | 247 | VAL | p.Val266Leu | P02749 |
| HOMO SAPIENS | -1 | 2205 | 1h7u | A | 18 | ILE | p.Ile18Val | P54278 |
| HOMO SAPIENS | -1 | 2205 | 1h7u | A | 20 | ARG | p.Arg20Gln | P54278 |
| HOMO SAPIENS | 9 | 2205 | 1h7u | A | 46 | SER | p.Ser46Ile | P54278 |
| HOMO SAPIENS | 0 | 2205 | 1h7u | A | 60 | ASP | p.Asp60Glu | P54278 |
| HOMO SAPIENS | 0 | 2205 | 1h7u | A | 80 | PHE | p.Phe80Leu | P54278 |
| HOMO SAPIENS | 0 | 2205 | 1h7u | A | 118 | SER | p.Ser118Arg | P54278 |
| HOMO SAPIENS | 0 | 2205 | 1h7u | A | 126 | HIS | p.His126Gln | P54278 |
| HOMO SAPIENS | 0 | 2205 | 1h7u | A | 128 | SER | p.Ser128Leu | P54278 |
| HOMO SAPIENS | 1 | 2205 | 1h7u | A | 134 | ARG | p.Arg134Ter | P54278 |
| HOMO SAPIENS | 0 | 2205 | 1h7u | A | 166 | LEU | p.Leu166Pro | P54278 |
| HOMO SAPIENS | 1 | 2205 | 1h7u | A | 191 | TYR | p.Tyr191Cys | P54278 |

| | | | | | | | | |
|---|---|---|---|---|---|---|---|---|
| HOMO SAPIENS | 0 | 2205 | 1h7u | A | 211 | ARG | p.Arg211Ter | P54278 |
| HOMO SAPIENS | 0 | 2205 | 1h7u | A | 233 | GLN | p.Gln233Ter | P54278 |
| HOMO SAPIENS | 0 | 2205 | 1h7u | A | 255 | TYR | p.Tyr255Ter | P54278 |
| HOMO SAPIENS | 0 | 2205 | 1h7u | A | 277 | THR | p.Thr277Lys | P54278 |
| HOMO SAPIENS | 0 | 2205 | 1h7u | A | 301 | LYS | p.Lys301Asn | P54278 |
| HOMO SAPIENS | 0 | 2205 | 1h7u | A | 312 | MET | p.Met312Ile | P54278 |
| HOMO SAPIENS | 0 | 2205 | 1h7u | A | 312 | MET | p.Met312Thr | P54278 |
| HOMO SAPIENS | 0 | 2205 | 1h7u | A | 315 | ARG | p.Arg315Ter | P54278 |
| HOMO SAPIENS | 0 | 2205 | 1h7u | A | 318 | TYR | p.Tyr318Cys | P54278 |
| HOMO SAPIENS | 0 | 2205 | 1h7u | A | 335 | ASN | p.Asn335Ile | P54278 |
| HOMO SAPIENS | 0 | 2205 | 1h7u | A | 360 | ILE | p.Ile360Met | P54278 |
| HOMO SAPIENS | 6 | 20754 | 5mpo | B | 7 | VAL | p.Val7Phe | O96033 |
| HOMO SAPIENS | 4 | 20754 | 5mpo | B | 64 | VAL | p.Val64Ile | O96033 |
| HOMO SAPIENS | 5 | 34944 | 4qam | B | 1114 | ASP | p.Asp1114Gly | Q96KN7 |
| HOMO SAPIENS | 5 | 34944 | 4qam | B | 1240 | GLY | p.Gly1240Glu | Q96KN7 |
| HOMO SAPIENS | 9 | 40496 | 3vnn | A | 278 | ARG | p.Arg278His | P49917 |
| HOMO SAPIENS | 6 | 30131 | 4dt7 | E | 40 | PRO | p.Pro210Leu | P04070 |
| HOMO SAPIENS | -1 | 30131 | 4dt7 | E | 50 | ARG | p.Arg220Trp | P04070 |
| HOMO SAPIENS | 7 | 15311 | 2pqt | A | 64 | ARG | p.Arg64Trp | P18440 |
| HOMO SAPIENS | 2 | 15311 | 2pqt | A | 149 | VAL | p.Val149Ile | P18440 |
| HOMO SAPIENS | -1 | 19581 | 3hy8 | A | 33 | ASP | p.Asp33Val | Q9NVS9 |
| HOMO SAPIENS | 5 | 19581 | 3hy8 | A | 116 | ARG | p.Arg116Gln | Q9NVS9 |
| HOMO SAPIENS | 4 | 19581 | 3hy8 | A | 225 | ARG | p.Arg225His | Q9NVS9 |
| HOMO SAPIENS | 2 | 19581 | 3hy8 | A | 229 | TRP | p.Arg229Trp | Q9NVS9 |
| HOMO SAPIENS | 2 | 19581 | 3hy8 | A | 229 | TRP | p.Arg229Gln | Q9NVS9 |
| HOMO SAPIENS | 0 | 10266 | 3t1i | A | 41 | ASP | p.Asp41His | P49959 |
| HOMO SAPIENS | 1 | 10266 | 3t1i | A | 305 | ARG | p.Arg305Trp | P49959 |
| HOMO SAPIENS | 0 | 10266 | 3t1i | A | 308 | PHE | p.Phe308Leu | P49959 |
| HOMO SAPIENS | 0 | 10266 | 3t1i | A | 334 | SER | p.Ser334Arg | P49959 |
| HOMO SAPIENS | 1 | 1600 | 3rbf | A | 61 | GLU | p.Glu61Asp | P20711 |
| HOMO SAPIENS | 0 | 1600 | 3rbf | A | 91 | ALA | p.Ala91Val | P20711 |
| HOMO SAPIENS | -1 | 1600 | 3rbf | A | 102 | GLY | p.Gly102Ser | P20711 |
| HOMO SAPIENS | 3 | 1600 | 3rbf | A | 147 | SER | p.Ser147Arg | P20711 |
| HOMO SAPIENS | 1 | 1600 | 3rbf | A | 210 | PRO | p.Pro210Leu | P20711 |
| HOMO SAPIENS | 0 | 1600 | 3rbf | A | 217 | MET | p.Met217Val | P20711 |
| HOMO SAPIENS | 4 | 1600 | 3rbf | A | 250 | SER | p.Ser250Phe | P20711 |
| HOMO SAPIENS | 2 | 1600 | 3rbf | A | 275 | ALA | p.Ala275Thr | P20711 |
| HOMO SAPIENS | 1 | 1600 | 3rbf | A | 309 | PHE | p.Phe309Leu | P20711 |
| HOMO SAPIENS | -1 | 1600 | 3rbf | A | 347 | ARG | p.Arg347Gln | P20711 |
| HOMO SAPIENS | 0 | 1600 | 3rbf | A | 462 | ARG | p.Arg462Gln | P20711 |
| HOMO SAPIENS | 6 | 20830 | 5y4f | A | 714 | THR | p.Thr715Ile | Q01484 |
| HOMO SAPIENS | 7 | 20830 | 5y4f | A | 719 | ASP | p.Asp720Glu | Q01484 |
| HOMO SAPIENS | 0 | 20830 | 5y4f | A | 814 | VAL | p.Val815Ile | Q01484 |
| HOMO SAPIENS | -1 | 302 | 2c9y | A | 9 | GLU | p.Glu9Ter | P54819 |
| HOMO SAPIENS | 0 | 302 | 2c9y | A | 103 | ARG | p.Arg103Trp | P54819 |
| HOMO SAPIENS | 3 | 302 | 2c9y | A | 165 | ASP | p.Asp165Gly | P54819 |
| HOMO SAPIENS | 0 | 302 | 2c9y | A | 182 | ALA | p.Ala182Asp | P54819 |
| HOMO SAPIENS | 2 | 302 | 2c9y | A | 186 | ARG | p.Arg186Cys | P54819 |
| HOMO SAPIENS | 0 | 302 | 2c9y | A | 209 | ALA | p.Ala209Thr | P54819 |
| HOMO SAPIENS | -1 | 16920 | 1jlj | B | 10 | ASN | p.Asn10Tyr | Q9NQX3 |
| HOMO SAPIENS | 9 | 8462 | 1zbq | B | 16 | GLY | p.Gly16Ser | P51659 |
| HOMO SAPIENS | 5 | 8462 | 1zbq | B | 19 | ALA | p.Ala19Gly | P51659 |
| HOMO SAPIENS | 1 | 8462 | 1zbq | B | 34 | ALA | p.Ala34Val | P51659 |
| HOMO SAPIENS | 1 | 8462 | 1zbq | B | 106 | ARG | p.Arg106His | P51659 |

| | | | | | | | | |
|---|---|---|---|---|---|---|---|---|
| HOMO SAPIENS | 0 | 8462 | 1zbq | B | 140 | LYS | p.Lys140Asn | P51659 |
| HOMO SAPIENS | 0 | 8462 | 1zbq | B | 292 | THR | p.Thr292Ser | P51659 |
| HOMO SAPIENS | 0 | 4375 | 4ndi | A | 199 | ARG | p.Arg213His | Q7Z2E3 |
| HOMO SAPIENS | 3 | 4375 | 4ndi | A | 201 | HIS | p.His215Arg | Q7Z2E3 |
| HOMO SAPIENS | 0 | 4375 | 4ndi | A | 206 | PRO | p.Pro220Leu | Q7Z2E3 |
| HOMO SAPIENS | 2 | 4375 | 4ndi | A | 223 | LEU | p.Leu237Phe | Q7Z2E3 |
| HOMO SAPIENS | 0 | 4375 | 4ndi | A | 245 | ARG | p.Arg259His | Q7Z2E3 |
| HOMO SAPIENS | 0 | 4375 | 4ndi | A | 263 | VAL | p.Val277Gly | Q7Z2E3 |
| HOMO SAPIENS | 0 | 4375 | 4ndi | A | 279 | TRP | p.Trp293Ter | Q7Z2E3 |
| HOMO SAPIENS | 0 | 27004 | 4ccg | Y | 321 | ASP | p.Asp321Glu | Q9NW38 |
| HOMO SAPIENS | -1 | 27004 | 4ccg | Y | 372 | GLY | p.Gly372Ala | Q9NW38 |
| HOMO SAPIENS | -1 | 6383 | 5fai | A | 34 | ALA | p.Ala34Gly | Q92979 |
| HOMO SAPIENS | 8 | 6383 | 5fai | A | 86 | ASP | p.Asp86Gly | Q92979 |
| HOMO SAPIENS | 2 | 24585 | 4mi5 | A | 684 | ARG | p.Arg679Cys | Q15910 |
| HOMO SAPIENS | 0 | 15967 | 3nhb | A | 723 | ARG | p.Arg723Gln | Q9NP58 |
| HOMO SAPIENS | 5 | 18152 | 1oey | A | 389 | HIS | p.His389Gln | P19878 |
| HOMO SAPIENS | 5 | 18152 | 1oey | A | 395 | ARG | p.Arg395Trp | P19878 |
| HOMO SAPIENS | -1 | 1044 | 1li1 | F | 2 | VAL | p.Val1486Ile | P08572 |
| HOMO SAPIENS | 6 | 1044 | 1li1 | F | 37 | GLU | p.Glu1521Lys | P08572 |
| HOMO SAPIENS | 5 | 1044 | 1li1 | F | 157 | SER | p.Ser1639Leu | P08572 |
| HOMO SAPIENS | 5 | 1044 | 1li1 | F | 179 | ARG | p.Arg1662His | P08572 |
| HOMO SAPIENS | 5 | 1044 | 1li1 | F | 180 | GLY | p.Gly1663Arg | P08572 |
| HOMO SAPIENS | 6 | 20076 | 4uyi | A | 671 | LEU | p.Leu671Ser | Q8IY92 |
| HOMO SAPIENS | 0 | 19588 | 1jmj | A | 189 | ARG | p.Arg208His | P05546 |
| HOMO SAPIENS | -1 | 38023 | 1ry7 | B | 65 | GLY | p.Gly65Arg | P22607 |
| HOMO SAPIENS | -1 | 38023 | 1ry7 | B | 139 | ASP | p.Asp139Glu | P22607 |
| HOMO SAPIENS | 0 | 38023 | 1ry7 | B | 248 | ARG | p.Arg248Cys | P22607 |
| HOMO SAPIENS | 0 | 38023 | 1ry7 | B | 249 | SER | p.Ser249Cys | P22607 |
| HOMO SAPIENS | 0 | 38023 | 1ry7 | B | 250 | PRO | p.Pro250Arg | P22607 |
| HOMO SAPIENS | 0 | 38023 | 1ry7 | B | 322 | GLU | p.Glu322Lys | P22607 |
| HOMO SAPIENS | 5 | 16884 | 5eli | A | 33 | GLN | p.Gln33Ter | Q9NZC2 |
| HOMO SAPIENS | 0 | 16884 | 5eli | A | 44 | TRP | p.Trp44Ter | Q9NZC2 |
| HOMO SAPIENS | 0 | 16884 | 5eli | A | 47 | ARG | p.Arg47His | Q9NZC2 |
| HOMO SAPIENS | 0 | 16884 | 5eli | A | 66 | THR | p.Thr66Met | Q9NZC2 |
| HOMO SAPIENS | 0 | 16884 | 5eli | A | 78 | TRP | p.Trp78Ter | Q9NZC2 |
| HOMO SAPIENS | 1 | 16884 | 5eli | A | 96 | THR | p.Thr96Lys | Q9NZC2 |
| HOMO SAPIENS | 9 | 16884 | 5eli | A | 126 | VAL | p.Val126Gly | Q9NZC2 |
| HOMO SAPIENS | -1 | 12267 | 3w5o | A | 3 | ALA | p.Ala3Val | P49917 |
| HOMO SAPIENS | 1 | 12267 | 3w5o | A | 9 | THR | p.Thr9Ile | P49917 |
| HOMO SAPIENS | 2 | 12267 | 3w5o | A | 188 | ARG | p.Arg188Gln | P49917 |
| HOMO SAPIENS | 0 | 12267 | 3w5o | A | 469 | GLY | p.Gly469Glu | P49917 |
| HOMO SAPIENS | 1 | 12267 | 3w5o | A | 580 | ARG | p.Arg580Ter | P49917 |
| HOMO SAPIENS | 0 | 12267 | 3w5o | A | 600 | GLU | p.Glu600Lys | P49917 |
| HOMO SAPIENS | 9 | 2839 | 2hw5 | B | 54 | ARG | p.Arg54Cys | P30084 |
| HOMO SAPIENS | 9 | 2839 | 2hw5 | B | 59 | ASN | p.Asn59Ser | P30084 |
| HOMO SAPIENS | 0 | 2839 | 2hw5 | B | 273 | LYS | p.Lys273Glu | P30084 |
| HOMO SAPIENS | 1 | 12266 | 2o2k | B | 939 | GLY | p.Gly939Cys | Q99707 |
| HOMO SAPIENS | 5 | 12266 | 2o2k | B | 1164 | ARG | p.Arg1164His | Q99707 |
| HOMO SAPIENS | 1 | 12266 | 2o2k | B | 1166 | LEU | p.Leu1166Met | Q99707 |
| HOMO SAPIENS | 5 | 12266 | 2o2k | B | 1173 | PRO | p.Pro1173Leu | Q99707 |
| HOMO SAPIENS | 5 | 12266 | 2o2k | B | 1222 | ASN | p.Asn1222Ser | Q99707 |
| HOMO SAPIENS | 0 | 30891 | 3ch4 | B | 138 | ARG | p.Arg138Ter | Q15126 |
| HOMO SAPIENS | -1 | 4433 | 1qlz | A | 54 | GLY | p.Gly54Ser | P04156 |
| HOMO SAPIENS | 0 | 22568 | 3cxl | A | 20 | LEU | p.Leu20Phe | P15882 |

| | | | | | | | | |
|---|---|---|---|---|---|---|---|---|
| HOMO SAPIENS | 0 | 22568 | 3cxl | A | 126 | ILE | p.Ile126Met | P15882 |
| HOMO SAPIENS | 0 | 22568 | 3cxl | A | 141 | PRO | p.Pro141Leu | P15882 |
| HOMO SAPIENS | 0 | 22568 | 3cxl | A | 143 | TYR | p.Tyr143His | P15882 |
| HOMO SAPIENS | 0 | 22568 | 3cxl | A | 223 | ALA | p.Ala223Val | P15882 |
| HOMO SAPIENS | 0 | 22568 | 3cxl | A | 228 | GLY | p.Gly228Ser | P15882 |
| HOMO SAPIENS | 0 | 22568 | 3cxl | A | 252 | PRO | p.Pro252Gln | P15882 |
| HOMO SAPIENS | 0 | 22568 | 3cxl | A | 313 | GLU | p.Glu313Lys | P15882 |
| HOMO SAPIENS | 3 | 2668 | 5eqg | A | 34 | ASN | p.Asn34Ile | P11166 |
| HOMO SAPIENS | 3 | 2668 | 5eqg | A | 77 | MET | p.Met77Val | P11166 |
| HOMO SAPIENS | 6 | 2668 | 5eqg | A | 91 | GLY | p.Gly91Asp | P11166 |
| HOMO SAPIENS | 6 | 2668 | 5eqg | A | 92 | ARG | p.Arg92Trp | P11166 |
| HOMO SAPIENS | 6 | 2668 | 5eqg | A | 93 | ARG | p.Arg93Trp | P11166 |
| HOMO SAPIENS | 0 | 2668 | 5eqg | A | 95 | SER | p.Ser95Ile | P11166 |
| HOMO SAPIENS | 3 | 2668 | 5eqg | A | 126 | ARG | p.Arg126His | P11166 |
| HOMO SAPIENS | 3 | 2668 | 5eqg | A | 126 | ARG | p.Arg126Cys | P11166 |
| HOMO SAPIENS | 4 | 2668 | 5eqg | A | 232 | ARG | p.Arg232Cys | P11166 |
| HOMO SAPIENS | 3 | 2668 | 5eqg | A | 249 | ARG | p.Arg249Gln | P11166 |
| HOMO SAPIENS | 3 | 2668 | 5eqg | A | 255 | LYS | p.Lys255Arg | P11166 |
| HOMO SAPIENS | 0 | 2668 | 5eqg | A | 275 | ALA | p.Ala275Thr | P11166 |
| HOMO SAPIENS | 4 | 2668 | 5eqg | A | 299 | GLU | p.Glu299Lys | P11166 |
| HOMO SAPIENS | 6 | 2668 | 5eqg | A | 314 | GLY | p.Gly314Ser | P11166 |
| HOMO SAPIENS | 7 | 2668 | 5eqg | A | 315 | ILE | p.Ile315Met | P11166 |
| HOMO SAPIENS | -1 | 2668 | 5eqg | A | 458 | ARG | p.Arg458Trp | P11166 |
| HOMO SAPIENS | -1 | 2668 | 5eqg | A | 468 | ARG | p.Arg468Trp | P11166 |
| HOMO SAPIENS | 3 | 2291 | 2amy | A | 31 | PHE | p.Phe31Leu | O15305 |
| HOMO SAPIENS | 7 | 2291 | 2amy | A | 32 | LEU | p.Leu32Arg | O15305 |
| HOMO SAPIENS | 7 | 2291 | 2amy | A | 65 | ASP | p.Asp65Tyr | O15305 |
| HOMO SAPIENS | 1 | 2291 | 2amy | A | 108 | ALA | p.Ala108Val | O15305 |
| HOMO SAPIENS | 3 | 2291 | 2amy | A | 113 | PRO | p.Pro113Leu | O15305 |
| HOMO SAPIENS | 3 | 2291 | 2amy | A | 113 | PRO | p.Pro113Ser | O15305 |
| HOMO SAPIENS | 6 | 2291 | 2amy | A | 117 | GLY | p.Gly117Arg | O15305 |
| HOMO SAPIENS | 6 | 2291 | 2amy | A | 119 | PHE | p.Phe119Leu | O15305 |
| HOMO SAPIENS | 6 | 2291 | 2amy | A | 123 | ARG | p.Arg123Gln | O15305 |
| HOMO SAPIENS | 0 | 2291 | 2amy | A | 129 | VAL | p.Val129Met | O15305 |
| HOMO SAPIENS | 7 | 2291 | 2amy | A | 139 | GLU | p.Glu139Lys | O15305 |
| HOMO SAPIENS | 5 | 2291 | 2amy | A | 141 | ARG | p.Arg141His | O15305 |
| HOMO SAPIENS | 7 | 2291 | 2amy | A | 144 | PHE | p.Phe144Leu | O15305 |
| HOMO SAPIENS | 6 | 2291 | 2amy | A | 148 | ASP | p.Asp148Asn | O15305 |
| HOMO SAPIENS | 6 | 2291 | 2amy | A | 153 | ILE | p.Ile153Thr | O15305 |
| HOMO SAPIENS | 2 | 2291 | 2amy | A | 157 | PHE | p.Phe157Ser | O15305 |
| HOMO SAPIENS | 1 | 2291 | 2amy | A | 162 | ARG | p.Arg162Trp | O15305 |
| HOMO SAPIENS | 6 | 2291 | 2amy | A | 188 | ASP | p.Asp188Gly | O15305 |
| HOMO SAPIENS | 5 | 2291 | 2amy | A | 194 | ARG | p.Arg194Ter | O15305 |
| HOMO SAPIENS | 0 | 2291 | 2amy | A | 207 | PHE | p.Phe207Ser | O15305 |
| HOMO SAPIENS | 5 | 2291 | 2amy | A | 208 | GLY | p.Gly208Ala | O15305 |
| HOMO SAPIENS | 1 | 2291 | 2amy | A | 216 | ASN | p.Asn216Ile | O15305 |
| HOMO SAPIENS | 5 | 2291 | 2amy | A | 218 | HIS | p.His218Leu | O15305 |
| HOMO SAPIENS | 6 | 2291 | 2amy | A | 223 | ASP | p.Asp223Glu | O15305 |
| HOMO SAPIENS | 6 | 2291 | 2amy | A | 226 | THR | p.Thr226Ser | O15305 |
| HOMO SAPIENS | 5 | 2291 | 2amy | A | 231 | VAL | p.Val231Met | O15305 |
| HOMO SAPIENS | 1 | 2291 | 2amy | A | 237 | THR | p.Thr237Met | O15305 |
| HOMO SAPIENS | 1 | 2291 | 2amy | A | 241 | CYS | p.Cys241Ser | O15305 |
| HOMO SAPIENS | 5 | 35907 | 1d6g | A | 21 | GLY | p.Gly21Arg | P32238 |
| HOMO SAPIENS | -1 | 33209 | 1x9d | A | 197 | PRO | p.Pro197Leu | Q9UKM7 |

| | | | | | | | | |
|---|---|---|---|---|---|---|---|---|
| HOMO SAPIENS | -1 | 33209 | 1x9d | A | 201 | PRO | p.Pro201Leu | Q9UKM7 |
| HOMO SAPIENS | 6 | 21305 | 4yep | B | 1326 | ARG | p.Arg1326Ter | P24043 |
| HOMO SAPIENS | 6 | 21305 | 4yep | B | 1350 | ARG | p.Arg1350Ter | P24043 |
| HOMO SAPIENS | 0 | 21553 | 3qyb | A | 1120 | VAL | p.Val1119Ala | O60343 |
| HOMO SAPIENS | 2 | 21553 | 3qyb | A | 1148 | THR | p.Thr1147Met | O60343 |
| HOMO SAPIENS | -1 | 30645 | 3m0c | C | -17 | TRP | p.Trp4Ter | P01130 |
| HOMO SAPIENS | -1 | 30645 | 3m0c | C | -17 | TRP | p.Trp4Ter | P01130 |
| HOMO SAPIENS | -1 | 30645 | 3m0c | C | 12 | GLN | p.Gln33Ter | P01130 |
| HOMO SAPIENS | -1 | 30645 | 3m0c | C | 15 | ASP | p.Asp36Glu | P01130 |
| HOMO SAPIENS | -1 | 30645 | 3m0c | C | 28 | SER | p.Ser49Arg | P01130 |
| HOMO SAPIENS | -1 | 30645 | 3m0c | C | 41 | THR | p.Thr62Met | P01130 |
| HOMO SAPIENS | -1 | 30645 | 3m0c | C | 69 | ASP | p.Asp90Asn | P01130 |
| HOMO SAPIENS | -1 | 30645 | 3m0c | C | 76 | ASN | p.Asn97Lys | P01130 |
| HOMO SAPIENS | -1 | 30645 | 3m0c | C | 78 | SER | p.Ser99Ter | P01130 |
| HOMO SAPIENS | -1 | 30645 | 3m0c | C | 81 | GLN | p.Gln102Ter | P01130 |
| HOMO SAPIENS | -1 | 30645 | 3m0c | C | 84 | PRO | p.Pro105Ser | P01130 |
| HOMO SAPIENS | -1 | 30645 | 3m0c | C | 194 | ARG | p.Arg215Cys | P01130 |
| HOMO SAPIENS | -1 | 30645 | 3m0c | C | 198 | GLY | p.Gly219Asp | P01130 |
| HOMO SAPIENS | -1 | 30645 | 3m0c | C | 201 | CYS | p.Cys222Tyr | P01130 |
| HOMO SAPIENS | -1 | 30645 | 3m0c | C | 201 | CYS | p.Cys222Ter | P01130 |
| HOMO SAPIENS | -1 | 30645 | 3m0c | C | 206 | ASP | p.Asp227Glu | P01130 |
| HOMO SAPIENS | -1 | 30645 | 3m0c | C | 249 | CYS | p.Cys270Ter | P01130 |
| HOMO SAPIENS | -1 | 30645 | 3m0c | C | 250 | VAL | p.Val271Ile | P01130 |
| HOMO SAPIENS | 0 | 30645 | 3m0c | C | 255 | CYS | p.Cys276Tyr | P01130 |
| HOMO SAPIENS | 0 | 30645 | 3m0c | C | 255 | CYS | p.Cys276Ter | P01130 |
| HOMO SAPIENS | 0 | 30645 | 3m0c | C | 256 | GLU | p.Glu277Lys | P01130 |
| HOMO SAPIENS | 0 | 30645 | 3m0c | C | 265 | SER | p.Ser286Arg | P01130 |
| HOMO SAPIENS | 0 | 30645 | 3m0c | C | 284 | TRP | p.Trp305Ter | P01130 |
| HOMO SAPIENS | 0 | 30645 | 3m0c | C | 292 | CYS | p.Cys313Ter | P01130 |
| HOMO SAPIENS | 1 | 30645 | 3m0c | C | 385 | ARG | p.Arg406Trp | P01130 |
| HOMO SAPIENS | 0 | 30645 | 3m0c | C | 395 | ARG | p.Arg416Gln | P01130 |
| HOMO SAPIENS | 0 | 30645 | 3m0c | C | 398 | TYR | p.Tyr419Ter | P01130 |
| HOMO SAPIENS | 1 | 30645 | 3m0c | C | 415 | VAL | p.Val436Ala | P01130 |
| HOMO SAPIENS | 1 | 30645 | 3m0c | C | 424 | ASP | p.Asp445Glu | P01130 |
| HOMO SAPIENS | 1 | 30645 | 3m0c | C | 425 | LEU | p.Leu446Val | P01130 |
| HOMO SAPIENS | -1 | 30645 | 3m0c | C | 440 | GLY | p.Gly461Cys | P01130 |
| HOMO SAPIENS | -1 | 30645 | 3m0c | C | 441 | VAL | p.Val462Ile | P01130 |
| HOMO SAPIENS | 1 | 30645 | 3m0c | C | 447 | VAL | p.Val468Ile | P01130 |
| HOMO SAPIENS | 1 | 30645 | 3m0c | C | 489 | THR | p.Thr510Met | P01130 |
| HOMO SAPIENS | 0 | 30645 | 3m0c | C | 524 | GLY | p.Gly545Glu | P01130 |
| HOMO SAPIENS | 1 | 30645 | 3m0c | C | 615 | GLY | p.Gly636Ser | P01130 |
| HOMO SAPIENS | 1 | 30645 | 3m0c | C | 625 | LEU | p.Leu646Ile | P01130 |
| HOMO SAPIENS | 1 | 30645 | 3m0c | C | 630 | ASP | p.Asp651Gly | P01130 |
| HOMO SAPIENS | 3 | 30645 | 3m0c | C | 646 | CYS | p.Cys667Tyr | P01130 |
| HOMO SAPIENS | 4 | 30645 | 3m0c | C | 654 | GLY | p.Gly675Ser | P01130 |
| HOMO SAPIENS | 6 | 30645 | 3m0c | C | 660 | CYS | p.Cys681Ter | P01130 |
| HOMO SAPIENS | 5 | 30645 | 3m0c | C | 676 | ALA | p.Ala697Pro | P01130 |
| HOMO SAPIENS | -1 | 30645 | 3m0c | C | 749 | GLN | p.Gln770Ter | P01130 |
| HOMO SAPIENS | -1 | 41323 | 4bxo | A | 1812 | PRO | p.Pro1812Ala | Q8IYD8 |
| HOMO SAPIENS | 5 | 8335 | 3h0t | C | 12 | GLY | p.Gly71Asp | P81172 |
| HOMO SAPIENS | 6 | 609 | 1bhg | B | 148 | PRO | p.Pro148Ser | P08236 |
| HOMO SAPIENS | 1 | 609 | 1bhg | B | 176 | LEU | p.Leu176Phe | P08236 |
| HOMO SAPIENS | 6 | 609 | 1bhg | B | 216 | ARG | p.Arg216Trp | P08236 |
| HOMO SAPIENS | 0 | 609 | 1bhg | B | 350 | LYS | p.Lys350Asn | P08236 |

| | | | | | | | | |
|---|---|---|---|---|---|---|---|---|
| HOMO SAPIENS | 1 | 609 | 1bhg | B | 354 | ALA | p.Ala354Val | P08236 |
| HOMO SAPIENS | 1 | 609 | 1bhg | B | 357 | ARG | p.Arg357Ter | P08236 |
| HOMO SAPIENS | 0 | 609 | 1bhg | B | 382 | ARG | p.Arg382Cys | P08236 |
| HOMO SAPIENS | 0 | 609 | 1bhg | B | 495 | TYR | p.Tyr495Cys | P08236 |
| HOMO SAPIENS | 2 | 609 | 1bhg | B | 507 | TRP | p.Trp507Ter | P08236 |
| HOMO SAPIENS | 0 | 609 | 1bhg | B | 577 | ARG | p.Arg577Leu | P08236 |
| HOMO SAPIENS | 1 | 609 | 1bhg | B | 611 | ARG | p.Arg611Trp | P08236 |
| HOMO SAPIENS | 0 | 609 | 1bhg | B | 619 | ALA | p.Ala619Val | P08236 |
| HOMO SAPIENS | 0 | 609 | 1bhg | B | 627 | TRP | p.Trp627Cys | P08236 |
| HOMO SAPIENS | 5 | 3422 | 1x9n | A | 566 | GLU | p.Glu566Lys | P18858 |
| HOMO SAPIENS | 2 | 3422 | 1x9n | A | 771 | ARG | p.Arg771Trp | P18858 |
| HOMO SAPIENS | 0 | 3279 | 5tby | B | 797 | ALA | p.Ala797Thr | P12883 |
| HOMO SAPIENS | -1 | 3279 | 5tby | B | 982 | MET | p.Met982Thr | P12883 |
| HOMO SAPIENS | -1 | 3279 | 5tby | B | 992 | LEU | p.Leu992Met | P12883 |
| HOMO SAPIENS | -1 | 3279 | 5tby | B | 1004 | GLN | p.Gln1004Glu | P12883 |
| HOMO SAPIENS | -1 | 3279 | 5tby | B | 1013 | GLU | p.Glu1013Lys | P12883 |
| HOMO SAPIENS | -1 | 3279 | 5tby | B | 1032 | ASP | p.Asp1032Asn | P12883 |
| HOMO SAPIENS | -1 | 3279 | 5tby | B | 1053 | ARG | p.Arg1053Gln | P12883 |
| HOMO SAPIENS | -1 | 3279 | 5tby | B | 1117 | GLU | p.Glu1117Ter | P12883 |
| HOMO SAPIENS | -1 | 3279 | 5tby | B | 1193 | ARG | p.Arg1193His | P12883 |
| HOMO SAPIENS | -1 | 3279 | 5tby | B | 1205 | GLU | p.Glu1205Lys | P12883 |
| HOMO SAPIENS | -1 | 3279 | 5tby | B | 1207 | ILE | p.Ile1207Met | P12883 |
| HOMO SAPIENS | -1 | 3279 | 5tby | B | 1250 | ARG | p.Arg1250Pro | P12883 |
| HOMO SAPIENS | -1 | 3279 | 5tby | B | 1259 | HIS | p.His1259Gln | P12883 |
| HOMO SAPIENS | -1 | 3279 | 5tby | B | 1272 | ASP | p.Asp1272Asn | P12883 |
| HOMO SAPIENS | -1 | 3279 | 5tby | B | 1295 | GLU | p.Glu1295Lys | P12883 |
| HOMO SAPIENS | -1 | 3279 | 5tby | B | 1335 | SER | p.Ser1335Leu | P12883 |
| HOMO SAPIENS | -1 | 3279 | 5tby | B | 1399 | GLU | p.Glu1399Asp | P12883 |
| HOMO SAPIENS | -1 | 3279 | 5tby | B | 1413 | SER | p.Ser1413Ter | P12883 |
| HOMO SAPIENS | -1 | 3279 | 5tby | B | 1459 | LYS | p.Lys1459Asn | P12883 |
| HOMO SAPIENS | -1 | 3279 | 5tby | B | 1467 | LEU | p.Leu1467Val | P12883 |
| HOMO SAPIENS | -1 | 3279 | 5tby | B | 1491 | SER | p.Ser1491Cys | P12883 |
| HOMO SAPIENS | -1 | 3279 | 5tby | B | 1496 | GLU | p.Glu1496Ala | P12883 |
| HOMO SAPIENS | -1 | 3279 | 5tby | B | 1500 | ARG | p.Arg1500Pro | P12883 |
| HOMO SAPIENS | -1 | 3279 | 5tby | B | 1606 | ARG | p.Arg1606His | P12883 |
| HOMO SAPIENS | -1 | 3279 | 5tby | B | 1664 | ASN | p.Asn1664Lys | P12883 |
| HOMO SAPIENS | -1 | 3279 | 5tby | B | 1712 | ARG | p.Arg1712Trp | P12883 |
| HOMO SAPIENS | -1 | 3279 | 5tby | B | 1769 | LEU | p.Leu1769Met | P12883 |
| HOMO SAPIENS | -1 | 3279 | 5tby | B | 1808 | GLY | p.Gly1808Ser | P12883 |
| HOMO SAPIENS | -1 | 3279 | 5tby | B | 1818 | ARG | p.Arg1818Trp | P12883 |
| HOMO SAPIENS | -1 | 3279 | 5tby | B | 1820 | ARG | p.Arg1820Trp | P12883 |
| HOMO SAPIENS | -1 | 3279 | 5tby | B | 1844 | GLU | p.Glu1844Lys | P12883 |
| HOMO SAPIENS | -1 | 3279 | 5tby | B | 1845 | ARG | p.Arg1845Trp | P12883 |
| HOMO SAPIENS | -1 | 3279 | 5tby | B | 1854 | THR | p.Thr1854Met | P12883 |
| HOMO SAPIENS | -1 | 3279 | 5tby | B | 1909 | ARG | p.Arg1909Trp | P12883 |
| HOMO SAPIENS | 2 | 2260 | 4kl1 | A | 553 | ASP | p.Asp553Asn | Q9Y3Q4 |
| HOMO SAPIENS | 0 | 40570 | 5lst | A | 522 | ARG | p.Arg522Cys | O94761 |
| HOMO SAPIENS | 0 | 40570 | 5lst | A | 522 | ARG | p.Arg522His | O94761 |
| HOMO SAPIENS | 0 | 40570 | 5lst | A | 591 | PRO | p.Pro591Leu | O94761 |
| HOMO SAPIENS | 0 | 40570 | 5lst | A | 793 | PRO | p.Pro793Ser | O94761 |
| HOMO SAPIENS | 4 | 40570 | 5lst | A | 799 | VAL | p.Val799Met | O94761 |
| HOMO SAPIENS | 0 | 40570 | 5lst | A | 1004 | ARG | p.Arg1004Trp | O94761 |
| HOMO SAPIENS | 0 | 40570 | 5lst | A | 1005 | ARG | p.Arg1005Gln | O94761 |
| HOMO SAPIENS | 0 | 40570 | 5lst | A | 1021 | ARG | p.Arg1021Gln | O94761 |

| | | | | | | | | |
|---|---|---|---|---|---|---|---|---|
| HOMO SAPIENS | 0 | 40570 | 5lst | A | 1043 | LEU | p.Leu1043Pro | O94761 |
| HOMO SAPIENS | 0 | 40570 | 5lst | A | 1045 | ALA | p.Ala1045Thr | O94761 |
| HOMO SAPIENS | 0 | 40570 | 5lst | A | 1105 | GLY | p.Gly1105Asp | O94761 |
| HOMO SAPIENS | 0 | 40570 | 5lst | A | 1105 | GLY | p.Gly1105Ser | O94761 |
| HOMO SAPIENS | -1 | 40570 | 5lst | A | 1113 | GLY | p.Gly1113Arg | O94761 |
| HOMO SAPIENS | -1 | 18865 | 1gxc | A | 64 | GLU | p.Glu64Lys | O96017 |
| HOMO SAPIENS | -1 | 11673 | 3dzu | D | 85 | PRO | p.Pro113Gln | P37231 |
| HOMO SAPIENS | 0 | 19383 | 3b96 | A | 63 | LYS | p.Lys103Arg | P49748 |
| HOMO SAPIENS | 4 | 19383 | 3b96 | A | 140 | ALA | p.Ala180Thr | P49748 |
| HOMO SAPIENS | 1 | 19383 | 3b96 | A | 145 | GLY | p.Gly185Ser | P49748 |
| HOMO SAPIENS | 0 | 19383 | 3b96 | A | 220 | THR | p.Thr260Met | P49748 |
| HOMO SAPIENS | 5 | 19383 | 3b96 | A | 243 | VAL | p.Val283Ala | P49748 |
| HOMO SAPIENS | 0 | 19383 | 3b96 | A | 249 | GLY | p.Gly289Arg | P49748 |
| HOMO SAPIENS | 0 | 19383 | 3b96 | A | 254 | GLY | p.Gly294Glu | P49748 |
| HOMO SAPIENS | 7 | 19383 | 3b96 | A | 326 | ARG | p.Arg366Cys | P49748 |
| HOMO SAPIENS | 7 | 19383 | 3b96 | A | 326 | ARG | p.Arg366His | P49748 |
| HOMO SAPIENS | 5 | 19383 | 3b96 | A | 342 | LYS | p.Lys382Gln | P49748 |
| HOMO SAPIENS | 0 | 19383 | 3b96 | A | 369 | THR | p.Thr409Met | P49748 |
| HOMO SAPIENS | 5 | 19383 | 3b96 | A | 376 | ALA | p.Ala416Ser | P49748 |
| HOMO SAPIENS | 0 | 19383 | 3b96 | A | 385 | ALA | p.Ala425Thr | P49748 |
| HOMO SAPIENS | 5 | 19383 | 3b96 | A | 399 | GLY | p.Gly439Asp | P49748 |
| HOMO SAPIENS | 7 | 19383 | 3b96 | A | 401 | GLY | p.Gly441Asp | P49748 |
| HOMO SAPIENS | 9 | 19383 | 3b96 | A | 413 | ARG | p.Arg453Gln | P49748 |
| HOMO SAPIENS | 2 | 19383 | 3b96 | A | 418 | PHE | p.Phe458Leu | P49748 |
| HOMO SAPIENS | 6 | 19383 | 3b96 | A | 419 | ARG | p.Arg459Trp | P49748 |
| HOMO SAPIENS | 6 | 19383 | 3b96 | A | 419 | ARG | p.Arg459Gln | P49748 |
| HOMO SAPIENS | -1 | 19383 | 3b96 | A | 471 | ARG | p.Arg511Gln | P49748 |
| HOMO SAPIENS | 0 | 19383 | 3b96 | A | 573 | ARG | p.Arg613Trp | P49748 |
| HOMO SAPIENS | 0 | 19383 | 3b96 | A | 575 | ARG | p.Arg615Gln | P49748 |
| HOMO SAPIENS | 0 | 22286 | 3k2s | A | 3 | PRO | p.Pro27His | P02647 |
| HOMO SAPIENS | 0 | 22286 | 3k2s | A | 4 | PRO | p.Pro28Arg | P02647 |
| HOMO SAPIENS | 0 | 22286 | 3k2s | A | 10 | ARG | p.Arg34Leu | P02647 |
| HOMO SAPIENS | 0 | 22286 | 3k2s | A | 26 | GLY | p.Gly50Arg | P02647 |
| HOMO SAPIENS | 0 | 22286 | 3k2s | A | 37 | ALA | p.Ala61Thr | P02647 |
| HOMO SAPIENS | 1 | 1622 | 2qrv | D | 631 | ARG | p.Arg635Trp | Q9Y6K1 |
| HOMO SAPIENS | 1 | 1622 | 2qrv | D | 900 | PRO | p.Pro904Leu | Q9Y6K1 |
| HOMO SAPIENS | 6 | 40796 | 1v18 | B | 1509 | LEU | p.Leu1509Val | P25054 |
| HOMO SAPIENS | 1 | 22375 | 4pki | G | 1214 | LYS | p.Asp214Asn | P28289 |
| HOMO SAPIENS | 1 | 16307 | 5mv8 | B | 483 | ALA | p.Ala483Ser | Q9Y6N9 |
| HOMO SAPIENS | 6 | 16307 | 5mv8 | B | 506 | ASN | p.Asn506Lys | Q9Y6N9 |
| HOMO SAPIENS | 4 | 16307 | 5mv8 | B | 519 | ASP | p.Glu519Asp | Q9Y6N9 |
| HOMO SAPIENS | 0 | 16307 | 5mv8 | B | 530 | GLY | p.Gly530Arg | Q9Y6N9 |
| HOMO SAPIENS | 4 | 36221 | 4cvo | A | 127 | VAL | p.Val127Ile | Q03468 |
| HOMO SAPIENS | 7 | 36221 | 4cvo | A | 134 | ARG | p.Arg134Trp | Q03468 |
| HOMO SAPIENS | 0 | 36221 | 4cvo | A | 156 | GLN | p.Gln156Ter | Q03468 |
| HOMO SAPIENS | 0 | 10829 | 4a11 | B | 13 | GLU | p.Glu13Ter | Q13216 |
| HOMO SAPIENS | 1 | 10829 | 4a11 | B | 100 | TYR | p.Tyr100Ter | Q13216 |
| HOMO SAPIENS | 0 | 10829 | 4a11 | B | 160 | ALA | p.Ala160Val | Q13216 |
| HOMO SAPIENS | 1 | 6759 | 5lev | A | 29 | ILE | p.Ile29Phe | Q9H3H5 |
| HOMO SAPIENS | 0 | 6759 | 5lev | A | 108 | MET | p.Met108Ile | Q9H3H5 |
| HOMO SAPIENS | 2 | 6759 | 5lev | A | 117 | VAL | p.Val117Ile | Q9H3H5 |
| HOMO SAPIENS | 1 | 6759 | 5lev | A | 170 | TYR | p.Tyr170Cys | Q9H3H5 |
| HOMO SAPIENS | 0 | 6759 | 5lev | A | 264 | GLY | p.Val264Gly | Q9H3H5 |
| HOMO SAPIENS | 0 | 6759 | 5lev | A | 332 | PHE | p.Phe332Val | Q9H3H5 |

| | | | | | | | | |
|---|---|---|---|---|---|---|---|---|
| HOMO SAPIENS | 7 | 6759 | 5lev | A | 393 | ILE | p.Ile393Val | Q9H3H5 |
| HOMO SAPIENS | -1 | 33473 | 5m5e | C | 9 | ASN | p.Asn31Ser | P31785 |
| HOMO SAPIENS | 0 | 26155 | 1ic8 | B | 98 | ALA | p.Ala98Val | P20823 |
| HOMO SAPIENS | 0 | 15329 | 4iqr | B | 127 | ARG | p.Arg136Trp | P41235 |
| HOMO SAPIENS | 0 | 20725 | 4bsj | A | 350 | GLU | p.Glu350Val | P35916 |
| HOMO SAPIENS | 0 | 20725 | 4bsj | A | 494 | THR | p.Thr494Ala | P35916 |
| HOMO SAPIENS | 0 | 20725 | 4bsj | A | 527 | ASN | p.Asn527Ser | P35916 |
| HOMO SAPIENS | 6 | 29959 | 1xiw | B | 47 | ARG | p.Arg68Ter | P04234 |
| HOMO SAPIENS | 2 | 29959 | 1xiw | B | 72 | CYS | p.Cys93Ter | P04234 |
| HOMO SAPIENS | 3 | 20242 | 5tjc | A | 102 | ARG | p.Arg102Ter | Q9GZU1 |
| HOMO SAPIENS | 3 | 20242 | 5tjc | A | 232 | THR | p.Thr232Pro | Q9GZU1 |
| HOMO SAPIENS | 7 | 9590 | 4ap8 | A | 50 | THR | p.Thr50Ala | O96007 |
| HOMO SAPIENS | 4 | 9590 | 4ap8 | A | 168 | GLU | p.Glu168Lys | O96007 |
| HOMO SAPIENS | -1 | 4297 | 5do7 | B | 19 | ASP | p.Asp19His | Q9H221 |
| HOMO SAPIENS | -1 | 4297 | 5do7 | B | 54 | TYR | p.Tyr54Cys | Q9H221 |
| HOMO SAPIENS | -1 | 4297 | 5do7 | B | 80 | CYS | p.Cys80Tyr | Q9H221 |
| HOMO SAPIENS | 5 | 4297 | 5do7 | B | 107 | SER | p.Ser107Ter | Q9H221 |
| HOMO SAPIENS | 1 | 4297 | 5do7 | B | 164 | ARG | p.Arg164Pro | Q9H221 |
| HOMO SAPIENS | 0 | 4297 | 5do7 | B | 210 | VAL | p.Val210Met | Q9H221 |
| HOMO SAPIENS | 0 | 4297 | 5do7 | B | 231 | PRO | p.Pro231Thr | Q9H221 |
| HOMO SAPIENS | 0 | 4297 | 5do7 | B | 263 | ARG | p.Arg263Gln | Q9H221 |
| HOMO SAPIENS | 0 | 4297 | 5do7 | B | 361 | TRP | p.Trp361Ter | Q9H221 |
| HOMO SAPIENS | 0 | 4297 | 5do7 | B | 400 | THR | p.Thr400Lys | Q9H221 |
| HOMO SAPIENS | 0 | 4297 | 5do7 | B | 549 | ALA | p.Ala549Pro | Q9H221 |
| HOMO SAPIENS | 0 | 4297 | 5do7 | B | 596 | LEU | p.Leu596Arg | Q9H221 |
| HOMO SAPIENS | 0 | 4297 | 5do7 | B | 632 | VAL | p.Val632Ala | Q9H221 |
| HOMO SAPIENS | 0 | 4297 | 5do7 | B | 658 | TYR | p.Tyr658Ter | Q9H221 |
| HOMO SAPIENS | 2 | 9356 | 4igg | B | 206 | GLN | p.Gln206His | P35221 |
| HOMO SAPIENS | 6 | 2179 | 2wk6 | A | 44 | ARG | p.Arg44Gln | P19971 |
| HOMO SAPIENS | 6 | 2179 | 2wk6 | A | 81 | ARG | p.Arg81Gln | P19971 |
| HOMO SAPIENS | 6 | 2179 | 2wk6 | A | 145 | GLY | p.Gly145Arg | P19971 |
| HOMO SAPIENS | 7 | 2179 | 2wk6 | A | 146 | ARG | p.Arg146His | P19971 |
| HOMO SAPIENS | 7 | 2179 | 2wk6 | A | 153 | GLY | p.Gly153Ser | P19971 |
| HOMO SAPIENS | 0 | 2179 | 2wk6 | A | 173 | MET | p.Met173Ile | P19971 |
| HOMO SAPIENS | 6 | 2179 | 2wk6 | A | 202 | ARG | p.Arg202Lys | P19971 |
| HOMO SAPIENS | 6 | 2179 | 2wk6 | A | 208 | VAL | p.Val208Met | P19971 |
| HOMO SAPIENS | 5 | 2179 | 2wk6 | A | 222 | LYS | p.Lys222Arg | P19971 |
| HOMO SAPIENS | 5 | 2179 | 2wk6 | A | 289 | GLU | p.Glu289Ala | P19971 |
| HOMO SAPIENS | 2 | 2179 | 2wk6 | A | 311 | GLY | p.Gly311Arg | P19971 |
| HOMO SAPIENS | 2 | 2179 | 2wk6 | A | 376 | ARG | p.Arg376Pro | P19971 |
| HOMO SAPIENS | 2 | 2179 | 2wk6 | A | 465 | ALA | p.Ala465Thr | P19971 |
| HOMO SAPIENS | -1 | 11856 | 3orh | A | 3 | ALA | p.Ala3Ser | Q14353 |
| HOMO SAPIENS | 1 | 11856 | 3orh | A | 20 | TRP | p.Trp20Ser | Q14353 |
| HOMO SAPIENS | 1 | 11856 | 3orh | A | 50 | MET | p.Met50Leu | Q14353 |
| HOMO SAPIENS | 1 | 11856 | 3orh | A | 110 | VAL | p.Val110Phe | Q14353 |
| HOMO SAPIENS | 8 | 11856 | 3orh | A | 164 | GLY | p.Gly164Asp | Q14353 |
| HOMO SAPIENS | 1 | 11856 | 3orh | A | 169 | CYS | p.Cys169Tyr | Q14353 |
| HOMO SAPIENS | 1 | 11856 | 3orh | A | 174 | TRP | p.Trp174Ter | Q14353 |
| HOMO SAPIENS | 2 | 11856 | 3orh | A | 196 | ALA | p.Ala196Val | Q14353 |
| HOMO SAPIENS | 1 | 11856 | 3orh | A | 209 | THR | p.Thr209Met | Q14353 |
| HOMO SAPIENS | 1 | 11856 | 3orh | A | 219 | ASP | p.Asp219Asn | Q14353 |
| HOMO SAPIENS | 1 | 11856 | 3orh | A | 224 | ALA | p.Ala224Thr | Q14353 |
| HOMO SAPIENS | 1 | 13244 | 2m6i | D | 218 | ARG | p.Arg246Leu | P23415 |
| HOMO SAPIENS | 1 | 13244 | 2m6i | D | 244 | ILE | p.Ile272Asn | P23415 |

| | | | | | | | | |
|---|---|---|---|---|---|---|---|---|
| HOMO SAPIENS | 5 | 13244 | 2m6i | D | 250 | PRO | p.Pro278Thr | P23415 |
| HOMO SAPIENS | 6 | 13244 | 2m6i | D | 266 | GLN | p.Gln294His | P23415 |
| HOMO SAPIENS | 1 | 13244 | 2m6i | D | 271 | ARG | p.Arg299Leu | P23415 |
| HOMO SAPIENS | 1 | 13244 | 2m6i | D | 276 | LYS | p.Lys304Glu | P23415 |
| HOMO SAPIENS | 1 | 13244 | 2m6i | D | 279 | TYR | p.Tyr307Cys | P23415 |
| HOMO SAPIENS | 0 | 13244 | 2m6i | D | 316 | GLY | p.Gly378Ser | P23415 |
| HOMO SAPIENS | 1 | 13244 | 2m6i | D | 400 | ARG | p.Arg428His | P23415 |
| HOMO SAPIENS | 3 | 1967 | 2i6b | A | 301 | ALA | p.Ala318Glu | P55263 |
| HOMO SAPIENS | 4 | 11096 | 1h8b | A | 61 | ALA | p.Ala882Thr | P35609 |
| HOMO SAPIENS | 0 | 15423 | 5ngz | A | 2 | GLN | p.Gln2Glu | Q9NPD8 |
| HOMO SAPIENS | 3 | 30475 | 3lpw | A | 21 | SER | p.Ser22894Tyr | Q8WZ42 |
| HOMO SAPIENS | 6 | 32687 | 2arf | A | 1041 | ARG | p.Arg1041Trp | P35670 |
| HOMO SAPIENS | 8 | 32687 | 2arf | A | 1061 | GLY | p.Gly1061Glu | P35670 |
| HOMO SAPIENS | 6 | 32687 | 2arf | A | 1064 | GLU | p.Glu1064Ala | P35670 |
| HOMO SAPIENS | 2 | 32687 | 2arf | A | 1069 | HIS | p.His1069Gln | P35670 |
| HOMO SAPIENS | 2 | 32687 | 2arf | A | 1088 | LEU | p.Leu1088Ter | P35670 |
| HOMO SAPIENS | 0 | 32687 | 2arf | A | 1099 | GLY | p.Gly1099Ser | P35670 |
| HOMO SAPIENS | 4 | 32687 | 2arf | A | 1106 | VAL | p.Val1106Asp | P35670 |
| HOMO SAPIENS | 2 | 32687 | 2arf | A | 1142 | GLN | p.Gln1142His | P35670 |
| HOMO SAPIENS | 8 | 32687 | 2arf | A | 1148 | ILE | p.Ile1148Thr | P35670 |
| HOMO SAPIENS | 6 | 32687 | 2arf | A | 1151 | ARG | p.Arg1151Cys | P35670 |
| HOMO SAPIENS | 4 | 32687 | 2arf | A | 1169 | MET | p.Met1169Val | P35670 |
| HOMO SAPIENS | 4 | 32687 | 2arf | A | 1173 | GLU | p.Glu1173Lys | P35670 |
| HOMO SAPIENS | 2 | 32687 | 2arf | A | 1176 | GLY | p.Gly1176Arg | P35670 |
| HOMO SAPIENS | 3 | 15544 | 5uq3 | C | 575 | PRO | p.Pro69Leu | P46527 |
| HOMO SAPIENS | 6 | 15544 | 5uq3 | C | 592 | GLU | p.Glu86Asp | P46527 |
| HOMO SAPIENS | 6 | 15544 | 5uq3 | C | 599 | ARG | p.Arg93Trp | P46527 |
| HOMO SAPIENS | -1 | 15544 | 5uq3 | C | 615 | VAL | p.Val109Gly | P46527 |
| HOMO SAPIENS | -1 | 15544 | 5uq3 | C | 631 | SER | p.Ser125Pro | P46527 |
| HOMO SAPIENS | -1 | 15544 | 5uq3 | C | 639 | PRO | p.Pro133Ser | P46527 |
| HOMO SAPIENS | 2 | 2703 | 4wxq | A | 245 | CYS | p.Cys245Tyr | Q99972 |
| HOMO SAPIENS | 0 | 2703 | 4wxq | A | 252 | GLY | p.Gly252Arg | Q99972 |
| HOMO SAPIENS | 1 | 2703 | 4wxq | A | 337 | GLN | p.Gln337Arg | Q99972 |
| HOMO SAPIENS | 5 | 2703 | 4wxq | A | 347 | TYR | p.Tyr347Ter | Q99972 |
| HOMO SAPIENS | 3 | 2703 | 4wxq | A | 364 | GLY | p.Gly364Val | Q99972 |
| HOMO SAPIENS | 2 | 2703 | 4wxq | A | 367 | GLY | p.Gly367Arg | Q99972 |
| HOMO SAPIENS | 2 | 2703 | 4wxq | A | 368 | GLN | p.Gln368Ter | Q99972 |
| HOMO SAPIENS | 6 | 2703 | 4wxq | A | 370 | PRO | p.Pro370Leu | Q99972 |
| HOMO SAPIENS | 1 | 2703 | 4wxq | A | 380 | ASP | p.Asp380His | Q99972 |
| HOMO SAPIENS | 8 | 2703 | 4wxq | A | 399 | GLY | p.Gly399Val | Q99972 |
| HOMO SAPIENS | 0 | 2703 | 4wxq | A | 423 | LYS | p.Lys423Glu | Q99972 |
| HOMO SAPIENS | 3 | 2703 | 4wxq | A | 433 | CYS | p.Cys433Arg | Q99972 |
| HOMO SAPIENS | 3 | 2703 | 4wxq | A | 437 | TYR | p.Tyr437His | Q99972 |
| HOMO SAPIENS | 2 | 2703 | 4wxq | A | 477 | ILE | p.Ile477Ser | Q99972 |
| HOMO SAPIENS | 3 | 2703 | 4wxq | A | 480 | ASN | p.Asn480Lys | Q99972 |
| HOMO SAPIENS | -1 | 34100 | 2fot | C | 1171 | GLN | p.Gln1173Pro | Q13813 |
| HOMO SAPIENS | 0 | 8773 | 1kw2 | A | 432 | ASP | p.Asp432Glu | P02774 |
| HOMO SAPIENS | 0 | 8773 | 1kw2 | A | 436 | LYS | p.Lys436Thr | P02774 |
| HOMO SAPIENS | 0 | 8402 | 4dzo | B | 649 | TYR | p.Tyr649Ter | Q9Y6D9 |
| HOMO SAPIENS | 0 | 39962 | 1nst | A | 611 | GLY | p.Gly611Ser | P52848 |
| HOMO SAPIENS | 2 | 22578 | 3dof | B | 45 | MET | p.Met45Arg | Q9Y2Y0 |
| HOMO SAPIENS | 1 | 23290 | 5j1s | A | 292 | ILE | p.Ile292Thr | O14656 |
| HOMO SAPIENS | 4 | 9505 | 4j1y | A | 534 | ARG | p.Arg534Trp | P09871 |
| HOMO SAPIENS | 9 | 260 | 2h57 | A | 31 | THR | p.Thr31Met | Q9H0F7 |

| | | | | | | | | |
|---|---|---|---|---|---|---|---|---|
| HOMO SAPIENS | 0 | 260 | 2h57 | A | 91 | ILE | p.Ile91Thr | Q9H0F7 |
| HOMO SAPIENS | 0 | 260 | 2h57 | A | 121 | ARG | p.Arg121His | Q9H0F7 |
| HOMO SAPIENS | 0 | 260 | 2h57 | A | 122 | ARG | p.Arg122Ter | Q9H0F7 |
| HOMO SAPIENS | 2 | 260 | 2h57 | A | 169 | GLY | p.Gly169Ala | Q9H0F7 |
| HOMO SAPIENS | 4 | 260 | 2h57 | A | 170 | LEU | p.Leu170Trp | Q9H0F7 |
| HOMO SAPIENS | 4 | 34023 | 2cr3 | A | 18 | GLY | p.Gly48Ser | P11362 |
| HOMO SAPIENS | 0 | 37667 | 3ii7 | A | 420 | ARG | p.Arg420Cys | Q8IXQ5 |
| HOMO SAPIENS | 0 | 41720 | 2k9u | A | 68 | ARG | p.Arg2364His | Q14315 |
| HOMO SAPIENS | 1 | 29250 | 5fmv | A | 568 | HIS | p.His568Gln | P08575 |
| HOMO SAPIENS | 1 | 1280 | 1fb1 | A | 69 | PRO | p.Pro69Leu | P30793 |
| HOMO SAPIENS | 3 | 1280 | 1fb1 | A | 88 | ARG | p.Arg88Trp | P30793 |
| HOMO SAPIENS | 2 | 1280 | 1fb1 | A | 108 | GLY | p.Gly108Asp | P30793 |
| HOMO SAPIENS | 4 | 1280 | 1fb1 | A | 134 | ASP | p.Asp134Val | P30793 |
| HOMO SAPIENS | 6 | 1280 | 1fb1 | A | 135 | ILE | p.Ile135Lys | P30793 |
| HOMO SAPIENS | 9 | 1280 | 1fb1 | A | 144 | HIS | p.His144Pro | P30793 |
| HOMO SAPIENS | 9 | 1280 | 1fb1 | A | 184 | ARG | p.Arg184His | P30793 |
| HOMO SAPIENS | 1 | 1280 | 1fb1 | A | 196 | ALA | p.Ala196Ser | P30793 |
| HOMO SAPIENS | 3 | 1280 | 1fb1 | A | 201 | GLY | p.Gly201Glu | P30793 |
| HOMO SAPIENS | 2 | 1280 | 1fb1 | A | 204 | VAL | p.Val204Ile | P30793 |
| HOMO SAPIENS | 2 | 1280 | 1fb1 | A | 211 | MET | p.Met211Ile | P30793 |
| HOMO SAPIENS | 0 | 1280 | 1fb1 | A | 221 | MET | p.Met221Thr | P30793 |
| HOMO SAPIENS | 3 | 1280 | 1fb1 | A | 224 | LYS | p.Lys224Arg | P30793 |
| HOMO SAPIENS | 2 | 1280 | 1fb1 | A | 249 | ARG | p.Arg249Ser | P30793 |
| HOMO SAPIENS | 8 | 41352 | 4k12 | A | 47 | ILE | p.Ile551Thr | P08603 |
| HOMO SAPIENS | 9 | 20703 | 5nir | A | 57 | CYS | p.Cys57Tyr | P02458 |
| HOMO SAPIENS | 9 | 20703 | 5nir | A | 64 | CYS | p.Cys64Ter | P02458 |
| HOMO SAPIENS | 0 | 20087 | 4cmn | A | 299 | MET | p.Met299Ile | Q01968 |
| HOMO SAPIENS | 1 | 20819 | 3aaf | A | 979 | ILE | p.Ile979Met | Q14191 |
| HOMO SAPIENS | 1 | 20819 | 3aaf | A | 984 | LEU | p.Leu984Ile | Q14191 |
| HOMO SAPIENS | 3 | 20819 | 3aaf | A | 987 | ARG | p.Arg987Ter | Q14191 |
| HOMO SAPIENS | -1 | 20819 | 3aaf | A | 1074 | PHE | p.Leu1074Phe | Q14191 |
| HOMO SAPIENS | 0 | 3619 | 3dem | A | 206 | ASN | p.Asn225Ser | P48740 |
| HOMO SAPIENS | 1 | 3619 | 3dem | A | 269 | ARG | p.Arg288Leu | P48740 |
| HOMO SAPIENS | 0 | 3105 | 3ooi | A | 164 | TYR | p.Tyr2015His | Q96L73 |
| HOMO SAPIENS | 6 | 15828 | 2lkx | A | 16 | LEU | p.Leu100Gln | Q99697 |
| HOMO SAPIENS | 8 | 15828 | 2lkx | A | 24 | ARG | p.Arg108His | Q99697 |
| HOMO SAPIENS | 2 | 15828 | 2lkx | A | 30 | THR | p.Thr114Pro | Q99697 |
| HOMO SAPIENS | 8 | 15828 | 2lkx | A | 45 | VAL | p.Val129Leu | Q99697 |
| HOMO SAPIENS | 8 | 15828 | 2lkx | A | 46 | ARG | p.Arg130Trp | Q99697 |
| HOMO SAPIENS | 9 | 15828 | 2lkx | A | 53 | ARG | p.Arg137Pro | Q99697 |
| HOMO SAPIENS | 9 | 13017 | 4irq | A | 141 | ARG | p.Arg141Trp | Q9UBV7 |
| HOMO SAPIENS | 0 | 13017 | 4irq | A | 186 | ALA | p.Ala186Asp | Q9UBV7 |
| HOMO SAPIENS | 0 | 13017 | 4irq | A | 206 | LEU | p.Leu206Pro | Q9UBV7 |
| HOMO SAPIENS | 1 | 13017 | 4irq | A | 214 | CYS | p.Cys214Tyr | Q9UBV7 |
| HOMO SAPIENS | 0 | 13017 | 4irq | A | 270 | ARG | p.Arg270Cys | Q9UBV7 |
| HOMO SAPIENS | -1 | 13053 | 4mqw | X | 307 | ALA | p.Ala307Thr | P23945 |
| HOMO SAPIENS | 0 | 19406 | 4b3g | A | 43 | ARG | p.Arg43Ter | P38935 |
| HOMO SAPIENS | 0 | 19406 | 4b3g | A | 46 | CYS | p.Cys46Ter | P38935 |
| HOMO SAPIENS | 0 | 19406 | 4b3g | A | 75 | ALA | p.Ala75Thr | P38935 |
| HOMO SAPIENS | 0 | 19406 | 4b3g | A | 201 | LEU | p.Leu201Ser | P38935 |
| HOMO SAPIENS | 3 | 19406 | 4b3g | A | 213 | HIS | p.His213Arg | P38935 |
| HOMO SAPIENS | 0 | 19406 | 4b3g | A | 275 | ILE | p.Ile275Val | P38935 |
| HOMO SAPIENS | -1 | 19406 | 4b3g | A | 320 | ARG | p.Arg320Ter | P38935 |
| HOMO SAPIENS | 0 | 19406 | 4b3g | A | 361 | LEU | p.Leu361Pro | P38935 |

| | | | | | | | | |
|---|---|---|---|---|---|---|---|---|
| HOMO SAPIENS | 0 | 19406 | 4b3g | A | 369 | PHE | p.Phe369Leu | P38935 |
| HOMO SAPIENS | 0 | 19406 | 4b3g | A | 493 | THR | p.Thr493Ile | P38935 |
| HOMO SAPIENS | 0 | 19406 | 4b3g | A | 496 | CYS | p.Cys496Ter | P38935 |
| HOMO SAPIENS | 0 | 19406 | 4b3g | A | 514 | GLU | p.Glu514Lys | P38935 |
| HOMO SAPIENS | 0 | 19406 | 4b3g | A | 528 | ALA | p.Ala528Thr | P38935 |
| HOMO SAPIENS | 0 | 19406 | 4b3g | A | 531 | PRO | p.Pro531Thr | P38935 |
| HOMO SAPIENS | 0 | 19406 | 4b3g | A | 580 | VAL | p.Val580Ile | P38935 |
| HOMO SAPIENS | 6 | 19406 | 4b3g | A | 603 | ARG | p.Arg603His | P38935 |
| HOMO SAPIENS | 0 | 21501 | 5jde | A | 40 | SER | p.Ser2831Asn | Q8WZ42 |
| HOMO SAPIENS | 5 | 21501 | 5jde | A | 189 | ALA | p.Ala2980Thr | Q8WZ42 |
| HOMO SAPIENS | 6 | 15008 | 3mop | H | 12 | ARG | p.Arg12Cys | Q9NWZ3 |
| HOMO SAPIENS | 8 | 17455 | 3jtc | C | 15 | ARG | p.Arg57Trp | P04070 |
| HOMO SAPIENS | -1 | 17455 | 3jtc | C | 20 | CGU | p.Glu62Ala | P04070 |
| HOMO SAPIENS | 0 | 27795 | 5gox | B | 625 | TYR | p.Tyr625Ter | Q92878 |
| HOMO SAPIENS | 0 | 27795 | 5gox | B | 653 | SER | p.Ser653Ter | Q92878 |
| HOMO SAPIENS | -1 | 6558 | 5nug | A | 137 | VAL | p.Val137Ile | Q14204 |
| HOMO SAPIENS | -1 | 6558 | 5nug | A | 567 | ARG | p.Arg567His | Q14204 |
| HOMO SAPIENS | -1 | 6558 | 5nug | A | 1140 | THR | p.Thr1140Met | Q14204 |
| HOMO SAPIENS | -1 | 6558 | 5nug | A | 1991 | ASP | p.Asp1991Asn | Q14204 |
| HOMO SAPIENS | -1 | 6558 | 5nug | A | 2401 | LYS | p.Lys2401Asn | Q14204 |
| HOMO SAPIENS | 0 | 6558 | 5nug | A | 2488 | ARG | p.Arg2488His | Q14204 |
| HOMO SAPIENS | 0 | 6558 | 5nug | A | 3907 | HIS | p.His3907Gln | Q14204 |
| HOMO SAPIENS | 0 | 6558 | 5nug | A | 3981 | THR | p.Thr3981Arg | Q14204 |
| HOMO SAPIENS | 0 | 6558 | 5nug | A | 4394 | THR | p.Thr4394Met | Q14204 |
| HOMO SAPIENS | 0 | 6558 | 5nug | A | 4573 | ASN | p.Asn4573Ser | Q14204 |
| HOMO SAPIENS | 0 | 6558 | 5nug | A | 4603 | SER | p.Ser4603Ile | Q14204 |
| HOMO SAPIENS | 1 | 4244 | 1umk | A | 57 | ARG | p.Arg58Gln | P00387 |
| HOMO SAPIENS | 1 | 4244 | 1umk | A | 72 | LEU | p.Leu73Pro | P00387 |
| HOMO SAPIENS | 1 | 4244 | 1umk | A | 105 | VAL | p.Val106Met | P00387 |
| HOMO SAPIENS | 0 | 4244 | 1umk | A | 116 | THR | p.Thr117Ser | P00387 |
| HOMO SAPIENS | 8 | 4244 | 1umk | A | 127 | SER | p.Ser128Pro | P00387 |
| HOMO SAPIENS | 0 | 4244 | 1umk | A | 148 | LEU | p.Leu149Pro | P00387 |
| HOMO SAPIENS | 0 | 4244 | 1umk | A | 159 | ARG | p.Arg160Ter | P00387 |
| HOMO SAPIENS | 0 | 4244 | 1umk | A | 203 | CYS | p.Cys204Arg | P00387 |
| HOMO SAPIENS | 0 | 4244 | 1umk | A | 203 | CYS | p.Cys204Tyr | P00387 |
| HOMO SAPIENS | 2 | 4244 | 1umk | A | 239 | ASP | p.Asp240Gly | P00387 |
| HOMO SAPIENS | 0 | 4244 | 1umk | A | 291 | GLY | p.Gly292Asp | P00387 |
| HOMO SAPIENS | 4 | 34495 | 1s35 | A | 1151 | ASP | p.Asn1151Asp | P11277 |
| HOMO SAPIENS | 3 | 30110 | 3zqs | A | 224 | THR | p.Thr224Ala | Q9NW38 |
| HOMO SAPIENS | 3 | 17588 | 5dse | B | 53 | LEU | p.Leu53Pro | Q9BYI3 |
| HOMO SAPIENS | 0 | 1564 | 4yqf | B | 72 | HIS | p.His347Gln | Q9UHD8 |
| HOMO SAPIENS | 0 | 17673 | 4jnw | A | 32 | ARG | p.Arg32186His | Q8WZ42 |
| HOMO SAPIENS | 3 | 17673 | 4jnw | A | 140 | ILE | p.Ile32294Val | Q8WZ42 |
| HOMO SAPIENS | 5 | 17673 | 4jnw | A | 269 | ALA | p.Ala32423Thr | Q8WZ42 |
| HOMO SAPIENS | 3 | 12858 | 3s97 | C | 302 | LEU | p.Leu302Pro | Q12860 |
| HOMO SAPIENS | 0 | 6147 | 4nn0 | A | 114 | ARG | p.Arg114Gln | Q9BXJ0 |
| HOMO SAPIENS | 0 | 7549 | 1tr2 | A | 277 | LEU | p.Leu277Met | P18206 |
| HOMO SAPIENS | 1 | 7549 | 1tr2 | A | 413 | ALA | p.Ala413Thr | P18206 |
| HOMO SAPIENS | 1 | 7549 | 1tr2 | A | 519 | ILE | p.Ile519Leu | P18206 |
| HOMO SAPIENS | 5 | 7549 | 1tr2 | A | 547 | ARG | p.Arg547Ter | P18206 |
| HOMO SAPIENS | 2 | 7549 | 1tr2 | A | 636 | HIS | p.His636Arg | P18206 |
| HOMO SAPIENS | 2 | 7549 | 1tr2 | A | 824 | ILE | p.Ile824Met | P18206 |
| HOMO SAPIENS | 2 | 13900 | 3gro | A | 45 | CYS | p.Cys45Tyr | P50897 |
| HOMO SAPIENS | 2 | 13900 | 3gro | A | 75 | THR | p.Thr75Pro | P50897 |

| | | | | | | | | |
|---|---|---|---|---|---|---|---|---|
| HOMO SAPIENS | 2 | 13900 | 3gro | A | 79 | ASP | p.Asp79Gly | P50897 |
| HOMO SAPIENS | 5 | 13900 | 3gro | A | 91 | GLN | p.Gln91Pro | P50897 |
| HOMO SAPIENS | 5 | 13900 | 3gro | A | 96 | CYS | p.Cys96Tyr | P50897 |
| HOMO SAPIENS | 0 | 13900 | 3gro | A | 104 | LYS | p.Lys104Ter | P50897 |
| HOMO SAPIENS | 2 | 13900 | 3gro | A | 108 | GLY | p.Gly108Arg | P50897 |
| HOMO SAPIENS | 2 | 13900 | 3gro | A | 109 | TYR | p.Tyr109Asp | P50897 |
| HOMO SAPIENS | 0 | 13900 | 3gro | A | 122 | ARG | p.Arg122Trp | P50897 |
| HOMO SAPIENS | 0 | 13900 | 3gro | A | 138 | SER | p.Ser138Leu | P50897 |
| HOMO SAPIENS | 5 | 13900 | 3gro | A | 151 | ARG | p.Arg151Ter | P50897 |
| HOMO SAPIENS | 7 | 13900 | 3gro | A | 164 | ARG | p.Arg164Ter | P50897 |
| HOMO SAPIENS | 0 | 13900 | 3gro | A | 177 | GLN | p.Gln177Glu | P50897 |
| HOMO SAPIENS | 0 | 13900 | 3gro | A | 181 | VAL | p.Val181Leu | P50897 |
| HOMO SAPIENS | 0 | 13900 | 3gro | A | 184 | GLU | p.Glu184Lys | P50897 |
| HOMO SAPIENS | 5 | 13900 | 3gro | A | 186 | TRP | p.Trp186Ter | P50897 |
| HOMO SAPIENS | 0 | 13900 | 3gro | A | 219 | LEU | p.Leu219Gln | P50897 |
| HOMO SAPIENS | 0 | 13900 | 3gro | A | 222 | LEU | p.Leu222Pro | P50897 |
| HOMO SAPIENS | 5 | 13900 | 3gro | A | 296 | TRP | p.Trp296Ter | P50897 |
| HOMO SAPIENS | 6 | 35028 | 5mwb | A | 444 | CYS | p.Cys444Tyr | Q04721 |
| HOMO SAPIENS | 0 | 35028 | 5mwb | A | 514 | ASN | p.Asn514Ser | Q04721 |
| HOMO SAPIENS | 5 | 1483 | 4gl2 | A | 822 | ARG | p.Arg822Gln | Q9BYX4 |
| HOMO SAPIENS | 0 | 1483 | 4gl2 | A | 843 | ARG | p.His843Arg | Q9BYX4 |
| HOMO SAPIENS | -1 | 28838 | 3cl3 | D | 153 | LEU | p.Leu153Arg | Q9Y6K9 |
| HOMO SAPIENS | 2 | 22563 | 4zel | B | 79 | ARG | p.Arg79Trp | P09172 |
| HOMO SAPIENS | 2 | 22563 | 4zel | B | 101 | VAL | p.Val101Met | P09172 |
| HOMO SAPIENS | 2 | 22563 | 4zel | B | 114 | ASP | p.Asp114Glu | P09172 |
| HOMO SAPIENS | 0 | 22563 | 4zel | B | 201 | ASN | p.Asn201Ser | P09172 |
| HOMO SAPIENS | 2 | 22563 | 4zel | B | 211 | ALA | p.Ala211Thr | P09172 |
| HOMO SAPIENS | -1 | 22563 | 4zel | B | 290 | ASP | p.Asp290Asn | P09172 |
| HOMO SAPIENS | 2 | 22563 | 4zel | B | 318 | ALA | p.Ala318Ser | P09172 |
| HOMO SAPIENS | 2 | 22563 | 4zel | B | 345 | ASP | p.Asp345Asn | P09172 |
| HOMO SAPIENS | 2 | 22563 | 4zel | B | 362 | ALA | p.Ala362Glu | P09172 |
| HOMO SAPIENS | 2 | 22563 | 4zel | B | 549 | ARG | p.Arg549Cys | P09172 |
| HOMO SAPIENS | 9 | 9780 | 1y6k | R | 36 | TYR | p.Tyr57Cys | Q13651 |
| HOMO SAPIENS | 1 | 9780 | 1y6k | R | 40 | LEU | p.Leu61Val | Q13651 |
| HOMO SAPIENS | 1 | 9780 | 1y6k | R | 63 | THR | p.Thr84Ile | Q13651 |
| HOMO SAPIENS | 9 | 9780 | 1y6k | R | 80 | ARG | p.Arg101Trp | Q13651 |
| HOMO SAPIENS | 0 | 9780 | 1y6k | R | 120 | GLY | p.Gly141Arg | Q13651 |
| HOMO SAPIENS | 0 | 9780 | 1y6k | R | 138 | SER | p.Ser159Gly | Q13651 |
| HOMO SAPIENS | 0 | 9780 | 1y6k | R | 203 | ILE | p.Ile224Val | Q13651 |
| HOMO SAPIENS | 0 | 16019 | 3rep | A | 211 | ARG | p.Arg211Cys | Q13418 |
| HOMO SAPIENS | 3 | 7879 | 5ec3 | A | 33 | ALA | p.Ala33Thr | P32754 |
| HOMO SAPIENS | 4 | 7879 | 5ec3 | A | 113 | ARG | p.Arg113Gln | P32754 |
| HOMO SAPIENS | 3 | 7879 | 5ec3 | A | 340 | VAL | p.Val340Leu | P32754 |
| HOMO SAPIENS | 0 | 8904 | 1qk9 | A | 6 | LYS | p.Lys82Arg | P51608 |
| HOMO SAPIENS | 4 | 8904 | 1qk9 | A | 24 | LEU | p.Leu100Val | P51608 |
| HOMO SAPIENS | 7 | 8904 | 1qk9 | A | 30 | ARG | p.Arg106Trp | P51608 |
| HOMO SAPIENS | 6 | 8904 | 1qk9 | A | 57 | ARG | p.Arg133Cys | P51608 |
| HOMO SAPIENS | 3 | 8904 | 1qk9 | A | 61 | GLU | p.Glu137Gly | P51608 |
| HOMO SAPIENS | 2 | 8904 | 1qk9 | A | 64 | ALA | p.Ala140Val | P51608 |
| HOMO SAPIENS | 5 | 8904 | 1qk9 | A | 79 | PHE | p.Phe155Ser | P51608 |
| HOMO SAPIENS | 3 | 8904 | 1qk9 | A | 82 | THR | p.Thr158Met | P51608 |
| HOMO SAPIENS | 1 | 23121 | 3e9k | A | 198 | THR | p.Thr198Ala | Q16719 |
| HOMO SAPIENS | -1 | 26090 | 3tje | F | 19 | LEU | p.Leu35Met | P25445 |
| HOMO SAPIENS | 8 | 26090 | 3tje | F | 106 | THR | p.Thr122Ile | P25445 |

| | | | | | | | | |
|---|---|---|---|---|---|---|---|---|
| HOMO SAPIENS | 2 | 36404 | 5cqr | A | 765 | GLY | p.Gly765Glu | O95163 |
| HOMO SAPIENS | 2 | 36404 | 5cqr | A | 816 | ILE | p.Ile816Leu | O95163 |
| HOMO SAPIENS | 2 | 36404 | 5cqr | A | 830 | ILE | p.Ile830Met | O95163 |
| HOMO SAPIENS | 5 | 36404 | 5cqr | A | 939 | TYR | p.Tyr939Ter | O95163 |
| HOMO SAPIENS | 2 | 36404 | 5cqr | A | 952 | LYS | p.Lys952Ile | O95163 |
| HOMO SAPIENS | 5 | 36404 | 5cqr | A | 1013 | GLY | p.Gly1013Ser | O95163 |
| HOMO SAPIENS | 5 | 36404 | 5cqr | A | 1072 | CYS | p.Cys1072Ser | O95163 |
| HOMO SAPIENS | -1 | 36404 | 5cqr | A | 1158 | PRO | p.Pro1158Leu | O95163 |
| HOMO SAPIENS | 1 | 34149 | 2k40 | A | 18 | ASN | p.Asn125Ser | Q9UBX0 |
| HOMO SAPIENS | 0 | 34149 | 2k40 | A | 42 | LEU | p.Glu149Lys | Q9UBX0 |
| HOMO SAPIENS | 9 | 34149 | 2k40 | A | 53 | ARG | p.Arg160Cys | Q9UBX0 |
| HOMO SAPIENS | 0 | 34149 | 2k40 | A | 63 | SER | p.Ser170Leu | Q9UBX0 |
| HOMO SAPIENS | 2 | 2468 | 3vp6 | A | 484 | LEU | p.Leu484Met | Q99259 |
| HOMO SAPIENS | 0 | 2468 | 3vp6 | A | 532 | ARG | p.Arg532Gln | Q99259 |
| HOMO SAPIENS | 5 | 1253 | 5xtd | S | 8 | GLY | p.Gly8Arg | O15239 |
| HOMO SAPIENS | 7 | 1253 | 5xtd | S | 32 | GLY | p.Gly32Arg | O15239 |
| HOMO SAPIENS | 7 | 1253 | 5xtd | S | 37 | ARG | p.Arg37Ser | O15239 |
| HOMO SAPIENS | -1 | 13292 | 3u1k | A | 121 | VAL | p.Ile121Val | Q8TCS8 |
| HOMO SAPIENS | 2 | 13292 | 3u1k | A | 165 | PRO | p.Pro165Ser | Q8TCS8 |
| HOMO SAPIENS | 2 | 13292 | 3u1k | A | 192 | ARG | p.Arg192Gln | Q8TCS8 |
| HOMO SAPIENS | 1 | 13292 | 3u1k | A | 315 | LYS | p.Lys315Arg | Q8TCS8 |
| HOMO SAPIENS | 4 | 13292 | 3u1k | A | 509 | VAL | p.Val509Ile | Q8TCS8 |
| HOMO SAPIENS | 7 | 13292 | 3u1k | A | 531 | THR | p.Thr531Arg | Q8TCS8 |
| HOMO SAPIENS | 0 | 13292 | 3u1k | A | 590 | ASN | p.Asn590Asp | Q8TCS8 |
| HOMO SAPIENS | 0 | 27048 | 4p5w | B | 184 | ARG | p.Arg184Ter | Q9UJS0 |
| HOMO SAPIENS | 0 | 27048 | 4p5w | B | 225 | SER | p.Ser225Ter | Q9UJS0 |
| HOMO SAPIENS | -1 | 27048 | 4p5w | B | 631 | ALA | p.Arg360Ter | Q9UJS0 |
| HOMO SAPIENS | 0 | 24240 | 4mdr | A | 318 | ARG | p.Arg318Ter | O00189 |
| HOMO SAPIENS | 0 | 24240 | 4mdr | A | 338 | ARG | p.Arg338Ter | O00189 |
| HOMO SAPIENS | 0 | 13429 | 4cg4 | A | 479 | PHE | p.Phe479Leu | O15553 |
| HOMO SAPIENS | 0 | 13429 | 4cg4 | A | 487 | VAL | p.Val487Leu | O15553 |
| HOMO SAPIENS | 3 | 13429 | 4cg4 | A | 653 | ARG | p.Arg653His | O15553 |
| HOMO SAPIENS | 1 | 13429 | 4cg4 | A | 680 | MET | p.Met680Ile | O15553 |
| HOMO SAPIENS | 1 | 13429 | 4cg4 | A | 688 | TYR | p.Tyr688Ter | O15553 |
| HOMO SAPIENS | 0 | 13429 | 4cg4 | A | 694 | MET | p.Met694Ile | O15553 |
| HOMO SAPIENS | 0 | 13429 | 4cg4 | A | 694 | MET | p.Met694Val | O15553 |
| HOMO SAPIENS | 1 | 13429 | 4cg4 | A | 726 | VAL | p.Val726Ala | O15553 |
| HOMO SAPIENS | 0 | 13429 | 4cg4 | A | 761 | ARG | p.Arg761His | O15553 |
| HOMO SAPIENS | 0 | 2079 | 4okh | A | 691 | GLY | p.Gly691Arg | P20807 |
| HOMO SAPIENS | 2 | 2079 | 4okh | A | 748 | ARG | p.Arg748Ter | P20807 |
| HOMO SAPIENS | 0 | 2079 | 4okh | A | 763 | TYR | p.Tyr763Cys | P20807 |
| HOMO SAPIENS | 1 | 2079 | 4okh | A | 769 | ARG | p.Arg769Gln | P20807 |
| HOMO SAPIENS | 9 | 2079 | 4okh | A | 777 | ILE | p.Ile777Val | P20807 |
| HOMO SAPIENS | 8 | 2079 | 4okh | A | 778 | ASP | p.Asp778His | P20807 |
| HOMO SAPIENS | 5 | 2079 | 4okh | A | 780 | ASP | p.Asp780His | P20807 |
| HOMO SAPIENS | 1 | 1200 | 2gi7 | A | 142 | ARG | p.Arg162Trp | Q9HCN6 |
| HOMO SAPIENS | 4 | 9199 | 3ezq | C | 260 | ASP | p.Asp260Val | P25445 |
| HOMO SAPIENS | 0 | 36522 | 5ms9 | A | 122 | ARG | p.Arg122Cys | P35555 |
| HOMO SAPIENS | 0 | 36522 | 5ms9 | A | 240 | ARG | p.Arg240Cys | P35555 |
| HOMO SAPIENS | 0 | 17448 | 3g2f | A | 225 | ARG | p.Arg225His | Q13873 |
| HOMO SAPIENS | 0 | 17448 | 3g2f | A | 347 | CYS | p.Cys347Tyr | Q13873 |
| HOMO SAPIENS | 3 | 17448 | 3g2f | A | 485 | ASP | p.Asp485Gly | Q13873 |
| HOMO SAPIENS | 7 | 17448 | 3g2f | A | 491 | ARG | p.Arg491Gln | Q13873 |
| HOMO SAPIENS | 7 | 17448 | 3g2f | A | 491 | ARG | p.Arg491Trp | Q13873 |

| | | | | | | | | |
|---|---|---|---|---|---|---|---|---|
| HOMO SAPIENS | 0 | 4861 | 4ux8 | A | 56 | LEU | p.Leu56Met | P07949 |
| HOMO SAPIENS | 1 | 4861 | 4ux8 | A | 88 | ILE | p.Ile88Leu | P07949 |
| HOMO SAPIENS | 0 | 4861 | 4ux8 | A | 114 | ARG | p.Arg114His | P07949 |
| HOMO SAPIENS | 5 | 4861 | 4ux8 | A | 180 | ARG | p.Arg180Gln | P07949 |
| HOMO SAPIENS | 0 | 4861 | 4ux8 | A | 208 | GLU | p.Glu208Asp | P07949 |
| HOMO SAPIENS | 0 | 4861 | 4ux8 | A | 373 | ALA | p.Ala373Val | P07949 |
| HOMO SAPIENS | 0 | 4861 | 4ux8 | A | 375 | LEU | p.Leu375Gln | P07949 |
| HOMO SAPIENS | -1 | 4861 | 4ux8 | A | 386 | ALA | p.Ala386Val | P07949 |
| HOMO SAPIENS | 0 | 4861 | 4ux8 | A | 446 | GLY | p.Gly446Arg | P07949 |
| HOMO SAPIENS | 0 | 4861 | 4ux8 | A | 480 | GLU | p.Glu480Lys | P07949 |
| HOMO SAPIENS | 0 | 4861 | 4ux8 | A | 489 | ASP | p.Asp489Asn | P07949 |
| HOMO SAPIENS | -1 | 4861 | 4ux8 | A | 510 | ALA | p.Ala510Val | P07949 |
| HOMO SAPIENS | -1 | 4861 | 4ux8 | A | 533 | GLY | p.Gly533Cys | P07949 |
| HOMO SAPIENS | -1 | 4861 | 4ux8 | A | 567 | ASP | p.Asp567Asn | P07949 |
| HOMO SAPIENS | -1 | 4861 | 4ux8 | A | 609 | CYS | p.Cys609Tyr | P07949 |
| HOMO SAPIENS | -1 | 4861 | 4ux8 | A | 609 | CYS | p.Cys609Gly | P07949 |
| HOMO SAPIENS | -1 | 4861 | 4ux8 | A | 611 | CYS | p.Cys611Trp | P07949 |
| HOMO SAPIENS | -1 | 4861 | 4ux8 | A | 618 | CYS | p.Cys618Gly | P07949 |
| HOMO SAPIENS | -1 | 4861 | 4ux8 | A | 618 | CYS | p.Cys618Phe | P07949 |
| HOMO SAPIENS | -1 | 4861 | 4ux8 | A | 620 | CYS | p.Cys620Gly | P07949 |
| HOMO SAPIENS | -1 | 4861 | 4ux8 | A | 620 | CYS | p.Cys620Ser | P07949 |
| HOMO SAPIENS | -1 | 4861 | 4ux8 | A | 620 | CYS | p.Cys620Trp | P07949 |
| HOMO SAPIENS | -1 | 4861 | 4ux8 | A | 623 | GLU | p.Glu623Lys | P07949 |
| HOMO SAPIENS | -1 | 4861 | 4ux8 | A | 631 | ASP | p.Asp631Gly | P07949 |
| HOMO SAPIENS | -1 | 4861 | 4ux8 | A | 631 | ASP | p.Asp631Glu | P07949 |
| HOMO SAPIENS | -1 | 4861 | 4ux8 | A | 634 | CYS | p.Cys634Gly | P07949 |
| HOMO SAPIENS | -1 | 4861 | 4ux8 | A | 634 | CYS | p.Cys634Phe | P07949 |
| HOMO SAPIENS | -1 | 4861 | 4ux8 | A | 634 | CYS | p.Cys634Trp | P07949 |
| HOMO SAPIENS | 0 | 36308 | 4nkq | B | 123 | PRO | p.Pro142Leu | P15509 |
| HOMO SAPIENS | 0 | 36308 | 4nkq | B | 145 | ARG | p.Arg164Gln | P15509 |
| HOMO SAPIENS | -1 | 36308 | 4nkq | B | 177 | GLY | p.Gly196Arg | P15509 |
| HOMO SAPIENS | 0 | 36006 | 1oqa | A | 107 | HIS | p.His1860Pro | P38398 |
| HOMO SAPIENS | 7 | 15394 | 2ozo | A | 465 | ARG | p.Arg465His | P43403 |
| HOMO SAPIENS | 7 | 15394 | 2ozo | A | 465 | ARG | p.Arg465Cys | P43403 |
| HOMO SAPIENS | 4 | 15394 | 2ozo | A | 518 | SER | p.Ser518Arg | P43403 |
| HOMO SAPIENS | 1 | 20813 | 3o5z | A | 19 | ARG | p.Arg19Trp | O00459 |
| HOMO SAPIENS | 2 | 4329 | 2bb5 | A | 5 | ILE | p.Ile23Val | P20062 |
| HOMO SAPIENS | 4 | 4329 | 2bb5 | A | 59 | LYS | p.Lys77Met | P20062 |
| HOMO SAPIENS | 4 | 4329 | 2bb5 | A | 197 | ARG | p.Arg215Trp | P20062 |
| HOMO SAPIENS | 1 | 4329 | 2bb5 | A | 241 | PRO | p.Arg259Pro | P20062 |
| HOMO SAPIENS | 2 | 4329 | 2bb5 | A | 315 | THR | p.Thr333Lys | P20062 |
| HOMO SAPIENS | 5 | 4329 | 2bb5 | A | 330 | SER | p.Ser348Phe | P20062 |
| HOMO SAPIENS | 4 | 4329 | 2bb5 | A | 358 | LEU | p.Leu376Ser | P20062 |
| HOMO SAPIENS | 5 | 4329 | 2bb5 | A | 381 | ARG | p.Arg399Gln | P20062 |
| HOMO SAPIENS | 4 | 18294 | 4r30 | A | 163 | ASN | p.Asn163Asp | O95278 |
| HOMO SAPIENS | 6 | 18294 | 4r30 | A | 227 | ALA | p.Ala227Val | O95278 |
| HOMO SAPIENS | 4 | 18294 | 4r30 | A | 241 | ARG | p.Arg241Ter | O95278 |
| HOMO SAPIENS | 4 | 18294 | 4r30 | A | 325 | ARG | p.Arg325His | O95278 |
| HOMO SAPIENS | 0 | 2632 | 4xem | A | 22 | GLU | p.Glu22Lys | P49588 |
| HOMO SAPIENS | 0 | 2632 | 4xem | A | 275 | GLY | p.Gly275Asp | P49588 |
| HOMO SAPIENS | 1 | 2632 | 4xem | A | 329 | ARG | p.Arg329His | P49588 |
| HOMO SAPIENS | 0 | 11431 | 5t4n | A | 284 | GLN | p.Gln305Glu | Q96QU1 |
| HOMO SAPIENS | 0 | 11431 | 5t4n | A | 315 | ARG | p.Arg336Ter | Q96QU1 |
| HOMO SAPIENS | 0 | 11431 | 5t4n | A | 322 | ARG | p.Arg343Lys | Q96QU1 |

| | | | | | | | | |
|---|---|---|---|---|---|---|---|---|
| HOMO SAPIENS | 1 | 11431 | 5t4n | A | 326 | LEU | p.Leu347Phe | Q96QU1 |
| HOMO SAPIENS | 0 | 11431 | 5t4n | A | 373 | TYR | p.Tyr394Cys | Q96QU1 |
| HOMO SAPIENS | 0 | 11431 | 5t4n | A | 398 | ASN | p.Asn419Ser | Q96QU1 |
| HOMO SAPIENS | 0 | 11431 | 5t4n | A | 414 | ALA | p.Asp435Ala | Q96QU1 |
| HOMO SAPIENS | 0 | 11431 | 5t4n | A | 433 | VAL | p.Val454Phe | Q96QU1 |
| HOMO SAPIENS | 1 | 11431 | 5t4n | A | 507 | VAL | p.Val528Asp | Q96QU1 |
| HOMO SAPIENS | 0 | 11431 | 5t4n | A | 547 | ALA | p.Ala568Ser | Q96QU1 |
| HOMO SAPIENS | 3 | 3194 | 5xtd | L | 84 | ARG | p.Arg84His | O43181 |
| HOMO SAPIENS | 1 | 3194 | 5xtd | L | 130 | THR | p.Thr130Ser | O43181 |
| HOMO SAPIENS | 8 | 3194 | 5xtd | L | 136 | SER | p.Ser136Ala | O43181 |
| HOMO SAPIENS | 0 | 40749 | 1nun | A | 106 | CYS | p.Cys106Phe | O15520 |
| HOMO SAPIENS | 9 | 40749 | 1nun | A | 156 | ILE | p.Ile156Arg | O15520 |
| HOMO SAPIENS | 1 | 40749 | 1nun | A | 193 | ARG | p.Arg193Ter | O15520 |
| HOMO SAPIENS | 0 | 40749 | 1nun | A | 207 | HIS | p.His207Pro | O15520 |
| HOMO SAPIENS | 0 | 7026 | 3n1f | D | 875 | SER | p.Ser875Gly | Q4KMG0 |
| HOMO SAPIENS | 6 | 23802 | 3rzv | A | 424 | ARG | p.Arg424Ter | O95630 |
| HOMO SAPIENS | 0 | 36720 | 1jm7 | A | 12 | GLN | p.Gln12Ter | P38398 |
| HOMO SAPIENS | 3 | 36720 | 1jm7 | A | 23 | GLU | p.Glu23Asp | P38398 |
| HOMO SAPIENS | 1 | 36720 | 1jm7 | A | 45 | LYS | p.Lys45Gln | P38398 |
| HOMO SAPIENS | 8 | 36720 | 1jm7 | A | 47 | CYS | p.Cys47Ter | P38398 |
| HOMO SAPIENS | 0 | 36720 | 1jm7 | A | 59 | SER | p.Ser59Ter | P38398 |
| HOMO SAPIENS | 8 | 36720 | 1jm7 | A | 61 | CYS | p.Cys61Gly | P38398 |
| HOMO SAPIENS | 8 | 36720 | 1jm7 | A | 63 | LEU | p.Leu63Ter | P38398 |
| HOMO SAPIENS | 7 | 36720 | 1jm7 | A | 64 | CYS | p.Cys64Gly | P38398 |
| HOMO SAPIENS | 0 | 36720 | 1jm7 | A | 101 | TYR | p.Tyr101Ter | P38398 |
| HOMO SAPIENS | 0 | 36720 | 1jm7 | A | null | TYR | p.Tyr105Cys | P38398 |
| HOMO SAPIENS | 0 | 28500 | 3kuz | A | 1217 | CYS | p.Cys1217Arg | O60486 |
| HOMO SAPIENS | 5 | 8097 | 3p56 | C | 69 | ARG | p.Arg69Trp | Q8TDP1 |
| HOMO SAPIENS | 5 | 8097 | 3p56 | C | 143 | LYS | p.Lys143Ile | Q8TDP1 |
| HOMO SAPIENS | 2 | 30073 | 3nmz | A | 380 | ARG | p.Arg380Gln | P25054 |
| HOMO SAPIENS | 1 | 22573 | 1cjy | A | 111 | SER | p.Ser111Pro | P47712 |
| HOMO SAPIENS | 0 | 22573 | 1cjy | A | 485 | ARG | p.Arg485His | P47712 |
| HOMO SAPIENS | 2 | 965 | 3E+077 | A | 99 | ALA | p.Ala99Val | Q9Y617 |
| HOMO SAPIENS | 2 | 965 | 3E+077 | A | 100 | ASP | p.Asp100Ala | Q9Y617 |
| HOMO SAPIENS | 4 | 1503 | 3kxc | A | 62 | ARG | p.Arg62Trp | O43617 |
| HOMO SAPIENS | 6 | 9418 | 1hdr | A | 22 | GLY | p.Gly23Asp | P09417 |
| HOMO SAPIENS | 2 | 9418 | 1hdr | A | 35 | TRP | p.Trp36Arg | P09417 |
| HOMO SAPIENS | 2 | 9418 | 1hdr | A | 107 | TRP | p.Trp108Gly | P09417 |
| HOMO SAPIENS | 6 | 9418 | 1hdr | A | 149 | TYR | p.Tyr150Cys | P09417 |
| HOMO SAPIENS | -1 | 12079 | 3tnu | B | 373 | GLY | p.Gly373Ala | P13647 |
| HOMO SAPIENS | 1 | 12079 | 3tnu | B | 463 | LEU | p.Leu463Pro | P13647 |
| HOMO SAPIENS | 1 | 12079 | 3tnu | B | 471 | ARG | p.Arg471Cys | P13647 |
| HOMO SAPIENS | 0 | 8135 | 3rbn | B | 89 | SER | p.Ser556Ile | P40692 |
| HOMO SAPIENS | 0 | 8135 | 3rbn | B | 111 | GLU | p.Glu578Gly | P40692 |
| HOMO SAPIENS | 2 | 8135 | 3rbn | B | 114 | PRO | p.Pro581Leu | P40692 |
| HOMO SAPIENS | 0 | 8135 | 3rbn | B | 133 | GLU | p.Glu600Gly | P40692 |
| HOMO SAPIENS | 2 | 8135 | 3rbn | B | 136 | PRO | p.Pro603Arg | P40692 |
| HOMO SAPIENS | 2 | 8135 | 3rbn | B | 136 | PRO | p.Pro603Leu | P40692 |
| HOMO SAPIENS | 7 | 8135 | 3rbn | B | 140 | LEU | p.Leu607His | P40692 |
| HOMO SAPIENS | 2 | 8135 | 3rbn | B | 151 | LYS | p.Lys618Ala | P40692 |
| HOMO SAPIENS | 2 | 8135 | 3rbn | B | 151 | LYS | p.Lys618Thr | P40692 |
| HOMO SAPIENS | 5 | 8135 | 3rbn | B | 181 | PRO | p.Pro648Ser | P40692 |
| HOMO SAPIENS | 2 | 8135 | 3rbn | B | 188 | ILE | p.Ile655Val | P40692 |
| HOMO SAPIENS | 2 | 8135 | 3rbn | B | 214 | ALA | p.Ala681Thr | P40692 |

| | | | | | | | | |
|---|---|---|---|---|---|---|---|---|
| HOMO SAPIENS | 2 | 8135 | 3rbn | B | 220 | ARG | p.Arg687Trp | P40692 |
| HOMO SAPIENS | 2 | 8135 | 3rbn | B | 222 | GLN | p.Gln689Arg | P40692 |
| HOMO SAPIENS | -1 | 8135 | 3rbn | B | 236 | GLN | p.Gln701Ter | P40692 |
| HOMO SAPIENS | 7 | 8135 | 3rbn | B | 251 | HIS | p.His718Tyr | P40692 |
| HOMO SAPIENS | 0 | 8135 | 3rbn | B | 284 | LYS | p.Lys751Arg | P40692 |
| HOMO SAPIENS | 1 | 21998 | 5d68 | A | 355 | LEU | p.Leu355Phe | O00522 |
| HOMO SAPIENS | 3 | 21998 | 5d68 | A | 380 | THR | p.Thr380Met | O00522 |
| HOMO SAPIENS | 0 | 21998 | 5d68 | A | 415 | ILE | p.Ile415Met | O00522 |
| HOMO SAPIENS | 0 | 33789 | 2m0c | A | 13 | ARG | p.Arg218Gln | Q9H161 |
| HOMO SAPIENS | 7 | 33789 | 2m0c | A | 52 | ARG | p.Arg257Thr | Q9H161 |
| HOMO SAPIENS | 0 | 33789 | 2m0c | A | 67 | ARG | p.Arg272Pro | Q9H161 |
| HOMO SAPIENS | 6 | 8263 | 2osg | A | 51 | GLY | p.Gly355Arg | Q9UDY2 |
| HOMO SAPIENS | 5 | 8263 | 2osg | A | 60 | THR | p.Thr364Met | Q9UDY2 |
| HOMO SAPIENS | 1 | 2475 | 5E+026 | B | 234 | THR | p.Thr234Ala | Q9BZ23 |
| HOMO SAPIENS | 2 | 2475 | 5E+026 | B | 264 | ARG | p.Arg264Trp | Q9BZ23 |
| HOMO SAPIENS | 4 | 2475 | 5E+026 | B | 286 | ARG | p.Arg286Cys | Q9BZ23 |
| HOMO SAPIENS | 0 | 2475 | 5E+026 | B | 357 | ARG | p.Arg357Trp | Q9BZ23 |
| HOMO SAPIENS | 2 | 2475 | 5E+026 | B | 481 | ARG | p.Arg481Ter | Q9BZ23 |
| HOMO SAPIENS | 8 | 2475 | 5E+026 | B | 521 | GLY | p.Gly521Arg | Q9BZ23 |
| HOMO SAPIENS | 2 | 2475 | 5E+026 | B | 528 | THR | p.Thr528Met | Q9BZ23 |
| HOMO SAPIENS | 1 | 3742 | 5xtd | E | 108 | HIS | p.His134Tyr | P56556 |
| HOMO SAPIENS | 1 | 16725 | 3g04 | C | 36 | ASP | p.Asp36His | P16473 |
| HOMO SAPIENS | 3 | 16725 | 3g04 | C | 52 | PRO | p.Pro52Thr | P16473 |
| HOMO SAPIENS | 1 | 16725 | 3g04 | C | 162 | PRO | p.Pro162Ala | P16473 |
| HOMO SAPIENS | -1 | 12366 | 5d2d | C | 750 | PRO | p.Pro750Leu | P13569 |
| HOMO SAPIENS | 0 | 12366 | 5d2d | C | 764 | ARG | p.Arg764Ter | P13569 |
| HOMO SAPIENS | 0 | 34845 | 2y23 | A | 1151 | ASP | p.Asp1151Glu | P52179 |
| HOMO SAPIENS | 0 | 34845 | 2y23 | A | 1180 | ASP | p.Asp1180Ala | P52179 |
| HOMO SAPIENS | 6 | 36740 | 3lbx | B | 2053 | ALA | p.Ala2053Pro | P11277 |
| HOMO SAPIENS | 0 | 19289 | 3ihj | A | 404 | ARG | p.Arg404Ter | Q8TD30 |
| HOMO SAPIENS | 5 | 1034 | 5xtd | Q | 228 | ARG | p.Arg228Gln | O75306 |
| HOMO SAPIENS | 5 | 1034 | 5xtd | Q | 229 | PRO | p.Pro229Gln | O75306 |
| HOMO SAPIENS | 4 | 1034 | 5xtd | Q | 323 | ARG | p.Arg323Gln | O75306 |
| HOMO SAPIENS | 4 | 1034 | 5xtd | Q | 352 | PRO | p.Pro352Ala | O75306 |
| HOMO SAPIENS | 5 | 1034 | 5xtd | Q | 413 | SER | p.Ser413Pro | O75306 |
| HOMO SAPIENS | 1 | 2772 | 2vaf | A | 1047 | THR | p.Thr66Ala | O14958 |
| HOMO SAPIENS | 5 | 2772 | 2vaf | A | 1057 | VAL | p.Val76Met | O14958 |
| HOMO SAPIENS | 0 | 2772 | 2vaf | A | 1148 | LEU | p.Leu167His | O14958 |
| HOMO SAPIENS | 1 | 2772 | 2vaf | A | 1149 | ILE | p.Ile168Thr | O14958 |
| HOMO SAPIENS | 8 | 2772 | 2vaf | A | 1225 | HIS | p.His244Arg | O14958 |
| HOMO SAPIENS | 8 | 2772 | 2vaf | A | 1225 | HIS | p.His244Tyr | O14958 |
| HOMO SAPIENS | 7 | 2772 | 2vaf | A | 1251 | ILE | p.Ile270Thr | O14958 |
| HOMO SAPIENS | 3 | 2772 | 2vaf | A | 1288 | ASP | p.Asp307His | O14958 |
| HOMO SAPIENS | 5 | 2772 | 2vaf | A | 1289 | PRO | p.Pro308Leu | O14958 |
| HOMO SAPIENS | 1 | 31885 | 2aex | A | 132 | ALA | p.Ala132Val | P36551 |
| HOMO SAPIENS | 1 | 31885 | 2aex | A | 272 | ASN | p.Asn272His | P36551 |
| HOMO SAPIENS | 8 | 31885 | 2aex | A | 279 | GLY | p.Gly279Arg | P36551 |
| HOMO SAPIENS | 5 | 31885 | 2aex | A | 294 | VAL | p.Val294Ile | P36551 |
| HOMO SAPIENS | 5 | 31885 | 2aex | A | 295 | HIS | p.His295Asp | P36551 |
| HOMO SAPIENS | 1 | 31885 | 2aex | A | 331 | ARG | p.Arg331Trp | P36551 |
| HOMO SAPIENS | 3 | 31885 | 2aex | A | 352 | ARG | p.Arg352Cys | P36551 |
| HOMO SAPIENS | 1 | 31885 | 2aex | A | 404 | LYS | p.Lys404Glu | P36551 |
| HOMO SAPIENS | 2 | 31885 | 2aex | A | 447 | ARG | p.Arg447Cys | P36551 |
| HOMO SAPIENS | 5 | 11721 | 2r3v | A | 20 | HIS | p.His20Pro | Q03426 |

| | | | | | | | | |
|---|---|---|---|---|---|---|---|---|
| HOMO SAPIENS | 1 | 11721 | 2r3v | A | 116 | TYR | p.Tyr116His | Q03426 |
| HOMO SAPIENS | 1 | 11721 | 2r3v | A | 148 | ALA | p.Ala148Thr | Q03426 |
| HOMO SAPIENS | 1 | 11721 | 2r3v | A | 167 | PRO | p.Pro167Leu | Q03426 |
| HOMO SAPIENS | 6 | 11721 | 2r3v | A | 203 | VAL | p.Val203Ala | Q03426 |
| HOMO SAPIENS | 2 | 11721 | 2r3v | A | 215 | ARG | p.Arg215Ter | Q03426 |
| HOMO SAPIENS | 2 | 11721 | 2r3v | A | 237 | THR | p.Thr237Ser | Q03426 |
| HOMO SAPIENS | 0 | 11721 | 2r3v | A | 268 | ILE | p.Ile268Thr | Q03426 |
| HOMO SAPIENS | 0 | 11721 | 2r3v | A | 268 | ILE | p.Ile268Val | Q03426 |
| HOMO SAPIENS | 0 | 11721 | 2r3v | A | 301 | ASN | p.Asn301Ser | Q03426 |
| HOMO SAPIENS | 6 | 11721 | 2r3v | A | 310 | VAL | p.Val310Met | Q03426 |
| HOMO SAPIENS | 5 | 11721 | 2r3v | A | 334 | ALA | p.Ala334Thr | Q03426 |
| HOMO SAPIENS | 2 | 11721 | 2r3v | A | 377 | VAL | p.Val377Ile | Q03426 |
| HOMO SAPIENS | 1 | 11721 | 2r3v | A | 386 | ASP | p.Asp386Asn | Q03426 |
| HOMO SAPIENS | 1 | 11721 | 2r3v | A | 388 | ARG | p.Arg388Ter | Q03426 |
| HOMO SAPIENS | 8 | 5010 | 2hgs | A | 125 | ARG | p.Arg125Cys | P48637 |
| HOMO SAPIENS | 1 | 5010 | 2hgs | A | 164 | ARG | p.Arg164Gln | P48637 |
| HOMO SAPIENS | 0 | 5010 | 2hgs | A | 219 | ASP | p.Asp219Gly | P48637 |
| HOMO SAPIENS | 7 | 5010 | 2hgs | A | 267 | ARG | p.Arg267Trp | P48637 |
| HOMO SAPIENS | 0 | 5010 | 2hgs | A | 283 | ARG | p.Arg283Cys | P48637 |
| HOMO SAPIENS | 0 | 6892 | 5erd | B | 7 | VAL | p.Val56Met | Q14126 |
| HOMO SAPIENS | 1 | 6892 | 5erd | B | 25 | HIS | p.His74Arg | Q14126 |
| HOMO SAPIENS | 0 | 6892 | 5erd | B | 40 | TYR | p.Tyr89Cys | Q14126 |
| HOMO SAPIENS | 0 | 6892 | 5erd | B | 97 | ARG | p.Arg146Leu | Q14126 |
| HOMO SAPIENS | 7 | 6892 | 5erd | B | 217 | ASN | p.Asn266Ser | Q14126 |
| HOMO SAPIENS | 0 | 6892 | 5erd | B | 243 | ARG | p.Arg292His | Q14126 |
| HOMO SAPIENS | 0 | 6892 | 5erd | B | 244 | ILE | p.Ile293Val | Q14126 |
| HOMO SAPIENS | 0 | 6892 | 5erd | B | 343 | VAL | p.Val392Ile | Q14126 |
| HOMO SAPIENS | 4 | 6892 | 5erd | B | 350 | GLU | p.Glu399Lys | Q14126 |
| HOMO SAPIENS | 0 | 6892 | 5erd | B | 410 | TYR | p.Tyr459Cys | Q14126 |
| HOMO SAPIENS | 0 | 6892 | 5erd | B | 466 | VAL | p.Val515Ile | Q14126 |
| HOMO SAPIENS | 0 | 6892 | 5erd | B | 482 | PHE | p.Phe531Cys | Q14126 |
| HOMO SAPIENS | 0 | 6892 | 5erd | B | 484 | VAL | p.Val533Ile | Q14126 |
| HOMO SAPIENS | 0 | 6892 | 5erd | B | 545 | LEU | p.Leu594Pro | Q14126 |
| HOMO SAPIENS | 0 | 1338 | 2ax4 | A | 58 | THR | p.Thr48Arg | O95340 |
| HOMO SAPIENS | 7 | 8675 | 2zkm | X | 385 | LYS | p.Lys385Arg | Q00722 |
| HOMO SAPIENS | 1 | 11769 | 2wya | B | 54 | VAL | p.Val54Met | P54868 |
| HOMO SAPIENS | 1 | 11769 | 2wya | B | 116 | ARG | p.Arg116Cys | P54868 |
| HOMO SAPIENS | 5 | 11769 | 2wya | B | 167 | TYR | p.Tyr167Cys | P54868 |
| HOMO SAPIENS | 1 | 11769 | 2wya | B | 174 | PHE | p.Phe174Leu | P54868 |
| HOMO SAPIENS | 0 | 11769 | 2wya | B | 212 | GLY | p.Gly212Arg | P54868 |
| HOMO SAPIENS | 1 | 11769 | 2wya | B | 288 | ARG | p.Arg288Ter | P54868 |
| HOMO SAPIENS | 0 | 11769 | 2wya | B | 375 | SER | p.Ser375Cys | P54868 |
| HOMO SAPIENS | 4 | 11769 | 2wya | B | 500 | ARG | p.Arg500His | P54868 |
| HOMO SAPIENS | 0 | 3102 | 2iwo | A | 1880 | ARG | p.Arg1908Lys | O75970 |
| HOMO SAPIENS | 6 | 14296 | 2m9w | A | 38 | GLY | p.Gly296Ser | P43694 |
| HOMO SAPIENS | 0 | 14296 | 2m9w | A | 58 | GLN | p.Gln316Glu | P43694 |
| HOMO SAPIENS | 2 | 704 | 5ekd | B | 239 | ARG | p.Arg239Ter | Q9UGM6 |
| HOMO SAPIENS | 2 | 704 | 5ekd | B | 264 | TYR | p.Tyr264Cys | Q9UGM6 |
| HOMO SAPIENS | 6 | 2228 | 1ey2 | A | 47 | SER | p.Ser47Ter | Q93099 |
| HOMO SAPIENS | 6 | 2228 | 1ey2 | A | 48 | ALA | p.Ala48Ser | Q93099 |
| HOMO SAPIENS | 6 | 2228 | 1ey2 | A | 53 | ARG | p.Arg53Gln | Q93099 |
| HOMO SAPIENS | 3 | 2228 | 1ey2 | A | 80 | HIS | p.Gln80His | Q93099 |
| HOMO SAPIENS | 0 | 2228 | 1ey2 | A | 120 | CYS | p.Cys120Trp | Q93099 |
| HOMO SAPIENS | 3 | 2228 | 1ey2 | A | 122 | ALA | p.Ala122Val | Q93099 |

| | | | | | | | | |
|---|---|---|---|---|---|---|---|---|
| HOMO SAPIENS | 3 | 2228 | 1ey2 | A | 161 | GLY | p.Gly161Arg | Q93099 |
| HOMO SAPIENS | 6 | 2228 | 1ey2 | A | 225 | ARG | p.Arg225His | Q93099 |
| HOMO SAPIENS | 0 | 2228 | 1ey2 | A | 230 | PRO | p.Pro230Ser | Q93099 |
| HOMO SAPIENS | 6 | 2228 | 1ey2 | A | 270 | GLY | p.Gly270Arg | Q93099 |
| HOMO SAPIENS | 6 | 2228 | 1ey2 | A | 330 | ARG | p.Arg330Ser | Q93099 |
| HOMO SAPIENS | -1 | 2228 | 1ey2 | A | 368 | MSE | p.Met368Val | Q93099 |
| HOMO SAPIENS | 6 | 2228 | 1ey2 | A | 371 | HIS | p.His371Arg | Q93099 |
| HOMO SAPIENS | 6 | 2228 | 1ey2 | A | 401 | GLU | p.Glu401Gln | Q93099 |
| HOMO SAPIENS | 5 | 34648 | 2k27 | A | 31 | ARG | p.Arg31His | Q06710 |
| HOMO SAPIENS | 7 | 34648 | 2k27 | A | 40 | GLN | p.Gln40Pro | Q06710 |
| HOMO SAPIENS | 5 | 34648 | 2k27 | A | 57 | CYS | p.Cys57Tyr | Q06710 |
| HOMO SAPIENS | 5 | 34648 | 2k27 | A | 62 | LEU | p.Leu62Arg | Q06710 |
| HOMO SAPIENS | 1 | 21399 | 3edu | A | 1819 | GLU | p.Glu1819Ter | P11277 |
| HOMO SAPIENS | 1 | 18153 | 1oey | J | 245 | TYR | p.Tyr245Ter | Q15080 |
| HOMO SAPIENS | 2 | 41567 | 3sw0 | X | 1050 | ASN | p.Asn1050Tyr | P08603 |
| HOMO SAPIENS | 5 | 41567 | 3sw0 | X | 1059 | ILE | p.Ile1059Thr | P08603 |
| HOMO SAPIENS | 0 | 36362 | 4wrm | A | 362 | HIS | p.His362Arg | P07333 |
| HOMO SAPIENS | -1 | 36362 | 4wrm | A | 413 | GLY | p.Gly413Ser | P07333 |
| HOMO SAPIENS | 0 | 31255 | 2w2j | A | 100 | SER | p.Ser100Pro | P35219 |
| HOMO SAPIENS | 0 | 31255 | 2w2j | A | 100 | SER | p.Ser100Leu | P35219 |
| HOMO SAPIENS | 5 | 31255 | 2w2j | A | 237 | ARG | p.Arg237Gln | P35219 |
| HOMO SAPIENS | -1 | 26450 | 1th1 | C | 1395 | SER | p.Ser1395Cys | P25054 |
| HOMO SAPIENS | -1 | 26450 | 1th1 | C | 1413 | MET | p.Met1413Val | P25054 |
| HOMO SAPIENS | -1 | 26450 | 1th1 | C | 1446 | ALA | p.Ala1446Thr | P25054 |
| HOMO SAPIENS | -1 | 26450 | 1th1 | C | 1454 | LYS | p.Lys1454Glu | P25054 |
| HOMO SAPIENS | 0 | 26450 | 1th1 | C | 1474 | ALA | p.Ala1474Thr | P25054 |
| HOMO SAPIENS | 1 | 981 | 5xtb | F | 14 | LEU | p.Leu14Val | O43678 |
| HOMO SAPIENS | 1 | 11069 | 3kq4 | A | 46 | SER | p.Ser46Leu | P27352 |
| HOMO SAPIENS | 1 | 11069 | 3kq4 | A | 97 | MET | p.Met97Thr | P27352 |
| HOMO SAPIENS | 1 | 11069 | 3kq4 | A | 255 | ASN | p.Asn255Ser | P27352 |
| HOMO SAPIENS | 1 | 11069 | 3kq4 | A | 330 | ASN | p.Asn330Lys | P27352 |
| HOMO SAPIENS | 0 | 41489 | 3erb | A | 53 | PRO | p.Pro73Leu | P06681 |
| HOMO SAPIENS | 6 | 8715 | 2bq8 | X | 23 | THR | p.Thr44Met | P13686 |
| HOMO SAPIENS | 0 | 8715 | 2bq8 | X | 88 | GLY | p.Gly109Arg | P13686 |
| HOMO SAPIENS | 6 | 8715 | 2bq8 | X | 127 | VAL | p.Val148Met | P13686 |
| HOMO SAPIENS | 2 | 8715 | 2bq8 | X | 179 | VAL | p.Val200Met | P13686 |
| HOMO SAPIENS | 6 | 8715 | 2bq8 | X | 251 | ARG | p.Arg272Cys | P13686 |
| HOMO SAPIENS | 7 | 8112 | 3p56 | A | 37 | GLY | p.Gly37Ser | O75792 |
| HOMO SAPIENS | -1 | 8112 | 3p56 | A | 202 | LEU | p.Leu202Ser | O75792 |
| HOMO SAPIENS | 1 | 8112 | 3p56 | A | 205 | ASP | p.Asp205Glu | O75792 |
| HOMO SAPIENS | 4 | 8112 | 3p56 | A | 212 | ASN | p.Asn212Ile | O75792 |
| HOMO SAPIENS | 0 | 8112 | 3p56 | A | 235 | ARG | p.Arg235Gln | O75792 |
| HOMO SAPIENS | 4 | 8112 | 3p56 | A | 240 | THR | p.Thr240Met | O75792 |
| HOMO SAPIENS | 0 | 8112 | 3p56 | A | 291 | ARG | p.Arg291His | O75792 |
| HOMO SAPIENS | 0 | 37603 | 2yqg | A | 10 | ARG | p.Arg49His | Q14126 |
| HOMO SAPIENS | 0 | 21178 | 3k1r | B | 458 | ASP | p.Asp458Val | Q495M9 |
| HOMO SAPIENS | -1 | 36828 | 4fq3 | B | 507 | GLY | p.Gly507Asp | P35637 |
| HOMO SAPIENS | 5 | 36828 | 4fq3 | B | 517 | HIS | p.His517Gln | P35637 |
| HOMO SAPIENS | 8 | 36828 | 4fq3 | B | 518 | ARG | p.Arg518Lys | P35637 |
| HOMO SAPIENS | 6 | 36828 | 4fq3 | B | 521 | ARG | p.Arg521Cys | P35637 |
| HOMO SAPIENS | 6 | 36828 | 4fq3 | B | 521 | ARG | p.Arg521His | P35637 |
| HOMO SAPIENS | 0 | 6363 | 3j7y | c | null | HIS | p.His13Arg | Q9H9J2 |
| HOMO SAPIENS | 2 | 6363 | 3j7y | c | 105 | ARG | p.Arg105Cys | Q9H9J2 |
| HOMO SAPIENS | 2 | 6363 | 3j7y | c | 138 | THR | p.Thr138Ile | Q9H9J2 |

| | | | | | | | | |
|---|---|---|---|---|---|---|---|---|
| HOMO SAPIENS | 1 | 6363 | 3j7y | c | 156 | LEU | p.Leu156Arg | Q9H9J2 |
| HOMO SAPIENS | 1 | 6782 | 5l46 | A | 109 | HIS | p.His109Arg | Q9UI17 |
| HOMO SAPIENS | 0 | 23951 | 5t76 | A | 175 | GLY | p.Gly931Ser | P49588 |
| HOMO SAPIENS | 8 | 12604 | 3p0l | A | 182 | ARG | p.Arg182Leu | P49675 |
| HOMO SAPIENS | 2 | 12604 | 3p0l | A | 187 | VAL | p.Val187Met | P49675 |
| HOMO SAPIENS | 2 | 12604 | 3p0l | A | 188 | ARG | p.Arg188Cys | P49675 |
| HOMO SAPIENS | 0 | 12604 | 3p0l | A | 193 | ARG | p.Arg193Ter | P49675 |
| HOMO SAPIENS | 9 | 12604 | 3p0l | A | 217 | ARG | p.Arg217Thr | P49675 |
| HOMO SAPIENS | 4 | 12604 | 3p0l | A | 218 | ALA | p.Ala218Val | P49675 |
| HOMO SAPIENS | 0 | 12604 | 3p0l | A | 250 | TRP | p.Trp250Ter | P49675 |
| HOMO SAPIENS | 1 | 12604 | 3p0l | A | 258 | GLN | p.Gln258Ter | P49675 |
| HOMO SAPIENS | -1 | 27765 | 4wkr | A | 4 | GLU | p.Glu4Lys | Q4G0J3 |
| HOMO SAPIENS | 2 | 3587 | 4kit | C | 2301 | PRO | p.Pro2301Thr | Q6P2Q9 |
| HOMO SAPIENS | 3 | 3587 | 4kit | C | 2304 | PHE | p.Phe2304Leu | Q6P2Q9 |
| HOMO SAPIENS | 3 | 3587 | 4kit | C | 2309 | HIS | p.His2309Pro | Q6P2Q9 |
| HOMO SAPIENS | 3 | 3587 | 4kit | C | 2310 | ARG | p.Arg2310Lys | Q6P2Q9 |
| HOMO SAPIENS | 0 | 9073 | 1n69 | B | 21 | ASN | p.Asn215His | P07602 |
| HOMO SAPIENS | 0 | 9073 | 1n69 | B | 23 | THR | p.Thr217Ile | P07602 |
| HOMO SAPIENS | 0 | 9073 | 1n69 | B | 47 | CYS | p.Cys241Ser | P07602 |
| HOMO SAPIENS | 0 | 475 | 2vd5 | A | 225 | THR | p.Thr225Met | Q09013 |
| HOMO SAPIENS | 1 | 9066 | 4mzv | A | 31 | ASN | p.Asn31Lys | P16422 |
| HOMO SAPIENS | 1 | 9066 | 4mzv | A | 47 | GLN | p.Gln47His | P16422 |
| HOMO SAPIENS | 3 | 9066 | 4mzv | A | 66 | CYS | p.Cys66Tyr | P16422 |
| HOMO SAPIENS | 1 | 9066 | 4mzv | A | 102 | SER | p.Ser102Gly | P16422 |
| HOMO SAPIENS | 5 | 9066 | 4mzv | A | 115 | THR | p.Met115Thr | P16422 |
| HOMO SAPIENS | 1 | 9066 | 4mzv | A | 138 | ARG | p.Arg138Ter | P16422 |
| HOMO SAPIENS | 1 | 9066 | 4mzv | A | 153 | ARG | p.Arg153Thr | P16422 |
| HOMO SAPIENS | 1 | 9066 | 4mzv | A | 165 | ALA | p.Ala165Thr | P16422 |
| HOMO SAPIENS | 8 | 1827 | 2nz2 | A | 14 | GLY | p.Gly14Ser | P00966 |
| HOMO SAPIENS | 5 | 1827 | 2nz2 | A | 18 | SER | p.Ser18Leu | P00966 |
| HOMO SAPIENS | 3 | 1827 | 2nz2 | A | 86 | ARG | p.Arg86Cys | P00966 |
| HOMO SAPIENS | 3 | 1827 | 2nz2 | A | 86 | ARG | p.Arg86His | P00966 |
| HOMO SAPIENS | 1 | 1827 | 2nz2 | A | 179 | TRP | p.Trp179Arg | P00966 |
| HOMO SAPIENS | 7 | 1827 | 2nz2 | A | 180 | SER | p.Ser180Asn | P00966 |
| HOMO SAPIENS | 7 | 1827 | 2nz2 | A | 191 | GLU | p.Glu191Lys | P00966 |
| HOMO SAPIENS | 4 | 1827 | 2nz2 | A | 256 | GLU | p.Glu256Lys | P00966 |
| HOMO SAPIENS | 4 | 1827 | 2nz2 | A | 263 | VAL | p.Val263Met | P00966 |
| HOMO SAPIENS | 5 | 1827 | 2nz2 | A | 265 | ARG | p.Arg265Cys | P00966 |
| HOMO SAPIENS | 5 | 1827 | 2nz2 | A | 268 | ILE | p.Ile268Met | P00966 |
| HOMO SAPIENS | 6 | 1827 | 2nz2 | A | 269 | VAL | p.Val269Met | P00966 |
| HOMO SAPIENS | 7 | 1827 | 2nz2 | A | 272 | ARG | p.Arg272Cys | P00966 |
| HOMO SAPIENS | 3 | 1827 | 2nz2 | A | 304 | ARG | p.Arg304Trp | P00966 |
| HOMO SAPIENS | 3 | 1827 | 2nz2 | A | 307 | ARG | p.Arg307Cys | P00966 |
| HOMO SAPIENS | 3 | 1827 | 2nz2 | A | 310 | LYS | p.Lys310Gln | P00966 |
| HOMO SAPIENS | 2 | 1827 | 2nz2 | A | 362 | GLY | p.Gly362Val | P00966 |
| HOMO SAPIENS | 2 | 1827 | 2nz2 | A | 363 | ARG | p.Arg363Trp | P00966 |
| HOMO SAPIENS | 2 | 1827 | 2nz2 | A | 363 | ARG | p.Arg363Gln | P00966 |
| HOMO SAPIENS | 1 | 1827 | 2nz2 | A | 390 | GLY | p.Gly390Arg | P00966 |
| HOMO SAPIENS | 0 | 29524 | 5vbn | B | 2165 | ARG | p.Arg2165His | Q07864 |
| HOMO SAPIENS | 2 | 29524 | 5vbn | B | 2255 | ILE | p.Ile2255Phe | Q07864 |
| HOMO SAPIENS | 2 | 29524 | 5vbn | B | 2274 | LEU | p.Leu2274Val | Q07864 |
| HOMO SAPIENS | 0 | 31897 | 3t0o | A | 184 | CYS | p.Cys184Arg | O00584 |
| HOMO SAPIENS | -1 | 31897 | 3t0o | A | 215 | GLU | p.Glu215Lys | O00584 |
| HOMO SAPIENS | 0 | 31897 | 3t0o | A | 236 | ARG | p.Arg236Trp | O00584 |

| | | | | | | | | |
|---|---|---|---|---|---|---|---|---|
| HOMO SAPIENS | 7 | 18490 | 4pcw | B | 31 | GLU | p.Glu32Ter | P20930 |
| HOMO SAPIENS | -1 | 39865 | 2h9e | L | 152 | GLY | p.Gly192Arg | P00742 |
| HOMO SAPIENS | 2 | 4309 | 2x0b | F | 174 | THR | p.Thr207Met | P01019 |
| HOMO SAPIENS | 3 | 4309 | 2x0b | F | 342 | ARG | p.Arg375Gln | P01019 |
| HOMO SAPIENS | 2 | 3094 | 1zll | B | 9 | ARG | p.Arg9Cys | P26678 |
| HOMO SAPIENS | 9 | 3094 | 1zll | B | 39 | LEU | p.Leu39Ter | P26678 |
| HOMO SAPIENS | 5 | 3094 | 1zll | B | 49 | VAL | p.Val49Met | P26678 |
| HOMO SAPIENS | -1 | 41498 | 3ffd | P | 8 | LEU | p.Leu44Pro | P12272 |
| HOMO SAPIENS | 1 | 20697 | 1igr | A | 407 | ARG | p.Arg437His | P08069 |
| HOMO SAPIENS | 0 | 20697 | 1igr | A | 416 | MET | p.Met446Val | P08069 |
| HOMO SAPIENS | -1 | 20697 | 1igr | A | 462 | VAL | p.Val492Ile | P08069 |
| HOMO SAPIENS | 0 | 3973 | 4cbz | A | 37 | LEU | p.Leu37Ser | P78504 |
| HOMO SAPIENS | 0 | 3973 | 4cbz | A | 45 | VAL | p.Val45Leu | P78504 |
| HOMO SAPIENS | 0 | 3973 | 4cbz | A | 134 | LEU | p.Leu134Met | P78504 |
| HOMO SAPIENS | -1 | 3973 | 4cbz | A | 146 | VAL | p.Val146Ile | P78504 |
| HOMO SAPIENS | 0 | 3973 | 4cbz | A | 176 | VAL | p.Val176Ile | P78504 |
| HOMO SAPIENS | 0 | 3973 | 4cbz | A | 184 | ARG | p.Arg184Cys | P78504 |
| HOMO SAPIENS | 6 | 1697 | 4hmy | S | 33 | ARG | p.Arg33Trp | Q96PC3 |
| HOMO SAPIENS | 2 | 23752 | 5j4o | A | 13 | ARG | p.Arg1611Ser | P02549 |
| HOMO SAPIENS | 0 | 23752 | 5j4o | A | 42 | ALA | p.Ala1640Gly | P02549 |
| HOMO SAPIENS | 0 | 23752 | 5j4o | A | 108 | HIS | p.His1706Gln | P02549 |
| HOMO SAPIENS | 6 | 23752 | 5j4o | A | 206 | LEU | p.Leu1804Phe | P02549 |
| HOMO SAPIENS | 8 | 531 | 2qz4 | A | 349 | GLY | p.Gly349Ser | Q9UQ90 |
| HOMO SAPIENS | 0 | 531 | 2qz4 | A | 486 | ARG | p.Arg486Gln | Q9UQ90 |
| HOMO SAPIENS | 1 | 531 | 2qz4 | A | 510 | ALA | p.Ala510Val | Q9UQ90 |
| HOMO SAPIENS | 0 | 531 | 2qz4 | A | 529 | ALA | p.Ala529Val | Q9UQ90 |
| HOMO SAPIENS | 0 | 531 | 2qz4 | A | 558 | LYS | p.Lys558Ter | Q9UQ90 |
| HOMO SAPIENS | 1 | 531 | 2qz4 | A | 559 | LYS | p.Lys559Ter | Q9UQ90 |
| HOMO SAPIENS | 2 | 7736 | 1fnh | A | 114 | TRP | p.Trp1834Arg | P02751 |
| HOMO SAPIENS | 0 | 7736 | 1fnh | A | 163 | LEU | p.Leu1883Arg | P02751 |
| HOMO SAPIENS | 3 | 20337 | 5wy5 | B | 264 | LEU | p.Leu264Phe | Q96MG7 |
| HOMO SAPIENS | 8 | 36096 | 2jku | A | 643 | GLY | p.Gly668Arg | P05165 |
| HOMO SAPIENS | 6 | 36096 | 2jku | A | 663 | CYS | p.Cys688Arg | P05165 |
| HOMO SAPIENS | 0 | 9850 | 3kbt | A | 1606 | TYR | p.Tyr1606Ter | P11277 |
| HOMO SAPIENS | 4 | 6876 | 4jkq | A | 217 | ILE | p.Ile217Val | Q92736 |
| HOMO SAPIENS | 5 | 6876 | 4jkq | A | 420 | ARG | p.Arg420Trp | Q92736 |
| HOMO SAPIENS | 2 | 27603 | 1itv | B | 62 | PRO | p.Arg574Pro | P14780 |
| HOMO SAPIENS | 4 | 27603 | 1itv | B | 156 | ARG | p.Arg668Gln | P14780 |
| HOMO SAPIENS | 2 | 22954 | 4f52 | E | 48 | ILE | p.Ile48Met | Q92990 |
| HOMO SAPIENS | 0 | 22954 | 4f52 | E | 91 | ASP | p.Asp91Asn | Q92990 |
| HOMO SAPIENS | 0 | 22954 | 4f52 | E | 146 | ALA | p.Ala146Thr | Q92990 |
| HOMO SAPIENS | 0 | 22954 | 4f52 | E | 181 | ILE | p.Ile181Lys | Q92990 |
| HOMO SAPIENS | 0 | 22954 | 4f52 | E | 194 | ASN | p.Asn194Ser | Q92990 |
| HOMO SAPIENS | 2 | 22954 | 4f52 | E | 254 | PRO | p.Pro254Arg | Q92990 |
| HOMO SAPIENS | 2 | 22954 | 4f52 | E | 300 | ILE | p.Ile300Thr | Q92990 |
| HOMO SAPIENS | 2 | 22954 | 4f52 | E | 336 | LEU | p.Leu336Ser | Q92990 |
| HOMO SAPIENS | 0 | 22954 | 4f52 | E | 353 | TYR | p.Tyr353His | Q92990 |
| HOMO SAPIENS | 0 | 22954 | 4f52 | E | 387 | MET | p.Met387Val | Q92990 |
| HOMO SAPIENS | 4 | 2946 | 5gl7 | A | 344 | TYR | p.Tyr344Cys | P56192 |
| HOMO SAPIENS | 0 | 2946 | 5gl7 | A | 370 | PHE | p.Phe370Leu | P56192 |
| HOMO SAPIENS | 0 | 2946 | 5gl7 | A | 393 | ALA | p.Ala393Thr | P56192 |
| HOMO SAPIENS | 0 | 2946 | 5gl7 | A | 523 | ILE | p.Ile523Thr | P56192 |
| HOMO SAPIENS | 0 | 2946 | 5gl7 | A | 567 | SER | p.Ser567Leu | P56192 |
| HOMO SAPIENS | 0 | 2946 | 5gl7 | A | 800 | PRO | p.Pro800Thr | P56192 |

| | | | | | | | | |
|---|---|---|---|---|---|---|---|---|
| HOMO SAPIENS | 3 | 34961 | 4wy4 | C | 44 | TYR | p.Tyr44His | O95721 |
| HOMO SAPIENS | 8 | 34961 | 4wy4 | C | 89 | GLU | p.Glu89Lys | O95721 |
| HOMO SAPIENS | 0 | 9427 | 4chl | B | 21 | ALA | p.Ala21Ser | O95571 |
| HOMO SAPIENS | 2 | 9427 | 4chl | B | 62 | ALA | p.Ala62Thr | O95571 |
| HOMO SAPIENS | 0 | 9427 | 4chl | B | 114 | ILE | p.Ile114Phe | O95571 |
| HOMO SAPIENS | 2 | 9427 | 4chl | B | 159 | ARG | p.Arg159His | O95571 |
| HOMO SAPIENS | 8 | 9427 | 4chl | B | 163 | ARG | p.Arg163Trp | O95571 |
| HOMO SAPIENS | 8 | 9427 | 4chl | B | 163 | ARG | p.Arg163Gln | O95571 |
| HOMO SAPIENS | 2 | 9427 | 4chl | B | 185 | LEU | p.Leu185Arg | O95571 |
| HOMO SAPIENS | 0 | 40917 | 2e8o | A | 11 | PRO | p.Pro26Leu | Q9Y3Z3 |
| HOMO SAPIENS | -1 | 39190 | 4gow | A | 556 | TYR | p.Tyr556Ter | P56696 |
| HOMO SAPIENS | 0 | 2399 | 2mkv | A | 8 | GLY | p.Gly10Ser | P54710 |
| HOMO SAPIENS | 1 | 2399 | 2mkv | A | 24 | VAL | p.Val26Ile | P54710 |
| HOMO SAPIENS | 4 | 2399 | 2mkv | A | 39 | GLY | p.Gly41Arg | P54710 |
| HOMO SAPIENS | 0 | 20801 | 2zej | B | 1441 | ARG | p.Arg1441Cys | Q5S007 |
| HOMO SAPIENS | 0 | 20801 | 2zej | B | 1441 | ARG | p.Arg1441His | Q5S007 |
| HOMO SAPIENS | 5 | 1019 | 5xtb | M | 62 | ARG | p.Arg62Gln | P28331 |
| HOMO SAPIENS | 1 | 1019 | 5xtb | M | 85 | ALA | p.Ala85Thr | P28331 |
| HOMO SAPIENS | 5 | 1019 | 5xtb | M | 231 | LEU | p.Leu231Val | P28331 |
| HOMO SAPIENS | 2 | 1019 | 5xtb | M | 241 | ARG | p.Arg241Trp | P28331 |
| HOMO SAPIENS | 1 | 1019 | 5xtb | M | 252 | ASP | p.Asp252Gly | P28331 |
| HOMO SAPIENS | 1 | 1019 | 5xtb | M | 408 | ARG | p.Arg408Cys | P28331 |
| HOMO SAPIENS | 2 | 1019 | 5xtb | M | 417 | ARG | p.Arg417Ser | P28331 |
| HOMO SAPIENS | 1 | 1019 | 5xtb | M | 557 | ARG | p.Arg557Ter | P28331 |
| HOMO SAPIENS | 0 | 1019 | 5xtb | M | 567 | ILE | p.Ile567Val | P28331 |
| HOMO SAPIENS | 1 | 1019 | 5xtb | M | 595 | THR | p.Thr595Ala | P28331 |
| HOMO SAPIENS | 1 | 1019 | 5xtb | M | 703 | ALA | p.Ala703Thr | P28331 |
| HOMO SAPIENS | -1 | 3437 | 5hyn | A | 185 | ASP | p.Asp185His | Q15910 |
| HOMO SAPIENS | -1 | 3437 | 5hyn | A | 482 | ALA | p.Ala482Thr | Q15910 |
| HOMO SAPIENS | 0 | 8201 | 2wfd | A | 373 | TYR | p.Tyr373Cys | Q9P2J5 |
| HOMO SAPIENS | 1 | 3590 | 3edi | A | 90 | VAL | p.Val238Ala | O43897 |
| HOMO SAPIENS | 6 | 4555 | 1ytq | A | 64 | ALA | p.Ala65Ser | P43320 |
| HOMO SAPIENS | 0 | 4555 | 1ytq | A | 194 | TRP | p.Trp195Gly | P43320 |
| HOMO SAPIENS | 0 | 1013 | 2nn6 | C | 272 | SER | p.Ser272Thr | Q96B26 |
| HOMO SAPIENS | -1 | 29026 | 5m73 | D | 530 | ILE | p.Ile530Asn | O76094 |
| HOMO SAPIENS | 8 | 2252 | 1d5v | A | 57 | SER | p.Ser125Leu | Q99958 |
| HOMO SAPIENS | 1 | 4337 | 3ecr | A | 29 | ARG | p.Arg26His | P08397 |
| HOMO SAPIENS | 2 | 4337 | 3ecr | A | 34 | ALA | p.Ala31Thr | P08397 |
| HOMO SAPIENS | 2 | 4337 | 3ecr | A | 37 | GLN | p.Gln34Lys | P08397 |
| HOMO SAPIENS | 1 | 4337 | 3ecr | A | 58 | ALA | p.Ala55Ser | P08397 |
| HOMO SAPIENS | 0 | 4337 | 3ecr | A | 84 | LEU | p.Leu81Pro | P08397 |
| HOMO SAPIENS | 0 | 4337 | 3ecr | A | 114 | GLY | p.Gly111Arg | P08397 |
| HOMO SAPIENS | 1 | 4337 | 3ecr | A | 119 | ARG | p.Arg116Trp | P08397 |
| HOMO SAPIENS | 1 | 4337 | 3ecr | A | 152 | ARG | p.Arg149Gln | P08397 |
| HOMO SAPIENS | 1 | 4337 | 3ecr | A | 170 | ARG | p.Arg167Trp | P08397 |
| HOMO SAPIENS | 1 | 4337 | 3ecr | A | 176 | ARG | p.Arg173Gln | P08397 |
| HOMO SAPIENS | 0 | 4337 | 3ecr | A | 180 | LEU | p.Leu177Arg | P08397 |
| HOMO SAPIENS | 0 | 4337 | 3ecr | A | 181 | ASP | p.Asp178Asn | P08397 |
| HOMO SAPIENS | 1 | 4337 | 3ecr | A | 198 | ARG | p.Arg195Cys | P08397 |
| HOMO SAPIENS | 0 | 4337 | 3ecr | A | 201 | TRP | p.Trp198Ter | P08397 |
| HOMO SAPIENS | 1 | 4337 | 3ecr | A | 204 | ARG | p.Arg201Trp | P08397 |
| HOMO SAPIENS | 1 | 4337 | 3ecr | A | 219 | GLY | p.Gly216Asp | P08397 |
| HOMO SAPIENS | 1 | 4337 | 3ecr | A | 226 | GLU | p.Glu223Lys | P08397 |
| HOMO SAPIENS | 0 | 4337 | 3ecr | A | 248 | LEU | p.Leu245Arg | P08397 |

| | | | | | | | | |
|---|---|---|---|---|---|---|---|---|
| HOMO SAPIENS | 1 | 4337 | 3ecr | A | 250 | CYS | p.Cys247Arg | P08397 |
| HOMO SAPIENS | 0 | 4337 | 3ecr | A | 253 | GLU | p.Glu250Lys | P08397 |
| HOMO SAPIENS | 0 | 4337 | 3ecr | A | 255 | ALA | p.Ala252Thr | P08397 |
| HOMO SAPIENS | 0 | 4337 | 3ecr | A | 255 | ALA | p.Ala252Val | P08397 |
| HOMO SAPIENS | 0 | 4337 | 3ecr | A | 259 | HIS | p.His256Asn | P08397 |
| HOMO SAPIENS | 0 | 4337 | 3ecr | A | 324 | ARG | p.Arg321His | P08397 |
| HOMO SAPIENS | -1 | 4337 | 3ecr | A | 362 | ASP | p.Asp359Asn | P08397 |
| HOMO SAPIENS | 1 | 1204 | 5lwv | A | 854 | ILE | p.Ala864Thr | O15294 |
| HOMO SAPIENS | 1 | 25016 | 1m4u | A | 35 | PRO | p.Pro35Arg | Q13253 |
| HOMO SAPIENS | 1 | 25016 | 1m4u | A | 35 | PRO | p.Pro35Ser | Q13253 |
| HOMO SAPIENS | 1 | 25016 | 1m4u | A | 167 | ARG | p.Arg167Gly | Q13253 |
| HOMO SAPIENS | 1 | 25016 | 1m4u | A | 184 | CYS | p.Cys184Tyr | Q13253 |
| HOMO SAPIENS | 0 | 25016 | 1m4u | A | 189 | GLY | p.Gly189Cys | Q13253 |
| HOMO SAPIENS | 1 | 25016 | 1m4u | A | 204 | ARG | p.Arg204Leu | Q13253 |
| HOMO SAPIENS | 1 | 25016 | 1m4u | A | 205 | TRP | p.Trp205Cys | Q13253 |
| HOMO SAPIENS | 1 | 25016 | 1m4u | A | 217 | TRP | p.Trp217Gly | Q13253 |
| HOMO SAPIENS | 0 | 25016 | 1m4u | A | 222 | TYR | p.Tyr222Cys | Q13253 |
| HOMO SAPIENS | 0 | 25016 | 1m4u | A | 222 | TYR | p.Tyr222Asp | Q13253 |
| HOMO SAPIENS | 1 | 25016 | 1m4u | A | 223 | PRO | p.Pro223Leu | Q13253 |
| HOMO SAPIENS | 0 | 1394 | 1um7 | A | 95 | ALA | p.Ala463Val | Q92796 |
| HOMO SAPIENS | 0 | 21294 | 4jpz | B | 1788 | GLU | p.Glu1788Asp | Q99250 |
| HOMO SAPIENS | 9 | 24068 | 2db6 | A | 47 | ARG | p.Arg119Cys | Q96MF2 |
| HOMO SAPIENS | 5 | 1869 | 4omt | A | 39 | ARG | p.Arg39Leu | P08237 |
| HOMO SAPIENS | 2 | 1869 | 4omt | A | 100 | ARG | p.Arg100Gln | P08237 |
| HOMO SAPIENS | 7 | 1869 | 4omt | A | 543 | ASP | p.Asp543Ala | P08237 |
| HOMO SAPIENS | 1 | 1869 | 4omt | A | 686 | TRP | p.Trp686Cys | P08237 |
| HOMO SAPIENS | 1 | 1869 | 4omt | A | 696 | ARG | p.Arg696His | P08237 |
| HOMO SAPIENS | 0 | 12163 | 4wxx | A | 495 | TYR | p.Tyr495Cys | P26358 |
| HOMO SAPIENS | 0 | 12163 | 4wxx | A | 528 | ILE | p.Ile528Met | P26358 |
| HOMO SAPIENS | 0 | 12391 | 1j1d | C | 36 | LYS | p.Lys36Gln | P19429 |
| HOMO SAPIENS | 0 | 12391 | 1j1d | C | 82 | PRO | p.Pro82Ser | P19429 |
| HOMO SAPIENS | 7 | 12391 | 1j1d | C | 102 | ALA | p.Ala102Thr | P19429 |
| HOMO SAPIENS | 1 | 5000 | 4cca | A | 17 | GLY | p.Gly17Arg | Q15833 |
| HOMO SAPIENS | 0 | 5000 | 4cca | A | 205 | VAL | p.Val205Ile | Q15833 |
| HOMO SAPIENS | 0 | 5000 | 4cca | A | 209 | LEU | p.Leu209Pro | Q15833 |
| HOMO SAPIENS | 1 | 5000 | 4cca | A | 459 | ARG | p.Arg459Trp | Q15833 |
| HOMO SAPIENS | 7 | 5000 | 4cca | A | 477 | PRO | p.Pro477Leu | Q15833 |
| HOMO SAPIENS | -1 | 5000 | 4cca | A | 526 | ILE | p.Ile526Val | Q15833 |
| HOMO SAPIENS | 0 | 5000 | 4cca | A | 541 | GLY | p.Gly541Ser | Q15833 |
| HOMO SAPIENS | 0 | 5000 | 4cca | A | 555 | ARG | p.Arg555Gly | Q15833 |
| HOMO SAPIENS | 5 | 28551 | 3vcm | P | 26 | ARG | p.Arg49Ter | P00797 |
| HOMO SAPIENS | 6 | 36453 | 5dzz | A | 2070 | ASP | p.Asp2070His | P15924 |
| HOMO SAPIENS | 0 | 36453 | 5dzz | A | 2166 | ARG | p.Arg2166Ter | P15924 |
| HOMO SAPIENS | 0 | 20268 | 2da6 | A | 96 | ASN | p.Asn321Ser | P35680 |
| HOMO SAPIENS | 1 | 1405 | 3iar | A | 8 | ASP | p.Asp8Asn | P00813 |
| HOMO SAPIENS | 8 | 1405 | 3iar | A | 15 | HIS | p.His15Asp | P00813 |
| HOMO SAPIENS | 6 | 1405 | 3iar | A | 74 | GLY | p.Gly74Val | P00813 |
| HOMO SAPIENS | 6 | 1405 | 3iar | A | 80 | LYS | p.Lys80Arg | P00813 |
| HOMO SAPIENS | 5 | 1405 | 3iar | A | 97 | TYR | p.Tyr97Cys | P00813 |
| HOMO SAPIENS | 5 | 1405 | 3iar | A | 101 | ARG | p.Arg101Leu | P00813 |
| HOMO SAPIENS | 5 | 1405 | 3iar | A | 101 | ARG | p.Arg101Trp | P00813 |
| HOMO SAPIENS | 5 | 1405 | 3iar | A | 106 | LEU | p.Leu106Val | P00813 |
| HOMO SAPIENS | 0 | 1405 | 3iar | A | 107 | LEU | p.Leu107Pro | P00813 |
| HOMO SAPIENS | 9 | 1405 | 3iar | A | 156 | ARG | p.Arg156Cys | P00813 |

| | | | | | | | | |
|---|---|---|---|---|---|---|---|---|
| HOMO SAPIENS | 6 | 1405 | 3iar | A | 211 | ARG | p.Arg211His | P00813 |
| HOMO SAPIENS | 0 | 1405 | 3iar | A | 216 | GLY | p.Gly216Arg | P00813 |
| HOMO SAPIENS | 0 | 1405 | 3iar | A | 291 | SER | p.Ser291Leu | P00813 |
| HOMO SAPIENS | 6 | 1405 | 3iar | A | 304 | LEU | p.Leu304Arg | P00813 |
| HOMO SAPIENS | 0 | 1405 | 3iar | A | 329 | ALA | p.Ala329Val | P00813 |
| HOMO SAPIENS | -1 | 31340 | 4atu | I | 186 | ARG | p.Arg186His | O43602 |
| HOMO SAPIENS | -1 | 31340 | 4atu | I | 192 | ARG | p.Arg192Trp | O43602 |
| HOMO SAPIENS | -1 | 31340 | 4atu | I | 196 | ARG | p.Arg196His | O43602 |
| HOMO SAPIENS | -1 | 31340 | 4atu | I | 203 | THR | p.Thr203Arg | O43602 |
| HOMO SAPIENS | -1 | 31340 | 4atu | I | 286 | ALA | p.Ala286Ser | O43602 |
| HOMO SAPIENS | 4 | 30734 | 1fsu | A | 95 | ARG | p.Arg95Gln | P15848 |
| HOMO SAPIENS | 0 | 30734 | 1fsu | A | 117 | CYS | p.Cys117Arg | P15848 |
| HOMO SAPIENS | 0 | 30734 | 1fsu | A | 137 | GLY | p.Gly137Val | P15848 |
| HOMO SAPIENS | 0 | 30734 | 1fsu | A | 210 | TYR | p.Tyr210Cys | P15848 |
| HOMO SAPIENS | 0 | 30734 | 1fsu | A | 236 | LEU | p.Leu236Pro | P15848 |
| HOMO SAPIENS | 1 | 30734 | 1fsu | A | 315 | ARG | p.Arg315Gln | P15848 |
| HOMO SAPIENS | 0 | 30734 | 1fsu | A | 358 | VAL | p.Val358Met | P15848 |
| HOMO SAPIENS | 2 | 30734 | 1fsu | A | 376 | VAL | p.Val376Met | P15848 |
| HOMO SAPIENS | 0 | 30734 | 1fsu | A | 384 | SER | p.Ser384Asn | P15848 |
| HOMO SAPIENS | 0 | 30734 | 1fsu | A | 393 | HIS | p.His393Pro | P15848 |
| HOMO SAPIENS | 0 | 30734 | 1fsu | A | 405 | CYS | p.Cys405Tyr | P15848 |
| HOMO SAPIENS | 0 | 30734 | 1fsu | A | 458 | ASN | p.Asn458Ser | P15848 |
| HOMO SAPIENS | 1 | 30734 | 1fsu | A | 484 | ARG | p.Arg484Gly | P15848 |
| HOMO SAPIENS | 0 | 38072 | 1wmv | A | 51 | PRO | p.Pro98Leu | Q9NZC7 |
| HOMO SAPIENS | -1 | 5674 | 1j1e | C | 185 | ASN | p.Asn185Lys | P19429 |
| HOMO SAPIENS | -1 | 5674 | 1j1e | C | 192 | ARG | p.Arg192His | P19429 |
| HOMO SAPIENS | -1 | 5674 | 1j1e | C | 196 | ASP | p.Asp196Asn | P19429 |
| HOMO SAPIENS | -1 | 5674 | 1j1e | C | 206 | LYS | p.Lys206Gln | P19429 |
| HOMO SAPIENS | 0 | 38067 | 1wlp | A | 13 | PRO | p.Pro156Gln | P13498 |
| HOMO SAPIENS | -1 | 20441 | 3vfd | A | 289 | HIS | p.His289Tyr | Q9UBP0 |
| HOMO SAPIENS | 2 | 20441 | 3vfd | A | 344 | ILE | p.Ile344Lys | Q9UBP0 |
| HOMO SAPIENS | 0 | 20441 | 3vfd | A | 362 | SER | p.Ser362Cys | Q9UBP0 |
| HOMO SAPIENS | 1 | 20441 | 3vfd | A | 386 | ASN | p.Asn386Ser | Q9UBP0 |
| HOMO SAPIENS | 9 | 20441 | 3vfd | A | 441 | ASP | p.Asp441Gly | Q9UBP0 |
| HOMO SAPIENS | -1 | 20441 | 3vfd | A | 448 | CYS | p.Cys448Tyr | Q9UBP0 |
| HOMO SAPIENS | 2 | 20441 | 3vfd | A | 500 | ARG | p.Arg499Cys | Q9UBP0 |
| HOMO SAPIENS | 1 | 20441 | 3vfd | A | 563 | ARG | p.Arg562Gly | Q9UBP0 |
| HOMO SAPIENS | 3 | 18471 | 4dx8 | J | 26 | ARG | p.Arg26Gln | O00522 |
| HOMO SAPIENS | -1 | 18471 | 4dx8 | J | 101 | GLY | p.Gly101Val | O00522 |
| HOMO SAPIENS | -1 | 34735 | 2ly4 | B | 83 | ALA | p.Ala83Glu | P04637 |
| HOMO SAPIENS | 0 | 31128 | 4cvh | A | 116 | ARG | p.Arg116Cys | A4D126 |
| HOMO SAPIENS | 8 | 31128 | 4cvh | A | 215 | GLN | p.Gln215Ter | A4D126 |
| HOMO SAPIENS | 0 | 31128 | 4cvh | A | 268 | ARG | p.Arg268Ter | A4D126 |
| HOMO SAPIENS | 0 | 31128 | 4cvh | A | 316 | THR | p.Thr316Lys | A4D126 |
| HOMO SAPIENS | 1 | 7164 | 3mt5 | A | 369 | ASP | p.Asp434Gly | Q12791 |
| HOMO SAPIENS | 2 | 26812 | 2wtk | F | 49 | TYR | p.Tyr49Asp | Q15831 |
| HOMO SAPIENS | 0 | 26812 | 2wtk | F | 67 | LEU | p.Leu67Pro | Q15831 |
| HOMO SAPIENS | 4 | 26812 | 2wtk | F | 104 | ARG | p.Arg104Ser | Q15831 |
| HOMO SAPIENS | 0 | 26812 | 2wtk | F | 135 | GLY | p.Gly135Arg | Q15831 |
| HOMO SAPIENS | 0 | 26812 | 2wtk | F | 163 | GLY | p.Gly163Asp | Q15831 |
| HOMO SAPIENS | 2 | 26812 | 2wtk | F | 194 | ALA | p.Asp194Asn | Q15831 |
| HOMO SAPIENS | 2 | 1357 | 5xtd | W | 57 | ARG | p.Arg57His | Q9P0J0 |
| HOMO SAPIENS | 0 | 28086 | 4nn2 | A | 229 | HIS | p.His229Arg | Q8IWS0 |
| HOMO SAPIENS | 0 | 28086 | 4nn2 | A | 234 | LYS | p.Lys234Glu | Q8IWS0 |

| | | | | | | | | |
|---|---|---|---|---|---|---|---|---|
| HOMO SAPIENS | 0 | 28086 | 4nn2 | A | 257 | ARG | p.Arg257Gly | Q8IWS0 |
| HOMO SAPIENS | 5 | 31182 | 2a1x | A | 176 | GLN | p.Gln176Lys | O14832 |
| HOMO SAPIENS | 1 | 31182 | 2a1x | A | 204 | GLY | p.Gly204Ser | O14832 |
| HOMO SAPIENS | 0 | 31182 | 2a1x | A | 269 | ASN | p.Asn269His | O14832 |
| HOMO SAPIENS | 3 | 31182 | 2a1x | A | 275 | ARG | p.Arg275Gln | O14832 |
| HOMO SAPIENS | 3 | 31182 | 2a1x | A | 275 | ARG | p.Arg275Trp | O14832 |
| HOMO SAPIENS | 4 | 2823 | 5xtb | P | 145 | THR | p.Thr145Ile | O75489 |
| HOMO SAPIENS | 5 | 2823 | 5xtb | P | 199 | ARG | p.Arg199Trp | O75489 |
| HOMO SAPIENS | 0 | 8645 | 2h5g | A | 372 | SER | p.Ser372Tyr | P54886 |
| HOMO SAPIENS | 3 | 8645 | 2h5g | A | 715 | ASP | p.Asp715His | P54886 |
| HOMO SAPIENS | 0 | 8645 | 2h5g | A | 765 | ARG | p.Arg765Gln | P54886 |
| HOMO SAPIENS | 0 | 8645 | 2h5g | A | 782 | TYR | p.Tyr782Cys | P54886 |
| HOMO SAPIENS | 0 | 8645 | 2h5g | A | 784 | HIS | p.His784Tyr | P54886 |
| HOMO SAPIENS | 1 | 13992 | 2cue | A | 39 | ARG | p.Arg242Thr | P26367 |
| HOMO SAPIENS | 9 | 13992 | 2cue | A | 55 | PHE | p.Phe258Ser | P26367 |
| HOMO SAPIENS | 0 | 30418 | 2xsw | A | 303 | VAL | p.Val303Met | Q9NRR6 |
| HOMO SAPIENS | 0 | 30418 | 2xsw | A | 315 | PRO | p.Pro315Leu | Q9NRR6 |
| HOMO SAPIENS | 0 | 30418 | 2xsw | A | 341 | GLY | p.Gly341Ser | Q9NRR6 |
| HOMO SAPIENS | 0 | 30418 | 2xsw | A | 378 | ARG | p.Arg378Cys | Q9NRR6 |
| HOMO SAPIENS | 0 | 30418 | 2xsw | A | 435 | ARG | p.Arg435Gln | Q9NRR6 |
| HOMO SAPIENS | 0 | 30418 | 2xsw | A | 515 | ARG | p.Arg515Trp | Q9NRR6 |
| HOMO SAPIENS | 1 | 30418 | 2xsw | A | 526 | PRO | p.Pro526Leu | Q9NRR6 |
| HOMO SAPIENS | 0 | 30418 | 2xsw | A | 563 | ARG | p.Arg563His | Q9NRR6 |
| HOMO SAPIENS | 2 | 30418 | 2xsw | A | 585 | ARG | p.Arg585His | Q9NRR6 |
| HOMO SAPIENS | 2 | 11167 | 4y99 | B | 205 | ARG | p.Arg215Leu | P45379 |
| HOMO SAPIENS | 5 | 11167 | 4y99 | B | 244 | GLU | p.Glu254Asp | P45379 |
| HOMO SAPIENS | 1 | 11167 | 4y99 | B | 253 | LYS | p.Lys263Arg | P45379 |
| HOMO SAPIENS | 8 | 11167 | 4y99 | B | 265 | ARG | p.Arg275Gln | P45379 |
| HOMO SAPIENS | 3 | 11167 | 4y99 | B | 270 | ASP | p.Asp280Asn | P45379 |
| HOMO SAPIENS | -1 | 11167 | 4y99 | B | 278 | ARG | p.Arg288Pro | P45379 |
| HOMO SAPIENS | -1 | 11167 | 4y99 | B | 286 | ARG | p.Arg296Cys | P45379 |
| HOMO SAPIENS | -1 | 11167 | 4y99 | B | 286 | ARG | p.Arg296His | P45379 |
| HOMO SAPIENS | 0 | 15993 | 4po6 | A | 197 | ARG | p.Arg197His | P29597 |
| HOMO SAPIENS | -1 | 15993 | 4po6 | A | 362 | VAL | p.Val362Phe | P29597 |
| HOMO SAPIENS | -1 | 15993 | 4po6 | A | 363 | GLY | p.Gly363Ser | P29597 |
| HOMO SAPIENS | 0 | 5646 | 5uak | A | 7 | GLU | p.Glu7Ter | P13569 |
| HOMO SAPIENS | 0 | 5646 | 5uak | A | 27 | GLY | p.Gly27Ter | P13569 |
| HOMO SAPIENS | 0 | 5646 | 5uak | A | 46 | ALA | p.Ala46Asp | P13569 |
| HOMO SAPIENS | 0 | 5646 | 5uak | A | 60 | GLU | p.Glu60Ter | P13569 |
| HOMO SAPIENS | 0 | 5646 | 5uak | A | 67 | PRO | p.Pro67Leu | P13569 |
| HOMO SAPIENS | 0 | 5646 | 5uak | A | 74 | ARG | p.Arg74Trp | P13569 |
| HOMO SAPIENS | 0 | 5646 | 5uak | A | 75 | ARG | p.Arg75Ter | P13569 |
| HOMO SAPIENS | 0 | 5646 | 5uak | A | 85 | GLY | p.Gly85Glu | P13569 |
| HOMO SAPIENS | 0 | 5646 | 5uak | A | 91 | GLY | p.Gly91Arg | P13569 |
| HOMO SAPIENS | 0 | 5646 | 5uak | A | 92 | GLU | p.Glu92Lys | P13569 |
| HOMO SAPIENS | 0 | 5646 | 5uak | A | 98 | GLN | p.Gln98Arg | P13569 |
| HOMO SAPIENS | 0 | 5646 | 5uak | A | 98 | GLN | p.Gln98Ter | P13569 |
| HOMO SAPIENS | 0 | 5646 | 5uak | A | 109 | TYR | p.Tyr109Cys | P13569 |
| HOMO SAPIENS | 0 | 5646 | 5uak | A | 110 | ASP | p.Asp110His | P13569 |
| HOMO SAPIENS | 0 | 5646 | 5uak | A | 117 | ARG | p.Arg117Cys | P13569 |
| HOMO SAPIENS | 0 | 5646 | 5uak | A | 117 | ARG | p.Arg117His | P13569 |
| HOMO SAPIENS | 0 | 5646 | 5uak | A | 125 | ILE | p.Ile125Thr | P13569 |
| HOMO SAPIENS | 0 | 5646 | 5uak | A | 148 | ILE | p.Ile148Thr | P13569 |
| HOMO SAPIENS | 0 | 5646 | 5uak | A | 170 | ARG | p.Arg170His | P13569 |

| | | | | | | | | |
|---|---|---|---|---|---|---|---|---|
| HOMO SAPIENS | 0 | 5646 | 5uak | A | 178 | GLY | p.Gly178Arg | P13569 |
| HOMO SAPIENS | 0 | 5646 | 5uak | A | 206 | LEU | p.Leu206Trp | P13569 |
| HOMO SAPIENS | 0 | 5646 | 5uak | A | 373 | ASP | p.Asp373Asn | P13569 |
| HOMO SAPIENS | -1 | 5646 | 5uak | A | 709 | ARG | p.Arg709Ter | P13569 |
| HOMO SAPIENS | -1 | 5646 | 5uak | A | 732 | LEU | p.Leu732Ter | P13569 |
| HOMO SAPIENS | -1 | 5646 | 5uak | A | 785 | ARG | p.Arg785Ter | P13569 |
| HOMO SAPIENS | -1 | 5646 | 5uak | A | 831 | GLU | p.Glu831Ter | P13569 |
| HOMO SAPIENS | -1 | 5646 | 5uak | A | 836 | ASP | p.Asp836Tyr | P13569 |
| HOMO SAPIENS | 0 | 5646 | 5uak | A | 851 | ARG | p.Arg851Ter | P13569 |
| HOMO SAPIENS | 0 | 5646 | 5uak | A | 913 | TYR | p.Tyr913Cys | P13569 |
| HOMO SAPIENS | 0 | 5646 | 5uak | A | 913 | TYR | p.Tyr913Ter | P13569 |
| HOMO SAPIENS | 0 | 5646 | 5uak | A | 945 | SER | p.Ser945Leu | P13569 |
| HOMO SAPIENS | 0 | 5646 | 5uak | A | 949 | HIS | p.His949Tyr | P13569 |
| HOMO SAPIENS | 0 | 5646 | 5uak | A | 970 | GLY | p.Gly970Asp | P13569 |
| HOMO SAPIENS | 0 | 5646 | 5uak | A | 1027 | ILE | p.Ile1027Thr | P13569 |
| HOMO SAPIENS | 0 | 5646 | 5uak | A | 1032 | TYR | p.Tyr1032Cys | P13569 |
| HOMO SAPIENS | 0 | 5646 | 5uak | A | 1061 | GLY | p.Gly1061Arg | P13569 |
| HOMO SAPIENS | 1 | 5646 | 5uak | A | 1065 | LEU | p.Leu1065Pro | P13569 |
| HOMO SAPIENS | 1 | 5646 | 5uak | A | 1066 | ARG | p.Arg1066Cys | P13569 |
| HOMO SAPIENS | 1 | 5646 | 5uak | A | 1066 | ARG | p.Arg1066His | P13569 |
| HOMO SAPIENS | 1 | 5646 | 5uak | A | 1067 | ALA | p.Ala1067Thr | P13569 |
| HOMO SAPIENS | 0 | 5646 | 5uak | A | 1069 | GLY | p.Gly1069Arg | P13569 |
| HOMO SAPIENS | 0 | 5646 | 5uak | A | 1070 | ARG | p.Arg1070Trp | P13569 |
| HOMO SAPIENS | 1 | 5646 | 5uak | A | 1071 | GLN | p.Gln1071Pro | P13569 |
| HOMO SAPIENS | 0 | 5646 | 5uak | A | 1074 | PHE | p.Phe1074Leu | P13569 |
| HOMO SAPIENS | 0 | 5646 | 5uak | A | 1077 | LEU | p.Leu1077Pro | P13569 |
| HOMO SAPIENS | 0 | 5646 | 5uak | A | 1089 | TRP | p.Trp1089Ter | P13569 |
| HOMO SAPIENS | 0 | 5646 | 5uak | A | 1092 | TYR | p.Tyr1092Ter | P13569 |
| HOMO SAPIENS | 0 | 5646 | 5uak | A | 1101 | MET | p.Met1101Lys | P13569 |
| HOMO SAPIENS | 0 | 5646 | 5uak | A | 1156 | LEU | p.Leu1156Phe | P13569 |
| HOMO SAPIENS | 0 | 5646 | 5uak | A | 1158 | ARG | p.Arg1158Ter | P13569 |
| HOMO SAPIENS | 0 | 5646 | 5uak | A | 1162 | ARG | p.Arg1162Leu | P13569 |
| HOMO SAPIENS | 0 | 5646 | 5uak | A | 1162 | ARG | p.Arg1162Ter | P13569 |
| HOMO SAPIENS | -1 | 5646 | 5uak | A | 1476 | GLN | p.Gln1476Ter | P13569 |
| HOMO SAPIENS | 0 | 22763 | 1imv | A | 52 | MET | p.Thr72Met | P36955 |
| HOMO SAPIENS | 0 | 22763 | 1imv | A | 301 | TYR | p.Tyr321Ter | P36955 |
| HOMO SAPIENS | 0 | 23179 | 5fql | A | 160 | PRO | p.Pro160Arg | P22304 |
| HOMO SAPIENS | 0 | 23179 | 5fql | A | 172 | ARG | p.Arg172Ter | P22304 |
| HOMO SAPIENS | 0 | 23179 | 5fql | A | 214 | THR | p.Thr214Met | P22304 |
| HOMO SAPIENS | 0 | 23179 | 5fql | A | 309 | THR | p.Thr309Ala | P22304 |
| HOMO SAPIENS | 0 | 23179 | 5fql | A | 333 | SER | p.Ser333Leu | P22304 |
| HOMO SAPIENS | 0 | 23179 | 5fql | A | 443 | ARG | p.Arg443Ter | P22304 |
| HOMO SAPIENS | 0 | 23179 | 5fql | A | 468 | ARG | p.Arg468Leu | P22304 |
| HOMO SAPIENS | 0 | 23179 | 5fql | A | 468 | ARG | p.Arg468Trp | P22304 |
| HOMO SAPIENS | 0 | 23179 | 5fql | A | 502 | TRP | p.Trp502Ser | P22304 |
| HOMO SAPIENS | 9 | 20989 | 1nfi | F | 185 | THR | p.Thr185Met | P25963 |
| HOMO SAPIENS | 4 | 20989 | 1nfi | F | 194 | GLY | p.Gly194Ala | P25963 |
| HOMO SAPIENS | 0 | 11538 | 3eab | A | 162 | VAL | p.Val162Ile | Q9UBP0 |
| HOMO SAPIENS | 0 | 33399 | 2p39 | A | 71 | SER | p.Ser71Gly | Q9GZV9 |
| HOMO SAPIENS | 3 | 33399 | 2p39 | A | 96 | MET | p.Met96Thr | Q9GZV9 |
| HOMO SAPIENS | 0 | 33399 | 2p39 | A | 129 | SER | p.Ser129Phe | Q9GZV9 |
| HOMO SAPIENS | -1 | 33399 | 2p39 | A | 172 | PRO | p.Pro172Leu | Q9GZV9 |
| HOMO SAPIENS | -1 | 33399 | 2p39 | A | 176 | ARG | p.Arg176Gln | Q9GZV9 |
| HOMO SAPIENS | -1 | 33399 | 2p39 | A | 179 | ARG | p.Arg179Trp | Q9GZV9 |

| | | | | | | | | |
|---|---|---|---|---|---|---|---|---|
| HOMO SAPIENS | -1 | 33399 | 2p39 | A | 179 | ARG | p.Arg179Gln | Q9GZV9 |
| HOMO SAPIENS | 0 | 16217 | 2dne | A | 79 | ILE | p.Ile163Leu | P10515 |
| HOMO SAPIENS | 8 | 7666 | 4kbq | A | 65 | ASN | p.Asn65Ser | Q9UNE7 |
| HOMO SAPIENS | 1 | 6407 | 1ikt | A | 71 | THR | p.Thr687Ile | P51659 |
| HOMO SAPIENS | 0 | 6407 | 1ikt | A | 112 | MET | p.Met728Val | P51659 |
| HOMO SAPIENS | 0 | 5849 | 4ihb | A | 67 | VAL | p.Val67Asp | O75923 |
| HOMO SAPIENS | 6 | 17703 | 5fcn | A | 20 | HIS | p.His99Gln | Q66GS9 |
| HOMO SAPIENS | 8 | 17703 | 5fcn | A | 33 | ARG | p.Arg112His | Q66GS9 |
| HOMO SAPIENS | 0 | 34095 | 2E+073 | A | 101 | HIS | p.His101Tyr | P05129 |
| HOMO SAPIENS | -1 | 41358 | 4liq | E | 507 | THR | p.Thr507Met | P07333 |
| HOMO SAPIENS | -1 | 21290 | 1fch | A | 235 | MET | p.Met272Arg | P50542 |
| HOMO SAPIENS | 5 | 23431 | 6pax | A | 23 | ARG | p.Arg26Gly | P26367 |
| HOMO SAPIENS | 1 | 23431 | 6pax | A | 61 | GLY | p.Gly64Val | P26367 |
| HOMO SAPIENS | 1 | 23431 | 6pax | A | 65 | PRO | p.Pro68Ser | P26367 |
| HOMO SAPIENS | 1 | 23431 | 6pax | A | 116 | SER | p.Ser119Arg | P26367 |
| HOMO SAPIENS | 9 | 23431 | 6pax | A | 123 | VAL | p.Val126Asp | P26367 |
| HOMO SAPIENS | 9 | 23431 | 6pax | A | 125 | ARG | p.Arg128Cys | P26367 |
| HOMO SAPIENS | 9 | 33003 | 4zrj | B | 535 | LEU | p.Leu535Pro | P35240 |
| HOMO SAPIENS | 7 | 33003 | 4zrj | B | 538 | GLN | p.Gln538Pro | P35240 |
| HOMO SAPIENS | 0 | 33003 | 4zrj | B | 585 | TRP | p.Ala585Thr | P35240 |
| HOMO SAPIENS | 0 | 3800 | 2nzj | A | 193 | VAL | p.Val193Met | O75628 |

| gene_name | max_cons | max_bs_id | max_pdb_id | max_chain_id |
|---|---|---|---|---|
| AKR1D1 | 0.306 | 22 | 3dop | A |
| AKR1D1 | 0.216 | 34 | 3cav | A |
| AKR1D1 | 0.276 | 21 | 3cot | A |
| AKR1D1 | 0.258 | 56 | 3cas | A |
| ALPP | None | None | None | None |
| BMPR1A | None | None | None | None |
| BMPR1A | None | None | None | None |
| FGFR2 | None | None | None | None |
| PROC | 0.587 | 3 | 1aut | C |
| PROC | None | None | None | None |
| PROC | None | None | None | None |
| PROC | 0.396 | 10 | 3f6u | H |
| PROC | 0.247 | 13 | 3f6u | H |
| PROC | None | None | None | None |
| PROC | None | None | None | None |
| PROC | None | None | None | None |
| PROC | None | None | None | None |
| PROC | 0.587 | 3 | 1aut | C |
| PROC | None | None | None | None |
| PROC | None | None | None | None |
| INS | 1 | 2 | 3zs2 | J |
| INS | 0.85 | 1 | 4ajx | D |
| INS | 1 | 1 | 3zs2 | J |
| INS | 1 | 1 | 3zs2 | J |
| FTL | 0.627 | 11 | 2fg8 | B |
| FTL | 0.313 | 62 | 2fg8 | G |
| FTL | 0.544 | 24 | 2fg8 | D |
| TNFSF11 | None | None | None | None |
| EHMT1 | None | None | None | None |
| EHMT1 | None | None | None | None |
| EHMT1 | None | None | None | None |
| CA2 | 0.939 | 1 | 4idr | X |
| CA2 | 0.939 | 1 | 4idr | X |
| CA2 | 0.902 | 5 | 4idr | X |
| CA2 | 0.902 | 5 | 4idr | X |
| SMAD4 | None | None | None | None |
| SMAD4 | None | None | None | None |
| TTR | 0.691 | 12 | 1ttb | B |
| TTR | 0.961 | 7 | 3grb | A |
| TTR | 0.891 | 4 | 3dk2 | B |
| TTR | 0.947 | 3 | 1tta | A |
| TTR | 0.923 | 1 | 3grb | B |
| TTR | 0.891 | 1 | 4y9b | B |
| TTR | 0.891 | 1 | 4y9b | B |
| TTR | 0.882 | 15 | 3grb | A |
| TTR | 0.909 | 5 | 4tl4 | A |
| TTR | 0.836 | 7 | 1z7j | B |
| TTR | 0.882 | 15 | 3grb | A |
| TTR | 0.891 | 5 | 3i9i | B |
| TTR | 0.902 | 9 | 3m1o | B |
| TTR | 0.902 | 9 | 3m1o | B |
| TTR | 0.902 | 9 | 3m1o | B |
| TTR | 0.636 | 16 | 1thc | A |

| | | | | |
|---|---|---|---|---|
| TTR | 0.902 | 9 | 3m1o | B |
| TTR | 0.836 | 7 | 1bze | A |
| TTR | 0.846 | 16 | 3gps | A |
| TTR | 0.846 | 16 | 3gps | A |
| TTR | 0.909 | 5 | 4tl4 | A |
| TTR | 0.909 | 5 | 4tl4 | A |
| TTR | 0.673 | 15 | 3dk2 | B |
| TTR | 0.945 | 4 | 4deu | A |
| TTR | 0.941 | 3 | 3did | B |
| TTR | 0.941 | 3 | 3did | B |
| TTR | 0.868 | 6 | 4l1t | A |
| TTR | 0.882 | 19 | 3gps | B |
| TTR | 0.891 | 4 | 3dk2 | B |
| TTR | 0.941 | 3 | 3did | B |
| TTR | 0.923 | 1 | 4hiq | A |
| TTR | 0.923 | 1 | 4hiq | A |
| TTR | 0.961 | 7 | 3grb | A |
| TTR | 0.839 | 7 | 1tsh | B |
| TTR | 0.891 | 6 | 1ttb | B |
| ALK | 0.682 | 1 | 4cd0 | A |
| ALK | None | None | None | None |
| THRB | None | None | None | None |
| THRB | None | None | None | None |
| THRB | None | None | None | None |
| THRB | None | None | None | None |
| THRB | None | None | None | None |
| THRB | None | None | None | None |
| THRB | None | None | None | None |
| THRB | None | None | None | None |
| THRB | None | None | None | None |
| THRB | None | None | None | None |
| THRB | None | None | None | None |
| THRB | None | None | None | None |
| THRB | None | None | None | None |
| THRB | None | None | None | None |
| THRB | None | None | None | None |
| THRB | None | None | None | None |
| THRB | None | None | None | None |
| THRB | None | None | None | None |
| THRB | 0.577 | 3 | 2j4a | A |
| IL2RG | None | None | None | None |
| IL2RG | None | None | None | None |
| IL2RG | None | None | None | None |
| IL2RG | 0.256 | 1 | 2b5i | C |
| IL2RG | None | None | None | None |
| ITGB3 | None | None | None | None |
| ITGB3 | None | None | None | None |
| ITGB3 | 0.294 | 1 | 1jv2 | B |
| ITGB3 | None | None | None | None |
| ITGB3 | None | None | None | None |
| ITGB3 | None | None | None | None |
| ITGB3 | None | None | None | None |
| ITGB3 | None | None | None | None |
| ITGB3 | None | None | None | None |

| | | | | |
|---|---|---|---|---|
| PKD2 | None | None | None | None |
| PKD2 | None | None | None | None |
| PKD2 | None | None | None | None |
| PKD2 | None | None | None | None |
| PKD2 | None | None | None | None |
| PKD2 | None | None | None | None |
| PKD2 | None | None | None | None |
| TP53 | 0.571 | 1 | 1saf | B |
| TP53 | 0.571 | 1 | 1saf | B |
| TP53 | None | None | None | None |
| TP53 | 0.585 | 15 | 4mzr | B |
| TP53 | None | None | None | None |
| TP53 | None | None | None | None |
| EGFR | None | None | None | None |
| EGFR | None | None | None | None |
| XRCC4 | None | None | None | None |
| XRCC4 | None | None | None | None |
| HBB | 0.438 | 2 | 2w6v | D |
| HBB | 0.467 | 2 | 3qje | D |
| HBB | 0.467 | 2 | 3qje | D |
| HBB | 0.303 | 15 | 2w72 | B |
| HBB | 0.303 | 21 | 2w72 | B |
| HBB | 0.332 | 7 | 1cbm | A |
| HBB | 0.421 | 11 | 3ic2 | B |
| HBB | 0.459 | 2 | 5ee4 | D |
| HBB | 0.459 | 2 | 5ee4 | D |
| HBB | 0.459 | 2 | 5ee4 | D |
| HBB | 0.535 | 3 | 3r5i | B |
| HBB | 0.383 | 3 | 1lfv | B |
| HBB | 0.383 | 3 | 1lfv | B |
| HBB | 0.402 | 11 | 1m9p | D |
| HBB | 0.256 | 24 | 4n7p | D |
| HBB | 0.459 | 2 | 5ee4 | D |
| HBB | 0.566 | 1 | 1cbm | B |
| HBB | 0.51 | 3 | 1abw | B |
| HBB | 0.392 | 2 | 1lfz | B |
| HBB | 0.474 | 1 | 2h35 | B |
| HBB | 0.474 | 1 | 2h35 | B |
| HBB | 0.38 | 2 | 1j7y | B |
| HBB | 0.508 | 7 | 3ic2 | B |
| HBB | 0.545 | 3 | 1bij | D |
| HBB | 0.6 | 1 | 2w72 | D |
| HBB | 0.332 | 4 | 1fn3 | D |
| HBB | 0.354 | 15 | 1bij | D |
| HBB | 0.308 | 15 | 4rol | D |
| HBB | 0.733 | 1 | 1yff | D |
| HBB | 0.692 | 2 | 1buw | B |
| RPS6KA3 | None | None | None | None |
| RPS6KA3 | 0.555 | 1 | 4nw5 | A |
| RPS6KA3 | 0.226 | 6 | 4nw5 | A |
| RPS6KA3 | None | None | None | None |
| RPS6KA3 | None | None | None | None |
| RPS6KA3 | None | None | None | None |
| GBA | 1 | 15 | 3gxd | B |

| | | | | | |
|---|---|---|---|---|---|
| GBA | | 1 | 55 | 3gxd | B |
| GBA | | 1 | 15 | 3gxd | B |
| GBA | | 0.75 | 45 | 3ril | A |
| GBA | | 0.25 | 33 | 2nt0 | A |
| GBA | | 0.25 | 33 | 2nt0 | A |
| GBA | | 1 | 12 | 1ogs | A |
| GBA | | 0.75 | 122 | 3gxi | A |
| GBA | | 1 | 2 | 1ogs | A |
| GBA | | 1 | 11 | 1ogs | A |
| GBA | | 0.5 | 59 | 3gxf | B |
| GBA | | 0.75 | 45 | 3ril | A |
| GBA | | 1 | 18 | 1ogs | A |
| GBA | | 1 | 3 | 1ogs | A |
| GBA | None | None | | None | None |
| GBA | | 1 | 2 | 1ogs | A |
| GBA | | 0.25 | 35 | 2nt0 | A |
| GBA | | 0.75 | 76 | 3gxm | C |
| GBA | | 1 | 3 | 1ogs | A |
| GBA | | 1 | 3 | 1ogs | A |
| GBA | | 0.75 | 62 | 3ril | A |
| GBA | | 1 | 28 | 1ogs | A |
| GBA | None | None | | None | None |
| GBA | | 0.5 | 78 | 3gxi | A |
| GBA | | 0.5 | 92 | 3rik | B |
| GBA | | 0.5 | 42 | 2xwe | A |
| GBA | | 0.333 | 117 | 1ogs | A |
| GBA | | 0.5 | 92 | 3rik | B |
| GBA | | 0.75 | 49 | 3ril | A |
| GBA | | 0.25 | 109 | 2f61 | A |
| GBA | | 0.667 | 60 | 1ogs | A |
| GBA | None | None | | None | None |
| GBA | | 0.5 | 11 | 2nt1 | A |
| GBA | | 0.75 | 21 | 3gxi | A |
| GBA | None | None | | None | None |
| GBA | | 0.5 | 35 | 2xwe | A |
| GBA | | 0.75 | 25 | 3keh | B |
| GBA | | 0.75 | 70 | 3ril | A |
| GBA | | 0.75 | 70 | 3ril | A |
| GBA | | 1 | 15 | 3gxd | B |
| GBA | | 1 | 44 | 3gxd | B |
| GBA | | 0.5 | 57 | 3gxf | D |
| GBA | | 1 | 44 | 3gxd | B |
| GBA | | 1 | 44 | 3gxd | B |
| GBA | None | None | | None | None |
| GBA | | 0.75 | 94 | 3gxi | A |
| GBA | | 0.75 | 94 | 3gxi | A |
| IFIH1 | None | None | | None | None |
| TGFBR2 | None | None | | None | None |
| TGFBR2 | None | None | | None | None |
| TGFBR2 | None | None | | None | None |
| TGFBR2 | None | None | | None | None |
| TGFBR2 | None | None | | None | None |
| TGFBR2 | None | None | | None | None |
| TGFBR2 | None | None | | None | None |

| | | | | |
|---|---|---|---|---|
| TGFBR2 | None | None | None | None |
| TGFBR2 | None | None | None | None |
| TGFBR2 | 0.742 | 1 | 5E+091 | A |
| TGFBR2 | None | None | None | None |
| TGFBR2 | None | None | None | None |
| TGFBR2 | 0.206 | 6 | 5E+092 | A |
| TGFBR2 | 0.676 | 1 | 5e8v | A |
| TGFBR2 | 0.676 | 1 | 5e8v | A |
| ACE | 0.684 | 17 | 3nxq | A |
| ACE | 0.474 | 25 | 4bxk | A |
| ACE | 0.263 | 31 | 2c6f | A |
| ACE | None | None | None | None |
| ACE | 0.211 | 93 | 4ca6 | A |
| ACE | 0.211 | 92 | 5ama | D |
| COL10A1 | None | None | None | None |
| COL10A1 | None | None | None | None |
| COL10A1 | None | None | None | None |
| COL10A1 | 0.409 | 1 | 1gr3 | A |
| COL10A1 | None | None | None | None |
| COL10A1 | 0.318 | 4 | 1gr3 | A |
| COL10A1 | None | None | None | None |
| COL10A1 | 0.273 | 6 | 1gr3 | A |
| COL10A1 | 0.318 | 4 | 1gr3 | A |
| KLF1 | None | None | None | None |
| C3 | None | None | None | None |
| CTSC | None | None | None | None |
| CTSC | 0.934 | 1 | 2djg | C |
| CTSC | None | None | None | None |
| LYZ | 0.877 | 3 | 1bb5 | A |
| LYZ | 0.923 | 1 | 1lz4 | A |
| LYZ | 0.852 | 2 | 1ubz | A |
| DHODH | None | None | None | None |
| DHODH | 0.207 | 18 | 5mut | A |
| DHODH | None | None | None | None |
| DHODH | None | None | None | None |
| DHODH | None | None | None | None |
| DHODH | None | None | None | None |
| DHODH | None | None | None | None |
| ADH1C | 0.527 | 11 | 1ht0 | A |
| GDNF | None | None | None | None |
| TNF | 0.5 | 8 | 2e7a | B |
| ALDH4A1 | 0.636 | 3 | 3v9h | A |
| ALDH4A1 | None | None | None | None |
| ALDH4A1 | None | None | None | None |
| BRAF | None | None | None | None |
| BRAF | None | None | None | None |
| BRAF | 0.752 | 2 | 3psb | A |
| BRAF | 0.752 | 2 | 3psb | A |
| BRAF | 0.737 | 1 | 3pri | A |
| BRAF | 0.737 | 1 | 3pri | A |
| BRAF | 0.737 | 1 | 3pri | A |
| BRAF | 0.737 | 1 | 3prf | A |
| BRAF | 0.737 | 1 | 3prf | A |
| BRAF | 0.752 | 2 | 3psb | A |

| | | | | |
|---|---|---|---|---|
| BRAF | 0.752 | 2 | 3psb | A |
| BRAF | 0.752 | 2 | 3psb | A |
| BRAF | 0.752 | 2 | 3psb | A |
| BRAF | 0.555 | 3 | 5vam | A |
| BRAF | 0.663 | 1 | 4ehe | B |
| BRAF | 0.734 | 1 | 5csx | A |
| TBX5 | None | None | None | None |
| TBX5 | None | None | None | None |
| TBX5 | None | None | None | None |
| TBX5 | None | None | None | None |
| TBX5 | None | None | None | None |
| TBX5 | None | None | None | None |
| TBX5 | None | None | None | None |
| TBX5 | None | None | None | None |
| TBX5 | None | None | None | None |
| RB1 | None | None | None | None |
| RB1 | None | None | None | None |
| RB1 | None | None | None | None |
| RB1 | None | None | None | None |
| RB1 | None | None | None | None |
| RB1 | None | None | None | None |
| CLIC2 | 0.5 | 3 | 2per | A |
| CLIC2 | None | None | None | None |
| ALDH7A1 | None | None | None | None |
| ALDH7A1 | None | None | None | None |
| ALDH7A1 | 0.405 | 4 | 2j6l | D |
| ALDH7A1 | 0.301 | 14 | 4zuk | A |
| ALDH7A1 | 0.409 | 3 | 4zvw | C |
| ALDH7A1 | None | None | None | None |
| ALDH7A1 | 0.534 | 3 | 2j6l | A |
| ALDH7A1 | None | None | None | None |
| ALDH7A1 | 0.333 | 6 | 4x0t | A |
| ALDH7A1 | None | None | None | None |
| ALDH7A1 | None | None | None | None |
| ALDH7A1 | 0.534 | 3 | 2j6l | A |
| ALDH7A1 | None | None | None | None |
| ALDH7A1 | None | None | None | None |
| ALDH7A1 | 0.632 | 1 | 4zuk | A |
| ALDH7A1 | None | None | None | None |
| KDR | 0.575 | 5 | 1y6a | A |
| MUT | None | None | None | None |
| MUT | None | None | None | None |
| MUT | None | None | None | None |
| MUT | None | None | None | None |
| MUT | None | None | None | None |
| MUT | None | None | None | None |
| MUT | None | None | None | None |
| MUT | None | None | None | None |
| MUT | None | None | None | None |
| MUT | None | None | None | None |
| MUT | None | None | None | None |
| MUT | None | None | None | None |
| MUT | None | None | None | None |
| MUT | None | None | None | None |

| | | | | |
|---|---|---|---|---|
| MUT | None | None | None | None |
| MUT | None | None | None | None |
| MUT | 0.6 | 1 | 3bic | A |
| MUT | 0.6 | 2 | 2xiq | A |
| MUT | 0.6 | 2 | 2xiq | A |
| MUT | None | None | None | None |
| MUT | None | None | None | None |
| MUT | 0.4 | 1 | 2xiq | A |
| MUT | None | None | None | None |
| MUT | None | None | None | None |
| MUT | None | None | None | None |
| MUT | None | None | None | None |
| MUT | None | None | None | None |
| MUT | None | None | None | None |
| MUT | None | None | None | None |
| MUT | None | None | None | None |
| MUT | None | None | None | None |
| MUT | None | None | None | None |
| MUT | None | None | None | None |
| MUT | None | None | None | None |
| SOD1 | 0.778 | 1 | 2gbu | A |
| SOD1 | 0.727 | 4 | 4xcr | A |
| SOD1 | 0.35 | 26 | 1n19 | A |
| SOD1 | 0.621 | 12 | 1hl5 | H |
| NECTIN3 | None | None | None | None |
| HFE | None | None | None | None |
| HFE | None | None | None | None |
| HFE | None | None | None | None |
| HFE | 0.5 | 1 | 1de4 | A |
| HFE | None | None | None | None |
| HFE | None | None | None | None |
| HFE | 0.25 | 15 | 1a6z | A |
| EIF2AK3 | None | None | None | None |
| EIF2AK3 | None | None | None | None |
| TERT | None | None | None | None |
| TERT | None | None | None | None |
| SPR | None | None | None | None |
| SPR | None | None | None | None |
| SPR | None | None | None | None |
| SPR | None | None | None | None |
| SPR | None | None | None | None |
| F7 | 0.625 | 2 | 1klj | L |
| UNG | None | None | None | None |
| MMP13 | 0.356 | 22 | 2ozr | E |
| MMP13 | 0.786 | 1 | 3tvc | A |
| MMP13 | 0.766 | 1 | 3zxh | B |
| MMP13 | 0.766 | 1 | 3zxh | B |
| CD207 | 0.621 | 1 | 3kqg | B |
| F2 | 1 | 2 | 1nrq | L |
| SPTBN2 | None | None | None | None |
| PKLR | 0.568 | 9 | 2vgb | A |
| PKLR | 0.486 | 17 | 2vgg | D |
| PKLR | None | None | None | None |
| PKLR | 0.486 | 8 | 2vgg | D |

| | | | | |
|---|---|---|---|---|
| PKLR | 0.568 | 3 | 4ip7 | D |
| PKLR | 0.432 | 13 | 2vgi | A |
| PKLR | 0.378 | 45 | 4ip7 | B |
| PKLR | 0.324 | 25 | 2vgf | B |
| SACS | None | None | None | None |
| HRAS | 0.529 | 3 | 6q21 | A |
| HRAS | 0.529 | 3 | 6q21 | A |
| HRAS | 0.59 | 1 | 1gnq | A |
| HRAS | 0.59 | 1 | 1gnq | A |
| HRAS | 0.664 | 1 | 2lwi | A |
| HRAS | 0.742 | 1 | 1ctq | A |
| HRAS | 0.69 | 1 | 1xd2 | A |
| HRAS | 0.415 | 3 | 6q21 | D |
| HRAS | 0.529 | 3 | 6q21 | A |
| HRAS | 0.403 | 3 | 2n46 | A |
| HRAS | 0.521 | 4 | 4nyj | R |
| HRAS | 0.664 | 1 | 2lwi | A |
| HRAS | 0.664 | 1 | 2lwi | A |
| VHL | None | None | None | None |
| VHL | None | None | None | None |
| VHL | None | None | None | None |
| VHL | None | None | None | None |
| VHL | 0.5 | 22 | 4w9l | C |
| VHL | 0.5 | 22 | 5nvy | C |
| VHL | None | None | None | None |
| VHL | None | None | None | None |
| VHL | None | None | None | None |
| VHL | 0.5 | 2 | 5nvy | C |
| VHL | 0.5 | 1 | 5nvy | C |
| VHL | 0.5 | 1 | 5nvy | C |
| VHL | 0.5 | 4 | 5nvy | C |
| VHL | None | None | None | None |
| VHL | 0.5 | 6 | 5nvy | C |
| VHL | 0.5 | 2 | 5nvy | C |
| VHL | 0.5 | 8 | 3ztc | C |
| VHL | 0.5 | 2 | 5nvy | C |
| VHL | 0.5 | 2 | 5nvy | C |
| VHL | None | None | None | None |
| VHL | 0.5 | 7 | 4w9h | C |
| VHL | 0.5 | 2 | 5nvy | C |
| VHL | 0.5 | 2 | 5nvy | C |
| VHL | 0.5 | 19 | 4w9l | C |
| VHL | None | None | None | None |
| TTBK1 | None | None | None | None |
| RAD51 | None | None | None | None |
| ALB | None | None | None | None |
| ALB | 0.5 | 2 | 4n0u | D |
| ALB | None | None | None | None |
| ALB | None | None | None | None |
| MET | None | None | None | None |
| MET | None | None | None | None |
| MET | None | None | None | None |
| MET | None | None | None | None |
| MET | None | None | None | None |

| | | | | |
|---|---|---|---|---|
| MET | None | None | None | None |
| MET | None | None | None | None |
| MET | None | None | None | None |
| MET | None | None | None | None |
| CP | None | None | None | None |
| CP | None | None | None | None |
| PGM1 | None | None | None | None |
| PGM1 | 0.409 | 2 | 5f9c | A |
| PGM1 | 0.409 | 2 | 5f9c | A |
| PGM1 | None | None | None | None |
| PGM1 | None | None | None | None |
| PGM1 | None | None | None | None |
| PGM1 | None | None | None | None |
| NR5A1 | None | None | None | None |
| NR5A1 | None | None | None | None |
| NR5A1 | None | None | None | None |
| F9 | None | None | None | None |
| F9 | None | None | None | None |
| HPRT1 | None | None | None | None |
| HPRT1 | None | None | None | None |
| HPRT1 | 0.25 | 7 | 3gep | B |
| HPRT1 | 0.375 | 15 | 3gep | B |
| HPRT1 | None | None | None | None |
| HPRT1 | 0.871 | 3 | 4rac | B |
| HPRT1 | 0.806 | 2 | 4ran | D |
| HPRT1 | None | None | None | None |
| HPRT1 | 0.25 | 4 | 3gep | B |
| HPRT1 | 0.44 | 3 | 1d6n | B |
| HPRT1 | None | None | None | None |
| HPRT1 | None | None | None | None |
| HPRT1 | 0.24 | 25 | 1d6n | B |
| HPRT1 | 0.703 | 4 | 5w8v | A |
| HPRT1 | None | None | None | None |
| HPRT1 | 0.321 | 8 | 4kn6 | A |
| HPRT1 | 0.312 | 27 | 3ggj | B |
| HPRT1 | 0.312 | 12 | 3gep | B |
| HPRT1 | 0.871 | 3 | 4rac | B |
| HPRT1 | 0.871 | 3 | 4rac | B |
| HPRT1 | 0.765 | 2 | 4rao | D |
| HPRT1 | 0.774 | 2 | 4rac | B |
| GPD1L | None | None | None | None |
| GPD1L | None | None | None | None |
| GPD1L | None | None | None | None |
| FGFR2 | None | None | None | None |
| FGFR2 | None | None | None | None |
| FGFR2 | None | None | None | None |
| FGFR2 | None | None | None | None |
| FGFR2 | None | None | None | None |
| FGFR2 | None | None | None | None |
| FGFR2 | None | None | None | None |
| FGFR2 | None | None | None | None |
| FGFR2 | None | None | None | None |
| FGFR2 | None | None | None | None |
| FGFR2 | None | None | None | None |

| | | | | |
|---|---|---|---|---|
| FGFR2 | None | None | None | None |
| FGFR2 | None | None | None | None |
| FGFR2 | None | None | None | None |
| FGFR2 | None | None | None | None |
| FGFR2 | None | None | None | None |
| FGFR2 | None | None | None | None |
| FGFR2 | None | None | None | None |
| FGFR2 | None | None | None | None |
| HSPD1 | None | None | None | None |
| HSPD1 | None | None | None | None |
| FGF8 | None | None | None | None |
| FGF8 | None | None | None | None |
| FGF8 | None | None | None | None |
| F9 | 0.438 | 1 | 5vyg | B |
| NCF4 | None | None | None | None |
| NCF4 | None | None | None | None |
| LDLR | None | None | None | None |
| LDLR | None | None | None | None |
| LDLR | 0.478 | 1 | 3gcx | E |
| LDLR | 0.478 | 1 | 3gcx | E |
| LDLR | 0.478 | 1 | 3gcx | E |
| LDLR | None | None | None | None |
| C3 | None | None | None | None |
| C3 | None | None | None | None |
| C3 | None | None | None | None |
| C3 | None | None | None | None |
| C3 | None | None | None | None |
| C3 | None | None | None | None |
| C3 | None | None | None | None |
| C3 | None | None | None | None |
| TULP1 | None | None | None | None |
| TULP1 | None | None | None | None |
| TULP1 | None | None | None | None |
| TULP1 | None | None | None | None |
| IKBKG | None | None | None | None |
| IKBKG | None | None | None | None |
| IKBKG | None | None | None | None |
| CD40 | None | None | None | None |
| CD40 | None | None | None | None |
| SLC9A3R1 | None | None | None | None |
| BCHE | 0.278 | 18 | 2xqg | A |
| BCHE | 0.439 | 5 | 2wil | B |
| BCHE | 0.544 | 11 | 1p0q | A |
| BCHE | None | None | None | None |
| BCHE | None | None | None | None |
| BCHE | None | None | None | None |
| BCHE | 0.63 | 4 | 2xqg | A |
| BCHE | None | None | None | None |
| BCHE | None | None | None | None |
| BCHE | 0.255 | 18 | 2xqj | A |
| BCHE | None | None | None | None |
| BCKDHB | 0.833 | 1 | 1v11 | B |
| BCKDHB | 0.5 | 18 | 2bfc | B |
| BCKDHB | None | None | None | None |

| | | | | |
|---|---|---|---|---|
| BCKDHB | 0.667 | 11 | 2bfd | B |
| BCKDHB | 0.833 | 10 | 1olx | B |
| BCKDHB | 0.833 | 1 | 1v11 | B |
| BCKDHB | 0.833 | 1 | 1v11 | B |
| BCKDHB | 0.625 | 2 | 1x7z | B |
| BCKDHB | 0.429 | 16 | 1ols | B |
| BCKDHB | 0.333 | 99 | 1olx | B |
| BCKDHB | 0.333 | 64 | 1x7y | B |
| BCKDHB | 0.857 | 1 | 2bff | B |
| CDH1 | 0.278 | 8 | 3ff7 | B |
| CDH1 | 0.575 | 1 | 3ff8 | A |
| CDH1 | 0.543 | 1 | 3ff7 | A |
| CDH1 | None | None | None | None |
| CDH1 | 0.2 | 20 | 2o72 | A |
| CTNNB1 | None | None | None | None |
| AIRE | None | None | None | None |
| EDA | 1 | 1 | 1rj8 | A |
| EDA | None | None | None | None |
| EDA | None | None | None | None |
| EDA | None | None | None | None |
| EDA | None | None | None | None |
| SH2D1A | 0.444 | 5 | 1ka7 | A |
| SH2D1A | None | None | None | None |
| SH2D1A | None | None | None | None |
| GCK | 0.556 | 37 | 3idh | A |
| GCK | 0.333 | 32 | 4isf | A |
| GCK | None | None | None | None |
| GCK | None | None | None | None |
| GCK | None | None | None | None |
| GCK | 0.611 | 13 | 3h1v | X |
| GCK | None | None | None | None |
| GCK | None | None | None | None |
| GCK | None | None | None | None |
| GCK | 0.778 | 5 | 3id8 | A |
| GCK | 0.833 | 1 | 4dhy | A |
| GCK | 0.389 | 6 | 4lc9 | B |
| GCK | 0.222 | 33 | 4mle | A |
| GCK | 0.611 | 6 | 3id8 | A |
| GCK | None | None | None | None |
| GCK | None | None | None | None |
| GCK | None | None | None | None |
| GCK | 0.5 | 5 | 3vey | A |
| GCK | None | None | None | None |
| GCK | None | None | None | None |
| AQP2 | None | None | None | None |
| AQP2 | None | None | None | None |
| AQP2 | None | None | None | None |
| AQP2 | None | None | None | None |
| AQP2 | None | None | None | None |
| AQP2 | None | None | None | None |
| AQP2 | None | None | None | None |
| AQP2 | None | None | None | None |
| AQP2 | None | None | None | None |
| AQP2 | None | None | None | None |

| | | | | |
|---|---|---|---|---|
| AQP2 | None | None | None | None |
| AQP2 | None | None | None | None |
| AQP2 | None | None | None | None |
| SUMF1 | 0.75 | 6 | 1z70 | X |
| SUMF1 | 0.556 | 30 | 1y1i | X |
| SUMF1 | 0.5 | 38 | 2aij | X |
| SUMF1 | 0.625 | 18 | 2aij | X |
| SUMF1 | 0.875 | 35 | 2afy | X |
| SUMF1 | 0.667 | 19 | 1y1i | X |
| SUMF1 | 0.75 | 17 | 1y1j | X |
| SUMF1 | 0.75 | 17 | 1y1j | X |
| SUMF1 | 0.625 | 27 | 1y1g | X |
| FAM20A | None | None | None | None |
| FAM20A | None | None | None | None |
| FAM20A | None | None | None | None |
| FAM20A | None | None | None | None |
| NT5E | None | None | None | None |
| NT5E | None | None | None | None |
| NOS3 | 0.564 | 54 | 1m9k | B |
| ACTG1 | None | None | None | None |
| ACTG1 | None | None | None | None |
| ACTG1 | 0.229 | 48 | 5jlh | E |
| ACTG1 | None | None | None | None |
| ACTG1 | 0.571 | 8 | 5jlh | E |
| ACTG1 | None | None | None | None |
| FCGR1A | None | None | None | None |
| GPI | 0.304 | 64 | 1iri | A |
| GPI | None | None | None | None |
| GPI | 1 | 3 | 1jiq | A |
| GPI | None | None | None | None |
| GPI | 0.696 | 22 | 1iat | A |
| GPI | 0.522 | 22 | 1nuh | A |
| GPI | 0.696 | 27 | 1iat | A |
| GPI | 0.304 | 87 | 1jlh | B |
| GPI | 0.391 | 141 | 1jlh | B |
| CFH | None | None | None | None |
| OGG1 | 0.895 | 13 | 1m3h | A |
| TPK1 | None | None | None | None |
| TPK1 | None | None | None | None |
| TPK1 | None | None | None | None |
| TPK1 | None | None | None | None |
| SEC23A | None | None | None | None |
| SEC23A | None | None | None | None |
| TYRP1 | None | None | None | None |
| TYRP1 | None | None | None | None |
| TYRP1 | None | None | None | None |
| TYRP1 | None | None | None | None |
| TYRP1 | 0.3 | 5 | 5m8m | A |
| TYRP1 | None | None | None | None |
| CSTB | None | None | None | None |
| CSTB | None | None | None | None |
| CSTB | None | None | None | None |
| ACP1 | None | None | None | None |
| IL17F | None | None | None | None |

| | | | | |
|---|---|---|---|---|
| IL17F | None | None | None | None |
| IL17F | None | None | None | None |
| IL17F | None | None | None | None |
| IL17F | None | None | None | None |
| IDH2 | 0.5 | 2 | 5svn | B |
| IDH2 | 0.5 | 2 | 5svn | B |
| IDH2 | 0.48 | 13 | 5i96 | A |
| IDH2 | None | None | None | None |
| TP53 | 0.925 | 11 | 2ocj | A |
| TP53 | 0.8 | 16 | 2ahi | D |
| TP53 | 0.65 | 49 | 4ibq | A |
| TP53 | 0.725 | 27 | 5g4o | A |
| TP53 | 0.455 | 21 | 3q06 | A |
| TP53 | 0.9 | 1 | 4agp | A |
| TP53 | 0.9 | 1 | 4agp | A |
| TP53 | 0.925 | 10 | 4hje | A |
| TP53 | 0.7 | 29 | 3d05 | A |
| TP53 | 0.875 | 7 | 3kmd | B |
| TP53 | 0.875 | 7 | 3kmd | B |
| TP53 | 0.925 | 6 | 3kz8 | B |
| TP53 | 0.925 | 6 | 3kz8 | B |
| TP53 | 0.609 | 22 | 3q01 | B |
| TP53 | 0.925 | 5 | 3kmd | B |
| TP53 | 0.925 | 5 | 3kmd | B |
| TP53 | 0.925 | 8 | 3kz8 | B |
| TP53 | 0.925 | 8 | 3kz8 | B |
| TP53 | 0.925 | 1 | 3kmd | B |
| TP53 | 0.9 | 11 | 2pcx | A |
| TP53 | 0.9 | 1 | 5aol | A |
| TP53 | 0.85 | 5 | 5lgy | A |
| TP53 | 0.9 | 1 | 3q06 | B |
| TP53 | 0.725 | 19 | 5lgy | A |
| TP53 | 0.775 | 14 | 4mzi | A |
| TP53 | 0.825 | 8 | 1kzy | A |
| TP53 | 0.825 | 8 | 1kzy | A |
| TP53 | 0.8 | 11 | 4ibt | A |
| TP53 | 0.8 | 11 | 4ibt | A |
| TP53 | 0.725 | 15 | 5lgy | A |
| TP53 | 0.725 | 29 | 2h1l | O |
| TP53 | 0.875 | 7 | 3kmd | B |
| TP53 | 0.725 | 15 | 5lgy | A |
| TP53 | 0.8 | 10 | 4agp | A |
| TP53 | 0.8 | 10 | 4agp | A |
| TP53 | 0.8 | 10 | 4agp | A |
| TP53 | 0.8 | 10 | 4agp | A |
| TP53 | 0.675 | 35 | 4ibq | A |
| TP53 | 0.575 | 37 | 4agm | A |
| TP53 | 0.675 | 17 | 3kz8 | B |
| TP53 | 0.356 | 63 | 4mzr | C |
| TP53 | None | None | None | None |
| LMAN1 | None | None | None | None |
| CTSK | 0.75 | 2 | 4n79 | A |
| CTSK | None | None | None | None |
| CTSK | 0.413 | 12 | 4x6j | A |

| | | | | |
|---|---|---|---|---|
| FGB | None | None | None | None |
| FGB | None | None | None | None |
| FGB | 0.688 | 1 | 1fzf | E |
| FGB | 0.281 | 10 | 2ffd | B |
| ACADM | None | None | None | None |
| ACADM | None | None | None | None |
| ACADM | 0.218 | 11 | 1t9g | C |
| ACADM | 0.471 | 3 | 4p13 | D |
| ACADM | None | None | None | None |
| ACADM | None | None | None | None |
| ACADM | 0.389 | 6 | 1t9g | D |
| ACADM | None | None | None | None |
| ACADM | 0.327 | 12 | 4p13 | C |
| ACADM | 0.389 | 6 | 1t9g | D |
| ACADM | None | None | None | None |
| ACADM | None | None | None | None |
| ACADM | None | None | None | None |
| ACADM | 0.291 | 14 | 1t9g | C |
| ACADM | None | None | None | None |
| ACADM | 0.481 | 2 | 4p13 | C |
| ACADM | 0.389 | 6 | 1t9g | D |
| ACADM | None | None | None | None |
| ACADM | None | None | None | None |
| ACADM | None | None | None | None |
| ACADM | 0.564 | 1 | 1t9g | C |
| ACADM | 0.255 | 10 | 1t9g | C |
| ACADM | None | None | None | None |
| ACADM | None | None | None | None |
| ACADM | 0.364 | 6 | 2a1t | A |
| ACADM | 0.312 | 12 | 4p13 | A |
| ACADM | 0.347 | 7 | 4p13 | B |
| PTS | None | None | None | None |
| PTS | 0.417 | 2 | 3i2b | E |
| PTS | None | None | None | None |
| PTS | 0.778 | 1 | 3i2b | D |
| PTS | None | None | None | None |
| PTS | None | None | None | None |
| PTS | None | None | None | None |
| PTS | None | None | None | None |
| PTS | None | None | None | None |
| INS | 0.75 | 2 | 3zqr | E |
| INS | 1 | 5 | 2l1z | A |
| INS | 1 | 5 | 2l1z | A |
| INS | 1 | 1 | 1his | A |
| FBP1 | 0.71 | 26 | 2y5l | C |
| FBP1 | None | None | None | None |
| FBP1 | 0.5 | 18 | 1fta | A |
| FBP1 | 0.7 | 10 | 2y5k | A |
| FBP1 | 0.516 | 38 | 3kc1 | B |
| FBP1 | 0.733 | 10 | 3kc1 | A |
| FBP1 | 0.633 | 43 | 2jjk | A |
| FBP1 | 0.8 | 11 | 2y5k | A |
| FBP1 | 0.548 | 41 | 2vt5 | D |
| FBP1 | 0.438 | 25 | 2fie | A |

| | | | | |
|---|---|---|---|---|
| FBP1 | 0.643 | 7 | 4mjo | B |
| KCNQ1 | None | None | None | None |
| KCNQ1 | None | None | None | None |
| KCNQ1 | None | None | None | None |
| UPP1 | 0.706 | 4 | 3euf | A |
| UMPS | 1 | 3 | 2qcl | A |
| UMPS | 0.556 | 59 | 2qcl | B |
| UMPS | None | None | None | None |
| DHFR | None | None | None | None |
| KHK | 0.667 | 2 | 2hw1 | A |
| TTPA | None | None | None | None |
| TTPA | None | None | None | None |
| TTPA | 0.5 | 1 | 1r5l | A |
| TTPA | None | None | None | None |
| MAP1LC3B | 0.387 | 8 | 3vtv | A |
| MAP1LC3B | None | None | None | None |
| ZAP70 | 0.317 | 1 | 2oq1 | A |
| PROC | 0.536 | 1 | 3f6u | L |
| SQSTM1 | None | None | None | None |
| AHCY | None | None | None | None |
| AHCY | None | None | None | None |
| PRNP | None | None | None | None |
| PRNP | None | None | None | None |
| PRNP | None | None | None | None |
| PRNP | None | None | None | None |
| PRNP | 0.277 | 1 | 1qlx | A |
| PRNP | None | None | None | None |
| PRNP | None | None | None | None |
| PRNP | None | None | None | None |
| PRNP | None | None | None | None |
| PRNP | 0.238 | 2 | 3hj5 | A |
| PRNP | 0.238 | 2 | 3hj5 | A |
| PRNP | None | None | None | None |
| PRNP | None | None | None | None |
| PRNP | None | None | None | None |
| PRNP | None | None | None | None |
| PRNP | 0.277 | 1 | 1qlx | A |
| REN | 0.431 | 2 | 2v13 | A |
| REN | 0.431 | 2 | 2v13 | A |
| HINT1 | 0.556 | 2 | 5ed6 | B |
| HINT1 | 0.897 | 1 | 5km4 | A |
| HINT1 | 0.227 | 60 | 5ed6 | A |
| HINT1 | 0.667 | 11 | 5ed6 | B |
| HINT1 | 0.923 | 1 | 5ipc | A |
| OAT | None | None | None | None |
| OAT | None | None | None | None |
| OAT | 0.381 | 5 | 2can | B |
| OAT | 0.381 | 5 | 2can | B |
| OAT | 0.381 | 7 | 2byj | B |
| OAT | None | None | None | None |
| OAT | None | None | None | None |
| OAT | None | None | None | None |
| OAT | 0.278 | 12 | 2byj | A |
| OAT | None | None | None | None |

| | | | | |
|---|---|---|---|---|
| OAT | 0.675 | 2 | 2byl | B |
| OAT | None | None | None | None |
| OAT | None | None | None | None |
| OAT | None | None | None | None |
| OAT | None | None | None | None |
| OAT | 0.758 | 1 | 2byl | A |
| OAT | 0.375 | 5 | 2oat | C |
| OAT | 0.525 | 4 | 2byl | B |
| OAT | 0.429 | 6 | 2byj | B |
| OAT | None | None | None | None |
| OAT | None | None | None | None |
| OAT | None | None | None | None |
| OAT | None | None | None | None |
| OAT | None | None | None | None |
| OAT | None | None | None | None |
| OAT | None | None | None | None |
| OAT | None | None | None | None |
| OAT | None | None | None | None |
| CTSA | None | None | None | None |
| CTSA | None | None | None | None |
| PNP | 0.375 | 24 | 1ulb | A |
| PNP | 0.29 | 111 | 1yry | E |
| PNP | 0.406 | 43 | 3phb | E |
| PNP | 0.706 | 5 | 4ear | C |
| PNP | 0.367 | 28 | 3ggs | C |
| PNP | 0.5 | 17 | 1pf7 | E |
| PNP | 0.933 | 1 | 3ggs | C |
| PNP | 0.833 | 1 | 3d1v | A |
| MYL2 | None | None | None | None |
| MYL2 | None | None | None | None |
| ANG | 0.548 | 4 | 1k5b | A |
| ANG | 0.419 | 8 | 4ahi | A |
| ANG | 0.419 | 8 | 4ahi | A |
| ANG | None | None | None | None |
| ANG | 0.761 | 1 | 5eop | A |
| ANG | 0.576 | 4 | 5m9v | A |
| ANG | 0.579 | 4 | 5m9q | A |
| ANG | None | None | None | None |
| ANG | None | None | None | None |
| ANG | 0.52 | 5 | 5m9g | A |
| POLH | 0.241 | 101 | 4dl5 | A |
| POLH | None | None | None | None |
| POLH | 0.208 | 9 | 3si8 | A |
| IL2RA | None | None | None | None |
| TGFB1 | None | None | None | None |
| TP53 | None | None | None | None |
| TP53 | None | None | None | None |
| PAH | None | None | None | None |
| PAH | None | None | None | None |
| PAH | 0.44 | 11 | 2pah | A |
| PAH | 0.44 | 11 | 2pah | A |
| PAH | 0.32 | 19 | 6pah | A |
| PAH | 0.32 | 19 | 6pah | A |
| PAH | 0.48 | 2 | 1pah | A |

| | | | | |
|---|---|---|---|---|
| PAH | 0.48 | 2 | 1pah | A |
| PAH | 0.4 | 11 | 3pah | A |
| PAH | 0.2 | 51 | 1j8u | A |
| PAH | 0.2 | 51 | 1j8u | A |
| PAH | 0.36 | 7 | 2pah | B |
| PAH | 0.44 | 9 | 1dmw | A |
| PAH | 0.48 | 1 | 1dmw | A |
| PAH | 0.48 | 1 | 1dmw | A |
| PAH | 0.44 | 8 | 2pah | B |
| PAH | 0.44 | 8 | 2pah | B |
| PAH | 0.417 | 3 | 1j8t | A |
| PAH | 0.48 | 2 | 4anp | A |
| PAH | 0.48 | 1 | 1dmw | A |
| PAH | 0.48 | 4 | 2pah | B |
| PAH | 0.44 | 11 | 2pah | A |
| PAH | 0.48 | 4 | 1mmt | A |
| PAH | 0.68 | 6 | 2pah | A |
| PAH | 0.76 | 1 | 1tdw | A |
| PAH | None | None | None | None |
| PAH | None | None | None | None |
| PAH | 0.4 | 14 | 1j8u | A |
| PAH | None | None | None | None |
| PAH | None | None | None | None |
| PAH | None | None | None | None |
| PAH | 0.2 | 43 | 3pah | A |
| PAH | 0.64 | 2 | 1lrm | A |
| PAH | 0.4 | 11 | 2pah | B |
| PAH | 0.28 | 31 | 4anp | A |
| PAH | 0.4 | 14 | 1j8u | A |
| PAH | 0.4 | 14 | 1j8u | A |
| PAH | None | None | None | None |
| PAH | 0.48 | 1 | 1dmw | A |
| PAH | 0.48 | 1 | 1dmw | A |
| PAH | None | None | None | None |
| HDAC8 | None | None | None | None |
| HDAC8 | 0.732 | 3 | 3rqd | A |
| VCP | 0.438 | 4 | 3tiw | A |
| VCP | 0.5 | 2 | 3qq8 | A |
| VCP | 0.5 | 2 | 3qq8 | A |
| VCP | None | None | None | None |
| VCP | None | None | None | None |
| VCP | None | None | None | None |
| VCP | None | None | None | None |
| B3GAT3 | None | None | None | None |
| B3GAT3 | None | None | None | None |
| CSF2RB | None | None | None | None |
| CSF2RB | None | None | None | None |
| CSF2RB | None | None | None | None |
| NF2 | 0.5 | 6 | 1h4r | B |
| NF2 | None | None | None | None |
| NF2 | None | None | None | None |
| PIK3CA | None | None | None | None |
| PHGDH | None | None | None | None |
| PHGDH | None | None | None | None |

| | | | | |
|---|---|---|---|---|
| PHGDH | None | None | None | None |
| PHGDH | 0.66 | 1 | 5n53 | A |
| POLH | None | None | None | None |
| XIAP | 0.5 | 7 | 5c7b | A |
| XIAP | None | None | None | None |
| PDHX | None | None | None | None |
| PDHX | None | None | None | None |
| PDHX | None | None | None | None |
| PDHX | None | None | None | None |
| PDHX | None | None | None | None |
| SAMHD1 | 0.5 | 12 | 5ao0 | A |
| SAMHD1 | 0.357 | 51 | 4zwe | D |
| SAMHD1 | 0.429 | 7 | 4tnr | D |
| SAMHD1 | None | None | None | None |
| SAMHD1 | 0.571 | 3 | 4qfy | D |
| SAMHD1 | None | None | None | None |
| SAMHD1 | 0.308 | 18 | 4qg2 | B |
| SAMHD1 | 0.286 | 57 | 5ao0 | B |
| CD36 | None | None | None | None |
| CD36 | None | None | None | None |
| CD36 | None | None | None | None |
| ASPH | None | None | None | None |
| CCT5 | None | None | None | None |
| CCT5 | None | None | None | None |
| CCT5 | None | None | None | None |
| SMN1 | None | None | None | None |
| SMN1 | None | None | None | None |
| SMN1 | None | None | None | None |
| SMN1 | None | None | None | None |
| ABCA1 | None | None | None | None |
| ABCA1 | None | None | None | None |
| ABCA1 | None | None | None | None |
| ABCA1 | None | None | None | None |
| ABCA1 | None | None | None | None |
| ABCA1 | None | None | None | None |
| ABCA1 | None | None | None | None |
| ABCA1 | None | None | None | None |
| ABCA1 | None | None | None | None |
| ABCA1 | None | None | None | None |
| ABCA1 | None | None | None | None |
| ABCA1 | None | None | None | None |
| ABCA1 | None | None | None | None |
| ABCA1 | None | None | None | None |
| ABCA1 | None | None | None | None |
| ABCA1 | None | None | None | None |
| ABCA1 | None | None | None | None |
| ABCA1 | None | None | None | None |
| ABCA1 | None | None | None | None |
| ABCA1 | None | None | None | None |
| ABCA1 | None | None | None | None |
| ABCA1 | None | None | None | None |
| ABCA1 | None | None | None | None |
| ABCA1 | None | None | None | None |

| | | | | |
|---|---|---|---|---|
| ABCA1 | None | None | None | None |
| ABCA1 | None | None | None | None |
| ABCA1 | None | None | None | None |
| ABCA1 | None | None | None | None |
| ABCA1 | None | None | None | None |
| ABCA1 | None | None | None | None |
| ABCA1 | None | None | None | None |
| ABCA1 | None | None | None | None |
| ABCA1 | None | None | None | None |
| ABCA1 | None | None | None | None |
| ABCA1 | None | None | None | None |
| ERCC1 | None | None | None | None |
| FECH | 0.524 | 27 | 2po5 | A |
| FECH | 0.6 | 60 | 2hre | A |
| FECH | 0.636 | 20 | 2po7 | A |
| FECH | 0.476 | 61 | 4klr | A |
| FECH | 0.5 | 84 | 2hrc | A |
| FECH | 0.667 | 10 | 2po5 | A |
| FECH | 0.455 | 30 | 4kla | B |
| PTEN | None | None | None | None |
| PTEN | None | None | None | None |
| PTEN | None | None | None | None |
| PTEN | None | None | None | None |
| PTEN | None | None | None | None |
| PTEN | None | None | None | None |
| PTEN | None | None | None | None |
| PTEN | None | None | None | None |
| PTEN | None | None | None | None |
| PTEN | None | None | None | None |
| PTEN | None | None | None | None |
| PTEN | None | None | None | None |
| PTEN | None | None | None | None |
| PTEN | None | None | None | None |
| PTEN | None | None | None | None |
| PTEN | None | None | None | None |
| PTEN | None | None | None | None |
| PTEN | None | None | None | None |
| PTEN | None | None | None | None |
| PTEN | None | None | None | None |
| ACAT1 | 0.667 | 1 | 2ib9 | C |
| ACAT1 | None | None | None | None |
| ACAT1 | None | None | None | None |
| ACAT1 | 0.667 | 1 | 2ib9 | C |
| ACAT1 | 0.667 | 1 | 2ib9 | C |
| ACAT1 | 0.472 | 11 | 2f2s | A |
| ACAT1 | 0.833 | 6 | 2ib7 | C |
| ACAT1 | 0.229 | 68 | 2ib8 | B |
| ACAT1 | None | None | None | None |
| ACAT1 | 0.528 | 9 | 2ibw | D |
| ACAT1 | 0.703 | 16 | 2ibu | A |
| CREBBP | 0.283 | 35 | 5mqg | A |
| ITGA2B | None | None | None | None |
| ITGA2B | None | None | None | None |
| ITGA2B | 0.214 | 20 | 2vdn | A |

| | | | | |
|---|---|---|---|---|
| ITGA2B | 0.538 | 1 | 3nig | A |
| TSC1 | None | None | None | None |
| ADA2 | None | None | None | None |
| ADA2 | None | None | None | None |
| ADA2 | 0.75 | 1 | 3lgd | A |
| ADA2 | None | None | None | None |
| ADA2 | None | None | None | None |
| ADA2 | None | None | None | None |
| ADA2 | None | None | None | None |
| ADA2 | None | None | None | None |
| ADA2 | 1 | 1 | 3lgd | B |
| ADA2 | None | None | None | None |
| PGK1 | None | None | None | None |
| PGK1 | 0.536 | 8 | 2xe8 | A |
| PGK1 | 0.25 | 38 | 4axx | A |
| PGK1 | 0.214 | 33 | 2y3i | A |
| PGK1 | 0.571 | 3 | 2ybe | A |
| TUBA1A | None | None | None | None |
| TUBA1A | 0.207 | 59 | 5jco | G |
| TUBA1A | 0.207 | 59 | 5jco | G |
| TUBA1A | None | None | None | None |
| TUBA1A | None | None | None | None |
| PTPN11 | None | None | None | None |
| PTPN11 | 0.237 | 25 | 4ohh | A |
| PTPN11 | 0.362 | 13 | 4ohl | A |
| PTPN11 | 0.327 | 15 | 4ohi | A |
| PTPN11 | None | None | None | None |
| PTPN11 | 0.22 | 4 | 4je4 | A |
| PTPN11 | 0.22 | 4 | 4je4 | A |
| PTPN11 | 0.22 | 4 | 4je4 | A |
| PTPN11 | None | None | None | None |
| PTPN11 | 0.404 | 5 | 3tl0 | A |
| PTPN11 | 0.404 | 5 | 3tl0 | A |
| PTPN11 | 0.404 | 5 | 3tl0 | A |
| PTPN11 | None | None | None | None |
| PTPN11 | 0.466 | 7 | 3zm1 | A |
| PTPN11 | 0.621 | 2 | 3b7o | A |
| PTPN11 | 0.776 | 1 | 3zm1 | A |
| PTPN11 | 0.453 | 5 | 4ohd | A |
| PTPN11 | 0.216 | 29 | 3b7o | A |
| PTPN11 | 0.216 | 29 | 3b7o | A |
| PTPN11 | None | None | None | None |
| PTPN11 | None | None | None | None |
| PTPN11 | 0.776 | 1 | 3zm1 | A |
| PTPN11 | 0.621 | 6 | 3zm1 | A |
| PTPN11 | None | None | None | None |
| PTPN11 | 0.216 | 29 | 3b7o | A |
| PTPN11 | 0.621 | 6 | 3zm1 | A |
| GLB1 | 0.333 | 34 | 3wez | A |
| GLB1 | 0.333 | 67 | 3wf4 | A |
| UQCRQ | None | None | None | None |
| VPS35 | None | None | None | None |
| NR3C1 | None | None | None | None |
| NR3C1 | None | None | None | None |

| | | | | |
|---|---|---|---|---|
| NR3C1 | None | None | None | None |
| NR3C1 | 0.359 | 21 | 1nhz | A |
| NR3C1 | 0.42 | 6 | 4mdd | A |
| NR3C1 | None | None | None | None |
| NR3C1 | None | None | None | None |
| NR3C1 | None | None | None | None |
| NR3C1 | 0.422 | 14 | 1nhz | A |
| APOA1 | None | None | None | None |
| APOA1 | None | None | None | None |
| APOA1 | None | None | None | None |
| APOA1 | None | None | None | None |
| APOA1 | None | None | None | None |
| APOA1 | None | None | None | None |
| APOA1 | None | None | None | None |
| BCKDHA | None | None | None | None |
| BCKDHA | 0.583 | 76 | 1x7x | A |
| BCKDHA | 0.963 | 1 | 1ols | A |
| BCKDHA | 0.696 | 3 | 1v1r | A |
| BCKDHA | 0.625 | 16 | 2beu | A |
| BCKDHA | None | None | None | None |
| BCKDHA | 0.652 | 11 | 1v1r | A |
| BCKDHA | 1 | 1 | 2beu | A |
| BCKDHA | 0.84 | 2 | 2j9f | A |
| BCKDHA | 0.556 | 42 | 1ols | A |
| BCKDHA | 0.609 | 90 | 1v1r | A |
| BCKDHA | 0.652 | 25 | 1v1r | A |
| BCKDHA | 0.346 | 5 | 1wci | A |
| BCKDHA | 0.625 | 60 | 2beu | A |
| BCKDHA | 0.625 | 28 | 2beu | A |
| TOP1 | None | None | None | None |
| ANTXR2 | None | None | None | None |
| ANTXR2 | None | None | None | None |
| ANTXR2 | None | None | None | None |
| TF | 0.625 | 35 | 5dyh | A |
| TF | 0.328 | 52 | 5dyh | A |
| FSHB | None | None | None | None |
| GALK1 | None | None | None | None |
| GALK1 | None | None | None | None |
| GALK1 | None | None | None | None |
| CYP17A1 | None | None | None | None |
| CYP17A1 | 0.353 | 2 | 4nkx | A |
| CYP17A1 | 0.353 | 2 | 4nkx | A |
| CYP17A1 | None | None | None | None |
| CYP17A1 | None | None | None | None |
| CYP17A1 | None | None | None | None |
| CYP17A1 | 0.244 | 5 | 4nkx | A |
| CYP17A1 | None | None | None | None |
| CYP17A1 | None | None | None | None |
| CYP17A1 | None | None | None | None |
| CYP17A1 | None | None | None | None |
| CYP17A1 | None | None | None | None |
| CYP17A1 | None | None | None | None |
| CYP17A1 | None | None | None | None |
| CYP17A1 | None | None | None | None |

| | | | | |
|---|---|---|---|---|
| CYP17A1 | None | None | None | None |
| CYP17A1 | None | None | None | None |
| CD44 | 0.667 | 24 | 1uuh | A |
| MDH2 | 0.426 | 5 | 4wlo | B |
| MDH2 | None | None | None | None |
| NDUFV1 | None | None | None | None |
| NDUFV1 | None | None | None | None |
| NDUFV1 | None | None | None | None |
| NDUFV1 | None | None | None | None |
| NDUFV1 | None | None | None | None |
| NDUFV1 | None | None | None | None |
| NDUFV1 | None | None | None | None |
| NDUFV1 | None | None | None | None |
| NDUFV1 | None | None | None | None |
| NDUFV1 | None | None | None | None |
| CTSC | None | None | None | None |
| CTSC | None | None | None | None |
| CTSC | None | None | None | None |
| CTSC | 0.342 | 13 | 3pdf | A |
| CTSC | 0.643 | 1 | 4oel | A |
| CTSC | None | None | None | None |
| CTSC | None | None | None | None |
| VWF | None | None | None | None |
| VWF | None | None | None | None |
| VWF | None | None | None | None |
| VWF | None | None | None | None |
| VWF | None | None | None | None |
| VWF | None | None | None | None |
| VWF | None | None | None | None |
| ORC4 | None | None | None | None |
| ORC4 | None | None | None | None |
| CRYAB | None | None | None | None |
| CRYAB | None | None | None | None |
| CRYAB | None | None | None | None |
| MAP2K2 | None | None | None | None |
| MAP2K2 | None | None | None | None |
| MAP2K2 | 0.299 | 4 | 1s9i | A |
| MAP2K2 | None | None | None | None |
| MAP2K2 | 0.272 | 4 | 1s9i | B |
| MAP2K2 | None | None | None | None |
| MAP2K2 | None | None | None | None |
| MAP2K2 | None | None | None | None |
| MAP2K2 | None | None | None | None |
| CBL | None | None | None | None |
| CBL | None | None | None | None |
| CBL | None | None | None | None |
| CBL | None | None | None | None |
| CBL | None | None | None | None |
| CBL | None | None | None | None |
| CBL | None | None | None | None |
| PFN1 | None | None | None | None |
| PFN1 | 0.75 | 2 | 1fik | A |
| ACSM2A | 0.224 | 4 | 3eq6 | A |
| HSPG2 | None | None | None | None |

| | | | | |
|---|---|---|---|---|
| HSPG2 | None | None | None | None |
| HSPG2 | None | None | None | None |
| HSPG2 | None | None | None | None |
| PARK7 | 0.444 | 48 | 3cyf | A |
| PARK7 | 0.769 | 4 | 1pe0 | A |
| PARK7 | 0.808 | 2 | 1pe0 | A |
| PARK7 | 0.667 | 8 | 4rky | A |
| PARK7 | 0.889 | 1 | 3cz9 | A |
| PARK7 | 0.75 | 2 | 1pdw | B |
| INSR | None | None | None | None |
| INSR | None | None | None | None |
| INSR | None | None | None | None |
| INSR | None | None | None | None |
| INSR | None | None | None | None |
| DSC2 | None | None | None | None |
| DSC2 | None | None | None | None |
| DSC2 | None | None | None | None |
| DSC2 | None | None | None | None |
| DSC2 | None | None | None | None |
| DSC2 | None | None | None | None |
| DSC2 | None | None | None | None |
| DSC2 | None | None | None | None |
| DSC2 | None | None | None | None |
| DSC2 | None | None | None | None |
| MICU1 | None | None | None | None |
| MICU1 | None | None | None | None |
| ADAMTS13 | None | None | None | None |
| ADAMTS13 | None | None | None | None |
| ADAMTS13 | None | None | None | None |
| ADAMTS13 | None | None | None | None |
| ADAMTS13 | None | None | None | None |
| ADAMTS13 | None | None | None | None |
| ADAMTS13 | None | None | None | None |
| IRAK4 | None | None | None | None |
| IRAK4 | None | None | None | None |
| RNASEH2B | None | None | None | None |
| RNASEH2B | None | None | None | None |
| RNASEH2B | None | None | None | None |
| NTRK1 | None | None | None | None |
| MPO | 0.75 | 3 | 1d5l | C |
| MPO | 0.6 | 25 | 1myp | D |
| MPO | 0.25 | 86 | 1d5l | C |
| PEX3 | None | None | None | None |
| SERPINA1 | None | None | None | None |
| SERPINA1 | None | None | None | None |
| SERPINA1 | None | None | None | None |
| SERPINA1 | None | None | None | None |
| SERPINA1 | None | None | None | None |
| SERPINA1 | None | None | None | None |
| SERPINA1 | None | None | None | None |
| SERPINA1 | 0.6 | 1 | 1ezx | A |
| SERPINA1 | None | None | None | None |
| SERPINA1 | None | None | None | None |
| YARS2 | None | None | None | None |

| | | | | |
|---|---|---|---|---|
| YARS2 | None | None | None | None |
| YARS2 | None | None | None | None |
| YARS2 | None | None | None | None |
| YARS2 | None | None | None | None |
| YARS2 | 0.781 | 1 | 2pid | B |
| YARS2 | None | None | None | None |
| YARS2 | None | None | None | None |
| YARS2 | None | None | None | None |
| PDHA1 | None | None | None | None |
| PDHA1 | None | None | None | None |
| PDHA1 | None | None | None | None |
| PDHA1 | None | None | None | None |
| PDHA1 | None | None | None | None |
| PDHA1 | None | None | None | None |
| PDHA1 | None | None | None | None |
| PDHA1 | 0.96 | 1 | 3exh | C |
| PDHA1 | 0.3 | 5 | 1ni4 | A |
| PDHA1 | None | None | None | None |
| PDHA1 | None | None | None | None |
| CUL4B | None | None | None | None |
| AGXT | None | None | None | None |
| AGXT | None | None | None | None |
| AGXT | None | None | None | None |
| AGXT | 0.692 | 3 | 2yob | A |
| AGXT | 0.786 | 7 | 4kxk | C |
| AGXT | 0.538 | 21 | 3r9a | A |
| AGXT | 0.353 | 33 | 3r9a | C |
| AGXT | None | None | None | None |
| AGXT | None | None | None | None |
| AGXT | 0.611 | 4 | 5hhy | B |
| AGXT | 0.333 | 40 | 4cbs | A |
| AGXT | 0.333 | 59 | 5f9s | A |
| AGXT | 0.308 | 45 | 3r9a | A |
| AGXT | 0.294 | 55 | 3r9a | C |
| AGXT | 0.429 | 28 | 4kxk | C |
| AGXT | 0.684 | 1 | 1h0c | A |
| AGXT | 0.5 | 2 | 4cbr | A |
| AGXT | 0.538 | 19 | 4kyo | A |
| AGXT | 0.538 | 19 | 4kyo | A |
| AGXT | 0.75 | 2 | 5luc | B |
| AGXT | 0.722 | 3 | 5hhy | B |
| AGXT | 0.571 | 3 | 4kxk | C |
| AGXT | 0.571 | 5 | 4kxk | C |
| AGXT | None | None | None | None |
| AGXT | 0.778 | 1 | 5hhy | B |
| AGXT | 0.778 | 1 | 5hhy | B |
| AGXT | 0.421 | 12 | 1h0c | A |
| AGXT | 0.692 | 7 | 4kyo | A |
| AGXT | 0.5 | 18 | 4cbr | A |
| AGXT | 0.5 | 18 | 4cbr | A |
| AGXT | 0.615 | 13 | 3r9a | A |
| AGXT | 0.786 | 7 | 4kxk | C |
| AGXT | 0.846 | 1 | 4kxk | A |
| AGXT | 0.615 | 36 | 3r9a | A |

| | | | | |
|---|---|---|---|---|
| AGXT | 0.571 | 10 | 4kxk | C |
| SFTPD | 0.509 | 2 | 3g83 | B |
| SFTPD | None | None | None | None |
| PIK3CA | None | None | None | None |
| PIK3CA | None | None | None | None |
| PIK3CA | None | None | None | None |
| AUH | 0.22 | 3 | 1hzd | E |
| AUH | None | None | None | None |
| AUH | None | None | None | None |
| AUH | 0.293 | 16 | 2zqr | D |
| F11 | None | None | None | None |
| F11 | None | None | None | None |
| F11 | None | None | None | None |
| F11 | None | None | None | None |
| MEN1 | 0.6 | 81 | 5db0 | A |
| MEN1 | 0.8 | 54 | 4i80 | A |
| MEN1 | 0.8 | 84 | 5db0 | A |
| MEN1 | 0.8 | 30 | 5ddb | A |
| MEN1 | 1 | 2 | 4i80 | A |
| MEN1 | 0.8 | 17 | 5db0 | A |
| MEN1 | 0.6 | 17 | 5ddb | A |
| MEN1 | 0.6 | 31 | 3u88 | A |
| MEN1 | None | None | None | None |
| MEN1 | 0.6 | 56 | 5ddb | A |
| MYBPC1 | None | None | None | None |
| MYBPC1 | None | None | None | None |
| MYBPC1 | None | None | None | None |
| PDCD10 | None | None | None | None |
| SCARB2 | None | None | None | None |
| SCARB2 | None | None | None | None |
| SCARB2 | None | None | None | None |
| SCARB2 | None | None | None | None |
| SCARB2 | None | None | None | None |
| TREX2 | None | None | None | None |
| TBK1 | None | None | None | None |
| TBK1 | None | None | None | None |
| DLD | None | None | None | None |
| DLD | None | None | None | None |
| DLD | 0.505 | 3 | 1zy8 | B |
| DLD | None | None | None | None |
| DLD | 0.24 | 11 | 1zy8 | A |
| DLD | None | None | None | None |
| LDHA | 0.802 | 1 | 4qt0 | A |
| TNNC1 | None | None | None | None |
| TNNC1 | 0.461 | 2 | 1wrl | A |
| TNNC1 | None | None | None | None |
| SMAD3 | None | None | None | None |
| HARS | None | None | None | None |
| HARS | None | None | None | None |
| TNNI3 | None | None | None | None |
| TNNI3 | None | None | None | None |
| TNNI3 | None | None | None | None |
| TNNI3 | None | None | None | None |
| TNNI3 | None | None | None | None |

| | | | | |
|---|---|---|---|---|
| TNNI3 | None | None | None | None |
| ACKR1 | None | None | None | None |
| BUB1B | None | None | None | None |
| BUB1B | None | None | None | None |
| BUB1B | None | None | None | None |
| AR | 0.522 | 6 | 1xj7 | A |
| AR | 0.273 | 23 | 2ao6 | A |
| AR | 0.305 | 55 | 4olm | A |
| AR | 0.305 | 55 | 4olm | A |
| AR | None | None | None | None |
| FTO | None | None | None | None |
| FTO | 0.333 | 1 | 4ie6 | A |
| TFAM | None | None | None | None |
| PYGL | None | None | None | None |
| PYGL | 0.778 | 3 | 1fa9 | A |
| PYGL | None | None | None | None |
| PYGL | 0.722 | 2 | 1fa9 | A |
| PYGL | 0.5 | 19 | 3dds | A |
| PYGL | 0.611 | 25 | 3ceh | B |
| PYGL | 0.611 | 32 | 1l7x | B |
| PYGL | None | None | None | None |
| PYGL | 0.722 | 2 | 1fc0 | B |
| PYGL | 0.778 | 3 | 1fa9 | A |
| PYGL | 0.278 | 167 | 1l7x | B |
| PYGL | 0.722 | 12 | 2ati | A |
| PYGL | 0.722 | 1 | 3ceh | B |
| PYGL | 0.611 | 11 | 3ceh | B |
| PYGL | 0.611 | 18 | 1fc0 | B |
| PYGL | 0.722 | 2 | 1fc0 | B |
| GALNS | None | None | None | None |
| GALNS | None | None | None | None |
| GALNS | None | None | None | None |
| GALNS | None | None | None | None |
| GALNS | None | None | None | None |
| GALNS | None | None | None | None |
| GALNS | None | None | None | None |
| GALNS | None | None | None | None |
| GALNS | None | None | None | None |
| GALNS | None | None | None | None |
| GALNS | None | None | None | None |
| GALNS | None | None | None | None |
| GALNS | None | None | None | None |
| GALNS | None | None | None | None |
| GALNS | None | None | None | None |
| GALNS | None | None | None | None |
| GALNS | None | None | None | None |
| GALNS | None | None | None | None |
| GALNS | None | None | None | None |
| GALNS | None | None | None | None |
| GALNS | None | None | None | None |
| GALNS | None | None | None | None |
| GALNS | None | None | None | None |
| GALNS | None | None | None | None |

| | | | | |
|---|---|---|---|---|
| GALNS | None | None | None | None |
| GALNS | None | None | None | None |
| GALNS | None | None | None | None |
| GALNS | None | None | None | None |
| GALNS | None | None | None | None |
| GALNS | None | None | None | None |
| GALNS | None | None | None | None |
| GALNS | None | None | None | None |
| F10 | 0.304 | 21 | 1z6e | L |
| GLUD1 | None | None | None | None |
| GLUD1 | None | None | None | None |
| GLUD1 | None | None | None | None |
| GLUD1 | None | None | None | None |
| GLUD1 | None | None | None | None |
| GLUD1 | None | None | None | None |
| GLUD1 | None | None | None | None |
| GLUD1 | None | None | None | None |
| GLUD1 | None | None | None | None |
| GLUD1 | None | None | None | None |
| NOTCH1 | 0.528 | 3 | 3v79 | K |
| POGLUT1 | None | None | None | None |
| POGLUT1 | None | None | None | None |
| NR3C1 | 0.455 | 1 | 4hn5 | B |
| CASP8 | 0.535 | 6 | 1f9e | C |
| CASP8 | 0.592 | 8 | 1qdu | C |
| TRAP1 | None | None | None | None |
| FANCM | None | None | None | None |
| EPOR | None | None | None | None |
| EPOR | None | None | None | None |
| NDP | None | None | None | None |
| NDP | None | None | None | None |
| NDP | None | None | None | None |
| NDP | None | None | None | None |
| NDP | None | None | None | None |
| NDP | None | None | None | None |
| NDP | None | None | None | None |
| NDP | None | None | None | None |
| NDP | 0.238 | 1 | 5bqc | A |
| NDP | 0.238 | 1 | 5bqc | A |
| NDP | None | None | None | None |
| NDP | None | None | None | None |
| NDP | None | None | None | None |
| NDP | None | None | None | None |
| NDP | None | None | None | None |
| NDP | None | None | None | None |
| NDP | None | None | None | None |
| ROGDI | None | None | None | None |
| CYP19A1 | None | None | None | None |
| CYP19A1 | None | None | None | None |
| CYP19A1 | None | None | None | None |
| CYP19A1 | None | None | None | None |
| CYP19A1 | None | None | None | None |
| CYP19A1 | 0.487 | 1 | 3eqm | A |
| CYP19A1 | 0.487 | 1 | 3eqm | A |

| | | | | |
|---|---|---|---|---|
| CYP19A1 | 0.257 | 5 | 4kq8 | A |
| UBA5 | None | None | None | None |
| PRPS1 | None | None | None | None |
| PRPS1 | 0.316 | 13 | 4f8e | A |
| PRPS1 | None | None | None | None |
| PRPS1 | None | None | None | None |
| PRPS1 | None | None | None | None |
| PRPS1 | None | None | None | None |
| PRPS1 | None | None | None | None |
| PRPS1 | None | None | None | None |
| F2 | None | None | None | None |
| F2 | None | None | None | None |
| NTF4 | None | None | None | None |
| NTF4 | None | None | None | None |
| PSAP | None | None | None | None |
| PSAP | None | None | None | None |
| TAP1 | None | None | None | None |
| TAP1 | None | None | None | None |
| TAP1 | None | None | None | None |
| ARG1 | None | None | None | None |
| ARG1 | 0.714 | 1 | 2pha | A |
| ARG1 | None | None | None | None |
| ARG1 | 0.471 | 10 | 3th7 | A |
| ARG1 | None | None | None | None |
| ARG1 | None | None | None | None |
| CRYM | None | None | None | None |
| ADSL | None | None | None | None |
| ADSL | None | None | None | None |
| ADSL | None | None | None | None |
| ADSL | None | None | None | None |
| ADSL | None | None | None | None |
| ADSL | None | None | None | None |
| ADSL | 0.312 | 8 | 2vd6 | B |
| ADSL | 0.312 | 8 | 2vd6 | B |
| ADSL | None | None | None | None |
| ADSL | 0.353 | 5 | 2vd6 | D |
| ADSL | None | None | None | None |
| ADSL | 0.294 | 11 | 2vd6 | D |
| ADSL | 0.647 | 1 | 2vd6 | D |
| ADSL | 0.435 | 1 | 4ffx | A |
| ADSL | None | None | None | None |
| ADSL | None | None | None | None |
| ADSL | None | None | None | None |
| ADSL | None | None | None | None |
| ADSL | None | None | None | None |
| ADSL | None | None | None | None |
| VWF | None | None | None | None |
| PLA2G7 | None | None | None | None |
| MMP13 | None | None | None | None |
| MMP13 | None | None | None | None |
| MMP13 | None | None | None | None |
| MMP13 | None | None | None | None |
| POMGNT1 | None | None | None | None |
| POMGNT1 | None | None | None | None |

| | | | | | |
|---|---|---|---|---|---|
| COL3A1 | None | None | None | None | |
| COL3A1 | None | None | None | None | |
| COL3A1 | None | None | None | None | |
| COL3A1 | None | None | None | None | |
| ACTN2 | None | None | None | None | |
| ACTN2 | None | None | None | None | |
| ACTN2 | None | None | None | None | |
| ACTN2 | None | None | None | None | |
| ACTN2 | None | None | None | None | |
| ACTN2 | None | None | None | None | |
| ACTN2 | None | None | None | None | |
| UBA5 | None | None | None | None | |
| ADAR | 0.75 | | 1 | 1qbj | B |
| UMPS | None | None | None | None | |
| UMPS | None | None | None | None | |
| UMPS | None | None | None | None | |
| TRNT1 | None | None | None | None | |
| TRNT1 | None | None | None | None | |
| NHEJ1 | None | None | None | None | |
| NHEJ1 | None | None | None | None | |
| NHEJ1 | None | None | None | None | |
| C8A | None | None | None | None | |
| GLUL | None | None | None | None | |
| GLUL | 0.375 | | 6 | 2qc8 | C |
| GLUL | 0.467 | | 9 | 2qc8 | B |
| GYG1 | 0.615 | | 45 | 3rmv | A |
| GYG1 | 0.923 | | 1 | 3u2u | B |
| FLNA | None | None | None | None | |
| FLNA | None | None | None | None | |
| FLNA | None | None | None | None | |
| FLNA | None | None | None | None | |
| FLNA | None | None | None | None | |
| FLNA | None | None | None | None | |
| FLNA | None | None | None | None | |
| TTN | None | None | None | None | |
| TTN | None | None | None | None | |
| TTN | None | None | None | None | |
| TTN | None | None | None | None | |
| SMS | None | None | None | None | |
| SMS | None | None | None | None | |
| SMS | None | None | None | None | |
| C5 | None | None | None | None | |
| SERPINC1 | None | None | None | None | |
| SERPINC1 | None | None | None | None | |
| SERPINC1 | None | None | None | None | |
| SERPINC1 | None | None | None | None | |
| SERPINC1 | None | None | None | None | |
| SERPINC1 | None | None | None | None | |
| SERPINC1 | None | None | None | None | |
| SERPINC1 | None | None | None | None | |
| SERPINC1 | None | None | None | None | |
| SERPINC1 | 0.273 | | 3 | 1ant | I |
| SERPINC1 | 0.262 | | 2 | 2znh | B |
| SERPINC1 | None | None | None | None | |

| | | | | |
|---|---|---|---|---|
| SERPINC1 | None | None | None | None |
| SERPINC1 | None | None | None | None |
| SERPINC1 | None | None | None | None |
| SERPINC1 | None | None | None | None |
| SERPINC1 | None | None | None | None |
| SERPINC1 | None | None | None | None |
| SERPINC1 | None | None | None | None |
| SERPINC1 | None | None | None | None |
| SERPINC1 | None | None | None | None |
| ETFB | None | None | None | None |
| ETFB | None | None | None | None |
| ETFB | None | None | None | None |
| ETFB | None | None | None | None |
| ETFB | None | None | None | None |
| ETFB | None | None | None | None |
| DIABLO | None | None | None | None |
| DIABLO | None | None | None | None |
| DPYS | 0.333 | 24 | 2vr2 | A |
| DPYS | None | None | None | None |
| DPYS | None | None | None | None |
| DPYS | 0.533 | 7 | 2vr2 | A |
| DPYS | None | None | None | None |
| GNE | None | None | None | None |
| GNE | None | None | None | None |
| GNE | None | None | None | None |
| GNE | None | None | None | None |
| GNE | None | None | None | None |
| GNE | None | None | None | None |
| GNE | None | None | None | None |
| GNE | None | None | None | None |
| GNE | None | None | None | None |
| GNMT | None | None | None | None |
| GNMT | 0.2 | 12 | 2azt | B |
| CTH | 0.324 | 20 | 3cog | D |
| CTH | 0.214 | 15 | 2nmp | B |
| CTH | None | None | None | None |
| USH1C | None | None | None | None |
| PSEN1 | None | None | None | None |
| CHEK2 | None | None | None | None |
| CHEK2 | None | None | None | None |
| CHEK2 | None | None | None | None |
| CHEK2 | None | None | None | None |
| CHEK2 | None | None | None | None |
| CHEK2 | None | None | None | None |
| CHEK2 | None | None | None | None |
| CHEK2 | None | None | None | None |
| CHEK2 | None | None | None | None |
| SNCA | None | None | None | None |
| SNCA | None | None | None | None |
| GAN | None | None | None | None |
| GAN | None | None | None | None |
| GAN | None | None | None | None |
| SBDS | None | None | None | None |
| SBDS | None | None | None | None |

| | | | | |
|---|---|---|---|---|
| SBDS | None | None | None | None |
| SBDS | None | None | None | None |
| SBDS | None | None | None | None |
| SBDS | None | None | None | None |
| MPO | 1 | 3 | 3f9p | A |
| MPO | 0.6 | 12 | 5mfa | A |
| KCNH2 | 0.292 | 2 | 4hp9 | A |
| KCNH2 | None | None | None | None |
| KCNH2 | 0.75 | 1 | 4hp9 | A |
| KCNH2 | None | None | None | None |
| TEK | None | None | None | None |
| TEK | None | None | None | None |
| TEK | None | None | None | None |
| TEK | None | None | None | None |
| NPC2 | None | None | None | None |
| NPC2 | None | None | None | None |
| NPC2 | None | None | None | None |
| NPC2 | None | None | None | None |
| NPC2 | None | None | None | None |
| NPC2 | None | None | None | None |
| NPC2 | None | None | None | None |
| PLG | None | None | None | None |
| PCSK9 | None | None | None | None |
| SFTPC | None | None | None | None |
| SFTPC | None | None | None | None |
| SFTPC | None | None | None | None |
| CD40LG | None | None | None | None |
| CD40LG | None | None | None | None |
| CD40LG | None | None | None | None |
| CD40LG | None | None | None | None |
| CD40LG | None | None | None | None |
| TAT | None | None | None | None |
| TAT | None | None | None | None |
| TAT | None | None | None | None |
| EP300 | None | None | None | None |
| FARS2 | None | None | None | None |
| FARS2 | None | None | None | None |
| FARS2 | None | None | None | None |
| FARS2 | None | None | None | None |
| FARS2 | None | None | None | None |
| FARS2 | None | None | None | None |
| FARS2 | None | None | None | None |
| FARS2 | None | None | None | None |
| FARS2 | 0.5 | 1 | 5mgw | A |
| FARS2 | None | None | None | None |
| FARS2 | None | None | None | None |
| FARS2 | None | None | None | None |
| FARS2 | None | None | None | None |
| MASP2 | None | None | None | None |
| MASP2 | None | None | None | None |
| MASP2 | None | None | None | None |
| MASP2 | None | None | None | None |
| STIM1 | None | None | None | None |
| CHRM2 | None | None | None | None |

| | | | | | |
|---|---|---|---|---|---|
| MCFD2 | | 0.206 | 9 | 3whu | B |
| MCFD2 | | 0.574 | 1 | 4yge | B |
| MCFD2 | | 0.623 | 1 | 3wnx | B |
| KITLG | None | None | | None | None |
| F12 | | 0.345 | 1 | 4bdx | A |
| F12 | | 0.345 | 1 | 4bdx | A |
| ERAP1 | | 0.381 | 5 | 3mdj | A |
| GTF2H5 | None | None | | None | None |
| GTF2H5 | None | None | | None | None |
| THBD | None | None | | None | None |
| MRPL28 | None | None | | None | None |
| MRPL28 | None | None | | None | None |
| HTRA1 | None | None | | None | None |
| HTRA1 | None | None | | None | None |
| HTRA1 | None | None | | None | None |
| HTRA1 | None | None | | None | None |
| HTRA1 | | 0.237 | 1 | 3nwu | B |
| HTRA1 | None | None | | None | None |
| KARS | None | None | | None | None |
| KARS | None | None | | None | None |
| KARS | None | None | | None | None |
| KARS | None | None | | None | None |
| KARS | None | None | | None | None |
| HK1 | | 0.722 | 11 | 4fpb | B |
| GARS | None | None | | None | None |
| GARS | None | None | | None | None |
| GARS | None | None | | None | None |
| GARS | None | None | | None | None |
| GARS | None | None | | None | None |
| GARS | None | None | | None | None |
| GARS | None | None | | None | None |
| IL12B | None | None | | None | None |
| HADH | None | None | | None | None |
| HADH | None | None | | None | None |
| HADH | | 0.52 | 4 | 1lsj | A |
| HADH | | 0.458 | 3 | 1m75 | A |
| HADH | | 0.52 | 4 | 1lsj | A |
| RRM2B | None | None | | None | None |
| RRM2B | None | None | | None | None |
| RRM2B | None | None | | None | None |
| RRM2B | None | None | | None | None |
| RRM2B | | 0.28 | 4 | 4djn | B |
| RRM2B | None | None | | None | None |
| RRM2B | | 0.333 | 2 | 3hf1 | B |
| RRM2B | None | None | | None | None |
| RRM2B | None | None | | None | None |
| RRM2B | | 0.286 | 8 | 2vux | A |
| RRM2B | None | None | | None | None |
| RRM2B | None | None | | None | None |
| RRM2B | None | None | | None | None |
| RRM2B | None | None | | None | None |
| RRM2B | None | None | | None | None |
| MARVELD2 | None | None | | None | None |
| TLR4 | None | None | | None | None |

| | | | | |
|---|---|---|---|---|
| TTN | None | None | None | None |
| TTN | None | None | None | None |
| TNFRSF13B | None | None | None | None |
| POLG | None | None | None | None |
| POLG | None | None | None | None |
| POLG | None | None | None | None |
| POLG | None | None | None | None |
| POLG | None | None | None | None |
| POLG | None | None | None | None |
| POLG | None | None | None | None |
| POLG | None | None | None | None |
| POLG | None | None | None | None |
| POLG | None | None | None | None |
| POLG | None | None | None | None |
| POLG | None | None | None | None |
| POLG | None | None | None | None |
| POLG | None | None | None | None |
| POLG | None | None | None | None |
| POLG | None | None | None | None |
| POLG | None | None | None | None |
| POLG | None | None | None | None |
| POLG | None | None | None | None |
| POLG | None | None | None | None |
| POLG | None | None | None | None |
| POLG | None | None | None | None |
| POLG | None | None | None | None |
| POLG | None | None | None | None |
| POLG | None | None | None | None |
| POLG | None | None | None | None |
| POLG | None | None | None | None |
| POLG | 0.467 | 4 | 5c53 | A |
| POLG | 0.533 | 1 | 4ztu | A |
| POLG | None | None | None | None |
| POLG | None | None | None | None |
| POLG | None | None | None | None |
| POLG | None | None | None | None |
| POLG | None | None | None | None |
| POLG | 0.533 | 1 | 4ztu | A |
| POLG | 0.2 | 8 | 4ztu | A |
| POLG | 0.2 | 8 | 4ztu | A |
| POLG | None | None | None | None |
| POLG | None | None | None | None |
| POLG | None | None | None | None |
| POLG | None | None | None | None |
| POLG | None | None | None | None |
| POLG | None | None | None | None |
| POLG | None | None | None | None |
| POLG | None | None | None | None |
| POLG | None | None | None | None |
| POLG | 0.267 | 6 | 4ztu | A |
| POLG | None | None | None | None |
| POLG | None | None | None | None |
| UROD | 0.632 | 24 | 1jpk | A |

| | | | | | |
|---|---|---|---|---|---|
| UROD | | 0.579 | 33 | 1r3v | A |
| UROD | None | None | | None | None |
| UROD | | 0.526 | 49 | 1jph | A |
| UROD | None | None | | None | None |
| UROD | | 0.632 | 13 | 3gvw | A |
| UROD | | 0.789 | 6 | 3gvw | A |
| UROD | | 0.842 | 2 | 3gvv | A |
| UROD | | 0.737 | 31 | 1r3q | A |
| UROD | | 0.667 | 14 | 2q71 | A |
| PCK1 | | 0.429 | 154 | 1khg | A |
| PCK1 | None | None | | None | None |
| PCK1 | None | None | | None | None |
| PCK1 | None | None | | None | None |
| PCK1 | | 0.524 | 60 | 1nhx | A |
| HEXA | None | None | | None | None |
| HEXA | None | None | | None | None |
| HEXA | None | None | | None | None |
| HEXA | None | None | | None | None |
| HEXA | None | None | | None | None |
| HEXA | | 0.364 | 2 | 2gjx | A |
| HEXA | | 0.364 | 2 | 2gjx | A |
| HEXA | None | None | | None | None |
| HEXA | None | None | | None | None |
| HEXA | None | None | | None | None |
| HEXA | None | None | | None | None |
| HEXA | | 0.364 | 2 | 2gjx | A |
| HEXA | None | None | | None | None |
| HEXA | None | None | | None | None |
| HEXA | None | None | | None | None |
| HEXA | | 0.455 | 1 | 2gjx | A |
| HEXA | None | None | | None | None |
| HEXA | None | None | | None | None |
| HEXA | None | None | | None | None |
| HEXA | None | None | | None | None |
| HEXA | None | None | | None | None |
| HEXA | None | None | | None | None |
| HEXA | None | None | | None | None |
| HEXA | None | None | | None | None |
| HEXA | | 0.273 | 6 | 2gjx | A |
| HEXA | None | None | | None | None |
| HEXA | None | None | | None | None |
| HEXA | | 0.455 | 3 | 2gjx | A |
| HEXA | | 0.455 | 3 | 2gjx | A |
| HEXA | None | None | | None | None |
| HEXA | None | None | | None | None |
| CFTR | None | None | | None | None |
| CFTR | None | None | | None | None |
| CFTR | None | None | | None | None |
| CFTR | None | None | | None | None |
| CFTR | None | None | | None | None |
| CFTR | None | None | | None | None |
| CFTR | None | None | | None | None |
| CFTR | None | None | | None | None |
| CFTR | None | None | | None | None |

| | | | | |
|---|---|---|---|---|
| CFTR | None | None | None | None |
| CFTR | 0.462 | 1 | 2pzg | A |
| CFTR | 0.321 | 1 | 1xmj | A |
| CFTR | 0.462 | 2 | 2pzg | A |
| CFTR | None | None | None | None |
| CFTR | None | None | None | None |
| CFTR | None | None | None | None |
| CFTR | None | None | None | None |
| CFTR | None | None | None | None |
| CFTR | None | None | None | None |
| CFTR | None | None | None | None |
| CFTR | None | None | None | None |
| CFTR | None | None | None | None |
| CFTR | None | None | None | None |
| CFTR | None | None | None | None |
| CFTR | None | None | None | None |
| CFTR | 0.462 | 1 | 2pzg | A |
| CFTR | None | None | None | None |
| CFTR | 0.462 | 2 | 2pzg | A |
| CFTR | None | None | None | None |
| CFTR | None | None | None | None |
| CFTR | None | None | None | None |
| CFTR | 0.346 | 3 | 2pzg | A |
| CFTR | None | None | None | None |
| CFTR | None | None | None | None |
| FAN1 | None | None | None | None |
| CBS | None | None | None | None |
| CBS | None | None | None | None |
| CBS | None | None | None | None |
| CBS | None | None | None | None |
| CBS | 0.406 | 6 | 4pcu | B |
| CBS | None | None | None | None |
| CBS | 0.469 | 6 | 4pcu | A |
| CBS | 0.597 | 3 | 5mms | C |
| CBS | 0.54 | 2 | 5mms | A |
| CBS | 0.54 | 2 | 5mms | A |
| CBS | 0.328 | 9 | 4pcu | A |
| CBS | None | None | None | None |
| CBS | None | None | None | None |
| CBS | 0.208 | 20 | 1m54 | B |
| CBS | 0.516 | 4 | 5mms | C |
| CBS | None | None | None | None |
| CBS | None | None | None | None |
| CBS | None | None | None | None |
| CBS | 0.698 | 2 | 1m54 | B |
| CBS | None | None | None | None |
| CBS | None | None | None | None |
| CBS | None | None | None | None |
| CBS | None | None | None | None |
| CBS | None | None | None | None |
| CBS | None | None | None | None |
| CBS | None | None | None | None |
| CBS | 0.424 | 6 | 4coo | B |
| CBS | None | None | None | None |

| | | | | |
|---|---|---|---|---|
| CBS | 0.603 | 3 | 4coo | A |
| CBS | 0.603 | 3 | 4coo | A |
| CBS | None | None | None | None |
| CBS | None | None | None | None |
| CBS | None | None | None | None |
| CBS | None | None | None | None |
| CBS | None | None | None | None |
| CBS | None | None | None | None |
| TLR3 | None | None | None | None |
| LIG4 | None | None | None | None |
| LIG4 | None | None | None | None |
| ASPA | 0.769 | 1 | 2i3c | B |
| ASPA | 0.769 | 1 | 2i3c | B |
| ASPA | None | None | None | None |
| ASPA | None | None | None | None |
| ASPA | None | None | None | None |
| ASPA | None | None | None | None |
| ASPA | None | None | None | None |
| ASPA | None | None | None | None |
| ASPA | None | None | None | None |
| ASPA | None | None | None | None |
| ASPA | 0.583 | 2 | 4tnu | A |
| ASPA | 0.583 | 2 | 4tnu | A |
| ASPA | None | None | None | None |
| CFH | None | None | None | None |
| FCGR3A | None | None | None | None |
| FCGR3A | None | None | None | None |
| GRIN2A | None | None | None | None |
| GRIN2A | 0.658 | 1 | 5i2n | A |
| GRIN2A | None | None | None | None |
| GLA | 0.467 | 10 | 1r46 | B |
| GLA | 0.375 | 19 | 3hg3 | B |
| GLA | 0.375 | 14 | 3hg5 | B |
| GLA | 0.312 | 13 | 3tv8 | B |
| GLA | 0.375 | 14 | 3hg5 | B |
| GLA | 0.375 | 14 | 3hg5 | B |
| GLA | 0.267 | 22 | 3lxc | B |
| GLA | 0.647 | 2 | 3s5y | A |
| GLA | 0.562 | 20 | 3gxt | B |
| GLA | 0.353 | 21 | 3s5y | A |
| GLA | None | None | None | None |
| GLA | 0.75 | 3 | 3gxn | B |
| GLA | 0.75 | 3 | 3gxn | B |
| GLA | 0.438 | 7 | 1r47 | B |
| GLA | 0.375 | 3 | 3hg4 | B |
| GLA | 0.375 | 11 | 3hg2 | A |
| GLA | None | None | None | None |
| GLA | 0.25 | 42 | 3hg5 | A |
| GLA | 0.25 | 42 | 3hg5 | A |
| GLA | 0.312 | 13 | 3hg2 | B |
| GLA | 0.375 | 19 | 3hg3 | B |
| GLA | 0.4 | 5 | 3lxa | A |
| GLA | 0.214 | 50 | 3lxb | A |
| GLA | 0.214 | 50 | 3lxb | A |

| | | | | |
|---|---|---|---|---|
| GLA | 0.214 | 61 | 3lxb | A |
| GLA | 0.467 | 22 | 3gxn | A |
| GLA | 0.214 | 40 | 3lxb | A |
| GLA | 0.214 | 40 | 3lxb | A |
| GLA | 0.2 | 58 | 3lxc | B |
| GLA | 0.214 | 69 | 3lxb | A |
| GLA | 0.286 | 36 | 3lxb | A |
| SERPINA3 | None | None | None | None |
| MTAP | None | None | None | None |
| MTAP | 0.462 | 25 | 3oze | B |
| MTAP | None | None | None | None |
| BLMH | None | None | None | None |
| TUSC3 | None | None | None | None |
| ISCU | None | None | None | None |
| CYP11B2 | None | None | None | None |
| CYP11B2 | None | None | None | None |
| CYP11B2 | None | None | None | None |
| CYP11B2 | None | None | None | None |
| CYP11B2 | None | None | None | None |
| CYP11B2 | None | None | None | None |
| CYP11B2 | None | None | None | None |
| CYP11B2 | None | None | None | None |
| CYP11B2 | None | None | None | None |
| CYP11B2 | None | None | None | None |
| CYP11B2 | None | None | None | None |
| CYP11B2 | None | None | None | None |
| CYP11B2 | None | None | None | None |
| CYP11B2 | None | None | None | None |
| CYP11B2 | None | None | None | None |
| CYP11B2 | None | None | None | None |
| CYP11B2 | None | None | None | None |
| CYP11B2 | None | None | None | None |
| CYP11B2 | None | None | None | None |
| CYP11B2 | None | None | None | None |
| CRADD | None | None | None | None |
| CRADD | None | None | None | None |
| FGA | None | None | None | None |
| FGA | None | None | None | None |
| FGA | None | None | None | None |
| FGA | None | None | None | None |
| FGA | None | None | None | None |
| FGA | None | None | None | None |
| FGA | None | None | None | None |
| FGA | None | None | None | None |
| IYD | 1 | 1 | 4ttb | A |
| IYD | None | None | None | None |
| IYD | None | None | None | None |
| IYD | None | None | None | None |
| TCAP | None | None | None | None |
| TCAP | None | None | None | None |
| TCAP | None | None | None | None |
| TCAP | None | None | None | None |
| TCAP | None | None | None | None |
| CYP7A1 | None | None | None | None |

| | | | | | |
|---|---|---|---|---|---|
| TEK | None | None | | None | None |
| TEK | None | None | | None | None |
| TEK | None | None | | None | None |
| TEK | None | None | | None | None |
| TEK | None | None | | None | None |
| TEK | None | None | | None | None |
| TEK | None | None | | None | None |
| CD8A | None | None | | None | None |
| HERC2 | None | None | | None | None |
| NR1H3 | 0.537 | | 3 | 3ipq | A |
| GRHPR | None | None | | None | None |
| GRHPR | None | None | | None | None |
| GRHPR | 0.458 | | 2 | 2q50 | A |
| GRHPR | 0.458 | | 2 | 2q50 | A |
| GRHPR | 0.697 | | 1 | 2wwr | D |
| GRHPR | 0.697 | | 1 | 2wwr | D |
| GRHPR | None | None | | None | None |
| GBE1 | None | None | | None | None |
| GBE1 | None | None | | None | None |
| GBE1 | None | None | | None | None |
| GBE1 | None | None | | None | None |
| GBE1 | None | None | | None | None |
| GBE1 | None | None | | None | None |
| GBE1 | None | None | | None | None |
| GBE1 | None | None | | None | None |
| GBE1 | None | None | | None | None |
| GBE1 | None | None | | None | None |
| GBE1 | None | None | | None | None |
| GBE1 | None | None | | None | None |
| GBE1 | None | None | | None | None |
| GBE1 | None | None | | None | None |
| GBE1 | None | None | | None | None |
| GBE1 | None | None | | None | None |
| GBE1 | None | None | | None | None |
| GBE1 | None | None | | None | None |
| GBE1 | None | None | | None | None |
| PC | None | None | | None | None |
| PC | 0.429 | | 1 | 3bg3 | A |
| PC | None | None | | None | None |
| PC | 0.4 | | 2 | 3bg9 | B |
| PC | None | None | | None | None |
| PC | None | None | | None | None |
| PC | 0.4 | | 6 | 3bg3 | B |
| PC | None | None | | None | None |
| MSH6 | None | None | | None | None |
| MSH6 | None | None | | None | None |
| MSH6 | None | None | | None | None |
| MSH6 | None | None | | None | None |
| MSH6 | None | None | | None | None |
| MSH6 | None | None | | None | None |
| MSH6 | None | None | | None | None |
| MSH6 | None | None | | None | None |
| MSH6 | None | None | | None | None |

| | | | | | |
|---|---|---|---|---|---|
| MSH6 | None | None | None | | None |
| MSH6 | None | None | None | | None |
| MSH6 | None | None | None | | None |
| MSH6 | None | None | None | | None |
| MSH6 | None | None | None | | None |
| MSH6 | None | None | None | | None |
| MSH6 | None | None | None | | None |
| MSH6 | None | None | None | | None |
| MSH6 | None | None | None | | None |
| MSH6 | None | None | None | | None |
| MSH6 | None | None | None | | None |
| MSH6 | None | None | None | | None |
| MSH6 | None | None | None | | None |
| MSH6 | None | None | None | | None |
| MSH6 | None | None | None | | None |
| MSH6 | None | None | None | | None |
| MSH6 | None | None | None | | None |
| MSH6 | None | None | None | | None |
| MSH6 | None | None | None | | None |
| MSH6 | None | None | None | | None |
| MSH6 | None | None | None | | None |
| RNASEL | 0.32 | 6 | | 4g8l | A |
| ORC1 | None | None | None | | None |
| ORC1 | None | None | None | | None |
| ORC1 | None | None | None | | None |
| KCNH2 | None | None | None | | None |
| KCNH2 | None | None | None | | None |
| KCNH2 | None | None | None | | None |
| KCNH2 | None | None | None | | None |
| KCNH2 | None | None | None | | None |
| KCNH2 | None | None | None | | None |
| KCNH2 | None | None | None | | None |
| KCNH2 | None | None | None | | None |
| KCNH2 | None | None | None | | None |
| KCNH2 | None | None | None | | None |
| KCNH2 | None | None | None | | None |
| KCNH2 | None | None | None | | None |
| KCNH2 | None | None | None | | None |
| KCNH2 | None | None | None | | None |
| KCNH2 | None | None | None | | None |
| KCNH2 | None | None | None | | None |
| KCNH2 | None | None | None | | None |
| KCNH2 | None | None | None | | None |
| KCNH2 | None | None | None | | None |
| KCNH2 | None | None | None | | None |
| KCNH2 | None | None | None | | None |
| KCNH2 | None | None | None | | None |
| KCNH2 | None | None | None | | None |
| KCNH2 | None | None | None | | None |
| MAT1A | 0.429 | 3 | | 2obv | A |
| MAT1A | 0.429 | 14 | | 2obv | A |
| MAT1A | 0.571 | 1 | | 2obv | A |
| MAT1A | 0.571 | 1 | | 2obv | A |
| MAT1A | None | None | None | | None |

| | | | | | |
|---|---|---|---|---|---|
| MAT1A | None | None | | None | None |
| MAT1A | None | None | | None | None |
| MAT1A | None | None | | None | None |
| INSR | None | None | | None | None |
| INSR | None | None | | None | None |
| INSR | None | None | | None | None |
| INSR | None | None | | None | None |
| AMT | 0.545 | | 5 | 1wsr | A |
| AMT | None | None | | None | None |
| AMT | None | None | | None | None |
| AMT | None | None | | None | None |
| AMT | None | None | | None | None |
| AMT | None | None | | None | None |
| AMT | None | None | | None | None |
| AMT | None | None | | None | None |
| AMT | None | None | | None | None |
| CSNK1D | None | None | | None | None |
| SF3B1 | None | None | | None | None |
| AGA | None | None | | None | None |
| AGA | None | None | | None | None |
| AGA | None | None | | None | None |
| AGA | None | None | | None | None |
| AGA | None | None | | None | None |
| AGA | None | None | | None | None |
| AGA | None | None | | None | None |
| NECTIN4 | None | None | | None | None |
| RS1 | None | None | | None | None |
| RS1 | None | None | | None | None |
| RS1 | None | None | | None | None |
| RS1 | None | None | | None | None |
| TBC1D7 | None | None | | None | None |
| CTSA | None | None | | None | None |
| CTSA | None | None | | None | None |
| TKT | 0.273 | | 20 | 4kxu | A |
| RP2 | None | None | | None | None |
| RP2 | None | None | | None | None |
| RP2 | None | None | | None | None |
| RP2 | None | None | | None | None |
| LYRM4 | None | None | | None | None |
| SOS1 | None | None | | None | None |
| SOS1 | None | None | | None | None |
| SOS1 | None | None | | None | None |
| SOS1 | None | None | | None | None |
| SOS1 | None | None | | None | None |
| SOS1 | None | None | | None | None |
| HUWE1 | None | None | | None | None |
| HUWE1 | None | None | | None | None |
| TPI1 | None | None | | None | None |
| TPI1 | 0.886 | | 1 | 4unk | B |
| TPI1 | None | None | | None | None |
| MSH2 | None | None | | None | None |
| MSH2 | None | None | | None | None |
| MSH2 | None | None | | None | None |
| MSH2 | None | None | | None | None |

| | | | | |
|---|---|---|---|---|
| MSH2 | None | None | None | None |
| MSH2 | None | None | None | None |
| MSH2 | None | None | None | None |
| MSH2 | None | None | None | None |
| MSH2 | None | None | None | None |
| MSH2 | None | None | None | None |
| MSH2 | None | None | None | None |
| MSH2 | None | None | None | None |
| MSH2 | None | None | None | None |
| MSH2 | None | None | None | None |
| MSH2 | None | None | None | None |
| MSH2 | None | None | None | None |
| MSH2 | None | None | None | None |
| MSH2 | None | None | None | None |
| MSH2 | None | None | None | None |
| MSH2 | None | None | None | None |
| MSH2 | None | None | None | None |
| MSH2 | None | None | None | None |
| MSH2 | None | None | None | None |
| MSH2 | None | None | None | None |
| MSH2 | None | None | None | None |
| MSH2 | None | None | None | None |
| MSH2 | None | None | None | None |
| MSH2 | None | None | None | None |
| MSH2 | None | None | None | None |
| MSH2 | None | None | None | None |
| MSH2 | None | None | None | None |
| MSH2 | None | None | None | None |
| MSH2 | None | None | None | None |
| MSH2 | None | None | None | None |
| MSH2 | None | None | None | None |
| MSH2 | None | None | None | None |
| GALE | None | None | None | None |
| GALE | 0.78 | 1 | 1i3n | A |
| GALE | None | None | None | None |
| GALE | None | None | None | None |
| GALE | None | None | None | None |
| GALE | None | None | None | None |
| GALE | None | None | None | None |
| GALE | 0.259 | 5 | 1i3m | A |
| NSD2 | None | None | None | None |
| SRP72 | None | None | None | None |
| SRP72 | None | None | None | None |
| SRP72 | None | None | None | None |
| POLK | None | None | None | None |
| GALT | None | None | None | None |
| GALT | None | None | None | None |
| GALT | None | None | None | None |
| GALT | None | None | None | None |
| GALT | None | None | None | None |
| GALT | None | None | None | None |
| GALT | None | None | None | None |
| GALT | None | None | None | None |
| GALT | None | None | None | None |

| | | | | |
|---|---|---|---|---|
| GALT | None | None | None | None |
| GALT | None | None | None | None |
| GALT | None | None | None | None |
| GALT | None | None | None | None |
| GALT | None | None | None | None |
| GALT | None | None | None | None |
| GALT | None | None | None | None |
| GALT | None | None | None | None |
| GALT | None | None | None | None |
| GALT | None | None | None | None |
| GALT | None | None | None | None |
| GALT | None | None | None | None |
| GALT | None | None | None | None |
| GALT | None | None | None | None |
| GALT | None | None | None | None |
| GALT | None | None | None | None |
| GALT | None | None | None | None |
| GALT | None | None | None | None |
| GALT | None | None | None | None |
| GALT | None | None | None | None |
| GALT | None | None | None | None |
| GALT | None | None | None | None |
| GALT | None | None | None | None |
| GALT | None | None | None | None |
| GALT | None | None | None | None |
| GALT | None | None | None | None |
| GALT | None | None | None | None |
| GALT | None | None | None | None |
| PYCR1 | 0.75 | 11 | 5uav | A |
| PYCR1 | 0.75 | 11 | 5uav | A |
| PYCR1 | 0.5 | 2 | 2izz | A |
| PYCR1 | 0.556 | 2 | 5uax | B |
| F8 | None | None | None | None |
| F8 | None | None | None | None |
| F8 | None | None | None | None |
| F8 | None | None | None | None |
| F8 | None | None | None | None |
| F8 | None | None | None | None |
| F8 | None | None | None | None |
| F8 | None | None | None | None |
| F8 | None | None | None | None |
| F8 | None | None | None | None |
| F8 | None | None | None | None |
| F8 | None | None | None | None |
| F8 | None | None | None | None |
| F8 | None | None | None | None |
| F8 | None | None | None | None |
| F8 | None | None | None | None |
| ANK1 | None | None | None | None |
| CHAT | None | None | None | None |
| CHAT | None | None | None | None |
| CHAT | 0.615 | 8 | 2fy5 | A |
| CHAT | 0.538 | 2 | 2fy5 | A |
| CHAT | None | None | None | None |

| | | | | | |
|---|---|---|---|---|---|
| CHAT | None | None | | None | None |
| CHAT | None | None | | None | None |
| CHAT | None | None | | None | None |
| CHAT | None | None | | None | None |
| CHAT | 0.538 | | 6 | 2fy5 | A |
| CHAT | None | None | | None | None |
| CHAT | None | None | | None | None |
| CHAT | None | None | | None | None |
| CHAT | None | None | | None | None |
| CHAT | None | None | | None | None |
| CHAT | None | None | | None | None |
| CHAT | None | None | | None | None |
| GHR | None | None | | None | None |
| GHR | 0.312 | | 2 | 3hhr | C |
| GHR | None | None | | None | None |
| GHR | 0.312 | | 1 | 3hhr | C |
| GHR | None | None | | None | None |
| GHR | None | None | | None | None |
| GHR | None | None | | None | None |
| GHR | None | None | | None | None |
| GHR | None | None | | None | None |
| GHR | None | None | | None | None |
| GHR | None | None | | None | None |
| GHR | None | None | | None | None |
| GHR | None | None | | None | None |
| GHR | None | None | | None | None |
| FZD4 | None | None | | None | None |
| FZD4 | None | None | | None | None |
| FZD4 | None | None | | None | None |
| MTR | None | None | | None | None |
| MTR | None | None | | None | None |
| MTR | None | None | | None | None |
| GM2A | None | None | | None | None |
| GM2A | None | None | | None | None |
| GM2A | None | None | | None | None |
| GM2A | None | None | | None | None |
| GM2A | None | None | | None | None |
| CRYGD | 0.444 | | 3 | 2g98 | A |
| CRYGD | None | None | | None | None |
| CRYGD | 0.522 | | 1 | 2kfb | A |
| CRYGD | 0.4 | | 3 | 4gr7 | X |
| CRYGD | 0.235 | | 9 | 1h4a | X |
| CRYGD | 0.333 | | 9 | 1hk0 | X |
| CRYGD | None | None | | None | None |
| CRYGD | None | None | | None | None |
| CRYGD | 0.556 | | 2 | 2g98 | A |
| TLR5 | None | None | | None | None |
| HSD17B10 | 0.855 | | 1 | 1u7t | B |
| HSD17B10 | None | None | | None | None |
| HSD17B10 | 0.542 | | 1 | 1so8 | A |
| HSD17B10 | 0.375 | | 3 | 1so8 | A |
| TBC1D20 | None | None | | None | None |
| FBN1 | None | None | | None | None |
| FBN1 | None | None | | None | None |

| | | | | |
|---|---|---|---|---|
| FBN1 | None | None | None | None |
| FBN1 | None | None | None | None |
| FBN1 | None | None | None | None |
| FBN1 | None | None | None | None |
| FBN1 | None | None | None | None |
| FXN | None | None | None | None |
| FXN | None | None | None | None |
| FXN | None | None | None | None |
| FXN | None | None | None | None |
| MYO5A | None | None | None | None |
| CTSK | None | None | None | None |
| CTSK | None | None | None | None |
| SCN5A | None | None | None | None |
| SCN5A | None | None | None | None |
| SOS1 | None | None | None | None |
| SOS1 | None | None | None | None |
| SOS1 | None | None | None | None |
| SOS1 | None | None | None | None |
| SOS1 | None | None | None | None |
| SOS1 | None | None | None | None |
| SOS1 | None | None | None | None |
| SOS1 | None | None | None | None |
| SOS1 | None | None | None | None |
| SOS1 | None | None | None | None |
| SOS1 | None | None | None | None |
| SOS1 | None | None | None | None |
| GATM | None | None | None | None |
| GATM | None | None | None | None |
| GATM | 0.75 | 4 | 4jdw | A |
| GATM | None | None | None | None |
| GATM | None | None | None | None |
| GATM | 0.75 | 6 | 6jdw | A |
| GATM | 0.75 | 11 | 6jdw | A |
| MMP2 | None | None | None | None |
| MMP2 | None | None | None | None |
| MMP2 | 0.407 | 7 | 1qib | A |
| MMP2 | None | None | None | None |
| NDUFB9 | None | None | None | None |
| NDUFB9 | None | None | None | None |
| GH1 | None | None | None | None |
| GH1 | None | None | None | None |
| GH1 | None | None | None | None |
| SLC6A4 | None | None | None | None |
| G6PD | None | None | None | None |
| G6PD | None | None | None | None |
| G6PD | None | None | None | None |
| G6PD | None | None | None | None |
| G6PD | None | None | None | None |
| G6PD | None | None | None | None |
| G6PD | None | None | None | None |
| G6PD | None | None | None | None |
| G6PD | 0.545 | 1 | 1qki | B |
| G6PD | None | None | None | None |
| G6PD | None | None | None | None |

| | | | | |
|---|---|---|---|---|
| G6PD | None | None | None | None |
| G6PD | None | None | None | None |
| G6PD | None | None | None | None |
| G6PD | None | None | None | None |
| G6PD | 0.455 | 1 | 1qki | D |
| G6PD | None | None | None | None |
| G6PD | None | None | None | None |
| G6PD | None | None | None | None |
| G6PD | None | None | None | None |
| CST3 | None | None | None | None |
| LMNA | None | None | None | None |
| LMNA | None | None | None | None |
| NDUFA9 | None | None | None | None |
| NDUFA9 | None | None | None | None |
| NDUFA9 | None | None | None | None |
| NDUFA9 | None | None | None | None |
| YARS | 0.581 | 1 | 5thl | A |
| YARS | None | None | None | None |
| YARS | None | None | None | None |
| YARS | None | None | None | None |
| EPHA2 | None | None | None | None |
| EPHA2 | None | None | None | None |
| EPHA2 | None | None | None | None |
| EPHA2 | None | None | None | None |
| DICER1 | None | None | None | None |
| DICER1 | None | None | None | None |
| SNX10 | None | None | None | None |
| SNX10 | None | None | None | None |
| TTN | None | None | None | None |
| TTN | None | None | None | None |
| GATA3 | None | None | None | None |
| C5 | None | None | None | None |
| THBD | None | None | None | None |
| BIN1 | None | None | None | None |
| BIN1 | None | None | None | None |
| SMARCAL1 | None | None | None | None |
| CFB | None | None | None | None |
| CYC1 | None | None | None | None |
| ANK2 | None | None | None | None |
| CHMP4B | None | None | None | None |
| CHMP4B | None | None | None | None |
| ERBB2 | None | None | None | None |
| ERBB2 | None | None | None | None |
| POLG2 | None | None | None | None |
| POLG2 | None | None | None | None |
| POLG2 | None | None | None | None |
| POLG2 | None | None | None | None |
| POLG2 | None | None | None | None |
| POLG2 | None | None | None | None |
| POLG2 | None | None | None | None |
| POLG2 | None | None | None | None |
| C3 | None | None | None | None |
| C3 | None | None | None | None |
| RPGR | None | None | None | None |

| | | | | |
|---|---|---|---|---|
| RPGR | None | None | None | None |
| RPGR | None | None | None | None |
| RPGR | None | None | None | None |
| RPGR | None | None | None | None |
| C1QA | None | None | None | None |
| PRKG1 | None | None | None | None |
| IGBP1 | None | None | None | None |
| ENO3 | None | None | None | None |
| ENO3 | None | None | None | None |
| ENO3 | 0.41 | 43 | 2xsx | A |
| ENO3 | 0.487 | 39 | 2xsx | A |
| ENO3 | 0.359 | 47 | 2xsx | B |
| ENO3 | 0.897 | 2 | 2xsx | B |
| MCPH1 | None | None | None | None |
| MCPH1 | None | None | None | None |
| MCPH1 | None | None | None | None |
| MCPH1 | None | None | None | None |
| MCPH1 | None | None | None | None |
| SNRNP200 | None | None | None | None |
| SNRNP200 | None | None | None | None |
| DNMT1 | None | None | None | None |
| DNMT1 | None | None | None | None |
| DNMT1 | None | None | None | None |
| AKT1 | None | None | None | None |
| NDUFS2 | None | None | None | None |
| ACVR2B | None | None | None | None |
| HMGCL | None | None | None | None |
| HMGCL | None | None | None | None |
| HMGCL | None | None | None | None |
| JUP | None | None | None | None |
| JUP | None | None | None | None |
| JUP | None | None | None | None |
| JUP | None | None | None | None |
| JUP | None | None | None | None |
| JUP | None | None | None | None |
| JUP | None | None | None | None |
| JUP | None | None | None | None |
| SPINK1 | None | None | None | None |
| SPINK1 | None | None | None | None |
| SPINK1 | None | None | None | None |
| VCL | None | None | None | None |
| VCL | None | None | None | None |
| NOTCH1 | None | None | None | None |
| NOTCH1 | None | None | None | None |
| NOTCH1 | None | None | None | None |
| NDUFB3 | None | None | None | None |
| IKBKG | None | None | None | None |
| GSN | 0.529 | 2 | 2fh1 | A |
| GSN | None | None | None | None |
| GSN | None | None | None | None |
| GSN | 0.5 | 3 | 1p8x | A |
| GSN | 0.263 | 13 | 2fh3 | A |
| GSN | None | None | None | None |
| GSN | None | None | None | None |

| | | | | |
|---|---|---|---|---|
| GSN | None | None | None | None |
| GSN | None | None | None | None |
| CLC | None | None | None | None |
| RAF1 | None | None | None | None |
| RAF1 | None | None | None | None |
| RAF1 | None | None | None | None |
| NDUFS8 | None | None | None | None |
| NDUFS8 | None | None | None | None |
| NDUFS8 | None | None | None | None |
| NDUFS8 | None | None | None | None |
| NDUFS8 | None | None | None | None |
| CASR | None | None | None | None |
| CASR | None | None | None | None |
| CASR | None | None | None | None |
| CASR | None | None | None | None |
| CASR | None | None | None | None |
| CASR | None | None | None | None |
| CASR | None | None | None | None |
| CASR | None | None | None | None |
| CASR | None | None | None | None |
| CASR | None | None | None | None |
| CASR | None | None | None | None |
| CASR | None | None | None | None |
| CASR | None | None | None | None |
| CASR | None | None | None | None |
| CASR | None | None | None | None |
| CASR | None | None | None | None |
| CASR | None | None | None | None |
| CASR | None | None | None | None |
| CASR | None | None | None | None |
| CASR | None | None | None | None |
| CASR | None | None | None | None |
| CASR | None | None | None | None |
| CASR | None | None | None | None |
| CASR | None | None | None | None |
| CASR | None | None | None | None |
| CASR | None | None | None | None |
| CASR | None | None | None | None |
| CASR | None | None | None | None |
| CASR | None | None | None | None |
| VCL | None | None | None | None |
| VCL | None | None | None | None |
| SMPD1 | None | None | None | None |
| SMPD1 | None | None | None | None |
| SMPD1 | None | None | None | None |
| KDM6A | None | None | None | None |
| KCNQ1 | None | None | None | None |
| KCNQ1 | None | None | None | None |
| KCNQ1 | None | None | None | None |
| KCNQ1 | None | None | None | None |
| KCNQ1 | None | None | None | None |
| KCNQ1 | None | None | None | None |
| NDUFA12 | None | None | None | None |
| GLRX5 | None | None | None | None |

| | | | | |
|---|---|---|---|---|
| PALB2 | None | None | None | None |
| PALB2 | None | None | None | None |
| PALB2 | None | None | None | None |
| PALB2 | None | None | None | None |
| PALB2 | None | None | None | None |
| PALB2 | None | None | None | None |
| PALB2 | None | None | None | None |
| PALB2 | None | None | None | None |
| PALB2 | None | None | None | None |
| PALB2 | None | None | None | None |
| ALDOB | 0.462 | 89 | 1qo5 | C |
| ALDOB | 0.731 | 2 | 1xdm | A |
| ALDOB | 0.731 | 1 | 1qo5 | B |
| ALDOB | 0.538 | 6 | 1xdm | A |
| ALDOB | 0.692 | 7 | 1qo5 | C |
| ALDOB | None | None | None | None |
| ALDOB | 0.577 | 25 | 1qo5 | B |
| GNE | None | None | None | None |
| GNE | None | None | None | None |
| GNE | None | None | None | None |
| GNE | None | None | None | None |
| GNE | None | None | None | None |
| GNE | None | None | None | None |
| GNE | None | None | None | None |
| GNE | None | None | None | None |
| GNE | None | None | None | None |
| GNE | None | None | None | None |
| GNE | None | None | None | None |
| GNE | None | None | None | None |
| CHRNB2 | None | None | None | None |
| CHRNB2 | None | None | None | None |
| ERCC3 | None | None | None | None |
| ERCC3 | None | None | None | None |
| ERCC3 | None | None | None | None |
| ERCC3 | None | None | None | None |
| ERCC3 | None | None | None | None |
| UNC119 | None | None | None | None |
| ATL1 | None | None | None | None |
| ATL1 | None | None | None | None |
| ATL1 | None | None | None | None |
| ATL1 | 0.8 | 1 | 3q5d | A |
| ATL1 | None | None | None | None |
| ATL1 | None | None | None | None |
| ATL1 | None | None | None | None |
| ATL1 | None | None | None | None |
| ATL1 | None | None | None | None |
| ATL1 | None | None | None | None |
| NR3C2 | 0.701 | 1 | 2oax | A |
| TBK1 | None | None | None | None |
| HYAL1 | None | None | None | None |
| HYAL1 | None | None | None | None |
| HYAL1 | None | None | None | None |
| HYAL1 | None | None | None | None |
| CTSD | 0.633 | 1 | 4obz | A |

| | | | | | |
|---|---|---|---|---|---|
| PRKCG | None | None | None | None | |
| TGFBR1 | None | None | None | None | |
| XDH | 0.308 | | 2 | 2e1q | A |
| XDH | None | None | None | None | |
| XDH | None | None | None | None | |
| XDH | None | None | None | None | |
| XDH | None | None | None | None | |
| XDH | None | None | None | None | |
| NMNAT1 | 0.833 | | 1 | 1kqn | E |
| NMNAT1 | None | None | None | None | |
| NMNAT1 | None | None | None | None | |
| NMNAT1 | None | None | None | None | |
| NMNAT1 | None | None | None | None | |
| NMNAT1 | None | None | None | None | |
| SERPING1 | None | None | None | None | |
| SERPING1 | None | None | None | None | |
| SERPING1 | None | None | None | None | |
| SERPING1 | None | None | None | None | |
| SERPING1 | None | None | None | None | |
| PDZRN3 | None | None | None | None | |
| FABP2 | None | None | None | None | |
| IFNGR1 | None | None | None | None | |
| IFNGR1 | None | None | None | None | |
| TDP1 | None | None | None | None | |
| TDP1 | 0.333 | | 11 | 1rg1 | A |
| INS | 0.263 | | 5 | 2lgb | B |
| INS | 0.769 | | 1 | 2r34 | C |
| INS | 0.769 | | 1 | 2r34 | C |
| PPOX | 0.857 | | 1 | 4ivm | B |
| PPOX | None | None | None | None | |
| PPOX | None | None | None | None | |
| PPOX | None | None | None | None | |
| PPOX | None | None | None | None | |
| PPOX | None | None | None | None | |
| PPOX | None | None | None | None | |
| PPOX | None | None | None | None | |
| PPOX | None | None | None | None | |
| GFER | None | None | None | None | |
| GFER | 0.769 | | 4 | 3tk0 | A |
| GFER | None | None | None | None | |
| GFER | None | None | None | None | |
| LIFR | None | None | None | None | |
| EGLN1 | 0.556 | | 21 | 5a3u | C |
| EGLN1 | 0.889 | | 6 | 5v18 | A |
| EGLN1 | 0.944 | | 2 | 5a3u | C |
| GP1BA | None | None | None | None | |
| GP1BA | None | None | None | None | |
| GP1BA | None | None | None | None | |
| GP1BA | None | None | None | None | |
| GP1BA | None | None | None | None | |
| MMADHC | None | None | None | None | |
| MMADHC | None | None | None | None | |
| MMADHC | None | None | None | None | |
| MMADHC | None | None | None | None | |

| | | | | |
|---|---|---|---|---|
| MMADHC | None | None | None | None |
| MMADHC | None | None | None | None |
| MMADHC | None | None | None | None |
| ACVRL1 | None | None | None | None |
| ACVRL1 | None | None | None | None |
| RFXANK | None | None | None | None |
| CHIT1 | None | None | None | None |
| UBR1 | None | None | None | None |
| BLM | None | None | None | None |
| BLM | None | None | None | None |
| BLM | None | None | None | None |
| BLM | None | None | None | None |
| BLM | None | None | None | None |
| BLM | None | None | None | None |
| BLM | None | None | None | None |
| BLM | None | None | None | None |
| BLM | None | None | None | None |
| BLM | None | None | None | None |
| BLM | None | None | None | None |
| BLM | None | None | None | None |
| BLM | None | None | None | None |
| VDR | 0.378 | 7 | 1kb4 | B |
| VDR | None | None | None | None |
| VDR | 0.378 | 4 | 1kb4 | A |
| VDR | 0.578 | 1 | 1kb2 | B |
| VDR | 0.578 | 1 | 1kb2 | B |
| VDR | 0.578 | 1 | 1kb2 | B |
| EPHA2 | None | None | None | None |
| EPHA2 | None | None | None | None |
| EPHA2 | None | None | None | None |
| EPHA2 | None | None | None | None |
| EPHA2 | None | None | None | None |
| EPHA2 | None | None | None | None |
| EPHA2 | None | None | None | None |
| EPHA2 | None | None | None | None |
| EPHA2 | None | None | None | None |
| EPHA2 | None | None | None | None |
| UBA1 | None | None | None | None |
| STAT1 | None | None | None | None |
| STAT1 | None | None | None | None |
| STAT1 | None | None | None | None |
| STAT1 | None | None | None | None |
| STAT1 | None | None | None | None |
| STAT1 | None | None | None | None |
| STAT1 | None | None | None | None |
| STAT1 | None | None | None | None |
| DNM1L | None | None | None | None |
| DNM1L | None | None | None | None |
| OTC | None | None | None | None |
| OTC | None | None | None | None |
| OTC | None | None | None | None |
| OTC | None | None | None | None |
| OTC | None | None | None | None |
| OTC | None | None | None | None |

| | | | | |
|---|---|---|---|---|
| OTC | None | None | None | None |
| OTC | None | None | None | None |
| OTC | None | None | None | None |
| OTC | 0.281 | 3 | 1fvo | A |
| OTC | None | None | None | None |
| OTC | None | None | None | None |
| OTC | None | None | None | None |
| OTC | None | None | None | None |
| OTC | None | None | None | None |
| OTC | None | None | None | None |
| OTC | None | None | None | None |
| RARS | None | None | None | None |
| RARS | None | None | None | None |
| RARS | None | None | None | None |
| SEPSECS | None | None | None | None |
| SEPSECS | None | None | None | None |
| SEPSECS | None | None | None | None |
| SEPSECS | None | None | None | None |
| SEPSECS | None | None | None | None |
| BRCA1 | None | None | None | None |
| BRCA1 | None | None | None | None |
| BRCA1 | 0.538 | 1 | 3pxc | X |
| BRCA1 | None | None | None | None |
| BRCA1 | None | None | None | None |
| BRCA1 | 0.462 | 1 | 1t2u | A |
| BRCA1 | None | None | None | None |
| BRCA1 | None | None | None | None |
| BRCA1 | None | None | None | None |
| BRCA1 | None | None | None | None |
| BRCA1 | None | None | None | None |
| BRCA1 | None | None | None | None |
| BRCA1 | None | None | None | None |
| BRCA1 | None | None | None | None |
| BRCA1 | None | None | None | None |
| BRCA1 | None | None | None | None |
| BRCA1 | None | None | None | None |
| BRCA1 | None | None | None | None |
| BRCA1 | None | None | None | None |
| BRCA1 | None | None | None | None |
| BRCA1 | None | None | None | None |
| BRCA1 | None | None | None | None |
| CUBN | None | None | None | None |
| ITGB3 | None | None | None | None |
| MTRR | None | None | None | None |
| MTRR | None | None | None | None |
| MTRR | None | None | None | None |
| MTRR | None | None | None | None |
| MTRR | None | None | None | None |
| MTRR | None | None | None | None |
| MTRR | None | None | None | None |
| MTRR | None | None | None | None |
| IL7R | None | None | None | None |
| IL7R | None | None | None | None |
| RPS24 | None | None | None | None |

| | | | | |
|---|---|---|---|---|
| RPS24 | None | None | None | None |
| HOGA1 | None | None | None | None |
| HOGA1 | None | None | None | None |
| HOGA1 | None | None | None | None |
| HOGA1 | None | None | None | None |
| HOGA1 | None | None | None | None |
| HOGA1 | None | None | None | None |
| HOGA1 | 0.791 | 1 | 3s5n | A |
| HOGA1 | None | None | None | None |
| HOGA1 | None | None | None | None |
| HOGA1 | None | None | None | None |
| HOGA1 | None | None | None | None |
| HOGA1 | None | None | None | None |
| HOGA1 | None | None | None | None |
| GJB2 | None | None | None | None |
| GJB2 | None | None | None | None |
| GJB2 | None | None | None | None |
| GJB2 | None | None | None | None |
| GJB2 | None | None | None | None |
| GJB2 | None | None | None | None |
| GJB2 | None | None | None | None |
| GJB2 | None | None | None | None |
| GJB2 | None | None | None | None |
| GJB2 | None | None | None | None |
| GJB2 | None | None | None | None |
| GJB2 | None | None | None | None |
| GJB2 | None | None | None | None |
| GJB2 | None | None | None | None |
| GJB2 | None | None | None | None |
| GJB2 | None | None | None | None |
| GJB2 | None | None | None | None |
| GJB2 | None | None | None | None |
| GJB2 | None | None | None | None |
| GJB2 | None | None | None | None |
| GJB2 | None | None | None | None |
| GJB2 | None | None | None | None |
| GJB2 | None | None | None | None |
| GJB2 | None | None | None | None |
| GJB2 | None | None | None | None |
| GJB2 | None | None | None | None |
| GJB2 | None | None | None | None |
| GJB2 | None | None | None | None |
| GJB2 | None | None | None | None |
| GJB2 | None | None | None | None |
| GJB2 | None | None | None | None |
| GJB2 | None | None | None | None |
| GJB2 | None | None | None | None |
| GJB2 | None | None | None | None |
| GJB2 | None | None | None | None |
| GJB2 | None | None | None | None |
| GJB2 | None | None | None | None |
| GJB2 | None | None | None | None |
| GJB2 | None | None | None | None |
| GJB2 | None | None | None | None |

| | | | | |
|---|---|---|---|---|
| GJB2 | None | None | None | None |
| GJB2 | None | None | None | None |
| GJB2 | None | None | None | None |
| GJB2 | None | None | None | None |
| GJB2 | None | None | None | None |
| GJB2 | None | None | None | None |
| GJB2 | None | None | None | None |
| GJB2 | None | None | None | None |
| GJB2 | None | None | None | None |
| GJB2 | None | None | None | None |
| GJB2 | None | None | None | None |
| GJB2 | None | None | None | None |
| GJB2 | None | None | None | None |
| GJB2 | None | None | None | None |
| CPS1 | None | None | None | None |
| CPS1 | 0.545 | 127 | 5dou | A |
| CPS1 | None | None | None | None |
| CPS1 | None | None | None | None |
| CPS1 | 0.364 | 19 | 5dou | A |
| CPS1 | 0.364 | 19 | 5dou | A |
| CPS1 | None | None | None | None |
| CPS1 | None | None | None | None |
| CPS1 | None | None | None | None |
| CPS1 | None | None | None | None |
| PSPH | None | None | None | None |
| PSPH | None | None | None | None |
| PSPH | None | None | None | None |
| PSPH | None | None | None | None |
| PSPH | None | None | None | None |
| PSPH | None | None | None | None |
| PEPD | 0.512 | 2 | 5m4l | B |
| PEPD | None | None | None | None |
| PEPD | None | None | None | None |
| PEPD | 0.643 | 2 | 5m4j | B |
| PEPD | 0.643 | 2 | 5m4j | B |
| PEPD | 0.419 | 5 | 5m4l | B |
| PEPD | None | None | None | None |
| PEPD | None | None | None | None |
| PEPD | None | None | None | None |
| KDM5C | None | None | None | None |
| KDM5C | 0.222 | 117 | 5fwj | A |
| KDM5C | 0.333 | 55 | 5fwj | A |
| ITGB6 | 0.312 | 2 | 5ffo | B |
| ITGB6 | None | None | None | None |
| ITGB6 | None | None | None | None |
| DARS | None | None | None | None |
| DARS | None | None | None | None |
| DARS | None | None | None | None |
| DARS | None | None | None | None |
| BTK | None | None | None | None |
| BTK | None | None | None | None |
| BTK | None | None | None | None |
| HDAC4 | None | None | None | None |
| ITGA2B | None | None | None | None |

| | | | | |
|---|---|---|---|---|
| ITGA2B | None | None | None | None |
| ITGA2B | None | None | None | None |
| ITGA2B | None | None | None | None |
| PCSK9 | None | None | None | None |
| PCSK9 | None | None | None | None |
| PCSK9 | None | None | None | None |
| PCSK9 | None | None | None | None |
| MUTYH | None | None | None | None |
| MUTYH | None | None | None | None |
| MUTYH | None | None | None | None |
| MUTYH | None | None | None | None |
| MUTYH | None | None | None | None |
| MUTYH | 0.375 | 2 | 3n5n | X |
| MUTYH | None | None | None | None |
| MUTYH | None | None | None | None |
| MUTYH | None | None | None | None |
| MUTYH | 0.375 | 2 | 3n5n | X |
| MUTYH | 0.375 | 2 | 3n5n | X |
| MUTYH | None | None | None | None |
| MUTYH | 0.375 | 2 | 3n5n | X |
| MUTYH | None | None | None | None |
| MUTYH | None | None | None | None |
| MUTYH | None | None | None | None |
| MUTYH | None | None | None | None |
| MUTYH | None | None | None | None |
| MUTYH | None | None | None | None |
| MUTYH | None | None | None | None |
| MUTYH | None | None | None | None |
| MUTYH | None | None | None | None |
| MUTYH | None | None | None | None |
| MUTYH | None | None | None | None |
| SLC25A12 | None | None | None | None |
| GSN | None | None | None | None |
| IHH | 0.909 | 1 | 3n1p | B |
| IHH | None | None | None | None |
| IHH | None | None | None | None |
| IHH | 0.818 | 2 | 3n1p | B |
| IHH | 0.727 | 12 | 3n1p | B |
| IHH | 0.636 | 15 | 3n1p | B |
| IHH | 0.818 | 7 | 3n1p | B |
| CD247 | None | None | None | None |
| EP300 | None | None | None | None |
| TTN | None | None | None | None |
| MRI1 | None | None | None | None |
| SGSH | None | None | None | None |
| SGSH | None | None | None | None |
| SGSH | None | None | None | None |
| SGSH | None | None | None | None |
| SGSH | None | None | None | None |
| SGSH | None | None | None | None |
| SGSH | None | None | None | None |
| SGSH | None | None | None | None |
| SGSH | None | None | None | None |
| SGSH | None | None | None | None |

| | | | | |
|---|---|---|---|---|
| SGSH | None | None | None | None |
| SGSH | None | None | None | None |
| SGSH | None | None | None | None |
| SGSH | None | None | None | None |
| SGSH | None | None | None | None |
| SGSH | None | None | None | None |
| SGSH | None | None | None | None |
| SGSH | None | None | None | None |
| SGSH | None | None | None | None |
| SGSH | None | None | None | None |
| SGSH | None | None | None | None |
| SGSH | None | None | None | None |
| CBL | None | None | None | None |
| CANT1 | None | None | None | None |
| CANT1 | None | None | None | None |
| CANT1 | None | None | None | None |
| CANT1 | None | None | None | None |
| CANT1 | None | None | None | None |
| CANT1 | None | None | None | None |
| CANT1 | None | None | None | None |
| SLC1A3 | None | None | None | None |
| SLC1A3 | None | None | None | None |
| SLC1A5 | None | None | None | None |
| SLC1A3 | None | None | None | None |
| SLC1A5 | None | None | None | None |
| SLC1A3 | None | None | None | None |
| SLC1A3 | None | None | None | None |
| CTSD | 0.423 | 15 | 4od9 | D |
| CTSD | None | None | None | None |
| CTSD | None | None | None | None |
| CTSD | None | None | None | None |
| CTSD | None | None | None | None |
| MTOR | None | None | None | None |
| ILK | None | None | None | None |
| PCDH15 | None | None | None | None |
| PCDH15 | None | None | None | None |
| PCDH15 | None | None | None | None |
| PCDH15 | None | None | None | None |
| PCDH15 | None | None | None | None |
| PCDH15 | None | None | None | None |
| KLHL40 | None | None | None | None |
| KLHL40 | None | None | None | None |
| KLHL40 | None | None | None | None |
| KLHL40 | None | None | None | None |
| CSTA | None | None | None | None |
| CSTA | None | None | None | None |
| HNMT | None | None | None | None |
| HNMT | None | None | None | None |
| IFNGR2 | None | None | None | None |
| IFNGR2 | None | None | None | None |
| PEX5 | None | None | None | None |
| KIF7 | None | None | None | None |
| KIF7 | None | None | None | None |
| MYL3 | None | None | None | None |

| | | | | |
|---|---|---|---|---|
| MYL3 | None | None | None | None |
| MYL3 | None | None | None | None |
| MYL3 | None | None | None | None |
| MYL3 | None | None | None | None |
| MYL3 | None | None | None | None |
| MYL3 | None | None | None | None |
| NAGLU | None | None | None | None |
| NAGLU | None | None | None | None |
| NAGLU | None | None | None | None |
| NAGLU | None | None | None | None |
| NAGLU | None | None | None | None |
| NAGLU | None | None | None | None |
| NAGLU | None | None | None | None |
| NAGLU | None | None | None | None |
| NAGLU | None | None | None | None |
| NAGLU | None | None | None | None |
| NAGLU | None | None | None | None |
| NAGLU | None | None | None | None |
| NAGLU | None | None | None | None |
| TNFRSF13C | None | None | None | None |
| KIF21A | None | None | None | None |
| KIF21A | None | None | None | None |
| KIF21A | None | None | None | None |
| KIF21A | None | None | None | None |
| KIF21A | None | None | None | None |
| DOCK2 | None | None | None | None |
| DSP | None | None | None | None |
| CDKN2A | None | None | None | None |
| CDKN2A | None | None | None | None |
| CDKN2A | None | None | None | None |
| CDKN2A | None | None | None | None |
| CDKN2A | None | None | None | None |
| CDKN2A | None | None | None | None |
| CDKN2A | None | None | None | None |
| CDKN2A | None | None | None | None |
| CDKN2A | None | None | None | None |
| CDKN2A | None | None | None | None |
| CDKN2A | None | None | None | None |
| CDKN2A | None | None | None | None |
| CDKN2A | None | None | None | None |
| CDKN2A | None | None | None | None |
| VCL | None | None | None | None |
| TPP1 | None | None | None | None |
| TPP1 | None | None | None | None |
| TPP1 | None | None | None | None |
| TPP1 | None | None | None | None |
| TPP1 | None | None | None | None |
| TPP1 | None | None | None | None |
| TPP1 | None | None | None | None |
| TPP1 | None | None | None | None |
| TPP1 | None | None | None | None |
| TPP1 | None | None | None | None |
| TPP1 | None | None | None | None |
| TPP1 | None | None | None | None |

| | | | | |
|---|---|---|---|---|
| TPP1 | None | None | None | None |
| TPP1 | None | None | None | None |
| TPP1 | None | None | None | None |
| TPP1 | None | None | None | None |
| TPP1 | None | None | None | None |
| TPP1 | None | None | None | None |
| TPP1 | None | None | None | None |
| TPP1 | None | None | None | None |
| TPP1 | None | None | None | None |
| TPP1 | None | None | None | None |
| IL10RB | None | None | None | None |
| IL10RB | None | None | None | None |
| CCM2 | None | None | None | None |
| CCM2 | None | None | None | None |
| CCM2 | None | None | None | None |
| DCX | None | None | None | None |
| DCX | None | None | None | None |
| DCX | None | None | None | None |
| DCX | None | None | None | None |
| DCX | None | None | None | None |
| DCX | None | None | None | None |
| DCX | None | None | None | None |
| DCX | None | None | None | None |
| BRCA2 | None | None | None | None |
| NNT | None | None | None | None |
| C2 | None | None | None | None |
| C2 | None | None | None | None |
| C2 | None | None | None | None |
| C2 | None | None | None | None |
| WRN | None | None | None | None |
| WRN | None | None | None | None |
| F13A1 | None | None | None | None |
| F13A1 | None | None | None | None |
| F13A1 | None | None | None | None |
| F13A1 | None | None | None | None |
| TF | None | None | None | None |
| TF | None | None | None | None |
| TF | 0.516 | 2 | 3ve1 | B |
| TF | 0.562 | 39 | 3skp | A |
| TF | None | None | None | None |
| ITPA | None | None | None | None |
| ITPA | None | None | None | None |
| SNCA | None | None | None | None |
| ITGB4 | None | None | None | None |
| ITGB4 | None | None | None | None |
| ITGB4 | None | None | None | None |
| LRP6 | None | None | None | None |
| LRP6 | None | None | None | None |
| LRP6 | None | None | None | None |
| PLG | None | None | None | None |
| PLG | 0.61 | 1 | 1i5k | A |
| TYROBP | None | None | None | None |
| UQCRC2 | None | None | None | None |
| GPSM2 | None | None | None | None |

| | | | | |
|---|---|---|---|---|
| GPSM2 | None | None | None | None |
| GPSM2 | None | None | None | None |
| SCN5A | None | None | None | None |
| SCN5A | None | None | None | None |
| SCN5A | None | None | None | None |
| SCN5A | None | None | None | None |
| SCN5A | None | None | None | None |
| ATIC | 0.6 | 6 | 1p4r | A |
| EZH2 | None | None | None | None |
| CPA1 | 0.387 | 33 | 4uez | A |
| SCN4B | None | None | None | None |
| SCN4B | None | None | None | None |
| MAN1B1 | None | None | None | None |
| MAN1B1 | 0.6 | 9 | 1fmi | A |
| MAN1B1 | None | None | None | None |
| MAN1B1 | None | None | None | None |
| CYCS | 0.74 | 1 | 2n9i | A |
| IDUA | None | None | None | None |
| IDUA | None | None | None | None |
| IDUA | None | None | None | None |
| IDUA | None | None | None | None |
| IDUA | None | None | None | None |
| IDUA | None | None | None | None |
| IDUA | None | None | None | None |
| IDUA | None | None | None | None |
| IDUA | None | None | None | None |
| IDUA | None | None | None | None |
| IDUA | None | None | None | None |
| IDUA | None | None | None | None |
| IDUA | None | None | None | None |
| IDUA | None | None | None | None |
| IDUA | None | None | None | None |
| IDUA | None | None | None | None |
| IDUA | None | None | None | None |
| IDUA | None | None | None | None |
| IDUA | None | None | None | None |
| IDUA | None | None | None | None |
| IDUA | None | None | None | None |
| IDUA | None | None | None | None |
| IDUA | None | None | None | None |
| IDUA | None | None | None | None |
| HIKESHI | None | None | None | None |
| APC | None | None | None | None |
| APC | None | None | None | None |
| APC | None | None | None | None |
| APP | None | None | None | None |
| CEP152 | None | None | None | None |
| GAA | 0.609 | 1 | 5kzx | A |
| GAA | 0.609 | 1 | 5kzx | A |
| GAA | None | None | None | None |
| GAA | None | None | None | None |
| GAA | 0.261 | 10 | 5kzx | A |
| GAA | 0.609 | 1 | 5kzx | A |

| | | | | |
|---|---|---|---|---|
| GAA | 0.609 | 1 | 5kzx | A |
| GAA | 0.609 | 1 | 5kzx | A |
| GAA | 0.609 | 1 | 5kzx | A |
| GAA | 0.609 | 1 | 5kzx | A |
| GAA | 0.609 | 1 | 5kzx | A |
| GAA | 0.609 | 1 | 5kzx | A |
| GAA | 0.609 | 1 | 5kzx | A |
| GAA | 0.609 | 1 | 5kzx | A |
| GAA | 0.609 | 1 | 5kzx | A |
| GAA | 0.609 | 1 | 5kzx | A |
| GAA | 0.609 | 1 | 5kzx | A |
| GAA | 0.609 | 1 | 5kzx | A |
| GAA | 0.609 | 1 | 5kzx | A |
| GAA | 0.609 | 1 | 5kzx | A |
| GAA | 0.609 | 1 | 5kzx | A |
| GAA | 0.609 | 1 | 5kzx | A |
| GAA | 0.609 | 1 | 5kzx | A |
| GAA | 0.609 | 1 | 5kzx | A |
| GAA | 0.609 | 1 | 5kzx | A |
| GAA | 0.609 | 1 | 5kzx | A |
| GAA | 0.609 | 1 | 5kzx | A |
| GAA | 0.609 | 1 | 5kzx | A |
| GAA | 0.609 | 1 | 5kzx | A |
| GAA | 0.609 | 1 | 5kzx | A |
| GAA | 0.609 | 1 | 5kzx | A |
| GAA | 0.609 | 1 | 5kzx | A |
| GAA | 0.609 | 1 | 5kzx | A |
| GAA | 0.609 | 1 | 5kzx | A |
| GAA | 0.609 | 1 | 5kzx | A |
| GAA | 0.609 | 1 | 5kzx | A |
| GAA | 0.609 | 1 | 5kzx | A |
| GAA | 0.609 | 1 | 5kzx | A |
| GAA | 0.609 | 1 | 5kzx | A |
| GAA | 0.609 | 1 | 5kzx | A |
| GAA | 0.609 | 1 | 5kzx | A |
| GAA | 0.609 | 1 | 5kzx | A |
| GAA | 0.609 | 1 | 5kzx | A |
| GAA | 0.609 | 1 | 5kzx | A |
| GAA | 0.609 | 1 | 5kzx | A |
| GAA | 0.609 | 1 | 5kzx | A |
| GAA | 0.609 | 1 | 5kzx | A |
| GAA | 0.609 | 1 | 5kzx | A |
| GAA | 0.261 | 10 | 5kzx | A |
| FOLR1 | None | None | None | None |
| DSP | None | None | None | None |
| DSP | None | None | None | None |
| DSP | None | None | None | None |
| DSP | None | None | None | None |
| DSP | None | None | None | None |
| CCM2 | None | None | None | None |
| FXN | None | None | None | None |
| FXN | None | None | None | None |
| COMP | None | None | None | None |
| COMP | None | None | None | None |

| | | | | | |
|---|---|---|---|---|---|
| COMP | None | None | | None | None |
| COMP | None | None | | None | None |
| COMP | None | None | | None | None |
| COMP | None | None | | None | None |
| COMP | None | None | | None | None |
| COMP | None | None | | None | None |
| COMP | None | None | | None | None |
| COMP | None | None | | None | None |
| COMP | None | None | | None | None |
| COMP | None | None | | None | None |
| AIFM1 | None | None | | None | None |
| AIFM1 | None | None | | None | None |
| AIFM1 | 0.229 | | 23 | 5fs8 | A |
| AIFM1 | None | None | | None | None |
| AIFM1 | 0.262 | | 40 | 5fs7 | A |
| AIFM1 | None | None | | None | None |
| AIFM1 | None | None | | None | None |
| MYH9 | None | None | | None | None |
| FHL2 | None | None | | None | None |
| FHL2 | None | None | | None | None |
| TNFRSF4 | None | None | | None | None |
| CFB | None | None | | None | None |
| CFB | None | None | | None | None |
| CFB | None | None | | None | None |
| CFB | None | None | | None | None |
| CFB | None | None | | None | None |
| CFB | None | None | | None | None |
| CFB | 0.258 | | 12 | 1dle | B |
| CFB | None | None | | None | None |
| CFB | None | None | | None | None |
| CFB | None | None | | None | None |
| MVD | None | None | | None | None |
| MVD | None | None | | None | None |
| HSD17B4 | None | None | | None | None |
| HSD17B4 | None | None | | None | None |
| HSD17B4 | None | None | | None | None |
| HSD17B4 | 0.429 | | 2 | 1s9c | A |
| HSD17B4 | None | None | | None | None |
| HSD17B4 | None | None | | None | None |
| F11 | None | None | | None | None |
| F11 | None | None | | None | None |
| F11 | None | None | | None | None |
| F11 | None | None | | None | None |
| F11 | None | None | | None | None |
| F11 | None | None | | None | None |
| F11 | None | None | | None | None |
| F11 | None | None | | None | None |
| F11 | None | None | | None | None |
| SPTA1 | None | None | | None | None |
| SPTA1 | None | None | | None | None |
| SPTA1 | None | None | | None | None |
| SPTA1 | None | None | | None | None |
| SPTA1 | None | None | | None | None |
| SPTA1 | None | None | | None | None |

| | | | | |
|---|---|---|---|---|
| GMPR | None | None | None | None |
| MITF | None | None | None | None |
| NPC1 | None | None | None | None |
| NPC1 | None | None | None | None |
| NPC1 | None | None | None | None |
| NPC1 | None | None | None | None |
| NPC1 | None | None | None | None |
| NPC1 | None | None | None | None |
| NPC1 | None | None | None | None |
| NPC1 | None | None | None | None |
| NPC1 | None | None | None | None |
| NPC1 | None | None | None | None |
| NPC1 | None | None | None | None |
| NPC1 | None | None | None | None |
| NPC1 | None | None | None | None |
| NPC1 | None | None | None | None |
| NPC1 | None | None | None | None |
| NPC1 | None | None | None | None |
| NPC1 | None | None | None | None |
| NPC1 | None | None | None | None |
| NPC1 | None | None | None | None |
| NPC1 | None | None | None | None |
| NPC1 | None | None | None | None |
| NPC1 | None | None | None | None |
| NPC1 | None | None | None | None |
| NPC1 | None | None | None | None |
| NPC1 | None | None | None | None |
| NPC1 | None | None | None | None |
| NPC1 | None | None | None | None |
| NPC1 | None | None | None | None |
| NPC1 | None | None | None | None |
| NPC1 | None | None | None | None |
| NPC1 | None | None | None | None |
| NPC1 | None | None | None | None |
| NPC1 | None | None | None | None |
| NPC1 | None | None | None | None |
| NPC1 | None | None | None | None |
| NPC1 | None | None | None | None |
| NPC1 | None | None | None | None |
| NPC1 | None | None | None | None |
| NPC1 | None | None | None | None |
| NPC1 | None | None | None | None |
| NPC1 | None | None | None | None |
| NPC1 | None | None | None | None |
| NPC1 | None | None | None | None |
| C3 | None | None | None | None |
| NLRX1 | None | None | None | None |
| IGLL1 | 0.244 | 2 | 2h3n | D |
| APRT | 0.769 | 4 | 1zn9 | B |
| APRT | 0.417 | 5 | 4x44 | A |
| APRT | None | None | None | None |
| APRT | 0.923 | 1 | 1zn8 | A |
| APRT | None | None | None | None |
| ZIC3 | None | None | None | None |

| | | | | |
|---|---|---|---|---|
| ZIC3 | None | None | None | None |
| KLHL3 | None | None | None | None |
| RPS6KA3 | None | None | None | None |
| RPS6KA3 | None | None | None | None |
| RPS6KA3 | None | None | None | None |
| F8 | None | None | None | None |
| F8 | None | None | None | None |
| F8 | None | None | None | None |
| F8 | None | None | None | None |
| F8 | None | None | None | None |
| F8 | None | None | None | None |
| F8 | None | None | None | None |
| F8 | None | None | None | None |
| F8 | None | None | None | None |
| F8 | None | None | None | None |
| F8 | None | None | None | None |
| F8 | None | None | None | None |
| F8 | None | None | None | None |
| F8 | None | None | None | None |
| F8 | None | None | None | None |
| F8 | None | None | None | None |
| F8 | None | None | None | None |
| ISG15 | None | None | None | None |
| EXOSC3 | None | None | None | None |
| EXOSC3 | None | None | None | None |
| EXOSC3 | None | None | None | None |
| NKX2-5 | 0.403 | 1 | 4s0h | B |
| NKX2-5 | None | None | None | None |
| NKX2-5 | 0.403 | 1 | 4s0h | B |
| NKX2-5 | 0.403 | 1 | 4s0h | B |
| FH | 0.207 | 44 | 30000 | A |
| FH | None | None | None | None |
| FH | None | None | None | None |
| FH | 0.304 | 30 | 30000 | D |
| FH | 0.621 | 4 | 30000 | A |
| FH | 0.462 | 6 | 30000 | C |
| FH | None | None | None | None |
| FH | None | None | None | None |
| FH | 0.64 | 6 | 5d6b | A |
| FH | 0.308 | 49 | 30000 | C |
| FH | 0.64 | 1 | 5d6b | A |
| FH | 0.64 | 1 | 5d6b | A |
| SNRNP200 | None | None | None | None |
| SNRNP200 | None | None | None | None |
| SNRNP200 | None | None | None | None |
| SNRNP200 | None | None | None | None |
| CHRNA1 | None | None | None | None |
| CHRNA1 | None | None | None | None |
| MYOM1 | None | None | None | None |
| MYOM1 | None | None | None | None |
| MYOM1 | None | None | None | None |
| MYOM1 | None | None | None | None |
| MYOM1 | None | None | None | None |
| MYOM1 | None | None | None | None |

| | | | | |
|---|---|---|---|---|
| LDLR | None | None | None | None |
| LDLR | None | None | None | None |
| LDLR | None | None | None | None |
| LDLR | None | None | None | None |
| LDLR | None | None | None | None |
| LDLR | None | None | None | None |
| EHMT1 | 0.286 | 8 | 3b7b | A |
| EHMT1 | 0.545 | 3 | 3b7b | A |
| EHMT1 | None | None | None | None |
| COL9A2 | None | None | None | None |
| HFE2 | None | None | None | None |
| HFE2 | None | None | None | None |
| CD3G | None | None | None | None |
| MMAB | None | None | None | None |
| MMAB | 0.5 | 3 | 2idx | C |
| MMAB | 0.45 | 7 | 2idx | C |
| MMAB | 0.65 | 3 | 2idx | A |
| MMAB | 0.65 | 3 | 2idx | A |
| MMAB | 0.7 | 2 | 2idx | B |
| MMAB | 0.7 | 2 | 2idx | B |
| MMAB | 0.35 | 4 | 2idx | A |
| MMAB | 0.4 | 11 | 2idx | C |
| MMAB | None | None | None | None |
| TAF1 | None | None | None | None |
| TAF1 | None | None | None | None |
| CD4 | None | None | None | None |
| QARS | None | None | None | None |
| QARS | None | None | None | None |
| QARS | None | None | None | None |
| QARS | None | None | None | None |
| QARS | None | None | None | None |
| FGFR3 | None | None | None | None |
| FGFR3 | None | None | None | None |
| FGFR3 | None | None | None | None |
| FGFR3 | None | None | None | None |
| FGFR3 | None | None | None | None |
| FGFR3 | None | None | None | None |
| FLNB | None | None | None | None |
| FLNB | None | None | None | None |
| FLNB | None | None | None | None |
| FLNB | None | None | None | None |
| FLNB | None | None | None | None |
| FLNB | None | None | None | None |
| FLNB | None | None | None | None |
| FLNB | None | None | None | None |
| IL21R | None | None | None | None |
| RB1 | None | None | None | None |
| RB1 | None | None | None | None |
| RB1 | None | None | None | None |
| NTRK1 | 0.533 | 1 | 1he7 | A |
| NOTCH1 | None | None | None | None |
| MYOM1 | None | None | None | None |
| MYOM1 | None | None | None | None |
| MYOM1 | None | None | None | None |

| | | | | |
|---|---|---|---|---|
| MYOM1 | None | None | None | None |
| MYOM1 | None | None | None | None |
| MYOM1 | None | None | None | None |
| COQ9 | None | None | None | None |
| COQ9 | None | None | None | None |
| CRBN | None | None | None | None |
| DCTN1 | None | None | None | None |
| DCTN1 | None | None | None | None |
| DCTN1 | None | None | None | None |
| DCTN1 | None | None | None | None |
| DCTN1 | None | None | None | None |
| DCTN1 | 0.333 | 1 | 2hkn | A |
| THAP1 | None | None | None | None |
| THAP1 | None | None | None | None |
| GNAS | None | None | None | None |
| GNAS | None | None | None | None |
| HNF4A | None | None | None | None |
| HNF4A | None | None | None | None |
| ALDH3A2 | None | None | None | None |
| ALDH3A2 | None | None | None | None |
| ALDH3A2 | 0.412 | 5 | 4qgk | B |
| ALDH3A2 | 0.557 | 2 | 4qgk | A |
| ALDH3A2 | None | None | None | None |
| ALDH3A2 | 0.569 | 3 | 4qgk | B |
| ALDH3A2 | None | None | None | None |
| ALDH3A2 | None | None | None | None |
| ALDH3A2 | None | None | None | None |
| HCN1 | None | None | None | None |
| PLEC | None | None | None | None |
| SCN5A | None | None | None | None |
| SCN5A | None | None | None | None |
| STAMBP | None | None | None | None |
| STAMBP | None | None | None | None |
| RNF125 | None | None | None | None |
| RNF125 | None | None | None | None |
| USH1C | None | None | None | None |
| OBSL1 | None | None | None | None |
| GRIN2B | None | None | None | None |
| GRIN2B | None | None | None | None |
| HCN4 | None | None | None | None |
| HCN4 | 0.222 | 5 | 3u11 | A |
| TALDO1 | None | None | None | None |
| ITGB3 | None | None | None | None |
| CHKB | None | None | None | None |
| CHKB | None | None | None | None |
| CHKB | 0.333 | 7 | 3lq3 | A |
| CHKB | None | None | None | None |
| ACTN4 | None | None | None | None |
| ACTN4 | None | None | None | None |
| ACTN4 | None | None | None | None |
| ACTN4 | None | None | None | None |
| CDT1 | None | None | None | None |
| CDT1 | None | None | None | None |
| CDT1 | None | None | None | None |

| | | | | |
|---|---|---|---|---|
| CDT1 | None | None | None | None |
| CDT1 | None | None | None | None |
| CDT1 | None | None | None | None |
| MAPT | None | None | None | None |
| F5 | 0.517 | 4 | 1czs | A |
| F5 | None | None | None | None |
| F5 | None | None | None | None |
| ESRRB | None | None | None | None |
| DGUOK | None | None | None | None |
| DGUOK | 0.471 | 8 | 2ocp | A |
| DGUOK | None | None | None | None |
| DGUOK | 0.375 | 6 | 2ocp | B |
| DGUOK | 0.765 | 1 | 2ocp | A |
| DGUOK | 0.438 | 11 | 2ocp | B |
| DGUOK | 0.294 | 19 | 2ocp | A |
| DGUOK | None | None | None | None |
| DGUOK | None | None | None | None |
| CENPJ | None | None | None | None |
| TTN | None | None | None | None |
| TTN | None | None | None | None |
| TTN | None | None | None | None |
| TTN | None | None | None | None |
| TNFRSF1A | None | None | None | None |
| TNFRSF1A | None | None | None | None |
| TNFRSF1A | None | None | None | None |
| TNFRSF1A | None | None | None | None |
| TNFRSF1A | None | None | None | None |
| TNFRSF1A | None | None | None | None |
| TNFRSF1A | None | None | None | None |
| TNFRSF1A | None | None | None | None |
| TNFRSF1A | None | None | None | None |
| TNFRSF1A | None | None | None | None |
| TNFRSF1A | None | None | None | None |
| TNFRSF1A | None | None | None | None |
| ASL | None | None | None | None |
| ASL | 0.625 | 25 | 1aos | A |
| ASL | None | None | None | None |
| ASL | 0.375 | 28 | 1k62 | A |
| ASL | 0.438 | 4 | 1k62 | B |
| ASL | 0.562 | 30 | 1k62 | A |
| ASL | 0.625 | 1 | 1aos | A |
| ASL | None | None | None | None |
| ASL | None | None | None | None |
| ASL | None | None | None | None |
| MYO7A | None | None | None | None |
| MYO7A | None | None | None | None |
| MYO7A | None | None | None | None |
| MYO7A | None | None | None | None |
| MYO7A | None | None | None | None |
| MYO7A | None | None | None | None |
| MYO7A | None | None | None | None |
| MYO7A | None | None | None | None |
| MYO7A | None | None | None | None |
| MYO7A | None | None | None | None |

| | | | | |
|---|---|---|---|---|
| MYO7A | None | None | None | None |
| MYO7A | None | None | None | None |
| MYO7A | None | None | None | None |
| MYO7A | None | None | None | None |
| MYO7A | None | None | None | None |
| MYO7A | None | None | None | None |
| MYO7A | None | None | None | None |
| MYO7A | None | None | None | None |
| MYO7A | None | None | None | None |
| MYO7A | None | None | None | None |
| APC | None | None | None | None |
| ALAD | None | None | None | None |
| ALAD | None | None | None | None |
| ALAD | None | None | None | None |
| ALAD | 0.542 | 1 | 1E+051 | A |
| ALAD | None | None | None | None |
| ALAD | None | None | None | None |
| POR | None | None | None | None |
| CSF1R | None | None | None | None |
| CSF1R | None | None | None | None |
| CSF1R | None | None | None | None |
| CSF1R | None | None | None | None |
| CSF1R | None | None | None | None |
| CSF1R | None | None | None | None |
| CSF1R | None | None | None | None |
| CSF1R | None | None | None | None |
| CSF1R | None | None | None | None |
| CSF1R | None | None | None | None |
| CSF1R | None | None | None | None |
| VWF | None | None | None | None |
| VWF | None | None | None | None |
| PHF8 | 0.333 | 6 | 3k3n | A |
| ACY1 | None | None | None | None |
| SMN1 | None | None | None | None |
| SMN1 | None | None | None | None |
| DLAT | None | None | None | None |
| DLAT | None | None | None | None |
| B2M | None | None | None | None |
| SOX9 | None | None | None | None |
| SOX9 | None | None | None | None |
| SOX9 | None | None | None | None |
| SOX9 | None | None | None | None |
| SOX9 | None | None | None | None |
| CFH | None | None | None | None |
| FLNC | None | None | None | None |
| FLNC | None | None | None | None |
| FLNC | None | None | None | None |
| ITGB2 | None | None | None | None |
| ITGB2 | 0.261 | 2 | 5e6r | B |
| ITGB2 | None | None | None | None |
| ITGB2 | None | None | None | None |
| ITGB2 | None | None | None | None |
| ITGB2 | None | None | None | None |
| ITGB2 | None | None | None | None |

| | | | | |
|---|---|---|---|---|
| ITGB2 | None | None | None | None |
| ITGB2 | None | None | None | None |
| PSEN1 | None | None | None | None |
| PSEN1 | None | None | None | None |
| PSEN1 | None | None | None | None |
| PSEN1 | None | None | None | None |
| PSEN1 | None | None | None | None |
| PSEN1 | None | None | None | None |
| OXCT1 | None | None | None | None |
| OXCT1 | 0.385 | 24 | 3dlx | A |
| OXCT1 | None | None | None | None |
| OXCT1 | None | None | None | None |
| OXCT1 | None | None | None | None |
| OXCT1 | None | None | None | None |
| OXCT1 | None | None | None | None |
| DHH | None | None | None | None |
| PRKN | None | None | None | None |
| PRKN | None | None | None | None |
| PRKN | None | None | None | None |
| PRKN | None | None | None | None |
| PRKN | None | None | None | None |
| PRKN | None | None | None | None |
| PRKN | None | None | None | None |
| PRKN | None | None | None | None |
| PRKN | None | None | None | None |
| PRKN | None | None | None | None |
| PRKN | None | None | None | None |
| LCAT | None | None | None | None |
| LCAT | None | None | None | None |
| LCAT | None | None | None | None |
| LCAT | None | None | None | None |
| LCAT | None | None | None | None |
| LCAT | None | None | None | None |
| LCAT | None | None | None | None |
| LCAT | None | None | None | None |
| LCAT | None | None | None | None |
| LCAT | None | None | None | None |
| LCAT | None | None | None | None |
| LCAT | None | None | None | None |
| FOXP3 | None | None | None | None |
| FOXP3 | None | None | None | None |
| FOXP3 | None | None | None | None |
| FOXP3 | None | None | None | None |
| SUFU | None | None | None | None |
| HSPA9 | 0.571 | 14 | 4kbo | A |
| HSPA9 | None | None | None | None |
| SQSTM1 | None | None | None | None |
| DARS2 | None | None | None | None |
| DARS2 | None | None | None | None |
| DARS2 | None | None | None | None |
| DARS2 | None | None | None | None |
| DARS2 | 0.412 | 2 | 4ah6 | A |
| DARS2 | None | None | None | None |
| DARS2 | None | None | None | None |

| | | | | |
|---|---|---|---|---|
| DARS2 | None | None | None | None |
| DARS2 | None | None | None | None |
| DARS2 | None | None | None | None |
| DARS2 | None | None | None | None |
| DARS2 | None | None | None | None |
| NT5C3A | None | None | None | None |
| NT5C3A | 0.4 | 18 | 2vkq | A |
| NT5C3A | 0.4 | 7 | 2vkq | A |
| NT5C3A | 0.6 | 14 | 2vkq | A |
| ETFA | None | None | None | None |
| ETFA | None | None | None | None |
| ETFA | None | None | None | None |
| ETFA | None | None | None | None |
| ETFA | None | None | None | None |
| ETFA | None | None | None | None |
| ETFA | None | None | None | None |
| TTN | None | None | None | None |
| TTN | None | None | None | None |
| FLT4 | None | None | None | None |
| FGB | None | None | None | None |
| HPCA | None | None | None | None |
| TAP2 | None | None | None | None |
| TAP2 | None | None | None | None |
| PTPRC | None | None | None | None |
| SLC4A1 | None | None | None | None |
| SLC4A1 | None | None | None | None |
| SLC4A1 | None | None | None | None |
| SLC4A1 | None | None | None | None |
| SLC4A1 | None | None | None | None |
| SLC4A1 | None | None | None | None |
| MYH7 | None | None | None | None |
| MYH7 | None | None | None | None |
| MYH7 | None | None | None | None |
| MYH7 | None | None | None | None |
| MYH7 | None | None | None | None |
| MYH7 | None | None | None | None |
| MYH7 | None | None | None | None |
| MYH7 | None | None | None | None |
| MYH7 | None | None | None | None |
| PARN | None | None | None | None |
| PARN | None | None | None | None |
| NDUFS6 | None | None | None | None |
| VRK1 | None | None | None | None |
| VRK1 | None | None | None | None |
| VRK1 | None | None | None | None |
| VRK1 | 0.29 | 8 | 3op5 | A |
| VRK1 | None | None | None | None |
| VRK1 | None | None | None | None |
| COLEC11 | 0.229 | 23 | 4ymd | B |
| VHL | None | None | None | None |
| TTN | None | None | None | None |
| NCF2 | None | None | None | None |
| NCF2 | None | None | None | None |
| NCF2 | None | None | None | None |

| | | | | |
|---|---|---|---|---|
| NCF2 | None | None | None | None |
| NCF2 | None | None | None | None |
| RPS26 | None | None | None | None |
| CETP | None | None | None | None |
| CETP | None | None | None | None |
| CETP | None | None | None | None |
| CETP | None | None | None | None |
| CETP | None | None | None | None |
| CETP | None | None | None | None |
| CETP | None | None | None | None |
| CETP | None | None | None | None |
| CETP | None | None | None | None |
| CETP | None | None | None | None |
| CETP | None | None | None | None |
| CETP | None | None | None | None |
| CETP | None | None | None | None |
| FLNA | None | None | None | None |
| FLNA | None | None | None | None |
| FLNA | None | None | None | None |
| FLNA | None | None | None | None |
| FLNA | None | None | None | None |
| FLNA | None | None | None | None |
| FLNA | None | None | None | None |
| MYBPC3 | None | None | None | None |
| MYBPC3 | None | None | None | None |
| MYBPC3 | None | None | None | None |
| MYBPC3 | None | None | None | None |
| MYBPC3 | None | None | None | None |
| MYBPC3 | None | None | None | None |
| MYBPC3 | None | None | None | None |
| MYBPC3 | None | None | None | None |
| MYBPC3 | None | None | None | None |
| MYBPC3 | None | None | None | None |
| LSS | None | None | None | None |
| C6 | None | None | None | None |
| C6 | None | None | None | None |
| RLBP1 | None | None | None | None |
| RLBP1 | None | None | None | None |
| RLBP1 | None | None | None | None |
| RLBP1 | None | None | None | None |
| RLBP1 | None | None | None | None |
| RLBP1 | None | None | None | None |
| RLBP1 | None | None | None | None |
| RLBP1 | None | None | None | None |
| MMAA | None | None | None | None |
| MMAA | 0.3 | 1 | 2www | C |
| MMAA | 0.286 | 4 | 2www | D |
| MMAA | None | None | None | None |
| MMAA | None | None | None | None |
| MMAA | None | None | None | None |
| IL1RN | None | None | None | None |
| MASTL | None | None | None | None |
| FHL1 | None | None | None | None |
| FHL1 | None | None | None | None |

| | | | | |
|---|---|---|---|---|
| FHL1 | None | None | None | None |
| FHL1 | None | None | None | None |
| FHL1 | None | None | None | None |
| FHL1 | None | None | None | None |
| FHL1 | None | None | None | None |
| FHL1 | None | None | None | None |
| FHL1 | None | None | None | None |
| EIF2B1 | 0.636 | 1 | 3ecs | H |
| EIF2B1 | 0.636 | 1 | 3ecs | H |
| COQ8A | None | None | None | None |
| COQ8A | None | None | None | None |
| COQ8A | None | None | None | None |
| COQ8A | None | None | None | None |
| COQ8A | None | None | None | None |
| COQ8A | None | None | None | None |
| COQ8A | None | None | None | None |
| COQ8A | None | None | None | None |
| COQ8A | None | None | None | None |
| COQ8A | None | None | None | None |
| COQ8A | None | None | None | None |
| COQ8A | None | None | None | None |
| COQ8A | None | None | None | None |
| COQ8A | None | None | None | None |
| COQ8A | None | None | None | None |
| COQ8A | None | None | None | None |
| UNC119 | None | None | None | None |
| IGHMBP2 | None | None | None | None |
| JAG1 | None | None | None | None |
| JAG1 | None | None | None | None |
| JAG1 | None | None | None | None |
| GNB1 | None | None | None | None |
| GNB1 | 0.3 | 3 | 5ukl | B |
| FLNA | None | None | None | None |
| FLNA | None | None | None | None |
| FLNA | None | None | None | None |
| FLNA | None | None | None | None |
| FLNA | None | None | None | None |
| CREBBP | None | None | None | None |
| CSF3R | None | None | None | None |
| CSF3R | None | None | None | None |
| HIBCH | None | None | None | None |
| HIBCH | None | None | None | None |
| HIBCH | None | None | None | None |
| MMACHC | None | None | None | None |
| MMACHC | None | None | None | None |
| MMACHC | None | None | None | None |
| MMACHC | None | None | None | None |
| MMACHC | None | None | None | None |
| MMACHC | None | None | None | None |
| MMACHC | None | None | None | None |
| MMACHC | None | None | None | None |
| MMACHC | None | None | None | None |
| MMACHC | None | None | None | None |
| MMACHC | None | None | None | None |

| | | | | |
|---|---|---|---|---|
| KLF1 | None | None | None | None |
| NAGS | None | None | None | None |
| NAGS | None | None | None | None |
| NAGS | None | None | None | None |
| PIK3R2 | None | None | None | None |
| EDNRB | None | None | None | None |
| EDNRB | None | None | None | None |
| EDNRB | None | None | None | None |
| EDNRB | None | None | None | None |
| E | None | None | None | None |
| RUNX1 | None | None | None | None |
| RUNX1 | None | None | None | None |
| CTNNB1 | None | None | None | None |
| CTNNB1 | None | None | None | None |
| CTNNB1 | None | None | None | None |
| CTNNB1 | None | None | None | None |
| CTNNB1 | None | None | None | None |
| CTNNB1 | None | None | None | None |
| TTN | None | None | None | None |
| TTN | None | None | None | None |
| TYK2 | None | None | None | None |
| TYK2 | None | None | None | None |
| TYK2 | None | None | None | None |
| ERCC4 | None | None | None | None |
| ERCC4 | None | None | None | None |
| ERCC4 | None | None | None | None |
| ERCC4 | None | None | None | None |
| CFI | None | None | None | None |
| CFI | None | None | None | None |
| CFI | None | None | None | None |
| CFI | None | None | None | None |
| CFI | None | None | None | None |
| CFI | None | None | None | None |
| CFI | None | None | None | None |
| CFI | None | None | None | None |
| CFI | None | None | None | None |
| CFI | 0.301 | 5 | 2xrc | A |
| CFI | None | None | None | None |
| CFI | 0.691 | 2 | 5o32 | J |
| CFI | 0.691 | 2 | 5o32 | J |
| CFI | 0.524 | 5 | 5o32 | I |
| NOTCH3 | None | None | None | None |
| NOTCH3 | None | None | None | None |
| NOTCH3 | None | None | None | None |
| NOTCH3 | None | None | None | None |
| TBX1 | None | None | None | None |
| TBX1 | None | None | None | None |
| DDB2 | None | None | None | None |
| DDB2 | None | None | None | None |
| TP53 | None | None | None | None |
| POMGNT1 | None | None | None | None |
| POMGNT1 | None | None | None | None |
| POMGNT1 | None | None | None | None |
| POMGNT1 | None | None | None | None |

| | | | | |
|---|---|---|---|---|
| POMGNT1 | None | None | None | None |
| POMGNT1 | None | None | None | None |
| POMGNT1 | None | None | None | None |
| POMGNT1 | None | None | None | None |
| POMGNT1 | None | None | None | None |
| POMGNT1 | None | None | None | None |
| POMGNT1 | None | None | None | None |
| POMGNT1 | None | None | None | None |
| POMGNT1 | None | None | None | None |
| FMR1 | None | None | None | None |
| FMR1 | None | None | None | None |
| SLC9A3R1 | None | None | None | None |
| HSPB1 | None | None | None | None |
| HSPB1 | None | None | None | None |
| HSPB1 | None | None | None | None |
| HSPB1 | None | None | None | None |
| HSPB1 | None | None | None | None |
| HSPB1 | None | None | None | None |
| HSPB1 | None | None | None | None |
| SOD3 | None | None | None | None |
| AGA | 1 | 2 | 1apz | D |
| AGA | None | None | None | None |
| AGA | None | None | None | None |
| NDUFS7 | None | None | None | None |
| NDUFS7 | None | None | None | None |
| SOX2 | None | None | None | None |
| CSRP3 | None | None | None | None |
| CSRP3 | None | None | None | None |
| PAH | None | None | None | None |
| PAH | None | None | None | None |
| PAH | None | None | None | None |
| PAH | None | None | None | None |
| CACNA1S | None | None | None | None |
| TRPV4 | None | None | None | None |
| TRPV4 | None | None | None | None |
| TRPV4 | None | None | None | None |
| TRPV4 | 0.58 | 3 | 4dx2 | A |
| TRPV4 | None | None | None | None |
| TRPV4 | None | None | None | None |
| TRPV4 | None | None | None | None |
| TRPV4 | None | None | None | None |
| TRPV4 | None | None | None | None |
| TRPV4 | None | None | None | None |
| TRPV4 | None | None | None | None |
| TRPV4 | None | None | None | None |
| TRPV4 | None | None | None | None |
| TNNI3 | None | None | None | None |
| MLYCD | None | None | None | None |
| MLYCD | None | None | None | None |
| MLYCD | None | None | None | None |
| MLYCD | None | None | None | None |
| AIP | None | None | None | None |
| AIP | None | None | None | None |
| ERCC2 | None | None | None | None |

| | | | | |
|---|---|---|---|---|
| ERCC2 | None | None | None | None |
| ERCC2 | None | None | None | None |
| ERCC2 | None | None | None | None |
| ERCC2 | None | None | None | None |
| ERCC2 | None | None | None | None |
| ERCC2 | None | None | None | None |
| ERCC2 | None | None | None | None |
| ERCC2 | None | None | None | None |
| ERCC2 | None | None | None | None |
| ERCC2 | None | None | None | None |
| ERCC2 | None | None | None | None |
| ERCC2 | None | None | None | None |
| ERCC2 | None | None | None | None |
| PUS1 | None | None | None | None |
| PUS1 | None | None | None | None |
| PUS1 | None | None | None | None |
| CASK | None | None | None | None |
| CASK | None | None | None | None |
| BLVRA | None | None | None | None |
| DICER1 | None | None | None | None |
| DICER1 | None | None | None | None |
| DICER1 | None | None | None | None |
| EIF2AK3 | None | None | None | None |
| EIF2AK3 | None | None | None | None |
| EIF2AK3 | None | None | None | None |
| KIT | None | None | None | None |
| KIT | None | None | None | None |
| KIT | None | None | None | None |
| KIT | None | None | None | None |
| KIT | None | None | None | None |
| KIT | None | None | None | None |
| DNMT3B | None | None | None | None |
| TGM1 | None | None | None | None |
| NDUFA10 | None | None | None | None |
| NDUFA10 | None | None | None | None |
| PRPF31 | None | None | None | None |
| PRPF31 | None | None | None | None |
| ECE1 | None | None | None | None |
| ECE1 | None | None | None | None |
| ECE1 | None | None | None | None |
| NDUFV2 | None | None | None | None |
| NDUFV2 | None | None | None | None |
| NDUFV2 | None | None | None | None |
| GHRHR | None | None | None | None |
| GHRHR | None | None | None | None |
| GHRHR | None | None | None | None |
| SLC4A1 | None | None | None | None |
| FGF9 | None | None | None | None |
| BARD1 | None | None | None | None |
| BARD1 | None | None | None | None |
| BARD1 | None | None | None | None |
| PTPN12 | 0.741 | 2 | 5j8r | D |
| MASP2 | None | None | None | None |
| MASP2 | None | None | None | None |

| | | | | |
|---|---|---|---|---|
| MASP2 | None | None | None | None |
| MASP2 | None | None | None | None |
| HOXB1 | None | None | None | None |
| SLC4A1 | None | None | None | None |
| SLC4A1 | None | None | None | None |
| SLC4A1 | None | None | None | None |
| SLC4A1 | None | None | None | None |
| SLC4A1 | None | None | None | None |
| SLC4A1 | None | None | None | None |
| SLC4A1 | None | None | None | None |
| SLC4A1 | None | None | None | None |
| SLC4A1 | None | None | None | None |
| SLC4A1 | None | None | None | None |
| SLC4A1 | None | None | None | None |
| SLC4A1 | None | None | None | None |
| SLC4A1 | None | None | None | None |
| SLC4A1 | None | None | None | None |
| SLC4A1 | None | None | None | None |
| SLC4A1 | None | None | None | None |
| SLC4A1 | None | None | None | None |
| CBL | None | None | None | None |
| FAS | None | None | None | None |
| MTMR2 | None | None | None | None |
| MTMR2 | None | None | None | None |
| MTMR2 | None | None | None | None |
| MTMR2 | None | None | None | None |
| MTMR2 | None | None | None | None |
| MTMR2 | None | None | None | None |
| MTMR2 | None | None | None | None |
| MTMR2 | None | None | None | None |
| KLHL3 | None | None | None | None |
| KLHL3 | None | None | None | None |
| KLHL3 | 0.667 | 1 | 4ch9 | B |
| KLHL3 | 0.733 | 1 | 5nkp | A |
| KLHL3 | None | None | None | None |
| KLHL3 | None | None | None | None |
| RNASEH1 | None | None | None | None |
| ACTN1 | None | None | None | None |
| PHKG2 | None | None | None | None |
| PHKG2 | None | None | None | None |
| PHKG2 | 0.472 | 2 | 2y7j | A |
| NTRK1 | None | None | None | None |
| APOH | None | None | None | None |
| PMS2 | None | None | None | None |
| PMS2 | None | None | None | None |
| PMS2 | 0.797 | 1 | 1ea6 | A |
| PMS2 | None | None | None | None |
| PMS2 | None | None | None | None |
| PMS2 | None | None | None | None |
| PMS2 | None | None | None | None |
| PMS2 | None | None | None | None |
| PMS2 | None | None | None | None |
| PMS2 | None | None | None | None |
| PMS2 | None | None | None | None |

| | | | | |
|---|---|---|---|---|
| PMS2 | None | None | None | None |
| PMS2 | None | None | None | None |
| PMS2 | None | None | None | None |
| PMS2 | None | None | None | None |
| PMS2 | None | None | None | None |
| PMS2 | None | None | None | None |
| PMS2 | None | None | None | None |
| PMS2 | None | None | None | None |
| PMS2 | None | None | None | None |
| PMS2 | None | None | None | None |
| PMS2 | None | None | None | None |
| MOCS2 | None | None | None | None |
| MOCS2 | None | None | None | None |
| RPGRIP1 | None | None | None | None |
| RPGRIP1 | None | None | None | None |
| LIG4 | None | None | None | None |
| PROC | None | None | None | None |
| PROC | None | None | None | None |
| NAT1 | None | None | None | None |
| NAT1 | None | None | None | None |
| PNPO | None | None | None | None |
| PNPO | 0.5 | 2 | 1nrg | A |
| PNPO | None | None | None | None |
| PNPO | None | None | None | None |
| PNPO | None | None | None | None |
| MRE11 | None | None | None | None |
| MRE11 | None | None | None | None |
| MRE11 | None | None | None | None |
| MRE11 | None | None | None | None |
| DDC | None | None | None | None |
| DDC | None | None | None | None |
| DDC | None | None | None | None |
| DDC | None | None | None | None |
| DDC | None | None | None | None |
| DDC | 0.909 | 7 | 3rch | B |
| DDC | 0.909 | 7 | 3rch | B |
| DDC | 0.643 | 4 | 3rbf | A |
| DDC | None | None | None | None |
| DDC | None | None | None | None |
| DDC | None | None | None | None |
| ANK2 | 0.2 | 7 | 4rlv | A |
| ANK2 | 0.242 | 5 | 4rlv | A |
| ANK2 | None | None | None | None |
| AK2 | None | None | None | None |
| AK2 | 0.241 | 12 | 2c9y | A |
| AK2 | 0.207 | 17 | 2c9y | A |
| AK2 | None | None | None | None |
| AK2 | None | None | None | None |
| AK2 | None | None | None | None |
| GPHN | None | None | None | None |
| HSD17B4 | 0.66 | 1 | 1zbq | A |
| HSD17B4 | 0.66 | 1 | 1zbq | A |
| HSD17B4 | None | None | None | None |
| HSD17B4 | 0.297 | 7 | 1zbq | B |

| | | | | |
|---|---|---|---|---|
| HSD17B4 | None | None | None | None |
| HSD17B4 | None | None | None | None |
| APTX | 0.6 | 5 | 4ndg | A |
| APTX | 0.6 | 5 | 4ndg | A |
| APTX | None | None | None | None |
| APTX | None | None | None | None |
| APTX | None | None | None | None |
| APTX | None | None | None | None |
| APTX | None | None | None | None |
| FANCL | None | None | None | None |
| FANCL | None | None | None | None |
| EMG1 | None | None | None | None |
| EMG1 | None | None | None | None |
| EZH2 | 0.205 | 1 | 4mi5 | A |
| ABCB6 | None | None | None | None |
| NCF2 | None | None | None | None |
| NCF2 | None | None | None | None |
| COL4A2 | None | None | None | None |
| COL4A2 | 1 | 1 | 1li1 | F |
| COL4A2 | None | None | None | None |
| COL4A2 | None | None | None | None |
| COL4A2 | 0.75 | 16 | 1li1 | C |
| SLX4 | None | None | None | None |
| SERPIND1 | None | None | None | None |
| FGFR3 | None | None | None | None |
| FGFR3 | None | None | None | None |
| FGFR3 | None | None | None | None |
| FGFR3 | None | None | None | None |
| FGFR3 | None | None | None | None |
| FGFR3 | None | None | None | None |
| TREM2 | None | None | None | None |
| TREM2 | None | None | None | None |
| TREM2 | None | None | None | None |
| TREM2 | None | None | None | None |
| TREM2 | None | None | None | None |
| TREM2 | None | None | None | None |
| TREM2 | None | None | None | None |
| LIG4 | None | None | None | None |
| LIG4 | None | None | None | None |
| LIG4 | None | None | None | None |
| LIG4 | None | None | None | None |
| LIG4 | None | None | None | None |
| LIG4 | None | None | None | None |
| ECHS1 | None | None | None | None |
| ECHS1 | 0.279 | 4 | 2hw5 | A |
| ECHS1 | None | None | None | None |
| MTR | None | None | None | None |
| MTR | None | None | None | None |
| MTR | None | None | None | None |
| MTR | None | None | None | None |
| MTR | None | None | None | None |
| PMVK | None | None | None | None |
| PRNP | None | None | None | None |
| CHN1 | None | None | None | None |

| | | | | |
|---|---|---|---|---|
| CHN1 | None | None | None | None |
| CHN1 | None | None | None | None |
| CHN1 | None | None | None | None |
| CHN1 | None | None | None | None |
| CHN1 | None | None | None | None |
| CHN1 | None | None | None | None |
| CHN1 | None | None | None | None |
| SLC2A1 | None | None | None | None |
| SLC2A1 | None | None | None | None |
| SLC2A1 | None | None | None | None |
| SLC2A1 | None | None | None | None |
| SLC2A1 | None | None | None | None |
| SLC2A1 | None | None | None | None |
| SLC2A1 | None | None | None | None |
| SLC2A1 | None | None | None | None |
| SLC2A1 | None | None | None | None |
| SLC2A1 | None | None | None | None |
| SLC2A1 | None | None | None | None |
| SLC2A1 | None | None | None | None |
| SLC2A1 | None | None | None | None |
| SLC2A1 | None | None | None | None |
| SLC2A1 | None | None | None | None |
| SLC2A1 | None | None | None | None |
| SLC2A1 | None | None | None | None |
| PMM2 | None | None | None | None |
| PMM2 | None | None | None | None |
| PMM2 | None | None | None | None |
| PMM2 | None | None | None | None |
| PMM2 | None | None | None | None |
| PMM2 | None | None | None | None |
| PMM2 | None | None | None | None |
| PMM2 | None | None | None | None |
| PMM2 | None | None | None | None |
| PMM2 | None | None | None | None |
| PMM2 | None | None | None | None |
| PMM2 | None | None | None | None |
| PMM2 | None | None | None | None |
| PMM2 | None | None | None | None |
| PMM2 | None | None | None | None |
| PMM2 | None | None | None | None |
| PMM2 | None | None | None | None |
| PMM2 | None | None | None | None |
| PMM2 | None | None | None | None |
| PMM2 | None | None | None | None |
| PMM2 | None | None | None | None |
| PMM2 | None | None | None | None |
| PMM2 | None | None | None | None |
| PMM2 | None | None | None | None |
| PMM2 | None | None | None | None |
| PMM2 | None | None | None | None |
| PMM2 | None | None | None | None |
| PMM2 | None | None | None | None |
| PMM2 | None | None | None | None |
| CCKAR | None | None | None | None |
| MAN1B1 | None | None | None | None |

| | | | | |
|---|---|---|---|---|
| MAN1B1 | None | None | None | None |
| LAMA2 | None | None | None | None |
| LAMA2 | None | None | None | None |
| TBC1D4 | None | None | None | None |
| TBC1D4 | None | None | None | None |
| LDLR | None | None | None | None |
| LDLR | None | None | None | None |
| LDLR | None | None | None | None |
| LDLR | None | None | None | None |
| LDLR | None | None | None | None |
| LDLR | None | None | None | None |
| LDLR | None | None | None | None |
| LDLR | None | None | None | None |
| LDLR | None | None | None | None |
| LDLR | None | None | None | None |
| LDLR | None | None | None | None |
| LDLR | None | None | None | None |
| LDLR | None | None | None | None |
| LDLR | None | None | None | None |
| LDLR | None | None | None | None |
| LDLR | None | None | None | None |
| LDLR | None | None | None | None |
| LDLR | None | None | None | None |
| LDLR | None | None | None | None |
| LDLR | None | None | None | None |
| LDLR | None | None | None | None |
| LDLR | None | None | None | None |
| LDLR | None | None | None | None |
| LDLR | None | None | None | None |
| LDLR | None | None | None | None |
| LDLR | None | None | None | None |
| LDLR | None | None | None | None |
| LDLR | None | None | None | None |
| LDLR | None | None | None | None |
| LDLR | None | None | None | None |
| LDLR | None | None | None | None |
| LDLR | None | None | None | None |
| LDLR | None | None | None | None |
| LDLR | None | None | None | None |
| LDLR | None | None | None | None |
| LDLR | None | None | None | None |
| LDLR | None | None | None | None |
| LDLR | None | None | None | None |
| LDLR | None | None | None | None |
| LDLR | None | None | None | None |
| LDLR | None | None | None | None |
| LDLR | 0.5 | 1 | 1ijq | A |
| LDLR | None | None | None | None |
| FANCM | None | None | None | None |
| HAMP | None | None | None | None |
| GUSB | 0.462 | 8 | 3hn3 | A |
| GUSB | None | None | None | None |
| GUSB | 0.45 | 16 | 3hn3 | B |
| GUSB | None | None | None | None |

| | | | | |
|---|---|---|---|---|
| GUSB | None | None | None | None |
| GUSB | None | None | None | None |
| GUSB | None | None | None | None |
| GUSB | None | None | None | None |
| GUSB | None | None | None | None |
| GUSB | None | None | None | None |
| GUSB | None | None | None | None |
| GUSB | None | None | None | None |
| GUSB | None | None | None | None |
| LIG1 | None | None | None | None |
| LIG1 | None | None | None | None |
| MYH7 | None | None | None | None |
| MYH7 | None | None | None | None |
| MYH7 | None | None | None | None |
| MYH7 | None | None | None | None |
| MYH7 | None | None | None | None |
| MYH7 | None | None | None | None |
| MYH7 | None | None | None | None |
| MYH7 | None | None | None | None |
| MYH7 | None | None | None | None |
| MYH7 | None | None | None | None |
| MYH7 | None | None | None | None |
| MYH7 | None | None | None | None |
| MYH7 | None | None | None | None |
| MYH7 | None | None | None | None |
| MYH7 | None | None | None | None |
| MYH7 | None | None | None | None |
| MYH7 | None | None | None | None |
| MYH7 | None | None | None | None |
| MYH7 | None | None | None | None |
| MYH7 | None | None | None | None |
| MYH7 | None | None | None | None |
| MYH7 | None | None | None | None |
| MYH7 | None | None | None | None |
| MYH7 | None | None | None | None |
| MYH7 | None | None | None | None |
| MYH7 | None | None | None | None |
| MYH7 | None | None | None | None |
| MYH7 | None | None | None | None |
| MYH7 | None | None | None | None |
| MYH7 | None | None | None | None |
| MYH7 | None | None | None | None |
| MYH7 | None | None | None | None |
| MYH7 | None | None | None | None |
| MYH7 | None | None | None | None |
| MYH7 | None | None | None | None |
| HCN4 | None | None | None | None |
| RECQL4 | None | None | None | None |
| RECQL4 | None | None | None | None |
| RECQL4 | None | None | None | None |
| RECQL4 | None | None | None | None |
| RECQL4 | None | None | None | None |
| RECQL4 | None | None | None | None |
| RECQL4 | None | None | None | None |
| RECQL4 | None | None | None | None |

| | | | | |
|---|---|---|---|---|
| RECQL4 | None | None | None | None |
| RECQL4 | None | None | None | None |
| RECQL4 | None | None | None | None |
| RECQL4 | None | None | None | None |
| RECQL4 | None | None | None | None |
| CHEK2 | None | None | None | None |
| PPARG | None | None | None | None |
| ACADVL | None | None | None | None |
| ACADVL | 0.385 | 4 | 3b96 | A |
| ACADVL | None | None | None | None |
| ACADVL | None | None | None | None |
| ACADVL | None | None | None | None |
| ACADVL | None | None | None | None |
| ACADVL | None | None | None | None |
| ACADVL | 0.346 | 14 | 3b96 | A |
| ACADVL | 0.346 | 14 | 3b96 | A |
| ACADVL | None | None | None | None |
| ACADVL | None | None | None | None |
| ACADVL | 0.231 | 10 | 3b96 | A |
| ACADVL | None | None | None | None |
| ACADVL | 0.346 | 14 | 3b96 | A |
| ACADVL | 0.269 | 12 | 3b96 | A |
| ACADVL | 0.423 | 2 | 3b96 | A |
| ACADVL | None | None | None | None |
| ACADVL | None | None | None | None |
| ACADVL | None | None | None | None |
| ACADVL | None | None | None | None |
| ACADVL | None | None | None | None |
| ACADVL | None | None | None | None |
| APOA1 | None | None | None | None |
| APOA1 | None | None | None | None |
| APOA1 | None | None | None | None |
| APOA1 | None | None | None | None |
| APOA1 | None | None | None | None |
| DNMT3A | None | None | None | None |
| DNMT3A | None | None | None | None |
| APC | None | None | None | None |
| TMOD1 | None | None | None | None |
| USH1C | None | None | None | None |
| USH1C | None | None | None | None |
| USH1C | None | None | None | None |
| USH1C | None | None | None | None |
| ERCC6 | None | None | None | None |
| ERCC6 | None | None | None | None |
| ERCC6 | None | None | None | None |
| ERCC8 | None | None | None | None |
| ERCC8 | None | None | None | None |
| ERCC8 | None | None | None | None |
| DPAGT1 | None | None | None | None |
| DPAGT1 | None | None | None | None |
| DPAGT1 | None | None | None | None |
| DPAGT1 | None | None | None | None |
| DPAGT1 | None | None | None | None |
| DPAGT1 | None | None | None | None |

| | | | | |
|---|---|---|---|---|
| DPAGT1 | None | None | None | None |
| IL2RG | None | None | None | None |
| HNF1A | None | None | None | None |
| HNF4A | None | None | None | None |
| FLT4 | None | None | None | None |
| FLT4 | None | None | None | None |
| FLT4 | None | None | None | None |
| CD3D | None | None | None | None |
| CD3D | None | None | None | None |
| MCOLN1 | None | None | None | None |
| MCOLN1 | None | None | None | None |
| MOCS2 | None | None | None | None |
| MOCS2 | None | None | None | None |
| ABCG8 | None | None | None | None |
| ABCG8 | None | None | None | None |
| ABCG8 | None | None | None | None |
| ABCG8 | None | None | None | None |
| ABCG8 | None | None | None | None |
| ABCG8 | None | None | None | None |
| ABCG8 | None | None | None | None |
| ABCG8 | None | None | None | None |
| ABCG8 | None | None | None | None |
| ABCG8 | None | None | None | None |
| ABCG8 | None | None | None | None |
| ABCG8 | None | None | None | None |
| ABCG8 | None | None | None | None |
| ABCG8 | None | None | None | None |
| CTNNA1 | None | None | None | None |
| TYMP | None | None | None | None |
| TYMP | None | None | None | None |
| TYMP | None | None | None | None |
| TYMP | None | None | None | None |
| TYMP | None | None | None | None |
| TYMP | None | None | None | None |
| TYMP | None | None | None | None |
| TYMP | None | None | None | None |
| TYMP | None | None | None | None |
| TYMP | None | None | None | None |
| TYMP | None | None | None | None |
| TYMP | None | None | None | None |
| TYMP | None | None | None | None |
| GAMT | None | None | None | None |
| GAMT | None | None | None | None |
| GAMT | None | None | None | None |
| GAMT | None | None | None | None |
| GAMT | None | None | None | None |
| GAMT | None | None | None | None |
| GAMT | None | None | None | None |
| GAMT | None | None | None | None |
| GAMT | None | None | None | None |
| GAMT | None | None | None | None |
| GAMT | None | None | None | None |
| GLRA1 | None | None | None | None |
| GLRA1 | None | None | None | None |

| | | | | |
|---|---|---|---|---|
| GLRA1 | None | None | None | None |
| GLRA1 | None | None | None | None |
| GLRA1 | None | None | None | None |
| GLRA1 | None | None | None | None |
| GLRA1 | None | None | None | None |
| GLRA1 | None | None | None | None |
| GLRA1 | None | None | None | None |
| ADK | None | None | None | None |
| ACTN2 | None | None | None | None |
| UBE2T | None | None | None | None |
| TTN | None | None | None | None |
| ATP7B | None | None | None | None |
| ATP7B | None | None | None | None |
| ATP7B | None | None | None | None |
| ATP7B | None | None | None | None |
| ATP7B | None | None | None | None |
| ATP7B | None | None | None | None |
| ATP7B | None | None | None | None |
| ATP7B | None | None | None | None |
| ATP7B | None | None | None | None |
| ATP7B | None | None | None | None |
| ATP7B | None | None | None | None |
| ATP7B | None | None | None | None |
| ATP7B | None | None | None | None |
| CDKN1B | None | None | None | None |
| CDKN1B | None | None | None | None |
| CDKN1B | None | None | None | None |
| CDKN1B | None | None | None | None |
| CDKN1B | None | None | None | None |
| CDKN1B | None | None | None | None |
| MYOC | None | None | None | None |
| MYOC | None | None | None | None |
| MYOC | None | None | None | None |
| MYOC | None | None | None | None |
| MYOC | None | None | None | None |
| MYOC | None | None | None | None |
| MYOC | None | None | None | None |
| MYOC | None | None | None | None |
| MYOC | None | None | None | None |
| MYOC | None | None | None | None |
| MYOC | None | None | None | None |
| MYOC | None | None | None | None |
| MYOC | None | None | None | None |
| MYOC | None | None | None | None |
| MYOC | None | None | None | None |
| SPTAN1 | None | None | None | None |
| GC | None | None | None | None |
| GC | None | None | None | None |
| MAD1L1 | None | None | None | None |
| NDST1 | None | None | None | None |
| ARL2BP | None | None | None | None |
| TOR1A | None | None | None | None |
| C1S | None | None | None | None |
| ARL6 | 0.508 | 1 | 2h57 | A |

| | | | | |
|---|---|---|---|---|
| ARL6 | None | None | None | None |
| ARL6 | None | None | None | None |
| ARL6 | None | None | None | None |
| ARL6 | None | None | None | None |
| ARL6 | None | None | None | None |
| FGFR1 | None | None | None | None |
| KLHL7 | None | None | None | None |
| FLNC | None | None | None | None |
| PTPRC | None | None | None | None |
| GCH1 | None | None | None | None |
| GCH1 | None | None | None | None |
| GCH1 | None | None | None | None |
| GCH1 | None | None | None | None |
| GCH1 | None | None | None | None |
| GCH1 | 0.2 | 17 | 1fb1 | A |
| GCH1 | None | None | None | None |
| GCH1 | None | None | None | None |
| GCH1 | None | None | None | None |
| GCH1 | None | None | None | None |
| GCH1 | 0.3 | 1 | 1fb1 | D |
| GCH1 | 0.2 | 3 | 1fb1 | D |
| GCH1 | 0.3 | 11 | 1fb1 | B |
| GCH1 | None | None | None | None |
| CFH | None | None | None | None |
| COL2A1 | None | None | None | None |
| COL2A1 | None | None | None | None |
| OCRL | None | None | None | None |
| WRN | None | None | None | None |
| WRN | None | None | None | None |
| WRN | None | None | None | None |
| WRN | None | None | None | None |
| MASP1 | None | None | None | None |
| MASP1 | None | None | None | None |
| NSD1 | None | None | None | None |
| PITX2 | None | None | None | None |
| PITX2 | None | None | None | None |
| PITX2 | None | None | None | None |
| PITX2 | 0.362 | 2 | 2lkx | A |
| PITX2 | 0.362 | 2 | 2lkx | A |
| PITX2 | 0.362 | 2 | 2lkx | A |
| B4GALT7 | 0.917 | 1 | 4irq | A |
| B4GALT7 | None | None | None | None |
| B4GALT7 | None | None | None | None |
| B4GALT7 | None | None | None | None |
| B4GALT7 | None | None | None | None |
| FSHR | None | None | None | None |
| IGHMBP2 | None | None | None | None |
| IGHMBP2 | None | None | None | None |
| IGHMBP2 | None | None | None | None |
| IGHMBP2 | None | None | None | None |
| IGHMBP2 | None | None | None | None |
| IGHMBP2 | None | None | None | None |
| IGHMBP2 | None | None | None | None |
| IGHMBP2 | None | None | None | None |

| | | | | |
|---|---|---|---|---|
| IGHMBP2 | None | None | None | None |
| IGHMBP2 | None | None | None | None |
| IGHMBP2 | None | None | None | None |
| IGHMBP2 | None | None | None | None |
| IGHMBP2 | None | None | None | None |
| IGHMBP2 | None | None | None | None |
| IGHMBP2 | None | None | None | None |
| IGHMBP2 | None | None | None | None |
| TTN | None | None | None | None |
| TTN | None | None | None | None |
| IRAK4 | None | None | None | None |
| PROC | None | None | None | None |
| PROC | None | None | None | None |
| RAD50 | None | None | None | None |
| RAD50 | None | None | None | None |
| DYNC1H1 | None | None | None | None |
| DYNC1H1 | None | None | None | None |
| DYNC1H1 | None | None | None | None |
| DYNC1H1 | None | None | None | None |
| DYNC1H1 | None | None | None | None |
| DYNC1H1 | None | None | None | None |
| DYNC1H1 | None | None | None | None |
| DYNC1H1 | None | None | None | None |
| DYNC1H1 | None | None | None | None |
| DYNC1H1 | None | None | None | None |
| DYNC1H1 | None | None | None | None |
| CYB5R3 | None | None | None | None |
| CYB5R3 | None | None | None | None |
| CYB5R3 | None | None | None | None |
| CYB5R3 | None | None | None | None |
| CYB5R3 | None | None | None | None |
| CYB5R3 | None | None | None | None |
| CYB5R3 | None | None | None | None |
| CYB5R3 | None | None | None | None |
| CYB5R3 | None | None | None | None |
| CYB5R3 | None | None | None | None |
| CYB5R3 | None | None | None | None |
| SPTB | None | None | None | None |
| FANCL | None | None | None | None |
| FAM126A | None | None | None | None |
| SEPT9 | None | None | None | None |
| TTN | None | None | None | None |
| TTN | None | None | None | None |
| TTN | 0.268 | 2 | 4jnw | A |
| CNTN1 | None | None | None | None |
| C1QTNF5 | None | None | None | None |
| VCL | None | None | None | None |
| VCL | None | None | None | None |
| VCL | None | None | None | None |
| VCL | None | None | None | None |
| VCL | None | None | None | None |
| VCL | None | None | None | None |
| PPT1 | None | None | None | None |
| PPT1 | None | None | None | None |

| | | | | |
|---|---|---|---|---|
| PPT1 | None | None | None | None |
| PPT1 | None | None | None | None |
| PPT1 | None | None | None | None |
| PPT1 | None | None | None | None |
| PPT1 | None | None | None | None |
| PPT1 | None | None | None | None |
| PPT1 | None | None | None | None |
| PPT1 | None | None | None | None |
| PPT1 | None | None | None | None |
| PPT1 | None | None | None | None |
| PPT1 | None | None | None | None |
| PPT1 | None | None | None | None |
| PPT1 | None | None | None | None |
| PPT1 | None | None | None | None |
| PPT1 | None | None | None | None |
| PPT1 | None | None | None | None |
| PPT1 | None | None | None | None |
| NOTCH2 | None | None | None | None |
| NOTCH2 | None | None | None | None |
| IFIH1 | None | None | None | None |
| IFIH1 | None | None | None | None |
| IKBKG | None | None | None | None |
| DBH | None | None | None | None |
| DBH | None | None | None | None |
| DBH | None | None | None | None |
| DBH | None | None | None | None |
| DBH | None | None | None | None |
| DBH | None | None | None | None |
| DBH | None | None | None | None |
| DBH | None | None | None | None |
| DBH | None | None | None | None |
| DBH | None | None | None | None |
| IL10RA | None | None | None | None |
| IL10RA | None | None | None | None |
| IL10RA | None | None | None | None |
| IL10RA | None | None | None | None |
| IL10RA | None | None | None | None |
| IL10RA | None | None | None | None |
| IL10RA | None | None | None | None |
| ILK | None | None | None | None |
| HPD | None | None | None | None |
| HPD | None | None | None | None |
| HPD | None | None | None | None |
| MECP2 | None | None | None | None |
| MECP2 | None | None | None | None |
| MECP2 | None | None | None | None |
| MECP2 | None | None | None | None |
| MECP2 | None | None | None | None |
| MECP2 | None | None | None | None |
| MECP2 | None | None | None | None |
| MECP2 | None | None | None | None |
| KYNU | None | None | None | None |
| FAS | None | None | None | None |
| FAS | None | None | None | None |

| | | | | |
|---|---|---|---|---|
| ELP1 | None | None | None | None |
| ELP1 | None | None | None | None |
| ELP1 | None | None | None | None |
| ELP1 | None | None | None | None |
| ELP1 | None | None | None | None |
| ELP1 | None | None | None | None |
| ELP1 | None | None | None | None |
| ELP1 | None | None | None | None |
| HESX1 | None | None | None | None |
| HESX1 | None | None | None | None |
| HESX1 | None | None | None | None |
| HESX1 | None | None | None | None |
| GAD1 | None | None | None | None |
| GAD1 | None | None | None | None |
| NDUFA1 | None | None | None | None |
| NDUFA1 | None | None | None | None |
| NDUFA1 | None | None | None | None |
| PNPT1 | None | None | None | None |
| PNPT1 | None | None | None | None |
| PNPT1 | None | None | None | None |
| PNPT1 | None | None | None | None |
| PNPT1 | 0.389 | 1 | 3u1k | A |
| PNPT1 | None | None | None | None |
| PNPT1 | None | None | None | None |
| SLC25A13 | None | None | None | None |
| SLC25A13 | None | None | None | None |
| SLC25A13 | None | None | None | None |
| AP4M1 | None | None | None | None |
| AP4M1 | None | None | None | None |
| MEFV | None | None | None | None |
| MEFV | None | None | None | None |
| MEFV | None | None | None | None |
| MEFV | None | None | None | None |
| MEFV | None | None | None | None |
| MEFV | None | None | None | None |
| MEFV | None | None | None | None |
| MEFV | None | None | None | None |
| MEFV | None | None | None | None |
| CAPN3 | None | None | None | None |
| CAPN3 | None | None | None | None |
| CAPN3 | None | None | None | None |
| CAPN3 | None | None | None | None |
| CAPN3 | 0.474 | 2 | 4okh | A |
| CAPN3 | 0.474 | 2 | 4okh | A |
| CAPN3 | None | None | None | None |
| GP6 | None | None | None | None |
| FAS | None | None | None | None |
| FBN1 | None | None | None | None |
| FBN1 | None | None | None | None |
| BMPR2 | None | None | None | None |
| BMPR2 | None | None | None | None |
| BMPR2 | None | None | None | None |
| BMPR2 | 0.327 | 6 | 3g2f | A |
| BMPR2 | 0.327 | 6 | 3g2f | A |

| | | | | |
|---|---|---|---|---|
| RET | None | None | None | None |
| RET | None | None | None | None |
| RET | None | None | None | None |
| RET | None | None | None | None |
| RET | None | None | None | None |
| RET | None | None | None | None |
| RET | None | None | None | None |
| RET | None | None | None | None |
| RET | None | None | None | None |
| RET | None | None | None | None |
| RET | None | None | None | None |
| RET | None | None | None | None |
| RET | None | None | None | None |
| RET | None | None | None | None |
| RET | None | None | None | None |
| RET | None | None | None | None |
| RET | None | None | None | None |
| RET | None | None | None | None |
| RET | None | None | None | None |
| RET | None | None | None | None |
| RET | None | None | None | None |
| RET | None | None | None | None |
| RET | None | None | None | None |
| RET | None | None | None | None |
| RET | None | None | None | None |
| RET | None | None | None | None |
| RET | None | None | None | None |
| RET | None | None | None | None |
| CSF2RA | None | None | None | None |
| CSF2RA | None | None | None | None |
| CSF2RA | None | None | None | None |
| BRCA1 | None | None | None | None |
| ZAP70 | 0.636 | 1 | 2ozo | A |
| ZAP70 | 0.636 | 1 | 2ozo | A |
| ZAP70 | 0.26 | 9 | 2ozo | A |
| PIK3R2 | None | None | None | None |
| TCN2 | None | None | None | None |
| TCN2 | None | None | None | None |
| TCN2 | None | None | None | None |
| TCN2 | None | None | None | None |
| TCN2 | None | None | None | None |
| TCN2 | None | None | None | None |
| TCN2 | None | None | None | None |
| TCN2 | None | None | None | None |
| EPM2A | None | None | None | None |
| EPM2A | None | None | None | None |
| EPM2A | None | None | None | None |
| EPM2A | None | None | None | None |
| AARS | None | None | None | None |
| AARS | None | None | None | None |
| AARS | None | None | None | None |
| PCDH15 | None | None | None | None |
| PCDH15 | None | None | None | None |
| PCDH15 | None | None | None | None |

| | | | | |
|---|---|---|---|---|
| PCDH15 | None | None | None | None |
| PCDH15 | None | None | None | None |
| PCDH15 | None | None | None | None |
| PCDH15 | None | None | None | None |
| PCDH15 | None | None | None | None |
| PCDH15 | None | None | None | None |
| PCDH15 | None | None | None | None |
| NDUFS4 | None | None | None | None |
| NDUFS4 | None | None | None | None |
| NDUFS4 | None | None | None | None |
| FGF10 | None | None | None | None |
| FGF10 | 0.558 | 3 | 1nun | A |
| FGF10 | None | None | None | None |
| FGF10 | None | None | None | None |
| CDON | None | None | None | None |
| STAMBP | 0.909 | 2 | 3rzv | A |
| BRCA1 | None | None | None | None |
| BRCA1 | None | None | None | None |
| BRCA1 | None | None | None | None |
| BRCA1 | None | None | None | None |
| BRCA1 | None | None | None | None |
| BRCA1 | None | None | None | None |
| BRCA1 | None | None | None | None |
| BRCA1 | None | None | None | None |
| BRCA1 | None | None | None | None |
| BRCA1 | None | None | None | None |
| PLXNC1 | None | None | None | None |
| RNASEH2C | None | None | None | None |
| RNASEH2C | None | None | None | None |
| APC | None | None | None | None |
| PLA2G4A | None | None | None | None |
| PLA2G4A | None | None | None | None |
| PSAT1 | None | None | None | None |
| PSAT1 | None | None | None | None |
| TRAPPC3 | None | None | None | None |
| QDPR | None | None | None | None |
| QDPR | None | None | None | None |
| QDPR | None | None | None | None |
| QDPR | None | None | None | None |
| KRT5 | None | None | None | None |
| KRT5 | None | None | None | None |
| KRT5 | None | None | None | None |
| MLH1 | None | None | None | None |
| MLH1 | None | None | None | None |
| MLH1 | None | None | None | None |
| MLH1 | None | None | None | None |
| MLH1 | None | None | None | None |
| MLH1 | None | None | None | None |
| MLH1 | None | None | None | None |
| MLH1 | None | None | None | None |
| MLH1 | None | None | None | None |
| MLH1 | None | None | None | None |
| MLH1 | None | None | None | None |
| MLH1 | None | None | None | None |

| | | | | |
|---|---|---|---|---|
| MLH1 | None | None | None | None |
| MLH1 | None | None | None | None |
| MLH1 | None | None | None | None |
| MLH1 | None | None | None | None |
| MLH1 | None | None | None | None |
| KRIT1 | None | None | None | None |
| KRIT1 | None | None | None | None |
| KRIT1 | None | None | None | None |
| ALX4 | None | None | None | None |
| ALX4 | None | None | None | None |
| ALX4 | None | None | None | None |
| TJP2 | None | None | None | None |
| TJP2 | None | None | None | None |
| PANK2 | None | None | None | None |
| PANK2 | None | None | None | None |
| PANK2 | None | None | None | None |
| PANK2 | None | None | None | None |
| PANK2 | None | None | None | None |
| PANK2 | 0.333 | 2 | 5E+026 | B |
| PANK2 | None | None | None | None |
| NDUFA6 | None | None | None | None |
| TSHR | None | None | None | None |
| TSHR | None | None | None | None |
| TSHR | None | None | None | None |
| CFTR | None | None | None | None |
| CFTR | None | None | None | None |
| MYOM1 | None | None | None | None |
| MYOM1 | None | None | None | None |
| SPTB | None | None | None | None |
| GPT2 | None | None | None | None |
| NDUFS2 | None | None | None | None |
| NDUFS2 | None | None | None | None |
| NDUFS2 | None | None | None | None |
| NDUFS2 | None | None | None | None |
| NDUFS2 | None | None | None | None |
| CASQ2 | None | None | None | None |
| CASQ2 | None | None | None | None |
| CASQ2 | None | None | None | None |
| CASQ2 | None | None | None | None |
| CASQ2 | None | None | None | None |
| CASQ2 | None | None | None | None |
| CASQ2 | None | None | None | None |
| CASQ2 | None | None | None | None |
| CASQ2 | None | None | None | None |
| CPOX | None | None | None | None |
| CPOX | None | None | None | None |
| CPOX | 0.857 | 2 | 2aex | A |
| CPOX | None | None | None | None |
| CPOX | None | None | None | None |
| CPOX | None | None | None | None |
| CPOX | None | None | None | None |
| CPOX | None | None | None | None |
| CPOX | None | None | None | None |
| MVK | None | None | None | None |

| | | | | |
|---|---|---|---|---|
| MVK | None | None | None | None |
| MVK | None | None | None | None |
| MVK | None | None | None | None |
| MVK | None | None | None | None |
| MVK | None | None | None | None |
| MVK | None | None | None | None |
| MVK | None | None | None | None |
| MVK | None | None | None | None |
| MVK | None | None | None | None |
| MVK | None | None | None | None |
| MVK | None | None | None | None |
| MVK | None | None | None | None |
| MVK | None | None | None | None |
| MVK | None | None | None | None |
| GSS | None | None | None | None |
| GSS | None | None | None | None |
| GSS | None | None | None | None |
| GSS | None | None | None | None |
| GSS | None | None | None | None |
| DSG2 | None | None | None | None |
| DSG2 | None | None | None | None |
| DSG2 | None | None | None | None |
| DSG2 | None | None | None | None |
| DSG2 | None | None | None | None |
| DSG2 | None | None | None | None |
| DSG2 | None | None | None | None |
| DSG2 | None | None | None | None |
| DSG2 | None | None | None | None |
| DSG2 | None | None | None | None |
| DSG2 | None | None | None | None |
| DSG2 | None | None | None | None |
| DSG2 | None | None | None | None |
| DSG2 | None | None | None | None |
| PAPSS2 | None | None | None | None |
| PLCB2 | None | None | None | None |
| HMGCS2 | None | None | None | None |
| HMGCS2 | None | None | None | None |
| HMGCS2 | 0.938 | 1 | 2wya | C |
| HMGCS2 | None | None | None | None |
| HMGCS2 | None | None | None | None |
| HMGCS2 | None | None | None | None |
| HMGCS2 | 0.562 | 4 | 2wya | C |
| HMGCS2 | None | None | None | None |
| MPDZ | None | None | None | None |
| GATA4 | None | None | None | None |
| GATA4 | None | None | None | None |
| WARS2 | None | None | None | None |
| WARS2 | None | None | None | None |
| HGD | 1 | 47 | 1eyb | A |
| HGD | 1 | 47 | 1eyb | A |
| HGD | 1 | 70 | 1eyb | A |
| HGD | None | None | None | None |
| HGD | None | None | None | None |
| HGD | None | None | None | None |

| | | | | | |
|---|---|---|---|---|---|
| HGD | None | None | None | | None |
| HGD | 0.5 | | 51 | 1ey2 | A |
| HGD | None | None | None | | None |
| HGD | 1 | | 1 | 1ey2 | A |
| HGD | 1 | | 9 | 1ey2 | A |
| HGD | None | None | None | | None |
| HGD | 1 | | 10 | 1eyb | A |
| HGD | 1 | | 7 | 1ey2 | A |
| PAX8 | None | None | None | | None |
| PAX8 | None | None | None | | None |
| PAX8 | None | None | None | | None |
| PAX8 | None | None | None | | None |
| SPTB | None | None | None | | None |
| NCF4 | None | None | None | | None |
| CFH | None | None | None | | None |
| CFH | None | None | None | | None |
| CSF1R | None | None | None | | None |
| CSF1R | None | None | None | | None |
| CA8 | None | None | None | | None |
| CA8 | None | None | None | | None |
| CA8 | 0.748 | | 7 | 2w2j | A |
| APC | None | None | None | | None |
| APC | None | None | None | | None |
| APC | None | None | None | | None |
| APC | None | None | None | | None |
| APC | None | None | None | | None |
| NDUFA2 | None | None | None | | None |
| GIF | None | None | None | | None |
| GIF | None | None | None | | None |
| GIF | None | None | None | | None |
| GIF | None | None | None | | None |
| C2 | None | None | None | | None |
| ACP5 | None | None | None | | None |
| ACP5 | None | None | None | | None |
| ACP5 | None | None | None | | None |
| ACP5 | None | None | None | | None |
| ACP5 | None | None | None | | None |
| RNASEH2A | None | None | None | | None |
| RNASEH2A | None | None | None | | None |
| RNASEH2A | None | None | None | | None |
| RNASEH2A | None | None | None | | None |
| RNASEH2A | None | None | None | | None |
| RNASEH2A | None | None | None | | None |
| RNASEH2A | None | None | None | | None |
| DSG2 | None | None | None | | None |
| USH1G | None | None | None | | None |
| FUS | None | None | None | | None |
| FUS | None | None | None | | None |
| FUS | None | None | None | | None |
| FUS | None | None | None | | None |
| FUS | None | None | None | | None |
| MRPL44 | None | None | None | | None |
| MRPL44 | None | None | None | | None |
| MRPL44 | None | None | None | | None |

| | | | | |
|---|---|---|---|---|
| MRPL44 | None | None | None | None |
| DMGDH | None | None | None | None |
| AARS | None | None | None | None |
| STAR | None | None | None | None |
| STAR | None | None | None | None |
| STAR | None | None | None | None |
| STAR | None | None | None | None |
| STAR | None | None | None | None |
| STAR | None | None | None | None |
| STAR | None | None | None | None |
| STAR | None | None | None | None |
| LARP7 | None | None | None | None |
| PRPF8 | None | None | None | None |
| PRPF8 | None | None | None | None |
| PRPF8 | None | None | None | None |
| PRPF8 | None | None | None | None |
| PSAP | None | None | None | None |
| PSAP | None | None | None | None |
| PSAP | None | None | None | None |
| DMPK | None | None | None | None |
| EPCAM | None | None | None | None |
| EPCAM | None | None | None | None |
| EPCAM | None | None | None | None |
| EPCAM | None | None | None | None |
| EPCAM | None | None | None | None |
| EPCAM | None | None | None | None |
| EPCAM | None | None | None | None |
| EPCAM | None | None | None | None |
| ASS1 | None | None | None | None |
| ASS1 | None | None | None | None |
| ASS1 | 0.571 | 3 | 2nz2 | A |
| ASS1 | 0.571 | 3 | 2nz2 | A |
| ASS1 | None | None | None | None |
| ASS1 | 0.857 | 20 | 2nz2 | A |
| ASS1 | 0.857 | 9 | 2nz2 | A |
| ASS1 | None | None | None | None |
| ASS1 | 0.429 | 13 | 2nz2 | A |
| ASS1 | 0.571 | 4 | 2nz2 | A |
| ASS1 | 0.857 | 9 | 2nz2 | A |
| ASS1 | 0.857 | 9 | 2nz2 | A |
| ASS1 | 0.857 | 9 | 2nz2 | A |
| ASS1 | None | None | None | None |
| ASS1 | None | None | None | None |
| ASS1 | None | None | None | None |
| ASS1 | None | None | None | None |
| ASS1 | None | None | None | None |
| ASS1 | None | None | None | None |
| ASS1 | None | None | None | None |
| POLE | None | None | None | None |
| POLE | None | None | None | None |
| POLE | None | None | None | None |
| RNASET2 | None | None | None | None |
| RNASET2 | None | None | None | None |
| RNASET2 | None | None | None | None |

| | | | | |
|---|---|---|---|---|
| FLG | 0.394 | | 2 4pcw | A |
| F10 | None | None | None | None |
| AGT | None | None | None | None |
| AGT | None | None | None | None |
| PLN | None | None | None | None |
| PLN | None | None | None | None |
| PLN | None | None | None | None |
| PTHLH | None | None | None | None |
| IGF1R | None | None | None | None |
| IGF1R | None | None | None | None |
| IGF1R | None | None | None | None |
| JAG1 | None | None | None | None |
| JAG1 | None | None | None | None |
| JAG1 | None | None | None | None |
| JAG1 | None | None | None | None |
| JAG1 | None | None | None | None |
| JAG1 | None | None | None | None |
| AP1S3 | None | None | None | None |
| SPTA1 | None | None | None | None |
| SPTA1 | None | None | None | None |
| SPTA1 | None | None | None | None |
| SPTA1 | None | None | None | None |
| SPG7 | None | None | None | None |
| SPG7 | None | None | None | None |
| SPG7 | None | None | None | None |
| SPG7 | None | None | None | None |
| SPG7 | None | None | None | None |
| SPG7 | None | None | None | None |
| FN1 | None | None | None | None |
| FN1 | None | None | None | None |
| NSMCE3 | None | None | None | None |
| PCCA | None | None | None | None |
| PCCA | None | None | None | None |
| SPTB | None | None | None | None |
| RYR2 | None | None | None | None |
| RYR2 | None | None | None | None |
| MMP9 | None | None | None | None |
| MMP9 | None | None | None | None |
| GLMN | None | None | None | None |
| GLMN | None | None | None | None |
| GLMN | None | None | None | None |
| GLMN | None | None | None | None |
| GLMN | None | None | None | None |
| GLMN | None | None | None | None |
| GLMN | None | None | None | None |
| GLMN | None | None | None | None |
| GLMN | None | None | None | None |
| GLMN | None | None | None | None |
| MARS | 0.333 | | 2 5goy | A |
| MARS | None | None | None | None |
| MARS | None | None | None | None |
| MARS | None | None | None | None |
| MARS | None | None | None | None |
| MARS | None | None | None | None |

| | | | | |
|---|---|---|---|---|
| SNAP29 | None | None | None | None |
| SNAP29 | None | None | None | None |
| ETHE1 | None | None | None | None |
| ETHE1 | None | None | None | None |
| ETHE1 | None | None | None | None |
| ETHE1 | None | None | None | None |
| ETHE1 | None | None | None | None |
| ETHE1 | None | None | None | None |
| ETHE1 | None | None | None | None |
| SAMHD1 | None | None | None | None |
| KCNQ4 | None | None | None | None |
| FXYD2 | None | None | None | None |
| FXYD2 | None | None | None | None |
| FXYD2 | None | None | None | None |
| LRRK2 | None | None | None | None |
| LRRK2 | None | None | None | None |
| NDUFS1 | None | None | None | None |
| NDUFS1 | None | None | None | None |
| NDUFS1 | None | None | None | None |
| NDUFS1 | None | None | None | None |
| NDUFS1 | None | None | None | None |
| NDUFS1 | None | None | None | None |
| NDUFS1 | None | None | None | None |
| NDUFS1 | None | None | None | None |
| NDUFS1 | None | None | None | None |
| NDUFS1 | None | None | None | None |
| NDUFS1 | None | None | None | None |
| EZH2 | None | None | None | None |
| EZH2 | None | None | None | None |
| LARS | None | None | None | None |
| TLL1 | None | None | None | None |
| CRYBB2 | None | None | None | None |
| CRYBB2 | None | None | None | None |
| EXOSC8 | None | None | None | None |
| SRP72 | None | None | None | None |
| FOXC2 | None | None | None | None |
| HMBS | None | None | None | None |
| HMBS | None | None | None | None |
| HMBS | None | None | None | None |
| HMBS | None | None | None | None |
| HMBS | None | None | None | None |
| HMBS | None | None | None | None |
| HMBS | None | None | None | None |
| HMBS | None | None | None | None |
| HMBS | None | None | None | None |
| HMBS | None | None | None | None |
| HMBS | None | None | None | None |
| HMBS | None | None | None | None |
| HMBS | None | None | None | None |
| HMBS | None | None | None | None |
| HMBS | None | None | None | None |
| HMBS | None | None | None | None |
| HMBS | None | None | None | None |
| HMBS | None | None | None | None |

| | | | | |
|---|---|---|---|---|
| HMBS | None | None | None | None |
| HMBS | None | None | None | None |
| HMBS | None | None | None | None |
| HMBS | None | None | None | None |
| HMBS | None | None | None | None |
| HMBS | None | None | None | None |
| HMBS | None | None | None | None |
| OGT | None | None | None | None |
| NOG | None | None | None | None |
| NOG | None | None | None | None |
| NOG | None | None | None | None |
| NOG | None | None | None | None |
| NOG | None | None | None | None |
| NOG | None | None | None | None |
| NOG | None | None | None | None |
| NOG | None | None | None | None |
| NOG | None | None | None | None |
| NOG | None | None | None | None |
| NOG | None | None | None | None |
| DLG3 | None | None | None | None |
| SCN2A | None | None | None | None |
| STAC3 | None | None | None | None |
| PFKM | None | None | None | None |
| PFKM | None | None | None | None |
| PFKM | None | None | None | None |
| PFKM | None | None | None | None |
| PFKM | None | None | None | None |
| DNMT1 | None | None | None | None |
| DNMT1 | None | None | None | None |
| TNNI3 | None | None | None | None |
| TNNI3 | None | None | None | None |
| TNNI3 | None | None | None | None |
| STXBP2 | None | None | None | None |
| STXBP2 | None | None | None | None |
| STXBP2 | None | None | None | None |
| STXBP2 | None | None | None | None |
| STXBP2 | None | None | None | None |
| STXBP2 | None | None | None | None |
| STXBP2 | None | None | None | None |
| STXBP2 | None | None | None | None |
| REN | None | None | None | None |
| DSP | None | None | None | None |
| DSP | None | None | None | None |
| HNF1B | None | None | None | None |
| ADA | None | None | None | None |
| ADA | None | None | None | None |
| ADA | 0.615 | 7 | 3iar | A |
| ADA | None | None | None | None |
| ADA | None | None | None | None |
| ADA | 0.538 | 5 | 3iar | A |
| ADA | 0.538 | 5 | 3iar | A |
| ADA | None | None | None | None |
| ADA | None | None | None | None |
| ADA | 0.615 | 1 | 3iar | A |

| | | | | |
|---|---|---|---|---|
| ADA | 0.462 | 6 | 3iar | A |
| ADA | 0.615 | 1 | 3iar | A |
| ADA | None | None | None | None |
| ADA | None | None | None | None |
| ADA | None | None | None | None |
| DCX | None | None | None | None |
| DCX | None | None | None | None |
| DCX | None | None | None | None |
| DCX | None | None | None | None |
| DCX | None | None | None | None |
| ARSB | None | None | None | None |
| ARSB | None | None | None | None |
| ARSB | None | None | None | None |
| ARSB | None | None | None | None |
| ARSB | None | None | None | None |
| ARSB | None | None | None | None |
| ARSB | None | None | None | None |
| ARSB | None | None | None | None |
| ARSB | None | None | None | None |
| ARSB | None | None | None | None |
| ARSB | None | None | None | None |
| ARSB | None | None | None | None |
| ARSB | None | None | None | None |
| WWOX | None | None | None | None |
| TNNI3 | None | None | None | None |
| TNNI3 | None | None | None | None |
| TNNI3 | None | None | None | None |
| TNNI3 | None | None | None | None |
| CYBA | None | None | None | None |
| SPAST | None | None | None | None |
| SPAST | None | None | None | None |
| SPAST | None | None | None | None |
| SPAST | None | None | None | None |
| SPAST | None | None | None | None |
| SPAST | None | None | None | None |
| SPAST | None | None | None | None |
| SPAST | None | None | None | None |
| KRIT1 | None | None | None | None |
| KRIT1 | None | None | None | None |
| TP53 | None | None | None | None |
| ISPD | None | None | None | None |
| ISPD | None | None | None | None |
| ISPD | None | None | None | None |
| ISPD | None | None | None | None |
| KCNMA1 | None | None | None | None |
| STK11 | None | None | None | None |
| STK11 | None | None | None | None |
| STK11 | None | None | None | None |
| STK11 | None | None | None | None |
| STK11 | None | None | None | None |
| STK11 | 0.412 | 2 | 2wtk | F |
| NDUFA13 | None | None | None | None |
| PHF6 | None | None | None | None |
| PHF6 | None | None | None | None |

| | | | | |
|---|---|---|---|---|
| PHF6 | None | None | None | None |
| PHYH | None | None | None | None |
| PHYH | None | None | None | None |
| PHYH | None | None | None | None |
| PHYH | None | None | None | None |
| PHYH | None | None | None | None |
| NDUFS3 | None | None | None | None |
| NDUFS3 | None | None | None | None |
| ALDH18A1 | None | None | None | None |
| ALDH18A1 | None | None | None | None |
| ALDH18A1 | None | None | None | None |
| ALDH18A1 | None | None | None | None |
| ALDH18A1 | None | None | None | None |
| PAX6 | None | None | None | None |
| PAX6 | 0.339 | 1 | 2cue | A |
| INPP5E | None | None | None | None |
| INPP5E | None | None | None | None |
| INPP5E | None | None | None | None |
| INPP5E | None | None | None | None |
| INPP5E | None | None | None | None |
| INPP5E | None | None | None | None |
| INPP5E | None | None | None | None |
| INPP5E | None | None | None | None |
| INPP5E | None | None | None | None |
| TNNT2 | None | None | None | None |
| TNNT2 | None | None | None | None |
| TNNT2 | None | None | None | None |
| TNNT2 | None | None | None | None |
| TNNT2 | None | None | None | None |
| TNNT2 | None | None | None | None |
| TNNT2 | None | None | None | None |
| TNNT2 | None | None | None | None |
| TYK2 | None | None | None | None |
| TYK2 | None | None | None | None |
| TYK2 | None | None | None | None |
| CFTR | None | None | None | None |
| CFTR | None | None | None | None |
| CFTR | None | None | None | None |
| CFTR | None | None | None | None |
| CFTR | None | None | None | None |
| CFTR | None | None | None | None |
| CFTR | None | None | None | None |
| CFTR | None | None | None | None |
| CFTR | None | None | None | None |
| CFTR | None | None | None | None |
| CFTR | None | None | None | None |
| CFTR | None | None | None | None |
| CFTR | None | None | None | None |
| CFTR | None | None | None | None |
| CFTR | None | None | None | None |
| CFTR | None | None | None | None |
| CFTR | None | None | None | None |
| CFTR | None | None | None | None |
| CFTR | None | None | None | None |

| | | | | |
|---|---|---|---|---|
| CFTR | None | None | None | None |
| CFTR | None | None | None | None |
| CFTR | None | None | None | None |
| CFTR | None | None | None | None |
| CFTR | None | None | None | None |
| CFTR | None | None | None | None |
| CFTR | None | None | None | None |
| CFTR | None | None | None | None |
| CFTR | None | None | None | None |
| CFTR | None | None | None | None |
| CFTR | None | None | None | None |
| CFTR | None | None | None | None |
| CFTR | None | None | None | None |
| CFTR | None | None | None | None |
| CFTR | None | None | None | None |
| CFTR | None | None | None | None |
| CFTR | None | None | None | None |
| CFTR | None | None | None | None |
| CFTR | None | None | None | None |
| CFTR | None | None | None | None |
| CFTR | None | None | None | None |
| CFTR | None | None | None | None |
| CFTR | None | None | None | None |
| CFTR | None | None | None | None |
| CFTR | None | None | None | None |
| CFTR | None | None | None | None |
| CFTR | None | None | None | None |
| CFTR | None | None | None | None |
| CFTR | None | None | None | None |
| CFTR | None | None | None | None |
| CFTR | None | None | None | None |
| CFTR | None | None | None | None |
| CFTR | None | None | None | None |
| CFTR | None | None | None | None |
| SERPINF1 | None | None | None | None |
| SERPINF1 | None | None | None | None |
| IDS | None | None | None | None |
| IDS | None | None | None | None |
| IDS | None | None | None | None |
| IDS | None | None | None | None |
| IDS | None | None | None | None |
| IDS | None | None | None | None |
| IDS | None | None | None | None |
| IDS | None | None | None | None |
| IDS | None | None | None | None |
| NFKBIA | 0.7 | 1 | 1nfi | F |
| NFKBIA | None | None | None | None |
| SPAST | None | None | None | None |
| FGF23 | None | None | None | None |
| FGF23 | None | None | None | None |
| FGF23 | None | None | None | None |
| FGF23 | None | None | None | None |
| FGF23 | None | None | None | None |
| FGF23 | None | None | None | None |

| | | | | |
|---|---|---|---|---|
| FGF23 | None | None | None | None |
| DLAT | None | None | None | None |
| STUB1 | None | None | None | None |
| HSD17B4 | None | None | None | None |
| HSD17B4 | None | None | None | None |
| DYSF | None | None | None | None |
| CEP135 | None | None | None | None |
| CEP135 | None | None | None | None |
| PRKCG | None | None | None | None |
| CSF1R | None | None | None | None |
| PEX5 | None | None | None | None |
| PAX6 | None | None | None | None |
| PAX6 | None | None | None | None |
| PAX6 | None | None | None | None |
| PAX6 | None | None | None | None |
| PAX6 | None | None | None | None |
| PAX6 | None | None | None | None |
| NF2 | None | None | None | None |
| NF2 | None | None | None | None |
| NF2 | None | None | None | None |
| REM1 | None | None | None | None |

| source_id | classification | binding_site | binding_site_single_water | protein_binding_site |
|---|---|---|---|---|
| rs121918343 | pathogenic | 1 | 1 | 0 |
| rs267606649 | pathogenic | 1 | 1 | 1 |
| rs121918342 | pathogenic | 0 | 1 | 0 |
| rs267606650 | pathogenic | 1 | 1 | 1 |
| rs1130335 | benign | 1 | 1 | 1 |
| rs199476087 | pathogenic | 1 | 1 | 1 |
| rs772163112 | benign | 0 | 0 | 0 |
| rs755793 | benign | 0 | 0 | 0 |
| rs121918155 | pathogenic | 1 | 1 | 0 |
| rs121918156 | pathogenic | 1 | 1 | 0 |
| rs121918154 | pathogenic | 1 | 1 | 1 |
| rs121918151 | pathogenic | 1 | 1 | 1 |
| rs121918144 | pathogenic | 0 | 1 | 0 |
| rs121918146 | pathogenic | 1 | 1 | 1 |
| rs121918160 | pathogenic | 1 | 1 | 1 |
| rs121918150 | pathogenic | 1 | 1 | 1 |
| rs121918141 | pathogenic | 1 | 1 | 1 |
| rs142742242 | pathogenic | 1 | 1 | 1 |
| rs121918142 | pathogenic | 1 | 1 | 1 |
| rs121918157 | pathogenic | 1 | 1 | 1 |
| rs121918101 | pathogenic | 1 | 1 | 1 |
| rs121908260 | pathogenic | 1 | 1 | 1 |
| rs80356667 | pathogenic | 1 | 1 | 1 |
| rs80356668 | pathogenic | 1 | 1 | 1 |
| rs397514540 | pathogenic | 1 | 1 | 1 |
| rs104894685 | pathogenic | 0 | 1 | 0 |
| rs199869995 | pathogenic | 1 | 1 | 1 |
| rs121909072 | pathogenic | 0 | 0 | 0 |
| rs143669310 | benign | 1 | 1 | 1 |
| rs747347498 | benign | 0 | 0 | 0 |
| rs138283222 | benign | 0 | 0 | 0 |
| rs118203931 | pathogenic | 1 | 1 | 1 |
| rs118203933 | pathogenic | 1 | 1 | 0 |
| rs118203932 | pathogenic | 1 | 1 | 0 |
| rs2228063 | benign | 1 | 1 | 1 |
| rs377767360 | pathogenic | 0 | 0 | 0 |
| rs397518413 | pathogenic | 1 | 1 | 1 |
| rs1800458 | benign | 1 | 1 | 0 |
| rs121918083 | pathogenic | 1 | 1 | 1 |
| rs121918094 | pathogenic | 1 | 1 | 1 |
| rs121918098 | pathogenic | 1 | 1 | 1 |
| rs121918093 | pathogenic | 1 | 1 | 1 |
| rs79977247 | pathogenic | 1 | 1 | 1 |
| rs28933979 | pathogenic | 1 | 1 | 1 |
| rs121918068 | pathogenic | 1 | 1 | 1 |
| rs121918077 | pathogenic | 1 | 1 | 1 |
| rs11541796 | pathogenic | 1 | 1 | 1 |
| rs104894665 | pathogenic | 1 | 1 | 1 |
| rs121918090 | pathogenic | 1 | 1 | 1 |
| rs121918081 | pathogenic | 1 | 1 | 1 |
| rs121918080 | pathogenic | 1 | 1 | 1 |
| rs121918076 | pathogenic | 1 | 1 | 1 |
| rs121918097 | pathogenic | 1 | 1 | 1 |

| | | | | |
|---|---|---|---|---|
| rs121918079 | pathogenic | 1 | 1 | 1 |
| rs121918069 | pathogenic | 1 | 1 | 1 |
| rs121918086 | pathogenic | 1 | 1 | 1 |
| rs121918091 | pathogenic | 1 | 1 | 1 |
| rs121918085 | pathogenic | 1 | 1 | 1 |
| rs121918100 | pathogenic | 1 | 1 | 1 |
| rs121918084 | pathogenic | 1 | 1 | 1 |
| rs121918071 | pathogenic | 1 | 1 | 1 |
| rs121918072 | pathogenic | 1 | 1 | 1 |
| rs121918082 | pathogenic | 1 | 1 | 1 |
| rs121918087 | pathogenic | 1 | 1 | 1 |
| rs76410435 | benign | 1 | 1 | 1 |
| rs121918089 | pathogenic | 1 | 1 | 1 |
| rs121918073 | pathogenic | 1 | 1 | 1 |
| rs121918075 | pathogenic | 1 | 1 | 1 |
| rs121918088 | pathogenic | 1 | 1 | 1 |
| rs76992529 | pathogenic | 1 | 1 | 1 |
| rs536294863 | benign | 1 | 1 | 1 |
| rs730881162 | benign | 1 | 1 | 1 |
| rs113994087 | pathogenic | 1 | 1 | 1 |
| rs74716434 | benign | 1 | 1 | 1 |
| rs121918694 | pathogenic | 1 | 1 | 1 |
| rs121918707 | pathogenic | 1 | 1 | 1 |
| rs121918695 | pathogenic | 1 | 1 | 0 |
| rs121918690 | pathogenic | 1 | 1 | 0 |
| rs121918696 | pathogenic | 1 | 1 | 0 |
| rs121918693 | pathogenic | 1 | 1 | 0 |
| rs28999969 | pathogenic | 1 | 1 | 1 |
| rs121918697 | pathogenic | 1 | 1 | 1 |
| rs121918688 | pathogenic | 1 | 1 | 0 |
| rs28999970 | pathogenic | 1 | 1 | 0 |
| rs121918686 | pathogenic | 1 | 1 | 0 |
| rs28999971 | pathogenic | 1 | 1 | 1 |
| rs121918698 | pathogenic | 1 | 1 | 1 |
| rs121918691 | pathogenic | 1 | 1 | 0 |
| rs121918692 | pathogenic | 1 | 1 | 1 |
| rs121918703 | pathogenic | 1 | 1 | 1 |
| rs121918687 | pathogenic | 1 | 1 | 1 |
| rs28933408 | pathogenic | 1 | 1 | 1 |
| rs121918702 | pathogenic | 1 | 1 | 1 |
| rs17875899 | benign | 1 | 1 | 1 |
| rs111033620 | pathogenic | 1 | 1 | 1 |
| rs111033622 | pathogenic | 1 | 1 | 1 |
| rs111033621 | pathogenic | 1 | 1 | 1 |
| rs111033618 | pathogenic | 1 | 1 | 1 |
| rs5918 | benign | 0 | 0 | 0 |
| rs121918452 | pathogenic | 0 | 0 | 0 |
| rs121918445 | pathogenic | 1 | 1 | 1 |
| rs5917 | benign | 1 | 1 | 1 |
| rs121918444 | pathogenic | 1 | 1 | 1 |
| rs121918446 | pathogenic | 1 | 1 | 1 |
| rs121918449 | pathogenic | 0 | 0 | 0 |
| rs121918448 | benign | 0 | 0 | 0 |
| rs13306487 | benign | 1 | 1 | 1 |

| | | | | |
|---|---|---|---|---|
| rs1805044 | benign | 0 | 0 | 0 |
| rs117078377 | benign | 0 | 0 | 0 |
| rs138132026 | benign | 1 | 1 | 1 |
| rs75762896 | benign | 1 | 1 | 0 |
| rs121918043 | pathogenic | 1 | 1 | 0 |
| rs143581690 | benign | 1 | 1 | 0 |
| rs147654263 | benign | 0 | 0 | 0 |
| rs587782529 | pathogenic | 1 | 1 | 1 |
| rs121912664 | pathogenic | 1 | 1 | 1 |
| rs150293825 | benign | 1 | 1 | 1 |
| rs17882252 | benign | 1 | 1 | 1 |
| rs121912662 | pathogenic | 1 | 1 | 1 |
| rs35993958 | benign | 1 | 1 | 1 |
| rs606231253 | pathogenic | 1 | 1 | 1 |
| rs2227983 | benign | 1 | 1 | 1 |
| rs587779351 | pathogenic | 0 | 0 | 0 |
| rs779773463 | pathogenic | 1 | 1 | 1 |
| rs713040 | benign | 1 | 1 | 1 |
| rs334 | pathogenic | 1 | 1 | 1 |
| rs33930165 | pathogenic | 1 | 1 | 1 |
| rs33918131 | benign | 1 | 1 | 1 |
| rs63750783 | pathogenic | 1 | 1 | 1 |
| rs33986703 | pathogenic | 1 | 1 | 1 |
| rs33972047 | pathogenic | 1 | 1 | 1 |
| rs33936254 | pathogenic | 1 | 1 | 1 |
| rs33959855 | benign | 1 | 1 | 1 |
| rs33950507 | pathogenic | 1 | 1 | 1 |
| rs33916412 | pathogenic | 1 | 1 | 0 |
| rs1135071 | pathogenic | 1 | 1 | 1 |
| rs33960103 | pathogenic | 1 | 1 | 1 |
| rs11549407 | pathogenic | 1 | 1 | 1 |
| rs33922842 | pathogenic | 1 | 1 | 1 |
| rs35278874 | benign | 1 | 1 | 1 |
| rs34621955 | benign | 1 | 1 | 1 |
| rs33995148 | pathogenic | 1 | 1 | 1 |
| rs33945705 | benign | 1 | 1 | 1 |
| rs33993568 | benign | 1 | 1 | 1 |
| rs35553496 | benign | 1 | 1 | 1 |
| rs33913712 | pathogenic | 1 | 1 | 1 |
| rs33917785 | pathogenic | 1 | 1 | 1 |
| rs34515413 | benign | 1 | 1 | 1 |
| rs35256489 | pathogenic | 1 | 1 | 0 |
| rs34484056 | pathogenic | 1 | 1 | 1 |
| rs33947020 | benign | 1 | 1 | 1 |
| rs33946267 | pathogenic | 1 | 1 | 1 |
| rs33966761 | pathogenic | 1 | 1 | 1 |
| rs36020563 | benign | 1 | 1 | 1 |
| rs122454124 | pathogenic | 1 | 1 | 1 |
| rs122454126 | pathogenic | 1 | 1 | 0 |
| rs122454127 | pathogenic | 1 | 1 | 1 |
| rs122454130 | pathogenic | 1 | 1 | 1 |
| rs122454125 | pathogenic | 1 | 1 | 1 |
| rs122454131 | pathogenic | 1 | 1 | 0 |
| rs121908302 | pathogenic | 0 | 1 | 0 |

| | | | | |
|---|---|---|---|---|
| rs77829017 | pathogenic | 0 | 1 | 0 |
| rs1141814 | pathogenic | 0 | 1 | 0 |
| rs121908312 | pathogenic | 0 | 1 | 0 |
| rs79653797 | pathogenic | 1 | 1 | 0 |
| rs439898 | pathogenic | 1 | 1 | 0 |
| rs121908299 | pathogenic | 0 | 1 | 0 |
| rs80356763 | pathogenic | 1 | 1 | 1 |
| rs781152868 | pathogenic | 0 | 1 | 0 |
| rs121908297 | pathogenic | 0 | 1 | 0 |
| rs398123531 | benign | 0 | 1 | 0 |
| rs398123532 | pathogenic | 0 | 1 | 0 |
| rs364897 | pathogenic | 1 | 1 | 1 |
| rs1064644 | pathogenic | 0 | 1 | 0 |
| rs409652 | pathogenic | 1 | 1 | 1 |
| rs121908300 | pathogenic | 0 | 1 | 0 |
| rs381737 | pathogenic | 0 | 1 | 0 |
| rs74500255 | pathogenic | 0 | 1 | 0 |
| rs121908313 | pathogenic | 0 | 1 | 0 |
| rs367968666 | pathogenic | 0 | 1 | 0 |
| rs78973108 | pathogenic | 1 | 1 | 1 |
| rs121908298 | pathogenic | 1 | 1 | 1 |
| rs78396650 | pathogenic | 0 | 0 | 0 |
| rs78198234 | pathogenic | 1 | 1 | 0 |
| rs121908304 | pathogenic | 1 | 1 | 1 |
| rs76539814 | pathogenic | 1 | 1 | 1 |
| rs121908305 | pathogenic | 0 | 1 | 0 |
| rs121908306 | pathogenic | 1 | 1 | 0 |
| rs121908308 | pathogenic | 1 | 1 | 1 |
| rs121908307 | pathogenic | 0 | 1 | 0 |
| rs75548401 | benign | 0 | 1 | 0 |
| rs76763715 | pathogenic | 0 | 0 | 0 |
| rs121908314 | pathogenic | 0 | 1 | 0 |
| rs398123528 | pathogenic | 0 | 1 | 0 |
| rs121908311 | pathogenic | 0 | 0 | 0 |
| rs121908310 | pathogenic | 1 | 1 | 0 |
| rs74598136 | pathogenic | 0 | 1 | 0 |
| rs1064651 | pathogenic | 1 | 1 | 1 |
| rs77369218 | pathogenic | 1 | 1 | 1 |
| rs121908295 | pathogenic | 0 | 1 | 0 |
| rs421016 | pathogenic | 0 | 1 | 0 |
| rs368060 | pathogenic | 0 | 1 | 0 |
| rs80356771 | pathogenic | 1 | 1 | 1 |
| rs80356772 | pathogenic | 1 | 1 | 1 |
| rs121908301 | pathogenic | 0 | 0 | 0 |
| rs747506979 | pathogenic | 0 | 1 | 0 |
| rs75822236 | pathogenic | 0 | 1 | 0 |
| rs1990760 | benign | 1 | 1 | 0 |
| rs28934568 | pathogenic | 1 | 1 | 0 |
| rs34833812 | benign | 1 | 1 | 0 |
| rs193922666 | benign | 1 | 1 | 0 |
| rs104893812 | pathogenic | 1 | 1 | 0 |
| rs144701411 | benign | 0 | 0 | 0 |
| rs104893813 | pathogenic | 0 | 1 | 0 |
| rs199660234 | pathogenic | 1 | 1 | 0 |

| | | | | |
|---|---|---|---|---|
| rs104893814 | pathogenic | 1 | 1 | 0 |
| rs143095746 | benign | 1 | 1 | 1 |
| rs104893807 | pathogenic | 1 | 1 | 0 |
| rs104893811 | pathogenic | 0 | 0 | 0 |
| rs104893816 | pathogenic | 0 | 0 | 0 |
| rs121918714 | pathogenic | 0 | 1 | 0 |
| rs104893810 | pathogenic | 1 | 1 | 0 |
| rs104893815 | pathogenic | 1 | 1 | 0 |
| rs4303 | benign | 0 | 1 | 0 |
| rs121912704 | pathogenic | 0 | 1 | 0 |
| rs56394458 | benign | 1 | 1 | 0 |
| rs397514688 | pathogenic | 0 | 0 | 0 |
| rs367797185 | pathogenic | 1 | 1 | 0 |
| rs12709426 | benign | 0 | 1 | 0 |
| rs2228547 | benign | 0 | 0 | 0 |
| rs111033546 | pathogenic | 0 | 0 | 0 |
| rs111033553 | pathogenic | 0 | 0 | 0 |
| rs111033554 | pathogenic | 1 | 1 | 1 |
| rs111033544 | pathogenic | 1 | 1 | 1 |
| rs111033555 | pathogenic | 1 | 1 | 1 |
| rs111033545 | pathogenic | 1 | 1 | 1 |
| rs111033549 | pathogenic | 0 | 1 | 0 |
| rs111033552 | pathogenic | 1 | 1 | 1 |
| rs752204035 | benign | 0 | 0 | 0 |
| rs11569534 | benign | 0 | 0 | 0 |
| rs28937571 | pathogenic | 1 | 1 | 1 |
| rs104894215 | pathogenic | 1 | 1 | 1 |
| rs3888798 | benign | 1 | 1 | 1 |
| rs764263034 | benign | 1 | 1 | 1 |
| rs1800973 | benign | 1 | 1 | 1 |
| rs367623154 | benign | 1 | 1 | 1 |
| rs201230446 | pathogenic | 1 | 1 | 1 |
| rs267606766 | pathogenic | 0 | 1 | 0 |
| rs267606769 | pathogenic | 0 | 1 | 0 |
| rs267606767 | pathogenic | 1 | 1 | 0 |
| rs267606768 | pathogenic | 0 | 1 | 0 |
| rs61733129 | benign | 0 | 1 | 0 |
| rs201947120 | pathogenic | 1 | 1 | 1 |
| rs283413 | benign | 0 | 1 | 0 |
| rs36119840 | benign | 0 | 0 | 0 |
| rs281865419 | pathogenic | 1 | 1 | 1 |
| rs137852937 | pathogenic | 0 | 1 | 0 |
| rs2230709 | benign | 0 | 1 | 0 |
| rs6695033 | benign | 0 | 1 | 0 |
| rs180177032 | pathogenic | 1 | 1 | 1 |
| rs180177033 | pathogenic | 1 | 1 | 1 |
| rs121913351 | pathogenic | 1 | 1 | 1 |
| rs397507473 | pathogenic | 1 | 1 | 1 |
| rs121913355 | pathogenic | 1 | 1 | 1 |
| rs121913357 | pathogenic | 1 | 1 | 1 |
| rs180177036 | pathogenic | 1 | 1 | 1 |
| rs180177039 | pathogenic | 1 | 1 | 1 |
| rs180177038 | pathogenic | 1 | 1 | 1 |
| rs180177040 | pathogenic | 1 | 1 | 1 |

| | | | | |
|---|---|---|---|---|
| rs121913338 | pathogenic | 1 | 1 | 1 |
| rs121913366 | pathogenic | 1 | 1 | 1 |
| rs121913369 | pathogenic | 1 | 1 | 1 |
| rs113488022 | pathogenic | 1 | 1 | 1 |
| rs121913364 | pathogenic | 1 | 1 | 1 |
| rs180177042 | pathogenic | 1 | 1 | 1 |
| rs200461617 | benign | 0 | 0 | 0 |
| rs141609745 | benign | 0 | 0 | 0 |
| rs104894383 | pathogenic | 1 | 1 | 1 |
| rs104894384 | pathogenic | 1 | 1 | 1 |
| rs104894381 | pathogenic | 1 | 1 | 0 |
| rs147710408 | benign | 1 | 1 | 1 |
| rs77357563 | benign | 1 | 1 | 1 |
| rs104894378 | pathogenic | 0 | 0 | 0 |
| rs104894382 | pathogenic | 0 | 0 | 0 |
| rs137853292 | pathogenic | 1 | 1 | 1 |
| rs587778834 | pathogenic | 0 | 0 | 0 |
| rs137853294 | pathogenic | 1 | 1 | 1 |
| rs551747882 | benign | 0 | 0 | 0 |
| rs3092903 | benign | 1 | 1 | 1 |
| rs137853296 | pathogenic | 0 | 0 | 0 |
| rs398122917 | pathogenic | 0 | 1 | 0 |
| rs41304992 | benign | 0 | 0 | 0 |
| rs121912708 | pathogenic | 1 | 1 | 1 |
| rs140102105 | pathogenic | 1 | 1 | 1 |
| rs376917645 | pathogenic | 1 | 1 | 0 |
| rs372660425 | pathogenic | 1 | 1 | 0 |
| rs555896752 | pathogenic | 1 | 1 | 1 |
| rs777829351 | pathogenic | 1 | 1 | 0 |
| rs121912711 | pathogenic | 1 | 1 | 0 |
| rs754449549 | pathogenic | 0 | 1 | 0 |
| rs141701364 | benign | 0 | 1 | 0 |
| rs761011399 | benign | 1 | 1 | 1 |
| rs121912710 | benign | 1 | 1 | 0 |
| rs121912707 | pathogenic | 1 | 1 | 0 |
| rs747597620 | pathogenic | 0 | 1 | 0 |
| rs186558364 | pathogenic | 1 | 1 | 1 |
| rs556400964 | pathogenic | 1 | 1 | 1 |
| rs61757684 | benign | 1 | 1 | 1 |
| rs121917766 | pathogenic | 1 | 1 | 1 |
| rs121918248 | pathogenic | 0 | 0 | 0 |
| rs398123278 | pathogenic | 0 | 0 | 0 |
| rs121918251 | pathogenic | 1 | 1 | 1 |
| rs727504022 | pathogenic | 0 | 0 | 0 |
| rs190834116 | pathogenic | 1 | 1 | 1 |
| rs121918249 | pathogenic | 1 | 1 | 0 |
| rs121918257 | pathogenic | 1 | 1 | 0 |
| rs142068898 | benign | 0 | 0 | 0 |
| rs757000253 | pathogenic | 0 | 0 | 0 |
| rs200908035 | pathogenic | 1 | 1 | 0 |
| rs760782399 | pathogenic | 0 | 0 | 0 |
| rs778702777 | pathogenic | 0 | 0 | 0 |
| rs121918258 | pathogenic | 1 | 1 | 0 |
| rs121918256 | pathogenic | 0 | 0 | 0 |

| | | | | |
|---|---|---|---|---|
| rs200596762 | pathogenic | 1 | 1 | 0 |
| rs547709692 | pathogenic | 0 | 0 | 0 |
| rs758577372 | pathogenic | 0 | 1 | 0 |
| rs564069299 | pathogenic | 1 | 1 | 1 |
| rs772552898 | pathogenic | 1 | 1 | 1 |
| rs368790885 | pathogenic | 1 | 1 | 1 |
| rs121918250 | pathogenic | 0 | 0 | 0 |
| rs533755473 | pathogenic | 1 | 1 | 1 |
| rs753288303 | pathogenic | 1 | 1 | 1 |
| rs2229385 | benign | 0 | 0 | 0 |
| rs1141321 | benign | 0 | 0 | 0 |
| rs753564352 | pathogenic | 0 | 0 | 0 |
| rs775593146 | pathogenic | 0 | 0 | 0 |
| rs121918254 | pathogenic | 0 | 0 | 0 |
| rs8589 | benign | 1 | 1 | 0 |
| rs777758903 | pathogenic | 0 | 0 | 0 |
| rs140600746 | pathogenic | 1 | 1 | 0 |
| rs121918255 | pathogenic | 1 | 1 | 0 |
| rs121918252 | pathogenic | 0 | 0 | 0 |
| rs779990936 | pathogenic | 1 | 1 | 0 |
| rs121912455 | pathogenic | 1 | 1 | 1 |
| rs121912459 | pathogenic | 1 | 1 | 1 |
| rs74315452 | pathogenic | 1 | 1 | 1 |
| rs121912451 | pathogenic | 1 | 1 | 1 |
| rs79006549 | pathogenic | 0 | 0 | 0 |
| rs28934889 | benign | 0 | 1 | 0 |
| rs111033557 | benign | 1 | 1 | 1 |
| rs28934597 | pathogenic | 1 | 1 | 1 |
| rs28934596 | pathogenic | 1 | 1 | 1 |
| rs28934595 | pathogenic | 1 | 1 | 1 |
| rs146519482 | pathogenic | 1 | 1 | 1 |
| rs111033563 | pathogenic | 1 | 1 | 1 |
| rs75385605 | benign | 1 | 1 | 1 |
| rs121908569 | pathogenic | 1 | 1 | 1 |
| rs35719940 | benign | 0 | 0 | 0 |
| rs121918664 | pathogenic | 0 | 0 | 0 |
| rs104893665 | pathogenic | 1 | 1 | 1 |
| rs104893666 | pathogenic | 1 | 1 | 1 |
| rs200845457 | benign | 1 | 1 | 1 |
| rs779204655 | pathogenic | 1 | 1 | 1 |
| rs121917747 | pathogenic | 1 | 1 | 1 |
| rs6042 | benign | 1 | 1 | 1 |
| rs104894380 | pathogenic | 0 | 0 | 0 |
| rs369083541 | pathogenic | 1 | 1 | 1 |
| rs145243532 | benign | 1 | 1 | 1 |
| rs121909500 | pathogenic | 1 | 1 | 1 |
| rs140993310 | benign | 1 | 1 | 1 |
| rs200837270 | pathogenic | 1 | 1 | 1 |
| rs121918484 | pathogenic | 1 | 1 | 1 |
| rs121918306 | pathogenic | 0 | 0 | 0 |
| rs118204089 | pathogenic | 1 | 1 | 0 |
| rs118204083 | pathogenic | 1 | 1 | 0 |
| rs201953584 | pathogenic | 1 | 1 | 1 |
| rs74315362 | pathogenic | 1 | 1 | 1 |

| | | | | |
|---|---|---|---|---|
| rs118204084 | pathogenic | 1 | 1 | 0 |
| rs118204085 | pathogenic | 1 | 1 | 1 |
| rs113403872 | pathogenic | 1 | 1 | 0 |
| rs1052176 | benign | 1 | 1 | 1 |
| rs2031640 | benign | 0 | 0 | 0 |
| rs104894230 | pathogenic | 1 | 1 | 1 |
| rs104894229 | pathogenic | 1 | 1 | 1 |
| rs104894228 | pathogenic | 1 | 1 | 1 |
| rs104894226 | pathogenic | 1 | 1 | 1 |
| rs121917757 | pathogenic | 1 | 1 | 1 |
| rs121917758 | pathogenic | 1 | 1 | 1 |
| rs28933406 | pathogenic | 1 | 1 | 1 |
| rs121917756 | pathogenic | 1 | 1 | 1 |
| rs104894227 | pathogenic | 1 | 1 | 1 |
| rs369106578 | pathogenic | 1 | 1 | 1 |
| rs397517142 | benign | 1 | 1 | 1 |
| rs104894231 | pathogenic | 1 | 1 | 1 |
| rs121917759 | pathogenic | 1 | 1 | 1 |
| rs104893827 | pathogenic | 0 | 0 | 0 |
| rs104893826 | pathogenic | 1 | 1 | 1 |
| rs5030802 | pathogenic | 1 | 1 | 1 |
| rs5030827 | pathogenic | 1 | 1 | 0 |
| rs119103277 | pathogenic | 1 | 1 | 1 |
| rs5030808 | pathogenic | 0 | 1 | 0 |
| rs5030809 | pathogenic | 1 | 1 | 1 |
| rs765978945 | pathogenic | 1 | 1 | 1 |
| rs104893824 | pathogenic | 1 | 1 | 1 |
| rs104893831 | pathogenic | 0 | 1 | 0 |
| rs104893830 | pathogenic | 0 | 1 | 0 |
| rs119103278 | pathogenic | 0 | 1 | 0 |
| rs5030818 | pathogenic | 1 | 1 | 1 |
| rs397516444 | pathogenic | 1 | 1 | 1 |
| rs28940297 | pathogenic | 1 | 1 | 1 |
| rs267607170 | pathogenic | 1 | 1 | 1 |
| rs104893825 | pathogenic | 1 | 1 | 1 |
| rs5030820 | pathogenic | 0 | 1 | 0 |
| rs5030821 | pathogenic | 0 | 1 | 0 |
| rs5030823 | pathogenic | 1 | 1 | 1 |
| rs5030824 | pathogenic | 1 | 1 | 1 |
| rs28940301 | pathogenic | 0 | 1 | 0 |
| rs28940300 | pathogenic | 0 | 1 | 0 |
| rs28940298 | pathogenic | 0 | 1 | 0 |
| rs200019083 | benign | 0 | 0 | 0 |
| rs759748655 | pathogenic | 0 | 0 | 0 |
| rs121917739 | pathogenic | 1 | 1 | 1 |
| rs75353611 | pathogenic | 1 | 1 | 1 |
| rs79744198 | benign | 1 | 1 | 1 |
| rs79228041 | pathogenic | 1 | 1 | 1 |
| rs80106970 | benign | 1 | 1 | 1 |
| rs375353223 | benign | 0 | 0 | 0 |
| rs56311081 | benign | 0 | 0 | 0 |
| rs55985569 | benign | 1 | 1 | 1 |
| rs45483396 | benign | 0 | 0 | 0 |
| rs35284565 | benign | 1 | 1 | 1 |

| | | | | |
|---|---|---|---|---|
| rs34349517 | benign | 0 | 0 | 0 |
| rs35776110 | benign | 0 | 0 | 0 |
| rs33917957 | benign | 1 | 1 | 1 |
| rs587778445 | benign | 0 | 0 | 0 |
| rs34624984 | benign | 0 | 0 | 0 |
| rs701753 | benign | 0 | 0 | 0 |
| rs855314 | benign | 0 | 0 | 0 |
| rs121918371 | pathogenic | 1 | 1 | 0 |
| rs398122912 | pathogenic | 0 | 1 | 0 |
| rs1126728 | benign | 0 | 0 | 0 |
| rs587777404 | pathogenic | 1 | 1 | 0 |
| rs11208257 | benign | 0 | 0 | 0 |
| rs6676290 | benign | 0 | 0 | 0 |
| rs104894118 | pathogenic | 0 | 0 | 0 |
| rs121918655 | pathogenic | 1 | 1 | 1 |
| rs104894120 | pathogenic | 1 | 1 | 1 |
| rs137852238 | pathogenic | 0 | 0 | 0 |
| rs137852237 | pathogenic | 0 | 0 | 0 |
| rs137852499 | pathogenic | 1 | 1 | 1 |
| rs137852480 | pathogenic | 1 | 1 | 1 |
| rs137852494 | pathogenic | 1 | 1 | 1 |
| rs137852502 | pathogenic | 1 | 1 | 1 |
| rs137852500 | pathogenic | 1 | 1 | 1 |
| rs137852487 | pathogenic | 1 | 1 | 1 |
| rs137852481 | pathogenic | 1 | 1 | 1 |
| rs137852501 | pathogenic | 0 | 0 | 0 |
| rs137852478 | pathogenic | 1 | 1 | 1 |
| rs137852485 | pathogenic | 1 | 1 | 1 |
| rs137852482 | pathogenic | 1 | 1 | 0 |
| rs137852483 | pathogenic | 0 | 1 | 0 |
| rs137852477 | pathogenic | 1 | 1 | 0 |
| rs137852496 | pathogenic | 1 | 1 | 0 |
| rs137852484 | pathogenic | 0 | 1 | 0 |
| rs137852498 | pathogenic | 1 | 1 | 0 |
| rs137852497 | pathogenic | 1 | 1 | 0 |
| rs137852493 | pathogenic | 1 | 1 | 0 |
| rs137852504 | pathogenic | 1 | 1 | 1 |
| rs267606863 | pathogenic | 1 | 1 | 1 |
| rs137852486 | pathogenic | 1 | 1 | 1 |
| rs137852479 | pathogenic | 1 | 1 | 1 |
| rs112122950 | benign | 0 | 0 | 0 |
| rs72546648 | benign | 1 | 1 | 1 |
| rs72552291 | pathogenic | 0 | 0 | 0 |
| rs79184941 | pathogenic | 1 | 1 | 1 |
| rs77543610 | pathogenic | 1 | 1 | 1 |
| rs121918505 | pathogenic | 1 | 1 | 1 |
| rs776587763 | pathogenic | 1 | 1 | 1 |
| rs121918497 | pathogenic | 1 | 1 | 1 |
| rs121918499 | pathogenic | 1 | 1 | 1 |
| rs121918501 | pathogenic | 1 | 1 | 1 |
| rs121918500 | pathogenic | 0 | 0 | 0 |
| rs121918504 | pathogenic | 1 | 1 | 1 |
| rs121918510 | pathogenic | 1 | 1 | 1 |
| rs121918493 | pathogenic | 1 | 1 | 1 |

| | | | | |
|---|---|---|---|---|
| rs121918489 | pathogenic | 0 | 0 | 0 |
| rs121918495 | pathogenic | 1 | 1 | 1 |
| rs121918488 | pathogenic | 0 | 0 | 0 |
| rs121918496 | pathogenic | 0 | 0 | 0 |
| rs121918492 | pathogenic | 1 | 1 | 1 |
| rs121918494 | pathogenic | 1 | 1 | 1 |
| rs121918502 | pathogenic | 1 | 1 | 1 |
| rs121918490 | pathogenic | 1 | 1 | 1 |
| rs72466451 | pathogenic | 1 | 1 | 1 |
| rs66468541 | pathogenic | 0 | 0 | 0 |
| rs137852662 | pathogenic | 0 | 0 | 0 |
| rs137852663 | pathogenic | 0 | 0 | 0 |
| rs606231407 | pathogenic | 0 | 0 | 0 |
| rs137852233 | pathogenic | 1 | 1 | 1 |
| rs112306225 | benign | 0 | 0 | 0 |
| rs387906808 | pathogenic | 1 | 1 | 0 |
| rs730882094 | pathogenic | 1 | 1 | 1 |
| rs373869746 | pathogenic | 1 | 1 | 1 |
| rs777524402 | pathogenic | 1 | 1 | 1 |
| rs769737896 | pathogenic | 0 | 1 | 0 |
| rs13306515 | benign | 1 | 1 | 0 |
| rs746982741 | pathogenic | 0 | 0 | 0 |
| rs2230200 | benign | 1 | 1 | 1 |
| rs1047286 | benign | 0 | 0 | 0 |
| rs141737564 | benign | 0 | 0 | 0 |
| rs774826179 | benign | 0 | 0 | 0 |
| rs11569422 | benign | 1 | 1 | 1 |
| rs146613648 | benign | 0 | 0 | 0 |
| rs144432231 | benign | 0 | 0 | 0 |
| rs140655115 | benign | 0 | 0 | 0 |
| rs387906837 | pathogenic | 1 | 1 | 0 |
| rs121909076 | pathogenic | 1 | 1 | 0 |
| rs121909077 | pathogenic | 0 | 0 | 0 |
| rs121909074 | pathogenic | 0 | 0 | 0 |
| rs137853330 | pathogenic | 1 | 1 | 1 |
| rs137853331 | pathogenic | 1 | 1 | 1 |
| rs137853332 | pathogenic | 1 | 1 | 1 |
| rs28931586 | pathogenic | 1 | 1 | 0 |
| rs11569321 | benign | 1 | 1 | 1 |
| rs119486097 | benign | 0 | 0 | 0 |
| rs1799807 | pathogenic | 1 | 1 | 1 |
| rs201820739 | pathogenic | 1 | 1 | 0 |
| rs114706984 | pathogenic | 1 | 1 | 0 |
| rs140080572 | pathogenic | 1 | 1 | 0 |
| rs28933389 | pathogenic | 0 | 1 | 0 |
| rs16849700 | benign | 1 | 1 | 0 |
| rs115624085 | pathogenic | 1 | 1 | 1 |
| rs747196387 | pathogenic | 1 | 1 | 1 |
| rs104893684 | pathogenic | 1 | 1 | 0 |
| rs121918557 | pathogenic | 1 | 1 | 0 |
| rs121918556 | pathogenic | 1 | 1 | 1 |
| rs398124571 | pathogenic | 1 | 1 | 1 |
| rs776631396 | pathogenic | 0 | 1 | 0 |
| rs398124576 | pathogenic | 1 | 1 | 1 |

| | | | | |
|---|---|---|---|---|
| rs398124581 | pathogenic | 1 | 1 | 1 |
| rs398124582 | pathogenic | 1 | 1 | 1 |
| rs149766077 | pathogenic | 0 | 1 | 0 |
| rs79761867 | pathogenic | 0 | 1 | 0 |
| rs148905512 | pathogenic | 0 | 1 | 0 |
| rs121965004 | pathogenic | 1 | 1 | 1 |
| rs386834233 | pathogenic | 0 | 1 | 0 |
| rs398124598 | pathogenic | 0 | 1 | 0 |
| rs398124603 | pathogenic | 1 | 1 | 1 |
| rs546716073 | benign | 1 | 1 | 1 |
| rs200310662 | benign | 1 | 1 | 1 |
| rs115494727 | benign | 1 | 1 | 1 |
| rs781513008 | benign | 0 | 0 | 0 |
| rs587776398 | benign | 1 | 1 | 0 |
| rs35288908 | benign | 1 | 1 | 1 |
| rs150634562 | pathogenic | 1 | 1 | 0 |
| rs142948132 | pathogenic | 1 | 1 | 1 |
| rs132630321 | pathogenic | 1 | 1 | 1 |
| rs132630317 | pathogenic | 1 | 1 | 1 |
| rs132630320 | pathogenic | 0 | 0 | 0 |
| rs781394318 | pathogenic | 1 | 1 | 1 |
| rs111033624 | pathogenic | 1 | 1 | 1 |
| rs111033630 | pathogenic | 1 | 1 | 1 |
| rs111033627 | pathogenic | 1 | 1 | 1 |
| rs193922287 | pathogenic | 1 | 1 | 1 |
| rs759072800 | pathogenic | 0 | 1 | 0 |
| rs104894010 | pathogenic | 0 | 0 | 0 |
| rs193922296 | pathogenic | 0 | 0 | 0 |
| rs104894006 | pathogenic | 1 | 1 | 1 |
| rs193922311 | pathogenic | 1 | 1 | 0 |
| rs80356654 | pathogenic | 1 | 1 | 0 |
| rs144723656 | benign | 1 | 1 | 1 |
| rs148311934 | pathogenic | 1 | 1 | 0 |
| rs80356655 | pathogenic | 1 | 1 | 1 |
| rs769268803 | pathogenic | 1 | 1 | 0 |
| rs104894008 | pathogenic | 0 | 1 | 0 |
| rs104894005 | pathogenic | 1 | 1 | 1 |
| rs104894009 | pathogenic | 1 | 1 | 0 |
| rs193922340 | pathogenic | 0 | 0 | 0 |
| rs104894016 | pathogenic | 0 | 1 | 0 |
| rs193921338 | pathogenic | 0 | 0 | 0 |
| rs193922272 | pathogenic | 1 | 1 | 1 |
| rs104894012 | pathogenic | 1 | 1 | 0 |
| rs193922285 | pathogenic | 1 | 1 | 1 |
| rs104894336 | pathogenic | 1 | 1 | 1 |
| rs28931580 | pathogenic | 1 | 1 | 1 |
| rs104894326 | pathogenic | 1 | 1 | 1 |
| rs104894331 | pathogenic | 0 | 0 | 0 |
| rs104894338 | pathogenic | 0 | 0 | 0 |
| rs104894333 | pathogenic | 1 | 1 | 1 |
| rs104894330 | pathogenic | 1 | 1 | 1 |
| rs143886391 | benign | 1 | 1 | 1 |
| rs104894335 | pathogenic | 1 | 1 | 1 |
| rs104894337 | pathogenic | 0 | 0 | 0 |

| | | | | |
|---|---|---|---|---|
| rs104894328 | pathogenic | 0 | 0 | 0 |
| rs104894341 | pathogenic | 1 | 1 | 0 |
| rs104894329 | pathogenic | 0 | 0 | 0 |
| rs137852850 | pathogenic | 1 | 1 | 1 |
| rs137852854 | pathogenic | 0 | 1 | 0 |
| rs137852849 | pathogenic | 0 | 1 | 0 |
| rs137852848 | pathogenic | 1 | 1 | 1 |
| rs137852852 | pathogenic | 0 | 1 | 0 |
| rs137852853 | pathogenic | 0 | 1 | 0 |
| rs137852847 | pathogenic | 0 | 1 | 0 |
| rs137852846 | pathogenic | 0 | 1 | 0 |
| rs137852844 | pathogenic | 0 | 1 | 0 |
| rs144411158 | pathogenic | 0 | 0 | 0 |
| rs2302234 | benign | 0 | 0 | 0 |
| rs139620139 | pathogenic | 0 | 0 | 0 |
| rs146689929 | benign | 0 | 0 | 0 |
| rs373328681 | pathogenic | 1 | 1 | 0 |
| rs387906620 | pathogenic | 1 | 1 | 1 |
| rs1799983 | benign | 0 | 1 | 0 |
| rs28999111 | pathogenic | 1 | 1 | 1 |
| rs104894544 | pathogenic | 1 | 1 | 1 |
| rs104894546 | pathogenic | 1 | 1 | 1 |
| rs28999112 | pathogenic | 0 | 1 | 0 |
| rs104894545 | pathogenic | 1 | 1 | 1 |
| rs104894547 | pathogenic | 1 | 1 | 1 |
| rs74315310 | pathogenic | 0 | 0 | 0 |
| rs267606852 | pathogenic | 0 | 1 | 0 |
| rs137853586 | pathogenic | 0 | 1 | 0 |
| rs137853582 | pathogenic | 1 | 1 | 1 |
| rs61754634 | pathogenic | 1 | 1 | 1 |
| rs137853587 | pathogenic | 0 | 1 | 0 |
| rs267606851 | pathogenic | 1 | 1 | 1 |
| rs267606853 | pathogenic | 1 | 1 | 1 |
| rs137853584 | pathogenic | 1 | 1 | 1 |
| rs137853585 | pathogenic | 1 | 1 | 1 |
| rs460184 | pathogenic | 1 | 1 | 1 |
| rs104893751 | pathogenic | 0 | 1 | 0 |
| rs77358162 | benign | 0 | 0 | 0 |
| rs387906936 | pathogenic | 0 | 0 | 0 |
| rs769525399 | pathogenic | 0 | 0 | 0 |
| rs371271054 | pathogenic | 1 | 1 | 0 |
| rs118204000 | pathogenic | 0 | 0 | 0 |
| rs138568622 | pathogenic | 1 | 1 | 1 |
| rs34509359 | benign | 0 | 0 | 0 |
| rs104894130 | pathogenic | 1 | 1 | 0 |
| rs281865424 | pathogenic | 0 | 0 | 0 |
| rs121912778 | pathogenic | 1 | 1 | 0 |
| rs771055145 | pathogenic | 1 | 1 | 0 |
| rs776174514 | pathogenic | 1 | 1 | 0 |
| rs545986367 | pathogenic | 1 | 1 | 1 |
| rs570768038 | benign | 1 | 1 | 1 |
| rs74315442 | pathogenic | 1 | 1 | 1 |
| rs79716074 | benign | 1 | 1 | 1 |
| rs748486078 | pathogenic | 1 | 1 | 1 |

| | | | | |
|---|---|---|---|---|
| rs2397084 | benign | 1 | 1 | 1 |
| rs11465553 | benign | 1 | 1 | 1 |
| rs144576902 | benign | 1 | 1 | 1 |
| rs763780 | benign | 1 | 1 | 1 |
| rs267606870 | pathogenic | 1 | 1 | 0 |
| rs121913502 | pathogenic | 1 | 1 | 0 |
| rs118101777 | benign | 1 | 1 | 1 |
| rs118053940 | benign | 0 | 0 | 0 |
| rs368771578 | benign | 1 | 1 | 1 |
| rs11540654 | pathogenic | 1 | 1 | 1 |
| rs121912658 | pathogenic | 1 | 1 | 1 |
| rs28934873 | pathogenic | 0 | 1 | 0 |
| rs780442292 | pathogenic | 1 | 1 | 1 |
| rs758781593 | benign | 1 | 1 | 1 |
| rs28934875 | pathogenic | 1 | 1 | 1 |
| rs28934874 | pathogenic | 1 | 1 | 0 |
| rs587782705 | pathogenic | 1 | 1 | 1 |
| rs587780068 | pathogenic | 1 | 1 | 1 |
| rs587782144 | pathogenic | 1 | 1 | 1 |
| rs138729528 | pathogenic | 1 | 1 | 1 |
| rs28934578 | pathogenic | 1 | 1 | 1 |
| rs397514495 | pathogenic | 1 | 1 | 1 |
| rs121912665 | pathogenic | 1 | 1 | 1 |
| rs397516435 | pathogenic | 1 | 1 | 1 |
| rs587778720 | pathogenic | 1 | 1 | 1 |
| rs397516436 | pathogenic | 1 | 1 | 1 |
| rs587781386 | benign | 1 | 1 | 1 |
| rs121912666 | pathogenic | 1 | 1 | 0 |
| rs587780073 | pathogenic | 1 | 1 | 1 |
| rs587782664 | pathogenic | 1 | 1 | 1 |
| rs730882005 | pathogenic | 1 | 1 | 1 |
| rs28934573 | pathogenic | 1 | 1 | 1 |
| rs121912655 | pathogenic | 1 | 1 | 1 |
| rs28934575 | pathogenic | 1 | 1 | 1 |
| rs121912656 | pathogenic | 1 | 1 | 1 |
| rs11540652 | pathogenic | 1 | 1 | 1 |
| rs121912651 | pathogenic | 1 | 1 | 1 |
| rs730882027 | pathogenic | 1 | 1 | 1 |
| rs121912653 | pathogenic | 1 | 1 | 1 |
| rs121912652 | pathogenic | 1 | 1 | 1 |
| rs121912657 | pathogenic | 0 | 1 | 0 |
| rs121913343 | pathogenic | 1 | 1 | 1 |
| rs28934576 | pathogenic | 1 | 1 | 1 |
| rs763098116 | pathogenic | 1 | 1 | 1 |
| rs121912660 | pathogenic | 1 | 1 | 1 |
| rs28934574 | pathogenic | 1 | 1 | 1 |
| rs371409680 | pathogenic | 1 | 1 | 1 |
| rs121912667 | pathogenic | 1 | 1 | 1 |
| rs121912663 | pathogenic | 1 | 1 | 1 |
| rs201744589 | pathogenic | 0 | 0 | 0 |
| rs33926449 | benign | 0 | 0 | 0 |
| rs74315302 | pathogenic | 0 | 1 | 0 |
| rs202040269 | pathogenic | 1 | 1 | 1 |
| rs375958814 | pathogenic | 1 | 1 | 1 |

| | | | | |
|---|---|---|---|---|
| rs121909624 | pathogenic | 1 | 1 | 1 |
| rs121909621 | pathogenic | 1 | 1 | 1 |
| rs121909622 | pathogenic | 1 | 1 | 0 |
| rs4220 | benign | 1 | 1 | 1 |
| rs147559466 | pathogenic | 1 | 1 | 1 |
| rs757434857 | pathogenic | 1 | 1 | 1 |
| rs398123072 | pathogenic | 1 | 1 | 1 |
| rs121434280 | pathogenic | 0 | 1 | 0 |
| rs398123074 | pathogenic | 0 | 1 | 0 |
| rs762114560 | pathogenic | 1 | 1 | 1 |
| rs121434283 | pathogenic | 1 | 1 | 0 |
| rs121434277 | pathogenic | 0 | 1 | 0 |
| rs745844469 | pathogenic | 1 | 1 | 1 |
| rs121434279 | pathogenic | 1 | 1 | 0 |
| rs121434278 | pathogenic | 1 | 1 | 1 |
| rs373715782 | pathogenic | 1 | 1 | 1 |
| rs200724875 | pathogenic | 1 | 1 | 1 |
| rs149678400 | pathogenic | 0 | 1 | 0 |
| rs770273135 | pathogenic | 1 | 1 | 1 |
| rs121434276 | pathogenic | 0 | 1 | 0 |
| rs121434281 | pathogenic | 1 | 1 | 0 |
| rs768884003 | pathogenic | 1 | 1 | 1 |
| rs201375579 | pathogenic | 0 | 0 | 0 |
| rs121434274 | pathogenic | 1 | 1 | 0 |
| rs121434282 | pathogenic | 1 | 1 | 0 |
| rs779759347 | pathogenic | 0 | 1 | 0 |
| rs77931234 | pathogenic | 1 | 1 | 1 |
| rs148207467 | pathogenic | 1 | 1 | 1 |
| rs766140986 | pathogenic | 1 | 1 | 1 |
| rs121434275 | pathogenic | 1 | 1 | 1 |
| rs748110745 | pathogenic | 1 | 1 | 1 |
| rs104894274 | pathogenic | 1 | 1 | 1 |
| rs104894273 | pathogenic | 0 | 1 | 0 |
| rs104894278 | pathogenic | 1 | 1 | 1 |
| rs104894275 | pathogenic | 1 | 1 | 1 |
| rs104894277 | pathogenic | 1 | 1 | 1 |
| rs104894276 | pathogenic | 0 | 0 | 0 |
| rs104894280 | pathogenic | 1 | 1 | 1 |
| rs145882709 | pathogenic | 1 | 1 | 1 |
| rs104894279 | pathogenic | 1 | 1 | 1 |
| rs80356670 | pathogenic | 1 | 1 | 1 |
| rs121918102 | pathogenic | 1 | 1 | 1 |
| rs80356671 | pathogenic | 1 | 1 | 1 |
| rs80356672 | pathogenic | 1 | 1 | 1 |
| rs758609113 | pathogenic | 1 | 1 | 0 |
| rs121918188 | pathogenic | 0 | 1 | 0 |
| rs121918189 | pathogenic | 1 | 1 | 0 |
| rs121918191 | pathogenic | 1 | 1 | 1 |
| rs1769259 | benign | 1 | 1 | 1 |
| rs140222720 | benign | 1 | 1 | 1 |
| rs2297085 | benign | 1 | 1 | 1 |
| rs200679026 | pathogenic | 1 | 1 | 1 |
| rs121918192 | pathogenic | 0 | 1 | 0 |
| rs145771944 | benign | 0 | 1 | 0 |

| | | | | |
|---|---|---|---|---|
| rs201508880 | pathogenic | 0 | 1 | 0 |
| rs17221854 | pathogenic | 0 | 0 | 0 |
| rs120074189 | pathogenic | 1 | 1 | 1 |
| rs120074190 | pathogenic | 1 | 1 | 1 |
| rs758600283 | benign | 1 | 1 | 1 |
| rs200305064 | pathogenic | 1 | 1 | 1 |
| rs121917891 | pathogenic | 1 | 1 | 1 |
| rs3772809 | benign | 0 | 1 | 0 |
| rs387906619 | pathogenic | 1 | 1 | 0 |
| rs2304681 | benign | 0 | 1 | 0 |
| rs121917849 | pathogenic | 0 | 0 | 0 |
| rs143010236 | pathogenic | 0 | 0 | 0 |
| rs121917850 | pathogenic | 1 | 1 | 0 |
| rs35916840 | pathogenic | 1 | 1 | 0 |
| rs28939688 | pathogenic | 1 | 1 | 1 |
| rs11258194 | benign | 1 | 1 | 1 |
| rs199840952 | pathogenic | 1 | 1 | 1 |
| rs121918159 | pathogenic | 1 | 1 | 1 |
| rs200396166 | pathogenic | 0 | 0 | 0 |
| rs757966746 | pathogenic | 0 | 1 | 0 |
| rs121918608 | pathogenic | 0 | 1 | 0 |
| rs74315401 | pathogenic | 1 | 1 | 1 |
| rs11538758 | pathogenic | 0 | 0 | 0 |
| rs74315402 | pathogenic | 1 | 1 | 1 |
| rs1799990 | pathogenic | 1 | 1 | 1 |
| rs74315410 | pathogenic | 1 | 1 | 1 |
| rs150351644 | benign | 1 | 1 | 1 |
| rs16990018 | benign | 1 | 1 | 1 |
| rs74315403 | pathogenic | 1 | 1 | 1 |
| rs74315408 | pathogenic | 1 | 1 | 1 |
| rs74315411 | pathogenic | 1 | 1 | 1 |
| rs74315413 | pathogenic | 1 | 1 | 1 |
| rs74315405 | pathogenic | 1 | 1 | 1 |
| rs28933385 | pathogenic | 1 | 1 | 1 |
| rs74315412 | pathogenic | 1 | 1 | 1 |
| rs74315407 | pathogenic | 1 | 1 | 1 |
| rs74315406 | pathogenic | 1 | 1 | 1 |
| rs11571117 | benign | 1 | 1 | 1 |
| rs121917742 | pathogenic | 1 | 1 | 1 |
| rs149782619 | pathogenic | 1 | 1 | 1 |
| rs397514492 | pathogenic | 1 | 1 | 1 |
| rs397514489 | pathogenic | 0 | 1 | 0 |
| rs397514493 | pathogenic | 1 | 1 | 1 |
| rs373849532 | pathogenic | 1 | 1 | 1 |
| rs121965048 | pathogenic | 1 | 1 | 1 |
| rs121965037 | pathogenic | 1 | 1 | 1 |
| rs121965060 | pathogenic | 0 | 1 | 0 |
| rs386833603 | pathogenic | 1 | 1 | 1 |
| rs121965038 | pathogenic | 1 | 1 | 1 |
| rs121965039 | pathogenic | 1 | 1 | 1 |
| rs121965040 | pathogenic | 1 | 1 | 1 |
| rs267606925 | pathogenic | 1 | 1 | 1 |
| rs121965056 | pathogenic | 0 | 1 | 0 |
| rs121965059 | pathogenic | 0 | 0 | 0 |

| | | | | |
|---|---|---|---|---|
| rs386833615 | pathogenic | 1 | 1 | 0 |
| rs121965051 | pathogenic | 1 | 1 | 0 |
| rs121965046 | pathogenic | 0 | 1 | 0 |
| rs121965052 | pathogenic | 0 | 1 | 0 |
| rs386833617 | pathogenic | 0 | 1 | 0 |
| rs386833618 | pathogenic | 1 | 1 | 0 |
| rs121965041 | pathogenic | 0 | 1 | 0 |
| rs121965042 | pathogenic | 0 | 1 | 0 |
| rs121965057 | pathogenic | 1 | 1 | 1 |
| rs386833621 | pathogenic | 1 | 1 | 1 |
| rs121965049 | pathogenic | 1 | 1 | 1 |
| rs386833623 | pathogenic | 1 | 1 | 1 |
| rs121965047 | pathogenic | 1 | 1 | 1 |
| rs121965053 | pathogenic | 0 | 1 | 0 |
| rs121965045 | pathogenic | 0 | 1 | 0 |
| rs121965054 | pathogenic | 0 | 1 | 0 |
| rs121965043 | pathogenic | 0 | 1 | 0 |
| rs386833598 | pathogenic | 0 | 0 | 0 |
| rs137854545 | pathogenic | 1 | 1 | 1 |
| rs137854544 | pathogenic | 1 | 1 | 1 |
| rs104894455 | pathogenic | 0 | 1 | 0 |
| rs1049564 | benign | 0 | 1 | 0 |
| rs104894460 | pathogenic | 1 | 1 | 1 |
| rs104894453 | pathogenic | 1 | 1 | 1 |
| rs104894450 | pathogenic | 0 | 1 | 0 |
| rs104894454 | pathogenic | 0 | 1 | 0 |
| rs104894452 | pathogenic | 1 | 1 | 1 |
| rs104894451 | pathogenic | 0 | 1 | 0 |
| rs104894368 | pathogenic | 1 | 1 | 1 |
| rs730880950 | pathogenic | 1 | 1 | 1 |
| rs121909535 | pathogenic | 1 | 1 | 1 |
| rs121909537 | pathogenic | 1 | 1 | 1 |
| rs121909536 | pathogenic | 1 | 1 | 1 |
| rs121909542 | pathogenic | 1 | 1 | 1 |
| rs121909538 | pathogenic | 1 | 1 | 1 |
| rs121909539 | pathogenic | 1 | 1 | 1 |
| rs121909540 | pathogenic | 1 | 1 | 1 |
| rs17560 | benign | 1 | 1 | 1 |
| rs121909543 | pathogenic | 0 | 1 | 0 |
| rs121909544 | pathogenic | 1 | 1 | 1 |
| rs2307456 | benign | 0 | 1 | 0 |
| rs745778317 | pathogenic | 0 | 1 | 0 |
| rs759607901 | pathogenic | 0 | 1 | 0 |
| rs201188114 | benign | 0 | 0 | 0 |
| rs1800472 | benign | 0 | 0 | 0 |
| rs587780728 | benign | 1 | 1 | 1 |
| rs144386518 | benign | 1 | 1 | 1 |
| rs281865439 | pathogenic | 1 | 1 | 1 |
| rs140175796 | pathogenic | 1 | 1 | 1 |
| rs5030843 | pathogenic | 1 | 1 | 0 |
| rs75166491 | pathogenic | 1 | 1 | 0 |
| rs199475679 | pathogenic | 1 | 1 | 1 |
| rs281865440 | pathogenic | 1 | 1 | 1 |
| rs74486803 | pathogenic | 0 | 1 | 0 |

| | | | | |
|---|---|---|---|---|
| rs199475602 | pathogenic | 0 | 1 | 0 |
| rs5030844 | pathogenic | 1 | 1 | 0 |
| rs62514927 | pathogenic | 1 | 1 | 1 |
| rs62514928 | pathogenic | 1 | 1 | 1 |
| rs62516152 | pathogenic | 0 | 1 | 0 |
| rs62507348 | pathogenic | 1 | 1 | 1 |
| rs76687508 | pathogenic | 1 | 1 | 1 |
| rs62508730 | pathogenic | 1 | 1 | 1 |
| rs62508588 | pathogenic | 0 | 1 | 0 |
| rs5030846 | pathogenic | 0 | 1 | 0 |
| rs74503222 | pathogenic | 1 | 1 | 0 |
| rs5030847 | pathogenic | 1 | 1 | 0 |
| rs5030850 | pathogenic | 1 | 1 | 1 |
| rs778154939 | pathogenic | 0 | 1 | 0 |
| rs62514952 | pathogenic | 1 | 1 | 1 |
| rs78655458 | pathogenic | 1 | 1 | 1 |
| rs62508698 | pathogenic | 1 | 1 | 0 |
| rs5030851 | pathogenic | 1 | 1 | 0 |
| rs62642933 | pathogenic | 1 | 1 | 1 |
| rs5030853 | pathogenic | 1 | 1 | 1 |
| rs62642935 | pathogenic | 1 | 1 | 1 |
| rs199475615 | pathogenic | 0 | 0 | 0 |
| rs62516062 | pathogenic | 1 | 1 | 0 |
| rs62516092 | pathogenic | 0 | 0 | 0 |
| rs62516095 | pathogenic | 1 | 1 | 1 |
| rs62642937 | pathogenic | 1 | 1 | 1 |
| rs62516141 | pathogenic | 0 | 1 | 0 |
| rs5030856 | pathogenic | 1 | 1 | 1 |
| rs5030857 | pathogenic | 1 | 1 | 1 |
| rs5030858 | pathogenic | 1 | 1 | 1 |
| rs79931499 | pathogenic | 1 | 1 | 1 |
| rs5030860 | pathogenic | 1 | 1 | 1 |
| rs281865437 | pathogenic | 1 | 1 | 1 |
| rs62644499 | pathogenic | 1 | 1 | 1 |
| rs147689487 | benign | 0 | 0 | 0 |
| rs397515417 | pathogenic | 1 | 1 | 1 |
| rs121909332 | pathogenic | 1 | 1 | 1 |
| rs121909330 | pathogenic | 1 | 1 | 0 |
| rs121909329 | pathogenic | 1 | 1 | 0 |
| rs121909335 | pathogenic | 1 | 1 | 1 |
| rs387906789 | pathogenic | 1 | 1 | 1 |
| rs121909334 | pathogenic | 1 | 1 | 1 |
| rs121909331 | pathogenic | 1 | 1 | 1 |
| rs372487178 | pathogenic | 1 | 1 | 1 |
| rs387906937 | pathogenic | 1 | 1 | 0 |
| rs16845 | benign | 1 | 1 | 1 |
| rs672601313 | pathogenic | 1 | 1 | 1 |
| rs117805308 | benign | 0 | 0 | 0 |
| rs121434261 | pathogenic | 1 | 1 | 1 |
| rs141629512 | benign | 0 | 0 | 0 |
| rs199957176 | benign | 1 | 1 | 1 |
| rs2230461 | benign | 1 | 1 | 1 |
| rs764618040 | pathogenic | 1 | 1 | 1 |
| rs267606949 | pathogenic | 1 | 1 | 1 |

| | | | | |
|---|---|---|---|---|
| rs587777483 | pathogenic | 1 | 1 | 1 |
| rs267606947 | pathogenic | 1 | 1 | 1 |
| rs6941583 | benign | 0 | 0 | 0 |
| rs749597959 | benign | 1 | 1 | 1 |
| rs182340753 | benign | 1 | 1 | 1 |
| rs146678221 | benign | 0 | 0 | 0 |
| rs11539202 | benign | 0 | 0 | 0 |
| rs148723565 | benign | 0 | 0 | 0 |
| rs61752925 | benign | 1 | 1 | 1 |
| rs113309941 | pathogenic | 0 | 0 | 0 |
| rs121434520 | pathogenic | 1 | 1 | 1 |
| rs121434517 | pathogenic | 1 | 1 | 1 |
| rs267607027 | pathogenic | 1 | 1 | 1 |
| rs138603088 | pathogenic | 0 | 0 | 0 |
| rs121434516 | pathogenic | 1 | 1 | 0 |
| rs121434521 | pathogenic | 0 | 0 | 0 |
| rs145735112 | benign | 1 | 1 | 1 |
| rs121434519 | pathogenic | 1 | 1 | 1 |
| rs75326924 | pathogenic | 1 | 1 | 0 |
| rs142186404 | pathogenic | 0 | 0 | 0 |
| rs121918035 | pathogenic | 0 | 0 | 0 |
| rs374385878 | pathogenic | 1 | 1 | 0 |
| rs11557652 | benign | 0 | 0 | 0 |
| rs118203986 | pathogenic | 1 | 1 | 1 |
| rs141675330 | benign | 1 | 1 | 1 |
| rs104893927 | pathogenic | 0 | 0 | 0 |
| rs104893935 | pathogenic | 0 | 0 | 0 |
| rs104893933 | pathogenic | 1 | 1 | 0 |
| rs104893934 | pathogenic | 0 | 0 | 0 |
| rs145183203 | benign | 1 | 1 | 0 |
| rs2230806 | benign | 0 | 0 | 0 |
| rs9282541 | benign | 0 | 0 | 0 |
| rs559110055 | benign | 0 | 0 | 0 |
| rs200030513 | benign | 0 | 0 | 0 |
| rs9282543 | benign | 0 | 0 | 0 |
| rs147675550 | benign | 0 | 0 | 0 |
| rs137854496 | pathogenic | 0 | 0 | 0 |
| rs374190304 | benign | 0 | 0 | 0 |
| rs114620717 | benign | 0 | 0 | 0 |
| rs2066718 | benign | 0 | 0 | 0 |
| rs35819696 | benign | 0 | 0 | 0 |
| rs138880920 | benign | 0 | 0 | 0 |
| rs2066715 | benign | 0 | 0 | 0 |
| rs35207495 | benign | 0 | 0 | 0 |
| rs2066714 | benign | 0 | 0 | 0 |
| rs28937313 | pathogenic | 0 | 0 | 0 |
| rs28937314 | pathogenic | 0 | 0 | 0 |
| rs137854495 | pathogenic | 0 | 0 | 0 |
| rs140365800 | benign | 0 | 0 | 0 |
| rs13306073 | benign | 0 | 0 | 0 |
| rs28933692 | pathogenic | 0 | 0 | 0 |
| rs33918808 | benign | 0 | 0 | 0 |
| rs76881554 | benign | 0 | 0 | 0 |
| rs143180998 | benign | 0 | 0 | 0 |

| | | | | |
|---|---|---|---|---|
| rs137854500 | pathogenic | 0 | 0 | 0 |
| rs538614702 | benign | 0 | 0 | 0 |
| rs201879964 | benign | 0 | 0 | 0 |
| rs2230808 | benign | 0 | 0 | 0 |
| rs138422574 | benign | 0 | 0 | 0 |
| rs137854498 | pathogenic | 0 | 0 | 0 |
| rs150125857 | benign | 0 | 0 | 0 |
| rs146292819 | pathogenic | 0 | 0 | 0 |
| rs142688906 | benign | 0 | 0 | 0 |
| rs137854499 | pathogenic | 0 | 0 | 0 |
| rs34879708 | benign | 0 | 0 | 0 |
| rs121913028 | pathogenic | 1 | 1 | 1 |
| rs1041951 | benign | 1 | 1 | 0 |
| rs138714094 | benign | 1 | 1 | 0 |
| rs200650502 | benign | 0 | 1 | 0 |
| rs370708663 | pathogenic | 1 | 1 | 1 |
| rs118204040 | pathogenic | 0 | 1 | 0 |
| rs146899669 | pathogenic | 1 | 1 | 0 |
| rs118204039 | pathogenic | 0 | 1 | 0 |
| rs121909233 | pathogenic | 0 | 0 | 0 |
| rs121909225 | pathogenic | 1 | 1 | 0 |
| rs121909236 | pathogenic | 0 | 0 | 0 |
| rs398123317 | pathogenic | 0 | 0 | 0 |
| rs121909226 | pathogenic | 1 | 1 | 0 |
| rs121909238 | pathogenic | 1 | 1 | 0 |
| rs121909230 | pathogenic | 0 | 0 | 0 |
| rs121909237 | pathogenic | 0 | 0 | 0 |
| rs121909222 | pathogenic | 1 | 1 | 0 |
| rs121909223 | pathogenic | 1 | 1 | 0 |
| rs121909218 | pathogenic | 1 | 1 | 0 |
| rs121909229 | pathogenic | 1 | 1 | 0 |
| rs121909224 | pathogenic | 1 | 1 | 0 |
| rs121909241 | pathogenic | 0 | 0 | 0 |
| rs121909221 | pathogenic | 0 | 0 | 0 |
| rs121913294 | pathogenic | 0 | 0 | 0 |
| rs121909234 | pathogenic | 0 | 0 | 0 |
| rs121909240 | pathogenic | 0 | 0 | 0 |
| rs767623493 | benign | 0 | 0 | 0 |
| rs121909239 | pathogenic | 0 | 0 | 0 |
| rs120074145 | pathogenic | 1 | 1 | 1 |
| rs120074148 | pathogenic | 1 | 1 | 1 |
| rs762991875 | pathogenic | 0 | 0 | 0 |
| rs148639841 | pathogenic | 1 | 1 | 1 |
| rs199524907 | pathogenic | 1 | 1 | 1 |
| rs120074141 | pathogenic | 1 | 1 | 1 |
| rs120074144 | pathogenic | 0 | 1 | 0 |
| rs120074146 | pathogenic | 1 | 1 | 1 |
| rs120074147 | pathogenic | 0 | 1 | 0 |
| rs120074143 | pathogenic | 0 | 1 | 0 |
| rs120074140 | pathogenic | 1 | 1 | 0 |
| rs28937315 | pathogenic | 1 | 1 | 1 |
| rs76066357 | benign | 1 | 1 | 1 |
| rs137852911 | pathogenic | 0 | 1 | 0 |
| rs137852907 | pathogenic | 1 | 1 | 0 |

| | | | | |
|---|---|---|---|---|
| rs137852910 | pathogenic | 1 | 1 | 1 |
| rs537585211 | benign | 0 | 0 | 0 |
| rs202134424 | pathogenic | 0 | 0 | 0 |
| rs200930463 | pathogenic | 0 | 0 | 0 |
| rs587777241 | pathogenic | 1 | 1 | 0 |
| rs77563738 | pathogenic | 0 | 0 | 0 |
| rs750868279 | pathogenic | 1 | 1 | 0 |
| rs148936893 | pathogenic | 0 | 0 | 0 |
| rs146597836 | pathogenic | 0 | 0 | 0 |
| rs775440641 | pathogenic | 1 | 1 | 1 |
| rs770689762 | pathogenic | 0 | 1 | 0 |
| rs376785840 | pathogenic | 0 | 0 | 0 |
| rs137852531 | pathogenic | 0 | 0 | 0 |
| rs137852538 | pathogenic | 1 | 1 | 0 |
| rs137852529 | pathogenic | 0 | 1 | 0 |
| rs137852528 | pathogenic | 0 | 1 | 0 |
| rs137852535 | pathogenic | 1 | 1 | 0 |
| rs137853045 | pathogenic | 0 | 1 | 0 |
| rs137853046 | pathogenic | 1 | 1 | 1 |
| rs137853043 | pathogenic | 1 | 1 | 1 |
| rs137853044 | pathogenic | 1 | 1 | 1 |
| rs137853047 | pathogenic | 1 | 1 | 1 |
| rs267606990 | pathogenic | 1 | 1 | 1 |
| rs751437780 | pathogenic | 1 | 1 | 0 |
| rs121918461 | pathogenic | 1 | 1 | 1 |
| rs121918460 | pathogenic | 1 | 1 | 1 |
| rs121918459 | pathogenic | 1 | 1 | 0 |
| rs397507511 | pathogenic | 1 | 1 | 1 |
| rs121918454 | pathogenic | 1 | 1 | 1 |
| rs121918453 | pathogenic | 1 | 1 | 1 |
| rs121918462 | pathogenic | 1 | 1 | 0 |
| rs121918465 | pathogenic | 1 | 1 | 0 |
| rs121918464 | pathogenic | 1 | 1 | 0 |
| rs121918466 | pathogenic | 1 | 1 | 1 |
| rs397507521 | pathogenic | 1 | 1 | 1 |
| rs397507527 | pathogenic | 1 | 1 | 1 |
| rs777603059 | pathogenic | 0 | 1 | 0 |
| rs121918456 | pathogenic | 1 | 1 | 1 |
| rs121918463 | pathogenic | 1 | 1 | 1 |
| rs121918455 | pathogenic | 0 | 1 | 0 |
| rs28933386 | pathogenic | 0 | 1 | 0 |
| rs774356443 | benign | 1 | 1 | 1 |
| rs121918467 | pathogenic | 1 | 1 | 1 |
| rs121918468 | pathogenic | 1 | 1 | 1 |
| rs121918469 | pathogenic | 1 | 1 | 1 |
| rs397507539 | pathogenic | 1 | 1 | 0 |
| rs121918458 | pathogenic | 1 | 1 | 1 |
| rs121918470 | pathogenic | 1 | 1 | 1 |
| rs35289681 | benign | 0 | 1 | 0 |
| rs4302331 | benign | 0 | 1 | 0 |
| rs11544803 | pathogenic | 1 | 1 | 1 |
| rs116254156 | benign | 0 | 0 | 0 |
| rs104893909 | pathogenic | 1 | 1 | 1 |
| rs104893911 | pathogenic | 1 | 1 | 1 |

| | | | | |
|---|---|---|---|---|
| rs104893908 | pathogenic | 1 | 1 | 0 |
| rs258751 | benign | 1 | 1 | 1 |
| rs104893914 | pathogenic | 1 | 1 | 0 |
| rs121909727 | pathogenic | 1 | 1 | 1 |
| rs104893910 | pathogenic | 1 | 1 | 1 |
| rs121909726 | pathogenic | 1 | 1 | 1 |
| rs104893912 | pathogenic | 1 | 1 | 1 |
| rs138407155 | benign | 1 | 1 | 1 |
| rs574061789 | benign | 1 | 1 | 1 |
| rs121912718 | pathogenic | 1 | 1 | 1 |
| rs121912719 | pathogenic | 1 | 1 | 1 |
| rs140770089 | benign | 1 | 1 | 1 |
| rs28931573 | pathogenic | 1 | 1 | 1 |
| rs121912717 | pathogenic | 1 | 1 | 1 |
| rs376456598 | pathogenic | 0 | 0 | 0 |
| rs398123499 | pathogenic | 1 | 1 | 1 |
| rs773048903 | pathogenic | 1 | 1 | 0 |
| rs398123503 | pathogenic | 1 | 1 | 1 |
| rs375785084 | pathogenic | 0 | 1 | 0 |
| rs137852874 | pathogenic | 1 | 1 | 1 |
| rs373713279 | pathogenic | 1 | 1 | 1 |
| rs137852873 | pathogenic | 1 | 1 | 0 |
| rs367823977 | pathogenic | 1 | 1 | 1 |
| rs398123508 | pathogenic | 1 | 1 | 1 |
| rs137852871 | pathogenic | 1 | 1 | 1 |
| rs753698250 | pathogenic | 0 | 1 | 0 |
| rs182923857 | pathogenic | 1 | 1 | 1 |
| rs148090804 | benign | 1 | 1 | 1 |
| rs137852870 | pathogenic | 1 | 1 | 1 |
| rs267607131 | pathogenic | 1 | 1 | 0 |
| rs137852902 | pathogenic | 0 | 0 | 0 |
| rs61741646 | benign | 1 | 1 | 1 |
| rs137852905 | pathogenic | 0 | 0 | 0 |
| rs121918681 | pathogenic | 0 | 1 | 0 |
| rs8177238 | benign | 1 | 1 | 0 |
| rs5030776 | pathogenic | 1 | 1 | 1 |
| rs104894572 | pathogenic | 0 | 0 | 0 |
| rs104894576 | pathogenic | 0 | 0 | 0 |
| rs80084721 | pathogenic | 0 | 0 | 0 |
| rs104894146 | pathogenic | 0 | 1 | 0 |
| rs104894138 | pathogenic | 1 | 1 | 0 |
| rs104894153 | pathogenic | 1 | 1 | 0 |
| rs104894135 | pathogenic | 1 | 1 | 0 |
| rs104894147 | pathogenic | 1 | 1 | 0 |
| rs104894148 | pathogenic | 1 | 1 | 0 |
| rs104894154 | pathogenic | 1 | 1 | 0 |
| rs104894150 | pathogenic | 1 | 1 | 0 |
| rs104894136 | pathogenic | 1 | 1 | 0 |
| rs104894144 | pathogenic | 0 | 0 | 0 |
| rs104894137 | pathogenic | 0 | 0 | 0 |
| rs61754278 | pathogenic | 1 | 1 | 0 |
| rs104894149 | pathogenic | 1 | 1 | 0 |
| rs104894139 | pathogenic | 1 | 1 | 0 |
| rs104894142 | pathogenic | 0 | 0 | 0 |

| | | | | |
|---|---|---|---|---|
| rs104894143 | pathogenic | 0 | 0 | 0 |
| rs104894145 | pathogenic | 0 | 0 | 0 |
| rs121909545 | benign | 0 | 1 | 0 |
| rs782308462 | pathogenic | 1 | 1 | 1 |
| rs375002796 | pathogenic | 0 | 1 | 0 |
| rs201727685 | pathogenic | 0 | 0 | 0 |
| rs768050261 | pathogenic | 1 | 1 | 1 |
| rs780851340 | benign | 0 | 0 | 0 |
| rs139824241 | benign | 1 | 1 | 0 |
| rs757486575 | pathogenic | 1 | 1 | 0 |
| rs750831299 | pathogenic | 1 | 1 | 1 |
| rs767602961 | pathogenic | 1 | 1 | 1 |
| rs121913661 | pathogenic | 1 | 1 | 1 |
| rs777692271 | pathogenic | 1 | 1 | 1 |
| rs121913659 | pathogenic | 1 | 1 | 1 |
| rs587777533 | pathogenic | 1 | 1 | 1 |
| rs104894210 | pathogenic | 0 | 0 | 0 |
| rs104894216 | pathogenic | 1 | 1 | 1 |
| rs104894207 | pathogenic | 1 | 1 | 1 |
| rs104894208 | pathogenic | 0 | 1 | 0 |
| rs104894214 | pathogenic | 1 | 1 | 1 |
| rs104894211 | pathogenic | 1 | 1 | 1 |
| rs61749384 | pathogenic | 0 | 0 | 0 |
| rs61750071 | pathogenic | 1 | 1 | 1 |
| rs61750074 | pathogenic | 0 | 0 | 0 |
| rs11063988 | benign | 0 | 0 | 0 |
| rs216311 | benign | 0 | 0 | 0 |
| rs1800382 | benign | 1 | 1 | 1 |
| rs11063987 | benign | 1 | 1 | 1 |
| rs387906847 | pathogenic | 1 | 1 | 1 |
| rs73003466 | benign | 0 | 0 | 0 |
| rs149787233 | benign | 1 | 1 | 1 |
| rs144451841 | pathogenic | 1 | 1 | 1 |
| rs141638421 | pathogenic | 1 | 1 | 1 |
| rs121434497 | pathogenic | 0 | 0 | 0 |
| rs121434498 | pathogenic | 0 | 0 | 0 |
| rs267607230 | pathogenic | 1 | 1 | 0 |
| rs121434499 | pathogenic | 1 | 1 | 0 |
| rs546749336 | benign | 1 | 1 | 1 |
| rs138873805 | benign | 0 | 0 | 0 |
| rs730880522 | pathogenic | 0 | 0 | 0 |
| rs200371894 | benign | 0 | 0 | 0 |
| rs730880509 | benign | 0 | 0 | 0 |
| rs267606704 | pathogenic | 1 | 1 | 1 |
| rs267606706 | pathogenic | 0 | 0 | 0 |
| rs267606705 | pathogenic | 1 | 1 | 1 |
| rs267606707 | pathogenic | 0 | 0 | 0 |
| rs387906665 | pathogenic | 1 | 1 | 1 |
| rs267606708 | pathogenic | 1 | 1 | 1 |
| rs148368481 | benign | 0 | 0 | 0 |
| rs13204 | benign | 0 | 0 | 0 |
| rs140547520 | pathogenic | 1 | 1 | 1 |
| rs755234990 | benign | 1 | 1 | 0 |
| rs151184052 | benign | 0 | 0 | 0 |

| | | | | |
|---|---|---|---|---|
| rs3736358 | benign | 0 | 0 | 0 |
| rs3736360 | benign | 0 | 0 | 0 |
| rs1138469 | benign | 0 | 0 | 0 |
| rs74315351 | pathogenic | 0 | 1 | 0 |
| rs137853051 | pathogenic | 1 | 1 | 0 |
| rs74315353 | pathogenic | 0 | 1 | 0 |
| rs74315352 | pathogenic | 0 | 1 | 0 |
| rs74315354 | pathogenic | 1 | 1 | 1 |
| rs28938172 | pathogenic | 1 | 1 | 1 |
| rs52836744 | pathogenic | 0 | 0 | 0 |
| rs121913159 | pathogenic | 1 | 1 | 1 |
| rs121913145 | pathogenic | 1 | 1 | 1 |
| rs891087 | benign | 1 | 1 | 1 |
| rs9282757 | benign | 1 | 1 | 1 |
| rs397517404 | pathogenic | 1 | 1 | 1 |
| rs199918720 | benign | 1 | 1 | 0 |
| rs368299411 | benign | 1 | 1 | 0 |
| rs139399951 | benign | 1 | 1 | 0 |
| rs763981974 | benign | 1 | 1 | 0 |
| rs144242114 | benign | 0 | 0 | 0 |
| rs750965801 | benign | 0 | 0 | 0 |
| rs561310777 | benign | 0 | 0 | 0 |
| rs148185335 | benign | 0 | 0 | 0 |
| rs147742157 | benign | 0 | 0 | 0 |
| rs375664373 | pathogenic | 1 | 1 | 1 |
| rs777327250 | pathogenic | 0 | 0 | 0 |
| rs121908471 | pathogenic | 1 | 1 | 0 |
| rs2301612 | pathogenic | 1 | 1 | 0 |
| rs121908476 | pathogenic | 0 | 0 | 0 |
| rs36220239 | benign | 0 | 0 | 0 |
| rs121908473 | pathogenic | 0 | 0 | 0 |
| rs28647808 | benign | 0 | 0 | 0 |
| rs36090624 | benign | 0 | 0 | 0 |
| rs121908002 | pathogenic | 1 | 1 | 1 |
| rs4251545 | benign | 1 | 1 | 1 |
| rs78705382 | pathogenic | 0 | 0 | 0 |
| rs75184679 | pathogenic | 1 | 1 | 1 |
| rs74555752 | pathogenic | 0 | 0 | 0 |
| rs56320207 | benign | 1 | 1 | 1 |
| rs28730837 | pathogenic | 1 | 1 | 1 |
| rs119469014 | pathogenic | 1 | 1 | 1 |
| rs119468010 | pathogenic | 1 | 1 | 1 |
| rs35220041 | benign | 0 | 0 | 0 |
| rs762321137 | pathogenic | 1 | 1 | 1 |
| rs28931570 | pathogenic | 1 | 1 | 1 |
| rs709932 | benign | 1 | 1 | 1 |
| rs746531546 | benign | 1 | 1 | 1 |
| rs6647 | benign | 1 | 1 | 1 |
| rs141095970 | benign | 1 | 1 | 1 |
| rs143370956 | benign | 1 | 1 | 1 |
| rs28929474 | pathogenic | 1 | 1 | 1 |
| rs61761869 | pathogenic | 1 | 1 | 1 |
| rs1303 | benign | 1 | 1 | 1 |
| rs149447502 | benign | 0 | 0 | 0 |

| | | | | |
|---|---|---|---|---|
| rs267607180 | pathogenic | 1 | 1 | 1 |
| rs150304121 | benign | 0 | 0 | 0 |
| rs147630375 | benign | 1 | 1 | 1 |
| rs11539445 | benign | 1 | 1 | 1 |
| rs372098364 | pathogenic | 1 | 1 | 0 |
| rs760072286 | benign | 0 | 0 | 0 |
| rs745595833 | pathogenic | 1 | 1 | 1 |
| rs587777214 | pathogenic | 0 | 0 | 0 |
| rs150318528 | benign | 0 | 0 | 0 |
| rs137853254 | pathogenic | 1 | 1 | 1 |
| rs137853255 | pathogenic | 0 | 0 | 0 |
| rs137853253 | pathogenic | 0 | 0 | 0 |
| rs202166915 | benign | 0 | 0 | 0 |
| rs137853259 | pathogenic | 0 | 0 | 0 |
| rs2229137 | benign | 0 | 0 | 0 |
| rs137853258 | pathogenic | 1 | 1 | 0 |
| rs137853252 | pathogenic | 1 | 1 | 1 |
| rs137853256 | pathogenic | 0 | 0 | 0 |
| rs137853250 | pathogenic | 1 | 1 | 1 |
| rs121434615 | pathogenic | 0 | 0 | 0 |
| rs115014558 | benign | 1 | 1 | 1 |
| rs34116584 | benign | 1 | 1 | 1 |
| rs34885252 | benign | 1 | 1 | 1 |
| rs180177262 | pathogenic | 1 | 1 | 1 |
| rs180177162 | pathogenic | 1 | 1 | 1 |
| rs121908523 | pathogenic | 1 | 1 | 1 |
| rs180177170 | pathogenic | 1 | 1 | 1 |
| rs180177180 | pathogenic | 1 | 1 | 1 |
| rs767586362 | pathogenic | 1 | 1 | 1 |
| rs121908521 | pathogenic | 1 | 1 | 1 |
| rs180177203 | pathogenic | 1 | 1 | 1 |
| rs138025751 | pathogenic | 1 | 1 | 1 |
| rs180177210 | pathogenic | 0 | 1 | 0 |
| rs180177211 | pathogenic | 0 | 1 | 0 |
| rs121908524 | pathogenic | 1 | 1 | 0 |
| rs121908530 | pathogenic | 1 | 1 | 0 |
| rs180177227 | pathogenic | 0 | 1 | 0 |
| rs180177230 | pathogenic | 1 | 1 | 1 |
| rs121908529 | pathogenic | 0 | 1 | 0 |
| rs117195882 | benign | 1 | 1 | 0 |
| rs180177238 | pathogenic | 1 | 1 | 0 |
| rs34664134 | benign | 1 | 1 | 1 |
| rs180177246 | pathogenic | 0 | 1 | 0 |
| rs536352238 | pathogenic | 0 | 0 | 0 |
| rs121908520 | pathogenic | 1 | 1 | 0 |
| rs180177248 | pathogenic | 1 | 1 | 0 |
| rs180177252 | pathogenic | 1 | 1 | 1 |
| rs121908526 | pathogenic | 1 | 1 | 1 |
| rs121908525 | pathogenic | 1 | 1 | 1 |
| rs147106773 | benign | 1 | 1 | 1 |
| rs180177264 | pathogenic | 0 | 1 | 0 |
| rs146525143 | pathogenic | 1 | 1 | 1 |
| rs140992177 | benign | 1 | 1 | 1 |
| rs13408961 | benign | 0 | 1 | 0 |

| | | | | |
|---|---|---|---|---|
| rs4426527 | benign | 1 | 1 | 1 |
| rs3088308 | benign | 0 | 1 | 0 |
| rs4469829 | benign | 1 | 1 | 0 |
| rs104886003 | pathogenic | 1 | 1 | 1 |
| rs121913286 | pathogenic | 1 | 1 | 1 |
| rs121913279 | pathogenic | 1 | 1 | 1 |
| rs121434636 | pathogenic | 1 | 1 | 1 |
| rs755299132 | pathogenic | 1 | 1 | 1 |
| rs748318386 | pathogenic | 1 | 1 | 1 |
| rs387906757 | pathogenic | 1 | 1 | 1 |
| rs121965064 | pathogenic | 1 | 1 | 1 |
| rs28934608 | pathogenic | 0 | 0 | 0 |
| rs5967 | benign | 1 | 1 | 1 |
| rs773905328 | pathogenic | 0 | 0 | 0 |
| rs104894256 | pathogenic | 0 | 1 | 0 |
| rs28931612 | pathogenic | 0 | 1 | 0 |
| rs104894263 | pathogenic | 1 | 1 | 1 |
| rs607969 | benign | 1 | 1 | 0 |
| rs104894262 | pathogenic | 0 | 1 | 0 |
| rs104894268 | pathogenic | 1 | 1 | 1 |
| rs104894264 | pathogenic | 0 | 1 | 0 |
| rs104894259 | pathogenic | 1 | 1 | 0 |
| rs2959656 | benign | 0 | 0 | 0 |
| rs121913035 | pathogenic | 1 | 1 | 0 |
| rs139605474 | benign | 0 | 0 | 0 |
| rs3817552 | benign | 0 | 0 | 0 |
| rs150541115 | benign | 0 | 0 | 0 |
| rs562763010 | benign | 1 | 1 | 1 |
| rs200053119 | pathogenic | 0 | 0 | 0 |
| rs73826386 | benign | 0 | 0 | 0 |
| rs143655258 | benign | 0 | 0 | 0 |
| rs758857853 | pathogenic | 1 | 1 | 0 |
| rs2228380 | benign | 0 | 0 | 0 |
| rs141078733 | benign | 1 | 1 | 1 |
| rs55824172 | pathogenic | 1 | 1 | 1 |
| rs756751089 | pathogenic | 1 | 1 | 1 |
| rs121964990 | pathogenic | 0 | 0 | 0 |
| rs143555186 | benign | 0 | 0 | 0 |
| rs121964993 | pathogenic | 1 | 1 | 0 |
| rs121964992 | pathogenic | 1 | 1 | 1 |
| rs121964991 | pathogenic | 1 | 1 | 0 |
| rs121964988 | pathogenic | 1 | 1 | 1 |
| rs4687 | benign | 1 | 1 | 1 |
| rs267607125 | pathogenic | 1 | 1 | 1 |
| rs751682878 | benign | 1 | 1 | 1 |
| rs267607126 | pathogenic | 1 | 1 | 1 |
| rs768713596 | pathogenic | 0 | 0 | 0 |
| rs147288996 | benign | 0 | 0 | 0 |
| rs387906639 | pathogenic | 1 | 1 | 1 |
| rs730881069 | pathogenic | 1 | 1 | 1 |
| rs730881071 | pathogenic | 1 | 1 | 1 |
| rs397516348 | pathogenic | 0 | 0 | 0 |
| rs121917760 | pathogenic | 0 | 0 | 0 |
| rs104894724 | pathogenic | 1 | 1 | 1 |

| | | | | |
|---|---|---|---|---|
| rs397516349 | pathogenic | 1 | 1 | 1 |
| rs12075 | benign | 0 | 0 | 0 |
| rs147549987 | benign | 0 | 0 | 0 |
| rs56079734 | benign | 0 | 0 | 0 |
| rs1017842 | benign | 1 | 1 | 1 |
| rs137852593 | pathogenic | 1 | 1 | 1 |
| rs137852571 | pathogenic | 1 | 1 | 1 |
| rs137852591 | pathogenic | 1 | 1 | 1 |
| rs9332971 | pathogenic | 1 | 1 | 1 |
| rs755226547 | pathogenic | 1 | 1 | 1 |
| rs79206939 | benign | 0 | 0 | 0 |
| rs121918214 | pathogenic | 1 | 1 | 0 |
| rs757075712 | pathogenic | 1 | 1 | 0 |
| rs150483902 | benign | 1 | 1 | 1 |
| rs113993973 | pathogenic | 0 | 1 | 0 |
| rs946616 | benign | 0 | 0 | 0 |
| rs113993976 | pathogenic | 1 | 1 | 0 |
| rs151022264 | benign | 0 | 1 | 0 |
| rs113993977 | pathogenic | 1 | 1 | 0 |
| rs113993978 | pathogenic | 1 | 1 | 0 |
| rs2228499 | benign | 0 | 1 | 0 |
| rs113993979 | pathogenic | 1 | 1 | 0 |
| rs113993980 | pathogenic | 1 | 1 | 0 |
| rs113993983 | pathogenic | 0 | 1 | 0 |
| rs150547274 | pathogenic | 1 | 1 | 0 |
| rs113993984 | pathogenic | 1 | 1 | 0 |
| rs113993986 | pathogenic | 1 | 1 | 0 |
| rs34313873 | benign | 0 | 1 | 0 |
| rs113993988 | pathogenic | 1 | 1 | 0 |
| rs199638097 | pathogenic | 1 | 1 | 1 |
| rs118204447 | pathogenic | 1 | 1 | 1 |
| rs11862754 | benign | 0 | 0 | 0 |
| rs118204445 | pathogenic | 0 | 0 | 0 |
| rs118204441 | pathogenic | 0 | 0 | 0 |
| rs118204438 | pathogenic | 0 | 0 | 0 |
| rs118204436 | pathogenic | 0 | 0 | 0 |
| rs146093755 | pathogenic | 0 | 0 | 0 |
| rs781439830 | pathogenic | 0 | 0 | 0 |
| rs398123438 | pathogenic | 0 | 0 | 0 |
| rs398123439 | pathogenic | 0 | 0 | 0 |
| rs118204444 | pathogenic | 0 | 0 | 0 |
| rs7187889 | benign | 0 | 0 | 0 |
| rs118204435 | pathogenic | 1 | 1 | 0 |
| rs34745339 | benign | 0 | 0 | 0 |
| rs398123440 | pathogenic | 0 | 0 | 0 |
| rs118204442 | pathogenic | 0 | 0 | 0 |
| rs770053354 | pathogenic | 0 | 0 | 0 |
| rs118204448 | pathogenic | 0 | 0 | 0 |
| rs118204443 | pathogenic | 0 | 0 | 0 |
| rs118204446 | pathogenic | 0 | 0 | 0 |
| rs746756997 | pathogenic | 0 | 0 | 0 |
| rs267606838 | pathogenic | 0 | 0 | 0 |
| rs118204437 | pathogenic | 0 | 0 | 0 |
| rs398123429 | pathogenic | 0 | 0 | 0 |

| | | | | |
|---|---|---|---|---|
| rs398123430 | pathogenic | 0 | 0 | 0 |
| rs2303269 | benign | 0 | 0 | 0 |
| rs114703967 | benign | 0 | 0 | 0 |
| rs118204439 | pathogenic | 0 | 0 | 0 |
| rs151296605 | benign | 0 | 0 | 0 |
| rs118204440 | pathogenic | 0 | 0 | 0 |
| rs78127134 | benign | 0 | 0 | 0 |
| rs372893383 | pathogenic | 0 | 0 | 0 |
| rs61753266 | pathogenic | 1 | 1 | 1 |
| rs139579928 | benign | 1 | 1 | 1 |
| rs56275071 | pathogenic | 1 | 1 | 1 |
| rs121909736 | pathogenic | 1 | 1 | 0 |
| rs121909737 | pathogenic | 1 | 1 | 0 |
| rs121909735 | pathogenic | 1 | 1 | 0 |
| rs121909731 | pathogenic | 1 | 1 | 1 |
| rs121909734 | pathogenic | 1 | 1 | 1 |
| rs121909733 | pathogenic | 1 | 1 | 1 |
| rs121909732 | pathogenic | 1 | 1 | 1 |
| rs121909730 | pathogenic | 1 | 1 | 1 |
| rs202198360 | benign | 1 | 1 | 1 |
| rs587777294 | pathogenic | 1 | 1 | 0 |
| rs587777297 | pathogenic | 1 | 1 | 0 |
| rs104893913 | pathogenic | 1 | 1 | 0 |
| rs17860424 | pathogenic | 1 | 1 | 1 |
| rs1045485 | benign | 1 | 1 | 1 |
| rs13926 | benign | 0 | 0 | 0 |
| rs79343837 | benign | 0 | 0 | 0 |
| rs45516306 | benign | 1 | 1 | 1 |
| rs139849355 | benign | 1 | 1 | 1 |
| rs104894874 | pathogenic | 1 | 1 | 1 |
| rs104894870 | pathogenic | 1 | 1 | 1 |
| rs137852221 | pathogenic | 1 | 1 | 1 |
| rs104894869 | pathogenic | 1 | 1 | 1 |
| rs104894880 | pathogenic | 1 | 1 | 1 |
| rs104894872 | pathogenic | 1 | 1 | 1 |
| rs104894868 | pathogenic | 1 | 1 | 1 |
| rs104894867 | pathogenic | 1 | 1 | 1 |
| rs104894871 | pathogenic | 1 | 1 | 1 |
| rs104894877 | pathogenic | 1 | 1 | 1 |
| rs104894883 | pathogenic | 1 | 1 | 1 |
| rs104894875 | pathogenic | 1 | 1 | 1 |
| rs104894876 | pathogenic | 1 | 1 | 1 |
| rs104894878 | pathogenic | 1 | 1 | 1 |
| rs137852220 | pathogenic | 1 | 1 | 1 |
| rs28933684 | pathogenic | 1 | 1 | 1 |
| rs104894873 | pathogenic | 1 | 1 | 1 |
| rs138409264 | benign | 0 | 0 | 0 |
| rs2236722 | benign | 0 | 0 | 0 |
| rs28757184 | benign | 0 | 0 | 0 |
| rs121434538 | pathogenic | 0 | 0 | 0 |
| rs700519 | benign | 0 | 0 | 0 |
| rs80051519 | pathogenic | 0 | 0 | 0 |
| rs121434536 | pathogenic | 1 | 1 | 0 |
| rs121434534 | pathogenic | 1 | 1 | 0 |

| | | | | |
|---|---|---|---|---|
| rs78310315 | pathogenic | 1 | 1 | 0 |
| rs540839115 | pathogenic | 1 | 1 | 1 |
| rs80338731 | pathogenic | 1 | 1 | 1 |
| rs180177151 | pathogenic | 1 | 1 | 1 |
| rs180177152 | pathogenic | 0 | 0 | 0 |
| rs80338732 | pathogenic | 1 | 1 | 1 |
| rs80338675 | pathogenic | 1 | 1 | 1 |
| rs80338676 | pathogenic | 1 | 1 | 1 |
| rs180177153 | pathogenic | 0 | 0 | 0 |
| rs180177154 | pathogenic | 1 | 1 | 1 |
| rs5896 | benign | 1 | 1 | 1 |
| rs121918477 | pathogenic | 0 | 0 | 0 |
| rs121918428 | pathogenic | 1 | 1 | 1 |
| rs121918427 | pathogenic | 1 | 1 | 1 |
| rs544300820 | benign | 1 | 1 | 1 |
| rs121918110 | pathogenic | 1 | 1 | 1 |
| rs1057141 | benign | 0 | 0 | 0 |
| rs1135216 | benign | 1 | 1 | 1 |
| rs121917702 | pathogenic | 1 | 1 | 1 |
| rs28941474 | pathogenic | 0 | 0 | 0 |
| rs104893944 | pathogenic | 1 | 1 | 1 |
| rs104893947 | pathogenic | 1 | 1 | 0 |
| rs104893943 | pathogenic | 1 | 1 | 1 |
| rs104893942 | pathogenic | 1 | 1 | 1 |
| rs104893940 | pathogenic | 1 | 1 | 1 |
| rs34045013 | benign | 0 | 0 | 0 |
| rs143083947 | pathogenic | 0 | 0 | 0 |
| rs145786986 | benign | 0 | 0 | 0 |
| rs119450942 | pathogenic | 0 | 0 | 0 |
| rs374259530 | pathogenic | 0 | 0 | 0 |
| rs756210458 | pathogenic | 0 | 0 | 0 |
| rs11089991 | benign | 0 | 0 | 0 |
| rs28941471 | pathogenic | 0 | 1 | 0 |
| rs750614500 | pathogenic | 0 | 1 | 0 |
| rs199761158 | benign | 1 | 1 | 1 |
| rs119450944 | pathogenic | 1 | 1 | 1 |
| rs373458753 | pathogenic | 1 | 1 | 1 |
| rs202064195 | pathogenic | 1 | 1 | 1 |
| rs776496275 | pathogenic | 1 | 1 | 1 |
| rs761493155 | pathogenic | 1 | 1 | 1 |
| rs763542069 | pathogenic | 0 | 0 | 0 |
| rs119450941 | pathogenic | 0 | 0 | 0 |
| rs119450940 | pathogenic | 0 | 0 | 0 |
| rs777821034 | pathogenic | 1 | 1 | 1 |
| rs771121666 | pathogenic | 1 | 1 | 1 |
| rs372895468 | pathogenic | 1 | 1 | 1 |
| rs61750603 | pathogenic | 0 | 0 | 0 |
| rs76863441 | pathogenic | 0 | 0 | 0 |
| rs142601143 | benign | 1 | 1 | 1 |
| rs185832993 | benign | 1 | 1 | 0 |
| rs17860568 | benign | 0 | 0 | 0 |
| rs142064825 | benign | 0 | 0 | 0 |
| rs564076183 | benign | 0 | 0 | 0 |
| rs386834034 | pathogenic | 0 | 0 | 0 |

| | | | | |
|---|---|---|---|---|
| rs147663769 | benign | 0 | 0 | 0 |
| rs776478974 | benign | 0 | 0 | 0 |
| rs111840783 | benign | 0 | 0 | 0 |
| rs1516446 | benign | 1 | 1 | 1 |
| rs143749154 | benign | 1 | 1 | 1 |
| rs80257412 | benign | 0 | 0 | 0 |
| rs142943120 | benign | 0 | 0 | 0 |
| rs144553482 | benign | 0 | 0 | 0 |
| rs35997569 | benign | 0 | 0 | 0 |
| rs772909106 | pathogenic | 0 | 0 | 0 |
| rs149433837 | benign | 1 | 1 | 1 |
| rs532178791 | pathogenic | 0 | 0 | 0 |
| rs145588689 | pathogenic | 1 | 1 | 0 |
| rs17843776 | benign | 1 | 1 | 0 |
| rs121917890 | pathogenic | 1 | 1 | 1 |
| rs121917892 | pathogenic | 0 | 0 | 0 |
| rs606231289 | pathogenic | 0 | 0 | 0 |
| rs370011798 | pathogenic | 0 | 0 | 0 |
| rs34689457 | benign | 0 | 0 | 0 |
| rs118204451 | pathogenic | 0 | 0 | 0 |
| rs118204452 | pathogenic | 1 | 1 | 1 |
| rs652785 | benign | 0 | 0 | 0 |
| rs35378126 | benign | 0 | 0 | 0 |
| rs80358214 | pathogenic | 1 | 1 | 1 |
| rs80358215 | pathogenic | 0 | 1 | 0 |
| rs267606858 | pathogenic | 1 | 1 | 1 |
| rs143137713 | pathogenic | 1 | 1 | 1 |
| rs782168634 | benign | 0 | 0 | 0 |
| rs143873938 | benign | 0 | 0 | 0 |
| rs782776511 | benign | 0 | 0 | 0 |
| rs201093148 | benign | 0 | 0 | 0 |
| rs267606815 | pathogenic | 0 | 0 | 0 |
| rs137853311 | pathogenic | 0 | 0 | 0 |
| rs267606817 | pathogenic | 0 | 0 | 0 |
| rs72650006 | benign | 0 | 0 | 0 |
| rs727503638 | benign | 1 | 1 | 0 |
| rs189286381 | benign | 0 | 0 | 0 |
| rs376644553 | benign | 0 | 0 | 0 |
| rs121434610 | pathogenic | 1 | 1 | 1 |
| rs267607076 | pathogenic | 1 | 1 | 1 |
| rs201275730 | benign | 0 | 0 | 0 |
| rs121909587 | pathogenic | 0 | 0 | 0 |
| rs121909558 | pathogenic | 1 | 1 | 0 |
| rs28929469 | pathogenic | 1 | 1 | 0 |
| rs121909547 | pathogenic | 1 | 1 | 0 |
| rs121909552 | pathogenic | 1 | 1 | 0 |
| rs121909573 | pathogenic | 1 | 1 | 1 |
| rs121909567 | pathogenic | 1 | 1 | 0 |
| rs121909569 | pathogenic | 0 | 0 | 0 |
| rs121909563 | pathogenic | 1 | 1 | 0 |
| rs121909570 | pathogenic | 1 | 1 | 1 |
| rs121909571 | pathogenic | 1 | 1 | 0 |
| rs121909572 | pathogenic | 1 | 1 | 1 |
| rs121909565 | pathogenic | 1 | 1 | 1 |

| | | | | |
|---|---|---|---|---|
| rs121909557 | pathogenic | 1 | 1 | 1 |
| rs121909568 | pathogenic | 1 | 1 | 1 |
| rs121909566 | pathogenic | 1 | 1 | 1 |
| rs121909549 | pathogenic | 1 | 1 | 1 |
| rs121909554 | pathogenic | 1 | 1 | 1 |
| rs121909550 | pathogenic | 1 | 1 | 1 |
| rs121909546 | pathogenic | 1 | 1 | 0 |
| rs121909555 | pathogenic | 1 | 1 | 1 |
| rs121909564 | pathogenic | 1 | 1 | 1 |
| rs548046212 | pathogenic | 0 | 0 | 0 |
| rs147353781 | benign | 0 | 0 | 0 |
| rs104894678 | pathogenic | 1 | 1 | 1 |
| rs1130426 | benign | 0 | 0 | 0 |
| rs104894677 | pathogenic | 1 | 1 | 1 |
| rs141917423 | benign | 1 | 1 | 1 |
| rs574777883 | benign | 0 | 0 | 0 |
| rs150199226 | benign | 1 | 1 | 1 |
| rs199618701 | benign | 0 | 1 | 0 |
| rs121964923 | pathogenic | 0 | 0 | 0 |
| rs121964924 | pathogenic | 0 | 0 | 0 |
| rs201258823 | pathogenic | 0 | 1 | 0 |
| rs267606773 | pathogenic | 0 | 0 | 0 |
| rs35638832 | benign | 0 | 0 | 0 |
| rs121908632 | pathogenic | 1 | 1 | 1 |
| rs121908625 | pathogenic | 1 | 1 | 1 |
| rs748949603 | pathogenic | 0 | 0 | 0 |
| rs757523840 | pathogenic | 0 | 0 | 0 |
| rs62541771 | pathogenic | 0 | 0 | 0 |
| rs776384541 | pathogenic | 1 | 1 | 0 |
| rs121908627 | pathogenic | 1 | 1 | 0 |
| rs28937594 | pathogenic | 0 | 0 | 0 |
| rs121907888 | pathogenic | 0 | 0 | 0 |
| rs121907889 | pathogenic | 0 | 1 | 0 |
| rs28941785 | pathogenic | 1 | 1 | 1 |
| rs28941786 | pathogenic | 1 | 1 | 1 |
| rs1021737 | benign | 0 | 0 | 0 |
| rs121908370 | pathogenic | 1 | 1 | 1 |
| rs63750083 | pathogenic | 0 | 0 | 0 |
| rs17883862 | benign | 0 | 0 | 0 |
| rs587782070 | pathogenic | 1 | 1 | 1 |
| rs587781269 | pathogenic | 1 | 1 | 1 |
| rs28909982 | pathogenic | 1 | 1 | 1 |
| rs373648967 | benign | 1 | 1 | 1 |
| rs368570187 | benign | 1 | 1 | 1 |
| rs730881701 | pathogenic | 1 | 1 | 1 |
| rs137853007 | pathogenic | 0 | 0 | 0 |
| rs575910805 | benign | 0 | 0 | 0 |
| rs104893875 | pathogenic | 1 | 1 | 1 |
| rs104893877 | pathogenic | 1 | 1 | 1 |
| rs119485093 | pathogenic | 1 | 1 | 1 |
| rs119485092 | pathogenic | 0 | 0 | 0 |
| rs119485090 | pathogenic | 0 | 0 | 0 |
| rs28942099 | pathogenic | 1 | 1 | 1 |
| rs373730800 | pathogenic | 1 | 1 | 1 |

| | | | | |
|---|---|---|---|---|
| rs113993995 | pathogenic | 0 | 0 | 0 |
| rs201070132 | pathogenic | 0 | 0 | 0 |
| rs113993996 | pathogenic | 0 | 0 | 0 |
| rs113993998 | pathogenic | 0 | 0 | 0 |
| rs78950939 | pathogenic | 1 | 1 | 1 |
| rs56378716 | pathogenic | 1 | 1 | 1 |
| rs754921704 | benign | 0 | 1 | 0 |
| rs766379103 | pathogenic | 1 | 1 | 0 |
| rs121912515 | pathogenic | 0 | 1 | 0 |
| rs764831888 | benign | 1 | 1 | 0 |
| rs572527340 | benign | 0 | 0 | 0 |
| rs35969327 | benign | 0 | 0 | 0 |
| rs35814893 | benign | 0 | 0 | 0 |
| rs34032300 | benign | 0 | 0 | 0 |
| rs80358260 | pathogenic | 0 | 0 | 0 |
| rs151220873 | benign | 0 | 0 | 0 |
| rs80358261 | pathogenic | 1 | 1 | 0 |
| rs11694 | pathogenic | 0 | 0 | 0 |
| rs80358264 | pathogenic | 0 | 0 | 0 |
| rs80358266 | pathogenic | 1 | 1 | 0 |
| rs104894458 | pathogenic | 1 | 1 | 1 |
| rs121918028 | pathogenic | 0 | 0 | 0 |
| rs11583680 | pathogenic | 0 | 0 | 0 |
| rs4715 | benign | 1 | 1 | 1 |
| rs34957318 | benign | 0 | 0 | 0 |
| rs1124 | benign | 1 | 1 | 1 |
| rs104894778 | pathogenic | 1 | 1 | 1 |
| rs104894777 | pathogenic | 1 | 1 | 1 |
| rs104894769 | pathogenic | 1 | 1 | 1 |
| rs104894768 | pathogenic | 1 | 1 | 1 |
| rs104894771 | pathogenic | 1 | 1 | 1 |
| rs575372249 | benign | 1 | 1 | 1 |
| rs118203914 | pathogenic | 0 | 0 | 0 |
| rs118203916 | pathogenic | 1 | 1 | 0 |
| rs140154690 | benign | 0 | 0 | 0 |
| rs34382405 | benign | 0 | 0 | 0 |
| rs73718082 | benign | 0 | 0 | 0 |
| rs764932724 | benign | 0 | 0 | 0 |
| rs554931092 | pathogenic | 1 | 1 | 0 |
| rs202060864 | pathogenic | 0 | 0 | 0 |
| rs116567033 | benign | 0 | 0 | 0 |
| rs11243011 | benign | 0 | 0 | 0 |
| rs142073519 | pathogenic | 0 | 0 | 0 |
| rs397514611 | pathogenic | 1 | 1 | 0 |
| rs397514612 | pathogenic | 0 | 0 | 0 |
| rs372054960 | pathogenic | 0 | 0 | 0 |
| rs775690041 | pathogenic | 0 | 0 | 0 |
| rs148921184 | pathogenic | 0 | 0 | 0 |
| rs12711521 | benign | 0 | 0 | 0 |
| rs2273346 | benign | 0 | 0 | 0 |
| rs12085877 | benign | 1 | 1 | 1 |
| rs144471433 | benign | 1 | 1 | 1 |
| rs397514671 | pathogenic | 1 | 1 | 1 |
| rs76394680 | benign | 0 | 0 | 0 |

| | | | | |
|---|---|---|---|---|
| rs78289603 | pathogenic | 1 | 1 | 1 |
| rs137852913 | pathogenic | 1 | 1 | 1 |
| rs137852914 | pathogenic | 1 | 1 | 1 |
| rs121918653 | pathogenic | 1 | 1 | 1 |
| rs35515200 | benign | 1 | 1 | 0 |
| rs17876030 | benign | 1 | 1 | 1 |
| rs556623443 | benign | 1 | 1 | 1 |
| rs121434365 | pathogenic | 1 | 1 | 1 |
| rs121434364 | pathogenic | 0 | 0 | 0 |
| rs41400249 | benign | 0 | 0 | 0 |
| rs149440376 | benign | 1 | 1 | 1 |
| rs181590179 | benign | 0 | 0 | 0 |
| rs113993968 | pathogenic | 0 | 0 | 0 |
| rs587776445 | pathogenic | 1 | 1 | 1 |
| rs113993969 | pathogenic | 1 | 1 | 1 |
| rs113993970 | pathogenic | 1 | 1 | 1 |
| rs587776449 | pathogenic | 1 | 1 | 1 |
| rs113993971 | pathogenic | 1 | 1 | 1 |
| rs267607194 | pathogenic | 1 | 1 | 1 |
| rs397514745 | pathogenic | 0 | 0 | 0 |
| rs201650281 | pathogenic | 0 | 0 | 0 |
| rs150529876 | benign | 1 | 1 | 1 |
| rs77573084 | benign | 0 | 0 | 0 |
| rs398122379 | pathogenic | 1 | 1 | 0 |
| rs765478968 | benign | 0 | 0 | 0 |
| rs201399681 | benign | 0 | 0 | 0 |
| rs17159287 | benign | 0 | 0 | 0 |
| rs538571144 | benign | 0 | 0 | 0 |
| rs137852647 | pathogenic | 0 | 0 | 0 |
| rs201358272 | pathogenic | 1 | 1 | 1 |
| rs181251337 | benign | 0 | 0 | 0 |
| rs3213119 | benign | 0 | 0 | 0 |
| rs137853102 | pathogenic | 1 | 1 | 0 |
| rs4956145 | benign | 0 | 0 | 0 |
| rs146036912 | pathogenic | 1 | 1 | 1 |
| rs375717077 | pathogenic | 1 | 1 | 1 |
| rs137853103 | pathogenic | 1 | 1 | 1 |
| rs776184830 | pathogenic | 1 | 1 | 1 |
| rs200273673 | pathogenic | 1 | 1 | 1 |
| rs267607025 | pathogenic | 1 | 1 | 1 |
| rs267607024 | pathogenic | 1 | 1 | 1 |
| rs515726188 | pathogenic | 1 | 1 | 1 |
| rs147315735 | benign | 0 | 0 | 0 |
| rs121918308 | pathogenic | 1 | 1 | 0 |
| rs515726195 | pathogenic | 0 | 0 | 0 |
| rs515726196 | pathogenic | 1 | 1 | 0 |
| rs121918311 | pathogenic | 0 | 1 | 0 |
| rs121918309 | pathogenic | 0 | 0 | 0 |
| rs387906891 | pathogenic | 0 | 0 | 0 |
| rs182614164 | pathogenic | 1 | 1 | 0 |
| rs121918307 | pathogenic | 0 | 0 | 0 |
| rs201440849 | benign | 0 | 0 | 0 |
| rs118203957 | pathogenic | 0 | 0 | 0 |
| rs4986790 | benign | 0 | 0 | 0 |

| | | | | |
|---|---|---|---|---|
| rs281864928 | pathogenic | 1 | 1 | 0 |
| rs267607156 | pathogenic | 0 | 0 | 0 |
| rs55916807 | benign | 0 | 0 | 0 |
| rs749750052 | benign | 0 | 0 | 0 |
| rs74382477 | benign | 0 | 0 | 0 |
| rs201016638 | benign | 0 | 0 | 0 |
| rs376683989 | benign | 0 | 0 | 0 |
| rs144439703 | pathogenic | 0 | 0 | 0 |
| rs779122022 | benign | 0 | 0 | 0 |
| rs121918056 | pathogenic | 0 | 0 | 0 |
| rs113994093 | pathogenic | 1 | 1 | 1 |
| rs141367015 | pathogenic | 0 | 0 | 0 |
| rs775538075 | pathogenic | 0 | 0 | 0 |
| rs121918055 | pathogenic | 0 | 0 | 0 |
| rs754246294 | benign | 1 | 1 | 1 |
| rs2307447 | benign | 1 | 1 | 1 |
| rs113994096 | pathogenic | 1 | 1 | 1 |
| rs367610201 | pathogenic | 1 | 1 | 1 |
| rs758438414 | pathogenic | 1 | 1 | 0 |
| rs121918046 | pathogenic | 0 | 0 | 0 |
| rs201871736 | benign | 0 | 0 | 0 |
| rs773994204 | benign | 0 | 0 | 0 |
| rs2307450 | benign | 1 | 1 | 1 |
| rs200257554 | benign | 0 | 0 | 0 |
| rs185645212 | benign | 0 | 0 | 0 |
| rs121918054 | pathogenic | 0 | 0 | 0 |
| rs113994097 | pathogenic | 0 | 0 | 0 |
| rs202037973 | pathogenic | 0 | 0 | 0 |
| rs769827124 | pathogenic | 1 | 1 | 0 |
| rs41549716 | benign | 1 | 1 | 1 |
| rs113994098 | pathogenic | 1 | 1 | 0 |
| rs144500145 | pathogenic | 1 | 1 | 0 |
| rs121918053 | pathogenic | 1 | 1 | 0 |
| rs121918050 | pathogenic | 1 | 1 | 0 |
| rs368587966 | pathogenic | 0 | 0 | 0 |
| rs139590686 | pathogenic | 0 | 0 | 0 |
| rs121918048 | pathogenic | 1 | 1 | 0 |
| rs768653086 | pathogenic | 1 | 1 | 0 |
| rs113994099 | pathogenic | 1 | 1 | 0 |
| rs121918051 | pathogenic | 1 | 1 | 0 |
| rs753160398 | pathogenic | 1 | 1 | 0 |
| rs201477273 | pathogenic | 1 | 1 | 0 |
| rs142347031 | pathogenic | 0 | 0 | 0 |
| rs121918049 | pathogenic | 0 | 0 | 0 |
| rs267606959 | pathogenic | 1 | 1 | 0 |
| rs201732356 | pathogenic | 0 | 0 | 0 |
| rs368435864 | pathogenic | 0 | 0 | 0 |
| rs764036283 | pathogenic | 0 | 0 | 0 |
| rs2307442 | benign | 0 | 0 | 0 |
| rs2307440 | benign | 0 | 0 | 0 |
| rs548076633 | pathogenic | 0 | 1 | 0 |
| rs750792237 | benign | 0 | 0 | 0 |
| rs3087374 | benign | 0 | 0 | 0 |
| rs121918060 | pathogenic | 0 | 1 | 0 |

| | | | | |
|---|---|---|---|---|
| rs397514765 | pathogenic | 1 | 1 | 1 |
| rs121918063 | pathogenic | 0 | 0 | 0 |
| rs121918058 | pathogenic | 1 | 1 | 1 |
| rs121918064 | pathogenic | 0 | 0 | 0 |
| rs36033115 | benign | 0 | 1 | 0 |
| rs121918057 | pathogenic | 0 | 1 | 0 |
| rs121918065 | pathogenic | 1 | 1 | 1 |
| rs121918061 | pathogenic | 1 | 1 | 1 |
| rs121918066 | pathogenic | 0 | 1 | 0 |
| rs707555 | benign | 0 | 1 | 0 |
| rs8192708 | benign | 0 | 1 | 0 |
| rs11552145 | benign | 0 | 0 | 0 |
| rs1804160 | benign | 0 | 1 | 0 |
| rs34103538 | benign | 0 | 1 | 0 |
| rs751393950 | pathogenic | 1 | 1 | 1 |
| rs121907979 | pathogenic | 1 | 1 | 1 |
| rs121907975 | pathogenic | 1 | 1 | 1 |
| rs121907962 | pathogenic | 1 | 1 | 1 |
| rs121907957 | pathogenic | 0 | 0 | 0 |
| rs121907953 | pathogenic | 1 | 1 | 1 |
| rs28941770 | pathogenic | 1 | 1 | 1 |
| rs28941771 | pathogenic | 1 | 1 | 1 |
| rs121907969 | pathogenic | 1 | 1 | 1 |
| rs121907973 | pathogenic | 1 | 1 | 1 |
| rs1800429 | pathogenic | 0 | 0 | 0 |
| rs121907976 | pathogenic | 1 | 1 | 0 |
| rs121907961 | pathogenic | 0 | 0 | 0 |
| rs121907974 | pathogenic | 0 | 0 | 0 |
| rs121907959 | pathogenic | 0 | 0 | 0 |
| rs121907971 | pathogenic | 1 | 1 | 0 |
| rs121907954 | pathogenic | 0 | 0 | 0 |
| rs121907977 | pathogenic | 0 | 0 | 0 |
| rs121907963 | pathogenic | 1 | 1 | 0 |
| rs370266293 | pathogenic | 1 | 1 | 0 |
| rs1800430 | benign | 0 | 0 | 0 |
| rs121907958 | pathogenic | 0 | 0 | 0 |
| rs202173526 | pathogenic | 0 | 0 | 0 |
| rs121907978 | pathogenic | 0 | 0 | 0 |
| rs121907981 | pathogenic | 0 | 1 | 0 |
| rs145012038 | benign | 1 | 1 | 1 |
| rs121907968 | pathogenic | 1 | 1 | 1 |
| rs121907966 | pathogenic | 1 | 1 | 1 |
| rs121907956 | pathogenic | 1 | 1 | 1 |
| rs28942071 | pathogenic | 1 | 1 | 1 |
| rs121907955 | pathogenic | 1 | 1 | 1 |
| rs4727853 | benign | 0 | 0 | 0 |
| rs201880593 | benign | 0 | 0 | 0 |
| rs201434579 | benign | 1 | 1 | 1 |
| rs74551128 | pathogenic | 1 | 1 | 0 |
| rs193922500 | pathogenic | 0 | 0 | 0 |
| rs121909009 | pathogenic | 1 | 1 | 0 |
| rs121908805 | pathogenic | 1 | 1 | 0 |
| rs139573311 | pathogenic | 0 | 0 | 0 |
| rs213950 | benign | 0 | 0 | 0 |

| | | | | |
|---|---|---|---|---|
| rs79282516 | pathogenic | 0 | 0 | 0 |
| rs121909017 | pathogenic | 1 | 1 | 1 |
| rs77101217 | pathogenic | 1 | 1 | 1 |
| rs1800092 | pathogenic | 0 | 1 | 0 |
| rs77646904 | pathogenic | 0 | 0 | 0 |
| rs1800095 | benign | 1 | 1 | 1 |
| rs387906368 | pathogenic | 1 | 1 | 1 |
| rs113993959 | pathogenic | 1 | 1 | 1 |
| rs121908755 | pathogenic | 1 | 1 | 1 |
| rs121909005 | pathogenic | 1 | 1 | 1 |
| rs121908757 | pathogenic | 1 | 1 | 1 |
| rs75527207 | pathogenic | 1 | 1 | 0 |
| rs121909013 | pathogenic | 1 | 1 | 0 |
| rs121909044 | pathogenic | 1 | 1 | 0 |
| rs74597325 | pathogenic | 1 | 1 | 0 |
| rs75549581 | pathogenic | 0 | 0 | 0 |
| rs80055610 | pathogenic | 1 | 1 | 1 |
| rs1800097 | pathogenic | 0 | 0 | 0 |
| rs121909006 | pathogenic | 0 | 1 | 0 |
| rs397508276 | pathogenic | 0 | 0 | 0 |
| rs397508296 | pathogenic | 1 | 1 | 1 |
| rs139468767 | pathogenic | 0 | 0 | 0 |
| rs121908759 | pathogenic | 1 | 1 | 0 |
| rs121909033 | pathogenic | 1 | 1 | 0 |
| rs397508331 | pathogenic | 0 | 0 | 0 |
| rs387907279 | pathogenic | 1 | 1 | 1 |
| rs546530618 | benign | 0 | 0 | 0 |
| rs558259739 | benign | 0 | 0 | 0 |
| rs201827340 | benign | 0 | 0 | 0 |
| rs148865119 | pathogenic | 1 | 1 | 0 |
| rs2229413 | benign | 1 | 1 | 1 |
| rs34040148 | benign | 1 | 1 | 1 |
| rs121964964 | pathogenic | 1 | 1 | 1 |
| rs760214620 | pathogenic | 1 | 1 | 1 |
| rs775992753 | pathogenic | 1 | 1 | 1 |
| rs770095972 | pathogenic | 1 | 1 | 1 |
| rs781444670 | pathogenic | 1 | 1 | 1 |
| rs147474549 | pathogenic | 0 | 0 | 0 |
| rs121964965 | pathogenic | 0 | 1 | 0 |
| rs121964963 | pathogenic | 1 | 1 | 1 |
| rs755952006 | pathogenic | 1 | 1 | 1 |
| rs121964970 | pathogenic | 1 | 1 | 1 |
| rs121964973 | pathogenic | 1 | 1 | 1 |
| rs773734233 | pathogenic | 1 | 1 | 1 |
| rs758236584 | pathogenic | 1 | 1 | 1 |
| rs149119723 | pathogenic | 1 | 1 | 0 |
| rs5742905 | pathogenic | 0 | 0 | 0 |
| rs121964962 | pathogenic | 1 | 1 | 1 |
| rs781567152 | pathogenic | 1 | 1 | 0 |
| rs398123151 | pathogenic | 1 | 1 | 1 |
| rs760417941 | pathogenic | 1 | 1 | 1 |
| rs774926464 | pathogenic | 0 | 0 | 0 |
| rs121964972 | pathogenic | 1 | 1 | 0 |
| rs372010465 | pathogenic | 1 | 1 | 1 |

| | | | | |
|---|---|---|---|---|
| rs763036586 | pathogenic | 1 | 1 | 1 |
| rs769080151 | pathogenic | 1 | 1 | 1 |
| rs121964967 | pathogenic | 1 | 1 | 1 |
| rs28934892 | pathogenic | 1 | 1 | 0 |
| rs28934891 | pathogenic | 1 | 1 | 1 |
| rs121964971 | pathogenic | 1 | 1 | 1 |
| rs121964968 | pathogenic | 1 | 1 | 1 |
| rs150828989 | benign | 1 | 1 | 1 |
| rs3775291 | benign | 1 | 1 | 0 |
| rs104894419 | pathogenic | 1 | 1 | 1 |
| rs2232642 | benign | 0 | 0 | 0 |
| rs104894551 | pathogenic | 1 | 1 | 0 |
| rs766328537 | pathogenic | 0 | 1 | 0 |
| rs104894553 | pathogenic | 1 | 1 | 0 |
| rs80099330 | benign | 0 | 0 | 0 |
| rs199565861 | pathogenic | 0 | 0 | 0 |
| rs104894548 | pathogenic | 0 | 0 | 0 |
| rs104894549 | pathogenic | 0 | 0 | 0 |
| rs104894550 | pathogenic | 1 | 1 | 1 |
| rs12948217 | pathogenic | 1 | 1 | 1 |
| rs761064915 | pathogenic | 0 | 0 | 0 |
| rs28940279 | pathogenic | 1 | 1 | 1 |
| rs774323189 | pathogenic | 1 | 1 | 1 |
| rs28940574 | pathogenic | 0 | 0 | 0 |
| rs515299 | benign | 1 | 1 | 1 |
| rs10127939 | benign | 1 | 1 | 1 |
| rs396991 | benign | 1 | 1 | 1 |
| rs145956175 | benign | 0 | 1 | 0 |
| rs747838255 | pathogenic | 1 | 1 | 1 |
| rs762659685 | pathogenic | 1 | 1 | 0 |
| rs104894831 | pathogenic | 0 | 1 | 0 |
| rs104894848 | pathogenic | 1 | 1 | 0 |
| rs104894833 | pathogenic | 0 | 1 | 0 |
| rs781838005 | pathogenic | 0 | 1 | 0 |
| rs104894834 | pathogenic | 1 | 1 | 0 |
| rs372966991 | pathogenic | 1 | 1 | 0 |
| rs138886989 | benign | 1 | 1 | 1 |
| rs28935195 | pathogenic | 0 | 1 | 0 |
| rs28935196 | pathogenic | 0 | 1 | 0 |
| rs372416832 | pathogenic | 1 | 1 | 1 |
| rs28935197 | pathogenic | 1 | 1 | 0 |
| rs28935486 | pathogenic | 1 | 1 | 0 |
| rs28935487 | pathogenic | 1 | 1 | 0 |
| rs28935488 | pathogenic | 1 | 1 | 1 |
| rs28935495 | pathogenic | 1 | 1 | 1 |
| rs28935485 | pathogenic | 1 | 1 | 1 |
| rs140329381 | pathogenic | 0 | 1 | 0 |
| rs104894830 | pathogenic | 0 | 1 | 0 |
| rs104894846 | pathogenic | 0 | 1 | 0 |
| rs28935489 | pathogenic | 0 | 1 | 0 |
| rs104894828 | pathogenic | 1 | 1 | 0 |
| rs28935491 | pathogenic | 1 | 1 | 0 |
| rs104894832 | pathogenic | 1 | 1 | 0 |
| rs28935492 | pathogenic | 1 | 1 | 0 |

| | | | | |
|---|---|---|---|---|
| rs730880437 | benign | 1 | 1 | 0 |
| rs28935493 | pathogenic | 0 | 1 | 0 |
| rs28935494 | pathogenic | 1 | 1 | 1 |
| rs111422676 | pathogenic | 1 | 1 | 1 |
| rs397515869 | benign | 1 | 1 | 0 |
| rs782449839 | pathogenic | 0 | 1 | 0 |
| rs104894852 | pathogenic | 1 | 1 | 1 |
| rs116929575 | pathogenic | 1 | 1 | 1 |
| rs7023954 | benign | 0 | 0 | 0 |
| rs184520335 | benign | 1 | 1 | 1 |
| rs143080527 | benign | 0 | 0 | 0 |
| rs1050565 | benign | 0 | 0 | 0 |
| rs11545035 | benign | 0 | 0 | 0 |
| rs267607190 | pathogenic | 1 | 1 | 1 |
| rs200283987 | benign | 0 | 0 | 0 |
| rs372556807 | benign | 1 | 1 | 0 |
| rs563073392 | benign | 0 | 0 | 0 |
| rs4539 | benign | 0 | 0 | 0 |
| rs28931609 | pathogenic | 1 | 1 | 1 |
| rs121912978 | pathogenic | 1 | 1 | 1 |
| rs104894072 | pathogenic | 1 | 1 | 1 |
| rs551933154 | benign | 1 | 1 | 1 |
| rs752962897 | benign | 1 | 1 | 0 |
| rs121912977 | pathogenic | 1 | 1 | 0 |
| rs4537 | benign | 1 | 1 | 1 |
| rs201830462 | benign | 0 | 0 | 0 |
| rs4544 | benign | 0 | 0 | 0 |
| rs562670189 | benign | 0 | 0 | 0 |
| rs4538 | benign | 0 | 1 | 0 |
| rs61757294 | pathogenic | 0 | 0 | 0 |
| rs5313 | benign | 0 | 0 | 0 |
| rs4545 | benign | 0 | 0 | 0 |
| rs72554627 | pathogenic | 1 | 1 | 1 |
| rs72554626 | pathogenic | 1 | 1 | 1 |
| rs387906861 | pathogenic | 1 | 1 | 1 |
| rs749655461 | pathogenic | 1 | 1 | 1 |
| rs121909606 | pathogenic | 1 | 1 | 1 |
| rs121909607 | pathogenic | 1 | 1 | 1 |
| rs200203992 | benign | 0 | 0 | 0 |
| rs6050 | benign | 0 | 0 | 0 |
| rs184635235 | benign | 0 | 0 | 0 |
| rs139146037 | benign | 0 | 0 | 0 |
| rs121909612 | pathogenic | 0 | 0 | 0 |
| rs78506343 | pathogenic | 0 | 0 | 0 |
| rs121918138 | pathogenic | 1 | 1 | 1 |
| rs121918139 | pathogenic | 1 | 1 | 1 |
| rs121918140 | pathogenic | 0 | 0 | 0 |
| rs36063028 | benign | 1 | 1 | 1 |
| rs45495192 | benign | 1 | 1 | 1 |
| rs45458802 | benign | 1 | 1 | 1 |
| rs552865793 | pathogenic | 1 | 1 | 1 |
| rs753744791 | pathogenic | 1 | 1 | 1 |
| rs777518512 | benign | 1 | 1 | 1 |
| rs142708991 | benign | 1 | 1 | 0 |

| | | | | |
|---|---|---|---|---|
| rs746069957 | benign | 0 | 0 | 0 |
| rs1334811 | benign | 0 | 0 | 0 |
| rs138894008 | benign | 1 | 1 | 0 |
| rs35030851 | benign | 1 | 1 | 0 |
| rs35378598 | benign | 0 | 0 | 0 |
| rs35818764 | benign | 1 | 1 | 1 |
| rs4631561 | benign | 1 | 1 | 1 |
| rs121918660 | pathogenic | 0 | 0 | 0 |
| rs373175587 | benign | 0 | 0 | 0 |
| rs61731956 | pathogenic | 1 | 1 | 1 |
| rs180177305 | pathogenic | 0 | 0 | 0 |
| rs119490108 | pathogenic | 0 | 0 | 0 |
| rs180177312 | pathogenic | 1 | 1 | 0 |
| rs180177314 | pathogenic | 1 | 1 | 0 |
| rs180177322 | pathogenic | 1 | 1 | 1 |
| rs180177323 | pathogenic | 1 | 1 | 1 |
| rs180177325 | pathogenic | 0 | 0 | 0 |
| rs35196441 | benign | 0 | 0 | 0 |
| rs28763904 | benign | 0 | 0 | 0 |
| rs2229519 | benign | 0 | 0 | 0 |
| rs137852886 | pathogenic | 0 | 0 | 0 |
| rs137852892 | pathogenic | 0 | 0 | 0 |
| rs770427750 | pathogenic | 0 | 0 | 0 |
| rs137852887 | pathogenic | 0 | 0 | 0 |
| rs137852893 | pathogenic | 0 | 0 | 0 |
| rs80338671 | pathogenic | 1 | 1 | 0 |
| rs2228389 | benign | 0 | 0 | 0 |
| rs80338672 | pathogenic | 0 | 0 | 0 |
| rs201958741 | pathogenic | 0 | 0 | 0 |
| rs80338673 | pathogenic | 0 | 0 | 0 |
| rs137852888 | pathogenic | 0 | 0 | 0 |
| rs137852889 | pathogenic | 0 | 0 | 0 |
| rs137852894 | pathogenic | 0 | 0 | 0 |
| rs137852890 | pathogenic | 0 | 0 | 0 |
| rs137852891 | pathogenic | 0 | 0 | 0 |
| rs766935302 | pathogenic | 0 | 0 | 0 |
| rs193074572 | benign | 0 | 0 | 0 |
| rs555210692 | benign | 1 | 1 | 0 |
| rs119103242 | pathogenic | 0 | 1 | 0 |
| rs143725101 | benign | 0 | 1 | 0 |
| rs28940589 | pathogenic | 1 | 1 | 1 |
| rs113994145 | pathogenic | 0 | 0 | 0 |
| rs773872994 | benign | 0 | 0 | 0 |
| rs28940590 | pathogenic | 1 | 1 | 0 |
| rs145400751 | benign | 1 | 1 | 1 |
| rs587782331 | benign | 0 | 0 | 0 |
| rs730881787 | benign | 0 | 0 | 0 |
| rs587778531 | benign | 0 | 0 | 0 |
| rs781652274 | benign | 0 | 0 | 0 |
| rs55882234 | benign | 0 | 0 | 0 |
| rs2020908 | benign | 0 | 0 | 0 |
| rs63750741 | pathogenic | 1 | 1 | 1 |
| rs41295268 | benign | 1 | 1 | 1 |
| rs63750897 | benign | 1 | 1 | 0 |

| | | | | |
|---|---|---|---|---|
| rs63751005 | benign | 1 | 1 | 0 |
| rs587779223 | benign | 0 | 0 | 0 |
| rs202127474 | benign | 0 | 0 | 0 |
| rs374230313 | benign | 0 | 0 | 0 |
| rs2020912 | benign | 0 | 0 | 0 |
| rs63751017 | pathogenic | 0 | 0 | 0 |
| rs587779246 | pathogenic | 0 | 0 | 0 |
| rs63749919 | pathogenic | 0 | 0 | 0 |
| rs63750563 | pathogenic | 1 | 1 | 1 |
| rs63749999 | pathogenic | 0 | 0 | 0 |
| rs200492211 | pathogenic | 0 | 0 | 0 |
| rs576269342 | benign | 0 | 0 | 0 |
| rs267608075 | benign | 0 | 0 | 0 |
| rs149605979 | benign | 0 | 0 | 0 |
| rs63750252 | benign | 0 | 0 | 0 |
| rs754289472 | benign | 0 | 0 | 0 |
| rs63750119 | pathogenic | 0 | 0 | 0 |
| rs587782109 | benign | 0 | 0 | 0 |
| rs34625968 | benign | 1 | 1 | 1 |
| rs764786814 | benign | 1 | 1 | 1 |
| rs199594809 | benign | 1 | 1 | 1 |
| rs74315364 | pathogenic | 1 | 1 | 1 |
| rs201253919 | pathogenic | 0 | 0 | 0 |
| rs387906828 | pathogenic | 1 | 1 | 1 |
| rs34521609 | benign | 1 | 1 | 1 |
| rs199472894 | pathogenic | 0 | 0 | 0 |
| rs199472902 | pathogenic | 0 | 0 | 0 |
| rs121912505 | pathogenic | 0 | 0 | 0 |
| rs28928905 | pathogenic | 0 | 0 | 0 |
| rs374376640 | benign | 0 | 0 | 0 |
| rs202102276 | benign | 0 | 0 | 0 |
| rs199472918 | pathogenic | 0 | 0 | 0 |
| rs121912516 | pathogenic | 1 | 1 | 1 |
| rs121912504 | pathogenic | 0 | 0 | 0 |
| rs9333649 | pathogenic | 0 | 0 | 0 |
| rs121912508 | pathogenic | 0 | 0 | 0 |
| rs199473428 | pathogenic | 1 | 1 | 1 |
| rs104894021 | pathogenic | 0 | 0 | 0 |
| rs28928904 | pathogenic | 1 | 1 | 1 |
| rs199473039 | benign | 1 | 1 | 1 |
| rs199472958 | pathogenic | 1 | 1 | 1 |
| rs1137617 | pathogenic | 1 | 1 | 1 |
| rs199473532 | pathogenic | 1 | 1 | 1 |
| rs199472988 | pathogenic | 0 | 0 | 0 |
| rs189014161 | pathogenic | 0 | 0 | 0 |
| rs12720441 | pathogenic | 0 | 0 | 0 |
| rs199473001 | pathogenic | 1 | 1 | 0 |
| rs121912506 | pathogenic | 1 | 1 | 0 |
| rs773724817 | pathogenic | 1 | 1 | 1 |
| rs118204002 | pathogenic | 1 | 1 | 1 |
| rs138556525 | pathogenic | 1 | 1 | 1 |
| rs72558181 | pathogenic | 1 | 1 | 1 |
| rs118204005 | pathogenic | 1 | 1 | 1 |
| rs118204004 | pathogenic | 0 | 0 | 0 |

| | | | | |
|---|---|---|---|---|
| rs118204001 | pathogenic | 1 | 1 | 1 |
| rs118204006 | pathogenic | 1 | 1 | 1 |
| rs118204003 | pathogenic | 0 | 0 | 0 |
| rs121913147 | pathogenic | 1 | 1 | 0 |
| rs2229429 | benign | 1 | 1 | 0 |
| rs182552223 | benign | 0 | 0 | 0 |
| rs146588336 | benign | 0 | 0 | 0 |
| rs121964983 | pathogenic | 0 | 1 | 0 |
| rs121964982 | pathogenic | 0 | 0 | 0 |
| rs386833679 | pathogenic | 0 | 0 | 0 |
| rs386833680 | pathogenic | 0 | 0 | 0 |
| rs121964981 | pathogenic | 1 | 1 | 0 |
| rs121964984 | pathogenic | 0 | 0 | 0 |
| rs144971200 | benign | 0 | 0 | 0 |
| rs121964985 | pathogenic | 1 | 1 | 0 |
| rs141246107 | benign | 0 | 0 | 0 |
| rs104894561 | pathogenic | 1 | 1 | 0 |
| rs377023736 | pathogenic | 0 | 0 | 0 |
| rs121964907 | pathogenic | 1 | 1 | 1 |
| rs121964909 | pathogenic | 1 | 1 | 1 |
| rs765070743 | pathogenic | 0 | 0 | 0 |
| rs2228119 | benign | 0 | 0 | 0 |
| rs745976989 | pathogenic | 1 | 1 | 1 |
| rs121964904 | pathogenic | 0 | 0 | 0 |
| rs386833430 | pathogenic | 1 | 1 | 1 |
| rs267606992 | pathogenic | 0 | 0 | 0 |
| rs61752063 | pathogenic | 0 | 0 | 0 |
| rs61752067 | pathogenic | 1 | 1 | 1 |
| rs104894934 | pathogenic | 0 | 0 | 0 |
| rs104894929 | pathogenic | 1 | 1 | 1 |
| rs80189640 | benign | 0 | 0 | 0 |
| rs137854547 | pathogenic | 1 | 1 | 1 |
| rs137854540 | pathogenic | 1 | 1 | 1 |
| rs782092363 | pathogenic | 1 | 1 | 1 |
| rs147783915 | benign | 0 | 0 | 0 |
| rs28933687 | pathogenic | 1 | 1 | 1 |
| rs149503319 | benign | 0 | 0 | 0 |
| rs1805147 | benign | 0 | 0 | 0 |
| rs587777218 | pathogenic | 1 | 1 | 1 |
| rs200786705 | benign | 0 | 0 | 0 |
| rs56219475 | benign | 0 | 0 | 0 |
| rs587781172 | benign | 0 | 0 | 0 |
| rs757094189 | pathogenic | 0 | 0 | 0 |
| rs140811086 | benign | 1 | 1 | 1 |
| rs574088829 | pathogenic | 1 | 1 | 1 |
| rs121918525 | pathogenic | 0 | 0 | 0 |
| rs121918527 | pathogenic | 0 | 0 | 0 |
| rs121964848 | pathogenic | 0 | 1 | 0 |
| rs121964845 | pathogenic | 1 | 1 | 1 |
| rs121964846 | pathogenic | 0 | 1 | 0 |
| rs17217716 | benign | 1 | 1 | 1 |
| rs141711342 | benign | 0 | 0 | 0 |
| rs372619120 | benign | 0 | 0 | 0 |
| rs63750002 | benign | 1 | 1 | 1 |

| | | | | |
|---|---|---|---|---|
| rs193922373 | benign | 0 | 0 | 0 |
| rs41295286 | benign | 0 | 0 | 0 |
| rs17217772 | benign | 0 | 0 | 0 |
| rs145649774 | benign | 0 | 0 | 0 |
| rs63750214 | pathogenic | 0 | 0 | 0 |
| rs63750716 | benign | 0 | 0 | 0 |
| rs202026056 | benign | 0 | 0 | 0 |
| rs4987188 | benign | 1 | 1 | 0 |
| rs267607938 | benign | 0 | 0 | 0 |
| rs17224367 | benign | 0 | 0 | 0 |
| rs63750006 | benign | 0 | 0 | 0 |
| rs587781373 | benign | 0 | 0 | 0 |
| rs63751207 | pathogenic | 1 | 1 | 0 |
| rs730881759 | benign | 0 | 0 | 0 |
| rs372350768 | benign | 1 | 1 | 1 |
| rs55778204 | benign | 1 | 1 | 0 |
| rs63749910 | benign | 0 | 0 | 0 |
| rs63750844 | benign | 0 | 0 | 0 |
| rs28929483 | pathogenic | 0 | 0 | 0 |
| rs61756468 | benign | 0 | 0 | 0 |
| rs63750875 | pathogenic | 1 | 1 | 0 |
| rs776528054 | benign | 1 | 1 | 0 |
| rs751939698 | benign | 0 | 0 | 0 |
| rs587780685 | benign | 0 | 0 | 0 |
| rs730881763 | benign | 1 | 1 | 0 |
| rs373226409 | benign | 0 | 0 | 0 |
| rs202145681 | benign | 1 | 1 | 1 |
| rs63749841 | benign | 1 | 1 | 1 |
| rs63750757 | benign | 1 | 1 | 1 |
| rs587778527 | benign | 1 | 1 | 1 |
| rs63750849 | benign | 0 | 0 | 0 |
| rs587779969 | benign | 0 | 0 | 0 |
| rs121908046 | pathogenic | 1 | 1 | 1 |
| rs28940882 | pathogenic | 1 | 1 | 0 |
| rs28940883 | pathogenic | 1 | 1 | 1 |
| rs727503943 | pathogenic | 1 | 1 | 1 |
| rs137853859 | pathogenic | 1 | 1 | 1 |
| rs121908045 | pathogenic | 1 | 1 | 0 |
| rs137853861 | pathogenic | 1 | 1 | 0 |
| rs3180383 | pathogenic | 0 | 1 | 0 |
| rs144431814 | benign | 0 | 0 | 0 |
| rs139502866 | benign | 0 | 0 | 0 |
| rs17524437 | benign | 0 | 0 | 0 |
| rs760960779 | benign | 0 | 0 | 0 |
| rs148960463 | pathogenic | 0 | 0 | 0 |
| rs111033647 | pathogenic | 1 | 1 | 1 |
| rs111033648 | pathogenic | 0 | 0 | 0 |
| rs1800461 | benign | 0 | 0 | 0 |
| rs111033656 | pathogenic | 0 | 0 | 0 |
| rs111033663 | pathogenic | 1 | 1 | 0 |
| rs111033669 | pathogenic | 1 | 1 | 1 |
| rs111033690 | pathogenic | 0 | 0 | 0 |
| rs111033686 | pathogenic | 0 | 0 | 0 |
| rs111033687 | pathogenic | 0 | 0 | 0 |

| | | | | |
|---|---|---|---|---|
| rs111033695 | pathogenic | 0 | 0 | 0 |
| rs111033697 | pathogenic | 0 | 0 | 0 |
| rs111033693 | pathogenic | 0 | 0 | 0 |
| rs111033694 | pathogenic | 0 | 0 | 0 |
| rs111033721 | pathogenic | 1 | 1 | 1 |
| rs111033722 | pathogenic | 0 | 0 | 0 |
| rs111033725 | pathogenic | 1 | 1 | 0 |
| rs75391579 | pathogenic | 1 | 1 | 1 |
| rs111033728 | pathogenic | 1 | 1 | 1 |
| rs111033735 | pathogenic | 1 | 1 | 1 |
| rs111033739 | pathogenic | 1 | 1 | 1 |
| rs111033736 | pathogenic | 0 | 0 | 0 |
| rs111033744 | pathogenic | 1 | 1 | 1 |
| rs111033750 | pathogenic | 1 | 1 | 1 |
| rs111033752 | pathogenic | 0 | 0 | 0 |
| rs111033751 | pathogenic | 0 | 0 | 0 |
| rs111033846 | pathogenic | 0 | 0 | 0 |
| rs368166217 | pathogenic | 0 | 0 | 0 |
| rs111033764 | pathogenic | 0 | 0 | 0 |
| rs111033847 | benign | 0 | 0 | 0 |
| rs111033778 | pathogenic | 0 | 0 | 0 |
| rs111033773 | pathogenic | 0 | 0 | 0 |
| rs111033792 | pathogenic | 0 | 0 | 0 |
| rs367543266 | pathogenic | 1 | 1 | 1 |
| rs111033794 | pathogenic | 0 | 0 | 0 |
| rs111033802 | pathogenic | 0 | 0 | 0 |
| rs111033803 | pathogenic | 0 | 0 | 0 |
| rs111033809 | pathogenic | 1 | 1 | 0 |
| rs121918376 | pathogenic | 0 | 1 | 0 |
| rs121918377 | pathogenic | 0 | 1 | 0 |
| rs121918378 | pathogenic | 1 | 1 | 1 |
| rs121918374 | pathogenic | 1 | 1 | 1 |
| rs137852379 | pathogenic | 0 | 0 | 0 |
| rs137852388 | pathogenic | 1 | 1 | 1 |
| rs137852368 | pathogenic | 0 | 0 | 0 |
| rs28935499 | pathogenic | 0 | 0 | 0 |
| rs28933668 | pathogenic | 0 | 0 | 0 |
| rs28933669 | pathogenic | 0 | 0 | 0 |
| rs28933670 | pathogenic | 0 | 0 | 0 |
| rs28933671 | pathogenic | 0 | 0 | 0 |
| rs28933672 | pathogenic | 0 | 0 | 0 |
| rs28937272 | pathogenic | 0 | 0 | 0 |
| rs35383156 | benign | 0 | 0 | 0 |
| rs137852418 | pathogenic | 0 | 0 | 0 |
| rs137852419 | pathogenic | 0 | 0 | 0 |
| rs137852428 | pathogenic | 0 | 0 | 0 |
| rs137852436 | pathogenic | 0 | 0 | 0 |
| rs28937285 | pathogenic | 0 | 0 | 0 |
| rs35213384 | benign | 1 | 1 | 1 |
| rs3810950 | benign | 0 | 0 | 0 |
| rs201479289 | pathogenic | 0 | 0 | 0 |
| rs121912820 | pathogenic | 0 | 1 | 0 |
| rs121912815 | pathogenic | 0 | 1 | 0 |
| rs115510708 | benign | 0 | 0 | 0 |

| | | | | |
|---|---|---|---|---|
| rs868749 | benign | 0 | 0 | 0 |
| rs121912823 | pathogenic | 0 | 0 | 0 |
| rs115212829 | benign | 0 | 0 | 0 |
| rs121912822 | pathogenic | 0 | 0 | 0 |
| rs121912816 | pathogenic | 0 | 1 | 0 |
| rs4838544 | benign | 0 | 0 | 0 |
| rs121912818 | pathogenic | 0 | 0 | 0 |
| rs121912821 | pathogenic | 0 | 0 | 0 |
| rs121912817 | pathogenic | 0 | 0 | 0 |
| rs8178992 | benign | 0 | 0 | 0 |
| rs121912819 | pathogenic | 1 | 1 | 0 |
| rs80097077 | benign | 1 | 1 | 0 |
| rs121909359 | pathogenic | 1 | 1 | 0 |
| rs121909358 | pathogenic | 1 | 1 | 1 |
| rs121909361 | pathogenic | 1 | 1 | 1 |
| rs121909357 | pathogenic | 0 | 1 | 0 |
| rs121909365 | pathogenic | 1 | 1 | 1 |
| rs6413484 | pathogenic | 1 | 1 | 1 |
| rs121909373 | pathogenic | 1 | 1 | 1 |
| rs121909366 | pathogenic | 1 | 1 | 1 |
| rs121909367 | pathogenic | 1 | 1 | 1 |
| rs121909368 | pathogenic | 1 | 1 | 1 |
| rs121909369 | pathogenic | 0 | 0 | 0 |
| rs121909363 | pathogenic | 1 | 1 | 1 |
| rs45588036 | pathogenic | 1 | 1 | 1 |
| rs121909364 | pathogenic | 1 | 1 | 1 |
| rs80358282 | benign | 0 | 0 | 0 |
| rs80358284 | pathogenic | 1 | 1 | 1 |
| rs61735303 | benign | 0 | 0 | 0 |
| rs2229274 | benign | 0 | 0 | 0 |
| rs145006491 | benign | 0 | 0 | 0 |
| rs121913580 | pathogenic | 1 | 1 | 0 |
| rs730882196 | pathogenic | 0 | 0 | 0 |
| rs153477 | benign | 0 | 0 | 0 |
| rs153478 | benign | 1 | 1 | 0 |
| rs61740602 | benign | 1 | 1 | 0 |
| rs104893892 | pathogenic | 1 | 1 | 0 |
| rs121909595 | pathogenic | 1 | 1 | 1 |
| rs28931605 | pathogenic | 1 | 1 | 1 |
| rs121909597 | pathogenic | 0 | 1 | 0 |
| rs202233735 | pathogenic | 0 | 1 | 0 |
| rs121909596 | pathogenic | 1 | 1 | 1 |
| rs75156162 | benign | 0 | 1 | 0 |
| rs2305430 | benign | 1 | 1 | 1 |
| rs150318966 | benign | 0 | 0 | 0 |
| rs121909598 | pathogenic | 0 | 1 | 0 |
| rs5744175 | benign | 0 | 0 | 0 |
| rs28935476 | pathogenic | 1 | 1 | 1 |
| rs28935475 | pathogenic | 1 | 1 | 1 |
| rs122461163 | pathogenic | 1 | 1 | 1 |
| rs62626305 | pathogenic | 1 | 1 | 1 |
| rs587777157 | pathogenic | 0 | 0 | 0 |
| rs137854478 | pathogenic | 1 | 1 | 0 |
| rs137854465 | pathogenic | 0 | 0 | 0 |

| | | | | |
|---|---|---|---|---|
| rs137854470 | pathogenic | 0 | 0 | 0 |
| rs137854482 | pathogenic | 0 | 0 | 0 |
| rs137854473 | pathogenic | 0 | 0 | 0 |
| rs137854456 | pathogenic | 0 | 0 | 0 |
| rs140598 | benign | 0 | 0 | 0 |
| rs104894105 | pathogenic | 1 | 1 | 1 |
| rs104894107 | pathogenic | 1 | 1 | 1 |
| rs104894106 | pathogenic | 1 | 1 | 1 |
| rs74621026 | benign | 1 | 1 | 1 |
| rs2290332 | benign | 0 | 0 | 0 |
| rs780202604 | pathogenic | 1 | 1 | 1 |
| rs74315305 | pathogenic | 1 | 1 | 0 |
| rs766751878 | pathogenic | 1 | 1 | 1 |
| rs137854603 | pathogenic | 1 | 1 | 1 |
| rs139592595 | benign | 1 | 1 | 0 |
| rs150565592 | benign | 1 | 1 | 1 |
| rs763520126 | benign | 0 | 0 | 0 |
| rs55980502 | benign | 0 | 0 | 0 |
| rs137852812 | pathogenic | 0 | 0 | 0 |
| rs137852813 | pathogenic | 0 | 0 | 0 |
| rs190222208 | benign | 0 | 0 | 0 |
| rs138459502 | pathogenic | 0 | 0 | 0 |
| rs730881041 | pathogenic | 0 | 0 | 0 |
| rs267607080 | pathogenic | 0 | 0 | 0 |
| rs137852814 | pathogenic | 0 | 0 | 0 |
| rs267607079 | pathogenic | 0 | 0 | 0 |
| rs192378417 | benign | 0 | 0 | 0 |
| rs1288775 | benign | 0 | 0 | 0 |
| rs80338737 | pathogenic | 1 | 1 | 1 |
| rs397514708 | pathogenic | 0 | 0 | 0 |
| rs775933965 | pathogenic | 0 | 0 | 0 |
| rs747608698 | benign | 0 | 1 | 0 |
| rs1145086 | benign | 0 | 1 | 0 |
| rs121912953 | pathogenic | 1 | 1 | 0 |
| rs243849 | benign | 0 | 0 | 0 |
| rs121912955 | pathogenic | 1 | 1 | 1 |
| rs17859943 | benign | 0 | 1 | 0 |
| rs755557988 | benign | 1 | 1 | 1 |
| rs10195 | benign | 0 | 0 | 0 |
| rs61735357 | benign | 1 | 1 | 1 |
| rs137882374 | benign | 0 | 0 | 0 |
| rs5388 | benign | 0 | 0 | 0 |
| rs6352 | benign | 0 | 0 | 0 |
| rs76645461 | pathogenic | 1 | 1 | 0 |
| rs1050828 | pathogenic | 0 | 0 | 0 |
| rs1050829 | pathogenic | 0 | 0 | 0 |
| rs137852314 | pathogenic | 0 | 0 | 0 |
| rs137852331 | pathogenic | 0 | 0 | 0 |
| rs5030872 | pathogenic | 0 | 0 | 0 |
| rs267606836 | pathogenic | 0 | 0 | 0 |
| rs5030868 | pathogenic | 0 | 0 | 0 |
| rs137852330 | pathogenic | 0 | 1 | 0 |
| rs137852326 | pathogenic | 1 | 1 | 1 |
| rs137852318 | pathogenic | 1 | 1 | 1 |

| | | | | |
|---|---|---|---|---|
| rs74575103 | benign | 0 | 0 | 0 |
| rs137852327 | pathogenic | 1 | 1 | 1 |
| rs76723693 | pathogenic | 0 | 0 | 0 |
| rs5030869 | pathogenic | 0 | 0 | 0 |
| rs34193178 | pathogenic | 1 | 1 | 1 |
| rs137852316 | pathogenic | 1 | 1 | 0 |
| rs398123546 | pathogenic | 1 | 1 | 1 |
| rs72554665 | pathogenic | 0 | 0 | 0 |
| rs72554664 | pathogenic | 0 | 0 | 0 |
| rs28939068 | pathogenic | 1 | 1 | 1 |
| rs59301204 | pathogenic | 1 | 1 | 1 |
| rs143715750 | benign | 1 | 1 | 1 |
| rs35263902 | pathogenic | 1 | 1 | 1 |
| rs202214518 | benign | 1 | 1 | 1 |
| rs34076756 | benign | 0 | 0 | 0 |
| rs199592341 | pathogenic | 1 | 1 | 0 |
| rs121908833 | pathogenic | 1 | 1 | 1 |
| rs121908834 | pathogenic | 1 | 1 | 0 |
| rs138454151 | benign | 0 | 0 | 0 |
| rs141323599 | benign | 0 | 0 | 0 |
| rs137853200 | pathogenic | 0 | 0 | 0 |
| rs137853199 | pathogenic | 0 | 0 | 0 |
| rs139787163 | benign | 0 | 0 | 0 |
| rs61731097 | benign | 1 | 1 | 1 |
| rs61729795 | benign | 0 | 0 | 0 |
| rs149242330 | benign | 0 | 0 | 0 |
| rs587777490 | pathogenic | 1 | 1 | 1 |
| rs398123011 | pathogenic | 1 | 1 | 0 |
| rs775186117 | pathogenic | 0 | 0 | 0 |
| rs72677237 | benign | 0 | 0 | 0 |
| rs104894163 | pathogenic | 1 | 1 | 1 |
| rs121909588 | pathogenic | 0 | 0 | 0 |
| rs1042579 | benign | 0 | 0 | 0 |
| rs121909273 | pathogenic | 0 | 0 | 0 |
| rs121909274 | pathogenic | 0 | 0 | 0 |
| rs119473034 | pathogenic | 1 | 1 | 1 |
| rs398123065 | pathogenic | 0 | 0 | 0 |
| rs587777042 | pathogenic | 1 | 1 | 0 |
| rs761413864 | pathogenic | 1 | 1 | 1 |
| rs118203965 | pathogenic | 0 | 0 | 0 |
| rs118203966 | pathogenic | 0 | 0 | 0 |
| rs1801201 | benign | 1 | 1 | 0 |
| rs1136201 | benign | 1 | 1 | 1 |
| rs144148008 | benign | 0 | 0 | 0 |
| rs200118378 | benign | 1 | 1 | 1 |
| rs112984118 | benign | 1 | 1 | 1 |
| rs149446102 | benign | 1 | 1 | 1 |
| rs1427463 | benign | 0 | 0 | 0 |
| rs201936720 | benign | 1 | 1 | 1 |
| rs61751983 | benign | 1 | 1 | 1 |
| rs104894632 | pathogenic | 1 | 1 | 1 |
| rs149202905 | benign | 0 | 0 | 0 |
| rs2230210 | benign | 1 | 1 | 1 |
| rs111631988 | benign | 0 | 0 | 0 |

| | | | | |
|---|---|---|---|---|
| rs137852550 | pathogenic | 0 | 0 | 0 |
| rs5963403 | benign | 0 | 0 | 0 |
| rs138018739 | benign | 0 | 0 | 0 |
| rs62640590 | benign | 0 | 0 | 0 |
| rs121909581 | pathogenic | 1 | 1 | 1 |
| rs202017913 | benign | 1 | 1 | 1 |
| rs143163513 | benign | 0 | 0 | 0 |
| rs238238 | benign | 0 | 1 | 0 |
| rs143945974 | benign | 0 | 0 | 0 |
| rs238239 | benign | 1 | 1 | 1 |
| rs121918403 | pathogenic | 0 | 1 | 0 |
| rs61735456 | benign | 0 | 1 | 0 |
| rs121918404 | pathogenic | 1 | 1 | 0 |
| rs12674488 | benign | 0 | 0 | 0 |
| rs201599657 | pathogenic | 0 | 0 | 0 |
| rs1057090 | benign | 0 | 0 | 0 |
| rs1057091 | benign | 0 | 0 | 0 |
| rs587783740 | benign | 0 | 0 | 0 |
| rs75956769 | benign | 0 | 0 | 0 |
| rs201691299 | benign | 0 | 0 | 0 |
| rs62621087 | benign | 0 | 0 | 0 |
| rs201308454 | benign | 0 | 0 | 0 |
| rs145657360 | benign | 0 | 0 | 0 |
| rs121434592 | pathogenic | 1 | 1 | 0 |
| rs375650413 | benign | 0 | 0 | 0 |
| rs121434437 | pathogenic | 1 | 1 | 0 |
| rs763494292 | pathogenic | 0 | 0 | 0 |
| rs121964997 | pathogenic | 1 | 1 | 0 |
| rs121964996 | pathogenic | 0 | 0 | 0 |
| rs41283425 | benign | 1 | 1 | 1 |
| rs376289528 | benign | 1 | 1 | 1 |
| rs144171604 | benign | 1 | 1 | 1 |
| rs782440692 | pathogenic | 1 | 1 | 1 |
| rs782058451 | pathogenic | 1 | 1 | 1 |
| rs78437817 | benign | 0 | 0 | 0 |
| rs200327969 | benign | 1 | 1 | 1 |
| rs143043662 | benign | 0 | 0 | 0 |
| rs111966833 | benign | 1 | 1 | 1 |
| rs35523678 | benign | 0 | 0 | 0 |
| rs515726208 | pathogenic | 0 | 0 | 0 |
| rs397517239 | pathogenic | 1 | 1 | 1 |
| rs148966602 | benign | 1 | 1 | 1 |
| rs76371972 | benign | 1 | 1 | 1 |
| rs2229968 | benign | 1 | 1 | 1 |
| rs61751538 | benign | 1 | 1 | 1 |
| rs200800978 | pathogenic | 1 | 1 | 1 |
| rs148695964 | benign | 1 | 1 | 1 |
| rs140042418 | benign | 1 | 1 | 1 |
| rs116185403 | benign | 0 | 0 | 0 |
| rs142828669 | benign | 1 | 1 | 1 |
| rs139832048 | benign | 0 | 1 | 0 |
| rs58750568 | benign | 0 | 1 | 0 |
| rs77681311 | benign | 0 | 0 | 0 |
| rs151208452 | benign | 0 | 0 | 0 |

| | | | | |
|---|---|---|---|---|
| rs76463933 | benign | 0 | 0 | 0 |
| rs144434647 | benign | 0 | 0 | 0 |
| rs117361485 | benign | 0 | 0 | 0 |
| rs587777588 | pathogenic | 0 | 0 | 0 |
| rs80338796 | pathogenic | 0 | 0 | 0 |
| rs121434594 | pathogenic | 0 | 0 | 0 |
| rs146766138 | pathogenic | 1 | 1 | 1 |
| rs28939679 | pathogenic | 1 | 1 | 1 |
| rs748754134 | pathogenic | 1 | 1 | 0 |
| rs764276946 | pathogenic | 1 | 1 | 1 |
| rs111033588 | pathogenic | 1 | 1 | 1 |
| rs201633414 | pathogenic | 1 | 1 | 1 |
| rs121909262 | pathogenic | 1 | 1 | 1 |
| rs104893702 | pathogenic | 1 | 1 | 1 |
| rs121909265 | pathogenic | 1 | 1 | 1 |
| rs121909266 | pathogenic | 1 | 1 | 1 |
| rs193922439 | pathogenic | 1 | 1 | 0 |
| rs104893691 | pathogenic | 1 | 1 | 1 |
| rs104893695 | pathogenic | 0 | 0 | 0 |
| rs104893708 | pathogenic | 1 | 1 | 1 |
| rs121909260 | pathogenic | 0 | 0 | 0 |
| rs104893696 | pathogenic | 0 | 0 | 0 |
| rs121909267 | pathogenic | 0 | 0 | 0 |
| rs121909263 | pathogenic | 0 | 0 | 0 |
| rs121909264 | pathogenic | 0 | 0 | 0 |
| rs104893694 | pathogenic | 1 | 1 | 1 |
| rs121909268 | pathogenic | 1 | 1 | 1 |
| rs104893689 | pathogenic | 1 | 1 | 0 |
| rs104893707 | pathogenic | 1 | 1 | 0 |
| rs104893697 | pathogenic | 1 | 1 | 1 |
| rs28936684 | pathogenic | 1 | 1 | 1 |
| rs62269092 | benign | 1 | 1 | 0 |
| rs121909259 | pathogenic | 1 | 1 | 0 |
| rs548403340 | benign | 1 | 1 | 0 |
| rs142818334 | benign | 0 | 0 | 0 |
| rs12493789 | benign | 1 | 1 | 1 |
| rs193922423 | pathogenic | 1 | 1 | 0 |
| rs104893719 | pathogenic | 1 | 1 | 1 |
| rs117375173 | benign | 1 | 1 | 0 |
| rs104893712 | pathogenic | 0 | 0 | 0 |
| rs121917776 | pathogenic | 1 | 1 | 1 |
| rs767325003 | benign | 1 | 1 | 1 |
| rs35824453 | benign | 0 | 0 | 0 |
| rs141641266 | benign | 0 | 0 | 0 |
| rs1050239 | benign | 0 | 0 | 0 |
| rs141303384 | benign | 0 | 0 | 0 |
| rs12720458 | pathogenic | 0 | 0 | 0 |
| rs368507376 | pathogenic | 1 | 1 | 1 |
| rs17215500 | pathogenic | 1 | 1 | 1 |
| rs120074188 | pathogenic | 1 | 1 | 1 |
| rs397508097 | pathogenic | 1 | 1 | 1 |
| rs138551008 | pathogenic | 0 | 0 | 0 |
| rs387907139 | pathogenic | 1 | 1 | 1 |
| rs765487627 | pathogenic | 0 | 0 | 0 |

| | | | | |
|---|---|---|---|---|
| rs587778587 | benign | 0 | 0 | 0 |
| rs45568339 | benign | 0 | 0 | 0 |
| rs78179744 | benign | 0 | 0 | 0 |
| rs45624036 | benign | 0 | 0 | 0 |
| rs61755166 | benign | 0 | 0 | 0 |
| rs180177132 | pathogenic | 1 | 1 | 1 |
| rs587776527 | pathogenic | 0 | 0 | 0 |
| rs373783514 | benign | 0 | 0 | 0 |
| rs730881896 | benign | 0 | 0 | 0 |
| rs118203998 | pathogenic | 0 | 0 | 0 |
| rs41281039 | pathogenic | 1 | 1 | 1 |
| rs118204430 | pathogenic | 1 | 1 | 1 |
| rs1800546 | pathogenic | 1 | 1 | 0 |
| rs76917243 | pathogenic | 0 | 1 | 0 |
| rs78340951 | pathogenic | 0 | 1 | 0 |
| rs77718928 | pathogenic | 0 | 0 | 0 |
| rs369586696 | pathogenic | 0 | 1 | 0 |
| rs745517517 | pathogenic | 1 | 1 | 0 |
| rs372872777 | pathogenic | 0 | 0 | 0 |
| rs748704459 | pathogenic | 0 | 0 | 0 |
| rs769215411 | pathogenic | 0 | 0 | 0 |
| rs139425890 | pathogenic | 0 | 0 | 0 |
| rs779694939 | pathogenic | 1 | 1 | 0 |
| rs773729410 | pathogenic | 0 | 0 | 0 |
| rs121908623 | pathogenic | 1 | 1 | 1 |
| rs121908622 | pathogenic | 1 | 1 | 1 |
| rs121908621 | pathogenic | 1 | 1 | 1 |
| rs762106720 | pathogenic | 1 | 1 | 0 |
| rs199877522 | pathogenic | 0 | 0 | 0 |
| rs74315291 | pathogenic | 1 | 1 | 1 |
| rs79137415 | benign | 0 | 0 | 0 |
| rs121913045 | pathogenic | 0 | 0 | 0 |
| rs34295337 | pathogenic | 0 | 0 | 0 |
| rs121913046 | pathogenic | 0 | 0 | 0 |
| rs121913047 | pathogenic | 0 | 0 | 0 |
| rs121913048 | pathogenic | 0 | 0 | 0 |
| rs267607166 | pathogenic | 0 | 0 | 0 |
| rs17850684 | benign | 1 | 1 | 1 |
| rs119476051 | pathogenic | 1 | 1 | 1 |
| rs17850683 | benign | 1 | 1 | 1 |
| rs119476049 | pathogenic | 1 | 1 | 1 |
| rs119476046 | pathogenic | 1 | 1 | 1 |
| rs119476048 | pathogenic | 0 | 0 | 0 |
| rs119476047 | pathogenic | 1 | 1 | 1 |
| rs28939094 | pathogenic | 1 | 1 | 1 |
| rs119476050 | pathogenic | 0 | 0 | 0 |
| rs387906941 | pathogenic | 1 | 1 | 1 |
| rs41511344 | pathogenic | 1 | 1 | 0 |
| rs748112833 | pathogenic | 1 | 1 | 1 |
| rs74342080 | benign | 1 | 1 | 0 |
| rs116482870 | benign | 0 | 0 | 0 |
| rs104893743 | pathogenic | 0 | 0 | 0 |
| rs117179004 | benign | 0 | 0 | 0 |
| rs587779409 | pathogenic | 1 | 1 | 1 |

| | | | | |
|---|---|---|---|---|
| rs3745406 | benign | 1 | 1 | 0 |
| rs387906696 | pathogenic | 0 | 0 | 0 |
| rs72549369 | pathogenic | 1 | 1 | 0 |
| rs45523133 | benign | 0 | 0 | 0 |
| rs119460972 | pathogenic | 0 | 1 | 0 |
| rs34929837 | benign | 0 | 1 | 0 |
| rs17323225 | benign | 0 | 1 | 0 |
| rs17011368 | benign | 0 | 0 | 0 |
| rs748902766 | pathogenic | 1 | 1 | 1 |
| rs387907292 | pathogenic | 0 | 0 | 0 |
| rs387907293 | pathogenic | 1 | 1 | 0 |
| rs371526758 | pathogenic | 1 | 1 | 1 |
| rs142968179 | pathogenic | 1 | 1 | 1 |
| rs387907291 | pathogenic | 1 | 1 | 0 |
| rs121907949 | pathogenic | 0 | 0 | 0 |
| rs121907947 | pathogenic | 0 | 0 | 0 |
| rs121907950 | pathogenic | 1 | 1 | 1 |
| rs121907948 | pathogenic | 1 | 1 | 1 |
| rs4926 | benign | 0 | 0 | 0 |
| rs748809996 | pathogenic | 0 | 0 | 0 |
| rs1799883 | benign | 1 | 1 | 0 |
| rs104893974 | pathogenic | 0 | 1 | 0 |
| rs104893973 | pathogenic | 0 | 1 | 0 |
| rs34452707 | benign | 0 | 0 | 0 |
| rs119467003 | pathogenic | 1 | 1 | 1 |
| rs121908261 | pathogenic | 0 | 1 | 0 |
| rs28933985 | pathogenic | 1 | 1 | 1 |
| rs80356669 | pathogenic | 1 | 1 | 1 |
| rs28936677 | pathogenic | 1 | 1 | 0 |
| rs121918326 | pathogenic | 0 | 0 | 0 |
| rs121918324 | pathogenic | 1 | 1 | 0 |
| rs121918325 | pathogenic | 1 | 1 | 0 |
| rs41270025 | pathogenic | 1 | 1 | 0 |
| rs121918323 | pathogenic | 0 | 0 | 0 |
| rs12735723 | pathogenic | 1 | 1 | 0 |
| rs36013429 | benign | 0 | 0 | 0 |
| rs28936676 | pathogenic | 1 | 1 | 0 |
| rs771809901 | pathogenic | 1 | 1 | 1 |
| rs758350837 | benign | 1 | 1 | 0 |
| rs121908192 | pathogenic | 1 | 1 | 1 |
| rs147568500 | pathogenic | 1 | 1 | 1 |
| rs3729734 | benign | 0 | 0 | 0 |
| rs371536347 | benign | 1 | 1 | 1 |
| rs80358193 | pathogenic | 1 | 1 | 1 |
| rs119476044 | pathogenic | 1 | 1 | 1 |
| rs111292798 | benign | 1 | 1 | 1 |
| rs121908063 | pathogenic | 1 | 1 | 0 |
| rs121908065 | pathogenic | 1 | 1 | 1 |
| rs121908062 | pathogenic | 1 | 1 | 1 |
| rs121908064 | pathogenic | 1 | 1 | 1 |
| rs61746421 | benign | 1 | 1 | 1 |
| rs34886916 | benign | 1 | 1 | 1 |
| rs118204045 | pathogenic | 0 | 0 | 0 |
| rs138607412 | benign | 1 | 1 | 1 |

| | | | | |
|---|---|---|---|---|
| rs118204046 | pathogenic | 0 | 0 | 0 |
| rs118204048 | pathogenic | 0 | 0 | 0 |
| rs118204044 | pathogenic | 0 | 0 | 0 |
| rs149664056 | benign | 0 | 0 | 0 |
| rs121909285 | pathogenic | 0 | 0 | 0 |
| rs1802498 | benign | 0 | 0 | 0 |
| rs1065761 | benign | 0 | 0 | 0 |
| rs119477054 | pathogenic | 0 | 0 | 0 |
| rs12720097 | benign | 0 | 0 | 0 |
| rs747281324 | pathogenic | 1 | 1 | 0 |
| rs367543028 | pathogenic | 1 | 1 | 0 |
| rs146077918 | benign | 0 | 0 | 0 |
| rs142551229 | benign | 0 | 0 | 0 |
| rs367543039 | pathogenic | 0 | 0 | 0 |
| rs587779884 | pathogenic | 1 | 1 | 0 |
| rs137853153 | pathogenic | 0 | 0 | 0 |
| rs367543029 | pathogenic | 0 | 0 | 0 |
| rs587783037 | pathogenic | 1 | 1 | 0 |
| rs142928725 | benign | 0 | 0 | 0 |
| rs28385141 | benign | 0 | 0 | 0 |
| rs587779888 | pathogenic | 0 | 0 | 0 |
| rs121909801 | pathogenic | 1 | 1 | 1 |
| rs121909790 | pathogenic | 1 | 1 | 0 |
| rs121909797 | pathogenic | 1 | 1 | 0 |
| rs121909794 | pathogenic | 1 | 1 | 1 |
| rs121909791 | pathogenic | 1 | 1 | 1 |
| rs121909793 | pathogenic | 1 | 1 | 1 |
| rs147977279 | benign | 1 | 1 | 1 |
| rs147352564 | benign | 0 | 0 | 0 |
| rs368758045 | benign | 1 | 1 | 1 |
| rs114498261 | benign | 1 | 1 | 0 |
| rs35484156 | benign | 0 | 0 | 0 |
| rs139176878 | benign | 0 | 0 | 0 |
| rs11543934 | benign | 0 | 0 | 0 |
| rs34192549 | benign | 0 | 0 | 0 |
| rs568736295 | benign | 0 | 0 | 0 |
| rs55747232 | benign | 0 | 0 | 0 |
| rs781887272 | benign | 0 | 0 | 0 |
| rs146273341 | benign | 0 | 0 | 0 |
| rs41473544 | benign | 0 | 0 | 0 |
| rs137852680 | pathogenic | 0 | 0 | 0 |
| rs55891000 | benign | 0 | 0 | 0 |
| rs140351189 | benign | 0 | 0 | 0 |
| rs137852679 | pathogenic | 1 | 1 | 0 |
| rs137852678 | pathogenic | 0 | 0 | 0 |
| rs137852677 | pathogenic | 1 | 1 | 1 |
| rs138620818 | benign | 0 | 0 | 0 |
| rs121908531 | pathogenic | 0 | 0 | 0 |
| rs72554307 | pathogenic | 0 | 0 | 0 |
| rs72554308 | pathogenic | 0 | 0 | 0 |
| rs72554312 | pathogenic | 1 | 1 | 0 |
| rs67939655 | pathogenic | 0 | 0 | 0 |
| rs67486158 | pathogenic | 0 | 0 | 0 |
| rs72554338 | pathogenic | 1 | 1 | 1 |

| | | | | |
|---|---|---|---|---|
| rs1800324 | pathogenic | 1 | 1 | 1 |
| rs72554356 | pathogenic | 1 | 1 | 1 |
| rs66656800 | pathogenic | 1 | 1 | 1 |
| rs68026851 | pathogenic | 1 | 1 | 1 |
| rs72558412 | pathogenic | 0 | 0 | 0 |
| rs72558415 | pathogenic | 0 | 0 | 0 |
| rs72558423 | pathogenic | 0 | 0 | 0 |
| rs67120076 | pathogenic | 1 | 1 | 1 |
| rs1800328 | pathogenic | 1 | 1 | 1 |
| rs72558451 | pathogenic | 1 | 1 | 1 |
| rs72558454 | pathogenic | 1 | 1 | 1 |
| rs672601372 | pathogenic | 0 | 0 | 0 |
| rs139644798 | pathogenic | 0 | 0 | 0 |
| rs369398935 | pathogenic | 0 | 0 | 0 |
| rs267607035 | pathogenic | 0 | 0 | 0 |
| rs267607036 | pathogenic | 1 | 1 | 1 |
| rs2302566 | benign | 1 | 1 | 0 |
| rs115959591 | benign | 0 | 0 | 0 |
| rs773876739 | pathogenic | 0 | 0 | 0 |
| rs1799967 | benign | 0 | 0 | 0 |
| rs80356968 | benign | 0 | 0 | 0 |
| rs28897695 | benign | 1 | 1 | 1 |
| rs80357314 | benign | 0 | 0 | 0 |
| rs80357169 | benign | 0 | 0 | 0 |
| rs397507239 | benign | 1 | 1 | 1 |
| rs80187739 | pathogenic | 1 | 1 | 1 |
| rs41293459 | pathogenic | 1 | 1 | 1 |
| rs55770810 | pathogenic | 1 | 1 | 1 |
| rs80356860 | pathogenic | 0 | 0 | 0 |
| rs56195342 | benign | 0 | 0 | 0 |
| rs80357442 | benign | 0 | 0 | 0 |
| rs41293463 | pathogenic | 1 | 1 | 1 |
| rs80357069 | pathogenic | 0 | 0 | 0 |
| rs397509275 | benign | 0 | 0 | 0 |
| rs80356920 | benign | 0 | 0 | 0 |
| rs397509284 | pathogenic | 0 | 0 | 0 |
| rs80357286 | benign | 0 | 0 | 0 |
| rs80357274 | benign | 0 | 0 | 0 |
| rs765656957 | benign | 0 | 0 | 0 |
| rs755427809 | benign | 0 | 0 | 0 |
| rs80357322 | benign | 0 | 0 | 0 |
| rs121434430 | pathogenic | 0 | 0 | 0 |
| rs121918450 | pathogenic | 1 | 1 | 1 |
| rs1532268 | benign | 0 | 0 | 0 |
| rs2303080 | benign | 0 | 0 | 0 |
| rs10064631 | benign | 0 | 0 | 0 |
| rs162036 | benign | 0 | 0 | 0 |
| rs2287780 | benign | 0 | 0 | 0 |
| rs16879334 | benign | 0 | 0 | 0 |
| rs137853061 | pathogenic | 1 | 1 | 0 |
| rs10380 | benign | 0 | 0 | 0 |
| rs1494558 | benign | 0 | 0 | 0 |
| rs1494555 | benign | 0 | 0 | 0 |
| rs192994060 | benign | 0 | 0 | 0 |

| | | | | |
|---|---|---|---|---|
| rs147128920 | benign | 1 | 1 | 0 |
| rs201803986 | pathogenic | 0 | 0 | 0 |
| rs746419489 | pathogenic | 0 | 1 | 0 |
| rs764396564 | pathogenic | 0 | 0 | 0 |
| rs758304537 | pathogenic | 0 | 0 | 0 |
| rs267606763 | pathogenic | 0 | 0 | 0 |
| rs267606762 | pathogenic | 0 | 0 | 0 |
| rs150702945 | pathogenic | 1 | 1 | 1 |
| rs202047589 | pathogenic | 0 | 0 | 0 |
| rs755562733 | pathogenic | 0 | 0 | 0 |
| rs267606764 | pathogenic | 1 | 1 | 1 |
| rs138207257 | pathogenic | 1 | 1 | 0 |
| rs149150736 | pathogenic | 1 | 1 | 1 |
| rs777046879 | pathogenic | 0 | 0 | 0 |
| rs104894408 | pathogenic | 1 | 1 | 1 |
| rs1801002 | pathogenic | 1 | 1 | 1 |
| rs28929485 | pathogenic | 1 | 1 | 1 |
| rs104894396 | pathogenic | 1 | 1 | 1 |
| rs371024165 | pathogenic | 0 | 0 | 0 |
| rs35887622 | pathogenic | 1 | 1 | 1 |
| rs587783644 | pathogenic | 1 | 1 | 1 |
| rs72474224 | pathogenic | 1 | 1 | 1 |
| rs111033296 | pathogenic | 1 | 1 | 1 |
| rs104894407 | pathogenic | 1 | 1 | 1 |
| rs72561723 | pathogenic | 1 | 1 | 1 |
| rs104894398 | pathogenic | 1 | 1 | 1 |
| rs28931594 | pathogenic | 1 | 1 | 1 |
| rs104894412 | pathogenic | 1 | 1 | 1 |
| rs111033297 | pathogenic | 1 | 1 | 1 |
| rs104894404 | pathogenic | 1 | 1 | 1 |
| rs104894410 | pathogenic | 1 | 1 | 1 |
| rs104894403 | pathogenic | 1 | 1 | 1 |
| rs397516870 | benign | 0 | 0 | 0 |
| rs200023879 | pathogenic | 1 | 1 | 1 |
| rs121912968 | pathogenic | 0 | 0 | 0 |
| rs104894402 | pathogenic | 1 | 1 | 1 |
| rs28931593 | pathogenic | 1 | 1 | 1 |
| rs104894397 | pathogenic | 0 | 0 | 0 |
| rs80338944 | pathogenic | 0 | 0 | 0 |
| rs199883710 | pathogenic | 0 | 0 | 0 |
| rs727504302 | pathogenic | 0 | 0 | 0 |
| rs145216882 | benign | 1 | 1 | 1 |
| rs781534323 | pathogenic | 1 | 1 | 1 |
| rs111033218 | benign | 1 | 1 | 1 |
| rs104894409 | pathogenic | 0 | 0 | 0 |
| rs80338945 | pathogenic | 1 | 1 | 1 |
| rs111033299 | pathogenic | 0 | 0 | 0 |
| rs143343083 | pathogenic | 1 | 1 | 1 |
| rs80338946 | pathogenic | 0 | 0 | 0 |
| rs150529554 | pathogenic | 0 | 0 | 0 |
| rs111033295 | pathogenic | 0 | 0 | 0 |
| rs111033188 | benign | 0 | 0 | 0 |
| rs397516874 | pathogenic | 0 | 0 | 0 |
| rs76434661 | pathogenic | 0 | 0 | 0 |

| | | | | |
|---|---|---|---|---|
| rs80338948 | pathogenic | 0 | 0 | 0 |
| rs104894401 | pathogenic | 0 | 0 | 0 |
| rs767178508 | pathogenic | 0 | 0 | 0 |
| rs772264564 | pathogenic | 0 | 0 | 0 |
| rs28931592 | pathogenic | 0 | 0 | 0 |
| rs80338949 | pathogenic | 1 | 1 | 1 |
| rs376898963 | benign | 1 | 1 | 1 |
| rs111033360 | benign | 0 | 0 | 0 |
| rs774518779 | pathogenic | 1 | 1 | 1 |
| rs28931595 | pathogenic | 1 | 1 | 1 |
| rs771748289 | pathogenic | 0 | 0 | 0 |
| rs104894406 | pathogenic | 0 | 0 | 0 |
| rs76838169 | benign | 0 | 0 | 0 |
| rs111033294 | pathogenic | 0 | 0 | 0 |
| rs200214298 | pathogenic | 0 | 0 | 0 |
| rs28940283 | pathogenic | 0 | 1 | 0 |
| rs1047883 | benign | 0 | 0 | 0 |
| rs138254425 | benign | 0 | 0 | 0 |
| rs121912592 | pathogenic | 1 | 1 | 0 |
| rs369061090 | pathogenic | 1 | 1 | 0 |
| rs758724746 | pathogenic | 0 | 1 | 0 |
| rs121912595 | pathogenic | 0 | 0 | 0 |
| rs76340296 | benign | 0 | 0 | 0 |
| rs140578009 | benign | 1 | 1 | 1 |
| rs74445297 | benign | 0 | 0 | 0 |
| rs78599516 | benign | 1 | 1 | 0 |
| rs104894035 | pathogenic | 1 | 1 | 0 |
| rs79451216 | benign | 0 | 0 | 0 |
| rs104894036 | pathogenic | 0 | 0 | 0 |
| rs73343757 | benign | 0 | 0 | 0 |
| rs121917722 | pathogenic | 1 | 1 | 1 |
| rs747700126 | pathogenic | 0 | 1 | 0 |
| rs121917725 | pathogenic | 1 | 1 | 1 |
| rs121917721 | pathogenic | 1 | 1 | 1 |
| rs121917723 | pathogenic | 1 | 1 | 0 |
| rs17569 | benign | 1 | 1 | 1 |
| rs2230062 | benign | 1 | 1 | 1 |
| rs17570 | benign | 0 | 0 | 0 |
| rs121917724 | pathogenic | 0 | 1 | 0 |
| rs199422235 | pathogenic | 0 | 0 | 0 |
| rs199422237 | pathogenic | 1 | 1 | 1 |
| rs199422234 | pathogenic | 1 | 1 | 0 |
| rs140015315 | pathogenic | 1 | 1 | 1 |
| rs730880298 | pathogenic | 1 | 1 | 1 |
| rs779692470 | pathogenic | 0 | 0 | 0 |
| rs369152939 | pathogenic | 0 | 0 | 0 |
| rs527236040 | pathogenic | 0 | 0 | 0 |
| rs587776985 | pathogenic | 1 | 1 | 1 |
| rs147077598 | pathogenic | 1 | 1 | 1 |
| rs128620185 | pathogenic | 1 | 1 | 1 |
| rs128620189 | pathogenic | 0 | 0 | 0 |
| rs128621190 | pathogenic | 0 | 0 | 0 |
| rs61754648 | benign | 0 | 0 | 0 |
| rs137852906 | pathogenic | 0 | 0 | 0 |

| | | | | |
|---|---|---|---|---|
| rs76811038 | pathogenic | 0 | 0 | 0 |
| rs149468422 | benign | 0 | 0 | 0 |
| rs5911 | benign | 0 | 0 | 0 |
| rs28942112 | pathogenic | 1 | 1 | 1 |
| rs143275858 | pathogenic | 1 | 1 | 0 |
| rs28362261 | benign | 0 | 0 | 0 |
| rs28362263 | benign | 1 | 1 | 1 |
| rs558707786 | pathogenic | 0 | 0 | 0 |
| rs138775799 | pathogenic | 0 | 0 | 0 |
| rs748170941 | pathogenic | 0 | 0 | 0 |
| rs121908380 | pathogenic | 0 | 0 | 0 |
| rs765123255 | pathogenic | 1 | 1 | 1 |
| rs730881832 | pathogenic | 0 | 1 | 0 |
| rs587782165 | benign | 1 | 1 | 1 |
| rs762307622 | pathogenic | 0 | 0 | 0 |
| rs34612342 | pathogenic | 1 | 1 | 0 |
| rs143353451 | pathogenic | 0 | 1 | 0 |
| rs747993448 | pathogenic | 0 | 1 | 0 |
| rs376561094 | pathogenic | 0 | 0 | 0 |
| rs776487884 | pathogenic | 1 | 1 | 0 |
| rs34126013 | pathogenic | 1 | 1 | 0 |
| rs200495564 | pathogenic | 1 | 1 | 0 |
| rs140342925 | pathogenic | 1 | 1 | 0 |
| rs587780749 | pathogenic | 1 | 1 | 0 |
| rs773087549 | pathogenic | 1 | 1 | 0 |
| rs730881833 | pathogenic | 0 | 0 | 0 |
| rs374950566 | pathogenic | 1 | 1 | 0 |
| rs587780752 | benign | 0 | 0 | 0 |
| rs3219489 | benign | 0 | 0 | 0 |
| rs199742231 | benign | 0 | 0 | 0 |
| rs587780082 | pathogenic | 0 | 0 | 0 |
| rs35881803 | benign | 0 | 0 | 0 |
| rs41305623 | benign | 1 | 1 | 1 |
| rs121917856 | pathogenic | 1 | 1 | 1 |
| rs576284122 | benign | 0 | 0 | 0 |
| rs142036701 | benign | 0 | 0 | 0 |
| rs121917852 | pathogenic | 1 | 1 | 1 |
| rs121917854 | pathogenic | 1 | 1 | 1 |
| rs121917853 | pathogenic | 1 | 1 | 1 |
| rs121917857 | pathogenic | 0 | 1 | 0 |
| rs193922740 | pathogenic | 1 | 1 | 1 |
| rs565779970 | pathogenic | 1 | 1 | 1 |
| rs6723526 | benign | 0 | 0 | 0 |
| rs141094096 | pathogenic | 1 | 1 | 1 |
| rs104894637 | pathogenic | 0 | 0 | 0 |
| rs104894636 | pathogenic | 0 | 0 | 0 |
| rs104894642 | pathogenic | 0 | 0 | 0 |
| rs104894638 | pathogenic | 0 | 0 | 0 |
| rs753666460 | pathogenic | 0 | 0 | 0 |
| rs104894643 | pathogenic | 1 | 1 | 0 |
| rs34520362 | pathogenic | 0 | 0 | 0 |
| rs374621913 | pathogenic | 0 | 0 | 0 |
| rs113641837 | benign | 0 | 0 | 0 |
| rs104894635 | pathogenic | 0 | 0 | 0 |

| | | | | |
|---|---|---|---|---|
| rs143947056 | pathogenic | 0 | 0 | 0 |
| rs138504221 | pathogenic | 0 | 0 | 0 |
| rs9894254 | benign | 0 | 0 | 0 |
| rs104894640 | pathogenic | 1 | 1 | 1 |
| rs58786455 | benign | 0 | 0 | 0 |
| rs144143780 | pathogenic | 0 | 0 | 0 |
| rs62620232 | benign | 0 | 0 | 0 |
| rs34297805 | benign | 1 | 1 | 1 |
| rs104894641 | pathogenic | 0 | 0 | 0 |
| rs148876719 | benign | 0 | 0 | 0 |
| rs104894639 | pathogenic | 0 | 0 | 0 |
| rs7503034 | benign | 0 | 0 | 0 |
| rs201631570 | benign | 0 | 0 | 0 |
| rs150181226 | pathogenic | 0 | 0 | 0 |
| rs377546036 | pathogenic | 0 | 0 | 0 |
| rs267606700 | pathogenic | 0 | 0 | 0 |
| rs267606701 | pathogenic | 1 | 1 | 0 |
| rs267606699 | pathogenic | 1 | 1 | 0 |
| rs9903215 | benign | 0 | 0 | 0 |
| rs139486406 | benign | 0 | 0 | 0 |
| rs143791614 | benign | 0 | 0 | 0 |
| rs573483474 | benign | 0 | 0 | 0 |
| rs2032892 | benign | 0 | 0 | 0 |
| rs137852619 | pathogenic | 0 | 0 | 0 |
| rs200243548 | benign | 0 | 0 | 0 |
| rs115702388 | benign | 0 | 0 | 0 |
| rs138085358 | pathogenic | 0 | 0 | 0 |
| rs121912789 | pathogenic | 1 | 1 | 1 |
| rs763407972 | pathogenic | 1 | 1 | 1 |
| rs147278302 | benign | 0 | 0 | 0 |
| rs374540411 | benign | 0 | 0 | 0 |
| rs121912790 | pathogenic | 1 | 1 | 0 |
| rs752049917 | benign | 0 | 0 | 0 |
| rs139640491 | benign | 1 | 1 | 1 |
| rs138010738 | benign | 1 | 1 | 0 |
| rs201328768 | pathogenic | 1 | 1 | 0 |
| rs2135720 | benign | 1 | 1 | 0 |
| rs201816080 | benign | 1 | 1 | 0 |
| rs754391973 | pathogenic | 0 | 0 | 0 |
| rs139087859 | benign | 0 | 0 | 0 |
| rs6805421 | benign | 0 | 0 | 0 |
| rs146266900 | benign | 0 | 0 | 0 |
| rs397509419 | pathogenic | 0 | 0 | 0 |
| rs149980411 | benign | 0 | 0 | 0 |
| rs747711488 | pathogenic | 1 | 1 | 1 |
| rs149474339 | pathogenic | 1 | 1 | 1 |
| rs758252808 | pathogenic | 1 | 1 | 0 |
| rs745756308 | pathogenic | 0 | 0 | 0 |
| rs4986958 | benign | 0 | 0 | 0 |
| rs74315444 | pathogenic | 0 | 0 | 0 |
| rs61752137 | pathogenic | 0 | 0 | 0 |
| rs8179065 | benign | 1 | 1 | 1 |
| rs758361736 | pathogenic | 1 | 1 | 0 |
| rs139794067 | pathogenic | 1 | 1 | 1 |

| | | | | |
|---|---|---|---|---|
| rs730880954 | pathogenic | 1 | 1 | 1 |
| rs730880955 | pathogenic | 1 | 1 | 1 |
| rs150634297 | pathogenic | 0 | 0 | 0 |
| rs104893750 | pathogenic | 1 | 1 | 1 |
| rs104893748 | pathogenic | 1 | 1 | 1 |
| rs104893749 | pathogenic | 1 | 1 | 1 |
| rs753520553 | pathogenic | 1 | 1 | 0 |
| rs61737294 | benign | 0 | 0 | 0 |
| rs104894601 | pathogenic | 0 | 0 | 0 |
| rs376090795 | pathogenic | 0 | 0 | 0 |
| rs104894596 | pathogenic | 0 | 0 | 0 |
| rs104894595 | pathogenic | 0 | 0 | 0 |
| rs104894598 | pathogenic | 0 | 0 | 0 |
| rs104894597 | pathogenic | 0 | 0 | 0 |
| rs104894591 | pathogenic | 0 | 0 | 0 |
| rs104894593 | pathogenic | 0 | 0 | 0 |
| rs104894594 | pathogenic | 0 | 0 | 0 |
| rs104894590 | pathogenic | 0 | 0 | 0 |
| rs86312 | benign | 0 | 0 | 0 |
| rs77874543 | benign | 0 | 0 | 0 |
| rs121912590 | pathogenic | 1 | 1 | 1 |
| rs121912589 | pathogenic | 1 | 1 | 1 |
| rs121912586 | pathogenic | 1 | 1 | 1 |
| rs121912585 | pathogenic | 1 | 1 | 1 |
| rs121912587 | pathogenic | 1 | 1 | 1 |
| rs762909359 | pathogenic | 0 | 0 | 0 |
| rs28931610 | pathogenic | 0 | 0 | 0 |
| rs104894097 | pathogenic | 1 | 1 | 1 |
| rs775176191 | benign | 0 | 0 | 0 |
| rs199907548 | pathogenic | 0 | 0 | 0 |
| rs104894095 | pathogenic | 1 | 1 | 1 |
| rs104894109 | pathogenic | 1 | 1 | 1 |
| rs372266620 | benign | 1 | 1 | 1 |
| rs104894099 | pathogenic | 0 | 0 | 0 |
| rs137854599 | pathogenic | 1 | 1 | 1 |
| rs104894094 | pathogenic | 0 | 0 | 0 |
| rs104894104 | pathogenic | 0 | 0 | 0 |
| rs6413463 | benign | 1 | 1 | 1 |
| rs104894098 | pathogenic | 0 | 0 | 0 |
| rs6413464 | benign | 0 | 0 | 0 |
| rs3731249 | benign | 0 | 0 | 0 |
| rs16931179 | benign | 0 | 0 | 0 |
| rs2734715 | benign | 0 | 0 | 0 |
| rs759080581 | pathogenic | 0 | 0 | 0 |
| rs1800746 | benign | 0 | 0 | 0 |
| rs202189057 | pathogenic | 0 | 0 | 0 |
| rs121908204 | pathogenic | 0 | 0 | 0 |
| rs756564767 | pathogenic | 0 | 0 | 0 |
| rs34758634 | benign | 1 | 1 | 1 |
| rs28940573 | pathogenic | 0 | 0 | 0 |
| rs119455955 | pathogenic | 1 | 1 | 0 |
| rs752164603 | pathogenic | 1 | 1 | 0 |
| rs763162812 | pathogenic | 0 | 0 | 0 |
| rs119455957 | pathogenic | 0 | 0 | 0 |

| | | | | |
|---|---|---|---|---|
| rs119455958 | pathogenic | 1 | 1 | 0 |
| rs750428882 | pathogenic | 0 | 0 | 0 |
| rs141482368 | benign | 0 | 0 | 0 |
| rs119455953 | pathogenic | 0 | 0 | 0 |
| rs119455954 | pathogenic | 0 | 0 | 0 |
| rs112812685 | benign | 1 | 1 | 0 |
| rs138744051 | benign | 1 | 1 | 0 |
| rs121908200 | pathogenic | 0 | 0 | 0 |
| rs119455956 | pathogenic | 0 | 0 | 0 |
| rs78970155 | benign | 0 | 0 | 0 |
| rs2834167 | benign | 0 | 0 | 0 |
| rs387907326 | pathogenic | 1 | 1 | 1 |
| rs2107732 | benign | 0 | 0 | 0 |
| rs11552377 | benign | 0 | 0 | 0 |
| rs137852843 | pathogenic | 0 | 0 | 0 |
| rs104894783 | pathogenic | 0 | 0 | 0 |
| rs122457137 | pathogenic | 1 | 1 | 1 |
| rs104894779 | pathogenic | 1 | 1 | 1 |
| rs104894786 | pathogenic | 0 | 0 | 0 |
| rs104894784 | pathogenic | 0 | 0 | 0 |
| rs104894785 | pathogenic | 0 | 0 | 0 |
| rs201870761 | pathogenic | 0 | 0 | 0 |
| rs104894781 | pathogenic | 0 | 0 | 0 |
| rs28897728 | benign | 0 | 0 | 0 |
| rs370273690 | pathogenic | 1 | 1 | 0 |
| rs9332739 | benign | 0 | 0 | 0 |
| rs150878060 | benign | 0 | 0 | 0 |
| rs573509224 | benign | 0 | 0 | 0 |
| rs36221133 | benign | 0 | 0 | 0 |
| rs2230009 | benign | 0 | 0 | 0 |
| rs267607008 | pathogenic | 0 | 0 | 0 |
| rs5985 | benign | 1 | 1 | 1 |
| rs5982 | benign | 1 | 1 | 1 |
| rs5987 | benign | 0 | 0 | 0 |
| rs5988 | benign | 0 | 0 | 0 |
| rs121918680 | pathogenic | 1 | 1 | 0 |
| rs121918679 | pathogenic | 1 | 1 | 0 |
| rs7845 | benign | 1 | 1 | 0 |
| rs1049296 | benign | 1 | 1 | 1 |
| rs121918677 | pathogenic | 1 | 1 | 0 |
| rs200086262 | pathogenic | 1 | 1 | 0 |
| rs746930990 | pathogenic | 1 | 1 | 0 |
| rs104893878 | pathogenic | 1 | 1 | 1 |
| rs121912468 | pathogenic | 1 | 1 | 1 |
| rs121912467 | pathogenic | 1 | 1 | 1 |
| rs145351926 | benign | 1 | 1 | 1 |
| rs141212743 | pathogenic | 1 | 1 | 1 |
| rs397515473 | pathogenic | 1 | 1 | 0 |
| rs121918313 | pathogenic | 0 | 1 | 0 |
| rs73015965 | pathogenic | 0 | 0 | 0 |
| rs121918030 | pathogenic | 0 | 1 | 0 |
| rs77782321 | benign | 0 | 0 | 0 |
| rs374661051 | pathogenic | 1 | 1 | 1 |
| rs267606854 | pathogenic | 0 | 0 | 0 |

| | | | | |
|---|---|---|---|---|
| rs41279678 | benign | 0 | 0 | 0 |
| rs61754640 | benign | 0 | 0 | 0 |
| rs199473314 | pathogenic | 0 | 0 | 0 |
| rs137854601 | pathogenic | 0 | 0 | 0 |
| rs137854614 | pathogenic | 0 | 0 | 0 |
| rs137854615 | pathogenic | 0 | 0 | 0 |
| rs199473635 | pathogenic | 0 | 0 | 0 |
| rs121434478 | pathogenic | 1 | 1 | 1 |
| rs775407864 | pathogenic | 1 | 1 | 1 |
| rs12706927 | benign | 0 | 1 | 0 |
| rs775614261 | benign | 0 | 0 | 0 |
| rs768926261 | benign | 0 | 0 | 0 |
| rs114057640 | benign | 0 | 0 | 0 |
| rs387906886 | pathogenic | 1 | 1 | 0 |
| rs138658585 | benign | 0 | 0 | 0 |
| rs4880091 | benign | 0 | 0 | 0 |
| rs121918552 | pathogenic | 1 | 1 | 1 |
| rs10794537 | benign | 0 | 0 | 0 |
| rs121965022 | pathogenic | 1 | 1 | 0 |
| rs121965020 | pathogenic | 0 | 0 | 0 |
| rs758452450 | pathogenic | 0 | 0 | 0 |
| rs747981483 | benign | 0 | 0 | 0 |
| rs58037052 | benign | 0 | 0 | 0 |
| rs121965029 | pathogenic | 1 | 1 | 0 |
| rs147490060 | benign | 0 | 0 | 0 |
| rs3755955 | benign | 0 | 0 | 0 |
| rs148789453 | pathogenic | 0 | 0 | 0 |
| rs121965030 | pathogenic | 0 | 0 | 0 |
| rs76722191 | benign | 0 | 0 | 0 |
| rs199801029 | pathogenic | 0 | 0 | 0 |
| rs121965033 | pathogenic | 0 | 0 | 0 |
| rs368454909 | pathogenic | 1 | 1 | 0 |
| rs6831280 | benign | 0 | 0 | 0 |
| rs121965024 | pathogenic | 0 | 0 | 0 |
| rs121965019 | pathogenic | 0 | 0 | 0 |
| rs11934801 | benign | 0 | 0 | 0 |
| rs73066479 | benign | 0 | 0 | 0 |
| rs121965027 | pathogenic | 0 | 0 | 0 |
| rs121965026 | pathogenic | 1 | 1 | 0 |
| rs121965021 | pathogenic | 0 | 0 | 0 |
| rs398123257 | benign | 0 | 0 | 0 |
| rs121965031 | pathogenic | 0 | 0 | 0 |
| rs202003795 | pathogenic | 0 | 0 | 0 |
| rs559510809 | pathogenic | 0 | 0 | 0 |
| rs762936223 | benign | 0 | 0 | 0 |
| rs137854580 | pathogenic | 1 | 1 | 1 |
| rs112263157 | benign | 0 | 0 | 0 |
| rs2289181 | benign | 0 | 0 | 0 |
| rs138034915 | benign | 0 | 1 | 0 |
| rs528367092 | pathogenic | 0 | 1 | 0 |
| rs1042393 | benign | 0 | 0 | 0 |
| rs370950728 | pathogenic | 0 | 1 | 0 |
| rs530478036 | benign | 0 | 1 | 0 |
| rs374569672 | benign | 0 | 1 | 0 |

| | | | | |
|---|---|---|---|---|
| rs121907944 | pathogenic | 0 | 1 | 0 |
| rs200856561 | pathogenic | 0 | 1 | 0 |
| rs577915581 | pathogenic | 0 | 1 | 0 |
| rs201896815 | pathogenic | 0 | 1 | 0 |
| rs121907945 | pathogenic | 0 | 1 | 0 |
| rs121907940 | pathogenic | 0 | 1 | 0 |
| rs543300039 | pathogenic | 0 | 1 | 0 |
| rs121907936 | pathogenic | 0 | 1 | 0 |
| rs766074609 | pathogenic | 0 | 1 | 0 |
| rs755253527 | pathogenic | 0 | 1 | 0 |
| rs560575383 | pathogenic | 0 | 1 | 0 |
| rs528369909 | benign | 0 | 1 | 0 |
| rs770610356 | pathogenic | 0 | 1 | 0 |
| rs747610090 | pathogenic | 0 | 1 | 0 |
| rs145712232 | benign | 0 | 1 | 0 |
| rs7215458 | benign | 1 | 1 | 0 |
| rs535644999 | pathogenic | 0 | 1 | 0 |
| rs772883420 | pathogenic | 1 | 1 | 0 |
| rs398123169 | pathogenic | 0 | 1 | 0 |
| rs121907937 | pathogenic | 0 | 1 | 0 |
| rs121907941 | pathogenic | 1 | 1 | 0 |
| rs121907942 | pathogenic | 0 | 1 | 0 |
| rs779556619 | pathogenic | 0 | 1 | 0 |
| rs1800307 | benign | 0 | 1 | 0 |
| rs377544304 | pathogenic | 1 | 1 | 0 |
| rs549029029 | pathogenic | 0 | 1 | 0 |
| rs753269119 | pathogenic | 1 | 1 | 1 |
| rs757617999 | pathogenic | 0 | 1 | 0 |
| rs28937909 | pathogenic | 0 | 1 | 0 |
| rs28940868 | pathogenic | 0 | 1 | 0 |
| rs368438393 | pathogenic | 0 | 1 | 0 |
| rs536906561 | pathogenic | 0 | 1 | 0 |
| rs374143224 | pathogenic | 0 | 1 | 0 |
| rs778418246 | pathogenic | 0 | 1 | 0 |
| rs398123172 | pathogenic | 0 | 1 | 0 |
| rs121907938 | pathogenic | 0 | 1 | 0 |
| rs1800312 | pathogenic | 0 | 1 | 0 |
| rs1126690 | benign | 0 | 1 | 0 |
| rs1800314 | benign | 0 | 1 | 0 |
| rs369532274 | pathogenic | 0 | 1 | 0 |
| rs121907943 | pathogenic | 0 | 1 | 0 |
| rs780321415 | pathogenic | 0 | 1 | 0 |
| rs377286472 | benign | 0 | 1 | 0 |
| rs121918406 | pathogenic | 1 | 1 | 1 |
| rs147484870 | benign | 0 | 0 | 0 |
| rs193922671 | pathogenic | 0 | 0 | 0 |
| rs201397978 | benign | 1 | 1 | 1 |
| rs34884895 | benign | 1 | 1 | 1 |
| rs138329459 | benign | 0 | 0 | 0 |
| rs2289366 | benign | 1 | 1 | 1 |
| rs141858334 | benign | 1 | 1 | 1 |
| rs59907886 | benign | 1 | 1 | 1 |
| rs137852653 | pathogenic | 0 | 0 | 0 |
| rs137852652 | pathogenic | 0 | 0 | 0 |

| | | | | |
|---|---|---|---|---|
| rs137852656 | pathogenic | 0 | 0 | 0 |
| rs61739916 | benign | 0 | 0 | 0 |
| rs28936668 | pathogenic | 0 | 0 | 0 |
| rs137852651 | pathogenic | 0 | 0 | 0 |
| rs137852650 | pathogenic | 0 | 0 | 0 |
| rs28936669 | pathogenic | 0 | 0 | 0 |
| rs137852654 | pathogenic | 0 | 0 | 0 |
| rs28936368 | pathogenic | 1 | 1 | 0 |
| rs137852655 | pathogenic | 0 | 0 | 0 |
| rs61752496 | benign | 0 | 0 | 0 |
| rs750098055 | benign | 0 | 0 | 0 |
| rs764149793 | benign | 0 | 0 | 0 |
| rs724160017 | pathogenic | 0 | 1 | 0 |
| rs724160019 | pathogenic | 0 | 0 | 0 |
| rs202219398 | pathogenic | 1 | 1 | 0 |
| rs724160023 | pathogenic | 1 | 1 | 0 |
| rs761953453 | pathogenic | 0 | 0 | 0 |
| rs80338835 | pathogenic | 1 | 1 | 1 |
| rs56284940 | benign | 0 | 0 | 0 |
| rs74575749 | benign | 0 | 0 | 0 |
| rs587777075 | pathogenic | 1 | 1 | 1 |
| rs641153 | benign | 0 | 0 | 0 |
| rs12614 | benign | 0 | 0 | 0 |
| rs117314762 | benign | 0 | 0 | 0 |
| rs537478097 | benign | 1 | 1 | 1 |
| rs144812066 | benign | 0 | 0 | 0 |
| rs4151651 | benign | 0 | 0 | 0 |
| rs138207668 | benign | 1 | 1 | 1 |
| rs149101394 | benign | 1 | 1 | 1 |
| rs4151659 | benign | 1 | 1 | 1 |
| rs45484591 | benign | 1 | 1 | 1 |
| rs761991070 | pathogenic | 0 | 0 | 0 |
| rs755948940 | pathogenic | 0 | 0 | 0 |
| rs137853097 | pathogenic | 0 | 0 | 0 |
| rs775297352 | benign | 0 | 0 | 0 |
| rs28943591 | benign | 0 | 0 | 0 |
| rs191468413 | pathogenic | 1 | 1 | 0 |
| rs11539471 | benign | 1 | 1 | 1 |
| rs11205 | benign | 0 | 0 | 0 |
| rs768409400 | pathogenic | 0 | 0 | 0 |
| rs762013077 | pathogenic | 0 | 0 | 0 |
| rs768474112 | pathogenic | 0 | 0 | 0 |
| rs121965063 | pathogenic | 1 | 1 | 1 |
| rs143648758 | pathogenic | 0 | 0 | 0 |
| rs121965066 | pathogenic | 0 | 0 | 0 |
| rs752907087 | pathogenic | 0 | 0 | 0 |
| rs747702749 | pathogenic | 0 | 0 | 0 |
| rs754087775 | pathogenic | 0 | 0 | 0 |
| rs121918641 | pathogenic | 1 | 1 | 1 |
| rs121918642 | pathogenic | 1 | 1 | 1 |
| rs121918640 | pathogenic | 1 | 1 | 1 |
| rs121918637 | pathogenic | 1 | 1 | 1 |
| rs121918644 | pathogenic | 1 | 1 | 1 |
| rs16840544 | benign | 0 | 0 | 0 |

| | | | | |
|---|---|---|---|---|
| rs1042391 | benign | 1 | 1 | 1 |
| rs104893744 | pathogenic | 1 | 1 | 1 |
| rs120074136 | pathogenic | 0 | 0 | 0 |
| rs1805081 | benign | 1 | 1 | 0 |
| rs80358251 | benign | 0 | 0 | 0 |
| rs774588125 | benign | 0 | 0 | 0 |
| rs120074134 | pathogenic | 0 | 0 | 0 |
| rs139751448 | pathogenic | 0 | 0 | 0 |
| rs77080672 | benign | 0 | 0 | 0 |
| rs372445155 | pathogenic | 1 | 1 | 0 |
| rs13381670 | benign | 0 | 0 | 0 |
| rs1788799 | benign | 0 | 0 | 0 |
| rs376213990 | pathogenic | 0 | 0 | 0 |
| rs757475924 | pathogenic | 0 | 0 | 0 |
| rs80358253 | pathogenic | 0 | 0 | 0 |
| rs1805082 | benign | 0 | 0 | 0 |
| rs120074130 | pathogenic | 0 | 0 | 0 |
| rs768999208 | pathogenic | 0 | 0 | 0 |
| rs34302553 | benign | 0 | 0 | 0 |
| rs28940897 | pathogenic | 0 | 0 | 0 |
| rs1140458 | benign | 0 | 0 | 0 |
| rs143124972 | pathogenic | 0 | 0 | 0 |
| rs120074135 | pathogenic | 0 | 0 | 0 |
| rs543206298 | pathogenic | 0 | 0 | 0 |
| rs120074132 | pathogenic | 0 | 0 | 0 |
| rs759826138 | pathogenic | 0 | 0 | 0 |
| rs28942108 | pathogenic | 0 | 0 | 0 |
| rs80358254 | pathogenic | 0 | 0 | 0 |
| rs80358257 | pathogenic | 0 | 0 | 0 |
| rs781261962 | pathogenic | 0 | 0 | 0 |
| rs28942107 | pathogenic | 0 | 0 | 0 |
| rs80358258 | pathogenic | 0 | 0 | 0 |
| rs80358259 | pathogenic | 1 | 1 | 0 |
| rs141440861 | benign | 0 | 0 | 0 |
| rs28942106 | pathogenic | 0 | 0 | 0 |
| rs778878523 | pathogenic | 0 | 0 | 0 |
| rs28942105 | pathogenic | 0 | 0 | 0 |
| rs61731969 | benign | 0 | 0 | 0 |
| rs200444084 | pathogenic | 0 | 0 | 0 |
| rs113371321 | pathogenic | 0 | 0 | 0 |
| rs369098773 | pathogenic | 0 | 0 | 0 |
| rs35248744 | benign | 0 | 0 | 0 |
| rs758902805 | pathogenic | 0 | 0 | 0 |
| rs120074131 | pathogenic | 0 | 0 | 0 |
| rs1805084 | benign | 0 | 0 | 0 |
| rs117793540 | benign | 0 | 0 | 0 |
| rs530058063 | benign | 0 | 0 | 0 |
| rs1064422 | pathogenic | 1 | 1 | 1 |
| rs104894506 | pathogenic | 1 | 1 | 1 |
| rs104894507 | pathogenic | 1 | 1 | 1 |
| rs104894508 | pathogenic | 1 | 1 | 0 |
| rs28999113 | pathogenic | 1 | 1 | 0 |
| rs281860266 | pathogenic | 0 | 0 | 0 |
| rs104894961 | pathogenic | 1 | 1 | 0 |

| | | | | |
|---|---|---|---|---|
| rs122463168 | pathogenic | 0 | 0 | 0 |
| rs199469624 | pathogenic | 1 | 1 | 1 |
| rs150107747 | benign | 0 | 0 | 0 |
| rs144984628 | pathogenic | 0 | 0 | 0 |
| rs28935171 | pathogenic | 0 | 0 | 0 |
| rs137852443 | pathogenic | 1 | 1 | 0 |
| rs28933674 | pathogenic | 1 | 1 | 1 |
| rs28933675 | pathogenic | 1 | 1 | 1 |
| rs28933676 | pathogenic | 1 | 1 | 1 |
| rs28933677 | pathogenic | 1 | 1 | 1 |
| rs28933678 | pathogenic | 0 | 0 | 0 |
| rs28933679 | pathogenic | 1 | 1 | 1 |
| rs28933680 | pathogenic | 0 | 0 | 0 |
| rs28933681 | pathogenic | 0 | 0 | 0 |
| rs28933682 | pathogenic | 0 | 0 | 0 |
| rs28937294 | pathogenic | 1 | 1 | 0 |
| rs137852459 | pathogenic | 0 | 0 | 0 |
| rs137852461 | pathogenic | 0 | 0 | 0 |
| rs137852464 | pathogenic | 1 | 1 | 1 |
| rs137852465 | pathogenic | 1 | 1 | 1 |
| rs1800297 | benign | 1 | 1 | 0 |
| rs137852374 | pathogenic | 1 | 1 | 1 |
| rs672601312 | pathogenic | 1 | 1 | 1 |
| rs387907196 | pathogenic | 1 | 1 | 1 |
| rs148348866 | benign | 0 | 0 | 0 |
| rs141138948 | pathogenic | 1 | 1 | 1 |
| rs137852685 | pathogenic | 1 | 1 | 0 |
| rs104893900 | pathogenic | 1 | 1 | 0 |
| rs137852686 | pathogenic | 1 | 1 | 1 |
| rs104893906 | pathogenic | 1 | 1 | 0 |
| rs587782207 | pathogenic | 1 | 1 | 1 |
| rs75086406 | pathogenic | 0 | 0 | 0 |
| rs199822819 | pathogenic | 0 | 0 | 0 |
| rs398123166 | pathogenic | 1 | 1 | 1 |
| rs121913123 | pathogenic | 1 | 1 | 1 |
| rs772190176 | pathogenic | 0 | 1 | 0 |
| rs587782216 | pathogenic | 1 | 1 | 1 |
| rs398123159 | pathogenic | 1 | 1 | 1 |
| rs121913122 | pathogenic | 1 | 1 | 1 |
| rs755436052 | pathogenic | 1 | 1 | 1 |
| rs200796606 | pathogenic | 1 | 1 | 1 |
| rs2070080 | pathogenic | 0 | 1 | 0 |
| rs149616320 | benign | 0 | 0 | 0 |
| rs143529458 | benign | 0 | 0 | 0 |
| rs267607077 | pathogenic | 1 | 1 | 1 |
| rs143898031 | benign | 1 | 1 | 0 |
| rs148304857 | benign | 1 | 1 | 1 |
| rs137852809 | pathogenic | 0 | 0 | 0 |
| rs3765623 | benign | 0 | 0 | 0 |
| rs16944397 | benign | 0 | 0 | 0 |
| rs730880167 | benign | 0 | 0 | 0 |
| rs187108957 | benign | 0 | 0 | 0 |
| rs117342470 | benign | 0 | 0 | 0 |
| rs751200138 | pathogenic | 1 | 1 | 1 |

| | | | | |
|---|---|---|---|---|
| rs755799528 | pathogenic | 0 | 0 | 0 |
| rs537484504 | benign | 0 | 0 | 0 |
| rs748944640 | pathogenic | 0 | 0 | 0 |
| rs199774121 | pathogenic | 0 | 0 | 0 |
| rs766094434 | pathogenic | 1 | 1 | 0 |
| rs777321035 | pathogenic | 1 | 1 | 1 |
| rs146269774 | benign | 0 | 1 | 0 |
| rs144085805 | benign | 0 | 1 | 0 |
| rs367643028 | benign | 0 | 0 | 0 |
| rs42524 | benign | 0 | 0 | 0 |
| rs28940586 | pathogenic | 0 | 0 | 0 |
| rs74315327 | pathogenic | 1 | 1 | 1 |
| rs199676861 | pathogenic | 0 | 0 | 0 |
| rs115802744 | benign | 1 | 1 | 0 |
| rs752866643 | pathogenic | 1 | 1 | 1 |
| rs28941784 | pathogenic | 1 | 1 | 1 |
| rs398124434 | pathogenic | 1 | 1 | 1 |
| rs756414548 | pathogenic | 1 | 1 | 1 |
| rs376128990 | pathogenic | 1 | 1 | 1 |
| rs746219370 | pathogenic | 1 | 1 | 1 |
| rs756195708 | pathogenic | 1 | 1 | 1 |
| rs369296618 | pathogenic | 1 | 1 | 1 |
| rs9593 | benign | 1 | 1 | 1 |
| rs139510844 | benign | 0 | 0 | 0 |
| rs28382158 | benign | 0 | 0 | 0 |
| rs28919570 | pathogenic | 0 | 0 | 0 |
| rs587777331 | pathogenic | 0 | 0 | 0 |
| rs587777333 | pathogenic | 0 | 0 | 0 |
| rs587777332 | pathogenic | 0 | 0 | 0 |
| rs774980346 | pathogenic | 0 | 0 | 0 |
| rs370934093 | pathogenic | 0 | 0 | 0 |
| rs121913479 | pathogenic | 1 | 1 | 1 |
| rs121913484 | pathogenic | 0 | 0 | 0 |
| rs121913485 | pathogenic | 1 | 1 | 1 |
| rs28931614 | pathogenic | 1 | 1 | 1 |
| rs17881656 | benign | 1 | 1 | 1 |
| rs28931615 | pathogenic | 1 | 1 | 1 |
| rs147846832 | benign | 0 | 0 | 0 |
| rs80356506 | pathogenic | 0 | 0 | 0 |
| rs80356494 | pathogenic | 0 | 0 | 0 |
| rs121908894 | pathogenic | 0 | 0 | 0 |
| rs121908895 | pathogenic | 0 | 0 | 0 |
| rs80356508 | pathogenic | 0 | 0 | 0 |
| rs80356507 | pathogenic | 0 | 0 | 0 |
| rs121908896 | pathogenic | 0 | 0 | 0 |
| rs745372589 | pathogenic | 1 | 1 | 0 |
| rs183898408 | benign | 0 | 0 | 0 |
| rs200844292 | benign | 0 | 0 | 0 |
| rs146897002 | benign | 0 | 0 | 0 |
| rs121964869 | pathogenic | 1 | 1 | 1 |
| rs201768800 | benign | 1 | 1 | 1 |
| rs9807556 | benign | 1 | 1 | 1 |
| rs184721031 | benign | 0 | 0 | 0 |
| rs201104206 | benign | 0 | 0 | 0 |

| | | | | |
|---|---|---|---|---|
| rs188677538 | benign | 0 | 0 | 0 |
| rs149528866 | benign | 0 | 0 | 0 |
| rs115382168 | benign | 0 | 0 | 0 |
| rs78846023 | benign | 1 | 1 | 1 |
| rs61730662 | benign | 0 | 0 | 0 |
| rs121918368 | pathogenic | 0 | 0 | 0 |
| rs566433112 | pathogenic | 1 | 1 | 1 |
| rs121909342 | pathogenic | 0 | 0 | 0 |
| rs67586389 | pathogenic | 0 | 0 | 0 |
| rs72466485 | pathogenic | 0 | 0 | 0 |
| rs72466486 | pathogenic | 1 | 1 | 1 |
| rs72466487 | pathogenic | 1 | 1 | 1 |
| rs267607112 | pathogenic | 0 | 0 | 0 |
| rs118204013 | pathogenic | 0 | 0 | 0 |
| rs121913495 | pathogenic | 0 | 0 | 0 |
| rs11554273 | pathogenic | 0 | 0 | 0 |
| rs193922477 | pathogenic | 1 | 1 | 1 |
| rs193922479 | pathogenic | 1 | 1 | 1 |
| rs72547554 | pathogenic | 0 | 0 | 0 |
| rs72547561 | pathogenic | 1 | 1 | 1 |
| rs72547562 | pathogenic | 1 | 1 | 0 |
| rs115977487 | benign | 1 | 1 | 0 |
| rs72547564 | pathogenic | 0 | 0 | 0 |
| rs72547568 | pathogenic | 1 | 1 | 0 |
| rs72547569 | pathogenic | 1 | 1 | 1 |
| rs72547571 | pathogenic | 0 | 0 | 0 |
| rs72547575 | pathogenic | 1 | 1 | 1 |
| rs544994462 | benign | 0 | 0 | 0 |
| rs190470017 | benign | 0 | 0 | 0 |
| rs199473266 | pathogenic | 1 | 1 | 1 |
| rs137854602 | pathogenic | 0 | 0 | 0 |
| rs143739249 | pathogenic | 0 | 0 | 0 |
| rs397514697 | pathogenic | 1 | 1 | 1 |
| rs373764886 | pathogenic | 0 | 0 | 0 |
| rs370242930 | pathogenic | 0 | 0 | 0 |
| rs377145777 | pathogenic | 1 | 1 | 1 |
| rs191256186 | benign | 0 | 0 | 0 |
| rs150070901 | benign | 1 | 1 | 1 |
| rs138098032 | benign | 0 | 0 | 0 |
| rs117731813 | benign | 1 | 1 | 1 |
| rs104894488 | pathogenic | 1 | 1 | 1 |
| rs11302 | benign | 0 | 1 | 0 |
| rs121918447 | pathogenic | 1 | 1 | 1 |
| rs138205828 | benign | 1 | 1 | 0 |
| rs149858290 | benign | 0 | 0 | 0 |
| rs750764003 | pathogenic | 0 | 1 | 0 |
| rs141381896 | benign | 0 | 0 | 0 |
| rs149027682 | benign | 0 | 0 | 0 |
| rs121908415 | pathogenic | 0 | 0 | 0 |
| rs121908416 | pathogenic | 0 | 0 | 0 |
| rs121908417 | pathogenic | 0 | 0 | 0 |
| rs587780306 | benign | 0 | 0 | 0 |
| rs3218725 | benign | 0 | 0 | 0 |
| rs3218729 | benign | 0 | 0 | 0 |

| | | | | |
|---|---|---|---|---|
| rs387906917 | pathogenic | 0 | 0 | 0 |
| rs200652608 | pathogenic | 0 | 0 | 0 |
| rs3218721 | benign | 0 | 0 | 0 |
| rs63750869 | pathogenic | 1 | 1 | 1 |
| rs118203910 | pathogenic | 0 | 1 | 0 |
| rs9332701 | benign | 1 | 1 | 0 |
| rs6027 | benign | 0 | 0 | 0 |
| rs121909110 | pathogenic | 0 | 1 | 0 |
| rs762550967 | pathogenic | 1 | 1 | 0 |
| rs763615602 | pathogenic | 1 | 1 | 0 |
| rs144907403 | benign | 0 | 0 | 0 |
| rs104893631 | pathogenic | 1 | 1 | 0 |
| rs144181978 | pathogenic | 1 | 1 | 0 |
| rs74874677 | benign | 1 | 1 | 1 |
| rs104893632 | pathogenic | 0 | 1 | 0 |
| rs371342464 | benign | 1 | 1 | 1 |
| rs104893633 | pathogenic | 1 | 1 | 0 |
| rs201088712 | pathogenic | 0 | 0 | 0 |
| rs201857541 | benign | 1 | 1 | 1 |
| rs139517732 | pathogenic | 1 | 1 | 0 |
| rs191057824 | benign | 1 | 1 | 0 |
| rs35683768 | benign | 1 | 1 | 0 |
| rs104895271 | pathogenic | 0 | 0 | 0 |
| rs104895217 | pathogenic | 1 | 1 | 1 |
| rs104895223 | pathogenic | 1 | 1 | 1 |
| rs104895218 | pathogenic | 1 | 1 | 1 |
| rs104895225 | pathogenic | 1 | 1 | 1 |
| rs4149637 | benign | 1 | 1 | 1 |
| rs104895219 | pathogenic | 1 | 1 | 1 |
| rs104895220 | pathogenic | 1 | 1 | 1 |
| rs104895228 | pathogenic | 1 | 1 | 1 |
| rs104895221 | pathogenic | 1 | 1 | 1 |
| rs104895222 | pathogenic | 1 | 1 | 1 |
| rs104895247 | pathogenic | 0 | 0 | 0 |
| rs28940585 | pathogenic | 0 | 0 | 0 |
| rs561367199 | pathogenic | 1 | 1 | 1 |
| rs143793815 | benign | 0 | 0 | 0 |
| rs28941473 | pathogenic | 1 | 1 | 1 |
| rs751590073 | pathogenic | 1 | 1 | 1 |
| rs111407265 | benign | 1 | 1 | 1 |
| rs373697663 | pathogenic | 1 | 1 | 1 |
| rs369879957 | pathogenic | 0 | 0 | 0 |
| rs28941472 | pathogenic | 1 | 1 | 1 |
| rs28940287 | pathogenic | 1 | 1 | 1 |
| rs77625410 | benign | 0 | 0 | 0 |
| rs111033477 | benign | 0 | 0 | 0 |
| rs115123584 | benign | 0 | 0 | 0 |
| rs374576916 | benign | 0 | 0 | 0 |
| rs397516317 | pathogenic | 0 | 0 | 0 |
| rs368657015 | pathogenic | 0 | 0 | 0 |
| rs397516321 | pathogenic | 0 | 0 | 0 |
| rs397516322 | pathogenic | 0 | 0 | 0 |
| rs111033215 | pathogenic | 0 | 0 | 0 |
| rs199606180 | pathogenic | 0 | 0 | 0 |

| | | | | |
|---|---|---|---|---|
| rs111033192 | pathogenic | 0 | 0 | 0 |
| rs948962 | benign | 1 | 1 | 1 |
| rs142293185 | benign | 0 | 0 | 0 |
| rs376764423 | pathogenic | 0 | 0 | 0 |
| rs773844428 | pathogenic | 0 | 0 | 0 |
| rs186644871 | benign | 0 | 0 | 0 |
| rs200313391 | benign | 0 | 0 | 0 |
| rs377670513 | pathogenic | 0 | 0 | 0 |
| rs1132036 | benign | 0 | 0 | 0 |
| rs111033231 | benign | 0 | 0 | 0 |
| rs145945630 | pathogenic | 0 | 0 | 0 |
| rs121912984 | pathogenic | 1 | 1 | 1 |
| rs1800435 | benign | 0 | 1 | 0 |
| rs121912980 | pathogenic | 0 | 0 | 0 |
| rs121912982 | pathogenic | 1 | 1 | 1 |
| rs121912983 | pathogenic | 0 | 0 | 0 |
| rs121912981 | pathogenic | 0 | 0 | 0 |
| rs1057868 | benign | 0 | 0 | 0 |
| rs56048668 | benign | 0 | 0 | 0 |
| rs55865465 | benign | 0 | 0 | 0 |
| rs748096324 | benign | 1 | 1 | 0 |
| rs150475750 | benign | 0 | 0 | 0 |
| rs143025739 | benign | 1 | 1 | 0 |
| rs79702016 | benign | 1 | 1 | 0 |
| rs41338945 | benign | 0 | 0 | 0 |
| rs146406037 | benign | 1 | 1 | 1 |
| rs143274491 | benign | 0 | 0 | 0 |
| rs3829986 | benign | 0 | 0 | 0 |
| rs149168939 | benign | 0 | 0 | 0 |
| rs1800385 | benign | 0 | 0 | 0 |
| rs1800386 | pathogenic | 0 | 0 | 0 |
| rs121918524 | pathogenic | 1 | 1 | 1 |
| rs121912700 | pathogenic | 0 | 0 | 0 |
| rs104893930 | pathogenic | 1 | 1 | 1 |
| rs104893931 | pathogenic | 0 | 0 | 0 |
| rs116125936 | benign | 0 | 0 | 0 |
| rs627441 | benign | 0 | 0 | 0 |
| rs104894481 | pathogenic | 0 | 0 | 0 |
| rs137853129 | pathogenic | 0 | 0 | 0 |
| rs137853130 | pathogenic | 1 | 1 | 0 |
| rs28940282 | pathogenic | 0 | 0 | 0 |
| rs2229989 | benign | 0 | 0 | 0 |
| rs104894647 | pathogenic | 0 | 0 | 0 |
| rs1061170 | benign | 1 | 1 | 1 |
| rs576402053 | benign | 0 | 0 | 0 |
| rs34972246 | benign | 0 | 0 | 0 |
| rs192725607 | benign | 0 | 0 | 0 |
| rs137852615 | pathogenic | 1 | 1 | 0 |
| rs137852617 | pathogenic | 1 | 1 | 1 |
| rs137852611 | pathogenic | 0 | 0 | 0 |
| rs137852612 | pathogenic | 0 | 0 | 0 |
| rs137852614 | pathogenic | 1 | 1 | 1 |
| rs137852610 | pathogenic | 1 | 1 | 1 |
| rs137852618 | pathogenic | 1 | 1 | 1 |

| | | | | |
|---|---|---|---|---|
| rs137852613 | pathogenic | 1 | 1 | 0 |
| rs137852609 | pathogenic | 0 | 0 | 0 |
| rs63750306 | pathogenic | 1 | 1 | 1 |
| rs63750265 | pathogenic | 1 | 1 | 1 |
| rs63751287 | pathogenic | 1 | 1 | 1 |
| rs199723282 | benign | 0 | 0 | 0 |
| rs121917807 | pathogenic | 0 | 0 | 0 |
| rs121917809 | pathogenic | 0 | 0 | 0 |
| rs75134564 | pathogenic | 0 | 0 | 0 |
| rs267606930 | pathogenic | 1 | 1 | 0 |
| rs121909302 | pathogenic | 0 | 0 | 0 |
| rs121909303 | pathogenic | 0 | 0 | 0 |
| rs121909299 | pathogenic | 0 | 0 | 0 |
| rs121909301 | pathogenic | 0 | 0 | 0 |
| rs121909300 | pathogenic | 1 | 1 | 0 |
| rs111033589 | pathogenic | 0 | 0 | 0 |
| rs137853059 | pathogenic | 0 | 0 | 0 |
| rs137853057 | pathogenic | 1 | 1 | 1 |
| rs1801474 | benign | 0 | 0 | 0 |
| rs137853060 | pathogenic | 1 | 1 | 1 |
| rs137853058 | pathogenic | 1 | 1 | 1 |
| rs34424986 | pathogenic | 0 | 0 | 0 |
| rs199657839 | benign | 1 | 1 | 1 |
| rs56092260 | benign | 0 | 0 | 0 |
| rs1801582 | benign | 1 | 1 | 1 |
| rs1801334 | benign | 1 | 1 | 1 |
| rs191486604 | pathogenic | 0 | 0 | 0 |
| rs121908051 | pathogenic | 0 | 0 | 0 |
| rs121908055 | pathogenic | 1 | 1 | 0 |
| rs28940886 | pathogenic | 0 | 0 | 0 |
| rs121908050 | pathogenic | 0 | 0 | 0 |
| rs28940887 | pathogenic | 0 | 0 | 0 |
| rs387906300 | pathogenic | 0 | 0 | 0 |
| rs28942087 | pathogenic | 0 | 0 | 0 |
| rs121908049 | pathogenic | 1 | 1 | 0 |
| rs121908054 | pathogenic | 1 | 1 | 0 |
| rs121908048 | pathogenic | 0 | 0 | 0 |
| rs28940888 | pathogenic | 1 | 1 | 1 |
| rs121908053 | pathogenic | 1 | 1 | 0 |
| rs122467169 | pathogenic | 1 | 1 | 1 |
| rs122467172 | pathogenic | 1 | 1 | 1 |
| rs122467170 | pathogenic | 1 | 1 | 1 |
| rs28935477 | pathogenic | 1 | 1 | 0 |
| rs28942088 | pathogenic | 0 | 0 | 0 |
| rs751478142 | pathogenic | 1 | 1 | 0 |
| rs765368797 | pathogenic | 0 | 0 | 0 |
| rs147810437 | benign | 0 | 0 | 0 |
| rs121918209 | pathogenic | 1 | 1 | 1 |
| rs121918208 | pathogenic | 1 | 1 | 1 |
| rs121918210 | pathogenic | 1 | 1 | 1 |
| rs35515638 | benign | 1 | 1 | 1 |
| rs121918207 | pathogenic | 1 | 1 | 1 |
| rs141298312 | benign | 0 | 0 | 0 |
| rs121918211 | pathogenic | 0 | 0 | 0 |

| | | | | |
|---|---|---|---|---|
| rs149660059 | benign | 0 | 0 | 0 |
| rs200670286 | pathogenic | 1 | 1 | 1 |
| rs121918212 | pathogenic | 1 | 1 | 1 |
| rs121918213 | pathogenic | 1 | 1 | 1 |
| rs121918205 | pathogenic | 1 | 1 | 1 |
| rs104894025 | pathogenic | 1 | 1 | 1 |
| rs104894026 | pathogenic | 1 | 1 | 1 |
| rs104894028 | pathogenic | 0 | 1 | 0 |
| rs104894029 | pathogenic | 0 | 1 | 0 |
| rs119458971 | pathogenic | 0 | 0 | 0 |
| rs119458969 | pathogenic | 1 | 1 | 1 |
| rs1801591 | benign | 1 | 1 | 1 |
| rs140169311 | benign | 0 | 0 | 0 |
| rs753363278 | benign | 1 | 1 | 1 |
| rs199763682 | pathogenic | 0 | 0 | 0 |
| rs119458970 | pathogenic | 1 | 1 | 1 |
| rs763722868 | pathogenic | 0 | 0 | 0 |
| rs56404770 | benign | 0 | 0 | 0 |
| rs34221241 | benign | 0 | 0 | 0 |
| rs121909625 | pathogenic | 0 | 0 | 0 |
| rs550921485 | pathogenic | 0 | 0 | 0 |
| rs1800454 | benign | 0 | 0 | 0 |
| rs241447 | benign | 0 | 0 | 0 |
| rs2230606 | benign | 1 | 1 | 0 |
| rs5036 | pathogenic | 1 | 1 | 1 |
| rs13306788 | benign | 0 | 0 | 0 |
| rs28929480 | pathogenic | 0 | 0 | 0 |
| rs121912749 | pathogenic | 0 | 0 | 0 |
| rs56361140 | pathogenic | 0 | 0 | 0 |
| rs28931583 | pathogenic | 1 | 1 | 1 |
| rs36211715 | pathogenic | 1 | 1 | 1 |
| rs138049878 | pathogenic | 1 | 1 | 1 |
| rs727503253 | pathogenic | 1 | 1 | 1 |
| rs267606908 | pathogenic | 0 | 0 | 0 |
| rs121913631 | pathogenic | 1 | 1 | 1 |
| rs36211716 | benign | 1 | 1 | 1 |
| rs121913639 | pathogenic | 1 | 1 | 1 |
| rs730880761 | pathogenic | 1 | 1 | 1 |
| rs121913629 | pathogenic | 1 | 1 | 1 |
| rs754368658 | pathogenic | 0 | 0 | 0 |
| rs777090017 | pathogenic | 0 | 0 | 0 |
| rs267606913 | pathogenic | 1 | 1 | 1 |
| rs773138218 | pathogenic | 0 | 0 | 0 |
| rs371295780 | pathogenic | 0 | 0 | 0 |
| rs387906830 | pathogenic | 1 | 1 | 0 |
| rs771364038 | pathogenic | 0 | 1 | 0 |
| rs772731615 | pathogenic | 0 | 0 | 0 |
| rs137853063 | pathogenic | 0 | 0 | 0 |
| rs387907076 | pathogenic | 0 | 1 | 0 |
| rs35460768 | benign | 0 | 0 | 0 |
| rs2278196 | benign | 1 | 1 | 1 |
| rs119103275 | pathogenic | 0 | 0 | 0 |
| rs119103276 | pathogenic | 1 | 1 | 1 |
| rs374402066 | pathogenic | 1 | 1 | 1 |

| | | | | |
|---|---|---|---|---|
| rs119103274 | pathogenic | 0 | 0 | 0 |
| rs2274064 | benign | 0 | 0 | 0 |
| rs267607023 | pathogenic | 1 | 1 | 0 |
| rs142117489 | benign | 1 | 1 | 0 |
| rs371258270 | benign | 0 | 0 | 0 |
| rs34716057 | benign | 0 | 0 | 0 |
| rs201778606 | benign | 0 | 0 | 0 |
| rs201438792 | benign | 0 | 0 | 0 |
| rs140547417 | benign | 0 | 0 | 0 |
| rs5881 | benign | 0 | 0 | 0 |
| rs752298084 | benign | 0 | 0 | 0 |
| rs34855278 | benign | 0 | 0 | 0 |
| rs5880 | benign | 1 | 1 | 0 |
| rs5882 | benign | 0 | 0 | 0 |
| rs1800777 | benign | 0 | 0 | 0 |
| rs577301331 | benign | 0 | 0 | 0 |
| rs137853313 | pathogenic | 0 | 0 | 0 |
| rs28935169 | pathogenic | 0 | 0 | 0 |
| rs137853315 | pathogenic | 0 | 0 | 0 |
| rs137853314 | pathogenic | 0 | 0 | 0 |
| rs28935469 | pathogenic | 0 | 0 | 0 |
| rs137853318 | pathogenic | 0 | 0 | 0 |
| rs28935470 | pathogenic | 0 | 0 | 0 |
| rs3729986 | benign | 0 | 0 | 0 |
| rs730880619 | pathogenic | 0 | 0 | 0 |
| rs201012766 | benign | 0 | 0 | 0 |
| rs727503218 | pathogenic | 0 | 0 | 0 |
| rs11570052 | benign | 0 | 0 | 0 |
| rs730880623 | pathogenic | 0 | 0 | 0 |
| rs138753870 | benign | 0 | 0 | 0 |
| rs397516068 | pathogenic | 0 | 0 | 0 |
| rs3729989 | benign | 1 | 1 | 1 |
| rs397516074 | pathogenic | 0 | 0 | 0 |
| rs561449819 | pathogenic | 0 | 0 | 0 |
| rs121917779 | benign | 0 | 0 | 0 |
| rs142881576 | pathogenic | 0 | 0 | 0 |
| rs77384282 | benign | 0 | 0 | 0 |
| rs201865787 | benign | 0 | 0 | 0 |
| rs142244640 | benign | 0 | 0 | 0 |
| rs200488706 | benign | 0 | 0 | 0 |
| rs137853291 | pathogenic | 1 | 1 | 0 |
| rs28933990 | pathogenic | 1 | 1 | 0 |
| rs140570189 | benign | 0 | 0 | 0 |
| rs201588178 | benign | 0 | 0 | 0 |
| rs104893846 | pathogenic | 0 | 0 | 0 |
| rs754545360 | pathogenic | 0 | 1 | 0 |
| rs104893851 | pathogenic | 0 | 1 | 0 |
| rs104893849 | pathogenic | 0 | 0 | 0 |
| rs571038432 | pathogenic | 1 | 1 | 0 |
| rs2270655 | benign | 1 | 1 | 1 |
| rs315952 | benign | 0 | 0 | 0 |
| rs151176548 | benign | 0 | 0 | 0 |
| rs122459147 | pathogenic | 1 | 1 | 0 |
| rs122458140 | pathogenic | 0 | 0 | 0 |

| | | | | |
|---|---|---|---|---|
| rs122458142 | pathogenic | 1 | 1 | 0 |
| rs267606812 | pathogenic | 1 | 1 | 0 |
| rs267606813 | pathogenic | 1 | 1 | 0 |
| rs122458143 | pathogenic | 0 | 0 | 0 |
| rs122459146 | pathogenic | 0 | 0 | 0 |
| rs122458144 | pathogenic | 0 | 0 | 0 |
| rs122458145 | pathogenic | 0 | 0 | 0 |
| rs113994007 | pathogenic | 1 | 1 | 0 |
| rs758746181 | pathogenic | 1 | 1 | 1 |
| rs145034527 | pathogenic | 1 | 1 | 0 |
| rs119468006 | pathogenic | 1 | 1 | 0 |
| rs201908721 | pathogenic | 0 | 0 | 0 |
| rs755933881 | pathogenic | 1 | 1 | 0 |
| rs771578775 | pathogenic | 0 | 0 | 0 |
| rs17849927 | benign | 0 | 0 | 0 |
| rs139133094 | benign | 0 | 0 | 0 |
| rs12593 | benign | 0 | 0 | 0 |
| rs578189699 | pathogenic | 1 | 1 | 0 |
| rs119468008 | pathogenic | 0 | 0 | 0 |
| rs768524626 | benign | 0 | 0 | 0 |
| rs119468009 | pathogenic | 0 | 0 | 0 |
| rs119468004 | pathogenic | 0 | 0 | 0 |
| rs199874519 | pathogenic | 0 | 0 | 0 |
| rs376478331 | pathogenic | 0 | 0 | 0 |
| rs61995958 | pathogenic | 0 | 0 | 0 |
| rs199714731 | benign | 0 | 0 | 0 |
| rs149185954 | benign | 0 | 0 | 0 |
| rs121918353 | pathogenic | 0 | 0 | 0 |
| rs1131695 | benign | 0 | 0 | 0 |
| rs28939668 | pathogenic | 0 | 0 | 0 |
| rs758432471 | pathogenic | 1 | 1 | 1 |
| rs752746786 | pathogenic | 1 | 1 | 1 |
| rs28935471 | pathogenic | 0 | 0 | 0 |
| rs80338837 | pathogenic | 0 | 0 | 0 |
| rs137853312 | pathogenic | 0 | 0 | 0 |
| rs28935472 | pathogenic | 0 | 0 | 0 |
| rs28935473 | pathogenic | 0 | 0 | 0 |
| rs267606752 | pathogenic | 0 | 0 | 0 |
| rs3917973 | benign | 0 | 0 | 0 |
| rs3918018 | benign | 0 | 0 | 0 |
| rs757976755 | pathogenic | 0 | 0 | 0 |
| rs121918329 | pathogenic | 1 | 1 | 0 |
| rs770114459 | pathogenic | 0 | 0 | 0 |
| rs6662272 | benign | 0 | 0 | 0 |
| rs121918242 | pathogenic | 1 | 1 | 0 |
| rs121918240 | pathogenic | 1 | 1 | 0 |
| rs200094982 | pathogenic | 0 | 0 | 0 |
| rs121918241 | pathogenic | 0 | 0 | 0 |
| rs140522266 | pathogenic | 1 | 1 | 0 |
| rs121918243 | pathogenic | 1 | 1 | 0 |
| rs370596113 | pathogenic | 1 | 1 | 0 |
| rs398124295 | pathogenic | 1 | 1 | 0 |
| rs587776889 | pathogenic | 1 | 1 | 0 |
| rs747527726 | pathogenic | 1 | 1 | 0 |

| | | | | |
|---|---|---|---|---|
| rs112631212 | benign | 0 | 0 | 0 |
| rs104894605 | pathogenic | 0 | 0 | 0 |
| rs104894607 | pathogenic | 0 | 0 | 0 |
| rs104894606 | pathogenic | 0 | 0 | 0 |
| rs143873873 | benign | 0 | 0 | 0 |
| rs104894388 | pathogenic | 1 | 1 | 0 |
| rs104894391 | pathogenic | 1 | 1 | 0 |
| rs5350 | benign | 1 | 1 | 1 |
| rs104894390 | pathogenic | 1 | 1 | 1 |
| rs5346 | benign | 1 | 1 | 1 |
| rs121912500 | pathogenic | 0 | 0 | 0 |
| rs74315450 | pathogenic | 1 | 1 | 0 |
| rs121913396 | pathogenic | 1 | 1 | 1 |
| rs28931588 | pathogenic | 1 | 1 | 1 |
| rs121913400 | pathogenic | 0 | 0 | 0 |
| rs28931589 | pathogenic | 1 | 1 | 1 |
| rs121913403 | pathogenic | 0 | 0 | 0 |
| rs121913413 | pathogenic | 0 | 0 | 0 |
| rs760107623 | pathogenic | 1 | 1 | 0 |
| rs16866378 | benign | 1 | 1 | 0 |
| rs201025290 | pathogenic | 1 | 1 | 0 |
| rs12720356 | benign | 1 | 1 | 0 |
| rs55882956 | benign | 0 | 1 | 0 |
| rs4986933 | benign | 1 | 1 | 0 |
| rs2020957 | benign | 1 | 1 | 1 |
| rs1800124 | benign | 1 | 1 | 1 |
| rs150077735 | benign | 1 | 1 | 1 |
| rs201419000 | benign | 0 | 0 | 0 |
| rs772044176 | pathogenic | 0 | 0 | 0 |
| rs138346388 | benign | 0 | 0 | 0 |
| rs121964916 | pathogenic | 0 | 0 | 0 |
| rs112534524 | pathogenic | 0 | 0 | 0 |
| rs182078921 | benign | 1 | 1 | 1 |
| rs113273712 | benign | 0 | 0 | 0 |
| rs74817407 | benign | 1 | 1 | 1 |
| rs61733901 | benign | 1 | 1 | 1 |
| rs121964912 | pathogenic | 1 | 1 | 1 |
| rs41278047 | benign | 1 | 1 | 1 |
| rs7437875 | benign | 1 | 1 | 1 |
| rs113460688 | benign | 1 | 1 | 1 |
| rs754572081 | benign | 1 | 1 | 1 |
| rs201082692 | benign | 1 | 1 | 1 |
| rs202085803 | benign | 1 | 1 | 1 |
| rs141320511 | benign | 0 | 0 | 0 |
| rs78501403 | benign | 0 | 0 | 0 |
| rs28939675 | pathogenic | 0 | 0 | 0 |
| rs74315522 | pathogenic | 0 | 0 | 0 |
| rs121434639 | pathogenic | 1 | 1 | 0 |
| rs121434640 | pathogenic | 1 | 1 | 1 |
| rs757282628 | benign | 1 | 1 | 1 |
| rs386834010 | pathogenic | 0 | 0 | 0 |
| rs200863680 | pathogenic | 0 | 0 | 0 |
| rs193919336 | pathogenic | 1 | 1 | 0 |
| rs386834011 | pathogenic | 0 | 0 | 0 |

| | | | | |
|---|---|---|---|---|
| rs28940869 | pathogenic | 0 | 0 | 0 |
| rs386834014 | pathogenic | 1 | 1 | 0 |
| rs267606960 | pathogenic | 0 | 0 | 0 |
| rs28942068 | pathogenic | 0 | 0 | 0 |
| rs193919335 | pathogenic | 0 | 0 | 0 |
| rs74374973 | pathogenic | 0 | 0 | 0 |
| rs386834018 | pathogenic | 0 | 0 | 0 |
| rs267606962 | pathogenic | 0 | 0 | 0 |
| rs6659553 | benign | 0 | 0 | 0 |
| rs29281 | benign | 0 | 0 | 0 |
| rs782265090 | benign | 0 | 0 | 0 |
| rs35910969 | pathogenic | 0 | 0 | 0 |
| rs121909113 | pathogenic | 1 | 1 | 1 |
| rs29001571 | pathogenic | 1 | 1 | 1 |
| rs587781250 | pathogenic | 1 | 1 | 1 |
| rs28939680 | pathogenic | 1 | 1 | 1 |
| rs28939681 | pathogenic | 1 | 1 | 1 |
| rs121909112 | pathogenic | 1 | 1 | 1 |
| rs28937568 | pathogenic | 1 | 1 | 1 |
| rs1799895 | pathogenic | 0 | 0 | 0 |
| rs386833431 | pathogenic | 1 | 1 | 1 |
| rs121964905 | pathogenic | 1 | 1 | 1 |
| rs121964906 | pathogenic | 1 | 1 | 1 |
| rs11551664 | pathogenic | 1 | 1 | 1 |
| rs104894705 | pathogenic | 1 | 1 | 1 |
| rs104893805 | pathogenic | 0 | 0 | 0 |
| rs104894205 | pathogenic | 0 | 0 | 0 |
| rs104894204 | pathogenic | 1 | 1 | 0 |
| rs62642926 | pathogenic | 0 | 0 | 0 |
| rs5030841 | pathogenic | 1 | 1 | 1 |
| rs75193786 | pathogenic | 1 | 1 | 1 |
| rs62508682 | pathogenic | 0 | 0 | 0 |
| rs3850625 | benign | 1 | 1 | 1 |
| rs141908793 | benign | 0 | 0 | 0 |
| rs187864727 | benign | 0 | 0 | 0 |
| rs548909101 | benign | 0 | 0 | 0 |
| rs3825394 | benign | 0 | 1 | 0 |
| rs143548402 | benign | 0 | 0 | 0 |
| rs267607146 | pathogenic | 0 | 0 | 0 |
| rs267607144 | pathogenic | 0 | 0 | 0 |
| rs267607148 | pathogenic | 1 | 1 | 1 |
| rs267607143 | pathogenic | 0 | 0 | 0 |
| rs267607145 | pathogenic | 0 | 0 | 0 |
| rs121912636 | pathogenic | 1 | 1 | 1 |
| rs121912634 | pathogenic | 1 | 1 | 1 |
| rs570675468 | benign | 0 | 0 | 0 |
| rs368861241 | pathogenic | 1 | 1 | 1 |
| rs104894528 | pathogenic | 1 | 1 | 1 |
| rs77493891 | benign | 0 | 0 | 0 |
| rs62640903 | benign | 0 | 0 | 0 |
| rs149926091 | benign | 1 | 1 | 1 |
| rs641081 | benign | 0 | 0 | 0 |
| rs4930199 | benign | 0 | 0 | 0 |
| rs121913020 | pathogenic | 0 | 0 | 0 |

| | | | | |
|---|---|---|---|---|
| rs1799793 | benign | 0 | 0 | 0 |
| rs121913016 | pathogenic | 0 | 0 | 0 |
| rs121913025 | pathogenic | 0 | 0 | 0 |
| rs121913019 | pathogenic | 0 | 0 | 0 |
| rs376556895 | pathogenic | 1 | 1 | 1 |
| rs121913024 | pathogenic | 1 | 1 | 1 |
| rs121913021 | pathogenic | 0 | 0 | 0 |
| rs121913023 | pathogenic | 0 | 0 | 0 |
| rs41556519 | pathogenic | 0 | 0 | 0 |
| rs121913022 | pathogenic | 0 | 0 | 0 |
| rs144564120 | pathogenic | 0 | 0 | 0 |
| rs121913026 | pathogenic | 1 | 1 | 1 |
| rs121913018 | pathogenic | 1 | 1 | 1 |
| rs142954643 | benign | 0 | 0 | 0 |
| rs76655496 | benign | 0 | 0 | 0 |
| rs104894371 | pathogenic | 1 | 1 | 0 |
| rs137964936 | benign | 0 | 0 | 0 |
| rs759736132 | pathogenic | 0 | 0 | 0 |
| rs387906595 | pathogenic | 1 | 1 | 0 |
| rs768248216 | pathogenic | 0 | 0 | 0 |
| rs548231008 | benign | 0 | 0 | 0 |
| rs757094384 | benign | 0 | 0 | 0 |
| rs867529 | benign | 0 | 0 | 0 |
| rs13045 | benign | 1 | 1 | 1 |
| rs121908570 | pathogenic | 0 | 0 | 0 |
| rs201872586 | benign | 1 | 1 | 0 |
| rs149092990 | benign | 0 | 0 | 0 |
| rs115585711 | benign | 0 | 0 | 0 |
| rs137909416 | benign | 1 | 1 | 0 |
| rs143388949 | benign | 1 | 1 | 1 |
| rs73137716 | benign | 1 | 1 | 1 |
| rs121908947 | pathogenic | 1 | 1 | 1 |
| rs398122904 | pathogenic | 1 | 1 | 1 |
| rs35715497 | benign | 1 | 1 | 0 |
| rs35462421 | benign | 0 | 0 | 0 |
| rs119475043 | pathogenic | 0 | 0 | 0 |
| rs119475042 | pathogenic | 0 | 0 | 0 |
| rs1076669 | benign | 1 | 1 | 0 |
| rs141146885 | benign | 0 | 0 | 0 |
| rs3026906 | pathogenic | 0 | 0 | 0 |
| rs75362221 | benign | 1 | 1 | 1 |
| rs138472116 | benign | 1 | 1 | 1 |
| rs371040282 | pathogenic | 1 | 1 | 1 |
| rs4988496 | benign | 0 | 0 | 0 |
| rs121918117 | pathogenic | 1 | 1 | 1 |
| rs4988498 | benign | 0 | 0 | 0 |
| rs5035 | benign | 0 | 0 | 0 |
| rs121918322 | pathogenic | 1 | 1 | 1 |
| rs370359540 | benign | 0 | 0 | 0 |
| rs758972589 | pathogenic | 0 | 0 | 0 |
| rs144856889 | benign | 0 | 0 | 0 |
| rs121434623 | pathogenic | 1 | 1 | 1 |
| rs61735600 | benign | 0 | 0 | 0 |
| rs72550870 | benign | 1 | 1 | 1 |

| | | | | |
|---|---|---|---|---|
| rs56392418 | benign | 0 | 0 | 0 |
| rs41307788 | benign | 1 | 1 | 1 |
| rs387907239 | pathogenic | 1 | 1 | 0 |
| rs28931584 | pathogenic | 0 | 0 | 0 |
| rs121912745 | pathogenic | 0 | 0 | 0 |
| rs121912744 | pathogenic | 0 | 0 | 0 |
| rs121912754 | pathogenic | 1 | 1 | 1 |
| rs121912746 | pathogenic | 0 | 0 | 0 |
| rs121912757 | pathogenic | 0 | 0 | 0 |
| rs121912758 | pathogenic | 0 | 0 | 0 |
| rs121912748 | pathogenic | 0 | 0 | 0 |
| rs121912755 | pathogenic | 0 | 0 | 0 |
| rs121912741 | pathogenic | 0 | 0 | 0 |
| rs121912753 | pathogenic | 0 | 0 | 0 |
| rs121912750 | pathogenic | 0 | 0 | 0 |
| rs2285644 | pathogenic | 0 | 0 | 0 |
| rs121912751 | pathogenic | 0 | 0 | 0 |
| rs5026 | benign | 0 | 0 | 0 |
| rs121912759 | pathogenic | 0 | 0 | 0 |
| rs28931585 | pathogenic | 0 | 0 | 0 |
| rs17122769 | benign | 0 | 0 | 0 |
| rs28362322 | benign | 1 | 1 | 1 |
| rs3824874 | benign | 0 | 0 | 0 |
| rs146004831 | benign | 0 | 0 | 0 |
| rs142155860 | benign | 0 | 0 | 0 |
| rs121434404 | pathogenic | 0 | 0 | 0 |
| rs61735578 | benign | 1 | 1 | 1 |
| rs558018 | benign | 0 | 0 | 0 |
| rs61735577 | benign | 1 | 1 | 1 |
| rs76784113 | benign | 0 | 0 | 0 |
| rs199469640 | pathogenic | 0 | 0 | 0 |
| rs199469628 | pathogenic | 1 | 1 | 0 |
| rs199469629 | pathogenic | 0 | 1 | 0 |
| rs387907156 | pathogenic | 0 | 1 | 0 |
| rs199469643 | pathogenic | 0 | 0 | 0 |
| rs199469646 | pathogenic | 0 | 0 | 0 |
| rs766294940 | pathogenic | 0 | 0 | 0 |
| rs387907347 | pathogenic | 0 | 0 | 0 |
| rs137853590 | pathogenic | 1 | 1 | 1 |
| rs137853589 | pathogenic | 1 | 1 | 1 |
| rs137853588 | pathogenic | 1 | 1 | 1 |
| rs55909005 | benign | 0 | 0 | 0 |
| rs4581 | benign | 0 | 0 | 0 |
| rs63750123 | benign | 0 | 0 | 0 |
| rs10254120 | benign | 0 | 0 | 0 |
| rs121434629 | pathogenic | 1 | 1 | 0 |
| rs200313585 | benign | 0 | 0 | 0 |
| rs143162541 | benign | 1 | 1 | 0 |
| rs760615315 | benign | 0 | 0 | 0 |
| rs768488890 | benign | 1 | 1 | 1 |
| rs116373169 | benign | 1 | 1 | 0 |
| rs63750871 | pathogenic | 1 | 1 | 0 |
| rs116349687 | benign | 0 | 0 | 0 |
| rs375289386 | benign | 0 | 0 | 0 |

| | | | | |
|---|---|---|---|---|
| rs760228510 | pathogenic | 1 | 1 | 1 |
| rs587779343 | pathogenic | 1 | 1 | 1 |
| rs573125799 | benign | 0 | 0 | 0 |
| rs1805322 | benign | 0 | 0 | 0 |
| rs267608153 | pathogenic | 0 | 0 | 0 |
| rs139194813 | benign | 0 | 0 | 0 |
| rs530021751 | benign | 0 | 0 | 0 |
| rs200640585 | pathogenic | 0 | 0 | 0 |
| rs139438201 | benign | 0 | 0 | 0 |
| rs200513014 | benign | 0 | 0 | 0 |
| rs567102013 | benign | 0 | 0 | 0 |
| rs121908608 | pathogenic | 0 | 0 | 0 |
| rs121908606 | pathogenic | 0 | 0 | 0 |
| rs17103671 | benign | 0 | 0 | 0 |
| rs34725281 | benign | 0 | 0 | 0 |
| rs104894421 | pathogenic | 1 | 1 | 0 |
| rs121918145 | pathogenic | 1 | 1 | 1 |
| rs121918152 | pathogenic | 1 | 1 | 1 |
| rs56379106 | benign | 0 | 0 | 0 |
| rs4987076 | benign | 0 | 0 | 0 |
| rs370243877 | pathogenic | 0 | 0 | 0 |
| rs17679445 | benign | 1 | 1 | 1 |
| rs550423482 | pathogenic | 1 | 1 | 1 |
| rs104894629 | pathogenic | 1 | 1 | 1 |
| rs773450573 | pathogenic | 1 | 1 | 1 |
| rs116679717 | benign | 0 | 0 | 0 |
| rs372000848 | pathogenic | 0 | 0 | 0 |
| rs138595973 | benign | 0 | 0 | 0 |
| rs115244417 | benign | 0 | 0 | 0 |
| rs11575292 | benign | 0 | 0 | 0 |
| rs137853211 | pathogenic | 1 | 1 | 1 |
| rs137853207 | pathogenic | 0 | 0 | 0 |
| rs137853210 | pathogenic | 1 | 1 | 1 |
| rs6262 | benign | 0 | 0 | 0 |
| rs6263 | benign | 0 | 1 | 0 |
| rs137853208 | pathogenic | 0 | 1 | 0 |
| rs137853212 | pathogenic | 1 | 1 | 0 |
| rs137853209 | pathogenic | 1 | 1 | 1 |
| rs201951824 | pathogenic | 0 | 0 | 0 |
| rs11575542 | benign | 0 | 0 | 0 |
| rs199681788 | benign | 0 | 1 | 0 |
| rs748835902 | benign | 1 | 1 | 1 |
| rs753719949 | benign | 0 | 0 | 0 |
| rs267606647 | pathogenic | 0 | 0 | 0 |
| rs267606648 | pathogenic | 1 | 1 | 1 |
| rs267606643 | pathogenic | 0 | 1 | 0 |
| rs559947967 | pathogenic | 1 | 1 | 0 |
| rs267606645 | pathogenic | 1 | 1 | 1 |
| rs12116440 | benign | 0 | 0 | 0 |
| rs121908539 | pathogenic | 0 | 0 | 0 |
| rs137853096 | pathogenic | 1 | 1 | 0 |
| rs148363262 | benign | 1 | 1 | 0 |
| rs587777442 | pathogenic | 0 | 0 | 0 |
| rs25640 | benign | 1 | 1 | 1 |

| | | | | |
|---|---|---|---|---|
| rs28943589 | benign | 0 | 0 | 0 |
| rs1143650 | benign | 1 | 1 | 1 |
| rs150886026 | benign | 0 | 1 | 0 |
| rs121908133 | pathogenic | 1 | 1 | 0 |
| rs121908131 | pathogenic | 0 | 0 | 0 |
| rs754477154 | pathogenic | 0 | 0 | 0 |
| rs747334007 | benign | 0 | 0 | 0 |
| rs121908132 | pathogenic | 0 | 0 | 0 |
| rs104894103 | pathogenic | 1 | 1 | 0 |
| rs140088149 | benign | 0 | 0 | 0 |
| rs149803148 | benign | 0 | 0 | 0 |
| rs11064480 | benign | 0 | 0 | 0 |
| rs74435397 | pathogenic | 1 | 1 | 1 |
| rs587783626 | pathogenic | 1 | 1 | 1 |
| rs148211042 | pathogenic | 1 | 1 | 1 |
| rs17849502 | benign | 0 | 0 | 0 |
| rs13306575 | pathogenic | 1 | 1 | 1 |
| rs201426733 | benign | 0 | 0 | 0 |
| rs182278784 | benign | 1 | 1 | 1 |
| rs77786415 | benign | 1 | 1 | 1 |
| rs200192119 | benign | 1 | 1 | 1 |
| rs12877501 | benign | 1 | 1 | 1 |
| rs77985244 | benign | 1 | 1 | 1 |
| rs5907 | pathogenic | 1 | 1 | 0 |
| rs2305178 | benign | 0 | 0 | 0 |
| rs3135867 | benign | 0 | 0 | 0 |
| rs121913482 | pathogenic | 1 | 1 | 1 |
| rs121913483 | pathogenic | 1 | 1 | 1 |
| rs4647924 | pathogenic | 1 | 1 | 1 |
| rs121913111 | pathogenic | 1 | 1 | 1 |
| rs104894002 | pathogenic | 0 | 0 | 0 |
| rs104894001 | pathogenic | 0 | 0 | 0 |
| rs75932628 | benign | 0 | 0 | 0 |
| rs201258663 | pathogenic | 0 | 0 | 0 |
| rs104893998 | pathogenic | 0 | 0 | 0 |
| rs2234253 | benign | 1 | 1 | 0 |
| rs121908402 | pathogenic | 1 | 1 | 0 |
| rs1805389 | benign | 0 | 0 | 0 |
| rs1805388 | benign | 0 | 0 | 0 |
| rs748385144 | pathogenic | 0 | 0 | 0 |
| rs104894420 | pathogenic | 0 | 0 | 0 |
| rs104894418 | pathogenic | 0 | 0 | 0 |
| rs61731910 | benign | 0 | 0 | 0 |
| rs758723288 | pathogenic | 1 | 1 | 0 |
| rs201865375 | pathogenic | 1 | 1 | 0 |
| rs565090080 | pathogenic | 1 | 1 | 1 |
| rs113042166 | benign | 0 | 0 | 0 |
| rs61736326 | benign | 1 | 1 | 1 |
| rs12030699 | benign | 1 | 1 | 1 |
| rs121913578 | pathogenic | 1 | 1 | 1 |
| rs61739582 | benign | 0 | 0 | 0 |
| rs745983207 | pathogenic | 0 | 0 | 0 |
| rs763524380 | benign | 0 | 0 | 0 |
| rs121912792 | pathogenic | 0 | 0 | 0 |

| | | | | |
|---|---|---|---|---|
| rs121912793 | pathogenic | 0 | 0 | 0 |
| rs387906599 | pathogenic | 0 | 0 | 0 |
| rs121912794 | pathogenic | 0 | 1 | 0 |
| rs121912795 | pathogenic | 0 | 0 | 0 |
| rs121912796 | pathogenic | 1 | 1 | 0 |
| rs121912797 | pathogenic | 0 | 1 | 0 |
| rs121912798 | pathogenic | 1 | 1 | 1 |
| rs80359812 | pathogenic | 1 | 1 | 0 |
| rs776583130 | benign | 1 | 1 | 0 |
| rs80359814 | pathogenic | 1 | 1 | 0 |
| rs202060209 | pathogenic | 1 | 1 | 0 |
| rs267607061 | pathogenic | 1 | 1 | 0 |
| rs267607060 | pathogenic | 0 | 0 | 0 |
| rs80359816 | pathogenic | 1 | 1 | 0 |
| rs80359818 | pathogenic | 1 | 1 | 0 |
| rs387907313 | pathogenic | 0 | 0 | 0 |
| rs587784395 | benign | 0 | 0 | 0 |
| rs5811 | benign | 0 | 0 | 0 |
| rs121909740 | pathogenic | 1 | 1 | 0 |
| rs148518827 | benign | 1 | 1 | 0 |
| rs121909739 | pathogenic | 1 | 1 | 0 |
| rs780528770 | benign | 1 | 1 | 0 |
| rs13306758 | pathogenic | 0 | 0 | 0 |
| rs267607059 | pathogenic | 0 | 0 | 0 |
| rs61730638 | benign | 0 | 0 | 0 |
| rs104894533 | pathogenic | 0 | 0 | 0 |
| rs104894527 | pathogenic | 0 | 0 | 0 |
| rs200503569 | pathogenic | 1 | 1 | 1 |
| rs80338700 | pathogenic | 1 | 1 | 1 |
| rs765433263 | pathogenic | 1 | 1 | 1 |
| rs104894530 | pathogenic | 1 | 1 | 1 |
| rs80338701 | pathogenic | 1 | 1 | 1 |
| rs141498002 | pathogenic | 1 | 1 | 1 |
| rs104894525 | pathogenic | 0 | 0 | 0 |
| rs80338703 | pathogenic | 0 | 0 | 0 |
| rs28936415 | pathogenic | 1 | 1 | 0 |
| rs150719105 | pathogenic | 0 | 0 | 0 |
| rs148032587 | pathogenic | 0 | 0 | 0 |
| rs150577656 | pathogenic | 0 | 0 | 0 |
| rs190521996 | pathogenic | 0 | 0 | 0 |
| rs104894526 | pathogenic | 1 | 1 | 1 |
| rs80338704 | pathogenic | 0 | 0 | 0 |
| rs199562225 | pathogenic | 1 | 1 | 1 |
| rs532870929 | pathogenic | 0 | 0 | 0 |
| rs398123309 | pathogenic | 0 | 0 | 0 |
| rs78290141 | pathogenic | 0 | 0 | 0 |
| rs80338705 | pathogenic | 0 | 0 | 0 |
| rs104894531 | pathogenic | 1 | 1 | 1 |
| rs80338706 | pathogenic | 0 | 0 | 0 |
| rs80338707 | pathogenic | 0 | 0 | 0 |
| rs80338708 | pathogenic | 0 | 0 | 0 |
| rs80338709 | pathogenic | 0 | 0 | 0 |
| rs104893833 | benign | 0 | 0 | 0 |
| rs61744585 | benign | 0 | 0 | 0 |

| | | | | |
|---|---|---|---|---|
| rs73569515 | benign | 0 | 0 | 0 |
| rs398123373 | pathogenic | 0 | 0 | 0 |
| rs756854513 | pathogenic | 0 | 0 | 0 |
| rs58232698 | benign | 0 | 0 | 0 |
| rs9600455 | benign | 0 | 0 | 0 |
| rs201016593 | pathogenic | 0 | 0 | 0 |
| rs756039188 | pathogenic | 0 | 0 | 0 |
| rs121908024 | pathogenic | 0 | 0 | 0 |
| rs373144619 | benign | 0 | 0 | 0 |
| rs138078086 | benign | 0 | 0 | 0 |
| rs376207800 | pathogenic | 0 | 0 | 0 |
| rs749038326 | pathogenic | 1 | 1 | 0 |
| rs372845091 | benign | 0 | 0 | 0 |
| rs377271627 | pathogenic | 0 | 0 | 0 |
| rs563390335 | pathogenic | 0 | 0 | 0 |
| rs13306510 | benign | 0 | 0 | 0 |
| rs764042910 | pathogenic | 0 | 0 | 0 |
| rs201384282 | benign | 1 | 1 | 1 |
| rs730882086 | pathogenic | 1 | 1 | 1 |
| rs756613387 | pathogenic | 1 | 1 | 1 |
| rs121908028 | benign | 1 | 1 | 0 |
| rs773328511 | benign | 0 | 0 | 0 |
| rs749220643 | pathogenic | 1 | 1 | 0 |
| rs730882089 | pathogenic | 0 | 0 | 0 |
| rs146651743 | benign | 0 | 0 | 0 |
| rs148698650 | pathogenic | 0 | 0 | 0 |
| rs140241383 | benign | 0 | 0 | 0 |
| rs757252110 | pathogenic | 0 | 0 | 0 |
| rs13306512 | pathogenic | 1 | 1 | 1 |
| rs121908043 | pathogenic | 1 | 1 | 1 |
| rs773658037 | pathogenic | 0 | 0 | 0 |
| rs774439908 | pathogenic | 0 | 0 | 0 |
| rs779732323 | pathogenic | 1 | 1 | 0 |
| rs749780672 | pathogenic | 0 | 0 | 0 |
| rs375651668 | benign | 1 | 1 | 1 |
| rs193922568 | pathogenic | 1 | 1 | 0 |
| rs750363970 | benign | 1 | 1 | 0 |
| rs5932 | benign | 0 | 0 | 0 |
| rs755154048 | pathogenic | 1 | 1 | 0 |
| rs759876319 | pathogenic | 0 | 0 | 0 |
| rs764550980 | pathogenic | 1 | 1 | 0 |
| rs779940524 | pathogenic | 0 | 0 | 0 |
| rs563382937 | pathogenic | 1 | 1 | 1 |
| rs28942083 | pathogenic | 0 | 1 | 0 |
| rs770744861 | pathogenic | 0 | 1 | 0 |
| rs121908031 | pathogenic | 0 | 1 | 0 |
| rs776217028 | pathogenic | 0 | 1 | 0 |
| rs763625913 | pathogenic | 0 | 0 | 0 |
| rs3736772 | benign | 0 | 0 | 0 |
| rs104894696 | benign | 1 | 1 | 1 |
| rs121918177 | pathogenic | 0 | 1 | 0 |
| rs121918181 | pathogenic | 1 | 1 | 0 |
| rs121918174 | pathogenic | 0 | 1 | 0 |
| rs121918182 | pathogenic | 0 | 0 | 0 |

| | | | | |
|---|---|---|---|---|
| rs121918175 | pathogenic | 1 | 1 | 1 |
| rs121918185 | pathogenic | 1 | 1 | 1 |
| rs121918173 | pathogenic | 1 | 1 | 0 |
| rs121918178 | pathogenic | 1 | 1 | 1 |
| rs121918179 | pathogenic | 1 | 1 | 1 |
| rs121918183 | pathogenic | 0 | 0 | 0 |
| rs121918176 | pathogenic | 1 | 1 | 1 |
| rs121918172 | pathogenic | 0 | 1 | 0 |
| rs121918184 | pathogenic | 0 | 0 | 0 |
| rs121434560 | pathogenic | 1 | 1 | 0 |
| rs121434561 | pathogenic | 1 | 1 | 0 |
| rs3218716 | pathogenic | 1 | 1 | 1 |
| rs145532615 | benign | 0 | 0 | 0 |
| rs149840927 | pathogenic | 0 | 0 | 0 |
| rs730880762 | pathogenic | 0 | 0 | 0 |
| rs730880764 | pathogenic | 0 | 0 | 0 |
| rs730880765 | pathogenic | 0 | 0 | 0 |
| rs587782962 | pathogenic | 0 | 0 | 0 |
| rs141735183 | pathogenic | 0 | 0 | 0 |
| rs397516187 | pathogenic | 0 | 0 | 0 |
| rs727505026 | pathogenic | 0 | 0 | 0 |
| rs529700838 | pathogenic | 0 | 0 | 0 |
| rs540263945 | pathogenic | 0 | 0 | 0 |
| rs149103761 | benign | 0 | 0 | 0 |
| rs730880906 | pathogenic | 0 | 0 | 0 |
| rs730880785 | pathogenic | 0 | 0 | 0 |
| rs397516199 | pathogenic | 0 | 0 | 0 |
| rs730880725 | benign | 0 | 0 | 0 |
| rs730880796 | pathogenic | 0 | 0 | 0 |
| rs201307101 | benign | 0 | 0 | 0 |
| rs397516214 | pathogenic | 0 | 0 | 0 |
| rs3729823 | benign | 0 | 0 | 0 |
| rs397516218 | pathogenic | 0 | 0 | 0 |
| rs121913647 | pathogenic | 0 | 0 | 0 |
| rs373514686 | pathogenic | 0 | 0 | 0 |
| rs763538103 | pathogenic | 0 | 0 | 0 |
| rs121913650 | pathogenic | 0 | 0 | 0 |
| rs139222507 | pathogenic | 0 | 0 | 0 |
| rs369940645 | pathogenic | 0 | 0 | 0 |
| rs763073072 | pathogenic | 0 | 0 | 0 |
| rs145734640 | pathogenic | 0 | 0 | 0 |
| rs730880821 | pathogenic | 0 | 0 | 0 |
| rs28933098 | pathogenic | 0 | 0 | 0 |
| rs372381770 | pathogenic | 0 | 0 | 0 |
| rs146796870 | benign | 0 | 0 | 0 |
| rs104894485 | pathogenic | 1 | 1 | 1 |
| rs35407712 | benign | 0 | 0 | 0 |
| rs35842750 | benign | 0 | 0 | 0 |
| rs2721191 | benign | 0 | 0 | 0 |
| rs35098923 | benign | 0 | 0 | 0 |
| rs34293591 | benign | 1 | 1 | 0 |
| rs36023964 | benign | 0 | 0 | 0 |
| rs4251691 | benign | 0 | 0 | 0 |
| rs34666647 | benign | 0 | 0 | 0 |

| | | | | |
|---|---|---|---|---|
| rs4925828 | benign | 0 | 0 | 0 |
| rs35348691 | benign | 0 | 0 | 0 |
| rs36078464 | benign | 0 | 0 | 0 |
| rs34915097 | benign | 0 | 0 | 0 |
| rs35101495 | benign | 0 | 0 | 0 |
| rs141568342 | pathogenic | 0 | 0 | 0 |
| rs1800571 | pathogenic | 0 | 0 | 0 |
| rs140566084 | benign | 1 | 1 | 1 |
| rs727503791 | pathogenic | 1 | 1 | 0 |
| rs545215807 | pathogenic | 0 | 0 | 0 |
| rs113994168 | pathogenic | 0 | 0 | 0 |
| rs113994167 | pathogenic | 0 | 0 | 0 |
| rs200788251 | pathogenic | 0 | 0 | 0 |
| rs200573371 | pathogenic | 0 | 0 | 0 |
| rs771874163 | pathogenic | 1 | 1 | 1 |
| rs112406105 | pathogenic | 1 | 1 | 1 |
| rs118204015 | pathogenic | 1 | 1 | 1 |
| rs113994169 | pathogenic | 0 | 0 | 0 |
| rs118204018 | pathogenic | 0 | 1 | 0 |
| rs138834083 | pathogenic | 0 | 0 | 0 |
| rs533055438 | pathogenic | 1 | 1 | 1 |
| rs2309689 | pathogenic | 1 | 1 | 1 |
| rs138058572 | pathogenic | 1 | 1 | 1 |
| rs118204017 | pathogenic | 1 | 1 | 0 |
| rs766742117 | pathogenic | 1 | 1 | 1 |
| rs751995154 | pathogenic | 1 | 1 | 1 |
| rs200771970 | pathogenic | 0 | 0 | 0 |
| rs118204014 | pathogenic | 0 | 0 | 0 |
| rs148584617 | pathogenic | 1 | 1 | 1 |
| rs121912720 | pathogenic | 0 | 0 | 0 |
| rs121912721 | pathogenic | 0 | 0 | 0 |
| rs28929476 | pathogenic | 0 | 0 | 0 |
| rs28931574 | pathogenic | 1 | 1 | 1 |
| rs12718465 | benign | 0 | 0 | 0 |
| rs144689354 | pathogenic | 0 | 0 | 0 |
| rs149095705 | pathogenic | 0 | 0 | 0 |
| rs572839648 | benign | 1 | 1 | 1 |
| rs121909715 | pathogenic | 1 | 1 | 0 |
| rs34077456 | benign | 1 | 1 | 1 |
| rs397517876 | benign | 0 | 0 | 0 |
| rs1064074 | benign | 0 | 0 | 0 |
| rs142751309 | benign | 0 | 0 | 0 |
| rs116275562 | benign | 0 | 0 | 0 |
| rs148095899 | benign | 0 | 0 | 0 |
| rs751838040 | pathogenic | 0 | 0 | 0 |
| rs121434324 | pathogenic | 1 | 1 | 1 |
| rs143367518 | pathogenic | 0 | 0 | 0 |
| rs121434325 | pathogenic | 0 | 0 | 0 |
| rs397515328 | pathogenic | 0 | 0 | 0 |
| rs376039938 | pathogenic | 0 | 0 | 0 |
| rs387907243 | pathogenic | 0 | 0 | 0 |
| rs28934876 | pathogenic | 0 | 0 | 0 |
| rs387907245 | pathogenic | 0 | 0 | 0 |
| rs138544311 | benign | 0 | 0 | 0 |

| | | | | |
|---|---|---|---|---|
| rs643788 | benign | 0 | 0 | 0 |
| rs749977756 | benign | 0 | 0 | 0 |
| rs1800574 | benign | 0 | 0 | 0 |
| rs137853336 | pathogenic | 1 | 1 | 1 |
| rs111916034 | benign | 0 | 0 | 0 |
| rs307826 | benign | 0 | 0 | 0 |
| rs35874891 | benign | 0 | 0 | 0 |
| rs111033580 | pathogenic | 1 | 1 | 1 |
| rs111033581 | pathogenic | 1 | 1 | 1 |
| rs121908373 | pathogenic | 0 | 0 | 0 |
| rs767122713 | pathogenic | 0 | 0 | 0 |
| rs2233213 | benign | 0 | 0 | 0 |
| rs121908605 | pathogenic | 1 | 1 | 0 |
| rs11887534 | benign | 0 | 0 | 0 |
| rs4148211 | benign | 0 | 0 | 0 |
| rs80025980 | benign | 0 | 0 | 0 |
| rs137854891 | pathogenic | 1 | 1 | 0 |
| rs74358901 | benign | 0 | 0 | 0 |
| rs9282574 | benign | 0 | 0 | 0 |
| rs137852993 | pathogenic | 0 | 0 | 0 |
| rs137852990 | pathogenic | 0 | 0 | 0 |
| rs137852987 | pathogenic | 0 | 0 | 0 |
| rs4148217 | benign | 0 | 0 | 0 |
| rs140690030 | benign | 0 | 0 | 0 |
| rs137852992 | pathogenic | 0 | 0 | 0 |
| rs6544718 | benign | 0 | 0 | 0 |
| rs137852989 | pathogenic | 0 | 0 | 0 |
| rs150893072 | benign | 1 | 1 | 1 |
| rs28931613 | pathogenic | 1 | 1 | 1 |
| rs143789597 | benign | 1 | 1 | 1 |
| rs121913037 | pathogenic | 1 | 1 | 0 |
| rs188802138 | pathogenic | 1 | 1 | 0 |
| rs121913038 | pathogenic | 1 | 1 | 0 |
| rs146835206 | benign | 0 | 0 | 0 |
| rs121913041 | pathogenic | 1 | 1 | 1 |
| rs121913039 | pathogenic | 1 | 1 | 1 |
| rs149977726 | pathogenic | 1 | 1 | 0 |
| rs121913036 | pathogenic | 0 | 0 | 0 |
| rs121913040 | pathogenic | 0 | 0 | 0 |
| rs765007158 | benign | 0 | 0 | 0 |
| rs112723255 | benign | 0 | 0 | 0 |
| rs574164748 | benign | 0 | 0 | 0 |
| rs80338734 | pathogenic | 1 | 1 | 0 |
| rs104894694 | pathogenic | 1 | 1 | 0 |
| rs753198836 | pathogenic | 0 | 0 | 0 |
| rs760101382 | pathogenic | 0 | 0 | 0 |
| rs121909272 | pathogenic | 1 | 1 | 0 |
| rs370421531 | pathogenic | 0 | 0 | 0 |
| rs565109128 | benign | 0 | 0 | 0 |
| rs17851582 | benign | 0 | 0 | 0 |
| rs753228876 | benign | 0 | 0 | 0 |
| rs141471799 | benign | 0 | 0 | 0 |
| rs281864916 | pathogenic | 1 | 1 | 0 |
| rs121918409 | pathogenic | 1 | 1 | 1 |

| | | | | |
|---|---|---|---|---|
| rs121918413 | pathogenic | 1 | 1 | 1 |
| rs121918411 | pathogenic | 1 | 1 | 1 |
| rs121918408 | pathogenic | 1 | 1 | 1 |
| rs121918412 | pathogenic | 1 | 1 | 1 |
| rs121918410 | pathogenic | 1 | 1 | 1 |
| rs116474260 | benign | 1 | 1 | 1 |
| rs281864919 | pathogenic | 1 | 1 | 1 |
| rs397514452 | pathogenic | 1 | 1 | 0 |
| rs747843638 | benign | 0 | 0 | 0 |
| rs774357609 | pathogenic | 1 | 1 | 1 |
| rs201804005 | benign | 0 | 0 | 0 |
| rs746485916 | pathogenic | 0 | 0 | 0 |
| rs764131178 | pathogenic | 0 | 0 | 0 |
| rs374094065 | pathogenic | 0 | 0 | 0 |
| rs76151636 | pathogenic | 1 | 1 | 0 |
| rs753250853 | pathogenic | 0 | 0 | 0 |
| rs761632029 | pathogenic | 0 | 0 | 0 |
| rs775541743 | pathogenic | 0 | 0 | 0 |
| rs778749563 | pathogenic | 0 | 0 | 0 |
| rs60431989 | pathogenic | 1 | 1 | 0 |
| rs755554442 | pathogenic | 1 | 1 | 0 |
| rs749085322 | pathogenic | 0 | 0 | 0 |
| rs756029120 | pathogenic | 1 | 1 | 0 |
| rs137853279 | pathogenic | 1 | 1 | 0 |
| rs777354267 | pathogenic | 1 | 1 | 1 |
| rs142940486 | benign | 1 | 1 | 1 |
| rs140167393 | benign | 1 | 1 | 1 |
| rs2066827 | benign | 0 | 0 | 0 |
| rs765681672 | benign | 0 | 0 | 0 |
| rs137985549 | benign | 0 | 0 | 0 |
| rs74315340 | pathogenic | 1 | 1 | 1 |
| rs74315341 | pathogenic | 0 | 0 | 0 |
| rs74315335 | pathogenic | 0 | 0 | 0 |
| rs61730974 | benign | 0 | 0 | 0 |
| rs121909193 | pathogenic | 0 | 0 | 0 |
| rs74315334 | pathogenic | 0 | 0 | 0 |
| rs74315329 | pathogenic | 0 | 0 | 0 |
| rs74315330 | pathogenic | 0 | 0 | 0 |
| rs121909194 | pathogenic | 0 | 0 | 0 |
| rs28936694 | pathogenic | 0 | 0 | 0 |
| rs74315336 | pathogenic | 0 | 0 | 0 |
| rs74315338 | pathogenic | 1 | 1 | 1 |
| rs74315328 | pathogenic | 0 | 0 | 0 |
| rs74315331 | pathogenic | 0 | 0 | 0 |
| rs74315332 | pathogenic | 0 | 0 | 0 |
| rs141696651 | benign | 0 | 0 | 0 |
| rs7041 | benign | 0 | 0 | 0 |
| rs4588 | benign | 0 | 0 | 0 |
| rs121908981 | pathogenic | 1 | 1 | 1 |
| rs606231459 | pathogenic | 0 | 0 | 0 |
| rs398123053 | pathogenic | 0 | 0 | 0 |
| rs774259802 | benign | 0 | 0 | 0 |
| rs121909582 | pathogenic | 1 | 1 | 1 |
| rs104893680 | pathogenic | 1 | 1 | 1 |

| | | | | |
|---|---|---|---|---|
| rs137854907 | pathogenic | 0 | 0 | 0 |
| rs765715798 | pathogenic | 0 | 0 | 0 |
| rs104893678 | pathogenic | 0 | 0 | 0 |
| rs104893679 | pathogenic | 0 | 1 | 0 |
| rs104893681 | pathogenic | 1 | 1 | 0 |
| rs121909640 | pathogenic | 1 | 1 | 0 |
| rs780705654 | pathogenic | 0 | 0 | 0 |
| rs201672146 | benign | 1 | 1 | 0 |
| rs12136658 | benign | 0 | 0 | 0 |
| rs56127440 | benign | 0 | 0 | 0 |
| rs104894433 | pathogenic | 1 | 1 | 1 |
| rs104894435 | pathogenic | 0 | 1 | 0 |
| rs104894437 | pathogenic | 1 | 1 | 1 |
| rs104894441 | pathogenic | 1 | 1 | 1 |
| rs104894440 | pathogenic | 1 | 1 | 1 |
| rs104894445 | pathogenic | 1 | 1 | 1 |
| rs104894436 | pathogenic | 0 | 0 | 0 |
| rs104894438 | pathogenic | 0 | 0 | 0 |
| rs200891969 | pathogenic | 1 | 1 | 0 |
| rs104894443 | pathogenic | 1 | 1 | 1 |
| rs104894434 | pathogenic | 1 | 1 | 1 |
| rs41298442 | pathogenic | 1 | 1 | 1 |
| rs104894442 | pathogenic | 1 | 1 | 1 |
| rs35453854 | benign | 0 | 0 | 0 |
| rs121912898 | pathogenic | 0 | 0 | 0 |
| rs121912897 | pathogenic | 1 | 1 | 0 |
| rs138260625 | benign | 0 | 0 | 0 |
| rs147802438 | benign | 1 | 1 | 0 |
| rs146443310 | benign | 1 | 1 | 0 |
| rs747319628 | pathogenic | 1 | 1 | 0 |
| rs1801195 | benign | 0 | 0 | 0 |
| rs145045341 | benign | 1 | 1 | 0 |
| rs116001173 | benign | 1 | 1 | 0 |
| rs587784175 | pathogenic | 0 | 0 | 0 |
| rs104893857 | pathogenic | 0 | 0 | 0 |
| rs104893862 | pathogenic | 1 | 1 | 0 |
| rs104893858 | pathogenic | 1 | 1 | 1 |
| rs121909249 | pathogenic | 0 | 1 | 0 |
| rs121909248 | pathogenic | 1 | 1 | 1 |
| rs104893859 | pathogenic | 1 | 1 | 0 |
| rs187063864 | pathogenic | 1 | 1 | 0 |
| rs121917817 | pathogenic | 1 | 1 | 0 |
| rs121917818 | pathogenic | 0 | 0 | 0 |
| rs753594601 | pathogenic | 0 | 0 | 0 |
| rs28937869 | pathogenic | 1 | 1 | 0 |
| rs6165 | benign | 0 | 0 | 0 |
| rs200089714 | pathogenic | 1 | 1 | 0 |
| rs372000714 | pathogenic | 0 | 0 | 0 |
| rs2228206 | benign | 0 | 0 | 0 |
| rs560096 | benign | 0 | 0 | 0 |
| rs137852666 | pathogenic | 0 | 0 | 0 |
| rs10896380 | benign | 0 | 0 | 0 |
| rs773690764 | pathogenic | 0 | 0 | 0 |
| rs201060167 | pathogenic | 0 | 0 | 0 |

| | | | | |
|---|---|---|---|---|
| rs137852670 | pathogenic | 0 | 0 | 0 |
| rs780594709 | pathogenic | 0 | 0 | 0 |
| rs145226920 | pathogenic | 0 | 0 | 0 |
| rs137852665 | pathogenic | 0 | 0 | 0 |
| rs724159960 | pathogenic | 0 | 0 | 0 |
| rs756985703 | pathogenic | 0 | 0 | 0 |
| rs137852667 | pathogenic | 0 | 0 | 0 |
| rs151079750 | pathogenic | 1 | 1 | 0 |
| rs2306636 | benign | 0 | 0 | 0 |
| rs72647885 | benign | 1 | 1 | 0 |
| rs377584435 | pathogenic | 1 | 1 | 1 |
| rs757583846 | pathogenic | 1 | 1 | 0 |
| rs121918148 | pathogenic | 0 | 0 | 0 |
| rs149201802 | benign | 0 | 0 | 0 |
| rs587781904 | pathogenic | 1 | 1 | 1 |
| rs143421325 | benign | 0 | 0 | 0 |
| rs587781253 | pathogenic | 0 | 0 | 0 |
| rs587780331 | benign | 0 | 0 | 0 |
| rs151001016 | benign | 0 | 0 | 0 |
| rs150888094 | benign | 0 | 0 | 0 |
| rs149581331 | benign | 0 | 0 | 0 |
| rs774840535 | benign | 0 | 0 | 0 |
| rs138428684 | benign | 0 | 0 | 0 |
| rs149300055 | benign | 0 | 0 | 0 |
| rs571005578 | benign | 0 | 0 | 0 |
| rs200731839 | benign | 0 | 0 | 0 |
| rs121965007 | pathogenic | 0 | 0 | 0 |
| rs121965013 | pathogenic | 0 | 1 | 0 |
| rs121965009 | pathogenic | 0 | 0 | 0 |
| rs1800457 | benign | 1 | 1 | 0 |
| rs121965006 | pathogenic | 1 | 1 | 0 |
| rs121965008 | pathogenic | 0 | 0 | 0 |
| rs61732609 | pathogenic | 0 | 0 | 0 |
| rs121965011 | pathogenic | 0 | 0 | 0 |
| rs121965015 | pathogenic | 0 | 0 | 0 |
| rs121965018 | pathogenic | 1 | 1 | 0 |
| rs121965016 | pathogenic | 0 | 0 | 0 |
| rs77806 | benign | 0 | 0 | 0 |
| rs149731356 | benign | 0 | 0 | 0 |
| rs72549407 | pathogenic | 0 | 0 | 0 |
| rs200363929 | benign | 0 | 0 | 0 |
| rs376403708 | benign | 1 | 1 | 0 |
| rs56376197 | benign | 1 | 1 | 1 |
| rs200237973 | benign | 1 | 1 | 1 |
| rs548330075 | benign | 0 | 0 | 0 |
| rs148291350 | benign | 0 | 0 | 0 |
| rs71579353 | pathogenic | 0 | 0 | 0 |
| rs146278697 | benign | 0 | 0 | 0 |
| rs141033098 | benign | 0 | 0 | 0 |
| rs397517234 | pathogenic | 0 | 0 | 0 |
| rs71579374 | benign | 0 | 0 | 0 |
| rs140381835 | benign | 0 | 0 | 0 |
| rs137852702 | pathogenic | 0 | 0 | 0 |
| rs137852696 | pathogenic | 1 | 1 | 1 |

| | | | | |
|---|---|---|---|---|
| rs137852697 | pathogenic | 0 | 0 | 0 |
| rs386833639 | pathogenic | 0 | 0 | 0 |
| rs386833640 | pathogenic | 0 | 0 | 0 |
| rs386833641 | benign | 0 | 0 | 0 |
| rs137852701 | pathogenic | 0 | 0 | 0 |
| rs386833642 | pathogenic | 0 | 0 | 0 |
| rs137852695 | pathogenic | 0 | 0 | 0 |
| rs386833646 | pathogenic | 0 | 0 | 0 |
| rs137852700 | pathogenic | 1 | 1 | 0 |
| rs386833649 | pathogenic | 1 | 1 | 0 |
| rs386833650 | pathogenic | 0 | 0 | 0 |
| rs148412181 | pathogenic | 0 | 0 | 0 |
| rs386833655 | pathogenic | 0 | 0 | 0 |
| rs386833656 | pathogenic | 1 | 1 | 0 |
| rs137852698 | pathogenic | 0 | 0 | 0 |
| rs386833661 | pathogenic | 0 | 0 | 0 |
| rs386833670 | pathogenic | 0 | 0 | 0 |
| rs111033632 | pathogenic | 1 | 1 | 1 |
| rs34308573 | benign | 1 | 1 | 1 |
| rs376048533 | pathogenic | 1 | 1 | 0 |
| rs3747517 | benign | 1 | 1 | 0 |
| rs137853328 | pathogenic | 0 | 0 | 0 |
| rs77273740 | benign | 0 | 0 | 0 |
| rs267606760 | pathogenic | 0 | 0 | 0 |
| rs77576840 | pathogenic | 0 | 0 | 0 |
| rs45465204 | benign | 1 | 1 | 0 |
| rs5320 | benign | 0 | 0 | 0 |
| rs5324 | benign | 0 | 0 | 0 |
| rs4531 | benign | 0 | 0 | 0 |
| rs267606761 | pathogenic | 0 | 0 | 0 |
| rs75215331 | pathogenic | 0 | 0 | 0 |
| rs6271 | benign | 0 | 0 | 0 |
| rs201643277 | pathogenic | 0 | 0 | 0 |
| rs4252250 | benign | 0 | 0 | 0 |
| rs137853580 | pathogenic | 0 | 0 | 0 |
| rs368287711 | pathogenic | 0 | 0 | 0 |
| rs137853579 | pathogenic | 0 | 0 | 0 |
| rs3135932 | benign | 0 | 0 | 0 |
| rs2228055 | benign | 0 | 0 | 0 |
| rs140322345 | benign | 1 | 1 | 1 |
| rs1154510 | pathogenic | 0 | 0 | 0 |
| rs11833399 | benign | 0 | 0 | 0 |
| rs36023382 | benign | 1 | 1 | 1 |
| rs61754444 | benign | 0 | 0 | 0 |
| rs28935168 | pathogenic | 0 | 0 | 0 |
| rs28934907 | pathogenic | 0 | 0 | 0 |
| rs28934904 | pathogenic | 1 | 1 | 0 |
| rs61748392 | pathogenic | 1 | 1 | 0 |
| rs28934908 | pathogenic | 0 | 0 | 0 |
| rs28934905 | pathogenic | 0 | 0 | 0 |
| rs28934906 | pathogenic | 1 | 1 | 0 |
| rs606231307 | pathogenic | 0 | 0 | 0 |
| rs9333296 | benign | 0 | 0 | 0 |
| rs3218614 | benign | 0 | 0 | 0 |

| | | | | |
|---|---|---|---|---|
| rs2230792 | benign | 0 | 0 | 0 |
| rs2230793 | benign | 0 | 0 | 0 |
| rs2230794 | benign | 0 | 0 | 0 |
| rs749052963 | pathogenic | 0 | 0 | 0 |
| rs2230798 | benign | 0 | 0 | 0 |
| rs2230795 | benign | 0 | 0 | 0 |
| rs3204145 | benign | 0 | 0 | 0 |
| rs1538660 | benign | 0 | 0 | 0 |
| rs9878928 | benign | 0 | 0 | 0 |
| rs104893742 | pathogenic | 0 | 0 | 0 |
| rs28936702 | pathogenic | 0 | 0 | 0 |
| rs28936703 | pathogenic | 0 | 0 | 0 |
| rs769392 | benign | 0 | 0 | 0 |
| rs769402 | benign | 1 | 1 | 1 |
| rs104894884 | pathogenic | 1 | 1 | 1 |
| rs1801316 | benign | 1 | 1 | 1 |
| rs104894885 | pathogenic | 1 | 1 | 1 |
| rs782572 | benign | 0 | 0 | 0 |
| rs151166046 | pathogenic | 0 | 0 | 0 |
| rs772153760 | pathogenic | 1 | 1 | 0 |
| rs35405862 | benign | 0 | 0 | 0 |
| rs146571352 | pathogenic | 0 | 1 | 0 |
| rs374698153 | pathogenic | 0 | 0 | 0 |
| rs7594497 | benign | 0 | 0 | 0 |
| rs80338716 | pathogenic | 0 | 0 | 0 |
| rs80338719 | pathogenic | 1 | 1 | 1 |
| rs80338721 | pathogenic | 0 | 0 | 0 |
| rs730882249 | pathogenic | 0 | 0 | 0 |
| rs146262009 | pathogenic | 0 | 0 | 0 |
| rs104895083 | pathogenic | 1 | 1 | 1 |
| rs104895100 | benign | 1 | 1 | 1 |
| rs104895085 | pathogenic | 0 | 0 | 0 |
| rs28940580 | pathogenic | 0 | 0 | 0 |
| rs104895098 | pathogenic | 0 | 0 | 0 |
| rs28940578 | pathogenic | 0 | 0 | 0 |
| rs61752717 | pathogenic | 0 | 0 | 0 |
| rs28940579 | pathogenic | 0 | 0 | 0 |
| rs104895097 | pathogenic | 0 | 0 | 0 |
| rs140425651 | benign | 1 | 1 | 0 |
| rs768090444 | pathogenic | 0 | 0 | 0 |
| rs764459544 | pathogenic | 0 | 0 | 0 |
| rs80338802 | pathogenic | 1 | 1 | 1 |
| rs149969786 | pathogenic | 1 | 1 | 0 |
| rs115311625 | pathogenic | 1 | 1 | 1 |
| rs778768583 | pathogenic | 1 | 1 | 1 |
| rs892090 | benign | 0 | 0 | 0 |
| rs28929498 | pathogenic | 1 | 1 | 1 |
| rs137854467 | pathogenic | 0 | 0 | 0 |
| rs137854480 | pathogenic | 0 | 0 | 0 |
| rs148682262 | benign | 0 | 0 | 0 |
| rs137852744 | pathogenic | 0 | 0 | 0 |
| rs137852745 | pathogenic | 0 | 1 | 0 |
| rs137852749 | pathogenic | 0 | 1 | 0 |
| rs137852746 | pathogenic | 0 | 1 | 0 |

| | | | | |
|---|---|---|---|---|
| rs145633958 | benign | 0 | 0 | 0 |
| rs141679950 | benign | 0 | 0 | 0 |
| rs76397662 | benign | 1 | 1 | 1 |
| rs370736139 | benign | 0 | 0 | 0 |
| rs781750106 | benign | 0 | 0 | 0 |
| rs546866208 | benign | 0 | 0 | 0 |
| rs142338976 | benign | 0 | 0 | 0 |
| rs115272158 | benign | 0 | 0 | 0 |
| rs115423919 | benign | 0 | 0 | 0 |
| rs537874538 | benign | 0 | 0 | 0 |
| rs9282834 | benign | 0 | 0 | 0 |
| rs201745826 | benign | 0 | 0 | 0 |
| rs75873440 | pathogenic | 0 | 0 | 0 |
| rs147219360 | benign | 0 | 0 | 0 |
| rs77939446 | pathogenic | 0 | 0 | 0 |
| rs77558292 | pathogenic | 0 | 0 | 0 |
| rs80069458 | pathogenic | 0 | 0 | 0 |
| rs76262710 | pathogenic | 0 | 0 | 0 |
| rs79781594 | pathogenic | 0 | 0 | 0 |
| rs77316810 | pathogenic | 0 | 0 | 0 |
| rs77503355 | pathogenic | 0 | 0 | 0 |
| rs79890926 | pathogenic | 0 | 0 | 0 |
| rs377767402 | benign | 0 | 0 | 0 |
| rs121913308 | pathogenic | 0 | 0 | 0 |
| rs55846256 | benign | 0 | 0 | 0 |
| rs75076352 | pathogenic | 0 | 0 | 0 |
| rs75996173 | pathogenic | 0 | 0 | 0 |
| rs77709286 | pathogenic | 0 | 0 | 0 |
| rs151058706 | benign | 0 | 0 | 0 |
| rs138964358 | benign | 0 | 0 | 0 |
| rs137852353 | pathogenic | 1 | 1 | 0 |
| rs201196020 | benign | 0 | 0 | 0 |
| rs137853201 | pathogenic | 1 | 1 | 1 |
| rs113994174 | pathogenic | 1 | 1 | 1 |
| rs104893674 | pathogenic | 1 | 1 | 1 |
| rs779519249 | benign | 1 | 1 | 1 |
| rs9606756 | benign | 0 | 0 | 0 |
| rs75680863 | benign | 1 | 1 | 1 |
| rs35838082 | benign | 0 | 0 | 0 |
| rs1801198 | benign | 0 | 0 | 0 |
| rs117458738 | benign | 1 | 1 | 0 |
| rs9621049 | benign | 1 | 1 | 0 |
| rs1131603 | benign | 1 | 1 | 0 |
| rs4820889 | benign | 0 | 0 | 0 |
| rs777767978 | pathogenic | 1 | 1 | 0 |
| rs147399860 | benign | 1 | 1 | 1 |
| rs104893950 | pathogenic | 0 | 0 | 0 |
| rs777308612 | benign | 0 | 0 | 0 |
| rs151091410 | benign | 0 | 0 | 0 |
| rs11537667 | benign | 0 | 0 | 0 |
| rs267606621 | pathogenic | 1 | 1 | 0 |
| rs143058902 | benign | 0 | 0 | 0 |
| rs370261904 | pathogenic | 0 | 0 | 0 |
| rs141169746 | benign | 0 | 0 | 0 |

| | | | | |
|---|---|---|---|---|
| rs111033436 | benign | 0 | 0 | 0 |
| rs148533341 | benign | 0 | 0 | 0 |
| rs143827620 | benign | 0 | 0 | 0 |
| rs4935502 | benign | 0 | 0 | 0 |
| rs61735473 | benign | 0 | 0 | 0 |
| rs267606932 | pathogenic | 1 | 1 | 0 |
| rs61730754 | benign | 0 | 0 | 0 |
| rs374491359 | pathogenic | 1 | 1 | 1 |
| rs761800077 | benign | 1 | 1 | 1 |
| rs760391381 | benign | 1 | 1 | 1 |
| rs104893885 | pathogenic | 0 | 0 | 0 |
| rs104893886 | pathogenic | 1 | 1 | 1 |
| rs104893884 | pathogenic | 0 | 0 | 0 |
| rs147715509 | benign | 0 | 0 | 0 |
| rs115533243 | benign | 0 | 0 | 0 |
| rs397509390 | pathogenic | 0 | 1 | 0 |
| rs80357134 | pathogenic | 0 | 0 | 0 |
| rs766004110 | benign | 1 | 1 | 1 |
| rs769650474 | benign | 0 | 0 | 0 |
| rs398122635 | benign | 1 | 1 | 0 |
| rs199522616 | pathogenic | 0 | 0 | 0 |
| rs28897672 | pathogenic | 1 | 1 | 0 |
| rs80357086 | pathogenic | 1 | 1 | 1 |
| rs80357064 | pathogenic | 1 | 1 | 0 |
| rs80356936 | benign | 0 | 0 | 0 |
| rs28897673 | benign | 0 | 0 | 0 |
| rs146747324 | benign | 0 | 0 | 0 |
| rs78635798 | pathogenic | 0 | 0 | 0 |
| rs75146158 | pathogenic | 1 | 1 | 1 |
| rs587782886 | benign | 0 | 0 | 0 |
| rs121434634 | pathogenic | 0 | 0 | 0 |
| rs121434635 | pathogenic | 0 | 0 | 0 |
| rs587777778 | pathogenic | 0 | 0 | 0 |
| rs118203967 | pathogenic | 0 | 0 | 0 |
| rs751375244 | pathogenic | 1 | 1 | 1 |
| rs104893863 | pathogenic | 1 | 1 | 0 |
| rs104893865 | pathogenic | 0 | 0 | 0 |
| rs104893864 | pathogenic | 1 | 1 | 1 |
| rs104893866 | pathogenic | 1 | 1 | 0 |
| rs561738497 | benign | 0 | 0 | 0 |
| rs57599352 | pathogenic | 1 | 1 | 1 |
| rs267607448 | pathogenic | 1 | 1 | 1 |
| rs63751596 | pathogenic | 0 | 0 | 0 |
| rs63751612 | benign | 0 | 0 | 0 |
| rs63751684 | benign | 0 | 0 | 0 |
| rs267607861 | benign | 0 | 0 | 0 |
| rs35831931 | benign | 0 | 0 | 0 |
| rs63750876 | benign | 0 | 0 | 0 |
| rs41295284 | benign | 0 | 0 | 0 |
| rs35502531 | benign | 0 | 0 | 0 |
| rs63750449 | benign | 0 | 0 | 0 |
| rs63750899 | pathogenic | 0 | 0 | 0 |
| rs55907433 | benign | 0 | 0 | 0 |
| rs63750217 | pathogenic | 0 | 0 | 0 |

| | | | | |
|---|---|---|---|---|
| rs63751275 | pathogenic | 0 | 0 | 0 |
| rs63750702 | benign | 1 | 1 | 1 |
| rs63750114 | benign | 0 | 0 | 0 |
| rs2020873 | benign | 0 | 0 | 0 |
| rs140195825 | benign | 1 | 1 | 1 |
| rs182763805 | benign | 0 | 0 | 0 |
| rs533696377 | benign | 0 | 0 | 0 |
| rs41278788 | benign | 0 | 0 | 0 |
| rs104894193 | pathogenic | 1 | 1 | 0 |
| rs3824915 | benign | 1 | 1 | 1 |
| rs104894196 | pathogenic | 0 | 0 | 0 |
| rs78681604 | benign | 1 | 1 | 1 |
| rs77321498 | benign | 1 | 1 | 1 |
| rs137852965 | pathogenic | 0 | 0 | 0 |
| rs137852961 | pathogenic | 0 | 0 | 0 |
| rs137852962 | pathogenic | 1 | 1 | 0 |
| rs753376100 | pathogenic | 0 | 0 | 0 |
| rs137852968 | pathogenic | 1 | 1 | 1 |
| rs137852959 | pathogenic | 1 | 1 | 0 |
| rs137852967 | pathogenic | 0 | 0 | 0 |
| rs113437301 | benign | 1 | 1 | 1 |
| rs61747482 | benign | 1 | 1 | 1 |
| rs2234919 | benign | 1 | 1 | 0 |
| rs121908863 | pathogenic | 0 | 0 | 0 |
| rs140455771 | pathogenic | 0 | 0 | 0 |
| rs121908810 | pathogenic | 1 | 1 | 1 |
| rs143879853 | benign | 0 | 0 | 0 |
| rs188319622 | benign | 0 | 0 | 0 |
| rs121918645 | pathogenic | 0 | 0 | 0 |
| rs115352435 | pathogenic | 0 | 0 | 0 |
| rs121434427 | pathogenic | 1 | 1 | 1 |
| rs121434428 | pathogenic | 0 | 0 | 0 |
| rs35086265 | benign | 1 | 1 | 1 |
| rs11576415 | benign | 0 | 0 | 0 |
| rs121434429 | pathogenic | 1 | 1 | 1 |
| rs4074536 | benign | 1 | 1 | 1 |
| rs10801999 | benign | 0 | 0 | 0 |
| rs121434550 | pathogenic | 0 | 0 | 0 |
| rs148057999 | pathogenic | 0 | 0 | 0 |
| rs28730716 | benign | 0 | 0 | 0 |
| rs142036299 | benign | 0 | 0 | 0 |
| rs762381137 | pathogenic | 0 | 0 | 0 |
| rs121434549 | pathogenic | 0 | 0 | 0 |
| rs139228801 | pathogenic | 0 | 0 | 0 |
| rs147219463 | benign | 0 | 0 | 0 |
| rs1131857 | benign | 0 | 0 | 0 |
| rs121917874 | pathogenic | 0 | 1 | 0 |
| rs2228056 | benign | 0 | 0 | 0 |
| rs121917870 | pathogenic | 0 | 0 | 0 |
| rs121917866 | pathogenic | 0 | 0 | 0 |
| rs11921054 | benign | 0 | 0 | 0 |
| rs121917868 | pathogenic | 0 | 0 | 0 |
| rs28931603 | pathogenic | 0 | 0 | 0 |
| rs104895295 | pathogenic | 1 | 1 | 0 |

| | | | | |
|---|---|---|---|---|
| rs104895382 | pathogenic | 0 | 0 | 0 |
| rs104895298 | pathogenic | 0 | 0 | 0 |
| rs104895300 | pathogenic | 0 | 0 | 0 |
| rs104895332 | pathogenic | 0 | 0 | 0 |
| rs758026399 | pathogenic | 1 | 1 | 1 |
| rs104895366 | pathogenic | 0 | 0 | 0 |
| rs104895304 | pathogenic | 0 | 0 | 0 |
| rs759997079 | pathogenic | 0 | 0 | 0 |
| rs121917789 | pathogenic | 0 | 0 | 0 |
| rs104895319 | pathogenic | 0 | 0 | 0 |
| rs104895317 | pathogenic | 1 | 1 | 0 |
| rs28934897 | pathogenic | 0 | 0 | 0 |
| rs104895380 | benign | 0 | 0 | 0 |
| rs104895360 | pathogenic | 0 | 0 | 0 |
| rs28936396 | pathogenic | 1 | 1 | 0 |
| rs121909307 | pathogenic | 1 | 1 | 1 |
| rs28938472 | pathogenic | 1 | 1 | 0 |
| rs121909308 | pathogenic | 1 | 1 | 0 |
| rs121909309 | pathogenic | 0 | 0 | 0 |
| rs121913013 | benign | 1 | 1 | 1 |
| rs201855245 | benign | 1 | 1 | 1 |
| rs2230232 | benign | 1 | 1 | 0 |
| rs113451409 | benign | 0 | 0 | 0 |
| rs121913011 | pathogenic | 1 | 1 | 0 |
| rs185821167 | pathogenic | 1 | 1 | 0 |
| rs2230234 | benign | 0 | 0 | 0 |
| rs193922639 | benign | 1 | 1 | 0 |
| rs774863785 | pathogenic | 1 | 1 | 0 |
| rs576404380 | pathogenic | 1 | 1 | 0 |
| rs2230235 | benign | 1 | 1 | 0 |
| rs200484060 | pathogenic | 1 | 1 | 0 |
| rs199761749 | benign | 0 | 0 | 0 |
| rs199681901 | benign | 0 | 0 | 0 |
| rs121908951 | pathogenic | 0 | 0 | 0 |
| rs769251460 | benign | 0 | 0 | 0 |
| rs28937320 | pathogenic | 0 | 0 | 0 |
| rs200607527 | pathogenic | 0 | 0 | 0 |
| rs137852640 | pathogenic | 1 | 1 | 1 |
| rs137852636 | pathogenic | 1 | 1 | 1 |
| rs137852638 | pathogenic | 0 | 0 | 0 |
| rs142637231 | pathogenic | 0 | 0 | 0 |
| rs151328418 | benign | 0 | 1 | 0 |
| rs137852639 | pathogenic | 1 | 1 | 1 |
| rs34605667 | benign | 0 | 0 | 0 |
| rs104894073 | pathogenic | 1 | 1 | 0 |
| rs56298569 | pathogenic | 0 | 0 | 0 |
| rs757600616 | pathogenic | 0 | 0 | 0 |
| rs139194636 | pathogenic | 0 | 0 | 0 |
| rs369517993 | pathogenic | 1 | 1 | 1 |
| rs138846036 | benign | 1 | 1 | 1 |
| rs200808744 | pathogenic | 1 | 1 | 1 |
| rs2255543 | benign | 0 | 0 | 0 |
| rs149165166 | pathogenic | 0 | 0 | 0 |
| rs544956641 | pathogenic | 1 | 1 | 1 |

| | | | | |
|---|---|---|---|---|
| rs28941783 | pathogenic | 0 | 0 | 0 |
| rs562853291 | pathogenic | 1 | 1 | 1 |
| rs28942100 | pathogenic | 0 | 0 | 0 |
| rs120074174 | pathogenic | 1 | 1 | 1 |
| rs120074171 | pathogenic | 1 | 1 | 1 |
| rs120074173 | pathogenic | 0 | 0 | 0 |
| rs120074172 | pathogenic | 1 | 1 | 1 |
| rs767159114 | pathogenic | 1 | 1 | 1 |
| rs104893657 | pathogenic | 0 | 0 | 0 |
| rs104893656 | pathogenic | 0 | 0 | 0 |
| rs104893659 | pathogenic | 0 | 0 | 0 |
| rs104893658 | pathogenic | 0 | 0 | 0 |
| rs200386310 | pathogenic | 0 | 0 | 0 |
| rs2072712 | benign | 0 | 0 | 0 |
| rs35274867 | benign | 0 | 0 | 0 |
| rs35343172 | benign | 0 | 0 | 0 |
| rs10079250 | benign | 0 | 0 | 0 |
| rs34951517 | benign | 0 | 0 | 0 |
| rs267606695 | pathogenic | 0 | 0 | 0 |
| rs149391728 | benign | 0 | 0 | 0 |
| rs387906598 | pathogenic | 1 | 1 | 0 |
| rs137854578 | pathogenic | 0 | 0 | 0 |
| rs141519952 | benign | 0 | 0 | 0 |
| rs146572883 | benign | 0 | 0 | 0 |
| rs111866410 | benign | 0 | 0 | 0 |
| rs139387758 | benign | 1 | 1 | 1 |
| rs35672074 | benign | 1 | 1 | 1 |
| rs121434322 | pathogenic | 0 | 0 | 0 |
| rs150884181 | benign | 0 | 0 | 0 |
| rs35867471 | benign | 0 | 0 | 0 |
| rs2867802 | benign | 0 | 0 | 0 |
| rs137902889 | benign | 0 | 0 | 0 |
| rs369804864 | pathogenic | 0 | 0 | 0 |
| rs781050795 | pathogenic | 0 | 0 | 0 |
| rs2305799 | benign | 0 | 0 | 0 |
| rs2229531 | benign | 0 | 0 | 0 |
| rs147025508 | benign | 0 | 0 | 0 |
| rs76857106 | pathogenic | 0 | 0 | 0 |
| rs7247284 | benign | 0 | 0 | 0 |
| rs62619782 | benign | 0 | 0 | 0 |
| rs377244188 | pathogenic | 0 | 0 | 0 |
| rs75718910 | pathogenic | 0 | 0 | 0 |
| rs79843600 | pathogenic | 0 | 0 | 0 |
| rs75037667 | pathogenic | 1 | 1 | 1 |
| rs121913006 | pathogenic | 0 | 0 | 0 |
| rs397517925 | pathogenic | 1 | 1 | 1 |
| rs267606831 | pathogenic | 1 | 1 | 1 |
| rs121909667 | pathogenic | 1 | 1 | 1 |
| rs121909669 | pathogenic | 1 | 1 | 1 |
| rs121909668 | pathogenic | 1 | 1 | 1 |
| rs121909671 | pathogenic | 1 | 1 | 1 |
| rs534596036 | benign | 0 | 0 | 0 |
| rs144037629 | pathogenic | 0 | 0 | 0 |
| rs11546406 | benign | 0 | 0 | 0 |

| | | | | |
|---|---|---|---|---|
| rs143697995 | pathogenic | 0 | 0 | 0 |
| rs121908331 | pathogenic | 0 | 0 | 0 |
| rs149377346 | benign | 0 | 0 | 0 |
| rs104894086 | pathogenic | 1 | 1 | 0 |
| rs104894089 | pathogenic | 0 | 0 | 0 |
| rs104894090 | pathogenic | 1 | 1 | 0 |
| rs387907235 | pathogenic | 0 | 0 | 0 |
| rs137852689 | pathogenic | 0 | 0 | 0 |
| rs137852690 | pathogenic | 1 | 1 | 0 |
| rs104894087 | pathogenic | 0 | 0 | 0 |
| rs104894085 | pathogenic | 0 | 0 | 0 |
| rs79383654 | benign | 0 | 0 | 0 |
| rs121434239 | pathogenic | 0 | 0 | 0 |
| rs121434240 | pathogenic | 0 | 0 | 0 |
| rs121434236 | pathogenic | 0 | 0 | 0 |
| rs121434238 | pathogenic | 1 | 1 | 1 |
| rs121918107 | pathogenic | 1 | 1 | 1 |
| rs121918103 | pathogenic | 1 | 1 | 1 |
| rs121918104 | pathogenic | 1 | 1 | 1 |
| rs61747614 | benign | 1 | 1 | 1 |
| rs555329870 | benign | 0 | 0 | 0 |
| rs186094931 | benign | 0 | 0 | 0 |
| rs267606785 | pathogenic | 0 | 0 | 0 |
| rs34474955 | benign | 0 | 0 | 0 |
| rs1126497 | benign | 1 | 1 | 1 |
| rs397514661 | pathogenic | 0 | 0 | 0 |
| rs189732445 | benign | 0 | 0 | 0 |
| rs146685071 | benign | 0 | 0 | 0 |
| rs121908636 | pathogenic | 1 | 1 | 0 |
| rs121908643 | pathogenic | 0 | 0 | 0 |
| rs121908644 | pathogenic | 1 | 1 | 1 |
| rs575001023 | pathogenic | 1 | 1 | 1 |
| rs121908646 | pathogenic | 1 | 1 | 0 |
| rs121908638 | pathogenic | 1 | 1 | 1 |
| rs777828000 | pathogenic | 1 | 1 | 1 |
| rs74923032 | benign | 0 | 0 | 0 |
| rs192838388 | pathogenic | 1 | 1 | 1 |
| rs148918985 | pathogenic | 1 | 1 | 1 |
| rs773540764 | benign | 1 | 1 | 1 |
| rs370595480 | pathogenic | 1 | 1 | 1 |
| rs762387914 | pathogenic | 1 | 1 | 1 |
| rs121908642 | pathogenic | 1 | 1 | 1 |
| rs183276875 | pathogenic | 1 | 1 | 1 |
| rs121908648 | pathogenic | 1 | 1 | 1 |
| rs121908647 | pathogenic | 1 | 1 | 1 |
| rs121908640 | pathogenic | 1 | 1 | 1 |
| rs771937610 | pathogenic | 1 | 1 | 1 |
| rs121908641 | pathogenic | 1 | 1 | 1 |
| rs5745068 | benign | 0 | 0 | 0 |
| rs73155056 | benign | 1 | 1 | 1 |
| rs148788180 | benign | 0 | 0 | 0 |
| rs121918137 | pathogenic | 0 | 0 | 0 |
| rs41269593 | benign | 0 | 0 | 0 |
| rs11159 | benign | 1 | 1 | 0 |

| | | | | |
|---|---|---|---|---|
| rs114733570 | pathogenic | 1 | 1 | 0 |
| rs3211783 | benign | 0 | 0 | 0 |
| rs4762 | benign | 0 | 0 | 0 |
| rs74315283 | pathogenic | 0 | 0 | 0 |
| rs111033559 | pathogenic | 0 | 0 | 0 |
| rs111033560 | pathogenic | 1 | 1 | 1 |
| rs749962743 | benign | 0 | 0 | 0 |
| rs267606986 | pathogenic | 0 | 0 | 0 |
| rs34516635 | benign | 0 | 0 | 0 |
| rs117440569 | benign | 0 | 0 | 0 |
| rs61740877 | benign | 0 | 0 | 0 |
| rs121918352 | pathogenic | 0 | 0 | 0 |
| rs183974372 | benign | 0 | 0 | 0 |
| rs140511204 | benign | 1 | 1 | 0 |
| rs6040067 | benign | 0 | 0 | 0 |
| rs199674138 | benign | 0 | 0 | 0 |
| rs121918350 | pathogenic | 0 | 0 | 0 |
| rs138292988 | benign | 1 | 1 | 1 |
| rs145288947 | benign | 0 | 0 | 0 |
| rs369714529 | benign | 0 | 0 | 0 |
| rs200945419 | benign | 0 | 0 | 0 |
| rs116959874 | benign | 0 | 0 | 0 |
| rs141659620 | pathogenic | 1 | 1 | 0 |
| rs111475461 | benign | 0 | 0 | 0 |
| rs61755320 | pathogenic | 0 | 0 | 0 |
| rs748600162 | pathogenic | 1 | 1 | 1 |
| rs369227537 | pathogenic | 1 | 1 | 1 |
| rs372981030 | pathogenic | 1 | 1 | 1 |
| rs137854486 | pathogenic | 0 | 0 | 0 |
| rs137854487 | pathogenic | 0 | 0 | 0 |
| rs199905054 | pathogenic | 0 | 0 | 0 |
| rs771438170 | pathogenic | 0 | 0 | 0 |
| rs774949844 | pathogenic | 0 | 0 | 0 |
| rs4899145 | benign | 0 | 0 | 0 |
| rs200642525 | benign | 0 | 0 | 0 |
| rs190140598 | pathogenic | 0 | 0 | 0 |
| rs2250889 | benign | 0 | 0 | 0 |
| rs17577 | benign | 0 | 0 | 0 |
| rs142032681 | benign | 0 | 0 | 0 |
| rs144577963 | benign | 0 | 0 | 0 |
| rs61754623 | benign | 0 | 0 | 0 |
| rs139873200 | benign | 0 | 0 | 0 |
| rs532530906 | benign | 0 | 0 | 0 |
| rs145762716 | benign | 0 | 0 | 0 |
| rs34062020 | benign | 0 | 0 | 0 |
| rs35258161 | benign | 0 | 0 | 0 |
| rs149792649 | benign | 0 | 0 | 0 |
| rs78101774 | benign | 0 | 0 | 0 |
| rs766466297 | pathogenic | 0 | 1 | 0 |
| rs140467171 | pathogenic | 0 | 0 | 0 |
| rs141340466 | pathogenic | 0 | 0 | 0 |
| rs201555303 | pathogenic | 0 | 0 | 0 |
| rs143592405 | pathogenic | 0 | 0 | 0 |
| rs781249411 | pathogenic | 0 | 0 | 0 |

| | | | | |
|---|---|---|---|---|
| rs116644127 | benign | 0 | 0 | 0 |
| rs151146863 | benign | 1 | 1 | 1 |
| rs116440799 | benign | 0 | 0 | 0 |
| rs138958351 | benign | 0 | 0 | 0 |
| rs143319354 | benign | 0 | 0 | 0 |
| rs768669208 | pathogenic | 1 | 1 | 1 |
| rs28940289 | pathogenic | 1 | 1 | 0 |
| rs745656120 | pathogenic | 1 | 1 | 0 |
| rs387906987 | pathogenic | 0 | 0 | 0 |
| rs147240777 | benign | 0 | 0 | 0 |
| rs140945833 | benign | 0 | 0 | 0 |
| rs146614981 | benign | 0 | 0 | 0 |
| rs149878562 | benign | 0 | 0 | 0 |
| rs28938168 | pathogenic | 0 | 0 | 0 |
| rs33939927 | pathogenic | 1 | 1 | 1 |
| rs34995376 | pathogenic | 1 | 1 | 1 |
| rs201034481 | pathogenic | 1 | 1 | 1 |
| rs774069983 | benign | 0 | 0 | 0 |
| rs199422226 | pathogenic | 1 | 1 | 0 |
| rs199422225 | pathogenic | 1 | 1 | 1 |
| rs199422224 | pathogenic | 1 | 1 | 0 |
| rs149271416 | pathogenic | 1 | 1 | 0 |
| rs1801318 | benign | 1 | 1 | 1 |
| rs372691318 | pathogenic | 1 | 1 | 1 |
| rs147685849 | benign | 0 | 0 | 0 |
| rs387907199 | pathogenic | 0 | 0 | 0 |
| rs767122069 | pathogenic | 0 | 0 | 0 |
| rs2302427 | benign | 0 | 0 | 0 |
| rs201135441 | benign | 0 | 0 | 0 |
| rs201861847 | pathogenic | 0 | 0 | 0 |
| rs137852952 | pathogenic | 0 | 0 | 0 |
| rs16986560 | benign | 0 | 0 | 0 |
| rs147344332 | pathogenic | 1 | 1 | 1 |
| rs36027220 | pathogenic | 0 | 0 | 0 |
| rs192401229 | benign | 0 | 0 | 0 |
| rs121909106 | pathogenic | 1 | 1 | 0 |
| rs118204103 | pathogenic | 1 | 1 | 0 |
| rs118204104 | pathogenic | 1 | 1 | 0 |
| rs118204105 | pathogenic | 1 | 1 | 0 |
| rs118204106 | pathogenic | 0 | 0 | 0 |
| rs118204119 | pathogenic | 0 | 0 | 0 |
| rs118204107 | pathogenic | 0 | 0 | 0 |
| rs118204094 | pathogenic | 0 | 0 | 0 |
| rs118204098 | pathogenic | 1 | 1 | 0 |
| rs118204101 | pathogenic | 1 | 1 | 0 |
| rs118204096 | pathogenic | 1 | 1 | 0 |
| rs118204108 | pathogenic | 0 | 0 | 0 |
| rs536814318 | pathogenic | 0 | 0 | 0 |
| rs34413634 | pathogenic | 1 | 1 | 0 |
| rs118204100 | pathogenic | 0 | 0 | 0 |
| rs118204109 | pathogenic | 0 | 0 | 0 |
| rs118204116 | pathogenic | 1 | 1 | 0 |
| rs118204110 | pathogenic | 0 | 0 | 0 |
| rs118204099 | pathogenic | 0 | 0 | 0 |

| | | | | |
|---|---|---|---|---|
| rs118204111 | pathogenic | 0 | 0 | 0 |
| rs118204112 | pathogenic | 0 | 0 | 0 |
| rs118204113 | pathogenic | 0 | 0 | 0 |
| rs118204114 | pathogenic | 0 | 0 | 0 |
| rs118204115 | pathogenic | 0 | 0 | 0 |
| rs150428209 | benign | 0 | 0 | 0 |
| rs144949995 | benign | 0 | 0 | 0 |
| rs190023981 | benign | 0 | 0 | 0 |
| rs104894611 | pathogenic | 1 | 1 | 1 |
| rs28937580 | pathogenic | 1 | 1 | 1 |
| rs121908949 | pathogenic | 0 | 0 | 0 |
| rs104894612 | pathogenic | 1 | 1 | 1 |
| rs104894609 | pathogenic | 1 | 1 | 1 |
| rs104894610 | pathogenic | 1 | 1 | 1 |
| rs104894615 | pathogenic | 0 | 0 | 0 |
| rs104894603 | pathogenic | 0 | 0 | 0 |
| rs104894602 | pathogenic | 1 | 1 | 1 |
| rs121908948 | pathogenic | 1 | 1 | 1 |
| rs104894608 | pathogenic | 1 | 1 | 1 |
| rs371545673 | benign | 0 | 0 | 0 |
| rs199925238 | benign | 0 | 0 | 0 |
| rs146313451 | benign | 0 | 0 | 0 |
| rs121918193 | pathogenic | 1 | 1 | 0 |
| rs2228500 | benign | 0 | 0 | 0 |
| rs121918194 | pathogenic | 1 | 1 | 0 |
| rs121918196 | pathogenic | 0 | 0 | 0 |
| rs41291971 | benign | 0 | 0 | 0 |
| rs199473690 | pathogenic | 0 | 0 | 0 |
| rs2228613 | benign | 0 | 0 | 0 |
| rs267607130 | pathogenic | 1 | 1 | 1 |
| rs77615401 | benign | 1 | 1 | 1 |
| rs374618872 | benign | 1 | 1 | 1 |
| rs146165014 | benign | 0 | 0 | 0 |
| rs144586070 | benign | 0 | 0 | 0 |
| rs121918541 | pathogenic | 0 | 0 | 0 |
| rs142105943 | benign | 0 | 0 | 0 |
| rs121918540 | pathogenic | 0 | 0 | 0 |
| rs6791 | benign | 0 | 0 | 0 |
| rs61736587 | pathogenic | 0 | 0 | 0 |
| rs61736586 | benign | 0 | 0 | 0 |
| rs121917741 | pathogenic | 1 | 1 | 1 |
| rs41302885 | benign | 0 | 0 | 0 |
| rs141026028 | benign | 0 | 0 | 0 |
| rs193922493 | pathogenic | 0 | 0 | 0 |
| rs73598374 | benign | 0 | 0 | 0 |
| rs121908725 | pathogenic | 1 | 1 | 0 |
| rs199422328 | pathogenic | 0 | 1 | 0 |
| rs11555566 | benign | 1 | 1 | 1 |
| rs267606634 | pathogenic | 0 | 0 | 0 |
| rs121908714 | pathogenic | 1 | 1 | 0 |
| rs121908717 | pathogenic | 1 | 1 | 0 |
| rs267606635 | pathogenic | 1 | 1 | 0 |
| rs121908739 | pathogenic | 0 | 0 | 0 |
| rs121908735 | pathogenic | 0 | 1 | 0 |

| | | | | |
|---|---|---|---|---|
| rs121908716 | pathogenic | 0 | 1 | 0 |
| rs121908723 | pathogenic | 0 | 1 | 0 |
| rs121908721 | pathogenic | 0 | 0 | 0 |
| rs199422327 | pathogenic | 0 | 0 | 0 |
| rs121908715 | pathogenic | 0 | 0 | 0 |
| rs587783563 | pathogenic | 0 | 0 | 0 |
| rs104894780 | pathogenic | 0 | 0 | 0 |
| rs56030372 | pathogenic | 1 | 1 | 1 |
| rs104894782 | pathogenic | 1 | 1 | 1 |
| rs149495971 | benign | 0 | 0 | 0 |
| rs118203942 | pathogenic | 1 | 1 | 0 |
| rs118203939 | pathogenic | 0 | 0 | 0 |
| rs118203938 | pathogenic | 0 | 0 | 0 |
| rs118203943 | pathogenic | 0 | 0 | 0 |
| rs118203940 | pathogenic | 0 | 0 | 0 |
| rs727503809 | pathogenic | 0 | 0 | 0 |
| rs1065757 | benign | 0 | 0 | 0 |
| rs1071598 | benign | 0 | 0 | 0 |
| rs25414 | benign | 0 | 0 | 0 |
| rs118203944 | pathogenic | 0 | 0 | 0 |
| rs118203941 | pathogenic | 0 | 0 | 0 |
| rs144222167 | benign | 0 | 0 | 0 |
| rs201101343 | pathogenic | 0 | 0 | 0 |
| rs144601717 | benign | 0 | 0 | 0 |
| rs267607129 | pathogenic | 0 | 0 | 0 |
| rs104894729 | pathogenic | 0 | 0 | 0 |
| rs104894727 | pathogenic | 0 | 0 | 0 |
| rs104894725 | pathogenic | 0 | 0 | 0 |
| rs104894515 | pathogenic | 1 | 1 | 1 |
| rs74688377 | benign | 0 | 0 | 0 |
| rs121908513 | pathogenic | 1 | 1 | 0 |
| rs121908509 | pathogenic | 0 | 0 | 0 |
| rs121908514 | pathogenic | 1 | 1 | 0 |
| rs121908512 | pathogenic | 1 | 1 | 0 |
| rs121908510 | pathogenic | 0 | 0 | 0 |
| rs121908511 | pathogenic | 1 | 1 | 1 |
| rs121908518 | pathogenic | 1 | 1 | 1 |
| rs34358665 | benign | 0 | 0 | 0 |
| rs147568834 | benign | 0 | 0 | 0 |
| rs201717599 | benign | 0 | 0 | 0 |
| rs61744487 | benign | 0 | 0 | 0 |
| rs370627877 | pathogenic | 1 | 1 | 1 |
| rs368593151 | pathogenic | 1 | 1 | 1 |
| rs114363936 | benign | 0 | 0 | 0 |
| rs137853333 | pathogenic | 0 | 0 | 0 |
| rs137853080 | pathogenic | 1 | 1 | 1 |
| rs137853077 | pathogenic | 1 | 1 | 1 |
| rs780749732 | benign | 1 | 1 | 1 |
| rs137853081 | pathogenic | 1 | 1 | 0 |
| rs137853078 | pathogenic | 0 | 0 | 0 |
| rs121913315 | pathogenic | 1 | 1 | 1 |
| rs752513525 | pathogenic | 1 | 1 | 1 |
| rs104894918 | pathogenic | 0 | 0 | 0 |
| rs104894917 | pathogenic | 0 | 0 | 0 |

| | | | | |
|---|---|---|---|---|
| rs104894919 | pathogenic | 0 | 0 | 0 |
| rs28939672 | pathogenic | 0 | 0 | 0 |
| rs104894173 | pathogenic | 0 | 0 | 0 |
| rs104894179 | pathogenic | 0 | 0 | 0 |
| rs104894174 | pathogenic | 1 | 1 | 0 |
| rs104894178 | pathogenic | 1 | 1 | 0 |
| rs28939714 | pathogenic | 1 | 1 | 1 |
| rs104894270 | pathogenic | 1 | 1 | 1 |
| rs3765571 | benign | 0 | 0 | 0 |
| rs752669339 | pathogenic | 1 | 1 | 1 |
| rs537043237 | pathogenic | 1 | 1 | 1 |
| rs774047299 | pathogenic | 1 | 1 | 1 |
| rs121434583 | pathogenic | 1 | 1 | 1 |
| rs121907927 | pathogenic | 0 | 0 | 0 |
| rs121907925 | pathogenic | 0 | 1 | 0 |
| rs746212325 | pathogenic | 0 | 0 | 0 |
| rs754637179 | pathogenic | 0 | 0 | 0 |
| rs780882740 | pathogenic | 0 | 0 | 0 |
| rs121918130 | pathogenic | 0 | 0 | 0 |
| rs121918129 | pathogenic | 0 | 0 | 0 |
| rs13297509 | pathogenic | 0 | 0 | 0 |
| rs746867724 | pathogenic | 0 | 0 | 0 |
| rs121918128 | pathogenic | 0 | 0 | 0 |
| rs752300607 | pathogenic | 0 | 0 | 0 |
| rs121964860 | pathogenic | 0 | 0 | 0 |
| rs45466197 | pathogenic | 1 | 1 | 1 |
| rs3730238 | benign | 1 | 1 | 1 |
| rs730881125 | pathogenic | 1 | 1 | 1 |
| rs121964861 | pathogenic | 1 | 1 | 1 |
| rs397516484 | pathogenic | 0 | 0 | 0 |
| rs367785431 | pathogenic | 0 | 0 | 0 |
| rs141121678 | pathogenic | 0 | 0 | 0 |
| rs12720263 | benign | 0 | 0 | 0 |
| rs2304256 | benign | 0 | 0 | 0 |
| rs2304255 | benign | 0 | 0 | 0 |
| rs121909045 | pathogenic | 0 | 0 | 0 |
| rs397508796 | pathogenic | 0 | 0 | 0 |
| rs151020603 | pathogenic | 0 | 0 | 0 |
| rs77284892 | pathogenic | 0 | 0 | 0 |
| rs368505753 | pathogenic | 0 | 0 | 0 |
| rs115545701 | pathogenic | 0 | 0 | 0 |
| rs121908749 | pathogenic | 0 | 0 | 0 |
| rs75961395 | pathogenic | 0 | 0 | 0 |
| rs121908750 | pathogenic | 0 | 0 | 0 |
| rs121908751 | pathogenic | 0 | 0 | 0 |
| rs397508464 | pathogenic | 0 | 0 | 0 |
| rs397508461 | pathogenic | 0 | 0 | 0 |
| rs121909031 | pathogenic | 0 | 0 | 0 |
| rs113993958 | pathogenic | 0 | 0 | 0 |
| rs77834169 | pathogenic | 0 | 0 | 0 |
| rs78655421 | pathogenic | 0 | 0 | 0 |
| rs141723617 | benign | 0 | 0 | 0 |
| rs35516286 | benign | 0 | 0 | 0 |
| rs1800079 | pathogenic | 1 | 1 | 0 |

| | | | | |
|---|---|---|---|---|
| rs80282562 | pathogenic | 0 | 0 | 0 |
| rs121908752 | pathogenic | 0 | 0 | 0 |
| rs556880586 | pathogenic | 0 | 0 | 0 |
| rs121908760 | pathogenic | 0 | 0 | 0 |
| rs397508350 | pathogenic | 0 | 0 | 0 |
| rs374946172 | pathogenic | 0 | 0 | 0 |
| rs397508387 | pathogenic | 0 | 0 | 0 |
| rs201386642 | pathogenic | 0 | 0 | 0 |
| rs121909012 | pathogenic | 0 | 0 | 0 |
| rs121909008 | pathogenic | 0 | 0 | 0 |
| rs149790377 | pathogenic | 0 | 0 | 0 |
| rs397508442 | pathogenic | 0 | 0 | 0 |
| rs121909035 | pathogenic | 0 | 0 | 0 |
| rs386134230 | pathogenic | 0 | 0 | 0 |
| rs1800112 | benign | 0 | 0 | 0 |
| rs144055758 | pathogenic | 0 | 0 | 0 |
| rs142394380 | pathogenic | 0 | 0 | 0 |
| rs121909036 | pathogenic | 0 | 0 | 0 |
| rs78194216 | pathogenic | 0 | 0 | 0 |
| rs121909019 | pathogenic | 0 | 0 | 0 |
| rs121909020 | pathogenic | 0 | 0 | 0 |
| rs200321110 | pathogenic | 0 | 0 | 0 |
| rs202179988 | pathogenic | 0 | 0 | 0 |
| rs121909037 | pathogenic | 0 | 0 | 0 |
| rs186045772 | pathogenic | 0 | 0 | 0 |
| rs139304906 | pathogenic | 0 | 0 | 0 |
| rs78802634 | pathogenic | 0 | 0 | 0 |
| rs121908761 | pathogenic | 0 | 0 | 0 |
| rs36210737 | pathogenic | 0 | 0 | 0 |
| rs139729994 | pathogenic | 0 | 0 | 0 |
| rs79850223 | pathogenic | 0 | 0 | 0 |
| rs1800120 | benign | 0 | 0 | 0 |
| rs74767530 | pathogenic | 0 | 0 | 0 |
| rs374705585 | pathogenic | 0 | 0 | 0 |
| rs1136287 | benign | 0 | 0 | 0 |
| rs6828 | benign | 0 | 0 | 0 |
| rs104894856 | pathogenic | 0 | 0 | 0 |
| rs104894860 | pathogenic | 0 | 0 | 0 |
| rs61736892 | pathogenic | 0 | 0 | 0 |
| rs145807417 | benign | 0 | 0 | 0 |
| rs104894853 | pathogenic | 0 | 0 | 0 |
| rs199422227 | pathogenic | 0 | 0 | 0 |
| rs113993946 | pathogenic | 0 | 0 | 0 |
| rs199422231 | pathogenic | 0 | 0 | 0 |
| rs199422228 | pathogenic | 0 | 0 | 0 |
| rs142195196 | benign | 0 | 1 | 0 |
| rs148656104 | benign | 1 | 1 | 1 |
| rs141944844 | benign | 0 | 0 | 0 |
| rs104894342 | pathogenic | 0 | 0 | 0 |
| rs104894343 | pathogenic | 0 | 0 | 0 |
| rs104894344 | pathogenic | 0 | 0 | 0 |
| rs573322878 | benign | 0 | 0 | 0 |
| rs104894347 | pathogenic | 0 | 0 | 0 |
| rs28937882 | pathogenic | 0 | 0 | 0 |

| | | | | |
|---|---|---|---|---|
| rs193922702 | pathogenic | 0 | 0 | 0 |
| rs200786104 | benign | 0 | 0 | 0 |
| rs690016544 | pathogenic | 1 | 1 | 1 |
| rs28943592 | benign | 0 | 0 | 0 |
| rs28943594 | benign | 0 | 0 | 0 |
| rs121908957 | pathogenic | 0 | 0 | 0 |
| rs115408010 | benign | 1 | 1 | 1 |
| rs77591659 | benign | 0 | 0 | 0 |
| rs121918511 | pathogenic | 0 | 0 | 0 |
| rs538359948 | benign | 0 | 0 | 0 |
| rs76708142 | benign | 0 | 0 | 0 |
| rs121907913 | pathogenic | 1 | 1 | 0 |
| rs121907920 | pathogenic | 0 | 0 | 0 |
| rs121907923 | pathogenic | 1 | 1 | 0 |
| rs121907928 | pathogenic | 1 | 1 | 0 |
| rs121907919 | pathogenic | 1 | 1 | 0 |
| rs121907918 | pathogenic | 1 | 1 | 0 |
| rs74315493 | pathogenic | 1 | 1 | 1 |
| rs74315494 | pathogenic | 1 | 1 | 1 |
| rs145446060 | benign | 1 | 1 | 1 |
| rs550409143 | benign | 1 | 1 | 0 |

| compound_binding_site | cofactor_binding_site | nucleic_binding_site | ion_binding_site |
|---|---|---|---|
| 1 | 0 | 0 | 0 |
| 1 | 0 | 0 | 0 |
| 0 | 0 | 0 | 0 |
| 1 | 0 | 0 | 0 |
| 0 | 0 | 0 | 0 |
| 0 | 0 | 0 | 0 |
| 0 | 0 | 0 | 0 |
| 0 | 0 | 0 | 0 |
| 1 | 0 | 0 | 1 |
| 1 | 0 | 0 | 0 |
| 0 | 0 | 0 | 1 |
| 1 | 0 | 1 | 0 |
| 0 | 0 | 0 | 0 |
| 0 | 0 | 0 | 0 |
| 0 | 0 | 0 | 0 |
| 0 | 0 | 0 | 0 |
| 0 | 0 | 0 | 0 |
| 1 | 0 | 0 | 1 |
| 1 | 0 | 1 | 0 |
| 0 | 0 | 0 | 0 |
| 1 | 0 | 0 | 1 |
| 1 | 0 | 0 | 0 |
| 1 | 0 | 0 | 0 |
| 1 | 0 | 0 | 0 |
| 0 | 0 | 0 | 0 |
| 0 | 0 | 0 | 0 |
| 1 | 0 | 0 | 1 |
| 0 | 0 | 0 | 0 |
| 0 | 0 | 0 | 0 |
| 0 | 0 | 0 | 0 |
| 0 | 0 | 0 | 0 |
| 1 | 0 | 0 | 0 |
| 1 | 0 | 0 | 1 |
| 1 | 0 | 0 | 0 |
| 0 | 0 | 0 | 0 |
| 0 | 0 | 0 | 0 |
| 0 | 0 | 0 | 0 |
| 1 | 0 | 0 | 1 |
| 1 | 0 | 0 | 1 |
| 1 | 0 | 0 | 1 |
| 1 | 0 | 0 | 0 |
| 1 | 0 | 0 | 0 |
| 1 | 0 | 0 | 0 |
| 1 | 0 | 0 | 0 |
| 1 | 0 | 0 | 0 |
| 1 | 0 | 0 | 0 |
| 1 | 0 | 0 | 0 |
| 1 | 0 | 0 | 0 |
| 1 | 0 | 0 | 0 |
| 1 | 0 | 0 | 0 |
| 1 | 0 | 0 | 0 |
| 1 | 0 | 0 | 0 |
| 1 | 0 | 0 | 0 |

| | | | |
|---|---|---|---|
| 1 | 0 | 0 | 1 |
| 1 | 0 | 0 | 0 |
| 1 | 0 | 0 | 0 |
| 1 | 0 | 0 | 0 |
| 0 | 0 | 0 | 0 |
| 1 | 0 | 0 | 0 |
| 0 | 0 | 0 | 0 |
| 1 | 0 | 0 | 0 |
| 1 | 0 | 0 | 0 |
| 1 | 0 | 0 | 1 |
| 1 | 0 | 0 | 0 |
| 1 | 0 | 0 | 0 |
| 1 | 0 | 0 | 0 |
| 1 | 0 | 0 | 0 |
| 1 | 0 | 0 | 0 |
| 1 | 0 | 0 | 0 |
| 1 | 0 | 0 | 0 |
| 1 | 0 | 0 | 1 |
| 1 | 0 | 0 | 1 |
| 1 | 1 | 0 | 1 |
| 1 | 0 | 0 | 0 |
| 1 | 0 | 0 | 0 |
| 1 | 0 | 0 | 0 |
| 1 | 0 | 0 | 0 |
| 1 | 1 | 0 | 0 |
| 1 | 0 | 0 | 0 |
| 1 | 0 | 0 | 0 |
| 1 | 0 | 0 | 0 |
| 1 | 0 | 0 | 0 |
| 1 | 0 | 0 | 0 |
| 1 | 1 | 0 | 0 |
| 1 | 1 | 0 | 0 |
| 1 | 1 | 0 | 0 |
| 1 | 1 | 0 | 0 |
| 1 | 1 | 0 | 0 |
| 1 | 0 | 0 | 0 |
| 1 | 1 | 0 | 0 |
| 1 | 0 | 0 | 0 |
| 1 | 0 | 0 | 0 |
| 1 | 1 | 0 | 0 |
| 0 | 0 | 0 | 0 |
| 0 | 0 | 0 | 0 |
| 0 | 0 | 0 | 0 |
| 0 | 0 | 0 | 0 |
| 0 | 0 | 0 | 0 |
| 0 | 0 | 0 | 0 |
| 0 | 0 | 0 | 0 |
| 1 | 0 | 0 | 1 |
| 0 | 0 | 0 | 0 |
| 1 | 0 | 0 | 0 |
| 1 | 0 | 0 | 0 |
| 0 | 0 | 0 | 0 |
| 0 | 0 | 0 | 0 |
| 0 | 0 | 0 | 0 |

| | | | |
|---|---|---|---|
| 0 | 0 | 0 | 0 |
| 0 | 0 | 0 | 0 |
| 0 | 0 | 0 | 0 |
| 1 | 0 | 0 | 0 |
| 1 | 0 | 0 | 0 |
| 1 | 0 | 0 | 0 |
| 0 | 0 | 0 | 0 |
| 0 | 0 | 1 | 0 |
| 0 | 0 | 1 | 0 |
| 0 | 0 | 1 | 0 |
| 0 | 0 | 1 | 0 |
| 0 | 0 | 1 | 0 |
| 0 | 0 | 0 | 0 |
| 0 | 0 | 0 | 0 |
| 1 | 0 | 0 | 0 |
| 0 | 0 | 0 | 0 |
| 0 | 0 | 0 | 0 |
| 1 | 0 | 0 | 0 |
| 1 | 0 | 0 | 0 |
| 1 | 0 | 0 | 0 |
| 1 | 0 | 0 | 0 |
| 1 | 1 | 0 | 0 |
| 1 | 0 | 0 | 0 |
| 1 | 0 | 0 | 0 |
| 0 | 0 | 0 | 0 |
| 0 | 0 | 0 | 0 |
| 1 | 0 | 0 | 0 |
| 1 | 1 | 0 | 0 |
| 1 | 0 | 0 | 0 |
| 1 | 0 | 0 | 0 |
| 1 | 1 | 0 | 0 |
| 1 | 1 | 0 | 0 |
| 1 | 1 | 0 | 0 |
| 1 | 1 | 0 | 0 |
| 1 | 1 | 0 | 0 |
| 1 | 1 | 0 | 0 |
| 1 | 1 | 0 | 0 |
| 1 | 1 | 0 | 0 |
| 1 | 1 | 0 | 0 |
| 1 | 1 | 0 | 0 |
| 1 | 1 | 0 | 0 |
| 1 | 1 | 0 | 0 |
| 1 | 0 | 0 | 0 |
| 1 | 0 | 0 | 0 |
| 1 | 0 | 0 | 0 |
| 1 | 1 | 0 | 0 |
| 1 | 0 | 0 | 0 |
| 1 | 1 | 1 | 0 |
| 1 | 1 | 1 | 0 |
| 1 | 1 | 0 | 0 |
| 1 | 1 | 0 | 0 |
| 1 | 0 | 0 | 0 |
| 1 | 0 | 0 | 0 |
| 0 | 0 | 0 | 0 |

| | | | |
|---|---|---|---|
| 0 | 0 | 0 | 0 |
| 0 | 0 | 0 | 0 |
| 0 | 0 | 0 | 0 |
| 1 | 0 | 0 | 0 |
| 1 | 0 | 0 | 0 |
| 0 | 0 | 0 | 0 |
| 0 | 0 | 0 | 0 |
| 0 | 0 | 0 | 0 |
| 0 | 0 | 0 | 0 |
| 0 | 0 | 0 | 0 |
| 0 | 0 | 0 | 0 |
| 0 | 0 | 0 | 0 |
| 0 | 0 | 0 | 0 |
| 0 | 0 | 0 | 0 |
| 0 | 0 | 0 | 0 |
| 0 | 0 | 0 | 0 |
| 0 | 0 | 0 | 0 |
| 0 | 0 | 0 | 0 |
| 0 | 0 | 0 | 0 |
| 0 | 0 | 0 | 0 |
| 0 | 0 | 0 | 0 |
| 0 | 0 | 0 | 0 |
| 1 | 0 | 0 | 0 |
| 1 | 0 | 0 | 0 |
| 0 | 0 | 0 | 0 |
| 0 | 0 | 0 | 0 |
| 1 | 0 | 0 | 0 |
| 0 | 0 | 0 | 0 |
| 0 | 0 | 0 | 0 |
| 0 | 0 | 0 | 0 |
| 0 | 0 | 0 | 0 |
| 0 | 0 | 0 | 0 |
| 0 | 0 | 0 | 0 |
| 0 | 0 | 0 | 0 |
| 1 | 0 | 0 | 0 |
| 0 | 0 | 0 | 0 |
| 0 | 0 | 0 | 0 |
| 0 | 0 | 0 | 0 |
| 0 | 0 | 0 | 0 |
| 0 | 0 | 0 | 0 |
| 0 | 0 | 0 | 0 |
| 0 | 0 | 0 | 0 |
| 0 | 0 | 0 | 0 |
| 0 | 0 | 0 | 0 |
| 0 | 0 | 0 | 0 |
| 0 | 0 | 0 | 0 |
| 0 | 0 | 1 | 0 |
| 1 | 0 | 0 | 0 |
| 1 | 0 | 0 | 0 |
| 1 | 1 | 0 | 0 |
| 1 | 0 | 0 | 0 |
| 0 | 0 | 0 | 0 |
| 0 | 0 | 0 | 0 |
| 1 | 0 | 0 | 0 |

| | | | |
|---|---|---|---|
| 1 | 0 | 0 | 0 |
| 1 | 0 | 0 | 0 |
| 1 | 0 | 0 | 0 |
| 0 | 0 | 0 | 0 |
| 0 | 0 | 0 | 0 |
| 0 | 0 | 0 | 0 |
| 1 | 0 | 0 | 0 |
| 1 | 0 | 0 | 0 |
| 0 | 0 | 0 | 0 |
| 0 | 0 | 0 | 0 |
| 1 | 0 | 0 | 0 |
| 0 | 0 | 0 | 0 |
| 1 | 0 | 0 | 0 |
| 0 | 0 | 0 | 0 |
| 0 | 0 | 0 | 0 |
| 0 | 0 | 0 | 0 |
| 0 | 0 | 0 | 0 |
| 0 | 0 | 0 | 0 |
| 0 | 0 | 0 | 0 |
| 0 | 0 | 0 | 0 |
| 0 | 0 | 0 | 0 |
| 0 | 0 | 0 | 0 |
| 0 | 0 | 0 | 0 |
| 0 | 0 | 0 | 0 |
| 0 | 0 | 0 | 0 |
| 0 | 0 | 0 | 0 |
| 1 | 0 | 0 | 0 |
| 1 | 0 | 0 | 0 |
| 1 | 0 | 0 | 0 |
| 1 | 0 | 0 | 0 |
| 1 | 0 | 0 | 0 |
| 1 | 0 | 0 | 0 |
| 0 | 0 | 0 | 0 |
| 0 | 0 | 0 | 0 |
| 1 | 0 | 0 | 0 |
| 0 | 0 | 0 | 0 |
| 0 | 0 | 0 | 0 |
| 0 | 0 | 0 | 0 |
| 0 | 0 | 0 | 0 |
| 0 | 0 | 0 | 0 |
| 0 | 0 | 0 | 0 |
| 0 | 0 | 0 | 0 |
| 0 | 0 | 0 | 0 |
| 0 | 0 | 0 | 0 |
| 1 | 1 | 0 | 0 |
| 1 | 1 | 0 | 0 |
| 1 | 1 | 0 | 1 |
| 1 | 1 | 0 | 1 |
| 1 | 1 | 0 | 1 |
| 1 | 1 | 0 | 1 |
| 1 | 1 | 0 | 0 |
| 1 | 1 | 0 | 1 |
| 1 | 1 | 0 | 1 |
| 1 | 1 | 0 | 1 |

| | | | |
|---|---|---|---|
| 1 | 1 | 0 | 1 |
| 1 | 1 | 0 | 1 |
| 1 | 1 | 0 | 1 |
| 1 | 1 | 0 | 1 |
| 1 | 1 | 0 | 1 |
| 1 | 0 | 0 | 0 |
| 0 | 0 | 0 | 0 |
| 0 | 0 | 0 | 0 |
| 0 | 0 | 0 | 0 |
| 1 | 0 | 0 | 0 |
| 0 | 0 | 1 | 0 |
| 0 | 0 | 0 | 0 |
| 0 | 0 | 0 | 0 |
| 0 | 0 | 0 | 0 |
| 0 | 0 | 0 | 0 |
| 0 | 0 | 0 | 0 |
| 0 | 0 | 0 | 0 |
| 0 | 0 | 0 | 0 |
| 0 | 0 | 0 | 0 |
| 0 | 0 | 0 | 0 |
| 0 | 0 | 0 | 0 |
| 0 | 0 | 0 | 0 |
| 0 | 0 | 0 | 0 |
| 0 | 0 | 0 | 0 |
| 0 | 0 | 0 | 0 |
| 1 | 1 | 0 | 0 |
| 1 | 1 | 0 | 0 |
| 1 | 1 | 0 | 0 |
| 0 | 1 | 0 | 0 |
| 1 | 0 | 0 | 0 |
| 0 | 0 | 0 | 0 |
| 0 | 0 | 0 | 0 |
| 0 | 0 | 0 | 0 |
| 1 | 0 | 0 | 0 |
| 1 | 1 | 0 | 0 |
| 0 | 0 | 0 | 0 |
| 0 | 0 | 0 | 0 |
| 0 | 0 | 0 | 0 |
| 0 | 0 | 0 | 0 |
| 1 | 0 | 0 | 0 |
| 0 | 0 | 0 | 0 |
| 0 | 0 | 0 | 0 |
| 0 | 0 | 0 | 0 |
| 0 | 0 | 0 | 0 |
| 0 | 0 | 0 | 0 |
| 0 | 1 | 0 | 0 |
| 0 | 1 | 0 | 0 |
| 0 | 0 | 0 | 0 |
| 0 | 0 | 0 | 0 |
| 0 | 1 | 0 | 0 |
| 0 | 0 | 0 | 0 |
| 0 | 0 | 0 | 0 |
| 0 | 1 | 0 | 0 |
| 0 | 0 | 0 | 0 |

| | | | |
|---|---|---|---|
| 1 | 1 | 0 | 0 |
| 0 | 0 | 0 | 0 |
| 0 | 0 | 0 | 0 |
| 0 | 0 | 0 | 0 |
| 0 | 0 | 0 | 0 |
| 0 | 0 | 0 | 0 |
| 0 | 0 | 0 | 0 |
| 0 | 0 | 0 | 0 |
| 0 | 0 | 0 | 0 |
| 0 | 0 | 0 | 0 |
| 0 | 0 | 0 | 0 |
| 0 | 0 | 0 | 0 |
| 0 | 0 | 0 | 0 |
| 0 | 0 | 0 | 0 |
| 0 | 1 | 0 | 0 |
| 0 | 0 | 0 | 0 |
| 0 | 1 | 0 | 0 |
| 0 | 1 | 0 | 0 |
| 0 | 0 | 0 | 0 |
| 0 | 1 | 0 | 0 |
| 1 | 0 | 0 | 1 |
| 1 | 1 | 0 | 0 |
| 1 | 0 | 0 | 0 |
| 1 | 0 | 0 | 1 |
| 0 | 0 | 0 | 0 |
| 0 | 0 | 0 | 0 |
| 0 | 0 | 0 | 0 |
| 0 | 0 | 0 | 0 |
| 1 | 0 | 0 | 0 |
| 0 | 0 | 0 | 0 |
| 1 | 0 | 0 | 0 |
| 0 | 0 | 0 | 0 |
| 0 | 0 | 0 | 0 |
| 0 | 0 | 0 | 0 |
| 0 | 0 | 0 | 0 |
| 0 | 0 | 0 | 0 |
| 0 | 0 | 0 | 0 |
| 1 | 1 | 0 | 0 |
| 1 | 1 | 0 | 0 |
| 1 | 1 | 0 | 0 |
| 0 | 0 | 0 | 0 |
| 0 | 0 | 0 | 0 |
| 0 | 0 | 0 | 0 |
| 1 | 0 | 0 | 0 |
| 1 | 0 | 0 | 1 |
| 1 | 0 | 0 | 1 |
| 1 | 0 | 0 | 0 |
| 1 | 0 | 0 | 1 |
| 1 | 0 | 0 | 0 |
| 0 | 0 | 0 | 0 |
| 1 | 1 | 0 | 0 |
| 1 | 1 | 0 | 0 |
| 0 | 0 | 0 | 0 |
| 0 | 0 | 0 | 0 |

| | | | |
|---|---|---|---|
| 1 | 0 | 0 | 0 |
| 1 | 0 | 0 | 0 |
| 1 | 0 | 0 | 0 |
| 0 | 0 | 0 | 0 |
| 0 | 0 | 0 | 0 |
| 1 | 1 | 0 | 1 |
| 1 | 1 | 0 | 1 |
| 1 | 1 | 0 | 1 |
| 1 | 1 | 0 | 1 |
| 1 | 1 | 0 | 1 |
| 1 | 1 | 0 | 1 |
| 1 | 1 | 0 | 1 |
| 1 | 1 | 0 | 0 |
| 1 | 1 | 0 | 0 |
| 1 | 1 | 1 | 1 |
| 0 | 0 | 0 | 0 |
| 1 | 1 | 0 | 0 |
| 1 | 1 | 0 | 0 |
| 0 | 0 | 0 | 0 |
| 0 | 0 | 0 | 0 |
| 0 | 0 | 0 | 0 |
| 1 | 0 | 0 | 0 |
| 1 | 0 | 0 | 0 |
| 0 | 0 | 0 | 0 |
| 1 | 0 | 0 | 0 |
| 1 | 0 | 0 | 0 |
| 1 | 0 | 0 | 0 |
| 0 | 0 | 0 | 0 |
| 0 | 0 | 0 | 0 |
| 0 | 0 | 0 | 0 |
| 0 | 0 | 0 | 0 |
| 0 | 0 | 0 | 0 |
| 0 | 0 | 0 | 0 |
| 0 | 0 | 0 | 0 |
| 0 | 0 | 0 | 0 |
| 0 | 0 | 0 | 0 |
| 0 | 0 | 0 | 0 |
| 0 | 0 | 0 | 0 |
| 0 | 0 | 0 | 0 |
| 0 | 0 | 0 | 0 |
| 0 | 0 | 0 | 0 |
| 0 | 0 | 0 | 0 |
| 0 | 0 | 0 | 0 |
| 0 | 0 | 0 | 0 |
| 0 | 0 | 0 | 0 |
| 1 | 0 | 0 | 0 |
| 1 | 0 | 0 | 0 |
| 1 | 0 | 0 | 0 |
| 1 | 0 | 0 | 0 |
| 0 | 0 | 0 | 0 |
| 0 | 0 | 0 | 0 |
| 0 | 0 | 0 | 0 |
| 0 | 0 | 0 | 0 |
| 0 | 0 | 0 | 0 |

| | | | |
|---|---|---|---|
| 0 | 0 | 0 | 0 |
| 0 | 0 | 0 | 0 |
| 0 | 0 | 0 | 0 |
| 0 | 0 | 0 | 0 |
| 0 | 0 | 0 | 0 |
| 0 | 0 | 0 | 0 |
| 0 | 0 | 0 | 0 |
| 1 | 0 | 0 | 1 |
| 0 | 0 | 0 | 0 |
| 0 | 0 | 0 | 0 |
| 1 | 0 | 0 | 0 |
| 0 | 0 | 0 | 0 |
| 0 | 0 | 0 | 0 |
| 0 | 0 | 0 | 0 |
| 0 | 0 | 0 | 0 |
| 1 | 0 | 0 | 0 |
| 0 | 0 | 0 | 0 |
| 0 | 0 | 0 | 0 |
| 0 | 0 | 0 | 0 |
| 0 | 0 | 0 | 0 |
| 0 | 0 | 0 | 0 |
| 0 | 0 | 0 | 0 |
| 0 | 0 | 0 | 0 |
| 1 | 1 | 0 | 1 |
| 0 | 0 | 0 | 0 |
| 0 | 0 | 0 | 0 |
| 0 | 0 | 0 | 0 |
| 1 | 1 | 0 | 1 |
| 1 | 1 | 0 | 0 |
| 0 | 0 | 0 | 0 |
| 1 | 0 | 0 | 0 |
| 1 | 1 | 0 | 0 |
| 0 | 0 | 0 | 0 |
| 1 | 0 | 0 | 0 |
| 1 | 1 | 0 | 0 |
| 1 | 0 | 0 | 0 |
| 1 | 1 | 0 | 1 |
| 1 | 1 | 0 | 1 |
| 1 | 0 | 0 | 0 |
| 0 | 0 | 0 | 0 |
| 0 | 0 | 0 | 0 |
| 0 | 0 | 0 | 0 |
| 0 | 0 | 0 | 0 |
| 0 | 0 | 0 | 0 |
| 0 | 0 | 0 | 0 |
| 0 | 0 | 0 | 0 |
| 0 | 0 | 0 | 0 |
| 0 | 0 | 0 | 0 |
| 0 | 0 | 0 | 0 |
| 0 | 0 | 0 | 0 |
| 0 | 0 | 0 | 0 |
| 0 | 0 | 0 | 0 |
| 0 | 0 | 0 | 0 |
| 0 | 0 | 0 | 0 |

| | | | |
|---|---|---|---|
| 0 | 0 | 0 | 0 |
| 0 | 0 | 0 | 0 |
| 0 | 0 | 0 | 0 |
| 0 | 0 | 0 | 0 |
| 0 | 0 | 0 | 0 |
| 0 | 0 | 0 | 0 |
| 0 | 0 | 0 | 0 |
| 0 | 0 | 0 | 0 |
| 0 | 0 | 0 | 0 |
| 0 | 0 | 0 | 0 |
| 0 | 0 | 0 | 0 |
| 0 | 0 | 0 | 0 |
| 0 | 0 | 0 | 0 |
| 1 | 1 | 0 | 0 |
| 0 | 0 | 0 | 0 |
| 1 | 0 | 0 | 0 |
| 0 | 0 | 0 | 1 |
| 0 | 0 | 0 | 0 |
| 0 | 0 | 0 | 0 |
| 0 | 0 | 0 | 0 |
| 0 | 0 | 0 | 0 |
| 0 | 0 | 0 | 0 |
| 0 | 0 | 0 | 0 |
| 0 | 0 | 0 | 0 |
| 0 | 0 | 0 | 0 |
| 0 | 0 | 0 | 0 |
| 0 | 0 | 0 | 0 |
| 0 | 0 | 0 | 0 |
| 0 | 0 | 0 | 0 |
| 0 | 0 | 0 | 0 |
| 1 | 0 | 0 | 0 |
| 1 | 0 | 0 | 0 |
| 0 | 0 | 0 | 0 |
| 0 | 0 | 0 | 0 |
| 0 | 0 | 0 | 0 |
| 0 | 0 | 0 | 0 |
| 0 | 0 | 0 | 0 |
| 0 | 0 | 0 | 0 |
| 0 | 0 | 0 | 0 |
| 0 | 0 | 0 | 0 |
| 1 | 0 | 0 | 0 |
| 1 | 0 | 0 | 0 |
| 1 | 0 | 0 | 0 |
| 1 | 0 | 0 | 0 |
| 0 | 0 | 0 | 0 |
| 1 | 0 | 0 | 0 |
| 0 | 0 | 0 | 0 |
| 0 | 0 | 0 | 0 |
| 1 | 0 | 0 | 0 |
| 1 | 0 | 0 | 0 |
| 0 | 0 | 0 | 0 |
| 0 | 0 | 0 | 0 |
| 0 | 0 | 0 | 0 |
| 0 | 0 | 0 | 0 |

| | | | |
|---|---|---|---|
| 0 | 0 | 0 | 0 |
| 0 | 0 | 0 | 0 |
| 0 | 0 | 0 | 0 |
| 0 | 0 | 0 | 0 |
| 0 | 0 | 0 | 0 |
| 0 | 0 | 0 | 0 |
| 0 | 0 | 0 | 0 |
| 0 | 0 | 0 | 0 |
| 0 | 0 | 0 | 0 |
| 0 | 0 | 0 | 0 |
| 0 | 0 | 0 | 0 |
| 0 | 0 | 0 | 0 |
| 0 | 0 | 0 | 0 |
| 0 | 0 | 0 | 0 |
| 0 | 0 | 0 | 0 |
| 1 | 0 | 0 | 1 |
| 0 | 0 | 0 | 0 |
| 0 | 0 | 0 | 0 |
| 0 | 0 | 0 | 0 |
| 0 | 0 | 0 | 0 |
| 0 | 0 | 0 | 0 |
| 1 | 1 | 0 | 0 |
| 1 | 1 | 0 | 0 |
| 1 | 0 | 0 | 0 |
| 1 | 0 | 0 | 0 |
| 0 | 0 | 0 | 0 |
| 0 | 0 | 0 | 0 |
| 0 | 0 | 0 | 0 |
| 0 | 0 | 0 | 0 |
| 1 | 0 | 0 | 0 |
| 1 | 0 | 0 | 0 |
| 1 | 0 | 0 | 0 |
| 1 | 1 | 0 | 0 |
| 1 | 1 | 0 | 0 |
| 1 | 0 | 0 | 0 |
| 0 | 0 | 0 | 0 |
| 0 | 0 | 0 | 0 |
| 0 | 1 | 0 | 0 |
| 0 | 0 | 0 | 0 |
| 0 | 0 | 0 | 0 |
| 0 | 0 | 0 | 0 |
| 1 | 1 | 0 | 0 |
| 1 | 0 | 0 | 0 |
| 1 | 0 | 0 | 0 |
| 0 | 0 | 0 | 0 |
| 0 | 0 | 0 | 0 |
| 0 | 0 | 0 | 0 |
| 0 | 0 | 0 | 0 |
| 0 | 0 | 0 | 0 |
| 0 | 0 | 0 | 0 |
| 0 | 0 | 0 | 0 |
| 0 | 0 | 0 | 0 |
| 0 | 0 | 0 | 0 |
| 0 | 0 | 0 | 0 |

| | | | |
|---|---|---|---|
| 0 | 0 | 0 | 0 |
| 1 | 0 | 0 | 0 |
| 0 | 0 | 0 | 0 |
| 0 | 0 | 0 | 0 |
| 0 | 0 | 0 | 0 |
| 0 | 0 | 0 | 0 |
| 1 | 0 | 0 | 0 |
| 0 | 0 | 0 | 0 |
| 0 | 0 | 0 | 0 |
| 0 | 0 | 0 | 0 |
| 0 | 0 | 0 | 0 |
| 0 | 0 | 0 | 0 |
| 0 | 0 | 0 | 0 |
| 0 | 0 | 0 | 0 |
| 0 | 0 | 0 | 0 |
| 0 | 0 | 0 | 0 |
| 0 | 1 | 0 | 0 |
| 0 | 0 | 0 | 0 |
| 0 | 0 | 0 | 0 |
| 0 | 0 | 0 | 0 |
| 1 | 0 | 0 | 0 |
| 0 | 0 | 0 | 0 |
| 0 | 0 | 0 | 0 |
| 1 | 1 | 0 | 0 |
| 0 | 0 | 0 | 0 |
| 0 | 0 | 0 | 0 |
| 0 | 0 | 0 | 0 |
| 0 | 0 | 0 | 0 |
| 1 | 0 | 0 | 0 |
| 0 | 0 | 0 | 0 |
| 0 | 0 | 0 | 0 |
| 0 | 0 | 0 | 0 |
| 0 | 0 | 0 | 0 |
| 0 | 0 | 0 | 0 |
| 0 | 0 | 0 | 0 |
| 1 | 0 | 0 | 0 |
| 0 | 0 | 0 | 0 |
| 0 | 0 | 0 | 0 |
| 0 | 0 | 0 | 0 |
| 0 | 0 | 0 | 0 |
| 1 | 1 | 0 | 0 |
| 0 | 0 | 0 | 0 |
| 0 | 0 | 0 | 0 |
| 0 | 0 | 0 | 0 |
| 0 | 0 | 0 | 0 |
| 0 | 0 | 0 | 0 |
| 1 | 0 | 0 | 0 |
| 1 | 0 | 0 | 1 |
| 1 | 0 | 0 | 0 |
| 0 | 0 | 0 | 0 |
| 0 | 0 | 0 | 0 |
| 0 | 0 | 0 | 0 |
| 1 | 0 | 0 | 0 |
| 1 | 0 | 0 | 0 |

| | | | |
|---|---|---|---|
| 1 | 0 | 0 | 0 |
| 0 | 0 | 0 | 0 |
| 0 | 0 | 0 | 0 |
| 0 | 0 | 0 | 0 |
| 1 | 1 | 0 | 0 |
| 1 | 1 | 0 | 0 |
| 0 | 0 | 0 | 0 |
| 0 | 0 | 0 | 0 |
| 1 | 0 | 0 | 0 |
| 1 | 0 | 0 | 0 |
| 1 | 0 | 1 | 0 |
| 0 | 0 | 0 | 0 |
| 0 | 0 | 1 | 0 |
| 0 | 0 | 1 | 0 |
| 0 | 0 | 1 | 0 |
| 1 | 0 | 0 | 0 |
| 1 | 0 | 0 | 0 |
| 0 | 0 | 0 | 0 |
| 0 | 0 | 0 | 0 |
| 1 | 0 | 0 | 1 |
| 1 | 0 | 0 | 1 |
| 1 | 0 | 0 | 0 |
| 0 | 0 | 0 | 0 |
| 0 | 0 | 0 | 0 |
| 0 | 0 | 0 | 0 |
| 0 | 0 | 0 | 0 |
| 0 | 0 | 0 | 0 |
| 1 | 0 | 0 | 0 |
| 0 | 0 | 0 | 0 |
| 0 | 0 | 0 | 1 |
| 0 | 0 | 0 | 1 |
| 0 | 0 | 1 | 1 |
| 0 | 0 | 1 | 1 |
| 0 | 0 | 1 | 1 |
| 0 | 0 | 1 | 1 |
| 0 | 0 | 1 | 0 |
| 0 | 0 | 1 | 0 |
| 0 | 0 | 0 | 0 |
| 0 | 0 | 0 | 0 |
| 0 | 0 | 0 | 0 |
| 0 | 0 | 0 | 0 |
| 0 | 0 | 1 | 0 |
| 0 | 0 | 1 | 0 |
| 0 | 0 | 1 | 0 |
| 0 | 0 | 1 | 0 |
| 0 | 0 | 1 | 0 |
| 0 | 0 | 1 | 0 |
| 0 | 0 | 1 | 0 |
| 0 | 0 | 1 | 0 |
| 0 | 0 | 0 | 0 |
| 0 | 0 | 0 | 0 |
| 0 | 0 | 0 | 0 |
| 0 | 0 | 0 | 0 |
| 1 | 0 | 0 | 0 |

| | | | |
|---|---|---|---|
| 0 | 0 | 0 | 0 |
| 0 | 0 | 0 | 0 |
| 0 | 0 | 0 | 0 |
| 1 | 0 | 0 | 0 |
| 0 | 0 | 0 | 0 |
| 0 | 0 | 0 | 0 |
| 0 | 0 | 0 | 0 |
| 0 | 0 | 0 | 0 |
| 0 | 0 | 0 | 0 |
| 0 | 0 | 0 | 0 |
| 1 | 1 | 0 | 0 |
| 0 | 0 | 0 | 0 |
| 1 | 1 | 0 | 0 |
| 1 | 1 | 0 | 0 |
| 0 | 1 | 0 | 0 |
| 0 | 0 | 0 | 0 |
| 0 | 0 | 0 | 0 |
| 0 | 0 | 0 | 0 |
| 0 | 0 | 0 | 0 |
| 0 | 0 | 0 | 0 |
| 1 | 1 | 0 | 0 |
| 0 | 1 | 0 | 0 |
| 0 | 0 | 0 | 0 |
| 0 | 1 | 0 | 0 |
| 1 | 1 | 0 | 0 |
| 0 | 0 | 0 | 0 |
| 0 | 0 | 0 | 0 |
| 1 | 1 | 0 | 0 |
| 1 | 1 | 0 | 0 |
| 0 | 1 | 0 | 0 |
| 0 | 0 | 0 | 0 |
| 0 | 0 | 0 | 0 |
| 0 | 0 | 0 | 0 |
| 0 | 0 | 0 | 0 |
| 0 | 0 | 0 | 0 |
| 0 | 0 | 0 | 0 |
| 0 | 0 | 0 | 0 |
| 0 | 0 | 0 | 0 |
| 0 | 0 | 0 | 0 |
| 0 | 0 | 0 | 0 |
| 1 | 0 | 0 | 0 |
| 1 | 0 | 0 | 0 |
| 1 | 0 | 0 | 0 |
| 1 | 0 | 0 | 0 |
| 1 | 1 | 0 | 1 |
| 0 | 0 | 0 | 0 |
| 1 | 0 | 0 | 0 |
| 0 | 0 | 0 | 0 |
| 1 | 0 | 0 | 0 |
| 0 | 0 | 0 | 0 |
| 1 | 0 | 0 | 0 |
| 1 | 0 | 0 | 0 |
| 0 | 0 | 0 | 0 |
| 0 | 0 | 0 | 0 |

| | | | |
|---|---|---|---|
| 0 | 0 | 0 | 0 |
| 0 | 0 | 0 | 0 |
| 0 | 0 | 0 | 0 |
| 0 | 0 | 0 | 0 |
| 1 | 0 | 0 | 0 |
| 1 | 0 | 0 | 0 |
| 1 | 0 | 0 | 0 |
| 0 | 0 | 0 | 0 |
| 1 | 1 | 0 | 0 |
| 0 | 0 | 0 | 0 |
| 0 | 0 | 0 | 0 |
| 0 | 0 | 0 | 0 |
| 1 | 1 | 0 | 0 |
| 1 | 0 | 0 | 0 |
| 0 | 0 | 0 | 0 |
| 0 | 0 | 0 | 0 |
| 1 | 1 | 0 | 0 |
| 0 | 0 | 0 | 0 |
| 0 | 0 | 0 | 0 |
| 0 | 0 | 0 | 0 |
| 0 | 0 | 0 | 0 |
| 0 | 0 | 0 | 0 |
| 0 | 0 | 0 | 0 |
| 0 | 0 | 0 | 0 |
| 1 | 0 | 0 | 0 |
| 0 | 0 | 0 | 0 |
| 1 | 0 | 0 | 0 |
| 0 | 0 | 0 | 0 |
| 0 | 0 | 0 | 0 |
| 0 | 0 | 0 | 0 |
| 1 | 0 | 0 | 0 |
| 1 | 0 | 0 | 0 |
| 0 | 0 | 0 | 0 |
| 0 | 0 | 0 | 0 |
| 1 | 0 | 0 | 0 |
| 0 | 0 | 0 | 0 |
| 0 | 0 | 0 | 0 |
| 1 | 0 | 0 | 0 |
| 0 | 0 | 0 | 0 |
| 1 | 0 | 0 | 0 |
| 1 | 0 | 0 | 0 |
| 0 | 0 | 0 | 0 |
| 0 | 0 | 0 | 0 |
| 1 | 1 | 0 | 0 |
| 1 | 0 | 0 | 0 |
| 1 | 1 | 0 | 0 |
| 0 | 0 | 0 | 0 |
| 0 | 0 | 0 | 0 |
| 0 | 0 | 0 | 0 |
| 0 | 0 | 0 | 0 |
| 1 | 1 | 0 | 0 |
| 0 | 0 | 0 | 0 |
| 0 | 0 | 0 | 0 |
| 0 | 0 | 0 | 0 |

| | | | |
|---|---|---|---|
| 0 | 1 | 0 | 0 |
| 1 | 0 | 0 | 0 |
| 0 | 0 | 0 | 0 |
| 0 | 0 | 0 | 0 |
| 0 | 0 | 0 | 0 |
| 1 | 1 | 0 | 0 |
| 0 | 0 | 0 | 0 |
| 0 | 0 | 0 | 0 |
| 0 | 0 | 0 | 0 |
| 1 | 0 | 0 | 0 |
| 1 | 0 | 0 | 0 |
| 0 | 0 | 0 | 0 |
| 0 | 0 | 0 | 0 |
| 0 | 0 | 0 | 0 |
| 0 | 0 | 0 | 0 |
| 0 | 0 | 0 | 0 |
| 0 | 0 | 0 | 0 |
| 0 | 0 | 0 | 0 |
| 0 | 0 | 0 | 0 |
| 0 | 0 | 0 | 0 |
| 0 | 0 | 0 | 0 |
| 0 | 0 | 0 | 0 |
| 0 | 0 | 0 | 0 |
| 1 | 1 | 0 | 0 |
| 0 | 0 | 0 | 0 |
| 0 | 0 | 0 | 0 |
| 1 | 1 | 1 | 0 |
| 0 | 0 | 0 | 0 |
| 0 | 0 | 0 | 0 |
| 0 | 0 | 0 | 0 |
| 1 | 1 | 1 | 0 |
| 1 | 0 | 0 | 0 |
| 1 | 0 | 0 | 0 |
| 1 | 0 | 0 | 0 |
| 1 | 0 | 0 | 0 |
| 1 | 1 | 1 | 0 |
| 1 | 1 | 1 | 0 |
| 0 | 0 | 0 | 0 |
| 0 | 0 | 0 | 0 |
| 1 | 1 | 1 | 0 |
| 0 | 0 | 0 | 0 |
| 0 | 0 | 0 | 0 |
| 0 | 0 | 0 | 0 |
| 0 | 0 | 0 | 0 |
| 0 | 0 | 0 | 0 |
| 0 | 0 | 0 | 0 |
| 0 | 0 | 0 | 0 |
| 0 | 0 | 0 | 0 |
| 1 | 0 | 0 | 0 |
| 1 | 0 | 0 | 0 |
| 1 | 0 | 0 | 0 |
| 0 | 0 | 0 | 0 |
| 0 | 0 | 0 | 0 |
| 0 | 0 | 0 | 0 |

| | | | |
|---|---|---|---|
| 0 | 0 | 0 | 0 |
| 1 | 0 | 0 | 0 |
| 0 | 0 | 0 | 0 |
| 0 | 0 | 0 | 0 |
| 0 | 0 | 0 | 0 |
| 0 | 0 | 0 | 0 |
| 0 | 0 | 0 | 0 |
| 0 | 0 | 0 | 0 |
| 0 | 0 | 0 | 0 |
| 0 | 0 | 0 | 0 |
| 1 | 1 | 0 | 0 |
| 1 | 1 | 0 | 0 |
| 0 | 0 | 0 | 0 |
| 0 | 0 | 0 | 0 |
| 0 | 0 | 0 | 0 |
| 1 | 0 | 0 | 0 |
| 1 | 0 | 0 | 0 |
| 1 | 1 | 0 | 1 |
| 0 | 0 | 0 | 0 |
| 0 | 0 | 0 | 0 |
| 0 | 0 | 0 | 0 |
| 0 | 0 | 0 | 0 |
| 1 | 0 | 0 | 1 |
| 0 | 0 | 0 | 0 |
| 0 | 0 | 0 | 0 |
| 1 | 0 | 0 | 0 |
| 0 | 0 | 0 | 0 |
| 0 | 0 | 0 | 0 |
| 0 | 0 | 0 | 0 |
| 0 | 0 | 0 | 0 |
| 0 | 0 | 0 | 0 |
| 0 | 0 | 0 | 0 |
| 0 | 0 | 0 | 0 |
| 0 | 0 | 0 | 0 |
| 0 | 0 | 0 | 0 |
| 1 | 0 | 0 | 0 |
| 1 | 0 | 0 | 0 |
| 1 | 1 | 0 | 0 |
| 1 | 1 | 0 | 0 |
| 0 | 0 | 0 | 0 |
| 0 | 0 | 0 | 0 |
| 1 | 0 | 0 | 0 |
| 0 | 0 | 0 | 0 |
| 1 | 0 | 0 | 0 |
| 1 | 1 | 0 | 0 |
| 0 | 0 | 0 | 0 |
| 0 | 0 | 0 | 0 |
| 0 | 0 | 0 | 0 |
| 0 | 0 | 0 | 0 |
| 0 | 0 | 0 | 0 |
| 0 | 0 | 0 | 0 |
| 1 | 0 | 0 | 0 |
| 0 | 0 | 0 | 0 |
| 0 | 0 | 0 | 0 |

| | | | |
|---|---|---|---|
| 0 | 0 | 0 | 0 |
| 1 | 1 | 0 | 0 |
| 0 | 0 | 0 | 0 |
| 1 | 0 | 0 | 0 |
| 1 | 0 | 0 | 0 |
| 0 | 0 | 0 | 0 |
| 0 | 0 | 0 | 0 |
| 0 | 0 | 0 | 0 |
| 0 | 0 | 0 | 0 |
| 0 | 0 | 0 | 0 |
| 0 | 0 | 0 | 0 |
| 1 | 1 | 0 | 0 |
| 1 | 1 | 0 | 1 |
| 0 | 0 | 0 | 0 |
| 0 | 1 | 0 | 0 |
| 0 | 0 | 0 | 0 |
| 0 | 0 | 0 | 0 |
| 0 | 0 | 0 | 0 |
| 0 | 0 | 0 | 0 |
| 0 | 0 | 0 | 0 |
| 0 | 0 | 0 | 0 |
| 1 | 0 | 0 | 0 |
| 0 | 0 | 0 | 0 |
| 0 | 0 | 0 | 0 |
| 0 | 0 | 0 | 0 |
| 0 | 0 | 0 | 0 |
| 0 | 0 | 0 | 0 |
| 1 | 0 | 0 | 0 |
| 0 | 0 | 0 | 0 |
| 0 | 0 | 0 | 0 |
| 0 | 0 | 0 | 0 |
| 0 | 0 | 0 | 0 |
| 0 | 0 | 0 | 0 |
| 0 | 0 | 0 | 0 |
| 0 | 0 | 0 | 0 |
| 0 | 0 | 0 | 0 |
| 0 | 0 | 0 | 0 |
| 0 | 0 | 0 | 0 |
| 0 | 0 | 0 | 0 |
| 0 | 0 | 0 | 0 |
| 0 | 0 | 0 | 0 |
| 0 | 0 | 0 | 0 |
| 0 | 0 | 0 | 0 |
| 0 | 0 | 0 | 0 |
| 0 | 0 | 0 | 0 |
| 0 | 0 | 0 | 0 |
| 0 | 0 | 0 | 0 |
| 0 | 0 | 0 | 0 |
| 0 | 0 | 0 | 0 |
| 0 | 0 | 0 | 0 |
| 0 | 0 | 0 | 0 |
| 0 | 0 | 0 | 0 |
| 0 | 0 | 0 | 0 |
| 0 | 0 | 0 | 0 |

| | | | |
|---|---|---|---|
| 0 | 0 | 0 | 0 |
| 0 | 0 | 0 | 0 |
| 0 | 0 | 0 | 0 |
| 0 | 0 | 0 | 0 |
| 0 | 0 | 0 | 0 |
| 0 | 0 | 0 | 0 |
| 0 | 0 | 0 | 0 |
| 0 | 0 | 0 | 0 |
| 0 | 0 | 0 | 0 |
| 0 | 0 | 0 | 0 |
| 0 | 0 | 0 | 0 |
| 0 | 0 | 0 | 0 |
| 1 | 0 | 0 | 0 |
| 1 | 1 | 0 | 0 |
| 0 | 0 | 0 | 0 |
| 1 | 0 | 0 | 0 |
| 0 | 0 | 0 | 0 |
| 1 | 0 | 0 | 0 |
| 0 | 0 | 0 | 0 |
| 0 | 0 | 0 | 0 |
| 1 | 0 | 0 | 0 |
| 0 | 0 | 0 | 0 |
| 0 | 0 | 0 | 0 |
| 1 | 0 | 0 | 0 |
| 1 | 0 | 0 | 0 |
| 0 | 0 | 0 | 0 |
| 0 | 0 | 0 | 0 |
| 1 | 0 | 0 | 0 |
| 1 | 0 | 0 | 0 |
| 1 | 0 | 0 | 0 |
| 1 | 0 | 0 | 0 |
| 1 | 0 | 0 | 0 |
| 0 | 0 | 0 | 0 |
| 0 | 0 | 0 | 0 |
| 0 | 0 | 0 | 0 |
| 0 | 0 | 0 | 0 |
| 0 | 0 | 0 | 0 |
| 0 | 0 | 0 | 0 |
| 0 | 0 | 0 | 0 |
| 0 | 0 | 0 | 0 |
| 0 | 0 | 0 | 0 |
| 0 | 0 | 0 | 0 |
| 0 | 0 | 0 | 0 |
| 0 | 0 | 0 | 0 |
| 1 | 1 | 0 | 0 |
| 0 | 0 | 0 | 0 |
| 0 | 0 | 0 | 0 |
| 0 | 0 | 0 | 0 |
| 0 | 0 | 0 | 0 |
| 1 | 1 | 0 | 0 |
| 1 | 0 | 0 | 0 |
| 0 | 0 | 0 | 0 |
| 0 | 0 | 0 | 0 |
| 0 | 0 | 0 | 1 |

| | | | |
|---|---|---|---|
| 0 | 0 | 0 | 0 |
| 0 | 0 | 0 | 0 |
| 0 | 0 | 0 | 0 |
| 0 | 0 | 0 | 0 |
| 1 | 0 | 0 | 1 |
| 0 | 0 | 0 | 0 |
| 1 | 0 | 0 | 0 |
| 0 | 0 | 0 | 0 |
| 0 | 0 | 0 | 0 |
| 0 | 0 | 0 | 0 |
| 0 | 0 | 0 | 0 |
| 0 | 0 | 0 | 0 |
| 0 | 0 | 0 | 0 |
| 1 | 0 | 0 | 0 |
| 0 | 0 | 0 | 0 |
| 0 | 0 | 0 | 0 |
| 1 | 1 | 0 | 0 |
| 0 | 0 | 0 | 0 |
| 1 | 0 | 0 | 0 |
| 1 | 0 | 0 | 0 |
| 0 | 0 | 0 | 0 |
| 0 | 0 | 0 | 0 |
| 0 | 0 | 0 | 0 |
| 1 | 0 | 0 | 0 |
| 1 | 0 | 0 | 0 |
| 1 | 0 | 0 | 0 |
| 1 | 0 | 0 | 0 |
| 1 | 0 | 0 | 0 |
| 1 | 0 | 0 | 0 |
| 1 | 0 | 0 | 0 |
| 1 | 0 | 0 | 0 |
| 1 | 0 | 0 | 0 |
| 1 | 0 | 0 | 0 |
| 1 | 0 | 0 | 0 |
| 0 | 0 | 0 | 0 |
| 0 | 0 | 0 | 0 |
| 0 | 0 | 0 | 0 |
| 1 | 0 | 0 | 0 |
| 1 | 0 | 0 | 0 |
| 0 | 0 | 0 | 0 |
| 0 | 0 | 0 | 0 |
| 1 | 0 | 0 | 0 |
| 0 | 0 | 0 | 0 |
| 1 | 0 | 0 | 0 |
| 1 | 0 | 0 | 0 |
| 1 | 0 | 0 | 0 |
| 1 | 0 | 0 | 0 |
| 1 | 0 | 0 | 0 |
| 0 | 0 | 0 | 0 |
| 0 | 0 | 0 | 0 |
| 1 | 0 | 0 | 0 |
| 0 | 0 | 0 | 0 |
| 1 | 0 | 0 | 0 |
| 1 | 0 | 0 | 0 |

| | | | |
|---|---|---|---|
| 1 | 0 | 0 | 0 |
| 1 | 0 | 0 | 0 |
| 1 | 0 | 0 | 0 |
| 1 | 0 | 0 | 0 |
| 1 | 0 | 0 | 0 |
| 1 | 0 | 0 | 0 |
| 0 | 0 | 0 | 0 |
| 1 | 0 | 0 | 0 |
| 1 | 0 | 0 | 0 |
| 1 | 0 | 0 | 0 |
| 1 | 0 | 0 | 0 |
| 1 | 0 | 0 | 0 |
| 1 | 0 | 0 | 0 |
| 1 | 0 | 0 | 0 |
| 0 | 0 | 0 | 0 |
| 0 | 0 | 0 | 0 |
| 1 | 1 | 0 | 0 |
| 1 | 1 | 0 | 0 |
| 0 | 0 | 0 | 0 |
| 0 | 0 | 0 | 0 |
| 0 | 0 | 0 | 0 |
| 1 | 1 | 0 | 1 |
| 0 | 0 | 0 | 0 |
| 0 | 0 | 0 | 0 |
| 0 | 0 | 0 | 0 |
| 0 | 0 | 0 | 0 |
| 0 | 0 | 0 | 0 |
| 0 | 0 | 0 | 0 |
| 0 | 0 | 0 | 0 |
| 1 | 0 | 1 | 0 |
| 0 | 0 | 0 | 0 |
| 0 | 0 | 0 | 0 |
| 0 | 0 | 0 | 0 |
| 0 | 0 | 0 | 0 |
| 0 | 0 | 0 | 0 |
| 0 | 0 | 0 | 0 |
| 0 | 0 | 0 | 0 |
| 0 | 0 | 0 | 0 |
| 0 | 0 | 0 | 0 |
| 0 | 0 | 0 | 0 |
| 1 | 1 | 0 | 0 |
| 1 | 1 | 0 | 0 |
| 1 | 0 | 0 | 0 |
| 1 | 1 | 0 | 0 |
| 1 | 0 | 0 | 0 |
| 1 | 1 | 0 | 0 |
| 1 | 0 | 0 | 0 |
| 1 | 0 | 0 | 0 |
| 0 | 0 | 0 | 0 |
| 0 | 0 | 0 | 0 |
| 1 | 0 | 0 | 0 |
| 1 | 0 | 0 | 0 |
| 1 | 0 | 0 | 0 |
| 0 | 0 | 0 | 0 |

| | | | |
|---|---|---|---|
| 0 | 0 | 0 | 0 |
| 0 | 0 | 0 | 0 |
| 0 | 0 | 0 | 0 |
| 1 | 1 | 1 | 0 |
| 0 | 0 | 0 | 0 |
| 0 | 0 | 0 | 0 |
| 0 | 0 | 0 | 0 |
| 0 | 0 | 0 | 0 |
| 0 | 1 | 0 | 0 |
| 0 | 1 | 0 | 0 |
| 0 | 1 | 0 | 0 |
| 0 | 0 | 0 | 0 |
| 0 | 0 | 0 | 0 |
| 0 | 0 | 0 | 0 |
| 1 | 0 | 0 | 0 |
| 1 | 0 | 0 | 0 |
| 0 | 0 | 0 | 0 |
| 0 | 0 | 0 | 0 |
| 1 | 0 | 0 | 0 |
| 0 | 0 | 0 | 0 |
| 1 | 0 | 0 | 0 |
| 1 | 0 | 0 | 0 |
| 0 | 0 | 0 | 0 |
| 0 | 0 | 0 | 0 |
| 0 | 0 | 0 | 0 |
| 0 | 0 | 0 | 0 |
| 0 | 0 | 0 | 0 |
| 0 | 0 | 1 | 0 |
| 0 | 0 | 0 | 0 |
| 0 | 1 | 0 | 0 |
| 0 | 0 | 0 | 0 |
| 0 | 0 | 0 | 0 |
| 0 | 0 | 0 | 0 |
| 0 | 0 | 0 | 0 |
| 0 | 0 | 0 | 0 |
| 0 | 0 | 0 | 0 |
| 1 | 0 | 0 | 0 |
| 1 | 0 | 0 | 0 |
| 1 | 0 | 0 | 0 |
| 0 | 0 | 0 | 0 |
| 0 | 0 | 0 | 0 |
| 0 | 0 | 0 | 0 |
| 0 | 0 | 0 | 0 |
| 0 | 0 | 0 | 0 |
| 0 | 0 | 0 | 0 |
| 0 | 0 | 0 | 0 |
| 0 | 0 | 0 | 0 |
| 0 | 0 | 0 | 1 |
| 0 | 0 | 0 | 0 |
| 0 | 0 | 0 | 0 |
| 0 | 0 | 0 | 0 |
| 0 | 0 | 0 | 0 |
| 1 | 1 | 0 | 0 |
| 0 | 0 | 0 | 0 |

| | | | |
|---|---|---|---|
| 0 | 0 | 0 | 0 |
| 0 | 0 | 0 | 0 |
| 0 | 0 | 0 | 0 |
| 0 | 0 | 0 | 0 |
| 1 | 0 | 0 | 0 |
| 0 | 0 | 0 | 0 |
| 0 | 0 | 0 | 0 |
| 0 | 0 | 0 | 0 |
| 0 | 0 | 0 | 0 |
| 0 | 0 | 0 | 0 |
| 0 | 0 | 0 | 0 |
| 0 | 0 | 0 | 0 |
| 0 | 0 | 0 | 0 |
| 0 | 0 | 0 | 0 |
| 0 | 0 | 0 | 0 |
| 0 | 0 | 0 | 0 |
| 0 | 0 | 0 | 0 |
| 0 | 0 | 0 | 0 |
| 0 | 0 | 0 | 0 |
| 0 | 0 | 0 | 0 |
| 0 | 0 | 0 | 0 |
| 0 | 0 | 0 | 0 |
| 0 | 0 | 0 | 0 |
| 0 | 0 | 0 | 0 |
| 0 | 0 | 0 | 0 |
| 0 | 0 | 0 | 0 |
| 0 | 0 | 0 | 0 |
| 0 | 0 | 0 | 0 |
| 0 | 0 | 0 | 0 |
| 0 | 0 | 0 | 0 |
| 0 | 0 | 0 | 0 |
| 0 | 0 | 0 | 0 |
| 0 | 0 | 0 | 0 |
| 0 | 0 | 0 | 0 |
| 0 | 0 | 0 | 0 |
| 0 | 0 | 0 | 0 |
| 0 | 0 | 0 | 0 |
| 0 | 0 | 0 | 0 |
| 0 | 0 | 0 | 0 |
| 0 | 0 | 0 | 0 |
| 1 | 1 | 0 | 0 |
| 0 | 0 | 0 | 0 |
| 0 | 0 | 0 | 0 |
| 0 | 0 | 0 | 0 |
| 0 | 0 | 0 | 0 |
| 1 | 0 | 0 | 0 |
| 1 | 0 | 0 | 0 |
| 0 | 0 | 0 | 0 |
| 1 | 0 | 0 | 0 |
| 0 | 0 | 0 | 0 |
| 0 | 0 | 0 | 0 |
| 1 | 0 | 0 | 0 |
| 1 | 0 | 0 | 0 |
| 0 | 0 | 0 | 0 |

| | | | |
|---|---|---|---|
| 0 | 0 | 0 | 0 |
| 0 | 0 | 0 | 0 |
| 0 | 0 | 0 | 0 |
| 0 | 0 | 0 | 0 |
| 1 | 0 | 0 | 0 |
| 0 | 0 | 0 | 0 |
| 0 | 0 | 0 | 0 |
| 0 | 0 | 0 | 0 |
| 0 | 0 | 0 | 0 |
| 0 | 0 | 0 | 0 |
| 0 | 0 | 0 | 0 |
| 0 | 0 | 0 | 0 |
| 0 | 0 | 0 | 0 |
| 0 | 0 | 0 | 0 |
| 0 | 0 | 0 | 0 |
| 1 | 1 | 0 | 1 |
| 0 | 0 | 0 | 0 |
| 0 | 0 | 0 | 0 |
| 0 | 0 | 0 | 0 |
| 0 | 0 | 0 | 0 |
| 0 | 0 | 0 | 0 |
| 0 | 0 | 0 | 0 |
| 0 | 0 | 0 | 0 |
| 0 | 0 | 0 | 0 |
| 0 | 0 | 0 | 0 |
| 0 | 0 | 0 | 0 |
| 0 | 0 | 0 | 0 |
| 1 | 0 | 0 | 0 |
| 1 | 0 | 0 | 0 |
| 0 | 0 | 0 | 0 |
| 1 | 1 | 0 | 0 |
| 0 | 0 | 0 | 0 |
| 0 | 0 | 0 | 0 |
| 0 | 0 | 0 | 0 |
| 1 | 1 | 0 | 0 |
| 1 | 1 | 0 | 0 |
| 0 | 0 | 0 | 0 |
| 1 | 0 | 0 | 0 |
| 0 | 0 | 0 | 0 |
| 1 | 1 | 0 | 0 |
| 0 | 1 | 0 | 0 |
| 0 | 0 | 0 | 0 |
| 0 | 0 | 0 | 0 |
| 0 | 0 | 0 | 0 |
| 1 | 1 | 0 | 0 |
| 1 | 1 | 0 | 0 |
| 1 | 1 | 0 | 0 |
| 0 | 0 | 0 | 0 |
| 0 | 0 | 0 | 0 |
| 0 | 0 | 0 | 0 |
| 0 | 0 | 0 | 0 |
| 0 | 0 | 0 | 0 |
| 0 | 0 | 0 | 0 |
| 0 | 0 | 0 | 0 |

| | | | |
|---|---|---|---|
| 0 | 0 | 0 | 0 |
| 0 | 0 | 0 | 0 |
| 1 | 0 | 0 | 0 |
| 1 | 0 | 0 | 0 |
| 0 | 0 | 0 | 0 |
| 1 | 0 | 0 | 0 |
| 0 | 0 | 0 | 0 |
| 0 | 0 | 0 | 0 |
| 0 | 0 | 0 | 0 |
| 0 | 1 | 0 | 0 |
| 0 | 0 | 0 | 0 |
| 0 | 0 | 0 | 0 |
| 0 | 0 | 0 | 0 |
| 0 | 0 | 0 | 0 |
| 0 | 0 | 0 | 0 |
| 0 | 0 | 0 | 0 |
| 0 | 0 | 0 | 0 |
| 1 | 0 | 0 | 0 |
| 0 | 0 | 0 | 0 |
| 1 | 0 | 0 | 0 |
| 0 | 0 | 0 | 0 |
| 1 | 0 | 0 | 0 |
| 0 | 0 | 0 | 0 |
| 1 | 0 | 0 | 0 |
| 0 | 0 | 0 | 0 |
| 0 | 0 | 0 | 0 |
| 0 | 0 | 0 | 0 |
| 0 | 0 | 0 | 0 |
| 0 | 0 | 0 | 0 |
| 0 | 0 | 0 | 0 |
| 0 | 0 | 0 | 0 |
| 1 | 0 | 0 | 0 |
| 0 | 0 | 0 | 0 |
| 0 | 0 | 0 | 0 |
| 0 | 0 | 0 | 0 |
| 0 | 0 | 0 | 0 |
| 0 | 0 | 0 | 0 |
| 0 | 0 | 0 | 0 |
| 1 | 1 | 0 | 0 |
| 1 | 1 | 0 | 0 |
| 1 | 1 | 0 | 0 |
| 1 | 1 | 0 | 0 |
| 1 | 1 | 0 | 0 |
| 1 | 0 | 0 | 0 |
| 1 | 0 | 0 | 0 |
| 1 | 0 | 0 | 0 |
| 0 | 0 | 0 | 0 |
| 0 | 0 | 0 | 0 |
| 0 | 0 | 0 | 0 |
| 0 | 0 | 0 | 0 |
| 0 | 0 | 0 | 0 |
| 0 | 0 | 0 | 0 |
| 0 | 0 | 0 | 0 |
| 0 | 0 | 0 | 0 |

| | | | |
|---|---|---|---|
| 0 | 0 | 0 | 0 |
| 0 | 0 | 0 | 0 |
| 0 | 0 | 0 | 0 |
| 0 | 0 | 0 | 0 |
| 0 | 0 | 0 | 0 |
| 1 | 0 | 0 | 0 |
| 1 | 0 | 0 | 0 |
| 0 | 0 | 0 | 0 |
| 0 | 0 | 0 | 0 |
| 0 | 0 | 0 | 0 |
| 0 | 0 | 0 | 0 |
| 1 | 0 | 0 | 0 |
| 0 | 0 | 1 | 0 |
| 0 | 0 | 0 | 0 |
| 0 | 0 | 0 | 0 |
| 0 | 0 | 0 | 0 |
| 1 | 0 | 0 | 0 |
| 0 | 0 | 0 | 0 |
| 1 | 0 | 0 | 0 |
| 1 | 0 | 0 | 0 |
| 0 | 0 | 0 | 0 |
| 1 | 0 | 0 | 0 |
| 1 | 1 | 0 | 0 |
| 0 | 0 | 0 | 0 |
| 1 | 1 | 0 | 0 |
| 1 | 1 | 0 | 0 |
| 1 | 1 | 0 | 0 |
| 0 | 0 | 0 | 0 |
| 1 | 0 | 0 | 0 |
| 0 | 0 | 0 | 0 |
| 0 | 0 | 0 | 0 |
| 0 | 0 | 0 | 0 |
| 0 | 0 | 0 | 0 |
| 0 | 0 | 0 | 0 |
| 0 | 0 | 0 | 0 |
| 0 | 0 | 0 | 0 |
| 0 | 0 | 0 | 0 |
| 0 | 0 | 0 | 0 |
| 0 | 0 | 0 | 0 |
| 0 | 0 | 0 | 0 |
| 0 | 0 | 0 | 0 |
| 0 | 0 | 0 | 0 |
| 0 | 0 | 0 | 0 |
| 0 | 0 | 0 | 0 |
| 0 | 0 | 0 | 0 |
| 0 | 0 | 0 | 0 |
| 0 | 0 | 0 | 0 |
| 0 | 0 | 0 | 0 |
| 0 | 0 | 0 | 0 |
| 0 | 0 | 0 | 0 |
| 0 | 0 | 0 | 0 |
| 0 | 0 | 0 | 0 |
| 0 | 0 | 0 | 0 |
| 0 | 0 | 0 | 0 |

| | | | |
|---|---|---|---|
| 0 | 0 | 0 | 0 |
| 0 | 0 | 0 | 0 |
| 0 | 0 | 0 | 0 |
| 0 | 0 | 0 | 0 |
| 0 | 0 | 0 | 0 |
| 0 | 0 | 0 | 0 |
| 0 | 0 | 0 | 0 |
| 0 | 0 | 0 | 0 |
| 0 | 0 | 0 | 0 |
| 0 | 0 | 0 | 0 |
| 0 | 1 | 0 | 0 |
| 0 | 1 | 0 | 0 |
| 0 | 1 | 0 | 0 |
| 0 | 1 | 0 | 0 |
| 0 | 0 | 0 | 0 |
| 0 | 1 | 0 | 0 |
| 0 | 1 | 0 | 0 |
| 1 | 1 | 0 | 0 |
| 0 | 1 | 0 | 0 |
| 1 | 0 | 0 | 0 |
| 1 | 1 | 0 | 0 |
| 1 | 1 | 0 | 0 |
| 0 | 0 | 1 | 1 |
| 0 | 0 | 0 | 0 |
| 1 | 0 | 0 | 0 |
| 0 | 0 | 0 | 0 |
| 0 | 0 | 0 | 0 |
| 0 | 0 | 0 | 0 |
| 0 | 0 | 0 | 0 |
| 0 | 0 | 0 | 0 |
| 0 | 0 | 0 | 0 |
| 0 | 0 | 0 | 0 |
| 0 | 0 | 0 | 0 |
| 0 | 0 | 0 | 0 |
| 0 | 0 | 0 | 0 |
| 0 | 0 | 0 | 0 |
| 0 | 0 | 0 | 0 |
| 0 | 0 | 0 | 0 |
| 0 | 0 | 0 | 0 |
| 0 | 0 | 0 | 0 |
| 1 | 0 | 0 | 0 |
| 0 | 0 | 0 | 0 |
| 1 | 0 | 0 | 0 |
| 1 | 0 | 0 | 0 |
| 0 | 0 | 0 | 0 |
| 1 | 0 | 0 | 0 |
| 0 | 0 | 0 | 0 |
| 0 | 0 | 0 | 0 |
| 0 | 0 | 0 | 0 |
| 0 | 0 | 0 | 0 |
| 0 | 0 | 0 | 0 |
| 0 | 0 | 0 | 0 |
| 1 | 1 | 0 | 0 |
| 1 | 1 | 0 | 0 |

| | | | |
|---|---|---|---|
| 1 | 1 | 0 | 0 |
| 0 | 0 | 0 | 0 |
| 0 | 0 | 0 | 0 |
| 0 | 0 | 0 | 0 |
| 0 | 0 | 0 | 0 |
| 0 | 0 | 0 | 0 |
| 0 | 1 | 0 | 0 |
| 0 | 0 | 0 | 0 |
| 0 | 0 | 0 | 0 |
| 0 | 1 | 0 | 0 |
| 1 | 0 | 0 | 0 |
| 0 | 0 | 0 | 0 |
| 0 | 0 | 1 | 0 |
| 0 | 0 | 1 | 0 |
| 0 | 0 | 0 | 0 |
| 0 | 0 | 0 | 0 |
| 0 | 0 | 0 | 0 |
| 0 | 1 | 0 | 0 |
| 0 | 0 | 0 | 0 |
| 0 | 0 | 0 | 0 |
| 1 | 0 | 0 | 0 |
| 0 | 0 | 0 | 1 |
| 0 | 0 | 0 | 0 |
| 0 | 0 | 0 | 0 |
| 0 | 0 | 0 | 0 |
| 0 | 0 | 0 | 0 |
| 0 | 0 | 0 | 0 |
| 0 | 0 | 0 | 0 |
| 0 | 0 | 0 | 0 |
| 0 | 0 | 0 | 0 |
| 0 | 0 | 0 | 0 |
| 0 | 0 | 0 | 0 |
| 0 | 0 | 0 | 0 |
| 0 | 0 | 0 | 0 |
| 0 | 0 | 0 | 0 |
| 0 | 0 | 0 | 0 |
| 1 | 0 | 0 | 0 |
| 0 | 0 | 0 | 0 |
| 1 | 0 | 0 | 0 |
| 0 | 0 | 0 | 0 |
| 0 | 0 | 0 | 0 |
| 0 | 0 | 0 | 0 |
| 0 | 0 | 0 | 0 |
| 0 | 0 | 0 | 0 |
| 0 | 0 | 0 | 0 |
| 0 | 0 | 0 | 0 |
| 0 | 0 | 0 | 0 |
| 0 | 0 | 0 | 0 |
| 0 | 0 | 0 | 0 |
| 1 | 0 | 0 | 0 |
| 0 | 0 | 0 | 0 |
| 0 | 0 | 0 | 0 |
| 0 | 0 | 0 | 0 |
| 0 | 0 | 0 | 0 |

| | | | |
|---|---|---|---|
| 0 | 0 | 0 | 0 |
| 0 | 0 | 0 | 0 |
| 0 | 0 | 0 | 0 |
| 0 | 0 | 0 | 0 |
| 0 | 0 | 0 | 0 |
| 0 | 0 | 0 | 0 |
| 0 | 0 | 0 | 0 |
| 0 | 0 | 0 | 0 |
| 0 | 0 | 0 | 0 |
| 0 | 0 | 0 | 0 |
| 0 | 0 | 0 | 0 |
| 0 | 0 | 0 | 0 |
| 0 | 0 | 1 | 0 |
| 1 | 0 | 0 | 0 |
| 1 | 0 | 0 | 0 |
| 0 | 0 | 0 | 0 |
| 0 | 0 | 0 | 0 |
| 0 | 0 | 0 | 0 |
| 0 | 0 | 0 | 0 |
| 0 | 0 | 0 | 0 |
| 0 | 0 | 0 | 0 |
| 0 | 0 | 0 | 0 |
| 0 | 0 | 0 | 0 |
| 1 | 1 | 0 | 1 |
| 0 | 0 | 0 | 0 |
| 1 | 1 | 0 | 0 |
| 1 | 1 | 0 | 1 |
| 0 | 0 | 0 | 0 |
| 0 | 0 | 0 | 0 |
| 0 | 0 | 0 | 0 |
| 0 | 0 | 0 | 0 |
| 0 | 0 | 0 | 0 |
| 0 | 0 | 0 | 0 |
| 0 | 0 | 0 | 0 |
| 0 | 0 | 0 | 0 |
| 1 | 0 | 0 | 0 |
| 0 | 0 | 0 | 0 |
| 0 | 0 | 0 | 0 |
| 0 | 0 | 0 | 0 |
| 0 | 0 | 0 | 0 |
| 0 | 0 | 0 | 0 |
| 0 | 0 | 0 | 0 |
| 1 | 0 | 0 | 0 |
| 1 | 0 | 0 | 0 |
| 1 | 0 | 0 | 0 |
| 1 | 0 | 0 | 0 |
| 1 | 0 | 0 | 0 |
| 0 | 0 | 0 | 0 |
| 0 | 0 | 0 | 0 |
| 1 | 0 | 0 | 0 |
| 1 | 0 | 0 | 0 |
| 0 | 0 | 0 | 0 |
| 1 | 0 | 0 | 0 |
| 1 | 0 | 0 | 0 |

| | | | |
|---|---|---|---|
| 0 | 0 | 0 | 0 |
| 1 | 0 | 0 | 0 |
| 0 | 0 | 0 | 0 |
| 0 | 0 | 0 | 0 |
| 0 | 0 | 0 | 0 |
| 0 | 0 | 0 | 0 |
| 1 | 0 | 0 | 0 |
| 0 | 0 | 0 | 0 |
| 0 | 0 | 0 | 0 |
| 0 | 0 | 0 | 0 |
| 0 | 0 | 0 | 0 |
| 1 | 1 | 0 | 0 |
| 0 | 0 | 0 | 0 |
| 0 | 0 | 0 | 0 |
| 0 | 0 | 0 | 0 |
| 0 | 0 | 0 | 0 |
| 0 | 0 | 0 | 0 |
| 0 | 0 | 0 | 0 |
| 0 | 0 | 0 | 0 |
| 0 | 0 | 0 | 0 |
| 0 | 0 | 0 | 0 |
| 0 | 0 | 0 | 0 |
| 0 | 0 | 0 | 0 |
| 0 | 0 | 0 | 0 |
| 0 | 0 | 0 | 0 |
| 0 | 0 | 0 | 0 |
| 0 | 0 | 0 | 0 |
| 0 | 0 | 0 | 0 |
| 1 | 1 | 0 | 0 |
| 1 | 0 | 0 | 0 |
| 0 | 0 | 0 | 0 |
| 0 | 0 | 0 | 0 |
| 0 | 0 | 0 | 0 |
| 0 | 0 | 0 | 0 |
| 0 | 0 | 0 | 0 |
| 0 | 0 | 0 | 0 |
| 0 | 0 | 0 | 0 |
| 0 | 0 | 0 | 0 |
| 0 | 0 | 0 | 0 |
| 0 | 0 | 0 | 0 |
| 0 | 0 | 0 | 0 |
| 0 | 0 | 0 | 0 |
| 0 | 0 | 0 | 0 |
| 0 | 0 | 0 | 0 |
| 0 | 0 | 0 | 0 |
| 0 | 0 | 0 | 0 |
| 0 | 0 | 0 | 0 |
| 0 | 0 | 0 | 0 |
| 0 | 0 | 0 | 0 |
| 0 | 0 | 0 | 0 |
| 0 | 0 | 0 | 0 |
| 0 | 0 | 0 | 0 |
| 0 | 0 | 1 | 0 |
| 0 | 0 | 0 | 0 |

| | | | |
|---|---|---|---|
| 0 | 0 | 0 | 0 |
| 0 | 0 | 0 | 0 |
| 0 | 0 | 0 | 0 |
| 0 | 0 | 0 | 0 |
| 1 | 0 | 0 | 0 |
| 0 | 0 | 0 | 0 |
| 0 | 0 | 0 | 0 |
| 1 | 1 | 0 | 0 |
| 0 | 0 | 0 | 0 |
| 1 | 1 | 0 | 0 |
| 0 | 0 | 0 | 0 |
| 0 | 0 | 0 | 0 |
| 0 | 0 | 0 | 0 |
| 0 | 0 | 0 | 0 |
| 0 | 0 | 0 | 0 |
| 0 | 0 | 0 | 0 |
| 1 | 0 | 0 | 0 |
| 0 | 0 | 0 | 0 |
| 0 | 0 | 0 | 0 |
| 1 | 0 | 0 | 0 |
| 1 | 0 | 0 | 0 |
| 0 | 0 | 0 | 0 |
| 0 | 0 | 0 | 0 |
| 0 | 0 | 0 | 0 |
| 0 | 0 | 0 | 0 |
| 0 | 0 | 0 | 0 |
| 1 | 0 | 0 | 0 |
| 0 | 0 | 0 | 0 |
| 0 | 0 | 0 | 0 |
| 1 | 0 | 0 | 0 |
| 0 | 0 | 0 | 0 |
| 0 | 0 | 0 | 0 |
| 0 | 0 | 0 | 0 |
| 1 | 1 | 0 | 0 |
| 0 | 0 | 0 | 0 |
| 0 | 0 | 0 | 0 |
| 0 | 0 | 0 | 0 |
| 0 | 0 | 0 | 0 |
| 0 | 1 | 0 | 0 |
| 0 | 0 | 0 | 0 |
| 0 | 0 | 0 | 0 |
| 0 | 0 | 0 | 0 |
| 0 | 0 | 0 | 0 |
| 1 | 1 | 0 | 0 |
| 0 | 0 | 0 | 0 |
| 0 | 0 | 0 | 0 |
| 0 | 0 | 0 | 0 |
| 0 | 0 | 0 | 0 |
| 0 | 0 | 0 | 0 |
| 0 | 0 | 0 | 0 |
| 0 | 0 | 0 | 0 |
| 1 | 0 | 0 | 0 |
| 0 | 0 | 0 | 0 |
| 0 | 0 | 0 | 0 |

| | | | |
|---|---|---|---|
| 1 | 0 | 0 | 1 |
| 1 | 0 | 0 | 1 |
| 1 | 0 | 0 | 1 |
| 0 | 0 | 0 | 0 |
| 0 | 0 | 0 | 0 |
| 0 | 0 | 0 | 0 |
| 1 | 0 | 0 | 0 |
| 0 | 0 | 0 | 0 |
| 0 | 0 | 0 | 0 |
| 0 | 0 | 0 | 0 |
| 0 | 0 | 1 | 0 |
| 0 | 0 | 0 | 0 |
| 0 | 0 | 0 | 0 |
| 0 | 0 | 0 | 0 |
| 0 | 0 | 0 | 0 |
| 0 | 0 | 0 | 0 |
| 1 | 0 | 0 | 0 |
| 0 | 0 | 0 | 0 |
| 0 | 0 | 0 | 0 |
| 0 | 0 | 0 | 0 |
| 0 | 0 | 0 | 0 |
| 0 | 0 | 0 | 0 |
| 0 | 0 | 0 | 0 |
| 1 | 1 | 0 | 0 |
| 0 | 0 | 0 | 0 |
| 0 | 0 | 0 | 0 |
| 0 | 0 | 0 | 0 |
| 0 | 0 | 0 | 0 |
| 0 | 0 | 0 | 0 |
| 0 | 0 | 0 | 0 |
| 0 | 0 | 0 | 0 |
| 0 | 0 | 0 | 0 |
| 0 | 1 | 0 | 0 |
| 0 | 0 | 0 | 0 |
| 1 | 1 | 0 | 0 |
| 0 | 0 | 0 | 0 |
| 0 | 0 | 0 | 0 |
| 0 | 0 | 0 | 0 |
| 0 | 0 | 0 | 0 |
| 0 | 0 | 0 | 0 |
| 0 | 0 | 0 | 0 |
| 1 | 0 | 0 | 1 |
| 0 | 0 | 0 | 0 |
| 1 | 0 | 0 | 1 |
| 0 | 0 | 0 | 0 |
| 1 | 0 | 0 | 1 |
| 0 | 0 | 0 | 0 |
| 0 | 0 | 0 | 0 |
| 0 | 0 | 0 | 0 |
| 1 | 0 | 0 | 0 |
| 0 | 0 | 0 | 0 |
| 0 | 0 | 0 | 0 |
| 0 | 0 | 0 | 0 |
| 0 | 0 | 0 | 0 |

| | | | |
|---|---|---|---|
| 0 | 0 | 0 | 0 |
| 0 | 0 | 0 | 0 |
| 0 | 0 | 0 | 0 |
| 0 | 0 | 0 | 0 |
| 0 | 0 | 0 | 0 |
| 0 | 0 | 0 | 0 |
| 0 | 0 | 0 | 0 |
| 0 | 0 | 0 | 0 |
| 0 | 0 | 0 | 0 |
| 0 | 0 | 0 | 0 |
| 0 | 0 | 0 | 0 |
| 0 | 0 | 0 | 0 |
| 0 | 0 | 0 | 0 |
| 0 | 0 | 0 | 0 |
| 0 | 0 | 0 | 0 |
| 0 | 0 | 0 | 0 |
| 0 | 0 | 0 | 0 |
| 0 | 0 | 0 | 0 |
| 0 | 0 | 1 | 0 |
| 0 | 0 | 0 | 0 |
| 0 | 0 | 0 | 0 |
| 0 | 0 | 0 | 0 |
| 0 | 0 | 0 | 0 |
| 0 | 0 | 0 | 0 |
| 0 | 0 | 0 | 0 |
| 0 | 0 | 0 | 0 |
| 0 | 0 | 0 | 0 |
| 0 | 0 | 0 | 0 |
| 0 | 0 | 1 | 0 |
| 0 | 0 | 0 | 0 |
| 0 | 0 | 1 | 0 |
| 1 | 0 | 0 | 0 |
| 1 | 1 | 1 | 0 |
| 1 | 1 | 1 | 0 |
| 0 | 0 | 0 | 0 |
| 0 | 0 | 0 | 0 |
| 0 | 1 | 0 | 0 |
| 1 | 1 | 1 | 0 |
| 1 | 1 | 1 | 0 |
| 0 | 0 | 1 | 0 |
| 0 | 0 | 1 | 0 |
| 1 | 0 | 0 | 0 |
| 0 | 0 | 0 | 0 |
| 0 | 0 | 0 | 0 |
| 1 | 1 | 0 | 0 |
| 0 | 0 | 0 | 0 |
| 0 | 0 | 0 | 0 |
| 0 | 0 | 0 | 0 |
| 0 | 0 | 0 | 0 |
| 0 | 0 | 0 | 0 |
| 0 | 0 | 0 | 0 |
| 0 | 0 | 0 | 0 |
| 0 | 0 | 0 | 0 |
| 0 | 0 | 0 | 0 |

| | | | |
|---|---|---|---|
| 0 | 0 | 0 | 0 |
| 0 | 0 | 0 | 0 |
| 0 | 0 | 0 | 0 |
| 0 | 0 | 0 | 0 |
| 0 | 0 | 0 | 0 |
| 0 | 0 | 0 | 0 |
| 0 | 0 | 0 | 0 |
| 0 | 0 | 0 | 0 |
| 0 | 0 | 0 | 0 |
| 0 | 0 | 0 | 0 |
| 0 | 0 | 0 | 0 |
| 0 | 0 | 0 | 0 |
| 0 | 0 | 0 | 0 |
| 0 | 0 | 0 | 0 |
| 0 | 0 | 0 | 0 |
| 0 | 0 | 0 | 0 |
| 0 | 0 | 0 | 0 |
| 0 | 0 | 0 | 0 |
| 0 | 0 | 0 | 0 |
| 1 | 0 | 0 | 0 |
| 1 | 0 | 0 | 0 |
| 0 | 0 | 0 | 0 |
| 0 | 0 | 0 | 0 |
| 0 | 0 | 0 | 0 |
| 0 | 0 | 0 | 0 |
| 1 | 0 | 0 | 0 |
| 0 | 0 | 0 | 0 |
| 0 | 0 | 0 | 0 |
| 0 | 0 | 0 | 0 |
| 1 | 0 | 0 | 0 |
| 0 | 0 | 0 | 0 |
| 0 | 0 | 0 | 0 |
| 1 | 0 | 0 | 0 |
| 1 | 0 | 0 | 0 |
| 0 | 0 | 0 | 0 |
| 0 | 0 | 0 | 0 |
| 0 | 0 | 0 | 0 |
| 0 | 0 | 0 | 0 |
| 0 | 0 | 0 | 0 |
| 0 | 0 | 0 | 0 |
| 0 | 0 | 0 | 0 |
| 0 | 0 | 0 | 0 |
| 0 | 0 | 0 | 0 |
| 0 | 0 | 0 | 0 |
| 0 | 0 | 0 | 0 |
| 0 | 0 | 0 | 0 |
| 0 | 0 | 0 | 0 |
| 0 | 1 | 0 | 0 |
| 0 | 1 | 0 | 0 |
| 0 | 0 | 0 | 0 |
| 0 | 1 | 0 | 0 |
| 0 | 1 | 0 | 0 |
| 0 | 0 | 0 | 0 |
| 0 | 0 | 0 | 0 |

| | | | |
|---|---|---|---|
| 0 | 0 | 0 | 0 |
| 0 | 0 | 0 | 0 |
| 1 | 1 | 0 | 1 |
| 0 | 0 | 0 | 0 |
| 0 | 0 | 0 | 0 |
| 0 | 0 | 0 | 0 |
| 0 | 0 | 0 | 0 |
| 0 | 1 | 0 | 0 |
| 1 | 1 | 0 | 0 |
| 1 | 1 | 0 | 0 |
| 1 | 1 | 0 | 0 |
| 1 | 1 | 0 | 0 |
| 1 | 1 | 0 | 0 |
| 0 | 1 | 0 | 0 |
| 0 | 1 | 0 | 0 |
| 0 | 0 | 0 | 0 |
| 0 | 0 | 0 | 0 |
| 0 | 0 | 0 | 0 |
| 0 | 0 | 0 | 0 |
| 0 | 0 | 0 | 0 |
| 0 | 0 | 0 | 0 |
| 0 | 0 | 0 | 0 |
| 0 | 1 | 0 | 0 |
| 0 | 1 | 0 | 0 |
| 0 | 0 | 0 | 0 |
| 0 | 0 | 1 | 0 |
| 0 | 0 | 0 | 0 |
| 0 | 0 | 0 | 0 |
| 0 | 0 | 0 | 0 |
| 0 | 1 | 0 | 0 |
| 0 | 0 | 0 | 0 |
| 0 | 0 | 0 | 0 |
| 0 | 0 | 0 | 0 |
| 0 | 0 | 0 | 0 |
| 0 | 0 | 0 | 0 |
| 0 | 0 | 0 | 0 |
| 0 | 0 | 0 | 0 |
| 0 | 0 | 0 | 0 |
| 0 | 0 | 0 | 0 |
| 1 | 0 | 0 | 0 |
| 1 | 1 | 0 | 0 |
| 1 | 0 | 0 | 0 |
| 0 | 0 | 0 | 0 |
| 0 | 1 | 0 | 0 |
| 1 | 1 | 0 | 0 |
| 0 | 1 | 0 | 0 |
| 0 | 0 | 0 | 0 |
| 1 | 0 | 0 | 0 |
| 0 | 1 | 0 | 0 |
| 0 | 0 | 0 | 0 |
| 0 | 0 | 0 | 0 |
| 0 | 0 | 0 | 0 |
| 1 | 1 | 0 | 0 |
| 0 | 0 | 0 | 0 |

| | | | |
|---|---|---|---|
| 0 | 0 | 0 | 0 |
| 0 | 0 | 0 | 0 |
| 1 | 0 | 0 | 0 |
| 1 | 1 | 0 | 0 |
| 1 | 1 | 0 | 0 |
| 0 | 0 | 0 | 0 |
| 0 | 0 | 0 | 0 |
| 0 | 0 | 0 | 0 |
| 1 | 0 | 0 | 0 |
| 0 | 0 | 0 | 0 |
| 0 | 0 | 0 | 0 |
| 1 | 0 | 0 | 1 |
| 0 | 0 | 0 | 0 |
| 1 | 0 | 0 | 0 |
| 0 | 0 | 0 | 0 |
| 0 | 0 | 0 | 0 |
| 0 | 0 | 0 | 0 |
| 0 | 0 | 0 | 0 |
| 0 | 0 | 0 | 0 |
| 0 | 0 | 0 | 0 |
| 0 | 0 | 0 | 0 |
| 1 | 0 | 0 | 1 |
| 1 | 0 | 0 | 0 |
| 0 | 0 | 0 | 0 |
| 0 | 0 | 0 | 0 |
| 0 | 0 | 0 | 0 |
| 0 | 0 | 0 | 0 |
| 0 | 0 | 0 | 0 |
| 1 | 0 | 0 | 0 |
| 1 | 0 | 0 | 0 |
| 0 | 0 | 0 | 0 |
| 0 | 0 | 0 | 0 |
| 0 | 0 | 0 | 0 |
| 0 | 0 | 0 | 0 |
| 0 | 0 | 0 | 0 |
| 0 | 0 | 0 | 0 |
| 1 | 0 | 0 | 0 |
| 0 | 0 | 0 | 0 |
| 0 | 0 | 0 | 0 |
| 1 | 0 | 0 | 0 |
| 1 | 0 | 0 | 0 |
| 1 | 0 | 0 | 0 |
| 1 | 0 | 0 | 0 |
| 0 | 0 | 0 | 0 |
| 0 | 0 | 0 | 0 |
| 1 | 0 | 0 | 0 |
| 0 | 0 | 0 | 0 |
| 0 | 0 | 0 | 0 |
| 0 | 0 | 0 | 0 |
| 0 | 0 | 0 | 0 |
| 0 | 0 | 0 | 0 |
| 1 | 0 | 0 | 0 |
| 1 | 0 | 0 | 0 |
| 1 | 0 | 0 | 0 |

| | | | |
|---|---|---|---|
| 1 | 0 | 0 | 0 |
| 0 | 0 | 0 | 0 |
| 0 | 0 | 0 | 0 |
| 0 | 0 | 0 | 0 |
| 1 | 0 | 0 | 0 |
| 0 | 0 | 0 | 0 |
| 1 | 0 | 0 | 0 |
| 0 | 0 | 0 | 0 |
| 0 | 0 | 0 | 0 |
| 0 | 0 | 0 | 0 |
| 0 | 0 | 0 | 0 |
| 0 | 0 | 0 | 0 |
| 0 | 0 | 0 | 0 |
| 0 | 0 | 0 | 0 |
| 0 | 0 | 0 | 0 |
| 1 | 0 | 0 | 0 |
| 0 | 0 | 0 | 0 |
| 0 | 0 | 0 | 0 |
| 0 | 0 | 0 | 0 |
| 0 | 0 | 0 | 0 |
| 0 | 0 | 0 | 0 |
| 0 | 0 | 0 | 0 |
| 1 | 0 | 0 | 0 |
| 1 | 0 | 0 | 0 |
| 0 | 0 | 0 | 0 |
| 0 | 0 | 0 | 0 |
| 0 | 0 | 0 | 0 |
| 0 | 0 | 0 | 0 |
| 0 | 0 | 0 | 0 |
| 0 | 0 | 0 | 0 |
| 0 | 0 | 0 | 0 |
| 0 | 0 | 0 | 0 |
| 1 | 0 | 0 | 0 |
| 0 | 0 | 0 | 0 |
| 0 | 0 | 0 | 0 |
| 0 | 0 | 0 | 0 |
| 0 | 0 | 0 | 0 |
| 0 | 0 | 0 | 0 |
| 0 | 0 | 0 | 0 |
| 0 | 0 | 0 | 0 |
| 0 | 0 | 0 | 0 |
| 0 | 0 | 0 | 0 |
| 0 | 0 | 0 | 0 |
| 0 | 0 | 0 | 0 |
| 0 | 1 | 0 | 0 |
| 0 | 0 | 0 | 0 |
| 0 | 0 | 0 | 0 |
| 0 | 0 | 0 | 0 |
| 0 | 0 | 0 | 0 |
| 0 | 0 | 0 | 0 |
| 0 | 0 | 0 | 0 |
| 0 | 0 | 0 | 0 |
| 0 | 0 | 0 | 0 |
| 1 | 0 | 0 | 0 |

| | | | |
|---|---|---|---|
| 0 | 0 | 0 | 0 |
| 0 | 0 | 0 | 0 |
| 0 | 0 | 0 | 0 |
| 0 | 0 | 0 | 0 |
| 0 | 0 | 0 | 0 |
| 0 | 0 | 0 | 0 |
| 0 | 0 | 0 | 0 |
| 0 | 0 | 0 | 0 |
| 0 | 0 | 0 | 0 |
| 1 | 0 | 0 | 0 |
| 0 | 0 | 0 | 0 |
| 0 | 0 | 0 | 0 |
| 1 | 1 | 0 | 0 |
| 1 | 1 | 0 | 0 |
| 1 | 1 | 0 | 0 |
| 1 | 1 | 0 | 0 |
| 0 | 0 | 0 | 0 |
| 0 | 0 | 0 | 0 |
| 0 | 0 | 0 | 0 |
| 0 | 0 | 0 | 0 |
| 0 | 0 | 0 | 0 |
| 0 | 0 | 0 | 0 |
| 0 | 0 | 0 | 0 |
| 0 | 0 | 0 | 0 |
| 0 | 0 | 0 | 0 |
| 0 | 0 | 0 | 1 |
| 0 | 0 | 0 | 0 |
| 0 | 0 | 0 | 0 |
| 0 | 0 | 0 | 0 |
| 0 | 0 | 0 | 0 |
| 0 | 0 | 0 | 0 |
| 0 | 0 | 0 | 0 |
| 0 | 0 | 0 | 0 |
| 0 | 0 | 0 | 0 |
| 0 | 0 | 0 | 0 |
| 0 | 0 | 0 | 0 |
| 0 | 0 | 0 | 0 |
| 1 | 1 | 0 | 0 |
| 0 | 0 | 0 | 0 |
| 0 | 0 | 0 | 0 |
| 1 | 1 | 0 | 0 |
| 0 | 0 | 0 | 0 |
| 0 | 0 | 0 | 0 |
| 1 | 0 | 0 | 1 |
| 0 | 0 | 0 | 0 |
| 0 | 0 | 0 | 0 |
| 0 | 0 | 0 | 0 |
| 0 | 0 | 0 | 0 |
| 0 | 0 | 0 | 0 |
| 0 | 0 | 0 | 0 |
| 0 | 0 | 0 | 0 |
| 0 | 0 | 1 | 0 |
| 0 | 0 | 1 | 0 |
| 0 | 0 | 1 | 0 |

| | | | |
|---|---|---|---|
| 0 | 0 | 1 | 0 |
| 0 | 0 | 0 | 0 |
| 0 | 0 | 0 | 0 |
| 0 | 0 | 0 | 0 |
| 0 | 0 | 0 | 0 |
| 0 | 0 | 0 | 0 |
| 0 | 0 | 0 | 0 |
| 0 | 0 | 0 | 0 |
| 0 | 0 | 0 | 0 |
| 0 | 0 | 0 | 0 |
| 0 | 0 | 0 | 0 |
| 0 | 0 | 0 | 0 |
| 0 | 0 | 0 | 0 |
| 0 | 0 | 0 | 0 |
| 0 | 0 | 0 | 0 |
| 0 | 0 | 0 | 0 |
| 0 | 0 | 0 | 0 |
| 0 | 0 | 0 | 0 |
| 0 | 0 | 0 | 0 |
| 0 | 0 | 0 | 0 |
| 0 | 0 | 0 | 0 |
| 0 | 0 | 0 | 0 |
| 0 | 0 | 0 | 0 |
| 0 | 1 | 0 | 1 |
| 0 | 0 | 0 | 0 |
| 0 | 0 | 0 | 0 |
| 0 | 0 | 0 | 0 |
| 0 | 0 | 0 | 0 |
| 0 | 0 | 0 | 0 |
| 0 | 0 | 0 | 0 |
| 0 | 0 | 0 | 0 |
| 0 | 0 | 0 | 0 |
| 0 | 0 | 0 | 0 |
| 0 | 0 | 0 | 0 |
| 0 | 0 | 0 | 0 |
| 0 | 0 | 0 | 0 |
| 0 | 0 | 0 | 0 |
| 0 | 0 | 0 | 0 |
| 0 | 0 | 0 | 0 |
| 0 | 0 | 0 | 0 |
| 0 | 0 | 0 | 0 |
| 0 | 0 | 0 | 0 |
| 0 | 0 | 0 | 0 |
| 0 | 0 | 0 | 0 |
| 0 | 0 | 0 | 0 |
| 0 | 0 | 0 | 0 |
| 1 | 0 | 0 | 0 |
| 1 | 0 | 0 | 0 |
| 0 | 0 | 0 | 0 |
| 0 | 1 | 0 | 0 |
| 1 | 1 | 0 | 1 |
| 1 | 1 | 0 | 1 |
| 1 | 1 | 0 | 1 |
| 0 | 0 | 0 | 0 |

| | | | |
|---|---|---|---|
| 1 | 1 | 0 | 0 |
| 0 | 0 | 0 | 0 |
| 0 | 0 | 0 | 0 |
| 0 | 0 | 0 | 0 |
| 0 | 0 | 0 | 0 |
| 0 | 0 | 0 | 0 |
| 0 | 0 | 0 | 0 |
| 0 | 0 | 0 | 0 |
| 0 | 0 | 0 | 0 |
| 0 | 0 | 0 | 0 |
| 0 | 0 | 0 | 0 |
| 0 | 1 | 0 | 0 |
| 0 | 0 | 0 | 0 |
| 0 | 0 | 0 | 0 |
| 0 | 1 | 0 | 0 |
| 0 | 0 | 0 | 0 |
| 1 | 1 | 0 | 0 |
| 0 | 0 | 0 | 0 |
| 0 | 0 | 0 | 0 |
| 1 | 0 | 0 | 0 |
| 0 | 0 | 0 | 0 |
| 0 | 0 | 0 | 0 |
| 0 | 0 | 0 | 0 |
| 0 | 0 | 0 | 0 |
| 0 | 0 | 0 | 0 |
| 0 | 0 | 0 | 0 |
| 0 | 0 | 0 | 0 |
| 0 | 0 | 0 | 0 |
| 0 | 0 | 0 | 0 |
| 0 | 0 | 0 | 0 |
| 0 | 0 | 0 | 0 |
| 0 | 0 | 0 | 0 |
| 0 | 0 | 0 | 0 |
| 1 | 0 | 0 | 0 |
| 0 | 0 | 0 | 0 |
| 0 | 1 | 0 | 0 |
| 0 | 0 | 0 | 0 |
| 0 | 0 | 0 | 0 |
| 0 | 0 | 0 | 0 |
| 0 | 0 | 0 | 0 |
| 0 | 0 | 0 | 0 |
| 0 | 0 | 0 | 0 |
| 0 | 0 | 0 | 0 |
| 0 | 0 | 0 | 0 |
| 0 | 0 | 0 | 0 |
| 0 | 0 | 0 | 0 |
| 0 | 0 | 0 | 0 |
| 0 | 0 | 0 | 0 |
| 1 | 1 | 0 | 0 |
| 0 | 0 | 0 | 0 |
| 0 | 0 | 0 | 0 |
| 0 | 0 | 0 | 0 |
| 0 | 0 | 0 | 0 |
| 0 | 0 | 0 | 0 |

| | | | |
|---|---|---|---|
| 0 | 0 | 0 | 0 |
| 0 | 0 | 0 | 0 |
| 0 | 0 | 0 | 0 |
| 0 | 0 | 0 | 0 |
| 0 | 0 | 0 | 0 |
| 0 | 0 | 0 | 0 |
| 0 | 0 | 0 | 0 |
| 0 | 1 | 0 | 0 |
| 0 | 0 | 0 | 0 |
| 0 | 0 | 0 | 0 |
| 0 | 0 | 0 | 0 |
| 0 | 0 | 0 | 0 |
| 0 | 0 | 1 | 0 |
| 0 | 0 | 0 | 0 |
| 0 | 0 | 1 | 0 |
| 0 | 0 | 1 | 0 |
| 0 | 0 | 0 | 0 |
| 0 | 0 | 0 | 0 |
| 0 | 0 | 0 | 0 |
| 0 | 0 | 0 | 0 |
| 0 | 1 | 0 | 0 |
| 0 | 1 | 0 | 0 |
| 0 | 0 | 0 | 0 |
| 0 | 0 | 0 | 0 |
| 0 | 1 | 0 | 0 |
| 0 | 0 | 0 | 0 |
| 0 | 0 | 0 | 0 |
| 0 | 1 | 0 | 0 |
| 0 | 0 | 0 | 0 |
| 0 | 0 | 0 | 0 |
| 0 | 0 | 0 | 0 |
| 0 | 0 | 0 | 0 |
| 1 | 1 | 0 | 0 |
| 1 | 1 | 0 | 0 |
| 1 | 1 | 0 | 0 |
| 0 | 1 | 0 | 0 |
| 1 | 0 | 0 | 0 |
| 0 | 1 | 0 | 0 |
| 1 | 1 | 0 | 0 |
| 0 | 0 | 0 | 0 |
| 0 | 0 | 0 | 0 |
| 0 | 0 | 0 | 0 |
| 0 | 0 | 0 | 0 |
| 0 | 0 | 0 | 0 |
| 0 | 0 | 0 | 0 |
| 0 | 0 | 0 | 0 |
| 0 | 0 | 0 | 0 |
| 0 | 0 | 0 | 0 |
| 0 | 0 | 0 | 0 |
| 1 | 1 | 0 | 0 |
| 1 | 1 | 0 | 0 |
| 0 | 0 | 0 | 0 |
| 0 | 0 | 0 | 0 |
| 0 | 0 | 0 | 0 |

| | | | |
|---|---|---|---|
| 0 | 0 | 0 | 0 |
| 0 | 0 | 0 | 0 |
| 0 | 0 | 0 | 0 |
| 0 | 0 | 0 | 0 |
| 0 | 0 | 0 | 0 |
| 0 | 0 | 0 | 0 |
| 1 | 1 | 0 | 0 |
| 1 | 1 | 0 | 0 |
| 0 | 0 | 0 | 0 |
| 0 | 0 | 0 | 0 |
| 0 | 0 | 0 | 0 |
| 0 | 0 | 0 | 0 |
| 0 | 0 | 0 | 0 |
| 0 | 0 | 0 | 0 |
| 0 | 0 | 0 | 0 |
| 0 | 0 | 0 | 0 |
| 0 | 0 | 0 | 0 |
| 0 | 0 | 0 | 0 |
| 0 | 0 | 0 | 0 |
| 0 | 0 | 0 | 0 |
| 0 | 0 | 0 | 0 |
| 0 | 0 | 0 | 0 |
| 0 | 0 | 0 | 0 |
| 0 | 0 | 0 | 0 |
| 0 | 0 | 0 | 0 |
| 0 | 0 | 0 | 0 |
| 0 | 0 | 0 | 0 |
| 1 | 1 | 0 | 0 |
| 0 | 0 | 0 | 0 |
| 0 | 0 | 0 | 0 |
| 0 | 0 | 0 | 0 |
| 0 | 0 | 0 | 0 |
| 0 | 0 | 0 | 0 |
| 0 | 0 | 0 | 0 |
| 0 | 0 | 0 | 0 |
| 0 | 0 | 0 | 0 |
| 0 | 0 | 0 | 0 |
| 0 | 0 | 0 | 0 |
| 0 | 0 | 0 | 0 |
| 0 | 0 | 0 | 0 |
| 0 | 0 | 0 | 0 |
| 0 | 0 | 0 | 0 |
| 0 | 0 | 0 | 0 |
| 0 | 0 | 0 | 0 |
| 0 | 0 | 0 | 0 |
| 0 | 0 | 0 | 0 |
| 0 | 0 | 0 | 0 |
| 0 | 0 | 0 | 0 |
| 0 | 0 | 0 | 0 |
| 0 | 0 | 0 | 0 |
| 0 | 0 | 0 | 0 |
| 0 | 0 | 0 | 0 |
| 0 | 0 | 0 | 0 |
| 0 | 0 | 0 | 0 |

| | | | |
|---|---|---|---|
| 0 | 0 | 0 | 0 |
| 0 | 0 | 0 | 0 |
| 0 | 0 | 0 | 0 |
| 0 | 0 | 0 | 0 |
| 0 | 0 | 0 | 0 |
| 0 | 0 | 0 | 0 |
| 0 | 0 | 0 | 0 |
| 0 | 0 | 0 | 0 |
| 0 | 0 | 0 | 0 |
| 0 | 0 | 0 | 0 |
| 1 | 0 | 0 | 0 |
| 1 | 1 | 0 | 0 |
| 1 | 0 | 0 | 0 |
| 0 | 0 | 0 | 0 |
| 0 | 0 | 0 | 0 |
| 0 | 0 | 0 | 0 |
| 0 | 0 | 0 | 0 |
| 0 | 0 | 0 | 0 |
| 0 | 0 | 0 | 0 |
| 0 | 0 | 0 | 0 |
| 0 | 0 | 0 | 0 |
| 0 | 0 | 0 | 0 |
| 0 | 0 | 0 | 0 |
| 0 | 0 | 0 | 0 |
| 0 | 0 | 0 | 0 |
| 0 | 0 | 0 | 0 |
| 0 | 0 | 0 | 0 |
| 0 | 0 | 0 | 0 |
| 0 | 0 | 0 | 0 |
| 0 | 0 | 0 | 0 |
| 0 | 0 | 0 | 0 |
| 0 | 1 | 0 | 0 |
| 0 | 0 | 0 | 0 |
| 0 | 0 | 0 | 0 |
| 1 | 0 | 0 | 0 |
| 1 | 0 | 0 | 0 |
| 1 | 0 | 0 | 0 |
| 0 | 0 | 0 | 0 |
| 0 | 0 | 0 | 0 |
| 0 | 0 | 0 | 0 |
| 0 | 0 | 0 | 0 |
| 0 | 0 | 0 | 0 |
| 0 | 0 | 0 | 0 |
| 0 | 0 | 0 | 0 |
| 0 | 0 | 0 | 0 |
| 0 | 0 | 0 | 0 |
| 0 | 0 | 0 | 0 |
| 0 | 0 | 0 | 0 |
| 0 | 0 | 0 | 0 |
| 0 | 0 | 0 | 0 |
| 0 | 0 | 0 | 0 |
| 0 | 0 | 0 | 0 |
| 0 | 0 | 0 | 1 |
| 0 | 0 | 0 | 0 |

| | | | |
|---|---|---|---|
| 0 | 0 | 0 | 0 |
| 0 | 0 | 0 | 0 |
| 0 | 0 | 0 | 0 |
| 0 | 0 | 0 | 0 |
| 0 | 0 | 0 | 0 |
| 0 | 0 | 0 | 0 |
| 0 | 0 | 0 | 0 |
| 0 | 0 | 0 | 0 |
| 0 | 0 | 0 | 0 |
| 0 | 0 | 0 | 0 |
| 0 | 0 | 0 | 0 |
| 1 | 0 | 0 | 0 |
| 0 | 0 | 0 | 0 |
| 0 | 0 | 0 | 0 |
| 0 | 0 | 1 | 0 |
| 0 | 0 | 0 | 0 |
| 0 | 0 | 0 | 0 |
| 0 | 0 | 0 | 0 |
| 0 | 0 | 0 | 0 |
| 0 | 0 | 0 | 0 |
| 0 | 0 | 0 | 0 |
| 0 | 0 | 0 | 0 |
| 0 | 0 | 0 | 0 |
| 0 | 0 | 0 | 0 |
| 0 | 0 | 0 | 0 |
| 0 | 0 | 0 | 0 |
| 0 | 0 | 0 | 0 |
| 0 | 0 | 0 | 0 |
| 0 | 0 | 0 | 0 |
| 0 | 0 | 0 | 0 |
| 0 | 0 | 0 | 0 |
| 0 | 0 | 0 | 0 |
| 0 | 0 | 0 | 0 |
| 1 | 0 | 0 | 0 |
| 0 | 0 | 0 | 0 |
| 1 | 0 | 0 | 1 |
| 0 | 0 | 0 | 0 |
| 0 | 0 | 0 | 0 |
| 0 | 0 | 0 | 0 |
| 0 | 0 | 0 | 0 |
| 0 | 0 | 0 | 0 |
| 0 | 0 | 0 | 0 |
| 0 | 0 | 0 | 0 |
| 0 | 1 | 0 | 0 |
| 0 | 0 | 0 | 0 |
| 0 | 0 | 0 | 0 |
| 0 | 0 | 0 | 0 |
| 0 | 0 | 0 | 0 |
| 0 | 0 | 0 | 0 |
| 0 | 0 | 0 | 0 |
| 0 | 0 | 0 | 0 |
| 0 | 0 | 0 | 0 |
| 0 | 0 | 0 | 0 |
| 0 | 0 | 0 | 0 |

| | | | |
|---|---|---|---|
| 0 | 0 | 0 | 0 |
| 0 | 0 | 0 | 0 |
| 0 | 0 | 0 | 0 |
| 0 | 0 | 0 | 0 |
| 0 | 0 | 0 | 0 |
| 0 | 1 | 0 | 0 |
| 0 | 0 | 0 | 0 |
| 0 | 0 | 0 | 0 |
| 0 | 0 | 0 | 0 |
| 0 | 0 | 0 | 0 |
| 0 | 0 | 0 | 0 |
| 0 | 0 | 0 | 0 |
| 0 | 0 | 0 | 0 |
| 0 | 0 | 0 | 0 |
| 0 | 0 | 0 | 0 |
| 0 | 1 | 0 | 0 |
| 1 | 1 | 0 | 0 |
| 0 | 0 | 1 | 0 |
| 0 | 0 | 0 | 0 |
| 0 | 0 | 0 | 0 |
| 0 | 0 | 0 | 0 |
| 0 | 0 | 0 | 0 |
| 0 | 0 | 0 | 0 |
| 0 | 0 | 0 | 0 |
| 0 | 0 | 0 | 0 |
| 0 | 0 | 0 | 0 |
| 0 | 0 | 0 | 0 |
| 1 | 0 | 0 | 0 |
| 0 | 0 | 0 | 0 |
| 0 | 0 | 0 | 0 |
| 0 | 0 | 1 | 0 |
| 0 | 0 | 0 | 0 |
| 0 | 0 | 0 | 0 |
| 0 | 0 | 0 | 0 |
| 0 | 0 | 0 | 0 |
| 0 | 0 | 0 | 0 |
| 0 | 0 | 0 | 0 |
| 1 | 1 | 0 | 0 |
| 0 | 0 | 0 | 0 |
| 0 | 0 | 0 | 0 |
| 0 | 0 | 0 | 0 |
| 1 | 0 | 0 | 0 |
| 1 | 0 | 0 | 0 |
| 0 | 0 | 0 | 0 |
| 0 | 0 | 0 | 0 |
| 0 | 0 | 0 | 0 |
| 0 | 0 | 0 | 0 |
| 0 | 0 | 0 | 0 |
| 0 | 0 | 0 | 0 |
| 0 | 0 | 0 | 0 |
| 0 | 0 | 0 | 0 |
| 0 | 0 | 0 | 0 |
| 0 | 0 | 0 | 0 |
| 0 | 0 | 0 | 0 |

| | | | |
|---|---|---|---|
| 0 | 0 | 0 | 0 |
| 0 | 0 | 0 | 0 |
| 0 | 0 | 0 | 0 |
| 0 | 0 | 0 | 0 |
| 0 | 0 | 0 | 0 |
| 0 | 0 | 0 | 0 |
| 0 | 0 | 0 | 0 |
| 0 | 0 | 0 | 0 |
| 0 | 0 | 0 | 0 |
| 0 | 0 | 0 | 0 |
| 0 | 0 | 0 | 0 |
| 0 | 0 | 0 | 0 |
| 1 | 0 | 0 | 0 |
| 0 | 0 | 0 | 0 |
| 0 | 0 | 0 | 0 |
| 0 | 0 | 0 | 0 |
| 0 | 0 | 0 | 0 |
| 0 | 0 | 0 | 0 |
| 0 | 0 | 0 | 0 |
| 0 | 0 | 0 | 0 |
| 0 | 0 | 0 | 0 |
| 0 | 0 | 0 | 0 |
| 0 | 0 | 0 | 0 |
| 1 | 0 | 0 | 0 |
| 0 | 0 | 0 | 0 |
| 0 | 0 | 0 | 0 |
| 0 | 0 | 0 | 0 |
| 1 | 1 | 0 | 1 |
| 0 | 0 | 0 | 0 |
| 0 | 0 | 0 | 0 |
| 0 | 0 | 0 | 0 |
| 0 | 0 | 0 | 0 |
| 0 | 0 | 0 | 0 |
| 0 | 0 | 0 | 0 |
| 0 | 0 | 0 | 0 |
| 0 | 0 | 0 | 0 |
| 0 | 0 | 0 | 0 |
| 0 | 0 | 0 | 0 |
| 0 | 0 | 0 | 0 |
| 0 | 0 | 0 | 0 |
| 0 | 0 | 0 | 0 |
| 0 | 0 | 0 | 0 |
| 0 | 0 | 0 | 0 |
| 0 | 0 | 0 | 0 |
| 0 | 0 | 0 | 0 |
| 0 | 0 | 0 | 0 |
| 0 | 0 | 0 | 0 |
| 0 | 0 | 0 | 0 |
| 0 | 0 | 0 | 0 |
| 0 | 0 | 0 | 0 |
| 0 | 0 | 0 | 0 |
| 0 | 0 | 0 | 0 |
| 0 | 0 | 0 | 0 |
| 0 | 0 | 0 | 0 |

| | | | |
|---|---|---|---|
| 0 | 0 | 0 | 0 |
| 0 | 0 | 0 | 0 |
| 0 | 0 | 0 | 0 |
| 0 | 0 | 0 | 0 |
| 0 | 0 | 0 | 0 |
| 0 | 0 | 0 | 0 |
| 0 | 0 | 0 | 0 |
| 0 | 0 | 0 | 0 |
| 1 | 0 | 0 | 0 |
| 1 | 0 | 0 | 0 |
| 1 | 0 | 0 | 0 |
| 0 | 0 | 0 | 0 |
| 1 | 0 | 0 | 0 |
| 0 | 0 | 0 | 0 |
| 0 | 0 | 0 | 0 |
| 1 | 0 | 0 | 0 |
| 0 | 0 | 0 | 0 |
| 0 | 0 | 0 | 0 |
| 0 | 0 | 0 | 0 |
| 0 | 0 | 0 | 0 |
| 0 | 0 | 0 | 0 |
| 0 | 0 | 0 | 0 |
| 0 | 0 | 0 | 0 |
| 0 | 0 | 0 | 0 |
| 0 | 0 | 0 | 0 |
| 0 | 0 | 0 | 0 |
| 0 | 0 | 0 | 0 |
| 1 | 0 | 0 | 0 |
| 1 | 0 | 0 | 0 |
| 0 | 0 | 0 | 0 |
| 0 | 0 | 0 | 0 |
| 0 | 0 | 0 | 0 |
| 1 | 0 | 0 | 0 |
| 0 | 0 | 0 | 0 |
| 0 | 0 | 0 | 0 |
| 0 | 0 | 0 | 0 |
| 0 | 0 | 0 | 0 |
| 0 | 0 | 0 | 0 |
| 0 | 0 | 0 | 0 |
| 0 | 0 | 0 | 0 |
| 1 | 0 | 0 | 0 |
| 1 | 0 | 0 | 0 |
| 0 | 0 | 0 | 0 |
| 0 | 0 | 0 | 0 |
| 0 | 0 | 0 | 0 |
| 0 | 0 | 0 | 0 |
| 0 | 0 | 0 | 0 |
| 0 | 0 | 0 | 0 |
| 0 | 0 | 0 | 0 |
| 0 | 0 | 0 | 0 |
| 0 | 0 | 0 | 0 |
| 0 | 0 | 0 | 0 |
| 0 | 0 | 0 | 0 |
| 0 | 0 | 0 | 0 |

| | | | |
|---|---|---|---|
| 0 | 0 | 0 | 0 |
| 0 | 0 | 0 | 0 |
| 0 | 0 | 0 | 0 |
| 0 | 0 | 0 | 0 |
| 0 | 0 | 0 | 0 |
| 0 | 0 | 0 | 0 |
| 0 | 0 | 0 | 0 |
| 0 | 0 | 0 | 0 |
| 0 | 0 | 0 | 0 |
| 0 | 0 | 0 | 0 |
| 1 | 0 | 0 | 0 |
| 0 | 0 | 0 | 0 |
| 1 | 0 | 0 | 0 |
| 0 | 0 | 0 | 0 |
| 0 | 0 | 0 | 0 |
| 0 | 0 | 0 | 0 |
| 0 | 0 | 0 | 0 |
| 0 | 1 | 0 | 0 |
| 0 | 0 | 0 | 0 |
| 0 | 0 | 0 | 0 |
| 0 | 0 | 0 | 0 |
| 0 | 0 | 0 | 0 |
| 1 | 0 | 0 | 0 |
| 0 | 0 | 0 | 0 |
| 0 | 0 | 0 | 0 |
| 1 | 0 | 0 | 0 |
| 1 | 0 | 0 | 0 |
| 1 | 0 | 0 | 0 |
| 0 | 0 | 0 | 0 |
| 0 | 0 | 0 | 0 |
| 0 | 0 | 0 | 0 |
| 0 | 0 | 0 | 0 |
| 0 | 0 | 0 | 0 |
| 0 | 0 | 0 | 0 |
| 0 | 0 | 0 | 0 |
| 0 | 0 | 0 | 0 |
| 0 | 0 | 0 | 0 |
| 0 | 0 | 0 | 0 |
| 0 | 0 | 0 | 0 |
| 0 | 0 | 0 | 0 |
| 1 | 1 | 0 | 0 |
| 0 | 0 | 0 | 0 |
| 0 | 0 | 0 | 0 |
| 0 | 0 | 0 | 0 |
| 0 | 0 | 0 | 0 |
| 0 | 0 | 0 | 0 |
| 0 | 0 | 0 | 0 |
| 1 | 0 | 0 | 0 |
| 0 | 0 | 0 | 0 |
| 0 | 0 | 0 | 0 |
| 0 | 0 | 0 | 0 |
| 0 | 0 | 0 | 0 |
| 0 | 0 | 0 | 0 |
| 0 | 0 | 0 | 0 |

| | | | |
|---|---|---|---|
| 1 | 0 | 0 | 0 |
| 0 | 0 | 0 | 0 |
| 1 | 0 | 0 | 0 |
| 0 | 0 | 0 | 0 |
| 0 | 0 | 0 | 0 |
| 0 | 0 | 0 | 0 |
| 0 | 0 | 0 | 0 |
| 0 | 0 | 0 | 0 |
| 1 | 1 | 0 | 0 |
| 0 | 0 | 0 | 0 |
| 1 | 1 | 0 | 0 |
| 1 | 1 | 0 | 0 |
| 0 | 0 | 0 | 0 |
| 1 | 1 | 0 | 0 |
| 0 | 0 | 0 | 0 |
| 0 | 0 | 0 | 0 |
| 0 | 0 | 0 | 0 |
| 0 | 0 | 0 | 0 |
| 0 | 0 | 0 | 0 |
| 0 | 0 | 0 | 0 |
| 1 | 0 | 0 | 0 |
| 0 | 0 | 0 | 0 |
| 0 | 0 | 0 | 0 |
| 0 | 0 | 0 | 0 |
| 1 | 0 | 0 | 0 |
| 0 | 0 | 0 | 0 |
| 1 | 0 | 0 | 0 |
| 1 | 0 | 0 | 0 |
| 1 | 1 | 0 | 0 |
| 0 | 0 | 0 | 0 |
| 1 | 1 | 0 | 0 |
| 1 | 1 | 0 | 0 |
| 1 | 1 | 0 | 0 |
| 0 | 0 | 0 | 0 |
| 1 | 1 | 0 | 0 |
| 0 | 0 | 0 | 0 |
| 1 | 0 | 0 | 0 |
| 0 | 0 | 0 | 0 |
| 1 | 1 | 0 | 0 |
| 1 | 1 | 0 | 0 |
| 1 | 0 | 0 | 0 |
| 0 | 0 | 0 | 0 |
| 0 | 0 | 0 | 0 |
| 1 | 0 | 0 | 0 |
| 0 | 0 | 0 | 0 |
| 0 | 0 | 0 | 0 |
| 1 | 0 | 0 | 0 |
| 0 | 0 | 0 | 0 |
| 1 | 0 | 0 | 0 |
| 0 | 0 | 0 | 0 |
| 0 | 0 | 0 | 0 |
| 0 | 0 | 0 | 0 |
| 0 | 0 | 0 | 0 |
| 0 | 0 | 0 | 0 |

| | | | |
|---|---|---|---|
| 0 | 0 | 0 | 0 |
| 0 | 0 | 0 | 0 |
| 0 | 0 | 0 | 0 |
| 0 | 0 | 0 | 0 |
| 0 | 0 | 0 | 0 |
| 0 | 0 | 0 | 0 |
| 0 | 0 | 0 | 0 |
| 0 | 0 | 0 | 0 |
| 0 | 0 | 0 | 0 |
| 1 | 1 | 0 | 0 |
| 1 | 1 | 0 | 0 |
| 0 | 0 | 0 | 0 |
| 0 | 0 | 0 | 0 |
| 0 | 0 | 0 | 0 |
| 1 | 0 | 1 | 0 |
| 0 | 0 | 0 | 0 |
| 0 | 0 | 0 | 0 |
| 1 | 0 | 1 | 0 |
| 0 | 0 | 0 | 0 |
| 0 | 0 | 0 | 0 |
| 0 | 0 | 0 | 0 |
| 0 | 0 | 1 | 0 |
| 1 | 0 | 1 | 0 |
| 0 | 0 | 1 | 0 |
| 0 | 0 | 1 | 0 |
| 0 | 0 | 1 | 1 |
| 0 | 0 | 1 | 1 |
| 0 | 0 | 0 | 0 |
| 0 | 0 | 0 | 0 |
| 0 | 0 | 0 | 0 |
| 0 | 0 | 0 | 0 |
| 0 | 0 | 0 | 0 |
| 0 | 0 | 0 | 0 |
| 0 | 0 | 0 | 0 |
| 0 | 0 | 0 | 0 |
| 0 | 0 | 0 | 0 |
| 0 | 0 | 0 | 0 |
| 0 | 0 | 0 | 0 |
| 0 | 0 | 0 | 0 |
| 0 | 0 | 0 | 0 |
| 0 | 0 | 0 | 0 |
| 0 | 0 | 0 | 0 |
| 0 | 0 | 0 | 0 |
| 0 | 0 | 1 | 0 |
| 0 | 0 | 0 | 0 |
| 0 | 0 | 0 | 0 |
| 0 | 0 | 0 | 0 |
| 0 | 0 | 0 | 0 |
| 0 | 0 | 0 | 0 |
| 0 | 0 | 0 | 0 |
| 1 | 0 | 0 | 0 |
| 0 | 0 | 0 | 0 |
| 0 | 0 | 0 | 0 |
| 0 | 0 | 0 | 0 |

| | | | |
|---|---|---|---|
| 0 | 0 | 0 | 0 |
| 0 | 0 | 0 | 0 |
| 0 | 0 | 0 | 0 |
| 1 | 0 | 0 | 0 |
| 0 | 0 | 0 | 0 |
| 0 | 0 | 0 | 0 |
| 0 | 0 | 0 | 0 |
| 0 | 0 | 0 | 0 |
| 1 | 0 | 0 | 0 |
| 1 | 0 | 0 | 0 |
| 0 | 0 | 0 | 0 |
| 0 | 0 | 0 | 0 |
| 0 | 0 | 0 | 0 |
| 0 | 0 | 0 | 0 |
| 0 | 0 | 0 | 0 |
| 0 | 0 | 0 | 0 |
| 0 | 0 | 1 | 0 |
| 0 | 0 | 0 | 0 |
| 0 | 0 | 0 | 0 |
| 0 | 0 | 0 | 0 |
| 0 | 0 | 0 | 0 |
| 0 | 0 | 0 | 0 |
| 0 | 0 | 0 | 0 |
| 0 | 0 | 0 | 0 |
| 0 | 0 | 0 | 0 |
| 0 | 0 | 0 | 0 |
| 0 | 0 | 0 | 0 |
| 0 | 0 | 0 | 0 |
| 0 | 0 | 0 | 0 |
| 0 | 0 | 0 | 0 |
| 0 | 0 | 0 | 0 |
| 0 | 0 | 0 | 0 |
| 0 | 0 | 0 | 0 |
| 0 | 0 | 0 | 0 |
| 0 | 0 | 0 | 0 |
| 0 | 0 | 0 | 0 |
| 0 | 0 | 0 | 0 |
| 0 | 0 | 0 | 0 |
| 0 | 0 | 0 | 0 |
| 0 | 0 | 0 | 0 |
| 0 | 0 | 0 | 0 |
| 0 | 0 | 0 | 0 |
| 0 | 0 | 0 | 0 |
| 0 | 0 | 0 | 0 |
| 0 | 0 | 0 | 0 |
| 0 | 0 | 0 | 0 |
| 0 | 0 | 0 | 0 |
| 0 | 0 | 0 | 0 |
| 0 | 0 | 0 | 0 |
| 0 | 1 | 0 | 0 |
| 0 | 0 | 0 | 0 |
| 0 | 0 | 0 | 0 |
| 0 | 0 | 0 | 0 |
| 0 | 0 | 0 | 0 |

| | | | |
|---|---|---|---|
| 0 | 0 | 1 | 0 |
| 0 | 0 | 0 | 0 |
| 0 | 0 | 0 | 0 |
| 0 | 0 | 0 | 0 |
| 0 | 0 | 0 | 0 |
| 0 | 0 | 0 | 0 |
| 0 | 0 | 0 | 0 |
| 1 | 0 | 0 | 0 |
| 0 | 0 | 0 | 0 |
| 0 | 0 | 0 | 0 |
| 0 | 0 | 0 | 0 |
| 1 | 0 | 0 | 0 |
| 0 | 0 | 0 | 0 |
| 0 | 0 | 0 | 0 |
| 0 | 0 | 0 | 0 |
| 0 | 0 | 0 | 0 |
| 0 | 0 | 0 | 0 |
| 0 | 0 | 0 | 0 |
| 0 | 0 | 0 | 0 |
| 0 | 0 | 0 | 0 |
| 0 | 0 | 0 | 0 |
| 0 | 0 | 0 | 0 |
| 0 | 0 | 0 | 0 |
| 0 | 0 | 0 | 0 |
| 0 | 0 | 0 | 0 |
| 0 | 0 | 0 | 0 |
| 0 | 0 | 0 | 0 |
| 0 | 0 | 0 | 0 |
| 0 | 0 | 0 | 0 |
| 0 | 0 | 0 | 0 |
| 0 | 0 | 0 | 0 |
| 0 | 0 | 0 | 0 |
| 0 | 0 | 0 | 0 |
| 0 | 0 | 0 | 0 |
| 0 | 0 | 0 | 0 |
| 0 | 0 | 0 | 0 |
| 0 | 0 | 0 | 0 |
| 0 | 0 | 0 | 0 |
| 0 | 0 | 0 | 0 |
| 0 | 0 | 0 | 0 |
| 0 | 0 | 0 | 0 |
| 0 | 0 | 0 | 0 |
| 0 | 0 | 0 | 0 |
| 0 | 0 | 0 | 0 |
| 0 | 0 | 0 | 0 |
| 0 | 0 | 0 | 0 |
| 0 | 0 | 0 | 0 |
| 0 | 0 | 0 | 0 |
| 0 | 0 | 0 | 0 |
| 0 | 0 | 0 | 0 |
| 0 | 0 | 0 | 0 |
| 0 | 0 | 0 | 0 |
| 0 | 0 | 0 | 0 |
| 0 | 0 | 0 | 0 |

| | | | |
|---|---|---|---|
| 0 | 0 | 0 | 0 |
| 0 | 0 | 0 | 0 |
| 0 | 0 | 0 | 0 |
| 0 | 0 | 0 | 0 |
| 0 | 0 | 0 | 0 |
| 0 | 0 | 0 | 0 |
| 0 | 0 | 0 | 0 |
| 0 | 0 | 0 | 0 |
| 0 | 0 | 0 | 0 |
| 0 | 0 | 0 | 0 |
| 0 | 0 | 0 | 0 |
| 0 | 0 | 0 | 0 |
| 0 | 0 | 0 | 0 |
| 0 | 0 | 0 | 0 |
| 0 | 0 | 0 | 0 |
| 0 | 0 | 0 | 0 |
| 0 | 0 | 0 | 0 |
| 0 | 0 | 0 | 0 |
| 0 | 0 | 0 | 1 |
| 0 | 1 | 0 | 1 |
| 0 | 0 | 0 | 0 |
| 0 | 0 | 0 | 0 |
| 0 | 0 | 0 | 0 |
| 0 | 0 | 0 | 0 |
| 0 | 0 | 0 | 0 |
| 1 | 0 | 0 | 0 |
| 1 | 0 | 0 | 0 |
| 0 | 0 | 0 | 0 |
| 0 | 0 | 0 | 0 |
| 0 | 0 | 0 | 0 |
| 0 | 0 | 0 | 0 |
| 0 | 0 | 0 | 0 |
| 0 | 0 | 0 | 0 |
| 1 | 0 | 0 | 1 |
| 1 | 0 | 0 | 0 |
| 1 | 0 | 0 | 1 |
| 1 | 0 | 0 | 0 |
| 0 | 0 | 0 | 0 |
| 0 | 0 | 0 | 0 |
| 0 | 0 | 0 | 0 |
| 0 | 0 | 0 | 0 |
| 1 | 0 | 0 | 0 |
| 1 | 0 | 0 | 1 |
| 1 | 0 | 0 | 0 |
| 0 | 0 | 0 | 0 |
| 0 | 0 | 0 | 0 |
| 0 | 0 | 0 | 0 |
| 0 | 0 | 0 | 0 |
| 0 | 0 | 0 | 0 |
| 1 | 0 | 0 | 0 |
| 0 | 0 | 0 | 0 |
| 0 | 0 | 0 | 0 |
| 0 | 0 | 0 | 0 |
| 0 | 0 | 0 | 0 |

| | | | |
|---|---|---|---|
| 0 | 0 | 0 | 0 |
| 0 | 0 | 0 | 0 |
| 0 | 0 | 0 | 0 |
| 1 | 0 | 0 | 0 |
| 1 | 0 | 0 | 0 |
| 0 | 0 | 0 | 0 |
| 1 | 0 | 0 | 0 |
| 0 | 0 | 0 | 0 |
| 0 | 0 | 0 | 0 |
| 0 | 0 | 0 | 0 |
| 0 | 0 | 0 | 0 |
| 1 | 0 | 1 | 0 |
| 0 | 0 | 0 | 0 |
| 0 | 0 | 0 | 0 |
| 0 | 0 | 0 | 0 |
| 0 | 0 | 1 | 0 |
| 0 | 0 | 0 | 0 |
| 0 | 0 | 0 | 0 |
| 0 | 0 | 0 | 0 |
| 0 | 0 | 1 | 0 |
| 1 | 0 | 1 | 0 |
| 1 | 0 | 0 | 0 |
| 1 | 0 | 0 | 0 |
| 1 | 0 | 0 | 0 |
| 0 | 0 | 1 | 0 |
| 0 | 0 | 0 | 0 |
| 1 | 0 | 0 | 0 |
| 0 | 0 | 0 | 0 |
| 0 | 0 | 0 | 0 |
| 0 | 0 | 0 | 0 |
| 0 | 0 | 0 | 0 |
| 0 | 0 | 0 | 0 |
| 0 | 0 | 0 | 0 |
| 0 | 0 | 0 | 0 |
| 0 | 0 | 0 | 0 |
| 0 | 0 | 0 | 0 |
| 0 | 0 | 0 | 1 |
| 0 | 0 | 0 | 1 |
| 0 | 0 | 0 | 1 |
| 0 | 0 | 0 | 0 |
| 0 | 0 | 0 | 0 |
| 0 | 0 | 0 | 0 |
| 0 | 0 | 0 | 0 |
| 0 | 0 | 0 | 0 |
| 0 | 0 | 0 | 0 |
| 0 | 0 | 0 | 0 |
| 0 | 0 | 0 | 0 |
| 0 | 0 | 0 | 0 |
| 0 | 0 | 0 | 0 |
| 0 | 0 | 0 | 0 |
| 0 | 0 | 0 | 0 |
| 0 | 0 | 0 | 0 |
| 0 | 0 | 0 | 0 |
| 0 | 0 | 0 | 0 |

| | | | |
|---|---|---|---|
| 0 | 0 | 0 | 0 |
| 0 | 0 | 0 | 0 |
| 0 | 0 | 0 | 0 |
| 0 | 0 | 0 | 0 |
| 0 | 0 | 0 | 0 |
| 0 | 0 | 0 | 0 |
| 0 | 0 | 0 | 0 |
| 0 | 0 | 0 | 0 |
| 0 | 0 | 0 | 0 |
| 0 | 0 | 0 | 0 |
| 0 | 0 | 0 | 0 |
| 0 | 0 | 0 | 0 |
| 0 | 0 | 0 | 0 |
| 0 | 0 | 0 | 0 |
| 0 | 0 | 0 | 0 |
| 0 | 0 | 0 | 0 |
| 0 | 1 | 0 | 0 |
| 0 | 1 | 0 | 0 |
| 0 | 0 | 0 | 0 |
| 0 | 0 | 0 | 0 |
| 0 | 0 | 0 | 0 |
| 0 | 0 | 0 | 0 |
| 0 | 0 | 0 | 0 |
| 0 | 0 | 0 | 0 |
| 0 | 0 | 0 | 0 |
| 0 | 0 | 0 | 0 |
| 0 | 0 | 0 | 0 |
| 0 | 0 | 0 | 0 |
| 0 | 0 | 0 | 0 |
| 0 | 0 | 0 | 0 |
| 0 | 0 | 0 | 0 |
| 1 | 0 | 0 | 0 |
| 0 | 0 | 0 | 0 |
| 0 | 0 | 0 | 0 |
| 0 | 0 | 0 | 0 |
| 0 | 0 | 0 | 0 |
| 0 | 0 | 0 | 0 |
| 0 | 0 | 0 | 0 |
| 0 | 0 | 0 | 0 |
| 0 | 0 | 0 | 0 |
| 0 | 0 | 0 | 0 |
| 0 | 0 | 0 | 0 |
| 0 | 0 | 0 | 0 |
| 0 | 0 | 0 | 0 |
| 0 | 0 | 0 | 0 |
| 0 | 0 | 0 | 0 |
| 1 | 1 | 0 | 0 |
| 0 | 0 | 0 | 0 |
| 0 | 0 | 0 | 0 |
| 0 | 0 | 0 | 0 |
| 0 | 0 | 0 | 0 |
| 0 | 0 | 0 | 0 |
| 1 | 0 | 0 | 0 |
| 0 | 0 | 0 | 0 |

| | | | |
|---|---|---|---|
| 0 | 0 | 0 | 0 |
| 0 | 0 | 0 | 0 |
| 0 | 0 | 0 | 0 |
| 0 | 0 | 0 | 0 |
| 0 | 0 | 0 | 0 |
| 0 | 0 | 0 | 0 |
| 1 | 0 | 0 | 0 |
| 0 | 0 | 0 | 0 |
| 0 | 0 | 0 | 0 |
| 0 | 0 | 0 | 0 |
| 0 | 0 | 0 | 0 |
| 0 | 0 | 0 | 0 |
| 0 | 0 | 0 | 0 |
| 0 | 0 | 0 | 0 |
| 0 | 0 | 0 | 0 |
| 0 | 0 | 0 | 0 |
| 0 | 0 | 0 | 0 |
| 0 | 0 | 0 | 0 |
| 0 | 0 | 0 | 0 |
| 0 | 0 | 0 | 0 |
| 0 | 0 | 0 | 0 |
| 0 | 0 | 0 | 0 |
| 0 | 0 | 0 | 0 |
| 0 | 0 | 0 | 0 |
| 0 | 0 | 0 | 0 |
| 0 | 0 | 0 | 0 |
| 0 | 0 | 0 | 0 |
| 0 | 0 | 0 | 0 |
| 0 | 0 | 0 | 0 |
| 0 | 0 | 0 | 0 |
| 0 | 0 | 0 | 0 |
| 0 | 0 | 0 | 0 |
| 0 | 0 | 0 | 0 |
| 0 | 0 | 0 | 0 |
| 0 | 0 | 0 | 0 |
| 0 | 0 | 0 | 0 |
| 0 | 0 | 0 | 0 |
| 0 | 0 | 0 | 0 |
| 0 | 0 | 0 | 0 |
| 0 | 0 | 0 | 0 |
| 0 | 0 | 0 | 0 |
| 0 | 0 | 0 | 0 |
| 0 | 0 | 0 | 0 |
| 0 | 0 | 0 | 0 |
| 0 | 0 | 0 | 0 |
| 0 | 0 | 0 | 0 |
| 0 | 0 | 0 | 0 |
| 0 | 0 | 0 | 0 |
| 0 | 0 | 0 | 0 |
| 0 | 0 | 0 | 0 |
| 0 | 0 | 0 | 0 |
| 0 | 0 | 0 | 0 |
| 0 | 0 | 0 | 0 |
| 0 | 0 | 0 | 0 |

| | | | |
|---|---|---|---|
| 0 | 0 | 0 | 0 |
| 0 | 0 | 0 | 0 |
| 0 | 0 | 0 | 0 |
| 0 | 0 | 0 | 0 |
| 0 | 0 | 0 | 0 |
| 0 | 0 | 0 | 0 |
| 0 | 0 | 0 | 0 |
| 0 | 0 | 0 | 0 |
| 0 | 0 | 0 | 0 |
| 0 | 0 | 0 | 0 |
| 0 | 0 | 0 | 0 |
| 0 | 0 | 0 | 0 |
| 0 | 0 | 0 | 0 |
| 0 | 0 | 0 | 0 |
| 0 | 0 | 0 | 0 |
| 0 | 0 | 0 | 0 |
| 0 | 0 | 0 | 0 |
| 0 | 0 | 0 | 0 |
| 0 | 0 | 0 | 0 |
| 0 | 0 | 0 | 0 |
| 0 | 0 | 0 | 0 |
| 0 | 0 | 0 | 0 |
| 0 | 0 | 0 | 0 |
| 0 | 0 | 0 | 0 |
| 1 | 1 | 0 | 0 |
| 0 | 0 | 0 | 0 |
| 0 | 0 | 0 | 0 |
| 0 | 0 | 0 | 0 |
| 0 | 0 | 0 | 0 |
| 0 | 0 | 0 | 0 |
| 0 | 0 | 0 | 0 |
| 0 | 0 | 0 | 0 |
| 0 | 0 | 0 | 0 |
| 0 | 0 | 0 | 0 |
| 0 | 0 | 0 | 0 |
| 1 | 0 | 0 | 0 |
| 1 | 0 | 0 | 1 |
| 1 | 0 | 0 | 0 |
| 0 | 0 | 0 | 0 |
| 1 | 0 | 0 | 0 |
| 1 | 1 | 0 | 0 |
| 1 | 1 | 0 | 0 |
| 0 | 0 | 0 | 0 |
| 0 | 0 | 0 | 0 |
| 0 | 0 | 0 | 0 |
| 0 | 0 | 0 | 0 |
| 0 | 0 | 0 | 0 |
| 0 | 0 | 0 | 0 |
| 0 | 0 | 0 | 0 |
| 0 | 0 | 0 | 0 |
| 0 | 0 | 0 | 0 |
| 0 | 0 | 0 | 0 |
| 0 | 0 | 0 | 0 |
| 0 | 0 | 0 | 0 |

| | | | |
|---|---|---|---|
| 0 | 0 | 0 | 0 |
| 0 | 0 | 0 | 0 |
| 0 | 0 | 0 | 0 |
| 0 | 0 | 0 | 0 |
| 0 | 0 | 0 | 0 |
| 0 | 0 | 0 | 0 |
| 0 | 0 | 0 | 0 |
| 0 | 0 | 0 | 0 |
| 0 | 0 | 0 | 0 |
| 0 | 0 | 0 | 0 |
| 0 | 0 | 0 | 0 |
| 0 | 0 | 0 | 0 |
| 0 | 0 | 0 | 0 |
| 1 | 0 | 0 | 0 |
| 0 | 0 | 0 | 0 |
| 0 | 0 | 0 | 0 |
| 1 | 1 | 0 | 0 |
| 0 | 0 | 0 | 0 |
| 0 | 0 | 0 | 0 |
| 0 | 0 | 0 | 0 |
| 0 | 0 | 0 | 0 |
| 0 | 0 | 0 | 0 |
| 0 | 0 | 0 | 0 |
| 1 | 0 | 0 | 0 |
| 0 | 0 | 0 | 0 |
| 0 | 0 | 0 | 0 |
| 0 | 0 | 0 | 0 |
| 0 | 0 | 0 | 0 |
| 0 | 0 | 0 | 0 |
| 0 | 0 | 0 | 0 |
| 0 | 0 | 0 | 0 |
| 1 | 0 | 0 | 0 |
| 0 | 0 | 0 | 0 |
| 0 | 0 | 0 | 0 |
| 0 | 0 | 0 | 0 |
| 0 | 0 | 0 | 0 |
| 0 | 0 | 0 | 0 |
| 0 | 0 | 0 | 0 |
| 0 | 0 | 0 | 0 |
| 0 | 0 | 0 | 0 |
| 0 | 0 | 0 | 0 |
| 0 | 0 | 0 | 0 |
| 0 | 0 | 0 | 0 |
| 0 | 0 | 0 | 0 |
| 0 | 0 | 0 | 0 |
| 0 | 0 | 0 | 0 |
| 0 | 0 | 0 | 0 |
| 0 | 0 | 0 | 0 |
| 0 | 0 | 0 | 0 |
| 0 | 0 | 0 | 0 |
| 0 | 0 | 0 | 0 |
| 0 | 0 | 0 | 0 |
| 0 | 0 | 0 | 0 |
| 0 | 0 | 0 | 0 |

| | | | |
|---|---|---|---|
| 0 | 0 | 0 | 0 |
| 0 | 0 | 0 | 0 |
| 0 | 0 | 0 | 0 |
| 0 | 0 | 0 | 0 |
| 0 | 0 | 0 | 0 |
| 0 | 0 | 0 | 0 |
| 0 | 0 | 0 | 0 |
| 0 | 0 | 0 | 0 |
| 0 | 0 | 0 | 0 |
| 0 | 0 | 0 | 0 |
| 0 | 0 | 0 | 0 |
| 0 | 0 | 0 | 0 |
| 0 | 0 | 0 | 0 |
| 0 | 0 | 0 | 0 |
| 0 | 0 | 0 | 0 |
| 0 | 0 | 0 | 0 |
| 0 | 0 | 0 | 0 |
| 1 | 0 | 0 | 0 |
| 0 | 0 | 0 | 0 |
| 0 | 0 | 0 | 0 |
| 0 | 0 | 0 | 0 |
| 0 | 0 | 0 | 0 |
| 0 | 0 | 0 | 0 |
| 0 | 0 | 0 | 0 |
| 1 | 0 | 0 | 0 |
| 0 | 0 | 0 | 0 |
| 0 | 0 | 0 | 0 |
| 0 | 0 | 0 | 0 |
| 0 | 0 | 0 | 0 |
| 0 | 0 | 0 | 0 |
| 0 | 0 | 0 | 0 |
| 0 | 0 | 0 | 0 |
| 0 | 0 | 0 | 0 |
| 0 | 0 | 0 | 0 |
| 0 | 0 | 0 | 0 |
| 0 | 0 | 0 | 0 |
| 0 | 0 | 0 | 0 |
| 0 | 0 | 0 | 0 |
| 0 | 0 | 0 | 0 |
| 0 | 0 | 0 | 0 |
| 0 | 0 | 0 | 0 |
| 0 | 0 | 0 | 0 |
| 0 | 0 | 0 | 0 |
| 0 | 0 | 0 | 0 |
| 0 | 0 | 0 | 0 |
| 0 | 0 | 0 | 0 |
| 0 | 0 | 0 | 0 |
| 0 | 0 | 0 | 0 |
| 0 | 0 | 0 | 0 |
| 0 | 0 | 0 | 0 |
| 0 | 0 | 0 | 0 |
| 0 | 0 | 0 | 0 |
| 0 | 0 | 0 | 0 |
| 0 | 0 | 0 | 0 |

| | | | |
|---|---|---|---|
| 0 | 0 | 0 | 0 |
| 0 | 0 | 0 | 0 |
| 0 | 0 | 0 | 0 |
| 0 | 0 | 0 | 0 |
| 0 | 0 | 0 | 0 |
| 0 | 0 | 0 | 0 |
| 0 | 0 | 0 | 0 |
| 0 | 0 | 0 | 0 |
| 0 | 0 | 0 | 0 |
| 0 | 0 | 0 | 0 |
| 0 | 0 | 0 | 0 |
| 0 | 0 | 0 | 0 |
| 0 | 0 | 0 | 0 |
| 0 | 0 | 0 | 0 |
| 1 | 1 | 0 | 0 |
| 1 | 1 | 0 | 0 |
| 0 | 0 | 0 | 0 |
| 0 | 0 | 0 | 0 |
| 0 | 0 | 0 | 0 |
| 0 | 0 | 0 | 0 |
| 0 | 0 | 0 | 0 |
| 0 | 0 | 0 | 0 |
| 0 | 0 | 0 | 0 |
| 0 | 0 | 0 | 0 |
| 0 | 0 | 0 | 0 |
| 0 | 0 | 0 | 0 |
| 0 | 0 | 0 | 0 |
| 1 | 0 | 0 | 0 |
| 0 | 0 | 0 | 0 |
| 1 | 0 | 0 | 0 |
| 0 | 0 | 0 | 0 |
| 0 | 0 | 0 | 0 |
| 0 | 0 | 0 | 0 |
| 0 | 0 | 0 | 0 |
| 0 | 0 | 0 | 0 |
| 0 | 0 | 0 | 0 |
| 1 | 0 | 0 | 0 |
| 0 | 0 | 0 | 0 |
| 0 | 0 | 0 | 0 |
| 0 | 0 | 0 | 0 |
| 0 | 0 | 0 | 0 |
| 0 | 0 | 0 | 0 |
| 0 | 0 | 0 | 0 |
| 0 | 0 | 0 | 0 |
| 0 | 0 | 0 | 0 |
| 0 | 0 | 0 | 0 |
| 0 | 0 | 0 | 0 |
| 0 | 0 | 0 | 0 |
| 0 | 0 | 0 | 0 |
| 0 | 0 | 0 | 0 |
| 0 | 0 | 0 | 0 |
| 0 | 0 | 0 | 0 |
| 0 | 0 | 0 | 0 |
| 0 | 0 | 0 | 0 |

| | | | |
|---|---|---|---|
| 0 | 0 | 0 | 0 |
| 0 | 0 | 0 | 0 |
| 0 | 0 | 0 | 0 |
| 0 | 0 | 0 | 0 |
| 0 | 0 | 0 | 0 |
| 0 | 0 | 0 | 0 |
| 0 | 0 | 0 | 0 |
| 0 | 0 | 0 | 0 |
| 0 | 0 | 0 | 0 |
| 0 | 0 | 0 | 0 |
| 0 | 0 | 0 | 0 |
| 0 | 0 | 0 | 0 |
| 0 | 0 | 0 | 0 |
| 0 | 0 | 0 | 0 |
| 0 | 0 | 0 | 0 |
| 0 | 0 | 0 | 0 |
| 0 | 0 | 0 | 0 |
| 0 | 0 | 0 | 0 |
| 0 | 0 | 0 | 0 |
| 0 | 0 | 0 | 0 |
| 0 | 0 | 0 | 0 |
| 0 | 0 | 0 | 0 |
| 0 | 0 | 0 | 0 |
| 0 | 0 | 0 | 0 |
| 0 | 0 | 0 | 0 |
| 0 | 0 | 0 | 0 |
| 0 | 0 | 0 | 0 |
| 0 | 0 | 0 | 0 |
| 0 | 0 | 0 | 0 |
| 0 | 0 | 0 | 0 |
| 0 | 0 | 0 | 0 |
| 0 | 0 | 0 | 0 |
| 0 | 0 | 0 | 0 |
| 0 | 0 | 0 | 0 |
| 0 | 0 | 0 | 0 |
| 0 | 0 | 0 | 0 |
| 0 | 0 | 0 | 0 |
| 0 | 0 | 0 | 0 |
| 0 | 0 | 0 | 0 |
| 0 | 0 | 0 | 0 |
| 0 | 0 | 0 | 0 |
| 0 | 0 | 0 | 0 |
| 0 | 0 | 0 | 0 |
| 0 | 0 | 0 | 0 |
| 0 | 0 | 0 | 0 |
| 0 | 0 | 0 | 0 |
| 0 | 0 | 0 | 0 |
| 0 | 0 | 0 | 0 |
| 1 | 0 | 0 | 1 |
| 0 | 0 | 0 | 0 |
| 1 | 0 | 0 | 0 |
| 1 | 0 | 0 | 0 |
| 0 | 0 | 0 | 0 |
| 0 | 0 | 0 | 1 |

| | | | |
|---|---|---|---|
| 0 | 0 | 0 | 0 |
| 0 | 0 | 0 | 0 |
| 0 | 0 | 0 | 0 |
| 0 | 0 | 0 | 0 |
| 0 | 0 | 0 | 0 |
| 0 | 0 | 0 | 0 |
| 0 | 0 | 0 | 0 |
| 0 | 0 | 0 | 0 |
| 0 | 0 | 0 | 0 |
| 0 | 0 | 0 | 0 |
| 0 | 0 | 0 | 0 |
| 0 | 0 | 0 | 0 |
| 0 | 0 | 0 | 0 |
| 0 | 0 | 0 | 0 |
| 0 | 0 | 0 | 0 |
| 0 | 0 | 0 | 0 |
| 0 | 0 | 0 | 0 |
| 0 | 0 | 0 | 0 |
| 0 | 0 | 0 | 0 |
| 0 | 0 | 0 | 0 |
| 0 | 0 | 0 | 0 |
| 0 | 0 | 0 | 0 |
| 0 | 0 | 0 | 0 |
| 0 | 0 | 0 | 0 |
| 0 | 0 | 0 | 0 |
| 0 | 0 | 0 | 0 |
| 0 | 0 | 1 | 0 |
| 0 | 0 | 1 | 0 |
| 0 | 0 | 1 | 0 |
| 0 | 0 | 1 | 0 |
| 0 | 0 | 0 | 0 |
| 0 | 0 | 0 | 0 |
| 0 | 0 | 0 | 0 |
| 1 | 0 | 0 | 0 |
| 1 | 0 | 0 | 0 |
| 0 | 0 | 0 | 0 |
| 0 | 0 | 0 | 0 |
| 0 | 0 | 0 | 0 |
| 0 | 0 | 0 | 0 |
| 1 | 0 | 0 | 0 |
| 0 | 0 | 0 | 0 |
| 0 | 0 | 0 | 0 |
| 0 | 0 | 0 | 0 |
| 0 | 0 | 0 | 0 |
| 0 | 0 | 0 | 0 |
| 0 | 1 | 0 | 0 |
| 0 | 0 | 0 | 0 |
| 0 | 0 | 0 | 0 |
| 0 | 0 | 0 | 0 |
| 0 | 0 | 0 | 0 |
| 0 | 0 | 0 | 0 |
| 0 | 0 | 0 | 0 |
| 0 | 0 | 0 | 0 |
| 0 | 0 | 0 | 0 |

| | | | |
|---|---|---|---|
| 0 | 0 | 0 | 0 |
| 0 | 0 | 0 | 0 |
| 0 | 0 | 0 | 0 |
| 0 | 0 | 0 | 0 |
| 0 | 0 | 0 | 0 |
| 0 | 0 | 0 | 1 |
| 0 | 0 | 0 | 0 |
| 0 | 0 | 0 | 0 |
| 0 | 0 | 0 | 0 |
| 0 | 0 | 0 | 0 |
| 0 | 0 | 0 | 0 |
| 0 | 0 | 0 | 0 |
| 0 | 0 | 0 | 0 |
| 0 | 1 | 0 | 0 |
| 0 | 1 | 0 | 0 |
| 0 | 1 | 0 | 0 |
| 0 | 1 | 0 | 0 |
| 0 | 1 | 0 | 0 |
| 0 | 0 | 0 | 0 |
| 0 | 0 | 0 | 0 |
| 0 | 0 | 0 | 0 |
| 0 | 0 | 0 | 0 |
| 0 | 0 | 0 | 0 |
| 0 | 0 | 0 | 0 |
| 0 | 0 | 0 | 0 |
| 0 | 0 | 0 | 0 |
| 0 | 0 | 0 | 0 |
| 0 | 0 | 0 | 0 |
| 0 | 0 | 0 | 0 |
| 0 | 0 | 0 | 0 |
| 0 | 0 | 0 | 0 |
| 0 | 0 | 0 | 0 |
| 0 | 0 | 0 | 0 |
| 0 | 0 | 0 | 0 |
| 0 | 0 | 0 | 0 |
| 0 | 0 | 0 | 0 |
| 0 | 0 | 0 | 0 |
| 0 | 0 | 0 | 0 |
| 0 | 0 | 0 | 0 |
| 0 | 0 | 0 | 0 |
| 0 | 0 | 0 | 0 |
| 0 | 0 | 0 | 0 |
| 0 | 0 | 0 | 0 |
| 0 | 0 | 0 | 0 |
| 0 | 0 | 0 | 0 |
| 0 | 0 | 0 | 0 |
| 0 | 0 | 0 | 0 |
| 0 | 0 | 0 | 0 |
| 0 | 0 | 0 | 0 |
| 0 | 0 | 0 | 0 |
| 0 | 0 | 0 | 0 |
| 0 | 0 | 0 | 0 |
| 0 | 0 | 0 | 0 |
| 0 | 0 | 0 | 0 |

| | | | |
|---|---|---|---|
| 0 | 0 | 0 | 0 |
| 0 | 0 | 0 | 0 |
| 0 | 0 | 0 | 0 |
| 1 | 0 | 0 | 0 |
| 0 | 0 | 0 | 0 |
| 0 | 0 | 0 | 0 |
| 0 | 0 | 0 | 0 |
| 0 | 0 | 0 | 0 |
| 0 | 0 | 0 | 0 |
| 0 | 0 | 0 | 0 |
| 0 | 0 | 0 | 0 |
| 0 | 0 | 0 | 0 |
| 0 | 0 | 0 | 0 |
| 0 | 0 | 0 | 0 |
| 0 | 0 | 0 | 0 |
| 0 | 0 | 0 | 0 |
| 0 | 0 | 0 | 0 |
| 0 | 0 | 0 | 0 |
| 0 | 0 | 0 | 0 |
| 0 | 0 | 0 | 0 |
| 1 | 1 | 0 | 0 |
| 1 | 1 | 0 | 0 |
| 0 | 0 | 0 | 0 |
| 0 | 1 | 0 | 0 |
| 0 | 0 | 0 | 0 |
| 0 | 0 | 0 | 0 |
| 0 | 0 | 0 | 0 |
| 0 | 0 | 0 | 0 |
| 0 | 0 | 0 | 0 |
| 0 | 0 | 0 | 0 |
| 0 | 0 | 0 | 0 |
| 0 | 0 | 0 | 0 |
| 0 | 0 | 0 | 0 |
| 0 | 0 | 0 | 0 |
| 0 | 0 | 0 | 0 |
| 0 | 0 | 0 | 0 |
| 0 | 0 | 0 | 0 |
| 0 | 0 | 0 | 0 |
| 0 | 0 | 0 | 0 |
| 1 | 0 | 0 | 0 |
| 1 | 0 | 0 | 0 |
| 0 | 0 | 0 | 0 |
| 0 | 0 | 0 | 0 |
| 1 | 0 | 0 | 0 |
| 0 | 0 | 0 | 0 |
| 0 | 0 | 0 | 0 |
| 0 | 0 | 0 | 0 |
| 0 | 0 | 0 | 0 |
| 0 | 0 | 0 | 0 |
| 0 | 0 | 0 | 0 |
| 0 | 0 | 0 | 0 |
| 0 | 0 | 0 | 0 |
| 0 | 0 | 0 | 0 |
| 0 | 0 | 0 | 0 |

| | | | |
|---|---|---|---|
| 0 | 0 | 0 | 0 |
| 0 | 0 | 0 | 0 |
| 0 | 0 | 0 | 0 |
| 0 | 0 | 0 | 0 |
| 0 | 0 | 0 | 0 |
| 1 | 0 | 0 | 0 |
| 0 | 0 | 0 | 0 |
| 0 | 0 | 0 | 0 |
| 0 | 1 | 0 | 0 |
| 1 | 1 | 0 | 0 |
| 0 | 0 | 0 | 0 |
| 1 | 1 | 0 | 0 |
| 1 | 0 | 0 | 0 |
| 0 | 0 | 0 | 0 |
| 0 | 0 | 0 | 0 |
| 0 | 0 | 0 | 0 |
| 1 | 1 | 0 | 0 |
| 0 | 0 | 0 | 0 |
| 0 | 0 | 0 | 0 |
| 1 | 0 | 0 | 0 |
| 0 | 0 | 0 | 0 |
| 0 | 0 | 0 | 0 |
| 0 | 0 | 0 | 0 |
| 0 | 0 | 0 | 0 |
| 0 | 0 | 0 | 0 |
| 0 | 0 | 0 | 0 |
| 0 | 0 | 0 | 0 |
| 0 | 0 | 0 | 0 |
| 0 | 0 | 0 | 0 |
| 0 | 0 | 0 | 0 |
| 1 | 0 | 0 | 0 |
| 0 | 0 | 0 | 0 |
| 0 | 0 | 0 | 0 |
| 0 | 0 | 0 | 0 |
| 0 | 0 | 0 | 0 |
| 1 | 0 | 0 | 0 |
| 0 | 0 | 0 | 0 |
| 0 | 0 | 0 | 0 |
| 0 | 0 | 0 | 0 |
| 0 | 0 | 0 | 0 |
| 0 | 0 | 0 | 0 |
| 0 | 0 | 0 | 0 |
| 0 | 0 | 0 | 0 |
| 0 | 0 | 0 | 0 |
| 0 | 0 | 0 | 0 |
| 0 | 0 | 0 | 0 |
| 0 | 0 | 0 | 0 |
| 0 | 0 | 0 | 0 |
| 0 | 0 | 0 | 0 |
| 0 | 0 | 0 | 0 |
| 0 | 0 | 0 | 0 |
| 0 | 0 | 0 | 0 |
| 0 | 0 | 0 | 0 |
| 0 | 0 | 0 | 0 |

| | | | |
|---|---|---|---|
| 0 | 0 | 0 | 0 |
| 0 | 0 | 0 | 0 |
| 0 | 0 | 0 | 0 |
| 0 | 0 | 0 | 0 |
| 0 | 0 | 0 | 0 |
| 0 | 0 | 0 | 0 |
| 0 | 0 | 0 | 0 |
| 0 | 0 | 0 | 0 |
| 0 | 0 | 0 | 0 |
| 0 | 0 | 0 | 0 |
| 0 | 0 | 0 | 0 |
| 0 | 0 | 0 | 0 |
| 0 | 0 | 0 | 0 |
| 0 | 0 | 0 | 0 |
| 0 | 0 | 0 | 1 |
| 0 | 0 | 0 | 0 |
| 0 | 0 | 0 | 0 |
| 0 | 0 | 0 | 0 |
| 0 | 0 | 0 | 0 |
| 0 | 0 | 0 | 0 |
| 0 | 0 | 0 | 0 |
| 0 | 0 | 0 | 0 |
| 0 | 0 | 0 | 0 |
| 0 | 0 | 0 | 0 |
| 0 | 0 | 0 | 0 |
| 0 | 0 | 0 | 0 |
| 0 | 0 | 0 | 0 |
| 0 | 0 | 0 | 0 |
| 0 | 0 | 0 | 0 |
| 0 | 0 | 0 | 0 |
| 0 | 0 | 0 | 0 |
| 0 | 0 | 0 | 0 |
| 0 | 0 | 0 | 0 |
| 0 | 0 | 0 | 0 |
| 0 | 0 | 0 | 0 |
| 0 | 0 | 0 | 0 |
| 0 | 0 | 0 | 0 |
| 0 | 0 | 0 | 0 |
| 0 | 0 | 0 | 0 |
| 0 | 0 | 1 | 0 |
| 0 | 0 | 0 | 0 |
| 0 | 0 | 0 | 0 |
| 0 | 0 | 0 | 0 |
| 1 | 0 | 0 | 0 |
| 0 | 0 | 0 | 0 |
| 0 | 0 | 0 | 0 |
| 0 | 0 | 0 | 0 |
| 0 | 0 | 0 | 0 |
| 1 | 0 | 0 | 1 |
| 0 | 0 | 0 | 0 |
| 0 | 0 | 0 | 0 |
| 0 | 0 | 0 | 0 |
| 0 | 0 | 0 | 0 |
| 0 | 0 | 0 | 0 |

| | | | |
|---|---|---|---|
| 0 | 0 | 0 | 0 |
| 0 | 0 | 0 | 0 |
| 0 | 0 | 0 | 0 |
| 0 | 0 | 0 | 0 |
| 0 | 0 | 0 | 0 |
| 0 | 0 | 0 | 0 |
| 0 | 0 | 0 | 0 |
| 0 | 0 | 0 | 0 |
| 0 | 0 | 0 | 0 |
| 1 | 0 | 0 | 0 |
| 0 | 0 | 0 | 0 |
| 0 | 0 | 0 | 0 |
| 0 | 0 | 0 | 0 |
| 0 | 0 | 0 | 0 |
| 1 | 0 | 0 | 0 |
| 0 | 0 | 0 | 0 |
| 0 | 0 | 0 | 0 |
| 0 | 0 | 0 | 0 |
| 0 | 0 | 0 | 0 |
| 0 | 0 | 0 | 0 |
| 0 | 0 | 0 | 0 |
| 0 | 0 | 0 | 0 |
| 0 | 0 | 0 | 0 |
| 0 | 0 | 0 | 0 |
| 1 | 0 | 0 | 0 |
| 0 | 0 | 0 | 0 |
| 0 | 0 | 0 | 0 |
| 0 | 0 | 0 | 0 |
| 0 | 0 | 0 | 0 |
| 0 | 0 | 0 | 0 |
| 0 | 0 | 0 | 0 |
| 0 | 0 | 0 | 0 |
| 0 | 0 | 0 | 0 |
| 0 | 0 | 0 | 0 |
| 1 | 0 | 0 | 0 |
| 1 | 0 | 0 | 0 |
| 0 | 0 | 0 | 0 |
| 0 | 0 | 0 | 0 |
| 1 | 0 | 0 | 0 |
| 0 | 0 | 1 | 0 |
| 0 | 0 | 1 | 0 |
| 0 | 0 | 1 | 0 |
| 0 | 0 | 1 | 0 |
| 0 | 0 | 0 | 0 |
| 1 | 0 | 0 | 0 |
| 0 | 0 | 0 | 0 |
| 0 | 0 | 0 | 0 |
| 0 | 0 | 0 | 0 |
| 0 | 0 | 1 | 0 |
| 0 | 0 | 0 | 0 |
| 0 | 0 | 0 | 0 |
| 0 | 0 | 0 | 0 |
| 0 | 0 | 0 | 0 |
| 0 | 0 | 0 | 0 |

| | | | |
|---|---|---|---|
| 0 | 0 | 0 | 0 |
| 0 | 0 | 0 | 0 |
| 0 | 0 | 0 | 0 |
| 0 | 0 | 0 | 0 |
| 0 | 0 | 0 | 0 |
| 0 | 0 | 0 | 0 |
| 0 | 0 | 0 | 0 |
| 0 | 0 | 0 | 0 |
| 0 | 0 | 0 | 0 |
| 0 | 0 | 0 | 0 |
| 0 | 1 | 0 | 0 |
| 0 | 0 | 0 | 0 |
| 0 | 0 | 0 | 0 |
| 0 | 0 | 0 | 0 |
| 0 | 0 | 0 | 0 |
| 0 | 1 | 0 | 0 |
| 0 | 0 | 0 | 0 |
| 0 | 0 | 0 | 0 |
| 0 | 0 | 0 | 0 |
| 0 | 0 | 0 | 0 |
| 0 | 0 | 0 | 0 |
| 0 | 0 | 0 | 0 |
| 0 | 0 | 0 | 0 |
| 0 | 0 | 0 | 0 |
| 0 | 0 | 0 | 0 |
| 0 | 0 | 0 | 0 |
| 0 | 0 | 0 | 0 |
| 0 | 0 | 0 | 0 |
| 0 | 0 | 0 | 0 |
| 0 | 0 | 0 | 0 |
| 0 | 0 | 0 | 0 |
| 0 | 0 | 0 | 0 |
| 0 | 0 | 0 | 0 |
| 0 | 0 | 0 | 0 |
| 0 | 0 | 0 | 0 |
| 0 | 0 | 0 | 0 |
| 0 | 0 | 0 | 0 |
| 0 | 0 | 0 | 0 |
| 0 | 0 | 0 | 0 |
| 0 | 0 | 0 | 0 |
| 0 | 0 | 0 | 0 |
| 0 | 0 | 0 | 0 |
| 0 | 0 | 0 | 0 |
| 0 | 0 | 0 | 0 |
| 1 | 1 | 0 | 0 |
| 0 | 0 | 0 | 0 |
| 0 | 0 | 0 | 0 |
| 0 | 0 | 0 | 0 |
| 0 | 0 | 0 | 0 |
| 0 | 0 | 0 | 0 |
| 0 | 0 | 0 | 0 |
| 0 | 0 | 0 | 0 |
| 0 | 0 | 0 | 0 |
| 0 | 0 | 0 | 0 |

| | | | |
|---|---|---|---|
| 0 | 0 | 0 | 0 |
| 0 | 0 | 0 | 0 |
| 0 | 0 | 1 | 0 |
| 1 | 0 | 0 | 0 |
| 0 | 0 | 0 | 0 |
| 0 | 0 | 0 | 0 |
| 0 | 0 | 0 | 0 |
| 0 | 0 | 0 | 0 |
| 0 | 0 | 0 | 0 |
| 0 | 0 | 0 | 0 |
| 0 | 0 | 0 | 0 |
| 0 | 0 | 0 | 0 |
| 1 | 0 | 0 | 0 |
| 0 | 0 | 0 | 0 |
| 0 | 0 | 0 | 0 |
| 0 | 0 | 0 | 0 |
| 0 | 0 | 0 | 0 |
| 0 | 0 | 0 | 0 |
| 0 | 0 | 0 | 0 |
| 0 | 0 | 0 | 0 |
| 0 | 0 | 0 | 0 |
| 0 | 0 | 0 | 0 |
| 0 | 0 | 0 | 0 |
| 0 | 0 | 0 | 0 |
| 0 | 0 | 0 | 0 |
| 0 | 0 | 0 | 0 |
| 0 | 0 | 0 | 0 |
| 0 | 0 | 0 | 0 |
| 0 | 0 | 0 | 0 |
| 0 | 0 | 0 | 0 |
| 0 | 0 | 0 | 0 |
| 0 | 0 | 0 | 0 |
| 0 | 0 | 0 | 0 |
| 0 | 0 | 0 | 0 |
| 0 | 0 | 0 | 0 |
| 0 | 0 | 0 | 0 |
| 0 | 0 | 0 | 0 |
| 0 | 0 | 0 | 0 |
| 0 | 0 | 0 | 0 |
| 0 | 0 | 0 | 0 |
| 0 | 1 | 0 | 0 |
| 1 | 1 | 0 | 0 |
| 0 | 0 | 0 | 0 |
| 0 | 0 | 0 | 0 |
| 0 | 0 | 0 | 0 |
| 0 | 0 | 0 | 0 |
| 0 | 0 | 0 | 0 |
| 0 | 0 | 0 | 0 |
| 0 | 1 | 0 | 0 |
| 0 | 0 | 0 | 0 |
| 0 | 0 | 0 | 0 |
| 0 | 0 | 0 | 0 |
| 0 | 0 | 0 | 1 |
| 0 | 0 | 0 | 0 |

| | | | |
|---|---|---|---|
| 0 | 0 | 0 | 1 |
| 0 | 0 | 0 | 1 |
| 0 | 0 | 0 | 1 |
| 0 | 0 | 0 | 0 |
| 0 | 0 | 0 | 0 |
| 0 | 0 | 0 | 0 |
| 0 | 0 | 0 | 0 |
| 1 | 0 | 0 | 0 |
| 0 | 0 | 0 | 0 |
| 1 | 1 | 0 | 0 |
| 0 | 1 | 0 | 0 |
| 0 | 0 | 0 | 0 |
| 1 | 1 | 0 | 0 |
| 0 | 0 | 0 | 0 |
| 0 | 0 | 0 | 0 |
| 0 | 0 | 0 | 0 |
| 0 | 0 | 0 | 0 |
| 1 | 0 | 0 | 0 |
| 0 | 0 | 0 | 0 |
| 0 | 0 | 0 | 0 |
| 0 | 0 | 0 | 0 |
| 0 | 0 | 0 | 0 |
| 0 | 0 | 0 | 0 |
| 0 | 0 | 0 | 0 |
| 0 | 0 | 0 | 0 |
| 0 | 0 | 0 | 0 |
| 0 | 0 | 0 | 0 |
| 0 | 0 | 0 | 0 |
| 0 | 0 | 0 | 0 |
| 0 | 0 | 0 | 0 |
| 0 | 0 | 0 | 0 |
| 1 | 0 | 0 | 0 |
| 0 | 0 | 0 | 0 |
| 0 | 0 | 0 | 0 |
| 0 | 0 | 0 | 0 |
| 0 | 0 | 0 | 0 |
| 0 | 0 | 0 | 0 |
| 0 | 0 | 0 | 0 |
| 0 | 0 | 0 | 0 |
| 0 | 0 | 0 | 0 |
| 0 | 0 | 0 | 0 |
| 1 | 1 | 0 | 0 |
| 0 | 0 | 0 | 0 |
| 0 | 0 | 0 | 0 |
| 1 | 0 | 0 | 0 |
| 0 | 1 | 0 | 0 |
| 0 | 0 | 0 | 0 |
| 0 | 0 | 0 | 0 |
| 0 | 1 | 0 | 0 |
| 1 | 1 | 0 | 0 |
| 1 | 1 | 0 | 0 |
| 1 | 1 | 0 | 0 |
| 1 | 1 | 0 | 0 |
| 1 | 1 | 0 | 0 |

| | | | |
|---|---|---|---|
| 0 | 0 | 0 | 0 |
| 0 | 0 | 0 | 0 |
| 0 | 0 | 0 | 0 |
| 0 | 0 | 0 | 0 |
| 0 | 0 | 0 | 0 |
| 1 | 0 | 0 | 0 |
| 1 | 0 | 0 | 0 |
| 0 | 0 | 0 | 0 |
| 0 | 0 | 0 | 0 |
| 1 | 0 | 0 | 0 |
| 0 | 0 | 0 | 0 |
| 0 | 0 | 1 | 0 |
| 0 | 0 | 0 | 0 |
| 0 | 0 | 0 | 0 |
| 0 | 0 | 0 | 0 |
| 0 | 0 | 0 | 0 |
| 0 | 0 | 0 | 0 |
| 0 | 0 | 0 | 0 |
| 0 | 0 | 0 | 0 |
| 0 | 0 | 0 | 0 |
| 1 | 1 | 0 | 0 |
| 1 | 0 | 0 | 0 |
| 0 | 0 | 0 | 0 |
| 0 | 0 | 1 | 0 |
| 0 | 0 | 0 | 0 |
| 0 | 0 | 1 | 0 |
| 0 | 0 | 0 | 0 |
| 0 | 0 | 0 | 0 |
| 0 | 0 | 0 | 0 |
| 0 | 0 | 0 | 0 |
| 0 | 0 | 0 | 0 |
| 0 | 0 | 0 | 0 |
| 0 | 0 | 0 | 0 |
| 0 | 0 | 0 | 0 |
| 0 | 0 | 0 | 0 |
| 0 | 0 | 0 | 0 |
| 1 | 0 | 0 | 0 |
| 0 | 0 | 0 | 0 |
| 1 | 0 | 0 | 0 |
| 1 | 0 | 0 | 0 |
| 1 | 0 | 0 | 0 |
| 0 | 0 | 0 | 0 |
| 0 | 0 | 0 | 0 |
| 0 | 0 | 0 | 0 |
| 0 | 0 | 0 | 0 |
| 0 | 0 | 0 | 0 |
| 0 | 0 | 0 | 0 |
| 1 | 0 | 1 | 0 |
| 0 | 0 | 0 | 0 |
| 0 | 0 | 0 | 0 |
| 0 | 0 | 0 | 0 |
| 0 | 0 | 0 | 0 |
| 1 | 1 | 0 | 0 |
| 0 | 0 | 0 | 0 |

| | | | |
|---|---|---|---|
| 0 | 0 | 0 | 0 |
| 0 | 1 | 0 | 0 |
| 0 | 0 | 0 | 0 |
| 0 | 0 | 0 | 0 |
| 0 | 0 | 0 | 0 |
| 0 | 0 | 0 | 0 |
| 0 | 0 | 0 | 0 |
| 0 | 0 | 0 | 0 |
| 0 | 0 | 0 | 0 |
| 0 | 0 | 0 | 0 |
| 0 | 0 | 0 | 0 |
| 0 | 0 | 0 | 0 |
| 0 | 0 | 0 | 0 |
| 0 | 0 | 0 | 0 |
| 0 | 0 | 0 | 0 |
| 0 | 0 | 0 | 0 |
| 0 | 0 | 0 | 0 |
| 0 | 0 | 0 | 0 |
| 0 | 0 | 0 | 0 |
| 0 | 0 | 0 | 0 |
| 1 | 0 | 0 | 0 |
| 0 | 0 | 0 | 0 |
| 0 | 0 | 0 | 0 |
| 0 | 0 | 0 | 0 |
| 1 | 0 | 0 | 0 |
| 0 | 0 | 0 | 0 |
| 0 | 0 | 0 | 0 |
| 0 | 0 | 0 | 1 |
| 0 | 0 | 0 | 0 |
| 0 | 0 | 0 | 0 |
| 0 | 0 | 0 | 0 |
| 0 | 0 | 0 | 0 |
| 0 | 0 | 0 | 0 |
| 0 | 0 | 0 | 0 |
| 0 | 0 | 0 | 0 |
| 0 | 0 | 0 | 0 |
| 0 | 0 | 0 | 0 |
| 0 | 0 | 0 | 0 |
| 0 | 0 | 0 | 0 |
| 0 | 0 | 0 | 0 |
| 0 | 0 | 0 | 0 |
| 0 | 0 | 0 | 0 |
| 0 | 0 | 0 | 0 |
| 0 | 0 | 0 | 0 |
| 0 | 0 | 0 | 0 |
| 0 | 0 | 0 | 0 |
| 0 | 0 | 0 | 0 |
| 0 | 0 | 0 | 0 |
| 0 | 0 | 0 | 0 |
| 0 | 0 | 0 | 0 |
| 0 | 0 | 0 | 0 |
| 0 | 0 | 0 | 0 |
| 0 | 0 | 0 | 0 |
| 0 | 0 | 0 | 0 |

| | | | |
|---|---|---|---|
| 0 | 0 | 0 | 0 |
| 0 | 0 | 0 | 0 |
| 0 | 0 | 0 | 0 |
| 0 | 0 | 0 | 0 |
| 0 | 0 | 0 | 0 |
| 0 | 0 | 0 | 0 |
| 0 | 0 | 0 | 0 |
| 0 | 0 | 0 | 0 |
| 0 | 0 | 0 | 0 |
| 0 | 0 | 0 | 0 |
| 0 | 0 | 0 | 0 |
| 0 | 0 | 0 | 0 |
| 0 | 0 | 0 | 0 |
| 0 | 0 | 0 | 0 |
| 0 | 0 | 0 | 0 |
| 1 | 0 | 0 | 0 |
| 0 | 0 | 0 | 0 |
| 0 | 0 | 0 | 0 |
| 0 | 1 | 0 | 0 |
| 0 | 0 | 0 | 0 |
| 0 | 0 | 0 | 0 |
| 0 | 0 | 0 | 0 |
| 0 | 0 | 0 | 0 |
| 0 | 0 | 0 | 0 |
| 0 | 0 | 0 | 0 |
| 1 | 0 | 0 | 0 |
| 0 | 0 | 0 | 0 |
| 0 | 0 | 0 | 0 |
| 0 | 0 | 0 | 0 |
| 0 | 0 | 0 | 0 |
| 0 | 0 | 0 | 0 |
| 0 | 0 | 0 | 0 |
| 0 | 0 | 0 | 0 |
| 1 | 1 | 0 | 0 |
| 0 | 0 | 0 | 0 |
| 0 | 0 | 0 | 0 |
| 0 | 0 | 0 | 0 |
| 0 | 0 | 0 | 0 |
| 0 | 0 | 0 | 0 |
| 0 | 0 | 0 | 0 |
| 0 | 0 | 0 | 0 |
| 0 | 0 | 0 | 0 |
| 0 | 0 | 0 | 0 |
| 0 | 0 | 0 | 0 |
| 0 | 0 | 0 | 0 |
| 0 | 0 | 0 | 0 |
| 0 | 0 | 0 | 0 |
| 1 | 0 | 0 | 0 |
| 0 | 0 | 0 | 0 |
| 0 | 0 | 0 | 0 |
| 0 | 0 | 0 | 0 |
| 0 | 0 | 0 | 0 |
| 0 | 0 | 0 | 0 |
| 1 | 0 | 0 | 1 |

| | | | |
|---|---|---|---|
| 0 | 0 | 0 | 0 |
| 0 | 0 | 0 | 0 |
| 0 | 0 | 1 | 0 |
| 0 | 0 | 0 | 0 |
| 0 | 0 | 0 | 0 |
| 0 | 0 | 0 | 0 |
| 0 | 0 | 0 | 0 |
| 0 | 0 | 0 | 0 |
| 0 | 0 | 0 | 0 |
| 0 | 0 | 0 | 0 |
| 0 | 0 | 0 | 0 |
| 0 | 0 | 0 | 0 |
| 0 | 0 | 0 | 0 |
| 0 | 0 | 0 | 0 |
| 0 | 0 | 0 | 0 |
| 0 | 0 | 0 | 0 |
| 0 | 0 | 0 | 0 |
| 0 | 0 | 0 | 0 |
| 0 | 0 | 0 | 0 |
| 0 | 0 | 0 | 0 |
| 0 | 0 | 0 | 0 |
| 0 | 0 | 0 | 0 |
| 0 | 0 | 0 | 0 |
| 0 | 0 | 0 | 0 |
| 0 | 0 | 0 | 0 |
| 0 | 0 | 0 | 0 |
| 0 | 0 | 0 | 0 |
| 0 | 0 | 0 | 0 |
| 0 | 0 | 0 | 0 |
| 0 | 0 | 0 | 0 |
| 0 | 0 | 0 | 0 |
| 1 | 0 | 0 | 0 |
| 0 | 0 | 0 | 0 |
| 0 | 0 | 0 | 0 |
| 0 | 0 | 0 | 0 |
| 0 | 0 | 0 | 0 |
| 0 | 0 | 0 | 0 |
| 0 | 0 | 0 | 0 |
| 0 | 0 | 0 | 0 |
| 0 | 0 | 0 | 0 |
| 1 | 1 | 0 | 0 |
| 0 | 0 | 0 | 0 |
| 0 | 0 | 0 | 0 |
| 0 | 0 | 0 | 0 |
| 0 | 0 | 0 | 0 |
| 1 | 1 | 0 | 0 |
| 0 | 0 | 0 | 0 |
| 1 | 1 | 0 | 0 |
| 0 | 0 | 0 | 0 |
| 1 | 0 | 0 | 0 |
| 1 | 0 | 0 | 0 |
| 1 | 0 | 0 | 0 |
| 0 | 0 | 0 | 0 |
| 0 | 0 | 0 | 0 |

| | | | |
|---|---|---|---|
| 1 | 0 | 0 | 0 |
| 0 | 0 | 0 | 0 |
| 0 | 0 | 0 | 0 |
| 0 | 0 | 0 | 0 |
| 0 | 0 | 0 | 0 |
| 0 | 0 | 0 | 0 |
| 0 | 0 | 0 | 0 |
| 0 | 0 | 0 | 0 |
| 0 | 0 | 0 | 0 |
| 0 | 0 | 0 | 0 |
| 0 | 0 | 0 | 0 |
| 0 | 0 | 0 | 0 |
| 0 | 0 | 0 | 0 |
| 0 | 0 | 0 | 0 |
| 0 | 0 | 0 | 0 |
| 1 | 1 | 1 | 0 |
| 0 | 0 | 0 | 0 |
| 1 | 0 | 0 | 0 |
| 0 | 0 | 0 | 0 |
| 0 | 0 | 0 | 0 |
| 0 | 0 | 0 | 0 |
| 0 | 1 | 0 | 0 |
| 0 | 0 | 0 | 0 |
| 0 | 1 | 0 | 0 |
| 0 | 1 | 0 | 0 |
| 0 | 0 | 0 | 0 |
| 0 | 0 | 0 | 0 |
| 0 | 0 | 0 | 0 |
| 0 | 0 | 0 | 0 |
| 0 | 0 | 0 | 0 |
| 0 | 0 | 0 | 0 |
| 0 | 0 | 0 | 0 |
| 1 | 1 | 0 | 0 |
| 0 | 0 | 0 | 0 |
| 0 | 0 | 0 | 0 |
| 0 | 0 | 0 | 0 |
| 0 | 1 | 0 | 0 |
| 1 | 1 | 0 | 0 |
| 0 | 0 | 0 | 0 |
| 0 | 0 | 0 | 0 |
| 0 | 0 | 0 | 0 |
| 0 | 0 | 0 | 0 |
| 0 | 0 | 0 | 0 |
| 0 | 0 | 0 | 0 |
| 1 | 1 | 0 | 0 |
| 0 | 0 | 0 | 0 |
| 1 | 0 | 0 | 0 |
| 1 | 1 | 0 | 0 |
| 0 | 0 | 0 | 0 |
| 0 | 0 | 0 | 0 |
| 1 | 1 | 0 | 0 |
| 1 | 1 | 0 | 0 |
| 0 | 0 | 0 | 0 |
| 1 | 1 | 0 | 0 |

| | | | |
|---|---|---|---|
| 0 | 0 | 0 | 0 |
| 0 | 0 | 0 | 0 |
| 0 | 0 | 0 | 0 |
| 1 | 1 | 0 | 0 |
| 0 | 0 | 0 | 0 |
| 0 | 0 | 0 | 0 |
| 0 | 0 | 0 | 0 |
| 0 | 0 | 0 | 0 |
| 1 | 0 | 0 | 0 |
| 0 | 0 | 0 | 0 |
| 0 | 0 | 0 | 0 |
| 0 | 0 | 0 | 0 |
| 1 | 0 | 1 | 0 |
| 1 | 0 | 0 | 0 |
| 0 | 0 | 0 | 0 |
| 0 | 0 | 0 | 0 |
| 0 | 0 | 0 | 0 |
| 0 | 0 | 0 | 0 |
| 0 | 0 | 0 | 0 |
| 0 | 0 | 0 | 0 |
| 0 | 0 | 0 | 0 |
| 0 | 0 | 0 | 0 |
| 0 | 0 | 0 | 0 |
| 1 | 0 | 0 | 0 |
| 0 | 0 | 0 | 0 |
| 0 | 0 | 0 | 0 |
| 0 | 0 | 0 | 0 |
| 0 | 0 | 0 | 0 |
| 0 | 0 | 0 | 0 |
| 0 | 0 | 0 | 0 |
| 0 | 0 | 0 | 0 |
| 0 | 0 | 0 | 0 |
| 0 | 0 | 0 | 0 |
| 0 | 0 | 0 | 0 |
| 0 | 0 | 0 | 0 |
| 0 | 0 | 0 | 0 |
| 0 | 0 | 0 | 0 |
| 0 | 0 | 0 | 0 |
| 0 | 0 | 0 | 0 |
| 0 | 0 | 0 | 0 |
| 0 | 0 | 0 | 0 |
| 0 | 0 | 0 | 0 |
| 0 | 0 | 0 | 0 |
| 1 | 1 | 0 | 0 |
| 1 | 1 | 0 | 0 |
| 0 | 0 | 0 | 0 |
| 0 | 0 | 0 | 0 |
| 0 | 1 | 0 | 0 |
| 0 | 1 | 0 | 0 |
| 0 | 1 | 0 | 0 |
| 0 | 0 | 0 | 0 |
| 0 | 0 | 0 | 0 |
| 0 | 0 | 0 | 0 |
| 0 | 0 | 0 | 0 |

| | | | |
|---|---|---|---|
| 0 | 0 | 0 | 0 |
| 0 | 0 | 0 | 0 |
| 0 | 0 | 0 | 0 |
| 0 | 0 | 0 | 0 |
| 1 | 0 | 0 | 0 |
| 0 | 0 | 0 | 0 |
| 0 | 0 | 0 | 0 |
| 1 | 0 | 0 | 0 |
| 1 | 0 | 0 | 0 |
| 1 | 0 | 0 | 0 |
| 1 | 0 | 0 | 0 |
| 1 | 0 | 0 | 0 |
| 0 | 0 | 0 | 0 |
| 1 | 0 | 0 | 0 |
| 1 | 0 | 0 | 0 |
| 0 | 0 | 0 | 0 |
| 0 | 0 | 0 | 0 |
| 0 | 0 | 0 | 0 |
| 1 | 0 | 0 | 0 |
| 1 | 0 | 0 | 0 |
| 1 | 0 | 0 | 0 |
| 1 | 0 | 0 | 0 |
| 0 | 0 | 0 | 0 |
| 0 | 0 | 0 | 0 |
| 0 | 0 | 0 | 0 |
| 0 | 0 | 0 | 0 |
| 0 | 0 | 0 | 0 |
| 0 | 0 | 0 | 0 |
| 0 | 0 | 0 | 0 |
| 0 | 0 | 0 | 0 |
| 0 | 0 | 0 | 0 |
| 0 | 0 | 0 | 0 |
| 1 | 0 | 0 | 0 |
| 0 | 0 | 0 | 0 |
| 0 | 0 | 0 | 0 |
| 1 | 0 | 0 | 0 |
| 0 | 0 | 0 | 0 |
| 0 | 0 | 0 | 0 |
| 0 | 0 | 0 | 0 |
| 0 | 0 | 0 | 0 |
| 0 | 0 | 0 | 0 |
| 0 | 0 | 0 | 0 |
| 0 | 0 | 0 | 0 |
| 0 | 0 | 0 | 0 |
| 0 | 0 | 0 | 0 |
| 0 | 0 | 0 | 0 |
| 0 | 0 | 0 | 0 |
| 0 | 0 | 0 | 0 |
| 0 | 0 | 0 | 0 |
| 0 | 0 | 0 | 0 |
| 0 | 0 | 0 | 0 |
| 0 | 0 | 0 | 0 |
| 0 | 0 | 0 | 0 |
| 0 | 0 | 0 | 0 |

| | | | |
|---|---|---|---|
| 0 | 0 | 0 | 0 |
| 0 | 0 | 0 | 0 |
| 0 | 0 | 0 | 0 |
| 0 | 0 | 0 | 0 |
| 0 | 0 | 0 | 0 |
| 0 | 0 | 0 | 0 |
| 0 | 0 | 0 | 0 |
| 0 | 0 | 0 | 0 |
| 0 | 0 | 0 | 0 |
| 0 | 0 | 0 | 0 |
| 0 | 0 | 0 | 0 |
| 0 | 0 | 0 | 1 |
| 0 | 0 | 0 | 0 |
| 0 | 0 | 0 | 0 |
| 0 | 0 | 0 | 0 |
| 0 | 0 | 0 | 0 |
| 0 | 0 | 0 | 0 |
| 0 | 0 | 0 | 1 |
| 0 | 0 | 0 | 0 |
| 0 | 0 | 0 | 0 |
| 0 | 0 | 0 | 1 |
| 0 | 0 | 0 | 0 |
| 0 | 0 | 0 | 0 |
| 0 | 0 | 0 | 0 |
| 0 | 0 | 0 | 0 |
| 0 | 0 | 0 | 0 |
| 0 | 0 | 0 | 0 |
| 0 | 0 | 0 | 0 |
| 0 | 0 | 0 | 1 |
| 0 | 0 | 0 | 0 |
| 0 | 0 | 0 | 0 |
| 0 | 0 | 0 | 0 |
| 0 | 0 | 0 | 0 |
| 0 | 0 | 0 | 0 |
| 0 | 0 | 0 | 0 |
| 0 | 0 | 0 | 0 |
| 0 | 0 | 0 | 0 |
| 0 | 0 | 0 | 0 |
| 0 | 0 | 0 | 0 |
| 0 | 0 | 0 | 0 |
| 0 | 0 | 0 | 0 |
| 0 | 0 | 0 | 0 |
| 0 | 0 | 0 | 0 |
| 0 | 0 | 0 | 0 |
| 0 | 0 | 0 | 0 |
| 0 | 0 | 0 | 0 |
| 0 | 0 | 0 | 0 |
| 0 | 0 | 0 | 0 |
| 0 | 0 | 0 | 0 |
| 0 | 0 | 0 | 0 |
| 0 | 0 | 0 | 0 |
| 0 | 0 | 0 | 0 |
| 0 | 0 | 0 | 0 |
| 0 | 0 | 0 | 0 |

| | | | |
|---|---|---|---|
| 0 | 0 | 0 | 0 |
| 0 | 0 | 0 | 0 |
| 1 | 0 | 0 | 0 |
| 0 | 0 | 0 | 0 |
| 1 | 0 | 0 | 0 |
| 0 | 0 | 0 | 0 |
| 0 | 0 | 0 | 0 |
| 0 | 0 | 0 | 0 |
| 0 | 0 | 0 | 0 |
| 1 | 1 | 0 | 0 |
| 0 | 0 | 1 | 0 |
| 0 | 0 | 0 | 0 |
| 0 | 0 | 0 | 0 |
| 0 | 0 | 0 | 0 |
| 0 | 0 | 0 | 0 |
| 0 | 0 | 0 | 0 |
| 0 | 0 | 0 | 0 |
| 0 | 0 | 0 | 0 |
| 0 | 0 | 0 | 0 |
| 0 | 0 | 0 | 0 |
| 0 | 0 | 0 | 0 |
| 0 | 0 | 0 | 0 |
| 0 | 0 | 0 | 0 |
| 0 | 0 | 0 | 0 |
| 0 | 0 | 0 | 0 |
| 0 | 0 | 0 | 0 |
| 0 | 0 | 0 | 0 |
| 0 | 0 | 0 | 0 |
| 0 | 0 | 0 | 0 |
| 0 | 0 | 0 | 0 |
| 0 | 0 | 0 | 0 |
| 0 | 0 | 0 | 0 |
| 0 | 0 | 0 | 0 |
| 0 | 0 | 0 | 0 |
| 0 | 0 | 0 | 0 |
| 0 | 0 | 0 | 0 |
| 0 | 0 | 0 | 0 |
| 0 | 0 | 0 | 0 |
| 0 | 0 | 0 | 0 |
| 0 | 0 | 0 | 0 |
| 0 | 0 | 0 | 0 |
| 0 | 0 | 0 | 0 |
| 0 | 0 | 0 | 0 |
| 0 | 0 | 0 | 0 |
| 0 | 0 | 0 | 0 |
| 1 | 0 | 0 | 0 |
| 0 | 0 | 0 | 0 |
| 0 | 0 | 0 | 0 |
| 0 | 0 | 0 | 0 |
| 0 | 0 | 0 | 0 |
| 1 | 0 | 0 | 0 |
| 0 | 0 | 0 | 0 |
| 0 | 0 | 0 | 0 |
| 0 | 0 | 0 | 0 |

| | | | |
|---|---|---|---|
| 0 | 0 | 0 | 0 |
| 0 | 0 | 0 | 0 |
| 0 | 0 | 0 | 0 |
| 0 | 0 | 0 | 0 |
| 0 | 0 | 0 | 0 |
| 0 | 0 | 0 | 0 |
| 0 | 0 | 0 | 0 |
| 0 | 0 | 0 | 0 |
| 1 | 1 | 0 | 0 |
| 0 | 0 | 0 | 0 |
| 0 | 0 | 0 | 0 |
| 0 | 0 | 0 | 0 |
| 0 | 0 | 0 | 0 |
| 0 | 0 | 0 | 0 |
| 0 | 1 | 0 | 0 |
| 0 | 1 | 0 | 0 |
| 0 | 1 | 0 | 0 |
| 0 | 0 | 0 | 0 |
| 0 | 0 | 0 | 0 |
| 0 | 0 | 0 | 0 |
| 0 | 1 | 0 | 0 |
| 0 | 0 | 0 | 0 |
| 0 | 0 | 0 | 0 |
| 1 | 1 | 0 | 0 |
| 0 | 1 | 0 | 0 |
| 0 | 1 | 0 | 0 |
| 0 | 0 | 0 | 0 |
| 0 | 0 | 0 | 0 |
| 0 | 0 | 0 | 0 |
| 0 | 0 | 0 | 0 |
| 0 | 0 | 0 | 0 |
| 0 | 0 | 0 | 0 |
| 0 | 0 | 0 | 0 |
| 0 | 0 | 0 | 0 |
| 0 | 0 | 0 | 0 |
| 0 | 0 | 0 | 0 |
| 0 | 0 | 0 | 0 |
| 1 | 0 | 0 | 0 |
| 0 | 0 | 0 | 0 |
| 0 | 0 | 0 | 0 |
| 0 | 0 | 0 | 0 |
| 0 | 0 | 0 | 0 |
| 0 | 0 | 0 | 0 |
| 0 | 0 | 0 | 0 |
| 0 | 0 | 0 | 0 |
| 0 | 0 | 0 | 0 |
| 0 | 0 | 0 | 0 |
| 0 | 0 | 0 | 0 |
| 0 | 0 | 0 | 0 |
| 0 | 0 | 0 | 0 |
| 0 | 0 | 0 | 0 |
| 0 | 0 | 0 | 0 |
| 0 | 0 | 0 | 0 |
| 0 | 0 | 0 | 0 |

| | | | |
|---|---|---|---|
| 0 | 0 | 0 | 0 |
| 0 | 0 | 0 | 0 |
| 0 | 0 | 0 | 0 |
| 0 | 0 | 1 | 0 |
| 0 | 0 | 0 | 0 |
| 0 | 0 | 0 | 0 |
| 0 | 0 | 0 | 0 |
| 0 | 0 | 0 | 0 |
| 0 | 0 | 0 | 0 |
| 0 | 0 | 0 | 0 |
| 0 | 0 | 0 | 0 |
| 0 | 0 | 0 | 0 |
| 1 | 0 | 0 | 0 |
| 0 | 0 | 0 | 0 |
| 0 | 0 | 0 | 0 |
| 0 | 0 | 0 | 0 |
| 0 | 1 | 0 | 0 |
| 0 | 0 | 0 | 0 |
| 0 | 0 | 0 | 0 |
| 0 | 0 | 0 | 0 |
| 0 | 0 | 0 | 0 |
| 0 | 0 | 0 | 0 |
| 0 | 0 | 0 | 0 |
| 0 | 0 | 0 | 0 |
| 0 | 0 | 0 | 0 |
| 0 | 0 | 0 | 0 |
| 0 | 0 | 0 | 0 |
| 0 | 0 | 0 | 0 |
| 0 | 0 | 0 | 0 |
| 0 | 0 | 0 | 0 |
| 1 | 0 | 0 | 0 |
| 1 | 0 | 0 | 0 |
| 1 | 0 | 0 | 0 |
| 0 | 0 | 0 | 0 |
| 1 | 0 | 0 | 0 |
| 1 | 0 | 0 | 0 |
| 1 | 0 | 0 | 0 |
| 0 | 0 | 0 | 0 |
| 0 | 0 | 0 | 0 |
| 0 | 0 | 0 | 0 |
| 0 | 0 | 0 | 0 |
| 0 | 0 | 0 | 0 |
| 0 | 1 | 0 | 0 |
| 0 | 1 | 0 | 0 |
| 0 | 0 | 0 | 0 |
| 0 | 0 | 0 | 0 |
| 1 | 0 | 0 | 0 |
| 0 | 0 | 0 | 0 |
| 0 | 0 | 0 | 0 |
| 0 | 0 | 0 | 0 |
| 0 | 0 | 0 | 0 |
| 0 | 0 | 0 | 0 |
| 1 | 0 | 0 | 0 |
| 0 | 0 | 0 | 0 |

| | | | |
|---|---|---|---|
| 0 | 0 | 0 | 0 |
| 1 | 0 | 0 | 0 |
| 1 | 0 | 0 | 0 |
| 0 | 0 | 0 | 0 |
| 0 | 0 | 0 | 0 |
| 0 | 0 | 0 | 0 |
| 0 | 0 | 0 | 0 |
| 0 | 1 | 0 | 0 |
| 0 | 0 | 0 | 0 |
| 0 | 0 | 0 | 0 |
| 0 | 0 | 0 | 0 |
| 0 | 0 | 0 | 0 |
| 0 | 0 | 0 | 0 |
| 0 | 0 | 0 | 0 |
| 1 | 0 | 0 | 0 |
| 0 | 0 | 0 | 0 |
| 0 | 0 | 0 | 0 |
| 0 | 0 | 0 | 0 |
| 0 | 0 | 0 | 0 |
| 1 | 0 | 0 | 0 |
| 1 | 0 | 0 | 0 |
| 0 | 0 | 0 | 0 |
| 1 | 0 | 0 | 0 |
| 1 | 0 | 0 | 0 |
| 0 | 0 | 0 | 0 |
| 0 | 0 | 0 | 0 |
| 0 | 0 | 0 | 0 |
| 0 | 0 | 0 | 0 |
| 0 | 0 | 0 | 0 |
| 0 | 0 | 0 | 0 |
| 0 | 0 | 0 | 0 |
| 0 | 0 | 0 | 0 |
| 0 | 0 | 0 | 0 |
| 0 | 0 | 0 | 0 |
| 0 | 0 | 0 | 0 |
| 0 | 0 | 0 | 0 |
| 0 | 0 | 0 | 0 |
| 0 | 0 | 0 | 0 |
| 0 | 0 | 0 | 0 |
| 0 | 0 | 0 | 0 |
| 0 | 0 | 0 | 0 |
| 0 | 0 | 0 | 0 |
| 0 | 0 | 0 | 0 |
| 0 | 0 | 0 | 0 |
| 0 | 0 | 0 | 0 |
| 0 | 0 | 0 | 0 |
| 0 | 0 | 0 | 0 |
| 0 | 0 | 0 | 0 |
| 0 | 0 | 0 | 0 |
| 0 | 0 | 0 | 0 |
| 0 | 0 | 0 | 0 |
| 0 | 0 | 0 | 0 |
| 0 | 0 | 0 | 0 |
| 1 | 1 | 0 | 1 |

| | | | |
|---|---|---|---|
| 0 | 0 | 0 | 0 |
| 0 | 0 | 0 | 0 |
| 0 | 0 | 0 | 0 |
| 0 | 0 | 0 | 0 |
| 0 | 1 | 0 | 0 |
| 0 | 0 | 0 | 0 |
| 0 | 0 | 0 | 0 |
| 0 | 1 | 0 | 0 |
| 0 | 0 | 0 | 0 |
| 0 | 0 | 0 | 0 |
| 0 | 0 | 0 | 0 |
| 0 | 0 | 0 | 0 |
| 0 | 0 | 0 | 0 |
| 0 | 0 | 0 | 0 |
| 0 | 1 | 0 | 1 |
| 0 | 1 | 0 | 0 |
| 0 | 0 | 0 | 0 |
| 0 | 0 | 0 | 0 |
| 0 | 1 | 0 | 0 |
| 0 | 1 | 0 | 1 |
| 0 | 0 | 0 | 0 |
| 0 | 0 | 0 | 0 |
| 0 | 0 | 0 | 0 |
| 0 | 0 | 0 | 0 |
| 0 | 0 | 0 | 0 |
| 1 | 0 | 0 | 0 |
| 0 | 0 | 0 | 0 |
| 0 | 0 | 1 | 0 |
| 0 | 0 | 1 | 0 |
| 0 | 0 | 1 | 0 |
| 0 | 0 | 0 | 0 |
| 0 | 0 | 0 | 0 |
| 0 | 0 | 0 | 0 |
| 0 | 0 | 0 | 0 |
| 0 | 0 | 0 | 0 |
| 0 | 0 | 1 | 0 |
| 0 | 0 | 0 | 0 |
| 0 | 0 | 0 | 0 |
| 1 | 0 | 1 | 0 |
| 0 | 0 | 1 | 0 |
| 1 | 1 | 0 | 0 |
| 0 | 1 | 0 | 0 |
| 0 | 0 | 0 | 0 |
| 0 | 0 | 0 | 0 |
| 1 | 0 | 0 | 0 |
| 0 | 0 | 0 | 0 |
| 0 | 0 | 1 | 0 |
| 0 | 0 | 0 | 0 |
| 0 | 0 | 0 | 0 |
| 0 | 0 | 0 | 0 |
| 0 | 0 | 0 | 0 |
| 0 | 0 | 0 | 0 |
| 0 | 0 | 0 | 0 |
| 0 | 0 | 0 | 0 |

| | | | |
|---|---|---|---|
| 0 | 0 | 0 | 0 |
| 0 | 0 | 0 | 0 |
| 0 | 0 | 0 | 0 |
| 0 | 0 | 0 | 0 |
| 0 | 0 | 0 | 0 |
| 0 | 0 | 0 | 0 |
| 0 | 0 | 0 | 0 |
| 1 | 1 | 0 | 0 |
| 0 | 0 | 0 | 0 |
| 0 | 0 | 0 | 0 |
| 0 | 0 | 0 | 0 |
| 0 | 0 | 0 | 1 |
| 0 | 0 | 0 | 0 |
| 0 | 0 | 0 | 0 |
| 0 | 0 | 0 | 0 |
| 0 | 0 | 0 | 0 |
| 0 | 0 | 0 | 0 |
| 0 | 0 | 0 | 0 |
| 0 | 0 | 0 | 0 |
| 0 | 0 | 0 | 0 |
| 0 | 0 | 0 | 0 |
| 0 | 0 | 0 | 0 |
| 0 | 0 | 0 | 0 |
| 0 | 0 | 0 | 0 |
| 0 | 0 | 0 | 0 |
| 0 | 0 | 0 | 0 |
| 0 | 0 | 0 | 0 |
| 0 | 0 | 0 | 0 |
| 0 | 0 | 0 | 0 |
| 0 | 1 | 0 | 0 |
| 0 | 1 | 0 | 0 |
| 0 | 0 | 0 | 0 |
| 0 | 0 | 0 | 0 |
| 0 | 0 | 0 | 0 |
| 0 | 0 | 0 | 0 |
| 0 | 1 | 0 | 0 |
| 0 | 0 | 0 | 0 |
| 0 | 0 | 0 | 0 |
| 0 | 0 | 0 | 0 |
| 0 | 0 | 0 | 0 |
| 0 | 0 | 0 | 0 |
| 1 | 1 | 0 | 0 |
| 1 | 0 | 0 | 0 |
| 0 | 0 | 0 | 0 |
| 0 | 0 | 0 | 0 |
| 0 | 0 | 0 | 0 |
| 0 | 0 | 0 | 0 |
| 0 | 0 | 0 | 0 |
| 0 | 0 | 0 | 0 |
| 0 | 0 | 0 | 0 |
| 0 | 0 | 0 | 0 |
| 0 | 0 | 0 | 0 |
| 0 | 0 | 0 | 0 |
| 0 | 0 | 0 | 0 |

| | | | |
|---|---|---|---|
| 0 | 0 | 0 | 0 |
| 0 | 0 | 0 | 0 |
| 0 | 0 | 0 | 0 |
| 0 | 0 | 0 | 0 |
| 0 | 0 | 0 | 0 |
| 0 | 0 | 0 | 0 |
| 0 | 0 | 0 | 0 |
| 0 | 0 | 0 | 0 |
| 1 | 0 | 0 | 0 |
| 1 | 0 | 0 | 0 |
| 0 | 0 | 0 | 0 |
| 0 | 0 | 0 | 0 |
| 0 | 0 | 0 | 0 |
| 1 | 0 | 0 | 0 |
| 0 | 0 | 0 | 0 |
| 0 | 0 | 0 | 0 |
| 0 | 0 | 0 | 0 |
| 0 | 0 | 0 | 0 |
| 0 | 0 | 0 | 0 |
| 1 | 1 | 0 | 0 |
| 0 | 0 | 1 | 0 |
| 0 | 0 | 0 | 0 |
| 0 | 0 | 0 | 0 |
| 0 | 0 | 0 | 0 |
| 0 | 0 | 0 | 0 |
| 0 | 0 | 0 | 0 |
| 0 | 0 | 0 | 0 |
| 0 | 0 | 0 | 0 |
| 0 | 0 | 0 | 0 |
| 0 | 0 | 0 | 0 |
| 0 | 0 | 0 | 0 |
| 0 | 0 | 0 | 0 |
| 0 | 0 | 0 | 0 |
| 0 | 0 | 0 | 0 |
| 0 | 0 | 0 | 0 |
| 0 | 0 | 0 | 0 |
| 0 | 0 | 0 | 0 |
| 0 | 0 | 0 | 0 |
| 0 | 0 | 0 | 0 |
| 0 | 0 | 0 | 0 |
| 0 | 0 | 0 | 0 |
| 0 | 0 | 0 | 0 |
| 0 | 0 | 0 | 0 |
| 0 | 0 | 0 | 0 |
| 0 | 0 | 0 | 0 |
| 1 | 0 | 1 | 0 |
| 1 | 0 | 0 | 0 |
| 0 | 0 | 0 | 0 |
| 0 | 0 | 0 | 0 |
| 0 | 0 | 1 | 0 |
| 0 | 0 | 0 | 0 |
| 0 | 0 | 0 | 0 |
| 0 | 0 | 0 | 0 |

| | | | |
|---|---|---|---|
| 0 | 0 | 0 | 0 |
| 0 | 0 | 0 | 0 |
| 0 | 0 | 0 | 0 |
| 0 | 0 | 0 | 0 |
| 0 | 0 | 0 | 0 |
| 0 | 0 | 0 | 0 |
| 0 | 0 | 0 | 0 |
| 0 | 0 | 0 | 0 |
| 0 | 0 | 0 | 0 |
| 0 | 0 | 0 | 0 |
| 0 | 0 | 0 | 0 |
| 0 | 0 | 0 | 0 |
| 0 | 0 | 0 | 0 |
| 0 | 0 | 0 | 0 |
| 0 | 0 | 0 | 0 |
| 0 | 0 | 0 | 0 |
| 0 | 0 | 0 | 0 |
| 0 | 0 | 0 | 0 |
| 0 | 0 | 0 | 0 |
| 1 | 0 | 0 | 0 |
| 0 | 0 | 0 | 0 |
| 0 | 0 | 0 | 0 |
| 0 | 0 | 0 | 0 |
| 0 | 0 | 0 | 0 |
| 0 | 0 | 0 | 0 |
| 0 | 0 | 0 | 0 |
| 0 | 0 | 0 | 0 |
| 0 | 0 | 0 | 0 |
| 0 | 0 | 0 | 0 |
| 0 | 0 | 0 | 0 |
| 0 | 0 | 0 | 0 |
| 0 | 0 | 0 | 0 |
| 0 | 0 | 0 | 0 |
| 0 | 0 | 0 | 0 |
| 0 | 0 | 0 | 0 |
| 0 | 0 | 0 | 0 |
| 0 | 0 | 0 | 0 |
| 0 | 0 | 0 | 0 |
| 1 | 0 | 0 | 1 |
| 0 | 0 | 0 | 0 |
| 0 | 0 | 0 | 0 |
| 0 | 0 | 0 | 0 |
| 0 | 0 | 0 | 1 |
| 0 | 0 | 0 | 1 |
| 0 | 0 | 0 | 1 |
| 0 | 0 | 0 | 0 |
| 0 | 0 | 0 | 0 |
| 0 | 0 | 0 | 0 |
| 0 | 0 | 0 | 0 |
| 0 | 0 | 0 | 0 |
| 0 | 0 | 0 | 0 |
| 0 | 0 | 0 | 0 |
| 0 | 0 | 0 | 0 |
| 0 | 0 | 0 | 0 |

| | | | |
|---|---|---|---|
| 0 | 0 | 0 | 0 |
| 0 | 0 | 0 | 0 |
| 0 | 0 | 0 | 0 |
| 0 | 0 | 0 | 0 |
| 0 | 0 | 0 | 0 |
| 0 | 0 | 0 | 0 |
| 0 | 0 | 0 | 0 |
| 0 | 0 | 0 | 0 |
| 0 | 0 | 0 | 0 |
| 0 | 0 | 0 | 0 |
| 0 | 0 | 0 | 0 |
| 0 | 0 | 0 | 0 |
| 0 | 0 | 0 | 0 |
| 0 | 0 | 0 | 0 |
| 0 | 0 | 0 | 0 |
| 0 | 0 | 0 | 0 |
| 0 | 0 | 0 | 0 |
| 0 | 0 | 0 | 0 |
| 0 | 0 | 0 | 0 |
| 0 | 0 | 0 | 0 |
| 0 | 0 | 0 | 0 |
| 0 | 0 | 0 | 0 |
| 0 | 0 | 0 | 0 |
| 0 | 0 | 0 | 0 |
| 0 | 0 | 0 | 0 |
| 0 | 0 | 0 | 0 |
| 0 | 0 | 0 | 0 |
| 0 | 0 | 0 | 0 |
| 0 | 0 | 0 | 0 |
| 0 | 0 | 0 | 0 |
| 0 | 0 | 0 | 0 |
| 0 | 0 | 0 | 0 |
| 1 | 1 | 0 | 1 |
| 1 | 1 | 0 | 1 |
| 1 | 0 | 0 | 0 |
| 0 | 0 | 0 | 0 |
| 0 | 0 | 0 | 0 |
| 0 | 0 | 0 | 0 |
| 0 | 0 | 0 | 0 |
| 0 | 0 | 0 | 0 |
| 0 | 0 | 0 | 0 |
| 0 | 0 | 0 | 0 |
| 1 | 1 | 0 | 0 |
| 0 | 0 | 0 | 0 |
| 1 | 0 | 0 | 0 |
| 0 | 0 | 0 | 0 |
| 0 | 0 | 0 | 0 |
| 0 | 0 | 0 | 0 |
| 0 | 0 | 0 | 0 |
| 0 | 0 | 0 | 0 |
| 1 | 0 | 0 | 0 |
| 0 | 0 | 0 | 0 |
| 0 | 0 | 0 | 0 |
| 0 | 0 | 0 | 0 |

| | | | |
|---|---|---|---|
| 0 | 0 | 0 | 0 |
| 0 | 0 | 0 | 0 |
| 0 | 0 | 0 | 0 |
| 0 | 0 | 0 | 0 |
| 0 | 0 | 0 | 0 |
| 0 | 0 | 0 | 0 |
| 0 | 0 | 0 | 0 |
| 0 | 0 | 0 | 0 |
| 0 | 0 | 0 | 0 |
| 0 | 0 | 0 | 0 |
| 0 | 0 | 0 | 0 |
| 0 | 0 | 0 | 0 |
| 0 | 0 | 0 | 0 |
| 0 | 0 | 0 | 0 |
| 0 | 0 | 0 | 0 |
| 0 | 0 | 0 | 0 |
| 0 | 0 | 0 | 0 |
| 0 | 0 | 0 | 0 |
| 0 | 0 | 0 | 0 |
| 0 | 0 | 0 | 1 |
| 0 | 0 | 0 | 0 |
| 0 | 0 | 0 | 1 |
| 0 | 0 | 0 | 1 |
| 0 | 0 | 0 | 1 |
| 0 | 0 | 0 | 0 |
| 0 | 0 | 0 | 0 |
| 0 | 0 | 0 | 0 |
| 0 | 0 | 0 | 0 |
| 0 | 0 | 0 | 0 |
| 0 | 0 | 0 | 0 |
| 0 | 0 | 0 | 0 |
| 0 | 0 | 0 | 0 |
| 0 | 0 | 0 | 0 |
| 0 | 0 | 0 | 0 |
| 0 | 0 | 0 | 0 |
| 0 | 1 | 0 | 0 |
| 0 | 0 | 0 | 0 |
| 0 | 0 | 0 | 0 |
| 0 | 1 | 0 | 0 |
| 0 | 0 | 0 | 0 |
| 0 | 0 | 0 | 0 |
| 0 | 0 | 0 | 0 |
| 0 | 0 | 0 | 0 |
| 0 | 0 | 0 | 0 |
| 0 | 0 | 0 | 0 |
| 0 | 0 | 0 | 0 |
| 0 | 0 | 0 | 0 |
| 0 | 0 | 0 | 0 |
| 0 | 0 | 0 | 0 |
| 0 | 0 | 0 | 0 |
| 0 | 0 | 0 | 0 |
| 0 | 0 | 0 | 0 |
| 0 | 0 | 0 | 0 |
| 0 | 0 | 0 | 0 |

| | | | |
|---|---|---|---|
| 0 | 0 | 0 | 0 |
| 0 | 0 | 0 | 0 |
| 0 | 0 | 0 | 0 |
| 0 | 0 | 0 | 0 |
| 0 | 0 | 0 | 0 |
| 0 | 0 | 0 | 0 |
| 0 | 0 | 0 | 0 |
| 0 | 0 | 0 | 0 |
| 0 | 0 | 1 | 0 |
| 0 | 0 | 1 | 0 |
| 0 | 0 | 0 | 0 |
| 0 | 0 | 0 | 0 |
| 0 | 0 | 0 | 0 |
| 0 | 0 | 0 | 0 |
| 0 | 0 | 0 | 0 |
| 0 | 1 | 0 | 0 |
| 0 | 0 | 0 | 0 |
| 0 | 0 | 0 | 0 |
| 1 | 1 | 0 | 0 |
| 0 | 0 | 0 | 0 |
| 0 | 0 | 0 | 0 |
| 0 | 0 | 0 | 0 |
| 0 | 0 | 0 | 0 |
| 0 | 0 | 0 | 0 |
| 0 | 0 | 0 | 0 |
| 0 | 0 | 0 | 0 |
| 0 | 0 | 0 | 0 |
| 0 | 0 | 0 | 0 |
| 0 | 0 | 0 | 0 |
| 0 | 0 | 0 | 0 |
| 0 | 0 | 0 | 0 |
| 0 | 0 | 0 | 0 |
| 0 | 0 | 0 | 0 |
| 0 | 0 | 0 | 0 |
| 0 | 0 | 0 | 0 |
| 0 | 0 | 0 | 0 |
| 0 | 0 | 0 | 0 |
| 0 | 0 | 0 | 0 |
| 0 | 0 | 0 | 0 |
| 0 | 0 | 0 | 0 |
| 0 | 0 | 0 | 0 |
| 0 | 0 | 0 | 0 |
| 0 | 0 | 0 | 0 |
| 0 | 0 | 0 | 0 |
| 0 | 0 | 0 | 0 |
| 0 | 0 | 0 | 0 |
| 0 | 0 | 0 | 0 |
| 0 | 0 | 0 | 0 |
| 0 | 0 | 0 | 0 |
| 0 | 0 | 0 | 0 |
| 0 | 0 | 0 | 0 |
| 0 | 0 | 0 | 0 |
| 0 | 0 | 0 | 0 |
| 1 | 0 | 0 | 0 |

| | | | |
|---|---|---|---|
| 0 | 0 | 0 | 0 |
| 0 | 0 | 0 | 0 |
| 0 | 0 | 0 | 0 |
| 0 | 0 | 0 | 0 |
| 0 | 0 | 0 | 0 |
| 0 | 0 | 0 | 0 |
| 0 | 0 | 0 | 0 |
| 0 | 0 | 0 | 0 |
| 0 | 0 | 0 | 0 |
| 0 | 0 | 0 | 0 |
| 1 | 0 | 0 | 0 |
| 0 | 0 | 0 | 0 |
| 0 | 0 | 0 | 0 |
| 0 | 0 | 0 | 0 |
| 1 | 1 | 0 | 1 |
| 0 | 0 | 0 | 0 |
| 1 | 0 | 0 | 0 |
| 1 | 1 | 0 | 0 |
| 0 | 0 | 0 | 0 |
| 0 | 0 | 0 | 0 |
| 0 | 0 | 0 | 0 |
| 0 | 0 | 0 | 0 |
| 0 | 0 | 0 | 0 |
| 0 | 0 | 0 | 1 |
| 0 | 0 | 0 | 0 |
| 0 | 0 | 0 | 0 |
| 0 | 0 | 0 | 0 |
| 0 | 0 | 0 | 1 |
| 0 | 0 | 0 | 0 |
| 0 | 0 | 0 | 0 |
| 0 | 0 | 0 | 0 |
| 0 | 0 | 0 | 0 |
| 0 | 0 | 0 | 0 |
| 0 | 0 | 0 | 0 |
| 0 | 0 | 0 | 0 |
| 0 | 0 | 0 | 0 |
| 0 | 0 | 0 | 0 |
| 0 | 0 | 0 | 0 |
| 0 | 0 | 0 | 0 |
| 0 | 0 | 0 | 0 |
| 0 | 0 | 0 | 0 |
| 0 | 0 | 0 | 0 |
| 0 | 0 | 0 | 0 |
| 0 | 0 | 0 | 0 |
| 0 | 0 | 1 | 0 |
| 0 | 0 | 0 | 0 |
| 0 | 0 | 0 | 0 |
| 0 | 0 | 0 | 0 |
| 0 | 0 | 0 | 0 |
| 0 | 0 | 0 | 0 |
| 0 | 0 | 0 | 0 |
| 0 | 0 | 0 | 0 |
| 0 | 0 | 0 | 0 |
| 0 | 0 | 0 | 0 |

| | | | |
|---|---|---|---|
| 0 | 0 | 0 | 0 |
| 0 | 0 | 0 | 0 |
| 0 | 0 | 0 | 0 |
| 0 | 0 | 0 | 0 |
| 0 | 0 | 0 | 0 |
| 0 | 0 | 0 | 0 |
| 1 | 0 | 0 | 1 |
| 1 | 0 | 0 | 0 |
| 0 | 0 | 0 | 0 |
| 0 | 0 | 0 | 0 |
| 0 | 0 | 0 | 0 |
| 0 | 0 | 0 | 0 |
| 0 | 0 | 0 | 0 |
| 0 | 0 | 0 | 0 |
| 0 | 0 | 0 | 0 |
| 0 | 0 | 0 | 0 |
| 0 | 0 | 0 | 0 |
| 0 | 0 | 0 | 0 |
| 0 | 0 | 0 | 0 |
| 0 | 0 | 0 | 0 |
| 1 | 0 | 0 | 0 |
| 0 | 0 | 0 | 0 |
| 0 | 0 | 0 | 0 |
| 0 | 0 | 0 | 0 |
| 0 | 0 | 0 | 0 |
| 0 | 0 | 0 | 0 |
| 0 | 0 | 0 | 0 |
| 0 | 0 | 0 | 0 |
| 0 | 0 | 0 | 0 |
| 0 | 0 | 0 | 0 |
| 0 | 0 | 0 | 0 |
| 0 | 0 | 0 | 0 |
| 0 | 0 | 0 | 0 |
| 0 | 0 | 0 | 0 |
| 0 | 0 | 0 | 0 |
| 0 | 0 | 0 | 0 |
| 0 | 0 | 0 | 0 |
| 0 | 0 | 0 | 0 |
| 0 | 0 | 0 | 0 |
| 0 | 0 | 0 | 0 |
| 0 | 0 | 0 | 0 |
| 0 | 0 | 0 | 0 |
| 0 | 0 | 0 | 0 |
| 0 | 0 | 0 | 0 |
| 0 | 0 | 0 | 0 |
| 0 | 0 | 0 | 0 |
| 0 | 0 | 0 | 0 |
| 0 | 0 | 0 | 0 |
| 0 | 0 | 0 | 0 |
| 0 | 0 | 0 | 0 |
| 0 | 0 | 0 | 0 |
| 0 | 0 | 0 | 0 |
| 0 | 0 | 0 | 0 |
| 0 | 0 | 0 | 0 |

| | | | |
|---|---|---|---|
| 0 | 0 | 0 | 0 |
| 0 | 0 | 0 | 0 |
| 0 | 0 | 0 | 0 |
| 1 | 0 | 0 | 0 |
| 0 | 0 | 0 | 0 |
| 1 | 0 | 0 | 0 |
| 0 | 0 | 0 | 0 |
| 0 | 0 | 0 | 0 |
| 1 | 0 | 0 | 0 |
| 0 | 0 | 0 | 0 |
| 0 | 0 | 0 | 0 |
| 0 | 0 | 0 | 0 |
| 0 | 0 | 0 | 0 |
| 0 | 0 | 0 | 0 |
| 0 | 0 | 0 | 0 |
| 0 | 0 | 0 | 0 |
| 0 | 0 | 0 | 0 |
| 0 | 0 | 0 | 0 |
| 1 | 0 | 0 | 0 |
| 0 | 0 | 0 | 0 |
| 0 | 0 | 0 | 0 |
| 0 | 0 | 0 | 0 |
| 0 | 0 | 0 | 0 |
| 0 | 0 | 0 | 0 |
| 0 | 0 | 0 | 0 |
| 0 | 0 | 0 | 0 |
| 0 | 0 | 0 | 0 |
| 0 | 0 | 0 | 0 |
| 0 | 1 | 0 | 0 |
| 0 | 0 | 0 | 0 |
| 0 | 0 | 0 | 0 |
| 0 | 0 | 0 | 0 |
| 0 | 1 | 0 | 0 |
| 1 | 1 | 0 | 0 |
| 1 | 0 | 0 | 0 |
| 0 | 0 | 0 | 0 |
| 0 | 0 | 0 | 0 |
| 0 | 0 | 0 | 0 |
| 0 | 0 | 0 | 0 |
| 0 | 0 | 0 | 0 |
| 1 | 0 | 0 | 0 |
| 0 | 0 | 0 | 0 |
| 0 | 0 | 0 | 0 |
| 0 | 0 | 0 | 0 |
| 0 | 0 | 0 | 0 |
| 0 | 0 | 0 | 0 |
| 0 | 0 | 0 | 0 |
| 0 | 0 | 0 | 0 |
| 0 | 0 | 0 | 0 |
| 0 | 0 | 0 | 0 |
| 0 | 0 | 0 | 0 |
| 0 | 0 | 0 | 0 |
| 0 | 0 | 0 | 0 |
| 1 | 0 | 0 | 0 |

| | | | |
|---|---|---|---|
| 1 | 0 | 0 | 1 |
| 0 | 0 | 0 | 0 |
| 0 | 0 | 0 | 0 |
| 0 | 0 | 0 | 0 |
| 0 | 0 | 0 | 0 |
| 0 | 0 | 0 | 0 |
| 0 | 0 | 0 | 0 |
| 0 | 0 | 0 | 0 |
| 0 | 0 | 0 | 0 |
| 0 | 0 | 0 | 0 |
| 0 | 0 | 0 | 0 |
| 0 | 0 | 0 | 0 |
| 0 | 0 | 0 | 0 |
| 0 | 0 | 0 | 0 |
| 0 | 0 | 0 | 0 |
| 0 | 0 | 0 | 0 |
| 0 | 0 | 0 | 0 |
| 0 | 0 | 0 | 0 |
| 0 | 0 | 0 | 0 |
| 0 | 0 | 0 | 0 |
| 0 | 0 | 0 | 0 |
| 0 | 0 | 0 | 0 |
| 0 | 1 | 0 | 0 |
| 0 | 0 | 0 | 0 |
| 0 | 0 | 0 | 0 |
| 0 | 0 | 0 | 0 |
| 0 | 0 | 0 | 0 |
| 0 | 0 | 0 | 0 |
| 0 | 0 | 0 | 0 |
| 0 | 0 | 0 | 0 |
| 0 | 0 | 0 | 0 |
| 0 | 0 | 0 | 0 |
| 0 | 0 | 0 | 0 |
| 0 | 0 | 0 | 0 |
| 0 | 0 | 0 | 0 |
| 0 | 0 | 0 | 0 |
| 0 | 0 | 0 | 0 |
| 0 | 0 | 0 | 0 |
| 0 | 0 | 0 | 0 |
| 0 | 0 | 0 | 0 |
| 0 | 0 | 0 | 0 |
| 0 | 0 | 0 | 0 |
| 0 | 0 | 0 | 0 |
| 0 | 0 | 0 | 0 |
| 0 | 0 | 0 | 0 |
| 0 | 0 | 0 | 0 |
| 0 | 0 | 0 | 0 |
| 0 | 0 | 0 | 0 |
| 0 | 0 | 0 | 0 |
| 0 | 0 | 0 | 0 |
| 0 | 0 | 0 | 0 |
| 0 | 0 | 0 | 0 |
| 0 | 0 | 0 | 0 |
| 0 | 0 | 0 | 0 |

| | | | |
|---|---|---|---|
| 0 | 0 | 0 | 0 |
| 0 | 0 | 0 | 0 |
| 0 | 0 | 0 | 0 |
| 0 | 0 | 0 | 0 |
| 0 | 0 | 0 | 0 |
| 0 | 0 | 0 | 0 |
| 0 | 1 | 0 | 0 |
| 0 | 1 | 0 | 0 |
| 0 | 0 | 0 | 0 |
| 0 | 0 | 0 | 0 |
| 0 | 0 | 0 | 0 |
| 0 | 0 | 0 | 0 |
| 0 | 0 | 0 | 0 |
| 0 | 0 | 0 | 0 |
| 0 | 0 | 0 | 0 |
| 0 | 0 | 0 | 0 |
| 1 | 0 | 0 | 0 |
| 0 | 0 | 0 | 0 |
| 1 | 0 | 0 | 0 |
| 1 | 0 | 0 | 0 |
| 1 | 0 | 0 | 0 |
| 1 | 0 | 0 | 0 |
| 0 | 0 | 0 | 0 |
| 0 | 0 | 0 | 0 |
| 0 | 0 | 0 | 0 |
| 0 | 0 | 0 | 0 |
| 0 | 0 | 0 | 0 |
| 0 | 0 | 0 | 0 |
| 0 | 0 | 0 | 0 |
| 0 | 0 | 0 | 0 |
| 0 | 0 | 0 | 0 |
| 0 | 0 | 0 | 0 |
| 0 | 0 | 0 | 0 |
| 0 | 0 | 0 | 0 |
| 0 | 0 | 0 | 0 |
| 0 | 0 | 1 | 0 |
| 1 | 0 | 0 | 0 |
| 1 | 0 | 0 | 0 |
| 1 | 0 | 0 | 0 |
| 0 | 0 | 0 | 0 |
| 0 | 0 | 0 | 0 |
| 0 | 0 | 0 | 0 |
| 0 | 0 | 0 | 0 |
| 1 | 0 | 0 | 0 |
| 1 | 0 | 0 | 0 |
| 1 | 0 | 0 | 0 |
| 0 | 0 | 0 | 0 |
| 0 | 0 | 0 | 0 |
| 1 | 0 | 0 | 0 |
| 0 | 0 | 0 | 0 |
| 0 | 0 | 0 | 0 |
| 1 | 0 | 0 | 0 |
| 0 | 0 | 0 | 0 |
| 0 | 0 | 0 | 0 |

| | | | |
|---|---|---|---|
| 0 | 0 | 0 | 0 |
| 0 | 0 | 0 | 0 |
| 0 | 0 | 0 | 0 |
| 0 | 0 | 0 | 0 |
| 0 | 0 | 0 | 0 |
| 0 | 0 | 0 | 0 |
| 0 | 0 | 0 | 0 |
| 0 | 0 | 0 | 0 |
| 0 | 0 | 0 | 0 |
| 0 | 0 | 0 | 0 |
| 0 | 0 | 0 | 0 |
| 0 | 0 | 0 | 0 |
| 0 | 0 | 0 | 0 |
| 0 | 0 | 0 | 0 |
| 0 | 0 | 0 | 0 |
| 0 | 0 | 0 | 0 |
| 0 | 0 | 0 | 0 |
| 0 | 0 | 0 | 0 |
| 0 | 0 | 0 | 0 |
| 0 | 0 | 0 | 0 |
| 0 | 0 | 0 | 0 |
| 0 | 0 | 0 | 0 |
| 1 | 1 | 0 | 0 |
| 0 | 0 | 0 | 0 |
| 0 | 1 | 0 | 0 |
| 0 | 0 | 0 | 0 |
| 0 | 0 | 0 | 0 |
| 0 | 0 | 0 | 0 |
| 0 | 0 | 0 | 0 |
| 0 | 0 | 0 | 0 |
| 0 | 0 | 0 | 0 |
| 0 | 0 | 0 | 0 |
| 0 | 0 | 0 | 0 |
| 0 | 0 | 0 | 0 |
| 0 | 0 | 0 | 0 |
| 0 | 0 | 0 | 0 |
| 0 | 0 | 0 | 0 |
| 0 | 0 | 0 | 0 |
| 0 | 0 | 0 | 0 |
| 0 | 0 | 0 | 0 |
| 0 | 0 | 0 | 0 |
| 0 | 0 | 0 | 0 |
| 0 | 0 | 0 | 0 |
| 0 | 0 | 0 | 0 |
| 0 | 0 | 0 | 0 |
| 1 | 0 | 0 | 1 |
| 0 | 0 | 0 | 0 |
| 0 | 0 | 0 | 0 |
| 0 | 0 | 0 | 0 |
| 1 | 0 | 0 | 0 |
| 1 | 0 | 0 | 0 |
| 1 | 0 | 0 | 0 |
| 0 | 0 | 0 | 0 |
| 0 | 0 | 0 | 0 |

| | | | |
|---|---|---|---|
| 0 | 0 | 0 | 0 |
| 0 | 0 | 0 | 0 |
| 0 | 0 | 0 | 0 |
| 0 | 0 | 0 | 0 |
| 0 | 0 | 0 | 0 |
| 0 | 0 | 0 | 0 |
| 0 | 0 | 0 | 0 |
| 0 | 0 | 0 | 0 |
| 0 | 0 | 0 | 0 |
| 0 | 0 | 0 | 0 |
| 0 | 0 | 0 | 1 |
| 0 | 0 | 0 | 0 |
| 0 | 0 | 0 | 0 |
| 0 | 0 | 0 | 0 |
| 0 | 0 | 0 | 0 |
| 0 | 0 | 0 | 0 |
| 0 | 0 | 0 | 0 |
| 0 | 0 | 0 | 0 |
| 0 | 0 | 0 | 0 |
| 0 | 0 | 0 | 0 |
| 0 | 0 | 0 | 0 |
| 0 | 0 | 0 | 0 |
| 0 | 0 | 0 | 0 |
| 0 | 0 | 0 | 0 |
| 0 | 0 | 0 | 0 |
| 0 | 0 | 0 | 0 |
| 0 | 0 | 0 | 0 |
| 0 | 0 | 0 | 0 |
| 0 | 0 | 0 | 0 |
| 0 | 0 | 0 | 0 |
| 0 | 1 | 0 | 0 |
| 0 | 0 | 0 | 0 |
| 1 | 1 | 0 | 0 |
| 1 | 1 | 0 | 0 |
| 0 | 0 | 0 | 0 |
| 0 | 0 | 0 | 0 |
| 0 | 0 | 0 | 0 |
| 0 | 0 | 0 | 0 |
| 0 | 0 | 0 | 0 |
| 0 | 0 | 0 | 0 |
| 0 | 0 | 0 | 0 |
| 0 | 1 | 0 | 0 |
| 0 | 0 | 0 | 0 |
| 0 | 0 | 0 | 0 |
| 0 | 0 | 0 | 0 |
| 1 | 0 | 0 | 0 |
| 1 | 0 | 0 | 0 |
| 1 | 0 | 0 | 0 |
| 1 | 1 | 1 | 0 |
| 0 | 0 | 0 | 0 |
| 1 | 1 | 1 | 1 |
| 0 | 0 | 0 | 0 |
| 0 | 0 | 0 | 0 |
| 0 | 0 | 0 | 0 |

| | | | |
|---|---|---|---|
| 0 | 0 | 0 | 0 |
| 0 | 0 | 0 | 0 |
| 0 | 0 | 0 | 0 |
| 0 | 0 | 0 | 0 |
| 1 | 0 | 0 | 0 |
| 1 | 0 | 0 | 0 |
| 0 | 0 | 0 | 0 |
| 0 | 0 | 0 | 0 |
| 0 | 0 | 0 | 0 |
| 0 | 0 | 0 | 0 |
| 0 | 0 | 0 | 0 |
| 0 | 0 | 0 | 0 |
| 0 | 0 | 0 | 0 |
| 0 | 0 | 0 | 0 |
| 0 | 0 | 0 | 0 |
| 0 | 0 | 0 | 0 |
| 0 | 0 | 0 | 0 |
| 0 | 0 | 0 | 0 |
| 0 | 0 | 0 | 0 |
| 0 | 0 | 0 | 0 |
| 0 | 0 | 0 | 0 |
| 0 | 0 | 0 | 0 |
| 0 | 0 | 0 | 0 |
| 0 | 0 | 0 | 0 |
| 0 | 0 | 0 | 0 |
| 0 | 0 | 0 | 0 |
| 0 | 0 | 0 | 0 |
| 0 | 0 | 0 | 0 |
| 0 | 0 | 0 | 0 |
| 0 | 0 | 0 | 0 |
| 0 | 0 | 0 | 0 |
| 0 | 0 | 0 | 0 |
| 0 | 0 | 0 | 0 |
| 0 | 0 | 0 | 0 |
| 0 | 0 | 0 | 0 |
| 0 | 0 | 0 | 0 |
| 0 | 0 | 0 | 0 |
| 0 | 0 | 0 | 0 |
| 0 | 0 | 0 | 0 |
| 0 | 0 | 0 | 0 |
| 0 | 0 | 0 | 0 |
| 0 | 0 | 0 | 0 |
| 0 | 0 | 0 | 0 |
| 0 | 0 | 0 | 0 |
| 0 | 0 | 0 | 0 |
| 0 | 0 | 0 | 0 |
| 0 | 0 | 0 | 0 |
| 0 | 0 | 0 | 0 |
| 0 | 0 | 0 | 0 |
| 0 | 0 | 0 | 0 |
| 0 | 0 | 0 | 0 |
| 0 | 0 | 0 | 0 |
| 0 | 0 | 0 | 0 |
| 0 | 1 | 0 | 0 |

| | | | |
|---|---|---|---|
| 0 | 0 | 0 | 0 |
| 0 | 0 | 0 | 0 |
| 0 | 0 | 0 | 0 |
| 0 | 0 | 0 | 0 |
| 0 | 0 | 0 | 0 |
| 0 | 0 | 0 | 0 |
| 0 | 0 | 0 | 0 |
| 0 | 0 | 0 | 0 |
| 0 | 0 | 0 | 0 |
| 0 | 0 | 0 | 0 |
| 0 | 0 | 0 | 0 |
| 0 | 0 | 0 | 0 |
| 0 | 0 | 0 | 0 |
| 0 | 0 | 0 | 0 |
| 0 | 0 | 0 | 0 |
| 0 | 0 | 0 | 0 |
| 0 | 0 | 0 | 0 |
| 0 | 0 | 0 | 0 |
| 0 | 0 | 0 | 0 |
| 0 | 0 | 0 | 0 |
| 0 | 0 | 0 | 0 |
| 0 | 0 | 0 | 0 |
| 0 | 0 | 0 | 0 |
| 0 | 0 | 0 | 0 |
| 0 | 0 | 0 | 0 |
| 0 | 0 | 0 | 0 |
| 0 | 0 | 0 | 0 |
| 0 | 0 | 0 | 0 |
| 0 | 0 | 0 | 0 |
| 0 | 0 | 0 | 0 |
| 0 | 0 | 0 | 0 |
| 0 | 0 | 0 | 0 |
| 0 | 0 | 0 | 0 |
| 0 | 0 | 0 | 0 |
| 0 | 0 | 0 | 0 |
| 0 | 0 | 0 | 0 |
| 0 | 0 | 0 | 0 |
| 0 | 0 | 0 | 0 |
| 0 | 0 | 0 | 0 |
| 0 | 0 | 0 | 0 |
| 0 | 0 | 0 | 0 |
| 0 | 0 | 0 | 0 |
| 0 | 0 | 0 | 0 |
| 0 | 0 | 0 | 0 |
| 0 | 0 | 0 | 0 |
| 0 | 0 | 0 | 0 |
| 0 | 0 | 0 | 0 |
| 0 | 0 | 0 | 0 |
| 0 | 0 | 0 | 0 |
| 0 | 0 | 0 | 0 |
| 0 | 0 | 0 | 0 |
| 0 | 0 | 0 | 0 |
| 0 | 0 | 0 | 0 |
| 0 | 0 | 0 | 0 |

| | | | |
|---|---|---|---|
| 0 | 0 | 0 | 0 |
| 0 | 0 | 0 | 0 |
| 0 | 0 | 0 | 0 |
| 0 | 0 | 0 | 0 |
| 0 | 0 | 0 | 0 |
| 0 | 0 | 0 | 0 |
| 0 | 0 | 0 | 0 |
| 0 | 0 | 0 | 0 |
| 0 | 0 | 0 | 0 |
| 0 | 0 | 0 | 0 |
| 0 | 0 | 0 | 0 |
| 0 | 0 | 1 | 0 |
| 0 | 0 | 0 | 0 |
| 0 | 0 | 1 | 0 |
| 0 | 0 | 1 | 0 |
| 0 | 0 | 1 | 0 |
| 0 | 0 | 1 | 0 |
| 0 | 0 | 0 | 0 |
| 0 | 0 | 0 | 0 |
| 0 | 0 | 0 | 0 |
| 1 | 0 | 0 | 0 |

| glycan_binding_site | water_binding_site | water_only_binding_site | non_binding_site |
|---|---|---|---|
| 0 | 1 | 0 | 0 |
| 0 | 0 | 0 | 0 |
| 0 | 0 | 0 | 1 |
| 0 | 0 | 0 | 0 |
| 0 | 0 | 0 | 0 |
| 0 | 0 | 0 | 0 |
| 0 | 0 | 0 | 1 |
| 0 | 0 | 0 | 1 |
| 0 | 1 | 0 | 0 |
| 0 | 0 | 0 | 0 |
| 0 | 0 | 0 | 0 |
| 0 | 1 | 0 | 0 |
| 0 | 0 | 0 | 1 |
| 1 | 0 | 0 | 0 |
| 0 | 0 | 0 | 0 |
| 0 | 0 | 0 | 0 |
| 0 | 0 | 0 | 0 |
| 0 | 1 | 0 | 0 |
| 0 | 0 | 0 | 0 |
| 0 | 0 | 0 | 0 |
| 0 | 1 | 0 | 0 |
| 1 | 1 | 0 | 0 |
| 1 | 1 | 0 | 0 |
| 0 | 1 | 0 | 0 |
| 0 | 1 | 0 | 0 |
| 0 | 1 | 1 | 1 |
| 0 | 1 | 0 | 0 |
| 0 | 0 | 0 | 1 |
| 0 | 0 | 0 | 0 |
| 0 | 0 | 0 | 1 |
| 0 | 0 | 0 | 1 |
| 0 | 1 | 0 | 0 |
| 0 | 1 | 0 | 0 |
| 0 | 1 | 0 | 0 |
| 0 | 1 | 0 | 0 |
| 0 | 0 | 0 | 1 |
| 0 | 0 | 0 | 0 |
| 0 | 1 | 0 | 0 |
| 0 | 1 | 0 | 0 |
| 0 | 1 | 0 | 0 |
| 0 | 1 | 0 | 0 |
| 0 | 1 | 0 | 0 |
| 0 | 1 | 0 | 0 |
| 0 | 1 | 0 | 0 |
| 0 | 1 | 0 | 0 |
| 0 | 1 | 0 | 0 |
| 0 | 1 | 0 | 0 |
| 0 | 1 | 0 | 0 |
| 0 | 1 | 0 | 0 |
| 0 | 1 | 0 | 0 |
| 0 | 1 | 0 | 0 |
| 0 | 1 | 0 | 0 |
| 0 | 1 | 0 | 0 |

| | | | |
|---|---|---|---|
| 0 | 1 | 0 | 0 |
| 0 | 1 | 0 | 0 |
| 0 | 1 | 0 | 0 |
| 0 | 1 | 0 | 0 |
| 0 | 1 | 0 | 0 |
| 0 | 1 | 0 | 0 |
| 0 | 1 | 0 | 0 |
| 0 | 1 | 0 | 0 |
| 0 | 1 | 0 | 0 |
| 0 | 1 | 0 | 0 |
| 0 | 1 | 0 | 0 |
| 0 | 1 | 0 | 0 |
| 0 | 1 | 0 | 0 |
| 0 | 1 | 0 | 0 |
| 0 | 1 | 0 | 0 |
| 0 | 1 | 0 | 0 |
| 0 | 1 | 0 | 0 |
| 0 | 1 | 0 | 0 |
| 0 | 1 | 0 | 0 |
| 0 | 1 | 0 | 0 |
| 0 | 0 | 0 | 0 |
| 0 | 0 | 0 | 0 |
| 0 | 0 | 0 | 0 |
| 0 | 0 | 0 | 0 |
| 0 | 0 | 0 | 0 |
| 0 | 0 | 0 | 0 |
| 0 | 0 | 0 | 0 |
| 0 | 0 | 0 | 0 |
| 0 | 0 | 0 | 0 |
| 0 | 0 | 0 | 0 |
| 0 | 0 | 0 | 0 |
| 0 | 0 | 0 | 0 |
| 0 | 0 | 0 | 0 |
| 0 | 0 | 0 | 0 |
| 0 | 0 | 0 | 0 |
| 0 | 0 | 0 | 0 |
| 0 | 0 | 0 | 0 |
| 0 | 0 | 0 | 0 |
| 0 | 0 | 0 | 0 |
| 0 | 1 | 0 | 0 |
| 1 | 0 | 0 | 0 |
| 0 | 0 | 0 | 0 |
| 1 | 0 | 0 | 0 |
| 1 | 0 | 0 | 0 |
| 0 | 0 | 0 | 0 |
| 0 | 0 | 0 | 1 |
| 0 | 0 | 0 | 1 |
| 0 | 0 | 0 | 0 |
| 1 | 0 | 0 | 0 |
| 0 | 0 | 0 | 0 |
| 0 | 0 | 0 | 0 |
| 0 | 0 | 0 | 1 |
| 0 | 0 | 0 | 1 |
| 0 | 0 | 0 | 0 |

| | | | |
|---|---|---|---|
| 0 | 0 | 0 | 1 |
| 0 | 0 | 0 | 1 |
| 0 | 0 | 0 | 0 |
| 0 | 0 | 0 | 0 |
| 0 | 0 | 0 | 0 |
| 0 | 0 | 0 | 0 |
| 0 | 0 | 0 | 1 |
| 0 | 1 | 0 | 0 |
| 0 | 1 | 0 | 0 |
| 0 | 0 | 0 | 0 |
| 0 | 1 | 0 | 0 |
| 0 | 0 | 0 | 0 |
| 0 | 0 | 0 | 0 |
| 1 | 0 | 0 | 0 |
| 1 | 0 | 0 | 0 |
| 0 | 0 | 0 | 1 |
| 0 | 0 | 0 | 0 |
| 0 | 1 | 0 | 0 |
| 0 | 1 | 0 | 0 |
| 0 | 1 | 0 | 0 |
| 0 | 1 | 0 | 0 |
| 0 | 1 | 0 | 0 |
| 0 | 1 | 0 | 0 |
| 0 | 1 | 0 | 0 |
| 0 | 1 | 0 | 0 |
| 0 | 1 | 0 | 0 |
| 0 | 1 | 0 | 0 |
| 0 | 1 | 0 | 0 |
| 0 | 1 | 0 | 0 |
| 0 | 1 | 0 | 0 |
| 0 | 1 | 0 | 0 |
| 0 | 0 | 0 | 0 |
| 0 | 1 | 0 | 0 |
| 0 | 1 | 0 | 0 |
| 0 | 1 | 0 | 0 |
| 0 | 1 | 0 | 0 |
| 0 | 1 | 0 | 0 |
| 0 | 1 | 0 | 0 |
| 0 | 1 | 0 | 0 |
| 0 | 1 | 0 | 0 |
| 0 | 1 | 0 | 0 |
| 0 | 1 | 0 | 0 |
| 0 | 1 | 0 | 0 |
| 0 | 1 | 0 | 0 |
| 0 | 1 | 0 | 0 |
| 0 | 1 | 0 | 0 |
| 1 | 1 | 0 | 0 |
| 0 | 0 | 0 | 0 |
| 0 | 1 | 0 | 0 |
| 0 | 0 | 0 | 0 |
| 0 | 0 | 0 | 0 |
| 0 | 0 | 0 | 0 |
| 0 | 0 | 0 | 0 |
| 0 | 1 | 1 | 1 |

| | | | |
|---|---|---|---|
| 0 | 1 | 1 | 1 |
| 0 | 1 | 1 | 1 |
| 0 | 1 | 1 | 1 |
| 0 | 0 | 0 | 0 |
| 0 | 0 | 0 | 0 |
| 0 | 1 | 1 | 1 |
| 0 | 1 | 0 | 0 |
| 0 | 1 | 1 | 1 |
| 0 | 1 | 1 | 1 |
| 0 | 1 | 1 | 1 |
| 0 | 1 | 1 | 1 |
| 0 | 1 | 0 | 0 |
| 0 | 1 | 1 | 1 |
| 0 | 0 | 0 | 0 |
| 0 | 1 | 1 | 1 |
| 0 | 0 | 0 | 1 |
| 0 | 1 | 1 | 1 |
| 0 | 1 | 1 | 1 |
| 0 | 1 | 1 | 1 |
| 0 | 1 | 0 | 0 |
| 0 | 1 | 0 | 0 |
| 0 | 0 | 0 | 1 |
| 0 | 1 | 0 | 0 |
| 0 | 1 | 0 | 0 |
| 0 | 1 | 0 | 0 |
| 0 | 1 | 1 | 1 |
| 0 | 1 | 0 | 0 |
| 0 | 1 | 0 | 0 |
| 0 | 0 | 0 | 1 |
| 0 | 1 | 1 | 1 |
| 0 | 0 | 0 | 1 |
| 0 | 1 | 1 | 1 |
| 0 | 1 | 1 | 1 |
| 0 | 0 | 0 | 1 |
| 0 | 1 | 0 | 0 |
| 0 | 1 | 1 | 1 |
| 1 | 1 | 0 | 0 |
| 1 | 1 | 0 | 0 |
| 0 | 1 | 1 | 1 |
| 0 | 1 | 1 | 1 |
| 0 | 1 | 1 | 1 |
| 0 | 1 | 0 | 0 |
| 0 | 1 | 0 | 0 |
| 0 | 0 | 0 | 1 |
| 0 | 1 | 1 | 1 |
| 0 | 1 | 1 | 1 |
| 0 | 0 | 0 | 0 |
| 0 | 0 | 0 | 0 |
| 0 | 0 | 0 | 0 |
| 0 | 0 | 0 | 0 |
| 0 | 0 | 0 | 0 |
| 0 | 0 | 0 | 1 |
| 0 | 0 | 0 | 1 |
| 0 | 0 | 0 | 0 |

| | | | |
|---|---|---|---|
| 0 | 0 | 0 | 0 |
| 0 | 0 | 0 | 0 |
| 0 | 1 | 0 | 0 |
| 0 | 0 | 0 | 1 |
| 0 | 0 | 0 | 1 |
| 0 | 0 | 0 | 1 |
| 0 | 1 | 0 | 0 |
| 0 | 1 | 0 | 0 |
| 0 | 1 | 1 | 1 |
| 0 | 1 | 1 | 1 |
| 1 | 0 | 0 | 0 |
| 0 | 0 | 0 | 1 |
| 1 | 0 | 0 | 0 |
| 0 | 0 | 0 | 1 |
| 0 | 0 | 0 | 1 |
| 0 | 0 | 0 | 1 |
| 0 | 0 | 0 | 1 |
| 0 | 1 | 0 | 0 |
| 0 | 0 | 0 | 0 |
| 0 | 1 | 0 | 0 |
| 0 | 0 | 0 | 0 |
| 0 | 0 | 0 | 1 |
| 0 | 1 | 0 | 0 |
| 0 | 0 | 0 | 1 |
| 0 | 0 | 0 | 1 |
| 0 | 0 | 0 | 0 |
| 0 | 1 | 0 | 0 |
| 0 | 0 | 0 | 0 |
| 0 | 1 | 0 | 0 |
| 0 | 1 | 0 | 0 |
| 0 | 1 | 0 | 0 |
| 0 | 0 | 0 | 0 |
| 0 | 0 | 0 | 1 |
| 0 | 0 | 0 | 1 |
| 0 | 0 | 0 | 0 |
| 0 | 0 | 0 | 1 |
| 0 | 0 | 0 | 1 |
| 0 | 0 | 0 | 0 |
| 0 | 1 | 1 | 1 |
| 0 | 0 | 0 | 1 |
| 0 | 1 | 0 | 0 |
| 0 | 1 | 1 | 1 |
| 0 | 0 | 0 | 1 |
| 0 | 0 | 0 | 1 |
| 0 | 0 | 0 | 0 |
| 0 | 0 | 0 | 0 |
| 0 | 1 | 0 | 0 |
| 0 | 1 | 0 | 0 |
| 0 | 1 | 0 | 0 |
| 0 | 1 | 0 | 0 |
| 0 | 1 | 0 | 0 |
| 0 | 1 | 0 | 0 |
| 0 | 1 | 0 | 0 |
| 0 | 1 | 0 | 0 |

| | | | |
|---|---|---|---|
| 0 | 1 | 0 | 0 |
| 0 | 1 | 0 | 0 |
| 0 | 1 | 0 | 0 |
| 0 | 1 | 0 | 0 |
| 0 | 1 | 0 | 0 |
| 0 | 1 | 0 | 0 |
| 0 | 0 | 0 | 1 |
| 0 | 0 | 0 | 1 |
| 0 | 0 | 0 | 0 |
| 0 | 0 | 0 | 0 |
| 0 | 0 | 0 | 0 |
| 0 | 0 | 0 | 0 |
| 0 | 0 | 0 | 0 |
| 0 | 0 | 0 | 1 |
| 0 | 0 | 0 | 1 |
| 0 | 0 | 0 | 0 |
| 0 | 0 | 0 | 1 |
| 0 | 0 | 0 | 0 |
| 0 | 0 | 0 | 1 |
| 0 | 0 | 0 | 0 |
| 0 | 0 | 0 | 1 |
| 0 | 1 | 1 | 1 |
| 0 | 0 | 0 | 1 |
| 0 | 0 | 0 | 0 |
| 0 | 0 | 0 | 0 |
| 0 | 1 | 0 | 0 |
| 0 | 1 | 0 | 0 |
| 0 | 1 | 0 | 0 |
| 0 | 0 | 0 | 0 |
| 0 | 1 | 0 | 0 |
| 0 | 0 | 0 | 1 |
| 0 | 1 | 1 | 1 |
| 0 | 0 | 0 | 0 |
| 0 | 0 | 0 | 0 |
| 0 | 1 | 0 | 0 |
| 0 | 0 | 0 | 1 |
| 0 | 0 | 0 | 0 |
| 0 | 1 | 0 | 0 |
| 0 | 0 | 0 | 0 |
| 0 | 1 | 0 | 0 |
| 0 | 0 | 0 | 1 |
| 0 | 0 | 0 | 1 |
| 0 | 0 | 0 | 0 |
| 0 | 0 | 0 | 1 |
| 0 | 0 | 0 | 0 |
| 0 | 0 | 0 | 0 |
| 0 | 0 | 0 | 0 |
| 0 | 0 | 0 | 1 |
| 0 | 0 | 0 | 1 |
| 0 | 0 | 0 | 0 |
| 0 | 0 | 0 | 1 |
| 0 | 0 | 0 | 1 |
| 0 | 0 | 0 | 0 |
| 0 | 0 | 0 | 1 |

| | | | |
|---|---|---|---|
| 0 | 0 | 0 | 0 |
| 0 | 0 | 0 | 1 |
| 0 | 1 | 1 | 1 |
| 0 | 1 | 0 | 0 |
| 0 | 1 | 0 | 0 |
| 0 | 0 | 0 | 0 |
| 0 | 0 | 0 | 1 |
| 0 | 1 | 0 | 0 |
| 0 | 0 | 0 | 0 |
| 0 | 0 | 0 | 1 |
| 0 | 0 | 0 | 1 |
| 0 | 0 | 0 | 1 |
| 0 | 0 | 0 | 1 |
| 0 | 0 | 0 | 1 |
| 0 | 0 | 0 | 0 |
| 0 | 0 | 0 | 1 |
| 0 | 0 | 0 | 0 |
| 0 | 0 | 0 | 0 |
| 0 | 0 | 0 | 1 |
| 0 | 0 | 0 | 0 |
| 0 | 1 | 0 | 0 |
| 0 | 1 | 0 | 0 |
| 0 | 1 | 0 | 0 |
| 0 | 1 | 0 | 0 |
| 0 | 0 | 0 | 1 |
| 0 | 0 | 0 | 1 |
| 0 | 0 | 0 | 0 |
| 0 | 0 | 0 | 0 |
| 0 | 1 | 0 | 0 |
| 0 | 0 | 0 | 0 |
| 0 | 0 | 0 | 0 |
| 0 | 0 | 0 | 0 |
| 0 | 0 | 0 | 0 |
| 0 | 0 | 0 | 0 |
| 0 | 0 | 0 | 1 |
| 0 | 0 | 0 | 1 |
| 0 | 0 | 0 | 0 |
| 0 | 0 | 0 | 0 |
| 0 | 0 | 0 | 0 |
| 0 | 0 | 0 | 0 |
| 0 | 0 | 0 | 0 |
| 0 | 1 | 0 | 0 |
| 0 | 0 | 0 | 1 |
| 0 | 1 | 0 | 0 |
| 0 | 1 | 0 | 0 |
| 0 | 1 | 0 | 0 |
| 0 | 1 | 0 | 0 |
| 0 | 1 | 0 | 0 |
| 1 | 1 | 0 | 0 |
| 0 | 0 | 0 | 1 |
| 0 | 1 | 0 | 0 |
| 0 | 1 | 0 | 0 |
| 0 | 0 | 0 | 0 |
| 0 | 1 | 0 | 0 |

| | | | |
|---|---|---|---|
| 0 | 1 | 0 | 0 |
| 0 | 1 | 0 | 0 |
| 0 | 1 | 0 | 0 |
| 0 | 1 | 0 | 0 |
| 0 | 0 | 0 | 1 |
| 0 | 1 | 0 | 0 |
| 0 | 1 | 0 | 0 |
| 0 | 1 | 0 | 0 |
| 0 | 1 | 0 | 0 |
| 0 | 1 | 0 | 0 |
| 0 | 1 | 0 | 0 |
| 0 | 1 | 0 | 0 |
| 0 | 1 | 0 | 0 |
| 0 | 1 | 0 | 0 |
| 0 | 1 | 0 | 0 |
| 0 | 1 | 0 | 0 |
| 0 | 1 | 0 | 0 |
| 0 | 1 | 0 | 0 |
| 0 | 0 | 0 | 1 |
| 0 | 0 | 0 | 0 |
| 0 | 0 | 0 | 0 |
| 0 | 0 | 0 | 0 |
| 0 | 1 | 0 | 0 |
| 0 | 1 | 1 | 1 |
| 0 | 0 | 0 | 0 |
| 0 | 0 | 0 | 0 |
| 0 | 0 | 0 | 0 |
| 0 | 1 | 1 | 1 |
| 0 | 1 | 1 | 1 |
| 0 | 1 | 1 | 1 |
| 0 | 1 | 0 | 0 |
| 0 | 0 | 0 | 0 |
| 0 | 1 | 0 | 0 |
| 0 | 1 | 0 | 0 |
| 0 | 1 | 0 | 0 |
| 0 | 1 | 1 | 1 |
| 0 | 1 | 1 | 1 |
| 0 | 0 | 0 | 0 |
| 0 | 1 | 0 | 0 |
| 0 | 1 | 1 | 1 |
| 0 | 1 | 1 | 1 |
| 0 | 1 | 1 | 1 |
| 0 | 0 | 0 | 1 |
| 0 | 0 | 0 | 1 |
| 0 | 0 | 0 | 0 |
| 0 | 0 | 0 | 0 |
| 0 | 1 | 0 | 0 |
| 0 | 0 | 0 | 0 |
| 0 | 0 | 0 | 0 |
| 0 | 0 | 0 | 1 |
| 0 | 0 | 0 | 1 |
| 0 | 0 | 0 | 0 |
| 0 | 0 | 0 | 1 |
| 0 | 0 | 0 | 0 |

| | | | |
|---|---|---|---|
| 0 | 0 | 0 | 1 |
| 0 | 0 | 0 | 1 |
| 0 | 0 | 0 | 0 |
| 0 | 0 | 0 | 1 |
| 0 | 0 | 0 | 1 |
| 0 | 0 | 0 | 1 |
| 0 | 0 | 0 | 1 |
| 0 | 1 | 0 | 0 |
| 0 | 1 | 1 | 1 |
| 0 | 0 | 0 | 1 |
| 0 | 0 | 0 | 0 |
| 0 | 0 | 0 | 1 |
| 0 | 0 | 0 | 1 |
| 0 | 0 | 0 | 1 |
| 0 | 0 | 0 | 0 |
| 0 | 0 | 0 | 0 |
| 0 | 0 | 0 | 1 |
| 0 | 0 | 0 | 1 |
| 0 | 0 | 0 | 0 |
| 0 | 0 | 0 | 0 |
| 0 | 0 | 0 | 0 |
| 0 | 1 | 0 | 0 |
| 0 | 0 | 0 | 0 |
| 0 | 1 | 0 | 0 |
| 0 | 1 | 0 | 0 |
| 0 | 0 | 0 | 1 |
| 0 | 0 | 0 | 0 |
| 0 | 1 | 0 | 0 |
| 0 | 0 | 0 | 0 |
| 0 | 0 | 0 | 1 |
| 0 | 0 | 0 | 0 |
| 0 | 1 | 0 | 0 |
| 0 | 0 | 0 | 1 |
| 0 | 1 | 0 | 0 |
| 0 | 1 | 0 | 0 |
| 0 | 1 | 0 | 0 |
| 0 | 1 | 0 | 0 |
| 0 | 1 | 0 | 0 |
| 0 | 1 | 0 | 0 |
| 0 | 1 | 0 | 0 |
| 0 | 0 | 0 | 1 |
| 0 | 0 | 0 | 0 |
| 0 | 0 | 0 | 1 |
| 0 | 0 | 0 | 0 |
| 0 | 0 | 0 | 0 |
| 1 | 0 | 0 | 0 |
| 0 | 0 | 0 | 0 |
| 1 | 0 | 0 | 0 |
| 0 | 0 | 0 | 0 |
| 0 | 0 | 0 | 0 |
| 0 | 0 | 0 | 1 |
| 0 | 0 | 0 | 0 |
| 0 | 0 | 0 | 0 |
| 0 | 0 | 0 | 0 |

| | | | |
|---|---|---|---|
| 0 | 0 | 0 | 1 |
| 0 | 0 | 0 | 0 |
| 0 | 0 | 0 | 1 |
| 0 | 0 | 0 | 1 |
| 1 | 0 | 0 | 0 |
| 0 | 0 | 0 | 0 |
| 1 | 0 | 0 | 0 |
| 1 | 0 | 0 | 0 |
| 0 | 0 | 0 | 0 |
| 0 | 0 | 0 | 1 |
| 0 | 0 | 0 | 1 |
| 0 | 0 | 0 | 1 |
| 0 | 0 | 0 | 1 |
| 1 | 1 | 0 | 0 |
| 0 | 0 | 0 | 1 |
| 0 | 0 | 0 | 0 |
| 0 | 0 | 0 | 0 |
| 0 | 0 | 0 | 0 |
| 1 | 1 | 0 | 0 |
| 0 | 1 | 1 | 1 |
| 1 | 1 | 0 | 0 |
| 0 | 0 | 0 | 1 |
| 0 | 0 | 0 | 0 |
| 0 | 0 | 0 | 1 |
| 0 | 0 | 0 | 1 |
| 0 | 0 | 0 | 1 |
| 0 | 0 | 0 | 0 |
| 0 | 0 | 0 | 1 |
| 0 | 0 | 0 | 1 |
| 0 | 0 | 0 | 1 |
| 0 | 0 | 0 | 0 |
| 0 | 0 | 0 | 0 |
| 0 | 0 | 0 | 1 |
| 0 | 0 | 0 | 1 |
| 0 | 0 | 0 | 0 |
| 0 | 0 | 0 | 0 |
| 0 | 0 | 0 | 0 |
| 1 | 0 | 0 | 0 |
| 0 | 0 | 0 | 0 |
| 0 | 0 | 0 | 1 |
| 0 | 0 | 0 | 0 |
| 0 | 1 | 0 | 0 |
| 0 | 1 | 0 | 0 |
| 0 | 0 | 0 | 0 |
| 0 | 0 | 0 | 1 |
| 1 | 0 | 0 | 0 |
| 0 | 1 | 0 | 0 |
| 0 | 0 | 0 | 0 |
| 0 | 0 | 0 | 0 |
| 1 | 0 | 0 | 0 |
| 0 | 0 | 0 | 0 |
| 0 | 1 | 0 | 0 |
| 0 | 1 | 1 | 1 |
| 0 | 0 | 0 | 0 |

| | | | |
|---|---|---|---|
| 0 | 1 | 0 | 0 |
| 0 | 1 | 0 | 0 |
| 0 | 1 | 1 | 1 |
| 0 | 1 | 1 | 1 |
| 0 | 1 | 1 | 1 |
| 0 | 1 | 0 | 0 |
| 0 | 1 | 1 | 1 |
| 0 | 1 | 1 | 1 |
| 0 | 1 | 0 | 0 |
| 0 | 0 | 0 | 0 |
| 1 | 1 | 0 | 0 |
| 1 | 1 | 0 | 0 |
| 0 | 0 | 0 | 1 |
| 1 | 0 | 0 | 0 |
| 0 | 0 | 0 | 0 |
| 0 | 0 | 0 | 0 |
| 0 | 1 | 0 | 0 |
| 0 | 0 | 0 | 0 |
| 0 | 0 | 0 | 0 |
| 0 | 0 | 0 | 1 |
| 0 | 0 | 0 | 0 |
| 0 | 1 | 0 | 0 |
| 0 | 0 | 0 | 0 |
| 0 | 0 | 0 | 0 |
| 0 | 1 | 0 | 0 |
| 0 | 1 | 1 | 1 |
| 0 | 0 | 0 | 1 |
| 0 | 0 | 0 | 1 |
| 0 | 0 | 0 | 0 |
| 0 | 1 | 0 | 0 |
| 0 | 0 | 0 | 0 |
| 0 | 0 | 0 | 0 |
| 0 | 0 | 0 | 0 |
| 0 | 1 | 0 | 0 |
| 0 | 1 | 0 | 0 |
| 0 | 1 | 1 | 1 |
| 0 | 0 | 0 | 0 |
| 0 | 1 | 0 | 0 |
| 0 | 0 | 0 | 1 |
| 0 | 0 | 0 | 1 |
| 0 | 0 | 0 | 1 |
| 0 | 1 | 0 | 0 |
| 0 | 0 | 0 | 0 |
| 0 | 0 | 0 | 0 |
| 0 | 0 | 0 | 0 |
| 0 | 0 | 0 | 0 |
| 0 | 0 | 0 | 0 |
| 0 | 0 | 0 | 1 |
| 0 | 0 | 0 | 1 |
| 0 | 0 | 0 | 0 |
| 0 | 0 | 0 | 0 |
| 0 | 0 | 0 | 0 |
| 0 | 0 | 0 | 0 |
| 0 | 0 | 0 | 1 |

| | | | |
|---|---|---|---|
| 0 | 0 | 0 | 1 |
| 0 | 0 | 0 | 0 |
| 0 | 0 | 0 | 1 |
| 0 | 1 | 0 | 0 |
| 0 | 1 | 1 | 1 |
| 0 | 1 | 1 | 1 |
| 0 | 1 | 0 | 0 |
| 0 | 1 | 1 | 1 |
| 0 | 1 | 1 | 1 |
| 0 | 1 | 1 | 1 |
| 0 | 1 | 1 | 1 |
| 0 | 1 | 1 | 1 |
| 0 | 0 | 0 | 1 |
| 0 | 0 | 0 | 1 |
| 0 | 0 | 0 | 1 |
| 0 | 0 | 0 | 1 |
| 0 | 0 | 0 | 0 |
| 0 | 0 | 0 | 0 |
| 0 | 1 | 1 | 1 |
| 0 | 0 | 0 | 0 |
| 0 | 0 | 0 | 0 |
| 0 | 0 | 0 | 0 |
| 0 | 0 | 0 | 1 |
| 0 | 1 | 0 | 0 |
| 0 | 0 | 0 | 0 |
| 0 | 0 | 0 | 1 |
| 0 | 1 | 1 | 1 |
| 0 | 0 | 0 | 1 |
| 0 | 1 | 0 | 0 |
| 0 | 0 | 0 | 0 |
| 0 | 1 | 1 | 1 |
| 0 | 1 | 0 | 0 |
| 0 | 1 | 0 | 0 |
| 0 | 1 | 0 | 0 |
| 0 | 1 | 0 | 0 |
| 0 | 0 | 0 | 0 |
| 0 | 1 | 1 | 1 |
| 0 | 0 | 0 | 1 |
| 0 | 0 | 0 | 1 |
| 0 | 0 | 0 | 1 |
| 0 | 0 | 0 | 0 |
| 0 | 0 | 0 | 1 |
| 0 | 0 | 0 | 0 |
| 0 | 0 | 0 | 1 |
| 1 | 0 | 0 | 0 |
| 0 | 0 | 0 | 1 |
| 0 | 0 | 0 | 0 |
| 0 | 1 | 0 | 0 |
| 1 | 0 | 0 | 0 |
| 0 | 0 | 0 | 0 |
| 0 | 0 | 0 | 0 |
| 0 | 0 | 0 | 0 |
| 0 | 0 | 0 | 0 |
| 0 | 0 | 0 | 0 |

| | | | |
|---|---|---|---|
| 0 | 0 | 0 | 0 |
| 0 | 0 | 0 | 0 |
| 0 | 0 | 0 | 0 |
| 0 | 0 | 0 | 0 |
| 0 | 1 | 0 | 0 |
| 0 | 1 | 0 | 0 |
| 0 | 1 | 0 | 0 |
| 0 | 0 | 0 | 1 |
| 0 | 1 | 0 | 0 |
| 0 | 1 | 0 | 0 |
| 0 | 1 | 0 | 0 |
| 0 | 1 | 1 | 1 |
| 0 | 1 | 0 | 0 |
| 0 | 1 | 0 | 0 |
| 0 | 1 | 0 | 0 |
| 0 | 1 | 0 | 0 |
| 0 | 1 | 0 | 0 |
| 0 | 1 | 0 | 0 |
| 0 | 1 | 0 | 0 |
| 0 | 1 | 0 | 0 |
| 0 | 1 | 0 | 0 |
| 0 | 1 | 0 | 0 |
| 0 | 1 | 0 | 0 |
| 0 | 1 | 0 | 0 |
| 0 | 1 | 0 | 0 |
| 0 | 1 | 0 | 0 |
| 0 | 1 | 0 | 0 |
| 0 | 1 | 0 | 0 |
| 0 | 1 | 0 | 0 |
| 0 | 1 | 0 | 0 |
| 0 | 1 | 0 | 0 |
| 0 | 1 | 0 | 0 |
| 0 | 1 | 0 | 0 |
| 0 | 1 | 0 | 0 |
| 0 | 1 | 0 | 0 |
| 0 | 1 | 0 | 0 |
| 0 | 1 | 0 | 0 |
| 0 | 1 | 0 | 0 |
| 0 | 1 | 0 | 0 |
| 0 | 1 | 0 | 0 |
| 0 | 1 | 1 | 1 |
| 0 | 1 | 0 | 0 |
| 0 | 1 | 0 | 0 |
| 0 | 1 | 0 | 0 |
| 0 | 1 | 0 | 0 |
| 0 | 1 | 0 | 0 |
| 0 | 1 | 0 | 0 |
| 0 | 1 | 0 | 0 |
| 0 | 1 | 0 | 0 |
| 0 | 0 | 0 | 1 |
| 0 | 0 | 0 | 1 |
| 0 | 1 | 1 | 1 |
| 1 | 0 | 0 | 0 |
| 0 | 1 | 0 | 0 |

| | | | |
|---|---|---|---|
| 0 | 0 | 0 | 0 |
| 0 | 0 | 0 | 0 |
| 1 | 1 | 0 | 0 |
| 1 | 0 | 0 | 0 |
| 0 | 0 | 0 | 0 |
| 0 | 0 | 0 | 0 |
| 0 | 0 | 0 | 0 |
| 0 | 1 | 1 | 1 |
| 0 | 0 | 0 | 1 |
| 0 | 0 | 0 | 0 |
| 0 | 1 | 0 | 0 |
| 0 | 0 | 0 | 1 |
| 0 | 1 | 0 | 0 |
| 0 | 1 | 0 | 0 |
| 0 | 0 | 0 | 0 |
| 0 | 0 | 0 | 0 |
| 0 | 0 | 0 | 0 |
| 0 | 0 | 0 | 1 |
| 0 | 0 | 0 | 0 |
| 0 | 1 | 1 | 1 |
| 0 | 1 | 0 | 0 |
| 0 | 0 | 0 | 0 |
| 0 | 0 | 0 | 1 |
| 0 | 0 | 0 | 0 |
| 0 | 1 | 0 | 0 |
| 0 | 0 | 0 | 1 |
| 0 | 0 | 0 | 0 |
| 0 | 0 | 0 | 0 |
| 0 | 1 | 0 | 0 |
| 0 | 1 | 0 | 0 |
| 0 | 1 | 0 | 0 |
| 0 | 0 | 0 | 0 |
| 0 | 1 | 1 | 1 |
| 0 | 0 | 0 | 0 |
| 0 | 1 | 0 | 0 |
| 0 | 0 | 0 | 0 |
| 0 | 0 | 0 | 1 |
| 0 | 0 | 0 | 0 |
| 0 | 0 | 0 | 0 |
| 0 | 0 | 0 | 0 |
| 0 | 1 | 0 | 0 |
| 0 | 1 | 0 | 0 |
| 0 | 1 | 0 | 0 |
| 0 | 1 | 0 | 0 |
| 0 | 1 | 0 | 0 |
| 0 | 0 | 0 | 1 |
| 0 | 1 | 0 | 0 |
| 0 | 1 | 0 | 0 |
| 0 | 1 | 0 | 0 |
| 0 | 1 | 0 | 0 |
| 0 | 1 | 0 | 0 |
| 0 | 1 | 0 | 0 |
| 0 | 1 | 1 | 1 |
| 0 | 1 | 1 | 1 |

| | | | |
|---|---|---|---|
| 0 | 1 | 1 | 1 |
| 0 | 0 | 0 | 1 |
| 0 | 0 | 0 | 0 |
| 0 | 0 | 0 | 0 |
| 0 | 1 | 0 | 0 |
| 0 | 1 | 0 | 0 |
| 0 | 1 | 0 | 0 |
| 0 | 0 | 0 | 1 |
| 0 | 0 | 0 | 0 |
| 0 | 1 | 1 | 1 |
| 0 | 0 | 0 | 1 |
| 0 | 0 | 0 | 1 |
| 0 | 1 | 0 | 0 |
| 0 | 0 | 0 | 0 |
| 0 | 1 | 0 | 0 |
| 0 | 0 | 0 | 0 |
| 0 | 1 | 0 | 0 |
| 1 | 1 | 0 | 0 |
| 0 | 0 | 0 | 1 |
| 0 | 0 | 0 | 1 |
| 0 | 0 | 0 | 1 |
| 0 | 0 | 0 | 0 |
| 0 | 0 | 0 | 1 |
| 0 | 0 | 0 | 0 |
| 0 | 0 | 0 | 0 |
| 0 | 0 | 0 | 0 |
| 0 | 0 | 0 | 0 |
| 0 | 0 | 0 | 0 |
| 0 | 0 | 0 | 0 |
| 0 | 0 | 0 | 0 |
| 0 | 0 | 0 | 0 |
| 0 | 0 | 0 | 0 |
| 0 | 0 | 0 | 0 |
| 0 | 0 | 0 | 0 |
| 0 | 0 | 0 | 0 |
| 0 | 0 | 0 | 0 |
| 0 | 0 | 0 | 0 |
| 0 | 1 | 0 | 0 |
| 1 | 1 | 0 | 0 |
| 0 | 1 | 0 | 0 |
| 0 | 1 | 0 | 0 |
| 0 | 0 | 0 | 1 |
| 0 | 1 | 0 | 0 |
| 0 | 1 | 0 | 0 |
| 0 | 0 | 0 | 0 |
| 0 | 0 | 0 | 0 |
| 0 | 1 | 1 | 1 |
| 0 | 1 | 0 | 0 |
| 0 | 1 | 0 | 0 |
| 0 | 0 | 0 | 0 |
| 0 | 0 | 0 | 0 |
| 0 | 0 | 0 | 0 |
| 0 | 0 | 0 | 1 |
| 0 | 0 | 0 | 1 |

| | | | |
|---|---|---|---|
| 0 | 1 | 0 | 0 |
| 0 | 0 | 0 | 0 |
| 0 | 0 | 0 | 1 |
| 0 | 0 | 0 | 1 |
| 0 | 0 | 0 | 1 |
| 0 | 1 | 0 | 0 |
| 0 | 1 | 1 | 1 |
| 0 | 1 | 1 | 1 |
| 0 | 1 | 0 | 0 |
| 0 | 0 | 0 | 0 |
| 0 | 0 | 0 | 0 |
| 0 | 0 | 0 | 0 |
| 0 | 0 | 0 | 0 |
| 0 | 0 | 0 | 1 |
| 0 | 0 | 0 | 1 |
| 0 | 0 | 0 | 1 |
| 0 | 0 | 0 | 1 |
| 0 | 0 | 0 | 1 |
| 0 | 0 | 0 | 0 |
| 0 | 0 | 0 | 0 |
| 0 | 1 | 1 | 1 |
| 0 | 0 | 0 | 1 |
| 0 | 1 | 0 | 0 |
| 0 | 1 | 0 | 0 |
| 0 | 1 | 1 | 1 |
| 0 | 1 | 1 | 1 |
| 0 | 1 | 0 | 0 |
| 0 | 1 | 1 | 1 |
| 0 | 0 | 0 | 0 |
| 0 | 0 | 0 | 0 |
| 0 | 1 | 0 | 0 |
| 0 | 1 | 0 | 0 |
| 0 | 1 | 0 | 0 |
| 1 | 0 | 0 | 0 |
| 1 | 1 | 0 | 0 |
| 0 | 1 | 0 | 0 |
| 0 | 1 | 0 | 0 |
| 0 | 0 | 0 | 0 |
| 0 | 0 | 0 | 1 |
| 0 | 1 | 0 | 0 |
| 0 | 0 | 0 | 1 |
| 0 | 0 | 0 | 1 |
| 0 | 0 | 0 | 1 |
| 0 | 0 | 0 | 1 |
| 0 | 0 | 0 | 1 |
| 0 | 0 | 0 | 0 |
| 0 | 0 | 0 | 0 |
| 0 | 0 | 0 | 0 |
| 0 | 0 | 0 | 0 |
| 0 | 1 | 0 | 0 |
| 0 | 1 | 0 | 0 |
| 0 | 1 | 0 | 0 |
| 0 | 1 | 0 | 0 |
| 0 | 1 | 1 | 1 |

| | | | |
|---|---|---|---|
| 0 | 1 | 1 | 1 |
| 0 | 1 | 0 | 0 |
| 0 | 0 | 0 | 0 |
| 0 | 0 | 0 | 0 |
| 0 | 1 | 1 | 1 |
| 0 | 1 | 0 | 0 |
| 0 | 1 | 0 | 0 |
| 0 | 1 | 0 | 0 |
| 0 | 1 | 1 | 1 |
| 0 | 1 | 1 | 1 |
| 0 | 1 | 0 | 0 |
| 0 | 1 | 0 | 0 |
| 0 | 1 | 0 | 0 |
| 0 | 1 | 1 | 1 |
| 0 | 1 | 0 | 0 |
| 0 | 1 | 0 | 0 |
| 0 | 1 | 0 | 0 |
| 0 | 1 | 0 | 0 |
| 0 | 0 | 0 | 0 |
| 0 | 0 | 0 | 0 |
| 0 | 1 | 0 | 0 |
| 0 | 0 | 0 | 1 |
| 0 | 0 | 0 | 0 |
| 0 | 0 | 0 | 1 |
| 0 | 0 | 0 | 0 |
| 0 | 1 | 0 | 0 |
| 0 | 1 | 1 | 1 |
| 0 | 0 | 0 | 0 |
| 0 | 1 | 0 | 0 |
| 0 | 1 | 0 | 0 |
| 0 | 0 | 0 | 0 |
| 0 | 1 | 0 | 0 |
| 0 | 1 | 0 | 0 |
| 0 | 0 | 0 | 0 |
| 0 | 0 | 0 | 1 |
| 0 | 1 | 0 | 0 |
| 0 | 1 | 0 | 0 |
| 0 | 1 | 0 | 0 |
| 0 | 1 | 0 | 0 |
| 0 | 0 | 0 | 0 |
| 0 | 0 | 0 | 0 |
| 0 | 0 | 0 | 0 |
| 0 | 0 | 0 | 0 |
| 0 | 0 | 0 | 0 |
| 0 | 0 | 0 | 0 |
| 0 | 0 | 0 | 0 |
| 0 | 0 | 0 | 0 |
| 0 | 0 | 0 | 1 |
| 0 | 1 | 0 | 0 |
| 0 | 0 | 0 | 1 |
| 0 | 0 | 0 | 0 |
| 0 | 0 | 0 | 0 |
| 0 | 0 | 0 | 0 |
| 0 | 0 | 0 | 0 |

| | | | |
|---|---|---|---|
| 0 | 0 | 0 | 0 |
| 0 | 1 | 0 | 0 |
| 0 | 0 | 0 | 1 |
| 0 | 1 | 0 | 0 |
| 0 | 0 | 0 | 0 |
| 0 | 0 | 0 | 1 |
| 0 | 0 | 0 | 1 |
| 0 | 0 | 0 | 1 |
| 0 | 0 | 0 | 0 |
| 0 | 0 | 0 | 1 |
| 0 | 1 | 0 | 0 |
| 0 | 1 | 0 | 0 |
| 0 | 1 | 0 | 0 |
| 0 | 0 | 0 | 1 |
| 0 | 1 | 0 | 0 |
| 0 | 0 | 0 | 1 |
| 0 | 1 | 0 | 0 |
| 0 | 0 | 0 | 0 |
| 1 | 0 | 0 | 0 |
| 0 | 0 | 0 | 1 |
| 0 | 0 | 0 | 1 |
| 0 | 0 | 0 | 0 |
| 0 | 0 | 0 | 1 |
| 0 | 0 | 0 | 0 |
| 0 | 0 | 0 | 0 |
| 0 | 0 | 0 | 1 |
| 0 | 0 | 0 | 1 |
| 0 | 0 | 0 | 0 |
| 0 | 0 | 0 | 1 |
| 1 | 0 | 0 | 0 |
| 0 | 0 | 0 | 1 |
| 0 | 0 | 0 | 1 |
| 0 | 0 | 0 | 1 |
| 0 | 0 | 0 | 1 |
| 0 | 0 | 0 | 1 |
| 0 | 0 | 0 | 1 |
| 0 | 0 | 0 | 1 |
| 0 | 0 | 0 | 1 |
| 0 | 0 | 0 | 1 |
| 0 | 0 | 0 | 1 |
| 0 | 0 | 0 | 1 |
| 0 | 0 | 0 | 1 |
| 0 | 0 | 0 | 1 |
| 0 | 0 | 0 | 1 |
| 0 | 0 | 0 | 1 |
| 0 | 0 | 0 | 1 |
| 0 | 0 | 0 | 1 |
| 0 | 0 | 0 | 1 |
| 0 | 0 | 0 | 1 |
| 0 | 0 | 0 | 1 |
| 0 | 0 | 0 | 1 |
| 0 | 0 | 0 | 1 |
| 0 | 0 | 0 | 1 |
| 0 | 0 | 0 | 1 |

| | | | |
|---|---|---|---|
| 0 | 0 | 0 | 1 |
| 0 | 0 | 0 | 1 |
| 0 | 0 | 0 | 1 |
| 0 | 0 | 0 | 1 |
| 0 | 0 | 0 | 1 |
| 0 | 0 | 0 | 1 |
| 0 | 0 | 0 | 1 |
| 0 | 0 | 0 | 1 |
| 0 | 0 | 0 | 1 |
| 0 | 0 | 0 | 1 |
| 0 | 0 | 0 | 1 |
| 0 | 0 | 0 | 0 |
| 0 | 1 | 0 | 0 |
| 0 | 1 | 0 | 0 |
| 0 | 1 | 1 | 1 |
| 0 | 1 | 0 | 0 |
| 0 | 1 | 1 | 1 |
| 0 | 1 | 0 | 0 |
| 0 | 1 | 1 | 1 |
| 0 | 0 | 0 | 1 |
| 0 | 0 | 0 | 0 |
| 0 | 0 | 0 | 1 |
| 0 | 0 | 0 | 1 |
| 0 | 0 | 0 | 0 |
| 0 | 0 | 0 | 0 |
| 0 | 0 | 0 | 1 |
| 0 | 0 | 0 | 1 |
| 0 | 0 | 0 | 0 |
| 0 | 0 | 0 | 0 |
| 0 | 0 | 0 | 0 |
| 0 | 0 | 0 | 0 |
| 0 | 0 | 0 | 0 |
| 0 | 0 | 0 | 1 |
| 0 | 0 | 0 | 1 |
| 0 | 0 | 0 | 1 |
| 0 | 0 | 0 | 1 |
| 0 | 0 | 0 | 1 |
| 0 | 0 | 0 | 1 |
| 0 | 0 | 0 | 1 |
| 0 | 1 | 0 | 0 |
| 0 | 0 | 0 | 0 |
| 0 | 0 | 0 | 1 |
| 0 | 1 | 0 | 0 |
| 0 | 1 | 0 | 0 |
| 0 | 1 | 0 | 0 |
| 0 | 1 | 1 | 1 |
| 0 | 0 | 0 | 0 |
| 0 | 0 | 0 | 1 |
| 0 | 1 | 1 | 1 |
| 0 | 1 | 0 | 0 |
| 0 | 0 | 0 | 0 |
| 0 | 0 | 0 | 0 |
| 0 | 0 | 0 | 1 |
| 0 | 0 | 0 | 0 |

| | | | |
|---|---|---|---|
| 0 | 1 | 0 | 0 |
| 0 | 0 | 0 | 1 |
| 0 | 0 | 0 | 1 |
| 0 | 0 | 0 | 1 |
| 0 | 1 | 0 | 0 |
| 0 | 0 | 0 | 1 |
| 0 | 0 | 0 | 0 |
| 0 | 0 | 0 | 1 |
| 0 | 0 | 0 | 1 |
| 0 | 0 | 0 | 0 |
| 0 | 1 | 1 | 1 |
| 0 | 0 | 0 | 1 |
| 0 | 0 | 0 | 1 |
| 0 | 1 | 0 | 0 |
| 0 | 0 | 0 | 1 |
| 0 | 0 | 0 | 1 |
| 0 | 1 | 0 | 0 |
| 0 | 0 | 0 | 1 |
| 0 | 0 | 0 | 0 |
| 0 | 0 | 0 | 0 |
| 0 | 0 | 0 | 0 |
| 0 | 0 | 0 | 0 |
| 0 | 0 | 0 | 0 |
| 0 | 0 | 0 | 0 |
| 0 | 1 | 0 | 0 |
| 0 | 1 | 0 | 0 |
| 0 | 0 | 0 | 0 |
| 0 | 0 | 0 | 0 |
| 0 | 0 | 0 | 0 |
| 0 | 0 | 0 | 0 |
| 0 | 0 | 0 | 0 |
| 0 | 1 | 0 | 0 |
| 0 | 1 | 0 | 0 |
| 0 | 1 | 0 | 0 |
| 0 | 0 | 0 | 0 |
| 0 | 1 | 0 | 0 |
| 0 | 1 | 1 | 1 |
| 0 | 1 | 0 | 0 |
| 0 | 1 | 0 | 0 |
| 0 | 0 | 0 | 1 |
| 0 | 0 | 0 | 1 |
| 0 | 0 | 0 | 0 |
| 0 | 0 | 0 | 0 |
| 0 | 1 | 0 | 0 |
| 0 | 1 | 0 | 0 |
| 0 | 0 | 0 | 0 |
| 0 | 0 | 0 | 0 |
| 0 | 1 | 0 | 0 |
| 0 | 1 | 1 | 1 |
| 0 | 1 | 1 | 1 |
| 0 | 0 | 0 | 0 |
| 0 | 0 | 0 | 1 |
| 0 | 0 | 0 | 0 |
| 0 | 0 | 0 | 0 |

| | | | |
|---|---|---|---|
| 0 | 0 | 0 | 0 |
| 0 | 1 | 0 | 0 |
| 0 | 1 | 0 | 0 |
| 0 | 0 | 0 | 0 |
| 0 | 0 | 0 | 0 |
| 0 | 0 | 0 | 0 |
| 0 | 1 | 0 | 0 |
| 0 | 0 | 0 | 0 |
| 0 | 0 | 0 | 0 |
| 0 | 0 | 0 | 0 |
| 0 | 0 | 0 | 0 |
| 0 | 0 | 0 | 0 |
| 0 | 0 | 0 | 0 |
| 0 | 0 | 0 | 0 |
| 0 | 0 | 0 | 1 |
| 0 | 1 | 0 | 0 |
| 0 | 1 | 0 | 0 |
| 0 | 1 | 0 | 0 |
| 0 | 1 | 1 | 1 |
| 0 | 0 | 0 | 0 |
| 0 | 1 | 0 | 0 |
| 0 | 1 | 0 | 0 |
| 0 | 1 | 0 | 0 |
| 0 | 1 | 0 | 0 |
| 0 | 1 | 0 | 0 |
| 0 | 1 | 1 | 1 |
| 0 | 1 | 0 | 0 |
| 0 | 1 | 0 | 0 |
| 0 | 1 | 0 | 0 |
| 0 | 0 | 0 | 0 |
| 0 | 0 | 0 | 1 |
| 0 | 0 | 0 | 0 |
| 0 | 0 | 0 | 1 |
| 0 | 1 | 1 | 1 |
| 1 | 1 | 0 | 0 |
| 0 | 0 | 0 | 0 |
| 0 | 0 | 0 | 1 |
| 0 | 0 | 0 | 1 |
| 0 | 0 | 0 | 1 |
| 0 | 0 | 0 | 1 |
| 0 | 1 | 0 | 0 |
| 0 | 1 | 0 | 0 |
| 0 | 0 | 0 | 0 |
| 0 | 0 | 0 | 0 |
| 0 | 0 | 0 | 0 |
| 0 | 0 | 0 | 0 |
| 0 | 0 | 0 | 0 |
| 0 | 0 | 0 | 0 |
| 0 | 0 | 0 | 1 |
| 0 | 0 | 0 | 1 |
| 0 | 0 | 0 | 0 |
| 0 | 0 | 0 | 0 |
| 0 | 0 | 0 | 0 |
| 0 | 0 | 0 | 1 |

| | | | |
|---|---|---|---|
| 0 | 0 | 0 | 1 |
| 0 | 0 | 0 | 1 |
| 0 | 1 | 1 | 1 |
| 0 | 1 | 0 | 0 |
| 0 | 0 | 0 | 1 |
| 0 | 0 | 0 | 1 |
| 0 | 0 | 0 | 0 |
| 0 | 0 | 0 | 1 |
| 0 | 0 | 0 | 0 |
| 0 | 0 | 0 | 0 |
| 0 | 0 | 0 | 0 |
| 0 | 0 | 0 | 0 |
| 0 | 0 | 0 | 0 |
| 0 | 0 | 0 | 0 |
| 0 | 0 | 0 | 0 |
| 0 | 0 | 0 | 0 |
| 0 | 0 | 0 | 1 |
| 0 | 0 | 0 | 0 |
| 0 | 1 | 0 | 0 |
| 0 | 1 | 1 | 1 |
| 0 | 0 | 0 | 0 |
| 0 | 0 | 0 | 0 |
| 0 | 0 | 0 | 1 |
| 0 | 0 | 0 | 0 |
| 0 | 0 | 0 | 1 |
| 0 | 0 | 0 | 1 |
| 0 | 0 | 0 | 1 |
| 0 | 0 | 0 | 0 |
| 0 | 0 | 0 | 0 |
| 0 | 0 | 0 | 0 |
| 0 | 0 | 0 | 1 |
| 0 | 0 | 0 | 0 |
| 0 | 0 | 0 | 0 |
| 0 | 0 | 0 | 0 |
| 0 | 0 | 0 | 1 |
| 0 | 0 | 0 | 1 |
| 0 | 0 | 0 | 0 |
| 0 | 0 | 0 | 0 |
| 0 | 0 | 0 | 0 |
| 0 | 0 | 0 | 1 |
| 0 | 0 | 0 | 1 |
| 0 | 0 | 0 | 1 |
| 0 | 0 | 0 | 1 |
| 0 | 0 | 0 | 0 |
| 0 | 0 | 0 | 1 |
| 0 | 0 | 0 | 0 |
| 0 | 0 | 0 | 1 |
| 0 | 0 | 0 | 0 |
| 0 | 0 | 0 | 0 |
| 0 | 0 | 0 | 1 |
| 0 | 0 | 0 | 1 |
| 0 | 1 | 0 | 0 |
| 0 | 0 | 0 | 0 |
| 0 | 0 | 0 | 1 |

| | | | |
|---|---|---|---|
| 0 | 0 | 0 | 1 |
| 0 | 0 | 0 | 1 |
| 0 | 0 | 0 | 1 |
| 0 | 1 | 1 | 1 |
| 0 | 1 | 0 | 0 |
| 0 | 1 | 1 | 1 |
| 0 | 1 | 1 | 1 |
| 0 | 1 | 0 | 0 |
| 0 | 1 | 0 | 0 |
| 0 | 0 | 0 | 1 |
| 0 | 0 | 0 | 0 |
| 0 | 0 | 0 | 0 |
| 1 | 0 | 0 | 0 |
| 1 | 0 | 0 | 0 |
| 0 | 0 | 0 | 0 |
| 1 | 0 | 0 | 0 |
| 1 | 0 | 0 | 0 |
| 1 | 0 | 0 | 0 |
| 1 | 0 | 0 | 0 |
| 0 | 0 | 0 | 1 |
| 0 | 0 | 0 | 1 |
| 0 | 0 | 0 | 1 |
| 0 | 0 | 0 | 1 |
| 0 | 0 | 0 | 1 |
| 0 | 0 | 0 | 0 |
| 0 | 0 | 0 | 1 |
| 1 | 0 | 0 | 0 |
| 1 | 0 | 0 | 0 |
| 0 | 0 | 0 | 1 |
| 0 | 0 | 0 | 1 |
| 0 | 0 | 0 | 1 |
| 0 | 0 | 0 | 1 |
| 0 | 0 | 0 | 1 |
| 0 | 0 | 0 | 0 |
| 0 | 0 | 0 | 0 |
| 0 | 0 | 0 | 1 |
| 0 | 0 | 0 | 0 |
| 0 | 0 | 0 | 1 |
| 0 | 0 | 0 | 0 |
| 0 | 1 | 0 | 0 |
| 0 | 1 | 0 | 0 |
| 0 | 0 | 0 | 0 |
| 0 | 0 | 0 | 1 |
| 0 | 0 | 0 | 0 |
| 0 | 0 | 0 | 0 |
| 1 | 0 | 0 | 0 |
| 1 | 0 | 0 | 0 |
| 1 | 0 | 0 | 0 |
| 0 | 0 | 0 | 0 |
| 0 | 0 | 0 | 0 |
| 0 | 1 | 0 | 0 |
| 1 | 0 | 0 | 0 |
| 1 | 0 | 0 | 0 |
| 0 | 0 | 0 | 1 |

| | | | |
|---|---|---|---|
| 0 | 0 | 0 | 0 |
| 0 | 0 | 0 | 1 |
| 0 | 0 | 0 | 0 |
| 0 | 0 | 0 | 0 |
| 0 | 1 | 0 | 0 |
| 0 | 0 | 0 | 1 |
| 0 | 0 | 0 | 0 |
| 0 | 0 | 0 | 1 |
| 0 | 0 | 0 | 1 |
| 0 | 0 | 0 | 0 |
| 0 | 0 | 0 | 1 |
| 0 | 0 | 0 | 1 |
| 0 | 0 | 0 | 1 |
| 0 | 0 | 0 | 1 |
| 0 | 0 | 0 | 1 |
| 0 | 1 | 0 | 0 |
| 0 | 1 | 0 | 0 |
| 0 | 0 | 0 | 1 |
| 0 | 0 | 0 | 0 |
| 0 | 0 | 0 | 1 |
| 0 | 0 | 0 | 0 |
| 0 | 0 | 0 | 0 |
| 0 | 0 | 0 | 0 |
| 0 | 1 | 0 | 0 |
| 0 | 1 | 0 | 0 |
| 0 | 1 | 0 | 0 |
| 0 | 1 | 0 | 0 |
| 0 | 0 | 0 | 0 |
| 0 | 0 | 0 | 0 |
| 0 | 1 | 0 | 0 |
| 0 | 1 | 0 | 0 |
| 0 | 1 | 0 | 0 |
| 0 | 1 | 1 | 1 |
| 0 | 0 | 0 | 1 |
| 0 | 1 | 0 | 0 |
| 0 | 1 | 0 | 0 |
| 0 | 1 | 1 | 1 |
| 0 | 1 | 0 | 0 |
| 0 | 1 | 1 | 1 |
| 0 | 1 | 0 | 0 |
| 0 | 1 | 0 | 0 |
| 0 | 1 | 0 | 0 |
| 0 | 1 | 1 | 1 |
| 0 | 0 | 0 | 1 |
| 0 | 1 | 0 | 0 |
| 0 | 1 | 0 | 0 |
| 0 | 1 | 0 | 0 |
| 0 | 1 | 0 | 0 |
| 0 | 1 | 0 | 0 |
| 0 | 1 | 0 | 0 |
| 0 | 1 | 1 | 1 |
| 0 | 1 | 0 | 0 |
| 0 | 1 | 0 | 0 |
| 0 | 1 | 1 | 1 |

| | | | |
|---|---|---|---|
| 0 | 1 | 0 | 0 |
| 0 | 1 | 1 | 1 |
| 0 | 0 | 0 | 0 |
| 0 | 0 | 0 | 0 |
| 0 | 0 | 0 | 0 |
| 0 | 0 | 0 | 0 |
| 0 | 0 | 0 | 0 |
| 0 | 0 | 0 | 0 |
| 0 | 0 | 0 | 0 |
| 0 | 0 | 0 | 0 |
| 0 | 0 | 0 | 0 |
| 0 | 0 | 0 | 1 |
| 0 | 0 | 0 | 0 |
| 0 | 0 | 0 | 1 |
| 0 | 1 | 1 | 1 |
| 0 | 1 | 1 | 1 |
| 0 | 1 | 0 | 0 |
| 0 | 1 | 0 | 0 |
| 0 | 1 | 1 | 1 |
| 0 | 1 | 0 | 0 |
| 0 | 1 | 1 | 1 |
| 0 | 1 | 0 | 0 |
| 0 | 0 | 0 | 1 |
| 0 | 1 | 0 | 0 |
| 0 | 0 | 0 | 1 |
| 0 | 0 | 0 | 1 |
| 0 | 0 | 0 | 1 |
| 0 | 0 | 0 | 0 |
| 0 | 0 | 0 | 1 |
| 0 | 0 | 0 | 1 |
| 0 | 0 | 0 | 1 |
| 1 | 0 | 0 | 0 |
| 0 | 0 | 0 | 1 |
| 0 | 0 | 0 | 0 |
| 0 | 0 | 0 | 0 |
| 0 | 0 | 0 | 0 |
| 0 | 0 | 0 | 1 |
| 0 | 0 | 0 | 1 |
| 0 | 1 | 0 | 0 |
| 0 | 0 | 0 | 0 |
| 0 | 0 | 0 | 0 |
| 0 | 0 | 0 | 0 |
| 0 | 1 | 0 | 0 |
| 0 | 0 | 0 | 0 |
| 0 | 1 | 0 | 0 |
| 0 | 0 | 0 | 0 |
| 0 | 0 | 0 | 1 |
| 0 | 0 | 0 | 1 |
| 0 | 0 | 0 | 0 |
| 0 | 0 | 0 | 0 |
| 0 | 0 | 0 | 0 |
| 0 | 0 | 0 | 1 |
| 0 | 0 | 0 | 1 |
| 0 | 0 | 0 | 0 |

| | | | |
|---|---|---|---|
| 0 | 0 | 0 | 0 |
| 0 | 0 | 0 | 1 |
| 0 | 0 | 0 | 1 |
| 0 | 0 | 0 | 1 |
| 0 | 0 | 0 | 0 |
| 0 | 1 | 0 | 0 |
| 0 | 0 | 0 | 0 |
| 0 | 1 | 0 | 0 |
| 0 | 1 | 0 | 0 |
| 0 | 0 | 0 | 0 |
| 0 | 0 | 0 | 1 |
| 0 | 1 | 0 | 0 |
| 0 | 0 | 0 | 0 |
| 0 | 0 | 0 | 0 |
| 0 | 1 | 1 | 1 |
| 0 | 0 | 0 | 1 |
| 0 | 1 | 0 | 0 |
| 0 | 1 | 1 | 1 |
| 0 | 1 | 0 | 0 |
| 0 | 1 | 0 | 0 |
| 0 | 0 | 0 | 1 |
| 0 | 1 | 0 | 0 |
| 0 | 1 | 0 | 0 |
| 0 | 0 | 0 | 1 |
| 0 | 1 | 0 | 0 |
| 0 | 1 | 0 | 0 |
| 0 | 1 | 0 | 0 |
| 0 | 1 | 1 | 1 |
| 0 | 1 | 0 | 0 |
| 0 | 0 | 0 | 0 |
| 0 | 0 | 0 | 0 |
| 0 | 0 | 0 | 1 |
| 0 | 0 | 0 | 1 |
| 0 | 0 | 0 | 1 |
| 0 | 0 | 0 | 1 |
| 0 | 0 | 0 | 1 |
| 0 | 0 | 0 | 1 |
| 0 | 0 | 0 | 1 |
| 0 | 0 | 0 | 1 |
| 0 | 0 | 0 | 1 |
| 0 | 0 | 0 | 1 |
| 0 | 0 | 0 | 1 |
| 1 | 0 | 0 | 0 |
| 0 | 0 | 0 | 1 |
| 0 | 0 | 0 | 1 |
| 0 | 0 | 0 | 1 |
| 0 | 0 | 0 | 1 |
| 0 | 0 | 0 | 1 |
| 0 | 0 | 0 | 1 |
| 0 | 0 | 0 | 1 |
| 0 | 0 | 0 | 1 |
| 0 | 0 | 0 | 1 |
| 0 | 0 | 0 | 1 |
| 0 | 0 | 0 | 1 |

| | | | |
|---|---|---|---|
| 0 | 0 | 0 | 1 |
| 0 | 0 | 0 | 1 |
| 0 | 0 | 0 | 1 |
| 0 | 0 | 0 | 1 |
| 0 | 0 | 0 | 1 |
| 0 | 0 | 0 | 1 |
| 0 | 0 | 0 | 1 |
| 0 | 0 | 0 | 1 |
| 0 | 1 | 0 | 0 |
| 0 | 0 | 0 | 0 |
| 0 | 0 | 0 | 0 |
| 0 | 0 | 0 | 0 |
| 0 | 0 | 0 | 0 |
| 0 | 0 | 0 | 0 |
| 0 | 0 | 0 | 0 |
| 0 | 0 | 0 | 0 |
| 0 | 0 | 0 | 0 |
| 0 | 0 | 0 | 0 |
| 0 | 0 | 0 | 0 |
| 0 | 1 | 0 | 0 |
| 1 | 0 | 0 | 0 |
| 1 | 0 | 0 | 0 |
| 0 | 1 | 0 | 0 |
| 0 | 1 | 0 | 0 |
| 0 | 1 | 0 | 0 |
| 0 | 0 | 0 | 1 |
| 0 | 0 | 0 | 1 |
| 0 | 0 | 0 | 0 |
| 0 | 0 | 0 | 0 |
| 1 | 0 | 0 | 0 |
| 1 | 0 | 0 | 0 |
| 0 | 0 | 0 | 0 |
| 1 | 0 | 0 | 0 |
| 1 | 0 | 0 | 0 |
| 0 | 0 | 0 | 0 |
| 0 | 0 | 0 | 0 |
| 0 | 0 | 0 | 0 |
| 0 | 0 | 0 | 0 |
| 0 | 0 | 0 | 0 |
| 0 | 0 | 0 | 0 |
| 0 | 0 | 0 | 0 |
| 0 | 0 | 0 | 0 |
| 1 | 0 | 0 | 0 |
| 1 | 0 | 0 | 0 |
| 1 | 0 | 0 | 0 |
| 0 | 0 | 0 | 0 |
| 0 | 0 | 0 | 1 |
| 0 | 0 | 0 | 1 |
| 0 | 0 | 0 | 1 |
| 0 | 0 | 0 | 1 |
| 0 | 0 | 0 | 1 |
| 0 | 0 | 0 | 1 |
| 0 | 1 | 0 | 0 |
| 0 | 1 | 0 | 0 |

| | | | |
|---|---|---|---|
| 0 | 0 | 0 | 0 |
| 0 | 0 | 0 | 0 |
| 0 | 0 | 0 | 0 |
| 0 | 1 | 0 | 0 |
| 0 | 0 | 0 | 1 |
| 0 | 0 | 0 | 0 |
| 0 | 0 | 0 | 0 |
| 0 | 0 | 0 | 0 |
| 0 | 0 | 0 | 1 |
| 0 | 0 | 0 | 0 |
| 1 | 0 | 0 | 0 |
| 0 | 0 | 0 | 1 |
| 0 | 0 | 0 | 0 |
| 0 | 0 | 0 | 0 |
| 0 | 0 | 0 | 0 |
| 0 | 0 | 0 | 0 |
| 0 | 0 | 0 | 1 |
| 0 | 0 | 0 | 0 |
| 0 | 0 | 0 | 0 |
| 0 | 0 | 0 | 1 |
| 0 | 1 | 0 | 0 |
| 0 | 0 | 0 | 0 |
| 0 | 1 | 0 | 0 |
| 0 | 0 | 0 | 0 |
| 0 | 0 | 0 | 0 |
| 0 | 0 | 0 | 1 |
| 0 | 0 | 0 | 1 |
| 0 | 0 | 0 | 1 |
| 0 | 0 | 0 | 1 |
| 0 | 0 | 0 | 1 |
| 0 | 0 | 0 | 1 |
| 0 | 0 | 0 | 1 |
| 0 | 1 | 1 | 1 |
| 0 | 1 | 1 | 1 |
| 0 | 0 | 0 | 0 |
| 0 | 1 | 0 | 0 |
| 0 | 0 | 0 | 0 |
| 0 | 0 | 0 | 0 |
| 0 | 1 | 0 | 0 |
| 0 | 1 | 0 | 0 |
| 0 | 0 | 0 | 1 |
| 0 | 0 | 0 | 1 |
| 0 | 0 | 0 | 1 |
| 0 | 0 | 0 | 0 |
| 0 | 0 | 0 | 0 |
| 0 | 0 | 0 | 0 |
| 0 | 0 | 0 | 1 |
| 0 | 0 | 0 | 1 |
| 0 | 0 | 0 | 0 |
| 0 | 0 | 0 | 0 |
| 0 | 0 | 0 | 1 |
| 0 | 0 | 0 | 1 |
| 0 | 0 | 0 | 1 |
| 0 | 0 | 0 | 1 |

| | | | |
|---|---|---|---|
| 0 | 0 | 0 | 1 |
| 0 | 0 | 0 | 1 |
| 0 | 0 | 0 | 1 |
| 0 | 0 | 0 | 0 |
| 0 | 0 | 0 | 0 |
| 0 | 0 | 0 | 1 |
| 0 | 0 | 0 | 1 |
| 0 | 0 | 0 | 1 |
| 0 | 0 | 0 | 1 |
| 0 | 0 | 0 | 1 |
| 0 | 0 | 0 | 0 |
| 0 | 0 | 0 | 1 |
| 0 | 1 | 0 | 0 |
| 0 | 0 | 0 | 0 |
| 0 | 0 | 0 | 0 |
| 0 | 0 | 0 | 1 |
| 0 | 0 | 0 | 1 |
| 0 | 0 | 0 | 1 |
| 0 | 0 | 0 | 1 |
| 0 | 0 | 0 | 1 |
| 0 | 0 | 0 | 0 |
| 0 | 0 | 0 | 1 |
| 0 | 0 | 0 | 1 |
| 0 | 1 | 0 | 0 |
| 0 | 1 | 1 | 1 |
| 0 | 1 | 0 | 0 |
| 0 | 1 | 0 | 0 |
| 0 | 0 | 0 | 1 |
| 0 | 0 | 0 | 1 |
| 0 | 0 | 0 | 1 |
| 0 | 0 | 0 | 1 |
| 0 | 0 | 0 | 1 |
| 0 | 0 | 0 | 1 |
| 0 | 0 | 0 | 1 |
| 0 | 0 | 0 | 1 |
| 0 | 0 | 0 | 0 |
| 0 | 0 | 0 | 1 |
| 0 | 0 | 0 | 1 |
| 0 | 0 | 0 | 0 |
| 0 | 0 | 0 | 0 |
| 0 | 0 | 0 | 1 |
| 0 | 0 | 0 | 1 |
| 1 | 0 | 0 | 0 |
| 1 | 0 | 0 | 0 |
| 0 | 0 | 0 | 0 |
| 0 | 0 | 0 | 0 |
| 1 | 0 | 0 | 0 |
| 1 | 0 | 0 | 0 |
| 0 | 0 | 0 | 1 |
| 1 | 0 | 0 | 0 |
| 1 | 0 | 0 | 0 |
| 1 | 0 | 0 | 0 |
| 1 | 0 | 0 | 0 |
| 1 | 0 | 0 | 0 |

| | | | |
|---|---|---|---|
| 0 | 0 | 0 | 0 |
| 0 | 0 | 0 | 0 |
| 1 | 0 | 0 | 0 |
| 1 | 0 | 0 | 0 |
| 1 | 0 | 0 | 0 |
| 1 | 0 | 0 | 0 |
| 0 | 0 | 0 | 0 |
| 0 | 0 | 0 | 0 |
| 0 | 0 | 0 | 0 |
| 0 | 0 | 0 | 1 |
| 0 | 0 | 0 | 1 |
| 0 | 0 | 0 | 0 |
| 0 | 0 | 0 | 1 |
| 0 | 0 | 0 | 0 |
| 0 | 0 | 0 | 0 |
| 0 | 0 | 0 | 1 |
| 0 | 0 | 0 | 0 |
| 0 | 1 | 1 | 1 |
| 0 | 0 | 0 | 1 |
| 0 | 0 | 0 | 1 |
| 0 | 1 | 1 | 1 |
| 0 | 0 | 0 | 1 |
| 0 | 0 | 0 | 1 |
| 0 | 0 | 0 | 0 |
| 0 | 0 | 0 | 0 |
| 0 | 0 | 0 | 1 |
| 0 | 0 | 0 | 1 |
| 0 | 0 | 0 | 1 |
| 0 | 0 | 0 | 0 |
| 0 | 0 | 0 | 0 |
| 0 | 0 | 0 | 1 |
| 0 | 0 | 0 | 1 |
| 0 | 0 | 0 | 1 |
| 0 | 1 | 0 | 0 |
| 0 | 0 | 0 | 0 |
| 0 | 0 | 0 | 1 |
| 0 | 0 | 0 | 0 |
| 0 | 0 | 0 | 1 |
| 0 | 0 | 0 | 1 |
| 0 | 0 | 0 | 0 |
| 0 | 0 | 0 | 0 |
| 0 | 0 | 0 | 0 |
| 0 | 0 | 0 | 0 |
| 0 | 0 | 0 | 0 |
| 0 | 0 | 0 | 0 |
| 0 | 0 | 0 | 1 |
| 0 | 0 | 0 | 1 |
| 0 | 0 | 0 | 0 |
| 0 | 0 | 0 | 0 |
| 0 | 0 | 0 | 0 |
| 0 | 0 | 0 | 1 |
| 0 | 0 | 0 | 1 |
| 0 | 0 | 0 | 0 |
| 0 | 0 | 0 | 0 |

| | | | |
|---|---|---|---|
| 0 | 0 | 0 | 1 |
| 0 | 0 | 0 | 1 |
| 0 | 0 | 0 | 1 |
| 0 | 0 | 0 | 1 |
| 0 | 1 | 0 | 0 |
| 0 | 1 | 0 | 0 |
| 0 | 0 | 0 | 1 |
| 0 | 0 | 0 | 0 |
| 0 | 1 | 1 | 1 |
| 0 | 0 | 0 | 0 |
| 0 | 0 | 0 | 1 |
| 0 | 0 | 0 | 1 |
| 0 | 0 | 0 | 1 |
| 0 | 0 | 0 | 1 |
| 0 | 0 | 0 | 1 |
| 0 | 0 | 0 | 1 |
| 0 | 0 | 0 | 0 |
| 0 | 0 | 0 | 1 |
| 0 | 0 | 0 | 1 |
| 0 | 0 | 0 | 0 |
| 0 | 0 | 0 | 0 |
| 0 | 0 | 0 | 1 |
| 0 | 0 | 0 | 1 |
| 0 | 0 | 0 | 0 |
| 0 | 0 | 0 | 1 |
| 0 | 0 | 0 | 0 |
| 0 | 0 | 0 | 0 |
| 0 | 0 | 0 | 0 |
| 0 | 0 | 0 | 0 |
| 0 | 0 | 0 | 0 |
| 0 | 0 | 0 | 0 |
| 0 | 0 | 0 | 0 |
| 0 | 0 | 0 | 1 |
| 0 | 0 | 0 | 0 |
| 0 | 0 | 0 | 1 |
| 0 | 0 | 0 | 1 |
| 0 | 0 | 0 | 1 |
| 0 | 0 | 0 | 1 |
| 0 | 0 | 0 | 0 |
| 0 | 0 | 0 | 1 |
| 0 | 0 | 0 | 1 |
| 0 | 0 | 0 | 1 |
| 0 | 0 | 0 | 1 |
| 0 | 1 | 0 | 0 |
| 0 | 0 | 0 | 1 |
| 0 | 0 | 0 | 1 |
| 0 | 0 | 0 | 1 |
| 0 | 0 | 0 | 1 |
| 0 | 0 | 0 | 1 |
| 0 | 0 | 0 | 1 |
| 0 | 0 | 0 | 0 |
| 0 | 0 | 0 | 0 |
| 0 | 0 | 0 | 0 |
| 0 | 0 | 0 | 1 |

| | | | |
|---|---|---|---|
| 0 | 0 | 0 | 0 |
| 0 | 1 | 0 | 0 |
| 0 | 1 | 0 | 0 |
| 0 | 0 | 0 | 0 |
| 1 | 1 | 0 | 0 |
| 1 | 1 | 0 | 0 |
| 0 | 1 | 0 | 0 |
| 0 | 0 | 0 | 0 |
| 0 | 0 | 0 | 1 |
| 0 | 0 | 0 | 1 |
| 0 | 0 | 0 | 0 |
| 0 | 0 | 0 | 1 |
| 0 | 0 | 0 | 1 |
| 0 | 0 | 0 | 0 |
| 0 | 0 | 0 | 0 |
| 0 | 0 | 0 | 0 |
| 0 | 0 | 0 | 0 |
| 0 | 0 | 0 | 0 |
| 0 | 0 | 0 | 0 |
| 0 | 0 | 0 | 1 |
| 0 | 0 | 0 | 1 |
| 0 | 0 | 0 | 0 |
| 0 | 0 | 0 | 1 |
| 0 | 1 | 0 | 0 |
| 0 | 0 | 0 | 1 |
| 0 | 0 | 0 | 1 |
| 0 | 0 | 0 | 1 |
| 0 | 0 | 0 | 1 |
| 0 | 0 | 0 | 1 |
| 0 | 0 | 0 | 0 |
| 0 | 0 | 0 | 1 |
| 0 | 0 | 0 | 1 |
| 0 | 0 | 0 | 0 |
| 0 | 0 | 0 | 1 |
| 0 | 1 | 0 | 0 |
| 0 | 1 | 0 | 0 |
| 0 | 1 | 0 | 0 |
| 0 | 0 | 0 | 0 |
| 0 | 0 | 0 | 0 |
| 0 | 0 | 0 | 0 |
| 0 | 0 | 0 | 0 |
| 0 | 0 | 0 | 0 |
| 0 | 0 | 0 | 1 |
| 0 | 1 | 0 | 0 |
| 0 | 0 | 0 | 1 |
| 0 | 0 | 0 | 0 |
| 0 | 0 | 0 | 1 |
| 0 | 0 | 0 | 1 |
| 0 | 0 | 0 | 1 |
| 0 | 0 | 0 | 0 |
| 0 | 0 | 0 | 1 |
| 0 | 0 | 0 | 1 |
| 0 | 0 | 0 | 1 |
| 0 | 0 | 0 | 1 |

| | | | |
|---|---|---|---|
| 1 | 0 | 0 | 0 |
| 0 | 0 | 0 | 1 |
| 0 | 0 | 0 | 1 |
| 0 | 0 | 0 | 1 |
| 0 | 0 | 0 | 1 |
| 0 | 0 | 0 | 1 |
| 0 | 0 | 0 | 1 |
| 0 | 0 | 0 | 1 |
| 0 | 0 | 0 | 1 |
| 0 | 0 | 0 | 1 |
| 0 | 0 | 0 | 0 |
| 0 | 0 | 0 | 1 |
| 0 | 0 | 0 | 1 |
| 0 | 0 | 0 | 1 |
| 0 | 0 | 0 | 0 |
| 0 | 0 | 0 | 0 |
| 0 | 0 | 0 | 0 |
| 0 | 0 | 0 | 0 |
| 0 | 0 | 0 | 0 |
| 0 | 0 | 0 | 1 |
| 0 | 0 | 0 | 1 |
| 0 | 0 | 0 | 1 |
| 0 | 0 | 0 | 0 |
| 0 | 0 | 0 | 1 |
| 0 | 0 | 0 | 1 |
| 0 | 0 | 0 | 1 |
| 0 | 0 | 0 | 1 |
| 0 | 0 | 0 | 1 |
| 0 | 0 | 0 | 0 |
| 0 | 0 | 0 | 0 |
| 0 | 0 | 0 | 0 |
| 0 | 1 | 0 | 0 |
| 0 | 1 | 0 | 0 |
| 0 | 0 | 0 | 0 |
| 0 | 0 | 0 | 1 |
| 0 | 0 | 0 | 1 |
| 0 | 0 | 0 | 0 |
| 0 | 0 | 0 | 0 |
| 0 | 1 | 0 | 0 |
| 0 | 0 | 0 | 0 |
| 0 | 0 | 0 | 0 |
| 0 | 0 | 0 | 0 |
| 0 | 0 | 0 | 1 |
| 0 | 0 | 0 | 1 |
| 0 | 0 | 0 | 0 |
| 0 | 0 | 0 | 1 |
| 0 | 0 | 0 | 1 |
| 0 | 0 | 0 | 1 |
| 0 | 0 | 0 | 1 |
| 0 | 0 | 0 | 1 |
| 0 | 0 | 0 | 1 |
| 0 | 0 | 0 | 1 |
| 0 | 0 | 0 | 1 |
| 0 | 1 | 1 | 1 |

| | | | |
|---|---|---|---|
| 0 | 1 | 0 | 0 |
| 0 | 0 | 0 | 1 |
| 0 | 1 | 0 | 0 |
| 0 | 0 | 0 | 1 |
| 0 | 1 | 1 | 1 |
| 0 | 1 | 1 | 1 |
| 0 | 1 | 0 | 0 |
| 0 | 1 | 0 | 0 |
| 0 | 1 | 1 | 1 |
| 0 | 1 | 1 | 1 |
| 0 | 0 | 0 | 1 |
| 0 | 0 | 0 | 1 |
| 0 | 0 | 0 | 1 |
| 0 | 1 | 1 | 1 |
| 0 | 0 | 0 | 0 |
| 0 | 0 | 0 | 0 |
| 0 | 0 | 0 | 0 |
| 0 | 0 | 0 | 0 |
| 0 | 0 | 0 | 1 |
| 0 | 1 | 0 | 0 |
| 0 | 1 | 0 | 0 |
| 0 | 0 | 0 | 0 |
| 0 | 0 | 0 | 0 |
| 0 | 0 | 0 | 0 |
| 0 | 0 | 0 | 1 |
| 0 | 1 | 0 | 0 |
| 0 | 0 | 0 | 1 |
| 0 | 0 | 0 | 1 |
| 0 | 0 | 0 | 1 |
| 0 | 1 | 0 | 0 |
| 0 | 0 | 0 | 1 |
| 0 | 0 | 0 | 1 |
| 0 | 0 | 0 | 0 |
| 0 | 0 | 0 | 0 |
| 0 | 0 | 0 | 1 |
| 0 | 0 | 0 | 1 |
| 0 | 0 | 0 | 1 |
| 0 | 0 | 0 | 1 |
| 0 | 0 | 0 | 1 |
| 0 | 0 | 0 | 0 |
| 0 | 0 | 0 | 0 |
| 0 | 1 | 0 | 0 |
| 0 | 1 | 0 | 0 |
| 0 | 0 | 0 | 0 |
| 0 | 0 | 0 | 0 |
| 0 | 0 | 0 | 1 |
| 0 | 0 | 0 | 1 |
| 0 | 0 | 0 | 0 |
| 0 | 0 | 0 | 0 |
| 0 | 0 | 0 | 1 |
| 0 | 0 | 0 | 0 |
| 0 | 0 | 0 | 0 |
| 0 | 0 | 0 | 1 |
| 0 | 0 | 0 | 1 |

| | | | |
|---|---|---|---|
| 0 | 0 | 0 | 1 |
| 0 | 1 | 0 | 0 |
| 0 | 1 | 0 | 0 |
| 0 | 1 | 1 | 1 |
| 0 | 0 | 0 | 1 |
| 0 | 0 | 0 | 0 |
| 0 | 0 | 0 | 0 |
| 0 | 0 | 0 | 0 |
| 0 | 0 | 0 | 0 |
| 0 | 0 | 0 | 0 |
| 0 | 0 | 0 | 0 |
| 0 | 0 | 0 | 0 |
| 0 | 0 | 0 | 0 |
| 0 | 0 | 0 | 0 |
| 0 | 0 | 0 | 0 |
| 0 | 0 | 0 | 1 |
| 0 | 1 | 0 | 0 |
| 0 | 0 | 0 | 1 |
| 0 | 1 | 1 | 1 |
| 0 | 0 | 0 | 1 |
| 0 | 0 | 0 | 0 |
| 0 | 0 | 0 | 1 |
| 0 | 1 | 0 | 0 |
| 0 | 0 | 0 | 0 |
| 0 | 0 | 0 | 1 |
| 0 | 0 | 0 | 0 |
| 0 | 0 | 0 | 1 |
| 0 | 0 | 0 | 1 |
| 0 | 0 | 0 | 1 |
| 0 | 0 | 0 | 0 |
| 0 | 1 | 0 | 0 |
| 0 | 0 | 0 | 0 |
| 0 | 1 | 0 | 0 |
| 0 | 1 | 0 | 0 |
| 0 | 1 | 0 | 0 |
| 0 | 1 | 0 | 0 |
| 0 | 1 | 0 | 0 |
| 0 | 0 | 0 | 1 |
| 0 | 0 | 0 | 1 |
| 0 | 0 | 0 | 0 |
| 0 | 1 | 0 | 0 |
| 0 | 0 | 0 | 0 |
| 0 | 0 | 0 | 0 |
| 0 | 0 | 0 | 0 |
| 0 | 1 | 0 | 0 |
| 0 | 0 | 0 | 0 |
| 0 | 0 | 0 | 1 |
| 0 | 0 | 0 | 0 |
| 0 | 0 | 0 | 0 |
| 0 | 0 | 0 | 0 |
| 0 | 0 | 0 | 0 |
| 0 | 0 | 0 | 1 |
| 0 | 1 | 0 | 0 |
| 0 | 0 | 0 | 0 |

| | | | |
|---|---|---|---|
| 0 | 1 | 0 | 0 |
| 0 | 1 | 0 | 0 |
| 0 | 0 | 0 | 0 |
| 0 | 0 | 0 | 0 |
| 0 | 0 | 0 | 0 |
| 0 | 0 | 0 | 0 |
| 0 | 0 | 0 | 0 |
| 0 | 0 | 0 | 0 |
| 1 | 0 | 0 | 0 |
| 0 | 0 | 0 | 0 |
| 0 | 0 | 0 | 1 |
| 0 | 1 | 0 | 0 |
| 0 | 1 | 1 | 1 |
| 0 | 0 | 0 | 0 |
| 0 | 0 | 0 | 1 |
| 0 | 0 | 0 | 1 |
| 0 | 0 | 0 | 1 |
| 0 | 0 | 0 | 1 |
| 0 | 0 | 0 | 0 |
| 0 | 0 | 0 | 0 |
| 0 | 0 | 0 | 1 |
| 0 | 1 | 0 | 0 |
| 0 | 1 | 0 | 0 |
| 0 | 0 | 0 | 1 |
| 0 | 0 | 0 | 0 |
| 0 | 0 | 0 | 0 |
| 0 | 0 | 0 | 0 |
| 0 | 0 | 0 | 1 |
| 0 | 1 | 0 | 0 |
| 0 | 0 | 0 | 0 |
| 0 | 1 | 1 | 1 |
| 1 | 1 | 0 | 0 |
| 0 | 1 | 1 | 1 |
| 0 | 1 | 1 | 1 |
| 1 | 1 | 0 | 0 |
| 1 | 1 | 0 | 0 |
| 1 | 0 | 0 | 0 |
| 0 | 1 | 1 | 1 |
| 0 | 1 | 1 | 1 |
| 0 | 1 | 0 | 0 |
| 1 | 0 | 0 | 0 |
| 0 | 1 | 0 | 0 |
| 0 | 1 | 0 | 0 |
| 1 | 1 | 0 | 0 |
| 1 | 1 | 0 | 0 |
| 0 | 1 | 0 | 0 |
| 0 | 0 | 0 | 1 |
| 0 | 0 | 0 | 1 |
| 0 | 0 | 0 | 1 |
| 0 | 1 | 1 | 1 |
| 1 | 1 | 0 | 0 |
| 0 | 1 | 0 | 0 |
| 0 | 0 | 0 | 0 |
| 0 | 0 | 0 | 0 |

| | | | |
|---|---|---|---|
| 0 | 0 | 0 | 0 |
| 0 | 1 | 1 | 1 |
| 0 | 0 | 0 | 0 |
| 0 | 0 | 0 | 0 |
| 0 | 0 | 0 | 0 |
| 0 | 0 | 0 | 1 |
| 0 | 0 | 0 | 0 |
| 0 | 0 | 0 | 0 |
| 0 | 0 | 0 | 1 |
| 0 | 1 | 0 | 0 |
| 0 | 0 | 0 | 1 |
| 0 | 0 | 0 | 1 |
| 0 | 0 | 0 | 1 |
| 0 | 0 | 0 | 0 |
| 0 | 0 | 0 | 1 |
| 0 | 0 | 0 | 0 |
| 0 | 0 | 0 | 1 |
| 0 | 0 | 0 | 1 |
| 0 | 0 | 0 | 0 |
| 0 | 0 | 0 | 0 |
| 0 | 0 | 0 | 0 |
| 0 | 0 | 0 | 0 |
| 0 | 0 | 0 | 0 |
| 0 | 0 | 0 | 0 |
| 0 | 0 | 0 | 0 |
| 0 | 0 | 0 | 1 |
| 0 | 0 | 0 | 1 |
| 0 | 0 | 0 | 1 |
| 0 | 0 | 0 | 1 |
| 0 | 0 | 0 | 1 |
| 0 | 0 | 0 | 1 |
| 0 | 0 | 0 | 1 |
| 0 | 0 | 0 | 0 |
| 0 | 0 | 0 | 0 |
| 0 | 0 | 0 | 0 |
| 0 | 0 | 0 | 0 |
| 0 | 0 | 0 | 0 |
| 0 | 0 | 0 | 0 |
| 0 | 0 | 0 | 1 |
| 0 | 0 | 0 | 1 |
| 0 | 0 | 0 | 1 |
| 0 | 0 | 0 | 1 |
| 0 | 0 | 0 | 1 |
| 0 | 0 | 0 | 1 |
| 0 | 1 | 0 | 0 |
| 0 | 0 | 0 | 0 |
| 0 | 0 | 0 | 1 |
| 0 | 0 | 0 | 0 |
| 0 | 0 | 0 | 0 |
| 0 | 0 | 0 | 0 |
| 0 | 0 | 0 | 0 |
| 0 | 0 | 0 | 0 |
| 0 | 0 | 0 | 0 |
| 0 | 0 | 0 | 0 |

| | | | |
|---|---|---|---|
| 0 | 0 | 0 | 1 |
| 0 | 0 | 0 | 1 |
| 1 | 0 | 0 | 0 |
| 1 | 0 | 0 | 0 |
| 0 | 0 | 0 | 1 |
| 0 | 0 | 0 | 0 |
| 0 | 0 | 0 | 0 |
| 0 | 0 | 0 | 1 |
| 0 | 0 | 0 | 1 |
| 0 | 1 | 0 | 0 |
| 0 | 0 | 0 | 1 |
| 0 | 0 | 0 | 1 |
| 0 | 1 | 0 | 0 |
| 0 | 1 | 0 | 0 |
| 0 | 1 | 0 | 0 |
| 0 | 1 | 0 | 0 |
| 0 | 0 | 0 | 1 |
| 0 | 0 | 0 | 1 |
| 0 | 0 | 0 | 1 |
| 0 | 0 | 0 | 1 |
| 0 | 0 | 0 | 1 |
| 0 | 0 | 0 | 1 |
| 0 | 0 | 0 | 1 |
| 0 | 0 | 0 | 1 |
| 0 | 0 | 0 | 1 |
| 0 | 0 | 0 | 0 |
| 0 | 0 | 0 | 1 |
| 0 | 0 | 0 | 1 |
| 0 | 0 | 0 | 1 |
| 0 | 0 | 0 | 1 |
| 0 | 0 | 0 | 1 |
| 0 | 0 | 0 | 1 |
| 0 | 0 | 0 | 1 |
| 0 | 0 | 0 | 1 |
| 0 | 0 | 0 | 1 |
| 0 | 0 | 0 | 1 |
| 0 | 0 | 0 | 1 |
| 0 | 0 | 0 | 0 |
| 0 | 1 | 1 | 1 |
| 0 | 0 | 0 | 1 |
| 0 | 1 | 0 | 0 |
| 0 | 0 | 0 | 1 |
| 0 | 0 | 0 | 1 |
| 0 | 1 | 0 | 0 |
| 0 | 0 | 0 | 0 |
| 0 | 0 | 0 | 1 |
| 0 | 0 | 0 | 1 |
| 0 | 0 | 0 | 1 |
| 0 | 0 | 0 | 1 |
| 0 | 0 | 0 | 1 |
| 0 | 0 | 0 | 1 |
| 0 | 0 | 0 | 0 |
| 0 | 0 | 0 | 0 |
| 0 | 0 | 0 | 0 |

| | | | |
|---|---|---|---|
| 0 | 0 | 0 | 0 |
| 0 | 0 | 0 | 1 |
| 0 | 0 | 0 | 1 |
| 0 | 0 | 0 | 1 |
| 0 | 0 | 0 | 1 |
| 0 | 0 | 0 | 1 |
| 0 | 0 | 0 | 1 |
| 0 | 0 | 0 | 1 |
| 0 | 0 | 0 | 0 |
| 0 | 0 | 0 | 1 |
| 0 | 0 | 0 | 1 |
| 0 | 0 | 0 | 1 |
| 0 | 0 | 0 | 1 |
| 0 | 0 | 0 | 1 |
| 0 | 0 | 0 | 1 |
| 0 | 0 | 0 | 1 |
| 0 | 0 | 0 | 1 |
| 0 | 0 | 0 | 1 |
| 0 | 0 | 0 | 0 |
| 0 | 0 | 0 | 0 |
| 0 | 0 | 0 | 0 |
| 0 | 1 | 0 | 0 |
| 0 | 0 | 0 | 1 |
| 0 | 0 | 0 | 0 |
| 0 | 0 | 0 | 0 |
| 0 | 0 | 0 | 1 |
| 0 | 0 | 0 | 1 |
| 0 | 0 | 0 | 1 |
| 0 | 0 | 0 | 1 |
| 0 | 0 | 0 | 1 |
| 0 | 0 | 0 | 1 |
| 0 | 0 | 0 | 1 |
| 0 | 0 | 0 | 0 |
| 0 | 0 | 0 | 1 |
| 0 | 0 | 0 | 1 |
| 0 | 0 | 0 | 1 |
| 0 | 0 | 0 | 0 |
| 0 | 0 | 0 | 1 |
| 0 | 0 | 0 | 0 |
| 0 | 0 | 0 | 0 |
| 0 | 0 | 0 | 0 |
| 0 | 0 | 0 | 0 |
| 0 | 0 | 0 | 0 |
| 0 | 0 | 0 | 1 |
| 0 | 0 | 0 | 1 |
| 0 | 0 | 0 | 1 |
| 0 | 0 | 0 | 0 |
| 0 | 0 | 0 | 0 |
| 0 | 0 | 0 | 0 |
| 0 | 1 | 0 | 0 |
| 0 | 1 | 0 | 0 |
| 0 | 1 | 0 | 0 |
| 0 | 1 | 0 | 0 |
| 0 | 0 | 0 | 1 |

| | | | |
|---|---|---|---|
| 0 | 0 | 0 | 0 |
| 0 | 0 | 0 | 0 |
| 0 | 0 | 0 | 1 |
| 1 | 0 | 0 | 0 |
| 1 | 0 | 0 | 0 |
| 0 | 0 | 0 | 1 |
| 0 | 0 | 0 | 1 |
| 0 | 1 | 1 | 1 |
| 0 | 0 | 0 | 1 |
| 0 | 0 | 0 | 1 |
| 0 | 0 | 0 | 1 |
| 0 | 0 | 0 | 0 |
| 0 | 0 | 0 | 1 |
| 0 | 0 | 0 | 1 |
| 0 | 0 | 0 | 0 |
| 0 | 0 | 0 | 1 |
| 0 | 0 | 0 | 0 |
| 0 | 0 | 0 | 1 |
| 0 | 0 | 0 | 0 |
| 1 | 0 | 0 | 0 |
| 0 | 0 | 0 | 1 |
| 0 | 0 | 0 | 1 |
| 0 | 0 | 0 | 0 |
| 0 | 0 | 0 | 1 |
| 0 | 0 | 0 | 0 |
| 0 | 0 | 0 | 1 |
| 0 | 0 | 0 | 1 |
| 0 | 0 | 0 | 0 |
| 0 | 0 | 0 | 1 |
| 0 | 0 | 0 | 0 |
| 0 | 0 | 0 | 1 |
| 0 | 0 | 0 | 0 |
| 0 | 0 | 0 | 0 |
| 0 | 0 | 0 | 0 |
| 0 | 0 | 0 | 1 |
| 0 | 0 | 0 | 0 |
| 0 | 0 | 0 | 1 |
| 0 | 0 | 0 | 1 |
| 0 | 0 | 0 | 0 |
| 0 | 0 | 0 | 1 |
| 0 | 0 | 0 | 1 |
| 0 | 0 | 0 | 1 |
| 0 | 0 | 0 | 1 |
| 0 | 0 | 0 | 0 |
| 0 | 0 | 0 | 0 |
| 0 | 0 | 0 | 1 |
| 0 | 0 | 0 | 1 |
| 0 | 0 | 0 | 1 |
| 0 | 1 | 0 | 0 |
| 0 | 0 | 0 | 1 |
| 0 | 0 | 0 | 0 |
| 0 | 0 | 0 | 1 |
| 0 | 0 | 0 | 1 |
| 0 | 0 | 0 | 0 |

| | | | |
|---|---|---|---|
| 0 | 0 | 0 | 1 |
| 0 | 0 | 0 | 1 |
| 0 | 0 | 0 | 1 |
| 0 | 0 | 0 | 1 |
| 0 | 0 | 0 | 1 |
| 0 | 0 | 0 | 1 |
| 0 | 0 | 0 | 1 |
| 0 | 0 | 0 | 0 |
| 0 | 0 | 0 | 1 |
| 0 | 0 | 0 | 1 |
| 0 | 0 | 0 | 1 |
| 0 | 0 | 0 | 1 |
| 0 | 0 | 0 | 0 |
| 0 | 0 | 0 | 1 |
| 0 | 0 | 0 | 0 |
| 0 | 0 | 0 | 0 |
| 0 | 0 | 0 | 1 |
| 0 | 0 | 0 | 1 |
| 0 | 0 | 0 | 1 |
| 0 | 0 | 0 | 1 |
| 0 | 0 | 0 | 0 |
| 0 | 0 | 0 | 0 |
| 0 | 0 | 0 | 1 |
| 0 | 0 | 0 | 1 |
| 0 | 0 | 0 | 0 |
| 0 | 0 | 0 | 1 |
| 0 | 0 | 0 | 0 |
| 0 | 0 | 0 | 0 |
| 0 | 0 | 0 | 0 |
| 0 | 0 | 0 | 0 |
| 0 | 0 | 0 | 1 |
| 0 | 0 | 0 | 1 |
| 0 | 0 | 0 | 0 |
| 0 | 1 | 0 | 0 |
| 0 | 0 | 0 | 0 |
| 0 | 0 | 0 | 0 |
| 0 | 0 | 0 | 0 |
| 0 | 0 | 0 | 0 |
| 0 | 0 | 0 | 0 |
| 0 | 0 | 0 | 1 |
| 0 | 0 | 0 | 1 |
| 0 | 0 | 0 | 1 |
| 0 | 0 | 0 | 1 |
| 0 | 0 | 0 | 1 |
| 0 | 0 | 0 | 1 |
| 0 | 0 | 0 | 0 |
| 0 | 0 | 0 | 1 |
| 0 | 0 | 0 | 1 |
| 0 | 0 | 0 | 1 |
| 0 | 0 | 0 | 0 |
| 0 | 0 | 0 | 0 |
| 0 | 0 | 0 | 1 |
| 0 | 0 | 0 | 1 |
| 0 | 0 | 0 | 1 |

| | | | |
|---|---|---|---|
| 0 | 0 | 0 | 1 |
| 0 | 0 | 0 | 1 |
| 0 | 0 | 0 | 1 |
| 0 | 0 | 0 | 1 |
| 0 | 0 | 0 | 0 |
| 0 | 0 | 0 | 1 |
| 0 | 0 | 0 | 0 |
| 0 | 0 | 0 | 0 |
| 0 | 0 | 0 | 0 |
| 0 | 0 | 0 | 0 |
| 0 | 0 | 0 | 0 |
| 0 | 0 | 0 | 1 |
| 0 | 0 | 0 | 0 |
| 0 | 0 | 0 | 0 |
| 0 | 0 | 0 | 1 |
| 0 | 0 | 0 | 1 |
| 0 | 0 | 0 | 1 |
| 0 | 0 | 0 | 1 |
| 0 | 0 | 0 | 1 |
| 0 | 0 | 0 | 1 |
| 0 | 0 | 0 | 1 |
| 0 | 0 | 0 | 1 |
| 0 | 0 | 0 | 1 |
| 0 | 0 | 0 | 0 |
| 0 | 0 | 0 | 1 |
| 0 | 0 | 0 | 1 |
| 0 | 0 | 0 | 1 |
| 0 | 0 | 0 | 0 |
| 0 | 1 | 1 | 1 |
| 0 | 1 | 1 | 1 |
| 0 | 1 | 0 | 0 |
| 0 | 1 | 0 | 0 |
| 0 | 0 | 0 | 1 |
| 0 | 0 | 0 | 0 |
| 0 | 0 | 0 | 1 |
| 0 | 0 | 0 | 1 |
| 0 | 0 | 0 | 1 |
| 0 | 0 | 0 | 1 |
| 0 | 0 | 0 | 1 |
| 0 | 0 | 0 | 1 |
| 0 | 0 | 0 | 1 |
| 0 | 0 | 0 | 1 |
| 0 | 0 | 0 | 1 |
| 0 | 0 | 0 | 1 |
| 0 | 0 | 0 | 1 |
| 0 | 0 | 0 | 1 |
| 0 | 0 | 0 | 1 |
| 0 | 0 | 0 | 1 |
| 0 | 0 | 0 | 0 |
| 0 | 0 | 0 | 1 |
| 0 | 0 | 0 | 1 |
| 0 | 1 | 1 | 1 |
| 0 | 1 | 1 | 1 |
| 0 | 0 | 0 | 1 |

| | | | |
|---|---|---|---|
| 0 | 0 | 0 | 1 |
| 0 | 0 | 0 | 1 |
| 0 | 0 | 0 | 1 |
| 0 | 0 | 0 | 1 |
| 0 | 1 | 1 | 1 |
| 0 | 0 | 0 | 1 |
| 0 | 0 | 0 | 1 |
| 0 | 0 | 0 | 1 |
| 0 | 0 | 0 | 1 |
| 0 | 0 | 0 | 1 |
| 0 | 0 | 0 | 0 |
| 0 | 0 | 0 | 0 |
| 0 | 0 | 0 | 0 |
| 1 | 1 | 0 | 0 |
| 1 | 0 | 0 | 0 |
| 0 | 1 | 1 | 1 |
| 1 | 0 | 0 | 0 |
| 0 | 0 | 0 | 0 |
| 0 | 0 | 0 | 0 |
| 0 | 0 | 0 | 0 |
| 0 | 0 | 0 | 0 |
| 1 | 0 | 0 | 0 |
| 0 | 0 | 0 | 1 |
| 0 | 0 | 0 | 0 |
| 1 | 0 | 0 | 0 |
| 1 | 0 | 0 | 0 |
| 0 | 0 | 0 | 1 |
| 0 | 0 | 0 | 0 |
| 0 | 0 | 0 | 1 |
| 0 | 0 | 0 | 1 |
| 0 | 0 | 0 | 1 |
| 0 | 0 | 0 | 0 |
| 0 | 0 | 0 | 1 |
| 0 | 0 | 0 | 1 |
| 0 | 0 | 0 | 0 |
| 0 | 0 | 0 | 0 |
| 0 | 0 | 0 | 0 |
| 0 | 1 | 0 | 0 |
| 0 | 0 | 0 | 0 |
| 0 | 1 | 1 | 1 |
| 0 | 1 | 1 | 1 |
| 0 | 0 | 0 | 0 |
| 0 | 1 | 1 | 1 |
| 0 | 0 | 0 | 0 |
| 0 | 0 | 0 | 1 |
| 0 | 1 | 1 | 1 |
| 0 | 0 | 0 | 1 |
| 0 | 1 | 0 | 0 |
| 0 | 0 | 0 | 0 |
| 0 | 1 | 0 | 0 |
| 0 | 1 | 0 | 0 |
| 0 | 0 | 0 | 1 |
| 0 | 0 | 0 | 0 |
| 0 | 0 | 0 | 1 |

| | | | |
|---|---|---|---|
| 0 | 0 | 0 | 1 |
| 0 | 0 | 0 | 1 |
| 0 | 0 | 0 | 1 |
| 0 | 0 | 0 | 1 |
| 0 | 0 | 0 | 1 |
| 0 | 0 | 0 | 0 |
| 0 | 0 | 0 | 0 |
| 0 | 0 | 0 | 0 |
| 0 | 0 | 0 | 0 |
| 0 | 0 | 0 | 1 |
| 1 | 0 | 0 | 0 |
| 1 | 0 | 0 | 0 |
| 0 | 0 | 0 | 0 |
| 0 | 0 | 0 | 0 |
| 0 | 0 | 0 | 0 |
| 0 | 0 | 0 | 0 |
| 0 | 0 | 0 | 1 |
| 0 | 0 | 0 | 1 |
| 0 | 0 | 0 | 1 |
| 0 | 0 | 0 | 1 |
| 0 | 0 | 0 | 1 |
| 0 | 0 | 0 | 1 |
| 0 | 0 | 0 | 1 |
| 0 | 0 | 0 | 1 |
| 0 | 0 | 0 | 1 |
| 0 | 0 | 0 | 1 |
| 0 | 0 | 0 | 1 |
| 0 | 0 | 0 | 1 |
| 0 | 1 | 0 | 0 |
| 0 | 0 | 0 | 1 |
| 0 | 0 | 0 | 1 |
| 0 | 1 | 1 | 1 |
| 0 | 1 | 1 | 1 |
| 0 | 0 | 0 | 0 |
| 0 | 0 | 0 | 1 |
| 0 | 1 | 0 | 0 |
| 0 | 0 | 0 | 1 |
| 0 | 0 | 0 | 0 |
| 0 | 0 | 0 | 1 |
| 0 | 0 | 0 | 0 |
| 0 | 0 | 0 | 1 |
| 0 | 0 | 0 | 1 |
| 0 | 0 | 0 | 1 |
| 0 | 0 | 0 | 0 |
| 0 | 0 | 0 | 1 |
| 0 | 0 | 0 | 1 |
| 0 | 0 | 0 | 1 |
| 0 | 0 | 0 | 1 |
| 0 | 0 | 0 | 1 |
| 0 | 0 | 0 | 1 |
| 0 | 0 | 0 | 1 |
| 0 | 1 | 1 | 1 |
| 0 | 0 | 0 | 0 |
| 0 | 0 | 0 | 0 |

| | | | |
|---|---|---|---|
| 0 | 0 | 0 | 1 |
| 0 | 0 | 0 | 0 |
| 0 | 0 | 0 | 1 |
| 0 | 0 | 0 | 1 |
| 0 | 1 | 0 | 0 |
| 0 | 0 | 0 | 0 |
| 0 | 0 | 0 | 0 |
| 0 | 0 | 0 | 1 |
| 0 | 0 | 0 | 1 |
| 0 | 0 | 0 | 0 |
| 0 | 0 | 0 | 0 |
| 0 | 0 | 0 | 0 |
| 0 | 0 | 0 | 0 |
| 0 | 0 | 0 | 0 |
| 0 | 0 | 0 | 1 |
| 0 | 0 | 0 | 0 |
| 0 | 1 | 0 | 0 |
| 0 | 0 | 0 | 0 |
| 0 | 0 | 0 | 1 |
| 0 | 0 | 0 | 1 |
| 0 | 0 | 0 | 1 |
| 0 | 0 | 0 | 1 |
| 0 | 0 | 0 | 1 |
| 0 | 0 | 0 | 0 |
| 0 | 0 | 0 | 1 |
| 0 | 0 | 0 | 1 |
| 0 | 0 | 0 | 0 |
| 0 | 0 | 0 | 0 |
| 0 | 0 | 0 | 1 |
| 0 | 0 | 0 | 1 |
| 0 | 0 | 0 | 0 |
| 0 | 0 | 0 | 1 |
| 0 | 0 | 0 | 1 |
| 0 | 0 | 0 | 1 |
| 0 | 0 | 0 | 1 |
| 0 | 0 | 0 | 0 |
| 0 | 0 | 0 | 1 |
| 0 | 0 | 0 | 0 |
| 0 | 0 | 0 | 0 |
| 0 | 0 | 0 | 1 |
| 0 | 0 | 0 | 1 |
| 0 | 0 | 0 | 0 |
| 0 | 0 | 0 | 0 |
| 0 | 0 | 0 | 1 |
| 0 | 0 | 0 | 0 |
| 0 | 0 | 0 | 0 |
| 0 | 0 | 0 | 0 |
| 0 | 0 | 0 | 1 |
| 0 | 0 | 0 | 0 |
| 0 | 0 | 0 | 0 |
| 0 | 0 | 0 | 0 |
| 0 | 0 | 0 | 1 |
| 0 | 0 | 0 | 0 |
| 0 | 0 | 0 | 1 |

| | | | |
|---|---|---|---|
| 0 | 0 | 0 | 1 |
| 0 | 0 | 0 | 1 |
| 0 | 0 | 0 | 1 |
| 0 | 0 | 0 | 1 |
| 0 | 0 | 0 | 0 |
| 0 | 0 | 0 | 0 |
| 0 | 0 | 0 | 1 |
| 0 | 0 | 0 | 1 |
| 0 | 0 | 0 | 1 |
| 0 | 1 | 0 | 0 |
| 0 | 1 | 1 | 1 |
| 0 | 1 | 1 | 1 |
| 0 | 1 | 0 | 0 |
| 0 | 0 | 0 | 1 |
| 0 | 0 | 0 | 1 |
| 0 | 0 | 0 | 1 |
| 0 | 0 | 0 | 1 |
| 0 | 0 | 0 | 1 |
| 0 | 0 | 0 | 1 |
| 0 | 0 | 0 | 1 |
| 0 | 0 | 0 | 1 |
| 0 | 0 | 0 | 1 |
| 0 | 0 | 0 | 1 |
| 0 | 0 | 0 | 0 |
| 0 | 0 | 0 | 1 |
| 1 | 0 | 0 | 0 |
| 0 | 0 | 0 | 1 |
| 0 | 0 | 0 | 0 |
| 0 | 0 | 0 | 1 |
| 0 | 0 | 0 | 0 |
| 0 | 0 | 0 | 0 |
| 0 | 0 | 0 | 0 |
| 0 | 0 | 0 | 0 |
| 0 | 0 | 0 | 0 |
| 0 | 0 | 0 | 1 |
| 0 | 0 | 0 | 0 |
| 0 | 0 | 0 | 1 |
| 0 | 0 | 0 | 0 |
| 0 | 0 | 0 | 1 |
| 0 | 0 | 0 | 1 |
| 0 | 0 | 0 | 0 |
| 0 | 0 | 0 | 0 |
| 0 | 0 | 0 | 0 |
| 0 | 0 | 0 | 0 |
| 0 | 0 | 0 | 0 |
| 0 | 0 | 0 | 0 |
| 0 | 0 | 0 | 0 |
| 0 | 1 | 0 | 0 |
| 0 | 0 | 0 | 1 |
| 0 | 0 | 0 | 0 |
| 0 | 1 | 1 | 1 |
| 0 | 0 | 0 | 1 |
| 0 | 0 | 0 | 1 |
| 0 | 0 | 0 | 1 |

| | | | |
|---|---|---|---|
| 0 | 0 | 0 | 1 |
| 0 | 0 | 0 | 1 |
| 0 | 0 | 0 | 1 |
| 0 | 0 | 0 | 1 |
| 0 | 0 | 0 | 1 |
| 0 | 0 | 0 | 1 |
| 0 | 0 | 0 | 0 |
| 0 | 0 | 0 | 0 |
| 0 | 0 | 0 | 0 |
| 0 | 0 | 0 | 0 |
| 0 | 0 | 0 | 0 |
| 0 | 0 | 0 | 0 |
| 0 | 0 | 0 | 0 |
| 0 | 0 | 0 | 0 |
| 0 | 0 | 0 | 0 |
| 0 | 0 | 0 | 0 |
| 1 | 0 | 0 | 0 |
| 0 | 0 | 0 | 0 |
| 0 | 0 | 0 | 1 |
| 0 | 0 | 0 | 0 |
| 0 | 0 | 0 | 1 |
| 0 | 0 | 0 | 1 |
| 0 | 0 | 0 | 1 |
| 0 | 0 | 0 | 1 |
| 0 | 0 | 0 | 1 |
| 0 | 0 | 0 | 0 |
| 0 | 0 | 0 | 0 |
| 0 | 0 | 0 | 0 |
| 0 | 0 | 0 | 0 |
| 0 | 0 | 0 | 0 |
| 0 | 0 | 0 | 0 |
| 1 | 0 | 0 | 0 |
| 0 | 0 | 0 | 0 |
| 1 | 0 | 0 | 0 |
| 0 | 0 | 0 | 1 |
| 1 | 0 | 0 | 0 |
| 1 | 0 | 0 | 0 |
| 0 | 0 | 0 | 0 |
| 1 | 0 | 0 | 0 |
| 0 | 0 | 0 | 1 |
| 0 | 0 | 0 | 0 |
| 0 | 0 | 0 | 0 |
| 0 | 0 | 0 | 1 |
| 0 | 0 | 0 | 1 |
| 0 | 0 | 0 | 1 |
| 0 | 0 | 0 | 1 |
| 0 | 0 | 0 | 1 |
| 0 | 0 | 0 | 0 |
| 0 | 0 | 0 | 0 |
| 0 | 0 | 0 | 0 |
| 0 | 0 | 0 | 0 |
| 0 | 0 | 0 | 1 |
| 0 | 0 | 0 | 0 |
| 0 | 0 | 0 | 1 |

| | | | |
|---|---|---|---|
| 0 | 0 | 0 | 1 |
| 0 | 0 | 0 | 1 |
| 0 | 0 | 0 | 1 |
| 0 | 0 | 0 | 1 |
| 0 | 0 | 0 | 1 |
| 0 | 0 | 0 | 0 |
| 0 | 0 | 0 | 1 |
| 0 | 0 | 0 | 1 |
| 0 | 0 | 0 | 1 |
| 0 | 0 | 0 | 1 |
| 0 | 1 | 0 | 0 |
| 0 | 1 | 0 | 0 |
| 0 | 1 | 0 | 0 |
| 0 | 1 | 1 | 1 |
| 0 | 1 | 1 | 1 |
| 0 | 0 | 0 | 1 |
| 0 | 1 | 1 | 1 |
| 0 | 0 | 0 | 0 |
| 0 | 0 | 0 | 1 |
| 0 | 0 | 0 | 1 |
| 0 | 0 | 0 | 1 |
| 0 | 0 | 0 | 1 |
| 0 | 0 | 0 | 0 |
| 0 | 0 | 0 | 1 |
| 0 | 0 | 0 | 0 |
| 0 | 0 | 0 | 0 |
| 0 | 0 | 0 | 0 |
| 0 | 0 | 0 | 0 |
| 0 | 0 | 0 | 1 |
| 0 | 0 | 0 | 0 |
| 0 | 0 | 0 | 1 |
| 0 | 0 | 0 | 1 |
| 0 | 0 | 0 | 1 |
| 0 | 0 | 0 | 1 |
| 0 | 0 | 0 | 1 |
| 0 | 0 | 0 | 1 |
| 0 | 0 | 0 | 1 |
| 0 | 0 | 0 | 0 |
| 0 | 0 | 0 | 0 |
| 0 | 0 | 0 | 0 |
| 0 | 1 | 0 | 0 |
| 0 | 0 | 0 | 0 |
| 0 | 0 | 0 | 1 |
| 0 | 0 | 0 | 0 |
| 0 | 0 | 0 | 0 |
| 0 | 0 | 0 | 1 |
| 0 | 0 | 0 | 0 |
| 0 | 1 | 0 | 0 |
| 0 | 0 | 0 | 0 |
| 1 | 0 | 0 | 0 |
| 0 | 0 | 0 | 1 |
| 0 | 0 | 0 | 1 |
| 0 | 0 | 0 | 1 |
| 0 | 1 | 0 | 0 |

| | | | |
|---|---|---|---|
| 0 | 0 | 0 | 0 |
| 0 | 0 | 0 | 1 |
| 0 | 1 | 0 | 0 |
| 0 | 0 | 0 | 1 |
| 0 | 0 | 0 | 1 |
| 0 | 0 | 0 | 1 |
| 0 | 0 | 0 | 1 |
| 0 | 0 | 0 | 1 |
| 0 | 1 | 0 | 0 |
| 0 | 0 | 0 | 1 |
| 0 | 0 | 0 | 0 |
| 0 | 0 | 0 | 0 |
| 0 | 0 | 0 | 0 |
| 0 | 0 | 0 | 0 |
| 0 | 0 | 0 | 1 |
| 0 | 0 | 0 | 1 |
| 0 | 0 | 0 | 0 |
| 0 | 0 | 0 | 0 |
| 0 | 0 | 0 | 1 |
| 0 | 0 | 0 | 1 |
| 0 | 0 | 0 | 0 |
| 0 | 0 | 0 | 1 |
| 0 | 0 | 0 | 1 |
| 0 | 0 | 0 | 1 |
| 0 | 1 | 0 | 0 |
| 0 | 0 | 0 | 1 |
| 0 | 1 | 0 | 0 |
| 0 | 1 | 0 | 0 |
| 0 | 1 | 0 | 0 |
| 0 | 0 | 0 | 1 |
| 0 | 0 | 0 | 0 |
| 0 | 0 | 0 | 0 |
| 0 | 0 | 0 | 0 |
| 0 | 0 | 0 | 1 |
| 0 | 0 | 0 | 0 |
| 0 | 0 | 0 | 1 |
| 0 | 0 | 0 | 0 |
| 0 | 0 | 0 | 0 |
| 0 | 1 | 0 | 0 |
| 0 | 0 | 0 | 0 |
| 0 | 0 | 0 | 0 |
| 0 | 0 | 0 | 1 |
| 0 | 1 | 0 | 0 |
| 0 | 1 | 0 | 0 |
| 0 | 1 | 0 | 0 |
| 1 | 0 | 0 | 0 |
| 0 | 0 | 0 | 0 |
| 1 | 0 | 0 | 0 |
| 1 | 0 | 0 | 0 |
| 1 | 0 | 0 | 0 |
| 0 | 0 | 0 | 0 |
| 0 | 0 | 0 | 0 |
| 0 | 0 | 0 | 1 |
| 0 | 0 | 0 | 0 |

| | | | |
|---|---|---|---|
| 0 | 0 | 0 | 1 |
| 0 | 0 | 0 | 1 |
| 0 | 0 | 0 | 1 |
| 0 | 0 | 0 | 1 |
| 0 | 0 | 0 | 1 |
| 0 | 0 | 0 | 1 |
| 0 | 0 | 0 | 1 |
| 0 | 0 | 0 | 1 |
| 0 | 0 | 0 | 1 |
| 0 | 0 | 0 | 0 |
| 0 | 0 | 0 | 0 |
| 0 | 0 | 0 | 1 |
| 0 | 0 | 0 | 1 |
| 0 | 0 | 0 | 1 |
| 0 | 0 | 0 | 0 |
| 0 | 0 | 0 | 1 |
| 0 | 0 | 0 | 1 |
| 0 | 0 | 0 | 0 |
| 0 | 0 | 0 | 1 |
| 0 | 0 | 0 | 1 |
| 0 | 0 | 0 | 1 |
| 0 | 1 | 0 | 0 |
| 0 | 0 | 0 | 0 |
| 0 | 1 | 0 | 0 |
| 0 | 1 | 0 | 0 |
| 0 | 1 | 0 | 0 |
| 0 | 1 | 0 | 0 |
| 0 | 0 | 0 | 0 |
| 0 | 0 | 0 | 1 |
| 0 | 0 | 0 | 0 |
| 1 | 0 | 0 | 0 |
| 0 | 0 | 0 | 1 |
| 0 | 0 | 0 | 1 |
| 0 | 0 | 0 | 1 |
| 0 | 0 | 0 | 1 |
| 0 | 0 | 0 | 1 |
| 0 | 0 | 0 | 1 |
| 0 | 0 | 0 | 1 |
| 0 | 0 | 0 | 1 |
| 0 | 0 | 0 | 1 |
| 0 | 0 | 0 | 1 |
| 0 | 0 | 0 | 1 |
| 0 | 0 | 0 | 1 |
| 0 | 0 | 0 | 0 |
| 0 | 0 | 0 | 1 |
| 0 | 0 | 0 | 0 |
| 0 | 0 | 0 | 1 |
| 0 | 0 | 0 | 1 |
| 0 | 0 | 0 | 1 |
| 0 | 0 | 0 | 1 |
| 0 | 0 | 0 | 0 |
| 0 | 0 | 0 | 1 |
| 0 | 0 | 0 | 1 |
| 0 | 0 | 0 | 0 |

| | | | |
|---|---|---|---|
| 0 | 0 | 0 | 0 |
| 0 | 0 | 0 | 0 |
| 0 | 0 | 0 | 0 |
| 0 | 0 | 0 | 0 |
| 0 | 0 | 0 | 1 |
| 0 | 0 | 0 | 1 |
| 0 | 0 | 0 | 1 |
| 0 | 0 | 0 | 0 |
| 0 | 0 | 0 | 0 |
| 0 | 0 | 0 | 0 |
| 0 | 0 | 0 | 0 |
| 0 | 0 | 0 | 1 |
| 0 | 0 | 0 | 1 |
| 0 | 0 | 0 | 1 |
| 0 | 0 | 0 | 1 |
| 0 | 0 | 0 | 0 |
| 0 | 0 | 0 | 0 |
| 0 | 0 | 0 | 1 |
| 0 | 0 | 0 | 1 |
| 0 | 0 | 0 | 1 |
| 0 | 0 | 0 | 1 |
| 0 | 1 | 0 | 0 |
| 0 | 0 | 0 | 1 |
| 0 | 0 | 0 | 1 |
| 0 | 1 | 0 | 0 |
| 0 | 0 | 0 | 0 |
| 0 | 0 | 0 | 0 |
| 0 | 0 | 0 | 0 |
| 0 | 0 | 0 | 1 |
| 0 | 0 | 0 | 1 |
| 0 | 0 | 0 | 1 |
| 0 | 0 | 0 | 0 |
| 0 | 0 | 0 | 1 |
| 0 | 0 | 0 | 1 |
| 0 | 0 | 0 | 1 |
| 0 | 0 | 0 | 1 |
| 0 | 0 | 0 | 1 |
| 0 | 0 | 0 | 1 |
| 0 | 0 | 0 | 1 |
| 0 | 0 | 0 | 1 |
| 0 | 0 | 0 | 1 |
| 0 | 0 | 0 | 1 |
| 0 | 0 | 0 | 0 |
| 0 | 0 | 0 | 1 |
| 0 | 0 | 0 | 1 |
| 0 | 0 | 0 | 1 |
| 0 | 0 | 0 | 1 |
| 0 | 0 | 0 | 1 |
| 0 | 0 | 0 | 1 |
| 0 | 0 | 0 | 0 |
| 0 | 0 | 0 | 1 |
| 0 | 0 | 0 | 1 |
| 0 | 0 | 0 | 1 |
| 0 | 0 | 0 | 1 |

| | | | |
|---|---|---|---|
| 0 | 0 | 0 | 0 |
| 0 | 0 | 0 | 1 |
| 0 | 0 | 0 | 1 |
| 0 | 0 | 0 | 1 |
| 0 | 0 | 0 | 1 |
| 0 | 0 | 0 | 1 |
| 0 | 0 | 0 | 1 |
| 0 | 1 | 0 | 0 |
| 0 | 0 | 0 | 1 |
| 0 | 0 | 0 | 1 |
| 0 | 0 | 0 | 0 |
| 0 | 0 | 0 | 0 |
| 0 | 0 | 0 | 0 |
| 0 | 0 | 0 | 1 |
| 0 | 0 | 0 | 0 |
| 0 | 0 | 0 | 0 |
| 0 | 0 | 0 | 0 |
| 0 | 0 | 0 | 0 |
| 0 | 0 | 0 | 1 |
| 0 | 0 | 0 | 0 |
| 0 | 0 | 0 | 0 |
| 0 | 0 | 0 | 0 |
| 0 | 0 | 0 | 0 |
| 0 | 0 | 0 | 0 |
| 0 | 0 | 0 | 0 |
| 0 | 0 | 0 | 0 |
| 0 | 0 | 0 | 0 |
| 0 | 0 | 0 | 0 |
| 0 | 0 | 0 | 0 |
| 0 | 0 | 0 | 0 |
| 0 | 0 | 0 | 0 |
| 0 | 0 | 0 | 0 |
| 0 | 0 | 0 | 1 |
| 0 | 0 | 0 | 0 |
| 0 | 0 | 0 | 1 |
| 0 | 0 | 0 | 0 |
| 0 | 0 | 0 | 0 |
| 0 | 0 | 0 | 1 |
| 0 | 0 | 0 | 1 |
| 0 | 0 | 0 | 1 |
| 0 | 0 | 0 | 1 |
| 0 | 0 | 0 | 0 |
| 0 | 0 | 0 | 0 |
| 0 | 0 | 0 | 0 |
| 0 | 0 | 0 | 1 |
| 0 | 0 | 0 | 0 |
| 0 | 0 | 0 | 1 |
| 0 | 0 | 0 | 0 |
| 0 | 0 | 0 | 1 |
| 0 | 0 | 0 | 1 |
| 0 | 0 | 0 | 1 |
| 0 | 0 | 0 | 1 |
| 0 | 0 | 0 | 1 |
| 0 | 0 | 0 | 1 |

| | | | |
|---|---|---|---|
| 0 | 0 | 0 | 1 |
| 0 | 0 | 0 | 1 |
| 0 | 0 | 0 | 1 |
| 0 | 0 | 0 | 1 |
| 0 | 0 | 0 | 1 |
| 0 | 0 | 0 | 0 |
| 0 | 0 | 0 | 0 |
| 0 | 0 | 0 | 1 |
| 0 | 0 | 0 | 0 |
| 0 | 0 | 0 | 0 |
| 0 | 0 | 0 | 1 |
| 0 | 0 | 0 | 1 |
| 0 | 0 | 0 | 1 |
| 0 | 0 | 0 | 1 |
| 0 | 0 | 0 | 1 |
| 0 | 1 | 1 | 1 |
| 0 | 0 | 0 | 1 |
| 0 | 0 | 0 | 1 |
| 0 | 1 | 0 | 0 |
| 0 | 1 | 0 | 0 |
| 0 | 0 | 0 | 1 |
| 0 | 0 | 0 | 1 |
| 0 | 0 | 0 | 1 |
| 0 | 0 | 0 | 0 |
| 0 | 0 | 0 | 1 |
| 0 | 0 | 0 | 0 |
| 0 | 0 | 0 | 0 |
| 0 | 0 | 0 | 1 |
| 0 | 0 | 0 | 1 |
| 0 | 0 | 0 | 1 |
| 0 | 1 | 0 | 0 |
| 0 | 0 | 0 | 1 |
| 0 | 0 | 0 | 0 |
| 0 | 1 | 0 | 0 |
| 0 | 1 | 0 | 0 |
| 0 | 1 | 0 | 0 |
| 0 | 0 | 0 | 0 |
| 0 | 0 | 0 | 1 |
| 0 | 0 | 0 | 1 |
| 0 | 0 | 0 | 1 |
| 0 | 0 | 0 | 0 |
| 0 | 1 | 0 | 0 |
| 0 | 1 | 0 | 0 |
| 0 | 0 | 0 | 0 |
| 0 | 0 | 0 | 1 |
| 0 | 0 | 0 | 1 |
| 0 | 0 | 0 | 1 |
| 0 | 0 | 0 | 0 |
| 0 | 0 | 0 | 0 |
| 0 | 0 | 0 | 0 |
| 0 | 0 | 0 | 1 |
| 0 | 0 | 0 | 1 |
| 0 | 0 | 0 | 1 |
| 0 | 0 | 0 | 1 |

| | | | |
|---|---|---|---|
| 0 | 0 | 0 | 1 |
| 0 | 0 | 0 | 1 |
| 0 | 0 | 0 | 1 |
| 0 | 0 | 0 | 0 |
| 0 | 0 | 0 | 0 |
| 0 | 0 | 0 | 1 |
| 0 | 0 | 0 | 0 |
| 0 | 0 | 0 | 1 |
| 0 | 0 | 0 | 1 |
| 0 | 0 | 0 | 1 |
| 0 | 0 | 0 | 1 |
| 0 | 0 | 0 | 0 |
| 0 | 1 | 1 | 1 |
| 0 | 0 | 0 | 0 |
| 0 | 0 | 0 | 1 |
| 0 | 0 | 0 | 0 |
| 0 | 1 | 1 | 1 |
| 0 | 1 | 1 | 1 |
| 0 | 0 | 0 | 1 |
| 0 | 1 | 0 | 0 |
| 0 | 0 | 0 | 0 |
| 0 | 0 | 0 | 0 |
| 0 | 0 | 0 | 0 |
| 0 | 0 | 0 | 0 |
| 0 | 0 | 0 | 0 |
| 0 | 0 | 0 | 1 |
| 0 | 0 | 0 | 0 |
| 0 | 0 | 0 | 1 |
| 0 | 0 | 0 | 1 |
| 0 | 0 | 0 | 1 |
| 0 | 0 | 0 | 1 |
| 0 | 0 | 0 | 1 |
| 0 | 0 | 0 | 0 |
| 0 | 1 | 0 | 0 |
| 0 | 0 | 0 | 1 |
| 0 | 0 | 0 | 1 |
| 0 | 1 | 0 | 0 |
| 0 | 1 | 0 | 0 |
| 0 | 1 | 0 | 0 |
| 0 | 1 | 1 | 1 |
| 0 | 0 | 0 | 0 |
| 0 | 0 | 0 | 0 |
| 0 | 0 | 0 | 1 |
| 0 | 0 | 0 | 0 |
| 0 | 0 | 0 | 1 |
| 0 | 0 | 0 | 1 |
| 0 | 0 | 0 | 1 |
| 0 | 0 | 0 | 1 |
| 0 | 0 | 0 | 1 |
| 1 | 0 | 0 | 0 |
| 0 | 0 | 0 | 1 |
| 0 | 0 | 0 | 1 |
| 0 | 0 | 0 | 1 |
| 0 | 0 | 0 | 1 |

| | | | |
|---|---|---|---|
| 0 | 0 | 0 | 1 |
| 0 | 0 | 0 | 1 |
| 0 | 0 | 0 | 1 |
| 0 | 0 | 0 | 0 |
| 0 | 0 | 0 | 1 |
| 0 | 0 | 0 | 1 |
| 0 | 0 | 0 | 1 |
| 0 | 0 | 0 | 0 |
| 0 | 0 | 0 | 1 |
| 0 | 0 | 0 | 1 |
| 0 | 0 | 0 | 1 |
| 0 | 0 | 0 | 1 |
| 0 | 0 | 0 | 1 |
| 0 | 0 | 0 | 1 |
| 0 | 0 | 0 | 1 |
| 0 | 0 | 0 | 1 |
| 0 | 0 | 0 | 0 |
| 0 | 0 | 0 | 0 |
| 0 | 0 | 0 | 1 |
| 0 | 0 | 0 | 1 |
| 0 | 0 | 0 | 1 |
| 0 | 0 | 0 | 1 |
| 0 | 0 | 0 | 1 |
| 0 | 0 | 0 | 1 |
| 0 | 0 | 0 | 1 |
| 0 | 0 | 0 | 1 |
| 0 | 0 | 0 | 1 |
| 0 | 1 | 0 | 0 |
| 0 | 0 | 0 | 0 |
| 0 | 0 | 0 | 1 |
| 0 | 0 | 0 | 1 |
| 0 | 0 | 0 | 0 |
| 0 | 0 | 0 | 1 |
| 0 | 0 | 0 | 0 |
| 1 | 0 | 0 | 0 |
| 1 | 0 | 0 | 0 |
| 1 | 0 | 0 | 0 |
| 1 | 0 | 0 | 0 |
| 0 | 0 | 0 | 1 |
| 0 | 0 | 0 | 1 |
| 0 | 0 | 0 | 1 |
| 0 | 0 | 0 | 1 |
| 0 | 0 | 0 | 1 |
| 0 | 0 | 0 | 1 |
| 0 | 0 | 0 | 0 |
| 0 | 0 | 0 | 0 |
| 0 | 0 | 0 | 0 |
| 0 | 0 | 0 | 1 |
| 0 | 0 | 0 | 1 |
| 0 | 0 | 0 | 1 |
| 0 | 0 | 0 | 1 |
| 0 | 0 | 0 | 0 |
| 0 | 0 | 0 | 0 |
| 0 | 0 | 0 | 0 |

| | | | |
|---|---|---|---|
| 0 | 0 | 0 | 0 |
| 0 | 0 | 0 | 0 |
| 0 | 0 | 0 | 1 |
| 0 | 0 | 0 | 0 |
| 0 | 0 | 0 | 0 |
| 0 | 0 | 0 | 0 |
| 0 | 0 | 0 | 0 |
| 0 | 0 | 0 | 1 |
| 0 | 0 | 0 | 1 |
| 0 | 0 | 0 | 1 |
| 0 | 0 | 0 | 1 |
| 0 | 0 | 0 | 1 |
| 0 | 0 | 0 | 1 |
| 0 | 0 | 0 | 1 |
| 0 | 0 | 0 | 1 |
| 0 | 0 | 0 | 1 |
| 0 | 0 | 0 | 1 |
| 0 | 0 | 0 | 1 |
| 0 | 0 | 0 | 1 |
| 0 | 0 | 0 | 1 |
| 0 | 0 | 0 | 0 |
| 0 | 0 | 0 | 0 |
| 0 | 0 | 0 | 0 |
| 0 | 0 | 0 | 0 |
| 0 | 0 | 0 | 0 |
| 0 | 0 | 0 | 1 |
| 0 | 0 | 0 | 1 |
| 0 | 0 | 0 | 0 |
| 0 | 0 | 0 | 1 |
| 0 | 0 | 0 | 1 |
| 0 | 0 | 0 | 0 |
| 0 | 0 | 0 | 0 |
| 0 | 0 | 0 | 0 |
| 0 | 0 | 0 | 1 |
| 0 | 0 | 0 | 0 |
| 0 | 0 | 0 | 1 |
| 0 | 0 | 0 | 1 |
| 0 | 0 | 0 | 0 |
| 0 | 0 | 0 | 1 |
| 0 | 0 | 0 | 1 |
| 0 | 0 | 0 | 1 |
| 0 | 0 | 0 | 1 |
| 0 | 0 | 0 | 1 |
| 0 | 0 | 0 | 1 |
| 0 | 0 | 0 | 1 |
| 0 | 0 | 0 | 1 |
| 0 | 0 | 0 | 1 |
| 0 | 0 | 0 | 1 |
| 0 | 0 | 0 | 0 |
| 0 | 0 | 0 | 1 |
| 1 | 0 | 0 | 0 |
| 1 | 0 | 0 | 0 |
| 0 | 0 | 0 | 1 |
| 0 | 0 | 0 | 1 |

| | | | |
|---|---|---|---|
| 1 | 0 | 0 | 0 |
| 0 | 0 | 0 | 1 |
| 0 | 0 | 0 | 1 |
| 0 | 0 | 0 | 1 |
| 0 | 0 | 0 | 1 |
| 1 | 0 | 0 | 0 |
| 1 | 0 | 0 | 0 |
| 0 | 0 | 0 | 1 |
| 0 | 0 | 0 | 1 |
| 0 | 0 | 0 | 1 |
| 0 | 0 | 0 | 1 |
| 0 | 0 | 0 | 0 |
| 0 | 0 | 0 | 1 |
| 0 | 0 | 0 | 1 |
| 0 | 0 | 0 | 1 |
| 0 | 0 | 0 | 1 |
| 0 | 0 | 0 | 0 |
| 0 | 0 | 0 | 0 |
| 0 | 0 | 0 | 1 |
| 0 | 0 | 0 | 1 |
| 0 | 0 | 0 | 1 |
| 0 | 0 | 0 | 1 |
| 0 | 0 | 0 | 1 |
| 0 | 0 | 0 | 1 |
| 0 | 0 | 0 | 0 |
| 0 | 0 | 0 | 1 |
| 0 | 0 | 0 | 1 |
| 0 | 0 | 0 | 1 |
| 0 | 0 | 0 | 1 |
| 0 | 0 | 0 | 1 |
| 0 | 0 | 0 | 1 |
| 0 | 0 | 0 | 0 |
| 0 | 0 | 0 | 0 |
| 0 | 0 | 0 | 1 |
| 0 | 0 | 0 | 1 |
| 0 | 0 | 0 | 0 |
| 1 | 0 | 0 | 0 |
| 0 | 1 | 0 | 0 |
| 1 | 1 | 0 | 0 |
| 0 | 0 | 0 | 0 |
| 0 | 0 | 0 | 0 |
| 0 | 0 | 0 | 0 |
| 0 | 0 | 0 | 0 |
| 0 | 0 | 0 | 0 |
| 0 | 0 | 0 | 0 |
| 0 | 0 | 0 | 0 |
| 0 | 0 | 0 | 0 |
| 1 | 0 | 0 | 0 |
| 0 | 0 | 0 | 1 |
| 0 | 0 | 0 | 1 |
| 0 | 1 | 1 | 1 |
| 0 | 0 | 0 | 1 |
| 0 | 0 | 0 | 0 |
| 0 | 0 | 0 | 1 |

| | | | |
|---|---|---|---|
| 0 | 0 | 0 | 1 |
| 0 | 0 | 0 | 1 |
| 0 | 0 | 0 | 1 |
| 0 | 0 | 0 | 1 |
| 0 | 0 | 0 | 1 |
| 0 | 0 | 0 | 1 |
| 0 | 0 | 0 | 1 |
| 0 | 1 | 0 | 0 |
| 0 | 0 | 0 | 0 |
| 0 | 1 | 1 | 1 |
| 0 | 0 | 0 | 1 |
| 0 | 0 | 0 | 1 |
| 0 | 0 | 0 | 1 |
| 0 | 1 | 0 | 0 |
| 0 | 0 | 0 | 1 |
| 0 | 0 | 0 | 1 |
| 0 | 1 | 0 | 0 |
| 0 | 0 | 0 | 1 |
| 1 | 0 | 0 | 0 |
| 0 | 0 | 0 | 1 |
| 0 | 0 | 0 | 1 |
| 0 | 0 | 0 | 1 |
| 0 | 0 | 0 | 1 |
| 0 | 0 | 0 | 0 |
| 0 | 0 | 0 | 1 |
| 0 | 0 | 0 | 1 |
| 0 | 0 | 0 | 1 |
| 0 | 0 | 0 | 1 |
| 0 | 0 | 0 | 1 |
| 0 | 0 | 0 | 1 |
| 0 | 0 | 0 | 1 |
| 1 | 0 | 0 | 0 |
| 0 | 0 | 0 | 1 |
| 0 | 0 | 0 | 1 |
| 0 | 0 | 0 | 1 |
| 0 | 0 | 0 | 1 |
| 0 | 0 | 0 | 1 |
| 0 | 0 | 0 | 1 |
| 1 | 0 | 0 | 0 |
| 0 | 0 | 0 | 1 |
| 0 | 0 | 0 | 1 |
| 0 | 0 | 0 | 1 |
| 0 | 0 | 0 | 1 |
| 0 | 0 | 0 | 1 |
| 0 | 0 | 0 | 1 |
| 0 | 0 | 0 | 0 |
| 0 | 0 | 0 | 1 |
| 0 | 0 | 0 | 1 |
| 0 | 1 | 1 | 1 |
| 0 | 1 | 1 | 1 |
| 0 | 0 | 0 | 1 |
| 0 | 0 | 0 | 1 |
| 0 | 0 | 0 | 1 |
| 0 | 1 | 1 | 1 |

| | | | |
|---|---|---|---|
| 0 | 1 | 1 | 1 |
| 0 | 1 | 1 | 1 |
| 0 | 1 | 1 | 1 |
| 0 | 1 | 1 | 1 |
| 0 | 1 | 1 | 1 |
| 0 | 1 | 1 | 1 |
| 0 | 1 | 1 | 1 |
| 0 | 1 | 1 | 1 |
| 0 | 1 | 1 | 1 |
| 0 | 1 | 1 | 1 |
| 0 | 1 | 1 | 1 |
| 0 | 1 | 1 | 1 |
| 0 | 1 | 1 | 1 |
| 0 | 1 | 1 | 1 |
| 0 | 1 | 1 | 1 |
| 1 | 1 | 0 | 0 |
| 0 | 1 | 1 | 1 |
| 0 | 1 | 0 | 0 |
| 0 | 1 | 1 | 1 |
| 0 | 1 | 1 | 1 |
| 1 | 1 | 0 | 0 |
| 0 | 1 | 1 | 1 |
| 0 | 1 | 1 | 1 |
| 0 | 1 | 1 | 1 |
| 0 | 1 | 0 | 0 |
| 0 | 1 | 1 | 1 |
| 0 | 1 | 0 | 0 |
| 0 | 1 | 1 | 1 |
| 0 | 1 | 1 | 1 |
| 0 | 1 | 1 | 1 |
| 0 | 1 | 1 | 1 |
| 0 | 1 | 1 | 1 |
| 0 | 1 | 1 | 1 |
| 0 | 1 | 1 | 1 |
| 0 | 1 | 1 | 1 |
| 0 | 1 | 1 | 1 |
| 0 | 1 | 1 | 1 |
| 0 | 1 | 1 | 1 |
| 0 | 1 | 1 | 1 |
| 0 | 1 | 1 | 1 |
| 0 | 1 | 1 | 1 |
| 0 | 1 | 1 | 1 |
| 0 | 0 | 0 | 1 |
| 0 | 0 | 0 | 0 |
| 0 | 0 | 0 | 1 |
| 0 | 0 | 0 | 1 |
| 0 | 0 | 0 | 0 |
| 0 | 0 | 0 | 0 |
| 0 | 0 | 0 | 1 |
| 0 | 0 | 0 | 0 |
| 0 | 0 | 0 | 0 |
| 0 | 0 | 0 | 0 |
| 0 | 0 | 0 | 1 |
| 0 | 0 | 0 | 1 |

| | | | |
|---|---|---|---|
| 0 | 0 | 0 | 1 |
| 0 | 0 | 0 | 1 |
| 0 | 0 | 0 | 1 |
| 0 | 0 | 0 | 1 |
| 0 | 0 | 0 | 1 |
| 0 | 0 | 0 | 1 |
| 0 | 0 | 0 | 1 |
| 1 | 0 | 0 | 0 |
| 0 | 0 | 0 | 1 |
| 0 | 0 | 0 | 1 |
| 0 | 0 | 0 | 1 |
| 0 | 0 | 0 | 1 |
| 0 | 0 | 0 | 1 |
| 0 | 0 | 0 | 1 |
| 0 | 0 | 0 | 0 |
| 0 | 0 | 0 | 0 |
| 0 | 0 | 0 | 1 |
| 0 | 0 | 0 | 0 |
| 0 | 0 | 0 | 1 |
| 0 | 0 | 0 | 1 |
| 0 | 0 | 0 | 0 |
| 0 | 0 | 0 | 1 |
| 0 | 0 | 0 | 1 |
| 0 | 0 | 0 | 1 |
| 0 | 0 | 0 | 0 |
| 0 | 0 | 0 | 1 |
| 0 | 0 | 0 | 1 |
| 1 | 0 | 0 | 0 |
| 0 | 0 | 0 | 0 |
| 1 | 0 | 0 | 0 |
| 1 | 0 | 0 | 0 |
| 0 | 0 | 0 | 1 |
| 0 | 0 | 0 | 1 |
| 0 | 0 | 0 | 1 |
| 0 | 0 | 0 | 1 |
| 0 | 0 | 0 | 1 |
| 0 | 1 | 0 | 0 |
| 0 | 0 | 0 | 0 |
| 0 | 0 | 0 | 1 |
| 0 | 0 | 0 | 1 |
| 0 | 0 | 0 | 1 |
| 0 | 0 | 0 | 1 |
| 0 | 0 | 0 | 0 |
| 0 | 0 | 0 | 1 |
| 0 | 0 | 0 | 1 |
| 0 | 0 | 0 | 1 |
| 0 | 0 | 0 | 1 |
| 0 | 0 | 0 | 1 |
| 0 | 0 | 0 | 0 |
| 0 | 0 | 0 | 0 |
| 0 | 0 | 0 | 0 |
| 0 | 0 | 0 | 0 |
| 0 | 0 | 0 | 0 |
| 0 | 0 | 0 | 1 |

| | | | |
|---|---|---|---|
| 0 | 0 | 0 | 0 |
| 0 | 0 | 0 | 0 |
| 0 | 0 | 0 | 1 |
| 1 | 0 | 0 | 0 |
| 0 | 0 | 0 | 1 |
| 0 | 0 | 0 | 1 |
| 0 | 0 | 0 | 1 |
| 0 | 0 | 0 | 1 |
| 0 | 0 | 0 | 1 |
| 1 | 0 | 0 | 0 |
| 0 | 0 | 0 | 1 |
| 0 | 0 | 0 | 1 |
| 0 | 0 | 0 | 1 |
| 0 | 0 | 0 | 1 |
| 0 | 0 | 0 | 1 |
| 0 | 0 | 0 | 1 |
| 0 | 0 | 0 | 1 |
| 0 | 0 | 0 | 1 |
| 0 | 0 | 0 | 1 |
| 0 | 0 | 0 | 1 |
| 0 | 0 | 0 | 1 |
| 0 | 0 | 0 | 1 |
| 0 | 0 | 0 | 1 |
| 0 | 0 | 0 | 1 |
| 0 | 0 | 0 | 1 |
| 0 | 0 | 0 | 1 |
| 0 | 0 | 0 | 1 |
| 0 | 0 | 0 | 1 |
| 0 | 0 | 0 | 1 |
| 0 | 0 | 0 | 1 |
| 0 | 0 | 0 | 1 |
| 0 | 0 | 0 | 1 |
| 1 | 0 | 0 | 0 |
| 0 | 0 | 0 | 1 |
| 0 | 0 | 0 | 1 |
| 0 | 0 | 0 | 1 |
| 0 | 0 | 0 | 1 |
| 0 | 0 | 0 | 1 |
| 0 | 0 | 0 | 1 |
| 0 | 0 | 0 | 1 |
| 0 | 0 | 0 | 1 |
| 0 | 0 | 0 | 1 |
| 0 | 0 | 0 | 1 |
| 0 | 0 | 0 | 1 |
| 0 | 0 | 0 | 1 |
| 0 | 0 | 0 | 1 |
| 0 | 0 | 0 | 1 |
| 0 | 0 | 0 | 0 |
| 0 | 1 | 0 | 0 |
| 0 | 1 | 0 | 0 |
| 0 | 0 | 0 | 0 |
| 0 | 1 | 0 | 0 |
| 0 | 0 | 0 | 1 |
| 0 | 0 | 0 | 0 |

| | | | |
|---|---|---|---|
| 0 | 0 | 0 | 1 |
| 0 | 0 | 0 | 0 |
| 0 | 0 | 0 | 1 |
| 0 | 0 | 0 | 1 |
| 0 | 0 | 0 | 1 |
| 1 | 0 | 0 | 0 |
| 0 | 0 | 0 | 0 |
| 0 | 0 | 0 | 0 |
| 1 | 0 | 0 | 0 |
| 0 | 0 | 0 | 0 |
| 0 | 0 | 0 | 1 |
| 0 | 0 | 0 | 0 |
| 0 | 0 | 0 | 1 |
| 0 | 0 | 0 | 1 |
| 0 | 0 | 0 | 1 |
| 1 | 0 | 0 | 0 |
| 0 | 0 | 0 | 1 |
| 0 | 0 | 0 | 1 |
| 0 | 0 | 0 | 0 |
| 0 | 0 | 0 | 0 |
| 1 | 0 | 0 | 0 |
| 1 | 0 | 0 | 0 |
| 0 | 0 | 0 | 0 |
| 0 | 0 | 0 | 0 |
| 0 | 0 | 0 | 1 |
| 0 | 0 | 0 | 0 |
| 0 | 1 | 0 | 0 |
| 0 | 0 | 0 | 0 |
| 0 | 1 | 0 | 0 |
| 0 | 1 | 0 | 0 |
| 0 | 0 | 0 | 0 |
| 0 | 0 | 0 | 1 |
| 0 | 0 | 0 | 1 |
| 0 | 1 | 0 | 0 |
| 0 | 1 | 0 | 0 |
| 0 | 1 | 1 | 1 |
| 0 | 0 | 0 | 0 |
| 0 | 0 | 0 | 0 |
| 0 | 1 | 0 | 0 |
| 0 | 1 | 0 | 0 |
| 0 | 1 | 0 | 0 |
| 0 | 1 | 1 | 1 |
| 0 | 0 | 0 | 1 |
| 0 | 0 | 0 | 1 |
| 0 | 0 | 0 | 0 |
| 0 | 0 | 0 | 0 |
| 0 | 0 | 0 | 0 |
| 0 | 0 | 0 | 1 |
| 0 | 0 | 0 | 1 |
| 0 | 0 | 0 | 1 |
| 0 | 0 | 0 | 1 |
| 0 | 0 | 0 | 1 |
| 0 | 0 | 0 | 1 |
| 0 | 0 | 0 | 0 |

| | | | |
|---|---|---|---|
| 0 | 0 | 0 | 1 |
| 0 | 0 | 0 | 1 |
| 0 | 0 | 0 | 1 |
| 0 | 0 | 0 | 1 |
| 1 | 0 | 0 | 0 |
| 0 | 0 | 0 | 0 |
| 0 | 0 | 0 | 1 |
| 0 | 1 | 1 | 1 |
| 0 | 0 | 0 | 1 |
| 0 | 0 | 0 | 1 |
| 0 | 0 | 0 | 1 |
| 0 | 0 | 0 | 0 |
| 0 | 0 | 0 | 1 |
| 0 | 0 | 0 | 0 |
| 0 | 1 | 0 | 0 |
| 0 | 1 | 0 | 0 |
| 0 | 1 | 0 | 0 |
| 0 | 1 | 0 | 0 |
| 0 | 1 | 0 | 0 |
| 0 | 1 | 0 | 0 |
| 0 | 1 | 0 | 0 |
| 0 | 1 | 0 | 0 |
| 0 | 0 | 0 | 0 |
| 0 | 0 | 0 | 1 |
| 0 | 0 | 0 | 1 |
| 0 | 0 | 0 | 1 |
| 0 | 0 | 0 | 1 |
| 0 | 0 | 0 | 1 |
| 0 | 0 | 0 | 1 |
| 0 | 0 | 0 | 1 |
| 0 | 0 | 0 | 1 |
| 0 | 0 | 0 | 0 |
| 0 | 0 | 0 | 1 |
| 0 | 0 | 0 | 0 |
| 0 | 0 | 0 | 0 |
| 0 | 0 | 0 | 0 |
| 0 | 0 | 0 | 0 |
| 0 | 0 | 0 | 1 |
| 0 | 0 | 0 | 1 |
| 0 | 0 | 0 | 1 |
| 0 | 0 | 0 | 1 |
| 0 | 0 | 0 | 1 |
| 0 | 0 | 0 | 1 |
| 0 | 0 | 0 | 1 |
| 0 | 0 | 0 | 1 |
| 1 | 0 | 0 | 0 |
| 0 | 0 | 0 | 1 |
| 0 | 0 | 0 | 1 |
| 0 | 0 | 0 | 1 |
| 1 | 1 | 0 | 0 |
| 1 | 0 | 0 | 0 |
| 0 | 0 | 0 | 0 |
| 0 | 0 | 0 | 1 |
| 0 | 0 | 0 | 1 |

| | | | |
|---|---|---|---|
| 0 | 0 | 0 | 1 |
| 0 | 0 | 0 | 1 |
| 0 | 0 | 0 | 1 |
| 0 | 0 | 0 | 0 |
| 0 | 0 | 0 | 1 |
| 0 | 0 | 0 | 1 |
| 0 | 0 | 0 | 0 |
| 0 | 0 | 0 | 1 |
| 0 | 0 | 0 | 1 |
| 0 | 0 | 0 | 1 |
| 0 | 0 | 0 | 0 |
| 0 | 1 | 0 | 0 |
| 0 | 0 | 0 | 1 |
| 0 | 0 | 0 | 1 |
| 0 | 0 | 0 | 1 |
| 0 | 0 | 0 | 1 |
| 0 | 0 | 0 | 0 |
| 0 | 0 | 0 | 0 |
| 0 | 0 | 0 | 1 |
| 0 | 0 | 0 | 0 |
| 0 | 1 | 0 | 0 |
| 0 | 1 | 0 | 0 |
| 0 | 0 | 0 | 1 |
| 0 | 1 | 0 | 0 |
| 0 | 0 | 0 | 0 |
| 0 | 0 | 0 | 1 |
| 0 | 0 | 0 | 0 |
| 0 | 0 | 0 | 1 |
| 0 | 0 | 0 | 1 |
| 0 | 0 | 0 | 0 |
| 0 | 0 | 0 | 1 |
| 0 | 0 | 0 | 1 |
| 0 | 0 | 0 | 0 |
| 0 | 0 | 0 | 1 |
| 0 | 0 | 0 | 1 |
| 0 | 0 | 0 | 0 |
| 0 | 0 | 0 | 1 |
| 0 | 0 | 0 | 0 |
| 0 | 0 | 0 | 1 |
| 0 | 0 | 0 | 0 |
| 0 | 0 | 0 | 0 |
| 0 | 0 | 0 | 1 |
| 0 | 0 | 0 | 0 |
| 0 | 0 | 0 | 0 |
| 0 | 0 | 0 | 1 |
| 0 | 1 | 1 | 1 |
| 0 | 0 | 0 | 1 |
| 0 | 0 | 0 | 1 |
| 0 | 0 | 0 | 1 |
| 0 | 0 | 0 | 1 |
| 0 | 0 | 0 | 1 |
| 0 | 0 | 0 | 1 |
| 0 | 0 | 0 | 1 |
| 0 | 0 | 0 | 1 |

| | | | |
|---|---|---|---|
| 0 | 0 | 0 | 1 |
| 0 | 0 | 0 | 1 |
| 0 | 0 | 0 | 1 |
| 0 | 0 | 0 | 0 |
| 0 | 1 | 1 | 1 |
| 0 | 0 | 0 | 0 |
| 0 | 0 | 0 | 1 |
| 0 | 0 | 0 | 1 |
| 0 | 0 | 0 | 0 |
| 0 | 1 | 0 | 0 |
| 0 | 0 | 0 | 1 |
| 0 | 1 | 0 | 0 |
| 0 | 1 | 0 | 0 |
| 0 | 1 | 0 | 0 |
| 0 | 0 | 0 | 1 |
| 0 | 0 | 0 | 0 |
| 0 | 0 | 0 | 0 |
| 0 | 0 | 0 | 1 |
| 1 | 0 | 0 | 0 |
| 1 | 0 | 0 | 0 |
| 1 | 0 | 0 | 0 |
| 1 | 0 | 0 | 0 |
| 0 | 0 | 0 | 1 |
| 0 | 0 | 0 | 0 |
| 0 | 0 | 0 | 0 |
| 0 | 0 | 0 | 0 |
| 0 | 0 | 0 | 0 |
| 0 | 0 | 0 | 0 |
| 0 | 0 | 0 | 0 |
| 0 | 0 | 0 | 0 |
| 0 | 0 | 0 | 0 |
| 0 | 0 | 0 | 0 |
| 0 | 0 | 0 | 0 |
| 0 | 0 | 0 | 1 |
| 0 | 0 | 0 | 1 |
| 0 | 1 | 0 | 0 |
| 0 | 0 | 0 | 1 |
| 0 | 1 | 0 | 0 |
| 0 | 1 | 0 | 0 |
| 0 | 1 | 0 | 0 |
| 0 | 1 | 0 | 0 |
| 0 | 0 | 0 | 1 |
| 0 | 0 | 0 | 0 |
| 0 | 0 | 0 | 0 |
| 0 | 0 | 0 | 1 |
| 0 | 0 | 0 | 1 |
| 0 | 0 | 0 | 1 |
| 0 | 0 | 0 | 1 |
| 0 | 0 | 0 | 1 |
| 0 | 0 | 0 | 1 |
| 0 | 0 | 0 | 1 |
| 0 | 0 | 0 | 1 |
| 0 | 0 | 0 | 1 |
| 0 | 0 | 0 | 1 |

| | | | |
|---|---|---|---|
| 0 | 0 | 0 | 1 |
| 0 | 0 | 0 | 0 |
| 0 | 0 | 0 | 1 |
| 0 | 0 | 0 | 1 |
| 0 | 0 | 0 | 1 |
| 0 | 0 | 0 | 1 |
| 0 | 0 | 0 | 1 |
| 0 | 0 | 0 | 1 |
| 0 | 0 | 0 | 1 |
| 0 | 0 | 0 | 1 |
| 0 | 0 | 0 | 1 |
| 0 | 0 | 0 | 0 |
| 0 | 0 | 0 | 1 |
| 0 | 0 | 0 | 1 |
| 0 | 1 | 0 | 0 |
| 0 | 0 | 0 | 1 |
| 0 | 0 | 0 | 1 |
| 0 | 0 | 0 | 1 |
| 0 | 0 | 0 | 1 |
| 0 | 0 | 0 | 1 |
| 1 | 0 | 0 | 0 |
| 0 | 0 | 0 | 1 |
| 1 | 0 | 0 | 0 |
| 1 | 0 | 0 | 0 |
| 0 | 0 | 0 | 1 |
| 0 | 0 | 0 | 0 |
| 0 | 0 | 0 | 1 |
| 0 | 0 | 0 | 1 |
| 0 | 0 | 0 | 1 |
| 0 | 0 | 0 | 1 |
| 0 | 0 | 0 | 1 |
| 0 | 1 | 0 | 0 |
| 0 | 0 | 0 | 1 |
| 0 | 0 | 0 | 0 |
| 0 | 0 | 0 | 1 |
| 0 | 0 | 0 | 1 |
| 0 | 0 | 0 | 1 |
| 0 | 0 | 0 | 1 |
| 0 | 0 | 0 | 1 |
| 0 | 0 | 0 | 0 |
| 0 | 0 | 0 | 1 |
| 0 | 0 | 0 | 1 |
| 0 | 0 | 0 | 1 |
| 0 | 0 | 0 | 0 |
| 0 | 0 | 0 | 1 |
| 0 | 0 | 0 | 1 |
| 0 | 0 | 0 | 1 |
| 1 | 0 | 0 | 0 |
| 0 | 0 | 0 | 0 |
| 0 | 0 | 0 | 1 |
| 0 | 0 | 0 | 1 |
| 0 | 0 | 0 | 0 |
| 1 | 0 | 0 | 0 |
| 0 | 0 | 0 | 0 |

| | | | |
|---|---|---|---|
| 1 | 0 | 0 | 0 |
| 0 | 0 | 0 | 1 |
| 0 | 0 | 0 | 0 |
| 0 | 0 | 0 | 0 |
| 0 | 0 | 0 | 0 |
| 0 | 0 | 0 | 1 |
| 0 | 0 | 0 | 1 |
| 0 | 0 | 0 | 1 |
| 0 | 0 | 0 | 1 |
| 0 | 1 | 0 | 0 |
| 0 | 0 | 0 | 1 |
| 0 | 0 | 0 | 1 |
| 0 | 0 | 0 | 1 |
| 0 | 0 | 0 | 1 |
| 0 | 0 | 0 | 0 |
| 0 | 0 | 0 | 1 |
| 0 | 0 | 0 | 1 |
| 0 | 0 | 0 | 0 |
| 0 | 0 | 0 | 1 |
| 0 | 0 | 0 | 0 |
| 0 | 0 | 0 | 0 |
| 0 | 0 | 0 | 1 |
| 0 | 0 | 0 | 0 |
| 0 | 0 | 0 | 1 |
| 0 | 0 | 0 | 0 |
| 0 | 0 | 0 | 0 |
| 0 | 0 | 0 | 1 |
| 0 | 0 | 0 | 1 |
| 1 | 0 | 0 | 0 |
| 0 | 0 | 0 | 1 |
| 0 | 0 | 0 | 1 |
| 0 | 0 | 0 | 1 |
| 0 | 0 | 0 | 1 |
| 0 | 0 | 0 | 1 |
| 0 | 0 | 0 | 0 |
| 0 | 0 | 0 | 0 |
| 0 | 0 | 0 | 1 |
| 0 | 0 | 0 | 0 |
| 0 | 0 | 0 | 0 |
| 0 | 0 | 0 | 0 |
| 0 | 0 | 0 | 0 |
| 0 | 0 | 0 | 0 |
| 0 | 0 | 0 | 0 |
| 0 | 0 | 0 | 1 |
| 0 | 1 | 0 | 0 |
| 0 | 0 | 0 | 1 |
| 0 | 0 | 0 | 1 |
| 0 | 0 | 0 | 0 |
| 0 | 0 | 0 | 0 |
| 0 | 0 | 0 | 0 |
| 0 | 0 | 0 | 0 |
| 0 | 1 | 0 | 0 |
| 0 | 0 | 0 | 1 |
| 0 | 0 | 0 | 1 |

| | | | |
|---|---|---|---|
| 0 | 0 | 0 | 1 |
| 0 | 0 | 0 | 0 |
| 0 | 0 | 0 | 0 |
| 0 | 0 | 0 | 0 |
| 0 | 0 | 0 | 0 |
| 0 | 0 | 0 | 0 |
| 0 | 1 | 0 | 0 |
| 0 | 1 | 1 | 1 |
| 0 | 1 | 1 | 1 |
| 0 | 0 | 0 | 1 |
| 0 | 0 | 0 | 0 |
| 0 | 0 | 0 | 0 |
| 0 | 0 | 0 | 1 |
| 0 | 0 | 0 | 0 |
| 0 | 0 | 0 | 1 |
| 0 | 0 | 0 | 0 |
| 0 | 0 | 0 | 1 |
| 0 | 0 | 0 | 1 |
| 0 | 0 | 0 | 1 |
| 0 | 0 | 0 | 1 |
| 0 | 0 | 0 | 1 |
| 0 | 0 | 0 | 1 |
| 0 | 0 | 0 | 1 |
| 1 | 0 | 0 | 0 |
| 0 | 0 | 0 | 0 |
| 0 | 0 | 0 | 1 |
| 0 | 0 | 0 | 1 |
| 0 | 0 | 0 | 1 |
| 0 | 0 | 0 | 1 |
| 0 | 0 | 0 | 0 |
| 0 | 0 | 0 | 0 |
| 0 | 0 | 0 | 0 |
| 0 | 0 | 0 | 0 |
| 0 | 0 | 0 | 1 |
| 0 | 0 | 0 | 0 |
| 0 | 0 | 0 | 0 |
| 0 | 0 | 0 | 0 |
| 0 | 0 | 0 | 0 |
| 0 | 0 | 0 | 0 |
| 0 | 0 | 0 | 1 |
| 0 | 0 | 0 | 1 |
| 0 | 0 | 0 | 0 |
| 0 | 0 | 0 | 1 |
| 0 | 0 | 0 | 1 |
| 0 | 0 | 0 | 0 |
| 0 | 0 | 0 | 1 |
| 0 | 0 | 0 | 1 |
| 0 | 0 | 0 | 1 |
| 0 | 0 | 0 | 1 |
| 0 | 0 | 0 | 1 |
| 0 | 0 | 0 | 0 |
| 0 | 0 | 0 | 1 |
| 0 | 0 | 0 | 0 |
| 0 | 0 | 0 | 0 |

| | | | |
|---|---|---|---|
| 0 | 0 | 0 | 1 |
| 0 | 0 | 0 | 1 |
| 0 | 0 | 0 | 0 |
| 0 | 0 | 0 | 0 |
| 0 | 0 | 0 | 1 |
| 0 | 0 | 0 | 1 |
| 0 | 0 | 0 | 1 |
| 0 | 0 | 0 | 1 |
| 0 | 0 | 0 | 1 |
| 0 | 0 | 0 | 1 |
| 0 | 0 | 0 | 1 |
| 0 | 0 | 0 | 1 |
| 0 | 0 | 0 | 0 |
| 0 | 0 | 0 | 1 |
| 0 | 0 | 0 | 1 |
| 0 | 0 | 0 | 1 |
| 0 | 0 | 0 | 1 |
| 0 | 0 | 0 | 1 |
| 0 | 0 | 0 | 1 |
| 0 | 0 | 0 | 1 |
| 0 | 0 | 0 | 1 |
| 0 | 0 | 0 | 1 |
| 0 | 0 | 0 | 1 |
| 0 | 0 | 0 | 1 |
| 0 | 0 | 0 | 1 |
| 0 | 0 | 0 | 1 |
| 0 | 0 | 0 | 1 |
| 0 | 0 | 0 | 1 |
| 0 | 0 | 0 | 1 |
| 0 | 0 | 0 | 1 |
| 0 | 0 | 0 | 1 |
| 0 | 0 | 0 | 0 |
| 0 | 0 | 0 | 1 |
| 0 | 0 | 0 | 1 |
| 0 | 0 | 0 | 1 |
| 0 | 0 | 0 | 1 |
| 0 | 0 | 0 | 1 |
| 0 | 0 | 0 | 1 |
| 0 | 0 | 0 | 1 |
| 0 | 0 | 0 | 1 |
| 0 | 0 | 0 | 0 |
| 0 | 0 | 0 | 0 |
| 0 | 0 | 0 | 1 |
| 0 | 0 | 0 | 1 |
| 0 | 0 | 0 | 1 |
| 0 | 1 | 1 | 1 |
| 0 | 0 | 0 | 1 |
| 0 | 0 | 0 | 1 |
| 0 | 0 | 0 | 0 |
| 0 | 0 | 0 | 0 |
| 0 | 0 | 0 | 1 |
| 0 | 0 | 0 | 1 |
| 0 | 0 | 0 | 0 |
| 0 | 0 | 0 | 1 |

| | | | |
|---|---|---|---|
| 0 | 0 | 0 | 0 |
| 0 | 0 | 0 | 0 |
| 0 | 0 | 0 | 0 |
| 0 | 0 | 0 | 1 |
| 0 | 0 | 0 | 1 |
| 0 | 0 | 0 | 1 |
| 0 | 0 | 0 | 1 |
| 0 | 1 | 0 | 0 |
| 0 | 1 | 0 | 0 |
| 0 | 0 | 0 | 0 |
| 0 | 0 | 0 | 0 |
| 0 | 0 | 0 | 1 |
| 0 | 0 | 0 | 0 |
| 0 | 0 | 0 | 1 |
| 0 | 0 | 0 | 1 |
| 0 | 0 | 0 | 1 |
| 0 | 0 | 0 | 1 |
| 0 | 0 | 0 | 0 |
| 0 | 0 | 0 | 1 |
| 0 | 0 | 0 | 1 |
| 0 | 0 | 0 | 1 |
| 0 | 0 | 0 | 1 |
| 0 | 0 | 0 | 1 |
| 0 | 0 | 0 | 1 |
| 0 | 0 | 0 | 1 |
| 0 | 0 | 0 | 1 |
| 0 | 0 | 0 | 1 |
| 0 | 0 | 0 | 1 |
| 0 | 0 | 0 | 1 |
| 0 | 0 | 0 | 1 |
| 0 | 0 | 0 | 0 |
| 0 | 1 | 0 | 0 |
| 0 | 0 | 0 | 1 |
| 0 | 0 | 0 | 1 |
| 0 | 0 | 0 | 1 |
| 0 | 0 | 0 | 1 |
| 0 | 0 | 0 | 1 |
| 0 | 0 | 0 | 1 |
| 0 | 0 | 0 | 1 |
| 0 | 0 | 0 | 1 |
| 0 | 0 | 0 | 1 |
| 0 | 0 | 0 | 0 |
| 0 | 0 | 0 | 1 |
| 0 | 0 | 0 | 1 |
| 0 | 0 | 0 | 0 |
| 0 | 0 | 0 | 0 |
| 0 | 0 | 0 | 1 |
| 0 | 0 | 0 | 1 |
| 0 | 0 | 0 | 0 |
| 0 | 0 | 0 | 0 |
| 0 | 0 | 0 | 0 |
| 0 | 0 | 0 | 0 |
| 0 | 0 | 0 | 0 |
| 0 | 0 | 0 | 0 |

| | | | |
|---|---|---|---|
| 0 | 0 | 0 | 1 |
| 0 | 0 | 0 | 1 |
| 0 | 0 | 0 | 1 |
| 0 | 0 | 0 | 1 |
| 0 | 0 | 0 | 1 |
| 0 | 0 | 0 | 0 |
| 0 | 0 | 0 | 0 |
| 0 | 0 | 0 | 0 |
| 0 | 0 | 0 | 0 |
| 0 | 0 | 0 | 0 |
| 0 | 0 | 0 | 1 |
| 0 | 0 | 0 | 0 |
| 0 | 0 | 0 | 0 |
| 0 | 0 | 0 | 0 |
| 0 | 0 | 0 | 1 |
| 0 | 0 | 0 | 0 |
| 0 | 0 | 0 | 1 |
| 0 | 0 | 0 | 1 |
| 1 | 0 | 0 | 0 |
| 1 | 0 | 0 | 0 |
| 0 | 0 | 0 | 0 |
| 0 | 0 | 0 | 0 |
| 0 | 0 | 0 | 1 |
| 0 | 0 | 0 | 0 |
| 0 | 0 | 0 | 0 |
| 0 | 0 | 0 | 0 |
| 0 | 0 | 0 | 0 |
| 0 | 0 | 0 | 1 |
| 0 | 0 | 0 | 1 |
| 0 | 0 | 0 | 1 |
| 0 | 0 | 0 | 1 |
| 0 | 0 | 0 | 1 |
| 0 | 0 | 0 | 0 |
| 0 | 0 | 0 | 1 |
| 0 | 0 | 0 | 0 |
| 0 | 0 | 0 | 0 |
| 0 | 1 | 0 | 0 |
| 0 | 0 | 0 | 0 |
| 0 | 1 | 0 | 0 |
| 0 | 1 | 0 | 0 |
| 0 | 1 | 0 | 0 |
| 1 | 0 | 0 | 0 |
| 0 | 0 | 0 | 0 |
| 0 | 0 | 0 | 1 |
| 0 | 0 | 0 | 1 |
| 0 | 0 | 0 | 1 |
| 0 | 0 | 0 | 1 |
| 0 | 0 | 0 | 0 |
| 0 | 0 | 0 | 0 |
| 0 | 0 | 0 | 0 |
| 0 | 0 | 0 | 1 |
| 0 | 0 | 0 | 1 |
| 0 | 0 | 0 | 0 |
| 0 | 0 | 0 | 1 |

| | | | |
|---|---|---|---|
| 0 | 0 | 0 | 1 |
| 0 | 0 | 0 | 0 |
| 0 | 0 | 0 | 1 |
| 0 | 0 | 0 | 1 |
| 0 | 0 | 0 | 1 |
| 0 | 0 | 0 | 1 |
| 0 | 0 | 0 | 1 |
| 0 | 0 | 0 | 1 |
| 0 | 0 | 0 | 1 |
| 0 | 0 | 0 | 1 |
| 0 | 0 | 0 | 1 |
| 0 | 0 | 0 | 1 |
| 0 | 0 | 0 | 0 |
| 0 | 0 | 0 | 0 |
| 0 | 0 | 0 | 0 |
| 0 | 0 | 0 | 0 |
| 0 | 0 | 0 | 0 |
| 0 | 0 | 0 | 0 |
| 0 | 0 | 0 | 0 |
| 0 | 0 | 0 | 1 |
| 1 | 1 | 0 | 0 |
| 0 | 0 | 0 | 0 |
| 0 | 0 | 0 | 0 |
| 0 | 0 | 0 | 0 |
| 0 | 0 | 0 | 0 |
| 0 | 0 | 0 | 1 |
| 0 | 0 | 0 | 1 |
| 0 | 0 | 0 | 0 |
| 0 | 0 | 0 | 1 |
| 0 | 0 | 0 | 0 |
| 0 | 0 | 0 | 0 |
| 0 | 0 | 0 | 1 |
| 0 | 0 | 0 | 0 |
| 0 | 0 | 0 | 1 |
| 0 | 0 | 0 | 1 |
| 0 | 0 | 0 | 1 |
| 0 | 1 | 1 | 1 |
| 0 | 0 | 0 | 1 |
| 0 | 0 | 0 | 1 |
| 0 | 0 | 0 | 1 |
| 0 | 0 | 0 | 0 |
| 0 | 0 | 0 | 1 |
| 0 | 0 | 0 | 1 |
| 0 | 0 | 0 | 0 |
| 0 | 0 | 0 | 0 |
| 0 | 0 | 0 | 1 |
| 0 | 0 | 0 | 0 |
| 0 | 0 | 0 | 0 |
| 0 | 0 | 0 | 1 |
| 0 | 0 | 0 | 1 |
| 0 | 0 | 0 | 0 |
| 0 | 0 | 0 | 1 |
| 0 | 0 | 0 | 1 |
| 0 | 0 | 0 | 1 |

| | | | |
|---|---|---|---|
| 0 | 0 | 0 | 1 |
| 0 | 0 | 0 | 1 |
| 0 | 0 | 0 | 1 |
| 0 | 0 | 0 | 1 |
| 0 | 0 | 0 | 0 |
| 0 | 0 | 0 | 0 |
| 0 | 0 | 0 | 1 |
| 0 | 0 | 0 | 1 |
| 0 | 0 | 0 | 1 |
| 0 | 0 | 0 | 1 |
| 0 | 0 | 0 | 1 |
| 0 | 0 | 0 | 0 |
| 0 | 0 | 0 | 0 |
| 0 | 0 | 0 | 1 |
| 0 | 0 | 0 | 1 |
| 0 | 0 | 0 | 0 |
| 0 | 0 | 0 | 1 |
| 0 | 0 | 0 | 1 |
| 0 | 0 | 0 | 0 |
| 0 | 0 | 0 | 1 |
| 0 | 0 | 0 | 1 |
| 0 | 0 | 0 | 1 |
| 0 | 0 | 0 | 1 |
| 0 | 0 | 0 | 0 |
| 0 | 0 | 0 | 1 |
| 0 | 0 | 0 | 0 |
| 0 | 0 | 0 | 1 |
| 0 | 0 | 0 | 1 |
| 1 | 0 | 0 | 0 |
| 1 | 0 | 0 | 0 |
| 0 | 0 | 0 | 0 |
| 0 | 0 | 0 | 0 |
| 0 | 0 | 0 | 0 |
| 0 | 0 | 0 | 0 |
| 0 | 0 | 0 | 1 |
| 0 | 0 | 0 | 1 |
| 0 | 0 | 0 | 1 |
| 1 | 0 | 0 | 0 |
| 0 | 0 | 0 | 1 |
| 0 | 0 | 0 | 1 |
| 0 | 0 | 0 | 0 |
| 0 | 0 | 0 | 0 |
| 0 | 0 | 0 | 0 |
| 0 | 0 | 0 | 1 |
| 0 | 0 | 0 | 0 |
| 0 | 0 | 0 | 1 |
| 0 | 0 | 0 | 1 |
| 0 | 0 | 0 | 0 |
| 0 | 0 | 0 | 1 |
| 0 | 0 | 0 | 1 |
| 0 | 0 | 0 | 1 |
| 0 | 1 | 0 | 0 |
| 0 | 0 | 0 | 1 |
| 0 | 0 | 0 | 0 |

| | | | |
|---|---|---|---|
| 0 | 0 | 0 | 1 |
| 0 | 0 | 0 | 0 |
| 0 | 0 | 0 | 0 |
| 0 | 0 | 0 | 1 |
| 0 | 0 | 0 | 1 |
| 0 | 0 | 0 | 1 |
| 0 | 0 | 0 | 0 |
| 0 | 0 | 0 | 1 |
| 0 | 0 | 0 | 1 |
| 0 | 0 | 0 | 1 |
| 0 | 0 | 0 | 1 |
| 0 | 0 | 0 | 1 |
| 0 | 0 | 0 | 1 |
| 0 | 0 | 0 | 1 |
| 0 | 0 | 0 | 1 |
| 0 | 0 | 0 | 1 |
| 0 | 0 | 0 | 1 |
| 0 | 0 | 0 | 1 |
| 0 | 0 | 0 | 1 |
| 0 | 0 | 0 | 1 |
| 0 | 0 | 0 | 1 |
| 0 | 0 | 0 | 0 |
| 0 | 0 | 0 | 1 |
| 0 | 0 | 0 | 1 |
| 0 | 0 | 0 | 1 |
| 0 | 0 | 0 | 1 |
| 0 | 0 | 0 | 0 |
| 0 | 0 | 0 | 1 |
| 0 | 0 | 0 | 0 |
| 0 | 0 | 0 | 1 |
| 0 | 0 | 0 | 1 |
| 0 | 0 | 0 | 0 |
| 0 | 1 | 1 | 1 |
| 0 | 1 | 1 | 1 |
| 0 | 0 | 0 | 1 |
| 0 | 0 | 0 | 1 |
| 0 | 0 | 0 | 1 |
| 0 | 0 | 0 | 1 |
| 0 | 0 | 0 | 0 |
| 0 | 0 | 0 | 0 |
| 0 | 1 | 0 | 0 |
| 0 | 0 | 0 | 1 |
| 0 | 0 | 0 | 1 |
| 0 | 0 | 0 | 1 |
| 0 | 0 | 0 | 1 |
| 0 | 1 | 0 | 0 |
| 0 | 0 | 0 | 1 |
| 0 | 0 | 0 | 0 |
| 0 | 0 | 0 | 1 |
| 0 | 0 | 0 | 0 |
| 0 | 0 | 0 | 0 |
| 0 | 0 | 0 | 0 |
| 0 | 0 | 0 | 1 |
| 0 | 0 | 0 | 1 |

| | | | |
|---|---|---|---|
| 0 | 0 | 0 | 0 |
| 0 | 0 | 0 | 0 |
| 0 | 0 | 0 | 1 |
| 0 | 0 | 0 | 1 |
| 0 | 0 | 0 | 1 |
| 0 | 0 | 0 | 1 |
| 0 | 0 | 0 | 1 |
| 0 | 0 | 0 | 1 |
| 0 | 0 | 0 | 1 |
| 0 | 0 | 0 | 1 |
| 0 | 0 | 0 | 1 |
| 0 | 0 | 0 | 1 |
| 0 | 0 | 0 | 1 |
| 0 | 0 | 0 | 1 |
| 0 | 0 | 0 | 1 |
| 0 | 0 | 0 | 0 |
| 0 | 0 | 0 | 0 |
| 0 | 0 | 0 | 0 |
| 0 | 0 | 0 | 1 |
| 0 | 0 | 0 | 1 |
| 0 | 0 | 0 | 1 |
| 0 | 1 | 0 | 0 |
| 0 | 0 | 0 | 0 |
| 0 | 0 | 0 | 0 |
| 0 | 0 | 0 | 0 |
| 0 | 0 | 0 | 1 |
| 0 | 0 | 0 | 1 |
| 0 | 0 | 0 | 1 |
| 0 | 0 | 0 | 1 |
| 0 | 0 | 0 | 1 |
| 0 | 0 | 0 | 0 |
| 0 | 0 | 0 | 1 |
| 0 | 0 | 0 | 0 |
| 0 | 0 | 0 | 1 |
| 0 | 1 | 1 | 1 |
| 0 | 1 | 1 | 1 |
| 0 | 1 | 0 | 0 |
| 0 | 0 | 0 | 0 |
| 0 | 0 | 0 | 1 |
| 0 | 0 | 0 | 1 |
| 0 | 0 | 0 | 1 |
| 0 | 0 | 0 | 0 |
| 0 | 0 | 0 | 1 |
| 0 | 0 | 0 | 1 |
| 0 | 0 | 0 | 0 |
| 0 | 0 | 0 | 1 |
| 0 | 0 | 0 | 0 |
| 0 | 0 | 0 | 0 |
| 0 | 0 | 0 | 1 |
| 0 | 0 | 0 | 1 |
| 0 | 1 | 0 | 0 |
| 0 | 1 | 0 | 0 |
| 0 | 0 | 0 | 1 |
| 0 | 0 | 0 | 0 |

| | | | |
|---|---|---|---|
| 0 | 0 | 0 | 1 |
| 0 | 0 | 0 | 0 |
| 0 | 1 | 1 | 1 |
| 0 | 1 | 0 | 0 |
| 0 | 0 | 0 | 1 |
| 0 | 0 | 0 | 1 |
| 0 | 0 | 0 | 1 |
| 0 | 0 | 0 | 1 |
| 0 | 0 | 0 | 0 |
| 0 | 0 | 0 | 1 |
| 0 | 0 | 0 | 1 |
| 0 | 0 | 0 | 1 |
| 0 | 0 | 0 | 0 |
| 0 | 0 | 0 | 0 |
| 0 | 0 | 0 | 0 |
| 0 | 0 | 0 | 1 |
| 0 | 0 | 0 | 0 |
| 0 | 0 | 0 | 1 |
| 0 | 1 | 0 | 0 |
| 0 | 0 | 0 | 0 |
| 0 | 0 | 0 | 0 |
| 0 | 1 | 0 | 0 |
| 0 | 0 | 0 | 0 |
| 1 | 0 | 0 | 0 |
| 0 | 0 | 0 | 1 |
| 0 | 0 | 0 | 1 |
| 0 | 0 | 0 | 0 |
| 0 | 0 | 0 | 0 |
| 0 | 0 | 0 | 0 |
| 0 | 0 | 0 | 0 |
| 0 | 0 | 0 | 1 |
| 0 | 0 | 0 | 1 |
| 0 | 0 | 0 | 1 |
| 0 | 0 | 0 | 1 |
| 0 | 0 | 0 | 1 |
| 1 | 0 | 0 | 0 |
| 1 | 0 | 0 | 0 |
| 0 | 0 | 0 | 1 |
| 0 | 0 | 0 | 1 |
| 0 | 0 | 0 | 1 |
| 0 | 0 | 0 | 1 |
| 0 | 0 | 0 | 1 |
| 0 | 0 | 0 | 1 |
| 0 | 0 | 0 | 0 |
| 0 | 0 | 0 | 0 |
| 0 | 0 | 0 | 0 |
| 0 | 0 | 0 | 1 |
| 0 | 0 | 0 | 0 |
| 0 | 0 | 0 | 0 |
| 0 | 0 | 0 | 0 |
| 0 | 0 | 0 | 1 |
| 0 | 0 | 0 | 1 |
| 0 | 0 | 0 | 1 |
| 0 | 0 | 0 | 1 |

| | | | |
|---|---|---|---|
| 0 | 0 | 0 | 1 |
| 0 | 0 | 0 | 1 |
| 0 | 0 | 0 | 1 |
| 0 | 0 | 0 | 1 |
| 0 | 0 | 0 | 0 |
| 0 | 0 | 0 | 1 |
| 0 | 0 | 0 | 0 |
| 0 | 0 | 0 | 0 |
| 0 | 0 | 0 | 0 |
| 0 | 0 | 0 | 0 |
| 0 | 0 | 0 | 0 |
| 0 | 0 | 0 | 0 |
| 0 | 0 | 0 | 1 |
| 0 | 0 | 0 | 0 |
| 0 | 0 | 0 | 0 |
| 0 | 0 | 0 | 1 |
| 0 | 0 | 0 | 1 |
| 0 | 0 | 0 | 1 |
| 0 | 0 | 0 | 0 |
| 0 | 0 | 0 | 0 |
| 0 | 0 | 0 | 0 |
| 0 | 0 | 0 | 0 |
| 0 | 0 | 0 | 1 |
| 0 | 0 | 0 | 1 |
| 0 | 0 | 0 | 1 |
| 0 | 0 | 0 | 1 |
| 0 | 0 | 0 | 1 |
| 0 | 0 | 0 | 0 |
| 0 | 0 | 0 | 0 |
| 0 | 0 | 0 | 0 |
| 0 | 0 | 0 | 0 |
| 0 | 0 | 0 | 0 |
| 0 | 0 | 0 | 0 |
| 0 | 0 | 0 | 1 |
| 0 | 0 | 0 | 1 |
| 0 | 0 | 0 | 0 |
| 0 | 0 | 0 | 1 |
| 0 | 0 | 0 | 1 |
| 0 | 0 | 0 | 1 |
| 0 | 0 | 0 | 1 |
| 0 | 0 | 0 | 0 |
| 0 | 0 | 0 | 1 |
| 0 | 0 | 0 | 0 |
| 0 | 0 | 0 | 1 |
| 0 | 0 | 0 | 1 |
| 0 | 0 | 0 | 1 |
| 0 | 0 | 0 | 1 |
| 0 | 0 | 0 | 0 |
| 0 | 0 | 0 | 1 |
| 0 | 0 | 0 | 1 |
| 0 | 0 | 0 | 1 |
| 0 | 0 | 0 | 1 |
| 0 | 0 | 0 | 1 |
| 0 | 0 | 0 | 1 |

| | | | |
|---|---|---|---|
| 0 | 0 | 0 | 1 |
| 0 | 0 | 0 | 1 |
| 0 | 0 | 0 | 1 |
| 0 | 0 | 0 | 1 |
| 0 | 0 | 0 | 1 |
| 0 | 0 | 0 | 1 |
| 0 | 0 | 0 | 1 |
| 0 | 0 | 0 | 1 |
| 0 | 0 | 0 | 1 |
| 0 | 0 | 0 | 1 |
| 0 | 0 | 0 | 1 |
| 0 | 0 | 0 | 0 |
| 0 | 0 | 0 | 1 |
| 0 | 0 | 0 | 1 |
| 0 | 0 | 0 | 1 |
| 0 | 0 | 0 | 1 |
| 0 | 0 | 0 | 1 |
| 0 | 0 | 0 | 0 |
| 0 | 0 | 0 | 0 |
| 0 | 0 | 0 | 0 |
| 0 | 0 | 0 | 0 |
| 0 | 0 | 0 | 1 |
| 1 | 0 | 0 | 0 |
| 0 | 0 | 0 | 1 |
| 0 | 0 | 0 | 1 |
| 0 | 0 | 0 | 1 |
| 0 | 0 | 0 | 1 |
| 0 | 0 | 0 | 1 |
| 0 | 0 | 0 | 0 |
| 0 | 0 | 0 | 0 |
| 0 | 0 | 0 | 1 |
| 0 | 0 | 0 | 1 |
| 1 | 0 | 0 | 0 |
| 0 | 0 | 0 | 1 |
| 0 | 0 | 0 | 0 |
| 1 | 0 | 0 | 0 |
| 1 | 0 | 0 | 0 |
| 0 | 0 | 0 | 1 |
| 1 | 0 | 0 | 0 |
| 0 | 0 | 0 | 1 |
| 1 | 0 | 0 | 0 |
| 0 | 0 | 0 | 1 |
| 0 | 0 | 0 | 0 |
| 0 | 0 | 0 | 1 |
| 0 | 0 | 0 | 1 |
| 0 | 0 | 0 | 1 |
| 0 | 1 | 1 | 1 |
| 0 | 0 | 0 | 1 |
| 0 | 0 | 0 | 1 |
| 0 | 0 | 0 | 0 |
| 0 | 1 | 1 | 1 |
| 1 | 0 | 0 | 0 |
| 0 | 1 | 1 | 1 |
| 0 | 0 | 0 | 1 |

| | | | |
|---|---|---|---|
| 0 | 0 | 0 | 0 |
| 0 | 0 | 0 | 0 |
| 0 | 0 | 0 | 0 |
| 0 | 0 | 0 | 0 |
| 0 | 0 | 0 | 0 |
| 0 | 0 | 0 | 1 |
| 0 | 0 | 0 | 0 |
| 0 | 0 | 0 | 1 |
| 0 | 0 | 0 | 1 |
| 0 | 0 | 0 | 0 |
| 0 | 0 | 0 | 0 |
| 0 | 0 | 0 | 0 |
| 0 | 0 | 0 | 1 |
| 0 | 0 | 0 | 1 |
| 0 | 0 | 0 | 1 |
| 0 | 0 | 0 | 1 |
| 0 | 0 | 0 | 1 |
| 0 | 0 | 0 | 1 |
| 0 | 0 | 0 | 1 |
| 0 | 0 | 0 | 1 |
| 0 | 0 | 0 | 1 |
| 0 | 0 | 0 | 1 |
| 0 | 0 | 0 | 1 |
| 0 | 0 | 0 | 1 |
| 0 | 0 | 0 | 1 |
| 0 | 0 | 0 | 1 |
| 0 | 0 | 0 | 1 |
| 0 | 0 | 0 | 1 |
| 0 | 0 | 0 | 1 |
| 0 | 0 | 0 | 1 |
| 0 | 0 | 0 | 1 |
| 0 | 0 | 0 | 1 |
| 0 | 0 | 0 | 1 |
| 0 | 0 | 0 | 1 |
| 0 | 0 | 0 | 1 |
| 0 | 0 | 0 | 1 |
| 0 | 0 | 0 | 1 |
| 0 | 0 | 0 | 1 |
| 0 | 0 | 0 | 1 |
| 0 | 0 | 0 | 1 |
| 0 | 0 | 0 | 1 |
| 0 | 0 | 0 | 1 |
| 0 | 0 | 0 | 1 |
| 0 | 0 | 0 | 1 |
| 0 | 0 | 0 | 1 |
| 0 | 0 | 0 | 1 |
| 0 | 0 | 0 | 0 |
| 0 | 0 | 0 | 1 |
| 0 | 0 | 0 | 1 |
| 0 | 0 | 0 | 1 |
| 0 | 0 | 0 | 1 |
| 0 | 0 | 0 | 0 |
| 0 | 0 | 0 | 1 |
| 0 | 0 | 0 | 1 |
| 0 | 0 | 0 | 1 |

| | | | |
|---|---|---|---|
| 0 | 0 | 0 | 1 |
| 0 | 0 | 0 | 1 |
| 0 | 0 | 0 | 1 |
| 0 | 0 | 0 | 1 |
| 0 | 0 | 0 | 1 |
| 0 | 0 | 0 | 1 |
| 0 | 0 | 0 | 1 |
| 0 | 0 | 0 | 0 |
| 0 | 1 | 0 | 0 |
| 0 | 0 | 0 | 1 |
| 0 | 0 | 0 | 1 |
| 0 | 0 | 0 | 1 |
| 0 | 0 | 0 | 1 |
| 0 | 0 | 0 | 1 |
| 0 | 1 | 0 | 0 |
| 0 | 1 | 0 | 0 |
| 0 | 0 | 0 | 0 |
| 0 | 0 | 0 | 1 |
| 0 | 0 | 0 | 1 |
| 0 | 0 | 0 | 1 |
| 0 | 1 | 0 | 0 |
| 0 | 0 | 0 | 0 |
| 0 | 1 | 0 | 0 |
| 0 | 0 | 0 | 0 |
| 0 | 0 | 0 | 0 |
| 0 | 0 | 0 | 0 |
| 0 | 0 | 0 | 1 |
| 0 | 0 | 0 | 1 |
| 0 | 0 | 0 | 0 |
| 0 | 0 | 0 | 1 |
| 0 | 0 | 0 | 1 |
| 0 | 0 | 0 | 1 |
| 0 | 0 | 0 | 0 |
| 0 | 0 | 0 | 1 |
| 0 | 0 | 0 | 1 |
| 0 | 0 | 0 | 1 |
| 0 | 0 | 0 | 0 |
| 0 | 0 | 0 | 0 |
| 0 | 0 | 0 | 0 |
| 0 | 0 | 0 | 1 |
| 0 | 0 | 0 | 1 |
| 0 | 0 | 0 | 1 |
| 0 | 0 | 0 | 1 |
| 0 | 0 | 0 | 1 |
| 0 | 0 | 0 | 1 |
| 0 | 0 | 0 | 0 |
| 0 | 0 | 0 | 1 |
| 0 | 0 | 0 | 1 |
| 0 | 0 | 0 | 1 |
| 0 | 0 | 0 | 1 |
| 0 | 0 | 0 | 1 |
| 0 | 0 | 0 | 1 |
| 0 | 0 | 0 | 1 |
| 0 | 0 | 0 | 1 |

| | | | |
|---|---|---|---|
| 0 | 0 | 0 | 1 |
| 0 | 0 | 0 | 1 |
| 0 | 0 | 0 | 1 |
| 0 | 0 | 0 | 0 |
| 0 | 0 | 0 | 1 |
| 0 | 0 | 0 | 1 |
| 0 | 0 | 0 | 1 |
| 0 | 0 | 0 | 0 |
| 0 | 0 | 0 | 0 |
| 0 | 0 | 0 | 1 |
| 0 | 0 | 0 | 1 |
| 0 | 0 | 0 | 1 |
| 0 | 0 | 0 | 0 |
| 0 | 0 | 0 | 1 |
| 0 | 0 | 0 | 1 |
| 0 | 0 | 0 | 1 |
| 0 | 0 | 0 | 0 |
| 0 | 0 | 0 | 1 |
| 0 | 0 | 0 | 1 |
| 0 | 0 | 0 | 1 |
| 0 | 0 | 0 | 1 |
| 0 | 0 | 0 | 1 |
| 0 | 0 | 0 | 1 |
| 0 | 0 | 0 | 1 |
| 0 | 0 | 0 | 1 |
| 0 | 0 | 0 | 1 |
| 0 | 0 | 0 | 1 |
| 0 | 0 | 0 | 0 |
| 0 | 0 | 0 | 0 |
| 0 | 0 | 0 | 0 |
| 0 | 0 | 0 | 0 |
| 0 | 0 | 0 | 0 |
| 0 | 0 | 0 | 0 |
| 0 | 0 | 0 | 1 |
| 0 | 0 | 0 | 0 |
| 0 | 0 | 0 | 0 |
| 0 | 0 | 0 | 0 |
| 0 | 0 | 0 | 1 |
| 0 | 0 | 0 | 1 |
| 0 | 0 | 0 | 1 |
| 0 | 0 | 0 | 1 |
| 0 | 0 | 0 | 1 |
| 0 | 0 | 0 | 0 |
| 0 | 0 | 0 | 0 |
| 0 | 0 | 0 | 1 |
| 0 | 0 | 0 | 1 |
| 0 | 0 | 0 | 0 |
| 0 | 0 | 0 | 1 |
| 0 | 0 | 0 | 1 |
| 0 | 0 | 0 | 1 |
| 0 | 0 | 0 | 1 |
| 0 | 0 | 0 | 1 |
| 0 | 0 | 0 | 0 |
| 0 | 0 | 0 | 0 |

| | | | |
|---|---|---|---|
| 0 | 0 | 0 | 0 |
| 0 | 0 | 0 | 0 |
| 0 | 0 | 0 | 0 |
| 0 | 0 | 0 | 0 |
| 0 | 0 | 0 | 0 |
| 0 | 0 | 0 | 0 |
| 0 | 0 | 0 | 0 |
| 0 | 0 | 0 | 0 |
| 0 | 0 | 0 | 1 |
| 0 | 0 | 0 | 0 |
| 0 | 0 | 0 | 1 |
| 0 | 0 | 0 | 1 |
| 0 | 0 | 0 | 1 |
| 0 | 0 | 0 | 1 |
| 0 | 0 | 0 | 0 |
| 0 | 0 | 0 | 1 |
| 0 | 0 | 0 | 1 |
| 0 | 0 | 0 | 1 |
| 0 | 0 | 0 | 1 |
| 0 | 0 | 0 | 0 |
| 0 | 0 | 0 | 0 |
| 0 | 0 | 0 | 1 |
| 0 | 0 | 0 | 0 |
| 0 | 0 | 0 | 0 |
| 0 | 0 | 0 | 0 |
| 0 | 0 | 0 | 0 |
| 0 | 0 | 0 | 0 |
| 0 | 0 | 0 | 1 |
| 0 | 0 | 0 | 1 |
| 0 | 0 | 0 | 1 |
| 0 | 0 | 0 | 0 |
| 0 | 0 | 0 | 1 |
| 0 | 0 | 0 | 1 |
| 0 | 0 | 0 | 1 |
| 0 | 0 | 0 | 1 |
| 0 | 0 | 0 | 1 |
| 0 | 0 | 0 | 1 |
| 0 | 0 | 0 | 1 |
| 0 | 0 | 0 | 1 |
| 0 | 0 | 0 | 1 |
| 0 | 0 | 0 | 1 |
| 0 | 0 | 0 | 0 |
| 0 | 0 | 0 | 1 |
| 0 | 0 | 0 | 1 |
| 0 | 0 | 0 | 1 |
| 0 | 0 | 0 | 1 |
| 0 | 0 | 0 | 1 |
| 0 | 0 | 0 | 1 |
| 0 | 0 | 0 | 0 |
| 0 | 0 | 0 | 1 |
| 0 | 0 | 0 | 1 |
| 0 | 0 | 0 | 1 |
| 0 | 0 | 0 | 0 |
| 0 | 1 | 0 | 0 |

| | | | |
|---|---|---|---|
| 0 | 0 | 0 | 1 |
| 0 | 0 | 0 | 1 |
| 0 | 0 | 0 | 1 |
| 0 | 0 | 0 | 1 |
| 0 | 0 | 0 | 0 |
| 1 | 0 | 0 | 0 |
| 0 | 0 | 0 | 1 |
| 0 | 0 | 0 | 0 |
| 0 | 0 | 0 | 1 |
| 0 | 0 | 0 | 1 |
| 0 | 0 | 0 | 0 |
| 0 | 0 | 0 | 1 |
| 0 | 0 | 0 | 0 |
| 0 | 0 | 0 | 0 |
| 0 | 0 | 0 | 0 |
| 0 | 0 | 0 | 0 |
| 0 | 0 | 0 | 1 |
| 0 | 0 | 0 | 1 |
| 0 | 0 | 0 | 0 |
| 0 | 1 | 0 | 0 |
| 0 | 0 | 0 | 0 |
| 0 | 1 | 0 | 0 |
| 0 | 0 | 0 | 0 |
| 0 | 0 | 0 | 1 |
| 0 | 0 | 0 | 1 |
| 0 | 0 | 0 | 0 |
| 0 | 0 | 0 | 1 |
| 0 | 0 | 0 | 0 |
| 0 | 0 | 0 | 0 |
| 0 | 0 | 0 | 0 |
| 0 | 0 | 0 | 1 |
| 1 | 0 | 0 | 0 |
| 1 | 0 | 0 | 0 |
| 0 | 0 | 0 | 1 |
| 0 | 0 | 0 | 1 |
| 0 | 0 | 0 | 0 |
| 0 | 0 | 0 | 0 |
| 0 | 1 | 1 | 1 |
| 0 | 1 | 0 | 0 |
| 0 | 1 | 0 | 0 |
| 0 | 1 | 0 | 0 |
| 0 | 0 | 0 | 0 |
| 0 | 0 | 0 | 1 |
| 0 | 0 | 0 | 1 |
| 0 | 0 | 0 | 0 |
| 0 | 0 | 0 | 1 |
| 0 | 0 | 0 | 0 |
| 0 | 0 | 0 | 1 |
| 0 | 0 | 0 | 1 |
| 0 | 0 | 0 | 1 |
| 0 | 0 | 0 | 1 |
| 0 | 0 | 0 | 1 |
| 0 | 0 | 0 | 1 |
| 0 | 0 | 0 | 1 |

| | | | |
|---|---|---|---|
| 0 | 0 | 0 | 1 |
| 0 | 0 | 0 | 1 |
| 0 | 0 | 0 | 1 |
| 0 | 0 | 0 | 1 |
| 0 | 0 | 0 | 1 |
| 0 | 0 | 0 | 1 |
| 0 | 0 | 0 | 1 |
| 0 | 0 | 0 | 0 |
| 0 | 0 | 0 | 1 |
| 1 | 0 | 0 | 0 |
| 0 | 0 | 0 | 0 |
| 0 | 0 | 0 | 0 |
| 0 | 0 | 0 | 1 |
| 0 | 0 | 0 | 1 |
| 0 | 0 | 0 | 0 |
| 0 | 0 | 0 | 1 |
| 0 | 0 | 0 | 1 |
| 0 | 0 | 0 | 1 |
| 0 | 0 | 0 | 1 |
| 0 | 0 | 0 | 1 |
| 0 | 0 | 0 | 1 |
| 0 | 0 | 0 | 1 |
| 0 | 0 | 0 | 1 |
| 0 | 0 | 0 | 1 |
| 0 | 0 | 0 | 1 |
| 0 | 0 | 0 | 1 |
| 0 | 0 | 0 | 1 |
| 0 | 0 | 0 | 1 |
| 0 | 0 | 0 | 1 |
| 0 | 0 | 0 | 0 |
| 0 | 0 | 0 | 0 |
| 0 | 0 | 0 | 1 |
| 0 | 0 | 0 | 1 |
| 0 | 0 | 0 | 1 |
| 0 | 0 | 0 | 1 |
| 0 | 0 | 0 | 0 |
| 0 | 0 | 0 | 1 |
| 0 | 0 | 0 | 1 |
| 0 | 0 | 0 | 1 |
| 0 | 0 | 0 | 1 |
| 0 | 0 | 0 | 1 |
| 0 | 0 | 0 | 0 |
| 0 | 0 | 0 | 0 |
| 0 | 0 | 0 | 0 |
| 0 | 0 | 0 | 1 |
| 0 | 0 | 0 | 1 |
| 0 | 0 | 0 | 1 |
| 0 | 0 | 0 | 1 |
| 0 | 0 | 0 | 1 |
| 0 | 0 | 0 | 1 |
| 0 | 0 | 0 | 1 |
| 0 | 0 | 0 | 1 |
| 0 | 0 | 0 | 1 |
| 0 | 0 | 0 | 0 |

| | | | |
|---|---|---|---|
| 0 | 0 | 0 | 1 |
| 0 | 0 | 0 | 1 |
| 0 | 0 | 0 | 1 |
| 0 | 0 | 0 | 1 |
| 0 | 0 | 0 | 1 |
| 0 | 0 | 0 | 1 |
| 0 | 0 | 0 | 1 |
| 0 | 0 | 0 | 1 |
| 0 | 0 | 0 | 0 |
| 0 | 0 | 0 | 0 |
| 0 | 0 | 0 | 1 |
| 0 | 0 | 0 | 1 |
| 0 | 0 | 0 | 1 |
| 0 | 0 | 0 | 0 |
| 0 | 0 | 0 | 1 |
| 0 | 0 | 0 | 1 |
| 0 | 0 | 0 | 1 |
| 1 | 0 | 0 | 0 |
| 1 | 0 | 0 | 0 |
| 0 | 0 | 0 | 0 |
| 0 | 0 | 0 | 0 |
| 0 | 0 | 0 | 1 |
| 0 | 0 | 0 | 1 |
| 0 | 0 | 0 | 1 |
| 0 | 0 | 0 | 1 |
| 1 | 0 | 0 | 0 |
| 0 | 0 | 0 | 1 |
| 0 | 0 | 0 | 1 |
| 0 | 0 | 0 | 1 |
| 0 | 0 | 0 | 1 |
| 0 | 0 | 0 | 1 |
| 0 | 0 | 0 | 1 |
| 0 | 0 | 0 | 1 |
| 0 | 0 | 0 | 1 |
| 0 | 0 | 0 | 1 |
| 0 | 0 | 0 | 1 |
| 0 | 0 | 0 | 1 |
| 0 | 0 | 0 | 1 |
| 0 | 0 | 0 | 1 |
| 0 | 0 | 0 | 0 |
| 0 | 0 | 0 | 1 |
| 0 | 0 | 0 | 1 |
| 0 | 0 | 0 | 0 |
| 0 | 0 | 0 | 1 |
| 0 | 0 | 0 | 1 |
| 0 | 0 | 0 | 1 |
| 0 | 0 | 0 | 0 |
| 0 | 0 | 0 | 0 |
| 0 | 0 | 0 | 1 |
| 0 | 0 | 0 | 1 |
| 0 | 0 | 0 | 0 |
| 0 | 0 | 0 | 1 |
| 0 | 0 | 0 | 1 |
| 0 | 0 | 0 | 1 |

| | | | |
|---|---|---|---|
| 0 | 0 | 0 | 1 |
| 0 | 0 | 0 | 1 |
| 0 | 0 | 0 | 1 |
| 0 | 0 | 0 | 1 |
| 0 | 0 | 0 | 1 |
| 0 | 0 | 0 | 1 |
| 0 | 0 | 0 | 1 |
| 0 | 0 | 0 | 1 |
| 0 | 0 | 0 | 1 |
| 0 | 0 | 0 | 1 |
| 0 | 0 | 0 | 1 |
| 0 | 0 | 0 | 1 |
| 0 | 0 | 0 | 1 |
| 0 | 0 | 0 | 0 |
| 0 | 0 | 0 | 0 |
| 0 | 0 | 0 | 0 |
| 0 | 0 | 0 | 0 |
| 0 | 0 | 0 | 1 |
| 0 | 0 | 0 | 1 |
| 0 | 0 | 0 | 0 |
| 0 | 0 | 0 | 1 |
| 0 | 1 | 1 | 1 |
| 0 | 0 | 0 | 1 |
| 0 | 0 | 0 | 1 |
| 0 | 0 | 0 | 1 |
| 0 | 0 | 0 | 0 |
| 0 | 0 | 0 | 1 |
| 0 | 0 | 0 | 1 |
| 0 | 0 | 0 | 1 |
| 0 | 0 | 0 | 0 |
| 0 | 0 | 0 | 0 |
| 0 | 0 | 0 | 1 |
| 0 | 0 | 0 | 1 |
| 0 | 0 | 0 | 1 |
| 0 | 0 | 0 | 1 |
| 0 | 0 | 0 | 1 |
| 0 | 0 | 0 | 1 |
| 0 | 0 | 0 | 1 |
| 0 | 0 | 0 | 0 |
| 0 | 0 | 0 | 1 |
| 0 | 0 | 0 | 1 |
| 0 | 0 | 0 | 0 |
| 0 | 1 | 0 | 0 |
| 0 | 1 | 0 | 0 |
| 0 | 0 | 0 | 0 |
| 0 | 0 | 0 | 1 |
| 0 | 0 | 0 | 0 |
| 0 | 0 | 0 | 1 |
| 0 | 0 | 0 | 1 |
| 0 | 0 | 0 | 1 |
| 0 | 0 | 0 | 1 |
| 0 | 0 | 0 | 1 |
| 0 | 1 | 1 | 1 |
| 0 | 1 | 1 | 1 |

| | | | |
|---|---|---|---|
| 0 | 0 | 0 | 1 |
| 0 | 0 | 0 | 1 |
| 0 | 0 | 0 | 0 |
| 0 | 0 | 0 | 1 |
| 0 | 0 | 0 | 1 |
| 0 | 0 | 0 | 1 |
| 0 | 0 | 0 | 1 |
| 0 | 0 | 0 | 1 |
| 0 | 0 | 0 | 1 |
| 0 | 0 | 0 | 1 |
| 0 | 0 | 0 | 1 |
| 0 | 0 | 0 | 1 |
| 0 | 0 | 0 | 1 |
| 0 | 0 | 0 | 1 |
| 0 | 0 | 0 | 1 |
| 0 | 0 | 0 | 1 |
| 0 | 0 | 0 | 1 |
| 0 | 0 | 0 | 1 |
| 0 | 0 | 0 | 1 |
| 0 | 0 | 0 | 1 |
| 0 | 0 | 0 | 1 |
| 0 | 0 | 0 | 1 |
| 0 | 0 | 0 | 1 |
| 0 | 0 | 0 | 1 |
| 0 | 0 | 0 | 1 |
| 0 | 0 | 0 | 1 |
| 0 | 0 | 0 | 1 |
| 0 | 0 | 0 | 1 |
| 0 | 0 | 0 | 1 |
| 0 | 0 | 0 | 1 |
| 1 | 0 | 0 | 0 |
| 0 | 0 | 0 | 1 |
| 0 | 1 | 0 | 0 |
| 0 | 1 | 0 | 0 |
| 0 | 0 | 0 | 0 |
| 0 | 0 | 0 | 0 |
| 0 | 0 | 0 | 1 |
| 0 | 0 | 0 | 0 |
| 0 | 0 | 0 | 1 |
| 0 | 0 | 0 | 1 |
| 1 | 0 | 0 | 0 |
| 1 | 0 | 0 | 0 |
| 0 | 0 | 0 | 0 |
| 0 | 0 | 0 | 1 |
| 0 | 0 | 0 | 0 |
| 0 | 0 | 0 | 0 |
| 0 | 0 | 0 | 1 |
| 0 | 0 | 0 | 1 |
| 0 | 0 | 0 | 1 |
| 0 | 0 | 0 | 1 |
| 0 | 0 | 0 | 0 |
| 0 | 0 | 0 | 1 |
| 0 | 0 | 0 | 1 |
| 0 | 0 | 0 | 1 |

| | | | |
|---|---|---|---|
| 0 | 0 | 0 | 1 |
| 0 | 0 | 0 | 1 |
| 0 | 0 | 0 | 1 |
| 0 | 0 | 0 | 1 |
| 0 | 0 | 0 | 1 |
| 1 | 0 | 0 | 0 |
| 0 | 0 | 0 | 1 |
| 0 | 0 | 0 | 0 |
| 0 | 0 | 0 | 0 |
| 0 | 0 | 0 | 0 |
| 0 | 0 | 0 | 1 |
| 0 | 1 | 0 | 0 |
| 0 | 0 | 0 | 1 |
| 0 | 0 | 0 | 1 |
| 0 | 0 | 0 | 1 |
| 0 | 1 | 1 | 1 |
| 0 | 0 | 0 | 1 |
| 0 | 0 | 0 | 0 |
| 0 | 0 | 0 | 1 |
| 0 | 0 | 0 | 0 |
| 0 | 0 | 0 | 1 |
| 0 | 0 | 0 | 0 |
| 0 | 0 | 0 | 0 |
| 0 | 0 | 0 | 0 |
| 0 | 0 | 0 | 1 |
| 0 | 0 | 0 | 1 |
| 0 | 0 | 0 | 1 |
| 0 | 0 | 0 | 1 |
| 0 | 0 | 0 | 0 |
| 0 | 0 | 0 | 1 |
| 0 | 0 | 0 | 1 |
| 0 | 0 | 0 | 1 |
| 0 | 0 | 0 | 1 |
| 0 | 0 | 0 | 1 |
| 0 | 0 | 0 | 0 |
| 0 | 0 | 0 | 0 |
| 0 | 0 | 0 | 1 |
| 0 | 0 | 0 | 0 |
| 0 | 0 | 0 | 0 |
| 0 | 0 | 0 | 1 |
| 0 | 0 | 0 | 0 |
| 0 | 0 | 0 | 0 |
| 0 | 0 | 0 | 1 |
| 0 | 0 | 0 | 1 |
| 0 | 0 | 0 | 1 |
| 0 | 0 | 0 | 1 |
| 0 | 0 | 0 | 1 |
| 0 | 0 | 0 | 1 |
| 0 | 0 | 0 | 1 |
| 0 | 0 | 0 | 1 |
| 0 | 0 | 0 | 1 |
| 0 | 0 | 0 | 1 |
| 0 | 0 | 0 | 1 |
| 0 | 0 | 0 | 1 |

| | | | |
|---|---|---|---|
| 0 | 0 | 0 | 1 |
| 0 | 0 | 0 | 0 |
| 0 | 0 | 0 | 1 |
| 0 | 0 | 0 | 1 |
| 0 | 0 | 0 | 0 |
| 0 | 0 | 0 | 1 |
| 0 | 0 | 0 | 1 |
| 0 | 0 | 0 | 1 |
| 0 | 0 | 0 | 0 |
| 0 | 0 | 0 | 0 |
| 0 | 0 | 0 | 1 |
| 0 | 0 | 0 | 0 |
| 0 | 0 | 0 | 0 |
| 0 | 0 | 0 | 1 |
| 0 | 0 | 0 | 1 |
| 0 | 0 | 0 | 0 |
| 0 | 0 | 0 | 1 |
| 0 | 0 | 0 | 0 |
| 0 | 1 | 0 | 0 |
| 0 | 0 | 0 | 1 |
| 0 | 0 | 0 | 0 |
| 0 | 0 | 0 | 0 |
| 1 | 0 | 0 | 0 |
| 0 | 0 | 0 | 1 |
| 0 | 0 | 0 | 1 |
| 0 | 0 | 0 | 0 |
| 0 | 0 | 0 | 1 |
| 0 | 0 | 0 | 1 |
| 0 | 0 | 0 | 1 |
| 0 | 0 | 0 | 1 |
| 0 | 0 | 0 | 0 |
| 0 | 0 | 0 | 1 |
| 0 | 0 | 0 | 0 |
| 0 | 0 | 0 | 1 |
| 0 | 0 | 0 | 0 |
| 0 | 0 | 0 | 0 |
| 0 | 0 | 0 | 1 |
| 0 | 0 | 0 | 1 |
| 0 | 0 | 0 | 1 |
| 0 | 0 | 0 | 1 |
| 0 | 0 | 0 | 1 |
| 0 | 0 | 0 | 1 |
| 0 | 0 | 0 | 1 |
| 0 | 0 | 0 | 1 |
| 0 | 0 | 0 | 1 |
| 0 | 0 | 0 | 1 |
| 0 | 1 | 1 | 1 |
| 0 | 0 | 0 | 1 |
| 0 | 0 | 0 | 1 |
| 0 | 0 | 0 | 1 |
| 0 | 0 | 0 | 1 |
| 0 | 0 | 0 | 1 |
| 0 | 0 | 0 | 1 |
| 0 | 0 | 0 | 0 |

| | | | |
|---|---|---|---|
| 0 | 0 | 0 | 1 |
| 0 | 0 | 0 | 1 |
| 0 | 0 | 0 | 1 |
| 0 | 0 | 0 | 1 |
| 0 | 0 | 0 | 0 |
| 0 | 0 | 0 | 1 |
| 0 | 0 | 0 | 1 |
| 0 | 0 | 0 | 1 |
| 0 | 0 | 0 | 1 |
| 0 | 0 | 0 | 1 |
| 0 | 0 | 0 | 0 |
| 0 | 0 | 0 | 1 |
| 0 | 0 | 0 | 1 |
| 0 | 0 | 0 | 1 |
| 0 | 0 | 0 | 0 |
| 0 | 0 | 0 | 0 |
| 0 | 0 | 0 | 0 |
| 0 | 0 | 0 | 0 |
| 0 | 0 | 0 | 1 |
| 0 | 0 | 0 | 0 |
| 0 | 0 | 0 | 0 |
| 1 | 0 | 0 | 0 |
| 0 | 0 | 0 | 1 |
| 0 | 0 | 0 | 0 |
| 1 | 0 | 0 | 0 |
| 0 | 0 | 0 | 1 |
| 1 | 0 | 0 | 0 |
| 0 | 0 | 0 | 0 |
| 1 | 0 | 0 | 0 |
| 1 | 0 | 0 | 0 |
| 1 | 0 | 0 | 0 |
| 0 | 0 | 0 | 1 |
| 0 | 0 | 0 | 1 |
| 0 | 0 | 0 | 1 |
| 0 | 0 | 0 | 1 |
| 0 | 0 | 0 | 1 |
| 0 | 0 | 0 | 1 |
| 0 | 1 | 0 | 0 |
| 0 | 0 | 0 | 0 |
| 0 | 0 | 0 | 1 |
| 0 | 0 | 0 | 1 |
| 0 | 1 | 1 | 1 |
| 0 | 0 | 0 | 0 |
| 0 | 0 | 0 | 1 |
| 0 | 0 | 0 | 0 |
| 0 | 0 | 0 | 1 |
| 0 | 0 | 0 | 1 |
| 0 | 0 | 0 | 1 |
| 0 | 1 | 0 | 0 |
| 0 | 1 | 0 | 0 |
| 0 | 1 | 0 | 0 |
| 0 | 0 | 0 | 1 |
| 0 | 0 | 0 | 1 |
| 0 | 0 | 0 | 0 |

| | | | |
|---|---|---|---|
| 0 | 0 | 0 | 1 |
| 0 | 1 | 0 | 0 |
| 0 | 0 | 0 | 1 |
| 0 | 1 | 0 | 0 |
| 0 | 1 | 0 | 0 |
| 0 | 0 | 0 | 1 |
| 0 | 1 | 0 | 0 |
| 0 | 1 | 0 | 0 |
| 0 | 0 | 0 | 1 |
| 0 | 0 | 0 | 1 |
| 0 | 0 | 0 | 1 |
| 0 | 0 | 0 | 1 |
| 0 | 0 | 0 | 1 |
| 0 | 0 | 0 | 1 |
| 0 | 0 | 0 | 1 |
| 0 | 0 | 0 | 1 |
| 0 | 0 | 0 | 1 |
| 0 | 0 | 0 | 1 |
| 0 | 0 | 0 | 1 |
| 0 | 0 | 0 | 1 |
| 1 | 1 | 0 | 0 |
| 0 | 0 | 0 | 1 |
| 0 | 0 | 0 | 1 |
| 0 | 0 | 0 | 1 |
| 0 | 0 | 0 | 1 |
| 0 | 0 | 0 | 0 |
| 0 | 0 | 0 | 0 |
| 0 | 0 | 0 | 1 |
| 0 | 0 | 0 | 1 |
| 0 | 0 | 0 | 1 |
| 0 | 0 | 0 | 1 |
| 0 | 0 | 0 | 1 |
| 0 | 0 | 0 | 1 |
| 0 | 0 | 0 | 1 |
| 0 | 0 | 0 | 1 |
| 0 | 0 | 0 | 1 |
| 0 | 0 | 0 | 1 |
| 0 | 0 | 0 | 1 |
| 0 | 0 | 0 | 1 |
| 0 | 0 | 0 | 1 |
| 0 | 0 | 0 | 1 |
| 0 | 0 | 0 | 1 |
| 0 | 0 | 0 | 1 |
| 0 | 0 | 0 | 0 |
| 0 | 0 | 0 | 1 |
| 0 | 0 | 0 | 0 |
| 0 | 0 | 0 | 0 |
| 0 | 0 | 0 | 0 |
| 0 | 0 | 0 | 0 |
| 0 | 0 | 0 | 0 |
| 0 | 0 | 0 | 0 |
| 0 | 0 | 0 | 1 |
| 0 | 0 | 0 | 1 |
| 0 | 0 | 0 | 1 |

| | | | |
|---|---|---|---|
| 0 | 0 | 0 | 1 |
| 0 | 0 | 0 | 1 |
| 0 | 0 | 0 | 1 |
| 0 | 0 | 0 | 0 |
| 0 | 0 | 0 | 1 |
| 0 | 0 | 0 | 0 |
| 0 | 0 | 0 | 1 |
| 0 | 0 | 0 | 1 |
| 0 | 0 | 0 | 0 |
| 0 | 0 | 0 | 1 |
| 0 | 0 | 0 | 1 |
| 0 | 0 | 0 | 1 |
| 0 | 0 | 0 | 1 |
| 0 | 0 | 0 | 1 |
| 0 | 0 | 0 | 1 |
| 0 | 0 | 0 | 0 |
| 0 | 0 | 0 | 0 |
| 0 | 0 | 0 | 0 |
| 0 | 0 | 0 | 0 |
| 0 | 0 | 0 | 0 |
| 0 | 0 | 0 | 1 |
| 0 | 0 | 0 | 1 |
| 0 | 0 | 0 | 1 |
| 0 | 0 | 0 | 1 |
| 0 | 0 | 0 | 0 |
| 0 | 0 | 0 | 1 |
| 0 | 0 | 0 | 1 |
| 0 | 0 | 0 | 1 |
| 0 | 0 | 0 | 0 |
| 0 | 0 | 0 | 1 |
| 0 | 1 | 0 | 0 |
| 0 | 1 | 0 | 0 |
| 0 | 0 | 0 | 0 |
| 0 | 1 | 0 | 0 |
| 0 | 1 | 0 | 0 |
| 0 | 0 | 0 | 1 |
| 0 | 1 | 0 | 0 |
| 0 | 1 | 0 | 0 |
| 0 | 1 | 0 | 0 |
| 0 | 1 | 0 | 0 |
| 0 | 1 | 0 | 0 |
| 0 | 0 | 0 | 0 |
| 0 | 0 | 0 | 0 |
| 0 | 0 | 0 | 0 |
| 0 | 0 | 0 | 0 |
| 0 | 0 | 0 | 0 |
| 0 | 0 | 0 | 0 |
| 0 | 0 | 0 | 0 |
| 0 | 0 | 0 | 1 |
| 0 | 0 | 0 | 0 |
| 0 | 0 | 0 | 1 |
| 0 | 0 | 0 | 1 |
| 0 | 0 | 0 | 1 |
| 0 | 0 | 0 | 0 |

| | | | |
|---|---|---|---|
| 0 | 1 | 0 | 0 |
| 0 | 0 | 0 | 1 |
| 0 | 0 | 0 | 1 |
| 0 | 0 | 0 | 1 |
| 0 | 0 | 0 | 1 |
| 0 | 0 | 0 | 0 |
| 0 | 0 | 0 | 1 |
| 0 | 0 | 0 | 1 |
| 0 | 0 | 0 | 1 |
| 0 | 0 | 0 | 1 |
| 0 | 0 | 0 | 1 |
| 0 | 0 | 0 | 1 |
| 0 | 0 | 0 | 1 |
| 1 | 0 | 0 | 0 |
| 0 | 0 | 0 | 1 |
| 0 | 0 | 0 | 1 |
| 0 | 0 | 0 | 1 |
| 0 | 0 | 0 | 0 |
| 0 | 0 | 0 | 1 |
| 0 | 0 | 0 | 1 |
| 0 | 0 | 0 | 1 |
| 0 | 0 | 0 | 1 |
| 0 | 0 | 0 | 0 |
| 0 | 0 | 0 | 1 |
| 0 | 0 | 0 | 1 |
| 0 | 0 | 0 | 0 |
| 0 | 0 | 0 | 0 |
| 0 | 0 | 0 | 0 |
| 0 | 0 | 0 | 1 |
| 0 | 0 | 0 | 1 |
| 0 | 0 | 0 | 1 |
| 0 | 0 | 0 | 1 |
| 0 | 0 | 0 | 1 |
| 0 | 0 | 0 | 1 |
| 0 | 0 | 0 | 1 |
| 0 | 0 | 0 | 1 |
| 0 | 0 | 0 | 1 |
| 0 | 0 | 0 | 1 |
| 0 | 0 | 0 | 1 |
| 0 | 0 | 0 | 1 |
| 0 | 0 | 0 | 1 |
| 0 | 0 | 0 | 1 |
| 0 | 0 | 0 | 1 |
| 0 | 0 | 0 | 1 |
| 0 | 0 | 0 | 1 |
| 0 | 0 | 0 | 1 |
| 0 | 0 | 0 | 1 |
| 0 | 0 | 0 | 1 |
| 0 | 1 | 1 | 1 |
| 0 | 0 | 0 | 1 |
| 0 | 0 | 0 | 1 |
| 0 | 0 | 0 | 1 |
| 0 | 0 | 0 | 1 |
| 0 | 0 | 0 | 1 |

| | | | |
|---|---|---|---|
| 0 | 0 | 0 | 1 |
| 0 | 0 | 0 | 0 |
| 0 | 0 | 0 | 1 |
| 0 | 0 | 0 | 1 |
| 0 | 0 | 0 | 1 |
| 0 | 0 | 0 | 0 |
| 0 | 0 | 0 | 0 |
| 0 | 0 | 0 | 0 |
| 0 | 0 | 0 | 1 |
| 0 | 0 | 0 | 1 |
| 0 | 0 | 0 | 1 |
| 0 | 0 | 0 | 1 |
| 0 | 0 | 0 | 1 |
| 0 | 0 | 0 | 1 |
| 0 | 0 | 0 | 0 |
| 0 | 0 | 0 | 0 |
| 0 | 0 | 0 | 0 |
| 0 | 0 | 0 | 1 |
| 0 | 0 | 0 | 0 |
| 0 | 0 | 0 | 0 |
| 0 | 0 | 0 | 0 |
| 0 | 0 | 0 | 0 |
| 0 | 0 | 0 | 0 |
| 0 | 0 | 0 | 0 |
| 0 | 0 | 0 | 1 |
| 0 | 0 | 0 | 1 |
| 0 | 0 | 0 | 1 |
| 0 | 0 | 0 | 1 |
| 0 | 0 | 0 | 1 |
| 0 | 0 | 0 | 1 |
| 0 | 0 | 0 | 1 |
| 0 | 0 | 0 | 1 |
| 0 | 0 | 0 | 0 |
| 0 | 0 | 0 | 1 |
| 0 | 0 | 0 | 1 |
| 0 | 0 | 0 | 0 |
| 0 | 0 | 0 | 0 |
| 0 | 0 | 0 | 0 |
| 0 | 0 | 0 | 0 |
| 0 | 0 | 0 | 1 |
| 0 | 0 | 0 | 1 |
| 0 | 0 | 0 | 1 |
| 0 | 0 | 0 | 1 |
| 0 | 0 | 0 | 0 |
| 0 | 0 | 0 | 0 |
| 0 | 0 | 0 | 0 |
| 0 | 0 | 0 | 1 |
| 0 | 0 | 0 | 1 |
| 0 | 0 | 0 | 0 |
| 0 | 0 | 0 | 1 |
| 0 | 0 | 0 | 1 |
| 0 | 0 | 0 | 0 |
| 0 | 0 | 0 | 1 |
| 0 | 0 | 0 | 1 |

| | | | |
|---|---|---|---|
| 0 | 0 | 0 | 1 |
| 0 | 0 | 0 | 1 |
| 0 | 0 | 0 | 1 |
| 0 | 0 | 0 | 1 |
| 0 | 0 | 0 | 1 |
| 0 | 0 | 0 | 1 |
| 0 | 0 | 0 | 1 |
| 0 | 0 | 0 | 1 |
| 0 | 0 | 0 | 0 |
| 0 | 0 | 0 | 0 |
| 0 | 0 | 0 | 1 |
| 0 | 0 | 0 | 0 |
| 0 | 0 | 0 | 0 |
| 0 | 0 | 0 | 0 |
| 0 | 0 | 0 | 1 |
| 0 | 0 | 0 | 1 |
| 0 | 0 | 0 | 0 |
| 0 | 0 | 0 | 0 |
| 0 | 0 | 0 | 0 |
| 0 | 0 | 0 | 1 |
| 0 | 0 | 0 | 1 |
| 0 | 0 | 0 | 1 |
| 0 | 0 | 0 | 0 |
| 0 | 0 | 0 | 1 |
| 0 | 0 | 0 | 0 |
| 0 | 0 | 0 | 1 |
| 0 | 0 | 0 | 1 |
| 0 | 0 | 0 | 1 |
| 0 | 0 | 0 | 1 |
| 0 | 0 | 0 | 0 |
| 0 | 0 | 0 | 0 |
| 0 | 0 | 0 | 0 |
| 0 | 0 | 0 | 1 |
| 0 | 0 | 0 | 1 |
| 0 | 0 | 0 | 1 |
| 0 | 0 | 0 | 1 |
| 0 | 0 | 0 | 1 |
| 0 | 0 | 0 | 1 |
| 0 | 0 | 0 | 1 |
| 0 | 0 | 0 | 1 |
| 0 | 0 | 0 | 0 |
| 0 | 0 | 0 | 1 |
| 0 | 0 | 0 | 1 |
| 0 | 0 | 0 | 1 |
| 0 | 0 | 0 | 1 |
| 0 | 0 | 0 | 0 |
| 0 | 1 | 1 | 1 |
| 0 | 0 | 0 | 0 |
| 0 | 0 | 0 | 1 |
| 0 | 1 | 0 | 0 |
| 0 | 1 | 0 | 0 |
| 0 | 0 | 0 | 0 |
| 0 | 0 | 0 | 1 |
| 0 | 1 | 1 | 1 |

| | | | |
|---|---|---|---|
| 0 | 1 | 1 | 1 |
| 0 | 1 | 1 | 1 |
| 0 | 0 | 0 | 1 |
| 0 | 0 | 0 | 1 |
| 0 | 0 | 0 | 1 |
| 0 | 0 | 0 | 1 |
| 0 | 0 | 0 | 1 |
| 0 | 0 | 0 | 0 |
| 0 | 0 | 0 | 0 |
| 0 | 0 | 0 | 1 |
| 0 | 0 | 0 | 0 |
| 0 | 0 | 0 | 1 |
| 0 | 0 | 0 | 1 |
| 0 | 0 | 0 | 1 |
| 0 | 0 | 0 | 1 |
| 0 | 0 | 0 | 1 |
| 0 | 0 | 0 | 1 |
| 0 | 0 | 0 | 1 |
| 0 | 0 | 0 | 1 |
| 0 | 0 | 0 | 1 |
| 0 | 0 | 0 | 1 |
| 0 | 0 | 0 | 1 |
| 0 | 0 | 0 | 1 |
| 0 | 0 | 0 | 1 |
| 0 | 0 | 0 | 1 |
| 0 | 0 | 0 | 1 |
| 0 | 0 | 0 | 1 |
| 0 | 0 | 0 | 1 |
| 0 | 0 | 0 | 0 |
| 0 | 0 | 0 | 1 |
| 0 | 0 | 0 | 0 |
| 0 | 0 | 0 | 1 |
| 0 | 0 | 0 | 0 |
| 0 | 0 | 0 | 0 |
| 0 | 0 | 0 | 1 |
| 0 | 0 | 0 | 0 |
| 0 | 0 | 0 | 0 |
| 0 | 0 | 0 | 1 |
| 0 | 0 | 0 | 1 |
| 0 | 0 | 0 | 1 |
| 0 | 0 | 0 | 1 |
| 0 | 0 | 0 | 0 |
| 0 | 0 | 0 | 0 |
| 0 | 0 | 0 | 1 |
| 0 | 0 | 0 | 1 |
| 0 | 0 | 0 | 0 |
| 0 | 0 | 0 | 0 |
| 0 | 0 | 0 | 0 |
| 0 | 0 | 0 | 0 |
| 0 | 0 | 0 | 1 |
| 0 | 1 | 0 | 0 |
| 0 | 0 | 0 | 0 |
| 0 | 0 | 0 | 1 |
| 0 | 0 | 0 | 1 |

| | | | |
|---|---|---|---|
| 0 | 0 | 0 | 1 |
| 0 | 0 | 0 | 1 |
| 0 | 0 | 0 | 1 |
| 0 | 0 | 0 | 1 |
| 0 | 0 | 0 | 0 |
| 0 | 0 | 0 | 0 |
| 0 | 0 | 0 | 0 |
| 0 | 0 | 0 | 0 |
| 0 | 0 | 0 | 1 |
| 0 | 0 | 0 | 0 |
| 0 | 0 | 0 | 0 |
| 0 | 0 | 0 | 0 |
| 0 | 0 | 0 | 0 |
| 0 | 0 | 0 | 1 |
| 0 | 1 | 1 | 1 |
| 0 | 0 | 0 | 1 |
| 0 | 0 | 0 | 1 |
| 0 | 0 | 0 | 1 |
| 0 | 0 | 0 | 1 |
| 0 | 0 | 0 | 1 |
| 0 | 0 | 0 | 1 |
| 0 | 0 | 0 | 1 |
| 0 | 0 | 0 | 1 |
| 0 | 0 | 0 | 1 |
| 0 | 0 | 0 | 1 |
| 0 | 0 | 0 | 0 |
| 0 | 0 | 0 | 0 |
| 0 | 0 | 0 | 0 |
| 0 | 0 | 0 | 0 |
| 0 | 0 | 0 | 1 |
| 0 | 0 | 0 | 1 |
| 0 | 0 | 0 | 1 |
| 0 | 0 | 0 | 1 |
| 0 | 0 | 0 | 1 |
| 0 | 0 | 0 | 1 |
| 0 | 0 | 0 | 1 |
| 0 | 0 | 0 | 1 |
| 0 | 0 | 0 | 1 |
| 0 | 0 | 0 | 1 |
| 0 | 0 | 0 | 1 |
| 0 | 0 | 0 | 1 |
| 0 | 0 | 0 | 1 |
| 0 | 0 | 0 | 1 |
| 0 | 0 | 0 | 1 |
| 0 | 0 | 0 | 1 |
| 0 | 0 | 0 | 1 |
| 0 | 0 | 0 | 1 |
| 0 | 0 | 0 | 1 |
| 0 | 0 | 0 | 1 |
| 0 | 0 | 0 | 1 |
| 0 | 0 | 0 | 1 |
| 0 | 0 | 0 | 1 |
| 0 | 0 | 0 | 1 |
| 0 | 0 | 0 | 0 |

| | | | |
|---|---|---|---|
| 0 | 0 | 0 | 1 |
| 0 | 0 | 0 | 1 |
| 0 | 0 | 0 | 1 |
| 0 | 0 | 0 | 1 |
| 0 | 0 | 0 | 1 |
| 0 | 0 | 0 | 1 |
| 0 | 0 | 0 | 1 |
| 0 | 0 | 0 | 1 |
| 0 | 0 | 0 | 1 |
| 0 | 0 | 0 | 1 |
| 0 | 0 | 0 | 1 |
| 0 | 0 | 0 | 1 |
| 0 | 0 | 0 | 1 |
| 0 | 0 | 0 | 1 |
| 0 | 0 | 0 | 1 |
| 0 | 0 | 0 | 1 |
| 0 | 0 | 0 | 1 |
| 0 | 0 | 0 | 1 |
| 0 | 0 | 0 | 1 |
| 0 | 0 | 0 | 1 |
| 0 | 0 | 0 | 1 |
| 0 | 0 | 0 | 1 |
| 0 | 0 | 0 | 1 |
| 0 | 0 | 0 | 1 |
| 0 | 0 | 0 | 1 |
| 0 | 0 | 0 | 1 |
| 0 | 0 | 0 | 1 |
| 0 | 0 | 0 | 1 |
| 0 | 0 | 0 | 1 |
| 0 | 0 | 0 | 1 |
| 0 | 0 | 0 | 1 |
| 0 | 0 | 0 | 1 |
| 0 | 0 | 0 | 1 |
| 0 | 0 | 0 | 1 |
| 0 | 0 | 0 | 1 |
| 0 | 0 | 0 | 1 |
| 0 | 0 | 0 | 1 |
| 0 | 0 | 0 | 1 |
| 0 | 0 | 0 | 1 |
| 0 | 0 | 0 | 1 |
| 0 | 0 | 0 | 1 |
| 0 | 0 | 0 | 1 |
| 0 | 0 | 0 | 1 |
| 0 | 0 | 0 | 1 |
| 0 | 0 | 0 | 1 |
| 0 | 0 | 0 | 1 |
| 0 | 1 | 1 | 1 |
| 0 | 0 | 0 | 0 |
| 0 | 0 | 0 | 1 |
| 0 | 0 | 0 | 1 |
| 0 | 0 | 0 | 1 |
| 0 | 0 | 0 | 1 |
| 0 | 0 | 0 | 1 |
| 0 | 0 | 0 | 1 |
| 0 | 0 | 0 | 1 |

| | | | |
|---|---|---|---|
| 0 | 0 | 0 | 1 |
| 0 | 0 | 0 | 1 |
| 0 | 0 | 0 | 0 |
| 0 | 0 | 0 | 1 |
| 0 | 0 | 0 | 1 |
| 0 | 0 | 0 | 1 |
| 0 | 0 | 0 | 0 |
| 0 | 0 | 0 | 1 |
| 0 | 0 | 0 | 1 |
| 0 | 0 | 0 | 1 |
| 0 | 0 | 0 | 1 |
| 0 | 0 | 0 | 0 |
| 0 | 0 | 0 | 1 |
| 0 | 0 | 0 | 0 |
| 0 | 0 | 0 | 0 |
| 0 | 0 | 0 | 0 |
| 0 | 0 | 0 | 0 |
| 0 | 0 | 0 | 0 |
| 0 | 0 | 0 | 0 |
| 0 | 0 | 0 | 0 |
| 0 | 0 | 0 | 0 |

| surface_non_binding_site | non_binding_site_single_water | surface_non_binding_site_single_water |
|---|---|---|
| 0 | 0 | 0 |
| 0 | 0 | 0 |
| 0 | 0 | 0 |
| 0 | 0 | 0 |
| 0 | 0 | 0 |
| 0 | 0 | 0 |
| 1 | 1 | 1 |
| 1 | 1 | 1 |
| 0 | 0 | 0 |
| 0 | 0 | 0 |
| 0 | 0 | 0 |
| 0 | 0 | 0 |
| 0 | 0 | 0 |
| 0 | 0 | 0 |
| 0 | 0 | 0 |
| 0 | 0 | 0 |
| 0 | 0 | 0 |
| 0 | 0 | 0 |
| 0 | 0 | 0 |
| 0 | 0 | 0 |
| 0 | 0 | 0 |
| 0 | 0 | 0 |
| 0 | 0 | 0 |
| 0 | 0 | 0 |
| 0 | 0 | 0 |
| 1 | 0 | 0 |
| 0 | 0 | 0 |
| 1 | 1 | 1 |
| 0 | 0 | 0 |
| 0 | 1 | 0 |
| 0 | 1 | 0 |
| 0 | 0 | 0 |
| 0 | 0 | 0 |
| 0 | 0 | 0 |
| 0 | 0 | 0 |
| 1 | 1 | 1 |
| 0 | 0 | 0 |
| 0 | 0 | 0 |
| 0 | 0 | 0 |
| 0 | 0 | 0 |
| 0 | 0 | 0 |
| 0 | 0 | 0 |
| 0 | 0 | 0 |
| 0 | 0 | 0 |
| 0 | 0 | 0 |
| 0 | 0 | 0 |
| 0 | 0 | 0 |
| 0 | 0 | 0 |
| 0 | 0 | 0 |
| 0 | 0 | 0 |
| 0 | 0 | 0 |
| 0 | 0 | 0 |
| 0 | 0 | 0 |

| | | |
|---|---|---|
| 0 | 0 | 0 |
| 0 | 0 | 0 |
| 0 | 0 | 0 |
| 0 | 0 | 0 |
| 0 | 0 | 0 |
| 0 | 0 | 0 |
| 0 | 0 | 0 |
| 0 | 0 | 0 |
| 0 | 0 | 0 |
| 0 | 0 | 0 |
| 0 | 0 | 0 |
| 0 | 0 | 0 |
| 0 | 0 | 0 |
| 0 | 0 | 0 |
| 0 | 0 | 0 |
| 0 | 0 | 0 |
| 0 | 0 | 0 |
| 0 | 0 | 0 |
| 0 | 0 | 0 |
| 0 | 0 | 0 |
| 0 | 0 | 0 |
| 0 | 0 | 0 |
| 0 | 0 | 0 |
| 0 | 0 | 0 |
| 0 | 0 | 0 |
| 0 | 0 | 0 |
| 0 | 0 | 0 |
| 0 | 0 | 0 |
| 0 | 0 | 0 |
| 0 | 0 | 0 |
| 0 | 0 | 0 |
| 0 | 0 | 0 |
| 0 | 0 | 0 |
| 0 | 0 | 0 |
| 0 | 0 | 0 |
| 0 | 0 | 0 |
| 0 | 0 | 0 |
| 0 | 0 | 0 |
| 0 | 0 | 0 |
| 0 | 0 | 0 |
| 0 | 0 | 0 |
| 0 | 0 | 0 |
| 0 | 0 | 0 |
| 0 | 0 | 0 |
| 0 | 0 | 0 |
| 1 | 1 | 1 |
| 0 | 1 | 0 |
| 0 | 0 | 0 |
| 0 | 0 | 0 |
| 0 | 0 | 0 |
| 0 | 0 | 0 |
| 1 | 1 | 1 |
| 1 | 1 | 1 |
| 0 | 0 | 0 |

| | | |
|---|---|---|
| 0 | 1 | 0 |
| 0 | 1 | 0 |
| 0 | 0 | 0 |
| 0 | 0 | 0 |
| 0 | 0 | 0 |
| 0 | 0 | 0 |
| 0 | 1 | 0 |
| 0 | 0 | 0 |
| 0 | 0 | 0 |
| 0 | 0 | 0 |
| 0 | 0 | 0 |
| 0 | 0 | 0 |
| 0 | 0 | 0 |
| 0 | 0 | 0 |
| 0 | 0 | 0 |
| 1 | 1 | 1 |
| 0 | 0 | 0 |
| 0 | 0 | 0 |
| 0 | 0 | 0 |
| 0 | 0 | 0 |
| 0 | 0 | 0 |
| 0 | 0 | 0 |
| 0 | 0 | 0 |
| 0 | 0 | 0 |
| 0 | 0 | 0 |
| 0 | 0 | 0 |
| 0 | 0 | 0 |
| 0 | 0 | 0 |
| 0 | 0 | 0 |
| 0 | 0 | 0 |
| 0 | 0 | 0 |
| 0 | 0 | 0 |
| 0 | 0 | 0 |
| 0 | 0 | 0 |
| 0 | 0 | 0 |
| 0 | 0 | 0 |
| 0 | 0 | 0 |
| 0 | 0 | 0 |
| 0 | 0 | 0 |
| 0 | 0 | 0 |
| 0 | 0 | 0 |
| 0 | 0 | 0 |
| 0 | 0 | 0 |
| 0 | 0 | 0 |
| 0 | 0 | 0 |
| 0 | 0 | 0 |
| 0 | 0 | 0 |
| 0 | 0 | 0 |
| 0 | 0 | 0 |
| 0 | 0 | 0 |
| 0 | 0 | 0 |
| 0 | 0 | 0 |
| 0 | 0 | 0 |
| 0 | 0 | 0 |

| | | |
|---|---|---|
| 1 | 0 | 0 |
| 1 | 0 | 0 |
| 1 | 0 | 0 |
| 0 | 0 | 0 |
| 0 | 0 | 0 |
| 0 | 0 | 0 |
| 0 | 0 | 0 |
| 1 | 0 | 0 |
| 0 | 0 | 0 |
| 1 | 0 | 0 |
| 1 | 0 | 0 |
| 0 | 0 | 0 |
| 1 | 0 | 0 |
| 0 | 0 | 0 |
| 0 | 0 | 0 |
| 1 | 0 | 0 |
| 1 | 0 | 0 |
| 1 | 0 | 0 |
| 1 | 0 | 0 |
| 0 | 0 | 0 |
| 0 | 0 | 0 |
| 0 | 1 | 0 |
| 0 | 0 | 0 |
| 0 | 0 | 0 |
| 0 | 0 | 0 |
| 1 | 0 | 0 |
| 0 | 0 | 0 |
| 0 | 0 | 0 |
| 1 | 0 | 0 |
| 1 | 0 | 0 |
| 1 | 1 | 1 |
| 0 | 0 | 0 |
| 0 | 0 | 0 |
| 0 | 1 | 0 |
| 0 | 0 | 0 |
| 0 | 0 | 0 |
| 0 | 0 | 0 |
| 0 | 0 | 0 |
| 0 | 0 | 0 |
| 1 | 0 | 0 |
| 1 | 0 | 0 |
| 0 | 0 | 0 |
| 0 | 0 | 0 |
| 1 | 1 | 1 |
| 1 | 0 | 0 |
| 1 | 0 | 0 |
| 0 | 0 | 0 |
| 0 | 0 | 0 |
| 0 | 0 | 0 |
| 0 | 0 | 0 |
| 0 | 0 | 0 |
| 0 | 1 | 0 |
| 1 | 0 | 0 |
| 0 | 0 | 0 |

| | | |
|---|---|---|
| 0 | 0 | 0 |
| 0 | 0 | 0 |
| 0 | 0 | 0 |
| 1 | 1 | 1 |
| 1 | 1 | 1 |
| 1 | 0 | 0 |
| 0 | 0 | 0 |
| 0 | 0 | 0 |
| 1 | 0 | 0 |
| 1 | 0 | 0 |
| 0 | 0 | 0 |
| 0 | 1 | 0 |
| 0 | 0 | 0 |
| 1 | 0 | 0 |
| 0 | 1 | 0 |
| 1 | 1 | 1 |
| 1 | 1 | 1 |
| 0 | 0 | 0 |
| 0 | 0 | 0 |
| 0 | 0 | 0 |
| 0 | 0 | 0 |
| 1 | 0 | 0 |
| 0 | 0 | 0 |
| 0 | 1 | 0 |
| 1 | 1 | 1 |
| 0 | 0 | 0 |
| 0 | 0 | 0 |
| 0 | 0 | 0 |
| 0 | 0 | 0 |
| 0 | 0 | 0 |
| 0 | 0 | 0 |
| 0 | 0 | 0 |
| 1 | 0 | 0 |
| 1 | 0 | 0 |
| 0 | 0 | 0 |
| 1 | 0 | 0 |
| 0 | 0 | 0 |
| 0 | 0 | 0 |
| 1 | 0 | 0 |
| 0 | 1 | 0 |
| 0 | 0 | 0 |
| 1 | 0 | 0 |
| 1 | 0 | 0 |
| 1 | 0 | 0 |
| 0 | 0 | 0 |
| 0 | 0 | 0 |
| 0 | 0 | 0 |
| 0 | 0 | 0 |
| 0 | 0 | 0 |
| 0 | 0 | 0 |
| 0 | 0 | 0 |
| 0 | 0 | 0 |
| 0 | 0 | 0 |
| 0 | 0 | 0 |

| | | |
|---|---|---|
| 0 | 0 | 0 |
| 0 | 0 | 0 |
| 0 | 0 | 0 |
| 0 | 0 | 0 |
| 0 | 0 | 0 |
| 0 | 0 | 0 |
| 0 | 1 | 0 |
| 0 | 1 | 0 |
| 0 | 0 | 0 |
| 0 | 0 | 0 |
| 0 | 0 | 0 |
| 0 | 0 | 0 |
| 0 | 0 | 0 |
| 0 | 1 | 0 |
| 0 | 1 | 0 |
| 0 | 0 | 0 |
| 0 | 1 | 0 |
| 0 | 0 | 0 |
| 1 | 1 | 1 |
| 0 | 0 | 0 |
| 1 | 1 | 1 |
| 1 | 0 | 0 |
| 1 | 1 | 1 |
| 0 | 0 | 0 |
| 0 | 0 | 0 |
| 0 | 0 | 0 |
| 0 | 0 | 0 |
| 0 | 0 | 0 |
| 0 | 0 | 0 |
| 0 | 0 | 0 |
| 0 | 0 | 0 |
| 1 | 0 | 0 |
| 0 | 0 | 0 |
| 0 | 0 | 0 |
| 0 | 0 | 0 |
| 0 | 0 | 0 |
| 0 | 0 | 0 |
| 0 | 0 | 0 |
| 0 | 0 | 0 |
| 0 | 0 | 0 |
| 0 | 1 | 0 |
| 0 | 1 | 0 |
| 0 | 0 | 0 |
| 0 | 1 | 0 |
| 0 | 0 | 0 |
| 0 | 0 | 0 |
| 0 | 0 | 0 |
| 1 | 1 | 1 |
| 1 | 1 | 1 |
| 0 | 0 | 0 |
| 0 | 1 | 0 |
| 1 | 1 | 1 |
| 0 | 0 | 0 |
| 0 | 1 | 0 |

| | | |
|---|---|---|
| 0 | 0 | 0 |
| 1 | 1 | 1 |
| 0 | 0 | 0 |
| 0 | 0 | 0 |
| 0 | 0 | 0 |
| 0 | 0 | 0 |
| 0 | 1 | 0 |
| 0 | 0 | 0 |
| 0 | 0 | 0 |
| 1 | 1 | 1 |
| 1 | 1 | 1 |
| 1 | 1 | 1 |
| 1 | 1 | 1 |
| 1 | 1 | 1 |
| 0 | 0 | 0 |
| 1 | 1 | 1 |
| 0 | 0 | 0 |
| 0 | 0 | 0 |
| 1 | 1 | 1 |
| 0 | 0 | 0 |
| 0 | 0 | 0 |
| 0 | 0 | 0 |
| 0 | 0 | 0 |
| 0 | 0 | 0 |
| 1 | 1 | 1 |
| 1 | 0 | 0 |
| 0 | 0 | 0 |
| 0 | 0 | 0 |
| 0 | 0 | 0 |
| 0 | 0 | 0 |
| 0 | 0 | 0 |
| 0 | 0 | 0 |
| 0 | 0 | 0 |
| 0 | 0 | 0 |
| 1 | 1 | 1 |
| 1 | 1 | 1 |
| 0 | 0 | 0 |
| 0 | 0 | 0 |
| 0 | 0 | 0 |
| 0 | 0 | 0 |
| 0 | 0 | 0 |
| 0 | 0 | 0 |
| 0 | 1 | 0 |
| 0 | 0 | 0 |
| 0 | 0 | 0 |
| 0 | 0 | 0 |
| 0 | 0 | 0 |
| 0 | 0 | 0 |
| 0 | 0 | 0 |
| 1 | 1 | 1 |
| 0 | 0 | 0 |
| 0 | 0 | 0 |
| 0 | 0 | 0 |
| 0 | 0 | 0 |

| | | |
|---|---|---|
| 0 | 0 | 0 |
| 0 | 0 | 0 |
| 0 | 0 | 0 |
| 0 | 0 | 0 |
| 1 | 1 | 1 |
| 0 | 0 | 0 |
| 0 | 0 | 0 |
| 0 | 0 | 0 |
| 0 | 0 | 0 |
| 0 | 0 | 0 |
| 0 | 0 | 0 |
| 0 | 0 | 0 |
| 0 | 0 | 0 |
| 0 | 0 | 0 |
| 0 | 0 | 0 |
| 0 | 0 | 0 |
| 0 | 0 | 0 |
| 0 | 0 | 0 |
| 1 | 1 | 1 |
| 0 | 0 | 0 |
| 0 | 0 | 0 |
| 0 | 0 | 0 |
| 0 | 0 | 0 |
| 1 | 0 | 0 |
| 0 | 0 | 0 |
| 0 | 0 | 0 |
| 0 | 0 | 0 |
| 1 | 0 | 0 |
| 1 | 0 | 0 |
| 1 | 0 | 0 |
| 0 | 0 | 0 |
| 0 | 0 | 0 |
| 0 | 0 | 0 |
| 0 | 0 | 0 |
| 0 | 0 | 0 |
| 1 | 0 | 0 |
| 1 | 0 | 0 |
| 0 | 0 | 0 |
| 0 | 0 | 0 |
| 1 | 0 | 0 |
| 1 | 0 | 0 |
| 1 | 0 | 0 |
| 0 | 1 | 0 |
| 1 | 1 | 1 |
| 0 | 0 | 0 |
| 0 | 0 | 0 |
| 0 | 0 | 0 |
| 0 | 0 | 0 |
| 0 | 0 | 0 |
| 0 | 1 | 0 |
| 1 | 1 | 1 |
| 0 | 0 | 0 |
| 1 | 1 | 1 |
| 0 | 0 | 0 |

| | | |
|---|---|---|
| 1 | 1 | 1 |
| 1 | 1 | 1 |
| 0 | 0 | 0 |
| 1 | 1 | 1 |
| 1 | 1 | 1 |
| 1 | 1 | 1 |
| 1 | 1 | 1 |
| 0 | 0 | 0 |
| 1 | 0 | 0 |
| 1 | 1 | 1 |
| 0 | 0 | 0 |
| 1 | 1 | 1 |
| 1 | 1 | 1 |
| 0 | 1 | 0 |
| 0 | 0 | 0 |
| 0 | 0 | 0 |
| 1 | 1 | 1 |
| 1 | 1 | 1 |
| 0 | 0 | 0 |
| 0 | 0 | 0 |
| 0 | 0 | 0 |
| 0 | 0 | 0 |
| 0 | 0 | 0 |
| 0 | 0 | 0 |
| 0 | 0 | 0 |
| 1 | 1 | 1 |
| 0 | 0 | 0 |
| 0 | 0 | 0 |
| 0 | 0 | 0 |
| 1 | 0 | 0 |
| 0 | 0 | 0 |
| 0 | 0 | 0 |
| 0 | 0 | 0 |
| 0 | 0 | 0 |
| 0 | 0 | 0 |
| 0 | 0 | 0 |
| 0 | 0 | 0 |
| 0 | 0 | 0 |
| 0 | 0 | 0 |
| 0 | 0 | 0 |
| 0 | 1 | 0 |
| 0 | 0 | 0 |
| 1 | 1 | 1 |
| 0 | 0 | 0 |
| 0 | 0 | 0 |
| 0 | 0 | 0 |
| 0 | 0 | 0 |
| 0 | 0 | 0 |
| 0 | 0 | 0 |
| 0 | 0 | 0 |
| 0 | 1 | 0 |
| 0 | 0 | 0 |
| 0 | 0 | 0 |
| 0 | 0 | 0 |

| | | |
|---|---|---|
| 1 | 1 | 1 |
| 0 | 0 | 0 |
| 0 | 1 | 0 |
| 0 | 1 | 0 |
| 0 | 0 | 0 |
| 0 | 0 | 0 |
| 0 | 0 | 0 |
| 0 | 0 | 0 |
| 0 | 0 | 0 |
| 0 | 1 | 0 |
| 1 | 1 | 1 |
| 1 | 1 | 1 |
| 1 | 1 | 1 |
| 0 | 0 | 0 |
| 1 | 1 | 1 |
| 0 | 0 | 0 |
| 0 | 0 | 0 |
| 0 | 0 | 0 |
| 0 | 0 | 0 |
| 1 | 0 | 0 |
| 0 | 0 | 0 |
| 1 | 1 | 1 |
| 0 | 0 | 0 |
| 0 | 1 | 0 |
| 1 | 1 | 1 |
| 1 | 1 | 1 |
| 0 | 0 | 0 |
| 1 | 1 | 1 |
| 1 | 1 | 1 |
| 1 | 1 | 1 |
| 0 | 0 | 0 |
| 0 | 0 | 0 |
| 1 | 1 | 1 |
| 1 | 1 | 1 |
| 0 | 0 | 0 |
| 0 | 0 | 0 |
| 0 | 0 | 0 |
| 0 | 0 | 0 |
| 0 | 0 | 0 |
| 1 | 1 | 1 |
| 0 | 0 | 0 |
| 0 | 0 | 0 |
| 0 | 0 | 0 |
| 0 | 0 | 0 |
| 1 | 0 | 0 |
| 0 | 0 | 0 |
| 0 | 0 | 0 |
| 0 | 0 | 0 |
| 0 | 0 | 0 |
| 0 | 0 | 0 |
| 0 | 0 | 0 |
| 0 | 0 | 0 |
| 1 | 0 | 0 |
| 0 | 0 | 0 |

| | | |
|---|---|---|
| 0 | 0 | 0 |
| 0 | 0 | 0 |
| 0 | 0 | 0 |
| 0 | 0 | 0 |
| 0 | 0 | 0 |
| 0 | 0 | 0 |
| 0 | 0 | 0 |
| 1 | 0 | 0 |
| 0 | 0 | 0 |
| 0 | 0 | 0 |
| 0 | 0 | 0 |
| 0 | 0 | 0 |
| 1 | 1 | 1 |
| 0 | 0 | 0 |
| 0 | 0 | 0 |
| 0 | 0 | 0 |
| 0 | 0 | 0 |
| 0 | 0 | 0 |
| 0 | 0 | 0 |
| 1 | 1 | 1 |
| 0 | 0 | 0 |
| 0 | 0 | 0 |
| 0 | 0 | 0 |
| 0 | 0 | 0 |
| 0 | 0 | 0 |
| 1 | 0 | 0 |
| 1 | 1 | 1 |
| 0 | 1 | 0 |
| 0 | 0 | 0 |
| 0 | 0 | 0 |
| 0 | 0 | 0 |
| 0 | 0 | 0 |
| 0 | 0 | 0 |
| 0 | 0 | 0 |
| 0 | 0 | 0 |
| 1 | 0 | 0 |
| 0 | 0 | 0 |
| 0 | 0 | 0 |
| 1 | 1 | 1 |
| 1 | 0 | 0 |
| 0 | 1 | 0 |
| 0 | 0 | 0 |
| 0 | 0 | 0 |
| 0 | 0 | 0 |
| 0 | 0 | 0 |
| 0 | 0 | 0 |
| 0 | 0 | 0 |
| 1 | 1 | 1 |
| 1 | 1 | 1 |
| 0 | 0 | 0 |
| 0 | 0 | 0 |
| 0 | 0 | 0 |
| 0 | 0 | 0 |
| 1 | 1 | 1 |

| | | |
|---|---|---|
| 1 | 1 | 1 |
| 0 | 0 | 0 |
| 0 | 1 | 0 |
| 0 | 0 | 0 |
| 1 | 0 | 0 |
| 1 | 0 | 0 |
| 0 | 0 | 0 |
| 1 | 0 | 0 |
| 1 | 0 | 0 |
| 1 | 0 | 0 |
| 1 | 0 | 0 |
| 1 | 0 | 0 |
| 1 | 1 | 1 |
| 1 | 1 | 1 |
| 1 | 1 | 1 |
| 1 | 1 | 1 |
| 0 | 0 | 0 |
| 0 | 0 | 0 |
| 1 | 0 | 0 |
| 0 | 0 | 0 |
| 0 | 0 | 0 |
| 0 | 0 | 0 |
| 1 | 0 | 0 |
| 0 | 0 | 0 |
| 0 | 0 | 0 |
| 1 | 1 | 1 |
| 1 | 0 | 0 |
| 1 | 0 | 0 |
| 0 | 0 | 0 |
| 0 | 0 | 0 |
| 0 | 0 | 0 |
| 0 | 0 | 0 |
| 0 | 0 | 0 |
| 0 | 0 | 0 |
| 0 | 0 | 0 |
| 0 | 0 | 0 |
| 1 | 0 | 0 |
| 1 | 1 | 1 |
| 1 | 1 | 1 |
| 1 | 1 | 1 |
| 0 | 0 | 0 |
| 1 | 1 | 1 |
| 0 | 0 | 0 |
| 1 | 1 | 1 |
| 0 | 0 | 0 |
| 0 | 1 | 0 |
| 0 | 0 | 0 |
| 0 | 0 | 0 |
| 0 | 0 | 0 |
| 0 | 0 | 0 |
| 0 | 0 | 0 |
| 0 | 0 | 0 |
| 0 | 0 | 0 |
| 0 | 0 | 0 |

| | | |
|---|---|---|
| 0 | 0 | 0 |
| 0 | 0 | 0 |
| 0 | 0 | 0 |
| 0 | 0 | 0 |
| 0 | 0 | 0 |
| 0 | 0 | 0 |
| 0 | 0 | 0 |
| 1 | 1 | 1 |
| 0 | 0 | 0 |
| 0 | 0 | 0 |
| 0 | 0 | 0 |
| 1 | 0 | 0 |
| 0 | 0 | 0 |
| 0 | 0 | 0 |
| 0 | 0 | 0 |
| 0 | 0 | 0 |
| 0 | 0 | 0 |
| 0 | 0 | 0 |
| 0 | 0 | 0 |
| 0 | 0 | 0 |
| 0 | 0 | 0 |
| 0 | 0 | 0 |
| 0 | 0 | 0 |
| 0 | 0 | 0 |
| 0 | 0 | 0 |
| 0 | 0 | 0 |
| 0 | 0 | 0 |
| 0 | 0 | 0 |
| 0 | 0 | 0 |
| 0 | 0 | 0 |
| 0 | 0 | 0 |
| 0 | 0 | 0 |
| 0 | 0 | 0 |
| 0 | 0 | 0 |
| 0 | 0 | 0 |
| 0 | 0 | 0 |
| 0 | 0 | 0 |
| 0 | 0 | 0 |
| 0 | 0 | 0 |
| 0 | 0 | 0 |
| 1 | 0 | 0 |
| 0 | 0 | 0 |
| 0 | 0 | 0 |
| 0 | 0 | 0 |
| 0 | 0 | 0 |
| 0 | 0 | 0 |
| 0 | 0 | 0 |
| 0 | 0 | 0 |
| 0 | 0 | 0 |
| 0 | 1 | 0 |
| 0 | 1 | 0 |
| 1 | 0 | 0 |
| 0 | 0 | 0 |
| 0 | 0 | 0 |

| | | |
|---|---|---|
| 0 | 0 | 0 |
| 0 | 0 | 0 |
| 0 | 0 | 0 |
| 0 | 0 | 0 |
| 0 | 0 | 0 |
| 0 | 0 | 0 |
| 0 | 0 | 0 |
| 1 | 0 | 0 |
| 1 | 0 | 0 |
| 0 | 0 | 0 |
| 0 | 0 | 0 |
| 1 | 0 | 0 |
| 0 | 0 | 0 |
| 0 | 0 | 0 |
| 0 | 0 | 0 |
| 0 | 0 | 0 |
| 0 | 0 | 0 |
| 1 | 0 | 0 |
| 0 | 0 | 0 |
| 1 | 0 | 0 |
| 0 | 0 | 0 |
| 0 | 0 | 0 |
| 1 | 1 | 1 |
| 0 | 0 | 0 |
| 0 | 0 | 0 |
| 1 | 0 | 0 |
| 0 | 0 | 0 |
| 0 | 0 | 0 |
| 0 | 0 | 0 |
| 0 | 0 | 0 |
| 0 | 0 | 0 |
| 0 | 0 | 0 |
| 1 | 0 | 0 |
| 0 | 0 | 0 |
| 0 | 0 | 0 |
| 0 | 0 | 0 |
| 1 | 1 | 1 |
| 0 | 0 | 0 |
| 0 | 0 | 0 |
| 0 | 0 | 0 |
| 0 | 0 | 0 |
| 0 | 0 | 0 |
| 0 | 0 | 0 |
| 0 | 0 | 0 |
| 0 | 0 | 0 |
| 0 | 0 | 0 |
| 0 | 0 | 0 |
| 0 | 0 | 0 |
| 0 | 0 | 0 |
| 0 | 0 | 0 |
| 0 | 0 | 0 |
| 0 | 0 | 0 |
| 0 | 0 | 0 |
| 0 | 0 | 0 |

| | | |
|---|---|---|
| 1 | 0 | 0 |
| 0 | 1 | 0 |
| 0 | 0 | 0 |
| 0 | 0 | 0 |
| 0 | 0 | 0 |
| 0 | 0 | 0 |
| 0 | 0 | 0 |
| 0 | 0 | 0 |
| 0 | 0 | 0 |
| 1 | 0 | 0 |
| 0 | 1 | 0 |
| 1 | 1 | 1 |
| 0 | 0 | 0 |
| 0 | 0 | 0 |
| 0 | 0 | 0 |
| 0 | 0 | 0 |
| 0 | 0 | 0 |
| 0 | 0 | 0 |
| 1 | 1 | 1 |
| 1 | 0 | 0 |
| 1 | 0 | 0 |
| 0 | 0 | 0 |
| 0 | 1 | 0 |
| 0 | 0 | 0 |
| 0 | 0 | 0 |
| 0 | 0 | 0 |
| 0 | 0 | 0 |
| 0 | 0 | 0 |
| 0 | 0 | 0 |
| 0 | 0 | 0 |
| 0 | 0 | 0 |
| 0 | 0 | 0 |
| 0 | 0 | 0 |
| 0 | 0 | 0 |
| 0 | 0 | 0 |
| 0 | 0 | 0 |
| 0 | 0 | 0 |
| 0 | 0 | 0 |
| 0 | 0 | 0 |
| 0 | 0 | 0 |
| 0 | 0 | 0 |
| 1 | 0 | 0 |
| 0 | 0 | 0 |
| 0 | 0 | 0 |
| 0 | 0 | 0 |
| 0 | 0 | 0 |
| 1 | 0 | 0 |
| 0 | 0 | 0 |
| 0 | 0 | 0 |
| 0 | 0 | 0 |
| 0 | 0 | 0 |
| 0 | 0 | 0 |
| 1 | 0 | 0 |
| 0 | 1 | 0 |

| | | |
|---|---|---|
| 0 | 0 | 0 |
| 0 | 0 | 0 |
| 1 | 0 | 0 |
| 1 | 0 | 0 |
| 1 | 0 | 0 |
| 0 | 0 | 0 |
| 1 | 0 | 0 |
| 0 | 0 | 0 |
| 0 | 0 | 0 |
| 0 | 0 | 0 |
| 0 | 0 | 0 |
| 0 | 0 | 0 |
| 0 | 0 | 0 |
| 1 | 0 | 0 |
| 1 | 0 | 0 |
| 1 | 0 | 0 |
| 0 | 0 | 0 |
| 1 | 1 | 1 |
| 0 | 0 | 0 |
| 0 | 0 | 0 |
| 1 | 0 | 0 |
| 1 | 0 | 0 |
| 0 | 0 | 0 |
| 0 | 0 | 0 |
| 1 | 0 | 0 |
| 0 | 0 | 0 |
| 0 | 0 | 0 |
| 1 | 0 | 0 |
| 0 | 0 | 0 |
| 0 | 0 | 0 |
| 0 | 0 | 0 |
| 0 | 0 | 0 |
| 0 | 0 | 0 |
| 0 | 0 | 0 |
| 0 | 0 | 0 |
| 0 | 0 | 0 |
| 0 | 0 | 0 |
| 0 | 0 | 0 |
| 0 | 0 | 0 |
| 0 | 0 | 0 |
| 1 | 0 | 0 |
| 1 | 0 | 0 |
| 1 | 0 | 0 |
| 1 | 1 | 1 |
| 1 | 1 | 1 |
| 0 | 0 | 0 |
| 0 | 0 | 0 |
| 0 | 0 | 0 |
| 0 | 0 | 0 |
| 0 | 0 | 0 |
| 0 | 0 | 0 |
| 0 | 0 | 0 |
| 0 | 0 | 0 |
| 1 | 0 | 0 |

| | | |
|---|---|---|
| 1 | 0 | 0 |
| 0 | 0 | 0 |
| 0 | 0 | 0 |
| 0 | 0 | 0 |
| 1 | 0 | 0 |
| 0 | 0 | 0 |
| 0 | 0 | 0 |
| 0 | 0 | 0 |
| 1 | 0 | 0 |
| 1 | 0 | 0 |
| 0 | 0 | 0 |
| 0 | 0 | 0 |
| 0 | 0 | 0 |
| 1 | 0 | 0 |
| 0 | 0 | 0 |
| 0 | 0 | 0 |
| 0 | 0 | 0 |
| 0 | 0 | 0 |
| 0 | 0 | 0 |
| 0 | 0 | 0 |
| 0 | 0 | 0 |
| 1 | 1 | 1 |
| 0 | 0 | 0 |
| 0 | 1 | 0 |
| 0 | 0 | 0 |
| 0 | 0 | 0 |
| 0 | 0 | 0 |
| 0 | 0 | 0 |
| 0 | 0 | 0 |
| 0 | 0 | 0 |
| 0 | 0 | 0 |
| 0 | 0 | 0 |
| 0 | 0 | 0 |
| 0 | 0 | 0 |
| 0 | 1 | 0 |
| 0 | 0 | 0 |
| 0 | 0 | 0 |
| 0 | 0 | 0 |
| 0 | 0 | 0 |
| 0 | 0 | 0 |
| 0 | 0 | 0 |
| 0 | 0 | 0 |
| 0 | 0 | 0 |
| 0 | 0 | 0 |
| 0 | 0 | 0 |
| 0 | 0 | 0 |
| 0 | 0 | 0 |
| 1 | 1 | 1 |
| 0 | 0 | 0 |
| 1 | 1 | 1 |
| 0 | 0 | 0 |
| 0 | 0 | 0 |
| 0 | 0 | 0 |
| 0 | 0 | 0 |

| | | |
|---|---|---|
| 0 | 0 | 0 |
| 0 | 0 | 0 |
| 1 | 1 | 1 |
| 0 | 0 | 0 |
| 0 | 0 | 0 |
| 0 | 1 | 0 |
| 0 | 1 | 0 |
| 0 | 1 | 0 |
| 0 | 0 | 0 |
| 0 | 1 | 0 |
| 0 | 0 | 0 |
| 0 | 0 | 0 |
| 0 | 0 | 0 |
| 1 | 1 | 1 |
| 0 | 0 | 0 |
| 0 | 1 | 0 |
| 0 | 0 | 0 |
| 0 | 0 | 0 |
| 0 | 0 | 0 |
| 1 | 1 | 1 |
| 1 | 1 | 1 |
| 0 | 0 | 0 |
| 1 | 1 | 1 |
| 0 | 0 | 0 |
| 0 | 0 | 0 |
| 1 | 1 | 1 |
| 1 | 1 | 1 |
| 0 | 0 | 0 |
| 1 | 1 | 1 |
| 0 | 0 | 0 |
| 0 | 1 | 0 |
| 0 | 1 | 0 |
| 1 | 1 | 1 |
| 0 | 1 | 0 |
| 1 | 1 | 1 |
| 1 | 1 | 1 |
| 1 | 1 | 1 |
| 1 | 1 | 1 |
| 1 | 1 | 1 |
| 1 | 1 | 1 |
| 1 | 1 | 1 |
| 1 | 1 | 1 |
| 0 | 1 | 0 |
| 0 | 1 | 0 |
| 0 | 1 | 0 |
| 0 | 1 | 0 |
| 0 | 1 | 0 |
| 0 | 1 | 0 |
| 0 | 1 | 0 |
| 0 | 1 | 0 |
| 0 | 1 | 0 |
| 0 | 1 | 0 |
| 0 | 1 | 0 |
| 0 | 1 | 0 |

| | | |
|---|---|---|
| 0 | 1 | 0 |
| 1 | 1 | 1 |
| 1 | 1 | 1 |
| 1 | 1 | 1 |
| 1 | 1 | 1 |
| 1 | 1 | 1 |
| 1 | 1 | 1 |
| 0 | 1 | 0 |
| 0 | 1 | 0 |
| 0 | 1 | 0 |
| 0 | 1 | 0 |
| 0 | 0 | 0 |
| 0 | 0 | 0 |
| 0 | 0 | 0 |
| 1 | 0 | 0 |
| 0 | 0 | 0 |
| 1 | 0 | 0 |
| 0 | 0 | 0 |
| 1 | 0 | 0 |
| 1 | 1 | 1 |
| 0 | 0 | 0 |
| 1 | 1 | 1 |
| 1 | 1 | 1 |
| 0 | 0 | 0 |
| 0 | 0 | 0 |
| 1 | 1 | 1 |
| 1 | 1 | 1 |
| 0 | 0 | 0 |
| 0 | 0 | 0 |
| 0 | 0 | 0 |
| 0 | 0 | 0 |
| 0 | 0 | 0 |
| 0 | 1 | 0 |
| 0 | 1 | 0 |
| 1 | 1 | 1 |
| 1 | 1 | 1 |
| 1 | 1 | 1 |
| 1 | 1 | 1 |
| 1 | 1 | 1 |
| 0 | 0 | 0 |
| 0 | 0 | 0 |
| 0 | 1 | 0 |
| 0 | 0 | 0 |
| 0 | 0 | 0 |
| 0 | 0 | 0 |
| 1 | 0 | 0 |
| 0 | 0 | 0 |
| 1 | 0 | 0 |
| 1 | 0 | 0 |
| 0 | 0 | 0 |
| 0 | 0 | 0 |
| 0 | 0 | 0 |
| 0 | 0 | 0 |
| 0 | 0 | 0 |

| | | |
|---|---|---|
| 0 | 0 | 0 |
| 0 | 1 | 0 |
| 1 | 1 | 1 |
| 1 | 1 | 1 |
| 0 | 0 | 0 |
| 1 | 1 | 1 |
| 0 | 0 | 0 |
| 1 | 1 | 1 |
| 1 | 1 | 1 |
| 0 | 0 | 0 |
| 1 | 0 | 0 |
| 1 | 1 | 1 |
| 1 | 1 | 1 |
| 0 | 0 | 0 |
| 1 | 0 | 0 |
| 1 | 0 | 0 |
| 0 | 0 | 0 |
| 1 | 0 | 0 |
| 0 | 0 | 0 |
| 0 | 0 | 0 |
| 0 | 0 | 0 |
| 0 | 0 | 0 |
| 0 | 0 | 0 |
| 0 | 0 | 0 |
| 0 | 0 | 0 |
| 0 | 0 | 0 |
| 0 | 0 | 0 |
| 0 | 0 | 0 |
| 0 | 0 | 0 |
| 0 | 0 | 0 |
| 0 | 0 | 0 |
| 0 | 0 | 0 |
| 0 | 0 | 0 |
| 0 | 0 | 0 |
| 0 | 0 | 0 |
| 0 | 0 | 0 |
| 1 | 0 | 0 |
| 0 | 0 | 0 |
| 0 | 0 | 0 |
| 0 | 0 | 0 |
| 0 | 0 | 0 |
| 0 | 0 | 0 |
| 0 | 0 | 0 |
| 0 | 0 | 0 |
| 0 | 0 | 0 |
| 0 | 0 | 0 |
| 0 | 0 | 0 |
| 0 | 0 | 0 |
| 1 | 0 | 0 |
| 1 | 0 | 0 |
| 0 | 0 | 0 |
| 1 | 1 | 1 |
| 0 | 0 | 0 |
| 0 | 0 | 0 |

| | | |
|---|---|---|
| 0 | 0 | 0 |
| 0 | 0 | 0 |
| 0 | 0 | 0 |
| 0 | 0 | 0 |
| 0 | 0 | 0 |
| 0 | 0 | 0 |
| 0 | 0 | 0 |
| 0 | 0 | 0 |
| 0 | 0 | 0 |
| 0 | 0 | 0 |
| 0 | 0 | 0 |
| 0 | 0 | 0 |
| 0 | 0 | 0 |
| 0 | 0 | 0 |
| 0 | 1 | 0 |
| 0 | 0 | 0 |
| 0 | 0 | 0 |
| 0 | 0 | 0 |
| 1 | 0 | 0 |
| 0 | 0 | 0 |
| 0 | 0 | 0 |
| 0 | 0 | 0 |
| 0 | 0 | 0 |
| 0 | 0 | 0 |
| 0 | 0 | 0 |
| 1 | 0 | 0 |
| 0 | 0 | 0 |
| 0 | 0 | 0 |
| 0 | 0 | 0 |
| 0 | 0 | 0 |
| 1 | 1 | 1 |
| 0 | 0 | 0 |
| 1 | 1 | 1 |
| 1 | 0 | 0 |
| 0 | 0 | 0 |
| 0 | 0 | 0 |
| 1 | 1 | 1 |
| 1 | 1 | 1 |
| 1 | 1 | 1 |
| 1 | 0 | 0 |
| 0 | 0 | 0 |
| 0 | 0 | 0 |
| 0 | 0 | 0 |
| 0 | 0 | 0 |
| 0 | 0 | 0 |
| 0 | 0 | 0 |
| 0 | 0 | 0 |
| 0 | 0 | 0 |
| 1 | 1 | 1 |
| 1 | 1 | 1 |
| 0 | 0 | 0 |
| 0 | 0 | 0 |
| 0 | 0 | 0 |
| 0 | 1 | 0 |

| | | |
|---|---|---|
| 1 | 1 | 1 |
| 1 | 1 | 1 |
| 1 | 0 | 0 |
| 0 | 0 | 0 |
| 1 | 0 | 0 |
| 1 | 1 | 1 |
| 0 | 0 | 0 |
| 1 | 1 | 1 |
| 0 | 0 | 0 |
| 0 | 0 | 0 |
| 0 | 0 | 0 |
| 0 | 0 | 0 |
| 0 | 0 | 0 |
| 0 | 0 | 0 |
| 0 | 0 | 0 |
| 0 | 0 | 0 |
| 0 | 1 | 0 |
| 0 | 0 | 0 |
| 0 | 0 | 0 |
| 1 | 0 | 0 |
| 0 | 0 | 0 |
| 0 | 0 | 0 |
| 1 | 1 | 1 |
| 0 | 0 | 0 |
| 1 | 1 | 1 |
| 1 | 1 | 1 |
| 0 | 1 | 0 |
| 0 | 0 | 0 |
| 0 | 0 | 0 |
| 0 | 0 | 0 |
| 1 | 1 | 1 |
| 0 | 0 | 0 |
| 0 | 0 | 0 |
| 0 | 0 | 0 |
| 0 | 1 | 0 |
| 0 | 1 | 0 |
| 0 | 0 | 0 |
| 0 | 0 | 0 |
| 0 | 0 | 0 |
| 1 | 1 | 1 |
| 1 | 1 | 1 |
| 0 | 1 | 0 |
| 0 | 1 | 0 |
| 0 | 0 | 0 |
| 1 | 1 | 1 |
| 0 | 0 | 0 |
| 1 | 1 | 1 |
| 0 | 0 | 0 |
| 0 | 0 | 0 |
| 1 | 1 | 1 |
| 0 | 1 | 0 |
| 0 | 0 | 0 |
| 0 | 0 | 0 |
| 1 | 1 | 1 |

| | | |
|---|---|---|
| 1 | 1 | 1 |
| 1 | 1 | 1 |
| 0 | 1 | 0 |
| 0 | 0 | 0 |
| 0 | 0 | 0 |
| 1 | 0 | 0 |
| 1 | 0 | 0 |
| 0 | 0 | 0 |
| 0 | 0 | 0 |
| 1 | 1 | 1 |
| 0 | 0 | 0 |
| 0 | 0 | 0 |
| 0 | 0 | 0 |
| 0 | 0 | 0 |
| 0 | 0 | 0 |
| 0 | 0 | 0 |
| 0 | 0 | 0 |
| 0 | 0 | 0 |
| 0 | 0 | 0 |
| 0 | 1 | 0 |
| 1 | 1 | 1 |
| 1 | 1 | 1 |
| 1 | 1 | 1 |
| 1 | 1 | 1 |
| 0 | 0 | 0 |
| 0 | 1 | 0 |
| 0 | 0 | 0 |
| 0 | 0 | 0 |
| 1 | 1 | 1 |
| 1 | 1 | 1 |
| 1 | 1 | 1 |
| 1 | 1 | 1 |
| 1 | 1 | 1 |
| 0 | 0 | 0 |
| 0 | 0 | 0 |
| 1 | 1 | 1 |
| 0 | 0 | 0 |
| 0 | 1 | 0 |
| 0 | 0 | 0 |
| 0 | 0 | 0 |
| 0 | 0 | 0 |
| 0 | 0 | 0 |
| 0 | 1 | 0 |
| 0 | 0 | 0 |
| 0 | 0 | 0 |
| 0 | 0 | 0 |
| 0 | 0 | 0 |
| 0 | 0 | 0 |
| 0 | 0 | 0 |
| 0 | 0 | 0 |
| 0 | 0 | 0 |
| 0 | 0 | 0 |
| 0 | 0 | 0 |
| 0 | 1 | 0 |

| | | |
|---|---|---|
| 0 | 0 | 0 |
| 0 | 1 | 0 |
| 0 | 0 | 0 |
| 0 | 0 | 0 |
| 0 | 0 | 0 |
| 1 | 1 | 1 |
| 0 | 0 | 0 |
| 1 | 1 | 1 |
| 0 | 1 | 0 |
| 0 | 0 | 0 |
| 0 | 1 | 0 |
| 0 | 1 | 0 |
| 0 | 1 | 0 |
| 0 | 1 | 0 |
| 0 | 1 | 0 |
| 0 | 0 | 0 |
| 0 | 0 | 0 |
| 0 | 1 | 0 |
| 0 | 0 | 0 |
| 1 | 1 | 1 |
| 0 | 0 | 0 |
| 0 | 0 | 0 |
| 0 | 0 | 0 |
| 0 | 0 | 0 |
| 0 | 0 | 0 |
| 0 | 0 | 0 |
| 0 | 0 | 0 |
| 0 | 0 | 0 |
| 0 | 0 | 0 |
| 0 | 0 | 0 |
| 0 | 0 | 0 |
| 0 | 0 | 0 |
| 1 | 0 | 0 |
| 1 | 0 | 0 |
| 0 | 0 | 0 |
| 0 | 0 | 0 |
| 0 | 0 | 0 |
| 0 | 0 | 0 |
| 1 | 0 | 0 |
| 0 | 0 | 0 |
| 0 | 0 | 0 |
| 0 | 0 | 0 |
| 1 | 0 | 0 |
| 0 | 1 | 0 |
| 0 | 0 | 0 |
| 0 | 0 | 0 |
| 0 | 0 | 0 |
| 0 | 0 | 0 |
| 0 | 0 | 0 |
| 0 | 0 | 0 |
| 1 | 0 | 0 |
| 0 | 0 | 0 |
| 0 | 0 | 0 |
| 1 | 0 | 0 |

| | | |
|---|---|---|
| 0 | 0 | 0 |
| 1 | 0 | 0 |
| 0 | 0 | 0 |
| 0 | 0 | 0 |
| 0 | 0 | 0 |
| 0 | 0 | 0 |
| 0 | 0 | 0 |
| 0 | 0 | 0 |
| 0 | 0 | 0 |
| 0 | 0 | 0 |
| 0 | 0 | 0 |
| 1 | 1 | 1 |
| 0 | 0 | 0 |
| 1 | 1 | 1 |
| 1 | 0 | 0 |
| 1 | 0 | 0 |
| 0 | 0 | 0 |
| 0 | 0 | 0 |
| 0 | 0 | 0 |
| 0 | 0 | 0 |
| 1 | 0 | 0 |
| 0 | 0 | 0 |
| 0 | 1 | 0 |
| 0 | 0 | 0 |
| 1 | 1 | 1 |
| 1 | 1 | 1 |
| 1 | 1 | 1 |
| 0 | 0 | 0 |
| 1 | 1 | 1 |
| 1 | 1 | 1 |
| 1 | 1 | 1 |
| 0 | 0 | 0 |
| 1 | 1 | 1 |
| 0 | 0 | 0 |
| 0 | 0 | 0 |
| 0 | 0 | 0 |
| 0 | 1 | 0 |
| 1 | 1 | 1 |
| 0 | 0 | 0 |
| 0 | 0 | 0 |
| 0 | 0 | 0 |
| 0 | 0 | 0 |
| 0 | 0 | 0 |
| 0 | 0 | 0 |
| 0 | 0 | 0 |
| 0 | 0 | 0 |
| 1 | 1 | 1 |
| 1 | 1 | 1 |
| 0 | 0 | 0 |
| 0 | 0 | 0 |
| 0 | 0 | 0 |
| 1 | 1 | 1 |
| 1 | 1 | 1 |
| 0 | 0 | 0 |

| | | |
|---|---|---|
| 0 | 0 | 0 |
| 0 | 1 | 0 |
| 0 | 1 | 0 |
| 1 | 1 | 1 |
| 0 | 0 | 0 |
| 0 | 0 | 0 |
| 0 | 0 | 0 |
| 0 | 0 | 0 |
| 0 | 0 | 0 |
| 0 | 0 | 0 |
| 1 | 1 | 1 |
| 0 | 0 | 0 |
| 0 | 0 | 0 |
| 0 | 0 | 0 |
| 1 | 0 | 0 |
| 1 | 1 | 1 |
| 0 | 0 | 0 |
| 1 | 0 | 0 |
| 0 | 0 | 0 |
| 0 | 0 | 0 |
| 1 | 0 | 0 |
| 0 | 0 | 0 |
| 0 | 0 | 0 |
| 1 | 0 | 0 |
| 0 | 0 | 0 |
| 0 | 0 | 0 |
| 0 | 0 | 0 |
| 1 | 0 | 0 |
| 0 | 0 | 0 |
| 0 | 0 | 0 |
| 0 | 0 | 0 |
| 1 | 1 | 1 |
| 1 | 1 | 1 |
| 1 | 1 | 1 |
| 0 | 1 | 0 |
| 0 | 1 | 0 |
| 0 | 1 | 0 |
| 1 | 1 | 1 |
| 1 | 1 | 1 |
| 1 | 1 | 1 |
| 0 | 1 | 0 |
| 1 | 1 | 1 |
| 0 | 0 | 0 |
| 0 | 1 | 0 |
| 1 | 1 | 1 |
| 1 | 1 | 1 |
| 0 | 1 | 0 |
| 1 | 1 | 1 |
| 1 | 1 | 1 |
| 1 | 1 | 1 |
| 1 | 1 | 1 |
| 1 | 1 | 1 |
| 1 | 1 | 1 |
| 1 | 1 | 1 |

| | | |
|---|---|---|
| 0 | 1 | 0 |
| 0 | 1 | 0 |
| 1 | 1 | 1 |
| 1 | 1 | 1 |
| 1 | 1 | 1 |
| 1 | 1 | 1 |
| 1 | 1 | 1 |
| 1 | 1 | 1 |
| 0 | 0 | 0 |
| 0 | 0 | 0 |
| 0 | 0 | 0 |
| 0 | 0 | 0 |
| 0 | 0 | 0 |
| 0 | 0 | 0 |
| 0 | 0 | 0 |
| 0 | 0 | 0 |
| 0 | 0 | 0 |
| 0 | 0 | 0 |
| 0 | 0 | 0 |
| 0 | 0 | 0 |
| 0 | 0 | 0 |
| 0 | 0 | 0 |
| 0 | 0 | 0 |
| 0 | 0 | 0 |
| 0 | 0 | 0 |
| 1 | 1 | 1 |
| 1 | 1 | 1 |
| 0 | 0 | 0 |
| 0 | 0 | 0 |
| 0 | 0 | 0 |
| 0 | 0 | 0 |
| 0 | 0 | 0 |
| 0 | 0 | 0 |
| 0 | 0 | 0 |
| 0 | 0 | 0 |
| 0 | 0 | 0 |
| 0 | 0 | 0 |
| 0 | 0 | 0 |
| 0 | 0 | 0 |
| 0 | 0 | 0 |
| 0 | 0 | 0 |
| 0 | 0 | 0 |
| 0 | 0 | 0 |
| 0 | 0 | 0 |
| 0 | 0 | 0 |
| 0 | 0 | 0 |
| 1 | 1 | 1 |
| 0 | 1 | 0 |
| 1 | 1 | 1 |
| 1 | 1 | 1 |
| 1 | 1 | 1 |
| 1 | 1 | 1 |
| 0 | 0 | 0 |
| 0 | 0 | 0 |

| | | |
|---|---|---|
| 0 | 0 | 0 |
| 0 | 0 | 0 |
| 0 | 0 | 0 |
| 0 | 0 | 0 |
| 0 | 1 | 0 |
| 0 | 0 | 0 |
| 0 | 0 | 0 |
| 0 | 0 | 0 |
| 1 | 1 | 1 |
| 0 | 0 | 0 |
| 0 | 0 | 0 |
| 0 | 1 | 0 |
| 0 | 0 | 0 |
| 0 | 0 | 0 |
| 0 | 0 | 0 |
| 0 | 0 | 0 |
| 1 | 1 | 1 |
| 0 | 0 | 0 |
| 0 | 0 | 0 |
| 1 | 1 | 1 |
| 0 | 0 | 0 |
| 0 | 0 | 0 |
| 0 | 0 | 0 |
| 0 | 0 | 0 |
| 0 | 0 | 0 |
| 0 | 1 | 0 |
| 0 | 1 | 0 |
| 1 | 1 | 1 |
| 1 | 1 | 1 |
| 0 | 1 | 0 |
| 1 | 1 | 1 |
| 1 | 1 | 1 |
| 1 | 0 | 0 |
| 1 | 0 | 0 |
| 0 | 0 | 0 |
| 0 | 0 | 0 |
| 0 | 0 | 0 |
| 0 | 0 | 0 |
| 0 | 0 | 0 |
| 0 | 0 | 0 |
| 1 | 1 | 1 |
| 1 | 1 | 1 |
| 1 | 1 | 1 |
| 0 | 0 | 0 |
| 0 | 0 | 0 |
| 0 | 0 | 0 |
| 1 | 1 | 1 |
| 0 | 1 | 0 |
| 0 | 0 | 0 |
| 0 | 0 | 0 |
| 1 | 1 | 1 |
| 1 | 1 | 1 |
| 1 | 1 | 1 |
| 1 | 1 | 1 |

| | | |
|---|---|---|
| 0 | 1 | 0 |
| 1 | 1 | 1 |
| 0 | 1 | 0 |
| 0 | 0 | 0 |
| 0 | 0 | 0 |
| 1 | 1 | 1 |
| 1 | 1 | 1 |
| 1 | 1 | 1 |
| 1 | 1 | 1 |
| 1 | 1 | 1 |
| 0 | 0 | 0 |
| 0 | 1 | 0 |
| 0 | 0 | 0 |
| 0 | 0 | 0 |
| 0 | 0 | 0 |
| 1 | 1 | 1 |
| 1 | 1 | 1 |
| 0 | 1 | 0 |
| 1 | 1 | 1 |
| 1 | 1 | 1 |
| 0 | 0 | 0 |
| 1 | 1 | 1 |
| 1 | 1 | 1 |
| 0 | 0 | 0 |
| 1 | 0 | 0 |
| 0 | 0 | 0 |
| 0 | 0 | 0 |
| 1 | 1 | 1 |
| 1 | 1 | 1 |
| 1 | 1 | 1 |
| 1 | 1 | 1 |
| 1 | 1 | 1 |
| 1 | 1 | 1 |
| 1 | 1 | 1 |
| 1 | 1 | 1 |
| 0 | 0 | 0 |
| 1 | 1 | 1 |
| 1 | 1 | 1 |
| 0 | 0 | 0 |
| 0 | 0 | 0 |
| 1 | 1 | 1 |
| 0 | 1 | 0 |
| 0 | 0 | 0 |
| 0 | 0 | 0 |
| 0 | 0 | 0 |
| 0 | 0 | 0 |
| 0 | 0 | 0 |
| 0 | 0 | 0 |
| 1 | 1 | 1 |
| 0 | 0 | 0 |
| 0 | 0 | 0 |
| 0 | 0 | 0 |
| 0 | 0 | 0 |
| 0 | 0 | 0 |

| | | |
|---|---|---|
| 0 | 0 | 0 |
| 0 | 0 | 0 |
| 0 | 0 | 0 |
| 0 | 0 | 0 |
| 0 | 0 | 0 |
| 0 | 0 | 0 |
| 0 | 0 | 0 |
| 0 | 0 | 0 |
| 0 | 0 | 0 |
| 0 | 1 | 0 |
| 0 | 1 | 0 |
| 0 | 0 | 0 |
| 1 | 1 | 1 |
| 0 | 0 | 0 |
| 0 | 0 | 0 |
| 0 | 1 | 0 |
| 0 | 0 | 0 |
| 1 | 0 | 0 |
| 1 | 1 | 1 |
| 1 | 1 | 1 |
| 0 | 0 | 0 |
| 0 | 1 | 0 |
| 0 | 1 | 0 |
| 0 | 0 | 0 |
| 0 | 0 | 0 |
| 0 | 1 | 0 |
| 0 | 1 | 0 |
| 1 | 1 | 1 |
| 0 | 0 | 0 |
| 0 | 0 | 0 |
| 1 | 1 | 1 |
| 1 | 1 | 1 |
| 0 | 0 | 0 |
| 0 | 0 | 0 |
| 0 | 0 | 0 |
| 0 | 1 | 0 |
| 0 | 0 | 0 |
| 0 | 1 | 0 |
| 0 | 1 | 0 |
| 0 | 0 | 0 |
| 0 | 0 | 0 |
| 0 | 0 | 0 |
| 0 | 0 | 0 |
| 0 | 0 | 0 |
| 0 | 0 | 0 |
| 1 | 1 | 1 |
| 0 | 1 | 0 |
| 0 | 0 | 0 |
| 0 | 0 | 0 |
| 0 | 0 | 0 |
| 1 | 1 | 1 |
| 1 | 1 | 1 |
| 0 | 0 | 0 |
| 0 | 0 | 0 |

| | | |
|---|---|---|
| 1 | 1 | 1 |
| 1 | 1 | 1 |
| 1 | 1 | 1 |
| 1 | 1 | 1 |
| 0 | 0 | 0 |
| 0 | 0 | 0 |
| 1 | 0 | 0 |
| 0 | 0 | 0 |
| 1 | 0 | 0 |
| 0 | 0 | 0 |
| 1 | 1 | 1 |
| 1 | 1 | 1 |
| 1 | 1 | 1 |
| 1 | 1 | 1 |
| 1 | 1 | 1 |
| 1 | 1 | 1 |
| 0 | 0 | 0 |
| 1 | 1 | 1 |
| 1 | 1 | 1 |
| 0 | 0 | 0 |
| 0 | 0 | 0 |
| 0 | 1 | 0 |
| 0 | 1 | 0 |
| 0 | 0 | 0 |
| 0 | 1 | 0 |
| 0 | 0 | 0 |
| 0 | 0 | 0 |
| 0 | 0 | 0 |
| 0 | 0 | 0 |
| 0 | 0 | 0 |
| 0 | 0 | 0 |
| 0 | 0 | 0 |
| 0 | 1 | 0 |
| 0 | 0 | 0 |
| 1 | 1 | 1 |
| 1 | 1 | 1 |
| 1 | 1 | 1 |
| 1 | 1 | 1 |
| 0 | 0 | 0 |
| 1 | 1 | 1 |
| 1 | 1 | 1 |
| 1 | 1 | 1 |
| 1 | 1 | 1 |
| 0 | 0 | 0 |
| 1 | 1 | 1 |
| 1 | 1 | 1 |
| 1 | 1 | 1 |
| 1 | 1 | 1 |
| 1 | 1 | 1 |
| 1 | 1 | 1 |
| 0 | 0 | 0 |
| 0 | 0 | 0 |
| 0 | 0 | 0 |
| 0 | 1 | 0 |

| | | |
|---|---|---|
| 0 | 0 | 0 |
| 0 | 0 | 0 |
| 0 | 0 | 0 |
| 0 | 0 | 0 |
| 0 | 0 | 0 |
| 0 | 0 | 0 |
| 0 | 0 | 0 |
| 0 | 0 | 0 |
| 1 | 1 | 1 |
| 1 | 1 | 1 |
| 0 | 0 | 0 |
| 1 | 1 | 1 |
| 1 | 1 | 1 |
| 0 | 0 | 0 |
| 0 | 0 | 0 |
| 0 | 0 | 0 |
| 0 | 0 | 0 |
| 0 | 0 | 0 |
| 0 | 0 | 0 |
| 1 | 1 | 1 |
| 1 | 1 | 1 |
| 0 | 0 | 0 |
| 1 | 1 | 1 |
| 0 | 0 | 0 |
| 1 | 1 | 1 |
| 1 | 1 | 1 |
| 1 | 1 | 1 |
| 0 | 1 | 0 |
| 0 | 1 | 0 |
| 0 | 0 | 0 |
| 0 | 1 | 0 |
| 1 | 1 | 1 |
| 0 | 0 | 0 |
| 1 | 1 | 1 |
| 0 | 0 | 0 |
| 0 | 0 | 0 |
| 0 | 0 | 0 |
| 0 | 0 | 0 |
| 0 | 0 | 0 |
| 0 | 0 | 0 |
| 0 | 0 | 0 |
| 0 | 0 | 0 |
| 1 | 1 | 1 |
| 0 | 0 | 0 |
| 1 | 1 | 1 |
| 0 | 0 | 0 |
| 1 | 0 | 0 |
| 1 | 1 | 1 |
| 1 | 1 | 1 |
| 0 | 0 | 0 |
| 1 | 1 | 1 |
| 1 | 1 | 1 |
| 0 | 1 | 0 |
| 1 | 1 | 1 |

| | | |
|---|---|---|
| 0 | 0 | 0 |
| 0 | 1 | 0 |
| 1 | 1 | 1 |
| 0 | 1 | 0 |
| 0 | 1 | 0 |
| 0 | 1 | 0 |
| 0 | 1 | 0 |
| 1 | 1 | 1 |
| 1 | 1 | 1 |
| 1 | 1 | 1 |
| 0 | 0 | 0 |
| 1 | 1 | 1 |
| 1 | 1 | 1 |
| 0 | 1 | 0 |
| 0 | 0 | 0 |
| 0 | 0 | 0 |
| 0 | 0 | 0 |
| 0 | 0 | 0 |
| 0 | 0 | 0 |
| 0 | 1 | 0 |
| 0 | 1 | 0 |
| 1 | 1 | 1 |
| 0 | 0 | 0 |
| 0 | 1 | 0 |
| 0 | 1 | 0 |
| 0 | 1 | 0 |
| 1 | 1 | 1 |
| 1 | 1 | 1 |
| 0 | 0 | 0 |
| 0 | 0 | 0 |
| 0 | 0 | 0 |
| 0 | 0 | 0 |
| 0 | 0 | 0 |
| 0 | 0 | 0 |
| 1 | 1 | 1 |
| 1 | 1 | 1 |
| 0 | 0 | 0 |
| 0 | 0 | 0 |
| 0 | 0 | 0 |
| 0 | 0 | 0 |
| 0 | 0 | 0 |
| 0 | 0 | 0 |
| 1 | 1 | 1 |
| 1 | 1 | 1 |
| 0 | 0 | 0 |
| 1 | 1 | 1 |
| 1 | 1 | 1 |
| 1 | 1 | 1 |
| 1 | 1 | 1 |
| 1 | 1 | 1 |
| 1 | 0 | 0 |
| 0 | 1 | 0 |
| 0 | 1 | 0 |
| 1 | 0 | 0 |

| | | |
|---|---|---|
| 0 | 0 | 0 |
| 1 | 1 | 1 |
| 0 | 0 | 0 |
| 0 | 1 | 0 |
| 1 | 0 | 0 |
| 0 | 0 | 0 |
| 0 | 0 | 0 |
| 0 | 0 | 0 |
| 1 | 0 | 0 |
| 1 | 0 | 0 |
| 0 | 0 | 0 |
| 1 | 1 | 1 |
| 1 | 0 | 0 |
| 1 | 0 | 0 |
| 0 | 0 | 0 |
| 0 | 0 | 0 |
| 0 | 0 | 0 |
| 0 | 0 | 0 |
| 1 | 1 | 1 |
| 0 | 0 | 0 |
| 0 | 0 | 0 |
| 0 | 0 | 0 |
| 0 | 0 | 0 |
| 0 | 0 | 0 |
| 1 | 1 | 1 |
| 0 | 0 | 0 |
| 1 | 1 | 1 |
| 0 | 1 | 0 |
| 1 | 1 | 1 |
| 0 | 0 | 0 |
| 1 | 1 | 1 |
| 1 | 1 | 1 |
| 0 | 0 | 0 |
| 0 | 0 | 0 |
| 1 | 1 | 1 |
| 1 | 1 | 1 |
| 1 | 1 | 1 |
| 0 | 1 | 0 |
| 1 | 0 | 0 |
| 0 | 0 | 0 |
| 0 | 0 | 0 |
| 0 | 0 | 0 |
| 0 | 0 | 0 |
| 0 | 0 | 0 |
| 0 | 0 | 0 |
| 0 | 1 | 0 |
| 0 | 1 | 0 |
| 0 | 0 | 0 |
| 0 | 0 | 0 |
| 1 | 1 | 1 |
| 0 | 0 | 0 |
| 0 | 0 | 0 |
| 0 | 1 | 0 |
| 1 | 1 | 1 |

| | | |
|---|---|---|
| 1 | 1 | 1 |
| 0 | 0 | 0 |
| 0 | 0 | 0 |
| 1 | 0 | 0 |
| 1 | 1 | 1 |
| 0 | 0 | 0 |
| 0 | 0 | 0 |
| 0 | 0 | 0 |
| 0 | 0 | 0 |
| 0 | 0 | 0 |
| 0 | 0 | 0 |
| 0 | 0 | 0 |
| 0 | 0 | 0 |
| 0 | 0 | 0 |
| 0 | 0 | 0 |
| 0 | 1 | 0 |
| 0 | 0 | 0 |
| 1 | 1 | 1 |
| 1 | 0 | 0 |
| 1 | 1 | 1 |
| 0 | 0 | 0 |
| 1 | 1 | 1 |
| 0 | 0 | 0 |
| 0 | 0 | 0 |
| 1 | 1 | 1 |
| 0 | 0 | 0 |
| 0 | 1 | 0 |
| 0 | 1 | 0 |
| 0 | 1 | 0 |
| 0 | 0 | 0 |
| 0 | 0 | 0 |
| 0 | 0 | 0 |
| 0 | 0 | 0 |
| 0 | 0 | 0 |
| 0 | 0 | 0 |
| 0 | 0 | 0 |
| 0 | 0 | 0 |
| 1 | 1 | 1 |
| 1 | 0 | 0 |
| 0 | 0 | 0 |
| 0 | 0 | 0 |
| 0 | 0 | 0 |
| 0 | 0 | 0 |
| 0 | 0 | 0 |
| 0 | 0 | 0 |
| 0 | 0 | 0 |
| 1 | 1 | 1 |
| 0 | 0 | 0 |
| 0 | 0 | 0 |
| 0 | 0 | 0 |
| 0 | 0 | 0 |
| 0 | 1 | 0 |
| 0 | 0 | 0 |
| 0 | 0 | 0 |

| | | |
|---|---|---|
| 0 | 0 | 0 |
| 0 | 0 | 0 |
| 0 | 0 | 0 |
| 0 | 0 | 0 |
| 0 | 0 | 0 |
| 0 | 0 | 0 |
| 0 | 0 | 0 |
| 0 | 0 | 0 |
| 0 | 0 | 0 |
| 0 | 0 | 0 |
| 1 | 1 | 1 |
| 0 | 0 | 0 |
| 1 | 0 | 0 |
| 0 | 0 | 0 |
| 0 | 1 | 0 |
| 1 | 1 | 1 |
| 1 | 1 | 1 |
| 1 | 1 | 1 |
| 0 | 0 | 0 |
| 0 | 0 | 0 |
| 1 | 1 | 1 |
| 0 | 0 | 0 |
| 0 | 0 | 0 |
| 1 | 1 | 1 |
| 0 | 0 | 0 |
| 0 | 0 | 0 |
| 0 | 0 | 0 |
| 1 | 0 | 0 |
| 0 | 0 | 0 |
| 0 | 0 | 0 |
| 1 | 0 | 0 |
| 0 | 0 | 0 |
| 1 | 0 | 0 |
| 1 | 0 | 0 |
| 0 | 0 | 0 |
| 0 | 0 | 0 |
| 0 | 0 | 0 |
| 1 | 0 | 0 |
| 1 | 0 | 0 |
| 0 | 0 | 0 |
| 0 | 0 | 0 |
| 0 | 0 | 0 |
| 0 | 0 | 0 |
| 0 | 0 | 0 |
| 0 | 0 | 0 |
| 0 | 0 | 0 |
| 0 | 0 | 0 |
| 0 | 0 | 0 |
| 0 | 0 | 0 |
| 0 | 0 | 0 |
| 0 | 0 | 0 |
| 0 | 0 | 0 |
| 0 | 0 | 0 |
| 0 | 0 | 0 |

| | | |
|---|---|---|
| 0 | 0 | 0 |
| 1 | 0 | 0 |
| 0 | 0 | 0 |
| 0 | 0 | 0 |
| 0 | 0 | 0 |
| 1 | 0 | 0 |
| 0 | 0 | 0 |
| 0 | 0 | 0 |
| 1 | 1 | 1 |
| 0 | 0 | 0 |
| 0 | 1 | 0 |
| 1 | 1 | 1 |
| 1 | 1 | 1 |
| 0 | 0 | 0 |
| 1 | 1 | 1 |
| 0 | 0 | 0 |
| 1 | 1 | 1 |
| 1 | 1 | 1 |
| 0 | 0 | 0 |
| 0 | 0 | 0 |
| 0 | 0 | 0 |
| 0 | 0 | 0 |
| 0 | 0 | 0 |
| 0 | 0 | 0 |
| 0 | 0 | 0 |
| 0 | 1 | 0 |
| 1 | 1 | 1 |
| 1 | 1 | 1 |
| 1 | 0 | 0 |
| 1 | 1 | 1 |
| 1 | 1 | 1 |
| 0 | 1 | 0 |
| 0 | 0 | 0 |
| 0 | 0 | 0 |
| 0 | 0 | 0 |
| 0 | 0 | 0 |
| 0 | 0 | 0 |
| 0 | 0 | 0 |
| 0 | 1 | 0 |
| 0 | 1 | 0 |
| 0 | 1 | 0 |
| 0 | 1 | 0 |
| 0 | 1 | 0 |
| 0 | 1 | 0 |
| 0 | 0 | 0 |
| 0 | 0 | 0 |
| 0 | 1 | 0 |
| 0 | 0 | 0 |
| 0 | 0 | 0 |
| 0 | 0 | 0 |
| 0 | 0 | 0 |
| 0 | 0 | 0 |
| 0 | 0 | 0 |
| 0 | 0 | 0 |

| | | |
|---|---|---|
| 1 | 1 | 1 |
| 1 | 1 | 1 |
| 0 | 0 | 0 |
| 0 | 0 | 0 |
| 1 | 1 | 1 |
| 0 | 0 | 0 |
| 0 | 0 | 0 |
| 1 | 1 | 1 |
| 1 | 1 | 1 |
| 0 | 0 | 0 |
| 1 | 1 | 1 |
| 1 | 1 | 1 |
| 0 | 0 | 0 |
| 0 | 0 | 0 |
| 0 | 0 | 0 |
| 0 | 0 | 0 |
| 0 | 1 | 0 |
| 0 | 1 | 0 |
| 1 | 1 | 1 |
| 1 | 1 | 1 |
| 1 | 1 | 1 |
| 0 | 1 | 0 |
| 1 | 1 | 1 |
| 1 | 1 | 1 |
| 1 | 1 | 1 |
| 0 | 0 | 0 |
| 1 | 1 | 1 |
| 1 | 1 | 1 |
| 1 | 1 | 1 |
| 0 | 1 | 0 |
| 0 | 1 | 0 |
| 1 | 1 | 1 |
| 1 | 1 | 1 |
| 1 | 1 | 1 |
| 1 | 1 | 1 |
| 1 | 1 | 1 |
| 1 | 1 | 1 |
| 0 | 0 | 0 |
| 0 | 0 | 0 |
| 1 | 0 | 0 |
| 0 | 0 | 0 |
| 1 | 1 | 1 |
| 1 | 1 | 1 |
| 0 | 0 | 0 |
| 0 | 0 | 0 |
| 0 | 1 | 0 |
| 0 | 1 | 0 |
| 0 | 1 | 0 |
| 1 | 1 | 1 |
| 1 | 1 | 1 |
| 1 | 1 | 1 |
| 0 | 0 | 0 |
| 0 | 0 | 0 |
| 0 | 0 | 0 |

| | | |
|---|---|---|
| 0 | 0 | 0 |
| 1 | 1 | 1 |
| 1 | 1 | 1 |
| 1 | 1 | 1 |
| 1 | 1 | 1 |
| 1 | 1 | 1 |
| 1 | 1 | 1 |
| 1 | 1 | 1 |
| 0 | 0 | 0 |
| 1 | 1 | 1 |
| 1 | 1 | 1 |
| 1 | 1 | 1 |
| 0 | 1 | 0 |
| 0 | 1 | 0 |
| 1 | 1 | 1 |
| 1 | 1 | 1 |
| 0 | 1 | 0 |
| 0 | 1 | 0 |
| 0 | 0 | 0 |
| 0 | 0 | 0 |
| 0 | 0 | 0 |
| 0 | 0 | 0 |
| 0 | 1 | 0 |
| 0 | 0 | 0 |
| 0 | 0 | 0 |
| 1 | 1 | 1 |
| 1 | 1 | 1 |
| 1 | 1 | 1 |
| 1 | 1 | 1 |
| 1 | 1 | 1 |
| 1 | 1 | 1 |
| 1 | 1 | 1 |
| 0 | 0 | 0 |
| 1 | 1 | 1 |
| 1 | 1 | 1 |
| 0 | 1 | 0 |
| 0 | 0 | 0 |
| 1 | 1 | 1 |
| 0 | 0 | 0 |
| 0 | 0 | 0 |
| 0 | 0 | 0 |
| 0 | 0 | 0 |
| 0 | 0 | 0 |
| 1 | 1 | 1 |
| 1 | 1 | 1 |
| 1 | 1 | 1 |
| 0 | 0 | 0 |
| 0 | 0 | 0 |
| 0 | 0 | 0 |
| 0 | 0 | 0 |
| 0 | 0 | 0 |
| 0 | 0 | 0 |
| 0 | 0 | 0 |
| 0 | 1 | 0 |

| | | |
|---|---|---|
| 0 | 0 | 0 |
| 0 | 0 | 0 |
| 0 | 1 | 0 |
| 0 | 0 | 0 |
| 0 | 0 | 0 |
| 1 | 1 | 1 |
| 1 | 1 | 1 |
| 1 | 0 | 0 |
| 1 | 1 | 1 |
| 1 | 1 | 1 |
| 1 | 1 | 1 |
| 0 | 0 | 0 |
| 1 | 1 | 1 |
| 1 | 1 | 1 |
| 0 | 0 | 0 |
| 1 | 1 | 1 |
| 0 | 0 | 0 |
| 1 | 1 | 1 |
| 0 | 0 | 0 |
| 0 | 0 | 0 |
| 1 | 1 | 1 |
| 1 | 1 | 1 |
| 0 | 0 | 0 |
| 1 | 1 | 1 |
| 0 | 0 | 0 |
| 1 | 1 | 1 |
| 1 | 1 | 1 |
| 0 | 0 | 0 |
| 1 | 1 | 1 |
| 0 | 0 | 0 |
| 0 | 1 | 0 |
| 0 | 0 | 0 |
| 0 | 0 | 0 |
| 0 | 0 | 0 |
| 1 | 1 | 1 |
| 0 | 0 | 0 |
| 1 | 1 | 1 |
| 1 | 1 | 1 |
| 0 | 0 | 0 |
| 1 | 1 | 1 |
| 1 | 1 | 1 |
| 1 | 1 | 1 |
| 1 | 1 | 1 |
| 0 | 0 | 0 |
| 0 | 0 | 0 |
| 1 | 1 | 1 |
| 1 | 1 | 1 |
| 1 | 0 | 0 |
| 0 | 0 | 0 |
| 1 | 0 | 0 |
| 0 | 0 | 0 |
| 1 | 1 | 1 |
| 1 | 1 | 1 |
| 0 | 0 | 0 |

| | | |
|---|---|---|
| 1 | 1 | 1 |
| 1 | 1 | 1 |
| 1 | 1 | 1 |
| 1 | 1 | 1 |
| 0 | 1 | 0 |
| 1 | 1 | 1 |
| 1 | 1 | 1 |
| 0 | 0 | 0 |
| 1 | 1 | 1 |
| 1 | 1 | 1 |
| 1 | 1 | 1 |
| 1 | 1 | 1 |
| 0 | 0 | 0 |
| 1 | 1 | 1 |
| 0 | 0 | 0 |
| 0 | 0 | 0 |
| 1 | 1 | 1 |
| 1 | 1 | 1 |
| 1 | 1 | 1 |
| 1 | 1 | 1 |
| 0 | 0 | 0 |
| 0 | 0 | 0 |
| 1 | 1 | 1 |
| 1 | 1 | 1 |
| 0 | 0 | 0 |
| 1 | 1 | 1 |
| 0 | 0 | 0 |
| 0 | 0 | 0 |
| 0 | 0 | 0 |
| 0 | 0 | 0 |
| 0 | 1 | 0 |
| 0 | 1 | 0 |
| 0 | 0 | 0 |
| 0 | 0 | 0 |
| 0 | 0 | 0 |
| 0 | 0 | 0 |
| 0 | 0 | 0 |
| 0 | 0 | 0 |
| 0 | 0 | 0 |
| 1 | 0 | 0 |
| 1 | 1 | 1 |
| 0 | 1 | 0 |
| 0 | 1 | 0 |
| 0 | 1 | 0 |
| 0 | 1 | 0 |
| 0 | 0 | 0 |
| 0 | 1 | 0 |
| 0 | 1 | 0 |
| 1 | 1 | 1 |
| 0 | 0 | 0 |
| 0 | 0 | 0 |
| 1 | 1 | 1 |
| 1 | 1 | 1 |
| 1 | 1 | 1 |

| | | |
|---|---|---|
| 1 | 1 | 1 |
| 1 | 1 | 1 |
| 1 | 1 | 1 |
| 1 | 1 | 1 |
| 0 | 0 | 0 |
| 1 | 1 | 1 |
| 0 | 0 | 0 |
| 0 | 0 | 0 |
| 0 | 0 | 0 |
| 0 | 0 | 0 |
| 0 | 0 | 0 |
| 1 | 1 | 1 |
| 0 | 0 | 0 |
| 0 | 0 | 0 |
| 1 | 1 | 1 |
| 1 | 1 | 1 |
| 1 | 1 | 1 |
| 1 | 1 | 1 |
| 1 | 1 | 1 |
| 1 | 1 | 1 |
| 1 | 1 | 1 |
| 1 | 1 | 1 |
| 1 | 1 | 1 |
| 0 | 0 | 0 |
| 1 | 1 | 1 |
| 1 | 1 | 1 |
| 1 | 1 | 1 |
| 0 | 0 | 0 |
| 1 | 0 | 0 |
| 1 | 0 | 0 |
| 0 | 0 | 0 |
| 0 | 0 | 0 |
| 1 | 1 | 1 |
| 0 | 0 | 0 |
| 0 | 1 | 0 |
| 1 | 1 | 1 |
| 1 | 1 | 1 |
| 1 | 1 | 1 |
| 1 | 1 | 1 |
| 1 | 1 | 1 |
| 1 | 1 | 1 |
| 1 | 1 | 1 |
| 1 | 1 | 1 |
| 1 | 1 | 1 |
| 1 | 1 | 1 |
| 1 | 1 | 1 |
| 1 | 1 | 1 |
| 1 | 1 | 1 |
| 0 | 0 | 0 |
| 0 | 1 | 0 |
| 1 | 1 | 1 |
| 1 | 0 | 0 |
| 1 | 0 | 0 |
| 1 | 1 | 1 |

| | | |
|---|---|---|
| 1 | 1 | 1 |
| 0 | 1 | 0 |
| 1 | 1 | 1 |
| 0 | 1 | 0 |
| 1 | 0 | 0 |
| 1 | 1 | 1 |
| 1 | 1 | 1 |
| 1 | 1 | 1 |
| 1 | 1 | 1 |
| 1 | 1 | 1 |
| 0 | 0 | 0 |
| 0 | 0 | 0 |
| 0 | 0 | 0 |
| 0 | 0 | 0 |
| 0 | 0 | 0 |
| 1 | 0 | 0 |
| 0 | 0 | 0 |
| 0 | 0 | 0 |
| 0 | 0 | 0 |
| 0 | 0 | 0 |
| 0 | 0 | 0 |
| 0 | 0 | 0 |
| 0 | 1 | 0 |
| 0 | 0 | 0 |
| 0 | 0 | 0 |
| 0 | 0 | 0 |
| 1 | 1 | 1 |
| 0 | 0 | 0 |
| 0 | 1 | 0 |
| 0 | 1 | 0 |
| 0 | 1 | 0 |
| 0 | 0 | 0 |
| 1 | 1 | 1 |
| 1 | 1 | 1 |
| 0 | 0 | 0 |
| 0 | 0 | 0 |
| 0 | 0 | 0 |
| 0 | 0 | 0 |
| 0 | 0 | 0 |
| 1 | 0 | 0 |
| 1 | 0 | 0 |
| 0 | 0 | 0 |
| 1 | 0 | 0 |
| 0 | 0 | 0 |
| 1 | 1 | 1 |
| 1 | 0 | 0 |
| 1 | 1 | 1 |
| 0 | 0 | 0 |
| 0 | 0 | 0 |
| 0 | 0 | 0 |
| 0 | 0 | 0 |
| 1 | 1 | 1 |
| 0 | 0 | 0 |
| 1 | 1 | 1 |

| | | |
|---|---|---|
| 1 | 1 | 1 |
| 1 | 1 | 1 |
| 1 | 1 | 1 |
| 1 | 1 | 1 |
| 1 | 1 | 1 |
| 0 | 0 | 0 |
| 0 | 0 | 0 |
| 0 | 0 | 0 |
| 0 | 0 | 0 |
| 0 | 1 | 0 |
| 0 | 0 | 0 |
| 0 | 0 | 0 |
| 0 | 0 | 0 |
| 0 | 0 | 0 |
| 0 | 0 | 0 |
| 0 | 0 | 0 |
| 1 | 1 | 1 |
| 0 | 1 | 0 |
| 1 | 1 | 1 |
| 1 | 1 | 1 |
| 1 | 1 | 1 |
| 1 | 1 | 1 |
| 1 | 1 | 1 |
| 1 | 1 | 1 |
| 1 | 1 | 1 |
| 1 | 1 | 1 |
| 1 | 1 | 1 |
| 1 | 1 | 1 |
| 0 | 0 | 0 |
| 0 | 1 | 0 |
| 0 | 1 | 0 |
| 1 | 0 | 0 |
| 1 | 0 | 0 |
| 0 | 0 | 0 |
| 1 | 1 | 1 |
| 0 | 0 | 0 |
| 1 | 0 | 0 |
| 0 | 0 | 0 |
| 1 | 1 | 1 |
| 0 | 0 | 0 |
| 1 | 1 | 1 |
| 1 | 1 | 1 |
| 0 | 1 | 0 |
| 0 | 0 | 0 |
| 1 | 1 | 1 |
| 1 | 1 | 1 |
| 1 | 1 | 1 |
| 1 | 1 | 1 |
| 1 | 1 | 1 |
| 1 | 1 | 1 |
| 1 | 1 | 1 |
| 1 | 0 | 0 |
| 0 | 0 | 0 |
| 0 | 0 | 0 |

| | | |
|---|---|---|
| 1 | 1 | 1 |
| 0 | 0 | 0 |
| 1 | 1 | 1 |
| 1 | 1 | 1 |
| 0 | 0 | 0 |
| 0 | 0 | 0 |
| 0 | 0 | 0 |
| 1 | 1 | 1 |
| 1 | 1 | 1 |
| 0 | 0 | 0 |
| 0 | 0 | 0 |
| 0 | 0 | 0 |
| 0 | 0 | 0 |
| 0 | 0 | 0 |
| 1 | 1 | 1 |
| 0 | 0 | 0 |
| 0 | 0 | 0 |
| 0 | 0 | 0 |
| 1 | 1 | 1 |
| 1 | 1 | 1 |
| 1 | 1 | 1 |
| 1 | 1 | 1 |
| 1 | 1 | 1 |
| 0 | 0 | 0 |
| 0 | 1 | 0 |
| 0 | 1 | 0 |
| 0 | 0 | 0 |
| 0 | 0 | 0 |
| 1 | 1 | 1 |
| 1 | 1 | 1 |
| 0 | 0 | 0 |
| 1 | 1 | 1 |
| 1 | 1 | 1 |
| 0 | 1 | 0 |
| 1 | 1 | 1 |
| 0 | 0 | 0 |
| 0 | 1 | 0 |
| 0 | 0 | 0 |
| 0 | 0 | 0 |
| 0 | 1 | 0 |
| 0 | 1 | 0 |
| 0 | 0 | 0 |
| 0 | 0 | 0 |
| 0 | 1 | 0 |
| 0 | 0 | 0 |
| 0 | 0 | 0 |
| 0 | 0 | 0 |
| 0 | 1 | 0 |
| 0 | 0 | 0 |
| 0 | 0 | 0 |
| 0 | 0 | 0 |
| 1 | 1 | 1 |
| 0 | 0 | 0 |
| 1 | 1 | 1 |

| | | |
|---|---|---|
| 1 | 1 | 1 |
| 1 | 1 | 1 |
| 1 | 1 | 1 |
| 1 | 1 | 1 |
| 0 | 0 | 0 |
| 0 | 0 | 0 |
| 1 | 1 | 1 |
| 1 | 0 | 0 |
| 1 | 1 | 1 |
| 0 | 0 | 0 |
| 1 | 0 | 0 |
| 1 | 0 | 0 |
| 0 | 0 | 0 |
| 1 | 1 | 1 |
| 1 | 1 | 1 |
| 1 | 1 | 1 |
| 1 | 1 | 1 |
| 1 | 1 | 1 |
| 1 | 1 | 1 |
| 1 | 1 | 1 |
| 1 | 1 | 1 |
| 0 | 1 | 0 |
| 1 | 1 | 1 |
| 0 | 0 | 0 |
| 1 | 1 | 1 |
| 0 | 0 | 0 |
| 1 | 1 | 1 |
| 0 | 0 | 0 |
| 1 | 1 | 1 |
| 0 | 0 | 0 |
| 0 | 0 | 0 |
| 0 | 0 | 0 |
| 0 | 0 | 0 |
| 0 | 0 | 0 |
| 0 | 1 | 0 |
| 0 | 0 | 0 |
| 1 | 1 | 1 |
| 0 | 0 | 0 |
| 1 | 1 | 1 |
| 1 | 1 | 1 |
| 0 | 0 | 0 |
| 0 | 0 | 0 |
| 0 | 0 | 0 |
| 0 | 0 | 0 |
| 0 | 0 | 0 |
| 0 | 0 | 0 |
| 0 | 0 | 0 |
| 0 | 0 | 0 |
| 1 | 1 | 1 |
| 0 | 0 | 0 |
| 1 | 0 | 0 |
| 1 | 0 | 0 |
| 1 | 1 | 1 |
| 1 | 1 | 1 |

| | | |
|---|---|---|
| 1 | 1 | 1 |
| 1 | 1 | 1 |
| 1 | 1 | 1 |
| 0 | 1 | 0 |
| 0 | 1 | 0 |
| 0 | 1 | 0 |
| 0 | 0 | 0 |
| 0 | 0 | 0 |
| 0 | 0 | 0 |
| 0 | 0 | 0 |
| 0 | 0 | 0 |
| 0 | 0 | 0 |
| 0 | 0 | 0 |
| 0 | 0 | 0 |
| 0 | 0 | 0 |
| 0 | 0 | 0 |
| 0 | 0 | 0 |
| 0 | 0 | 0 |
| 1 | 1 | 1 |
| 0 | 0 | 0 |
| 0 | 1 | 0 |
| 0 | 1 | 0 |
| 0 | 1 | 0 |
| 1 | 1 | 1 |
| 0 | 1 | 0 |
| 0 | 0 | 0 |
| 0 | 0 | 0 |
| 0 | 0 | 0 |
| 0 | 0 | 0 |
| 0 | 0 | 0 |
| 0 | 0 | 0 |
| 0 | 0 | 0 |
| 0 | 0 | 0 |
| 0 | 0 | 0 |
| 1 | 1 | 1 |
| 0 | 0 | 0 |
| 0 | 0 | 0 |
| 0 | 0 | 0 |
| 0 | 0 | 0 |
| 0 | 1 | 0 |
| 0 | 0 | 0 |
| 0 | 0 | 0 |
| 1 | 1 | 1 |
| 0 | 1 | 0 |
| 1 | 1 | 1 |
| 0 | 1 | 0 |
| 0 | 1 | 0 |
| 0 | 0 | 0 |
| 0 | 0 | 0 |
| 0 | 0 | 0 |
| 0 | 0 | 0 |
| 0 | 1 | 0 |
| 0 | 0 | 0 |
| 1 | 1 | 1 |

| | | |
|---|---|---|
| 0 | 1 | 0 |
| 1 | 1 | 1 |
| 1 | 1 | 1 |
| 1 | 1 | 1 |
| 1 | 1 | 1 |
| 0 | 0 | 0 |
| 0 | 1 | 0 |
| 1 | 1 | 1 |
| 1 | 1 | 1 |
| 1 | 1 | 1 |
| 0 | 0 | 0 |
| 0 | 0 | 0 |
| 0 | 0 | 0 |
| 0 | 0 | 0 |
| 0 | 0 | 0 |
| 0 | 1 | 0 |
| 1 | 0 | 0 |
| 0 | 0 | 0 |
| 1 | 1 | 1 |
| 1 | 1 | 1 |
| 1 | 1 | 1 |
| 1 | 1 | 1 |
| 0 | 0 | 0 |
| 1 | 1 | 1 |
| 0 | 0 | 0 |
| 0 | 0 | 0 |
| 0 | 0 | 0 |
| 0 | 0 | 0 |
| 1 | 1 | 1 |
| 0 | 0 | 0 |
| 1 | 1 | 1 |
| 0 | 1 | 0 |
| 0 | 1 | 0 |
| 0 | 1 | 0 |
| 1 | 1 | 1 |
| 1 | 1 | 1 |
| 0 | 1 | 0 |
| 0 | 0 | 0 |
| 0 | 0 | 0 |
| 0 | 0 | 0 |
| 0 | 0 | 0 |
| 0 | 0 | 0 |
| 1 | 1 | 1 |
| 0 | 0 | 0 |
| 0 | 0 | 0 |
| 1 | 1 | 1 |
| 0 | 0 | 0 |
| 0 | 0 | 0 |
| 0 | 0 | 0 |
| 0 | 0 | 0 |
| 1 | 1 | 1 |
| 1 | 1 | 1 |
| 1 | 1 | 1 |
| 0 | 0 | 0 |

| | | |
|---|---|---|
| 0 | 0 | 0 |
| 1 | 1 | 1 |
| 0 | 0 | 0 |
| 0 | 1 | 0 |
| 1 | 0 | 0 |
| 1 | 0 | 0 |
| 1 | 0 | 0 |
| 1 | 1 | 1 |
| 0 | 0 | 0 |
| 1 | 1 | 1 |
| 0 | 0 | 0 |
| 0 | 0 | 0 |
| 0 | 0 | 0 |
| 0 | 0 | 0 |
| 1 | 1 | 1 |
| 1 | 1 | 1 |
| 0 | 0 | 0 |
| 0 | 0 | 0 |
| 0 | 1 | 0 |
| 1 | 1 | 1 |
| 0 | 0 | 0 |
| 1 | 0 | 0 |
| 1 | 0 | 0 |
| 1 | 1 | 1 |
| 0 | 0 | 0 |
| 1 | 0 | 0 |
| 0 | 0 | 0 |
| 0 | 0 | 0 |
| 0 | 0 | 0 |
| 1 | 1 | 1 |
| 0 | 0 | 0 |
| 0 | 0 | 0 |
| 0 | 0 | 0 |
| 1 | 1 | 1 |
| 0 | 0 | 0 |
| 1 | 1 | 1 |
| 0 | 0 | 0 |
| 0 | 0 | 0 |
| 0 | 0 | 0 |
| 0 | 0 | 0 |
| 0 | 0 | 0 |
| 1 | 1 | 1 |
| 0 | 0 | 0 |
| 0 | 0 | 0 |
| 0 | 0 | 0 |
| 0 | 0 | 0 |
| 0 | 0 | 0 |
| 0 | 0 | 0 |
| 0 | 0 | 0 |
| 0 | 0 | 0 |
| 0 | 0 | 0 |
| 0 | 0 | 0 |
| 1 | 1 | 1 |
| 0 | 0 | 0 |

| | | |
|---|---|---|
| 0 | 1 | 0 |
| 1 | 1 | 1 |
| 1 | 1 | 1 |
| 1 | 1 | 1 |
| 1 | 1 | 1 |
| 1 | 1 | 1 |
| 1 | 1 | 1 |
| 1 | 1 | 1 |
| 1 | 1 | 1 |
| 0 | 0 | 0 |
| 0 | 0 | 0 |
| 1 | 1 | 1 |
| 1 | 1 | 1 |
| 1 | 1 | 1 |
| 0 | 0 | 0 |
| 1 | 1 | 1 |
| 1 | 1 | 1 |
| 0 | 0 | 0 |
| 0 | 1 | 0 |
| 0 | 1 | 0 |
| 0 | 1 | 0 |
| 0 | 0 | 0 |
| 0 | 0 | 0 |
| 0 | 0 | 0 |
| 0 | 0 | 0 |
| 0 | 0 | 0 |
| 0 | 0 | 0 |
| 0 | 0 | 0 |
| 1 | 1 | 1 |
| 0 | 0 | 0 |
| 0 | 0 | 0 |
| 1 | 1 | 1 |
| 1 | 1 | 1 |
| 1 | 1 | 1 |
| 1 | 1 | 1 |
| 1 | 1 | 1 |
| 1 | 1 | 1 |
| 1 | 1 | 1 |
| 1 | 1 | 1 |
| 1 | 1 | 1 |
| 1 | 1 | 1 |
| 1 | 1 | 1 |
| 1 | 1 | 1 |
| 0 | 0 | 0 |
| 1 | 1 | 1 |
| 0 | 0 | 0 |
| 1 | 1 | 1 |
| 1 | 1 | 1 |
| 1 | 1 | 1 |
| 1 | 1 | 1 |
| 0 | 0 | 0 |
| 1 | 1 | 1 |
| 1 | 1 | 1 |
| 0 | 0 | 0 |

| | | |
|---|---|---|
| 0 | 0 | 0 |
| 0 | 0 | 0 |
| 0 | 0 | 0 |
| 0 | 0 | 0 |
| 1 | 1 | 1 |
| 0 | 1 | 0 |
| 1 | 1 | 1 |
| 0 | 0 | 0 |
| 0 | 0 | 0 |
| 0 | 0 | 0 |
| 0 | 0 | 0 |
| 1 | 1 | 1 |
| 1 | 1 | 1 |
| 1 | 1 | 1 |
| 1 | 1 | 1 |
| 0 | 0 | 0 |
| 0 | 0 | 0 |
| 0 | 1 | 0 |
| 0 | 1 | 0 |
| 1 | 1 | 1 |
| 1 | 1 | 1 |
| 0 | 0 | 0 |
| 1 | 1 | 1 |
| 0 | 1 | 0 |
| 0 | 0 | 0 |
| 0 | 0 | 0 |
| 0 | 0 | 0 |
| 0 | 0 | 0 |
| 1 | 1 | 1 |
| 1 | 1 | 1 |
| 1 | 1 | 1 |
| 0 | 0 | 0 |
| 1 | 1 | 1 |
| 1 | 1 | 1 |
| 1 | 1 | 1 |
| 1 | 1 | 1 |
| 1 | 1 | 1 |
| 1 | 1 | 1 |
| 1 | 1 | 1 |
| 1 | 1 | 1 |
| 1 | 1 | 1 |
| 1 | 1 | 1 |
| 0 | 0 | 0 |
| 0 | 1 | 0 |
| 0 | 1 | 0 |
| 1 | 1 | 1 |
| 0 | 1 | 0 |
| 1 | 1 | 1 |
| 1 | 1 | 1 |
| 0 | 0 | 0 |
| 1 | 1 | 1 |
| 1 | 1 | 1 |
| 1 | 1 | 1 |
| 1 | 1 | 1 |

| | | |
|---|---|---|
| 0 | 0 | 0 |
| 1 | 1 | 1 |
| 0 | 0 | 0 |
| 0 | 1 | 0 |
| 1 | 1 | 1 |
| 1 | 1 | 1 |
| 1 | 1 | 1 |
| 0 | 0 | 0 |
| 1 | 1 | 1 |
| 1 | 1 | 1 |
| 0 | 0 | 0 |
| 0 | 0 | 0 |
| 0 | 0 | 0 |
| 1 | 1 | 1 |
| 0 | 0 | 0 |
| 0 | 0 | 0 |
| 0 | 0 | 0 |
| 0 | 0 | 0 |
| 0 | 1 | 0 |
| 0 | 0 | 0 |
| 0 | 0 | 0 |
| 0 | 0 | 0 |
| 0 | 0 | 0 |
| 0 | 0 | 0 |
| 0 | 0 | 0 |
| 0 | 0 | 0 |
| 0 | 0 | 0 |
| 0 | 0 | 0 |
| 0 | 0 | 0 |
| 0 | 0 | 0 |
| 0 | 0 | 0 |
| 0 | 0 | 0 |
| 1 | 1 | 1 |
| 0 | 0 | 0 |
| 1 | 1 | 1 |
| 0 | 0 | 0 |
| 0 | 0 | 0 |
| 1 | 1 | 1 |
| 1 | 1 | 1 |
| 0 | 1 | 0 |
| 0 | 1 | 0 |
| 0 | 0 | 0 |
| 0 | 0 | 0 |
| 0 | 0 | 0 |
| 1 | 1 | 1 |
| 0 | 0 | 0 |
| 1 | 1 | 1 |
| 0 | 0 | 0 |
| 0 | 1 | 0 |
| 0 | 1 | 0 |
| 0 | 1 | 0 |
| 0 | 1 | 0 |
| 0 | 1 | 0 |
| 1 | 1 | 1 |

| | | |
|---|---|---|
| 1 | 1 | 1 |
| 1 | 1 | 1 |
| 0 | 1 | 0 |
| 1 | 1 | 1 |
| 1 | 1 | 1 |
| 0 | 0 | 0 |
| 0 | 0 | 0 |
| 1 | 1 | 1 |
| 0 | 0 | 0 |
| 0 | 0 | 0 |
| 1 | 1 | 1 |
| 0 | 1 | 0 |
| 1 | 1 | 1 |
| 1 | 1 | 1 |
| 1 | 1 | 1 |
| 0 | 0 | 0 |
| 1 | 1 | 1 |
| 1 | 1 | 1 |
| 0 | 0 | 0 |
| 0 | 0 | 0 |
| 0 | 0 | 0 |
| 1 | 1 | 1 |
| 1 | 1 | 1 |
| 0 | 0 | 0 |
| 1 | 1 | 1 |
| 0 | 0 | 0 |
| 0 | 0 | 0 |
| 1 | 1 | 1 |
| 1 | 1 | 1 |
| 1 | 1 | 1 |
| 0 | 0 | 0 |
| 1 | 0 | 0 |
| 0 | 0 | 0 |
| 0 | 0 | 0 |
| 0 | 0 | 0 |
| 0 | 0 | 0 |
| 0 | 0 | 0 |
| 1 | 1 | 1 |
| 0 | 0 | 0 |
| 0 | 1 | 0 |
| 0 | 0 | 0 |
| 0 | 0 | 0 |
| 0 | 0 | 0 |
| 0 | 0 | 0 |
| 1 | 1 | 1 |
| 0 | 1 | 0 |
| 0 | 1 | 0 |
| 0 | 0 | 0 |
| 0 | 0 | 0 |
| 0 | 0 | 0 |
| 1 | 1 | 1 |
| 0 | 1 | 0 |
| 1 | 1 | 1 |
| 1 | 1 | 1 |

| | | |
|---|---|---|
| 1 | 1 | 1 |
| 0 | 1 | 0 |
| 0 | 1 | 0 |
| 0 | 0 | 0 |
| 0 | 0 | 0 |
| 1 | 1 | 1 |
| 0 | 0 | 0 |
| 1 | 1 | 1 |
| 1 | 1 | 1 |
| 1 | 1 | 1 |
| 1 | 1 | 1 |
| 0 | 0 | 0 |
| 1 | 0 | 0 |
| 0 | 0 | 0 |
| 1 | 1 | 1 |
| 0 | 0 | 0 |
| 0 | 0 | 0 |
| 0 | 0 | 0 |
| 1 | 1 | 1 |
| 0 | 0 | 0 |
| 0 | 0 | 0 |
| 0 | 0 | 0 |
| 0 | 0 | 0 |
| 0 | 0 | 0 |
| 0 | 0 | 0 |
| 1 | 1 | 1 |
| 0 | 0 | 0 |
| 0 | 1 | 0 |
| 0 | 1 | 0 |
| 0 | 1 | 0 |
| 0 | 1 | 0 |
| 0 | 1 | 0 |
| 0 | 0 | 0 |
| 0 | 0 | 0 |
| 1 | 1 | 1 |
| 1 | 1 | 1 |
| 0 | 0 | 0 |
| 0 | 0 | 0 |
| 0 | 0 | 0 |
| 1 | 0 | 0 |
| 0 | 0 | 0 |
| 0 | 0 | 0 |
| 1 | 1 | 1 |
| 0 | 0 | 0 |
| 1 | 1 | 1 |
| 1 | 1 | 1 |
| 1 | 1 | 1 |
| 1 | 1 | 1 |
| 1 | 1 | 1 |
| 0 | 0 | 0 |
| 1 | 1 | 1 |
| 1 | 1 | 1 |
| 1 | 1 | 1 |
| 0 | 1 | 0 |

| | | |
|---|---|---|
| 1 | 1 | 1 |
| 1 | 1 | 1 |
| 1 | 1 | 1 |
| 0 | 0 | 0 |
| 1 | 1 | 1 |
| 1 | 1 | 1 |
| 0 | 1 | 0 |
| 0 | 0 | 0 |
| 1 | 1 | 1 |
| 1 | 1 | 1 |
| 1 | 1 | 1 |
| 1 | 1 | 1 |
| 1 | 1 | 1 |
| 0 | 1 | 0 |
| 0 | 1 | 0 |
| 0 | 1 | 0 |
| 0 | 0 | 0 |
| 0 | 0 | 0 |
| 0 | 1 | 0 |
| 1 | 1 | 1 |
| 0 | 1 | 0 |
| 1 | 1 | 1 |
| 0 | 1 | 0 |
| 1 | 1 | 1 |
| 1 | 1 | 1 |
| 1 | 1 | 1 |
| 1 | 1 | 1 |
| 0 | 0 | 0 |
| 0 | 0 | 0 |
| 1 | 1 | 1 |
| 1 | 1 | 1 |
| 0 | 0 | 0 |
| 0 | 1 | 0 |
| 0 | 0 | 0 |
| 0 | 0 | 0 |
| 0 | 0 | 0 |
| 0 | 0 | 0 |
| 0 | 0 | 0 |
| 1 | 1 | 1 |
| 1 | 1 | 1 |
| 1 | 1 | 1 |
| 0 | 1 | 0 |
| 1 | 1 | 1 |
| 1 | 1 | 1 |
| 0 | 0 | 0 |
| 0 | 0 | 0 |
| 0 | 0 | 0 |
| 1 | 1 | 1 |
| 1 | 1 | 1 |
| 1 | 1 | 1 |
| 1 | 1 | 1 |
| 0 | 0 | 0 |
| 0 | 0 | 0 |
| 0 | 0 | 0 |

| | | |
|---|---|---|
| 0 | 0 | 0 |
| 0 | 0 | 0 |
| 1 | 1 | 1 |
| 0 | 0 | 0 |
| 0 | 0 | 0 |
| 0 | 0 | 0 |
| 0 | 0 | 0 |
| 1 | 1 | 1 |
| 1 | 1 | 1 |
| 1 | 1 | 1 |
| 1 | 1 | 1 |
| 1 | 1 | 1 |
| 0 | 1 | 0 |
| 0 | 1 | 0 |
| 1 | 1 | 1 |
| 1 | 1 | 1 |
| 1 | 1 | 1 |
| 1 | 1 | 1 |
| 1 | 1 | 1 |
| 1 | 1 | 1 |
| 0 | 0 | 0 |
| 0 | 0 | 0 |
| 0 | 0 | 0 |
| 0 | 0 | 0 |
| 0 | 0 | 0 |
| 1 | 1 | 1 |
| 1 | 1 | 1 |
| 0 | 0 | 0 |
| 0 | 1 | 0 |
| 0 | 1 | 0 |
| 0 | 0 | 0 |
| 0 | 0 | 0 |
| 0 | 0 | 0 |
| 1 | 1 | 1 |
| 0 | 0 | 0 |
| 1 | 1 | 1 |
| 0 | 1 | 0 |
| 0 | 0 | 0 |
| 1 | 1 | 1 |
| 1 | 1 | 1 |
| 0 | 1 | 0 |
| 0 | 1 | 0 |
| 1 | 1 | 1 |
| 1 | 1 | 1 |
| 1 | 1 | 1 |
| 1 | 1 | 1 |
| 1 | 1 | 1 |
| 1 | 1 | 1 |
| 0 | 0 | 0 |
| 1 | 1 | 1 |
| 0 | 0 | 0 |
| 0 | 0 | 0 |
| 1 | 1 | 1 |
| 0 | 1 | 0 |

| | | |
|---|---|---|
| 0 | 0 | 0 |
| 1 | 1 | 1 |
| 1 | 1 | 1 |
| 1 | 1 | 1 |
| 1 | 1 | 1 |
| 0 | 0 | 0 |
| 0 | 0 | 0 |
| 1 | 1 | 1 |
| 0 | 1 | 0 |
| 1 | 1 | 1 |
| 1 | 1 | 1 |
| 0 | 0 | 0 |
| 0 | 1 | 0 |
| 1 | 1 | 1 |
| 0 | 1 | 0 |
| 1 | 1 | 1 |
| 0 | 0 | 0 |
| 0 | 0 | 0 |
| 1 | 1 | 1 |
| 1 | 1 | 1 |
| 1 | 1 | 1 |
| 1 | 1 | 1 |
| 1 | 1 | 1 |
| 1 | 1 | 1 |
| 0 | 0 | 0 |
| 1 | 1 | 1 |
| 1 | 1 | 1 |
| 1 | 1 | 1 |
| 1 | 1 | 1 |
| 1 | 1 | 1 |
| 0 | 1 | 0 |
| 0 | 0 | 0 |
| 0 | 0 | 0 |
| 0 | 1 | 0 |
| 1 | 1 | 1 |
| 0 | 0 | 0 |
| 0 | 0 | 0 |
| 0 | 0 | 0 |
| 0 | 0 | 0 |
| 0 | 0 | 0 |
| 0 | 0 | 0 |
| 0 | 0 | 0 |
| 0 | 0 | 0 |
| 0 | 0 | 0 |
| 0 | 0 | 0 |
| 0 | 0 | 0 |
| 0 | 0 | 0 |
| 0 | 0 | 0 |
| 1 | 0 | 0 |
| 1 | 1 | 1 |
| 1 | 0 | 0 |
| 1 | 1 | 1 |
| 0 | 0 | 0 |
| 1 | 1 | 1 |

| | | |
|---|---|---|
| 1 | 1 | 1 |
| 1 | 1 | 1 |
| 1 | 1 | 1 |
| 1 | 1 | 1 |
| 1 | 1 | 1 |
| 1 | 1 | 1 |
| 1 | 1 | 1 |
| 0 | 0 | 0 |
| 0 | 0 | 0 |
| 1 | 0 | 0 |
| 1 | 1 | 1 |
| 1 | 1 | 1 |
| 1 | 1 | 1 |
| 0 | 0 | 0 |
| 1 | 1 | 1 |
| 1 | 1 | 1 |
| 0 | 0 | 0 |
| 1 | 1 | 1 |
| 0 | 0 | 0 |
| 0 | 1 | 0 |
| 1 | 1 | 1 |
| 1 | 1 | 1 |
| 1 | 1 | 1 |
| 0 | 0 | 0 |
| 1 | 1 | 1 |
| 1 | 1 | 1 |
| 0 | 1 | 0 |
| 0 | 1 | 0 |
| 0 | 1 | 0 |
| 0 | 1 | 0 |
| 1 | 1 | 1 |
| 0 | 0 | 0 |
| 1 | 1 | 1 |
| 0 | 1 | 0 |
| 1 | 1 | 1 |
| 1 | 1 | 1 |
| 1 | 1 | 1 |
| 1 | 1 | 1 |
| 0 | 0 | 0 |
| 1 | 1 | 1 |
| 1 | 1 | 1 |
| 1 | 1 | 1 |
| 0 | 1 | 0 |
| 1 | 1 | 1 |
| 1 | 1 | 1 |
| 0 | 0 | 0 |
| 0 | 1 | 0 |
| 0 | 1 | 0 |
| 1 | 0 | 0 |
| 1 | 0 | 0 |
| 0 | 1 | 0 |
| 1 | 0 | 0 |
| 0 | 0 | 0 |
| 1 | 0 | 0 |

| | | |
|---|---|---|
| 1 | 0 | 0 |
| 1 | 0 | 0 |
| 1 | 0 | 0 |
| 1 | 0 | 0 |
| 0 | 0 | 0 |
| 0 | 0 | 0 |
| 0 | 0 | 0 |
| 0 | 0 | 0 |
| 1 | 0 | 0 |
| 1 | 0 | 0 |
| 1 | 0 | 0 |
| 1 | 0 | 0 |
| 1 | 0 | 0 |
| 0 | 0 | 0 |
| 1 | 0 | 0 |
| 0 | 0 | 0 |
| 1 | 0 | 0 |
| 0 | 0 | 0 |
| 1 | 0 | 0 |
| 0 | 0 | 0 |
| 0 | 0 | 0 |
| 1 | 0 | 0 |
| 1 | 0 | 0 |
| 0 | 0 | 0 |
| 0 | 0 | 0 |
| 1 | 0 | 0 |
| 0 | 0 | 0 |
| 0 | 0 | 0 |
| 0 | 0 | 0 |
| 1 | 0 | 0 |
| 1 | 0 | 0 |
| 0 | 0 | 0 |
| 1 | 0 | 0 |
| 1 | 0 | 0 |
| 1 | 0 | 0 |
| 1 | 0 | 0 |
| 1 | 0 | 0 |
| 1 | 0 | 0 |
| 1 | 0 | 0 |
| 1 | 0 | 0 |
| 1 | 0 | 0 |
| 1 | 0 | 0 |
| 1 | 0 | 0 |
| 0 | 0 | 0 |
| 1 | 1 | 1 |
| 1 | 1 | 1 |
| 0 | 0 | 0 |
| 0 | 0 | 0 |
| 0 | 1 | 0 |
| 0 | 0 | 0 |
| 0 | 0 | 0 |
| 0 | 0 | 0 |
| 1 | 1 | 1 |
| 1 | 1 | 1 |

| | | |
|---|---|---|
| 1 | 1 | 1 |
| 1 | 1 | 1 |
| 1 | 1 | 1 |
| 1 | 1 | 1 |
| 1 | 1 | 1 |
| 1 | 1 | 1 |
| 1 | 1 | 1 |
| 0 | 0 | 0 |
| 1 | 1 | 1 |
| 1 | 1 | 1 |
| 0 | 1 | 0 |
| 0 | 1 | 0 |
| 1 | 0 | 0 |
| 1 | 1 | 1 |
| 0 | 0 | 0 |
| 0 | 0 | 0 |
| 0 | 1 | 0 |
| 0 | 0 | 0 |
| 1 | 1 | 1 |
| 1 | 1 | 1 |
| 0 | 0 | 0 |
| 0 | 1 | 0 |
| 0 | 1 | 0 |
| 1 | 1 | 1 |
| 0 | 0 | 0 |
| 1 | 1 | 1 |
| 0 | 1 | 0 |
| 0 | 0 | 0 |
| 0 | 0 | 0 |
| 0 | 0 | 0 |
| 0 | 0 | 0 |
| 1 | 1 | 1 |
| 1 | 1 | 1 |
| 0 | 1 | 0 |
| 1 | 1 | 1 |
| 1 | 1 | 1 |
| 0 | 0 | 0 |
| 0 | 0 | 0 |
| 1 | 1 | 1 |
| 1 | 1 | 1 |
| 1 | 1 | 1 |
| 1 | 1 | 1 |
| 0 | 0 | 0 |
| 0 | 1 | 0 |
| 1 | 1 | 1 |
| 1 | 1 | 1 |
| 1 | 1 | 1 |
| 1 | 1 | 1 |
| 0 | 0 | 0 |
| 0 | 0 | 0 |
| 0 | 0 | 0 |
| 0 | 0 | 0 |
| 0 | 0 | 0 |
| 1 | 1 | 1 |

| | | |
|---|---|---|
| 0 | 0 | 0 |
| 0 | 0 | 0 |
| 0 | 1 | 0 |
| 0 | 0 | 0 |
| 1 | 1 | 1 |
| 0 | 1 | 0 |
| 0 | 1 | 0 |
| 1 | 1 | 1 |
| 1 | 1 | 1 |
| 0 | 0 | 0 |
| 1 | 1 | 1 |
| 0 | 1 | 0 |
| 1 | 1 | 1 |
| 0 | 1 | 0 |
| 0 | 1 | 0 |
| 0 | 1 | 0 |
| 0 | 1 | 0 |
| 1 | 1 | 1 |
| 1 | 1 | 1 |
| 0 | 1 | 0 |
| 0 | 1 | 0 |
| 0 | 1 | 0 |
| 0 | 1 | 0 |
| 0 | 1 | 0 |
| 0 | 1 | 0 |
| 0 | 1 | 0 |
| 0 | 1 | 0 |
| 0 | 1 | 0 |
| 0 | 1 | 0 |
| 0 | 1 | 0 |
| 1 | 1 | 1 |
| 1 | 1 | 1 |
| 0 | 0 | 0 |
| 1 | 1 | 1 |
| 1 | 1 | 1 |
| 1 | 1 | 1 |
| 1 | 1 | 1 |
| 0 | 1 | 0 |
| 1 | 1 | 1 |
| 1 | 1 | 1 |
| 1 | 1 | 1 |
| 1 | 1 | 1 |
| 1 | 1 | 1 |
| 0 | 1 | 0 |
| 0 | 1 | 0 |
| 1 | 1 | 1 |
| 1 | 1 | 1 |
| 0 | 0 | 0 |
| 0 | 0 | 0 |
| 0 | 0 | 0 |
| 0 | 0 | 0 |
| 0 | 0 | 0 |
| 1 | 1 | 1 |
| 0 | 0 | 0 |

| | | |
|---|---|---|
| 1 | 1 | 1 |
| 0 | 0 | 0 |
| 1 | 1 | 1 |
| 0 | 1 | 0 |
| 0 | 1 | 0 |
| 0 | 0 | 0 |
| 0 | 0 | 0 |
| 0 | 0 | 0 |
| 0 | 0 | 0 |
| 0 | 0 | 0 |
| 0 | 1 | 0 |
| 0 | 0 | 0 |
| 0 | 1 | 0 |
| 1 | 1 | 1 |
| 0 | 1 | 0 |
| 0 | 0 | 0 |
| 1 | 1 | 1 |
| 1 | 1 | 1 |
| 0 | 0 | 0 |
| 0 | 0 | 0 |
| 0 | 0 | 0 |
| 0 | 0 | 0 |
| 0 | 0 | 0 |
| 0 | 0 | 0 |
| 0 | 1 | 0 |
| 0 | 0 | 0 |
| 0 | 0 | 0 |
| 0 | 0 | 0 |
| 0 | 0 | 0 |
| 0 | 0 | 0 |
| 0 | 0 | 0 |
| 1 | 1 | 1 |
| 1 | 1 | 1 |
| 0 | 0 | 0 |
| 0 | 0 | 0 |
| 1 | 0 | 0 |
| 0 | 0 | 0 |
| 0 | 0 | 0 |
| 0 | 0 | 0 |
| 0 | 0 | 0 |
| 0 | 0 | 0 |
| 1 | 0 | 0 |
| 1 | 1 | 1 |
| 1 | 1 | 1 |
| 0 | 0 | 0 |
| 0 | 0 | 0 |
| 0 | 0 | 0 |
| 1 | 1 | 1 |
| 1 | 1 | 1 |
| 1 | 1 | 1 |
| 1 | 1 | 1 |
| 1 | 1 | 1 |
| 1 | 1 | 1 |
| 0 | 0 | 0 |

| | | |
|---|---|---|
| 1 | 1 | 1 |
| 1 | 1 | 1 |
| 1 | 1 | 1 |
| 1 | 1 | 1 |
| 0 | 0 | 0 |
| 0 | 0 | 0 |
| 1 | 0 | 0 |
| 1 | 0 | 0 |
| 0 | 1 | 0 |
| 1 | 1 | 1 |
| 1 | 1 | 1 |
| 0 | 0 | 0 |
| 1 | 1 | 1 |
| 0 | 0 | 0 |
| 0 | 0 | 0 |
| 0 | 0 | 0 |
| 0 | 0 | 0 |
| 0 | 0 | 0 |
| 0 | 0 | 0 |
| 0 | 0 | 0 |
| 0 | 0 | 0 |
| 0 | 0 | 0 |
| 0 | 0 | 0 |
| 0 | 1 | 0 |
| 0 | 1 | 0 |
| 0 | 1 | 0 |
| 1 | 1 | 1 |
| 1 | 1 | 1 |
| 1 | 1 | 1 |
| 1 | 1 | 1 |
| 1 | 1 | 1 |
| 0 | 0 | 0 |
| 1 | 1 | 1 |
| 0 | 0 | 0 |
| 0 | 0 | 0 |
| 0 | 0 | 0 |
| 0 | 0 | 0 |
| 1 | 1 | 1 |
| 1 | 1 | 1 |
| 0 | 1 | 0 |
| 1 | 1 | 1 |
| 1 | 1 | 1 |
| 1 | 1 | 1 |
| 0 | 1 | 0 |
| 1 | 1 | 1 |
| 0 | 0 | 0 |
| 1 | 1 | 1 |
| 1 | 1 | 1 |
| 1 | 1 | 1 |
| 0 | 0 | 0 |
| 0 | 0 | 0 |
| 0 | 0 | 0 |
| 0 | 1 | 0 |
| 0 | 1 | 0 |

| | | |
|---|---|---|
| 0 | 1 | 0 |
| 0 | 1 | 0 |
| 0 | 1 | 0 |
| 0 | 0 | 0 |
| 0 | 1 | 0 |
| 1 | 1 | 1 |
| 0 | 0 | 0 |
| 0 | 1 | 0 |
| 1 | 1 | 1 |
| 1 | 1 | 1 |
| 0 | 0 | 0 |
| 0 | 0 | 0 |
| 1 | 1 | 1 |
| 1 | 1 | 1 |
| 0 | 1 | 0 |
| 0 | 1 | 0 |
| 0 | 0 | 0 |
| 0 | 0 | 0 |
| 1 | 1 | 1 |
| 0 | 0 | 0 |
| 0 | 0 | 0 |
| 0 | 0 | 0 |
| 0 | 1 | 0 |
| 0 | 0 | 0 |
| 0 | 0 | 0 |
| 1 | 1 | 1 |
| 0 | 0 | 0 |
| 0 | 1 | 0 |
| 1 | 1 | 1 |
| 0 | 0 | 0 |
| 0 | 1 | 0 |
| 1 | 1 | 1 |
| 0 | 0 | 0 |
| 0 | 1 | 0 |
| 0 | 1 | 0 |
| 0 | 0 | 0 |
| 1 | 1 | 1 |
| 0 | 0 | 0 |
| 1 | 1 | 1 |
| 0 | 0 | 0 |
| 0 | 0 | 0 |
| 1 | 0 | 0 |
| 0 | 0 | 0 |
| 0 | 0 | 0 |
| 1 | 1 | 1 |
| 1 | 0 | 0 |
| 0 | 1 | 0 |
| 1 | 1 | 1 |
| 1 | 1 | 1 |
| 1 | 1 | 1 |
| 1 | 1 | 1 |
| 0 | 1 | 0 |
| 0 | 1 | 0 |
| 0 | 1 | 0 |

| | | |
|---|---|---|
| 0 | 1 | 0 |
| 0 | 1 | 0 |
| 0 | 1 | 0 |
| 0 | 0 | 0 |
| 1 | 0 | 0 |
| 0 | 0 | 0 |
| 1 | 1 | 1 |
| 1 | 0 | 0 |
| 0 | 0 | 0 |
| 0 | 0 | 0 |
| 1 | 1 | 1 |
| 0 | 0 | 0 |
| 0 | 0 | 0 |
| 0 | 0 | 0 |
| 1 | 0 | 0 |
| 0 | 0 | 0 |
| 0 | 0 | 0 |
| 0 | 1 | 0 |
| 0 | 0 | 0 |
| 0 | 0 | 0 |
| 0 | 0 | 0 |
| 0 | 0 | 0 |
| 1 | 1 | 1 |
| 0 | 0 | 0 |
| 0 | 0 | 0 |
| 0 | 0 | 0 |
| 0 | 0 | 0 |
| 0 | 0 | 0 |
| 0 | 0 | 0 |
| 0 | 0 | 0 |
| 0 | 0 | 0 |
| 0 | 0 | 0 |
| 0 | 0 | 0 |
| 0 | 1 | 0 |
| 0 | 1 | 0 |
| 0 | 0 | 0 |
| 1 | 1 | 1 |
| 0 | 0 | 0 |
| 0 | 0 | 0 |
| 0 | 0 | 0 |
| 0 | 0 | 0 |
| 1 | 1 | 1 |
| 0 | 0 | 0 |
| 0 | 0 | 0 |
| 1 | 1 | 1 |
| 0 | 1 | 0 |
| 1 | 1 | 1 |
| 1 | 1 | 1 |
| 1 | 1 | 1 |
| 0 | 1 | 0 |
| 1 | 1 | 1 |
| 1 | 1 | 1 |
| 1 | 1 | 1 |
| 1 | 1 | 1 |

| | | |
|---|---|---|
| 1 | 1 | 1 |
| 0 | 0 | 0 |
| 1 | 1 | 1 |
| 1 | 1 | 1 |
| 1 | 1 | 1 |
| 1 | 1 | 1 |
| 1 | 1 | 1 |
| 1 | 1 | 1 |
| 1 | 1 | 1 |
| 0 | 1 | 0 |
| 1 | 1 | 1 |
| 0 | 0 | 0 |
| 1 | 0 | 0 |
| 0 | 1 | 0 |
| 0 | 0 | 0 |
| 0 | 1 | 0 |
| 0 | 1 | 0 |
| 1 | 1 | 1 |
| 1 | 1 | 1 |
| 1 | 1 | 1 |
| 0 | 0 | 0 |
| 0 | 1 | 0 |
| 0 | 0 | 0 |
| 0 | 0 | 0 |
| 1 | 1 | 1 |
| 0 | 0 | 0 |
| 1 | 1 | 1 |
| 1 | 1 | 1 |
| 1 | 1 | 1 |
| 1 | 1 | 1 |
| 1 | 1 | 1 |
| 0 | 0 | 0 |
| 1 | 1 | 1 |
| 0 | 0 | 0 |
| 1 | 1 | 1 |
| 0 | 1 | 0 |
| 0 | 1 | 0 |
| 0 | 1 | 0 |
| 1 | 1 | 1 |
| 0 | 0 | 0 |
| 1 | 1 | 1 |
| 1 | 1 | 1 |
| 0 | 1 | 0 |
| 0 | 0 | 0 |
| 1 | 1 | 1 |
| 1 | 1 | 1 |
| 1 | 1 | 1 |
| 0 | 0 | 0 |
| 0 | 0 | 0 |
| 1 | 1 | 1 |
| 1 | 1 | 1 |
| 0 | 0 | 0 |
| 0 | 0 | 0 |
| 0 | 0 | 0 |

| | | |
|---|---|---|
| 0 | 0 | 0 |
| 1 | 1 | 1 |
| 0 | 0 | 0 |
| 0 | 0 | 0 |
| 0 | 0 | 0 |
| 1 | 1 | 1 |
| 1 | 1 | 1 |
| 0 | 1 | 0 |
| 1 | 1 | 1 |
| 0 | 0 | 0 |
| 1 | 1 | 1 |
| 0 | 1 | 0 |
| 1 | 1 | 1 |
| 1 | 1 | 1 |
| 0 | 0 | 0 |
| 0 | 1 | 0 |
| 1 | 1 | 1 |
| 0 | 0 | 0 |
| 1 | 1 | 1 |
| 0 | 0 | 0 |
| 0 | 0 | 0 |
| 1 | 1 | 1 |
| 0 | 0 | 0 |
| 1 | 1 | 1 |
| 0 | 0 | 0 |
| 0 | 0 | 0 |
| 1 | 1 | 1 |
| 0 | 1 | 0 |
| 0 | 0 | 0 |
| 1 | 1 | 1 |
| 1 | 1 | 1 |
| 1 | 1 | 1 |
| 1 | 1 | 1 |
| 0 | 1 | 0 |
| 0 | 0 | 0 |
| 0 | 0 | 0 |
| 1 | 1 | 1 |
| 0 | 0 | 0 |
| 0 | 0 | 0 |
| 0 | 0 | 0 |
| 0 | 0 | 0 |
| 0 | 0 | 0 |
| 0 | 0 | 0 |
| 0 | 1 | 0 |
| 0 | 0 | 0 |
| 1 | 1 | 1 |
| 0 | 1 | 0 |
| 0 | 0 | 0 |
| 0 | 0 | 0 |
| 0 | 0 | 0 |
| 0 | 0 | 0 |
| 0 | 0 | 0 |
| 1 | 1 | 1 |
| 1 | 1 | 1 |

| | | |
|---|---|---|
| 1 | 1 | 1 |
| 0 | 0 | 0 |
| 0 | 0 | 0 |
| 0 | 0 | 0 |
| 0 | 0 | 0 |
| 0 | 0 | 0 |
| 0 | 0 | 0 |
| 1 | 0 | 0 |
| 1 | 0 | 0 |
| 1 | 1 | 1 |
| 0 | 0 | 0 |
| 0 | 0 | 0 |
| 1 | 1 | 1 |
| 0 | 0 | 0 |
| 1 | 1 | 1 |
| 0 | 0 | 0 |
| 1 | 1 | 1 |
| 1 | 1 | 1 |
| 0 | 1 | 0 |
| 0 | 1 | 0 |
| 0 | 1 | 0 |
| 1 | 1 | 1 |
| 0 | 1 | 0 |
| 0 | 0 | 0 |
| 0 | 0 | 0 |
| 1 | 1 | 1 |
| 1 | 1 | 1 |
| 1 | 1 | 1 |
| 1 | 1 | 1 |
| 0 | 0 | 0 |
| 0 | 0 | 0 |
| 0 | 0 | 0 |
| 0 | 0 | 0 |
| 1 | 1 | 1 |
| 0 | 0 | 0 |
| 0 | 0 | 0 |
| 0 | 0 | 0 |
| 0 | 0 | 0 |
| 0 | 0 | 0 |
| 1 | 1 | 1 |
| 1 | 1 | 1 |
| 0 | 0 | 0 |
| 1 | 1 | 1 |
| 1 | 1 | 1 |
| 0 | 0 | 0 |
| 1 | 0 | 0 |
| 1 | 1 | 1 |
| 0 | 1 | 0 |
| 1 | 0 | 0 |
| 0 | 1 | 0 |
| 0 | 0 | 0 |
| 1 | 1 | 1 |
| 0 | 0 | 0 |
| 0 | 0 | 0 |

| | | |
|---|---|---|
| 0 | 1 | 0 |
| 1 | 1 | 1 |
| 0 | 0 | 0 |
| 0 | 0 | 0 |
| 1 | 1 | 1 |
| 1 | 1 | 1 |
| 1 | 1 | 1 |
| 1 | 1 | 1 |
| 1 | 1 | 1 |
| 1 | 1 | 1 |
| 1 | 1 | 1 |
| 1 | 1 | 1 |
| 0 | 0 | 0 |
| 1 | 1 | 1 |
| 1 | 1 | 1 |
| 1 | 1 | 1 |
| 1 | 1 | 1 |
| 1 | 1 | 1 |
| 1 | 1 | 1 |
| 1 | 1 | 1 |
| 1 | 1 | 1 |
| 1 | 1 | 1 |
| 1 | 1 | 1 |
| 1 | 1 | 1 |
| 1 | 1 | 1 |
| 1 | 1 | 1 |
| 0 | 1 | 0 |
| 1 | 1 | 1 |
| 1 | 1 | 1 |
| 1 | 1 | 1 |
| 1 | 1 | 1 |
| 0 | 0 | 0 |
| 1 | 1 | 1 |
| 0 | 1 | 0 |
| 1 | 1 | 1 |
| 1 | 1 | 1 |
| 0 | 1 | 0 |
| 1 | 1 | 1 |
| 1 | 1 | 1 |
| 1 | 1 | 1 |
| 0 | 0 | 0 |
| 0 | 0 | 0 |
| 1 | 1 | 1 |
| 1 | 1 | 1 |
| 1 | 1 | 1 |
| 1 | 0 | 0 |
| 1 | 0 | 0 |
| 1 | 1 | 1 |
| 0 | 0 | 0 |
| 0 | 0 | 0 |
| 1 | 1 | 1 |
| 0 | 1 | 0 |
| 0 | 0 | 0 |
| 1 | 1 | 1 |

| | | |
|---|---|---|
| 0 | 0 | 0 |
| 0 | 0 | 0 |
| 0 | 0 | 0 |
| 1 | 1 | 1 |
| 1 | 1 | 1 |
| 1 | 1 | 1 |
| 1 | 1 | 1 |
| 0 | 0 | 0 |
| 0 | 0 | 0 |
| 0 | 0 | 0 |
| 0 | 0 | 0 |
| 1 | 1 | 1 |
| 0 | 0 | 0 |
| 1 | 1 | 1 |
| 1 | 1 | 1 |
| 1 | 1 | 1 |
| 1 | 1 | 1 |
| 0 | 0 | 0 |
| 1 | 1 | 1 |
| 1 | 1 | 1 |
| 1 | 1 | 1 |
| 1 | 1 | 1 |
| 0 | 1 | 0 |
| 1 | 1 | 1 |
| 1 | 1 | 1 |
| 0 | 1 | 0 |
| 1 | 1 | 1 |
| 1 | 1 | 1 |
| 1 | 1 | 1 |
| 1 | 1 | 1 |
| 0 | 0 | 0 |
| 0 | 0 | 0 |
| 1 | 1 | 1 |
| 1 | 1 | 1 |
| 1 | 1 | 1 |
| 1 | 1 | 1 |
| 1 | 1 | 1 |
| 1 | 1 | 1 |
| 1 | 1 | 1 |
| 1 | 1 | 1 |
| 1 | 1 | 1 |
| 0 | 0 | 0 |
| 1 | 1 | 1 |
| 1 | 1 | 1 |
| 0 | 0 | 0 |
| 0 | 0 | 0 |
| 1 | 1 | 1 |
| 1 | 1 | 1 |
| 0 | 0 | 0 |
| 0 | 0 | 0 |
| 0 | 0 | 0 |
| 0 | 0 | 0 |
| 0 | 0 | 0 |
| 0 | 0 | 0 |

| | | |
|---|---|---|
| 1 | 1 | 1 |
| 1 | 1 | 1 |
| 1 | 1 | 1 |
| 1 | 1 | 1 |
| 1 | 1 | 1 |
| 0 | 0 | 0 |
| 0 | 0 | 0 |
| 0 | 0 | 0 |
| 0 | 0 | 0 |
| 0 | 0 | 0 |
| 0 | 1 | 0 |
| 0 | 0 | 0 |
| 0 | 0 | 0 |
| 0 | 0 | 0 |
| 0 | 1 | 0 |
| 0 | 0 | 0 |
| 0 | 1 | 0 |
| 0 | 1 | 0 |
| 0 | 0 | 0 |
| 0 | 0 | 0 |
| 0 | 0 | 0 |
| 0 | 0 | 0 |
| 1 | 0 | 0 |
| 0 | 0 | 0 |
| 0 | 0 | 0 |
| 0 | 0 | 0 |
| 0 | 0 | 0 |
| 1 | 1 | 1 |
| 1 | 1 | 1 |
| 1 | 1 | 1 |
| 1 | 1 | 1 |
| 1 | 1 | 1 |
| 0 | 0 | 0 |
| 0 | 1 | 0 |
| 0 | 0 | 0 |
| 0 | 0 | 0 |
| 0 | 0 | 0 |
| 0 | 0 | 0 |
| 0 | 0 | 0 |
| 0 | 0 | 0 |
| 0 | 0 | 0 |
| 0 | 0 | 0 |
| 0 | 0 | 0 |
| 1 | 1 | 1 |
| 1 | 1 | 1 |
| 1 | 1 | 1 |
| 1 | 1 | 1 |
| 0 | 0 | 0 |
| 0 | 0 | 0 |
| 0 | 0 | 0 |
| 1 | 1 | 1 |
| 1 | 1 | 1 |
| 0 | 0 | 0 |
| 1 | 1 | 1 |

| | | |
|---|---|---|
| 1 | 1 | 1 |
| 0 | 0 | 0 |
| 0 | 1 | 0 |
| 0 | 1 | 0 |
| 1 | 1 | 1 |
| 1 | 1 | 1 |
| 0 | 1 | 0 |
| 1 | 1 | 1 |
| 0 | 1 | 0 |
| 1 | 1 | 1 |
| 1 | 1 | 1 |
| 1 | 1 | 1 |
| 0 | 0 | 0 |
| 0 | 0 | 0 |
| 0 | 0 | 0 |
| 0 | 0 | 0 |
| 0 | 0 | 0 |
| 0 | 0 | 0 |
| 0 | 0 | 0 |
| 0 | 1 | 0 |
| 0 | 0 | 0 |
| 0 | 0 | 0 |
| 0 | 0 | 0 |
| 0 | 0 | 0 |
| 0 | 0 | 0 |
| 1 | 1 | 1 |
| 1 | 1 | 1 |
| 0 | 0 | 0 |
| 0 | 1 | 0 |
| 0 | 0 | 0 |
| 0 | 0 | 0 |
| 1 | 1 | 1 |
| 0 | 0 | 0 |
| 1 | 1 | 1 |
| 1 | 1 | 1 |
| 1 | 1 | 1 |
| 1 | 0 | 0 |
| 1 | 1 | 1 |
| 1 | 1 | 1 |
| 1 | 1 | 1 |
| 0 | 0 | 0 |
| 1 | 1 | 1 |
| 1 | 1 | 1 |
| 0 | 0 | 0 |
| 0 | 0 | 0 |
| 1 | 1 | 1 |
| 0 | 0 | 0 |
| 0 | 0 | 0 |
| 1 | 1 | 1 |
| 1 | 1 | 1 |
| 0 | 0 | 0 |
| 1 | 1 | 1 |
| 1 | 1 | 1 |
| 1 | 1 | 1 |

| | | |
|---|---|---|
| 1 | 1 | 1 |
| 1 | 1 | 1 |
| 1 | 1 | 1 |
| 1 | 1 | 1 |
| 0 | 0 | 0 |
| 0 | 0 | 0 |
| 1 | 1 | 1 |
| 1 | 1 | 1 |
| 1 | 1 | 1 |
| 0 | 1 | 0 |
| 1 | 1 | 1 |
| 0 | 0 | 0 |
| 0 | 0 | 0 |
| 1 | 1 | 1 |
| 1 | 1 | 1 |
| 0 | 0 | 0 |
| 0 | 1 | 0 |
| 1 | 1 | 1 |
| 0 | 0 | 0 |
| 1 | 1 | 1 |
| 1 | 1 | 1 |
| 1 | 1 | 1 |
| 1 | 1 | 1 |
| 0 | 0 | 0 |
| 1 | 1 | 1 |
| 0 | 0 | 0 |
| 1 | 1 | 1 |
| 1 | 1 | 1 |
| 0 | 0 | 0 |
| 0 | 0 | 0 |
| 0 | 0 | 0 |
| 0 | 0 | 0 |
| 0 | 0 | 0 |
| 0 | 0 | 0 |
| 1 | 1 | 1 |
| 1 | 1 | 1 |
| 1 | 1 | 1 |
| 0 | 0 | 0 |
| 1 | 1 | 1 |
| 1 | 1 | 1 |
| 0 | 0 | 0 |
| 0 | 0 | 0 |
| 0 | 0 | 0 |
| 1 | 1 | 1 |
| 0 | 0 | 0 |
| 0 | 1 | 0 |
| 0 | 1 | 0 |
| 0 | 0 | 0 |
| 1 | 1 | 1 |
| 1 | 1 | 1 |
| 1 | 1 | 1 |
| 0 | 0 | 0 |
| 1 | 1 | 1 |
| 0 | 0 | 0 |

| | | |
|---|---|---|
| 1 | 1 | 1 |
| 0 | 0 | 0 |
| 0 | 0 | 0 |
| 0 | 1 | 0 |
| 1 | 1 | 1 |
| 1 | 1 | 1 |
| 0 | 0 | 0 |
| 1 | 1 | 1 |
| 0 | 1 | 0 |
| 0 | 1 | 0 |
| 1 | 1 | 1 |
| 1 | 1 | 1 |
| 1 | 1 | 1 |
| 1 | 1 | 1 |
| 1 | 1 | 1 |
| 1 | 1 | 1 |
| 1 | 1 | 1 |
| 1 | 1 | 1 |
| 1 | 1 | 1 |
| 1 | 1 | 1 |
| 1 | 1 | 1 |
| 0 | 0 | 0 |
| 0 | 1 | 0 |
| 0 | 1 | 0 |
| 1 | 1 | 1 |
| 1 | 1 | 1 |
| 0 | 0 | 0 |
| 1 | 1 | 1 |
| 0 | 0 | 0 |
| 0 | 1 | 0 |
| 1 | 1 | 1 |
| 0 | 0 | 0 |
| 0 | 0 | 0 |
| 1 | 0 | 0 |
| 1 | 1 | 1 |
| 1 | 1 | 1 |
| 1 | 1 | 1 |
| 1 | 1 | 1 |
| 0 | 0 | 0 |
| 0 | 0 | 0 |
| 0 | 0 | 0 |
| 1 | 1 | 1 |
| 1 | 1 | 1 |
| 0 | 1 | 0 |
| 0 | 1 | 0 |
| 0 | 0 | 0 |
| 1 | 1 | 1 |
| 0 | 0 | 0 |
| 1 | 1 | 1 |
| 0 | 0 | 0 |
| 0 | 0 | 0 |
| 0 | 0 | 0 |
| 1 | 1 | 1 |
| 1 | 1 | 1 |

| | | |
|---|---|---|
| 0 | 0 | 0 |
| 0 | 0 | 0 |
| 1 | 1 | 1 |
| 1 | 1 | 1 |
| 1 | 1 | 1 |
| 1 | 1 | 1 |
| 1 | 1 | 1 |
| 1 | 1 | 1 |
| 1 | 1 | 1 |
| 1 | 1 | 1 |
| 1 | 1 | 1 |
| 1 | 1 | 1 |
| 1 | 1 | 1 |
| 1 | 1 | 1 |
| 1 | 1 | 1 |
| 0 | 0 | 0 |
| 0 | 0 | 0 |
| 0 | 0 | 0 |
| 1 | 1 | 1 |
| 1 | 1 | 1 |
| 0 | 1 | 0 |
| 0 | 0 | 0 |
| 0 | 0 | 0 |
| 0 | 0 | 0 |
| 0 | 0 | 0 |
| 1 | 1 | 1 |
| 1 | 1 | 1 |
| 1 | 1 | 1 |
| 1 | 1 | 1 |
| 1 | 1 | 1 |
| 0 | 0 | 0 |
| 0 | 1 | 0 |
| 0 | 0 | 0 |
| 1 | 1 | 1 |
| 1 | 0 | 0 |
| 1 | 0 | 0 |
| 0 | 0 | 0 |
| 0 | 0 | 0 |
| 0 | 1 | 0 |
| 1 | 1 | 1 |
| 1 | 0 | 0 |
| 0 | 0 | 0 |
| 1 | 1 | 1 |
| 0 | 1 | 0 |
| 0 | 0 | 0 |
| 1 | 0 | 0 |
| 0 | 0 | 0 |
| 0 | 0 | 0 |
| 1 | 1 | 1 |
| 0 | 1 | 0 |
| 0 | 0 | 0 |
| 0 | 0 | 0 |
| 1 | 1 | 1 |
| 0 | 0 | 0 |

| | | |
|---|---|---|
| 0 | 1 | 0 |
| 0 | 0 | 0 |
| 1 | 0 | 0 |
| 0 | 0 | 0 |
| 1 | 1 | 1 |
| 1 | 1 | 1 |
| 1 | 1 | 1 |
| 0 | 1 | 0 |
| 0 | 0 | 0 |
| 1 | 1 | 1 |
| 0 | 1 | 0 |
| 0 | 1 | 0 |
| 0 | 0 | 0 |
| 0 | 0 | 0 |
| 0 | 0 | 0 |
| 1 | 1 | 1 |
| 0 | 0 | 0 |
| 0 | 1 | 0 |
| 0 | 0 | 0 |
| 0 | 0 | 0 |
| 0 | 0 | 0 |
| 0 | 0 | 0 |
| 0 | 0 | 0 |
| 0 | 0 | 0 |
| 0 | 1 | 0 |
| 0 | 1 | 0 |
| 0 | 0 | 0 |
| 0 | 0 | 0 |
| 0 | 0 | 0 |
| 0 | 0 | 0 |
| 1 | 1 | 1 |
| 1 | 1 | 1 |
| 1 | 1 | 1 |
| 1 | 1 | 1 |
| 1 | 1 | 1 |
| 0 | 0 | 0 |
| 0 | 0 | 0 |
| 0 | 1 | 0 |
| 1 | 1 | 1 |
| 1 | 1 | 1 |
| 0 | 1 | 0 |
| 1 | 1 | 1 |
| 1 | 1 | 1 |
| 0 | 0 | 0 |
| 0 | 0 | 0 |
| 0 | 0 | 0 |
| 1 | 1 | 1 |
| 0 | 0 | 0 |
| 0 | 0 | 0 |
| 0 | 0 | 0 |
| 1 | 1 | 1 |
| 1 | 1 | 1 |
| 0 | 1 | 0 |
| 0 | 1 | 0 |

| | | |
|---|---|---|
| 0 | 1 | 0 |
| 1 | 1 | 1 |
| 1 | 0 | 0 |
| 1 | 1 | 1 |
| 0 | 0 | 0 |
| 1 | 0 | 0 |
| 0 | 0 | 0 |
| 0 | 0 | 0 |
| 0 | 0 | 0 |
| 0 | 0 | 0 |
| 0 | 0 | 0 |
| 0 | 0 | 0 |
| 0 | 1 | 0 |
| 0 | 0 | 0 |
| 0 | 0 | 0 |
| 1 | 1 | 1 |
| 1 | 1 | 1 |
| 1 | 1 | 1 |
| 0 | 0 | 0 |
| 0 | 0 | 0 |
| 0 | 0 | 0 |
| 0 | 0 | 0 |
| 0 | 1 | 0 |
| 0 | 1 | 0 |
| 1 | 1 | 1 |
| 1 | 1 | 1 |
| 1 | 1 | 1 |
| 0 | 0 | 0 |
| 0 | 0 | 0 |
| 0 | 0 | 0 |
| 0 | 0 | 0 |
| 0 | 0 | 0 |
| 0 | 0 | 0 |
| 0 | 1 | 0 |
| 1 | 1 | 1 |
| 0 | 0 | 0 |
| 1 | 1 | 1 |
| 1 | 1 | 1 |
| 1 | 1 | 1 |
| 1 | 1 | 1 |
| 0 | 0 | 0 |
| 1 | 1 | 1 |
| 0 | 0 | 0 |
| 0 | 1 | 0 |
| 1 | 1 | 1 |
| 1 | 1 | 1 |
| 1 | 1 | 1 |
| 0 | 0 | 0 |
| 1 | 1 | 1 |
| 1 | 1 | 1 |
| 1 | 1 | 1 |
| 1 | 1 | 1 |
| 1 | 1 | 1 |
| 0 | 1 | 0 |

| | | |
|---|---|---|
| 0 | 1 | 0 |
| 1 | 1 | 1 |
| 1 | 1 | 1 |
| 0 | 1 | 0 |
| 1 | 1 | 1 |
| 0 | 1 | 0 |
| 0 | 1 | 0 |
| 0 | 1 | 0 |
| 0 | 1 | 0 |
| 0 | 1 | 0 |
| 0 | 1 | 0 |
| 0 | 0 | 0 |
| 0 | 1 | 0 |
| 0 | 1 | 0 |
| 0 | 1 | 0 |
| 0 | 1 | 0 |
| 0 | 1 | 0 |
| 0 | 0 | 0 |
| 0 | 0 | 0 |
| 0 | 0 | 0 |
| 0 | 0 | 0 |
| 0 | 1 | 0 |
| 0 | 0 | 0 |
| 1 | 1 | 1 |
| 1 | 1 | 1 |
| 1 | 1 | 1 |
| 1 | 1 | 1 |
| 0 | 1 | 0 |
| 0 | 0 | 0 |
| 0 | 0 | 0 |
| 1 | 1 | 1 |
| 1 | 1 | 1 |
| 0 | 0 | 0 |
| 1 | 1 | 1 |
| 0 | 0 | 0 |
| 0 | 0 | 0 |
| 0 | 0 | 0 |
| 1 | 1 | 1 |
| 0 | 0 | 0 |
| 1 | 1 | 1 |
| 0 | 0 | 0 |
| 1 | 1 | 1 |
| 0 | 0 | 0 |
| 1 | 0 | 0 |
| 1 | 0 | 0 |
| 1 | 0 | 0 |
| 1 | 0 | 0 |
| 0 | 1 | 0 |
| 0 | 1 | 0 |
| 0 | 0 | 0 |
| 1 | 0 | 0 |
| 0 | 0 | 0 |
| 1 | 0 | 0 |
| 1 | 1 | 1 |

| | | |
|---|---|---|
| 0 | 0 | 0 |
| 0 | 0 | 0 |
| 0 | 0 | 0 |
| 0 | 0 | 0 |
| 0 | 0 | 0 |
| 1 | 1 | 1 |
| 0 | 0 | 0 |
| 0 | 0 | 0 |
| 1 | 1 | 1 |
| 0 | 0 | 0 |
| 0 | 0 | 0 |
| 0 | 0 | 0 |
| 0 | 1 | 0 |
| 0 | 1 | 0 |
| 0 | 1 | 0 |
| 0 | 1 | 0 |
| 0 | 1 | 0 |
| 0 | 1 | 0 |
| 0 | 1 | 0 |
| 0 | 1 | 0 |
| 0 | 1 | 0 |
| 0 | 1 | 0 |
| 0 | 1 | 0 |
| 0 | 1 | 0 |
| 0 | 1 | 0 |
| 0 | 1 | 0 |
| 0 | 1 | 0 |
| 0 | 1 | 0 |
| 0 | 1 | 0 |
| 0 | 1 | 0 |
| 0 | 1 | 0 |
| 0 | 1 | 0 |
| 0 | 1 | 0 |
| 0 | 1 | 0 |
| 0 | 1 | 0 |
| 0 | 1 | 0 |
| 0 | 1 | 0 |
| 0 | 1 | 0 |
| 0 | 1 | 0 |
| 0 | 1 | 0 |
| 0 | 1 | 0 |
| 0 | 1 | 0 |
| 0 | 1 | 0 |
| 0 | 1 | 0 |
| 0 | 1 | 0 |
| 0 | 0 | 0 |
| 1 | 1 | 1 |
| 1 | 1 | 1 |
| 1 | 1 | 1 |
| 1 | 1 | 1 |
| 0 | 0 | 0 |
| 1 | 1 | 1 |
| 1 | 1 | 1 |
| 0 | 1 | 0 |

| | | |
|---|---|---|
| 1 | 1 | 1 |
| 1 | 1 | 1 |
| 1 | 1 | 1 |
| 1 | 1 | 1 |
| 0 | 1 | 0 |
| 0 | 1 | 0 |
| 0 | 1 | 0 |
| 0 | 0 | 0 |
| 0 | 0 | 0 |
| 1 | 1 | 1 |
| 0 | 1 | 0 |
| 1 | 1 | 1 |
| 1 | 1 | 1 |
| 1 | 1 | 1 |
| 0 | 0 | 0 |
| 0 | 0 | 0 |
| 0 | 0 | 0 |
| 1 | 1 | 1 |
| 1 | 0 | 0 |
| 1 | 1 | 1 |
| 0 | 0 | 0 |
| 0 | 0 | 0 |
| 0 | 0 | 0 |
| 0 | 0 | 0 |
| 0 | 0 | 0 |
| 0 | 0 | 0 |
| 0 | 1 | 0 |
| 1 | 1 | 1 |
| 0 | 0 | 0 |
| 1 | 1 | 1 |
| 1 | 1 | 1 |
| 1 | 1 | 1 |
| 0 | 0 | 0 |
| 1 | 1 | 1 |
| 1 | 1 | 1 |
| 1 | 1 | 1 |
| 0 | 0 | 0 |
| 0 | 0 | 0 |
| 0 | 0 | 0 |
| 1 | 1 | 1 |
| 1 | 1 | 1 |
| 1 | 1 | 1 |
| 1 | 1 | 1 |
| 1 | 1 | 1 |
| 0 | 1 | 0 |
| 0 | 0 | 0 |
| 1 | 1 | 1 |
| 0 | 1 | 0 |
| 1 | 1 | 1 |
| 0 | 1 | 0 |
| 1 | 1 | 1 |
| 1 | 1 | 1 |
| 0 | 1 | 0 |
| 1 | 1 | 1 |

| | | |
|---|---|---|
| 1 | 1 | 1 |
| 0 | 1 | 0 |
| 1 | 1 | 1 |
| 0 | 0 | 0 |
| 1 | 1 | 1 |
| 1 | 1 | 1 |
| 1 | 1 | 1 |
| 0 | 0 | 0 |
| 0 | 0 | 0 |
| 1 | 1 | 1 |
| 1 | 1 | 1 |
| 1 | 1 | 1 |
| 0 | 0 | 0 |
| 0 | 1 | 0 |
| 0 | 1 | 0 |
| 0 | 1 | 0 |
| 0 | 0 | 0 |
| 1 | 1 | 1 |
| 1 | 1 | 1 |
| 1 | 1 | 1 |
| 1 | 1 | 1 |
| 1 | 1 | 1 |
| 1 | 1 | 1 |
| 1 | 1 | 1 |
| 0 | 1 | 0 |
| 1 | 1 | 1 |
| 0 | 1 | 0 |
| 0 | 0 | 0 |
| 0 | 0 | 0 |
| 0 | 0 | 0 |
| 0 | 0 | 0 |
| 0 | 0 | 0 |
| 0 | 0 | 0 |
| 0 | 1 | 0 |
| 0 | 0 | 0 |
| 0 | 0 | 0 |
| 0 | 0 | 0 |
| 1 | 1 | 1 |
| 1 | 1 | 1 |
| 1 | 1 | 1 |
| 1 | 1 | 1 |
| 0 | 1 | 0 |
| 0 | 0 | 0 |
| 0 | 0 | 0 |
| 1 | 1 | 1 |
| 1 | 1 | 1 |
| 0 | 0 | 0 |
| 1 | 1 | 1 |
| 1 | 1 | 1 |
| 1 | 1 | 1 |
| 1 | 1 | 1 |
| 1 | 1 | 1 |
| 0 | 0 | 0 |
| 0 | 0 | 0 |

| | | |
|---|---|---|
| 0 | 0 | 0 |
| 0 | 0 | 0 |
| 0 | 0 | 0 |
| 0 | 0 | 0 |
| 0 | 0 | 0 |
| 0 | 0 | 0 |
| 0 | 0 | 0 |
| 0 | 0 | 0 |
| 1 | 1 | 1 |
| 0 | 0 | 0 |
| 1 | 1 | 1 |
| 1 | 1 | 1 |
| 1 | 1 | 1 |
| 1 | 1 | 1 |
| 0 | 0 | 0 |
| 1 | 1 | 1 |
| 1 | 1 | 1 |
| 1 | 1 | 1 |
| 1 | 1 | 1 |
| 0 | 0 | 0 |
| 0 | 0 | 0 |
| 1 | 1 | 1 |
| 0 | 0 | 0 |
| 0 | 0 | 0 |
| 0 | 0 | 0 |
| 0 | 0 | 0 |
| 0 | 0 | 0 |
| 0 | 1 | 0 |
| 0 | 1 | 0 |
| 0 | 1 | 0 |
| 0 | 0 | 0 |
| 1 | 1 | 1 |
| 1 | 1 | 1 |
| 1 | 1 | 1 |
| 1 | 1 | 1 |
| 1 | 1 | 1 |
| 1 | 1 | 1 |
| 1 | 1 | 1 |
| 1 | 1 | 1 |
| 1 | 1 | 1 |
| 0 | 1 | 0 |
| 0 | 0 | 0 |
| 1 | 1 | 1 |
| 1 | 1 | 1 |
| 1 | 1 | 1 |
| 0 | 1 | 0 |
| 1 | 1 | 1 |
| 1 | 1 | 1 |
| 0 | 0 | 0 |
| 0 | 1 | 0 |
| 1 | 1 | 1 |
| 1 | 1 | 1 |
| 0 | 0 | 0 |
| 0 | 0 | 0 |

| | | |
|---|---|---|
| 1 | 1 | 1 |
| 1 | 1 | 1 |
| 0 | 1 | 0 |
| 1 | 0 | 0 |
| 0 | 0 | 0 |
| 0 | 0 | 0 |
| 0 | 1 | 0 |
| 0 | 0 | 0 |
| 1 | 1 | 1 |
| 1 | 1 | 1 |
| 0 | 0 | 0 |
| 1 | 0 | 0 |
| 0 | 0 | 0 |
| 0 | 0 | 0 |
| 0 | 0 | 0 |
| 0 | 0 | 0 |
| 1 | 1 | 1 |
| 1 | 1 | 1 |
| 0 | 0 | 0 |
| 0 | 0 | 0 |
| 0 | 0 | 0 |
| 0 | 0 | 0 |
| 0 | 0 | 0 |
| 1 | 1 | 1 |
| 1 | 1 | 1 |
| 0 | 0 | 0 |
| 1 | 1 | 1 |
| 0 | 0 | 0 |
| 0 | 0 | 0 |
| 0 | 0 | 0 |
| 0 | 1 | 0 |
| 0 | 0 | 0 |
| 0 | 0 | 0 |
| 1 | 1 | 1 |
| 1 | 1 | 1 |
| 0 | 0 | 0 |
| 0 | 0 | 0 |
| 1 | 0 | 0 |
| 0 | 0 | 0 |
| 0 | 0 | 0 |
| 0 | 0 | 0 |
| 0 | 0 | 0 |
| 0 | 1 | 0 |
| 1 | 1 | 1 |
| 0 | 0 | 0 |
| 0 | 1 | 0 |
| 0 | 0 | 0 |
| 1 | 1 | 1 |
| 1 | 1 | 1 |
| 1 | 1 | 1 |
| 1 | 1 | 1 |
| 1 | 1 | 1 |
| 0 | 1 | 0 |
| 1 | 1 | 1 |

| | | |
|---|---|---|
| 1 | 1 | 1 |
| 1 | 1 | 1 |
| 1 | 1 | 1 |
| 0 | 1 | 0 |
| 1 | 1 | 1 |
| 1 | 1 | 1 |
| 0 | 1 | 0 |
| 0 | 0 | 0 |
| 1 | 1 | 1 |
| 0 | 0 | 0 |
| 0 | 0 | 0 |
| 0 | 0 | 0 |
| 0 | 1 | 0 |
| 1 | 1 | 1 |
| 0 | 0 | 0 |
| 0 | 1 | 0 |
| 0 | 1 | 0 |
| 0 | 1 | 0 |
| 0 | 1 | 0 |
| 0 | 1 | 0 |
| 1 | 1 | 1 |
| 1 | 1 | 1 |
| 1 | 1 | 1 |
| 0 | 1 | 0 |
| 1 | 1 | 1 |
| 1 | 1 | 1 |
| 1 | 1 | 1 |
| 1 | 0 | 0 |
| 1 | 1 | 1 |
| 0 | 0 | 0 |
| 0 | 0 | 0 |
| 1 | 1 | 1 |
| 1 | 1 | 1 |
| 0 | 1 | 0 |
| 0 | 1 | 0 |
| 0 | 0 | 0 |
| 1 | 1 | 1 |
| 1 | 1 | 1 |
| 1 | 1 | 1 |
| 1 | 1 | 1 |
| 1 | 1 | 1 |
| 0 | 0 | 0 |
| 0 | 0 | 0 |
| 0 | 0 | 0 |
| 1 | 1 | 1 |
| 1 | 1 | 1 |
| 1 | 1 | 1 |
| 1 | 1 | 1 |
| 1 | 1 | 1 |
| 1 | 1 | 1 |
| 1 | 1 | 1 |
| 1 | 1 | 1 |
| 1 | 1 | 1 |
| 0 | 0 | 0 |

| | | |
|---|---|---|
| 1 | 1 | 1 |
| 1 | 1 | 1 |
| 1 | 1 | 1 |
| 1 | 1 | 1 |
| 1 | 1 | 1 |
| 1 | 1 | 1 |
| 1 | 1 | 1 |
| 0 | 1 | 0 |
| 0 | 0 | 0 |
| 0 | 0 | 0 |
| 1 | 1 | 1 |
| 0 | 1 | 0 |
| 0 | 1 | 0 |
| 0 | 0 | 0 |
| 1 | 1 | 1 |
| 1 | 1 | 1 |
| 1 | 1 | 1 |
| 0 | 0 | 0 |
| 0 | 0 | 0 |
| 0 | 0 | 0 |
| 0 | 0 | 0 |
| 0 | 1 | 0 |
| 1 | 1 | 1 |
| 1 | 1 | 1 |
| 1 | 1 | 1 |
| 0 | 0 | 0 |
| 1 | 1 | 1 |
| 0 | 1 | 0 |
| 1 | 1 | 1 |
| 1 | 1 | 1 |
| 1 | 1 | 1 |
| 1 | 1 | 1 |
| 1 | 1 | 1 |
| 1 | 1 | 1 |
| 1 | 1 | 1 |
| 1 | 1 | 1 |
| 1 | 1 | 1 |
| 1 | 1 | 1 |
| 1 | 1 | 1 |
| 0 | 0 | 0 |
| 1 | 1 | 1 |
| 1 | 1 | 1 |
| 0 | 0 | 0 |
| 1 | 1 | 1 |
| 1 | 1 | 1 |
| 1 | 1 | 1 |
| 0 | 0 | 0 |
| 0 | 0 | 0 |
| 1 | 1 | 1 |
| 1 | 1 | 1 |
| 0 | 0 | 0 |
| 1 | 1 | 1 |
| 0 | 1 | 0 |
| 1 | 1 | 1 |

| | | |
|---|---|---|
| 1 | 1 | 1 |
| 1 | 1 | 1 |
| 1 | 1 | 1 |
| 1 | 1 | 1 |
| 1 | 1 | 1 |
| 1 | 1 | 1 |
| 1 | 1 | 1 |
| 0 | 1 | 0 |
| 1 | 1 | 1 |
| 1 | 1 | 1 |
| 1 | 1 | 1 |
| 1 | 1 | 1 |
| 1 | 1 | 1 |
| 0 | 0 | 0 |
| 0 | 0 | 0 |
| 0 | 0 | 0 |
| 0 | 0 | 0 |
| 0 | 1 | 0 |
| 1 | 1 | 1 |
| 0 | 0 | 0 |
| 1 | 1 | 1 |
| 1 | 0 | 0 |
| 1 | 1 | 1 |
| 1 | 1 | 1 |
| 1 | 1 | 1 |
| 0 | 0 | 0 |
| 0 | 1 | 0 |
| 1 | 1 | 1 |
| 1 | 1 | 1 |
| 0 | 0 | 0 |
| 0 | 0 | 0 |
| 1 | 1 | 1 |
| 1 | 1 | 1 |
| 1 | 1 | 1 |
| 1 | 1 | 1 |
| 1 | 1 | 1 |
| 1 | 1 | 1 |
| 1 | 1 | 1 |
| 0 | 0 | 0 |
| 1 | 1 | 1 |
| 1 | 1 | 1 |
| 0 | 0 | 0 |
| 0 | 0 | 0 |
| 0 | 0 | 0 |
| 0 | 0 | 0 |
| 1 | 1 | 1 |
| 0 | 0 | 0 |
| 1 | 1 | 1 |
| 1 | 1 | 1 |
| 1 | 1 | 1 |
| 0 | 1 | 0 |
| 1 | 0 | 0 |
| 1 | 0 | 0 |
| 1 | 0 | 0 |

| | | |
|---|---|---|
| 1 | 1 | 1 |
| 1 | 1 | 1 |
| 0 | 0 | 0 |
| 1 | 1 | 1 |
| 1 | 1 | 1 |
| 1 | 1 | 1 |
| 1 | 1 | 1 |
| 0 | 1 | 0 |
| 1 | 1 | 1 |
| 1 | 1 | 1 |
| 1 | 1 | 1 |
| 0 | 1 | 0 |
| 0 | 1 | 0 |
| 0 | 1 | 0 |
| 0 | 1 | 0 |
| 0 | 1 | 0 |
| 0 | 1 | 0 |
| 0 | 1 | 0 |
| 0 | 1 | 0 |
| 0 | 1 | 0 |
| 0 | 1 | 0 |
| 0 | 1 | 0 |
| 0 | 1 | 0 |
| 0 | 1 | 0 |
| 0 | 1 | 0 |
| 0 | 1 | 0 |
| 0 | 1 | 0 |
| 0 | 1 | 0 |
| 1 | 1 | 1 |
| 1 | 1 | 1 |
| 0 | 0 | 0 |
| 1 | 1 | 1 |
| 0 | 0 | 0 |
| 0 | 0 | 0 |
| 0 | 0 | 0 |
| 0 | 0 | 0 |
| 1 | 1 | 1 |
| 0 | 0 | 0 |
| 1 | 1 | 1 |
| 1 | 1 | 1 |
| 0 | 0 | 0 |
| 0 | 0 | 0 |
| 0 | 0 | 0 |
| 1 | 1 | 1 |
| 0 | 0 | 0 |
| 0 | 0 | 0 |
| 1 | 1 | 1 |
| 1 | 1 | 1 |
| 1 | 1 | 1 |
| 1 | 1 | 1 |
| 0 | 0 | 0 |
| 1 | 1 | 1 |
| 1 | 1 | 1 |
| 1 | 1 | 1 |

| | | |
|---|---|---|
| 1 | 1 | 1 |
| 1 | 1 | 1 |
| 1 | 1 | 1 |
| 1 | 1 | 1 |
| 1 | 1 | 1 |
| 0 | 0 | 0 |
| 1 | 1 | 1 |
| 0 | 0 | 0 |
| 0 | 0 | 0 |
| 0 | 0 | 0 |
| 1 | 1 | 1 |
| 0 | 0 | 0 |
| 1 | 1 | 1 |
| 1 | 1 | 1 |
| 1 | 1 | 1 |
| 1 | 0 | 0 |
| 1 | 1 | 1 |
| 0 | 0 | 0 |
| 1 | 1 | 1 |
| 0 | 0 | 0 |
| 1 | 1 | 1 |
| 0 | 0 | 0 |
| 0 | 0 | 0 |
| 0 | 0 | 0 |
| 1 | 1 | 1 |
| 0 | 1 | 0 |
| 1 | 1 | 1 |
| 1 | 1 | 1 |
| 0 | 0 | 0 |
| 1 | 1 | 1 |
| 1 | 1 | 1 |
| 1 | 1 | 1 |
| 1 | 1 | 1 |
| 1 | 1 | 1 |
| 0 | 0 | 0 |
| 0 | 0 | 0 |
| 1 | 1 | 1 |
| 0 | 0 | 0 |
| 0 | 0 | 0 |
| 0 | 1 | 0 |
| 0 | 0 | 0 |
| 0 | 0 | 0 |
| 0 | 1 | 0 |
| 0 | 1 | 0 |
| 1 | 1 | 1 |
| 0 | 1 | 0 |
| 1 | 1 | 1 |
| 1 | 1 | 1 |
| 1 | 1 | 1 |
| 1 | 1 | 1 |
| 1 | 1 | 1 |
| 1 | 1 | 1 |
| 1 | 1 | 1 |
| 1 | 1 | 1 |

| | | |
|---|---|---|
| 1 | 1 | 1 |
| 0 | 0 | 0 |
| 0 | 1 | 0 |
| 1 | 1 | 1 |
| 0 | 0 | 0 |
| 1 | 1 | 1 |
| 1 | 1 | 1 |
| 1 | 1 | 1 |
| 0 | 0 | 0 |
| 0 | 0 | 0 |
| 1 | 1 | 1 |
| 0 | 0 | 0 |
| 0 | 0 | 0 |
| 1 | 1 | 1 |
| 1 | 1 | 1 |
| 0 | 0 | 0 |
| 0 | 1 | 0 |
| 0 | 0 | 0 |
| 0 | 0 | 0 |
| 1 | 1 | 1 |
| 0 | 0 | 0 |
| 0 | 0 | 0 |
| 0 | 0 | 0 |
| 1 | 1 | 1 |
| 0 | 1 | 0 |
| 0 | 0 | 0 |
| 1 | 1 | 1 |
| 1 | 1 | 1 |
| 1 | 1 | 1 |
| 1 | 1 | 1 |
| 0 | 0 | 0 |
| 1 | 1 | 1 |
| 0 | 0 | 0 |
| 1 | 1 | 1 |
| 0 | 0 | 0 |
| 0 | 0 | 0 |
| 1 | 1 | 1 |
| 1 | 1 | 1 |
| 1 | 1 | 1 |
| 1 | 1 | 1 |
| 1 | 1 | 1 |
| 1 | 1 | 1 |
| 1 | 1 | 1 |
| 1 | 1 | 1 |
| 1 | 1 | 1 |
| 1 | 1 | 1 |
| 1 | 0 | 0 |
| 1 | 1 | 1 |
| 1 | 1 | 1 |
| 1 | 1 | 1 |
| 1 | 1 | 1 |
| 1 | 1 | 1 |
| 1 | 1 | 1 |
| 0 | 0 | 0 |

| | | |
|---|---|---|
| 1 | 1 | 1 |
| 1 | 1 | 1 |
| 1 | 1 | 1 |
| 1 | 1 | 1 |
| 0 | 0 | 0 |
| 1 | 1 | 1 |
| 0 | 1 | 0 |
| 0 | 1 | 0 |
| 0 | 1 | 0 |
| 1 | 1 | 1 |
| 0 | 0 | 0 |
| 1 | 1 | 1 |
| 1 | 1 | 1 |
| 1 | 1 | 1 |
| 0 | 0 | 0 |
| 0 | 0 | 0 |
| 0 | 0 | 0 |
| 0 | 0 | 0 |
| 0 | 1 | 0 |
| 0 | 0 | 0 |
| 0 | 0 | 0 |
| 0 | 0 | 0 |
| 1 | 1 | 1 |
| 0 | 0 | 0 |
| 0 | 0 | 0 |
| 1 | 1 | 1 |
| 0 | 0 | 0 |
| 0 | 0 | 0 |
| 0 | 0 | 0 |
| 0 | 0 | 0 |
| 0 | 0 | 0 |
| 1 | 1 | 1 |
| 1 | 1 | 1 |
| 0 | 1 | 0 |
| 1 | 1 | 1 |
| 1 | 1 | 1 |
| 1 | 1 | 1 |
| 0 | 0 | 0 |
| 0 | 0 | 0 |
| 0 | 1 | 0 |
| 1 | 1 | 1 |
| 0 | 0 | 0 |
| 0 | 0 | 0 |
| 0 | 1 | 0 |
| 0 | 0 | 0 |
| 1 | 1 | 1 |
| 1 | 1 | 1 |
| 1 | 1 | 1 |
| 0 | 0 | 0 |
| 0 | 0 | 0 |
| 0 | 0 | 0 |
| 1 | 1 | 1 |
| 0 | 1 | 0 |
| 0 | 0 | 0 |

| | | |
|---|---|---|
| 1 | 1 | 1 |
| 0 | 0 | 0 |
| 1 | 1 | 1 |
| 0 | 0 | 0 |
| 0 | 0 | 0 |
| 0 | 1 | 0 |
| 0 | 0 | 0 |
| 0 | 0 | 0 |
| 1 | 1 | 1 |
| 1 | 1 | 1 |
| 1 | 1 | 1 |
| 1 | 1 | 1 |
| 1 | 1 | 1 |
| 1 | 1 | 1 |
| 1 | 1 | 1 |
| 1 | 1 | 1 |
| 1 | 1 | 1 |
| 0 | 1 | 0 |
| 1 | 1 | 1 |
| 1 | 1 | 1 |
| 0 | 0 | 0 |
| 0 | 1 | 0 |
| 0 | 1 | 0 |
| 0 | 1 | 0 |
| 0 | 1 | 0 |
| 0 | 0 | 0 |
| 0 | 0 | 0 |
| 1 | 1 | 1 |
| 1 | 1 | 1 |
| 1 | 1 | 1 |
| 1 | 1 | 1 |
| 1 | 1 | 1 |
| 1 | 1 | 1 |
| 0 | 1 | 0 |
| 1 | 1 | 1 |
| 1 | 1 | 1 |
| 1 | 1 | 1 |
| 1 | 1 | 1 |
| 0 | 1 | 0 |
| 1 | 1 | 1 |
| 1 | 1 | 1 |
| 0 | 1 | 0 |
| 1 | 1 | 1 |
| 0 | 0 | 0 |
| 1 | 1 | 1 |
| 0 | 0 | 0 |
| 0 | 0 | 0 |
| 0 | 0 | 0 |
| 0 | 0 | 0 |
| 0 | 0 | 0 |
| 0 | 0 | 0 |
| 0 | 1 | 0 |
| 1 | 1 | 1 |
| 1 | 1 | 1 |

| | | |
|---|---|---|
| 1 | 1 | 1 |
| 1 | 1 | 1 |
| 1 | 1 | 1 |
| 0 | 0 | 0 |
| 1 | 1 | 1 |
| 0 | 0 | 0 |
| 1 | 1 | 1 |
| 1 | 1 | 1 |
| 0 | 0 | 0 |
| 1 | 1 | 1 |
| 1 | 1 | 1 |
| 0 | 1 | 0 |
| 1 | 1 | 1 |
| 1 | 1 | 1 |
| 1 | 1 | 1 |
| 0 | 0 | 0 |
| 0 | 0 | 0 |
| 0 | 0 | 0 |
| 0 | 0 | 0 |
| 0 | 0 | 0 |
| 1 | 1 | 1 |
| 1 | 1 | 1 |
| 1 | 1 | 1 |
| 1 | 1 | 1 |
| 0 | 0 | 0 |
| 1 | 1 | 1 |
| 1 | 1 | 1 |
| 1 | 1 | 1 |
| 0 | 0 | 0 |
| 1 | 1 | 1 |
| 0 | 0 | 0 |
| 0 | 0 | 0 |
| 0 | 0 | 0 |
| 0 | 0 | 0 |
| 0 | 0 | 0 |
| 1 | 1 | 1 |
| 0 | 0 | 0 |
| 0 | 0 | 0 |
| 0 | 0 | 0 |
| 0 | 0 | 0 |
| 0 | 0 | 0 |
| 0 | 0 | 0 |
| 0 | 0 | 0 |
| 0 | 0 | 0 |
| 0 | 0 | 0 |
| 0 | 0 | 0 |
| 0 | 0 | 0 |
| 0 | 0 | 0 |
| 1 | 1 | 1 |
| 0 | 0 | 0 |
| 1 | 1 | 1 |
| 1 | 1 | 1 |
| 0 | 1 | 0 |
| 0 | 0 | 0 |

| | | |
|---|---|---|
| 0 | 0 | 0 |
| 0 | 1 | 0 |
| 1 | 1 | 1 |
| 1 | 1 | 1 |
| 1 | 1 | 1 |
| 0 | 0 | 0 |
| 1 | 1 | 1 |
| 0 | 1 | 0 |
| 1 | 1 | 1 |
| 0 | 1 | 0 |
| 0 | 1 | 0 |
| 0 | 1 | 0 |
| 1 | 1 | 1 |
| 0 | 0 | 0 |
| 0 | 1 | 0 |
| 1 | 1 | 1 |
| 1 | 1 | 1 |
| 0 | 0 | 0 |
| 1 | 1 | 1 |
| 1 | 1 | 1 |
| 1 | 1 | 1 |
| 1 | 1 | 1 |
| 0 | 0 | 0 |
| 1 | 1 | 1 |
| 1 | 1 | 1 |
| 0 | 0 | 0 |
| 0 | 0 | 0 |
| 0 | 0 | 0 |
| 1 | 1 | 1 |
| 0 | 1 | 0 |
| 1 | 1 | 1 |
| 1 | 1 | 1 |
| 1 | 1 | 1 |
| 1 | 1 | 1 |
| 1 | 1 | 1 |
| 1 | 1 | 1 |
| 1 | 1 | 1 |
| 1 | 1 | 1 |
| 1 | 1 | 1 |
| 1 | 1 | 1 |
| 1 | 1 | 1 |
| 1 | 1 | 1 |
| 1 | 1 | 1 |
| 1 | 1 | 1 |
| 1 | 1 | 1 |
| 1 | 1 | 1 |
| 1 | 1 | 1 |
| 1 | 1 | 1 |
| 1 | 0 | 0 |
| 1 | 1 | 1 |
| 1 | 1 | 1 |
| 1 | 1 | 1 |
| 1 | 1 | 1 |
| 1 | 1 | 1 |

| | | |
|---|---|---|
| 1 | 1 | 1 |
| 0 | 0 | 0 |
| 1 | 1 | 1 |
| 1 | 1 | 1 |
| 1 | 1 | 1 |
| 0 | 0 | 0 |
| 0 | 0 | 0 |
| 0 | 0 | 0 |
| 1 | 1 | 1 |
| 1 | 1 | 1 |
| 0 | 1 | 0 |
| 1 | 1 | 1 |
| 1 | 1 | 1 |
| 1 | 1 | 1 |
| 0 | 0 | 0 |
| 0 | 0 | 0 |
| 0 | 0 | 0 |
| 1 | 1 | 1 |
| 0 | 0 | 0 |
| 0 | 0 | 0 |
| 0 | 0 | 0 |
| 0 | 0 | 0 |
| 0 | 0 | 0 |
| 0 | 0 | 0 |
| 0 | 1 | 0 |
| 1 | 1 | 1 |
| 1 | 1 | 1 |
| 0 | 1 | 0 |
| 0 | 1 | 0 |
| 1 | 1 | 1 |
| 1 | 1 | 1 |
| 1 | 1 | 1 |
| 0 | 0 | 0 |
| 1 | 1 | 1 |
| 0 | 1 | 0 |
| 0 | 0 | 0 |
| 0 | 0 | 0 |
| 0 | 0 | 0 |
| 0 | 0 | 0 |
| 1 | 1 | 1 |
| 0 | 1 | 0 |
| 1 | 1 | 1 |
| 1 | 1 | 1 |
| 0 | 0 | 0 |
| 0 | 0 | 0 |
| 0 | 0 | 0 |
| 0 | 1 | 0 |
| 1 | 1 | 1 |
| 0 | 0 | 0 |
| 1 | 1 | 1 |
| 1 | 1 | 1 |
| 0 | 0 | 0 |
| 1 | 1 | 1 |
| 0 | 1 | 0 |

| | | |
|---|---|---|
| 1 | 1 | 1 |
| 0 | 1 | 0 |
| 1 | 1 | 1 |
| 1 | 1 | 1 |
| 1 | 1 | 1 |
| 1 | 1 | 1 |
| 0 | 1 | 0 |
| 1 | 1 | 1 |
| 0 | 0 | 0 |
| 0 | 0 | 0 |
| 1 | 1 | 1 |
| 0 | 0 | 0 |
| 0 | 0 | 0 |
| 0 | 0 | 0 |
| 1 | 1 | 1 |
| 1 | 1 | 1 |
| 0 | 0 | 0 |
| 0 | 0 | 0 |
| 0 | 0 | 0 |
| 1 | 1 | 1 |
| 1 | 1 | 1 |
| 1 | 1 | 1 |
| 0 | 0 | 0 |
| 1 | 1 | 1 |
| 0 | 0 | 0 |
| 1 | 1 | 1 |
| 1 | 1 | 1 |
| 1 | 1 | 1 |
| 1 | 1 | 1 |
| 0 | 0 | 0 |
| 0 | 0 | 0 |
| 0 | 0 | 0 |
| 1 | 1 | 1 |
| 0 | 1 | 0 |
| 1 | 1 | 1 |
| 1 | 1 | 1 |
| 1 | 1 | 1 |
| 0 | 1 | 0 |
| 0 | 1 | 0 |
| 0 | 1 | 0 |
| 0 | 0 | 0 |
| 1 | 1 | 1 |
| 1 | 1 | 1 |
| 1 | 1 | 1 |
| 1 | 1 | 1 |
| 0 | 0 | 0 |
| 1 | 0 | 0 |
| 0 | 0 | 0 |
| 1 | 1 | 1 |
| 0 | 0 | 0 |
| 0 | 0 | 0 |
| 0 | 0 | 0 |
| 0 | 1 | 0 |
| 1 | 0 | 0 |

| | | |
|---|---|---|
| 1 | 0 | 0 |
| 0 | 0 | 0 |
| 0 | 1 | 0 |
| 1 | 1 | 1 |
| 0 | 1 | 0 |
| 0 | 1 | 0 |
| 0 | 1 | 0 |
| 0 | 0 | 0 |
| 0 | 0 | 0 |
| 0 | 1 | 0 |
| 0 | 0 | 0 |
| 1 | 1 | 1 |
| 1 | 1 | 1 |
| 1 | 1 | 1 |
| 0 | 1 | 0 |
| 1 | 1 | 1 |
| 1 | 1 | 1 |
| 1 | 1 | 1 |
| 1 | 1 | 1 |
| 0 | 1 | 0 |
| 1 | 1 | 1 |
| 1 | 1 | 1 |
| 1 | 1 | 1 |
| 1 | 1 | 1 |
| 0 | 1 | 0 |
| 0 | 1 | 0 |
| 0 | 1 | 0 |
| 0 | 1 | 0 |
| 0 | 0 | 0 |
| 0 | 1 | 0 |
| 0 | 0 | 0 |
| 1 | 1 | 1 |
| 0 | 0 | 0 |
| 0 | 0 | 0 |
| 0 | 1 | 0 |
| 0 | 0 | 0 |
| 0 | 0 | 0 |
| 1 | 1 | 1 |
| 0 | 1 | 0 |
| 0 | 1 | 0 |
| 1 | 1 | 1 |
| 0 | 0 | 0 |
| 0 | 0 | 0 |
| 1 | 1 | 1 |
| 1 | 1 | 1 |
| 0 | 0 | 0 |
| 0 | 0 | 0 |
| 0 | 0 | 0 |
| 0 | 0 | 0 |
| 0 | 1 | 0 |
| 0 | 0 | 0 |
| 0 | 0 | 0 |
| 1 | 1 | 1 |
| 1 | 1 | 1 |

| | | |
|---|---|---|
| 1 | 1 | 1 |
| 1 | 1 | 1 |
| 1 | 1 | 1 |
| 1 | 1 | 1 |
| 0 | 0 | 0 |
| 0 | 0 | 0 |
| 0 | 0 | 0 |
| 0 | 0 | 0 |
| 1 | 1 | 1 |
| 0 | 0 | 0 |
| 0 | 0 | 0 |
| 0 | 0 | 0 |
| 0 | 0 | 0 |
| 1 | 1 | 1 |
| 1 | 0 | 0 |
| 0 | 1 | 0 |
| 1 | 1 | 1 |
| 1 | 1 | 1 |
| 1 | 1 | 1 |
| 0 | 1 | 0 |
| 1 | 1 | 1 |
| 1 | 1 | 1 |
| 1 | 1 | 1 |
| 1 | 1 | 1 |
| 1 | 1 | 1 |
| 0 | 0 | 0 |
| 0 | 0 | 0 |
| 0 | 0 | 0 |
| 0 | 0 | 0 |
| 0 | 1 | 0 |
| 0 | 1 | 0 |
| 0 | 1 | 0 |
| 1 | 1 | 1 |
| 0 | 1 | 0 |
| 0 | 1 | 0 |
| 1 | 1 | 1 |
| 1 | 1 | 1 |
| 1 | 1 | 1 |
| 1 | 1 | 1 |
| 1 | 1 | 1 |
| 1 | 1 | 1 |
| 1 | 1 | 1 |
| 1 | 1 | 1 |
| 1 | 1 | 1 |
| 1 | 1 | 1 |
| 1 | 1 | 1 |
| 1 | 1 | 1 |
| 1 | 1 | 1 |
| 1 | 1 | 1 |
| 1 | 1 | 1 |
| 1 | 1 | 1 |
| 1 | 1 | 1 |
| 1 | 1 | 1 |
| 0 | 0 | 0 |

| | | |
|---|---|---|
| 1 | 1 | 1 |
| 1 | 1 | 1 |
| 1 | 1 | 1 |
| 0 | 1 | 0 |
| 0 | 1 | 0 |
| 0 | 1 | 0 |
| 0 | 1 | 0 |
| 0 | 1 | 0 |
| 1 | 1 | 1 |
| 1 | 1 | 1 |
| 1 | 1 | 1 |
| 1 | 1 | 1 |
| 1 | 1 | 1 |
| 1 | 1 | 1 |
| 1 | 1 | 1 |
| 1 | 1 | 1 |
| 1 | 1 | 1 |
| 1 | 1 | 1 |
| 1 | 1 | 1 |
| 1 | 1 | 1 |
| 1 | 1 | 1 |
| 1 | 1 | 1 |
| 1 | 1 | 1 |
| 1 | 1 | 1 |
| 1 | 1 | 1 |
| 1 | 1 | 1 |
| 1 | 1 | 1 |
| 1 | 1 | 1 |
| 1 | 1 | 1 |
| 1 | 1 | 1 |
| 1 | 1 | 1 |
| 1 | 1 | 1 |
| 1 | 1 | 1 |
| 0 | 1 | 0 |
| 1 | 1 | 1 |
| 1 | 1 | 1 |
| 1 | 1 | 1 |
| 1 | 1 | 1 |
| 1 | 1 | 1 |
| 1 | 1 | 1 |
| 0 | 1 | 0 |
| 1 | 1 | 1 |
| 1 | 1 | 1 |
| 1 | 1 | 1 |
| 0 | 1 | 0 |
| 1 | 0 | 0 |
| 0 | 0 | 0 |
| 1 | 1 | 1 |
| 0 | 1 | 0 |
| 1 | 1 | 1 |
| 1 | 1 | 1 |
| 0 | 1 | 0 |
| 0 | 1 | 0 |
| 0 | 1 | 0 |

| | | |
|---|---|---|
| 0 | 1 | 0 |
| 1 | 1 | 1 |
| 0 | 0 | 0 |
| 1 | 1 | 1 |
| 1 | 1 | 1 |
| 1 | 1 | 1 |
| 0 | 0 | 0 |
| 1 | 1 | 1 |
| 1 | 1 | 1 |
| 0 | 1 | 0 |
| 0 | 1 | 0 |
| 0 | 0 | 0 |
| 1 | 1 | 1 |
| 0 | 0 | 0 |
| 0 | 0 | 0 |
| 0 | 0 | 0 |
| 0 | 0 | 0 |
| 0 | 0 | 0 |
| 0 | 0 | 0 |
| 0 | 0 | 0 |
| 0 | 0 | 0 |

| organism_scientific | evo_cons | cluster_id | pdb_id | chain_id | resi | resn | change | uniprot_id |
|---|---|---|---|---|---|---|---|---|
| HOMO SAPIENS | 5 | 87 | 3uzw | A | 106 | LEU | p.Leu106Phe | P51857 |
| HOMO SAPIENS | 3 | 87 | 3uzw | A | 133 | PRO | p.Pro133Arg | P51857 |
| HOMO SAPIENS | 0 | 87 | 3uzw | A | 198 | PRO | p.Pro198Leu | P51857 |
| HOMO SAPIENS | 7 | 87 | 3uzw | A | 261 | ARG | p.Arg261Cys | P51857 |
| HOMO SAPIENS | 0 | 4693 | 3mk1 | A | 3 | PRO | p.Pro25Leu | P05187 |
| HOMO SAPIENS | 8 | 4830 | 2h64 | B | 101 | CYS | p.Cys124Arg | P36894 |
| HOMO SAPIENS | 2 | 4830 | 2h64 | B | 116 | PRO | p.Pro139Ser | P36894 |
| HOMO SAPIENS | 0 | 6937 | 3dar | A | 186 | MET | p.Met186Thr | P21802 |
| HOMO SAPIENS | 3 | 5 | 1aut | C | 30 | GLN | p.Gln226His | P04070 |
| HOMO SAPIENS | 1 | 5 | 1aut | C | 68 | LEU | p.Leu265Phe | P04070 |
| HOMO SAPIENS | 0 | 5 | 1aut | C | 75 | ARG | p.Arg272Cys | P04070 |
| HOMO SAPIENS | 6 | 5 | 1aut | C | 92 | PRO | p.Pro289Leu | P04070 |
| HOMO SAPIENS | 0 | 5 | 1aut | C | 104 | ALA | p.Ala301Val | P04070 |
| HOMO SAPIENS | 1 | 5 | 1aut | C | 112 | ALA | p.Ala309Thr | P04070 |
| HOMO SAPIENS | 2 | 5 | 1aut | C | 115 | SER | p.Ser312Leu | P04070 |
| HOMO SAPIENS | 1 | 5 | 1aut | C | 133 | GLY | p.Gly334Ser | P04070 |
| HOMO SAPIENS | 0 | 5 | 1aut | C | 147 | ARG | p.Arg348Ter | P04070 |
| HOMO SAPIENS | 8 | 5 | 1aut | C | 194 | ASP | p.Asp401Asn | P04070 |
| HOMO SAPIENS | 7 | 5 | 1aut | C | 237 | TRP | p.Trp444Cys | P04070 |
| HOMO SAPIENS | 0 | 5 | 1aut | C | 238 | ILE | p.Ile445Met | P04070 |
| HOMO SAPIENS | 8 | 43 | 1aiy | F | 10 | HIS | p.His34Asp | P01308 |
| HOMO SAPIENS | 2 | 43 | 1aiy | F | 22 | ARG | p.Arg46Gln | P01308 |
| HOMO SAPIENS | 4 | 43 | 1aiy | F | 23 | GLY | p.Gly47Val | P01308 |
| HOMO SAPIENS | 7 | 43 | 1aiy | F | 24 | PHE | p.Phe48Ser | P01308 |
| HOMO SAPIENS | 5 | 24 | 2fg8 | A | 33 | THR | p.Thr30Ile | P02792 |
| HOMO SAPIENS | 4 | 24 | 2fg8 | A | 99 | ALA | p.Ala96Thr | P02792 |
| HOMO SAPIENS | 5 | 24 | 2fg8 | A | 107 | GLU | p.Glu104Ter | P02792 |
| HOMO SAPIENS | 3 | 5409 | 5bnq | A | 199 | MET | p.Met199Lys | O14788 |
| HOMO SAPIENS | -1 | 839 | 3swc | A | 959 | GLN | p.Gln990His | Q9H9B1 |
| HOMO SAPIENS | -1 | 839 | 3swc | A | 968 | ALA | p.Ala999Val | Q9H9B1 |
| HOMO SAPIENS | -1 | 839 | 3swc | A | 973 | SER | p.Ser1004Ile | Q9H9B1 |
| HOMO SAPIENS | 8 | 17 | 1eou | A | 18 | LYS | p.Lys18Glu | P00918 |
| HOMO SAPIENS | 2 | 17 | 1eou | A | 107 | HIS | p.His107Tyr | P00918 |
| HOMO SAPIENS | 1 | 17 | 1eou | A | 236 | PRO | p.Pro236His | P00918 |
| HOMO SAPIENS | 6 | 17 | 1eou | A | 252 | ASN | p.Asn252Asp | P00918 |
| HOMO SAPIENS | 2 | 9140 | 5c4v | A | 445 | ARG | p.Arg445Ter | Q13485 |
| HOMO SAPIENS | 7 | 9140 | 5c4v | A | 496 | ARG | p.Arg496Cys | Q13485 |
| HOMO SAPIENS | -1 | 42 | 4pwi | A | 6 | GLY | p.Gly26Ser | P02766 |
| HOMO SAPIENS | 6 | 42 | 4pwi | A | 10 | CYS | p.Cys30Arg | P02766 |
| HOMO SAPIENS | 8 | 42 | 4pwi | A | 12 | LEU | p.Leu32Pro | P02766 |
| HOMO SAPIENS | 9 | 42 | 4pwi | A | 18 | ASP | p.Asp38Gly | P02766 |
| HOMO SAPIENS | 8 | 42 | 4pwi | A | 20 | VAL | p.Val40Ile | P02766 |
| HOMO SAPIENS | 4 | 42 | 4pwi | A | 30 | MET | p.Val50Ala | P02766 |
| HOMO SAPIENS | 4 | 42 | 4pwi | A | 30 | MET | p.Val50Leu | P02766 |
| HOMO SAPIENS | 6 | 42 | 4pwi | A | 33 | PHE | p.Phe53Ile | P02766 |
| HOMO SAPIENS | 7 | 42 | 4pwi | A | 36 | ALA | p.Ala56Pro | P02766 |
| HOMO SAPIENS | 7 | 42 | 4pwi | A | 42 | GLU | p.Glu62Gly | P02766 |
| HOMO SAPIENS | 8 | 42 | 4pwi | A | 44 | PHE | p.Phe64Ser | P02766 |
| HOMO SAPIENS | 4 | 42 | 4pwi | A | 47 | GLY | p.Gly67Ala | P02766 |
| HOMO SAPIENS | 6 | 42 | 4pwi | A | 49 | THR | p.Thr69Ala | P02766 |
| HOMO SAPIENS | 6 | 42 | 4pwi | A | 50 | SER | p.Ser70Ile | P02766 |
| HOMO SAPIENS | 6 | 42 | 4pwi | A | 50 | SER | p.Ser70Arg | P02766 |
| HOMO SAPIENS | 0 | 42 | 4pwi | A | 53 | GLY | p.Gly73Glu | P02766 |

| | | | | | | | | |
|---|---|---|---|---|---|---|---|---|
| HOMO SAPIENS | 7 | 42 | 4pwi | A | 55 | LEU | p.Leu75Pro | P02766 |
| HOMO SAPIENS | 6 | 42 | 4pwi | A | 58 | LEU | p.Leu78His | P02766 |
| HOMO SAPIENS | 7 | 42 | 4pwi | A | 61 | GLU | p.Glu81Lys | P02766 |
| HOMO SAPIENS | 8 | 42 | 4pwi | A | 64 | PHE | p.Phe84Leu | P02766 |
| HOMO SAPIENS | 8 | 42 | 4pwi | A | 68 | ILE | p.Ile88Leu | P02766 |
| HOMO SAPIENS | 8 | 42 | 4pwi | A | 69 | TYR | p.Tyr89His | P02766 |
| HOMO SAPIENS | 4 | 42 | 4pwi | A | 71 | VAL | p.Val91Ala | P02766 |
| HOMO SAPIENS | 6 | 42 | 4pwi | A | 77 | SER | p.Ser97Tyr | P02766 |
| HOMO SAPIENS | 4 | 42 | 4pwi | A | 84 | ILE | p.Ile104Ser | P02766 |
| HOMO SAPIENS | 8 | 42 | 4pwi | A | 89 | GLU | p.Glu109Gln | P02766 |
| HOMO SAPIENS | 8 | 42 | 4pwi | A | 97 | ALA | p.Ala117Gly | P02766 |
| HOMO SAPIENS | 7 | 42 | 4pwi | A | 99 | ASP | p.Asp119Asn | P02766 |
| HOMO SAPIENS | 9 | 42 | 4pwi | A | 107 | ILE | p.Ile127Val | P02766 |
| HOMO SAPIENS | 9 | 42 | 4pwi | A | 111 | LEU | p.Leu131Met | P02766 |
| HOMO SAPIENS | 7 | 42 | 4pwi | A | 114 | TYR | p.Tyr134Cys | P02766 |
| HOMO SAPIENS | 7 | 42 | 4pwi | A | 114 | TYR | p.Tyr134His | P02766 |
| HOMO SAPIENS | 7 | 42 | 4pwi | A | 122 | VAL | p.Val142Ile | P02766 |
| HOMO SAPIENS | -1 | 42 | 4pwi | A | 126 | LYS | p.Lys146Arg | P02766 |
| HOMO SAPIENS | -1 | 42 | 4pwi | A | 127 | GLU | p.Glu147Ter | P02766 |
| HOMO SAPIENS | -1 | 1312 | 4dce | A | 1275 | ARG | p.Arg1275Gln | Q9UM73 |
| HOMO SAPIENS | -1 | 1312 | 4dce | A | 1280 | ALA | p.Ala1280Val | Q9UM73 |
| HOMO SAPIENS | 1 | 20 | 1y0x | X | 234 | ALA | p.Ala234Thr | P10828 |
| HOMO SAPIENS | 0 | 20 | 1y0x | X | 243 | ARG | p.Arg243Trp | P10828 |
| HOMO SAPIENS | 3 | 20 | 1y0x | X | 316 | ARG | p.Arg316His | P10828 |
| HOMO SAPIENS | 1 | 20 | 1y0x | X | 317 | ALA | p.Ala317Thr | P10828 |
| HOMO SAPIENS | 2 | 20 | 1y0x | X | 320 | ARG | p.Arg320Cys | P10828 |
| HOMO SAPIENS | 2 | 20 | 1y0x | X | 320 | ARG | p.Arg320His | P10828 |
| HOMO SAPIENS | 0 | 20 | 1y0x | X | 332 | GLY | p.Gly332Arg | P10828 |
| HOMO SAPIENS | 1 | 20 | 1y0x | X | 338 | ARG | p.Arg338Trp | P10828 |
| HOMO SAPIENS | 0 | 20 | 1y0x | X | 340 | GLN | p.Gln340His | P10828 |
| HOMO SAPIENS | 0 | 20 | 1y0x | X | 345 | GLY | p.Gly345Val | P10828 |
| HOMO SAPIENS | 0 | 20 | 1y0x | X | 345 | GLY | p.Gly345Ser | P10828 |
| HOMO SAPIENS | 0 | 20 | 1y0x | X | 347 | GLY | p.Gly347Glu | P10828 |
| HOMO SAPIENS | 1 | 20 | 1y0x | X | 438 | ARG | p.Arg438His | P10828 |
| HOMO SAPIENS | 0 | 20 | 1y0x | X | 442 | MET | p.Met442Val | P10828 |
| HOMO SAPIENS | 0 | 20 | 1y0x | X | 443 | LYS | p.Lys443Glu | P10828 |
| HOMO SAPIENS | 0 | 20 | 1y0x | X | 446 | CYS | p.Cys446Arg | P10828 |
| HOMO SAPIENS | 2 | 20 | 1y0x | X | 453 | PRO | p.Pro453His | P10828 |
| HOMO SAPIENS | 2 | 20 | 1y0x | X | 453 | PRO | p.Pro453Thr | P10828 |
| HOMO SAPIENS | 3 | 20 | 1y0x | X | 459 | PHE | p.Phe459Cys | P10828 |
| HOMO SAPIENS | 0 | 4409 | 3qb7 | D | 87 | GLU | p.Glu109Lys | P31785 |
| HOMO SAPIENS | 1 | 4409 | 3qb7 | D | 92 | GLY | p.Gly114Asp | P31785 |
| HOMO SAPIENS | 1 | 4409 | 3qb7 | D | 93 | CYS | p.Cys115Arg | P31785 |
| HOMO SAPIENS | 0 | 4409 | 3qb7 | D | 131 | ILE | p.Ile153Asn | P31785 |
| HOMO SAPIENS | 9 | 4409 | 3qb7 | D | 200 | ARG | p.Arg222Cys | P31785 |
| HOMO SAPIENS | 0 | 3891 | 3ije | B | 33 | LEU | p.Leu59Pro | P05106 |
| HOMO SAPIENS | 0 | 3891 | 3ije | B | 117 | LEU | p.Leu143Trp | P05106 |
| HOMO SAPIENS | 3 | 3891 | 3ije | B | 119 | ASP | p.Asp145Tyr | P05106 |
| HOMO SAPIENS | 2 | 3891 | 3ije | B | 143 | ARG | p.Arg169Gln | P05106 |
| HOMO SAPIENS | 2 | 3891 | 3ije | B | 214 | ARG | p.Arg240Gln | P05106 |
| HOMO SAPIENS | 2 | 3891 | 3ije | B | 214 | ARG | p.Arg240Trp | P05106 |
| HOMO SAPIENS | 2 | 3891 | 3ije | B | 374 | CYS | p.Cys400Tyr | P05106 |
| HOMO SAPIENS | 1 | 3891 | 3ije | B | 407 | PRO | p.Pro433Ala | P05106 |
| HOMO SAPIENS | 0 | 3891 | 3ije | B | 489 | ARG | p.Arg515Gln | P05106 |

| | | | | | | | | |
|---|---|---|---|---|---|---|---|---|
| HOMO SAPIENS | -1 | 9613 | 5mke | B | 28 | ARG | p.Arg28Pro | Q13563 |
| HOMO SAPIENS | -1 | 9613 | 5mke | B | 190 | ALA | p.Ala190Thr | Q13563 |
| HOMO SAPIENS | 4 | 9613 | 5mke | B | 262 | VAL | p.Val262Met | Q13563 |
| HOMO SAPIENS | 0 | 9613 | 5mke | B | 482 | PHE | p.Phe482Cys | Q13563 |
| HOMO SAPIENS | 4 | 9613 | 5mke | B | 511 | ASP | p.Asp511Val | Q13563 |
| HOMO SAPIENS | 5 | 9613 | 5mke | B | 516 | VAL | p.Val516Leu | Q13563 |
| HOMO SAPIENS | -1 | 9613 | 5mke | B | 807 | ARG | p.Arg807Gln | Q13563 |
| HOMO SAPIENS | 4 | 2428 | 1olh | C | 337 | ARG | p.Arg337Cys | P04637 |
| HOMO SAPIENS | 4 | 2428 | 1olh | C | 337 | ARG | p.Arg337His | P04637 |
| HOMO SAPIENS | 8 | 2428 | 1olh | C | 338 | PHE | p.Phe338Leu | P04637 |
| HOMO SAPIENS | 8 | 2428 | 1olh | C | 339 | GLU | p.Glu339Lys | P04637 |
| HOMO SAPIENS | 7 | 2428 | 1olh | C | 344 | LEU | p.Leu344Pro | P04637 |
| HOMO SAPIENS | 0 | 2428 | 1olh | C | 360 | GLY | p.Gly360Ala | P04637 |
| HOMO SAPIENS | 7 | 4867 | 1mox | B | 404 | GLY | p.Gly428Asp | P00533 |
| HOMO SAPIENS | 5 | 4867 | 1mox | B | 497 | ARG | p.Arg521Lys | P00533 |
| HOMO SAPIENS | 1 | 4630 | 1fu1 | A | 43 | TRP | p.Trp43Arg | Q13426 |
| HOMO SAPIENS | 2 | 4630 | 1fu1 | A | 161 | ARG | p.Arg161Ter | Q13426 |
| HOMO SAPIENS | 2 | 15 | 1lfy | B | 2 | HIS | p.His3Gln | P68871 |
| HOMO SAPIENS | 2 | 15 | 1lfy | B | 6 | GLU | p.Glu7Ala | P68871 |
| HOMO SAPIENS | 2 | 15 | 1lfy | B | 6 | GLU | p.Glu7Lys | P68871 |
| HOMO SAPIENS | 4 | 15 | 1lfy | B | 9 | SER | p.Ser10Cys | P68871 |
| HOMO SAPIENS | 6 | 15 | 1lfy | B | 15 | TRP | p.Trp16Ter | P68871 |
| HOMO SAPIENS | 3 | 15 | 1lfy | B | 17 | LYS | p.Lys18Glu | P68871 |
| HOMO SAPIENS | 6 | 15 | 1lfy | B | 19 | ASN | p.Asn20Ser | P68871 |
| HOMO SAPIENS | 6 | 15 | 1lfy | B | 22 | GLU | p.Glu23Ala | P68871 |
| HOMO SAPIENS | 6 | 15 | 1lfy | B | 22 | GLU | p.Glu23Lys | P68871 |
| HOMO SAPIENS | 8 | 15 | 1lfy | B | 26 | GLU | p.Glu27Lys | P68871 |
| HOMO SAPIENS | 8 | 15 | 1lfy | B | 28 | LEU | p.Leu29Pro | P68871 |
| HOMO SAPIENS | 8 | 15 | 1lfy | B | 30 | ARG | p.Arg31Ser | P68871 |
| HOMO SAPIENS | 8 | 15 | 1lfy | B | 30 | ARG | p.Arg31Lys | P68871 |
| HOMO SAPIENS | 6 | 15 | 1lfy | B | 39 | GLN | p.Gln40Ter | P68871 |
| HOMO SAPIENS | 2 | 15 | 1lfy | B | 43 | GLU | p.Glu44Gln | P68871 |
| HOMO SAPIENS | 6 | 15 | 1lfy | B | 57 | ASN | p.Asn58Lys | P68871 |
| HOMO SAPIENS | 6 | 15 | 1lfy | B | 59 | LYS | p.Lys60Asn | P68871 |
| HOMO SAPIENS | 8 | 15 | 1lfy | B | 61 | LYS | p.Lys62Glu | P68871 |
| HOMO SAPIENS | 4 | 15 | 1lfy | B | 73 | ASP | p.Asp74Tyr | P68871 |
| HOMO SAPIENS | 4 | 15 | 1lfy | B | 87 | THR | p.Thr88Ile | P68871 |
| HOMO SAPIENS | 4 | 15 | 1lfy | B | 87 | THR | p.Thr88Pro | P68871 |
| HOMO SAPIENS | 5 | 15 | 1lfy | B | 90 | GLU | p.Glu91Lys | P68871 |
| HOMO SAPIENS | 6 | 15 | 1lfy | B | 91 | LEU | p.Leu92Pro | P68871 |
| HOMO SAPIENS | 5 | 15 | 1lfy | B | 97 | HIS | p.His98Gln | P68871 |
| HOMO SAPIENS | 6 | 15 | 1lfy | B | 110 | LEU | p.Leu111Pro | P68871 |
| HOMO SAPIENS | 6 | 15 | 1lfy | B | 113 | VAL | p.Val114Glu | P68871 |
| HOMO SAPIENS | 2 | 15 | 1lfy | B | 119 | GLY | p.Gly120Ala | P68871 |
| HOMO SAPIENS | 4 | 15 | 1lfy | B | 121 | GLU | p.Glu122Gln | P68871 |
| HOMO SAPIENS | 5 | 15 | 1lfy | B | 134 | VAL | p.Val135Glu | P68871 |
| HOMO SAPIENS | 3 | 15 | 1lfy | B | 143 | HIS | p.His144Gln | P68871 |
| HOMO SAPIENS | 4 | 91 | 5d9k | A | 75 | GLY | p.Gly75Val | P51812 |
| HOMO SAPIENS | 5 | 91 | 5d9k | A | 82 | VAL | p.Val82Phe | P51812 |
| HOMO SAPIENS | 0 | 91 | 5d9k | A | 114 | ARG | p.Arg114Trp | P51812 |
| HOMO SAPIENS | 4 | 91 | 5d9k | A | 189 | ILE | p.Ile189Lys | P51812 |
| HOMO SAPIENS | -1 | 91 | 5d9k | A | 227 | SER | p.Ser227Ala | P51812 |
| HOMO SAPIENS | 1 | 91 | 5d9k | A | 268 | PHE | p.Phe268Ser | P51812 |
| HOMO SAPIENS | 0 | 954 | 2vt0 | A | 15 | VAL | p.Val54Leu | P04062 |

| | | | | | | | | |
|---|---|---|---|---|---|---|---|---|
| HOMO SAPIENS | 5 | 954 | 2vt0 | A | 46 | GLY | p.Gly85Glu | P04062 |
| HOMO SAPIENS | 5 | 954 | 2vt0 | A | 48 | ARG | p.Arg87Trp | P04062 |
| HOMO SAPIENS | 6 | 954 | 2vt0 | A | 79 | LYS | p.Lys118Asn | P04062 |
| HOMO SAPIENS | 0 | 954 | 2vt0 | A | 120 | ARG | p.Arg159Gln | P04062 |
| HOMO SAPIENS | 0 | 954 | 2vt0 | A | 120 | ARG | p.Arg159Trp | P04062 |
| HOMO SAPIENS | 0 | 954 | 2vt0 | A | 122 | PRO | p.Pro161Ser | P04062 |
| HOMO SAPIENS | 5 | 954 | 2vt0 | A | 131 | ARG | p.Arg170Leu | P04062 |
| HOMO SAPIENS | 5 | 954 | 2vt0 | A | 135 | TYR | p.Tyr174Cys | P04062 |
| HOMO SAPIENS | 0 | 954 | 2vt0 | A | 157 | LYS | p.Lys196Gln | P04062 |
| HOMO SAPIENS | 5 | 954 | 2vt0 | A | 163 | ARG | p.Arg202Gln | P04062 |
| HOMO SAPIENS | 5 | 954 | 2vt0 | A | 170 | ARG | p.Arg209Cys | P04062 |
| HOMO SAPIENS | 5 | 954 | 2vt0 | A | 188 | ASN | p.Asn227Ser | P04062 |
| HOMO SAPIENS | 3 | 954 | 2vt0 | A | 196 | SER | p.Ser235Pro | P04062 |
| HOMO SAPIENS | 5 | 954 | 2vt0 | A | 202 | GLY | p.Gly241Arg | P04062 |
| HOMO SAPIENS | 0 | 954 | 2vt0 | A | 212 | TYR | p.Tyr251His | P04062 |
| HOMO SAPIENS | 0 | 954 | 2vt0 | A | 213 | PHE | p.Phe252Ile | P04062 |
| HOMO SAPIENS | 5 | 954 | 2vt0 | A | 216 | PHE | p.Phe255Tyr | P04062 |
| HOMO SAPIENS | 6 | 954 | 2vt0 | A | 251 | PHE | p.Phe290Leu | P04062 |
| HOMO SAPIENS | 5 | 954 | 2vt0 | A | 255 | HIS | p.His294Gln | P04062 |
| HOMO SAPIENS | 6 | 954 | 2vt0 | A | 257 | ARG | p.Arg296Gln | P04062 |
| HOMO SAPIENS | 5 | 954 | 2vt0 | A | 289 | PRO | p.Pro328Leu | P04062 |
| HOMO SAPIENS | 0 | 954 | 2vt0 | A | 309 | ALA | p.Ala348Val | P04062 |
| HOMO SAPIENS | 0 | 954 | 2vt0 | A | 311 | HIS | p.His350Arg | P04062 |
| HOMO SAPIENS | 6 | 954 | 2vt0 | A | 312 | TRP | p.Trp351Cys | P04062 |
| HOMO SAPIENS | 5 | 954 | 2vt0 | A | 323 | THR | p.Thr362Ile | P04062 |
| HOMO SAPIENS | 5 | 954 | 2vt0 | A | 325 | GLY | p.Gly364Arg | P04062 |
| HOMO SAPIENS | 6 | 954 | 2vt0 | A | 342 | CYS | p.Cys381Gly | P04062 |
| HOMO SAPIENS | 5 | 954 | 2vt0 | A | 353 | ARG | p.Arg392Gly | P04062 |
| HOMO SAPIENS | 3 | 954 | 2vt0 | A | 364 | SER | p.Ser403Thr | P04062 |
| HOMO SAPIENS | 5 | 954 | 2vt0 | A | 369 | THR | p.Thr408Met | P04062 |
| HOMO SAPIENS | 3 | 954 | 2vt0 | A | 370 | ASN | p.Asn409Ser | P04062 |
| HOMO SAPIENS | 0 | 954 | 2vt0 | A | 371 | LEU | p.Leu410Val | P04062 |
| HOMO SAPIENS | 0 | 954 | 2vt0 | A | 375 | VAL | p.Val414Leu | P04062 |
| HOMO SAPIENS | 0 | 954 | 2vt0 | A | 377 | GLY | p.Gly416Ser | P04062 |
| HOMO SAPIENS | 8 | 954 | 2vt0 | A | 398 | VAL | p.Val437Phe | P04062 |
| HOMO SAPIENS | 0 | 954 | 2vt0 | A | 401 | PRO | p.Pro440Leu | P04062 |
| HOMO SAPIENS | 3 | 954 | 2vt0 | A | 409 | ASP | p.Asp448His | P04062 |
| HOMO SAPIENS | 3 | 954 | 2vt0 | A | 409 | ASP | p.Asp448Val | P04062 |
| HOMO SAPIENS | 0 | 954 | 2vt0 | A | 415 | PRO | p.Pro454Arg | P04062 |
| HOMO SAPIENS | 5 | 954 | 2vt0 | A | 444 | LEU | p.Leu483Arg | P04062 |
| HOMO SAPIENS | 5 | 954 | 2vt0 | A | 456 | ALA | p.Ala495Pro | P04062 |
| HOMO SAPIENS | 5 | 954 | 2vt0 | A | 463 | ARG | p.Arg502Cys | P04062 |
| HOMO SAPIENS | 5 | 954 | 2vt0 | A | 463 | ARG | p.Arg502His | P04062 |
| HOMO SAPIENS | 5 | 954 | 2vt0 | A | 478 | GLY | p.Gly517Ser | P04062 |
| HOMO SAPIENS | 5 | 954 | 2vt0 | A | 496 | ARG | p.Arg535Cys | P04062 |
| HOMO SAPIENS | 5 | 954 | 2vt0 | A | 496 | ARG | p.Arg535His | P04062 |
| HOMO SAPIENS | 1 | 9037 | 2rqb | A | 946 | ALA | p.Ala946Thr | Q9BYX4 |
| HOMO SAPIENS | 0 | 253 | 5E+092 | A | 308 | LEU | p.Leu308Pro | P37173 |
| HOMO SAPIENS | 0 | 253 | 5E+092 | A | 315 | THR | p.Thr315Met | P37173 |
| HOMO SAPIENS | 0 | 253 | 5E+092 | A | 328 | HIS | p.His328Gln | P37173 |
| HOMO SAPIENS | 0 | 253 | 5E+092 | A | 336 | TYR | p.Tyr336Asn | P37173 |
| HOMO SAPIENS | 0 | 253 | 5E+092 | A | 348 | ARG | p.Arg348Cys | P37173 |
| HOMO SAPIENS | 0 | 253 | 5E+092 | A | 355 | ALA | p.Ala355Pro | P37173 |
| HOMO SAPIENS | 1 | 253 | 5E+092 | A | 356 | ARG | p.Arg356Gly | P37173 |

| | | | | | | | | |
|---|---|---|---|---|---|---|---|---|
| HOMO SAPIENS | 0 | 253 | 5E+092 | A | 357 | GLY | p.Gly357Trp | P37173 |
| HOMO SAPIENS | 1 | 253 | 5E+092 | A | 403 | ARG | p.Arg403His | P37173 |
| HOMO SAPIENS | 5 | 253 | 5E+092 | A | 449 | SER | p.Ser449Phe | P37173 |
| HOMO SAPIENS | 0 | 253 | 5E+092 | A | 460 | ARG | p.Arg460Cys | P37173 |
| HOMO SAPIENS | 0 | 253 | 5E+092 | A | 460 | ARG | p.Arg460His | P37173 |
| HOMO SAPIENS | 2 | 253 | 5E+092 | A | 526 | GLU | p.Glu526Gln | P37173 |
| HOMO SAPIENS | 5 | 253 | 5E+092 | A | 528 | ARG | p.Arg528Cys | P37173 |
| HOMO SAPIENS | 5 | 253 | 5E+092 | A | 528 | ARG | p.Arg528His | P37173 |
| HOMO SAPIENS | 6 | 158 | 5amc | A | 231 | ARG | p.Arg260Leu | P12821 |
| HOMO SAPIENS | 7 | 158 | 5amc | A | 237 | TYR | p.Tyr266Ter | P12821 |
| HOMO SAPIENS | 6 | 158 | 5amc | A | 325 | GLY | p.Gly354Arg | P12821 |
| HOMO SAPIENS | 0 | 158 | 5amc | A | 467 | ARG | p.Arg496Ter | P12821 |
| HOMO SAPIENS | 4 | 158 | 5amc | A | 479 | ARG | p.Arg508Ter | P12821 |
| HOMO SAPIENS | 0 | 158 | 5amc | A | 563 | ASP | p.Asp592Gly | P12821 |
| HOMO SAPIENS | -1 | 33062 | 1gr3 | A | 545 | GLY | p.Gly545Arg | Q03692 |
| HOMO SAPIENS | 5 | 33062 | 1gr3 | A | 591 | CYS | p.Cys591Arg | Q03692 |
| HOMO SAPIENS | 2 | 33062 | 1gr3 | A | 595 | GLY | p.Gly595Glu | Q03692 |
| HOMO SAPIENS | 0 | 33062 | 1gr3 | A | 597 | TYR | p.Tyr597Cys | Q03692 |
| HOMO SAPIENS | 5 | 33062 | 1gr3 | A | 598 | TYR | p.Tyr598Asp | Q03692 |
| HOMO SAPIENS | 7 | 33062 | 1gr3 | A | 600 | SER | p.Ser600Pro | Q03692 |
| HOMO SAPIENS | 1 | 33062 | 1gr3 | A | 614 | LEU | p.Leu614Pro | Q03692 |
| HOMO SAPIENS | 6 | 33062 | 1gr3 | A | 651 | TRP | p.Trp651Arg | Q03692 |
| HOMO SAPIENS | 7 | 33062 | 1gr3 | A | 671 | SER | p.Ser671Pro | Q03692 |
| HOMO SAPIENS | -1 | 38257 | 2l2i | B | 87 | PRO | p.Pro87Ala | Q13351 |
| HOMO SAPIENS | 2 | 2036 | 1ghq | A | 231 | GLY | p.Gly1224Asp | P01024 |
| HOMO SAPIENS | 5 | 3594 | 4cde | F | 388 | TYR | p.Tyr412Cys | P53634 |
| HOMO SAPIENS | 9 | 3594 | 4cde | F | 405 | TRP | p.Trp429Cys | P53634 |
| HOMO SAPIENS | 0 | 3594 | 4cde | F | 429 | ILE | p.Ile453Val | P53634 |
| HOMO SAPIENS | 7 | 18 | 1jkb | A | 34 | TRP | p.Trp52Ter | P61626 |
| HOMO SAPIENS | 9 | 18 | 1jkb | A | 70 | THR | p.Thr88Asn | P61626 |
| HOMO SAPIENS | 8 | 18 | 1jkb | A | 121 | VAL | p.Val139Ile | P61626 |
| HOMO SAPIENS | 2 | 414 | 3fj6 | A | 136 | ARG | p.Arg135Cys | Q02127 |
| HOMO SAPIENS | 1 | 414 | 3fj6 | A | 153 | GLY | p.Gly152Arg | Q02127 |
| HOMO SAPIENS | 0 | 414 | 3fj6 | A | 200 | ARG | p.Arg199Cys | Q02127 |
| HOMO SAPIENS | 0 | 414 | 3fj6 | A | 203 | GLY | p.Gly202Ala | Q02127 |
| HOMO SAPIENS | 0 | 414 | 3fj6 | A | 245 | ARG | p.Arg244Trp | Q02127 |
| HOMO SAPIENS | 0 | 414 | 3fj6 | A | 342 | ALA | p.Ala341Val | Q02127 |
| HOMO SAPIENS | 1 | 414 | 3fj6 | A | 347 | ARG | p.Arg346Trp | Q02127 |
| HOMO SAPIENS | 6 | 103 | 1u3w | A | 77 | GLY | p.Gly78Ter | P00326 |
| HOMO SAPIENS | -1 | 8369 | 4ux8 | F | 16 | ARG | p.Arg93Trp | P39905 |
| HOMO SAPIENS | 0 | 366 | 2zjc | A | 32 | GLY | p.Arg108Trp | P01375 |
| HOMO SAPIENS | 0 | 1457 | 3v9i | A | 352 | LEU | p.Ser352Leu | P30038 |
| HOMO SAPIENS | 0 | 1457 | 3v9i | A | 470 | VAL | p.Val470Ile | P30038 |
| HOMO SAPIENS | 0 | 1457 | 3v9i | A | 473 | THR | p.Thr473Ala | P30038 |
| HOMO SAPIENS | 1 | 2 | 4wo5 | A | 462 | ARG | p.Arg462Ile | P15056 |
| HOMO SAPIENS | 3 | 2 | 4wo5 | A | 463 | ILE | p.Ile463Ser | P15056 |
| HOMO SAPIENS | 1 | 2 | 4wo5 | A | 466 | GLY | p.Gly466Val | P15056 |
| HOMO SAPIENS | 1 | 2 | 4wo5 | A | 468 | PHE | p.Phe468Ser | P15056 |
| HOMO SAPIENS | 2 | 2 | 4wo5 | A | 469 | GLY | p.Gly469Ala | P15056 |
| HOMO SAPIENS | 2 | 2 | 4wo5 | A | 469 | GLY | p.Gly469Arg | P15056 |
| HOMO SAPIENS | 2 | 2 | 4wo5 | A | 485 | LEU | p.Leu485Phe | P15056 |
| HOMO SAPIENS | 0 | 2 | 4wo5 | A | 501 | GLU | p.Glu501Gly | P15056 |
| HOMO SAPIENS | 0 | 2 | 4wo5 | A | 501 | GLU | p.Glu501Lys | P15056 |
| HOMO SAPIENS | 7 | 2 | 4wo5 | A | 581 | ASN | p.Asn581Asp | P15056 |

| | | | | | | | | |
|---|---|---|---|---|---|---|---|---|
| HOMO SAPIENS | 8 | 2 | 4wo5 | A | 594 | ASP | p.Asp594Gly | P15056 |
| HOMO SAPIENS | 0 | 2 | 4wo5 | A | 597 | LEU | p.Leu597Arg | P15056 |
| HOMO SAPIENS | 0 | 2 | 4wo5 | A | 597 | LEU | p.Leu597Val | P15056 |
| HOMO SAPIENS | 0 | 2 | 4wo5 | A | 600 | VAL | p.Val600Glu | P15056 |
| HOMO SAPIENS | 2 | 2 | 4wo5 | A | 601 | LYS | p.Lys601Glu | P15056 |
| HOMO SAPIENS | 8 | 2 | 4wo5 | A | 638 | ASP | p.Asp638Glu | P15056 |
| HOMO SAPIENS | -1 | 30366 | 5bqd | B | 19 | ALA | p.Ala19Ser | Q99593 |
| HOMO SAPIENS | -1 | 30366 | 5bqd | B | 23 | PRO | p.Pro23His | Q99593 |
| HOMO SAPIENS | 1 | 30366 | 5bqd | B | 49 | GLN | p.Gln49Lys | Q99593 |
| HOMO SAPIENS | 2 | 30366 | 5bqd | B | 54 | ILE | p.Ile54Thr | Q99593 |
| HOMO SAPIENS | 8 | 30366 | 5bqd | B | 80 | GLY | p.Gly80Arg | Q99593 |
| HOMO SAPIENS | 6 | 30366 | 5bqd | B | 106 | ILE | p.Ile106Val | Q99593 |
| HOMO SAPIENS | 7 | 30366 | 5bqd | B | 111 | ASP | p.Asp111Tyr | Q99593 |
| HOMO SAPIENS | -1 | 30366 | 5bqd | B | 237 | ARG | p.Arg237Gln | Q99593 |
| HOMO SAPIENS | -1 | 30366 | 5bqd | B | 237 | ARG | p.Arg237Trp | Q99593 |
| HOMO SAPIENS | 5 | 4789 | 1n4m | A | 567 | SER | p.Ser567Leu | P06400 |
| HOMO SAPIENS | 0 | 4789 | 1n4m | A | 658 | ALA | p.Ala658Asp | P06400 |
| HOMO SAPIENS | 7 | 4789 | 1n4m | A | 661 | ARG | p.Arg661Trp | P06400 |
| HOMO SAPIENS | 6 | 4789 | 1n4m | A | 668 | ARG | p.Arg668His | P06400 |
| HOMO SAPIENS | 8 | 4789 | 1n4m | A | 697 | ASP | p.Asp697Glu | P06400 |
| HOMO SAPIENS | 6 | 4789 | 1n4m | A | 712 | CYS | p.Cys712Arg | P06400 |
| HOMO SAPIENS | 1 | 1798 | 2r5g | A | 101 | HIS | p.His101Gln | O15247 |
| HOMO SAPIENS | 0 | 1798 | 2r5g | A | 160 | PRO | p.Pro160Ala | O15247 |
| HOMO SAPIENS | 2 | 22 | 4x0t | A | 82 | ARG | p.Arg110Ter | P49419 |
| HOMO SAPIENS | 0 | 22 | 4x0t | A | 149 | ALA | p.Ala177Pro | P49419 |
| HOMO SAPIENS | 4 | 22 | 4x0t | A | 164 | THR | p.Thr192Met | P49419 |
| HOMO SAPIENS | 9 | 22 | 4x0t | A | 167 | ASN | p.Asn195Ser | P49419 |
| HOMO SAPIENS | 0 | 22 | 4x0t | A | 175 | TRP | p.Trp203Gly | P49419 |
| HOMO SAPIENS | 0 | 22 | 4x0t | A | 194 | THR | p.Thr222Ala | P49419 |
| HOMO SAPIENS | 0 | 22 | 4x0t | A | 273 | ASN | p.Asn301Ile | P49419 |
| HOMO SAPIENS | 0 | 22 | 4x0t | A | 307 | ARG | p.Arg335Gln | P49419 |
| HOMO SAPIENS | 3 | 22 | 4x0t | A | 313 | SER | p.Ser341Asn | P49419 |
| HOMO SAPIENS | 0 | 22 | 4x0t | A | 329 | ILE | p.Ile357Val | P49419 |
| HOMO SAPIENS | 1 | 22 | 4x0t | A | 380 | TYR | p.Tyr408Ter | P49419 |
| HOMO SAPIENS | 9 | 22 | 4x0t | A | 399 | GLU | p.Glu427Gln | P49419 |
| HOMO SAPIENS | 0 | 22 | 4x0t | A | 406 | TYR | p.Tyr434Cys | P49419 |
| HOMO SAPIENS | 0 | 22 | 4x0t | A | 431 | ILE | p.Ile459Phe | P49419 |
| HOMO SAPIENS | 4 | 22 | 4x0t | A | 477 | GLY | p.Gly505Arg | P49419 |
| HOMO SAPIENS | 0 | 22 | 4x0t | A | 495 | THR | p.Thr523Ala | P49419 |
| HOMO SAPIENS | 8 | 11042 | 1y6b | A | 1145 | PRO | p.Pro1147Ser | P35968 |
| HOMO SAPIENS | -1 | 3867 | 2xij | A | 18 | GLN | p.Gln18Ter | P22033 |
| HOMO SAPIENS | -1 | 3867 | 2xij | A | 31 | ARG | p.Arg31Ter | P22033 |
| HOMO SAPIENS | 2 | 3867 | 2xij | A | 93 | ARG | p.Arg93His | P22033 |
| HOMO SAPIENS | 0 | 3867 | 2xij | A | 94 | GLY | p.Gly94Arg | P22033 |
| HOMO SAPIENS | 2 | 3867 | 2xij | A | 95 | PRO | p.Pro95Leu | P22033 |
| HOMO SAPIENS | 2 | 3867 | 2xij | A | 105 | TRP | p.Trp105Arg | P22033 |
| HOMO SAPIENS | 2 | 3867 | 2xij | A | 108 | ARG | p.Arg108Cys | P22033 |
| HOMO SAPIENS | 3 | 3867 | 2xij | A | 154 | ARG | p.Arg154His | P22033 |
| HOMO SAPIENS | 3 | 3867 | 2xij | A | 156 | ASP | p.Asp156Val | P22033 |
| HOMO SAPIENS | 0 | 3867 | 2xij | A | 189 | ASN | p.Asn189Ile | P22033 |
| HOMO SAPIENS | 0 | 3867 | 2xij | A | 191 | ALA | p.Ala191Glu | P22033 |
| HOMO SAPIENS | 0 | 3867 | 2xij | A | 203 | GLY | p.Gly203Arg | P22033 |
| HOMO SAPIENS | 2 | 3867 | 2xij | A | 215 | GLY | p.Gly215Ser | P22033 |
| HOMO SAPIENS | 0 | 3867 | 2xij | A | 219 | ASN | p.Asn219Tyr | P22033 |

| | | | | | | | | |
|---|---|---|---|---|---|---|---|---|
| HOMO SAPIENS | 3 | 3867 | 2xij | A | 228 | ARG | p.Arg228Ter | P22033 |
| HOMO SAPIENS | 1 | 3867 | 2xij | A | 297 | THR | p.Thr297Ile | P22033 |
| HOMO SAPIENS | 0 | 3867 | 2xij | A | 326 | ARG | p.Arg326Lys | P22033 |
| HOMO SAPIENS | 3 | 3867 | 2xij | A | 369 | ARG | p.Arg369His | P22033 |
| HOMO SAPIENS | 3 | 3867 | 2xij | A | 369 | ARG | p.Arg369Cys | P22033 |
| HOMO SAPIENS | 0 | 3867 | 2xij | A | 370 | THR | p.Thr370Pro | P22033 |
| HOMO SAPIENS | 0 | 3867 | 2xij | A | 377 | ALA | p.Ala377Glu | P22033 |
| HOMO SAPIENS | 2 | 3867 | 2xij | A | 426 | GLY | p.Gly426Ala | P22033 |
| HOMO SAPIENS | 2 | 3867 | 2xij | A | 427 | GLY | p.Gly427Asp | P22033 |
| HOMO SAPIENS | 1 | 3867 | 2xij | A | 499 | THR | p.Ala499Thr | P22033 |
| HOMO SAPIENS | 2 | 3867 | 2xij | A | 532 | ARG | p.Arg532His | P22033 |
| HOMO SAPIENS | 3 | 3867 | 2xij | A | 555 | ALA | p.Ala555Thr | P22033 |
| HOMO SAPIENS | 0 | 3867 | 2xij | A | 573 | PHE | p.Phe573Ser | P22033 |
| HOMO SAPIENS | 3 | 3867 | 2xij | A | 623 | GLY | p.Gly623Arg | P22033 |
| HOMO SAPIENS | 5 | 3867 | 2xij | A | 671 | VAL | p.Ile671Val | P22033 |
| HOMO SAPIENS | 4 | 3867 | 2xij | A | 694 | ARG | p.Arg694Trp | P22033 |
| HOMO SAPIENS | 1 | 3867 | 2xij | A | 700 | MET | p.Met700Lys | P22033 |
| HOMO SAPIENS | 8 | 3867 | 2xij | A | 703 | GLY | p.Gly703Arg | P22033 |
| HOMO SAPIENS | 0 | 3867 | 2xij | A | 717 | GLY | p.Gly717Val | P22033 |
| HOMO SAPIENS | 1 | 3867 | 2xij | A | 727 | ARG | p.Arg727Ter | P22033 |
| HOMO SAPIENS | -1 | 52 | 3k91 | B | 72 | GLY | p.Gly73Ser | P00441 |
| HOMO SAPIENS | 6 | 52 | 3k91 | B | 96 | ASP | p.Asp97Asn | P00441 |
| HOMO SAPIENS | 5 | 52 | 3k91 | B | 112 | ILE | p.Ile113Thr | P00441 |
| HOMO SAPIENS | -1 | 52 | 3k91 | B | 134 | SER | p.Ser135Asn | P00441 |
| HOMO SAPIENS | 0 | 3950 | 4fom | A | 296 | ASN | p.Asn296His | Q9NQS3 |
| HOMO SAPIENS | 1 | 10 | 1de4 | A | 31 | VAL | p.Val53Met | Q30201 |
| HOMO SAPIENS | 0 | 10 | 1de4 | A | 37 | VAL | p.Val59Met | Q30201 |
| HOMO SAPIENS | 0 | 10 | 1de4 | A | 71 | GLY | p.Gly93Arg | Q30201 |
| HOMO SAPIENS | 0 | 10 | 1de4 | A | 83 | ILE | p.Ile105Thr | Q30201 |
| HOMO SAPIENS | 0 | 10 | 1de4 | A | 105 | GLN | p.Gln127His | Q30201 |
| HOMO SAPIENS | 1 | 10 | 1de4 | A | 146 | GLU | p.Glu168Ter | Q30201 |
| HOMO SAPIENS | 2 | 10 | 1de4 | A | 261 | GLN | p.Gln283Pro | Q30201 |
| HOMO SAPIENS | 0 | 8038 | 4x7n | A | 586 | ILE | p.Ile586Leu | Q9NZJ5 |
| HOMO SAPIENS | 0 | 8038 | 4x7n | A | 588 | ARG | p.Arg588Gln | Q9NZJ5 |
| HOMO SAPIENS | 0 | 35175 | 5ugw | A | 1062 | ALA | p.Ala1062Thr | O14746 |
| HOMO SAPIENS | 2 | 35175 | 5ugw | A | 1090 | VAL | p.Val1090Met | O14746 |
| HOMO SAPIENS | 0 | 1722 | 4z3k | C | 147 | ARG | p.Arg150Gly | P35270 |
| HOMO SAPIENS | 0 | 1722 | 4z3k | C | 160 | PRO | p.Pro163Leu | P35270 |
| HOMO SAPIENS | 0 | 1722 | 4z3k | C | 207 | ARG | p.Arg210Trp | P35270 |
| HOMO SAPIENS | 0 | 1722 | 4z3k | C | 216 | ARG | p.Arg219Ter | P35270 |
| HOMO SAPIENS | 0 | 1722 | 4z3k | C | 248 | LYS | p.Lys251Ter | P35270 |
| HOMO SAPIENS | 1 | 2832 | 4x8v | L | 115 | HIS | p.His175Gln | P08709 |
| HOMO SAPIENS | 0 | 825 | 1emh | A | 242 | PHE | p.Phe251Ser | P13051 |
| HOMO SAPIENS | 0 | 122 | 5uwk | A | 109 | ARG | p.Arg109Ter | P45452 |
| HOMO SAPIENS | 1 | 122 | 5uwk | A | 170 | LYS | p.Lys170Thr | P45452 |
| HOMO SAPIENS | 9 | 122 | 5uwk | A | 232 | HIS | p.His232Asn | P45452 |
| HOMO SAPIENS | 5 | 122 | 5uwk | A | 257 | ASP | p.Asp257Val | P45452 |
| HOMO SAPIENS | 9 | 1074 | 3p5i | A | 264 | TRP | p.Trp264Arg | Q9UJ71 |
| HOMO SAPIENS | 0 | 146 | 3r3g | A | 14C | GLU | p.Glu352Lys | P00734 |
| HOMO SAPIENS | 3 | 9522 | 1wyq | A | 83 | LEU | p.Leu253Pro | O15020 |
| HOMO SAPIENS | 8 | 67 | 2vgb | A | 130 | SER | p.Ser130Tyr | P30613 |
| HOMO SAPIENS | 8 | 67 | 2vgb | A | 163 | ARG | p.Arg163Cys | P30613 |
| HOMO SAPIENS | 4 | 67 | 2vgb | A | 241 | GLU | p.Glu241Ter | P30613 |
| HOMO SAPIENS | 9 | 67 | 2vgb | A | 384 | THR | p.Thr384Met | P30613 |

| | | | | | | | | |
|---|---|---|---|---|---|---|---|---|
| HOMO SAPIENS | 0 | 67 | 2vgb | A | 421 | GLN | p.Gln421Lys | P30613 |
| HOMO SAPIENS | 6 | 67 | 2vgb | A | 479 | ARG | p.Arg479His | P30613 |
| HOMO SAPIENS | 8 | 67 | 2vgb | A | 510 | ARG | p.Arg510Gln | P30613 |
| HOMO SAPIENS | 4 | 67 | 2vgb | A | 569 | ARG | p.Arg569Trp | P30613 |
| HOMO SAPIENS | 0 | 9088 | 5v44 | A | 232 | ASN | p.Asn232Lys | Q9NZJ4 |
| HOMO SAPIENS | 4 | 16 | 3lbn | A | 12 | GLY | p.Gly12Val | P01112 |
| HOMO SAPIENS | 4 | 16 | 3lbn | A | 12 | GLY | p.Gly12Ser | P01112 |
| HOMO SAPIENS | 4 | 16 | 3lbn | A | 13 | GLY | p.Gly13Cys | P01112 |
| HOMO SAPIENS | 4 | 16 | 3lbn | A | 13 | GLY | p.Gly13Asp | P01112 |
| HOMO SAPIENS | 5 | 16 | 3lbn | A | 22 | GLN | p.Gln22Lys | P01112 |
| HOMO SAPIENS | 5 | 16 | 3lbn | A | 58 | THR | p.Thr58Ile | P01112 |
| HOMO SAPIENS | 3 | 16 | 3lbn | A | 61 | GLN | p.Gln61Lys | P01112 |
| HOMO SAPIENS | 1 | 16 | 3lbn | A | 63 | GLU | p.Glu63Lys | P01112 |
| HOMO SAPIENS | 9 | 16 | 3lbn | A | 117 | LYS | p.Lys117Arg | P01112 |
| HOMO SAPIENS | 6 | 16 | 3lbn | A | 123 | ARG | p.Arg123Cys | P01112 |
| HOMO SAPIENS | 1 | 16 | 3lbn | A | 138 | GLY | p.Gly138Ser | P01112 |
| HOMO SAPIENS | 9 | 16 | 3lbn | A | 146 | ALA | p.Ala146Thr | P01112 |
| HOMO SAPIENS | 9 | 16 | 3lbn | A | 146 | ALA | p.Ala146Val | P01112 |
| HOMO SAPIENS | 1 | 359 | 5nvy | C | 63 | LEU | p.Leu63Pro | P40337 |
| HOMO SAPIENS | 1 | 359 | 5nvy | C | 64 | ARG | p.Arg64Pro | P40337 |
| HOMO SAPIENS | 1 | 359 | 5nvy | C | 70 | GLU | p.Glu70Lys | P40337 |
| HOMO SAPIENS | 0 | 359 | 5nvy | C | 84 | VAL | p.Val84Leu | P40337 |
| HOMO SAPIENS | 1 | 359 | 5nvy | C | 88 | TRP | p.Trp88Ser | P40337 |
| HOMO SAPIENS | 1 | 359 | 5nvy | C | 93 | GLY | p.Gly93Cys | P40337 |
| HOMO SAPIENS | 1 | 359 | 5nvy | C | 98 | TYR | p.Tyr98His | P40337 |
| HOMO SAPIENS | 1 | 359 | 5nvy | C | 111 | SER | p.Ser111Arg | P40337 |
| HOMO SAPIENS | 1 | 359 | 5nvy | C | 112 | TYR | p.Tyr112His | P40337 |
| HOMO SAPIENS | 1 | 359 | 5nvy | C | 126 | ASP | p.Asp126Tyr | P40337 |
| HOMO SAPIENS | 1 | 359 | 5nvy | C | 130 | VAL | p.Val130Leu | P40337 |
| HOMO SAPIENS | 1 | 359 | 5nvy | C | 135 | LEU | p.Leu135Phe | P40337 |
| HOMO SAPIENS | 9 | 359 | 5nvy | C | 161 | ARG | p.Arg161Gly | P40337 |
| HOMO SAPIENS | 7 | 359 | 5nvy | C | 162 | CYS | p.Cys162Phe | P40337 |
| HOMO SAPIENS | 4 | 359 | 5nvy | C | 163 | LEU | p.Leu163Pro | P40337 |
| HOMO SAPIENS | 3 | 359 | 5nvy | C | 164 | GLN | p.Gln164Arg | P40337 |
| HOMO SAPIENS | 9 | 359 | 5nvy | C | 166 | VAL | p.Val166Phe | P40337 |
| HOMO SAPIENS | 7 | 359 | 5nvy | C | 167 | ARG | p.Arg167Gly | P40337 |
| HOMO SAPIENS | 7 | 359 | 5nvy | C | 167 | ARG | p.Arg167Gln | P40337 |
| HOMO SAPIENS | 1 | 359 | 5nvy | C | 183 | SER | p.Ser183Ter | P40337 |
| HOMO SAPIENS | 2 | 359 | 5nvy | C | 188 | LEU | p.Leu188Val | P40337 |
| HOMO SAPIENS | 1 | 359 | 5nvy | C | 191 | HIS | p.His191Asp | P40337 |
| HOMO SAPIENS | 1 | 359 | 5nvy | C | 192 | PRO | p.Pro192Ser | P40337 |
| HOMO SAPIENS | 1 | 359 | 5nvy | C | 200 | ARG | p.Arg200Trp | P40337 |
| HOMO SAPIENS | -1 | 359 | 5nvy | C | 211 | MET | p.Met211Leu | P40337 |
| HOMO SAPIENS | 0 | 14099 | 4btm | A | 282 | ARG | p.Arg258Gln | Q5TCY1 |
| HOMO SAPIENS | 0 | 696 | 5h1b | A | 150 | ARG | p.Arg150Gln | Q06609 |
| HOMO SAPIENS | -1 | 208 | 2bxl | A | 1 | ASP | p.Asp25Val | P02768 |
| HOMO SAPIENS | 0 | 208 | 2bxl | A | 269 | ASP | p.Asp293Gly | P02768 |
| HOMO SAPIENS | 0 | 208 | 2bxl | A | 570 | GLU | p.Glu594Lys | P02768 |
| HOMO SAPIENS | 0 | 208 | 2bxl | A | 573 | LYS | p.Lys597Glu | P02768 |
| HOMO SAPIENS | -1 | 12813 | 2uzx | B | 35 | MET | p.Met35Leu | P08581 |
| HOMO SAPIENS | 1 | 12813 | 2uzx | B | 156 | SER | p.Ser156Leu | P08581 |
| HOMO SAPIENS | 0 | 12813 | 2uzx | B | 168 | GLU | p.Glu168Asp | P08581 |
| HOMO SAPIENS | 2 | 12813 | 2uzx | B | 211 | LEU | p.Leu211Trp | P08581 |
| HOMO SAPIENS | 2 | 12813 | 2uzx | B | 218 | ARG | p.Arg218Ser | P08581 |

| | | | | | | | | |
|---|---|---|---|---|---|---|---|---|
| HOMO SAPIENS | 1 | 12813 | 2uzx | B | 238 | LEU | p.Leu238Ser | P08581 |
| HOMO SAPIENS | 4 | 12813 | 2uzx | B | 320 | ALA | p.Ala320Val | P08581 |
| HOMO SAPIENS | 1 | 12813 | 2uzx | B | 375 | ASN | p.Asn375Ser | P08581 |
| HOMO SAPIENS | 0 | 12813 | 2uzx | B | 575 | LEU | p.Leu575Ile | P08581 |
| HOMO SAPIENS | 1 | 10780 | 4ejx | A | 348 | ARG | p.Arg367Cys | P00450 |
| HOMO SAPIENS | 2 | 10780 | 4ejx | A | 525 | ASP | p.Asp544Glu | P00450 |
| HOMO SAPIENS | 1 | 1407 | 5jn5 | A | 88 | ILE | p.Ile88Val | P36871 |
| HOMO SAPIENS | 7 | 1407 | 5jn5 | A | 115 | THR | p.Thr115Ala | P36871 |
| HOMO SAPIENS | 3 | 1407 | 5jn5 | A | 121 | GLY | p.Gly121Arg | P36871 |
| HOMO SAPIENS | 2 | 1407 | 5jn5 | A | 221 | ARG | p.Arg221Cys | P36871 |
| HOMO SAPIENS | 6 | 1407 | 5jn5 | A | 263 | TYR | p.Asp263Tyr | P36871 |
| HOMO SAPIENS | 3 | 1407 | 5jn5 | A | 420 | TYR | p.Tyr420His | P36871 |
| HOMO SAPIENS | 0 | 1407 | 5jn5 | A | 501 | VAL | p.Val501Ile | P36871 |
| HOMO SAPIENS | -1 | 1878 | 1yow | A | 255 | ARG | p.Arg255Leu | Q13285 |
| HOMO SAPIENS | 6 | 1878 | 1yow | A | 293 | ASP | p.Asp293Asn | Q13285 |
| HOMO SAPIENS | 1 | 1878 | 1yow | A | 437 | LEU | p.Leu437Gln | Q13285 |
| HOMO SAPIENS | 0 | 3071 | 2wpm | E | 145 | ARG | p.Arg191His | P00740 |
| HOMO SAPIENS | 0 | 3071 | 2wpm | E | 145 | ARG | p.Arg191Cys | P00740 |
| HOMO SAPIENS | 3 | 589 | 3ggc | B | 15 | GLY | p.Gly16Ser | P00492 |
| HOMO SAPIENS | 4 | 589 | 3ggc | B | 40 | LEU | p.Leu41Pro | P00492 |
| HOMO SAPIENS | 2 | 589 | 3ggc | B | 50 | ARG | p.Arg51Gly | P00492 |
| HOMO SAPIENS | 5 | 589 | 3ggc | B | 51 | ASP | p.Asp52Gly | P00492 |
| HOMO SAPIENS | 0 | 589 | 3ggc | B | 57 | GLY | p.Gly58Arg | P00492 |
| HOMO SAPIENS | 6 | 589 | 3ggc | B | 69 | GLY | p.Gly70Glu | P00492 |
| HOMO SAPIENS | 1 | 589 | 3ggc | B | 73 | PHE | p.Phe74Leu | P00492 |
| HOMO SAPIENS | 0 | 589 | 3ggc | B | 77 | LEU | p.Leu78Val | P00492 |
| HOMO SAPIENS | 3 | 589 | 3ggc | B | 79 | ASP | p.Asp80Val | P00492 |
| HOMO SAPIENS | 0 | 589 | 3ggc | B | 103 | SER | p.Ser104Arg | P00492 |
| HOMO SAPIENS | 0 | 589 | 3ggc | B | 109 | SER | p.Ser110Leu | P00492 |
| HOMO SAPIENS | 1 | 589 | 3ggc | B | 129 | VAL | p.Val130Asp | P00492 |
| HOMO SAPIENS | 4 | 589 | 3ggc | B | 131 | ILE | p.Ile132Met | P00492 |
| HOMO SAPIENS | 7 | 589 | 3ggc | B | 142 | MET | p.Met143Lys | P00492 |
| HOMO SAPIENS | 0 | 589 | 3ggc | B | 160 | ALA | p.Ala161Ser | P00492 |
| HOMO SAPIENS | 6 | 589 | 3ggc | B | 167 | THR | p.Thr168Ile | P00492 |
| HOMO SAPIENS | 0 | 589 | 3ggc | B | 169 | ARG | p.Arg170Ter | P00492 |
| HOMO SAPIENS | 5 | 589 | 3ggc | B | 175 | PRO | p.Pro176Leu | P00492 |
| HOMO SAPIENS | 8 | 589 | 3ggc | B | 193 | ASP | p.Asp194Glu | P00492 |
| HOMO SAPIENS | 8 | 589 | 3ggc | B | 193 | ASP | p.Asp194Asn | P00492 |
| HOMO SAPIENS | 8 | 589 | 3ggc | B | 198 | PHE | p.Phe199Val | P00492 |
| HOMO SAPIENS | 8 | 589 | 3ggc | B | 200 | ASP | p.Asp201Gly | P00492 |
| HOMO SAPIENS | 0 | 4269 | 2pla | B | 174 | GLU | p.Glu174Lys | Q8N335 |
| HOMO SAPIENS | 1 | 4269 | 2pla | B | 220 | ARG | p.Arg220His | Q8N335 |
| HOMO SAPIENS | 4 | 4269 | 2pla | B | 280 | ALA | p.Ala280Val | Q8N335 |
| HOMO SAPIENS | 0 | 2712 | 1ii4 | E | 252 | TRP | p.Ser252Leu | P21802 |
| HOMO SAPIENS | 0 | 2712 | 1ii4 | E | 253 | PRO | p.Pro253Arg | P21802 |
| HOMO SAPIENS | 5 | 2712 | 1ii4 | E | 267 | SER | p.Ser267Pro | P21802 |
| HOMO SAPIENS | 0 | 2712 | 1ii4 | E | 278 | CYS | p.Cys278Phe | P21802 |
| HOMO SAPIENS | 0 | 2712 | 1ii4 | E | 289 | GLN | p.Gln289Pro | P21802 |
| HOMO SAPIENS | 0 | 2712 | 1ii4 | E | 290 | TRP | p.Trp290Cys | P21802 |
| HOMO SAPIENS | 0 | 2712 | 1ii4 | E | 290 | TRP | p.Trp290Arg | P21802 |
| HOMO SAPIENS | 0 | 2712 | 1ii4 | E | 292 | LYS | p.Lys292Glu | P21802 |
| HOMO SAPIENS | 0 | 2712 | 1ii4 | E | 315 | ALA | p.Ala315Ser | P21802 |
| HOMO SAPIENS | 0 | 2712 | 1ii4 | E | 321 | ASP | p.Asp321Ala | P21802 |
| HOMO SAPIENS | 1 | 2712 | 1ii4 | E | 328 | TYR | p.Tyr328Cys | P21802 |

| | | | | | | | | |
|---|---|---|---|---|---|---|---|---|
| HOMO SAPIENS | 9 | 2712 | 1ii4 | E | 340 | TYR | p.Tyr340His | P21802 |
| HOMO SAPIENS | 1 | 2712 | 1ii4 | E | 341 | THR | p.Thr341Pro | P21802 |
| HOMO SAPIENS | 0 | 2712 | 1ii4 | E | 342 | CYS | p.Cys342Arg | P21802 |
| HOMO SAPIENS | 0 | 2712 | 1ii4 | E | 342 | CYS | p.Cys342Trp | P21802 |
| HOMO SAPIENS | 0 | 2712 | 1ii4 | E | 344 | ALA | p.Ala344Gly | P21802 |
| HOMO SAPIENS | 0 | 2712 | 1ii4 | E | 347 | SER | p.Ser347Cys | P21802 |
| HOMO SAPIENS | 0 | 2712 | 1ii4 | E | 351 | SER | p.Ser351Cys | P21802 |
| HOMO SAPIENS | 1 | 2712 | 1ii4 | E | 354 | SER | p.Ser354Cys | P21802 |
| HOMO SAPIENS | 2 | 53 | 4pj1 | H | 5 | ASP | p.Asp29Gly | P10809 |
| HOMO SAPIENS | 0 | 53 | 4pj1 | H | 74 | VAL | p.Val98Ile | P10809 |
| HOMO SAPIENS | 0 | 29968 | 2fdb | M | 71 | LYS | p.Lys89Glu | P55075 |
| HOMO SAPIENS | 2 | 29968 | 2fdb | M | 98 | ARG | p.Arg116Gly | P55075 |
| HOMO SAPIENS | 2 | 29968 | 2fdb | M | 122 | GLY | p.Gly140Ser | P55075 |
| HOMO SAPIENS | 8 | 5061 | 5f85 | B | 60 | GLY | p.Gly106Ser | P00740 |
| HOMO SAPIENS | 0 | 35944 | 1h6h | A | 85 | THR | p.Thr85Asn | Q15080 |
| HOMO SAPIENS | 8 | 35944 | 1h6h | A | 105 | ARG | p.Arg105Gln | Q15080 |
| HOMO SAPIENS | 7 | 12016 | 1hj7 | A | 295 | ASN | p.Asn316Ser | P01130 |
| HOMO SAPIENS | 5 | 12016 | 1hj7 | A | 302 | GLY | p.Gly323Ser | P01130 |
| HOMO SAPIENS | 6 | 12016 | 1hj7 | A | 319 | CYS | p.Cys340Trp | P01130 |
| HOMO SAPIENS | 7 | 12016 | 1hj7 | A | 329 | ARG | p.Arg350Ter | P01130 |
| HOMO SAPIENS | 5 | 12016 | 1hj7 | A | 331 | CYS | p.Cys352Ter | P01130 |
| HOMO SAPIENS | 2 | 12016 | 1hj7 | A | 345 | GLN | p.Gln366Arg | P01130 |
| HOMO SAPIENS | 2 | 1481 | 2i07 | A | 239 | TYR | p.Tyr261Ter | P01024 |
| HOMO SAPIENS | -1 | 1481 | 2i07 | A | 292 | LEU | p.Pro314Leu | P01024 |
| HOMO SAPIENS | 2 | 1481 | 2i07 | A | 326 | ILE | p.Ile348Val | P01024 |
| HOMO SAPIENS | 2 | 1481 | 2i07 | A | 413 | GLU | p.Glu435Lys | P01024 |
| HOMO SAPIENS | 4 | 1481 | 2i07 | A | 447 | GLU | p.Glu469Asp | P01024 |
| HOMO SAPIENS | 2 | 1481 | 2i07 | A | 597 | VAL | p.Val619Met | P01024 |
| HOMO SAPIENS | 5 | 1481 | 2i07 | A | 603 | ILE | p.Ile625Phe | P01024 |
| HOMO SAPIENS | 0 | 1481 | 2i07 | A | 611 | LYS | p.Lys633Arg | P01024 |
| HOMO SAPIENS | 0 | 11123 | 3c5n | A | 315 | GLY | p.Gly368Trp | O00294 |
| HOMO SAPIENS | 8 | 11123 | 3c5n | A | 329 | PHE | p.Phe382Ser | O00294 |
| HOMO SAPIENS | 5 | 11123 | 3c5n | A | 429 | ARG | p.Arg482Trp | O00294 |
| HOMO SAPIENS | 8 | 11123 | 3c5n | A | 438 | PHE | p.Phe491Leu | O00294 |
| HOMO SAPIENS | 3 | 4398 | 3fx0 | B | 1288 | ALA | p.Ala288Gly | Q9Y6K9 |
| HOMO SAPIENS | 7 | 4398 | 3fx0 | B | 1315 | GLU | p.Glu315Ala | Q9Y6K9 |
| HOMO SAPIENS | 8 | 4398 | 3fx0 | B | 1319 | ARG | p.Arg319Gln | Q9Y6K9 |
| HOMO SAPIENS | 6 | 3989 | 3qd6 | S | 83 | CYS | p.Cys83Arg | P25942 |
| HOMO SAPIENS | 1 | 3989 | 3qd6 | S | 124 | SER | p.Ser124Leu | P25942 |
| HOMO SAPIENS | 2 | 24003 | 2m0v | A | 225 | GLU | p.Glu225Lys | O14745 |
| HOMO SAPIENS | 2 | 46 | 4aqd | A | 70 | ASP | p.Asp98Gly | P06276 |
| HOMO SAPIENS | 4 | 46 | 4aqd | A | 115 | GLY | p.Gly143Asp | P06276 |
| HOMO SAPIENS | 3 | 46 | 4aqd | A | 184 | ALA | p.Ala212Val | P06276 |
| HOMO SAPIENS | 0 | 46 | 4aqd | A | 225 | GLY | p.Gly253Ter | P06276 |
| HOMO SAPIENS | 2 | 46 | 4aqd | A | 243 | THR | p.Thr271Met | P06276 |
| HOMO SAPIENS | 2 | 46 | 4aqd | A | 255 | GLU | p.Glu283Asp | P06276 |
| HOMO SAPIENS | 5 | 46 | 4aqd | A | 267 | LYS | p.Lys295Arg | P06276 |
| HOMO SAPIENS | 3 | 46 | 4aqd | A | 271 | GLU | p.Glu299Ter | P06276 |
| HOMO SAPIENS | 2 | 46 | 4aqd | A | 307 | LEU | p.Leu335Pro | P06276 |
| HOMO SAPIENS | 1 | 46 | 4aqd | A | 330 | LEU | p.Leu358Ile | P06276 |
| HOMO SAPIENS | 0 | 46 | 4aqd | A | 497 | GLU | p.Glu525Val | P06276 |
| HOMO SAPIENS | 3 | 786 | 1ols | B | 51 | GLY | p.Gly101Asp | P21953 |
| HOMO SAPIENS | 2 | 786 | 1ols | B | 87 | ALA | p.Ala137Val | P21953 |
| HOMO SAPIENS | 0 | 786 | 1ols | B | 110 | ILE | p.Ile160Ser | P21953 |

| | | | | | | | | |
|---|---|---|---|---|---|---|---|---|
| HOMO SAPIENS | 4 | 786 | 1ols | B | 120 | ARG | p.Arg170Cys | P21953 |
| HOMO SAPIENS | 2 | 786 | 1ols | B | 126 | ASN | p.Asn176Tyr | P21953 |
| HOMO SAPIENS | 0 | 786 | 1ols | B | 133 | ARG | p.Arg183Gly | P21953 |
| HOMO SAPIENS | 0 | 786 | 1ols | B | 133 | ARG | p.Arg183Pro | P21953 |
| HOMO SAPIENS | 0 | 786 | 1ols | B | 135 | PRO | p.Pro185Leu | P21953 |
| HOMO SAPIENS | 3 | 786 | 1ols | B | 156 | HIS | p.His206Tyr | P21953 |
| HOMO SAPIENS | 0 | 786 | 1ols | B | 228 | GLY | p.Gly278Ser | P21953 |
| HOMO SAPIENS | 1 | 786 | 1ols | B | 235 | ARG | p.Arg285Ter | P21953 |
| HOMO SAPIENS | 6 | 786 | 1ols | B | 274 | ARG | p.Arg324Ter | P21953 |
| HOMO SAPIENS | 1 | 831 | 4zt1 | B | 48 | VAL | p.Val202Ile | P12830 |
| HOMO SAPIENS | 6 | 831 | 4zt1 | B | 70 | ARG | p.Arg224Cys | P12830 |
| HOMO SAPIENS | 5 | 831 | 4zt1 | B | 79 | HIS | p.His233Gln | P12830 |
| HOMO SAPIENS | 4 | 831 | 4zt1 | B | 120 | GLY | p.Gly274Ser | P12830 |
| HOMO SAPIENS | 7 | 831 | 4zt1 | B | 187 | TYR | p.Tyr341Ter | P12830 |
| HOMO SAPIENS | 0 | 8497 | 3sla | A | 287 | ASN | p.Asn287Ser | P35222 |
| HOMO SAPIENS | 7 | 23986 | 2ke1 | A | 301 | VAL | p.Val301Met | O43918 |
| HOMO SAPIENS | 0 | 4967 | 1rj7 | B | 334 | ARG | p.Arg334His | Q92838 |
| HOMO SAPIENS | 0 | 4967 | 1rj7 | B | 338 | THR | p.Thr338Met | Q92838 |
| HOMO SAPIENS | 3 | 4967 | 1rj7 | B | 349 | ALA | p.Ala349Thr | Q92838 |
| HOMO SAPIENS | 2 | 4967 | 1rj7 | B | 358 | GLN | p.Gln358Glu | Q92838 |
| HOMO SAPIENS | 1 | 4967 | 1rj7 | B | 375 | LYS | p.Lys375Glu | Q92838 |
| HOMO SAPIENS | 9 | 7623 | 1d4t | A | 32 | ARG | p.Arg32Thr | O60880 |
| HOMO SAPIENS | 6 | 7623 | 1d4t | A | 55 | ARG | p.Arg55Leu | O60880 |
| HOMO SAPIENS | 0 | 7623 | 1d4t | A | 68 | THR | p.Thr68Ile | O60880 |
| HOMO SAPIENS | 6 | 705 | 3idh | A | 59 | PRO | p.Pro59Ser | P35557 |
| HOMO SAPIENS | 5 | 705 | 3idh | A | 124 | ASP | p.Asp124Asn | P35557 |
| HOMO SAPIENS | 4 | 705 | 3idh | A | 131 | SER | p.Ser131Pro | P35557 |
| HOMO SAPIENS | 0 | 705 | 3idh | A | 147 | GLY | p.Gly147Asp | P35557 |
| HOMO SAPIENS | 5 | 705 | 3idh | A | 186 | ARG | p.Arg186Ter | P35557 |
| HOMO SAPIENS | 0 | 705 | 3idh | A | 202 | MET | p.Met202Thr | P35557 |
| HOMO SAPIENS | 2 | 705 | 3idh | A | 210 | MET | p.Met210Lys | P35557 |
| HOMO SAPIENS | 1 | 705 | 3idh | A | 215 | TYR | p.Tyr215Ter | P35557 |
| HOMO SAPIENS | 0 | 705 | 3idh | A | 226 | VAL | p.Val226Met | P35557 |
| HOMO SAPIENS | 9 | 705 | 3idh | A | 228 | THR | p.Thr228Met | P35557 |
| HOMO SAPIENS | 9 | 705 | 3idh | A | 256 | GLU | p.Glu256Lys | P35557 |
| HOMO SAPIENS | 2 | 705 | 3idh | A | 261 | GLY | p.Gly261Arg | P35557 |
| HOMO SAPIENS | 6 | 705 | 3idh | A | 279 | GLU | p.Glu279Gln | P35557 |
| HOMO SAPIENS | 7 | 705 | 3idh | A | 299 | GLY | p.Gly299Arg | P35557 |
| HOMO SAPIENS | 2 | 705 | 3idh | A | 318 | GLY | p.Gly318Trp | P35557 |
| HOMO SAPIENS | 4 | 705 | 3idh | A | 378 | ALA | p.Ala378Thr | P35557 |
| HOMO SAPIENS | 0 | 705 | 3idh | A | 387 | ALA | p.Ala387Glu | P35557 |
| HOMO SAPIENS | 5 | 705 | 3idh | A | 414 | LYS | p.Lys414Glu | P35557 |
| HOMO SAPIENS | 0 | 705 | 3idh | A | 455 | VAL | p.Val455Met | P35557 |
| HOMO SAPIENS | -1 | 705 | 3idh | A | 462 | MET | p.Met462Ile | P35557 |
| HOMO SAPIENS | 1 | 509 | 4nef | A | 22 | LEU | p.Leu22Val | P41181 |
| HOMO SAPIENS | 4 | 509 | 4nef | A | 57 | GLN | p.Gln57Pro | P41181 |
| HOMO SAPIENS | 8 | 509 | 4nef | A | 64 | GLY | p.Gly64Arg | P41181 |
| HOMO SAPIENS | 7 | 509 | 4nef | A | 68 | ASN | p.Asn68Ser | P41181 |
| HOMO SAPIENS | 1 | 509 | 4nef | A | 100 | GLY | p.Gly100Val | P41181 |
| HOMO SAPIENS | 0 | 509 | 4nef | A | 125 | THR | p.Thr125Met | P41181 |
| HOMO SAPIENS | 0 | 509 | 4nef | A | 126 | THR | p.Thr126Met | P41181 |
| HOMO SAPIENS | 4 | 509 | 4nef | A | 146 | PHE | p.Phe146Leu | P41181 |
| HOMO SAPIENS | 0 | 509 | 4nef | A | 175 | GLY | p.Gly175Arg | P41181 |
| HOMO SAPIENS | 5 | 509 | 4nef | A | 181 | CYS | p.Cys181Trp | P41181 |

| | | | | | | | | | |
|---|---|---|---|---|---|---|---|---|---|
| HOMO SAPIENS | 9 | 509 | 4nef | A | | 187 | ARG | p.Arg187Cys | P41181 |
| HOMO SAPIENS | 0 | 509 | 4nef | A | | 190 | ALA | p.Ala190Thr | P41181 |
| HOMO SAPIENS | 0 | 509 | 4nef | A | | 216 | SER | p.Ser216Pro | P41181 |
| HOMO SAPIENS | 4 | 2242 | 1z70 | X | | 1155 | SER | p.Ser155Pro | Q8NBK3 |
| HOMO SAPIENS | 3 | 2242 | 1z70 | X | | 1218 | CYS | p.Cys218Tyr | Q8NBK3 |
| HOMO SAPIENS | 0 | 2242 | 1z70 | X | | 1279 | ALA | p.Ala279Val | Q8NBK3 |
| HOMO SAPIENS | -1 | 2242 | 1z70 | X | | 1336 | OCS | p.Cys336Arg | Q8NBK3 |
| HOMO SAPIENS | 4 | 2242 | 1z70 | X | | 1345 | ARG | p.Arg345Cys | Q8NBK3 |
| HOMO SAPIENS | 2 | 2242 | 1z70 | X | | 1348 | ALA | p.Ala348Pro | Q8NBK3 |
| HOMO SAPIENS | 5 | 2242 | 1z70 | X | | 1349 | ARG | p.Arg349Gln | Q8NBK3 |
| HOMO SAPIENS | 5 | 2242 | 1z70 | X | | 1349 | ARG | p.Arg349Trp | Q8NBK3 |
| HOMO SAPIENS | 3 | 2242 | 1z70 | X | | 1359 | SER | p.Ser359Ter | Q8NBK3 |
| HOMO SAPIENS | 5 | 8269 | 5wrr | B | | 136 | ARG | p.Arg136Ter | Q96MK3 |
| HOMO SAPIENS | 1 | 8269 | 5wrr | B | | 332 | LYS | p.Asn332Lys | Q96MK3 |
| HOMO SAPIENS | 3 | 8269 | 5wrr | B | | 478 | ARG | p.Arg478Ter | Q96MK3 |
| HOMO SAPIENS | 5 | 8269 | 5wrr | B | | 504 | THR | p.Thr504Ile | Q96MK3 |
| HOMO SAPIENS | 5 | 1411 | 4h1s | B | | 221 | SER | p.Ser221Ter | P21589 |
| HOMO SAPIENS | 1 | 1411 | 4h1s | B | | 358 | CYS | p.Cys358Tyr | P21589 |
| HOMO SAPIENS | 2 | 23 | 5vvd | B | | 298 | GLU | p.Glu298Asp | P29474 |
| HOMO SAPIENS | 0 | 19 | 5jlh | E | | 88 | THR | p.Thr89Ile | P63261 |
| HOMO SAPIENS | 6 | 19 | 5jlh | E | | 117 | LYS | p.Lys118Met | P63261 |
| HOMO SAPIENS | 5 | 19 | 5jlh | E | | 263 | PRO | p.Pro264Leu | P63261 |
| HOMO SAPIENS | 1 | 19 | 5jlh | E | | 277 | THR | p.Thr278Ile | P63261 |
| HOMO SAPIENS | 6 | 19 | 5jlh | E | | 331 | PRO | p.Pro332Ala | P63261 |
| HOMO SAPIENS | 5 | 19 | 5jlh | E | | 369 | VAL | p.Val370Ala | P63261 |
| HOMO SAPIENS | 0 | 14306 | 4w4o | C | | 92 | ARG | p.Arg92Ter | P12314 |
| HOMO SAPIENS | 3 | 603 | 1nuh | A | | 4 | THR | p.Thr5Ile | P06744 |
| HOMO SAPIENS | 2 | 603 | 1nuh | A | | 19 | HIS | p.His20Pro | P06744 |
| HOMO SAPIENS | 8 | 603 | 1nuh | A | | 158 | GLY | p.Gly159Ser | P06744 |
| HOMO SAPIENS | 6 | 603 | 1nuh | A | | 223 | THR | p.Thr224Met | P06744 |
| HOMO SAPIENS | 0 | 603 | 1nuh | A | | 338 | LEU | p.Leu339Pro | P06744 |
| HOMO SAPIENS | 6 | 603 | 1nuh | A | | 342 | GLN | p.Gln343Arg | P06744 |
| HOMO SAPIENS | 6 | 603 | 1nuh | A | | 374 | THR | p.Thr375Arg | P06744 |
| HOMO SAPIENS | 4 | 603 | 1nuh | A | | 524 | ILE | p.Ile525Thr | P06744 |
| HOMO SAPIENS | 4 | 603 | 1nuh | A | | 538 | ASP | p.Asp539Asn | P06744 |
| HOMO SAPIENS | 1 | 3107 | 3kzj | A | | 1197 | VAL | p.Val1197Ala | P08603 |
| HOMO SAPIENS | 9 | 1748 | 2noh | A | | 46 | ARG | p.Arg46Gln | O15527 |
| HOMO SAPIENS | 2 | 11750 | 3s4y | A | | 33 | ARG | p.Arg33His | Q9H3S4 |
| HOMO SAPIENS | 2 | 11750 | 3s4y | A | | 40 | LEU | p.Leu40Pro | Q9H3S4 |
| HOMO SAPIENS | 4 | 11750 | 3s4y | A | | 142 | LEU | p.Leu142Phe | Q9H3S4 |
| HOMO SAPIENS | 5 | 11750 | 3s4y | A | | 219 | ASN | p.Asn219Ser | Q9H3S4 |
| HOMO SAPIENS | 1 | 2591 | 5vnj | A | | 382 | PHE | p.Phe382Leu | Q15436 |
| HOMO SAPIENS | 2 | 2591 | 5vnj | A | | 702 | MET | p.Met702Val | Q15436 |
| HOMO SAPIENS | 1 | 2426 | 5m8t | A | | 87 | ARG | p.Arg87Trp | P17643 |
| HOMO SAPIENS | 1 | 2426 | 5m8t | A | | 166 | SER | p.Ser166Ter | P17643 |
| HOMO SAPIENS | 0 | 2426 | 5m8t | A | | 356 | ARG | p.Arg356Gln | P17643 |
| HOMO SAPIENS | 2 | 2426 | 5m8t | A | | 374 | SER | p.Arg374Ter | P17643 |
| HOMO SAPIENS | 6 | 2426 | 5m8t | A | | 378 | ASN | p.Asn378Ser | P17643 |
| HOMO SAPIENS | 6 | 2426 | 5m8t | A | | 382 | LEU | p.Leu382Pro | P17643 |
| HOMO SAPIENS | 7 | 10097 | 4n6v | | 1 | 46 | GLN | p.Gln46Ter | P04080 |
| HOMO SAPIENS | 3 | 10097 | 4n6v | | 1 | 65 | VAL | p.Val65Ile | P04080 |
| HOMO SAPIENS | 8 | 10097 | 4n6v | | 1 | 68 | ARG | p.Arg68Ter | P04080 |
| HOMO SAPIENS | 2 | 859 | 4z9a | A | | 105 | GLN | p.Gln106Arg | P24666 |
| HOMO SAPIENS | 8 | 1420 | 5nan | F | | 95 | SER | p.Ser95Leu | Q96PD4 |

| | | | | | | | | |
|---|---|---|---|---|---|---|---|---|
| HOMO SAPIENS | 6 | 1420 | 5nan | F | 126 | GLU | p.Glu126Gly | Q96PD4 |
| HOMO SAPIENS | 8 | 1420 | 5nan | F | 155 | VAL | p.Val155Ile | Q96PD4 |
| HOMO SAPIENS | 6 | 1420 | 5nan | F | 157 | PRO | p.Pro157Ser | Q96PD4 |
| HOMO SAPIENS | -1 | 1420 | 5nan | F | 161 | HIS | p.His161Arg | Q96PD4 |
| HOMO SAPIENS | 3 | 288 | 5i96 | B | 140 | GLN | p.Arg140Gly | P48735 |
| HOMO SAPIENS | 3 | 288 | 5i96 | B | 140 | GLN | p.Arg140Gln | P48735 |
| HOMO SAPIENS | 2 | 288 | 5i96 | B | 261 | ARG | p.Arg261His | P48735 |
| HOMO SAPIENS | 2 | 288 | 5i96 | B | 435 | THR | p.Thr435Met | P48735 |
| HOMO SAPIENS | 9 | 120 | 1tup | B | 107 | TYR | p.Tyr107His | P04637 |
| HOMO SAPIENS | 8 | 120 | 1tup | B | 110 | ARG | p.Arg110Leu | P04637 |
| HOMO SAPIENS | 5 | 120 | 1tup | B | 120 | LYS | p.Lys120Glu | P04637 |
| HOMO SAPIENS | 4 | 120 | 1tup | B | 133 | MET | p.Met133Thr | P04637 |
| HOMO SAPIENS | 3 | 120 | 1tup | B | 134 | PHE | p.Phe134Cys | P04637 |
| HOMO SAPIENS | 8 | 120 | 1tup | B | 136 | GLN | p.Gln136His | P04637 |
| HOMO SAPIENS | 9 | 120 | 1tup | B | 138 | ALA | p.Ala138Pro | P04637 |
| HOMO SAPIENS | 9 | 120 | 1tup | B | 151 | PRO | p.Pro151Ser | P04637 |
| HOMO SAPIENS | 9 | 120 | 1tup | B | 152 | PRO | p.Pro152Leu | P04637 |
| HOMO SAPIENS | 9 | 120 | 1tup | B | 158 | ARG | p.Arg158Cys | P04637 |
| HOMO SAPIENS | 9 | 120 | 1tup | B | 158 | ARG | p.Arg158His | P04637 |
| HOMO SAPIENS | 9 | 120 | 1tup | B | 175 | ARG | p.Arg175Cys | P04637 |
| HOMO SAPIENS | 9 | 120 | 1tup | B | 175 | ARG | p.Arg175Leu | P04637 |
| HOMO SAPIENS | 8 | 120 | 1tup | B | 181 | ARG | p.Arg181Leu | P04637 |
| HOMO SAPIENS | 9 | 120 | 1tup | B | 189 | ALA | p.Ala189Val | P04637 |
| HOMO SAPIENS | 9 | 120 | 1tup | B | 196 | ARG | p.Arg196Gly | P04637 |
| HOMO SAPIENS | 9 | 120 | 1tup | B | 213 | ARG | p.Arg213Gln | P04637 |
| HOMO SAPIENS | 9 | 120 | 1tup | B | 213 | ARG | p.Arg213Ter | P04637 |
| HOMO SAPIENS | 9 | 120 | 1tup | B | 214 | HIS | p.His214Gln | P04637 |
| HOMO SAPIENS | 9 | 120 | 1tup | B | 220 | TYR | p.Tyr220Cys | P04637 |
| HOMO SAPIENS | 0 | 120 | 1tup | B | 234 | TYR | p.Tyr234Cys | P04637 |
| HOMO SAPIENS | 9 | 120 | 1tup | B | 237 | MET | p.Met237Ile | P04637 |
| HOMO SAPIENS | 0 | 120 | 1tup | B | 238 | CYS | p.Cys238Tyr | P04637 |
| HOMO SAPIENS | 9 | 120 | 1tup | B | 241 | SER | p.Ser241Phe | P04637 |
| HOMO SAPIENS | 9 | 120 | 1tup | B | 242 | CYS | p.Cys242Ser | P04637 |
| HOMO SAPIENS | 9 | 120 | 1tup | B | 245 | GLY | p.Gly245Cys | P04637 |
| HOMO SAPIENS | 9 | 120 | 1tup | B | 245 | GLY | p.Gly245Asp | P04637 |
| HOMO SAPIENS | 9 | 120 | 1tup | B | 248 | ARG | p.Arg248Leu | P04637 |
| HOMO SAPIENS | 9 | 120 | 1tup | B | 248 | ARG | p.Arg248Trp | P04637 |
| HOMO SAPIENS | 9 | 120 | 1tup | B | 251 | ILE | p.Ile251Ser | P04637 |
| HOMO SAPIENS | 9 | 120 | 1tup | B | 252 | LEU | p.Leu252Pro | P04637 |
| HOMO SAPIENS | 9 | 120 | 1tup | B | 258 | GLU | p.Glu258Lys | P04637 |
| HOMO SAPIENS | 8 | 120 | 1tup | B | 272 | VAL | p.Val272Leu | P04637 |
| HOMO SAPIENS | 9 | 120 | 1tup | B | 273 | ARG | p.Arg273Cys | P04637 |
| HOMO SAPIENS | 9 | 120 | 1tup | B | 273 | ARG | p.Arg273His | P04637 |
| HOMO SAPIENS | 9 | 120 | 1tup | B | 277 | CYS | p.Cys277Phe | P04637 |
| HOMO SAPIENS | 9 | 120 | 1tup | B | 280 | ARG | p.Arg280Ile | P04637 |
| HOMO SAPIENS | 9 | 120 | 1tup | B | 282 | ARG | p.Arg282Trp | P04637 |
| HOMO SAPIENS | 9 | 120 | 1tup | B | 283 | ARG | p.Arg283His | P04637 |
| HOMO SAPIENS | 9 | 120 | 1tup | B | 285 | GLU | p.Glu285Val | P04637 |
| HOMO SAPIENS | -1 | 120 | 1tup | B | 292 | LYS | p.Lys292Ile | P04637 |
| HOMO SAPIENS | -1 | 120 | 1tup | B | 298 | GLU | p.Glu298Lys | P04637 |
| HOMO SAPIENS | -1 | 2791 | 4ygb | A | 39 | VAL | p.Val39Ala | P49257 |
| HOMO SAPIENS | 2 | 106 | 4x6j | A | 32 | GLY | p.Gly146Arg | P43235 |
| HOMO SAPIENS | 0 | 106 | 4x6j | A | 76 | GLN | p.Gln190Ter | P43235 |
| HOMO SAPIENS | 7 | 106 | 4x6j | A | 198 | ARG | p.Arg312Ter | P43235 |

| | | | | | | | | |
|---|---|---|---|---|---|---|---|---|
| HOMO SAPIENS | 5 | 490 | 2hlo | B | 172 | LEU | p.Leu202Gln | P02675 |
| HOMO SAPIENS | 0 | 490 | 2hlo | B | 353 | LEU | p.Leu383Arg | P02675 |
| HOMO SAPIENS | 4 | 490 | 2hlo | B | 400 | GLY | p.Gly430Asp | P02675 |
| HOMO SAPIENS | 7 | 490 | 2hlo | B | 448 | ARG | p.Arg478Lys | P02675 |
| HOMO SAPIENS | 1 | 289 | 4p13 | B | 18 | GLU | p.Glu43Lys | P11310 |
| HOMO SAPIENS | 2 | 289 | 4p13 | B | 20 | GLN | p.Gln45Arg | P11310 |
| HOMO SAPIENS | 5 | 289 | 4p13 | B | 28 | ARG | p.Arg53Cys | P11310 |
| HOMO SAPIENS | 1 | 289 | 4p13 | B | 42 | TYR | p.Tyr67His | P11310 |
| HOMO SAPIENS | 1 | 289 | 4p13 | B | 53 | ILE | p.Ile78Thr | P11310 |
| HOMO SAPIENS | 1 | 289 | 4p13 | B | 59 | LEU | p.Leu84Phe | P11310 |
| HOMO SAPIENS | 2 | 289 | 4p13 | B | 96 | THR | p.Thr121Ile | P11310 |
| HOMO SAPIENS | 0 | 289 | 4p13 | B | 124 | MET | p.Met149Ile | P11310 |
| HOMO SAPIENS | 5 | 289 | 4p13 | B | 143 | ASP | p.Asp168Ala | P11310 |
| HOMO SAPIENS | 7 | 289 | 4p13 | B | 168 | THR | p.Thr193Ala | P11310 |
| HOMO SAPIENS | 1 | 289 | 4p13 | B | 170 | GLY | p.Gly195Arg | P11310 |
| HOMO SAPIENS | 2 | 289 | 4p13 | B | 181 | ARG | p.Arg206Cys | P11310 |
| HOMO SAPIENS | 2 | 289 | 4p13 | B | 181 | ARG | p.Arg206His | P11310 |
| HOMO SAPIENS | 8 | 289 | 4p13 | B | 203 | THR | p.Thr228Ile | P11310 |
| HOMO SAPIENS | 2 | 289 | 4p13 | B | 208 | ILE | p.Ile233Asn | P11310 |
| HOMO SAPIENS | 1 | 289 | 4p13 | B | 219 | CYS | p.Cys244Arg | P11310 |
| HOMO SAPIENS | 6 | 289 | 4p13 | B | 220 | SER | p.Ser245Leu | P11310 |
| HOMO SAPIENS | 5 | 289 | 4p13 | B | 228 | GLU | p.Glu253Lys | P11310 |
| HOMO SAPIENS | 2 | 289 | 4p13 | B | 241 | ASP | p.Asp266Gly | P11310 |
| HOMO SAPIENS | 1 | 289 | 4p13 | B | 242 | GLY | p.Gly267Arg | P11310 |
| HOMO SAPIENS | 7 | 289 | 4p13 | B | 256 | ARG | p.Arg281Thr | P11310 |
| HOMO SAPIENS | 4 | 289 | 4p13 | B | 269 | ARG | p.Arg294Thr | P11310 |
| HOMO SAPIENS | 4 | 289 | 4p13 | B | 304 | GLU | p.Lys329Glu | P11310 |
| HOMO SAPIENS | 1 | 289 | 4p13 | B | 324 | ARG | p.Arg349Ter | P11310 |
| HOMO SAPIENS | 3 | 289 | 4p13 | B | 326 | THR | p.Thr351Ile | P11310 |
| HOMO SAPIENS | 7 | 289 | 4p13 | B | 350 | ILE | p.Ile375Thr | P11310 |
| HOMO SAPIENS | 3 | 289 | 4p13 | B | 359 | GLU | p.Glu384Ter | P11310 |
| HOMO SAPIENS | 1 | 4563 | 3i2b | A | 16 | ARG | p.Arg16Cys | Q03393 |
| HOMO SAPIENS | 5 | 4563 | 3i2b | A | 25 | ARG | p.Arg25Gln | Q03393 |
| HOMO SAPIENS | 1 | 4563 | 3i2b | A | 47 | ASN | p.Asn47Asp | Q03393 |
| HOMO SAPIENS | 8 | 4563 | 3i2b | A | 52 | ASN | p.Asn52Ser | Q03393 |
| HOMO SAPIENS | 1 | 4563 | 3i2b | A | 56 | VAL | p.Val56Met | Q03393 |
| HOMO SAPIENS | 1 | 4563 | 3i2b | A | 87 | PRO | p.Pro87Ser | Q03393 |
| HOMO SAPIENS | 1 | 4563 | 3i2b | A | 96 | ASP | p.Asp96Asn | Q03393 |
| HOMO SAPIENS | 0 | 4563 | 3i2b | A | 99 | TYR | p.Tyr99Ter | Q03393 |
| HOMO SAPIENS | 2 | 4563 | 3i2b | A | 116 | ASP | p.Asp116Gly | Q03393 |
| HOMO SAPIENS | 2 | 41 | 3exx | A | 1 | GLY | p.Gly90Cys | P01308 |
| HOMO SAPIENS | 5 | 41 | 3exx | A | 3 | VAL | p.Val92Leu | P01308 |
| HOMO SAPIENS | 4 | 41 | 3exx | A | 7 | CYS | p.Cys96Ser | P01308 |
| HOMO SAPIENS | 9 | 41 | 3exx | A | 19 | TYR | p.Tyr108Cys | P01308 |
| HOMO SAPIENS | 9 | 131 | 4mjo | D | 118 | ASP | p.Asp119Asn | P09467 |
| HOMO SAPIENS | 0 | 131 | 4mjo | D | 163 | GLY | p.Gly164Ser | P09467 |
| HOMO SAPIENS | 0 | 131 | 4mjo | D | 176 | ALA | p.Ala177Asp | P09467 |
| HOMO SAPIENS | 5 | 131 | 4mjo | D | 193 | PHE | p.Phe194Ser | P09467 |
| HOMO SAPIENS | 4 | 131 | 4mjo | D | 217 | ARG | p.Arg218Lys | P09467 |
| HOMO SAPIENS | 4 | 131 | 4mjo | D | 223 | VAL | p.Val224Ile | P09467 |
| HOMO SAPIENS | 4 | 131 | 4mjo | D | 232 | PHE | p.Phe233Ile | P09467 |
| HOMO SAPIENS | 5 | 131 | 4mjo | D | 243 | ARG | p.Arg244Trp | P09467 |
| HOMO SAPIENS | 0 | 131 | 4mjo | D | 283 | PRO | p.Pro284Arg | P09467 |
| HOMO SAPIENS | 0 | 131 | 4mjo | D | 288 | MET | p.Met289Leu | P09467 |

| | | | | | | | | |
|---|---|---|---|---|---|---|---|---|
| HOMO SAPIENS | 0 | 131 | 4mjo | D | 323 | ASP | p.Asp324Asn | P09467 |
| HOMO SAPIENS | -1 | 13122 | 3hfc | A | 583 | ARG | p.Arg583Cys | P51787 |
| HOMO SAPIENS | 1 | 13122 | 3hfc | A | 587 | THR | p.Thr587Met | P51787 |
| HOMO SAPIENS | 0 | 13122 | 3hfc | A | 589 | GLY | p.Gly589Asp | P51787 |
| HOMO SAPIENS | 4 | 3002 | 3nbq | B | 94 | ARG | p.Arg94Cys | Q16831 |
| HOMO SAPIENS | 1 | 3616 | 3dbp | A | 97 | ILE | p.Ile286Asn | P11172 |
| HOMO SAPIENS | 0 | 3616 | 3dbp | A | 240 | GLY | p.Gly429Arg | P11172 |
| HOMO SAPIENS | 0 | 3616 | 3dbp | A | 257 | ILE | p.Ile446Val | P11172 |
| HOMO SAPIENS | 1 | 584 | 1drf | A | 79 | LEU | p.Leu80Phe | P00374 |
| HOMO SAPIENS | 3 | 1609 | 3qa2 | B | 49 | VAL | p.Val49Ile | P50053 |
| HOMO SAPIENS | 0 | 20560 | 5mug | A | 101 | HIS | p.His101Gln | P49638 |
| HOMO SAPIENS | 2 | 20560 | 5mug | A | 120 | ALA | p.Ala120Thr | P49638 |
| HOMO SAPIENS | 3 | 20560 | 5mug | A | 192 | ARG | p.Arg192His | P49638 |
| HOMO SAPIENS | 3 | 20560 | 5mug | A | 221 | ARG | p.Arg221Trp | P49638 |
| HOMO SAPIENS | 4 | 256 | 3vtv | A | 65 | THR | p.Glu50Lys | Q9GZQ8 |
| HOMO SAPIENS | 4 | 256 | 3vtv | A | 113 | VAL | p.Met98Lys | Q9GZQ8 |
| HOMO SAPIENS | 4 | 5397 | 1m61 | A | 192 | ARG | p.Arg192Trp | P43403 |
| HOMO SAPIENS | 5 | 39881 | 1aut | L | 107 | HIS | p.His149Pro | P04070 |
| HOMO SAPIENS | 5 | 4010 | 4uf8 | I | 33 | ALA | p.Ala33Val | Q13501 |
| HOMO SAPIENS | 3 | 313 | 5w49 | A | 98 | PRO | p.Pro98Leu | P23526 |
| HOMO SAPIENS | 2 | 313 | 5w49 | A | 143 | TYR | p.Tyr143Cys | P23526 |
| HOMO SAPIENS | -1 | 1643 | 2m8t | A | 102 | PRO | p.Pro102Leu | P04156 |
| HOMO SAPIENS | -1 | 1643 | 2m8t | A | 105 | PRO | p.Pro105Leu | P04156 |
| HOMO SAPIENS | -1 | 1643 | 2m8t | A | 117 | ALA | p.Ala117Val | P04156 |
| HOMO SAPIENS | 5 | 1643 | 2m8t | A | 129 | MET | p.Met129Val | P04156 |
| HOMO SAPIENS | 5 | 1643 | 2m8t | A | 131 | GLY | p.Gly131Val | P04156 |
| HOMO SAPIENS | 2 | 1643 | 2m8t | A | 142 | GLY | p.Gly142Ser | P04156 |
| HOMO SAPIENS | 3 | 1643 | 2m8t | A | 171 | ASN | p.Asn171Ser | P04156 |
| HOMO SAPIENS | 9 | 1643 | 2m8t | A | 178 | ASP | p.Asp178Asn | P04156 |
| HOMO SAPIENS | 9 | 1643 | 2m8t | A | 180 | VAL | p.Val180Ile | P04156 |
| HOMO SAPIENS | 9 | 1643 | 2m8t | A | 183 | THR | p.Thr183Ala | P04156 |
| HOMO SAPIENS | 9 | 1643 | 2m8t | A | 187 | HIS | p.His187Arg | P04156 |
| HOMO SAPIENS | 8 | 1643 | 2m8t | A | 198 | PHE | p.Phe198Ser | P04156 |
| HOMO SAPIENS | 9 | 1643 | 2m8t | A | 200 | GLU | p.Glu200Lys | P04156 |
| HOMO SAPIENS | 9 | 1643 | 2m8t | A | 208 | ARG | p.Arg208His | P04156 |
| HOMO SAPIENS | 3 | 1643 | 2m8t | A | 210 | VAL | p.Val210Ile | P04156 |
| HOMO SAPIENS | 6 | 1643 | 2m8t | A | 217 | GLN | p.Gln217Arg | P04156 |
| HOMO SAPIENS | 1 | 29 | 5kos | A | 151 | GLY | p.Gly217Arg | P00797 |
| HOMO SAPIENS | 1 | 29 | 5kos | A | 164 | ARG | p.Arg230Lys | P00797 |
| HOMO SAPIENS | 3 | 504 | 4zkv | B | 37 | ARG | p.Arg37Pro | P49773 |
| HOMO SAPIENS | 2 | 504 | 4zkv | B | 62 | GLN | p.Gln62Ter | P49773 |
| HOMO SAPIENS | 2 | 504 | 4zkv | B | 84 | CYS | p.Cys84Arg | P49773 |
| HOMO SAPIENS | 4 | 504 | 4zkv | B | 93 | GLY | p.Gly93Asp | P49773 |
| HOMO SAPIENS | 9 | 504 | 4zkv | B | 112 | HIS | p.His112Asn | P49773 |
| HOMO SAPIENS | 0 | 74 | 5vwo | C | 54 | ASN | p.Asn54Lys | P04181 |
| HOMO SAPIENS | 0 | 74 | 5vwo | C | 55 | TYR | p.Tyr55His | P04181 |
| HOMO SAPIENS | 2 | 74 | 5vwo | C | 90 | GLN | p.Gln90Glu | P04181 |
| HOMO SAPIENS | 6 | 74 | 5vwo | C | 91 | GLY | p.Gly91Glu | P04181 |
| HOMO SAPIENS | 2 | 74 | 5vwo | C | 93 | CYS | p.Cys93Phe | P04181 |
| HOMO SAPIENS | 1 | 74 | 5vwo | C | 154 | ARG | p.Arg154Leu | P04181 |
| HOMO SAPIENS | 0 | 74 | 5vwo | C | 180 | ARG | p.Arg180Thr | P04181 |
| HOMO SAPIENS | 0 | 74 | 5vwo | C | 199 | PRO | p.Pro199Gln | P04181 |
| HOMO SAPIENS | 4 | 74 | 5vwo | C | 209 | TYR | p.Tyr209Ter | P04181 |
| HOMO SAPIENS | 0 | 74 | 5vwo | C | 226 | ALA | p.Ala226Val | P04181 |

| | | | | | | | | |
|---|---|---|---|---|---|---|---|---|
| HOMO SAPIENS | 5 | 74 | 5vwo | C | 233 | GLN | p.Gln233Arg | P04181 |
| HOMO SAPIENS | 5 | 74 | 5vwo | C | 241 | PRO | p.Pro241Leu | P04181 |
| HOMO SAPIENS | 2 | 74 | 5vwo | C | 245 | TYR | p.Tyr245Cys | P04181 |
| HOMO SAPIENS | 6 | 74 | 5vwo | C | 250 | ARG | p.Arg250Pro | P04181 |
| HOMO SAPIENS | 6 | 74 | 5vwo | C | 250 | ARG | p.Arg250Ter | P04181 |
| HOMO SAPIENS | 8 | 74 | 5vwo | C | 267 | THR | p.Thr267Ile | P04181 |
| HOMO SAPIENS | 0 | 74 | 5vwo | C | 270 | ALA | p.Ala270Pro | P04181 |
| HOMO SAPIENS | 0 | 74 | 5vwo | C | 271 | ARG | p.Arg271Lys | P04181 |
| HOMO SAPIENS | 1 | 74 | 5vwo | C | 299 | TYR | p.Tyr299Ter | P04181 |
| HOMO SAPIENS | 0 | 74 | 5vwo | C | 318 | GLU | p.Glu318Lys | P04181 |
| HOMO SAPIENS | 1 | 74 | 5vwo | C | 319 | HIS | p.His319Tyr | P04181 |
| HOMO SAPIENS | 1 | 74 | 5vwo | C | 331 | ARG | p.Arg331Ter | P04181 |
| HOMO SAPIENS | 0 | 74 | 5vwo | C | 332 | VAL | p.Val332Met | P04181 |
| HOMO SAPIENS | 1 | 74 | 5vwo | C | 353 | GLY | p.Gly353Asp | P04181 |
| HOMO SAPIENS | 2 | 74 | 5vwo | C | 375 | GLY | p.Gly375Ala | P04181 |
| HOMO SAPIENS | 0 | 74 | 5vwo | C | 394 | CYS | p.Cys394Arg | P04181 |
| HOMO SAPIENS | 0 | 74 | 5vwo | C | 402 | LEU | p.Leu402Pro | P04181 |
| HOMO SAPIENS | 0 | 74 | 5vwo | C | 436 | ILE | p.Ile436Asn | P04181 |
| HOMO SAPIENS | 7 | 24356 | 4az0 | A | 104 | VAL | p.Val132Met | P10619 |
| HOMO SAPIENS | 4 | 24356 | 4az0 | A | 221 | TYR | p.Tyr249Asn | P10619 |
| HOMO SAPIENS | 3 | 245 | 1rsz | A | 24 | ARG | p.Arg24Ter | P00491 |
| HOMO SAPIENS | 3 | 245 | 1rsz | A | 51 | SER | p.Gly51Ser | P00491 |
| HOMO SAPIENS | 1 | 245 | 1rsz | A | 58 | ARG | p.Arg58Ter | P00491 |
| HOMO SAPIENS | 4 | 245 | 1rsz | A | 89 | GLU | p.Glu89Lys | P00491 |
| HOMO SAPIENS | 6 | 245 | 1rsz | A | 128 | ASP | p.Asp128Gly | P00491 |
| HOMO SAPIENS | 0 | 245 | 1rsz | A | 174 | ALA | p.Ala174Pro | P00491 |
| HOMO SAPIENS | 8 | 245 | 1rsz | A | 192 | TYR | p.Tyr192Cys | P00491 |
| HOMO SAPIENS | 8 | 245 | 1rsz | A | 234 | ARG | p.Arg234Pro | P00491 |
| HOMO SAPIENS | 0 | 37 | 5tby | E | 22 | GLU | p.Glu22Lys | P10916 |
| HOMO SAPIENS | 1 | 37 | 5tby | E | 86 | PHE | p.Phe86Ser | P10916 |
| HOMO SAPIENS | 9 | 71 | 5m9j | A | 12 | GLN | p.Gln36Leu | P03950 |
| HOMO SAPIENS | 3 | 71 | 5m9j | A | 17 | LYS | p.Lys41Glu | P03950 |
| HOMO SAPIENS | 3 | 71 | 5m9j | A | 17 | LYS | p.Lys41Ile | P03950 |
| HOMO SAPIENS | 4 | 71 | 5m9j | A | 28 | SER | p.Ser52Asn | P03950 |
| HOMO SAPIENS | 7 | 71 | 5m9j | A | 31 | ARG | p.Arg55Lys | P03950 |
| HOMO SAPIENS | 9 | 71 | 5m9j | A | 39 | CYS | p.Cys63Trp | P03950 |
| HOMO SAPIENS | 9 | 71 | 5m9j | A | 40 | LYS | p.Lys64Ile | P03950 |
| HOMO SAPIENS | 2 | 71 | 5m9j | A | 60 | GLU | p.Lys84Glu | P03950 |
| HOMO SAPIENS | 0 | 71 | 5m9j | A | 112 | PRO | p.Pro136Leu | P03950 |
| HOMO SAPIENS | 9 | 71 | 5m9j | A | 113 | VAL | p.Val137Ile | P03950 |
| HOMO SAPIENS | 3 | 326 | 5dqi | A | 209 | GLY | p.Gly209Val | Q9Y253 |
| HOMO SAPIENS | 1 | 326 | 5dqi | A | 242 | SER | p.Ser242Ter | Q9Y253 |
| HOMO SAPIENS | 1 | 326 | 5dqi | A | 303 | ARG | p.Arg303Ter | Q9Y253 |
| HOMO SAPIENS | 1 | 8203 | 3iu3 | K | 132 | VAL | p.Val153Ile | P01589 |
| HOMO SAPIENS | 0 | 4995 | 5ffo | H | 234 | THR | p.Thr263Ile | P01137 |
| HOMO SAPIENS | 5 | 38609 | 2ruk | A | 49 | ASP | p.Asp49His | P04637 |
| HOMO SAPIENS | 5 | 38609 | 2ruk | A | 58 | PRO | p.Pro58Arg | P04637 |
| HOMO SAPIENS | 0 | 1607 | 6pah | A | 117 | THR | p.Thr117Ile | P00439 |
| HOMO SAPIENS | 3 | 1607 | 6pah | A | 145 | ASP | p.Asp145Val | P00439 |
| HOMO SAPIENS | 5 | 1607 | 6pah | A | 158 | ARG | p.Arg158Gln | P00439 |
| HOMO SAPIENS | 5 | 1607 | 6pah | A | 158 | ARG | p.Arg158Trp | P00439 |
| HOMO SAPIENS | 5 | 1607 | 6pah | A | 169 | ARG | p.Arg169His | P00439 |
| HOMO SAPIENS | 5 | 1607 | 6pah | A | 169 | ARG | p.Arg169Cys | P00439 |
| HOMO SAPIENS | 3 | 1607 | 6pah | A | 176 | ARG | p.Arg176Leu | P00439 |

| | | | | | | | | |
|---|---|---|---|---|---|---|---|---|
| HOMO SAPIENS | 3 | 1607 | 6pah | A | 177 | VAL | p.Val177Leu | P00439 |
| HOMO SAPIENS | 1 | 1607 | 6pah | A | 194 | LEU | p.Leu194Pro | P00439 |
| HOMO SAPIENS | 1 | 1607 | 6pah | A | 204 | TYR | p.Tyr204Cys | P00439 |
| HOMO SAPIENS | 1 | 1607 | 6pah | A | 204 | TYR | p.Tyr204Ter | P00439 |
| HOMO SAPIENS | 2 | 1607 | 6pah | A | 230 | VAL | p.Val230Ile | P00439 |
| HOMO SAPIENS | 4 | 1607 | 6pah | A | 232 | GLN | p.Gln232Ter | P00439 |
| HOMO SAPIENS | 4 | 1607 | 6pah | A | 241 | ARG | p.Arg241Cys | P00439 |
| HOMO SAPIENS | 4 | 1607 | 6pah | A | 241 | ARG | p.Arg241His | P00439 |
| HOMO SAPIENS | 5 | 1607 | 6pah | A | 243 | ARG | p.Arg243Gln | P00439 |
| HOMO SAPIENS | 5 | 1607 | 6pah | A | 243 | ARG | p.Arg243Ter | P00439 |
| HOMO SAPIENS | 7 | 1607 | 6pah | A | 249 | LEU | p.Leu249Phe | P00439 |
| HOMO SAPIENS | 5 | 1607 | 6pah | A | 252 | ARG | p.Arg252Gly | P00439 |
| HOMO SAPIENS | 5 | 1607 | 6pah | A | 261 | ARG | p.Arg261Ter | P00439 |
| HOMO SAPIENS | 4 | 1607 | 6pah | A | 267 | GLN | p.Gln267Leu | P00439 |
| HOMO SAPIENS | 1 | 1607 | 6pah | A | 272 | GLY | p.Gly272Ter | P00439 |
| HOMO SAPIENS | 2 | 1607 | 6pah | A | 277 | TYR | p.Tyr277Asp | P00439 |
| HOMO SAPIENS | 7 | 1607 | 6pah | A | 280 | GLU | p.Glu280Lys | P00439 |
| HOMO SAPIENS | 8 | 1607 | 6pah | A | 281 | PRO | p.Pro281Leu | P00439 |
| HOMO SAPIENS | 0 | 1607 | 6pah | A | 299 | PHE | p.Phe299Cys | P00439 |
| HOMO SAPIENS | 5 | 1607 | 6pah | A | 300 | ALA | p.Ala300Ser | P00439 |
| HOMO SAPIENS | 5 | 1607 | 6pah | A | 309 | ALA | p.Ala309Asp | P00439 |
| HOMO SAPIENS | 4 | 1607 | 6pah | A | 320 | LYS | p.Lys320Asn | P00439 |
| HOMO SAPIENS | 0 | 1607 | 6pah | A | 345 | ALA | p.Ala345Ser | P00439 |
| HOMO SAPIENS | 0 | 1607 | 6pah | A | 348 | LEU | p.Leu348Val | P00439 |
| HOMO SAPIENS | 6 | 1607 | 6pah | A | 356 | TYR | p.Tyr356Ter | P00439 |
| HOMO SAPIENS | 2 | 1607 | 6pah | A | 380 | THR | p.Thr380Met | P00439 |
| HOMO SAPIENS | 0 | 1607 | 6pah | A | 386 | TYR | p.Tyr386Cys | P00439 |
| HOMO SAPIENS | 5 | 1607 | 6pah | A | 390 | GLU | p.Glu390Gly | P00439 |
| HOMO SAPIENS | 5 | 1607 | 6pah | A | 403 | ALA | p.Ala403Val | P00439 |
| HOMO SAPIENS | 4 | 1607 | 6pah | A | 408 | ARG | p.Arg408Trp | P00439 |
| HOMO SAPIENS | 3 | 1607 | 6pah | A | 413 | ARG | p.Arg413Pro | P00439 |
| HOMO SAPIENS | 4 | 1607 | 6pah | A | 414 | TYR | p.Tyr414Cys | P00439 |
| HOMO SAPIENS | 4 | 1607 | 6pah | A | 414 | TYR | p.Tyr414His | P00439 |
| HOMO SAPIENS | 4 | 1607 | 6pah | A | 415 | ASP | p.Asp415Asn | P00439 |
| HOMO SAPIENS | -1 | 676 | 5dc5 | A | 7 | PRO | p.Pro7Leu | Q9BY41 |
| HOMO SAPIENS | 0 | 676 | 5dc5 | A | 311 | THR | p.Thr311Met | Q9BY41 |
| HOMO SAPIENS | 1 | 124 | 5ftk | E | 95 | ARG | p.Arg95Gly | P55072 |
| HOMO SAPIENS | 1 | 124 | 5ftk | E | 155 | ARG | p.Arg155Cys | P55072 |
| HOMO SAPIENS | 1 | 124 | 5ftk | E | 155 | ARG | p.Arg155His | P55072 |
| HOMO SAPIENS | 1 | 124 | 5ftk | E | 159 | ARG | p.Arg159His | P55072 |
| HOMO SAPIENS | 1 | 124 | 5ftk | E | 159 | ARG | p.Arg159Gly | P55072 |
| HOMO SAPIENS | 0 | 124 | 5ftk | E | 191 | ARG | p.Arg191Gln | P55072 |
| HOMO SAPIENS | 0 | 124 | 5ftk | E | 232 | ALA | p.Ala232Glu | P55072 |
| HOMO SAPIENS | 5 | 5442 | 1fgg | B | 223 | GLY | p.Gly223Ser | O94766 |
| HOMO SAPIENS | 0 | 5442 | 1fgg | B | 277 | ARG | p.Arg277Gln | O94766 |
| HOMO SAPIENS | 0 | 9072 | 2gys | B | 225 | GLU | p.Glu249Gln | P32927 |
| HOMO SAPIENS | 0 | 9072 | 2gys | B | 247 | SER | p.Ser271Ter | P32927 |
| HOMO SAPIENS | 0 | 9072 | 2gys | B | 294 | ASP | p.Asp312Asn | P32927 |
| HOMO SAPIENS | 0 | 1484 | 4zrj | A | 62 | PHE | p.Phe62Ser | P35240 |
| HOMO SAPIENS | 3 | 1484 | 4zrj | A | 205 | MET | p.Met205Val | P35240 |
| HOMO SAPIENS | 3 | 1484 | 4zrj | A | 301 | ILE | p.Ile301Met | P35240 |
| HOMO SAPIENS | 4 | 40923 | 2enq | A | 68 | ILE | p.Ile391Met | P42336 |
| HOMO SAPIENS | 1 | 6179 | 5ofv | A | 125 | THR | p.Thr125Met | O43175 |
| HOMO SAPIENS | 0 | 6179 | 5ofv | A | 135 | ARG | p.Arg135Trp | O43175 |

| | | | | | | | | |
|---|---|---|---|---|---|---|---|---|
| HOMO SAPIENS | 1 | 6179 | 5ofv | A | 163 | ARG | p.Arg163Gln | O43175 |
| HOMO SAPIENS | 9 | 6179 | 5ofv | A | 261 | VAL | p.Val261Met | O43175 |
| HOMO SAPIENS | 7 | 34118 | 2i5o | A | 24 | MET | p.Met647Leu | Q9Y253 |
| HOMO SAPIENS | 8 | 3234 | 5c84 | A | 311 | LYS | p.Lys311Asn | P98170 |
| HOMO SAPIENS | 3 | 3234 | 5c84 | A | 321 | ALA | p.Ala321Gly | P98170 |
| HOMO SAPIENS | -1 | 15129 | 1zy8 | K | 1 | GLY | p.Gly54Ser | O00330 |
| HOMO SAPIENS | -1 | 15129 | 1zy8 | K | 48 | THR | p.Thr101Ala | O00330 |
| HOMO SAPIENS | -1 | 15129 | 1zy8 | K | 105 | GLU | p.Glu158Asp | O00330 |
| HOMO SAPIENS | 8 | 15129 | 1zy8 | K | 136 | ARG | p.Arg189His | O00330 |
| HOMO SAPIENS | -1 | 15129 | 1zy8 | K | 195 | GLN | p.Gln248Ter | O00330 |
| HOMO SAPIENS | 8 | 202 | 4bzb | A | 123 | HIS | p.His123Pro | Q9Y3Z3 |
| HOMO SAPIENS | 4 | 202 | 4bzb | A | 145 | ARG | p.Arg145Ter | Q9Y3Z3 |
| HOMO SAPIENS | 6 | 202 | 4bzb | A | 164 | ARG | p.Arg164Ter | Q9Y3Z3 |
| HOMO SAPIENS | 3 | 202 | 4bzb | A | 201 | ILE | p.Ile201Asn | Q9Y3Z3 |
| HOMO SAPIENS | 0 | 202 | 4bzb | A | 209 | GLY | p.Gly209Ser | Q9Y3Z3 |
| HOMO SAPIENS | 0 | 202 | 4bzb | A | 254 | MET | p.Met254Val | Q9Y3Z3 |
| HOMO SAPIENS | 5 | 202 | 4bzb | A | 531 | ARG | p.Arg531Ser | Q9Y3Z3 |
| HOMO SAPIENS | 6 | 202 | 4bzb | A | 548 | GLN | p.Gln548Ter | Q9Y3Z3 |
| HOMO SAPIENS | 3 | 30947 | 5lgd | A | 90 | PRO | p.Pro90Ser | P16671 |
| HOMO SAPIENS | 6 | 30947 | 5lgd | A | 254 | PHE | p.Phe254Leu | P16671 |
| HOMO SAPIENS | 0 | 30947 | 5lgd | A | 413 | ILE | p.Ile413Leu | P16671 |
| HOMO SAPIENS | 2 | 3906 | 5jtc | A | 735 | ARG | p.Arg735Trp | Q12797 |
| HOMO SAPIENS | 1 | 45 | 5uyx | C | 146 | GLU | p.Glu146Val | P48643 |
| HOMO SAPIENS | 1 | 45 | 5uyx | C | 147 | HIS | p.His147Arg | P48643 |
| HOMO SAPIENS | 0 | 45 | 5uyx | C | 362 | ILE | p.Ile362Met | P48643 |
| HOMO SAPIENS | 8 | 39596 | 1mhn | A | 95 | GLY | p.Gly95Arg | Q16637 |
| HOMO SAPIENS | 4 | 39596 | 1mhn | A | 111 | ALA | p.Ala111Gly | Q16637 |
| HOMO SAPIENS | 9 | 39596 | 1mhn | A | 116 | ILE | p.Ile116Phe | Q16637 |
| HOMO SAPIENS | 1 | 39596 | 1mhn | A | 136 | GLN | p.Gln136Glu | Q16637 |
| HOMO SAPIENS | 1 | 6408 | 5xjy | A | 85 | PRO | p.Pro85Leu | O95477 |
| HOMO SAPIENS | 0 | 6408 | 5xjy | A | 219 | ARG | p.Arg219Lys | O95477 |
| HOMO SAPIENS | 0 | 6408 | 5xjy | A | 230 | ARG | p.Arg230Cys | O95477 |
| HOMO SAPIENS | 1 | 6408 | 5xjy | A | 281 | MET | p.Met281Thr | O95477 |
| HOMO SAPIENS | -1 | 6408 | 5xjy | A | 343 | ALA | p.Ala343Gly | O95477 |
| HOMO SAPIENS | 1 | 6408 | 5xjy | A | 399 | VAL | p.Val399Ala | O95477 |
| HOMO SAPIENS | 1 | 6408 | 5xjy | A | 496 | ARG | p.Arg496Trp | O95477 |
| HOMO SAPIENS | 1 | 6408 | 5xjy | A | 590 | TRP | p.Trp590Ser | O95477 |
| HOMO SAPIENS | 1 | 6408 | 5xjy | A | 638 | ARG | p.Arg638Gln | O95477 |
| HOMO SAPIENS | 4 | 6408 | 5xjy | A | 697 | ALA | p.Ala697Thr | O95477 |
| HOMO SAPIENS | 1 | 6408 | 5xjy | A | 771 | VAL | p.Val771Met | O95477 |
| HOMO SAPIENS | 1 | 6408 | 5xjy | A | 774 | THR | p.Thr774Pro | O95477 |
| HOMO SAPIENS | 1 | 6408 | 5xjy | A | 776 | LYS | p.Lys776Asn | O95477 |
| HOMO SAPIENS | 0 | 6408 | 5xjy | A | 825 | VAL | p.Val825Ile | O95477 |
| HOMO SAPIENS | -1 | 6408 | 5xjy | A | 868 | GLU | p.Glu868Lys | O95477 |
| HOMO SAPIENS | -1 | 6408 | 5xjy | A | 883 | ILE | p.Ile883Met | O95477 |
| HOMO SAPIENS | 0 | 6408 | 5xjy | A | 935 | ASN | p.Asn935Ser | O95477 |
| HOMO SAPIENS | 0 | 6408 | 5xjy | A | 935 | ASN | p.Asn935His | O95477 |
| HOMO SAPIENS | 0 | 6408 | 5xjy | A | 937 | ALA | p.Ala937Val | O95477 |
| HOMO SAPIENS | 0 | 6408 | 5xjy | A | 1018 | ASP | p.Asp1018Gly | O95477 |
| HOMO SAPIENS | 0 | 6408 | 5xjy | A | 1096 | VAL | p.Val1096Ile | O95477 |
| HOMO SAPIENS | 0 | 6408 | 5xjy | A | 1099 | ASP | p.Asp1099Tyr | O95477 |
| HOMO SAPIENS | 0 | 6408 | 5xjy | A | 1172 | GLU | p.Glu1172Asp | O95477 |
| HOMO SAPIENS | 0 | 6408 | 5xjy | A | 1181 | SER | p.Ser1181Phe | O95477 |
| HOMO SAPIENS | 0 | 6408 | 5xjy | A | 1182 | ALA | p.Ala1182Thr | O95477 |

| | | | | | | | | |
|---|---|---|---|---|---|---|---|---|
| HOMO SAPIENS | -1 | 6408 | 5xjy | A | 1289 | ASP | p.Asp1289Asn | O95477 |
| HOMO SAPIENS | 0 | 6408 | 5xjy | A | 1399 | THR | p.Thr1399Ser | O95477 |
| HOMO SAPIENS | 1 | 6408 | 5xjy | A | 1408 | LEU | p.Leu1408Phe | O95477 |
| HOMO SAPIENS | 0 | 6408 | 5xjy | A | 1587 | LYS | p.Lys1587Arg | O95477 |
| HOMO SAPIENS | 1 | 6408 | 5xjy | A | 1674 | VAL | p.Val1674Ile | O95477 |
| HOMO SAPIENS | 1 | 6408 | 5xjy | A | 1680 | ARG | p.Arg1680Trp | O95477 |
| HOMO SAPIENS | 1 | 6408 | 5xjy | A | 1680 | ARG | p.Arg1680Gln | O95477 |
| HOMO SAPIENS | 0 | 6408 | 5xjy | A | 1800 | ASN | p.Asn1800His | O95477 |
| HOMO SAPIENS | -1 | 6408 | 5xjy | A | 1925 | ARG | p.Arg1925Gln | O95477 |
| HOMO SAPIENS | 0 | 6408 | 5xjy | A | 2009 | PHE | p.Phe2009Ser | O95477 |
| HOMO SAPIENS | -1 | 6408 | 5xjy | A | 2243 | ASP | p.Asp2243Glu | O95477 |
| HOMO SAPIENS | 4 | 23760 | 2mut | A | 231 | LEU | p.Phe231Leu | P07992 |
| HOMO SAPIENS | 6 | 1460 | 2hre | B | 96 | ARG | p.Arg96Gln | P22830 |
| HOMO SAPIENS | 8 | 1460 | 2hre | B | 129 | GLY | p.Gly129Arg | P22830 |
| HOMO SAPIENS | 9 | 1460 | 2hre | B | 172 | GLU | p.Glu172Lys | P22830 |
| HOMO SAPIENS | 4 | 1460 | 2hre | B | 285 | GLN | p.Gln285Arg | P22830 |
| HOMO SAPIENS | 5 | 1460 | 2hre | B | 362 | VAL | p.Val362Gly | P22830 |
| HOMO SAPIENS | 6 | 1460 | 2hre | B | 411 | CYS | p.Cys411Gly | P22830 |
| HOMO SAPIENS | 6 | 1460 | 2hre | B | 417 | PHE | p.Phe417Ser | P22830 |
| HOMO SAPIENS | 0 | 2210 | 5bug | A | 19 | ASP | p.Asp19Asn | P60484 |
| HOMO SAPIENS | 0 | 2210 | 5bug | A | 35 | MET | p.Met35Arg | P60484 |
| HOMO SAPIENS | 1 | 2210 | 5bug | A | 61 | HIS | p.His61Asp | P60484 |
| HOMO SAPIENS | 0 | 2210 | 5bug | A | 68 | TYR | p.Tyr68His | P60484 |
| HOMO SAPIENS | 8 | 2210 | 5bug | A | 70 | LEU | p.Leu70Pro | P60484 |
| HOMO SAPIENS | 6 | 2210 | 5bug | A | 93 | HIS | p.His93Arg | P60484 |
| HOMO SAPIENS | 1 | 2210 | 5bug | A | 112 | LEU | p.Leu112Pro | P60484 |
| HOMO SAPIENS | 0 | 2210 | 5bug | A | 121 | ALA | p.Ala121Gly | P60484 |
| HOMO SAPIENS | 3 | 2210 | 5bug | A | 123 | HIS | p.His123Arg | P60484 |
| HOMO SAPIENS | 7 | 2210 | 5bug | A | 124 | CYS | p.Cys124Arg | P60484 |
| HOMO SAPIENS | 9 | 2210 | 5bug | A | 129 | GLY | p.Gly129Glu | P60484 |
| HOMO SAPIENS | 9 | 2210 | 5bug | A | 130 | ARG | p.Arg130Gln | P60484 |
| HOMO SAPIENS | 9 | 2210 | 5bug | A | 130 | ARG | p.Arg130Gly | P60484 |
| HOMO SAPIENS | 0 | 2210 | 5bug | A | 132 | GLY | p.Gly132Val | P60484 |
| HOMO SAPIENS | 0 | 2210 | 5bug | A | 170 | SER | p.Ser170Arg | P60484 |
| HOMO SAPIENS | 1 | 2210 | 5bug | A | 173 | ARG | p.Arg173His | P60484 |
| HOMO SAPIENS | 0 | 2210 | 5bug | A | 217 | VAL | p.Val217Ile | P60484 |
| HOMO SAPIENS | 0 | 2210 | 5bug | A | 241 | PHE | p.Phe241Ser | P60484 |
| HOMO SAPIENS | 0 | 2210 | 5bug | A | 250 | CYS | p.Cys250Ter | P60484 |
| HOMO SAPIENS | 1 | 2210 | 5bug | A | 252 | ASP | p.Asp252Gly | P60484 |
| HOMO SAPIENS | 3 | 123 | 2ibu | A | 93 | ASN | p.Asn93Ser | P24752 |
| HOMO SAPIENS | 0 | 123 | 2ibu | A | 145 | GLN | p.Gln145Glu | P24752 |
| HOMO SAPIENS | 0 | 123 | 2ibu | A | 152 | GLY | p.Gly152Ala | P24752 |
| HOMO SAPIENS | 2 | 123 | 2ibu | A | 158 | ASN | p.Asn158Asp | P24752 |
| HOMO SAPIENS | 2 | 123 | 2ibu | A | 158 | ASN | p.Asn158Ser | P24752 |
| HOMO SAPIENS | 5 | 123 | 2ibu | A | 183 | GLY | p.Gly183Arg | P24752 |
| HOMO SAPIENS | 3 | 123 | 2ibu | A | 272 | GLN | p.Gln272Ter | P24752 |
| HOMO SAPIENS | 3 | 123 | 2ibu | A | 312 | ILE | p.Ile312Thr | P24752 |
| HOMO SAPIENS | 1 | 123 | 2ibu | A | 333 | ALA | p.Ala333Pro | P24752 |
| HOMO SAPIENS | 5 | 123 | 2ibu | A | 379 | GLY | p.Gly379Val | P24752 |
| HOMO SAPIENS | 0 | 123 | 2ibu | A | 380 | ALA | p.Ala380Thr | P24752 |
| HOMO SAPIENS | 2 | 369 | 5nrw | A | 1175 | TYR | p.Tyr1175Cys | Q92793 |
| HOMO SAPIENS | 2 | 830 | 2vc2 | A | 116 | LEU | p.Leu147Val | P08514 |
| HOMO SAPIENS | 0 | 830 | 2vc2 | A | 183 | LEU | p.Leu214Pro | P08514 |
| HOMO SAPIENS | 4 | 830 | 2vc2 | A | 242 | GLY | p.Gly273Asp | P08514 |

| | | | | | | | | |
|---|---|---|---|---|---|---|---|---|
| HOMO SAPIENS | 6 | 830 | 2vc2 | A | 324 | GLU | p.Glu355Lys | P08514 |
| HOMO SAPIENS | -1 | 10353 | 5ejc | E | 989 | ALA | p.Ala989Ser | Q92574 |
| HOMO SAPIENS | 0 | 15448 | 3lgg | A | 21 | GLY | p.Gly47Arg | Q9NZK5 |
| HOMO SAPIENS | 0 | 15448 | 3lgg | A | 21 | GLY | p.Gly47Val | Q9NZK5 |
| HOMO SAPIENS | 9 | 15448 | 3lgg | A | 86 | HIS | p.His112Gln | Q9NZK5 |
| HOMO SAPIENS | 0 | 15448 | 3lgg | A | 143 | ARG | p.Arg169Gln | Q9NZK5 |
| HOMO SAPIENS | 3 | 15448 | 3lgg | A | 221 | ALA | p.Ala247Val | Q9NZK5 |
| HOMO SAPIENS | 0 | 15448 | 3lgg | A | 225 | PRO | p.Pro251Leu | Q9NZK5 |
| HOMO SAPIENS | 0 | 15448 | 3lgg | A | 283 | MET | p.Met309Ile | Q9NZK5 |
| HOMO SAPIENS | 2 | 15448 | 3lgg | A | 334 | THR | p.Thr360Ala | Q9NZK5 |
| HOMO SAPIENS | 7 | 15448 | 3lgg | A | 357 | GLY | p.Gly383Ser | Q9NZK5 |
| HOMO SAPIENS | 0 | 15448 | 3lgg | A | 427 | TYR | p.Tyr453Cys | Q9NZK5 |
| HOMO SAPIENS | 0 | 872 | 2y3i | A | 87 | LEU | p.Leu88Pro | P00558 |
| HOMO SAPIENS | 1 | 872 | 2y3i | A | 163 | ASP | p.Asp164Val | P00558 |
| HOMO SAPIENS | 4 | 872 | 2y3i | A | 205 | ARG | p.Arg206Pro | P00558 |
| HOMO SAPIENS | 4 | 872 | 2y3i | A | 267 | ASP | p.Asp268Asn | P00558 |
| HOMO SAPIENS | 8 | 872 | 2y3i | A | 284 | ASP | p.Asp285Val | P00558 |
| HOMO SAPIENS | 4 | 8 | 5jco | B | 188 | ILE | p.Ile188Leu | Q71U36 |
| HOMO SAPIENS | 7 | 8 | 5jco | B | 263 | PRO | p.Pro263Thr | Q71U36 |
| HOMO SAPIENS | 6 | 8 | 5jco | B | 264 | ARG | p.Arg264Cys | Q71U36 |
| HOMO SAPIENS | 7 | 8 | 5jco | B | 402 | ARG | p.Arg402His | Q71U36 |
| HOMO SAPIENS | 7 | 8 | 5jco | B | 419 | SER | p.Ser419Leu | Q71U36 |
| HOMO SAPIENS | -1 | 880 | 4nwf | A | 2 | THR | p.Thr2Ile | Q06124 |
| HOMO SAPIENS | 2 | 880 | 4nwf | A | 58 | ASN | p.Asn58Ser | Q06124 |
| HOMO SAPIENS | 4 | 880 | 4nwf | A | 61 | ASP | p.Asp61Gly | Q06124 |
| HOMO SAPIENS | 2 | 880 | 4nwf | A | 62 | TYR | p.Tyr62Asp | Q06124 |
| HOMO SAPIENS | 3 | 880 | 4nwf | A | 63 | TYR | p.Tyr63Cys | Q06124 |
| HOMO SAPIENS | 2 | 880 | 4nwf | A | 69 | GLU | p.Glu69Gln | Q06124 |
| HOMO SAPIENS | 2 | 880 | 4nwf | A | 72 | ALA | p.Ala72Gly | Q06124 |
| HOMO SAPIENS | 2 | 880 | 4nwf | A | 72 | ALA | p.Ala72Ser | Q06124 |
| HOMO SAPIENS | 3 | 880 | 4nwf | A | 73 | THR | p.Thr73Ile | Q06124 |
| HOMO SAPIENS | 5 | 880 | 4nwf | A | 76 | GLU | p.Glu76Ala | Q06124 |
| HOMO SAPIENS | 5 | 880 | 4nwf | A | 76 | GLU | p.Glu76Lys | Q06124 |
| HOMO SAPIENS | 3 | 880 | 4nwf | A | 79 | GLN | p.Gln79Arg | Q06124 |
| HOMO SAPIENS | 5 | 880 | 4nwf | A | 152 | ARG | p.Arg152His | Q06124 |
| HOMO SAPIENS | 2 | 880 | 4nwf | A | 268 | GLY | p.Gly268Cys | Q06124 |
| HOMO SAPIENS | 9 | 880 | 4nwf | A | 276 | LYS | p.Lys276Arg | Q06124 |
| HOMO SAPIENS | 9 | 880 | 4nwf | A | 279 | TYR | p.Tyr279Cys | Q06124 |
| HOMO SAPIENS | 5 | 880 | 4nwf | A | 285 | PHE | p.Phe285Ser | Q06124 |
| HOMO SAPIENS | 0 | 880 | 4nwf | A | 308 | ASP | p.Asn308Ser | Q06124 |
| HOMO SAPIENS | 0 | 880 | 4nwf | A | 308 | ASP | p.Asn308Asp | Q06124 |
| HOMO SAPIENS | 5 | 880 | 4nwf | A | 392 | ALA | p.Ala392Thr | Q06124 |
| HOMO SAPIENS | 1 | 880 | 4nwf | A | 411 | THR | p.Thr415Met | Q06124 |
| HOMO SAPIENS | 0 | 880 | 4nwf | A | 461 | ALA | p.Ala465Thr | Q06124 |
| HOMO SAPIENS | 0 | 880 | 4nwf | A | 464 | GLY | p.Gly468Ala | Q06124 |
| HOMO SAPIENS | 1 | 880 | 4nwf | A | 491 | PRO | p.Pro495Ser | Q06124 |
| HOMO SAPIENS | 1 | 880 | 4nwf | A | 502 | SER | p.Ser506Thr | Q06124 |
| HOMO SAPIENS | 8 | 880 | 4nwf | A | 510 | GLN | p.Gln514Pro | Q06124 |
| HOMO SAPIENS | 4 | 1872 | 3wf1 | D | 109 | ARG | p.Arg109Trp | P16278 |
| HOMO SAPIENS | 0 | 1872 | 3wf1 | D | 521 | ARG | p.Arg521Cys | P16278 |
| HOMO SAPIENS | 5 | 910 | 5xte | A | 45 | SER | p.Ser45Phe | O14949 |
| HOMO SAPIENS | 2 | 9017 | 5f0k | A | 382 | LYS | p.Lys382Arg | Q96QK1 |
| HOMO SAPIENS | 1 | 710 | 4lsj | A | 559 | ILE | p.Ile559Asn | P04150 |
| HOMO SAPIENS | 5 | 710 | 4lsj | A | 571 | VAL | p.Val571Ala | P04150 |

| | | | | | | | | |
|---|---|---|---|---|---|---|---|---|
| HOMO SAPIENS | 0 | 710 | 4lsj | A | 641 | ASP | p.Asp641Val | P04150 |
| HOMO SAPIENS | 1 | 710 | 4lsj | A | 678 | ASP | p.Asp678Glu | P04150 |
| HOMO SAPIENS | 2 | 710 | 4lsj | A | 679 | GLY | p.Gly679Ser | P04150 |
| HOMO SAPIENS | 1 | 710 | 4lsj | A | 737 | PHE | p.Phe737Leu | P04150 |
| HOMO SAPIENS | 1 | 710 | 4lsj | A | 747 | ILE | p.Ile747Met | P04150 |
| HOMO SAPIENS | 1 | 710 | 4lsj | A | 753 | LEU | p.Leu753Phe | P04150 |
| HOMO SAPIENS | 1 | 710 | 4lsj | A | 773 | LEU | p.Leu773Pro | P04150 |
| HOMO SAPIENS | 1 | 2145 | 1av1 | B | 71 | PHE | p.Phe95Tyr | P02647 |
| HOMO SAPIENS | 1 | 2145 | 1av1 | B | 128 | GLU | p.Glu152Lys | P02647 |
| HOMO SAPIENS | 1 | 2145 | 1av1 | B | 136 | GLU | p.Glu160Lys | P02647 |
| HOMO SAPIENS | 2 | 2145 | 1av1 | B | 143 | PRO | p.Pro167Arg | P02647 |
| HOMO SAPIENS | 6 | 2145 | 1av1 | B | 164 | ALA | p.Ala188Ser | P02647 |
| HOMO SAPIENS | 7 | 2145 | 1av1 | B | 173 | ARG | p.Arg197Cys | P02647 |
| HOMO SAPIENS | 2 | 2145 | 1av1 | B | 198 | GLU | p.Glu222Lys | P02647 |
| HOMO SAPIENS | -1 | 3432 | 2beu | A | 1 | SER | p.Ser46Ter | P12694 |
| HOMO SAPIENS | 5 | 3432 | 2beu | A | 79 | ARG | p.Arg124Trp | P12694 |
| HOMO SAPIENS | 7 | 3432 | 2beu | A | 114 | ARG | p.Arg159Gln | P12694 |
| HOMO SAPIENS | 6 | 3432 | 2beu | A | 166 | THR | p.Thr211Met | P12694 |
| HOMO SAPIENS | 4 | 3432 | 2beu | A | 175 | ALA | p.Ala220Val | P12694 |
| HOMO SAPIENS | 5 | 3432 | 2beu | A | 204 | GLY | p.Gly249Ser | P12694 |
| HOMO SAPIENS | 4 | 3432 | 2beu | A | 209 | ALA | p.Ala254Asp | P12694 |
| HOMO SAPIENS | 6 | 3432 | 2beu | A | 220 | ARG | p.Arg265Trp | P12694 |
| HOMO SAPIENS | 6 | 3432 | 2beu | A | 234 | TYR | p.Tyr279Cys | P12694 |
| HOMO SAPIENS | 5 | 3432 | 2beu | A | 240 | ALA | p.Ala285Pro | P12694 |
| HOMO SAPIENS | 4 | 3432 | 2beu | A | 245 | GLY | p.Gly290Arg | P12694 |
| HOMO SAPIENS | 5 | 3432 | 2beu | A | 269 | ARG | p.Arg314Ter | P12694 |
| HOMO SAPIENS | 3 | 3432 | 2beu | A | 301 | ARG | p.Arg346Cys | P12694 |
| HOMO SAPIENS | 5 | 3432 | 2beu | A | 359 | ASN | p.Asn404Ser | P12694 |
| HOMO SAPIENS | 4 | 3432 | 2beu | A | 393 | TYR | p.Tyr438Asn | P12694 |
| HOMO SAPIENS | 8 | 4541 | 1rr8 | C | 533 | ASP | p.Asp533Gly | P11387 |
| HOMO SAPIENS | 2 | 3247 | 1tzn | d | 1105 | GLY | p.Gly105Asp | P58335 |
| HOMO SAPIENS | 3 | 3247 | 1tzn | d | 1111 | ARG | p.Arg111Cys | P58335 |
| HOMO SAPIENS | 2 | 3247 | 1tzn | d | 1189 | ILE | p.Ile189Thr | P58335 |
| HOMO SAPIENS | 9 | 467 | 1suv | C | 58 | ASP | p.Asp77Asn | P02787 |
| HOMO SAPIENS | 2 | 467 | 1suv | C | 277 | ASP | p.Asp296Gly | P02787 |
| HOMO SAPIENS | 9 | 6485 | 1xwd | B | 51 | CYS | p.Cys69Gly | P01225 |
| HOMO SAPIENS | 1 | 694 | 1wuu | A | 28 | PRO | p.Pro28Thr | P51570 |
| HOMO SAPIENS | 0 | 694 | 1wuu | A | 32 | VAL | p.Val32Met | P51570 |
| HOMO SAPIENS | 0 | 694 | 1wuu | A | 198 | ALA | p.Ala198Val | P51570 |
| HOMO SAPIENS | 0 | 143 | 4nkx | A | 93 | PHE | p.Phe93Cys | P05093 |
| HOMO SAPIENS | 3 | 143 | 4nkx | A | 96 | ARG | p.Arg96Trp | P05093 |
| HOMO SAPIENS | 3 | 143 | 4nkx | A | 96 | ARG | p.Arg96Gln | P05093 |
| HOMO SAPIENS | 0 | 143 | 4nkx | A | 106 | SER | p.Ser106Pro | P05093 |
| HOMO SAPIENS | 1 | 143 | 4nkx | A | 114 | PHE | p.Phe114Val | P05093 |
| HOMO SAPIENS | 1 | 143 | 4nkx | A | 116 | ASP | p.Asp116Val | P05093 |
| HOMO SAPIENS | 3 | 143 | 4nkx | A | 125 | ARG | p.Arg125Gln | P05093 |
| HOMO SAPIENS | 0 | 143 | 4nkx | A | 201 | TYR | p.Tyr201Asp | P05093 |
| HOMO SAPIENS | 0 | 143 | 4nkx | A | 239 | ARG | p.Arg239Ter | P05093 |
| HOMO SAPIENS | 0 | 143 | 4nkx | A | 329 | TYR | p.Tyr329Asp | P05093 |
| HOMO SAPIENS | 1 | 143 | 4nkx | A | 342 | PRO | p.Pro342Thr | P05093 |
| HOMO SAPIENS | 2 | 143 | 4nkx | A | 347 | ARG | p.Arg347His | P05093 |
| HOMO SAPIENS | 2 | 143 | 4nkx | A | 347 | ARG | p.Arg347Cys | P05093 |
| HOMO SAPIENS | 1 | 143 | 4nkx | A | 358 | ARG | p.Arg358Gln | P05093 |
| HOMO SAPIENS | 0 | 143 | 4nkx | A | 362 | ARG | p.Arg362Cys | P05093 |

| | | | | | | | | |
|---|---|---|---|---|---|---|---|---|
| HOMO SAPIENS | 1 | 143 | 4nkx | A | 406 | TRP | p.Trp406Arg | P05093 |
| HOMO SAPIENS | 0 | 143 | 4nkx | A | 428 | PRO | p.Pro428Leu | P05093 |
| HOMO SAPIENS | 6 | 1441 | 1uuh | A | 46 | ARG | p.Arg46Gly | P16070 |
| HOMO SAPIENS | 3 | 82 | 4wlv | D | 37 | GLY | p.Gly37Arg | P40926 |
| HOMO SAPIENS | 0 | 82 | 4wlv | D | 133 | PRO | p.Pro133Leu | P40926 |
| HOMO SAPIENS | 2 | 392 | 5xtd | A | 56 | SER | p.Ser56Pro | P49821 |
| HOMO SAPIENS | 2 | 392 | 5xtd | A | 59 | ARG | p.Arg59Ter | P49821 |
| HOMO SAPIENS | 2 | 392 | 5xtd | A | 74 | ASP | p.Asp74Asn | P49821 |
| HOMO SAPIENS | 4 | 392 | 5xtd | A | 116 | ASN | p.Asn116Lys | P49821 |
| HOMO SAPIENS | 4 | 392 | 5xtd | A | 117 | ALA | p.Ala117Thr | P49821 |
| HOMO SAPIENS | 5 | 392 | 5xtd | A | 122 | PRO | p.Pro122Leu | P49821 |
| HOMO SAPIENS | 3 | 392 | 5xtd | A | 160 | GLY | p.Gly160Glu | P49821 |
| HOMO SAPIENS | 4 | 392 | 5xtd | A | 214 | GLU | p.Glu214Lys | P49821 |
| HOMO SAPIENS | 2 | 392 | 5xtd | A | 357 | MET | p.Met357Thr | P49821 |
| HOMO SAPIENS | 4 | 392 | 5xtd | A | 423 | THR | p.Thr423Met | P49821 |
| HOMO SAPIENS | 0 | 22593 | 3pdf | A | 8 | TYR | p.Tyr32Ter | P53634 |
| HOMO SAPIENS | 0 | 22593 | 3pdf | A | 15 | TRP | p.Trp39Ser | P53634 |
| HOMO SAPIENS | 0 | 22593 | 3pdf | A | 103 | HIS | p.His127Pro | P53634 |
| HOMO SAPIENS | 9 | 22593 | 3pdf | A | 228 | GLN | p.Gln252Leu | P53634 |
| HOMO SAPIENS | 3 | 22593 | 3pdf | A | 262 | GLN | p.Gln286Arg | P53634 |
| HOMO SAPIENS | 4 | 22593 | 3pdf | A | 277 | GLY | p.Gly301Ser | P53634 |
| HOMO SAPIENS | 0 | 22593 | 3pdf | A | 323 | TYR | p.Tyr347Cys | P53634 |
| HOMO SAPIENS | 5 | 3461 | 4c2a | A | 1306 | GLN | p.Arg1306Trp | P04275 |
| HOMO SAPIENS | 5 | 3461 | 4c2a | A | 1374 | ARG | p.Arg1374Cys | P04275 |
| HOMO SAPIENS | 5 | 3461 | 4c2a | A | 1379 | ARG | p.Arg1379Cys | P04275 |
| HOMO SAPIENS | 4 | 3461 | 4c2a | A | 1380 | ILE | p.Ile1380Val | P04275 |
| HOMO SAPIENS | 0 | 3461 | 4c2a | A | 1381 | ALA | p.Thr1381Ala | P04275 |
| HOMO SAPIENS | 5 | 3461 | 4c2a | A | 1399 | ARG | p.Arg1399His | P04275 |
| HOMO SAPIENS | 5 | 3461 | 4c2a | A | 1435 | ASN | p.Asn1435Ser | P04275 |
| HOMO SAPIENS | 0 | 4533 | 5uj7 | C | 174 | TYR | p.Tyr174Cys | O43929 |
| HOMO SAPIENS | 0 | 4533 | 5uj7 | C | 202 | LEU | p.Leu202Val | O43929 |
| HOMO SAPIENS | 0 | 1374 | 3j07 | O | 39 | PRO | p.Pro39Leu | P02511 |
| HOMO SAPIENS | 0 | 1374 | 3j07 | O | 107 | ARG | p.Arg107His | P02511 |
| HOMO SAPIENS | 0 | 1374 | 3j07 | O | 157 | ARG | p.Arg157His | P02511 |
| HOMO SAPIENS | -1 | 942 | 1s9i | B | 57 | PHE | p.Phe57Cys | P36507 |
| HOMO SAPIENS | -1 | 942 | 1s9i | B | 57 | PHE | p.Phe57Val | P36507 |
| HOMO SAPIENS | 2 | 942 | 1s9i | B | 128 | PRO | p.Pro128Gln | P36507 |
| HOMO SAPIENS | 1 | 942 | 1s9i | B | 134 | TYR | p.Tyr134His | P36507 |
| HOMO SAPIENS | 1 | 942 | 1s9i | B | 188 | HIS | p.His188Gln | P36507 |
| HOMO SAPIENS | 0 | 942 | 1s9i | B | 262 | VAL | p.Val262Phe | P36507 |
| HOMO SAPIENS | 0 | 942 | 1s9i | B | 263 | GLY | p.Gly263Arg | P36507 |
| HOMO SAPIENS | -1 | 942 | 1s9i | B | 298 | PRO | p.Pro298Leu | P36507 |
| HOMO SAPIENS | -1 | 942 | 1s9i | B | 305 | VAL | p.Val305Ile | P36507 |
| HOMO SAPIENS | 0 | 3245 | 1fbv | A | 367 | GLN | p.Gln367Pro | P22681 |
| HOMO SAPIENS | 0 | 3245 | 1fbv | A | 371 | TYR | p.Tyr371His | P22681 |
| HOMO SAPIENS | 1 | 3245 | 1fbv | A | 382 | LYS | p.Lys382Glu | P22681 |
| HOMO SAPIENS | 2 | 3245 | 1fbv | A | 390 | ASP | p.Asp390Tyr | P22681 |
| HOMO SAPIENS | 9 | 3245 | 1fbv | A | 396 | CYS | p.Cys396Arg | P22681 |
| HOMO SAPIENS | 7 | 3245 | 1fbv | A | 420 | ARG | p.Arg420Gln | P22681 |
| HOMO SAPIENS | 1 | 3245 | 1fbv | A | 429 | ILE | p.Ile429Met | P22681 |
| HOMO SAPIENS | 0 | 2604 | 2pbd | P | 111 | LEU | p.Leu112Met | P07737 |
| HOMO SAPIENS | 3 | 2604 | 2pbd | P | 116 | GLU | p.Glu117Gly | P07737 |
| HOMO SAPIENS | 4 | 2013 | 3b7w | A | 472 | ARG | p.Arg472Gly | Q08AH3 |
| HOMO SAPIENS | 0 | 24346 | 3sh5 | A | 85 | ARG | p.Arg4281His | P98160 |

| | | | | | | | | |
|---|---|---|---|---|---|---|---|---|
| HOMO SAPIENS | 0 | 24346 | 3sh5 | A | 122 | ARG | p.Arg4318Trp | P98160 |
| HOMO SAPIENS | 0 | 24346 | 3sh5 | A | 135 | SER | p.Ser4331Asn | P98160 |
| HOMO SAPIENS | 0 | 24346 | 3sh5 | A | 136 | VAL | p.Val4332Ile | P98160 |
| HOMO SAPIENS | 0 | 707 | 4p2g | A | 26 | MET | p.Met26Ile | Q99497 |
| HOMO SAPIENS | 5 | 707 | 4p2g | A | 39 | ALA | p.Ala39Ser | Q99497 |
| HOMO SAPIENS | 5 | 707 | 4p2g | A | 64 | GLU | p.Glu64Asp | Q99497 |
| HOMO SAPIENS | 4 | 707 | 4p2g | A | 149 | ASP | p.Asp149Ala | Q99497 |
| HOMO SAPIENS | 6 | 707 | 4p2g | A | 163 | GLU | p.Glu163Lys | Q99497 |
| HOMO SAPIENS | 6 | 707 | 4p2g | A | 166 | LEU | p.Leu166Pro | Q99497 |
| HOMO SAPIENS | 3 | 5121 | 4xss | E | 31 | GLY | p.Gly58Arg | P06213 |
| HOMO SAPIENS | 2 | 5121 | 4xss | E | 119 | ILE | p.Ile146Met | P06213 |
| HOMO SAPIENS | 8 | 5121 | 4xss | E | 209 | HIS | p.His236Arg | P06213 |
| HOMO SAPIENS | 2 | 5121 | 4xss | E | 234 | ASP | p.Asp261Glu | P06213 |
| HOMO SAPIENS | 1 | 5121 | 4xss | E | 276 | GLN | p.Gln303His | P06213 |
| HOMO SAPIENS | 1 | 27790 | 5erp | B | 140 | THR | p.Thr275Met | Q02487 |
| HOMO SAPIENS | 0 | 27790 | 5erp | B | 150 | ILE | p.Ile285Thr | Q02487 |
| HOMO SAPIENS | 1 | 27790 | 5erp | B | 205 | THR | p.Thr340Ala | Q02487 |
| HOMO SAPIENS | 1 | 27790 | 5erp | B | 223 | THR | p.Thr358Ile | Q02487 |
| HOMO SAPIENS | 0 | 27790 | 5erp | B | 301 | GLY | p.Gly436Ala | Q02487 |
| HOMO SAPIENS | -1 | 27790 | 5erp | B | 315 | ARG | p.Arg450Ser | Q02487 |
| HOMO SAPIENS | 0 | 27790 | 5erp | B | 369 | SER | p.Ser504Gly | Q02487 |
| HOMO SAPIENS | 0 | 27790 | 5erp | B | 385 | ILE | p.Ile520Thr | Q02487 |
| HOMO SAPIENS | 0 | 27790 | 5erp | B | 461 | ALA | p.Ala596Val | Q02487 |
| HOMO SAPIENS | 0 | 27790 | 5erp | B | 503 | GLN | p.Gln638His | Q02487 |
| HOMO SAPIENS | 5 | 6599 | 4nsc | A | 129 | ARG | p.Arg129Leu | Q9BPX6 |
| HOMO SAPIENS | -1 | 6599 | 4nsc | A | 183 | GLN | p.Gln183Ter | Q9BPX6 |
| HOMO SAPIENS | 0 | 20420 | 3ghm | A | 398 | ARG | p.Arg398His | Q76LX8 |
| HOMO SAPIENS | 0 | 20420 | 3ghm | A | 448 | GLN | p.Gln448Glu | Q76LX8 |
| HOMO SAPIENS | 1 | 20420 | 3ghm | A | 449 | GLN | p.Gln449Ter | Q76LX8 |
| HOMO SAPIENS | 0 | 20420 | 3ghm | A | 456 | GLN | p.Gln456His | Q76LX8 |
| HOMO SAPIENS | 1 | 20420 | 3ghm | A | 528 | ARG | p.Arg528Gly | Q76LX8 |
| HOMO SAPIENS | 0 | 20420 | 3ghm | A | 618 | PRO | p.Pro618Ala | Q76LX8 |
| HOMO SAPIENS | 0 | 20420 | 3ghm | A | 625 | ARG | p.Arg625His | Q76LX8 |
| HOMO SAPIENS | 0 | 275 | 4rmz | A | 293 | GLN | p.Gln293Ter | Q9NWZ3 |
| HOMO SAPIENS | 0 | 275 | 4rmz | A | 428 | ALA | p.Ala428Thr | Q9NWZ3 |
| HOMO SAPIENS | 2 | 10458 | 3puf | B | 138 | LEU | p.Leu138Phe | Q5TBB1 |
| HOMO SAPIENS | 6 | 10458 | 3puf | B | 177 | ALA | p.Ala177Thr | Q5TBB1 |
| HOMO SAPIENS | -1 | 10458 | 3puf | B | 185 | VAL | p.Val185Gly | Q5TBB1 |
| HOMO SAPIENS | 0 | 26648 | 2n90 | A | 36 | ARG | p.Arg444Gln | P04629 |
| HOMO SAPIENS | 2 | 977 | 1dnw | D | 166 | ALA | p.Ala332Val | P05164 |
| HOMO SAPIENS | 4 | 977 | 1dnw | D | 333 | ARG | p.Arg499Cys | P05164 |
| HOMO SAPIENS | 1 | 977 | 1dnw | D | 403 | ARG | p.Arg569Trp | P05164 |
| HOMO SAPIENS | 0 | 25704 | 3ajb | A | 82 | GLN | p.Gln82Arg | P56589 |
| HOMO SAPIENS | 3 | 255 | 3t1p | A | 38 | TYR | p.Tyr62Ter | P01009 |
| HOMO SAPIENS | 3 | 255 | 3t1p | A | 39 | ARG | p.Arg63Cys | P01009 |
| HOMO SAPIENS | 0 | 255 | 3t1p | A | 101 | ARG | p.Arg125His | P01009 |
| HOMO SAPIENS | 0 | 255 | 3t1p | A | 212 | GLN | p.Gln236Arg | P01009 |
| HOMO SAPIENS | 0 | 255 | 3t1p | A | 213 | VAL | p.Val237Ala | P01009 |
| HOMO SAPIENS | 1 | 255 | 3t1p | A | 269 | HIS | p.His293Gln | P01009 |
| HOMO SAPIENS | 8 | 255 | 3t1p | A | 341 | ASP | p.Asp365Asn | P01009 |
| HOMO SAPIENS | 9 | 255 | 3t1p | A | 342 | GLU | p.Glu366Lys | P01009 |
| HOMO SAPIENS | 0 | 255 | 3t1p | A | 369 | PRO | p.Pro393Ser | P01009 |
| HOMO SAPIENS | 0 | 255 | 3t1p | A | 376 | GLU | p.Glu400Asp | P01009 |
| HOMO SAPIENS | -1 | 5175 | 3zxi | A | 35 | ALA | p.Ala35Asp | Q9Y2Z4 |

| | | | | | | | | | |
|---|---|---|---|---|---|---|---|---|---|
| HOMO SAPIENS | 0 | 5175 | 3zxi | A | | 52 | PHE | p.Phe52Leu | Q9Y2Z4 |
| HOMO SAPIENS | -1 | 5175 | 3zxi | A | | 68 | GLY | p.Gly68Ser | Q9Y2Z4 |
| HOMO SAPIENS | 0 | 5175 | 3zxi | A | | 179 | LYS | p.Lys179Gln | Q9Y2Z4 |
| HOMO SAPIENS | 0 | 5175 | 3zxi | A | | 191 | GLY | p.Gly191Val | Q9Y2Z4 |
| HOMO SAPIENS | 0 | 5175 | 3zxi | A | | 251 | ILE | p.Ile251Val | Q9Y2Z4 |
| HOMO SAPIENS | 2 | 5175 | 3zxi | A | | 295 | ASP | p.Asp295Asn | Q9Y2Z4 |
| HOMO SAPIENS | 0 | 5175 | 3zxi | A | | 311 | ASP | p.Asp311Glu | Q9Y2Z4 |
| HOMO SAPIENS | 0 | 5175 | 3zxi | A | | 360 | ARG | p.Arg360Ter | Q9Y2Z4 |
| HOMO SAPIENS | 1 | 3016 | 3exg | | 3 | 4 | ASP | p.Asp33His | P08559 |
| HOMO SAPIENS | 1 | 3016 | 3exg | | 3 | 176 | PHE | p.Phe205Leu | P08559 |
| HOMO SAPIENS | 1 | 3016 | 3exg | | 3 | 214 | TYR | p.Tyr243Asn | P08559 |
| HOMO SAPIENS | 1 | 3016 | 3exg | | 3 | 229 | ASP | p.Asp258Ala | P08559 |
| HOMO SAPIENS | 0 | 3016 | 3exg | | 3 | 233 | VAL | p.Val262Ile | P08559 |
| HOMO SAPIENS | 1 | 3016 | 3exg | | 3 | 234 | ARG | p.Arg263Gly | P08559 |
| HOMO SAPIENS | 0 | 3016 | 3exg | | 3 | 253 | MET | p.Met282Leu | P08559 |
| HOMO SAPIENS | 7 | 3016 | 3exg | | 3 | 259 | ARG | p.Arg288His | P08559 |
| HOMO SAPIENS | -1 | 3016 | 3exg | | 3 | 273 | ARG | p.Arg302Cys | P08559 |
| HOMO SAPIENS | 2 | 3016 | 3exg | | 3 | 286 | ASP | p.Asp315Asn | P08559 |
| HOMO SAPIENS | 1 | 3016 | 3exg | | 3 | 349 | ARG | p.Arg378His | P08559 |
| HOMO SAPIENS | 5 | 6729 | 4a0l | E | | 572 | ARG | p.Arg572Cys | Q13620 |
| HOMO SAPIENS | 0 | 1067 | 4kyo | A | | 9 | THR | p.Thr9Asn | P21549 |
| HOMO SAPIENS | 0 | 1067 | 4kyo | A | | 11 | PRO | p.Pro11Leu | P21549 |
| HOMO SAPIENS | 0 | 1067 | 4kyo | A | | 22 | ASN | p.Asn22Ser | P21549 |
| HOMO SAPIENS | 4 | 1067 | 4kyo | A | | 25 | LEU | p.Leu25Arg | P21549 |
| HOMO SAPIENS | 7 | 1067 | 4kyo | A | | 36 | ARG | p.Arg36His | P21549 |
| HOMO SAPIENS | 3 | 1067 | 4kyo | A | | 41 | GLY | p.Gly41Arg | P21549 |
| HOMO SAPIENS | 0 | 1067 | 4kyo | A | | 42 | GLY | p.Gly42Glu | P21549 |
| HOMO SAPIENS | 3 | 1067 | 4kyo | A | | 56 | ILE | p.Ile56Asn | P21549 |
| HOMO SAPIENS | 3 | 1067 | 4kyo | A | | 59 | GLU | p.Glu59Lys | P21549 |
| HOMO SAPIENS | 3 | 1067 | 4kyo | A | | 66 | TYR | p.Tyr66Ter | P21549 |
| HOMO SAPIENS | 3 | 1067 | 4kyo | A | | 111 | ARG | p.Arg111Leu | P21549 |
| HOMO SAPIENS | 7 | 1067 | 4kyo | A | | 118 | ARG | p.Arg118His | P21549 |
| HOMO SAPIENS | 3 | 1067 | 4kyo | A | | 122 | ARG | p.Arg122Ter | P21549 |
| HOMO SAPIENS | 3 | 1067 | 4kyo | A | | 124 | HIS | p.His124Pro | P21549 |
| HOMO SAPIENS | 0 | 1067 | 4kyo | A | | 152 | PHE | p.Phe152Ile | P21549 |
| HOMO SAPIENS | 1 | 1067 | 4kyo | A | | 156 | GLY | p.Gly156Arg | P21549 |
| HOMO SAPIENS | 0 | 1067 | 4kyo | A | | 161 | GLY | p.Gly161Arg | P21549 |
| HOMO SAPIENS | 3 | 1067 | 4kyo | A | | 166 | LEU | p.Leu166Pro | P21549 |
| HOMO SAPIENS | 5 | 1067 | 4kyo | A | | 170 | GLY | p.Gly170Arg | P21549 |
| HOMO SAPIENS | 5 | 1067 | 4kyo | A | | 186 | ALA | p.Ala186Val | P21549 |
| HOMO SAPIENS | 3 | 1067 | 4kyo | A | | 187 | SER | p.Ser187Phe | P21549 |
| HOMO SAPIENS | 3 | 1067 | 4kyo | A | | 197 | ARG | p.Arg197Pro | P21549 |
| HOMO SAPIENS | 5 | 1067 | 4kyo | A | | 201 | ASP | p.Asp201Glu | P21549 |
| HOMO SAPIENS | 0 | 1067 | 4kyo | A | | 202 | ILE | p.Ile202Asn | P21549 |
| HOMO SAPIENS | 1 | 1067 | 4kyo | A | | 205 | SER | p.Ser205Pro | P21549 |
| HOMO SAPIENS | 1 | 1067 | 4kyo | A | | 205 | SER | p.Ser205Ter | P21549 |
| HOMO SAPIENS | 2 | 1067 | 4kyo | A | | 216 | GLY | p.Gly216Arg | P21549 |
| HOMO SAPIENS | 5 | 1067 | 4kyo | A | | 233 | ARG | p.Arg233Cys | P21549 |
| HOMO SAPIENS | 5 | 1067 | 4kyo | A | | 244 | ILE | p.Ile244Thr | P21549 |
| HOMO SAPIENS | 5 | 1067 | 4kyo | A | | 244 | ILE | p.Ile244Met | P21549 |
| HOMO SAPIENS | 4 | 1067 | 4kyo | A | | 253 | CYS | p.Cys253Arg | P21549 |
| HOMO SAPIENS | 7 | 1067 | 4kyo | A | | 274 | GLU | p.Glu274Asp | P21549 |
| HOMO SAPIENS | 7 | 1067 | 4kyo | A | | 279 | ILE | p.Ile279Thr | P21549 |
| HOMO SAPIENS | 6 | 1067 | 4kyo | A | | 295 | ALA | p.Ala295Thr | P21549 |

| | | | | | | | | |
|---|---|---|---|---|---|---|---|---|
| HOMO SAPIENS | 5 | 1067 | 4kyo | A | 340 | ILE | p.Ile340Met | P21549 |
| HOMO SAPIENS | 1 | 790 | 2ric | C | 270 | SER | p.Ser290Thr | P35247 |
| HOMO SAPIENS | 0 | 790 | 2ric | C | 289 | GLU | p.Glu309Lys | P35247 |
| HOMO SAPIENS | 1 | 212 | 5sxe | A | 545 | GLU | p.Glu545Lys | P42336 |
| HOMO SAPIENS | 3 | 212 | 5sxe | A | 546 | GLN | p.Gln546Glu | P42336 |
| HOMO SAPIENS | 2 | 212 | 5sxe | A | 1047 | HIS | p.His1047Leu | P42336 |
| HOMO SAPIENS | 7 | 152 | 2zqq | F | 197 | ARG | p.Arg197Ter | Q13825 |
| HOMO SAPIENS | 0 | 152 | 2zqq | F | 248 | ALA | p.Ala248Thr | Q13825 |
| HOMO SAPIENS | 0 | 152 | 2zqq | F | 275 | ALA | p.Ala275Val | Q13825 |
| HOMO SAPIENS | 5 | 152 | 2zqq | F | 331 | LYS | p.Lys331Ter | Q13825 |
| HOMO SAPIENS | 6 | 17801 | 2j8l | A | 283 | PHE | p.Phe301Leu | P03951 |
| HOMO SAPIENS | 3 | 17801 | 2j8l | A | 308 | ARG | p.Arg326Cys | P03951 |
| HOMO SAPIENS | 0 | 17801 | 2j8l | A | 321 | CYS | p.Cys339Phe | P03951 |
| HOMO SAPIENS | 3 | 17801 | 2j8l | A | 351 | TYR | p.Tyr369Ter | P03951 |
| HOMO SAPIENS | 7 | 1802 | 5db0 | A | 22 | LEU | p.Leu22Arg | O00255 |
| HOMO SAPIENS | 7 | 1802 | 5db0 | A | 26 | GLU | p.Glu26Lys | O00255 |
| HOMO SAPIENS | 7 | 1802 | 5db0 | A | 139 | HIS | p.His139Asp | O00255 |
| HOMO SAPIENS | 0 | 1802 | 5db0 | A | 171 | ARG | p.Arg176Gln | O00255 |
| HOMO SAPIENS | 0 | 1802 | 5db0 | A | 184 | VAL | p.Val189Glu | O00255 |
| HOMO SAPIENS | 7 | 1802 | 5db0 | A | 255 | GLU | p.Glu260Lys | O00255 |
| HOMO SAPIENS | 4 | 1802 | 5db0 | A | 418 | ASP | p.Asp423Asn | O00255 |
| HOMO SAPIENS | 0 | 1802 | 5db0 | A | 436 | TRP | p.Trp441Arg | O00255 |
| HOMO SAPIENS | -1 | 1802 | 5db0 | A | 541 | ALA | p.Thr546Ala | O00255 |
| HOMO SAPIENS | 4 | 1802 | 5db0 | A | 552 | THR | p.Thr557Ser | O00255 |
| HOMO SAPIENS | 0 | 3450 | 2yuz | A | 36 | PRO | p.Pro461Gln | Q00872 |
| HOMO SAPIENS | 1 | 3450 | 2yuz | A | 56 | GLN | p.His481Gln | Q00872 |
| HOMO SAPIENS | 2 | 3450 | 2yuz | A | 68 | ALA | p.Ala493Val | Q00872 |
| HOMO SAPIENS | 0 | 3211 | 3rqg | A | 72 | VAL | p.Val72Leu | Q9BUL8 |
| HOMO SAPIENS | 2 | 4168 | 4q4f | A | 121 | ARG | p.Arg121Ter | Q14108 |
| HOMO SAPIENS | 2 | 4168 | 4q4f | A | 121 | ARG | p.Arg121Gln | Q14108 |
| HOMO SAPIENS | 2 | 4168 | 4q4f | A | 159 | MET | p.Met159Val | Q14108 |
| HOMO SAPIENS | 4 | 4168 | 4q4f | A | 363 | HIS | p.His363Asn | Q14108 |
| HOMO SAPIENS | 2 | 4168 | 4q4f | A | 396 | VAL | p.Val396Ile | Q14108 |
| HOMO SAPIENS | 0 | 1448 | 1y97 | A | 87 | ARG | p.Arg130Ter | Q9BQ50 |
| HOMO SAPIENS | 0 | 3444 | 4iwp | A | 151 | SER | p.Ser151Phe | Q9UHD2 |
| HOMO SAPIENS | 0 | 3444 | 4iwp | A | 401 | LYS | p.Lys401Glu | Q9UHD2 |
| HOMO SAPIENS | 0 | 116 | 1zy8 | B | 194 | GLY | p.Gly229Cys | P09622 |
| HOMO SAPIENS | 0 | 116 | 1zy8 | B | 255 | ASP | p.Asp290Gly | P09622 |
| HOMO SAPIENS | 2 | 116 | 1zy8 | B | 326 | MET | p.Met361Val | P09622 |
| HOMO SAPIENS | 0 | 116 | 1zy8 | B | 340 | GLU | p.Glu375Lys | P09622 |
| HOMO SAPIENS | 2 | 116 | 1zy8 | B | 358 | ILE | p.Ile393Thr | P09622 |
| HOMO SAPIENS | 1 | 116 | 1zy8 | B | 453 | PRO | p.Pro488Leu | P09622 |
| HOMO SAPIENS | 7 | 127 | 4qo7 | B | 160 | SER | p.Ser161Arg | P00338 |
| HOMO SAPIENS | 0 | 2025 | 1wrl | A | 8 | ALA | p.Ala8Val | P63316 |
| HOMO SAPIENS | 8 | 2025 | 1wrl | A | 24 | PHE | p.Phe24Leu | P63316 |
| HOMO SAPIENS | 0 | 2025 | 1wrl | A | 84 | SER | p.Cys84Tyr | P63316 |
| HOMO SAPIENS | 8 | 3791 | 1mhd | B | 93 | ARG | p.Arg93Ter | P84022 |
| HOMO SAPIENS | 0 | 1575 | 4phc | A | 205 | GLY | p.Gly205Asp | P12081 |
| HOMO SAPIENS | 0 | 1575 | 4phc | A | 454 | TYR | p.Tyr454Ser | P12081 |
| HOMO SAPIENS | 0 | 39620 | 1ozs | B | 135 | ARG | p.Arg136Gln | P19429 |
| HOMO SAPIENS | 5 | 39620 | 1ozs | B | 140 | ARG | p.Arg141Trp | P19429 |
| HOMO SAPIENS | 5 | 39620 | 1ozs | B | 142 | THR | p.Thr143Asn | P19429 |
| HOMO SAPIENS | 5 | 39620 | 1ozs | B | 143 | LEU | p.Leu144Gln | P19429 |
| HOMO SAPIENS | 5 | 39620 | 1ozs | B | 144 | ARG | p.Arg145Gly | P19429 |

| | | | | | | | | |
|---|---|---|---|---|---|---|---|---|
| HOMO SAPIENS | 5 | 39620 | 1ozs | B | 144 | ARG | p.Arg145Gln | P19429 |
| HOMO SAPIENS | -1 | 20934 | 4nuv | D | 42 | GLY | p.Gly42Asp | Q16570 |
| HOMO SAPIENS | -1 | 40565 | 5lcw | S | 4 | VAL | p.Val4Met | O60566 |
| HOMO SAPIENS | 0 | 40565 | 5lcw | S | 40 | THR | p.Thr40Met | O60566 |
| HOMO SAPIENS | 4 | 40565 | 5lcw | S | 198 | ALA | p.Glu390Asp | O60566 |
| HOMO SAPIENS | 4 | 758 | 4oiu | A | 726 | ARG | p.Arg727Leu | P10275 |
| HOMO SAPIENS | 4 | 758 | 4oiu | A | 730 | VAL | p.Val731Met | P10275 |
| HOMO SAPIENS | 2 | 758 | 4oiu | A | 798 | GLN | p.Gln799Glu | P10275 |
| HOMO SAPIENS | 3 | 758 | 4oiu | A | 855 | ARG | p.Arg856His | P10275 |
| HOMO SAPIENS | 1 | 758 | 4oiu | A | 886 | MET | p.Met887Val | P10275 |
| HOMO SAPIENS | 0 | 3303 | 4ie0 | A | 134 | ALA | p.Ala134Thr | Q9C0B1 |
| HOMO SAPIENS | 9 | 3303 | 4ie0 | A | 316 | ARG | p.Arg316Gln | Q9C0B1 |
| HOMO SAPIENS | 1 | 41502 | 3fgh | A | 136 | PRO | p.Pro178Leu | Q00059 |
| HOMO SAPIENS | 5 | 175 | 3ceh | B | 58 | THR | p.Thr59Lys | P06737 |
| HOMO SAPIENS | 8 | 175 | 3ceh | B | 93 | ARG | p.Arg94Ter | P06737 |
| HOMO SAPIENS | 5 | 175 | 3ceh | B | 221 | VAL | p.Val222Ile | P06737 |
| HOMO SAPIENS | 0 | 175 | 3ceh | B | 338 | ASN | p.Asn339Ser | P06737 |
| HOMO SAPIENS | 1 | 175 | 3ceh | B | 372 | PHE | p.Phe373Leu | P06737 |
| HOMO SAPIENS | 9 | 175 | 3ceh | B | 376 | ASN | p.Asn377Lys | P06737 |
| HOMO SAPIENS | 8 | 175 | 3ceh | B | 398 | ARG | p.Arg399Ter | P06737 |
| HOMO SAPIENS | 5 | 175 | 3ceh | B | 424 | ARG | p.Arg425Pro | P06737 |
| HOMO SAPIENS | 9 | 175 | 3ceh | B | 455 | VAL | p.Val456Met | P06737 |
| HOMO SAPIENS | 8 | 175 | 3ceh | B | 490 | ARG | p.Arg491Cys | P06737 |
| HOMO SAPIENS | 8 | 175 | 3ceh | B | 631 | ASN | p.Asn632Ile | P06737 |
| HOMO SAPIENS | 9 | 175 | 3ceh | B | 648 | TYR | p.Tyr649Ter | P06737 |
| HOMO SAPIENS | 9 | 175 | 3ceh | B | 672 | GLU | p.Glu673Lys | P06737 |
| HOMO SAPIENS | 9 | 175 | 3ceh | B | 674 | SER | p.Ser675Leu | P06737 |
| HOMO SAPIENS | 0 | 175 | 3ceh | B | 805 | ILE | p.Ile806Leu | P06737 |
| HOMO SAPIENS | 0 | 175 | 3ceh | B | 820 | TYR | p.Tyr821His | P06737 |
| HOMO SAPIENS | 1 | 6750 | 4fdj | A | 47 | GLY | p.Gly47Arg | P34059 |
| HOMO SAPIENS | 9 | 6750 | 4fdj | A | 60 | ASP | p.Asp60Asn | P34059 |
| HOMO SAPIENS | 5 | 6750 | 4fdj | A | 67 | LEU | p.Leu67Met | P34059 |
| HOMO SAPIENS | 0 | 6750 | 4fdj | A | 69 | PHE | p.Phe69Val | P34059 |
| HOMO SAPIENS | 3 | 6750 | 4fdj | A | 94 | ARG | p.Arg94Gly | P34059 |
| HOMO SAPIENS | 0 | 6750 | 4fdj | A | 113 | ILE | p.Ile113Phe | P34059 |
| HOMO SAPIENS | 0 | 6750 | 4fdj | A | 138 | VAL | p.Val138Ala | P34059 |
| HOMO SAPIENS | 0 | 6750 | 4fdj | A | 139 | GLY | p.Gly139Ser | P34059 |
| HOMO SAPIENS | 0 | 6750 | 4fdj | A | 151 | PRO | p.Pro151Thr | P34059 |
| HOMO SAPIENS | 0 | 6750 | 4fdj | A | 155 | GLY | p.Gly155Arg | P34059 |
| HOMO SAPIENS | 0 | 6750 | 4fdj | A | 159 | TRP | p.Trp159Ter | P34059 |
| HOMO SAPIENS | 0 | 6750 | 4fdj | A | 162 | SER | p.Ser162Phe | P34059 |
| HOMO SAPIENS | 0 | 6750 | 4fdj | A | 200 | THR | p.Thr200Met | P34059 |
| HOMO SAPIENS | 0 | 6750 | 4fdj | A | 204 | ASN | p.Asn204Lys | P34059 |
| HOMO SAPIENS | 0 | 6750 | 4fdj | A | 231 | ALA | p.Ala231Gly | P34059 |
| HOMO SAPIENS | 2 | 6750 | 4fdj | A | 235 | THR | p.Thr235Lys | P34059 |
| HOMO SAPIENS | 1 | 6750 | 4fdj | A | 259 | ARG | p.Arg259Gln | P34059 |
| HOMO SAPIENS | 0 | 6750 | 4fdj | A | 287 | SER | p.Ser287Leu | P34059 |
| HOMO SAPIENS | 0 | 6750 | 4fdj | A | 291 | ALA | p.Ala291Thr | P34059 |
| HOMO SAPIENS | 3 | 6750 | 4fdj | A | 301 | GLY | p.Gly301Cys | P34059 |
| HOMO SAPIENS | 0 | 6750 | 4fdj | A | 312 | THR | p.Thr312Ser | P34059 |
| HOMO SAPIENS | 0 | 6750 | 4fdj | A | 318 | MET | p.Met318Arg | P34059 |
| HOMO SAPIENS | 0 | 6750 | 4fdj | A | 340 | GLY | p.Gly340Asp | P34059 |
| HOMO SAPIENS | 0 | 6750 | 4fdj | A | 386 | ARG | p.Arg386Cys | P34059 |
| HOMO SAPIENS | 0 | 6750 | 4fdj | A | 391 | MET | p.Met391Val | P34059 |

| | | | | | | | | |
|---|---|---|---|---|---|---|---|---|
| HOMO SAPIENS | 0 | 6750 | 4fdj | A | 392 | ALA | p.Ala392Val | P34059 |
| HOMO SAPIENS | 0 | 6750 | 4fdj | A | 393 | ALA | p.Ala393Ser | P34059 |
| HOMO SAPIENS | 0 | 6750 | 4fdj | A | 459 | ALA | p.Ala459Val | P34059 |
| HOMO SAPIENS | 0 | 6750 | 4fdj | A | 473 | GLN | p.Gln473Ter | P34059 |
| HOMO SAPIENS | 2 | 6750 | 4fdj | A | 480 | VAL | p.Val480Phe | P34059 |
| HOMO SAPIENS | 0 | 6750 | 4fdj | A | 487 | ASN | p.Asn487Ser | P34059 |
| HOMO SAPIENS | 0 | 6750 | 4fdj | A | 488 | VAL | p.Val488Met | P34059 |
| HOMO SAPIENS | 0 | 6750 | 4fdj | A | 520 | TRP | p.Trp520Ter | P34059 |
| HOMO SAPIENS | 4 | 1088 | 2p3u | A | 102 | GLU | p.Glu142Lys | P00742 |
| HOMO SAPIENS | 6 | 370 | 1nr1 | D | 73 | ASP | p.Asp126His | P00367 |
| HOMO SAPIENS | 3 | 370 | 1nr1 | D | 221 | ARG | p.Arg274Cys | P00367 |
| HOMO SAPIENS | 2 | 370 | 1nr1 | D | 265 | ARG | p.Arg318Lys | P00367 |
| HOMO SAPIENS | 1 | 370 | 1nr1 | D | 269 | ARG | p.Arg322His | P00367 |
| HOMO SAPIENS | 1 | 370 | 1nr1 | D | 296 | GLU | p.Glu349Ala | P00367 |
| HOMO SAPIENS | 1 | 370 | 1nr1 | D | 445 | SER | p.Ser498Leu | P00367 |
| HOMO SAPIENS | 1 | 370 | 1nr1 | D | 446 | GLY | p.Gly499Asp | P00367 |
| HOMO SAPIENS | 1 | 370 | 1nr1 | D | 446 | GLY | p.Gly499Ser | P00367 |
| HOMO SAPIENS | 1 | 370 | 1nr1 | D | 448 | SER | p.Ser501Pro | P00367 |
| HOMO SAPIENS | 2 | 370 | 1nr1 | D | 454 | HIS | p.His507Tyr | P00367 |
| HOMO SAPIENS | 1 | 7720 | 2he0 | A | 147 | HIS | p.His2018Gln | P46531 |
| HOMO SAPIENS | 2 | 11844 | 5l0v | A | 218 | ARG | p.Arg218Ter | Q8NBL1 |
| HOMO SAPIENS | 2 | 11844 | 5l0v | A | 279 | ARG | p.Arg279Trp | Q8NBL1 |
| HOMO SAPIENS | 9 | 1485 | 5va0 | A | 477 | ARG | p.Arg477His | P04150 |
| HOMO SAPIENS | 0 | 235 | 2fun | B | 2248 | ARG | p.Arg248Trp | Q14790 |
| HOMO SAPIENS | 4 | 235 | 2fun | B | 2285 | HIS | p.Asp285His | Q14790 |
| HOMO SAPIENS | 0 | 4822 | 5y3n | A | 307 | ARG | p.Arg307Gly | Q12931 |
| HOMO SAPIENS | 8 | 32928 | 4m6w | A | 1886 | HIS | p.His1886Tyr | Q8IYD8 |
| HOMO SAPIENS | 0 | 2711 | 4y5v | F | 22 | GLY | p.Gly46Glu | P19235 |
| HOMO SAPIENS | -1 | 2711 | 4y5v | F | 166 | ALA | p.Ala190Ser | P19235 |
| HOMO SAPIENS | 4 | 5161 | 5bpu | A | 42 | HIS | p.His42Arg | Q00604 |
| HOMO SAPIENS | 0 | 5161 | 5bpu | A | 44 | TYR | p.Tyr44Cys | Q00604 |
| HOMO SAPIENS | 0 | 5161 | 5bpu | A | 45 | VAL | p.Val45Glu | Q00604 |
| HOMO SAPIENS | 0 | 5161 | 5bpu | A | 60 | VAL | p.Val60Glu | Q00604 |
| HOMO SAPIENS | 0 | 5161 | 5bpu | A | 61 | LEU | p.Leu61Phe | Q00604 |
| HOMO SAPIENS | 7 | 5161 | 5bpu | A | 69 | CYS | p.Cys69Ser | Q00604 |
| HOMO SAPIENS | 0 | 5161 | 5bpu | A | 75 | SER | p.Ser75Cys | Q00604 |
| HOMO SAPIENS | 0 | 5161 | 5bpu | A | 90 | ARG | p.Arg90Pro | Q00604 |
| HOMO SAPIENS | 9 | 5161 | 5bpu | A | 96 | CYS | p.Cys96Tyr | Q00604 |
| HOMO SAPIENS | 9 | 5161 | 5bpu | A | 96 | CYS | p.Cys96Trp | Q00604 |
| HOMO SAPIENS | 0 | 5161 | 5bpu | A | 101 | SER | p.Ser101Phe | Q00604 |
| HOMO SAPIENS | 0 | 5161 | 5bpu | A | 105 | ALA | p.Ala105Thr | Q00604 |
| HOMO SAPIENS | 0 | 5161 | 5bpu | A | 110 | CYS | p.Cys110Gly | Q00604 |
| HOMO SAPIENS | 0 | 5161 | 5bpu | A | 121 | ARG | p.Arg121Trp | Q00604 |
| HOMO SAPIENS | 0 | 5161 | 5bpu | A | 121 | ARG | p.Arg121Leu | Q00604 |
| HOMO SAPIENS | 0 | 5161 | 5bpu | A | 124 | LEU | p.Leu124Phe | Q00604 |
| HOMO SAPIENS | 8 | 5161 | 5bpu | A | 128 | CYS | p.Cys128Ter | Q00604 |
| HOMO SAPIENS | 1 | 13944 | 5xqh | A | 238 | ARG | p.Arg238His | Q9GZN7 |
| HOMO SAPIENS | -1 | 7508 | 5jl6 | A | 39 | TRP | p.Trp39Arg | P11511 |
| HOMO SAPIENS | 0 | 7508 | 5jl6 | A | 201 | THR | p.Thr201Met | P11511 |
| HOMO SAPIENS | 0 | 7508 | 5jl6 | A | 210 | GLU | p.Glu210Lys | P11511 |
| HOMO SAPIENS | 1 | 7508 | 5jl6 | A | 264 | ARG | p.Arg264Cys | P11511 |
| HOMO SAPIENS | 2 | 7508 | 5jl6 | A | 365 | ARG | p.Arg365Gln | P11511 |
| HOMO SAPIENS | 7 | 7508 | 5jl6 | A | 375 | ARG | p.Arg375Cys | P11511 |
| HOMO SAPIENS | 8 | 7508 | 5jl6 | A | 435 | ARG | p.Arg435Cys | P11511 |

| | | | | | | | | |
|---|---|---|---|---|---|---|---|---|
| HOMO SAPIENS | 9 | 7508 | 5jl6 | A | 437 | CYS | p.Cys437Tyr | P11511 |
| HOMO SAPIENS | -1 | 30206 | 5l95 | B | 246 | ARG | p.Arg246Ter | Q9GZZ9 |
| HOMO SAPIENS | 4 | 875 | 3efh | B | 43 | GLU | p.Glu43Asp | P60891 |
| HOMO SAPIENS | 5 | 875 | 3efh | B | 65 | ASP | p.Asp65Asn | P60891 |
| HOMO SAPIENS | 0 | 875 | 3efh | B | 87 | ALA | p.Ala87Thr | P60891 |
| HOMO SAPIENS | 2 | 875 | 3efh | B | 115 | MET | p.Met115Thr | P60891 |
| HOMO SAPIENS | 8 | 875 | 3efh | B | 133 | GLN | p.Gln133Pro | P60891 |
| HOMO SAPIENS | 8 | 875 | 3efh | B | 152 | LEU | p.Leu152Pro | P60891 |
| HOMO SAPIENS | 7 | 875 | 3efh | B | 290 | ILE | p.Ile290Thr | P60891 |
| HOMO SAPIENS | 6 | 875 | 3efh | B | 306 | GLY | p.Gly306Arg | P60891 |
| HOMO SAPIENS | 0 | 7978 | 5edm | A | 122 | MET | p.Thr165Met | P00734 |
| HOMO SAPIENS | -1 | 7978 | 5edm | A | 257 | ARG | p.Arg314Cys | P00734 |
| HOMO SAPIENS | 0 | 1716 | 1b98 | M | 126 | ARG | p.Arg206Gln | P34130 |
| HOMO SAPIENS | 0 | 1716 | 1b98 | M | 126 | ARG | p.Arg206Trp | P34130 |
| HOMO SAPIENS | 9 | 3960 | 2qyp | B | 28 | ILE | p.Ile338Val | P07602 |
| HOMO SAPIENS | 1 | 3960 | 2qyp | B | 39 | LEU | p.Leu349Pro | P07602 |
| HOMO SAPIENS | 0 | 31414 | 5u1d | A | 333 | ILE | p.Ile393Val | Q03518 |
| HOMO SAPIENS | 0 | 31414 | 5u1d | A | 637 | ASP | p.Asp697Gly | Q03518 |
| HOMO SAPIENS | 0 | 31414 | 5u1d | A | 659 | ARG | p.Arg719Gln | Q03518 |
| HOMO SAPIENS | 3 | 119 | 2pho | A | 11 | ILE | p.Ile11Thr | P05089 |
| HOMO SAPIENS | 8 | 119 | 2pho | A | 21 | ARG | p.Arg21Ter | P05089 |
| HOMO SAPIENS | 0 | 119 | 2pho | A | 122 | TRP | p.Trp122Ter | P05089 |
| HOMO SAPIENS | 6 | 119 | 2pho | A | 138 | GLY | p.Gly138Val | P05089 |
| HOMO SAPIENS | 5 | 119 | 2pho | A | 290 | THR | p.Thr290Ser | P05089 |
| HOMO SAPIENS | 5 | 119 | 2pho | A | 291 | ARG | p.Arg291Ter | P05089 |
| HOMO SAPIENS | 0 | 3732 | 2i99 | A | 247 | TYR | p.Tyr247Ter | Q14894 |
| HOMO SAPIENS | -1 | 1382 | 4ffx | A | 2 | ALA | p.Ala2Val | P30566 |
| HOMO SAPIENS | 1 | 1382 | 4ffx | A | 72 | ILE | p.Ile72Met | P30566 |
| HOMO SAPIENS | 1 | 1382 | 4ffx | A | 100 | PRO | p.Pro100Ala | P30566 |
| HOMO SAPIENS | 0 | 1382 | 4ffx | A | 114 | TYR | p.Tyr114His | P30566 |
| HOMO SAPIENS | 1 | 1382 | 4ffx | A | 141 | ARG | p.Arg141Trp | P30566 |
| HOMO SAPIENS | 0 | 1382 | 4ffx | A | 147 | LYS | p.Lys147Met | P30566 |
| HOMO SAPIENS | 3 | 1382 | 4ffx | A | 190 | ARG | p.Arg190Gln | P30566 |
| HOMO SAPIENS | 3 | 1382 | 4ffx | A | 190 | ARG | p.Arg190Ter | P30566 |
| HOMO SAPIENS | 0 | 1382 | 4ffx | A | 217 | HIS | p.His217Asp | P30566 |
| HOMO SAPIENS | 2 | 1382 | 4ffx | A | 246 | LYS | p.Lys246Glu | P30566 |
| HOMO SAPIENS | 5 | 1382 | 4ffx | A | 303 | ARG | p.Arg303Cys | P30566 |
| HOMO SAPIENS | 1 | 1382 | 4ffx | A | 318 | PRO | p.Pro318Leu | P30566 |
| HOMO SAPIENS | 6 | 1382 | 4ffx | A | 332 | ASP | p.Asp332His | P30566 |
| HOMO SAPIENS | 6 | 1382 | 4ffx | A | 337 | ARG | p.Arg337Ter | P30566 |
| HOMO SAPIENS | 1 | 1382 | 4ffx | A | 396 | ARG | p.Arg396His | P30566 |
| HOMO SAPIENS | 1 | 1382 | 4ffx | A | 426 | ARG | p.Arg426His | P30566 |
| HOMO SAPIENS | 0 | 1382 | 4ffx | A | 438 | SER | p.Ser438Pro | P30566 |
| HOMO SAPIENS | 1 | 1382 | 4ffx | A | 447 | SER | p.Ser447Pro | P30566 |
| HOMO SAPIENS | 2 | 1382 | 4ffx | A | 448 | SER | p.Ser448Pro | P30566 |
| HOMO SAPIENS | 2 | 1382 | 4ffx | A | 450 | THR | p.Thr450Ser | P30566 |
| HOMO SAPIENS | 0 | 6212 | 1ao3 | A | 46 | SER | p.Ser1731Thr | P04275 |
| HOMO SAPIENS | 0 | 1711 | 3f9c | A | 279 | VAL | p.Val279Phe | Q13093 |
| HOMO SAPIENS | 0 | 6112 | 4fvl | A | 312 | HIS | p.His312Arg | P45452 |
| HOMO SAPIENS | 0 | 6112 | 4fvl | A | 323 | THR | p.Thr323Met | P45452 |
| HOMO SAPIENS | 0 | 6112 | 4fvl | A | 390 | ASP | p.Asp390Gly | P45452 |
| HOMO SAPIENS | 0 | 6112 | 4fvl | A | 458 | ARG | p.Arg458Cys | P45452 |
| HOMO SAPIENS | 2 | 6180 | 5ggn | A | 97 | ARG | p.Arg97Trp | Q8WZA1 |
| HOMO SAPIENS | 2 | 6180 | 5ggn | A | 215 | ARG | p.Arg215Ter | Q8WZA1 |

| | | | | | | | | |
|---|---|---|---|---|---|---|---|---|
| HOMO SAPIENS | -1 | 5526 | 4ak3 | A | 11 | ASP | p.Asp1232Asn | P02461 |
| HOMO SAPIENS | 6 | 5526 | 4ak3 | A | 38 | ALA | p.Ala1259Thr | P02461 |
| HOMO SAPIENS | 0 | 5526 | 4ak3 | A | 92 | LYS | p.Lys1313Arg | P02461 |
| HOMO SAPIENS | 0 | 5526 | 4ak3 | A | 132 | GLN | p.His1353Gln | P02461 |
| HOMO SAPIENS | 4 | 16616 | 1hci | B | 433 | SER | p.Ser433Leu | P35609 |
| HOMO SAPIENS | 1 | 16616 | 1hci | B | 475 | ASP | p.Asp475Asn | P35609 |
| HOMO SAPIENS | 1 | 16616 | 1hci | B | 476 | ALA | p.Ala476Ser | P35609 |
| HOMO SAPIENS | 2 | 16616 | 1hci | B | 507 | MET | p.Met507Val | P35609 |
| HOMO SAPIENS | 1 | 16616 | 1hci | B | 604 | MET | p.Met604Val | P35609 |
| HOMO SAPIENS | 2 | 16616 | 1hci | B | 640 | ARG | p.Arg640His | P35609 |
| HOMO SAPIENS | 1 | 16616 | 1hci | B | 721 | ARG | p.Arg721Cys | P35609 |
| HOMO SAPIENS | -1 | 3402 | 5iaa | B | 57 | MET | p.Met57Val | Q9GZZ9 |
| HOMO SAPIENS | 7 | 3250 | 3irq | B | 193 | PRO | p.Pro193Ala | P55265 |
| HOMO SAPIENS | 2 | 15200 | 2wns | A | 30 | SER | p.Ser30Gly | P11172 |
| HOMO SAPIENS | 0 | 15200 | 2wns | A | 96 | ARG | p.Arg96Gly | P11172 |
| HOMO SAPIENS | 0 | 15200 | 2wns | A | 109 | VAL | p.Val109Gly | P11172 |
| HOMO SAPIENS | 2 | 15759 | 4x4w | B | 140 | LEU | p.Leu166Ser | Q96Q11 |
| HOMO SAPIENS | 0 | 15759 | 4x4w | B | 197 | ILE | p.Ile223Thr | Q96Q11 |
| HOMO SAPIENS | 0 | 3156 | 3w03 | B | 14 | ALA | p.Ala14Thr | Q9H9Q4 |
| HOMO SAPIENS | 0 | 3156 | 3w03 | B | 57 | ARG | p.Arg57Gly | Q9H9Q4 |
| HOMO SAPIENS | 0 | 3156 | 3w03 | B | 123 | CYS | p.Cys123Arg | Q9H9Q4 |
| HOMO SAPIENS | 0 | 11693 | 3ojy | A | 63 | GLN | p.Gln93Lys | P07357 |
| HOMO SAPIENS | 0 | 1421 | 2qc8 | E | 275 | GLU | p.Glu275Asp | P15104 |
| HOMO SAPIENS | 9 | 1421 | 2qc8 | E | 324 | ARG | p.Arg324Cys | P15104 |
| HOMO SAPIENS | 6 | 1421 | 2qc8 | E | 341 | ARG | p.Arg341Cys | P15104 |
| HOMO SAPIENS | 6 | 2850 | 3qvb | A | 83 | THR | p.Thr83Met | P46976 |
| HOMO SAPIENS | 8 | 2850 | 3qvb | A | 102 | ASP | p.Asp102His | P46976 |
| HOMO SAPIENS | 1 | 24314 | 4m9p | A | 480 | PRO | p.Pro480Leu | P21333 |
| HOMO SAPIENS | 0 | 24314 | 4m9p | A | 528 | VAL | p.Val528Met | P21333 |
| HOMO SAPIENS | 0 | 24314 | 4m9p | A | 590 | VAL | p.Val590Ile | P21333 |
| HOMO SAPIENS | 0 | 24314 | 4m9p | A | 605 | ASP | p.Asp605Asn | P21333 |
| HOMO SAPIENS | 1 | 24314 | 4m9p | A | 637 | PRO | p.Pro637Gln | P21333 |
| HOMO SAPIENS | 0 | 24314 | 4m9p | A | 656 | LEU | p.Leu656Phe | P21333 |
| HOMO SAPIENS | 2 | 24314 | 4m9p | A | 711 | VAL | p.Val711Asp | P21333 |
| HOMO SAPIENS | 0 | 4951 | 5joe | A | 40 | THR | p.Thr9621Ser | Q8WZ42 |
| HOMO SAPIENS | 1 | 4951 | 5joe | A | 41 | GLU | p.Glu9622Gln | Q8WZ42 |
| HOMO SAPIENS | 0 | 4951 | 5joe | A | 82 | ALA | p.Ala9663Ser | Q8WZ42 |
| HOMO SAPIENS | 4 | 4951 | 5joe | A | 89 | ILE | p.Ile9670Asn | Q8WZ42 |
| HOMO SAPIENS | 0 | 9664 | 3c6m | A | 56 | GLY | p.Gly56Ser | P52788 |
| HOMO SAPIENS | 0 | 9664 | 3c6m | A | 132 | VAL | p.Val132Gly | P52788 |
| HOMO SAPIENS | 1 | 9664 | 3c6m | A | 190 | THR | p.Thr190Ile | P52788 |
| HOMO SAPIENS | -1 | 12118 | 5hcd | B | 19 | GLN | p.Gln19Ter | P01031 |
| HOMO SAPIENS | 0 | 1583 | 2b4x | L | 7 | ILE | p.Ile39Asn | P01008 |
| HOMO SAPIENS | 0 | 1583 | 2b4x | L | 24 | ARG | p.Arg56Cys | P01008 |
| HOMO SAPIENS | 0 | 1583 | 2b4x | L | 47 | ARG | p.Arg79Cys | P01008 |
| HOMO SAPIENS | 0 | 1583 | 2b4x | L | 47 | ARG | p.Arg79His | P01008 |
| HOMO SAPIENS | 1 | 1583 | 2b4x | L | 95 | CYS | p.Cys127Arg | P01008 |
| HOMO SAPIENS | 1 | 1583 | 2b4x | L | 99 | LEU | p.Leu131Phe | P01008 |
| HOMO SAPIENS | 0 | 1583 | 2b4x | L | 116 | SER | p.Ser148Pro | P01008 |
| HOMO SAPIENS | 0 | 1583 | 2b4x | L | 129 | ARG | p.Arg161Gln | P01008 |
| HOMO SAPIENS | 0 | 1583 | 2b4x | L | 135 | ASN | p.Asn167Thr | P01008 |
| HOMO SAPIENS | 2 | 1583 | 2b4x | L | 187 | ASN | p.Asn219Asp | P01008 |
| HOMO SAPIENS | 0 | 1583 | 2b4x | L | 191 | SER | p.Ser223Pro | P01008 |
| HOMO SAPIENS | 2 | 1583 | 2b4x | L | 349 | SER | p.Ser381Pro | P01008 |

| | | | | | | | | |
|---|---|---|---|---|---|---|---|---|
| HOMO SAPIENS | 0 | 1583 | 2b4x | L | 382 | ALA | p.Ala414Thr | P01008 |
| HOMO SAPIENS | 0 | 1583 | 2b4x | L | 387 | ALA | p.Ala419Val | P01008 |
| HOMO SAPIENS | 0 | 1583 | 2b4x | L | 392 | GLY | p.Gly424Asp | P01008 |
| HOMO SAPIENS | 0 | 1583 | 2b4x | L | 393 | ARG | p.Arg425His | P01008 |
| HOMO SAPIENS | 0 | 1583 | 2b4x | L | 393 | ARG | p.Arg425Cys | P01008 |
| HOMO SAPIENS | 0 | 1583 | 2b4x | L | 394 | SER | p.Ser426Leu | P01008 |
| HOMO SAPIENS | 0 | 1583 | 2b4x | L | 404 | ALA | p.Ala436Thr | P01008 |
| HOMO SAPIENS | 2 | 1583 | 2b4x | L | 407 | PRO | p.Pro439Leu | P01008 |
| HOMO SAPIENS | 2 | 1583 | 2b4x | L | 429 | PRO | p.Pro461Leu | P01008 |
| HOMO SAPIENS | 0 | 11647 | 2a1t | S | 78 | ALA | p.Ala78Ser | P38117 |
| HOMO SAPIENS | 0 | 11647 | 2a1t | S | 98 | ARG | p.Arg98Cys | P38117 |
| HOMO SAPIENS | 8 | 11647 | 2a1t | S | 128 | ASP | p.Asp128Asn | P38117 |
| HOMO SAPIENS | 5 | 11647 | 2a1t | S | 154 | THR | p.Thr154Met | P38117 |
| HOMO SAPIENS | 6 | 11647 | 2a1t | S | 164 | ARG | p.Arg164Gln | P38117 |
| HOMO SAPIENS | 5 | 11647 | 2a1t | S | 174 | ARG | p.Arg174His | P38117 |
| HOMO SAPIENS | -1 | 11200 | 4tx5 | A | 4 | ILE | p.Ile59Val | Q9NR28 |
| HOMO SAPIENS | 0 | 11200 | 4tx5 | A | 169 | GLY | p.Gly224Arg | Q9NR28 |
| HOMO SAPIENS | 3 | 417 | 2vr2 | A | 6 | ARG | p.Arg6Gln | Q14117 |
| HOMO SAPIENS | 1 | 417 | 2vr2 | A | 334 | GLN | p.Gln334Arg | Q14117 |
| HOMO SAPIENS | 2 | 417 | 2vr2 | A | 360 | TRP | p.Trp360Arg | Q14117 |
| HOMO SAPIENS | 0 | 417 | 2vr2 | A | 379 | SER | p.Ser379Arg | Q14117 |
| HOMO SAPIENS | 0 | 417 | 2vr2 | A | 435 | GLY | p.Gly435Arg | Q14117 |
| HOMO SAPIENS | 0 | 12498 | 3eo3 | B | 423 | ILE | p.Ile423Val | Q9Y223 |
| HOMO SAPIENS | 5 | 12498 | 3eo3 | B | 572 | VAL | p.Val572Leu | Q9Y223 |
| HOMO SAPIENS | 0 | 12498 | 3eo3 | B | 576 | GLY | p.Gly576Glu | Q9Y223 |
| HOMO SAPIENS | 0 | 12498 | 3eo3 | B | 587 | ILE | p.Ile587Thr | Q9Y223 |
| HOMO SAPIENS | 0 | 12498 | 3eo3 | B | 615 | SER | p.Ser615Ter | Q9Y223 |
| HOMO SAPIENS | 1 | 12498 | 3eo3 | B | 631 | ALA | p.Ala631Val | Q9Y223 |
| HOMO SAPIENS | 1 | 12498 | 3eo3 | B | 669 | GLY | p.Gly669Arg | Q9Y223 |
| HOMO SAPIENS | 1 | 12498 | 3eo3 | B | 696 | VAL | p.Val696Met | Q9Y223 |
| HOMO SAPIENS | 2 | 12498 | 3eo3 | B | 712 | MET | p.Met712Thr | Q9Y223 |
| HOMO SAPIENS | 0 | 1568 | 1r74 | A | 49 | LEU | p.Leu50Pro | Q14749 |
| HOMO SAPIENS | 0 | 1568 | 1r74 | A | 176 | HIS | p.His177Asn | Q14749 |
| HOMO SAPIENS | 3 | 147 | 5eig | A | 67 | THR | p.Thr67Ile | P32929 |
| HOMO SAPIENS | 3 | 147 | 5eig | A | 240 | GLN | p.Gln240Glu | P32929 |
| HOMO SAPIENS | -1 | 147 | 5eig | A | 403 | SER | p.Ser403Ile | P32929 |
| HOMO SAPIENS | 8 | 20036 | 2lsr | A | 31 | ARG | p.Arg31Ter | Q9Y6N9 |
| HOMO SAPIENS | -1 | 41382 | 4uis | B | 431 | UNK | p.Ala431Glu | P49768 |
| HOMO SAPIENS | -1 | 8536 | 3i6w | B | 85 | PRO | p.Pro85Leu | O96017 |
| HOMO SAPIENS | 0 | 8536 | 3i6w | B | 93 | TRP | p.Trp93Ter | O96017 |
| HOMO SAPIENS | 0 | 8536 | 3i6w | B | 95 | ARG | p.Arg95Ter | O96017 |
| HOMO SAPIENS | 0 | 8536 | 3i6w | B | 117 | ARG | p.Arg117Gly | O96017 |
| HOMO SAPIENS | 0 | 8536 | 3i6w | B | 119 | LYS | p.Lys119Arg | O96017 |
| HOMO SAPIENS | 0 | 8536 | 3i6w | B | 137 | ARG | p.Arg137Gln | O96017 |
| HOMO SAPIENS | 0 | 8536 | 3i6w | B | 137 | ARG | p.Arg137Ter | O96017 |
| HOMO SAPIENS | 0 | 8536 | 3i6w | B | 145 | ARG | p.Arg145Trp | O96017 |
| HOMO SAPIENS | 0 | 8536 | 3i6w | B | 160 | ILE | p.Ile160Met | O96017 |
| HOMO SAPIENS | 0 | 41322 | 4bxl | C | 46 | GLU | p.Glu46Lys | P37840 |
| HOMO SAPIENS | 5 | 41322 | 4bxl | C | 53 | ALA | p.Ala53Thr | P37840 |
| HOMO SAPIENS | 2 | 24966 | 3hve | A | 15 | ARG | p.Arg15Ser | Q9H2C0 |
| HOMO SAPIENS | 1 | 24966 | 3hve | A | 138 | ARG | p.Arg138His | Q9H2C0 |
| HOMO SAPIENS | 0 | 24966 | 3hve | A | 201 | ARG | p.Arg201Ter | Q9H2C0 |
| HOMO SAPIENS | 0 | 11680 | 5an9 | J | 8 | ASN | p.Asn8Lys | Q9Y3A5 |
| HOMO SAPIENS | 1 | 11680 | 5an9 | J | 33 | LYS | p.Lys33Arg | Q9Y3A5 |

| | | | | | | | | |
|---|---|---|---|---|---|---|---|---|
| HOMO SAPIENS | 1 | 11680 | 5an9 | J | 126 | ARG | p.Arg126Thr | Q9Y3A5 |
| HOMO SAPIENS | 1 | 11680 | 5an9 | J | 130 | VAL | p.Val130Met | Q9Y3A5 |
| HOMO SAPIENS | 1 | 11680 | 5an9 | J | 169 | ARG | p.Arg169Cys | Q9Y3A5 |
| HOMO SAPIENS | 1 | 11680 | 5an9 | J | 218 | ARG | p.Arg218Ter | Q9Y3A5 |
| HOMO SAPIENS | 4 | 978 | 3zs0 | B | 7 | TYR | p.Tyr173Cys | P05164 |
| HOMO SAPIENS | 1 | 978 | 3zs0 | B | 85 | MET | p.Met251Thr | P05164 |
| HOMO SAPIENS | 4 | 5140 | 2l4r | A | 52 | CYS | p.Cys52Ter | Q12809 |
| HOMO SAPIENS | 4 | 5140 | 2l4r | A | 63 | PRO | p.Pro63His | Q12809 |
| HOMO SAPIENS | 0 | 5140 | 2l4r | A | 100 | ARG | p.Arg100Gly | Q12809 |
| HOMO SAPIENS | 4 | 5140 | 2l4r | A | 129 | PHE | p.Phe129Leu | Q12809 |
| HOMO SAPIENS | 0 | 25836 | 2gy7 | B | 103 | GLU | p.Glu103Asp | Q02763 |
| HOMO SAPIENS | 0 | 25836 | 2gy7 | B | 148 | ILE | p.Ile148Thr | Q02763 |
| HOMO SAPIENS | 0 | 25836 | 2gy7 | B | 226 | ALA | p.Ala226Val | Q02763 |
| HOMO SAPIENS | 0 | 25836 | 2gy7 | B | 391 | THR | p.Thr391Ile | Q02763 |
| HOMO SAPIENS | 4 | 12563 | 5kwy | D | 20 | GLU | p.Glu20Ter | P61916 |
| HOMO SAPIENS | 2 | 12563 | 5kwy | D | 30 | VAL | p.Val30Met | P61916 |
| HOMO SAPIENS | 4 | 12563 | 5kwy | D | 39 | VAL | p.Val39Met | P61916 |
| HOMO SAPIENS | 8 | 12563 | 5kwy | D | 67 | SER | p.Ser67Pro | P61916 |
| HOMO SAPIENS | 8 | 12563 | 5kwy | D | 99 | CYS | p.Cys99Arg | P61916 |
| HOMO SAPIENS | 2 | 12563 | 5kwy | D | 118 | GLU | p.Glu118Ter | P61916 |
| HOMO SAPIENS | 2 | 12563 | 5kwy | D | 120 | PRO | p.Pro120Ser | P61916 |
| HOMO SAPIENS | -1 | 1679 | 1pmk | A | -2 | VAL | p.Val374Phe | P00747 |
| HOMO SAPIENS | -1 | 5315 | 2w2n | P | 53 | ALA | p.Ala53Val | Q8NBP7 |
| HOMO SAPIENS | 8 | 13496 | 2yad | A | 138 | THR | p.Asn138Thr | P11686 |
| HOMO SAPIENS | -1 | 13496 | 2yad | A | 173 | ARG | p.Arg167Gln | P11686 |
| HOMO SAPIENS | 2 | 13496 | 2yad | A | 186 | SER | p.Asn186Ser | P11686 |
| HOMO SAPIENS | 0 | 4426 | 3qd6 | C | 123 | ALA | p.Ala123Glu | P29965 |
| HOMO SAPIENS | 1 | 4426 | 3qd6 | C | 140 | TRP | p.Trp140Gly | P29965 |
| HOMO SAPIENS | 0 | 4426 | 3qd6 | C | 155 | LEU | p.Leu155Pro | P29965 |
| HOMO SAPIENS | 6 | 4426 | 3qd6 | C | 227 | GLY | p.Gly227Val | P29965 |
| HOMO SAPIENS | 0 | 4426 | 3qd6 | C | 235 | ALA | p.Ala235Pro | P29965 |
| HOMO SAPIENS | 0 | 10734 | 3dyd | B | 45 | PRO | p.Pro45Ser | P17735 |
| HOMO SAPIENS | -1 | 10734 | 3dyd | B | 57 | ARG | p.Arg57Ter | P17735 |
| HOMO SAPIENS | 8 | 10734 | 3dyd | B | 417 | ARG | p.Arg417Ter | P17735 |
| HOMO SAPIENS | 0 | 6956 | 4pzs | A | 1600 | LEU | p.Leu1600Phe | Q09472 |
| HOMO SAPIENS | 0 | 9370 | 3hfv | A | 21 | SER | p.Ser57Cys | O95363 |
| HOMO SAPIENS | 1 | 9370 | 3hfv | A | 25 | ASP | p.Asp61Glu | O95363 |
| HOMO SAPIENS | 1 | 9370 | 3hfv | A | 130 | ALA | p.Ala166Thr | O95363 |
| HOMO SAPIENS | 5 | 9370 | 3hfv | A | 148 | ASP | p.Asp184Asn | O95363 |
| HOMO SAPIENS | 0 | 9370 | 3hfv | A | 187 | ARG | p.Arg223Cys | O95363 |
| HOMO SAPIENS | 0 | 9370 | 3hfv | A | 210 | THR | p.Thr246Met | O95363 |
| HOMO SAPIENS | 1 | 9370 | 3hfv | A | 244 | ASN | p.Asn280Ser | O95363 |
| HOMO SAPIENS | 2 | 9370 | 3hfv | A | 288 | TYR | p.Tyr324Cys | O95363 |
| HOMO SAPIENS | 5 | 9370 | 3hfv | A | 293 | ILE | p.Ile329Thr | O95363 |
| HOMO SAPIENS | 1 | 9370 | 3hfv | A | 355 | ASP | p.Asp391Val | O95363 |
| HOMO SAPIENS | 1 | 9370 | 3hfv | A | 371 | THR | p.Thr407Met | O95363 |
| HOMO SAPIENS | 1 | 9370 | 3hfv | A | 383 | ARG | p.Arg419Cys | O95363 |
| HOMO SAPIENS | 1 | 9370 | 3hfv | A | 387 | ARG | p.Arg423Gln | O95363 |
| HOMO SAPIENS | 0 | 14401 | 1q3x | A | 371 | ASP | p.Asp371Tyr | O00187 |
| HOMO SAPIENS | 0 | 14401 | 1q3x | A | 377 | VAL | p.Val377Ala | O00187 |
| HOMO SAPIENS | 0 | 14401 | 1q3x | A | 439 | ARG | p.Arg439His | O00187 |
| HOMO SAPIENS | 0 | 14401 | 1q3x | A | 577 | GLN | p.Gln577His | O00187 |
| HOMO SAPIENS | 2 | 18270 | 3teq | A | 429 | ARG | p.Arg429Cys | Q13586 |
| HOMO SAPIENS | -1 | 16081 | 4mqt | A | 231 | VAL | p.Val231Ile | P08172 |

| | | | | | | | | |
|---|---|---|---|---|---|---|---|---|
| HOMO SAPIENS | 9 | 6607 | 3whu | B | 81 | ASP | p.Asp81His | Q8NI22 |
| HOMO SAPIENS | 9 | 6607 | 3whu | B | 129 | ASP | p.Asp129Glu | Q8NI22 |
| HOMO SAPIENS | 9 | 6607 | 3whu | B | 136 | ILE | p.Ile136Thr | Q8NI22 |
| HOMO SAPIENS | 0 | 7208 | 2e9w | D | 11 | ASN | p.Asn36Ser | P21583 |
| HOMO SAPIENS | 0 | 24566 | 4bdx | A | 10 | LEU | p.Leu140Val | P00748 |
| HOMO SAPIENS | 5 | 24566 | 4bdx | A | 77 | PRO | p.Ala207Pro | P00748 |
| HOMO SAPIENS | 7 | 606 | 3qnf | A | 319 | MET | p.Met319Val | Q9NZ08 |
| HOMO SAPIENS | 8 | 4097 | 1ydl | A | 21 | LEU | p.Leu21Pro | Q6ZYL4 |
| HOMO SAPIENS | 6 | 4097 | 1ydl | A | 56 | ARG | p.Arg56Ter | Q6ZYL4 |
| HOMO SAPIENS | 8 | 36691 | 1dqb | A | 42 | ARG | p.Arg403Lys | P07204 |
| HOMO SAPIENS | 0 | 5642 | 3j7y | X | 59 | ARG | p.Arg59Pro | Q13084 |
| HOMO SAPIENS | 1 | 5642 | 3j7y | X | 204 | VAL | p.Val204Met | Q13084 |
| HOMO SAPIENS | 1 | 1119 | 3nwu | B | 252 | ALA | p.Ala252Thr | Q92743 |
| HOMO SAPIENS | 3 | 1119 | 3nwu | B | 274 | ARG | p.Arg274Gln | Q92743 |
| HOMO SAPIENS | 9 | 1119 | 3nwu | B | 297 | VAL | p.Val297Met | Q92743 |
| HOMO SAPIENS | -1 | 1119 | 3nwu | B | 302 | ARG | p.Arg302Ter | Q92743 |
| HOMO SAPIENS | 9 | 1119 | 3nwu | B | 321 | ALA | p.Ala321Thr | Q92743 |
| HOMO SAPIENS | -1 | 1119 | 3nwu | B | 370 | ARG | p.Arg370Ter | Q92743 |
| HOMO SAPIENS | 0 | 416 | 4ycu | B | 105 | LEU | p.Leu105His | Q15046 |
| HOMO SAPIENS | 0 | 416 | 4ycu | B | 145 | TYR | p.Tyr145His | Q15046 |
| HOMO SAPIENS | 0 | 416 | 4ycu | B | 200 | PRO | p.Pro200Leu | Q15046 |
| HOMO SAPIENS | 0 | 416 | 4ycu | B | 201 | TYR | p.Tyr201His | Q15046 |
| HOMO SAPIENS | 0 | 416 | 4ycu | B | 420 | ARG | p.Arg420Cys | Q15046 |
| HOMO SAPIENS | 9 | 3297 | 4fpb | A | 680 | THR | p.Thr680Ser | P19367 |
| HOMO SAPIENS | 0 | 6771 | 2zt6 | A | 201 | ALA | p.Ala255Val | P41250 |
| HOMO SAPIENS | 0 | 6771 | 2zt6 | A | 202 | ASP | p.Asp256His | P41250 |
| HOMO SAPIENS | 0 | 6771 | 2zt6 | A | 334 | ARG | p.Arg388Gln | P41250 |
| HOMO SAPIENS | -1 | 6771 | 2zt6 | A | 439 | ASN | p.Asn493Ser | P41250 |
| HOMO SAPIENS | -1 | 6771 | 2zt6 | A | 500 | ASP | p.Asp554Asn | P41250 |
| HOMO SAPIENS | 1 | 6771 | 2zt6 | A | 581 | SER | p.Ser635Leu | P41250 |
| HOMO SAPIENS | -1 | 6771 | 2zt6 | A | 684 | GLU | p.Glu738Lys | P41250 |
| HOMO SAPIENS | 0 | 2687 | 4grw | B | 276 | VAL | p.Val298Phe | P29460 |
| HOMO SAPIENS | 6 | 913 | 1m75 | B | 45 | ASP | p.Asp57Glu | Q16836 |
| HOMO SAPIENS | 2 | 913 | 1m75 | B | 74 | PRO | p.Leu86Pro | Q16836 |
| HOMO SAPIENS | 3 | 913 | 1m75 | B | 214 | TYR | p.Tyr226His | Q16836 |
| HOMO SAPIENS | 3 | 913 | 1m75 | B | 224 | ARG | p.Arg236Ter | Q16836 |
| HOMO SAPIENS | 4 | 913 | 1m75 | B | 246 | PRO | p.Pro258Leu | Q16836 |
| HOMO SAPIENS | 2 | 4713 | 4djn | B | 40 | ARG | p.Arg40Cys | Q7LG56 |
| HOMO SAPIENS | 1 | 4713 | 4djn | B | 41 | ARG | p.Arg41Pro | Q7LG56 |
| HOMO SAPIENS | 2 | 4713 | 4djn | B | 110 | ARG | p.Arg110His | Q7LG56 |
| HOMO SAPIENS | 2 | 4713 | 4djn | B | 121 | ARG | p.Arg121His | Q7LG56 |
| HOMO SAPIENS | 9 | 4713 | 4djn | B | 131 | GLU | p.Glu131Lys | Q7LG56 |
| HOMO SAPIENS | 2 | 4713 | 4djn | B | 182 | THR | p.Thr182Ala | Q7LG56 |
| HOMO SAPIENS | 6 | 4713 | 4djn | B | 194 | GLU | p.Glu194Lys | Q7LG56 |
| HOMO SAPIENS | 2 | 4713 | 4djn | B | 211 | ARG | p.Arg211Lys | Q7LG56 |
| HOMO SAPIENS | 0 | 4713 | 4djn | B | 224 | ILE | p.Ile224Ser | Q7LG56 |
| HOMO SAPIENS | 3 | 4713 | 4djn | B | 229 | GLY | p.Gly229Val | Q7LG56 |
| HOMO SAPIENS | 2 | 4713 | 4djn | B | 236 | CYS | p.Cys236Phe | Q7LG56 |
| HOMO SAPIENS | 0 | 4713 | 4djn | B | 273 | GLY | p.Gly273Ser | Q7LG56 |
| HOMO SAPIENS | -1 | 4713 | 4djn | B | 282 | MSE | p.Met282Ile | Q7LG56 |
| HOMO SAPIENS | 3 | 4713 | 4djn | B | 284 | GLN | p.Gln284Ter | Q7LG56 |
| HOMO SAPIENS | 3 | 4713 | 4djn | B | 305 | ALA | p.Ala305Ser | Q7LG56 |
| HOMO SAPIENS | -1 | 8974 | 5n7i | A | 500 | ARG | p.Arg500Ter | Q8N4S9 |
| HOMO SAPIENS | 0 | 17072 | 3fxi | B | 299 | ASP | p.Asp299Gly | O00206 |

| | | | | | | | | |
|---|---|---|---|---|---|---|---|---|
| HOMO SAPIENS | 1 | 14355 | 2wwk | T | 55 | ILE | p.Ile34306Asn | Q8WZ42 |
| HOMO SAPIENS | 0 | 14355 | 2wwk | T | 64 | LEU | p.Leu34315Pro | Q8WZ42 |
| HOMO SAPIENS | 2 | 17797 | 1xu1 | S | 72 | ARG | p.Arg72His | O14836 |
| HOMO SAPIENS | -1 | 10836 | 4ztu | A | 39 | GLN | p.Gln39Arg | P54098 |
| HOMO SAPIENS | -1 | 10836 | 4ztu | A | 42 | ARG | p.Arg42Gln | P54098 |
| HOMO SAPIENS | -1 | 10836 | 4ztu | A | 45 | GLN | p.Gln45Arg | P54098 |
| HOMO SAPIENS | -1 | 10836 | 4ztu | A | 52 | GLN | p.Gln52Glu | P54098 |
| HOMO SAPIENS | 0 | 10836 | 4ztu | A | 88 | PHE | p.Phe88Leu | P54098 |
| HOMO SAPIENS | 1 | 10836 | 4ztu | A | 194 | ALA | p.Ala194Val | P54098 |
| HOMO SAPIENS | 1 | 10836 | 4ztu | A | 227 | ARG | p.Arg227Trp | P54098 |
| HOMO SAPIENS | 0 | 10836 | 4ztu | A | 232 | ARG | p.Arg232His | P54098 |
| HOMO SAPIENS | 1 | 10836 | 4ztu | A | 285 | GLN | p.Gln285His | P54098 |
| HOMO SAPIENS | 1 | 10836 | 4ztu | A | 434 | TYR | p.Tyr434His | P54098 |
| HOMO SAPIENS | -1 | 10836 | 4ztu | A | 511 | SER | p.Ser511Asn | P54098 |
| HOMO SAPIENS | 0 | 10836 | 4ztu | A | 533 | CYS | p.Cys533Trp | P54098 |
| HOMO SAPIENS | 1 | 10836 | 4ztu | A | 546 | ARG | p.Arg546Cys | P54098 |
| HOMO SAPIENS | 1 | 10836 | 4ztu | A | 587 | PRO | p.Pro587Leu | P54098 |
| HOMO SAPIENS | 0 | 10836 | 4ztu | A | 603 | MET | p.Met603Thr | P54098 |
| HOMO SAPIENS | 0 | 10836 | 4ztu | A | 623 | LEU | p.Leu623Trp | P54098 |
| HOMO SAPIENS | -1 | 10836 | 4ztu | A | 627 | ARG | p.Arg627Trp | P54098 |
| HOMO SAPIENS | -1 | 10836 | 4ztu | A | 628 | ARG | p.Arg628Pro | P54098 |
| HOMO SAPIENS | 0 | 10836 | 4ztu | A | 635 | PRO | p.Pro635Leu | P54098 |
| HOMO SAPIENS | 0 | 10836 | 4ztu | A | 662 | GLU | p.Glu662Lys | P54098 |
| HOMO SAPIENS | -1 | 10836 | 4ztu | A | 674 | GLY | p.Gly674Asp | P54098 |
| HOMO SAPIENS | -1 | 10836 | 4ztu | A | 722 | ARG | p.Arg722His | P54098 |
| HOMO SAPIENS | -1 | 10836 | 4ztu | A | 737 | GLY | p.Gly737Arg | P54098 |
| HOMO SAPIENS | 0 | 10836 | 4ztu | A | 748 | TRP | p.Trp748Ser | P54098 |
| HOMO SAPIENS | 0 | 10836 | 4ztu | A | 749 | PHE | p.Phe749Ser | P54098 |
| HOMO SAPIENS | 1 | 10836 | 4ztu | A | 807 | ARG | p.Arg807Cys | P54098 |
| HOMO SAPIENS | 1 | 10836 | 4ztu | A | 831 | TYR | p.Tyr831Cys | P54098 |
| HOMO SAPIENS | 8 | 10836 | 4ztu | A | 848 | GLY | p.Gly848Ser | P54098 |
| HOMO SAPIENS | 1 | 10836 | 4ztu | A | 852 | ARG | p.Arg852Cys | P54098 |
| HOMO SAPIENS | 9 | 10836 | 4ztu | A | 853 | ARG | p.Arg853Trp | P54098 |
| HOMO SAPIENS | 1 | 10836 | 4ztu | A | 864 | ASN | p.Asn864Ser | P54098 |
| HOMO SAPIENS | 1 | 10836 | 4ztu | A | 879 | GLN | p.Gln879Arg | P54098 |
| HOMO SAPIENS | 1 | 10836 | 4ztu | A | 914 | THR | p.Thr914Pro | P54098 |
| HOMO SAPIENS | 6 | 10836 | 4ztu | A | 932 | HIS | p.His932Tyr | P54098 |
| HOMO SAPIENS | 2 | 10836 | 4ztu | A | 944 | GLU | p.Glu944Lys | P54098 |
| HOMO SAPIENS | 6 | 10836 | 4ztu | A | 955 | TYR | p.Tyr955Cys | P54098 |
| HOMO SAPIENS | 1 | 10836 | 4ztu | A | 957 | ALA | p.Ala957Ser | P54098 |
| HOMO SAPIENS | 1 | 10836 | 4ztu | A | 957 | ALA | p.Ala957Val | P54098 |
| HOMO SAPIENS | 1 | 10836 | 4ztu | A | 964 | ARG | p.Arg964Cys | P54098 |
| HOMO SAPIENS | 1 | 10836 | 4ztu | A | 966 | LEU | p.Leu966Arg | P54098 |
| HOMO SAPIENS | 0 | 10836 | 4ztu | A | 1051 | GLY | p.Gly1051Arg | P54098 |
| HOMO SAPIENS | 1 | 10836 | 4ztu | A | 1073 | PRO | p.Pro1073Leu | P54098 |
| HOMO SAPIENS | 1 | 10836 | 4ztu | A | 1096 | ARG | p.Arg1096Cys | P54098 |
| HOMO SAPIENS | 1 | 10836 | 4ztu | A | 1096 | ARG | p.Arg1096His | P54098 |
| HOMO SAPIENS | 0 | 10836 | 4ztu | A | 1116 | MET | p.Met1116Thr | P54098 |
| HOMO SAPIENS | 1 | 10836 | 4ztu | A | 1142 | ARG | p.Arg1142Trp | P54098 |
| HOMO SAPIENS | 1 | 10836 | 4ztu | A | 1146 | ARG | p.Arg1146Cys | P54098 |
| HOMO SAPIENS | 1 | 10836 | 4ztu | A | 1157 | ASN | p.Asn1157Ser | P54098 |
| HOMO SAPIENS | -1 | 10836 | 4ztu | A | 1234 | ARG | p.Arg1234Gln | P54098 |
| HOMO SAPIENS | -1 | 10836 | 4ztu | A | 1236 | GLN | p.Gln1236His | P54098 |
| HOMO SAPIENS | 5 | 2666 | 1jph | A | 62 | PRO | p.Pro62Leu | P06132 |

| | | | | | | | | |
|---|---|---|---|---|---|---|---|---|
| HOMO SAPIENS | 5 | 2666 | 1jph | A | 116 | GLN | p.Gln116Ter | P06132 |
| HOMO SAPIENS | 0 | 2666 | 1jph | A | 165 | MET | p.Met165Arg | P06132 |
| HOMO SAPIENS | 6 | 2666 | 1jph | A | 167 | GLU | p.Glu167Lys | P06132 |
| HOMO SAPIENS | 0 | 2666 | 1jph | A | 195 | LEU | p.Leu195Phe | P06132 |
| HOMO SAPIENS | 7 | 2666 | 1jph | A | 253 | LEU | p.Leu253Gln | P06132 |
| HOMO SAPIENS | 0 | 2666 | 1jph | A | 281 | GLY | p.Gly281Glu | P06132 |
| HOMO SAPIENS | 9 | 2666 | 1jph | A | 304 | ASN | p.Asn304Lys | P06132 |
| HOMO SAPIENS | 8 | 2666 | 1jph | A | 311 | TYR | p.Tyr311Cys | P06132 |
| HOMO SAPIENS | 5 | 2666 | 1jph | A | 332 | ARG | p.Arg332His | P06132 |
| HOMO SAPIENS | 1 | 889 | 1m51 | A | 184 | LEU | p.Val184Leu | P35558 |
| HOMO SAPIENS | 0 | 889 | 1m51 | A | 267 | VAL | p.Ile267Val | P35558 |
| HOMO SAPIENS | 2 | 889 | 1m51 | A | 276 | GLU | p.Glu276Lys | P35558 |
| HOMO SAPIENS | 6 | 889 | 1m51 | A | 368 | VAL | p.Val368Ile | P35558 |
| HOMO SAPIENS | 0 | 889 | 1m51 | A | 579 | SER | p.Ser579Arg | P35558 |
| HOMO SAPIENS | 1 | 4084 | 2gjx | A | 28 | GLN | p.Gln28Ter | P06865 |
| HOMO SAPIENS | 0 | 4084 | 2gjx | A | 39 | LEU | p.Leu39Arg | P06865 |
| HOMO SAPIENS | 0 | 4084 | 2gjx | A | 127 | LEU | p.Leu127Arg | P06865 |
| HOMO SAPIENS | 6 | 4084 | 2gjx | A | 137 | ARG | p.Arg137Ter | P06865 |
| HOMO SAPIENS | 6 | 4084 | 2gjx | A | 170 | ARG | p.Arg170Gln | P06865 |
| HOMO SAPIENS | 9 | 4084 | 2gjx | A | 178 | ARG | p.Arg178Cys | P06865 |
| HOMO SAPIENS | 9 | 4084 | 2gjx | A | 178 | ARG | p.Arg178His | P06865 |
| HOMO SAPIENS | 5 | 4084 | 2gjx | A | 180 | TYR | p.Tyr180His | P06865 |
| HOMO SAPIENS | 5 | 4084 | 2gjx | A | 180 | TYR | p.Tyr180Ter | P06865 |
| HOMO SAPIENS | 1 | 4084 | 2gjx | A | 197 | LYS | p.Lys197Thr | P06865 |
| HOMO SAPIENS | 0 | 4084 | 2gjx | A | 200 | VAL | p.Val200Met | P06865 |
| HOMO SAPIENS | 9 | 4084 | 2gjx | A | 204 | HIS | p.His204Arg | P06865 |
| HOMO SAPIENS | 5 | 4084 | 2gjx | A | 210 | SER | p.Ser210Phe | P06865 |
| HOMO SAPIENS | 0 | 4084 | 2gjx | A | 211 | PHE | p.Phe211Ser | P06865 |
| HOMO SAPIENS | 2 | 4084 | 2gjx | A | 250 | GLY | p.Gly250Asp | P06865 |
| HOMO SAPIENS | 9 | 4084 | 2gjx | A | 258 | ASP | p.Asp258His | P06865 |
| HOMO SAPIENS | 0 | 4084 | 2gjx | A | 269 | GLY | p.Gly269Ser | P06865 |
| HOMO SAPIENS | 0 | 4084 | 2gjx | A | 301 | MET | p.Met301Arg | P06865 |
| HOMO SAPIENS | 3 | 4084 | 2gjx | A | 393 | ARG | p.Arg393Ter | P06865 |
| HOMO SAPIENS | 3 | 4084 | 2gjx | A | 393 | ARG | p.Arg393Pro | P06865 |
| HOMO SAPIENS | 0 | 4084 | 2gjx | A | 399 | ASN | p.Asn399Asp | P06865 |
| HOMO SAPIENS | 2 | 4084 | 2gjx | A | 420 | TRP | p.Trp420Cys | P06865 |
| HOMO SAPIENS | 1 | 4084 | 2gjx | A | 434 | PHE | p.Phe434Leu | P06865 |
| HOMO SAPIENS | 0 | 4084 | 2gjx | A | 454 | GLY | p.Gly454Ser | P06865 |
| HOMO SAPIENS | 1 | 4084 | 2gjx | A | 474 | TRP | p.Trp474Cys | P06865 |
| HOMO SAPIENS | 0 | 4084 | 2gjx | A | 479 | ALA | p.Ala479Thr | P06865 |
| HOMO SAPIENS | 5 | 4084 | 2gjx | A | 485 | TRP | p.Trp485Arg | P06865 |
| HOMO SAPIENS | 4 | 4084 | 2gjx | A | 499 | ARG | p.Arg499Cys | P06865 |
| HOMO SAPIENS | 4 | 4084 | 2gjx | A | 499 | ARG | p.Arg499His | P06865 |
| HOMO SAPIENS | 2 | 4084 | 2gjx | A | 504 | ARG | p.Arg504Cys | P06865 |
| HOMO SAPIENS | 2 | 4084 | 2gjx | A | 504 | ARG | p.Arg504His | P06865 |
| HOMO SAPIENS | -1 | 1943 | 1xmj | A | 417 | ASN | p.Asn417Lys | P13569 |
| HOMO SAPIENS | -1 | 1943 | 1xmj | A | 422 | SER | p.Ser422Phe | P13569 |
| HOMO SAPIENS | 1 | 1943 | 1xmj | A | 438 | THR | p.Thr438Ala | P13569 |
| HOMO SAPIENS | 0 | 1943 | 1xmj | A | 455 | ALA | p.Ala455Glu | P13569 |
| HOMO SAPIENS | 4 | 1943 | 1xmj | A | 456 | VAL | p.Val456Ala | P13569 |
| HOMO SAPIENS | 9 | 1943 | 1xmj | A | 458 | GLY | p.Gly458Val | P13569 |
| HOMO SAPIENS | 8 | 1943 | 1xmj | A | 466 | SER | p.Ser466Ter | P13569 |
| HOMO SAPIENS | 0 | 1943 | 1xmj | A | 467 | LEU | p.Leu467Pro | P13569 |
| HOMO SAPIENS | 1 | 1943 | 1xmj | A | 470 | MET | p.Val470Met | P13569 |

| | | | | | | | | |
|---|---|---|---|---|---|---|---|---|
| HOMO SAPIENS | 5 | 1943 | 1xmj | A | 480 | GLY | p.Gly480Cys | P13569 |
| HOMO SAPIENS | 3 | 1943 | 1xmj | A | 492 | SER | p.Ser492Phe | P13569 |
| HOMO SAPIENS | 8 | 1943 | 1xmj | A | 493 | GLN | p.Gln493Ter | P13569 |
| HOMO SAPIENS | 5 | 1943 | 1xmj | A | 506 | ILE | p.Ile506Met | P13569 |
| HOMO SAPIENS | 3 | 1943 | 1xmj | A | 520 | VAL | p.Val520Phe | P13569 |
| HOMO SAPIENS | 1 | 1943 | 1xmj | A | 528 | GLU | p.Glu528Asp | P13569 |
| HOMO SAPIENS | 1 | 1943 | 1xmj | A | 534 | ALA | p.Ala534Glu | P13569 |
| HOMO SAPIENS | 4 | 1943 | 1xmj | A | 542 | GLY | p.Gly542Ter | P13569 |
| HOMO SAPIENS | 6 | 1943 | 1xmj | A | 549 | SER | p.Ser549Asn | P13569 |
| HOMO SAPIENS | 6 | 1943 | 1xmj | A | 549 | SER | p.Ser549Arg | P13569 |
| HOMO SAPIENS | 6 | 1943 | 1xmj | A | 549 | SER | p.Ser549Arg | P13569 |
| HOMO SAPIENS | 6 | 1943 | 1xmj | A | 551 | GLY | p.Gly551Asp | P13569 |
| HOMO SAPIENS | 6 | 1943 | 1xmj | A | 551 | GLY | p.Gly551Ser | P13569 |
| HOMO SAPIENS | 6 | 1943 | 1xmj | A | 553 | GLN | p.Arg553Gln | P13569 |
| HOMO SAPIENS | 6 | 1943 | 1xmj | A | 553 | GLN | p.Arg553Ter | P13569 |
| HOMO SAPIENS | 0 | 1943 | 1xmj | A | 559 | ALA | p.Ala559Thr | P13569 |
| HOMO SAPIENS | 7 | 1943 | 1xmj | A | 560 | ARG | p.Arg560Lys | P13569 |
| HOMO SAPIENS | 4 | 1943 | 1xmj | A | 562 | VAL | p.Val562Ile | P13569 |
| HOMO SAPIENS | 3 | 1943 | 1xmj | A | 563 | TYR | p.Tyr563Asn | P13569 |
| HOMO SAPIENS | 3 | 1943 | 1xmj | A | 569 | TYR | p.Tyr569Asp | P13569 |
| HOMO SAPIENS | 3 | 1943 | 1xmj | A | 585 | GLU | p.Glu585Ter | P13569 |
| HOMO SAPIENS | 1 | 1943 | 1xmj | A | 618 | ILE | p.Ile618Thr | P13569 |
| HOMO SAPIENS | 9 | 1943 | 1xmj | A | 622 | GLY | p.Gly622Asp | P13569 |
| HOMO SAPIENS | -1 | 1943 | 1xmj | A | 648 | ASP | p.Asp648Val | P13569 |
| HOMO SAPIENS | 0 | 1943 | 1xmj | A | 673 | GLY | p.Gly673Ter | P13569 |
| HOMO SAPIENS | 5 | 5111 | 4rec | A | 749 | ARG | p.Arg749Ter | Q9Y2M0 |
| HOMO SAPIENS | -1 | 4580 | 4l0d | B | 2 | GLY | p.Pro2Leu | P35520 |
| HOMO SAPIENS | -1 | 4580 | 4l0d | B | 12 | PRO | p.Pro12Ser | P35520 |
| HOMO SAPIENS | -1 | 4580 | 4l0d | B | 18 | ARG | p.Arg18Cys | P35520 |
| HOMO SAPIENS | 1 | 4580 | 4l0d | B | 49 | PRO | p.Pro49Leu | P35520 |
| HOMO SAPIENS | 2 | 4580 | 4l0d | B | 70 | PRO | p.Pro70Leu | P35520 |
| HOMO SAPIENS | 1 | 4580 | 4l0d | B | 102 | LYS | p.Lys102Gln | P35520 |
| HOMO SAPIENS | 7 | 4580 | 4l0d | B | 114 | ALA | p.Ala114Val | P35520 |
| HOMO SAPIENS | 5 | 4580 | 4l0d | B | 116 | GLY | p.Gly116Arg | P35520 |
| HOMO SAPIENS | 8 | 4580 | 4l0d | B | 121 | ARG | p.Arg121Cys | P35520 |
| HOMO SAPIENS | 8 | 4580 | 4l0d | B | 121 | ARG | p.Arg121His | P35520 |
| HOMO SAPIENS | 5 | 4580 | 4l0d | B | 125 | ARG | p.Arg125Gln | P35520 |
| HOMO SAPIENS | 4 | 4580 | 4l0d | B | 134 | GLY | p.Gly134Arg | P35520 |
| HOMO SAPIENS | 4 | 4580 | 4l0d | B | 139 | GLY | p.Gly139Arg | P35520 |
| HOMO SAPIENS | 0 | 4580 | 4l0d | B | 145 | PRO | p.Pro145Leu | P35520 |
| HOMO SAPIENS | 0 | 4580 | 4l0d | B | 148 | GLY | p.Gly148Arg | P35520 |
| HOMO SAPIENS | 0 | 4580 | 4l0d | B | 168 | VAL | p.Val168Met | P35520 |
| HOMO SAPIENS | 2 | 4580 | 4l0d | B | 191 | THR | p.Thr191Met | P35520 |
| HOMO SAPIENS | 5 | 4580 | 4l0d | B | 234 | ASP | p.Asp234Asn | P35520 |
| HOMO SAPIENS | 0 | 4580 | 4l0d | B | 257 | THR | p.Thr257Met | P35520 |
| HOMO SAPIENS | 4 | 4580 | 4l0d | B | 262 | THR | p.Thr262Met | P35520 |
| HOMO SAPIENS | 0 | 4580 | 4l0d | B | 278 | ILE | p.Ile278Thr | P35520 |
| HOMO SAPIENS | 2 | 4580 | 4l0d | B | 307 | GLY | p.Gly307Ser | P35520 |
| HOMO SAPIENS | 4 | 4580 | 4l0d | B | 320 | VAL | p.Val320Ala | P35520 |
| HOMO SAPIENS | 2 | 4580 | 4l0d | B | 336 | ARG | p.Arg336Cys | P35520 |
| HOMO SAPIENS | 2 | 4580 | 4l0d | B | 336 | ARG | p.Arg336His | P35520 |
| HOMO SAPIENS | 0 | 4580 | 4l0d | B | 351 | GLY | p.Gly351Arg | P35520 |
| HOMO SAPIENS | 0 | 4580 | 4l0d | B | 353 | THR | p.Thr353Met | P35520 |
| HOMO SAPIENS | 0 | 4580 | 4l0d | B | 371 | VAL | p.Val371Met | P35520 |

| | | | | | | | | |
|---|---|---|---|---|---|---|---|---|
| HOMO SAPIENS | 2 | 4580 | 4l0d | B | 379 | ARG | p.Arg379Gln | P35520 |
| HOMO SAPIENS | 2 | 4580 | 4l0d | B | 379 | ARG | p.Arg379Trp | P35520 |
| HOMO SAPIENS | 2 | 4580 | 4l0d | B | 384 | LYS | p.Lys384Glu | P35520 |
| HOMO SAPIENS | 0 | 4580 | 4l0d | B | 422 | PRO | p.Pro422Leu | P35520 |
| HOMO SAPIENS | 0 | 4580 | 4l0d | B | 444 | ASP | p.Asp444Asn | P35520 |
| HOMO SAPIENS | 0 | 4580 | 4l0d | B | 466 | SER | p.Ser466Leu | P35520 |
| HOMO SAPIENS | 0 | 4580 | 4l0d | B | 539 | LEU | p.Leu539Ser | P35520 |
| HOMO SAPIENS | 0 | 4580 | 4l0d | B | 548 | ARG | p.Arg548Gln | P35520 |
| HOMO SAPIENS | 1 | 6536 | 3ulu | A | 412 | LEU | p.Leu412Phe | O15455 |
| HOMO SAPIENS | 1 | 28493 | 3ii6 | X | 814 | ARG | p.Arg814Ter | P49917 |
| HOMO SAPIENS | 1 | 28493 | 3ii6 | X | 857 | ALA | p.Ala857Thr | P49917 |
| HOMO SAPIENS | 9 | 3543 | 2q51 | A | 24 | GLU | p.Glu24Gly | P45381 |
| HOMO SAPIENS | 1 | 3543 | 2q51 | A | 27 | GLY | p.Gly27Arg | P45381 |
| HOMO SAPIENS | 9 | 3543 | 2q51 | A | 71 | ARG | p.Arg71His | P45381 |
| HOMO SAPIENS | -1 | 3543 | 2q51 | A | 82 | MSE | p.Met82Thr | P45381 |
| HOMO SAPIENS | 0 | 3543 | 2q51 | A | 143 | ILE | p.Ile143Phe | P45381 |
| HOMO SAPIENS | 1 | 3543 | 2q51 | A | 152 | CYS | p.Cys152Arg | P45381 |
| HOMO SAPIENS | 2 | 3543 | 2q51 | A | 218 | CYS | p.Cys218Ter | P45381 |
| HOMO SAPIENS | 1 | 3543 | 2q51 | A | 231 | TYR | p.Tyr231Cys | P45381 |
| HOMO SAPIENS | 1 | 3543 | 2q51 | A | 231 | TYR | p.Tyr231Ter | P45381 |
| HOMO SAPIENS | 2 | 3543 | 2q51 | A | 274 | GLY | p.Gly274Arg | P45381 |
| HOMO SAPIENS | 6 | 3543 | 2q51 | A | 285 | GLU | p.Glu285Ala | P45381 |
| HOMO SAPIENS | 5 | 3543 | 2q51 | A | 287 | ALA | p.Ala287Thr | P45381 |
| HOMO SAPIENS | 2 | 3543 | 2q51 | A | 305 | ALA | p.Ala305Glu | P45381 |
| HOMO SAPIENS | 2 | 24687 | 1hfi | A | 25 | SER | p.Ser890Ile | P08603 |
| HOMO SAPIENS | 0 | 1508 | 3wn5 | C | 45 | LEU | p.Leu66Arg | P08637 |
| HOMO SAPIENS | 0 | 1508 | 3wn5 | C | 155 | VAL | p.Phe176Val | P08637 |
| HOMO SAPIENS | 0 | 750 | 5tpa | A | 54 | VAL | p.Val452Met | Q12879 |
| HOMO SAPIENS | 9 | 750 | 5tpa | A | 120 | ARG | p.Arg518Cys | Q12879 |
| HOMO SAPIENS | 0 | 750 | 5tpa | A | 199 | ALA | p.Ala716Thr | Q12879 |
| HOMO SAPIENS | 4 | 1363 | 3hg2 | B | 40 | PRO | p.Pro40Ser | P06280 |
| HOMO SAPIENS | 3 | 1363 | 3hg2 | B | 65 | SER | p.Ser65Thr | P06280 |
| HOMO SAPIENS | 4 | 1363 | 3hg2 | B | 66 | GLU | p.Glu66Gln | P06280 |
| HOMO SAPIENS | 2 | 1363 | 3hg2 | B | 80 | GLY | p.Gly80Asp | P06280 |
| HOMO SAPIENS | 3 | 1363 | 3hg2 | B | 112 | ARG | p.Arg112Cys | P06280 |
| HOMO SAPIENS | 3 | 1363 | 3hg2 | B | 112 | ARG | p.Arg112His | P06280 |
| HOMO SAPIENS | 0 | 1363 | 3hg2 | B | 139 | ASN | p.Asn139Ser | P06280 |
| HOMO SAPIENS | 7 | 1363 | 3hg2 | B | 156 | ALA | p.Ala156Thr | P06280 |
| HOMO SAPIENS | 5 | 1363 | 3hg2 | B | 162 | TRP | p.Trp162Arg | P06280 |
| HOMO SAPIENS | 5 | 1363 | 3hg2 | B | 207 | TYR | p.Tyr207His | P06280 |
| HOMO SAPIENS | 0 | 1363 | 3hg2 | B | 215 | ASN | p.Asn215Ser | P06280 |
| HOMO SAPIENS | 0 | 1363 | 3hg2 | B | 264 | ASP | p.Asp264Val | P06280 |
| HOMO SAPIENS | 5 | 1363 | 3hg2 | B | 266 | ASP | p.Asp266Val | P06280 |
| HOMO SAPIENS | 2 | 1363 | 3hg2 | B | 269 | VAL | p.Val269Ala | P06280 |
| HOMO SAPIENS | 3 | 1363 | 3hg2 | B | 272 | ASN | p.Asn272Ser | P06280 |
| HOMO SAPIENS | 3 | 1363 | 3hg2 | B | 279 | GLN | p.Gln279Glu | P06280 |
| HOMO SAPIENS | 0 | 1363 | 3hg2 | B | 289 | ILE | p.Ile289Val | P06280 |
| HOMO SAPIENS | 0 | 1363 | 3hg2 | B | 296 | MET | p.Met296Val | P06280 |
| HOMO SAPIENS | 0 | 1363 | 3hg2 | B | 296 | MET | p.Met296Ile | P06280 |
| HOMO SAPIENS | 0 | 1363 | 3hg2 | B | 297 | SER | p.Ser297Phe | P06280 |
| HOMO SAPIENS | 2 | 1363 | 3hg2 | B | 301 | ARG | p.Arg301Gln | P06280 |
| HOMO SAPIENS | 3 | 1363 | 3hg2 | B | 327 | GLN | p.Gln327Lys | P06280 |
| HOMO SAPIENS | 0 | 1363 | 3hg2 | B | 328 | GLY | p.Gly328Arg | P06280 |
| HOMO SAPIENS | 0 | 1363 | 3hg2 | B | 328 | GLY | p.Gly328Ala | P06280 |

| | | | | | | | | |
|---|---|---|---|---|---|---|---|---|
| HOMO SAPIENS | 2 | 1363 | 3hg2 | B | 331 | LEU | p.Leu331Phe | P06280 |
| HOMO SAPIENS | 0 | 1363 | 3hg2 | B | 342 | ARG | p.Arg342Gln | P06280 |
| HOMO SAPIENS | 1 | 1363 | 3hg2 | B | 361 | GLY | p.Gly361Arg | P06280 |
| HOMO SAPIENS | 1 | 1363 | 3hg2 | B | 363 | ARG | p.Arg363His | P06280 |
| HOMO SAPIENS | 1 | 1363 | 3hg2 | B | 385 | THR | p.Thr385Ala | P06280 |
| HOMO SAPIENS | 1 | 1363 | 3hg2 | B | 399 | TRP | p.Trp399Ser | P06280 |
| HOMO SAPIENS | 1 | 1363 | 3hg2 | B | 410 | THR | p.Thr410Ala | P06280 |
| HOMO SAPIENS | 2 | 3904 | 3caa | B | 385 | MET | p.Met414Val | P01011 |
| HOMO SAPIENS | 0 | 558 | 5eub | A | 56 | VAL | p.Val56Ile | Q13126 |
| HOMO SAPIENS | 0 | 558 | 5eub | A | 189 | TRP | p.Trp189Leu | Q13126 |
| HOMO SAPIENS | 0 | 558 | 5eub | A | 218 | ALA | p.Ala218Gly | Q13126 |
| HOMO SAPIENS | 1 | 3865 | 1cb5 | B | 443 | ILE | p.Ile443Val | Q13867 |
| HOMO SAPIENS | 2 | 14329 | 4m92 | A | 24 | ILE | p.Ile65Val | Q13454 |
| HOMO SAPIENS | 1 | 7381 | 5kz5 | k | 50 | GLY | p.Gly50Glu | Q9H1K1 |
| HOMO SAPIENS | 0 | 1102 | 4fdh | E | 86 | PRO | p.Pro86Ala | P19099 |
| HOMO SAPIENS | 0 | 1102 | 4fdh | E | 118 | ALA | p.Ala118Thr | P19099 |
| HOMO SAPIENS | 0 | 1102 | 4fdh | E | 159 | PRO | p.Pro159Leu | P19099 |
| HOMO SAPIENS | 0 | 1102 | 4fdh | E | 173 | LYS | p.Lys173Arg | P19099 |
| HOMO SAPIENS | 0 | 1102 | 4fdh | E | 181 | ARG | p.Arg181Trp | P19099 |
| HOMO SAPIENS | 0 | 1102 | 4fdh | E | 185 | THR | p.Thr185Ile | P19099 |
| HOMO SAPIENS | 0 | 1102 | 4fdh | E | 198 | GLU | p.Glu198Asp | P19099 |
| HOMO SAPIENS | 0 | 1102 | 4fdh | E | 214 | HIS | p.His214Asp | P19099 |
| HOMO SAPIENS | 0 | 1102 | 4fdh | E | 251 | LYS | p.Lys251Arg | P19099 |
| HOMO SAPIENS | 0 | 1102 | 4fdh | E | 255 | GLU | p.Glu255Ter | P19099 |
| HOMO SAPIENS | 0 | 1102 | 4fdh | E | 281 | ASN | p.Asn281Ser | P19099 |
| HOMO SAPIENS | 0 | 1102 | 4fdh | E | 320 | ALA | p.Ala320Val | P19099 |
| HOMO SAPIENS | 0 | 1102 | 4fdh | E | 339 | ILE | p.Ile339Thr | P19099 |
| HOMO SAPIENS | 0 | 1102 | 4fdh | E | 341 | ARG | p.Arg341Cys | P19099 |
| HOMO SAPIENS | 2 | 1102 | 4fdh | E | 374 | ARG | p.Arg374Trp | P19099 |
| HOMO SAPIENS | 0 | 1102 | 4fdh | E | 386 | VAL | p.Val386Ala | P19099 |
| HOMO SAPIENS | 0 | 1102 | 4fdh | E | 390 | LEU | p.Leu390Phe | P19099 |
| HOMO SAPIENS | -1 | 1102 | 4fdh | E | 435 | GLY | p.Gly435Ser | P19099 |
| HOMO SAPIENS | 0 | 1102 | 4fdh | E | 461 | LEU | p.Leu461Pro | P19099 |
| HOMO SAPIENS | 0 | 1102 | 4fdh | E | 498 | THR | p.Thr498Ala | P19099 |
| HOMO SAPIENS | 6 | 10292 | 2of5 | F | 128 | GLY | p.Gly128Arg | P78560 |
| HOMO SAPIENS | 6 | 10292 | 2of5 | F | 170 | ARG | p.Arg170Cys | P78560 |
| HOMO SAPIENS | -1 | 18965 | 3ghg | A | 16 | ARG | p.Arg35Cys | P02671 |
| HOMO SAPIENS | -1 | 18965 | 3ghg | A | 16 | ARG | p.Arg35His | P02671 |
| HOMO SAPIENS | -1 | 18965 | 3ghg | A | 283 | PRO | p.Pro302Ala | P02671 |
| HOMO SAPIENS | -1 | 18965 | 3ghg | A | 312 | THR | p.Thr331Ala | P02671 |
| HOMO SAPIENS | -1 | 18965 | 3ghg | A | 381 | SER | p.Ser400Phe | P02671 |
| HOMO SAPIENS | -1 | 18965 | 3ghg | A | 463 | VAL | p.Val482Met | P02671 |
| HOMO SAPIENS | -1 | 18965 | 3ghg | A | 526 | GLU | p.Glu545Val | P02671 |
| HOMO SAPIENS | -1 | 18965 | 3ghg | A | 554 | ARG | p.Arg573Leu | P02671 |
| HOMO SAPIENS | 8 | 3023 | 4ttb | A | 101 | ARG | p.Arg101Trp | Q6PHW0 |
| HOMO SAPIENS | 2 | 3023 | 4ttb | A | 116 | ILE | p.Ile116Thr | Q6PHW0 |
| HOMO SAPIENS | 0 | 3023 | 4ttb | A | 220 | ALA | p.Ala220Thr | Q6PHW0 |
| HOMO SAPIENS | 2 | 3023 | 4ttb | A | 271 | GLU | p.Glu271Lys | Q6PHW0 |
| HOMO SAPIENS | 3 | 33226 | 1ya5 | T | 11 | SER | p.Ser11Ter | O15273 |
| HOMO SAPIENS | 6 | 33226 | 1ya5 | T | 64 | SER | p.Ser64Leu | O15273 |
| HOMO SAPIENS | 6 | 33226 | 1ya5 | T | 70 | ARG | p.Arg70Gln | O15273 |
| HOMO SAPIENS | 3 | 33226 | 1ya5 | T | 75 | GLY | p.Gly75Ser | O15273 |
| HOMO SAPIENS | 0 | 33226 | 1ya5 | T | 87 | ARG | p.Arg87Trp | O15273 |
| HOMO SAPIENS | 2 | 5667 | 3dax | A | 398 | PRO | p.Pro398Ala | P22680 |

| | | | | | | | | |
|---|---|---|---|---|---|---|---|---|
| HOMO SAPIENS | 0 | 12861 | 5mya | A | 464 | ASN | p.Asn464His | Q02763 |
| HOMO SAPIENS | 2 | 12861 | 5mya | A | 486 | VAL | p.Val486Ile | Q02763 |
| HOMO SAPIENS | 0 | 12861 | 5mya | A | 522 | ARG | p.Arg522His | Q02763 |
| HOMO SAPIENS | 2 | 12861 | 5mya | A | 600 | VAL | p.Val600Leu | Q02763 |
| HOMO SAPIENS | 0 | 12861 | 5mya | A | 634 | LEU | p.Leu634Phe | Q02763 |
| HOMO SAPIENS | 0 | 12861 | 5mya | A | 684 | HIS | p.His684Gln | Q02763 |
| HOMO SAPIENS | 0 | 12861 | 5mya | A | 724 | ALA | p.Ala724Thr | Q02763 |
| HOMO SAPIENS | 0 | 7743 | 3qzw | G | 90 | GLY | p.Gly111Ser | P01732 |
| HOMO SAPIENS | 2 | 9498 | 3kci | A | 4267 | ASP | p.Asp4267Glu | O95714 |
| HOMO SAPIENS | 5 | 201 | 3ips | B | 415 | ARG | p.Arg415Gln | Q13133 |
| HOMO SAPIENS | 3 | 229 | 2q50 | A | 68 | LEU | p.Leu68Pro | Q9UBQ7 |
| HOMO SAPIENS | 2 | 229 | 2q50 | A | 99 | ARG | p.Arg99Ter | Q9UBQ7 |
| HOMO SAPIENS | 3 | 229 | 2q50 | A | 160 | GLY | p.Gly160Arg | Q9UBQ7 |
| HOMO SAPIENS | 5 | 229 | 2q50 | A | 165 | GLY | p.Gly165Asp | Q9UBQ7 |
| HOMO SAPIENS | 4 | 229 | 2q50 | A | 302 | ARG | p.Arg302Cys | Q9UBQ7 |
| HOMO SAPIENS | 4 | 229 | 2q50 | A | 302 | ARG | p.Arg302His | Q9UBQ7 |
| HOMO SAPIENS | -1 | 229 | 2q50 | A | 322 | MSE | p.Met322Arg | Q9UBQ7 |
| HOMO SAPIENS | -1 | 4519 | 5clw | A | 40 | PRO | p.Pro40Thr | Q04446 |
| HOMO SAPIENS | 1 | 4519 | 5clw | A | 59 | ILE | p.Ile59Thr | Q04446 |
| HOMO SAPIENS | 0 | 4519 | 5clw | A | 190 | ARG | p.Arg190Gly | Q04446 |
| HOMO SAPIENS | 1 | 4519 | 5clw | A | 224 | LEU | p.Leu224Pro | Q04446 |
| HOMO SAPIENS | 0 | 4519 | 5clw | A | 236 | GLN | p.Gln236His | Q04446 |
| HOMO SAPIENS | 1 | 4519 | 5clw | A | 254 | THR | p.Thr254Ala | Q04446 |
| HOMO SAPIENS | 0 | 4519 | 5clw | A | 257 | PHE | p.Phe257Leu | Q04446 |
| HOMO SAPIENS | 1 | 4519 | 5clw | A | 262 | ARG | p.Arg262Cys | Q04446 |
| HOMO SAPIENS | 3 | 4519 | 5clw | A | 329 | TYR | p.Tyr329Ser | Q04446 |
| HOMO SAPIENS | 0 | 4519 | 5clw | A | 507 | THR | p.Thr507Ala | Q04446 |
| HOMO SAPIENS | 0 | 4519 | 5clw | A | 515 | ARG | p.Arg515Cys | Q04446 |
| HOMO SAPIENS | 0 | 4519 | 5clw | A | 515 | ARG | p.Arg515His | Q04446 |
| HOMO SAPIENS | 0 | 4519 | 5clw | A | 524 | ARG | p.Arg524Gln | Q04446 |
| HOMO SAPIENS | 0 | 4519 | 5clw | A | 524 | ARG | p.Arg524Ter | Q04446 |
| HOMO SAPIENS | 0 | 4519 | 5clw | A | 545 | HIS | p.His545Arg | Q04446 |
| HOMO SAPIENS | 2 | 4519 | 5clw | A | 548 | TRP | p.Trp548Ter | Q04446 |
| HOMO SAPIENS | 0 | 4519 | 5clw | A | 592 | GLU | p.Glu592Ter | Q04446 |
| HOMO SAPIENS | 0 | 4519 | 5clw | A | 628 | HIS | p.His628Arg | Q04446 |
| HOMO SAPIENS | 0 | 4519 | 5clw | A | 637 | ARG | p.Arg637Ter | Q04446 |
| HOMO SAPIENS | 0 | 4519 | 5clw | A | 673 | ALA | p.Ala673Thr | Q04446 |
| HOMO SAPIENS | 1 | 10405 | 3bg3 | D | 507 | VAL | p.Val507Ile | P11498 |
| HOMO SAPIENS | 0 | 10405 | 3bg3 | D | 583 | ARG | p.Arg583Leu | P11498 |
| HOMO SAPIENS | 1 | 10405 | 3bg3 | D | 589 | LYS | p.Lys589Asn | P11498 |
| HOMO SAPIENS | 6 | 10405 | 3bg3 | D | 610 | ALA | p.Ala610Thr | P11498 |
| HOMO SAPIENS | 2 | 10405 | 3bg3 | D | 631 | ARG | p.Arg631Gln | P11498 |
| HOMO SAPIENS | 1 | 10405 | 3bg3 | D | 732 | ARG | p.Arg732Gln | P11498 |
| HOMO SAPIENS | 6 | 10405 | 3bg3 | D | 743 | MET | p.Met743Ile | P11498 |
| HOMO SAPIENS | 0 | 10405 | 3bg3 | D | 750 | THR | p.Thr750Met | P11498 |
| HOMO SAPIENS | -1 | 13751 | 2o8d | B | 350 | ALA | p.Ala350Val | P52701 |
| HOMO SAPIENS | -1 | 13751 | 2o8d | B | 352 | VAL | p.Val352Ile | P52701 |
| HOMO SAPIENS | -1 | 13751 | 2o8d | B | 355 | GLY | p.Gly355Ser | P52701 |
| HOMO SAPIENS | 0 | 13751 | 2o8d | B | 385 | ARG | p.Arg385Trp | P52701 |
| HOMO SAPIENS | 0 | 13751 | 2o8d | B | 390 | ASP | p.Asp390Glu | P52701 |
| HOMO SAPIENS | 0 | 13751 | 2o8d | B | 396 | LEU | p.Leu396Val | P52701 |
| HOMO SAPIENS | 0 | 13751 | 2o8d | B | 449 | LEU | p.Leu449Pro | P52701 |
| HOMO SAPIENS | 0 | 13751 | 2o8d | B | 468 | ARG | p.Arg468His | P52701 |
| HOMO SAPIENS | 0 | 13751 | 2o8d | B | 503 | SER | p.Ser503Cys | P52701 |

| | | | | | | | | |
|---|---|---|---|---|---|---|---|---|
| HOMO SAPIENS | 1 | 13751 | 2o8d | B | 509 | VAL | p.Val509Ala | P52701 |
| HOMO SAPIENS | 0 | 13751 | 2o8d | B | 675 | GLU | p.Glu675Asp | P52701 |
| HOMO SAPIENS | 0 | 13751 | 2o8d | B | 795 | ILE | p.Ile795Thr | P52701 |
| HOMO SAPIENS | 0 | 13751 | 2o8d | B | 850 | TYR | p.Tyr850Ter | P52701 |
| HOMO SAPIENS | 0 | 13751 | 2o8d | B | 878 | VAL | p.Val878Ala | P52701 |
| HOMO SAPIENS | 0 | 13751 | 2o8d | B | 911 | ARG | p.Arg911Ter | P52701 |
| HOMO SAPIENS | 0 | 13751 | 2o8d | B | 922 | ARG | p.Arg922Ter | P52701 |
| HOMO SAPIENS | 0 | 13751 | 2o8d | B | 969 | TYR | p.Tyr969Cys | P52701 |
| HOMO SAPIENS | 0 | 13751 | 2o8d | B | 1005 | ARG | p.Arg1005Ter | P52701 |
| HOMO SAPIENS | 0 | 13751 | 2o8d | B | 1035 | ARG | p.Arg1035Ter | P52701 |
| HOMO SAPIENS | 1 | 13751 | 2o8d | B | 1048 | GLN | p.Gln1048Ter | P52701 |
| HOMO SAPIENS | 1 | 13751 | 2o8d | B | 1051 | VAL | p.Val1051Ile | P52701 |
| HOMO SAPIENS | 0 | 13751 | 2o8d | B | 1054 | ILE | p.Ile1054Phe | P52701 |
| HOMO SAPIENS | 0 | 13751 | 2o8d | B | 1054 | ILE | p.Ile1054Met | P52701 |
| HOMO SAPIENS | 0 | 13751 | 2o8d | B | 1163 | GLU | p.Glu1163Val | P52701 |
| HOMO SAPIENS | 0 | 13751 | 2o8d | B | 1237 | GLU | p.Glu1237Asp | P52701 |
| HOMO SAPIENS | 0 | 13751 | 2o8d | B | 1242 | ARG | p.Arg1242His | P52701 |
| HOMO SAPIENS | -1 | 13751 | 2o8d | B | 1278 | PRO | p.Pro1278Thr | P52701 |
| HOMO SAPIENS | 0 | 13751 | 2o8d | B | 1304 | ARG | p.Arg1304Lys | P52701 |
| HOMO SAPIENS | 0 | 13751 | 2o8d | B | 1317 | HIS | p.His1317Gln | P52701 |
| HOMO SAPIENS | 0 | 13751 | 2o8d | B | 1329 | SER | p.Ser1329Leu | P52701 |
| HOMO SAPIENS | 3 | 13273 | 4g8k | A | 265 | GLU | p.Glu265Ter | Q05823 |
| HOMO SAPIENS | -1 | 11088 | 5uj7 | A | 666 | ARG | p.Arg666Trp | Q13415 |
| HOMO SAPIENS | 4 | 11088 | 5uj7 | A | 720 | ARG | p.Arg720Gln | Q13415 |
| HOMO SAPIENS | 0 | 11088 | 5uj7 | A | 816 | MET | p.Met816Thr | Q13415 |
| HOMO SAPIENS | 1 | 4352 | 5va1 | A | 421 | THR | p.Thr421Met | Q12809 |
| HOMO SAPIENS | 0 | 4352 | 5va1 | A | 451 | PRO | p.Pro451Leu | Q12809 |
| HOMO SAPIENS | 0 | 4352 | 5va1 | A | 470 | ASN | p.Asn470Asp | Q12809 |
| HOMO SAPIENS | 0 | 4352 | 5va1 | A | 490 | ALA | p.Ala490Thr | Q12809 |
| HOMO SAPIENS | 0 | 4352 | 5va1 | A | 491 | VAL | p.Val491Ile | Q12809 |
| HOMO SAPIENS | 1 | 4352 | 5va1 | A | 533 | VAL | p.Val533Gly | Q12809 |
| HOMO SAPIENS | 0 | 4352 | 5va1 | A | 552 | LEU | p.Leu552Ser | Q12809 |
| HOMO SAPIENS | 0 | 4352 | 5va1 | A | 558 | ALA | p.Ala558Pro | Q12809 |
| HOMO SAPIENS | 1 | 4352 | 5va1 | A | 561 | ALA | p.Ala561Val | Q12809 |
| HOMO SAPIENS | 0 | 4352 | 5va1 | A | 572 | GLY | p.Gly572Arg | Q12809 |
| HOMO SAPIENS | -1 | 4352 | 5va1 | A | 582 | ARG | p.Arg582Cys | Q12809 |
| HOMO SAPIENS | 0 | 4352 | 5va1 | A | 584 | GLY | p.Gly584Ser | Q12809 |
| HOMO SAPIENS | 0 | 4352 | 5va1 | A | 588 | ASN | p.Asn588Lys | Q12809 |
| HOMO SAPIENS | 0 | 4352 | 5va1 | A | 593 | ILE | p.Ile593Arg | Q12809 |
| HOMO SAPIENS | 1 | 4352 | 5va1 | A | 627 | PHE | p.Phe627Leu | Q12809 |
| HOMO SAPIENS | 0 | 4352 | 5va1 | A | 630 | VAL | p.Val630Leu | Q12809 |
| HOMO SAPIENS | 0 | 4352 | 5va1 | A | 652 | TYR | p.Tyr652Ter | Q12809 |
| HOMO SAPIENS | 0 | 4352 | 5va1 | A | 711 | ILE | p.Ile711Val | Q12809 |
| HOMO SAPIENS | 0 | 4352 | 5va1 | A | 735 | SER | p.Ser735Leu | Q12809 |
| HOMO SAPIENS | 0 | 4352 | 5va1 | A | 744 | ARG | p.Arg744Ter | Q12809 |
| HOMO SAPIENS | 0 | 4352 | 5va1 | A | 784 | ARG | p.Arg784Trp | Q12809 |
| HOMO SAPIENS | 1 | 4352 | 5va1 | A | 820 | GLY | p.Gly820Arg | Q12809 |
| HOMO SAPIENS | 5 | 4352 | 5va1 | A | 822 | VAL | p.Val822Met | Q12809 |
| HOMO SAPIENS | 1 | 4352 | 5va1 | A | 863 | ARG | p.Arg863Ter | Q12809 |
| HOMO SAPIENS | 5 | 303 | 2obv | A | 55 | ALA | p.Ala55Asp | Q00266 |
| HOMO SAPIENS | 7 | 303 | 2obv | A | 259 | ALA | p.Ala259Gly | Q00266 |
| HOMO SAPIENS | 9 | 303 | 2obv | A | 264 | ARG | p.Arg264His | Q00266 |
| HOMO SAPIENS | 9 | 303 | 2obv | A | 264 | ARG | p.Arg264Cys | Q00266 |
| HOMO SAPIENS | 0 | 303 | 2obv | A | 305 | LEU | p.Leu305Pro | Q00266 |

| | | | | | | | | |
|---|---|---|---|---|---|---|---|---|
| HOMO SAPIENS | 7 | 303 | 2obv | A | 322 | ILE | p.Ile322Met | Q00266 |
| HOMO SAPIENS | 0 | 303 | 2obv | A | 336 | GLY | p.Gly336Arg | Q00266 |
| HOMO SAPIENS | 0 | 303 | 2obv | A | 357 | PRO | p.Pro357Leu | Q00266 |
| HOMO SAPIENS | 0 | 2077 | 4zxb | E | 462 | ASN | p.Asn489Ser | P06213 |
| HOMO SAPIENS | -1 | 2077 | 4zxb | E | 519 | ASP | p.Asp546Glu | P06213 |
| HOMO SAPIENS | 2 | 2077 | 4zxb | E | 819 | THR | p.Thr858Ala | P06213 |
| HOMO SAPIENS | 0 | 2077 | 4zxb | E | 907 | ASP | p.Asp946Glu | P06213 |
| HOMO SAPIENS | 7 | 3274 | 1wsv | B | 14 | HIS | p.His42Arg | P48728 |
| HOMO SAPIENS | 0 | 3274 | 1wsv | B | 19 | GLY | p.Gly47Arg | P48728 |
| HOMO SAPIENS | 4 | 3274 | 1wsv | B | 45 | ARG | p.Arg73Cys | P48728 |
| HOMO SAPIENS | 1 | 3274 | 1wsv | B | 49 | SER | p.Ser77Leu | P48728 |
| HOMO SAPIENS | 3 | 3274 | 1wsv | B | 241 | GLY | p.Gly269Asp | P48728 |
| HOMO SAPIENS | 7 | 3274 | 1wsv | B | 248 | ASP | p.Asp276His | P48728 |
| HOMO SAPIENS | 0 | 3274 | 1wsv | B | 272 | MET | p.Met300Val | P48728 |
| HOMO SAPIENS | 4 | 3274 | 1wsv | B | 292 | ARG | p.Arg320His | P48728 |
| HOMO SAPIENS | 0 | 3274 | 1wsv | B | 354 | ARG | p.Arg382Gln | P48728 |
| HOMO SAPIENS | -1 | 435 | 3uyt | A | 44 | THR | p.Thr44Ala | P48730 |
| HOMO SAPIENS | 1 | 4698 | 5ife | C | 666 | LYS | p.Lys666Asn | O75533 |
| HOMO SAPIENS | 1 | 16917 | 1apy | C | 37 | GLY | p.Gly60Asp | P20933 |
| HOMO SAPIENS | 0 | 16917 | 1apy | C | 49 | SER | p.Ser72Pro | P20933 |
| HOMO SAPIENS | 0 | 16917 | 1apy | C | 84 | ARG | p.Arg107Ter | P20933 |
| HOMO SAPIENS | 0 | 16917 | 1apy | C | 126 | SER | p.Thr149Ser | P20933 |
| HOMO SAPIENS | 0 | 16917 | 1apy | C | 135 | TRP | p.Trp158Ter | P20933 |
| HOMO SAPIENS | 0 | 16917 | 1apy | C | 140 | CYS | p.Cys163Ser | P20933 |
| HOMO SAPIENS | 0 | 16917 | 1apy | C | 145 | TRP | p.Trp168Ter | P20933 |
| HOMO SAPIENS | 0 | 10413 | 4frw | A | 185 | THR | p.Thr185Met | Q96NY8 |
| HOMO SAPIENS | 0 | 5217 | 5n6w | G | 96 | TRP | p.Trp96Arg | O15537 |
| HOMO SAPIENS | 1 | 5217 | 5n6w | G | 102 | ARG | p.Arg102Trp | O15537 |
| HOMO SAPIENS | 0 | 5217 | 5n6w | G | 109 | GLY | p.Gly109Arg | O15537 |
| HOMO SAPIENS | -1 | 5217 | 5n6w | G | 223 | CYS | p.Cys223Arg | O15537 |
| HOMO SAPIENS | 0 | 9485 | 4z6y | E | 138 | ALA | p.Ala138Asp | Q9P0N9 |
| HOMO SAPIENS | 3 | 4504 | 4mwt | B | 411 | GLY | p.Gly439Ser | P10619 |
| HOMO SAPIENS | 3 | 4504 | 4mwt | B | 412 | PHE | p.Phe440Val | P10619 |
| HOMO SAPIENS | 6 | 6391 | 4kxy | A | 318 | ARG | p.Arg318Cys | P29401 |
| HOMO SAPIENS | 6 | 19546 | 2bx6 | A | 87 | THR | p.Thr87Ile | O75695 |
| HOMO SAPIENS | 1 | 19546 | 2bx6 | A | 118 | ARG | p.Arg118His | O75695 |
| HOMO SAPIENS | 1 | 19546 | 2bx6 | A | 201 | ILE | p.Ile201Val | O75695 |
| HOMO SAPIENS | 1 | 19546 | 2bx6 | A | 282 | ARG | p.Arg282Trp | O75695 |
| HOMO SAPIENS | 1 | 5265 | 5usr | H | 68 | ARG | p.Arg68Leu | Q9HD34 |
| HOMO SAPIENS | 2 | 1012 | 4nyi | S | 569 | LEU | p.Leu569Val | Q07889 |
| HOMO SAPIENS | 2 | 1012 | 4nyi | S | 655 | PRO | p.Pro655Leu | Q07889 |
| HOMO SAPIENS | 4 | 1012 | 4nyi | S | 663 | ILE | p.Ile663Met | Q07889 |
| HOMO SAPIENS | 2 | 1012 | 4nyi | S | 702 | TYR | p.Tyr702Cys | Q07889 |
| HOMO SAPIENS | 2 | 1012 | 4nyi | S | 708 | ALA | p.Ala708Thr | Q07889 |
| HOMO SAPIENS | 4 | 1012 | 4nyi | S | 733 | ILE | p.Ile733Phe | Q07889 |
| HOMO SAPIENS | 1 | 1848 | 3g1n | A | 4013 | ARG | p.Arg4013Trp | Q7Z6Z7 |
| HOMO SAPIENS | 1 | 1848 | 3g1n | A | 4187 | ARG | p.Arg4187Cys | Q7Z6Z7 |
| HOMO SAPIENS | 1 | 77 | 4pod | A | 41 | CYS | p.Cys79Tyr | P60174 |
| HOMO SAPIENS | 9 | 77 | 4pod | A | 104 | GLU | p.Glu142Asp | P60174 |
| HOMO SAPIENS | 1 | 77 | 4pod | A | 122 | GLY | p.Gly160Arg | P60174 |
| HOMO SAPIENS | -1 | 8672 | 3thz | A | 8 | THR | p.Thr8Met | P43246 |
| HOMO SAPIENS | 0 | 8672 | 3thz | A | 19 | PHE | p.Phe19Leu | P43246 |
| HOMO SAPIENS | 0 | 8672 | 3thz | A | 23 | PHE | p.Phe23Leu | P43246 |
| HOMO SAPIENS | 0 | 8672 | 3thz | A | 96 | ARG | p.Arg96His | P43246 |

| | | | | | | | | |
|---|---|---|---|---|---|---|---|---|
| HOMO SAPIENS | 0 | 8672 | 3thz | A | 102 | VAL | p.Val102Ile | P43246 |
| HOMO SAPIENS | 0 | 8672 | 3thz | A | 106 | ARG | p.Arg106Lys | P43246 |
| HOMO SAPIENS | 0 | 8672 | 3thz | A | 127 | ASN | p.Asn127Ser | P43246 |
| HOMO SAPIENS | 0 | 8672 | 3thz | A | 128 | LEU | p.Leu128Val | P43246 |
| HOMO SAPIENS | 0 | 8672 | 3thz | A | 163 | VAL | p.Val163Gly | P43246 |
| HOMO SAPIENS | 0 | 8672 | 3thz | A | 169 | ILE | p.Ile169Val | P43246 |
| HOMO SAPIENS | 0 | 8672 | 3thz | A | 315 | GLY | p.Gly315Asp | P43246 |
| HOMO SAPIENS | -1 | 8672 | 3thz | A | 322 | GLY | p.Gly322Asp | P43246 |
| HOMO SAPIENS | 0 | 8672 | 3thz | A | 331 | ASN | p.Asn331Asp | P43246 |
| HOMO SAPIENS | 1 | 8672 | 3thz | A | 390 | LEU | p.Leu390Phe | P43246 |
| HOMO SAPIENS | 0 | 8672 | 3thz | A | 419 | GLN | p.Gln419Lys | P43246 |
| HOMO SAPIENS | 0 | 8672 | 3thz | A | 447 | PHE | p.Phe447Leu | P43246 |
| HOMO SAPIENS | 0 | 8672 | 3thz | A | 524 | ARG | p.Arg524Pro | P43246 |
| HOMO SAPIENS | 0 | 8672 | 3thz | A | 539 | PHE | p.Phe539Leu | P43246 |
| HOMO SAPIENS | -1 | 8672 | 3thz | A | 546 | LYS | p.Lys546Asn | P43246 |
| HOMO SAPIENS | 0 | 8672 | 3thz | A | 564 | THR | p.Thr564Ala | P43246 |
| HOMO SAPIENS | 0 | 8672 | 3thz | A | 577 | ILE | p.Ile577Thr | P43246 |
| HOMO SAPIENS | 0 | 8672 | 3thz | A | 588 | TYR | p.Tyr588Ter | P43246 |
| HOMO SAPIENS | 0 | 8672 | 3thz | A | 622 | PRO | p.Pro622Leu | P43246 |
| HOMO SAPIENS | 0 | 8672 | 3thz | A | 629 | GLN | p.Gln629Arg | P43246 |
| HOMO SAPIENS | 0 | 8672 | 3thz | A | 636 | ALA | p.Ala636Pro | P43246 |
| HOMO SAPIENS | 2 | 8672 | 3thz | A | 642 | VAL | p.Val642Ile | P43246 |
| HOMO SAPIENS | 1 | 8672 | 3thz | A | 654 | ASP | p.Asp654Glu | P43246 |
| HOMO SAPIENS | 0 | 8672 | 3thz | A | 662 | GLN | p.Gln662His | P43246 |
| HOMO SAPIENS | 2 | 8672 | 3thz | A | 681 | GLN | p.Gln681His | P43246 |
| HOMO SAPIENS | 0 | 8672 | 3thz | A | 707 | CYS | p.Cys707Ser | P43246 |
| HOMO SAPIENS | 0 | 8672 | 3thz | A | 809 | GLU | p.Glu809Lys | P43246 |
| HOMO SAPIENS | 0 | 8672 | 3thz | A | 813 | MET | p.Met813Val | P43246 |
| HOMO SAPIENS | 1 | 8672 | 3thz | A | 834 | ALA | p.Ala834Thr | P43246 |
| HOMO SAPIENS | 0 | 8672 | 3thz | A | 849 | LEU | p.Leu849Val | P43246 |
| HOMO SAPIENS | -1 | 8672 | 3thz | A | 860 | SER | p.Ser860Leu | P43246 |
| HOMO SAPIENS | -1 | 8672 | 3thz | A | 934 | THR | p.Thr934Lys | P43246 |
| HOMO SAPIENS | 2 | 292 | 1i3k | A | 34 | ASN | p.Asn34Ser | Q14376 |
| HOMO SAPIENS | 7 | 292 | 1i3k | A | 90 | GLY | p.Gly90Glu | Q14376 |
| HOMO SAPIENS | 2 | 292 | 1i3k | A | 103 | ASP | p.Asp103Gly | Q14376 |
| HOMO SAPIENS | 2 | 292 | 1i3k | A | 136 | TYR | p.Tyr136Ter | Q14376 |
| HOMO SAPIENS | 0 | 292 | 1i3k | A | 169 | ARG | p.Arg169Trp | Q14376 |
| HOMO SAPIENS | 0 | 292 | 1i3k | A | 183 | LEU | p.Leu183Pro | Q14376 |
| HOMO SAPIENS | 0 | 292 | 1i3k | A | 302 | GLY | p.Gly302Asp | Q14376 |
| HOMO SAPIENS | 0 | 292 | 1i3k | A | 313 | LEU | p.Leu313Met | Q14376 |
| HOMO SAPIENS | 1 | 2084 | 5vc8 | A | 295 | GLN | p.Gln295Leu | O96028 |
| HOMO SAPIENS | -1 | 10157 | 5wrw | A | 7 | GLY | p.Gly7Ala | O76094 |
| HOMO SAPIENS | -1 | 10157 | 5wrw | A | 7 | GLY | p.Gly7Arg | O76094 |
| HOMO SAPIENS | -1 | 10157 | 5wrw | A | 8 | GLY | p.Gly8Ala | O76094 |
| HOMO SAPIENS | 0 | 3113 | 3in5 | A | 29 | GLU | p.Glu29Lys | Q9UBT6 |
| HOMO SAPIENS | 5 | 4546 | 5in3 | B | 44 | VAL | p.Val44Leu | P07902 |
| HOMO SAPIENS | -1 | 4546 | 5in3 | B | 51 | ARG | p.Arg51Leu | P07902 |
| HOMO SAPIENS | -1 | 4546 | 5in3 | B | 62 | LEU | p.Leu62Met | P07902 |
| HOMO SAPIENS | 4 | 4546 | 5in3 | B | 66 | PRO | p.Pro66His | P07902 |
| HOMO SAPIENS | 2 | 4546 | 5in3 | B | 74 | LEU | p.Leu74Pro | P07902 |
| HOMO SAPIENS | 5 | 4546 | 5in3 | B | 97 | ASN | p.Asn97Ser | P07902 |
| HOMO SAPIENS | 1 | 4546 | 5in3 | B | 135 | SER | p.Ser135Leu | P07902 |
| HOMO SAPIENS | 5 | 4546 | 5in3 | B | 138 | THR | p.Thr138Met | P07902 |
| HOMO SAPIENS | 7 | 4546 | 5in3 | B | 139 | LEU | p.Leu139Pro | P07902 |

| | | | | | | | | |
|---|---|---|---|---|---|---|---|---|
| HOMO SAPIENS | 2 | 4546 | 5in3 | B | 142 | MET | p.Met142Lys | P07902 |
| HOMO SAPIENS | 1 | 4546 | 5in3 | B | 143 | SER | p.Ser143Leu | P07902 |
| HOMO SAPIENS | 2 | 4546 | 5in3 | B | 148 | ARG | p.Arg148Gly | P07902 |
| HOMO SAPIENS | 2 | 4546 | 5in3 | B | 148 | ARG | p.Arg148Gln | P07902 |
| HOMO SAPIENS | 5 | 4546 | 5in3 | B | 183 | PRO | p.Pro183Thr | P07902 |
| HOMO SAPIENS | 5 | 4546 | 5in3 | B | 185 | PRO | p.Pro185His | P07902 |
| HOMO SAPIENS | 5 | 4546 | 5in3 | B | 186 | HIS | p.His186Tyr | P07902 |
| HOMO SAPIENS | 7 | 4546 | 5in3 | B | 188 | GLN | p.Gln188Arg | P07902 |
| HOMO SAPIENS | 5 | 4546 | 5in3 | B | 195 | LEU | p.Leu195Pro | P07902 |
| HOMO SAPIENS | 4 | 4546 | 5in3 | B | 201 | ARG | p.Arg201His | P07902 |
| HOMO SAPIENS | 4 | 4546 | 5in3 | B | 201 | ARG | p.Arg201Cys | P07902 |
| HOMO SAPIENS | 4 | 4546 | 5in3 | B | 203 | GLU | p.Glu203Lys | P07902 |
| HOMO SAPIENS | 2 | 4546 | 5in3 | B | 209 | TYR | p.Tyr209Cys | P07902 |
| HOMO SAPIENS | 5 | 4546 | 5in3 | B | 223 | ARG | p.Arg223Ser | P07902 |
| HOMO SAPIENS | 5 | 4546 | 5in3 | B | 226 | LEU | p.Leu226Pro | P07902 |
| HOMO SAPIENS | 5 | 4546 | 5in3 | B | 226 | LEU | p.Leu226Val | P07902 |
| HOMO SAPIENS | 2 | 4546 | 5in3 | B | 227 | LEU | p.Leu227Pro | P07902 |
| HOMO SAPIENS | 8 | 4546 | 5in3 | B | 258 | ARG | p.Arg258Cys | P07902 |
| HOMO SAPIENS | 1 | 4546 | 5in3 | B | 265 | PRO | p.Pro265Ala | P07902 |
| HOMO SAPIENS | 8 | 4546 | 5in3 | B | 268 | THR | p.Thr268Asn | P07902 |
| HOMO SAPIENS | 7 | 4546 | 5in3 | B | 278 | ILE | p.Ile278Asn | P07902 |
| HOMO SAPIENS | 5 | 4546 | 5in3 | B | 285 | LYS | p.Lys285Asn | P07902 |
| HOMO SAPIENS | 1 | 4546 | 5in3 | B | 319 | HIS | p.His319Gln | P07902 |
| HOMO SAPIENS | 4 | 4546 | 5in3 | B | 321 | HIS | p.His321Tyr | P07902 |
| HOMO SAPIENS | 4 | 4546 | 5in3 | B | 325 | PRO | p.Pro325Leu | P07902 |
| HOMO SAPIENS | 2 | 4546 | 5in3 | B | 328 | ARG | p.Arg328His | P07902 |
| HOMO SAPIENS | 2 | 4546 | 5in3 | B | 329 | SER | p.Ser329Phe | P07902 |
| HOMO SAPIENS | 5 | 4546 | 5in3 | B | 334 | LYS | p.Lys334Arg | P07902 |
| HOMO SAPIENS | 6 | 1290 | 5uav | C | 119 | ARG | p.Arg119Gly | P32322 |
| HOMO SAPIENS | 6 | 1290 | 5uav | C | 119 | ARG | p.Arg119His | P32322 |
| HOMO SAPIENS | 3 | 1290 | 5uav | C | 251 | ARG | p.Arg251His | P32322 |
| HOMO SAPIENS | 7 | 1290 | 5uav | C | 266 | ARG | p.Arg266Gln | P32322 |
| HOMO SAPIENS | 0 | 9618 | 2r7e | A | 22 | GLY | p.Gly41Cys | P00451 |
| HOMO SAPIENS | 2 | 9618 | 2r7e | A | 113 | GLU | p.Glu132Asp | P00451 |
| HOMO SAPIENS | -1 | 9618 | 2r7e | A | 336 | ARG | p.Arg355Ter | P00451 |
| HOMO SAPIENS | 0 | 9618 | 2r7e | A | 372 | ARG | p.Arg391His | P00451 |
| HOMO SAPIENS | 0 | 9618 | 2r7e | A | 373 | SER | p.Ser392Leu | P00451 |
| HOMO SAPIENS | 0 | 9618 | 2r7e | A | 373 | SER | p.Ser392Pro | P00451 |
| HOMO SAPIENS | 3 | 9618 | 2r7e | A | 386 | ILE | p.Ile405Ser | P00451 |
| HOMO SAPIENS | 6 | 9618 | 2r7e | A | 390 | GLU | p.Glu409Gly | P00451 |
| HOMO SAPIENS | 2 | 9618 | 2r7e | A | 412 | LEU | p.Leu431Phe | P00451 |
| HOMO SAPIENS | 5 | 9618 | 2r7e | A | 425 | LYS | p.Lys444Arg | P00451 |
| HOMO SAPIENS | 0 | 9618 | 2r7e | A | 484 | ARG | p.Arg503His | P00451 |
| HOMO SAPIENS | 2 | 9618 | 2r7e | A | 531 | ARG | p.Arg550His | P00451 |
| HOMO SAPIENS | 3 | 9618 | 2r7e | A | 535 | SER | p.Ser554Gly | P00451 |
| HOMO SAPIENS | 2 | 9618 | 2r7e | A | 593 | ARG | p.Arg612Cys | P00451 |
| HOMO SAPIENS | 0 | 9618 | 2r7e | A | 704 | ALA | p.Ala723Thr | P00451 |
| HOMO SAPIENS | 0 | 9618 | 2r7e | A | 720 | GLU | p.Glu739Lys | P00451 |
| HOMO SAPIENS | 1 | 13085 | 3ud1 | A | 1075 | ILE | p.Thr1075Ile | P16157 |
| HOMO SAPIENS | -1 | 3289 | 2fy4 | A | 2 | ALA | p.Ala120Thr | P28329 |
| HOMO SAPIENS | 7 | 3289 | 2fy4 | A | 18 | VAL | p.Val136Met | P28329 |
| HOMO SAPIENS | 5 | 3289 | 2fy4 | A | 92 | LEU | p.Leu210Pro | P28329 |
| HOMO SAPIENS | 2 | 3289 | 2fy4 | A | 93 | PRO | p.Pro211Ala | P28329 |
| HOMO SAPIENS | 4 | 3289 | 2fy4 | A | 131 | LEU | p.Leu249Val | P28329 |

| | | | | | | | | |
|---|---|---|---|---|---|---|---|---|
| HOMO SAPIENS | 2 | 3289 | 2fy4 | A | 181 | PRO | p.Pro299Leu | P28329 |
| HOMO SAPIENS | 0 | 3289 | 2fy4 | A | 218 | ILE | p.Ile336Thr | P28329 |
| HOMO SAPIENS | 2 | 3289 | 2fy4 | A | 261 | ASP | p.Asp379His | P28329 |
| HOMO SAPIENS | 0 | 3289 | 2fy4 | A | 302 | ARG | p.Arg420Cys | P28329 |
| HOMO SAPIENS | 5 | 3289 | 2fy4 | A | 323 | GLU | p.Glu441Lys | P28329 |
| HOMO SAPIENS | 0 | 3289 | 2fy4 | A | 343 | MET | p.Val461Met | P28329 |
| HOMO SAPIENS | 2 | 3289 | 2fy4 | A | 364 | ARG | p.Arg482Gly | P28329 |
| HOMO SAPIENS | 3 | 3289 | 2fy4 | A | 380 | SER | p.Ser498Leu | P28329 |
| HOMO SAPIENS | 6 | 3289 | 2fy4 | A | 388 | VAL | p.Val506Leu | P28329 |
| HOMO SAPIENS | 7 | 3289 | 2fy4 | A | 429 | HIS | p.His547Gln | P28329 |
| HOMO SAPIENS | 8 | 3289 | 2fy4 | A | 442 | ARG | p.Arg560His | P28329 |
| HOMO SAPIENS | 3 | 3289 | 2fy4 | A | 443 | ARG | p.Arg561Gln | P28329 |
| HOMO SAPIENS | 0 | 4137 | 3hhr | B | 38 | CYS | p.Cys56Ter | P10912 |
| HOMO SAPIENS | 0 | 4137 | 3hhr | B | 43 | ARG | p.Arg61Ter | P10912 |
| HOMO SAPIENS | 0 | 4137 | 3hhr | B | 44 | GLU | p.Glu62Lys | P10912 |
| HOMO SAPIENS | 0 | 4137 | 3hhr | B | 96 | PHE | p.Phe114Ser | P10912 |
| HOMO SAPIENS | 5 | 4137 | 3hhr | B | 131 | PRO | p.Pro149Gln | P10912 |
| HOMO SAPIENS | 0 | 4137 | 3hhr | B | 144 | VAL | p.Val162Ile | P10912 |
| HOMO SAPIENS | 0 | 4137 | 3hhr | B | 150 | HIS | p.His168Gln | P10912 |
| HOMO SAPIENS | 0 | 4137 | 3hhr | B | 152 | ASP | p.Asp170His | P10912 |
| HOMO SAPIENS | 0 | 4137 | 3hhr | B | 153 | ILE | p.Ile171Thr | P10912 |
| HOMO SAPIENS | 0 | 4137 | 3hhr | B | 154 | GLN | p.Gln172Pro | P10912 |
| HOMO SAPIENS | 0 | 4137 | 3hhr | B | 155 | VAL | p.Val173Gly | P10912 |
| HOMO SAPIENS | 0 | 4137 | 3hhr | B | 217 | ARG | p.Arg235Ter | P10912 |
| HOMO SAPIENS | 5 | 4137 | 3hhr | B | 224 | GLU | p.Glu242Asp | P10912 |
| HOMO SAPIENS | 5 | 4137 | 3hhr | B | 224 | GLU | p.Glu242Ter | P10912 |
| HOMO SAPIENS | 6 | 2493 | 5bpb | A | 69 | HIS | p.His69Tyr | Q9ULV1 |
| HOMO SAPIENS | 6 | 2493 | 5bpb | A | 105 | MET | p.Met105Val | Q9ULV1 |
| HOMO SAPIENS | -1 | 2493 | 5bpb | A | 168 | PRO | p.Pro168Ser | Q9ULV1 |
| HOMO SAPIENS | 0 | 32123 | 4ccz | A | 314 | ASP | p.Asp314Asn | Q99707 |
| HOMO SAPIENS | 0 | 32123 | 4ccz | A | 345 | VAL | p.Val345Phe | Q99707 |
| HOMO SAPIENS | 1 | 32123 | 4ccz | A | 585 | ARG | p.Arg585Ter | Q99707 |
| HOMO SAPIENS | 4 | 4618 | 2ag4 | A | 26 | PRO | p.Pro55Leu | P17900 |
| HOMO SAPIENS | 4 | 4618 | 2ag4 | A | 30 | VAL | p.Ile59Val | P17900 |
| HOMO SAPIENS | 2 | 4618 | 2ag4 | A | 40 | VAL | p.Met69Val | P17900 |
| HOMO SAPIENS | 2 | 4618 | 2ag4 | A | 124 | VAL | p.Val153Ala | P17900 |
| HOMO SAPIENS | 8 | 4618 | 2ag4 | A | 140 | ARG | p.Arg169Pro | P17900 |
| HOMO SAPIENS | 5 | 1082 | 1hk0 | X | 14 | ARG | p.Arg15Cys | P07320 |
| HOMO SAPIENS | 7 | 1082 | 1hk0 | X | 23 | PRO | p.Pro24Thr | P07320 |
| HOMO SAPIENS | 6 | 1082 | 1hk0 | X | 36 | ARG | p.Arg37Ser | P07320 |
| HOMO SAPIENS | 5 | 1082 | 1hk0 | X | 55 | TYR | p.Tyr56Ter | P07320 |
| HOMO SAPIENS | 3 | 1082 | 1hk0 | X | 58 | ARG | p.Arg59His | P07320 |
| HOMO SAPIENS | 7 | 1082 | 1hk0 | X | 88 | HIS | p.His88Gln | P07320 |
| HOMO SAPIENS | 6 | 1082 | 1hk0 | X | 95 | ARG | p.Arg95Ser | P07320 |
| HOMO SAPIENS | 5 | 1082 | 1hk0 | X | 126 | VAL | p.Val126Met | P07320 |
| HOMO SAPIENS | 8 | 1082 | 1hk0 | X | 157 | TRP | p.Trp157Ter | P07320 |
| HOMO SAPIENS | 0 | 22879 | 3j0a | B | 644 | ILE | p.Ile644Phe | O60602 |
| HOMO SAPIENS | 8 | 6 | 2o23 | B | 122 | LEU | p.Leu122Val | Q99714 |
| HOMO SAPIENS | 1 | 6 | 2o23 | B | 130 | ARG | p.Arg130Cys | Q99714 |
| HOMO SAPIENS | 1 | 6 | 2o23 | B | 247 | ASN | p.Asn247Ser | Q99714 |
| HOMO SAPIENS | 0 | 6 | 2o23 | B | 249 | GLU | p.Glu249Gln | Q99714 |
| HOMO SAPIENS | 1 | 12910 | 4hl4 | A | 67 | ARG | p.Arg67Ter | Q96BZ9 |
| HOMO SAPIENS | 8 | 16105 | 1lmj | A | 7 | GLU | p.Glu1073Lys | P35555 |
| HOMO SAPIENS | 8 | 16105 | 1lmj | A | 8 | CYS | p.Cys1074Arg | P35555 |

| | | | | | | | | |
|---|---|---|---|---|---|---|---|---|
| HOMO SAPIENS | 9 | 16105 | 1lmj | A | 51 | CYS | p.Cys1117Tyr | P35555 |
| HOMO SAPIENS | 9 | 16105 | 1lmj | A | 63 | CYS | p.Cys1129Tyr | P35555 |
| HOMO SAPIENS | 9 | 16105 | 1lmj | A | 65 | ASN | p.Asn1131Tyr | P35555 |
| HOMO SAPIENS | 4 | 16105 | 1lmj | A | 71 | ARG | p.Arg1137Pro | P35555 |
| HOMO SAPIENS | 1 | 16105 | 1lmj | A | 82 | PRO | p.Pro1148Ala | P35555 |
| HOMO SAPIENS | 0 | 4795 | 3s5f | A | 106 | LEU | p.Leu106Ser | Q16595 |
| HOMO SAPIENS | 0 | 4795 | 3s5f | A | 130 | GLY | p.Gly130Val | Q16595 |
| HOMO SAPIENS | 0 | 4795 | 3s5f | A | 154 | ILE | p.Ile154Phe | Q16595 |
| HOMO SAPIENS | -1 | 4795 | 3s5f | A | 209 | ASP | p.Asp209Gly | Q16595 |
| HOMO SAPIENS | -1 | 2884 | 4lli | A | 1655 | ASP | p.Asp1655Glu | Q9Y4I1 |
| HOMO SAPIENS | 0 | 1786 | 7pck | C | 56P | TYR | p.Tyr71Ter | P43235 |
| HOMO SAPIENS | 0 | 1786 | 7pck | C | 64P | GLY | p.Gly79Glu | P43235 |
| HOMO SAPIENS | 7 | 38286 | 2l53 | B | 1914 | ARG | p.Arg1914Gly | Q14524 |
| HOMO SAPIENS | 1 | 38286 | 2l53 | B | 1924 | ALA | p.Ala1924Thr | Q14524 |
| HOMO SAPIENS | 0 | 31747 | 3ksy | A | 25 | PRO | p.Pro25Ser | Q07889 |
| HOMO SAPIENS | 3 | 31747 | 3ksy | A | 37 | THR | p.Thr37Ala | Q07889 |
| HOMO SAPIENS | 0 | 31747 | 3ksy | A | 54 | GLN | p.Gln54His | Q07889 |
| HOMO SAPIENS | -1 | 31747 | 3ksy | A | 190 | ASP | p.Asp190Glu | Q07889 |
| HOMO SAPIENS | 2 | 31747 | 3ksy | A | 266 | THR | p.Thr266Lys | Q07889 |
| HOMO SAPIENS | 1 | 31747 | 3ksy | A | 269 | MET | p.Met269Arg | Q07889 |
| HOMO SAPIENS | 1 | 31747 | 3ksy | A | 340 | PRO | p.Pro340Ser | Q07889 |
| HOMO SAPIENS | 1 | 31747 | 3ksy | A | 423 | ASN | p.Asn423Lys | Q07889 |
| HOMO SAPIENS | 1 | 31747 | 3ksy | A | 424 | GLU | p.Glu424Lys | Q07889 |
| HOMO SAPIENS | 1 | 31747 | 3ksy | A | 432 | TRP | p.Trp432Arg | Q07889 |
| HOMO SAPIENS | 1 | 31747 | 3ksy | A | 552 | ARG | p.Arg552Gly | Q07889 |
| HOMO SAPIENS | 1 | 31747 | 3ksy | A | 552 | ARG | p.Arg552Ser | Q07889 |
| HOMO SAPIENS | 3 | 7529 | 6jdw | A | 93 | ILE | p.Ile93Met | P50440 |
| HOMO SAPIENS | 2 | 7529 | 6jdw | A | 110 | GLN | p.Gln110His | P50440 |
| HOMO SAPIENS | 5 | 7529 | 6jdw | A | 149 | TRP | p.Trp149Ter | P50440 |
| HOMO SAPIENS | 0 | 7529 | 6jdw | A | 169 | ARG | p.Arg169Ter | P50440 |
| HOMO SAPIENS | 0 | 7529 | 6jdw | A | 260 | ARG | p.Arg260Ter | P50440 |
| HOMO SAPIENS | 3 | 7529 | 6jdw | A | 344 | ILE | p.Ile344Val | P50440 |
| HOMO SAPIENS | 5 | 7529 | 6jdw | A | 418 | LEU | p.Leu418Ile | P50440 |
| HOMO SAPIENS | 1 | 14788 | 1eak | A | 101 | ARG | p.Arg101His | P08253 |
| HOMO SAPIENS | 0 | 14788 | 1eak | A | 383 | ASP | p.Asp383Glu | P08253 |
| HOMO SAPIENS | 7 | 14788 | 1eak | A | 404 | GLN | p.Glu404Lys | P08253 |
| HOMO SAPIENS | 2 | 14788 | 1eak | A | 447 | ALA | p.Ala447Val | P08253 |
| HOMO SAPIENS | 0 | 1255 | 5xtc | p | 8 | PRO | p.Pro8His | Q9Y6M9 |
| HOMO SAPIENS | 0 | 1255 | 5xtc | p | 146 | PRO | p.Pro146Ser | Q9Y6M9 |
| HOMO SAPIENS | 3 | 6721 | 1a22 | A | 25 | PHE | p.Phe51Tyr | P01241 |
| HOMO SAPIENS | 2 | 6721 | 1a22 | A | 95 | SER | p.Ser121Arg | P01241 |
| HOMO SAPIENS | 2 | 6721 | 1a22 | A | 110 | VAL | p.Val136Ile | P01241 |
| HOMO SAPIENS | 0 | 1380 | 5i6z | A | 605 | LYS | p.Lys605Asn | P31645 |
| HOMO SAPIENS | 0 | 941 | 1qki | D | 48 | ILE | p.Ile48Thr | P11413 |
| HOMO SAPIENS | 3 | 941 | 1qki | D | 68 | VAL | p.Val68Met | P11413 |
| HOMO SAPIENS | 2 | 941 | 1qki | D | 126 | ASN | p.Asn126Asp | P11413 |
| HOMO SAPIENS | 8 | 941 | 1qki | D | 163 | GLY | p.Gly163Ser | P11413 |
| HOMO SAPIENS | 6 | 941 | 1qki | D | 165 | ASN | p.Asn165Asp | P11413 |
| HOMO SAPIENS | 4 | 941 | 1qki | D | 181 | ASP | p.Asp181Val | P11413 |
| HOMO SAPIENS | 4 | 941 | 1qki | D | 182 | ARG | p.Arg182Trp | P11413 |
| HOMO SAPIENS | 5 | 941 | 1qki | D | 188 | SER | p.Ser188Phe | P11413 |
| HOMO SAPIENS | 9 | 941 | 1qki | D | 198 | ARG | p.Arg198Cys | P11413 |
| HOMO SAPIENS | 8 | 941 | 1qki | D | 213 | VAL | p.Val213Leu | P11413 |
| HOMO SAPIENS | 5 | 941 | 1qki | D | 282 | ASP | p.Asp282His | P11413 |

| | | | | | | | | |
|---|---|---|---|---|---|---|---|---|
| HOMO SAPIENS | 7 | 941 | 1qki | D | 285 | ARG | p.Arg285His | P11413 |
| HOMO SAPIENS | 5 | 941 | 1qki | D | 291 | VAL | p.Val291Met | P11413 |
| HOMO SAPIENS | 8 | 941 | 1qki | D | 323 | LEU | p.Leu323Pro | P11413 |
| HOMO SAPIENS | 3 | 941 | 1qki | D | 335 | ALA | p.Ala335Thr | P11413 |
| HOMO SAPIENS | 6 | 941 | 1qki | D | 350 | ASP | p.Asp350His | P11413 |
| HOMO SAPIENS | 5 | 941 | 1qki | D | 393 | ARG | p.Arg393His | P11413 |
| HOMO SAPIENS | 5 | 941 | 1qki | D | 454 | ARG | p.Arg454Cys | P11413 |
| HOMO SAPIENS | 2 | 941 | 1qki | D | 459 | LEU | p.Arg459Leu | P11413 |
| HOMO SAPIENS | 4 | 941 | 1qki | D | 463 | ARG | p.Arg463His | P11413 |
| HOMO SAPIENS | 4 | 2483 | 1tij | A | 68 | LEU | p.Leu94Gln | P01034 |
| HOMO SAPIENS | 2 | 20632 | 3v5b | A | 331 | ARG | p.Arg331Pro | P02545 |
| HOMO SAPIENS | 2 | 20632 | 3v5b | A | 374 | HIS | p.His374Gln | P02545 |
| HOMO SAPIENS | 4 | 13929 | 5xtd | J | 75 | ARG | p.Arg75His | Q16795 |
| HOMO SAPIENS | 3 | 13929 | 5xtd | J | 243 | VAL | p.Val243Ala | Q16795 |
| HOMO SAPIENS | 1 | 13929 | 5xtd | J | 294 | PRO | p.Pro294Leu | Q16795 |
| HOMO SAPIENS | 2 | 13929 | 5xtd | J | 321 | ARG | p.Arg321Pro | Q16795 |
| HOMO SAPIENS | 0 | 9411 | 5thl | A | 41 | ARG | p.Gly41Arg | P54577 |
| HOMO SAPIENS | 0 | 9411 | 5thl | A | 196 | GLU | p.Glu196Lys | P54577 |
| HOMO SAPIENS | 0 | 9411 | 5thl | A | 316 | ALA | p.Ala316Ser | P54577 |
| HOMO SAPIENS | 0 | 9411 | 5thl | A | 339 | ALA | p.Ala339Thr | P54577 |
| HOMO SAPIENS | 4 | 16273 | 2e8n | A | 46 | THR | p.Thr940Ile | P29317 |
| HOMO SAPIENS | 7 | 16273 | 2e8n | A | 54 | GLY | p.Gly948Trp | P29317 |
| HOMO SAPIENS | 6 | 16273 | 2e8n | A | 65 | ALA | p.Ala959Thr | P29317 |
| HOMO SAPIENS | 7 | 30387 | 2k9y | A | 541 | VAL | p.Val541Met | P29317 |
| HOMO SAPIENS | 0 | 6961 | 4nha | A | 843 | ILE | p.Ile853Val | Q9UPY3 |
| HOMO SAPIENS | -1 | 6961 | 4nha | A | 862 | ALA | p.Ala872Thr | Q9UPY3 |
| HOMO SAPIENS | 3 | 15708 | 4pzg | A | 16 | ARG | p.Arg16Ter | Q9Y5X0 |
| HOMO SAPIENS | 4 | 15708 | 4pzg | A | 51 | ARG | p.Arg51Gln | Q9Y5X0 |
| HOMO SAPIENS | 3 | 30166 | 4o00 | A | 44 | ARG | p.Arg14257Ter | Q8WZ42 |
| HOMO SAPIENS | 2 | 30166 | 4o00 | A | 54 | ARG | p.Arg14267His | Q8WZ42 |
| HOMO SAPIENS | 8 | 11158 | 4hca | A | 275 | TRP | p.Trp274Arg | P23771 |
| HOMO SAPIENS | 2 | 4328 | 4e0s | A | 1476 | ARG | p.Arg1476Ter | P01031 |
| HOMO SAPIENS | 0 | 11462 | 1dx5 | I | 455 | ALA | p.Ala473Val | P07204 |
| HOMO SAPIENS | -1 | 11150 | 2fic | A | 45 | LYS | p.Lys35Asn | O00499 |
| HOMO SAPIENS | 6 | 11150 | 2fic | A | 161 | ASP | p.Asp151Asn | O00499 |
| HOMO SAPIENS | 8 | 27382 | 4mqv | B | 17 | ARG | p.Arg17Ter | Q9NZC9 |
| HOMO SAPIENS | -1 | 37991 | 1q0p | A | 231 | GLN | p.Gln256Ter | P00751 |
| HOMO SAPIENS | 1 | 443 | 5xte | U | 215 | LEU | p.Leu215Phe | P08574 |
| HOMO SAPIENS | 8 | 21471 | 5y4e | A | 1461 | ASN | p.Thr692Ala | Q01484 |
| HOMO SAPIENS | -1 | 35443 | 3um3 | B | 129 | ASP | p.Asp129Val | Q9H444 |
| HOMO SAPIENS | -1 | 35443 | 3um3 | B | 161 | GLU | p.Glu161Lys | Q9H444 |
| HOMO SAPIENS | 6 | 28781 | 2n2a | B | 111 | ILE | p.Ile654Val | P04626 |
| HOMO SAPIENS | 7 | 28781 | 2n2a | B | 112 | ILE | p.Ile655Val | P04626 |
| HOMO SAPIENS | -1 | 3346 | 4ztz | C | 61 | GLY | p.Gly61Arg | Q9UHN1 |
| HOMO SAPIENS | 4 | 3346 | 4ztz | C | 103 | GLY | p.Gly103Ser | Q9UHN1 |
| HOMO SAPIENS | 4 | 3346 | 4ztz | C | 133 | HIS | p.His133Tyr | Q9UHN1 |
| HOMO SAPIENS | -1 | 3346 | 4ztz | C | 153 | LEU | p.Leu153Val | Q9UHN1 |
| HOMO SAPIENS | -1 | 3346 | 4ztz | C | 169 | ALA | p.Ala169Thr | Q9UHN1 |
| HOMO SAPIENS | 7 | 3346 | 4ztz | C | 369 | ARG | p.Arg369Gly | Q9UHN1 |
| HOMO SAPIENS | 4 | 3346 | 4ztz | C | 386 | ASP | p.Asp386Glu | Q9UHN1 |
| HOMO SAPIENS | 4 | 3346 | 4ztz | C | 451 | GLY | p.Gly451Glu | Q9UHN1 |
| HOMO SAPIENS | 4 | 2131 | 5fo9 | E | 1549 | LEU | p.Leu1549Met | P01024 |
| HOMO SAPIENS | 2 | 2131 | 5fo9 | E | 1619 | SER | p.Ser1619Arg | P01024 |
| HOMO SAPIENS | 3 | 12662 | 4jhp | C | 75 | ILE | p.Ile75Val | Q92834 |

| | | | | | | | | |
|---|---|---|---|---|---|---|---|---|
| HOMO SAPIENS | 1 | 12662 | 4jhp | C | 173 | GLY | p.Gly173Arg | Q92834 |
| HOMO SAPIENS | 0 | 12662 | 4jhp | C | 184 | GLN | p.Gln184His | Q92834 |
| HOMO SAPIENS | 3 | 12662 | 4jhp | C | 262 | ALA | p.Ala262Gly | Q92834 |
| HOMO SAPIENS | 3 | 12662 | 4jhp | C | 302 | CYS | p.Cys302Tyr | Q92834 |
| HOMO SAPIENS | 5 | 1093 | 2jg9 | A | 186 | GLN | p.Gln208Ter | P02745 |
| HOMO SAPIENS | 0 | 13181 | 4r4l | A | 12 | MET | p.Met13Ile | Q13976 |
| HOMO SAPIENS | 8 | 25939 | 4iyp | A | 106 | ARG | p.Arg106Gln | P78318 |
| HOMO SAPIENS | 1 | 217 | 2xsx | A | 71 | SER | p.Asn71Ser | P13929 |
| HOMO SAPIENS | 0 | 217 | 2xsx | A | 76 | ALA | p.Ala76Pro | P13929 |
| HOMO SAPIENS | 1 | 217 | 2xsx | A | 85 | ALA | p.Val85Ala | P13929 |
| HOMO SAPIENS | 8 | 217 | 2xsx | A | 156 | GLY | p.Gly156Asp | P13929 |
| HOMO SAPIENS | 5 | 217 | 2xsx | A | 332 | VAL | p.Val332Ile | P13929 |
| HOMO SAPIENS | 7 | 217 | 2xsx | A | 374 | GLY | p.Gly374Glu | P13929 |
| HOMO SAPIENS | 1 | 5508 | 3szm | A | 682 | THR | p.Thr682Asn | Q8NEM0 |
| HOMO SAPIENS | 0 | 5508 | 3szm | A | 715 | TRP | p.Trp715Ter | Q8NEM0 |
| HOMO SAPIENS | 1 | 5508 | 3szm | A | 761 | VAL | p.Ala761Val | Q8NEM0 |
| HOMO SAPIENS | 1 | 5508 | 3szm | A | 828 | PRO | p.Pro828Ser | Q8NEM0 |
| HOMO SAPIENS | 4 | 5508 | 3szm | A | 829 | GLU | p.Glu829Asp | Q8NEM0 |
| HOMO SAPIENS | 2 | 16353 | 2q0z | X | 171 | GLY | p.Gly1978Ala | O75643 |
| HOMO SAPIENS | 2 | 16353 | 2q0z | X | 188 | ALA | p.Ala1995Thr | O75643 |
| HOMO SAPIENS | 0 | 4127 | 3swr | A | 860 | GLY | p.Gly860Arg | P26358 |
| HOMO SAPIENS | -1 | 4127 | 3swr | A | 1122 | SER | p.Ser1122Cys | P26358 |
| HOMO SAPIENS | 0 | 4127 | 3swr | A | 1216 | ASP | p.Asp1216Glu | P26358 |
| HOMO SAPIENS | 0 | 6382 | 3o96 | A | 17 | GLU | p.Glu17Lys | P31749 |
| HOMO SAPIENS | 6 | 14314 | 5xtc | Q | 71 | PRO | p.Pro71Leu | O75306 |
| HOMO SAPIENS | 2 | 5886 | 4fao | E | 22 | ARG | p.Arg40His | Q13705 |
| HOMO SAPIENS | 1 | 2534 | 3mp4 | A | 37 | GLU | p.Glu37Ter | P35914 |
| HOMO SAPIENS | 3 | 2534 | 3mp4 | A | 41 | MET | p.Arg41Gln | P35914 |
| HOMO SAPIENS | 1 | 2534 | 3mp4 | A | 70 | VAL | p.Val70Leu | P35914 |
| HOMO SAPIENS | 2 | 2196 | 3ifq | A | 142 | ARG | p.Arg142His | P14923 |
| HOMO SAPIENS | 2 | 2196 | 3ifq | A | 153 | ASP | p.Asp153Glu | P14923 |
| HOMO SAPIENS | 3 | 2196 | 3ifq | A | 176 | ARG | p.Arg176Gln | P14923 |
| HOMO SAPIENS | 3 | 2196 | 3ifq | A | 265 | ARG | p.Arg265His | P14923 |
| HOMO SAPIENS | 2 | 2196 | 3ifq | A | 301 | GLU | p.Glu301Gly | P14923 |
| HOMO SAPIENS | 0 | 2196 | 3ifq | A | 456 | VAL | p.Val456Ile | P14923 |
| HOMO SAPIENS | 3 | 2196 | 3ifq | A | 603 | VAL | p.Val603Leu | P14923 |
| HOMO SAPIENS | 0 | 2196 | 3ifq | A | 648 | VAL | p.Val648Ile | P14923 |
| HOMO SAPIENS | 1 | 8746 | 1cgj | I | 32 | PRO | p.Pro55Ser | P00995 |
| HOMO SAPIENS | 1 | 8746 | 1cgj | I | 44 | ARG | p.Arg67His | P00995 |
| HOMO SAPIENS | 1 | 8746 | 1cgj | I | 44 | ARG | p.Arg67Cys | P00995 |
| HOMO SAPIENS | 3 | 3212 | 1syq | A | 105 | ARG | p.Arg105Ter | P18206 |
| HOMO SAPIENS | 3 | 3212 | 1syq | A | 126 | PHE | p.Phe126Leu | P18206 |
| HOMO SAPIENS | 4 | 10028 | 3eto | A | 1609 | ARG | p.Arg1608His | P46531 |
| HOMO SAPIENS | 1 | 10028 | 3eto | A | 1672 | VAL | p.Val1671Ile | P46531 |
| HOMO SAPIENS | 5 | 10028 | 3eto | A | 1692 | GLN | p.Gln1691His | P46531 |
| HOMO SAPIENS | 3 | 3749 | 5xtc | Z | 70 | GLY | p.Gly70Ter | O43676 |
| HOMO SAPIENS | 2 | 17039 | 3brv | B | 57 | GLU | p.Glu57Lys | Q9Y6K9 |
| HOMO SAPIENS | 8 | 3589 | 2fh3 | A | 433 | VAL | p.Val460Leu | P06396 |
| HOMO SAPIENS | 1 | 3589 | 2fh3 | A | 454 | ARG | p.Arg481Cys | P06396 |
| HOMO SAPIENS | 4 | 3589 | 2fh3 | A | 498 | ARG | p.Arg525His | P06396 |
| HOMO SAPIENS | 2 | 3589 | 2fh3 | A | 520 | TYR | p.Tyr547Cys | P06396 |
| HOMO SAPIENS | 1 | 3589 | 2fh3 | A | 529 | GLY | p.Gly556Arg | P06396 |
| HOMO SAPIENS | 2 | 3589 | 2fh3 | A | 536 | THR | p.Thr563Ser | P06396 |
| HOMO SAPIENS | 2 | 3589 | 2fh3 | A | 579 | VAL | p.Val606Met | P06396 |

| | | | | | | | | |
|---|---|---|---|---|---|---|---|---|
| HOMO SAPIENS | 1 | 3589 | 2fh3 | A | 589 | THR | p.Thr616Met | P06396 |
| HOMO SAPIENS | 5 | 3589 | 2fh3 | A | 668 | THR | p.Thr695Met | P06396 |
| HOMO SAPIENS | 0 | 148 | 1lcl | A | 130 | ILE | p.Ile130Ser | Q05315 |
| HOMO SAPIENS | -1 | 24727 | 4fj3 | P | 237 | ALA | p.Ala237Thr | P04049 |
| HOMO SAPIENS | 0 | 24727 | 4fj3 | P | 257 | SER | p.Ser257Leu | P04049 |
| HOMO SAPIENS | 0 | 24727 | 4fj3 | P | 261 | PRO | p.Pro261Ala | P04049 |
| HOMO SAPIENS | 0 | 2949 | 5xtb | B | 77 | ARG | p.Arg77Gly | O00217 |
| HOMO SAPIENS | 0 | 2949 | 5xtb | B | 79 | PRO | p.Pro79Leu | O00217 |
| HOMO SAPIENS | 0 | 2949 | 5xtb | B | 100 | ALA | p.Ala100Val | O00217 |
| HOMO SAPIENS | 1 | 2949 | 5xtb | B | 115 | LYS | p.Lys115Glu | O00217 |
| HOMO SAPIENS | 0 | 2949 | 5xtb | B | 138 | ARG | p.Arg138His | O00217 |
| HOMO SAPIENS | 1 | 3959 | 5k5s | B | 25 | ARG | p.Arg25Ter | P41180 |
| HOMO SAPIENS | 4 | 3959 | 5k5s | B | 39 | PRO | p.Pro39Ala | P41180 |
| HOMO SAPIENS | 0 | 3959 | 5k5s | B | 47 | LYS | p.Lys47Asn | P41180 |
| HOMO SAPIENS | 2 | 3959 | 5k5s | B | 62 | ARG | p.Arg62Met | P41180 |
| HOMO SAPIENS | 2 | 3959 | 5k5s | B | 66 | ARG | p.Arg66Cys | P41180 |
| HOMO SAPIENS | 4 | 3959 | 5k5s | B | 90 | ASN | p.Asn90Thr | P41180 |
| HOMO SAPIENS | 1 | 3959 | 5k5s | B | 116 | ALA | p.Ala116Thr | P41180 |
| HOMO SAPIENS | 1 | 3959 | 5k5s | B | 118 | ASN | p.Asn118Lys | P41180 |
| HOMO SAPIENS | -1 | 3959 | 5k5s | B | 125 | LEU | p.Leu125Pro | P41180 |
| HOMO SAPIENS | -1 | 3959 | 5k5s | B | 127 | GLU | p.Glu127Ala | P41180 |
| HOMO SAPIENS | -1 | 3959 | 5k5s | B | 128 | PHE | p.Phe128Leu | P41180 |
| HOMO SAPIENS | -1 | 3959 | 5k5s | B | 131 | CYS | p.Cys131Trp | P41180 |
| HOMO SAPIENS | 1 | 3959 | 5k5s | B | 138 | THR | p.Thr138Met | P41180 |
| HOMO SAPIENS | 0 | 3959 | 5k5s | B | 143 | GLY | p.Gly143Glu | P41180 |
| HOMO SAPIENS | 5 | 3959 | 5k5s | B | 151 | THR | p.Thr151Met | P41180 |
| HOMO SAPIENS | 4 | 3959 | 5k5s | B | 180 | PHE | p.Phe180Cys | P41180 |
| HOMO SAPIENS | 0 | 3959 | 5k5s | B | 185 | ARG | p.Arg185Gln | P41180 |
| HOMO SAPIENS | 0 | 3959 | 5k5s | B | 185 | ARG | p.Arg185Ter | P41180 |
| HOMO SAPIENS | 2 | 3959 | 5k5s | B | 191 | GLU | p.Glu191Lys | P41180 |
| HOMO SAPIENS | 2 | 3959 | 5k5s | B | 227 | ARG | p.Arg227Leu | P41180 |
| HOMO SAPIENS | 2 | 3959 | 5k5s | B | 250 | GLU | p.Glu250Lys | P41180 |
| HOMO SAPIENS | 6 | 3959 | 5k5s | B | 297 | GLU | p.Glu297Lys | P41180 |
| HOMO SAPIENS | 1 | 3959 | 5k5s | B | 336 | LYS | p.Lys336Asn | P41180 |
| HOMO SAPIENS | 6 | 3959 | 5k5s | B | 429 | HIS | p.His429Tyr | P41180 |
| HOMO SAPIENS | 0 | 3959 | 5k5s | B | 445 | THR | p.Thr445Ala | P41180 |
| HOMO SAPIENS | 3 | 3959 | 5k5s | B | 509 | GLY | p.Gly509Arg | P41180 |
| HOMO SAPIENS | 1 | 3959 | 5k5s | B | 553 | GLY | p.Gly553Arg | P41180 |
| HOMO SAPIENS | 3 | 3959 | 5k5s | B | 592 | ASN | p.Asn592Ser | P41180 |
| HOMO SAPIENS | -1 | 3959 | 5k5s | B | 604 | GLU | p.Glu604Lys | P41180 |
| HOMO SAPIENS | 4 | 2564 | 5l0c | C | 975 | ARG | p.Arg975Trp | P18206 |
| HOMO SAPIENS | 4 | 2564 | 5l0c | C | 975 | ARG | p.Arg975Leu | P18206 |
| HOMO SAPIENS | 1 | 5695 | 5i8r | A | 294 | ARG | p.Arg294Gln | P17405 |
| HOMO SAPIENS | 0 | 5695 | 5i8r | A | 485 | ALA | p.Ala485Val | P17405 |
| HOMO SAPIENS | 1 | 5695 | 5i8r | A | 506 | GLY | p.Gly506Arg | P17405 |
| HOMO SAPIENS | -1 | 1978 | 3avs | A | 1060 | HIS | p.His1060Leu | O15550 |
| HOMO SAPIENS | 0 | 11173 | 4v0c | B | 362 | LYS | p.Lys362Arg | P51787 |
| HOMO SAPIENS | 3 | 11173 | 4v0c | B | 381 | CYS | p.Cys381Tyr | P51787 |
| HOMO SAPIENS | 7 | 11173 | 4v0c | B | 518 | ARG | p.Arg518Gly | P51787 |
| HOMO SAPIENS | 2 | 11173 | 4v0c | B | 525 | ALA | p.Ala525Thr | P51787 |
| HOMO SAPIENS | 0 | 11173 | 4v0c | B | 530 | GLN | p.Gln530Ter | P51787 |
| HOMO SAPIENS | -1 | 11173 | 4v0c | B | 536 | TYR | p.Tyr536Ter | P51787 |
| HOMO SAPIENS | 2 | 1113 | 5xtd | N | 60 | ARG | p.Arg60Ter | Q9UI09 |
| HOMO SAPIENS | 3 | 6061 | 2mmz | A | 148 | LEU | p.Leu148Ser | Q86SX6 |

| | | | | | | | | |
|---|---|---|---|---|---|---|---|---|
| HOMO SAPIENS | -1 | 26081 | 3eu7 | A | 837 | GLU | p.Glu837Ter | Q86YC2 |
| HOMO SAPIENS | 0 | 26081 | 3eu7 | A | 864 | PRO | p.Pro864Ser | Q86YC2 |
| HOMO SAPIENS | 5 | 26081 | 3eu7 | A | 891 | CYS | p.Cys891Trp | Q86YC2 |
| HOMO SAPIENS | 0 | 26081 | 3eu7 | A | 932 | VAL | p.Val932Met | Q86YC2 |
| HOMO SAPIENS | 5 | 26081 | 3eu7 | A | 961 | LEU | p.Leu961Met | Q86YC2 |
| HOMO SAPIENS | 5 | 26081 | 3eu7 | A | 1038 | TRP | p.Trp1038Ter | Q86YC2 |
| HOMO SAPIENS | -1 | 26081 | 3eu7 | A | 1086 | ARG | p.Arg1086Ter | Q86YC2 |
| HOMO SAPIENS | 0 | 26081 | 3eu7 | A | 1122 | ASP | p.Asp1122Glu | Q86YC2 |
| HOMO SAPIENS | 5 | 26081 | 3eu7 | A | 1158 | HIS | p.His1158Gln | Q86YC2 |
| HOMO SAPIENS | 5 | 26081 | 3eu7 | A | 1183 | TYR | p.Tyr1183Ter | Q86YC2 |
| HOMO SAPIENS | 4 | 199 | 1qo5 | C | 45 | ARG | p.Arg46Trp | P05062 |
| HOMO SAPIENS | 5 | 199 | 1qo5 | C | 147 | TRP | p.Trp148Arg | P05062 |
| HOMO SAPIENS | 0 | 199 | 1qo5 | C | 149 | ALA | p.Ala150Pro | P05062 |
| HOMO SAPIENS | 0 | 199 | 1qo5 | C | 174 | ALA | p.Ala175Asp | P05062 |
| HOMO SAPIENS | 0 | 199 | 1qo5 | C | 334 | ASN | p.Asn335Lys | P05062 |
| HOMO SAPIENS | 0 | 199 | 1qo5 | C | 337 | ALA | p.Ala338Val | P05062 |
| HOMO SAPIENS | 5 | 199 | 1qo5 | C | 342 | TYR | p.Tyr343His | P05062 |
| HOMO SAPIENS | 0 | 6910 | 4zht | B | 59 | ARG | p.Arg59Ter | Q9Y223 |
| HOMO SAPIENS | 4 | 6910 | 4zht | B | 129 | ARG | p.Arg129Ter | Q9Y223 |
| HOMO SAPIENS | 4 | 6910 | 4zht | B | 129 | ARG | p.Arg129Gln | Q9Y223 |
| HOMO SAPIENS | 1 | 6910 | 4zht | B | 162 | ARG | p.Arg162Cys | Q9Y223 |
| HOMO SAPIENS | 0 | 6910 | 4zht | B | 176 | ASP | p.Asp176Val | Q9Y223 |
| HOMO SAPIENS | 0 | 6910 | 4zht | B | 216 | VAL | p.Val216Ala | Q9Y223 |
| HOMO SAPIENS | 0 | 6910 | 4zht | B | 246 | ARG | p.Arg246Trp | Q9Y223 |
| HOMO SAPIENS | 0 | 6910 | 4zht | B | 263 | ARG | p.Arg263Leu | Q9Y223 |
| HOMO SAPIENS | 0 | 6910 | 4zht | B | 266 | ARG | p.Arg266Gln | Q9Y223 |
| HOMO SAPIENS | 0 | 6910 | 4zht | B | 266 | ARG | p.Arg266Trp | Q9Y223 |
| HOMO SAPIENS | 0 | 6910 | 4zht | B | 277 | ARG | p.Arg277Cys | Q9Y223 |
| HOMO SAPIENS | 0 | 6910 | 4zht | B | 378 | ASP | p.Asp378Tyr | Q9Y223 |
| HOMO SAPIENS | 3 | 211 | 5kxi | C | 262 | VAL | p.Val287Leu | P17787 |
| HOMO SAPIENS | 0 | 211 | 5kxi | C | 369 | PHE | p.Phe478Leu | P17787 |
| HOMO SAPIENS | 0 | 2454 | 5iy9 | V | null | PHE | p.Phe99Ser | P19447 |
| HOMO SAPIENS | 0 | 2454 | 5iy9 | V | null | ARG | p.Arg109Ter | P19447 |
| HOMO SAPIENS | 0 | 2454 | 5iy9 | V | null | THR | p.Thr119Pro | P19447 |
| HOMO SAPIENS | 0 | 2454 | 5iy9 | V | 425 | ARG | p.Arg425Ter | P19447 |
| HOMO SAPIENS | 1 | 2454 | 5iy9 | V | 545 | GLN | p.Gln545Ter | P19447 |
| HOMO SAPIENS | -1 | 3805 | 3rbq | A | 57 | LYS | p.Lys57Ter | Q13432 |
| HOMO SAPIENS | 2 | 2649 | 4ido | A | 43 | ASP | p.Asp43Glu | Q8WXF7 |
| HOMO SAPIENS | 2 | 2649 | 4ido | A | 157 | LEU | p.Leu157Trp | Q8WXF7 |
| HOMO SAPIENS | 2 | 2649 | 4ido | A | 193 | PHE | p.Phe193Cys | Q8WXF7 |
| HOMO SAPIENS | 7 | 2649 | 4ido | A | 217 | ARG | p.Arg217Gln | Q8WXF7 |
| HOMO SAPIENS | 4 | 2649 | 4ido | A | 239 | ARG | p.Arg239Cys | Q8WXF7 |
| HOMO SAPIENS | 3 | 2649 | 4ido | A | 258 | HIS | p.His258Arg | Q8WXF7 |
| HOMO SAPIENS | 4 | 2649 | 4ido | A | 259 | SER | p.Ser259Tyr | Q8WXF7 |
| HOMO SAPIENS | 4 | 2649 | 4ido | A | 408 | MET | p.Met408Val | Q8WXF7 |
| HOMO SAPIENS | 2 | 2649 | 4ido | A | 415 | ARG | p.Arg415Trp | Q8WXF7 |
| HOMO SAPIENS | 3 | 2649 | 4ido | A | 416 | ARG | p.Arg416Cys | Q8WXF7 |
| HOMO SAPIENS | 5 | 2203 | 3wff | A | 810 | LEU | p.Ser810Leu | P08235 |
| HOMO SAPIENS | 4 | 12648 | 5ep6 | B | 696 | GLU | p.Glu696Lys | Q9UHD2 |
| HOMO SAPIENS | 0 | 30622 | 2pe4 | A | 90 | GLU | p.Glu90Asp | Q12794 |
| HOMO SAPIENS | 0 | 30622 | 2pe4 | A | 256 | GLY | p.Gly256Arg | Q12794 |
| HOMO SAPIENS | 1 | 30622 | 2pe4 | A | 268 | GLU | p.Glu268Lys | Q12794 |
| HOMO SAPIENS | 0 | 30622 | 2pe4 | A | 392 | THR | p.Thr392Met | Q12794 |
| HOMO SAPIENS | 5 | 5411 | 1lya | A | 93 | SER | p.Ser157Leu | P07339 |

| | | | | | | | | |
|---|---|---|---|---|---|---|---|---|
| HOMO SAPIENS | 3 | 5743 | 2uzp | A | 189 | ASN | p.Asn189Lys | P05129 |
| HOMO SAPIENS | 1 | 12529 | 3kfd | J | 21 | ASN | p.Asn45Ser | P36897 |
| HOMO SAPIENS | 1 | 1063 | 2e1q | B | 149 | ARG | p.Arg149Cys | P47989 |
| HOMO SAPIENS | -1 | 1063 | 2e1q | B | 172 | GLY | p.Gly172Arg | P47989 |
| HOMO SAPIENS | 4 | 1063 | 2e1q | B | 228 | ARG | p.Arg228Ter | P47989 |
| HOMO SAPIENS | 5 | 1063 | 2e1q | B | 395 | LYS | p.Lys395Met | P47989 |
| HOMO SAPIENS | 2 | 1063 | 2e1q | B | 646 | ILE | p.Ile646Val | P47989 |
| HOMO SAPIENS | 4 | 1063 | 2e1q | B | 703 | ILE | p.Ile703Val | P47989 |
| HOMO SAPIENS | 8 | 2525 | 1kqn | F | 18 | ASN | p.Asn18Ser | Q9HAN9 |
| HOMO SAPIENS | 0 | 2525 | 1kqn | F | 151 | VAL | p.Val151Phe | Q9HAN9 |
| HOMO SAPIENS | 0 | 2525 | 1kqn | F | 153 | LEU | p.Leu153Val | Q9HAN9 |
| HOMO SAPIENS | 0 | 2525 | 1kqn | F | 169 | TRP | p.Trp169Ter | Q9HAN9 |
| HOMO SAPIENS | 0 | 2525 | 1kqn | F | 207 | ARG | p.Arg207Trp | Q9HAN9 |
| HOMO SAPIENS | 0 | 2525 | 1kqn | F | 273 | ASN | p.Asn273Asp | Q9HAN9 |
| HOMO SAPIENS | 0 | 12299 | 5duq | A | 432 | VAL | p.Val454Glu | P05155 |
| HOMO SAPIENS | 0 | 12299 | 5duq | A | 436 | ALA | p.Ala458Thr | P05155 |
| HOMO SAPIENS | 0 | 12299 | 5duq | A | 443 | ALA | p.Ala465Val | P05155 |
| HOMO SAPIENS | 0 | 12299 | 5duq | A | 444 | ARG | p.Arg466His | P05155 |
| HOMO SAPIENS | 0 | 12299 | 5duq | A | 458 | MET | p.Val480Met | P05155 |
| HOMO SAPIENS | 0 | 327 | 1uhp | A | 36 | ASP | p.Asp274Asn | Q9UPQ7 |
| HOMO SAPIENS | 0 | 64 | 2mo5 | A | 54 | ALA | p.Ala55Thr | P12104 |
| HOMO SAPIENS | 2 | 35969 | 1jrh | I | 60 | CYS | p.Cys77Tyr | P15260 |
| HOMO SAPIENS | 4 | 35969 | 1jrh | I | 70 | ILE | p.Ile87Thr | P15260 |
| HOMO SAPIENS | 2 | 1068 | 1mu7 | A | 304 | ARG | p.Arg304Gln | Q9NUW8 |
| HOMO SAPIENS | 8 | 1068 | 1mu7 | A | 493 | HIS | p.His493Arg | Q9NUW8 |
| HOMO SAPIENS | 0 | 32360 | 2kqp | A | 31 | ARG | p.Arg55Cys | P01308 |
| HOMO SAPIENS | 0 | 32360 | 2kqp | A | 65 | ARG | p.Arg89His | P01308 |
| HOMO SAPIENS | 0 | 32360 | 2kqp | A | 65 | ARG | p.Arg89Cys | P01308 |
| HOMO SAPIENS | 6 | 10805 | 3nks | A | 12 | ILE | p.Ile12Thr | P50336 |
| HOMO SAPIENS | 0 | 10805 | 3nks | A | 20 | HIS | p.His20Pro | P50336 |
| HOMO SAPIENS | 0 | 10805 | 3nks | A | 59 | ARG | p.Arg59Trp | P50336 |
| HOMO SAPIENS | 0 | 10805 | 3nks | A | 168 | ARG | p.Arg168Cys | P50336 |
| HOMO SAPIENS | 0 | 10805 | 3nks | A | 168 | ARG | p.Arg168His | P50336 |
| HOMO SAPIENS | 2 | 10805 | 3nks | A | 232 | GLY | p.Gly232Arg | P50336 |
| HOMO SAPIENS | 3 | 10805 | 3nks | A | 256 | PRO | p.Pro256Arg | P50336 |
| HOMO SAPIENS | 0 | 10805 | 3nks | A | 304 | ARG | p.Arg304His | P50336 |
| HOMO SAPIENS | 0 | 10805 | 3nks | A | 349 | ASP | p.Asp349Ala | P50336 |
| HOMO SAPIENS | 9 | 7008 | 3u2l | A | 125 | GLN | p.Gln125Ter | P55789 |
| HOMO SAPIENS | 4 | 7008 | 3u2l | A | 172 | HIS | p.His172Asn | P55789 |
| HOMO SAPIENS | 4 | 7008 | 3u2l | A | 194 | ARG | p.Arg194His | P55789 |
| HOMO SAPIENS | 3 | 7008 | 3u2l | A | 198 | GLY | p.Gly198Ser | P55789 |
| HOMO SAPIENS | 0 | 20821 | 3e0g | A | 65 | HIS | p.His116Tyr | P42702 |
| HOMO SAPIENS | 3 | 1151 | 5l9v | A | 278 | ASP | p.Asp278Asn | Q9GZT9 |
| HOMO SAPIENS | 4 | 1151 | 5l9v | A | 317 | CYS | p.Pro317Arg | Q9GZT9 |
| HOMO SAPIENS | 5 | 1151 | 5l9v | A | 371 | ARG | p.Arg371His | Q9GZT9 |
| HOMO SAPIENS | 2 | 3935 | 1gwb | A | 36 | ARG | p.Arg36Gly | P07359 |
| HOMO SAPIENS | 1 | 3935 | 1gwb | A | 73 | LEU | p.Leu73Phe | P07359 |
| HOMO SAPIENS | 1 | 3935 | 1gwb | A | 172 | ALA | p.Ala172Val | P07359 |
| HOMO SAPIENS | 0 | 3935 | 1gwb | A | 249 | GLY | p.Gly249Val | P07359 |
| HOMO SAPIENS | 0 | 3935 | 1gwb | A | 255 | MET | p.Met255Val | P07359 |
| HOMO SAPIENS | 2 | 14541 | 5cuz | A | 138 | GLU | p.Glu138Lys | Q9H3L0 |
| HOMO SAPIENS | 2 | 14541 | 5cuz | A | 143 | SER | p.Ser143Ile | Q9H3L0 |
| HOMO SAPIENS | 4 | 14541 | 5cuz | A | 182 | THR | p.Thr182Asn | Q9H3L0 |
| HOMO SAPIENS | 6 | 14541 | 5cuz | A | 206 | ASN | p.Asn206Ile | Q9H3L0 |

| | | | | | | | | |
|---|---|---|---|---|---|---|---|---|
| HOMO SAPIENS | 0 | 14541 | 5cuz | A | 249 | TYR | p.Tyr249Cys | Q9H3L0 |
| HOMO SAPIENS | 4 | 14541 | 5cuz | A | 250 | ARG | p.Arg250Ter | Q9H3L0 |
| HOMO SAPIENS | 2 | 14541 | 5cuz | A | 259 | LEU | p.Leu259Pro | Q9H3L0 |
| HOMO SAPIENS | 0 | 4206 | 4fao | D | 30 | PRO | p.Pro30Ser | P37023 |
| HOMO SAPIENS | 1 | 4206 | 4fao | D | 50 | TRP | p.Trp50Cys | P37023 |
| HOMO SAPIENS | 0 | 16553 | 3v30 | A | 251 | GLN | p.Gln251Glu | O14593 |
| HOMO SAPIENS | 0 | 5012 | 5hbf | A | 442 | ALA | p.Ala442Gly | Q13231 |
| HOMO SAPIENS | 5 | 4844 | 5tdc | A | 136 | HIS | p.His136Arg | Q8IWV7 |
| HOMO SAPIENS | 0 | 15435 | 4cgz | A | 643 | ARG | p.Arg643His | P54132 |
| HOMO SAPIENS | 1 | 15435 | 4cgz | A | 672 | GLN | p.Gln672Arg | P54132 |
| HOMO SAPIENS | 1 | 15435 | 4cgz | A | 700 | GLN | p.Gln700Ter | P54132 |
| HOMO SAPIENS | 0 | 15435 | 4cgz | A | 707 | PRO | p.Pro707Ser | P54132 |
| HOMO SAPIENS | 0 | 15435 | 4cgz | A | 755 | LYS | p.Lys755Glu | P54132 |
| HOMO SAPIENS | 0 | 15435 | 4cgz | A | 881 | TRP | p.Trp881Ter | P54132 |
| HOMO SAPIENS | 2 | 15435 | 4cgz | A | 899 | ARG | p.Arg899Ter | P54132 |
| HOMO SAPIENS | 0 | 15435 | 4cgz | A | 1036 | CYS | p.Cys1036Phe | P54132 |
| HOMO SAPIENS | 0 | 15435 | 4cgz | A | 1055 | CYS | p.Cys1055Ser | P54132 |
| HOMO SAPIENS | 0 | 15435 | 4cgz | A | 1139 | ARG | p.Arg1139Ter | P54132 |
| HOMO SAPIENS | -1 | 15435 | 4cgz | A | 1198 | VAL | p.Val1198Met | P54132 |
| HOMO SAPIENS | -1 | 15435 | 4cgz | A | 1205 | VAL | p.Val1205Ile | P54132 |
| HOMO SAPIENS | 0 | 15435 | 4cgz | A | 1251 | GLU | p.Glu1251Gln | P54132 |
| HOMO SAPIENS | 0 | 11185 | 1kb4 | A | 30 | ARG | p.Arg30Ter | P11473 |
| HOMO SAPIENS | 5 | 11185 | 1kb4 | A | 33 | GLY | p.Gly33Asp | P11473 |
| HOMO SAPIENS | 5 | 11185 | 1kb4 | A | 46 | GLY | p.Gly46Asp | P11473 |
| HOMO SAPIENS | 6 | 11185 | 1kb4 | A | 50 | ARG | p.Arg50Gln | P11473 |
| HOMO SAPIENS | 9 | 11185 | 1kb4 | A | 73 | ARG | p.Arg73Gln | P11473 |
| HOMO SAPIENS | 9 | 11185 | 1kb4 | A | 80 | ARG | p.Arg80Gln | P11473 |
| HOMO SAPIENS | 0 | 5382 | 3fl7 | A | 41 | LEU | p.Leu41Val | P29317 |
| HOMO SAPIENS | 2 | 5382 | 3fl7 | A | 175 | ARG | p.Arg175Cys | P29317 |
| HOMO SAPIENS | 1 | 5382 | 3fl7 | A | 195 | ARG | p.Arg195Cys | P29317 |
| HOMO SAPIENS | 0 | 5382 | 3fl7 | A | 232 | ASP | p.Asp232Gly | P29317 |
| HOMO SAPIENS | 0 | 5382 | 3fl7 | A | 277 | SER | p.Ser277Leu | P29317 |
| HOMO SAPIENS | 0 | 5382 | 3fl7 | A | 315 | ARG | p.Arg315Gln | P29317 |
| HOMO SAPIENS | 7 | 5382 | 3fl7 | A | 350 | PRO | p.Pro350Thr | P29317 |
| HOMO SAPIENS | 0 | 5382 | 3fl7 | A | 391 | GLY | p.Gly391Arg | P29317 |
| HOMO SAPIENS | 0 | 5382 | 3fl7 | A | 418 | VAL | p.Val418Ile | P29317 |
| HOMO SAPIENS | 0 | 5382 | 3fl7 | A | 511 | THR | p.Thr511Met | P29317 |
| HOMO SAPIENS | 2 | 29168 | 4p22 | B | 144 | VAL | p.Val144Ile | P22314 |
| HOMO SAPIENS | 5 | 4348 | 1bf5 | A | 241 | ARG | p.Arg241Gln | P42224 |
| HOMO SAPIENS | 2 | 4348 | 1bf5 | A | 266 | VAL | p.Val266Ile | P42224 |
| HOMO SAPIENS | 4 | 4348 | 1bf5 | A | 320 | GLU | p.Glu320Gln | P42224 |
| HOMO SAPIENS | 1 | 4348 | 1bf5 | A | 372 | ASN | p.Asn372Lys | P42224 |
| HOMO SAPIENS | 1 | 4348 | 1bf5 | A | 447 | ASP | p.Asp447Glu | P42224 |
| HOMO SAPIENS | 2 | 4348 | 1bf5 | A | 463 | GLN | p.Gln463His | P42224 |
| HOMO SAPIENS | 4 | 4348 | 1bf5 | A | 600 | LEU | p.Leu600Pro | P42224 |
| HOMO SAPIENS | 0 | 4348 | 1bf5 | A | 706 | LEU | p.Leu706Ser | P42224 |
| HOMO SAPIENS | 4 | 4991 | 4bej | B | 141 | ASN | p.Asn141Ser | O00429 |
| HOMO SAPIENS | 0 | 4991 | 4bej | B | 395 | ALA | p.Ala395Asp | O00429 |
| HOMO SAPIENS | 2 | 11792 | 1fvo | A | 40 | ARG | p.Arg40Cys | P00480 |
| HOMO SAPIENS | 2 | 11792 | 1fvo | A | 40 | ARG | p.Arg40His | P00480 |
| HOMO SAPIENS | 0 | 11792 | 1fvo | A | 45 | LEU | p.Leu45Pro | P00480 |
| HOMO SAPIENS | 1 | 11792 | 1fvo | A | 47 | ASN | p.Asn47Ile | P00480 |
| HOMO SAPIENS | 0 | 11792 | 1fvo | A | 50 | GLY | p.Gly50Arg | P00480 |
| HOMO SAPIENS | 5 | 11792 | 1fvo | A | 87 | GLU | p.Glu87Lys | P00480 |

| | | | | | | | | |
|---|---|---|---|---|---|---|---|---|
| HOMO SAPIENS | 3 | 11792 | 1fvo | A | 111 | LEU | p.Leu111Pro | P00480 |
| HOMO SAPIENS | 1 | 11792 | 1fvo | A | 125 | THR | p.Thr125Met | P00480 |
| HOMO SAPIENS | 3 | 11792 | 1fvo | A | 129 | ARG | p.Arg129His | P00480 |
| HOMO SAPIENS | 9 | 11792 | 1fvo | A | 141 | ARG | p.Arg141Pro | P00480 |
| HOMO SAPIENS | 0 | 11792 | 1fvo | A | 206 | MET | p.Met206Arg | P00480 |
| HOMO SAPIENS | 0 | 11792 | 1fvo | A | 207 | SER | p.Ser207Arg | P00480 |
| HOMO SAPIENS | 0 | 11792 | 1fvo | A | 216 | GLN | p.Gln216Glu | P00480 |
| HOMO SAPIENS | 1 | 11792 | 1fvo | A | 225 | PRO | p.Pro225Leu | P00480 |
| HOMO SAPIENS | 0 | 11792 | 1fvo | A | 270 | ARG | p.Gln270Arg | P00480 |
| HOMO SAPIENS | 0 | 11792 | 1fvo | A | 270 | ARG | p.Gln270Ter | P00480 |
| HOMO SAPIENS | 2 | 11792 | 1fvo | A | 277 | ARG | p.Arg277Trp | P00480 |
| HOMO SAPIENS | 0 | 3721 | 4r3z | B | 2 | ASP | p.Asp2Gly | P54136 |
| HOMO SAPIENS | 0 | 3721 | 4r3z | B | 456 | SER | p.Ser456Leu | P54136 |
| HOMO SAPIENS | 5 | 3721 | 4r3z | B | 512 | ARG | p.Arg512Gln | P54136 |
| HOMO SAPIENS | 3 | 5749 | 3hl2 | A | 239 | ALA | p.Ala239Thr | Q9HD40 |
| HOMO SAPIENS | 1 | 5749 | 3hl2 | A | 334 | TYR | p.Tyr334Cys | Q9HD40 |
| HOMO SAPIENS | 1 | 5749 | 3hl2 | A | 452 | LYS | p.Lys452Asn | Q9HD40 |
| HOMO SAPIENS | -1 | 5749 | 3hl2 | A | 465 | ARG | p.Arg465Leu | Q9HD40 |
| HOMO SAPIENS | -1 | 5749 | 3hl2 | A | 489 | ASP | p.Asp489Gly | Q9HD40 |
| HOMO SAPIENS | 5 | 1721 | 3k15 | A | 1652 | MET | p.Met1652Ile | P38398 |
| HOMO SAPIENS | 5 | 1721 | 3k15 | A | 1652 | MET | p.Met1652Lys | P38398 |
| HOMO SAPIENS | 5 | 1721 | 3k15 | A | 1662 | PHE | p.Phe1662Cys | P38398 |
| HOMO SAPIENS | 5 | 1721 | 3k15 | A | 1664 | LEU | p.Leu1664Pro | P38398 |
| HOMO SAPIENS | 0 | 1721 | 3k15 | A | 1665 | VAL | p.Val1665Met | P38398 |
| HOMO SAPIENS | 5 | 1721 | 3k15 | A | 1690 | LYS | p.Lys1690Gln | P38398 |
| HOMO SAPIENS | 5 | 1721 | 3k15 | A | 1692 | ASP | p.Asp1692Asn | P38398 |
| HOMO SAPIENS | 5 | 1721 | 3k15 | A | 1699 | ARG | p.Arg1699Gln | P38398 |
| HOMO SAPIENS | 5 | 1721 | 3k15 | A | 1699 | ARG | p.Arg1699Trp | P38398 |
| HOMO SAPIENS | 3 | 1721 | 3k15 | A | 1706 | GLY | p.Gly1706Glu | P38398 |
| HOMO SAPIENS | 5 | 1721 | 3k15 | A | 1720 | THR | p.Thr1720Ala | P38398 |
| HOMO SAPIENS | 5 | 1721 | 3k15 | A | 1751 | ARG | p.Arg1751Pro | P38398 |
| HOMO SAPIENS | 4 | 1721 | 3k15 | A | 1775 | MET | p.Met1775Arg | P38398 |
| HOMO SAPIENS | 2 | 1721 | 3k15 | A | 1788 | GLY | p.Gly1788Val | P38398 |
| HOMO SAPIENS | 5 | 1721 | 3k15 | A | 1794 | GLU | p.Glu1794Asp | P38398 |
| HOMO SAPIENS | 5 | 1721 | 3k15 | A | 1804 | VAL | p.Val1804Asp | P38398 |
| HOMO SAPIENS | 3 | 1721 | 3k15 | A | 1815 | TRP | p.Trp1815Ter | P38398 |
| HOMO SAPIENS | 1 | 1721 | 3k15 | A | 1819 | ASN | p.Asn1819Ser | P38398 |
| HOMO SAPIENS | 5 | 1721 | 3k15 | A | 1856 | PRO | p.Pro1856Ser | P38398 |
| HOMO SAPIENS | 5 | 1721 | 3k15 | A | 1858 | ILE | p.Ile1858Leu | P38398 |
| HOMO SAPIENS | 5 | 1721 | 3k15 | A | 1858 | ILE | p.Ile1858Thr | P38398 |
| HOMO SAPIENS | 5 | 1721 | 3k15 | A | 1859 | PRO | p.Pro1859Arg | P38398 |
| HOMO SAPIENS | 2 | 22086 | 3kq4 | B | 1297 | PRO | p.Pro1297Leu | O60494 |
| HOMO SAPIENS | 1 | 12135 | 2k9j | B | 724 | ARG | p.Arg750Ter | P05106 |
| HOMO SAPIENS | -1 | 24027 | 2qtl | A | 175 | LEU | p.Ser202Leu | Q9UBK8 |
| HOMO SAPIENS | 0 | 24027 | 2qtl | A | 257 | SER | p.Ser284Thr | Q9UBK8 |
| HOMO SAPIENS | 0 | 24027 | 2qtl | A | 333 | LEU | p.Leu360Val | Q9UBK8 |
| HOMO SAPIENS | 0 | 24027 | 2qtl | A | 350 | LYS | p.Lys377Arg | Q9UBK8 |
| HOMO SAPIENS | 0 | 24027 | 2qtl | A | 415 | ARG | p.Arg442Cys | Q9UBK8 |
| HOMO SAPIENS | 2 | 24027 | 2qtl | A | 450 | PRO | p.Pro477Arg | Q9UBK8 |
| HOMO SAPIENS | 4 | 24027 | 2qtl | A | 487 | GLY | p.Gly514Arg | Q9UBK8 |
| HOMO SAPIENS | 1 | 24027 | 2qtl | A | 595 | HIS | p.His622Tyr | Q9UBK8 |
| HOMO SAPIENS | 0 | 5758 | 3di2 | B | 46 | THR | p.Thr66Ile | P16871 |
| HOMO SAPIENS | 1 | 5758 | 3di2 | B | 118 | VAL | p.Ile138Val | P16871 |
| HOMO SAPIENS | 4 | 848 | 5oa3 | Y | 79 | LEU | p.Leu79Val | P62847 |

| | | | | | | | | |
|---|---|---|---|---|---|---|---|---|
| HOMO SAPIENS | 2 | 848 | 5oa3 | Y | 124 | ASN | p.Asn124Ser | P62847 |
| HOMO SAPIENS | 0 | 81 | 3s5o | A | 36 | ALA | p.Ala36Glu | Q86XE5 |
| HOMO SAPIENS | 0 | 81 | 3s5o | A | 39 | TYR | p.Tyr39Ter | Q86XE5 |
| HOMO SAPIENS | 0 | 81 | 3s5o | A | 45 | PRO | p.Pro45Leu | Q86XE5 |
| HOMO SAPIENS | 0 | 81 | 3s5o | A | 70 | ARG | p.Arg70Ter | Q86XE5 |
| HOMO SAPIENS | 0 | 81 | 3s5o | A | 70 | ARG | p.Arg70Pro | Q86XE5 |
| HOMO SAPIENS | 1 | 81 | 3s5o | A | 97 | ARG | p.Arg97Cys | Q86XE5 |
| HOMO SAPIENS | 4 | 81 | 3s5o | A | 113 | GLU | p.Glu113Lys | Q86XE5 |
| HOMO SAPIENS | 1 | 81 | 3s5o | A | 190 | PRO | p.Pro190Leu | Q86XE5 |
| HOMO SAPIENS | 0 | 81 | 3s5o | A | 245 | VAL | p.Val245Phe | Q86XE5 |
| HOMO SAPIENS | 0 | 81 | 3s5o | A | 257 | CYS | p.Cys257Gly | Q86XE5 |
| HOMO SAPIENS | 0 | 81 | 3s5o | A | 287 | GLY | p.Gly287Val | Q86XE5 |
| HOMO SAPIENS | 7 | 81 | 3s5o | A | 303 | ARG | p.Arg303Cys | Q86XE5 |
| HOMO SAPIENS | 0 | 81 | 3s5o | A | 325 | GLY | p.Gly325Ser | Q86XE5 |
| HOMO SAPIENS | -1 | 622 | 5era | A | 12 | GLY | p.Gly12Arg | P29033 |
| HOMO SAPIENS | -1 | 622 | 5era | A | 12 | GLY | p.Gly12Asp | P29033 |
| HOMO SAPIENS | -1 | 622 | 5era | A | 17 | SER | p.Ser17Phe | P29033 |
| HOMO SAPIENS | 5 | 622 | 5era | A | 24 | TRP | p.Trp24Ter | P29033 |
| HOMO SAPIENS | 0 | 622 | 5era | A | 32 | ARG | p.Arg32Cys | P29033 |
| HOMO SAPIENS | 5 | 622 | 5era | A | 34 | MET | p.Met34Thr | P29033 |
| HOMO SAPIENS | 5 | 622 | 5era | A | 36 | LEU | p.Leu36Pro | P29033 |
| HOMO SAPIENS | 5 | 622 | 5era | A | 37 | VAL | p.Val37Ile | P29033 |
| HOMO SAPIENS | 5 | 622 | 5era | A | 40 | ALA | p.Ala40Glu | P29033 |
| HOMO SAPIENS | 5 | 622 | 5era | A | 44 | TRP | p.Trp44Cys | P29033 |
| HOMO SAPIENS | 5 | 622 | 5era | A | 45 | GLY | p.Gly45Glu | P29033 |
| HOMO SAPIENS | 5 | 622 | 5era | A | 47 | GLU | p.Glu47Ter | P29033 |
| HOMO SAPIENS | 5 | 622 | 5era | A | 50 | ASP | p.Asp50Asn | P29033 |
| HOMO SAPIENS | 5 | 622 | 5era | A | 54 | ASN | p.Asn54Lys | P29033 |
| HOMO SAPIENS | 5 | 622 | 5era | A | 57 | GLN | p.Gln57Ter | P29033 |
| HOMO SAPIENS | 5 | 622 | 5era | A | 59 | GLY | p.Gly59Ala | P29033 |
| HOMO SAPIENS | 5 | 622 | 5era | A | 59 | GLY | p.Gly59Ser | P29033 |
| HOMO SAPIENS | 5 | 622 | 5era | A | 66 | ASP | p.Asp66His | P29033 |
| HOMO SAPIENS | 5 | 622 | 5era | A | 68 | TYR | p.Tyr68Cys | P29033 |
| HOMO SAPIENS | 5 | 622 | 5era | A | 70 | PRO | p.Pro70Ala | P29033 |
| HOMO SAPIENS | 5 | 622 | 5era | A | 73 | HIS | p.His73Arg | P29033 |
| HOMO SAPIENS | 5 | 622 | 5era | A | 75 | ARG | p.Arg75Trp | P29033 |
| HOMO SAPIENS | 5 | 622 | 5era | A | 75 | ARG | p.Arg75Gln | P29033 |
| HOMO SAPIENS | 2 | 622 | 5era | A | 77 | TRP | p.Trp77Arg | P29033 |
| HOMO SAPIENS | 2 | 622 | 5era | A | 77 | TRP | p.Trp77Ter | P29033 |
| HOMO SAPIENS | 0 | 622 | 5era | A | 80 | GLN | p.Gln80Ter | P29033 |
| HOMO SAPIENS | 0 | 622 | 5era | A | 80 | GLN | p.Gln80Pro | P29033 |
| HOMO SAPIENS | 5 | 622 | 5era | A | 81 | LEU | p.Leu81Val | P29033 |
| HOMO SAPIENS | 5 | 622 | 5era | A | 82 | ILE | p.Ile82Met | P29033 |
| HOMO SAPIENS | 5 | 622 | 5era | A | 83 | PHE | p.Phe83Leu | P29033 |
| HOMO SAPIENS | 5 | 622 | 5era | A | 84 | VAL | p.Val84Leu | P29033 |
| HOMO SAPIENS | 5 | 622 | 5era | A | 90 | LEU | p.Leu90Pro | P29033 |
| HOMO SAPIENS | 5 | 622 | 5era | A | 95 | VAL | p.Val95Met | P29033 |
| HOMO SAPIENS | -1 | 622 | 5era | A | 100 | HIS | p.His100Tyr | P29033 |
| HOMO SAPIENS | -1 | 622 | 5era | A | 113 | SER | p.Ser113Arg | P29033 |
| HOMO SAPIENS | -1 | 622 | 5era | A | 119 | GLU | p.Glu119Lys | P29033 |
| HOMO SAPIENS | -1 | 622 | 5era | A | 122 | LYS | p.Lys122Ile | P29033 |
| HOMO SAPIENS | -1 | 622 | 5era | A | 123 | THR | p.Thr123Asn | P29033 |
| HOMO SAPIENS | -1 | 622 | 5era | A | 124 | GLN | p.Gln124Ter | P29033 |
| HOMO SAPIENS | 5 | 622 | 5era | A | 139 | SER | p.Ser139Asn | P29033 |

| | | | | | | | | |
|---|---|---|---|---|---|---|---|---|
| HOMO SAPIENS | 5 | 622 | 5era | A | 143 | ARG | p.Arg143Trp | P29033 |
| HOMO SAPIENS | 5 | 622 | 5era | A | 143 | ARG | p.Arg143Gln | P29033 |
| HOMO SAPIENS | 0 | 622 | 5era | A | 147 | GLU | p.Glu147Lys | P29033 |
| HOMO SAPIENS | 7 | 622 | 5era | A | 155 | TYR | p.Tyr155Ter | P29033 |
| HOMO SAPIENS | 2 | 622 | 5era | A | 159 | ASP | p.Asp159Val | P29033 |
| HOMO SAPIENS | 5 | 622 | 5era | A | 163 | MET | p.Met163Val | P29033 |
| HOMO SAPIENS | 5 | 622 | 5era | A | 165 | ARG | p.Arg165Trp | P29033 |
| HOMO SAPIENS | 5 | 622 | 5era | A | 167 | VAL | p.Val167Met | P29033 |
| HOMO SAPIENS | 5 | 622 | 5era | A | 169 | CYS | p.Cys169Tyr | P29033 |
| HOMO SAPIENS | 5 | 622 | 5era | A | 179 | ASP | p.Asp179Asn | P29033 |
| HOMO SAPIENS | 0 | 622 | 5era | A | 199 | SER | p.Ser199Phe | P29033 |
| HOMO SAPIENS | 0 | 622 | 5era | A | 202 | CYS | p.Cys202Phe | P29033 |
| HOMO SAPIENS | 5 | 622 | 5era | A | 203 | ILE | p.Ile203Thr | P29033 |
| HOMO SAPIENS | 5 | 622 | 5era | A | 206 | ASN | p.Asn206Ser | P29033 |
| HOMO SAPIENS | 0 | 8103 | 5dot | A | 163 | GLY | p.Gly163Ter | P31327 |
| HOMO SAPIENS | 0 | 8103 | 5dot | A | 337 | HIS | p.His337Arg | P31327 |
| HOMO SAPIENS | 3 | 8103 | 5dot | A | 344 | THR | p.Thr344Ala | P31327 |
| HOMO SAPIENS | 1 | 8103 | 5dot | A | 405 | THR | p.Thr405Asn | P31327 |
| HOMO SAPIENS | 5 | 8103 | 5dot | A | 544 | THR | p.Thr544Met | P31327 |
| HOMO SAPIENS | 5 | 8103 | 5dot | A | 716 | ASN | p.Asn716Lys | P31327 |
| HOMO SAPIENS | -1 | 8103 | 5dot | A | 780 | ARG | p.Arg780His | P31327 |
| HOMO SAPIENS | 0 | 8103 | 5dot | A | 982 | GLY | p.Gly982Asp | P31327 |
| HOMO SAPIENS | 3 | 8103 | 5dot | A | 1119 | ALA | p.Ala1119Ser | P31327 |
| HOMO SAPIENS | 1 | 8103 | 5dot | A | 1376 | GLY | p.Gly1376Ser | P31327 |
| HOMO SAPIENS | 4 | 6833 | 1l8l | B | 27 | ARG | p.Arg27Ser | P78330 |
| HOMO SAPIENS | 3 | 6833 | 1l8l | B | 32 | ASP | p.Asp32Gly | P78330 |
| HOMO SAPIENS | 3 | 6833 | 1l8l | B | 32 | ASP | p.Asp32Asn | P78330 |
| HOMO SAPIENS | 1 | 6833 | 1l8l | B | 49 | ARG | p.Arg49Trp | P78330 |
| HOMO SAPIENS | 0 | 6833 | 1l8l | B | 52 | MET | p.Met52Thr | P78330 |
| HOMO SAPIENS | 0 | 6833 | 1l8l | B | 83 | GLN | p.Gln83His | P78330 |
| HOMO SAPIENS | 6 | 810 | 2iw2 | A | 185 | ARG | p.Arg184Gln | P12955 |
| HOMO SAPIENS | 0 | 810 | 2iw2 | A | 213 | ALA | p.Ala212Pro | P12955 |
| HOMO SAPIENS | 1 | 810 | 2iw2 | A | 266 | ARG | p.Arg265Ter | P12955 |
| HOMO SAPIENS | 9 | 810 | 2iw2 | A | 277 | ASP | p.Asp276Asn | P12955 |
| HOMO SAPIENS | 0 | 810 | 2iw2 | A | 279 | GLY | p.Gly278Asp | P12955 |
| HOMO SAPIENS | 9 | 810 | 2iw2 | A | 378 | HIS | p.His377Gln | P12955 |
| HOMO SAPIENS | 0 | 810 | 2iw2 | A | 389 | ARG | p.Arg388His | P12955 |
| HOMO SAPIENS | 0 | 810 | 2iw2 | A | 436 | PHE | p.Leu435Phe | P12955 |
| HOMO SAPIENS | 0 | 810 | 2iw2 | A | 449 | GLY | p.Gly448Arg | P12955 |
| HOMO SAPIENS | -1 | 1530 | 5fwj | A | 388 | ALA | p.Ala388Pro | P41229 |
| HOMO SAPIENS | 4 | 1530 | 5fwj | A | 451 | SER | p.Ser451Arg | P41229 |
| HOMO SAPIENS | 3 | 1530 | 5fwj | A | 731 | LEU | p.Leu731Phe | P41229 |
| HOMO SAPIENS | 3 | 578 | 5net | B | 123 | ALA | p.Ala143Thr | P18564 |
| HOMO SAPIENS | 6 | 578 | 5net | B | 176 | PRO | p.Pro196Thr | P18564 |
| HOMO SAPIENS | 3 | 578 | 5net | B | 280 | GLU | p.Glu300Lys | P18564 |
| HOMO SAPIENS | -1 | 5247 | 4j15 | A | 274 | ALA | p.Ala274Val | P14868 |
| HOMO SAPIENS | -1 | 5247 | 4j15 | A | 280 | HIS | p.His280Leu | P14868 |
| HOMO SAPIENS | 2 | 5247 | 4j15 | A | 460 | ARG | p.Arg460His | P14868 |
| HOMO SAPIENS | 5 | 5247 | 4j15 | A | 494 | ARG | p.Arg494Cys | P14868 |
| HOMO SAPIENS | 7 | 2566 | 1b55 | A | 28 | ARG | p.Arg28His | Q06187 |
| HOMO SAPIENS | 5 | 2566 | 1b55 | A | 33 | THR | p.Thr33Pro | Q06187 |
| HOMO SAPIENS | 0 | 2566 | 1b55 | A | 113 | VAL | p.Val113Asp | Q06187 |
| HOMO SAPIENS | 0 | 356 | 5a2s | A | 786 | ALA | p.Ala786Thr | P56524 |
| HOMO SAPIENS | 5 | 2960 | 4cak | A | 553 | ARG | p.Arg584Ter | P08514 |

| | | | | | | | | |
|---|---|---|---|---|---|---|---|---|
| HOMO SAPIENS | 5 | 2960 | 4cak | A | 565 | ILE | p.Ile596Thr | P08514 |
| HOMO SAPIENS | -1 | 2960 | 4cak | A | 841 | LEU | p.Leu872Met | P08514 |
| HOMO SAPIENS | -1 | 2960 | 4cak | A | 843 | ILE | p.Ile874Ser | P08514 |
| HOMO SAPIENS | 0 | 5054 | 4k8r | B | 216 | PHE | p.Phe216Leu | Q8NBP7 |
| HOMO SAPIENS | 1 | 5054 | 4k8r | B | 417 | HIS | p.His417Gln | Q8NBP7 |
| HOMO SAPIENS | 1 | 5054 | 4k8r | B | 425 | ASN | p.Asn425Ser | Q8NBP7 |
| HOMO SAPIENS | 1 | 5054 | 4k8r | B | 443 | ALA | p.Ala443Thr | Q8NBP7 |
| HOMO SAPIENS | 0 | 27283 | 3n5n | X | 71 | HIS | p.His82Arg | Q9UIF7 |
| HOMO SAPIENS | 1 | 27283 | 3n5n | X | 83 | ARG | p.Arg94Ter | Q9UIF7 |
| HOMO SAPIENS | 8 | 27283 | 3n5n | X | 89 | TRP | p.Trp100Ter | Q9UIF7 |
| HOMO SAPIENS | 8 | 27283 | 3n5n | X | 90 | TYR | p.Tyr101Ter | Q9UIF7 |
| HOMO SAPIENS | 7 | 27283 | 3n5n | X | 95 | ARG | p.Arg106Trp | Q9UIF7 |
| HOMO SAPIENS | 7 | 27283 | 3n5n | X | 117 | TRP | p.Trp128Arg | Q9UIF7 |
| HOMO SAPIENS | 0 | 27283 | 3n5n | X | 137 | GLY | p.Gly148Arg | Q9UIF7 |
| HOMO SAPIENS | 5 | 27283 | 3n5n | X | 142 | TRP | p.Trp153Ter | Q9UIF7 |
| HOMO SAPIENS | 0 | 27283 | 3n5n | X | 165 | TYR | p.Tyr176Cys | Q9UIF7 |
| HOMO SAPIENS | 0 | 27283 | 3n5n | X | 168 | ARG | p.Arg179His | Q9UIF7 |
| HOMO SAPIENS | 0 | 27283 | 3n5n | X | 168 | ARG | p.Arg179Cys | Q9UIF7 |
| HOMO SAPIENS | 0 | 27283 | 3n5n | X | 196 | GLN | p.Gln207Ter | Q9UIF7 |
| HOMO SAPIENS | 8 | 27283 | 3n5n | X | 201 | VAL | p.Val212Met | Q9UIF7 |
| HOMO SAPIENS | 9 | 27283 | 3n5n | X | 227 | ARG | p.Arg238Trp | Q9UIF7 |
| HOMO SAPIENS | 9 | 27283 | 3n5n | X | 231 | ARG | p.Arg242Cys | Q9UIF7 |
| HOMO SAPIENS | 9 | 27283 | 3n5n | X | 231 | ARG | p.Arg242His | Q9UIF7 |
| HOMO SAPIENS | 3 | 27283 | 3n5n | X | 232 | VAL | p.Val243Phe | Q9UIF7 |
| HOMO SAPIENS | 3 | 27283 | 3n5n | X | 246 | GLN | p.Gln257Ter | Q9UIF7 |
| HOMO SAPIENS | 9 | 27283 | 3n5n | X | 272 | GLY | p.Gly283Glu | Q9UIF7 |
| HOMO SAPIENS | 9 | 27283 | 3n5n | X | 281 | PRO | p.Pro292Leu | Q9UIF7 |
| HOMO SAPIENS | -1 | 27283 | 3n5n | X | 314 | ASP | p.Asp325Glu | Q9UIF7 |
| HOMO SAPIENS | 0 | 27283 | 3n5n | X | 324 | GLN | p.Gln335His | Q9UIF7 |
| HOMO SAPIENS | 0 | 27283 | 3n5n | X | 324 | GLN | p.Gln335Arg | Q9UIF7 |
| HOMO SAPIENS | 0 | 27283 | 3n5n | X | 324 | GLN | p.Gln335Ter | Q9UIF7 |
| HOMO SAPIENS | -1 | 20852 | 4p60 | B | 243 | ARG | p.Arg243Lys | O75746 |
| HOMO SAPIENS | -1 | 1875 | 5dd2 | G | 152 | VAL | p.Val179Leu | P06396 |
| HOMO SAPIENS | 9 | 2126 | 3n1p | B | 46 | PRO | p.Pro46Leu | Q14623 |
| HOMO SAPIENS | 8 | 2126 | 3n1p | B | 76 | GLU | p.Glu76Ala | Q14623 |
| HOMO SAPIENS | 9 | 2126 | 3n1p | B | 77 | ARG | p.Arg77Ser | Q14623 |
| HOMO SAPIENS | 6 | 2126 | 3n1p | B | 95 | GLU | p.Glu95Lys | Q14623 |
| HOMO SAPIENS | 9 | 2126 | 3n1p | B | 100 | ASP | p.Asp100Glu | Q14623 |
| HOMO SAPIENS | 9 | 2126 | 3n1p | B | 131 | GLU | p.Glu131Lys | Q14623 |
| HOMO SAPIENS | 0 | 2126 | 3n1p | B | 190 | VAL | p.Val190Ala | Q14623 |
| HOMO SAPIENS | 5 | 40410 | 3ioz | B | 70 | GLN | p.Gln70Leu | P20963 |
| HOMO SAPIENS | 1 | 5754 | 5lkz | A | 1108 | THR | p.Tyr1191Ter | Q09472 |
| HOMO SAPIENS | 3 | 8159 | 1tiu | A | 3 | GLU | p.Lys12679Glu | Q8WZ42 |
| HOMO SAPIENS | 4 | 3275 | 4ldq | A | 210 | ARG | p.Arg210Gln | Q9BV20 |
| HOMO SAPIENS | 0 | 8143 | 4miv | B | 66 | SER | p.Ser66Trp | P51688 |
| HOMO SAPIENS | 1 | 8143 | 4miv | B | 74 | ARG | p.Arg74Cys | P51688 |
| HOMO SAPIENS | 0 | 8143 | 4miv | B | 128 | PRO | p.Pro128Leu | P51688 |
| HOMO SAPIENS | 1 | 8143 | 4miv | B | 150 | ARG | p.Arg150Gln | P51688 |
| HOMO SAPIENS | 0 | 8143 | 4miv | B | 191 | GLY | p.Gly191Arg | P51688 |
| HOMO SAPIENS | 0 | 8143 | 4miv | B | 206 | ARG | p.Arg206Pro | P51688 |
| HOMO SAPIENS | 0 | 8143 | 4miv | B | 225 | PHE | p.Phe225Leu | P51688 |
| HOMO SAPIENS | 0 | 8143 | 4miv | B | 233 | ARG | p.Arg233Ter | P51688 |
| HOMO SAPIENS | 0 | 8143 | 4miv | B | 234 | ALA | p.Ala234Gly | P51688 |
| HOMO SAPIENS | 0 | 8143 | 4miv | B | 245 | ARG | p.Arg245His | P51688 |

| | | | | | | | | |
|---|---|---|---|---|---|---|---|---|
| HOMO SAPIENS | 0 | 8143 | 4miv | B | 293 | PRO | p.Pro293Ser | P51688 |
| HOMO SAPIENS | 1 | 8143 | 4miv | B | 298 | SER | p.Ser298Pro | P51688 |
| HOMO SAPIENS | 0 | 8143 | 4miv | B | 361 | VAL | p.Val361Ile | P51688 |
| HOMO SAPIENS | 0 | 8143 | 4miv | B | 369 | GLU | p.Glu369Lys | P51688 |
| HOMO SAPIENS | 0 | 8143 | 4miv | B | 372 | MET | p.Met372Ile | P51688 |
| HOMO SAPIENS | 1 | 8143 | 4miv | B | 380 | GLN | p.Gln380Arg | P51688 |
| HOMO SAPIENS | 0 | 8143 | 4miv | B | 387 | VAL | p.Val387Met | P51688 |
| HOMO SAPIENS | 0 | 8143 | 4miv | B | 394 | MET | p.Met394Ile | P51688 |
| HOMO SAPIENS | 0 | 8143 | 4miv | B | 433 | ARG | p.Arg433Gln | P51688 |
| HOMO SAPIENS | 1 | 8143 | 4miv | B | 446 | HIS | p.His446Arg | P51688 |
| HOMO SAPIENS | 5 | 8143 | 4miv | B | 447 | GLU | p.Glu447Lys | P51688 |
| HOMO SAPIENS | 2 | 8143 | 4miv | B | 456 | ARG | p.Arg456His | P51688 |
| HOMO SAPIENS | 1 | 21880 | 2oo9 | B | 857 | LEU | p.Leu857Phe | P22681 |
| HOMO SAPIENS | 0 | 9467 | 1s1d | A | 154 | LEU | p.Leu224Pro | Q8WVQ1 |
| HOMO SAPIENS | 0 | 9467 | 1s1d | A | 156 | VAL | p.Val226Met | Q8WVQ1 |
| HOMO SAPIENS | 0 | 9467 | 1s1d | A | 229 | PRO | p.Pro299Leu | Q8WVQ1 |
| HOMO SAPIENS | 4 | 9467 | 1s1d | A | 230 | ARG | p.Arg300Cys | Q8WVQ1 |
| HOMO SAPIENS | 4 | 9467 | 1s1d | A | 230 | ARG | p.Arg300His | Q8WVQ1 |
| HOMO SAPIENS | 0 | 9467 | 1s1d | A | 253 | ALA | p.Ala323Thr | Q8WVQ1 |
| HOMO SAPIENS | 2 | 9467 | 1s1d | A | 305 | MET | p.Met375Leu | Q8WVQ1 |
| HOMO SAPIENS | -1 | 2361 | 5llm | A | 5 | ASN | p.Asn5Lys | P43003 |
| HOMO SAPIENS | 0 | 2361 | 5llm | A | 71 | TYR | p.Tyr71Cys | P43003 |
| HOMO SAPIENS | -1 | 2361 | 5llm | A | 211 | LYS | p.Glu219Asp | Q15758 |
| HOMO SAPIENS | 8 | 2361 | 5llm | A | 270 | PRO | p.Pro290Arg | P43003 |
| HOMO SAPIENS | 1 | 2361 | 5llm | A | 321 | THR | p.Ala329Thr | Q15758 |
| HOMO SAPIENS | 1 | 2361 | 5llm | A | 365 | ARG | p.Arg385His | P43003 |
| HOMO SAPIENS | 1 | 2361 | 5llm | A | 479 | ARG | p.Arg499Gln | P43003 |
| HOMO SAPIENS | 3 | 5412 | 4obz | D | 165 | PHE | p.Phe229Ile | P07339 |
| HOMO SAPIENS | 2 | 5412 | 4obz | D | 187 | ASP | p.Asp251Asn | P07339 |
| HOMO SAPIENS | 0 | 5412 | 4obz | D | 218 | GLY | p.Gly282Arg | P07339 |
| HOMO SAPIENS | 0 | 5412 | 4obz | D | 245 | ARG | p.Arg309His | P07339 |
| HOMO SAPIENS | 2 | 5412 | 4obz | D | 319 | TRP | p.Trp383Cys | P07339 |
| HOMO SAPIENS | 0 | 3544 | 5h64 | A | 277 | MET | p.Met277Ile | P42345 |
| HOMO SAPIENS | 0 | 14230 | 2kbx | A | 145 | ARG | p.Arg145Ser | Q13418 |
| HOMO SAPIENS | 2 | 34970 | 4xhz | A | 834 | ARG | p.Arg855Trp | Q96QU1 |
| HOMO SAPIENS | 4 | 34970 | 4xhz | A | 854 | SER | p.Ser875Ter | Q96QU1 |
| HOMO SAPIENS | 1 | 34970 | 4xhz | A | 908 | ARG | p.Arg929Gln | Q96QU1 |
| HOMO SAPIENS | 1 | 34970 | 4xhz | A | 941 | ARG | p.Arg962Cys | Q96QU1 |
| HOMO SAPIENS | 0 | 34970 | 4xhz | A | 970 | ARG | p.Arg991Ter | Q96QU1 |
| HOMO SAPIENS | 0 | 34970 | 4xhz | A | 976 | GLU | p.Glu997Gly | Q96QU1 |
| HOMO SAPIENS | 0 | 25734 | 4asc | A | 345 | SER | p.Asn345Ser | Q2TBA0 |
| HOMO SAPIENS | 0 | 25734 | 4asc | A | 514 | VAL | p.Val514Met | Q2TBA0 |
| HOMO SAPIENS | 6 | 25734 | 4asc | A | 528 | GLU | p.Glu528Lys | Q2TBA0 |
| HOMO SAPIENS | 0 | 25734 | 4asc | A | 580 | LEU | p.Leu580Ile | Q2TBA0 |
| HOMO SAPIENS | 8 | 2356 | 3kse | D | 22 | LYS | p.Lys22Ter | P01040 |
| HOMO SAPIENS | 1 | 2356 | 3kse | D | 58 | ARG | p.Arg58Ter | P01040 |
| HOMO SAPIENS | 2 | 6000 | 1jqd | A | 60 | GLY | p.Gly60Asp | P50135 |
| HOMO SAPIENS | 9 | 6000 | 1jqd | A | 208 | LEU | p.Leu208Pro | P50135 |
| HOMO SAPIENS | 0 | 14741 | 5eh1 | A | 58 | THR | p.Thr58Arg | P38484 |
| HOMO SAPIENS | 0 | 14741 | 5eh1 | A | 168 | THR | p.Thr168Asn | P38484 |
| HOMO SAPIENS | 1 | 4814 | 4kxk | D | 427 | ARG | p.Arg427Ter | P50542 |
| HOMO SAPIENS | 0 | 188 | 2xt3 | A | 49 | ASP | p.Asp52Asn | Q2M1P5 |
| HOMO SAPIENS | 3 | 188 | 2xt3 | A | 142 | TYR | p.Tyr145Ser | Q2M1P5 |
| HOMO SAPIENS | 9 | 8681 | 5tby | D | 57 | ALA | p.Ala57Asp | P08590 |

| | | | | | | | | |
|---|---|---|---|---|---|---|---|---|
| HOMO SAPIENS | 3 | 8681 | 5tby | D | 62 | ASP | p.Asp62Asn | P08590 |
| HOMO SAPIENS | 0 | 8681 | 5tby | D | 65 | PRO | p.Pro65Arg | P08590 |
| HOMO SAPIENS | 7 | 8681 | 5tby | D | 79 | VAL | p.Val79Ile | P08590 |
| HOMO SAPIENS | 0 | 8681 | 5tby | D | 143 | GLU | p.Glu143Lys | P08590 |
| HOMO SAPIENS | 0 | 8681 | 5tby | D | 149 | MET | p.Met149Val | P08590 |
| HOMO SAPIENS | 3 | 8681 | 5tby | D | 154 | ARG | p.Arg154His | P08590 |
| HOMO SAPIENS | 6 | 30984 | 4xwh | A | 140 | TYR | p.Tyr140Cys | P54802 |
| HOMO SAPIENS | 4 | 30984 | 4xwh | A | 141 | SER | p.Ser141Thr | P54802 |
| HOMO SAPIENS | 2 | 30984 | 4xwh | A | 234 | ARG | p.Arg234Cys | P54802 |
| HOMO SAPIENS | 2 | 30984 | 4xwh | A | 336 | GLU | p.Glu336Ter | P54802 |
| HOMO SAPIENS | 6 | 30984 | 4xwh | A | 482 | ARG | p.Arg482Trp | P54802 |
| HOMO SAPIENS | 2 | 30984 | 4xwh | A | 521 | PRO | p.Pro521Leu | P54802 |
| HOMO SAPIENS | 0 | 30984 | 4xwh | A | 565 | ARG | p.Arg565Gln | P54802 |
| HOMO SAPIENS | 0 | 30984 | 4xwh | A | 565 | ARG | p.Arg565Trp | P54802 |
| HOMO SAPIENS | 6 | 30984 | 4xwh | A | 626 | ARG | p.Arg626Ter | P54802 |
| HOMO SAPIENS | 6 | 30984 | 4xwh | A | 643 | ARG | p.Arg643His | P54802 |
| HOMO SAPIENS | 6 | 30984 | 4xwh | A | 643 | ARG | p.Arg643Cys | P54802 |
| HOMO SAPIENS | 2 | 30984 | 4xwh | A | 674 | ARG | p.Arg674His | P54802 |
| HOMO SAPIENS | 2 | 30984 | 4xwh | A | 737 | ARG | p.Arg737Gly | P54802 |
| HOMO SAPIENS | 5 | 10545 | 1oqe | N | 6 | PRO | p.Pro21Arg | Q96RJ3 |
| HOMO SAPIENS | 1 | 16995 | 5d3a | B | 12 | MET | p.Met947Arg | Q7Z4S6 |
| HOMO SAPIENS | 1 | 16995 | 5d3a | B | 12 | MET | p.Met947Val | Q7Z4S6 |
| HOMO SAPIENS | 2 | 16995 | 5d3a | B | 19 | ARG | p.Arg954Gln | Q7Z4S6 |
| HOMO SAPIENS | 2 | 16995 | 5d3a | B | 19 | ARG | p.Arg954Trp | Q7Z4S6 |
| HOMO SAPIENS | 0 | 16995 | 5d3a | B | 75 | ILE | p.Ile1010Thr | Q7Z4S6 |
| HOMO SAPIENS | 0 | 17091 | 2yin | B | 1324 | GLN | p.Gln1324Ter | Q92608 |
| HOMO SAPIENS | 5 | 27606 | 1lm7 | B | 2366 | ARG | p.Arg2366Cys | P15924 |
| HOMO SAPIENS | 2 | 15731 | 1bi7 | B | 24 | ARG | p.Arg24Pro | P42771 |
| HOMO SAPIENS | 0 | 15731 | 1bi7 | B | 28 | VAL | p.Val28Gly | P42771 |
| HOMO SAPIENS | 0 | 15731 | 1bi7 | B | 49 | ILE | p.Ile49Thr | P42771 |
| HOMO SAPIENS | 0 | 15731 | 1bi7 | B | 53 | MET | p.Met53Ile | P42771 |
| HOMO SAPIENS | 2 | 15731 | 1bi7 | B | 56 | SER | p.Ser56Ile | P42771 |
| HOMO SAPIENS | 1 | 15731 | 1bi7 | B | 57 | ALA | p.Ala57Val | P42771 |
| HOMO SAPIENS | 4 | 15731 | 1bi7 | B | 59 | VAL | p.Val59Gly | P42771 |
| HOMO SAPIENS | 9 | 15731 | 1bi7 | B | 89 | GLY | p.Gly89Asp | P42771 |
| HOMO SAPIENS | 0 | 15731 | 1bi7 | B | 101 | GLY | p.Gly101Trp | P42771 |
| HOMO SAPIENS | 0 | 15731 | 1bi7 | B | 114 | PRO | p.Pro114Ser | P42771 |
| HOMO SAPIENS | 9 | 15731 | 1bi7 | B | 123 | HIS | p.His123Gln | P42771 |
| HOMO SAPIENS | 8 | 15731 | 1bi7 | B | 126 | VAL | p.Val126Asp | P42771 |
| HOMO SAPIENS | 7 | 15731 | 1bi7 | B | 127 | ALA | p.Ala127Ser | P42771 |
| HOMO SAPIENS | -1 | 15731 | 1bi7 | B | 148 | ALA | p.Ala148Thr | P42771 |
| HOMO SAPIENS | -1 | 19858 | 3jbk | M | 866 | ALA | p.Ala934Val | P18206 |
| HOMO SAPIENS | 0 | 19484 | 3ee6 | B | 62 | SER | p.Ser62Leu | O14773 |
| HOMO SAPIENS | 1 | 19484 | 3ee6 | B | 66 | GLN | p.Gln66Ter | O14773 |
| HOMO SAPIENS | 0 | 19484 | 3ee6 | B | 100 | GLN | p.Gln100Arg | O14773 |
| HOMO SAPIENS | 0 | 19484 | 3ee6 | B | 104 | LEU | p.Leu104Ter | O14773 |
| HOMO SAPIENS | 0 | 19484 | 3ee6 | B | 127 | ARG | p.Arg127Gln | O14773 |
| HOMO SAPIENS | 0 | 19484 | 3ee6 | B | 127 | ARG | p.Arg127Ter | O14773 |
| HOMO SAPIENS | -1 | 19484 | 3ee6 | B | 185 | ARG | p.Arg185Cys | O14773 |
| HOMO SAPIENS | 0 | 19484 | 3ee6 | B | 206 | ARG | p.Arg206Cys | O14773 |
| HOMO SAPIENS | 0 | 19484 | 3ee6 | B | 208 | ARG | p.Arg208Ter | O14773 |
| HOMO SAPIENS | 0 | 19484 | 3ee6 | B | 214 | GLN | p.Gln214Ter | O14773 |
| HOMO SAPIENS | 1 | 19484 | 3ee6 | B | 276 | ASP | p.Asp276Val | O14773 |
| HOMO SAPIENS | 0 | 19484 | 3ee6 | B | 284 | GLY | p.Gly284Val | O14773 |

| | | | | | | | | |
|---|---|---|---|---|---|---|---|---|
| HOMO SAPIENS | 0 | 19484 | 3ee6 | B | 286 | ASN | p.Asn286Ser | O14773 |
| HOMO SAPIENS | 0 | 19484 | 3ee6 | B | 339 | ARG | p.Arg339Gly | O14773 |
| HOMO SAPIENS | 2 | 19484 | 3ee6 | B | 345 | MET | p.Met345Leu | O14773 |
| HOMO SAPIENS | 0 | 19484 | 3ee6 | B | 365 | CYS | p.Cys365Arg | O14773 |
| HOMO SAPIENS | 0 | 19484 | 3ee6 | B | 365 | CYS | p.Cys365Tyr | O14773 |
| HOMO SAPIENS | 0 | 19484 | 3ee6 | B | 373 | GLN | p.Gln373Glu | O14773 |
| HOMO SAPIENS | 6 | 19484 | 3ee6 | B | 418 | ARG | p.Arg418Gln | O14773 |
| HOMO SAPIENS | 7 | 19484 | 3ee6 | B | 422 | GLN | p.Gln422His | O14773 |
| HOMO SAPIENS | 0 | 19484 | 3ee6 | B | 447 | ARG | p.Arg447His | O14773 |
| HOMO SAPIENS | 0 | 19484 | 3ee6 | B | 466 | VAL | p.Val466Met | O14773 |
| HOMO SAPIENS | 0 | 4654 | 5t5w | A | 47 | LYS | p.Lys47Glu | Q08334 |
| HOMO SAPIENS | 0 | 4654 | 5t5w | A | 141 | GLU | p.Glu141Ter | Q08334 |
| HOMO SAPIENS | -1 | 18815 | 4wj7 | A | 53 | VAL | p.Val53Ile | Q9BSQ5 |
| HOMO SAPIENS | 0 | 18815 | 4wj7 | A | 120 | VAL | p.Val120Ile | Q9BSQ5 |
| HOMO SAPIENS | 0 | 18815 | 4wj7 | A | 198 | LEU | p.Leu198Arg | Q9BSQ5 |
| HOMO SAPIENS | 0 | 4602 | 1mjd | A | 47 | SER | p.Ser47Arg | O43602 |
| HOMO SAPIENS | 6 | 4602 | 1mjd | A | 59 | ARG | p.Arg59Leu | O43602 |
| HOMO SAPIENS | 7 | 4602 | 1mjd | A | 62 | ASP | p.Asp62Asn | O43602 |
| HOMO SAPIENS | 7 | 4602 | 1mjd | A | 71 | ALA | p.Ala71Ser | O43602 |
| HOMO SAPIENS | 7 | 4602 | 1mjd | A | 78 | ARG | p.Arg78Leu | O43602 |
| HOMO SAPIENS | 7 | 4602 | 1mjd | A | 89 | ARG | p.Arg89Gly | O43602 |
| HOMO SAPIENS | 8 | 4602 | 1mjd | A | 104 | ILE | p.Ile104Phe | O43602 |
| HOMO SAPIENS | 7 | 4602 | 1mjd | A | 125 | TYR | p.Tyr125His | O43602 |
| HOMO SAPIENS | 5 | 34467 | 1n0w | B | 1529 | GLY | p.Gly1529Arg | P51587 |
| HOMO SAPIENS | 8 | 5738 | 1u31 | A | 130 | ASN | p.Asn1009Lys | Q13423 |
| HOMO SAPIENS | 0 | 5760 | 2i6s | A | 298 | GLU | p.Glu318Asp | P06681 |
| HOMO SAPIENS | 0 | 5760 | 2i6s | A | 573 | ARG | p.Arg593Gln | P06681 |
| HOMO SAPIENS | 0 | 5760 | 2i6s | A | 592 | SER | p.Ser612Asn | P06681 |
| HOMO SAPIENS | 0 | 5760 | 2i6s | A | 621 | VAL | p.Val641Ala | P06681 |
| HOMO SAPIENS | 1 | 10977 | 2fby | A | 114 | VAL | p.Val114Ile | Q14191 |
| HOMO SAPIENS | 0 | 10977 | 2fby | A | 135 | LYS | p.Lys135Glu | Q14191 |
| HOMO SAPIENS | -1 | 1175 | 1qrk | A | 34 | VAL | p.Val35Leu | P00488 |
| HOMO SAPIENS | 4 | 1175 | 1qrk | A | 564 | PRO | p.Pro565Leu | P00488 |
| HOMO SAPIENS | 0 | 1175 | 1qrk | A | 650 | VAL | p.Val651Ile | P00488 |
| HOMO SAPIENS | 0 | 1175 | 1qrk | A | 651 | GLU | p.Gln652Glu | P00488 |
| HOMO SAPIENS | 3 | 909 | 4h0w | A | 375 | GLU | p.Glu394Lys | P02787 |
| HOMO SAPIENS | 0 | 909 | 4h0w | A | 458 | ALA | p.Ala477Pro | P02787 |
| HOMO SAPIENS | 6 | 909 | 4h0w | A | 515 | TYR | p.Tyr534Ter | P02787 |
| HOMO SAPIENS | 7 | 909 | 4h0w | A | 570 | PRO | p.Pro589Ser | P02787 |
| HOMO SAPIENS | 3 | 909 | 4h0w | A | 652 | GLY | p.Gly671Glu | P02787 |
| HOMO SAPIENS | 5 | 1964 | 2car | A | 151 | TRP | p.Trp151Ter | Q9BY32 |
| HOMO SAPIENS | 7 | 1964 | 2car | A | 178 | ARG | p.Arg178Cys | Q9BY32 |
| HOMO SAPIENS | 0 | 144 | 2n0a | J | 30 | ALA | p.Ala30Pro | P37840 |
| HOMO SAPIENS | 1 | 12288 | 4q58 | D | 1225 | ARG | p.Arg1225His | P16144 |
| HOMO SAPIENS | 2 | 12288 | 4q58 | D | 1281 | ARG | p.Arg1281Trp | P16144 |
| HOMO SAPIENS | 2 | 12288 | 4q58 | D | 1287 | ASN | p.Asn1287Lys | P16144 |
| HOMO SAPIENS | 0 | 5085 | 3s94 | A | 360 | ARG | p.Arg360His | O75581 |
| HOMO SAPIENS | 0 | 5085 | 3s94 | A | 433 | ASN | p.Asn433Ser | O75581 |
| HOMO SAPIENS | 2 | 5085 | 3s94 | A | 611 | ARG | p.Arg611Cys | O75581 |
| HOMO SAPIENS | 0 | 15452 | 4dur | A | 19 | LYS | p.Lys38Glu | P00747 |
| HOMO SAPIENS | 2 | 15452 | 4dur | A | 216 | ARG | p.Arg235His | P00747 |
| HOMO SAPIENS | 7 | 8406 | 4wo1 | D | 28 | VAL | p.Val55Leu | O43914 |
| HOMO SAPIENS | 2 | 895 | 5xte | K | 183 | ARG | p.Arg183Trp | P22695 |
| HOMO SAPIENS | 2 | 4191 | 4wnd | A | 120 | ARG | p.Arg127Ter | P81274 |

| | | | | | | | | |
|---|---|---|---|---|---|---|---|---|
| HOMO SAPIENS | 2 | 4191 | 4wnd | A | 120 | ARG | p.Arg127Gln | P81274 |
| HOMO SAPIENS | 1 | 4191 | 4wnd | A | 349 | GLY | p.Gly356Arg | P81274 |
| HOMO SAPIENS | 0 | 41734 | 2kbi | A | 1777 | VAL | p.Val1777Met | Q14524 |
| HOMO SAPIENS | 1 | 41734 | 2kbi | A | 1784 | GLU | p.Glu1784Lys | Q14524 |
| HOMO SAPIENS | 5 | 41734 | 2kbi | A | 1795 | TYR | p.Tyr1795Cys | Q14524 |
| HOMO SAPIENS | 5 | 41734 | 2kbi | A | 1795 | TYR | p.Tyr1795His | Q14524 |
| HOMO SAPIENS | 7 | 41734 | 2kbi | A | 1826 | ARG | p.Arg1826Cys | Q14524 |
| HOMO SAPIENS | 3 | 2119 | 1pl0 | A | 426 | LYS | p.Lys426Arg | P31939 |
| HOMO SAPIENS | 5 | 3475 | 5u62 | C | 50 | LEU | p.Leu50Ser | Q15910 |
| HOMO SAPIENS | 3 | 379 | 4uez | A | 200 | TYR | p.Tyr200Ter | P15085 |
| HOMO SAPIENS | 0 | 8195 | 5xaw | B | 67 | ARG | p.Arg67Trp | Q8IWT1 |
| HOMO SAPIENS | 0 | 8195 | 5xaw | B | 94 | THR | p.Thr94Met | Q8IWT1 |
| HOMO SAPIENS | 1 | 5711 | 1fo3 | A | 265 | ARG | p.Arg265His | Q9UKM7 |
| HOMO SAPIENS | 4 | 5711 | 1fo3 | A | 334 | ARG | p.Arg334Cys | Q9UKM7 |
| HOMO SAPIENS | 1 | 5711 | 1fo3 | A | 478 | LYS | p.Lys478Arg | Q9UKM7 |
| HOMO SAPIENS | 1 | 5711 | 1fo3 | A | 682 | ASP | p.Asp682Glu | Q9UKM7 |
| HOMO SAPIENS | 0 | 170 | 2n3y | A | 41 | GLY | p.Gly42Ser | P99999 |
| HOMO SAPIENS | 0 | 4122 | 3w81 | A | 33 | HIS | p.His33Gln | P35475 |
| HOMO SAPIENS | 0 | 4122 | 3w81 | A | 64 | TYR | p.Tyr64Ter | P35475 |
| HOMO SAPIENS | 0 | 4122 | 3w81 | A | 70 | GLN | p.Gln70Ter | P35475 |
| HOMO SAPIENS | 0 | 4122 | 3w81 | A | 75 | ALA | p.Ala75Thr | P35475 |
| HOMO SAPIENS | 0 | 4122 | 3w81 | A | 79 | ALA | p.Ala79Val | P35475 |
| HOMO SAPIENS | 0 | 4122 | 3w81 | A | 79 | ALA | p.Ala79Thr | P35475 |
| HOMO SAPIENS | 0 | 4122 | 3w81 | A | 89 | ARG | p.Arg89Gln | P35475 |
| HOMO SAPIENS | 0 | 4122 | 3w81 | A | 99 | THR | p.Thr99Ile | P35475 |
| HOMO SAPIENS | 0 | 4122 | 3w81 | A | 105 | ARG | p.Arg105Gln | P35475 |
| HOMO SAPIENS | 0 | 4122 | 3w81 | A | 238 | LEU | p.Leu238Gln | P35475 |
| HOMO SAPIENS | 0 | 4122 | 3w81 | A | 300 | ALA | p.Ala300Thr | P35475 |
| HOMO SAPIENS | 0 | 4122 | 3w81 | A | 322 | VAL | p.Val322Glu | P35475 |
| HOMO SAPIENS | 0 | 4122 | 3w81 | A | 327 | ALA | p.Ala327Pro | P35475 |
| HOMO SAPIENS | 0 | 4122 | 3w81 | A | 346 | LEU | p.Leu346Arg | P35475 |
| HOMO SAPIENS | 0 | 4122 | 3w81 | A | 349 | ASP | p.Asp349Asn | P35475 |
| HOMO SAPIENS | 0 | 4122 | 3w81 | A | 361 | ALA | p.Ala361Thr | P35475 |
| HOMO SAPIENS | 0 | 4122 | 3w81 | A | 366 | THR | p.Thr366Pro | P35475 |
| HOMO SAPIENS | 0 | 4122 | 3w81 | A | 402 | TRP | p.Trp402Ter | P35475 |
| HOMO SAPIENS | 0 | 4122 | 3w81 | A | 409 | GLY | p.Gly409Arg | P35475 |
| HOMO SAPIENS | 0 | 4122 | 3w81 | A | 454 | VAL | p.Val454Ile | P35475 |
| HOMO SAPIENS | 1 | 4122 | 3w81 | A | 490 | LEU | p.Leu490Pro | P35475 |
| HOMO SAPIENS | 1 | 4122 | 3w81 | A | 492 | ARG | p.Arg492Pro | P35475 |
| HOMO SAPIENS | 0 | 4122 | 3w81 | A | 533 | PRO | p.Pro533Arg | P35475 |
| HOMO SAPIENS | 0 | 4122 | 3w81 | A | 591 | ALA | p.Ala591Thr | P35475 |
| HOMO SAPIENS | 7 | 4122 | 3w81 | A | 619 | ARG | p.Arg619Gly | P35475 |
| HOMO SAPIENS | 0 | 15325 | 3wvz | B | 54 | VAL | p.Val54Leu | Q53FT3 |
| HOMO SAPIENS | 3 | 3228 | 5iz8 | A | 421 | TRP | p.Trp421Ter | P25054 |
| HOMO SAPIENS | 2 | 3228 | 5iz8 | A | 437 | PRO | p.Pro437Arg | P25054 |
| HOMO SAPIENS | 3 | 3228 | 5iz8 | A | 499 | ARG | p.Arg499Ter | P25054 |
| HOMO SAPIENS | -1 | 29229 | 5buo | A | 539 | SER | p.Ser614Gly | P05067 |
| HOMO SAPIENS | -1 | 34931 | 4n7v | C | 54 | SER | p.Ser54Leu | O94986 |
| HOMO SAPIENS | 0 | 2450 | 5kzx | A | 123 | GLY | p.Gly123Glu | P10253 |
| HOMO SAPIENS | 1 | 2450 | 5kzx | A | 190 | ARG | p.Arg190His | P10253 |
| HOMO SAPIENS | -1 | 2450 | 5kzx | A | 199 | HIS | p.His199Arg | P10253 |
| HOMO SAPIENS | 0 | 2450 | 5kzx | A | 219 | GLY | p.Gly219Arg | P10253 |
| HOMO SAPIENS | 0 | 2450 | 5kzx | A | 220 | VAL | p.Val220Leu | P10253 |
| HOMO SAPIENS | 0 | 2450 | 5kzx | A | 222 | VAL | p.Val222Met | P10253 |

| | | | | | | | | |
|---|---|---|---|---|---|---|---|---|
| HOMO SAPIENS | 0 | 2450 | 5kzx | A | 237 | ALA | p.Ala237Val | P10253 |
| HOMO SAPIENS | 0 | 2450 | 5kzx | A | 251 | SER | p.Ser251Leu | P10253 |
| HOMO SAPIENS | 0 | 2450 | 5kzx | A | 254 | SER | p.Ser254Leu | P10253 |
| HOMO SAPIENS | 0 | 2450 | 5kzx | A | 262 | GLU | p.Glu262Lys | P10253 |
| HOMO SAPIENS | 0 | 2450 | 5kzx | A | 293 | GLY | p.Gly293Arg | P10253 |
| HOMO SAPIENS | 0 | 2450 | 5kzx | A | 299 | LEU | p.Leu299Arg | P10253 |
| HOMO SAPIENS | 0 | 2450 | 5kzx | A | 309 | GLY | p.Gly309Arg | P10253 |
| HOMO SAPIENS | 0 | 2450 | 5kzx | A | 318 | MET | p.Met318Thr | P10253 |
| HOMO SAPIENS | 0 | 2450 | 5kzx | A | 355 | LEU | p.Leu355Pro | P10253 |
| HOMO SAPIENS | 1 | 2450 | 5kzx | A | 361 | PRO | p.Pro361Leu | P10253 |
| HOMO SAPIENS | 4 | 2450 | 5kzx | A | 408 | MET | p.Met408Val | P10253 |
| HOMO SAPIENS | 1 | 2450 | 5kzx | A | 429 | GLN | p.Gln429Glu | P10253 |
| HOMO SAPIENS | 4 | 2450 | 5kzx | A | 437 | ARG | p.Arg437Cys | P10253 |
| HOMO SAPIENS | 0 | 2450 | 5kzx | A | 439 | MET | p.Met439Lys | P10253 |
| HOMO SAPIENS | 1 | 2450 | 5kzx | A | 448 | SER | p.Ser448Thr | P10253 |
| HOMO SAPIENS | 0 | 2450 | 5kzx | A | 451 | PRO | p.Pro451Arg | P10253 |
| HOMO SAPIENS | 1 | 2450 | 5kzx | A | 459 | ASP | p.Asp459His | P10253 |
| HOMO SAPIENS | 8 | 2450 | 5kzx | A | 481 | TRP | p.Trp481Arg | P10253 |
| HOMO SAPIENS | 1 | 2450 | 5kzx | A | 489 | ASP | p.Asp489Asn | P10253 |
| HOMO SAPIENS | 0 | 2450 | 5kzx | A | 521 | GLU | p.Glu521Lys | P10253 |
| HOMO SAPIENS | 0 | 2450 | 5kzx | A | 529 | SER | p.Ser529Val | P10253 |
| HOMO SAPIENS | 0 | 2450 | 5kzx | A | 545 | PRO | p.Pro545Leu | P10253 |
| HOMO SAPIENS | 0 | 2450 | 5kzx | A | 552 | LEU | p.Leu552Pro | P10253 |
| HOMO SAPIENS | 0 | 2450 | 5kzx | A | 576 | GLY | p.Gly576Ser | P10253 |
| HOMO SAPIENS | 9 | 2450 | 5kzx | A | 600 | ARG | p.Arg600His | P10253 |
| HOMO SAPIENS | 8 | 2450 | 5kzx | A | 615 | GLY | p.Gly615Arg | P10253 |
| HOMO SAPIENS | 3 | 2450 | 5kzx | A | 619 | SER | p.Ser619Asn | P10253 |
| HOMO SAPIENS | 0 | 2450 | 5kzx | A | 638 | GLY | p.Gly638Trp | P10253 |
| HOMO SAPIENS | 0 | 2450 | 5kzx | A | 643 | GLY | p.Gly643Arg | P10253 |
| HOMO SAPIENS | 9 | 2450 | 5kzx | A | 645 | ASP | p.Asp645Glu | P10253 |
| HOMO SAPIENS | 9 | 2450 | 5kzx | A | 645 | ASP | p.Asp645His | P10253 |
| HOMO SAPIENS | 0 | 2450 | 5kzx | A | 648 | GLY | p.Gly648Ser | P10253 |
| HOMO SAPIENS | 5 | 2450 | 5kzx | A | 660 | ARG | p.Arg660His | P10253 |
| HOMO SAPIENS | 8 | 2450 | 5kzx | A | 672 | ARG | p.Arg672Gln | P10253 |
| HOMO SAPIENS | 0 | 2450 | 5kzx | A | 702 | ARG | p.Arg702Leu | P10253 |
| HOMO SAPIENS | 3 | 2450 | 5kzx | A | 725 | ARG | p.Arg725Trp | P10253 |
| HOMO SAPIENS | 0 | 2450 | 5kzx | A | 746 | TRP | p.Trp746Cys | P10253 |
| HOMO SAPIENS | 0 | 2450 | 5kzx | A | 780 | ILE | p.Val780Ile | P10253 |
| HOMO SAPIENS | 1 | 2450 | 5kzx | A | 816 | VAL | p.Val816Ile | P10253 |
| HOMO SAPIENS | 0 | 2450 | 5kzx | A | 838 | GLN | p.Gln838Ter | P10253 |
| HOMO SAPIENS | 2 | 2450 | 5kzx | A | 854 | ARG | p.Arg854Ter | P10253 |
| HOMO SAPIENS | 3 | 2450 | 5kzx | A | 870 | ARG | p.Arg870Ter | P10253 |
| HOMO SAPIENS | 0 | 2450 | 5kzx | A | 890 | VAL | p.Val890Leu | P10253 |
| HOMO SAPIENS | 6 | 1882 | 4km6 | A | 175 | CYS | p.Cys175Ter | P15328 |
| HOMO SAPIENS | 3 | 14767 | 1lm5 | A | 2628 | LEU | p.Leu2628Pro | P15924 |
| HOMO SAPIENS | 5 | 14767 | 1lm5 | A | 2655 | ALA | p.Ala2655Gly | P15924 |
| HOMO SAPIENS | 3 | 14767 | 1lm5 | A | 2731 | TYR | p.Tyr2731His | P15924 |
| HOMO SAPIENS | 1 | 14767 | 1lm5 | A | 2767 | THR | p.Thr2767Asn | P15924 |
| HOMO SAPIENS | -1 | 14767 | 1lm5 | A | 2819 | MET | p.Met2819Leu | P15924 |
| HOMO SAPIENS | -1 | 11246 | 4y5o | A | 289 | SER | p.Ser289Asn | Q9BSQ5 |
| HOMO SAPIENS | 0 | 7380 | 5kz5 | I | 60 | ARG | p.Arg60His | Q16595 |
| HOMO SAPIENS | 0 | 7380 | 5kz5 | I | 76 | MET | p.Met76Val | Q16595 |
| HOMO SAPIENS | 2 | 22187 | 3fby | A | 328 | CYS | p.Cys328Arg | P49747 |
| HOMO SAPIENS | 2 | 22187 | 3fby | A | 342 | ASP | p.Asp342Tyr | P49747 |

| | | | | | | | | |
|---|---|---|---|---|---|---|---|---|
| HOMO SAPIENS | 2 | 22187 | 3fby | A | 348 | CYS | p.Cys348Arg | P49747 |
| HOMO SAPIENS | 2 | 22187 | 3fby | A | 386 | ASN | p.Asn386Asp | P49747 |
| HOMO SAPIENS | 2 | 22187 | 3fby | A | 453 | ASN | p.Asn453Ser | P49747 |
| HOMO SAPIENS | 2 | 22187 | 3fby | A | 468 | CYS | p.Cys468Tyr | P49747 |
| HOMO SAPIENS | 3 | 22187 | 3fby | A | 472 | ASP | p.Asp472Tyr | P49747 |
| HOMO SAPIENS | 3 | 22187 | 3fby | A | 473 | ASP | p.Asp473Gly | P49747 |
| HOMO SAPIENS | 2 | 22187 | 3fby | A | 523 | ASN | p.Asn523Lys | P49747 |
| HOMO SAPIENS | 1 | 22187 | 3fby | A | 718 | ARG | p.Arg718Trp | P49747 |
| HOMO SAPIENS | 1 | 22187 | 3fby | A | 719 | GLY | p.Gly719Asp | P49747 |
| HOMO SAPIENS | 0 | 22187 | 3fby | A | 756 | GLN | p.Gln756Arg | P49747 |
| HOMO SAPIENS | -1 | 2784 | 5kvh | A | 88 | MET | p.Met88Val | O95831 |
| HOMO SAPIENS | -1 | 2784 | 5kvh | A | 96 | ASN | p.Asn96Ser | O95831 |
| HOMO SAPIENS | 0 | 2784 | 5kvh | A | 282 | THR | p.Thr282Met | O95831 |
| HOMO SAPIENS | 1 | 2784 | 5kvh | A | 366 | ASN | p.Asn366Ser | O95831 |
| HOMO SAPIENS | 3 | 2784 | 5kvh | A | 463 | ARG | p.Arg463Ile | O95831 |
| HOMO SAPIENS | 0 | 2784 | 5kvh | A | 498 | VAL | p.Val498Met | O95831 |
| HOMO SAPIENS | -1 | 2784 | 5kvh | A | 549 | ALA | p.Ala549Val | O95831 |
| HOMO SAPIENS | 0 | 38316 | 2lnk | C | 1933 | ARG | p.Arg1933Ter | P35579 |
| HOMO SAPIENS | 0 | 33808 | 2miu | A | 29 | VAL | p.Val29Met | Q14192 |
| HOMO SAPIENS | 1 | 33808 | 2miu | A | 37 | ALA | p.Ala37Ser | Q14192 |
| HOMO SAPIENS | 2 | 20994 | 2hey | R | 65 | ARG | p.Arg65Cys | P43489 |
| HOMO SAPIENS | -1 | 5687 | 2xwj | I | 7 | ARG | p.Arg32Gln | P00751 |
| HOMO SAPIENS | -1 | 5687 | 2xwj | I | 7 | ARG | p.Arg32Trp | P00751 |
| HOMO SAPIENS | 0 | 5687 | 2xwj | I | 49 | ARG | p.Arg74His | P00751 |
| HOMO SAPIENS | 0 | 5687 | 2xwj | I | 177 | ARG | p.Arg202Gly | P00751 |
| HOMO SAPIENS | 0 | 5687 | 2xwj | I | 217 | ILE | p.Ile242Leu | P00751 |
| HOMO SAPIENS | -1 | 5687 | 2xwj | I | 227 | GLY | p.Gly252Ser | P00751 |
| HOMO SAPIENS | 0 | 5687 | 2xwj | I | 483 | HIS | p.His508Gln | P00751 |
| HOMO SAPIENS | 0 | 5687 | 2xwj | I | 508 | LYS | p.Lys533Arg | P00751 |
| HOMO SAPIENS | 0 | 5687 | 2xwj | I | 540 | LYS | p.Lys565Glu | P00751 |
| HOMO SAPIENS | 0 | 5687 | 2xwj | I | 541 | GLU | p.Glu566Ala | P00751 |
| HOMO SAPIENS | 2 | 953 | 3d4j | A | 249 | PHE | p.Phe249Ser | P53602 |
| HOMO SAPIENS | 1 | 953 | 3d4j | A | 292 | ASN | p.Asn292Ser | P53602 |
| HOMO SAPIENS | 0 | 3300 | 1s9c | A | 140 | ASN | p.Asn457Tyr | P51659 |
| HOMO SAPIENS | 2 | 3300 | 1s9c | A | 167 | ILE | p.Ile484Val | P51659 |
| HOMO SAPIENS | 0 | 3300 | 1s9c | A | 174 | ALA | p.Ala491Thr | P51659 |
| HOMO SAPIENS | 8 | 3300 | 1s9c | A | 193 | ASP | p.Asp510Asn | P51659 |
| HOMO SAPIENS | 3 | 3300 | 1s9c | A | 194 | TRP | p.Trp511Arg | P51659 |
| HOMO SAPIENS | 0 | 3300 | 1s9c | A | 242 | VAL | p.Ile559Val | P51659 |
| HOMO SAPIENS | 0 | 10810 | 5i25 | A | 5 | GLN | p.Gln23Ter | P03951 |
| HOMO SAPIENS | 0 | 10810 | 5i25 | A | 55 | TRP | p.Trp73Ter | P03951 |
| HOMO SAPIENS | 0 | 10810 | 5i25 | A | 91 | ALA | p.Ala109Thr | P03951 |
| HOMO SAPIENS | 0 | 10810 | 5i25 | A | 117 | GLU | p.Glu135Ter | P03951 |
| HOMO SAPIENS | 0 | 10810 | 5i25 | A | 118 | CYS | p.Cys136Ter | P03951 |
| HOMO SAPIENS | 0 | 10810 | 5i25 | A | 128 | CYS | p.Cys146Ter | P03951 |
| HOMO SAPIENS | 0 | 10810 | 5i25 | A | 210 | ARG | p.Arg228Ter | P03951 |
| HOMO SAPIENS | 0 | 10810 | 5i25 | A | 226 | GLN | p.Gln244Ter | P03951 |
| HOMO SAPIENS | 0 | 10810 | 5i25 | A | 233 | GLN | p.Gln251Ter | P03951 |
| HOMO SAPIENS | 0 | 16668 | 3lbx | A | 28 | ARG | p.Arg28Leu | P02549 |
| HOMO SAPIENS | 0 | 16668 | 3lbx | A | 28 | ARG | p.Arg28Cys | P02549 |
| HOMO SAPIENS | 1 | 16668 | 3lbx | A | 41 | ARG | p.Arg41Trp | P02549 |
| HOMO SAPIENS | 1 | 16668 | 3lbx | A | 45 | ARG | p.Arg45Ser | P02549 |
| HOMO SAPIENS | 2 | 16668 | 3lbx | A | 48 | LYS | p.Lys48Arg | P02549 |
| HOMO SAPIENS | 1 | 16668 | 3lbx | A | 152 | ASP | p.Asp152Asn | P02549 |

| | | | | | | | | |
|---|---|---|---|---|---|---|---|---|
| HOMO SAPIENS | 0 | 226 | 2ble | A | 256 | ILE | p.Phe256Ile | P36959 |
| HOMO SAPIENS | 2 | 36808 | 4c7n | A | 3 | SER | p.Ser357Pro | O75030 |
| HOMO SAPIENS | 0 | 4092 | 5jnx | A | 113 | CYS | p.Cys113Arg | O15118 |
| HOMO SAPIENS | 4 | 4092 | 5jnx | A | 215 | HIS | p.His215Arg | O15118 |
| HOMO SAPIENS | 6 | 4092 | 5jnx | A | 237 | PRO | p.Pro237Ser | O15118 |
| HOMO SAPIENS | 0 | 4092 | 5jnx | A | 363 | ALA | p.Ala363Val | O15118 |
| HOMO SAPIENS | 0 | 4092 | 5jnx | A | 378 | VAL | p.Val378Ala | O15118 |
| HOMO SAPIENS | 8 | 4092 | 5jnx | A | 404 | ARG | p.Arg404Gln | O15118 |
| HOMO SAPIENS | 8 | 4092 | 5jnx | A | 411 | ARG | p.Arg411Gln | O15118 |
| HOMO SAPIENS | 2 | 4092 | 5jnx | A | 474 | PRO | p.Pro474Leu | O15118 |
| HOMO SAPIENS | 8 | 4092 | 5jnx | A | 511 | THR | p.Thr511Met | O15118 |
| HOMO SAPIENS | 0 | 4092 | 5jnx | A | 642 | MET | p.Met642Ile | O15118 |
| HOMO SAPIENS | 2 | 4092 | 5jnx | A | 664 | VAL | p.Val664Met | O15118 |
| HOMO SAPIENS | 0 | 4092 | 5jnx | A | 734 | SER | p.Ser734Ile | O15118 |
| HOMO SAPIENS | -1 | 4092 | 5jnx | A | 775 | GLN | p.Gln775Pro | O15118 |
| HOMO SAPIENS | 0 | 4092 | 5jnx | A | 858 | ILE | p.Ile858Val | O15118 |
| HOMO SAPIENS | 0 | 4092 | 5jnx | A | 889 | VAL | p.Val889Met | O15118 |
| HOMO SAPIENS | 0 | 4092 | 5jnx | A | 910 | GLY | p.Gly910Ser | O15118 |
| HOMO SAPIENS | 0 | 4092 | 5jnx | A | 911 | GLY | p.Gly911Ser | O15118 |
| HOMO SAPIENS | 0 | 4092 | 5jnx | A | 928 | GLN | p.Gln928Pro | O15118 |
| HOMO SAPIENS | 0 | 4092 | 5jnx | A | 931 | ASN | p.Asn931Lys | O15118 |
| HOMO SAPIENS | 0 | 4092 | 5jnx | A | 940 | SER | p.Ser940Leu | O15118 |
| HOMO SAPIENS | 0 | 4092 | 5jnx | A | 950 | VAL | p.Val950Met | O15118 |
| HOMO SAPIENS | 0 | 4092 | 5jnx | A | 954 | SER | p.Ser954Leu | O15118 |
| HOMO SAPIENS | 0 | 4092 | 5jnx | A | 958 | ARG | p.Arg958Gln | O15118 |
| HOMO SAPIENS | 0 | 4092 | 5jnx | A | 958 | ARG | p.Arg958Ter | O15118 |
| HOMO SAPIENS | -1 | 4092 | 5jnx | A | 978 | ARG | p.Arg978Cys | O15118 |
| HOMO SAPIENS | 0 | 4092 | 5jnx | A | 992 | GLY | p.Gly992Trp | O15118 |
| HOMO SAPIENS | 0 | 4092 | 5jnx | A | 1007 | PRO | p.Pro1007Ala | O15118 |
| HOMO SAPIENS | 0 | 4092 | 5jnx | A | 1019 | TYR | p.Tyr1019Cys | O15118 |
| HOMO SAPIENS | 2 | 4092 | 5jnx | A | 1035 | ALA | p.Ala1035Val | O15118 |
| HOMO SAPIENS | 2 | 4092 | 5jnx | A | 1054 | ALA | p.Ala1054Thr | O15118 |
| HOMO SAPIENS | 2 | 4092 | 5jnx | A | 1061 | ILE | p.Ile1061Thr | O15118 |
| HOMO SAPIENS | 0 | 4092 | 5jnx | A | 1073 | GLY | p.Gly1073Ser | O15118 |
| HOMO SAPIENS | 2 | 4092 | 5jnx | A | 1088 | TYR | p.Tyr1088Cys | O15118 |
| HOMO SAPIENS | 2 | 4092 | 5jnx | A | 1142 | MET | p.Met1142Thr | O15118 |
| HOMO SAPIENS | 2 | 4092 | 5jnx | A | 1156 | ASN | p.Asn1156Ser | O15118 |
| HOMO SAPIENS | 0 | 4092 | 5jnx | A | 1179 | MET | p.Met1179Val | O15118 |
| HOMO SAPIENS | 0 | 4092 | 5jnx | A | 1186 | ARG | p.Arg1186His | O15118 |
| HOMO SAPIENS | 0 | 4092 | 5jnx | A | 1187 | ALA | p.Ala1187Gly | O15118 |
| HOMO SAPIENS | 2 | 4092 | 5jnx | A | 1189 | GLU | p.Glu1189Gly | O15118 |
| HOMO SAPIENS | 2 | 4092 | 5jnx | A | 1200 | SER | p.Ser1200Gly | O15118 |
| HOMO SAPIENS | 2 | 4092 | 5jnx | A | 1205 | THR | p.Thr1205Lys | O15118 |
| HOMO SAPIENS | 0 | 4092 | 5jnx | A | 1213 | LEU | p.Leu1213Phe | O15118 |
| HOMO SAPIENS | -1 | 4092 | 5jnx | A | 1266 | ARG | p.Arg1266Gln | O15118 |
| HOMO SAPIENS | 9 | 16498 | 4i6o | A | 64 | ARG | p.Arg735Trp | P01024 |
| HOMO SAPIENS | 1 | 21902 | 3un9 | C | 751 | ARG | p.Arg751His | Q86UT6 |
| HOMO SAPIENS | 7 | 20391 | 2h32 | B | 105 | PRO | p.Pro142Leu | P15814 |
| HOMO SAPIENS | 6 | 1635 | 4x45 | A | 65 | ASP | p.Asp65Val | P07741 |
| HOMO SAPIENS | 1 | 1635 | 4x45 | A | 98 | TRP | p.Trp98Ter | P07741 |
| HOMO SAPIENS | 0 | 1635 | 4x45 | A | 110 | LEU | p.Leu110Pro | P07741 |
| HOMO SAPIENS | 7 | 1635 | 4x45 | A | 136 | MET | p.Met136Thr | P07741 |
| HOMO SAPIENS | 1 | 1635 | 4x45 | A | 150 | VAL | p.Val150Phe | P07741 |
| HOMO SAPIENS | 4 | 34810 | 2rpc | A | 16 | CYS | p.Cys253Ser | O60481 |

| | | | | | | | | |
|---|---|---|---|---|---|---|---|---|
| HOMO SAPIENS | 1 | 34810 | 2rpc | A | 18 | TRP | p.Trp255Gly | O60481 |
| HOMO SAPIENS | 1 | 14359 | 4hxi | A | 78 | MET | p.Met78Val | Q9UH77 |
| HOMO SAPIENS | 0 | 5342 | 4d9t | A | 628 | GLU | p.Glu628Asp | P51812 |
| HOMO SAPIENS | -1 | 5342 | 4d9t | A | 723 | ARG | p.Arg723His | P51812 |
| HOMO SAPIENS | -1 | 5342 | 4d9t | A | 729 | ARG | p.Arg729Gln | P51812 |
| HOMO SAPIENS | 1 | 9619 | 3j2s | B | 1781 | ARG | p.Arg1800Cys | P00451 |
| HOMO SAPIENS | 0 | 9619 | 3j2s | B | 1823 | MET | p.Met1842Ile | P00451 |
| HOMO SAPIENS | 1 | 9619 | 3j2s | B | 1825 | PRO | p.Pro1844Ser | P00451 |
| HOMO SAPIENS | 0 | 9619 | 3j2s | B | 1826 | THR | p.Thr1845Pro | P00451 |
| HOMO SAPIENS | 0 | 9619 | 3j2s | B | 1834 | ALA | p.Ala1853Val | P00451 |
| HOMO SAPIENS | 0 | 9619 | 3j2s | B | 1846 | ASP | p.Asp1865Asn | P00451 |
| HOMO SAPIENS | 1 | 9619 | 3j2s | B | 1848 | HIS | p.His1867Arg | P00451 |
| HOMO SAPIENS | 0 | 9619 | 3j2s | B | 1854 | PRO | p.Pro1873Arg | P00451 |
| HOMO SAPIENS | 1 | 9619 | 3j2s | B | 1885 | GLU | p.Glu1904Lys | P00451 |
| HOMO SAPIENS | 0 | 9619 | 3j2s | B | 1922 | ASN | p.Asn1941Ser | P00451 |
| HOMO SAPIENS | 1 | 9619 | 3j2s | B | 1941 | ARG | p.Arg1960Gln | P00451 |
| HOMO SAPIENS | 4 | 9619 | 3j2s | B | 2105 | TYR | p.Tyr2124Cys | P00451 |
| HOMO SAPIENS | 6 | 9619 | 3j2s | B | 2150 | ARG | p.Arg2169His | P00451 |
| HOMO SAPIENS | 0 | 9619 | 3j2s | B | 2159 | ARG | p.Arg2178Cys | P00451 |
| HOMO SAPIENS | 0 | 9619 | 3j2s | B | 2159 | ARG | p.Arg2178His | P00451 |
| HOMO SAPIENS | 2 | 9619 | 3j2s | B | 2238 | MET | p.Met2257Val | P00451 |
| HOMO SAPIENS | 2 | 9619 | 3j2s | B | 2300 | PRO | p.Pro2319Ser | P00451 |
| HOMO SAPIENS | 1 | 667 | 1z2m | A | 127 | GLU | p.Glu127Ter | P05161 |
| HOMO SAPIENS | 0 | 15717 | 2nn6 | G | 30 | GLY | p.Gly31Ala | Q9NQT5 |
| HOMO SAPIENS | -1 | 15717 | 2nn6 | G | 55 | ASN | p.Asn56His | Q9NQT5 |
| HOMO SAPIENS | 7 | 15717 | 2nn6 | G | 131 | ASP | p.Asp132Ala | Q9NQT5 |
| HOMO SAPIENS | 7 | 784 | 3rkq | A | 161 | ARG | p.Arg161Pro | P52952 |
| HOMO SAPIENS | 6 | 784 | 3rkq | A | 178 | THR | p.Thr178Met | P52952 |
| HOMO SAPIENS | 7 | 784 | 3rkq | A | 183 | LYS | p.Lys183Glu | P52952 |
| HOMO SAPIENS | 8 | 784 | 3rkq | A | 190 | ARG | p.Arg190Cys | P52952 |
| HOMO SAPIENS | 6 | 385 | 30000 | C | 70 | ALA | p.Ala70Thr | P07954 |
| HOMO SAPIENS | 2 | 385 | 30000 | C | 101 | ARG | p.Arg101Pro | P07954 |
| HOMO SAPIENS | 2 | 385 | 30000 | C | 174 | PRO | p.Pro174Arg | P07954 |
| HOMO SAPIENS | 6 | 385 | 30000 | C | 187 | SER | p.Ser187Ter | P07954 |
| HOMO SAPIENS | 9 | 385 | 30000 | C | 233 | ARG | p.Arg233His | P07954 |
| HOMO SAPIENS | 3 | 385 | 30000 | C | 273 | ALA | p.Ala273Thr | P07954 |
| HOMO SAPIENS | 4 | 385 | 30000 | C | 334 | SER | p.Ser334Arg | P07954 |
| HOMO SAPIENS | 6 | 385 | 30000 | C | 340 | ASN | p.Asn340Lys | P07954 |
| HOMO SAPIENS | 8 | 385 | 30000 | C | 343 | ARG | p.Arg343Ter | P07954 |
| HOMO SAPIENS | 3 | 385 | 30000 | C | 350 | ARG | p.Arg350Trp | P07954 |
| HOMO SAPIENS | 4 | 385 | 30000 | C | 376 | GLN | p.Gln376Pro | P07954 |
| HOMO SAPIENS | 6 | 385 | 30000 | C | 434 | CYS | p.Cys434Ter | P07954 |
| HOMO SAPIENS | 0 | 3316 | 4f91 | B | 934 | THR | p.Thr934Ala | O75643 |
| HOMO SAPIENS | 0 | 3316 | 4f91 | B | 1002 | ASN | p.Asn1002Ser | O75643 |
| HOMO SAPIENS | 0 | 3316 | 4f91 | B | 1087 | SER | p.Ser1087Leu | O75643 |
| HOMO SAPIENS | 4 | 3316 | 4f91 | B | 1389 | VAL | p.Val1389Ile | O75643 |
| HOMO SAPIENS | 0 | 619 | 5hbt | B | 195 | ASP | p.Asp240His | P02708 |
| HOMO SAPIENS | 1 | 619 | 5hbt | B | 209 | ARG | p.Arg254Leu | P02708 |
| HOMO SAPIENS | 0 | 18171 | 2y25 | B | 1408 | ASP | p.Asp1408Asn | P52179 |
| HOMO SAPIENS | 0 | 18171 | 2y25 | B | 1453 | MET | p.Met1453Thr | P52179 |
| HOMO SAPIENS | 0 | 18171 | 2y25 | B | 1459 | ILE | p.Ile1459Lys | P52179 |
| HOMO SAPIENS | 0 | 18171 | 2y25 | B | 1558 | PHE | p.Phe1558Leu | P52179 |
| HOMO SAPIENS | 0 | 18171 | 2y25 | B | 1573 | ARG | p.Arg1573Gln | P52179 |
| HOMO SAPIENS | 7 | 18171 | 2y25 | B | 1663 | VAL | p.Val1663Met | P52179 |

| | | | | | | | | |
|---|---|---|---|---|---|---|---|---|
| HOMO SAPIENS | 8 | 14262 | 2fcw | B | 117 | SER | p.Ser138Ter | P01130 |
| HOMO SAPIENS | 8 | 14262 | 2fcw | B | 118 | ASP | p.Asp139Glu | P01130 |
| HOMO SAPIENS | 9 | 14262 | 2fcw | B | 119 | GLU | p.Glu140Lys | P01130 |
| HOMO SAPIENS | 7 | 14262 | 2fcw | B | 122 | CYS | p.Cys143Ter | P01130 |
| HOMO SAPIENS | 6 | 14262 | 2fcw | B | 134 | CYS | p.Cys155Ter | P01130 |
| HOMO SAPIENS | 7 | 14262 | 2fcw | B | 147 | ASP | p.Asp168Glu | P01130 |
| HOMO SAPIENS | 4 | 2624 | 3b7b | A | 855 | VAL | p.Val886Met | Q9H9B1 |
| HOMO SAPIENS | 7 | 2624 | 3b7b | A | 868 | ILE | p.Ile899Val | Q9H9B1 |
| HOMO SAPIENS | 0 | 2624 | 3b7b | A | 892 | GLU | p.Glu923Lys | Q9H9B1 |
| HOMO SAPIENS | 0 | 4791 | 5ctd | B | 68 | LEU | p.Pro549Ala | Q14055 |
| HOMO SAPIENS | 5 | 460 | 4ui1 | D | 80 | CYS | p.Cys80Arg | Q6ZVN8 |
| HOMO SAPIENS | 3 | 460 | 4ui1 | D | 101 | LEU | p.Leu101Pro | Q6ZVN8 |
| HOMO SAPIENS | 0 | 40782 | 1sy6 | A | 47 | LYS | p.Lys69Ter | P09693 |
| HOMO SAPIENS | -1 | 1092 | 2idx | A | 62 | THR | p.Thr62Met | Q96EY8 |
| HOMO SAPIENS | 6 | 1092 | 2idx | A | 183 | HIS | p.His183Leu | Q96EY8 |
| HOMO SAPIENS | 7 | 1092 | 2idx | A | 186 | ARG | p.Arg186Trp | Q96EY8 |
| HOMO SAPIENS | 6 | 1092 | 2idx | A | 190 | ARG | p.Arg190Cys | Q96EY8 |
| HOMO SAPIENS | 6 | 1092 | 2idx | A | 190 | ARG | p.Arg190His | Q96EY8 |
| HOMO SAPIENS | 7 | 1092 | 2idx | A | 191 | ARG | p.Arg191Trp | Q96EY8 |
| HOMO SAPIENS | 7 | 1092 | 2idx | A | 191 | ARG | p.Arg191Gln | Q96EY8 |
| HOMO SAPIENS | 6 | 1092 | 2idx | A | 195 | ARG | p.Arg195His | Q96EY8 |
| HOMO SAPIENS | 1 | 1092 | 2idx | A | 234 | GLN | p.Gln234Ter | Q96EY8 |
| HOMO SAPIENS | 0 | 1092 | 2idx | A | 239 | MET | p.Met239Lys | Q96EY8 |
| HOMO SAPIENS | -1 | 6418 | 5fur | G | 11 | THR | p.Thr11Ile | P21675 |
| HOMO SAPIENS | -1 | 6418 | 5fur | G | 290 | LEU | p.Leu269Val | P21675 |
| HOMO SAPIENS | 0 | 6100 | 1wiq | A | 240 | ARG | p.Arg265Trp | P01730 |
| HOMO SAPIENS | 0 | 13757 | 4ye6 | A | 45 | GLY | p.Gly45Val | P47897 |
| HOMO SAPIENS | 1 | 13757 | 4ye6 | A | 57 | TYR | p.Tyr57His | P47897 |
| HOMO SAPIENS | 8 | 13757 | 4ye6 | A | 403 | ARG | p.Arg403Trp | P47897 |
| HOMO SAPIENS | 7 | 13757 | 4ye6 | A | 524 | ARG | p.Arg524Trp | P47897 |
| HOMO SAPIENS | 6 | 13757 | 4ye6 | A | 690 | ARG | p.Arg690Cys | P47897 |
| HOMO SAPIENS | 0 | 26958 | 2lzl | A | 370 | GLY | p.Gly370Cys | P22607 |
| HOMO SAPIENS | 0 | 26958 | 2lzl | A | 371 | SER | p.Ser371Cys | P22607 |
| HOMO SAPIENS | 0 | 26958 | 2lzl | A | 373 | TYR | p.Tyr373Cys | P22607 |
| HOMO SAPIENS | 0 | 26958 | 2lzl | A | 380 | GLY | p.Gly380Arg | P22607 |
| HOMO SAPIENS | 0 | 26958 | 2lzl | A | 384 | PHE | p.Phe384Leu | P22607 |
| HOMO SAPIENS | 5 | 26958 | 2lzl | A | 391 | ALA | p.Ala391Glu | P22607 |
| HOMO SAPIENS | 6 | 11325 | 2wa6 | A | 121 | TYR | p.Tyr121Phe | O75369 |
| HOMO SAPIENS | 2 | 11325 | 2wa6 | A | 161 | PHE | p.Phe161Cys | O75369 |
| HOMO SAPIENS | 0 | 11325 | 2wa6 | A | 171 | LEU | p.Leu171Arg | O75369 |
| HOMO SAPIENS | 2 | 11325 | 2wa6 | A | 173 | ALA | p.Ala173Val | O75369 |
| HOMO SAPIENS | 0 | 11325 | 2wa6 | A | 202 | MET | p.Met202Val | O75369 |
| HOMO SAPIENS | 3 | 11325 | 2wa6 | A | 227 | GLU | p.Glu227Lys | O75369 |
| HOMO SAPIENS | 0 | 11325 | 2wa6 | A | 234 | LEU | p.Leu234Val | O75369 |
| HOMO SAPIENS | 2 | 11325 | 2wa6 | A | 235 | SER | p.Ser235Pro | O75369 |
| HOMO SAPIENS | 9 | 7240 | 3tgx | A | 137 | TYR | p.Tyr156Cys | Q9HBE5 |
| HOMO SAPIENS | 0 | 35479 | 4elj | A | 307 | THR | p.Thr307Ile | P06400 |
| HOMO SAPIENS | 0 | 35479 | 4elj | A | 310 | GLY | p.Gly310Glu | P06400 |
| HOMO SAPIENS | 0 | 35479 | 4elj | A | 377 | THR | p.Thr377Ser | P06400 |
| HOMO SAPIENS | 9 | 36706 | 1he7 | A | 359 | TYR | p.Tyr359Cys | P04629 |
| HOMO SAPIENS | 4 | 6538 | 4d0e | A | 504 | ARG | p.Arg504His | P46531 |
| HOMO SAPIENS | 1 | 26518 | 5fm5 | M | 600 | GLU | p.Glu600Val | P52179 |
| HOMO SAPIENS | -1 | 26518 | 5fm5 | M | 651 | ARG | p.Arg651Gln | P52179 |
| HOMO SAPIENS | -1 | 26518 | 5fm5 | M | 660 | PRO | p.Pro660Ala | P52179 |

| | | | | | | | | |
|---|---|---|---|---|---|---|---|---|
| HOMO SAPIENS | -1 | 26518 | 5fm5 | M | 688 | THR | p.Thr688Ser | P52179 |
| HOMO SAPIENS | -1 | 26518 | 5fm5 | M | 704 | GLU | p.Glu704Lys | P52179 |
| HOMO SAPIENS | -1 | 26518 | 5fm5 | M | 727 | THR | p.Thr727Ala | P52179 |
| HOMO SAPIENS | 2 | 27054 | 4rhp | B | 209 | LEU | p.Leu209Val | O75208 |
| HOMO SAPIENS | -1 | 27054 | 4rhp | B | 288 | LYS | p.Lys288Asn | O75208 |
| HOMO SAPIENS | 5 | 2384 | 5fqd | B | 419 | ARG | p.Arg419Ter | Q96SW2 |
| HOMO SAPIENS | 4 | 5154 | 2hl3 | A | 56 | LYS | p.Lys56Arg | Q14203 |
| HOMO SAPIENS | 0 | 5154 | 2hl3 | A | 59 | GLY | p.Gly59Ser | Q14203 |
| HOMO SAPIENS | 0 | 5154 | 2hl3 | A | 71 | GLY | p.Gly71Ala | Q14203 |
| HOMO SAPIENS | 0 | 5154 | 2hl3 | A | 71 | GLY | p.Gly71Arg | Q14203 |
| HOMO SAPIENS | 7 | 5154 | 2hl3 | A | 72 | THR | p.Thr72Pro | Q14203 |
| HOMO SAPIENS | 2 | 5154 | 2hl3 | A | 74 | GLN | p.Gln74Pro | Q14203 |
| HOMO SAPIENS | 2 | 20039 | 2l1g | A | 9 | GLY | p.Gly9Cys | Q9NVV9 |
| HOMO SAPIENS | 5 | 20039 | 2l1g | A | 81 | PHE | p.Phe81Leu | Q9NVV9 |
| HOMO SAPIENS | -1 | 97 | 5uz7 | A | 201 | ARG | p.Arg201His | P63092 |
| HOMO SAPIENS | -1 | 97 | 5uz7 | A | 201 | ARG | p.Arg201Cys | P63092 |
| HOMO SAPIENS | 0 | 3257 | 4b7w | B | 278 | GLU | p.Glu278Asp | P41235 |
| HOMO SAPIENS | 1 | 3257 | 4b7w | B | 331 | ARG | p.Arg331Cys | P41235 |
| HOMO SAPIENS | 0 | 2814 | 4qgk | B | 10 | GLN | p.Gln10Ter | P51648 |
| HOMO SAPIENS | 0 | 2814 | 4qgk | B | 177 | ARG | p.Arg177Ter | P51648 |
| HOMO SAPIENS | 9 | 2814 | 4qgk | B | 184 | THR | p.Thr184Met | P51648 |
| HOMO SAPIENS | 1 | 2814 | 4qgk | B | 188 | ALA | p.Ala188Val | P51648 |
| HOMO SAPIENS | 0 | 2814 | 4qgk | B | 214 | CYS | p.Cys214Tyr | P51648 |
| HOMO SAPIENS | 0 | 2814 | 4qgk | B | 245 | ASP | p.Asp245Asn | P51648 |
| HOMO SAPIENS | 0 | 2814 | 4qgk | B | 266 | LYS | p.Lys266Asn | P51648 |
| HOMO SAPIENS | 0 | 2814 | 4qgk | B | 315 | PRO | p.Pro315Ser | P51648 |
| HOMO SAPIENS | 0 | 2814 | 4qgk | B | 386 | ASN | p.Asn386Ser | P51648 |
| HOMO SAPIENS | -1 | 5486 | 5u6o | B | 47 | GLY | p.Gly47Val | O60741 |
| HOMO SAPIENS | 1 | 21475 | 3pdy | A | 679 | THR | p.Thr789Ile | Q15149 |
| HOMO SAPIENS | 0 | 32890 | 4djc | B | 1501 | LEU | p.Leu1501Val | Q14524 |
| HOMO SAPIENS | -1 | 32890 | 4djc | B | 1512 | ARG | p.Arg1512Trp | Q14524 |
| HOMO SAPIENS | 0 | 26683 | 2xze | B | 38 | ARG | p.Arg38Cys | O95630 |
| HOMO SAPIENS | 0 | 26683 | 2xze | B | 100 | PHE | p.Phe100Tyr | O95630 |
| HOMO SAPIENS | -1 | 23175 | 5dka | A | 163 | SER | p.Ser163Leu | Q96EQ8 |
| HOMO SAPIENS | -1 | 23175 | 5dka | A | 174 | ARG | p.Arg174Cys | Q96EQ8 |
| HOMO SAPIENS | 6 | 21177 | 3k1r | A | 155 | ARG | p.Arg155Ter | Q9Y6N9 |
| HOMO SAPIENS | 0 | 1237 | 2wwm | O | 40 | PRO | p.Pro40Arg | O75147 |
| HOMO SAPIENS | 1 | 2543 | 5ewl | B | 64 | VAL | p.Val64Leu | Q13224 |
| HOMO SAPIENS | 2 | 2543 | 5ewl | B | 271 | ALA | p.Ala271Glu | Q13224 |
| HOMO SAPIENS | 3 | 34262 | 2mng | A | 613 | PHE | p.Phe613Leu | Q9Y3Q4 |
| HOMO SAPIENS | 7 | 34262 | 2mng | A | 672 | SER | p.Ser672Arg | Q9Y3Q4 |
| HOMO SAPIENS | 4 | 1667 | 1f05 | A | 321 | LYS | p.Lys321Met | P37837 |
| HOMO SAPIENS | 0 | 25179 | 2knc | B | 752 | SER | p.Ser778Pro | P05106 |
| HOMO SAPIENS | 0 | 2185 | 2ig7 | A | 50 | TYR | p.Tyr50Cys | Q9Y259 |
| HOMO SAPIENS | 0 | 2185 | 2ig7 | A | 224 | ASN | p.Asn224His | Q9Y259 |
| HOMO SAPIENS | 0 | 2185 | 2ig7 | A | 270 | TYR | p.Tyr270Ter | Q9Y259 |
| HOMO SAPIENS | 0 | 2185 | 2ig7 | A | 328 | GLN | p.Gln328Arg | Q9Y259 |
| HOMO SAPIENS | 3 | 2201 | 2r0o | A | 179 | PRO | p.Pro179Leu | O43707 |
| HOMO SAPIENS | 3 | 2201 | 2r0o | A | 255 | GLU | p.Lys255Glu | O43707 |
| HOMO SAPIENS | 3 | 2201 | 2r0o | A | 259 | THR | p.Thr259Ile | O43707 |
| HOMO SAPIENS | 4 | 2201 | 2r0o | A | 262 | SER | p.Ser262Pro | O43707 |
| HOMO SAPIENS | -1 | 31221 | 2wvr | C | 44 | THR | p.Thr44Pro | Q9H211 |
| HOMO SAPIENS | -1 | 31221 | 2wvr | C | 135 | ALA | p.Ala135Val | Q9H211 |
| HOMO SAPIENS | -1 | 31221 | 2wvr | C | 456 | GLU | p.Glu456Ala | Q9H211 |

| | | | | | | | | |
|---|---|---|---|---|---|---|---|---|
| HOMO SAPIENS | -1 | 31221 | 2wvr | C | 462 | ARG | p.Arg462Gln | Q9H211 |
| HOMO SAPIENS | -1 | 31221 | 2wvr | C | 468 | GLU | p.Glu468Lys | Q9H211 |
| HOMO SAPIENS | -1 | 31221 | 2wvr | C | 537 | ALA | p.Ala537Val | Q9H211 |
| HOMO SAPIENS | 5 | 2576 | 5o3l | C | 363 | VAL | p.Val680Ile | P10636 |
| HOMO SAPIENS | 5 | 1345 | 1czt | A | 37 | ARG | p.Arg2102Cys | P12259 |
| HOMO SAPIENS | 1 | 1345 | 1czt | A | 83 | MET | p.Met2148Thr | P12259 |
| HOMO SAPIENS | 1 | 1345 | 1czt | A | 157 | ASP | p.Asp2222Gly | P12259 |
| HOMO SAPIENS | 8 | 35985 | 1lo1 | A | 110 | ALA | p.Ala110Val | O95718 |
| HOMO SAPIENS | 1 | 617 | 2ocp | B | 44 | GLU | p.Glu44Lys | Q16854 |
| HOMO SAPIENS | 5 | 617 | 2ocp | B | 46 | ASN | p.Asn46Ser | Q16854 |
| HOMO SAPIENS | 0 | 617 | 2ocp | B | 59 | THR | p.Thr59Met | Q16854 |
| HOMO SAPIENS | 7 | 617 | 2ocp | B | 142 | ARG | p.Arg142Lys | Q16854 |
| HOMO SAPIENS | 3 | 617 | 2ocp | B | 154 | ASN | p.Asn154Lys | Q16854 |
| HOMO SAPIENS | 3 | 617 | 2ocp | B | 170 | GLN | p.Gln170Arg | Q16854 |
| HOMO SAPIENS | 3 | 617 | 2ocp | B | 227 | GLU | p.Glu227Lys | Q16854 |
| HOMO SAPIENS | 3 | 617 | 2ocp | B | 233 | LYS | p.Lys233Glu | Q16854 |
| HOMO SAPIENS | 3 | 617 | 2ocp | B | 255 | ASP | p.Asp255Tyr | Q16854 |
| HOMO SAPIENS | -1 | 38975 | 5itz | D | 421 | GLN | p.Gln421His | Q9HC77 |
| HOMO SAPIENS | 2 | 6235 | 2f8v | A | 15 | VAL | p.Val15Ile | Q8WZ42 |
| HOMO SAPIENS | 0 | 6235 | 2f8v | A | 54 | VAL | p.Val54Met | Q8WZ42 |
| HOMO SAPIENS | 0 | 6235 | 2f8v | A | 59 | SER | p.Ser59Arg | Q8WZ42 |
| HOMO SAPIENS | 0 | 6235 | 2f8v | A | 60 | ASP | p.Asp60Tyr | Q8WZ42 |
| HOMO SAPIENS | 0 | 12630 | 1ext | B | 12 | ASP | p.Asp41Glu | P19438 |
| HOMO SAPIENS | 3 | 12630 | 1ext | B | 30 | CYS | p.Cys59Arg | P19438 |
| HOMO SAPIENS | 3 | 12630 | 1ext | B | 30 | CYS | p.Cys59Ser | P19438 |
| HOMO SAPIENS | 5 | 12630 | 1ext | B | 33 | CYS | p.Cys62Tyr | P19438 |
| HOMO SAPIENS | 5 | 12630 | 1ext | B | 33 | CYS | p.Cys62Gly | P19438 |
| HOMO SAPIENS | 1 | 12630 | 1ext | B | 46 | PRO | p.Pro75Leu | P19438 |
| HOMO SAPIENS | 3 | 12630 | 1ext | B | 50 | THR | p.Thr79Met | P19438 |
| HOMO SAPIENS | 6 | 12630 | 1ext | B | 52 | CYS | p.Cys81Phe | P19438 |
| HOMO SAPIENS | 6 | 12630 | 1ext | B | 70 | CYS | p.Cys99Ser | P19438 |
| HOMO SAPIENS | 5 | 12630 | 1ext | B | 88 | CYS | p.Cys117Arg | P19438 |
| HOMO SAPIENS | 5 | 12630 | 1ext | B | 88 | CYS | p.Cys117Tyr | P19438 |
| HOMO SAPIENS | -1 | 12630 | 1ext | B | 170 | ILE | p.Ile199Asn | P19438 |
| HOMO SAPIENS | 0 | 994 | 1aos | B | 95 | ARG | p.Arg95Cys | P04424 |
| HOMO SAPIENS | 0 | 994 | 1aos | B | 111 | ARG | p.Arg111Gln | P04424 |
| HOMO SAPIENS | 0 | 994 | 1aos | B | 131 | THR | p.Thr131Met | P04424 |
| HOMO SAPIENS | 6 | 994 | 1aos | B | 178 | VAL | p.Val178Met | P04424 |
| HOMO SAPIENS | 9 | 994 | 1aos | B | 182 | ARG | p.Arg182Gln | P04424 |
| HOMO SAPIENS | 9 | 994 | 1aos | B | 186 | ARG | p.Arg186Trp | P04424 |
| HOMO SAPIENS | 8 | 994 | 1aos | B | 193 | ARG | p.Arg193Gln | P04424 |
| HOMO SAPIENS | 5 | 994 | 1aos | B | 217 | ARG | p.Arg217Ter | P04424 |
| HOMO SAPIENS | 0 | 994 | 1aos | B | 286 | GLN | p.Gln286Arg | P04424 |
| HOMO SAPIENS | 5 | 994 | 1aos | B | 379 | ARG | p.Arg379Cys | P04424 |
| HOMO SAPIENS | 4 | 16066 | 5mv9 | A | 1719 | TYR | p.Tyr1719Cys | Q13402 |
| HOMO SAPIENS | -1 | 16066 | 5mv9 | A | 1739 | ARG | p.Arg1739Ter | Q13402 |
| HOMO SAPIENS | 4 | 16066 | 5mv9 | A | 1775 | ILE | p.Ile1775Thr | Q13402 |
| HOMO SAPIENS | 2 | 16066 | 5mv9 | A | 1786 | SER | p.Ser1786Thr | Q13402 |
| HOMO SAPIENS | 4 | 16066 | 5mv9 | A | 1798 | GLN | p.Gln1798Ter | Q13402 |
| HOMO SAPIENS | 0 | 16066 | 5mv9 | A | 1858 | LEU | p.Leu1858Pro | Q13402 |
| HOMO SAPIENS | 6 | 16066 | 5mv9 | A | 1873 | ARG | p.Arg1873Trp | Q13402 |
| HOMO SAPIENS | 6 | 16066 | 5mv9 | A | 1873 | ARG | p.Arg1873Leu | Q13402 |
| HOMO SAPIENS | 6 | 16066 | 5mv9 | A | 1883 | ARG | p.Arg1883Gln | Q13402 |
| HOMO SAPIENS | 4 | 16066 | 5mv9 | A | 1887 | PRO | p.Pro1887Leu | Q13402 |

| | | | | | | | | |
|---|---|---|---|---|---|---|---|---|
| HOMO SAPIENS | 5 | 16066 | 5mv9 | A | 1942 | GLY | p.Gly1942Ter | Q13402 |
| HOMO SAPIENS | 5 | 16066 | 5mv9 | A | 1954 | LEU | p.Leu1954Ile | Q13402 |
| HOMO SAPIENS | 5 | 16066 | 5mv9 | A | 1956 | VAL | p.Val1956Ile | Q13402 |
| HOMO SAPIENS | 5 | 16066 | 5mv9 | A | 1967 | ARG | p.Arg1967Ter | Q13402 |
| HOMO SAPIENS | 5 | 16066 | 5mv9 | A | 1990 | GLN | p.Gln1990Ter | Q13402 |
| HOMO SAPIENS | 2 | 16066 | 5mv9 | A | 2005 | LYS | p.Lys2005Glu | Q13402 |
| HOMO SAPIENS | 1 | 16066 | 5mv9 | A | 2072 | VAL | p.Val2072Ile | Q13402 |
| HOMO SAPIENS | 4 | 16066 | 5mv9 | A | 2109 | THR | p.Thr2109Ile | Q13402 |
| HOMO SAPIENS | 4 | 16066 | 5mv9 | A | 2142 | ASP | p.Asp2142Asn | Q13402 |
| HOMO SAPIENS | -1 | 16066 | 5mv9 | A | 2214 | GLY | p.Gly2214Ser | Q13402 |
| HOMO SAPIENS | 7 | 27942 | 1deb | A | 24 | ARG | p.Arg24Ter | P25054 |
| HOMO SAPIENS | 0 | 876 | 5hnr | B | 12 | PHE | p.Phe12Leu | P13716 |
| HOMO SAPIENS | 0 | 876 | 5hnr | B | 59 | LYS | p.Lys59Asn | P13716 |
| HOMO SAPIENS | -1 | 876 | 5hnr | B | 133 | GLY | p.Gly133Arg | P13716 |
| HOMO SAPIENS | 2 | 876 | 5hnr | B | 240 | ARG | p.Arg240Trp | P13716 |
| HOMO SAPIENS | 0 | 876 | 5hnr | B | 274 | ALA | p.Ala274Thr | P13716 |
| HOMO SAPIENS | 0 | 876 | 5hnr | B | 275 | VAL | p.Val275Met | P13716 |
| HOMO SAPIENS | 0 | 1142 | 3qe2 | A | 503 | VAL | p.Ala500Val | P16435 |
| HOMO SAPIENS | 8 | 14358 | 4dkd | C | 32 | VAL | p.Val32Gly | P07333 |
| HOMO SAPIENS | 0 | 14358 | 4dkd | C | 60 | LEU | p.Leu60Val | P07333 |
| HOMO SAPIENS | 7 | 14358 | 4dkd | C | 75 | THR | p.Thr75Ile | P07333 |
| HOMO SAPIENS | -1 | 14358 | 4dkd | C | 90 | PRO | p.Pro90Thr | P07333 |
| HOMO SAPIENS | 0 | 14358 | 4dkd | C | 106 | ARG | p.Arg106Trp | P07333 |
| HOMO SAPIENS | 0 | 14358 | 4dkd | C | 241 | ASN | p.Asn241Asp | P07333 |
| HOMO SAPIENS | 0 | 14358 | 4dkd | C | 245 | ALA | p.Ala245Ser | P07333 |
| HOMO SAPIENS | 0 | 14358 | 4dkd | C | 255 | ASN | p.Asn255Ile | P07333 |
| HOMO SAPIENS | 0 | 14358 | 4dkd | C | 268 | VAL | p.Val268Ile | P07333 |
| HOMO SAPIENS | 0 | 14358 | 4dkd | C | 279 | VAL | p.Val279Met | P07333 |
| HOMO SAPIENS | 0 | 14358 | 4dkd | C | 294 | ARG | p.Arg294Gln | P07333 |
| HOMO SAPIENS | 5 | 8968 | 3ppy | A | 1565 | VAL | p.Val1565Leu | P04275 |
| HOMO SAPIENS | 4 | 8968 | 3ppy | A | 1584 | TYR | p.Tyr1584Cys | P04275 |
| HOMO SAPIENS | 5 | 5341 | 3k3n | A | 279 | PHE | p.Phe315Ser | Q9UPP1 |
| HOMO SAPIENS | 0 | 29463 | 1q7l | A | 197 | ARG | p.Arg197Trp | Q03154 |
| HOMO SAPIENS | 0 | 33761 | 2leh | B | 30 | ASP | p.Asp30Asn | Q16637 |
| HOMO SAPIENS | 6 | 33761 | 2leh | B | 44 | ASP | p.Asp44Val | Q16637 |
| HOMO SAPIENS | -1 | 12249 | 2pnr | C | 224 | GLU | p.Glu310Lys | P10515 |
| HOMO SAPIENS | -1 | 12249 | 2pnr | C | 232 | VAL | p.Val318Ala | P10515 |
| HOMO SAPIENS | -1 | 7 | 5whk | B | -9 | ALA | p.Ala11Pro | P61769 |
| HOMO SAPIENS | 8 | 20298 | 4euw | A | 154 | PHE | p.Phe154Leu | P48436 |
| HOMO SAPIENS | 9 | 20298 | 4euw | A | 158 | ALA | p.Ala158Thr | P48436 |
| HOMO SAPIENS | 5 | 20298 | 4euw | A | 165 | HIS | p.His165Tyr | P48436 |
| HOMO SAPIENS | 3 | 20298 | 4euw | A | 169 | HIS | p.His169Gln | P48436 |
| HOMO SAPIENS | 0 | 20298 | 4euw | A | 173 | LYS | p.Lys173Glu | P48436 |
| HOMO SAPIENS | 1 | 16465 | 2jgx | A | 402 | TYR | p.Tyr402His | P08603 |
| HOMO SAPIENS | 0 | 8993 | 3v8o | A | 659 | LEU | p.Leu659Arg | Q14315 |
| HOMO SAPIENS | 6 | 8993 | 3v8o | A | 693 | ASP | p.Asp693Ala | Q14315 |
| HOMO SAPIENS | 0 | 8993 | 3v8o | A | 709 | ALA | p.Ala709Thr | Q14315 |
| HOMO SAPIENS | 2 | 3852 | 5es4 | B | 106 | ASP | p.Asp128Asn | P05107 |
| HOMO SAPIENS | 2 | 3852 | 5es4 | B | 116 | SER | p.Ser138Pro | P05107 |
| HOMO SAPIENS | 1 | 3852 | 5es4 | B | 127 | LEU | p.Leu149Pro | P05107 |
| HOMO SAPIENS | 0 | 3852 | 5es4 | B | 147 | GLY | p.Gly169Arg | P05107 |
| HOMO SAPIENS | 3 | 3852 | 5es4 | B | 156 | PRO | p.Pro178Leu | P05107 |
| HOMO SAPIENS | 0 | 3852 | 5es4 | B | 174 | LYS | p.Lys196Thr | P05107 |
| HOMO SAPIENS | 2 | 3852 | 5es4 | B | 251 | GLY | p.Gly273Arg | P05107 |

| | | | | | | | | |
|---|---|---|---|---|---|---|---|---|
| HOMO SAPIENS | 2 | 3852 | 5es4 | B | 329 | ASN | p.Asn351Ser | P05107 |
| HOMO SAPIENS | 2 | 3852 | 5es4 | B | 571 | ARG | p.Arg593Cys | P05107 |
| HOMO SAPIENS | 7 | 5962 | 5fn2 | B | 146 | MET | p.Met146Leu | P49768 |
| HOMO SAPIENS | 3 | 5962 | 5fn2 | B | 166 | LEU | p.Leu166Pro | P49768 |
| HOMO SAPIENS | 1 | 5962 | 5fn2 | B | 233 | MET | p.Met233Leu | P49768 |
| HOMO SAPIENS | 1 | 5962 | 5fn2 | B | 261 | VAL | p.Val261Gly | P49768 |
| HOMO SAPIENS | 1 | 5962 | 5fn2 | B | 266 | GLY | p.Gly266Ser | P49768 |
| HOMO SAPIENS | -1 | 5962 | 5fn2 | B | 333 | ASP | p.Asp333Gly | P49768 |
| HOMO SAPIENS | 6 | 1918 | 3dlx | A | 58 | THR | p.Thr58Met | P55809 |
| HOMO SAPIENS | 5 | 1918 | 3dlx | A | 133 | VAL | p.Val133Glu | P55809 |
| HOMO SAPIENS | 5 | 1918 | 3dlx | A | 219 | GLY | p.Gly219Glu | P55809 |
| HOMO SAPIENS | 0 | 1918 | 3dlx | A | 221 | VAL | p.Val221Met | P55809 |
| HOMO SAPIENS | 5 | 1918 | 3dlx | A | 283 | SER | p.Ser283Ter | P55809 |
| HOMO SAPIENS | 4 | 1918 | 3dlx | A | 324 | GLY | p.Gly324Glu | P55809 |
| HOMO SAPIENS | 0 | 1918 | 3dlx | A | 456 | CYS | p.Cys456Phe | P55809 |
| HOMO SAPIENS | 0 | 9010 | 2wfr | A | 162 | LEU | p.Leu162Pro | O43323 |
| HOMO SAPIENS | 5 | 3770 | 5n38 | A | 56 | VAL | p.Val56Glu | O60260 |
| HOMO SAPIENS | 1 | 3770 | 5n38 | A | 161 | LYS | p.Lys161Asn | O60260 |
| HOMO SAPIENS | 1 | 3770 | 5n38 | A | 167 | SER | p.Ser167Asn | O60260 |
| HOMO SAPIENS | 1 | 3770 | 5n38 | A | 211 | LYS | p.Lys211Asn | O60260 |
| HOMO SAPIENS | 1 | 3770 | 5n38 | A | 212 | CYS | p.Cys212Tyr | O60260 |
| HOMO SAPIENS | 1 | 3770 | 5n38 | A | 275 | ARG | p.Arg275Trp | O60260 |
| HOMO SAPIENS | 2 | 3770 | 5n38 | A | 334 | ARG | p.Arg334Cys | O60260 |
| HOMO SAPIENS | 1 | 3770 | 5n38 | A | 366 | ARG | p.Arg366Trp | O60260 |
| HOMO SAPIENS | -1 | 3770 | 5n38 | A | 380 | VAL | p.Val380Leu | O60260 |
| HOMO SAPIENS | 1 | 3770 | 5n38 | A | 394 | ASP | p.Asp394Asn | O60260 |
| HOMO SAPIENS | 1 | 3770 | 5n38 | A | 430 | GLY | p.Gly430Asp | O60260 |
| HOMO SAPIENS | -1 | 2467 | 4xx1 | A | 10 | PRO | p.Pro34Leu | P04180 |
| HOMO SAPIENS | 4 | 2467 | 4xx1 | A | 83 | TYR | p.Tyr107Ter | P04180 |
| HOMO SAPIENS | 4 | 2467 | 4xx1 | A | 93 | ALA | p.Ala117Thr | P04180 |
| HOMO SAPIENS | 4 | 2467 | 4xx1 | A | 123 | THR | p.Thr147Ile | P04180 |
| HOMO SAPIENS | 4 | 2467 | 4xx1 | A | 135 | ARG | p.Arg159Trp | P04180 |
| HOMO SAPIENS | 3 | 2467 | 4xx1 | A | 158 | ARG | p.Arg182Cys | P04180 |
| HOMO SAPIENS | 0 | 2467 | 4xx1 | A | 209 | LEU | p.Leu233Pro | P04180 |
| HOMO SAPIENS | 1 | 2467 | 4xx1 | A | 228 | ASN | p.Asn252Lys | P04180 |
| HOMO SAPIENS | 4 | 2467 | 4xx1 | A | 252 | MET | p.Met276Lys | P04180 |
| HOMO SAPIENS | 3 | 2467 | 4xx1 | A | 293 | MET | p.Met317Ile | P04180 |
| HOMO SAPIENS | 5 | 2467 | 4xx1 | A | 321 | THR | p.Thr345Met | P04180 |
| HOMO SAPIENS | 9 | 2467 | 4xx1 | A | 347 | THR | p.Thr371Met | P04180 |
| HOMO SAPIENS | 1 | 5834 | 3qrf | G | 371 | PHE | p.Phe371Cys | Q9BZS1 |
| HOMO SAPIENS | 1 | 5834 | 3qrf | G | 373 | PHE | p.Phe373Ala | Q9BZS1 |
| HOMO SAPIENS | 9 | 5834 | 3qrf | G | 384 | ALA | p.Ala384Thr | Q9BZS1 |
| HOMO SAPIENS | 6 | 5834 | 3qrf | G | 397 | ARG | p.Arg397Trp | Q9BZS1 |
| HOMO SAPIENS | -1 | 11869 | 4km8 | A | 15 | PRO | p.Pro15Leu | Q9UMX1 |
| HOMO SAPIENS | 5 | 262 | 4kbo | A | 126 | ARG | p.Arg126Trp | P38646 |
| HOMO SAPIENS | 3 | 262 | 4kbo | A | 128 | TYR | p.Tyr128Cys | P38646 |
| HOMO SAPIENS | -1 | 11347 | 4uf9 | D | 117 | ALA | p.Ala117Val | Q13501 |
| HOMO SAPIENS | 0 | 2173 | 4ah6 | A | 45 | SER | p.Ser45Gly | Q6PI48 |
| HOMO SAPIENS | 2 | 2173 | 4ah6 | A | 152 | CYS | p.Cys152Phe | Q6PI48 |
| HOMO SAPIENS | 4 | 2173 | 4ah6 | A | 179 | ARG | p.Arg179His | Q6PI48 |
| HOMO SAPIENS | 0 | 2173 | 4ah6 | A | 196 | LYS | p.Lys196Arg | Q6PI48 |
| HOMO SAPIENS | 8 | 2173 | 4ah6 | A | 263 | ARG | p.Arg263Gln | Q6PI48 |
| HOMO SAPIENS | 1 | 2173 | 4ah6 | A | 338 | GLY | p.Gly338Glu | Q6PI48 |
| HOMO SAPIENS | 0 | 2173 | 4ah6 | A | 425 | GLU | p.Glu425Ter | Q6PI48 |

| | | | | | | | | |
|---|---|---|---|---|---|---|---|---|
| HOMO SAPIENS | 1 | 2173 | 4ah6 | A | 548 | LEU | p.Leu548Met | Q6PI48 |
| HOMO SAPIENS | 0 | 2173 | 4ah6 | A | 609 | ARG | p.Arg609Trp | Q6PI48 |
| HOMO SAPIENS | 2 | 2173 | 4ah6 | A | 613 | LEU | p.Leu613Phe | Q6PI48 |
| HOMO SAPIENS | 0 | 2173 | 4ah6 | A | 626 | LEU | p.Leu626Gln | Q6PI48 |
| HOMO SAPIENS | 0 | 2173 | 4ah6 | A | 626 | LEU | p.Leu626Val | Q6PI48 |
| HOMO SAPIENS | 7 | 3203 | 2cn1 | A | 87 | ASP | p.Asp137Val | Q9H0P0 |
| HOMO SAPIENS | 8 | 3203 | 2cn1 | A | 166 | GLN | p.Gln216Ter | Q9H0P0 |
| HOMO SAPIENS | 8 | 3203 | 2cn1 | A | 179 | ASN | p.Asn229Ser | Q9H0P0 |
| HOMO SAPIENS | 7 | 3203 | 2cn1 | A | 230 | GLY | p.Gly280Arg | Q9H0P0 |
| HOMO SAPIENS | 2 | 3424 | 2a1t | R | 116 | GLY | p.Gly116Arg | P13804 |
| HOMO SAPIENS | 2 | 3424 | 2a1t | R | 157 | VAL | p.Val157Gly | P13804 |
| HOMO SAPIENS | 2 | 3424 | 2a1t | R | 171 | THR | p.Thr171Ile | P13804 |
| HOMO SAPIENS | 0 | 3424 | 2a1t | R | 178 | THR | p.Thr178Arg | P13804 |
| HOMO SAPIENS | 0 | 3424 | 2a1t | R | 194 | VAL | p.Val194Met | P13804 |
| HOMO SAPIENS | 1 | 3424 | 2a1t | R | 209 | ARG | p.Arg209Ter | P13804 |
| HOMO SAPIENS | 6 | 3424 | 2a1t | R | 266 | THR | p.Thr266Met | P13804 |
| HOMO SAPIENS | 0 | 27598 | 1g1c | A | 26 | ARG | p.Arg2098Gly | Q8WZ42 |
| HOMO SAPIENS | 1 | 27598 | 1g1c | A | 46 | ILE | p.Ile2118Thr | Q8WZ42 |
| HOMO SAPIENS | 0 | 32885 | 4bsk | A | 149 | ASN | p.Asn149Asp | P35916 |
| HOMO SAPIENS | -1 | 8074 | 3ghg | B | 17 | ARG | p.Arg47Ter | P02675 |
| HOMO SAPIENS | -1 | 447 | 5g4p | A | 190 | ALA | p.Ala190Thr | P84074 |
| HOMO SAPIENS | 0 | 10761 | 5u1d | B | 379 | VAL | p.Val379Ile | Q03519 |
| HOMO SAPIENS | 0 | 10761 | 5u1d | B | 665 | THR | p.Thr665Ala | Q03519 |
| HOMO SAPIENS | 5 | 29251 | 5fn7 | A | 79 | ILE | p.Ile294Leu | P08575 |
| HOMO SAPIENS | -1 | 27727 | 4ky9 | A | 56 | LYS | p.Lys56Glu | P02730 |
| HOMO SAPIENS | 2 | 27727 | 4ky9 | A | 72 | GLU | p.Glu72Asp | P02730 |
| HOMO SAPIENS | 2 | 27727 | 4ky9 | A | 90 | GLU | p.Glu90Lys | P02730 |
| HOMO SAPIENS | 1 | 27727 | 4ky9 | A | 130 | GLY | p.Gly130Arg | P02730 |
| HOMO SAPIENS | 2 | 27727 | 4ky9 | A | 150 | ARG | p.Arg150Ter | P02730 |
| HOMO SAPIENS | 0 | 27727 | 4ky9 | A | 327 | PRO | p.Pro327Arg | P02730 |
| HOMO SAPIENS | 2 | 13202 | 2fxo | B | 870 | ARG | p.Arg870His | P12883 |
| HOMO SAPIENS | 2 | 13202 | 2fxo | B | 870 | ARG | p.Arg870Cys | P12883 |
| HOMO SAPIENS | 4 | 13202 | 2fxo | B | 904 | ARG | p.Arg904Cys | P12883 |
| HOMO SAPIENS | 4 | 13202 | 2fxo | B | 906 | ASP | p.Asp906Gly | P12883 |
| HOMO SAPIENS | 2 | 13202 | 2fxo | B | 908 | LEU | p.Leu908Val | P12883 |
| HOMO SAPIENS | 2 | 13202 | 2fxo | B | 923 | ASN | p.Asn923Lys | P12883 |
| HOMO SAPIENS | 7 | 13202 | 2fxo | B | 935 | GLU | p.Glu935Lys | P12883 |
| HOMO SAPIENS | 7 | 13202 | 2fxo | B | 935 | GLU | p.Glu935Val | P12883 |
| HOMO SAPIENS | 8 | 13202 | 2fxo | B | 949 | GLU | p.Glu949Lys | P12883 |
| HOMO SAPIENS | 0 | 13200 | 2a1s | A | 349 | ARG | p.Arg349Trp | O95453 |
| HOMO SAPIENS | 0 | 13200 | 2a1s | A | 421 | LYS | p.Lys421Arg | O95453 |
| HOMO SAPIENS | 6 | 3196 | 5xtb | T | 115 | CYS | p.Cys115Tyr | O75380 |
| HOMO SAPIENS | 0 | 1108 | 5ukf | A | 89 | ARG | p.Arg89Gln | Q99986 |
| HOMO SAPIENS | 0 | 1108 | 5ukf | A | 119 | HIS | p.His119Arg | Q99986 |
| HOMO SAPIENS | 0 | 1108 | 5ukf | A | 133 | ARG | p.Arg133Cys | Q99986 |
| HOMO SAPIENS | 0 | 1108 | 5ukf | A | 236 | VAL | p.Val236Leu | Q99986 |
| HOMO SAPIENS | 0 | 1108 | 5ukf | A | 321 | ARG | p.Arg321Cys | Q99986 |
| HOMO SAPIENS | -1 | 1108 | 5ukf | A | 358 | ARG | p.Arg358Ter | Q99986 |
| HOMO SAPIENS | 0 | 5842 | 4yli | C | 204 | GLY | p.Gly204Ser | Q9BWP8 |
| HOMO SAPIENS | -1 | 31362 | 4wqo | A | 25 | PRO | p.Pro25Leu | P40337 |
| HOMO SAPIENS | 2 | 10107 | 2j8h | A | 34 | VAL | p.Val31886Ile | Q8WZ42 |
| HOMO SAPIENS | 8 | 20190 | 1wm5 | A | 77 | ARG | p.Arg77Gln | P19878 |
| HOMO SAPIENS | 1 | 20190 | 1wm5 | A | 100 | GLN | p.Gln100Ter | P19878 |
| HOMO SAPIENS | 0 | 20190 | 1wm5 | A | 102 | ARG | p.Arg102Ter | P19878 |

| | | | | | | | | | |
|---|---|---|---|---|---|---|---|---|---|
| HOMO SAPIENS | 0 | 20190 | 1wm5 | A | | 128 | ALA | p.Ala128Val | P19878 |
| HOMO SAPIENS | 0 | 20190 | 1wm5 | A | | 181 | LYS | p.Lys181Arg | P19878 |
| HOMO SAPIENS | 7 | 1278 | 5a2q | a | | 33 | ASP | p.Asp33Asn | P62854 |
| HOMO SAPIENS | 3 | 19515 | 2obd | A | | 11 | ILE | p.Ile28Met | P11597 |
| HOMO SAPIENS | 3 | 19515 | 2obd | A | | 89 | VAL | p.Val106Met | P11597 |
| HOMO SAPIENS | 3 | 19515 | 2obd | A | | 137 | ARG | p.Arg154Trp | P11597 |
| HOMO SAPIENS | 3 | 19515 | 2obd | A | | 139 | ASP | p.Asp156Asn | P11597 |
| HOMO SAPIENS | 3 | 19515 | 2obd | A | | 204 | SER | p.Ser221Arg | P11597 |
| HOMO SAPIENS | 0 | 19515 | 2obd | A | | 297 | GLU | p.Glu314Lys | P11597 |
| HOMO SAPIENS | 0 | 19515 | 2obd | A | | 314 | GLY | p.Gly331Ser | P11597 |
| HOMO SAPIENS | 0 | 19515 | 2obd | A | | 332 | SER | p.Ser349Tyr | P11597 |
| HOMO SAPIENS | 0 | 19515 | 2obd | A | | 368 | VAL | p.Val385Met | P11597 |
| HOMO SAPIENS | 0 | 19515 | 2obd | A | | 373 | ALA | p.Ala390Pro | P11597 |
| HOMO SAPIENS | 3 | 19515 | 2obd | A | | 405 | ILE | p.Val422Ile | P11597 |
| HOMO SAPIENS | 0 | 19515 | 2obd | A | | 451 | ARG | p.Arg468Gln | P11597 |
| HOMO SAPIENS | 0 | 19515 | 2obd | A | | 451 | ARG | p.Arg468Ter | P11597 |
| HOMO SAPIENS | 0 | 6626 | 3hop | A | | 39 | ALA | p.Ala39Gly | P21333 |
| HOMO SAPIENS | 1 | 6626 | 3hop | A | | 82 | GLU | p.Glu82Val | P21333 |
| HOMO SAPIENS | 1 | 6626 | 3hop | A | | 128 | ALA | p.Ala128Val | P21333 |
| HOMO SAPIENS | 2 | 6626 | 3hop | A | | 203 | ASP | p.Asp203Tyr | P21333 |
| HOMO SAPIENS | 1 | 6626 | 3hop | A | | 207 | PRO | p.Pro207Leu | P21333 |
| HOMO SAPIENS | 2 | 6626 | 3hop | A | | 210 | CYS | p.Cys210Phe | P21333 |
| HOMO SAPIENS | 7 | 6626 | 3hop | A | | 254 | GLU | p.Glu254Lys | P21333 |
| HOMO SAPIENS | 4 | 4210 | 2avg | A | | 8 | VAL | p.Val158Met | Q14896 |
| HOMO SAPIENS | 6 | 4210 | 2avg | A | | 15 | GLU | p.Glu165Asp | Q14896 |
| HOMO SAPIENS | 4 | 4210 | 2avg | A | | 27 | ARG | p.Arg177His | Q14896 |
| HOMO SAPIENS | 0 | 4210 | 2avg | A | | 28 | VAL | p.Val178Met | Q14896 |
| HOMO SAPIENS | 6 | 4210 | 2avg | A | | 39 | VAL | p.Val189Ile | Q14896 |
| HOMO SAPIENS | 2 | 4210 | 2avg | A | | 52 | LYS | p.Lys202Gln | Q14896 |
| HOMO SAPIENS | 4 | 4210 | 2avg | A | | 67 | SER | p.Ser217Gly | Q14896 |
| HOMO SAPIENS | 6 | 4210 | 2avg | A | | 69 | VAL | p.Val219Leu | Q14896 |
| HOMO SAPIENS | 2 | 4210 | 2avg | A | | 86 | GLY | p.Ser236Gly | Q14896 |
| HOMO SAPIENS | 4 | 4210 | 2avg | A | | 108 | GLU | p.Glu258Lys | Q14896 |
| HOMO SAPIENS | 0 | 22237 | 1w6k | A | | 588 | GLY | p.Gly588Ser | P48449 |
| HOMO SAPIENS | 0 | 10765 | 4a5w | B | | 98 | GLU | p.Glu98Ala | P13671 |
| HOMO SAPIENS | 0 | 10765 | 4a5w | B | | 596 | ARG | p.Arg596Ter | P13671 |
| HOMO SAPIENS | -1 | 15515 | 3hx3 | A | | 11 | MSE | p.Met10Lys | P12271 |
| HOMO SAPIENS | 8 | 15515 | 3hx3 | A | | 64 | ARG | p.Arg64Gln | P12271 |
| HOMO SAPIENS | 0 | 15515 | 3hx3 | A | | 182 | PHE | p.Phe182Cys | P12271 |
| HOMO SAPIENS | 0 | 15515 | 3hx3 | A | | 216 | ARG | p.Arg216Gln | P12271 |
| HOMO SAPIENS | -1 | 15515 | 3hx3 | A | | 226 | MSE | p.Met226Lys | P12271 |
| HOMO SAPIENS | 0 | 15515 | 3hx3 | A | | 234 | TRP | p.Arg234Trp | P12271 |
| HOMO SAPIENS | 0 | 15515 | 3hx3 | A | | 273 | LEU | p.Leu273Phe | P12271 |
| HOMO SAPIENS | 0 | 15515 | 3hx3 | A | | 292 | THR | p.Thr292Met | P12271 |
| HOMO SAPIENS | 0 | 5152 | 2www | D | | 95 | GLN | p.Gln95Ter | Q8IVH4 |
| HOMO SAPIENS | 0 | 5152 | 2www | D | | 133 | GLN | p.Gln133Ter | Q8IVH4 |
| HOMO SAPIENS | 0 | 5152 | 2www | D | | 145 | ARG | p.Arg145Ter | Q8IVH4 |
| HOMO SAPIENS | 0 | 5152 | 2www | D | | 207 | TYR | p.Tyr207Cys | Q8IVH4 |
| HOMO SAPIENS | 8 | 5152 | 2www | D | | 330 | ARG | p.Arg330Ter | Q8IVH4 |
| HOMO SAPIENS | 0 | 5152 | 2www | D | | 363 | GLN | p.Gln363His | Q8IVH4 |
| HOMO SAPIENS | 1 | 1626 | 1ilr | | 1 | 105 | SER | p.Ser130Arg | P18510 |
| HOMO SAPIENS | -1 | 929 | 5loh | A | | 12 | GLY | p.Gly12Arg | Q96GX5 |
| HOMO SAPIENS | 9 | 3206 | 1x63 | A | | 21 | CYS | p.Cys104Arg | Q13642 |
| HOMO SAPIENS | 8 | 3206 | 1x63 | A | | 39 | TRP | p.Trp122Ser | Q13642 |

| | | | | | | | | |
|---|---|---|---|---|---|---|---|---|
| HOMO SAPIENS | 9 | 3206 | 1x63 | A | 40 | HIS | p.His123Tyr | Q13642 |
| HOMO SAPIENS | 9 | 3206 | 1x63 | A | 40 | HIS | p.His123Leu | Q13642 |
| HOMO SAPIENS | 9 | 3206 | 1x63 | A | 40 | HIS | p.His123Gln | Q13642 |
| HOMO SAPIENS | 3 | 3206 | 1x63 | A | 49 | CYS | p.Cys132Phe | Q13642 |
| HOMO SAPIENS | 3 | 3206 | 1x63 | A | 67 | CYS | p.Cys150Tyr | Q13642 |
| HOMO SAPIENS | 3 | 3206 | 1x63 | A | 70 | CYS | p.Cys153Arg | Q13642 |
| HOMO SAPIENS | 3 | 3206 | 1x63 | A | 70 | CYS | p.Cys153Tyr | Q13642 |
| HOMO SAPIENS | 5 | 1052 | 3ecs | E | 208 | ASN | p.Asn208Tyr | Q14232 |
| HOMO SAPIENS | 2 | 1052 | 3ecs | E | 275 | TYR | p.Tyr275Cys | Q14232 |
| HOMO SAPIENS | 0 | 24253 | 5i35 | A | 271 | ARG | p.Arg271Cys | Q8NI60 |
| HOMO SAPIENS | 0 | 24253 | 5i35 | A | 272 | GLY | p.Gly272Asp | Q8NI60 |
| HOMO SAPIENS | 0 | 24253 | 5i35 | A | 299 | ARG | p.Arg299Trp | Q8NI60 |
| HOMO SAPIENS | 1 | 24253 | 5i35 | A | 339 | ALA | p.Ala339Thr | Q8NI60 |
| HOMO SAPIENS | 0 | 24253 | 5i35 | A | 348 | ARG | p.Arg348Ter | Q8NI60 |
| HOMO SAPIENS | 0 | 24253 | 5i35 | A | 395 | ILE | p.Ile395Met | Q8NI60 |
| HOMO SAPIENS | 0 | 24253 | 5i35 | A | 396 | ASP | p.Asp396Glu | Q8NI60 |
| HOMO SAPIENS | 0 | 24253 | 5i35 | A | 480 | PHE | p.Phe480Leu | Q8NI60 |
| HOMO SAPIENS | 2 | 24253 | 5i35 | A | 511 | THR | p.Thr511Met | Q8NI60 |
| HOMO SAPIENS | 0 | 24253 | 5i35 | A | 514 | TYR | p.Tyr514Cys | Q8NI60 |
| HOMO SAPIENS | 0 | 24253 | 5i35 | A | 538 | ARG | p.Arg538Gln | Q8NI60 |
| HOMO SAPIENS | 0 | 24253 | 5i35 | A | 549 | GLY | p.Gly549Ser | Q8NI60 |
| HOMO SAPIENS | 0 | 24253 | 5i35 | A | 551 | GLU | p.Glu551Lys | Q8NI60 |
| HOMO SAPIENS | -1 | 24253 | 5i35 | A | 555 | MSE | p.Met555Ile | Q8NI60 |
| HOMO SAPIENS | 0 | 24253 | 5i35 | A | 568 | GLU | p.Glu568Ter | Q8NI60 |
| HOMO SAPIENS | 0 | 24253 | 5i35 | A | 602 | PRO | p.Pro602Arg | Q8NI60 |
| HOMO SAPIENS | -1 | 15505 | 4goj | C | 22 | GLY | p.Gly22Val | Q13432 |
| HOMO SAPIENS | 5 | 23902 | 2lrr | A | 765 | HIS | p.His765Gln | P38935 |
| HOMO SAPIENS | 1 | 29650 | 2vj2 | A | 234 | CYS | p.Cys234Tyr | P78504 |
| HOMO SAPIENS | 0 | 29650 | 2vj2 | A | 255 | TYR | p.Tyr255Ter | P78504 |
| HOMO SAPIENS | 3 | 29650 | 2vj2 | A | 274 | GLY | p.Gly274Asp | P78504 |
| HOMO SAPIENS | 1 | 105 | 5he0 | B | 77 | GLY | p.Gly77Ser | P62873 |
| HOMO SAPIENS | 2 | 105 | 5he0 | B | 80 | ILE | p.Ile80Thr | P62873 |
| HOMO SAPIENS | 3 | 4587 | 3rgh | A | 159 | ASP | p.Asp1159Ala | P21333 |
| HOMO SAPIENS | 3 | 4587 | 3rgh | A | 184 | ASP | p.Asp1184Glu | P21333 |
| HOMO SAPIENS | 2 | 4587 | 3rgh | A | 186 | SER | p.Ser1186Leu | P21333 |
| HOMO SAPIENS | 4 | 4587 | 3rgh | A | 188 | ALA | p.Ala1188Thr | P21333 |
| HOMO SAPIENS | 2 | 4587 | 3rgh | A | 199 | SER | p.Ser1199Leu | P21333 |
| HOMO SAPIENS | 0 | 20283 | 4n3w | A | 1278 | GLU | p.Glu1278Lys | Q92793 |
| HOMO SAPIENS | 1 | 14294 | 2d9q | B | 208 | MET | p.Met231Thr | Q99062 |
| HOMO SAPIENS | 6 | 14294 | 2d9q | B | 297 | ASP | p.Asp320Asn | Q99062 |
| HOMO SAPIENS | 0 | 11642 | 3bpt | A | 66 | ARG | p.Arg66Trp | Q6NVY1 |
| HOMO SAPIENS | 0 | 11642 | 3bpt | A | 122 | TYR | p.Tyr122Cys | Q6NVY1 |
| HOMO SAPIENS | 0 | 11642 | 3bpt | A | 345 | GLY | p.Gly345Ser | Q6NVY1 |
| HOMO SAPIENS | 1 | 3942 | 3sby | A | 60 | ASP | p.Asp60His | Q9Y4U1 |
| HOMO SAPIENS | 0 | 3942 | 3sby | A | 111 | ARG | p.Arg111Ter | Q9Y4U1 |
| HOMO SAPIENS | 1 | 3942 | 3sby | A | 116 | LEU | p.Leu116Pro | Q9Y4U1 |
| HOMO SAPIENS | 1 | 3942 | 3sby | A | 130 | TYR | p.Tyr130Cys | Q9Y4U1 |
| HOMO SAPIENS | 1 | 3942 | 3sby | A | 132 | ARG | p.Arg132Ter | Q9Y4U1 |
| HOMO SAPIENS | 1 | 3942 | 3sby | A | 147 | GLY | p.Gly147Ala | Q9Y4U1 |
| HOMO SAPIENS | 1 | 3942 | 3sby | A | 161 | ARG | p.Arg161Gln | Q9Y4U1 |
| HOMO SAPIENS | 1 | 3942 | 3sby | A | 161 | ARG | p.Arg161Ter | Q9Y4U1 |
| HOMO SAPIENS | 1 | 3942 | 3sby | A | 203 | TRP | p.Trp203Ter | Q9Y4U1 |
| HOMO SAPIENS | 1 | 3942 | 3sby | A | 203 | TRP | p.Trp203Ter | Q9Y4U1 |
| HOMO SAPIENS | 1 | 3942 | 3sby | A | 205 | TYR | p.Tyr205Ter | Q9Y4U1 |

| | | | | | | | | | |
|---|---|---|---|---|---|---|---|---|---|
| HOMO SAPIENS | 0 | 38340 | 2mbh | B | | 39 | MET | p.Met39Leu | Q13351 |
| HOMO SAPIENS | 2 | 7360 | 4k30 | A | | 430 | LEU | p.Leu430Pro | Q8N159 |
| HOMO SAPIENS | 2 | 7360 | 4k30 | A | | 433 | GLU | p.Glu433Asp | Q8N159 |
| HOMO SAPIENS | 7 | 7360 | 4k30 | A | | 484 | TRP | p.Trp484Arg | Q8N159 |
| HOMO SAPIENS | 1 | 41903 | 2xs6 | A | | 151 | PRO | p.Pro151Ser | O00459 |
| HOMO SAPIENS | 0 | 172 | 5glh | A | | 183 | ALA | p.Ala183Gly | P24530 |
| HOMO SAPIENS | 0 | 172 | 5glh | A | | 201 | ARG | p.Arg201Ter | P24530 |
| HOMO SAPIENS | 0 | 172 | 5glh | A | | 244 | THR | p.Thr244Met | P24530 |
| HOMO SAPIENS | 0 | 172 | 5glh | A | | 253 | ARG | p.Arg253Ter | P24530 |
| HOMO SAPIENS | 9 | 172 | 5glh | A | | 1017 | ILE | p.Leu17Phe | P00720 |
| HOMO SAPIENS | -1 | 3342 | 1ljm | A | | 1058 | HIS | p.His58Asn | Q01196 |
| HOMO SAPIENS | 4 | 3342 | 1ljm | A | | 1174 | ARG | p.Arg174Gln | Q01196 |
| HOMO SAPIENS | 5 | 25607 | 1p22 | C | | 32 | ASP | p.Asp32Ala | P35222 |
| HOMO SAPIENS | 5 | 25607 | 1p22 | C | | 32 | ASP | p.Asp32Tyr | P35222 |
| HOMO SAPIENS | -1 | 25607 | 1p22 | C | | 33 | SEP | p.Ser33Phe | P35222 |
| HOMO SAPIENS | 5 | 25607 | 1p22 | C | | 34 | GLY | p.Gly34Glu | P35222 |
| HOMO SAPIENS | -1 | 25607 | 1p22 | C | | 37 | SEP | p.Ser37Cys | P35222 |
| HOMO SAPIENS | -1 | 25607 | 1p22 | C | | 41 | THR | p.Thr41Ile | P35222 |
| HOMO SAPIENS | 1 | 48 | 4uow | | 9 | 60 | GLN | p.Arg34111His | Q8WZ42 |
| HOMO SAPIENS | 0 | 48 | 4uow | | 9 | 64 | LEU | p.Val34115Ala | Q8WZ42 |
| HOMO SAPIENS | 0 | 1127 | 5c01 | A | | 638 | ARG | p.Arg638Ter | P29597 |
| HOMO SAPIENS | 0 | 1127 | 5c01 | A | | 684 | ILE | p.Ile684Ser | P29597 |
| HOMO SAPIENS | 0 | 1127 | 5c01 | A | | 703 | ARG | p.Arg703Trp | P29597 |
| HOMO SAPIENS | 5 | 16389 | 2aq0 | B | | 128 | ALA | p.Ala860Asp | Q92889 |
| HOMO SAPIENS | 2 | 16389 | 2aq0 | B | | 141 | ILE | p.Ile873Val | Q92889 |
| HOMO SAPIENS | 5 | 16389 | 2aq0 | B | | 143 | GLU | p.Glu875Gly | Q92889 |
| HOMO SAPIENS | 0 | 16389 | 2aq0 | B | | 180 | GLY | p.Gly912Arg | Q92889 |
| HOMO SAPIENS | 0 | 15816 | 2xrc | A | | 89 | THR | p.Thr107Ala | P05156 |
| HOMO SAPIENS | 0 | 15816 | 2xrc | A | | 133 | ASN | p.Asn151Ser | P05156 |
| HOMO SAPIENS | 0 | 15816 | 2xrc | A | | 185 | THR | p.Thr203Ile | P05156 |
| HOMO SAPIENS | 0 | 15816 | 2xrc | A | | 225 | GLY | p.Gly243Asp | P05156 |
| HOMO SAPIENS | 0 | 15816 | 2xrc | A | | 243 | GLY | p.Gly261Ala | P05156 |
| HOMO SAPIENS | 0 | 15816 | 2xrc | A | | 269 | GLY | p.Gly287Arg | P05156 |
| HOMO SAPIENS | -1 | 15816 | 2xrc | A | | 288 | ILE | p.Ile306Val | P05156 |
| HOMO SAPIENS | -1 | 15816 | 2xrc | A | | 388 | ARG | p.Arg406His | P05156 |
| HOMO SAPIENS | 7 | 15816 | 2xrc | A | | 398 | ILE | p.Ile416Leu | P05156 |
| HOMO SAPIENS | 9 | 15816 | 2xrc | A | | 400 | HIS | p.His418Leu | P05156 |
| HOMO SAPIENS | 0 | 15816 | 2xrc | A | | 423 | LYS | p.Lys441Arg | P05156 |
| HOMO SAPIENS | -1 | 15816 | 2xrc | A | | 530 | GLU | p.Glu548Gln | P05156 |
| HOMO SAPIENS | 0 | 15816 | 2xrc | A | | 535 | PRO | p.Pro553Ala | P05156 |
| HOMO SAPIENS | 0 | 15816 | 2xrc | A | | 536 | GLU | p.Glu554Gly | P05156 |
| HOMO SAPIENS | 1 | 12385 | 4zlp | B | | 1450 | ALA | p.Ala1450Pro | Q9UM47 |
| HOMO SAPIENS | 7 | 12385 | 4zlp | B | | 1482 | GLY | p.Gly1482Arg | Q9UM47 |
| HOMO SAPIENS | 2 | 12385 | 4zlp | B | | 1518 | LEU | p.Leu1518Met | Q9UM47 |
| HOMO SAPIENS | 0 | 12385 | 4zlp | B | | 1560 | ARG | p.Arg1560Pro | Q9UM47 |
| HOMO SAPIENS | 0 | 1227 | 4a04 | A | | 148 | PHE | p.Phe148Tyr | O43435 |
| HOMO SAPIENS | 4 | 1227 | 4a04 | A | | 194 | HIS | p.His194Gln | O43435 |
| HOMO SAPIENS | 2 | 8017 | 3ei4 | B | | 244 | LYS | p.Lys244Glu | Q92466 |
| HOMO SAPIENS | 1 | 8017 | 3ei4 | B | | 273 | ARG | p.Arg273His | Q92466 |
| HOMO SAPIENS | 3 | 36112 | 2k8f | B | | 9 | SER | p.Ser9Arg | P04637 |
| HOMO SAPIENS | 2 | 12880 | 5ggf | A | | 265 | ARG | p.Arg265His | Q8WZA1 |
| HOMO SAPIENS | 2 | 12880 | 5ggf | A | | 287 | ILE | p.Ile287Ser | Q8WZA1 |
| HOMO SAPIENS | 2 | 12880 | 5ggf | A | | 311 | ARG | p.Arg311Gln | Q8WZA1 |
| HOMO SAPIENS | 2 | 12880 | 5ggf | A | | 425 | TRP | p.Trp425Ser | Q8WZA1 |

| | | | | | | | | |
|---|---|---|---|---|---|---|---|---|
| HOMO SAPIENS | 2 | 12880 | 5ggf | A | 442 | ARG | p.Arg442Cys | Q8WZA1 |
| HOMO SAPIENS | 2 | 12880 | 5ggf | A | 448 | GLY | p.Gly448Arg | Q8WZA1 |
| HOMO SAPIENS | 0 | 12880 | 5ggf | A | 490 | CYS | p.Cys490Tyr | Q8WZA1 |
| HOMO SAPIENS | 0 | 12880 | 5ggf | A | 493 | PRO | p.Pro493Arg | Q8WZA1 |
| HOMO SAPIENS | 3 | 12880 | 5ggf | A | 550 | SER | p.Ser550Asn | Q8WZA1 |
| HOMO SAPIENS | 1 | 12880 | 5ggf | A | 556 | ASP | p.Asp556Asn | Q8WZA1 |
| HOMO SAPIENS | 0 | 12880 | 5ggf | A | 580 | ARG | p.Arg580Ter | Q8WZA1 |
| HOMO SAPIENS | 2 | 12880 | 5ggf | A | 605 | ARG | p.Arg605Pro | Q8WZA1 |
| HOMO SAPIENS | 0 | 12880 | 5ggf | A | 623 | VAL | p.Met623Val | Q8WZA1 |
| HOMO SAPIENS | 1 | 10187 | 4qw2 | A | 145 | ALA | p.Ala145Ser | Q06787 |
| HOMO SAPIENS | 1 | 10187 | 4qw2 | A | 174 | ASN | p.Asn174Lys | Q06787 |
| HOMO SAPIENS | 0 | 24002 | 2m0t | A | 110 | LEU | p.Leu110Val | O14745 |
| HOMO SAPIENS | 9 | 3126 | 4mjh | A | 99 | LEU | p.Leu99Met | P04792 |
| HOMO SAPIENS | 1 | 3126 | 4mjh | A | 127 | ARG | p.Arg127Trp | P04792 |
| HOMO SAPIENS | 1 | 3126 | 4mjh | A | 127 | ARG | p.Arg127Leu | P04792 |
| HOMO SAPIENS | 1 | 3126 | 4mjh | A | 135 | SER | p.Ser135Phe | P04792 |
| HOMO SAPIENS | 2 | 3126 | 4mjh | A | 136 | ARG | p.Arg136Trp | P04792 |
| HOMO SAPIENS | 1 | 3126 | 4mjh | A | 140 | ARG | p.Arg140Gly | P04792 |
| HOMO SAPIENS | 0 | 3126 | 4mjh | A | 151 | THR | p.Thr151Ile | P04792 |
| HOMO SAPIENS | -1 | 15780 | 2jlp | A | 213 | ARG | p.Arg231Gly | P08294 |
| HOMO SAPIENS | 8 | 3578 | 1apz | D | 203 | GLY | p.Gly226Asp | P20933 |
| HOMO SAPIENS | 1 | 3578 | 1apz | D | 279 | GLY | p.Gly302Arg | P20933 |
| HOMO SAPIENS | 0 | 3578 | 1apz | D | 283 | CYS | p.Cys306Arg | P20933 |
| HOMO SAPIENS | 5 | 1335 | 5xtd | C | 115 | ARG | p.Arg115Cys | O75251 |
| HOMO SAPIENS | 0 | 1335 | 5xtd | C | 122 | VAL | p.Val122Met | O75251 |
| HOMO SAPIENS | 2 | 5063 | 2le4 | A | 37 | ARG | p.Arg74Pro | P48431 |
| HOMO SAPIENS | 1 | 25870 | 2o10 | A | 44 | LEU | p.Leu44Pro | P50461 |
| HOMO SAPIENS | 4 | 25870 | 2o10 | A | 58 | CYS | p.Cys58Gly | P50461 |
| HOMO SAPIENS | 0 | 18748 | 5fii | C | 39 | PHE | p.Phe39Leu | P00439 |
| HOMO SAPIENS | 6 | 18748 | 5fii | C | 48 | LEU | p.Leu48Ser | P00439 |
| HOMO SAPIENS | 2 | 18748 | 5fii | C | 65 | ILE | p.Ile65Asn | P00439 |
| HOMO SAPIENS | 3 | 18748 | 5fii | C | 95 | ILE | p.Ile95Phe | P00439 |
| HOMO SAPIENS | 1 | 14261 | 2vay | B | 1539 | ARG | p.Arg1539Cys | Q13698 |
| HOMO SAPIENS | 0 | 2494 | 4dx1 | A | 183 | GLU | p.Glu183Asp | Q9HBA0 |
| HOMO SAPIENS | 0 | 2494 | 4dx1 | A | 217 | ALA | p.Ala217Ser | Q9HBA0 |
| HOMO SAPIENS | 0 | 2494 | 4dx1 | A | 217 | ALA | p.Ala217Val | Q9HBA0 |
| HOMO SAPIENS | 4 | 2494 | 4dx1 | A | 224 | ARG | p.Arg224Trp | Q9HBA0 |
| HOMO SAPIENS | 4 | 2494 | 4dx1 | A | 254 | VAL | p.Val254Leu | Q9HBA0 |
| HOMO SAPIENS | 0 | 2494 | 4dx1 | A | 269 | ARG | p.Arg269Cys | Q9HBA0 |
| HOMO SAPIENS | 0 | 2494 | 4dx1 | A | 269 | ARG | p.Arg269His | Q9HBA0 |
| HOMO SAPIENS | 0 | 2494 | 4dx1 | A | 278 | GLU | p.Glu278Lys | Q9HBA0 |
| HOMO SAPIENS | 0 | 2494 | 4dx1 | A | 315 | ARG | p.Arg315Trp | Q9HBA0 |
| HOMO SAPIENS | 0 | 2494 | 4dx1 | A | 316 | ARG | p.Arg316Cys | Q9HBA0 |
| HOMO SAPIENS | 0 | 2494 | 4dx1 | A | 331 | ILE | p.Ile331Phe | Q9HBA0 |
| HOMO SAPIENS | 0 | 2494 | 4dx1 | A | 333 | ASP | p.Asp333Gly | Q9HBA0 |
| HOMO SAPIENS | 1 | 2494 | 4dx1 | A | 365 | VAL | p.Val365Met | Q9HBA0 |
| HOMO SAPIENS | 0 | 23998 | 2l1r | B | 161 | ARG | p.Arg162Trp | P19429 |
| HOMO SAPIENS | 0 | 8354 | 4f0x | D | 187 | SER | p.Ser187Ter | O95822 |
| HOMO SAPIENS | 2 | 8354 | 4f0x | D | 259 | GLY | p.Gly259Ala | O95822 |
| HOMO SAPIENS | 2 | 8354 | 4f0x | D | 284 | THR | p.Thr284Ala | O95822 |
| HOMO SAPIENS | 3 | 8354 | 4f0x | D | 309 | LYS | p.Lys309Gln | O95822 |
| HOMO SAPIENS | 0 | 21182 | 4apo | A | 228 | GLN | p.Gln228Lys | O00170 |
| HOMO SAPIENS | 1 | 21182 | 4apo | A | 307 | GLN | p.Gln307Arg | O00170 |
| HOMO SAPIENS | 0 | 2287 | 5of4 | B | 112 | ARG | p.Arg112His | P18074 |

| | | | | | | | | |
|---|---|---|---|---|---|---|---|---|
| HOMO SAPIENS | 0 | 2287 | 5of4 | B | 312 | ASP | p.Asp312Asn | P18074 |
| HOMO SAPIENS | 3 | 2287 | 5of4 | B | 461 | LEU | p.Leu461Val | P18074 |
| HOMO SAPIENS | 0 | 2287 | 5of4 | B | 485 | LEU | p.Leu485Pro | P18074 |
| HOMO SAPIENS | 1 | 2287 | 5of4 | B | 541 | SER | p.Ser541Arg | P18074 |
| HOMO SAPIENS | 0 | 2287 | 5of4 | B | 616 | ARG | p.Arg616Pro | P18074 |
| HOMO SAPIENS | 0 | 2287 | 5of4 | B | 616 | ARG | p.Arg616Trp | P18074 |
| HOMO SAPIENS | 1 | 2287 | 5of4 | B | 658 | ARG | p.Arg658Cys | P18074 |
| HOMO SAPIENS | 1 | 2287 | 5of4 | B | 681 | ASP | p.Asp681Asn | P18074 |
| HOMO SAPIENS | 1 | 2287 | 5of4 | B | 683 | ARG | p.Arg683Trp | P18074 |
| HOMO SAPIENS | 0 | 2287 | 5of4 | B | 713 | GLY | p.Gly713Arg | P18074 |
| HOMO SAPIENS | 0 | 2287 | 5of4 | B | 717 | ALA | p.Ala717Gly | P18074 |
| HOMO SAPIENS | 0 | 2287 | 5of4 | B | 722 | ARG | p.Arg722Trp | P18074 |
| HOMO SAPIENS | 0 | 2287 | 5of4 | B | 725 | ALA | p.Ala725Pro | P18074 |
| HOMO SAPIENS | 4 | 12188 | 4its | A | 122 | ARG | p.Arg122Gly | Q9Y606 |
| HOMO SAPIENS | 4 | 12188 | 4its | A | 133 | ASP | p.Asp133Asn | Q9Y606 |
| HOMO SAPIENS | 4 | 12188 | 4its | A | 144 | ARG | p.Arg144Trp | Q9Y606 |
| HOMO SAPIENS | 0 | 31529 | 1kgd | A | 761 | ARG | p.Arg766Gln | O14936 |
| HOMO SAPIENS | 1 | 31529 | 1kgd | A | 824 | GLN | p.Gln829Arg | O14936 |
| HOMO SAPIENS | 0 | 10906 | 2h63 | C | 18 | ARG | p.Arg18Ter | P53004 |
| HOMO SAPIENS | 9 | 27767 | 4wyq | A | 280 | LEU | p.Leu278Phe | Q9UPY3 |
| HOMO SAPIENS | 0 | 27767 | 4wyq | A | 297 | SER | p.Ser295Cys | Q9UPY3 |
| HOMO SAPIENS | 1 | 27767 | 4wyq | A | 334 | TYR | p.Tyr332Ter | Q9UPY3 |
| HOMO SAPIENS | 7 | 7775 | 5sv7 | C | 136 | SER | p.Ser136Cys | Q9NZJ5 |
| HOMO SAPIENS | 2 | 7775 | 5sv7 | C | 166 | GLN | p.Gln166Arg | Q9NZJ5 |
| HOMO SAPIENS | 2 | 7775 | 5sv7 | C | 332 | GLU | p.Glu332Ter | Q9NZJ5 |
| HOMO SAPIENS | 0 | 21449 | 2e9w | B | 84 | THR | p.Thr84Met | P10721 |
| HOMO SAPIENS | 0 | 21449 | 2e9w | B | 168 | ALA | p.Ala168Glu | P10721 |
| HOMO SAPIENS | 0 | 21449 | 2e9w | B | 178 | ALA | p.Ala178Thr | P10721 |
| HOMO SAPIENS | 0 | 21449 | 2e9w | B | 293 | ASN | p.Asn293Ser | P10721 |
| HOMO SAPIENS | -1 | 21449 | 2e9w | B | 318 | MET | p.Met318Leu | P10721 |
| HOMO SAPIENS | 0 | 21449 | 2e9w | B | 374 | VAL | p.Val374Ile | P10721 |
| HOMO SAPIENS | 6 | 3010 | 3flg | A | 66 | SER | p.Ser270Pro | Q9UBC3 |
| HOMO SAPIENS | 1 | 33918 | 2xzz | A | 760 | ARG | p.Arg760Ter | P22735 |
| HOMO SAPIENS | 2 | 1254 | 5xtc | w | 65 | ASN | p.Asn65Ser | O95299 |
| HOMO SAPIENS | 1 | 1254 | 5xtc | w | 238 | GLU | p.Glu238Lys | O95299 |
| HOMO SAPIENS | 3 | 10245 | 3siv | B | 194 | ALA | p.Ala194Glu | Q8WWY3 |
| HOMO SAPIENS | 4 | 10245 | 3siv | B | 216 | ALA | p.Ala216Pro | Q8WWY3 |
| HOMO SAPIENS | 0 | 2670 | 3dwb | A | 341 | THR | p.Thr341Ile | P42892 |
| HOMO SAPIENS | 1 | 2670 | 3dwb | A | 462 | SER | p.Ser462Asn | P42892 |
| HOMO SAPIENS | 1 | 2670 | 3dwb | A | 754 | ARG | p.Arg754Cys | P42892 |
| HOMO SAPIENS | 1 | 2961 | 5xtb | O | 40 | VAL | p.Val40Met | P19404 |
| HOMO SAPIENS | 5 | 2961 | 5xtb | O | 172 | ILE | p.Ile172Thr | P19404 |
| HOMO SAPIENS | 5 | 2961 | 5xtb | O | 190 | ASP | p.Asp190Gly | P19404 |
| HOMO SAPIENS | 5 | 29655 | 2xdg | A | 57 | ALA | p.Ala57Thr | Q02643 |
| HOMO SAPIENS | 4 | 29655 | 2xdg | A | 72 | GLU | p.Glu72Ter | Q02643 |
| HOMO SAPIENS | 0 | 29655 | 2xdg | A | 121 | GLU | p.Glu121Asp | Q02643 |
| HOMO SAPIENS | -1 | 17935 | 1hyn | P | 38 | ASP | p.Asp38Ala | P02730 |
| HOMO SAPIENS | 7 | 12825 | 1g82 | D | 99 | SER | p.Ser99Asn | P31371 |
| HOMO SAPIENS | 6 | 24927 | 1jm7 | B | 85 | VAL | p.Val85Leu | Q99728 |
| HOMO SAPIENS | 1 | 24927 | 1jm7 | B | 112 | ARG | p.Arg112Ter | Q99728 |
| HOMO SAPIENS | 0 | 24927 | 1jm7 | B | 116 | HIS | p.His116Tyr | Q99728 |
| HOMO SAPIENS | 7 | 92 | 5hde | A | 61 | LYS | p.Lys61Arg | Q05209 |
| HOMO SAPIENS | 0 | 17246 | 1szb | A | 84 | ARG | p.Arg99Gln | O00187 |
| HOMO SAPIENS | 1 | 17246 | 1szb | A | 105 | ASP | p.Asp120Gly | O00187 |

| | | | | | | | | |
|---|---|---|---|---|---|---|---|---|
| HOMO SAPIENS | 0 | 17246 | 1szb | A | 111 | PRO | p.Pro126Leu | O00187 |
| HOMO SAPIENS | 9 | 17246 | 1szb | A | 141 | CYS | p.Cys156Tyr | O00187 |
| HOMO SAPIENS | 0 | 32473 | 1b72 | A | 207 | ARG | p.Arg207Cys | P14653 |
| HOMO SAPIENS | 0 | 1920 | 4yzf | B | 488 | VAL | p.Val488Met | P02730 |
| HOMO SAPIENS | 0 | 1920 | 4yzf | B | 589 | ARG | p.Arg589Cys | P02730 |
| HOMO SAPIENS | 0 | 1920 | 4yzf | B | 589 | ARG | p.Arg589His | P02730 |
| HOMO SAPIENS | 0 | 1920 | 4yzf | B | 602 | ARG | p.Arg602Pro | P02730 |
| HOMO SAPIENS | 0 | 1920 | 4yzf | B | 613 | SER | p.Ser613Phe | P02730 |
| HOMO SAPIENS | -1 | 1920 | 4yzf | B | 646 | ARG | p.Arg646Gln | P02730 |
| HOMO SAPIENS | -1 | 1920 | 4yzf | B | 646 | ARG | p.Arg646Trp | P02730 |
| HOMO SAPIENS | 0 | 1920 | 4yzf | B | 701 | GLY | p.Gly701Asp | P02730 |
| HOMO SAPIENS | 0 | 1920 | 4yzf | B | 760 | ARG | p.Arg760Gln | P02730 |
| HOMO SAPIENS | 0 | 1920 | 4yzf | B | 771 | GLY | p.Gly771Asp | P02730 |
| HOMO SAPIENS | 0 | 1920 | 4yzf | B | 773 | SER | p.Ser773Pro | P02730 |
| HOMO SAPIENS | 0 | 1920 | 4yzf | B | 837 | THR | p.Thr837Ala | P02730 |
| HOMO SAPIENS | 0 | 1920 | 4yzf | B | 854 | PRO | p.Pro854Leu | P02730 |
| HOMO SAPIENS | 0 | 1920 | 4yzf | B | 858 | ALA | p.Ala858Asp | P02730 |
| HOMO SAPIENS | 0 | 1920 | 4yzf | B | 862 | VAL | p.Val862Ile | P02730 |
| HOMO SAPIENS | 0 | 1920 | 4yzf | B | 868 | PRO | p.Pro868Leu | P02730 |
| HOMO SAPIENS | 0 | 1920 | 4yzf | B | 870 | ARG | p.Arg870Trp | P02730 |
| HOMO SAPIENS | 0 | 9863 | 2juj | A | 54 | VAL | p.Val904Ile | P22681 |
| HOMO SAPIENS | 0 | 21934 | 2na7 | A | 184 | ILE | p.Ile184Val | P25445 |
| HOMO SAPIENS | -1 | 6746 | 1lw3 | A | 3 | THR | p.Lys3Thr | Q13614 |
| HOMO SAPIENS | -1 | 6746 | 1lw3 | A | 44 | VAL | p.Val44Ile | Q13614 |
| HOMO SAPIENS | 3 | 6746 | 1lw3 | A | 183 | ALA | p.Ala183Ser | Q13614 |
| HOMO SAPIENS | 4 | 6746 | 1lw3 | A | 482 | GLN | p.Gln482Ter | Q13614 |
| HOMO SAPIENS | 4 | 6746 | 1lw3 | A | 502 | GLU | p.Glu502Gln | Q13614 |
| HOMO SAPIENS | 3 | 6746 | 1lw3 | A | 545 | ASN | p.Asn545Ser | Q13614 |
| HOMO SAPIENS | 0 | 6746 | 1lw3 | A | 586 | ARG | p.Arg586Trp | Q13614 |
| HOMO SAPIENS | -1 | 6746 | 1lw3 | A | 602 | ALA | p.Ala602Gly | Q13614 |
| HOMO SAPIENS | 1 | 319 | 4ch9 | B | 336 | ARG | p.Arg336Ile | Q9UH77 |
| HOMO SAPIENS | 3 | 319 | 4ch9 | B | 340 | ALA | p.Ala340Val | Q9UH77 |
| HOMO SAPIENS | 0 | 319 | 4ch9 | B | 384 | ARG | p.Arg384Gln | Q9UH77 |
| HOMO SAPIENS | 5 | 319 | 4ch9 | B | 426 | PRO | p.Pro426Leu | Q9UH77 |
| HOMO SAPIENS | 3 | 319 | 4ch9 | B | 431 | ARG | p.Arg431Gln | Q9UH77 |
| HOMO SAPIENS | 2 | 319 | 4ch9 | B | 575 | ARG | p.Arg575Trp | Q9UH77 |
| HOMO SAPIENS | 0 | 574 | 2qkk | A | 142 | VAL | p.Val142Ile | O60930 |
| HOMO SAPIENS | 0 | 24021 | 2n8y | A | 752 | ARG | p.Arg752Gln | P12814 |
| HOMO SAPIENS | 0 | 18693 | 2y7j | B | 44 | ARG | p.Arg44Ter | P15735 |
| HOMO SAPIENS | 0 | 18693 | 2y7j | B | 106 | VAL | p.Val106Glu | P15735 |
| HOMO SAPIENS | 3 | 18693 | 2y7j | B | 189 | GLY | p.Gly189Glu | P15735 |
| HOMO SAPIENS | 0 | 17087 | 2ifg | A | 237 | THR | p.Thr237Met | P04629 |
| HOMO SAPIENS | 0 | 7980 | 1qub | A | 247 | VAL | p.Val266Leu | P02749 |
| HOMO SAPIENS | -1 | 2205 | 1h7u | A | 18 | ILE | p.Ile18Val | P54278 |
| HOMO SAPIENS | -1 | 2205 | 1h7u | A | 20 | ARG | p.Arg20Gln | P54278 |
| HOMO SAPIENS | 9 | 2205 | 1h7u | A | 46 | SER | p.Ser46Ile | P54278 |
| HOMO SAPIENS | 0 | 2205 | 1h7u | A | 60 | ASP | p.Asp60Glu | P54278 |
| HOMO SAPIENS | 0 | 2205 | 1h7u | A | 80 | PHE | p.Phe80Leu | P54278 |
| HOMO SAPIENS | 0 | 2205 | 1h7u | A | 118 | SER | p.Ser118Arg | P54278 |
| HOMO SAPIENS | 0 | 2205 | 1h7u | A | 126 | HIS | p.His126Gln | P54278 |
| HOMO SAPIENS | 0 | 2205 | 1h7u | A | 128 | SER | p.Ser128Leu | P54278 |
| HOMO SAPIENS | 1 | 2205 | 1h7u | A | 134 | ARG | p.Arg134Ter | P54278 |
| HOMO SAPIENS | 0 | 2205 | 1h7u | A | 166 | LEU | p.Leu166Pro | P54278 |
| HOMO SAPIENS | 1 | 2205 | 1h7u | A | 191 | TYR | p.Tyr191Cys | P54278 |

| | | | | | | | | |
|---|---|---|---|---|---|---|---|---|
| HOMO SAPIENS | 0 | 2205 | 1h7u | A | 211 | ARG | p.Arg211Ter | P54278 |
| HOMO SAPIENS | 0 | 2205 | 1h7u | A | 233 | GLN | p.Gln233Ter | P54278 |
| HOMO SAPIENS | 0 | 2205 | 1h7u | A | 255 | TYR | p.Tyr255Ter | P54278 |
| HOMO SAPIENS | 0 | 2205 | 1h7u | A | 277 | THR | p.Thr277Lys | P54278 |
| HOMO SAPIENS | 0 | 2205 | 1h7u | A | 301 | LYS | p.Lys301Asn | P54278 |
| HOMO SAPIENS | 0 | 2205 | 1h7u | A | 312 | MET | p.Met312Ile | P54278 |
| HOMO SAPIENS | 0 | 2205 | 1h7u | A | 312 | MET | p.Met312Thr | P54278 |
| HOMO SAPIENS | 0 | 2205 | 1h7u | A | 315 | ARG | p.Arg315Ter | P54278 |
| HOMO SAPIENS | 0 | 2205 | 1h7u | A | 318 | TYR | p.Tyr318Cys | P54278 |
| HOMO SAPIENS | 0 | 2205 | 1h7u | A | 335 | ASN | p.Asn335Ile | P54278 |
| HOMO SAPIENS | 0 | 2205 | 1h7u | A | 360 | ILE | p.Ile360Met | P54278 |
| HOMO SAPIENS | 6 | 20754 | 5mpo | B | 7 | VAL | p.Val7Phe | O96033 |
| HOMO SAPIENS | 4 | 20754 | 5mpo | B | 64 | VAL | p.Val64Ile | O96033 |
| HOMO SAPIENS | 5 | 34944 | 4qam | B | 1114 | ASP | p.Asp1114Gly | Q96KN7 |
| HOMO SAPIENS | 5 | 34944 | 4qam | B | 1240 | GLY | p.Gly1240Glu | Q96KN7 |
| HOMO SAPIENS | 9 | 40496 | 3vnn | A | 278 | ARG | p.Arg278His | P49917 |
| HOMO SAPIENS | 6 | 30131 | 4dt7 | E | 40 | PRO | p.Pro210Leu | P04070 |
| HOMO SAPIENS | -1 | 30131 | 4dt7 | E | 50 | ARG | p.Arg220Trp | P04070 |
| HOMO SAPIENS | 7 | 15311 | 2pqt | A | 64 | ARG | p.Arg64Trp | P18440 |
| HOMO SAPIENS | 2 | 15311 | 2pqt | A | 149 | VAL | p.Val149Ile | P18440 |
| HOMO SAPIENS | -1 | 19581 | 3hy8 | A | 33 | ASP | p.Asp33Val | Q9NVS9 |
| HOMO SAPIENS | 5 | 19581 | 3hy8 | A | 116 | ARG | p.Arg116Gln | Q9NVS9 |
| HOMO SAPIENS | 4 | 19581 | 3hy8 | A | 225 | ARG | p.Arg225His | Q9NVS9 |
| HOMO SAPIENS | 2 | 19581 | 3hy8 | A | 229 | TRP | p.Arg229Trp | Q9NVS9 |
| HOMO SAPIENS | 2 | 19581 | 3hy8 | A | 229 | TRP | p.Arg229Gln | Q9NVS9 |
| HOMO SAPIENS | 0 | 10266 | 3t1i | A | 41 | ASP | p.Asp41His | P49959 |
| HOMO SAPIENS | 1 | 10266 | 3t1i | A | 305 | ARG | p.Arg305Trp | P49959 |
| HOMO SAPIENS | 0 | 10266 | 3t1i | A | 308 | PHE | p.Phe308Leu | P49959 |
| HOMO SAPIENS | 0 | 10266 | 3t1i | A | 334 | SER | p.Ser334Arg | P49959 |
| HOMO SAPIENS | 1 | 1600 | 3rbf | A | 61 | GLU | p.Glu61Asp | P20711 |
| HOMO SAPIENS | 0 | 1600 | 3rbf | A | 91 | ALA | p.Ala91Val | P20711 |
| HOMO SAPIENS | -1 | 1600 | 3rbf | A | 102 | GLY | p.Gly102Ser | P20711 |
| HOMO SAPIENS | 3 | 1600 | 3rbf | A | 147 | SER | p.Ser147Arg | P20711 |
| HOMO SAPIENS | 1 | 1600 | 3rbf | A | 210 | PRO | p.Pro210Leu | P20711 |
| HOMO SAPIENS | 0 | 1600 | 3rbf | A | 217 | MET | p.Met217Val | P20711 |
| HOMO SAPIENS | 4 | 1600 | 3rbf | A | 250 | SER | p.Ser250Phe | P20711 |
| HOMO SAPIENS | 2 | 1600 | 3rbf | A | 275 | ALA | p.Ala275Thr | P20711 |
| HOMO SAPIENS | 1 | 1600 | 3rbf | A | 309 | PHE | p.Phe309Leu | P20711 |
| HOMO SAPIENS | -1 | 1600 | 3rbf | A | 347 | ARG | p.Arg347Gln | P20711 |
| HOMO SAPIENS | 0 | 1600 | 3rbf | A | 462 | ARG | p.Arg462Gln | P20711 |
| HOMO SAPIENS | 6 | 20830 | 5y4f | A | 714 | THR | p.Thr715Ile | Q01484 |
| HOMO SAPIENS | 7 | 20830 | 5y4f | A | 719 | ASP | p.Asp720Glu | Q01484 |
| HOMO SAPIENS | 0 | 20830 | 5y4f | A | 814 | VAL | p.Val815Ile | Q01484 |
| HOMO SAPIENS | -1 | 302 | 2c9y | A | 9 | GLU | p.Glu9Ter | P54819 |
| HOMO SAPIENS | 0 | 302 | 2c9y | A | 103 | ARG | p.Arg103Trp | P54819 |
| HOMO SAPIENS | 3 | 302 | 2c9y | A | 165 | ASP | p.Asp165Gly | P54819 |
| HOMO SAPIENS | 0 | 302 | 2c9y | A | 182 | ALA | p.Ala182Asp | P54819 |
| HOMO SAPIENS | 2 | 302 | 2c9y | A | 186 | ARG | p.Arg186Cys | P54819 |
| HOMO SAPIENS | 0 | 302 | 2c9y | A | 209 | ALA | p.Ala209Thr | P54819 |
| HOMO SAPIENS | -1 | 16920 | 1jlj | B | 10 | ASN | p.Asn10Tyr | Q9NQX3 |
| HOMO SAPIENS | 9 | 8462 | 1zbq | B | 16 | GLY | p.Gly16Ser | P51659 |
| HOMO SAPIENS | 5 | 8462 | 1zbq | B | 19 | ALA | p.Ala19Gly | P51659 |
| HOMO SAPIENS | 1 | 8462 | 1zbq | B | 34 | ALA | p.Ala34Val | P51659 |
| HOMO SAPIENS | 1 | 8462 | 1zbq | B | 106 | ARG | p.Arg106His | P51659 |

| | | | | | | | | |
|---|---|---|---|---|---|---|---|---|
| HOMO SAPIENS | 0 | 8462 | 1zbq | B | 140 | LYS | p.Lys140Asn | P51659 |
| HOMO SAPIENS | 0 | 8462 | 1zbq | B | 292 | THR | p.Thr292Ser | P51659 |
| HOMO SAPIENS | 0 | 4375 | 4ndi | A | 199 | ARG | p.Arg213His | Q7Z2E3 |
| HOMO SAPIENS | 3 | 4375 | 4ndi | A | 201 | HIS | p.His215Arg | Q7Z2E3 |
| HOMO SAPIENS | 0 | 4375 | 4ndi | A | 206 | PRO | p.Pro220Leu | Q7Z2E3 |
| HOMO SAPIENS | 2 | 4375 | 4ndi | A | 223 | LEU | p.Leu237Phe | Q7Z2E3 |
| HOMO SAPIENS | 0 | 4375 | 4ndi | A | 245 | ARG | p.Arg259His | Q7Z2E3 |
| HOMO SAPIENS | 0 | 4375 | 4ndi | A | 263 | VAL | p.Val277Gly | Q7Z2E3 |
| HOMO SAPIENS | 0 | 4375 | 4ndi | A | 279 | TRP | p.Trp293Ter | Q7Z2E3 |
| HOMO SAPIENS | 0 | 27004 | 4ccg | Y | 321 | ASP | p.Asp321Glu | Q9NW38 |
| HOMO SAPIENS | -1 | 27004 | 4ccg | Y | 372 | GLY | p.Gly372Ala | Q9NW38 |
| HOMO SAPIENS | -1 | 6383 | 5fai | A | 34 | ALA | p.Ala34Gly | Q92979 |
| HOMO SAPIENS | 8 | 6383 | 5fai | A | 86 | ASP | p.Asp86Gly | Q92979 |
| HOMO SAPIENS | 2 | 24585 | 4mi5 | A | 684 | ARG | p.Arg679Cys | Q15910 |
| HOMO SAPIENS | 0 | 15967 | 3nhb | A | 723 | ARG | p.Arg723Gln | Q9NP58 |
| HOMO SAPIENS | 5 | 18152 | 1oey | A | 389 | HIS | p.His389Gln | P19878 |
| HOMO SAPIENS | 5 | 18152 | 1oey | A | 395 | ARG | p.Arg395Trp | P19878 |
| HOMO SAPIENS | -1 | 1044 | 1li1 | F | 2 | VAL | p.Val1486Ile | P08572 |
| HOMO SAPIENS | 6 | 1044 | 1li1 | F | 37 | GLU | p.Glu1521Lys | P08572 |
| HOMO SAPIENS | 5 | 1044 | 1li1 | F | 157 | SER | p.Ser1639Leu | P08572 |
| HOMO SAPIENS | 5 | 1044 | 1li1 | F | 179 | ARG | p.Arg1662His | P08572 |
| HOMO SAPIENS | 5 | 1044 | 1li1 | F | 180 | GLY | p.Gly1663Arg | P08572 |
| HOMO SAPIENS | 6 | 20076 | 4uyi | A | 671 | LEU | p.Leu671Ser | Q8IY92 |
| HOMO SAPIENS | 0 | 19588 | 1jmj | A | 189 | ARG | p.Arg208His | P05546 |
| HOMO SAPIENS | -1 | 38023 | 1ry7 | B | 65 | GLY | p.Gly65Arg | P22607 |
| HOMO SAPIENS | -1 | 38023 | 1ry7 | B | 139 | ASP | p.Asp139Glu | P22607 |
| HOMO SAPIENS | 0 | 38023 | 1ry7 | B | 248 | ARG | p.Arg248Cys | P22607 |
| HOMO SAPIENS | 0 | 38023 | 1ry7 | B | 249 | SER | p.Ser249Cys | P22607 |
| HOMO SAPIENS | 0 | 38023 | 1ry7 | B | 250 | PRO | p.Pro250Arg | P22607 |
| HOMO SAPIENS | 0 | 38023 | 1ry7 | B | 322 | GLU | p.Glu322Lys | P22607 |
| HOMO SAPIENS | 5 | 16884 | 5eli | A | 33 | GLN | p.Gln33Ter | Q9NZC2 |
| HOMO SAPIENS | 0 | 16884 | 5eli | A | 44 | TRP | p.Trp44Ter | Q9NZC2 |
| HOMO SAPIENS | 0 | 16884 | 5eli | A | 47 | ARG | p.Arg47His | Q9NZC2 |
| HOMO SAPIENS | 0 | 16884 | 5eli | A | 66 | THR | p.Thr66Met | Q9NZC2 |
| HOMO SAPIENS | 0 | 16884 | 5eli | A | 78 | TRP | p.Trp78Ter | Q9NZC2 |
| HOMO SAPIENS | 1 | 16884 | 5eli | A | 96 | THR | p.Thr96Lys | Q9NZC2 |
| HOMO SAPIENS | 9 | 16884 | 5eli | A | 126 | VAL | p.Val126Gly | Q9NZC2 |
| HOMO SAPIENS | -1 | 12267 | 3w5o | A | 3 | ALA | p.Ala3Val | P49917 |
| HOMO SAPIENS | 1 | 12267 | 3w5o | A | 9 | THR | p.Thr9Ile | P49917 |
| HOMO SAPIENS | 2 | 12267 | 3w5o | A | 188 | ARG | p.Arg188Gln | P49917 |
| HOMO SAPIENS | 0 | 12267 | 3w5o | A | 469 | GLY | p.Gly469Glu | P49917 |
| HOMO SAPIENS | 1 | 12267 | 3w5o | A | 580 | ARG | p.Arg580Ter | P49917 |
| HOMO SAPIENS | 0 | 12267 | 3w5o | A | 600 | GLU | p.Glu600Lys | P49917 |
| HOMO SAPIENS | 9 | 2839 | 2hw5 | B | 54 | ARG | p.Arg54Cys | P30084 |
| HOMO SAPIENS | 9 | 2839 | 2hw5 | B | 59 | ASN | p.Asn59Ser | P30084 |
| HOMO SAPIENS | 0 | 2839 | 2hw5 | B | 273 | LYS | p.Lys273Glu | P30084 |
| HOMO SAPIENS | 1 | 12266 | 2o2k | B | 939 | GLY | p.Gly939Cys | Q99707 |
| HOMO SAPIENS | 5 | 12266 | 2o2k | B | 1164 | ARG | p.Arg1164His | Q99707 |
| HOMO SAPIENS | 1 | 12266 | 2o2k | B | 1166 | LEU | p.Leu1166Met | Q99707 |
| HOMO SAPIENS | 5 | 12266 | 2o2k | B | 1173 | PRO | p.Pro1173Leu | Q99707 |
| HOMO SAPIENS | 5 | 12266 | 2o2k | B | 1222 | ASN | p.Asn1222Ser | Q99707 |
| HOMO SAPIENS | 0 | 30891 | 3ch4 | B | 138 | ARG | p.Arg138Ter | Q15126 |
| HOMO SAPIENS | -1 | 4433 | 1qlz | A | 54 | GLY | p.Gly54Ser | P04156 |
| HOMO SAPIENS | 0 | 22568 | 3cxl | A | 20 | LEU | p.Leu20Phe | P15882 |

| | | | | | | | | |
|---|---|---|---|---|---|---|---|---|
| HOMO SAPIENS | 0 | 22568 | 3cxl | A | 126 | ILE | p.Ile126Met | P15882 |
| HOMO SAPIENS | 0 | 22568 | 3cxl | A | 141 | PRO | p.Pro141Leu | P15882 |
| HOMO SAPIENS | 0 | 22568 | 3cxl | A | 143 | TYR | p.Tyr143His | P15882 |
| HOMO SAPIENS | 0 | 22568 | 3cxl | A | 223 | ALA | p.Ala223Val | P15882 |
| HOMO SAPIENS | 0 | 22568 | 3cxl | A | 228 | GLY | p.Gly228Ser | P15882 |
| HOMO SAPIENS | 0 | 22568 | 3cxl | A | 252 | PRO | p.Pro252Gln | P15882 |
| HOMO SAPIENS | 0 | 22568 | 3cxl | A | 313 | GLU | p.Glu313Lys | P15882 |
| HOMO SAPIENS | 3 | 2668 | 5eqg | A | 34 | ASN | p.Asn34Ile | P11166 |
| HOMO SAPIENS | 3 | 2668 | 5eqg | A | 77 | MET | p.Met77Val | P11166 |
| HOMO SAPIENS | 6 | 2668 | 5eqg | A | 91 | GLY | p.Gly91Asp | P11166 |
| HOMO SAPIENS | 6 | 2668 | 5eqg | A | 92 | ARG | p.Arg92Trp | P11166 |
| HOMO SAPIENS | 6 | 2668 | 5eqg | A | 93 | ARG | p.Arg93Trp | P11166 |
| HOMO SAPIENS | 0 | 2668 | 5eqg | A | 95 | SER | p.Ser95Ile | P11166 |
| HOMO SAPIENS | 3 | 2668 | 5eqg | A | 126 | ARG | p.Arg126His | P11166 |
| HOMO SAPIENS | 3 | 2668 | 5eqg | A | 126 | ARG | p.Arg126Cys | P11166 |
| HOMO SAPIENS | 4 | 2668 | 5eqg | A | 232 | ARG | p.Arg232Cys | P11166 |
| HOMO SAPIENS | 3 | 2668 | 5eqg | A | 249 | ARG | p.Arg249Gln | P11166 |
| HOMO SAPIENS | 3 | 2668 | 5eqg | A | 255 | LYS | p.Lys255Arg | P11166 |
| HOMO SAPIENS | 0 | 2668 | 5eqg | A | 275 | ALA | p.Ala275Thr | P11166 |
| HOMO SAPIENS | 4 | 2668 | 5eqg | A | 299 | GLU | p.Glu299Lys | P11166 |
| HOMO SAPIENS | 6 | 2668 | 5eqg | A | 314 | GLY | p.Gly314Ser | P11166 |
| HOMO SAPIENS | 7 | 2668 | 5eqg | A | 315 | ILE | p.Ile315Met | P11166 |
| HOMO SAPIENS | -1 | 2668 | 5eqg | A | 458 | ARG | p.Arg458Trp | P11166 |
| HOMO SAPIENS | -1 | 2668 | 5eqg | A | 468 | ARG | p.Arg468Trp | P11166 |
| HOMO SAPIENS | 3 | 2291 | 2amy | A | 31 | PHE | p.Phe31Leu | O15305 |
| HOMO SAPIENS | 7 | 2291 | 2amy | A | 32 | LEU | p.Leu32Arg | O15305 |
| HOMO SAPIENS | 7 | 2291 | 2amy | A | 65 | ASP | p.Asp65Tyr | O15305 |
| HOMO SAPIENS | 1 | 2291 | 2amy | A | 108 | ALA | p.Ala108Val | O15305 |
| HOMO SAPIENS | 3 | 2291 | 2amy | A | 113 | PRO | p.Pro113Leu | O15305 |
| HOMO SAPIENS | 3 | 2291 | 2amy | A | 113 | PRO | p.Pro113Ser | O15305 |
| HOMO SAPIENS | 6 | 2291 | 2amy | A | 117 | GLY | p.Gly117Arg | O15305 |
| HOMO SAPIENS | 6 | 2291 | 2amy | A | 119 | PHE | p.Phe119Leu | O15305 |
| HOMO SAPIENS | 6 | 2291 | 2amy | A | 123 | ARG | p.Arg123Gln | O15305 |
| HOMO SAPIENS | 0 | 2291 | 2amy | A | 129 | VAL | p.Val129Met | O15305 |
| HOMO SAPIENS | 7 | 2291 | 2amy | A | 139 | GLU | p.Glu139Lys | O15305 |
| HOMO SAPIENS | 5 | 2291 | 2amy | A | 141 | ARG | p.Arg141His | O15305 |
| HOMO SAPIENS | 7 | 2291 | 2amy | A | 144 | PHE | p.Phe144Leu | O15305 |
| HOMO SAPIENS | 6 | 2291 | 2amy | A | 148 | ASP | p.Asp148Asn | O15305 |
| HOMO SAPIENS | 6 | 2291 | 2amy | A | 153 | ILE | p.Ile153Thr | O15305 |
| HOMO SAPIENS | 2 | 2291 | 2amy | A | 157 | PHE | p.Phe157Ser | O15305 |
| HOMO SAPIENS | 1 | 2291 | 2amy | A | 162 | ARG | p.Arg162Trp | O15305 |
| HOMO SAPIENS | 6 | 2291 | 2amy | A | 188 | ASP | p.Asp188Gly | O15305 |
| HOMO SAPIENS | 5 | 2291 | 2amy | A | 194 | ARG | p.Arg194Ter | O15305 |
| HOMO SAPIENS | 0 | 2291 | 2amy | A | 207 | PHE | p.Phe207Ser | O15305 |
| HOMO SAPIENS | 5 | 2291 | 2amy | A | 208 | GLY | p.Gly208Ala | O15305 |
| HOMO SAPIENS | 1 | 2291 | 2amy | A | 216 | ASN | p.Asn216Ile | O15305 |
| HOMO SAPIENS | 5 | 2291 | 2amy | A | 218 | HIS | p.His218Leu | O15305 |
| HOMO SAPIENS | 6 | 2291 | 2amy | A | 223 | ASP | p.Asp223Glu | O15305 |
| HOMO SAPIENS | 6 | 2291 | 2amy | A | 226 | THR | p.Thr226Ser | O15305 |
| HOMO SAPIENS | 5 | 2291 | 2amy | A | 231 | VAL | p.Val231Met | O15305 |
| HOMO SAPIENS | 1 | 2291 | 2amy | A | 237 | THR | p.Thr237Met | O15305 |
| HOMO SAPIENS | 1 | 2291 | 2amy | A | 241 | CYS | p.Cys241Ser | O15305 |
| HOMO SAPIENS | 5 | 35907 | 1d6g | A | 21 | GLY | p.Gly21Arg | P32238 |
| HOMO SAPIENS | -1 | 33209 | 1x9d | A | 197 | PRO | p.Pro197Leu | Q9UKM7 |

| | | | | | | | | |
|---|---|---|---|---|---|---|---|---|
| HOMO SAPIENS | -1 | 33209 | 1x9d | A | 201 | PRO | p.Pro201Leu | Q9UKM7 |
| HOMO SAPIENS | 6 | 21305 | 4yep | B | 1326 | ARG | p.Arg1326Ter | P24043 |
| HOMO SAPIENS | 6 | 21305 | 4yep | B | 1350 | ARG | p.Arg1350Ter | P24043 |
| HOMO SAPIENS | 0 | 21553 | 3qyb | A | 1120 | VAL | p.Val1119Ala | O60343 |
| HOMO SAPIENS | 2 | 21553 | 3qyb | A | 1148 | THR | p.Thr1147Met | O60343 |
| HOMO SAPIENS | -1 | 30645 | 3m0c | C | -17 | TRP | p.Trp4Ter | P01130 |
| HOMO SAPIENS | -1 | 30645 | 3m0c | C | -17 | TRP | p.Trp4Ter | P01130 |
| HOMO SAPIENS | -1 | 30645 | 3m0c | C | 12 | GLN | p.Gln33Ter | P01130 |
| HOMO SAPIENS | -1 | 30645 | 3m0c | C | 15 | ASP | p.Asp36Glu | P01130 |
| HOMO SAPIENS | -1 | 30645 | 3m0c | C | 28 | SER | p.Ser49Arg | P01130 |
| HOMO SAPIENS | -1 | 30645 | 3m0c | C | 41 | THR | p.Thr62Met | P01130 |
| HOMO SAPIENS | -1 | 30645 | 3m0c | C | 69 | ASP | p.Asp90Asn | P01130 |
| HOMO SAPIENS | -1 | 30645 | 3m0c | C | 76 | ASN | p.Asn97Lys | P01130 |
| HOMO SAPIENS | -1 | 30645 | 3m0c | C | 78 | SER | p.Ser99Ter | P01130 |
| HOMO SAPIENS | -1 | 30645 | 3m0c | C | 81 | GLN | p.Gln102Ter | P01130 |
| HOMO SAPIENS | -1 | 30645 | 3m0c | C | 84 | PRO | p.Pro105Ser | P01130 |
| HOMO SAPIENS | -1 | 30645 | 3m0c | C | 194 | ARG | p.Arg215Cys | P01130 |
| HOMO SAPIENS | -1 | 30645 | 3m0c | C | 198 | GLY | p.Gly219Asp | P01130 |
| HOMO SAPIENS | -1 | 30645 | 3m0c | C | 201 | CYS | p.Cys222Tyr | P01130 |
| HOMO SAPIENS | -1 | 30645 | 3m0c | C | 201 | CYS | p.Cys222Ter | P01130 |
| HOMO SAPIENS | -1 | 30645 | 3m0c | C | 206 | ASP | p.Asp227Glu | P01130 |
| HOMO SAPIENS | -1 | 30645 | 3m0c | C | 249 | CYS | p.Cys270Ter | P01130 |
| HOMO SAPIENS | -1 | 30645 | 3m0c | C | 250 | VAL | p.Val271Ile | P01130 |
| HOMO SAPIENS | 0 | 30645 | 3m0c | C | 255 | CYS | p.Cys276Tyr | P01130 |
| HOMO SAPIENS | 0 | 30645 | 3m0c | C | 255 | CYS | p.Cys276Ter | P01130 |
| HOMO SAPIENS | 0 | 30645 | 3m0c | C | 256 | GLU | p.Glu277Lys | P01130 |
| HOMO SAPIENS | 0 | 30645 | 3m0c | C | 265 | SER | p.Ser286Arg | P01130 |
| HOMO SAPIENS | 0 | 30645 | 3m0c | C | 284 | TRP | p.Trp305Ter | P01130 |
| HOMO SAPIENS | 0 | 30645 | 3m0c | C | 292 | CYS | p.Cys313Ter | P01130 |
| HOMO SAPIENS | 1 | 30645 | 3m0c | C | 385 | ARG | p.Arg406Trp | P01130 |
| HOMO SAPIENS | 0 | 30645 | 3m0c | C | 395 | ARG | p.Arg416Gln | P01130 |
| HOMO SAPIENS | 0 | 30645 | 3m0c | C | 398 | TYR | p.Tyr419Ter | P01130 |
| HOMO SAPIENS | 1 | 30645 | 3m0c | C | 415 | VAL | p.Val436Ala | P01130 |
| HOMO SAPIENS | 1 | 30645 | 3m0c | C | 424 | ASP | p.Asp445Glu | P01130 |
| HOMO SAPIENS | 1 | 30645 | 3m0c | C | 425 | LEU | p.Leu446Val | P01130 |
| HOMO SAPIENS | -1 | 30645 | 3m0c | C | 440 | GLY | p.Gly461Cys | P01130 |
| HOMO SAPIENS | -1 | 30645 | 3m0c | C | 441 | VAL | p.Val462Ile | P01130 |
| HOMO SAPIENS | 1 | 30645 | 3m0c | C | 447 | VAL | p.Val468Ile | P01130 |
| HOMO SAPIENS | 1 | 30645 | 3m0c | C | 489 | THR | p.Thr510Met | P01130 |
| HOMO SAPIENS | 0 | 30645 | 3m0c | C | 524 | GLY | p.Gly545Glu | P01130 |
| HOMO SAPIENS | 1 | 30645 | 3m0c | C | 615 | GLY | p.Gly636Ser | P01130 |
| HOMO SAPIENS | 1 | 30645 | 3m0c | C | 625 | LEU | p.Leu646Ile | P01130 |
| HOMO SAPIENS | 1 | 30645 | 3m0c | C | 630 | ASP | p.Asp651Gly | P01130 |
| HOMO SAPIENS | 3 | 30645 | 3m0c | C | 646 | CYS | p.Cys667Tyr | P01130 |
| HOMO SAPIENS | 4 | 30645 | 3m0c | C | 654 | GLY | p.Gly675Ser | P01130 |
| HOMO SAPIENS | 6 | 30645 | 3m0c | C | 660 | CYS | p.Cys681Ter | P01130 |
| HOMO SAPIENS | 5 | 30645 | 3m0c | C | 676 | ALA | p.Ala697Pro | P01130 |
| HOMO SAPIENS | -1 | 30645 | 3m0c | C | 749 | GLN | p.Gln770Ter | P01130 |
| HOMO SAPIENS | -1 | 41323 | 4bxo | A | 1812 | PRO | p.Pro1812Ala | Q8IYD8 |
| HOMO SAPIENS | 5 | 8335 | 3h0t | C | 12 | GLY | p.Gly71Asp | P81172 |
| HOMO SAPIENS | 6 | 609 | 1bhg | B | 148 | PRO | p.Pro148Ser | P08236 |
| HOMO SAPIENS | 1 | 609 | 1bhg | B | 176 | LEU | p.Leu176Phe | P08236 |
| HOMO SAPIENS | 6 | 609 | 1bhg | B | 216 | ARG | p.Arg216Trp | P08236 |
| HOMO SAPIENS | 0 | 609 | 1bhg | B | 350 | LYS | p.Lys350Asn | P08236 |

| | | | | | | | | |
|---|---|---|---|---|---|---|---|---|
| HOMO SAPIENS | 1 | 609 | 1bhg | B | 354 | ALA | p.Ala354Val | P08236 |
| HOMO SAPIENS | 1 | 609 | 1bhg | B | 357 | ARG | p.Arg357Ter | P08236 |
| HOMO SAPIENS | 0 | 609 | 1bhg | B | 382 | ARG | p.Arg382Cys | P08236 |
| HOMO SAPIENS | 0 | 609 | 1bhg | B | 495 | TYR | p.Tyr495Cys | P08236 |
| HOMO SAPIENS | 2 | 609 | 1bhg | B | 507 | TRP | p.Trp507Ter | P08236 |
| HOMO SAPIENS | 0 | 609 | 1bhg | B | 577 | ARG | p.Arg577Leu | P08236 |
| HOMO SAPIENS | 1 | 609 | 1bhg | B | 611 | ARG | p.Arg611Trp | P08236 |
| HOMO SAPIENS | 0 | 609 | 1bhg | B | 619 | ALA | p.Ala619Val | P08236 |
| HOMO SAPIENS | 0 | 609 | 1bhg | B | 627 | TRP | p.Trp627Cys | P08236 |
| HOMO SAPIENS | 5 | 3422 | 1x9n | A | 566 | GLU | p.Glu566Lys | P18858 |
| HOMO SAPIENS | 2 | 3422 | 1x9n | A | 771 | ARG | p.Arg771Trp | P18858 |
| HOMO SAPIENS | 0 | 3279 | 5tby | B | 797 | ALA | p.Ala797Thr | P12883 |
| HOMO SAPIENS | -1 | 3279 | 5tby | B | 982 | MET | p.Met982Thr | P12883 |
| HOMO SAPIENS | -1 | 3279 | 5tby | B | 992 | LEU | p.Leu992Met | P12883 |
| HOMO SAPIENS | -1 | 3279 | 5tby | B | 1004 | GLN | p.Gln1004Glu | P12883 |
| HOMO SAPIENS | -1 | 3279 | 5tby | B | 1013 | GLU | p.Glu1013Lys | P12883 |
| HOMO SAPIENS | -1 | 3279 | 5tby | B | 1032 | ASP | p.Asp1032Asn | P12883 |
| HOMO SAPIENS | -1 | 3279 | 5tby | B | 1053 | ARG | p.Arg1053Gln | P12883 |
| HOMO SAPIENS | -1 | 3279 | 5tby | B | 1117 | GLU | p.Glu1117Ter | P12883 |
| HOMO SAPIENS | -1 | 3279 | 5tby | B | 1193 | ARG | p.Arg1193His | P12883 |
| HOMO SAPIENS | -1 | 3279 | 5tby | B | 1205 | GLU | p.Glu1205Lys | P12883 |
| HOMO SAPIENS | -1 | 3279 | 5tby | B | 1207 | ILE | p.Ile1207Met | P12883 |
| HOMO SAPIENS | -1 | 3279 | 5tby | B | 1250 | ARG | p.Arg1250Pro | P12883 |
| HOMO SAPIENS | -1 | 3279 | 5tby | B | 1259 | HIS | p.His1259Gln | P12883 |
| HOMO SAPIENS | -1 | 3279 | 5tby | B | 1272 | ASP | p.Asp1272Asn | P12883 |
| HOMO SAPIENS | -1 | 3279 | 5tby | B | 1295 | GLU | p.Glu1295Lys | P12883 |
| HOMO SAPIENS | -1 | 3279 | 5tby | B | 1335 | SER | p.Ser1335Leu | P12883 |
| HOMO SAPIENS | -1 | 3279 | 5tby | B | 1399 | GLU | p.Glu1399Asp | P12883 |
| HOMO SAPIENS | -1 | 3279 | 5tby | B | 1413 | SER | p.Ser1413Ter | P12883 |
| HOMO SAPIENS | -1 | 3279 | 5tby | B | 1459 | LYS | p.Lys1459Asn | P12883 |
| HOMO SAPIENS | -1 | 3279 | 5tby | B | 1467 | LEU | p.Leu1467Val | P12883 |
| HOMO SAPIENS | -1 | 3279 | 5tby | B | 1491 | SER | p.Ser1491Cys | P12883 |
| HOMO SAPIENS | -1 | 3279 | 5tby | B | 1496 | GLU | p.Glu1496Ala | P12883 |
| HOMO SAPIENS | -1 | 3279 | 5tby | B | 1500 | ARG | p.Arg1500Pro | P12883 |
| HOMO SAPIENS | -1 | 3279 | 5tby | B | 1606 | ARG | p.Arg1606His | P12883 |
| HOMO SAPIENS | -1 | 3279 | 5tby | B | 1664 | ASN | p.Asn1664Lys | P12883 |
| HOMO SAPIENS | -1 | 3279 | 5tby | B | 1712 | ARG | p.Arg1712Trp | P12883 |
| HOMO SAPIENS | -1 | 3279 | 5tby | B | 1769 | LEU | p.Leu1769Met | P12883 |
| HOMO SAPIENS | -1 | 3279 | 5tby | B | 1808 | GLY | p.Gly1808Ser | P12883 |
| HOMO SAPIENS | -1 | 3279 | 5tby | B | 1818 | ARG | p.Arg1818Trp | P12883 |
| HOMO SAPIENS | -1 | 3279 | 5tby | B | 1820 | ARG | p.Arg1820Trp | P12883 |
| HOMO SAPIENS | -1 | 3279 | 5tby | B | 1844 | GLU | p.Glu1844Lys | P12883 |
| HOMO SAPIENS | -1 | 3279 | 5tby | B | 1845 | ARG | p.Arg1845Trp | P12883 |
| HOMO SAPIENS | -1 | 3279 | 5tby | B | 1854 | THR | p.Thr1854Met | P12883 |
| HOMO SAPIENS | -1 | 3279 | 5tby | B | 1909 | ARG | p.Arg1909Trp | P12883 |
| HOMO SAPIENS | 2 | 2260 | 4kl1 | A | 553 | ASP | p.Asp553Asn | Q9Y3Q4 |
| HOMO SAPIENS | 0 | 40570 | 5lst | A | 522 | ARG | p.Arg522Cys | O94761 |
| HOMO SAPIENS | 0 | 40570 | 5lst | A | 522 | ARG | p.Arg522His | O94761 |
| HOMO SAPIENS | 0 | 40570 | 5lst | A | 591 | PRO | p.Pro591Leu | O94761 |
| HOMO SAPIENS | 0 | 40570 | 5lst | A | 793 | PRO | p.Pro793Ser | O94761 |
| HOMO SAPIENS | 4 | 40570 | 5lst | A | 799 | VAL | p.Val799Met | O94761 |
| HOMO SAPIENS | 0 | 40570 | 5lst | A | 1004 | ARG | p.Arg1004Trp | O94761 |
| HOMO SAPIENS | 0 | 40570 | 5lst | A | 1005 | ARG | p.Arg1005Gln | O94761 |
| HOMO SAPIENS | 0 | 40570 | 5lst | A | 1021 | ARG | p.Arg1021Gln | O94761 |

| | | | | | | | | |
|---|---|---|---|---|---|---|---|---|
| HOMO SAPIENS | 0 | 40570 | 5lst | A | 1043 | LEU | p.Leu1043Pro | O94761 |
| HOMO SAPIENS | 0 | 40570 | 5lst | A | 1045 | ALA | p.Ala1045Thr | O94761 |
| HOMO SAPIENS | 0 | 40570 | 5lst | A | 1105 | GLY | p.Gly1105Asp | O94761 |
| HOMO SAPIENS | 0 | 40570 | 5lst | A | 1105 | GLY | p.Gly1105Ser | O94761 |
| HOMO SAPIENS | -1 | 40570 | 5lst | A | 1113 | GLY | p.Gly1113Arg | O94761 |
| HOMO SAPIENS | -1 | 18865 | 1gxc | A | 64 | GLU | p.Glu64Lys | O96017 |
| HOMO SAPIENS | -1 | 11673 | 3dzu | D | 85 | PRO | p.Pro113Gln | P37231 |
| HOMO SAPIENS | 0 | 19383 | 3b96 | A | 63 | LYS | p.Lys103Arg | P49748 |
| HOMO SAPIENS | 4 | 19383 | 3b96 | A | 140 | ALA | p.Ala180Thr | P49748 |
| HOMO SAPIENS | 1 | 19383 | 3b96 | A | 145 | GLY | p.Gly185Ser | P49748 |
| HOMO SAPIENS | 0 | 19383 | 3b96 | A | 220 | THR | p.Thr260Met | P49748 |
| HOMO SAPIENS | 5 | 19383 | 3b96 | A | 243 | VAL | p.Val283Ala | P49748 |
| HOMO SAPIENS | 0 | 19383 | 3b96 | A | 249 | GLY | p.Gly289Arg | P49748 |
| HOMO SAPIENS | 0 | 19383 | 3b96 | A | 254 | GLY | p.Gly294Glu | P49748 |
| HOMO SAPIENS | 7 | 19383 | 3b96 | A | 326 | ARG | p.Arg366Cys | P49748 |
| HOMO SAPIENS | 7 | 19383 | 3b96 | A | 326 | ARG | p.Arg366His | P49748 |
| HOMO SAPIENS | 5 | 19383 | 3b96 | A | 342 | LYS | p.Lys382Gln | P49748 |
| HOMO SAPIENS | 0 | 19383 | 3b96 | A | 369 | THR | p.Thr409Met | P49748 |
| HOMO SAPIENS | 5 | 19383 | 3b96 | A | 376 | ALA | p.Ala416Ser | P49748 |
| HOMO SAPIENS | 0 | 19383 | 3b96 | A | 385 | ALA | p.Ala425Thr | P49748 |
| HOMO SAPIENS | 5 | 19383 | 3b96 | A | 399 | GLY | p.Gly439Asp | P49748 |
| HOMO SAPIENS | 7 | 19383 | 3b96 | A | 401 | GLY | p.Gly441Asp | P49748 |
| HOMO SAPIENS | 9 | 19383 | 3b96 | A | 413 | ARG | p.Arg453Gln | P49748 |
| HOMO SAPIENS | 2 | 19383 | 3b96 | A | 418 | PHE | p.Phe458Leu | P49748 |
| HOMO SAPIENS | 6 | 19383 | 3b96 | A | 419 | ARG | p.Arg459Trp | P49748 |
| HOMO SAPIENS | 6 | 19383 | 3b96 | A | 419 | ARG | p.Arg459Gln | P49748 |
| HOMO SAPIENS | -1 | 19383 | 3b96 | A | 471 | ARG | p.Arg511Gln | P49748 |
| HOMO SAPIENS | 0 | 19383 | 3b96 | A | 573 | ARG | p.Arg613Trp | P49748 |
| HOMO SAPIENS | 0 | 19383 | 3b96 | A | 575 | ARG | p.Arg615Gln | P49748 |
| HOMO SAPIENS | 0 | 22286 | 3k2s | A | 3 | PRO | p.Pro27His | P02647 |
| HOMO SAPIENS | 0 | 22286 | 3k2s | A | 4 | PRO | p.Pro28Arg | P02647 |
| HOMO SAPIENS | 0 | 22286 | 3k2s | A | 10 | ARG | p.Arg34Leu | P02647 |
| HOMO SAPIENS | 0 | 22286 | 3k2s | A | 26 | GLY | p.Gly50Arg | P02647 |
| HOMO SAPIENS | 0 | 22286 | 3k2s | A | 37 | ALA | p.Ala61Thr | P02647 |
| HOMO SAPIENS | 1 | 1622 | 2qrv | D | 631 | ARG | p.Arg635Trp | Q9Y6K1 |
| HOMO SAPIENS | 1 | 1622 | 2qrv | D | 900 | PRO | p.Pro904Leu | Q9Y6K1 |
| HOMO SAPIENS | 6 | 40796 | 1v18 | B | 1509 | LEU | p.Leu1509Val | P25054 |
| HOMO SAPIENS | 1 | 22375 | 4pki | G | 1214 | LYS | p.Asp214Asn | P28289 |
| HOMO SAPIENS | 1 | 16307 | 5mv8 | B | 483 | ALA | p.Ala483Ser | Q9Y6N9 |
| HOMO SAPIENS | 6 | 16307 | 5mv8 | B | 506 | ASN | p.Asn506Lys | Q9Y6N9 |
| HOMO SAPIENS | 4 | 16307 | 5mv8 | B | 519 | ASP | p.Glu519Asp | Q9Y6N9 |
| HOMO SAPIENS | 0 | 16307 | 5mv8 | B | 530 | GLY | p.Gly530Arg | Q9Y6N9 |
| HOMO SAPIENS | 4 | 36221 | 4cvo | A | 127 | VAL | p.Val127Ile | Q03468 |
| HOMO SAPIENS | 7 | 36221 | 4cvo | A | 134 | ARG | p.Arg134Trp | Q03468 |
| HOMO SAPIENS | 0 | 36221 | 4cvo | A | 156 | GLN | p.Gln156Ter | Q03468 |
| HOMO SAPIENS | 0 | 10829 | 4a11 | B | 13 | GLU | p.Glu13Ter | Q13216 |
| HOMO SAPIENS | 1 | 10829 | 4a11 | B | 100 | TYR | p.Tyr100Ter | Q13216 |
| HOMO SAPIENS | 0 | 10829 | 4a11 | B | 160 | ALA | p.Ala160Val | Q13216 |
| HOMO SAPIENS | 1 | 6759 | 5lev | A | 29 | ILE | p.Ile29Phe | Q9H3H5 |
| HOMO SAPIENS | 0 | 6759 | 5lev | A | 108 | MET | p.Met108Ile | Q9H3H5 |
| HOMO SAPIENS | 2 | 6759 | 5lev | A | 117 | VAL | p.Val117Ile | Q9H3H5 |
| HOMO SAPIENS | 1 | 6759 | 5lev | A | 170 | TYR | p.Tyr170Cys | Q9H3H5 |
| HOMO SAPIENS | 0 | 6759 | 5lev | A | 264 | GLY | p.Val264Gly | Q9H3H5 |
| HOMO SAPIENS | 0 | 6759 | 5lev | A | 332 | PHE | p.Phe332Val | Q9H3H5 |

| | | | | | | | | |
|---|---|---|---|---|---|---|---|---|
| HOMO SAPIENS | 7 | 6759 | 5lev | A | 393 | ILE | p.Ile393Val | Q9H3H5 |
| HOMO SAPIENS | -1 | 33473 | 5m5e | C | 9 | ASN | p.Asn31Ser | P31785 |
| HOMO SAPIENS | 0 | 26155 | 1ic8 | B | 98 | ALA | p.Ala98Val | P20823 |
| HOMO SAPIENS | 0 | 15329 | 4iqr | B | 127 | ARG | p.Arg136Trp | P41235 |
| HOMO SAPIENS | 0 | 20725 | 4bsj | A | 350 | GLU | p.Glu350Val | P35916 |
| HOMO SAPIENS | 0 | 20725 | 4bsj | A | 494 | THR | p.Thr494Ala | P35916 |
| HOMO SAPIENS | 0 | 20725 | 4bsj | A | 527 | ASN | p.Asn527Ser | P35916 |
| HOMO SAPIENS | 6 | 29959 | 1xiw | B | 47 | ARG | p.Arg68Ter | P04234 |
| HOMO SAPIENS | 2 | 29959 | 1xiw | B | 72 | CYS | p.Cys93Ter | P04234 |
| HOMO SAPIENS | 3 | 20242 | 5tjc | A | 102 | ARG | p.Arg102Ter | Q9GZU1 |
| HOMO SAPIENS | 3 | 20242 | 5tjc | A | 232 | THR | p.Thr232Pro | Q9GZU1 |
| HOMO SAPIENS | 7 | 9590 | 4ap8 | A | 50 | THR | p.Thr50Ala | O96007 |
| HOMO SAPIENS | 4 | 9590 | 4ap8 | A | 168 | GLU | p.Glu168Lys | O96007 |
| HOMO SAPIENS | -1 | 4297 | 5do7 | B | 19 | ASP | p.Asp19His | Q9H221 |
| HOMO SAPIENS | -1 | 4297 | 5do7 | B | 54 | TYR | p.Tyr54Cys | Q9H221 |
| HOMO SAPIENS | -1 | 4297 | 5do7 | B | 80 | CYS | p.Cys80Tyr | Q9H221 |
| HOMO SAPIENS | 5 | 4297 | 5do7 | B | 107 | SER | p.Ser107Ter | Q9H221 |
| HOMO SAPIENS | 1 | 4297 | 5do7 | B | 164 | ARG | p.Arg164Pro | Q9H221 |
| HOMO SAPIENS | 0 | 4297 | 5do7 | B | 210 | VAL | p.Val210Met | Q9H221 |
| HOMO SAPIENS | 0 | 4297 | 5do7 | B | 231 | PRO | p.Pro231Thr | Q9H221 |
| HOMO SAPIENS | 0 | 4297 | 5do7 | B | 263 | ARG | p.Arg263Gln | Q9H221 |
| HOMO SAPIENS | 0 | 4297 | 5do7 | B | 361 | TRP | p.Trp361Ter | Q9H221 |
| HOMO SAPIENS | 0 | 4297 | 5do7 | B | 400 | THR | p.Thr400Lys | Q9H221 |
| HOMO SAPIENS | 0 | 4297 | 5do7 | B | 549 | ALA | p.Ala549Pro | Q9H221 |
| HOMO SAPIENS | 0 | 4297 | 5do7 | B | 596 | LEU | p.Leu596Arg | Q9H221 |
| HOMO SAPIENS | 0 | 4297 | 5do7 | B | 632 | VAL | p.Val632Ala | Q9H221 |
| HOMO SAPIENS | 0 | 4297 | 5do7 | B | 658 | TYR | p.Tyr658Ter | Q9H221 |
| HOMO SAPIENS | 2 | 9356 | 4igg | B | 206 | GLN | p.Gln206His | P35221 |
| HOMO SAPIENS | 6 | 2179 | 2wk6 | A | 44 | ARG | p.Arg44Gln | P19971 |
| HOMO SAPIENS | 6 | 2179 | 2wk6 | A | 81 | ARG | p.Arg81Gln | P19971 |
| HOMO SAPIENS | 6 | 2179 | 2wk6 | A | 145 | GLY | p.Gly145Arg | P19971 |
| HOMO SAPIENS | 7 | 2179 | 2wk6 | A | 146 | ARG | p.Arg146His | P19971 |
| HOMO SAPIENS | 7 | 2179 | 2wk6 | A | 153 | GLY | p.Gly153Ser | P19971 |
| HOMO SAPIENS | 0 | 2179 | 2wk6 | A | 173 | MET | p.Met173Ile | P19971 |
| HOMO SAPIENS | 6 | 2179 | 2wk6 | A | 202 | ARG | p.Arg202Lys | P19971 |
| HOMO SAPIENS | 6 | 2179 | 2wk6 | A | 208 | VAL | p.Val208Met | P19971 |
| HOMO SAPIENS | 5 | 2179 | 2wk6 | A | 222 | LYS | p.Lys222Arg | P19971 |
| HOMO SAPIENS | 5 | 2179 | 2wk6 | A | 289 | GLU | p.Glu289Ala | P19971 |
| HOMO SAPIENS | 2 | 2179 | 2wk6 | A | 311 | GLY | p.Gly311Arg | P19971 |
| HOMO SAPIENS | 2 | 2179 | 2wk6 | A | 376 | ARG | p.Arg376Pro | P19971 |
| HOMO SAPIENS | 2 | 2179 | 2wk6 | A | 465 | ALA | p.Ala465Thr | P19971 |
| HOMO SAPIENS | -1 | 11856 | 3orh | A | 3 | ALA | p.Ala3Ser | Q14353 |
| HOMO SAPIENS | 1 | 11856 | 3orh | A | 20 | TRP | p.Trp20Ser | Q14353 |
| HOMO SAPIENS | 1 | 11856 | 3orh | A | 50 | MET | p.Met50Leu | Q14353 |
| HOMO SAPIENS | 1 | 11856 | 3orh | A | 110 | VAL | p.Val110Phe | Q14353 |
| HOMO SAPIENS | 8 | 11856 | 3orh | A | 164 | GLY | p.Gly164Asp | Q14353 |
| HOMO SAPIENS | 1 | 11856 | 3orh | A | 169 | CYS | p.Cys169Tyr | Q14353 |
| HOMO SAPIENS | 1 | 11856 | 3orh | A | 174 | TRP | p.Trp174Ter | Q14353 |
| HOMO SAPIENS | 2 | 11856 | 3orh | A | 196 | ALA | p.Ala196Val | Q14353 |
| HOMO SAPIENS | 1 | 11856 | 3orh | A | 209 | THR | p.Thr209Met | Q14353 |
| HOMO SAPIENS | 1 | 11856 | 3orh | A | 219 | ASP | p.Asp219Asn | Q14353 |
| HOMO SAPIENS | 1 | 11856 | 3orh | A | 224 | ALA | p.Ala224Thr | Q14353 |
| HOMO SAPIENS | 1 | 13244 | 2m6i | D | 218 | ARG | p.Arg246Leu | P23415 |
| HOMO SAPIENS | 1 | 13244 | 2m6i | D | 244 | ILE | p.Ile272Asn | P23415 |

| | | | | | | | | |
|---|---|---|---|---|---|---|---|---|
| HOMO SAPIENS | 5 | 13244 | 2m6i | D | 250 | PRO | p.Pro278Thr | P23415 |
| HOMO SAPIENS | 6 | 13244 | 2m6i | D | 266 | GLN | p.Gln294His | P23415 |
| HOMO SAPIENS | 1 | 13244 | 2m6i | D | 271 | ARG | p.Arg299Leu | P23415 |
| HOMO SAPIENS | 1 | 13244 | 2m6i | D | 276 | LYS | p.Lys304Glu | P23415 |
| HOMO SAPIENS | 1 | 13244 | 2m6i | D | 279 | TYR | p.Tyr307Cys | P23415 |
| HOMO SAPIENS | 0 | 13244 | 2m6i | D | 316 | GLY | p.Gly378Ser | P23415 |
| HOMO SAPIENS | 1 | 13244 | 2m6i | D | 400 | ARG | p.Arg428His | P23415 |
| HOMO SAPIENS | 3 | 1967 | 2i6b | A | 301 | ALA | p.Ala318Glu | P55263 |
| HOMO SAPIENS | 4 | 11096 | 1h8b | A | 61 | ALA | p.Ala882Thr | P35609 |
| HOMO SAPIENS | 0 | 15423 | 5ngz | A | 2 | GLN | p.Gln2Glu | Q9NPD8 |
| HOMO SAPIENS | 3 | 30475 | 3lpw | A | 21 | SER | p.Ser22894Tyr | Q8WZ42 |
| HOMO SAPIENS | 6 | 32687 | 2arf | A | 1041 | ARG | p.Arg1041Trp | P35670 |
| HOMO SAPIENS | 8 | 32687 | 2arf | A | 1061 | GLY | p.Gly1061Glu | P35670 |
| HOMO SAPIENS | 6 | 32687 | 2arf | A | 1064 | GLU | p.Glu1064Ala | P35670 |
| HOMO SAPIENS | 2 | 32687 | 2arf | A | 1069 | HIS | p.His1069Gln | P35670 |
| HOMO SAPIENS | 2 | 32687 | 2arf | A | 1088 | LEU | p.Leu1088Ter | P35670 |
| HOMO SAPIENS | 0 | 32687 | 2arf | A | 1099 | GLY | p.Gly1099Ser | P35670 |
| HOMO SAPIENS | 4 | 32687 | 2arf | A | 1106 | VAL | p.Val1106Asp | P35670 |
| HOMO SAPIENS | 2 | 32687 | 2arf | A | 1142 | GLN | p.Gln1142His | P35670 |
| HOMO SAPIENS | 8 | 32687 | 2arf | A | 1148 | ILE | p.Ile1148Thr | P35670 |
| HOMO SAPIENS | 6 | 32687 | 2arf | A | 1151 | ARG | p.Arg1151Cys | P35670 |
| HOMO SAPIENS | 4 | 32687 | 2arf | A | 1169 | MET | p.Met1169Val | P35670 |
| HOMO SAPIENS | 4 | 32687 | 2arf | A | 1173 | GLU | p.Glu1173Lys | P35670 |
| HOMO SAPIENS | 2 | 32687 | 2arf | A | 1176 | GLY | p.Gly1176Arg | P35670 |
| HOMO SAPIENS | 3 | 15544 | 5uq3 | C | 575 | PRO | p.Pro69Leu | P46527 |
| HOMO SAPIENS | 6 | 15544 | 5uq3 | C | 592 | GLU | p.Glu86Asp | P46527 |
| HOMO SAPIENS | 6 | 15544 | 5uq3 | C | 599 | ARG | p.Arg93Trp | P46527 |
| HOMO SAPIENS | -1 | 15544 | 5uq3 | C | 615 | VAL | p.Val109Gly | P46527 |
| HOMO SAPIENS | -1 | 15544 | 5uq3 | C | 631 | SER | p.Ser125Pro | P46527 |
| HOMO SAPIENS | -1 | 15544 | 5uq3 | C | 639 | PRO | p.Pro133Ser | P46527 |
| HOMO SAPIENS | 2 | 2703 | 4wxq | A | 245 | CYS | p.Cys245Tyr | Q99972 |
| HOMO SAPIENS | 0 | 2703 | 4wxq | A | 252 | GLY | p.Gly252Arg | Q99972 |
| HOMO SAPIENS | 1 | 2703 | 4wxq | A | 337 | GLN | p.Gln337Arg | Q99972 |
| HOMO SAPIENS | 5 | 2703 | 4wxq | A | 347 | TYR | p.Tyr347Ter | Q99972 |
| HOMO SAPIENS | 3 | 2703 | 4wxq | A | 364 | GLY | p.Gly364Val | Q99972 |
| HOMO SAPIENS | 2 | 2703 | 4wxq | A | 367 | GLY | p.Gly367Arg | Q99972 |
| HOMO SAPIENS | 2 | 2703 | 4wxq | A | 368 | GLN | p.Gln368Ter | Q99972 |
| HOMO SAPIENS | 6 | 2703 | 4wxq | A | 370 | PRO | p.Pro370Leu | Q99972 |
| HOMO SAPIENS | 1 | 2703 | 4wxq | A | 380 | ASP | p.Asp380His | Q99972 |
| HOMO SAPIENS | 8 | 2703 | 4wxq | A | 399 | GLY | p.Gly399Val | Q99972 |
| HOMO SAPIENS | 0 | 2703 | 4wxq | A | 423 | LYS | p.Lys423Glu | Q99972 |
| HOMO SAPIENS | 3 | 2703 | 4wxq | A | 433 | CYS | p.Cys433Arg | Q99972 |
| HOMO SAPIENS | 3 | 2703 | 4wxq | A | 437 | TYR | p.Tyr437His | Q99972 |
| HOMO SAPIENS | 2 | 2703 | 4wxq | A | 477 | ILE | p.Ile477Ser | Q99972 |
| HOMO SAPIENS | 3 | 2703 | 4wxq | A | 480 | ASN | p.Asn480Lys | Q99972 |
| HOMO SAPIENS | -1 | 34100 | 2fot | C | 1171 | GLN | p.Gln1173Pro | Q13813 |
| HOMO SAPIENS | 0 | 8773 | 1kw2 | A | 432 | ASP | p.Asp432Glu | P02774 |
| HOMO SAPIENS | 0 | 8773 | 1kw2 | A | 436 | LYS | p.Lys436Thr | P02774 |
| HOMO SAPIENS | 0 | 8402 | 4dzo | B | 649 | TYR | p.Tyr649Ter | Q9Y6D9 |
| HOMO SAPIENS | 0 | 39962 | 1nst | A | 611 | GLY | p.Gly611Ser | P52848 |
| HOMO SAPIENS | 2 | 22578 | 3dof | B | 45 | MET | p.Met45Arg | Q9Y2Y0 |
| HOMO SAPIENS | 1 | 23290 | 5j1s | A | 292 | ILE | p.Ile292Thr | O14656 |
| HOMO SAPIENS | 4 | 9505 | 4j1y | A | 534 | ARG | p.Arg534Trp | P09871 |
| HOMO SAPIENS | 9 | 260 | 2h57 | A | 31 | THR | p.Thr31Met | Q9H0F7 |

| | | | | | | | | |
|---|---|---|---|---|---|---|---|---|
| HOMO SAPIENS | 0 | 260 | 2h57 | A | 91 | ILE | p.Ile91Thr | Q9H0F7 |
| HOMO SAPIENS | 0 | 260 | 2h57 | A | 121 | ARG | p.Arg121His | Q9H0F7 |
| HOMO SAPIENS | 0 | 260 | 2h57 | A | 122 | ARG | p.Arg122Ter | Q9H0F7 |
| HOMO SAPIENS | 2 | 260 | 2h57 | A | 169 | GLY | p.Gly169Ala | Q9H0F7 |
| HOMO SAPIENS | 4 | 260 | 2h57 | A | 170 | LEU | p.Leu170Trp | Q9H0F7 |
| HOMO SAPIENS | 4 | 34023 | 2cr3 | A | 18 | GLY | p.Gly48Ser | P11362 |
| HOMO SAPIENS | 0 | 37667 | 3ii7 | A | 420 | ARG | p.Arg420Cys | Q8IXQ5 |
| HOMO SAPIENS | 0 | 41720 | 2k9u | A | 68 | ARG | p.Arg2364His | Q14315 |
| HOMO SAPIENS | 1 | 29250 | 5fmv | A | 568 | HIS | p.His568Gln | P08575 |
| HOMO SAPIENS | 1 | 1280 | 1fb1 | A | 69 | PRO | p.Pro69Leu | P30793 |
| HOMO SAPIENS | 3 | 1280 | 1fb1 | A | 88 | ARG | p.Arg88Trp | P30793 |
| HOMO SAPIENS | 2 | 1280 | 1fb1 | A | 108 | GLY | p.Gly108Asp | P30793 |
| HOMO SAPIENS | 4 | 1280 | 1fb1 | A | 134 | ASP | p.Asp134Val | P30793 |
| HOMO SAPIENS | 6 | 1280 | 1fb1 | A | 135 | ILE | p.Ile135Lys | P30793 |
| HOMO SAPIENS | 9 | 1280 | 1fb1 | A | 144 | HIS | p.His144Pro | P30793 |
| HOMO SAPIENS | 9 | 1280 | 1fb1 | A | 184 | ARG | p.Arg184His | P30793 |
| HOMO SAPIENS | 1 | 1280 | 1fb1 | A | 196 | ALA | p.Ala196Ser | P30793 |
| HOMO SAPIENS | 3 | 1280 | 1fb1 | A | 201 | GLY | p.Gly201Glu | P30793 |
| HOMO SAPIENS | 2 | 1280 | 1fb1 | A | 204 | VAL | p.Val204Ile | P30793 |
| HOMO SAPIENS | 2 | 1280 | 1fb1 | A | 211 | MET | p.Met211Ile | P30793 |
| HOMO SAPIENS | 0 | 1280 | 1fb1 | A | 221 | MET | p.Met221Thr | P30793 |
| HOMO SAPIENS | 3 | 1280 | 1fb1 | A | 224 | LYS | p.Lys224Arg | P30793 |
| HOMO SAPIENS | 2 | 1280 | 1fb1 | A | 249 | ARG | p.Arg249Ser | P30793 |
| HOMO SAPIENS | 8 | 41352 | 4k12 | A | 47 | ILE | p.Ile551Thr | P08603 |
| HOMO SAPIENS | 9 | 20703 | 5nir | A | 57 | CYS | p.Cys57Tyr | P02458 |
| HOMO SAPIENS | 9 | 20703 | 5nir | A | 64 | CYS | p.Cys64Ter | P02458 |
| HOMO SAPIENS | 0 | 20087 | 4cmn | A | 299 | MET | p.Met299Ile | Q01968 |
| HOMO SAPIENS | 1 | 20819 | 3aaf | A | 979 | ILE | p.Ile979Met | Q14191 |
| HOMO SAPIENS | 1 | 20819 | 3aaf | A | 984 | LEU | p.Leu984Ile | Q14191 |
| HOMO SAPIENS | 3 | 20819 | 3aaf | A | 987 | ARG | p.Arg987Ter | Q14191 |
| HOMO SAPIENS | -1 | 20819 | 3aaf | A | 1074 | PHE | p.Leu1074Phe | Q14191 |
| HOMO SAPIENS | 0 | 3619 | 3dem | A | 206 | ASN | p.Asn225Ser | P48740 |
| HOMO SAPIENS | 1 | 3619 | 3dem | A | 269 | ARG | p.Arg288Leu | P48740 |
| HOMO SAPIENS | 0 | 3105 | 3ooi | A | 164 | TYR | p.Tyr2015His | Q96L73 |
| HOMO SAPIENS | 6 | 15828 | 2lkx | A | 16 | LEU | p.Leu100Gln | Q99697 |
| HOMO SAPIENS | 8 | 15828 | 2lkx | A | 24 | ARG | p.Arg108His | Q99697 |
| HOMO SAPIENS | 2 | 15828 | 2lkx | A | 30 | THR | p.Thr114Pro | Q99697 |
| HOMO SAPIENS | 8 | 15828 | 2lkx | A | 45 | VAL | p.Val129Leu | Q99697 |
| HOMO SAPIENS | 8 | 15828 | 2lkx | A | 46 | ARG | p.Arg130Trp | Q99697 |
| HOMO SAPIENS | 9 | 15828 | 2lkx | A | 53 | ARG | p.Arg137Pro | Q99697 |
| HOMO SAPIENS | 9 | 13017 | 4irq | A | 141 | ARG | p.Arg141Trp | Q9UBV7 |
| HOMO SAPIENS | 0 | 13017 | 4irq | A | 186 | ALA | p.Ala186Asp | Q9UBV7 |
| HOMO SAPIENS | 0 | 13017 | 4irq | A | 206 | LEU | p.Leu206Pro | Q9UBV7 |
| HOMO SAPIENS | 1 | 13017 | 4irq | A | 214 | CYS | p.Cys214Tyr | Q9UBV7 |
| HOMO SAPIENS | 0 | 13017 | 4irq | A | 270 | ARG | p.Arg270Cys | Q9UBV7 |
| HOMO SAPIENS | -1 | 13053 | 4mqw | X | 307 | ALA | p.Ala307Thr | P23945 |
| HOMO SAPIENS | 0 | 19406 | 4b3g | A | 43 | ARG | p.Arg43Ter | P38935 |
| HOMO SAPIENS | 0 | 19406 | 4b3g | A | 46 | CYS | p.Cys46Ter | P38935 |
| HOMO SAPIENS | 0 | 19406 | 4b3g | A | 75 | ALA | p.Ala75Thr | P38935 |
| HOMO SAPIENS | 0 | 19406 | 4b3g | A | 201 | LEU | p.Leu201Ser | P38935 |
| HOMO SAPIENS | 3 | 19406 | 4b3g | A | 213 | HIS | p.His213Arg | P38935 |
| HOMO SAPIENS | 0 | 19406 | 4b3g | A | 275 | ILE | p.Ile275Val | P38935 |
| HOMO SAPIENS | -1 | 19406 | 4b3g | A | 320 | ARG | p.Arg320Ter | P38935 |
| HOMO SAPIENS | 0 | 19406 | 4b3g | A | 361 | LEU | p.Leu361Pro | P38935 |

| | | | | | | | | |
|---|---|---|---|---|---|---|---|---|
| HOMO SAPIENS | 0 | 19406 | 4b3g | A | 369 | PHE | p.Phe369Leu | P38935 |
| HOMO SAPIENS | 0 | 19406 | 4b3g | A | 493 | THR | p.Thr493Ile | P38935 |
| HOMO SAPIENS | 0 | 19406 | 4b3g | A | 496 | CYS | p.Cys496Ter | P38935 |
| HOMO SAPIENS | 0 | 19406 | 4b3g | A | 514 | GLU | p.Glu514Lys | P38935 |
| HOMO SAPIENS | 0 | 19406 | 4b3g | A | 528 | ALA | p.Ala528Thr | P38935 |
| HOMO SAPIENS | 0 | 19406 | 4b3g | A | 531 | PRO | p.Pro531Thr | P38935 |
| HOMO SAPIENS | 0 | 19406 | 4b3g | A | 580 | VAL | p.Val580Ile | P38935 |
| HOMO SAPIENS | 6 | 19406 | 4b3g | A | 603 | ARG | p.Arg603His | P38935 |
| HOMO SAPIENS | 0 | 21501 | 5jde | A | 40 | SER | p.Ser2831Asn | Q8WZ42 |
| HOMO SAPIENS | 5 | 21501 | 5jde | A | 189 | ALA | p.Ala2980Thr | Q8WZ42 |
| HOMO SAPIENS | 6 | 15008 | 3mop | H | 12 | ARG | p.Arg12Cys | Q9NWZ3 |
| HOMO SAPIENS | 8 | 17455 | 3jtc | C | 15 | ARG | p.Arg57Trp | P04070 |
| HOMO SAPIENS | -1 | 17455 | 3jtc | C | 20 | CGU | p.Glu62Ala | P04070 |
| HOMO SAPIENS | 0 | 27795 | 5gox | B | 625 | TYR | p.Tyr625Ter | Q92878 |
| HOMO SAPIENS | 0 | 27795 | 5gox | B | 653 | SER | p.Ser653Ter | Q92878 |
| HOMO SAPIENS | -1 | 6558 | 5nug | A | 137 | VAL | p.Val137Ile | Q14204 |
| HOMO SAPIENS | -1 | 6558 | 5nug | A | 567 | ARG | p.Arg567His | Q14204 |
| HOMO SAPIENS | -1 | 6558 | 5nug | A | 1140 | THR | p.Thr1140Met | Q14204 |
| HOMO SAPIENS | -1 | 6558 | 5nug | A | 1991 | ASP | p.Asp1991Asn | Q14204 |
| HOMO SAPIENS | -1 | 6558 | 5nug | A | 2401 | LYS | p.Lys2401Asn | Q14204 |
| HOMO SAPIENS | 0 | 6558 | 5nug | A | 2488 | ARG | p.Arg2488His | Q14204 |
| HOMO SAPIENS | 0 | 6558 | 5nug | A | 3907 | HIS | p.His3907Gln | Q14204 |
| HOMO SAPIENS | 0 | 6558 | 5nug | A | 3981 | THR | p.Thr3981Arg | Q14204 |
| HOMO SAPIENS | 0 | 6558 | 5nug | A | 4394 | THR | p.Thr4394Met | Q14204 |
| HOMO SAPIENS | 0 | 6558 | 5nug | A | 4573 | ASN | p.Asn4573Ser | Q14204 |
| HOMO SAPIENS | 0 | 6558 | 5nug | A | 4603 | SER | p.Ser4603Ile | Q14204 |
| HOMO SAPIENS | 1 | 4244 | 1umk | A | 57 | ARG | p.Arg58Gln | P00387 |
| HOMO SAPIENS | 1 | 4244 | 1umk | A | 72 | LEU | p.Leu73Pro | P00387 |
| HOMO SAPIENS | 1 | 4244 | 1umk | A | 105 | VAL | p.Val106Met | P00387 |
| HOMO SAPIENS | 0 | 4244 | 1umk | A | 116 | THR | p.Thr117Ser | P00387 |
| HOMO SAPIENS | 8 | 4244 | 1umk | A | 127 | SER | p.Ser128Pro | P00387 |
| HOMO SAPIENS | 0 | 4244 | 1umk | A | 148 | LEU | p.Leu149Pro | P00387 |
| HOMO SAPIENS | 0 | 4244 | 1umk | A | 159 | ARG | p.Arg160Ter | P00387 |
| HOMO SAPIENS | 0 | 4244 | 1umk | A | 203 | CYS | p.Cys204Arg | P00387 |
| HOMO SAPIENS | 0 | 4244 | 1umk | A | 203 | CYS | p.Cys204Tyr | P00387 |
| HOMO SAPIENS | 2 | 4244 | 1umk | A | 239 | ASP | p.Asp240Gly | P00387 |
| HOMO SAPIENS | 0 | 4244 | 1umk | A | 291 | GLY | p.Gly292Asp | P00387 |
| HOMO SAPIENS | 4 | 34495 | 1s35 | A | 1151 | ASP | p.Asn1151Asp | P11277 |
| HOMO SAPIENS | 3 | 30110 | 3zqs | A | 224 | THR | p.Thr224Ala | Q9NW38 |
| HOMO SAPIENS | 3 | 17588 | 5dse | B | 53 | LEU | p.Leu53Pro | Q9BYI3 |
| HOMO SAPIENS | 0 | 1564 | 4yqf | B | 72 | HIS | p.His347Gln | Q9UHD8 |
| HOMO SAPIENS | 0 | 17673 | 4jnw | A | 32 | ARG | p.Arg32186His | Q8WZ42 |
| HOMO SAPIENS | 3 | 17673 | 4jnw | A | 140 | ILE | p.Ile32294Val | Q8WZ42 |
| HOMO SAPIENS | 5 | 17673 | 4jnw | A | 269 | ALA | p.Ala32423Thr | Q8WZ42 |
| HOMO SAPIENS | 3 | 12858 | 3s97 | C | 302 | LEU | p.Leu302Pro | Q12860 |
| HOMO SAPIENS | 0 | 6147 | 4nn0 | A | 114 | ARG | p.Arg114Gln | Q9BXJ0 |
| HOMO SAPIENS | 0 | 7549 | 1tr2 | A | 277 | LEU | p.Leu277Met | P18206 |
| HOMO SAPIENS | 1 | 7549 | 1tr2 | A | 413 | ALA | p.Ala413Thr | P18206 |
| HOMO SAPIENS | 1 | 7549 | 1tr2 | A | 519 | ILE | p.Ile519Leu | P18206 |
| HOMO SAPIENS | 5 | 7549 | 1tr2 | A | 547 | ARG | p.Arg547Ter | P18206 |
| HOMO SAPIENS | 2 | 7549 | 1tr2 | A | 636 | HIS | p.His636Arg | P18206 |
| HOMO SAPIENS | 2 | 7549 | 1tr2 | A | 824 | ILE | p.Ile824Met | P18206 |
| HOMO SAPIENS | 2 | 13900 | 3gro | A | 45 | CYS | p.Cys45Tyr | P50897 |
| HOMO SAPIENS | 2 | 13900 | 3gro | A | 75 | THR | p.Thr75Pro | P50897 |

| | | | | | | | | |
|---|---|---|---|---|---|---|---|---|
| HOMO SAPIENS | 2 | 13900 | 3gro | A | 79 | ASP | p.Asp79Gly | P50897 |
| HOMO SAPIENS | 5 | 13900 | 3gro | A | 91 | GLN | p.Gln91Pro | P50897 |
| HOMO SAPIENS | 5 | 13900 | 3gro | A | 96 | CYS | p.Cys96Tyr | P50897 |
| HOMO SAPIENS | 0 | 13900 | 3gro | A | 104 | LYS | p.Lys104Ter | P50897 |
| HOMO SAPIENS | 2 | 13900 | 3gro | A | 108 | GLY | p.Gly108Arg | P50897 |
| HOMO SAPIENS | 2 | 13900 | 3gro | A | 109 | TYR | p.Tyr109Asp | P50897 |
| HOMO SAPIENS | 0 | 13900 | 3gro | A | 122 | ARG | p.Arg122Trp | P50897 |
| HOMO SAPIENS | 0 | 13900 | 3gro | A | 138 | SER | p.Ser138Leu | P50897 |
| HOMO SAPIENS | 5 | 13900 | 3gro | A | 151 | ARG | p.Arg151Ter | P50897 |
| HOMO SAPIENS | 7 | 13900 | 3gro | A | 164 | ARG | p.Arg164Ter | P50897 |
| HOMO SAPIENS | 0 | 13900 | 3gro | A | 177 | GLN | p.Gln177Glu | P50897 |
| HOMO SAPIENS | 0 | 13900 | 3gro | A | 181 | VAL | p.Val181Leu | P50897 |
| HOMO SAPIENS | 0 | 13900 | 3gro | A | 184 | GLU | p.Glu184Lys | P50897 |
| HOMO SAPIENS | 5 | 13900 | 3gro | A | 186 | TRP | p.Trp186Ter | P50897 |
| HOMO SAPIENS | 0 | 13900 | 3gro | A | 219 | LEU | p.Leu219Gln | P50897 |
| HOMO SAPIENS | 0 | 13900 | 3gro | A | 222 | LEU | p.Leu222Pro | P50897 |
| HOMO SAPIENS | 5 | 13900 | 3gro | A | 296 | TRP | p.Trp296Ter | P50897 |
| HOMO SAPIENS | 6 | 35028 | 5mwb | A | 444 | CYS | p.Cys444Tyr | Q04721 |
| HOMO SAPIENS | 0 | 35028 | 5mwb | A | 514 | ASN | p.Asn514Ser | Q04721 |
| HOMO SAPIENS | 5 | 1483 | 4gl2 | A | 822 | ARG | p.Arg822Gln | Q9BYX4 |
| HOMO SAPIENS | 0 | 1483 | 4gl2 | A | 843 | ARG | p.His843Arg | Q9BYX4 |
| HOMO SAPIENS | -1 | 28838 | 3cl3 | D | 153 | LEU | p.Leu153Arg | Q9Y6K9 |
| HOMO SAPIENS | 2 | 22563 | 4zel | B | 79 | ARG | p.Arg79Trp | P09172 |
| HOMO SAPIENS | 2 | 22563 | 4zel | B | 101 | VAL | p.Val101Met | P09172 |
| HOMO SAPIENS | 2 | 22563 | 4zel | B | 114 | ASP | p.Asp114Glu | P09172 |
| HOMO SAPIENS | 0 | 22563 | 4zel | B | 201 | ASN | p.Asn201Ser | P09172 |
| HOMO SAPIENS | 2 | 22563 | 4zel | B | 211 | ALA | p.Ala211Thr | P09172 |
| HOMO SAPIENS | -1 | 22563 | 4zel | B | 290 | ASP | p.Asp290Asn | P09172 |
| HOMO SAPIENS | 2 | 22563 | 4zel | B | 318 | ALA | p.Ala318Ser | P09172 |
| HOMO SAPIENS | 2 | 22563 | 4zel | B | 345 | ASP | p.Asp345Asn | P09172 |
| HOMO SAPIENS | 2 | 22563 | 4zel | B | 362 | ALA | p.Ala362Glu | P09172 |
| HOMO SAPIENS | 2 | 22563 | 4zel | B | 549 | ARG | p.Arg549Cys | P09172 |
| HOMO SAPIENS | 9 | 9780 | 1y6k | R | 36 | TYR | p.Tyr57Cys | Q13651 |
| HOMO SAPIENS | 1 | 9780 | 1y6k | R | 40 | LEU | p.Leu61Val | Q13651 |
| HOMO SAPIENS | 1 | 9780 | 1y6k | R | 63 | THR | p.Thr84Ile | Q13651 |
| HOMO SAPIENS | 9 | 9780 | 1y6k | R | 80 | ARG | p.Arg101Trp | Q13651 |
| HOMO SAPIENS | 0 | 9780 | 1y6k | R | 120 | GLY | p.Gly141Arg | Q13651 |
| HOMO SAPIENS | 0 | 9780 | 1y6k | R | 138 | SER | p.Ser159Gly | Q13651 |
| HOMO SAPIENS | 0 | 9780 | 1y6k | R | 203 | ILE | p.Ile224Val | Q13651 |
| HOMO SAPIENS | 0 | 16019 | 3rep | A | 211 | ARG | p.Arg211Cys | Q13418 |
| HOMO SAPIENS | 3 | 7879 | 5ec3 | A | 33 | ALA | p.Ala33Thr | P32754 |
| HOMO SAPIENS | 4 | 7879 | 5ec3 | A | 113 | ARG | p.Arg113Gln | P32754 |
| HOMO SAPIENS | 3 | 7879 | 5ec3 | A | 340 | VAL | p.Val340Leu | P32754 |
| HOMO SAPIENS | 0 | 8904 | 1qk9 | A | 6 | LYS | p.Lys82Arg | P51608 |
| HOMO SAPIENS | 4 | 8904 | 1qk9 | A | 24 | LEU | p.Leu100Val | P51608 |
| HOMO SAPIENS | 7 | 8904 | 1qk9 | A | 30 | ARG | p.Arg106Trp | P51608 |
| HOMO SAPIENS | 6 | 8904 | 1qk9 | A | 57 | ARG | p.Arg133Cys | P51608 |
| HOMO SAPIENS | 3 | 8904 | 1qk9 | A | 61 | GLU | p.Glu137Gly | P51608 |
| HOMO SAPIENS | 2 | 8904 | 1qk9 | A | 64 | ALA | p.Ala140Val | P51608 |
| HOMO SAPIENS | 5 | 8904 | 1qk9 | A | 79 | PHE | p.Phe155Ser | P51608 |
| HOMO SAPIENS | 3 | 8904 | 1qk9 | A | 82 | THR | p.Thr158Met | P51608 |
| HOMO SAPIENS | 1 | 23121 | 3e9k | A | 198 | THR | p.Thr198Ala | Q16719 |
| HOMO SAPIENS | -1 | 26090 | 3tje | F | 19 | LEU | p.Leu35Met | P25445 |
| HOMO SAPIENS | 8 | 26090 | 3tje | F | 106 | THR | p.Thr122Ile | P25445 |

| | | | | | | | | |
|---|---|---|---|---|---|---|---|---|
| HOMO SAPIENS | 2 | 36404 | 5cqr | A | 765 | GLY | p.Gly765Glu | O95163 |
| HOMO SAPIENS | 2 | 36404 | 5cqr | A | 816 | ILE | p.Ile816Leu | O95163 |
| HOMO SAPIENS | 2 | 36404 | 5cqr | A | 830 | ILE | p.Ile830Met | O95163 |
| HOMO SAPIENS | 5 | 36404 | 5cqr | A | 939 | TYR | p.Tyr939Ter | O95163 |
| HOMO SAPIENS | 2 | 36404 | 5cqr | A | 952 | LYS | p.Lys952Ile | O95163 |
| HOMO SAPIENS | 5 | 36404 | 5cqr | A | 1013 | GLY | p.Gly1013Ser | O95163 |
| HOMO SAPIENS | 5 | 36404 | 5cqr | A | 1072 | CYS | p.Cys1072Ser | O95163 |
| HOMO SAPIENS | -1 | 36404 | 5cqr | A | 1158 | PRO | p.Pro1158Leu | O95163 |
| HOMO SAPIENS | 1 | 34149 | 2k40 | A | 18 | ASN | p.Asn125Ser | Q9UBX0 |
| HOMO SAPIENS | 0 | 34149 | 2k40 | A | 42 | LEU | p.Glu149Lys | Q9UBX0 |
| HOMO SAPIENS | 9 | 34149 | 2k40 | A | 53 | ARG | p.Arg160Cys | Q9UBX0 |
| HOMO SAPIENS | 0 | 34149 | 2k40 | A | 63 | SER | p.Ser170Leu | Q9UBX0 |
| HOMO SAPIENS | 2 | 2468 | 3vp6 | A | 484 | LEU | p.Leu484Met | Q99259 |
| HOMO SAPIENS | 0 | 2468 | 3vp6 | A | 532 | ARG | p.Arg532Gln | Q99259 |
| HOMO SAPIENS | 5 | 1253 | 5xtd | S | 8 | GLY | p.Gly8Arg | O15239 |
| HOMO SAPIENS | 7 | 1253 | 5xtd | S | 32 | GLY | p.Gly32Arg | O15239 |
| HOMO SAPIENS | 7 | 1253 | 5xtd | S | 37 | ARG | p.Arg37Ser | O15239 |
| HOMO SAPIENS | -1 | 13292 | 3u1k | A | 121 | VAL | p.Ile121Val | Q8TCS8 |
| HOMO SAPIENS | 2 | 13292 | 3u1k | A | 165 | PRO | p.Pro165Ser | Q8TCS8 |
| HOMO SAPIENS | 2 | 13292 | 3u1k | A | 192 | ARG | p.Arg192Gln | Q8TCS8 |
| HOMO SAPIENS | 1 | 13292 | 3u1k | A | 315 | LYS | p.Lys315Arg | Q8TCS8 |
| HOMO SAPIENS | 4 | 13292 | 3u1k | A | 509 | VAL | p.Val509Ile | Q8TCS8 |
| HOMO SAPIENS | 7 | 13292 | 3u1k | A | 531 | THR | p.Thr531Arg | Q8TCS8 |
| HOMO SAPIENS | 0 | 13292 | 3u1k | A | 590 | ASN | p.Asn590Asp | Q8TCS8 |
| HOMO SAPIENS | 0 | 27048 | 4p5w | B | 184 | ARG | p.Arg184Ter | Q9UJS0 |
| HOMO SAPIENS | 0 | 27048 | 4p5w | B | 225 | SER | p.Ser225Ter | Q9UJS0 |
| HOMO SAPIENS | -1 | 27048 | 4p5w | B | 631 | ALA | p.Arg360Ter | Q9UJS0 |
| HOMO SAPIENS | 0 | 24240 | 4mdr | A | 318 | ARG | p.Arg318Ter | O00189 |
| HOMO SAPIENS | 0 | 24240 | 4mdr | A | 338 | ARG | p.Arg338Ter | O00189 |
| HOMO SAPIENS | 0 | 13429 | 4cg4 | A | 479 | PHE | p.Phe479Leu | O15553 |
| HOMO SAPIENS | 0 | 13429 | 4cg4 | A | 487 | VAL | p.Val487Leu | O15553 |
| HOMO SAPIENS | 3 | 13429 | 4cg4 | A | 653 | ARG | p.Arg653His | O15553 |
| HOMO SAPIENS | 1 | 13429 | 4cg4 | A | 680 | MET | p.Met680Ile | O15553 |
| HOMO SAPIENS | 1 | 13429 | 4cg4 | A | 688 | TYR | p.Tyr688Ter | O15553 |
| HOMO SAPIENS | 0 | 13429 | 4cg4 | A | 694 | MET | p.Met694Ile | O15553 |
| HOMO SAPIENS | 0 | 13429 | 4cg4 | A | 694 | MET | p.Met694Val | O15553 |
| HOMO SAPIENS | 1 | 13429 | 4cg4 | A | 726 | VAL | p.Val726Ala | O15553 |
| HOMO SAPIENS | 0 | 13429 | 4cg4 | A | 761 | ARG | p.Arg761His | O15553 |
| HOMO SAPIENS | 0 | 2079 | 4okh | A | 691 | GLY | p.Gly691Arg | P20807 |
| HOMO SAPIENS | 2 | 2079 | 4okh | A | 748 | ARG | p.Arg748Ter | P20807 |
| HOMO SAPIENS | 0 | 2079 | 4okh | A | 763 | TYR | p.Tyr763Cys | P20807 |
| HOMO SAPIENS | 1 | 2079 | 4okh | A | 769 | ARG | p.Arg769Gln | P20807 |
| HOMO SAPIENS | 9 | 2079 | 4okh | A | 777 | ILE | p.Ile777Val | P20807 |
| HOMO SAPIENS | 8 | 2079 | 4okh | A | 778 | ASP | p.Asp778His | P20807 |
| HOMO SAPIENS | 5 | 2079 | 4okh | A | 780 | ASP | p.Asp780His | P20807 |
| HOMO SAPIENS | 1 | 1200 | 2gi7 | A | 142 | ARG | p.Arg162Trp | Q9HCN6 |
| HOMO SAPIENS | 4 | 9199 | 3ezq | C | 260 | ASP | p.Asp260Val | P25445 |
| HOMO SAPIENS | 0 | 36522 | 5ms9 | A | 122 | ARG | p.Arg122Cys | P35555 |
| HOMO SAPIENS | 0 | 36522 | 5ms9 | A | 240 | ARG | p.Arg240Cys | P35555 |
| HOMO SAPIENS | 0 | 17448 | 3g2f | A | 225 | ARG | p.Arg225His | Q13873 |
| HOMO SAPIENS | 0 | 17448 | 3g2f | A | 347 | CYS | p.Cys347Tyr | Q13873 |
| HOMO SAPIENS | 3 | 17448 | 3g2f | A | 485 | ASP | p.Asp485Gly | Q13873 |
| HOMO SAPIENS | 7 | 17448 | 3g2f | A | 491 | ARG | p.Arg491Gln | Q13873 |
| HOMO SAPIENS | 7 | 17448 | 3g2f | A | 491 | ARG | p.Arg491Trp | Q13873 |

| | | | | | | | | |
|---|---|---|---|---|---|---|---|---|
| HOMO SAPIENS | 0 | 4861 | 4ux8 | A | 56 | LEU | p.Leu56Met | P07949 |
| HOMO SAPIENS | 1 | 4861 | 4ux8 | A | 88 | ILE | p.Ile88Leu | P07949 |
| HOMO SAPIENS | 0 | 4861 | 4ux8 | A | 114 | ARG | p.Arg114His | P07949 |
| HOMO SAPIENS | 5 | 4861 | 4ux8 | A | 180 | ARG | p.Arg180Gln | P07949 |
| HOMO SAPIENS | 0 | 4861 | 4ux8 | A | 208 | GLU | p.Glu208Asp | P07949 |
| HOMO SAPIENS | 0 | 4861 | 4ux8 | A | 373 | ALA | p.Ala373Val | P07949 |
| HOMO SAPIENS | 0 | 4861 | 4ux8 | A | 375 | LEU | p.Leu375Gln | P07949 |
| HOMO SAPIENS | -1 | 4861 | 4ux8 | A | 386 | ALA | p.Ala386Val | P07949 |
| HOMO SAPIENS | 0 | 4861 | 4ux8 | A | 446 | GLY | p.Gly446Arg | P07949 |
| HOMO SAPIENS | 0 | 4861 | 4ux8 | A | 480 | GLU | p.Glu480Lys | P07949 |
| HOMO SAPIENS | 0 | 4861 | 4ux8 | A | 489 | ASP | p.Asp489Asn | P07949 |
| HOMO SAPIENS | -1 | 4861 | 4ux8 | A | 510 | ALA | p.Ala510Val | P07949 |
| HOMO SAPIENS | -1 | 4861 | 4ux8 | A | 533 | GLY | p.Gly533Cys | P07949 |
| HOMO SAPIENS | -1 | 4861 | 4ux8 | A | 567 | ASP | p.Asp567Asn | P07949 |
| HOMO SAPIENS | -1 | 4861 | 4ux8 | A | 609 | CYS | p.Cys609Tyr | P07949 |
| HOMO SAPIENS | -1 | 4861 | 4ux8 | A | 609 | CYS | p.Cys609Gly | P07949 |
| HOMO SAPIENS | -1 | 4861 | 4ux8 | A | 611 | CYS | p.Cys611Trp | P07949 |
| HOMO SAPIENS | -1 | 4861 | 4ux8 | A | 618 | CYS | p.Cys618Gly | P07949 |
| HOMO SAPIENS | -1 | 4861 | 4ux8 | A | 618 | CYS | p.Cys618Phe | P07949 |
| HOMO SAPIENS | -1 | 4861 | 4ux8 | A | 620 | CYS | p.Cys620Gly | P07949 |
| HOMO SAPIENS | -1 | 4861 | 4ux8 | A | 620 | CYS | p.Cys620Ser | P07949 |
| HOMO SAPIENS | -1 | 4861 | 4ux8 | A | 620 | CYS | p.Cys620Trp | P07949 |
| HOMO SAPIENS | -1 | 4861 | 4ux8 | A | 623 | GLU | p.Glu623Lys | P07949 |
| HOMO SAPIENS | -1 | 4861 | 4ux8 | A | 631 | ASP | p.Asp631Gly | P07949 |
| HOMO SAPIENS | -1 | 4861 | 4ux8 | A | 631 | ASP | p.Asp631Glu | P07949 |
| HOMO SAPIENS | -1 | 4861 | 4ux8 | A | 634 | CYS | p.Cys634Gly | P07949 |
| HOMO SAPIENS | -1 | 4861 | 4ux8 | A | 634 | CYS | p.Cys634Phe | P07949 |
| HOMO SAPIENS | -1 | 4861 | 4ux8 | A | 634 | CYS | p.Cys634Trp | P07949 |
| HOMO SAPIENS | 0 | 36308 | 4nkq | B | 123 | PRO | p.Pro142Leu | P15509 |
| HOMO SAPIENS | 0 | 36308 | 4nkq | B | 145 | ARG | p.Arg164Gln | P15509 |
| HOMO SAPIENS | -1 | 36308 | 4nkq | B | 177 | GLY | p.Gly196Arg | P15509 |
| HOMO SAPIENS | 0 | 36006 | 1oqa | A | 107 | HIS | p.His1860Pro | P38398 |
| HOMO SAPIENS | 7 | 15394 | 2ozo | A | 465 | ARG | p.Arg465His | P43403 |
| HOMO SAPIENS | 7 | 15394 | 2ozo | A | 465 | ARG | p.Arg465Cys | P43403 |
| HOMO SAPIENS | 4 | 15394 | 2ozo | A | 518 | SER | p.Ser518Arg | P43403 |
| HOMO SAPIENS | 1 | 20813 | 3o5z | A | 19 | ARG | p.Arg19Trp | O00459 |
| HOMO SAPIENS | 2 | 4329 | 2bb5 | A | 5 | ILE | p.Ile23Val | P20062 |
| HOMO SAPIENS | 4 | 4329 | 2bb5 | A | 59 | LYS | p.Lys77Met | P20062 |
| HOMO SAPIENS | 4 | 4329 | 2bb5 | A | 197 | ARG | p.Arg215Trp | P20062 |
| HOMO SAPIENS | 1 | 4329 | 2bb5 | A | 241 | PRO | p.Arg259Pro | P20062 |
| HOMO SAPIENS | 2 | 4329 | 2bb5 | A | 315 | THR | p.Thr333Lys | P20062 |
| HOMO SAPIENS | 5 | 4329 | 2bb5 | A | 330 | SER | p.Ser348Phe | P20062 |
| HOMO SAPIENS | 4 | 4329 | 2bb5 | A | 358 | LEU | p.Leu376Ser | P20062 |
| HOMO SAPIENS | 5 | 4329 | 2bb5 | A | 381 | ARG | p.Arg399Gln | P20062 |
| HOMO SAPIENS | 4 | 18294 | 4r30 | A | 163 | ASN | p.Asn163Asp | O95278 |
| HOMO SAPIENS | 6 | 18294 | 4r30 | A | 227 | ALA | p.Ala227Val | O95278 |
| HOMO SAPIENS | 4 | 18294 | 4r30 | A | 241 | ARG | p.Arg241Ter | O95278 |
| HOMO SAPIENS | 4 | 18294 | 4r30 | A | 325 | ARG | p.Arg325His | O95278 |
| HOMO SAPIENS | 0 | 2632 | 4xem | A | 22 | GLU | p.Glu22Lys | P49588 |
| HOMO SAPIENS | 0 | 2632 | 4xem | A | 275 | GLY | p.Gly275Asp | P49588 |
| HOMO SAPIENS | 1 | 2632 | 4xem | A | 329 | ARG | p.Arg329His | P49588 |
| HOMO SAPIENS | 0 | 11431 | 5t4n | A | 284 | GLN | p.Gln305Glu | Q96QU1 |
| HOMO SAPIENS | 0 | 11431 | 5t4n | A | 315 | ARG | p.Arg336Ter | Q96QU1 |
| HOMO SAPIENS | 0 | 11431 | 5t4n | A | 322 | ARG | p.Arg343Lys | Q96QU1 |

| | | | | | | | | |
|---|---|---|---|---|---|---|---|---|
| HOMO SAPIENS | 1 | 11431 | 5t4n | A | 326 | LEU | p.Leu347Phe | Q96QU1 |
| HOMO SAPIENS | 0 | 11431 | 5t4n | A | 373 | TYR | p.Tyr394Cys | Q96QU1 |
| HOMO SAPIENS | 0 | 11431 | 5t4n | A | 398 | ASN | p.Asn419Ser | Q96QU1 |
| HOMO SAPIENS | 0 | 11431 | 5t4n | A | 414 | ALA | p.Asp435Ala | Q96QU1 |
| HOMO SAPIENS | 0 | 11431 | 5t4n | A | 433 | VAL | p.Val454Phe | Q96QU1 |
| HOMO SAPIENS | 1 | 11431 | 5t4n | A | 507 | VAL | p.Val528Asp | Q96QU1 |
| HOMO SAPIENS | 0 | 11431 | 5t4n | A | 547 | ALA | p.Ala568Ser | Q96QU1 |
| HOMO SAPIENS | 3 | 3194 | 5xtd | L | 84 | ARG | p.Arg84His | O43181 |
| HOMO SAPIENS | 1 | 3194 | 5xtd | L | 130 | THR | p.Thr130Ser | O43181 |
| HOMO SAPIENS | 8 | 3194 | 5xtd | L | 136 | SER | p.Ser136Ala | O43181 |
| HOMO SAPIENS | 0 | 40749 | 1nun | A | 106 | CYS | p.Cys106Phe | O15520 |
| HOMO SAPIENS | 9 | 40749 | 1nun | A | 156 | ILE | p.Ile156Arg | O15520 |
| HOMO SAPIENS | 1 | 40749 | 1nun | A | 193 | ARG | p.Arg193Ter | O15520 |
| HOMO SAPIENS | 0 | 40749 | 1nun | A | 207 | HIS | p.His207Pro | O15520 |
| HOMO SAPIENS | 0 | 7026 | 3n1f | D | 875 | SER | p.Ser875Gly | Q4KMG0 |
| HOMO SAPIENS | 6 | 23802 | 3rzv | A | 424 | ARG | p.Arg424Ter | O95630 |
| HOMO SAPIENS | 0 | 36720 | 1jm7 | A | 12 | GLN | p.Gln12Ter | P38398 |
| HOMO SAPIENS | 3 | 36720 | 1jm7 | A | 23 | GLU | p.Glu23Asp | P38398 |
| HOMO SAPIENS | 1 | 36720 | 1jm7 | A | 45 | LYS | p.Lys45Gln | P38398 |
| HOMO SAPIENS | 8 | 36720 | 1jm7 | A | 47 | CYS | p.Cys47Ter | P38398 |
| HOMO SAPIENS | 0 | 36720 | 1jm7 | A | 59 | SER | p.Ser59Ter | P38398 |
| HOMO SAPIENS | 8 | 36720 | 1jm7 | A | 61 | CYS | p.Cys61Gly | P38398 |
| HOMO SAPIENS | 8 | 36720 | 1jm7 | A | 63 | LEU | p.Leu63Ter | P38398 |
| HOMO SAPIENS | 7 | 36720 | 1jm7 | A | 64 | CYS | p.Cys64Gly | P38398 |
| HOMO SAPIENS | 0 | 36720 | 1jm7 | A | 101 | TYR | p.Tyr101Ter | P38398 |
| HOMO SAPIENS | 0 | 36720 | 1jm7 | A | null | TYR | p.Tyr105Cys | P38398 |
| HOMO SAPIENS | 0 | 28500 | 3kuz | A | 1217 | CYS | p.Cys1217Arg | O60486 |
| HOMO SAPIENS | 5 | 8097 | 3p56 | C | 69 | ARG | p.Arg69Trp | Q8TDP1 |
| HOMO SAPIENS | 5 | 8097 | 3p56 | C | 143 | LYS | p.Lys143Ile | Q8TDP1 |
| HOMO SAPIENS | 2 | 30073 | 3nmz | A | 380 | ARG | p.Arg380Gln | P25054 |
| HOMO SAPIENS | 1 | 22573 | 1cjy | A | 111 | SER | p.Ser111Pro | P47712 |
| HOMO SAPIENS | 0 | 22573 | 1cjy | A | 485 | ARG | p.Arg485His | P47712 |
| HOMO SAPIENS | 2 | 965 | 3E+077 | A | 99 | ALA | p.Ala99Val | Q9Y617 |
| HOMO SAPIENS | 2 | 965 | 3E+077 | A | 100 | ASP | p.Asp100Ala | Q9Y617 |
| HOMO SAPIENS | 4 | 1503 | 3kxc | A | 62 | ARG | p.Arg62Trp | O43617 |
| HOMO SAPIENS | 6 | 9418 | 1hdr | A | 22 | GLY | p.Gly23Asp | P09417 |
| HOMO SAPIENS | 2 | 9418 | 1hdr | A | 35 | TRP | p.Trp36Arg | P09417 |
| HOMO SAPIENS | 2 | 9418 | 1hdr | A | 107 | TRP | p.Trp108Gly | P09417 |
| HOMO SAPIENS | 6 | 9418 | 1hdr | A | 149 | TYR | p.Tyr150Cys | P09417 |
| HOMO SAPIENS | -1 | 12079 | 3tnu | B | 373 | GLY | p.Gly373Ala | P13647 |
| HOMO SAPIENS | 1 | 12079 | 3tnu | B | 463 | LEU | p.Leu463Pro | P13647 |
| HOMO SAPIENS | 1 | 12079 | 3tnu | B | 471 | ARG | p.Arg471Cys | P13647 |
| HOMO SAPIENS | 0 | 8135 | 3rbn | B | 89 | SER | p.Ser556Ile | P40692 |
| HOMO SAPIENS | 0 | 8135 | 3rbn | B | 111 | GLU | p.Glu578Gly | P40692 |
| HOMO SAPIENS | 2 | 8135 | 3rbn | B | 114 | PRO | p.Pro581Leu | P40692 |
| HOMO SAPIENS | 0 | 8135 | 3rbn | B | 133 | GLU | p.Glu600Gly | P40692 |
| HOMO SAPIENS | 2 | 8135 | 3rbn | B | 136 | PRO | p.Pro603Arg | P40692 |
| HOMO SAPIENS | 2 | 8135 | 3rbn | B | 136 | PRO | p.Pro603Leu | P40692 |
| HOMO SAPIENS | 7 | 8135 | 3rbn | B | 140 | LEU | p.Leu607His | P40692 |
| HOMO SAPIENS | 2 | 8135 | 3rbn | B | 151 | LYS | p.Lys618Ala | P40692 |
| HOMO SAPIENS | 2 | 8135 | 3rbn | B | 151 | LYS | p.Lys618Thr | P40692 |
| HOMO SAPIENS | 5 | 8135 | 3rbn | B | 181 | PRO | p.Pro648Ser | P40692 |
| HOMO SAPIENS | 2 | 8135 | 3rbn | B | 188 | ILE | p.Ile655Val | P40692 |
| HOMO SAPIENS | 2 | 8135 | 3rbn | B | 214 | ALA | p.Ala681Thr | P40692 |

| | | | | | | | | |
|---|---|---|---|---|---|---|---|---|
| HOMO SAPIENS | 2 | 8135 | 3rbn | B | 220 | ARG | p.Arg687Trp | P40692 |
| HOMO SAPIENS | 2 | 8135 | 3rbn | B | 222 | GLN | p.Gln689Arg | P40692 |
| HOMO SAPIENS | -1 | 8135 | 3rbn | B | 236 | GLN | p.Gln701Ter | P40692 |
| HOMO SAPIENS | 7 | 8135 | 3rbn | B | 251 | HIS | p.His718Tyr | P40692 |
| HOMO SAPIENS | 0 | 8135 | 3rbn | B | 284 | LYS | p.Lys751Arg | P40692 |
| HOMO SAPIENS | 1 | 21998 | 5d68 | A | 355 | LEU | p.Leu355Phe | O00522 |
| HOMO SAPIENS | 3 | 21998 | 5d68 | A | 380 | THR | p.Thr380Met | O00522 |
| HOMO SAPIENS | 0 | 21998 | 5d68 | A | 415 | ILE | p.Ile415Met | O00522 |
| HOMO SAPIENS | 0 | 33789 | 2m0c | A | 13 | ARG | p.Arg218Gln | Q9H161 |
| HOMO SAPIENS | 7 | 33789 | 2m0c | A | 52 | ARG | p.Arg257Thr | Q9H161 |
| HOMO SAPIENS | 0 | 33789 | 2m0c | A | 67 | ARG | p.Arg272Pro | Q9H161 |
| HOMO SAPIENS | 6 | 8263 | 2osg | A | 51 | GLY | p.Gly355Arg | Q9UDY2 |
| HOMO SAPIENS | 5 | 8263 | 2osg | A | 60 | THR | p.Thr364Met | Q9UDY2 |
| HOMO SAPIENS | 1 | 2475 | 5E+026 | B | 234 | THR | p.Thr234Ala | Q9BZ23 |
| HOMO SAPIENS | 2 | 2475 | 5E+026 | B | 264 | ARG | p.Arg264Trp | Q9BZ23 |
| HOMO SAPIENS | 4 | 2475 | 5E+026 | B | 286 | ARG | p.Arg286Cys | Q9BZ23 |
| HOMO SAPIENS | 0 | 2475 | 5E+026 | B | 357 | ARG | p.Arg357Trp | Q9BZ23 |
| HOMO SAPIENS | 2 | 2475 | 5E+026 | B | 481 | ARG | p.Arg481Ter | Q9BZ23 |
| HOMO SAPIENS | 8 | 2475 | 5E+026 | B | 521 | GLY | p.Gly521Arg | Q9BZ23 |
| HOMO SAPIENS | 2 | 2475 | 5E+026 | B | 528 | THR | p.Thr528Met | Q9BZ23 |
| HOMO SAPIENS | 1 | 3742 | 5xtd | E | 108 | HIS | p.His134Tyr | P56556 |
| HOMO SAPIENS | 1 | 16725 | 3g04 | C | 36 | ASP | p.Asp36His | P16473 |
| HOMO SAPIENS | 3 | 16725 | 3g04 | C | 52 | PRO | p.Pro52Thr | P16473 |
| HOMO SAPIENS | 1 | 16725 | 3g04 | C | 162 | PRO | p.Pro162Ala | P16473 |
| HOMO SAPIENS | -1 | 12366 | 5d2d | C | 750 | PRO | p.Pro750Leu | P13569 |
| HOMO SAPIENS | 0 | 12366 | 5d2d | C | 764 | ARG | p.Arg764Ter | P13569 |
| HOMO SAPIENS | 0 | 34845 | 2y23 | A | 1151 | ASP | p.Asp1151Glu | P52179 |
| HOMO SAPIENS | 0 | 34845 | 2y23 | A | 1180 | ASP | p.Asp1180Ala | P52179 |
| HOMO SAPIENS | 6 | 36740 | 3lbx | B | 2053 | ALA | p.Ala2053Pro | P11277 |
| HOMO SAPIENS | 0 | 19289 | 3ihj | A | 404 | ARG | p.Arg404Ter | Q8TD30 |
| HOMO SAPIENS | 5 | 1034 | 5xtd | Q | 228 | ARG | p.Arg228Gln | O75306 |
| HOMO SAPIENS | 5 | 1034 | 5xtd | Q | 229 | PRO | p.Pro229Gln | O75306 |
| HOMO SAPIENS | 4 | 1034 | 5xtd | Q | 323 | ARG | p.Arg323Gln | O75306 |
| HOMO SAPIENS | 4 | 1034 | 5xtd | Q | 352 | PRO | p.Pro352Ala | O75306 |
| HOMO SAPIENS | 5 | 1034 | 5xtd | Q | 413 | SER | p.Ser413Pro | O75306 |
| HOMO SAPIENS | 1 | 2772 | 2vaf | A | 1047 | THR | p.Thr66Ala | O14958 |
| HOMO SAPIENS | 5 | 2772 | 2vaf | A | 1057 | VAL | p.Val76Met | O14958 |
| HOMO SAPIENS | 0 | 2772 | 2vaf | A | 1148 | LEU | p.Leu167His | O14958 |
| HOMO SAPIENS | 1 | 2772 | 2vaf | A | 1149 | ILE | p.Ile168Thr | O14958 |
| HOMO SAPIENS | 8 | 2772 | 2vaf | A | 1225 | HIS | p.His244Arg | O14958 |
| HOMO SAPIENS | 8 | 2772 | 2vaf | A | 1225 | HIS | p.His244Tyr | O14958 |
| HOMO SAPIENS | 7 | 2772 | 2vaf | A | 1251 | ILE | p.Ile270Thr | O14958 |
| HOMO SAPIENS | 3 | 2772 | 2vaf | A | 1288 | ASP | p.Asp307His | O14958 |
| HOMO SAPIENS | 5 | 2772 | 2vaf | A | 1289 | PRO | p.Pro308Leu | O14958 |
| HOMO SAPIENS | 1 | 31885 | 2aex | A | 132 | ALA | p.Ala132Val | P36551 |
| HOMO SAPIENS | 1 | 31885 | 2aex | A | 272 | ASN | p.Asn272His | P36551 |
| HOMO SAPIENS | 8 | 31885 | 2aex | A | 279 | GLY | p.Gly279Arg | P36551 |
| HOMO SAPIENS | 5 | 31885 | 2aex | A | 294 | VAL | p.Val294Ile | P36551 |
| HOMO SAPIENS | 5 | 31885 | 2aex | A | 295 | HIS | p.His295Asp | P36551 |
| HOMO SAPIENS | 1 | 31885 | 2aex | A | 331 | ARG | p.Arg331Trp | P36551 |
| HOMO SAPIENS | 3 | 31885 | 2aex | A | 352 | ARG | p.Arg352Cys | P36551 |
| HOMO SAPIENS | 1 | 31885 | 2aex | A | 404 | LYS | p.Lys404Glu | P36551 |
| HOMO SAPIENS | 2 | 31885 | 2aex | A | 447 | ARG | p.Arg447Cys | P36551 |
| HOMO SAPIENS | 5 | 11721 | 2r3v | A | 20 | HIS | p.His20Pro | Q03426 |

| | | | | | | | | |
|---|---|---|---|---|---|---|---|---|
| HOMO SAPIENS | 1 | 11721 | 2r3v | A | 116 | TYR | p.Tyr116His | Q03426 |
| HOMO SAPIENS | 1 | 11721 | 2r3v | A | 148 | ALA | p.Ala148Thr | Q03426 |
| HOMO SAPIENS | 1 | 11721 | 2r3v | A | 167 | PRO | p.Pro167Leu | Q03426 |
| HOMO SAPIENS | 6 | 11721 | 2r3v | A | 203 | VAL | p.Val203Ala | Q03426 |
| HOMO SAPIENS | 2 | 11721 | 2r3v | A | 215 | ARG | p.Arg215Ter | Q03426 |
| HOMO SAPIENS | 2 | 11721 | 2r3v | A | 237 | THR | p.Thr237Ser | Q03426 |
| HOMO SAPIENS | 0 | 11721 | 2r3v | A | 268 | ILE | p.Ile268Thr | Q03426 |
| HOMO SAPIENS | 0 | 11721 | 2r3v | A | 268 | ILE | p.Ile268Val | Q03426 |
| HOMO SAPIENS | 0 | 11721 | 2r3v | A | 301 | ASN | p.Asn301Ser | Q03426 |
| HOMO SAPIENS | 6 | 11721 | 2r3v | A | 310 | VAL | p.Val310Met | Q03426 |
| HOMO SAPIENS | 5 | 11721 | 2r3v | A | 334 | ALA | p.Ala334Thr | Q03426 |
| HOMO SAPIENS | 2 | 11721 | 2r3v | A | 377 | VAL | p.Val377Ile | Q03426 |
| HOMO SAPIENS | 1 | 11721 | 2r3v | A | 386 | ASP | p.Asp386Asn | Q03426 |
| HOMO SAPIENS | 1 | 11721 | 2r3v | A | 388 | ARG | p.Arg388Ter | Q03426 |
| HOMO SAPIENS | 8 | 5010 | 2hgs | A | 125 | ARG | p.Arg125Cys | P48637 |
| HOMO SAPIENS | 1 | 5010 | 2hgs | A | 164 | ARG | p.Arg164Gln | P48637 |
| HOMO SAPIENS | 0 | 5010 | 2hgs | A | 219 | ASP | p.Asp219Gly | P48637 |
| HOMO SAPIENS | 7 | 5010 | 2hgs | A | 267 | ARG | p.Arg267Trp | P48637 |
| HOMO SAPIENS | 0 | 5010 | 2hgs | A | 283 | ARG | p.Arg283Cys | P48637 |
| HOMO SAPIENS | 0 | 6892 | 5erd | B | 7 | VAL | p.Val56Met | Q14126 |
| HOMO SAPIENS | 1 | 6892 | 5erd | B | 25 | HIS | p.His74Arg | Q14126 |
| HOMO SAPIENS | 0 | 6892 | 5erd | B | 40 | TYR | p.Tyr89Cys | Q14126 |
| HOMO SAPIENS | 0 | 6892 | 5erd | B | 97 | ARG | p.Arg146Leu | Q14126 |
| HOMO SAPIENS | 7 | 6892 | 5erd | B | 217 | ASN | p.Asn266Ser | Q14126 |
| HOMO SAPIENS | 0 | 6892 | 5erd | B | 243 | ARG | p.Arg292His | Q14126 |
| HOMO SAPIENS | 0 | 6892 | 5erd | B | 244 | ILE | p.Ile293Val | Q14126 |
| HOMO SAPIENS | 0 | 6892 | 5erd | B | 343 | VAL | p.Val392Ile | Q14126 |
| HOMO SAPIENS | 4 | 6892 | 5erd | B | 350 | GLU | p.Glu399Lys | Q14126 |
| HOMO SAPIENS | 0 | 6892 | 5erd | B | 410 | TYR | p.Tyr459Cys | Q14126 |
| HOMO SAPIENS | 0 | 6892 | 5erd | B | 466 | VAL | p.Val515Ile | Q14126 |
| HOMO SAPIENS | 0 | 6892 | 5erd | B | 482 | PHE | p.Phe531Cys | Q14126 |
| HOMO SAPIENS | 0 | 6892 | 5erd | B | 484 | VAL | p.Val533Ile | Q14126 |
| HOMO SAPIENS | 0 | 6892 | 5erd | B | 545 | LEU | p.Leu594Pro | Q14126 |
| HOMO SAPIENS | 0 | 1338 | 2ax4 | A | 58 | THR | p.Thr48Arg | O95340 |
| HOMO SAPIENS | 7 | 8675 | 2zkm | X | 385 | LYS | p.Lys385Arg | Q00722 |
| HOMO SAPIENS | 1 | 11769 | 2wya | B | 54 | VAL | p.Val54Met | P54868 |
| HOMO SAPIENS | 1 | 11769 | 2wya | B | 116 | ARG | p.Arg116Cys | P54868 |
| HOMO SAPIENS | 5 | 11769 | 2wya | B | 167 | TYR | p.Tyr167Cys | P54868 |
| HOMO SAPIENS | 1 | 11769 | 2wya | B | 174 | PHE | p.Phe174Leu | P54868 |
| HOMO SAPIENS | 0 | 11769 | 2wya | B | 212 | GLY | p.Gly212Arg | P54868 |
| HOMO SAPIENS | 1 | 11769 | 2wya | B | 288 | ARG | p.Arg288Ter | P54868 |
| HOMO SAPIENS | 0 | 11769 | 2wya | B | 375 | SER | p.Ser375Cys | P54868 |
| HOMO SAPIENS | 4 | 11769 | 2wya | B | 500 | ARG | p.Arg500His | P54868 |
| HOMO SAPIENS | 0 | 3102 | 2iwo | A | 1880 | ARG | p.Arg1908Lys | O75970 |
| HOMO SAPIENS | 6 | 14296 | 2m9w | A | 38 | GLY | p.Gly296Ser | P43694 |
| HOMO SAPIENS | 0 | 14296 | 2m9w | A | 58 | GLN | p.Gln316Glu | P43694 |
| HOMO SAPIENS | 2 | 704 | 5ekd | B | 239 | ARG | p.Arg239Ter | Q9UGM6 |
| HOMO SAPIENS | 2 | 704 | 5ekd | B | 264 | TYR | p.Tyr264Cys | Q9UGM6 |
| HOMO SAPIENS | 6 | 2228 | 1ey2 | A | 47 | SER | p.Ser47Ter | Q93099 |
| HOMO SAPIENS | 6 | 2228 | 1ey2 | A | 48 | ALA | p.Ala48Ser | Q93099 |
| HOMO SAPIENS | 6 | 2228 | 1ey2 | A | 53 | ARG | p.Arg53Gln | Q93099 |
| HOMO SAPIENS | 3 | 2228 | 1ey2 | A | 80 | HIS | p.Gln80His | Q93099 |
| HOMO SAPIENS | 0 | 2228 | 1ey2 | A | 120 | CYS | p.Cys120Trp | Q93099 |
| HOMO SAPIENS | 3 | 2228 | 1ey2 | A | 122 | ALA | p.Ala122Val | Q93099 |

| | | | | | | | | |
|---|---|---|---|---|---|---|---|---|
| HOMO SAPIENS | 3 | 2228 | 1ey2 | A | 161 | GLY | p.Gly161Arg | Q93099 |
| HOMO SAPIENS | 6 | 2228 | 1ey2 | A | 225 | ARG | p.Arg225His | Q93099 |
| HOMO SAPIENS | 0 | 2228 | 1ey2 | A | 230 | PRO | p.Pro230Ser | Q93099 |
| HOMO SAPIENS | 6 | 2228 | 1ey2 | A | 270 | GLY | p.Gly270Arg | Q93099 |
| HOMO SAPIENS | 6 | 2228 | 1ey2 | A | 330 | ARG | p.Arg330Ser | Q93099 |
| HOMO SAPIENS | -1 | 2228 | 1ey2 | A | 368 | MSE | p.Met368Val | Q93099 |
| HOMO SAPIENS | 6 | 2228 | 1ey2 | A | 371 | HIS | p.His371Arg | Q93099 |
| HOMO SAPIENS | 6 | 2228 | 1ey2 | A | 401 | GLU | p.Glu401Gln | Q93099 |
| HOMO SAPIENS | 5 | 34648 | 2k27 | A | 31 | ARG | p.Arg31His | Q06710 |
| HOMO SAPIENS | 7 | 34648 | 2k27 | A | 40 | GLN | p.Gln40Pro | Q06710 |
| HOMO SAPIENS | 5 | 34648 | 2k27 | A | 57 | CYS | p.Cys57Tyr | Q06710 |
| HOMO SAPIENS | 5 | 34648 | 2k27 | A | 62 | LEU | p.Leu62Arg | Q06710 |
| HOMO SAPIENS | 1 | 21399 | 3edu | A | 1819 | GLU | p.Glu1819Ter | P11277 |
| HOMO SAPIENS | 1 | 18153 | 1oey | J | 245 | TYR | p.Tyr245Ter | Q15080 |
| HOMO SAPIENS | 2 | 41567 | 3sw0 | X | 1050 | ASN | p.Asn1050Tyr | P08603 |
| HOMO SAPIENS | 5 | 41567 | 3sw0 | X | 1059 | ILE | p.Ile1059Thr | P08603 |
| HOMO SAPIENS | 0 | 36362 | 4wrm | A | 362 | HIS | p.His362Arg | P07333 |
| HOMO SAPIENS | -1 | 36362 | 4wrm | A | 413 | GLY | p.Gly413Ser | P07333 |
| HOMO SAPIENS | 0 | 31255 | 2w2j | A | 100 | SER | p.Ser100Pro | P35219 |
| HOMO SAPIENS | 0 | 31255 | 2w2j | A | 100 | SER | p.Ser100Leu | P35219 |
| HOMO SAPIENS | 5 | 31255 | 2w2j | A | 237 | ARG | p.Arg237Gln | P35219 |
| HOMO SAPIENS | -1 | 26450 | 1th1 | C | 1395 | SER | p.Ser1395Cys | P25054 |
| HOMO SAPIENS | -1 | 26450 | 1th1 | C | 1413 | MET | p.Met1413Val | P25054 |
| HOMO SAPIENS | -1 | 26450 | 1th1 | C | 1446 | ALA | p.Ala1446Thr | P25054 |
| HOMO SAPIENS | -1 | 26450 | 1th1 | C | 1454 | LYS | p.Lys1454Glu | P25054 |
| HOMO SAPIENS | 0 | 26450 | 1th1 | C | 1474 | ALA | p.Ala1474Thr | P25054 |
| HOMO SAPIENS | 1 | 981 | 5xtb | F | 14 | LEU | p.Leu14Val | O43678 |
| HOMO SAPIENS | 1 | 11069 | 3kq4 | A | 46 | SER | p.Ser46Leu | P27352 |
| HOMO SAPIENS | 1 | 11069 | 3kq4 | A | 97 | MET | p.Met97Thr | P27352 |
| HOMO SAPIENS | 1 | 11069 | 3kq4 | A | 255 | ASN | p.Asn255Ser | P27352 |
| HOMO SAPIENS | 1 | 11069 | 3kq4 | A | 330 | ASN | p.Asn330Lys | P27352 |
| HOMO SAPIENS | 0 | 41489 | 3erb | A | 53 | PRO | p.Pro73Leu | P06681 |
| HOMO SAPIENS | 6 | 8715 | 2bq8 | X | 23 | THR | p.Thr44Met | P13686 |
| HOMO SAPIENS | 0 | 8715 | 2bq8 | X | 88 | GLY | p.Gly109Arg | P13686 |
| HOMO SAPIENS | 6 | 8715 | 2bq8 | X | 127 | VAL | p.Val148Met | P13686 |
| HOMO SAPIENS | 2 | 8715 | 2bq8 | X | 179 | VAL | p.Val200Met | P13686 |
| HOMO SAPIENS | 6 | 8715 | 2bq8 | X | 251 | ARG | p.Arg272Cys | P13686 |
| HOMO SAPIENS | 7 | 8112 | 3p56 | A | 37 | GLY | p.Gly37Ser | O75792 |
| HOMO SAPIENS | -1 | 8112 | 3p56 | A | 202 | LEU | p.Leu202Ser | O75792 |
| HOMO SAPIENS | 1 | 8112 | 3p56 | A | 205 | ASP | p.Asp205Glu | O75792 |
| HOMO SAPIENS | 4 | 8112 | 3p56 | A | 212 | ASN | p.Asn212Ile | O75792 |
| HOMO SAPIENS | 0 | 8112 | 3p56 | A | 235 | ARG | p.Arg235Gln | O75792 |
| HOMO SAPIENS | 4 | 8112 | 3p56 | A | 240 | THR | p.Thr240Met | O75792 |
| HOMO SAPIENS | 0 | 8112 | 3p56 | A | 291 | ARG | p.Arg291His | O75792 |
| HOMO SAPIENS | 0 | 37603 | 2yqg | A | 10 | ARG | p.Arg49His | Q14126 |
| HOMO SAPIENS | 0 | 21178 | 3k1r | B | 458 | ASP | p.Asp458Val | Q495M9 |
| HOMO SAPIENS | -1 | 36828 | 4fq3 | B | 507 | GLY | p.Gly507Asp | P35637 |
| HOMO SAPIENS | 5 | 36828 | 4fq3 | B | 517 | HIS | p.His517Gln | P35637 |
| HOMO SAPIENS | 8 | 36828 | 4fq3 | B | 518 | ARG | p.Arg518Lys | P35637 |
| HOMO SAPIENS | 6 | 36828 | 4fq3 | B | 521 | ARG | p.Arg521Cys | P35637 |
| HOMO SAPIENS | 6 | 36828 | 4fq3 | B | 521 | ARG | p.Arg521His | P35637 |
| HOMO SAPIENS | 0 | 6363 | 3j7y | c | null | HIS | p.His13Arg | Q9H9J2 |
| HOMO SAPIENS | 2 | 6363 | 3j7y | c | 105 | ARG | p.Arg105Cys | Q9H9J2 |
| HOMO SAPIENS | 2 | 6363 | 3j7y | c | 138 | THR | p.Thr138Ile | Q9H9J2 |

| | | | | | | | | |
|---|---|---|---|---|---|---|---|---|
| HOMO SAPIENS | 1 | 6363 | 3j7y | c | 156 | LEU | p.Leu156Arg | Q9H9J2 |
| HOMO SAPIENS | 1 | 6782 | 5l46 | A | 109 | HIS | p.His109Arg | Q9UI17 |
| HOMO SAPIENS | 0 | 23951 | 5t76 | A | 175 | GLY | p.Gly931Ser | P49588 |
| HOMO SAPIENS | 8 | 12604 | 3p0l | A | 182 | ARG | p.Arg182Leu | P49675 |
| HOMO SAPIENS | 2 | 12604 | 3p0l | A | 187 | VAL | p.Val187Met | P49675 |
| HOMO SAPIENS | 2 | 12604 | 3p0l | A | 188 | ARG | p.Arg188Cys | P49675 |
| HOMO SAPIENS | 0 | 12604 | 3p0l | A | 193 | ARG | p.Arg193Ter | P49675 |
| HOMO SAPIENS | 9 | 12604 | 3p0l | A | 217 | ARG | p.Arg217Thr | P49675 |
| HOMO SAPIENS | 4 | 12604 | 3p0l | A | 218 | ALA | p.Ala218Val | P49675 |
| HOMO SAPIENS | 0 | 12604 | 3p0l | A | 250 | TRP | p.Trp250Ter | P49675 |
| HOMO SAPIENS | 1 | 12604 | 3p0l | A | 258 | GLN | p.Gln258Ter | P49675 |
| HOMO SAPIENS | -1 | 27765 | 4wkr | A | 4 | GLU | p.Glu4Lys | Q4G0J3 |
| HOMO SAPIENS | 2 | 3587 | 4kit | C | 2301 | PRO | p.Pro2301Thr | Q6P2Q9 |
| HOMO SAPIENS | 3 | 3587 | 4kit | C | 2304 | PHE | p.Phe2304Leu | Q6P2Q9 |
| HOMO SAPIENS | 3 | 3587 | 4kit | C | 2309 | HIS | p.His2309Pro | Q6P2Q9 |
| HOMO SAPIENS | 3 | 3587 | 4kit | C | 2310 | ARG | p.Arg2310Lys | Q6P2Q9 |
| HOMO SAPIENS | 0 | 9073 | 1n69 | B | 21 | ASN | p.Asn215His | P07602 |
| HOMO SAPIENS | 0 | 9073 | 1n69 | B | 23 | THR | p.Thr217Ile | P07602 |
| HOMO SAPIENS | 0 | 9073 | 1n69 | B | 47 | CYS | p.Cys241Ser | P07602 |
| HOMO SAPIENS | 0 | 475 | 2vd5 | A | 225 | THR | p.Thr225Met | Q09013 |
| HOMO SAPIENS | 1 | 9066 | 4mzv | A | 31 | ASN | p.Asn31Lys | P16422 |
| HOMO SAPIENS | 1 | 9066 | 4mzv | A | 47 | GLN | p.Gln47His | P16422 |
| HOMO SAPIENS | 3 | 9066 | 4mzv | A | 66 | CYS | p.Cys66Tyr | P16422 |
| HOMO SAPIENS | 1 | 9066 | 4mzv | A | 102 | SER | p.Ser102Gly | P16422 |
| HOMO SAPIENS | 5 | 9066 | 4mzv | A | 115 | THR | p.Met115Thr | P16422 |
| HOMO SAPIENS | 1 | 9066 | 4mzv | A | 138 | ARG | p.Arg138Ter | P16422 |
| HOMO SAPIENS | 1 | 9066 | 4mzv | A | 153 | ARG | p.Arg153Thr | P16422 |
| HOMO SAPIENS | 1 | 9066 | 4mzv | A | 165 | ALA | p.Ala165Thr | P16422 |
| HOMO SAPIENS | 8 | 1827 | 2nz2 | A | 14 | GLY | p.Gly14Ser | P00966 |
| HOMO SAPIENS | 5 | 1827 | 2nz2 | A | 18 | SER | p.Ser18Leu | P00966 |
| HOMO SAPIENS | 3 | 1827 | 2nz2 | A | 86 | ARG | p.Arg86Cys | P00966 |
| HOMO SAPIENS | 3 | 1827 | 2nz2 | A | 86 | ARG | p.Arg86His | P00966 |
| HOMO SAPIENS | 1 | 1827 | 2nz2 | A | 179 | TRP | p.Trp179Arg | P00966 |
| HOMO SAPIENS | 7 | 1827 | 2nz2 | A | 180 | SER | p.Ser180Asn | P00966 |
| HOMO SAPIENS | 7 | 1827 | 2nz2 | A | 191 | GLU | p.Glu191Lys | P00966 |
| HOMO SAPIENS | 4 | 1827 | 2nz2 | A | 256 | GLU | p.Glu256Lys | P00966 |
| HOMO SAPIENS | 4 | 1827 | 2nz2 | A | 263 | VAL | p.Val263Met | P00966 |
| HOMO SAPIENS | 5 | 1827 | 2nz2 | A | 265 | ARG | p.Arg265Cys | P00966 |
| HOMO SAPIENS | 5 | 1827 | 2nz2 | A | 268 | ILE | p.Ile268Met | P00966 |
| HOMO SAPIENS | 6 | 1827 | 2nz2 | A | 269 | VAL | p.Val269Met | P00966 |
| HOMO SAPIENS | 7 | 1827 | 2nz2 | A | 272 | ARG | p.Arg272Cys | P00966 |
| HOMO SAPIENS | 3 | 1827 | 2nz2 | A | 304 | ARG | p.Arg304Trp | P00966 |
| HOMO SAPIENS | 3 | 1827 | 2nz2 | A | 307 | ARG | p.Arg307Cys | P00966 |
| HOMO SAPIENS | 3 | 1827 | 2nz2 | A | 310 | LYS | p.Lys310Gln | P00966 |
| HOMO SAPIENS | 2 | 1827 | 2nz2 | A | 362 | GLY | p.Gly362Val | P00966 |
| HOMO SAPIENS | 2 | 1827 | 2nz2 | A | 363 | ARG | p.Arg363Trp | P00966 |
| HOMO SAPIENS | 2 | 1827 | 2nz2 | A | 363 | ARG | p.Arg363Gln | P00966 |
| HOMO SAPIENS | 1 | 1827 | 2nz2 | A | 390 | GLY | p.Gly390Arg | P00966 |
| HOMO SAPIENS | 0 | 29524 | 5vbn | B | 2165 | ARG | p.Arg2165His | Q07864 |
| HOMO SAPIENS | 2 | 29524 | 5vbn | B | 2255 | ILE | p.Ile2255Phe | Q07864 |
| HOMO SAPIENS | 2 | 29524 | 5vbn | B | 2274 | LEU | p.Leu2274Val | Q07864 |
| HOMO SAPIENS | 0 | 31897 | 3t0o | A | 184 | CYS | p.Cys184Arg | O00584 |
| HOMO SAPIENS | -1 | 31897 | 3t0o | A | 215 | GLU | p.Glu215Lys | O00584 |
| HOMO SAPIENS | 0 | 31897 | 3t0o | A | 236 | ARG | p.Arg236Trp | O00584 |

| | | | | | | | | |
|---|---|---|---|---|---|---|---|---|
| HOMO SAPIENS | 7 | 18490 | 4pcw | B | 31 | GLU | p.Glu32Ter | P20930 |
| HOMO SAPIENS | -1 | 39865 | 2h9e | L | 152 | GLY | p.Gly192Arg | P00742 |
| HOMO SAPIENS | 2 | 4309 | 2x0b | F | 174 | THR | p.Thr207Met | P01019 |
| HOMO SAPIENS | 3 | 4309 | 2x0b | F | 342 | ARG | p.Arg375Gln | P01019 |
| HOMO SAPIENS | 2 | 3094 | 1zll | B | 9 | ARG | p.Arg9Cys | P26678 |
| HOMO SAPIENS | 9 | 3094 | 1zll | B | 39 | LEU | p.Leu39Ter | P26678 |
| HOMO SAPIENS | 5 | 3094 | 1zll | B | 49 | VAL | p.Val49Met | P26678 |
| HOMO SAPIENS | -1 | 41498 | 3ffd | P | 8 | LEU | p.Leu44Pro | P12272 |
| HOMO SAPIENS | 1 | 20697 | 1igr | A | 407 | ARG | p.Arg437His | P08069 |
| HOMO SAPIENS | 0 | 20697 | 1igr | A | 416 | MET | p.Met446Val | P08069 |
| HOMO SAPIENS | -1 | 20697 | 1igr | A | 462 | VAL | p.Val492Ile | P08069 |
| HOMO SAPIENS | 0 | 3973 | 4cbz | A | 37 | LEU | p.Leu37Ser | P78504 |
| HOMO SAPIENS | 0 | 3973 | 4cbz | A | 45 | VAL | p.Val45Leu | P78504 |
| HOMO SAPIENS | 0 | 3973 | 4cbz | A | 134 | LEU | p.Leu134Met | P78504 |
| HOMO SAPIENS | -1 | 3973 | 4cbz | A | 146 | VAL | p.Val146Ile | P78504 |
| HOMO SAPIENS | 0 | 3973 | 4cbz | A | 176 | VAL | p.Val176Ile | P78504 |
| HOMO SAPIENS | 0 | 3973 | 4cbz | A | 184 | ARG | p.Arg184Cys | P78504 |
| HOMO SAPIENS | 6 | 1697 | 4hmy | S | 33 | ARG | p.Arg33Trp | Q96PC3 |
| HOMO SAPIENS | 2 | 23752 | 5j4o | A | 13 | ARG | p.Arg1611Ser | P02549 |
| HOMO SAPIENS | 0 | 23752 | 5j4o | A | 42 | ALA | p.Ala1640Gly | P02549 |
| HOMO SAPIENS | 0 | 23752 | 5j4o | A | 108 | HIS | p.His1706Gln | P02549 |
| HOMO SAPIENS | 6 | 23752 | 5j4o | A | 206 | LEU | p.Leu1804Phe | P02549 |
| HOMO SAPIENS | 8 | 531 | 2qz4 | A | 349 | GLY | p.Gly349Ser | Q9UQ90 |
| HOMO SAPIENS | 0 | 531 | 2qz4 | A | 486 | ARG | p.Arg486Gln | Q9UQ90 |
| HOMO SAPIENS | 1 | 531 | 2qz4 | A | 510 | ALA | p.Ala510Val | Q9UQ90 |
| HOMO SAPIENS | 0 | 531 | 2qz4 | A | 529 | ALA | p.Ala529Val | Q9UQ90 |
| HOMO SAPIENS | 0 | 531 | 2qz4 | A | 558 | LYS | p.Lys558Ter | Q9UQ90 |
| HOMO SAPIENS | 1 | 531 | 2qz4 | A | 559 | LYS | p.Lys559Ter | Q9UQ90 |
| HOMO SAPIENS | 2 | 7736 | 1fnh | A | 114 | TRP | p.Trp1834Arg | P02751 |
| HOMO SAPIENS | 0 | 7736 | 1fnh | A | 163 | LEU | p.Leu1883Arg | P02751 |
| HOMO SAPIENS | 3 | 20337 | 5wy5 | B | 264 | LEU | p.Leu264Phe | Q96MG7 |
| HOMO SAPIENS | 8 | 36096 | 2jku | A | 643 | GLY | p.Gly668Arg | P05165 |
| HOMO SAPIENS | 6 | 36096 | 2jku | A | 663 | CYS | p.Cys688Arg | P05165 |
| HOMO SAPIENS | 0 | 9850 | 3kbt | A | 1606 | TYR | p.Tyr1606Ter | P11277 |
| HOMO SAPIENS | 4 | 6876 | 4jkq | A | 217 | ILE | p.Ile217Val | Q92736 |
| HOMO SAPIENS | 5 | 6876 | 4jkq | A | 420 | ARG | p.Arg420Trp | Q92736 |
| HOMO SAPIENS | 2 | 27603 | 1itv | B | 62 | PRO | p.Arg574Pro | P14780 |
| HOMO SAPIENS | 4 | 27603 | 1itv | B | 156 | ARG | p.Arg668Gln | P14780 |
| HOMO SAPIENS | 2 | 22954 | 4f52 | E | 48 | ILE | p.Ile48Met | Q92990 |
| HOMO SAPIENS | 0 | 22954 | 4f52 | E | 91 | ASP | p.Asp91Asn | Q92990 |
| HOMO SAPIENS | 0 | 22954 | 4f52 | E | 146 | ALA | p.Ala146Thr | Q92990 |
| HOMO SAPIENS | 0 | 22954 | 4f52 | E | 181 | ILE | p.Ile181Lys | Q92990 |
| HOMO SAPIENS | 0 | 22954 | 4f52 | E | 194 | ASN | p.Asn194Ser | Q92990 |
| HOMO SAPIENS | 2 | 22954 | 4f52 | E | 254 | PRO | p.Pro254Arg | Q92990 |
| HOMO SAPIENS | 2 | 22954 | 4f52 | E | 300 | ILE | p.Ile300Thr | Q92990 |
| HOMO SAPIENS | 2 | 22954 | 4f52 | E | 336 | LEU | p.Leu336Ser | Q92990 |
| HOMO SAPIENS | 0 | 22954 | 4f52 | E | 353 | TYR | p.Tyr353His | Q92990 |
| HOMO SAPIENS | 0 | 22954 | 4f52 | E | 387 | MET | p.Met387Val | Q92990 |
| HOMO SAPIENS | 4 | 2946 | 5gl7 | A | 344 | TYR | p.Tyr344Cys | P56192 |
| HOMO SAPIENS | 0 | 2946 | 5gl7 | A | 370 | PHE | p.Phe370Leu | P56192 |
| HOMO SAPIENS | 0 | 2946 | 5gl7 | A | 393 | ALA | p.Ala393Thr | P56192 |
| HOMO SAPIENS | 0 | 2946 | 5gl7 | A | 523 | ILE | p.Ile523Thr | P56192 |
| HOMO SAPIENS | 0 | 2946 | 5gl7 | A | 567 | SER | p.Ser567Leu | P56192 |
| HOMO SAPIENS | 0 | 2946 | 5gl7 | A | 800 | PRO | p.Pro800Thr | P56192 |

| | | | | | | | | |
|---|---|---|---|---|---|---|---|---|
| HOMO SAPIENS | 3 | 34961 | 4wy4 | C | 44 | TYR | p.Tyr44His | O95721 |
| HOMO SAPIENS | 8 | 34961 | 4wy4 | C | 89 | GLU | p.Glu89Lys | O95721 |
| HOMO SAPIENS | 0 | 9427 | 4chl | B | 21 | ALA | p.Ala21Ser | O95571 |
| HOMO SAPIENS | 2 | 9427 | 4chl | B | 62 | ALA | p.Ala62Thr | O95571 |
| HOMO SAPIENS | 0 | 9427 | 4chl | B | 114 | ILE | p.Ile114Phe | O95571 |
| HOMO SAPIENS | 2 | 9427 | 4chl | B | 159 | ARG | p.Arg159His | O95571 |
| HOMO SAPIENS | 8 | 9427 | 4chl | B | 163 | ARG | p.Arg163Trp | O95571 |
| HOMO SAPIENS | 8 | 9427 | 4chl | B | 163 | ARG | p.Arg163Gln | O95571 |
| HOMO SAPIENS | 2 | 9427 | 4chl | B | 185 | LEU | p.Leu185Arg | O95571 |
| HOMO SAPIENS | 0 | 40917 | 2e8o | A | 11 | PRO | p.Pro26Leu | Q9Y3Z3 |
| HOMO SAPIENS | -1 | 39190 | 4gow | A | 556 | TYR | p.Tyr556Ter | P56696 |
| HOMO SAPIENS | 0 | 2399 | 2mkv | A | 8 | GLY | p.Gly10Ser | P54710 |
| HOMO SAPIENS | 1 | 2399 | 2mkv | A | 24 | VAL | p.Val26Ile | P54710 |
| HOMO SAPIENS | 4 | 2399 | 2mkv | A | 39 | GLY | p.Gly41Arg | P54710 |
| HOMO SAPIENS | 0 | 20801 | 2zej | B | 1441 | ARG | p.Arg1441Cys | Q5S007 |
| HOMO SAPIENS | 0 | 20801 | 2zej | B | 1441 | ARG | p.Arg1441His | Q5S007 |
| HOMO SAPIENS | 5 | 1019 | 5xtb | M | 62 | ARG | p.Arg62Gln | P28331 |
| HOMO SAPIENS | 1 | 1019 | 5xtb | M | 85 | ALA | p.Ala85Thr | P28331 |
| HOMO SAPIENS | 5 | 1019 | 5xtb | M | 231 | LEU | p.Leu231Val | P28331 |
| HOMO SAPIENS | 2 | 1019 | 5xtb | M | 241 | ARG | p.Arg241Trp | P28331 |
| HOMO SAPIENS | 1 | 1019 | 5xtb | M | 252 | ASP | p.Asp252Gly | P28331 |
| HOMO SAPIENS | 1 | 1019 | 5xtb | M | 408 | ARG | p.Arg408Cys | P28331 |
| HOMO SAPIENS | 2 | 1019 | 5xtb | M | 417 | ARG | p.Arg417Ser | P28331 |
| HOMO SAPIENS | 1 | 1019 | 5xtb | M | 557 | ARG | p.Arg557Ter | P28331 |
| HOMO SAPIENS | 0 | 1019 | 5xtb | M | 567 | ILE | p.Ile567Val | P28331 |
| HOMO SAPIENS | 1 | 1019 | 5xtb | M | 595 | THR | p.Thr595Ala | P28331 |
| HOMO SAPIENS | 1 | 1019 | 5xtb | M | 703 | ALA | p.Ala703Thr | P28331 |
| HOMO SAPIENS | -1 | 3437 | 5hyn | A | 185 | ASP | p.Asp185His | Q15910 |
| HOMO SAPIENS | -1 | 3437 | 5hyn | A | 482 | ALA | p.Ala482Thr | Q15910 |
| HOMO SAPIENS | 0 | 8201 | 2wfd | A | 373 | TYR | p.Tyr373Cys | Q9P2J5 |
| HOMO SAPIENS | 1 | 3590 | 3edi | A | 90 | VAL | p.Val238Ala | O43897 |
| HOMO SAPIENS | 6 | 4555 | 1ytq | A | 64 | ALA | p.Ala65Ser | P43320 |
| HOMO SAPIENS | 0 | 4555 | 1ytq | A | 194 | TRP | p.Trp195Gly | P43320 |
| HOMO SAPIENS | 0 | 1013 | 2nn6 | C | 272 | SER | p.Ser272Thr | Q96B26 |
| HOMO SAPIENS | -1 | 29026 | 5m73 | D | 530 | ILE | p.Ile530Asn | O76094 |
| HOMO SAPIENS | 8 | 2252 | 1d5v | A | 57 | SER | p.Ser125Leu | Q99958 |
| HOMO SAPIENS | 1 | 4337 | 3ecr | A | 29 | ARG | p.Arg26His | P08397 |
| HOMO SAPIENS | 2 | 4337 | 3ecr | A | 34 | ALA | p.Ala31Thr | P08397 |
| HOMO SAPIENS | 2 | 4337 | 3ecr | A | 37 | GLN | p.Gln34Lys | P08397 |
| HOMO SAPIENS | 1 | 4337 | 3ecr | A | 58 | ALA | p.Ala55Ser | P08397 |
| HOMO SAPIENS | 0 | 4337 | 3ecr | A | 84 | LEU | p.Leu81Pro | P08397 |
| HOMO SAPIENS | 0 | 4337 | 3ecr | A | 114 | GLY | p.Gly111Arg | P08397 |
| HOMO SAPIENS | 1 | 4337 | 3ecr | A | 119 | ARG | p.Arg116Trp | P08397 |
| HOMO SAPIENS | 1 | 4337 | 3ecr | A | 152 | ARG | p.Arg149Gln | P08397 |
| HOMO SAPIENS | 1 | 4337 | 3ecr | A | 170 | ARG | p.Arg167Trp | P08397 |
| HOMO SAPIENS | 1 | 4337 | 3ecr | A | 176 | ARG | p.Arg173Gln | P08397 |
| HOMO SAPIENS | 0 | 4337 | 3ecr | A | 180 | LEU | p.Leu177Arg | P08397 |
| HOMO SAPIENS | 0 | 4337 | 3ecr | A | 181 | ASP | p.Asp178Asn | P08397 |
| HOMO SAPIENS | 1 | 4337 | 3ecr | A | 198 | ARG | p.Arg195Cys | P08397 |
| HOMO SAPIENS | 0 | 4337 | 3ecr | A | 201 | TRP | p.Trp198Ter | P08397 |
| HOMO SAPIENS | 1 | 4337 | 3ecr | A | 204 | ARG | p.Arg201Trp | P08397 |
| HOMO SAPIENS | 1 | 4337 | 3ecr | A | 219 | GLY | p.Gly216Asp | P08397 |
| HOMO SAPIENS | 1 | 4337 | 3ecr | A | 226 | GLU | p.Glu223Lys | P08397 |
| HOMO SAPIENS | 0 | 4337 | 3ecr | A | 248 | LEU | p.Leu245Arg | P08397 |

| | | | | | | | | |
|---|---|---|---|---|---|---|---|---|
| HOMO SAPIENS | 1 | 4337 | 3ecr | A | 250 | CYS | p.Cys247Arg | P08397 |
| HOMO SAPIENS | 0 | 4337 | 3ecr | A | 253 | GLU | p.Glu250Lys | P08397 |
| HOMO SAPIENS | 0 | 4337 | 3ecr | A | 255 | ALA | p.Ala252Thr | P08397 |
| HOMO SAPIENS | 0 | 4337 | 3ecr | A | 255 | ALA | p.Ala252Val | P08397 |
| HOMO SAPIENS | 0 | 4337 | 3ecr | A | 259 | HIS | p.His256Asn | P08397 |
| HOMO SAPIENS | 0 | 4337 | 3ecr | A | 324 | ARG | p.Arg321His | P08397 |
| HOMO SAPIENS | -1 | 4337 | 3ecr | A | 362 | ASP | p.Asp359Asn | P08397 |
| HOMO SAPIENS | 1 | 1204 | 5lwv | A | 854 | ILE | p.Ala864Thr | O15294 |
| HOMO SAPIENS | 1 | 25016 | 1m4u | A | 35 | PRO | p.Pro35Arg | Q13253 |
| HOMO SAPIENS | 1 | 25016 | 1m4u | A | 35 | PRO | p.Pro35Ser | Q13253 |
| HOMO SAPIENS | 1 | 25016 | 1m4u | A | 167 | ARG | p.Arg167Gly | Q13253 |
| HOMO SAPIENS | 1 | 25016 | 1m4u | A | 184 | CYS | p.Cys184Tyr | Q13253 |
| HOMO SAPIENS | 0 | 25016 | 1m4u | A | 189 | GLY | p.Gly189Cys | Q13253 |
| HOMO SAPIENS | 1 | 25016 | 1m4u | A | 204 | ARG | p.Arg204Leu | Q13253 |
| HOMO SAPIENS | 1 | 25016 | 1m4u | A | 205 | TRP | p.Trp205Cys | Q13253 |
| HOMO SAPIENS | 1 | 25016 | 1m4u | A | 217 | TRP | p.Trp217Gly | Q13253 |
| HOMO SAPIENS | 0 | 25016 | 1m4u | A | 222 | TYR | p.Tyr222Cys | Q13253 |
| HOMO SAPIENS | 0 | 25016 | 1m4u | A | 222 | TYR | p.Tyr222Asp | Q13253 |
| HOMO SAPIENS | 1 | 25016 | 1m4u | A | 223 | PRO | p.Pro223Leu | Q13253 |
| HOMO SAPIENS | 0 | 1394 | 1um7 | A | 95 | ALA | p.Ala463Val | Q92796 |
| HOMO SAPIENS | 0 | 21294 | 4jpz | B | 1788 | GLU | p.Glu1788Asp | Q99250 |
| HOMO SAPIENS | 9 | 24068 | 2db6 | A | 47 | ARG | p.Arg119Cys | Q96MF2 |
| HOMO SAPIENS | 5 | 1869 | 4omt | A | 39 | ARG | p.Arg39Leu | P08237 |
| HOMO SAPIENS | 2 | 1869 | 4omt | A | 100 | ARG | p.Arg100Gln | P08237 |
| HOMO SAPIENS | 7 | 1869 | 4omt | A | 543 | ASP | p.Asp543Ala | P08237 |
| HOMO SAPIENS | 1 | 1869 | 4omt | A | 686 | TRP | p.Trp686Cys | P08237 |
| HOMO SAPIENS | 1 | 1869 | 4omt | A | 696 | ARG | p.Arg696His | P08237 |
| HOMO SAPIENS | 0 | 12163 | 4wxx | A | 495 | TYR | p.Tyr495Cys | P26358 |
| HOMO SAPIENS | 0 | 12163 | 4wxx | A | 528 | ILE | p.Ile528Met | P26358 |
| HOMO SAPIENS | 0 | 12391 | 1j1d | C | 36 | LYS | p.Lys36Gln | P19429 |
| HOMO SAPIENS | 0 | 12391 | 1j1d | C | 82 | PRO | p.Pro82Ser | P19429 |
| HOMO SAPIENS | 7 | 12391 | 1j1d | C | 102 | ALA | p.Ala102Thr | P19429 |
| HOMO SAPIENS | 1 | 5000 | 4cca | A | 17 | GLY | p.Gly17Arg | Q15833 |
| HOMO SAPIENS | 0 | 5000 | 4cca | A | 205 | VAL | p.Val205Ile | Q15833 |
| HOMO SAPIENS | 0 | 5000 | 4cca | A | 209 | LEU | p.Leu209Pro | Q15833 |
| HOMO SAPIENS | 1 | 5000 | 4cca | A | 459 | ARG | p.Arg459Trp | Q15833 |
| HOMO SAPIENS | 7 | 5000 | 4cca | A | 477 | PRO | p.Pro477Leu | Q15833 |
| HOMO SAPIENS | -1 | 5000 | 4cca | A | 526 | ILE | p.Ile526Val | Q15833 |
| HOMO SAPIENS | 0 | 5000 | 4cca | A | 541 | GLY | p.Gly541Ser | Q15833 |
| HOMO SAPIENS | 0 | 5000 | 4cca | A | 555 | ARG | p.Arg555Gly | Q15833 |
| HOMO SAPIENS | 5 | 28551 | 3vcm | P | 26 | ARG | p.Arg49Ter | P00797 |
| HOMO SAPIENS | 6 | 36453 | 5dzz | A | 2070 | ASP | p.Asp2070His | P15924 |
| HOMO SAPIENS | 0 | 36453 | 5dzz | A | 2166 | ARG | p.Arg2166Ter | P15924 |
| HOMO SAPIENS | 0 | 20268 | 2da6 | A | 96 | ASN | p.Asn321Ser | P35680 |
| HOMO SAPIENS | 1 | 1405 | 3iar | A | 8 | ASP | p.Asp8Asn | P00813 |
| HOMO SAPIENS | 8 | 1405 | 3iar | A | 15 | HIS | p.His15Asp | P00813 |
| HOMO SAPIENS | 6 | 1405 | 3iar | A | 74 | GLY | p.Gly74Val | P00813 |
| HOMO SAPIENS | 6 | 1405 | 3iar | A | 80 | LYS | p.Lys80Arg | P00813 |
| HOMO SAPIENS | 5 | 1405 | 3iar | A | 97 | TYR | p.Tyr97Cys | P00813 |
| HOMO SAPIENS | 5 | 1405 | 3iar | A | 101 | ARG | p.Arg101Leu | P00813 |
| HOMO SAPIENS | 5 | 1405 | 3iar | A | 101 | ARG | p.Arg101Trp | P00813 |
| HOMO SAPIENS | 5 | 1405 | 3iar | A | 106 | LEU | p.Leu106Val | P00813 |
| HOMO SAPIENS | 0 | 1405 | 3iar | A | 107 | LEU | p.Leu107Pro | P00813 |
| HOMO SAPIENS | 9 | 1405 | 3iar | A | 156 | ARG | p.Arg156Cys | P00813 |

| | | | | | | | | |
|---|---|---|---|---|---|---|---|---|
| HOMO SAPIENS | 6 | 1405 | 3iar | A | 211 | ARG | p.Arg211His | P00813 |
| HOMO SAPIENS | 0 | 1405 | 3iar | A | 216 | GLY | p.Gly216Arg | P00813 |
| HOMO SAPIENS | 0 | 1405 | 3iar | A | 291 | SER | p.Ser291Leu | P00813 |
| HOMO SAPIENS | 6 | 1405 | 3iar | A | 304 | LEU | p.Leu304Arg | P00813 |
| HOMO SAPIENS | 0 | 1405 | 3iar | A | 329 | ALA | p.Ala329Val | P00813 |
| HOMO SAPIENS | -1 | 31340 | 4atu | I | 186 | ARG | p.Arg186His | O43602 |
| HOMO SAPIENS | -1 | 31340 | 4atu | I | 192 | ARG | p.Arg192Trp | O43602 |
| HOMO SAPIENS | -1 | 31340 | 4atu | I | 196 | ARG | p.Arg196His | O43602 |
| HOMO SAPIENS | -1 | 31340 | 4atu | I | 203 | THR | p.Thr203Arg | O43602 |
| HOMO SAPIENS | -1 | 31340 | 4atu | I | 286 | ALA | p.Ala286Ser | O43602 |
| HOMO SAPIENS | 4 | 30734 | 1fsu | A | 95 | ARG | p.Arg95Gln | P15848 |
| HOMO SAPIENS | 0 | 30734 | 1fsu | A | 117 | CYS | p.Cys117Arg | P15848 |
| HOMO SAPIENS | 0 | 30734 | 1fsu | A | 137 | GLY | p.Gly137Val | P15848 |
| HOMO SAPIENS | 0 | 30734 | 1fsu | A | 210 | TYR | p.Tyr210Cys | P15848 |
| HOMO SAPIENS | 0 | 30734 | 1fsu | A | 236 | LEU | p.Leu236Pro | P15848 |
| HOMO SAPIENS | 1 | 30734 | 1fsu | A | 315 | ARG | p.Arg315Gln | P15848 |
| HOMO SAPIENS | 0 | 30734 | 1fsu | A | 358 | VAL | p.Val358Met | P15848 |
| HOMO SAPIENS | 2 | 30734 | 1fsu | A | 376 | VAL | p.Val376Met | P15848 |
| HOMO SAPIENS | 0 | 30734 | 1fsu | A | 384 | SER | p.Ser384Asn | P15848 |
| HOMO SAPIENS | 0 | 30734 | 1fsu | A | 393 | HIS | p.His393Pro | P15848 |
| HOMO SAPIENS | 0 | 30734 | 1fsu | A | 405 | CYS | p.Cys405Tyr | P15848 |
| HOMO SAPIENS | 0 | 30734 | 1fsu | A | 458 | ASN | p.Asn458Ser | P15848 |
| HOMO SAPIENS | 1 | 30734 | 1fsu | A | 484 | ARG | p.Arg484Gly | P15848 |
| HOMO SAPIENS | 0 | 38072 | 1wmv | A | 51 | PRO | p.Pro98Leu | Q9NZC7 |
| HOMO SAPIENS | -1 | 5674 | 1j1e | C | 185 | ASN | p.Asn185Lys | P19429 |
| HOMO SAPIENS | -1 | 5674 | 1j1e | C | 192 | ARG | p.Arg192His | P19429 |
| HOMO SAPIENS | -1 | 5674 | 1j1e | C | 196 | ASP | p.Asp196Asn | P19429 |
| HOMO SAPIENS | -1 | 5674 | 1j1e | C | 206 | LYS | p.Lys206Gln | P19429 |
| HOMO SAPIENS | 0 | 38067 | 1wlp | A | 13 | PRO | p.Pro156Gln | P13498 |
| HOMO SAPIENS | -1 | 20441 | 3vfd | A | 289 | HIS | p.His289Tyr | Q9UBP0 |
| HOMO SAPIENS | 2 | 20441 | 3vfd | A | 344 | ILE | p.Ile344Lys | Q9UBP0 |
| HOMO SAPIENS | 0 | 20441 | 3vfd | A | 362 | SER | p.Ser362Cys | Q9UBP0 |
| HOMO SAPIENS | 1 | 20441 | 3vfd | A | 386 | ASN | p.Asn386Ser | Q9UBP0 |
| HOMO SAPIENS | 9 | 20441 | 3vfd | A | 441 | ASP | p.Asp441Gly | Q9UBP0 |
| HOMO SAPIENS | -1 | 20441 | 3vfd | A | 448 | CYS | p.Cys448Tyr | Q9UBP0 |
| HOMO SAPIENS | 2 | 20441 | 3vfd | A | 500 | ARG | p.Arg499Cys | Q9UBP0 |
| HOMO SAPIENS | 1 | 20441 | 3vfd | A | 563 | ARG | p.Arg562Gly | Q9UBP0 |
| HOMO SAPIENS | 3 | 18471 | 4dx8 | J | 26 | ARG | p.Arg26Gln | O00522 |
| HOMO SAPIENS | -1 | 18471 | 4dx8 | J | 101 | GLY | p.Gly101Val | O00522 |
| HOMO SAPIENS | -1 | 34735 | 2ly4 | B | 83 | ALA | p.Ala83Glu | P04637 |
| HOMO SAPIENS | 0 | 31128 | 4cvh | A | 116 | ARG | p.Arg116Cys | A4D126 |
| HOMO SAPIENS | 8 | 31128 | 4cvh | A | 215 | GLN | p.Gln215Ter | A4D126 |
| HOMO SAPIENS | 0 | 31128 | 4cvh | A | 268 | ARG | p.Arg268Ter | A4D126 |
| HOMO SAPIENS | 0 | 31128 | 4cvh | A | 316 | THR | p.Thr316Lys | A4D126 |
| HOMO SAPIENS | 1 | 7164 | 3mt5 | A | 369 | ASP | p.Asp434Gly | Q12791 |
| HOMO SAPIENS | 2 | 26812 | 2wtk | F | 49 | TYR | p.Tyr49Asp | Q15831 |
| HOMO SAPIENS | 0 | 26812 | 2wtk | F | 67 | LEU | p.Leu67Pro | Q15831 |
| HOMO SAPIENS | 4 | 26812 | 2wtk | F | 104 | ARG | p.Arg104Ser | Q15831 |
| HOMO SAPIENS | 0 | 26812 | 2wtk | F | 135 | GLY | p.Gly135Arg | Q15831 |
| HOMO SAPIENS | 0 | 26812 | 2wtk | F | 163 | GLY | p.Gly163Asp | Q15831 |
| HOMO SAPIENS | 2 | 26812 | 2wtk | F | 194 | ALA | p.Asp194Asn | Q15831 |
| HOMO SAPIENS | 2 | 1357 | 5xtd | W | 57 | ARG | p.Arg57His | Q9P0J0 |
| HOMO SAPIENS | 0 | 28086 | 4nn2 | A | 229 | HIS | p.His229Arg | Q8IWS0 |
| HOMO SAPIENS | 0 | 28086 | 4nn2 | A | 234 | LYS | p.Lys234Glu | Q8IWS0 |

| | | | | | | | | |
|---|---|---|---|---|---|---|---|---|
| HOMO SAPIENS | 0 | 28086 | 4nn2 | A | 257 | ARG | p.Arg257Gly | Q8IWS0 |
| HOMO SAPIENS | 5 | 31182 | 2a1x | A | 176 | GLN | p.Gln176Lys | O14832 |
| HOMO SAPIENS | 1 | 31182 | 2a1x | A | 204 | GLY | p.Gly204Ser | O14832 |
| HOMO SAPIENS | 0 | 31182 | 2a1x | A | 269 | ASN | p.Asn269His | O14832 |
| HOMO SAPIENS | 3 | 31182 | 2a1x | A | 275 | ARG | p.Arg275Gln | O14832 |
| HOMO SAPIENS | 3 | 31182 | 2a1x | A | 275 | ARG | p.Arg275Trp | O14832 |
| HOMO SAPIENS | 4 | 2823 | 5xtb | P | 145 | THR | p.Thr145Ile | O75489 |
| HOMO SAPIENS | 5 | 2823 | 5xtb | P | 199 | ARG | p.Arg199Trp | O75489 |
| HOMO SAPIENS | 0 | 8645 | 2h5g | A | 372 | SER | p.Ser372Tyr | P54886 |
| HOMO SAPIENS | 3 | 8645 | 2h5g | A | 715 | ASP | p.Asp715His | P54886 |
| HOMO SAPIENS | 0 | 8645 | 2h5g | A | 765 | ARG | p.Arg765Gln | P54886 |
| HOMO SAPIENS | 0 | 8645 | 2h5g | A | 782 | TYR | p.Tyr782Cys | P54886 |
| HOMO SAPIENS | 0 | 8645 | 2h5g | A | 784 | HIS | p.His784Tyr | P54886 |
| HOMO SAPIENS | 1 | 13992 | 2cue | A | 39 | ARG | p.Arg242Thr | P26367 |
| HOMO SAPIENS | 9 | 13992 | 2cue | A | 55 | PHE | p.Phe258Ser | P26367 |
| HOMO SAPIENS | 0 | 30418 | 2xsw | A | 303 | VAL | p.Val303Met | Q9NRR6 |
| HOMO SAPIENS | 0 | 30418 | 2xsw | A | 315 | PRO | p.Pro315Leu | Q9NRR6 |
| HOMO SAPIENS | 0 | 30418 | 2xsw | A | 341 | GLY | p.Gly341Ser | Q9NRR6 |
| HOMO SAPIENS | 0 | 30418 | 2xsw | A | 378 | ARG | p.Arg378Cys | Q9NRR6 |
| HOMO SAPIENS | 0 | 30418 | 2xsw | A | 435 | ARG | p.Arg435Gln | Q9NRR6 |
| HOMO SAPIENS | 0 | 30418 | 2xsw | A | 515 | ARG | p.Arg515Trp | Q9NRR6 |
| HOMO SAPIENS | 1 | 30418 | 2xsw | A | 526 | PRO | p.Pro526Leu | Q9NRR6 |
| HOMO SAPIENS | 0 | 30418 | 2xsw | A | 563 | ARG | p.Arg563His | Q9NRR6 |
| HOMO SAPIENS | 2 | 30418 | 2xsw | A | 585 | ARG | p.Arg585His | Q9NRR6 |
| HOMO SAPIENS | 2 | 11167 | 4y99 | B | 205 | ARG | p.Arg215Leu | P45379 |
| HOMO SAPIENS | 5 | 11167 | 4y99 | B | 244 | GLU | p.Glu254Asp | P45379 |
| HOMO SAPIENS | 1 | 11167 | 4y99 | B | 253 | LYS | p.Lys263Arg | P45379 |
| HOMO SAPIENS | 8 | 11167 | 4y99 | B | 265 | ARG | p.Arg275Gln | P45379 |
| HOMO SAPIENS | 3 | 11167 | 4y99 | B | 270 | ASP | p.Asp280Asn | P45379 |
| HOMO SAPIENS | -1 | 11167 | 4y99 | B | 278 | ARG | p.Arg288Pro | P45379 |
| HOMO SAPIENS | -1 | 11167 | 4y99 | B | 286 | ARG | p.Arg296Cys | P45379 |
| HOMO SAPIENS | -1 | 11167 | 4y99 | B | 286 | ARG | p.Arg296His | P45379 |
| HOMO SAPIENS | 0 | 15993 | 4po6 | A | 197 | ARG | p.Arg197His | P29597 |
| HOMO SAPIENS | -1 | 15993 | 4po6 | A | 362 | VAL | p.Val362Phe | P29597 |
| HOMO SAPIENS | -1 | 15993 | 4po6 | A | 363 | GLY | p.Gly363Ser | P29597 |
| HOMO SAPIENS | 0 | 5646 | 5uak | A | 7 | GLU | p.Glu7Ter | P13569 |
| HOMO SAPIENS | 0 | 5646 | 5uak | A | 27 | GLY | p.Gly27Ter | P13569 |
| HOMO SAPIENS | 0 | 5646 | 5uak | A | 46 | ALA | p.Ala46Asp | P13569 |
| HOMO SAPIENS | 0 | 5646 | 5uak | A | 60 | GLU | p.Glu60Ter | P13569 |
| HOMO SAPIENS | 0 | 5646 | 5uak | A | 67 | PRO | p.Pro67Leu | P13569 |
| HOMO SAPIENS | 0 | 5646 | 5uak | A | 74 | ARG | p.Arg74Trp | P13569 |
| HOMO SAPIENS | 0 | 5646 | 5uak | A | 75 | ARG | p.Arg75Ter | P13569 |
| HOMO SAPIENS | 0 | 5646 | 5uak | A | 85 | GLY | p.Gly85Glu | P13569 |
| HOMO SAPIENS | 0 | 5646 | 5uak | A | 91 | GLY | p.Gly91Arg | P13569 |
| HOMO SAPIENS | 0 | 5646 | 5uak | A | 92 | GLU | p.Glu92Lys | P13569 |
| HOMO SAPIENS | 0 | 5646 | 5uak | A | 98 | GLN | p.Gln98Arg | P13569 |
| HOMO SAPIENS | 0 | 5646 | 5uak | A | 98 | GLN | p.Gln98Ter | P13569 |
| HOMO SAPIENS | 0 | 5646 | 5uak | A | 109 | TYR | p.Tyr109Cys | P13569 |
| HOMO SAPIENS | 0 | 5646 | 5uak | A | 110 | ASP | p.Asp110His | P13569 |
| HOMO SAPIENS | 0 | 5646 | 5uak | A | 117 | ARG | p.Arg117Cys | P13569 |
| HOMO SAPIENS | 0 | 5646 | 5uak | A | 117 | ARG | p.Arg117His | P13569 |
| HOMO SAPIENS | 0 | 5646 | 5uak | A | 125 | ILE | p.Ile125Thr | P13569 |
| HOMO SAPIENS | 0 | 5646 | 5uak | A | 148 | ILE | p.Ile148Thr | P13569 |
| HOMO SAPIENS | 0 | 5646 | 5uak | A | 170 | ARG | p.Arg170His | P13569 |

| | | | | | | | | |
|---|---|---|---|---|---|---|---|---|
| HOMO SAPIENS | 0 | 5646 | 5uak | A | 178 | GLY | p.Gly178Arg | P13569 |
| HOMO SAPIENS | 0 | 5646 | 5uak | A | 206 | LEU | p.Leu206Trp | P13569 |
| HOMO SAPIENS | 0 | 5646 | 5uak | A | 373 | ASP | p.Asp373Asn | P13569 |
| HOMO SAPIENS | -1 | 5646 | 5uak | A | 709 | ARG | p.Arg709Ter | P13569 |
| HOMO SAPIENS | -1 | 5646 | 5uak | A | 732 | LEU | p.Leu732Ter | P13569 |
| HOMO SAPIENS | -1 | 5646 | 5uak | A | 785 | ARG | p.Arg785Ter | P13569 |
| HOMO SAPIENS | -1 | 5646 | 5uak | A | 831 | GLU | p.Glu831Ter | P13569 |
| HOMO SAPIENS | -1 | 5646 | 5uak | A | 836 | ASP | p.Asp836Tyr | P13569 |
| HOMO SAPIENS | 0 | 5646 | 5uak | A | 851 | ARG | p.Arg851Ter | P13569 |
| HOMO SAPIENS | 0 | 5646 | 5uak | A | 913 | TYR | p.Tyr913Cys | P13569 |
| HOMO SAPIENS | 0 | 5646 | 5uak | A | 913 | TYR | p.Tyr913Ter | P13569 |
| HOMO SAPIENS | 0 | 5646 | 5uak | A | 945 | SER | p.Ser945Leu | P13569 |
| HOMO SAPIENS | 0 | 5646 | 5uak | A | 949 | HIS | p.His949Tyr | P13569 |
| HOMO SAPIENS | 0 | 5646 | 5uak | A | 970 | GLY | p.Gly970Asp | P13569 |
| HOMO SAPIENS | 0 | 5646 | 5uak | A | 1027 | ILE | p.Ile1027Thr | P13569 |
| HOMO SAPIENS | 0 | 5646 | 5uak | A | 1032 | TYR | p.Tyr1032Cys | P13569 |
| HOMO SAPIENS | 0 | 5646 | 5uak | A | 1061 | GLY | p.Gly1061Arg | P13569 |
| HOMO SAPIENS | 1 | 5646 | 5uak | A | 1065 | LEU | p.Leu1065Pro | P13569 |
| HOMO SAPIENS | 1 | 5646 | 5uak | A | 1066 | ARG | p.Arg1066Cys | P13569 |
| HOMO SAPIENS | 1 | 5646 | 5uak | A | 1066 | ARG | p.Arg1066His | P13569 |
| HOMO SAPIENS | 1 | 5646 | 5uak | A | 1067 | ALA | p.Ala1067Thr | P13569 |
| HOMO SAPIENS | 0 | 5646 | 5uak | A | 1069 | GLY | p.Gly1069Arg | P13569 |
| HOMO SAPIENS | 0 | 5646 | 5uak | A | 1070 | ARG | p.Arg1070Trp | P13569 |
| HOMO SAPIENS | 1 | 5646 | 5uak | A | 1071 | GLN | p.Gln1071Pro | P13569 |
| HOMO SAPIENS | 0 | 5646 | 5uak | A | 1074 | PHE | p.Phe1074Leu | P13569 |
| HOMO SAPIENS | 0 | 5646 | 5uak | A | 1077 | LEU | p.Leu1077Pro | P13569 |
| HOMO SAPIENS | 0 | 5646 | 5uak | A | 1089 | TRP | p.Trp1089Ter | P13569 |
| HOMO SAPIENS | 0 | 5646 | 5uak | A | 1092 | TYR | p.Tyr1092Ter | P13569 |
| HOMO SAPIENS | 0 | 5646 | 5uak | A | 1101 | MET | p.Met1101Lys | P13569 |
| HOMO SAPIENS | 0 | 5646 | 5uak | A | 1156 | LEU | p.Leu1156Phe | P13569 |
| HOMO SAPIENS | 0 | 5646 | 5uak | A | 1158 | ARG | p.Arg1158Ter | P13569 |
| HOMO SAPIENS | 0 | 5646 | 5uak | A | 1162 | ARG | p.Arg1162Leu | P13569 |
| HOMO SAPIENS | 0 | 5646 | 5uak | A | 1162 | ARG | p.Arg1162Ter | P13569 |
| HOMO SAPIENS | -1 | 5646 | 5uak | A | 1476 | GLN | p.Gln1476Ter | P13569 |
| HOMO SAPIENS | 0 | 22763 | 1imv | A | 52 | MET | p.Thr72Met | P36955 |
| HOMO SAPIENS | 0 | 22763 | 1imv | A | 301 | TYR | p.Tyr321Ter | P36955 |
| HOMO SAPIENS | 0 | 23179 | 5fql | A | 160 | PRO | p.Pro160Arg | P22304 |
| HOMO SAPIENS | 0 | 23179 | 5fql | A | 172 | ARG | p.Arg172Ter | P22304 |
| HOMO SAPIENS | 0 | 23179 | 5fql | A | 214 | THR | p.Thr214Met | P22304 |
| HOMO SAPIENS | 0 | 23179 | 5fql | A | 309 | THR | p.Thr309Ala | P22304 |
| HOMO SAPIENS | 0 | 23179 | 5fql | A | 333 | SER | p.Ser333Leu | P22304 |
| HOMO SAPIENS | 0 | 23179 | 5fql | A | 443 | ARG | p.Arg443Ter | P22304 |
| HOMO SAPIENS | 0 | 23179 | 5fql | A | 468 | ARG | p.Arg468Leu | P22304 |
| HOMO SAPIENS | 0 | 23179 | 5fql | A | 468 | ARG | p.Arg468Trp | P22304 |
| HOMO SAPIENS | 0 | 23179 | 5fql | A | 502 | TRP | p.Trp502Ser | P22304 |
| HOMO SAPIENS | 9 | 20989 | 1nfi | F | 185 | THR | p.Thr185Met | P25963 |
| HOMO SAPIENS | 4 | 20989 | 1nfi | F | 194 | GLY | p.Gly194Ala | P25963 |
| HOMO SAPIENS | 0 | 11538 | 3eab | A | 162 | VAL | p.Val162Ile | Q9UBP0 |
| HOMO SAPIENS | 0 | 33399 | 2p39 | A | 71 | SER | p.Ser71Gly | Q9GZV9 |
| HOMO SAPIENS | 3 | 33399 | 2p39 | A | 96 | MET | p.Met96Thr | Q9GZV9 |
| HOMO SAPIENS | 0 | 33399 | 2p39 | A | 129 | SER | p.Ser129Phe | Q9GZV9 |
| HOMO SAPIENS | -1 | 33399 | 2p39 | A | 172 | PRO | p.Pro172Leu | Q9GZV9 |
| HOMO SAPIENS | -1 | 33399 | 2p39 | A | 176 | ARG | p.Arg176Gln | Q9GZV9 |
| HOMO SAPIENS | -1 | 33399 | 2p39 | A | 179 | ARG | p.Arg179Trp | Q9GZV9 |

| | | | | | | | | |
|---|---|---|---|---|---|---|---|---|
| HOMO SAPIENS | -1 | 33399 | 2p39 | A | 179 | ARG | p.Arg179Gln | Q9GZV9 |
| HOMO SAPIENS | 0 | 16217 | 2dne | A | 79 | ILE | p.Ile163Leu | P10515 |
| HOMO SAPIENS | 8 | 7666 | 4kbq | A | 65 | ASN | p.Asn65Ser | Q9UNE7 |
| HOMO SAPIENS | 1 | 6407 | 1ikt | A | 71 | THR | p.Thr687Ile | P51659 |
| HOMO SAPIENS | 0 | 6407 | 1ikt | A | 112 | MET | p.Met728Val | P51659 |
| HOMO SAPIENS | 0 | 5849 | 4ihb | A | 67 | VAL | p.Val67Asp | O75923 |
| HOMO SAPIENS | 6 | 17703 | 5fcn | A | 20 | HIS | p.His99Gln | Q66GS9 |
| HOMO SAPIENS | 8 | 17703 | 5fcn | A | 33 | ARG | p.Arg112His | Q66GS9 |
| HOMO SAPIENS | 0 | 34095 | 2E+073 | A | 101 | HIS | p.His101Tyr | P05129 |
| HOMO SAPIENS | -1 | 41358 | 4liq | E | 507 | THR | p.Thr507Met | P07333 |
| HOMO SAPIENS | -1 | 21290 | 1fch | A | 235 | MET | p.Met272Arg | P50542 |
| HOMO SAPIENS | 5 | 23431 | 6pax | A | 23 | ARG | p.Arg26Gly | P26367 |
| HOMO SAPIENS | 1 | 23431 | 6pax | A | 61 | GLY | p.Gly64Val | P26367 |
| HOMO SAPIENS | 1 | 23431 | 6pax | A | 65 | PRO | p.Pro68Ser | P26367 |
| HOMO SAPIENS | 1 | 23431 | 6pax | A | 116 | SER | p.Ser119Arg | P26367 |
| HOMO SAPIENS | 9 | 23431 | 6pax | A | 123 | VAL | p.Val126Asp | P26367 |
| HOMO SAPIENS | 9 | 23431 | 6pax | A | 125 | ARG | p.Arg128Cys | P26367 |
| HOMO SAPIENS | 9 | 33003 | 4zrj | B | 535 | LEU | p.Leu535Pro | P35240 |
| HOMO SAPIENS | 7 | 33003 | 4zrj | B | 538 | GLN | p.Gln538Pro | P35240 |
| HOMO SAPIENS | 0 | 33003 | 4zrj | B | 585 | TRP | p.Ala585Thr | P35240 |
| HOMO SAPIENS | 0 | 3800 | 2nzj | A | 193 | VAL | p.Val193Met | O75628 |

| gene_name | max_cons | max_bs_id | max_pdb_id | max_chain_id |
|---|---|---|---|---|
| AKR1D1 | 0.306 | 22 | 3dop | A |
| AKR1D1 | None | None | None | None |
| AKR1D1 | None | None | None | None |
| AKR1D1 | None | None | None | None |
| ALPP | None | None | None | None |
| BMPR1A | None | None | None | None |
| BMPR1A | None | None | None | None |
| FGFR2 | None | None | None | None |
| PROC | 0.587 | 3 | 1aut | C |
| PROC | None | None | None | None |
| PROC | None | None | None | None |
| PROC | 0.396 | 10 | 3f6u | H |
| PROC | None | None | None | None |
| PROC | None | None | None | None |
| PROC | None | None | None | None |
| PROC | None | None | None | None |
| PROC | None | None | None | None |
| PROC | 0.587 | 3 | 1aut | C |
| PROC | None | None | None | None |
| PROC | None | None | None | None |
| INS | 1 | 2 | 3zs2 | J |
| INS | 0.85 | 1 | 4ajx | D |
| INS | 1 | 1 | 3zs2 | J |
| INS | 1 | 1 | 3zs2 | J |
| FTL | 0.627 | 11 | 2fg8 | B |
| FTL | 0.313 | 62 | 2fg8 | G |
| FTL | 0.544 | 24 | 2fg8 | D |
| TNFSF11 | None | None | None | None |
| EHMT1 | None | None | None | None |
| EHMT1 | None | None | None | None |
| EHMT1 | None | None | None | None |
| CA2 | 0.939 | 1 | 4idr | X |
| CA2 | 0.939 | 1 | 4idr | X |
| CA2 | 0.902 | 5 | 4idr | X |
| CA2 | 0.902 | 5 | 4idr | X |
| SMAD4 | None | None | None | None |
| SMAD4 | None | None | None | None |
| TTR | 0.691 | 12 | 1ttb | B |
| TTR | 0.961 | 7 | 3grb | A |
| TTR | 0.891 | 4 | 3dk2 | B |
| TTR | 0.947 | 3 | 1tta | A |
| TTR | 0.923 | 1 | 3grb | B |
| TTR | 0.891 | 1 | 4y9b | B |
| TTR | 0.891 | 1 | 4y9b | B |
| TTR | 0.882 | 15 | 3grb | A |
| TTR | 0.909 | 5 | 4tl4 | A |
| TTR | 0.836 | 7 | 1z7j | B |
| TTR | 0.882 | 15 | 3grb | A |
| TTR | 0.891 | 5 | 3i9i | B |
| TTR | 0.902 | 9 | 3m1o | B |
| TTR | 0.902 | 9 | 3m1o | B |
| TTR | 0.902 | 9 | 3m1o | B |
| TTR | 0.636 | 16 | 1thc | A |

| | | | | | |
|---|---|---|---|---|---|
| TTR | | 0.902 | 9 | 3m1o | B |
| TTR | | 0.836 | 7 | 1bze | A |
| TTR | | 0.846 | 16 | 3gps | A |
| TTR | | 0.846 | 16 | 3gps | A |
| TTR | | 0.909 | 5 | 4tl4 | A |
| TTR | | 0.909 | 5 | 4tl4 | A |
| TTR | | 0.673 | 15 | 3dk2 | B |
| TTR | | 0.945 | 4 | 4deu | A |
| TTR | | 0.941 | 3 | 3did | B |
| TTR | | 0.941 | 3 | 3did | B |
| TTR | | 0.868 | 6 | 4l1t | A |
| TTR | | 0.882 | 19 | 3gps | B |
| TTR | | 0.891 | 4 | 3dk2 | B |
| TTR | | 0.941 | 3 | 3did | B |
| TTR | | 0.923 | 1 | 4hiq | A |
| TTR | | 0.923 | 1 | 4hiq | A |
| TTR | | 0.961 | 7 | 3grb | A |
| TTR | | 0.839 | 7 | 1tsh | B |
| TTR | | 0.891 | 6 | 1ttb | B |
| ALK | | 0.682 | 1 | 4cd0 | A |
| ALK | None | None | | None | None |
| THRB | None | None | | None | None |
| THRB | None | None | | None | None |
| THRB | None | None | | None | None |
| THRB | None | None | | None | None |
| THRB | None | None | | None | None |
| THRB | None | None | | None | None |
| THRB | None | None | | None | None |
| THRB | None | None | | None | None |
| THRB | None | None | | None | None |
| THRB | None | None | | None | None |
| THRB | None | None | | None | None |
| THRB | None | None | | None | None |
| THRB | None | None | | None | None |
| THRB | None | None | | None | None |
| THRB | None | None | | None | None |
| THRB | None | None | | None | None |
| THRB | None | None | | None | None |
| THRB | None | None | | None | None |
| THRB | | 0.577 | 3 | 2j4a | A |
| IL2RG | None | None | | None | None |
| IL2RG | None | None | | None | None |
| IL2RG | None | None | | None | None |
| IL2RG | None | None | | None | None |
| IL2RG | None | None | | None | None |
| ITGB3 | None | None | | None | None |
| ITGB3 | None | None | | None | None |
| ITGB3 | None | None | | None | None |
| ITGB3 | None | None | | None | None |
| ITGB3 | None | None | | None | None |
| ITGB3 | None | None | | None | None |
| ITGB3 | None | None | | None | None |
| ITGB3 | None | None | | None | None |
| ITGB3 | None | None | | None | None |

| | | | | |
|---|---|---|---|---|
| PKD2 | None | None | None | None |
| PKD2 | None | None | None | None |
| PKD2 | None | None | None | None |
| PKD2 | None | None | None | None |
| PKD2 | None | None | None | None |
| PKD2 | None | None | None | None |
| PKD2 | None | None | None | None |
| TP53 | 0.571 | 1 | 1saf | B |
| TP53 | 0.571 | 1 | 1saf | B |
| TP53 | None | None | None | None |
| TP53 | 0.585 | 15 | 4mzr | B |
| TP53 | None | None | None | None |
| TP53 | None | None | None | None |
| EGFR | None | None | None | None |
| EGFR | None | None | None | None |
| XRCC4 | None | None | None | None |
| XRCC4 | None | None | None | None |
| HBB | 0.438 | 2 | 2w6v | D |
| HBB | 0.467 | 2 | 3qje | D |
| HBB | 0.467 | 2 | 3qje | D |
| HBB | 0.303 | 15 | 2w72 | B |
| HBB | 0.303 | 21 | 2w72 | B |
| HBB | 0.332 | 7 | 1cbm | A |
| HBB | 0.421 | 11 | 3ic2 | B |
| HBB | 0.459 | 2 | 5ee4 | D |
| HBB | 0.459 | 2 | 5ee4 | D |
| HBB | 0.459 | 2 | 5ee4 | D |
| HBB | 0.535 | 3 | 3r5i | B |
| HBB | 0.383 | 3 | 1lfv | B |
| HBB | 0.383 | 3 | 1lfv | B |
| HBB | 0.402 | 11 | 1m9p | D |
| HBB | None | None | None | None |
| HBB | 0.459 | 2 | 5ee4 | D |
| HBB | 0.566 | 1 | 1cbm | B |
| HBB | 0.51 | 3 | 1abw | B |
| HBB | 0.392 | 2 | 1lfz | B |
| HBB | 0.474 | 1 | 2h35 | B |
| HBB | 0.474 | 1 | 2h35 | B |
| HBB | 0.38 | 2 | 1j7y | B |
| HBB | 0.508 | 7 | 3ic2 | B |
| HBB | 0.545 | 3 | 1bij | D |
| HBB | 0.6 | 1 | 2w72 | D |
| HBB | 0.332 | 4 | 1fn3 | D |
| HBB | 0.354 | 15 | 1bij | D |
| HBB | 0.308 | 15 | 4rol | D |
| HBB | 0.733 | 1 | 1yff | D |
| HBB | 0.692 | 2 | 1buw | B |
| RPS6KA3 | None | None | None | None |
| RPS6KA3 | 0.555 | 1 | 4nw5 | A |
| RPS6KA3 | None | None | None | None |
| RPS6KA3 | None | None | None | None |
| RPS6KA3 | None | None | None | None |
| RPS6KA3 | None | None | None | None |
| GBA | 1 | 15 | 3gxd | B |

| | | | | | |
|---|---|---|---|---|---|
| GBA | | 1 | 55 | 3gxd | B |
| GBA | | 1 | 15 | 3gxd | B |
| GBA | | 0.75 | 45 | 3ril | A |
| GBA | None | None | | None | None |
| GBA | None | None | | None | None |
| GBA | | 1 | 12 | 1ogs | A |
| GBA | | 0.75 | 122 | 3gxi | A |
| GBA | | 1 | 2 | 1ogs | A |
| GBA | | 1 | 11 | 1ogs | A |
| GBA | | 0.5 | 59 | 3gxf | B |
| GBA | | 0.75 | 45 | 3ril | A |
| GBA | | 1 | 18 | 1ogs | A |
| GBA | | 1 | 3 | 1ogs | A |
| GBA | None | None | | None | None |
| GBA | | 1 | 2 | 1ogs | A |
| GBA | None | None | | None | None |
| GBA | | 0.75 | 76 | 3gxm | C |
| GBA | | 1 | 3 | 1ogs | A |
| GBA | | 1 | 3 | 1ogs | A |
| GBA | | 0.75 | 62 | 3ril | A |
| GBA | | 1 | 28 | 1ogs | A |
| GBA | None | None | | None | None |
| GBA | | 0.5 | 78 | 3gxi | A |
| GBA | | 0.5 | 92 | 3rik | B |
| GBA | | 0.5 | 42 | 2xwe | A |
| GBA | | 0.333 | 117 | 1ogs | A |
| GBA | | 0.5 | 92 | 3rik | B |
| GBA | | 0.75 | 49 | 3ril | A |
| GBA | None | None | | None | None |
| GBA | | 0.667 | 60 | 1ogs | A |
| GBA | None | None | | None | None |
| GBA | | 0.5 | 11 | 2nt1 | A |
| GBA | | 0.75 | 21 | 3gxi | A |
| GBA | None | None | | None | None |
| GBA | | 0.5 | 35 | 2xwe | A |
| GBA | | 0.75 | 25 | 3keh | B |
| GBA | | 0.75 | 70 | 3ril | A |
| GBA | | 0.75 | 70 | 3ril | A |
| GBA | | 1 | 15 | 3gxd | B |
| GBA | | 1 | 44 | 3gxd | B |
| GBA | | 0.5 | 57 | 3gxf | D |
| GBA | | 1 | 44 | 3gxd | B |
| GBA | | 1 | 44 | 3gxd | B |
| GBA | None | None | | None | None |
| GBA | | 0.75 | 94 | 3gxi | A |
| GBA | | 0.75 | 94 | 3gxi | A |
| IFIH1 | None | None | | None | None |
| TGFBR2 | None | None | | None | None |
| TGFBR2 | None | None | | None | None |
| TGFBR2 | None | None | | None | None |
| TGFBR2 | None | None | | None | None |
| TGFBR2 | None | None | | None | None |
| TGFBR2 | None | None | | None | None |
| TGFBR2 | None | None | | None | None |

| | | | | |
|---|---|---|---|---|
| TGFBR2 | None | None | None | None |
| TGFBR2 | None | None | None | None |
| TGFBR2 | 0.742 | 1 | 5E+091 | A |
| TGFBR2 | None | None | None | None |
| TGFBR2 | None | None | None | None |
| TGFBR2 | None | None | None | None |
| TGFBR2 | 0.676 | 1 | 5e8v | A |
| TGFBR2 | 0.676 | 1 | 5e8v | A |
| ACE | 0.684 | 17 | 3nxq | A |
| ACE | 0.474 | 25 | 4bxk | A |
| ACE | None | None | None | None |
| ACE | None | None | None | None |
| ACE | None | None | None | None |
| ACE | None | None | None | None |
| COL10A1 | None | None | None | None |
| COL10A1 | None | None | None | None |
| COL10A1 | None | None | None | None |
| COL10A1 | 0.409 | 1 | 1gr3 | A |
| COL10A1 | None | None | None | None |
| COL10A1 | 0.318 | 4 | 1gr3 | A |
| COL10A1 | None | None | None | None |
| COL10A1 | None | None | None | None |
| COL10A1 | 0.318 | 4 | 1gr3 | A |
| KLF1 | None | None | None | None |
| C3 | None | None | None | None |
| CTSC | None | None | None | None |
| CTSC | 0.934 | 1 | 2djg | C |
| CTSC | None | None | None | None |
| LYZ | 0.877 | 3 | 1bb5 | A |
| LYZ | 0.923 | 1 | 1lz4 | A |
| LYZ | 0.852 | 2 | 1ubz | A |
| DHODH | None | None | None | None |
| DHODH | None | None | None | None |
| DHODH | None | None | None | None |
| DHODH | None | None | None | None |
| DHODH | None | None | None | None |
| DHODH | None | None | None | None |
| DHODH | None | None | None | None |
| ADH1C | 0.527 | 11 | 1ht0 | A |
| GDNF | None | None | None | None |
| TNF | 0.5 | 8 | 2e7a | B |
| ALDH4A1 | 0.636 | 3 | 3v9h | A |
| ALDH4A1 | None | None | None | None |
| ALDH4A1 | None | None | None | None |
| BRAF | None | None | None | None |
| BRAF | None | None | None | None |
| BRAF | 0.752 | 2 | 3psb | A |
| BRAF | 0.752 | 2 | 3psb | A |
| BRAF | 0.737 | 1 | 3pri | A |
| BRAF | 0.737 | 1 | 3pri | A |
| BRAF | 0.737 | 1 | 3pri | A |
| BRAF | 0.737 | 1 | 3prf | A |
| BRAF | 0.737 | 1 | 3prf | A |
| BRAF | 0.752 | 2 | 3psb | A |

| | | | | |
|---|---|---|---|---|
| BRAF | 0.752 | 2 | 3psb | A |
| BRAF | 0.752 | 2 | 3psb | A |
| BRAF | 0.752 | 2 | 3psb | A |
| BRAF | 0.555 | 3 | 5vam | A |
| BRAF | 0.663 | 1 | 4ehe | B |
| BRAF | 0.734 | 1 | 5csx | A |
| TBX5 | None | None | None | None |
| TBX5 | None | None | None | None |
| TBX5 | None | None | None | None |
| TBX5 | None | None | None | None |
| TBX5 | None | None | None | None |
| TBX5 | None | None | None | None |
| TBX5 | None | None | None | None |
| TBX5 | None | None | None | None |
| TBX5 | None | None | None | None |
| RB1 | None | None | None | None |
| RB1 | None | None | None | None |
| RB1 | None | None | None | None |
| RB1 | None | None | None | None |
| RB1 | None | None | None | None |
| RB1 | None | None | None | None |
| CLIC2 | 0.5 | 3 | 2per | A |
| CLIC2 | None | None | None | None |
| ALDH7A1 | None | None | None | None |
| ALDH7A1 | None | None | None | None |
| ALDH7A1 | 0.405 | 4 | 2j6l | D |
| ALDH7A1 | 0.301 | 14 | 4zuk | A |
| ALDH7A1 | 0.409 | 3 | 4zvw | C |
| ALDH7A1 | None | None | None | None |
| ALDH7A1 | 0.534 | 3 | 2j6l | A |
| ALDH7A1 | None | None | None | None |
| ALDH7A1 | 0.333 | 6 | 4x0t | A |
| ALDH7A1 | None | None | None | None |
| ALDH7A1 | None | None | None | None |
| ALDH7A1 | 0.534 | 3 | 2j6l | A |
| ALDH7A1 | None | None | None | None |
| ALDH7A1 | None | None | None | None |
| ALDH7A1 | 0.632 | 1 | 4zuk | A |
| ALDH7A1 | None | None | None | None |
| KDR | 0.575 | 5 | 1y6a | A |
| MUT | None | None | None | None |
| MUT | None | None | None | None |
| MUT | None | None | None | None |
| MUT | None | None | None | None |
| MUT | None | None | None | None |
| MUT | None | None | None | None |
| MUT | None | None | None | None |
| MUT | None | None | None | None |
| MUT | None | None | None | None |
| MUT | None | None | None | None |
| MUT | None | None | None | None |
| MUT | None | None | None | None |
| MUT | None | None | None | None |
| MUT | None | None | None | None |

| | | | | |
|---|---|---|---|---|
| MUT | None | None | None | None |
| MUT | None | None | None | None |
| MUT | 0.6 | 1 | 3bic | A |
| MUT | 0.6 | 2 | 2xiq | A |
| MUT | 0.6 | 2 | 2xiq | A |
| MUT | None | None | None | None |
| MUT | None | None | None | None |
| MUT | 0.4 | 1 | 2xiq | A |
| MUT | None | None | None | None |
| MUT | None | None | None | None |
| MUT | None | None | None | None |
| MUT | None | None | None | None |
| MUT | None | None | None | None |
| MUT | None | None | None | None |
| MUT | None | None | None | None |
| MUT | None | None | None | None |
| MUT | None | None | None | None |
| MUT | None | None | None | None |
| MUT | None | None | None | None |
| MUT | None | None | None | None |
| SOD1 | 0.778 | 1 | 2gbu | A |
| SOD1 | 0.727 | 4 | 4xcr | A |
| SOD1 | 0.35 | 26 | 1n19 | A |
| SOD1 | 0.621 | 12 | 1hl5 | H |
| NECTIN3 | None | None | None | None |
| HFE | None | None | None | None |
| HFE | None | None | None | None |
| HFE | None | None | None | None |
| HFE | 0.5 | 1 | 1de4 | A |
| HFE | None | None | None | None |
| HFE | None | None | None | None |
| HFE | None | None | None | None |
| EIF2AK3 | None | None | None | None |
| EIF2AK3 | None | None | None | None |
| TERT | None | None | None | None |
| TERT | None | None | None | None |
| SPR | None | None | None | None |
| SPR | None | None | None | None |
| SPR | None | None | None | None |
| SPR | None | None | None | None |
| SPR | None | None | None | None |
| F7 | 0.625 | 2 | 1klj | L |
| UNG | None | None | None | None |
| MMP13 | 0.356 | 22 | 2ozr | E |
| MMP13 | 0.786 | 1 | 3tvc | A |
| MMP13 | 0.766 | 1 | 3zxh | B |
| MMP13 | 0.766 | 1 | 3zxh | B |
| CD207 | 0.621 | 1 | 3kqg | B |
| F2 | 1 | 2 | 1nrq | L |
| SPTBN2 | None | None | None | None |
| PKLR | 0.568 | 9 | 2vgb | A |
| PKLR | 0.486 | 17 | 2vgg | D |
| PKLR | None | None | None | None |
| PKLR | 0.486 | 8 | 2vgg | D |

| | | | | |
|---|---|---|---|---|
| PKLR | 0.568 | 3 | 4ip7 | D |
| PKLR | 0.432 | 13 | 2vgi | A |
| PKLR | 0.378 | 45 | 4ip7 | B |
| PKLR | 0.324 | 25 | 2vgf | B |
| SACS | None | None | None | None |
| HRAS | 0.529 | 3 | 6q21 | A |
| HRAS | 0.529 | 3 | 6q21 | A |
| HRAS | 0.59 | 1 | 1gnq | A |
| HRAS | 0.59 | 1 | 1gnq | A |
| HRAS | 0.664 | 1 | 2lwi | A |
| HRAS | 0.742 | 1 | 1ctq | A |
| HRAS | 0.69 | 1 | 1xd2 | A |
| HRAS | 0.415 | 3 | 6q21 | D |
| HRAS | 0.529 | 3 | 6q21 | A |
| HRAS | 0.403 | 3 | 2n46 | A |
| HRAS | 0.521 | 4 | 4nyj | R |
| HRAS | 0.664 | 1 | 2lwi | A |
| HRAS | 0.664 | 1 | 2lwi | A |
| VHL | None | None | None | None |
| VHL | None | None | None | None |
| VHL | None | None | None | None |
| VHL | None | None | None | None |
| VHL | 0.5 | 22 | 4w9l | C |
| VHL | 0.5 | 22 | 5nvy | C |
| VHL | None | None | None | None |
| VHL | None | None | None | None |
| VHL | None | None | None | None |
| VHL | 0.5 | 2 | 5nvy | C |
| VHL | 0.5 | 1 | 5nvy | C |
| VHL | 0.5 | 1 | 5nvy | C |
| VHL | 0.5 | 4 | 5nvy | C |
| VHL | None | None | None | None |
| VHL | 0.5 | 6 | 5nvy | C |
| VHL | 0.5 | 2 | 5nvy | C |
| VHL | 0.5 | 8 | 3ztc | C |
| VHL | 0.5 | 2 | 5nvy | C |
| VHL | 0.5 | 2 | 5nvy | C |
| VHL | None | None | None | None |
| VHL | 0.5 | 7 | 4w9h | C |
| VHL | 0.5 | 2 | 5nvy | C |
| VHL | 0.5 | 2 | 5nvy | C |
| VHL | 0.5 | 19 | 4w9l | C |
| VHL | None | None | None | None |
| TTBK1 | None | None | None | None |
| RAD51 | None | None | None | None |
| ALB | None | None | None | None |
| ALB | 0.5 | 2 | 4n0u | D |
| ALB | None | None | None | None |
| ALB | None | None | None | None |
| MET | None | None | None | None |
| MET | None | None | None | None |
| MET | None | None | None | None |
| MET | None | None | None | None |
| MET | None | None | None | None |

| | | | | |
|---|---|---|---|---|
| MET | None | None | None | None |
| MET | None | None | None | None |
| MET | None | None | None | None |
| MET | None | None | None | None |
| CP | None | None | None | None |
| CP | None | None | None | None |
| PGM1 | None | None | None | None |
| PGM1 | 0.409 | 2 | 5f9c | A |
| PGM1 | 0.409 | 2 | 5f9c | A |
| PGM1 | None | None | None | None |
| PGM1 | None | None | None | None |
| PGM1 | None | None | None | None |
| PGM1 | None | None | None | None |
| NR5A1 | None | None | None | None |
| NR5A1 | None | None | None | None |
| NR5A1 | None | None | None | None |
| F9 | None | None | None | None |
| F9 | None | None | None | None |
| HPRT1 | None | None | None | None |
| HPRT1 | None | None | None | None |
| HPRT1 | None | None | None | None |
| HPRT1 | 0.375 | 15 | 3gep | B |
| HPRT1 | None | None | None | None |
| HPRT1 | 0.871 | 3 | 4rac | B |
| HPRT1 | 0.806 | 2 | 4ran | D |
| HPRT1 | None | None | None | None |
| HPRT1 | None | None | None | None |
| HPRT1 | 0.44 | 3 | 1d6n | B |
| HPRT1 | None | None | None | None |
| HPRT1 | None | None | None | None |
| HPRT1 | None | None | None | None |
| HPRT1 | 0.703 | 4 | 5w8v | A |
| HPRT1 | None | None | None | None |
| HPRT1 | 0.321 | 8 | 4kn6 | A |
| HPRT1 | 0.312 | 27 | 3ggj | B |
| HPRT1 | 0.312 | 12 | 3gep | B |
| HPRT1 | 0.871 | 3 | 4rac | B |
| HPRT1 | 0.871 | 3 | 4rac | B |
| HPRT1 | 0.765 | 2 | 4rao | D |
| HPRT1 | 0.774 | 2 | 4rac | B |
| GPD1L | None | None | None | None |
| GPD1L | None | None | None | None |
| GPD1L | None | None | None | None |
| FGFR2 | None | None | None | None |
| FGFR2 | None | None | None | None |
| FGFR2 | None | None | None | None |
| FGFR2 | None | None | None | None |
| FGFR2 | None | None | None | None |
| FGFR2 | None | None | None | None |
| FGFR2 | None | None | None | None |
| FGFR2 | None | None | None | None |
| FGFR2 | None | None | None | None |
| FGFR2 | None | None | None | None |
| FGFR2 | None | None | None | None |

| | | | | | |
|---|---|---|---|---|---|
| FGFR2 | None | None | None | | None |
| FGFR2 | None | None | None | | None |
| FGFR2 | None | None | None | | None |
| FGFR2 | None | None | None | | None |
| FGFR2 | None | None | None | | None |
| FGFR2 | None | None | None | | None |
| FGFR2 | None | None | None | | None |
| FGFR2 | None | None | None | | None |
| HSPD1 | None | None | None | | None |
| HSPD1 | None | None | None | | None |
| FGF8 | None | None | None | | None |
| FGF8 | None | None | None | | None |
| FGF8 | None | None | None | | None |
| F9 | 0.438 | 1 | 5vyg | | B |
| NCF4 | None | None | None | | None |
| NCF4 | None | None | None | | None |
| LDLR | None | None | None | | None |
| LDLR | None | None | None | | None |
| LDLR | 0.478 | 1 | 3gcx | | E |
| LDLR | 0.478 | 1 | 3gcx | | E |
| LDLR | 0.478 | 1 | 3gcx | | E |
| LDLR | None | None | None | | None |
| C3 | None | None | None | | None |
| C3 | None | None | None | | None |
| C3 | None | None | None | | None |
| C3 | None | None | None | | None |
| C3 | None | None | None | | None |
| C3 | None | None | None | | None |
| C3 | None | None | None | | None |
| C3 | None | None | None | | None |
| TULP1 | None | None | None | | None |
| TULP1 | None | None | None | | None |
| TULP1 | None | None | None | | None |
| TULP1 | None | None | None | | None |
| IKBKG | None | None | None | | None |
| IKBKG | None | None | None | | None |
| IKBKG | None | None | None | | None |
| CD40 | None | None | None | | None |
| CD40 | None | None | None | | None |
| SLC9A3R1 | None | None | None | | None |
| BCHE | None | None | None | | None |
| BCHE | 0.439 | 5 | 2wil | | B |
| BCHE | 0.544 | 11 | 1p0q | | A |
| BCHE | None | None | None | | None |
| BCHE | None | None | None | | None |
| BCHE | None | None | None | | None |
| BCHE | 0.63 | 4 | 2xqg | | A |
| BCHE | None | None | None | | None |
| BCHE | None | None | None | | None |
| BCHE | None | None | None | | None |
| BCHE | None | None | None | | None |
| BCKDHB | 0.833 | 1 | 1v11 | | B |
| BCKDHB | 0.5 | 18 | 2bfc | | B |
| BCKDHB | None | None | None | | None |

| | | | | |
|---|---|---|---|---|
| BCKDHB | 0.667 | 11 | 2bfd | B |
| BCKDHB | 0.833 | 10 | 1olx | B |
| BCKDHB | 0.833 | 1 | 1v11 | B |
| BCKDHB | 0.833 | 1 | 1v11 | B |
| BCKDHB | 0.625 | 2 | 1x7z | B |
| BCKDHB | 0.429 | 16 | 1ols | B |
| BCKDHB | 0.333 | 99 | 1olx | B |
| BCKDHB | 0.333 | 64 | 1x7y | B |
| BCKDHB | 0.857 | 1 | 2bff | B |
| CDH1 | None | None | None | None |
| CDH1 | 0.575 | 1 | 3ff8 | A |
| CDH1 | 0.543 | 1 | 3ff7 | A |
| CDH1 | None | None | None | None |
| CDH1 | None | None | None | None |
| CTNNB1 | None | None | None | None |
| AIRE | None | None | None | None |
| EDA | 1 | 1 | 1rj8 | A |
| EDA | None | None | None | None |
| EDA | None | None | None | None |
| EDA | None | None | None | None |
| EDA | None | None | None | None |
| SH2D1A | 0.444 | 5 | 1ka7 | A |
| SH2D1A | None | None | None | None |
| SH2D1A | None | None | None | None |
| GCK | 0.556 | 37 | 3idh | A |
| GCK | 0.333 | 32 | 4isf | A |
| GCK | None | None | None | None |
| GCK | None | None | None | None |
| GCK | None | None | None | None |
| GCK | 0.611 | 13 | 3h1v | X |
| GCK | None | None | None | None |
| GCK | None | None | None | None |
| GCK | None | None | None | None |
| GCK | 0.778 | 5 | 3id8 | A |
| GCK | 0.833 | 1 | 4dhy | A |
| GCK | 0.389 | 6 | 4lc9 | B |
| GCK | None | None | None | None |
| GCK | 0.611 | 6 | 3id8 | A |
| GCK | None | None | None | None |
| GCK | None | None | None | None |
| GCK | None | None | None | None |
| GCK | 0.5 | 5 | 3vey | A |
| GCK | None | None | None | None |
| GCK | None | None | None | None |
| AQP2 | None | None | None | None |
| AQP2 | None | None | None | None |
| AQP2 | None | None | None | None |
| AQP2 | None | None | None | None |
| AQP2 | None | None | None | None |
| AQP2 | None | None | None | None |
| AQP2 | None | None | None | None |
| AQP2 | None | None | None | None |
| AQP2 | None | None | None | None |
| AQP2 | None | None | None | None |

| | | | | |
|---|---|---|---|---|
| AQP2 | None | None | None | None |
| AQP2 | None | None | None | None |
| AQP2 | None | None | None | None |
| SUMF1 | 0.75 | 6 | 1z70 | X |
| SUMF1 | 0.556 | 30 | 1y1i | X |
| SUMF1 | 0.5 | 38 | 2aij | X |
| SUMF1 | 0.625 | 18 | 2aij | X |
| SUMF1 | 0.875 | 35 | 2afy | X |
| SUMF1 | 0.667 | 19 | 1y1i | X |
| SUMF1 | 0.75 | 17 | 1y1j | X |
| SUMF1 | 0.75 | 17 | 1y1j | X |
| SUMF1 | 0.625 | 27 | 1y1g | X |
| FAM20A | None | None | None | None |
| FAM20A | None | None | None | None |
| FAM20A | None | None | None | None |
| FAM20A | None | None | None | None |
| NT5E | None | None | None | None |
| NT5E | None | None | None | None |
| NOS3 | 0.564 | 54 | 1m9k | B |
| ACTG1 | None | None | None | None |
| ACTG1 | None | None | None | None |
| ACTG1 | None | None | None | None |
| ACTG1 | None | None | None | None |
| ACTG1 | 0.571 | 8 | 5jlh | E |
| ACTG1 | None | None | None | None |
| FCGR1A | None | None | None | None |
| GPI | 0.304 | 64 | 1iri | A |
| GPI | None | None | None | None |
| GPI | 1 | 3 | 1jiq | A |
| GPI | None | None | None | None |
| GPI | 0.696 | 22 | 1iat | A |
| GPI | 0.522 | 22 | 1nuh | A |
| GPI | 0.696 | 27 | 1iat | A |
| GPI | 0.304 | 87 | 1jlh | B |
| GPI | 0.391 | 141 | 1jlh | B |
| CFH | None | None | None | None |
| OGG1 | 0.895 | 13 | 1m3h | A |
| TPK1 | None | None | None | None |
| TPK1 | None | None | None | None |
| TPK1 | None | None | None | None |
| TPK1 | None | None | None | None |
| SEC23A | None | None | None | None |
| SEC23A | None | None | None | None |
| TYRP1 | None | None | None | None |
| TYRP1 | None | None | None | None |
| TYRP1 | None | None | None | None |
| TYRP1 | None | None | None | None |
| TYRP1 | 0.3 | 5 | 5m8m | A |
| TYRP1 | None | None | None | None |
| CSTB | None | None | None | None |
| CSTB | None | None | None | None |
| CSTB | None | None | None | None |
| ACP1 | None | None | None | None |
| IL17F | None | None | None | None |

| | | | | |
|---|---|---|---|---|
| IL17F | None | None | None | None |
| IL17F | None | None | None | None |
| IL17F | None | None | None | None |
| IL17F | None | None | None | None |
| IDH2 | 0.5 | 2 | 5svn | B |
| IDH2 | 0.5 | 2 | 5svn | B |
| IDH2 | 0.48 | 13 | 5i96 | A |
| IDH2 | None | None | None | None |
| TP53 | 0.925 | 11 | 2ocj | A |
| TP53 | 0.8 | 16 | 2ahi | D |
| TP53 | 0.65 | 49 | 4ibq | A |
| TP53 | 0.725 | 27 | 5g4o | A |
| TP53 | 0.455 | 21 | 3q06 | A |
| TP53 | 0.9 | 1 | 4agp | A |
| TP53 | 0.9 | 1 | 4agp | A |
| TP53 | 0.925 | 10 | 4hje | A |
| TP53 | 0.7 | 29 | 3d05 | A |
| TP53 | 0.875 | 7 | 3kmd | B |
| TP53 | 0.875 | 7 | 3kmd | B |
| TP53 | 0.925 | 6 | 3kz8 | B |
| TP53 | 0.925 | 6 | 3kz8 | B |
| TP53 | 0.609 | 22 | 3q01 | B |
| TP53 | 0.925 | 5 | 3kmd | B |
| TP53 | 0.925 | 5 | 3kmd | B |
| TP53 | 0.925 | 8 | 3kz8 | B |
| TP53 | 0.925 | 8 | 3kz8 | B |
| TP53 | 0.925 | 1 | 3kmd | B |
| TP53 | 0.9 | 11 | 2pcx | A |
| TP53 | 0.9 | 1 | 5aol | A |
| TP53 | 0.85 | 5 | 5lgy | A |
| TP53 | 0.9 | 1 | 3q06 | B |
| TP53 | 0.725 | 19 | 5lgy | A |
| TP53 | 0.775 | 14 | 4mzi | A |
| TP53 | 0.825 | 8 | 1kzy | A |
| TP53 | 0.825 | 8 | 1kzy | A |
| TP53 | 0.8 | 11 | 4ibt | A |
| TP53 | 0.8 | 11 | 4ibt | A |
| TP53 | 0.725 | 15 | 5lgy | A |
| TP53 | 0.725 | 29 | 2h1l | O |
| TP53 | 0.875 | 7 | 3kmd | B |
| TP53 | 0.725 | 15 | 5lgy | A |
| TP53 | 0.8 | 10 | 4agp | A |
| TP53 | 0.8 | 10 | 4agp | A |
| TP53 | 0.8 | 10 | 4agp | A |
| TP53 | 0.8 | 10 | 4agp | A |
| TP53 | 0.675 | 35 | 4ibq | A |
| TP53 | 0.575 | 37 | 4agm | A |
| TP53 | 0.675 | 17 | 3kz8 | B |
| TP53 | 0.356 | 63 | 4mzr | C |
| TP53 | None | None | None | None |
| LMAN1 | None | None | None | None |
| CTSK | 0.75 | 2 | 4n79 | A |
| CTSK | None | None | None | None |
| CTSK | 0.413 | 12 | 4x6j | A |

| | | | | |
|---|---|---|---|---|
| FGB | None | None | None | None |
| FGB | None | None | None | None |
| FGB | 0.688 | 1 | 1fzf | E |
| FGB | None | None | None | None |
| ACADM | None | None | None | None |
| ACADM | None | None | None | None |
| ACADM | None | None | None | None |
| ACADM | 0.471 | 3 | 4p13 | D |
| ACADM | None | None | None | None |
| ACADM | None | None | None | None |
| ACADM | 0.389 | 6 | 1t9g | D |
| ACADM | None | None | None | None |
| ACADM | 0.327 | 12 | 4p13 | C |
| ACADM | 0.389 | 6 | 1t9g | D |
| ACADM | None | None | None | None |
| ACADM | None | None | None | None |
| ACADM | None | None | None | None |
| ACADM | None | None | None | None |
| ACADM | None | None | None | None |
| ACADM | 0.481 | 2 | 4p13 | C |
| ACADM | 0.389 | 6 | 1t9g | D |
| ACADM | None | None | None | None |
| ACADM | None | None | None | None |
| ACADM | None | None | None | None |
| ACADM | 0.564 | 1 | 1t9g | C |
| ACADM | None | None | None | None |
| ACADM | None | None | None | None |
| ACADM | None | None | None | None |
| ACADM | 0.364 | 6 | 2a1t | A |
| ACADM | 0.312 | 12 | 4p13 | A |
| ACADM | 0.347 | 7 | 4p13 | B |
| PTS | None | None | None | None |
| PTS | 0.417 | 2 | 3i2b | E |
| PTS | None | None | None | None |
| PTS | 0.778 | 1 | 3i2b | D |
| PTS | None | None | None | None |
| PTS | None | None | None | None |
| PTS | None | None | None | None |
| PTS | None | None | None | None |
| PTS | None | None | None | None |
| INS | 0.75 | 2 | 3zqr | E |
| INS | 1 | 5 | 2l1z | A |
| INS | 1 | 5 | 2l1z | A |
| INS | 1 | 1 | 1his | A |
| FBP1 | 0.71 | 26 | 2y5l | C |
| FBP1 | None | None | None | None |
| FBP1 | 0.5 | 18 | 1fta | A |
| FBP1 | 0.7 | 10 | 2y5k | A |
| FBP1 | 0.516 | 38 | 3kc1 | B |
| FBP1 | 0.733 | 10 | 3kc1 | A |
| FBP1 | 0.633 | 43 | 2jjk | A |
| FBP1 | 0.8 | 11 | 2y5k | A |
| FBP1 | 0.548 | 41 | 2vt5 | D |
| FBP1 | 0.438 | 25 | 2fie | A |

| | | | | | |
|---|---|---|---|---|---|
| FBP1 | | 0.643 | 7 | 4mjo | B |
| KCNQ1 | None | None | | None | None |
| KCNQ1 | None | None | | None | None |
| KCNQ1 | None | None | | None | None |
| UPP1 | | 0.706 | 4 | 3euf | A |
| UMPS | | 1 | 3 | 2qcl | A |
| UMPS | | 0.556 | 59 | 2qcl | B |
| UMPS | None | None | | None | None |
| DHFR | None | None | | None | None |
| KHK | | 0.667 | 2 | 2hw1 | A |
| TTPA | None | None | | None | None |
| TTPA | None | None | | None | None |
| TTPA | | 0.5 | 1 | 1r5l | A |
| TTPA | None | None | | None | None |
| MAP1LC3B | | 0.387 | 8 | 3vtv | A |
| MAP1LC3B | None | None | | None | None |
| ZAP70 | | 0.317 | 1 | 2oq1 | A |
| PROC | | 0.536 | 1 | 3f6u | L |
| SQSTM1 | None | None | | None | None |
| AHCY | None | None | | None | None |
| AHCY | None | None | | None | None |
| PRNP | None | None | | None | None |
| PRNP | None | None | | None | None |
| PRNP | None | None | | None | None |
| PRNP | None | None | | None | None |
| PRNP | None | None | | None | None |
| PRNP | None | None | | None | None |
| PRNP | None | None | | None | None |
| PRNP | None | None | | None | None |
| PRNP | None | None | | None | None |
| PRNP | None | None | | None | None |
| PRNP | None | None | | None | None |
| PRNP | None | None | | None | None |
| PRNP | None | None | | None | None |
| PRNP | None | None | | None | None |
| PRNP | None | None | | None | None |
| PRNP | None | None | | None | None |
| REN | | 0.431 | 2 | 2v13 | A |
| REN | | 0.431 | 2 | 2v13 | A |
| HINT1 | | 0.556 | 2 | 5ed6 | B |
| HINT1 | | 0.897 | 1 | 5km4 | A |
| HINT1 | None | None | | None | None |
| HINT1 | | 0.667 | 11 | 5ed6 | B |
| HINT1 | | 0.923 | 1 | 5ipc | A |
| OAT | None | None | | None | None |
| OAT | None | None | | None | None |
| OAT | | 0.381 | 5 | 2can | B |
| OAT | | 0.381 | 5 | 2can | B |
| OAT | | 0.381 | 7 | 2byj | B |
| OAT | None | None | | None | None |
| OAT | None | None | | None | None |
| OAT | None | None | | None | None |
| OAT | None | None | | None | None |
| OAT | None | None | | None | None |

| | | | | |
|---|---|---|---|---|
| OAT | 0.675 | 2 | 2byl | B |
| OAT | None | None | None | None |
| OAT | None | None | None | None |
| OAT | None | None | None | None |
| OAT | None | None | None | None |
| OAT | 0.758 | 1 | 2byl | A |
| OAT | 0.375 | 5 | 2oat | C |
| OAT | 0.525 | 4 | 2byl | B |
| OAT | 0.429 | 6 | 2byj | B |
| OAT | None | None | None | None |
| OAT | None | None | None | None |
| OAT | None | None | None | None |
| OAT | None | None | None | None |
| OAT | None | None | None | None |
| OAT | None | None | None | None |
| OAT | None | None | None | None |
| OAT | None | None | None | None |
| OAT | None | None | None | None |
| CTSA | None | None | None | None |
| CTSA | None | None | None | None |
| PNP | 0.375 | 24 | 1ulb | A |
| PNP | None | None | None | None |
| PNP | 0.406 | 43 | 3phb | E |
| PNP | 0.706 | 5 | 4ear | C |
| PNP | 0.367 | 28 | 3ggs | C |
| PNP | 0.5 | 17 | 1pf7 | E |
| PNP | 0.933 | 1 | 3ggs | C |
| PNP | 0.833 | 1 | 3d1v | A |
| MYL2 | None | None | None | None |
| MYL2 | None | None | None | None |
| ANG | 0.548 | 4 | 1k5b | A |
| ANG | 0.419 | 8 | 4ahi | A |
| ANG | 0.419 | 8 | 4ahi | A |
| ANG | None | None | None | None |
| ANG | 0.761 | 1 | 5eop | A |
| ANG | 0.576 | 4 | 5m9v | A |
| ANG | 0.579 | 4 | 5m9q | A |
| ANG | None | None | None | None |
| ANG | None | None | None | None |
| ANG | 0.52 | 5 | 5m9g | A |
| POLH | None | None | None | None |
| POLH | None | None | None | None |
| POLH | None | None | None | None |
| IL2RA | None | None | None | None |
| TGFB1 | None | None | None | None |
| TP53 | None | None | None | None |
| TP53 | None | None | None | None |
| PAH | None | None | None | None |
| PAH | None | None | None | None |
| PAH | 0.44 | 11 | 2pah | A |
| PAH | 0.44 | 11 | 2pah | A |
| PAH | 0.32 | 19 | 6pah | A |
| PAH | 0.32 | 19 | 6pah | A |
| PAH | 0.48 | 2 | 1pah | A |

| | | | | |
|---|---|---|---|---|
| PAH | 0.48 | 2 | 1pah | A |
| PAH | 0.4 | 11 | 3pah | A |
| PAH | None | None | None | None |
| PAH | None | None | None | None |
| PAH | 0.36 | 7 | 2pah | B |
| PAH | 0.44 | 9 | 1dmw | A |
| PAH | 0.48 | 1 | 1dmw | A |
| PAH | 0.48 | 1 | 1dmw | A |
| PAH | 0.44 | 8 | 2pah | B |
| PAH | 0.44 | 8 | 2pah | B |
| PAH | 0.417 | 3 | 1j8t | A |
| PAH | 0.48 | 2 | 4anp | A |
| PAH | 0.48 | 1 | 1dmw | A |
| PAH | 0.48 | 4 | 2pah | B |
| PAH | 0.44 | 11 | 2pah | A |
| PAH | 0.48 | 4 | 1mmt | A |
| PAH | 0.68 | 6 | 2pah | A |
| PAH | 0.76 | 1 | 1tdw | A |
| PAH | None | None | None | None |
| PAH | None | None | None | None |
| PAH | 0.4 | 14 | 1j8u | A |
| PAH | None | None | None | None |
| PAH | None | None | None | None |
| PAH | None | None | None | None |
| PAH | None | None | None | None |
| PAH | 0.64 | 2 | 1lrm | A |
| PAH | 0.4 | 11 | 2pah | B |
| PAH | None | None | None | None |
| PAH | 0.4 | 14 | 1j8u | A |
| PAH | 0.4 | 14 | 1j8u | A |
| PAH | None | None | None | None |
| PAH | 0.48 | 1 | 1dmw | A |
| PAH | 0.48 | 1 | 1dmw | A |
| PAH | None | None | None | None |
| HDAC8 | None | None | None | None |
| HDAC8 | 0.732 | 3 | 3rqd | A |
| VCP | 0.438 | 4 | 3tiw | A |
| VCP | 0.5 | 2 | 3qq8 | A |
| VCP | 0.5 | 2 | 3qq8 | A |
| VCP | None | None | None | None |
| VCP | None | None | None | None |
| VCP | None | None | None | None |
| VCP | None | None | None | None |
| B3GAT3 | None | None | None | None |
| B3GAT3 | None | None | None | None |
| CSF2RB | None | None | None | None |
| CSF2RB | None | None | None | None |
| CSF2RB | None | None | None | None |
| NF2 | 0.5 | 6 | 1h4r | B |
| NF2 | None | None | None | None |
| NF2 | None | None | None | None |
| PIK3CA | None | None | None | None |
| PHGDH | None | None | None | None |
| PHGDH | None | None | None | None |

| | | | | |
|---|---|---|---|---|
| PHGDH | None | None | None | None |
| PHGDH | 0.66 | 1 | 5n53 | A |
| POLH | None | None | None | None |
| XIAP | 0.5 | 7 | 5c7b | A |
| XIAP | None | None | None | None |
| PDHX | None | None | None | None |
| PDHX | None | None | None | None |
| PDHX | None | None | None | None |
| PDHX | None | None | None | None |
| PDHX | None | None | None | None |
| SAMHD1 | 0.5 | 12 | 5ao0 | A |
| SAMHD1 | 0.357 | 51 | 4zwe | D |
| SAMHD1 | 0.429 | 7 | 4tnr | D |
| SAMHD1 | None | None | None | None |
| SAMHD1 | 0.571 | 3 | 4qfy | D |
| SAMHD1 | None | None | None | None |
| SAMHD1 | 0.308 | 18 | 4qg2 | B |
| SAMHD1 | None | None | None | None |
| CD36 | None | None | None | None |
| CD36 | None | None | None | None |
| CD36 | None | None | None | None |
| ASPH | None | None | None | None |
| CCT5 | None | None | None | None |
| CCT5 | None | None | None | None |
| CCT5 | None | None | None | None |
| SMN1 | None | None | None | None |
| SMN1 | None | None | None | None |
| SMN1 | None | None | None | None |
| SMN1 | None | None | None | None |
| ABCA1 | None | None | None | None |
| ABCA1 | None | None | None | None |
| ABCA1 | None | None | None | None |
| ABCA1 | None | None | None | None |
| ABCA1 | None | None | None | None |
| ABCA1 | None | None | None | None |
| ABCA1 | None | None | None | None |
| ABCA1 | None | None | None | None |
| ABCA1 | None | None | None | None |
| ABCA1 | None | None | None | None |
| ABCA1 | None | None | None | None |
| ABCA1 | None | None | None | None |
| ABCA1 | None | None | None | None |
| ABCA1 | None | None | None | None |
| ABCA1 | None | None | None | None |
| ABCA1 | None | None | None | None |
| ABCA1 | None | None | None | None |
| ABCA1 | None | None | None | None |
| ABCA1 | None | None | None | None |
| ABCA1 | None | None | None | None |
| ABCA1 | None | None | None | None |
| ABCA1 | None | None | None | None |
| ABCA1 | None | None | None | None |
| ABCA1 | None | None | None | None |

| | | | | |
|---|---|---|---|---|
| ABCA1 | None | None | None | None |
| ABCA1 | None | None | None | None |
| ABCA1 | None | None | None | None |
| ABCA1 | None | None | None | None |
| ABCA1 | None | None | None | None |
| ABCA1 | None | None | None | None |
| ABCA1 | None | None | None | None |
| ABCA1 | None | None | None | None |
| ABCA1 | None | None | None | None |
| ABCA1 | None | None | None | None |
| ABCA1 | None | None | None | None |
| ERCC1 | None | None | None | None |
| FECH | 0.524 | 27 | 2po5 | A |
| FECH | 0.6 | 60 | 2hre | A |
| FECH | 0.636 | 20 | 2po7 | A |
| FECH | 0.476 | 61 | 4klr | A |
| FECH | 0.5 | 84 | 2hrc | A |
| FECH | 0.667 | 10 | 2po5 | A |
| FECH | 0.455 | 30 | 4kla | B |
| PTEN | None | None | None | None |
| PTEN | None | None | None | None |
| PTEN | None | None | None | None |
| PTEN | None | None | None | None |
| PTEN | None | None | None | None |
| PTEN | None | None | None | None |
| PTEN | None | None | None | None |
| PTEN | None | None | None | None |
| PTEN | None | None | None | None |
| PTEN | None | None | None | None |
| PTEN | None | None | None | None |
| PTEN | None | None | None | None |
| PTEN | None | None | None | None |
| PTEN | None | None | None | None |
| PTEN | None | None | None | None |
| PTEN | None | None | None | None |
| PTEN | None | None | None | None |
| PTEN | None | None | None | None |
| PTEN | None | None | None | None |
| PTEN | None | None | None | None |
| ACAT1 | 0.667 | 1 | 2ib9 | C |
| ACAT1 | None | None | None | None |
| ACAT1 | None | None | None | None |
| ACAT1 | 0.667 | 1 | 2ib9 | C |
| ACAT1 | 0.667 | 1 | 2ib9 | C |
| ACAT1 | 0.472 | 11 | 2f2s | A |
| ACAT1 | 0.833 | 6 | 2ib7 | C |
| ACAT1 | None | None | None | None |
| ACAT1 | None | None | None | None |
| ACAT1 | 0.528 | 9 | 2ibw | D |
| ACAT1 | 0.703 | 16 | 2ibu | A |
| CREBBP | None | None | None | None |
| ITGA2B | None | None | None | None |
| ITGA2B | None | None | None | None |
| ITGA2B | None | None | None | None |

| | | | | |
|---|---|---|---|---|
| ITGA2B | 0.538 | 1 | 3nig | A |
| TSC1 | None | None | None | None |
| ADA2 | None | None | None | None |
| ADA2 | None | None | None | None |
| ADA2 | 0.75 | 1 | 3lgd | A |
| ADA2 | None | None | None | None |
| ADA2 | None | None | None | None |
| ADA2 | None | None | None | None |
| ADA2 | None | None | None | None |
| ADA2 | None | None | None | None |
| ADA2 | 1 | 1 | 3lgd | B |
| ADA2 | None | None | None | None |
| PGK1 | None | None | None | None |
| PGK1 | 0.536 | 8 | 2xe8 | A |
| PGK1 | None | None | None | None |
| PGK1 | None | None | None | None |
| PGK1 | 0.571 | 3 | 2ybe | A |
| TUBA1A | None | None | None | None |
| TUBA1A | None | None | None | None |
| TUBA1A | None | None | None | None |
| TUBA1A | None | None | None | None |
| TUBA1A | None | None | None | None |
| PTPN11 | None | None | None | None |
| PTPN11 | None | None | None | None |
| PTPN11 | 0.362 | 13 | 4ohl | A |
| PTPN11 | 0.327 | 15 | 4ohi | A |
| PTPN11 | None | None | None | None |
| PTPN11 | None | None | None | None |
| PTPN11 | None | None | None | None |
| PTPN11 | None | None | None | None |
| PTPN11 | None | None | None | None |
| PTPN11 | 0.404 | 5 | 3tl0 | A |
| PTPN11 | 0.404 | 5 | 3tl0 | A |
| PTPN11 | 0.404 | 5 | 3tl0 | A |
| PTPN11 | None | None | None | None |
| PTPN11 | 0.466 | 7 | 3zm1 | A |
| PTPN11 | 0.621 | 2 | 3b7o | A |
| PTPN11 | 0.776 | 1 | 3zm1 | A |
| PTPN11 | 0.453 | 5 | 4ohd | A |
| PTPN11 | None | None | None | None |
| PTPN11 | None | None | None | None |
| PTPN11 | None | None | None | None |
| PTPN11 | None | None | None | None |
| PTPN11 | 0.776 | 1 | 3zm1 | A |
| PTPN11 | 0.621 | 6 | 3zm1 | A |
| PTPN11 | None | None | None | None |
| PTPN11 | None | None | None | None |
| PTPN11 | 0.621 | 6 | 3zm1 | A |
| GLB1 | 0.333 | 34 | 3wez | A |
| GLB1 | 0.333 | 67 | 3wf4 | A |
| UQCRQ | None | None | None | None |
| VPS35 | None | None | None | None |
| NR3C1 | None | None | None | None |
| NR3C1 | None | None | None | None |

| | | | | |
|---|---|---|---|---|
| NR3C1 | None | None | None | None |
| NR3C1 | 0.359 | 21 | 1nhz | A |
| NR3C1 | 0.42 | 6 | 4mdd | A |
| NR3C1 | None | None | None | None |
| NR3C1 | None | None | None | None |
| NR3C1 | None | None | None | None |
| NR3C1 | 0.422 | 14 | 1nhz | A |
| APOA1 | None | None | None | None |
| APOA1 | None | None | None | None |
| APOA1 | None | None | None | None |
| APOA1 | None | None | None | None |
| APOA1 | None | None | None | None |
| APOA1 | None | None | None | None |
| APOA1 | None | None | None | None |
| BCKDHA | None | None | None | None |
| BCKDHA | 0.583 | 76 | 1x7x | A |
| BCKDHA | 0.963 | 1 | 1ols | A |
| BCKDHA | 0.696 | 3 | 1v1r | A |
| BCKDHA | 0.625 | 16 | 2beu | A |
| BCKDHA | None | None | None | None |
| BCKDHA | 0.652 | 11 | 1v1r | A |
| BCKDHA | 1 | 1 | 2beu | A |
| BCKDHA | 0.84 | 2 | 2j9f | A |
| BCKDHA | 0.556 | 42 | 1ols | A |
| BCKDHA | 0.609 | 90 | 1v1r | A |
| BCKDHA | 0.652 | 25 | 1v1r | A |
| BCKDHA | 0.346 | 5 | 1wci | A |
| BCKDHA | 0.625 | 60 | 2beu | A |
| BCKDHA | 0.625 | 28 | 2beu | A |
| TOP1 | None | None | None | None |
| ANTXR2 | None | None | None | None |
| ANTXR2 | None | None | None | None |
| ANTXR2 | None | None | None | None |
| TF | 0.625 | 35 | 5dyh | A |
| TF | 0.328 | 52 | 5dyh | A |
| FSHB | None | None | None | None |
| GALK1 | None | None | None | None |
| GALK1 | None | None | None | None |
| GALK1 | None | None | None | None |
| CYP17A1 | None | None | None | None |
| CYP17A1 | 0.353 | 2 | 4nkx | A |
| CYP17A1 | 0.353 | 2 | 4nkx | A |
| CYP17A1 | None | None | None | None |
| CYP17A1 | None | None | None | None |
| CYP17A1 | None | None | None | None |
| CYP17A1 | None | None | None | None |
| CYP17A1 | None | None | None | None |
| CYP17A1 | None | None | None | None |
| CYP17A1 | None | None | None | None |
| CYP17A1 | None | None | None | None |
| CYP17A1 | None | None | None | None |
| CYP17A1 | None | None | None | None |
| CYP17A1 | None | None | None | None |
| CYP17A1 | None | None | None | None |

| | | | | |
|---|---|---|---|---|
| CYP17A1 | None | None | None | None |
| CYP17A1 | None | None | None | None |
| CD44 | 0.667 | 24 | 1uuh | A |
| MDH2 | 0.426 | 5 | 4wlo | B |
| MDH2 | None | None | None | None |
| NDUFV1 | None | None | None | None |
| NDUFV1 | None | None | None | None |
| NDUFV1 | None | None | None | None |
| NDUFV1 | None | None | None | None |
| NDUFV1 | None | None | None | None |
| NDUFV1 | None | None | None | None |
| NDUFV1 | None | None | None | None |
| NDUFV1 | None | None | None | None |
| NDUFV1 | None | None | None | None |
| NDUFV1 | None | None | None | None |
| CTSC | None | None | None | None |
| CTSC | None | None | None | None |
| CTSC | None | None | None | None |
| CTSC | 0.342 | 13 | 3pdf | A |
| CTSC | 0.643 | 1 | 4oel | A |
| CTSC | None | None | None | None |
| CTSC | None | None | None | None |
| VWF | None | None | None | None |
| VWF | None | None | None | None |
| VWF | None | None | None | None |
| VWF | None | None | None | None |
| VWF | None | None | None | None |
| VWF | None | None | None | None |
| VWF | None | None | None | None |
| ORC4 | None | None | None | None |
| ORC4 | None | None | None | None |
| CRYAB | None | None | None | None |
| CRYAB | None | None | None | None |
| CRYAB | None | None | None | None |
| MAP2K2 | None | None | None | None |
| MAP2K2 | None | None | None | None |
| MAP2K2 | None | None | None | None |
| MAP2K2 | None | None | None | None |
| MAP2K2 | None | None | None | None |
| MAP2K2 | None | None | None | None |
| MAP2K2 | None | None | None | None |
| MAP2K2 | None | None | None | None |
| MAP2K2 | None | None | None | None |
| CBL | None | None | None | None |
| CBL | None | None | None | None |
| CBL | None | None | None | None |
| CBL | None | None | None | None |
| CBL | None | None | None | None |
| CBL | None | None | None | None |
| CBL | None | None | None | None |
| PFN1 | None | None | None | None |
| PFN1 | 0.75 | 2 | 1fik | A |
| ACSM2A | None | None | None | None |
| HSPG2 | None | None | None | None |

| | | | | |
|---|---|---|---|---|
| HSPG2 | None | None | None | None |
| HSPG2 | None | None | None | None |
| HSPG2 | None | None | None | None |
| PARK7 | 0.444 | 48 | 3cyf | A |
| PARK7 | 0.769 | 4 | 1pe0 | A |
| PARK7 | 0.808 | 2 | 1pe0 | A |
| PARK7 | 0.667 | 8 | 4rky | A |
| PARK7 | 0.889 | 1 | 3cz9 | A |
| PARK7 | 0.75 | 2 | 1pdw | B |
| INSR | None | None | None | None |
| INSR | None | None | None | None |
| INSR | None | None | None | None |
| INSR | None | None | None | None |
| INSR | None | None | None | None |
| DSC2 | None | None | None | None |
| DSC2 | None | None | None | None |
| DSC2 | None | None | None | None |
| DSC2 | None | None | None | None |
| DSC2 | None | None | None | None |
| DSC2 | None | None | None | None |
| DSC2 | None | None | None | None |
| DSC2 | None | None | None | None |
| DSC2 | None | None | None | None |
| DSC2 | None | None | None | None |
| MICU1 | None | None | None | None |
| MICU1 | None | None | None | None |
| ADAMTS13 | None | None | None | None |
| ADAMTS13 | None | None | None | None |
| ADAMTS13 | None | None | None | None |
| ADAMTS13 | None | None | None | None |
| ADAMTS13 | None | None | None | None |
| ADAMTS13 | None | None | None | None |
| ADAMTS13 | None | None | None | None |
| IRAK4 | None | None | None | None |
| IRAK4 | None | None | None | None |
| RNASEH2B | None | None | None | None |
| RNASEH2B | None | None | None | None |
| RNASEH2B | None | None | None | None |
| NTRK1 | None | None | None | None |
| MPO | 0.75 | 3 | 1d5l | C |
| MPO | 0.6 | 25 | 1myp | D |
| MPO | None | None | None | None |
| PEX3 | None | None | None | None |
| SERPINA1 | None | None | None | None |
| SERPINA1 | None | None | None | None |
| SERPINA1 | None | None | None | None |
| SERPINA1 | None | None | None | None |
| SERPINA1 | None | None | None | None |
| SERPINA1 | None | None | None | None |
| SERPINA1 | None | None | None | None |
| SERPINA1 | 0.6 | 1 | 1ezx | A |
| SERPINA1 | None | None | None | None |
| SERPINA1 | None | None | None | None |
| YARS2 | None | None | None | None |

| | | | | |
|---|---|---|---|---|
| YARS2 | None | None | None | None |
| YARS2 | None | None | None | None |
| YARS2 | None | None | None | None |
| YARS2 | None | None | None | None |
| YARS2 | 0.781 | 1 | 2pid | B |
| YARS2 | None | None | None | None |
| YARS2 | None | None | None | None |
| YARS2 | None | None | None | None |
| PDHA1 | None | None | None | None |
| PDHA1 | None | None | None | None |
| PDHA1 | None | None | None | None |
| PDHA1 | None | None | None | None |
| PDHA1 | None | None | None | None |
| PDHA1 | None | None | None | None |
| PDHA1 | None | None | None | None |
| PDHA1 | 0.96 | 1 | 3exh | C |
| PDHA1 | 0.3 | 5 | 1ni4 | A |
| PDHA1 | None | None | None | None |
| PDHA1 | None | None | None | None |
| CUL4B | None | None | None | None |
| AGXT | None | None | None | None |
| AGXT | None | None | None | None |
| AGXT | None | None | None | None |
| AGXT | 0.692 | 3 | 2yob | A |
| AGXT | 0.786 | 7 | 4kxk | C |
| AGXT | 0.538 | 21 | 3r9a | A |
| AGXT | 0.353 | 33 | 3r9a | C |
| AGXT | None | None | None | None |
| AGXT | None | None | None | None |
| AGXT | 0.611 | 4 | 5hhy | B |
| AGXT | 0.333 | 40 | 4cbs | A |
| AGXT | 0.333 | 59 | 5f9s | A |
| AGXT | 0.308 | 45 | 3r9a | A |
| AGXT | None | None | None | None |
| AGXT | 0.429 | 28 | 4kxk | C |
| AGXT | 0.684 | 1 | 1h0c | A |
| AGXT | 0.5 | 2 | 4cbr | A |
| AGXT | 0.538 | 19 | 4kyo | A |
| AGXT | 0.538 | 19 | 4kyo | A |
| AGXT | 0.75 | 2 | 5luc | B |
| AGXT | 0.722 | 3 | 5hhy | B |
| AGXT | 0.571 | 3 | 4kxk | C |
| AGXT | 0.571 | 5 | 4kxk | C |
| AGXT | None | None | None | None |
| AGXT | 0.778 | 1 | 5hhy | B |
| AGXT | 0.778 | 1 | 5hhy | B |
| AGXT | 0.421 | 12 | 1h0c | A |
| AGXT | 0.692 | 7 | 4kyo | A |
| AGXT | 0.5 | 18 | 4cbr | A |
| AGXT | 0.5 | 18 | 4cbr | A |
| AGXT | 0.615 | 13 | 3r9a | A |
| AGXT | 0.786 | 7 | 4kxk | C |
| AGXT | 0.846 | 1 | 4kxk | A |
| AGXT | 0.615 | 36 | 3r9a | A |

| | | | | |
|---|---|---|---|---|
| AGXT | 0.571 | 10 | 4kxk | C |
| SFTPD | 0.509 | 2 | 3g83 | B |
| SFTPD | None | None | None | None |
| PIK3CA | None | None | None | None |
| PIK3CA | None | None | None | None |
| PIK3CA | None | None | None | None |
| AUH | None | None | None | None |
| AUH | None | None | None | None |
| AUH | None | None | None | None |
| AUH | None | None | None | None |
| F11 | None | None | None | None |
| F11 | None | None | None | None |
| F11 | None | None | None | None |
| F11 | None | None | None | None |
| MEN1 | 0.6 | 81 | 5db0 | A |
| MEN1 | 0.8 | 54 | 4i80 | A |
| MEN1 | 0.8 | 84 | 5db0 | A |
| MEN1 | 0.8 | 30 | 5ddb | A |
| MEN1 | 1 | 2 | 4i80 | A |
| MEN1 | 0.8 | 17 | 5db0 | A |
| MEN1 | 0.6 | 17 | 5ddb | A |
| MEN1 | 0.6 | 31 | 3u88 | A |
| MEN1 | None | None | None | None |
| MEN1 | 0.6 | 56 | 5ddb | A |
| MYBPC1 | None | None | None | None |
| MYBPC1 | None | None | None | None |
| MYBPC1 | None | None | None | None |
| PDCD10 | None | None | None | None |
| SCARB2 | None | None | None | None |
| SCARB2 | None | None | None | None |
| SCARB2 | None | None | None | None |
| SCARB2 | None | None | None | None |
| SCARB2 | None | None | None | None |
| TREX2 | None | None | None | None |
| TBK1 | None | None | None | None |
| TBK1 | None | None | None | None |
| DLD | None | None | None | None |
| DLD | None | None | None | None |
| DLD | 0.505 | 3 | 1zy8 | B |
| DLD | None | None | None | None |
| DLD | None | None | None | None |
| DLD | None | None | None | None |
| LDHA | 0.802 | 1 | 4qt0 | A |
| TNNC1 | None | None | None | None |
| TNNC1 | 0.461 | 2 | 1wrl | A |
| TNNC1 | None | None | None | None |
| SMAD3 | None | None | None | None |
| HARS | None | None | None | None |
| HARS | None | None | None | None |
| TNNI3 | None | None | None | None |
| TNNI3 | None | None | None | None |
| TNNI3 | None | None | None | None |
| TNNI3 | None | None | None | None |
| TNNI3 | None | None | None | None |

| | | | | |
|---|---|---|---|---|
| TNNI3 | None | None | None | None |
| ACKR1 | None | None | None | None |
| BUB1B | None | None | None | None |
| BUB1B | None | None | None | None |
| BUB1B | None | None | None | None |
| AR | 0.522 | 6 | 1xj7 | A |
| AR | None | None | None | None |
| AR | 0.305 | 55 | 4olm | A |
| AR | 0.305 | 55 | 4olm | A |
| AR | None | None | None | None |
| FTO | None | None | None | None |
| FTO | 0.333 | 1 | 4ie6 | A |
| TFAM | None | None | None | None |
| PYGL | None | None | None | None |
| PYGL | 0.778 | 3 | 1fa9 | A |
| PYGL | None | None | None | None |
| PYGL | 0.722 | 2 | 1fa9 | A |
| PYGL | 0.5 | 19 | 3dds | A |
| PYGL | 0.611 | 25 | 3ceh | B |
| PYGL | 0.611 | 32 | 1l7x | B |
| PYGL | None | None | None | None |
| PYGL | 0.722 | 2 | 1fc0 | B |
| PYGL | 0.778 | 3 | 1fa9 | A |
| PYGL | None | None | None | None |
| PYGL | 0.722 | 12 | 2ati | A |
| PYGL | 0.722 | 1 | 3ceh | B |
| PYGL | 0.611 | 11 | 3ceh | B |
| PYGL | 0.611 | 18 | 1fc0 | B |
| PYGL | 0.722 | 2 | 1fc0 | B |
| GALNS | None | None | None | None |
| GALNS | None | None | None | None |
| GALNS | None | None | None | None |
| GALNS | None | None | None | None |
| GALNS | None | None | None | None |
| GALNS | None | None | None | None |
| GALNS | None | None | None | None |
| GALNS | None | None | None | None |
| GALNS | None | None | None | None |
| GALNS | None | None | None | None |
| GALNS | None | None | None | None |
| GALNS | None | None | None | None |
| GALNS | None | None | None | None |
| GALNS | None | None | None | None |
| GALNS | None | None | None | None |
| GALNS | None | None | None | None |
| GALNS | None | None | None | None |
| GALNS | None | None | None | None |
| GALNS | None | None | None | None |
| GALNS | None | None | None | None |
| GALNS | None | None | None | None |
| GALNS | None | None | None | None |
| GALNS | None | None | None | None |
| GALNS | None | None | None | None |
| GALNS | None | None | None | None |

| | | | | | |
|---|---|---|---|---|---|
| GALNS | None | None | | None | None |
| GALNS | None | None | | None | None |
| GALNS | None | None | | None | None |
| GALNS | None | None | | None | None |
| GALNS | None | None | | None | None |
| GALNS | None | None | | None | None |
| GALNS | None | None | | None | None |
| GALNS | None | None | | None | None |
| F10 | 0.304 | | 21 | 1z6e | L |
| GLUD1 | None | None | | None | None |
| GLUD1 | None | None | | None | None |
| GLUD1 | None | None | | None | None |
| GLUD1 | None | None | | None | None |
| GLUD1 | None | None | | None | None |
| GLUD1 | None | None | | None | None |
| GLUD1 | None | None | | None | None |
| GLUD1 | None | None | | None | None |
| GLUD1 | None | None | | None | None |
| GLUD1 | None | None | | None | None |
| NOTCH1 | 0.528 | | 3 | 3v79 | K |
| POGLUT1 | None | None | | None | None |
| POGLUT1 | None | None | | None | None |
| NR3C1 | 0.455 | | 1 | 4hn5 | B |
| CASP8 | 0.535 | | 6 | 1f9e | C |
| CASP8 | 0.592 | | 8 | 1qdu | C |
| TRAP1 | None | None | | None | None |
| FANCM | None | None | | None | None |
| EPOR | None | None | | None | None |
| EPOR | None | None | | None | None |
| NDP | None | None | | None | None |
| NDP | None | None | | None | None |
| NDP | None | None | | None | None |
| NDP | None | None | | None | None |
| NDP | None | None | | None | None |
| NDP | None | None | | None | None |
| NDP | None | None | | None | None |
| NDP | None | None | | None | None |
| NDP | None | None | | None | None |
| NDP | None | None | | None | None |
| NDP | None | None | | None | None |
| NDP | None | None | | None | None |
| NDP | None | None | | None | None |
| NDP | None | None | | None | None |
| NDP | None | None | | None | None |
| NDP | None | None | | None | None |
| NDP | None | None | | None | None |
| ROGDI | None | None | | None | None |
| CYP19A1 | None | None | | None | None |
| CYP19A1 | None | None | | None | None |
| CYP19A1 | None | None | | None | None |
| CYP19A1 | None | None | | None | None |
| CYP19A1 | None | None | | None | None |
| CYP19A1 | 0.487 | | 1 | 3eqm | A |
| CYP19A1 | 0.487 | | 1 | 3eqm | A |

| | | | | |
|---|---|---|---|---|
| CYP19A1 | None | None | None | None |
| UBA5 | None | None | None | None |
| PRPS1 | None | None | None | None |
| PRPS1 | 0.316 | 13 | 4f8e | A |
| PRPS1 | None | None | None | None |
| PRPS1 | None | None | None | None |
| PRPS1 | None | None | None | None |
| PRPS1 | None | None | None | None |
| PRPS1 | None | None | None | None |
| PRPS1 | None | None | None | None |
| F2 | None | None | None | None |
| F2 | None | None | None | None |
| NTF4 | None | None | None | None |
| NTF4 | None | None | None | None |
| PSAP | None | None | None | None |
| PSAP | None | None | None | None |
| TAP1 | None | None | None | None |
| TAP1 | None | None | None | None |
| TAP1 | None | None | None | None |
| ARG1 | None | None | None | None |
| ARG1 | 0.714 | 1 | 2pha | A |
| ARG1 | None | None | None | None |
| ARG1 | 0.471 | 10 | 3th7 | A |
| ARG1 | None | None | None | None |
| ARG1 | None | None | None | None |
| CRYM | None | None | None | None |
| ADSL | None | None | None | None |
| ADSL | None | None | None | None |
| ADSL | None | None | None | None |
| ADSL | None | None | None | None |
| ADSL | None | None | None | None |
| ADSL | None | None | None | None |
| ADSL | 0.312 | 8 | 2vd6 | B |
| ADSL | 0.312 | 8 | 2vd6 | B |
| ADSL | None | None | None | None |
| ADSL | 0.353 | 5 | 2vd6 | D |
| ADSL | None | None | None | None |
| ADSL | None | None | None | None |
| ADSL | 0.647 | 1 | 2vd6 | D |
| ADSL | 0.435 | 1 | 4ffx | A |
| ADSL | None | None | None | None |
| ADSL | None | None | None | None |
| ADSL | None | None | None | None |
| ADSL | None | None | None | None |
| ADSL | None | None | None | None |
| ADSL | None | None | None | None |
| VWF | None | None | None | None |
| PLA2G7 | None | None | None | None |
| MMP13 | None | None | None | None |
| MMP13 | None | None | None | None |
| MMP13 | None | None | None | None |
| MMP13 | None | None | None | None |
| POMGNT1 | None | None | None | None |
| POMGNT1 | None | None | None | None |

| | | | | |
|---|---|---|---|---|
| COL3A1 | None | None | None | None |
| COL3A1 | None | None | None | None |
| COL3A1 | None | None | None | None |
| COL3A1 | None | None | None | None |
| ACTN2 | None | None | None | None |
| ACTN2 | None | None | None | None |
| ACTN2 | None | None | None | None |
| ACTN2 | None | None | None | None |
| ACTN2 | None | None | None | None |
| ACTN2 | None | None | None | None |
| ACTN2 | None | None | None | None |
| UBA5 | None | None | None | None |
| ADAR | 0.75 | 1 | 1qbj | B |
| UMPS | None | None | None | None |
| UMPS | None | None | None | None |
| UMPS | None | None | None | None |
| TRNT1 | None | None | None | None |
| TRNT1 | None | None | None | None |
| NHEJ1 | None | None | None | None |
| NHEJ1 | None | None | None | None |
| NHEJ1 | None | None | None | None |
| C8A | None | None | None | None |
| GLUL | None | None | None | None |
| GLUL | 0.375 | 6 | 2qc8 | C |
| GLUL | 0.467 | 9 | 2qc8 | B |
| GYG1 | 0.615 | 45 | 3rmv | A |
| GYG1 | 0.923 | 1 | 3u2u | B |
| FLNA | None | None | None | None |
| FLNA | None | None | None | None |
| FLNA | None | None | None | None |
| FLNA | None | None | None | None |
| FLNA | None | None | None | None |
| FLNA | None | None | None | None |
| FLNA | None | None | None | None |
| TTN | None | None | None | None |
| TTN | None | None | None | None |
| TTN | None | None | None | None |
| TTN | None | None | None | None |
| SMS | None | None | None | None |
| SMS | None | None | None | None |
| SMS | None | None | None | None |
| C5 | None | None | None | None |
| SERPINC1 | None | None | None | None |
| SERPINC1 | None | None | None | None |
| SERPINC1 | None | None | None | None |
| SERPINC1 | None | None | None | None |
| SERPINC1 | None | None | None | None |
| SERPINC1 | None | None | None | None |
| SERPINC1 | None | None | None | None |
| SERPINC1 | None | None | None | None |
| SERPINC1 | None | None | None | None |
| SERPINC1 | None | None | None | None |
| SERPINC1 | None | None | None | None |
| SERPINC1 | None | None | None | None |

| | | | | |
|---|---|---|---|---|
| SERPINC1 | None | None | None | None |
| SERPINC1 | None | None | None | None |
| SERPINC1 | None | None | None | None |
| SERPINC1 | None | None | None | None |
| SERPINC1 | None | None | None | None |
| SERPINC1 | None | None | None | None |
| SERPINC1 | None | None | None | None |
| SERPINC1 | None | None | None | None |
| SERPINC1 | None | None | None | None |
| ETFB | None | None | None | None |
| ETFB | None | None | None | None |
| ETFB | None | None | None | None |
| ETFB | None | None | None | None |
| ETFB | None | None | None | None |
| ETFB | None | None | None | None |
| DIABLO | None | None | None | None |
| DIABLO | None | None | None | None |
| DPYS | 0.333 | 24 | 2vr2 | A |
| DPYS | None | None | None | None |
| DPYS | None | None | None | None |
| DPYS | 0.533 | 7 | 2vr2 | A |
| DPYS | None | None | None | None |
| GNE | None | None | None | None |
| GNE | None | None | None | None |
| GNE | None | None | None | None |
| GNE | None | None | None | None |
| GNE | None | None | None | None |
| GNE | None | None | None | None |
| GNE | None | None | None | None |
| GNE | None | None | None | None |
| GNE | None | None | None | None |
| GNMT | None | None | None | None |
| GNMT | None | None | None | None |
| CTH | 0.324 | 20 | 3cog | D |
| CTH | None | None | None | None |
| CTH | None | None | None | None |
| USH1C | None | None | None | None |
| PSEN1 | None | None | None | None |
| CHEK2 | None | None | None | None |
| CHEK2 | None | None | None | None |
| CHEK2 | None | None | None | None |
| CHEK2 | None | None | None | None |
| CHEK2 | None | None | None | None |
| CHEK2 | None | None | None | None |
| CHEK2 | None | None | None | None |
| CHEK2 | None | None | None | None |
| CHEK2 | None | None | None | None |
| SNCA | None | None | None | None |
| SNCA | None | None | None | None |
| GAN | None | None | None | None |
| GAN | None | None | None | None |
| GAN | None | None | None | None |
| SBDS | None | None | None | None |
| SBDS | None | None | None | None |

| | | | | |
|---|---|---|---|---|
| SBDS | None | None | None | None |
| SBDS | None | None | None | None |
| SBDS | None | None | None | None |
| SBDS | None | None | None | None |
| MPO | | 1 | 3 3f9p | A |
| MPO | | 0.6 | 12 5mfa | A |
| KCNH2 | None | None | None | None |
| KCNH2 | None | None | None | None |
| KCNH2 | | 0.75 | 1 4hp9 | A |
| KCNH2 | None | None | None | None |
| TEK | None | None | None | None |
| TEK | None | None | None | None |
| TEK | None | None | None | None |
| TEK | None | None | None | None |
| NPC2 | None | None | None | None |
| NPC2 | None | None | None | None |
| NPC2 | None | None | None | None |
| NPC2 | None | None | None | None |
| NPC2 | None | None | None | None |
| NPC2 | None | None | None | None |
| NPC2 | None | None | None | None |
| PLG | None | None | None | None |
| PCSK9 | None | None | None | None |
| SFTPC | None | None | None | None |
| SFTPC | None | None | None | None |
| SFTPC | None | None | None | None |
| CD40LG | None | None | None | None |
| CD40LG | None | None | None | None |
| CD40LG | None | None | None | None |
| CD40LG | None | None | None | None |
| CD40LG | None | None | None | None |
| TAT | None | None | None | None |
| TAT | None | None | None | None |
| TAT | None | None | None | None |
| EP300 | None | None | None | None |
| FARS2 | None | None | None | None |
| FARS2 | None | None | None | None |
| FARS2 | None | None | None | None |
| FARS2 | None | None | None | None |
| FARS2 | None | None | None | None |
| FARS2 | None | None | None | None |
| FARS2 | None | None | None | None |
| FARS2 | None | None | None | None |
| FARS2 | | 0.5 | 1 5mgw | A |
| FARS2 | None | None | None | None |
| FARS2 | None | None | None | None |
| FARS2 | None | None | None | None |
| FARS2 | None | None | None | None |
| MASP2 | None | None | None | None |
| MASP2 | None | None | None | None |
| MASP2 | None | None | None | None |
| MASP2 | None | None | None | None |
| STIM1 | None | None | None | None |
| CHRM2 | None | None | None | None |

| | | | | |
|---|---|---|---|---|
| MCFD2 | None | None | None | None |
| MCFD2 | 0.574 | 1 | 4yge | B |
| MCFD2 | 0.623 | 1 | 3wnx | B |
| KITLG | None | None | None | None |
| F12 | 0.345 | 1 | 4bdx | A |
| F12 | 0.345 | 1 | 4bdx | A |
| ERAP1 | 0.381 | 5 | 3mdj | A |
| GTF2H5 | None | None | None | None |
| GTF2H5 | None | None | None | None |
| THBD | None | None | None | None |
| MRPL28 | None | None | None | None |
| MRPL28 | None | None | None | None |
| HTRA1 | None | None | None | None |
| HTRA1 | None | None | None | None |
| HTRA1 | None | None | None | None |
| HTRA1 | None | None | None | None |
| HTRA1 | None | None | None | None |
| HTRA1 | None | None | None | None |
| KARS | None | None | None | None |
| KARS | None | None | None | None |
| KARS | None | None | None | None |
| KARS | None | None | None | None |
| KARS | None | None | None | None |
| HK1 | 0.722 | 11 | 4fpb | B |
| GARS | None | None | None | None |
| GARS | None | None | None | None |
| GARS | None | None | None | None |
| GARS | None | None | None | None |
| GARS | None | None | None | None |
| GARS | None | None | None | None |
| GARS | None | None | None | None |
| IL12B | None | None | None | None |
| HADH | None | None | None | None |
| HADH | None | None | None | None |
| HADH | 0.52 | 4 | 1lsj | A |
| HADH | 0.458 | 3 | 1m75 | A |
| HADH | 0.52 | 4 | 1lsj | A |
| RRM2B | None | None | None | None |
| RRM2B | None | None | None | None |
| RRM2B | None | None | None | None |
| RRM2B | None | None | None | None |
| RRM2B | None | None | None | None |
| RRM2B | None | None | None | None |
| RRM2B | 0.333 | 2 | 3hf1 | B |
| RRM2B | None | None | None | None |
| RRM2B | None | None | None | None |
| RRM2B | None | None | None | None |
| RRM2B | None | None | None | None |
| RRM2B | None | None | None | None |
| RRM2B | None | None | None | None |
| RRM2B | None | None | None | None |
| RRM2B | None | None | None | None |
| MARVELD2 | None | None | None | None |
| TLR4 | None | None | None | None |

| | | | | |
|---|---|---|---|---|
| TTN | None | None | None | None |
| TTN | None | None | None | None |
| TNFRSF13B | None | None | None | None |
| POLG | None | None | None | None |
| POLG | None | None | None | None |
| POLG | None | None | None | None |
| POLG | None | None | None | None |
| POLG | None | None | None | None |
| POLG | None | None | None | None |
| POLG | None | None | None | None |
| POLG | None | None | None | None |
| POLG | None | None | None | None |
| POLG | None | None | None | None |
| POLG | None | None | None | None |
| POLG | None | None | None | None |
| POLG | None | None | None | None |
| POLG | None | None | None | None |
| POLG | None | None | None | None |
| POLG | None | None | None | None |
| POLG | None | None | None | None |
| POLG | None | None | None | None |
| POLG | None | None | None | None |
| POLG | None | None | None | None |
| POLG | None | None | None | None |
| POLG | None | None | None | None |
| POLG | None | None | None | None |
| POLG | None | None | None | None |
| POLG | None | None | None | None |
| POLG | None | None | None | None |
| POLG | None | None | None | None |
| POLG | None | None | None | None |
| POLG | 0.467 | 4 | 5c53 | A |
| POLG | 0.533 | 1 | 4ztu | A |
| POLG | None | None | None | None |
| POLG | None | None | None | None |
| POLG | None | None | None | None |
| POLG | None | None | None | None |
| POLG | None | None | None | None |
| POLG | 0.533 | 1 | 4ztu | A |
| POLG | None | None | None | None |
| POLG | None | None | None | None |
| POLG | None | None | None | None |
| POLG | None | None | None | None |
| POLG | None | None | None | None |
| POLG | None | None | None | None |
| POLG | None | None | None | None |
| POLG | None | None | None | None |
| POLG | None | None | None | None |
| POLG | None | None | None | None |
| POLG | None | None | None | None |
| POLG | None | None | None | None |
| POLG | None | None | None | None |
| POLG | None | None | None | None |
| UROD | 0.632 | 24 | 1jpk | A |

| | | | | | |
|---|---|---|---|---|---|
| UROD | | 0.579 | 33 | 1r3v | A |
| UROD | None | None | | None | None |
| UROD | | 0.526 | 49 | 1jph | A |
| UROD | None | None | | None | None |
| UROD | | 0.632 | 13 | 3gvw | A |
| UROD | | 0.789 | 6 | 3gvw | A |
| UROD | | 0.842 | 2 | 3gvv | A |
| UROD | | 0.737 | 31 | 1r3q | A |
| UROD | | 0.667 | 14 | 2q71 | A |
| PCK1 | | 0.429 | 154 | 1khg | A |
| PCK1 | None | None | | None | None |
| PCK1 | None | None | | None | None |
| PCK1 | None | None | | None | None |
| PCK1 | | 0.524 | 60 | 1nhx | A |
| HEXA | None | None | | None | None |
| HEXA | None | None | | None | None |
| HEXA | None | None | | None | None |
| HEXA | None | None | | None | None |
| HEXA | None | None | | None | None |
| HEXA | | 0.364 | 2 | 2gjx | A |
| HEXA | | 0.364 | 2 | 2gjx | A |
| HEXA | None | None | | None | None |
| HEXA | None | None | | None | None |
| HEXA | None | None | | None | None |
| HEXA | None | None | | None | None |
| HEXA | | 0.364 | 2 | 2gjx | A |
| HEXA | None | None | | None | None |
| HEXA | None | None | | None | None |
| HEXA | None | None | | None | None |
| HEXA | | 0.455 | 1 | 2gjx | A |
| HEXA | None | None | | None | None |
| HEXA | None | None | | None | None |
| HEXA | None | None | | None | None |
| HEXA | None | None | | None | None |
| HEXA | None | None | | None | None |
| HEXA | None | None | | None | None |
| HEXA | None | None | | None | None |
| HEXA | None | None | | None | None |
| HEXA | None | None | | None | None |
| HEXA | None | None | | None | None |
| HEXA | None | None | | None | None |
| HEXA | | 0.455 | 3 | 2gjx | A |
| HEXA | | 0.455 | 3 | 2gjx | A |
| HEXA | None | None | | None | None |
| HEXA | None | None | | None | None |
| CFTR | None | None | | None | None |
| CFTR | None | None | | None | None |
| CFTR | None | None | | None | None |
| CFTR | None | None | | None | None |
| CFTR | None | None | | None | None |
| CFTR | None | None | | None | None |
| CFTR | None | None | | None | None |
| CFTR | None | None | | None | None |
| CFTR | None | None | | None | None |

| | | | | |
|---|---|---|---|---|
| CFTR | None | None | None | None |
| CFTR | 0.462 | 1 | 2pzg | A |
| CFTR | 0.321 | 1 | 1xmj | A |
| CFTR | 0.462 | 2 | 2pzg | A |
| CFTR | None | None | None | None |
| CFTR | None | None | None | None |
| CFTR | None | None | None | None |
| CFTR | None | None | None | None |
| CFTR | None | None | None | None |
| CFTR | None | None | None | None |
| CFTR | None | None | None | None |
| CFTR | None | None | None | None |
| CFTR | None | None | None | None |
| CFTR | None | None | None | None |
| CFTR | None | None | None | None |
| CFTR | None | None | None | None |
| CFTR | 0.462 | 1 | 2pzg | A |
| CFTR | None | None | None | None |
| CFTR | 0.462 | 2 | 2pzg | A |
| CFTR | None | None | None | None |
| CFTR | None | None | None | None |
| CFTR | None | None | None | None |
| CFTR | 0.346 | 3 | 2pzg | A |
| CFTR | None | None | None | None |
| CFTR | None | None | None | None |
| FAN1 | None | None | None | None |
| CBS | None | None | None | None |
| CBS | None | None | None | None |
| CBS | None | None | None | None |
| CBS | None | None | None | None |
| CBS | 0.406 | 6 | 4pcu | B |
| CBS | None | None | None | None |
| CBS | 0.469 | 6 | 4pcu | A |
| CBS | 0.597 | 3 | 5mms | C |
| CBS | 0.54 | 2 | 5mms | A |
| CBS | 0.54 | 2 | 5mms | A |
| CBS | 0.328 | 9 | 4pcu | A |
| CBS | None | None | None | None |
| CBS | None | None | None | None |
| CBS | None | None | None | None |
| CBS | 0.516 | 4 | 5mms | C |
| CBS | None | None | None | None |
| CBS | None | None | None | None |
| CBS | None | None | None | None |
| CBS | 0.698 | 2 | 1m54 | B |
| CBS | None | None | None | None |
| CBS | None | None | None | None |
| CBS | None | None | None | None |
| CBS | None | None | None | None |
| CBS | None | None | None | None |
| CBS | None | None | None | None |
| CBS | None | None | None | None |
| CBS | 0.424 | 6 | 4coo | B |
| CBS | None | None | None | None |

| | | | | |
|---|---|---|---|---|
| CBS | 0.603 | 3 | 4coo | A |
| CBS | 0.603 | 3 | 4coo | A |
| CBS | None | None | None | None |
| CBS | None | None | None | None |
| CBS | None | None | None | None |
| CBS | None | None | None | None |
| CBS | None | None | None | None |
| CBS | None | None | None | None |
| TLR3 | None | None | None | None |
| LIG4 | None | None | None | None |
| LIG4 | None | None | None | None |
| ASPA | 0.769 | 1 | 2i3c | B |
| ASPA | 0.769 | 1 | 2i3c | B |
| ASPA | None | None | None | None |
| ASPA | None | None | None | None |
| ASPA | None | None | None | None |
| ASPA | None | None | None | None |
| ASPA | None | None | None | None |
| ASPA | None | None | None | None |
| ASPA | None | None | None | None |
| ASPA | None | None | None | None |
| ASPA | 0.583 | 2 | 4tnu | A |
| ASPA | 0.583 | 2 | 4tnu | A |
| ASPA | None | None | None | None |
| CFH | None | None | None | None |
| FCGR3A | None | None | None | None |
| FCGR3A | None | None | None | None |
| GRIN2A | None | None | None | None |
| GRIN2A | 0.658 | 1 | 5i2n | A |
| GRIN2A | None | None | None | None |
| GLA | 0.467 | 10 | 1r46 | B |
| GLA | 0.375 | 19 | 3hg3 | B |
| GLA | 0.375 | 14 | 3hg5 | B |
| GLA | 0.312 | 13 | 3tv8 | B |
| GLA | 0.375 | 14 | 3hg5 | B |
| GLA | 0.375 | 14 | 3hg5 | B |
| GLA | None | None | None | None |
| GLA | 0.647 | 2 | 3s5y | A |
| GLA | 0.562 | 20 | 3gxt | B |
| GLA | 0.353 | 21 | 3s5y | A |
| GLA | None | None | None | None |
| GLA | 0.75 | 3 | 3gxn | B |
| GLA | 0.75 | 3 | 3gxn | B |
| GLA | 0.438 | 7 | 1r47 | B |
| GLA | 0.375 | 3 | 3hg4 | B |
| GLA | 0.375 | 11 | 3hg2 | A |
| GLA | None | None | None | None |
| GLA | None | None | None | None |
| GLA | None | None | None | None |
| GLA | 0.312 | 13 | 3hg2 | B |
| GLA | 0.375 | 19 | 3hg3 | B |
| GLA | 0.4 | 5 | 3lxa | A |
| GLA | None | None | None | None |
| GLA | None | None | None | None |

| | | | | |
|---|---|---|---|---|
| GLA | None | None | None | None |
| GLA | 0.467 | 22 | 3gxn | A |
| GLA | None | None | None | None |
| GLA | None | None | None | None |
| GLA | None | None | None | None |
| GLA | None | None | None | None |
| GLA | None | None | None | None |
| SERPINA3 | None | None | None | None |
| MTAP | None | None | None | None |
| MTAP | 0.462 | 25 | 3oze | B |
| MTAP | None | None | None | None |
| BLMH | None | None | None | None |
| TUSC3 | None | None | None | None |
| ISCU | None | None | None | None |
| CYP11B2 | None | None | None | None |
| CYP11B2 | None | None | None | None |
| CYP11B2 | None | None | None | None |
| CYP11B2 | None | None | None | None |
| CYP11B2 | None | None | None | None |
| CYP11B2 | None | None | None | None |
| CYP11B2 | None | None | None | None |
| CYP11B2 | None | None | None | None |
| CYP11B2 | None | None | None | None |
| CYP11B2 | None | None | None | None |
| CYP11B2 | None | None | None | None |
| CYP11B2 | None | None | None | None |
| CYP11B2 | None | None | None | None |
| CYP11B2 | None | None | None | None |
| CYP11B2 | None | None | None | None |
| CYP11B2 | None | None | None | None |
| CYP11B2 | None | None | None | None |
| CYP11B2 | None | None | None | None |
| CYP11B2 | None | None | None | None |
| CYP11B2 | None | None | None | None |
| CRADD | None | None | None | None |
| CRADD | None | None | None | None |
| FGA | None | None | None | None |
| FGA | None | None | None | None |
| FGA | None | None | None | None |
| FGA | None | None | None | None |
| FGA | None | None | None | None |
| FGA | None | None | None | None |
| FGA | None | None | None | None |
| FGA | None | None | None | None |
| IYD | 1 | 1 | 4ttb | A |
| IYD | None | None | None | None |
| IYD | None | None | None | None |
| IYD | None | None | None | None |
| TCAP | None | None | None | None |
| TCAP | None | None | None | None |
| TCAP | None | None | None | None |
| TCAP | None | None | None | None |
| TCAP | None | None | None | None |
| CYP7A1 | None | None | None | None |

| | | | | | |
|---|---|---|---|---|---|
| TEK | None | None | | None | None |
| TEK | None | None | | None | None |
| TEK | None | None | | None | None |
| TEK | None | None | | None | None |
| TEK | None | None | | None | None |
| TEK | None | None | | None | None |
| TEK | None | None | | None | None |
| CD8A | None | None | | None | None |
| HERC2 | None | None | | None | None |
| NR1H3 | 0.537 | | 3 | 3ipq | A |
| GRHPR | None | None | | None | None |
| GRHPR | None | None | | None | None |
| GRHPR | 0.458 | | 2 | 2q50 | A |
| GRHPR | 0.458 | | 2 | 2q50 | A |
| GRHPR | 0.697 | | 1 | 2wwr | D |
| GRHPR | 0.697 | | 1 | 2wwr | D |
| GRHPR | None | None | | None | None |
| GBE1 | None | None | | None | None |
| GBE1 | None | None | | None | None |
| GBE1 | None | None | | None | None |
| GBE1 | None | None | | None | None |
| GBE1 | None | None | | None | None |
| GBE1 | None | None | | None | None |
| GBE1 | None | None | | None | None |
| GBE1 | None | None | | None | None |
| GBE1 | None | None | | None | None |
| GBE1 | None | None | | None | None |
| GBE1 | None | None | | None | None |
| GBE1 | None | None | | None | None |
| GBE1 | None | None | | None | None |
| GBE1 | None | None | | None | None |
| GBE1 | None | None | | None | None |
| GBE1 | None | None | | None | None |
| GBE1 | None | None | | None | None |
| GBE1 | None | None | | None | None |
| GBE1 | None | None | | None | None |
| PC | None | None | | None | None |
| PC | 0.429 | | 1 | 3bg3 | A |
| PC | None | None | | None | None |
| PC | 0.4 | | 2 | 3bg9 | B |
| PC | None | None | | None | None |
| PC | None | None | | None | None |
| PC | 0.4 | | 6 | 3bg3 | B |
| PC | None | None | | None | None |
| MSH6 | None | None | | None | None |
| MSH6 | None | None | | None | None |
| MSH6 | None | None | | None | None |
| MSH6 | None | None | | None | None |
| MSH6 | None | None | | None | None |
| MSH6 | None | None | | None | None |
| MSH6 | None | None | | None | None |
| MSH6 | None | None | | None | None |
| MSH6 | None | None | | None | None |

| | | | | | |
|---|---|---|---|---|---|
| MSH6 | None | None | None | | None |
| MSH6 | None | None | None | | None |
| MSH6 | None | None | None | | None |
| MSH6 | None | None | None | | None |
| MSH6 | None | None | None | | None |
| MSH6 | None | None | None | | None |
| MSH6 | None | None | None | | None |
| MSH6 | None | None | None | | None |
| MSH6 | None | None | None | | None |
| MSH6 | None | None | None | | None |
| MSH6 | None | None | None | | None |
| MSH6 | None | None | None | | None |
| MSH6 | None | None | None | | None |
| MSH6 | None | None | None | | None |
| MSH6 | None | None | None | | None |
| MSH6 | None | None | None | | None |
| MSH6 | None | None | None | | None |
| MSH6 | None | None | None | | None |
| MSH6 | None | None | None | | None |
| MSH6 | None | None | None | | None |
| MSH6 | None | None | None | | None |
| RNASEL | 0.32 | | 6 | 4g8l | A |
| ORC1 | None | None | None | | None |
| ORC1 | None | None | None | | None |
| ORC1 | None | None | None | | None |
| KCNH2 | None | None | None | | None |
| KCNH2 | None | None | None | | None |
| KCNH2 | None | None | None | | None |
| KCNH2 | None | None | None | | None |
| KCNH2 | None | None | None | | None |
| KCNH2 | None | None | None | | None |
| KCNH2 | None | None | None | | None |
| KCNH2 | None | None | None | | None |
| KCNH2 | None | None | None | | None |
| KCNH2 | None | None | None | | None |
| KCNH2 | None | None | None | | None |
| KCNH2 | None | None | None | | None |
| KCNH2 | None | None | None | | None |
| KCNH2 | None | None | None | | None |
| KCNH2 | None | None | None | | None |
| KCNH2 | None | None | None | | None |
| KCNH2 | None | None | None | | None |
| KCNH2 | None | None | None | | None |
| KCNH2 | None | None | None | | None |
| KCNH2 | None | None | None | | None |
| KCNH2 | None | None | None | | None |
| KCNH2 | None | None | None | | None |
| KCNH2 | None | None | None | | None |
| KCNH2 | None | None | None | | None |
| MAT1A | 0.429 | | 3 | 2obv | A |
| MAT1A | 0.429 | | 14 | 2obv | A |
| MAT1A | 0.571 | | 1 | 2obv | A |
| MAT1A | 0.571 | | 1 | 2obv | A |
| MAT1A | None | None | None | | None |

| | | | | |
|---|---|---|---|---|
| MAT1A | None | None | None | None |
| MAT1A | None | None | None | None |
| MAT1A | None | None | None | None |
| INSR | None | None | None | None |
| INSR | None | None | None | None |
| INSR | None | None | None | None |
| INSR | None | None | None | None |
| AMT | 0.545 | 5 | 1wsr | A |
| AMT | None | None | None | None |
| AMT | None | None | None | None |
| AMT | None | None | None | None |
| AMT | None | None | None | None |
| AMT | None | None | None | None |
| AMT | None | None | None | None |
| AMT | None | None | None | None |
| AMT | None | None | None | None |
| CSNK1D | None | None | None | None |
| SF3B1 | None | None | None | None |
| AGA | None | None | None | None |
| AGA | None | None | None | None |
| AGA | None | None | None | None |
| AGA | None | None | None | None |
| AGA | None | None | None | None |
| AGA | None | None | None | None |
| AGA | None | None | None | None |
| NECTIN4 | None | None | None | None |
| RS1 | None | None | None | None |
| RS1 | None | None | None | None |
| RS1 | None | None | None | None |
| RS1 | None | None | None | None |
| TBC1D7 | None | None | None | None |
| CTSA | None | None | None | None |
| CTSA | None | None | None | None |
| TKT | None | None | None | None |
| RP2 | None | None | None | None |
| RP2 | None | None | None | None |
| RP2 | None | None | None | None |
| RP2 | None | None | None | None |
| LYRM4 | None | None | None | None |
| SOS1 | None | None | None | None |
| SOS1 | None | None | None | None |
| SOS1 | None | None | None | None |
| SOS1 | None | None | None | None |
| SOS1 | None | None | None | None |
| SOS1 | None | None | None | None |
| HUWE1 | None | None | None | None |
| HUWE1 | None | None | None | None |
| TPI1 | None | None | None | None |
| TPI1 | 0.886 | 1 | 4unk | B |
| TPI1 | None | None | None | None |
| MSH2 | None | None | None | None |
| MSH2 | None | None | None | None |
| MSH2 | None | None | None | None |
| MSH2 | None | None | None | None |

| | | | | |
|---|---|---|---|---|
| MSH2 | None | None | None | None |
| MSH2 | None | None | None | None |
| MSH2 | None | None | None | None |
| MSH2 | None | None | None | None |
| MSH2 | None | None | None | None |
| MSH2 | None | None | None | None |
| MSH2 | None | None | None | None |
| MSH2 | None | None | None | None |
| MSH2 | None | None | None | None |
| MSH2 | None | None | None | None |
| MSH2 | None | None | None | None |
| MSH2 | None | None | None | None |
| MSH2 | None | None | None | None |
| MSH2 | None | None | None | None |
| MSH2 | None | None | None | None |
| MSH2 | None | None | None | None |
| MSH2 | None | None | None | None |
| MSH2 | None | None | None | None |
| MSH2 | None | None | None | None |
| MSH2 | None | None | None | None |
| MSH2 | None | None | None | None |
| MSH2 | None | None | None | None |
| MSH2 | None | None | None | None |
| MSH2 | None | None | None | None |
| MSH2 | None | None | None | None |
| MSH2 | None | None | None | None |
| MSH2 | None | None | None | None |
| MSH2 | None | None | None | None |
| MSH2 | None | None | None | None |
| MSH2 | None | None | None | None |
| MSH2 | None | None | None | None |
| MSH2 | None | None | None | None |
| GALE | None | None | None | None |
| GALE | 0.78 | 1 | 1i3n | A |
| GALE | None | None | None | None |
| GALE | None | None | None | None |
| GALE | None | None | None | None |
| GALE | None | None | None | None |
| GALE | None | None | None | None |
| GALE | None | None | None | None |
| NSD2 | None | None | None | None |
| SRP72 | None | None | None | None |
| SRP72 | None | None | None | None |
| SRP72 | None | None | None | None |
| POLK | None | None | None | None |
| GALT | None | None | None | None |
| GALT | None | None | None | None |
| GALT | None | None | None | None |
| GALT | None | None | None | None |
| GALT | None | None | None | None |
| GALT | None | None | None | None |
| GALT | None | None | None | None |
| GALT | None | None | None | None |
| GALT | None | None | None | None |

| | | | | |
|---|---|---|---|---|
| GALT | None | None | None | None |
| GALT | None | None | None | None |
| GALT | None | None | None | None |
| GALT | None | None | None | None |
| GALT | None | None | None | None |
| GALT | None | None | None | None |
| GALT | None | None | None | None |
| GALT | None | None | None | None |
| GALT | None | None | None | None |
| GALT | None | None | None | None |
| GALT | None | None | None | None |
| GALT | None | None | None | None |
| GALT | None | None | None | None |
| GALT | None | None | None | None |
| GALT | None | None | None | None |
| GALT | None | None | None | None |
| GALT | None | None | None | None |
| GALT | None | None | None | None |
| GALT | None | None | None | None |
| GALT | None | None | None | None |
| GALT | None | None | None | None |
| GALT | None | None | None | None |
| GALT | None | None | None | None |
| GALT | None | None | None | None |
| GALT | None | None | None | None |
| GALT | None | None | None | None |
| GALT | None | None | None | None |
| GALT | None | None | None | None |
| PYCR1 | 0.75 | 11 | 5uav | A |
| PYCR1 | 0.75 | 11 | 5uav | A |
| PYCR1 | 0.5 | 2 | 2izz | A |
| PYCR1 | 0.556 | 2 | 5uax | B |
| F8 | None | None | None | None |
| F8 | None | None | None | None |
| F8 | None | None | None | None |
| F8 | None | None | None | None |
| F8 | None | None | None | None |
| F8 | None | None | None | None |
| F8 | None | None | None | None |
| F8 | None | None | None | None |
| F8 | None | None | None | None |
| F8 | None | None | None | None |
| F8 | None | None | None | None |
| F8 | None | None | None | None |
| F8 | None | None | None | None |
| F8 | None | None | None | None |
| F8 | None | None | None | None |
| F8 | None | None | None | None |
| ANK1 | None | None | None | None |
| CHAT | None | None | None | None |
| CHAT | None | None | None | None |
| CHAT | 0.615 | 8 | 2fy5 | A |
| CHAT | 0.538 | 2 | 2fy5 | A |
| CHAT | None | None | None | None |

| | | | | |
|---|---|---|---|---|
| CHAT | None | None | None | None |
| CHAT | None | None | None | None |
| CHAT | None | None | None | None |
| CHAT | None | None | None | None |
| CHAT | 0.538 | 6 | 2fy5 | A |
| CHAT | None | None | None | None |
| CHAT | None | None | None | None |
| CHAT | None | None | None | None |
| CHAT | None | None | None | None |
| CHAT | None | None | None | None |
| CHAT | None | None | None | None |
| CHAT | None | None | None | None |
| GHR | None | None | None | None |
| GHR | 0.312 | 2 | 3hhr | C |
| GHR | None | None | None | None |
| GHR | 0.312 | 1 | 3hhr | C |
| GHR | None | None | None | None |
| GHR | None | None | None | None |
| GHR | None | None | None | None |
| GHR | None | None | None | None |
| GHR | None | None | None | None |
| GHR | None | None | None | None |
| GHR | None | None | None | None |
| GHR | None | None | None | None |
| GHR | None | None | None | None |
| GHR | None | None | None | None |
| FZD4 | None | None | None | None |
| FZD4 | None | None | None | None |
| FZD4 | None | None | None | None |
| MTR | None | None | None | None |
| MTR | None | None | None | None |
| MTR | None | None | None | None |
| GM2A | None | None | None | None |
| GM2A | None | None | None | None |
| GM2A | None | None | None | None |
| GM2A | None | None | None | None |
| GM2A | None | None | None | None |
| CRYGD | 0.444 | 3 | 2g98 | A |
| CRYGD | None | None | None | None |
| CRYGD | 0.522 | 1 | 2kfb | A |
| CRYGD | 0.4 | 3 | 4gr7 | X |
| CRYGD | None | None | None | None |
| CRYGD | 0.333 | 9 | 1hk0 | X |
| CRYGD | None | None | None | None |
| CRYGD | None | None | None | None |
| CRYGD | 0.556 | 2 | 2g98 | A |
| TLR5 | None | None | None | None |
| HSD17B10 | 0.855 | 1 | 1u7t | B |
| HSD17B10 | None | None | None | None |
| HSD17B10 | 0.542 | 1 | 1so8 | A |
| HSD17B10 | 0.375 | 3 | 1so8 | A |
| TBC1D20 | None | None | None | None |
| FBN1 | None | None | None | None |
| FBN1 | None | None | None | None |

| | | | | |
|---|---|---|---|---|
| FBN1 | None | None | None | None |
| FBN1 | None | None | None | None |
| FBN1 | None | None | None | None |
| FBN1 | None | None | None | None |
| FBN1 | None | None | None | None |
| FXN | None | None | None | None |
| FXN | None | None | None | None |
| FXN | None | None | None | None |
| FXN | None | None | None | None |
| MYO5A | None | None | None | None |
| CTSK | None | None | None | None |
| CTSK | None | None | None | None |
| SCN5A | None | None | None | None |
| SCN5A | None | None | None | None |
| SOS1 | None | None | None | None |
| SOS1 | None | None | None | None |
| SOS1 | None | None | None | None |
| SOS1 | None | None | None | None |
| SOS1 | None | None | None | None |
| SOS1 | None | None | None | None |
| SOS1 | None | None | None | None |
| SOS1 | None | None | None | None |
| SOS1 | None | None | None | None |
| SOS1 | None | None | None | None |
| SOS1 | None | None | None | None |
| SOS1 | None | None | None | None |
| GATM | None | None | None | None |
| GATM | None | None | None | None |
| GATM | 0.75 | 4 | 4jdw | A |
| GATM | None | None | None | None |
| GATM | None | None | None | None |
| GATM | 0.75 | 6 | 6jdw | A |
| GATM | 0.75 | 11 | 6jdw | A |
| MMP2 | None | None | None | None |
| MMP2 | None | None | None | None |
| MMP2 | 0.407 | 7 | 1qib | A |
| MMP2 | None | None | None | None |
| NDUFB9 | None | None | None | None |
| NDUFB9 | None | None | None | None |
| GH1 | None | None | None | None |
| GH1 | None | None | None | None |
| GH1 | None | None | None | None |
| SLC6A4 | None | None | None | None |
| G6PD | None | None | None | None |
| G6PD | None | None | None | None |
| G6PD | None | None | None | None |
| G6PD | None | None | None | None |
| G6PD | None | None | None | None |
| G6PD | None | None | None | None |
| G6PD | None | None | None | None |
| G6PD | None | None | None | None |
| G6PD | 0.545 | 1 | 1qki | B |
| G6PD | None | None | None | None |
| G6PD | None | None | None | None |

| | | | | |
|---|---|---|---|---|
| G6PD | None | None | None | None |
| G6PD | None | None | None | None |
| G6PD | None | None | None | None |
| G6PD | None | None | None | None |
| G6PD | 0.455 | 1 | 1qki | D |
| G6PD | None | None | None | None |
| G6PD | None | None | None | None |
| G6PD | None | None | None | None |
| G6PD | None | None | None | None |
| CST3 | None | None | None | None |
| LMNA | None | None | None | None |
| LMNA | None | None | None | None |
| NDUFA9 | None | None | None | None |
| NDUFA9 | None | None | None | None |
| NDUFA9 | None | None | None | None |
| NDUFA9 | None | None | None | None |
| YARS | 0.581 | 1 | 5thl | A |
| YARS | None | None | None | None |
| YARS | None | None | None | None |
| YARS | None | None | None | None |
| EPHA2 | None | None | None | None |
| EPHA2 | None | None | None | None |
| EPHA2 | None | None | None | None |
| EPHA2 | None | None | None | None |
| DICER1 | None | None | None | None |
| DICER1 | None | None | None | None |
| SNX10 | None | None | None | None |
| SNX10 | None | None | None | None |
| TTN | None | None | None | None |
| TTN | None | None | None | None |
| GATA3 | None | None | None | None |
| C5 | None | None | None | None |
| THBD | None | None | None | None |
| BIN1 | None | None | None | None |
| BIN1 | None | None | None | None |
| SMARCAL1 | None | None | None | None |
| CFB | None | None | None | None |
| CYC1 | None | None | None | None |
| ANK2 | None | None | None | None |
| CHMP4B | None | None | None | None |
| CHMP4B | None | None | None | None |
| ERBB2 | None | None | None | None |
| ERBB2 | None | None | None | None |
| POLG2 | None | None | None | None |
| POLG2 | None | None | None | None |
| POLG2 | None | None | None | None |
| POLG2 | None | None | None | None |
| POLG2 | None | None | None | None |
| POLG2 | None | None | None | None |
| POLG2 | None | None | None | None |
| POLG2 | None | None | None | None |
| C3 | None | None | None | None |
| C3 | None | None | None | None |
| RPGR | None | None | None | None |

| | | | | |
|---|---|---|---|---|
| RPGR | None | None | None | None |
| RPGR | None | None | None | None |
| RPGR | None | None | None | None |
| RPGR | None | None | None | None |
| C1QA | None | None | None | None |
| PRKG1 | None | None | None | None |
| IGBP1 | None | None | None | None |
| ENO3 | None | None | None | None |
| ENO3 | None | None | None | None |
| ENO3 | 0.41 | 43 | 2xsx | A |
| ENO3 | 0.487 | 39 | 2xsx | A |
| ENO3 | 0.359 | 47 | 2xsx | B |
| ENO3 | 0.897 | 2 | 2xsx | B |
| MCPH1 | None | None | None | None |
| MCPH1 | None | None | None | None |
| MCPH1 | None | None | None | None |
| MCPH1 | None | None | None | None |
| MCPH1 | None | None | None | None |
| SNRNP200 | None | None | None | None |
| SNRNP200 | None | None | None | None |
| DNMT1 | None | None | None | None |
| DNMT1 | None | None | None | None |
| DNMT1 | None | None | None | None |
| AKT1 | None | None | None | None |
| NDUFS2 | None | None | None | None |
| ACVR2B | None | None | None | None |
| HMGCL | None | None | None | None |
| HMGCL | None | None | None | None |
| HMGCL | None | None | None | None |
| JUP | None | None | None | None |
| JUP | None | None | None | None |
| JUP | None | None | None | None |
| JUP | None | None | None | None |
| JUP | None | None | None | None |
| JUP | None | None | None | None |
| JUP | None | None | None | None |
| JUP | None | None | None | None |
| SPINK1 | None | None | None | None |
| SPINK1 | None | None | None | None |
| SPINK1 | None | None | None | None |
| VCL | None | None | None | None |
| VCL | None | None | None | None |
| NOTCH1 | None | None | None | None |
| NOTCH1 | None | None | None | None |
| NOTCH1 | None | None | None | None |
| NDUFB3 | None | None | None | None |
| IKBKG | None | None | None | None |
| GSN | 0.529 | 2 | 2fh1 | A |
| GSN | None | None | None | None |
| GSN | None | None | None | None |
| GSN | 0.5 | 3 | 1p8x | A |
| GSN | None | None | None | None |
| GSN | None | None | None | None |
| GSN | None | None | None | None |

| | | | | |
|---|---|---|---|---|
| GSN | None | None | None | None |
| GSN | None | None | None | None |
| CLC | None | None | None | None |
| RAF1 | None | None | None | None |
| RAF1 | None | None | None | None |
| RAF1 | None | None | None | None |
| NDUFS8 | None | None | None | None |
| NDUFS8 | None | None | None | None |
| NDUFS8 | None | None | None | None |
| NDUFS8 | None | None | None | None |
| NDUFS8 | None | None | None | None |
| CASR | None | None | None | None |
| CASR | None | None | None | None |
| CASR | None | None | None | None |
| CASR | None | None | None | None |
| CASR | None | None | None | None |
| CASR | None | None | None | None |
| CASR | None | None | None | None |
| CASR | None | None | None | None |
| CASR | None | None | None | None |
| CASR | None | None | None | None |
| CASR | None | None | None | None |
| CASR | None | None | None | None |
| CASR | None | None | None | None |
| CASR | None | None | None | None |
| CASR | None | None | None | None |
| CASR | None | None | None | None |
| CASR | None | None | None | None |
| CASR | None | None | None | None |
| CASR | None | None | None | None |
| CASR | None | None | None | None |
| CASR | None | None | None | None |
| CASR | None | None | None | None |
| CASR | None | None | None | None |
| CASR | None | None | None | None |
| CASR | None | None | None | None |
| CASR | None | None | None | None |
| CASR | None | None | None | None |
| CASR | None | None | None | None |
| CASR | None | None | None | None |
| CASR | None | None | None | None |
| VCL | None | None | None | None |
| VCL | None | None | None | None |
| SMPD1 | None | None | None | None |
| SMPD1 | None | None | None | None |
| SMPD1 | None | None | None | None |
| KDM6A | None | None | None | None |
| KCNQ1 | None | None | None | None |
| KCNQ1 | None | None | None | None |
| KCNQ1 | None | None | None | None |
| KCNQ1 | None | None | None | None |
| KCNQ1 | None | None | None | None |
| KCNQ1 | None | None | None | None |
| NDUFA12 | None | None | None | None |
| GLRX5 | None | None | None | None |

| | | | | |
|---|---|---|---|---|
| PALB2 | None | None | None | None |
| PALB2 | None | None | None | None |
| PALB2 | None | None | None | None |
| PALB2 | None | None | None | None |
| PALB2 | None | None | None | None |
| PALB2 | None | None | None | None |
| PALB2 | None | None | None | None |
| PALB2 | None | None | None | None |
| PALB2 | None | None | None | None |
| PALB2 | None | None | None | None |
| ALDOB | 0.462 | 89 | 1qo5 | C |
| ALDOB | 0.731 | 2 | 1xdm | A |
| ALDOB | 0.731 | 1 | 1qo5 | B |
| ALDOB | 0.538 | 6 | 1xdm | A |
| ALDOB | 0.692 | 7 | 1qo5 | C |
| ALDOB | None | None | None | None |
| ALDOB | 0.577 | 25 | 1qo5 | B |
| GNE | None | None | None | None |
| GNE | None | None | None | None |
| GNE | None | None | None | None |
| GNE | None | None | None | None |
| GNE | None | None | None | None |
| GNE | None | None | None | None |
| GNE | None | None | None | None |
| GNE | None | None | None | None |
| GNE | None | None | None | None |
| GNE | None | None | None | None |
| GNE | None | None | None | None |
| GNE | None | None | None | None |
| CHRNB2 | None | None | None | None |
| CHRNB2 | None | None | None | None |
| ERCC3 | None | None | None | None |
| ERCC3 | None | None | None | None |
| ERCC3 | None | None | None | None |
| ERCC3 | None | None | None | None |
| ERCC3 | None | None | None | None |
| UNC119 | None | None | None | None |
| ATL1 | None | None | None | None |
| ATL1 | None | None | None | None |
| ATL1 | None | None | None | None |
| ATL1 | 0.8 | 1 | 3q5d | A |
| ATL1 | None | None | None | None |
| ATL1 | None | None | None | None |
| ATL1 | None | None | None | None |
| ATL1 | None | None | None | None |
| ATL1 | None | None | None | None |
| ATL1 | None | None | None | None |
| NR3C2 | 0.701 | 1 | 2oax | A |
| TBK1 | None | None | None | None |
| HYAL1 | None | None | None | None |
| HYAL1 | None | None | None | None |
| HYAL1 | None | None | None | None |
| HYAL1 | None | None | None | None |
| CTSD | 0.633 | 1 | 4obz | A |

| | | | | |
|---|---|---|---|---|
| PRKCG | None | None | None | None |
| TGFBR1 | None | None | None | None |
| XDH | 0.308 | 2 | 2e1q | A |
| XDH | None | None | None | None |
| XDH | None | None | None | None |
| XDH | None | None | None | None |
| XDH | None | None | None | None |
| XDH | None | None | None | None |
| NMNAT1 | 0.833 | 1 | 1kqn | E |
| NMNAT1 | None | None | None | None |
| NMNAT1 | None | None | None | None |
| NMNAT1 | None | None | None | None |
| NMNAT1 | None | None | None | None |
| NMNAT1 | None | None | None | None |
| SERPING1 | None | None | None | None |
| SERPING1 | None | None | None | None |
| SERPING1 | None | None | None | None |
| SERPING1 | None | None | None | None |
| SERPING1 | None | None | None | None |
| PDZRN3 | None | None | None | None |
| FABP2 | None | None | None | None |
| IFNGR1 | None | None | None | None |
| IFNGR1 | None | None | None | None |
| TDP1 | None | None | None | None |
| TDP1 | 0.333 | 11 | 1rg1 | A |
| INS | None | None | None | None |
| INS | 0.769 | 1 | 2r34 | C |
| INS | 0.769 | 1 | 2r34 | C |
| PPOX | 0.857 | 1 | 4ivm | B |
| PPOX | None | None | None | None |
| PPOX | None | None | None | None |
| PPOX | None | None | None | None |
| PPOX | None | None | None | None |
| PPOX | None | None | None | None |
| PPOX | None | None | None | None |
| PPOX | None | None | None | None |
| PPOX | None | None | None | None |
| GFER | None | None | None | None |
| GFER | 0.769 | 4 | 3tk0 | A |
| GFER | None | None | None | None |
| GFER | None | None | None | None |
| LIFR | None | None | None | None |
| EGLN1 | 0.556 | 21 | 5a3u | C |
| EGLN1 | 0.889 | 6 | 5v18 | A |
| EGLN1 | 0.944 | 2 | 5a3u | C |
| GP1BA | None | None | None | None |
| GP1BA | None | None | None | None |
| GP1BA | None | None | None | None |
| GP1BA | None | None | None | None |
| GP1BA | None | None | None | None |
| MMADHC | None | None | None | None |
| MMADHC | None | None | None | None |
| MMADHC | None | None | None | None |
| MMADHC | None | None | None | None |

| | | | | |
|---|---|---|---|---|
| MMADHC | None | None | None | None |
| MMADHC | None | None | None | None |
| MMADHC | None | None | None | None |
| ACVRL1 | None | None | None | None |
| ACVRL1 | None | None | None | None |
| RFXANK | None | None | None | None |
| CHIT1 | None | None | None | None |
| UBR1 | None | None | None | None |
| BLM | None | None | None | None |
| BLM | None | None | None | None |
| BLM | None | None | None | None |
| BLM | None | None | None | None |
| BLM | None | None | None | None |
| BLM | None | None | None | None |
| BLM | None | None | None | None |
| BLM | None | None | None | None |
| BLM | None | None | None | None |
| BLM | None | None | None | None |
| BLM | None | None | None | None |
| BLM | None | None | None | None |
| BLM | None | None | None | None |
| VDR | 0.378 | 7 | 1kb4 | B |
| VDR | None | None | None | None |
| VDR | 0.378 | 4 | 1kb4 | A |
| VDR | 0.578 | 1 | 1kb2 | B |
| VDR | 0.578 | 1 | 1kb2 | B |
| VDR | 0.578 | 1 | 1kb2 | B |
| EPHA2 | None | None | None | None |
| EPHA2 | None | None | None | None |
| EPHA2 | None | None | None | None |
| EPHA2 | None | None | None | None |
| EPHA2 | None | None | None | None |
| EPHA2 | None | None | None | None |
| EPHA2 | None | None | None | None |
| EPHA2 | None | None | None | None |
| EPHA2 | None | None | None | None |
| EPHA2 | None | None | None | None |
| UBA1 | None | None | None | None |
| STAT1 | None | None | None | None |
| STAT1 | None | None | None | None |
| STAT1 | None | None | None | None |
| STAT1 | None | None | None | None |
| STAT1 | None | None | None | None |
| STAT1 | None | None | None | None |
| STAT1 | None | None | None | None |
| STAT1 | None | None | None | None |
| DNM1L | None | None | None | None |
| DNM1L | None | None | None | None |
| OTC | None | None | None | None |
| OTC | None | None | None | None |
| OTC | None | None | None | None |
| OTC | None | None | None | None |
| OTC | None | None | None | None |
| OTC | None | None | None | None |

| | | | | |
|---|---|---|---|---|
| OTC | None | None | None | None |
| OTC | None | None | None | None |
| OTC | None | None | None | None |
| OTC | None | None | None | None |
| OTC | None | None | None | None |
| OTC | None | None | None | None |
| OTC | None | None | None | None |
| OTC | None | None | None | None |
| OTC | None | None | None | None |
| OTC | None | None | None | None |
| OTC | None | None | None | None |
| RARS | None | None | None | None |
| RARS | None | None | None | None |
| RARS | None | None | None | None |
| SEPSECS | None | None | None | None |
| SEPSECS | None | None | None | None |
| SEPSECS | None | None | None | None |
| SEPSECS | None | None | None | None |
| SEPSECS | None | None | None | None |
| BRCA1 | None | None | None | None |
| BRCA1 | None | None | None | None |
| BRCA1 | 0.538 | 1 | 3pxc | X |
| BRCA1 | None | None | None | None |
| BRCA1 | None | None | None | None |
| BRCA1 | 0.462 | 1 | 1t2u | A |
| BRCA1 | None | None | None | None |
| BRCA1 | None | None | None | None |
| BRCA1 | None | None | None | None |
| BRCA1 | None | None | None | None |
| BRCA1 | None | None | None | None |
| BRCA1 | None | None | None | None |
| BRCA1 | None | None | None | None |
| BRCA1 | None | None | None | None |
| BRCA1 | None | None | None | None |
| BRCA1 | None | None | None | None |
| BRCA1 | None | None | None | None |
| BRCA1 | None | None | None | None |
| BRCA1 | None | None | None | None |
| BRCA1 | None | None | None | None |
| BRCA1 | None | None | None | None |
| BRCA1 | None | None | None | None |
| CUBN | None | None | None | None |
| ITGB3 | None | None | None | None |
| MTRR | None | None | None | None |
| MTRR | None | None | None | None |
| MTRR | None | None | None | None |
| MTRR | None | None | None | None |
| MTRR | None | None | None | None |
| MTRR | None | None | None | None |
| MTRR | None | None | None | None |
| MTRR | None | None | None | None |
| IL7R | None | None | None | None |
| IL7R | None | None | None | None |
| RPS24 | None | None | None | None |

| | | | | |
|---|---|---|---|---|
| RPS24 | None | None | None | None |
| HOGA1 | None | None | None | None |
| HOGA1 | None | None | None | None |
| HOGA1 | None | None | None | None |
| HOGA1 | None | None | None | None |
| HOGA1 | None | None | None | None |
| HOGA1 | None | None | None | None |
| HOGA1 | 0.791 | 1 | 3s5n | A |
| HOGA1 | None | None | None | None |
| HOGA1 | None | None | None | None |
| HOGA1 | None | None | None | None |
| HOGA1 | None | None | None | None |
| HOGA1 | None | None | None | None |
| HOGA1 | None | None | None | None |
| GJB2 | None | None | None | None |
| GJB2 | None | None | None | None |
| GJB2 | None | None | None | None |
| GJB2 | None | None | None | None |
| GJB2 | None | None | None | None |
| GJB2 | None | None | None | None |
| GJB2 | None | None | None | None |
| GJB2 | None | None | None | None |
| GJB2 | None | None | None | None |
| GJB2 | None | None | None | None |
| GJB2 | None | None | None | None |
| GJB2 | None | None | None | None |
| GJB2 | None | None | None | None |
| GJB2 | None | None | None | None |
| GJB2 | None | None | None | None |
| GJB2 | None | None | None | None |
| GJB2 | None | None | None | None |
| GJB2 | None | None | None | None |
| GJB2 | None | None | None | None |
| GJB2 | None | None | None | None |
| GJB2 | None | None | None | None |
| GJB2 | None | None | None | None |
| GJB2 | None | None | None | None |
| GJB2 | None | None | None | None |
| GJB2 | None | None | None | None |
| GJB2 | None | None | None | None |
| GJB2 | None | None | None | None |
| GJB2 | None | None | None | None |
| GJB2 | None | None | None | None |
| GJB2 | None | None | None | None |
| GJB2 | None | None | None | None |
| GJB2 | None | None | None | None |
| GJB2 | None | None | None | None |
| GJB2 | None | None | None | None |
| GJB2 | None | None | None | None |
| GJB2 | None | None | None | None |
| GJB2 | None | None | None | None |
| GJB2 | None | None | None | None |
| GJB2 | None | None | None | None |
| GJB2 | None | None | None | None |

| | | | | |
|---|---|---|---|---|
| GJB2 | None | None | None | None |
| GJB2 | None | None | None | None |
| GJB2 | None | None | None | None |
| GJB2 | None | None | None | None |
| GJB2 | None | None | None | None |
| GJB2 | None | None | None | None |
| GJB2 | None | None | None | None |
| GJB2 | None | None | None | None |
| GJB2 | None | None | None | None |
| GJB2 | None | None | None | None |
| GJB2 | None | None | None | None |
| GJB2 | None | None | None | None |
| GJB2 | None | None | None | None |
| GJB2 | None | None | None | None |
| CPS1 | None | None | None | None |
| CPS1 | 0.545 | 127 | 5dou | A |
| CPS1 | None | None | None | None |
| CPS1 | None | None | None | None |
| CPS1 | 0.364 | 19 | 5dou | A |
| CPS1 | 0.364 | 19 | 5dou | A |
| CPS1 | None | None | None | None |
| CPS1 | None | None | None | None |
| CPS1 | None | None | None | None |
| CPS1 | None | None | None | None |
| PSPH | None | None | None | None |
| PSPH | None | None | None | None |
| PSPH | None | None | None | None |
| PSPH | None | None | None | None |
| PSPH | None | None | None | None |
| PSPH | None | None | None | None |
| PEPD | 0.512 | 2 | 5m4l | B |
| PEPD | None | None | None | None |
| PEPD | None | None | None | None |
| PEPD | 0.643 | 2 | 5m4j | B |
| PEPD | 0.643 | 2 | 5m4j | B |
| PEPD | 0.419 | 5 | 5m4l | B |
| PEPD | None | None | None | None |
| PEPD | None | None | None | None |
| PEPD | None | None | None | None |
| KDM5C | None | None | None | None |
| KDM5C | None | None | None | None |
| KDM5C | 0.333 | 55 | 5fwj | A |
| ITGB6 | 0.312 | 2 | 5ffo | B |
| ITGB6 | None | None | None | None |
| ITGB6 | None | None | None | None |
| DARS | None | None | None | None |
| DARS | None | None | None | None |
| DARS | None | None | None | None |
| DARS | None | None | None | None |
| BTK | None | None | None | None |
| BTK | None | None | None | None |
| BTK | None | None | None | None |
| HDAC4 | None | None | None | None |
| ITGA2B | None | None | None | None |

| | | | | |
|---|---|---|---|---|
| ITGA2B | None | None | None | None |
| ITGA2B | None | None | None | None |
| ITGA2B | None | None | None | None |
| PCSK9 | None | None | None | None |
| PCSK9 | None | None | None | None |
| PCSK9 | None | None | None | None |
| PCSK9 | None | None | None | None |
| MUTYH | None | None | None | None |
| MUTYH | None | None | None | None |
| MUTYH | None | None | None | None |
| MUTYH | None | None | None | None |
| MUTYH | None | None | None | None |
| MUTYH | 0.375 | 2 | 3n5n | X |
| MUTYH | None | None | None | None |
| MUTYH | None | None | None | None |
| MUTYH | None | None | None | None |
| MUTYH | 0.375 | 2 | 3n5n | X |
| MUTYH | 0.375 | 2 | 3n5n | X |
| MUTYH | None | None | None | None |
| MUTYH | 0.375 | 2 | 3n5n | X |
| MUTYH | None | None | None | None |
| MUTYH | None | None | None | None |
| MUTYH | None | None | None | None |
| MUTYH | None | None | None | None |
| MUTYH | None | None | None | None |
| MUTYH | None | None | None | None |
| MUTYH | None | None | None | None |
| MUTYH | None | None | None | None |
| MUTYH | None | None | None | None |
| MUTYH | None | None | None | None |
| MUTYH | None | None | None | None |
| SLC25A12 | None | None | None | None |
| GSN | None | None | None | None |
| IHH | 0.909 | 1 | 3n1p | B |
| IHH | None | None | None | None |
| IHH | None | None | None | None |
| IHH | 0.818 | 2 | 3n1p | B |
| IHH | 0.727 | 12 | 3n1p | B |
| IHH | 0.636 | 15 | 3n1p | B |
| IHH | 0.818 | 7 | 3n1p | B |
| CD247 | None | None | None | None |
| EP300 | None | None | None | None |
| TTN | None | None | None | None |
| MRI1 | None | None | None | None |
| SGSH | None | None | None | None |
| SGSH | None | None | None | None |
| SGSH | None | None | None | None |
| SGSH | None | None | None | None |
| SGSH | None | None | None | None |
| SGSH | None | None | None | None |
| SGSH | None | None | None | None |
| SGSH | None | None | None | None |
| SGSH | None | None | None | None |
| SGSH | None | None | None | None |

| | | | | |
|---|---|---|---|---|
| SGSH | None | None | None | None |
| SGSH | None | None | None | None |
| SGSH | None | None | None | None |
| SGSH | None | None | None | None |
| SGSH | None | None | None | None |
| SGSH | None | None | None | None |
| SGSH | None | None | None | None |
| SGSH | None | None | None | None |
| SGSH | None | None | None | None |
| SGSH | None | None | None | None |
| SGSH | None | None | None | None |
| SGSH | None | None | None | None |
| CBL | None | None | None | None |
| CANT1 | None | None | None | None |
| CANT1 | None | None | None | None |
| CANT1 | None | None | None | None |
| CANT1 | None | None | None | None |
| CANT1 | None | None | None | None |
| CANT1 | None | None | None | None |
| CANT1 | None | None | None | None |
| SLC1A3 | None | None | None | None |
| SLC1A3 | None | None | None | None |
| SLC1A5 | None | None | None | None |
| SLC1A3 | None | None | None | None |
| SLC1A5 | None | None | None | None |
| SLC1A3 | None | None | None | None |
| SLC1A3 | None | None | None | None |
| CTSD | 0.423 | 15 | 4od9 | D |
| CTSD | None | None | None | None |
| CTSD | None | None | None | None |
| CTSD | None | None | None | None |
| CTSD | None | None | None | None |
| MTOR | None | None | None | None |
| ILK | None | None | None | None |
| PCDH15 | None | None | None | None |
| PCDH15 | None | None | None | None |
| PCDH15 | None | None | None | None |
| PCDH15 | None | None | None | None |
| PCDH15 | None | None | None | None |
| PCDH15 | None | None | None | None |
| KLHL40 | None | None | None | None |
| KLHL40 | None | None | None | None |
| KLHL40 | None | None | None | None |
| KLHL40 | None | None | None | None |
| CSTA | None | None | None | None |
| CSTA | None | None | None | None |
| HNMT | None | None | None | None |
| HNMT | None | None | None | None |
| IFNGR2 | None | None | None | None |
| IFNGR2 | None | None | None | None |
| PEX5 | None | None | None | None |
| KIF7 | None | None | None | None |
| KIF7 | None | None | None | None |
| MYL3 | None | None | None | None |

| | | | | |
|---|---|---|---|---|
| MYL3 | None | None | None | None |
| MYL3 | None | None | None | None |
| MYL3 | None | None | None | None |
| MYL3 | None | None | None | None |
| MYL3 | None | None | None | None |
| MYL3 | None | None | None | None |
| NAGLU | None | None | None | None |
| NAGLU | None | None | None | None |
| NAGLU | None | None | None | None |
| NAGLU | None | None | None | None |
| NAGLU | None | None | None | None |
| NAGLU | None | None | None | None |
| NAGLU | None | None | None | None |
| NAGLU | None | None | None | None |
| NAGLU | None | None | None | None |
| NAGLU | None | None | None | None |
| NAGLU | None | None | None | None |
| NAGLU | None | None | None | None |
| NAGLU | None | None | None | None |
| TNFRSF13C | None | None | None | None |
| KIF21A | None | None | None | None |
| KIF21A | None | None | None | None |
| KIF21A | None | None | None | None |
| KIF21A | None | None | None | None |
| KIF21A | None | None | None | None |
| DOCK2 | None | None | None | None |
| DSP | None | None | None | None |
| CDKN2A | None | None | None | None |
| CDKN2A | None | None | None | None |
| CDKN2A | None | None | None | None |
| CDKN2A | None | None | None | None |
| CDKN2A | None | None | None | None |
| CDKN2A | None | None | None | None |
| CDKN2A | None | None | None | None |
| CDKN2A | None | None | None | None |
| CDKN2A | None | None | None | None |
| CDKN2A | None | None | None | None |
| CDKN2A | None | None | None | None |
| CDKN2A | None | None | None | None |
| CDKN2A | None | None | None | None |
| CDKN2A | None | None | None | None |
| VCL | None | None | None | None |
| TPP1 | None | None | None | None |
| TPP1 | None | None | None | None |
| TPP1 | None | None | None | None |
| TPP1 | None | None | None | None |
| TPP1 | None | None | None | None |
| TPP1 | None | None | None | None |
| TPP1 | None | None | None | None |
| TPP1 | None | None | None | None |
| TPP1 | None | None | None | None |
| TPP1 | None | None | None | None |
| TPP1 | None | None | None | None |
| TPP1 | None | None | None | None |

| | | | | |
|---|---|---|---|---|
| TPP1 | None | None | None | None |
| TPP1 | None | None | None | None |
| TPP1 | None | None | None | None |
| TPP1 | None | None | None | None |
| TPP1 | None | None | None | None |
| TPP1 | None | None | None | None |
| TPP1 | None | None | None | None |
| TPP1 | None | None | None | None |
| TPP1 | None | None | None | None |
| TPP1 | None | None | None | None |
| IL10RB | None | None | None | None |
| IL10RB | None | None | None | None |
| CCM2 | None | None | None | None |
| CCM2 | None | None | None | None |
| CCM2 | None | None | None | None |
| DCX | None | None | None | None |
| DCX | None | None | None | None |
| DCX | None | None | None | None |
| DCX | None | None | None | None |
| DCX | None | None | None | None |
| DCX | None | None | None | None |
| DCX | None | None | None | None |
| DCX | None | None | None | None |
| BRCA2 | None | None | None | None |
| NNT | None | None | None | None |
| C2 | None | None | None | None |
| C2 | None | None | None | None |
| C2 | None | None | None | None |
| C2 | None | None | None | None |
| WRN | None | None | None | None |
| WRN | None | None | None | None |
| F13A1 | None | None | None | None |
| F13A1 | None | None | None | None |
| F13A1 | None | None | None | None |
| F13A1 | None | None | None | None |
| TF | None | None | None | None |
| TF | None | None | None | None |
| TF | 0.516 | 2 | 3ve1 | B |
| TF | 0.562 | 39 | 3skp | A |
| TF | None | None | None | None |
| ITPA | None | None | None | None |
| ITPA | None | None | None | None |
| SNCA | None | None | None | None |
| ITGB4 | None | None | None | None |
| ITGB4 | None | None | None | None |
| ITGB4 | None | None | None | None |
| LRP6 | None | None | None | None |
| LRP6 | None | None | None | None |
| LRP6 | None | None | None | None |
| PLG | None | None | None | None |
| PLG | 0.61 | 1 | 1i5k | A |
| TYROBP | None | None | None | None |
| UQCRC2 | None | None | None | None |
| GPSM2 | None | None | None | None |

| | | | | |
|---|---|---|---|---|
| GPSM2 | None | None | None | None |
| GPSM2 | None | None | None | None |
| SCN5A | None | None | None | None |
| SCN5A | None | None | None | None |
| SCN5A | None | None | None | None |
| SCN5A | None | None | None | None |
| SCN5A | None | None | None | None |
| ATIC | 0.6 | 6 | 1p4r | A |
| EZH2 | None | None | None | None |
| CPA1 | 0.387 | 33 | 4uez | A |
| SCN4B | None | None | None | None |
| SCN4B | None | None | None | None |
| MAN1B1 | None | None | None | None |
| MAN1B1 | 0.6 | 9 | 1fmi | A |
| MAN1B1 | None | None | None | None |
| MAN1B1 | None | None | None | None |
| CYCS | 0.74 | 1 | 2n9i | A |
| IDUA | None | None | None | None |
| IDUA | None | None | None | None |
| IDUA | None | None | None | None |
| IDUA | None | None | None | None |
| IDUA | None | None | None | None |
| IDUA | None | None | None | None |
| IDUA | None | None | None | None |
| IDUA | None | None | None | None |
| IDUA | None | None | None | None |
| IDUA | None | None | None | None |
| IDUA | None | None | None | None |
| IDUA | None | None | None | None |
| IDUA | None | None | None | None |
| IDUA | None | None | None | None |
| IDUA | None | None | None | None |
| IDUA | None | None | None | None |
| IDUA | None | None | None | None |
| IDUA | None | None | None | None |
| IDUA | None | None | None | None |
| IDUA | None | None | None | None |
| IDUA | None | None | None | None |
| IDUA | None | None | None | None |
| IDUA | None | None | None | None |
| IDUA | None | None | None | None |
| IDUA | None | None | None | None |
| HIKESHI | None | None | None | None |
| APC | None | None | None | None |
| APC | None | None | None | None |
| APC | None | None | None | None |
| APP | None | None | None | None |
| CEP152 | None | None | None | None |
| GAA | 0.609 | 1 | 5kzx | A |
| GAA | 0.609 | 1 | 5kzx | A |
| GAA | None | None | None | None |
| GAA | None | None | None | None |
| GAA | None | None | None | None |
| GAA | 0.609 | 1 | 5kzx | A |

| | | | | |
|---|---|---|---|---|
| GAA | 0.609 | 1 | 5kzx | A |
| GAA | 0.609 | 1 | 5kzx | A |
| GAA | 0.609 | 1 | 5kzx | A |
| GAA | 0.609 | 1 | 5kzx | A |
| GAA | 0.609 | 1 | 5kzx | A |
| GAA | 0.609 | 1 | 5kzx | A |
| GAA | 0.609 | 1 | 5kzx | A |
| GAA | 0.609 | 1 | 5kzx | A |
| GAA | 0.609 | 1 | 5kzx | A |
| GAA | 0.609 | 1 | 5kzx | A |
| GAA | 0.609 | 1 | 5kzx | A |
| GAA | 0.609 | 1 | 5kzx | A |
| GAA | 0.609 | 1 | 5kzx | A |
| GAA | 0.609 | 1 | 5kzx | A |
| GAA | 0.609 | 1 | 5kzx | A |
| GAA | 0.609 | 1 | 5kzx | A |
| GAA | 0.609 | 1 | 5kzx | A |
| GAA | 0.609 | 1 | 5kzx | A |
| GAA | 0.609 | 1 | 5kzx | A |
| GAA | 0.609 | 1 | 5kzx | A |
| GAA | 0.609 | 1 | 5kzx | A |
| GAA | 0.609 | 1 | 5kzx | A |
| GAA | 0.609 | 1 | 5kzx | A |
| GAA | 0.609 | 1 | 5kzx | A |
| GAA | 0.609 | 1 | 5kzx | A |
| GAA | 0.609 | 1 | 5kzx | A |
| GAA | 0.609 | 1 | 5kzx | A |
| GAA | 0.609 | 1 | 5kzx | A |
| GAA | 0.609 | 1 | 5kzx | A |
| GAA | 0.609 | 1 | 5kzx | A |
| GAA | 0.609 | 1 | 5kzx | A |
| GAA | 0.609 | 1 | 5kzx | A |
| GAA | 0.609 | 1 | 5kzx | A |
| GAA | 0.609 | 1 | 5kzx | A |
| GAA | 0.609 | 1 | 5kzx | A |
| GAA | 0.609 | 1 | 5kzx | A |
| GAA | 0.609 | 1 | 5kzx | A |
| GAA | 0.609 | 1 | 5kzx | A |
| GAA | 0.609 | 1 | 5kzx | A |
| GAA | 0.609 | 1 | 5kzx | A |
| GAA | 0.609 | 1 | 5kzx | A |
| GAA | 0.609 | 1 | 5kzx | A |
| GAA | None | None | None | None |
| FOLR1 | None | None | None | None |
| DSP | None | None | None | None |
| DSP | None | None | None | None |
| DSP | None | None | None | None |
| DSP | None | None | None | None |
| DSP | None | None | None | None |
| CCM2 | None | None | None | None |
| FXN | None | None | None | None |
| FXN | None | None | None | None |
| COMP | None | None | None | None |
| COMP | None | None | None | None |

| | | | | |
|---|---|---|---|---|
| COMP | None | None | None | None |
| COMP | None | None | None | None |
| COMP | None | None | None | None |
| COMP | None | None | None | None |
| COMP | None | None | None | None |
| COMP | None | None | None | None |
| COMP | None | None | None | None |
| COMP | None | None | None | None |
| COMP | None | None | None | None |
| COMP | None | None | None | None |
| AIFM1 | None | None | None | None |
| AIFM1 | None | None | None | None |
| AIFM1 | None | None | None | None |
| AIFM1 | None | None | None | None |
| AIFM1 | None | None | None | None |
| AIFM1 | None | None | None | None |
| AIFM1 | None | None | None | None |
| MYH9 | None | None | None | None |
| FHL2 | None | None | None | None |
| FHL2 | None | None | None | None |
| TNFRSF4 | None | None | None | None |
| CFB | None | None | None | None |
| CFB | None | None | None | None |
| CFB | None | None | None | None |
| CFB | None | None | None | None |
| CFB | None | None | None | None |
| CFB | None | None | None | None |
| CFB | None | None | None | None |
| CFB | None | None | None | None |
| CFB | None | None | None | None |
| CFB | None | None | None | None |
| MVD | None | None | None | None |
| MVD | None | None | None | None |
| HSD17B4 | None | None | None | None |
| HSD17B4 | None | None | None | None |
| HSD17B4 | None | None | None | None |
| HSD17B4 | 0.429 | 2 | 1s9c | A |
| HSD17B4 | None | None | None | None |
| HSD17B4 | None | None | None | None |
| F11 | None | None | None | None |
| F11 | None | None | None | None |
| F11 | None | None | None | None |
| F11 | None | None | None | None |
| F11 | None | None | None | None |
| F11 | None | None | None | None |
| F11 | None | None | None | None |
| F11 | None | None | None | None |
| F11 | None | None | None | None |
| SPTA1 | None | None | None | None |
| SPTA1 | None | None | None | None |
| SPTA1 | None | None | None | None |
| SPTA1 | None | None | None | None |
| SPTA1 | None | None | None | None |
| SPTA1 | None | None | None | None |

| | | | | |
|---|---|---|---|---|
| GMPR | None | None | None | None |
| MITF | None | None | None | None |
| NPC1 | None | None | None | None |
| NPC1 | None | None | None | None |
| NPC1 | None | None | None | None |
| NPC1 | None | None | None | None |
| NPC1 | None | None | None | None |
| NPC1 | None | None | None | None |
| NPC1 | None | None | None | None |
| NPC1 | None | None | None | None |
| NPC1 | None | None | None | None |
| NPC1 | None | None | None | None |
| NPC1 | None | None | None | None |
| NPC1 | None | None | None | None |
| NPC1 | None | None | None | None |
| NPC1 | None | None | None | None |
| NPC1 | None | None | None | None |
| NPC1 | None | None | None | None |
| NPC1 | None | None | None | None |
| NPC1 | None | None | None | None |
| NPC1 | None | None | None | None |
| NPC1 | None | None | None | None |
| NPC1 | None | None | None | None |
| NPC1 | None | None | None | None |
| NPC1 | None | None | None | None |
| NPC1 | None | None | None | None |
| NPC1 | None | None | None | None |
| NPC1 | None | None | None | None |
| NPC1 | None | None | None | None |
| NPC1 | None | None | None | None |
| NPC1 | None | None | None | None |
| NPC1 | None | None | None | None |
| NPC1 | None | None | None | None |
| NPC1 | None | None | None | None |
| NPC1 | None | None | None | None |
| NPC1 | None | None | None | None |
| NPC1 | None | None | None | None |
| NPC1 | None | None | None | None |
| NPC1 | None | None | None | None |
| NPC1 | None | None | None | None |
| NPC1 | None | None | None | None |
| NPC1 | None | None | None | None |
| NPC1 | None | None | None | None |
| NPC1 | None | None | None | None |
| NPC1 | None | None | None | None |
| NPC1 | None | None | None | None |
| C3 | None | None | None | None |
| NLRX1 | None | None | None | None |
| IGLL1 | None | None | None | None |
| APRT | 0.769 | 4 | 1zn9 | B |
| APRT | 0.417 | 5 | 4x44 | A |
| APRT | None | None | None | None |
| APRT | 0.923 | 1 | 1zn8 | A |
| APRT | None | None | None | None |
| ZIC3 | None | None | None | None |

| | | | | | |
|---|---|---|---|---|---|
| ZIC3 | None | None | None | | None |
| KLHL3 | None | None | None | | None |
| RPS6KA3 | None | None | None | | None |
| RPS6KA3 | None | None | None | | None |
| RPS6KA3 | None | None | None | | None |
| F8 | None | None | None | | None |
| F8 | None | None | None | | None |
| F8 | None | None | None | | None |
| F8 | None | None | None | | None |
| F8 | None | None | None | | None |
| F8 | None | None | None | | None |
| F8 | None | None | None | | None |
| F8 | None | None | None | | None |
| F8 | None | None | None | | None |
| F8 | None | None | None | | None |
| F8 | None | None | None | | None |
| F8 | None | None | None | | None |
| F8 | None | None | None | | None |
| F8 | None | None | None | | None |
| F8 | None | None | None | | None |
| F8 | None | None | None | | None |
| F8 | None | None | None | | None |
| ISG15 | None | None | None | | None |
| EXOSC3 | None | None | None | | None |
| EXOSC3 | None | None | None | | None |
| EXOSC3 | None | None | None | | None |
| NKX2-5 | 0.403 | 1 | 4s0h | | B |
| NKX2-5 | None | None | None | | None |
| NKX2-5 | 0.403 | 1 | 4s0h | | B |
| NKX2-5 | 0.403 | 1 | 4s0h | | B |
| FH | None | None | None | | None |
| FH | None | None | None | | None |
| FH | None | None | None | | None |
| FH | 0.304 | 30 | | 30000 | D |
| FH | 0.621 | 4 | | 30000 | A |
| FH | 0.462 | 6 | | 30000 | C |
| FH | None | None | None | | None |
| FH | None | None | None | | None |
| FH | 0.64 | 6 | 5d6b | | A |
| FH | 0.308 | 49 | | 30000 | C |
| FH | 0.64 | 1 | 5d6b | | A |
| FH | 0.64 | 1 | 5d6b | | A |
| SNRNP200 | None | None | None | | None |
| SNRNP200 | None | None | None | | None |
| SNRNP200 | None | None | None | | None |
| SNRNP200 | None | None | None | | None |
| CHRNA1 | None | None | None | | None |
| CHRNA1 | None | None | None | | None |
| MYOM1 | None | None | None | | None |
| MYOM1 | None | None | None | | None |
| MYOM1 | None | None | None | | None |
| MYOM1 | None | None | None | | None |
| MYOM1 | None | None | None | | None |
| MYOM1 | None | None | None | | None |

| | | | | |
|---|---|---|---|---|
| LDLR | None | None | None | None |
| LDLR | None | None | None | None |
| LDLR | None | None | None | None |
| LDLR | None | None | None | None |
| LDLR | None | None | None | None |
| LDLR | None | None | None | None |
| EHMT1 | None | None | None | None |
| EHMT1 | 0.545 | 3 | 3b7b | A |
| EHMT1 | None | None | None | None |
| COL9A2 | None | None | None | None |
| HFE2 | None | None | None | None |
| HFE2 | None | None | None | None |
| CD3G | None | None | None | None |
| MMAB | None | None | None | None |
| MMAB | 0.5 | 3 | 2idx | C |
| MMAB | 0.45 | 7 | 2idx | C |
| MMAB | 0.65 | 3 | 2idx | A |
| MMAB | 0.65 | 3 | 2idx | A |
| MMAB | 0.7 | 2 | 2idx | B |
| MMAB | 0.7 | 2 | 2idx | B |
| MMAB | 0.35 | 4 | 2idx | A |
| MMAB | 0.4 | 11 | 2idx | C |
| MMAB | None | None | None | None |
| TAF1 | None | None | None | None |
| TAF1 | None | None | None | None |
| CD4 | None | None | None | None |
| QARS | None | None | None | None |
| QARS | None | None | None | None |
| QARS | None | None | None | None |
| QARS | None | None | None | None |
| QARS | None | None | None | None |
| FGFR3 | None | None | None | None |
| FGFR3 | None | None | None | None |
| FGFR3 | None | None | None | None |
| FGFR3 | None | None | None | None |
| FGFR3 | None | None | None | None |
| FGFR3 | None | None | None | None |
| FLNB | None | None | None | None |
| FLNB | None | None | None | None |
| FLNB | None | None | None | None |
| FLNB | None | None | None | None |
| FLNB | None | None | None | None |
| FLNB | None | None | None | None |
| FLNB | None | None | None | None |
| FLNB | None | None | None | None |
| IL21R | None | None | None | None |
| RB1 | None | None | None | None |
| RB1 | None | None | None | None |
| RB1 | None | None | None | None |
| NTRK1 | 0.533 | 1 | 1he7 | A |
| NOTCH1 | None | None | None | None |
| MYOM1 | None | None | None | None |
| MYOM1 | None | None | None | None |
| MYOM1 | None | None | None | None |

| | | | | |
|---|---|---|---|---|
| MYOM1 | None | None | None | None |
| MYOM1 | None | None | None | None |
| MYOM1 | None | None | None | None |
| COQ9 | None | None | None | None |
| COQ9 | None | None | None | None |
| CRBN | None | None | None | None |
| DCTN1 | None | None | None | None |
| DCTN1 | None | None | None | None |
| DCTN1 | None | None | None | None |
| DCTN1 | None | None | None | None |
| DCTN1 | None | None | None | None |
| DCTN1 | 0.333 | 1 | 2hkn | A |
| THAP1 | None | None | None | None |
| THAP1 | None | None | None | None |
| GNAS | None | None | None | None |
| GNAS | None | None | None | None |
| HNF4A | None | None | None | None |
| HNF4A | None | None | None | None |
| ALDH3A2 | None | None | None | None |
| ALDH3A2 | None | None | None | None |
| ALDH3A2 | 0.412 | 5 | 4qgk | B |
| ALDH3A2 | 0.557 | 2 | 4qgk | A |
| ALDH3A2 | None | None | None | None |
| ALDH3A2 | 0.569 | 3 | 4qgk | B |
| ALDH3A2 | None | None | None | None |
| ALDH3A2 | None | None | None | None |
| ALDH3A2 | None | None | None | None |
| HCN1 | None | None | None | None |
| PLEC | None | None | None | None |
| SCN5A | None | None | None | None |
| SCN5A | None | None | None | None |
| STAMBP | None | None | None | None |
| STAMBP | None | None | None | None |
| RNF125 | None | None | None | None |
| RNF125 | None | None | None | None |
| USH1C | None | None | None | None |
| OBSL1 | None | None | None | None |
| GRIN2B | None | None | None | None |
| GRIN2B | None | None | None | None |
| HCN4 | None | None | None | None |
| HCN4 | None | None | None | None |
| TALDO1 | None | None | None | None |
| ITGB3 | None | None | None | None |
| CHKB | None | None | None | None |
| CHKB | None | None | None | None |
| CHKB | 0.333 | 7 | 3lq3 | A |
| CHKB | None | None | None | None |
| ACTN4 | None | None | None | None |
| ACTN4 | None | None | None | None |
| ACTN4 | None | None | None | None |
| ACTN4 | None | None | None | None |
| CDT1 | None | None | None | None |
| CDT1 | None | None | None | None |
| CDT1 | None | None | None | None |

| | | | | |
|---|---|---|---|---|
| CDT1 | None | None | None | None |
| CDT1 | None | None | None | None |
| CDT1 | None | None | None | None |
| MAPT | None | None | None | None |
| F5 | 0.517 | 4 | 1czs | A |
| F5 | None | None | None | None |
| F5 | None | None | None | None |
| ESRRB | None | None | None | None |
| DGUOK | None | None | None | None |
| DGUOK | 0.471 | 8 | 2ocp | A |
| DGUOK | None | None | None | None |
| DGUOK | 0.375 | 6 | 2ocp | B |
| DGUOK | 0.765 | 1 | 2ocp | A |
| DGUOK | 0.438 | 11 | 2ocp | B |
| DGUOK | None | None | None | None |
| DGUOK | None | None | None | None |
| DGUOK | None | None | None | None |
| CENPJ | None | None | None | None |
| TTN | None | None | None | None |
| TTN | None | None | None | None |
| TTN | None | None | None | None |
| TTN | None | None | None | None |
| TNFRSF1A | None | None | None | None |
| TNFRSF1A | None | None | None | None |
| TNFRSF1A | None | None | None | None |
| TNFRSF1A | None | None | None | None |
| TNFRSF1A | None | None | None | None |
| TNFRSF1A | None | None | None | None |
| TNFRSF1A | None | None | None | None |
| TNFRSF1A | None | None | None | None |
| TNFRSF1A | None | None | None | None |
| TNFRSF1A | None | None | None | None |
| TNFRSF1A | None | None | None | None |
| TNFRSF1A | None | None | None | None |
| ASL | None | None | None | None |
| ASL | 0.625 | 25 | 1aos | A |
| ASL | None | None | None | None |
| ASL | 0.375 | 28 | 1k62 | A |
| ASL | 0.438 | 4 | 1k62 | B |
| ASL | 0.562 | 30 | 1k62 | A |
| ASL | 0.625 | 1 | 1aos | A |
| ASL | None | None | None | None |
| ASL | None | None | None | None |
| ASL | None | None | None | None |
| MYO7A | None | None | None | None |
| MYO7A | None | None | None | None |
| MYO7A | None | None | None | None |
| MYO7A | None | None | None | None |
| MYO7A | None | None | None | None |
| MYO7A | None | None | None | None |
| MYO7A | None | None | None | None |
| MYO7A | None | None | None | None |
| MYO7A | None | None | None | None |
| MYO7A | None | None | None | None |

| | | | | |
|---|---|---|---|---|
| MYO7A | None | None | None | None |
| MYO7A | None | None | None | None |
| MYO7A | None | None | None | None |
| MYO7A | None | None | None | None |
| MYO7A | None | None | None | None |
| MYO7A | None | None | None | None |
| MYO7A | None | None | None | None |
| MYO7A | None | None | None | None |
| MYO7A | None | None | None | None |
| MYO7A | None | None | None | None |
| APC | None | None | None | None |
| ALAD | None | None | None | None |
| ALAD | None | None | None | None |
| ALAD | None | None | None | None |
| ALAD | 0.542 | 1 | 1E+051 | A |
| ALAD | None | None | None | None |
| ALAD | None | None | None | None |
| POR | None | None | None | None |
| CSF1R | None | None | None | None |
| CSF1R | None | None | None | None |
| CSF1R | None | None | None | None |
| CSF1R | None | None | None | None |
| CSF1R | None | None | None | None |
| CSF1R | None | None | None | None |
| CSF1R | None | None | None | None |
| CSF1R | None | None | None | None |
| CSF1R | None | None | None | None |
| CSF1R | None | None | None | None |
| CSF1R | None | None | None | None |
| VWF | None | None | None | None |
| VWF | None | None | None | None |
| PHF8 | 0.333 | 6 | 3k3n | A |
| ACY1 | None | None | None | None |
| SMN1 | None | None | None | None |
| SMN1 | None | None | None | None |
| DLAT | None | None | None | None |
| DLAT | None | None | None | None |
| B2M | None | None | None | None |
| SOX9 | None | None | None | None |
| SOX9 | None | None | None | None |
| SOX9 | None | None | None | None |
| SOX9 | None | None | None | None |
| SOX9 | None | None | None | None |
| CFH | None | None | None | None |
| FLNC | None | None | None | None |
| FLNC | None | None | None | None |
| FLNC | None | None | None | None |
| ITGB2 | None | None | None | None |
| ITGB2 | None | None | None | None |
| ITGB2 | None | None | None | None |
| ITGB2 | None | None | None | None |
| ITGB2 | None | None | None | None |
| ITGB2 | None | None | None | None |
| ITGB2 | None | None | None | None |

| | | | | |
|---|---|---|---|---|
| ITGB2 | None | None | None | None |
| ITGB2 | None | None | None | None |
| PSEN1 | None | None | None | None |
| PSEN1 | None | None | None | None |
| PSEN1 | None | None | None | None |
| PSEN1 | None | None | None | None |
| PSEN1 | None | None | None | None |
| PSEN1 | None | None | None | None |
| OXCT1 | None | None | None | None |
| OXCT1 | 0.385 | 24 | 3dlx | A |
| OXCT1 | None | None | None | None |
| OXCT1 | None | None | None | None |
| OXCT1 | None | None | None | None |
| OXCT1 | None | None | None | None |
| OXCT1 | None | None | None | None |
| DHH | None | None | None | None |
| PRKN | None | None | None | None |
| PRKN | None | None | None | None |
| PRKN | None | None | None | None |
| PRKN | None | None | None | None |
| PRKN | None | None | None | None |
| PRKN | None | None | None | None |
| PRKN | None | None | None | None |
| PRKN | None | None | None | None |
| PRKN | None | None | None | None |
| PRKN | None | None | None | None |
| PRKN | None | None | None | None |
| LCAT | None | None | None | None |
| LCAT | None | None | None | None |
| LCAT | None | None | None | None |
| LCAT | None | None | None | None |
| LCAT | None | None | None | None |
| LCAT | None | None | None | None |
| LCAT | None | None | None | None |
| LCAT | None | None | None | None |
| LCAT | None | None | None | None |
| LCAT | None | None | None | None |
| LCAT | None | None | None | None |
| LCAT | None | None | None | None |
| FOXP3 | None | None | None | None |
| FOXP3 | None | None | None | None |
| FOXP3 | None | None | None | None |
| FOXP3 | None | None | None | None |
| SUFU | None | None | None | None |
| HSPA9 | 0.571 | 14 | 4kbo | A |
| HSPA9 | None | None | None | None |
| SQSTM1 | None | None | None | None |
| DARS2 | None | None | None | None |
| DARS2 | None | None | None | None |
| DARS2 | None | None | None | None |
| DARS2 | None | None | None | None |
| DARS2 | 0.412 | 2 | 4ah6 | A |
| DARS2 | None | None | None | None |
| DARS2 | None | None | None | None |

| | | | | |
|---|---|---|---|---|
| DARS2 | None | None | None | None |
| DARS2 | None | None | None | None |
| DARS2 | None | None | None | None |
| DARS2 | None | None | None | None |
| DARS2 | None | None | None | None |
| NT5C3A | None | None | None | None |
| NT5C3A | 0.4 | 18 | 2vkq | A |
| NT5C3A | 0.4 | 7 | 2vkq | A |
| NT5C3A | 0.6 | 14 | 2vkq | A |
| ETFA | None | None | None | None |
| ETFA | None | None | None | None |
| ETFA | None | None | None | None |
| ETFA | None | None | None | None |
| ETFA | None | None | None | None |
| ETFA | None | None | None | None |
| ETFA | None | None | None | None |
| TTN | None | None | None | None |
| TTN | None | None | None | None |
| FLT4 | None | None | None | None |
| FGB | None | None | None | None |
| HPCA | None | None | None | None |
| TAP2 | None | None | None | None |
| TAP2 | None | None | None | None |
| PTPRC | None | None | None | None |
| SLC4A1 | None | None | None | None |
| SLC4A1 | None | None | None | None |
| SLC4A1 | None | None | None | None |
| SLC4A1 | None | None | None | None |
| SLC4A1 | None | None | None | None |
| SLC4A1 | None | None | None | None |
| MYH7 | None | None | None | None |
| MYH7 | None | None | None | None |
| MYH7 | None | None | None | None |
| MYH7 | None | None | None | None |
| MYH7 | None | None | None | None |
| MYH7 | None | None | None | None |
| MYH7 | None | None | None | None |
| MYH7 | None | None | None | None |
| MYH7 | None | None | None | None |
| PARN | None | None | None | None |
| PARN | None | None | None | None |
| NDUFS6 | None | None | None | None |
| VRK1 | None | None | None | None |
| VRK1 | None | None | None | None |
| VRK1 | None | None | None | None |
| VRK1 | None | None | None | None |
| VRK1 | None | None | None | None |
| VRK1 | None | None | None | None |
| COLEC11 | None | None | None | None |
| VHL | None | None | None | None |
| TTN | None | None | None | None |
| NCF2 | None | None | None | None |
| NCF2 | None | None | None | None |
| NCF2 | None | None | None | None |

| | | | | |
|---|---|---|---|---|
| NCF2 | None | None | None | None |
| NCF2 | None | None | None | None |
| RPS26 | None | None | None | None |
| CETP | None | None | None | None |
| CETP | None | None | None | None |
| CETP | None | None | None | None |
| CETP | None | None | None | None |
| CETP | None | None | None | None |
| CETP | None | None | None | None |
| CETP | None | None | None | None |
| CETP | None | None | None | None |
| CETP | None | None | None | None |
| CETP | None | None | None | None |
| CETP | None | None | None | None |
| CETP | None | None | None | None |
| CETP | None | None | None | None |
| FLNA | None | None | None | None |
| FLNA | None | None | None | None |
| FLNA | None | None | None | None |
| FLNA | None | None | None | None |
| FLNA | None | None | None | None |
| FLNA | None | None | None | None |
| FLNA | None | None | None | None |
| MYBPC3 | None | None | None | None |
| MYBPC3 | None | None | None | None |
| MYBPC3 | None | None | None | None |
| MYBPC3 | None | None | None | None |
| MYBPC3 | None | None | None | None |
| MYBPC3 | None | None | None | None |
| MYBPC3 | None | None | None | None |
| MYBPC3 | None | None | None | None |
| MYBPC3 | None | None | None | None |
| MYBPC3 | None | None | None | None |
| LSS | None | None | None | None |
| C6 | None | None | None | None |
| C6 | None | None | None | None |
| RLBP1 | None | None | None | None |
| RLBP1 | None | None | None | None |
| RLBP1 | None | None | None | None |
| RLBP1 | None | None | None | None |
| RLBP1 | None | None | None | None |
| RLBP1 | None | None | None | None |
| RLBP1 | None | None | None | None |
| RLBP1 | None | None | None | None |
| MMAA | None | None | None | None |
| MMAA | 0.3 | 1 | 2www | C |
| MMAA | None | None | None | None |
| MMAA | None | None | None | None |
| MMAA | None | None | None | None |
| MMAA | None | None | None | None |
| IL1RN | None | None | None | None |
| MASTL | None | None | None | None |
| FHL1 | None | None | None | None |
| FHL1 | None | None | None | None |

| | | | | |
|---|---|---|---|---|
| FHL1 | None | None | None | None |
| FHL1 | None | None | None | None |
| FHL1 | None | None | None | None |
| FHL1 | None | None | None | None |
| FHL1 | None | None | None | None |
| FHL1 | None | None | None | None |
| FHL1 | None | None | None | None |
| EIF2B1 | 0.636 | 1 | 3ecs | H |
| EIF2B1 | 0.636 | 1 | 3ecs | H |
| COQ8A | None | None | None | None |
| COQ8A | None | None | None | None |
| COQ8A | None | None | None | None |
| COQ8A | None | None | None | None |
| COQ8A | None | None | None | None |
| COQ8A | None | None | None | None |
| COQ8A | None | None | None | None |
| COQ8A | None | None | None | None |
| COQ8A | None | None | None | None |
| COQ8A | None | None | None | None |
| COQ8A | None | None | None | None |
| COQ8A | None | None | None | None |
| COQ8A | None | None | None | None |
| COQ8A | None | None | None | None |
| COQ8A | None | None | None | None |
| COQ8A | None | None | None | None |
| UNC119 | None | None | None | None |
| IGHMBP2 | None | None | None | None |
| JAG1 | None | None | None | None |
| JAG1 | None | None | None | None |
| JAG1 | None | None | None | None |
| GNB1 | None | None | None | None |
| GNB1 | 0.3 | 3 | 5ukl | B |
| FLNA | None | None | None | None |
| FLNA | None | None | None | None |
| FLNA | None | None | None | None |
| FLNA | None | None | None | None |
| FLNA | None | None | None | None |
| CREBBP | None | None | None | None |
| CSF3R | None | None | None | None |
| CSF3R | None | None | None | None |
| HIBCH | None | None | None | None |
| HIBCH | None | None | None | None |
| HIBCH | None | None | None | None |
| MMACHC | None | None | None | None |
| MMACHC | None | None | None | None |
| MMACHC | None | None | None | None |
| MMACHC | None | None | None | None |
| MMACHC | None | None | None | None |
| MMACHC | None | None | None | None |
| MMACHC | None | None | None | None |
| MMACHC | None | None | None | None |
| MMACHC | None | None | None | None |
| MMACHC | None | None | None | None |
| MMACHC | None | None | None | None |

| | | | | |
|---|---|---|---|---|
| KLF1 | None | None | None | None |
| NAGS | None | None | None | None |
| NAGS | None | None | None | None |
| NAGS | None | None | None | None |
| PIK3R2 | None | None | None | None |
| EDNRB | None | None | None | None |
| EDNRB | None | None | None | None |
| EDNRB | None | None | None | None |
| EDNRB | None | None | None | None |
| E | None | None | None | None |
| RUNX1 | None | None | None | None |
| RUNX1 | None | None | None | None |
| CTNNB1 | None | None | None | None |
| CTNNB1 | None | None | None | None |
| CTNNB1 | None | None | None | None |
| CTNNB1 | None | None | None | None |
| CTNNB1 | None | None | None | None |
| CTNNB1 | None | None | None | None |
| TTN | None | None | None | None |
| TTN | None | None | None | None |
| TYK2 | None | None | None | None |
| TYK2 | None | None | None | None |
| TYK2 | None | None | None | None |
| ERCC4 | None | None | None | None |
| ERCC4 | None | None | None | None |
| ERCC4 | None | None | None | None |
| ERCC4 | None | None | None | None |
| CFI | None | None | None | None |
| CFI | None | None | None | None |
| CFI | None | None | None | None |
| CFI | None | None | None | None |
| CFI | None | None | None | None |
| CFI | None | None | None | None |
| CFI | None | None | None | None |
| CFI | None | None | None | None |
| CFI | None | None | None | None |
| CFI | 0.301 | 5 | 2xrc | A |
| CFI | None | None | None | None |
| CFI | 0.691 | 2 | 5o32 | J |
| CFI | 0.691 | 2 | 5o32 | J |
| CFI | 0.524 | 5 | 5o32 | I |
| NOTCH3 | None | None | None | None |
| NOTCH3 | None | None | None | None |
| NOTCH3 | None | None | None | None |
| NOTCH3 | None | None | None | None |
| TBX1 | None | None | None | None |
| TBX1 | None | None | None | None |
| DDB2 | None | None | None | None |
| DDB2 | None | None | None | None |
| TP53 | None | None | None | None |
| POMGNT1 | None | None | None | None |
| POMGNT1 | None | None | None | None |
| POMGNT1 | None | None | None | None |
| POMGNT1 | None | None | None | None |

| | | | | |
|---|---|---|---|---|
| POMGNT1 | None | None | None | None |
| POMGNT1 | None | None | None | None |
| POMGNT1 | None | None | None | None |
| POMGNT1 | None | None | None | None |
| POMGNT1 | None | None | None | None |
| POMGNT1 | None | None | None | None |
| POMGNT1 | None | None | None | None |
| POMGNT1 | None | None | None | None |
| POMGNT1 | None | None | None | None |
| FMR1 | None | None | None | None |
| FMR1 | None | None | None | None |
| SLC9A3R1 | None | None | None | None |
| HSPB1 | None | None | None | None |
| HSPB1 | None | None | None | None |
| HSPB1 | None | None | None | None |
| HSPB1 | None | None | None | None |
| HSPB1 | None | None | None | None |
| HSPB1 | None | None | None | None |
| HSPB1 | None | None | None | None |
| SOD3 | None | None | None | None |
| AGA | 1 | 2 | 1apz | D |
| AGA | None | None | None | None |
| AGA | None | None | None | None |
| NDUFS7 | None | None | None | None |
| NDUFS7 | None | None | None | None |
| SOX2 | None | None | None | None |
| CSRP3 | None | None | None | None |
| CSRP3 | None | None | None | None |
| PAH | None | None | None | None |
| PAH | None | None | None | None |
| PAH | None | None | None | None |
| PAH | None | None | None | None |
| CACNA1S | None | None | None | None |
| TRPV4 | None | None | None | None |
| TRPV4 | None | None | None | None |
| TRPV4 | None | None | None | None |
| TRPV4 | 0.58 | 3 | 4dx2 | A |
| TRPV4 | None | None | None | None |
| TRPV4 | None | None | None | None |
| TRPV4 | None | None | None | None |
| TRPV4 | None | None | None | None |
| TRPV4 | None | None | None | None |
| TRPV4 | None | None | None | None |
| TRPV4 | None | None | None | None |
| TRPV4 | None | None | None | None |
| TRPV4 | None | None | None | None |
| TNNI3 | None | None | None | None |
| MLYCD | None | None | None | None |
| MLYCD | None | None | None | None |
| MLYCD | None | None | None | None |
| MLYCD | None | None | None | None |
| AIP | None | None | None | None |
| AIP | None | None | None | None |
| ERCC2 | None | None | None | None |

| | | | | |
|---|---|---|---|---|
| ERCC2 | None | None | None | None |
| ERCC2 | None | None | None | None |
| ERCC2 | None | None | None | None |
| ERCC2 | None | None | None | None |
| ERCC2 | None | None | None | None |
| ERCC2 | None | None | None | None |
| ERCC2 | None | None | None | None |
| ERCC2 | None | None | None | None |
| ERCC2 | None | None | None | None |
| ERCC2 | None | None | None | None |
| ERCC2 | None | None | None | None |
| ERCC2 | None | None | None | None |
| ERCC2 | None | None | None | None |
| PUS1 | None | None | None | None |
| PUS1 | None | None | None | None |
| PUS1 | None | None | None | None |
| CASK | None | None | None | None |
| CASK | None | None | None | None |
| BLVRA | None | None | None | None |
| DICER1 | None | None | None | None |
| DICER1 | None | None | None | None |
| DICER1 | None | None | None | None |
| EIF2AK3 | None | None | None | None |
| EIF2AK3 | None | None | None | None |
| EIF2AK3 | None | None | None | None |
| KIT | None | None | None | None |
| KIT | None | None | None | None |
| KIT | None | None | None | None |
| KIT | None | None | None | None |
| KIT | None | None | None | None |
| KIT | None | None | None | None |
| DNMT3B | None | None | None | None |
| TGM1 | None | None | None | None |
| NDUFA10 | None | None | None | None |
| NDUFA10 | None | None | None | None |
| PRPF31 | None | None | None | None |
| PRPF31 | None | None | None | None |
| ECE1 | None | None | None | None |
| ECE1 | None | None | None | None |
| ECE1 | None | None | None | None |
| NDUFV2 | None | None | None | None |
| NDUFV2 | None | None | None | None |
| NDUFV2 | None | None | None | None |
| GHRHR | None | None | None | None |
| GHRHR | None | None | None | None |
| GHRHR | None | None | None | None |
| SLC4A1 | None | None | None | None |
| FGF9 | None | None | None | None |
| BARD1 | None | None | None | None |
| BARD1 | None | None | None | None |
| BARD1 | None | None | None | None |
| PTPN12 | 0.741 | 2 | 5j8r | D |
| MASP2 | None | None | None | None |
| MASP2 | None | None | None | None |

| | | | | |
|---|---|---|---|---|
| MASP2 | None | None | None | None |
| MASP2 | None | None | None | None |
| HOXB1 | None | None | None | None |
| SLC4A1 | None | None | None | None |
| SLC4A1 | None | None | None | None |
| SLC4A1 | None | None | None | None |
| SLC4A1 | None | None | None | None |
| SLC4A1 | None | None | None | None |
| SLC4A1 | None | None | None | None |
| SLC4A1 | None | None | None | None |
| SLC4A1 | None | None | None | None |
| SLC4A1 | None | None | None | None |
| SLC4A1 | None | None | None | None |
| SLC4A1 | None | None | None | None |
| SLC4A1 | None | None | None | None |
| SLC4A1 | None | None | None | None |
| SLC4A1 | None | None | None | None |
| SLC4A1 | None | None | None | None |
| SLC4A1 | None | None | None | None |
| SLC4A1 | None | None | None | None |
| CBL | None | None | None | None |
| FAS | None | None | None | None |
| MTMR2 | None | None | None | None |
| MTMR2 | None | None | None | None |
| MTMR2 | None | None | None | None |
| MTMR2 | None | None | None | None |
| MTMR2 | None | None | None | None |
| MTMR2 | None | None | None | None |
| MTMR2 | None | None | None | None |
| MTMR2 | None | None | None | None |
| KLHL3 | None | None | None | None |
| KLHL3 | None | None | None | None |
| KLHL3 | 0.667 | 1 | 4ch9 | B |
| KLHL3 | 0.733 | 1 | 5nkp | A |
| KLHL3 | None | None | None | None |
| KLHL3 | None | None | None | None |
| RNASEH1 | None | None | None | None |
| ACTN1 | None | None | None | None |
| PHKG2 | None | None | None | None |
| PHKG2 | None | None | None | None |
| PHKG2 | 0.472 | 2 | 2y7j | A |
| NTRK1 | None | None | None | None |
| APOH | None | None | None | None |
| PMS2 | None | None | None | None |
| PMS2 | None | None | None | None |
| PMS2 | 0.797 | 1 | 1ea6 | A |
| PMS2 | None | None | None | None |
| PMS2 | None | None | None | None |
| PMS2 | None | None | None | None |
| PMS2 | None | None | None | None |
| PMS2 | None | None | None | None |
| PMS2 | None | None | None | None |
| PMS2 | None | None | None | None |
| PMS2 | None | None | None | None |

| | | | | |
|---|---|---|---|---|
| PMS2 | None | None | None | None |
| PMS2 | None | None | None | None |
| PMS2 | None | None | None | None |
| PMS2 | None | None | None | None |
| PMS2 | None | None | None | None |
| PMS2 | None | None | None | None |
| PMS2 | None | None | None | None |
| PMS2 | None | None | None | None |
| PMS2 | None | None | None | None |
| PMS2 | None | None | None | None |
| PMS2 | None | None | None | None |
| MOCS2 | None | None | None | None |
| MOCS2 | None | None | None | None |
| RPGRIP1 | None | None | None | None |
| RPGRIP1 | None | None | None | None |
| LIG4 | None | None | None | None |
| PROC | None | None | None | None |
| PROC | None | None | None | None |
| NAT1 | None | None | None | None |
| NAT1 | None | None | None | None |
| PNPO | None | None | None | None |
| PNPO | 0.5 | 2 | 1nrg | A |
| PNPO | None | None | None | None |
| PNPO | None | None | None | None |
| PNPO | None | None | None | None |
| MRE11 | None | None | None | None |
| MRE11 | None | None | None | None |
| MRE11 | None | None | None | None |
| MRE11 | None | None | None | None |
| DDC | None | None | None | None |
| DDC | None | None | None | None |
| DDC | None | None | None | None |
| DDC | None | None | None | None |
| DDC | None | None | None | None |
| DDC | 0.909 | 7 | 3rch | B |
| DDC | 0.909 | 7 | 3rch | B |
| DDC | 0.643 | 4 | 3rbf | A |
| DDC | None | None | None | None |
| DDC | None | None | None | None |
| DDC | None | None | None | None |
| ANK2 | None | None | None | None |
| ANK2 | None | None | None | None |
| ANK2 | None | None | None | None |
| AK2 | None | None | None | None |
| AK2 | None | None | None | None |
| AK2 | None | None | None | None |
| AK2 | None | None | None | None |
| AK2 | None | None | None | None |
| AK2 | None | None | None | None |
| GPHN | None | None | None | None |
| HSD17B4 | 0.66 | 1 | 1zbq | A |
| HSD17B4 | 0.66 | 1 | 1zbq | A |
| HSD17B4 | None | None | None | None |
| HSD17B4 | None | None | None | None |

| | | | | |
|---|---|---|---|---|
| HSD17B4 | None | None | None | None |
| HSD17B4 | None | None | None | None |
| APTX | 0.6 | 5 | 4ndg | A |
| APTX | 0.6 | 5 | 4ndg | A |
| APTX | None | None | None | None |
| APTX | None | None | None | None |
| APTX | None | None | None | None |
| APTX | None | None | None | None |
| APTX | None | None | None | None |
| FANCL | None | None | None | None |
| FANCL | None | None | None | None |
| EMG1 | None | None | None | None |
| EMG1 | None | None | None | None |
| EZH2 | None | None | None | None |
| ABCB6 | None | None | None | None |
| NCF2 | None | None | None | None |
| NCF2 | None | None | None | None |
| COL4A2 | None | None | None | None |
| COL4A2 | 1 | 1 | 1li1 | F |
| COL4A2 | None | None | None | None |
| COL4A2 | None | None | None | None |
| COL4A2 | 0.75 | 16 | 1li1 | C |
| SLX4 | None | None | None | None |
| SERPIND1 | None | None | None | None |
| FGFR3 | None | None | None | None |
| FGFR3 | None | None | None | None |
| FGFR3 | None | None | None | None |
| FGFR3 | None | None | None | None |
| FGFR3 | None | None | None | None |
| FGFR3 | None | None | None | None |
| TREM2 | None | None | None | None |
| TREM2 | None | None | None | None |
| TREM2 | None | None | None | None |
| TREM2 | None | None | None | None |
| TREM2 | None | None | None | None |
| TREM2 | None | None | None | None |
| TREM2 | None | None | None | None |
| LIG4 | None | None | None | None |
| LIG4 | None | None | None | None |
| LIG4 | None | None | None | None |
| LIG4 | None | None | None | None |
| LIG4 | None | None | None | None |
| LIG4 | None | None | None | None |
| ECHS1 | None | None | None | None |
| ECHS1 | None | None | None | None |
| ECHS1 | None | None | None | None |
| MTR | None | None | None | None |
| MTR | None | None | None | None |
| MTR | None | None | None | None |
| MTR | None | None | None | None |
| MTR | None | None | None | None |
| PMVK | None | None | None | None |
| PRNP | None | None | None | None |
| CHN1 | None | None | None | None |

| | | | | |
|---|---|---|---|---|
| CHN1 | None | None | None | None |
| CHN1 | None | None | None | None |
| CHN1 | None | None | None | None |
| CHN1 | None | None | None | None |
| CHN1 | None | None | None | None |
| CHN1 | None | None | None | None |
| CHN1 | None | None | None | None |
| SLC2A1 | None | None | None | None |
| SLC2A1 | None | None | None | None |
| SLC2A1 | None | None | None | None |
| SLC2A1 | None | None | None | None |
| SLC2A1 | None | None | None | None |
| SLC2A1 | None | None | None | None |
| SLC2A1 | None | None | None | None |
| SLC2A1 | None | None | None | None |
| SLC2A1 | None | None | None | None |
| SLC2A1 | None | None | None | None |
| SLC2A1 | None | None | None | None |
| SLC2A1 | None | None | None | None |
| SLC2A1 | None | None | None | None |
| SLC2A1 | None | None | None | None |
| SLC2A1 | None | None | None | None |
| SLC2A1 | None | None | None | None |
| SLC2A1 | None | None | None | None |
| PMM2 | None | None | None | None |
| PMM2 | None | None | None | None |
| PMM2 | None | None | None | None |
| PMM2 | None | None | None | None |
| PMM2 | None | None | None | None |
| PMM2 | None | None | None | None |
| PMM2 | None | None | None | None |
| PMM2 | None | None | None | None |
| PMM2 | None | None | None | None |
| PMM2 | None | None | None | None |
| PMM2 | None | None | None | None |
| PMM2 | None | None | None | None |
| PMM2 | None | None | None | None |
| PMM2 | None | None | None | None |
| PMM2 | None | None | None | None |
| PMM2 | None | None | None | None |
| PMM2 | None | None | None | None |
| PMM2 | None | None | None | None |
| PMM2 | None | None | None | None |
| PMM2 | None | None | None | None |
| PMM2 | None | None | None | None |
| PMM2 | None | None | None | None |
| PMM2 | None | None | None | None |
| PMM2 | None | None | None | None |
| PMM2 | None | None | None | None |
| PMM2 | None | None | None | None |
| PMM2 | None | None | None | None |
| PMM2 | None | None | None | None |
| CCKAR | None | None | None | None |
| MAN1B1 | None | None | None | None |

| | | | | |
|---|---|---|---|---|
| MAN1B1 | None | None | None | None |
| LAMA2 | None | None | None | None |
| LAMA2 | None | None | None | None |
| TBC1D4 | None | None | None | None |
| TBC1D4 | None | None | None | None |
| LDLR | None | None | None | None |
| LDLR | None | None | None | None |
| LDLR | None | None | None | None |
| LDLR | None | None | None | None |
| LDLR | None | None | None | None |
| LDLR | None | None | None | None |
| LDLR | None | None | None | None |
| LDLR | None | None | None | None |
| LDLR | None | None | None | None |
| LDLR | None | None | None | None |
| LDLR | None | None | None | None |
| LDLR | None | None | None | None |
| LDLR | None | None | None | None |
| LDLR | None | None | None | None |
| LDLR | None | None | None | None |
| LDLR | None | None | None | None |
| LDLR | None | None | None | None |
| LDLR | None | None | None | None |
| LDLR | None | None | None | None |
| LDLR | None | None | None | None |
| LDLR | None | None | None | None |
| LDLR | None | None | None | None |
| LDLR | None | None | None | None |
| LDLR | None | None | None | None |
| LDLR | None | None | None | None |
| LDLR | None | None | None | None |
| LDLR | None | None | None | None |
| LDLR | None | None | None | None |
| LDLR | None | None | None | None |
| LDLR | None | None | None | None |
| LDLR | None | None | None | None |
| LDLR | None | None | None | None |
| LDLR | None | None | None | None |
| LDLR | None | None | None | None |
| LDLR | None | None | None | None |
| LDLR | None | None | None | None |
| LDLR | None | None | None | None |
| LDLR | None | None | None | None |
| LDLR | None | None | None | None |
| LDLR | None | None | None | None |
| LDLR | | 0.5 | 1 | 1ijq | A |
| LDLR | None | None | None | None |
| FANCM | None | None | None | None |
| HAMP | None | None | None | None |
| GUSB | | 0.462 | 8 | 3hn3 | A |
| GUSB | None | None | None | None |
| GUSB | | 0.45 | 16 | 3hn3 | B |
| GUSB | None | None | None | None |

| | | | | |
|---|---|---|---|---|
| GUSB | None | None | None | None |
| GUSB | None | None | None | None |
| GUSB | None | None | None | None |
| GUSB | None | None | None | None |
| GUSB | None | None | None | None |
| GUSB | None | None | None | None |
| GUSB | None | None | None | None |
| GUSB | None | None | None | None |
| GUSB | None | None | None | None |
| LIG1 | None | None | None | None |
| LIG1 | None | None | None | None |
| MYH7 | None | None | None | None |
| MYH7 | None | None | None | None |
| MYH7 | None | None | None | None |
| MYH7 | None | None | None | None |
| MYH7 | None | None | None | None |
| MYH7 | None | None | None | None |
| MYH7 | None | None | None | None |
| MYH7 | None | None | None | None |
| MYH7 | None | None | None | None |
| MYH7 | None | None | None | None |
| MYH7 | None | None | None | None |
| MYH7 | None | None | None | None |
| MYH7 | None | None | None | None |
| MYH7 | None | None | None | None |
| MYH7 | None | None | None | None |
| MYH7 | None | None | None | None |
| MYH7 | None | None | None | None |
| MYH7 | None | None | None | None |
| MYH7 | None | None | None | None |
| MYH7 | None | None | None | None |
| MYH7 | None | None | None | None |
| MYH7 | None | None | None | None |
| MYH7 | None | None | None | None |
| MYH7 | None | None | None | None |
| MYH7 | None | None | None | None |
| MYH7 | None | None | None | None |
| MYH7 | None | None | None | None |
| MYH7 | None | None | None | None |
| MYH7 | None | None | None | None |
| MYH7 | None | None | None | None |
| MYH7 | None | None | None | None |
| MYH7 | None | None | None | None |
| MYH7 | None | None | None | None |
| MYH7 | None | None | None | None |
| HCN4 | None | None | None | None |
| RECQL4 | None | None | None | None |
| RECQL4 | None | None | None | None |
| RECQL4 | None | None | None | None |
| RECQL4 | None | None | None | None |
| RECQL4 | None | None | None | None |
| RECQL4 | None | None | None | None |
| RECQL4 | None | None | None | None |
| RECQL4 | None | None | None | None |

| | | | | |
|---|---|---|---|---|
| RECQL4 | None | None | None | None |
| RECQL4 | None | None | None | None |
| RECQL4 | None | None | None | None |
| RECQL4 | None | None | None | None |
| RECQL4 | None | None | None | None |
| CHEK2 | None | None | None | None |
| PPARG | None | None | None | None |
| ACADVL | None | None | None | None |
| ACADVL | 0.385 | 4 | 3b96 | A |
| ACADVL | None | None | None | None |
| ACADVL | None | None | None | None |
| ACADVL | None | None | None | None |
| ACADVL | None | None | None | None |
| ACADVL | None | None | None | None |
| ACADVL | 0.346 | 14 | 3b96 | A |
| ACADVL | 0.346 | 14 | 3b96 | A |
| ACADVL | None | None | None | None |
| ACADVL | None | None | None | None |
| ACADVL | None | None | None | None |
| ACADVL | None | None | None | None |
| ACADVL | 0.346 | 14 | 3b96 | A |
| ACADVL | None | None | None | None |
| ACADVL | 0.423 | 2 | 3b96 | A |
| ACADVL | None | None | None | None |
| ACADVL | None | None | None | None |
| ACADVL | None | None | None | None |
| ACADVL | None | None | None | None |
| ACADVL | None | None | None | None |
| ACADVL | None | None | None | None |
| APOA1 | None | None | None | None |
| APOA1 | None | None | None | None |
| APOA1 | None | None | None | None |
| APOA1 | None | None | None | None |
| APOA1 | None | None | None | None |
| DNMT3A | None | None | None | None |
| DNMT3A | None | None | None | None |
| APC | None | None | None | None |
| TMOD1 | None | None | None | None |
| USH1C | None | None | None | None |
| USH1C | None | None | None | None |
| USH1C | None | None | None | None |
| USH1C | None | None | None | None |
| ERCC6 | None | None | None | None |
| ERCC6 | None | None | None | None |
| ERCC6 | None | None | None | None |
| ERCC8 | None | None | None | None |
| ERCC8 | None | None | None | None |
| ERCC8 | None | None | None | None |
| DPAGT1 | None | None | None | None |
| DPAGT1 | None | None | None | None |
| DPAGT1 | None | None | None | None |
| DPAGT1 | None | None | None | None |
| DPAGT1 | None | None | None | None |
| DPAGT1 | None | None | None | None |

| | | | | |
|---|---|---|---|---|
| DPAGT1 | None | None | None | None |
| IL2RG | None | None | None | None |
| HNF1A | None | None | None | None |
| HNF4A | None | None | None | None |
| FLT4 | None | None | None | None |
| FLT4 | None | None | None | None |
| FLT4 | None | None | None | None |
| CD3D | None | None | None | None |
| CD3D | None | None | None | None |
| MCOLN1 | None | None | None | None |
| MCOLN1 | None | None | None | None |
| MOCS2 | None | None | None | None |
| MOCS2 | None | None | None | None |
| ABCG8 | None | None | None | None |
| ABCG8 | None | None | None | None |
| ABCG8 | None | None | None | None |
| ABCG8 | None | None | None | None |
| ABCG8 | None | None | None | None |
| ABCG8 | None | None | None | None |
| ABCG8 | None | None | None | None |
| ABCG8 | None | None | None | None |
| ABCG8 | None | None | None | None |
| ABCG8 | None | None | None | None |
| ABCG8 | None | None | None | None |
| ABCG8 | None | None | None | None |
| ABCG8 | None | None | None | None |
| ABCG8 | None | None | None | None |
| CTNNA1 | None | None | None | None |
| TYMP | None | None | None | None |
| TYMP | None | None | None | None |
| TYMP | None | None | None | None |
| TYMP | None | None | None | None |
| TYMP | None | None | None | None |
| TYMP | None | None | None | None |
| TYMP | None | None | None | None |
| TYMP | None | None | None | None |
| TYMP | None | None | None | None |
| TYMP | None | None | None | None |
| TYMP | None | None | None | None |
| TYMP | None | None | None | None |
| TYMP | None | None | None | None |
| GAMT | None | None | None | None |
| GAMT | None | None | None | None |
| GAMT | None | None | None | None |
| GAMT | None | None | None | None |
| GAMT | None | None | None | None |
| GAMT | None | None | None | None |
| GAMT | None | None | None | None |
| GAMT | None | None | None | None |
| GAMT | None | None | None | None |
| GAMT | None | None | None | None |
| GAMT | None | None | None | None |
| GLRA1 | None | None | None | None |
| GLRA1 | None | None | None | None |

| | | | | |
|---|---|---|---|---|
| GLRA1 | None | None | None | None |
| GLRA1 | None | None | None | None |
| GLRA1 | None | None | None | None |
| GLRA1 | None | None | None | None |
| GLRA1 | None | None | None | None |
| GLRA1 | None | None | None | None |
| GLRA1 | None | None | None | None |
| ADK | None | None | None | None |
| ACTN2 | None | None | None | None |
| UBE2T | None | None | None | None |
| TTN | None | None | None | None |
| ATP7B | None | None | None | None |
| ATP7B | None | None | None | None |
| ATP7B | None | None | None | None |
| ATP7B | None | None | None | None |
| ATP7B | None | None | None | None |
| ATP7B | None | None | None | None |
| ATP7B | None | None | None | None |
| ATP7B | None | None | None | None |
| ATP7B | None | None | None | None |
| ATP7B | None | None | None | None |
| ATP7B | None | None | None | None |
| ATP7B | None | None | None | None |
| ATP7B | None | None | None | None |
| CDKN1B | None | None | None | None |
| CDKN1B | None | None | None | None |
| CDKN1B | None | None | None | None |
| CDKN1B | None | None | None | None |
| CDKN1B | None | None | None | None |
| CDKN1B | None | None | None | None |
| MYOC | None | None | None | None |
| MYOC | None | None | None | None |
| MYOC | None | None | None | None |
| MYOC | None | None | None | None |
| MYOC | None | None | None | None |
| MYOC | None | None | None | None |
| MYOC | None | None | None | None |
| MYOC | None | None | None | None |
| MYOC | None | None | None | None |
| MYOC | None | None | None | None |
| MYOC | None | None | None | None |
| MYOC | None | None | None | None |
| MYOC | None | None | None | None |
| MYOC | None | None | None | None |
| MYOC | None | None | None | None |
| SPTAN1 | None | None | None | None |
| GC | None | None | None | None |
| GC | None | None | None | None |
| MAD1L1 | None | None | None | None |
| NDST1 | None | None | None | None |
| ARL2BP | None | None | None | None |
| TOR1A | None | None | None | None |
| C1S | None | None | None | None |
| ARL6 | 0.508 | 1 | 2h57 | A |

| | | | | |
|---|---|---|---|---|
| ARL6 | None | None | None | None |
| ARL6 | None | None | None | None |
| ARL6 | None | None | None | None |
| ARL6 | None | None | None | None |
| ARL6 | None | None | None | None |
| FGFR1 | None | None | None | None |
| KLHL7 | None | None | None | None |
| FLNC | None | None | None | None |
| PTPRC | None | None | None | None |
| GCH1 | None | None | None | None |
| GCH1 | None | None | None | None |
| GCH1 | None | None | None | None |
| GCH1 | None | None | None | None |
| GCH1 | None | None | None | None |
| GCH1 | None | None | None | None |
| GCH1 | None | None | None | None |
| GCH1 | None | None | None | None |
| GCH1 | None | None | None | None |
| GCH1 | None | None | None | None |
| GCH1 | 0.3 | 1 | 1fb1 | D |
| GCH1 | None | None | None | None |
| GCH1 | 0.3 | 11 | 1fb1 | B |
| GCH1 | None | None | None | None |
| CFH | None | None | None | None |
| COL2A1 | None | None | None | None |
| COL2A1 | None | None | None | None |
| OCRL | None | None | None | None |
| WRN | None | None | None | None |
| WRN | None | None | None | None |
| WRN | None | None | None | None |
| WRN | None | None | None | None |
| MASP1 | None | None | None | None |
| MASP1 | None | None | None | None |
| NSD1 | None | None | None | None |
| PITX2 | None | None | None | None |
| PITX2 | None | None | None | None |
| PITX2 | None | None | None | None |
| PITX2 | 0.362 | 2 | 2lkx | A |
| PITX2 | 0.362 | 2 | 2lkx | A |
| PITX2 | 0.362 | 2 | 2lkx | A |
| B4GALT7 | 0.917 | 1 | 4irq | A |
| B4GALT7 | None | None | None | None |
| B4GALT7 | None | None | None | None |
| B4GALT7 | None | None | None | None |
| B4GALT7 | None | None | None | None |
| FSHR | None | None | None | None |
| IGHMBP2 | None | None | None | None |
| IGHMBP2 | None | None | None | None |
| IGHMBP2 | None | None | None | None |
| IGHMBP2 | None | None | None | None |
| IGHMBP2 | None | None | None | None |
| IGHMBP2 | None | None | None | None |
| IGHMBP2 | None | None | None | None |
| IGHMBP2 | None | None | None | None |

| | | | | |
|---|---|---|---|---|
| IGHMBP2 | None | None | None | None |
| IGHMBP2 | None | None | None | None |
| IGHMBP2 | None | None | None | None |
| IGHMBP2 | None | None | None | None |
| IGHMBP2 | None | None | None | None |
| IGHMBP2 | None | None | None | None |
| IGHMBP2 | None | None | None | None |
| IGHMBP2 | None | None | None | None |
| TTN | None | None | None | None |
| TTN | None | None | None | None |
| IRAK4 | None | None | None | None |
| PROC | None | None | None | None |
| PROC | None | None | None | None |
| RAD50 | None | None | None | None |
| RAD50 | None | None | None | None |
| DYNC1H1 | None | None | None | None |
| DYNC1H1 | None | None | None | None |
| DYNC1H1 | None | None | None | None |
| DYNC1H1 | None | None | None | None |
| DYNC1H1 | None | None | None | None |
| DYNC1H1 | None | None | None | None |
| DYNC1H1 | None | None | None | None |
| DYNC1H1 | None | None | None | None |
| DYNC1H1 | None | None | None | None |
| DYNC1H1 | None | None | None | None |
| DYNC1H1 | None | None | None | None |
| CYB5R3 | None | None | None | None |
| CYB5R3 | None | None | None | None |
| CYB5R3 | None | None | None | None |
| CYB5R3 | None | None | None | None |
| CYB5R3 | None | None | None | None |
| CYB5R3 | None | None | None | None |
| CYB5R3 | None | None | None | None |
| CYB5R3 | None | None | None | None |
| CYB5R3 | None | None | None | None |
| CYB5R3 | None | None | None | None |
| CYB5R3 | None | None | None | None |
| SPTB | None | None | None | None |
| FANCL | None | None | None | None |
| FAM126A | None | None | None | None |
| SEPT9 | None | None | None | None |
| TTN | None | None | None | None |
| TTN | None | None | None | None |
| TTN | None | None | None | None |
| CNTN1 | None | None | None | None |
| C1QTNF5 | None | None | None | None |
| VCL | None | None | None | None |
| VCL | None | None | None | None |
| VCL | None | None | None | None |
| VCL | None | None | None | None |
| VCL | None | None | None | None |
| VCL | None | None | None | None |
| PPT1 | None | None | None | None |
| PPT1 | None | None | None | None |

| | | | | |
|---|---|---|---|---|
| PPT1 | None | None | None | None |
| PPT1 | None | None | None | None |
| PPT1 | None | None | None | None |
| PPT1 | None | None | None | None |
| PPT1 | None | None | None | None |
| PPT1 | None | None | None | None |
| PPT1 | None | None | None | None |
| PPT1 | None | None | None | None |
| PPT1 | None | None | None | None |
| PPT1 | None | None | None | None |
| PPT1 | None | None | None | None |
| PPT1 | None | None | None | None |
| PPT1 | None | None | None | None |
| PPT1 | None | None | None | None |
| PPT1 | None | None | None | None |
| PPT1 | None | None | None | None |
| PPT1 | None | None | None | None |
| NOTCH2 | None | None | None | None |
| NOTCH2 | None | None | None | None |
| IFIH1 | None | None | None | None |
| IFIH1 | None | None | None | None |
| IKBKG | None | None | None | None |
| DBH | None | None | None | None |
| DBH | None | None | None | None |
| DBH | None | None | None | None |
| DBH | None | None | None | None |
| DBH | None | None | None | None |
| DBH | None | None | None | None |
| DBH | None | None | None | None |
| DBH | None | None | None | None |
| DBH | None | None | None | None |
| DBH | None | None | None | None |
| IL10RA | None | None | None | None |
| IL10RA | None | None | None | None |
| IL10RA | None | None | None | None |
| IL10RA | None | None | None | None |
| IL10RA | None | None | None | None |
| IL10RA | None | None | None | None |
| IL10RA | None | None | None | None |
| ILK | None | None | None | None |
| HPD | None | None | None | None |
| HPD | None | None | None | None |
| HPD | None | None | None | None |
| MECP2 | None | None | None | None |
| MECP2 | None | None | None | None |
| MECP2 | None | None | None | None |
| MECP2 | None | None | None | None |
| MECP2 | None | None | None | None |
| MECP2 | None | None | None | None |
| MECP2 | None | None | None | None |
| MECP2 | None | None | None | None |
| KYNU | None | None | None | None |
| FAS | None | None | None | None |
| FAS | None | None | None | None |

| | | | | | |
|---|---|---|---|---|---|
| ELP1 | None | None | | None | None |
| ELP1 | None | None | | None | None |
| ELP1 | None | None | | None | None |
| ELP1 | None | None | | None | None |
| ELP1 | None | None | | None | None |
| ELP1 | None | None | | None | None |
| ELP1 | None | None | | None | None |
| ELP1 | None | None | | None | None |
| HESX1 | None | None | | None | None |
| HESX1 | None | None | | None | None |
| HESX1 | None | None | | None | None |
| HESX1 | None | None | | None | None |
| GAD1 | None | None | | None | None |
| GAD1 | None | None | | None | None |
| NDUFA1 | None | None | | None | None |
| NDUFA1 | None | None | | None | None |
| NDUFA1 | None | None | | None | None |
| PNPT1 | None | None | | None | None |
| PNPT1 | None | None | | None | None |
| PNPT1 | None | None | | None | None |
| PNPT1 | None | None | | None | None |
| PNPT1 | 0.389 | | 1 | 3u1k | A |
| PNPT1 | None | None | | None | None |
| PNPT1 | None | None | | None | None |
| SLC25A13 | None | None | | None | None |
| SLC25A13 | None | None | | None | None |
| SLC25A13 | None | None | | None | None |
| AP4M1 | None | None | | None | None |
| AP4M1 | None | None | | None | None |
| MEFV | None | None | | None | None |
| MEFV | None | None | | None | None |
| MEFV | None | None | | None | None |
| MEFV | None | None | | None | None |
| MEFV | None | None | | None | None |
| MEFV | None | None | | None | None |
| MEFV | None | None | | None | None |
| MEFV | None | None | | None | None |
| MEFV | None | None | | None | None |
| CAPN3 | None | None | | None | None |
| CAPN3 | None | None | | None | None |
| CAPN3 | None | None | | None | None |
| CAPN3 | None | None | | None | None |
| CAPN3 | 0.474 | | 2 | 4okh | A |
| CAPN3 | 0.474 | | 2 | 4okh | A |
| CAPN3 | None | None | | None | None |
| GP6 | None | None | | None | None |
| FAS | None | None | | None | None |
| FBN1 | None | None | | None | None |
| FBN1 | None | None | | None | None |
| BMPR2 | None | None | | None | None |
| BMPR2 | None | None | | None | None |
| BMPR2 | None | None | | None | None |
| BMPR2 | 0.327 | | 6 | 3g2f | A |
| BMPR2 | 0.327 | | 6 | 3g2f | A |

| | | | | |
|---|---|---|---|---|
| RET | None | None | None | None |
| RET | None | None | None | None |
| RET | None | None | None | None |
| RET | None | None | None | None |
| RET | None | None | None | None |
| RET | None | None | None | None |
| RET | None | None | None | None |
| RET | None | None | None | None |
| RET | None | None | None | None |
| RET | None | None | None | None |
| RET | None | None | None | None |
| RET | None | None | None | None |
| RET | None | None | None | None |
| RET | None | None | None | None |
| RET | None | None | None | None |
| RET | None | None | None | None |
| RET | None | None | None | None |
| RET | None | None | None | None |
| RET | None | None | None | None |
| RET | None | None | None | None |
| RET | None | None | None | None |
| RET | None | None | None | None |
| RET | None | None | None | None |
| RET | None | None | None | None |
| RET | None | None | None | None |
| RET | None | None | None | None |
| RET | None | None | None | None |
| CSF2RA | None | None | None | None |
| CSF2RA | None | None | None | None |
| CSF2RA | None | None | None | None |
| BRCA1 | None | None | None | None |
| ZAP70 | 0.636 | 1 | 2ozo | A |
| ZAP70 | 0.636 | 1 | 2ozo | A |
| ZAP70 | None | None | None | None |
| PIK3R2 | None | None | None | None |
| TCN2 | None | None | None | None |
| TCN2 | None | None | None | None |
| TCN2 | None | None | None | None |
| TCN2 | None | None | None | None |
| TCN2 | None | None | None | None |
| TCN2 | None | None | None | None |
| TCN2 | None | None | None | None |
| TCN2 | None | None | None | None |
| EPM2A | None | None | None | None |
| EPM2A | None | None | None | None |
| EPM2A | None | None | None | None |
| EPM2A | None | None | None | None |
| AARS | None | None | None | None |
| AARS | None | None | None | None |
| AARS | None | None | None | None |
| PCDH15 | None | None | None | None |
| PCDH15 | None | None | None | None |
| PCDH15 | None | None | None | None |

| | | | | |
|---|---|---|---|---|
| PCDH15 | None | None | None | None |
| PCDH15 | None | None | None | None |
| PCDH15 | None | None | None | None |
| PCDH15 | None | None | None | None |
| PCDH15 | None | None | None | None |
| PCDH15 | None | None | None | None |
| PCDH15 | None | None | None | None |
| NDUFS4 | None | None | None | None |
| NDUFS4 | None | None | None | None |
| NDUFS4 | None | None | None | None |
| FGF10 | None | None | None | None |
| FGF10 | 0.558 | 3 | 1nun | A |
| FGF10 | None | None | None | None |
| FGF10 | None | None | None | None |
| CDON | None | None | None | None |
| STAMBP | 0.909 | 2 | 3rzv | A |
| BRCA1 | None | None | None | None |
| BRCA1 | None | None | None | None |
| BRCA1 | None | None | None | None |
| BRCA1 | None | None | None | None |
| BRCA1 | None | None | None | None |
| BRCA1 | None | None | None | None |
| BRCA1 | None | None | None | None |
| BRCA1 | None | None | None | None |
| BRCA1 | None | None | None | None |
| BRCA1 | None | None | None | None |
| PLXNC1 | None | None | None | None |
| RNASEH2C | None | None | None | None |
| RNASEH2C | None | None | None | None |
| APC | None | None | None | None |
| PLA2G4A | None | None | None | None |
| PLA2G4A | None | None | None | None |
| PSAT1 | None | None | None | None |
| PSAT1 | None | None | None | None |
| TRAPPC3 | None | None | None | None |
| QDPR | None | None | None | None |
| QDPR | None | None | None | None |
| QDPR | None | None | None | None |
| QDPR | None | None | None | None |
| KRT5 | None | None | None | None |
| KRT5 | None | None | None | None |
| KRT5 | None | None | None | None |
| MLH1 | None | None | None | None |
| MLH1 | None | None | None | None |
| MLH1 | None | None | None | None |
| MLH1 | None | None | None | None |
| MLH1 | None | None | None | None |
| MLH1 | None | None | None | None |
| MLH1 | None | None | None | None |
| MLH1 | None | None | None | None |
| MLH1 | None | None | None | None |
| MLH1 | None | None | None | None |
| MLH1 | None | None | None | None |
| MLH1 | None | None | None | None |

| | | | | |
|---|---|---|---|---|
| MLH1 | None | None | None | None |
| MLH1 | None | None | None | None |
| MLH1 | None | None | None | None |
| MLH1 | None | None | None | None |
| MLH1 | None | None | None | None |
| KRIT1 | None | None | None | None |
| KRIT1 | None | None | None | None |
| KRIT1 | None | None | None | None |
| ALX4 | None | None | None | None |
| ALX4 | None | None | None | None |
| ALX4 | None | None | None | None |
| TJP2 | None | None | None | None |
| TJP2 | None | None | None | None |
| PANK2 | None | None | None | None |
| PANK2 | None | None | None | None |
| PANK2 | None | None | None | None |
| PANK2 | None | None | None | None |
| PANK2 | None | None | None | None |
| PANK2 | 0.333 | 2 | 5E+026 | B |
| PANK2 | None | None | None | None |
| NDUFA6 | None | None | None | None |
| TSHR | None | None | None | None |
| TSHR | None | None | None | None |
| TSHR | None | None | None | None |
| CFTR | None | None | None | None |
| CFTR | None | None | None | None |
| MYOM1 | None | None | None | None |
| MYOM1 | None | None | None | None |
| SPTB | None | None | None | None |
| GPT2 | None | None | None | None |
| NDUFS2 | None | None | None | None |
| NDUFS2 | None | None | None | None |
| NDUFS2 | None | None | None | None |
| NDUFS2 | None | None | None | None |
| NDUFS2 | None | None | None | None |
| CASQ2 | None | None | None | None |
| CASQ2 | None | None | None | None |
| CASQ2 | None | None | None | None |
| CASQ2 | None | None | None | None |
| CASQ2 | None | None | None | None |
| CASQ2 | None | None | None | None |
| CASQ2 | None | None | None | None |
| CASQ2 | None | None | None | None |
| CASQ2 | None | None | None | None |
| CPOX | None | None | None | None |
| CPOX | None | None | None | None |
| CPOX | 0.857 | 2 | 2aex | A |
| CPOX | None | None | None | None |
| CPOX | None | None | None | None |
| CPOX | None | None | None | None |
| CPOX | None | None | None | None |
| CPOX | None | None | None | None |
| CPOX | None | None | None | None |
| MVK | None | None | None | None |

| | | | | |
|---|---|---|---|---|
| MVK | None | None | None | None |
| MVK | None | None | None | None |
| MVK | None | None | None | None |
| MVK | None | None | None | None |
| MVK | None | None | None | None |
| MVK | None | None | None | None |
| MVK | None | None | None | None |
| MVK | None | None | None | None |
| MVK | None | None | None | None |
| MVK | None | None | None | None |
| MVK | None | None | None | None |
| MVK | None | None | None | None |
| MVK | None | None | None | None |
| MVK | None | None | None | None |
| GSS | None | None | None | None |
| GSS | None | None | None | None |
| GSS | None | None | None | None |
| GSS | None | None | None | None |
| GSS | None | None | None | None |
| DSG2 | None | None | None | None |
| DSG2 | None | None | None | None |
| DSG2 | None | None | None | None |
| DSG2 | None | None | None | None |
| DSG2 | None | None | None | None |
| DSG2 | None | None | None | None |
| DSG2 | None | None | None | None |
| DSG2 | None | None | None | None |
| DSG2 | None | None | None | None |
| DSG2 | None | None | None | None |
| DSG2 | None | None | None | None |
| DSG2 | None | None | None | None |
| DSG2 | None | None | None | None |
| DSG2 | None | None | None | None |
| PAPSS2 | None | None | None | None |
| PLCB2 | None | None | None | None |
| HMGCS2 | None | None | None | None |
| HMGCS2 | None | None | None | None |
| HMGCS2 | 0.938 | 1 | 2wya | C |
| HMGCS2 | None | None | None | None |
| HMGCS2 | None | None | None | None |
| HMGCS2 | None | None | None | None |
| HMGCS2 | 0.562 | 4 | 2wya | C |
| HMGCS2 | None | None | None | None |
| MPDZ | None | None | None | None |
| GATA4 | None | None | None | None |
| GATA4 | None | None | None | None |
| WARS2 | None | None | None | None |
| WARS2 | None | None | None | None |
| HGD | 1 | 47 | 1eyb | A |
| HGD | 1 | 47 | 1eyb | A |
| HGD | 1 | 70 | 1eyb | A |
| HGD | None | None | None | None |
| HGD | None | None | None | None |
| HGD | None | None | None | None |

| | | | | |
|---|---|---|---|---|
| HGD | None | None | None | None |
| HGD | 0.5 | 51 | 1ey2 | A |
| HGD | None | None | None | None |
| HGD | 1 | 1 | 1ey2 | A |
| HGD | 1 | 9 | 1ey2 | A |
| HGD | None | None | None | None |
| HGD | 1 | 10 | 1eyb | A |
| HGD | 1 | 7 | 1ey2 | A |
| PAX8 | None | None | None | None |
| PAX8 | None | None | None | None |
| PAX8 | None | None | None | None |
| PAX8 | None | None | None | None |
| SPTB | None | None | None | None |
| NCF4 | None | None | None | None |
| CFH | None | None | None | None |
| CFH | None | None | None | None |
| CSF1R | None | None | None | None |
| CSF1R | None | None | None | None |
| CA8 | None | None | None | None |
| CA8 | None | None | None | None |
| CA8 | 0.748 | 7 | 2w2j | A |
| APC | None | None | None | None |
| APC | None | None | None | None |
| APC | None | None | None | None |
| APC | None | None | None | None |
| APC | None | None | None | None |
| NDUFA2 | None | None | None | None |
| GIF | None | None | None | None |
| GIF | None | None | None | None |
| GIF | None | None | None | None |
| GIF | None | None | None | None |
| C2 | None | None | None | None |
| ACP5 | None | None | None | None |
| ACP5 | None | None | None | None |
| ACP5 | None | None | None | None |
| ACP5 | None | None | None | None |
| ACP5 | None | None | None | None |
| RNASEH2A | None | None | None | None |
| RNASEH2A | None | None | None | None |
| RNASEH2A | None | None | None | None |
| RNASEH2A | None | None | None | None |
| RNASEH2A | None | None | None | None |
| RNASEH2A | None | None | None | None |
| RNASEH2A | None | None | None | None |
| DSG2 | None | None | None | None |
| USH1G | None | None | None | None |
| FUS | None | None | None | None |
| FUS | None | None | None | None |
| FUS | None | None | None | None |
| FUS | None | None | None | None |
| FUS | None | None | None | None |
| MRPL44 | None | None | None | None |
| MRPL44 | None | None | None | None |
| MRPL44 | None | None | None | None |

| | | | | |
|---|---|---|---|---|
| MRPL44 | None | None | None | None |
| DMGDH | None | None | None | None |
| AARS | None | None | None | None |
| STAR | None | None | None | None |
| STAR | None | None | None | None |
| STAR | None | None | None | None |
| STAR | None | None | None | None |
| STAR | None | None | None | None |
| STAR | None | None | None | None |
| STAR | None | None | None | None |
| STAR | None | None | None | None |
| LARP7 | None | None | None | None |
| PRPF8 | None | None | None | None |
| PRPF8 | None | None | None | None |
| PRPF8 | None | None | None | None |
| PRPF8 | None | None | None | None |
| PSAP | None | None | None | None |
| PSAP | None | None | None | None |
| PSAP | None | None | None | None |
| DMPK | None | None | None | None |
| EPCAM | None | None | None | None |
| EPCAM | None | None | None | None |
| EPCAM | None | None | None | None |
| EPCAM | None | None | None | None |
| EPCAM | None | None | None | None |
| EPCAM | None | None | None | None |
| EPCAM | None | None | None | None |
| EPCAM | None | None | None | None |
| ASS1 | None | None | None | None |
| ASS1 | None | None | None | None |
| ASS1 | 0.571 | 3 | 2nz2 | A |
| ASS1 | 0.571 | 3 | 2nz2 | A |
| ASS1 | None | None | None | None |
| ASS1 | 0.857 | 20 | 2nz2 | A |
| ASS1 | 0.857 | 9 | 2nz2 | A |
| ASS1 | None | None | None | None |
| ASS1 | 0.429 | 13 | 2nz2 | A |
| ASS1 | 0.571 | 4 | 2nz2 | A |
| ASS1 | 0.857 | 9 | 2nz2 | A |
| ASS1 | 0.857 | 9 | 2nz2 | A |
| ASS1 | 0.857 | 9 | 2nz2 | A |
| ASS1 | None | None | None | None |
| ASS1 | None | None | None | None |
| ASS1 | None | None | None | None |
| ASS1 | None | None | None | None |
| ASS1 | None | None | None | None |
| ASS1 | None | None | None | None |
| ASS1 | None | None | None | None |
| POLE | None | None | None | None |
| POLE | None | None | None | None |
| POLE | None | None | None | None |
| RNASET2 | None | None | None | None |
| RNASET2 | None | None | None | None |
| RNASET2 | None | None | None | None |

| | | | | |
|---|---|---|---|---|
| FLG | 0.394 | | 2 4pcw | A |
| F10 | None | None | None | None |
| AGT | None | None | None | None |
| AGT | None | None | None | None |
| PLN | None | None | None | None |
| PLN | None | None | None | None |
| PLN | None | None | None | None |
| PTHLH | None | None | None | None |
| IGF1R | None | None | None | None |
| IGF1R | None | None | None | None |
| IGF1R | None | None | None | None |
| JAG1 | None | None | None | None |
| JAG1 | None | None | None | None |
| JAG1 | None | None | None | None |
| JAG1 | None | None | None | None |
| JAG1 | None | None | None | None |
| JAG1 | None | None | None | None |
| AP1S3 | None | None | None | None |
| SPTA1 | None | None | None | None |
| SPTA1 | None | None | None | None |
| SPTA1 | None | None | None | None |
| SPTA1 | None | None | None | None |
| SPG7 | None | None | None | None |
| SPG7 | None | None | None | None |
| SPG7 | None | None | None | None |
| SPG7 | None | None | None | None |
| SPG7 | None | None | None | None |
| SPG7 | None | None | None | None |
| FN1 | None | None | None | None |
| FN1 | None | None | None | None |
| NSMCE3 | None | None | None | None |
| PCCA | None | None | None | None |
| PCCA | None | None | None | None |
| SPTB | None | None | None | None |
| RYR2 | None | None | None | None |
| RYR2 | None | None | None | None |
| MMP9 | None | None | None | None |
| MMP9 | None | None | None | None |
| GLMN | None | None | None | None |
| GLMN | None | None | None | None |
| GLMN | None | None | None | None |
| GLMN | None | None | None | None |
| GLMN | None | None | None | None |
| GLMN | None | None | None | None |
| GLMN | None | None | None | None |
| GLMN | None | None | None | None |
| GLMN | None | None | None | None |
| GLMN | None | None | None | None |
| MARS | 0.333 | | 2 5goy | A |
| MARS | None | None | None | None |
| MARS | None | None | None | None |
| MARS | None | None | None | None |
| MARS | None | None | None | None |
| MARS | None | None | None | None |

| | | | | |
|---|---|---|---|---|
| SNAP29 | None | None | None | None |
| SNAP29 | None | None | None | None |
| ETHE1 | None | None | None | None |
| ETHE1 | None | None | None | None |
| ETHE1 | None | None | None | None |
| ETHE1 | None | None | None | None |
| ETHE1 | None | None | None | None |
| ETHE1 | None | None | None | None |
| ETHE1 | None | None | None | None |
| SAMHD1 | None | None | None | None |
| KCNQ4 | None | None | None | None |
| FXYD2 | None | None | None | None |
| FXYD2 | None | None | None | None |
| FXYD2 | None | None | None | None |
| LRRK2 | None | None | None | None |
| LRRK2 | None | None | None | None |
| NDUFS1 | None | None | None | None |
| NDUFS1 | None | None | None | None |
| NDUFS1 | None | None | None | None |
| NDUFS1 | None | None | None | None |
| NDUFS1 | None | None | None | None |
| NDUFS1 | None | None | None | None |
| NDUFS1 | None | None | None | None |
| NDUFS1 | None | None | None | None |
| NDUFS1 | None | None | None | None |
| NDUFS1 | None | None | None | None |
| NDUFS1 | None | None | None | None |
| EZH2 | None | None | None | None |
| EZH2 | None | None | None | None |
| LARS | None | None | None | None |
| TLL1 | None | None | None | None |
| CRYBB2 | None | None | None | None |
| CRYBB2 | None | None | None | None |
| EXOSC8 | None | None | None | None |
| SRP72 | None | None | None | None |
| FOXC2 | None | None | None | None |
| HMBS | None | None | None | None |
| HMBS | None | None | None | None |
| HMBS | None | None | None | None |
| HMBS | None | None | None | None |
| HMBS | None | None | None | None |
| HMBS | None | None | None | None |
| HMBS | None | None | None | None |
| HMBS | None | None | None | None |
| HMBS | None | None | None | None |
| HMBS | None | None | None | None |
| HMBS | None | None | None | None |
| HMBS | None | None | None | None |
| HMBS | None | None | None | None |
| HMBS | None | None | None | None |
| HMBS | None | None | None | None |
| HMBS | None | None | None | None |
| HMBS | None | None | None | None |
| HMBS | None | None | None | None |

| | | | | |
|---|---|---|---|---|
| HMBS | None | None | None | None |
| HMBS | None | None | None | None |
| HMBS | None | None | None | None |
| HMBS | None | None | None | None |
| HMBS | None | None | None | None |
| HMBS | None | None | None | None |
| HMBS | None | None | None | None |
| OGT | None | None | None | None |
| NOG | None | None | None | None |
| NOG | None | None | None | None |
| NOG | None | None | None | None |
| NOG | None | None | None | None |
| NOG | None | None | None | None |
| NOG | None | None | None | None |
| NOG | None | None | None | None |
| NOG | None | None | None | None |
| NOG | None | None | None | None |
| NOG | None | None | None | None |
| NOG | None | None | None | None |
| DLG3 | None | None | None | None |
| SCN2A | None | None | None | None |
| STAC3 | None | None | None | None |
| PFKM | None | None | None | None |
| PFKM | None | None | None | None |
| PFKM | None | None | None | None |
| PFKM | None | None | None | None |
| PFKM | None | None | None | None |
| DNMT1 | None | None | None | None |
| DNMT1 | None | None | None | None |
| TNNI3 | None | None | None | None |
| TNNI3 | None | None | None | None |
| TNNI3 | None | None | None | None |
| STXBP2 | None | None | None | None |
| STXBP2 | None | None | None | None |
| STXBP2 | None | None | None | None |
| STXBP2 | None | None | None | None |
| STXBP2 | None | None | None | None |
| STXBP2 | None | None | None | None |
| STXBP2 | None | None | None | None |
| STXBP2 | None | None | None | None |
| REN | None | None | None | None |
| DSP | None | None | None | None |
| DSP | None | None | None | None |
| HNF1B | None | None | None | None |
| ADA | None | None | None | None |
| ADA | None | None | None | None |
| ADA | 0.615 | 7 | 3iar | A |
| ADA | None | None | None | None |
| ADA | None | None | None | None |
| ADA | 0.538 | 5 | 3iar | A |
| ADA | 0.538 | 5 | 3iar | A |
| ADA | None | None | None | None |
| ADA | None | None | None | None |
| ADA | 0.615 | 1 | 3iar | A |

| | | | | |
|---|---|---|---|---|
| ADA | 0.462 | 6 | 3iar | A |
| ADA | 0.615 | 1 | 3iar | A |
| ADA | None | None | None | None |
| ADA | None | None | None | None |
| ADA | None | None | None | None |
| DCX | None | None | None | None |
| DCX | None | None | None | None |
| DCX | None | None | None | None |
| DCX | None | None | None | None |
| DCX | None | None | None | None |
| ARSB | None | None | None | None |
| ARSB | None | None | None | None |
| ARSB | None | None | None | None |
| ARSB | None | None | None | None |
| ARSB | None | None | None | None |
| ARSB | None | None | None | None |
| ARSB | None | None | None | None |
| ARSB | None | None | None | None |
| ARSB | None | None | None | None |
| ARSB | None | None | None | None |
| ARSB | None | None | None | None |
| ARSB | None | None | None | None |
| ARSB | None | None | None | None |
| WWOX | None | None | None | None |
| TNNI3 | None | None | None | None |
| TNNI3 | None | None | None | None |
| TNNI3 | None | None | None | None |
| TNNI3 | None | None | None | None |
| CYBA | None | None | None | None |
| SPAST | None | None | None | None |
| SPAST | None | None | None | None |
| SPAST | None | None | None | None |
| SPAST | None | None | None | None |
| SPAST | None | None | None | None |
| SPAST | None | None | None | None |
| SPAST | None | None | None | None |
| SPAST | None | None | None | None |
| KRIT1 | None | None | None | None |
| KRIT1 | None | None | None | None |
| TP53 | None | None | None | None |
| ISPD | None | None | None | None |
| ISPD | None | None | None | None |
| ISPD | None | None | None | None |
| ISPD | None | None | None | None |
| KCNMA1 | None | None | None | None |
| STK11 | None | None | None | None |
| STK11 | None | None | None | None |
| STK11 | None | None | None | None |
| STK11 | None | None | None | None |
| STK11 | None | None | None | None |
| STK11 | 0.412 | 2 | 2wtk | F |
| NDUFA13 | None | None | None | None |
| PHF6 | None | None | None | None |
| PHF6 | None | None | None | None |

| | | | | |
|---|---|---|---|---|
| PHF6 | None | None | None | None |
| PHYH | None | None | None | None |
| PHYH | None | None | None | None |
| PHYH | None | None | None | None |
| PHYH | None | None | None | None |
| PHYH | None | None | None | None |
| NDUFS3 | None | None | None | None |
| NDUFS3 | None | None | None | None |
| ALDH18A1 | None | None | None | None |
| ALDH18A1 | None | None | None | None |
| ALDH18A1 | None | None | None | None |
| ALDH18A1 | None | None | None | None |
| ALDH18A1 | None | None | None | None |
| PAX6 | None | None | None | None |
| PAX6 | 0.339 | 1 | 2cue | A |
| INPP5E | None | None | None | None |
| INPP5E | None | None | None | None |
| INPP5E | None | None | None | None |
| INPP5E | None | None | None | None |
| INPP5E | None | None | None | None |
| INPP5E | None | None | None | None |
| INPP5E | None | None | None | None |
| INPP5E | None | None | None | None |
| INPP5E | None | None | None | None |
| TNNT2 | None | None | None | None |
| TNNT2 | None | None | None | None |
| TNNT2 | None | None | None | None |
| TNNT2 | None | None | None | None |
| TNNT2 | None | None | None | None |
| TNNT2 | None | None | None | None |
| TNNT2 | None | None | None | None |
| TNNT2 | None | None | None | None |
| TYK2 | None | None | None | None |
| TYK2 | None | None | None | None |
| TYK2 | None | None | None | None |
| CFTR | None | None | None | None |
| CFTR | None | None | None | None |
| CFTR | None | None | None | None |
| CFTR | None | None | None | None |
| CFTR | None | None | None | None |
| CFTR | None | None | None | None |
| CFTR | None | None | None | None |
| CFTR | None | None | None | None |
| CFTR | None | None | None | None |
| CFTR | None | None | None | None |
| CFTR | None | None | None | None |
| CFTR | None | None | None | None |
| CFTR | None | None | None | None |
| CFTR | None | None | None | None |
| CFTR | None | None | None | None |
| CFTR | None | None | None | None |
| CFTR | None | None | None | None |
| CFTR | None | None | None | None |
| CFTR | None | None | None | None |

| | | | | |
|---|---|---|---|---|
| CFTR | None | None | None | None |
| CFTR | None | None | None | None |
| CFTR | None | None | None | None |
| CFTR | None | None | None | None |
| CFTR | None | None | None | None |
| CFTR | None | None | None | None |
| CFTR | None | None | None | None |
| CFTR | None | None | None | None |
| CFTR | None | None | None | None |
| CFTR | None | None | None | None |
| CFTR | None | None | None | None |
| CFTR | None | None | None | None |
| CFTR | None | None | None | None |
| CFTR | None | None | None | None |
| CFTR | None | None | None | None |
| CFTR | None | None | None | None |
| CFTR | None | None | None | None |
| CFTR | None | None | None | None |
| CFTR | None | None | None | None |
| CFTR | None | None | None | None |
| CFTR | None | None | None | None |
| CFTR | None | None | None | None |
| CFTR | None | None | None | None |
| CFTR | None | None | None | None |
| CFTR | None | None | None | None |
| CFTR | None | None | None | None |
| CFTR | None | None | None | None |
| CFTR | None | None | None | None |
| CFTR | None | None | None | None |
| CFTR | None | None | None | None |
| CFTR | None | None | None | None |
| CFTR | None | None | None | None |
| CFTR | None | None | None | None |
| CFTR | None | None | None | None |
| SERPINF1 | None | None | None | None |
| SERPINF1 | None | None | None | None |
| IDS | None | None | None | None |
| IDS | None | None | None | None |
| IDS | None | None | None | None |
| IDS | None | None | None | None |
| IDS | None | None | None | None |
| IDS | None | None | None | None |
| IDS | None | None | None | None |
| IDS | None | None | None | None |
| IDS | None | None | None | None |
| NFKBIA | 0.7 | 1 | 1nfi | F |
| NFKBIA | None | None | None | None |
| SPAST | None | None | None | None |
| FGF23 | None | None | None | None |
| FGF23 | None | None | None | None |
| FGF23 | None | None | None | None |
| FGF23 | None | None | None | None |
| FGF23 | None | None | None | None |
| FGF23 | None | None | None | None |

| | | | | |
|---|---|---|---|---|
| FGF23 | None | None | None | None |
| DLAT | None | None | None | None |
| STUB1 | None | None | None | None |
| HSD17B4 | None | None | None | None |
| HSD17B4 | None | None | None | None |
| DYSF | None | None | None | None |
| CEP135 | None | None | None | None |
| CEP135 | None | None | None | None |
| PRKCG | None | None | None | None |
| CSF1R | None | None | None | None |
| PEX5 | None | None | None | None |
| PAX6 | None | None | None | None |
| PAX6 | None | None | None | None |
| PAX6 | None | None | None | None |
| PAX6 | None | None | None | None |
| PAX6 | None | None | None | None |
| PAX6 | None | None | None | None |
| NF2 | None | None | None | None |
| NF2 | None | None | None | None |
| NF2 | None | None | None | None |
| REM1 | None | None | None | None |

| source_id | classification | binding_site | binding_site_single_water | protein_binding_site |
|---|---|---|---|---|
| rs121918343 | pathogenic | 1 | 1 | 0 |
| rs267606649 | pathogenic | 1 | 1 | 1 |
| rs121918342 | pathogenic | 0 | 1 | 0 |
| rs267606650 | pathogenic | 1 | 1 | 1 |
| rs1130335 | benign | 1 | 1 | 1 |
| rs199476087 | pathogenic | 1 | 1 | 1 |
| rs772163112 | benign | 0 | 0 | 0 |
| rs755793 | benign | 0 | 0 | 0 |
| rs121918155 | pathogenic | 1 | 1 | 0 |
| rs121918156 | pathogenic | 1 | 1 | 0 |
| rs121918154 | pathogenic | 1 | 1 | 1 |
| rs121918151 | pathogenic | 1 | 1 | 1 |
| rs121918144 | pathogenic | 0 | 1 | 0 |
| rs121918146 | pathogenic | 1 | 1 | 1 |
| rs121918160 | pathogenic | 1 | 1 | 1 |
| rs121918150 | pathogenic | 1 | 1 | 1 |
| rs121918141 | pathogenic | 1 | 1 | 1 |
| rs142742242 | pathogenic | 1 | 1 | 1 |
| rs121918142 | pathogenic | 1 | 1 | 1 |
| rs121918157 | pathogenic | 1 | 1 | 1 |
| rs121918101 | pathogenic | 1 | 1 | 1 |
| rs121908260 | pathogenic | 1 | 1 | 1 |
| rs80356667 | pathogenic | 1 | 1 | 1 |
| rs80356668 | pathogenic | 1 | 1 | 1 |
| rs397514540 | pathogenic | 1 | 1 | 1 |
| rs104894685 | pathogenic | 0 | 1 | 0 |
| rs199869995 | pathogenic | 1 | 1 | 1 |
| rs121909072 | pathogenic | 0 | 0 | 0 |
| rs143669310 | benign | 1 | 1 | 1 |
| rs747347498 | benign | 0 | 0 | 0 |
| rs138283222 | benign | 0 | 0 | 0 |
| rs118203931 | pathogenic | 1 | 1 | 1 |
| rs118203933 | pathogenic | 1 | 1 | 0 |
| rs118203932 | pathogenic | 1 | 1 | 0 |
| rs2228063 | benign | 1 | 1 | 1 |
| rs377767360 | pathogenic | 0 | 0 | 0 |
| rs397518413 | pathogenic | 1 | 1 | 1 |
| rs1800458 | benign | 1 | 1 | 0 |
| rs121918083 | pathogenic | 1 | 1 | 1 |
| rs121918094 | pathogenic | 1 | 1 | 1 |
| rs121918098 | pathogenic | 1 | 1 | 1 |
| rs121918093 | pathogenic | 1 | 1 | 1 |
| rs79977247 | pathogenic | 1 | 1 | 1 |
| rs28933979 | pathogenic | 1 | 1 | 1 |
| rs121918068 | pathogenic | 1 | 1 | 1 |
| rs121918077 | pathogenic | 1 | 1 | 1 |
| rs11541796 | pathogenic | 1 | 1 | 1 |
| rs104894665 | pathogenic | 1 | 1 | 1 |
| rs121918090 | pathogenic | 1 | 1 | 1 |
| rs121918081 | pathogenic | 1 | 1 | 1 |
| rs121918080 | pathogenic | 1 | 1 | 1 |
| rs121918076 | pathogenic | 1 | 1 | 1 |
| rs121918097 | pathogenic | 1 | 1 | 1 |

| | | | | |
|---|---|---|---|---|
| rs121918079 | pathogenic | 1 | 1 | 1 |
| rs121918069 | pathogenic | 1 | 1 | 1 |
| rs121918086 | pathogenic | 1 | 1 | 1 |
| rs121918091 | pathogenic | 1 | 1 | 1 |
| rs121918085 | pathogenic | 1 | 1 | 1 |
| rs121918100 | pathogenic | 1 | 1 | 1 |
| rs121918084 | pathogenic | 1 | 1 | 1 |
| rs121918071 | pathogenic | 1 | 1 | 1 |
| rs121918072 | pathogenic | 1 | 1 | 1 |
| rs121918082 | pathogenic | 1 | 1 | 1 |
| rs121918087 | pathogenic | 1 | 1 | 1 |
| rs76410435 | benign | 1 | 1 | 1 |
| rs121918089 | pathogenic | 1 | 1 | 1 |
| rs121918073 | pathogenic | 1 | 1 | 1 |
| rs121918075 | pathogenic | 1 | 1 | 1 |
| rs121918088 | pathogenic | 1 | 1 | 1 |
| rs76992529 | pathogenic | 1 | 1 | 1 |
| rs536294863 | benign | 1 | 1 | 1 |
| rs730881162 | benign | 1 | 1 | 1 |
| rs113994087 | pathogenic | 1 | 1 | 1 |
| rs74716434 | benign | 1 | 1 | 1 |
| rs121918694 | pathogenic | 1 | 1 | 1 |
| rs121918707 | pathogenic | 1 | 1 | 1 |
| rs121918695 | pathogenic | 1 | 1 | 0 |
| rs121918690 | pathogenic | 1 | 1 | 0 |
| rs121918696 | pathogenic | 1 | 1 | 0 |
| rs121918693 | pathogenic | 1 | 1 | 0 |
| rs28999969 | pathogenic | 1 | 1 | 1 |
| rs121918697 | pathogenic | 1 | 1 | 1 |
| rs121918688 | pathogenic | 1 | 1 | 0 |
| rs28999970 | pathogenic | 1 | 1 | 0 |
| rs121918686 | pathogenic | 1 | 1 | 0 |
| rs28999971 | pathogenic | 1 | 1 | 1 |
| rs121918698 | pathogenic | 1 | 1 | 1 |
| rs121918691 | pathogenic | 1 | 1 | 0 |
| rs121918692 | pathogenic | 1 | 1 | 1 |
| rs121918703 | pathogenic | 1 | 1 | 1 |
| rs121918687 | pathogenic | 1 | 1 | 1 |
| rs28933408 | pathogenic | 1 | 1 | 1 |
| rs121918702 | pathogenic | 1 | 1 | 1 |
| rs17875899 | benign | 1 | 1 | 1 |
| rs111033620 | pathogenic | 1 | 1 | 1 |
| rs111033622 | pathogenic | 1 | 1 | 1 |
| rs111033621 | pathogenic | 1 | 1 | 1 |
| rs111033618 | pathogenic | 1 | 1 | 1 |
| rs5918 | benign | 0 | 0 | 0 |
| rs121918452 | pathogenic | 0 | 0 | 0 |
| rs121918445 | pathogenic | 1 | 1 | 1 |
| rs5917 | benign | 1 | 1 | 1 |
| rs121918444 | pathogenic | 1 | 1 | 1 |
| rs121918446 | pathogenic | 1 | 1 | 1 |
| rs121918449 | pathogenic | 0 | 0 | 0 |
| rs121918448 | benign | 0 | 0 | 0 |
| rs13306487 | benign | 1 | 1 | 1 |

| | | | | |
|---|---|---|---|---|
| rs1805044 | benign | 0 | 0 | 0 |
| rs117078377 | benign | 0 | 0 | 0 |
| rs138132026 | benign | 1 | 1 | 1 |
| rs75762896 | benign | 1 | 1 | 0 |
| rs121918043 | pathogenic | 1 | 1 | 0 |
| rs143581690 | benign | 1 | 1 | 0 |
| rs147654263 | benign | 0 | 0 | 0 |
| rs587782529 | pathogenic | 1 | 1 | 1 |
| rs121912664 | pathogenic | 1 | 1 | 1 |
| rs150293825 | benign | 1 | 1 | 1 |
| rs17882252 | benign | 1 | 1 | 1 |
| rs121912662 | pathogenic | 1 | 1 | 1 |
| rs35993958 | benign | 1 | 1 | 1 |
| rs606231253 | pathogenic | 1 | 1 | 1 |
| rs2227983 | benign | 1 | 1 | 1 |
| rs587779351 | pathogenic | 0 | 0 | 0 |
| rs779773463 | pathogenic | 1 | 1 | 1 |
| rs713040 | benign | 1 | 1 | 1 |
| rs334 | pathogenic | 1 | 1 | 1 |
| rs33930165 | pathogenic | 1 | 1 | 1 |
| rs33918131 | benign | 1 | 1 | 1 |
| rs63750783 | pathogenic | 1 | 1 | 1 |
| rs33986703 | pathogenic | 1 | 1 | 1 |
| rs33972047 | pathogenic | 1 | 1 | 1 |
| rs33936254 | pathogenic | 1 | 1 | 1 |
| rs33959855 | benign | 1 | 1 | 1 |
| rs33950507 | pathogenic | 1 | 1 | 1 |
| rs33916412 | pathogenic | 1 | 1 | 0 |
| rs1135071 | pathogenic | 1 | 1 | 1 |
| rs33960103 | pathogenic | 1 | 1 | 1 |
| rs11549407 | pathogenic | 1 | 1 | 1 |
| rs33922842 | pathogenic | 1 | 1 | 1 |
| rs35278874 | benign | 1 | 1 | 1 |
| rs34621955 | benign | 1 | 1 | 1 |
| rs33995148 | pathogenic | 1 | 1 | 1 |
| rs33945705 | benign | 1 | 1 | 1 |
| rs33993568 | benign | 1 | 1 | 1 |
| rs35553496 | benign | 1 | 1 | 1 |
| rs33913712 | pathogenic | 1 | 1 | 1 |
| rs33917785 | pathogenic | 1 | 1 | 1 |
| rs34515413 | benign | 1 | 1 | 1 |
| rs35256489 | pathogenic | 1 | 1 | 0 |
| rs34484056 | pathogenic | 1 | 1 | 1 |
| rs33947020 | benign | 1 | 1 | 1 |
| rs33946267 | pathogenic | 1 | 1 | 1 |
| rs33966761 | pathogenic | 1 | 1 | 1 |
| rs36020563 | benign | 1 | 1 | 1 |
| rs122454124 | pathogenic | 1 | 1 | 1 |
| rs122454126 | pathogenic | 1 | 1 | 0 |
| rs122454127 | pathogenic | 1 | 1 | 1 |
| rs122454130 | pathogenic | 1 | 1 | 1 |
| rs122454125 | pathogenic | 1 | 1 | 1 |
| rs122454131 | pathogenic | 1 | 1 | 0 |
| rs121908302 | pathogenic | 0 | 1 | 0 |

| | | | | |
|---|---|---|---|---|
| rs77829017 | pathogenic | 0 | 1 | 0 |
| rs1141814 | pathogenic | 0 | 1 | 0 |
| rs121908312 | pathogenic | 0 | 1 | 0 |
| rs79653797 | pathogenic | 1 | 1 | 0 |
| rs439898 | pathogenic | 1 | 1 | 0 |
| rs121908299 | pathogenic | 0 | 1 | 0 |
| rs80356763 | pathogenic | 1 | 1 | 1 |
| rs781152868 | pathogenic | 0 | 1 | 0 |
| rs121908297 | pathogenic | 0 | 1 | 0 |
| rs398123531 | benign | 0 | 1 | 0 |
| rs398123532 | pathogenic | 0 | 1 | 0 |
| rs364897 | pathogenic | 1 | 1 | 1 |
| rs1064644 | pathogenic | 0 | 1 | 0 |
| rs409652 | pathogenic | 1 | 1 | 1 |
| rs121908300 | pathogenic | 0 | 1 | 0 |
| rs381737 | pathogenic | 0 | 1 | 0 |
| rs74500255 | pathogenic | 0 | 1 | 0 |
| rs121908313 | pathogenic | 0 | 1 | 0 |
| rs367968666 | pathogenic | 0 | 1 | 0 |
| rs78973108 | pathogenic | 1 | 1 | 1 |
| rs121908298 | pathogenic | 1 | 1 | 1 |
| rs78396650 | pathogenic | 0 | 0 | 0 |
| rs78198234 | pathogenic | 1 | 1 | 0 |
| rs121908304 | pathogenic | 1 | 1 | 1 |
| rs76539814 | pathogenic | 1 | 1 | 1 |
| rs121908305 | pathogenic | 0 | 1 | 0 |
| rs121908306 | pathogenic | 1 | 1 | 0 |
| rs121908308 | pathogenic | 1 | 1 | 1 |
| rs121908307 | pathogenic | 0 | 1 | 0 |
| rs75548401 | benign | 0 | 1 | 0 |
| rs76763715 | pathogenic | 0 | 0 | 0 |
| rs121908314 | pathogenic | 0 | 1 | 0 |
| rs398123528 | pathogenic | 0 | 1 | 0 |
| rs121908311 | pathogenic | 0 | 0 | 0 |
| rs121908310 | pathogenic | 1 | 1 | 0 |
| rs74598136 | pathogenic | 0 | 1 | 0 |
| rs1064651 | pathogenic | 1 | 1 | 1 |
| rs77369218 | pathogenic | 1 | 1 | 1 |
| rs121908295 | pathogenic | 0 | 1 | 0 |
| rs421016 | pathogenic | 0 | 1 | 0 |
| rs368060 | pathogenic | 0 | 1 | 0 |
| rs80356771 | pathogenic | 1 | 1 | 1 |
| rs80356772 | pathogenic | 1 | 1 | 1 |
| rs121908301 | pathogenic | 0 | 0 | 0 |
| rs747506979 | pathogenic | 0 | 1 | 0 |
| rs75822236 | pathogenic | 0 | 1 | 0 |
| rs1990760 | benign | 1 | 1 | 0 |
| rs28934568 | pathogenic | 1 | 1 | 0 |
| rs34833812 | benign | 1 | 1 | 0 |
| rs193922666 | benign | 1 | 1 | 0 |
| rs104893812 | pathogenic | 1 | 1 | 0 |
| rs144701411 | benign | 0 | 0 | 0 |
| rs104893813 | pathogenic | 0 | 1 | 0 |
| rs199660234 | pathogenic | 1 | 1 | 0 |

| | | | | |
|---|---|---|---|---|
| rs104893814 | pathogenic | 1 | 1 | 0 |
| rs143095746 | benign | 1 | 1 | 1 |
| rs104893807 | pathogenic | 1 | 1 | 0 |
| rs104893811 | pathogenic | 0 | 0 | 0 |
| rs104893816 | pathogenic | 0 | 0 | 0 |
| rs121918714 | pathogenic | 0 | 1 | 0 |
| rs104893810 | pathogenic | 1 | 1 | 0 |
| rs104893815 | pathogenic | 1 | 1 | 0 |
| rs4303 | benign | 0 | 1 | 0 |
| rs121912704 | pathogenic | 0 | 1 | 0 |
| rs56394458 | benign | 1 | 1 | 0 |
| rs397514688 | pathogenic | 0 | 0 | 0 |
| rs367797185 | pathogenic | 1 | 1 | 0 |
| rs12709426 | benign | 0 | 1 | 0 |
| rs2228547 | benign | 0 | 0 | 0 |
| rs111033546 | pathogenic | 0 | 0 | 0 |
| rs111033553 | pathogenic | 0 | 0 | 0 |
| rs111033554 | pathogenic | 1 | 1 | 1 |
| rs111033544 | pathogenic | 1 | 1 | 1 |
| rs111033555 | pathogenic | 1 | 1 | 1 |
| rs111033545 | pathogenic | 1 | 1 | 1 |
| rs111033549 | pathogenic | 0 | 1 | 0 |
| rs111033552 | pathogenic | 1 | 1 | 1 |
| rs752204035 | benign | 0 | 0 | 0 |
| rs11569534 | benign | 0 | 0 | 0 |
| rs28937571 | pathogenic | 1 | 1 | 1 |
| rs104894215 | pathogenic | 1 | 1 | 1 |
| rs3888798 | benign | 1 | 1 | 1 |
| rs764263034 | benign | 1 | 1 | 1 |
| rs1800973 | benign | 1 | 1 | 1 |
| rs367623154 | benign | 1 | 1 | 1 |
| rs201230446 | pathogenic | 1 | 1 | 1 |
| rs267606766 | pathogenic | 0 | 1 | 0 |
| rs267606769 | pathogenic | 0 | 1 | 0 |
| rs267606767 | pathogenic | 1 | 1 | 0 |
| rs267606768 | pathogenic | 0 | 1 | 0 |
| rs61733129 | benign | 0 | 1 | 0 |
| rs201947120 | pathogenic | 1 | 1 | 1 |
| rs283413 | benign | 0 | 1 | 0 |
| rs36119840 | benign | 0 | 0 | 0 |
| rs281865419 | pathogenic | 1 | 1 | 1 |
| rs137852937 | pathogenic | 0 | 1 | 0 |
| rs2230709 | benign | 0 | 1 | 0 |
| rs6695033 | benign | 0 | 1 | 0 |
| rs180177032 | pathogenic | 1 | 1 | 1 |
| rs180177033 | pathogenic | 1 | 1 | 1 |
| rs121913351 | pathogenic | 1 | 1 | 1 |
| rs397507473 | pathogenic | 1 | 1 | 1 |
| rs121913355 | pathogenic | 1 | 1 | 1 |
| rs121913357 | pathogenic | 1 | 1 | 1 |
| rs180177036 | pathogenic | 1 | 1 | 1 |
| rs180177039 | pathogenic | 1 | 1 | 1 |
| rs180177038 | pathogenic | 1 | 1 | 1 |
| rs180177040 | pathogenic | 1 | 1 | 1 |

| | | | | |
|---|---|---|---|---|
| rs121913338 | pathogenic | 1 | 1 | 1 |
| rs121913366 | pathogenic | 1 | 1 | 1 |
| rs121913369 | pathogenic | 1 | 1 | 1 |
| rs113488022 | pathogenic | 1 | 1 | 1 |
| rs121913364 | pathogenic | 1 | 1 | 1 |
| rs180177042 | pathogenic | 1 | 1 | 1 |
| rs200461617 | benign | 0 | 0 | 0 |
| rs141609745 | benign | 0 | 0 | 0 |
| rs104894383 | pathogenic | 1 | 1 | 1 |
| rs104894384 | pathogenic | 1 | 1 | 1 |
| rs104894381 | pathogenic | 1 | 1 | 0 |
| rs147710408 | benign | 1 | 1 | 1 |
| rs77357563 | benign | 1 | 1 | 1 |
| rs104894378 | pathogenic | 0 | 0 | 0 |
| rs104894382 | pathogenic | 0 | 0 | 0 |
| rs137853292 | pathogenic | 1 | 1 | 1 |
| rs587778834 | pathogenic | 0 | 0 | 0 |
| rs137853294 | pathogenic | 1 | 1 | 1 |
| rs551747882 | benign | 0 | 0 | 0 |
| rs3092903 | benign | 1 | 1 | 1 |
| rs137853296 | pathogenic | 0 | 0 | 0 |
| rs398122917 | pathogenic | 0 | 1 | 0 |
| rs41304992 | benign | 0 | 0 | 0 |
| rs121912708 | pathogenic | 1 | 1 | 1 |
| rs140102105 | pathogenic | 1 | 1 | 1 |
| rs376917645 | pathogenic | 1 | 1 | 0 |
| rs372660425 | pathogenic | 1 | 1 | 0 |
| rs555896752 | pathogenic | 1 | 1 | 1 |
| rs777829351 | pathogenic | 1 | 1 | 0 |
| rs121912711 | pathogenic | 1 | 1 | 0 |
| rs754449549 | pathogenic | 0 | 1 | 0 |
| rs141701364 | benign | 0 | 1 | 0 |
| rs761011399 | benign | 1 | 1 | 1 |
| rs121912710 | benign | 1 | 1 | 0 |
| rs121912707 | pathogenic | 1 | 1 | 0 |
| rs747597620 | pathogenic | 0 | 1 | 0 |
| rs186558364 | pathogenic | 1 | 1 | 1 |
| rs556400964 | pathogenic | 1 | 1 | 1 |
| rs61757684 | benign | 1 | 1 | 1 |
| rs121917766 | pathogenic | 1 | 1 | 1 |
| rs121918248 | pathogenic | 0 | 0 | 0 |
| rs398123278 | pathogenic | 0 | 0 | 0 |
| rs121918251 | pathogenic | 1 | 1 | 1 |
| rs727504022 | pathogenic | 0 | 0 | 0 |
| rs190834116 | pathogenic | 1 | 1 | 1 |
| rs121918249 | pathogenic | 1 | 1 | 0 |
| rs121918257 | pathogenic | 1 | 1 | 0 |
| rs142068898 | benign | 0 | 0 | 0 |
| rs757000253 | pathogenic | 0 | 0 | 0 |
| rs200908035 | pathogenic | 1 | 1 | 0 |
| rs760782399 | pathogenic | 0 | 0 | 0 |
| rs778702777 | pathogenic | 0 | 0 | 0 |
| rs121918258 | pathogenic | 1 | 1 | 0 |
| rs121918256 | pathogenic | 0 | 0 | 0 |

| | | | | |
|---|---|---|---|---|
| rs200596762 | pathogenic | 1 | 1 | 0 |
| rs547709692 | pathogenic | 0 | 0 | 0 |
| rs758577372 | pathogenic | 0 | 1 | 0 |
| rs564069299 | pathogenic | 1 | 1 | 1 |
| rs772552898 | pathogenic | 1 | 1 | 1 |
| rs368790885 | pathogenic | 1 | 1 | 1 |
| rs121918250 | pathogenic | 0 | 0 | 0 |
| rs533755473 | pathogenic | 1 | 1 | 1 |
| rs753288303 | pathogenic | 1 | 1 | 1 |
| rs2229385 | benign | 0 | 0 | 0 |
| rs1141321 | benign | 0 | 0 | 0 |
| rs753564352 | pathogenic | 0 | 0 | 0 |
| rs775593146 | pathogenic | 0 | 0 | 0 |
| rs121918254 | pathogenic | 0 | 0 | 0 |
| rs8589 | benign | 1 | 1 | 0 |
| rs777758903 | pathogenic | 0 | 0 | 0 |
| rs140600746 | pathogenic | 1 | 1 | 0 |
| rs121918255 | pathogenic | 1 | 1 | 0 |
| rs121918252 | pathogenic | 0 | 0 | 0 |
| rs779990936 | pathogenic | 1 | 1 | 0 |
| rs121912455 | pathogenic | 1 | 1 | 1 |
| rs121912459 | pathogenic | 1 | 1 | 1 |
| rs74315452 | pathogenic | 1 | 1 | 1 |
| rs121912451 | pathogenic | 1 | 1 | 1 |
| rs79006549 | pathogenic | 0 | 0 | 0 |
| rs28934889 | benign | 0 | 1 | 0 |
| rs111033557 | benign | 1 | 1 | 1 |
| rs28934597 | pathogenic | 1 | 1 | 1 |
| rs28934596 | pathogenic | 1 | 1 | 1 |
| rs28934595 | pathogenic | 1 | 1 | 1 |
| rs146519482 | pathogenic | 1 | 1 | 1 |
| rs111033563 | pathogenic | 1 | 1 | 1 |
| rs75385605 | benign | 1 | 1 | 1 |
| rs121908569 | pathogenic | 1 | 1 | 1 |
| rs35719940 | benign | 0 | 0 | 0 |
| rs121918664 | pathogenic | 0 | 0 | 0 |
| rs104893665 | pathogenic | 1 | 1 | 1 |
| rs104893666 | pathogenic | 1 | 1 | 1 |
| rs200845457 | benign | 1 | 1 | 1 |
| rs779204655 | pathogenic | 1 | 1 | 1 |
| rs121917747 | pathogenic | 1 | 1 | 1 |
| rs6042 | benign | 1 | 1 | 1 |
| rs104894380 | pathogenic | 0 | 0 | 0 |
| rs369083541 | pathogenic | 1 | 1 | 1 |
| rs145243532 | benign | 1 | 1 | 1 |
| rs121909500 | pathogenic | 1 | 1 | 1 |
| rs140993310 | benign | 1 | 1 | 1 |
| rs200837270 | pathogenic | 1 | 1 | 1 |
| rs121918484 | pathogenic | 1 | 1 | 1 |
| rs121918306 | pathogenic | 0 | 0 | 0 |
| rs118204089 | pathogenic | 1 | 1 | 0 |
| rs118204083 | pathogenic | 1 | 1 | 0 |
| rs201953584 | pathogenic | 1 | 1 | 1 |
| rs74315362 | pathogenic | 1 | 1 | 1 |

| | | | | |
|---|---|---|---|---|
| rs118204084 | pathogenic | 1 | 1 | 0 |
| rs118204085 | pathogenic | 1 | 1 | 1 |
| rs113403872 | pathogenic | 1 | 1 | 0 |
| rs1052176 | benign | 1 | 1 | 1 |
| rs2031640 | benign | 0 | 0 | 0 |
| rs104894230 | pathogenic | 1 | 1 | 1 |
| rs104894229 | pathogenic | 1 | 1 | 1 |
| rs104894228 | pathogenic | 1 | 1 | 1 |
| rs104894226 | pathogenic | 1 | 1 | 1 |
| rs121917757 | pathogenic | 1 | 1 | 1 |
| rs121917758 | pathogenic | 1 | 1 | 1 |
| rs28933406 | pathogenic | 1 | 1 | 1 |
| rs121917756 | pathogenic | 1 | 1 | 1 |
| rs104894227 | pathogenic | 1 | 1 | 1 |
| rs369106578 | pathogenic | 1 | 1 | 1 |
| rs397517142 | benign | 1 | 1 | 1 |
| rs104894231 | pathogenic | 1 | 1 | 1 |
| rs121917759 | pathogenic | 1 | 1 | 1 |
| rs104893827 | pathogenic | 0 | 0 | 0 |
| rs104893826 | pathogenic | 1 | 1 | 1 |
| rs5030802 | pathogenic | 1 | 1 | 1 |
| rs5030827 | pathogenic | 1 | 1 | 0 |
| rs119103277 | pathogenic | 1 | 1 | 1 |
| rs5030808 | pathogenic | 0 | 1 | 0 |
| rs5030809 | pathogenic | 1 | 1 | 1 |
| rs765978945 | pathogenic | 1 | 1 | 1 |
| rs104893824 | pathogenic | 1 | 1 | 1 |
| rs104893831 | pathogenic | 0 | 1 | 0 |
| rs104893830 | pathogenic | 0 | 1 | 0 |
| rs119103278 | pathogenic | 0 | 1 | 0 |
| rs5030818 | pathogenic | 1 | 1 | 1 |
| rs397516444 | pathogenic | 1 | 1 | 1 |
| rs28940297 | pathogenic | 1 | 1 | 1 |
| rs267607170 | pathogenic | 1 | 1 | 1 |
| rs104893825 | pathogenic | 1 | 1 | 1 |
| rs5030820 | pathogenic | 0 | 1 | 0 |
| rs5030821 | pathogenic | 0 | 1 | 0 |
| rs5030823 | pathogenic | 1 | 1 | 1 |
| rs5030824 | pathogenic | 1 | 1 | 1 |
| rs28940301 | pathogenic | 0 | 1 | 0 |
| rs28940300 | pathogenic | 0 | 1 | 0 |
| rs28940298 | pathogenic | 0 | 1 | 0 |
| rs200019083 | benign | 0 | 0 | 0 |
| rs759748655 | pathogenic | 0 | 0 | 0 |
| rs121917739 | pathogenic | 1 | 1 | 1 |
| rs75353611 | pathogenic | 1 | 1 | 1 |
| rs79744198 | benign | 1 | 1 | 1 |
| rs79228041 | pathogenic | 1 | 1 | 1 |
| rs80106970 | benign | 1 | 1 | 1 |
| rs375353223 | benign | 0 | 0 | 0 |
| rs56311081 | benign | 0 | 0 | 0 |
| rs55985569 | benign | 1 | 1 | 1 |
| rs45483396 | benign | 0 | 0 | 0 |
| rs35284565 | benign | 1 | 1 | 1 |

| | | | | |
|---|---|---|---|---|
| rs34349517 | benign | 0 | 0 | 0 |
| rs35776110 | benign | 0 | 0 | 0 |
| rs33917957 | benign | 1 | 1 | 1 |
| rs587778445 | benign | 0 | 0 | 0 |
| rs34624984 | benign | 0 | 0 | 0 |
| rs701753 | benign | 0 | 0 | 0 |
| rs855314 | benign | 0 | 0 | 0 |
| rs121918371 | pathogenic | 1 | 1 | 0 |
| rs398122912 | pathogenic | 0 | 1 | 0 |
| rs1126728 | benign | 0 | 0 | 0 |
| rs587777404 | pathogenic | 1 | 1 | 0 |
| rs11208257 | benign | 0 | 0 | 0 |
| rs6676290 | benign | 0 | 0 | 0 |
| rs104894118 | pathogenic | 0 | 0 | 0 |
| rs121918655 | pathogenic | 1 | 1 | 1 |
| rs104894120 | pathogenic | 1 | 1 | 1 |
| rs137852238 | pathogenic | 0 | 0 | 0 |
| rs137852237 | pathogenic | 0 | 0 | 0 |
| rs137852499 | pathogenic | 1 | 1 | 1 |
| rs137852480 | pathogenic | 1 | 1 | 1 |
| rs137852494 | pathogenic | 1 | 1 | 1 |
| rs137852502 | pathogenic | 1 | 1 | 1 |
| rs137852500 | pathogenic | 1 | 1 | 1 |
| rs137852487 | pathogenic | 1 | 1 | 1 |
| rs137852481 | pathogenic | 1 | 1 | 1 |
| rs137852501 | pathogenic | 0 | 0 | 0 |
| rs137852478 | pathogenic | 1 | 1 | 1 |
| rs137852485 | pathogenic | 1 | 1 | 1 |
| rs137852482 | pathogenic | 1 | 1 | 0 |
| rs137852483 | pathogenic | 0 | 1 | 0 |
| rs137852477 | pathogenic | 1 | 1 | 0 |
| rs137852496 | pathogenic | 1 | 1 | 0 |
| rs137852484 | pathogenic | 0 | 1 | 0 |
| rs137852498 | pathogenic | 1 | 1 | 0 |
| rs137852497 | pathogenic | 1 | 1 | 0 |
| rs137852493 | pathogenic | 1 | 1 | 0 |
| rs137852504 | pathogenic | 1 | 1 | 1 |
| rs267606863 | pathogenic | 1 | 1 | 1 |
| rs137852486 | pathogenic | 1 | 1 | 1 |
| rs137852479 | pathogenic | 1 | 1 | 1 |
| rs112122950 | benign | 0 | 0 | 0 |
| rs72546648 | benign | 1 | 1 | 1 |
| rs72552291 | pathogenic | 0 | 0 | 0 |
| rs79184941 | pathogenic | 1 | 1 | 1 |
| rs77543610 | pathogenic | 1 | 1 | 1 |
| rs121918505 | pathogenic | 1 | 1 | 1 |
| rs776587763 | pathogenic | 1 | 1 | 1 |
| rs121918497 | pathogenic | 1 | 1 | 1 |
| rs121918499 | pathogenic | 1 | 1 | 1 |
| rs121918501 | pathogenic | 1 | 1 | 1 |
| rs121918500 | pathogenic | 0 | 0 | 0 |
| rs121918504 | pathogenic | 1 | 1 | 1 |
| rs121918510 | pathogenic | 1 | 1 | 1 |
| rs121918493 | pathogenic | 1 | 1 | 1 |

| | | | | |
|---|---|---|---|---|
| rs121918489 | pathogenic | 0 | 0 | 0 |
| rs121918495 | pathogenic | 1 | 1 | 1 |
| rs121918488 | pathogenic | 0 | 0 | 0 |
| rs121918496 | pathogenic | 0 | 0 | 0 |
| rs121918492 | pathogenic | 1 | 1 | 1 |
| rs121918494 | pathogenic | 1 | 1 | 1 |
| rs121918502 | pathogenic | 1 | 1 | 1 |
| rs121918490 | pathogenic | 1 | 1 | 1 |
| rs72466451 | pathogenic | 1 | 1 | 1 |
| rs66468541 | pathogenic | 0 | 0 | 0 |
| rs137852662 | pathogenic | 0 | 0 | 0 |
| rs137852663 | pathogenic | 0 | 0 | 0 |
| rs606231407 | pathogenic | 0 | 0 | 0 |
| rs137852233 | pathogenic | 1 | 1 | 1 |
| rs112306225 | benign | 0 | 0 | 0 |
| rs387906808 | pathogenic | 1 | 1 | 0 |
| rs730882094 | pathogenic | 1 | 1 | 1 |
| rs373869746 | pathogenic | 1 | 1 | 1 |
| rs777524402 | pathogenic | 1 | 1 | 1 |
| rs769737896 | pathogenic | 0 | 1 | 0 |
| rs13306515 | benign | 1 | 1 | 0 |
| rs746982741 | pathogenic | 0 | 0 | 0 |
| rs2230200 | benign | 1 | 1 | 1 |
| rs1047286 | benign | 0 | 0 | 0 |
| rs141737564 | benign | 0 | 0 | 0 |
| rs774826179 | benign | 0 | 0 | 0 |
| rs11569422 | benign | 1 | 1 | 1 |
| rs146613648 | benign | 0 | 0 | 0 |
| rs144432231 | benign | 0 | 0 | 0 |
| rs140655115 | benign | 0 | 0 | 0 |
| rs387906837 | pathogenic | 1 | 1 | 0 |
| rs121909076 | pathogenic | 1 | 1 | 0 |
| rs121909077 | pathogenic | 0 | 0 | 0 |
| rs121909074 | pathogenic | 0 | 0 | 0 |
| rs137853330 | pathogenic | 1 | 1 | 1 |
| rs137853331 | pathogenic | 1 | 1 | 1 |
| rs137853332 | pathogenic | 1 | 1 | 1 |
| rs28931586 | pathogenic | 1 | 1 | 0 |
| rs11569321 | benign | 1 | 1 | 1 |
| rs119486097 | benign | 0 | 0 | 0 |
| rs1799807 | pathogenic | 1 | 1 | 1 |
| rs201820739 | pathogenic | 1 | 1 | 0 |
| rs114706984 | pathogenic | 1 | 1 | 0 |
| rs140080572 | pathogenic | 1 | 1 | 0 |
| rs28933389 | pathogenic | 0 | 1 | 0 |
| rs16849700 | benign | 1 | 1 | 0 |
| rs115624085 | pathogenic | 1 | 1 | 1 |
| rs747196387 | pathogenic | 1 | 1 | 1 |
| rs104893684 | pathogenic | 1 | 1 | 0 |
| rs121918557 | pathogenic | 1 | 1 | 0 |
| rs121918556 | pathogenic | 1 | 1 | 1 |
| rs398124571 | pathogenic | 1 | 1 | 1 |
| rs776631396 | pathogenic | 0 | 1 | 0 |
| rs398124576 | pathogenic | 1 | 1 | 1 |

| | | | | |
|---|---|---|---|---|
| rs398124581 | pathogenic | 1 | 1 | 1 |
| rs398124582 | pathogenic | 1 | 1 | 1 |
| rs149766077 | pathogenic | 0 | 1 | 0 |
| rs79761867 | pathogenic | 0 | 1 | 0 |
| rs148905512 | pathogenic | 0 | 1 | 0 |
| rs121965004 | pathogenic | 1 | 1 | 1 |
| rs386834233 | pathogenic | 0 | 1 | 0 |
| rs398124598 | pathogenic | 0 | 1 | 0 |
| rs398124603 | pathogenic | 1 | 1 | 1 |
| rs546716073 | benign | 1 | 1 | 1 |
| rs200310662 | benign | 1 | 1 | 1 |
| rs115494727 | benign | 1 | 1 | 1 |
| rs781513008 | benign | 0 | 0 | 0 |
| rs587776398 | benign | 1 | 1 | 0 |
| rs35288908 | benign | 1 | 1 | 1 |
| rs150634562 | pathogenic | 1 | 1 | 0 |
| rs142948132 | pathogenic | 1 | 1 | 1 |
| rs132630321 | pathogenic | 1 | 1 | 1 |
| rs132630317 | pathogenic | 1 | 1 | 1 |
| rs132630320 | pathogenic | 0 | 0 | 0 |
| rs781394318 | pathogenic | 1 | 1 | 1 |
| rs111033624 | pathogenic | 1 | 1 | 1 |
| rs111033630 | pathogenic | 1 | 1 | 1 |
| rs111033627 | pathogenic | 1 | 1 | 1 |
| rs193922287 | pathogenic | 1 | 1 | 1 |
| rs759072800 | pathogenic | 0 | 1 | 0 |
| rs104894010 | pathogenic | 0 | 0 | 0 |
| rs193922296 | pathogenic | 0 | 0 | 0 |
| rs104894006 | pathogenic | 1 | 1 | 1 |
| rs193922311 | pathogenic | 1 | 1 | 0 |
| rs80356654 | pathogenic | 1 | 1 | 0 |
| rs144723656 | benign | 1 | 1 | 1 |
| rs148311934 | pathogenic | 1 | 1 | 0 |
| rs80356655 | pathogenic | 1 | 1 | 1 |
| rs769268803 | pathogenic | 1 | 1 | 0 |
| rs104894008 | pathogenic | 0 | 1 | 0 |
| rs104894005 | pathogenic | 1 | 1 | 1 |
| rs104894009 | pathogenic | 1 | 1 | 0 |
| rs193922340 | pathogenic | 0 | 0 | 0 |
| rs104894016 | pathogenic | 0 | 1 | 0 |
| rs193921338 | pathogenic | 0 | 0 | 0 |
| rs193922272 | pathogenic | 1 | 1 | 1 |
| rs104894012 | pathogenic | 1 | 1 | 0 |
| rs193922285 | pathogenic | 1 | 1 | 1 |
| rs104894336 | pathogenic | 1 | 1 | 1 |
| rs28931580 | pathogenic | 1 | 1 | 1 |
| rs104894326 | pathogenic | 1 | 1 | 1 |
| rs104894331 | pathogenic | 0 | 0 | 0 |
| rs104894338 | pathogenic | 0 | 0 | 0 |
| rs104894333 | pathogenic | 1 | 1 | 1 |
| rs104894330 | pathogenic | 1 | 1 | 1 |
| rs143886391 | benign | 1 | 1 | 1 |
| rs104894335 | pathogenic | 1 | 1 | 1 |
| rs104894337 | pathogenic | 0 | 0 | 0 |

| | | | | |
|---|---|---|---|---|
| rs104894328 | pathogenic | 0 | 0 | 0 |
| rs104894341 | pathogenic | 1 | 1 | 0 |
| rs104894329 | pathogenic | 0 | 0 | 0 |
| rs137852850 | pathogenic | 1 | 1 | 1 |
| rs137852854 | pathogenic | 0 | 1 | 0 |
| rs137852849 | pathogenic | 0 | 1 | 0 |
| rs137852848 | pathogenic | 1 | 1 | 1 |
| rs137852852 | pathogenic | 0 | 1 | 0 |
| rs137852853 | pathogenic | 0 | 1 | 0 |
| rs137852847 | pathogenic | 0 | 1 | 0 |
| rs137852846 | pathogenic | 0 | 1 | 0 |
| rs137852844 | pathogenic | 0 | 1 | 0 |
| rs144411158 | pathogenic | 0 | 0 | 0 |
| rs2302234 | benign | 0 | 0 | 0 |
| rs139620139 | pathogenic | 0 | 0 | 0 |
| rs146689929 | benign | 0 | 0 | 0 |
| rs373328681 | pathogenic | 1 | 1 | 0 |
| rs387906620 | pathogenic | 1 | 1 | 1 |
| rs1799983 | benign | 0 | 1 | 0 |
| rs28999111 | pathogenic | 1 | 1 | 1 |
| rs104894544 | pathogenic | 1 | 1 | 1 |
| rs104894546 | pathogenic | 1 | 1 | 1 |
| rs28999112 | pathogenic | 0 | 1 | 0 |
| rs104894545 | pathogenic | 1 | 1 | 1 |
| rs104894547 | pathogenic | 1 | 1 | 1 |
| rs74315310 | pathogenic | 0 | 0 | 0 |
| rs267606852 | pathogenic | 0 | 1 | 0 |
| rs137853586 | pathogenic | 0 | 1 | 0 |
| rs137853582 | pathogenic | 1 | 1 | 1 |
| rs61754634 | pathogenic | 1 | 1 | 1 |
| rs137853587 | pathogenic | 0 | 1 | 0 |
| rs267606851 | pathogenic | 1 | 1 | 1 |
| rs267606853 | pathogenic | 1 | 1 | 1 |
| rs137853584 | pathogenic | 1 | 1 | 1 |
| rs137853585 | pathogenic | 1 | 1 | 1 |
| rs460184 | pathogenic | 1 | 1 | 1 |
| rs104893751 | pathogenic | 0 | 1 | 0 |
| rs77358162 | benign | 0 | 0 | 0 |
| rs387906936 | pathogenic | 0 | 0 | 0 |
| rs769525399 | pathogenic | 0 | 0 | 0 |
| rs371271054 | pathogenic | 1 | 1 | 0 |
| rs118204000 | pathogenic | 0 | 0 | 0 |
| rs138568622 | pathogenic | 1 | 1 | 1 |
| rs34509359 | benign | 0 | 0 | 0 |
| rs104894130 | pathogenic | 1 | 1 | 0 |
| rs281865424 | pathogenic | 0 | 0 | 0 |
| rs121912778 | pathogenic | 1 | 1 | 0 |
| rs771055145 | pathogenic | 1 | 1 | 0 |
| rs776174514 | pathogenic | 1 | 1 | 0 |
| rs545986367 | pathogenic | 1 | 1 | 1 |
| rs570768038 | benign | 1 | 1 | 1 |
| rs74315442 | pathogenic | 1 | 1 | 1 |
| rs79716074 | benign | 1 | 1 | 1 |
| rs748486078 | pathogenic | 1 | 1 | 1 |

| | | | | |
|---|---|---|---|---|
| rs2397084 | benign | 1 | 1 | 1 |
| rs11465553 | benign | 1 | 1 | 1 |
| rs144576902 | benign | 1 | 1 | 1 |
| rs763780 | benign | 1 | 1 | 1 |
| rs267606870 | pathogenic | 1 | 1 | 0 |
| rs121913502 | pathogenic | 1 | 1 | 0 |
| rs118101777 | benign | 1 | 1 | 1 |
| rs118053940 | benign | 0 | 0 | 0 |
| rs368771578 | benign | 1 | 1 | 1 |
| rs11540654 | pathogenic | 1 | 1 | 1 |
| rs121912658 | pathogenic | 1 | 1 | 1 |
| rs28934873 | pathogenic | 0 | 1 | 0 |
| rs780442292 | pathogenic | 1 | 1 | 1 |
| rs758781593 | benign | 1 | 1 | 1 |
| rs28934875 | pathogenic | 1 | 1 | 1 |
| rs28934874 | pathogenic | 1 | 1 | 0 |
| rs587782705 | pathogenic | 1 | 1 | 1 |
| rs587780068 | pathogenic | 1 | 1 | 1 |
| rs587782144 | pathogenic | 1 | 1 | 1 |
| rs138729528 | pathogenic | 1 | 1 | 1 |
| rs28934578 | pathogenic | 1 | 1 | 1 |
| rs397514495 | pathogenic | 1 | 1 | 1 |
| rs121912665 | pathogenic | 1 | 1 | 1 |
| rs397516435 | pathogenic | 1 | 1 | 1 |
| rs587778720 | pathogenic | 1 | 1 | 1 |
| rs397516436 | pathogenic | 1 | 1 | 1 |
| rs587781386 | benign | 1 | 1 | 1 |
| rs121912666 | pathogenic | 1 | 1 | 0 |
| rs587780073 | pathogenic | 1 | 1 | 1 |
| rs587782664 | pathogenic | 1 | 1 | 1 |
| rs730882005 | pathogenic | 1 | 1 | 1 |
| rs28934573 | pathogenic | 1 | 1 | 1 |
| rs121912655 | pathogenic | 1 | 1 | 1 |
| rs28934575 | pathogenic | 1 | 1 | 1 |
| rs121912656 | pathogenic | 1 | 1 | 1 |
| rs11540652 | pathogenic | 1 | 1 | 1 |
| rs121912651 | pathogenic | 1 | 1 | 1 |
| rs730882027 | pathogenic | 1 | 1 | 1 |
| rs121912653 | pathogenic | 1 | 1 | 1 |
| rs121912652 | pathogenic | 1 | 1 | 1 |
| rs121912657 | pathogenic | 0 | 1 | 0 |
| rs121913343 | pathogenic | 1 | 1 | 1 |
| rs28934576 | pathogenic | 1 | 1 | 1 |
| rs763098116 | pathogenic | 1 | 1 | 1 |
| rs121912660 | pathogenic | 1 | 1 | 1 |
| rs28934574 | pathogenic | 1 | 1 | 1 |
| rs371409680 | pathogenic | 1 | 1 | 1 |
| rs121912667 | pathogenic | 1 | 1 | 1 |
| rs121912663 | pathogenic | 1 | 1 | 1 |
| rs201744589 | pathogenic | 0 | 0 | 0 |
| rs33926449 | benign | 0 | 0 | 0 |
| rs74315302 | pathogenic | 0 | 1 | 0 |
| rs202040269 | pathogenic | 1 | 1 | 1 |
| rs375958814 | pathogenic | 1 | 1 | 1 |

| | | | | |
|---|---|---|---|---|
| rs121909624 | pathogenic | 1 | 1 | 1 |
| rs121909621 | pathogenic | 1 | 1 | 1 |
| rs121909622 | pathogenic | 1 | 1 | 0 |
| rs4220 | benign | 1 | 1 | 1 |
| rs147559466 | pathogenic | 1 | 1 | 1 |
| rs757434857 | pathogenic | 1 | 1 | 1 |
| rs398123072 | pathogenic | 1 | 1 | 1 |
| rs121434280 | pathogenic | 0 | 1 | 0 |
| rs398123074 | pathogenic | 0 | 1 | 0 |
| rs762114560 | pathogenic | 1 | 1 | 1 |
| rs121434283 | pathogenic | 1 | 1 | 0 |
| rs121434277 | pathogenic | 0 | 1 | 0 |
| rs745844469 | pathogenic | 1 | 1 | 1 |
| rs121434279 | pathogenic | 1 | 1 | 0 |
| rs121434278 | pathogenic | 1 | 1 | 1 |
| rs373715782 | pathogenic | 1 | 1 | 1 |
| rs200724875 | pathogenic | 1 | 1 | 1 |
| rs149678400 | pathogenic | 0 | 1 | 0 |
| rs770273135 | pathogenic | 1 | 1 | 1 |
| rs121434276 | pathogenic | 0 | 1 | 0 |
| rs121434281 | pathogenic | 1 | 1 | 0 |
| rs768884003 | pathogenic | 1 | 1 | 1 |
| rs201375579 | pathogenic | 0 | 0 | 0 |
| rs121434274 | pathogenic | 1 | 1 | 0 |
| rs121434282 | pathogenic | 1 | 1 | 0 |
| rs779759347 | pathogenic | 0 | 1 | 0 |
| rs77931234 | pathogenic | 1 | 1 | 1 |
| rs148207467 | pathogenic | 1 | 1 | 1 |
| rs766140986 | pathogenic | 1 | 1 | 1 |
| rs121434275 | pathogenic | 1 | 1 | 1 |
| rs748110745 | pathogenic | 1 | 1 | 1 |
| rs104894274 | pathogenic | 1 | 1 | 1 |
| rs104894273 | pathogenic | 0 | 1 | 0 |
| rs104894278 | pathogenic | 1 | 1 | 1 |
| rs104894275 | pathogenic | 1 | 1 | 1 |
| rs104894277 | pathogenic | 1 | 1 | 1 |
| rs104894276 | pathogenic | 0 | 0 | 0 |
| rs104894280 | pathogenic | 1 | 1 | 1 |
| rs145882709 | pathogenic | 1 | 1 | 1 |
| rs104894279 | pathogenic | 1 | 1 | 1 |
| rs80356670 | pathogenic | 1 | 1 | 1 |
| rs121918102 | pathogenic | 1 | 1 | 1 |
| rs80356671 | pathogenic | 1 | 1 | 1 |
| rs80356672 | pathogenic | 1 | 1 | 1 |
| rs758609113 | pathogenic | 1 | 1 | 0 |
| rs121918188 | pathogenic | 0 | 1 | 0 |
| rs121918189 | pathogenic | 1 | 1 | 0 |
| rs121918191 | pathogenic | 1 | 1 | 1 |
| rs1769259 | benign | 1 | 1 | 1 |
| rs140222720 | benign | 1 | 1 | 1 |
| rs2297085 | benign | 1 | 1 | 1 |
| rs200679026 | pathogenic | 1 | 1 | 1 |
| rs121918192 | pathogenic | 0 | 1 | 0 |
| rs145771944 | benign | 0 | 1 | 0 |

| | | | | |
|---|---|---|---|---|
| rs201508880 | pathogenic | 0 | 1 | 0 |
| rs17221854 | pathogenic | 0 | 0 | 0 |
| rs120074189 | pathogenic | 1 | 1 | 1 |
| rs120074190 | pathogenic | 1 | 1 | 1 |
| rs758600283 | benign | 1 | 1 | 1 |
| rs200305064 | pathogenic | 1 | 1 | 1 |
| rs121917891 | pathogenic | 1 | 1 | 1 |
| rs3772809 | benign | 0 | 1 | 0 |
| rs387906619 | pathogenic | 1 | 1 | 0 |
| rs2304681 | benign | 0 | 1 | 0 |
| rs121917849 | pathogenic | 0 | 0 | 0 |
| rs143010236 | pathogenic | 0 | 0 | 0 |
| rs121917850 | pathogenic | 1 | 1 | 0 |
| rs35916840 | pathogenic | 1 | 1 | 0 |
| rs28939688 | pathogenic | 1 | 1 | 1 |
| rs11258194 | benign | 1 | 1 | 1 |
| rs199840952 | pathogenic | 1 | 1 | 1 |
| rs121918159 | pathogenic | 1 | 1 | 1 |
| rs200396166 | pathogenic | 0 | 0 | 0 |
| rs757966746 | pathogenic | 0 | 1 | 0 |
| rs121918608 | pathogenic | 0 | 1 | 0 |
| rs74315401 | pathogenic | 1 | 1 | 1 |
| rs11538758 | pathogenic | 0 | 0 | 0 |
| rs74315402 | pathogenic | 1 | 1 | 1 |
| rs1799990 | pathogenic | 1 | 1 | 1 |
| rs74315410 | pathogenic | 1 | 1 | 1 |
| rs150351644 | benign | 1 | 1 | 1 |
| rs16990018 | benign | 1 | 1 | 1 |
| rs74315403 | pathogenic | 1 | 1 | 1 |
| rs74315408 | pathogenic | 1 | 1 | 1 |
| rs74315411 | pathogenic | 1 | 1 | 1 |
| rs74315413 | pathogenic | 1 | 1 | 1 |
| rs74315405 | pathogenic | 1 | 1 | 1 |
| rs28933385 | pathogenic | 1 | 1 | 1 |
| rs74315412 | pathogenic | 1 | 1 | 1 |
| rs74315407 | pathogenic | 1 | 1 | 1 |
| rs74315406 | pathogenic | 1 | 1 | 1 |
| rs11571117 | benign | 1 | 1 | 1 |
| rs121917742 | pathogenic | 1 | 1 | 1 |
| rs149782619 | pathogenic | 1 | 1 | 1 |
| rs397514492 | pathogenic | 1 | 1 | 1 |
| rs397514489 | pathogenic | 0 | 1 | 0 |
| rs397514493 | pathogenic | 1 | 1 | 1 |
| rs373849532 | pathogenic | 1 | 1 | 1 |
| rs121965048 | pathogenic | 1 | 1 | 1 |
| rs121965037 | pathogenic | 1 | 1 | 1 |
| rs121965060 | pathogenic | 0 | 1 | 0 |
| rs386833603 | pathogenic | 1 | 1 | 1 |
| rs121965038 | pathogenic | 1 | 1 | 1 |
| rs121965039 | pathogenic | 1 | 1 | 1 |
| rs121965040 | pathogenic | 1 | 1 | 1 |
| rs267606925 | pathogenic | 1 | 1 | 1 |
| rs121965056 | pathogenic | 0 | 1 | 0 |
| rs121965059 | pathogenic | 0 | 0 | 0 |

| | | | | |
|---|---|---|---|---|
| rs386833615 | pathogenic | 1 | 1 | 0 |
| rs121965051 | pathogenic | 1 | 1 | 0 |
| rs121965046 | pathogenic | 0 | 1 | 0 |
| rs121965052 | pathogenic | 0 | 1 | 0 |
| rs386833617 | pathogenic | 0 | 1 | 0 |
| rs386833618 | pathogenic | 1 | 1 | 0 |
| rs121965041 | pathogenic | 0 | 1 | 0 |
| rs121965042 | pathogenic | 0 | 1 | 0 |
| rs121965057 | pathogenic | 1 | 1 | 1 |
| rs386833621 | pathogenic | 1 | 1 | 1 |
| rs121965049 | pathogenic | 1 | 1 | 1 |
| rs386833623 | pathogenic | 1 | 1 | 1 |
| rs121965047 | pathogenic | 1 | 1 | 1 |
| rs121965053 | pathogenic | 0 | 1 | 0 |
| rs121965045 | pathogenic | 0 | 1 | 0 |
| rs121965054 | pathogenic | 0 | 1 | 0 |
| rs121965043 | pathogenic | 0 | 1 | 0 |
| rs386833598 | pathogenic | 0 | 0 | 0 |
| rs137854545 | pathogenic | 1 | 1 | 1 |
| rs137854544 | pathogenic | 1 | 1 | 1 |
| rs104894455 | pathogenic | 0 | 1 | 0 |
| rs1049564 | benign | 0 | 1 | 0 |
| rs104894460 | pathogenic | 1 | 1 | 1 |
| rs104894453 | pathogenic | 1 | 1 | 1 |
| rs104894450 | pathogenic | 0 | 1 | 0 |
| rs104894454 | pathogenic | 0 | 1 | 0 |
| rs104894452 | pathogenic | 1 | 1 | 1 |
| rs104894451 | pathogenic | 0 | 1 | 0 |
| rs104894368 | pathogenic | 1 | 1 | 1 |
| rs730880950 | pathogenic | 1 | 1 | 1 |
| rs121909535 | pathogenic | 1 | 1 | 1 |
| rs121909537 | pathogenic | 1 | 1 | 1 |
| rs121909536 | pathogenic | 1 | 1 | 1 |
| rs121909542 | pathogenic | 1 | 1 | 1 |
| rs121909538 | pathogenic | 1 | 1 | 1 |
| rs121909539 | pathogenic | 1 | 1 | 1 |
| rs121909540 | pathogenic | 1 | 1 | 1 |
| rs17560 | benign | 1 | 1 | 1 |
| rs121909543 | pathogenic | 0 | 1 | 0 |
| rs121909544 | pathogenic | 1 | 1 | 1 |
| rs2307456 | benign | 0 | 1 | 0 |
| rs745778317 | pathogenic | 0 | 1 | 0 |
| rs759607901 | pathogenic | 0 | 1 | 0 |
| rs201188114 | benign | 0 | 0 | 0 |
| rs1800472 | benign | 0 | 0 | 0 |
| rs587780728 | benign | 1 | 1 | 1 |
| rs144386518 | benign | 1 | 1 | 1 |
| rs281865439 | pathogenic | 1 | 1 | 1 |
| rs140175796 | pathogenic | 1 | 1 | 1 |
| rs5030843 | pathogenic | 1 | 1 | 0 |
| rs75166491 | pathogenic | 1 | 1 | 0 |
| rs199475679 | pathogenic | 1 | 1 | 1 |
| rs281865440 | pathogenic | 1 | 1 | 1 |
| rs74486803 | pathogenic | 0 | 1 | 0 |

| | | | | |
|---|---|---|---|---|
| rs199475602 | pathogenic | 0 | 1 | 0 |
| rs5030844 | pathogenic | 1 | 1 | 0 |
| rs62514927 | pathogenic | 1 | 1 | 1 |
| rs62514928 | pathogenic | 1 | 1 | 1 |
| rs62516152 | pathogenic | 0 | 1 | 0 |
| rs62507348 | pathogenic | 1 | 1 | 1 |
| rs76687508 | pathogenic | 1 | 1 | 1 |
| rs62508730 | pathogenic | 1 | 1 | 1 |
| rs62508588 | pathogenic | 0 | 1 | 0 |
| rs5030846 | pathogenic | 0 | 1 | 0 |
| rs74503222 | pathogenic | 1 | 1 | 0 |
| rs5030847 | pathogenic | 1 | 1 | 0 |
| rs5030850 | pathogenic | 1 | 1 | 1 |
| rs778154939 | pathogenic | 0 | 1 | 0 |
| rs62514952 | pathogenic | 1 | 1 | 1 |
| rs78655458 | pathogenic | 1 | 1 | 1 |
| rs62508698 | pathogenic | 1 | 1 | 0 |
| rs5030851 | pathogenic | 1 | 1 | 0 |
| rs62642933 | pathogenic | 1 | 1 | 1 |
| rs5030853 | pathogenic | 1 | 1 | 1 |
| rs62642935 | pathogenic | 1 | 1 | 1 |
| rs199475615 | pathogenic | 0 | 0 | 0 |
| rs62516062 | pathogenic | 1 | 1 | 0 |
| rs62516092 | pathogenic | 0 | 0 | 0 |
| rs62516095 | pathogenic | 1 | 1 | 1 |
| rs62642937 | pathogenic | 1 | 1 | 1 |
| rs62516141 | pathogenic | 0 | 1 | 0 |
| rs5030856 | pathogenic | 1 | 1 | 1 |
| rs5030857 | pathogenic | 1 | 1 | 1 |
| rs5030858 | pathogenic | 1 | 1 | 1 |
| rs79931499 | pathogenic | 1 | 1 | 1 |
| rs5030860 | pathogenic | 1 | 1 | 1 |
| rs281865437 | pathogenic | 1 | 1 | 1 |
| rs62644499 | pathogenic | 1 | 1 | 1 |
| rs147689487 | benign | 0 | 0 | 0 |
| rs397515417 | pathogenic | 1 | 1 | 1 |
| rs121909332 | pathogenic | 1 | 1 | 1 |
| rs121909330 | pathogenic | 1 | 1 | 0 |
| rs121909329 | pathogenic | 1 | 1 | 0 |
| rs121909335 | pathogenic | 1 | 1 | 1 |
| rs387906789 | pathogenic | 1 | 1 | 1 |
| rs121909334 | pathogenic | 1 | 1 | 1 |
| rs121909331 | pathogenic | 1 | 1 | 1 |
| rs372487178 | pathogenic | 1 | 1 | 1 |
| rs387906937 | pathogenic | 1 | 1 | 0 |
| rs16845 | benign | 1 | 1 | 1 |
| rs672601313 | pathogenic | 1 | 1 | 1 |
| rs117805308 | benign | 0 | 0 | 0 |
| rs121434261 | pathogenic | 1 | 1 | 1 |
| rs141629512 | benign | 0 | 0 | 0 |
| rs199957176 | benign | 1 | 1 | 1 |
| rs2230461 | benign | 1 | 1 | 1 |
| rs764618040 | pathogenic | 1 | 1 | 1 |
| rs267606949 | pathogenic | 1 | 1 | 1 |

| | | | | |
|---|---|---|---|---|
| rs587777483 | pathogenic | 1 | 1 | 1 |
| rs267606947 | pathogenic | 1 | 1 | 1 |
| rs6941583 | benign | 0 | 0 | 0 |
| rs749597959 | benign | 1 | 1 | 1 |
| rs182340753 | benign | 1 | 1 | 1 |
| rs146678221 | benign | 0 | 0 | 0 |
| rs11539202 | benign | 0 | 0 | 0 |
| rs148723565 | benign | 0 | 0 | 0 |
| rs61752925 | benign | 1 | 1 | 1 |
| rs113309941 | pathogenic | 0 | 0 | 0 |
| rs121434520 | pathogenic | 1 | 1 | 1 |
| rs121434517 | pathogenic | 1 | 1 | 1 |
| rs267607027 | pathogenic | 1 | 1 | 1 |
| rs138603088 | pathogenic | 0 | 0 | 0 |
| rs121434516 | pathogenic | 1 | 1 | 0 |
| rs121434521 | pathogenic | 0 | 0 | 0 |
| rs145735112 | benign | 1 | 1 | 1 |
| rs121434519 | pathogenic | 1 | 1 | 1 |
| rs75326924 | pathogenic | 1 | 1 | 0 |
| rs142186404 | pathogenic | 0 | 0 | 0 |
| rs121918035 | pathogenic | 0 | 0 | 0 |
| rs374385878 | pathogenic | 1 | 1 | 0 |
| rs11557652 | benign | 0 | 0 | 0 |
| rs118203986 | pathogenic | 1 | 1 | 1 |
| rs141675330 | benign | 1 | 1 | 1 |
| rs104893927 | pathogenic | 0 | 0 | 0 |
| rs104893935 | pathogenic | 0 | 0 | 0 |
| rs104893933 | pathogenic | 1 | 1 | 0 |
| rs104893934 | pathogenic | 0 | 0 | 0 |
| rs145183203 | benign | 1 | 1 | 0 |
| rs2230806 | benign | 0 | 0 | 0 |
| rs9282541 | benign | 0 | 0 | 0 |
| rs559110055 | benign | 0 | 0 | 0 |
| rs200030513 | benign | 0 | 0 | 0 |
| rs9282543 | benign | 0 | 0 | 0 |
| rs147675550 | benign | 0 | 0 | 0 |
| rs137854496 | pathogenic | 0 | 0 | 0 |
| rs374190304 | benign | 0 | 0 | 0 |
| rs114620717 | benign | 0 | 0 | 0 |
| rs2066718 | benign | 0 | 0 | 0 |
| rs35819696 | benign | 0 | 0 | 0 |
| rs138880920 | benign | 0 | 0 | 0 |
| rs2066715 | benign | 0 | 0 | 0 |
| rs35207495 | benign | 0 | 0 | 0 |
| rs2066714 | benign | 0 | 0 | 0 |
| rs28937313 | pathogenic | 0 | 0 | 0 |
| rs28937314 | pathogenic | 0 | 0 | 0 |
| rs137854495 | pathogenic | 0 | 0 | 0 |
| rs140365800 | benign | 0 | 0 | 0 |
| rs13306073 | benign | 0 | 0 | 0 |
| rs28933692 | pathogenic | 0 | 0 | 0 |
| rs33918808 | benign | 0 | 0 | 0 |
| rs76881554 | benign | 0 | 0 | 0 |
| rs143180998 | benign | 0 | 0 | 0 |

| | | | | |
|---|---|---|---|---|
| rs137854500 | pathogenic | 0 | 0 | 0 |
| rs538614702 | benign | 0 | 0 | 0 |
| rs201879964 | benign | 0 | 0 | 0 |
| rs2230808 | benign | 0 | 0 | 0 |
| rs138422574 | benign | 0 | 0 | 0 |
| rs137854498 | pathogenic | 0 | 0 | 0 |
| rs150125857 | benign | 0 | 0 | 0 |
| rs146292819 | pathogenic | 0 | 0 | 0 |
| rs142688906 | benign | 0 | 0 | 0 |
| rs137854499 | pathogenic | 0 | 0 | 0 |
| rs34879708 | benign | 0 | 0 | 0 |
| rs121913028 | pathogenic | 1 | 1 | 1 |
| rs1041951 | benign | 1 | 1 | 0 |
| rs138714094 | benign | 1 | 1 | 0 |
| rs200650502 | benign | 0 | 1 | 0 |
| rs370708663 | pathogenic | 1 | 1 | 1 |
| rs118204040 | pathogenic | 0 | 1 | 0 |
| rs146899669 | pathogenic | 1 | 1 | 0 |
| rs118204039 | pathogenic | 0 | 1 | 0 |
| rs121909233 | pathogenic | 0 | 0 | 0 |
| rs121909225 | pathogenic | 1 | 1 | 0 |
| rs121909236 | pathogenic | 0 | 0 | 0 |
| rs398123317 | pathogenic | 0 | 0 | 0 |
| rs121909226 | pathogenic | 1 | 1 | 0 |
| rs121909238 | pathogenic | 1 | 1 | 0 |
| rs121909230 | pathogenic | 0 | 0 | 0 |
| rs121909237 | pathogenic | 0 | 0 | 0 |
| rs121909222 | pathogenic | 1 | 1 | 0 |
| rs121909223 | pathogenic | 1 | 1 | 0 |
| rs121909218 | pathogenic | 1 | 1 | 0 |
| rs121909229 | pathogenic | 1 | 1 | 0 |
| rs121909224 | pathogenic | 1 | 1 | 0 |
| rs121909241 | pathogenic | 0 | 0 | 0 |
| rs121909221 | pathogenic | 0 | 0 | 0 |
| rs121913294 | pathogenic | 0 | 0 | 0 |
| rs121909234 | pathogenic | 0 | 0 | 0 |
| rs121909240 | pathogenic | 0 | 0 | 0 |
| rs767623493 | benign | 0 | 0 | 0 |
| rs121909239 | pathogenic | 0 | 0 | 0 |
| rs120074145 | pathogenic | 1 | 1 | 1 |
| rs120074148 | pathogenic | 1 | 1 | 1 |
| rs762991875 | pathogenic | 0 | 0 | 0 |
| rs148639841 | pathogenic | 1 | 1 | 1 |
| rs199524907 | pathogenic | 1 | 1 | 1 |
| rs120074141 | pathogenic | 1 | 1 | 1 |
| rs120074144 | pathogenic | 0 | 1 | 0 |
| rs120074146 | pathogenic | 1 | 1 | 1 |
| rs120074147 | pathogenic | 0 | 1 | 0 |
| rs120074143 | pathogenic | 0 | 1 | 0 |
| rs120074140 | pathogenic | 1 | 1 | 0 |
| rs28937315 | pathogenic | 1 | 1 | 1 |
| rs76066357 | benign | 1 | 1 | 1 |
| rs137852911 | pathogenic | 0 | 1 | 0 |
| rs137852907 | pathogenic | 1 | 1 | 0 |

| | | | | |
|---|---|---|---|---|
| rs137852910 | pathogenic | 1 | 1 | 1 |
| rs537585211 | benign | 0 | 0 | 0 |
| rs202134424 | pathogenic | 0 | 0 | 0 |
| rs200930463 | pathogenic | 0 | 0 | 0 |
| rs587777241 | pathogenic | 1 | 1 | 0 |
| rs77563738 | pathogenic | 0 | 0 | 0 |
| rs750868279 | pathogenic | 1 | 1 | 0 |
| rs148936893 | pathogenic | 0 | 0 | 0 |
| rs146597836 | pathogenic | 0 | 0 | 0 |
| rs775440641 | pathogenic | 1 | 1 | 1 |
| rs770689762 | pathogenic | 0 | 1 | 0 |
| rs376785840 | pathogenic | 0 | 0 | 0 |
| rs137852531 | pathogenic | 0 | 0 | 0 |
| rs137852538 | pathogenic | 1 | 1 | 0 |
| rs137852529 | pathogenic | 0 | 1 | 0 |
| rs137852528 | pathogenic | 0 | 1 | 0 |
| rs137852535 | pathogenic | 1 | 1 | 0 |
| rs137853045 | pathogenic | 0 | 1 | 0 |
| rs137853046 | pathogenic | 1 | 1 | 1 |
| rs137853043 | pathogenic | 1 | 1 | 1 |
| rs137853044 | pathogenic | 1 | 1 | 1 |
| rs137853047 | pathogenic | 1 | 1 | 1 |
| rs267606990 | pathogenic | 1 | 1 | 1 |
| rs751437780 | pathogenic | 1 | 1 | 0 |
| rs121918461 | pathogenic | 1 | 1 | 1 |
| rs121918460 | pathogenic | 1 | 1 | 1 |
| rs121918459 | pathogenic | 1 | 1 | 0 |
| rs397507511 | pathogenic | 1 | 1 | 1 |
| rs121918454 | pathogenic | 1 | 1 | 1 |
| rs121918453 | pathogenic | 1 | 1 | 1 |
| rs121918462 | pathogenic | 1 | 1 | 0 |
| rs121918465 | pathogenic | 1 | 1 | 0 |
| rs121918464 | pathogenic | 1 | 1 | 0 |
| rs121918466 | pathogenic | 1 | 1 | 1 |
| rs397507521 | pathogenic | 1 | 1 | 1 |
| rs397507527 | pathogenic | 1 | 1 | 1 |
| rs777603059 | pathogenic | 0 | 1 | 0 |
| rs121918456 | pathogenic | 1 | 1 | 1 |
| rs121918463 | pathogenic | 1 | 1 | 1 |
| rs121918455 | pathogenic | 0 | 1 | 0 |
| rs28933386 | pathogenic | 0 | 1 | 0 |
| rs774356443 | benign | 1 | 1 | 1 |
| rs121918467 | pathogenic | 1 | 1 | 1 |
| rs121918468 | pathogenic | 1 | 1 | 1 |
| rs121918469 | pathogenic | 1 | 1 | 1 |
| rs397507539 | pathogenic | 1 | 1 | 0 |
| rs121918458 | pathogenic | 1 | 1 | 1 |
| rs121918470 | pathogenic | 1 | 1 | 1 |
| rs35289681 | benign | 0 | 1 | 0 |
| rs4302331 | benign | 0 | 1 | 0 |
| rs11544803 | pathogenic | 1 | 1 | 1 |
| rs116254156 | benign | 0 | 0 | 0 |
| rs104893909 | pathogenic | 1 | 1 | 1 |
| rs104893911 | pathogenic | 1 | 1 | 1 |

| | | | | |
|---|---|---|---|---|
| rs104893908 | pathogenic | 1 | 1 | 0 |
| rs258751 | benign | 1 | 1 | 1 |
| rs104893914 | pathogenic | 1 | 1 | 0 |
| rs121909727 | pathogenic | 1 | 1 | 1 |
| rs104893910 | pathogenic | 1 | 1 | 1 |
| rs121909726 | pathogenic | 1 | 1 | 1 |
| rs104893912 | pathogenic | 1 | 1 | 1 |
| rs138407155 | benign | 1 | 1 | 1 |
| rs574061789 | benign | 1 | 1 | 1 |
| rs121912718 | pathogenic | 1 | 1 | 1 |
| rs121912719 | pathogenic | 1 | 1 | 1 |
| rs140770089 | benign | 1 | 1 | 1 |
| rs28931573 | pathogenic | 1 | 1 | 1 |
| rs121912717 | pathogenic | 1 | 1 | 1 |
| rs376456598 | pathogenic | 0 | 0 | 0 |
| rs398123499 | pathogenic | 1 | 1 | 1 |
| rs773048903 | pathogenic | 1 | 1 | 0 |
| rs398123503 | pathogenic | 1 | 1 | 1 |
| rs375785084 | pathogenic | 0 | 1 | 0 |
| rs137852874 | pathogenic | 1 | 1 | 1 |
| rs373713279 | pathogenic | 1 | 1 | 1 |
| rs137852873 | pathogenic | 1 | 1 | 0 |
| rs367823977 | pathogenic | 1 | 1 | 1 |
| rs398123508 | pathogenic | 1 | 1 | 1 |
| rs137852871 | pathogenic | 1 | 1 | 1 |
| rs753698250 | pathogenic | 0 | 1 | 0 |
| rs182923857 | pathogenic | 1 | 1 | 1 |
| rs148090804 | benign | 1 | 1 | 1 |
| rs137852870 | pathogenic | 1 | 1 | 1 |
| rs267607131 | pathogenic | 1 | 1 | 0 |
| rs137852902 | pathogenic | 0 | 0 | 0 |
| rs61741646 | benign | 1 | 1 | 1 |
| rs137852905 | pathogenic | 0 | 0 | 0 |
| rs121918681 | pathogenic | 0 | 1 | 0 |
| rs8177238 | benign | 1 | 1 | 0 |
| rs5030776 | pathogenic | 1 | 1 | 1 |
| rs104894572 | pathogenic | 0 | 0 | 0 |
| rs104894576 | pathogenic | 0 | 0 | 0 |
| rs80084721 | pathogenic | 0 | 0 | 0 |
| rs104894146 | pathogenic | 0 | 1 | 0 |
| rs104894138 | pathogenic | 1 | 1 | 0 |
| rs104894153 | pathogenic | 1 | 1 | 0 |
| rs104894135 | pathogenic | 1 | 1 | 0 |
| rs104894147 | pathogenic | 1 | 1 | 0 |
| rs104894148 | pathogenic | 1 | 1 | 0 |
| rs104894154 | pathogenic | 1 | 1 | 0 |
| rs104894150 | pathogenic | 1 | 1 | 0 |
| rs104894136 | pathogenic | 1 | 1 | 0 |
| rs104894144 | pathogenic | 0 | 0 | 0 |
| rs104894137 | pathogenic | 0 | 0 | 0 |
| rs61754278 | pathogenic | 1 | 1 | 0 |
| rs104894149 | pathogenic | 1 | 1 | 0 |
| rs104894139 | pathogenic | 1 | 1 | 0 |
| rs104894142 | pathogenic | 0 | 0 | 0 |

| | | | | |
|---|---|---|---|---|
| rs104894143 | pathogenic | 0 | 0 | 0 |
| rs104894145 | pathogenic | 0 | 0 | 0 |
| rs121909545 | benign | 0 | 1 | 0 |
| rs782308462 | pathogenic | 1 | 1 | 1 |
| rs375002796 | pathogenic | 0 | 1 | 0 |
| rs201727685 | pathogenic | 0 | 0 | 0 |
| rs768050261 | pathogenic | 1 | 1 | 1 |
| rs780851340 | benign | 0 | 0 | 0 |
| rs139824241 | benign | 1 | 1 | 0 |
| rs757486575 | pathogenic | 1 | 1 | 0 |
| rs750831299 | pathogenic | 1 | 1 | 1 |
| rs767602961 | pathogenic | 1 | 1 | 1 |
| rs121913661 | pathogenic | 1 | 1 | 1 |
| rs777692271 | pathogenic | 1 | 1 | 1 |
| rs121913659 | pathogenic | 1 | 1 | 1 |
| rs587777533 | pathogenic | 1 | 1 | 1 |
| rs104894210 | pathogenic | 0 | 0 | 0 |
| rs104894216 | pathogenic | 1 | 1 | 1 |
| rs104894207 | pathogenic | 1 | 1 | 1 |
| rs104894208 | pathogenic | 0 | 1 | 0 |
| rs104894214 | pathogenic | 1 | 1 | 1 |
| rs104894211 | pathogenic | 1 | 1 | 1 |
| rs61749384 | pathogenic | 0 | 0 | 0 |
| rs61750071 | pathogenic | 1 | 1 | 1 |
| rs61750074 | pathogenic | 0 | 0 | 0 |
| rs11063988 | benign | 0 | 0 | 0 |
| rs216311 | benign | 0 | 0 | 0 |
| rs1800382 | benign | 1 | 1 | 1 |
| rs11063987 | benign | 1 | 1 | 1 |
| rs387906847 | pathogenic | 1 | 1 | 1 |
| rs73003466 | benign | 0 | 0 | 0 |
| rs149787233 | benign | 1 | 1 | 1 |
| rs144451841 | pathogenic | 1 | 1 | 1 |
| rs141638421 | pathogenic | 1 | 1 | 1 |
| rs121434497 | pathogenic | 0 | 0 | 0 |
| rs121434498 | pathogenic | 0 | 0 | 0 |
| rs267607230 | pathogenic | 1 | 1 | 0 |
| rs121434499 | pathogenic | 1 | 1 | 0 |
| rs546749336 | benign | 1 | 1 | 1 |
| rs138873805 | benign | 0 | 0 | 0 |
| rs730880522 | pathogenic | 0 | 0 | 0 |
| rs200371894 | benign | 0 | 0 | 0 |
| rs730880509 | benign | 0 | 0 | 0 |
| rs267606704 | pathogenic | 1 | 1 | 1 |
| rs267606706 | pathogenic | 0 | 0 | 0 |
| rs267606705 | pathogenic | 1 | 1 | 1 |
| rs267606707 | pathogenic | 0 | 0 | 0 |
| rs387906665 | pathogenic | 1 | 1 | 1 |
| rs267606708 | pathogenic | 1 | 1 | 1 |
| rs148368481 | benign | 0 | 0 | 0 |
| rs13204 | benign | 0 | 0 | 0 |
| rs140547520 | pathogenic | 1 | 1 | 1 |
| rs755234990 | benign | 1 | 1 | 0 |
| rs151184052 | benign | 0 | 0 | 0 |

| | | | | |
|---|---|---|---|---|
| rs3736358 | benign | 0 | 0 | 0 |
| rs3736360 | benign | 0 | 0 | 0 |
| rs1138469 | benign | 0 | 0 | 0 |
| rs74315351 | pathogenic | 0 | 1 | 0 |
| rs137853051 | pathogenic | 1 | 1 | 0 |
| rs74315353 | pathogenic | 0 | 1 | 0 |
| rs74315352 | pathogenic | 0 | 1 | 0 |
| rs74315354 | pathogenic | 1 | 1 | 1 |
| rs28938172 | pathogenic | 1 | 1 | 1 |
| rs52836744 | pathogenic | 0 | 0 | 0 |
| rs121913159 | pathogenic | 1 | 1 | 1 |
| rs121913145 | pathogenic | 1 | 1 | 1 |
| rs891087 | benign | 1 | 1 | 1 |
| rs9282757 | benign | 1 | 1 | 1 |
| rs397517404 | pathogenic | 1 | 1 | 1 |
| rs199918720 | benign | 1 | 1 | 0 |
| rs368299411 | benign | 1 | 1 | 0 |
| rs139399951 | benign | 1 | 1 | 0 |
| rs763981974 | benign | 1 | 1 | 0 |
| rs144242114 | benign | 0 | 0 | 0 |
| rs750965801 | benign | 0 | 0 | 0 |
| rs561310777 | benign | 0 | 0 | 0 |
| rs148185335 | benign | 0 | 0 | 0 |
| rs147742157 | benign | 0 | 0 | 0 |
| rs375664373 | pathogenic | 1 | 1 | 1 |
| rs777327250 | pathogenic | 0 | 0 | 0 |
| rs121908471 | pathogenic | 1 | 1 | 0 |
| rs2301612 | pathogenic | 1 | 1 | 0 |
| rs121908476 | pathogenic | 0 | 0 | 0 |
| rs36220239 | benign | 0 | 0 | 0 |
| rs121908473 | pathogenic | 0 | 0 | 0 |
| rs28647808 | benign | 0 | 0 | 0 |
| rs36090624 | benign | 0 | 0 | 0 |
| rs121908002 | pathogenic | 1 | 1 | 1 |
| rs4251545 | benign | 1 | 1 | 1 |
| rs78705382 | pathogenic | 0 | 0 | 0 |
| rs75184679 | pathogenic | 1 | 1 | 1 |
| rs74555752 | pathogenic | 0 | 0 | 0 |
| rs56320207 | benign | 1 | 1 | 1 |
| rs28730837 | pathogenic | 1 | 1 | 1 |
| rs119469014 | pathogenic | 1 | 1 | 1 |
| rs119468010 | pathogenic | 1 | 1 | 1 |
| rs35220041 | benign | 0 | 0 | 0 |
| rs762321137 | pathogenic | 1 | 1 | 1 |
| rs28931570 | pathogenic | 1 | 1 | 1 |
| rs709932 | benign | 1 | 1 | 1 |
| rs746531546 | benign | 1 | 1 | 1 |
| rs6647 | benign | 1 | 1 | 1 |
| rs141095970 | benign | 1 | 1 | 1 |
| rs143370956 | benign | 1 | 1 | 1 |
| rs28929474 | pathogenic | 1 | 1 | 1 |
| rs61761869 | pathogenic | 1 | 1 | 1 |
| rs1303 | benign | 1 | 1 | 1 |
| rs149447502 | benign | 0 | 0 | 0 |

| | | | | |
|---|---|---|---|---|
| rs267607180 | pathogenic | 1 | 1 | 1 |
| rs150304121 | benign | 0 | 0 | 0 |
| rs147630375 | benign | 1 | 1 | 1 |
| rs11539445 | benign | 1 | 1 | 1 |
| rs372098364 | pathogenic | 1 | 1 | 0 |
| rs760072286 | benign | 0 | 0 | 0 |
| rs745595833 | pathogenic | 1 | 1 | 1 |
| rs587777214 | pathogenic | 0 | 0 | 0 |
| rs150318528 | benign | 0 | 0 | 0 |
| rs137853254 | pathogenic | 1 | 1 | 1 |
| rs137853255 | pathogenic | 0 | 0 | 0 |
| rs137853253 | pathogenic | 0 | 0 | 0 |
| rs202166915 | benign | 0 | 0 | 0 |
| rs137853259 | pathogenic | 0 | 0 | 0 |
| rs2229137 | benign | 0 | 0 | 0 |
| rs137853258 | pathogenic | 1 | 1 | 0 |
| rs137853252 | pathogenic | 1 | 1 | 1 |
| rs137853256 | pathogenic | 0 | 0 | 0 |
| rs137853250 | pathogenic | 1 | 1 | 1 |
| rs121434615 | pathogenic | 0 | 0 | 0 |
| rs115014558 | benign | 1 | 1 | 1 |
| rs34116584 | benign | 1 | 1 | 1 |
| rs34885252 | benign | 1 | 1 | 1 |
| rs180177262 | pathogenic | 1 | 1 | 1 |
| rs180177162 | pathogenic | 1 | 1 | 1 |
| rs121908523 | pathogenic | 1 | 1 | 1 |
| rs180177170 | pathogenic | 1 | 1 | 1 |
| rs180177180 | pathogenic | 1 | 1 | 1 |
| rs767586362 | pathogenic | 1 | 1 | 1 |
| rs121908521 | pathogenic | 1 | 1 | 1 |
| rs180177203 | pathogenic | 1 | 1 | 1 |
| rs138025751 | pathogenic | 1 | 1 | 1 |
| rs180177210 | pathogenic | 0 | 1 | 0 |
| rs180177211 | pathogenic | 0 | 1 | 0 |
| rs121908524 | pathogenic | 1 | 1 | 0 |
| rs121908530 | pathogenic | 1 | 1 | 0 |
| rs180177227 | pathogenic | 0 | 1 | 0 |
| rs180177230 | pathogenic | 1 | 1 | 1 |
| rs121908529 | pathogenic | 0 | 1 | 0 |
| rs117195882 | benign | 1 | 1 | 0 |
| rs180177238 | pathogenic | 1 | 1 | 0 |
| rs34664134 | benign | 1 | 1 | 1 |
| rs180177246 | pathogenic | 0 | 1 | 0 |
| rs536352238 | pathogenic | 0 | 0 | 0 |
| rs121908520 | pathogenic | 1 | 1 | 0 |
| rs180177248 | pathogenic | 1 | 1 | 0 |
| rs180177252 | pathogenic | 1 | 1 | 1 |
| rs121908526 | pathogenic | 1 | 1 | 1 |
| rs121908525 | pathogenic | 1 | 1 | 1 |
| rs147106773 | benign | 1 | 1 | 1 |
| rs180177264 | pathogenic | 0 | 1 | 0 |
| rs146525143 | pathogenic | 1 | 1 | 1 |
| rs140992177 | benign | 1 | 1 | 1 |
| rs13408961 | benign | 0 | 1 | 0 |

| | | | | |
|---|---|---|---|---|
| rs4426527 | benign | 1 | 1 | 1 |
| rs3088308 | benign | 0 | 1 | 0 |
| rs4469829 | benign | 1 | 1 | 0 |
| rs104886003 | pathogenic | 1 | 1 | 1 |
| rs121913286 | pathogenic | 1 | 1 | 1 |
| rs121913279 | pathogenic | 1 | 1 | 1 |
| rs121434636 | pathogenic | 1 | 1 | 1 |
| rs755299132 | pathogenic | 1 | 1 | 1 |
| rs748318386 | pathogenic | 1 | 1 | 1 |
| rs387906757 | pathogenic | 1 | 1 | 1 |
| rs121965064 | pathogenic | 1 | 1 | 1 |
| rs28934608 | pathogenic | 0 | 0 | 0 |
| rs5967 | benign | 1 | 1 | 1 |
| rs773905328 | pathogenic | 0 | 0 | 0 |
| rs104894256 | pathogenic | 0 | 1 | 0 |
| rs28931612 | pathogenic | 0 | 1 | 0 |
| rs104894263 | pathogenic | 1 | 1 | 1 |
| rs607969 | benign | 1 | 1 | 0 |
| rs104894262 | pathogenic | 0 | 1 | 0 |
| rs104894268 | pathogenic | 1 | 1 | 1 |
| rs104894264 | pathogenic | 0 | 1 | 0 |
| rs104894259 | pathogenic | 1 | 1 | 0 |
| rs2959656 | benign | 0 | 0 | 0 |
| rs121913035 | pathogenic | 1 | 1 | 0 |
| rs139605474 | benign | 0 | 0 | 0 |
| rs3817552 | benign | 0 | 0 | 0 |
| rs150541115 | benign | 0 | 0 | 0 |
| rs562763010 | benign | 1 | 1 | 1 |
| rs200053119 | pathogenic | 0 | 0 | 0 |
| rs73826386 | benign | 0 | 0 | 0 |
| rs143655258 | benign | 0 | 0 | 0 |
| rs758857853 | pathogenic | 1 | 1 | 0 |
| rs2228380 | benign | 0 | 0 | 0 |
| rs141078733 | benign | 1 | 1 | 1 |
| rs55824172 | pathogenic | 1 | 1 | 1 |
| rs756751089 | pathogenic | 1 | 1 | 1 |
| rs121964990 | pathogenic | 0 | 0 | 0 |
| rs143555186 | benign | 0 | 0 | 0 |
| rs121964993 | pathogenic | 1 | 1 | 0 |
| rs121964992 | pathogenic | 1 | 1 | 1 |
| rs121964991 | pathogenic | 1 | 1 | 0 |
| rs121964988 | pathogenic | 1 | 1 | 1 |
| rs4687 | benign | 1 | 1 | 1 |
| rs267607125 | pathogenic | 1 | 1 | 1 |
| rs751682878 | benign | 1 | 1 | 1 |
| rs267607126 | pathogenic | 1 | 1 | 1 |
| rs768713596 | pathogenic | 0 | 0 | 0 |
| rs147288996 | benign | 0 | 0 | 0 |
| rs387906639 | pathogenic | 1 | 1 | 1 |
| rs730881069 | pathogenic | 1 | 1 | 1 |
| rs730881071 | pathogenic | 1 | 1 | 1 |
| rs397516348 | pathogenic | 0 | 0 | 0 |
| rs121917760 | pathogenic | 0 | 0 | 0 |
| rs104894724 | pathogenic | 1 | 1 | 1 |

| | | | | |
|---|---|---|---|---|
| rs397516349 | pathogenic | 1 | 1 | 1 |
| rs12075 | benign | 0 | 0 | 0 |
| rs147549987 | benign | 0 | 0 | 0 |
| rs56079734 | benign | 0 | 0 | 0 |
| rs1017842 | benign | 1 | 1 | 1 |
| rs137852593 | pathogenic | 1 | 1 | 1 |
| rs137852571 | pathogenic | 1 | 1 | 1 |
| rs137852591 | pathogenic | 1 | 1 | 1 |
| rs9332971 | pathogenic | 1 | 1 | 1 |
| rs755226547 | pathogenic | 1 | 1 | 1 |
| rs79206939 | benign | 0 | 0 | 0 |
| rs121918214 | pathogenic | 1 | 1 | 0 |
| rs757075712 | pathogenic | 1 | 1 | 0 |
| rs150483902 | benign | 1 | 1 | 1 |
| rs113993973 | pathogenic | 0 | 1 | 0 |
| rs946616 | benign | 0 | 0 | 0 |
| rs113993976 | pathogenic | 1 | 1 | 0 |
| rs151022264 | benign | 0 | 1 | 0 |
| rs113993977 | pathogenic | 1 | 1 | 0 |
| rs113993978 | pathogenic | 1 | 1 | 0 |
| rs2228499 | benign | 0 | 1 | 0 |
| rs113993979 | pathogenic | 1 | 1 | 0 |
| rs113993980 | pathogenic | 1 | 1 | 0 |
| rs113993983 | pathogenic | 0 | 1 | 0 |
| rs150547274 | pathogenic | 1 | 1 | 0 |
| rs113993984 | pathogenic | 1 | 1 | 0 |
| rs113993986 | pathogenic | 1 | 1 | 0 |
| rs34313873 | benign | 0 | 1 | 0 |
| rs113993988 | pathogenic | 1 | 1 | 0 |
| rs199638097 | pathogenic | 1 | 1 | 1 |
| rs118204447 | pathogenic | 1 | 1 | 1 |
| rs11862754 | benign | 0 | 0 | 0 |
| rs118204445 | pathogenic | 0 | 0 | 0 |
| rs118204441 | pathogenic | 0 | 0 | 0 |
| rs118204438 | pathogenic | 0 | 0 | 0 |
| rs118204436 | pathogenic | 0 | 0 | 0 |
| rs146093755 | pathogenic | 0 | 0 | 0 |
| rs781439830 | pathogenic | 0 | 0 | 0 |
| rs398123438 | pathogenic | 0 | 0 | 0 |
| rs398123439 | pathogenic | 0 | 0 | 0 |
| rs118204444 | pathogenic | 0 | 0 | 0 |
| rs7187889 | benign | 0 | 0 | 0 |
| rs118204435 | pathogenic | 1 | 1 | 0 |
| rs34745339 | benign | 0 | 0 | 0 |
| rs398123440 | pathogenic | 0 | 0 | 0 |
| rs118204442 | pathogenic | 0 | 0 | 0 |
| rs770053354 | pathogenic | 0 | 0 | 0 |
| rs118204448 | pathogenic | 0 | 0 | 0 |
| rs118204443 | pathogenic | 0 | 0 | 0 |
| rs118204446 | pathogenic | 0 | 0 | 0 |
| rs746756997 | pathogenic | 0 | 0 | 0 |
| rs267606838 | pathogenic | 0 | 0 | 0 |
| rs118204437 | pathogenic | 0 | 0 | 0 |
| rs398123429 | pathogenic | 0 | 0 | 0 |

| | | | | |
|---|---|---|---|---|
| rs398123430 | pathogenic | 0 | 0 | 0 |
| rs2303269 | benign | 0 | 0 | 0 |
| rs114703967 | benign | 0 | 0 | 0 |
| rs118204439 | pathogenic | 0 | 0 | 0 |
| rs151296605 | benign | 0 | 0 | 0 |
| rs118204440 | pathogenic | 0 | 0 | 0 |
| rs78127134 | benign | 0 | 0 | 0 |
| rs372893383 | pathogenic | 0 | 0 | 0 |
| rs61753266 | pathogenic | 1 | 1 | 1 |
| rs139579928 | benign | 1 | 1 | 1 |
| rs56275071 | pathogenic | 1 | 1 | 1 |
| rs121909736 | pathogenic | 1 | 1 | 0 |
| rs121909737 | pathogenic | 1 | 1 | 0 |
| rs121909735 | pathogenic | 1 | 1 | 0 |
| rs121909731 | pathogenic | 1 | 1 | 1 |
| rs121909734 | pathogenic | 1 | 1 | 1 |
| rs121909733 | pathogenic | 1 | 1 | 1 |
| rs121909732 | pathogenic | 1 | 1 | 1 |
| rs121909730 | pathogenic | 1 | 1 | 1 |
| rs202198360 | benign | 1 | 1 | 1 |
| rs587777294 | pathogenic | 1 | 1 | 0 |
| rs587777297 | pathogenic | 1 | 1 | 0 |
| rs104893913 | pathogenic | 1 | 1 | 0 |
| rs17860424 | pathogenic | 1 | 1 | 1 |
| rs1045485 | benign | 1 | 1 | 1 |
| rs13926 | benign | 0 | 0 | 0 |
| rs79343837 | benign | 0 | 0 | 0 |
| rs45516306 | benign | 1 | 1 | 1 |
| rs139849355 | benign | 1 | 1 | 1 |
| rs104894874 | pathogenic | 1 | 1 | 1 |
| rs104894870 | pathogenic | 1 | 1 | 1 |
| rs137852221 | pathogenic | 1 | 1 | 1 |
| rs104894869 | pathogenic | 1 | 1 | 1 |
| rs104894880 | pathogenic | 1 | 1 | 1 |
| rs104894872 | pathogenic | 1 | 1 | 1 |
| rs104894868 | pathogenic | 1 | 1 | 1 |
| rs104894867 | pathogenic | 1 | 1 | 1 |
| rs104894871 | pathogenic | 1 | 1 | 1 |
| rs104894877 | pathogenic | 1 | 1 | 1 |
| rs104894883 | pathogenic | 1 | 1 | 1 |
| rs104894875 | pathogenic | 1 | 1 | 1 |
| rs104894876 | pathogenic | 1 | 1 | 1 |
| rs104894878 | pathogenic | 1 | 1 | 1 |
| rs137852220 | pathogenic | 1 | 1 | 1 |
| rs28933684 | pathogenic | 1 | 1 | 1 |
| rs104894873 | pathogenic | 1 | 1 | 1 |
| rs138409264 | benign | 0 | 0 | 0 |
| rs2236722 | benign | 0 | 0 | 0 |
| rs28757184 | benign | 0 | 0 | 0 |
| rs121434538 | pathogenic | 0 | 0 | 0 |
| rs700519 | benign | 0 | 0 | 0 |
| rs80051519 | pathogenic | 0 | 0 | 0 |
| rs121434536 | pathogenic | 1 | 1 | 0 |
| rs121434534 | pathogenic | 1 | 1 | 0 |

| | | | | |
|---|---|---|---|---|
| rs78310315 | pathogenic | 1 | 1 | 0 |
| rs540839115 | pathogenic | 1 | 1 | 1 |
| rs80338731 | pathogenic | 1 | 1 | 1 |
| rs180177151 | pathogenic | 1 | 1 | 1 |
| rs180177152 | pathogenic | 0 | 0 | 0 |
| rs80338732 | pathogenic | 1 | 1 | 1 |
| rs80338675 | pathogenic | 1 | 1 | 1 |
| rs80338676 | pathogenic | 1 | 1 | 1 |
| rs180177153 | pathogenic | 0 | 0 | 0 |
| rs180177154 | pathogenic | 1 | 1 | 1 |
| rs5896 | benign | 1 | 1 | 1 |
| rs121918477 | pathogenic | 0 | 0 | 0 |
| rs121918428 | pathogenic | 1 | 1 | 1 |
| rs121918427 | pathogenic | 1 | 1 | 1 |
| rs544300820 | benign | 1 | 1 | 1 |
| rs121918110 | pathogenic | 1 | 1 | 1 |
| rs1057141 | benign | 0 | 0 | 0 |
| rs1135216 | benign | 1 | 1 | 1 |
| rs121917702 | pathogenic | 1 | 1 | 1 |
| rs28941474 | pathogenic | 0 | 0 | 0 |
| rs104893944 | pathogenic | 1 | 1 | 1 |
| rs104893947 | pathogenic | 1 | 1 | 0 |
| rs104893943 | pathogenic | 1 | 1 | 1 |
| rs104893942 | pathogenic | 1 | 1 | 1 |
| rs104893940 | pathogenic | 1 | 1 | 1 |
| rs34045013 | benign | 0 | 0 | 0 |
| rs143083947 | pathogenic | 0 | 0 | 0 |
| rs145786986 | benign | 0 | 0 | 0 |
| rs119450942 | pathogenic | 0 | 0 | 0 |
| rs374259530 | pathogenic | 0 | 0 | 0 |
| rs756210458 | pathogenic | 0 | 0 | 0 |
| rs11089991 | benign | 0 | 0 | 0 |
| rs28941471 | pathogenic | 0 | 1 | 0 |
| rs750614500 | pathogenic | 0 | 1 | 0 |
| rs199761158 | benign | 1 | 1 | 1 |
| rs119450944 | pathogenic | 1 | 1 | 1 |
| rs373458753 | pathogenic | 1 | 1 | 1 |
| rs202064195 | pathogenic | 1 | 1 | 1 |
| rs776496275 | pathogenic | 1 | 1 | 1 |
| rs761493155 | pathogenic | 1 | 1 | 1 |
| rs763542069 | pathogenic | 0 | 0 | 0 |
| rs119450941 | pathogenic | 0 | 0 | 0 |
| rs119450940 | pathogenic | 0 | 0 | 0 |
| rs777821034 | pathogenic | 1 | 1 | 1 |
| rs771121666 | pathogenic | 1 | 1 | 1 |
| rs372895468 | pathogenic | 1 | 1 | 1 |
| rs61750603 | pathogenic | 0 | 0 | 0 |
| rs76863441 | pathogenic | 0 | 0 | 0 |
| rs142601143 | benign | 1 | 1 | 1 |
| rs185832993 | benign | 1 | 1 | 0 |
| rs17860568 | benign | 0 | 0 | 0 |
| rs142064825 | benign | 0 | 0 | 0 |
| rs564076183 | benign | 0 | 0 | 0 |
| rs386834034 | pathogenic | 0 | 0 | 0 |

| | | | | |
|---|---|---|---|---|
| rs147663769 | benign | 0 | 0 | 0 |
| rs776478974 | benign | 0 | 0 | 0 |
| rs111840783 | benign | 0 | 0 | 0 |
| rs1516446 | benign | 1 | 1 | 1 |
| rs143749154 | benign | 1 | 1 | 1 |
| rs80257412 | benign | 0 | 0 | 0 |
| rs142943120 | benign | 0 | 0 | 0 |
| rs144553482 | benign | 0 | 0 | 0 |
| rs35997569 | benign | 0 | 0 | 0 |
| rs772909106 | pathogenic | 0 | 0 | 0 |
| rs149433837 | benign | 1 | 1 | 1 |
| rs532178791 | pathogenic | 0 | 0 | 0 |
| rs145588689 | pathogenic | 1 | 1 | 0 |
| rs17843776 | benign | 1 | 1 | 0 |
| rs121917890 | pathogenic | 1 | 1 | 1 |
| rs121917892 | pathogenic | 0 | 0 | 0 |
| rs606231289 | pathogenic | 0 | 0 | 0 |
| rs370011798 | pathogenic | 0 | 0 | 0 |
| rs34689457 | benign | 0 | 0 | 0 |
| rs118204451 | pathogenic | 0 | 0 | 0 |
| rs118204452 | pathogenic | 1 | 1 | 1 |
| rs652785 | benign | 0 | 0 | 0 |
| rs35378126 | benign | 0 | 0 | 0 |
| rs80358214 | pathogenic | 1 | 1 | 1 |
| rs80358215 | pathogenic | 0 | 1 | 0 |
| rs267606858 | pathogenic | 1 | 1 | 1 |
| rs143137713 | pathogenic | 1 | 1 | 1 |
| rs782168634 | benign | 0 | 0 | 0 |
| rs143873938 | benign | 0 | 0 | 0 |
| rs782776511 | benign | 0 | 0 | 0 |
| rs201093148 | benign | 0 | 0 | 0 |
| rs267606815 | pathogenic | 0 | 0 | 0 |
| rs137853311 | pathogenic | 0 | 0 | 0 |
| rs267606817 | pathogenic | 0 | 0 | 0 |
| rs72650006 | benign | 0 | 0 | 0 |
| rs727503638 | benign | 1 | 1 | 0 |
| rs189286381 | benign | 0 | 0 | 0 |
| rs376644553 | benign | 0 | 0 | 0 |
| rs121434610 | pathogenic | 1 | 1 | 1 |
| rs267607076 | pathogenic | 1 | 1 | 1 |
| rs201275730 | benign | 0 | 0 | 0 |
| rs121909587 | pathogenic | 0 | 0 | 0 |
| rs121909558 | pathogenic | 1 | 1 | 0 |
| rs28929469 | pathogenic | 1 | 1 | 0 |
| rs121909547 | pathogenic | 1 | 1 | 0 |
| rs121909552 | pathogenic | 1 | 1 | 0 |
| rs121909573 | pathogenic | 1 | 1 | 1 |
| rs121909567 | pathogenic | 1 | 1 | 0 |
| rs121909569 | pathogenic | 0 | 0 | 0 |
| rs121909563 | pathogenic | 1 | 1 | 0 |
| rs121909570 | pathogenic | 1 | 1 | 1 |
| rs121909571 | pathogenic | 1 | 1 | 0 |
| rs121909572 | pathogenic | 1 | 1 | 1 |
| rs121909565 | pathogenic | 1 | 1 | 1 |

| | | | | |
|---|---|---|---|---|
| rs121909557 | pathogenic | 1 | 1 | 1 |
| rs121909568 | pathogenic | 1 | 1 | 1 |
| rs121909566 | pathogenic | 1 | 1 | 1 |
| rs121909549 | pathogenic | 1 | 1 | 1 |
| rs121909554 | pathogenic | 1 | 1 | 1 |
| rs121909550 | pathogenic | 1 | 1 | 1 |
| rs121909546 | pathogenic | 1 | 1 | 0 |
| rs121909555 | pathogenic | 1 | 1 | 1 |
| rs121909564 | pathogenic | 1 | 1 | 1 |
| rs548046212 | pathogenic | 0 | 0 | 0 |
| rs147353781 | benign | 0 | 0 | 0 |
| rs104894678 | pathogenic | 1 | 1 | 1 |
| rs1130426 | benign | 0 | 0 | 0 |
| rs104894677 | pathogenic | 1 | 1 | 1 |
| rs141917423 | benign | 1 | 1 | 1 |
| rs574777883 | benign | 0 | 0 | 0 |
| rs150199226 | benign | 1 | 1 | 1 |
| rs199618701 | benign | 0 | 1 | 0 |
| rs121964923 | pathogenic | 0 | 0 | 0 |
| rs121964924 | pathogenic | 0 | 0 | 0 |
| rs201258823 | pathogenic | 0 | 1 | 0 |
| rs267606773 | pathogenic | 0 | 0 | 0 |
| rs35638832 | benign | 0 | 0 | 0 |
| rs121908632 | pathogenic | 1 | 1 | 1 |
| rs121908625 | pathogenic | 1 | 1 | 1 |
| rs748949603 | pathogenic | 0 | 0 | 0 |
| rs757523840 | pathogenic | 0 | 0 | 0 |
| rs62541771 | pathogenic | 0 | 0 | 0 |
| rs776384541 | pathogenic | 1 | 1 | 0 |
| rs121908627 | pathogenic | 1 | 1 | 0 |
| rs28937594 | pathogenic | 0 | 0 | 0 |
| rs121907888 | pathogenic | 0 | 0 | 0 |
| rs121907889 | pathogenic | 0 | 1 | 0 |
| rs28941785 | pathogenic | 1 | 1 | 1 |
| rs28941786 | pathogenic | 1 | 1 | 1 |
| rs1021737 | benign | 0 | 0 | 0 |
| rs121908370 | pathogenic | 1 | 1 | 1 |
| rs63750083 | pathogenic | 0 | 0 | 0 |
| rs17883862 | benign | 0 | 0 | 0 |
| rs587782070 | pathogenic | 1 | 1 | 1 |
| rs587781269 | pathogenic | 1 | 1 | 1 |
| rs28909982 | pathogenic | 1 | 1 | 1 |
| rs373648967 | benign | 1 | 1 | 1 |
| rs368570187 | benign | 1 | 1 | 1 |
| rs730881701 | pathogenic | 1 | 1 | 1 |
| rs137853007 | pathogenic | 0 | 0 | 0 |
| rs575910805 | benign | 0 | 0 | 0 |
| rs104893875 | pathogenic | 1 | 1 | 1 |
| rs104893877 | pathogenic | 1 | 1 | 1 |
| rs119485093 | pathogenic | 1 | 1 | 1 |
| rs119485092 | pathogenic | 0 | 0 | 0 |
| rs119485090 | pathogenic | 0 | 0 | 0 |
| rs28942099 | pathogenic | 1 | 1 | 1 |
| rs373730800 | pathogenic | 1 | 1 | 1 |

| | | | | |
|---|---|---|---|---|
| rs113993995 | pathogenic | 0 | 0 | 0 |
| rs201070132 | pathogenic | 0 | 0 | 0 |
| rs113993996 | pathogenic | 0 | 0 | 0 |
| rs113993998 | pathogenic | 0 | 0 | 0 |
| rs78950939 | pathogenic | 1 | 1 | 1 |
| rs56378716 | pathogenic | 1 | 1 | 1 |
| rs754921704 | benign | 0 | 1 | 0 |
| rs766379103 | pathogenic | 1 | 1 | 0 |
| rs121912515 | pathogenic | 0 | 1 | 0 |
| rs764831888 | benign | 1 | 1 | 0 |
| rs572527340 | benign | 0 | 0 | 0 |
| rs35969327 | benign | 0 | 0 | 0 |
| rs35814893 | benign | 0 | 0 | 0 |
| rs34032300 | benign | 0 | 0 | 0 |
| rs80358260 | pathogenic | 0 | 0 | 0 |
| rs151220873 | benign | 0 | 0 | 0 |
| rs80358261 | pathogenic | 1 | 1 | 0 |
| rs11694 | pathogenic | 0 | 0 | 0 |
| rs80358264 | pathogenic | 0 | 0 | 0 |
| rs80358266 | pathogenic | 1 | 1 | 0 |
| rs104894458 | pathogenic | 1 | 1 | 1 |
| rs121918028 | pathogenic | 0 | 0 | 0 |
| rs11583680 | pathogenic | 0 | 0 | 0 |
| rs4715 | benign | 1 | 1 | 1 |
| rs34957318 | benign | 0 | 0 | 0 |
| rs1124 | benign | 1 | 1 | 1 |
| rs104894778 | pathogenic | 1 | 1 | 1 |
| rs104894777 | pathogenic | 1 | 1 | 1 |
| rs104894769 | pathogenic | 1 | 1 | 1 |
| rs104894768 | pathogenic | 1 | 1 | 1 |
| rs104894771 | pathogenic | 1 | 1 | 1 |
| rs575372249 | benign | 1 | 1 | 1 |
| rs118203914 | pathogenic | 0 | 0 | 0 |
| rs118203916 | pathogenic | 1 | 1 | 0 |
| rs140154690 | benign | 0 | 0 | 0 |
| rs34382405 | benign | 0 | 0 | 0 |
| rs73718082 | benign | 0 | 0 | 0 |
| rs764932724 | benign | 0 | 0 | 0 |
| rs554931092 | pathogenic | 1 | 1 | 0 |
| rs202060864 | pathogenic | 0 | 0 | 0 |
| rs116567033 | benign | 0 | 0 | 0 |
| rs11243011 | benign | 0 | 0 | 0 |
| rs142073519 | pathogenic | 0 | 0 | 0 |
| rs397514611 | pathogenic | 1 | 1 | 0 |
| rs397514612 | pathogenic | 0 | 0 | 0 |
| rs372054960 | pathogenic | 0 | 0 | 0 |
| rs775690041 | pathogenic | 0 | 0 | 0 |
| rs148921184 | pathogenic | 0 | 0 | 0 |
| rs12711521 | benign | 0 | 0 | 0 |
| rs2273346 | benign | 0 | 0 | 0 |
| rs12085877 | benign | 1 | 1 | 1 |
| rs144471433 | benign | 1 | 1 | 1 |
| rs397514671 | pathogenic | 1 | 1 | 1 |
| rs76394680 | benign | 0 | 0 | 0 |

| | | | | |
|---|---|---|---|---|
| rs78289603 | pathogenic | 1 | 1 | 1 |
| rs137852913 | pathogenic | 1 | 1 | 1 |
| rs137852914 | pathogenic | 1 | 1 | 1 |
| rs121918653 | pathogenic | 1 | 1 | 1 |
| rs35515200 | benign | 1 | 1 | 0 |
| rs17876030 | benign | 1 | 1 | 1 |
| rs556623443 | benign | 1 | 1 | 1 |
| rs121434365 | pathogenic | 1 | 1 | 1 |
| rs121434364 | pathogenic | 0 | 0 | 0 |
| rs41400249 | benign | 0 | 0 | 0 |
| rs149440376 | benign | 1 | 1 | 1 |
| rs181590179 | benign | 0 | 0 | 0 |
| rs113993968 | pathogenic | 0 | 0 | 0 |
| rs587776445 | pathogenic | 1 | 1 | 1 |
| rs113993969 | pathogenic | 1 | 1 | 1 |
| rs113993970 | pathogenic | 1 | 1 | 1 |
| rs587776449 | pathogenic | 1 | 1 | 1 |
| rs113993971 | pathogenic | 1 | 1 | 1 |
| rs267607194 | pathogenic | 1 | 1 | 1 |
| rs397514745 | pathogenic | 0 | 0 | 0 |
| rs201650281 | pathogenic | 0 | 0 | 0 |
| rs150529876 | benign | 1 | 1 | 1 |
| rs77573084 | benign | 0 | 0 | 0 |
| rs398122379 | pathogenic | 1 | 1 | 0 |
| rs765478968 | benign | 0 | 0 | 0 |
| rs201399681 | benign | 0 | 0 | 0 |
| rs17159287 | benign | 0 | 0 | 0 |
| rs538571144 | benign | 0 | 0 | 0 |
| rs137852647 | pathogenic | 0 | 0 | 0 |
| rs201358272 | pathogenic | 1 | 1 | 1 |
| rs181251337 | benign | 0 | 0 | 0 |
| rs3213119 | benign | 0 | 0 | 0 |
| rs137853102 | pathogenic | 1 | 1 | 0 |
| rs4956145 | benign | 0 | 0 | 0 |
| rs146036912 | pathogenic | 1 | 1 | 1 |
| rs375717077 | pathogenic | 1 | 1 | 1 |
| rs137853103 | pathogenic | 1 | 1 | 1 |
| rs776184830 | pathogenic | 1 | 1 | 1 |
| rs200273673 | pathogenic | 1 | 1 | 1 |
| rs267607025 | pathogenic | 1 | 1 | 1 |
| rs267607024 | pathogenic | 1 | 1 | 1 |
| rs515726188 | pathogenic | 1 | 1 | 1 |
| rs147315735 | benign | 0 | 0 | 0 |
| rs121918308 | pathogenic | 1 | 1 | 0 |
| rs515726195 | pathogenic | 0 | 0 | 0 |
| rs515726196 | pathogenic | 1 | 1 | 0 |
| rs121918311 | pathogenic | 0 | 1 | 0 |
| rs121918309 | pathogenic | 0 | 0 | 0 |
| rs387906891 | pathogenic | 0 | 0 | 0 |
| rs182614164 | pathogenic | 1 | 1 | 0 |
| rs121918307 | pathogenic | 0 | 0 | 0 |
| rs201440849 | benign | 0 | 0 | 0 |
| rs118203957 | pathogenic | 0 | 0 | 0 |
| rs4986790 | benign | 0 | 0 | 0 |

| | | | | |
|---|---|---|---|---|
| rs281864928 | pathogenic | 1 | 1 | 0 |
| rs267607156 | pathogenic | 0 | 0 | 0 |
| rs55916807 | benign | 0 | 0 | 0 |
| rs749750052 | benign | 0 | 0 | 0 |
| rs74382477 | benign | 0 | 0 | 0 |
| rs201016638 | benign | 0 | 0 | 0 |
| rs376683989 | benign | 0 | 0 | 0 |
| rs144439703 | pathogenic | 0 | 0 | 0 |
| rs779122022 | benign | 0 | 0 | 0 |
| rs121918056 | pathogenic | 0 | 0 | 0 |
| rs113994093 | pathogenic | 1 | 1 | 1 |
| rs141367015 | pathogenic | 0 | 0 | 0 |
| rs775538075 | pathogenic | 0 | 0 | 0 |
| rs121918055 | pathogenic | 0 | 0 | 0 |
| rs754246294 | benign | 1 | 1 | 1 |
| rs2307447 | benign | 1 | 1 | 1 |
| rs113994096 | pathogenic | 1 | 1 | 1 |
| rs367610201 | pathogenic | 1 | 1 | 1 |
| rs758438414 | pathogenic | 1 | 1 | 0 |
| rs121918046 | pathogenic | 0 | 0 | 0 |
| rs201871736 | benign | 0 | 0 | 0 |
| rs773994204 | benign | 0 | 0 | 0 |
| rs2307450 | benign | 1 | 1 | 1 |
| rs200257554 | benign | 0 | 0 | 0 |
| rs185645212 | benign | 0 | 0 | 0 |
| rs121918054 | pathogenic | 0 | 0 | 0 |
| rs113994097 | pathogenic | 0 | 0 | 0 |
| rs202037973 | pathogenic | 0 | 0 | 0 |
| rs769827124 | pathogenic | 1 | 1 | 0 |
| rs41549716 | benign | 1 | 1 | 1 |
| rs113994098 | pathogenic | 1 | 1 | 0 |
| rs144500145 | pathogenic | 1 | 1 | 0 |
| rs121918053 | pathogenic | 1 | 1 | 0 |
| rs121918050 | pathogenic | 1 | 1 | 0 |
| rs368587966 | pathogenic | 0 | 0 | 0 |
| rs139590686 | pathogenic | 0 | 0 | 0 |
| rs121918048 | pathogenic | 1 | 1 | 0 |
| rs768653086 | pathogenic | 1 | 1 | 0 |
| rs113994099 | pathogenic | 1 | 1 | 0 |
| rs121918051 | pathogenic | 1 | 1 | 0 |
| rs753160398 | pathogenic | 1 | 1 | 0 |
| rs201477273 | pathogenic | 1 | 1 | 0 |
| rs142347031 | pathogenic | 0 | 0 | 0 |
| rs121918049 | pathogenic | 0 | 0 | 0 |
| rs267606959 | pathogenic | 1 | 1 | 0 |
| rs201732356 | pathogenic | 0 | 0 | 0 |
| rs368435864 | pathogenic | 0 | 0 | 0 |
| rs764036283 | pathogenic | 0 | 0 | 0 |
| rs2307442 | benign | 0 | 0 | 0 |
| rs2307440 | benign | 0 | 0 | 0 |
| rs548076633 | pathogenic | 0 | 1 | 0 |
| rs750792237 | benign | 0 | 0 | 0 |
| rs3087374 | benign | 0 | 0 | 0 |
| rs121918060 | pathogenic | 0 | 1 | 0 |

| | | | | |
|---|---|---|---|---|
| rs397514765 | pathogenic | 1 | 1 | 1 |
| rs121918063 | pathogenic | 0 | 0 | 0 |
| rs121918058 | pathogenic | 1 | 1 | 1 |
| rs121918064 | pathogenic | 0 | 0 | 0 |
| rs36033115 | benign | 0 | 1 | 0 |
| rs121918057 | pathogenic | 0 | 1 | 0 |
| rs121918065 | pathogenic | 1 | 1 | 1 |
| rs121918061 | pathogenic | 1 | 1 | 1 |
| rs121918066 | pathogenic | 0 | 1 | 0 |
| rs707555 | benign | 0 | 1 | 0 |
| rs8192708 | benign | 0 | 1 | 0 |
| rs11552145 | benign | 0 | 0 | 0 |
| rs1804160 | benign | 0 | 1 | 0 |
| rs34103538 | benign | 0 | 1 | 0 |
| rs751393950 | pathogenic | 1 | 1 | 1 |
| rs121907979 | pathogenic | 1 | 1 | 1 |
| rs121907975 | pathogenic | 1 | 1 | 1 |
| rs121907962 | pathogenic | 1 | 1 | 1 |
| rs121907957 | pathogenic | 0 | 0 | 0 |
| rs121907953 | pathogenic | 1 | 1 | 1 |
| rs28941770 | pathogenic | 1 | 1 | 1 |
| rs28941771 | pathogenic | 1 | 1 | 1 |
| rs121907969 | pathogenic | 1 | 1 | 1 |
| rs121907973 | pathogenic | 1 | 1 | 1 |
| rs1800429 | pathogenic | 0 | 0 | 0 |
| rs121907976 | pathogenic | 1 | 1 | 0 |
| rs121907961 | pathogenic | 0 | 0 | 0 |
| rs121907974 | pathogenic | 0 | 0 | 0 |
| rs121907959 | pathogenic | 0 | 0 | 0 |
| rs121907971 | pathogenic | 1 | 1 | 0 |
| rs121907954 | pathogenic | 0 | 0 | 0 |
| rs121907977 | pathogenic | 0 | 0 | 0 |
| rs121907963 | pathogenic | 1 | 1 | 0 |
| rs370266293 | pathogenic | 1 | 1 | 0 |
| rs1800430 | benign | 0 | 0 | 0 |
| rs121907958 | pathogenic | 0 | 0 | 0 |
| rs202173526 | pathogenic | 0 | 0 | 0 |
| rs121907978 | pathogenic | 0 | 0 | 0 |
| rs121907981 | pathogenic | 0 | 1 | 0 |
| rs145012038 | benign | 1 | 1 | 1 |
| rs121907968 | pathogenic | 1 | 1 | 1 |
| rs121907966 | pathogenic | 1 | 1 | 1 |
| rs121907956 | pathogenic | 1 | 1 | 1 |
| rs28942071 | pathogenic | 1 | 1 | 1 |
| rs121907955 | pathogenic | 1 | 1 | 1 |
| rs4727853 | benign | 0 | 0 | 0 |
| rs201880593 | benign | 0 | 0 | 0 |
| rs201434579 | benign | 1 | 1 | 1 |
| rs74551128 | pathogenic | 1 | 1 | 0 |
| rs193922500 | pathogenic | 0 | 0 | 0 |
| rs121909009 | pathogenic | 1 | 1 | 0 |
| rs121908805 | pathogenic | 1 | 1 | 0 |
| rs139573311 | pathogenic | 0 | 0 | 0 |
| rs213950 | benign | 0 | 0 | 0 |

| | | | | |
|---|---|---|---|---|
| rs79282516 | pathogenic | 0 | 0 | 0 |
| rs121909017 | pathogenic | 1 | 1 | 1 |
| rs77101217 | pathogenic | 1 | 1 | 1 |
| rs1800092 | pathogenic | 0 | 1 | 0 |
| rs77646904 | pathogenic | 0 | 0 | 0 |
| rs1800095 | benign | 1 | 1 | 1 |
| rs387906368 | pathogenic | 1 | 1 | 1 |
| rs113993959 | pathogenic | 1 | 1 | 1 |
| rs121908755 | pathogenic | 1 | 1 | 1 |
| rs121909005 | pathogenic | 1 | 1 | 1 |
| rs121908757 | pathogenic | 1 | 1 | 1 |
| rs75527207 | pathogenic | 1 | 1 | 0 |
| rs121909013 | pathogenic | 1 | 1 | 0 |
| rs121909044 | pathogenic | 1 | 1 | 0 |
| rs74597325 | pathogenic | 1 | 1 | 0 |
| rs75549581 | pathogenic | 0 | 0 | 0 |
| rs80055610 | pathogenic | 1 | 1 | 1 |
| rs1800097 | pathogenic | 0 | 0 | 0 |
| rs121909006 | pathogenic | 0 | 1 | 0 |
| rs397508276 | pathogenic | 0 | 0 | 0 |
| rs397508296 | pathogenic | 1 | 1 | 1 |
| rs139468767 | pathogenic | 0 | 0 | 0 |
| rs121908759 | pathogenic | 1 | 1 | 0 |
| rs121909033 | pathogenic | 1 | 1 | 0 |
| rs397508331 | pathogenic | 0 | 0 | 0 |
| rs387907279 | pathogenic | 1 | 1 | 1 |
| rs546530618 | benign | 0 | 0 | 0 |
| rs558259739 | benign | 0 | 0 | 0 |
| rs201827340 | benign | 0 | 0 | 0 |
| rs148865119 | pathogenic | 1 | 1 | 0 |
| rs2229413 | benign | 1 | 1 | 1 |
| rs34040148 | benign | 1 | 1 | 1 |
| rs121964964 | pathogenic | 1 | 1 | 1 |
| rs760214620 | pathogenic | 1 | 1 | 1 |
| rs775992753 | pathogenic | 1 | 1 | 1 |
| rs770095972 | pathogenic | 1 | 1 | 1 |
| rs781444670 | pathogenic | 1 | 1 | 1 |
| rs147474549 | pathogenic | 0 | 0 | 0 |
| rs121964965 | pathogenic | 0 | 1 | 0 |
| rs121964963 | pathogenic | 1 | 1 | 1 |
| rs755952006 | pathogenic | 1 | 1 | 1 |
| rs121964970 | pathogenic | 1 | 1 | 1 |
| rs121964973 | pathogenic | 1 | 1 | 1 |
| rs773734233 | pathogenic | 1 | 1 | 1 |
| rs758236584 | pathogenic | 1 | 1 | 1 |
| rs149119723 | pathogenic | 1 | 1 | 0 |
| rs5742905 | pathogenic | 0 | 0 | 0 |
| rs121964962 | pathogenic | 1 | 1 | 1 |
| rs781567152 | pathogenic | 1 | 1 | 0 |
| rs398123151 | pathogenic | 1 | 1 | 1 |
| rs760417941 | pathogenic | 1 | 1 | 1 |
| rs774926464 | pathogenic | 0 | 0 | 0 |
| rs121964972 | pathogenic | 1 | 1 | 0 |
| rs372010465 | pathogenic | 1 | 1 | 1 |

| | | | | |
|---|---|---|---|---|
| rs763036586 | pathogenic | 1 | 1 | 1 |
| rs769080151 | pathogenic | 1 | 1 | 1 |
| rs121964967 | pathogenic | 1 | 1 | 1 |
| rs28934892 | pathogenic | 1 | 1 | 0 |
| rs28934891 | pathogenic | 1 | 1 | 1 |
| rs121964971 | pathogenic | 1 | 1 | 1 |
| rs121964968 | pathogenic | 1 | 1 | 1 |
| rs150828989 | benign | 1 | 1 | 1 |
| rs3775291 | benign | 1 | 1 | 0 |
| rs104894419 | pathogenic | 1 | 1 | 1 |
| rs2232642 | benign | 0 | 0 | 0 |
| rs104894551 | pathogenic | 1 | 1 | 0 |
| rs766328537 | pathogenic | 0 | 1 | 0 |
| rs104894553 | pathogenic | 1 | 1 | 0 |
| rs80099330 | benign | 0 | 0 | 0 |
| rs199565861 | pathogenic | 0 | 0 | 0 |
| rs104894548 | pathogenic | 0 | 0 | 0 |
| rs104894549 | pathogenic | 0 | 0 | 0 |
| rs104894550 | pathogenic | 1 | 1 | 1 |
| rs12948217 | pathogenic | 1 | 1 | 1 |
| rs761064915 | pathogenic | 0 | 0 | 0 |
| rs28940279 | pathogenic | 1 | 1 | 1 |
| rs774323189 | pathogenic | 1 | 1 | 1 |
| rs28940574 | pathogenic | 0 | 0 | 0 |
| rs515299 | benign | 1 | 1 | 1 |
| rs10127939 | benign | 1 | 1 | 1 |
| rs396991 | benign | 1 | 1 | 1 |
| rs145956175 | benign | 0 | 1 | 0 |
| rs747838255 | pathogenic | 1 | 1 | 1 |
| rs762659685 | pathogenic | 1 | 1 | 0 |
| rs104894831 | pathogenic | 0 | 1 | 0 |
| rs104894848 | pathogenic | 1 | 1 | 0 |
| rs104894833 | pathogenic | 0 | 1 | 0 |
| rs781838005 | pathogenic | 0 | 1 | 0 |
| rs104894834 | pathogenic | 1 | 1 | 0 |
| rs372966991 | pathogenic | 1 | 1 | 0 |
| rs138886989 | benign | 1 | 1 | 1 |
| rs28935195 | pathogenic | 0 | 1 | 0 |
| rs28935196 | pathogenic | 0 | 1 | 0 |
| rs372416832 | pathogenic | 1 | 1 | 1 |
| rs28935197 | pathogenic | 1 | 1 | 0 |
| rs28935486 | pathogenic | 1 | 1 | 0 |
| rs28935487 | pathogenic | 1 | 1 | 0 |
| rs28935488 | pathogenic | 1 | 1 | 1 |
| rs28935495 | pathogenic | 1 | 1 | 1 |
| rs28935485 | pathogenic | 1 | 1 | 1 |
| rs140329381 | pathogenic | 0 | 1 | 0 |
| rs104894830 | pathogenic | 0 | 1 | 0 |
| rs104894846 | pathogenic | 0 | 1 | 0 |
| rs28935489 | pathogenic | 0 | 1 | 0 |
| rs104894828 | pathogenic | 1 | 1 | 0 |
| rs28935491 | pathogenic | 1 | 1 | 0 |
| rs104894832 | pathogenic | 1 | 1 | 0 |
| rs28935492 | pathogenic | 1 | 1 | 0 |

| | | | | |
|---|---|---|---|---|
| rs730880437 | benign | 1 | 1 | 0 |
| rs28935493 | pathogenic | 0 | 1 | 0 |
| rs28935494 | pathogenic | 1 | 1 | 1 |
| rs111422676 | pathogenic | 1 | 1 | 1 |
| rs397515869 | benign | 1 | 1 | 0 |
| rs782449839 | pathogenic | 0 | 1 | 0 |
| rs104894852 | pathogenic | 1 | 1 | 1 |
| rs116929575 | pathogenic | 1 | 1 | 1 |
| rs7023954 | benign | 0 | 0 | 0 |
| rs184520335 | benign | 1 | 1 | 1 |
| rs143080527 | benign | 0 | 0 | 0 |
| rs1050565 | benign | 0 | 0 | 0 |
| rs11545035 | benign | 0 | 0 | 0 |
| rs267607190 | pathogenic | 1 | 1 | 1 |
| rs200283987 | benign | 0 | 0 | 0 |
| rs372556807 | benign | 1 | 1 | 0 |
| rs563073392 | benign | 0 | 0 | 0 |
| rs4539 | benign | 0 | 0 | 0 |
| rs28931609 | pathogenic | 1 | 1 | 1 |
| rs121912978 | pathogenic | 1 | 1 | 1 |
| rs104894072 | pathogenic | 1 | 1 | 1 |
| rs551933154 | benign | 1 | 1 | 1 |
| rs752962897 | benign | 1 | 1 | 0 |
| rs121912977 | pathogenic | 1 | 1 | 0 |
| rs4537 | benign | 1 | 1 | 1 |
| rs201830462 | benign | 0 | 0 | 0 |
| rs4544 | benign | 0 | 0 | 0 |
| rs562670189 | benign | 0 | 0 | 0 |
| rs4538 | benign | 0 | 1 | 0 |
| rs61757294 | pathogenic | 0 | 0 | 0 |
| rs5313 | benign | 0 | 0 | 0 |
| rs4545 | benign | 0 | 0 | 0 |
| rs72554627 | pathogenic | 1 | 1 | 1 |
| rs72554626 | pathogenic | 1 | 1 | 1 |
| rs387906861 | pathogenic | 1 | 1 | 1 |
| rs749655461 | pathogenic | 1 | 1 | 1 |
| rs121909606 | pathogenic | 1 | 1 | 1 |
| rs121909607 | pathogenic | 1 | 1 | 1 |
| rs200203992 | benign | 0 | 0 | 0 |
| rs6050 | benign | 0 | 0 | 0 |
| rs184635235 | benign | 0 | 0 | 0 |
| rs139146037 | benign | 0 | 0 | 0 |
| rs121909612 | pathogenic | 0 | 0 | 0 |
| rs78506343 | pathogenic | 0 | 0 | 0 |
| rs121918138 | pathogenic | 1 | 1 | 1 |
| rs121918139 | pathogenic | 1 | 1 | 1 |
| rs121918140 | pathogenic | 0 | 0 | 0 |
| rs36063028 | benign | 1 | 1 | 1 |
| rs45495192 | benign | 1 | 1 | 1 |
| rs45458802 | benign | 1 | 1 | 1 |
| rs552865793 | pathogenic | 1 | 1 | 1 |
| rs753744791 | pathogenic | 1 | 1 | 1 |
| rs777518512 | benign | 1 | 1 | 1 |
| rs142708991 | benign | 1 | 1 | 0 |

| | | | | |
|---|---|---|---|---|
| rs746069957 | benign | 0 | 0 | 0 |
| rs1334811 | benign | 0 | 0 | 0 |
| rs138894008 | benign | 1 | 1 | 0 |
| rs35030851 | benign | 1 | 1 | 0 |
| rs35378598 | benign | 0 | 0 | 0 |
| rs35818764 | benign | 1 | 1 | 1 |
| rs4631561 | benign | 1 | 1 | 1 |
| rs121918660 | pathogenic | 0 | 0 | 0 |
| rs373175587 | benign | 0 | 0 | 0 |
| rs61731956 | pathogenic | 1 | 1 | 1 |
| rs180177305 | pathogenic | 0 | 0 | 0 |
| rs119490108 | pathogenic | 0 | 0 | 0 |
| rs180177312 | pathogenic | 1 | 1 | 0 |
| rs180177314 | pathogenic | 1 | 1 | 0 |
| rs180177322 | pathogenic | 1 | 1 | 1 |
| rs180177323 | pathogenic | 1 | 1 | 1 |
| rs180177325 | pathogenic | 0 | 0 | 0 |
| rs35196441 | benign | 0 | 0 | 0 |
| rs28763904 | benign | 0 | 0 | 0 |
| rs2229519 | benign | 0 | 0 | 0 |
| rs137852886 | pathogenic | 0 | 0 | 0 |
| rs137852892 | pathogenic | 0 | 0 | 0 |
| rs770427750 | pathogenic | 0 | 0 | 0 |
| rs137852887 | pathogenic | 0 | 0 | 0 |
| rs137852893 | pathogenic | 0 | 0 | 0 |
| rs80338671 | pathogenic | 1 | 1 | 0 |
| rs2228389 | benign | 0 | 0 | 0 |
| rs80338672 | pathogenic | 0 | 0 | 0 |
| rs201958741 | pathogenic | 0 | 0 | 0 |
| rs80338673 | pathogenic | 0 | 0 | 0 |
| rs137852888 | pathogenic | 0 | 0 | 0 |
| rs137852889 | pathogenic | 0 | 0 | 0 |
| rs137852894 | pathogenic | 0 | 0 | 0 |
| rs137852890 | pathogenic | 0 | 0 | 0 |
| rs137852891 | pathogenic | 0 | 0 | 0 |
| rs766935302 | pathogenic | 0 | 0 | 0 |
| rs193074572 | benign | 0 | 0 | 0 |
| rs555210692 | benign | 1 | 1 | 0 |
| rs119103242 | pathogenic | 0 | 1 | 0 |
| rs143725101 | benign | 0 | 1 | 0 |
| rs28940589 | pathogenic | 1 | 1 | 1 |
| rs113994145 | pathogenic | 0 | 0 | 0 |
| rs773872994 | benign | 0 | 0 | 0 |
| rs28940590 | pathogenic | 1 | 1 | 0 |
| rs145400751 | benign | 1 | 1 | 1 |
| rs587782331 | benign | 0 | 0 | 0 |
| rs730881787 | benign | 0 | 0 | 0 |
| rs587778531 | benign | 0 | 0 | 0 |
| rs781652274 | benign | 0 | 0 | 0 |
| rs55882234 | benign | 0 | 0 | 0 |
| rs2020908 | benign | 0 | 0 | 0 |
| rs63750741 | pathogenic | 1 | 1 | 1 |
| rs41295268 | benign | 1 | 1 | 1 |
| rs63750897 | benign | 1 | 1 | 0 |

| | | | | |
|---|---|---|---|---|
| rs63751005 | benign | 1 | 1 | 0 |
| rs587779223 | benign | 0 | 0 | 0 |
| rs202127474 | benign | 0 | 0 | 0 |
| rs374230313 | benign | 0 | 0 | 0 |
| rs2020912 | benign | 0 | 0 | 0 |
| rs63751017 | pathogenic | 0 | 0 | 0 |
| rs587779246 | pathogenic | 0 | 0 | 0 |
| rs63749919 | pathogenic | 0 | 0 | 0 |
| rs63750563 | pathogenic | 1 | 1 | 1 |
| rs63749999 | pathogenic | 0 | 0 | 0 |
| rs200492211 | pathogenic | 0 | 0 | 0 |
| rs576269342 | benign | 0 | 0 | 0 |
| rs267608075 | benign | 0 | 0 | 0 |
| rs149605979 | benign | 0 | 0 | 0 |
| rs63750252 | benign | 0 | 0 | 0 |
| rs754289472 | benign | 0 | 0 | 0 |
| rs63750119 | pathogenic | 0 | 0 | 0 |
| rs587782109 | benign | 0 | 0 | 0 |
| rs34625968 | benign | 1 | 1 | 1 |
| rs764786814 | benign | 1 | 1 | 1 |
| rs199594809 | benign | 1 | 1 | 1 |
| rs74315364 | pathogenic | 1 | 1 | 1 |
| rs201253919 | pathogenic | 0 | 0 | 0 |
| rs387906828 | pathogenic | 1 | 1 | 1 |
| rs34521609 | benign | 1 | 1 | 1 |
| rs199472894 | pathogenic | 0 | 0 | 0 |
| rs199472902 | pathogenic | 0 | 0 | 0 |
| rs121912505 | pathogenic | 0 | 0 | 0 |
| rs28928905 | pathogenic | 0 | 0 | 0 |
| rs374376640 | benign | 0 | 0 | 0 |
| rs202102276 | benign | 0 | 0 | 0 |
| rs199472918 | pathogenic | 0 | 0 | 0 |
| rs121912516 | pathogenic | 1 | 1 | 1 |
| rs121912504 | pathogenic | 0 | 0 | 0 |
| rs9333649 | pathogenic | 0 | 0 | 0 |
| rs121912508 | pathogenic | 0 | 0 | 0 |
| rs199473428 | pathogenic | 1 | 1 | 1 |
| rs104894021 | pathogenic | 0 | 0 | 0 |
| rs28928904 | pathogenic | 1 | 1 | 1 |
| rs199473039 | benign | 1 | 1 | 1 |
| rs199472958 | pathogenic | 1 | 1 | 1 |
| rs1137617 | pathogenic | 1 | 1 | 1 |
| rs199473532 | pathogenic | 1 | 1 | 1 |
| rs199472988 | pathogenic | 0 | 0 | 0 |
| rs189014161 | pathogenic | 0 | 0 | 0 |
| rs12720441 | pathogenic | 0 | 0 | 0 |
| rs199473001 | pathogenic | 1 | 1 | 0 |
| rs121912506 | pathogenic | 1 | 1 | 0 |
| rs773724817 | pathogenic | 1 | 1 | 1 |
| rs118204002 | pathogenic | 1 | 1 | 1 |
| rs138556525 | pathogenic | 1 | 1 | 1 |
| rs72558181 | pathogenic | 1 | 1 | 1 |
| rs118204005 | pathogenic | 1 | 1 | 1 |
| rs118204004 | pathogenic | 0 | 0 | 0 |

| | | | | |
|---|---|---|---|---|
| rs118204001 | pathogenic | 1 | 1 | 1 |
| rs118204006 | pathogenic | 1 | 1 | 1 |
| rs118204003 | pathogenic | 0 | 0 | 0 |
| rs121913147 | pathogenic | 1 | 1 | 0 |
| rs2229429 | benign | 1 | 1 | 0 |
| rs182552223 | benign | 0 | 0 | 0 |
| rs146588336 | benign | 0 | 0 | 0 |
| rs121964983 | pathogenic | 0 | 1 | 0 |
| rs121964982 | pathogenic | 0 | 0 | 0 |
| rs386833679 | pathogenic | 0 | 0 | 0 |
| rs386833680 | pathogenic | 0 | 0 | 0 |
| rs121964981 | pathogenic | 1 | 1 | 0 |
| rs121964984 | pathogenic | 0 | 0 | 0 |
| rs144971200 | benign | 0 | 0 | 0 |
| rs121964985 | pathogenic | 1 | 1 | 0 |
| rs141246107 | benign | 0 | 0 | 0 |
| rs104894561 | pathogenic | 1 | 1 | 0 |
| rs377023736 | pathogenic | 0 | 0 | 0 |
| rs121964907 | pathogenic | 1 | 1 | 1 |
| rs121964909 | pathogenic | 1 | 1 | 1 |
| rs765070743 | pathogenic | 0 | 0 | 0 |
| rs2228119 | benign | 0 | 0 | 0 |
| rs745976989 | pathogenic | 1 | 1 | 1 |
| rs121964904 | pathogenic | 0 | 0 | 0 |
| rs386833430 | pathogenic | 1 | 1 | 1 |
| rs267606992 | pathogenic | 0 | 0 | 0 |
| rs61752063 | pathogenic | 0 | 0 | 0 |
| rs61752067 | pathogenic | 1 | 1 | 1 |
| rs104894934 | pathogenic | 0 | 0 | 0 |
| rs104894929 | pathogenic | 1 | 1 | 1 |
| rs80189640 | benign | 0 | 0 | 0 |
| rs137854547 | pathogenic | 1 | 1 | 1 |
| rs137854540 | pathogenic | 1 | 1 | 1 |
| rs782092363 | pathogenic | 1 | 1 | 1 |
| rs147783915 | benign | 0 | 0 | 0 |
| rs28933687 | pathogenic | 1 | 1 | 1 |
| rs149503319 | benign | 0 | 0 | 0 |
| rs1805147 | benign | 0 | 0 | 0 |
| rs587777218 | pathogenic | 1 | 1 | 1 |
| rs200786705 | benign | 0 | 0 | 0 |
| rs56219475 | benign | 0 | 0 | 0 |
| rs587781172 | benign | 0 | 0 | 0 |
| rs757094189 | pathogenic | 0 | 0 | 0 |
| rs140811086 | benign | 1 | 1 | 1 |
| rs574088829 | pathogenic | 1 | 1 | 1 |
| rs121918525 | pathogenic | 0 | 0 | 0 |
| rs121918527 | pathogenic | 0 | 0 | 0 |
| rs121964848 | pathogenic | 0 | 1 | 0 |
| rs121964845 | pathogenic | 1 | 1 | 1 |
| rs121964846 | pathogenic | 0 | 1 | 0 |
| rs17217716 | benign | 1 | 1 | 1 |
| rs141711342 | benign | 0 | 0 | 0 |
| rs372619120 | benign | 0 | 0 | 0 |
| rs63750002 | benign | 1 | 1 | 1 |

| | | | | |
|---|---|---|---|---|
| rs193922373 | benign | 0 | 0 | 0 |
| rs41295286 | benign | 0 | 0 | 0 |
| rs17217772 | benign | 0 | 0 | 0 |
| rs145649774 | benign | 0 | 0 | 0 |
| rs63750214 | pathogenic | 0 | 0 | 0 |
| rs63750716 | benign | 0 | 0 | 0 |
| rs202026056 | benign | 0 | 0 | 0 |
| rs4987188 | benign | 1 | 1 | 0 |
| rs267607938 | benign | 0 | 0 | 0 |
| rs17224367 | benign | 0 | 0 | 0 |
| rs63750006 | benign | 0 | 0 | 0 |
| rs587781373 | benign | 0 | 0 | 0 |
| rs63751207 | pathogenic | 1 | 1 | 0 |
| rs730881759 | benign | 0 | 0 | 0 |
| rs372350768 | benign | 1 | 1 | 1 |
| rs55778204 | benign | 1 | 1 | 0 |
| rs63749910 | benign | 0 | 0 | 0 |
| rs63750844 | benign | 0 | 0 | 0 |
| rs28929483 | pathogenic | 0 | 0 | 0 |
| rs61756468 | benign | 0 | 0 | 0 |
| rs63750875 | pathogenic | 1 | 1 | 0 |
| rs776528054 | benign | 1 | 1 | 0 |
| rs751939698 | benign | 0 | 0 | 0 |
| rs587780685 | benign | 0 | 0 | 0 |
| rs730881763 | benign | 1 | 1 | 0 |
| rs373226409 | benign | 0 | 0 | 0 |
| rs202145681 | benign | 1 | 1 | 1 |
| rs63749841 | benign | 1 | 1 | 1 |
| rs63750757 | benign | 1 | 1 | 1 |
| rs587778527 | benign | 1 | 1 | 1 |
| rs63750849 | benign | 0 | 0 | 0 |
| rs587779969 | benign | 0 | 0 | 0 |
| rs121908046 | pathogenic | 1 | 1 | 1 |
| rs28940882 | pathogenic | 1 | 1 | 0 |
| rs28940883 | pathogenic | 1 | 1 | 1 |
| rs727503943 | pathogenic | 1 | 1 | 1 |
| rs137853859 | pathogenic | 1 | 1 | 1 |
| rs121908045 | pathogenic | 1 | 1 | 0 |
| rs137853861 | pathogenic | 1 | 1 | 0 |
| rs3180383 | pathogenic | 0 | 1 | 0 |
| rs144431814 | benign | 0 | 0 | 0 |
| rs139502866 | benign | 0 | 0 | 0 |
| rs17524437 | benign | 0 | 0 | 0 |
| rs760960779 | benign | 0 | 0 | 0 |
| rs148960463 | pathogenic | 0 | 0 | 0 |
| rs111033647 | pathogenic | 1 | 1 | 1 |
| rs111033648 | pathogenic | 0 | 0 | 0 |
| rs1800461 | benign | 0 | 0 | 0 |
| rs111033656 | pathogenic | 0 | 0 | 0 |
| rs111033663 | pathogenic | 1 | 1 | 0 |
| rs111033669 | pathogenic | 1 | 1 | 1 |
| rs111033690 | pathogenic | 0 | 0 | 0 |
| rs111033686 | pathogenic | 0 | 0 | 0 |
| rs111033687 | pathogenic | 0 | 0 | 0 |

| | | | | |
|---|---|---|---|---|
| rs111033695 | pathogenic | 0 | 0 | 0 |
| rs111033697 | pathogenic | 0 | 0 | 0 |
| rs111033693 | pathogenic | 0 | 0 | 0 |
| rs111033694 | pathogenic | 0 | 0 | 0 |
| rs111033721 | pathogenic | 1 | 1 | 1 |
| rs111033722 | pathogenic | 0 | 0 | 0 |
| rs111033725 | pathogenic | 1 | 1 | 0 |
| rs75391579 | pathogenic | 1 | 1 | 1 |
| rs111033728 | pathogenic | 1 | 1 | 1 |
| rs111033735 | pathogenic | 1 | 1 | 1 |
| rs111033739 | pathogenic | 1 | 1 | 1 |
| rs111033736 | pathogenic | 0 | 0 | 0 |
| rs111033744 | pathogenic | 1 | 1 | 1 |
| rs111033750 | pathogenic | 1 | 1 | 1 |
| rs111033752 | pathogenic | 0 | 0 | 0 |
| rs111033751 | pathogenic | 0 | 0 | 0 |
| rs111033846 | pathogenic | 0 | 0 | 0 |
| rs368166217 | pathogenic | 0 | 0 | 0 |
| rs111033764 | pathogenic | 0 | 0 | 0 |
| rs111033847 | benign | 0 | 0 | 0 |
| rs111033778 | pathogenic | 0 | 0 | 0 |
| rs111033773 | pathogenic | 0 | 0 | 0 |
| rs111033792 | pathogenic | 0 | 0 | 0 |
| rs367543266 | pathogenic | 1 | 1 | 1 |
| rs111033794 | pathogenic | 0 | 0 | 0 |
| rs111033802 | pathogenic | 0 | 0 | 0 |
| rs111033803 | pathogenic | 0 | 0 | 0 |
| rs111033809 | pathogenic | 1 | 1 | 0 |
| rs121918376 | pathogenic | 0 | 1 | 0 |
| rs121918377 | pathogenic | 0 | 1 | 0 |
| rs121918378 | pathogenic | 1 | 1 | 1 |
| rs121918374 | pathogenic | 1 | 1 | 1 |
| rs137852379 | pathogenic | 0 | 0 | 0 |
| rs137852388 | pathogenic | 1 | 1 | 1 |
| rs137852368 | pathogenic | 0 | 0 | 0 |
| rs28935499 | pathogenic | 0 | 0 | 0 |
| rs28933668 | pathogenic | 0 | 0 | 0 |
| rs28933669 | pathogenic | 0 | 0 | 0 |
| rs28933670 | pathogenic | 0 | 0 | 0 |
| rs28933671 | pathogenic | 0 | 0 | 0 |
| rs28933672 | pathogenic | 0 | 0 | 0 |
| rs28937272 | pathogenic | 0 | 0 | 0 |
| rs35383156 | benign | 0 | 0 | 0 |
| rs137852418 | pathogenic | 0 | 0 | 0 |
| rs137852419 | pathogenic | 0 | 0 | 0 |
| rs137852428 | pathogenic | 0 | 0 | 0 |
| rs137852436 | pathogenic | 0 | 0 | 0 |
| rs28937285 | pathogenic | 0 | 0 | 0 |
| rs35213384 | benign | 1 | 1 | 1 |
| rs3810950 | benign | 0 | 0 | 0 |
| rs201479289 | pathogenic | 0 | 0 | 0 |
| rs121912820 | pathogenic | 0 | 1 | 0 |
| rs121912815 | pathogenic | 0 | 1 | 0 |
| rs115510708 | benign | 0 | 0 | 0 |

| | | | | |
|---|---|---|---|---|
| rs868749 | benign | 0 | 0 | 0 |
| rs121912823 | pathogenic | 0 | 0 | 0 |
| rs115212829 | benign | 0 | 0 | 0 |
| rs121912822 | pathogenic | 0 | 0 | 0 |
| rs121912816 | pathogenic | 0 | 1 | 0 |
| rs4838544 | benign | 0 | 0 | 0 |
| rs121912818 | pathogenic | 0 | 0 | 0 |
| rs121912821 | pathogenic | 0 | 0 | 0 |
| rs121912817 | pathogenic | 0 | 0 | 0 |
| rs8178992 | benign | 0 | 0 | 0 |
| rs121912819 | pathogenic | 1 | 1 | 0 |
| rs80097077 | benign | 1 | 1 | 0 |
| rs121909359 | pathogenic | 1 | 1 | 0 |
| rs121909358 | pathogenic | 1 | 1 | 1 |
| rs121909361 | pathogenic | 1 | 1 | 1 |
| rs121909357 | pathogenic | 0 | 1 | 0 |
| rs121909365 | pathogenic | 1 | 1 | 1 |
| rs6413484 | pathogenic | 1 | 1 | 1 |
| rs121909373 | pathogenic | 1 | 1 | 1 |
| rs121909366 | pathogenic | 1 | 1 | 1 |
| rs121909367 | pathogenic | 1 | 1 | 1 |
| rs121909368 | pathogenic | 1 | 1 | 1 |
| rs121909369 | pathogenic | 0 | 0 | 0 |
| rs121909363 | pathogenic | 1 | 1 | 1 |
| rs45588036 | pathogenic | 1 | 1 | 1 |
| rs121909364 | pathogenic | 1 | 1 | 1 |
| rs80358282 | benign | 0 | 0 | 0 |
| rs80358284 | pathogenic | 1 | 1 | 1 |
| rs61735303 | benign | 0 | 0 | 0 |
| rs2229274 | benign | 0 | 0 | 0 |
| rs145006491 | benign | 0 | 0 | 0 |
| rs121913580 | pathogenic | 1 | 1 | 0 |
| rs730882196 | pathogenic | 0 | 0 | 0 |
| rs153477 | benign | 0 | 0 | 0 |
| rs153478 | benign | 1 | 1 | 0 |
| rs61740602 | benign | 1 | 1 | 0 |
| rs104893892 | pathogenic | 1 | 1 | 0 |
| rs121909595 | pathogenic | 1 | 1 | 1 |
| rs28931605 | pathogenic | 1 | 1 | 1 |
| rs121909597 | pathogenic | 0 | 1 | 0 |
| rs202233735 | pathogenic | 0 | 1 | 0 |
| rs121909596 | pathogenic | 1 | 1 | 1 |
| rs75156162 | benign | 0 | 1 | 0 |
| rs2305430 | benign | 1 | 1 | 1 |
| rs150318966 | benign | 0 | 0 | 0 |
| rs121909598 | pathogenic | 0 | 1 | 0 |
| rs5744175 | benign | 0 | 0 | 0 |
| rs28935476 | pathogenic | 1 | 1 | 1 |
| rs28935475 | pathogenic | 1 | 1 | 1 |
| rs122461163 | pathogenic | 1 | 1 | 1 |
| rs62626305 | pathogenic | 1 | 1 | 1 |
| rs587777157 | pathogenic | 0 | 0 | 0 |
| rs137854478 | pathogenic | 1 | 1 | 0 |
| rs137854465 | pathogenic | 0 | 0 | 0 |

| | | | | |
|---|---|---|---|---|
| rs137854470 | pathogenic | 0 | 0 | 0 |
| rs137854482 | pathogenic | 0 | 0 | 0 |
| rs137854473 | pathogenic | 0 | 0 | 0 |
| rs137854456 | pathogenic | 0 | 0 | 0 |
| rs140598 | benign | 0 | 0 | 0 |
| rs104894105 | pathogenic | 1 | 1 | 1 |
| rs104894107 | pathogenic | 1 | 1 | 1 |
| rs104894106 | pathogenic | 1 | 1 | 1 |
| rs74621026 | benign | 1 | 1 | 1 |
| rs2290332 | benign | 0 | 0 | 0 |
| rs780202604 | pathogenic | 1 | 1 | 1 |
| rs74315305 | pathogenic | 1 | 1 | 0 |
| rs766751878 | pathogenic | 1 | 1 | 1 |
| rs137854603 | pathogenic | 1 | 1 | 1 |
| rs139592595 | benign | 1 | 1 | 0 |
| rs150565592 | benign | 1 | 1 | 1 |
| rs763520126 | benign | 0 | 0 | 0 |
| rs55980502 | benign | 0 | 0 | 0 |
| rs137852812 | pathogenic | 0 | 0 | 0 |
| rs137852813 | pathogenic | 0 | 0 | 0 |
| rs190222208 | benign | 0 | 0 | 0 |
| rs138459502 | pathogenic | 0 | 0 | 0 |
| rs730881041 | pathogenic | 0 | 0 | 0 |
| rs267607080 | pathogenic | 0 | 0 | 0 |
| rs137852814 | pathogenic | 0 | 0 | 0 |
| rs267607079 | pathogenic | 0 | 0 | 0 |
| rs192378417 | benign | 0 | 0 | 0 |
| rs1288775 | benign | 0 | 0 | 0 |
| rs80338737 | pathogenic | 1 | 1 | 1 |
| rs397514708 | pathogenic | 0 | 0 | 0 |
| rs775933965 | pathogenic | 0 | 0 | 0 |
| rs747608698 | benign | 0 | 1 | 0 |
| rs1145086 | benign | 0 | 1 | 0 |
| rs121912953 | pathogenic | 1 | 1 | 0 |
| rs243849 | benign | 0 | 0 | 0 |
| rs121912955 | pathogenic | 1 | 1 | 1 |
| rs17859943 | benign | 0 | 1 | 0 |
| rs755557988 | benign | 1 | 1 | 1 |
| rs10195 | benign | 0 | 0 | 0 |
| rs61735357 | benign | 1 | 1 | 1 |
| rs137882374 | benign | 0 | 0 | 0 |
| rs5388 | benign | 0 | 0 | 0 |
| rs6352 | benign | 0 | 0 | 0 |
| rs76645461 | pathogenic | 1 | 1 | 0 |
| rs1050828 | pathogenic | 0 | 0 | 0 |
| rs1050829 | pathogenic | 0 | 0 | 0 |
| rs137852314 | pathogenic | 0 | 0 | 0 |
| rs137852331 | pathogenic | 0 | 0 | 0 |
| rs5030872 | pathogenic | 0 | 0 | 0 |
| rs267606836 | pathogenic | 0 | 0 | 0 |
| rs5030868 | pathogenic | 0 | 0 | 0 |
| rs137852330 | pathogenic | 0 | 1 | 0 |
| rs137852326 | pathogenic | 1 | 1 | 1 |
| rs137852318 | pathogenic | 1 | 1 | 1 |

| | | | | |
|---|---|---|---|---|
| rs74575103 | benign | 0 | 0 | 0 |
| rs137852327 | pathogenic | 1 | 1 | 1 |
| rs76723693 | pathogenic | 0 | 0 | 0 |
| rs5030869 | pathogenic | 0 | 0 | 0 |
| rs34193178 | pathogenic | 1 | 1 | 1 |
| rs137852316 | pathogenic | 1 | 1 | 0 |
| rs398123546 | pathogenic | 1 | 1 | 1 |
| rs72554665 | pathogenic | 0 | 0 | 0 |
| rs72554664 | pathogenic | 0 | 0 | 0 |
| rs28939068 | pathogenic | 1 | 1 | 1 |
| rs59301204 | pathogenic | 1 | 1 | 1 |
| rs143715750 | benign | 1 | 1 | 1 |
| rs35263902 | pathogenic | 1 | 1 | 1 |
| rs202214518 | benign | 1 | 1 | 1 |
| rs34076756 | benign | 0 | 0 | 0 |
| rs199592341 | pathogenic | 1 | 1 | 0 |
| rs121908833 | pathogenic | 1 | 1 | 1 |
| rs121908834 | pathogenic | 1 | 1 | 0 |
| rs138454151 | benign | 0 | 0 | 0 |
| rs141323599 | benign | 0 | 0 | 0 |
| rs137853200 | pathogenic | 0 | 0 | 0 |
| rs137853199 | pathogenic | 0 | 0 | 0 |
| rs139787163 | benign | 0 | 0 | 0 |
| rs61731097 | benign | 1 | 1 | 1 |
| rs61729795 | benign | 0 | 0 | 0 |
| rs149242330 | benign | 0 | 0 | 0 |
| rs587777490 | pathogenic | 1 | 1 | 1 |
| rs398123011 | pathogenic | 1 | 1 | 0 |
| rs775186117 | pathogenic | 0 | 0 | 0 |
| rs72677237 | benign | 0 | 0 | 0 |
| rs104894163 | pathogenic | 1 | 1 | 1 |
| rs121909588 | pathogenic | 0 | 0 | 0 |
| rs1042579 | benign | 0 | 0 | 0 |
| rs121909273 | pathogenic | 0 | 0 | 0 |
| rs121909274 | pathogenic | 0 | 0 | 0 |
| rs119473034 | pathogenic | 1 | 1 | 1 |
| rs398123065 | pathogenic | 0 | 0 | 0 |
| rs587777042 | pathogenic | 1 | 1 | 0 |
| rs761413864 | pathogenic | 1 | 1 | 1 |
| rs118203965 | pathogenic | 0 | 0 | 0 |
| rs118203966 | pathogenic | 0 | 0 | 0 |
| rs1801201 | benign | 1 | 1 | 0 |
| rs1136201 | benign | 1 | 1 | 1 |
| rs144148008 | benign | 0 | 0 | 0 |
| rs200118378 | benign | 1 | 1 | 1 |
| rs112984118 | benign | 1 | 1 | 1 |
| rs149446102 | benign | 1 | 1 | 1 |
| rs1427463 | benign | 0 | 0 | 0 |
| rs201936720 | benign | 1 | 1 | 1 |
| rs61751983 | benign | 1 | 1 | 1 |
| rs104894632 | pathogenic | 1 | 1 | 1 |
| rs149202905 | benign | 0 | 0 | 0 |
| rs2230210 | benign | 1 | 1 | 1 |
| rs111631988 | benign | 0 | 0 | 0 |

| | | | | |
|---|---|---|---|---|
| rs137852550 | pathogenic | 0 | 0 | 0 |
| rs5963403 | benign | 0 | 0 | 0 |
| rs138018739 | benign | 0 | 0 | 0 |
| rs62640590 | benign | 0 | 0 | 0 |
| rs121909581 | pathogenic | 1 | 1 | 1 |
| rs202017913 | benign | 1 | 1 | 1 |
| rs143163513 | benign | 0 | 0 | 0 |
| rs238238 | benign | 0 | 1 | 0 |
| rs143945974 | benign | 0 | 0 | 0 |
| rs238239 | benign | 1 | 1 | 1 |
| rs121918403 | pathogenic | 0 | 1 | 0 |
| rs61735456 | benign | 0 | 1 | 0 |
| rs121918404 | pathogenic | 1 | 1 | 0 |
| rs12674488 | benign | 0 | 0 | 0 |
| rs201599657 | pathogenic | 0 | 0 | 0 |
| rs1057090 | benign | 0 | 0 | 0 |
| rs1057091 | benign | 0 | 0 | 0 |
| rs587783740 | benign | 0 | 0 | 0 |
| rs75956769 | benign | 0 | 0 | 0 |
| rs201691299 | benign | 0 | 0 | 0 |
| rs62621087 | benign | 0 | 0 | 0 |
| rs201308454 | benign | 0 | 0 | 0 |
| rs145657360 | benign | 0 | 0 | 0 |
| rs121434592 | pathogenic | 1 | 1 | 0 |
| rs375650413 | benign | 0 | 0 | 0 |
| rs121434437 | pathogenic | 1 | 1 | 0 |
| rs763494292 | pathogenic | 0 | 0 | 0 |
| rs121964997 | pathogenic | 1 | 1 | 0 |
| rs121964996 | pathogenic | 0 | 0 | 0 |
| rs41283425 | benign | 1 | 1 | 1 |
| rs376289528 | benign | 1 | 1 | 1 |
| rs144171604 | benign | 1 | 1 | 1 |
| rs782440692 | pathogenic | 1 | 1 | 1 |
| rs782058451 | pathogenic | 1 | 1 | 1 |
| rs78437817 | benign | 0 | 0 | 0 |
| rs200327969 | benign | 1 | 1 | 1 |
| rs143043662 | benign | 0 | 0 | 0 |
| rs111966833 | benign | 1 | 1 | 1 |
| rs35523678 | benign | 0 | 0 | 0 |
| rs515726208 | pathogenic | 0 | 0 | 0 |
| rs397517239 | pathogenic | 1 | 1 | 1 |
| rs148966602 | benign | 1 | 1 | 1 |
| rs76371972 | benign | 1 | 1 | 1 |
| rs2229968 | benign | 1 | 1 | 1 |
| rs61751538 | benign | 1 | 1 | 1 |
| rs200800978 | pathogenic | 1 | 1 | 1 |
| rs148695964 | benign | 1 | 1 | 1 |
| rs140042418 | benign | 1 | 1 | 1 |
| rs116185403 | benign | 0 | 0 | 0 |
| rs142828669 | benign | 1 | 1 | 1 |
| rs139832048 | benign | 0 | 1 | 0 |
| rs58750568 | benign | 0 | 1 | 0 |
| rs77681311 | benign | 0 | 0 | 0 |
| rs151208452 | benign | 0 | 0 | 0 |

| | | | | |
|---|---|---|---|---|
| rs76463933 | benign | 0 | 0 | 0 |
| rs144434647 | benign | 0 | 0 | 0 |
| rs117361485 | benign | 0 | 0 | 0 |
| rs587777588 | pathogenic | 0 | 0 | 0 |
| rs80338796 | pathogenic | 0 | 0 | 0 |
| rs121434594 | pathogenic | 0 | 0 | 0 |
| rs146766138 | pathogenic | 1 | 1 | 1 |
| rs28939679 | pathogenic | 1 | 1 | 1 |
| rs748754134 | pathogenic | 1 | 1 | 0 |
| rs764276946 | pathogenic | 1 | 1 | 1 |
| rs111033588 | pathogenic | 1 | 1 | 1 |
| rs201633414 | pathogenic | 1 | 1 | 1 |
| rs121909262 | pathogenic | 1 | 1 | 1 |
| rs104893702 | pathogenic | 1 | 1 | 1 |
| rs121909265 | pathogenic | 1 | 1 | 1 |
| rs121909266 | pathogenic | 1 | 1 | 1 |
| rs193922439 | pathogenic | 1 | 1 | 0 |
| rs104893691 | pathogenic | 1 | 1 | 1 |
| rs104893695 | pathogenic | 0 | 0 | 0 |
| rs104893708 | pathogenic | 1 | 1 | 1 |
| rs121909260 | pathogenic | 0 | 0 | 0 |
| rs104893696 | pathogenic | 0 | 0 | 0 |
| rs121909267 | pathogenic | 0 | 0 | 0 |
| rs121909263 | pathogenic | 0 | 0 | 0 |
| rs121909264 | pathogenic | 0 | 0 | 0 |
| rs104893694 | pathogenic | 1 | 1 | 1 |
| rs121909268 | pathogenic | 1 | 1 | 1 |
| rs104893689 | pathogenic | 1 | 1 | 0 |
| rs104893707 | pathogenic | 1 | 1 | 0 |
| rs104893697 | pathogenic | 1 | 1 | 1 |
| rs28936684 | pathogenic | 1 | 1 | 1 |
| rs62269092 | benign | 1 | 1 | 0 |
| rs121909259 | pathogenic | 1 | 1 | 0 |
| rs548403340 | benign | 1 | 1 | 0 |
| rs142818334 | benign | 0 | 0 | 0 |
| rs12493789 | benign | 1 | 1 | 1 |
| rs193922423 | pathogenic | 1 | 1 | 0 |
| rs104893719 | pathogenic | 1 | 1 | 1 |
| rs117375173 | benign | 1 | 1 | 0 |
| rs104893712 | pathogenic | 0 | 0 | 0 |
| rs121917776 | pathogenic | 1 | 1 | 1 |
| rs767325003 | benign | 1 | 1 | 1 |
| rs35824453 | benign | 0 | 0 | 0 |
| rs141641266 | benign | 0 | 0 | 0 |
| rs1050239 | benign | 0 | 0 | 0 |
| rs141303384 | benign | 0 | 0 | 0 |
| rs12720458 | pathogenic | 0 | 0 | 0 |
| rs368507376 | pathogenic | 1 | 1 | 1 |
| rs17215500 | pathogenic | 1 | 1 | 1 |
| rs120074188 | pathogenic | 1 | 1 | 1 |
| rs397508097 | pathogenic | 1 | 1 | 1 |
| rs138551008 | pathogenic | 0 | 0 | 0 |
| rs387907139 | pathogenic | 1 | 1 | 1 |
| rs765487627 | pathogenic | 0 | 0 | 0 |

| | | | | |
|---|---|---|---|---|
| rs587778587 | benign | 0 | 0 | 0 |
| rs45568339 | benign | 0 | 0 | 0 |
| rs78179744 | benign | 0 | 0 | 0 |
| rs45624036 | benign | 0 | 0 | 0 |
| rs61755166 | benign | 0 | 0 | 0 |
| rs180177132 | pathogenic | 1 | 1 | 1 |
| rs587776527 | pathogenic | 0 | 0 | 0 |
| rs373783514 | benign | 0 | 0 | 0 |
| rs730881896 | benign | 0 | 0 | 0 |
| rs118203998 | pathogenic | 0 | 0 | 0 |
| rs41281039 | pathogenic | 1 | 1 | 1 |
| rs118204430 | pathogenic | 1 | 1 | 1 |
| rs1800546 | pathogenic | 1 | 1 | 0 |
| rs76917243 | pathogenic | 0 | 1 | 0 |
| rs78340951 | pathogenic | 0 | 1 | 0 |
| rs77718928 | pathogenic | 0 | 0 | 0 |
| rs369586696 | pathogenic | 0 | 1 | 0 |
| rs745517517 | pathogenic | 1 | 1 | 0 |
| rs372872777 | pathogenic | 0 | 0 | 0 |
| rs748704459 | pathogenic | 0 | 0 | 0 |
| rs769215411 | pathogenic | 0 | 0 | 0 |
| rs139425890 | pathogenic | 0 | 0 | 0 |
| rs779694939 | pathogenic | 1 | 1 | 0 |
| rs773729410 | pathogenic | 0 | 0 | 0 |
| rs121908623 | pathogenic | 1 | 1 | 1 |
| rs121908622 | pathogenic | 1 | 1 | 1 |
| rs121908621 | pathogenic | 1 | 1 | 1 |
| rs762106720 | pathogenic | 1 | 1 | 0 |
| rs199877522 | pathogenic | 0 | 0 | 0 |
| rs74315291 | pathogenic | 1 | 1 | 1 |
| rs79137415 | benign | 0 | 0 | 0 |
| rs121913045 | pathogenic | 0 | 0 | 0 |
| rs34295337 | pathogenic | 0 | 0 | 0 |
| rs121913046 | pathogenic | 0 | 0 | 0 |
| rs121913047 | pathogenic | 0 | 0 | 0 |
| rs121913048 | pathogenic | 0 | 0 | 0 |
| rs267607166 | pathogenic | 0 | 0 | 0 |
| rs17850684 | benign | 1 | 1 | 1 |
| rs119476051 | pathogenic | 1 | 1 | 1 |
| rs17850683 | benign | 1 | 1 | 1 |
| rs119476049 | pathogenic | 1 | 1 | 1 |
| rs119476046 | pathogenic | 1 | 1 | 1 |
| rs119476048 | pathogenic | 0 | 0 | 0 |
| rs119476047 | pathogenic | 1 | 1 | 1 |
| rs28939094 | pathogenic | 1 | 1 | 1 |
| rs119476050 | pathogenic | 0 | 0 | 0 |
| rs387906941 | pathogenic | 1 | 1 | 1 |
| rs41511344 | pathogenic | 1 | 1 | 0 |
| rs748112833 | pathogenic | 1 | 1 | 1 |
| rs74342080 | benign | 1 | 1 | 0 |
| rs116482870 | benign | 0 | 0 | 0 |
| rs104893743 | pathogenic | 0 | 0 | 0 |
| rs117179004 | benign | 0 | 0 | 0 |
| rs587779409 | pathogenic | 1 | 1 | 1 |

| | | | | |
|---|---|---|---|---|
| rs3745406 | benign | 1 | 1 | 0 |
| rs387906696 | pathogenic | 0 | 0 | 0 |
| rs72549369 | pathogenic | 1 | 1 | 0 |
| rs45523133 | benign | 0 | 0 | 0 |
| rs119460972 | pathogenic | 0 | 1 | 0 |
| rs34929837 | benign | 0 | 1 | 0 |
| rs17323225 | benign | 0 | 1 | 0 |
| rs17011368 | benign | 0 | 0 | 0 |
| rs748902766 | pathogenic | 1 | 1 | 1 |
| rs387907292 | pathogenic | 0 | 0 | 0 |
| rs387907293 | pathogenic | 1 | 1 | 0 |
| rs371526758 | pathogenic | 1 | 1 | 1 |
| rs142968179 | pathogenic | 1 | 1 | 1 |
| rs387907291 | pathogenic | 1 | 1 | 0 |
| rs121907949 | pathogenic | 0 | 0 | 0 |
| rs121907947 | pathogenic | 0 | 0 | 0 |
| rs121907950 | pathogenic | 1 | 1 | 1 |
| rs121907948 | pathogenic | 1 | 1 | 1 |
| rs4926 | benign | 0 | 0 | 0 |
| rs748809996 | pathogenic | 0 | 0 | 0 |
| rs1799883 | benign | 1 | 1 | 0 |
| rs104893974 | pathogenic | 0 | 1 | 0 |
| rs104893973 | pathogenic | 0 | 1 | 0 |
| rs34452707 | benign | 0 | 0 | 0 |
| rs119467003 | pathogenic | 1 | 1 | 1 |
| rs121908261 | pathogenic | 0 | 1 | 0 |
| rs28933985 | pathogenic | 1 | 1 | 1 |
| rs80356669 | pathogenic | 1 | 1 | 1 |
| rs28936677 | pathogenic | 1 | 1 | 0 |
| rs121918326 | pathogenic | 0 | 0 | 0 |
| rs121918324 | pathogenic | 1 | 1 | 0 |
| rs121918325 | pathogenic | 1 | 1 | 0 |
| rs41270025 | pathogenic | 1 | 1 | 0 |
| rs121918323 | pathogenic | 0 | 0 | 0 |
| rs12735723 | pathogenic | 1 | 1 | 0 |
| rs36013429 | benign | 0 | 0 | 0 |
| rs28936676 | pathogenic | 1 | 1 | 0 |
| rs771809901 | pathogenic | 1 | 1 | 1 |
| rs758350837 | benign | 1 | 1 | 0 |
| rs121908192 | pathogenic | 1 | 1 | 1 |
| rs147568500 | pathogenic | 1 | 1 | 1 |
| rs3729734 | benign | 0 | 0 | 0 |
| rs371536347 | benign | 1 | 1 | 1 |
| rs80358193 | pathogenic | 1 | 1 | 1 |
| rs119476044 | pathogenic | 1 | 1 | 1 |
| rs111292798 | benign | 1 | 1 | 1 |
| rs121908063 | pathogenic | 1 | 1 | 0 |
| rs121908065 | pathogenic | 1 | 1 | 1 |
| rs121908062 | pathogenic | 1 | 1 | 1 |
| rs121908064 | pathogenic | 1 | 1 | 1 |
| rs61746421 | benign | 1 | 1 | 1 |
| rs34886916 | benign | 1 | 1 | 1 |
| rs118204045 | pathogenic | 0 | 0 | 0 |
| rs138607412 | benign | 1 | 1 | 1 |

| | | | | |
|---|---|---|---|---|
| rs118204046 | pathogenic | 0 | 0 | 0 |
| rs118204048 | pathogenic | 0 | 0 | 0 |
| rs118204044 | pathogenic | 0 | 0 | 0 |
| rs149664056 | benign | 0 | 0 | 0 |
| rs121909285 | pathogenic | 0 | 0 | 0 |
| rs1802498 | benign | 0 | 0 | 0 |
| rs1065761 | benign | 0 | 0 | 0 |
| rs119477054 | pathogenic | 0 | 0 | 0 |
| rs12720097 | benign | 0 | 0 | 0 |
| rs747281324 | pathogenic | 1 | 1 | 0 |
| rs367543028 | pathogenic | 1 | 1 | 0 |
| rs146077918 | benign | 0 | 0 | 0 |
| rs142551229 | benign | 0 | 0 | 0 |
| rs367543039 | pathogenic | 0 | 0 | 0 |
| rs587779884 | pathogenic | 1 | 1 | 0 |
| rs137853153 | pathogenic | 0 | 0 | 0 |
| rs367543029 | pathogenic | 0 | 0 | 0 |
| rs587783037 | pathogenic | 1 | 1 | 0 |
| rs142928725 | benign | 0 | 0 | 0 |
| rs28385141 | benign | 0 | 0 | 0 |
| rs587779888 | pathogenic | 0 | 0 | 0 |
| rs121909801 | pathogenic | 1 | 1 | 1 |
| rs121909790 | pathogenic | 1 | 1 | 0 |
| rs121909797 | pathogenic | 1 | 1 | 0 |
| rs121909794 | pathogenic | 1 | 1 | 1 |
| rs121909791 | pathogenic | 1 | 1 | 1 |
| rs121909793 | pathogenic | 1 | 1 | 1 |
| rs147977279 | benign | 1 | 1 | 1 |
| rs147352564 | benign | 0 | 0 | 0 |
| rs368758045 | benign | 1 | 1 | 1 |
| rs114498261 | benign | 1 | 1 | 0 |
| rs35484156 | benign | 0 | 0 | 0 |
| rs139176878 | benign | 0 | 0 | 0 |
| rs11543934 | benign | 0 | 0 | 0 |
| rs34192549 | benign | 0 | 0 | 0 |
| rs568736295 | benign | 0 | 0 | 0 |
| rs55747232 | benign | 0 | 0 | 0 |
| rs781887272 | benign | 0 | 0 | 0 |
| rs146273341 | benign | 0 | 0 | 0 |
| rs41473544 | benign | 0 | 0 | 0 |
| rs137852680 | pathogenic | 0 | 0 | 0 |
| rs55891000 | benign | 0 | 0 | 0 |
| rs140351189 | benign | 0 | 0 | 0 |
| rs137852679 | pathogenic | 1 | 1 | 0 |
| rs137852678 | pathogenic | 0 | 0 | 0 |
| rs137852677 | pathogenic | 1 | 1 | 1 |
| rs138620818 | benign | 0 | 0 | 0 |
| rs121908531 | pathogenic | 0 | 0 | 0 |
| rs72554307 | pathogenic | 0 | 0 | 0 |
| rs72554308 | pathogenic | 0 | 0 | 0 |
| rs72554312 | pathogenic | 1 | 1 | 0 |
| rs67939655 | pathogenic | 0 | 0 | 0 |
| rs67486158 | pathogenic | 0 | 0 | 0 |
| rs72554338 | pathogenic | 1 | 1 | 1 |

| | | | | |
|---|---|---|---|---|
| rs1800324 | pathogenic | 1 | 1 | 1 |
| rs72554356 | pathogenic | 1 | 1 | 1 |
| rs66656800 | pathogenic | 1 | 1 | 1 |
| rs68026851 | pathogenic | 1 | 1 | 1 |
| rs72558412 | pathogenic | 0 | 0 | 0 |
| rs72558415 | pathogenic | 0 | 0 | 0 |
| rs72558423 | pathogenic | 0 | 0 | 0 |
| rs67120076 | pathogenic | 1 | 1 | 1 |
| rs1800328 | pathogenic | 1 | 1 | 1 |
| rs72558451 | pathogenic | 1 | 1 | 1 |
| rs72558454 | pathogenic | 1 | 1 | 1 |
| rs672601372 | pathogenic | 0 | 0 | 0 |
| rs139644798 | pathogenic | 0 | 0 | 0 |
| rs369398935 | pathogenic | 0 | 0 | 0 |
| rs267607035 | pathogenic | 0 | 0 | 0 |
| rs267607036 | pathogenic | 1 | 1 | 1 |
| rs2302566 | benign | 1 | 1 | 0 |
| rs115959591 | benign | 0 | 0 | 0 |
| rs773876739 | pathogenic | 0 | 0 | 0 |
| rs1799967 | benign | 0 | 0 | 0 |
| rs80356968 | benign | 0 | 0 | 0 |
| rs28897695 | benign | 1 | 1 | 1 |
| rs80357314 | benign | 0 | 0 | 0 |
| rs80357169 | benign | 0 | 0 | 0 |
| rs397507239 | benign | 1 | 1 | 1 |
| rs80187739 | pathogenic | 1 | 1 | 1 |
| rs41293459 | pathogenic | 1 | 1 | 1 |
| rs55770810 | pathogenic | 1 | 1 | 1 |
| rs80356860 | pathogenic | 0 | 0 | 0 |
| rs56195342 | benign | 0 | 0 | 0 |
| rs80357442 | benign | 0 | 0 | 0 |
| rs41293463 | pathogenic | 1 | 1 | 1 |
| rs80357069 | pathogenic | 0 | 0 | 0 |
| rs397509275 | benign | 0 | 0 | 0 |
| rs80356920 | benign | 0 | 0 | 0 |
| rs397509284 | pathogenic | 0 | 0 | 0 |
| rs80357286 | benign | 0 | 0 | 0 |
| rs80357274 | benign | 0 | 0 | 0 |
| rs765656957 | benign | 0 | 0 | 0 |
| rs755427809 | benign | 0 | 0 | 0 |
| rs80357322 | benign | 0 | 0 | 0 |
| rs121434430 | pathogenic | 0 | 0 | 0 |
| rs121918450 | pathogenic | 1 | 1 | 1 |
| rs1532268 | benign | 0 | 0 | 0 |
| rs2303080 | benign | 0 | 0 | 0 |
| rs10064631 | benign | 0 | 0 | 0 |
| rs162036 | benign | 0 | 0 | 0 |
| rs2287780 | benign | 0 | 0 | 0 |
| rs16879334 | benign | 0 | 0 | 0 |
| rs137853061 | pathogenic | 1 | 1 | 0 |
| rs10380 | benign | 0 | 0 | 0 |
| rs1494558 | benign | 0 | 0 | 0 |
| rs1494555 | benign | 0 | 0 | 0 |
| rs192994060 | benign | 0 | 0 | 0 |

| | | | | |
|---|---|---|---|---|
| rs147128920 | benign | 1 | 1 | 0 |
| rs201803986 | pathogenic | 0 | 0 | 0 |
| rs746419489 | pathogenic | 0 | 1 | 0 |
| rs764396564 | pathogenic | 0 | 0 | 0 |
| rs758304537 | pathogenic | 0 | 0 | 0 |
| rs267606763 | pathogenic | 0 | 0 | 0 |
| rs267606762 | pathogenic | 0 | 0 | 0 |
| rs150702945 | pathogenic | 1 | 1 | 1 |
| rs202047589 | pathogenic | 0 | 0 | 0 |
| rs755562733 | pathogenic | 0 | 0 | 0 |
| rs267606764 | pathogenic | 1 | 1 | 1 |
| rs138207257 | pathogenic | 1 | 1 | 0 |
| rs149150736 | pathogenic | 1 | 1 | 1 |
| rs777046879 | pathogenic | 0 | 0 | 0 |
| rs104894408 | pathogenic | 1 | 1 | 1 |
| rs1801002 | pathogenic | 1 | 1 | 1 |
| rs28929485 | pathogenic | 1 | 1 | 1 |
| rs104894396 | pathogenic | 1 | 1 | 1 |
| rs371024165 | pathogenic | 0 | 0 | 0 |
| rs35887622 | pathogenic | 1 | 1 | 1 |
| rs587783644 | pathogenic | 1 | 1 | 1 |
| rs72474224 | pathogenic | 1 | 1 | 1 |
| rs111033296 | pathogenic | 1 | 1 | 1 |
| rs104894407 | pathogenic | 1 | 1 | 1 |
| rs72561723 | pathogenic | 1 | 1 | 1 |
| rs104894398 | pathogenic | 1 | 1 | 1 |
| rs28931594 | pathogenic | 1 | 1 | 1 |
| rs104894412 | pathogenic | 1 | 1 | 1 |
| rs111033297 | pathogenic | 1 | 1 | 1 |
| rs104894404 | pathogenic | 1 | 1 | 1 |
| rs104894410 | pathogenic | 1 | 1 | 1 |
| rs104894403 | pathogenic | 1 | 1 | 1 |
| rs397516870 | benign | 0 | 0 | 0 |
| rs200023879 | pathogenic | 1 | 1 | 1 |
| rs121912968 | pathogenic | 0 | 0 | 0 |
| rs104894402 | pathogenic | 1 | 1 | 1 |
| rs28931593 | pathogenic | 1 | 1 | 1 |
| rs104894397 | pathogenic | 0 | 0 | 0 |
| rs80338944 | pathogenic | 0 | 0 | 0 |
| rs199883710 | pathogenic | 0 | 0 | 0 |
| rs727504302 | pathogenic | 0 | 0 | 0 |
| rs145216882 | benign | 1 | 1 | 1 |
| rs781534323 | pathogenic | 1 | 1 | 1 |
| rs111033218 | benign | 1 | 1 | 1 |
| rs104894409 | pathogenic | 0 | 0 | 0 |
| rs80338945 | pathogenic | 1 | 1 | 1 |
| rs111033299 | pathogenic | 0 | 0 | 0 |
| rs143343083 | pathogenic | 1 | 1 | 1 |
| rs80338946 | pathogenic | 0 | 0 | 0 |
| rs150529554 | pathogenic | 0 | 0 | 0 |
| rs111033295 | pathogenic | 0 | 0 | 0 |
| rs111033188 | benign | 0 | 0 | 0 |
| rs397516874 | pathogenic | 0 | 0 | 0 |
| rs76434661 | pathogenic | 0 | 0 | 0 |

| | | | | |
|---|---|---|---|---|
| rs80338948 | pathogenic | 0 | 0 | 0 |
| rs104894401 | pathogenic | 0 | 0 | 0 |
| rs767178508 | pathogenic | 0 | 0 | 0 |
| rs772264564 | pathogenic | 0 | 0 | 0 |
| rs28931592 | pathogenic | 0 | 0 | 0 |
| rs80338949 | pathogenic | 1 | 1 | 1 |
| rs376898963 | benign | 1 | 1 | 1 |
| rs111033360 | benign | 0 | 0 | 0 |
| rs774518779 | pathogenic | 1 | 1 | 1 |
| rs28931595 | pathogenic | 1 | 1 | 1 |
| rs771748289 | pathogenic | 0 | 0 | 0 |
| rs104894406 | pathogenic | 0 | 0 | 0 |
| rs76838169 | benign | 0 | 0 | 0 |
| rs111033294 | pathogenic | 0 | 0 | 0 |
| rs200214298 | pathogenic | 0 | 0 | 0 |
| rs28940283 | pathogenic | 0 | 1 | 0 |
| rs1047883 | benign | 0 | 0 | 0 |
| rs138254425 | benign | 0 | 0 | 0 |
| rs121912592 | pathogenic | 1 | 1 | 0 |
| rs369061090 | pathogenic | 1 | 1 | 0 |
| rs758724746 | pathogenic | 0 | 1 | 0 |
| rs121912595 | pathogenic | 0 | 0 | 0 |
| rs76340296 | benign | 0 | 0 | 0 |
| rs140578009 | benign | 1 | 1 | 1 |
| rs74445297 | benign | 0 | 0 | 0 |
| rs78599516 | benign | 1 | 1 | 0 |
| rs104894035 | pathogenic | 1 | 1 | 0 |
| rs79451216 | benign | 0 | 0 | 0 |
| rs104894036 | pathogenic | 0 | 0 | 0 |
| rs73343757 | benign | 0 | 0 | 0 |
| rs121917722 | pathogenic | 1 | 1 | 1 |
| rs747700126 | pathogenic | 0 | 1 | 0 |
| rs121917725 | pathogenic | 1 | 1 | 1 |
| rs121917721 | pathogenic | 1 | 1 | 1 |
| rs121917723 | pathogenic | 1 | 1 | 0 |
| rs17569 | benign | 1 | 1 | 1 |
| rs2230062 | benign | 1 | 1 | 1 |
| rs17570 | benign | 0 | 0 | 0 |
| rs121917724 | pathogenic | 0 | 1 | 0 |
| rs199422235 | pathogenic | 0 | 0 | 0 |
| rs199422237 | pathogenic | 1 | 1 | 1 |
| rs199422234 | pathogenic | 1 | 1 | 0 |
| rs140015315 | pathogenic | 1 | 1 | 1 |
| rs730880298 | pathogenic | 1 | 1 | 1 |
| rs779692470 | pathogenic | 0 | 0 | 0 |
| rs369152939 | pathogenic | 0 | 0 | 0 |
| rs527236040 | pathogenic | 0 | 0 | 0 |
| rs587776985 | pathogenic | 1 | 1 | 1 |
| rs147077598 | pathogenic | 1 | 1 | 1 |
| rs128620185 | pathogenic | 1 | 1 | 1 |
| rs128620189 | pathogenic | 0 | 0 | 0 |
| rs128621190 | pathogenic | 0 | 0 | 0 |
| rs61754648 | benign | 0 | 0 | 0 |
| rs137852906 | pathogenic | 0 | 0 | 0 |

| | | | | |
|---|---|---|---|---|
| rs76811038 | pathogenic | 0 | 0 | 0 |
| rs149468422 | benign | 0 | 0 | 0 |
| rs5911 | benign | 0 | 0 | 0 |
| rs28942112 | pathogenic | 1 | 1 | 1 |
| rs143275858 | pathogenic | 1 | 1 | 0 |
| rs28362261 | benign | 0 | 0 | 0 |
| rs28362263 | benign | 1 | 1 | 1 |
| rs558707786 | pathogenic | 0 | 0 | 0 |
| rs138775799 | pathogenic | 0 | 0 | 0 |
| rs748170941 | pathogenic | 0 | 0 | 0 |
| rs121908380 | pathogenic | 0 | 0 | 0 |
| rs765123255 | pathogenic | 1 | 1 | 1 |
| rs730881832 | pathogenic | 0 | 1 | 0 |
| rs587782165 | benign | 1 | 1 | 1 |
| rs762307622 | pathogenic | 0 | 0 | 0 |
| rs34612342 | pathogenic | 1 | 1 | 0 |
| rs143353451 | pathogenic | 0 | 1 | 0 |
| rs747993448 | pathogenic | 0 | 1 | 0 |
| rs376561094 | pathogenic | 0 | 0 | 0 |
| rs776487884 | pathogenic | 1 | 1 | 0 |
| rs34126013 | pathogenic | 1 | 1 | 0 |
| rs200495564 | pathogenic | 1 | 1 | 0 |
| rs140342925 | pathogenic | 1 | 1 | 0 |
| rs587780749 | pathogenic | 1 | 1 | 0 |
| rs773087549 | pathogenic | 1 | 1 | 0 |
| rs730881833 | pathogenic | 0 | 0 | 0 |
| rs374950566 | pathogenic | 1 | 1 | 0 |
| rs587780752 | benign | 0 | 0 | 0 |
| rs3219489 | benign | 0 | 0 | 0 |
| rs199742231 | benign | 0 | 0 | 0 |
| rs587780082 | pathogenic | 0 | 0 | 0 |
| rs35881803 | benign | 0 | 0 | 0 |
| rs41305623 | benign | 1 | 1 | 1 |
| rs121917856 | pathogenic | 1 | 1 | 1 |
| rs576284122 | benign | 0 | 0 | 0 |
| rs142036701 | benign | 0 | 0 | 0 |
| rs121917852 | pathogenic | 1 | 1 | 1 |
| rs121917854 | pathogenic | 1 | 1 | 1 |
| rs121917853 | pathogenic | 1 | 1 | 1 |
| rs121917857 | pathogenic | 0 | 1 | 0 |
| rs193922740 | pathogenic | 1 | 1 | 1 |
| rs565779970 | pathogenic | 1 | 1 | 1 |
| rs6723526 | benign | 0 | 0 | 0 |
| rs141094096 | pathogenic | 1 | 1 | 1 |
| rs104894637 | pathogenic | 0 | 0 | 0 |
| rs104894636 | pathogenic | 0 | 0 | 0 |
| rs104894642 | pathogenic | 0 | 0 | 0 |
| rs104894638 | pathogenic | 0 | 0 | 0 |
| rs753666460 | pathogenic | 0 | 0 | 0 |
| rs104894643 | pathogenic | 1 | 1 | 0 |
| rs34520362 | pathogenic | 0 | 0 | 0 |
| rs374621913 | pathogenic | 0 | 0 | 0 |
| rs113641837 | benign | 0 | 0 | 0 |
| rs104894635 | pathogenic | 0 | 0 | 0 |

| | | | | |
|---|---|---|---|---|
| rs143947056 | pathogenic | 0 | 0 | 0 |
| rs138504221 | pathogenic | 0 | 0 | 0 |
| rs9894254 | benign | 0 | 0 | 0 |
| rs104894640 | pathogenic | 1 | 1 | 1 |
| rs58786455 | benign | 0 | 0 | 0 |
| rs144143780 | pathogenic | 0 | 0 | 0 |
| rs62620232 | benign | 0 | 0 | 0 |
| rs34297805 | benign | 1 | 1 | 1 |
| rs104894641 | pathogenic | 0 | 0 | 0 |
| rs148876719 | benign | 0 | 0 | 0 |
| rs104894639 | pathogenic | 0 | 0 | 0 |
| rs7503034 | benign | 0 | 0 | 0 |
| rs201631570 | benign | 0 | 0 | 0 |
| rs150181226 | pathogenic | 0 | 0 | 0 |
| rs377546036 | pathogenic | 0 | 0 | 0 |
| rs267606700 | pathogenic | 0 | 0 | 0 |
| rs267606701 | pathogenic | 1 | 1 | 0 |
| rs267606699 | pathogenic | 1 | 1 | 0 |
| rs9903215 | benign | 0 | 0 | 0 |
| rs139486406 | benign | 0 | 0 | 0 |
| rs143791614 | benign | 0 | 0 | 0 |
| rs573483474 | benign | 0 | 0 | 0 |
| rs2032892 | benign | 0 | 0 | 0 |
| rs137852619 | pathogenic | 0 | 0 | 0 |
| rs200243548 | benign | 0 | 0 | 0 |
| rs115702388 | benign | 0 | 0 | 0 |
| rs138085358 | pathogenic | 0 | 0 | 0 |
| rs121912789 | pathogenic | 1 | 1 | 1 |
| rs763407972 | pathogenic | 1 | 1 | 1 |
| rs147278302 | benign | 0 | 0 | 0 |
| rs374540411 | benign | 0 | 0 | 0 |
| rs121912790 | pathogenic | 1 | 1 | 0 |
| rs752049917 | benign | 0 | 0 | 0 |
| rs139640491 | benign | 1 | 1 | 1 |
| rs138010738 | benign | 1 | 1 | 0 |
| rs201328768 | pathogenic | 1 | 1 | 0 |
| rs2135720 | benign | 1 | 1 | 0 |
| rs201816080 | benign | 1 | 1 | 0 |
| rs754391973 | pathogenic | 0 | 0 | 0 |
| rs139087859 | benign | 0 | 0 | 0 |
| rs6805421 | benign | 0 | 0 | 0 |
| rs146266900 | benign | 0 | 0 | 0 |
| rs397509419 | pathogenic | 0 | 0 | 0 |
| rs149980411 | benign | 0 | 0 | 0 |
| rs747711488 | pathogenic | 1 | 1 | 1 |
| rs149474339 | pathogenic | 1 | 1 | 1 |
| rs758252808 | pathogenic | 1 | 1 | 0 |
| rs745756308 | pathogenic | 0 | 0 | 0 |
| rs4986958 | benign | 0 | 0 | 0 |
| rs74315444 | pathogenic | 0 | 0 | 0 |
| rs61752137 | pathogenic | 0 | 0 | 0 |
| rs8179065 | benign | 1 | 1 | 1 |
| rs758361736 | pathogenic | 1 | 1 | 0 |
| rs139794067 | pathogenic | 1 | 1 | 1 |

| | | | | |
|---|---|---|---|---|
| rs730880954 | pathogenic | 1 | 1 | 1 |
| rs730880955 | pathogenic | 1 | 1 | 1 |
| rs150634297 | pathogenic | 0 | 0 | 0 |
| rs104893750 | pathogenic | 1 | 1 | 1 |
| rs104893748 | pathogenic | 1 | 1 | 1 |
| rs104893749 | pathogenic | 1 | 1 | 1 |
| rs753520553 | pathogenic | 1 | 1 | 0 |
| rs61737294 | benign | 0 | 0 | 0 |
| rs104894601 | pathogenic | 0 | 0 | 0 |
| rs376090795 | pathogenic | 0 | 0 | 0 |
| rs104894596 | pathogenic | 0 | 0 | 0 |
| rs104894595 | pathogenic | 0 | 0 | 0 |
| rs104894598 | pathogenic | 0 | 0 | 0 |
| rs104894597 | pathogenic | 0 | 0 | 0 |
| rs104894591 | pathogenic | 0 | 0 | 0 |
| rs104894593 | pathogenic | 0 | 0 | 0 |
| rs104894594 | pathogenic | 0 | 0 | 0 |
| rs104894590 | pathogenic | 0 | 0 | 0 |
| rs86312 | benign | 0 | 0 | 0 |
| rs77874543 | benign | 0 | 0 | 0 |
| rs121912590 | pathogenic | 1 | 1 | 1 |
| rs121912589 | pathogenic | 1 | 1 | 1 |
| rs121912586 | pathogenic | 1 | 1 | 1 |
| rs121912585 | pathogenic | 1 | 1 | 1 |
| rs121912587 | pathogenic | 1 | 1 | 1 |
| rs762909359 | pathogenic | 0 | 0 | 0 |
| rs28931610 | pathogenic | 0 | 0 | 0 |
| rs104894097 | pathogenic | 1 | 1 | 1 |
| rs775176191 | benign | 0 | 0 | 0 |
| rs199907548 | pathogenic | 0 | 0 | 0 |
| rs104894095 | pathogenic | 1 | 1 | 1 |
| rs104894109 | pathogenic | 1 | 1 | 1 |
| rs372266620 | benign | 1 | 1 | 1 |
| rs104894099 | pathogenic | 0 | 0 | 0 |
| rs137854599 | pathogenic | 1 | 1 | 1 |
| rs104894094 | pathogenic | 0 | 0 | 0 |
| rs104894104 | pathogenic | 0 | 0 | 0 |
| rs6413463 | benign | 1 | 1 | 1 |
| rs104894098 | pathogenic | 0 | 0 | 0 |
| rs6413464 | benign | 0 | 0 | 0 |
| rs3731249 | benign | 0 | 0 | 0 |
| rs16931179 | benign | 0 | 0 | 0 |
| rs2734715 | benign | 0 | 0 | 0 |
| rs759080581 | pathogenic | 0 | 0 | 0 |
| rs1800746 | benign | 0 | 0 | 0 |
| rs202189057 | pathogenic | 0 | 0 | 0 |
| rs121908204 | pathogenic | 0 | 0 | 0 |
| rs756564767 | pathogenic | 0 | 0 | 0 |
| rs34758634 | benign | 1 | 1 | 1 |
| rs28940573 | pathogenic | 0 | 0 | 0 |
| rs119455955 | pathogenic | 1 | 1 | 0 |
| rs752164603 | pathogenic | 1 | 1 | 0 |
| rs763162812 | pathogenic | 0 | 0 | 0 |
| rs119455957 | pathogenic | 0 | 0 | 0 |

| | | | | |
|---|---|---|---|---|
| rs119455958 | pathogenic | 1 | 1 | 0 |
| rs750428882 | pathogenic | 0 | 0 | 0 |
| rs141482368 | benign | 0 | 0 | 0 |
| rs119455953 | pathogenic | 0 | 0 | 0 |
| rs119455954 | pathogenic | 0 | 0 | 0 |
| rs112812685 | benign | 1 | 1 | 0 |
| rs138744051 | benign | 1 | 1 | 0 |
| rs121908200 | pathogenic | 0 | 0 | 0 |
| rs119455956 | pathogenic | 0 | 0 | 0 |
| rs78970155 | benign | 0 | 0 | 0 |
| rs2834167 | benign | 0 | 0 | 0 |
| rs387907326 | pathogenic | 1 | 1 | 1 |
| rs2107732 | benign | 0 | 0 | 0 |
| rs11552377 | benign | 0 | 0 | 0 |
| rs137852843 | pathogenic | 0 | 0 | 0 |
| rs104894783 | pathogenic | 0 | 0 | 0 |
| rs122457137 | pathogenic | 1 | 1 | 1 |
| rs104894779 | pathogenic | 1 | 1 | 1 |
| rs104894786 | pathogenic | 0 | 0 | 0 |
| rs104894784 | pathogenic | 0 | 0 | 0 |
| rs104894785 | pathogenic | 0 | 0 | 0 |
| rs201870761 | pathogenic | 0 | 0 | 0 |
| rs104894781 | pathogenic | 0 | 0 | 0 |
| rs28897728 | benign | 0 | 0 | 0 |
| rs370273690 | pathogenic | 1 | 1 | 0 |
| rs9332739 | benign | 0 | 0 | 0 |
| rs150878060 | benign | 0 | 0 | 0 |
| rs573509224 | benign | 0 | 0 | 0 |
| rs36221133 | benign | 0 | 0 | 0 |
| rs2230009 | benign | 0 | 0 | 0 |
| rs267607008 | pathogenic | 0 | 0 | 0 |
| rs5985 | benign | 1 | 1 | 1 |
| rs5982 | benign | 1 | 1 | 1 |
| rs5987 | benign | 0 | 0 | 0 |
| rs5988 | benign | 0 | 0 | 0 |
| rs121918680 | pathogenic | 1 | 1 | 0 |
| rs121918679 | pathogenic | 1 | 1 | 0 |
| rs7845 | benign | 1 | 1 | 0 |
| rs1049296 | benign | 1 | 1 | 1 |
| rs121918677 | pathogenic | 1 | 1 | 0 |
| rs200086262 | pathogenic | 1 | 1 | 0 |
| rs746930990 | pathogenic | 1 | 1 | 0 |
| rs104893878 | pathogenic | 1 | 1 | 1 |
| rs121912468 | pathogenic | 1 | 1 | 1 |
| rs121912467 | pathogenic | 1 | 1 | 1 |
| rs145351926 | benign | 1 | 1 | 1 |
| rs141212743 | pathogenic | 1 | 1 | 1 |
| rs397515473 | pathogenic | 1 | 1 | 0 |
| rs121918313 | pathogenic | 0 | 1 | 0 |
| rs73015965 | pathogenic | 0 | 0 | 0 |
| rs121918030 | pathogenic | 0 | 1 | 0 |
| rs77782321 | benign | 0 | 0 | 0 |
| rs374661051 | pathogenic | 1 | 1 | 1 |
| rs267606854 | pathogenic | 0 | 0 | 0 |

| | | | | |
|---|---|---|---|---|
| rs41279678 | benign | 0 | 0 | 0 |
| rs61754640 | benign | 0 | 0 | 0 |
| rs199473314 | pathogenic | 0 | 0 | 0 |
| rs137854601 | pathogenic | 0 | 0 | 0 |
| rs137854614 | pathogenic | 0 | 0 | 0 |
| rs137854615 | pathogenic | 0 | 0 | 0 |
| rs199473635 | pathogenic | 0 | 0 | 0 |
| rs121434478 | pathogenic | 1 | 1 | 1 |
| rs775407864 | pathogenic | 1 | 1 | 1 |
| rs12706927 | benign | 0 | 1 | 0 |
| rs775614261 | benign | 0 | 0 | 0 |
| rs768926261 | benign | 0 | 0 | 0 |
| rs114057640 | benign | 0 | 0 | 0 |
| rs387906886 | pathogenic | 1 | 1 | 0 |
| rs138658585 | benign | 0 | 0 | 0 |
| rs4880091 | benign | 0 | 0 | 0 |
| rs121918552 | pathogenic | 1 | 1 | 1 |
| rs10794537 | benign | 0 | 0 | 0 |
| rs121965022 | pathogenic | 1 | 1 | 0 |
| rs121965020 | pathogenic | 0 | 0 | 0 |
| rs758452450 | pathogenic | 0 | 0 | 0 |
| rs747981483 | benign | 0 | 0 | 0 |
| rs58037052 | benign | 0 | 0 | 0 |
| rs121965029 | pathogenic | 1 | 1 | 0 |
| rs147490060 | benign | 0 | 0 | 0 |
| rs3755955 | benign | 0 | 0 | 0 |
| rs148789453 | pathogenic | 0 | 0 | 0 |
| rs121965030 | pathogenic | 0 | 0 | 0 |
| rs76722191 | benign | 0 | 0 | 0 |
| rs199801029 | pathogenic | 0 | 0 | 0 |
| rs121965033 | pathogenic | 0 | 0 | 0 |
| rs368454909 | pathogenic | 1 | 1 | 0 |
| rs6831280 | benign | 0 | 0 | 0 |
| rs121965024 | pathogenic | 0 | 0 | 0 |
| rs121965019 | pathogenic | 0 | 0 | 0 |
| rs11934801 | benign | 0 | 0 | 0 |
| rs73066479 | benign | 0 | 0 | 0 |
| rs121965027 | pathogenic | 0 | 0 | 0 |
| rs121965026 | pathogenic | 1 | 1 | 0 |
| rs121965021 | pathogenic | 0 | 0 | 0 |
| rs398123257 | benign | 0 | 0 | 0 |
| rs121965031 | pathogenic | 0 | 0 | 0 |
| rs202003795 | pathogenic | 0 | 0 | 0 |
| rs559510809 | pathogenic | 0 | 0 | 0 |
| rs762936223 | benign | 0 | 0 | 0 |
| rs137854580 | pathogenic | 1 | 1 | 1 |
| rs112263157 | benign | 0 | 0 | 0 |
| rs2289181 | benign | 0 | 0 | 0 |
| rs138034915 | benign | 0 | 1 | 0 |
| rs528367092 | pathogenic | 0 | 1 | 0 |
| rs1042393 | benign | 0 | 0 | 0 |
| rs370950728 | pathogenic | 0 | 1 | 0 |
| rs530478036 | benign | 0 | 1 | 0 |
| rs374569672 | benign | 0 | 1 | 0 |

| | | | | |
|---|---|---|---|---|
| rs121907944 | pathogenic | 0 | 1 | 0 |
| rs200856561 | pathogenic | 0 | 1 | 0 |
| rs577915581 | pathogenic | 0 | 1 | 0 |
| rs201896815 | pathogenic | 0 | 1 | 0 |
| rs121907945 | pathogenic | 0 | 1 | 0 |
| rs121907940 | pathogenic | 0 | 1 | 0 |
| rs543300039 | pathogenic | 0 | 1 | 0 |
| rs121907936 | pathogenic | 0 | 1 | 0 |
| rs766074609 | pathogenic | 0 | 1 | 0 |
| rs755253527 | pathogenic | 0 | 1 | 0 |
| rs560575383 | pathogenic | 0 | 1 | 0 |
| rs528369909 | benign | 0 | 1 | 0 |
| rs770610356 | pathogenic | 0 | 1 | 0 |
| rs747610090 | pathogenic | 0 | 1 | 0 |
| rs145712232 | benign | 0 | 1 | 0 |
| rs7215458 | benign | 1 | 1 | 0 |
| rs535644999 | pathogenic | 0 | 1 | 0 |
| rs772883420 | pathogenic | 1 | 1 | 0 |
| rs398123169 | pathogenic | 0 | 1 | 0 |
| rs121907937 | pathogenic | 0 | 1 | 0 |
| rs121907941 | pathogenic | 1 | 1 | 0 |
| rs121907942 | pathogenic | 0 | 1 | 0 |
| rs779556619 | pathogenic | 0 | 1 | 0 |
| rs1800307 | benign | 0 | 1 | 0 |
| rs377544304 | pathogenic | 1 | 1 | 0 |
| rs549029029 | pathogenic | 0 | 1 | 0 |
| rs753269119 | pathogenic | 1 | 1 | 1 |
| rs757617999 | pathogenic | 0 | 1 | 0 |
| rs28937909 | pathogenic | 0 | 1 | 0 |
| rs28940868 | pathogenic | 0 | 1 | 0 |
| rs368438393 | pathogenic | 0 | 1 | 0 |
| rs536906561 | pathogenic | 0 | 1 | 0 |
| rs374143224 | pathogenic | 0 | 1 | 0 |
| rs778418246 | pathogenic | 0 | 1 | 0 |
| rs398123172 | pathogenic | 0 | 1 | 0 |
| rs121907938 | pathogenic | 0 | 1 | 0 |
| rs1800312 | pathogenic | 0 | 1 | 0 |
| rs1126690 | benign | 0 | 1 | 0 |
| rs1800314 | benign | 0 | 1 | 0 |
| rs369532274 | pathogenic | 0 | 1 | 0 |
| rs121907943 | pathogenic | 0 | 1 | 0 |
| rs780321415 | pathogenic | 0 | 1 | 0 |
| rs377286472 | benign | 0 | 1 | 0 |
| rs121918406 | pathogenic | 1 | 1 | 1 |
| rs147484870 | benign | 0 | 0 | 0 |
| rs193922671 | pathogenic | 0 | 0 | 0 |
| rs201397978 | benign | 1 | 1 | 1 |
| rs34884895 | benign | 1 | 1 | 1 |
| rs138329459 | benign | 0 | 0 | 0 |
| rs2289366 | benign | 1 | 1 | 1 |
| rs141858334 | benign | 1 | 1 | 1 |
| rs59907886 | benign | 1 | 1 | 1 |
| rs137852653 | pathogenic | 0 | 0 | 0 |
| rs137852652 | pathogenic | 0 | 0 | 0 |

| | | | | |
|---|---|---|---|---|
| rs137852656 | pathogenic | 0 | 0 | 0 |
| rs61739916 | benign | 0 | 0 | 0 |
| rs28936668 | pathogenic | 0 | 0 | 0 |
| rs137852651 | pathogenic | 0 | 0 | 0 |
| rs137852650 | pathogenic | 0 | 0 | 0 |
| rs28936669 | pathogenic | 0 | 0 | 0 |
| rs137852654 | pathogenic | 0 | 0 | 0 |
| rs28936368 | pathogenic | 1 | 1 | 0 |
| rs137852655 | pathogenic | 0 | 0 | 0 |
| rs61752496 | benign | 0 | 0 | 0 |
| rs750098055 | benign | 0 | 0 | 0 |
| rs764149793 | benign | 0 | 0 | 0 |
| rs724160017 | pathogenic | 0 | 1 | 0 |
| rs724160019 | pathogenic | 0 | 0 | 0 |
| rs202219398 | pathogenic | 1 | 1 | 0 |
| rs724160023 | pathogenic | 1 | 1 | 0 |
| rs761953453 | pathogenic | 0 | 0 | 0 |
| rs80338835 | pathogenic | 1 | 1 | 1 |
| rs56284940 | benign | 0 | 0 | 0 |
| rs74575749 | benign | 0 | 0 | 0 |
| rs587777075 | pathogenic | 1 | 1 | 1 |
| rs641153 | benign | 0 | 0 | 0 |
| rs12614 | benign | 0 | 0 | 0 |
| rs117314762 | benign | 0 | 0 | 0 |
| rs537478097 | benign | 1 | 1 | 1 |
| rs144812066 | benign | 0 | 0 | 0 |
| rs4151651 | benign | 0 | 0 | 0 |
| rs138207668 | benign | 1 | 1 | 1 |
| rs149101394 | benign | 1 | 1 | 1 |
| rs4151659 | benign | 1 | 1 | 1 |
| rs45484591 | benign | 1 | 1 | 1 |
| rs761991070 | pathogenic | 0 | 0 | 0 |
| rs755948940 | pathogenic | 0 | 0 | 0 |
| rs137853097 | pathogenic | 0 | 0 | 0 |
| rs775297352 | benign | 0 | 0 | 0 |
| rs28943591 | benign | 0 | 0 | 0 |
| rs191468413 | pathogenic | 1 | 1 | 0 |
| rs11539471 | benign | 1 | 1 | 1 |
| rs11205 | benign | 0 | 0 | 0 |
| rs768409400 | pathogenic | 0 | 0 | 0 |
| rs762013077 | pathogenic | 0 | 0 | 0 |
| rs768474112 | pathogenic | 0 | 0 | 0 |
| rs121965063 | pathogenic | 1 | 1 | 1 |
| rs143648758 | pathogenic | 0 | 0 | 0 |
| rs121965066 | pathogenic | 0 | 0 | 0 |
| rs752907087 | pathogenic | 0 | 0 | 0 |
| rs747702749 | pathogenic | 0 | 0 | 0 |
| rs754087775 | pathogenic | 0 | 0 | 0 |
| rs121918641 | pathogenic | 1 | 1 | 1 |
| rs121918642 | pathogenic | 1 | 1 | 1 |
| rs121918640 | pathogenic | 1 | 1 | 1 |
| rs121918637 | pathogenic | 1 | 1 | 1 |
| rs121918644 | pathogenic | 1 | 1 | 1 |
| rs16840544 | benign | 0 | 0 | 0 |

| | | | | |
|---|---|---|---|---|
| rs1042391 | benign | 1 | 1 | 1 |
| rs104893744 | pathogenic | 1 | 1 | 1 |
| rs120074136 | pathogenic | 0 | 0 | 0 |
| rs1805081 | benign | 1 | 1 | 0 |
| rs80358251 | benign | 0 | 0 | 0 |
| rs774588125 | benign | 0 | 0 | 0 |
| rs120074134 | pathogenic | 0 | 0 | 0 |
| rs139751448 | pathogenic | 0 | 0 | 0 |
| rs77080672 | benign | 0 | 0 | 0 |
| rs372445155 | pathogenic | 1 | 1 | 0 |
| rs13381670 | benign | 0 | 0 | 0 |
| rs1788799 | benign | 0 | 0 | 0 |
| rs376213990 | pathogenic | 0 | 0 | 0 |
| rs757475924 | pathogenic | 0 | 0 | 0 |
| rs80358253 | pathogenic | 0 | 0 | 0 |
| rs1805082 | benign | 0 | 0 | 0 |
| rs120074130 | pathogenic | 0 | 0 | 0 |
| rs768999208 | pathogenic | 0 | 0 | 0 |
| rs34302553 | benign | 0 | 0 | 0 |
| rs28940897 | pathogenic | 0 | 0 | 0 |
| rs1140458 | benign | 0 | 0 | 0 |
| rs143124972 | pathogenic | 0 | 0 | 0 |
| rs120074135 | pathogenic | 0 | 0 | 0 |
| rs543206298 | pathogenic | 0 | 0 | 0 |
| rs120074132 | pathogenic | 0 | 0 | 0 |
| rs759826138 | pathogenic | 0 | 0 | 0 |
| rs28942108 | pathogenic | 0 | 0 | 0 |
| rs80358254 | pathogenic | 0 | 0 | 0 |
| rs80358257 | pathogenic | 0 | 0 | 0 |
| rs781261962 | pathogenic | 0 | 0 | 0 |
| rs28942107 | pathogenic | 0 | 0 | 0 |
| rs80358258 | pathogenic | 0 | 0 | 0 |
| rs80358259 | pathogenic | 1 | 1 | 0 |
| rs141440861 | benign | 0 | 0 | 0 |
| rs28942106 | pathogenic | 0 | 0 | 0 |
| rs778878523 | pathogenic | 0 | 0 | 0 |
| rs28942105 | pathogenic | 0 | 0 | 0 |
| rs61731969 | benign | 0 | 0 | 0 |
| rs200444084 | pathogenic | 0 | 0 | 0 |
| rs113371321 | pathogenic | 0 | 0 | 0 |
| rs369098773 | pathogenic | 0 | 0 | 0 |
| rs35248744 | benign | 0 | 0 | 0 |
| rs758902805 | pathogenic | 0 | 0 | 0 |
| rs120074131 | pathogenic | 0 | 0 | 0 |
| rs1805084 | benign | 0 | 0 | 0 |
| rs117793540 | benign | 0 | 0 | 0 |
| rs530058063 | benign | 0 | 0 | 0 |
| rs1064422 | pathogenic | 1 | 1 | 1 |
| rs104894506 | pathogenic | 1 | 1 | 1 |
| rs104894507 | pathogenic | 1 | 1 | 1 |
| rs104894508 | pathogenic | 1 | 1 | 0 |
| rs28999113 | pathogenic | 1 | 1 | 0 |
| rs281860266 | pathogenic | 0 | 0 | 0 |
| rs104894961 | pathogenic | 1 | 1 | 0 |

| | | | | |
|---|---|---|---|---|
| rs122463168 | pathogenic | 0 | 0 | 0 |
| rs199469624 | pathogenic | 1 | 1 | 1 |
| rs150107747 | benign | 0 | 0 | 0 |
| rs144984628 | pathogenic | 0 | 0 | 0 |
| rs28935171 | pathogenic | 0 | 0 | 0 |
| rs137852443 | pathogenic | 1 | 1 | 0 |
| rs28933674 | pathogenic | 1 | 1 | 1 |
| rs28933675 | pathogenic | 1 | 1 | 1 |
| rs28933676 | pathogenic | 1 | 1 | 1 |
| rs28933677 | pathogenic | 1 | 1 | 1 |
| rs28933678 | pathogenic | 0 | 0 | 0 |
| rs28933679 | pathogenic | 1 | 1 | 1 |
| rs28933680 | pathogenic | 0 | 0 | 0 |
| rs28933681 | pathogenic | 0 | 0 | 0 |
| rs28933682 | pathogenic | 0 | 0 | 0 |
| rs28937294 | pathogenic | 1 | 1 | 0 |
| rs137852459 | pathogenic | 0 | 0 | 0 |
| rs137852461 | pathogenic | 0 | 0 | 0 |
| rs137852464 | pathogenic | 1 | 1 | 1 |
| rs137852465 | pathogenic | 1 | 1 | 1 |
| rs1800297 | benign | 1 | 1 | 0 |
| rs137852374 | pathogenic | 1 | 1 | 1 |
| rs672601312 | pathogenic | 1 | 1 | 1 |
| rs387907196 | pathogenic | 1 | 1 | 1 |
| rs148348866 | benign | 0 | 0 | 0 |
| rs141138948 | pathogenic | 1 | 1 | 1 |
| rs137852685 | pathogenic | 1 | 1 | 0 |
| rs104893900 | pathogenic | 1 | 1 | 0 |
| rs137852686 | pathogenic | 1 | 1 | 1 |
| rs104893906 | pathogenic | 1 | 1 | 0 |
| rs587782207 | pathogenic | 1 | 1 | 1 |
| rs75086406 | pathogenic | 0 | 0 | 0 |
| rs199822819 | pathogenic | 0 | 0 | 0 |
| rs398123166 | pathogenic | 1 | 1 | 1 |
| rs121913123 | pathogenic | 1 | 1 | 1 |
| rs772190176 | pathogenic | 0 | 1 | 0 |
| rs587782216 | pathogenic | 1 | 1 | 1 |
| rs398123159 | pathogenic | 1 | 1 | 1 |
| rs121913122 | pathogenic | 1 | 1 | 1 |
| rs755436052 | pathogenic | 1 | 1 | 1 |
| rs200796606 | pathogenic | 1 | 1 | 1 |
| rs2070080 | pathogenic | 0 | 1 | 0 |
| rs149616320 | benign | 0 | 0 | 0 |
| rs143529458 | benign | 0 | 0 | 0 |
| rs267607077 | pathogenic | 1 | 1 | 1 |
| rs143898031 | benign | 1 | 1 | 0 |
| rs148304857 | benign | 1 | 1 | 1 |
| rs137852809 | pathogenic | 0 | 0 | 0 |
| rs3765623 | benign | 0 | 0 | 0 |
| rs16944397 | benign | 0 | 0 | 0 |
| rs730880167 | benign | 0 | 0 | 0 |
| rs187108957 | benign | 0 | 0 | 0 |
| rs117342470 | benign | 0 | 0 | 0 |
| rs751200138 | pathogenic | 1 | 1 | 1 |

| | | | | |
|---|---|---|---|---|
| rs755799528 | pathogenic | 0 | 0 | 0 |
| rs537484504 | benign | 0 | 0 | 0 |
| rs748944640 | pathogenic | 0 | 0 | 0 |
| rs199774121 | pathogenic | 0 | 0 | 0 |
| rs766094434 | pathogenic | 1 | 1 | 0 |
| rs777321035 | pathogenic | 1 | 1 | 1 |
| rs146269774 | benign | 0 | 1 | 0 |
| rs144085805 | benign | 0 | 1 | 0 |
| rs367643028 | benign | 0 | 0 | 0 |
| rs42524 | benign | 0 | 0 | 0 |
| rs28940586 | pathogenic | 0 | 0 | 0 |
| rs74315327 | pathogenic | 1 | 1 | 1 |
| rs199676861 | pathogenic | 0 | 0 | 0 |
| rs115802744 | benign | 1 | 1 | 0 |
| rs752866643 | pathogenic | 1 | 1 | 1 |
| rs28941784 | pathogenic | 1 | 1 | 1 |
| rs398124434 | pathogenic | 1 | 1 | 1 |
| rs756414548 | pathogenic | 1 | 1 | 1 |
| rs376128990 | pathogenic | 1 | 1 | 1 |
| rs746219370 | pathogenic | 1 | 1 | 1 |
| rs756195708 | pathogenic | 1 | 1 | 1 |
| rs369296618 | pathogenic | 1 | 1 | 1 |
| rs9593 | benign | 1 | 1 | 1 |
| rs139510844 | benign | 0 | 0 | 0 |
| rs28382158 | benign | 0 | 0 | 0 |
| rs28919570 | pathogenic | 0 | 0 | 0 |
| rs587777331 | pathogenic | 0 | 0 | 0 |
| rs587777333 | pathogenic | 0 | 0 | 0 |
| rs587777332 | pathogenic | 0 | 0 | 0 |
| rs774980346 | pathogenic | 0 | 0 | 0 |
| rs370934093 | pathogenic | 0 | 0 | 0 |
| rs121913479 | pathogenic | 1 | 1 | 1 |
| rs121913484 | pathogenic | 0 | 0 | 0 |
| rs121913485 | pathogenic | 1 | 1 | 1 |
| rs28931614 | pathogenic | 1 | 1 | 1 |
| rs17881656 | benign | 1 | 1 | 1 |
| rs28931615 | pathogenic | 1 | 1 | 1 |
| rs147846832 | benign | 0 | 0 | 0 |
| rs80356506 | pathogenic | 0 | 0 | 0 |
| rs80356494 | pathogenic | 0 | 0 | 0 |
| rs121908894 | pathogenic | 0 | 0 | 0 |
| rs121908895 | pathogenic | 0 | 0 | 0 |
| rs80356508 | pathogenic | 0 | 0 | 0 |
| rs80356507 | pathogenic | 0 | 0 | 0 |
| rs121908896 | pathogenic | 0 | 0 | 0 |
| rs745372589 | pathogenic | 1 | 1 | 0 |
| rs183898408 | benign | 0 | 0 | 0 |
| rs200844292 | benign | 0 | 0 | 0 |
| rs146897002 | benign | 0 | 0 | 0 |
| rs121964869 | pathogenic | 1 | 1 | 1 |
| rs201768800 | benign | 1 | 1 | 1 |
| rs9807556 | benign | 1 | 1 | 1 |
| rs184721031 | benign | 0 | 0 | 0 |
| rs201104206 | benign | 0 | 0 | 0 |

| | | | | |
|---|---|---|---|---|
| rs188677538 | benign | 0 | 0 | 0 |
| rs149528866 | benign | 0 | 0 | 0 |
| rs115382168 | benign | 0 | 0 | 0 |
| rs78846023 | benign | 1 | 1 | 1 |
| rs61730662 | benign | 0 | 0 | 0 |
| rs121918368 | pathogenic | 0 | 0 | 0 |
| rs566433112 | pathogenic | 1 | 1 | 1 |
| rs121909342 | pathogenic | 0 | 0 | 0 |
| rs67586389 | pathogenic | 0 | 0 | 0 |
| rs72466485 | pathogenic | 0 | 0 | 0 |
| rs72466486 | pathogenic | 1 | 1 | 1 |
| rs72466487 | pathogenic | 1 | 1 | 1 |
| rs267607112 | pathogenic | 0 | 0 | 0 |
| rs118204013 | pathogenic | 0 | 0 | 0 |
| rs121913495 | pathogenic | 0 | 0 | 0 |
| rs11554273 | pathogenic | 0 | 0 | 0 |
| rs193922477 | pathogenic | 1 | 1 | 1 |
| rs193922479 | pathogenic | 1 | 1 | 1 |
| rs72547554 | pathogenic | 0 | 0 | 0 |
| rs72547561 | pathogenic | 1 | 1 | 1 |
| rs72547562 | pathogenic | 1 | 1 | 0 |
| rs115977487 | benign | 1 | 1 | 0 |
| rs72547564 | pathogenic | 0 | 0 | 0 |
| rs72547568 | pathogenic | 1 | 1 | 0 |
| rs72547569 | pathogenic | 1 | 1 | 1 |
| rs72547571 | pathogenic | 0 | 0 | 0 |
| rs72547575 | pathogenic | 1 | 1 | 1 |
| rs544994462 | benign | 0 | 0 | 0 |
| rs190470017 | benign | 0 | 0 | 0 |
| rs199473266 | pathogenic | 1 | 1 | 1 |
| rs137854602 | pathogenic | 0 | 0 | 0 |
| rs143739249 | pathogenic | 0 | 0 | 0 |
| rs397514697 | pathogenic | 1 | 1 | 1 |
| rs373764886 | pathogenic | 0 | 0 | 0 |
| rs370242930 | pathogenic | 0 | 0 | 0 |
| rs377145777 | pathogenic | 1 | 1 | 1 |
| rs191256186 | benign | 0 | 0 | 0 |
| rs150070901 | benign | 1 | 1 | 1 |
| rs138098032 | benign | 0 | 0 | 0 |
| rs117731813 | benign | 1 | 1 | 1 |
| rs104894488 | pathogenic | 1 | 1 | 1 |
| rs11302 | benign | 0 | 1 | 0 |
| rs121918447 | pathogenic | 1 | 1 | 1 |
| rs138205828 | benign | 1 | 1 | 0 |
| rs149858290 | benign | 0 | 0 | 0 |
| rs750764003 | pathogenic | 0 | 1 | 0 |
| rs141381896 | benign | 0 | 0 | 0 |
| rs149027682 | benign | 0 | 0 | 0 |
| rs121908415 | pathogenic | 0 | 0 | 0 |
| rs121908416 | pathogenic | 0 | 0 | 0 |
| rs121908417 | pathogenic | 0 | 0 | 0 |
| rs587780306 | benign | 0 | 0 | 0 |
| rs3218725 | benign | 0 | 0 | 0 |
| rs3218729 | benign | 0 | 0 | 0 |

| | | | | |
|---|---|---|---|---|
| rs387906917 | pathogenic | 0 | 0 | 0 |
| rs200652608 | pathogenic | 0 | 0 | 0 |
| rs3218721 | benign | 0 | 0 | 0 |
| rs63750869 | pathogenic | 1 | 1 | 1 |
| rs118203910 | pathogenic | 0 | 1 | 0 |
| rs9332701 | benign | 1 | 1 | 0 |
| rs6027 | benign | 0 | 0 | 0 |
| rs121909110 | pathogenic | 0 | 1 | 0 |
| rs762550967 | pathogenic | 1 | 1 | 0 |
| rs763615602 | pathogenic | 1 | 1 | 0 |
| rs144907403 | benign | 0 | 0 | 0 |
| rs104893631 | pathogenic | 1 | 1 | 0 |
| rs144181978 | pathogenic | 1 | 1 | 0 |
| rs74874677 | benign | 1 | 1 | 1 |
| rs104893632 | pathogenic | 0 | 1 | 0 |
| rs371342464 | benign | 1 | 1 | 1 |
| rs104893633 | pathogenic | 1 | 1 | 0 |
| rs201088712 | pathogenic | 0 | 0 | 0 |
| rs201857541 | benign | 1 | 1 | 1 |
| rs139517732 | pathogenic | 1 | 1 | 0 |
| rs191057824 | benign | 1 | 1 | 0 |
| rs35683768 | benign | 1 | 1 | 0 |
| rs104895271 | pathogenic | 0 | 0 | 0 |
| rs104895217 | pathogenic | 1 | 1 | 1 |
| rs104895223 | pathogenic | 1 | 1 | 1 |
| rs104895218 | pathogenic | 1 | 1 | 1 |
| rs104895225 | pathogenic | 1 | 1 | 1 |
| rs4149637 | benign | 1 | 1 | 1 |
| rs104895219 | pathogenic | 1 | 1 | 1 |
| rs104895220 | pathogenic | 1 | 1 | 1 |
| rs104895228 | pathogenic | 1 | 1 | 1 |
| rs104895221 | pathogenic | 1 | 1 | 1 |
| rs104895222 | pathogenic | 1 | 1 | 1 |
| rs104895247 | pathogenic | 0 | 0 | 0 |
| rs28940585 | pathogenic | 0 | 0 | 0 |
| rs561367199 | pathogenic | 1 | 1 | 1 |
| rs143793815 | benign | 0 | 0 | 0 |
| rs28941473 | pathogenic | 1 | 1 | 1 |
| rs751590073 | pathogenic | 1 | 1 | 1 |
| rs111407265 | benign | 1 | 1 | 1 |
| rs373697663 | pathogenic | 1 | 1 | 1 |
| rs369879957 | pathogenic | 0 | 0 | 0 |
| rs28941472 | pathogenic | 1 | 1 | 1 |
| rs28940287 | pathogenic | 1 | 1 | 1 |
| rs77625410 | benign | 0 | 0 | 0 |
| rs111033477 | benign | 0 | 0 | 0 |
| rs115123584 | benign | 0 | 0 | 0 |
| rs374576916 | benign | 0 | 0 | 0 |
| rs397516317 | pathogenic | 0 | 0 | 0 |
| rs368657015 | pathogenic | 0 | 0 | 0 |
| rs397516321 | pathogenic | 0 | 0 | 0 |
| rs397516322 | pathogenic | 0 | 0 | 0 |
| rs111033215 | pathogenic | 0 | 0 | 0 |
| rs199606180 | pathogenic | 0 | 0 | 0 |

| | | | | |
|---|---|---|---|---|
| rs111033192 | pathogenic | 0 | 0 | 0 |
| rs948962 | benign | 1 | 1 | 1 |
| rs142293185 | benign | 0 | 0 | 0 |
| rs376764423 | pathogenic | 0 | 0 | 0 |
| rs773844428 | pathogenic | 0 | 0 | 0 |
| rs186644871 | benign | 0 | 0 | 0 |
| rs200313391 | benign | 0 | 0 | 0 |
| rs377670513 | pathogenic | 0 | 0 | 0 |
| rs1132036 | benign | 0 | 0 | 0 |
| rs111033231 | benign | 0 | 0 | 0 |
| rs145945630 | pathogenic | 0 | 0 | 0 |
| rs121912984 | pathogenic | 1 | 1 | 1 |
| rs1800435 | benign | 0 | 1 | 0 |
| rs121912980 | pathogenic | 0 | 0 | 0 |
| rs121912982 | pathogenic | 1 | 1 | 1 |
| rs121912983 | pathogenic | 0 | 0 | 0 |
| rs121912981 | pathogenic | 0 | 0 | 0 |
| rs1057868 | benign | 0 | 0 | 0 |
| rs56048668 | benign | 0 | 0 | 0 |
| rs55865465 | benign | 0 | 0 | 0 |
| rs748096324 | benign | 1 | 1 | 0 |
| rs150475750 | benign | 0 | 0 | 0 |
| rs143025739 | benign | 1 | 1 | 0 |
| rs79702016 | benign | 1 | 1 | 0 |
| rs41338945 | benign | 0 | 0 | 0 |
| rs146406037 | benign | 1 | 1 | 1 |
| rs143274491 | benign | 0 | 0 | 0 |
| rs3829986 | benign | 0 | 0 | 0 |
| rs149168939 | benign | 0 | 0 | 0 |
| rs1800385 | benign | 0 | 0 | 0 |
| rs1800386 | pathogenic | 0 | 0 | 0 |
| rs121918524 | pathogenic | 1 | 1 | 1 |
| rs121912700 | pathogenic | 0 | 0 | 0 |
| rs104893930 | pathogenic | 1 | 1 | 1 |
| rs104893931 | pathogenic | 0 | 0 | 0 |
| rs116125936 | benign | 0 | 0 | 0 |
| rs627441 | benign | 0 | 0 | 0 |
| rs104894481 | pathogenic | 0 | 0 | 0 |
| rs137853129 | pathogenic | 0 | 0 | 0 |
| rs137853130 | pathogenic | 1 | 1 | 0 |
| rs28940282 | pathogenic | 0 | 0 | 0 |
| rs2229989 | benign | 0 | 0 | 0 |
| rs104894647 | pathogenic | 0 | 0 | 0 |
| rs1061170 | benign | 1 | 1 | 1 |
| rs576402053 | benign | 0 | 0 | 0 |
| rs34972246 | benign | 0 | 0 | 0 |
| rs192725607 | benign | 0 | 0 | 0 |
| rs137852615 | pathogenic | 1 | 1 | 0 |
| rs137852617 | pathogenic | 1 | 1 | 1 |
| rs137852611 | pathogenic | 0 | 0 | 0 |
| rs137852612 | pathogenic | 0 | 0 | 0 |
| rs137852614 | pathogenic | 1 | 1 | 1 |
| rs137852610 | pathogenic | 1 | 1 | 1 |
| rs137852618 | pathogenic | 1 | 1 | 1 |

| | | | | |
|---|---|---|---|---|
| rs137852613 | pathogenic | 1 | 1 | 0 |
| rs137852609 | pathogenic | 0 | 0 | 0 |
| rs63750306 | pathogenic | 1 | 1 | 1 |
| rs63750265 | pathogenic | 1 | 1 | 1 |
| rs63751287 | pathogenic | 1 | 1 | 1 |
| rs199723282 | benign | 0 | 0 | 0 |
| rs121917807 | pathogenic | 0 | 0 | 0 |
| rs121917809 | pathogenic | 0 | 0 | 0 |
| rs75134564 | pathogenic | 0 | 0 | 0 |
| rs267606930 | pathogenic | 1 | 1 | 0 |
| rs121909302 | pathogenic | 0 | 0 | 0 |
| rs121909303 | pathogenic | 0 | 0 | 0 |
| rs121909299 | pathogenic | 0 | 0 | 0 |
| rs121909301 | pathogenic | 0 | 0 | 0 |
| rs121909300 | pathogenic | 1 | 1 | 0 |
| rs111033589 | pathogenic | 0 | 0 | 0 |
| rs137853059 | pathogenic | 0 | 0 | 0 |
| rs137853057 | pathogenic | 1 | 1 | 1 |
| rs1801474 | benign | 0 | 0 | 0 |
| rs137853060 | pathogenic | 1 | 1 | 1 |
| rs137853058 | pathogenic | 1 | 1 | 1 |
| rs34424986 | pathogenic | 0 | 0 | 0 |
| rs199657839 | benign | 1 | 1 | 1 |
| rs56092260 | benign | 0 | 0 | 0 |
| rs1801582 | benign | 1 | 1 | 1 |
| rs1801334 | benign | 1 | 1 | 1 |
| rs191486604 | pathogenic | 0 | 0 | 0 |
| rs121908051 | pathogenic | 0 | 0 | 0 |
| rs121908055 | pathogenic | 1 | 1 | 0 |
| rs28940886 | pathogenic | 0 | 0 | 0 |
| rs121908050 | pathogenic | 0 | 0 | 0 |
| rs28940887 | pathogenic | 0 | 0 | 0 |
| rs387906300 | pathogenic | 0 | 0 | 0 |
| rs28942087 | pathogenic | 0 | 0 | 0 |
| rs121908049 | pathogenic | 1 | 1 | 0 |
| rs121908054 | pathogenic | 1 | 1 | 0 |
| rs121908048 | pathogenic | 0 | 0 | 0 |
| rs28940888 | pathogenic | 1 | 1 | 1 |
| rs121908053 | pathogenic | 1 | 1 | 0 |
| rs122467169 | pathogenic | 1 | 1 | 1 |
| rs122467172 | pathogenic | 1 | 1 | 1 |
| rs122467170 | pathogenic | 1 | 1 | 1 |
| rs28935477 | pathogenic | 1 | 1 | 0 |
| rs28942088 | pathogenic | 0 | 0 | 0 |
| rs751478142 | pathogenic | 1 | 1 | 0 |
| rs765368797 | pathogenic | 0 | 0 | 0 |
| rs147810437 | benign | 0 | 0 | 0 |
| rs121918209 | pathogenic | 1 | 1 | 1 |
| rs121918208 | pathogenic | 1 | 1 | 1 |
| rs121918210 | pathogenic | 1 | 1 | 1 |
| rs35515638 | benign | 1 | 1 | 1 |
| rs121918207 | pathogenic | 1 | 1 | 1 |
| rs141298312 | benign | 0 | 0 | 0 |
| rs121918211 | pathogenic | 0 | 0 | 0 |

| | | | | |
|---|---|---|---|---|
| rs149660059 | benign | 0 | 0 | 0 |
| rs200670286 | pathogenic | 1 | 1 | 1 |
| rs121918212 | pathogenic | 1 | 1 | 1 |
| rs121918213 | pathogenic | 1 | 1 | 1 |
| rs121918205 | pathogenic | 1 | 1 | 1 |
| rs104894025 | pathogenic | 1 | 1 | 1 |
| rs104894026 | pathogenic | 1 | 1 | 1 |
| rs104894028 | pathogenic | 0 | 1 | 0 |
| rs104894029 | pathogenic | 0 | 1 | 0 |
| rs119458971 | pathogenic | 0 | 0 | 0 |
| rs119458969 | pathogenic | 1 | 1 | 1 |
| rs1801591 | benign | 1 | 1 | 1 |
| rs140169311 | benign | 0 | 0 | 0 |
| rs753363278 | benign | 1 | 1 | 1 |
| rs199763682 | pathogenic | 0 | 0 | 0 |
| rs119458970 | pathogenic | 1 | 1 | 1 |
| rs763722868 | pathogenic | 0 | 0 | 0 |
| rs56404770 | benign | 0 | 0 | 0 |
| rs34221241 | benign | 0 | 0 | 0 |
| rs121909625 | pathogenic | 0 | 0 | 0 |
| rs550921485 | pathogenic | 0 | 0 | 0 |
| rs1800454 | benign | 0 | 0 | 0 |
| rs241447 | benign | 0 | 0 | 0 |
| rs2230606 | benign | 1 | 1 | 0 |
| rs5036 | pathogenic | 1 | 1 | 1 |
| rs13306788 | benign | 0 | 0 | 0 |
| rs28929480 | pathogenic | 0 | 0 | 0 |
| rs121912749 | pathogenic | 0 | 0 | 0 |
| rs56361140 | pathogenic | 0 | 0 | 0 |
| rs28931583 | pathogenic | 1 | 1 | 1 |
| rs36211715 | pathogenic | 1 | 1 | 1 |
| rs138049878 | pathogenic | 1 | 1 | 1 |
| rs727503253 | pathogenic | 1 | 1 | 1 |
| rs267606908 | pathogenic | 0 | 0 | 0 |
| rs121913631 | pathogenic | 1 | 1 | 1 |
| rs36211716 | benign | 1 | 1 | 1 |
| rs121913639 | pathogenic | 1 | 1 | 1 |
| rs730880761 | pathogenic | 1 | 1 | 1 |
| rs121913629 | pathogenic | 1 | 1 | 1 |
| rs754368658 | pathogenic | 0 | 0 | 0 |
| rs777090017 | pathogenic | 0 | 0 | 0 |
| rs267606913 | pathogenic | 1 | 1 | 1 |
| rs773138218 | pathogenic | 0 | 0 | 0 |
| rs371295780 | pathogenic | 0 | 0 | 0 |
| rs387906830 | pathogenic | 1 | 1 | 0 |
| rs771364038 | pathogenic | 0 | 1 | 0 |
| rs772731615 | pathogenic | 0 | 0 | 0 |
| rs137853063 | pathogenic | 0 | 0 | 0 |
| rs387907076 | pathogenic | 0 | 1 | 0 |
| rs35460768 | benign | 0 | 0 | 0 |
| rs2278196 | benign | 1 | 1 | 1 |
| rs119103275 | pathogenic | 0 | 0 | 0 |
| rs119103276 | pathogenic | 1 | 1 | 1 |
| rs374402066 | pathogenic | 1 | 1 | 1 |

| | | | | |
|---|---|---|---|---|
| rs119103274 | pathogenic | 0 | 0 | 0 |
| rs2274064 | benign | 0 | 0 | 0 |
| rs267607023 | pathogenic | 1 | 1 | 0 |
| rs142117489 | benign | 1 | 1 | 0 |
| rs371258270 | benign | 0 | 0 | 0 |
| rs34716057 | benign | 0 | 0 | 0 |
| rs201778606 | benign | 0 | 0 | 0 |
| rs201438792 | benign | 0 | 0 | 0 |
| rs140547417 | benign | 0 | 0 | 0 |
| rs5881 | benign | 0 | 0 | 0 |
| rs752298084 | benign | 0 | 0 | 0 |
| rs34855278 | benign | 0 | 0 | 0 |
| rs5880 | benign | 1 | 1 | 0 |
| rs5882 | benign | 0 | 0 | 0 |
| rs1800777 | benign | 0 | 0 | 0 |
| rs577301331 | benign | 0 | 0 | 0 |
| rs137853313 | pathogenic | 0 | 0 | 0 |
| rs28935169 | pathogenic | 0 | 0 | 0 |
| rs137853315 | pathogenic | 0 | 0 | 0 |
| rs137853314 | pathogenic | 0 | 0 | 0 |
| rs28935469 | pathogenic | 0 | 0 | 0 |
| rs137853318 | pathogenic | 0 | 0 | 0 |
| rs28935470 | pathogenic | 0 | 0 | 0 |
| rs3729986 | benign | 0 | 0 | 0 |
| rs730880619 | pathogenic | 0 | 0 | 0 |
| rs201012766 | benign | 0 | 0 | 0 |
| rs727503218 | pathogenic | 0 | 0 | 0 |
| rs11570052 | benign | 0 | 0 | 0 |
| rs730880623 | pathogenic | 0 | 0 | 0 |
| rs138753870 | benign | 0 | 0 | 0 |
| rs397516068 | pathogenic | 0 | 0 | 0 |
| rs3729989 | benign | 1 | 1 | 1 |
| rs397516074 | pathogenic | 0 | 0 | 0 |
| rs561449819 | pathogenic | 0 | 0 | 0 |
| rs121917779 | benign | 0 | 0 | 0 |
| rs142881576 | pathogenic | 0 | 0 | 0 |
| rs77384282 | benign | 0 | 0 | 0 |
| rs201865787 | benign | 0 | 0 | 0 |
| rs142244640 | benign | 0 | 0 | 0 |
| rs200488706 | benign | 0 | 0 | 0 |
| rs137853291 | pathogenic | 1 | 1 | 0 |
| rs28933990 | pathogenic | 1 | 1 | 0 |
| rs140570189 | benign | 0 | 0 | 0 |
| rs201588178 | benign | 0 | 0 | 0 |
| rs104893846 | pathogenic | 0 | 0 | 0 |
| rs754545360 | pathogenic | 0 | 1 | 0 |
| rs104893851 | pathogenic | 0 | 1 | 0 |
| rs104893849 | pathogenic | 0 | 0 | 0 |
| rs571038432 | pathogenic | 1 | 1 | 0 |
| rs2270655 | benign | 1 | 1 | 1 |
| rs315952 | benign | 0 | 0 | 0 |
| rs151176548 | benign | 0 | 0 | 0 |
| rs122459147 | pathogenic | 1 | 1 | 0 |
| rs122458140 | pathogenic | 0 | 0 | 0 |

| | | | | |
|---|---|---|---|---|
| rs122458142 | pathogenic | 1 | 1 | 0 |
| rs267606812 | pathogenic | 1 | 1 | 0 |
| rs267606813 | pathogenic | 1 | 1 | 0 |
| rs122458143 | pathogenic | 0 | 0 | 0 |
| rs122459146 | pathogenic | 0 | 0 | 0 |
| rs122458144 | pathogenic | 0 | 0 | 0 |
| rs122458145 | pathogenic | 0 | 0 | 0 |
| rs113994007 | pathogenic | 1 | 1 | 0 |
| rs758746181 | pathogenic | 1 | 1 | 1 |
| rs145034527 | pathogenic | 1 | 1 | 0 |
| rs119468006 | pathogenic | 1 | 1 | 0 |
| rs201908721 | pathogenic | 0 | 0 | 0 |
| rs755933881 | pathogenic | 1 | 1 | 0 |
| rs771578775 | pathogenic | 0 | 0 | 0 |
| rs17849927 | benign | 0 | 0 | 0 |
| rs139133094 | benign | 0 | 0 | 0 |
| rs12593 | benign | 0 | 0 | 0 |
| rs578189699 | pathogenic | 1 | 1 | 0 |
| rs119468008 | pathogenic | 0 | 0 | 0 |
| rs768524626 | benign | 0 | 0 | 0 |
| rs119468009 | pathogenic | 0 | 0 | 0 |
| rs119468004 | pathogenic | 0 | 0 | 0 |
| rs199874519 | pathogenic | 0 | 0 | 0 |
| rs376478331 | pathogenic | 0 | 0 | 0 |
| rs61995958 | pathogenic | 0 | 0 | 0 |
| rs199714731 | benign | 0 | 0 | 0 |
| rs149185954 | benign | 0 | 0 | 0 |
| rs121918353 | pathogenic | 0 | 0 | 0 |
| rs1131695 | benign | 0 | 0 | 0 |
| rs28939668 | pathogenic | 0 | 0 | 0 |
| rs758432471 | pathogenic | 1 | 1 | 1 |
| rs752746786 | pathogenic | 1 | 1 | 1 |
| rs28935471 | pathogenic | 0 | 0 | 0 |
| rs80338837 | pathogenic | 0 | 0 | 0 |
| rs137853312 | pathogenic | 0 | 0 | 0 |
| rs28935472 | pathogenic | 0 | 0 | 0 |
| rs28935473 | pathogenic | 0 | 0 | 0 |
| rs267606752 | pathogenic | 0 | 0 | 0 |
| rs3917973 | benign | 0 | 0 | 0 |
| rs3918018 | benign | 0 | 0 | 0 |
| rs757976755 | pathogenic | 0 | 0 | 0 |
| rs121918329 | pathogenic | 1 | 1 | 0 |
| rs770114459 | pathogenic | 0 | 0 | 0 |
| rs6662272 | benign | 0 | 0 | 0 |
| rs121918242 | pathogenic | 1 | 1 | 0 |
| rs121918240 | pathogenic | 1 | 1 | 0 |
| rs200094982 | pathogenic | 0 | 0 | 0 |
| rs121918241 | pathogenic | 0 | 0 | 0 |
| rs140522266 | pathogenic | 1 | 1 | 0 |
| rs121918243 | pathogenic | 1 | 1 | 0 |
| rs370596113 | pathogenic | 1 | 1 | 0 |
| rs398124295 | pathogenic | 1 | 1 | 0 |
| rs587776889 | pathogenic | 1 | 1 | 0 |
| rs747527726 | pathogenic | 1 | 1 | 0 |

| | | | | |
|---|---|---|---|---|
| rs112631212 | benign | 0 | 0 | 0 |
| rs104894605 | pathogenic | 0 | 0 | 0 |
| rs104894607 | pathogenic | 0 | 0 | 0 |
| rs104894606 | pathogenic | 0 | 0 | 0 |
| rs143873873 | benign | 0 | 0 | 0 |
| rs104894388 | pathogenic | 1 | 1 | 0 |
| rs104894391 | pathogenic | 1 | 1 | 0 |
| rs5350 | benign | 1 | 1 | 1 |
| rs104894390 | pathogenic | 1 | 1 | 1 |
| rs5346 | benign | 1 | 1 | 1 |
| rs121912500 | pathogenic | 0 | 0 | 0 |
| rs74315450 | pathogenic | 1 | 1 | 0 |
| rs121913396 | pathogenic | 1 | 1 | 1 |
| rs28931588 | pathogenic | 1 | 1 | 1 |
| rs121913400 | pathogenic | 0 | 0 | 0 |
| rs28931589 | pathogenic | 1 | 1 | 1 |
| rs121913403 | pathogenic | 0 | 0 | 0 |
| rs121913413 | pathogenic | 0 | 0 | 0 |
| rs760107623 | pathogenic | 1 | 1 | 0 |
| rs16866378 | benign | 1 | 1 | 0 |
| rs201025290 | pathogenic | 1 | 1 | 0 |
| rs12720356 | benign | 1 | 1 | 0 |
| rs55882956 | benign | 0 | 1 | 0 |
| rs4986933 | benign | 1 | 1 | 0 |
| rs2020957 | benign | 1 | 1 | 1 |
| rs1800124 | benign | 1 | 1 | 1 |
| rs150077735 | benign | 1 | 1 | 1 |
| rs201419000 | benign | 0 | 0 | 0 |
| rs772044176 | pathogenic | 0 | 0 | 0 |
| rs138346388 | benign | 0 | 0 | 0 |
| rs121964916 | pathogenic | 0 | 0 | 0 |
| rs112534524 | pathogenic | 0 | 0 | 0 |
| rs182078921 | benign | 1 | 1 | 1 |
| rs113273712 | benign | 0 | 0 | 0 |
| rs74817407 | benign | 1 | 1 | 1 |
| rs61733901 | benign | 1 | 1 | 1 |
| rs121964912 | pathogenic | 1 | 1 | 1 |
| rs41278047 | benign | 1 | 1 | 1 |
| rs7437875 | benign | 1 | 1 | 1 |
| rs113460688 | benign | 1 | 1 | 1 |
| rs754572081 | benign | 1 | 1 | 1 |
| rs201082692 | benign | 1 | 1 | 1 |
| rs202085803 | benign | 1 | 1 | 1 |
| rs141320511 | benign | 0 | 0 | 0 |
| rs78501403 | benign | 0 | 0 | 0 |
| rs28939675 | pathogenic | 0 | 0 | 0 |
| rs74315522 | pathogenic | 0 | 0 | 0 |
| rs121434639 | pathogenic | 1 | 1 | 0 |
| rs121434640 | pathogenic | 1 | 1 | 1 |
| rs757282628 | benign | 1 | 1 | 1 |
| rs386834010 | pathogenic | 0 | 0 | 0 |
| rs200863680 | pathogenic | 0 | 0 | 0 |
| rs193919336 | pathogenic | 1 | 1 | 0 |
| rs386834011 | pathogenic | 0 | 0 | 0 |

| | | | | |
|---|---|---|---|---|
| rs28940869 | pathogenic | 0 | 0 | 0 |
| rs386834014 | pathogenic | 1 | 1 | 0 |
| rs267606960 | pathogenic | 0 | 0 | 0 |
| rs28942068 | pathogenic | 0 | 0 | 0 |
| rs193919335 | pathogenic | 0 | 0 | 0 |
| rs74374973 | pathogenic | 0 | 0 | 0 |
| rs386834018 | pathogenic | 0 | 0 | 0 |
| rs267606962 | pathogenic | 0 | 0 | 0 |
| rs6659553 | benign | 0 | 0 | 0 |
| rs29281 | benign | 0 | 0 | 0 |
| rs782265090 | benign | 0 | 0 | 0 |
| rs35910969 | pathogenic | 0 | 0 | 0 |
| rs121909113 | pathogenic | 1 | 1 | 1 |
| rs29001571 | pathogenic | 1 | 1 | 1 |
| rs587781250 | pathogenic | 1 | 1 | 1 |
| rs28939680 | pathogenic | 1 | 1 | 1 |
| rs28939681 | pathogenic | 1 | 1 | 1 |
| rs121909112 | pathogenic | 1 | 1 | 1 |
| rs28937568 | pathogenic | 1 | 1 | 1 |
| rs1799895 | pathogenic | 0 | 0 | 0 |
| rs386833431 | pathogenic | 1 | 1 | 1 |
| rs121964905 | pathogenic | 1 | 1 | 1 |
| rs121964906 | pathogenic | 1 | 1 | 1 |
| rs11551664 | pathogenic | 1 | 1 | 1 |
| rs104894705 | pathogenic | 1 | 1 | 1 |
| rs104893805 | pathogenic | 0 | 0 | 0 |
| rs104894205 | pathogenic | 0 | 0 | 0 |
| rs104894204 | pathogenic | 1 | 1 | 0 |
| rs62642926 | pathogenic | 0 | 0 | 0 |
| rs5030841 | pathogenic | 1 | 1 | 1 |
| rs75193786 | pathogenic | 1 | 1 | 1 |
| rs62508682 | pathogenic | 0 | 0 | 0 |
| rs3850625 | benign | 1 | 1 | 1 |
| rs141908793 | benign | 0 | 0 | 0 |
| rs187864727 | benign | 0 | 0 | 0 |
| rs548909101 | benign | 0 | 0 | 0 |
| rs3825394 | benign | 0 | 1 | 0 |
| rs143548402 | benign | 0 | 0 | 0 |
| rs267607146 | pathogenic | 0 | 0 | 0 |
| rs267607144 | pathogenic | 0 | 0 | 0 |
| rs267607148 | pathogenic | 1 | 1 | 1 |
| rs267607143 | pathogenic | 0 | 0 | 0 |
| rs267607145 | pathogenic | 0 | 0 | 0 |
| rs121912636 | pathogenic | 1 | 1 | 1 |
| rs121912634 | pathogenic | 1 | 1 | 1 |
| rs570675468 | benign | 0 | 0 | 0 |
| rs368861241 | pathogenic | 1 | 1 | 1 |
| rs104894528 | pathogenic | 1 | 1 | 1 |
| rs77493891 | benign | 0 | 0 | 0 |
| rs62640903 | benign | 0 | 0 | 0 |
| rs149926091 | benign | 1 | 1 | 1 |
| rs641081 | benign | 0 | 0 | 0 |
| rs4930199 | benign | 0 | 0 | 0 |
| rs121913020 | pathogenic | 0 | 0 | 0 |

| | | | | |
|---|---|---|---|---|
| rs1799793 | benign | 0 | 0 | 0 |
| rs121913016 | pathogenic | 0 | 0 | 0 |
| rs121913025 | pathogenic | 0 | 0 | 0 |
| rs121913019 | pathogenic | 0 | 0 | 0 |
| rs376556895 | pathogenic | 1 | 1 | 1 |
| rs121913024 | pathogenic | 1 | 1 | 1 |
| rs121913021 | pathogenic | 0 | 0 | 0 |
| rs121913023 | pathogenic | 0 | 0 | 0 |
| rs41556519 | pathogenic | 0 | 0 | 0 |
| rs121913022 | pathogenic | 0 | 0 | 0 |
| rs144564120 | pathogenic | 0 | 0 | 0 |
| rs121913026 | pathogenic | 1 | 1 | 1 |
| rs121913018 | pathogenic | 1 | 1 | 1 |
| rs142954643 | benign | 0 | 0 | 0 |
| rs76655496 | benign | 0 | 0 | 0 |
| rs104894371 | pathogenic | 1 | 1 | 0 |
| rs137964936 | benign | 0 | 0 | 0 |
| rs759736132 | pathogenic | 0 | 0 | 0 |
| rs387906595 | pathogenic | 1 | 1 | 0 |
| rs768248216 | pathogenic | 0 | 0 | 0 |
| rs548231008 | benign | 0 | 0 | 0 |
| rs757094384 | benign | 0 | 0 | 0 |
| rs867529 | benign | 0 | 0 | 0 |
| rs13045 | benign | 1 | 1 | 1 |
| rs121908570 | pathogenic | 0 | 0 | 0 |
| rs201872586 | benign | 1 | 1 | 0 |
| rs149092990 | benign | 0 | 0 | 0 |
| rs115585711 | benign | 0 | 0 | 0 |
| rs137909416 | benign | 1 | 1 | 0 |
| rs143388949 | benign | 1 | 1 | 1 |
| rs73137716 | benign | 1 | 1 | 1 |
| rs121908947 | pathogenic | 1 | 1 | 1 |
| rs398122904 | pathogenic | 1 | 1 | 1 |
| rs35715497 | benign | 1 | 1 | 0 |
| rs35462421 | benign | 0 | 0 | 0 |
| rs119475043 | pathogenic | 0 | 0 | 0 |
| rs119475042 | pathogenic | 0 | 0 | 0 |
| rs1076669 | benign | 1 | 1 | 0 |
| rs141146885 | benign | 0 | 0 | 0 |
| rs3026906 | pathogenic | 0 | 0 | 0 |
| rs75362221 | benign | 1 | 1 | 1 |
| rs138472116 | benign | 1 | 1 | 1 |
| rs371040282 | pathogenic | 1 | 1 | 1 |
| rs4988496 | benign | 0 | 0 | 0 |
| rs121918117 | pathogenic | 1 | 1 | 1 |
| rs4988498 | benign | 0 | 0 | 0 |
| rs5035 | benign | 0 | 0 | 0 |
| rs121918322 | pathogenic | 1 | 1 | 1 |
| rs370359540 | benign | 0 | 0 | 0 |
| rs758972589 | pathogenic | 0 | 0 | 0 |
| rs144856889 | benign | 0 | 0 | 0 |
| rs121434623 | pathogenic | 1 | 1 | 1 |
| rs61735600 | benign | 0 | 0 | 0 |
| rs72550870 | benign | 1 | 1 | 1 |

| | | | | |
|---|---|---|---|---|
| rs56392418 | benign | 0 | 0 | 0 |
| rs41307788 | benign | 1 | 1 | 1 |
| rs387907239 | pathogenic | 1 | 1 | 0 |
| rs28931584 | pathogenic | 0 | 0 | 0 |
| rs121912745 | pathogenic | 0 | 0 | 0 |
| rs121912744 | pathogenic | 0 | 0 | 0 |
| rs121912754 | pathogenic | 1 | 1 | 1 |
| rs121912746 | pathogenic | 0 | 0 | 0 |
| rs121912757 | pathogenic | 0 | 0 | 0 |
| rs121912758 | pathogenic | 0 | 0 | 0 |
| rs121912748 | pathogenic | 0 | 0 | 0 |
| rs121912755 | pathogenic | 0 | 0 | 0 |
| rs121912741 | pathogenic | 0 | 0 | 0 |
| rs121912753 | pathogenic | 0 | 0 | 0 |
| rs121912750 | pathogenic | 0 | 0 | 0 |
| rs2285644 | pathogenic | 0 | 0 | 0 |
| rs121912751 | pathogenic | 0 | 0 | 0 |
| rs5026 | benign | 0 | 0 | 0 |
| rs121912759 | pathogenic | 0 | 0 | 0 |
| rs28931585 | pathogenic | 0 | 0 | 0 |
| rs17122769 | benign | 0 | 0 | 0 |
| rs28362322 | benign | 1 | 1 | 1 |
| rs3824874 | benign | 0 | 0 | 0 |
| rs146004831 | benign | 0 | 0 | 0 |
| rs142155860 | benign | 0 | 0 | 0 |
| rs121434404 | pathogenic | 0 | 0 | 0 |
| rs61735578 | benign | 1 | 1 | 1 |
| rs558018 | benign | 0 | 0 | 0 |
| rs61735577 | benign | 1 | 1 | 1 |
| rs76784113 | benign | 0 | 0 | 0 |
| rs199469640 | pathogenic | 0 | 0 | 0 |
| rs199469628 | pathogenic | 1 | 1 | 0 |
| rs199469629 | pathogenic | 0 | 1 | 0 |
| rs387907156 | pathogenic | 0 | 1 | 0 |
| rs199469643 | pathogenic | 0 | 0 | 0 |
| rs199469646 | pathogenic | 0 | 0 | 0 |
| rs766294940 | pathogenic | 0 | 0 | 0 |
| rs387907347 | pathogenic | 0 | 0 | 0 |
| rs137853590 | pathogenic | 1 | 1 | 1 |
| rs137853589 | pathogenic | 1 | 1 | 1 |
| rs137853588 | pathogenic | 1 | 1 | 1 |
| rs55909005 | benign | 0 | 0 | 0 |
| rs4581 | benign | 0 | 0 | 0 |
| rs63750123 | benign | 0 | 0 | 0 |
| rs10254120 | benign | 0 | 0 | 0 |
| rs121434629 | pathogenic | 1 | 1 | 0 |
| rs200313585 | benign | 0 | 0 | 0 |
| rs143162541 | benign | 1 | 1 | 0 |
| rs760615315 | benign | 0 | 0 | 0 |
| rs768488890 | benign | 1 | 1 | 1 |
| rs116373169 | benign | 1 | 1 | 0 |
| rs63750871 | pathogenic | 1 | 1 | 0 |
| rs116349687 | benign | 0 | 0 | 0 |
| rs375289386 | benign | 0 | 0 | 0 |

| | | | | |
|---|---|---|---|---|
| rs760228510 | pathogenic | 1 | 1 | 1 |
| rs587779343 | pathogenic | 1 | 1 | 1 |
| rs573125799 | benign | 0 | 0 | 0 |
| rs1805322 | benign | 0 | 0 | 0 |
| rs267608153 | pathogenic | 0 | 0 | 0 |
| rs139194813 | benign | 0 | 0 | 0 |
| rs530021751 | benign | 0 | 0 | 0 |
| rs200640585 | pathogenic | 0 | 0 | 0 |
| rs139438201 | benign | 0 | 0 | 0 |
| rs200513014 | benign | 0 | 0 | 0 |
| rs567102013 | benign | 0 | 0 | 0 |
| rs121908608 | pathogenic | 0 | 0 | 0 |
| rs121908606 | pathogenic | 0 | 0 | 0 |
| rs17103671 | benign | 0 | 0 | 0 |
| rs34725281 | benign | 0 | 0 | 0 |
| rs104894421 | pathogenic | 1 | 1 | 0 |
| rs121918145 | pathogenic | 1 | 1 | 1 |
| rs121918152 | pathogenic | 1 | 1 | 1 |
| rs56379106 | benign | 0 | 0 | 0 |
| rs4987076 | benign | 0 | 0 | 0 |
| rs370243877 | pathogenic | 0 | 0 | 0 |
| rs17679445 | benign | 1 | 1 | 1 |
| rs550423482 | pathogenic | 1 | 1 | 1 |
| rs104894629 | pathogenic | 1 | 1 | 1 |
| rs773450573 | pathogenic | 1 | 1 | 1 |
| rs116679717 | benign | 0 | 0 | 0 |
| rs372000848 | pathogenic | 0 | 0 | 0 |
| rs138595973 | benign | 0 | 0 | 0 |
| rs115244417 | benign | 0 | 0 | 0 |
| rs11575292 | benign | 0 | 0 | 0 |
| rs137853211 | pathogenic | 1 | 1 | 1 |
| rs137853207 | pathogenic | 0 | 0 | 0 |
| rs137853210 | pathogenic | 1 | 1 | 1 |
| rs6262 | benign | 0 | 0 | 0 |
| rs6263 | benign | 0 | 1 | 0 |
| rs137853208 | pathogenic | 0 | 1 | 0 |
| rs137853212 | pathogenic | 1 | 1 | 0 |
| rs137853209 | pathogenic | 1 | 1 | 1 |
| rs201951824 | pathogenic | 0 | 0 | 0 |
| rs11575542 | benign | 0 | 0 | 0 |
| rs199681788 | benign | 0 | 1 | 0 |
| rs748835902 | benign | 1 | 1 | 1 |
| rs753719949 | benign | 0 | 0 | 0 |
| rs267606647 | pathogenic | 0 | 0 | 0 |
| rs267606648 | pathogenic | 1 | 1 | 1 |
| rs267606643 | pathogenic | 0 | 1 | 0 |
| rs559947967 | pathogenic | 1 | 1 | 0 |
| rs267606645 | pathogenic | 1 | 1 | 1 |
| rs12116440 | benign | 0 | 0 | 0 |
| rs121908539 | pathogenic | 0 | 0 | 0 |
| rs137853096 | pathogenic | 1 | 1 | 0 |
| rs148363262 | benign | 1 | 1 | 0 |
| rs587777442 | pathogenic | 0 | 0 | 0 |
| rs25640 | benign | 1 | 1 | 1 |

| | | | | |
|---|---|---|---|---|
| rs28943589 | benign | 0 | 0 | 0 |
| rs1143650 | benign | 1 | 1 | 1 |
| rs150886026 | benign | 0 | 1 | 0 |
| rs121908133 | pathogenic | 1 | 1 | 0 |
| rs121908131 | pathogenic | 0 | 0 | 0 |
| rs754477154 | pathogenic | 0 | 0 | 0 |
| rs747334007 | benign | 0 | 0 | 0 |
| rs121908132 | pathogenic | 0 | 0 | 0 |
| rs104894103 | pathogenic | 1 | 1 | 0 |
| rs140088149 | benign | 0 | 0 | 0 |
| rs149803148 | benign | 0 | 0 | 0 |
| rs11064480 | benign | 0 | 0 | 0 |
| rs74435397 | pathogenic | 1 | 1 | 1 |
| rs587783626 | pathogenic | 1 | 1 | 1 |
| rs148211042 | pathogenic | 1 | 1 | 1 |
| rs17849502 | benign | 0 | 0 | 0 |
| rs13306575 | pathogenic | 1 | 1 | 1 |
| rs201426733 | benign | 0 | 0 | 0 |
| rs182278784 | benign | 1 | 1 | 1 |
| rs77786415 | benign | 1 | 1 | 1 |
| rs200192119 | benign | 1 | 1 | 1 |
| rs12877501 | benign | 1 | 1 | 1 |
| rs77985244 | benign | 1 | 1 | 1 |
| rs5907 | pathogenic | 1 | 1 | 0 |
| rs2305178 | benign | 0 | 0 | 0 |
| rs3135867 | benign | 0 | 0 | 0 |
| rs121913482 | pathogenic | 1 | 1 | 1 |
| rs121913483 | pathogenic | 1 | 1 | 1 |
| rs4647924 | pathogenic | 1 | 1 | 1 |
| rs121913111 | pathogenic | 1 | 1 | 1 |
| rs104894002 | pathogenic | 0 | 0 | 0 |
| rs104894001 | pathogenic | 0 | 0 | 0 |
| rs75932628 | benign | 0 | 0 | 0 |
| rs201258663 | pathogenic | 0 | 0 | 0 |
| rs104893998 | pathogenic | 0 | 0 | 0 |
| rs2234253 | benign | 1 | 1 | 0 |
| rs121908402 | pathogenic | 1 | 1 | 0 |
| rs1805389 | benign | 0 | 0 | 0 |
| rs1805388 | benign | 0 | 0 | 0 |
| rs748385144 | pathogenic | 0 | 0 | 0 |
| rs104894420 | pathogenic | 0 | 0 | 0 |
| rs104894418 | pathogenic | 0 | 0 | 0 |
| rs61731910 | benign | 0 | 0 | 0 |
| rs758723288 | pathogenic | 1 | 1 | 0 |
| rs201865375 | pathogenic | 1 | 1 | 0 |
| rs565090080 | pathogenic | 1 | 1 | 1 |
| rs113042166 | benign | 0 | 0 | 0 |
| rs61736326 | benign | 1 | 1 | 1 |
| rs12030699 | benign | 1 | 1 | 1 |
| rs121913578 | pathogenic | 1 | 1 | 1 |
| rs61739582 | benign | 0 | 0 | 0 |
| rs745983207 | pathogenic | 0 | 0 | 0 |
| rs763524380 | benign | 0 | 0 | 0 |
| rs121912792 | pathogenic | 0 | 0 | 0 |

| | | | | |
|---|---|---|---|---|
| rs121912793 | pathogenic | 0 | 0 | 0 |
| rs387906599 | pathogenic | 0 | 0 | 0 |
| rs121912794 | pathogenic | 0 | 1 | 0 |
| rs121912795 | pathogenic | 0 | 0 | 0 |
| rs121912796 | pathogenic | 1 | 1 | 0 |
| rs121912797 | pathogenic | 0 | 1 | 0 |
| rs121912798 | pathogenic | 1 | 1 | 1 |
| rs80359812 | pathogenic | 1 | 1 | 0 |
| rs776583130 | benign | 1 | 1 | 0 |
| rs80359814 | pathogenic | 1 | 1 | 0 |
| rs202060209 | pathogenic | 1 | 1 | 0 |
| rs267607061 | pathogenic | 1 | 1 | 0 |
| rs267607060 | pathogenic | 0 | 0 | 0 |
| rs80359816 | pathogenic | 1 | 1 | 0 |
| rs80359818 | pathogenic | 1 | 1 | 0 |
| rs387907313 | pathogenic | 0 | 0 | 0 |
| rs587784395 | benign | 0 | 0 | 0 |
| rs5811 | benign | 0 | 0 | 0 |
| rs121909740 | pathogenic | 1 | 1 | 0 |
| rs148518827 | benign | 1 | 1 | 0 |
| rs121909739 | pathogenic | 1 | 1 | 0 |
| rs780528770 | benign | 1 | 1 | 0 |
| rs13306758 | pathogenic | 0 | 0 | 0 |
| rs267607059 | pathogenic | 0 | 0 | 0 |
| rs61730638 | benign | 0 | 0 | 0 |
| rs104894533 | pathogenic | 0 | 0 | 0 |
| rs104894527 | pathogenic | 0 | 0 | 0 |
| rs200503569 | pathogenic | 1 | 1 | 1 |
| rs80338700 | pathogenic | 1 | 1 | 1 |
| rs765433263 | pathogenic | 1 | 1 | 1 |
| rs104894530 | pathogenic | 1 | 1 | 1 |
| rs80338701 | pathogenic | 1 | 1 | 1 |
| rs141498002 | pathogenic | 1 | 1 | 1 |
| rs104894525 | pathogenic | 0 | 0 | 0 |
| rs80338703 | pathogenic | 0 | 0 | 0 |
| rs28936415 | pathogenic | 1 | 1 | 0 |
| rs150719105 | pathogenic | 0 | 0 | 0 |
| rs148032587 | pathogenic | 0 | 0 | 0 |
| rs150577656 | pathogenic | 0 | 0 | 0 |
| rs190521996 | pathogenic | 0 | 0 | 0 |
| rs104894526 | pathogenic | 1 | 1 | 1 |
| rs80338704 | pathogenic | 0 | 0 | 0 |
| rs199562225 | pathogenic | 1 | 1 | 1 |
| rs532870929 | pathogenic | 0 | 0 | 0 |
| rs398123309 | pathogenic | 0 | 0 | 0 |
| rs78290141 | pathogenic | 0 | 0 | 0 |
| rs80338705 | pathogenic | 0 | 0 | 0 |
| rs104894531 | pathogenic | 1 | 1 | 1 |
| rs80338706 | pathogenic | 0 | 0 | 0 |
| rs80338707 | pathogenic | 0 | 0 | 0 |
| rs80338708 | pathogenic | 0 | 0 | 0 |
| rs80338709 | pathogenic | 0 | 0 | 0 |
| rs104893833 | benign | 0 | 0 | 0 |
| rs61744585 | benign | 0 | 0 | 0 |

| | | | | |
|---|---|---|---|---|
| rs73569515 | benign | 0 | 0 | 0 |
| rs398123373 | pathogenic | 0 | 0 | 0 |
| rs756854513 | pathogenic | 0 | 0 | 0 |
| rs58232698 | benign | 0 | 0 | 0 |
| rs9600455 | benign | 0 | 0 | 0 |
| rs201016593 | pathogenic | 0 | 0 | 0 |
| rs756039188 | pathogenic | 0 | 0 | 0 |
| rs121908024 | pathogenic | 0 | 0 | 0 |
| rs373144619 | benign | 0 | 0 | 0 |
| rs138078086 | benign | 0 | 0 | 0 |
| rs376207800 | pathogenic | 0 | 0 | 0 |
| rs749038326 | pathogenic | 1 | 1 | 0 |
| rs372845091 | benign | 0 | 0 | 0 |
| rs377271627 | pathogenic | 0 | 0 | 0 |
| rs563390335 | pathogenic | 0 | 0 | 0 |
| rs13306510 | benign | 0 | 0 | 0 |
| rs764042910 | pathogenic | 0 | 0 | 0 |
| rs201384282 | benign | 1 | 1 | 1 |
| rs730882086 | pathogenic | 1 | 1 | 1 |
| rs756613387 | pathogenic | 1 | 1 | 1 |
| rs121908028 | benign | 1 | 1 | 0 |
| rs773328511 | benign | 0 | 0 | 0 |
| rs749220643 | pathogenic | 1 | 1 | 0 |
| rs730882089 | pathogenic | 0 | 0 | 0 |
| rs146651743 | benign | 0 | 0 | 0 |
| rs148698650 | pathogenic | 0 | 0 | 0 |
| rs140241383 | benign | 0 | 0 | 0 |
| rs757252110 | pathogenic | 0 | 0 | 0 |
| rs13306512 | pathogenic | 1 | 1 | 1 |
| rs121908043 | pathogenic | 1 | 1 | 1 |
| rs773658037 | pathogenic | 0 | 0 | 0 |
| rs774439908 | pathogenic | 0 | 0 | 0 |
| rs779732323 | pathogenic | 1 | 1 | 0 |
| rs749780672 | pathogenic | 0 | 0 | 0 |
| rs375651668 | benign | 1 | 1 | 1 |
| rs193922568 | pathogenic | 1 | 1 | 0 |
| rs750363970 | benign | 1 | 1 | 0 |
| rs5932 | benign | 0 | 0 | 0 |
| rs755154048 | pathogenic | 1 | 1 | 0 |
| rs759876319 | pathogenic | 0 | 0 | 0 |
| rs764550980 | pathogenic | 1 | 1 | 0 |
| rs779940524 | pathogenic | 0 | 0 | 0 |
| rs563382937 | pathogenic | 1 | 1 | 1 |
| rs28942083 | pathogenic | 0 | 1 | 0 |
| rs770744861 | pathogenic | 0 | 1 | 0 |
| rs121908031 | pathogenic | 0 | 1 | 0 |
| rs776217028 | pathogenic | 0 | 1 | 0 |
| rs763625913 | pathogenic | 0 | 0 | 0 |
| rs3736772 | benign | 0 | 0 | 0 |
| rs104894696 | benign | 1 | 1 | 1 |
| rs121918177 | pathogenic | 0 | 1 | 0 |
| rs121918181 | pathogenic | 1 | 1 | 0 |
| rs121918174 | pathogenic | 0 | 1 | 0 |
| rs121918182 | pathogenic | 0 | 0 | 0 |

| | | | | |
|---|---|---|---|---|
| rs121918175 | pathogenic | 1 | 1 | 1 |
| rs121918185 | pathogenic | 1 | 1 | 1 |
| rs121918173 | pathogenic | 1 | 1 | 0 |
| rs121918178 | pathogenic | 1 | 1 | 1 |
| rs121918179 | pathogenic | 1 | 1 | 1 |
| rs121918183 | pathogenic | 0 | 0 | 0 |
| rs121918176 | pathogenic | 1 | 1 | 1 |
| rs121918172 | pathogenic | 0 | 1 | 0 |
| rs121918184 | pathogenic | 0 | 0 | 0 |
| rs121434560 | pathogenic | 1 | 1 | 0 |
| rs121434561 | pathogenic | 1 | 1 | 0 |
| rs3218716 | pathogenic | 1 | 1 | 1 |
| rs145532615 | benign | 0 | 0 | 0 |
| rs149840927 | pathogenic | 0 | 0 | 0 |
| rs730880762 | pathogenic | 0 | 0 | 0 |
| rs730880764 | pathogenic | 0 | 0 | 0 |
| rs730880765 | pathogenic | 0 | 0 | 0 |
| rs587782962 | pathogenic | 0 | 0 | 0 |
| rs141735183 | pathogenic | 0 | 0 | 0 |
| rs397516187 | pathogenic | 0 | 0 | 0 |
| rs727505026 | pathogenic | 0 | 0 | 0 |
| rs529700838 | pathogenic | 0 | 0 | 0 |
| rs540263945 | pathogenic | 0 | 0 | 0 |
| rs149103761 | benign | 0 | 0 | 0 |
| rs730880906 | pathogenic | 0 | 0 | 0 |
| rs730880785 | pathogenic | 0 | 0 | 0 |
| rs397516199 | pathogenic | 0 | 0 | 0 |
| rs730880725 | benign | 0 | 0 | 0 |
| rs730880796 | pathogenic | 0 | 0 | 0 |
| rs201307101 | benign | 0 | 0 | 0 |
| rs397516214 | pathogenic | 0 | 0 | 0 |
| rs3729823 | benign | 0 | 0 | 0 |
| rs397516218 | pathogenic | 0 | 0 | 0 |
| rs121913647 | pathogenic | 0 | 0 | 0 |
| rs373514686 | pathogenic | 0 | 0 | 0 |
| rs763538103 | pathogenic | 0 | 0 | 0 |
| rs121913650 | pathogenic | 0 | 0 | 0 |
| rs139222507 | pathogenic | 0 | 0 | 0 |
| rs369940645 | pathogenic | 0 | 0 | 0 |
| rs763073072 | pathogenic | 0 | 0 | 0 |
| rs145734640 | pathogenic | 0 | 0 | 0 |
| rs730880821 | pathogenic | 0 | 0 | 0 |
| rs28933098 | pathogenic | 0 | 0 | 0 |
| rs372381770 | pathogenic | 0 | 0 | 0 |
| rs146796870 | benign | 0 | 0 | 0 |
| rs104894485 | pathogenic | 1 | 1 | 1 |
| rs35407712 | benign | 0 | 0 | 0 |
| rs35842750 | benign | 0 | 0 | 0 |
| rs2721191 | benign | 0 | 0 | 0 |
| rs35098923 | benign | 0 | 0 | 0 |
| rs34293591 | benign | 1 | 1 | 0 |
| rs36023964 | benign | 0 | 0 | 0 |
| rs4251691 | benign | 0 | 0 | 0 |
| rs34666647 | benign | 0 | 0 | 0 |

| | | | | |
|---|---|---|---|---|
| rs4925828 | benign | 0 | 0 | 0 |
| rs35348691 | benign | 0 | 0 | 0 |
| rs36078464 | benign | 0 | 0 | 0 |
| rs34915097 | benign | 0 | 0 | 0 |
| rs35101495 | benign | 0 | 0 | 0 |
| rs141568342 | pathogenic | 0 | 0 | 0 |
| rs1800571 | pathogenic | 0 | 0 | 0 |
| rs140566084 | benign | 1 | 1 | 1 |
| rs727503791 | pathogenic | 1 | 1 | 0 |
| rs545215807 | pathogenic | 0 | 0 | 0 |
| rs113994168 | pathogenic | 0 | 0 | 0 |
| rs113994167 | pathogenic | 0 | 0 | 0 |
| rs200788251 | pathogenic | 0 | 0 | 0 |
| rs200573371 | pathogenic | 0 | 0 | 0 |
| rs771874163 | pathogenic | 1 | 1 | 1 |
| rs112406105 | pathogenic | 1 | 1 | 1 |
| rs118204015 | pathogenic | 1 | 1 | 1 |
| rs113994169 | pathogenic | 0 | 0 | 0 |
| rs118204018 | pathogenic | 0 | 1 | 0 |
| rs138834083 | pathogenic | 0 | 0 | 0 |
| rs533055438 | pathogenic | 1 | 1 | 1 |
| rs2309689 | pathogenic | 1 | 1 | 1 |
| rs138058572 | pathogenic | 1 | 1 | 1 |
| rs118204017 | pathogenic | 1 | 1 | 0 |
| rs766742117 | pathogenic | 1 | 1 | 1 |
| rs751995154 | pathogenic | 1 | 1 | 1 |
| rs200771970 | pathogenic | 0 | 0 | 0 |
| rs118204014 | pathogenic | 0 | 0 | 0 |
| rs148584617 | pathogenic | 1 | 1 | 1 |
| rs121912720 | pathogenic | 0 | 0 | 0 |
| rs121912721 | pathogenic | 0 | 0 | 0 |
| rs28929476 | pathogenic | 0 | 0 | 0 |
| rs28931574 | pathogenic | 1 | 1 | 1 |
| rs12718465 | benign | 0 | 0 | 0 |
| rs144689354 | pathogenic | 0 | 0 | 0 |
| rs149095705 | pathogenic | 0 | 0 | 0 |
| rs572839648 | benign | 1 | 1 | 1 |
| rs121909715 | pathogenic | 1 | 1 | 0 |
| rs34077456 | benign | 1 | 1 | 1 |
| rs397517876 | benign | 0 | 0 | 0 |
| rs1064074 | benign | 0 | 0 | 0 |
| rs142751309 | benign | 0 | 0 | 0 |
| rs116275562 | benign | 0 | 0 | 0 |
| rs148095899 | benign | 0 | 0 | 0 |
| rs751838040 | pathogenic | 0 | 0 | 0 |
| rs121434324 | pathogenic | 1 | 1 | 1 |
| rs143367518 | pathogenic | 0 | 0 | 0 |
| rs121434325 | pathogenic | 0 | 0 | 0 |
| rs397515328 | pathogenic | 0 | 0 | 0 |
| rs376039938 | pathogenic | 0 | 0 | 0 |
| rs387907243 | pathogenic | 0 | 0 | 0 |
| rs28934876 | pathogenic | 0 | 0 | 0 |
| rs387907245 | pathogenic | 0 | 0 | 0 |
| rs138544311 | benign | 0 | 0 | 0 |

| | | | | |
|---|---|---|---|---|
| rs643788 | benign | 0 | 0 | 0 |
| rs749977756 | benign | 0 | 0 | 0 |
| rs1800574 | benign | 0 | 0 | 0 |
| rs137853336 | pathogenic | 1 | 1 | 1 |
| rs111916034 | benign | 0 | 0 | 0 |
| rs307826 | benign | 0 | 0 | 0 |
| rs35874891 | benign | 0 | 0 | 0 |
| rs111033580 | pathogenic | 1 | 1 | 1 |
| rs111033581 | pathogenic | 1 | 1 | 1 |
| rs121908373 | pathogenic | 0 | 0 | 0 |
| rs767122713 | pathogenic | 0 | 0 | 0 |
| rs2233213 | benign | 0 | 0 | 0 |
| rs121908605 | pathogenic | 1 | 1 | 0 |
| rs11887534 | benign | 0 | 0 | 0 |
| rs4148211 | benign | 0 | 0 | 0 |
| rs80025980 | benign | 0 | 0 | 0 |
| rs137854891 | pathogenic | 1 | 1 | 0 |
| rs74358901 | benign | 0 | 0 | 0 |
| rs9282574 | benign | 0 | 0 | 0 |
| rs137852993 | pathogenic | 0 | 0 | 0 |
| rs137852990 | pathogenic | 0 | 0 | 0 |
| rs137852987 | pathogenic | 0 | 0 | 0 |
| rs4148217 | benign | 0 | 0 | 0 |
| rs140690030 | benign | 0 | 0 | 0 |
| rs137852992 | pathogenic | 0 | 0 | 0 |
| rs6544718 | benign | 0 | 0 | 0 |
| rs137852989 | pathogenic | 0 | 0 | 0 |
| rs150893072 | benign | 1 | 1 | 1 |
| rs28931613 | pathogenic | 1 | 1 | 1 |
| rs143789597 | benign | 1 | 1 | 1 |
| rs121913037 | pathogenic | 1 | 1 | 0 |
| rs188802138 | pathogenic | 1 | 1 | 0 |
| rs121913038 | pathogenic | 1 | 1 | 0 |
| rs146835206 | benign | 0 | 0 | 0 |
| rs121913041 | pathogenic | 1 | 1 | 1 |
| rs121913039 | pathogenic | 1 | 1 | 1 |
| rs149977726 | pathogenic | 1 | 1 | 0 |
| rs121913036 | pathogenic | 0 | 0 | 0 |
| rs121913040 | pathogenic | 0 | 0 | 0 |
| rs765007158 | benign | 0 | 0 | 0 |
| rs112723255 | benign | 0 | 0 | 0 |
| rs574164748 | benign | 0 | 0 | 0 |
| rs80338734 | pathogenic | 1 | 1 | 0 |
| rs104894694 | pathogenic | 1 | 1 | 0 |
| rs753198836 | pathogenic | 0 | 0 | 0 |
| rs760101382 | pathogenic | 0 | 0 | 0 |
| rs121909272 | pathogenic | 1 | 1 | 0 |
| rs370421531 | pathogenic | 0 | 0 | 0 |
| rs565109128 | benign | 0 | 0 | 0 |
| rs17851582 | benign | 0 | 0 | 0 |
| rs753228876 | benign | 0 | 0 | 0 |
| rs141471799 | benign | 0 | 0 | 0 |
| rs281864916 | pathogenic | 1 | 1 | 0 |
| rs121918409 | pathogenic | 1 | 1 | 1 |

| | | | | |
|---|---|---|---|---|
| rs121918413 | pathogenic | 1 | 1 | 1 |
| rs121918411 | pathogenic | 1 | 1 | 1 |
| rs121918408 | pathogenic | 1 | 1 | 1 |
| rs121918412 | pathogenic | 1 | 1 | 1 |
| rs121918410 | pathogenic | 1 | 1 | 1 |
| rs116474260 | benign | 1 | 1 | 1 |
| rs281864919 | pathogenic | 1 | 1 | 1 |
| rs397514452 | pathogenic | 1 | 1 | 0 |
| rs747843638 | benign | 0 | 0 | 0 |
| rs774357609 | pathogenic | 1 | 1 | 1 |
| rs201804005 | benign | 0 | 0 | 0 |
| rs746485916 | pathogenic | 0 | 0 | 0 |
| rs764131178 | pathogenic | 0 | 0 | 0 |
| rs374094065 | pathogenic | 0 | 0 | 0 |
| rs76151636 | pathogenic | 1 | 1 | 0 |
| rs753250853 | pathogenic | 0 | 0 | 0 |
| rs761632029 | pathogenic | 0 | 0 | 0 |
| rs775541743 | pathogenic | 0 | 0 | 0 |
| rs778749563 | pathogenic | 0 | 0 | 0 |
| rs60431989 | pathogenic | 1 | 1 | 0 |
| rs755554442 | pathogenic | 1 | 1 | 0 |
| rs749085322 | pathogenic | 0 | 0 | 0 |
| rs756029120 | pathogenic | 1 | 1 | 0 |
| rs137853279 | pathogenic | 1 | 1 | 0 |
| rs777354267 | pathogenic | 1 | 1 | 1 |
| rs142940486 | benign | 1 | 1 | 1 |
| rs140167393 | benign | 1 | 1 | 1 |
| rs2066827 | benign | 0 | 0 | 0 |
| rs765681672 | benign | 0 | 0 | 0 |
| rs137985549 | benign | 0 | 0 | 0 |
| rs74315340 | pathogenic | 1 | 1 | 1 |
| rs74315341 | pathogenic | 0 | 0 | 0 |
| rs74315335 | pathogenic | 0 | 0 | 0 |
| rs61730974 | benign | 0 | 0 | 0 |
| rs121909193 | pathogenic | 0 | 0 | 0 |
| rs74315334 | pathogenic | 0 | 0 | 0 |
| rs74315329 | pathogenic | 0 | 0 | 0 |
| rs74315330 | pathogenic | 0 | 0 | 0 |
| rs121909194 | pathogenic | 0 | 0 | 0 |
| rs28936694 | pathogenic | 0 | 0 | 0 |
| rs74315336 | pathogenic | 0 | 0 | 0 |
| rs74315338 | pathogenic | 1 | 1 | 1 |
| rs74315328 | pathogenic | 0 | 0 | 0 |
| rs74315331 | pathogenic | 0 | 0 | 0 |
| rs74315332 | pathogenic | 0 | 0 | 0 |
| rs141696651 | benign | 0 | 0 | 0 |
| rs7041 | benign | 0 | 0 | 0 |
| rs4588 | benign | 0 | 0 | 0 |
| rs121908981 | pathogenic | 1 | 1 | 1 |
| rs606231459 | pathogenic | 0 | 0 | 0 |
| rs398123053 | pathogenic | 0 | 0 | 0 |
| rs774259802 | benign | 0 | 0 | 0 |
| rs121909582 | pathogenic | 1 | 1 | 1 |
| rs104893680 | pathogenic | 1 | 1 | 1 |

| | | | | |
|---|---|---|---|---|
| rs137854907 | pathogenic | 0 | 0 | 0 |
| rs765715798 | pathogenic | 0 | 0 | 0 |
| rs104893678 | pathogenic | 0 | 0 | 0 |
| rs104893679 | pathogenic | 0 | 1 | 0 |
| rs104893681 | pathogenic | 1 | 1 | 0 |
| rs121909640 | pathogenic | 1 | 1 | 0 |
| rs780705654 | pathogenic | 0 | 0 | 0 |
| rs201672146 | benign | 1 | 1 | 0 |
| rs12136658 | benign | 0 | 0 | 0 |
| rs56127440 | benign | 0 | 0 | 0 |
| rs104894433 | pathogenic | 1 | 1 | 1 |
| rs104894435 | pathogenic | 0 | 1 | 0 |
| rs104894437 | pathogenic | 1 | 1 | 1 |
| rs104894441 | pathogenic | 1 | 1 | 1 |
| rs104894440 | pathogenic | 1 | 1 | 1 |
| rs104894445 | pathogenic | 1 | 1 | 1 |
| rs104894436 | pathogenic | 0 | 0 | 0 |
| rs104894438 | pathogenic | 0 | 0 | 0 |
| rs200891969 | pathogenic | 1 | 1 | 0 |
| rs104894443 | pathogenic | 1 | 1 | 1 |
| rs104894434 | pathogenic | 1 | 1 | 1 |
| rs41298442 | pathogenic | 1 | 1 | 1 |
| rs104894442 | pathogenic | 1 | 1 | 1 |
| rs35453854 | benign | 0 | 0 | 0 |
| rs121912898 | pathogenic | 0 | 0 | 0 |
| rs121912897 | pathogenic | 1 | 1 | 0 |
| rs138260625 | benign | 0 | 0 | 0 |
| rs147802438 | benign | 1 | 1 | 0 |
| rs146443310 | benign | 1 | 1 | 0 |
| rs747319628 | pathogenic | 1 | 1 | 0 |
| rs1801195 | benign | 0 | 0 | 0 |
| rs145045341 | benign | 1 | 1 | 0 |
| rs116001173 | benign | 1 | 1 | 0 |
| rs587784175 | pathogenic | 0 | 0 | 0 |
| rs104893857 | pathogenic | 0 | 0 | 0 |
| rs104893862 | pathogenic | 1 | 1 | 0 |
| rs104893858 | pathogenic | 1 | 1 | 1 |
| rs121909249 | pathogenic | 0 | 1 | 0 |
| rs121909248 | pathogenic | 1 | 1 | 1 |
| rs104893859 | pathogenic | 1 | 1 | 0 |
| rs187063864 | pathogenic | 1 | 1 | 0 |
| rs121917817 | pathogenic | 1 | 1 | 0 |
| rs121917818 | pathogenic | 0 | 0 | 0 |
| rs753594601 | pathogenic | 0 | 0 | 0 |
| rs28937869 | pathogenic | 1 | 1 | 0 |
| rs6165 | benign | 0 | 0 | 0 |
| rs200089714 | pathogenic | 1 | 1 | 0 |
| rs372000714 | pathogenic | 0 | 0 | 0 |
| rs2228206 | benign | 0 | 0 | 0 |
| rs560096 | benign | 0 | 0 | 0 |
| rs137852666 | pathogenic | 0 | 0 | 0 |
| rs10896380 | benign | 0 | 0 | 0 |
| rs773690764 | pathogenic | 0 | 0 | 0 |
| rs201060167 | pathogenic | 0 | 0 | 0 |

| | | | | |
|---|---|---|---|---|
| rs137852670 | pathogenic | 0 | 0 | 0 |
| rs780594709 | pathogenic | 0 | 0 | 0 |
| rs145226920 | pathogenic | 0 | 0 | 0 |
| rs137852665 | pathogenic | 0 | 0 | 0 |
| rs724159960 | pathogenic | 0 | 0 | 0 |
| rs756985703 | pathogenic | 0 | 0 | 0 |
| rs137852667 | pathogenic | 0 | 0 | 0 |
| rs151079750 | pathogenic | 1 | 1 | 0 |
| rs2306636 | benign | 0 | 0 | 0 |
| rs72647885 | benign | 1 | 1 | 0 |
| rs377584435 | pathogenic | 1 | 1 | 1 |
| rs757583846 | pathogenic | 1 | 1 | 0 |
| rs121918148 | pathogenic | 0 | 0 | 0 |
| rs149201802 | benign | 0 | 0 | 0 |
| rs587781904 | pathogenic | 1 | 1 | 1 |
| rs143421325 | benign | 0 | 0 | 0 |
| rs587781253 | pathogenic | 0 | 0 | 0 |
| rs587780331 | benign | 0 | 0 | 0 |
| rs151001016 | benign | 0 | 0 | 0 |
| rs150888094 | benign | 0 | 0 | 0 |
| rs149581331 | benign | 0 | 0 | 0 |
| rs774840535 | benign | 0 | 0 | 0 |
| rs138428684 | benign | 0 | 0 | 0 |
| rs149300055 | benign | 0 | 0 | 0 |
| rs571005578 | benign | 0 | 0 | 0 |
| rs200731839 | benign | 0 | 0 | 0 |
| rs121965007 | pathogenic | 0 | 0 | 0 |
| rs121965013 | pathogenic | 0 | 1 | 0 |
| rs121965009 | pathogenic | 0 | 0 | 0 |
| rs1800457 | benign | 1 | 1 | 0 |
| rs121965006 | pathogenic | 1 | 1 | 0 |
| rs121965008 | pathogenic | 0 | 0 | 0 |
| rs61732609 | pathogenic | 0 | 0 | 0 |
| rs121965011 | pathogenic | 0 | 0 | 0 |
| rs121965015 | pathogenic | 0 | 0 | 0 |
| rs121965018 | pathogenic | 1 | 1 | 0 |
| rs121965016 | pathogenic | 0 | 0 | 0 |
| rs77806 | benign | 0 | 0 | 0 |
| rs149731356 | benign | 0 | 0 | 0 |
| rs72549407 | pathogenic | 0 | 0 | 0 |
| rs200363929 | benign | 0 | 0 | 0 |
| rs376403708 | benign | 1 | 1 | 0 |
| rs56376197 | benign | 1 | 1 | 1 |
| rs200237973 | benign | 1 | 1 | 1 |
| rs548330075 | benign | 0 | 0 | 0 |
| rs148291350 | benign | 0 | 0 | 0 |
| rs71579353 | pathogenic | 0 | 0 | 0 |
| rs146278697 | benign | 0 | 0 | 0 |
| rs141033098 | benign | 0 | 0 | 0 |
| rs397517234 | pathogenic | 0 | 0 | 0 |
| rs71579374 | benign | 0 | 0 | 0 |
| rs140381835 | benign | 0 | 0 | 0 |
| rs137852702 | pathogenic | 0 | 0 | 0 |
| rs137852696 | pathogenic | 1 | 1 | 1 |

| | | | | |
|---|---|---|---|---|
| rs137852697 | pathogenic | 0 | 0 | 0 |
| rs386833639 | pathogenic | 0 | 0 | 0 |
| rs386833640 | pathogenic | 0 | 0 | 0 |
| rs386833641 | benign | 0 | 0 | 0 |
| rs137852701 | pathogenic | 0 | 0 | 0 |
| rs386833642 | pathogenic | 0 | 0 | 0 |
| rs137852695 | pathogenic | 0 | 0 | 0 |
| rs386833646 | pathogenic | 0 | 0 | 0 |
| rs137852700 | pathogenic | 1 | 1 | 0 |
| rs386833649 | pathogenic | 1 | 1 | 0 |
| rs386833650 | pathogenic | 0 | 0 | 0 |
| rs148412181 | pathogenic | 0 | 0 | 0 |
| rs386833655 | pathogenic | 0 | 0 | 0 |
| rs386833656 | pathogenic | 1 | 1 | 0 |
| rs137852698 | pathogenic | 0 | 0 | 0 |
| rs386833661 | pathogenic | 0 | 0 | 0 |
| rs386833670 | pathogenic | 0 | 0 | 0 |
| rs111033632 | pathogenic | 1 | 1 | 1 |
| rs34308573 | benign | 1 | 1 | 1 |
| rs376048533 | pathogenic | 1 | 1 | 0 |
| rs3747517 | benign | 1 | 1 | 0 |
| rs137853328 | pathogenic | 0 | 0 | 0 |
| rs77273740 | benign | 0 | 0 | 0 |
| rs267606760 | pathogenic | 0 | 0 | 0 |
| rs77576840 | pathogenic | 0 | 0 | 0 |
| rs45465204 | benign | 1 | 1 | 0 |
| rs5320 | benign | 0 | 0 | 0 |
| rs5324 | benign | 0 | 0 | 0 |
| rs4531 | benign | 0 | 0 | 0 |
| rs267606761 | pathogenic | 0 | 0 | 0 |
| rs75215331 | pathogenic | 0 | 0 | 0 |
| rs6271 | benign | 0 | 0 | 0 |
| rs201643277 | pathogenic | 0 | 0 | 0 |
| rs4252250 | benign | 0 | 0 | 0 |
| rs137853580 | pathogenic | 0 | 0 | 0 |
| rs368287711 | pathogenic | 0 | 0 | 0 |
| rs137853579 | pathogenic | 0 | 0 | 0 |
| rs3135932 | benign | 0 | 0 | 0 |
| rs2228055 | benign | 0 | 0 | 0 |
| rs140322345 | benign | 1 | 1 | 1 |
| rs1154510 | pathogenic | 0 | 0 | 0 |
| rs11833399 | benign | 0 | 0 | 0 |
| rs36023382 | benign | 1 | 1 | 1 |
| rs61754444 | benign | 0 | 0 | 0 |
| rs28935168 | pathogenic | 0 | 0 | 0 |
| rs28934907 | pathogenic | 0 | 0 | 0 |
| rs28934904 | pathogenic | 1 | 1 | 0 |
| rs61748392 | pathogenic | 1 | 1 | 0 |
| rs28934908 | pathogenic | 0 | 0 | 0 |
| rs28934905 | pathogenic | 0 | 0 | 0 |
| rs28934906 | pathogenic | 1 | 1 | 0 |
| rs606231307 | pathogenic | 0 | 0 | 0 |
| rs9333296 | benign | 0 | 0 | 0 |
| rs3218614 | benign | 0 | 0 | 0 |

| | | | | |
|---|---|---|---|---|
| rs2230792 | benign | 0 | 0 | 0 |
| rs2230793 | benign | 0 | 0 | 0 |
| rs2230794 | benign | 0 | 0 | 0 |
| rs749052963 | pathogenic | 0 | 0 | 0 |
| rs2230798 | benign | 0 | 0 | 0 |
| rs2230795 | benign | 0 | 0 | 0 |
| rs3204145 | benign | 0 | 0 | 0 |
| rs1538660 | benign | 0 | 0 | 0 |
| rs9878928 | benign | 0 | 0 | 0 |
| rs104893742 | pathogenic | 0 | 0 | 0 |
| rs28936702 | pathogenic | 0 | 0 | 0 |
| rs28936703 | pathogenic | 0 | 0 | 0 |
| rs769392 | benign | 0 | 0 | 0 |
| rs769402 | benign | 1 | 1 | 1 |
| rs104894884 | pathogenic | 1 | 1 | 1 |
| rs1801316 | benign | 1 | 1 | 1 |
| rs104894885 | pathogenic | 1 | 1 | 1 |
| rs782572 | benign | 0 | 0 | 0 |
| rs151166046 | pathogenic | 0 | 0 | 0 |
| rs772153760 | pathogenic | 1 | 1 | 0 |
| rs35405862 | benign | 0 | 0 | 0 |
| rs146571352 | pathogenic | 0 | 1 | 0 |
| rs374698153 | pathogenic | 0 | 0 | 0 |
| rs7594497 | benign | 0 | 0 | 0 |
| rs80338716 | pathogenic | 0 | 0 | 0 |
| rs80338719 | pathogenic | 1 | 1 | 1 |
| rs80338721 | pathogenic | 0 | 0 | 0 |
| rs730882249 | pathogenic | 0 | 0 | 0 |
| rs146262009 | pathogenic | 0 | 0 | 0 |
| rs104895083 | pathogenic | 1 | 1 | 1 |
| rs104895100 | benign | 1 | 1 | 1 |
| rs104895085 | pathogenic | 0 | 0 | 0 |
| rs28940580 | pathogenic | 0 | 0 | 0 |
| rs104895098 | pathogenic | 0 | 0 | 0 |
| rs28940578 | pathogenic | 0 | 0 | 0 |
| rs61752717 | pathogenic | 0 | 0 | 0 |
| rs28940579 | pathogenic | 0 | 0 | 0 |
| rs104895097 | pathogenic | 0 | 0 | 0 |
| rs140425651 | benign | 1 | 1 | 0 |
| rs768090444 | pathogenic | 0 | 0 | 0 |
| rs764459544 | pathogenic | 0 | 0 | 0 |
| rs80338802 | pathogenic | 1 | 1 | 1 |
| rs149969786 | pathogenic | 1 | 1 | 0 |
| rs115311625 | pathogenic | 1 | 1 | 1 |
| rs778768583 | pathogenic | 1 | 1 | 1 |
| rs892090 | benign | 0 | 0 | 0 |
| rs28929498 | pathogenic | 1 | 1 | 1 |
| rs137854467 | pathogenic | 0 | 0 | 0 |
| rs137854480 | pathogenic | 0 | 0 | 0 |
| rs148682262 | benign | 0 | 0 | 0 |
| rs137852744 | pathogenic | 0 | 0 | 0 |
| rs137852745 | pathogenic | 0 | 1 | 0 |
| rs137852749 | pathogenic | 0 | 1 | 0 |
| rs137852746 | pathogenic | 0 | 1 | 0 |

| | | | | |
|---|---|---|---|---|
| rs145633958 | benign | 0 | 0 | 0 |
| rs141679950 | benign | 0 | 0 | 0 |
| rs76397662 | benign | 1 | 1 | 1 |
| rs370736139 | benign | 0 | 0 | 0 |
| rs781750106 | benign | 0 | 0 | 0 |
| rs546866208 | benign | 0 | 0 | 0 |
| rs142338976 | benign | 0 | 0 | 0 |
| rs115272158 | benign | 0 | 0 | 0 |
| rs115423919 | benign | 0 | 0 | 0 |
| rs537874538 | benign | 0 | 0 | 0 |
| rs9282834 | benign | 0 | 0 | 0 |
| rs201745826 | benign | 0 | 0 | 0 |
| rs75873440 | pathogenic | 0 | 0 | 0 |
| rs147219360 | benign | 0 | 0 | 0 |
| rs77939446 | pathogenic | 0 | 0 | 0 |
| rs77558292 | pathogenic | 0 | 0 | 0 |
| rs80069458 | pathogenic | 0 | 0 | 0 |
| rs76262710 | pathogenic | 0 | 0 | 0 |
| rs79781594 | pathogenic | 0 | 0 | 0 |
| rs77316810 | pathogenic | 0 | 0 | 0 |
| rs77503355 | pathogenic | 0 | 0 | 0 |
| rs79890926 | pathogenic | 0 | 0 | 0 |
| rs377767402 | benign | 0 | 0 | 0 |
| rs121913308 | pathogenic | 0 | 0 | 0 |
| rs55846256 | benign | 0 | 0 | 0 |
| rs75076352 | pathogenic | 0 | 0 | 0 |
| rs75996173 | pathogenic | 0 | 0 | 0 |
| rs77709286 | pathogenic | 0 | 0 | 0 |
| rs151058706 | benign | 0 | 0 | 0 |
| rs138964358 | benign | 0 | 0 | 0 |
| rs137852353 | pathogenic | 1 | 1 | 0 |
| rs201196020 | benign | 0 | 0 | 0 |
| rs137853201 | pathogenic | 1 | 1 | 1 |
| rs113994174 | pathogenic | 1 | 1 | 1 |
| rs104893674 | pathogenic | 1 | 1 | 1 |
| rs779519249 | benign | 1 | 1 | 1 |
| rs9606756 | benign | 0 | 0 | 0 |
| rs75680863 | benign | 1 | 1 | 1 |
| rs35838082 | benign | 0 | 0 | 0 |
| rs1801198 | benign | 0 | 0 | 0 |
| rs117458738 | benign | 1 | 1 | 0 |
| rs9621049 | benign | 1 | 1 | 0 |
| rs1131603 | benign | 1 | 1 | 0 |
| rs4820889 | benign | 0 | 0 | 0 |
| rs777767978 | pathogenic | 1 | 1 | 0 |
| rs147399860 | benign | 1 | 1 | 1 |
| rs104893950 | pathogenic | 0 | 0 | 0 |
| rs777308612 | benign | 0 | 0 | 0 |
| rs151091410 | benign | 0 | 0 | 0 |
| rs11537667 | benign | 0 | 0 | 0 |
| rs267606621 | pathogenic | 1 | 1 | 0 |
| rs143058902 | benign | 0 | 0 | 0 |
| rs370261904 | pathogenic | 0 | 0 | 0 |
| rs141169746 | benign | 0 | 0 | 0 |

| | | | | |
|---|---|---|---|---|
| rs111033436 | benign | 0 | 0 | 0 |
| rs148533341 | benign | 0 | 0 | 0 |
| rs143827620 | benign | 0 | 0 | 0 |
| rs4935502 | benign | 0 | 0 | 0 |
| rs61735473 | benign | 0 | 0 | 0 |
| rs267606932 | pathogenic | 1 | 1 | 0 |
| rs61730754 | benign | 0 | 0 | 0 |
| rs374491359 | pathogenic | 1 | 1 | 1 |
| rs761800077 | benign | 1 | 1 | 1 |
| rs760391381 | benign | 1 | 1 | 1 |
| rs104893885 | pathogenic | 0 | 0 | 0 |
| rs104893886 | pathogenic | 1 | 1 | 1 |
| rs104893884 | pathogenic | 0 | 0 | 0 |
| rs147715509 | benign | 0 | 0 | 0 |
| rs115533243 | benign | 0 | 0 | 0 |
| rs397509390 | pathogenic | 0 | 1 | 0 |
| rs80357134 | pathogenic | 0 | 0 | 0 |
| rs766004110 | benign | 1 | 1 | 1 |
| rs769650474 | benign | 0 | 0 | 0 |
| rs398122635 | benign | 1 | 1 | 0 |
| rs199522616 | pathogenic | 0 | 0 | 0 |
| rs28897672 | pathogenic | 1 | 1 | 0 |
| rs80357086 | pathogenic | 1 | 1 | 1 |
| rs80357064 | pathogenic | 1 | 1 | 0 |
| rs80356936 | benign | 0 | 0 | 0 |
| rs28897673 | benign | 0 | 0 | 0 |
| rs146747324 | benign | 0 | 0 | 0 |
| rs78635798 | pathogenic | 0 | 0 | 0 |
| rs75146158 | pathogenic | 1 | 1 | 1 |
| rs587782886 | benign | 0 | 0 | 0 |
| rs121434634 | pathogenic | 0 | 0 | 0 |
| rs121434635 | pathogenic | 0 | 0 | 0 |
| rs587777778 | pathogenic | 0 | 0 | 0 |
| rs118203967 | pathogenic | 0 | 0 | 0 |
| rs751375244 | pathogenic | 1 | 1 | 1 |
| rs104893863 | pathogenic | 1 | 1 | 0 |
| rs104893865 | pathogenic | 0 | 0 | 0 |
| rs104893864 | pathogenic | 1 | 1 | 1 |
| rs104893866 | pathogenic | 1 | 1 | 0 |
| rs561738497 | benign | 0 | 0 | 0 |
| rs57599352 | pathogenic | 1 | 1 | 1 |
| rs267607448 | pathogenic | 1 | 1 | 1 |
| rs63751596 | pathogenic | 0 | 0 | 0 |
| rs63751612 | benign | 0 | 0 | 0 |
| rs63751684 | benign | 0 | 0 | 0 |
| rs267607861 | benign | 0 | 0 | 0 |
| rs35831931 | benign | 0 | 0 | 0 |
| rs63750876 | benign | 0 | 0 | 0 |
| rs41295284 | benign | 0 | 0 | 0 |
| rs35502531 | benign | 0 | 0 | 0 |
| rs63750449 | benign | 0 | 0 | 0 |
| rs63750899 | pathogenic | 0 | 0 | 0 |
| rs55907433 | benign | 0 | 0 | 0 |
| rs63750217 | pathogenic | 0 | 0 | 0 |

| | | | | |
|---|---|---|---|---|
| rs63751275 | pathogenic | 0 | 0 | 0 |
| rs63750702 | benign | 1 | 1 | 1 |
| rs63750114 | benign | 0 | 0 | 0 |
| rs2020873 | benign | 0 | 0 | 0 |
| rs140195825 | benign | 1 | 1 | 1 |
| rs182763805 | benign | 0 | 0 | 0 |
| rs533696377 | benign | 0 | 0 | 0 |
| rs41278788 | benign | 0 | 0 | 0 |
| rs104894193 | pathogenic | 1 | 1 | 0 |
| rs3824915 | benign | 1 | 1 | 1 |
| rs104894196 | pathogenic | 0 | 0 | 0 |
| rs78681604 | benign | 1 | 1 | 1 |
| rs77321498 | benign | 1 | 1 | 1 |
| rs137852965 | pathogenic | 0 | 0 | 0 |
| rs137852961 | pathogenic | 0 | 0 | 0 |
| rs137852962 | pathogenic | 1 | 1 | 0 |
| rs753376100 | pathogenic | 0 | 0 | 0 |
| rs137852968 | pathogenic | 1 | 1 | 1 |
| rs137852959 | pathogenic | 1 | 1 | 0 |
| rs137852967 | pathogenic | 0 | 0 | 0 |
| rs113437301 | benign | 1 | 1 | 1 |
| rs61747482 | benign | 1 | 1 | 1 |
| rs2234919 | benign | 1 | 1 | 0 |
| rs121908863 | pathogenic | 0 | 0 | 0 |
| rs140455771 | pathogenic | 0 | 0 | 0 |
| rs121908810 | pathogenic | 1 | 1 | 1 |
| rs143879853 | benign | 0 | 0 | 0 |
| rs188319622 | benign | 0 | 0 | 0 |
| rs121918645 | pathogenic | 0 | 0 | 0 |
| rs115352435 | pathogenic | 0 | 0 | 0 |
| rs121434427 | pathogenic | 1 | 1 | 1 |
| rs121434428 | pathogenic | 0 | 0 | 0 |
| rs35086265 | benign | 1 | 1 | 1 |
| rs11576415 | benign | 0 | 0 | 0 |
| rs121434429 | pathogenic | 1 | 1 | 1 |
| rs4074536 | benign | 1 | 1 | 1 |
| rs10801999 | benign | 0 | 0 | 0 |
| rs121434550 | pathogenic | 0 | 0 | 0 |
| rs148057999 | pathogenic | 0 | 0 | 0 |
| rs28730716 | benign | 0 | 0 | 0 |
| rs142036299 | benign | 0 | 0 | 0 |
| rs762381137 | pathogenic | 0 | 0 | 0 |
| rs121434549 | pathogenic | 0 | 0 | 0 |
| rs139228801 | pathogenic | 0 | 0 | 0 |
| rs147219463 | benign | 0 | 0 | 0 |
| rs1131857 | benign | 0 | 0 | 0 |
| rs121917874 | pathogenic | 0 | 1 | 0 |
| rs2228056 | benign | 0 | 0 | 0 |
| rs121917870 | pathogenic | 0 | 0 | 0 |
| rs121917866 | pathogenic | 0 | 0 | 0 |
| rs11921054 | benign | 0 | 0 | 0 |
| rs121917868 | pathogenic | 0 | 0 | 0 |
| rs28931603 | pathogenic | 0 | 0 | 0 |
| rs104895295 | pathogenic | 1 | 1 | 0 |

| | | | | |
|---|---|---|---|---|
| rs104895382 | pathogenic | 0 | 0 | 0 |
| rs104895298 | pathogenic | 0 | 0 | 0 |
| rs104895300 | pathogenic | 0 | 0 | 0 |
| rs104895332 | pathogenic | 0 | 0 | 0 |
| rs758026399 | pathogenic | 1 | 1 | 1 |
| rs104895366 | pathogenic | 0 | 0 | 0 |
| rs104895304 | pathogenic | 0 | 0 | 0 |
| rs759997079 | pathogenic | 0 | 0 | 0 |
| rs121917789 | pathogenic | 0 | 0 | 0 |
| rs104895319 | pathogenic | 0 | 0 | 0 |
| rs104895317 | pathogenic | 1 | 1 | 0 |
| rs28934897 | pathogenic | 0 | 0 | 0 |
| rs104895380 | benign | 0 | 0 | 0 |
| rs104895360 | pathogenic | 0 | 0 | 0 |
| rs28936396 | pathogenic | 1 | 1 | 0 |
| rs121909307 | pathogenic | 1 | 1 | 1 |
| rs28938472 | pathogenic | 1 | 1 | 0 |
| rs121909308 | pathogenic | 1 | 1 | 0 |
| rs121909309 | pathogenic | 0 | 0 | 0 |
| rs121913013 | benign | 1 | 1 | 1 |
| rs201855245 | benign | 1 | 1 | 1 |
| rs2230232 | benign | 1 | 1 | 0 |
| rs113451409 | benign | 0 | 0 | 0 |
| rs121913011 | pathogenic | 1 | 1 | 0 |
| rs185821167 | pathogenic | 1 | 1 | 0 |
| rs2230234 | benign | 0 | 0 | 0 |
| rs193922639 | benign | 1 | 1 | 0 |
| rs774863785 | pathogenic | 1 | 1 | 0 |
| rs576404380 | pathogenic | 1 | 1 | 0 |
| rs2230235 | benign | 1 | 1 | 0 |
| rs200484060 | pathogenic | 1 | 1 | 0 |
| rs199761749 | benign | 0 | 0 | 0 |
| rs199681901 | benign | 0 | 0 | 0 |
| rs121908951 | pathogenic | 0 | 0 | 0 |
| rs769251460 | benign | 0 | 0 | 0 |
| rs28937320 | pathogenic | 0 | 0 | 0 |
| rs200607527 | pathogenic | 0 | 0 | 0 |
| rs137852640 | pathogenic | 1 | 1 | 1 |
| rs137852636 | pathogenic | 1 | 1 | 1 |
| rs137852638 | pathogenic | 0 | 0 | 0 |
| rs142637231 | pathogenic | 0 | 0 | 0 |
| rs151328418 | benign | 0 | 1 | 0 |
| rs137852639 | pathogenic | 1 | 1 | 1 |
| rs34605667 | benign | 0 | 0 | 0 |
| rs104894073 | pathogenic | 1 | 1 | 0 |
| rs56298569 | pathogenic | 0 | 0 | 0 |
| rs757600616 | pathogenic | 0 | 0 | 0 |
| rs139194636 | pathogenic | 0 | 0 | 0 |
| rs369517993 | pathogenic | 1 | 1 | 1 |
| rs138846036 | benign | 1 | 1 | 1 |
| rs200808744 | pathogenic | 1 | 1 | 1 |
| rs2255543 | benign | 0 | 0 | 0 |
| rs149165166 | pathogenic | 0 | 0 | 0 |
| rs544956641 | pathogenic | 1 | 1 | 1 |

| | | | | |
|---|---|---|---|---|
| rs28941783 | pathogenic | 0 | 0 | 0 |
| rs562853291 | pathogenic | 1 | 1 | 1 |
| rs28942100 | pathogenic | 0 | 0 | 0 |
| rs120074174 | pathogenic | 1 | 1 | 1 |
| rs120074171 | pathogenic | 1 | 1 | 1 |
| rs120074173 | pathogenic | 0 | 0 | 0 |
| rs120074172 | pathogenic | 1 | 1 | 1 |
| rs767159114 | pathogenic | 1 | 1 | 1 |
| rs104893657 | pathogenic | 0 | 0 | 0 |
| rs104893656 | pathogenic | 0 | 0 | 0 |
| rs104893659 | pathogenic | 0 | 0 | 0 |
| rs104893658 | pathogenic | 0 | 0 | 0 |
| rs200386310 | pathogenic | 0 | 0 | 0 |
| rs2072712 | benign | 0 | 0 | 0 |
| rs35274867 | benign | 0 | 0 | 0 |
| rs35343172 | benign | 0 | 0 | 0 |
| rs10079250 | benign | 0 | 0 | 0 |
| rs34951517 | benign | 0 | 0 | 0 |
| rs267606695 | pathogenic | 0 | 0 | 0 |
| rs149391728 | benign | 0 | 0 | 0 |
| rs387906598 | pathogenic | 1 | 1 | 0 |
| rs137854578 | pathogenic | 0 | 0 | 0 |
| rs141519952 | benign | 0 | 0 | 0 |
| rs146572883 | benign | 0 | 0 | 0 |
| rs111866410 | benign | 0 | 0 | 0 |
| rs139387758 | benign | 1 | 1 | 1 |
| rs35672074 | benign | 1 | 1 | 1 |
| rs121434322 | pathogenic | 0 | 0 | 0 |
| rs150884181 | benign | 0 | 0 | 0 |
| rs35867471 | benign | 0 | 0 | 0 |
| rs2867802 | benign | 0 | 0 | 0 |
| rs137902889 | benign | 0 | 0 | 0 |
| rs369804864 | pathogenic | 0 | 0 | 0 |
| rs781050795 | pathogenic | 0 | 0 | 0 |
| rs2305799 | benign | 0 | 0 | 0 |
| rs2229531 | benign | 0 | 0 | 0 |
| rs147025508 | benign | 0 | 0 | 0 |
| rs76857106 | pathogenic | 0 | 0 | 0 |
| rs7247284 | benign | 0 | 0 | 0 |
| rs62619782 | benign | 0 | 0 | 0 |
| rs377244188 | pathogenic | 0 | 0 | 0 |
| rs75718910 | pathogenic | 0 | 0 | 0 |
| rs79843600 | pathogenic | 0 | 0 | 0 |
| rs75037667 | pathogenic | 1 | 1 | 1 |
| rs121913006 | pathogenic | 0 | 0 | 0 |
| rs397517925 | pathogenic | 1 | 1 | 1 |
| rs267606831 | pathogenic | 1 | 1 | 1 |
| rs121909667 | pathogenic | 1 | 1 | 1 |
| rs121909669 | pathogenic | 1 | 1 | 1 |
| rs121909668 | pathogenic | 1 | 1 | 1 |
| rs121909671 | pathogenic | 1 | 1 | 1 |
| rs534596036 | benign | 0 | 0 | 0 |
| rs144037629 | pathogenic | 0 | 0 | 0 |
| rs11546406 | benign | 0 | 0 | 0 |

| | | | | |
|---|---|---|---|---|
| rs143697995 | pathogenic | 0 | 0 | 0 |
| rs121908331 | pathogenic | 0 | 0 | 0 |
| rs149377346 | benign | 0 | 0 | 0 |
| rs104894086 | pathogenic | 1 | 1 | 0 |
| rs104894089 | pathogenic | 0 | 0 | 0 |
| rs104894090 | pathogenic | 1 | 1 | 0 |
| rs387907235 | pathogenic | 0 | 0 | 0 |
| rs137852689 | pathogenic | 0 | 0 | 0 |
| rs137852690 | pathogenic | 1 | 1 | 0 |
| rs104894087 | pathogenic | 0 | 0 | 0 |
| rs104894085 | pathogenic | 0 | 0 | 0 |
| rs79383654 | benign | 0 | 0 | 0 |
| rs121434239 | pathogenic | 0 | 0 | 0 |
| rs121434240 | pathogenic | 0 | 0 | 0 |
| rs121434236 | pathogenic | 0 | 0 | 0 |
| rs121434238 | pathogenic | 1 | 1 | 1 |
| rs121918107 | pathogenic | 1 | 1 | 1 |
| rs121918103 | pathogenic | 1 | 1 | 1 |
| rs121918104 | pathogenic | 1 | 1 | 1 |
| rs61747614 | benign | 1 | 1 | 1 |
| rs555329870 | benign | 0 | 0 | 0 |
| rs186094931 | benign | 0 | 0 | 0 |
| rs267606785 | pathogenic | 0 | 0 | 0 |
| rs34474955 | benign | 0 | 0 | 0 |
| rs1126497 | benign | 1 | 1 | 1 |
| rs397514661 | pathogenic | 0 | 0 | 0 |
| rs189732445 | benign | 0 | 0 | 0 |
| rs146685071 | benign | 0 | 0 | 0 |
| rs121908636 | pathogenic | 1 | 1 | 0 |
| rs121908643 | pathogenic | 0 | 0 | 0 |
| rs121908644 | pathogenic | 1 | 1 | 1 |
| rs575001023 | pathogenic | 1 | 1 | 1 |
| rs121908646 | pathogenic | 1 | 1 | 0 |
| rs121908638 | pathogenic | 1 | 1 | 1 |
| rs777828000 | pathogenic | 1 | 1 | 1 |
| rs74923032 | benign | 0 | 0 | 0 |
| rs192838388 | pathogenic | 1 | 1 | 1 |
| rs148918985 | pathogenic | 1 | 1 | 1 |
| rs773540764 | benign | 1 | 1 | 1 |
| rs370595480 | pathogenic | 1 | 1 | 1 |
| rs762387914 | pathogenic | 1 | 1 | 1 |
| rs121908642 | pathogenic | 1 | 1 | 1 |
| rs183276875 | pathogenic | 1 | 1 | 1 |
| rs121908648 | pathogenic | 1 | 1 | 1 |
| rs121908647 | pathogenic | 1 | 1 | 1 |
| rs121908640 | pathogenic | 1 | 1 | 1 |
| rs771937610 | pathogenic | 1 | 1 | 1 |
| rs121908641 | pathogenic | 1 | 1 | 1 |
| rs5745068 | benign | 0 | 0 | 0 |
| rs73155056 | benign | 1 | 1 | 1 |
| rs148788180 | benign | 0 | 0 | 0 |
| rs121918137 | pathogenic | 0 | 0 | 0 |
| rs41269593 | benign | 0 | 0 | 0 |
| rs11159 | benign | 1 | 1 | 0 |

| | | | | |
|---|---|---|---|---|
| rs114733570 | pathogenic | 1 | 1 | 0 |
| rs3211783 | benign | 0 | 0 | 0 |
| rs4762 | benign | 0 | 0 | 0 |
| rs74315283 | pathogenic | 0 | 0 | 0 |
| rs111033559 | pathogenic | 0 | 0 | 0 |
| rs111033560 | pathogenic | 1 | 1 | 1 |
| rs749962743 | benign | 0 | 0 | 0 |
| rs267606986 | pathogenic | 0 | 0 | 0 |
| rs34516635 | benign | 0 | 0 | 0 |
| rs117440569 | benign | 0 | 0 | 0 |
| rs61740877 | benign | 0 | 0 | 0 |
| rs121918352 | pathogenic | 0 | 0 | 0 |
| rs183974372 | benign | 0 | 0 | 0 |
| rs140511204 | benign | 1 | 1 | 0 |
| rs6040067 | benign | 0 | 0 | 0 |
| rs199674138 | benign | 0 | 0 | 0 |
| rs121918350 | pathogenic | 0 | 0 | 0 |
| rs138292988 | benign | 1 | 1 | 1 |
| rs145288947 | benign | 0 | 0 | 0 |
| rs369714529 | benign | 0 | 0 | 0 |
| rs200945419 | benign | 0 | 0 | 0 |
| rs116959874 | benign | 0 | 0 | 0 |
| rs141659620 | pathogenic | 1 | 1 | 0 |
| rs111475461 | benign | 0 | 0 | 0 |
| rs61755320 | pathogenic | 0 | 0 | 0 |
| rs748600162 | pathogenic | 1 | 1 | 1 |
| rs369227537 | pathogenic | 1 | 1 | 1 |
| rs372981030 | pathogenic | 1 | 1 | 1 |
| rs137854486 | pathogenic | 0 | 0 | 0 |
| rs137854487 | pathogenic | 0 | 0 | 0 |
| rs199905054 | pathogenic | 0 | 0 | 0 |
| rs771438170 | pathogenic | 0 | 0 | 0 |
| rs774949844 | pathogenic | 0 | 0 | 0 |
| rs4899145 | benign | 0 | 0 | 0 |
| rs200642525 | benign | 0 | 0 | 0 |
| rs190140598 | pathogenic | 0 | 0 | 0 |
| rs2250889 | benign | 0 | 0 | 0 |
| rs17577 | benign | 0 | 0 | 0 |
| rs142032681 | benign | 0 | 0 | 0 |
| rs144577963 | benign | 0 | 0 | 0 |
| rs61754623 | benign | 0 | 0 | 0 |
| rs139873200 | benign | 0 | 0 | 0 |
| rs532530906 | benign | 0 | 0 | 0 |
| rs145762716 | benign | 0 | 0 | 0 |
| rs34062020 | benign | 0 | 0 | 0 |
| rs35258161 | benign | 0 | 0 | 0 |
| rs149792649 | benign | 0 | 0 | 0 |
| rs78101774 | benign | 0 | 0 | 0 |
| rs766466297 | pathogenic | 0 | 1 | 0 |
| rs140467171 | pathogenic | 0 | 0 | 0 |
| rs141340466 | pathogenic | 0 | 0 | 0 |
| rs201555303 | pathogenic | 0 | 0 | 0 |
| rs143592405 | pathogenic | 0 | 0 | 0 |
| rs781249411 | pathogenic | 0 | 0 | 0 |

| | | | | |
|---|---|---|---|---|
| rs116644127 | benign | 0 | 0 | 0 |
| rs151146863 | benign | 1 | 1 | 1 |
| rs116440799 | benign | 0 | 0 | 0 |
| rs138958351 | benign | 0 | 0 | 0 |
| rs143319354 | benign | 0 | 0 | 0 |
| rs768669208 | pathogenic | 1 | 1 | 1 |
| rs28940289 | pathogenic | 1 | 1 | 0 |
| rs745656120 | pathogenic | 1 | 1 | 0 |
| rs387906987 | pathogenic | 0 | 0 | 0 |
| rs147240777 | benign | 0 | 0 | 0 |
| rs140945833 | benign | 0 | 0 | 0 |
| rs146614981 | benign | 0 | 0 | 0 |
| rs149878562 | benign | 0 | 0 | 0 |
| rs28938168 | pathogenic | 0 | 0 | 0 |
| rs33939927 | pathogenic | 1 | 1 | 1 |
| rs34995376 | pathogenic | 1 | 1 | 1 |
| rs201034481 | pathogenic | 1 | 1 | 1 |
| rs774069983 | benign | 0 | 0 | 0 |
| rs199422226 | pathogenic | 1 | 1 | 0 |
| rs199422225 | pathogenic | 1 | 1 | 1 |
| rs199422224 | pathogenic | 1 | 1 | 0 |
| rs149271416 | pathogenic | 1 | 1 | 0 |
| rs1801318 | benign | 1 | 1 | 1 |
| rs372691318 | pathogenic | 1 | 1 | 1 |
| rs147685849 | benign | 0 | 0 | 0 |
| rs387907199 | pathogenic | 0 | 0 | 0 |
| rs767122069 | pathogenic | 0 | 0 | 0 |
| rs2302427 | benign | 0 | 0 | 0 |
| rs201135441 | benign | 0 | 0 | 0 |
| rs201861847 | pathogenic | 0 | 0 | 0 |
| rs137852952 | pathogenic | 0 | 0 | 0 |
| rs16986560 | benign | 0 | 0 | 0 |
| rs147344332 | pathogenic | 1 | 1 | 1 |
| rs36027220 | pathogenic | 0 | 0 | 0 |
| rs192401229 | benign | 0 | 0 | 0 |
| rs121909106 | pathogenic | 1 | 1 | 0 |
| rs118204103 | pathogenic | 1 | 1 | 0 |
| rs118204104 | pathogenic | 1 | 1 | 0 |
| rs118204105 | pathogenic | 1 | 1 | 0 |
| rs118204106 | pathogenic | 0 | 0 | 0 |
| rs118204119 | pathogenic | 0 | 0 | 0 |
| rs118204107 | pathogenic | 0 | 0 | 0 |
| rs118204094 | pathogenic | 0 | 0 | 0 |
| rs118204098 | pathogenic | 1 | 1 | 0 |
| rs118204101 | pathogenic | 1 | 1 | 0 |
| rs118204096 | pathogenic | 1 | 1 | 0 |
| rs118204108 | pathogenic | 0 | 0 | 0 |
| rs536814318 | pathogenic | 0 | 0 | 0 |
| rs34413634 | pathogenic | 1 | 1 | 0 |
| rs118204100 | pathogenic | 0 | 0 | 0 |
| rs118204109 | pathogenic | 0 | 0 | 0 |
| rs118204116 | pathogenic | 1 | 1 | 0 |
| rs118204110 | pathogenic | 0 | 0 | 0 |
| rs118204099 | pathogenic | 0 | 0 | 0 |

| | | | | |
|---|---|---|---|---|
| rs118204111 | pathogenic | 0 | 0 | 0 |
| rs118204112 | pathogenic | 0 | 0 | 0 |
| rs118204113 | pathogenic | 0 | 0 | 0 |
| rs118204114 | pathogenic | 0 | 0 | 0 |
| rs118204115 | pathogenic | 0 | 0 | 0 |
| rs150428209 | benign | 0 | 0 | 0 |
| rs144949995 | benign | 0 | 0 | 0 |
| rs190023981 | benign | 0 | 0 | 0 |
| rs104894611 | pathogenic | 1 | 1 | 1 |
| rs28937580 | pathogenic | 1 | 1 | 1 |
| rs121908949 | pathogenic | 0 | 0 | 0 |
| rs104894612 | pathogenic | 1 | 1 | 1 |
| rs104894609 | pathogenic | 1 | 1 | 1 |
| rs104894610 | pathogenic | 1 | 1 | 1 |
| rs104894615 | pathogenic | 0 | 0 | 0 |
| rs104894603 | pathogenic | 0 | 0 | 0 |
| rs104894602 | pathogenic | 1 | 1 | 1 |
| rs121908948 | pathogenic | 1 | 1 | 1 |
| rs104894608 | pathogenic | 1 | 1 | 1 |
| rs371545673 | benign | 0 | 0 | 0 |
| rs199925238 | benign | 0 | 0 | 0 |
| rs146313451 | benign | 0 | 0 | 0 |
| rs121918193 | pathogenic | 1 | 1 | 0 |
| rs2228500 | benign | 0 | 0 | 0 |
| rs121918194 | pathogenic | 1 | 1 | 0 |
| rs121918196 | pathogenic | 0 | 0 | 0 |
| rs41291971 | benign | 0 | 0 | 0 |
| rs199473690 | pathogenic | 0 | 0 | 0 |
| rs2228613 | benign | 0 | 0 | 0 |
| rs267607130 | pathogenic | 1 | 1 | 1 |
| rs77615401 | benign | 1 | 1 | 1 |
| rs374618872 | benign | 1 | 1 | 1 |
| rs146165014 | benign | 0 | 0 | 0 |
| rs144586070 | benign | 0 | 0 | 0 |
| rs121918541 | pathogenic | 0 | 0 | 0 |
| rs142105943 | benign | 0 | 0 | 0 |
| rs121918540 | pathogenic | 0 | 0 | 0 |
| rs6791 | benign | 0 | 0 | 0 |
| rs61736587 | pathogenic | 0 | 0 | 0 |
| rs61736586 | benign | 0 | 0 | 0 |
| rs121917741 | pathogenic | 1 | 1 | 1 |
| rs41302885 | benign | 0 | 0 | 0 |
| rs141026028 | benign | 0 | 0 | 0 |
| rs193922493 | pathogenic | 0 | 0 | 0 |
| rs73598374 | benign | 0 | 0 | 0 |
| rs121908725 | pathogenic | 1 | 1 | 0 |
| rs199422328 | pathogenic | 0 | 1 | 0 |
| rs11555566 | benign | 1 | 1 | 1 |
| rs267606634 | pathogenic | 0 | 0 | 0 |
| rs121908714 | pathogenic | 1 | 1 | 0 |
| rs121908717 | pathogenic | 1 | 1 | 0 |
| rs267606635 | pathogenic | 1 | 1 | 0 |
| rs121908739 | pathogenic | 0 | 0 | 0 |
| rs121908735 | pathogenic | 0 | 1 | 0 |

| | | | | |
|---|---|---|---|---|
| rs121908716 | pathogenic | 0 | 1 | 0 |
| rs121908723 | pathogenic | 0 | 1 | 0 |
| rs121908721 | pathogenic | 0 | 0 | 0 |
| rs199422327 | pathogenic | 0 | 0 | 0 |
| rs121908715 | pathogenic | 0 | 0 | 0 |
| rs587783563 | pathogenic | 0 | 0 | 0 |
| rs104894780 | pathogenic | 0 | 0 | 0 |
| rs56030372 | pathogenic | 1 | 1 | 1 |
| rs104894782 | pathogenic | 1 | 1 | 1 |
| rs149495971 | benign | 0 | 0 | 0 |
| rs118203942 | pathogenic | 1 | 1 | 0 |
| rs118203939 | pathogenic | 0 | 0 | 0 |
| rs118203938 | pathogenic | 0 | 0 | 0 |
| rs118203943 | pathogenic | 0 | 0 | 0 |
| rs118203940 | pathogenic | 0 | 0 | 0 |
| rs727503809 | pathogenic | 0 | 0 | 0 |
| rs1065757 | benign | 0 | 0 | 0 |
| rs1071598 | benign | 0 | 0 | 0 |
| rs25414 | benign | 0 | 0 | 0 |
| rs118203944 | pathogenic | 0 | 0 | 0 |
| rs118203941 | pathogenic | 0 | 0 | 0 |
| rs144222167 | benign | 0 | 0 | 0 |
| rs201101343 | pathogenic | 0 | 0 | 0 |
| rs144601717 | benign | 0 | 0 | 0 |
| rs267607129 | pathogenic | 0 | 0 | 0 |
| rs104894729 | pathogenic | 0 | 0 | 0 |
| rs104894727 | pathogenic | 0 | 0 | 0 |
| rs104894725 | pathogenic | 0 | 0 | 0 |
| rs104894515 | pathogenic | 1 | 1 | 1 |
| rs74688377 | benign | 0 | 0 | 0 |
| rs121908513 | pathogenic | 1 | 1 | 0 |
| rs121908509 | pathogenic | 0 | 0 | 0 |
| rs121908514 | pathogenic | 1 | 1 | 0 |
| rs121908512 | pathogenic | 1 | 1 | 0 |
| rs121908510 | pathogenic | 0 | 0 | 0 |
| rs121908511 | pathogenic | 1 | 1 | 1 |
| rs121908518 | pathogenic | 1 | 1 | 1 |
| rs34358665 | benign | 0 | 0 | 0 |
| rs147568834 | benign | 0 | 0 | 0 |
| rs201717599 | benign | 0 | 0 | 0 |
| rs61744487 | benign | 0 | 0 | 0 |
| rs370627877 | pathogenic | 1 | 1 | 1 |
| rs368593151 | pathogenic | 1 | 1 | 1 |
| rs114363936 | benign | 0 | 0 | 0 |
| rs137853333 | pathogenic | 0 | 0 | 0 |
| rs137853080 | pathogenic | 1 | 1 | 1 |
| rs137853077 | pathogenic | 1 | 1 | 1 |
| rs780749732 | benign | 1 | 1 | 1 |
| rs137853081 | pathogenic | 1 | 1 | 0 |
| rs137853078 | pathogenic | 0 | 0 | 0 |
| rs121913315 | pathogenic | 1 | 1 | 1 |
| rs752513525 | pathogenic | 1 | 1 | 1 |
| rs104894918 | pathogenic | 0 | 0 | 0 |
| rs104894917 | pathogenic | 0 | 0 | 0 |

| | | | | |
|---|---|---|---|---|
| rs104894919 | pathogenic | 0 | 0 | 0 |
| rs28939672 | pathogenic | 0 | 0 | 0 |
| rs104894173 | pathogenic | 0 | 0 | 0 |
| rs104894179 | pathogenic | 0 | 0 | 0 |
| rs104894174 | pathogenic | 1 | 1 | 0 |
| rs104894178 | pathogenic | 1 | 1 | 0 |
| rs28939714 | pathogenic | 1 | 1 | 1 |
| rs104894270 | pathogenic | 1 | 1 | 1 |
| rs3765571 | benign | 0 | 0 | 0 |
| rs752669339 | pathogenic | 1 | 1 | 1 |
| rs537043237 | pathogenic | 1 | 1 | 1 |
| rs774047299 | pathogenic | 1 | 1 | 1 |
| rs121434583 | pathogenic | 1 | 1 | 1 |
| rs121907927 | pathogenic | 0 | 0 | 0 |
| rs121907925 | pathogenic | 0 | 1 | 0 |
| rs746212325 | pathogenic | 0 | 0 | 0 |
| rs754637179 | pathogenic | 0 | 0 | 0 |
| rs780882740 | pathogenic | 0 | 0 | 0 |
| rs121918130 | pathogenic | 0 | 0 | 0 |
| rs121918129 | pathogenic | 0 | 0 | 0 |
| rs13297509 | pathogenic | 0 | 0 | 0 |
| rs746867724 | pathogenic | 0 | 0 | 0 |
| rs121918128 | pathogenic | 0 | 0 | 0 |
| rs752300607 | pathogenic | 0 | 0 | 0 |
| rs121964860 | pathogenic | 0 | 0 | 0 |
| rs45466197 | pathogenic | 1 | 1 | 1 |
| rs3730238 | benign | 1 | 1 | 1 |
| rs730881125 | pathogenic | 1 | 1 | 1 |
| rs121964861 | pathogenic | 1 | 1 | 1 |
| rs397516484 | pathogenic | 0 | 0 | 0 |
| rs367785431 | pathogenic | 0 | 0 | 0 |
| rs141121678 | pathogenic | 0 | 0 | 0 |
| rs12720263 | benign | 0 | 0 | 0 |
| rs2304256 | benign | 0 | 0 | 0 |
| rs2304255 | benign | 0 | 0 | 0 |
| rs121909045 | pathogenic | 0 | 0 | 0 |
| rs397508796 | pathogenic | 0 | 0 | 0 |
| rs151020603 | pathogenic | 0 | 0 | 0 |
| rs77284892 | pathogenic | 0 | 0 | 0 |
| rs368505753 | pathogenic | 0 | 0 | 0 |
| rs115545701 | pathogenic | 0 | 0 | 0 |
| rs121908749 | pathogenic | 0 | 0 | 0 |
| rs75961395 | pathogenic | 0 | 0 | 0 |
| rs121908750 | pathogenic | 0 | 0 | 0 |
| rs121908751 | pathogenic | 0 | 0 | 0 |
| rs397508464 | pathogenic | 0 | 0 | 0 |
| rs397508461 | pathogenic | 0 | 0 | 0 |
| rs121909031 | pathogenic | 0 | 0 | 0 |
| rs113993958 | pathogenic | 0 | 0 | 0 |
| rs77834169 | pathogenic | 0 | 0 | 0 |
| rs78655421 | pathogenic | 0 | 0 | 0 |
| rs141723617 | benign | 0 | 0 | 0 |
| rs35516286 | benign | 0 | 0 | 0 |
| rs1800079 | pathogenic | 1 | 1 | 0 |

| | | | | |
|---|---|---|---|---|
| rs80282562 | pathogenic | 0 | 0 | 0 |
| rs121908752 | pathogenic | 0 | 0 | 0 |
| rs556880586 | pathogenic | 0 | 0 | 0 |
| rs121908760 | pathogenic | 0 | 0 | 0 |
| rs397508350 | pathogenic | 0 | 0 | 0 |
| rs374946172 | pathogenic | 0 | 0 | 0 |
| rs397508387 | pathogenic | 0 | 0 | 0 |
| rs201386642 | pathogenic | 0 | 0 | 0 |
| rs121909012 | pathogenic | 0 | 0 | 0 |
| rs121909008 | pathogenic | 0 | 0 | 0 |
| rs149790377 | pathogenic | 0 | 0 | 0 |
| rs397508442 | pathogenic | 0 | 0 | 0 |
| rs121909035 | pathogenic | 0 | 0 | 0 |
| rs386134230 | pathogenic | 0 | 0 | 0 |
| rs1800112 | benign | 0 | 0 | 0 |
| rs144055758 | pathogenic | 0 | 0 | 0 |
| rs142394380 | pathogenic | 0 | 0 | 0 |
| rs121909036 | pathogenic | 0 | 0 | 0 |
| rs78194216 | pathogenic | 0 | 0 | 0 |
| rs121909019 | pathogenic | 0 | 0 | 0 |
| rs121909020 | pathogenic | 0 | 0 | 0 |
| rs200321110 | pathogenic | 0 | 0 | 0 |
| rs202179988 | pathogenic | 0 | 0 | 0 |
| rs121909037 | pathogenic | 0 | 0 | 0 |
| rs186045772 | pathogenic | 0 | 0 | 0 |
| rs139304906 | pathogenic | 0 | 0 | 0 |
| rs78802634 | pathogenic | 0 | 0 | 0 |
| rs121908761 | pathogenic | 0 | 0 | 0 |
| rs36210737 | pathogenic | 0 | 0 | 0 |
| rs139729994 | pathogenic | 0 | 0 | 0 |
| rs79850223 | pathogenic | 0 | 0 | 0 |
| rs1800120 | benign | 0 | 0 | 0 |
| rs74767530 | pathogenic | 0 | 0 | 0 |
| rs374705585 | pathogenic | 0 | 0 | 0 |
| rs1136287 | benign | 0 | 0 | 0 |
| rs6828 | benign | 0 | 0 | 0 |
| rs104894856 | pathogenic | 0 | 0 | 0 |
| rs104894860 | pathogenic | 0 | 0 | 0 |
| rs61736892 | pathogenic | 0 | 0 | 0 |
| rs145807417 | benign | 0 | 0 | 0 |
| rs104894853 | pathogenic | 0 | 0 | 0 |
| rs199422227 | pathogenic | 0 | 0 | 0 |
| rs113993946 | pathogenic | 0 | 0 | 0 |
| rs199422231 | pathogenic | 0 | 0 | 0 |
| rs199422228 | pathogenic | 0 | 0 | 0 |
| rs142195196 | benign | 0 | 1 | 0 |
| rs148656104 | benign | 1 | 1 | 1 |
| rs141944844 | benign | 0 | 0 | 0 |
| rs104894342 | pathogenic | 0 | 0 | 0 |
| rs104894343 | pathogenic | 0 | 0 | 0 |
| rs104894344 | pathogenic | 0 | 0 | 0 |
| rs573322878 | benign | 0 | 0 | 0 |
| rs104894347 | pathogenic | 0 | 0 | 0 |
| rs28937882 | pathogenic | 0 | 0 | 0 |

| | | | | |
|---|---|---|---|---|
| rs193922702 | pathogenic | 0 | 0 | 0 |
| rs200786104 | benign | 0 | 0 | 0 |
| rs690016544 | pathogenic | 1 | 1 | 1 |
| rs28943592 | benign | 0 | 0 | 0 |
| rs28943594 | benign | 0 | 0 | 0 |
| rs121908957 | pathogenic | 0 | 0 | 0 |
| rs115408010 | benign | 1 | 1 | 1 |
| rs77591659 | benign | 0 | 0 | 0 |
| rs121918511 | pathogenic | 0 | 0 | 0 |
| rs538359948 | benign | 0 | 0 | 0 |
| rs76708142 | benign | 0 | 0 | 0 |
| rs121907913 | pathogenic | 1 | 1 | 0 |
| rs121907920 | pathogenic | 0 | 0 | 0 |
| rs121907923 | pathogenic | 1 | 1 | 0 |
| rs121907928 | pathogenic | 1 | 1 | 0 |
| rs121907919 | pathogenic | 1 | 1 | 0 |
| rs121907918 | pathogenic | 1 | 1 | 0 |
| rs74315493 | pathogenic | 1 | 1 | 1 |
| rs74315494 | pathogenic | 1 | 1 | 1 |
| rs145446060 | benign | 1 | 1 | 1 |
| rs550409143 | benign | 1 | 1 | 0 |

| compound_binding_site | cofactor_binding_site | nucleic_binding_site | ion_binding_site |
|---|---|---|---|
| 1 | 0 | 0 | 0 |
| 1 | 0 | 0 | 0 |
| 0 | 0 | 0 | 0 |
| 1 | 0 | 0 | 0 |
| 0 | 0 | 0 | 0 |
| 0 | 0 | 0 | 0 |
| 0 | 0 | 0 | 0 |
| 0 | 0 | 0 | 0 |
| 1 | 0 | 0 | 1 |
| 1 | 0 | 0 | 0 |
| 0 | 0 | 0 | 1 |
| 1 | 0 | 1 | 0 |
| 0 | 0 | 0 | 0 |
| 0 | 0 | 0 | 0 |
| 0 | 0 | 0 | 0 |
| 0 | 0 | 0 | 0 |
| 0 | 0 | 0 | 0 |
| 1 | 0 | 0 | 1 |
| 1 | 0 | 1 | 0 |
| 0 | 0 | 0 | 0 |
| 1 | 0 | 0 | 1 |
| 1 | 0 | 0 | 0 |
| 1 | 0 | 0 | 0 |
| 1 | 0 | 0 | 0 |
| 0 | 0 | 0 | 0 |
| 0 | 0 | 0 | 0 |
| 1 | 0 | 0 | 1 |
| 0 | 0 | 0 | 0 |
| 0 | 0 | 0 | 0 |
| 0 | 0 | 0 | 0 |
| 0 | 0 | 0 | 0 |
| 1 | 0 | 0 | 0 |
| 1 | 0 | 0 | 1 |
| 1 | 0 | 0 | 0 |
| 0 | 0 | 0 | 0 |
| 0 | 0 | 0 | 0 |
| 0 | 0 | 0 | 0 |
| 1 | 0 | 0 | 1 |
| 1 | 0 | 0 | 1 |
| 1 | 0 | 0 | 1 |
| 1 | 0 | 0 | 0 |
| 1 | 0 | 0 | 0 |
| 1 | 0 | 0 | 0 |
| 1 | 0 | 0 | 0 |
| 1 | 0 | 0 | 0 |
| 1 | 0 | 0 | 0 |
| 1 | 0 | 0 | 0 |
| 1 | 0 | 0 | 0 |
| 1 | 0 | 0 | 0 |
| 1 | 0 | 0 | 0 |
| 1 | 0 | 0 | 0 |
| 1 | 0 | 0 | 0 |
| 1 | 0 | 0 | 0 |

| | | | |
|---|---|---|---|
| 1 | 0 | 0 | 1 |
| 1 | 0 | 0 | 0 |
| 1 | 0 | 0 | 0 |
| 1 | 0 | 0 | 0 |
| 0 | 0 | 0 | 0 |
| 1 | 0 | 0 | 0 |
| 0 | 0 | 0 | 0 |
| 1 | 0 | 0 | 0 |
| 1 | 0 | 0 | 0 |
| 1 | 0 | 0 | 1 |
| 1 | 0 | 0 | 0 |
| 1 | 0 | 0 | 0 |
| 1 | 0 | 0 | 0 |
| 1 | 0 | 0 | 0 |
| 1 | 0 | 0 | 0 |
| 1 | 0 | 0 | 0 |
| 1 | 0 | 0 | 0 |
| 1 | 0 | 0 | 1 |
| 1 | 0 | 0 | 1 |
| 1 | 1 | 0 | 1 |
| 1 | 0 | 0 | 0 |
| 1 | 0 | 0 | 0 |
| 1 | 0 | 0 | 0 |
| 1 | 0 | 0 | 0 |
| 1 | 1 | 0 | 0 |
| 1 | 0 | 0 | 0 |
| 1 | 0 | 0 | 0 |
| 1 | 0 | 0 | 0 |
| 1 | 0 | 0 | 0 |
| 1 | 0 | 0 | 0 |
| 1 | 1 | 0 | 0 |
| 1 | 1 | 0 | 0 |
| 1 | 1 | 0 | 0 |
| 1 | 1 | 0 | 0 |
| 1 | 1 | 0 | 0 |
| 1 | 0 | 0 | 0 |
| 1 | 1 | 0 | 0 |
| 1 | 0 | 0 | 0 |
| 1 | 0 | 0 | 0 |
| 1 | 1 | 0 | 0 |
| 0 | 0 | 0 | 0 |
| 0 | 0 | 0 | 0 |
| 0 | 0 | 0 | 0 |
| 0 | 0 | 0 | 0 |
| 0 | 0 | 0 | 0 |
| 0 | 0 | 0 | 0 |
| 0 | 0 | 0 | 0 |
| 1 | 0 | 0 | 1 |
| 0 | 0 | 0 | 0 |
| 1 | 0 | 0 | 0 |
| 1 | 0 | 0 | 0 |
| 0 | 0 | 0 | 0 |
| 0 | 0 | 0 | 0 |
| 0 | 0 | 0 | 0 |

| | | | |
|---|---|---|---|
| 0 | 0 | 0 | 0 |
| 0 | 0 | 0 | 0 |
| 0 | 0 | 0 | 0 |
| 1 | 0 | 0 | 0 |
| 1 | 0 | 0 | 0 |
| 1 | 0 | 0 | 0 |
| 0 | 0 | 0 | 0 |
| 0 | 0 | 1 | 0 |
| 0 | 0 | 1 | 0 |
| 0 | 0 | 1 | 0 |
| 0 | 0 | 1 | 0 |
| 0 | 0 | 1 | 0 |
| 0 | 0 | 0 | 0 |
| 0 | 0 | 0 | 0 |
| 1 | 0 | 0 | 0 |
| 0 | 0 | 0 | 0 |
| 0 | 0 | 0 | 0 |
| 1 | 0 | 0 | 0 |
| 1 | 0 | 0 | 0 |
| 1 | 0 | 0 | 0 |
| 1 | 0 | 0 | 0 |
| 1 | 1 | 0 | 0 |
| 1 | 0 | 0 | 0 |
| 1 | 0 | 0 | 0 |
| 0 | 0 | 0 | 0 |
| 0 | 0 | 0 | 0 |
| 1 | 0 | 0 | 0 |
| 1 | 1 | 0 | 0 |
| 1 | 0 | 0 | 0 |
| 1 | 0 | 0 | 0 |
| 1 | 1 | 0 | 0 |
| 1 | 1 | 0 | 0 |
| 1 | 1 | 0 | 0 |
| 1 | 1 | 0 | 0 |
| 1 | 1 | 0 | 0 |
| 1 | 1 | 0 | 0 |
| 1 | 1 | 0 | 0 |
| 1 | 1 | 0 | 0 |
| 1 | 1 | 0 | 0 |
| 1 | 1 | 0 | 0 |
| 1 | 1 | 0 | 0 |
| 1 | 1 | 0 | 0 |
| 1 | 0 | 0 | 0 |
| 1 | 0 | 0 | 0 |
| 1 | 0 | 0 | 0 |
| 1 | 1 | 0 | 0 |
| 1 | 0 | 0 | 0 |
| 1 | 1 | 1 | 0 |
| 1 | 1 | 1 | 0 |
| 1 | 1 | 0 | 0 |
| 1 | 1 | 0 | 0 |
| 1 | 0 | 0 | 0 |
| 1 | 0 | 0 | 0 |
| 0 | 0 | 0 | 0 |

| | | | |
|---|---|---|---|
| 0 | 0 | 0 | 0 |
| 0 | 0 | 0 | 0 |
| 0 | 0 | 0 | 0 |
| 1 | 0 | 0 | 0 |
| 1 | 0 | 0 | 0 |
| 0 | 0 | 0 | 0 |
| 0 | 0 | 0 | 0 |
| 0 | 0 | 0 | 0 |
| 0 | 0 | 0 | 0 |
| 0 | 0 | 0 | 0 |
| 0 | 0 | 0 | 0 |
| 0 | 0 | 0 | 0 |
| 0 | 0 | 0 | 0 |
| 0 | 0 | 0 | 0 |
| 0 | 0 | 0 | 0 |
| 0 | 0 | 0 | 0 |
| 0 | 0 | 0 | 0 |
| 0 | 0 | 0 | 0 |
| 0 | 0 | 0 | 0 |
| 0 | 0 | 0 | 0 |
| 0 | 0 | 0 | 0 |
| 0 | 0 | 0 | 0 |
| 1 | 0 | 0 | 0 |
| 1 | 0 | 0 | 0 |
| 0 | 0 | 0 | 0 |
| 0 | 0 | 0 | 0 |
| 1 | 0 | 0 | 0 |
| 0 | 0 | 0 | 0 |
| 0 | 0 | 0 | 0 |
| 0 | 0 | 0 | 0 |
| 0 | 0 | 0 | 0 |
| 0 | 0 | 0 | 0 |
| 0 | 0 | 0 | 0 |
| 0 | 0 | 0 | 0 |
| 1 | 0 | 0 | 0 |
| 0 | 0 | 0 | 0 |
| 0 | 0 | 0 | 0 |
| 0 | 0 | 0 | 0 |
| 0 | 0 | 0 | 0 |
| 0 | 0 | 0 | 0 |
| 0 | 0 | 0 | 0 |
| 0 | 0 | 0 | 0 |
| 0 | 0 | 0 | 0 |
| 0 | 0 | 0 | 0 |
| 0 | 0 | 0 | 0 |
| 0 | 0 | 0 | 0 |
| 0 | 0 | 1 | 0 |
| 1 | 0 | 0 | 0 |
| 1 | 0 | 0 | 0 |
| 1 | 1 | 0 | 0 |
| 1 | 0 | 0 | 0 |
| 0 | 0 | 0 | 0 |
| 0 | 0 | 0 | 0 |
| 1 | 0 | 0 | 0 |

| | | | |
|---|---|---|---|
| 1 | 0 | 0 | 0 |
| 1 | 0 | 0 | 0 |
| 1 | 0 | 0 | 0 |
| 0 | 0 | 0 | 0 |
| 0 | 0 | 0 | 0 |
| 0 | 0 | 0 | 0 |
| 1 | 0 | 0 | 0 |
| 1 | 0 | 0 | 0 |
| 0 | 0 | 0 | 0 |
| 0 | 0 | 0 | 0 |
| 1 | 0 | 0 | 0 |
| 0 | 0 | 0 | 0 |
| 1 | 0 | 0 | 0 |
| 0 | 0 | 0 | 0 |
| 0 | 0 | 0 | 0 |
| 0 | 0 | 0 | 0 |
| 0 | 0 | 0 | 0 |
| 0 | 0 | 0 | 0 |
| 0 | 0 | 0 | 0 |
| 0 | 0 | 0 | 0 |
| 0 | 0 | 0 | 0 |
| 0 | 0 | 0 | 0 |
| 0 | 0 | 0 | 0 |
| 0 | 0 | 0 | 0 |
| 0 | 0 | 0 | 0 |
| 0 | 0 | 0 | 0 |
| 1 | 0 | 0 | 0 |
| 1 | 0 | 0 | 0 |
| 1 | 0 | 0 | 0 |
| 1 | 0 | 0 | 0 |
| 1 | 0 | 0 | 0 |
| 1 | 0 | 0 | 0 |
| 0 | 0 | 0 | 0 |
| 0 | 0 | 0 | 0 |
| 1 | 0 | 0 | 0 |
| 0 | 0 | 0 | 0 |
| 0 | 0 | 0 | 0 |
| 0 | 0 | 0 | 0 |
| 0 | 0 | 0 | 0 |
| 0 | 0 | 0 | 0 |
| 0 | 0 | 0 | 0 |
| 0 | 0 | 0 | 0 |
| 0 | 0 | 0 | 0 |
| 0 | 0 | 0 | 0 |
| 1 | 1 | 0 | 0 |
| 1 | 1 | 0 | 0 |
| 1 | 1 | 0 | 1 |
| 1 | 1 | 0 | 1 |
| 1 | 1 | 0 | 1 |
| 1 | 1 | 0 | 1 |
| 1 | 1 | 0 | 0 |
| 1 | 1 | 0 | 1 |
| 1 | 1 | 0 | 1 |
| 1 | 1 | 0 | 1 |

| | | | |
|---|---|---|---|
| 1 | 1 | 0 | 1 |
| 1 | 1 | 0 | 1 |
| 1 | 1 | 0 | 1 |
| 1 | 1 | 0 | 1 |
| 1 | 1 | 0 | 1 |
| 1 | 0 | 0 | 0 |
| 0 | 0 | 0 | 0 |
| 0 | 0 | 0 | 0 |
| 0 | 0 | 0 | 0 |
| 1 | 0 | 0 | 0 |
| 0 | 0 | 1 | 0 |
| 0 | 0 | 0 | 0 |
| 0 | 0 | 0 | 0 |
| 0 | 0 | 0 | 0 |
| 0 | 0 | 0 | 0 |
| 0 | 0 | 0 | 0 |
| 0 | 0 | 0 | 0 |
| 0 | 0 | 0 | 0 |
| 0 | 0 | 0 | 0 |
| 0 | 0 | 0 | 0 |
| 0 | 0 | 0 | 0 |
| 0 | 0 | 0 | 0 |
| 0 | 0 | 0 | 0 |
| 0 | 0 | 0 | 0 |
| 0 | 0 | 0 | 0 |
| 1 | 1 | 0 | 0 |
| 1 | 1 | 0 | 0 |
| 1 | 1 | 0 | 0 |
| 0 | 1 | 0 | 0 |
| 1 | 0 | 0 | 0 |
| 0 | 0 | 0 | 0 |
| 0 | 0 | 0 | 0 |
| 0 | 0 | 0 | 0 |
| 1 | 0 | 0 | 0 |
| 1 | 1 | 0 | 0 |
| 0 | 0 | 0 | 0 |
| 0 | 0 | 0 | 0 |
| 0 | 0 | 0 | 0 |
| 0 | 0 | 0 | 0 |
| 1 | 0 | 0 | 0 |
| 0 | 0 | 0 | 0 |
| 0 | 0 | 0 | 0 |
| 0 | 0 | 0 | 0 |
| 0 | 0 | 0 | 0 |
| 0 | 0 | 0 | 0 |
| 0 | 1 | 0 | 0 |
| 0 | 1 | 0 | 0 |
| 0 | 0 | 0 | 0 |
| 0 | 0 | 0 | 0 |
| 0 | 1 | 0 | 0 |
| 0 | 0 | 0 | 0 |
| 0 | 0 | 0 | 0 |
| 0 | 1 | 0 | 0 |
| 0 | 0 | 0 | 0 |

| | | | |
|---|---|---|---|
| 1 | 1 | 0 | 0 |
| 0 | 0 | 0 | 0 |
| 0 | 0 | 0 | 0 |
| 0 | 0 | 0 | 0 |
| 0 | 0 | 0 | 0 |
| 0 | 0 | 0 | 0 |
| 0 | 0 | 0 | 0 |
| 0 | 0 | 0 | 0 |
| 0 | 0 | 0 | 0 |
| 0 | 0 | 0 | 0 |
| 0 | 0 | 0 | 0 |
| 0 | 0 | 0 | 0 |
| 0 | 0 | 0 | 0 |
| 0 | 0 | 0 | 0 |
| 0 | 1 | 0 | 0 |
| 0 | 0 | 0 | 0 |
| 0 | 1 | 0 | 0 |
| 0 | 1 | 0 | 0 |
| 0 | 0 | 0 | 0 |
| 0 | 1 | 0 | 0 |
| 1 | 0 | 0 | 1 |
| 1 | 1 | 0 | 0 |
| 1 | 0 | 0 | 0 |
| 1 | 0 | 0 | 1 |
| 0 | 0 | 0 | 0 |
| 0 | 0 | 0 | 0 |
| 0 | 0 | 0 | 0 |
| 0 | 0 | 0 | 0 |
| 1 | 0 | 0 | 0 |
| 0 | 0 | 0 | 0 |
| 1 | 0 | 0 | 0 |
| 0 | 0 | 0 | 0 |
| 0 | 0 | 0 | 0 |
| 0 | 0 | 0 | 0 |
| 0 | 0 | 0 | 0 |
| 0 | 0 | 0 | 0 |
| 0 | 0 | 0 | 0 |
| 1 | 1 | 0 | 0 |
| 1 | 1 | 0 | 0 |
| 1 | 1 | 0 | 0 |
| 0 | 0 | 0 | 0 |
| 0 | 0 | 0 | 0 |
| 0 | 0 | 0 | 0 |
| 1 | 0 | 0 | 0 |
| 1 | 0 | 0 | 1 |
| 1 | 0 | 0 | 1 |
| 1 | 0 | 0 | 0 |
| 1 | 0 | 0 | 1 |
| 1 | 0 | 0 | 0 |
| 0 | 0 | 0 | 0 |
| 1 | 1 | 0 | 0 |
| 1 | 1 | 0 | 0 |
| 0 | 0 | 0 | 0 |
| 0 | 0 | 0 | 0 |

| | | | |
|---|---|---|---|
| 1 | 0 | 0 | 0 |
| 1 | 0 | 0 | 0 |
| 1 | 0 | 0 | 0 |
| 0 | 0 | 0 | 0 |
| 0 | 0 | 0 | 0 |
| 1 | 1 | 0 | 1 |
| 1 | 1 | 0 | 1 |
| 1 | 1 | 0 | 1 |
| 1 | 1 | 0 | 1 |
| 1 | 1 | 0 | 1 |
| 1 | 1 | 0 | 1 |
| 1 | 1 | 0 | 1 |
| 1 | 1 | 0 | 0 |
| 1 | 1 | 0 | 0 |
| 1 | 1 | 1 | 1 |
| 0 | 0 | 0 | 0 |
| 1 | 1 | 0 | 0 |
| 1 | 1 | 0 | 0 |
| 0 | 0 | 0 | 0 |
| 0 | 0 | 0 | 0 |
| 0 | 0 | 0 | 0 |
| 1 | 0 | 0 | 0 |
| 1 | 0 | 0 | 0 |
| 0 | 0 | 0 | 0 |
| 1 | 0 | 0 | 0 |
| 1 | 0 | 0 | 0 |
| 1 | 0 | 0 | 0 |
| 0 | 0 | 0 | 0 |
| 0 | 0 | 0 | 0 |
| 0 | 0 | 0 | 0 |
| 0 | 0 | 0 | 0 |
| 0 | 0 | 0 | 0 |
| 0 | 0 | 0 | 0 |
| 0 | 0 | 0 | 0 |
| 0 | 0 | 0 | 0 |
| 0 | 0 | 0 | 0 |
| 0 | 0 | 0 | 0 |
| 0 | 0 | 0 | 0 |
| 0 | 0 | 0 | 0 |
| 0 | 0 | 0 | 0 |
| 0 | 0 | 0 | 0 |
| 0 | 0 | 0 | 0 |
| 0 | 0 | 0 | 0 |
| 0 | 0 | 0 | 0 |
| 0 | 0 | 0 | 0 |
| 1 | 0 | 0 | 0 |
| 1 | 0 | 0 | 0 |
| 1 | 0 | 0 | 0 |
| 1 | 0 | 0 | 0 |
| 0 | 0 | 0 | 0 |
| 0 | 0 | 0 | 0 |
| 0 | 0 | 0 | 0 |
| 0 | 0 | 0 | 0 |
| 0 | 0 | 0 | 0 |

| | | | |
|---|---|---|---|
| 0 | 0 | 0 | 0 |
| 0 | 0 | 0 | 0 |
| 0 | 0 | 0 | 0 |
| 0 | 0 | 0 | 0 |
| 0 | 0 | 0 | 0 |
| 0 | 0 | 0 | 0 |
| 0 | 0 | 0 | 0 |
| 1 | 0 | 0 | 1 |
| 0 | 0 | 0 | 0 |
| 0 | 0 | 0 | 0 |
| 1 | 0 | 0 | 0 |
| 0 | 0 | 0 | 0 |
| 0 | 0 | 0 | 0 |
| 0 | 0 | 0 | 0 |
| 0 | 0 | 0 | 0 |
| 1 | 0 | 0 | 0 |
| 0 | 0 | 0 | 0 |
| 0 | 0 | 0 | 0 |
| 0 | 0 | 0 | 0 |
| 0 | 0 | 0 | 0 |
| 0 | 0 | 0 | 0 |
| 0 | 0 | 0 | 0 |
| 0 | 0 | 0 | 0 |
| 1 | 1 | 0 | 1 |
| 0 | 0 | 0 | 0 |
| 0 | 0 | 0 | 0 |
| 0 | 0 | 0 | 0 |
| 1 | 1 | 0 | 1 |
| 1 | 1 | 0 | 0 |
| 0 | 0 | 0 | 0 |
| 1 | 0 | 0 | 0 |
| 1 | 1 | 0 | 0 |
| 0 | 0 | 0 | 0 |
| 1 | 0 | 0 | 0 |
| 1 | 1 | 0 | 0 |
| 1 | 0 | 0 | 0 |
| 1 | 1 | 0 | 1 |
| 1 | 1 | 0 | 1 |
| 1 | 0 | 0 | 0 |
| 0 | 0 | 0 | 0 |
| 0 | 0 | 0 | 0 |
| 0 | 0 | 0 | 0 |
| 0 | 0 | 0 | 0 |
| 0 | 0 | 0 | 0 |
| 0 | 0 | 0 | 0 |
| 0 | 0 | 0 | 0 |
| 0 | 0 | 0 | 0 |
| 0 | 0 | 0 | 0 |
| 0 | 0 | 0 | 0 |
| 0 | 0 | 0 | 0 |
| 0 | 0 | 0 | 0 |
| 0 | 0 | 0 | 0 |
| 0 | 0 | 0 | 0 |
| 0 | 0 | 0 | 0 |

| | | | |
|---|---|---|---|
| 0 | 0 | 0 | 0 |
| 0 | 0 | 0 | 0 |
| 0 | 0 | 0 | 0 |
| 0 | 0 | 0 | 0 |
| 0 | 0 | 0 | 0 |
| 0 | 0 | 0 | 0 |
| 0 | 0 | 0 | 0 |
| 0 | 0 | 0 | 0 |
| 0 | 0 | 0 | 0 |
| 0 | 0 | 0 | 0 |
| 0 | 0 | 0 | 0 |
| 0 | 0 | 0 | 0 |
| 0 | 0 | 0 | 0 |
| 1 | 1 | 0 | 0 |
| 0 | 0 | 0 | 0 |
| 1 | 0 | 0 | 0 |
| 0 | 0 | 0 | 1 |
| 0 | 0 | 0 | 0 |
| 0 | 0 | 0 | 0 |
| 0 | 0 | 0 | 0 |
| 0 | 0 | 0 | 0 |
| 0 | 0 | 0 | 0 |
| 0 | 0 | 0 | 0 |
| 0 | 0 | 0 | 0 |
| 0 | 0 | 0 | 0 |
| 0 | 0 | 0 | 0 |
| 0 | 0 | 0 | 0 |
| 0 | 0 | 0 | 0 |
| 0 | 0 | 0 | 0 |
| 0 | 0 | 0 | 0 |
| 1 | 0 | 0 | 0 |
| 1 | 0 | 0 | 0 |
| 0 | 0 | 0 | 0 |
| 0 | 0 | 0 | 0 |
| 0 | 0 | 0 | 0 |
| 0 | 0 | 0 | 0 |
| 0 | 0 | 0 | 0 |
| 0 | 0 | 0 | 0 |
| 0 | 0 | 0 | 0 |
| 0 | 0 | 0 | 0 |
| 1 | 0 | 0 | 0 |
| 1 | 0 | 0 | 0 |
| 1 | 0 | 0 | 0 |
| 1 | 0 | 0 | 0 |
| 0 | 0 | 0 | 0 |
| 1 | 0 | 0 | 0 |
| 0 | 0 | 0 | 0 |
| 0 | 0 | 0 | 0 |
| 1 | 0 | 0 | 0 |
| 1 | 0 | 0 | 0 |
| 0 | 0 | 0 | 0 |
| 0 | 0 | 0 | 0 |
| 0 | 0 | 0 | 0 |
| 0 | 0 | 0 | 0 |

| | | | |
|---|---|---|---|
| 0 | 0 | 0 | 0 |
| 0 | 0 | 0 | 0 |
| 0 | 0 | 0 | 0 |
| 0 | 0 | 0 | 0 |
| 0 | 0 | 0 | 0 |
| 0 | 0 | 0 | 0 |
| 0 | 0 | 0 | 0 |
| 0 | 0 | 0 | 0 |
| 0 | 0 | 0 | 0 |
| 0 | 0 | 0 | 0 |
| 0 | 0 | 0 | 0 |
| 0 | 0 | 0 | 0 |
| 0 | 0 | 0 | 0 |
| 0 | 0 | 0 | 0 |
| 0 | 0 | 0 | 0 |
| 1 | 0 | 0 | 1 |
| 0 | 0 | 0 | 0 |
| 0 | 0 | 0 | 0 |
| 0 | 0 | 0 | 0 |
| 0 | 0 | 0 | 0 |
| 0 | 0 | 0 | 0 |
| 1 | 1 | 0 | 0 |
| 1 | 1 | 0 | 0 |
| 1 | 0 | 0 | 0 |
| 1 | 0 | 0 | 0 |
| 0 | 0 | 0 | 0 |
| 0 | 0 | 0 | 0 |
| 0 | 0 | 0 | 0 |
| 0 | 0 | 0 | 0 |
| 1 | 0 | 0 | 0 |
| 1 | 0 | 0 | 0 |
| 1 | 0 | 0 | 0 |
| 1 | 1 | 0 | 0 |
| 1 | 1 | 0 | 0 |
| 1 | 0 | 0 | 0 |
| 0 | 0 | 0 | 0 |
| 0 | 0 | 0 | 0 |
| 0 | 1 | 0 | 0 |
| 0 | 0 | 0 | 0 |
| 0 | 0 | 0 | 0 |
| 0 | 0 | 0 | 0 |
| 1 | 1 | 0 | 0 |
| 1 | 0 | 0 | 0 |
| 1 | 0 | 0 | 0 |
| 0 | 0 | 0 | 0 |
| 0 | 0 | 0 | 0 |
| 0 | 0 | 0 | 0 |
| 0 | 0 | 0 | 0 |
| 0 | 0 | 0 | 0 |
| 0 | 0 | 0 | 0 |
| 0 | 0 | 0 | 0 |
| 0 | 0 | 0 | 0 |
| 0 | 0 | 0 | 0 |
| 0 | 0 | 0 | 0 |

| | | | |
|---|---|---|---|
| 0 | 0 | 0 | 0 |
| 1 | 0 | 0 | 0 |
| 0 | 0 | 0 | 0 |
| 0 | 0 | 0 | 0 |
| 0 | 0 | 0 | 0 |
| 0 | 0 | 0 | 0 |
| 1 | 0 | 0 | 0 |
| 0 | 0 | 0 | 0 |
| 0 | 0 | 0 | 0 |
| 0 | 0 | 0 | 0 |
| 0 | 0 | 0 | 0 |
| 0 | 0 | 0 | 0 |
| 0 | 0 | 0 | 0 |
| 0 | 0 | 0 | 0 |
| 0 | 0 | 0 | 0 |
| 0 | 0 | 0 | 0 |
| 0 | 1 | 0 | 0 |
| 0 | 0 | 0 | 0 |
| 0 | 0 | 0 | 0 |
| 0 | 0 | 0 | 0 |
| 1 | 0 | 0 | 0 |
| 0 | 0 | 0 | 0 |
| 0 | 0 | 0 | 0 |
| 1 | 1 | 0 | 0 |
| 0 | 0 | 0 | 0 |
| 0 | 0 | 0 | 0 |
| 0 | 0 | 0 | 0 |
| 0 | 0 | 0 | 0 |
| 1 | 0 | 0 | 0 |
| 0 | 0 | 0 | 0 |
| 0 | 0 | 0 | 0 |
| 0 | 0 | 0 | 0 |
| 0 | 0 | 0 | 0 |
| 0 | 0 | 0 | 0 |
| 0 | 0 | 0 | 0 |
| 1 | 0 | 0 | 0 |
| 0 | 0 | 0 | 0 |
| 0 | 0 | 0 | 0 |
| 0 | 0 | 0 | 0 |
| 0 | 0 | 0 | 0 |
| 1 | 1 | 0 | 0 |
| 0 | 0 | 0 | 0 |
| 0 | 0 | 0 | 0 |
| 0 | 0 | 0 | 0 |
| 0 | 0 | 0 | 0 |
| 0 | 0 | 0 | 0 |
| 1 | 0 | 0 | 0 |
| 1 | 0 | 0 | 1 |
| 1 | 0 | 0 | 0 |
| 0 | 0 | 0 | 0 |
| 0 | 0 | 0 | 0 |
| 0 | 0 | 0 | 0 |
| 1 | 0 | 0 | 0 |
| 1 | 0 | 0 | 0 |

| | | | |
|---|---|---|---|
| 1 | 0 | 0 | 0 |
| 0 | 0 | 0 | 0 |
| 0 | 0 | 0 | 0 |
| 0 | 0 | 0 | 0 |
| 1 | 1 | 0 | 0 |
| 1 | 1 | 0 | 0 |
| 0 | 0 | 0 | 0 |
| 0 | 0 | 0 | 0 |
| 1 | 0 | 0 | 0 |
| 1 | 0 | 0 | 0 |
| 1 | 0 | 1 | 0 |
| 0 | 0 | 0 | 0 |
| 0 | 0 | 1 | 0 |
| 0 | 0 | 1 | 0 |
| 0 | 0 | 1 | 0 |
| 1 | 0 | 0 | 0 |
| 1 | 0 | 0 | 0 |
| 0 | 0 | 0 | 0 |
| 0 | 0 | 0 | 0 |
| 1 | 0 | 0 | 1 |
| 1 | 0 | 0 | 1 |
| 1 | 0 | 0 | 0 |
| 0 | 0 | 0 | 0 |
| 0 | 0 | 0 | 0 |
| 0 | 0 | 0 | 0 |
| 0 | 0 | 0 | 0 |
| 0 | 0 | 0 | 0 |
| 1 | 0 | 0 | 0 |
| 0 | 0 | 0 | 0 |
| 0 | 0 | 0 | 1 |
| 0 | 0 | 0 | 1 |
| 0 | 0 | 1 | 1 |
| 0 | 0 | 1 | 1 |
| 0 | 0 | 1 | 1 |
| 0 | 0 | 1 | 1 |
| 0 | 0 | 1 | 0 |
| 0 | 0 | 1 | 0 |
| 0 | 0 | 0 | 0 |
| 0 | 0 | 0 | 0 |
| 0 | 0 | 0 | 0 |
| 0 | 0 | 0 | 0 |
| 0 | 0 | 1 | 0 |
| 0 | 0 | 1 | 0 |
| 0 | 0 | 1 | 0 |
| 0 | 0 | 1 | 0 |
| 0 | 0 | 1 | 0 |
| 0 | 0 | 1 | 0 |
| 0 | 0 | 1 | 0 |
| 0 | 0 | 1 | 0 |
| 0 | 0 | 0 | 0 |
| 0 | 0 | 0 | 0 |
| 0 | 0 | 0 | 0 |
| 0 | 0 | 0 | 0 |
| 1 | 0 | 0 | 0 |

| | | | |
|---|---|---|---|
| 0 | 0 | 0 | 0 |
| 0 | 0 | 0 | 0 |
| 0 | 0 | 0 | 0 |
| 1 | 0 | 0 | 0 |
| 0 | 0 | 0 | 0 |
| 0 | 0 | 0 | 0 |
| 0 | 0 | 0 | 0 |
| 0 | 0 | 0 | 0 |
| 0 | 0 | 0 | 0 |
| 0 | 0 | 0 | 0 |
| 1 | 1 | 0 | 0 |
| 0 | 0 | 0 | 0 |
| 1 | 1 | 0 | 0 |
| 1 | 1 | 0 | 0 |
| 0 | 1 | 0 | 0 |
| 0 | 0 | 0 | 0 |
| 0 | 0 | 0 | 0 |
| 0 | 0 | 0 | 0 |
| 0 | 0 | 0 | 0 |
| 0 | 0 | 0 | 0 |
| 1 | 1 | 0 | 0 |
| 0 | 1 | 0 | 0 |
| 0 | 0 | 0 | 0 |
| 0 | 1 | 0 | 0 |
| 1 | 1 | 0 | 0 |
| 0 | 0 | 0 | 0 |
| 0 | 0 | 0 | 0 |
| 1 | 1 | 0 | 0 |
| 1 | 1 | 0 | 0 |
| 0 | 1 | 0 | 0 |
| 0 | 0 | 0 | 0 |
| 0 | 0 | 0 | 0 |
| 0 | 0 | 0 | 0 |
| 0 | 0 | 0 | 0 |
| 0 | 0 | 0 | 0 |
| 0 | 0 | 0 | 0 |
| 0 | 0 | 0 | 0 |
| 0 | 0 | 0 | 0 |
| 0 | 0 | 0 | 0 |
| 0 | 0 | 0 | 0 |
| 1 | 0 | 0 | 0 |
| 1 | 0 | 0 | 0 |
| 1 | 0 | 0 | 0 |
| 1 | 0 | 0 | 0 |
| 1 | 1 | 0 | 1 |
| 0 | 0 | 0 | 0 |
| 1 | 0 | 0 | 0 |
| 0 | 0 | 0 | 0 |
| 1 | 0 | 0 | 0 |
| 0 | 0 | 0 | 0 |
| 1 | 0 | 0 | 0 |
| 1 | 0 | 0 | 0 |
| 0 | 0 | 0 | 0 |
| 0 | 0 | 0 | 0 |

| | | | |
|---|---|---|---|
| 0 | 0 | 0 | 0 |
| 0 | 0 | 0 | 0 |
| 0 | 0 | 0 | 0 |
| 0 | 0 | 0 | 0 |
| 1 | 0 | 0 | 0 |
| 1 | 0 | 0 | 0 |
| 1 | 0 | 0 | 0 |
| 0 | 0 | 0 | 0 |
| 1 | 1 | 0 | 0 |
| 0 | 0 | 0 | 0 |
| 0 | 0 | 0 | 0 |
| 0 | 0 | 0 | 0 |
| 1 | 1 | 0 | 0 |
| 1 | 0 | 0 | 0 |
| 0 | 0 | 0 | 0 |
| 0 | 0 | 0 | 0 |
| 1 | 1 | 0 | 0 |
| 0 | 0 | 0 | 0 |
| 0 | 0 | 0 | 0 |
| 0 | 0 | 0 | 0 |
| 0 | 0 | 0 | 0 |
| 0 | 0 | 0 | 0 |
| 0 | 0 | 0 | 0 |
| 0 | 0 | 0 | 0 |
| 1 | 0 | 0 | 0 |
| 0 | 0 | 0 | 0 |
| 1 | 0 | 0 | 0 |
| 0 | 0 | 0 | 0 |
| 0 | 0 | 0 | 0 |
| 0 | 0 | 0 | 0 |
| 1 | 0 | 0 | 0 |
| 1 | 0 | 0 | 0 |
| 0 | 0 | 0 | 0 |
| 0 | 0 | 0 | 0 |
| 1 | 0 | 0 | 0 |
| 0 | 0 | 0 | 0 |
| 0 | 0 | 0 | 0 |
| 1 | 0 | 0 | 0 |
| 0 | 0 | 0 | 0 |
| 1 | 0 | 0 | 0 |
| 1 | 0 | 0 | 0 |
| 0 | 0 | 0 | 0 |
| 0 | 0 | 0 | 0 |
| 1 | 1 | 0 | 0 |
| 1 | 0 | 0 | 0 |
| 1 | 1 | 0 | 0 |
| 0 | 0 | 0 | 0 |
| 0 | 0 | 0 | 0 |
| 0 | 0 | 0 | 0 |
| 0 | 0 | 0 | 0 |
| 1 | 1 | 0 | 0 |
| 0 | 0 | 0 | 0 |
| 0 | 0 | 0 | 0 |
| 0 | 0 | 0 | 0 |

| | | | |
|---|---|---|---|
| 0 | 1 | 0 | 0 |
| 1 | 0 | 0 | 0 |
| 0 | 0 | 0 | 0 |
| 0 | 0 | 0 | 0 |
| 0 | 0 | 0 | 0 |
| 1 | 1 | 0 | 0 |
| 0 | 0 | 0 | 0 |
| 0 | 0 | 0 | 0 |
| 0 | 0 | 0 | 0 |
| 1 | 0 | 0 | 0 |
| 1 | 0 | 0 | 0 |
| 0 | 0 | 0 | 0 |
| 0 | 0 | 0 | 0 |
| 0 | 0 | 0 | 0 |
| 0 | 0 | 0 | 0 |
| 0 | 0 | 0 | 0 |
| 0 | 0 | 0 | 0 |
| 0 | 0 | 0 | 0 |
| 0 | 0 | 0 | 0 |
| 0 | 0 | 0 | 0 |
| 0 | 0 | 0 | 0 |
| 0 | 0 | 0 | 0 |
| 0 | 0 | 0 | 0 |
| 1 | 1 | 0 | 0 |
| 0 | 0 | 0 | 0 |
| 0 | 0 | 0 | 0 |
| 1 | 1 | 1 | 0 |
| 0 | 0 | 0 | 0 |
| 0 | 0 | 0 | 0 |
| 0 | 0 | 0 | 0 |
| 1 | 1 | 1 | 0 |
| 1 | 0 | 0 | 0 |
| 1 | 0 | 0 | 0 |
| 1 | 0 | 0 | 0 |
| 1 | 0 | 0 | 0 |
| 1 | 1 | 1 | 0 |
| 1 | 1 | 1 | 0 |
| 0 | 0 | 0 | 0 |
| 0 | 0 | 0 | 0 |
| 1 | 1 | 1 | 0 |
| 0 | 0 | 0 | 0 |
| 0 | 0 | 0 | 0 |
| 0 | 0 | 0 | 0 |
| 0 | 0 | 0 | 0 |
| 0 | 0 | 0 | 0 |
| 0 | 0 | 0 | 0 |
| 0 | 0 | 0 | 0 |
| 0 | 0 | 0 | 0 |
| 1 | 0 | 0 | 0 |
| 1 | 0 | 0 | 0 |
| 1 | 0 | 0 | 0 |
| 0 | 0 | 0 | 0 |
| 0 | 0 | 0 | 0 |
| 0 | 0 | 0 | 0 |

| | | | |
|---|---|---|---|
| 0 | 0 | 0 | 0 |
| 1 | 0 | 0 | 0 |
| 0 | 0 | 0 | 0 |
| 0 | 0 | 0 | 0 |
| 0 | 0 | 0 | 0 |
| 0 | 0 | 0 | 0 |
| 0 | 0 | 0 | 0 |
| 0 | 0 | 0 | 0 |
| 0 | 0 | 0 | 0 |
| 0 | 0 | 0 | 0 |
| 1 | 1 | 0 | 0 |
| 1 | 1 | 0 | 0 |
| 0 | 0 | 0 | 0 |
| 0 | 0 | 0 | 0 |
| 0 | 0 | 0 | 0 |
| 1 | 0 | 0 | 0 |
| 1 | 0 | 0 | 0 |
| 1 | 1 | 0 | 1 |
| 0 | 0 | 0 | 0 |
| 0 | 0 | 0 | 0 |
| 0 | 0 | 0 | 0 |
| 0 | 0 | 0 | 0 |
| 1 | 0 | 0 | 1 |
| 0 | 0 | 0 | 0 |
| 0 | 0 | 0 | 0 |
| 1 | 0 | 0 | 0 |
| 0 | 0 | 0 | 0 |
| 0 | 0 | 0 | 0 |
| 0 | 0 | 0 | 0 |
| 0 | 0 | 0 | 0 |
| 0 | 0 | 0 | 0 |
| 0 | 0 | 0 | 0 |
| 0 | 0 | 0 | 0 |
| 0 | 0 | 0 | 0 |
| 0 | 0 | 0 | 0 |
| 1 | 0 | 0 | 0 |
| 1 | 0 | 0 | 0 |
| 1 | 1 | 0 | 0 |
| 1 | 1 | 0 | 0 |
| 0 | 0 | 0 | 0 |
| 0 | 0 | 0 | 0 |
| 1 | 0 | 0 | 0 |
| 0 | 0 | 0 | 0 |
| 1 | 0 | 0 | 0 |
| 1 | 1 | 0 | 0 |
| 0 | 0 | 0 | 0 |
| 0 | 0 | 0 | 0 |
| 0 | 0 | 0 | 0 |
| 0 | 0 | 0 | 0 |
| 0 | 0 | 0 | 0 |
| 0 | 0 | 0 | 0 |
| 1 | 0 | 0 | 0 |
| 0 | 0 | 0 | 0 |
| 0 | 0 | 0 | 0 |

| | | | |
|---|---|---|---|
| 0 | 0 | 0 | 0 |
| 1 | 1 | 0 | 0 |
| 0 | 0 | 0 | 0 |
| 1 | 0 | 0 | 0 |
| 1 | 0 | 0 | 0 |
| 0 | 0 | 0 | 0 |
| 0 | 0 | 0 | 0 |
| 0 | 0 | 0 | 0 |
| 0 | 0 | 0 | 0 |
| 0 | 0 | 0 | 0 |
| 0 | 0 | 0 | 0 |
| 1 | 1 | 0 | 0 |
| 1 | 1 | 0 | 1 |
| 0 | 0 | 0 | 0 |
| 0 | 1 | 0 | 0 |
| 0 | 0 | 0 | 0 |
| 0 | 0 | 0 | 0 |
| 0 | 0 | 0 | 0 |
| 0 | 0 | 0 | 0 |
| 0 | 0 | 0 | 0 |
| 0 | 0 | 0 | 0 |
| 1 | 0 | 0 | 0 |
| 0 | 0 | 0 | 0 |
| 0 | 0 | 0 | 0 |
| 0 | 0 | 0 | 0 |
| 0 | 0 | 0 | 0 |
| 0 | 0 | 0 | 0 |
| 1 | 0 | 0 | 0 |
| 0 | 0 | 0 | 0 |
| 0 | 0 | 0 | 0 |
| 0 | 0 | 0 | 0 |
| 0 | 0 | 0 | 0 |
| 0 | 0 | 0 | 0 |
| 0 | 0 | 0 | 0 |
| 0 | 0 | 0 | 0 |
| 0 | 0 | 0 | 0 |
| 0 | 0 | 0 | 0 |
| 0 | 0 | 0 | 0 |
| 0 | 0 | 0 | 0 |
| 0 | 0 | 0 | 0 |
| 0 | 0 | 0 | 0 |
| 0 | 0 | 0 | 0 |
| 0 | 0 | 0 | 0 |
| 0 | 0 | 0 | 0 |
| 0 | 0 | 0 | 0 |
| 0 | 0 | 0 | 0 |
| 0 | 0 | 0 | 0 |
| 0 | 0 | 0 | 0 |
| 0 | 0 | 0 | 0 |
| 0 | 0 | 0 | 0 |
| 0 | 0 | 0 | 0 |
| 0 | 0 | 0 | 0 |
| 0 | 0 | 0 | 0 |
| 0 | 0 | 0 | 0 |

| | | | |
|---|---|---|---|
| 0 | 0 | 0 | 0 |
| 0 | 0 | 0 | 0 |
| 0 | 0 | 0 | 0 |
| 0 | 0 | 0 | 0 |
| 0 | 0 | 0 | 0 |
| 0 | 0 | 0 | 0 |
| 0 | 0 | 0 | 0 |
| 0 | 0 | 0 | 0 |
| 0 | 0 | 0 | 0 |
| 0 | 0 | 0 | 0 |
| 0 | 0 | 0 | 0 |
| 0 | 0 | 0 | 0 |
| 1 | 0 | 0 | 0 |
| 1 | 1 | 0 | 0 |
| 0 | 0 | 0 | 0 |
| 1 | 0 | 0 | 0 |
| 0 | 0 | 0 | 0 |
| 1 | 0 | 0 | 0 |
| 0 | 0 | 0 | 0 |
| 0 | 0 | 0 | 0 |
| 1 | 0 | 0 | 0 |
| 0 | 0 | 0 | 0 |
| 0 | 0 | 0 | 0 |
| 1 | 0 | 0 | 0 |
| 1 | 0 | 0 | 0 |
| 0 | 0 | 0 | 0 |
| 0 | 0 | 0 | 0 |
| 1 | 0 | 0 | 0 |
| 1 | 0 | 0 | 0 |
| 1 | 0 | 0 | 0 |
| 1 | 0 | 0 | 0 |
| 1 | 0 | 0 | 0 |
| 0 | 0 | 0 | 0 |
| 0 | 0 | 0 | 0 |
| 0 | 0 | 0 | 0 |
| 0 | 0 | 0 | 0 |
| 0 | 0 | 0 | 0 |
| 0 | 0 | 0 | 0 |
| 0 | 0 | 0 | 0 |
| 0 | 0 | 0 | 0 |
| 0 | 0 | 0 | 0 |
| 0 | 0 | 0 | 0 |
| 0 | 0 | 0 | 0 |
| 0 | 0 | 0 | 0 |
| 1 | 1 | 0 | 0 |
| 0 | 0 | 0 | 0 |
| 0 | 0 | 0 | 0 |
| 0 | 0 | 0 | 0 |
| 0 | 0 | 0 | 0 |
| 1 | 1 | 0 | 0 |
| 1 | 0 | 0 | 0 |
| 0 | 0 | 0 | 0 |
| 0 | 0 | 0 | 0 |
| 0 | 0 | 0 | 1 |

| | | | |
|---|---|---|---|
| 0 | 0 | 0 | 0 |
| 0 | 0 | 0 | 0 |
| 0 | 0 | 0 | 0 |
| 0 | 0 | 0 | 0 |
| 1 | 0 | 0 | 1 |
| 0 | 0 | 0 | 0 |
| 1 | 0 | 0 | 0 |
| 0 | 0 | 0 | 0 |
| 0 | 0 | 0 | 0 |
| 0 | 0 | 0 | 0 |
| 0 | 0 | 0 | 0 |
| 0 | 0 | 0 | 0 |
| 0 | 0 | 0 | 0 |
| 1 | 0 | 0 | 0 |
| 0 | 0 | 0 | 0 |
| 0 | 0 | 0 | 0 |
| 1 | 1 | 0 | 0 |
| 0 | 0 | 0 | 0 |
| 1 | 0 | 0 | 0 |
| 1 | 0 | 0 | 0 |
| 0 | 0 | 0 | 0 |
| 0 | 0 | 0 | 0 |
| 0 | 0 | 0 | 0 |
| 1 | 0 | 0 | 0 |
| 1 | 0 | 0 | 0 |
| 1 | 0 | 0 | 0 |
| 1 | 0 | 0 | 0 |
| 1 | 0 | 0 | 0 |
| 1 | 0 | 0 | 0 |
| 1 | 0 | 0 | 0 |
| 1 | 0 | 0 | 0 |
| 1 | 0 | 0 | 0 |
| 1 | 0 | 0 | 0 |
| 1 | 0 | 0 | 0 |
| 0 | 0 | 0 | 0 |
| 0 | 0 | 0 | 0 |
| 0 | 0 | 0 | 0 |
| 1 | 0 | 0 | 0 |
| 1 | 0 | 0 | 0 |
| 0 | 0 | 0 | 0 |
| 0 | 0 | 0 | 0 |
| 1 | 0 | 0 | 0 |
| 0 | 0 | 0 | 0 |
| 1 | 0 | 0 | 0 |
| 1 | 0 | 0 | 0 |
| 1 | 0 | 0 | 0 |
| 1 | 0 | 0 | 0 |
| 1 | 0 | 0 | 0 |
| 0 | 0 | 0 | 0 |
| 0 | 0 | 0 | 0 |
| 1 | 0 | 0 | 0 |
| 0 | 0 | 0 | 0 |
| 1 | 0 | 0 | 0 |
| 1 | 0 | 0 | 0 |

| | | | |
|---|---|---|---|
| 1 | 0 | 0 | 0 |
| 1 | 0 | 0 | 0 |
| 1 | 0 | 0 | 0 |
| 1 | 0 | 0 | 0 |
| 1 | 0 | 0 | 0 |
| 1 | 0 | 0 | 0 |
| 0 | 0 | 0 | 0 |
| 1 | 0 | 0 | 0 |
| 1 | 0 | 0 | 0 |
| 1 | 0 | 0 | 0 |
| 1 | 0 | 0 | 0 |
| 1 | 0 | 0 | 0 |
| 1 | 0 | 0 | 0 |
| 1 | 0 | 0 | 0 |
| 0 | 0 | 0 | 0 |
| 0 | 0 | 0 | 0 |
| 1 | 1 | 0 | 0 |
| 1 | 1 | 0 | 0 |
| 0 | 0 | 0 | 0 |
| 0 | 0 | 0 | 0 |
| 0 | 0 | 0 | 0 |
| 1 | 1 | 0 | 1 |
| 0 | 0 | 0 | 0 |
| 0 | 0 | 0 | 0 |
| 0 | 0 | 0 | 0 |
| 0 | 0 | 0 | 0 |
| 0 | 0 | 0 | 0 |
| 0 | 0 | 0 | 0 |
| 0 | 0 | 0 | 0 |
| 1 | 0 | 1 | 0 |
| 0 | 0 | 0 | 0 |
| 0 | 0 | 0 | 0 |
| 0 | 0 | 0 | 0 |
| 0 | 0 | 0 | 0 |
| 0 | 0 | 0 | 0 |
| 0 | 0 | 0 | 0 |
| 0 | 0 | 0 | 0 |
| 0 | 0 | 0 | 0 |
| 0 | 0 | 0 | 0 |
| 0 | 0 | 0 | 0 |
| 1 | 1 | 0 | 0 |
| 1 | 1 | 0 | 0 |
| 1 | 0 | 0 | 0 |
| 1 | 1 | 0 | 0 |
| 1 | 0 | 0 | 0 |
| 1 | 1 | 0 | 0 |
| 1 | 0 | 0 | 0 |
| 1 | 0 | 0 | 0 |
| 0 | 0 | 0 | 0 |
| 0 | 0 | 0 | 0 |
| 1 | 0 | 0 | 0 |
| 1 | 0 | 0 | 0 |
| 1 | 0 | 0 | 0 |
| 0 | 0 | 0 | 0 |

| | | | |
|---|---|---|---|
| 0 | 0 | 0 | 0 |
| 0 | 0 | 0 | 0 |
| 0 | 0 | 0 | 0 |
| 1 | 1 | 1 | 0 |
| 0 | 0 | 0 | 0 |
| 0 | 0 | 0 | 0 |
| 0 | 0 | 0 | 0 |
| 0 | 0 | 0 | 0 |
| 0 | 1 | 0 | 0 |
| 0 | 1 | 0 | 0 |
| 0 | 1 | 0 | 0 |
| 0 | 0 | 0 | 0 |
| 0 | 0 | 0 | 0 |
| 0 | 0 | 0 | 0 |
| 1 | 0 | 0 | 0 |
| 1 | 0 | 0 | 0 |
| 0 | 0 | 0 | 0 |
| 0 | 0 | 0 | 0 |
| 1 | 0 | 0 | 0 |
| 0 | 0 | 0 | 0 |
| 1 | 0 | 0 | 0 |
| 1 | 0 | 0 | 0 |
| 0 | 0 | 0 | 0 |
| 0 | 0 | 0 | 0 |
| 0 | 0 | 0 | 0 |
| 0 | 0 | 0 | 0 |
| 0 | 0 | 0 | 0 |
| 0 | 0 | 1 | 0 |
| 0 | 0 | 0 | 0 |
| 0 | 1 | 0 | 0 |
| 0 | 0 | 0 | 0 |
| 0 | 0 | 0 | 0 |
| 0 | 0 | 0 | 0 |
| 0 | 0 | 0 | 0 |
| 0 | 0 | 0 | 0 |
| 0 | 0 | 0 | 0 |
| 1 | 0 | 0 | 0 |
| 1 | 0 | 0 | 0 |
| 1 | 0 | 0 | 0 |
| 0 | 0 | 0 | 0 |
| 0 | 0 | 0 | 0 |
| 0 | 0 | 0 | 0 |
| 0 | 0 | 0 | 0 |
| 0 | 0 | 0 | 0 |
| 0 | 0 | 0 | 0 |
| 0 | 0 | 0 | 0 |
| 0 | 0 | 0 | 0 |
| 0 | 0 | 0 | 1 |
| 0 | 0 | 0 | 0 |
| 0 | 0 | 0 | 0 |
| 0 | 0 | 0 | 0 |
| 0 | 0 | 0 | 0 |
| 1 | 1 | 0 | 0 |
| 0 | 0 | 0 | 0 |

| | | | |
|---|---|---|---|
| 0 | 0 | 0 | 0 |
| 0 | 0 | 0 | 0 |
| 0 | 0 | 0 | 0 |
| 0 | 0 | 0 | 0 |
| 1 | 0 | 0 | 0 |
| 0 | 0 | 0 | 0 |
| 0 | 0 | 0 | 0 |
| 0 | 0 | 0 | 0 |
| 0 | 0 | 0 | 0 |
| 0 | 0 | 0 | 0 |
| 0 | 0 | 0 | 0 |
| 0 | 0 | 0 | 0 |
| 0 | 0 | 0 | 0 |
| 0 | 0 | 0 | 0 |
| 0 | 0 | 0 | 0 |
| 0 | 0 | 0 | 0 |
| 0 | 0 | 0 | 0 |
| 0 | 0 | 0 | 0 |
| 0 | 0 | 0 | 0 |
| 0 | 0 | 0 | 0 |
| 0 | 0 | 0 | 0 |
| 0 | 0 | 0 | 0 |
| 0 | 0 | 0 | 0 |
| 0 | 0 | 0 | 0 |
| 0 | 0 | 0 | 0 |
| 0 | 0 | 0 | 0 |
| 0 | 0 | 0 | 0 |
| 0 | 0 | 0 | 0 |
| 0 | 0 | 0 | 0 |
| 0 | 0 | 0 | 0 |
| 0 | 0 | 0 | 0 |
| 0 | 0 | 0 | 0 |
| 0 | 0 | 0 | 0 |
| 0 | 0 | 0 | 0 |
| 0 | 0 | 0 | 0 |
| 0 | 0 | 0 | 0 |
| 0 | 0 | 0 | 0 |
| 0 | 0 | 0 | 0 |
| 0 | 0 | 0 | 0 |
| 0 | 0 | 0 | 0 |
| 1 | 1 | 0 | 0 |
| 0 | 0 | 0 | 0 |
| 0 | 0 | 0 | 0 |
| 0 | 0 | 0 | 0 |
| 0 | 0 | 0 | 0 |
| 1 | 0 | 0 | 0 |
| 1 | 0 | 0 | 0 |
| 0 | 0 | 0 | 0 |
| 1 | 0 | 0 | 0 |
| 0 | 0 | 0 | 0 |
| 0 | 0 | 0 | 0 |
| 1 | 0 | 0 | 0 |
| 1 | 0 | 0 | 0 |
| 0 | 0 | 0 | 0 |

| | | | |
|---|---|---|---|
| 0 | 0 | 0 | 0 |
| 0 | 0 | 0 | 0 |
| 0 | 0 | 0 | 0 |
| 0 | 0 | 0 | 0 |
| 1 | 0 | 0 | 0 |
| 0 | 0 | 0 | 0 |
| 0 | 0 | 0 | 0 |
| 0 | 0 | 0 | 0 |
| 0 | 0 | 0 | 0 |
| 0 | 0 | 0 | 0 |
| 0 | 0 | 0 | 0 |
| 0 | 0 | 0 | 0 |
| 0 | 0 | 0 | 0 |
| 0 | 0 | 0 | 0 |
| 0 | 0 | 0 | 0 |
| 1 | 1 | 0 | 1 |
| 0 | 0 | 0 | 0 |
| 0 | 0 | 0 | 0 |
| 0 | 0 | 0 | 0 |
| 0 | 0 | 0 | 0 |
| 0 | 0 | 0 | 0 |
| 0 | 0 | 0 | 0 |
| 0 | 0 | 0 | 0 |
| 0 | 0 | 0 | 0 |
| 0 | 0 | 0 | 0 |
| 0 | 0 | 0 | 0 |
| 0 | 0 | 0 | 0 |
| 1 | 0 | 0 | 0 |
| 1 | 0 | 0 | 0 |
| 0 | 0 | 0 | 0 |
| 1 | 1 | 0 | 0 |
| 0 | 0 | 0 | 0 |
| 0 | 0 | 0 | 0 |
| 0 | 0 | 0 | 0 |
| 1 | 1 | 0 | 0 |
| 1 | 1 | 0 | 0 |
| 0 | 0 | 0 | 0 |
| 1 | 0 | 0 | 0 |
| 0 | 0 | 0 | 0 |
| 1 | 1 | 0 | 0 |
| 0 | 1 | 0 | 0 |
| 0 | 0 | 0 | 0 |
| 0 | 0 | 0 | 0 |
| 0 | 0 | 0 | 0 |
| 1 | 1 | 0 | 0 |
| 1 | 1 | 0 | 0 |
| 1 | 1 | 0 | 0 |
| 0 | 0 | 0 | 0 |
| 0 | 0 | 0 | 0 |
| 0 | 0 | 0 | 0 |
| 0 | 0 | 0 | 0 |
| 0 | 0 | 0 | 0 |
| 0 | 0 | 0 | 0 |
| 0 | 0 | 0 | 0 |

| | | | |
|---|---|---|---|
| 0 | 0 | 0 | 0 |
| 0 | 0 | 0 | 0 |
| 1 | 0 | 0 | 0 |
| 1 | 0 | 0 | 0 |
| 0 | 0 | 0 | 0 |
| 1 | 0 | 0 | 0 |
| 0 | 0 | 0 | 0 |
| 0 | 0 | 0 | 0 |
| 0 | 0 | 0 | 0 |
| 0 | 1 | 0 | 0 |
| 0 | 0 | 0 | 0 |
| 0 | 0 | 0 | 0 |
| 0 | 0 | 0 | 0 |
| 0 | 0 | 0 | 0 |
| 0 | 0 | 0 | 0 |
| 0 | 0 | 0 | 0 |
| 0 | 0 | 0 | 0 |
| 1 | 0 | 0 | 0 |
| 0 | 0 | 0 | 0 |
| 1 | 0 | 0 | 0 |
| 0 | 0 | 0 | 0 |
| 1 | 0 | 0 | 0 |
| 0 | 0 | 0 | 0 |
| 1 | 0 | 0 | 0 |
| 0 | 0 | 0 | 0 |
| 0 | 0 | 0 | 0 |
| 0 | 0 | 0 | 0 |
| 0 | 0 | 0 | 0 |
| 0 | 0 | 0 | 0 |
| 0 | 0 | 0 | 0 |
| 0 | 0 | 0 | 0 |
| 1 | 0 | 0 | 0 |
| 0 | 0 | 0 | 0 |
| 0 | 0 | 0 | 0 |
| 0 | 0 | 0 | 0 |
| 0 | 0 | 0 | 0 |
| 0 | 0 | 0 | 0 |
| 0 | 0 | 0 | 0 |
| 1 | 1 | 0 | 0 |
| 1 | 1 | 0 | 0 |
| 1 | 1 | 0 | 0 |
| 1 | 1 | 0 | 0 |
| 1 | 1 | 0 | 0 |
| 1 | 0 | 0 | 0 |
| 1 | 0 | 0 | 0 |
| 1 | 0 | 0 | 0 |
| 0 | 0 | 0 | 0 |
| 0 | 0 | 0 | 0 |
| 0 | 0 | 0 | 0 |
| 0 | 0 | 0 | 0 |
| 0 | 0 | 0 | 0 |
| 0 | 0 | 0 | 0 |
| 0 | 0 | 0 | 0 |
| 0 | 0 | 0 | 0 |

| | | | |
|---|---|---|---|
| 0 | 0 | 0 | 0 |
| 0 | 0 | 0 | 0 |
| 0 | 0 | 0 | 0 |
| 0 | 0 | 0 | 0 |
| 0 | 0 | 0 | 0 |
| 1 | 0 | 0 | 0 |
| 1 | 0 | 0 | 0 |
| 0 | 0 | 0 | 0 |
| 0 | 0 | 0 | 0 |
| 0 | 0 | 0 | 0 |
| 0 | 0 | 0 | 0 |
| 1 | 0 | 0 | 0 |
| 0 | 0 | 1 | 0 |
| 0 | 0 | 0 | 0 |
| 0 | 0 | 0 | 0 |
| 0 | 0 | 0 | 0 |
| 1 | 0 | 0 | 0 |
| 0 | 0 | 0 | 0 |
| 1 | 0 | 0 | 0 |
| 1 | 0 | 0 | 0 |
| 0 | 0 | 0 | 0 |
| 1 | 0 | 0 | 0 |
| 1 | 1 | 0 | 0 |
| 0 | 0 | 0 | 0 |
| 1 | 1 | 0 | 0 |
| 1 | 1 | 0 | 0 |
| 1 | 1 | 0 | 0 |
| 0 | 0 | 0 | 0 |
| 1 | 0 | 0 | 0 |
| 0 | 0 | 0 | 0 |
| 0 | 0 | 0 | 0 |
| 0 | 0 | 0 | 0 |
| 0 | 0 | 0 | 0 |
| 0 | 0 | 0 | 0 |
| 0 | 0 | 0 | 0 |
| 0 | 0 | 0 | 0 |
| 0 | 0 | 0 | 0 |
| 0 | 0 | 0 | 0 |
| 0 | 0 | 0 | 0 |
| 0 | 0 | 0 | 0 |
| 0 | 0 | 0 | 0 |
| 0 | 0 | 0 | 0 |
| 0 | 0 | 0 | 0 |
| 0 | 0 | 0 | 0 |
| 0 | 0 | 0 | 0 |
| 0 | 0 | 0 | 0 |
| 0 | 0 | 0 | 0 |
| 0 | 0 | 0 | 0 |
| 0 | 0 | 0 | 0 |
| 0 | 0 | 0 | 0 |
| 0 | 0 | 0 | 0 |
| 0 | 0 | 0 | 0 |
| 0 | 0 | 0 | 0 |

| | | | |
|---|---|---|---|
| 0 | 0 | 0 | 0 |
| 0 | 0 | 0 | 0 |
| 0 | 0 | 0 | 0 |
| 0 | 0 | 0 | 0 |
| 0 | 0 | 0 | 0 |
| 0 | 0 | 0 | 0 |
| 0 | 0 | 0 | 0 |
| 0 | 0 | 0 | 0 |
| 0 | 0 | 0 | 0 |
| 0 | 0 | 0 | 0 |
| 0 | 1 | 0 | 0 |
| 0 | 1 | 0 | 0 |
| 0 | 1 | 0 | 0 |
| 0 | 1 | 0 | 0 |
| 0 | 0 | 0 | 0 |
| 0 | 1 | 0 | 0 |
| 0 | 1 | 0 | 0 |
| 1 | 1 | 0 | 0 |
| 0 | 1 | 0 | 0 |
| 1 | 0 | 0 | 0 |
| 1 | 1 | 0 | 0 |
| 1 | 1 | 0 | 0 |
| 0 | 0 | 1 | 1 |
| 0 | 0 | 0 | 0 |
| 1 | 0 | 0 | 0 |
| 0 | 0 | 0 | 0 |
| 0 | 0 | 0 | 0 |
| 0 | 0 | 0 | 0 |
| 0 | 0 | 0 | 0 |
| 0 | 0 | 0 | 0 |
| 0 | 0 | 0 | 0 |
| 0 | 0 | 0 | 0 |
| 0 | 0 | 0 | 0 |
| 0 | 0 | 0 | 0 |
| 0 | 0 | 0 | 0 |
| 0 | 0 | 0 | 0 |
| 0 | 0 | 0 | 0 |
| 0 | 0 | 0 | 0 |
| 0 | 0 | 0 | 0 |
| 0 | 0 | 0 | 0 |
| 1 | 0 | 0 | 0 |
| 0 | 0 | 0 | 0 |
| 1 | 0 | 0 | 0 |
| 1 | 0 | 0 | 0 |
| 0 | 0 | 0 | 0 |
| 1 | 0 | 0 | 0 |
| 0 | 0 | 0 | 0 |
| 0 | 0 | 0 | 0 |
| 0 | 0 | 0 | 0 |
| 0 | 0 | 0 | 0 |
| 0 | 0 | 0 | 0 |
| 0 | 0 | 0 | 0 |
| 1 | 1 | 0 | 0 |
| 1 | 1 | 0 | 0 |

| | | | |
|---|---|---|---|
| 1 | 1 | 0 | 0 |
| 0 | 0 | 0 | 0 |
| 0 | 0 | 0 | 0 |
| 0 | 0 | 0 | 0 |
| 0 | 0 | 0 | 0 |
| 0 | 0 | 0 | 0 |
| 0 | 1 | 0 | 0 |
| 0 | 0 | 0 | 0 |
| 0 | 0 | 0 | 0 |
| 0 | 1 | 0 | 0 |
| 1 | 0 | 0 | 0 |
| 0 | 0 | 0 | 0 |
| 0 | 0 | 1 | 0 |
| 0 | 0 | 1 | 0 |
| 0 | 0 | 0 | 0 |
| 0 | 0 | 0 | 0 |
| 0 | 0 | 0 | 0 |
| 0 | 1 | 0 | 0 |
| 0 | 0 | 0 | 0 |
| 0 | 0 | 0 | 0 |
| 1 | 0 | 0 | 0 |
| 0 | 0 | 0 | 1 |
| 0 | 0 | 0 | 0 |
| 0 | 0 | 0 | 0 |
| 0 | 0 | 0 | 0 |
| 0 | 0 | 0 | 0 |
| 0 | 0 | 0 | 0 |
| 0 | 0 | 0 | 0 |
| 0 | 0 | 0 | 0 |
| 0 | 0 | 0 | 0 |
| 0 | 0 | 0 | 0 |
| 0 | 0 | 0 | 0 |
| 0 | 0 | 0 | 0 |
| 0 | 0 | 0 | 0 |
| 0 | 0 | 0 | 0 |
| 0 | 0 | 0 | 0 |
| 1 | 0 | 0 | 0 |
| 0 | 0 | 0 | 0 |
| 1 | 0 | 0 | 0 |
| 0 | 0 | 0 | 0 |
| 0 | 0 | 0 | 0 |
| 0 | 0 | 0 | 0 |
| 0 | 0 | 0 | 0 |
| 0 | 0 | 0 | 0 |
| 0 | 0 | 0 | 0 |
| 0 | 0 | 0 | 0 |
| 0 | 0 | 0 | 0 |
| 0 | 0 | 0 | 0 |
| 0 | 0 | 0 | 0 |
| 1 | 0 | 0 | 0 |
| 0 | 0 | 0 | 0 |
| 0 | 0 | 0 | 0 |
| 0 | 0 | 0 | 0 |
| 0 | 0 | 0 | 0 |

| | | | |
|---|---|---|---|
| 0 | 0 | 0 | 0 |
| 0 | 0 | 0 | 0 |
| 0 | 0 | 0 | 0 |
| 0 | 0 | 0 | 0 |
| 0 | 0 | 0 | 0 |
| 0 | 0 | 0 | 0 |
| 0 | 0 | 0 | 0 |
| 0 | 0 | 0 | 0 |
| 0 | 0 | 0 | 0 |
| 0 | 0 | 0 | 0 |
| 0 | 0 | 0 | 0 |
| 0 | 0 | 0 | 0 |
| 0 | 0 | 1 | 0 |
| 1 | 0 | 0 | 0 |
| 1 | 0 | 0 | 0 |
| 0 | 0 | 0 | 0 |
| 0 | 0 | 0 | 0 |
| 0 | 0 | 0 | 0 |
| 0 | 0 | 0 | 0 |
| 0 | 0 | 0 | 0 |
| 0 | 0 | 0 | 0 |
| 0 | 0 | 0 | 0 |
| 0 | 0 | 0 | 0 |
| 1 | 1 | 0 | 1 |
| 0 | 0 | 0 | 0 |
| 1 | 1 | 0 | 0 |
| 1 | 1 | 0 | 1 |
| 0 | 0 | 0 | 0 |
| 0 | 0 | 0 | 0 |
| 0 | 0 | 0 | 0 |
| 0 | 0 | 0 | 0 |
| 0 | 0 | 0 | 0 |
| 0 | 0 | 0 | 0 |
| 0 | 0 | 0 | 0 |
| 0 | 0 | 0 | 0 |
| 1 | 0 | 0 | 0 |
| 0 | 0 | 0 | 0 |
| 0 | 0 | 0 | 0 |
| 0 | 0 | 0 | 0 |
| 0 | 0 | 0 | 0 |
| 0 | 0 | 0 | 0 |
| 0 | 0 | 0 | 0 |
| 1 | 0 | 0 | 0 |
| 1 | 0 | 0 | 0 |
| 1 | 0 | 0 | 0 |
| 1 | 0 | 0 | 0 |
| 1 | 0 | 0 | 0 |
| 0 | 0 | 0 | 0 |
| 0 | 0 | 0 | 0 |
| 1 | 0 | 0 | 0 |
| 1 | 0 | 0 | 0 |
| 0 | 0 | 0 | 0 |
| 1 | 0 | 0 | 0 |
| 1 | 0 | 0 | 0 |

| | | | |
|---|---|---|---|
| 0 | 0 | 0 | 0 |
| 1 | 0 | 0 | 0 |
| 0 | 0 | 0 | 0 |
| 0 | 0 | 0 | 0 |
| 0 | 0 | 0 | 0 |
| 0 | 0 | 0 | 0 |
| 1 | 0 | 0 | 0 |
| 0 | 0 | 0 | 0 |
| 0 | 0 | 0 | 0 |
| 0 | 0 | 0 | 0 |
| 0 | 0 | 0 | 0 |
| 1 | 1 | 0 | 0 |
| 0 | 0 | 0 | 0 |
| 0 | 0 | 0 | 0 |
| 0 | 0 | 0 | 0 |
| 0 | 0 | 0 | 0 |
| 0 | 0 | 0 | 0 |
| 0 | 0 | 0 | 0 |
| 0 | 0 | 0 | 0 |
| 0 | 0 | 0 | 0 |
| 0 | 0 | 0 | 0 |
| 0 | 0 | 0 | 0 |
| 0 | 0 | 0 | 0 |
| 0 | 0 | 0 | 0 |
| 0 | 0 | 0 | 0 |
| 0 | 0 | 0 | 0 |
| 0 | 0 | 0 | 0 |
| 0 | 0 | 0 | 0 |
| 1 | 1 | 0 | 0 |
| 1 | 0 | 0 | 0 |
| 0 | 0 | 0 | 0 |
| 0 | 0 | 0 | 0 |
| 0 | 0 | 0 | 0 |
| 0 | 0 | 0 | 0 |
| 0 | 0 | 0 | 0 |
| 0 | 0 | 0 | 0 |
| 0 | 0 | 0 | 0 |
| 0 | 0 | 0 | 0 |
| 0 | 0 | 0 | 0 |
| 0 | 0 | 0 | 0 |
| 0 | 0 | 0 | 0 |
| 0 | 0 | 0 | 0 |
| 0 | 0 | 0 | 0 |
| 0 | 0 | 0 | 0 |
| 0 | 0 | 0 | 0 |
| 0 | 0 | 0 | 0 |
| 0 | 0 | 0 | 0 |
| 0 | 0 | 0 | 0 |
| 0 | 0 | 0 | 0 |
| 0 | 0 | 0 | 0 |
| 0 | 0 | 0 | 0 |
| 0 | 0 | 0 | 0 |
| 0 | 0 | 1 | 0 |
| 0 | 0 | 0 | 0 |

| | | | |
|---|---|---|---|
| 0 | 0 | 0 | 0 |
| 0 | 0 | 0 | 0 |
| 0 | 0 | 0 | 0 |
| 0 | 0 | 0 | 0 |
| 1 | 0 | 0 | 0 |
| 0 | 0 | 0 | 0 |
| 0 | 0 | 0 | 0 |
| 1 | 1 | 0 | 0 |
| 0 | 0 | 0 | 0 |
| 1 | 1 | 0 | 0 |
| 0 | 0 | 0 | 0 |
| 0 | 0 | 0 | 0 |
| 0 | 0 | 0 | 0 |
| 0 | 0 | 0 | 0 |
| 0 | 0 | 0 | 0 |
| 0 | 0 | 0 | 0 |
| 1 | 0 | 0 | 0 |
| 0 | 0 | 0 | 0 |
| 0 | 0 | 0 | 0 |
| 1 | 0 | 0 | 0 |
| 1 | 0 | 0 | 0 |
| 0 | 0 | 0 | 0 |
| 0 | 0 | 0 | 0 |
| 0 | 0 | 0 | 0 |
| 0 | 0 | 0 | 0 |
| 0 | 0 | 0 | 0 |
| 1 | 0 | 0 | 0 |
| 0 | 0 | 0 | 0 |
| 0 | 0 | 0 | 0 |
| 1 | 0 | 0 | 0 |
| 0 | 0 | 0 | 0 |
| 0 | 0 | 0 | 0 |
| 0 | 0 | 0 | 0 |
| 1 | 1 | 0 | 0 |
| 0 | 0 | 0 | 0 |
| 0 | 0 | 0 | 0 |
| 0 | 0 | 0 | 0 |
| 0 | 0 | 0 | 0 |
| 0 | 1 | 0 | 0 |
| 0 | 0 | 0 | 0 |
| 0 | 0 | 0 | 0 |
| 0 | 0 | 0 | 0 |
| 0 | 0 | 0 | 0 |
| 1 | 1 | 0 | 0 |
| 0 | 0 | 0 | 0 |
| 0 | 0 | 0 | 0 |
| 0 | 0 | 0 | 0 |
| 0 | 0 | 0 | 0 |
| 0 | 0 | 0 | 0 |
| 0 | 0 | 0 | 0 |
| 0 | 0 | 0 | 0 |
| 1 | 0 | 0 | 0 |
| 0 | 0 | 0 | 0 |
| 0 | 0 | 0 | 0 |

| | | | |
|---|---|---|---|
| 1 | 0 | 0 | 1 |
| 1 | 0 | 0 | 1 |
| 1 | 0 | 0 | 1 |
| 0 | 0 | 0 | 0 |
| 0 | 0 | 0 | 0 |
| 0 | 0 | 0 | 0 |
| 1 | 0 | 0 | 0 |
| 0 | 0 | 0 | 0 |
| 0 | 0 | 0 | 0 |
| 0 | 0 | 0 | 0 |
| 0 | 0 | 1 | 0 |
| 0 | 0 | 0 | 0 |
| 0 | 0 | 0 | 0 |
| 0 | 0 | 0 | 0 |
| 0 | 0 | 0 | 0 |
| 0 | 0 | 0 | 0 |
| 1 | 0 | 0 | 0 |
| 0 | 0 | 0 | 0 |
| 0 | 0 | 0 | 0 |
| 0 | 0 | 0 | 0 |
| 0 | 0 | 0 | 0 |
| 0 | 0 | 0 | 0 |
| 0 | 0 | 0 | 0 |
| 1 | 1 | 0 | 0 |
| 0 | 0 | 0 | 0 |
| 0 | 0 | 0 | 0 |
| 0 | 0 | 0 | 0 |
| 0 | 0 | 0 | 0 |
| 0 | 0 | 0 | 0 |
| 0 | 0 | 0 | 0 |
| 0 | 0 | 0 | 0 |
| 0 | 0 | 0 | 0 |
| 0 | 1 | 0 | 0 |
| 0 | 0 | 0 | 0 |
| 1 | 1 | 0 | 0 |
| 0 | 0 | 0 | 0 |
| 0 | 0 | 0 | 0 |
| 0 | 0 | 0 | 0 |
| 0 | 0 | 0 | 0 |
| 0 | 0 | 0 | 0 |
| 0 | 0 | 0 | 0 |
| 1 | 0 | 0 | 1 |
| 0 | 0 | 0 | 0 |
| 1 | 0 | 0 | 1 |
| 0 | 0 | 0 | 0 |
| 1 | 0 | 0 | 1 |
| 0 | 0 | 0 | 0 |
| 0 | 0 | 0 | 0 |
| 0 | 0 | 0 | 0 |
| 1 | 0 | 0 | 0 |
| 0 | 0 | 0 | 0 |
| 0 | 0 | 0 | 0 |
| 0 | 0 | 0 | 0 |
| 0 | 0 | 0 | 0 |

| | | | |
|---|---|---|---|
| 0 | 0 | 0 | 0 |
| 0 | 0 | 0 | 0 |
| 0 | 0 | 0 | 0 |
| 0 | 0 | 0 | 0 |
| 0 | 0 | 0 | 0 |
| 0 | 0 | 0 | 0 |
| 0 | 0 | 0 | 0 |
| 0 | 0 | 0 | 0 |
| 0 | 0 | 0 | 0 |
| 0 | 0 | 0 | 0 |
| 0 | 0 | 0 | 0 |
| 0 | 0 | 0 | 0 |
| 0 | 0 | 0 | 0 |
| 0 | 0 | 0 | 0 |
| 0 | 0 | 0 | 0 |
| 0 | 0 | 0 | 0 |
| 0 | 0 | 0 | 0 |
| 0 | 0 | 0 | 0 |
| 0 | 0 | 1 | 0 |
| 0 | 0 | 0 | 0 |
| 0 | 0 | 0 | 0 |
| 0 | 0 | 0 | 0 |
| 0 | 0 | 0 | 0 |
| 0 | 0 | 0 | 0 |
| 0 | 0 | 0 | 0 |
| 0 | 0 | 0 | 0 |
| 0 | 0 | 0 | 0 |
| 0 | 0 | 0 | 0 |
| 0 | 0 | 1 | 0 |
| 0 | 0 | 0 | 0 |
| 0 | 0 | 1 | 0 |
| 1 | 0 | 0 | 0 |
| 1 | 1 | 1 | 0 |
| 1 | 1 | 1 | 0 |
| 0 | 0 | 0 | 0 |
| 0 | 0 | 0 | 0 |
| 0 | 1 | 0 | 0 |
| 1 | 1 | 1 | 0 |
| 1 | 1 | 1 | 0 |
| 0 | 0 | 1 | 0 |
| 0 | 0 | 1 | 0 |
| 1 | 0 | 0 | 0 |
| 0 | 0 | 0 | 0 |
| 0 | 0 | 0 | 0 |
| 1 | 1 | 0 | 0 |
| 0 | 0 | 0 | 0 |
| 0 | 0 | 0 | 0 |
| 0 | 0 | 0 | 0 |
| 0 | 0 | 0 | 0 |
| 0 | 0 | 0 | 0 |
| 0 | 0 | 0 | 0 |
| 0 | 0 | 0 | 0 |
| 0 | 0 | 0 | 0 |
| 0 | 0 | 0 | 0 |

| | | | |
|---|---|---|---|
| 0 | 0 | 0 | 0 |
| 0 | 0 | 0 | 0 |
| 0 | 0 | 0 | 0 |
| 0 | 0 | 0 | 0 |
| 0 | 0 | 0 | 0 |
| 0 | 0 | 0 | 0 |
| 0 | 0 | 0 | 0 |
| 0 | 0 | 0 | 0 |
| 0 | 0 | 0 | 0 |
| 0 | 0 | 0 | 0 |
| 0 | 0 | 0 | 0 |
| 0 | 0 | 0 | 0 |
| 0 | 0 | 0 | 0 |
| 0 | 0 | 0 | 0 |
| 0 | 0 | 0 | 0 |
| 0 | 0 | 0 | 0 |
| 0 | 0 | 0 | 0 |
| 0 | 0 | 0 | 0 |
| 0 | 0 | 0 | 0 |
| 1 | 0 | 0 | 0 |
| 1 | 0 | 0 | 0 |
| 0 | 0 | 0 | 0 |
| 0 | 0 | 0 | 0 |
| 0 | 0 | 0 | 0 |
| 0 | 0 | 0 | 0 |
| 1 | 0 | 0 | 0 |
| 0 | 0 | 0 | 0 |
| 0 | 0 | 0 | 0 |
| 0 | 0 | 0 | 0 |
| 1 | 0 | 0 | 0 |
| 0 | 0 | 0 | 0 |
| 0 | 0 | 0 | 0 |
| 1 | 0 | 0 | 0 |
| 1 | 0 | 0 | 0 |
| 0 | 0 | 0 | 0 |
| 0 | 0 | 0 | 0 |
| 0 | 0 | 0 | 0 |
| 0 | 0 | 0 | 0 |
| 0 | 0 | 0 | 0 |
| 0 | 0 | 0 | 0 |
| 0 | 0 | 0 | 0 |
| 0 | 0 | 0 | 0 |
| 0 | 0 | 0 | 0 |
| 0 | 0 | 0 | 0 |
| 0 | 0 | 0 | 0 |
| 0 | 0 | 0 | 0 |
| 0 | 0 | 0 | 0 |
| 0 | 1 | 0 | 0 |
| 0 | 1 | 0 | 0 |
| 0 | 0 | 0 | 0 |
| 0 | 1 | 0 | 0 |
| 0 | 1 | 0 | 0 |
| 0 | 0 | 0 | 0 |
| 0 | 0 | 0 | 0 |

| | | | |
|---|---|---|---|
| 0 | 0 | 0 | 0 |
| 0 | 0 | 0 | 0 |
| 1 | 1 | 0 | 1 |
| 0 | 0 | 0 | 0 |
| 0 | 0 | 0 | 0 |
| 0 | 0 | 0 | 0 |
| 0 | 0 | 0 | 0 |
| 0 | 1 | 0 | 0 |
| 1 | 1 | 0 | 0 |
| 1 | 1 | 0 | 0 |
| 1 | 1 | 0 | 0 |
| 1 | 1 | 0 | 0 |
| 1 | 1 | 0 | 0 |
| 0 | 1 | 0 | 0 |
| 0 | 1 | 0 | 0 |
| 0 | 0 | 0 | 0 |
| 0 | 0 | 0 | 0 |
| 0 | 0 | 0 | 0 |
| 0 | 0 | 0 | 0 |
| 0 | 0 | 0 | 0 |
| 0 | 0 | 0 | 0 |
| 0 | 0 | 0 | 0 |
| 0 | 1 | 0 | 0 |
| 0 | 1 | 0 | 0 |
| 0 | 0 | 0 | 0 |
| 0 | 0 | 1 | 0 |
| 0 | 0 | 0 | 0 |
| 0 | 0 | 0 | 0 |
| 0 | 0 | 0 | 0 |
| 0 | 1 | 0 | 0 |
| 0 | 0 | 0 | 0 |
| 0 | 0 | 0 | 0 |
| 0 | 0 | 0 | 0 |
| 0 | 0 | 0 | 0 |
| 0 | 0 | 0 | 0 |
| 0 | 0 | 0 | 0 |
| 0 | 0 | 0 | 0 |
| 0 | 0 | 0 | 0 |
| 0 | 0 | 0 | 0 |
| 1 | 0 | 0 | 0 |
| 1 | 1 | 0 | 0 |
| 1 | 0 | 0 | 0 |
| 0 | 0 | 0 | 0 |
| 0 | 1 | 0 | 0 |
| 1 | 1 | 0 | 0 |
| 0 | 1 | 0 | 0 |
| 0 | 0 | 0 | 0 |
| 1 | 0 | 0 | 0 |
| 0 | 1 | 0 | 0 |
| 0 | 0 | 0 | 0 |
| 0 | 0 | 0 | 0 |
| 0 | 0 | 0 | 0 |
| 1 | 1 | 0 | 0 |
| 0 | 0 | 0 | 0 |

| | | | |
|---|---|---|---|
| 0 | 0 | 0 | 0 |
| 0 | 0 | 0 | 0 |
| 1 | 0 | 0 | 0 |
| 1 | 1 | 0 | 0 |
| 1 | 1 | 0 | 0 |
| 0 | 0 | 0 | 0 |
| 0 | 0 | 0 | 0 |
| 0 | 0 | 0 | 0 |
| 1 | 0 | 0 | 0 |
| 0 | 0 | 0 | 0 |
| 0 | 0 | 0 | 0 |
| 1 | 0 | 0 | 1 |
| 0 | 0 | 0 | 0 |
| 1 | 0 | 0 | 0 |
| 0 | 0 | 0 | 0 |
| 0 | 0 | 0 | 0 |
| 0 | 0 | 0 | 0 |
| 0 | 0 | 0 | 0 |
| 0 | 0 | 0 | 0 |
| 0 | 0 | 0 | 0 |
| 0 | 0 | 0 | 0 |
| 1 | 0 | 0 | 1 |
| 1 | 0 | 0 | 0 |
| 0 | 0 | 0 | 0 |
| 0 | 0 | 0 | 0 |
| 0 | 0 | 0 | 0 |
| 0 | 0 | 0 | 0 |
| 0 | 0 | 0 | 0 |
| 1 | 0 | 0 | 0 |
| 1 | 0 | 0 | 0 |
| 0 | 0 | 0 | 0 |
| 0 | 0 | 0 | 0 |
| 0 | 0 | 0 | 0 |
| 0 | 0 | 0 | 0 |
| 0 | 0 | 0 | 0 |
| 0 | 0 | 0 | 0 |
| 1 | 0 | 0 | 0 |
| 0 | 0 | 0 | 0 |
| 0 | 0 | 0 | 0 |
| 1 | 0 | 0 | 0 |
| 1 | 0 | 0 | 0 |
| 1 | 0 | 0 | 0 |
| 1 | 0 | 0 | 0 |
| 0 | 0 | 0 | 0 |
| 0 | 0 | 0 | 0 |
| 1 | 0 | 0 | 0 |
| 0 | 0 | 0 | 0 |
| 0 | 0 | 0 | 0 |
| 0 | 0 | 0 | 0 |
| 0 | 0 | 0 | 0 |
| 0 | 0 | 0 | 0 |
| 1 | 0 | 0 | 0 |
| 1 | 0 | 0 | 0 |
| 1 | 0 | 0 | 0 |

| | | | |
|---|---|---|---|
| 1 | 0 | 0 | 0 |
| 0 | 0 | 0 | 0 |
| 0 | 0 | 0 | 0 |
| 0 | 0 | 0 | 0 |
| 1 | 0 | 0 | 0 |
| 0 | 0 | 0 | 0 |
| 1 | 0 | 0 | 0 |
| 0 | 0 | 0 | 0 |
| 0 | 0 | 0 | 0 |
| 0 | 0 | 0 | 0 |
| 0 | 0 | 0 | 0 |
| 0 | 0 | 0 | 0 |
| 0 | 0 | 0 | 0 |
| 0 | 0 | 0 | 0 |
| 0 | 0 | 0 | 0 |
| 1 | 0 | 0 | 0 |
| 0 | 0 | 0 | 0 |
| 0 | 0 | 0 | 0 |
| 0 | 0 | 0 | 0 |
| 0 | 0 | 0 | 0 |
| 0 | 0 | 0 | 0 |
| 0 | 0 | 0 | 0 |
| 1 | 0 | 0 | 0 |
| 1 | 0 | 0 | 0 |
| 0 | 0 | 0 | 0 |
| 0 | 0 | 0 | 0 |
| 0 | 0 | 0 | 0 |
| 0 | 0 | 0 | 0 |
| 0 | 0 | 0 | 0 |
| 0 | 0 | 0 | 0 |
| 0 | 0 | 0 | 0 |
| 0 | 0 | 0 | 0 |
| 1 | 0 | 0 | 0 |
| 0 | 0 | 0 | 0 |
| 0 | 0 | 0 | 0 |
| 0 | 0 | 0 | 0 |
| 0 | 0 | 0 | 0 |
| 0 | 0 | 0 | 0 |
| 0 | 0 | 0 | 0 |
| 0 | 0 | 0 | 0 |
| 0 | 0 | 0 | 0 |
| 0 | 0 | 0 | 0 |
| 0 | 0 | 0 | 0 |
| 0 | 0 | 0 | 0 |
| 0 | 1 | 0 | 0 |
| 0 | 0 | 0 | 0 |
| 0 | 0 | 0 | 0 |
| 0 | 0 | 0 | 0 |
| 0 | 0 | 0 | 0 |
| 0 | 0 | 0 | 0 |
| 0 | 0 | 0 | 0 |
| 0 | 0 | 0 | 0 |
| 0 | 0 | 0 | 0 |
| 1 | 0 | 0 | 0 |

| | | | |
|---|---|---|---|
| 0 | 0 | 0 | 0 |
| 0 | 0 | 0 | 0 |
| 0 | 0 | 0 | 0 |
| 0 | 0 | 0 | 0 |
| 0 | 0 | 0 | 0 |
| 0 | 0 | 0 | 0 |
| 0 | 0 | 0 | 0 |
| 0 | 0 | 0 | 0 |
| 0 | 0 | 0 | 0 |
| 1 | 0 | 0 | 0 |
| 0 | 0 | 0 | 0 |
| 0 | 0 | 0 | 0 |
| 1 | 1 | 0 | 0 |
| 1 | 1 | 0 | 0 |
| 1 | 1 | 0 | 0 |
| 1 | 1 | 0 | 0 |
| 0 | 0 | 0 | 0 |
| 0 | 0 | 0 | 0 |
| 0 | 0 | 0 | 0 |
| 0 | 0 | 0 | 0 |
| 0 | 0 | 0 | 0 |
| 0 | 0 | 0 | 0 |
| 0 | 0 | 0 | 0 |
| 0 | 0 | 0 | 0 |
| 0 | 0 | 0 | 0 |
| 0 | 0 | 0 | 1 |
| 0 | 0 | 0 | 0 |
| 0 | 0 | 0 | 0 |
| 0 | 0 | 0 | 0 |
| 0 | 0 | 0 | 0 |
| 0 | 0 | 0 | 0 |
| 0 | 0 | 0 | 0 |
| 0 | 0 | 0 | 0 |
| 0 | 0 | 0 | 0 |
| 0 | 0 | 0 | 0 |
| 0 | 0 | 0 | 0 |
| 0 | 0 | 0 | 0 |
| 1 | 1 | 0 | 0 |
| 0 | 0 | 0 | 0 |
| 0 | 0 | 0 | 0 |
| 1 | 1 | 0 | 0 |
| 0 | 0 | 0 | 0 |
| 0 | 0 | 0 | 0 |
| 1 | 0 | 0 | 1 |
| 0 | 0 | 0 | 0 |
| 0 | 0 | 0 | 0 |
| 0 | 0 | 0 | 0 |
| 0 | 0 | 0 | 0 |
| 0 | 0 | 0 | 0 |
| 0 | 0 | 0 | 0 |
| 0 | 0 | 0 | 0 |
| 0 | 0 | 1 | 0 |
| 0 | 0 | 1 | 0 |
| 0 | 0 | 1 | 0 |

| | | | |
|---|---|---|---|
| 0 | 0 | 1 | 0 |
| 0 | 0 | 0 | 0 |
| 0 | 0 | 0 | 0 |
| 0 | 0 | 0 | 0 |
| 0 | 0 | 0 | 0 |
| 0 | 0 | 0 | 0 |
| 0 | 0 | 0 | 0 |
| 0 | 0 | 0 | 0 |
| 0 | 0 | 0 | 0 |
| 0 | 0 | 0 | 0 |
| 0 | 0 | 0 | 0 |
| 0 | 0 | 0 | 0 |
| 0 | 0 | 0 | 0 |
| 0 | 0 | 0 | 0 |
| 0 | 0 | 0 | 0 |
| 0 | 0 | 0 | 0 |
| 0 | 0 | 0 | 0 |
| 0 | 0 | 0 | 0 |
| 0 | 0 | 0 | 0 |
| 0 | 0 | 0 | 0 |
| 0 | 0 | 0 | 0 |
| 0 | 0 | 0 | 0 |
| 0 | 0 | 0 | 0 |
| 0 | 1 | 0 | 1 |
| 0 | 0 | 0 | 0 |
| 0 | 0 | 0 | 0 |
| 0 | 0 | 0 | 0 |
| 0 | 0 | 0 | 0 |
| 0 | 0 | 0 | 0 |
| 0 | 0 | 0 | 0 |
| 0 | 0 | 0 | 0 |
| 0 | 0 | 0 | 0 |
| 0 | 0 | 0 | 0 |
| 0 | 0 | 0 | 0 |
| 0 | 0 | 0 | 0 |
| 0 | 0 | 0 | 0 |
| 0 | 0 | 0 | 0 |
| 0 | 0 | 0 | 0 |
| 0 | 0 | 0 | 0 |
| 0 | 0 | 0 | 0 |
| 0 | 0 | 0 | 0 |
| 0 | 0 | 0 | 0 |
| 0 | 0 | 0 | 0 |
| 0 | 0 | 0 | 0 |
| 0 | 0 | 0 | 0 |
| 0 | 0 | 0 | 0 |
| 1 | 0 | 0 | 0 |
| 1 | 0 | 0 | 0 |
| 0 | 0 | 0 | 0 |
| 0 | 1 | 0 | 0 |
| 1 | 1 | 0 | 1 |
| 1 | 1 | 0 | 1 |
| 1 | 1 | 0 | 1 |
| 0 | 0 | 0 | 0 |

| | | | |
|---|---|---|---|
| 1 | 1 | 0 | 0 |
| 0 | 0 | 0 | 0 |
| 0 | 0 | 0 | 0 |
| 0 | 0 | 0 | 0 |
| 0 | 0 | 0 | 0 |
| 0 | 0 | 0 | 0 |
| 0 | 0 | 0 | 0 |
| 0 | 0 | 0 | 0 |
| 0 | 0 | 0 | 0 |
| 0 | 0 | 0 | 0 |
| 0 | 0 | 0 | 0 |
| 0 | 1 | 0 | 0 |
| 0 | 0 | 0 | 0 |
| 0 | 0 | 0 | 0 |
| 0 | 1 | 0 | 0 |
| 0 | 0 | 0 | 0 |
| 1 | 1 | 0 | 0 |
| 0 | 0 | 0 | 0 |
| 0 | 0 | 0 | 0 |
| 1 | 0 | 0 | 0 |
| 0 | 0 | 0 | 0 |
| 0 | 0 | 0 | 0 |
| 0 | 0 | 0 | 0 |
| 0 | 0 | 0 | 0 |
| 0 | 0 | 0 | 0 |
| 0 | 0 | 0 | 0 |
| 0 | 0 | 0 | 0 |
| 0 | 0 | 0 | 0 |
| 0 | 0 | 0 | 0 |
| 0 | 0 | 0 | 0 |
| 0 | 0 | 0 | 0 |
| 0 | 0 | 0 | 0 |
| 0 | 0 | 0 | 0 |
| 1 | 0 | 0 | 0 |
| 0 | 0 | 0 | 0 |
| 0 | 1 | 0 | 0 |
| 0 | 0 | 0 | 0 |
| 0 | 0 | 0 | 0 |
| 0 | 0 | 0 | 0 |
| 0 | 0 | 0 | 0 |
| 0 | 0 | 0 | 0 |
| 0 | 0 | 0 | 0 |
| 0 | 0 | 0 | 0 |
| 0 | 0 | 0 | 0 |
| 0 | 0 | 0 | 0 |
| 0 | 0 | 0 | 0 |
| 0 | 0 | 0 | 0 |
| 0 | 0 | 0 | 0 |
| 1 | 1 | 0 | 0 |
| 0 | 0 | 0 | 0 |
| 0 | 0 | 0 | 0 |
| 0 | 0 | 0 | 0 |
| 0 | 0 | 0 | 0 |
| 0 | 0 | 0 | 0 |

| | | | |
|---|---|---|---|
| 0 | 0 | 0 | 0 |
| 0 | 0 | 0 | 0 |
| 0 | 0 | 0 | 0 |
| 0 | 0 | 0 | 0 |
| 0 | 0 | 0 | 0 |
| 0 | 0 | 0 | 0 |
| 0 | 0 | 0 | 0 |
| 0 | 1 | 0 | 0 |
| 0 | 0 | 0 | 0 |
| 0 | 0 | 0 | 0 |
| 0 | 0 | 0 | 0 |
| 0 | 0 | 0 | 0 |
| 0 | 0 | 1 | 0 |
| 0 | 0 | 0 | 0 |
| 0 | 0 | 1 | 0 |
| 0 | 0 | 1 | 0 |
| 0 | 0 | 0 | 0 |
| 0 | 0 | 0 | 0 |
| 0 | 0 | 0 | 0 |
| 0 | 0 | 0 | 0 |
| 0 | 1 | 0 | 0 |
| 0 | 1 | 0 | 0 |
| 0 | 0 | 0 | 0 |
| 0 | 0 | 0 | 0 |
| 0 | 1 | 0 | 0 |
| 0 | 0 | 0 | 0 |
| 0 | 0 | 0 | 0 |
| 0 | 1 | 0 | 0 |
| 0 | 0 | 0 | 0 |
| 0 | 0 | 0 | 0 |
| 0 | 0 | 0 | 0 |
| 0 | 0 | 0 | 0 |
| 1 | 1 | 0 | 0 |
| 1 | 1 | 0 | 0 |
| 1 | 1 | 0 | 0 |
| 0 | 1 | 0 | 0 |
| 1 | 0 | 0 | 0 |
| 0 | 1 | 0 | 0 |
| 1 | 1 | 0 | 0 |
| 0 | 0 | 0 | 0 |
| 0 | 0 | 0 | 0 |
| 0 | 0 | 0 | 0 |
| 0 | 0 | 0 | 0 |
| 0 | 0 | 0 | 0 |
| 0 | 0 | 0 | 0 |
| 0 | 0 | 0 | 0 |
| 0 | 0 | 0 | 0 |
| 0 | 0 | 0 | 0 |
| 0 | 0 | 0 | 0 |
| 1 | 1 | 0 | 0 |
| 1 | 1 | 0 | 0 |
| 0 | 0 | 0 | 0 |
| 0 | 0 | 0 | 0 |
| 0 | 0 | 0 | 0 |

| | | | |
|---|---|---|---|
| 0 | 0 | 0 | 0 |
| 0 | 0 | 0 | 0 |
| 0 | 0 | 0 | 0 |
| 0 | 0 | 0 | 0 |
| 0 | 0 | 0 | 0 |
| 0 | 0 | 0 | 0 |
| 1 | 1 | 0 | 0 |
| 1 | 1 | 0 | 0 |
| 0 | 0 | 0 | 0 |
| 0 | 0 | 0 | 0 |
| 0 | 0 | 0 | 0 |
| 0 | 0 | 0 | 0 |
| 0 | 0 | 0 | 0 |
| 0 | 0 | 0 | 0 |
| 0 | 0 | 0 | 0 |
| 0 | 0 | 0 | 0 |
| 0 | 0 | 0 | 0 |
| 0 | 0 | 0 | 0 |
| 0 | 0 | 0 | 0 |
| 0 | 0 | 0 | 0 |
| 0 | 0 | 0 | 0 |
| 0 | 0 | 0 | 0 |
| 0 | 0 | 0 | 0 |
| 0 | 0 | 0 | 0 |
| 0 | 0 | 0 | 0 |
| 0 | 0 | 0 | 0 |
| 0 | 0 | 0 | 0 |
| 1 | 1 | 0 | 0 |
| 0 | 0 | 0 | 0 |
| 0 | 0 | 0 | 0 |
| 0 | 0 | 0 | 0 |
| 0 | 0 | 0 | 0 |
| 0 | 0 | 0 | 0 |
| 0 | 0 | 0 | 0 |
| 0 | 0 | 0 | 0 |
| 0 | 0 | 0 | 0 |
| 0 | 0 | 0 | 0 |
| 0 | 0 | 0 | 0 |
| 0 | 0 | 0 | 0 |
| 0 | 0 | 0 | 0 |
| 0 | 0 | 0 | 0 |
| 0 | 0 | 0 | 0 |
| 0 | 0 | 0 | 0 |
| 0 | 0 | 0 | 0 |
| 0 | 0 | 0 | 0 |
| 0 | 0 | 0 | 0 |
| 0 | 0 | 0 | 0 |
| 0 | 0 | 0 | 0 |
| 0 | 0 | 0 | 0 |
| 0 | 0 | 0 | 0 |
| 0 | 0 | 0 | 0 |
| 0 | 0 | 0 | 0 |
| 0 | 0 | 0 | 0 |
| 0 | 0 | 0 | 0 |

| | | | |
|---|---|---|---|
| 0 | 0 | 0 | 0 |
| 0 | 0 | 0 | 0 |
| 0 | 0 | 0 | 0 |
| 0 | 0 | 0 | 0 |
| 0 | 0 | 0 | 0 |
| 0 | 0 | 0 | 0 |
| 0 | 0 | 0 | 0 |
| 0 | 0 | 0 | 0 |
| 0 | 0 | 0 | 0 |
| 0 | 0 | 0 | 0 |
| 1 | 0 | 0 | 0 |
| 1 | 1 | 0 | 0 |
| 1 | 0 | 0 | 0 |
| 0 | 0 | 0 | 0 |
| 0 | 0 | 0 | 0 |
| 0 | 0 | 0 | 0 |
| 0 | 0 | 0 | 0 |
| 0 | 0 | 0 | 0 |
| 0 | 0 | 0 | 0 |
| 0 | 0 | 0 | 0 |
| 0 | 0 | 0 | 0 |
| 0 | 0 | 0 | 0 |
| 0 | 0 | 0 | 0 |
| 0 | 0 | 0 | 0 |
| 0 | 0 | 0 | 0 |
| 0 | 0 | 0 | 0 |
| 0 | 0 | 0 | 0 |
| 0 | 0 | 0 | 0 |
| 0 | 0 | 0 | 0 |
| 0 | 0 | 0 | 0 |
| 0 | 0 | 0 | 0 |
| 0 | 1 | 0 | 0 |
| 0 | 0 | 0 | 0 |
| 0 | 0 | 0 | 0 |
| 1 | 0 | 0 | 0 |
| 1 | 0 | 0 | 0 |
| 1 | 0 | 0 | 0 |
| 0 | 0 | 0 | 0 |
| 0 | 0 | 0 | 0 |
| 0 | 0 | 0 | 0 |
| 0 | 0 | 0 | 0 |
| 0 | 0 | 0 | 0 |
| 0 | 0 | 0 | 0 |
| 0 | 0 | 0 | 0 |
| 0 | 0 | 0 | 0 |
| 0 | 0 | 0 | 0 |
| 0 | 0 | 0 | 0 |
| 0 | 0 | 0 | 0 |
| 0 | 0 | 0 | 0 |
| 0 | 0 | 0 | 0 |
| 0 | 0 | 0 | 0 |
| 0 | 0 | 0 | 0 |
| 0 | 0 | 0 | 1 |
| 0 | 0 | 0 | 0 |

| | | | |
|---|---|---|---|
| 0 | 0 | 0 | 0 |
| 0 | 0 | 0 | 0 |
| 0 | 0 | 0 | 0 |
| 0 | 0 | 0 | 0 |
| 0 | 0 | 0 | 0 |
| 0 | 0 | 0 | 0 |
| 0 | 0 | 0 | 0 |
| 0 | 0 | 0 | 0 |
| 0 | 0 | 0 | 0 |
| 0 | 0 | 0 | 0 |
| 0 | 0 | 0 | 0 |
| 1 | 0 | 0 | 0 |
| 0 | 0 | 0 | 0 |
| 0 | 0 | 0 | 0 |
| 0 | 0 | 1 | 0 |
| 0 | 0 | 0 | 0 |
| 0 | 0 | 0 | 0 |
| 0 | 0 | 0 | 0 |
| 0 | 0 | 0 | 0 |
| 0 | 0 | 0 | 0 |
| 0 | 0 | 0 | 0 |
| 0 | 0 | 0 | 0 |
| 0 | 0 | 0 | 0 |
| 0 | 0 | 0 | 0 |
| 0 | 0 | 0 | 0 |
| 0 | 0 | 0 | 0 |
| 0 | 0 | 0 | 0 |
| 0 | 0 | 0 | 0 |
| 0 | 0 | 0 | 0 |
| 0 | 0 | 0 | 0 |
| 0 | 0 | 0 | 0 |
| 0 | 0 | 0 | 0 |
| 0 | 0 | 0 | 0 |
| 1 | 0 | 0 | 0 |
| 0 | 0 | 0 | 0 |
| 1 | 0 | 0 | 1 |
| 0 | 0 | 0 | 0 |
| 0 | 0 | 0 | 0 |
| 0 | 0 | 0 | 0 |
| 0 | 0 | 0 | 0 |
| 0 | 0 | 0 | 0 |
| 0 | 0 | 0 | 0 |
| 0 | 0 | 0 | 0 |
| 0 | 1 | 0 | 0 |
| 0 | 0 | 0 | 0 |
| 0 | 0 | 0 | 0 |
| 0 | 0 | 0 | 0 |
| 0 | 0 | 0 | 0 |
| 0 | 0 | 0 | 0 |
| 0 | 0 | 0 | 0 |
| 0 | 0 | 0 | 0 |
| 0 | 0 | 0 | 0 |
| 0 | 0 | 0 | 0 |
| 0 | 0 | 0 | 0 |

| | | | |
|---|---|---|---|
| 0 | 0 | 0 | 0 |
| 0 | 0 | 0 | 0 |
| 0 | 0 | 0 | 0 |
| 0 | 0 | 0 | 0 |
| 0 | 0 | 0 | 0 |
| 0 | 1 | 0 | 0 |
| 0 | 0 | 0 | 0 |
| 0 | 0 | 0 | 0 |
| 0 | 0 | 0 | 0 |
| 0 | 0 | 0 | 0 |
| 0 | 0 | 0 | 0 |
| 0 | 0 | 0 | 0 |
| 0 | 0 | 0 | 0 |
| 0 | 0 | 0 | 0 |
| 0 | 0 | 0 | 0 |
| 0 | 1 | 0 | 0 |
| 1 | 1 | 0 | 0 |
| 0 | 0 | 1 | 0 |
| 0 | 0 | 0 | 0 |
| 0 | 0 | 0 | 0 |
| 0 | 0 | 0 | 0 |
| 0 | 0 | 0 | 0 |
| 0 | 0 | 0 | 0 |
| 0 | 0 | 0 | 0 |
| 0 | 0 | 0 | 0 |
| 0 | 0 | 0 | 0 |
| 0 | 0 | 0 | 0 |
| 1 | 0 | 0 | 0 |
| 0 | 0 | 0 | 0 |
| 0 | 0 | 0 | 0 |
| 0 | 0 | 1 | 0 |
| 0 | 0 | 0 | 0 |
| 0 | 0 | 0 | 0 |
| 0 | 0 | 0 | 0 |
| 0 | 0 | 0 | 0 |
| 0 | 0 | 0 | 0 |
| 0 | 0 | 0 | 0 |
| 1 | 1 | 0 | 0 |
| 0 | 0 | 0 | 0 |
| 0 | 0 | 0 | 0 |
| 0 | 0 | 0 | 0 |
| 1 | 0 | 0 | 0 |
| 1 | 0 | 0 | 0 |
| 0 | 0 | 0 | 0 |
| 0 | 0 | 0 | 0 |
| 0 | 0 | 0 | 0 |
| 0 | 0 | 0 | 0 |
| 0 | 0 | 0 | 0 |
| 0 | 0 | 0 | 0 |
| 0 | 0 | 0 | 0 |
| 0 | 0 | 0 | 0 |
| 0 | 0 | 0 | 0 |
| 0 | 0 | 0 | 0 |
| 0 | 0 | 0 | 0 |

| | | | |
|---|---|---|---|
| 0 | 0 | 0 | 0 |
| 0 | 0 | 0 | 0 |
| 0 | 0 | 0 | 0 |
| 0 | 0 | 0 | 0 |
| 0 | 0 | 0 | 0 |
| 0 | 0 | 0 | 0 |
| 0 | 0 | 0 | 0 |
| 0 | 0 | 0 | 0 |
| 0 | 0 | 0 | 0 |
| 0 | 0 | 0 | 0 |
| 0 | 0 | 0 | 0 |
| 0 | 0 | 0 | 0 |
| 1 | 0 | 0 | 0 |
| 0 | 0 | 0 | 0 |
| 0 | 0 | 0 | 0 |
| 0 | 0 | 0 | 0 |
| 0 | 0 | 0 | 0 |
| 0 | 0 | 0 | 0 |
| 0 | 0 | 0 | 0 |
| 0 | 0 | 0 | 0 |
| 0 | 0 | 0 | 0 |
| 0 | 0 | 0 | 0 |
| 0 | 0 | 0 | 0 |
| 1 | 0 | 0 | 0 |
| 0 | 0 | 0 | 0 |
| 0 | 0 | 0 | 0 |
| 0 | 0 | 0 | 0 |
| 1 | 1 | 0 | 1 |
| 0 | 0 | 0 | 0 |
| 0 | 0 | 0 | 0 |
| 0 | 0 | 0 | 0 |
| 0 | 0 | 0 | 0 |
| 0 | 0 | 0 | 0 |
| 0 | 0 | 0 | 0 |
| 0 | 0 | 0 | 0 |
| 0 | 0 | 0 | 0 |
| 0 | 0 | 0 | 0 |
| 0 | 0 | 0 | 0 |
| 0 | 0 | 0 | 0 |
| 0 | 0 | 0 | 0 |
| 0 | 0 | 0 | 0 |
| 0 | 0 | 0 | 0 |
| 0 | 0 | 0 | 0 |
| 0 | 0 | 0 | 0 |
| 0 | 0 | 0 | 0 |
| 0 | 0 | 0 | 0 |
| 0 | 0 | 0 | 0 |
| 0 | 0 | 0 | 0 |
| 0 | 0 | 0 | 0 |
| 0 | 0 | 0 | 0 |
| 0 | 0 | 0 | 0 |
| 0 | 0 | 0 | 0 |
| 0 | 0 | 0 | 0 |
| 0 | 0 | 0 | 0 |

| | | | |
|---|---|---|---|
| 0 | 0 | 0 | 0 |
| 0 | 0 | 0 | 0 |
| 0 | 0 | 0 | 0 |
| 0 | 0 | 0 | 0 |
| 0 | 0 | 0 | 0 |
| 0 | 0 | 0 | 0 |
| 0 | 0 | 0 | 0 |
| 0 | 0 | 0 | 0 |
| 1 | 0 | 0 | 0 |
| 1 | 0 | 0 | 0 |
| 1 | 0 | 0 | 0 |
| 0 | 0 | 0 | 0 |
| 1 | 0 | 0 | 0 |
| 0 | 0 | 0 | 0 |
| 0 | 0 | 0 | 0 |
| 1 | 0 | 0 | 0 |
| 0 | 0 | 0 | 0 |
| 0 | 0 | 0 | 0 |
| 0 | 0 | 0 | 0 |
| 0 | 0 | 0 | 0 |
| 0 | 0 | 0 | 0 |
| 0 | 0 | 0 | 0 |
| 0 | 0 | 0 | 0 |
| 0 | 0 | 0 | 0 |
| 0 | 0 | 0 | 0 |
| 0 | 0 | 0 | 0 |
| 0 | 0 | 0 | 0 |
| 1 | 0 | 0 | 0 |
| 1 | 0 | 0 | 0 |
| 0 | 0 | 0 | 0 |
| 0 | 0 | 0 | 0 |
| 0 | 0 | 0 | 0 |
| 1 | 0 | 0 | 0 |
| 0 | 0 | 0 | 0 |
| 0 | 0 | 0 | 0 |
| 0 | 0 | 0 | 0 |
| 0 | 0 | 0 | 0 |
| 0 | 0 | 0 | 0 |
| 0 | 0 | 0 | 0 |
| 0 | 0 | 0 | 0 |
| 1 | 0 | 0 | 0 |
| 1 | 0 | 0 | 0 |
| 0 | 0 | 0 | 0 |
| 0 | 0 | 0 | 0 |
| 0 | 0 | 0 | 0 |
| 0 | 0 | 0 | 0 |
| 0 | 0 | 0 | 0 |
| 0 | 0 | 0 | 0 |
| 0 | 0 | 0 | 0 |
| 0 | 0 | 0 | 0 |
| 0 | 0 | 0 | 0 |
| 0 | 0 | 0 | 0 |
| 0 | 0 | 0 | 0 |
| 0 | 0 | 0 | 0 |

| | | | |
|---|---|---|---|
| 0 | 0 | 0 | 0 |
| 0 | 0 | 0 | 0 |
| 0 | 0 | 0 | 0 |
| 0 | 0 | 0 | 0 |
| 0 | 0 | 0 | 0 |
| 0 | 0 | 0 | 0 |
| 0 | 0 | 0 | 0 |
| 0 | 0 | 0 | 0 |
| 0 | 0 | 0 | 0 |
| 0 | 0 | 0 | 0 |
| 1 | 0 | 0 | 0 |
| 0 | 0 | 0 | 0 |
| 1 | 0 | 0 | 0 |
| 0 | 0 | 0 | 0 |
| 0 | 0 | 0 | 0 |
| 0 | 0 | 0 | 0 |
| 0 | 0 | 0 | 0 |
| 0 | 1 | 0 | 0 |
| 0 | 0 | 0 | 0 |
| 0 | 0 | 0 | 0 |
| 0 | 0 | 0 | 0 |
| 0 | 0 | 0 | 0 |
| 1 | 0 | 0 | 0 |
| 0 | 0 | 0 | 0 |
| 0 | 0 | 0 | 0 |
| 1 | 0 | 0 | 0 |
| 1 | 0 | 0 | 0 |
| 1 | 0 | 0 | 0 |
| 0 | 0 | 0 | 0 |
| 0 | 0 | 0 | 0 |
| 0 | 0 | 0 | 0 |
| 0 | 0 | 0 | 0 |
| 0 | 0 | 0 | 0 |
| 0 | 0 | 0 | 0 |
| 0 | 0 | 0 | 0 |
| 0 | 0 | 0 | 0 |
| 0 | 0 | 0 | 0 |
| 0 | 0 | 0 | 0 |
| 0 | 0 | 0 | 0 |
| 0 | 0 | 0 | 0 |
| 1 | 1 | 0 | 0 |
| 0 | 0 | 0 | 0 |
| 0 | 0 | 0 | 0 |
| 0 | 0 | 0 | 0 |
| 0 | 0 | 0 | 0 |
| 0 | 0 | 0 | 0 |
| 0 | 0 | 0 | 0 |
| 1 | 0 | 0 | 0 |
| 0 | 0 | 0 | 0 |
| 0 | 0 | 0 | 0 |
| 0 | 0 | 0 | 0 |
| 0 | 0 | 0 | 0 |
| 0 | 0 | 0 | 0 |
| 0 | 0 | 0 | 0 |

| | | | |
|---|---|---|---|
| 1 | 0 | 0 | 0 |
| 0 | 0 | 0 | 0 |
| 1 | 0 | 0 | 0 |
| 0 | 0 | 0 | 0 |
| 0 | 0 | 0 | 0 |
| 0 | 0 | 0 | 0 |
| 0 | 0 | 0 | 0 |
| 0 | 0 | 0 | 0 |
| 1 | 1 | 0 | 0 |
| 0 | 0 | 0 | 0 |
| 1 | 1 | 0 | 0 |
| 1 | 1 | 0 | 0 |
| 0 | 0 | 0 | 0 |
| 1 | 1 | 0 | 0 |
| 0 | 0 | 0 | 0 |
| 0 | 0 | 0 | 0 |
| 0 | 0 | 0 | 0 |
| 0 | 0 | 0 | 0 |
| 0 | 0 | 0 | 0 |
| 0 | 0 | 0 | 0 |
| 1 | 0 | 0 | 0 |
| 0 | 0 | 0 | 0 |
| 0 | 0 | 0 | 0 |
| 0 | 0 | 0 | 0 |
| 1 | 0 | 0 | 0 |
| 0 | 0 | 0 | 0 |
| 1 | 0 | 0 | 0 |
| 1 | 0 | 0 | 0 |
| 1 | 1 | 0 | 0 |
| 0 | 0 | 0 | 0 |
| 1 | 1 | 0 | 0 |
| 1 | 1 | 0 | 0 |
| 1 | 1 | 0 | 0 |
| 0 | 0 | 0 | 0 |
| 1 | 1 | 0 | 0 |
| 0 | 0 | 0 | 0 |
| 1 | 0 | 0 | 0 |
| 0 | 0 | 0 | 0 |
| 1 | 1 | 0 | 0 |
| 1 | 1 | 0 | 0 |
| 1 | 0 | 0 | 0 |
| 0 | 0 | 0 | 0 |
| 0 | 0 | 0 | 0 |
| 1 | 0 | 0 | 0 |
| 0 | 0 | 0 | 0 |
| 0 | 0 | 0 | 0 |
| 1 | 0 | 0 | 0 |
| 0 | 0 | 0 | 0 |
| 1 | 0 | 0 | 0 |
| 0 | 0 | 0 | 0 |
| 0 | 0 | 0 | 0 |
| 0 | 0 | 0 | 0 |
| 0 | 0 | 0 | 0 |
| 0 | 0 | 0 | 0 |

| | | | |
|---|---|---|---|
| 0 | 0 | 0 | 0 |
| 0 | 0 | 0 | 0 |
| 0 | 0 | 0 | 0 |
| 0 | 0 | 0 | 0 |
| 0 | 0 | 0 | 0 |
| 0 | 0 | 0 | 0 |
| 0 | 0 | 0 | 0 |
| 0 | 0 | 0 | 0 |
| 0 | 0 | 0 | 0 |
| 1 | 1 | 0 | 0 |
| 1 | 1 | 0 | 0 |
| 0 | 0 | 0 | 0 |
| 0 | 0 | 0 | 0 |
| 0 | 0 | 0 | 0 |
| 1 | 0 | 1 | 0 |
| 0 | 0 | 0 | 0 |
| 0 | 0 | 0 | 0 |
| 1 | 0 | 1 | 0 |
| 0 | 0 | 0 | 0 |
| 0 | 0 | 0 | 0 |
| 0 | 0 | 0 | 0 |
| 0 | 0 | 1 | 0 |
| 1 | 0 | 1 | 0 |
| 0 | 0 | 1 | 0 |
| 0 | 0 | 1 | 0 |
| 0 | 0 | 1 | 1 |
| 0 | 0 | 1 | 1 |
| 0 | 0 | 0 | 0 |
| 0 | 0 | 0 | 0 |
| 0 | 0 | 0 | 0 |
| 0 | 0 | 0 | 0 |
| 0 | 0 | 0 | 0 |
| 0 | 0 | 0 | 0 |
| 0 | 0 | 0 | 0 |
| 0 | 0 | 0 | 0 |
| 0 | 0 | 0 | 0 |
| 0 | 0 | 0 | 0 |
| 0 | 0 | 0 | 0 |
| 0 | 0 | 0 | 0 |
| 0 | 0 | 0 | 0 |
| 0 | 0 | 0 | 0 |
| 0 | 0 | 0 | 0 |
| 0 | 0 | 0 | 0 |
| 0 | 0 | 1 | 0 |
| 0 | 0 | 0 | 0 |
| 0 | 0 | 0 | 0 |
| 0 | 0 | 0 | 0 |
| 0 | 0 | 0 | 0 |
| 0 | 0 | 0 | 0 |
| 0 | 0 | 0 | 0 |
| 1 | 0 | 0 | 0 |
| 0 | 0 | 0 | 0 |
| 0 | 0 | 0 | 0 |
| 0 | 0 | 0 | 0 |

| | | | |
|---|---|---|---|
| 0 | 0 | 0 | 0 |
| 0 | 0 | 0 | 0 |
| 0 | 0 | 0 | 0 |
| 1 | 0 | 0 | 0 |
| 0 | 0 | 0 | 0 |
| 0 | 0 | 0 | 0 |
| 0 | 0 | 0 | 0 |
| 0 | 0 | 0 | 0 |
| 1 | 0 | 0 | 0 |
| 1 | 0 | 0 | 0 |
| 0 | 0 | 0 | 0 |
| 0 | 0 | 0 | 0 |
| 0 | 0 | 0 | 0 |
| 0 | 0 | 0 | 0 |
| 0 | 0 | 0 | 0 |
| 0 | 0 | 0 | 0 |
| 0 | 0 | 1 | 0 |
| 0 | 0 | 0 | 0 |
| 0 | 0 | 0 | 0 |
| 0 | 0 | 0 | 0 |
| 0 | 0 | 0 | 0 |
| 0 | 0 | 0 | 0 |
| 0 | 0 | 0 | 0 |
| 0 | 0 | 0 | 0 |
| 0 | 0 | 0 | 0 |
| 0 | 0 | 0 | 0 |
| 0 | 0 | 0 | 0 |
| 0 | 0 | 0 | 0 |
| 0 | 0 | 0 | 0 |
| 0 | 0 | 0 | 0 |
| 0 | 0 | 0 | 0 |
| 0 | 0 | 0 | 0 |
| 0 | 0 | 0 | 0 |
| 0 | 0 | 0 | 0 |
| 0 | 0 | 0 | 0 |
| 0 | 0 | 0 | 0 |
| 0 | 0 | 0 | 0 |
| 0 | 0 | 0 | 0 |
| 0 | 0 | 0 | 0 |
| 0 | 0 | 0 | 0 |
| 0 | 0 | 0 | 0 |
| 0 | 0 | 0 | 0 |
| 0 | 0 | 0 | 0 |
| 0 | 0 | 0 | 0 |
| 0 | 0 | 0 | 0 |
| 0 | 0 | 0 | 0 |
| 0 | 0 | 0 | 0 |
| 0 | 0 | 0 | 0 |
| 0 | 0 | 0 | 0 |
| 0 | 1 | 0 | 0 |
| 0 | 0 | 0 | 0 |
| 0 | 0 | 0 | 0 |
| 0 | 0 | 0 | 0 |
| 0 | 0 | 0 | 0 |

| | | | |
|---|---|---|---|
| 0 | 0 | 1 | 0 |
| 0 | 0 | 0 | 0 |
| 0 | 0 | 0 | 0 |
| 0 | 0 | 0 | 0 |
| 0 | 0 | 0 | 0 |
| 0 | 0 | 0 | 0 |
| 0 | 0 | 0 | 0 |
| 1 | 0 | 0 | 0 |
| 0 | 0 | 0 | 0 |
| 0 | 0 | 0 | 0 |
| 0 | 0 | 0 | 0 |
| 1 | 0 | 0 | 0 |
| 0 | 0 | 0 | 0 |
| 0 | 0 | 0 | 0 |
| 0 | 0 | 0 | 0 |
| 0 | 0 | 0 | 0 |
| 0 | 0 | 0 | 0 |
| 0 | 0 | 0 | 0 |
| 0 | 0 | 0 | 0 |
| 0 | 0 | 0 | 0 |
| 0 | 0 | 0 | 0 |
| 0 | 0 | 0 | 0 |
| 0 | 0 | 0 | 0 |
| 0 | 0 | 0 | 0 |
| 0 | 0 | 0 | 0 |
| 0 | 0 | 0 | 0 |
| 0 | 0 | 0 | 0 |
| 0 | 0 | 0 | 0 |
| 0 | 0 | 0 | 0 |
| 0 | 0 | 0 | 0 |
| 0 | 0 | 0 | 0 |
| 0 | 0 | 0 | 0 |
| 0 | 0 | 0 | 0 |
| 0 | 0 | 0 | 0 |
| 0 | 0 | 0 | 0 |
| 0 | 0 | 0 | 0 |
| 0 | 0 | 0 | 0 |
| 0 | 0 | 0 | 0 |
| 0 | 0 | 0 | 0 |
| 0 | 0 | 0 | 0 |
| 0 | 0 | 0 | 0 |
| 0 | 0 | 0 | 0 |
| 0 | 0 | 0 | 0 |
| 0 | 0 | 0 | 0 |
| 0 | 0 | 0 | 0 |
| 0 | 0 | 0 | 0 |
| 0 | 0 | 0 | 0 |
| 0 | 0 | 0 | 0 |
| 0 | 0 | 0 | 0 |
| 0 | 0 | 0 | 0 |
| 0 | 0 | 0 | 0 |
| 0 | 0 | 0 | 0 |
| 0 | 0 | 0 | 0 |
| 0 | 0 | 0 | 0 |

| | | | |
|---|---|---|---|
| 0 | 0 | 0 | 0 |
| 0 | 0 | 0 | 0 |
| 0 | 0 | 0 | 0 |
| 0 | 0 | 0 | 0 |
| 0 | 0 | 0 | 0 |
| 0 | 0 | 0 | 0 |
| 0 | 0 | 0 | 0 |
| 0 | 0 | 0 | 0 |
| 0 | 0 | 0 | 0 |
| 0 | 0 | 0 | 0 |
| 0 | 0 | 0 | 0 |
| 0 | 0 | 0 | 0 |
| 0 | 0 | 0 | 0 |
| 0 | 0 | 0 | 0 |
| 0 | 0 | 0 | 0 |
| 0 | 0 | 0 | 0 |
| 0 | 0 | 0 | 0 |
| 0 | 0 | 0 | 0 |
| 0 | 0 | 0 | 1 |
| 0 | 1 | 0 | 1 |
| 0 | 0 | 0 | 0 |
| 0 | 0 | 0 | 0 |
| 0 | 0 | 0 | 0 |
| 0 | 0 | 0 | 0 |
| 0 | 0 | 0 | 0 |
| 1 | 0 | 0 | 0 |
| 1 | 0 | 0 | 0 |
| 0 | 0 | 0 | 0 |
| 0 | 0 | 0 | 0 |
| 0 | 0 | 0 | 0 |
| 0 | 0 | 0 | 0 |
| 0 | 0 | 0 | 0 |
| 0 | 0 | 0 | 0 |
| 1 | 0 | 0 | 1 |
| 1 | 0 | 0 | 0 |
| 1 | 0 | 0 | 1 |
| 1 | 0 | 0 | 0 |
| 0 | 0 | 0 | 0 |
| 0 | 0 | 0 | 0 |
| 0 | 0 | 0 | 0 |
| 0 | 0 | 0 | 0 |
| 1 | 0 | 0 | 0 |
| 1 | 0 | 0 | 1 |
| 1 | 0 | 0 | 0 |
| 0 | 0 | 0 | 0 |
| 0 | 0 | 0 | 0 |
| 0 | 0 | 0 | 0 |
| 0 | 0 | 0 | 0 |
| 0 | 0 | 0 | 0 |
| 1 | 0 | 0 | 0 |
| 0 | 0 | 0 | 0 |
| 0 | 0 | 0 | 0 |
| 0 | 0 | 0 | 0 |
| 0 | 0 | 0 | 0 |

| | | | |
|---|---|---|---|
| 0 | 0 | 0 | 0 |
| 0 | 0 | 0 | 0 |
| 0 | 0 | 0 | 0 |
| 1 | 0 | 0 | 0 |
| 1 | 0 | 0 | 0 |
| 0 | 0 | 0 | 0 |
| 1 | 0 | 0 | 0 |
| 0 | 0 | 0 | 0 |
| 0 | 0 | 0 | 0 |
| 0 | 0 | 0 | 0 |
| 0 | 0 | 0 | 0 |
| 1 | 0 | 1 | 0 |
| 0 | 0 | 0 | 0 |
| 0 | 0 | 0 | 0 |
| 0 | 0 | 0 | 0 |
| 0 | 0 | 1 | 0 |
| 0 | 0 | 0 | 0 |
| 0 | 0 | 0 | 0 |
| 0 | 0 | 0 | 0 |
| 0 | 0 | 1 | 0 |
| 1 | 0 | 1 | 0 |
| 1 | 0 | 0 | 0 |
| 1 | 0 | 0 | 0 |
| 1 | 0 | 0 | 0 |
| 0 | 0 | 1 | 0 |
| 0 | 0 | 0 | 0 |
| 1 | 0 | 0 | 0 |
| 0 | 0 | 0 | 0 |
| 0 | 0 | 0 | 0 |
| 0 | 0 | 0 | 0 |
| 0 | 0 | 0 | 0 |
| 0 | 0 | 0 | 0 |
| 0 | 0 | 0 | 0 |
| 0 | 0 | 0 | 0 |
| 0 | 0 | 0 | 0 |
| 0 | 0 | 0 | 0 |
| 0 | 0 | 0 | 1 |
| 0 | 0 | 0 | 1 |
| 0 | 0 | 0 | 1 |
| 0 | 0 | 0 | 0 |
| 0 | 0 | 0 | 0 |
| 0 | 0 | 0 | 0 |
| 0 | 0 | 0 | 0 |
| 0 | 0 | 0 | 0 |
| 0 | 0 | 0 | 0 |
| 0 | 0 | 0 | 0 |
| 0 | 0 | 0 | 0 |
| 0 | 0 | 0 | 0 |
| 0 | 0 | 0 | 0 |
| 0 | 0 | 0 | 0 |
| 0 | 0 | 0 | 0 |
| 0 | 0 | 0 | 0 |
| 0 | 0 | 0 | 0 |
| 0 | 0 | 0 | 0 |

| | | | |
|---|---|---|---|
| 0 | 0 | 0 | 0 |
| 0 | 0 | 0 | 0 |
| 0 | 0 | 0 | 0 |
| 0 | 0 | 0 | 0 |
| 0 | 0 | 0 | 0 |
| 0 | 0 | 0 | 0 |
| 0 | 0 | 0 | 0 |
| 0 | 0 | 0 | 0 |
| 0 | 0 | 0 | 0 |
| 0 | 0 | 0 | 0 |
| 0 | 0 | 0 | 0 |
| 0 | 0 | 0 | 0 |
| 0 | 0 | 0 | 0 |
| 0 | 0 | 0 | 0 |
| 0 | 0 | 0 | 0 |
| 0 | 0 | 0 | 0 |
| 0 | 1 | 0 | 0 |
| 0 | 1 | 0 | 0 |
| 0 | 0 | 0 | 0 |
| 0 | 0 | 0 | 0 |
| 0 | 0 | 0 | 0 |
| 0 | 0 | 0 | 0 |
| 0 | 0 | 0 | 0 |
| 0 | 0 | 0 | 0 |
| 0 | 0 | 0 | 0 |
| 0 | 0 | 0 | 0 |
| 0 | 0 | 0 | 0 |
| 0 | 0 | 0 | 0 |
| 0 | 0 | 0 | 0 |
| 0 | 0 | 0 | 0 |
| 0 | 0 | 0 | 0 |
| 1 | 0 | 0 | 0 |
| 0 | 0 | 0 | 0 |
| 0 | 0 | 0 | 0 |
| 0 | 0 | 0 | 0 |
| 0 | 0 | 0 | 0 |
| 0 | 0 | 0 | 0 |
| 0 | 0 | 0 | 0 |
| 0 | 0 | 0 | 0 |
| 0 | 0 | 0 | 0 |
| 0 | 0 | 0 | 0 |
| 0 | 0 | 0 | 0 |
| 0 | 0 | 0 | 0 |
| 0 | 0 | 0 | 0 |
| 0 | 0 | 0 | 0 |
| 0 | 0 | 0 | 0 |
| 1 | 1 | 0 | 0 |
| 0 | 0 | 0 | 0 |
| 0 | 0 | 0 | 0 |
| 0 | 0 | 0 | 0 |
| 0 | 0 | 0 | 0 |
| 0 | 0 | 0 | 0 |
| 1 | 0 | 0 | 0 |
| 0 | 0 | 0 | 0 |

| | | | |
|---|---|---|---|
| 0 | 0 | 0 | 0 |
| 0 | 0 | 0 | 0 |
| 0 | 0 | 0 | 0 |
| 0 | 0 | 0 | 0 |
| 0 | 0 | 0 | 0 |
| 0 | 0 | 0 | 0 |
| 1 | 0 | 0 | 0 |
| 0 | 0 | 0 | 0 |
| 0 | 0 | 0 | 0 |
| 0 | 0 | 0 | 0 |
| 0 | 0 | 0 | 0 |
| 0 | 0 | 0 | 0 |
| 0 | 0 | 0 | 0 |
| 0 | 0 | 0 | 0 |
| 0 | 0 | 0 | 0 |
| 0 | 0 | 0 | 0 |
| 0 | 0 | 0 | 0 |
| 0 | 0 | 0 | 0 |
| 0 | 0 | 0 | 0 |
| 0 | 0 | 0 | 0 |
| 0 | 0 | 0 | 0 |
| 0 | 0 | 0 | 0 |
| 0 | 0 | 0 | 0 |
| 0 | 0 | 0 | 0 |
| 0 | 0 | 0 | 0 |
| 0 | 0 | 0 | 0 |
| 0 | 0 | 0 | 0 |
| 0 | 0 | 0 | 0 |
| 0 | 0 | 0 | 0 |
| 0 | 0 | 0 | 0 |
| 0 | 0 | 0 | 0 |
| 0 | 0 | 0 | 0 |
| 0 | 0 | 0 | 0 |
| 0 | 0 | 0 | 0 |
| 0 | 0 | 0 | 0 |
| 0 | 0 | 0 | 0 |
| 0 | 0 | 0 | 0 |
| 0 | 0 | 0 | 0 |
| 0 | 0 | 0 | 0 |
| 0 | 0 | 0 | 0 |
| 0 | 0 | 0 | 0 |
| 0 | 0 | 0 | 0 |
| 0 | 0 | 0 | 0 |
| 0 | 0 | 0 | 0 |
| 0 | 0 | 0 | 0 |
| 0 | 0 | 0 | 0 |
| 0 | 0 | 0 | 0 |
| 0 | 0 | 0 | 0 |
| 0 | 0 | 0 | 0 |
| 0 | 0 | 0 | 0 |
| 0 | 0 | 0 | 0 |
| 0 | 0 | 0 | 0 |
| 0 | 0 | 0 | 0 |
| 0 | 0 | 0 | 0 |

| | | | |
|---|---|---|---|
| 0 | 0 | 0 | 0 |
| 0 | 0 | 0 | 0 |
| 0 | 0 | 0 | 0 |
| 0 | 0 | 0 | 0 |
| 0 | 0 | 0 | 0 |
| 0 | 0 | 0 | 0 |
| 0 | 0 | 0 | 0 |
| 0 | 0 | 0 | 0 |
| 0 | 0 | 0 | 0 |
| 0 | 0 | 0 | 0 |
| 0 | 0 | 0 | 0 |
| 0 | 0 | 0 | 0 |
| 0 | 0 | 0 | 0 |
| 0 | 0 | 0 | 0 |
| 0 | 0 | 0 | 0 |
| 0 | 0 | 0 | 0 |
| 0 | 0 | 0 | 0 |
| 0 | 0 | 0 | 0 |
| 0 | 0 | 0 | 0 |
| 0 | 0 | 0 | 0 |
| 0 | 0 | 0 | 0 |
| 0 | 0 | 0 | 0 |
| 0 | 0 | 0 | 0 |
| 0 | 0 | 0 | 0 |
| 1 | 1 | 0 | 0 |
| 0 | 0 | 0 | 0 |
| 0 | 0 | 0 | 0 |
| 0 | 0 | 0 | 0 |
| 0 | 0 | 0 | 0 |
| 0 | 0 | 0 | 0 |
| 0 | 0 | 0 | 0 |
| 0 | 0 | 0 | 0 |
| 0 | 0 | 0 | 0 |
| 0 | 0 | 0 | 0 |
| 0 | 0 | 0 | 0 |
| 1 | 0 | 0 | 0 |
| 1 | 0 | 0 | 1 |
| 1 | 0 | 0 | 0 |
| 0 | 0 | 0 | 0 |
| 1 | 0 | 0 | 0 |
| 1 | 1 | 0 | 0 |
| 1 | 1 | 0 | 0 |
| 0 | 0 | 0 | 0 |
| 0 | 0 | 0 | 0 |
| 0 | 0 | 0 | 0 |
| 0 | 0 | 0 | 0 |
| 0 | 0 | 0 | 0 |
| 0 | 0 | 0 | 0 |
| 0 | 0 | 0 | 0 |
| 0 | 0 | 0 | 0 |
| 0 | 0 | 0 | 0 |
| 0 | 0 | 0 | 0 |
| 0 | 0 | 0 | 0 |
| 0 | 0 | 0 | 0 |

| | | | |
|---|---|---|---|
| 0 | 0 | 0 | 0 |
| 0 | 0 | 0 | 0 |
| 0 | 0 | 0 | 0 |
| 0 | 0 | 0 | 0 |
| 0 | 0 | 0 | 0 |
| 0 | 0 | 0 | 0 |
| 0 | 0 | 0 | 0 |
| 0 | 0 | 0 | 0 |
| 0 | 0 | 0 | 0 |
| 0 | 0 | 0 | 0 |
| 0 | 0 | 0 | 0 |
| 0 | 0 | 0 | 0 |
| 0 | 0 | 0 | 0 |
| 1 | 0 | 0 | 0 |
| 0 | 0 | 0 | 0 |
| 0 | 0 | 0 | 0 |
| 1 | 1 | 0 | 0 |
| 0 | 0 | 0 | 0 |
| 0 | 0 | 0 | 0 |
| 0 | 0 | 0 | 0 |
| 0 | 0 | 0 | 0 |
| 0 | 0 | 0 | 0 |
| 0 | 0 | 0 | 0 |
| 1 | 0 | 0 | 0 |
| 0 | 0 | 0 | 0 |
| 0 | 0 | 0 | 0 |
| 0 | 0 | 0 | 0 |
| 0 | 0 | 0 | 0 |
| 0 | 0 | 0 | 0 |
| 0 | 0 | 0 | 0 |
| 0 | 0 | 0 | 0 |
| 1 | 0 | 0 | 0 |
| 0 | 0 | 0 | 0 |
| 0 | 0 | 0 | 0 |
| 0 | 0 | 0 | 0 |
| 0 | 0 | 0 | 0 |
| 0 | 0 | 0 | 0 |
| 0 | 0 | 0 | 0 |
| 0 | 0 | 0 | 0 |
| 0 | 0 | 0 | 0 |
| 0 | 0 | 0 | 0 |
| 0 | 0 | 0 | 0 |
| 0 | 0 | 0 | 0 |
| 0 | 0 | 0 | 0 |
| 0 | 0 | 0 | 0 |
| 0 | 0 | 0 | 0 |
| 0 | 0 | 0 | 0 |
| 0 | 0 | 0 | 0 |
| 0 | 0 | 0 | 0 |
| 0 | 0 | 0 | 0 |
| 0 | 0 | 0 | 0 |
| 0 | 0 | 0 | 0 |
| 0 | 0 | 0 | 0 |

| | | | |
|---|---|---|---|
| 0 | 0 | 0 | 0 |
| 0 | 0 | 0 | 0 |
| 0 | 0 | 0 | 0 |
| 0 | 0 | 0 | 0 |
| 0 | 0 | 0 | 0 |
| 0 | 0 | 0 | 0 |
| 0 | 0 | 0 | 0 |
| 0 | 0 | 0 | 0 |
| 0 | 0 | 0 | 0 |
| 0 | 0 | 0 | 0 |
| 0 | 0 | 0 | 0 |
| 0 | 0 | 0 | 0 |
| 0 | 0 | 0 | 0 |
| 0 | 0 | 0 | 0 |
| 0 | 0 | 0 | 0 |
| 0 | 0 | 0 | 0 |
| 0 | 0 | 0 | 0 |
| 1 | 0 | 0 | 0 |
| 0 | 0 | 0 | 0 |
| 0 | 0 | 0 | 0 |
| 0 | 0 | 0 | 0 |
| 0 | 0 | 0 | 0 |
| 0 | 0 | 0 | 0 |
| 0 | 0 | 0 | 0 |
| 1 | 0 | 0 | 0 |
| 0 | 0 | 0 | 0 |
| 0 | 0 | 0 | 0 |
| 0 | 0 | 0 | 0 |
| 0 | 0 | 0 | 0 |
| 0 | 0 | 0 | 0 |
| 0 | 0 | 0 | 0 |
| 0 | 0 | 0 | 0 |
| 0 | 0 | 0 | 0 |
| 0 | 0 | 0 | 0 |
| 0 | 0 | 0 | 0 |
| 0 | 0 | 0 | 0 |
| 0 | 0 | 0 | 0 |
| 0 | 0 | 0 | 0 |
| 0 | 0 | 0 | 0 |
| 0 | 0 | 0 | 0 |
| 0 | 0 | 0 | 0 |
| 0 | 0 | 0 | 0 |
| 0 | 0 | 0 | 0 |
| 0 | 0 | 0 | 0 |
| 0 | 0 | 0 | 0 |
| 0 | 0 | 0 | 0 |
| 0 | 0 | 0 | 0 |
| 0 | 0 | 0 | 0 |
| 0 | 0 | 0 | 0 |
| 0 | 0 | 0 | 0 |
| 0 | 0 | 0 | 0 |
| 0 | 0 | 0 | 0 |
| 0 | 0 | 0 | 0 |
| 0 | 0 | 0 | 0 |

| | | | |
|---|---|---|---|
| 0 | 0 | 0 | 0 |
| 0 | 0 | 0 | 0 |
| 0 | 0 | 0 | 0 |
| 0 | 0 | 0 | 0 |
| 0 | 0 | 0 | 0 |
| 0 | 0 | 0 | 0 |
| 0 | 0 | 0 | 0 |
| 0 | 0 | 0 | 0 |
| 0 | 0 | 0 | 0 |
| 0 | 0 | 0 | 0 |
| 0 | 0 | 0 | 0 |
| 0 | 0 | 0 | 0 |
| 0 | 0 | 0 | 0 |
| 0 | 0 | 0 | 0 |
| 1 | 1 | 0 | 0 |
| 1 | 1 | 0 | 0 |
| 0 | 0 | 0 | 0 |
| 0 | 0 | 0 | 0 |
| 0 | 0 | 0 | 0 |
| 0 | 0 | 0 | 0 |
| 0 | 0 | 0 | 0 |
| 0 | 0 | 0 | 0 |
| 0 | 0 | 0 | 0 |
| 0 | 0 | 0 | 0 |
| 0 | 0 | 0 | 0 |
| 0 | 0 | 0 | 0 |
| 0 | 0 | 0 | 0 |
| 1 | 0 | 0 | 0 |
| 0 | 0 | 0 | 0 |
| 1 | 0 | 0 | 0 |
| 0 | 0 | 0 | 0 |
| 0 | 0 | 0 | 0 |
| 0 | 0 | 0 | 0 |
| 0 | 0 | 0 | 0 |
| 0 | 0 | 0 | 0 |
| 0 | 0 | 0 | 0 |
| 1 | 0 | 0 | 0 |
| 0 | 0 | 0 | 0 |
| 0 | 0 | 0 | 0 |
| 0 | 0 | 0 | 0 |
| 0 | 0 | 0 | 0 |
| 0 | 0 | 0 | 0 |
| 0 | 0 | 0 | 0 |
| 0 | 0 | 0 | 0 |
| 0 | 0 | 0 | 0 |
| 0 | 0 | 0 | 0 |
| 0 | 0 | 0 | 0 |
| 0 | 0 | 0 | 0 |
| 0 | 0 | 0 | 0 |
| 0 | 0 | 0 | 0 |
| 0 | 0 | 0 | 0 |
| 0 | 0 | 0 | 0 |
| 0 | 0 | 0 | 0 |
| 0 | 0 | 0 | 0 |

| | | | |
|---|---|---|---|
| 0 | 0 | 0 | 0 |
| 0 | 0 | 0 | 0 |
| 0 | 0 | 0 | 0 |
| 0 | 0 | 0 | 0 |
| 0 | 0 | 0 | 0 |
| 0 | 0 | 0 | 0 |
| 0 | 0 | 0 | 0 |
| 0 | 0 | 0 | 0 |
| 0 | 0 | 0 | 0 |
| 0 | 0 | 0 | 0 |
| 0 | 0 | 0 | 0 |
| 0 | 0 | 0 | 0 |
| 0 | 0 | 0 | 0 |
| 0 | 0 | 0 | 0 |
| 0 | 0 | 0 | 0 |
| 0 | 0 | 0 | 0 |
| 0 | 0 | 0 | 0 |
| 0 | 0 | 0 | 0 |
| 0 | 0 | 0 | 0 |
| 0 | 0 | 0 | 0 |
| 0 | 0 | 0 | 0 |
| 0 | 0 | 0 | 0 |
| 0 | 0 | 0 | 0 |
| 0 | 0 | 0 | 0 |
| 0 | 0 | 0 | 0 |
| 0 | 0 | 0 | 0 |
| 0 | 0 | 0 | 0 |
| 0 | 0 | 0 | 0 |
| 0 | 0 | 0 | 0 |
| 0 | 0 | 0 | 0 |
| 0 | 0 | 0 | 0 |
| 0 | 0 | 0 | 0 |
| 0 | 0 | 0 | 0 |
| 0 | 0 | 0 | 0 |
| 0 | 0 | 0 | 0 |
| 0 | 0 | 0 | 0 |
| 0 | 0 | 0 | 0 |
| 0 | 0 | 0 | 0 |
| 0 | 0 | 0 | 0 |
| 0 | 0 | 0 | 0 |
| 0 | 0 | 0 | 0 |
| 0 | 0 | 0 | 0 |
| 0 | 0 | 0 | 0 |
| 0 | 0 | 0 | 0 |
| 0 | 0 | 0 | 0 |
| 0 | 0 | 0 | 0 |
| 0 | 0 | 0 | 0 |
| 0 | 0 | 0 | 0 |
| 1 | 0 | 0 | 1 |
| 0 | 0 | 0 | 0 |
| 1 | 0 | 0 | 0 |
| 1 | 0 | 0 | 0 |
| 0 | 0 | 0 | 0 |
| 0 | 0 | 0 | 1 |

| | | | |
|---|---|---|---|
| 0 | 0 | 0 | 0 |
| 0 | 0 | 0 | 0 |
| 0 | 0 | 0 | 0 |
| 0 | 0 | 0 | 0 |
| 0 | 0 | 0 | 0 |
| 0 | 0 | 0 | 0 |
| 0 | 0 | 0 | 0 |
| 0 | 0 | 0 | 0 |
| 0 | 0 | 0 | 0 |
| 0 | 0 | 0 | 0 |
| 0 | 0 | 0 | 0 |
| 0 | 0 | 0 | 0 |
| 0 | 0 | 0 | 0 |
| 0 | 0 | 0 | 0 |
| 0 | 0 | 0 | 0 |
| 0 | 0 | 0 | 0 |
| 0 | 0 | 0 | 0 |
| 0 | 0 | 0 | 0 |
| 0 | 0 | 0 | 0 |
| 0 | 0 | 0 | 0 |
| 0 | 0 | 0 | 0 |
| 0 | 0 | 0 | 0 |
| 0 | 0 | 0 | 0 |
| 0 | 0 | 0 | 0 |
| 0 | 0 | 0 | 0 |
| 0 | 0 | 0 | 0 |
| 0 | 0 | 1 | 0 |
| 0 | 0 | 1 | 0 |
| 0 | 0 | 1 | 0 |
| 0 | 0 | 1 | 0 |
| 0 | 0 | 0 | 0 |
| 0 | 0 | 0 | 0 |
| 0 | 0 | 0 | 0 |
| 1 | 0 | 0 | 0 |
| 1 | 0 | 0 | 0 |
| 0 | 0 | 0 | 0 |
| 0 | 0 | 0 | 0 |
| 0 | 0 | 0 | 0 |
| 0 | 0 | 0 | 0 |
| 1 | 0 | 0 | 0 |
| 0 | 0 | 0 | 0 |
| 0 | 0 | 0 | 0 |
| 0 | 0 | 0 | 0 |
| 0 | 0 | 0 | 0 |
| 0 | 0 | 0 | 0 |
| 0 | 1 | 0 | 0 |
| 0 | 0 | 0 | 0 |
| 0 | 0 | 0 | 0 |
| 0 | 0 | 0 | 0 |
| 0 | 0 | 0 | 0 |
| 0 | 0 | 0 | 0 |
| 0 | 0 | 0 | 0 |
| 0 | 0 | 0 | 0 |
| 0 | 0 | 0 | 0 |

| | | | |
|---|---|---|---|
| 0 | 0 | 0 | 0 |
| 0 | 0 | 0 | 0 |
| 0 | 0 | 0 | 0 |
| 0 | 0 | 0 | 0 |
| 0 | 0 | 0 | 0 |
| 0 | 0 | 0 | 1 |
| 0 | 0 | 0 | 0 |
| 0 | 0 | 0 | 0 |
| 0 | 0 | 0 | 0 |
| 0 | 0 | 0 | 0 |
| 0 | 0 | 0 | 0 |
| 0 | 0 | 0 | 0 |
| 0 | 0 | 0 | 0 |
| 0 | 1 | 0 | 0 |
| 0 | 1 | 0 | 0 |
| 0 | 1 | 0 | 0 |
| 0 | 1 | 0 | 0 |
| 0 | 1 | 0 | 0 |
| 0 | 0 | 0 | 0 |
| 0 | 0 | 0 | 0 |
| 0 | 0 | 0 | 0 |
| 0 | 0 | 0 | 0 |
| 0 | 0 | 0 | 0 |
| 0 | 0 | 0 | 0 |
| 0 | 0 | 0 | 0 |
| 0 | 0 | 0 | 0 |
| 0 | 0 | 0 | 0 |
| 0 | 0 | 0 | 0 |
| 0 | 0 | 0 | 0 |
| 0 | 0 | 0 | 0 |
| 0 | 0 | 0 | 0 |
| 0 | 0 | 0 | 0 |
| 0 | 0 | 0 | 0 |
| 0 | 0 | 0 | 0 |
| 0 | 0 | 0 | 0 |
| 0 | 0 | 0 | 0 |
| 0 | 0 | 0 | 0 |
| 0 | 0 | 0 | 0 |
| 0 | 0 | 0 | 0 |
| 0 | 0 | 0 | 0 |
| 0 | 0 | 0 | 0 |
| 0 | 0 | 0 | 0 |
| 0 | 0 | 0 | 0 |
| 0 | 0 | 0 | 0 |
| 0 | 0 | 0 | 0 |
| 0 | 0 | 0 | 0 |
| 0 | 0 | 0 | 0 |
| 0 | 0 | 0 | 0 |
| 0 | 0 | 0 | 0 |
| 0 | 0 | 0 | 0 |
| 0 | 0 | 0 | 0 |
| 0 | 0 | 0 | 0 |
| 0 | 0 | 0 | 0 |

| | | | |
|---|---|---|---|
| 0 | 0 | 0 | 0 |
| 0 | 0 | 0 | 0 |
| 0 | 0 | 0 | 0 |
| 1 | 0 | 0 | 0 |
| 0 | 0 | 0 | 0 |
| 0 | 0 | 0 | 0 |
| 0 | 0 | 0 | 0 |
| 0 | 0 | 0 | 0 |
| 0 | 0 | 0 | 0 |
| 0 | 0 | 0 | 0 |
| 0 | 0 | 0 | 0 |
| 0 | 0 | 0 | 0 |
| 0 | 0 | 0 | 0 |
| 0 | 0 | 0 | 0 |
| 0 | 0 | 0 | 0 |
| 0 | 0 | 0 | 0 |
| 0 | 0 | 0 | 0 |
| 0 | 0 | 0 | 0 |
| 0 | 0 | 0 | 0 |
| 0 | 0 | 0 | 0 |
| 1 | 1 | 0 | 0 |
| 1 | 1 | 0 | 0 |
| 0 | 0 | 0 | 0 |
| 0 | 1 | 0 | 0 |
| 0 | 0 | 0 | 0 |
| 0 | 0 | 0 | 0 |
| 0 | 0 | 0 | 0 |
| 0 | 0 | 0 | 0 |
| 0 | 0 | 0 | 0 |
| 0 | 0 | 0 | 0 |
| 0 | 0 | 0 | 0 |
| 0 | 0 | 0 | 0 |
| 0 | 0 | 0 | 0 |
| 0 | 0 | 0 | 0 |
| 0 | 0 | 0 | 0 |
| 0 | 0 | 0 | 0 |
| 0 | 0 | 0 | 0 |
| 0 | 0 | 0 | 0 |
| 0 | 0 | 0 | 0 |
| 1 | 0 | 0 | 0 |
| 1 | 0 | 0 | 0 |
| 0 | 0 | 0 | 0 |
| 0 | 0 | 0 | 0 |
| 1 | 0 | 0 | 0 |
| 0 | 0 | 0 | 0 |
| 0 | 0 | 0 | 0 |
| 0 | 0 | 0 | 0 |
| 0 | 0 | 0 | 0 |
| 0 | 0 | 0 | 0 |
| 0 | 0 | 0 | 0 |
| 0 | 0 | 0 | 0 |
| 0 | 0 | 0 | 0 |
| 0 | 0 | 0 | 0 |
| 0 | 0 | 0 | 0 |

| | | | |
|---|---|---|---|
| 0 | 0 | 0 | 0 |
| 0 | 0 | 0 | 0 |
| 0 | 0 | 0 | 0 |
| 0 | 0 | 0 | 0 |
| 0 | 0 | 0 | 0 |
| 1 | 0 | 0 | 0 |
| 0 | 0 | 0 | 0 |
| 0 | 0 | 0 | 0 |
| 0 | 1 | 0 | 0 |
| 1 | 1 | 0 | 0 |
| 0 | 0 | 0 | 0 |
| 1 | 1 | 0 | 0 |
| 1 | 0 | 0 | 0 |
| 0 | 0 | 0 | 0 |
| 0 | 0 | 0 | 0 |
| 0 | 0 | 0 | 0 |
| 1 | 1 | 0 | 0 |
| 0 | 0 | 0 | 0 |
| 0 | 0 | 0 | 0 |
| 1 | 0 | 0 | 0 |
| 0 | 0 | 0 | 0 |
| 0 | 0 | 0 | 0 |
| 0 | 0 | 0 | 0 |
| 0 | 0 | 0 | 0 |
| 0 | 0 | 0 | 0 |
| 0 | 0 | 0 | 0 |
| 0 | 0 | 0 | 0 |
| 0 | 0 | 0 | 0 |
| 0 | 0 | 0 | 0 |
| 0 | 0 | 0 | 0 |
| 1 | 0 | 0 | 0 |
| 0 | 0 | 0 | 0 |
| 0 | 0 | 0 | 0 |
| 0 | 0 | 0 | 0 |
| 0 | 0 | 0 | 0 |
| 1 | 0 | 0 | 0 |
| 0 | 0 | 0 | 0 |
| 0 | 0 | 0 | 0 |
| 0 | 0 | 0 | 0 |
| 0 | 0 | 0 | 0 |
| 0 | 0 | 0 | 0 |
| 0 | 0 | 0 | 0 |
| 0 | 0 | 0 | 0 |
| 0 | 0 | 0 | 0 |
| 0 | 0 | 0 | 0 |
| 0 | 0 | 0 | 0 |
| 0 | 0 | 0 | 0 |
| 0 | 0 | 0 | 0 |
| 0 | 0 | 0 | 0 |
| 0 | 0 | 0 | 0 |
| 0 | 0 | 0 | 0 |
| 0 | 0 | 0 | 0 |
| 0 | 0 | 0 | 0 |
| 0 | 0 | 0 | 0 |

| | | | |
|---|---|---|---|
| 0 | 0 | 0 | 0 |
| 0 | 0 | 0 | 0 |
| 0 | 0 | 0 | 0 |
| 0 | 0 | 0 | 0 |
| 0 | 0 | 0 | 0 |
| 0 | 0 | 0 | 0 |
| 0 | 0 | 0 | 0 |
| 0 | 0 | 0 | 0 |
| 0 | 0 | 0 | 0 |
| 0 | 0 | 0 | 0 |
| 0 | 0 | 0 | 0 |
| 0 | 0 | 0 | 0 |
| 0 | 0 | 0 | 0 |
| 0 | 0 | 0 | 0 |
| 0 | 0 | 0 | 1 |
| 0 | 0 | 0 | 0 |
| 0 | 0 | 0 | 0 |
| 0 | 0 | 0 | 0 |
| 0 | 0 | 0 | 0 |
| 0 | 0 | 0 | 0 |
| 0 | 0 | 0 | 0 |
| 0 | 0 | 0 | 0 |
| 0 | 0 | 0 | 0 |
| 0 | 0 | 0 | 0 |
| 0 | 0 | 0 | 0 |
| 0 | 0 | 0 | 0 |
| 0 | 0 | 0 | 0 |
| 0 | 0 | 0 | 0 |
| 0 | 0 | 0 | 0 |
| 0 | 0 | 0 | 0 |
| 0 | 0 | 0 | 0 |
| 0 | 0 | 0 | 0 |
| 0 | 0 | 0 | 0 |
| 0 | 0 | 0 | 0 |
| 0 | 0 | 0 | 0 |
| 0 | 0 | 0 | 0 |
| 0 | 0 | 0 | 0 |
| 0 | 0 | 0 | 0 |
| 0 | 0 | 0 | 0 |
| 0 | 0 | 1 | 0 |
| 0 | 0 | 0 | 0 |
| 0 | 0 | 0 | 0 |
| 0 | 0 | 0 | 0 |
| 1 | 0 | 0 | 0 |
| 0 | 0 | 0 | 0 |
| 0 | 0 | 0 | 0 |
| 0 | 0 | 0 | 0 |
| 0 | 0 | 0 | 0 |
| 1 | 0 | 0 | 1 |
| 0 | 0 | 0 | 0 |
| 0 | 0 | 0 | 0 |
| 0 | 0 | 0 | 0 |
| 0 | 0 | 0 | 0 |
| 0 | 0 | 0 | 0 |

| | | | |
|---|---|---|---|
| 0 | 0 | 0 | 0 |
| 0 | 0 | 0 | 0 |
| 0 | 0 | 0 | 0 |
| 0 | 0 | 0 | 0 |
| 0 | 0 | 0 | 0 |
| 0 | 0 | 0 | 0 |
| 0 | 0 | 0 | 0 |
| 0 | 0 | 0 | 0 |
| 0 | 0 | 0 | 0 |
| 1 | 0 | 0 | 0 |
| 0 | 0 | 0 | 0 |
| 0 | 0 | 0 | 0 |
| 0 | 0 | 0 | 0 |
| 0 | 0 | 0 | 0 |
| 1 | 0 | 0 | 0 |
| 0 | 0 | 0 | 0 |
| 0 | 0 | 0 | 0 |
| 0 | 0 | 0 | 0 |
| 0 | 0 | 0 | 0 |
| 0 | 0 | 0 | 0 |
| 0 | 0 | 0 | 0 |
| 0 | 0 | 0 | 0 |
| 0 | 0 | 0 | 0 |
| 0 | 0 | 0 | 0 |
| 1 | 0 | 0 | 0 |
| 0 | 0 | 0 | 0 |
| 0 | 0 | 0 | 0 |
| 0 | 0 | 0 | 0 |
| 0 | 0 | 0 | 0 |
| 0 | 0 | 0 | 0 |
| 0 | 0 | 0 | 0 |
| 0 | 0 | 0 | 0 |
| 0 | 0 | 0 | 0 |
| 0 | 0 | 0 | 0 |
| 1 | 0 | 0 | 0 |
| 1 | 0 | 0 | 0 |
| 0 | 0 | 0 | 0 |
| 0 | 0 | 0 | 0 |
| 1 | 0 | 0 | 0 |
| 0 | 0 | 1 | 0 |
| 0 | 0 | 1 | 0 |
| 0 | 0 | 1 | 0 |
| 0 | 0 | 1 | 0 |
| 0 | 0 | 0 | 0 |
| 1 | 0 | 0 | 0 |
| 0 | 0 | 0 | 0 |
| 0 | 0 | 0 | 0 |
| 0 | 0 | 0 | 0 |
| 0 | 0 | 1 | 0 |
| 0 | 0 | 0 | 0 |
| 0 | 0 | 0 | 0 |
| 0 | 0 | 0 | 0 |
| 0 | 0 | 0 | 0 |
| 0 | 0 | 0 | 0 |

| | | | |
|---|---|---|---|
| 0 | 0 | 0 | 0 |
| 0 | 0 | 0 | 0 |
| 0 | 0 | 0 | 0 |
| 0 | 0 | 0 | 0 |
| 0 | 0 | 0 | 0 |
| 0 | 0 | 0 | 0 |
| 0 | 0 | 0 | 0 |
| 0 | 0 | 0 | 0 |
| 0 | 0 | 0 | 0 |
| 0 | 0 | 0 | 0 |
| 0 | 1 | 0 | 0 |
| 0 | 0 | 0 | 0 |
| 0 | 0 | 0 | 0 |
| 0 | 0 | 0 | 0 |
| 0 | 0 | 0 | 0 |
| 0 | 1 | 0 | 0 |
| 0 | 0 | 0 | 0 |
| 0 | 0 | 0 | 0 |
| 0 | 0 | 0 | 0 |
| 0 | 0 | 0 | 0 |
| 0 | 0 | 0 | 0 |
| 0 | 0 | 0 | 0 |
| 0 | 0 | 0 | 0 |
| 0 | 0 | 0 | 0 |
| 0 | 0 | 0 | 0 |
| 0 | 0 | 0 | 0 |
| 0 | 0 | 0 | 0 |
| 0 | 0 | 0 | 0 |
| 0 | 0 | 0 | 0 |
| 0 | 0 | 0 | 0 |
| 0 | 0 | 0 | 0 |
| 0 | 0 | 0 | 0 |
| 0 | 0 | 0 | 0 |
| 0 | 0 | 0 | 0 |
| 0 | 0 | 0 | 0 |
| 0 | 0 | 0 | 0 |
| 0 | 0 | 0 | 0 |
| 0 | 0 | 0 | 0 |
| 0 | 0 | 0 | 0 |
| 0 | 0 | 0 | 0 |
| 0 | 0 | 0 | 0 |
| 0 | 0 | 0 | 0 |
| 0 | 0 | 0 | 0 |
| 0 | 0 | 0 | 0 |
| 1 | 1 | 0 | 0 |
| 0 | 0 | 0 | 0 |
| 0 | 0 | 0 | 0 |
| 0 | 0 | 0 | 0 |
| 0 | 0 | 0 | 0 |
| 0 | 0 | 0 | 0 |
| 0 | 0 | 0 | 0 |
| 0 | 0 | 0 | 0 |
| 0 | 0 | 0 | 0 |
| 0 | 0 | 0 | 0 |

| | | | |
|---|---|---|---|
| 0 | 0 | 0 | 0 |
| 0 | 0 | 0 | 0 |
| 0 | 0 | 1 | 0 |
| 1 | 0 | 0 | 0 |
| 0 | 0 | 0 | 0 |
| 0 | 0 | 0 | 0 |
| 0 | 0 | 0 | 0 |
| 0 | 0 | 0 | 0 |
| 0 | 0 | 0 | 0 |
| 0 | 0 | 0 | 0 |
| 0 | 0 | 0 | 0 |
| 0 | 0 | 0 | 0 |
| 1 | 0 | 0 | 0 |
| 0 | 0 | 0 | 0 |
| 0 | 0 | 0 | 0 |
| 0 | 0 | 0 | 0 |
| 0 | 0 | 0 | 0 |
| 0 | 0 | 0 | 0 |
| 0 | 0 | 0 | 0 |
| 0 | 0 | 0 | 0 |
| 0 | 0 | 0 | 0 |
| 0 | 0 | 0 | 0 |
| 0 | 0 | 0 | 0 |
| 0 | 0 | 0 | 0 |
| 0 | 0 | 0 | 0 |
| 0 | 0 | 0 | 0 |
| 0 | 0 | 0 | 0 |
| 0 | 0 | 0 | 0 |
| 0 | 0 | 0 | 0 |
| 0 | 0 | 0 | 0 |
| 0 | 0 | 0 | 0 |
| 0 | 0 | 0 | 0 |
| 0 | 0 | 0 | 0 |
| 0 | 0 | 0 | 0 |
| 0 | 0 | 0 | 0 |
| 0 | 0 | 0 | 0 |
| 0 | 0 | 0 | 0 |
| 0 | 0 | 0 | 0 |
| 0 | 0 | 0 | 0 |
| 0 | 0 | 0 | 0 |
| 0 | 1 | 0 | 0 |
| 1 | 1 | 0 | 0 |
| 0 | 0 | 0 | 0 |
| 0 | 0 | 0 | 0 |
| 0 | 0 | 0 | 0 |
| 0 | 0 | 0 | 0 |
| 0 | 0 | 0 | 0 |
| 0 | 0 | 0 | 0 |
| 0 | 1 | 0 | 0 |
| 0 | 0 | 0 | 0 |
| 0 | 0 | 0 | 0 |
| 0 | 0 | 0 | 0 |
| 0 | 0 | 0 | 1 |
| 0 | 0 | 0 | 0 |

| | | | |
|---|---|---|---|
| 0 | 0 | 0 | 1 |
| 0 | 0 | 0 | 1 |
| 0 | 0 | 0 | 1 |
| 0 | 0 | 0 | 0 |
| 0 | 0 | 0 | 0 |
| 0 | 0 | 0 | 0 |
| 0 | 0 | 0 | 0 |
| 1 | 0 | 0 | 0 |
| 0 | 0 | 0 | 0 |
| 1 | 1 | 0 | 0 |
| 0 | 1 | 0 | 0 |
| 0 | 0 | 0 | 0 |
| 1 | 1 | 0 | 0 |
| 0 | 0 | 0 | 0 |
| 0 | 0 | 0 | 0 |
| 0 | 0 | 0 | 0 |
| 0 | 0 | 0 | 0 |
| 1 | 0 | 0 | 0 |
| 0 | 0 | 0 | 0 |
| 0 | 0 | 0 | 0 |
| 0 | 0 | 0 | 0 |
| 0 | 0 | 0 | 0 |
| 0 | 0 | 0 | 0 |
| 0 | 0 | 0 | 0 |
| 0 | 0 | 0 | 0 |
| 0 | 0 | 0 | 0 |
| 0 | 0 | 0 | 0 |
| 0 | 0 | 0 | 0 |
| 0 | 0 | 0 | 0 |
| 0 | 0 | 0 | 0 |
| 0 | 0 | 0 | 0 |
| 1 | 0 | 0 | 0 |
| 0 | 0 | 0 | 0 |
| 0 | 0 | 0 | 0 |
| 0 | 0 | 0 | 0 |
| 0 | 0 | 0 | 0 |
| 0 | 0 | 0 | 0 |
| 0 | 0 | 0 | 0 |
| 0 | 0 | 0 | 0 |
| 0 | 0 | 0 | 0 |
| 0 | 0 | 0 | 0 |
| 1 | 1 | 0 | 0 |
| 0 | 0 | 0 | 0 |
| 0 | 0 | 0 | 0 |
| 1 | 0 | 0 | 0 |
| 0 | 1 | 0 | 0 |
| 0 | 0 | 0 | 0 |
| 0 | 0 | 0 | 0 |
| 0 | 1 | 0 | 0 |
| 1 | 1 | 0 | 0 |
| 1 | 1 | 0 | 0 |
| 1 | 1 | 0 | 0 |
| 1 | 1 | 0 | 0 |
| 1 | 1 | 0 | 0 |

| | | | |
|---|---|---|---|
| 0 | 0 | 0 | 0 |
| 0 | 0 | 0 | 0 |
| 0 | 0 | 0 | 0 |
| 0 | 0 | 0 | 0 |
| 0 | 0 | 0 | 0 |
| 1 | 0 | 0 | 0 |
| 1 | 0 | 0 | 0 |
| 0 | 0 | 0 | 0 |
| 0 | 0 | 0 | 0 |
| 1 | 0 | 0 | 0 |
| 0 | 0 | 0 | 0 |
| 0 | 0 | 1 | 0 |
| 0 | 0 | 0 | 0 |
| 0 | 0 | 0 | 0 |
| 0 | 0 | 0 | 0 |
| 0 | 0 | 0 | 0 |
| 0 | 0 | 0 | 0 |
| 0 | 0 | 0 | 0 |
| 0 | 0 | 0 | 0 |
| 0 | 0 | 0 | 0 |
| 1 | 1 | 0 | 0 |
| 1 | 0 | 0 | 0 |
| 0 | 0 | 0 | 0 |
| 0 | 0 | 1 | 0 |
| 0 | 0 | 0 | 0 |
| 0 | 0 | 1 | 0 |
| 0 | 0 | 0 | 0 |
| 0 | 0 | 0 | 0 |
| 0 | 0 | 0 | 0 |
| 0 | 0 | 0 | 0 |
| 0 | 0 | 0 | 0 |
| 0 | 0 | 0 | 0 |
| 0 | 0 | 0 | 0 |
| 0 | 0 | 0 | 0 |
| 0 | 0 | 0 | 0 |
| 0 | 0 | 0 | 0 |
| 1 | 0 | 0 | 0 |
| 0 | 0 | 0 | 0 |
| 1 | 0 | 0 | 0 |
| 1 | 0 | 0 | 0 |
| 1 | 0 | 0 | 0 |
| 0 | 0 | 0 | 0 |
| 0 | 0 | 0 | 0 |
| 0 | 0 | 0 | 0 |
| 0 | 0 | 0 | 0 |
| 0 | 0 | 0 | 0 |
| 0 | 0 | 0 | 0 |
| 1 | 0 | 1 | 0 |
| 0 | 0 | 0 | 0 |
| 0 | 0 | 0 | 0 |
| 0 | 0 | 0 | 0 |
| 0 | 0 | 0 | 0 |
| 1 | 1 | 0 | 0 |
| 0 | 0 | 0 | 0 |

| | | | |
|---|---|---|---|
| 0 | 0 | 0 | 0 |
| 0 | 1 | 0 | 0 |
| 0 | 0 | 0 | 0 |
| 0 | 0 | 0 | 0 |
| 0 | 0 | 0 | 0 |
| 0 | 0 | 0 | 0 |
| 0 | 0 | 0 | 0 |
| 0 | 0 | 0 | 0 |
| 0 | 0 | 0 | 0 |
| 0 | 0 | 0 | 0 |
| 0 | 0 | 0 | 0 |
| 0 | 0 | 0 | 0 |
| 0 | 0 | 0 | 0 |
| 0 | 0 | 0 | 0 |
| 0 | 0 | 0 | 0 |
| 0 | 0 | 0 | 0 |
| 0 | 0 | 0 | 0 |
| 0 | 0 | 0 | 0 |
| 0 | 0 | 0 | 0 |
| 0 | 0 | 0 | 0 |
| 1 | 0 | 0 | 0 |
| 0 | 0 | 0 | 0 |
| 0 | 0 | 0 | 0 |
| 0 | 0 | 0 | 0 |
| 1 | 0 | 0 | 0 |
| 0 | 0 | 0 | 0 |
| 0 | 0 | 0 | 0 |
| 0 | 0 | 0 | 1 |
| 0 | 0 | 0 | 0 |
| 0 | 0 | 0 | 0 |
| 0 | 0 | 0 | 0 |
| 0 | 0 | 0 | 0 |
| 0 | 0 | 0 | 0 |
| 0 | 0 | 0 | 0 |
| 0 | 0 | 0 | 0 |
| 0 | 0 | 0 | 0 |
| 0 | 0 | 0 | 0 |
| 0 | 0 | 0 | 0 |
| 0 | 0 | 0 | 0 |
| 0 | 0 | 0 | 0 |
| 0 | 0 | 0 | 0 |
| 0 | 0 | 0 | 0 |
| 0 | 0 | 0 | 0 |
| 0 | 0 | 0 | 0 |
| 0 | 0 | 0 | 0 |
| 0 | 0 | 0 | 0 |
| 0 | 0 | 0 | 0 |
| 0 | 0 | 0 | 0 |
| 0 | 0 | 0 | 0 |
| 0 | 0 | 0 | 0 |
| 0 | 0 | 0 | 0 |
| 0 | 0 | 0 | 0 |
| 0 | 0 | 0 | 0 |
| 0 | 0 | 0 | 0 |

| | | | |
|---|---|---|---|
| 0 | 0 | 0 | 0 |
| 0 | 0 | 0 | 0 |
| 0 | 0 | 0 | 0 |
| 0 | 0 | 0 | 0 |
| 0 | 0 | 0 | 0 |
| 0 | 0 | 0 | 0 |
| 0 | 0 | 0 | 0 |
| 0 | 0 | 0 | 0 |
| 0 | 0 | 0 | 0 |
| 0 | 0 | 0 | 0 |
| 0 | 0 | 0 | 0 |
| 0 | 0 | 0 | 0 |
| 0 | 0 | 0 | 0 |
| 0 | 0 | 0 | 0 |
| 0 | 0 | 0 | 0 |
| 1 | 0 | 0 | 0 |
| 0 | 0 | 0 | 0 |
| 0 | 0 | 0 | 0 |
| 0 | 1 | 0 | 0 |
| 0 | 0 | 0 | 0 |
| 0 | 0 | 0 | 0 |
| 0 | 0 | 0 | 0 |
| 0 | 0 | 0 | 0 |
| 0 | 0 | 0 | 0 |
| 0 | 0 | 0 | 0 |
| 1 | 0 | 0 | 0 |
| 0 | 0 | 0 | 0 |
| 0 | 0 | 0 | 0 |
| 0 | 0 | 0 | 0 |
| 0 | 0 | 0 | 0 |
| 0 | 0 | 0 | 0 |
| 0 | 0 | 0 | 0 |
| 0 | 0 | 0 | 0 |
| 1 | 1 | 0 | 0 |
| 0 | 0 | 0 | 0 |
| 0 | 0 | 0 | 0 |
| 0 | 0 | 0 | 0 |
| 0 | 0 | 0 | 0 |
| 0 | 0 | 0 | 0 |
| 0 | 0 | 0 | 0 |
| 0 | 0 | 0 | 0 |
| 0 | 0 | 0 | 0 |
| 0 | 0 | 0 | 0 |
| 0 | 0 | 0 | 0 |
| 0 | 0 | 0 | 0 |
| 0 | 0 | 0 | 0 |
| 0 | 0 | 0 | 0 |
| 1 | 0 | 0 | 0 |
| 0 | 0 | 0 | 0 |
| 0 | 0 | 0 | 0 |
| 0 | 0 | 0 | 0 |
| 0 | 0 | 0 | 0 |
| 0 | 0 | 0 | 0 |
| 1 | 0 | 0 | 1 |

| | | | |
|---|---|---|---|
| 0 | 0 | 0 | 0 |
| 0 | 0 | 0 | 0 |
| 0 | 0 | 1 | 0 |
| 0 | 0 | 0 | 0 |
| 0 | 0 | 0 | 0 |
| 0 | 0 | 0 | 0 |
| 0 | 0 | 0 | 0 |
| 0 | 0 | 0 | 0 |
| 0 | 0 | 0 | 0 |
| 0 | 0 | 0 | 0 |
| 0 | 0 | 0 | 0 |
| 0 | 0 | 0 | 0 |
| 0 | 0 | 0 | 0 |
| 0 | 0 | 0 | 0 |
| 0 | 0 | 0 | 0 |
| 0 | 0 | 0 | 0 |
| 0 | 0 | 0 | 0 |
| 0 | 0 | 0 | 0 |
| 0 | 0 | 0 | 0 |
| 0 | 0 | 0 | 0 |
| 0 | 0 | 0 | 0 |
| 0 | 0 | 0 | 0 |
| 0 | 0 | 0 | 0 |
| 0 | 0 | 0 | 0 |
| 0 | 0 | 0 | 0 |
| 0 | 0 | 0 | 0 |
| 0 | 0 | 0 | 0 |
| 0 | 0 | 0 | 0 |
| 0 | 0 | 0 | 0 |
| 0 | 0 | 0 | 0 |
| 0 | 0 | 0 | 0 |
| 1 | 0 | 0 | 0 |
| 0 | 0 | 0 | 0 |
| 0 | 0 | 0 | 0 |
| 0 | 0 | 0 | 0 |
| 0 | 0 | 0 | 0 |
| 0 | 0 | 0 | 0 |
| 0 | 0 | 0 | 0 |
| 0 | 0 | 0 | 0 |
| 0 | 0 | 0 | 0 |
| 1 | 1 | 0 | 0 |
| 0 | 0 | 0 | 0 |
| 0 | 0 | 0 | 0 |
| 0 | 0 | 0 | 0 |
| 0 | 0 | 0 | 0 |
| 1 | 1 | 0 | 0 |
| 0 | 0 | 0 | 0 |
| 1 | 1 | 0 | 0 |
| 0 | 0 | 0 | 0 |
| 1 | 0 | 0 | 0 |
| 1 | 0 | 0 | 0 |
| 1 | 0 | 0 | 0 |
| 0 | 0 | 0 | 0 |
| 0 | 0 | 0 | 0 |

| | | | |
|---|---|---|---|
| 1 | 0 | 0 | 0 |
| 0 | 0 | 0 | 0 |
| 0 | 0 | 0 | 0 |
| 0 | 0 | 0 | 0 |
| 0 | 0 | 0 | 0 |
| 0 | 0 | 0 | 0 |
| 0 | 0 | 0 | 0 |
| 0 | 0 | 0 | 0 |
| 0 | 0 | 0 | 0 |
| 0 | 0 | 0 | 0 |
| 0 | 0 | 0 | 0 |
| 0 | 0 | 0 | 0 |
| 0 | 0 | 0 | 0 |
| 0 | 0 | 0 | 0 |
| 0 | 0 | 0 | 0 |
| 1 | 1 | 1 | 0 |
| 0 | 0 | 0 | 0 |
| 1 | 0 | 0 | 0 |
| 0 | 0 | 0 | 0 |
| 0 | 0 | 0 | 0 |
| 0 | 0 | 0 | 0 |
| 0 | 1 | 0 | 0 |
| 0 | 0 | 0 | 0 |
| 0 | 1 | 0 | 0 |
| 0 | 1 | 0 | 0 |
| 0 | 0 | 0 | 0 |
| 0 | 0 | 0 | 0 |
| 0 | 0 | 0 | 0 |
| 0 | 0 | 0 | 0 |
| 0 | 0 | 0 | 0 |
| 0 | 0 | 0 | 0 |
| 0 | 0 | 0 | 0 |
| 1 | 1 | 0 | 0 |
| 0 | 0 | 0 | 0 |
| 0 | 0 | 0 | 0 |
| 0 | 0 | 0 | 0 |
| 0 | 1 | 0 | 0 |
| 1 | 1 | 0 | 0 |
| 0 | 0 | 0 | 0 |
| 0 | 0 | 0 | 0 |
| 0 | 0 | 0 | 0 |
| 0 | 0 | 0 | 0 |
| 0 | 0 | 0 | 0 |
| 0 | 0 | 0 | 0 |
| 1 | 1 | 0 | 0 |
| 0 | 0 | 0 | 0 |
| 1 | 0 | 0 | 0 |
| 1 | 1 | 0 | 0 |
| 0 | 0 | 0 | 0 |
| 0 | 0 | 0 | 0 |
| 1 | 1 | 0 | 0 |
| 1 | 1 | 0 | 0 |
| 0 | 0 | 0 | 0 |
| 1 | 1 | 0 | 0 |

| | | | |
|---|---|---|---|
| 0 | 0 | 0 | 0 |
| 0 | 0 | 0 | 0 |
| 0 | 0 | 0 | 0 |
| 1 | 1 | 0 | 0 |
| 0 | 0 | 0 | 0 |
| 0 | 0 | 0 | 0 |
| 0 | 0 | 0 | 0 |
| 0 | 0 | 0 | 0 |
| 1 | 0 | 0 | 0 |
| 0 | 0 | 0 | 0 |
| 0 | 0 | 0 | 0 |
| 0 | 0 | 0 | 0 |
| 1 | 0 | 1 | 0 |
| 1 | 0 | 0 | 0 |
| 0 | 0 | 0 | 0 |
| 0 | 0 | 0 | 0 |
| 0 | 0 | 0 | 0 |
| 0 | 0 | 0 | 0 |
| 0 | 0 | 0 | 0 |
| 0 | 0 | 0 | 0 |
| 0 | 0 | 0 | 0 |
| 0 | 0 | 0 | 0 |
| 0 | 0 | 0 | 0 |
| 1 | 0 | 0 | 0 |
| 0 | 0 | 0 | 0 |
| 0 | 0 | 0 | 0 |
| 0 | 0 | 0 | 0 |
| 0 | 0 | 0 | 0 |
| 0 | 0 | 0 | 0 |
| 0 | 0 | 0 | 0 |
| 0 | 0 | 0 | 0 |
| 0 | 0 | 0 | 0 |
| 0 | 0 | 0 | 0 |
| 0 | 0 | 0 | 0 |
| 0 | 0 | 0 | 0 |
| 0 | 0 | 0 | 0 |
| 0 | 0 | 0 | 0 |
| 0 | 0 | 0 | 0 |
| 0 | 0 | 0 | 0 |
| 0 | 0 | 0 | 0 |
| 0 | 0 | 0 | 0 |
| 0 | 0 | 0 | 0 |
| 0 | 0 | 0 | 0 |
| 1 | 1 | 0 | 0 |
| 1 | 1 | 0 | 0 |
| 0 | 0 | 0 | 0 |
| 0 | 0 | 0 | 0 |
| 0 | 1 | 0 | 0 |
| 0 | 1 | 0 | 0 |
| 0 | 1 | 0 | 0 |
| 0 | 0 | 0 | 0 |
| 0 | 0 | 0 | 0 |
| 0 | 0 | 0 | 0 |
| 0 | 0 | 0 | 0 |

| | | | |
|---|---|---|---|
| 0 | 0 | 0 | 0 |
| 0 | 0 | 0 | 0 |
| 0 | 0 | 0 | 0 |
| 0 | 0 | 0 | 0 |
| 1 | 0 | 0 | 0 |
| 0 | 0 | 0 | 0 |
| 0 | 0 | 0 | 0 |
| 1 | 0 | 0 | 0 |
| 1 | 0 | 0 | 0 |
| 1 | 0 | 0 | 0 |
| 1 | 0 | 0 | 0 |
| 1 | 0 | 0 | 0 |
| 0 | 0 | 0 | 0 |
| 1 | 0 | 0 | 0 |
| 1 | 0 | 0 | 0 |
| 0 | 0 | 0 | 0 |
| 0 | 0 | 0 | 0 |
| 0 | 0 | 0 | 0 |
| 1 | 0 | 0 | 0 |
| 1 | 0 | 0 | 0 |
| 1 | 0 | 0 | 0 |
| 1 | 0 | 0 | 0 |
| 0 | 0 | 0 | 0 |
| 0 | 0 | 0 | 0 |
| 0 | 0 | 0 | 0 |
| 0 | 0 | 0 | 0 |
| 0 | 0 | 0 | 0 |
| 0 | 0 | 0 | 0 |
| 0 | 0 | 0 | 0 |
| 0 | 0 | 0 | 0 |
| 0 | 0 | 0 | 0 |
| 0 | 0 | 0 | 0 |
| 1 | 0 | 0 | 0 |
| 0 | 0 | 0 | 0 |
| 0 | 0 | 0 | 0 |
| 1 | 0 | 0 | 0 |
| 0 | 0 | 0 | 0 |
| 0 | 0 | 0 | 0 |
| 0 | 0 | 0 | 0 |
| 0 | 0 | 0 | 0 |
| 0 | 0 | 0 | 0 |
| 0 | 0 | 0 | 0 |
| 0 | 0 | 0 | 0 |
| 0 | 0 | 0 | 0 |
| 0 | 0 | 0 | 0 |
| 0 | 0 | 0 | 0 |
| 0 | 0 | 0 | 0 |
| 0 | 0 | 0 | 0 |
| 0 | 0 | 0 | 0 |
| 0 | 0 | 0 | 0 |
| 0 | 0 | 0 | 0 |
| 0 | 0 | 0 | 0 |
| 0 | 0 | 0 | 0 |
| 0 | 0 | 0 | 0 |

| | | | |
|---|---|---|---|
| 0 | 0 | 0 | 0 |
| 0 | 0 | 0 | 0 |
| 0 | 0 | 0 | 0 |
| 0 | 0 | 0 | 0 |
| 0 | 0 | 0 | 0 |
| 0 | 0 | 0 | 0 |
| 0 | 0 | 0 | 0 |
| 0 | 0 | 0 | 0 |
| 0 | 0 | 0 | 0 |
| 0 | 0 | 0 | 0 |
| 0 | 0 | 0 | 0 |
| 0 | 0 | 0 | 1 |
| 0 | 0 | 0 | 0 |
| 0 | 0 | 0 | 0 |
| 0 | 0 | 0 | 0 |
| 0 | 0 | 0 | 0 |
| 0 | 0 | 0 | 0 |
| 0 | 0 | 0 | 1 |
| 0 | 0 | 0 | 0 |
| 0 | 0 | 0 | 0 |
| 0 | 0 | 0 | 1 |
| 0 | 0 | 0 | 0 |
| 0 | 0 | 0 | 0 |
| 0 | 0 | 0 | 0 |
| 0 | 0 | 0 | 0 |
| 0 | 0 | 0 | 0 |
| 0 | 0 | 0 | 0 |
| 0 | 0 | 0 | 0 |
| 0 | 0 | 0 | 1 |
| 0 | 0 | 0 | 0 |
| 0 | 0 | 0 | 0 |
| 0 | 0 | 0 | 0 |
| 0 | 0 | 0 | 0 |
| 0 | 0 | 0 | 0 |
| 0 | 0 | 0 | 0 |
| 0 | 0 | 0 | 0 |
| 0 | 0 | 0 | 0 |
| 0 | 0 | 0 | 0 |
| 0 | 0 | 0 | 0 |
| 0 | 0 | 0 | 0 |
| 0 | 0 | 0 | 0 |
| 0 | 0 | 0 | 0 |
| 0 | 0 | 0 | 0 |
| 0 | 0 | 0 | 0 |
| 0 | 0 | 0 | 0 |
| 0 | 0 | 0 | 0 |
| 0 | 0 | 0 | 0 |
| 0 | 0 | 0 | 0 |
| 0 | 0 | 0 | 0 |
| 0 | 0 | 0 | 0 |
| 0 | 0 | 0 | 0 |
| 0 | 0 | 0 | 0 |
| 0 | 0 | 0 | 0 |
| 0 | 0 | 0 | 0 |

| | | | |
|---|---|---|---|
| 0 | 0 | 0 | 0 |
| 0 | 0 | 0 | 0 |
| 1 | 0 | 0 | 0 |
| 0 | 0 | 0 | 0 |
| 1 | 0 | 0 | 0 |
| 0 | 0 | 0 | 0 |
| 0 | 0 | 0 | 0 |
| 0 | 0 | 0 | 0 |
| 0 | 0 | 0 | 0 |
| 1 | 1 | 0 | 0 |
| 0 | 0 | 1 | 0 |
| 0 | 0 | 0 | 0 |
| 0 | 0 | 0 | 0 |
| 0 | 0 | 0 | 0 |
| 0 | 0 | 0 | 0 |
| 0 | 0 | 0 | 0 |
| 0 | 0 | 0 | 0 |
| 0 | 0 | 0 | 0 |
| 0 | 0 | 0 | 0 |
| 0 | 0 | 0 | 0 |
| 0 | 0 | 0 | 0 |
| 0 | 0 | 0 | 0 |
| 0 | 0 | 0 | 0 |
| 0 | 0 | 0 | 0 |
| 0 | 0 | 0 | 0 |
| 0 | 0 | 0 | 0 |
| 0 | 0 | 0 | 0 |
| 0 | 0 | 0 | 0 |
| 0 | 0 | 0 | 0 |
| 0 | 0 | 0 | 0 |
| 0 | 0 | 0 | 0 |
| 0 | 0 | 0 | 0 |
| 0 | 0 | 0 | 0 |
| 0 | 0 | 0 | 0 |
| 0 | 0 | 0 | 0 |
| 0 | 0 | 0 | 0 |
| 0 | 0 | 0 | 0 |
| 0 | 0 | 0 | 0 |
| 0 | 0 | 0 | 0 |
| 0 | 0 | 0 | 0 |
| 0 | 0 | 0 | 0 |
| 0 | 0 | 0 | 0 |
| 0 | 0 | 0 | 0 |
| 0 | 0 | 0 | 0 |
| 0 | 0 | 0 | 0 |
| 1 | 0 | 0 | 0 |
| 0 | 0 | 0 | 0 |
| 0 | 0 | 0 | 0 |
| 0 | 0 | 0 | 0 |
| 0 | 0 | 0 | 0 |
| 1 | 0 | 0 | 0 |
| 0 | 0 | 0 | 0 |
| 0 | 0 | 0 | 0 |
| 0 | 0 | 0 | 0 |

| | | | |
|---|---|---|---|
| 0 | 0 | 0 | 0 |
| 0 | 0 | 0 | 0 |
| 0 | 0 | 0 | 0 |
| 0 | 0 | 0 | 0 |
| 0 | 0 | 0 | 0 |
| 0 | 0 | 0 | 0 |
| 0 | 0 | 0 | 0 |
| 0 | 0 | 0 | 0 |
| 1 | 1 | 0 | 0 |
| 0 | 0 | 0 | 0 |
| 0 | 0 | 0 | 0 |
| 0 | 0 | 0 | 0 |
| 0 | 0 | 0 | 0 |
| 0 | 0 | 0 | 0 |
| 0 | 1 | 0 | 0 |
| 0 | 1 | 0 | 0 |
| 0 | 1 | 0 | 0 |
| 0 | 0 | 0 | 0 |
| 0 | 0 | 0 | 0 |
| 0 | 0 | 0 | 0 |
| 0 | 1 | 0 | 0 |
| 0 | 0 | 0 | 0 |
| 0 | 0 | 0 | 0 |
| 1 | 1 | 0 | 0 |
| 0 | 1 | 0 | 0 |
| 0 | 1 | 0 | 0 |
| 0 | 0 | 0 | 0 |
| 0 | 0 | 0 | 0 |
| 0 | 0 | 0 | 0 |
| 0 | 0 | 0 | 0 |
| 0 | 0 | 0 | 0 |
| 0 | 0 | 0 | 0 |
| 0 | 0 | 0 | 0 |
| 0 | 0 | 0 | 0 |
| 0 | 0 | 0 | 0 |
| 0 | 0 | 0 | 0 |
| 0 | 0 | 0 | 0 |
| 1 | 0 | 0 | 0 |
| 0 | 0 | 0 | 0 |
| 0 | 0 | 0 | 0 |
| 0 | 0 | 0 | 0 |
| 0 | 0 | 0 | 0 |
| 0 | 0 | 0 | 0 |
| 0 | 0 | 0 | 0 |
| 0 | 0 | 0 | 0 |
| 0 | 0 | 0 | 0 |
| 0 | 0 | 0 | 0 |
| 0 | 0 | 0 | 0 |
| 0 | 0 | 0 | 0 |
| 0 | 0 | 0 | 0 |
| 0 | 0 | 0 | 0 |
| 0 | 0 | 0 | 0 |
| 0 | 0 | 0 | 0 |

| | | | |
|---|---|---|---|
| 0 | 0 | 0 | 0 |
| 0 | 0 | 0 | 0 |
| 0 | 0 | 0 | 0 |
| 0 | 0 | 1 | 0 |
| 0 | 0 | 0 | 0 |
| 0 | 0 | 0 | 0 |
| 0 | 0 | 0 | 0 |
| 0 | 0 | 0 | 0 |
| 0 | 0 | 0 | 0 |
| 0 | 0 | 0 | 0 |
| 0 | 0 | 0 | 0 |
| 0 | 0 | 0 | 0 |
| 1 | 0 | 0 | 0 |
| 0 | 0 | 0 | 0 |
| 0 | 0 | 0 | 0 |
| 0 | 0 | 0 | 0 |
| 0 | 1 | 0 | 0 |
| 0 | 0 | 0 | 0 |
| 0 | 0 | 0 | 0 |
| 0 | 0 | 0 | 0 |
| 0 | 0 | 0 | 0 |
| 0 | 0 | 0 | 0 |
| 0 | 0 | 0 | 0 |
| 0 | 0 | 0 | 0 |
| 0 | 0 | 0 | 0 |
| 0 | 0 | 0 | 0 |
| 0 | 0 | 0 | 0 |
| 0 | 0 | 0 | 0 |
| 0 | 0 | 0 | 0 |
| 0 | 0 | 0 | 0 |
| 1 | 0 | 0 | 0 |
| 1 | 0 | 0 | 0 |
| 1 | 0 | 0 | 0 |
| 0 | 0 | 0 | 0 |
| 1 | 0 | 0 | 0 |
| 1 | 0 | 0 | 0 |
| 1 | 0 | 0 | 0 |
| 0 | 0 | 0 | 0 |
| 0 | 0 | 0 | 0 |
| 0 | 0 | 0 | 0 |
| 0 | 0 | 0 | 0 |
| 0 | 0 | 0 | 0 |
| 0 | 1 | 0 | 0 |
| 0 | 1 | 0 | 0 |
| 0 | 0 | 0 | 0 |
| 0 | 0 | 0 | 0 |
| 1 | 0 | 0 | 0 |
| 0 | 0 | 0 | 0 |
| 0 | 0 | 0 | 0 |
| 0 | 0 | 0 | 0 |
| 0 | 0 | 0 | 0 |
| 0 | 0 | 0 | 0 |
| 1 | 0 | 0 | 0 |
| 0 | 0 | 0 | 0 |

| | | | |
|---|---|---|---|
| 0 | 0 | 0 | 0 |
| 1 | 0 | 0 | 0 |
| 1 | 0 | 0 | 0 |
| 0 | 0 | 0 | 0 |
| 0 | 0 | 0 | 0 |
| 0 | 0 | 0 | 0 |
| 0 | 0 | 0 | 0 |
| 0 | 1 | 0 | 0 |
| 0 | 0 | 0 | 0 |
| 0 | 0 | 0 | 0 |
| 0 | 0 | 0 | 0 |
| 0 | 0 | 0 | 0 |
| 0 | 0 | 0 | 0 |
| 0 | 0 | 0 | 0 |
| 1 | 0 | 0 | 0 |
| 0 | 0 | 0 | 0 |
| 0 | 0 | 0 | 0 |
| 0 | 0 | 0 | 0 |
| 0 | 0 | 0 | 0 |
| 1 | 0 | 0 | 0 |
| 1 | 0 | 0 | 0 |
| 0 | 0 | 0 | 0 |
| 1 | 0 | 0 | 0 |
| 1 | 0 | 0 | 0 |
| 0 | 0 | 0 | 0 |
| 0 | 0 | 0 | 0 |
| 0 | 0 | 0 | 0 |
| 0 | 0 | 0 | 0 |
| 0 | 0 | 0 | 0 |
| 0 | 0 | 0 | 0 |
| 0 | 0 | 0 | 0 |
| 0 | 0 | 0 | 0 |
| 0 | 0 | 0 | 0 |
| 0 | 0 | 0 | 0 |
| 0 | 0 | 0 | 0 |
| 0 | 0 | 0 | 0 |
| 0 | 0 | 0 | 0 |
| 0 | 0 | 0 | 0 |
| 0 | 0 | 0 | 0 |
| 0 | 0 | 0 | 0 |
| 0 | 0 | 0 | 0 |
| 0 | 0 | 0 | 0 |
| 0 | 0 | 0 | 0 |
| 0 | 0 | 0 | 0 |
| 0 | 0 | 0 | 0 |
| 0 | 0 | 0 | 0 |
| 0 | 0 | 0 | 0 |
| 0 | 0 | 0 | 0 |
| 0 | 0 | 0 | 0 |
| 0 | 0 | 0 | 0 |
| 0 | 0 | 0 | 0 |
| 0 | 0 | 0 | 0 |
| 0 | 0 | 0 | 0 |
| 1 | 1 | 0 | 1 |

| | | | |
|---|---|---|---|
| 0 | 0 | 0 | 0 |
| 0 | 0 | 0 | 0 |
| 0 | 0 | 0 | 0 |
| 0 | 0 | 0 | 0 |
| 0 | 1 | 0 | 0 |
| 0 | 0 | 0 | 0 |
| 0 | 0 | 0 | 0 |
| 0 | 1 | 0 | 0 |
| 0 | 0 | 0 | 0 |
| 0 | 0 | 0 | 0 |
| 0 | 0 | 0 | 0 |
| 0 | 0 | 0 | 0 |
| 0 | 0 | 0 | 0 |
| 0 | 0 | 0 | 0 |
| 0 | 1 | 0 | 1 |
| 0 | 1 | 0 | 0 |
| 0 | 0 | 0 | 0 |
| 0 | 0 | 0 | 0 |
| 0 | 1 | 0 | 0 |
| 0 | 1 | 0 | 1 |
| 0 | 0 | 0 | 0 |
| 0 | 0 | 0 | 0 |
| 0 | 0 | 0 | 0 |
| 0 | 0 | 0 | 0 |
| 0 | 0 | 0 | 0 |
| 1 | 0 | 0 | 0 |
| 0 | 0 | 0 | 0 |
| 0 | 0 | 1 | 0 |
| 0 | 0 | 1 | 0 |
| 0 | 0 | 1 | 0 |
| 0 | 0 | 0 | 0 |
| 0 | 0 | 0 | 0 |
| 0 | 0 | 0 | 0 |
| 0 | 0 | 0 | 0 |
| 0 | 0 | 0 | 0 |
| 0 | 0 | 1 | 0 |
| 0 | 0 | 0 | 0 |
| 0 | 0 | 0 | 0 |
| 1 | 0 | 1 | 0 |
| 0 | 0 | 1 | 0 |
| 1 | 1 | 0 | 0 |
| 0 | 1 | 0 | 0 |
| 0 | 0 | 0 | 0 |
| 0 | 0 | 0 | 0 |
| 1 | 0 | 0 | 0 |
| 0 | 0 | 0 | 0 |
| 0 | 0 | 1 | 0 |
| 0 | 0 | 0 | 0 |
| 0 | 0 | 0 | 0 |
| 0 | 0 | 0 | 0 |
| 0 | 0 | 0 | 0 |
| 0 | 0 | 0 | 0 |
| 0 | 0 | 0 | 0 |
| 0 | 0 | 0 | 0 |

| | | | |
|---|---|---|---|
| 0 | 0 | 0 | 0 |
| 0 | 0 | 0 | 0 |
| 0 | 0 | 0 | 0 |
| 0 | 0 | 0 | 0 |
| 0 | 0 | 0 | 0 |
| 0 | 0 | 0 | 0 |
| 0 | 0 | 0 | 0 |
| 1 | 1 | 0 | 0 |
| 0 | 0 | 0 | 0 |
| 0 | 0 | 0 | 0 |
| 0 | 0 | 0 | 0 |
| 0 | 0 | 0 | 1 |
| 0 | 0 | 0 | 0 |
| 0 | 0 | 0 | 0 |
| 0 | 0 | 0 | 0 |
| 0 | 0 | 0 | 0 |
| 0 | 0 | 0 | 0 |
| 0 | 0 | 0 | 0 |
| 0 | 0 | 0 | 0 |
| 0 | 0 | 0 | 0 |
| 0 | 0 | 0 | 0 |
| 0 | 0 | 0 | 0 |
| 0 | 0 | 0 | 0 |
| 0 | 0 | 0 | 0 |
| 0 | 0 | 0 | 0 |
| 0 | 0 | 0 | 0 |
| 0 | 0 | 0 | 0 |
| 0 | 0 | 0 | 0 |
| 0 | 0 | 0 | 0 |
| 0 | 1 | 0 | 0 |
| 0 | 1 | 0 | 0 |
| 0 | 0 | 0 | 0 |
| 0 | 0 | 0 | 0 |
| 0 | 0 | 0 | 0 |
| 0 | 0 | 0 | 0 |
| 0 | 1 | 0 | 0 |
| 0 | 0 | 0 | 0 |
| 0 | 0 | 0 | 0 |
| 0 | 0 | 0 | 0 |
| 0 | 0 | 0 | 0 |
| 0 | 0 | 0 | 0 |
| 1 | 1 | 0 | 0 |
| 1 | 0 | 0 | 0 |
| 0 | 0 | 0 | 0 |
| 0 | 0 | 0 | 0 |
| 0 | 0 | 0 | 0 |
| 0 | 0 | 0 | 0 |
| 0 | 0 | 0 | 0 |
| 0 | 0 | 0 | 0 |
| 0 | 0 | 0 | 0 |
| 0 | 0 | 0 | 0 |
| 0 | 0 | 0 | 0 |
| 0 | 0 | 0 | 0 |
| 0 | 0 | 0 | 0 |

| | | | |
|---|---|---|---|
| 0 | 0 | 0 | 0 |
| 0 | 0 | 0 | 0 |
| 0 | 0 | 0 | 0 |
| 0 | 0 | 0 | 0 |
| 0 | 0 | 0 | 0 |
| 0 | 0 | 0 | 0 |
| 0 | 0 | 0 | 0 |
| 0 | 0 | 0 | 0 |
| 1 | 0 | 0 | 0 |
| 1 | 0 | 0 | 0 |
| 0 | 0 | 0 | 0 |
| 0 | 0 | 0 | 0 |
| 0 | 0 | 0 | 0 |
| 1 | 0 | 0 | 0 |
| 0 | 0 | 0 | 0 |
| 0 | 0 | 0 | 0 |
| 0 | 0 | 0 | 0 |
| 0 | 0 | 0 | 0 |
| 0 | 0 | 0 | 0 |
| 1 | 1 | 0 | 0 |
| 0 | 0 | 1 | 0 |
| 0 | 0 | 0 | 0 |
| 0 | 0 | 0 | 0 |
| 0 | 0 | 0 | 0 |
| 0 | 0 | 0 | 0 |
| 0 | 0 | 0 | 0 |
| 0 | 0 | 0 | 0 |
| 0 | 0 | 0 | 0 |
| 0 | 0 | 0 | 0 |
| 0 | 0 | 0 | 0 |
| 0 | 0 | 0 | 0 |
| 0 | 0 | 0 | 0 |
| 0 | 0 | 0 | 0 |
| 0 | 0 | 0 | 0 |
| 0 | 0 | 0 | 0 |
| 0 | 0 | 0 | 0 |
| 0 | 0 | 0 | 0 |
| 0 | 0 | 0 | 0 |
| 0 | 0 | 0 | 0 |
| 0 | 0 | 0 | 0 |
| 0 | 0 | 0 | 0 |
| 0 | 0 | 0 | 0 |
| 0 | 0 | 0 | 0 |
| 0 | 0 | 0 | 0 |
| 0 | 0 | 0 | 0 |
| 0 | 0 | 0 | 0 |
| 1 | 0 | 1 | 0 |
| 1 | 0 | 0 | 0 |
| 0 | 0 | 0 | 0 |
| 0 | 0 | 0 | 0 |
| 0 | 0 | 1 | 0 |
| 0 | 0 | 0 | 0 |
| 0 | 0 | 0 | 0 |
| 0 | 0 | 0 | 0 |

| | | | |
|---|---|---|---|
| 0 | 0 | 0 | 0 |
| 0 | 0 | 0 | 0 |
| 0 | 0 | 0 | 0 |
| 0 | 0 | 0 | 0 |
| 0 | 0 | 0 | 0 |
| 0 | 0 | 0 | 0 |
| 0 | 0 | 0 | 0 |
| 0 | 0 | 0 | 0 |
| 0 | 0 | 0 | 0 |
| 0 | 0 | 0 | 0 |
| 0 | 0 | 0 | 0 |
| 0 | 0 | 0 | 0 |
| 0 | 0 | 0 | 0 |
| 0 | 0 | 0 | 0 |
| 0 | 0 | 0 | 0 |
| 0 | 0 | 0 | 0 |
| 0 | 0 | 0 | 0 |
| 0 | 0 | 0 | 0 |
| 0 | 0 | 0 | 0 |
| 1 | 0 | 0 | 0 |
| 0 | 0 | 0 | 0 |
| 0 | 0 | 0 | 0 |
| 0 | 0 | 0 | 0 |
| 0 | 0 | 0 | 0 |
| 0 | 0 | 0 | 0 |
| 0 | 0 | 0 | 0 |
| 0 | 0 | 0 | 0 |
| 0 | 0 | 0 | 0 |
| 0 | 0 | 0 | 0 |
| 0 | 0 | 0 | 0 |
| 0 | 0 | 0 | 0 |
| 0 | 0 | 0 | 0 |
| 0 | 0 | 0 | 0 |
| 0 | 0 | 0 | 0 |
| 0 | 0 | 0 | 0 |
| 0 | 0 | 0 | 0 |
| 0 | 0 | 0 | 0 |
| 0 | 0 | 0 | 0 |
| 1 | 0 | 0 | 1 |
| 0 | 0 | 0 | 0 |
| 0 | 0 | 0 | 0 |
| 0 | 0 | 0 | 0 |
| 0 | 0 | 0 | 1 |
| 0 | 0 | 0 | 1 |
| 0 | 0 | 0 | 1 |
| 0 | 0 | 0 | 0 |
| 0 | 0 | 0 | 0 |
| 0 | 0 | 0 | 0 |
| 0 | 0 | 0 | 0 |
| 0 | 0 | 0 | 0 |
| 0 | 0 | 0 | 0 |
| 0 | 0 | 0 | 0 |
| 0 | 0 | 0 | 0 |
| 0 | 0 | 0 | 0 |

| | | | |
|---|---|---|---|
| 0 | 0 | 0 | 0 |
| 0 | 0 | 0 | 0 |
| 0 | 0 | 0 | 0 |
| 0 | 0 | 0 | 0 |
| 0 | 0 | 0 | 0 |
| 0 | 0 | 0 | 0 |
| 0 | 0 | 0 | 0 |
| 0 | 0 | 0 | 0 |
| 0 | 0 | 0 | 0 |
| 0 | 0 | 0 | 0 |
| 0 | 0 | 0 | 0 |
| 0 | 0 | 0 | 0 |
| 0 | 0 | 0 | 0 |
| 0 | 0 | 0 | 0 |
| 0 | 0 | 0 | 0 |
| 0 | 0 | 0 | 0 |
| 0 | 0 | 0 | 0 |
| 0 | 0 | 0 | 0 |
| 0 | 0 | 0 | 0 |
| 0 | 0 | 0 | 0 |
| 0 | 0 | 0 | 0 |
| 0 | 0 | 0 | 0 |
| 0 | 0 | 0 | 0 |
| 0 | 0 | 0 | 0 |
| 0 | 0 | 0 | 0 |
| 0 | 0 | 0 | 0 |
| 0 | 0 | 0 | 0 |
| 0 | 0 | 0 | 0 |
| 0 | 0 | 0 | 0 |
| 0 | 0 | 0 | 0 |
| 0 | 0 | 0 | 0 |
| 0 | 0 | 0 | 0 |
| 1 | 1 | 0 | 1 |
| 1 | 1 | 0 | 1 |
| 1 | 0 | 0 | 0 |
| 0 | 0 | 0 | 0 |
| 0 | 0 | 0 | 0 |
| 0 | 0 | 0 | 0 |
| 0 | 0 | 0 | 0 |
| 0 | 0 | 0 | 0 |
| 0 | 0 | 0 | 0 |
| 0 | 0 | 0 | 0 |
| 1 | 1 | 0 | 0 |
| 0 | 0 | 0 | 0 |
| 1 | 0 | 0 | 0 |
| 0 | 0 | 0 | 0 |
| 0 | 0 | 0 | 0 |
| 0 | 0 | 0 | 0 |
| 0 | 0 | 0 | 0 |
| 0 | 0 | 0 | 0 |
| 1 | 0 | 0 | 0 |
| 0 | 0 | 0 | 0 |
| 0 | 0 | 0 | 0 |
| 0 | 0 | 0 | 0 |

| | | | |
|---|---|---|---|
| 0 | 0 | 0 | 0 |
| 0 | 0 | 0 | 0 |
| 0 | 0 | 0 | 0 |
| 0 | 0 | 0 | 0 |
| 0 | 0 | 0 | 0 |
| 0 | 0 | 0 | 0 |
| 0 | 0 | 0 | 0 |
| 0 | 0 | 0 | 0 |
| 0 | 0 | 0 | 0 |
| 0 | 0 | 0 | 0 |
| 0 | 0 | 0 | 0 |
| 0 | 0 | 0 | 0 |
| 0 | 0 | 0 | 0 |
| 0 | 0 | 0 | 0 |
| 0 | 0 | 0 | 0 |
| 0 | 0 | 0 | 0 |
| 0 | 0 | 0 | 0 |
| 0 | 0 | 0 | 0 |
| 0 | 0 | 0 | 0 |
| 0 | 0 | 0 | 1 |
| 0 | 0 | 0 | 0 |
| 0 | 0 | 0 | 1 |
| 0 | 0 | 0 | 1 |
| 0 | 0 | 0 | 1 |
| 0 | 0 | 0 | 0 |
| 0 | 0 | 0 | 0 |
| 0 | 0 | 0 | 0 |
| 0 | 0 | 0 | 0 |
| 0 | 0 | 0 | 0 |
| 0 | 0 | 0 | 0 |
| 0 | 0 | 0 | 0 |
| 0 | 0 | 0 | 0 |
| 0 | 0 | 0 | 0 |
| 0 | 0 | 0 | 0 |
| 0 | 0 | 0 | 0 |
| 0 | 1 | 0 | 0 |
| 0 | 0 | 0 | 0 |
| 0 | 0 | 0 | 0 |
| 0 | 1 | 0 | 0 |
| 0 | 0 | 0 | 0 |
| 0 | 0 | 0 | 0 |
| 0 | 0 | 0 | 0 |
| 0 | 0 | 0 | 0 |
| 0 | 0 | 0 | 0 |
| 0 | 0 | 0 | 0 |
| 0 | 0 | 0 | 0 |
| 0 | 0 | 0 | 0 |
| 0 | 0 | 0 | 0 |
| 0 | 0 | 0 | 0 |
| 0 | 0 | 0 | 0 |
| 0 | 0 | 0 | 0 |
| 0 | 0 | 0 | 0 |
| 0 | 0 | 0 | 0 |
| 0 | 0 | 0 | 0 |

| | | | |
|---|---|---|---|
| 0 | 0 | 0 | 0 |
| 0 | 0 | 0 | 0 |
| 0 | 0 | 0 | 0 |
| 0 | 0 | 0 | 0 |
| 0 | 0 | 0 | 0 |
| 0 | 0 | 0 | 0 |
| 0 | 0 | 0 | 0 |
| 0 | 0 | 0 | 0 |
| 0 | 0 | 1 | 0 |
| 0 | 0 | 1 | 0 |
| 0 | 0 | 0 | 0 |
| 0 | 0 | 0 | 0 |
| 0 | 0 | 0 | 0 |
| 0 | 0 | 0 | 0 |
| 0 | 0 | 0 | 0 |
| 0 | 1 | 0 | 0 |
| 0 | 0 | 0 | 0 |
| 0 | 0 | 0 | 0 |
| 1 | 1 | 0 | 0 |
| 0 | 0 | 0 | 0 |
| 0 | 0 | 0 | 0 |
| 0 | 0 | 0 | 0 |
| 0 | 0 | 0 | 0 |
| 0 | 0 | 0 | 0 |
| 0 | 0 | 0 | 0 |
| 0 | 0 | 0 | 0 |
| 0 | 0 | 0 | 0 |
| 0 | 0 | 0 | 0 |
| 0 | 0 | 0 | 0 |
| 0 | 0 | 0 | 0 |
| 0 | 0 | 0 | 0 |
| 0 | 0 | 0 | 0 |
| 0 | 0 | 0 | 0 |
| 0 | 0 | 0 | 0 |
| 0 | 0 | 0 | 0 |
| 0 | 0 | 0 | 0 |
| 0 | 0 | 0 | 0 |
| 0 | 0 | 0 | 0 |
| 0 | 0 | 0 | 0 |
| 0 | 0 | 0 | 0 |
| 0 | 0 | 0 | 0 |
| 0 | 0 | 0 | 0 |
| 0 | 0 | 0 | 0 |
| 0 | 0 | 0 | 0 |
| 0 | 0 | 0 | 0 |
| 0 | 0 | 0 | 0 |
| 0 | 0 | 0 | 0 |
| 0 | 0 | 0 | 0 |
| 0 | 0 | 0 | 0 |
| 0 | 0 | 0 | 0 |
| 0 | 0 | 0 | 0 |
| 0 | 0 | 0 | 0 |
| 0 | 0 | 0 | 0 |
| 1 | 0 | 0 | 0 |

| | | | |
|---|---|---|---|
| 0 | 0 | 0 | 0 |
| 0 | 0 | 0 | 0 |
| 0 | 0 | 0 | 0 |
| 0 | 0 | 0 | 0 |
| 0 | 0 | 0 | 0 |
| 0 | 0 | 0 | 0 |
| 0 | 0 | 0 | 0 |
| 0 | 0 | 0 | 0 |
| 0 | 0 | 0 | 0 |
| 0 | 0 | 0 | 0 |
| 1 | 0 | 0 | 0 |
| 0 | 0 | 0 | 0 |
| 0 | 0 | 0 | 0 |
| 0 | 0 | 0 | 0 |
| 1 | 1 | 0 | 1 |
| 0 | 0 | 0 | 0 |
| 1 | 0 | 0 | 0 |
| 1 | 1 | 0 | 0 |
| 0 | 0 | 0 | 0 |
| 0 | 0 | 0 | 0 |
| 0 | 0 | 0 | 0 |
| 0 | 0 | 0 | 0 |
| 0 | 0 | 0 | 0 |
| 0 | 0 | 0 | 1 |
| 0 | 0 | 0 | 0 |
| 0 | 0 | 0 | 0 |
| 0 | 0 | 0 | 0 |
| 0 | 0 | 0 | 1 |
| 0 | 0 | 0 | 0 |
| 0 | 0 | 0 | 0 |
| 0 | 0 | 0 | 0 |
| 0 | 0 | 0 | 0 |
| 0 | 0 | 0 | 0 |
| 0 | 0 | 0 | 0 |
| 0 | 0 | 0 | 0 |
| 0 | 0 | 0 | 0 |
| 0 | 0 | 0 | 0 |
| 0 | 0 | 0 | 0 |
| 0 | 0 | 0 | 0 |
| 0 | 0 | 0 | 0 |
| 0 | 0 | 0 | 0 |
| 0 | 0 | 0 | 0 |
| 0 | 0 | 0 | 0 |
| 0 | 0 | 0 | 0 |
| 0 | 0 | 1 | 0 |
| 0 | 0 | 0 | 0 |
| 0 | 0 | 0 | 0 |
| 0 | 0 | 0 | 0 |
| 0 | 0 | 0 | 0 |
| 0 | 0 | 0 | 0 |
| 0 | 0 | 0 | 0 |
| 0 | 0 | 0 | 0 |
| 0 | 0 | 0 | 0 |
| 0 | 0 | 0 | 0 |

| | | | |
|---|---|---|---|
| 0 | 0 | 0 | 0 |
| 0 | 0 | 0 | 0 |
| 0 | 0 | 0 | 0 |
| 0 | 0 | 0 | 0 |
| 0 | 0 | 0 | 0 |
| 0 | 0 | 0 | 0 |
| 1 | 0 | 0 | 1 |
| 1 | 0 | 0 | 0 |
| 0 | 0 | 0 | 0 |
| 0 | 0 | 0 | 0 |
| 0 | 0 | 0 | 0 |
| 0 | 0 | 0 | 0 |
| 0 | 0 | 0 | 0 |
| 0 | 0 | 0 | 0 |
| 0 | 0 | 0 | 0 |
| 0 | 0 | 0 | 0 |
| 0 | 0 | 0 | 0 |
| 0 | 0 | 0 | 0 |
| 0 | 0 | 0 | 0 |
| 0 | 0 | 0 | 0 |
| 1 | 0 | 0 | 0 |
| 0 | 0 | 0 | 0 |
| 0 | 0 | 0 | 0 |
| 0 | 0 | 0 | 0 |
| 0 | 0 | 0 | 0 |
| 0 | 0 | 0 | 0 |
| 0 | 0 | 0 | 0 |
| 0 | 0 | 0 | 0 |
| 0 | 0 | 0 | 0 |
| 0 | 0 | 0 | 0 |
| 0 | 0 | 0 | 0 |
| 0 | 0 | 0 | 0 |
| 0 | 0 | 0 | 0 |
| 0 | 0 | 0 | 0 |
| 0 | 0 | 0 | 0 |
| 0 | 0 | 0 | 0 |
| 0 | 0 | 0 | 0 |
| 0 | 0 | 0 | 0 |
| 0 | 0 | 0 | 0 |
| 0 | 0 | 0 | 0 |
| 0 | 0 | 0 | 0 |
| 0 | 0 | 0 | 0 |
| 0 | 0 | 0 | 0 |
| 0 | 0 | 0 | 0 |
| 0 | 0 | 0 | 0 |
| 0 | 0 | 0 | 0 |
| 0 | 0 | 0 | 0 |
| 0 | 0 | 0 | 0 |
| 0 | 0 | 0 | 0 |
| 0 | 0 | 0 | 0 |
| 0 | 0 | 0 | 0 |
| 0 | 0 | 0 | 0 |
| 0 | 0 | 0 | 0 |
| 0 | 0 | 0 | 0 |

| | | | |
|---|---|---|---|
| 0 | 0 | 0 | 0 |
| 0 | 0 | 0 | 0 |
| 0 | 0 | 0 | 0 |
| 1 | 0 | 0 | 0 |
| 0 | 0 | 0 | 0 |
| 1 | 0 | 0 | 0 |
| 0 | 0 | 0 | 0 |
| 0 | 0 | 0 | 0 |
| 1 | 0 | 0 | 0 |
| 0 | 0 | 0 | 0 |
| 0 | 0 | 0 | 0 |
| 0 | 0 | 0 | 0 |
| 0 | 0 | 0 | 0 |
| 0 | 0 | 0 | 0 |
| 0 | 0 | 0 | 0 |
| 0 | 0 | 0 | 0 |
| 0 | 0 | 0 | 0 |
| 0 | 0 | 0 | 0 |
| 1 | 0 | 0 | 0 |
| 0 | 0 | 0 | 0 |
| 0 | 0 | 0 | 0 |
| 0 | 0 | 0 | 0 |
| 0 | 0 | 0 | 0 |
| 0 | 0 | 0 | 0 |
| 0 | 0 | 0 | 0 |
| 0 | 0 | 0 | 0 |
| 0 | 0 | 0 | 0 |
| 0 | 0 | 0 | 0 |
| 0 | 1 | 0 | 0 |
| 0 | 0 | 0 | 0 |
| 0 | 0 | 0 | 0 |
| 0 | 0 | 0 | 0 |
| 0 | 1 | 0 | 0 |
| 1 | 1 | 0 | 0 |
| 1 | 0 | 0 | 0 |
| 0 | 0 | 0 | 0 |
| 0 | 0 | 0 | 0 |
| 0 | 0 | 0 | 0 |
| 0 | 0 | 0 | 0 |
| 0 | 0 | 0 | 0 |
| 1 | 0 | 0 | 0 |
| 0 | 0 | 0 | 0 |
| 0 | 0 | 0 | 0 |
| 0 | 0 | 0 | 0 |
| 0 | 0 | 0 | 0 |
| 0 | 0 | 0 | 0 |
| 0 | 0 | 0 | 0 |
| 0 | 0 | 0 | 0 |
| 0 | 0 | 0 | 0 |
| 0 | 0 | 0 | 0 |
| 0 | 0 | 0 | 0 |
| 0 | 0 | 0 | 0 |
| 0 | 0 | 0 | 0 |
| 1 | 0 | 0 | 0 |

| | | | |
|---|---|---|---|
| 1 | 0 | 0 | 1 |
| 0 | 0 | 0 | 0 |
| 0 | 0 | 0 | 0 |
| 0 | 0 | 0 | 0 |
| 0 | 0 | 0 | 0 |
| 0 | 0 | 0 | 0 |
| 0 | 0 | 0 | 0 |
| 0 | 0 | 0 | 0 |
| 0 | 0 | 0 | 0 |
| 0 | 0 | 0 | 0 |
| 0 | 0 | 0 | 0 |
| 0 | 0 | 0 | 0 |
| 0 | 0 | 0 | 0 |
| 0 | 0 | 0 | 0 |
| 0 | 0 | 0 | 0 |
| 0 | 0 | 0 | 0 |
| 0 | 0 | 0 | 0 |
| 0 | 0 | 0 | 0 |
| 0 | 0 | 0 | 0 |
| 0 | 0 | 0 | 0 |
| 0 | 0 | 0 | 0 |
| 0 | 0 | 0 | 0 |
| 0 | 1 | 0 | 0 |
| 0 | 0 | 0 | 0 |
| 0 | 0 | 0 | 0 |
| 0 | 0 | 0 | 0 |
| 0 | 0 | 0 | 0 |
| 0 | 0 | 0 | 0 |
| 0 | 0 | 0 | 0 |
| 0 | 0 | 0 | 0 |
| 0 | 0 | 0 | 0 |
| 0 | 0 | 0 | 0 |
| 0 | 0 | 0 | 0 |
| 0 | 0 | 0 | 0 |
| 0 | 0 | 0 | 0 |
| 0 | 0 | 0 | 0 |
| 0 | 0 | 0 | 0 |
| 0 | 0 | 0 | 0 |
| 0 | 0 | 0 | 0 |
| 0 | 0 | 0 | 0 |
| 0 | 0 | 0 | 0 |
| 0 | 0 | 0 | 0 |
| 0 | 0 | 0 | 0 |
| 0 | 0 | 0 | 0 |
| 0 | 0 | 0 | 0 |
| 0 | 0 | 0 | 0 |
| 0 | 0 | 0 | 0 |
| 0 | 0 | 0 | 0 |
| 0 | 0 | 0 | 0 |
| 0 | 0 | 0 | 0 |
| 0 | 0 | 0 | 0 |
| 0 | 0 | 0 | 0 |
| 0 | 0 | 0 | 0 |
| 0 | 0 | 0 | 0 |

| | | | |
|---|---|---|---|
| 0 | 0 | 0 | 0 |
| 0 | 0 | 0 | 0 |
| 0 | 0 | 0 | 0 |
| 0 | 0 | 0 | 0 |
| 0 | 0 | 0 | 0 |
| 0 | 0 | 0 | 0 |
| 0 | 1 | 0 | 0 |
| 0 | 1 | 0 | 0 |
| 0 | 0 | 0 | 0 |
| 0 | 0 | 0 | 0 |
| 0 | 0 | 0 | 0 |
| 0 | 0 | 0 | 0 |
| 0 | 0 | 0 | 0 |
| 0 | 0 | 0 | 0 |
| 0 | 0 | 0 | 0 |
| 0 | 0 | 0 | 0 |
| 1 | 0 | 0 | 0 |
| 0 | 0 | 0 | 0 |
| 1 | 0 | 0 | 0 |
| 1 | 0 | 0 | 0 |
| 1 | 0 | 0 | 0 |
| 1 | 0 | 0 | 0 |
| 0 | 0 | 0 | 0 |
| 0 | 0 | 0 | 0 |
| 0 | 0 | 0 | 0 |
| 0 | 0 | 0 | 0 |
| 0 | 0 | 0 | 0 |
| 0 | 0 | 0 | 0 |
| 0 | 0 | 0 | 0 |
| 0 | 0 | 0 | 0 |
| 0 | 0 | 0 | 0 |
| 0 | 0 | 0 | 0 |
| 0 | 0 | 0 | 0 |
| 0 | 0 | 0 | 0 |
| 0 | 0 | 0 | 0 |
| 0 | 0 | 1 | 0 |
| 1 | 0 | 0 | 0 |
| 1 | 0 | 0 | 0 |
| 1 | 0 | 0 | 0 |
| 0 | 0 | 0 | 0 |
| 0 | 0 | 0 | 0 |
| 0 | 0 | 0 | 0 |
| 0 | 0 | 0 | 0 |
| 1 | 0 | 0 | 0 |
| 1 | 0 | 0 | 0 |
| 1 | 0 | 0 | 0 |
| 0 | 0 | 0 | 0 |
| 0 | 0 | 0 | 0 |
| 1 | 0 | 0 | 0 |
| 0 | 0 | 0 | 0 |
| 0 | 0 | 0 | 0 |
| 1 | 0 | 0 | 0 |
| 0 | 0 | 0 | 0 |
| 0 | 0 | 0 | 0 |

| | | | |
|---|---|---|---|
| 0 | 0 | 0 | 0 |
| 0 | 0 | 0 | 0 |
| 0 | 0 | 0 | 0 |
| 0 | 0 | 0 | 0 |
| 0 | 0 | 0 | 0 |
| 0 | 0 | 0 | 0 |
| 0 | 0 | 0 | 0 |
| 0 | 0 | 0 | 0 |
| 0 | 0 | 0 | 0 |
| 0 | 0 | 0 | 0 |
| 0 | 0 | 0 | 0 |
| 0 | 0 | 0 | 0 |
| 0 | 0 | 0 | 0 |
| 0 | 0 | 0 | 0 |
| 0 | 0 | 0 | 0 |
| 0 | 0 | 0 | 0 |
| 0 | 0 | 0 | 0 |
| 0 | 0 | 0 | 0 |
| 0 | 0 | 0 | 0 |
| 0 | 0 | 0 | 0 |
| 0 | 0 | 0 | 0 |
| 0 | 0 | 0 | 0 |
| 1 | 1 | 0 | 0 |
| 0 | 0 | 0 | 0 |
| 0 | 1 | 0 | 0 |
| 0 | 0 | 0 | 0 |
| 0 | 0 | 0 | 0 |
| 0 | 0 | 0 | 0 |
| 0 | 0 | 0 | 0 |
| 0 | 0 | 0 | 0 |
| 0 | 0 | 0 | 0 |
| 0 | 0 | 0 | 0 |
| 0 | 0 | 0 | 0 |
| 0 | 0 | 0 | 0 |
| 0 | 0 | 0 | 0 |
| 0 | 0 | 0 | 0 |
| 0 | 0 | 0 | 0 |
| 0 | 0 | 0 | 0 |
| 0 | 0 | 0 | 0 |
| 0 | 0 | 0 | 0 |
| 0 | 0 | 0 | 0 |
| 0 | 0 | 0 | 0 |
| 0 | 0 | 0 | 0 |
| 0 | 0 | 0 | 0 |
| 0 | 0 | 0 | 0 |
| 1 | 0 | 0 | 1 |
| 0 | 0 | 0 | 0 |
| 0 | 0 | 0 | 0 |
| 0 | 0 | 0 | 0 |
| 1 | 0 | 0 | 0 |
| 1 | 0 | 0 | 0 |
| 1 | 0 | 0 | 0 |
| 0 | 0 | 0 | 0 |
| 0 | 0 | 0 | 0 |

| | | | |
|---|---|---|---|
| 0 | 0 | 0 | 0 |
| 0 | 0 | 0 | 0 |
| 0 | 0 | 0 | 0 |
| 0 | 0 | 0 | 0 |
| 0 | 0 | 0 | 0 |
| 0 | 0 | 0 | 0 |
| 0 | 0 | 0 | 0 |
| 0 | 0 | 0 | 0 |
| 0 | 0 | 0 | 0 |
| 0 | 0 | 0 | 0 |
| 0 | 0 | 0 | 1 |
| 0 | 0 | 0 | 0 |
| 0 | 0 | 0 | 0 |
| 0 | 0 | 0 | 0 |
| 0 | 0 | 0 | 0 |
| 0 | 0 | 0 | 0 |
| 0 | 0 | 0 | 0 |
| 0 | 0 | 0 | 0 |
| 0 | 0 | 0 | 0 |
| 0 | 0 | 0 | 0 |
| 0 | 0 | 0 | 0 |
| 0 | 0 | 0 | 0 |
| 0 | 0 | 0 | 0 |
| 0 | 0 | 0 | 0 |
| 0 | 0 | 0 | 0 |
| 0 | 0 | 0 | 0 |
| 0 | 0 | 0 | 0 |
| 0 | 0 | 0 | 0 |
| 0 | 0 | 0 | 0 |
| 0 | 0 | 0 | 0 |
| 0 | 1 | 0 | 0 |
| 0 | 0 | 0 | 0 |
| 1 | 1 | 0 | 0 |
| 1 | 1 | 0 | 0 |
| 0 | 0 | 0 | 0 |
| 0 | 0 | 0 | 0 |
| 0 | 0 | 0 | 0 |
| 0 | 0 | 0 | 0 |
| 0 | 0 | 0 | 0 |
| 0 | 0 | 0 | 0 |
| 0 | 0 | 0 | 0 |
| 0 | 1 | 0 | 0 |
| 0 | 0 | 0 | 0 |
| 0 | 0 | 0 | 0 |
| 0 | 0 | 0 | 0 |
| 1 | 0 | 0 | 0 |
| 1 | 0 | 0 | 0 |
| 1 | 0 | 0 | 0 |
| 1 | 1 | 1 | 0 |
| 0 | 0 | 0 | 0 |
| 1 | 1 | 1 | 1 |
| 0 | 0 | 0 | 0 |
| 0 | 0 | 0 | 0 |
| 0 | 0 | 0 | 0 |

| | | | |
|---|---|---|---|
| 0 | 0 | 0 | 0 |
| 0 | 0 | 0 | 0 |
| 0 | 0 | 0 | 0 |
| 0 | 0 | 0 | 0 |
| 1 | 0 | 0 | 0 |
| 1 | 0 | 0 | 0 |
| 0 | 0 | 0 | 0 |
| 0 | 0 | 0 | 0 |
| 0 | 0 | 0 | 0 |
| 0 | 0 | 0 | 0 |
| 0 | 0 | 0 | 0 |
| 0 | 0 | 0 | 0 |
| 0 | 0 | 0 | 0 |
| 0 | 0 | 0 | 0 |
| 0 | 0 | 0 | 0 |
| 0 | 0 | 0 | 0 |
| 0 | 0 | 0 | 0 |
| 0 | 0 | 0 | 0 |
| 0 | 0 | 0 | 0 |
| 0 | 0 | 0 | 0 |
| 0 | 0 | 0 | 0 |
| 0 | 0 | 0 | 0 |
| 0 | 0 | 0 | 0 |
| 0 | 0 | 0 | 0 |
| 0 | 0 | 0 | 0 |
| 0 | 0 | 0 | 0 |
| 0 | 0 | 0 | 0 |
| 0 | 0 | 0 | 0 |
| 0 | 0 | 0 | 0 |
| 0 | 0 | 0 | 0 |
| 0 | 0 | 0 | 0 |
| 0 | 0 | 0 | 0 |
| 0 | 0 | 0 | 0 |
| 0 | 0 | 0 | 0 |
| 0 | 0 | 0 | 0 |
| 0 | 0 | 0 | 0 |
| 0 | 0 | 0 | 0 |
| 0 | 0 | 0 | 0 |
| 0 | 0 | 0 | 0 |
| 0 | 0 | 0 | 0 |
| 0 | 0 | 0 | 0 |
| 0 | 0 | 0 | 0 |
| 0 | 0 | 0 | 0 |
| 0 | 0 | 0 | 0 |
| 0 | 0 | 0 | 0 |
| 0 | 0 | 0 | 0 |
| 0 | 0 | 0 | 0 |
| 0 | 0 | 0 | 0 |
| 0 | 0 | 0 | 0 |
| 0 | 0 | 0 | 0 |
| 0 | 0 | 0 | 0 |
| 0 | 0 | 0 | 0 |
| 0 | 0 | 0 | 0 |
| 0 | 1 | 0 | 0 |

| | | | |
|---|---|---|---|
| 0 | 0 | 0 | 0 |
| 0 | 0 | 0 | 0 |
| 0 | 0 | 0 | 0 |
| 0 | 0 | 0 | 0 |
| 0 | 0 | 0 | 0 |
| 0 | 0 | 0 | 0 |
| 0 | 0 | 0 | 0 |
| 0 | 0 | 0 | 0 |
| 0 | 0 | 0 | 0 |
| 0 | 0 | 0 | 0 |
| 0 | 0 | 0 | 0 |
| 0 | 0 | 0 | 0 |
| 0 | 0 | 0 | 0 |
| 0 | 0 | 0 | 0 |
| 0 | 0 | 0 | 0 |
| 0 | 0 | 0 | 0 |
| 0 | 0 | 0 | 0 |
| 0 | 0 | 0 | 0 |
| 0 | 0 | 0 | 0 |
| 0 | 0 | 0 | 0 |
| 0 | 0 | 0 | 0 |
| 0 | 0 | 0 | 0 |
| 0 | 0 | 0 | 0 |
| 0 | 0 | 0 | 0 |
| 0 | 0 | 0 | 0 |
| 0 | 0 | 0 | 0 |
| 0 | 0 | 0 | 0 |
| 0 | 0 | 0 | 0 |
| 0 | 0 | 0 | 0 |
| 0 | 0 | 0 | 0 |
| 0 | 0 | 0 | 0 |
| 0 | 0 | 0 | 0 |
| 0 | 0 | 0 | 0 |
| 0 | 0 | 0 | 0 |
| 0 | 0 | 0 | 0 |
| 0 | 0 | 0 | 0 |
| 0 | 0 | 0 | 0 |
| 0 | 0 | 0 | 0 |
| 0 | 0 | 0 | 0 |
| 0 | 0 | 0 | 0 |
| 0 | 0 | 0 | 0 |
| 0 | 0 | 0 | 0 |
| 0 | 0 | 0 | 0 |
| 0 | 0 | 0 | 0 |
| 0 | 0 | 0 | 0 |
| 0 | 0 | 0 | 0 |
| 0 | 0 | 0 | 0 |
| 0 | 0 | 0 | 0 |
| 0 | 0 | 0 | 0 |
| 0 | 0 | 0 | 0 |
| 0 | 0 | 0 | 0 |
| 0 | 0 | 0 | 0 |
| 0 | 0 | 0 | 0 |
| 0 | 0 | 0 | 0 |

| | | | |
|---|---|---|---|
| 0 | 0 | 0 | 0 |
| 0 | 0 | 0 | 0 |
| 0 | 0 | 0 | 0 |
| 0 | 0 | 0 | 0 |
| 0 | 0 | 0 | 0 |
| 0 | 0 | 0 | 0 |
| 0 | 0 | 0 | 0 |
| 0 | 0 | 0 | 0 |
| 0 | 0 | 0 | 0 |
| 0 | 0 | 0 | 0 |
| 0 | 0 | 0 | 0 |
| 0 | 0 | 1 | 0 |
| 0 | 0 | 0 | 0 |
| 0 | 0 | 1 | 0 |
| 0 | 0 | 1 | 0 |
| 0 | 0 | 1 | 0 |
| 0 | 0 | 1 | 0 |
| 0 | 0 | 0 | 0 |
| 0 | 0 | 0 | 0 |
| 0 | 0 | 0 | 0 |
| 1 | 0 | 0 | 0 |

| glycan_binding_site | water_binding_site | water_only_binding_site | non_binding_site |
|---|---|---|---|
| 0 | 0 | 0 | 0 |
| 0 | 0 | 0 | 0 |
| 0 | 0 | 0 | 1 |
| 0 | 0 | 0 | 0 |
| 0 | 0 | 0 | 0 |
| 0 | 0 | 0 | 0 |
| 0 | 0 | 0 | 1 |
| 0 | 0 | 0 | 1 |
| 0 | 1 | 0 | 0 |
| 0 | 0 | 0 | 0 |
| 0 | 0 | 0 | 0 |
| 0 | 0 | 0 | 0 |
| 0 | 0 | 0 | 1 |
| 1 | 0 | 0 | 0 |
| 0 | 0 | 0 | 0 |
| 0 | 0 | 0 | 0 |
| 0 | 0 | 0 | 0 |
| 0 | 1 | 0 | 0 |
| 0 | 0 | 0 | 0 |
| 0 | 0 | 0 | 0 |
| 0 | 1 | 0 | 0 |
| 1 | 1 | 0 | 0 |
| 1 | 1 | 0 | 0 |
| 0 | 1 | 0 | 0 |
| 0 | 1 | 0 | 0 |
| 0 | 0 | 0 | 1 |
| 0 | 1 | 0 | 0 |
| 0 | 0 | 0 | 1 |
| 0 | 0 | 0 | 0 |
| 0 | 0 | 0 | 1 |
| 0 | 0 | 0 | 1 |
| 0 | 1 | 0 | 0 |
| 0 | 1 | 0 | 0 |
| 0 | 1 | 0 | 0 |
| 0 | 1 | 0 | 0 |
| 0 | 0 | 0 | 1 |
| 0 | 0 | 0 | 0 |
| 0 | 1 | 0 | 0 |
| 0 | 1 | 0 | 0 |
| 0 | 1 | 0 | 0 |
| 0 | 1 | 0 | 0 |
| 0 | 1 | 0 | 0 |
| 0 | 1 | 0 | 0 |
| 0 | 1 | 0 | 0 |
| 0 | 1 | 0 | 0 |
| 0 | 1 | 0 | 0 |
| 0 | 1 | 0 | 0 |
| 0 | 1 | 0 | 0 |
| 0 | 1 | 0 | 0 |
| 0 | 1 | 0 | 0 |
| 0 | 1 | 0 | 0 |
| 0 | 1 | 0 | 0 |
| 0 | 1 | 0 | 0 |

| | | | |
|---|---|---|---|
| 0 | 1 | 0 | 0 |
| 0 | 1 | 0 | 0 |
| 0 | 1 | 0 | 0 |
| 0 | 1 | 0 | 0 |
| 0 | 1 | 0 | 0 |
| 0 | 1 | 0 | 0 |
| 0 | 1 | 0 | 0 |
| 0 | 1 | 0 | 0 |
| 0 | 1 | 0 | 0 |
| 0 | 1 | 0 | 0 |
| 0 | 1 | 0 | 0 |
| 0 | 1 | 0 | 0 |
| 0 | 1 | 0 | 0 |
| 0 | 1 | 0 | 0 |
| 0 | 1 | 0 | 0 |
| 0 | 1 | 0 | 0 |
| 0 | 1 | 0 | 0 |
| 0 | 1 | 0 | 0 |
| 0 | 1 | 0 | 0 |
| 0 | 1 | 0 | 0 |
| 0 | 0 | 0 | 0 |
| 0 | 0 | 0 | 0 |
| 0 | 0 | 0 | 0 |
| 0 | 0 | 0 | 0 |
| 0 | 0 | 0 | 0 |
| 0 | 0 | 0 | 0 |
| 0 | 0 | 0 | 0 |
| 0 | 0 | 0 | 0 |
| 0 | 0 | 0 | 0 |
| 0 | 0 | 0 | 0 |
| 0 | 0 | 0 | 0 |
| 0 | 0 | 0 | 0 |
| 0 | 0 | 0 | 0 |
| 0 | 0 | 0 | 0 |
| 0 | 0 | 0 | 0 |
| 0 | 0 | 0 | 0 |
| 0 | 0 | 0 | 0 |
| 0 | 0 | 0 | 0 |
| 0 | 0 | 0 | 0 |
| 0 | 1 | 0 | 0 |
| 1 | 0 | 0 | 0 |
| 0 | 0 | 0 | 0 |
| 1 | 0 | 0 | 0 |
| 1 | 0 | 0 | 0 |
| 0 | 0 | 0 | 0 |
| 0 | 0 | 0 | 1 |
| 0 | 0 | 0 | 1 |
| 0 | 0 | 0 | 0 |
| 1 | 0 | 0 | 0 |
| 0 | 0 | 0 | 0 |
| 0 | 0 | 0 | 0 |
| 0 | 0 | 0 | 1 |
| 0 | 0 | 0 | 1 |
| 0 | 0 | 0 | 0 |

| | | | |
|---|---|---|---|
| 0 | 0 | 0 | 1 |
| 0 | 0 | 0 | 1 |
| 0 | 0 | 0 | 0 |
| 0 | 0 | 0 | 0 |
| 0 | 0 | 0 | 0 |
| 0 | 0 | 0 | 0 |
| 0 | 0 | 0 | 1 |
| 0 | 1 | 0 | 0 |
| 0 | 1 | 0 | 0 |
| 0 | 0 | 0 | 0 |
| 0 | 1 | 0 | 0 |
| 0 | 0 | 0 | 0 |
| 0 | 0 | 0 | 0 |
| 1 | 0 | 0 | 0 |
| 1 | 0 | 0 | 0 |
| 0 | 0 | 0 | 1 |
| 0 | 0 | 0 | 0 |
| 0 | 1 | 0 | 0 |
| 0 | 1 | 0 | 0 |
| 0 | 1 | 0 | 0 |
| 0 | 0 | 0 | 0 |
| 0 | 0 | 0 | 0 |
| 0 | 0 | 0 | 0 |
| 0 | 1 | 0 | 0 |
| 0 | 1 | 0 | 0 |
| 0 | 1 | 0 | 0 |
| 0 | 1 | 0 | 0 |
| 0 | 1 | 0 | 0 |
| 0 | 0 | 0 | 0 |
| 0 | 0 | 0 | 0 |
| 0 | 1 | 0 | 0 |
| 0 | 0 | 0 | 0 |
| 0 | 1 | 0 | 0 |
| 0 | 1 | 0 | 0 |
| 0 | 1 | 0 | 0 |
| 0 | 0 | 0 | 0 |
| 0 | 1 | 0 | 0 |
| 0 | 1 | 0 | 0 |
| 0 | 0 | 0 | 0 |
| 0 | 1 | 0 | 0 |
| 0 | 1 | 0 | 0 |
| 0 | 1 | 0 | 0 |
| 0 | 0 | 0 | 0 |
| 0 | 0 | 0 | 0 |
| 0 | 0 | 0 | 0 |
| 0 | 1 | 0 | 0 |
| 1 | 1 | 0 | 0 |
| 0 | 0 | 0 | 0 |
| 0 | 1 | 0 | 0 |
| 0 | 0 | 0 | 0 |
| 0 | 0 | 0 | 0 |
| 0 | 0 | 0 | 0 |
| 0 | 0 | 0 | 0 |
| 0 | 1 | 1 | 1 |

| | | | |
|---|---|---|---|
| 0 | 1 | 1 | 1 |
| 0 | 1 | 1 | 1 |
| 0 | 1 | 1 | 1 |
| 0 | 0 | 0 | 0 |
| 0 | 0 | 0 | 0 |
| 0 | 1 | 1 | 1 |
| 0 | 1 | 0 | 0 |
| 0 | 1 | 1 | 1 |
| 0 | 1 | 1 | 1 |
| 0 | 1 | 1 | 1 |
| 0 | 1 | 1 | 1 |
| 0 | 1 | 0 | 0 |
| 0 | 1 | 1 | 1 |
| 0 | 0 | 0 | 0 |
| 0 | 1 | 1 | 1 |
| 0 | 0 | 0 | 1 |
| 0 | 1 | 1 | 1 |
| 0 | 1 | 1 | 1 |
| 0 | 1 | 1 | 1 |
| 0 | 1 | 0 | 0 |
| 0 | 1 | 0 | 0 |
| 0 | 0 | 0 | 1 |
| 0 | 1 | 0 | 0 |
| 0 | 1 | 0 | 0 |
| 0 | 1 | 0 | 0 |
| 0 | 0 | 0 | 1 |
| 0 | 1 | 0 | 0 |
| 0 | 1 | 0 | 0 |
| 0 | 0 | 0 | 1 |
| 0 | 1 | 1 | 1 |
| 0 | 0 | 0 | 1 |
| 0 | 1 | 1 | 1 |
| 0 | 1 | 1 | 1 |
| 0 | 0 | 0 | 1 |
| 0 | 1 | 0 | 0 |
| 0 | 1 | 1 | 1 |
| 1 | 1 | 0 | 0 |
| 1 | 1 | 0 | 0 |
| 0 | 1 | 1 | 1 |
| 0 | 1 | 1 | 1 |
| 0 | 1 | 1 | 1 |
| 0 | 1 | 0 | 0 |
| 0 | 1 | 0 | 0 |
| 0 | 0 | 0 | 1 |
| 0 | 1 | 1 | 1 |
| 0 | 1 | 1 | 1 |
| 0 | 0 | 0 | 0 |
| 0 | 0 | 0 | 0 |
| 0 | 0 | 0 | 0 |
| 0 | 0 | 0 | 0 |
| 0 | 0 | 0 | 0 |
| 0 | 0 | 0 | 1 |
| 0 | 0 | 0 | 1 |
| 0 | 0 | 0 | 0 |

| | | | |
|---|---|---|---|
| 0 | 0 | 0 | 0 |
| 0 | 0 | 0 | 0 |
| 0 | 1 | 0 | 0 |
| 0 | 0 | 0 | 1 |
| 0 | 0 | 0 | 1 |
| 0 | 0 | 0 | 1 |
| 0 | 1 | 0 | 0 |
| 0 | 1 | 0 | 0 |
| 0 | 1 | 1 | 1 |
| 0 | 1 | 1 | 1 |
| 1 | 0 | 0 | 0 |
| 0 | 0 | 0 | 1 |
| 1 | 0 | 0 | 0 |
| 0 | 0 | 0 | 1 |
| 0 | 0 | 0 | 1 |
| 0 | 0 | 0 | 1 |
| 0 | 0 | 0 | 1 |
| 0 | 1 | 0 | 0 |
| 0 | 0 | 0 | 0 |
| 0 | 0 | 0 | 0 |
| 0 | 0 | 0 | 0 |
| 0 | 0 | 0 | 1 |
| 0 | 0 | 0 | 0 |
| 0 | 0 | 0 | 1 |
| 0 | 0 | 0 | 1 |
| 0 | 0 | 0 | 0 |
| 0 | 1 | 0 | 0 |
| 0 | 0 | 0 | 0 |
| 0 | 1 | 0 | 0 |
| 0 | 1 | 0 | 0 |
| 0 | 1 | 0 | 0 |
| 0 | 0 | 0 | 0 |
| 0 | 0 | 0 | 1 |
| 0 | 0 | 0 | 1 |
| 0 | 0 | 0 | 0 |
| 0 | 0 | 0 | 1 |
| 0 | 0 | 0 | 1 |
| 0 | 0 | 0 | 0 |
| 0 | 1 | 1 | 1 |
| 0 | 0 | 0 | 1 |
| 0 | 1 | 0 | 0 |
| 0 | 1 | 1 | 1 |
| 0 | 0 | 0 | 1 |
| 0 | 0 | 0 | 1 |
| 0 | 0 | 0 | 0 |
| 0 | 0 | 0 | 0 |
| 0 | 1 | 0 | 0 |
| 0 | 1 | 0 | 0 |
| 0 | 1 | 0 | 0 |
| 0 | 1 | 0 | 0 |
| 0 | 1 | 0 | 0 |
| 0 | 1 | 0 | 0 |
| 0 | 1 | 0 | 0 |
| 0 | 1 | 0 | 0 |

| | | | |
|---|---|---|---|
| 0 | 1 | 0 | 0 |
| 0 | 1 | 0 | 0 |
| 0 | 1 | 0 | 0 |
| 0 | 1 | 0 | 0 |
| 0 | 1 | 0 | 0 |
| 0 | 1 | 0 | 0 |
| 0 | 0 | 0 | 1 |
| 0 | 0 | 0 | 1 |
| 0 | 0 | 0 | 0 |
| 0 | 0 | 0 | 0 |
| 0 | 0 | 0 | 0 |
| 0 | 0 | 0 | 0 |
| 0 | 0 | 0 | 0 |
| 0 | 0 | 0 | 1 |
| 0 | 0 | 0 | 1 |
| 0 | 0 | 0 | 0 |
| 0 | 0 | 0 | 1 |
| 0 | 0 | 0 | 0 |
| 0 | 0 | 0 | 1 |
| 0 | 0 | 0 | 0 |
| 0 | 0 | 0 | 1 |
| 0 | 1 | 1 | 1 |
| 0 | 0 | 0 | 1 |
| 0 | 0 | 0 | 0 |
| 0 | 0 | 0 | 0 |
| 0 | 1 | 0 | 0 |
| 0 | 0 | 0 | 0 |
| 0 | 1 | 0 | 0 |
| 0 | 0 | 0 | 0 |
| 0 | 1 | 0 | 0 |
| 0 | 0 | 0 | 1 |
| 0 | 0 | 0 | 1 |
| 0 | 0 | 0 | 0 |
| 0 | 0 | 0 | 0 |
| 0 | 1 | 0 | 0 |
| 0 | 0 | 0 | 1 |
| 0 | 0 | 0 | 0 |
| 0 | 1 | 0 | 0 |
| 0 | 0 | 0 | 0 |
| 0 | 1 | 0 | 0 |
| 0 | 0 | 0 | 1 |
| 0 | 0 | 0 | 1 |
| 0 | 0 | 0 | 0 |
| 0 | 0 | 0 | 1 |
| 0 | 0 | 0 | 0 |
| 0 | 0 | 0 | 0 |
| 0 | 0 | 0 | 0 |
| 0 | 0 | 0 | 1 |
| 0 | 0 | 0 | 1 |
| 0 | 0 | 0 | 0 |
| 0 | 0 | 0 | 1 |
| 0 | 0 | 0 | 1 |
| 0 | 0 | 0 | 0 |
| 0 | 0 | 0 | 1 |

| | | | |
|---|---|---|---|
| 0 | 0 | 0 | 0 |
| 0 | 0 | 0 | 1 |
| 0 | 1 | 1 | 1 |
| 0 | 1 | 0 | 0 |
| 0 | 1 | 0 | 0 |
| 0 | 0 | 0 | 0 |
| 0 | 0 | 0 | 1 |
| 0 | 1 | 0 | 0 |
| 0 | 0 | 0 | 0 |
| 0 | 0 | 0 | 1 |
| 0 | 0 | 0 | 1 |
| 0 | 0 | 0 | 1 |
| 0 | 0 | 0 | 1 |
| 0 | 0 | 0 | 1 |
| 0 | 0 | 0 | 0 |
| 0 | 0 | 0 | 1 |
| 0 | 0 | 0 | 0 |
| 0 | 0 | 0 | 0 |
| 0 | 0 | 0 | 1 |
| 0 | 0 | 0 | 0 |
| 0 | 1 | 0 | 0 |
| 0 | 1 | 0 | 0 |
| 0 | 0 | 0 | 0 |
| 0 | 1 | 0 | 0 |
| 0 | 0 | 0 | 1 |
| 0 | 0 | 0 | 1 |
| 0 | 0 | 0 | 0 |
| 0 | 0 | 0 | 0 |
| 0 | 1 | 0 | 0 |
| 0 | 0 | 0 | 0 |
| 0 | 0 | 0 | 0 |
| 0 | 0 | 0 | 0 |
| 0 | 0 | 0 | 0 |
| 0 | 0 | 0 | 0 |
| 0 | 0 | 0 | 1 |
| 0 | 0 | 0 | 1 |
| 0 | 0 | 0 | 0 |
| 0 | 0 | 0 | 0 |
| 0 | 0 | 0 | 0 |
| 0 | 0 | 0 | 0 |
| 0 | 0 | 0 | 0 |
| 0 | 1 | 0 | 0 |
| 0 | 0 | 0 | 1 |
| 0 | 0 | 0 | 0 |
| 0 | 1 | 0 | 0 |
| 0 | 1 | 0 | 0 |
| 0 | 1 | 0 | 0 |
| 0 | 1 | 0 | 0 |
| 1 | 1 | 0 | 0 |
| 0 | 0 | 0 | 1 |
| 0 | 1 | 0 | 0 |
| 0 | 1 | 0 | 0 |
| 0 | 0 | 0 | 0 |
| 0 | 1 | 0 | 0 |

| | | | |
|---|---|---|---|
| 0 | 1 | 0 | 0 |
| 0 | 1 | 0 | 0 |
| 0 | 0 | 0 | 0 |
| 0 | 0 | 0 | 0 |
| 0 | 0 | 0 | 1 |
| 0 | 1 | 0 | 0 |
| 0 | 1 | 0 | 0 |
| 0 | 1 | 0 | 0 |
| 0 | 1 | 0 | 0 |
| 0 | 1 | 0 | 0 |
| 0 | 1 | 0 | 0 |
| 0 | 1 | 0 | 0 |
| 0 | 1 | 0 | 0 |
| 0 | 1 | 0 | 0 |
| 0 | 1 | 0 | 0 |
| 0 | 1 | 0 | 0 |
| 0 | 1 | 0 | 0 |
| 0 | 1 | 0 | 0 |
| 0 | 0 | 0 | 1 |
| 0 | 0 | 0 | 0 |
| 0 | 0 | 0 | 0 |
| 0 | 0 | 0 | 0 |
| 0 | 1 | 0 | 0 |
| 0 | 1 | 1 | 1 |
| 0 | 0 | 0 | 0 |
| 0 | 0 | 0 | 0 |
| 0 | 0 | 0 | 0 |
| 0 | 1 | 1 | 1 |
| 0 | 1 | 1 | 1 |
| 0 | 1 | 1 | 1 |
| 0 | 1 | 0 | 0 |
| 0 | 0 | 0 | 0 |
| 0 | 1 | 0 | 0 |
| 0 | 1 | 0 | 0 |
| 0 | 1 | 0 | 0 |
| 0 | 1 | 1 | 1 |
| 0 | 1 | 1 | 1 |
| 0 | 0 | 0 | 0 |
| 0 | 1 | 0 | 0 |
| 0 | 1 | 1 | 1 |
| 0 | 1 | 1 | 1 |
| 0 | 1 | 1 | 1 |
| 0 | 0 | 0 | 1 |
| 0 | 0 | 0 | 1 |
| 0 | 0 | 0 | 0 |
| 0 | 0 | 0 | 0 |
| 0 | 1 | 0 | 0 |
| 0 | 0 | 0 | 0 |
| 0 | 0 | 0 | 0 |
| 0 | 0 | 0 | 1 |
| 0 | 0 | 0 | 1 |
| 0 | 0 | 0 | 0 |
| 0 | 0 | 0 | 1 |
| 0 | 0 | 0 | 0 |

| | | | |
|---|---|---|---|
| 0 | 0 | 0 | 1 |
| 0 | 0 | 0 | 1 |
| 0 | 0 | 0 | 0 |
| 0 | 0 | 0 | 1 |
| 0 | 0 | 0 | 1 |
| 0 | 0 | 0 | 1 |
| 0 | 0 | 0 | 1 |
| 0 | 1 | 0 | 0 |
| 0 | 1 | 1 | 1 |
| 0 | 0 | 0 | 1 |
| 0 | 0 | 0 | 0 |
| 0 | 0 | 0 | 1 |
| 0 | 0 | 0 | 1 |
| 0 | 0 | 0 | 1 |
| 0 | 0 | 0 | 0 |
| 0 | 0 | 0 | 0 |
| 0 | 0 | 0 | 1 |
| 0 | 0 | 0 | 1 |
| 0 | 0 | 0 | 0 |
| 0 | 0 | 0 | 0 |
| 0 | 0 | 0 | 0 |
| 0 | 0 | 0 | 0 |
| 0 | 0 | 0 | 0 |
| 0 | 1 | 0 | 0 |
| 0 | 1 | 0 | 0 |
| 0 | 0 | 0 | 1 |
| 0 | 0 | 0 | 0 |
| 0 | 1 | 0 | 0 |
| 0 | 0 | 0 | 0 |
| 0 | 0 | 0 | 1 |
| 0 | 0 | 0 | 0 |
| 0 | 1 | 0 | 0 |
| 0 | 0 | 0 | 1 |
| 0 | 0 | 0 | 0 |
| 0 | 0 | 0 | 0 |
| 0 | 0 | 0 | 0 |
| 0 | 1 | 0 | 0 |
| 0 | 1 | 0 | 0 |
| 0 | 1 | 0 | 0 |
| 0 | 1 | 0 | 0 |
| 0 | 0 | 0 | 1 |
| 0 | 0 | 0 | 0 |
| 0 | 0 | 0 | 1 |
| 0 | 0 | 0 | 0 |
| 0 | 0 | 0 | 0 |
| 1 | 0 | 0 | 0 |
| 0 | 0 | 0 | 0 |
| 1 | 0 | 0 | 0 |
| 0 | 0 | 0 | 0 |
| 0 | 0 | 0 | 0 |
| 0 | 0 | 0 | 1 |
| 0 | 0 | 0 | 0 |
| 0 | 0 | 0 | 0 |
| 0 | 0 | 0 | 0 |

| | | | |
|---|---|---|---|
| 0 | 0 | 0 | 1 |
| 0 | 0 | 0 | 0 |
| 0 | 0 | 0 | 1 |
| 0 | 0 | 0 | 1 |
| 1 | 0 | 0 | 0 |
| 0 | 0 | 0 | 0 |
| 1 | 0 | 0 | 0 |
| 1 | 0 | 0 | 0 |
| 0 | 0 | 0 | 0 |
| 0 | 0 | 0 | 1 |
| 0 | 0 | 0 | 1 |
| 0 | 0 | 0 | 1 |
| 0 | 0 | 0 | 1 |
| 1 | 1 | 0 | 0 |
| 0 | 0 | 0 | 1 |
| 0 | 0 | 0 | 0 |
| 0 | 0 | 0 | 0 |
| 0 | 0 | 0 | 0 |
| 1 | 1 | 0 | 0 |
| 0 | 1 | 1 | 1 |
| 1 | 1 | 0 | 0 |
| 0 | 0 | 0 | 1 |
| 0 | 0 | 0 | 0 |
| 0 | 0 | 0 | 1 |
| 0 | 0 | 0 | 1 |
| 0 | 0 | 0 | 1 |
| 0 | 0 | 0 | 0 |
| 0 | 0 | 0 | 1 |
| 0 | 0 | 0 | 1 |
| 0 | 0 | 0 | 1 |
| 0 | 0 | 0 | 0 |
| 0 | 0 | 0 | 0 |
| 0 | 0 | 0 | 1 |
| 0 | 0 | 0 | 1 |
| 0 | 0 | 0 | 0 |
| 0 | 0 | 0 | 0 |
| 0 | 0 | 0 | 0 |
| 1 | 0 | 0 | 0 |
| 0 | 0 | 0 | 0 |
| 0 | 0 | 0 | 1 |
| 0 | 0 | 0 | 0 |
| 0 | 1 | 0 | 0 |
| 0 | 1 | 0 | 0 |
| 0 | 0 | 0 | 0 |
| 0 | 0 | 0 | 1 |
| 1 | 0 | 0 | 0 |
| 0 | 1 | 0 | 0 |
| 0 | 0 | 0 | 0 |
| 0 | 0 | 0 | 0 |
| 1 | 0 | 0 | 0 |
| 0 | 0 | 0 | 0 |
| 0 | 1 | 0 | 0 |
| 0 | 1 | 1 | 1 |
| 0 | 0 | 0 | 0 |

| | | | |
|---|---|---|---|
| 0 | 1 | 0 | 0 |
| 0 | 1 | 0 | 0 |
| 0 | 1 | 1 | 1 |
| 0 | 1 | 1 | 1 |
| 0 | 1 | 1 | 1 |
| 0 | 1 | 0 | 0 |
| 0 | 0 | 0 | 1 |
| 0 | 0 | 0 | 1 |
| 0 | 1 | 0 | 0 |
| 0 | 0 | 0 | 0 |
| 1 | 1 | 0 | 0 |
| 1 | 1 | 0 | 0 |
| 0 | 0 | 0 | 1 |
| 1 | 0 | 0 | 0 |
| 0 | 0 | 0 | 0 |
| 0 | 0 | 0 | 0 |
| 0 | 1 | 0 | 0 |
| 0 | 0 | 0 | 0 |
| 0 | 0 | 0 | 0 |
| 0 | 0 | 0 | 1 |
| 0 | 0 | 0 | 0 |
| 0 | 1 | 0 | 0 |
| 0 | 0 | 0 | 0 |
| 0 | 0 | 0 | 0 |
| 0 | 1 | 0 | 0 |
| 0 | 0 | 0 | 1 |
| 0 | 0 | 0 | 1 |
| 0 | 0 | 0 | 1 |
| 0 | 0 | 0 | 0 |
| 0 | 1 | 0 | 0 |
| 0 | 0 | 0 | 0 |
| 0 | 0 | 0 | 0 |
| 0 | 0 | 0 | 0 |
| 0 | 1 | 0 | 0 |
| 0 | 1 | 0 | 0 |
| 0 | 0 | 0 | 1 |
| 0 | 0 | 0 | 0 |
| 0 | 1 | 0 | 0 |
| 0 | 0 | 0 | 1 |
| 0 | 0 | 0 | 1 |
| 0 | 0 | 0 | 1 |
| 0 | 1 | 0 | 0 |
| 0 | 0 | 0 | 0 |
| 0 | 0 | 0 | 0 |
| 0 | 0 | 0 | 0 |
| 0 | 0 | 0 | 0 |
| 0 | 0 | 0 | 0 |
| 0 | 0 | 0 | 1 |
| 0 | 0 | 0 | 1 |
| 0 | 0 | 0 | 0 |
| 0 | 0 | 0 | 0 |
| 0 | 0 | 0 | 0 |
| 0 | 0 | 0 | 0 |
| 0 | 0 | 0 | 1 |

| | | | |
|---|---|---|---|
| 0 | 0 | 0 | 1 |
| 0 | 0 | 0 | 0 |
| 0 | 0 | 0 | 1 |
| 0 | 1 | 0 | 0 |
| 0 | 1 | 1 | 1 |
| 0 | 1 | 1 | 1 |
| 0 | 1 | 0 | 0 |
| 0 | 1 | 1 | 1 |
| 0 | 1 | 1 | 1 |
| 0 | 1 | 1 | 1 |
| 0 | 1 | 1 | 1 |
| 0 | 1 | 1 | 1 |
| 0 | 0 | 0 | 1 |
| 0 | 0 | 0 | 1 |
| 0 | 0 | 0 | 1 |
| 0 | 0 | 0 | 1 |
| 0 | 0 | 0 | 0 |
| 0 | 0 | 0 | 0 |
| 0 | 1 | 1 | 1 |
| 0 | 0 | 0 | 0 |
| 0 | 0 | 0 | 0 |
| 0 | 0 | 0 | 0 |
| 0 | 0 | 0 | 1 |
| 0 | 1 | 0 | 0 |
| 0 | 0 | 0 | 0 |
| 0 | 0 | 0 | 1 |
| 0 | 0 | 0 | 1 |
| 0 | 0 | 0 | 1 |
| 0 | 1 | 0 | 0 |
| 0 | 0 | 0 | 0 |
| 0 | 1 | 1 | 1 |
| 0 | 1 | 0 | 0 |
| 0 | 1 | 0 | 0 |
| 0 | 0 | 0 | 0 |
| 0 | 0 | 0 | 0 |
| 0 | 0 | 0 | 0 |
| 0 | 1 | 1 | 1 |
| 0 | 0 | 0 | 1 |
| 0 | 0 | 0 | 1 |
| 0 | 0 | 0 | 1 |
| 0 | 0 | 0 | 0 |
| 0 | 0 | 0 | 1 |
| 0 | 0 | 0 | 0 |
| 0 | 0 | 0 | 1 |
| 1 | 0 | 0 | 0 |
| 0 | 0 | 0 | 1 |
| 0 | 0 | 0 | 0 |
| 0 | 0 | 0 | 0 |
| 1 | 0 | 0 | 0 |
| 0 | 0 | 0 | 0 |
| 0 | 0 | 0 | 0 |
| 0 | 0 | 0 | 0 |
| 0 | 0 | 0 | 0 |
| 0 | 0 | 0 | 0 |

| | | | |
|---|---|---|---|
| 0 | 0 | 0 | 0 |
| 0 | 0 | 0 | 0 |
| 0 | 0 | 0 | 0 |
| 0 | 0 | 0 | 0 |
| 0 | 1 | 0 | 0 |
| 0 | 1 | 0 | 0 |
| 0 | 1 | 0 | 0 |
| 0 | 0 | 0 | 1 |
| 0 | 1 | 0 | 0 |
| 0 | 1 | 0 | 0 |
| 0 | 1 | 0 | 0 |
| 0 | 1 | 1 | 1 |
| 0 | 1 | 0 | 0 |
| 0 | 1 | 0 | 0 |
| 0 | 1 | 0 | 0 |
| 0 | 1 | 0 | 0 |
| 0 | 1 | 0 | 0 |
| 0 | 1 | 0 | 0 |
| 0 | 1 | 0 | 0 |
| 0 | 1 | 0 | 0 |
| 0 | 1 | 0 | 0 |
| 0 | 1 | 0 | 0 |
| 0 | 1 | 0 | 0 |
| 0 | 1 | 0 | 0 |
| 0 | 1 | 0 | 0 |
| 0 | 1 | 0 | 0 |
| 0 | 1 | 0 | 0 |
| 0 | 1 | 0 | 0 |
| 0 | 1 | 0 | 0 |
| 0 | 1 | 0 | 0 |
| 0 | 1 | 0 | 0 |
| 0 | 1 | 0 | 0 |
| 0 | 1 | 0 | 0 |
| 0 | 1 | 0 | 0 |
| 0 | 1 | 0 | 0 |
| 0 | 1 | 0 | 0 |
| 0 | 1 | 0 | 0 |
| 0 | 1 | 0 | 0 |
| 0 | 1 | 0 | 0 |
| 0 | 1 | 0 | 0 |
| 0 | 1 | 1 | 1 |
| 0 | 1 | 0 | 0 |
| 0 | 1 | 0 | 0 |
| 0 | 1 | 0 | 0 |
| 0 | 1 | 0 | 0 |
| 0 | 1 | 0 | 0 |
| 0 | 1 | 0 | 0 |
| 0 | 1 | 0 | 0 |
| 0 | 0 | 0 | 0 |
| 0 | 0 | 0 | 1 |
| 0 | 0 | 0 | 1 |
| 0 | 1 | 1 | 1 |
| 1 | 0 | 0 | 0 |
| 0 | 1 | 0 | 0 |

| | | | |
|---|---|---|---|
| 0 | 0 | 0 | 0 |
| 0 | 0 | 0 | 0 |
| 1 | 1 | 0 | 0 |
| 1 | 0 | 0 | 0 |
| 0 | 0 | 0 | 0 |
| 0 | 0 | 0 | 0 |
| 0 | 0 | 0 | 0 |
| 0 | 1 | 1 | 1 |
| 0 | 0 | 0 | 1 |
| 0 | 0 | 0 | 0 |
| 0 | 0 | 0 | 0 |
| 0 | 0 | 0 | 1 |
| 0 | 0 | 0 | 0 |
| 0 | 0 | 0 | 0 |
| 0 | 0 | 0 | 0 |
| 0 | 0 | 0 | 0 |
| 0 | 0 | 0 | 0 |
| 0 | 0 | 0 | 1 |
| 0 | 0 | 0 | 0 |
| 0 | 1 | 1 | 1 |
| 0 | 0 | 0 | 0 |
| 0 | 0 | 0 | 0 |
| 0 | 0 | 0 | 1 |
| 0 | 0 | 0 | 0 |
| 0 | 1 | 0 | 0 |
| 0 | 0 | 0 | 1 |
| 0 | 0 | 0 | 0 |
| 0 | 0 | 0 | 0 |
| 0 | 0 | 0 | 0 |
| 0 | 0 | 0 | 0 |
| 0 | 0 | 0 | 0 |
| 0 | 0 | 0 | 0 |
| 0 | 1 | 1 | 1 |
| 0 | 0 | 0 | 0 |
| 0 | 1 | 0 | 0 |
| 0 | 0 | 0 | 0 |
| 0 | 0 | 0 | 1 |
| 0 | 0 | 0 | 0 |
| 0 | 0 | 0 | 0 |
| 0 | 0 | 0 | 0 |
| 0 | 1 | 0 | 0 |
| 0 | 1 | 0 | 0 |
| 0 | 1 | 0 | 0 |
| 0 | 1 | 0 | 0 |
| 0 | 1 | 0 | 0 |
| 0 | 0 | 0 | 1 |
| 0 | 1 | 0 | 0 |
| 0 | 1 | 0 | 0 |
| 0 | 1 | 0 | 0 |
| 0 | 1 | 0 | 0 |
| 0 | 1 | 0 | 0 |
| 0 | 1 | 0 | 0 |
| 0 | 1 | 1 | 1 |
| 0 | 1 | 1 | 1 |

| | | | |
|---|---|---|---|
| 0 | 1 | 1 | 1 |
| 0 | 0 | 0 | 1 |
| 0 | 0 | 0 | 0 |
| 0 | 0 | 0 | 0 |
| 0 | 1 | 0 | 0 |
| 0 | 1 | 0 | 0 |
| 0 | 1 | 0 | 0 |
| 0 | 0 | 0 | 1 |
| 0 | 0 | 0 | 0 |
| 0 | 1 | 1 | 1 |
| 0 | 0 | 0 | 1 |
| 0 | 0 | 0 | 1 |
| 0 | 1 | 0 | 0 |
| 0 | 0 | 0 | 0 |
| 0 | 0 | 0 | 0 |
| 0 | 0 | 0 | 0 |
| 0 | 0 | 0 | 0 |
| 1 | 1 | 0 | 0 |
| 0 | 0 | 0 | 1 |
| 0 | 0 | 0 | 1 |
| 0 | 0 | 0 | 1 |
| 0 | 0 | 0 | 0 |
| 0 | 0 | 0 | 1 |
| 0 | 0 | 0 | 0 |
| 0 | 0 | 0 | 0 |
| 0 | 0 | 0 | 0 |
| 0 | 0 | 0 | 0 |
| 0 | 0 | 0 | 0 |
| 0 | 0 | 0 | 0 |
| 0 | 0 | 0 | 0 |
| 0 | 0 | 0 | 0 |
| 0 | 0 | 0 | 0 |
| 0 | 0 | 0 | 0 |
| 0 | 0 | 0 | 0 |
| 0 | 0 | 0 | 0 |
| 0 | 0 | 0 | 0 |
| 0 | 0 | 0 | 0 |
| 0 | 1 | 0 | 0 |
| 1 | 1 | 0 | 0 |
| 0 | 1 | 0 | 0 |
| 0 | 1 | 0 | 0 |
| 0 | 0 | 0 | 1 |
| 0 | 1 | 0 | 0 |
| 0 | 1 | 0 | 0 |
| 0 | 0 | 0 | 0 |
| 0 | 0 | 0 | 0 |
| 0 | 0 | 0 | 1 |
| 0 | 0 | 0 | 0 |
| 0 | 0 | 0 | 0 |
| 0 | 0 | 0 | 0 |
| 0 | 0 | 0 | 0 |
| 0 | 0 | 0 | 0 |
| 0 | 0 | 0 | 1 |
| 0 | 0 | 0 | 1 |

| | | | |
|---|---|---|---|
| 0 | 1 | 0 | 0 |
| 0 | 0 | 0 | 0 |
| 0 | 0 | 0 | 1 |
| 0 | 0 | 0 | 1 |
| 0 | 0 | 0 | 1 |
| 0 | 1 | 0 | 0 |
| 0 | 0 | 0 | 1 |
| 0 | 1 | 1 | 1 |
| 0 | 1 | 0 | 0 |
| 0 | 0 | 0 | 0 |
| 0 | 0 | 0 | 0 |
| 0 | 0 | 0 | 0 |
| 0 | 0 | 0 | 0 |
| 0 | 0 | 0 | 1 |
| 0 | 0 | 0 | 1 |
| 0 | 0 | 0 | 1 |
| 0 | 0 | 0 | 1 |
| 0 | 0 | 0 | 1 |
| 0 | 0 | 0 | 0 |
| 0 | 0 | 0 | 0 |
| 0 | 0 | 0 | 1 |
| 0 | 0 | 0 | 1 |
| 0 | 1 | 0 | 0 |
| 0 | 1 | 0 | 0 |
| 0 | 0 | 0 | 1 |
| 0 | 1 | 1 | 1 |
| 0 | 1 | 0 | 0 |
| 0 | 1 | 1 | 1 |
| 0 | 0 | 0 | 0 |
| 0 | 0 | 0 | 0 |
| 0 | 1 | 0 | 0 |
| 0 | 1 | 0 | 0 |
| 0 | 1 | 0 | 0 |
| 1 | 0 | 0 | 0 |
| 1 | 1 | 0 | 0 |
| 0 | 1 | 0 | 0 |
| 0 | 1 | 0 | 0 |
| 0 | 0 | 0 | 0 |
| 0 | 0 | 0 | 1 |
| 0 | 1 | 0 | 0 |
| 0 | 0 | 0 | 1 |
| 0 | 0 | 0 | 1 |
| 0 | 0 | 0 | 1 |
| 0 | 0 | 0 | 1 |
| 0 | 0 | 0 | 1 |
| 0 | 0 | 0 | 0 |
| 0 | 0 | 0 | 0 |
| 0 | 0 | 0 | 0 |
| 0 | 0 | 0 | 0 |
| 0 | 1 | 0 | 0 |
| 0 | 1 | 0 | 0 |
| 0 | 0 | 0 | 0 |
| 0 | 0 | 0 | 0 |
| 0 | 1 | 1 | 1 |

| | | | |
|---|---|---|---|
| 0 | 1 | 1 | 1 |
| 0 | 1 | 0 | 0 |
| 0 | 0 | 0 | 0 |
| 0 | 0 | 0 | 0 |
| 0 | 0 | 0 | 1 |
| 0 | 1 | 0 | 0 |
| 0 | 1 | 0 | 0 |
| 0 | 1 | 0 | 0 |
| 0 | 1 | 1 | 1 |
| 0 | 1 | 1 | 1 |
| 0 | 1 | 0 | 0 |
| 0 | 1 | 0 | 0 |
| 0 | 1 | 0 | 0 |
| 0 | 1 | 1 | 1 |
| 0 | 1 | 0 | 0 |
| 0 | 1 | 0 | 0 |
| 0 | 1 | 0 | 0 |
| 0 | 1 | 0 | 0 |
| 0 | 0 | 0 | 0 |
| 0 | 0 | 0 | 0 |
| 0 | 1 | 0 | 0 |
| 0 | 0 | 0 | 1 |
| 0 | 0 | 0 | 0 |
| 0 | 0 | 0 | 1 |
| 0 | 0 | 0 | 0 |
| 0 | 1 | 0 | 0 |
| 0 | 1 | 1 | 1 |
| 0 | 0 | 0 | 0 |
| 0 | 1 | 0 | 0 |
| 0 | 1 | 0 | 0 |
| 0 | 0 | 0 | 0 |
| 0 | 1 | 0 | 0 |
| 0 | 1 | 0 | 0 |
| 0 | 0 | 0 | 0 |
| 0 | 0 | 0 | 1 |
| 0 | 1 | 0 | 0 |
| 0 | 1 | 0 | 0 |
| 0 | 1 | 0 | 0 |
| 0 | 1 | 0 | 0 |
| 0 | 0 | 0 | 0 |
| 0 | 0 | 0 | 0 |
| 0 | 0 | 0 | 0 |
| 0 | 0 | 0 | 0 |
| 0 | 0 | 0 | 0 |
| 0 | 0 | 0 | 0 |
| 0 | 0 | 0 | 0 |
| 0 | 0 | 0 | 0 |
| 0 | 0 | 0 | 1 |
| 0 | 1 | 0 | 0 |
| 0 | 0 | 0 | 1 |
| 0 | 0 | 0 | 0 |
| 0 | 0 | 0 | 0 |
| 0 | 0 | 0 | 0 |
| 0 | 0 | 0 | 0 |

| | | | |
|---|---|---|---|
| 0 | 0 | 0 | 0 |
| 0 | 1 | 0 | 0 |
| 0 | 0 | 0 | 1 |
| 0 | 1 | 0 | 0 |
| 0 | 0 | 0 | 0 |
| 0 | 0 | 0 | 1 |
| 0 | 0 | 0 | 1 |
| 0 | 0 | 0 | 1 |
| 0 | 0 | 0 | 0 |
| 0 | 0 | 0 | 1 |
| 0 | 1 | 0 | 0 |
| 0 | 0 | 0 | 0 |
| 0 | 1 | 0 | 0 |
| 0 | 0 | 0 | 1 |
| 0 | 1 | 0 | 0 |
| 0 | 0 | 0 | 1 |
| 0 | 0 | 0 | 0 |
| 0 | 0 | 0 | 0 |
| 1 | 0 | 0 | 0 |
| 0 | 0 | 0 | 1 |
| 0 | 0 | 0 | 1 |
| 0 | 0 | 0 | 0 |
| 0 | 0 | 0 | 1 |
| 0 | 0 | 0 | 0 |
| 0 | 0 | 0 | 0 |
| 0 | 0 | 0 | 1 |
| 0 | 0 | 0 | 1 |
| 0 | 0 | 0 | 0 |
| 0 | 0 | 0 | 1 |
| 1 | 0 | 0 | 0 |
| 0 | 0 | 0 | 1 |
| 0 | 0 | 0 | 1 |
| 0 | 0 | 0 | 1 |
| 0 | 0 | 0 | 1 |
| 0 | 0 | 0 | 1 |
| 0 | 0 | 0 | 1 |
| 0 | 0 | 0 | 1 |
| 0 | 0 | 0 | 1 |
| 0 | 0 | 0 | 1 |
| 0 | 0 | 0 | 1 |
| 0 | 0 | 0 | 1 |
| 0 | 0 | 0 | 1 |
| 0 | 0 | 0 | 1 |
| 0 | 0 | 0 | 1 |
| 0 | 0 | 0 | 1 |
| 0 | 0 | 0 | 1 |
| 0 | 0 | 0 | 1 |
| 0 | 0 | 0 | 1 |
| 0 | 0 | 0 | 1 |
| 0 | 0 | 0 | 1 |
| 0 | 0 | 0 | 1 |
| 0 | 0 | 0 | 1 |
| 0 | 0 | 0 | 1 |
| 0 | 0 | 0 | 1 |

| | | | |
|---|---|---|---|
| 0 | 0 | 0 | 1 |
| 0 | 0 | 0 | 1 |
| 0 | 0 | 0 | 1 |
| 0 | 0 | 0 | 1 |
| 0 | 0 | 0 | 1 |
| 0 | 0 | 0 | 1 |
| 0 | 0 | 0 | 1 |
| 0 | 0 | 0 | 1 |
| 0 | 0 | 0 | 1 |
| 0 | 0 | 0 | 1 |
| 0 | 0 | 0 | 1 |
| 0 | 0 | 0 | 0 |
| 0 | 1 | 0 | 0 |
| 0 | 1 | 0 | 0 |
| 0 | 1 | 1 | 1 |
| 0 | 1 | 0 | 0 |
| 0 | 1 | 1 | 1 |
| 0 | 1 | 0 | 0 |
| 0 | 1 | 1 | 1 |
| 0 | 0 | 0 | 1 |
| 0 | 0 | 0 | 0 |
| 0 | 0 | 0 | 1 |
| 0 | 0 | 0 | 1 |
| 0 | 0 | 0 | 0 |
| 0 | 0 | 0 | 0 |
| 0 | 0 | 0 | 1 |
| 0 | 0 | 0 | 1 |
| 0 | 0 | 0 | 0 |
| 0 | 0 | 0 | 0 |
| 0 | 0 | 0 | 0 |
| 0 | 0 | 0 | 0 |
| 0 | 0 | 0 | 0 |
| 0 | 0 | 0 | 1 |
| 0 | 0 | 0 | 1 |
| 0 | 0 | 0 | 1 |
| 0 | 0 | 0 | 1 |
| 0 | 0 | 0 | 1 |
| 0 | 0 | 0 | 1 |
| 0 | 0 | 0 | 1 |
| 0 | 1 | 0 | 0 |
| 0 | 0 | 0 | 0 |
| 0 | 0 | 0 | 1 |
| 0 | 1 | 0 | 0 |
| 0 | 1 | 0 | 0 |
| 0 | 1 | 0 | 0 |
| 0 | 1 | 1 | 1 |
| 0 | 0 | 0 | 0 |
| 0 | 0 | 0 | 1 |
| 0 | 1 | 1 | 1 |
| 0 | 1 | 0 | 0 |
| 0 | 0 | 0 | 0 |
| 0 | 0 | 0 | 0 |
| 0 | 0 | 0 | 1 |
| 0 | 0 | 0 | 0 |

| | | | |
|---|---|---|---|
| 0 | 1 | 0 | 0 |
| 0 | 0 | 0 | 1 |
| 0 | 0 | 0 | 1 |
| 0 | 0 | 0 | 1 |
| 0 | 1 | 0 | 0 |
| 0 | 0 | 0 | 1 |
| 0 | 0 | 0 | 0 |
| 0 | 0 | 0 | 1 |
| 0 | 0 | 0 | 1 |
| 0 | 0 | 0 | 0 |
| 0 | 1 | 1 | 1 |
| 0 | 0 | 0 | 1 |
| 0 | 0 | 0 | 1 |
| 0 | 1 | 0 | 0 |
| 0 | 0 | 0 | 1 |
| 0 | 0 | 0 | 1 |
| 0 | 1 | 0 | 0 |
| 0 | 0 | 0 | 1 |
| 0 | 0 | 0 | 0 |
| 0 | 0 | 0 | 0 |
| 0 | 0 | 0 | 0 |
| 0 | 0 | 0 | 0 |
| 0 | 0 | 0 | 0 |
| 0 | 0 | 0 | 0 |
| 0 | 0 | 0 | 0 |
| 0 | 0 | 0 | 0 |
| 0 | 0 | 0 | 0 |
| 0 | 0 | 0 | 0 |
| 0 | 0 | 0 | 0 |
| 0 | 0 | 0 | 0 |
| 0 | 0 | 0 | 0 |
| 0 | 1 | 0 | 0 |
| 0 | 1 | 0 | 0 |
| 0 | 1 | 0 | 0 |
| 0 | 0 | 0 | 0 |
| 0 | 1 | 0 | 0 |
| 0 | 1 | 1 | 1 |
| 0 | 1 | 0 | 0 |
| 0 | 1 | 0 | 0 |
| 0 | 0 | 0 | 1 |
| 0 | 0 | 0 | 1 |
| 0 | 0 | 0 | 0 |
| 0 | 0 | 0 | 0 |
| 0 | 1 | 0 | 0 |
| 0 | 1 | 0 | 0 |
| 0 | 0 | 0 | 0 |
| 0 | 0 | 0 | 0 |
| 0 | 1 | 0 | 0 |
| 0 | 0 | 0 | 1 |
| 0 | 0 | 0 | 1 |
| 0 | 0 | 0 | 0 |
| 0 | 0 | 0 | 1 |
| 0 | 0 | 0 | 0 |
| 0 | 0 | 0 | 0 |

| | | | |
|---|---|---|---|
| 0 | 0 | 0 | 0 |
| 0 | 0 | 0 | 0 |
| 0 | 1 | 0 | 0 |
| 0 | 0 | 0 | 0 |
| 0 | 0 | 0 | 0 |
| 0 | 0 | 0 | 0 |
| 0 | 1 | 0 | 0 |
| 0 | 0 | 0 | 0 |
| 0 | 0 | 0 | 0 |
| 0 | 0 | 0 | 0 |
| 0 | 0 | 0 | 0 |
| 0 | 0 | 0 | 0 |
| 0 | 0 | 0 | 0 |
| 0 | 0 | 0 | 0 |
| 0 | 0 | 0 | 1 |
| 0 | 1 | 0 | 0 |
| 0 | 1 | 0 | 0 |
| 0 | 1 | 0 | 0 |
| 0 | 1 | 1 | 1 |
| 0 | 0 | 0 | 0 |
| 0 | 1 | 0 | 0 |
| 0 | 1 | 0 | 0 |
| 0 | 1 | 0 | 0 |
| 0 | 1 | 0 | 0 |
| 0 | 1 | 0 | 0 |
| 0 | 1 | 1 | 1 |
| 0 | 0 | 0 | 0 |
| 0 | 1 | 0 | 0 |
| 0 | 1 | 0 | 0 |
| 0 | 0 | 0 | 0 |
| 0 | 0 | 0 | 1 |
| 0 | 0 | 0 | 0 |
| 0 | 0 | 0 | 1 |
| 0 | 1 | 1 | 1 |
| 1 | 0 | 0 | 0 |
| 0 | 0 | 0 | 0 |
| 0 | 0 | 0 | 1 |
| 0 | 0 | 0 | 1 |
| 0 | 0 | 0 | 1 |
| 0 | 0 | 0 | 1 |
| 0 | 0 | 0 | 0 |
| 0 | 0 | 0 | 0 |
| 0 | 0 | 0 | 0 |
| 0 | 0 | 0 | 0 |
| 0 | 0 | 0 | 0 |
| 0 | 0 | 0 | 0 |
| 0 | 0 | 0 | 0 |
| 0 | 0 | 0 | 0 |
| 0 | 0 | 0 | 1 |
| 0 | 0 | 0 | 1 |
| 0 | 0 | 0 | 0 |
| 0 | 0 | 0 | 0 |
| 0 | 0 | 0 | 0 |
| 0 | 0 | 0 | 1 |

| | | | |
|---|---|---|---|
| 0 | 0 | 0 | 1 |
| 0 | 0 | 0 | 1 |
| 0 | 1 | 1 | 1 |
| 0 | 1 | 0 | 0 |
| 0 | 0 | 0 | 1 |
| 0 | 0 | 0 | 1 |
| 0 | 0 | 0 | 0 |
| 0 | 0 | 0 | 1 |
| 0 | 0 | 0 | 0 |
| 0 | 0 | 0 | 0 |
| 0 | 0 | 0 | 0 |
| 0 | 0 | 0 | 0 |
| 0 | 0 | 0 | 0 |
| 0 | 0 | 0 | 0 |
| 0 | 0 | 0 | 0 |
| 0 | 0 | 0 | 0 |
| 0 | 0 | 0 | 1 |
| 0 | 0 | 0 | 0 |
| 0 | 0 | 0 | 0 |
| 0 | 1 | 1 | 1 |
| 0 | 0 | 0 | 0 |
| 0 | 0 | 0 | 0 |
| 0 | 0 | 0 | 1 |
| 0 | 0 | 0 | 0 |
| 0 | 0 | 0 | 1 |
| 0 | 0 | 0 | 1 |
| 0 | 0 | 0 | 1 |
| 0 | 0 | 0 | 0 |
| 0 | 0 | 0 | 0 |
| 0 | 0 | 0 | 0 |
| 0 | 0 | 0 | 1 |
| 0 | 0 | 0 | 0 |
| 0 | 0 | 0 | 0 |
| 0 | 0 | 0 | 0 |
| 0 | 0 | 0 | 1 |
| 0 | 0 | 0 | 1 |
| 0 | 0 | 0 | 0 |
| 0 | 0 | 0 | 0 |
| 0 | 0 | 0 | 0 |
| 0 | 0 | 0 | 1 |
| 0 | 0 | 0 | 1 |
| 0 | 0 | 0 | 1 |
| 0 | 0 | 0 | 1 |
| 0 | 0 | 0 | 0 |
| 0 | 0 | 0 | 1 |
| 0 | 0 | 0 | 0 |
| 0 | 0 | 0 | 1 |
| 0 | 0 | 0 | 0 |
| 0 | 0 | 0 | 0 |
| 0 | 0 | 0 | 1 |
| 0 | 0 | 0 | 1 |
| 0 | 1 | 0 | 0 |
| 0 | 0 | 0 | 0 |
| 0 | 0 | 0 | 1 |

| | | | |
|---|---|---|---|
| 0 | 0 | 0 | 1 |
| 0 | 0 | 0 | 1 |
| 0 | 0 | 0 | 1 |
| 0 | 1 | 1 | 1 |
| 0 | 1 | 0 | 0 |
| 0 | 1 | 1 | 1 |
| 0 | 1 | 1 | 1 |
| 0 | 1 | 0 | 0 |
| 0 | 1 | 0 | 0 |
| 0 | 0 | 0 | 1 |
| 0 | 0 | 0 | 0 |
| 0 | 0 | 0 | 0 |
| 1 | 0 | 0 | 0 |
| 1 | 0 | 0 | 0 |
| 0 | 0 | 0 | 0 |
| 1 | 0 | 0 | 0 |
| 1 | 0 | 0 | 0 |
| 1 | 0 | 0 | 0 |
| 1 | 0 | 0 | 0 |
| 0 | 0 | 0 | 1 |
| 0 | 0 | 0 | 1 |
| 0 | 0 | 0 | 1 |
| 0 | 0 | 0 | 1 |
| 0 | 0 | 0 | 1 |
| 0 | 0 | 0 | 0 |
| 0 | 0 | 0 | 1 |
| 1 | 0 | 0 | 0 |
| 1 | 0 | 0 | 0 |
| 0 | 0 | 0 | 1 |
| 0 | 0 | 0 | 1 |
| 0 | 0 | 0 | 1 |
| 0 | 0 | 0 | 1 |
| 0 | 0 | 0 | 1 |
| 0 | 0 | 0 | 0 |
| 0 | 0 | 0 | 0 |
| 0 | 0 | 0 | 1 |
| 0 | 0 | 0 | 0 |
| 0 | 0 | 0 | 1 |
| 0 | 0 | 0 | 0 |
| 0 | 1 | 0 | 0 |
| 0 | 1 | 0 | 0 |
| 0 | 0 | 0 | 0 |
| 0 | 0 | 0 | 1 |
| 0 | 0 | 0 | 0 |
| 0 | 0 | 0 | 0 |
| 1 | 0 | 0 | 0 |
| 1 | 0 | 0 | 0 |
| 1 | 0 | 0 | 0 |
| 0 | 0 | 0 | 0 |
| 0 | 0 | 0 | 0 |
| 0 | 1 | 0 | 0 |
| 1 | 0 | 0 | 0 |
| 1 | 0 | 0 | 0 |
| 0 | 0 | 0 | 1 |

| | | | |
|---|---|---|---|
| 0 | 0 | 0 | 0 |
| 0 | 0 | 0 | 1 |
| 0 | 0 | 0 | 0 |
| 0 | 0 | 0 | 0 |
| 0 | 1 | 0 | 0 |
| 0 | 0 | 0 | 1 |
| 0 | 0 | 0 | 0 |
| 0 | 0 | 0 | 1 |
| 0 | 0 | 0 | 1 |
| 0 | 0 | 0 | 0 |
| 0 | 0 | 0 | 1 |
| 0 | 0 | 0 | 1 |
| 0 | 0 | 0 | 1 |
| 0 | 0 | 0 | 1 |
| 0 | 0 | 0 | 1 |
| 0 | 1 | 0 | 0 |
| 0 | 0 | 0 | 0 |
| 0 | 0 | 0 | 1 |
| 0 | 0 | 0 | 0 |
| 0 | 0 | 0 | 1 |
| 0 | 0 | 0 | 0 |
| 0 | 0 | 0 | 0 |
| 0 | 0 | 0 | 0 |
| 0 | 1 | 0 | 0 |
| 0 | 1 | 0 | 0 |
| 0 | 1 | 0 | 0 |
| 0 | 0 | 0 | 0 |
| 0 | 0 | 0 | 0 |
| 0 | 0 | 0 | 0 |
| 0 | 1 | 0 | 0 |
| 0 | 0 | 0 | 0 |
| 0 | 0 | 0 | 0 |
| 0 | 0 | 0 | 1 |
| 0 | 0 | 0 | 1 |
| 0 | 1 | 0 | 0 |
| 0 | 1 | 0 | 0 |
| 0 | 1 | 1 | 1 |
| 0 | 1 | 0 | 0 |
| 0 | 1 | 1 | 1 |
| 0 | 1 | 0 | 0 |
| 0 | 1 | 0 | 0 |
| 0 | 1 | 0 | 0 |
| 0 | 1 | 1 | 1 |
| 0 | 0 | 0 | 1 |
| 0 | 1 | 0 | 0 |
| 0 | 1 | 0 | 0 |
| 0 | 1 | 0 | 0 |
| 0 | 1 | 0 | 0 |
| 0 | 1 | 0 | 0 |
| 0 | 1 | 0 | 0 |
| 0 | 1 | 1 | 1 |
| 0 | 1 | 0 | 0 |
| 0 | 1 | 0 | 0 |
| 0 | 1 | 1 | 1 |

| | | | |
|---|---|---|---|
| 0 | 1 | 0 | 0 |
| 0 | 1 | 1 | 1 |
| 0 | 0 | 0 | 0 |
| 0 | 0 | 0 | 0 |
| 0 | 0 | 0 | 0 |
| 0 | 0 | 0 | 0 |
| 0 | 0 | 0 | 0 |
| 0 | 0 | 0 | 0 |
| 0 | 0 | 0 | 0 |
| 0 | 0 | 0 | 0 |
| 0 | 0 | 0 | 0 |
| 0 | 0 | 0 | 1 |
| 0 | 0 | 0 | 0 |
| 0 | 0 | 0 | 1 |
| 0 | 1 | 1 | 1 |
| 0 | 1 | 1 | 1 |
| 0 | 1 | 0 | 0 |
| 0 | 1 | 0 | 0 |
| 0 | 1 | 1 | 1 |
| 0 | 1 | 0 | 0 |
| 0 | 1 | 1 | 1 |
| 0 | 1 | 0 | 0 |
| 0 | 0 | 0 | 1 |
| 0 | 1 | 0 | 0 |
| 0 | 0 | 0 | 1 |
| 0 | 0 | 0 | 1 |
| 0 | 0 | 0 | 1 |
| 0 | 0 | 0 | 0 |
| 0 | 0 | 0 | 1 |
| 0 | 0 | 0 | 1 |
| 0 | 0 | 0 | 1 |
| 1 | 0 | 0 | 0 |
| 0 | 0 | 0 | 1 |
| 0 | 0 | 0 | 0 |
| 0 | 0 | 0 | 0 |
| 0 | 0 | 0 | 0 |
| 0 | 0 | 0 | 1 |
| 0 | 0 | 0 | 1 |
| 0 | 1 | 0 | 0 |
| 0 | 0 | 0 | 0 |
| 0 | 0 | 0 | 0 |
| 0 | 0 | 0 | 0 |
| 0 | 1 | 0 | 0 |
| 0 | 0 | 0 | 0 |
| 0 | 1 | 0 | 0 |
| 0 | 0 | 0 | 0 |
| 0 | 0 | 0 | 1 |
| 0 | 0 | 0 | 1 |
| 0 | 0 | 0 | 0 |
| 0 | 0 | 0 | 0 |
| 0 | 0 | 0 | 0 |
| 0 | 0 | 0 | 1 |
| 0 | 0 | 0 | 1 |
| 0 | 0 | 0 | 0 |

| | | | |
|---|---|---|---|
| 0 | 0 | 0 | 0 |
| 0 | 0 | 0 | 1 |
| 0 | 0 | 0 | 1 |
| 0 | 0 | 0 | 1 |
| 0 | 0 | 0 | 0 |
| 0 | 1 | 0 | 0 |
| 0 | 0 | 0 | 0 |
| 0 | 0 | 0 | 0 |
| 0 | 0 | 0 | 0 |
| 0 | 0 | 0 | 0 |
| 0 | 0 | 0 | 1 |
| 0 | 0 | 0 | 0 |
| 0 | 0 | 0 | 0 |
| 0 | 0 | 0 | 0 |
| 0 | 1 | 1 | 1 |
| 0 | 0 | 0 | 1 |
| 0 | 1 | 0 | 0 |
| 0 | 1 | 1 | 1 |
| 0 | 1 | 0 | 0 |
| 0 | 1 | 0 | 0 |
| 0 | 0 | 0 | 1 |
| 0 | 1 | 0 | 0 |
| 0 | 1 | 0 | 0 |
| 0 | 0 | 0 | 1 |
| 0 | 1 | 0 | 0 |
| 0 | 1 | 0 | 0 |
| 0 | 1 | 0 | 0 |
| 0 | 1 | 1 | 1 |
| 0 | 1 | 0 | 0 |
| 0 | 0 | 0 | 0 |
| 0 | 0 | 0 | 0 |
| 0 | 0 | 0 | 1 |
| 0 | 0 | 0 | 1 |
| 0 | 0 | 0 | 1 |
| 0 | 0 | 0 | 1 |
| 0 | 0 | 0 | 1 |
| 0 | 0 | 0 | 1 |
| 0 | 0 | 0 | 1 |
| 0 | 0 | 0 | 1 |
| 0 | 0 | 0 | 1 |
| 0 | 0 | 0 | 1 |
| 0 | 0 | 0 | 1 |
| 1 | 0 | 0 | 0 |
| 0 | 0 | 0 | 1 |
| 0 | 0 | 0 | 1 |
| 0 | 0 | 0 | 1 |
| 0 | 0 | 0 | 1 |
| 0 | 0 | 0 | 1 |
| 0 | 0 | 0 | 1 |
| 0 | 0 | 0 | 1 |
| 0 | 0 | 0 | 1 |
| 0 | 0 | 0 | 1 |
| 0 | 0 | 0 | 1 |
| 0 | 0 | 0 | 1 |

| | | | |
|---|---|---|---|
| 0 | 0 | 0 | 1 |
| 0 | 0 | 0 | 1 |
| 0 | 0 | 0 | 1 |
| 0 | 0 | 0 | 1 |
| 0 | 0 | 0 | 1 |
| 0 | 0 | 0 | 1 |
| 0 | 0 | 0 | 1 |
| 0 | 0 | 0 | 1 |
| 0 | 0 | 0 | 0 |
| 0 | 0 | 0 | 0 |
| 0 | 0 | 0 | 0 |
| 0 | 0 | 0 | 0 |
| 0 | 0 | 0 | 0 |
| 0 | 0 | 0 | 0 |
| 0 | 0 | 0 | 0 |
| 0 | 0 | 0 | 0 |
| 0 | 0 | 0 | 0 |
| 0 | 0 | 0 | 0 |
| 0 | 0 | 0 | 0 |
| 0 | 1 | 0 | 0 |
| 1 | 0 | 0 | 0 |
| 1 | 0 | 0 | 0 |
| 0 | 1 | 0 | 0 |
| 0 | 1 | 0 | 0 |
| 0 | 1 | 0 | 0 |
| 0 | 0 | 0 | 1 |
| 0 | 0 | 0 | 1 |
| 0 | 0 | 0 | 0 |
| 0 | 0 | 0 | 0 |
| 1 | 0 | 0 | 0 |
| 1 | 0 | 0 | 0 |
| 0 | 0 | 0 | 0 |
| 1 | 0 | 0 | 0 |
| 1 | 0 | 0 | 0 |
| 0 | 0 | 0 | 0 |
| 0 | 0 | 0 | 0 |
| 0 | 0 | 0 | 0 |
| 0 | 0 | 0 | 0 |
| 0 | 0 | 0 | 0 |
| 0 | 0 | 0 | 0 |
| 0 | 0 | 0 | 0 |
| 0 | 0 | 0 | 0 |
| 1 | 0 | 0 | 0 |
| 1 | 0 | 0 | 0 |
| 1 | 0 | 0 | 0 |
| 0 | 0 | 0 | 0 |
| 0 | 0 | 0 | 1 |
| 0 | 0 | 0 | 1 |
| 0 | 0 | 0 | 1 |
| 0 | 0 | 0 | 1 |
| 0 | 0 | 0 | 1 |
| 0 | 0 | 0 | 1 |
| 0 | 1 | 0 | 0 |
| 0 | 1 | 0 | 0 |

| | | | |
|---|---|---|---|
| 0 | 0 | 0 | 0 |
| 0 | 0 | 0 | 0 |
| 0 | 0 | 0 | 0 |
| 0 | 0 | 0 | 0 |
| 0 | 0 | 0 | 1 |
| 0 | 0 | 0 | 0 |
| 0 | 0 | 0 | 0 |
| 0 | 0 | 0 | 0 |
| 0 | 0 | 0 | 1 |
| 0 | 0 | 0 | 0 |
| 1 | 0 | 0 | 0 |
| 0 | 0 | 0 | 1 |
| 0 | 0 | 0 | 0 |
| 0 | 0 | 0 | 0 |
| 0 | 0 | 0 | 0 |
| 0 | 0 | 0 | 0 |
| 0 | 0 | 0 | 1 |
| 0 | 0 | 0 | 0 |
| 0 | 0 | 0 | 0 |
| 0 | 0 | 0 | 1 |
| 0 | 1 | 0 | 0 |
| 0 | 0 | 0 | 0 |
| 0 | 1 | 0 | 0 |
| 0 | 0 | 0 | 0 |
| 0 | 0 | 0 | 0 |
| 0 | 0 | 0 | 1 |
| 0 | 0 | 0 | 1 |
| 0 | 0 | 0 | 1 |
| 0 | 0 | 0 | 1 |
| 0 | 0 | 0 | 1 |
| 0 | 0 | 0 | 1 |
| 0 | 0 | 0 | 1 |
| 0 | 0 | 0 | 1 |
| 0 | 0 | 0 | 1 |
| 0 | 0 | 0 | 0 |
| 0 | 0 | 0 | 0 |
| 0 | 0 | 0 | 0 |
| 0 | 0 | 0 | 0 |
| 0 | 1 | 0 | 0 |
| 0 | 1 | 0 | 0 |
| 0 | 0 | 0 | 1 |
| 0 | 0 | 0 | 1 |
| 0 | 0 | 0 | 1 |
| 0 | 0 | 0 | 0 |
| 0 | 0 | 0 | 0 |
| 0 | 0 | 0 | 0 |
| 0 | 0 | 0 | 1 |
| 0 | 0 | 0 | 1 |
| 0 | 0 | 0 | 0 |
| 0 | 0 | 0 | 0 |
| 0 | 0 | 0 | 1 |
| 0 | 0 | 0 | 1 |
| 0 | 0 | 0 | 1 |
| 0 | 0 | 0 | 1 |

| | | | |
|---|---|---|---|
| 0 | 0 | 0 | 1 |
| 0 | 0 | 0 | 1 |
| 0 | 0 | 0 | 1 |
| 0 | 0 | 0 | 0 |
| 0 | 0 | 0 | 0 |
| 0 | 0 | 0 | 1 |
| 0 | 0 | 0 | 1 |
| 0 | 0 | 0 | 1 |
| 0 | 0 | 0 | 1 |
| 0 | 0 | 0 | 1 |
| 0 | 0 | 0 | 0 |
| 0 | 0 | 0 | 1 |
| 0 | 1 | 0 | 0 |
| 0 | 0 | 0 | 0 |
| 0 | 0 | 0 | 0 |
| 0 | 0 | 0 | 1 |
| 0 | 0 | 0 | 1 |
| 0 | 0 | 0 | 1 |
| 0 | 0 | 0 | 1 |
| 0 | 0 | 0 | 1 |
| 0 | 0 | 0 | 0 |
| 0 | 0 | 0 | 1 |
| 0 | 0 | 0 | 1 |
| 0 | 0 | 0 | 0 |
| 0 | 1 | 1 | 1 |
| 0 | 1 | 0 | 0 |
| 0 | 1 | 0 | 0 |
| 0 | 0 | 0 | 1 |
| 0 | 0 | 0 | 1 |
| 0 | 0 | 0 | 1 |
| 0 | 0 | 0 | 1 |
| 0 | 0 | 0 | 1 |
| 0 | 0 | 0 | 1 |
| 0 | 0 | 0 | 1 |
| 0 | 0 | 0 | 1 |
| 0 | 0 | 0 | 0 |
| 0 | 0 | 0 | 1 |
| 0 | 0 | 0 | 1 |
| 0 | 0 | 0 | 0 |
| 0 | 0 | 0 | 0 |
| 0 | 0 | 0 | 1 |
| 0 | 0 | 0 | 1 |
| 1 | 0 | 0 | 0 |
| 1 | 0 | 0 | 0 |
| 0 | 0 | 0 | 0 |
| 0 | 0 | 0 | 0 |
| 1 | 0 | 0 | 0 |
| 1 | 0 | 0 | 0 |
| 0 | 0 | 0 | 1 |
| 1 | 0 | 0 | 0 |
| 1 | 0 | 0 | 0 |
| 1 | 0 | 0 | 0 |
| 1 | 0 | 0 | 0 |
| 1 | 0 | 0 | 0 |

| | | | |
|---|---|---|---|
| 0 | 0 | 0 | 0 |
| 0 | 0 | 0 | 0 |
| 1 | 0 | 0 | 0 |
| 1 | 0 | 0 | 0 |
| 1 | 0 | 0 | 0 |
| 1 | 0 | 0 | 0 |
| 0 | 0 | 0 | 0 |
| 0 | 0 | 0 | 0 |
| 0 | 0 | 0 | 0 |
| 0 | 0 | 0 | 1 |
| 0 | 0 | 0 | 1 |
| 0 | 0 | 0 | 0 |
| 0 | 0 | 0 | 1 |
| 0 | 0 | 0 | 0 |
| 0 | 0 | 0 | 0 |
| 0 | 0 | 0 | 1 |
| 0 | 0 | 0 | 0 |
| 0 | 0 | 0 | 1 |
| 0 | 0 | 0 | 1 |
| 0 | 0 | 0 | 1 |
| 0 | 1 | 1 | 1 |
| 0 | 0 | 0 | 1 |
| 0 | 0 | 0 | 1 |
| 0 | 0 | 0 | 0 |
| 0 | 0 | 0 | 0 |
| 0 | 0 | 0 | 1 |
| 0 | 0 | 0 | 1 |
| 0 | 0 | 0 | 1 |
| 0 | 0 | 0 | 0 |
| 0 | 0 | 0 | 0 |
| 0 | 0 | 0 | 1 |
| 0 | 0 | 0 | 1 |
| 0 | 0 | 0 | 1 |
| 0 | 0 | 0 | 0 |
| 0 | 0 | 0 | 0 |
| 0 | 0 | 0 | 1 |
| 0 | 0 | 0 | 0 |
| 0 | 0 | 0 | 1 |
| 0 | 0 | 0 | 1 |
| 0 | 0 | 0 | 0 |
| 0 | 0 | 0 | 0 |
| 0 | 0 | 0 | 0 |
| 0 | 0 | 0 | 0 |
| 0 | 0 | 0 | 0 |
| 0 | 0 | 0 | 0 |
| 0 | 0 | 0 | 1 |
| 0 | 0 | 0 | 1 |
| 0 | 0 | 0 | 0 |
| 0 | 0 | 0 | 0 |
| 0 | 0 | 0 | 0 |
| 0 | 0 | 0 | 1 |
| 0 | 0 | 0 | 1 |
| 0 | 0 | 0 | 0 |
| 0 | 0 | 0 | 0 |

| | | | |
|---|---|---|---|
| 0 | 0 | 0 | 1 |
| 0 | 0 | 0 | 1 |
| 0 | 0 | 0 | 1 |
| 0 | 0 | 0 | 1 |
| 0 | 1 | 0 | 0 |
| 0 | 1 | 0 | 0 |
| 0 | 0 | 0 | 1 |
| 0 | 0 | 0 | 0 |
| 0 | 1 | 1 | 1 |
| 0 | 0 | 0 | 0 |
| 0 | 0 | 0 | 1 |
| 0 | 0 | 0 | 1 |
| 0 | 0 | 0 | 1 |
| 0 | 0 | 0 | 1 |
| 0 | 0 | 0 | 1 |
| 0 | 0 | 0 | 1 |
| 0 | 0 | 0 | 0 |
| 0 | 0 | 0 | 1 |
| 0 | 0 | 0 | 1 |
| 0 | 0 | 0 | 0 |
| 0 | 0 | 0 | 0 |
| 0 | 0 | 0 | 1 |
| 0 | 0 | 0 | 1 |
| 0 | 0 | 0 | 0 |
| 0 | 0 | 0 | 1 |
| 0 | 0 | 0 | 0 |
| 0 | 0 | 0 | 0 |
| 0 | 0 | 0 | 0 |
| 0 | 0 | 0 | 0 |
| 0 | 0 | 0 | 0 |
| 0 | 0 | 0 | 0 |
| 0 | 0 | 0 | 0 |
| 0 | 0 | 0 | 1 |
| 0 | 0 | 0 | 0 |
| 0 | 0 | 0 | 1 |
| 0 | 0 | 0 | 1 |
| 0 | 0 | 0 | 1 |
| 0 | 0 | 0 | 1 |
| 0 | 0 | 0 | 0 |
| 0 | 0 | 0 | 1 |
| 0 | 0 | 0 | 1 |
| 0 | 0 | 0 | 1 |
| 0 | 0 | 0 | 1 |
| 0 | 1 | 0 | 0 |
| 0 | 0 | 0 | 1 |
| 0 | 0 | 0 | 1 |
| 0 | 0 | 0 | 1 |
| 0 | 0 | 0 | 1 |
| 0 | 0 | 0 | 1 |
| 0 | 0 | 0 | 1 |
| 0 | 0 | 0 | 0 |
| 0 | 0 | 0 | 0 |
| 0 | 0 | 0 | 0 |
| 0 | 0 | 0 | 1 |

| | | | |
|---|---|---|---|
| 0 | 0 | 0 | 0 |
| 0 | 1 | 0 | 0 |
| 0 | 1 | 0 | 0 |
| 0 | 0 | 0 | 0 |
| 1 | 0 | 0 | 0 |
| 1 | 0 | 0 | 0 |
| 0 | 0 | 0 | 0 |
| 0 | 0 | 0 | 0 |
| 0 | 0 | 0 | 1 |
| 0 | 0 | 0 | 1 |
| 0 | 0 | 0 | 0 |
| 0 | 0 | 0 | 1 |
| 0 | 0 | 0 | 1 |
| 0 | 0 | 0 | 0 |
| 0 | 0 | 0 | 0 |
| 0 | 0 | 0 | 0 |
| 0 | 0 | 0 | 0 |
| 0 | 0 | 0 | 0 |
| 0 | 0 | 0 | 0 |
| 0 | 0 | 0 | 1 |
| 0 | 0 | 0 | 1 |
| 0 | 0 | 0 | 0 |
| 0 | 0 | 0 | 1 |
| 0 | 1 | 0 | 0 |
| 0 | 0 | 0 | 1 |
| 0 | 0 | 0 | 1 |
| 0 | 0 | 0 | 1 |
| 0 | 0 | 0 | 1 |
| 0 | 0 | 0 | 1 |
| 0 | 0 | 0 | 0 |
| 0 | 0 | 0 | 1 |
| 0 | 0 | 0 | 1 |
| 0 | 0 | 0 | 0 |
| 0 | 0 | 0 | 1 |
| 0 | 1 | 0 | 0 |
| 0 | 1 | 0 | 0 |
| 0 | 1 | 0 | 0 |
| 0 | 0 | 0 | 0 |
| 0 | 0 | 0 | 0 |
| 0 | 0 | 0 | 0 |
| 0 | 0 | 0 | 0 |
| 0 | 0 | 0 | 0 |
| 0 | 0 | 0 | 1 |
| 0 | 0 | 0 | 0 |
| 0 | 0 | 0 | 1 |
| 0 | 0 | 0 | 0 |
| 0 | 0 | 0 | 1 |
| 0 | 0 | 0 | 1 |
| 0 | 0 | 0 | 1 |
| 0 | 0 | 0 | 0 |
| 0 | 0 | 0 | 1 |
| 0 | 0 | 0 | 1 |
| 0 | 0 | 0 | 1 |
| 0 | 0 | 0 | 1 |

| | | | |
|---|---|---|---|
| 1 | 0 | 0 | 0 |
| 0 | 0 | 0 | 1 |
| 0 | 0 | 0 | 1 |
| 0 | 0 | 0 | 1 |
| 0 | 0 | 0 | 1 |
| 0 | 0 | 0 | 1 |
| 0 | 0 | 0 | 1 |
| 0 | 0 | 0 | 1 |
| 0 | 0 | 0 | 1 |
| 0 | 0 | 0 | 1 |
| 0 | 0 | 0 | 0 |
| 0 | 0 | 0 | 1 |
| 0 | 0 | 0 | 1 |
| 0 | 0 | 0 | 1 |
| 0 | 0 | 0 | 0 |
| 0 | 0 | 0 | 0 |
| 0 | 0 | 0 | 0 |
| 0 | 0 | 0 | 0 |
| 0 | 0 | 0 | 0 |
| 0 | 0 | 0 | 1 |
| 0 | 0 | 0 | 1 |
| 0 | 0 | 0 | 1 |
| 0 | 0 | 0 | 0 |
| 0 | 0 | 0 | 1 |
| 0 | 0 | 0 | 1 |
| 0 | 0 | 0 | 1 |
| 0 | 0 | 0 | 1 |
| 0 | 0 | 0 | 1 |
| 0 | 0 | 0 | 0 |
| 0 | 0 | 0 | 0 |
| 0 | 0 | 0 | 0 |
| 0 | 1 | 0 | 0 |
| 0 | 1 | 0 | 0 |
| 0 | 0 | 0 | 0 |
| 0 | 0 | 0 | 1 |
| 0 | 0 | 0 | 1 |
| 0 | 0 | 0 | 0 |
| 0 | 0 | 0 | 0 |
| 0 | 1 | 0 | 0 |
| 0 | 0 | 0 | 0 |
| 0 | 0 | 0 | 0 |
| 0 | 0 | 0 | 0 |
| 0 | 0 | 0 | 1 |
| 0 | 0 | 0 | 1 |
| 0 | 0 | 0 | 0 |
| 0 | 0 | 0 | 1 |
| 0 | 0 | 0 | 1 |
| 0 | 0 | 0 | 1 |
| 0 | 0 | 0 | 1 |
| 0 | 0 | 0 | 1 |
| 0 | 0 | 0 | 1 |
| 0 | 0 | 0 | 1 |
| 0 | 0 | 0 | 1 |
| 0 | 1 | 1 | 1 |

| | | | |
|---|---|---|---|
| 0 | 1 | 0 | 0 |
| 0 | 0 | 0 | 1 |
| 0 | 1 | 0 | 0 |
| 0 | 0 | 0 | 1 |
| 0 | 1 | 1 | 1 |
| 0 | 1 | 1 | 1 |
| 0 | 1 | 0 | 0 |
| 0 | 1 | 0 | 0 |
| 0 | 1 | 1 | 1 |
| 0 | 1 | 1 | 1 |
| 0 | 0 | 0 | 1 |
| 0 | 0 | 0 | 1 |
| 0 | 0 | 0 | 1 |
| 0 | 1 | 1 | 1 |
| 0 | 0 | 0 | 0 |
| 0 | 0 | 0 | 0 |
| 0 | 0 | 0 | 0 |
| 0 | 0 | 0 | 0 |
| 0 | 0 | 0 | 1 |
| 0 | 0 | 0 | 0 |
| 0 | 0 | 0 | 0 |
| 0 | 0 | 0 | 0 |
| 0 | 0 | 0 | 0 |
| 0 | 0 | 0 | 0 |
| 0 | 0 | 0 | 1 |
| 0 | 0 | 0 | 0 |
| 0 | 0 | 0 | 1 |
| 0 | 0 | 0 | 1 |
| 0 | 0 | 0 | 1 |
| 0 | 1 | 0 | 0 |
| 0 | 0 | 0 | 1 |
| 0 | 0 | 0 | 1 |
| 0 | 0 | 0 | 0 |
| 0 | 0 | 0 | 0 |
| 0 | 0 | 0 | 1 |
| 0 | 0 | 0 | 1 |
| 0 | 0 | 0 | 1 |
| 0 | 0 | 0 | 1 |
| 0 | 0 | 0 | 1 |
| 0 | 0 | 0 | 0 |
| 0 | 0 | 0 | 0 |
| 0 | 1 | 0 | 0 |
| 0 | 1 | 0 | 0 |
| 0 | 0 | 0 | 0 |
| 0 | 0 | 0 | 0 |
| 0 | 0 | 0 | 1 |
| 0 | 0 | 0 | 1 |
| 0 | 0 | 0 | 0 |
| 0 | 0 | 0 | 0 |
| 0 | 0 | 0 | 1 |
| 0 | 0 | 0 | 0 |
| 0 | 0 | 0 | 0 |
| 0 | 0 | 0 | 1 |
| 0 | 0 | 0 | 1 |

| | | | |
|---|---|---|---|
| 0 | 0 | 0 | 1 |
| 0 | 1 | 0 | 0 |
| 0 | 0 | 0 | 0 |
| 0 | 1 | 1 | 1 |
| 0 | 0 | 0 | 1 |
| 0 | 0 | 0 | 0 |
| 0 | 0 | 0 | 0 |
| 0 | 0 | 0 | 0 |
| 0 | 0 | 0 | 0 |
| 0 | 0 | 0 | 0 |
| 0 | 0 | 0 | 0 |
| 0 | 0 | 0 | 0 |
| 0 | 0 | 0 | 0 |
| 0 | 0 | 0 | 0 |
| 0 | 0 | 0 | 0 |
| 0 | 0 | 0 | 1 |
| 0 | 1 | 0 | 0 |
| 0 | 0 | 0 | 1 |
| 0 | 1 | 1 | 1 |
| 0 | 0 | 0 | 1 |
| 0 | 0 | 0 | 0 |
| 0 | 0 | 0 | 1 |
| 0 | 0 | 0 | 0 |
| 0 | 0 | 0 | 0 |
| 0 | 0 | 0 | 1 |
| 0 | 0 | 0 | 0 |
| 0 | 0 | 0 | 1 |
| 0 | 0 | 0 | 1 |
| 0 | 0 | 0 | 1 |
| 0 | 0 | 0 | 0 |
| 0 | 1 | 0 | 0 |
| 0 | 0 | 0 | 0 |
| 0 | 1 | 0 | 0 |
| 0 | 1 | 0 | 0 |
| 0 | 1 | 0 | 0 |
| 0 | 1 | 0 | 0 |
| 0 | 0 | 0 | 0 |
| 0 | 0 | 0 | 1 |
| 0 | 0 | 0 | 1 |
| 0 | 0 | 0 | 0 |
| 0 | 1 | 0 | 0 |
| 0 | 0 | 0 | 0 |
| 0 | 0 | 0 | 0 |
| 0 | 0 | 0 | 0 |
| 0 | 1 | 0 | 0 |
| 0 | 0 | 0 | 0 |
| 0 | 0 | 0 | 1 |
| 0 | 0 | 0 | 0 |
| 0 | 0 | 0 | 0 |
| 0 | 0 | 0 | 0 |
| 0 | 0 | 0 | 0 |
| 0 | 0 | 0 | 1 |
| 0 | 1 | 0 | 0 |
| 0 | 0 | 0 | 0 |

| | | | |
|---|---|---|---|
| 0 | 1 | 0 | 0 |
| 0 | 1 | 0 | 0 |
| 0 | 0 | 0 | 0 |
| 0 | 0 | 0 | 0 |
| 0 | 0 | 0 | 0 |
| 0 | 0 | 0 | 0 |
| 0 | 0 | 0 | 0 |
| 0 | 0 | 0 | 0 |
| 1 | 0 | 0 | 0 |
| 0 | 0 | 0 | 0 |
| 0 | 0 | 0 | 1 |
| 0 | 1 | 0 | 0 |
| 0 | 1 | 1 | 1 |
| 0 | 0 | 0 | 0 |
| 0 | 0 | 0 | 1 |
| 0 | 0 | 0 | 1 |
| 0 | 0 | 0 | 1 |
| 0 | 0 | 0 | 1 |
| 0 | 0 | 0 | 0 |
| 0 | 0 | 0 | 0 |
| 0 | 0 | 0 | 1 |
| 0 | 1 | 0 | 0 |
| 0 | 1 | 0 | 0 |
| 0 | 0 | 0 | 1 |
| 0 | 0 | 0 | 0 |
| 0 | 0 | 0 | 0 |
| 0 | 0 | 0 | 0 |
| 0 | 0 | 0 | 1 |
| 0 | 1 | 0 | 0 |
| 0 | 0 | 0 | 0 |
| 0 | 1 | 1 | 1 |
| 1 | 0 | 0 | 0 |
| 0 | 0 | 0 | 1 |
| 0 | 0 | 0 | 1 |
| 1 | 0 | 0 | 0 |
| 1 | 0 | 0 | 0 |
| 1 | 0 | 0 | 0 |
| 0 | 1 | 1 | 1 |
| 0 | 1 | 1 | 1 |
| 0 | 0 | 0 | 0 |
| 1 | 0 | 0 | 0 |
| 0 | 1 | 0 | 0 |
| 0 | 1 | 0 | 0 |
| 1 | 1 | 0 | 0 |
| 1 | 0 | 0 | 0 |
| 0 | 0 | 0 | 0 |
| 0 | 0 | 0 | 1 |
| 0 | 0 | 0 | 1 |
| 0 | 0 | 0 | 1 |
| 0 | 0 | 0 | 1 |
| 1 | 0 | 0 | 0 |
| 0 | 1 | 0 | 0 |
| 0 | 0 | 0 | 0 |
| 0 | 0 | 0 | 0 |

| | | | |
|---|---|---|---|
| 0 | 0 | 0 | 0 |
| 0 | 1 | 1 | 1 |
| 0 | 0 | 0 | 0 |
| 0 | 0 | 0 | 0 |
| 0 | 0 | 0 | 0 |
| 0 | 0 | 0 | 1 |
| 0 | 0 | 0 | 0 |
| 0 | 0 | 0 | 0 |
| 0 | 0 | 0 | 1 |
| 0 | 1 | 0 | 0 |
| 0 | 0 | 0 | 1 |
| 0 | 0 | 0 | 1 |
| 0 | 0 | 0 | 1 |
| 0 | 0 | 0 | 0 |
| 0 | 0 | 0 | 1 |
| 0 | 0 | 0 | 0 |
| 0 | 0 | 0 | 1 |
| 0 | 0 | 0 | 1 |
| 0 | 0 | 0 | 0 |
| 0 | 0 | 0 | 0 |
| 0 | 0 | 0 | 0 |
| 0 | 0 | 0 | 0 |
| 0 | 0 | 0 | 0 |
| 0 | 0 | 0 | 0 |
| 0 | 0 | 0 | 0 |
| 0 | 0 | 0 | 1 |
| 0 | 0 | 0 | 1 |
| 0 | 0 | 0 | 1 |
| 0 | 0 | 0 | 1 |
| 0 | 0 | 0 | 1 |
| 0 | 0 | 0 | 1 |
| 0 | 0 | 0 | 1 |
| 0 | 0 | 0 | 0 |
| 0 | 0 | 0 | 0 |
| 0 | 0 | 0 | 0 |
| 0 | 0 | 0 | 0 |
| 0 | 0 | 0 | 0 |
| 0 | 0 | 0 | 0 |
| 0 | 0 | 0 | 1 |
| 0 | 0 | 0 | 1 |
| 0 | 0 | 0 | 1 |
| 0 | 0 | 0 | 1 |
| 0 | 0 | 0 | 1 |
| 0 | 0 | 0 | 1 |
| 0 | 1 | 0 | 0 |
| 0 | 0 | 0 | 0 |
| 0 | 0 | 0 | 1 |
| 0 | 0 | 0 | 0 |
| 0 | 0 | 0 | 0 |
| 0 | 0 | 0 | 0 |
| 0 | 0 | 0 | 0 |
| 0 | 0 | 0 | 0 |
| 0 | 0 | 0 | 0 |
| 0 | 0 | 0 | 0 |

| | | | |
|---|---|---|---|
| 0 | 0 | 0 | 1 |
| 0 | 0 | 0 | 1 |
| 1 | 0 | 0 | 0 |
| 1 | 0 | 0 | 0 |
| 0 | 0 | 0 | 1 |
| 0 | 0 | 0 | 0 |
| 0 | 0 | 0 | 0 |
| 0 | 0 | 0 | 1 |
| 0 | 0 | 0 | 1 |
| 0 | 1 | 0 | 0 |
| 0 | 0 | 0 | 1 |
| 0 | 0 | 0 | 1 |
| 0 | 1 | 0 | 0 |
| 0 | 1 | 0 | 0 |
| 0 | 1 | 0 | 0 |
| 0 | 1 | 0 | 0 |
| 0 | 0 | 0 | 1 |
| 0 | 0 | 0 | 1 |
| 0 | 0 | 0 | 1 |
| 0 | 0 | 0 | 1 |
| 0 | 0 | 0 | 1 |
| 0 | 0 | 0 | 1 |
| 0 | 0 | 0 | 1 |
| 0 | 0 | 0 | 1 |
| 0 | 0 | 0 | 1 |
| 0 | 0 | 0 | 0 |
| 0 | 0 | 0 | 1 |
| 0 | 0 | 0 | 1 |
| 0 | 0 | 0 | 1 |
| 0 | 0 | 0 | 1 |
| 0 | 0 | 0 | 1 |
| 0 | 0 | 0 | 1 |
| 0 | 0 | 0 | 1 |
| 0 | 0 | 0 | 1 |
| 0 | 0 | 0 | 1 |
| 0 | 0 | 0 | 1 |
| 0 | 0 | 0 | 1 |
| 0 | 0 | 0 | 0 |
| 0 | 1 | 1 | 1 |
| 0 | 0 | 0 | 1 |
| 0 | 1 | 0 | 0 |
| 0 | 0 | 0 | 1 |
| 0 | 0 | 0 | 1 |
| 0 | 1 | 0 | 0 |
| 0 | 0 | 0 | 0 |
| 0 | 0 | 0 | 1 |
| 0 | 0 | 0 | 1 |
| 0 | 0 | 0 | 1 |
| 0 | 0 | 0 | 1 |
| 0 | 0 | 0 | 1 |
| 0 | 0 | 0 | 1 |
| 0 | 0 | 0 | 0 |
| 0 | 0 | 0 | 0 |
| 0 | 0 | 0 | 0 |

| | | | |
|---|---|---|---|
| 0 | 0 | 0 | 0 |
| 0 | 0 | 0 | 1 |
| 0 | 0 | 0 | 1 |
| 0 | 0 | 0 | 1 |
| 0 | 0 | 0 | 1 |
| 0 | 0 | 0 | 1 |
| 0 | 0 | 0 | 1 |
| 0 | 0 | 0 | 1 |
| 0 | 0 | 0 | 0 |
| 0 | 0 | 0 | 1 |
| 0 | 0 | 0 | 1 |
| 0 | 0 | 0 | 1 |
| 0 | 0 | 0 | 1 |
| 0 | 0 | 0 | 1 |
| 0 | 0 | 0 | 1 |
| 0 | 0 | 0 | 1 |
| 0 | 0 | 0 | 1 |
| 0 | 0 | 0 | 1 |
| 0 | 0 | 0 | 0 |
| 0 | 0 | 0 | 0 |
| 0 | 0 | 0 | 0 |
| 0 | 0 | 0 | 0 |
| 0 | 0 | 0 | 1 |
| 0 | 0 | 0 | 0 |
| 0 | 0 | 0 | 0 |
| 0 | 0 | 0 | 1 |
| 0 | 0 | 0 | 1 |
| 0 | 0 | 0 | 1 |
| 0 | 0 | 0 | 1 |
| 0 | 0 | 0 | 1 |
| 0 | 0 | 0 | 1 |
| 0 | 0 | 0 | 1 |
| 0 | 0 | 0 | 0 |
| 0 | 0 | 0 | 1 |
| 0 | 0 | 0 | 1 |
| 0 | 0 | 0 | 1 |
| 0 | 0 | 0 | 0 |
| 0 | 0 | 0 | 1 |
| 0 | 0 | 0 | 0 |
| 0 | 0 | 0 | 0 |
| 0 | 0 | 0 | 0 |
| 0 | 0 | 0 | 0 |
| 0 | 0 | 0 | 0 |
| 0 | 0 | 0 | 1 |
| 0 | 0 | 0 | 1 |
| 0 | 0 | 0 | 1 |
| 0 | 0 | 0 | 0 |
| 0 | 0 | 0 | 0 |
| 0 | 0 | 0 | 0 |
| 0 | 1 | 0 | 0 |
| 0 | 1 | 0 | 0 |
| 0 | 1 | 0 | 0 |
| 0 | 1 | 0 | 0 |
| 0 | 0 | 0 | 1 |

| | | | |
|---|---|---|---|
| 0 | 0 | 0 | 0 |
| 0 | 0 | 0 | 0 |
| 0 | 0 | 0 | 1 |
| 1 | 0 | 0 | 0 |
| 1 | 0 | 0 | 0 |
| 0 | 0 | 0 | 1 |
| 0 | 0 | 0 | 1 |
| 0 | 1 | 1 | 1 |
| 0 | 0 | 0 | 1 |
| 0 | 0 | 0 | 1 |
| 0 | 0 | 0 | 1 |
| 0 | 0 | 0 | 0 |
| 0 | 0 | 0 | 1 |
| 0 | 0 | 0 | 1 |
| 0 | 0 | 0 | 0 |
| 0 | 0 | 0 | 1 |
| 0 | 0 | 0 | 0 |
| 0 | 0 | 0 | 1 |
| 0 | 0 | 0 | 0 |
| 1 | 0 | 0 | 0 |
| 0 | 0 | 0 | 1 |
| 0 | 0 | 0 | 1 |
| 0 | 0 | 0 | 0 |
| 0 | 0 | 0 | 1 |
| 0 | 0 | 0 | 0 |
| 0 | 0 | 0 | 1 |
| 0 | 0 | 0 | 1 |
| 0 | 0 | 0 | 0 |
| 0 | 0 | 0 | 1 |
| 0 | 0 | 0 | 0 |
| 0 | 0 | 0 | 1 |
| 0 | 0 | 0 | 0 |
| 0 | 0 | 0 | 0 |
| 0 | 0 | 0 | 0 |
| 0 | 0 | 0 | 1 |
| 0 | 0 | 0 | 0 |
| 0 | 0 | 0 | 1 |
| 0 | 0 | 0 | 1 |
| 0 | 0 | 0 | 0 |
| 0 | 0 | 0 | 1 |
| 0 | 0 | 0 | 1 |
| 0 | 0 | 0 | 1 |
| 0 | 0 | 0 | 1 |
| 0 | 0 | 0 | 0 |
| 0 | 0 | 0 | 0 |
| 0 | 0 | 0 | 1 |
| 0 | 0 | 0 | 1 |
| 0 | 0 | 0 | 1 |
| 0 | 1 | 0 | 0 |
| 0 | 0 | 0 | 1 |
| 0 | 0 | 0 | 0 |
| 0 | 0 | 0 | 1 |
| 0 | 0 | 0 | 1 |
| 0 | 0 | 0 | 0 |

| | | | |
|---|---|---|---|
| 0 | 0 | 0 | 1 |
| 0 | 0 | 0 | 1 |
| 0 | 0 | 0 | 1 |
| 0 | 0 | 0 | 1 |
| 0 | 0 | 0 | 1 |
| 0 | 0 | 0 | 1 |
| 0 | 0 | 0 | 1 |
| 0 | 0 | 0 | 0 |
| 0 | 0 | 0 | 1 |
| 0 | 0 | 0 | 1 |
| 0 | 0 | 0 | 1 |
| 0 | 0 | 0 | 1 |
| 0 | 0 | 0 | 0 |
| 0 | 0 | 0 | 1 |
| 0 | 0 | 0 | 0 |
| 0 | 0 | 0 | 0 |
| 0 | 0 | 0 | 1 |
| 0 | 0 | 0 | 1 |
| 0 | 0 | 0 | 1 |
| 0 | 0 | 0 | 1 |
| 0 | 0 | 0 | 0 |
| 0 | 0 | 0 | 0 |
| 0 | 0 | 0 | 1 |
| 0 | 0 | 0 | 1 |
| 0 | 0 | 0 | 0 |
| 0 | 0 | 0 | 1 |
| 0 | 0 | 0 | 0 |
| 0 | 0 | 0 | 0 |
| 0 | 0 | 0 | 0 |
| 0 | 0 | 0 | 0 |
| 0 | 0 | 0 | 1 |
| 0 | 0 | 0 | 1 |
| 0 | 0 | 0 | 0 |
| 0 | 1 | 0 | 0 |
| 0 | 0 | 0 | 0 |
| 0 | 0 | 0 | 0 |
| 0 | 0 | 0 | 0 |
| 0 | 0 | 0 | 0 |
| 0 | 0 | 0 | 0 |
| 0 | 0 | 0 | 1 |
| 0 | 0 | 0 | 1 |
| 0 | 0 | 0 | 1 |
| 0 | 0 | 0 | 1 |
| 0 | 0 | 0 | 1 |
| 0 | 0 | 0 | 1 |
| 0 | 0 | 0 | 0 |
| 0 | 0 | 0 | 1 |
| 0 | 0 | 0 | 1 |
| 0 | 0 | 0 | 1 |
| 0 | 0 | 0 | 0 |
| 0 | 0 | 0 | 0 |
| 0 | 0 | 0 | 1 |
| 0 | 0 | 0 | 1 |
| 0 | 0 | 0 | 1 |

| | | | |
|---|---|---|---|
| 0 | 0 | 0 | 1 |
| 0 | 0 | 0 | 1 |
| 0 | 0 | 0 | 1 |
| 0 | 0 | 0 | 1 |
| 0 | 0 | 0 | 0 |
| 0 | 0 | 0 | 1 |
| 0 | 0 | 0 | 0 |
| 0 | 0 | 0 | 0 |
| 0 | 0 | 0 | 0 |
| 0 | 0 | 0 | 0 |
| 0 | 0 | 0 | 0 |
| 0 | 0 | 0 | 1 |
| 0 | 0 | 0 | 0 |
| 0 | 0 | 0 | 0 |
| 0 | 0 | 0 | 1 |
| 0 | 0 | 0 | 1 |
| 0 | 0 | 0 | 1 |
| 0 | 0 | 0 | 1 |
| 0 | 0 | 0 | 1 |
| 0 | 0 | 0 | 1 |
| 0 | 0 | 0 | 1 |
| 0 | 0 | 0 | 1 |
| 0 | 0 | 0 | 1 |
| 0 | 0 | 0 | 0 |
| 0 | 0 | 0 | 1 |
| 0 | 0 | 0 | 1 |
| 0 | 0 | 0 | 1 |
| 0 | 0 | 0 | 0 |
| 0 | 1 | 1 | 1 |
| 0 | 1 | 1 | 1 |
| 0 | 1 | 0 | 0 |
| 0 | 1 | 0 | 0 |
| 0 | 0 | 0 | 1 |
| 0 | 0 | 0 | 0 |
| 0 | 0 | 0 | 1 |
| 0 | 0 | 0 | 1 |
| 0 | 0 | 0 | 1 |
| 0 | 0 | 0 | 1 |
| 0 | 0 | 0 | 1 |
| 0 | 0 | 0 | 1 |
| 0 | 0 | 0 | 1 |
| 0 | 0 | 0 | 1 |
| 0 | 0 | 0 | 1 |
| 0 | 0 | 0 | 1 |
| 0 | 0 | 0 | 1 |
| 0 | 0 | 0 | 1 |
| 0 | 0 | 0 | 1 |
| 0 | 0 | 0 | 1 |
| 0 | 0 | 0 | 0 |
| 0 | 0 | 0 | 1 |
| 0 | 0 | 0 | 1 |
| 0 | 1 | 1 | 1 |
| 0 | 1 | 1 | 1 |
| 0 | 0 | 0 | 1 |

| | | | |
|---|---|---|---|
| 0 | 0 | 0 | 1 |
| 0 | 0 | 0 | 1 |
| 0 | 0 | 0 | 1 |
| 0 | 0 | 0 | 1 |
| 0 | 1 | 1 | 1 |
| 0 | 0 | 0 | 1 |
| 0 | 0 | 0 | 1 |
| 0 | 0 | 0 | 1 |
| 0 | 0 | 0 | 1 |
| 0 | 0 | 0 | 1 |
| 0 | 0 | 0 | 0 |
| 0 | 0 | 0 | 0 |
| 0 | 0 | 0 | 0 |
| 1 | 0 | 0 | 0 |
| 1 | 0 | 0 | 0 |
| 0 | 0 | 0 | 1 |
| 1 | 0 | 0 | 0 |
| 0 | 0 | 0 | 0 |
| 0 | 0 | 0 | 0 |
| 0 | 0 | 0 | 0 |
| 0 | 0 | 0 | 0 |
| 1 | 0 | 0 | 0 |
| 0 | 0 | 0 | 1 |
| 0 | 0 | 0 | 0 |
| 1 | 0 | 0 | 0 |
| 1 | 0 | 0 | 0 |
| 0 | 0 | 0 | 1 |
| 0 | 0 | 0 | 0 |
| 0 | 0 | 0 | 1 |
| 0 | 0 | 0 | 1 |
| 0 | 0 | 0 | 1 |
| 0 | 0 | 0 | 0 |
| 0 | 0 | 0 | 1 |
| 0 | 0 | 0 | 1 |
| 0 | 0 | 0 | 0 |
| 0 | 0 | 0 | 0 |
| 0 | 0 | 0 | 0 |
| 0 | 1 | 0 | 0 |
| 0 | 0 | 0 | 0 |
| 0 | 1 | 1 | 1 |
| 0 | 1 | 1 | 1 |
| 0 | 0 | 0 | 0 |
| 0 | 0 | 0 | 1 |
| 0 | 0 | 0 | 0 |
| 0 | 0 | 0 | 1 |
| 0 | 1 | 1 | 1 |
| 0 | 0 | 0 | 1 |
| 0 | 1 | 0 | 0 |
| 0 | 0 | 0 | 0 |
| 0 | 1 | 0 | 0 |
| 0 | 0 | 0 | 0 |
| 0 | 0 | 0 | 1 |
| 0 | 0 | 0 | 0 |
| 0 | 0 | 0 | 1 |

| | | | |
|---|---|---|---|
| 0 | 0 | 0 | 1 |
| 0 | 0 | 0 | 1 |
| 0 | 0 | 0 | 1 |
| 0 | 0 | 0 | 1 |
| 0 | 0 | 0 | 1 |
| 0 | 0 | 0 | 0 |
| 0 | 0 | 0 | 0 |
| 0 | 0 | 0 | 0 |
| 0 | 0 | 0 | 0 |
| 0 | 0 | 0 | 1 |
| 1 | 0 | 0 | 0 |
| 1 | 0 | 0 | 0 |
| 0 | 0 | 0 | 0 |
| 0 | 0 | 0 | 0 |
| 0 | 0 | 0 | 0 |
| 0 | 0 | 0 | 0 |
| 0 | 0 | 0 | 1 |
| 0 | 0 | 0 | 1 |
| 0 | 0 | 0 | 1 |
| 0 | 0 | 0 | 1 |
| 0 | 0 | 0 | 1 |
| 0 | 0 | 0 | 1 |
| 0 | 0 | 0 | 1 |
| 0 | 0 | 0 | 1 |
| 0 | 0 | 0 | 1 |
| 0 | 0 | 0 | 1 |
| 0 | 0 | 0 | 1 |
| 0 | 0 | 0 | 1 |
| 0 | 1 | 0 | 0 |
| 0 | 0 | 0 | 1 |
| 0 | 0 | 0 | 1 |
| 0 | 1 | 1 | 1 |
| 0 | 1 | 1 | 1 |
| 0 | 0 | 0 | 0 |
| 0 | 0 | 0 | 1 |
| 0 | 1 | 0 | 0 |
| 0 | 0 | 0 | 1 |
| 0 | 0 | 0 | 0 |
| 0 | 0 | 0 | 1 |
| 0 | 0 | 0 | 0 |
| 0 | 0 | 0 | 1 |
| 0 | 0 | 0 | 1 |
| 0 | 0 | 0 | 1 |
| 0 | 0 | 0 | 0 |
| 0 | 0 | 0 | 1 |
| 0 | 0 | 0 | 1 |
| 0 | 0 | 0 | 1 |
| 0 | 0 | 0 | 1 |
| 0 | 0 | 0 | 1 |
| 0 | 0 | 0 | 1 |
| 0 | 0 | 0 | 1 |
| 0 | 1 | 1 | 1 |
| 0 | 0 | 0 | 0 |
| 0 | 0 | 0 | 0 |

| | | | |
|---|---|---|---|
| 0 | 0 | 0 | 1 |
| 0 | 0 | 0 | 0 |
| 0 | 0 | 0 | 1 |
| 0 | 0 | 0 | 1 |
| 0 | 1 | 0 | 0 |
| 0 | 0 | 0 | 0 |
| 0 | 0 | 0 | 0 |
| 0 | 0 | 0 | 1 |
| 0 | 0 | 0 | 1 |
| 0 | 0 | 0 | 0 |
| 0 | 0 | 0 | 0 |
| 0 | 0 | 0 | 0 |
| 0 | 0 | 0 | 0 |
| 0 | 0 | 0 | 0 |
| 0 | 0 | 0 | 1 |
| 0 | 0 | 0 | 0 |
| 0 | 1 | 0 | 0 |
| 0 | 0 | 0 | 0 |
| 0 | 0 | 0 | 1 |
| 0 | 0 | 0 | 1 |
| 0 | 0 | 0 | 1 |
| 0 | 0 | 0 | 1 |
| 0 | 0 | 0 | 1 |
| 0 | 0 | 0 | 0 |
| 0 | 0 | 0 | 1 |
| 0 | 0 | 0 | 1 |
| 0 | 0 | 0 | 0 |
| 0 | 0 | 0 | 0 |
| 0 | 0 | 0 | 1 |
| 0 | 0 | 0 | 1 |
| 0 | 0 | 0 | 0 |
| 0 | 0 | 0 | 1 |
| 0 | 0 | 0 | 1 |
| 0 | 0 | 0 | 1 |
| 0 | 0 | 0 | 1 |
| 0 | 0 | 0 | 0 |
| 0 | 0 | 0 | 1 |
| 0 | 0 | 0 | 0 |
| 0 | 0 | 0 | 0 |
| 0 | 0 | 0 | 1 |
| 0 | 0 | 0 | 1 |
| 0 | 0 | 0 | 0 |
| 0 | 0 | 0 | 0 |
| 0 | 0 | 0 | 1 |
| 0 | 0 | 0 | 0 |
| 0 | 0 | 0 | 0 |
| 0 | 0 | 0 | 0 |
| 0 | 0 | 0 | 1 |
| 0 | 0 | 0 | 0 |
| 0 | 0 | 0 | 0 |
| 0 | 0 | 0 | 0 |
| 0 | 0 | 0 | 1 |
| 0 | 0 | 0 | 0 |
| 0 | 0 | 0 | 1 |

| | | | |
|---|---|---|---|
| 0 | 0 | 0 | 1 |
| 0 | 0 | 0 | 1 |
| 0 | 0 | 0 | 1 |
| 0 | 0 | 0 | 1 |
| 0 | 0 | 0 | 0 |
| 0 | 0 | 0 | 0 |
| 0 | 0 | 0 | 1 |
| 0 | 0 | 0 | 1 |
| 0 | 0 | 0 | 1 |
| 0 | 1 | 0 | 0 |
| 0 | 1 | 1 | 1 |
| 0 | 0 | 0 | 1 |
| 0 | 1 | 0 | 0 |
| 0 | 0 | 0 | 1 |
| 0 | 0 | 0 | 1 |
| 0 | 0 | 0 | 1 |
| 0 | 0 | 0 | 1 |
| 0 | 0 | 0 | 1 |
| 0 | 0 | 0 | 1 |
| 0 | 0 | 0 | 1 |
| 0 | 0 | 0 | 1 |
| 0 | 0 | 0 | 1 |
| 0 | 0 | 0 | 1 |
| 0 | 0 | 0 | 0 |
| 0 | 0 | 0 | 1 |
| 1 | 0 | 0 | 0 |
| 0 | 0 | 0 | 1 |
| 0 | 0 | 0 | 0 |
| 0 | 0 | 0 | 1 |
| 0 | 0 | 0 | 0 |
| 0 | 0 | 0 | 0 |
| 0 | 0 | 0 | 0 |
| 0 | 0 | 0 | 0 |
| 0 | 0 | 0 | 0 |
| 0 | 0 | 0 | 1 |
| 0 | 0 | 0 | 0 |
| 0 | 0 | 0 | 1 |
| 0 | 0 | 0 | 0 |
| 0 | 0 | 0 | 1 |
| 0 | 0 | 0 | 1 |
| 0 | 0 | 0 | 0 |
| 0 | 0 | 0 | 0 |
| 0 | 0 | 0 | 0 |
| 0 | 0 | 0 | 0 |
| 0 | 0 | 0 | 0 |
| 0 | 0 | 0 | 0 |
| 0 | 0 | 0 | 0 |
| 0 | 1 | 0 | 0 |
| 0 | 0 | 0 | 1 |
| 0 | 0 | 0 | 0 |
| 0 | 1 | 1 | 1 |
| 0 | 0 | 0 | 1 |
| 0 | 0 | 0 | 1 |
| 0 | 0 | 0 | 1 |

| | | | |
|---|---|---|---|
| 0 | 0 | 0 | 1 |
| 0 | 0 | 0 | 1 |
| 0 | 0 | 0 | 1 |
| 0 | 0 | 0 | 1 |
| 0 | 0 | 0 | 1 |
| 0 | 0 | 0 | 1 |
| 0 | 0 | 0 | 0 |
| 0 | 0 | 0 | 0 |
| 0 | 0 | 0 | 0 |
| 0 | 0 | 0 | 0 |
| 0 | 0 | 0 | 0 |
| 0 | 0 | 0 | 0 |
| 0 | 0 | 0 | 0 |
| 0 | 0 | 0 | 0 |
| 0 | 0 | 0 | 0 |
| 0 | 0 | 0 | 0 |
| 1 | 0 | 0 | 0 |
| 0 | 0 | 0 | 0 |
| 0 | 0 | 0 | 1 |
| 0 | 0 | 0 | 0 |
| 0 | 0 | 0 | 1 |
| 0 | 0 | 0 | 1 |
| 0 | 0 | 0 | 1 |
| 0 | 0 | 0 | 1 |
| 0 | 0 | 0 | 1 |
| 0 | 0 | 0 | 0 |
| 0 | 0 | 0 | 0 |
| 0 | 0 | 0 | 0 |
| 0 | 0 | 0 | 0 |
| 0 | 0 | 0 | 0 |
| 0 | 0 | 0 | 0 |
| 1 | 0 | 0 | 0 |
| 0 | 0 | 0 | 0 |
| 1 | 0 | 0 | 0 |
| 0 | 0 | 0 | 1 |
| 1 | 0 | 0 | 0 |
| 1 | 0 | 0 | 0 |
| 0 | 0 | 0 | 0 |
| 1 | 0 | 0 | 0 |
| 0 | 0 | 0 | 1 |
| 0 | 0 | 0 | 0 |
| 0 | 0 | 0 | 0 |
| 0 | 0 | 0 | 1 |
| 0 | 0 | 0 | 1 |
| 0 | 0 | 0 | 1 |
| 0 | 0 | 0 | 1 |
| 0 | 0 | 0 | 1 |
| 0 | 0 | 0 | 0 |
| 0 | 0 | 0 | 0 |
| 0 | 0 | 0 | 0 |
| 0 | 0 | 0 | 0 |
| 0 | 0 | 0 | 1 |
| 0 | 0 | 0 | 0 |
| 0 | 0 | 0 | 1 |

| | | | |
|---|---|---|---|
| 0 | 0 | 0 | 1 |
| 0 | 0 | 0 | 1 |
| 0 | 0 | 0 | 1 |
| 0 | 0 | 0 | 1 |
| 0 | 0 | 0 | 1 |
| 0 | 0 | 0 | 0 |
| 0 | 0 | 0 | 1 |
| 0 | 0 | 0 | 1 |
| 0 | 0 | 0 | 1 |
| 0 | 0 | 0 | 1 |
| 0 | 1 | 0 | 0 |
| 0 | 1 | 0 | 0 |
| 0 | 1 | 0 | 0 |
| 0 | 1 | 1 | 1 |
| 0 | 1 | 1 | 1 |
| 0 | 0 | 0 | 1 |
| 0 | 1 | 1 | 1 |
| 0 | 0 | 0 | 0 |
| 0 | 0 | 0 | 1 |
| 0 | 0 | 0 | 1 |
| 0 | 0 | 0 | 1 |
| 0 | 0 | 0 | 1 |
| 0 | 0 | 0 | 0 |
| 0 | 0 | 0 | 1 |
| 0 | 0 | 0 | 0 |
| 0 | 0 | 0 | 0 |
| 0 | 0 | 0 | 0 |
| 0 | 0 | 0 | 0 |
| 0 | 0 | 0 | 1 |
| 0 | 0 | 0 | 0 |
| 0 | 0 | 0 | 1 |
| 0 | 0 | 0 | 1 |
| 0 | 0 | 0 | 1 |
| 0 | 0 | 0 | 1 |
| 0 | 0 | 0 | 1 |
| 0 | 0 | 0 | 1 |
| 0 | 0 | 0 | 1 |
| 0 | 0 | 0 | 0 |
| 0 | 0 | 0 | 0 |
| 0 | 0 | 0 | 0 |
| 0 | 1 | 0 | 0 |
| 0 | 0 | 0 | 0 |
| 0 | 0 | 0 | 1 |
| 0 | 0 | 0 | 0 |
| 0 | 0 | 0 | 0 |
| 0 | 0 | 0 | 1 |
| 0 | 0 | 0 | 0 |
| 0 | 1 | 0 | 0 |
| 0 | 0 | 0 | 0 |
| 1 | 0 | 0 | 0 |
| 0 | 0 | 0 | 1 |
| 0 | 0 | 0 | 1 |
| 0 | 0 | 0 | 1 |
| 0 | 1 | 0 | 0 |

| | | | |
|---|---|---|---|
| 0 | 0 | 0 | 0 |
| 0 | 0 | 0 | 1 |
| 0 | 0 | 0 | 0 |
| 0 | 0 | 0 | 1 |
| 0 | 0 | 0 | 1 |
| 0 | 0 | 0 | 1 |
| 0 | 0 | 0 | 1 |
| 0 | 0 | 0 | 1 |
| 0 | 1 | 0 | 0 |
| 0 | 0 | 0 | 1 |
| 0 | 0 | 0 | 0 |
| 0 | 0 | 0 | 0 |
| 0 | 0 | 0 | 0 |
| 0 | 0 | 0 | 0 |
| 0 | 0 | 0 | 1 |
| 0 | 0 | 0 | 1 |
| 0 | 0 | 0 | 0 |
| 0 | 0 | 0 | 0 |
| 0 | 0 | 0 | 1 |
| 0 | 0 | 0 | 1 |
| 0 | 0 | 0 | 0 |
| 0 | 0 | 0 | 1 |
| 0 | 0 | 0 | 1 |
| 0 | 0 | 0 | 1 |
| 0 | 0 | 0 | 0 |
| 0 | 0 | 0 | 1 |
| 0 | 1 | 0 | 0 |
| 0 | 1 | 0 | 0 |
| 0 | 1 | 0 | 0 |
| 0 | 0 | 0 | 1 |
| 0 | 0 | 0 | 0 |
| 0 | 0 | 0 | 0 |
| 0 | 0 | 0 | 0 |
| 0 | 0 | 0 | 1 |
| 0 | 0 | 0 | 0 |
| 0 | 0 | 0 | 1 |
| 0 | 0 | 0 | 0 |
| 0 | 0 | 0 | 0 |
| 0 | 1 | 0 | 0 |
| 0 | 0 | 0 | 0 |
| 0 | 0 | 0 | 0 |
| 0 | 0 | 0 | 1 |
| 0 | 1 | 0 | 0 |
| 0 | 1 | 0 | 0 |
| 0 | 1 | 0 | 0 |
| 1 | 0 | 0 | 0 |
| 0 | 0 | 0 | 0 |
| 1 | 0 | 0 | 0 |
| 1 | 0 | 0 | 0 |
| 1 | 0 | 0 | 0 |
| 0 | 0 | 0 | 0 |
| 0 | 0 | 0 | 0 |
| 0 | 0 | 0 | 1 |
| 0 | 0 | 0 | 0 |

| | | | |
|---|---|---|---|
| 0 | 0 | 0 | 1 |
| 0 | 0 | 0 | 1 |
| 0 | 0 | 0 | 1 |
| 0 | 0 | 0 | 1 |
| 0 | 0 | 0 | 1 |
| 0 | 0 | 0 | 1 |
| 0 | 0 | 0 | 1 |
| 0 | 0 | 0 | 1 |
| 0 | 0 | 0 | 1 |
| 0 | 0 | 0 | 0 |
| 0 | 0 | 0 | 0 |
| 0 | 0 | 0 | 1 |
| 0 | 0 | 0 | 1 |
| 0 | 0 | 0 | 1 |
| 0 | 0 | 0 | 0 |
| 0 | 0 | 0 | 1 |
| 0 | 0 | 0 | 1 |
| 0 | 0 | 0 | 0 |
| 0 | 0 | 0 | 1 |
| 0 | 0 | 0 | 1 |
| 0 | 0 | 0 | 1 |
| 0 | 0 | 0 | 0 |
| 0 | 0 | 0 | 0 |
| 0 | 0 | 0 | 0 |
| 0 | 1 | 0 | 0 |
| 0 | 1 | 0 | 0 |
| 0 | 1 | 0 | 0 |
| 0 | 0 | 0 | 0 |
| 0 | 0 | 0 | 1 |
| 0 | 0 | 0 | 0 |
| 1 | 0 | 0 | 0 |
| 0 | 0 | 0 | 1 |
| 0 | 0 | 0 | 1 |
| 0 | 0 | 0 | 1 |
| 0 | 0 | 0 | 1 |
| 0 | 0 | 0 | 1 |
| 0 | 0 | 0 | 1 |
| 0 | 0 | 0 | 1 |
| 0 | 0 | 0 | 1 |
| 0 | 0 | 0 | 1 |
| 0 | 0 | 0 | 1 |
| 0 | 0 | 0 | 1 |
| 0 | 0 | 0 | 1 |
| 0 | 0 | 0 | 0 |
| 0 | 0 | 0 | 1 |
| 0 | 0 | 0 | 0 |
| 0 | 0 | 0 | 1 |
| 0 | 0 | 0 | 1 |
| 0 | 0 | 0 | 1 |
| 0 | 0 | 0 | 1 |
| 0 | 0 | 0 | 0 |
| 0 | 0 | 0 | 1 |
| 0 | 0 | 0 | 1 |
| 0 | 0 | 0 | 0 |

| | | | |
|---|---|---|---|
| 0 | 0 | 0 | 0 |
| 0 | 0 | 0 | 0 |
| 0 | 0 | 0 | 0 |
| 0 | 0 | 0 | 0 |
| 0 | 0 | 0 | 1 |
| 0 | 0 | 0 | 1 |
| 0 | 0 | 0 | 1 |
| 0 | 0 | 0 | 0 |
| 0 | 0 | 0 | 0 |
| 0 | 0 | 0 | 0 |
| 0 | 0 | 0 | 0 |
| 0 | 0 | 0 | 1 |
| 0 | 0 | 0 | 1 |
| 0 | 0 | 0 | 1 |
| 0 | 0 | 0 | 1 |
| 0 | 0 | 0 | 0 |
| 0 | 0 | 0 | 0 |
| 0 | 0 | 0 | 1 |
| 0 | 0 | 0 | 1 |
| 0 | 0 | 0 | 1 |
| 0 | 0 | 0 | 1 |
| 0 | 1 | 0 | 0 |
| 0 | 0 | 0 | 1 |
| 0 | 0 | 0 | 1 |
| 0 | 1 | 0 | 0 |
| 0 | 0 | 0 | 0 |
| 0 | 0 | 0 | 0 |
| 0 | 0 | 0 | 0 |
| 0 | 0 | 0 | 1 |
| 0 | 0 | 0 | 1 |
| 0 | 0 | 0 | 1 |
| 0 | 0 | 0 | 0 |
| 0 | 0 | 0 | 1 |
| 0 | 0 | 0 | 1 |
| 0 | 0 | 0 | 1 |
| 0 | 0 | 0 | 1 |
| 0 | 0 | 0 | 1 |
| 0 | 0 | 0 | 1 |
| 0 | 0 | 0 | 1 |
| 0 | 0 | 0 | 1 |
| 0 | 0 | 0 | 1 |
| 0 | 0 | 0 | 1 |
| 0 | 0 | 0 | 0 |
| 0 | 0 | 0 | 1 |
| 0 | 0 | 0 | 1 |
| 0 | 0 | 0 | 1 |
| 0 | 0 | 0 | 1 |
| 0 | 0 | 0 | 1 |
| 0 | 0 | 0 | 1 |
| 0 | 0 | 0 | 0 |
| 0 | 0 | 0 | 1 |
| 0 | 0 | 0 | 1 |
| 0 | 0 | 0 | 1 |
| 0 | 0 | 0 | 1 |

| | | | |
|---|---|---|---|
| 0 | 0 | 0 | 0 |
| 0 | 0 | 0 | 1 |
| 0 | 0 | 0 | 1 |
| 0 | 0 | 0 | 1 |
| 0 | 0 | 0 | 1 |
| 0 | 0 | 0 | 1 |
| 0 | 0 | 0 | 1 |
| 0 | 1 | 0 | 0 |
| 0 | 0 | 0 | 1 |
| 0 | 0 | 0 | 1 |
| 0 | 0 | 0 | 0 |
| 0 | 0 | 0 | 0 |
| 0 | 0 | 0 | 0 |
| 0 | 0 | 0 | 1 |
| 0 | 0 | 0 | 0 |
| 0 | 0 | 0 | 0 |
| 0 | 0 | 0 | 0 |
| 0 | 0 | 0 | 0 |
| 0 | 0 | 0 | 1 |
| 0 | 0 | 0 | 0 |
| 0 | 0 | 0 | 0 |
| 0 | 0 | 0 | 0 |
| 0 | 0 | 0 | 0 |
| 0 | 0 | 0 | 0 |
| 0 | 0 | 0 | 0 |
| 0 | 0 | 0 | 0 |
| 0 | 0 | 0 | 0 |
| 0 | 0 | 0 | 0 |
| 0 | 0 | 0 | 0 |
| 0 | 0 | 0 | 0 |
| 0 | 0 | 0 | 0 |
| 0 | 0 | 0 | 0 |
| 0 | 0 | 0 | 1 |
| 0 | 0 | 0 | 0 |
| 0 | 0 | 0 | 1 |
| 0 | 0 | 0 | 0 |
| 0 | 0 | 0 | 0 |
| 0 | 0 | 0 | 1 |
| 0 | 0 | 0 | 1 |
| 0 | 0 | 0 | 1 |
| 0 | 0 | 0 | 1 |
| 0 | 0 | 0 | 0 |
| 0 | 0 | 0 | 0 |
| 0 | 0 | 0 | 0 |
| 0 | 0 | 0 | 1 |
| 0 | 0 | 0 | 0 |
| 0 | 0 | 0 | 1 |
| 0 | 0 | 0 | 0 |
| 0 | 0 | 0 | 1 |
| 0 | 0 | 0 | 1 |
| 0 | 0 | 0 | 1 |
| 0 | 0 | 0 | 1 |
| 0 | 0 | 0 | 1 |
| 0 | 0 | 0 | 1 |

| | | | |
|---|---|---|---|
| 0 | 0 | 0 | 1 |
| 0 | 0 | 0 | 1 |
| 0 | 0 | 0 | 1 |
| 0 | 0 | 0 | 1 |
| 0 | 0 | 0 | 1 |
| 0 | 0 | 0 | 0 |
| 0 | 0 | 0 | 0 |
| 0 | 0 | 0 | 1 |
| 0 | 0 | 0 | 0 |
| 0 | 0 | 0 | 0 |
| 0 | 0 | 0 | 1 |
| 0 | 0 | 0 | 1 |
| 0 | 0 | 0 | 1 |
| 0 | 0 | 0 | 1 |
| 0 | 0 | 0 | 1 |
| 0 | 1 | 1 | 1 |
| 0 | 0 | 0 | 1 |
| 0 | 0 | 0 | 1 |
| 0 | 0 | 0 | 0 |
| 0 | 0 | 0 | 0 |
| 0 | 0 | 0 | 1 |
| 0 | 0 | 0 | 1 |
| 0 | 0 | 0 | 1 |
| 0 | 0 | 0 | 0 |
| 0 | 0 | 0 | 1 |
| 0 | 0 | 0 | 0 |
| 0 | 0 | 0 | 0 |
| 0 | 0 | 0 | 1 |
| 0 | 0 | 0 | 1 |
| 0 | 0 | 0 | 1 |
| 0 | 1 | 0 | 0 |
| 0 | 0 | 0 | 1 |
| 0 | 0 | 0 | 0 |
| 0 | 1 | 0 | 0 |
| 0 | 1 | 0 | 0 |
| 0 | 1 | 0 | 0 |
| 0 | 0 | 0 | 0 |
| 0 | 0 | 0 | 1 |
| 0 | 0 | 0 | 1 |
| 0 | 0 | 0 | 1 |
| 0 | 0 | 0 | 0 |
| 0 | 0 | 0 | 0 |
| 0 | 0 | 0 | 0 |
| 0 | 0 | 0 | 0 |
| 0 | 0 | 0 | 1 |
| 0 | 0 | 0 | 1 |
| 0 | 0 | 0 | 1 |
| 0 | 0 | 0 | 0 |
| 0 | 0 | 0 | 0 |
| 0 | 0 | 0 | 0 |
| 0 | 0 | 0 | 1 |
| 0 | 0 | 0 | 1 |
| 0 | 0 | 0 | 1 |
| 0 | 0 | 0 | 1 |

| | | | |
|---|---|---|---|
| 0 | 0 | 0 | 1 |
| 0 | 0 | 0 | 1 |
| 0 | 0 | 0 | 1 |
| 0 | 0 | 0 | 0 |
| 0 | 0 | 0 | 0 |
| 0 | 0 | 0 | 1 |
| 0 | 0 | 0 | 0 |
| 0 | 0 | 0 | 1 |
| 0 | 0 | 0 | 1 |
| 0 | 0 | 0 | 1 |
| 0 | 0 | 0 | 1 |
| 0 | 0 | 0 | 0 |
| 0 | 0 | 0 | 1 |
| 0 | 0 | 0 | 0 |
| 0 | 0 | 0 | 1 |
| 0 | 0 | 0 | 0 |
| 0 | 0 | 0 | 1 |
| 0 | 0 | 0 | 1 |
| 0 | 0 | 0 | 1 |
| 0 | 0 | 0 | 0 |
| 0 | 0 | 0 | 0 |
| 0 | 0 | 0 | 0 |
| 0 | 0 | 0 | 0 |
| 0 | 0 | 0 | 0 |
| 0 | 0 | 0 | 0 |
| 0 | 0 | 0 | 1 |
| 0 | 0 | 0 | 0 |
| 0 | 0 | 0 | 1 |
| 0 | 0 | 0 | 1 |
| 0 | 0 | 0 | 1 |
| 0 | 0 | 0 | 1 |
| 0 | 0 | 0 | 1 |
| 0 | 0 | 0 | 0 |
| 0 | 1 | 0 | 0 |
| 0 | 0 | 0 | 1 |
| 0 | 0 | 0 | 1 |
| 0 | 1 | 0 | 0 |
| 0 | 1 | 0 | 0 |
| 0 | 1 | 0 | 0 |
| 0 | 1 | 1 | 1 |
| 0 | 0 | 0 | 0 |
| 0 | 0 | 0 | 0 |
| 0 | 0 | 0 | 1 |
| 0 | 0 | 0 | 0 |
| 0 | 0 | 0 | 1 |
| 0 | 0 | 0 | 1 |
| 0 | 0 | 0 | 1 |
| 0 | 0 | 0 | 1 |
| 0 | 0 | 0 | 1 |
| 1 | 0 | 0 | 0 |
| 0 | 0 | 0 | 1 |
| 0 | 0 | 0 | 1 |
| 0 | 0 | 0 | 1 |
| 0 | 0 | 0 | 1 |

| | | | |
|---|---|---|---|
| 0 | 0 | 0 | 1 |
| 0 | 0 | 0 | 1 |
| 0 | 0 | 0 | 1 |
| 0 | 0 | 0 | 0 |
| 0 | 0 | 0 | 1 |
| 0 | 0 | 0 | 1 |
| 0 | 0 | 0 | 1 |
| 0 | 0 | 0 | 0 |
| 0 | 0 | 0 | 1 |
| 0 | 0 | 0 | 1 |
| 0 | 0 | 0 | 1 |
| 0 | 0 | 0 | 1 |
| 0 | 0 | 0 | 1 |
| 0 | 0 | 0 | 1 |
| 0 | 0 | 0 | 1 |
| 0 | 0 | 0 | 1 |
| 0 | 0 | 0 | 0 |
| 0 | 0 | 0 | 0 |
| 0 | 0 | 0 | 1 |
| 0 | 0 | 0 | 1 |
| 0 | 0 | 0 | 1 |
| 0 | 0 | 0 | 1 |
| 0 | 0 | 0 | 1 |
| 0 | 0 | 0 | 1 |
| 0 | 0 | 0 | 1 |
| 0 | 0 | 0 | 1 |
| 0 | 0 | 0 | 1 |
| 0 | 1 | 0 | 0 |
| 0 | 0 | 0 | 0 |
| 0 | 0 | 0 | 1 |
| 0 | 0 | 0 | 1 |
| 0 | 0 | 0 | 0 |
| 0 | 0 | 0 | 1 |
| 0 | 0 | 0 | 0 |
| 1 | 0 | 0 | 0 |
| 1 | 0 | 0 | 0 |
| 1 | 0 | 0 | 0 |
| 1 | 0 | 0 | 0 |
| 0 | 0 | 0 | 1 |
| 0 | 0 | 0 | 1 |
| 0 | 0 | 0 | 1 |
| 0 | 0 | 0 | 1 |
| 0 | 0 | 0 | 1 |
| 0 | 0 | 0 | 1 |
| 0 | 0 | 0 | 0 |
| 0 | 0 | 0 | 0 |
| 0 | 0 | 0 | 0 |
| 0 | 0 | 0 | 1 |
| 0 | 0 | 0 | 1 |
| 0 | 0 | 0 | 1 |
| 0 | 0 | 0 | 1 |
| 0 | 0 | 0 | 0 |
| 0 | 0 | 0 | 0 |
| 0 | 0 | 0 | 0 |

| | | | |
|---|---|---|---|
| 0 | 0 | 0 | 0 |
| 0 | 0 | 0 | 0 |
| 0 | 0 | 0 | 1 |
| 0 | 0 | 0 | 0 |
| 0 | 0 | 0 | 0 |
| 0 | 0 | 0 | 0 |
| 0 | 0 | 0 | 0 |
| 0 | 0 | 0 | 1 |
| 0 | 0 | 0 | 1 |
| 0 | 0 | 0 | 1 |
| 0 | 0 | 0 | 1 |
| 0 | 0 | 0 | 1 |
| 0 | 0 | 0 | 1 |
| 0 | 0 | 0 | 1 |
| 0 | 0 | 0 | 1 |
| 0 | 0 | 0 | 1 |
| 0 | 0 | 0 | 1 |
| 0 | 0 | 0 | 1 |
| 0 | 0 | 0 | 1 |
| 0 | 0 | 0 | 1 |
| 0 | 0 | 0 | 0 |
| 0 | 0 | 0 | 0 |
| 0 | 0 | 0 | 0 |
| 0 | 0 | 0 | 0 |
| 0 | 0 | 0 | 0 |
| 0 | 0 | 0 | 1 |
| 0 | 0 | 0 | 1 |
| 0 | 0 | 0 | 0 |
| 0 | 0 | 0 | 1 |
| 0 | 0 | 0 | 1 |
| 0 | 0 | 0 | 0 |
| 0 | 0 | 0 | 0 |
| 0 | 0 | 0 | 0 |
| 0 | 0 | 0 | 1 |
| 0 | 0 | 0 | 0 |
| 0 | 0 | 0 | 1 |
| 0 | 0 | 0 | 1 |
| 0 | 0 | 0 | 0 |
| 0 | 0 | 0 | 1 |
| 0 | 0 | 0 | 1 |
| 0 | 0 | 0 | 1 |
| 0 | 0 | 0 | 1 |
| 0 | 0 | 0 | 1 |
| 0 | 0 | 0 | 1 |
| 0 | 0 | 0 | 1 |
| 0 | 0 | 0 | 1 |
| 0 | 0 | 0 | 1 |
| 0 | 0 | 0 | 1 |
| 0 | 0 | 0 | 0 |
| 0 | 0 | 0 | 1 |
| 1 | 0 | 0 | 0 |
| 1 | 0 | 0 | 0 |
| 0 | 0 | 0 | 1 |
| 0 | 0 | 0 | 1 |

| | | | |
|---|---|---|---|
| 1 | 0 | 0 | 0 |
| 0 | 0 | 0 | 1 |
| 0 | 0 | 0 | 1 |
| 0 | 0 | 0 | 1 |
| 0 | 0 | 0 | 1 |
| 1 | 0 | 0 | 0 |
| 1 | 0 | 0 | 0 |
| 0 | 0 | 0 | 1 |
| 0 | 0 | 0 | 1 |
| 0 | 0 | 0 | 1 |
| 0 | 0 | 0 | 1 |
| 0 | 0 | 0 | 0 |
| 0 | 0 | 0 | 1 |
| 0 | 0 | 0 | 1 |
| 0 | 0 | 0 | 1 |
| 0 | 0 | 0 | 1 |
| 0 | 0 | 0 | 0 |
| 0 | 0 | 0 | 0 |
| 0 | 0 | 0 | 1 |
| 0 | 0 | 0 | 1 |
| 0 | 0 | 0 | 1 |
| 0 | 0 | 0 | 1 |
| 0 | 0 | 0 | 1 |
| 0 | 0 | 0 | 1 |
| 0 | 0 | 0 | 0 |
| 0 | 0 | 0 | 1 |
| 0 | 0 | 0 | 1 |
| 0 | 0 | 0 | 1 |
| 0 | 0 | 0 | 1 |
| 0 | 0 | 0 | 1 |
| 0 | 0 | 0 | 1 |
| 0 | 0 | 0 | 0 |
| 0 | 0 | 0 | 0 |
| 0 | 0 | 0 | 1 |
| 0 | 0 | 0 | 1 |
| 0 | 0 | 0 | 0 |
| 1 | 0 | 0 | 0 |
| 0 | 1 | 0 | 0 |
| 1 | 1 | 0 | 0 |
| 0 | 0 | 0 | 0 |
| 0 | 0 | 0 | 0 |
| 0 | 0 | 0 | 0 |
| 0 | 0 | 0 | 0 |
| 0 | 0 | 0 | 0 |
| 0 | 0 | 0 | 0 |
| 0 | 0 | 0 | 0 |
| 0 | 0 | 0 | 0 |
| 1 | 0 | 0 | 0 |
| 0 | 0 | 0 | 1 |
| 0 | 0 | 0 | 1 |
| 0 | 1 | 1 | 1 |
| 0 | 0 | 0 | 1 |
| 0 | 0 | 0 | 0 |
| 0 | 0 | 0 | 1 |

| | | | |
|---|---|---|---|
| 0 | 0 | 0 | 1 |
| 0 | 0 | 0 | 1 |
| 0 | 0 | 0 | 1 |
| 0 | 0 | 0 | 1 |
| 0 | 0 | 0 | 1 |
| 0 | 0 | 0 | 1 |
| 0 | 0 | 0 | 1 |
| 0 | 1 | 0 | 0 |
| 0 | 0 | 0 | 0 |
| 0 | 0 | 0 | 1 |
| 0 | 0 | 0 | 1 |
| 0 | 0 | 0 | 1 |
| 0 | 0 | 0 | 1 |
| 0 | 1 | 0 | 0 |
| 0 | 0 | 0 | 1 |
| 0 | 0 | 0 | 1 |
| 0 | 1 | 0 | 0 |
| 0 | 0 | 0 | 1 |
| 1 | 0 | 0 | 0 |
| 0 | 0 | 0 | 1 |
| 0 | 0 | 0 | 1 |
| 0 | 0 | 0 | 1 |
| 0 | 0 | 0 | 1 |
| 0 | 0 | 0 | 0 |
| 0 | 0 | 0 | 1 |
| 0 | 0 | 0 | 1 |
| 0 | 0 | 0 | 1 |
| 0 | 0 | 0 | 1 |
| 0 | 0 | 0 | 1 |
| 0 | 0 | 0 | 1 |
| 0 | 0 | 0 | 1 |
| 1 | 0 | 0 | 0 |
| 0 | 0 | 0 | 1 |
| 0 | 0 | 0 | 1 |
| 0 | 0 | 0 | 1 |
| 0 | 0 | 0 | 1 |
| 0 | 0 | 0 | 1 |
| 0 | 0 | 0 | 1 |
| 1 | 0 | 0 | 0 |
| 0 | 0 | 0 | 1 |
| 0 | 0 | 0 | 1 |
| 0 | 0 | 0 | 1 |
| 0 | 0 | 0 | 1 |
| 0 | 0 | 0 | 1 |
| 0 | 0 | 0 | 1 |
| 0 | 0 | 0 | 0 |
| 0 | 0 | 0 | 1 |
| 0 | 0 | 0 | 1 |
| 0 | 1 | 1 | 1 |
| 0 | 1 | 1 | 1 |
| 0 | 0 | 0 | 1 |
| 0 | 0 | 0 | 1 |
| 0 | 0 | 0 | 1 |
| 0 | 1 | 1 | 1 |

| | | | |
|---|---|---|---|
| 0 | 1 | 1 | 1 |
| 0 | 1 | 1 | 1 |
| 0 | 1 | 1 | 1 |
| 0 | 1 | 1 | 1 |
| 0 | 1 | 1 | 1 |
| 0 | 1 | 1 | 1 |
| 0 | 1 | 1 | 1 |
| 0 | 1 | 1 | 1 |
| 0 | 1 | 1 | 1 |
| 0 | 1 | 1 | 1 |
| 0 | 1 | 1 | 1 |
| 0 | 1 | 1 | 1 |
| 0 | 1 | 1 | 1 |
| 0 | 1 | 1 | 1 |
| 0 | 1 | 1 | 1 |
| 1 | 1 | 0 | 0 |
| 0 | 1 | 1 | 1 |
| 0 | 1 | 0 | 0 |
| 0 | 1 | 1 | 1 |
| 0 | 1 | 1 | 1 |
| 1 | 1 | 0 | 0 |
| 0 | 1 | 1 | 1 |
| 0 | 1 | 1 | 1 |
| 0 | 1 | 1 | 1 |
| 0 | 1 | 0 | 0 |
| 0 | 1 | 1 | 1 |
| 0 | 1 | 0 | 0 |
| 0 | 1 | 1 | 1 |
| 0 | 1 | 1 | 1 |
| 0 | 1 | 1 | 1 |
| 0 | 1 | 1 | 1 |
| 0 | 1 | 1 | 1 |
| 0 | 1 | 1 | 1 |
| 0 | 1 | 1 | 1 |
| 0 | 1 | 1 | 1 |
| 0 | 1 | 1 | 1 |
| 0 | 1 | 1 | 1 |
| 0 | 1 | 1 | 1 |
| 0 | 1 | 1 | 1 |
| 0 | 1 | 1 | 1 |
| 0 | 1 | 1 | 1 |
| 0 | 1 | 1 | 1 |
| 0 | 0 | 0 | 1 |
| 0 | 0 | 0 | 0 |
| 0 | 0 | 0 | 1 |
| 0 | 0 | 0 | 1 |
| 0 | 0 | 0 | 0 |
| 0 | 0 | 0 | 0 |
| 0 | 0 | 0 | 1 |
| 0 | 0 | 0 | 0 |
| 0 | 0 | 0 | 0 |
| 0 | 0 | 0 | 0 |
| 0 | 0 | 0 | 1 |
| 0 | 0 | 0 | 1 |

| | | | |
|---|---|---|---|
| 0 | 0 | 0 | 1 |
| 0 | 0 | 0 | 1 |
| 0 | 0 | 0 | 1 |
| 0 | 0 | 0 | 1 |
| 0 | 0 | 0 | 1 |
| 0 | 0 | 0 | 1 |
| 0 | 0 | 0 | 1 |
| 1 | 0 | 0 | 0 |
| 0 | 0 | 0 | 1 |
| 0 | 0 | 0 | 1 |
| 0 | 0 | 0 | 1 |
| 0 | 0 | 0 | 1 |
| 0 | 0 | 0 | 1 |
| 0 | 0 | 0 | 1 |
| 0 | 0 | 0 | 0 |
| 0 | 0 | 0 | 0 |
| 0 | 0 | 0 | 1 |
| 0 | 0 | 0 | 0 |
| 0 | 0 | 0 | 1 |
| 0 | 0 | 0 | 1 |
| 0 | 0 | 0 | 0 |
| 0 | 0 | 0 | 1 |
| 0 | 0 | 0 | 1 |
| 0 | 0 | 0 | 1 |
| 0 | 0 | 0 | 0 |
| 0 | 0 | 0 | 1 |
| 0 | 0 | 0 | 1 |
| 1 | 0 | 0 | 0 |
| 0 | 0 | 0 | 0 |
| 1 | 0 | 0 | 0 |
| 1 | 0 | 0 | 0 |
| 0 | 0 | 0 | 1 |
| 0 | 0 | 0 | 1 |
| 0 | 0 | 0 | 1 |
| 0 | 0 | 0 | 1 |
| 0 | 0 | 0 | 1 |
| 0 | 1 | 0 | 0 |
| 0 | 0 | 0 | 0 |
| 0 | 0 | 0 | 1 |
| 0 | 0 | 0 | 1 |
| 0 | 0 | 0 | 1 |
| 0 | 0 | 0 | 1 |
| 0 | 0 | 0 | 0 |
| 0 | 0 | 0 | 1 |
| 0 | 0 | 0 | 1 |
| 0 | 0 | 0 | 1 |
| 0 | 0 | 0 | 1 |
| 0 | 0 | 0 | 1 |
| 0 | 0 | 0 | 0 |
| 0 | 0 | 0 | 0 |
| 0 | 0 | 0 | 0 |
| 0 | 0 | 0 | 0 |
| 0 | 0 | 0 | 0 |
| 0 | 0 | 0 | 1 |

| | | | |
|---|---|---|---|
| 0 | 0 | 0 | 0 |
| 0 | 0 | 0 | 0 |
| 0 | 0 | 0 | 1 |
| 1 | 0 | 0 | 0 |
| 0 | 0 | 0 | 1 |
| 0 | 0 | 0 | 1 |
| 0 | 0 | 0 | 1 |
| 0 | 0 | 0 | 1 |
| 0 | 0 | 0 | 1 |
| 1 | 0 | 0 | 0 |
| 0 | 0 | 0 | 1 |
| 0 | 0 | 0 | 1 |
| 0 | 0 | 0 | 1 |
| 0 | 0 | 0 | 1 |
| 0 | 0 | 0 | 1 |
| 0 | 0 | 0 | 1 |
| 0 | 0 | 0 | 1 |
| 0 | 0 | 0 | 1 |
| 0 | 0 | 0 | 1 |
| 0 | 0 | 0 | 1 |
| 0 | 0 | 0 | 1 |
| 0 | 0 | 0 | 1 |
| 0 | 0 | 0 | 1 |
| 0 | 0 | 0 | 1 |
| 0 | 0 | 0 | 1 |
| 0 | 0 | 0 | 1 |
| 0 | 0 | 0 | 1 |
| 0 | 0 | 0 | 1 |
| 0 | 0 | 0 | 1 |
| 0 | 0 | 0 | 1 |
| 0 | 0 | 0 | 1 |
| 0 | 0 | 0 | 1 |
| 1 | 0 | 0 | 0 |
| 0 | 0 | 0 | 1 |
| 0 | 0 | 0 | 1 |
| 0 | 0 | 0 | 1 |
| 0 | 0 | 0 | 1 |
| 0 | 0 | 0 | 1 |
| 0 | 0 | 0 | 1 |
| 0 | 0 | 0 | 1 |
| 0 | 0 | 0 | 1 |
| 0 | 0 | 0 | 1 |
| 0 | 0 | 0 | 1 |
| 0 | 0 | 0 | 1 |
| 0 | 0 | 0 | 1 |
| 0 | 0 | 0 | 1 |
| 0 | 0 | 0 | 1 |
| 0 | 0 | 0 | 0 |
| 0 | 1 | 0 | 0 |
| 0 | 1 | 0 | 0 |
| 0 | 0 | 0 | 0 |
| 0 | 1 | 0 | 0 |
| 0 | 0 | 0 | 1 |
| 0 | 0 | 0 | 0 |

| | | | |
|---|---|---|---|
| 0 | 0 | 0 | 1 |
| 0 | 0 | 0 | 0 |
| 0 | 0 | 0 | 1 |
| 0 | 0 | 0 | 1 |
| 0 | 0 | 0 | 1 |
| 1 | 0 | 0 | 0 |
| 0 | 0 | 0 | 0 |
| 0 | 0 | 0 | 0 |
| 1 | 0 | 0 | 0 |
| 0 | 0 | 0 | 0 |
| 0 | 0 | 0 | 1 |
| 0 | 0 | 0 | 0 |
| 0 | 0 | 0 | 1 |
| 0 | 0 | 0 | 1 |
| 0 | 0 | 0 | 1 |
| 1 | 0 | 0 | 0 |
| 0 | 0 | 0 | 1 |
| 0 | 0 | 0 | 1 |
| 0 | 0 | 0 | 0 |
| 0 | 0 | 0 | 0 |
| 1 | 0 | 0 | 0 |
| 1 | 0 | 0 | 0 |
| 0 | 0 | 0 | 0 |
| 0 | 0 | 0 | 0 |
| 0 | 0 | 0 | 1 |
| 0 | 0 | 0 | 0 |
| 0 | 1 | 0 | 0 |
| 0 | 0 | 0 | 0 |
| 0 | 1 | 0 | 0 |
| 0 | 1 | 0 | 0 |
| 0 | 0 | 0 | 0 |
| 0 | 0 | 0 | 1 |
| 0 | 0 | 0 | 1 |
| 0 | 0 | 0 | 0 |
| 0 | 1 | 0 | 0 |
| 0 | 1 | 1 | 1 |
| 0 | 0 | 0 | 0 |
| 0 | 0 | 0 | 0 |
| 0 | 1 | 0 | 0 |
| 0 | 0 | 0 | 0 |
| 0 | 1 | 0 | 0 |
| 0 | 1 | 1 | 1 |
| 0 | 0 | 0 | 1 |
| 0 | 0 | 0 | 1 |
| 0 | 0 | 0 | 0 |
| 0 | 0 | 0 | 0 |
| 0 | 0 | 0 | 0 |
| 0 | 0 | 0 | 1 |
| 0 | 0 | 0 | 1 |
| 0 | 0 | 0 | 1 |
| 0 | 0 | 0 | 1 |
| 0 | 0 | 0 | 1 |
| 0 | 0 | 0 | 1 |
| 0 | 0 | 0 | 0 |

| | | | |
|---|---|---|---|
| 0 | 0 | 0 | 1 |
| 0 | 0 | 0 | 1 |
| 0 | 0 | 0 | 1 |
| 0 | 0 | 0 | 1 |
| 1 | 0 | 0 | 0 |
| 0 | 0 | 0 | 0 |
| 0 | 0 | 0 | 1 |
| 0 | 1 | 1 | 1 |
| 0 | 0 | 0 | 1 |
| 0 | 0 | 0 | 1 |
| 0 | 0 | 0 | 1 |
| 0 | 0 | 0 | 0 |
| 0 | 0 | 0 | 1 |
| 0 | 0 | 0 | 0 |
| 0 | 1 | 0 | 0 |
| 0 | 1 | 0 | 0 |
| 0 | 1 | 0 | 0 |
| 0 | 1 | 0 | 0 |
| 0 | 1 | 0 | 0 |
| 0 | 1 | 0 | 0 |
| 0 | 0 | 0 | 0 |
| 0 | 1 | 0 | 0 |
| 0 | 0 | 0 | 0 |
| 0 | 0 | 0 | 1 |
| 0 | 0 | 0 | 1 |
| 0 | 0 | 0 | 1 |
| 0 | 0 | 0 | 1 |
| 0 | 0 | 0 | 1 |
| 0 | 0 | 0 | 1 |
| 0 | 0 | 0 | 1 |
| 0 | 0 | 0 | 1 |
| 0 | 0 | 0 | 0 |
| 0 | 0 | 0 | 1 |
| 0 | 0 | 0 | 0 |
| 0 | 0 | 0 | 0 |
| 0 | 0 | 0 | 0 |
| 0 | 0 | 0 | 0 |
| 0 | 0 | 0 | 1 |
| 0 | 0 | 0 | 1 |
| 0 | 0 | 0 | 1 |
| 0 | 0 | 0 | 1 |
| 0 | 0 | 0 | 1 |
| 0 | 0 | 0 | 1 |
| 0 | 0 | 0 | 1 |
| 0 | 0 | 0 | 1 |
| 1 | 0 | 0 | 0 |
| 0 | 0 | 0 | 1 |
| 0 | 0 | 0 | 1 |
| 0 | 0 | 0 | 1 |
| 1 | 1 | 0 | 0 |
| 1 | 0 | 0 | 0 |
| 0 | 0 | 0 | 0 |
| 0 | 0 | 0 | 1 |
| 0 | 0 | 0 | 1 |

| | | | |
|---|---|---|---|
| 0 | 0 | 0 | 1 |
| 0 | 0 | 0 | 1 |
| 0 | 0 | 0 | 1 |
| 0 | 0 | 0 | 0 |
| 0 | 0 | 0 | 1 |
| 0 | 0 | 0 | 1 |
| 0 | 0 | 0 | 0 |
| 0 | 0 | 0 | 1 |
| 0 | 0 | 0 | 1 |
| 0 | 0 | 0 | 1 |
| 0 | 0 | 0 | 0 |
| 0 | 0 | 0 | 0 |
| 0 | 0 | 0 | 1 |
| 0 | 0 | 0 | 1 |
| 0 | 0 | 0 | 1 |
| 0 | 0 | 0 | 1 |
| 0 | 0 | 0 | 0 |
| 0 | 0 | 0 | 0 |
| 0 | 0 | 0 | 1 |
| 0 | 0 | 0 | 0 |
| 0 | 1 | 0 | 0 |
| 0 | 1 | 0 | 0 |
| 0 | 0 | 0 | 1 |
| 0 | 1 | 0 | 0 |
| 0 | 0 | 0 | 0 |
| 0 | 0 | 0 | 1 |
| 0 | 0 | 0 | 0 |
| 0 | 0 | 0 | 1 |
| 0 | 0 | 0 | 1 |
| 0 | 0 | 0 | 0 |
| 0 | 0 | 0 | 1 |
| 0 | 0 | 0 | 1 |
| 0 | 0 | 0 | 0 |
| 0 | 0 | 0 | 1 |
| 0 | 0 | 0 | 1 |
| 0 | 0 | 0 | 0 |
| 0 | 0 | 0 | 1 |
| 0 | 0 | 0 | 0 |
| 0 | 0 | 0 | 1 |
| 0 | 0 | 0 | 0 |
| 0 | 0 | 0 | 0 |
| 0 | 0 | 0 | 1 |
| 0 | 0 | 0 | 0 |
| 0 | 0 | 0 | 0 |
| 0 | 0 | 0 | 1 |
| 0 | 0 | 0 | 1 |
| 0 | 0 | 0 | 1 |
| 0 | 0 | 0 | 1 |
| 0 | 0 | 0 | 1 |
| 0 | 0 | 0 | 1 |
| 0 | 0 | 0 | 1 |
| 0 | 0 | 0 | 1 |
| 0 | 0 | 0 | 1 |
| 0 | 0 | 0 | 1 |

| | | | |
|---|---|---|---|
| 0 | 0 | 0 | 1 |
| 0 | 0 | 0 | 1 |
| 0 | 0 | 0 | 1 |
| 0 | 0 | 0 | 0 |
| 0 | 1 | 1 | 1 |
| 0 | 0 | 0 | 0 |
| 0 | 0 | 0 | 1 |
| 0 | 0 | 0 | 1 |
| 0 | 0 | 0 | 0 |
| 0 | 1 | 0 | 0 |
| 0 | 0 | 0 | 1 |
| 0 | 0 | 0 | 0 |
| 0 | 1 | 0 | 0 |
| 0 | 1 | 0 | 0 |
| 0 | 0 | 0 | 1 |
| 0 | 0 | 0 | 0 |
| 0 | 0 | 0 | 0 |
| 0 | 0 | 0 | 1 |
| 1 | 0 | 0 | 0 |
| 1 | 0 | 0 | 0 |
| 1 | 0 | 0 | 0 |
| 1 | 0 | 0 | 0 |
| 0 | 0 | 0 | 1 |
| 0 | 0 | 0 | 0 |
| 0 | 0 | 0 | 0 |
| 0 | 0 | 0 | 0 |
| 0 | 0 | 0 | 0 |
| 0 | 0 | 0 | 0 |
| 0 | 0 | 0 | 0 |
| 0 | 0 | 0 | 0 |
| 0 | 0 | 0 | 0 |
| 0 | 0 | 0 | 0 |
| 0 | 0 | 0 | 0 |
| 0 | 0 | 0 | 1 |
| 0 | 0 | 0 | 1 |
| 0 | 1 | 0 | 0 |
| 0 | 0 | 0 | 1 |
| 0 | 0 | 0 | 0 |
| 0 | 1 | 0 | 0 |
| 0 | 1 | 0 | 0 |
| 0 | 1 | 0 | 0 |
| 0 | 0 | 0 | 1 |
| 0 | 0 | 0 | 0 |
| 0 | 0 | 0 | 0 |
| 0 | 0 | 0 | 1 |
| 0 | 0 | 0 | 1 |
| 0 | 0 | 0 | 1 |
| 0 | 0 | 0 | 1 |
| 0 | 0 | 0 | 1 |
| 0 | 0 | 0 | 1 |
| 0 | 0 | 0 | 1 |
| 0 | 0 | 0 | 1 |
| 0 | 0 | 0 | 1 |
| 0 | 0 | 0 | 1 |

| | | | |
|---|---|---|---|
| 0 | 0 | 0 | 1 |
| 0 | 0 | 0 | 0 |
| 0 | 0 | 0 | 1 |
| 0 | 0 | 0 | 1 |
| 0 | 0 | 0 | 1 |
| 0 | 0 | 0 | 1 |
| 0 | 0 | 0 | 1 |
| 0 | 0 | 0 | 1 |
| 0 | 0 | 0 | 1 |
| 0 | 0 | 0 | 1 |
| 0 | 0 | 0 | 1 |
| 0 | 0 | 0 | 0 |
| 0 | 0 | 0 | 1 |
| 0 | 0 | 0 | 1 |
| 0 | 1 | 0 | 0 |
| 0 | 0 | 0 | 1 |
| 0 | 0 | 0 | 1 |
| 0 | 0 | 0 | 1 |
| 0 | 0 | 0 | 1 |
| 0 | 0 | 0 | 1 |
| 1 | 0 | 0 | 0 |
| 0 | 0 | 0 | 1 |
| 1 | 0 | 0 | 0 |
| 1 | 0 | 0 | 0 |
| 0 | 0 | 0 | 1 |
| 0 | 0 | 0 | 0 |
| 0 | 0 | 0 | 1 |
| 0 | 0 | 0 | 1 |
| 0 | 0 | 0 | 1 |
| 0 | 0 | 0 | 1 |
| 0 | 0 | 0 | 1 |
| 0 | 0 | 0 | 0 |
| 0 | 0 | 0 | 1 |
| 0 | 0 | 0 | 0 |
| 0 | 0 | 0 | 1 |
| 0 | 0 | 0 | 1 |
| 0 | 0 | 0 | 1 |
| 0 | 0 | 0 | 1 |
| 0 | 0 | 0 | 1 |
| 0 | 0 | 0 | 0 |
| 0 | 0 | 0 | 1 |
| 0 | 0 | 0 | 1 |
| 0 | 0 | 0 | 1 |
| 0 | 0 | 0 | 0 |
| 0 | 0 | 0 | 1 |
| 0 | 0 | 0 | 1 |
| 0 | 0 | 0 | 1 |
| 1 | 0 | 0 | 0 |
| 0 | 0 | 0 | 0 |
| 0 | 0 | 0 | 1 |
| 0 | 0 | 0 | 1 |
| 0 | 0 | 0 | 0 |
| 1 | 0 | 0 | 0 |
| 0 | 0 | 0 | 0 |

| | | | |
|---|---|---|---|
| 1 | 0 | 0 | 0 |
| 0 | 0 | 0 | 1 |
| 0 | 0 | 0 | 0 |
| 0 | 0 | 0 | 0 |
| 0 | 0 | 0 | 0 |
| 0 | 0 | 0 | 1 |
| 0 | 0 | 0 | 1 |
| 0 | 0 | 0 | 1 |
| 0 | 0 | 0 | 1 |
| 0 | 0 | 0 | 0 |
| 0 | 0 | 0 | 1 |
| 0 | 0 | 0 | 1 |
| 0 | 0 | 0 | 1 |
| 0 | 0 | 0 | 1 |
| 0 | 0 | 0 | 0 |
| 0 | 0 | 0 | 1 |
| 0 | 0 | 0 | 1 |
| 0 | 0 | 0 | 0 |
| 0 | 0 | 0 | 1 |
| 0 | 0 | 0 | 0 |
| 0 | 0 | 0 | 0 |
| 0 | 0 | 0 | 1 |
| 0 | 0 | 0 | 0 |
| 0 | 0 | 0 | 1 |
| 0 | 0 | 0 | 0 |
| 0 | 0 | 0 | 0 |
| 0 | 0 | 0 | 1 |
| 0 | 0 | 0 | 1 |
| 1 | 0 | 0 | 0 |
| 0 | 0 | 0 | 1 |
| 0 | 0 | 0 | 1 |
| 0 | 0 | 0 | 1 |
| 0 | 0 | 0 | 1 |
| 0 | 0 | 0 | 1 |
| 0 | 0 | 0 | 0 |
| 0 | 0 | 0 | 0 |
| 0 | 0 | 0 | 1 |
| 0 | 0 | 0 | 0 |
| 0 | 0 | 0 | 0 |
| 0 | 0 | 0 | 0 |
| 0 | 0 | 0 | 0 |
| 0 | 0 | 0 | 0 |
| 0 | 0 | 0 | 0 |
| 0 | 0 | 0 | 1 |
| 0 | 1 | 0 | 0 |
| 0 | 0 | 0 | 1 |
| 0 | 0 | 0 | 1 |
| 0 | 0 | 0 | 0 |
| 0 | 0 | 0 | 0 |
| 0 | 0 | 0 | 0 |
| 0 | 0 | 0 | 0 |
| 0 | 1 | 0 | 0 |
| 0 | 0 | 0 | 1 |
| 0 | 0 | 0 | 1 |

| | | | |
|---|---|---|---|
| 0 | 0 | 0 | 1 |
| 0 | 0 | 0 | 0 |
| 0 | 0 | 0 | 0 |
| 0 | 0 | 0 | 0 |
| 0 | 0 | 0 | 0 |
| 0 | 0 | 0 | 0 |
| 0 | 1 | 0 | 0 |
| 0 | 1 | 1 | 1 |
| 0 | 1 | 1 | 1 |
| 0 | 0 | 0 | 1 |
| 0 | 0 | 0 | 0 |
| 0 | 0 | 0 | 0 |
| 0 | 0 | 0 | 1 |
| 0 | 0 | 0 | 0 |
| 0 | 0 | 0 | 1 |
| 0 | 0 | 0 | 0 |
| 0 | 0 | 0 | 1 |
| 0 | 0 | 0 | 1 |
| 0 | 0 | 0 | 1 |
| 0 | 0 | 0 | 1 |
| 0 | 0 | 0 | 1 |
| 0 | 0 | 0 | 1 |
| 0 | 0 | 0 | 1 |
| 1 | 0 | 0 | 0 |
| 0 | 0 | 0 | 0 |
| 0 | 0 | 0 | 1 |
| 0 | 0 | 0 | 1 |
| 0 | 0 | 0 | 1 |
| 0 | 0 | 0 | 1 |
| 0 | 0 | 0 | 0 |
| 0 | 0 | 0 | 0 |
| 0 | 0 | 0 | 0 |
| 0 | 0 | 0 | 0 |
| 0 | 0 | 0 | 1 |
| 0 | 0 | 0 | 0 |
| 0 | 0 | 0 | 0 |
| 0 | 0 | 0 | 0 |
| 0 | 0 | 0 | 0 |
| 0 | 0 | 0 | 0 |
| 0 | 0 | 0 | 1 |
| 0 | 0 | 0 | 1 |
| 0 | 0 | 0 | 0 |
| 0 | 0 | 0 | 1 |
| 0 | 0 | 0 | 1 |
| 0 | 0 | 0 | 0 |
| 0 | 0 | 0 | 1 |
| 0 | 0 | 0 | 1 |
| 0 | 0 | 0 | 1 |
| 0 | 0 | 0 | 1 |
| 0 | 0 | 0 | 1 |
| 0 | 0 | 0 | 0 |
| 0 | 0 | 0 | 1 |
| 0 | 0 | 0 | 0 |
| 0 | 0 | 0 | 0 |

| | | | |
|---|---|---|---|
| 0 | 0 | 0 | 1 |
| 0 | 0 | 0 | 1 |
| 0 | 0 | 0 | 0 |
| 0 | 0 | 0 | 0 |
| 0 | 0 | 0 | 1 |
| 0 | 0 | 0 | 1 |
| 0 | 0 | 0 | 1 |
| 0 | 0 | 0 | 1 |
| 0 | 0 | 0 | 1 |
| 0 | 0 | 0 | 1 |
| 0 | 0 | 0 | 1 |
| 0 | 0 | 0 | 1 |
| 0 | 0 | 0 | 0 |
| 0 | 0 | 0 | 1 |
| 0 | 0 | 0 | 1 |
| 0 | 0 | 0 | 1 |
| 0 | 0 | 0 | 1 |
| 0 | 0 | 0 | 1 |
| 0 | 0 | 0 | 1 |
| 0 | 0 | 0 | 1 |
| 0 | 0 | 0 | 1 |
| 0 | 0 | 0 | 1 |
| 0 | 0 | 0 | 1 |
| 0 | 0 | 0 | 1 |
| 0 | 0 | 0 | 1 |
| 0 | 0 | 0 | 1 |
| 0 | 0 | 0 | 1 |
| 0 | 0 | 0 | 1 |
| 0 | 0 | 0 | 1 |
| 0 | 0 | 0 | 1 |
| 0 | 0 | 0 | 1 |
| 0 | 0 | 0 | 0 |
| 0 | 0 | 0 | 1 |
| 0 | 0 | 0 | 1 |
| 0 | 0 | 0 | 1 |
| 0 | 0 | 0 | 1 |
| 0 | 0 | 0 | 1 |
| 0 | 0 | 0 | 1 |
| 0 | 0 | 0 | 1 |
| 0 | 0 | 0 | 1 |
| 0 | 0 | 0 | 0 |
| 0 | 0 | 0 | 0 |
| 0 | 0 | 0 | 1 |
| 0 | 0 | 0 | 1 |
| 0 | 0 | 0 | 1 |
| 0 | 0 | 0 | 1 |
| 0 | 0 | 0 | 1 |
| 0 | 0 | 0 | 1 |
| 0 | 0 | 0 | 0 |
| 0 | 0 | 0 | 0 |
| 0 | 0 | 0 | 1 |
| 0 | 0 | 0 | 1 |
| 0 | 0 | 0 | 0 |
| 0 | 0 | 0 | 1 |

| | | | |
|---|---|---|---|
| 0 | 0 | 0 | 0 |
| 0 | 0 | 0 | 0 |
| 0 | 0 | 0 | 0 |
| 0 | 0 | 0 | 1 |
| 0 | 0 | 0 | 1 |
| 0 | 0 | 0 | 1 |
| 0 | 0 | 0 | 1 |
| 0 | 1 | 0 | 0 |
| 0 | 1 | 0 | 0 |
| 0 | 0 | 0 | 0 |
| 0 | 0 | 0 | 0 |
| 0 | 0 | 0 | 1 |
| 0 | 0 | 0 | 0 |
| 0 | 0 | 0 | 1 |
| 0 | 0 | 0 | 1 |
| 0 | 0 | 0 | 1 |
| 0 | 0 | 0 | 1 |
| 0 | 0 | 0 | 0 |
| 0 | 0 | 0 | 1 |
| 0 | 0 | 0 | 1 |
| 0 | 0 | 0 | 1 |
| 0 | 0 | 0 | 1 |
| 0 | 0 | 0 | 1 |
| 0 | 0 | 0 | 1 |
| 0 | 0 | 0 | 1 |
| 0 | 0 | 0 | 1 |
| 0 | 0 | 0 | 1 |
| 0 | 0 | 0 | 1 |
| 0 | 0 | 0 | 1 |
| 0 | 0 | 0 | 1 |
| 0 | 0 | 0 | 0 |
| 0 | 0 | 0 | 0 |
| 0 | 0 | 0 | 1 |
| 0 | 0 | 0 | 1 |
| 0 | 0 | 0 | 1 |
| 0 | 0 | 0 | 1 |
| 0 | 0 | 0 | 1 |
| 0 | 0 | 0 | 1 |
| 0 | 0 | 0 | 1 |
| 0 | 0 | 0 | 1 |
| 0 | 0 | 0 | 1 |
| 0 | 0 | 0 | 0 |
| 0 | 0 | 0 | 1 |
| 0 | 0 | 0 | 1 |
| 0 | 0 | 0 | 0 |
| 0 | 0 | 0 | 0 |
| 0 | 0 | 0 | 1 |
| 0 | 0 | 0 | 1 |
| 0 | 0 | 0 | 0 |
| 0 | 0 | 0 | 0 |
| 0 | 0 | 0 | 0 |
| 0 | 0 | 0 | 0 |
| 0 | 0 | 0 | 0 |
| 0 | 0 | 0 | 0 |

| | | | |
|---|---|---|---|
| 0 | 0 | 0 | 1 |
| 0 | 0 | 0 | 1 |
| 0 | 0 | 0 | 1 |
| 0 | 0 | 0 | 1 |
| 0 | 0 | 0 | 1 |
| 0 | 0 | 0 | 0 |
| 0 | 0 | 0 | 0 |
| 0 | 0 | 0 | 0 |
| 0 | 0 | 0 | 0 |
| 0 | 0 | 0 | 0 |
| 0 | 0 | 0 | 1 |
| 0 | 0 | 0 | 0 |
| 0 | 0 | 0 | 0 |
| 0 | 0 | 0 | 0 |
| 0 | 0 | 0 | 1 |
| 0 | 0 | 0 | 0 |
| 0 | 0 | 0 | 1 |
| 0 | 0 | 0 | 1 |
| 1 | 0 | 0 | 0 |
| 1 | 0 | 0 | 0 |
| 0 | 0 | 0 | 0 |
| 0 | 0 | 0 | 0 |
| 0 | 0 | 0 | 1 |
| 0 | 0 | 0 | 0 |
| 0 | 0 | 0 | 0 |
| 0 | 0 | 0 | 0 |
| 0 | 0 | 0 | 0 |
| 0 | 0 | 0 | 1 |
| 0 | 0 | 0 | 1 |
| 0 | 0 | 0 | 1 |
| 0 | 0 | 0 | 1 |
| 0 | 0 | 0 | 1 |
| 0 | 0 | 0 | 0 |
| 0 | 0 | 0 | 1 |
| 0 | 0 | 0 | 0 |
| 0 | 0 | 0 | 0 |
| 0 | 0 | 0 | 0 |
| 0 | 0 | 0 | 0 |
| 0 | 1 | 0 | 0 |
| 0 | 1 | 0 | 0 |
| 0 | 1 | 0 | 0 |
| 1 | 0 | 0 | 0 |
| 0 | 0 | 0 | 0 |
| 0 | 0 | 0 | 1 |
| 0 | 0 | 0 | 1 |
| 0 | 0 | 0 | 1 |
| 0 | 0 | 0 | 1 |
| 0 | 0 | 0 | 0 |
| 0 | 0 | 0 | 0 |
| 0 | 0 | 0 | 0 |
| 0 | 0 | 0 | 1 |
| 0 | 0 | 0 | 1 |
| 0 | 0 | 0 | 0 |
| 0 | 0 | 0 | 1 |

| | | | |
|---|---|---|---|
| 0 | 0 | 0 | 1 |
| 0 | 0 | 0 | 0 |
| 0 | 0 | 0 | 1 |
| 0 | 0 | 0 | 1 |
| 0 | 0 | 0 | 1 |
| 0 | 0 | 0 | 1 |
| 0 | 0 | 0 | 1 |
| 0 | 0 | 0 | 1 |
| 0 | 0 | 0 | 1 |
| 0 | 0 | 0 | 1 |
| 0 | 0 | 0 | 1 |
| 0 | 0 | 0 | 1 |
| 0 | 0 | 0 | 0 |
| 0 | 0 | 0 | 0 |
| 0 | 0 | 0 | 0 |
| 0 | 0 | 0 | 0 |
| 0 | 0 | 0 | 0 |
| 0 | 0 | 0 | 0 |
| 0 | 0 | 0 | 0 |
| 0 | 0 | 0 | 1 |
| 1 | 1 | 0 | 0 |
| 0 | 0 | 0 | 0 |
| 0 | 0 | 0 | 0 |
| 0 | 0 | 0 | 0 |
| 0 | 0 | 0 | 0 |
| 0 | 0 | 0 | 1 |
| 0 | 0 | 0 | 1 |
| 0 | 0 | 0 | 0 |
| 0 | 0 | 0 | 1 |
| 0 | 0 | 0 | 0 |
| 0 | 0 | 0 | 0 |
| 0 | 0 | 0 | 1 |
| 0 | 0 | 0 | 0 |
| 0 | 0 | 0 | 1 |
| 0 | 0 | 0 | 1 |
| 0 | 0 | 0 | 1 |
| 0 | 1 | 1 | 1 |
| 0 | 0 | 0 | 1 |
| 0 | 0 | 0 | 1 |
| 0 | 0 | 0 | 1 |
| 0 | 0 | 0 | 0 |
| 0 | 0 | 0 | 1 |
| 0 | 0 | 0 | 1 |
| 0 | 0 | 0 | 0 |
| 0 | 0 | 0 | 0 |
| 0 | 0 | 0 | 1 |
| 0 | 0 | 0 | 0 |
| 0 | 0 | 0 | 0 |
| 0 | 0 | 0 | 1 |
| 0 | 0 | 0 | 1 |
| 0 | 0 | 0 | 0 |
| 0 | 0 | 0 | 1 |
| 0 | 0 | 0 | 1 |
| 0 | 0 | 0 | 1 |

| | | | |
|---|---|---|---|
| 0 | 0 | 0 | 1 |
| 0 | 0 | 0 | 1 |
| 0 | 0 | 0 | 1 |
| 0 | 0 | 0 | 1 |
| 0 | 0 | 0 | 0 |
| 0 | 0 | 0 | 0 |
| 0 | 0 | 0 | 1 |
| 0 | 0 | 0 | 1 |
| 0 | 0 | 0 | 1 |
| 0 | 0 | 0 | 1 |
| 0 | 0 | 0 | 1 |
| 0 | 0 | 0 | 0 |
| 0 | 0 | 0 | 0 |
| 0 | 0 | 0 | 1 |
| 0 | 0 | 0 | 1 |
| 0 | 0 | 0 | 0 |
| 0 | 0 | 0 | 1 |
| 0 | 0 | 0 | 1 |
| 0 | 0 | 0 | 0 |
| 0 | 0 | 0 | 1 |
| 0 | 0 | 0 | 1 |
| 0 | 0 | 0 | 1 |
| 0 | 0 | 0 | 1 |
| 0 | 0 | 0 | 0 |
| 0 | 0 | 0 | 1 |
| 0 | 0 | 0 | 0 |
| 0 | 0 | 0 | 1 |
| 0 | 0 | 0 | 1 |
| 1 | 0 | 0 | 0 |
| 1 | 0 | 0 | 0 |
| 0 | 0 | 0 | 0 |
| 0 | 0 | 0 | 0 |
| 0 | 0 | 0 | 0 |
| 0 | 0 | 0 | 0 |
| 0 | 0 | 0 | 1 |
| 0 | 0 | 0 | 1 |
| 0 | 0 | 0 | 1 |
| 1 | 0 | 0 | 0 |
| 0 | 0 | 0 | 1 |
| 0 | 0 | 0 | 1 |
| 0 | 0 | 0 | 0 |
| 0 | 0 | 0 | 0 |
| 0 | 0 | 0 | 0 |
| 0 | 0 | 0 | 1 |
| 0 | 0 | 0 | 0 |
| 0 | 0 | 0 | 1 |
| 0 | 0 | 0 | 1 |
| 0 | 0 | 0 | 0 |
| 0 | 0 | 0 | 1 |
| 0 | 0 | 0 | 1 |
| 0 | 0 | 0 | 1 |
| 0 | 1 | 0 | 0 |
| 0 | 0 | 0 | 1 |
| 0 | 0 | 0 | 0 |

| | | | |
|---|---|---|---|
| 0 | 0 | 0 | 1 |
| 0 | 0 | 0 | 0 |
| 0 | 0 | 0 | 0 |
| 0 | 0 | 0 | 1 |
| 0 | 0 | 0 | 1 |
| 0 | 0 | 0 | 1 |
| 0 | 0 | 0 | 0 |
| 0 | 0 | 0 | 1 |
| 0 | 0 | 0 | 1 |
| 0 | 0 | 0 | 1 |
| 0 | 0 | 0 | 1 |
| 0 | 0 | 0 | 1 |
| 0 | 0 | 0 | 1 |
| 0 | 0 | 0 | 1 |
| 0 | 0 | 0 | 1 |
| 0 | 0 | 0 | 1 |
| 0 | 0 | 0 | 1 |
| 0 | 0 | 0 | 1 |
| 0 | 0 | 0 | 1 |
| 0 | 0 | 0 | 1 |
| 0 | 0 | 0 | 1 |
| 0 | 0 | 0 | 0 |
| 0 | 0 | 0 | 1 |
| 0 | 0 | 0 | 1 |
| 0 | 0 | 0 | 1 |
| 0 | 0 | 0 | 1 |
| 0 | 0 | 0 | 0 |
| 0 | 0 | 0 | 1 |
| 0 | 0 | 0 | 0 |
| 0 | 0 | 0 | 1 |
| 0 | 0 | 0 | 1 |
| 0 | 0 | 0 | 0 |
| 0 | 1 | 1 | 1 |
| 0 | 1 | 1 | 1 |
| 0 | 0 | 0 | 1 |
| 0 | 0 | 0 | 1 |
| 0 | 0 | 0 | 1 |
| 0 | 0 | 0 | 1 |
| 0 | 0 | 0 | 0 |
| 0 | 0 | 0 | 0 |
| 0 | 1 | 0 | 0 |
| 0 | 0 | 0 | 1 |
| 0 | 0 | 0 | 1 |
| 0 | 0 | 0 | 1 |
| 0 | 0 | 0 | 1 |
| 0 | 1 | 0 | 0 |
| 0 | 0 | 0 | 1 |
| 0 | 0 | 0 | 0 |
| 0 | 0 | 0 | 1 |
| 0 | 0 | 0 | 0 |
| 0 | 0 | 0 | 0 |
| 0 | 0 | 0 | 0 |
| 0 | 0 | 0 | 1 |
| 0 | 0 | 0 | 1 |

| | | | |
|---|---|---|---|
| 0 | 0 | 0 | 0 |
| 0 | 0 | 0 | 0 |
| 0 | 0 | 0 | 1 |
| 0 | 0 | 0 | 1 |
| 0 | 0 | 0 | 1 |
| 0 | 0 | 0 | 1 |
| 0 | 0 | 0 | 1 |
| 0 | 0 | 0 | 1 |
| 0 | 0 | 0 | 1 |
| 0 | 0 | 0 | 1 |
| 0 | 0 | 0 | 1 |
| 0 | 0 | 0 | 1 |
| 0 | 0 | 0 | 1 |
| 0 | 0 | 0 | 1 |
| 0 | 0 | 0 | 1 |
| 0 | 0 | 0 | 0 |
| 0 | 0 | 0 | 0 |
| 0 | 0 | 0 | 0 |
| 0 | 0 | 0 | 1 |
| 0 | 0 | 0 | 1 |
| 0 | 0 | 0 | 1 |
| 0 | 1 | 0 | 0 |
| 0 | 0 | 0 | 0 |
| 0 | 0 | 0 | 0 |
| 0 | 0 | 0 | 0 |
| 0 | 0 | 0 | 1 |
| 0 | 0 | 0 | 1 |
| 0 | 0 | 0 | 1 |
| 0 | 0 | 0 | 1 |
| 0 | 0 | 0 | 1 |
| 0 | 0 | 0 | 0 |
| 0 | 0 | 0 | 1 |
| 0 | 0 | 0 | 0 |
| 0 | 0 | 0 | 1 |
| 0 | 1 | 1 | 1 |
| 0 | 1 | 1 | 1 |
| 0 | 1 | 0 | 0 |
| 0 | 0 | 0 | 0 |
| 0 | 0 | 0 | 1 |
| 0 | 0 | 0 | 1 |
| 0 | 0 | 0 | 1 |
| 0 | 0 | 0 | 0 |
| 0 | 0 | 0 | 1 |
| 0 | 0 | 0 | 1 |
| 0 | 0 | 0 | 0 |
| 0 | 0 | 0 | 1 |
| 0 | 0 | 0 | 0 |
| 0 | 0 | 0 | 0 |
| 0 | 0 | 0 | 1 |
| 0 | 0 | 0 | 1 |
| 0 | 1 | 0 | 0 |
| 0 | 1 | 0 | 0 |
| 0 | 0 | 0 | 1 |
| 0 | 0 | 0 | 0 |

| | | | |
|---|---|---|---|
| 0 | 0 | 0 | 1 |
| 0 | 0 | 0 | 0 |
| 0 | 1 | 1 | 1 |
| 0 | 1 | 0 | 0 |
| 0 | 0 | 0 | 1 |
| 0 | 0 | 0 | 1 |
| 0 | 0 | 0 | 1 |
| 0 | 0 | 0 | 1 |
| 0 | 0 | 0 | 0 |
| 0 | 0 | 0 | 1 |
| 0 | 0 | 0 | 1 |
| 0 | 0 | 0 | 1 |
| 0 | 0 | 0 | 0 |
| 0 | 0 | 0 | 0 |
| 0 | 0 | 0 | 0 |
| 0 | 0 | 0 | 1 |
| 0 | 0 | 0 | 0 |
| 0 | 0 | 0 | 1 |
| 0 | 1 | 0 | 0 |
| 0 | 0 | 0 | 0 |
| 0 | 0 | 0 | 0 |
| 0 | 1 | 0 | 0 |
| 0 | 0 | 0 | 0 |
| 1 | 0 | 0 | 0 |
| 0 | 0 | 0 | 1 |
| 0 | 0 | 0 | 1 |
| 0 | 0 | 0 | 0 |
| 0 | 0 | 0 | 0 |
| 0 | 0 | 0 | 0 |
| 0 | 0 | 0 | 0 |
| 0 | 0 | 0 | 1 |
| 0 | 0 | 0 | 1 |
| 0 | 0 | 0 | 1 |
| 0 | 0 | 0 | 1 |
| 0 | 0 | 0 | 1 |
| 1 | 0 | 0 | 0 |
| 1 | 0 | 0 | 0 |
| 0 | 0 | 0 | 1 |
| 0 | 0 | 0 | 1 |
| 0 | 0 | 0 | 1 |
| 0 | 0 | 0 | 1 |
| 0 | 0 | 0 | 1 |
| 0 | 0 | 0 | 1 |
| 0 | 0 | 0 | 0 |
| 0 | 0 | 0 | 0 |
| 0 | 0 | 0 | 0 |
| 0 | 0 | 0 | 1 |
| 0 | 0 | 0 | 0 |
| 0 | 0 | 0 | 0 |
| 0 | 0 | 0 | 0 |
| 0 | 0 | 0 | 1 |
| 0 | 0 | 0 | 1 |
| 0 | 0 | 0 | 1 |
| 0 | 0 | 0 | 1 |

| | | | |
|---|---|---|---|
| 0 | 0 | 0 | 1 |
| 0 | 0 | 0 | 1 |
| 0 | 0 | 0 | 1 |
| 0 | 0 | 0 | 1 |
| 0 | 0 | 0 | 0 |
| 0 | 0 | 0 | 1 |
| 0 | 0 | 0 | 0 |
| 0 | 0 | 0 | 0 |
| 0 | 0 | 0 | 0 |
| 0 | 0 | 0 | 0 |
| 0 | 0 | 0 | 0 |
| 0 | 0 | 0 | 0 |
| 0 | 0 | 0 | 1 |
| 0 | 0 | 0 | 0 |
| 0 | 0 | 0 | 0 |
| 0 | 0 | 0 | 1 |
| 0 | 0 | 0 | 1 |
| 0 | 0 | 0 | 1 |
| 0 | 0 | 0 | 0 |
| 0 | 0 | 0 | 0 |
| 0 | 0 | 0 | 0 |
| 0 | 0 | 0 | 0 |
| 0 | 0 | 0 | 1 |
| 0 | 0 | 0 | 1 |
| 0 | 0 | 0 | 1 |
| 0 | 0 | 0 | 1 |
| 0 | 0 | 0 | 1 |
| 0 | 0 | 0 | 0 |
| 0 | 0 | 0 | 0 |
| 0 | 0 | 0 | 0 |
| 0 | 0 | 0 | 0 |
| 0 | 0 | 0 | 0 |
| 0 | 0 | 0 | 0 |
| 0 | 0 | 0 | 1 |
| 0 | 0 | 0 | 1 |
| 0 | 0 | 0 | 0 |
| 0 | 0 | 0 | 1 |
| 0 | 0 | 0 | 1 |
| 0 | 0 | 0 | 1 |
| 0 | 0 | 0 | 1 |
| 0 | 0 | 0 | 0 |
| 0 | 0 | 0 | 1 |
| 0 | 0 | 0 | 0 |
| 0 | 0 | 0 | 1 |
| 0 | 0 | 0 | 1 |
| 0 | 0 | 0 | 1 |
| 0 | 0 | 0 | 1 |
| 0 | 0 | 0 | 0 |
| 0 | 0 | 0 | 1 |
| 0 | 0 | 0 | 1 |
| 0 | 0 | 0 | 1 |
| 0 | 0 | 0 | 1 |
| 0 | 0 | 0 | 1 |
| 0 | 0 | 0 | 1 |

| | | | |
|---|---|---|---|
| 0 | 0 | 0 | 1 |
| 0 | 0 | 0 | 1 |
| 0 | 0 | 0 | 1 |
| 0 | 0 | 0 | 1 |
| 0 | 0 | 0 | 1 |
| 0 | 0 | 0 | 1 |
| 0 | 0 | 0 | 1 |
| 0 | 0 | 0 | 1 |
| 0 | 0 | 0 | 1 |
| 0 | 0 | 0 | 1 |
| 0 | 0 | 0 | 1 |
| 0 | 0 | 0 | 0 |
| 0 | 0 | 0 | 1 |
| 0 | 0 | 0 | 1 |
| 0 | 0 | 0 | 1 |
| 0 | 0 | 0 | 1 |
| 0 | 0 | 0 | 1 |
| 0 | 0 | 0 | 0 |
| 0 | 0 | 0 | 0 |
| 0 | 0 | 0 | 0 |
| 0 | 0 | 0 | 0 |
| 0 | 0 | 0 | 1 |
| 1 | 0 | 0 | 0 |
| 0 | 0 | 0 | 1 |
| 0 | 0 | 0 | 1 |
| 0 | 0 | 0 | 1 |
| 0 | 0 | 0 | 1 |
| 0 | 0 | 0 | 1 |
| 0 | 0 | 0 | 0 |
| 0 | 0 | 0 | 0 |
| 0 | 0 | 0 | 1 |
| 0 | 0 | 0 | 1 |
| 1 | 0 | 0 | 0 |
| 0 | 0 | 0 | 1 |
| 0 | 0 | 0 | 0 |
| 1 | 0 | 0 | 0 |
| 1 | 0 | 0 | 0 |
| 0 | 0 | 0 | 1 |
| 1 | 0 | 0 | 0 |
| 0 | 0 | 0 | 1 |
| 1 | 0 | 0 | 0 |
| 0 | 0 | 0 | 1 |
| 0 | 0 | 0 | 0 |
| 0 | 0 | 0 | 1 |
| 0 | 0 | 0 | 1 |
| 0 | 0 | 0 | 1 |
| 0 | 1 | 1 | 1 |
| 0 | 0 | 0 | 1 |
| 0 | 0 | 0 | 1 |
| 0 | 0 | 0 | 0 |
| 0 | 1 | 1 | 1 |
| 1 | 0 | 0 | 0 |
| 0 | 1 | 1 | 1 |
| 0 | 0 | 0 | 1 |

| | | | |
|---|---|---|---|
| 0 | 0 | 0 | 0 |
| 0 | 0 | 0 | 0 |
| 0 | 0 | 0 | 0 |
| 0 | 0 | 0 | 0 |
| 0 | 0 | 0 | 0 |
| 0 | 0 | 0 | 1 |
| 0 | 0 | 0 | 0 |
| 0 | 0 | 0 | 1 |
| 0 | 0 | 0 | 1 |
| 0 | 0 | 0 | 0 |
| 0 | 0 | 0 | 0 |
| 0 | 0 | 0 | 0 |
| 0 | 0 | 0 | 1 |
| 0 | 0 | 0 | 1 |
| 0 | 0 | 0 | 1 |
| 0 | 0 | 0 | 1 |
| 0 | 0 | 0 | 1 |
| 0 | 0 | 0 | 1 |
| 0 | 0 | 0 | 1 |
| 0 | 0 | 0 | 1 |
| 0 | 0 | 0 | 1 |
| 0 | 0 | 0 | 1 |
| 0 | 0 | 0 | 1 |
| 0 | 0 | 0 | 1 |
| 0 | 0 | 0 | 1 |
| 0 | 0 | 0 | 1 |
| 0 | 0 | 0 | 1 |
| 0 | 0 | 0 | 1 |
| 0 | 0 | 0 | 1 |
| 0 | 0 | 0 | 1 |
| 0 | 0 | 0 | 1 |
| 0 | 0 | 0 | 1 |
| 0 | 0 | 0 | 1 |
| 0 | 0 | 0 | 1 |
| 0 | 0 | 0 | 1 |
| 0 | 0 | 0 | 1 |
| 0 | 0 | 0 | 1 |
| 0 | 0 | 0 | 1 |
| 0 | 0 | 0 | 1 |
| 0 | 0 | 0 | 1 |
| 0 | 0 | 0 | 1 |
| 0 | 0 | 0 | 1 |
| 0 | 0 | 0 | 1 |
| 0 | 0 | 0 | 1 |
| 0 | 0 | 0 | 1 |
| 0 | 0 | 0 | 0 |
| 0 | 0 | 0 | 1 |
| 0 | 0 | 0 | 1 |
| 0 | 0 | 0 | 1 |
| 0 | 0 | 0 | 1 |
| 0 | 0 | 0 | 0 |
| 0 | 0 | 0 | 1 |
| 0 | 0 | 0 | 1 |
| 0 | 0 | 0 | 1 |

| | | | |
|---|---|---|---|
| 0 | 0 | 0 | 1 |
| 0 | 0 | 0 | 1 |
| 0 | 0 | 0 | 1 |
| 0 | 0 | 0 | 1 |
| 0 | 0 | 0 | 1 |
| 0 | 0 | 0 | 1 |
| 0 | 0 | 0 | 1 |
| 0 | 0 | 0 | 0 |
| 0 | 0 | 0 | 0 |
| 0 | 0 | 0 | 1 |
| 0 | 0 | 0 | 1 |
| 0 | 0 | 0 | 1 |
| 0 | 0 | 0 | 1 |
| 0 | 0 | 0 | 1 |
| 0 | 0 | 0 | 0 |
| 0 | 0 | 0 | 0 |
| 0 | 0 | 0 | 0 |
| 0 | 0 | 0 | 1 |
| 0 | 0 | 0 | 1 |
| 0 | 0 | 0 | 1 |
| 0 | 0 | 0 | 0 |
| 0 | 0 | 0 | 0 |
| 0 | 1 | 0 | 0 |
| 0 | 0 | 0 | 0 |
| 0 | 0 | 0 | 0 |
| 0 | 0 | 0 | 0 |
| 0 | 0 | 0 | 1 |
| 0 | 0 | 0 | 1 |
| 0 | 0 | 0 | 0 |
| 0 | 0 | 0 | 1 |
| 0 | 0 | 0 | 1 |
| 0 | 0 | 0 | 1 |
| 0 | 0 | 0 | 0 |
| 0 | 0 | 0 | 1 |
| 0 | 0 | 0 | 1 |
| 0 | 0 | 0 | 1 |
| 0 | 0 | 0 | 0 |
| 0 | 0 | 0 | 0 |
| 0 | 0 | 0 | 0 |
| 0 | 0 | 0 | 1 |
| 0 | 0 | 0 | 1 |
| 0 | 0 | 0 | 1 |
| 0 | 0 | 0 | 1 |
| 0 | 0 | 0 | 1 |
| 0 | 0 | 0 | 1 |
| 0 | 0 | 0 | 0 |
| 0 | 0 | 0 | 1 |
| 0 | 0 | 0 | 1 |
| 0 | 0 | 0 | 1 |
| 0 | 0 | 0 | 1 |
| 0 | 0 | 0 | 1 |
| 0 | 0 | 0 | 1 |
| 0 | 0 | 0 | 1 |
| 0 | 0 | 0 | 1 |

| | | | |
|---|---|---|---|
| 0 | 0 | 0 | 1 |
| 0 | 0 | 0 | 1 |
| 0 | 0 | 0 | 1 |
| 0 | 0 | 0 | 0 |
| 0 | 0 | 0 | 1 |
| 0 | 0 | 0 | 1 |
| 0 | 0 | 0 | 1 |
| 0 | 0 | 0 | 0 |
| 0 | 0 | 0 | 0 |
| 0 | 0 | 0 | 1 |
| 0 | 0 | 0 | 1 |
| 0 | 0 | 0 | 1 |
| 0 | 0 | 0 | 0 |
| 0 | 0 | 0 | 1 |
| 0 | 0 | 0 | 1 |
| 0 | 0 | 0 | 1 |
| 0 | 0 | 0 | 0 |
| 0 | 0 | 0 | 1 |
| 0 | 0 | 0 | 1 |
| 0 | 0 | 0 | 1 |
| 0 | 0 | 0 | 1 |
| 0 | 0 | 0 | 1 |
| 0 | 0 | 0 | 1 |
| 0 | 0 | 0 | 1 |
| 0 | 0 | 0 | 1 |
| 0 | 0 | 0 | 1 |
| 0 | 0 | 0 | 1 |
| 0 | 0 | 0 | 0 |
| 0 | 0 | 0 | 0 |
| 0 | 0 | 0 | 0 |
| 0 | 0 | 0 | 0 |
| 0 | 0 | 0 | 0 |
| 0 | 0 | 0 | 0 |
| 0 | 0 | 0 | 1 |
| 0 | 0 | 0 | 0 |
| 0 | 0 | 0 | 0 |
| 0 | 0 | 0 | 0 |
| 0 | 0 | 0 | 1 |
| 0 | 0 | 0 | 1 |
| 0 | 0 | 0 | 1 |
| 0 | 0 | 0 | 1 |
| 0 | 0 | 0 | 1 |
| 0 | 0 | 0 | 0 |
| 0 | 0 | 0 | 0 |
| 0 | 0 | 0 | 1 |
| 0 | 0 | 0 | 1 |
| 0 | 0 | 0 | 0 |
| 0 | 0 | 0 | 1 |
| 0 | 0 | 0 | 1 |
| 0 | 0 | 0 | 1 |
| 0 | 0 | 0 | 1 |
| 0 | 0 | 0 | 1 |
| 0 | 0 | 0 | 0 |
| 0 | 0 | 0 | 0 |

| | | | |
|---|---|---|---|
| 0 | 0 | 0 | 0 |
| 0 | 0 | 0 | 0 |
| 0 | 0 | 0 | 0 |
| 0 | 0 | 0 | 0 |
| 0 | 0 | 0 | 0 |
| 0 | 0 | 0 | 0 |
| 0 | 0 | 0 | 0 |
| 0 | 0 | 0 | 0 |
| 0 | 0 | 0 | 1 |
| 0 | 0 | 0 | 0 |
| 0 | 0 | 0 | 1 |
| 0 | 0 | 0 | 1 |
| 0 | 0 | 0 | 1 |
| 0 | 0 | 0 | 1 |
| 0 | 0 | 0 | 0 |
| 0 | 0 | 0 | 1 |
| 0 | 0 | 0 | 1 |
| 0 | 0 | 0 | 1 |
| 0 | 0 | 0 | 1 |
| 0 | 0 | 0 | 0 |
| 0 | 0 | 0 | 0 |
| 0 | 0 | 0 | 1 |
| 0 | 0 | 0 | 0 |
| 0 | 0 | 0 | 0 |
| 0 | 0 | 0 | 0 |
| 0 | 0 | 0 | 0 |
| 0 | 0 | 0 | 0 |
| 0 | 0 | 0 | 1 |
| 0 | 0 | 0 | 1 |
| 0 | 0 | 0 | 1 |
| 0 | 0 | 0 | 0 |
| 0 | 0 | 0 | 1 |
| 0 | 0 | 0 | 1 |
| 0 | 0 | 0 | 1 |
| 0 | 0 | 0 | 1 |
| 0 | 0 | 0 | 1 |
| 0 | 0 | 0 | 1 |
| 0 | 0 | 0 | 1 |
| 0 | 0 | 0 | 1 |
| 0 | 0 | 0 | 1 |
| 0 | 0 | 0 | 1 |
| 0 | 0 | 0 | 0 |
| 0 | 0 | 0 | 1 |
| 0 | 0 | 0 | 1 |
| 0 | 0 | 0 | 1 |
| 0 | 0 | 0 | 1 |
| 0 | 0 | 0 | 1 |
| 0 | 0 | 0 | 1 |
| 0 | 0 | 0 | 0 |
| 0 | 0 | 0 | 1 |
| 0 | 0 | 0 | 1 |
| 0 | 0 | 0 | 1 |
| 0 | 0 | 0 | 0 |
| 0 | 1 | 0 | 0 |

| | | | |
|---|---|---|---|
| 0 | 0 | 0 | 1 |
| 0 | 0 | 0 | 1 |
| 0 | 0 | 0 | 1 |
| 0 | 0 | 0 | 1 |
| 0 | 0 | 0 | 0 |
| 1 | 0 | 0 | 0 |
| 0 | 0 | 0 | 1 |
| 0 | 0 | 0 | 0 |
| 0 | 0 | 0 | 1 |
| 0 | 0 | 0 | 1 |
| 0 | 0 | 0 | 0 |
| 0 | 0 | 0 | 1 |
| 0 | 0 | 0 | 0 |
| 0 | 0 | 0 | 0 |
| 0 | 0 | 0 | 0 |
| 0 | 0 | 0 | 0 |
| 0 | 0 | 0 | 1 |
| 0 | 0 | 0 | 1 |
| 0 | 0 | 0 | 0 |
| 0 | 0 | 0 | 0 |
| 0 | 0 | 0 | 0 |
| 0 | 0 | 0 | 0 |
| 0 | 0 | 0 | 0 |
| 0 | 0 | 0 | 1 |
| 0 | 0 | 0 | 1 |
| 0 | 0 | 0 | 0 |
| 0 | 0 | 0 | 1 |
| 0 | 0 | 0 | 0 |
| 0 | 0 | 0 | 0 |
| 0 | 0 | 0 | 0 |
| 0 | 0 | 0 | 1 |
| 1 | 0 | 0 | 0 |
| 1 | 0 | 0 | 0 |
| 0 | 0 | 0 | 1 |
| 0 | 0 | 0 | 1 |
| 0 | 0 | 0 | 0 |
| 0 | 0 | 0 | 0 |
| 0 | 0 | 0 | 1 |
| 0 | 0 | 0 | 0 |
| 0 | 0 | 0 | 0 |
| 0 | 1 | 0 | 0 |
| 0 | 0 | 0 | 0 |
| 0 | 0 | 0 | 1 |
| 0 | 0 | 0 | 1 |
| 0 | 0 | 0 | 0 |
| 0 | 0 | 0 | 1 |
| 0 | 0 | 0 | 0 |
| 0 | 0 | 0 | 1 |
| 0 | 0 | 0 | 1 |
| 0 | 0 | 0 | 1 |
| 0 | 0 | 0 | 1 |
| 0 | 0 | 0 | 1 |
| 0 | 0 | 0 | 1 |
| 0 | 0 | 0 | 1 |

| | | | |
|---|---|---|---|
| 0 | 0 | 0 | 1 |
| 0 | 0 | 0 | 1 |
| 0 | 0 | 0 | 1 |
| 0 | 0 | 0 | 1 |
| 0 | 0 | 0 | 1 |
| 0 | 0 | 0 | 1 |
| 0 | 0 | 0 | 1 |
| 0 | 0 | 0 | 0 |
| 0 | 0 | 0 | 1 |
| 1 | 0 | 0 | 0 |
| 0 | 0 | 0 | 0 |
| 0 | 0 | 0 | 0 |
| 0 | 0 | 0 | 1 |
| 0 | 0 | 0 | 1 |
| 0 | 0 | 0 | 0 |
| 0 | 0 | 0 | 1 |
| 0 | 0 | 0 | 1 |
| 0 | 0 | 0 | 1 |
| 0 | 0 | 0 | 1 |
| 0 | 0 | 0 | 1 |
| 0 | 0 | 0 | 1 |
| 0 | 0 | 0 | 1 |
| 0 | 0 | 0 | 1 |
| 0 | 0 | 0 | 1 |
| 0 | 0 | 0 | 1 |
| 0 | 0 | 0 | 1 |
| 0 | 0 | 0 | 1 |
| 0 | 0 | 0 | 1 |
| 0 | 0 | 0 | 1 |
| 0 | 0 | 0 | 0 |
| 0 | 0 | 0 | 0 |
| 0 | 0 | 0 | 1 |
| 0 | 0 | 0 | 1 |
| 0 | 0 | 0 | 1 |
| 0 | 0 | 0 | 1 |
| 0 | 0 | 0 | 0 |
| 0 | 0 | 0 | 1 |
| 0 | 0 | 0 | 1 |
| 0 | 0 | 0 | 1 |
| 0 | 0 | 0 | 1 |
| 0 | 0 | 0 | 1 |
| 0 | 0 | 0 | 0 |
| 0 | 0 | 0 | 0 |
| 0 | 0 | 0 | 0 |
| 0 | 0 | 0 | 1 |
| 0 | 0 | 0 | 1 |
| 0 | 0 | 0 | 1 |
| 0 | 0 | 0 | 1 |
| 0 | 0 | 0 | 1 |
| 0 | 0 | 0 | 1 |
| 0 | 0 | 0 | 1 |
| 0 | 0 | 0 | 1 |
| 0 | 0 | 0 | 1 |
| 0 | 0 | 0 | 0 |

| | | | |
|---|---|---|---|
| 0 | 0 | 0 | 1 |
| 0 | 0 | 0 | 1 |
| 0 | 0 | 0 | 1 |
| 0 | 0 | 0 | 1 |
| 0 | 0 | 0 | 1 |
| 0 | 0 | 0 | 1 |
| 0 | 0 | 0 | 1 |
| 0 | 0 | 0 | 1 |
| 0 | 0 | 0 | 0 |
| 0 | 0 | 0 | 0 |
| 0 | 0 | 0 | 1 |
| 0 | 0 | 0 | 1 |
| 0 | 0 | 0 | 1 |
| 0 | 0 | 0 | 0 |
| 0 | 0 | 0 | 1 |
| 0 | 0 | 0 | 1 |
| 0 | 0 | 0 | 1 |
| 1 | 0 | 0 | 0 |
| 1 | 0 | 0 | 0 |
| 0 | 0 | 0 | 0 |
| 0 | 0 | 0 | 0 |
| 0 | 0 | 0 | 1 |
| 0 | 0 | 0 | 1 |
| 0 | 0 | 0 | 1 |
| 0 | 0 | 0 | 1 |
| 1 | 0 | 0 | 0 |
| 0 | 0 | 0 | 1 |
| 0 | 0 | 0 | 1 |
| 0 | 0 | 0 | 1 |
| 0 | 0 | 0 | 1 |
| 0 | 0 | 0 | 1 |
| 0 | 0 | 0 | 1 |
| 0 | 0 | 0 | 1 |
| 0 | 0 | 0 | 1 |
| 0 | 0 | 0 | 1 |
| 0 | 0 | 0 | 1 |
| 0 | 0 | 0 | 1 |
| 0 | 0 | 0 | 1 |
| 0 | 0 | 0 | 1 |
| 0 | 0 | 0 | 0 |
| 0 | 0 | 0 | 1 |
| 0 | 0 | 0 | 1 |
| 0 | 0 | 0 | 0 |
| 0 | 0 | 0 | 1 |
| 0 | 0 | 0 | 1 |
| 0 | 0 | 0 | 1 |
| 0 | 0 | 0 | 0 |
| 0 | 0 | 0 | 0 |
| 0 | 0 | 0 | 1 |
| 0 | 0 | 0 | 1 |
| 0 | 0 | 0 | 0 |
| 0 | 0 | 0 | 1 |
| 0 | 0 | 0 | 1 |
| 0 | 0 | 0 | 1 |

| | | | |
|---|---|---|---|
| 0 | 0 | 0 | 1 |
| 0 | 0 | 0 | 1 |
| 0 | 0 | 0 | 1 |
| 0 | 0 | 0 | 1 |
| 0 | 0 | 0 | 1 |
| 0 | 0 | 0 | 1 |
| 0 | 0 | 0 | 1 |
| 0 | 0 | 0 | 1 |
| 0 | 0 | 0 | 1 |
| 0 | 0 | 0 | 1 |
| 0 | 0 | 0 | 1 |
| 0 | 0 | 0 | 1 |
| 0 | 0 | 0 | 1 |
| 0 | 0 | 0 | 0 |
| 0 | 0 | 0 | 0 |
| 0 | 0 | 0 | 0 |
| 0 | 0 | 0 | 0 |
| 0 | 0 | 0 | 1 |
| 0 | 0 | 0 | 1 |
| 0 | 0 | 0 | 0 |
| 0 | 0 | 0 | 1 |
| 0 | 0 | 0 | 1 |
| 0 | 0 | 0 | 1 |
| 0 | 0 | 0 | 1 |
| 0 | 0 | 0 | 1 |
| 0 | 0 | 0 | 0 |
| 0 | 0 | 0 | 1 |
| 0 | 0 | 0 | 1 |
| 0 | 0 | 0 | 1 |
| 0 | 0 | 0 | 0 |
| 0 | 0 | 0 | 0 |
| 0 | 0 | 0 | 1 |
| 0 | 0 | 0 | 1 |
| 0 | 0 | 0 | 1 |
| 0 | 0 | 0 | 1 |
| 0 | 0 | 0 | 1 |
| 0 | 0 | 0 | 1 |
| 0 | 0 | 0 | 1 |
| 0 | 0 | 0 | 0 |
| 0 | 0 | 0 | 1 |
| 0 | 0 | 0 | 1 |
| 0 | 0 | 0 | 0 |
| 0 | 1 | 0 | 0 |
| 0 | 1 | 0 | 0 |
| 0 | 0 | 0 | 0 |
| 0 | 0 | 0 | 1 |
| 0 | 0 | 0 | 0 |
| 0 | 0 | 0 | 1 |
| 0 | 0 | 0 | 1 |
| 0 | 0 | 0 | 1 |
| 0 | 0 | 0 | 1 |
| 0 | 0 | 0 | 1 |
| 0 | 0 | 0 | 1 |
| 0 | 0 | 0 | 1 |

| | | | |
|---|---|---|---|
| 0 | 0 | 0 | 1 |
| 0 | 0 | 0 | 1 |
| 0 | 0 | 0 | 0 |
| 0 | 0 | 0 | 1 |
| 0 | 0 | 0 | 1 |
| 0 | 0 | 0 | 1 |
| 0 | 0 | 0 | 1 |
| 0 | 0 | 0 | 1 |
| 0 | 0 | 0 | 1 |
| 0 | 0 | 0 | 1 |
| 0 | 0 | 0 | 1 |
| 0 | 0 | 0 | 1 |
| 0 | 0 | 0 | 1 |
| 0 | 0 | 0 | 1 |
| 0 | 0 | 0 | 1 |
| 0 | 0 | 0 | 1 |
| 0 | 0 | 0 | 1 |
| 0 | 0 | 0 | 1 |
| 0 | 0 | 0 | 1 |
| 0 | 0 | 0 | 1 |
| 0 | 0 | 0 | 1 |
| 0 | 0 | 0 | 1 |
| 0 | 0 | 0 | 1 |
| 0 | 0 | 0 | 1 |
| 0 | 0 | 0 | 1 |
| 0 | 0 | 0 | 1 |
| 0 | 0 | 0 | 1 |
| 0 | 0 | 0 | 1 |
| 0 | 0 | 0 | 1 |
| 0 | 0 | 0 | 1 |
| 1 | 0 | 0 | 0 |
| 0 | 0 | 0 | 1 |
| 0 | 1 | 0 | 0 |
| 0 | 1 | 0 | 0 |
| 0 | 0 | 0 | 0 |
| 0 | 0 | 0 | 0 |
| 0 | 0 | 0 | 1 |
| 0 | 0 | 0 | 0 |
| 0 | 0 | 0 | 1 |
| 0 | 0 | 0 | 1 |
| 1 | 0 | 0 | 0 |
| 1 | 0 | 0 | 0 |
| 0 | 0 | 0 | 0 |
| 0 | 0 | 0 | 1 |
| 0 | 0 | 0 | 0 |
| 0 | 0 | 0 | 0 |
| 0 | 0 | 0 | 1 |
| 0 | 0 | 0 | 1 |
| 0 | 0 | 0 | 1 |
| 0 | 0 | 0 | 1 |
| 0 | 0 | 0 | 0 |
| 0 | 0 | 0 | 1 |
| 0 | 0 | 0 | 1 |
| 0 | 0 | 0 | 1 |

| | | | |
|---|---|---|---|
| 0 | 0 | 0 | 1 |
| 0 | 0 | 0 | 1 |
| 0 | 0 | 0 | 1 |
| 0 | 0 | 0 | 1 |
| 0 | 0 | 0 | 1 |
| 1 | 0 | 0 | 0 |
| 0 | 0 | 0 | 1 |
| 0 | 0 | 0 | 0 |
| 0 | 0 | 0 | 0 |
| 0 | 0 | 0 | 0 |
| 0 | 0 | 0 | 1 |
| 0 | 1 | 0 | 0 |
| 0 | 0 | 0 | 1 |
| 0 | 0 | 0 | 1 |
| 0 | 0 | 0 | 1 |
| 0 | 1 | 1 | 1 |
| 0 | 0 | 0 | 1 |
| 0 | 0 | 0 | 0 |
| 0 | 0 | 0 | 1 |
| 0 | 0 | 0 | 0 |
| 0 | 0 | 0 | 1 |
| 0 | 0 | 0 | 0 |
| 0 | 0 | 0 | 0 |
| 0 | 0 | 0 | 0 |
| 0 | 0 | 0 | 1 |
| 0 | 0 | 0 | 1 |
| 0 | 0 | 0 | 1 |
| 0 | 0 | 0 | 1 |
| 0 | 0 | 0 | 0 |
| 0 | 0 | 0 | 1 |
| 0 | 0 | 0 | 1 |
| 0 | 0 | 0 | 1 |
| 0 | 0 | 0 | 1 |
| 0 | 0 | 0 | 1 |
| 0 | 0 | 0 | 0 |
| 0 | 0 | 0 | 0 |
| 0 | 0 | 0 | 1 |
| 0 | 0 | 0 | 0 |
| 0 | 0 | 0 | 0 |
| 0 | 0 | 0 | 1 |
| 0 | 0 | 0 | 0 |
| 0 | 0 | 0 | 0 |
| 0 | 0 | 0 | 1 |
| 0 | 0 | 0 | 1 |
| 0 | 0 | 0 | 1 |
| 0 | 0 | 0 | 1 |
| 0 | 0 | 0 | 1 |
| 0 | 0 | 0 | 1 |
| 0 | 0 | 0 | 1 |
| 0 | 0 | 0 | 1 |
| 0 | 0 | 0 | 1 |
| 0 | 0 | 0 | 1 |
| 0 | 0 | 0 | 1 |
| 0 | 0 | 0 | 1 |

| | | | |
|---|---|---|---|
| 0 | 0 | 0 | 1 |
| 0 | 0 | 0 | 0 |
| 0 | 0 | 0 | 1 |
| 0 | 0 | 0 | 1 |
| 0 | 0 | 0 | 0 |
| 0 | 0 | 0 | 1 |
| 0 | 0 | 0 | 1 |
| 0 | 0 | 0 | 1 |
| 0 | 0 | 0 | 0 |
| 0 | 0 | 0 | 0 |
| 0 | 0 | 0 | 1 |
| 0 | 0 | 0 | 0 |
| 0 | 0 | 0 | 0 |
| 0 | 0 | 0 | 1 |
| 0 | 0 | 0 | 1 |
| 0 | 0 | 0 | 0 |
| 0 | 0 | 0 | 1 |
| 0 | 0 | 0 | 0 |
| 0 | 0 | 0 | 0 |
| 0 | 0 | 0 | 1 |
| 0 | 0 | 0 | 0 |
| 0 | 0 | 0 | 0 |
| 1 | 0 | 0 | 0 |
| 0 | 0 | 0 | 1 |
| 0 | 0 | 0 | 1 |
| 0 | 0 | 0 | 0 |
| 0 | 0 | 0 | 1 |
| 0 | 0 | 0 | 1 |
| 0 | 0 | 0 | 1 |
| 0 | 0 | 0 | 1 |
| 0 | 0 | 0 | 0 |
| 0 | 0 | 0 | 1 |
| 0 | 0 | 0 | 0 |
| 0 | 0 | 0 | 1 |
| 0 | 0 | 0 | 0 |
| 0 | 0 | 0 | 0 |
| 0 | 0 | 0 | 1 |
| 0 | 0 | 0 | 1 |
| 0 | 0 | 0 | 1 |
| 0 | 0 | 0 | 1 |
| 0 | 0 | 0 | 1 |
| 0 | 0 | 0 | 1 |
| 0 | 0 | 0 | 1 |
| 0 | 0 | 0 | 1 |
| 0 | 0 | 0 | 1 |
| 0 | 0 | 0 | 1 |
| 0 | 1 | 1 | 1 |
| 0 | 0 | 0 | 1 |
| 0 | 0 | 0 | 1 |
| 0 | 0 | 0 | 1 |
| 0 | 0 | 0 | 1 |
| 0 | 0 | 0 | 1 |
| 0 | 0 | 0 | 1 |
| 0 | 0 | 0 | 0 |

| | | | |
|---|---|---|---|
| 0 | 0 | 0 | 1 |
| 0 | 0 | 0 | 1 |
| 0 | 0 | 0 | 1 |
| 0 | 0 | 0 | 1 |
| 0 | 0 | 0 | 0 |
| 0 | 0 | 0 | 1 |
| 0 | 0 | 0 | 1 |
| 0 | 0 | 0 | 1 |
| 0 | 0 | 0 | 1 |
| 0 | 0 | 0 | 1 |
| 0 | 0 | 0 | 0 |
| 0 | 0 | 0 | 1 |
| 0 | 0 | 0 | 1 |
| 0 | 0 | 0 | 1 |
| 0 | 0 | 0 | 0 |
| 0 | 0 | 0 | 0 |
| 0 | 0 | 0 | 0 |
| 0 | 0 | 0 | 0 |
| 0 | 0 | 0 | 1 |
| 0 | 0 | 0 | 0 |
| 0 | 0 | 0 | 0 |
| 1 | 0 | 0 | 0 |
| 0 | 0 | 0 | 1 |
| 0 | 0 | 0 | 0 |
| 1 | 0 | 0 | 0 |
| 0 | 0 | 0 | 1 |
| 1 | 0 | 0 | 0 |
| 0 | 0 | 0 | 0 |
| 1 | 0 | 0 | 0 |
| 1 | 0 | 0 | 0 |
| 1 | 0 | 0 | 0 |
| 0 | 0 | 0 | 1 |
| 0 | 0 | 0 | 1 |
| 0 | 0 | 0 | 1 |
| 0 | 0 | 0 | 1 |
| 0 | 0 | 0 | 1 |
| 0 | 0 | 0 | 1 |
| 0 | 1 | 0 | 0 |
| 0 | 0 | 0 | 0 |
| 0 | 0 | 0 | 1 |
| 0 | 0 | 0 | 1 |
| 0 | 1 | 1 | 1 |
| 0 | 0 | 0 | 0 |
| 0 | 0 | 0 | 1 |
| 0 | 0 | 0 | 0 |
| 0 | 0 | 0 | 1 |
| 0 | 0 | 0 | 1 |
| 0 | 0 | 0 | 1 |
| 0 | 1 | 0 | 0 |
| 0 | 1 | 0 | 0 |
| 0 | 1 | 0 | 0 |
| 0 | 0 | 0 | 1 |
| 0 | 0 | 0 | 1 |
| 0 | 0 | 0 | 0 |

| | | | |
|---|---|---|---|
| 0 | 0 | 0 | 1 |
| 0 | 1 | 0 | 0 |
| 0 | 0 | 0 | 1 |
| 0 | 1 | 0 | 0 |
| 0 | 1 | 0 | 0 |
| 0 | 0 | 0 | 1 |
| 0 | 1 | 0 | 0 |
| 0 | 1 | 0 | 0 |
| 0 | 0 | 0 | 1 |
| 0 | 0 | 0 | 1 |
| 0 | 0 | 0 | 1 |
| 0 | 0 | 0 | 1 |
| 0 | 0 | 0 | 1 |
| 0 | 0 | 0 | 1 |
| 0 | 0 | 0 | 1 |
| 0 | 0 | 0 | 1 |
| 0 | 0 | 0 | 1 |
| 0 | 0 | 0 | 1 |
| 0 | 0 | 0 | 1 |
| 0 | 0 | 0 | 1 |
| 1 | 1 | 0 | 0 |
| 0 | 0 | 0 | 1 |
| 0 | 0 | 0 | 1 |
| 0 | 0 | 0 | 1 |
| 0 | 0 | 0 | 1 |
| 0 | 0 | 0 | 0 |
| 0 | 0 | 0 | 0 |
| 0 | 0 | 0 | 1 |
| 0 | 0 | 0 | 1 |
| 0 | 0 | 0 | 1 |
| 0 | 0 | 0 | 1 |
| 0 | 0 | 0 | 1 |
| 0 | 0 | 0 | 1 |
| 0 | 0 | 0 | 1 |
| 0 | 0 | 0 | 1 |
| 0 | 0 | 0 | 1 |
| 0 | 0 | 0 | 1 |
| 0 | 0 | 0 | 1 |
| 0 | 0 | 0 | 1 |
| 0 | 0 | 0 | 1 |
| 0 | 0 | 0 | 1 |
| 0 | 0 | 0 | 1 |
| 0 | 0 | 0 | 1 |
| 0 | 0 | 0 | 0 |
| 0 | 0 | 0 | 1 |
| 0 | 0 | 0 | 0 |
| 0 | 0 | 0 | 0 |
| 0 | 0 | 0 | 0 |
| 0 | 0 | 0 | 0 |
| 0 | 0 | 0 | 0 |
| 0 | 0 | 0 | 0 |
| 0 | 0 | 0 | 1 |
| 0 | 0 | 0 | 1 |
| 0 | 0 | 0 | 1 |

| | | | |
|---|---|---|---|
| 0 | 0 | 0 | 1 |
| 0 | 0 | 0 | 1 |
| 0 | 0 | 0 | 1 |
| 0 | 0 | 0 | 0 |
| 0 | 0 | 0 | 1 |
| 0 | 0 | 0 | 0 |
| 0 | 0 | 0 | 1 |
| 0 | 0 | 0 | 1 |
| 0 | 0 | 0 | 0 |
| 0 | 0 | 0 | 1 |
| 0 | 0 | 0 | 1 |
| 0 | 0 | 0 | 1 |
| 0 | 0 | 0 | 1 |
| 0 | 0 | 0 | 1 |
| 0 | 0 | 0 | 1 |
| 0 | 0 | 0 | 0 |
| 0 | 0 | 0 | 0 |
| 0 | 0 | 0 | 0 |
| 0 | 0 | 0 | 0 |
| 0 | 0 | 0 | 0 |
| 0 | 0 | 0 | 1 |
| 0 | 0 | 0 | 1 |
| 0 | 0 | 0 | 1 |
| 0 | 0 | 0 | 1 |
| 0 | 0 | 0 | 0 |
| 0 | 0 | 0 | 1 |
| 0 | 0 | 0 | 1 |
| 0 | 0 | 0 | 1 |
| 0 | 0 | 0 | 0 |
| 0 | 0 | 0 | 1 |
| 0 | 1 | 0 | 0 |
| 0 | 1 | 0 | 0 |
| 0 | 0 | 0 | 0 |
| 0 | 1 | 0 | 0 |
| 0 | 1 | 0 | 0 |
| 0 | 0 | 0 | 1 |
| 0 | 1 | 0 | 0 |
| 0 | 1 | 0 | 0 |
| 0 | 1 | 0 | 0 |
| 0 | 1 | 0 | 0 |
| 0 | 1 | 0 | 0 |
| 0 | 0 | 0 | 0 |
| 0 | 0 | 0 | 0 |
| 0 | 0 | 0 | 0 |
| 0 | 0 | 0 | 0 |
| 0 | 0 | 0 | 0 |
| 0 | 0 | 0 | 0 |
| 0 | 0 | 0 | 0 |
| 0 | 0 | 0 | 1 |
| 0 | 0 | 0 | 0 |
| 0 | 0 | 0 | 1 |
| 0 | 0 | 0 | 1 |
| 0 | 0 | 0 | 1 |
| 0 | 0 | 0 | 0 |

| | | | |
|---|---|---|---|
| 0 | 0 | 0 | 0 |
| 0 | 0 | 0 | 1 |
| 0 | 0 | 0 | 1 |
| 0 | 0 | 0 | 1 |
| 0 | 0 | 0 | 1 |
| 0 | 0 | 0 | 0 |
| 0 | 0 | 0 | 1 |
| 0 | 0 | 0 | 1 |
| 0 | 0 | 0 | 1 |
| 0 | 0 | 0 | 1 |
| 0 | 0 | 0 | 1 |
| 0 | 0 | 0 | 1 |
| 0 | 0 | 0 | 1 |
| 1 | 0 | 0 | 0 |
| 0 | 0 | 0 | 1 |
| 0 | 0 | 0 | 1 |
| 0 | 0 | 0 | 1 |
| 0 | 0 | 0 | 0 |
| 0 | 0 | 0 | 1 |
| 0 | 0 | 0 | 1 |
| 0 | 0 | 0 | 1 |
| 0 | 0 | 0 | 1 |
| 0 | 0 | 0 | 0 |
| 0 | 0 | 0 | 1 |
| 0 | 0 | 0 | 1 |
| 0 | 0 | 0 | 0 |
| 0 | 0 | 0 | 0 |
| 0 | 0 | 0 | 0 |
| 0 | 0 | 0 | 1 |
| 0 | 0 | 0 | 1 |
| 0 | 0 | 0 | 1 |
| 0 | 0 | 0 | 1 |
| 0 | 0 | 0 | 1 |
| 0 | 0 | 0 | 1 |
| 0 | 0 | 0 | 1 |
| 0 | 0 | 0 | 1 |
| 0 | 0 | 0 | 1 |
| 0 | 0 | 0 | 1 |
| 0 | 0 | 0 | 1 |
| 0 | 0 | 0 | 1 |
| 0 | 0 | 0 | 1 |
| 0 | 0 | 0 | 1 |
| 0 | 0 | 0 | 1 |
| 0 | 0 | 0 | 1 |
| 0 | 0 | 0 | 1 |
| 0 | 0 | 0 | 1 |
| 0 | 0 | 0 | 1 |
| 0 | 0 | 0 | 1 |
| 0 | 0 | 0 | 1 |
| 0 | 0 | 0 | 1 |
| 0 | 0 | 0 | 1 |
| 0 | 0 | 0 | 1 |
| 0 | 0 | 0 | 1 |

| | | | |
|---|---|---|---|
| 0 | 0 | 0 | 1 |
| 0 | 0 | 0 | 0 |
| 0 | 0 | 0 | 1 |
| 0 | 0 | 0 | 1 |
| 0 | 0 | 0 | 1 |
| 0 | 0 | 0 | 0 |
| 0 | 0 | 0 | 0 |
| 0 | 0 | 0 | 0 |
| 0 | 0 | 0 | 1 |
| 0 | 0 | 0 | 1 |
| 0 | 0 | 0 | 1 |
| 0 | 0 | 0 | 1 |
| 0 | 0 | 0 | 1 |
| 0 | 0 | 0 | 1 |
| 0 | 0 | 0 | 0 |
| 0 | 0 | 0 | 0 |
| 0 | 0 | 0 | 0 |
| 0 | 0 | 0 | 1 |
| 0 | 0 | 0 | 0 |
| 0 | 0 | 0 | 0 |
| 0 | 0 | 0 | 0 |
| 0 | 0 | 0 | 0 |
| 0 | 0 | 0 | 0 |
| 0 | 0 | 0 | 0 |
| 0 | 0 | 0 | 1 |
| 0 | 0 | 0 | 1 |
| 0 | 0 | 0 | 1 |
| 0 | 0 | 0 | 1 |
| 0 | 0 | 0 | 1 |
| 0 | 0 | 0 | 1 |
| 0 | 0 | 0 | 1 |
| 0 | 0 | 0 | 1 |
| 0 | 0 | 0 | 0 |
| 0 | 0 | 0 | 1 |
| 0 | 0 | 0 | 1 |
| 0 | 0 | 0 | 0 |
| 0 | 0 | 0 | 0 |
| 0 | 0 | 0 | 0 |
| 0 | 0 | 0 | 0 |
| 0 | 0 | 0 | 1 |
| 0 | 0 | 0 | 1 |
| 0 | 0 | 0 | 1 |
| 0 | 0 | 0 | 1 |
| 0 | 0 | 0 | 0 |
| 0 | 0 | 0 | 0 |
| 0 | 0 | 0 | 0 |
| 0 | 0 | 0 | 1 |
| 0 | 0 | 0 | 1 |
| 0 | 0 | 0 | 0 |
| 0 | 0 | 0 | 1 |
| 0 | 0 | 0 | 1 |
| 0 | 0 | 0 | 0 |
| 0 | 0 | 0 | 1 |
| 0 | 0 | 0 | 1 |

| | | | |
|---|---|---|---|
| 0 | 0 | 0 | 1 |
| 0 | 0 | 0 | 1 |
| 0 | 0 | 0 | 1 |
| 0 | 0 | 0 | 1 |
| 0 | 0 | 0 | 1 |
| 0 | 0 | 0 | 1 |
| 0 | 0 | 0 | 1 |
| 0 | 0 | 0 | 1 |
| 0 | 0 | 0 | 0 |
| 0 | 0 | 0 | 0 |
| 0 | 0 | 0 | 1 |
| 0 | 0 | 0 | 0 |
| 0 | 0 | 0 | 0 |
| 0 | 0 | 0 | 0 |
| 0 | 0 | 0 | 1 |
| 0 | 0 | 0 | 1 |
| 0 | 0 | 0 | 0 |
| 0 | 0 | 0 | 0 |
| 0 | 0 | 0 | 0 |
| 0 | 0 | 0 | 1 |
| 0 | 0 | 0 | 1 |
| 0 | 0 | 0 | 1 |
| 0 | 0 | 0 | 0 |
| 0 | 0 | 0 | 1 |
| 0 | 0 | 0 | 0 |
| 0 | 0 | 0 | 1 |
| 0 | 0 | 0 | 1 |
| 0 | 0 | 0 | 1 |
| 0 | 0 | 0 | 1 |
| 0 | 0 | 0 | 0 |
| 0 | 0 | 0 | 0 |
| 0 | 0 | 0 | 0 |
| 0 | 0 | 0 | 1 |
| 0 | 0 | 0 | 1 |
| 0 | 0 | 0 | 1 |
| 0 | 0 | 0 | 1 |
| 0 | 0 | 0 | 1 |
| 0 | 0 | 0 | 1 |
| 0 | 0 | 0 | 1 |
| 0 | 0 | 0 | 1 |
| 0 | 0 | 0 | 0 |
| 0 | 0 | 0 | 1 |
| 0 | 0 | 0 | 1 |
| 0 | 0 | 0 | 1 |
| 0 | 0 | 0 | 1 |
| 0 | 0 | 0 | 0 |
| 0 | 1 | 1 | 1 |
| 0 | 0 | 0 | 0 |
| 0 | 0 | 0 | 1 |
| 0 | 1 | 0 | 0 |
| 0 | 1 | 0 | 0 |
| 0 | 0 | 0 | 0 |
| 0 | 0 | 0 | 1 |
| 0 | 1 | 1 | 1 |

| | | | |
|---|---|---|---|
| 0 | 1 | 1 | 1 |
| 0 | 1 | 1 | 1 |
| 0 | 0 | 0 | 1 |
| 0 | 0 | 0 | 1 |
| 0 | 0 | 0 | 1 |
| 0 | 0 | 0 | 1 |
| 0 | 0 | 0 | 1 |
| 0 | 0 | 0 | 0 |
| 0 | 0 | 0 | 0 |
| 0 | 0 | 0 | 1 |
| 0 | 0 | 0 | 0 |
| 0 | 0 | 0 | 1 |
| 0 | 0 | 0 | 1 |
| 0 | 0 | 0 | 1 |
| 0 | 0 | 0 | 1 |
| 0 | 0 | 0 | 1 |
| 0 | 0 | 0 | 1 |
| 0 | 0 | 0 | 1 |
| 0 | 0 | 0 | 1 |
| 0 | 0 | 0 | 1 |
| 0 | 0 | 0 | 1 |
| 0 | 0 | 0 | 1 |
| 0 | 0 | 0 | 1 |
| 0 | 0 | 0 | 1 |
| 0 | 0 | 0 | 1 |
| 0 | 0 | 0 | 1 |
| 0 | 0 | 0 | 1 |
| 0 | 0 | 0 | 1 |
| 0 | 0 | 0 | 0 |
| 0 | 0 | 0 | 1 |
| 0 | 0 | 0 | 0 |
| 0 | 0 | 0 | 1 |
| 0 | 0 | 0 | 0 |
| 0 | 0 | 0 | 0 |
| 0 | 0 | 0 | 1 |
| 0 | 0 | 0 | 0 |
| 0 | 0 | 0 | 0 |
| 0 | 0 | 0 | 1 |
| 0 | 0 | 0 | 1 |
| 0 | 0 | 0 | 1 |
| 0 | 0 | 0 | 1 |
| 0 | 0 | 0 | 0 |
| 0 | 0 | 0 | 0 |
| 0 | 0 | 0 | 1 |
| 0 | 0 | 0 | 1 |
| 0 | 0 | 0 | 0 |
| 0 | 0 | 0 | 0 |
| 0 | 0 | 0 | 0 |
| 0 | 0 | 0 | 0 |
| 0 | 0 | 0 | 1 |
| 0 | 1 | 0 | 0 |
| 0 | 0 | 0 | 0 |
| 0 | 0 | 0 | 1 |
| 0 | 0 | 0 | 1 |

| | | | |
|---|---|---|---|
| 0 | 0 | 0 | 1 |
| 0 | 0 | 0 | 1 |
| 0 | 0 | 0 | 1 |
| 0 | 0 | 0 | 1 |
| 0 | 0 | 0 | 0 |
| 0 | 0 | 0 | 0 |
| 0 | 0 | 0 | 0 |
| 0 | 0 | 0 | 0 |
| 0 | 0 | 0 | 1 |
| 0 | 0 | 0 | 0 |
| 0 | 0 | 0 | 0 |
| 0 | 0 | 0 | 0 |
| 0 | 0 | 0 | 0 |
| 0 | 0 | 0 | 1 |
| 0 | 0 | 0 | 1 |
| 0 | 0 | 0 | 1 |
| 0 | 0 | 0 | 1 |
| 0 | 0 | 0 | 1 |
| 0 | 0 | 0 | 1 |
| 0 | 0 | 0 | 1 |
| 0 | 0 | 0 | 1 |
| 0 | 0 | 0 | 1 |
| 0 | 0 | 0 | 1 |
| 0 | 0 | 0 | 1 |
| 0 | 0 | 0 | 1 |
| 0 | 0 | 0 | 0 |
| 0 | 0 | 0 | 0 |
| 0 | 0 | 0 | 0 |
| 0 | 0 | 0 | 0 |
| 0 | 0 | 0 | 1 |
| 0 | 0 | 0 | 1 |
| 0 | 0 | 0 | 1 |
| 0 | 0 | 0 | 1 |
| 0 | 0 | 0 | 1 |
| 0 | 0 | 0 | 1 |
| 0 | 0 | 0 | 1 |
| 0 | 0 | 0 | 1 |
| 0 | 0 | 0 | 1 |
| 0 | 0 | 0 | 1 |
| 0 | 0 | 0 | 1 |
| 0 | 0 | 0 | 1 |
| 0 | 0 | 0 | 1 |
| 0 | 0 | 0 | 1 |
| 0 | 0 | 0 | 1 |
| 0 | 0 | 0 | 1 |
| 0 | 0 | 0 | 1 |
| 0 | 0 | 0 | 1 |
| 0 | 0 | 0 | 1 |
| 0 | 0 | 0 | 1 |
| 0 | 0 | 0 | 1 |
| 0 | 0 | 0 | 1 |
| 0 | 0 | 0 | 1 |
| 0 | 0 | 0 | 1 |
| 0 | 0 | 0 | 0 |

| | | | |
|---|---|---|---|
| 0 | 0 | 0 | 1 |
| 0 | 0 | 0 | 1 |
| 0 | 0 | 0 | 1 |
| 0 | 0 | 0 | 1 |
| 0 | 0 | 0 | 1 |
| 0 | 0 | 0 | 1 |
| 0 | 0 | 0 | 1 |
| 0 | 0 | 0 | 1 |
| 0 | 0 | 0 | 1 |
| 0 | 0 | 0 | 1 |
| 0 | 0 | 0 | 1 |
| 0 | 0 | 0 | 1 |
| 0 | 0 | 0 | 1 |
| 0 | 0 | 0 | 1 |
| 0 | 0 | 0 | 1 |
| 0 | 0 | 0 | 1 |
| 0 | 0 | 0 | 1 |
| 0 | 0 | 0 | 1 |
| 0 | 0 | 0 | 1 |
| 0 | 0 | 0 | 1 |
| 0 | 0 | 0 | 1 |
| 0 | 0 | 0 | 1 |
| 0 | 0 | 0 | 1 |
| 0 | 0 | 0 | 1 |
| 0 | 0 | 0 | 1 |
| 0 | 0 | 0 | 1 |
| 0 | 0 | 0 | 1 |
| 0 | 0 | 0 | 1 |
| 0 | 0 | 0 | 1 |
| 0 | 0 | 0 | 1 |
| 0 | 0 | 0 | 1 |
| 0 | 0 | 0 | 1 |
| 0 | 0 | 0 | 1 |
| 0 | 0 | 0 | 1 |
| 0 | 0 | 0 | 1 |
| 0 | 0 | 0 | 1 |
| 0 | 0 | 0 | 1 |
| 0 | 0 | 0 | 1 |
| 0 | 0 | 0 | 1 |
| 0 | 0 | 0 | 1 |
| 0 | 0 | 0 | 1 |
| 0 | 0 | 0 | 1 |
| 0 | 0 | 0 | 1 |
| 0 | 0 | 0 | 1 |
| 0 | 0 | 0 | 1 |
| 0 | 1 | 1 | 1 |
| 0 | 0 | 0 | 0 |
| 0 | 0 | 0 | 1 |
| 0 | 0 | 0 | 1 |
| 0 | 0 | 0 | 1 |
| 0 | 0 | 0 | 1 |
| 0 | 0 | 0 | 1 |
| 0 | 0 | 0 | 1 |
| 0 | 0 | 0 | 1 |

| | | | |
|---|---|---|---|
| 0 | 0 | 0 | 1 |
| 0 | 0 | 0 | 1 |
| 0 | 0 | 0 | 0 |
| 0 | 0 | 0 | 1 |
| 0 | 0 | 0 | 1 |
| 0 | 0 | 0 | 1 |
| 0 | 0 | 0 | 0 |
| 0 | 0 | 0 | 1 |
| 0 | 0 | 0 | 1 |
| 0 | 0 | 0 | 1 |
| 0 | 0 | 0 | 1 |
| 0 | 0 | 0 | 0 |
| 0 | 0 | 0 | 1 |
| 0 | 0 | 0 | 0 |
| 0 | 0 | 0 | 0 |
| 0 | 0 | 0 | 0 |
| 0 | 0 | 0 | 0 |
| 0 | 0 | 0 | 0 |
| 0 | 0 | 0 | 0 |
| 0 | 0 | 0 | 0 |
| 0 | 0 | 0 | 0 |

| surface_non_binding_site | non_binding_site_single_water | surface_non_binding_site_single_water |
|---|---|---|
| 0 | 0 | 0 |
| 0 | 0 | 0 |
| 0 | 0 | 0 |
| 0 | 0 | 0 |
| 0 | 0 | 0 |
| 0 | 0 | 0 |
| 1 | 1 | 1 |
| 1 | 1 | 1 |
| 0 | 0 | 0 |
| 0 | 0 | 0 |
| 0 | 0 | 0 |
| 0 | 0 | 0 |
| 0 | 0 | 0 |
| 0 | 0 | 0 |
| 0 | 0 | 0 |
| 0 | 0 | 0 |
| 0 | 0 | 0 |
| 0 | 0 | 0 |
| 0 | 0 | 0 |
| 0 | 0 | 0 |
| 0 | 0 | 0 |
| 0 | 0 | 0 |
| 0 | 0 | 0 |
| 0 | 0 | 0 |
| 0 | 0 | 0 |
| 1 | 0 | 0 |
| 0 | 0 | 0 |
| 1 | 1 | 1 |
| 0 | 0 | 0 |
| 0 | 1 | 0 |
| 0 | 1 | 0 |
| 0 | 0 | 0 |
| 0 | 0 | 0 |
| 0 | 0 | 0 |
| 0 | 0 | 0 |
| 1 | 1 | 1 |
| 0 | 0 | 0 |
| 0 | 0 | 0 |
| 0 | 0 | 0 |
| 0 | 0 | 0 |
| 0 | 0 | 0 |
| 0 | 0 | 0 |
| 0 | 0 | 0 |
| 0 | 0 | 0 |
| 0 | 0 | 0 |
| 0 | 0 | 0 |
| 0 | 0 | 0 |
| 0 | 0 | 0 |
| 0 | 0 | 0 |
| 0 | 0 | 0 |
| 0 | 0 | 0 |
| 0 | 0 | 0 |
| 0 | 0 | 0 |

| | | |
|---|---|---|
| 0 | 0 | 0 |
| 0 | 0 | 0 |
| 0 | 0 | 0 |
| 0 | 0 | 0 |
| 0 | 0 | 0 |
| 0 | 0 | 0 |
| 0 | 0 | 0 |
| 0 | 0 | 0 |
| 0 | 0 | 0 |
| 0 | 0 | 0 |
| 0 | 0 | 0 |
| 0 | 0 | 0 |
| 0 | 0 | 0 |
| 0 | 0 | 0 |
| 0 | 0 | 0 |
| 0 | 0 | 0 |
| 0 | 0 | 0 |
| 0 | 0 | 0 |
| 0 | 0 | 0 |
| 0 | 0 | 0 |
| 0 | 0 | 0 |
| 0 | 0 | 0 |
| 0 | 0 | 0 |
| 0 | 0 | 0 |
| 0 | 0 | 0 |
| 0 | 0 | 0 |
| 0 | 0 | 0 |
| 0 | 0 | 0 |
| 0 | 0 | 0 |
| 0 | 0 | 0 |
| 0 | 0 | 0 |
| 0 | 0 | 0 |
| 0 | 0 | 0 |
| 0 | 0 | 0 |
| 0 | 0 | 0 |
| 0 | 0 | 0 |
| 0 | 0 | 0 |
| 0 | 0 | 0 |
| 0 | 0 | 0 |
| 0 | 0 | 0 |
| 0 | 0 | 0 |
| 0 | 0 | 0 |
| 0 | 0 | 0 |
| 0 | 0 | 0 |
| 0 | 0 | 0 |
| 1 | 1 | 1 |
| 0 | 1 | 0 |
| 0 | 0 | 0 |
| 0 | 0 | 0 |
| 0 | 0 | 0 |
| 0 | 0 | 0 |
| 1 | 1 | 1 |
| 1 | 1 | 1 |
| 0 | 0 | 0 |

| | | |
|---|---|---|
| 0 | 1 | 0 |
| 0 | 1 | 0 |
| 0 | 0 | 0 |
| 0 | 0 | 0 |
| 0 | 0 | 0 |
| 0 | 0 | 0 |
| 0 | 1 | 0 |
| 0 | 0 | 0 |
| 0 | 0 | 0 |
| 0 | 0 | 0 |
| 0 | 0 | 0 |
| 0 | 0 | 0 |
| 0 | 0 | 0 |
| 0 | 0 | 0 |
| 0 | 0 | 0 |
| 1 | 1 | 1 |
| 0 | 0 | 0 |
| 0 | 0 | 0 |
| 0 | 0 | 0 |
| 0 | 0 | 0 |
| 0 | 0 | 0 |
| 0 | 0 | 0 |
| 0 | 0 | 0 |
| 0 | 0 | 0 |
| 0 | 0 | 0 |
| 0 | 0 | 0 |
| 0 | 0 | 0 |
| 0 | 0 | 0 |
| 0 | 0 | 0 |
| 0 | 0 | 0 |
| 0 | 0 | 0 |
| 0 | 0 | 0 |
| 0 | 0 | 0 |
| 0 | 0 | 0 |
| 0 | 0 | 0 |
| 0 | 0 | 0 |
| 0 | 0 | 0 |
| 0 | 0 | 0 |
| 0 | 0 | 0 |
| 0 | 0 | 0 |
| 0 | 0 | 0 |
| 0 | 0 | 0 |
| 0 | 0 | 0 |
| 0 | 0 | 0 |
| 0 | 0 | 0 |
| 0 | 0 | 0 |
| 0 | 0 | 0 |
| 0 | 0 | 0 |
| 0 | 0 | 0 |
| 0 | 0 | 0 |
| 0 | 0 | 0 |
| 0 | 0 | 0 |
| 0 | 0 | 0 |

| | | |
|---|---|---|
| 1 | 0 | 0 |
| 1 | 0 | 0 |
| 1 | 0 | 0 |
| 0 | 0 | 0 |
| 0 | 0 | 0 |
| 0 | 0 | 0 |
| 0 | 0 | 0 |
| 1 | 0 | 0 |
| 0 | 0 | 0 |
| 1 | 0 | 0 |
| 1 | 0 | 0 |
| 0 | 0 | 0 |
| 1 | 0 | 0 |
| 0 | 0 | 0 |
| 0 | 0 | 0 |
| 1 | 0 | 0 |
| 1 | 0 | 0 |
| 1 | 0 | 0 |
| 1 | 0 | 0 |
| 0 | 0 | 0 |
| 0 | 0 | 0 |
| 0 | 1 | 0 |
| 0 | 0 | 0 |
| 0 | 0 | 0 |
| 0 | 0 | 0 |
| 1 | 0 | 0 |
| 0 | 0 | 0 |
| 0 | 0 | 0 |
| 1 | 0 | 0 |
| 1 | 0 | 0 |
| 1 | 1 | 1 |
| 0 | 0 | 0 |
| 0 | 0 | 0 |
| 0 | 1 | 0 |
| 0 | 0 | 0 |
| 0 | 0 | 0 |
| 0 | 0 | 0 |
| 0 | 0 | 0 |
| 0 | 0 | 0 |
| 1 | 0 | 0 |
| 1 | 0 | 0 |
| 0 | 0 | 0 |
| 0 | 0 | 0 |
| 1 | 1 | 1 |
| 1 | 0 | 0 |
| 1 | 0 | 0 |
| 0 | 0 | 0 |
| 0 | 0 | 0 |
| 0 | 0 | 0 |
| 0 | 0 | 0 |
| 0 | 0 | 0 |
| 0 | 1 | 0 |
| 1 | 0 | 0 |
| 0 | 0 | 0 |

| | | |
|---|---|---|
| 0 | 0 | 0 |
| 0 | 0 | 0 |
| 0 | 0 | 0 |
| 1 | 1 | 1 |
| 1 | 1 | 1 |
| 1 | 0 | 0 |
| 0 | 0 | 0 |
| 0 | 0 | 0 |
| 1 | 0 | 0 |
| 1 | 0 | 0 |
| 0 | 0 | 0 |
| 0 | 1 | 0 |
| 0 | 0 | 0 |
| 1 | 0 | 0 |
| 0 | 1 | 0 |
| 1 | 1 | 1 |
| 1 | 1 | 1 |
| 0 | 0 | 0 |
| 0 | 0 | 0 |
| 0 | 0 | 0 |
| 0 | 0 | 0 |
| 1 | 0 | 0 |
| 0 | 0 | 0 |
| 0 | 1 | 0 |
| 1 | 1 | 1 |
| 0 | 0 | 0 |
| 0 | 0 | 0 |
| 0 | 0 | 0 |
| 0 | 0 | 0 |
| 0 | 0 | 0 |
| 0 | 0 | 0 |
| 0 | 0 | 0 |
| 1 | 0 | 0 |
| 1 | 0 | 0 |
| 0 | 0 | 0 |
| 1 | 0 | 0 |
| 0 | 0 | 0 |
| 0 | 0 | 0 |
| 1 | 0 | 0 |
| 0 | 1 | 0 |
| 0 | 0 | 0 |
| 1 | 0 | 0 |
| 1 | 0 | 0 |
| 1 | 0 | 0 |
| 0 | 0 | 0 |
| 0 | 0 | 0 |
| 0 | 0 | 0 |
| 0 | 0 | 0 |
| 0 | 0 | 0 |
| 0 | 0 | 0 |
| 0 | 0 | 0 |
| 0 | 0 | 0 |
| 0 | 0 | 0 |
| 0 | 0 | 0 |

| | | |
|---|---|---|
| 0 | 0 | 0 |
| 0 | 0 | 0 |
| 0 | 0 | 0 |
| 0 | 0 | 0 |
| 0 | 0 | 0 |
| 0 | 0 | 0 |
| 0 | 1 | 0 |
| 0 | 1 | 0 |
| 0 | 0 | 0 |
| 0 | 0 | 0 |
| 0 | 0 | 0 |
| 0 | 0 | 0 |
| 0 | 0 | 0 |
| 0 | 1 | 0 |
| 0 | 1 | 0 |
| 0 | 0 | 0 |
| 0 | 1 | 0 |
| 0 | 0 | 0 |
| 1 | 1 | 1 |
| 0 | 0 | 0 |
| 1 | 1 | 1 |
| 1 | 0 | 0 |
| 1 | 1 | 1 |
| 0 | 0 | 0 |
| 0 | 0 | 0 |
| 0 | 0 | 0 |
| 0 | 0 | 0 |
| 0 | 0 | 0 |
| 0 | 0 | 0 |
| 0 | 0 | 0 |
| 0 | 0 | 0 |
| 1 | 0 | 0 |
| 0 | 0 | 0 |
| 0 | 0 | 0 |
| 0 | 0 | 0 |
| 0 | 0 | 0 |
| 0 | 0 | 0 |
| 0 | 0 | 0 |
| 0 | 0 | 0 |
| 0 | 0 | 0 |
| 0 | 1 | 0 |
| 0 | 1 | 0 |
| 0 | 0 | 0 |
| 0 | 1 | 0 |
| 0 | 0 | 0 |
| 0 | 0 | 0 |
| 0 | 0 | 0 |
| 1 | 1 | 1 |
| 1 | 1 | 1 |
| 0 | 0 | 0 |
| 0 | 1 | 0 |
| 1 | 1 | 1 |
| 0 | 0 | 0 |
| 0 | 1 | 0 |

| | | |
|---|---|---|
| 0 | 0 | 0 |
| 1 | 1 | 1 |
| 0 | 0 | 0 |
| 0 | 0 | 0 |
| 0 | 0 | 0 |
| 0 | 0 | 0 |
| 0 | 1 | 0 |
| 0 | 0 | 0 |
| 0 | 0 | 0 |
| 1 | 1 | 1 |
| 1 | 1 | 1 |
| 1 | 1 | 1 |
| 1 | 1 | 1 |
| 1 | 1 | 1 |
| 0 | 0 | 0 |
| 1 | 1 | 1 |
| 0 | 0 | 0 |
| 0 | 0 | 0 |
| 1 | 1 | 1 |
| 0 | 0 | 0 |
| 0 | 0 | 0 |
| 0 | 0 | 0 |
| 0 | 0 | 0 |
| 0 | 0 | 0 |
| 1 | 1 | 1 |
| 1 | 0 | 0 |
| 0 | 0 | 0 |
| 0 | 0 | 0 |
| 0 | 0 | 0 |
| 0 | 0 | 0 |
| 0 | 0 | 0 |
| 0 | 0 | 0 |
| 0 | 0 | 0 |
| 0 | 0 | 0 |
| 1 | 1 | 1 |
| 1 | 1 | 1 |
| 0 | 0 | 0 |
| 0 | 0 | 0 |
| 0 | 0 | 0 |
| 0 | 0 | 0 |
| 0 | 0 | 0 |
| 0 | 0 | 0 |
| 0 | 1 | 0 |
| 0 | 0 | 0 |
| 0 | 0 | 0 |
| 0 | 0 | 0 |
| 0 | 0 | 0 |
| 0 | 0 | 0 |
| 0 | 0 | 0 |
| 1 | 1 | 1 |
| 0 | 0 | 0 |
| 0 | 0 | 0 |
| 0 | 0 | 0 |
| 0 | 0 | 0 |

| | | |
|---|---|---|
| 0 | 0 | 0 |
| 0 | 0 | 0 |
| 0 | 0 | 0 |
| 0 | 0 | 0 |
| 1 | 1 | 1 |
| 0 | 0 | 0 |
| 0 | 0 | 0 |
| 0 | 0 | 0 |
| 0 | 0 | 0 |
| 0 | 0 | 0 |
| 0 | 0 | 0 |
| 0 | 0 | 0 |
| 0 | 0 | 0 |
| 0 | 0 | 0 |
| 0 | 0 | 0 |
| 0 | 0 | 0 |
| 0 | 0 | 0 |
| 0 | 0 | 0 |
| 1 | 1 | 1 |
| 0 | 0 | 0 |
| 0 | 0 | 0 |
| 0 | 0 | 0 |
| 0 | 0 | 0 |
| 1 | 0 | 0 |
| 0 | 0 | 0 |
| 0 | 0 | 0 |
| 0 | 0 | 0 |
| 1 | 0 | 0 |
| 1 | 0 | 0 |
| 1 | 0 | 0 |
| 0 | 0 | 0 |
| 0 | 0 | 0 |
| 0 | 0 | 0 |
| 0 | 0 | 0 |
| 0 | 0 | 0 |
| 1 | 0 | 0 |
| 1 | 0 | 0 |
| 0 | 0 | 0 |
| 0 | 0 | 0 |
| 1 | 0 | 0 |
| 1 | 0 | 0 |
| 1 | 0 | 0 |
| 0 | 1 | 0 |
| 1 | 1 | 1 |
| 0 | 0 | 0 |
| 0 | 0 | 0 |
| 0 | 0 | 0 |
| 0 | 0 | 0 |
| 0 | 0 | 0 |
| 0 | 1 | 0 |
| 1 | 1 | 1 |
| 0 | 0 | 0 |
| 1 | 1 | 1 |
| 0 | 0 | 0 |

| | | |
|---|---|---|
| 1 | 1 | 1 |
| 1 | 1 | 1 |
| 0 | 0 | 0 |
| 1 | 1 | 1 |
| 1 | 1 | 1 |
| 1 | 1 | 1 |
| 1 | 1 | 1 |
| 0 | 0 | 0 |
| 1 | 0 | 0 |
| 1 | 1 | 1 |
| 0 | 0 | 0 |
| 1 | 1 | 1 |
| 1 | 1 | 1 |
| 0 | 1 | 0 |
| 0 | 0 | 0 |
| 0 | 0 | 0 |
| 1 | 1 | 1 |
| 1 | 1 | 1 |
| 0 | 0 | 0 |
| 0 | 0 | 0 |
| 0 | 0 | 0 |
| 0 | 0 | 0 |
| 0 | 0 | 0 |
| 0 | 0 | 0 |
| 0 | 0 | 0 |
| 1 | 1 | 1 |
| 0 | 0 | 0 |
| 0 | 0 | 0 |
| 0 | 0 | 0 |
| 1 | 0 | 0 |
| 0 | 0 | 0 |
| 0 | 0 | 0 |
| 0 | 0 | 0 |
| 0 | 0 | 0 |
| 0 | 0 | 0 |
| 0 | 0 | 0 |
| 0 | 0 | 0 |
| 0 | 0 | 0 |
| 0 | 0 | 0 |
| 0 | 0 | 0 |
| 0 | 1 | 0 |
| 0 | 0 | 0 |
| 1 | 1 | 1 |
| 0 | 0 | 0 |
| 0 | 0 | 0 |
| 0 | 0 | 0 |
| 0 | 0 | 0 |
| 0 | 0 | 0 |
| 0 | 0 | 0 |
| 0 | 0 | 0 |
| 0 | 1 | 0 |
| 0 | 0 | 0 |
| 0 | 0 | 0 |
| 0 | 0 | 0 |

| | | |
|---|---|---|
| 1 | 1 | 1 |
| 0 | 0 | 0 |
| 0 | 1 | 0 |
| 0 | 1 | 0 |
| 0 | 0 | 0 |
| 0 | 0 | 0 |
| 0 | 0 | 0 |
| 0 | 0 | 0 |
| 0 | 0 | 0 |
| 0 | 1 | 0 |
| 1 | 1 | 1 |
| 1 | 1 | 1 |
| 1 | 1 | 1 |
| 0 | 0 | 0 |
| 1 | 1 | 1 |
| 0 | 0 | 0 |
| 0 | 0 | 0 |
| 0 | 0 | 0 |
| 0 | 0 | 0 |
| 1 | 0 | 0 |
| 0 | 0 | 0 |
| 1 | 1 | 1 |
| 0 | 0 | 0 |
| 0 | 1 | 0 |
| 1 | 1 | 1 |
| 1 | 1 | 1 |
| 0 | 0 | 0 |
| 1 | 1 | 1 |
| 1 | 1 | 1 |
| 1 | 1 | 1 |
| 0 | 0 | 0 |
| 0 | 0 | 0 |
| 1 | 1 | 1 |
| 1 | 1 | 1 |
| 0 | 0 | 0 |
| 0 | 0 | 0 |
| 0 | 0 | 0 |
| 0 | 0 | 0 |
| 0 | 0 | 0 |
| 1 | 1 | 1 |
| 0 | 0 | 0 |
| 0 | 0 | 0 |
| 0 | 0 | 0 |
| 0 | 0 | 0 |
| 1 | 0 | 0 |
| 0 | 0 | 0 |
| 0 | 0 | 0 |
| 0 | 0 | 0 |
| 0 | 0 | 0 |
| 0 | 0 | 0 |
| 0 | 0 | 0 |
| 0 | 0 | 0 |
| 1 | 0 | 0 |
| 0 | 0 | 0 |

| | | |
|---|---|---|
| 0 | 0 | 0 |
| 0 | 0 | 0 |
| 0 | 0 | 0 |
| 0 | 0 | 0 |
| 0 | 0 | 0 |
| 0 | 0 | 0 |
| 0 | 0 | 0 |
| 1 | 0 | 0 |
| 0 | 0 | 0 |
| 0 | 0 | 0 |
| 0 | 0 | 0 |
| 0 | 0 | 0 |
| 1 | 1 | 1 |
| 0 | 0 | 0 |
| 0 | 0 | 0 |
| 0 | 0 | 0 |
| 0 | 0 | 0 |
| 0 | 0 | 0 |
| 0 | 0 | 0 |
| 1 | 1 | 1 |
| 0 | 0 | 0 |
| 0 | 0 | 0 |
| 0 | 0 | 0 |
| 0 | 0 | 0 |
| 0 | 0 | 0 |
| 1 | 0 | 0 |
| 1 | 1 | 1 |
| 0 | 1 | 0 |
| 0 | 0 | 0 |
| 0 | 0 | 0 |
| 0 | 0 | 0 |
| 0 | 0 | 0 |
| 0 | 0 | 0 |
| 0 | 0 | 0 |
| 0 | 0 | 0 |
| 1 | 0 | 0 |
| 0 | 0 | 0 |
| 0 | 0 | 0 |
| 1 | 1 | 1 |
| 1 | 0 | 0 |
| 0 | 1 | 0 |
| 0 | 0 | 0 |
| 0 | 0 | 0 |
| 0 | 0 | 0 |
| 0 | 0 | 0 |
| 0 | 0 | 0 |
| 0 | 0 | 0 |
| 1 | 1 | 1 |
| 1 | 1 | 1 |
| 0 | 0 | 0 |
| 0 | 0 | 0 |
| 0 | 0 | 0 |
| 0 | 0 | 0 |
| 1 | 1 | 1 |

| | | |
|---|---|---|
| 1 | 1 | 1 |
| 0 | 0 | 0 |
| 0 | 1 | 0 |
| 0 | 0 | 0 |
| 1 | 0 | 0 |
| 1 | 0 | 0 |
| 0 | 0 | 0 |
| 1 | 0 | 0 |
| 1 | 0 | 0 |
| 1 | 0 | 0 |
| 1 | 0 | 0 |
| 1 | 0 | 0 |
| 1 | 1 | 1 |
| 1 | 1 | 1 |
| 1 | 1 | 1 |
| 1 | 1 | 1 |
| 0 | 0 | 0 |
| 0 | 0 | 0 |
| 1 | 0 | 0 |
| 0 | 0 | 0 |
| 0 | 0 | 0 |
| 0 | 0 | 0 |
| 1 | 0 | 0 |
| 0 | 0 | 0 |
| 0 | 0 | 0 |
| 1 | 1 | 1 |
| 1 | 0 | 0 |
| 1 | 0 | 0 |
| 0 | 0 | 0 |
| 0 | 0 | 0 |
| 0 | 0 | 0 |
| 0 | 0 | 0 |
| 0 | 0 | 0 |
| 0 | 0 | 0 |
| 0 | 0 | 0 |
| 0 | 0 | 0 |
| 1 | 0 | 0 |
| 1 | 1 | 1 |
| 1 | 1 | 1 |
| 1 | 1 | 1 |
| 0 | 0 | 0 |
| 1 | 1 | 1 |
| 0 | 0 | 0 |
| 1 | 1 | 1 |
| 0 | 0 | 0 |
| 0 | 1 | 0 |
| 0 | 0 | 0 |
| 0 | 0 | 0 |
| 0 | 0 | 0 |
| 0 | 0 | 0 |
| 0 | 0 | 0 |
| 0 | 0 | 0 |
| 0 | 0 | 0 |
| 0 | 0 | 0 |

| | | |
|---|---|---|
| 0 | 0 | 0 |
| 0 | 0 | 0 |
| 0 | 0 | 0 |
| 0 | 0 | 0 |
| 0 | 0 | 0 |
| 0 | 0 | 0 |
| 0 | 0 | 0 |
| 1 | 1 | 1 |
| 0 | 0 | 0 |
| 0 | 0 | 0 |
| 0 | 0 | 0 |
| 1 | 0 | 0 |
| 0 | 0 | 0 |
| 0 | 0 | 0 |
| 0 | 0 | 0 |
| 0 | 0 | 0 |
| 0 | 0 | 0 |
| 0 | 0 | 0 |
| 0 | 0 | 0 |
| 0 | 0 | 0 |
| 0 | 0 | 0 |
| 0 | 0 | 0 |
| 0 | 0 | 0 |
| 0 | 0 | 0 |
| 0 | 0 | 0 |
| 0 | 0 | 0 |
| 0 | 0 | 0 |
| 0 | 0 | 0 |
| 0 | 0 | 0 |
| 0 | 0 | 0 |
| 0 | 0 | 0 |
| 0 | 0 | 0 |
| 0 | 0 | 0 |
| 0 | 0 | 0 |
| 0 | 0 | 0 |
| 0 | 0 | 0 |
| 0 | 0 | 0 |
| 0 | 0 | 0 |
| 0 | 0 | 0 |
| 0 | 0 | 0 |
| 1 | 0 | 0 |
| 0 | 0 | 0 |
| 0 | 0 | 0 |
| 0 | 0 | 0 |
| 0 | 0 | 0 |
| 0 | 0 | 0 |
| 0 | 0 | 0 |
| 0 | 0 | 0 |
| 0 | 0 | 0 |
| 0 | 1 | 0 |
| 0 | 1 | 0 |
| 1 | 0 | 0 |
| 0 | 0 | 0 |
| 0 | 0 | 0 |

| | | |
|---|---|---|
| 0 | 0 | 0 |
| 0 | 0 | 0 |
| 0 | 0 | 0 |
| 0 | 0 | 0 |
| 0 | 0 | 0 |
| 0 | 0 | 0 |
| 0 | 0 | 0 |
| 1 | 0 | 0 |
| 1 | 0 | 0 |
| 0 | 0 | 0 |
| 0 | 0 | 0 |
| 1 | 0 | 0 |
| 0 | 0 | 0 |
| 0 | 0 | 0 |
| 0 | 0 | 0 |
| 0 | 0 | 0 |
| 0 | 0 | 0 |
| 1 | 0 | 0 |
| 0 | 0 | 0 |
| 1 | 0 | 0 |
| 0 | 0 | 0 |
| 0 | 0 | 0 |
| 1 | 1 | 1 |
| 0 | 0 | 0 |
| 0 | 0 | 0 |
| 1 | 0 | 0 |
| 0 | 0 | 0 |
| 0 | 0 | 0 |
| 0 | 0 | 0 |
| 0 | 0 | 0 |
| 0 | 0 | 0 |
| 0 | 0 | 0 |
| 1 | 0 | 0 |
| 0 | 0 | 0 |
| 0 | 0 | 0 |
| 0 | 0 | 0 |
| 1 | 1 | 1 |
| 0 | 0 | 0 |
| 0 | 0 | 0 |
| 0 | 0 | 0 |
| 0 | 0 | 0 |
| 0 | 0 | 0 |
| 0 | 0 | 0 |
| 0 | 0 | 0 |
| 0 | 0 | 0 |
| 0 | 0 | 0 |
| 0 | 0 | 0 |
| 0 | 0 | 0 |
| 0 | 0 | 0 |
| 0 | 0 | 0 |
| 0 | 0 | 0 |
| 0 | 0 | 0 |
| 0 | 0 | 0 |
| 0 | 0 | 0 |

| | | |
|---|---|---|
| 1 | 0 | 0 |
| 0 | 1 | 0 |
| 0 | 0 | 0 |
| 0 | 0 | 0 |
| 0 | 0 | 0 |
| 0 | 0 | 0 |
| 0 | 0 | 0 |
| 0 | 0 | 0 |
| 0 | 0 | 0 |
| 1 | 0 | 0 |
| 0 | 1 | 0 |
| 1 | 1 | 1 |
| 0 | 0 | 0 |
| 0 | 0 | 0 |
| 0 | 0 | 0 |
| 0 | 0 | 0 |
| 0 | 0 | 0 |
| 0 | 0 | 0 |
| 1 | 1 | 1 |
| 1 | 0 | 0 |
| 1 | 0 | 0 |
| 0 | 0 | 0 |
| 0 | 1 | 0 |
| 0 | 0 | 0 |
| 0 | 0 | 0 |
| 0 | 0 | 0 |
| 0 | 0 | 0 |
| 0 | 0 | 0 |
| 0 | 0 | 0 |
| 0 | 0 | 0 |
| 0 | 0 | 0 |
| 0 | 0 | 0 |
| 0 | 0 | 0 |
| 0 | 0 | 0 |
| 0 | 0 | 0 |
| 0 | 0 | 0 |
| 0 | 0 | 0 |
| 0 | 0 | 0 |
| 0 | 0 | 0 |
| 0 | 0 | 0 |
| 0 | 0 | 0 |
| 1 | 0 | 0 |
| 0 | 0 | 0 |
| 0 | 0 | 0 |
| 0 | 0 | 0 |
| 0 | 0 | 0 |
| 1 | 0 | 0 |
| 0 | 0 | 0 |
| 0 | 0 | 0 |
| 0 | 0 | 0 |
| 0 | 0 | 0 |
| 0 | 0 | 0 |
| 1 | 0 | 0 |
| 0 | 1 | 0 |

| | | |
|---|---|---|
| 0 | 0 | 0 |
| 0 | 0 | 0 |
| 1 | 0 | 0 |
| 1 | 0 | 0 |
| 1 | 0 | 0 |
| 0 | 0 | 0 |
| 1 | 0 | 0 |
| 0 | 0 | 0 |
| 0 | 0 | 0 |
| 0 | 0 | 0 |
| 0 | 0 | 0 |
| 0 | 0 | 0 |
| 0 | 0 | 0 |
| 1 | 0 | 0 |
| 1 | 0 | 0 |
| 1 | 0 | 0 |
| 0 | 0 | 0 |
| 1 | 1 | 1 |
| 0 | 0 | 0 |
| 0 | 0 | 0 |
| 1 | 0 | 0 |
| 1 | 0 | 0 |
| 0 | 0 | 0 |
| 0 | 0 | 0 |
| 1 | 0 | 0 |
| 0 | 0 | 0 |
| 0 | 0 | 0 |
| 1 | 0 | 0 |
| 0 | 0 | 0 |
| 0 | 0 | 0 |
| 0 | 0 | 0 |
| 0 | 0 | 0 |
| 0 | 0 | 0 |
| 0 | 0 | 0 |
| 0 | 0 | 0 |
| 0 | 0 | 0 |
| 0 | 0 | 0 |
| 0 | 0 | 0 |
| 0 | 0 | 0 |
| 0 | 0 | 0 |
| 1 | 0 | 0 |
| 1 | 0 | 0 |
| 1 | 0 | 0 |
| 1 | 1 | 1 |
| 1 | 1 | 1 |
| 0 | 0 | 0 |
| 0 | 0 | 0 |
| 0 | 0 | 0 |
| 0 | 0 | 0 |
| 0 | 0 | 0 |
| 0 | 0 | 0 |
| 0 | 0 | 0 |
| 0 | 0 | 0 |
| 1 | 0 | 0 |

| | | |
|---|---|---|
| 1 | 0 | 0 |
| 0 | 0 | 0 |
| 0 | 0 | 0 |
| 0 | 0 | 0 |
| 1 | 0 | 0 |
| 0 | 0 | 0 |
| 0 | 0 | 0 |
| 0 | 0 | 0 |
| 1 | 0 | 0 |
| 1 | 0 | 0 |
| 0 | 0 | 0 |
| 0 | 0 | 0 |
| 0 | 0 | 0 |
| 1 | 0 | 0 |
| 0 | 0 | 0 |
| 0 | 0 | 0 |
| 0 | 0 | 0 |
| 0 | 0 | 0 |
| 0 | 0 | 0 |
| 0 | 0 | 0 |
| 0 | 0 | 0 |
| 1 | 1 | 1 |
| 0 | 0 | 0 |
| 0 | 1 | 0 |
| 0 | 0 | 0 |
| 0 | 0 | 0 |
| 0 | 0 | 0 |
| 0 | 0 | 0 |
| 0 | 0 | 0 |
| 0 | 0 | 0 |
| 0 | 0 | 0 |
| 0 | 0 | 0 |
| 0 | 0 | 0 |
| 0 | 0 | 0 |
| 0 | 1 | 0 |
| 0 | 0 | 0 |
| 0 | 0 | 0 |
| 0 | 0 | 0 |
| 0 | 0 | 0 |
| 0 | 0 | 0 |
| 0 | 0 | 0 |
| 0 | 0 | 0 |
| 0 | 0 | 0 |
| 0 | 0 | 0 |
| 0 | 0 | 0 |
| 0 | 0 | 0 |
| 0 | 0 | 0 |
| 1 | 1 | 1 |
| 0 | 0 | 0 |
| 1 | 1 | 1 |
| 0 | 0 | 0 |
| 0 | 0 | 0 |
| 0 | 0 | 0 |
| 0 | 0 | 0 |

| | | |
|---|---|---|
| 0 | 0 | 0 |
| 0 | 0 | 0 |
| 1 | 1 | 1 |
| 0 | 0 | 0 |
| 0 | 0 | 0 |
| 0 | 1 | 0 |
| 0 | 1 | 0 |
| 0 | 1 | 0 |
| 0 | 0 | 0 |
| 0 | 1 | 0 |
| 0 | 0 | 0 |
| 0 | 0 | 0 |
| 0 | 0 | 0 |
| 1 | 1 | 1 |
| 0 | 0 | 0 |
| 0 | 1 | 0 |
| 0 | 0 | 0 |
| 0 | 0 | 0 |
| 0 | 0 | 0 |
| 1 | 1 | 1 |
| 1 | 1 | 1 |
| 0 | 0 | 0 |
| 1 | 1 | 1 |
| 0 | 0 | 0 |
| 0 | 0 | 0 |
| 1 | 1 | 1 |
| 1 | 1 | 1 |
| 0 | 0 | 0 |
| 1 | 1 | 1 |
| 0 | 0 | 0 |
| 0 | 1 | 0 |
| 0 | 1 | 0 |
| 1 | 1 | 1 |
| 0 | 1 | 0 |
| 1 | 1 | 1 |
| 1 | 1 | 1 |
| 1 | 1 | 1 |
| 1 | 1 | 1 |
| 1 | 1 | 1 |
| 1 | 1 | 1 |
| 1 | 1 | 1 |
| 1 | 1 | 1 |
| 0 | 1 | 0 |
| 0 | 1 | 0 |
| 0 | 1 | 0 |
| 0 | 1 | 0 |
| 0 | 1 | 0 |
| 0 | 1 | 0 |
| 0 | 1 | 0 |
| 0 | 1 | 0 |
| 0 | 1 | 0 |
| 0 | 1 | 0 |
| 0 | 1 | 0 |
| 0 | 1 | 0 |

| | | |
|---|---|---|
| 0 | 1 | 0 |
| 1 | 1 | 1 |
| 1 | 1 | 1 |
| 1 | 1 | 1 |
| 1 | 1 | 1 |
| 1 | 1 | 1 |
| 1 | 1 | 1 |
| 0 | 1 | 0 |
| 0 | 1 | 0 |
| 0 | 1 | 0 |
| 0 | 1 | 0 |
| 0 | 0 | 0 |
| 0 | 0 | 0 |
| 0 | 0 | 0 |
| 1 | 0 | 0 |
| 0 | 0 | 0 |
| 1 | 0 | 0 |
| 0 | 0 | 0 |
| 1 | 0 | 0 |
| 1 | 1 | 1 |
| 0 | 0 | 0 |
| 1 | 1 | 1 |
| 1 | 1 | 1 |
| 0 | 0 | 0 |
| 0 | 0 | 0 |
| 1 | 1 | 1 |
| 1 | 1 | 1 |
| 0 | 0 | 0 |
| 0 | 0 | 0 |
| 0 | 0 | 0 |
| 0 | 0 | 0 |
| 0 | 0 | 0 |
| 0 | 1 | 0 |
| 0 | 1 | 0 |
| 1 | 1 | 1 |
| 1 | 1 | 1 |
| 1 | 1 | 1 |
| 1 | 1 | 1 |
| 1 | 1 | 1 |
| 0 | 0 | 0 |
| 0 | 0 | 0 |
| 0 | 1 | 0 |
| 0 | 0 | 0 |
| 0 | 0 | 0 |
| 0 | 0 | 0 |
| 1 | 0 | 0 |
| 0 | 0 | 0 |
| 1 | 0 | 0 |
| 1 | 0 | 0 |
| 0 | 0 | 0 |
| 0 | 0 | 0 |
| 0 | 0 | 0 |
| 0 | 0 | 0 |
| 0 | 0 | 0 |

| | | |
|---|---|---|
| 0 | 0 | 0 |
| 0 | 1 | 0 |
| 1 | 1 | 1 |
| 1 | 1 | 1 |
| 0 | 0 | 0 |
| 1 | 1 | 1 |
| 0 | 0 | 0 |
| 1 | 1 | 1 |
| 1 | 1 | 1 |
| 0 | 0 | 0 |
| 1 | 0 | 0 |
| 1 | 1 | 1 |
| 1 | 1 | 1 |
| 0 | 0 | 0 |
| 1 | 0 | 0 |
| 1 | 0 | 0 |
| 0 | 0 | 0 |
| 1 | 0 | 0 |
| 0 | 0 | 0 |
| 0 | 0 | 0 |
| 0 | 0 | 0 |
| 0 | 0 | 0 |
| 0 | 0 | 0 |
| 0 | 0 | 0 |
| 0 | 0 | 0 |
| 0 | 0 | 0 |
| 0 | 0 | 0 |
| 0 | 0 | 0 |
| 0 | 0 | 0 |
| 0 | 0 | 0 |
| 0 | 0 | 0 |
| 0 | 0 | 0 |
| 0 | 0 | 0 |
| 0 | 0 | 0 |
| 0 | 0 | 0 |
| 0 | 0 | 0 |
| 1 | 0 | 0 |
| 0 | 0 | 0 |
| 0 | 0 | 0 |
| 0 | 0 | 0 |
| 0 | 0 | 0 |
| 0 | 0 | 0 |
| 0 | 0 | 0 |
| 0 | 0 | 0 |
| 0 | 0 | 0 |
| 0 | 0 | 0 |
| 0 | 0 | 0 |
| 0 | 0 | 0 |
| 1 | 0 | 0 |
| 1 | 0 | 0 |
| 0 | 0 | 0 |
| 1 | 1 | 1 |
| 0 | 0 | 0 |
| 0 | 0 | 0 |

| | | |
|---|---|---|
| 0 | 0 | 0 |
| 0 | 0 | 0 |
| 0 | 0 | 0 |
| 0 | 0 | 0 |
| 0 | 0 | 0 |
| 0 | 0 | 0 |
| 0 | 0 | 0 |
| 0 | 0 | 0 |
| 0 | 0 | 0 |
| 0 | 0 | 0 |
| 0 | 0 | 0 |
| 0 | 0 | 0 |
| 0 | 0 | 0 |
| 0 | 0 | 0 |
| 0 | 1 | 0 |
| 0 | 0 | 0 |
| 0 | 0 | 0 |
| 0 | 0 | 0 |
| 1 | 0 | 0 |
| 0 | 0 | 0 |
| 0 | 0 | 0 |
| 0 | 0 | 0 |
| 0 | 0 | 0 |
| 0 | 0 | 0 |
| 0 | 0 | 0 |
| 1 | 0 | 0 |
| 0 | 0 | 0 |
| 0 | 0 | 0 |
| 0 | 0 | 0 |
| 0 | 0 | 0 |
| 1 | 1 | 1 |
| 0 | 0 | 0 |
| 1 | 1 | 1 |
| 1 | 0 | 0 |
| 0 | 0 | 0 |
| 0 | 0 | 0 |
| 1 | 1 | 1 |
| 1 | 1 | 1 |
| 1 | 1 | 1 |
| 1 | 0 | 0 |
| 0 | 0 | 0 |
| 0 | 0 | 0 |
| 0 | 0 | 0 |
| 0 | 0 | 0 |
| 0 | 0 | 0 |
| 0 | 0 | 0 |
| 0 | 0 | 0 |
| 0 | 0 | 0 |
| 1 | 1 | 1 |
| 1 | 1 | 1 |
| 0 | 0 | 0 |
| 0 | 0 | 0 |
| 0 | 0 | 0 |
| 0 | 1 | 0 |

| | | |
|---|---|---|
| 1 | 1 | 1 |
| 1 | 1 | 1 |
| 1 | 0 | 0 |
| 0 | 0 | 0 |
| 1 | 0 | 0 |
| 1 | 1 | 1 |
| 0 | 0 | 0 |
| 1 | 1 | 1 |
| 0 | 0 | 0 |
| 0 | 0 | 0 |
| 0 | 0 | 0 |
| 0 | 0 | 0 |
| 0 | 0 | 0 |
| 0 | 0 | 0 |
| 0 | 0 | 0 |
| 0 | 0 | 0 |
| 0 | 1 | 0 |
| 0 | 0 | 0 |
| 0 | 0 | 0 |
| 1 | 0 | 0 |
| 0 | 0 | 0 |
| 0 | 0 | 0 |
| 1 | 1 | 1 |
| 0 | 0 | 0 |
| 1 | 1 | 1 |
| 1 | 1 | 1 |
| 0 | 1 | 0 |
| 0 | 0 | 0 |
| 0 | 0 | 0 |
| 0 | 0 | 0 |
| 1 | 1 | 1 |
| 0 | 0 | 0 |
| 0 | 0 | 0 |
| 0 | 0 | 0 |
| 0 | 1 | 0 |
| 0 | 1 | 0 |
| 0 | 0 | 0 |
| 0 | 0 | 0 |
| 0 | 0 | 0 |
| 1 | 1 | 1 |
| 1 | 1 | 1 |
| 0 | 1 | 0 |
| 0 | 1 | 0 |
| 0 | 0 | 0 |
| 1 | 1 | 1 |
| 0 | 0 | 0 |
| 1 | 1 | 1 |
| 0 | 0 | 0 |
| 0 | 0 | 0 |
| 1 | 1 | 1 |
| 0 | 1 | 0 |
| 0 | 0 | 0 |
| 0 | 0 | 0 |
| 1 | 1 | 1 |

| | | |
|---|---|---|
| 1 | 1 | 1 |
| 1 | 1 | 1 |
| 0 | 1 | 0 |
| 0 | 0 | 0 |
| 0 | 0 | 0 |
| 1 | 0 | 0 |
| 1 | 0 | 0 |
| 0 | 0 | 0 |
| 0 | 0 | 0 |
| 1 | 1 | 1 |
| 0 | 0 | 0 |
| 0 | 0 | 0 |
| 0 | 0 | 0 |
| 0 | 0 | 0 |
| 0 | 0 | 0 |
| 0 | 0 | 0 |
| 0 | 0 | 0 |
| 0 | 0 | 0 |
| 0 | 0 | 0 |
| 0 | 1 | 0 |
| 1 | 1 | 1 |
| 1 | 1 | 1 |
| 1 | 1 | 1 |
| 1 | 1 | 1 |
| 0 | 0 | 0 |
| 0 | 1 | 0 |
| 0 | 0 | 0 |
| 0 | 0 | 0 |
| 1 | 1 | 1 |
| 1 | 1 | 1 |
| 1 | 1 | 1 |
| 1 | 1 | 1 |
| 1 | 1 | 1 |
| 0 | 0 | 0 |
| 0 | 0 | 0 |
| 1 | 1 | 1 |
| 0 | 0 | 0 |
| 0 | 1 | 0 |
| 0 | 0 | 0 |
| 0 | 0 | 0 |
| 0 | 0 | 0 |
| 0 | 0 | 0 |
| 0 | 1 | 0 |
| 0 | 0 | 0 |
| 0 | 0 | 0 |
| 0 | 0 | 0 |
| 0 | 0 | 0 |
| 0 | 0 | 0 |
| 0 | 0 | 0 |
| 0 | 0 | 0 |
| 0 | 0 | 0 |
| 0 | 0 | 0 |
| 0 | 0 | 0 |
| 0 | 1 | 0 |

| | | |
|---|---|---|
| 0 | 0 | 0 |
| 0 | 1 | 0 |
| 0 | 0 | 0 |
| 0 | 0 | 0 |
| 0 | 0 | 0 |
| 1 | 1 | 1 |
| 0 | 0 | 0 |
| 1 | 1 | 1 |
| 0 | 1 | 0 |
| 0 | 0 | 0 |
| 0 | 1 | 0 |
| 0 | 1 | 0 |
| 0 | 1 | 0 |
| 0 | 1 | 0 |
| 0 | 1 | 0 |
| 0 | 0 | 0 |
| 0 | 0 | 0 |
| 0 | 1 | 0 |
| 0 | 0 | 0 |
| 1 | 1 | 1 |
| 0 | 0 | 0 |
| 0 | 0 | 0 |
| 0 | 0 | 0 |
| 0 | 0 | 0 |
| 0 | 0 | 0 |
| 0 | 0 | 0 |
| 0 | 0 | 0 |
| 0 | 0 | 0 |
| 0 | 0 | 0 |
| 0 | 0 | 0 |
| 0 | 0 | 0 |
| 0 | 0 | 0 |
| 1 | 0 | 0 |
| 1 | 0 | 0 |
| 0 | 0 | 0 |
| 0 | 0 | 0 |
| 0 | 0 | 0 |
| 0 | 0 | 0 |
| 1 | 0 | 0 |
| 0 | 0 | 0 |
| 0 | 0 | 0 |
| 0 | 0 | 0 |
| 1 | 0 | 0 |
| 0 | 1 | 0 |
| 0 | 0 | 0 |
| 0 | 0 | 0 |
| 0 | 0 | 0 |
| 0 | 0 | 0 |
| 0 | 0 | 0 |
| 0 | 0 | 0 |
| 1 | 0 | 0 |
| 0 | 0 | 0 |
| 0 | 0 | 0 |
| 1 | 0 | 0 |

| | | |
|---|---|---|
| 0 | 0 | 0 |
| 1 | 0 | 0 |
| 0 | 0 | 0 |
| 0 | 0 | 0 |
| 0 | 0 | 0 |
| 0 | 0 | 0 |
| 0 | 0 | 0 |
| 0 | 0 | 0 |
| 0 | 0 | 0 |
| 0 | 0 | 0 |
| 0 | 0 | 0 |
| 1 | 1 | 1 |
| 0 | 0 | 0 |
| 1 | 1 | 1 |
| 1 | 0 | 0 |
| 1 | 0 | 0 |
| 0 | 0 | 0 |
| 0 | 0 | 0 |
| 0 | 0 | 0 |
| 0 | 0 | 0 |
| 1 | 0 | 0 |
| 0 | 0 | 0 |
| 0 | 1 | 0 |
| 0 | 0 | 0 |
| 1 | 1 | 1 |
| 1 | 1 | 1 |
| 1 | 1 | 1 |
| 0 | 0 | 0 |
| 1 | 1 | 1 |
| 1 | 1 | 1 |
| 1 | 1 | 1 |
| 0 | 0 | 0 |
| 1 | 1 | 1 |
| 0 | 0 | 0 |
| 0 | 0 | 0 |
| 0 | 0 | 0 |
| 0 | 1 | 0 |
| 1 | 1 | 1 |
| 0 | 0 | 0 |
| 0 | 0 | 0 |
| 0 | 0 | 0 |
| 0 | 0 | 0 |
| 0 | 0 | 0 |
| 0 | 0 | 0 |
| 0 | 0 | 0 |
| 0 | 0 | 0 |
| 1 | 1 | 1 |
| 1 | 1 | 1 |
| 0 | 0 | 0 |
| 0 | 0 | 0 |
| 0 | 0 | 0 |
| 1 | 1 | 1 |
| 1 | 1 | 1 |
| 0 | 0 | 0 |

| | | |
|---|---|---|
| 0 | 0 | 0 |
| 0 | 1 | 0 |
| 0 | 1 | 0 |
| 1 | 1 | 1 |
| 0 | 0 | 0 |
| 0 | 0 | 0 |
| 0 | 0 | 0 |
| 0 | 0 | 0 |
| 0 | 0 | 0 |
| 0 | 0 | 0 |
| 1 | 1 | 1 |
| 0 | 0 | 0 |
| 0 | 0 | 0 |
| 0 | 0 | 0 |
| 1 | 0 | 0 |
| 1 | 1 | 1 |
| 0 | 0 | 0 |
| 1 | 0 | 0 |
| 0 | 0 | 0 |
| 0 | 0 | 0 |
| 1 | 0 | 0 |
| 0 | 0 | 0 |
| 0 | 0 | 0 |
| 1 | 0 | 0 |
| 0 | 0 | 0 |
| 0 | 0 | 0 |
| 0 | 0 | 0 |
| 1 | 0 | 0 |
| 0 | 0 | 0 |
| 0 | 0 | 0 |
| 0 | 0 | 0 |
| 1 | 1 | 1 |
| 1 | 1 | 1 |
| 1 | 1 | 1 |
| 0 | 1 | 0 |
| 0 | 1 | 0 |
| 0 | 1 | 0 |
| 1 | 1 | 1 |
| 1 | 1 | 1 |
| 1 | 1 | 1 |
| 0 | 1 | 0 |
| 1 | 1 | 1 |
| 0 | 0 | 0 |
| 0 | 1 | 0 |
| 1 | 1 | 1 |
| 1 | 1 | 1 |
| 0 | 1 | 0 |
| 1 | 1 | 1 |
| 1 | 1 | 1 |
| 1 | 1 | 1 |
| 1 | 1 | 1 |
| 1 | 1 | 1 |
| 1 | 1 | 1 |
| 1 | 1 | 1 |

| | | |
|---|---|---|
| 0 | 1 | 0 |
| 0 | 1 | 0 |
| 1 | 1 | 1 |
| 1 | 1 | 1 |
| 1 | 1 | 1 |
| 1 | 1 | 1 |
| 1 | 1 | 1 |
| 1 | 1 | 1 |
| 0 | 0 | 0 |
| 0 | 0 | 0 |
| 0 | 0 | 0 |
| 0 | 0 | 0 |
| 0 | 0 | 0 |
| 0 | 0 | 0 |
| 0 | 0 | 0 |
| 0 | 0 | 0 |
| 0 | 0 | 0 |
| 0 | 0 | 0 |
| 0 | 0 | 0 |
| 0 | 0 | 0 |
| 0 | 0 | 0 |
| 0 | 0 | 0 |
| 0 | 0 | 0 |
| 0 | 0 | 0 |
| 0 | 0 | 0 |
| 1 | 1 | 1 |
| 1 | 1 | 1 |
| 0 | 0 | 0 |
| 0 | 0 | 0 |
| 0 | 0 | 0 |
| 0 | 0 | 0 |
| 0 | 0 | 0 |
| 0 | 0 | 0 |
| 0 | 0 | 0 |
| 0 | 0 | 0 |
| 0 | 0 | 0 |
| 0 | 0 | 0 |
| 0 | 0 | 0 |
| 0 | 0 | 0 |
| 0 | 0 | 0 |
| 0 | 0 | 0 |
| 0 | 0 | 0 |
| 0 | 0 | 0 |
| 0 | 0 | 0 |
| 0 | 0 | 0 |
| 0 | 0 | 0 |
| 1 | 1 | 1 |
| 0 | 1 | 0 |
| 1 | 1 | 1 |
| 1 | 1 | 1 |
| 1 | 1 | 1 |
| 1 | 1 | 1 |
| 0 | 0 | 0 |
| 0 | 0 | 0 |

| | | |
|---|---|---|
| 0 | 0 | 0 |
| 0 | 0 | 0 |
| 0 | 0 | 0 |
| 0 | 0 | 0 |
| 0 | 1 | 0 |
| 0 | 0 | 0 |
| 0 | 0 | 0 |
| 0 | 0 | 0 |
| 1 | 1 | 1 |
| 0 | 0 | 0 |
| 0 | 0 | 0 |
| 0 | 1 | 0 |
| 0 | 0 | 0 |
| 0 | 0 | 0 |
| 0 | 0 | 0 |
| 0 | 0 | 0 |
| 1 | 1 | 1 |
| 0 | 0 | 0 |
| 0 | 0 | 0 |
| 1 | 1 | 1 |
| 0 | 0 | 0 |
| 0 | 0 | 0 |
| 0 | 0 | 0 |
| 0 | 0 | 0 |
| 0 | 0 | 0 |
| 0 | 1 | 0 |
| 0 | 1 | 0 |
| 1 | 1 | 1 |
| 1 | 1 | 1 |
| 0 | 1 | 0 |
| 1 | 1 | 1 |
| 1 | 1 | 1 |
| 1 | 0 | 0 |
| 1 | 0 | 0 |
| 0 | 0 | 0 |
| 0 | 0 | 0 |
| 0 | 0 | 0 |
| 0 | 0 | 0 |
| 0 | 0 | 0 |
| 0 | 0 | 0 |
| 1 | 1 | 1 |
| 1 | 1 | 1 |
| 1 | 1 | 1 |
| 0 | 0 | 0 |
| 0 | 0 | 0 |
| 0 | 0 | 0 |
| 1 | 1 | 1 |
| 0 | 1 | 0 |
| 0 | 0 | 0 |
| 0 | 0 | 0 |
| 1 | 1 | 1 |
| 1 | 1 | 1 |
| 1 | 1 | 1 |
| 1 | 1 | 1 |

| | | |
|---|---|---|
| 0 | 1 | 0 |
| 1 | 1 | 1 |
| 0 | 1 | 0 |
| 0 | 0 | 0 |
| 0 | 0 | 0 |
| 1 | 1 | 1 |
| 1 | 1 | 1 |
| 1 | 1 | 1 |
| 1 | 1 | 1 |
| 1 | 1 | 1 |
| 0 | 0 | 0 |
| 0 | 1 | 0 |
| 0 | 0 | 0 |
| 0 | 0 | 0 |
| 0 | 0 | 0 |
| 1 | 1 | 1 |
| 1 | 1 | 1 |
| 0 | 1 | 0 |
| 1 | 1 | 1 |
| 1 | 1 | 1 |
| 0 | 0 | 0 |
| 1 | 1 | 1 |
| 1 | 1 | 1 |
| 0 | 0 | 0 |
| 1 | 0 | 0 |
| 0 | 0 | 0 |
| 0 | 0 | 0 |
| 1 | 1 | 1 |
| 1 | 1 | 1 |
| 1 | 1 | 1 |
| 1 | 1 | 1 |
| 1 | 1 | 1 |
| 1 | 1 | 1 |
| 1 | 1 | 1 |
| 1 | 1 | 1 |
| 0 | 0 | 0 |
| 1 | 1 | 1 |
| 1 | 1 | 1 |
| 0 | 0 | 0 |
| 0 | 0 | 0 |
| 1 | 1 | 1 |
| 0 | 1 | 0 |
| 0 | 0 | 0 |
| 0 | 0 | 0 |
| 0 | 0 | 0 |
| 0 | 0 | 0 |
| 0 | 0 | 0 |
| 0 | 0 | 0 |
| 1 | 1 | 1 |
| 0 | 0 | 0 |
| 0 | 0 | 0 |
| 0 | 0 | 0 |
| 0 | 0 | 0 |
| 0 | 0 | 0 |

| | | |
|---|---|---|
| 0 | 0 | 0 |
| 0 | 0 | 0 |
| 0 | 0 | 0 |
| 0 | 0 | 0 |
| 0 | 0 | 0 |
| 0 | 0 | 0 |
| 0 | 0 | 0 |
| 0 | 0 | 0 |
| 0 | 0 | 0 |
| 0 | 1 | 0 |
| 0 | 1 | 0 |
| 0 | 0 | 0 |
| 1 | 1 | 1 |
| 0 | 0 | 0 |
| 0 | 0 | 0 |
| 0 | 1 | 0 |
| 0 | 0 | 0 |
| 1 | 0 | 0 |
| 1 | 1 | 1 |
| 1 | 1 | 1 |
| 0 | 0 | 0 |
| 0 | 1 | 0 |
| 0 | 1 | 0 |
| 0 | 0 | 0 |
| 0 | 0 | 0 |
| 0 | 1 | 0 |
| 0 | 1 | 0 |
| 1 | 1 | 1 |
| 0 | 0 | 0 |
| 0 | 0 | 0 |
| 1 | 1 | 1 |
| 1 | 1 | 1 |
| 0 | 0 | 0 |
| 0 | 0 | 0 |
| 0 | 0 | 0 |
| 0 | 1 | 0 |
| 0 | 0 | 0 |
| 0 | 1 | 0 |
| 0 | 1 | 0 |
| 0 | 0 | 0 |
| 0 | 0 | 0 |
| 0 | 0 | 0 |
| 0 | 0 | 0 |
| 0 | 0 | 0 |
| 0 | 0 | 0 |
| 1 | 1 | 1 |
| 0 | 1 | 0 |
| 0 | 0 | 0 |
| 0 | 0 | 0 |
| 0 | 0 | 0 |
| 1 | 1 | 1 |
| 1 | 1 | 1 |
| 0 | 0 | 0 |
| 0 | 0 | 0 |

| | | |
|---|---|---|
| 1 | 1 | 1 |
| 1 | 1 | 1 |
| 1 | 1 | 1 |
| 1 | 1 | 1 |
| 0 | 0 | 0 |
| 0 | 0 | 0 |
| 1 | 0 | 0 |
| 0 | 0 | 0 |
| 1 | 0 | 0 |
| 0 | 0 | 0 |
| 1 | 1 | 1 |
| 1 | 1 | 1 |
| 1 | 1 | 1 |
| 1 | 1 | 1 |
| 1 | 1 | 1 |
| 1 | 1 | 1 |
| 0 | 0 | 0 |
| 1 | 1 | 1 |
| 1 | 1 | 1 |
| 0 | 0 | 0 |
| 0 | 0 | 0 |
| 0 | 1 | 0 |
| 0 | 1 | 0 |
| 0 | 0 | 0 |
| 0 | 1 | 0 |
| 0 | 0 | 0 |
| 0 | 0 | 0 |
| 0 | 0 | 0 |
| 0 | 0 | 0 |
| 0 | 0 | 0 |
| 0 | 0 | 0 |
| 0 | 0 | 0 |
| 0 | 1 | 0 |
| 0 | 0 | 0 |
| 1 | 1 | 1 |
| 1 | 1 | 1 |
| 1 | 1 | 1 |
| 1 | 1 | 1 |
| 0 | 0 | 0 |
| 1 | 1 | 1 |
| 1 | 1 | 1 |
| 1 | 1 | 1 |
| 1 | 1 | 1 |
| 0 | 0 | 0 |
| 1 | 1 | 1 |
| 1 | 1 | 1 |
| 1 | 1 | 1 |
| 1 | 1 | 1 |
| 1 | 1 | 1 |
| 1 | 1 | 1 |
| 0 | 0 | 0 |
| 0 | 0 | 0 |
| 0 | 0 | 0 |
| 0 | 1 | 0 |

| | | |
|---|---|---|
| 0 | 0 | 0 |
| 0 | 0 | 0 |
| 0 | 0 | 0 |
| 0 | 0 | 0 |
| 0 | 0 | 0 |
| 0 | 0 | 0 |
| 0 | 0 | 0 |
| 0 | 0 | 0 |
| 1 | 1 | 1 |
| 1 | 1 | 1 |
| 0 | 0 | 0 |
| 1 | 1 | 1 |
| 1 | 1 | 1 |
| 0 | 0 | 0 |
| 0 | 0 | 0 |
| 0 | 0 | 0 |
| 0 | 0 | 0 |
| 0 | 0 | 0 |
| 0 | 0 | 0 |
| 1 | 1 | 1 |
| 1 | 1 | 1 |
| 0 | 0 | 0 |
| 1 | 1 | 1 |
| 0 | 0 | 0 |
| 1 | 1 | 1 |
| 1 | 1 | 1 |
| 1 | 1 | 1 |
| 0 | 1 | 0 |
| 0 | 1 | 0 |
| 0 | 0 | 0 |
| 0 | 1 | 0 |
| 1 | 1 | 1 |
| 0 | 0 | 0 |
| 1 | 1 | 1 |
| 0 | 0 | 0 |
| 0 | 0 | 0 |
| 0 | 0 | 0 |
| 0 | 0 | 0 |
| 0 | 0 | 0 |
| 0 | 0 | 0 |
| 0 | 0 | 0 |
| 0 | 0 | 0 |
| 1 | 1 | 1 |
| 0 | 0 | 0 |
| 1 | 1 | 1 |
| 0 | 0 | 0 |
| 1 | 0 | 0 |
| 1 | 1 | 1 |
| 1 | 1 | 1 |
| 0 | 0 | 0 |
| 1 | 1 | 1 |
| 1 | 1 | 1 |
| 0 | 1 | 0 |
| 1 | 1 | 1 |

| | | |
|---|---|---|
| 0 | 0 | 0 |
| 0 | 1 | 0 |
| 1 | 1 | 1 |
| 0 | 1 | 0 |
| 0 | 1 | 0 |
| 0 | 1 | 0 |
| 0 | 1 | 0 |
| 1 | 1 | 1 |
| 1 | 1 | 1 |
| 1 | 1 | 1 |
| 0 | 0 | 0 |
| 1 | 1 | 1 |
| 1 | 1 | 1 |
| 0 | 1 | 0 |
| 0 | 0 | 0 |
| 0 | 0 | 0 |
| 0 | 0 | 0 |
| 0 | 0 | 0 |
| 0 | 0 | 0 |
| 0 | 1 | 0 |
| 0 | 1 | 0 |
| 1 | 1 | 1 |
| 0 | 0 | 0 |
| 0 | 1 | 0 |
| 0 | 1 | 0 |
| 0 | 1 | 0 |
| 1 | 1 | 1 |
| 1 | 1 | 1 |
| 0 | 0 | 0 |
| 0 | 0 | 0 |
| 0 | 0 | 0 |
| 0 | 0 | 0 |
| 0 | 0 | 0 |
| 0 | 0 | 0 |
| 1 | 1 | 1 |
| 1 | 1 | 1 |
| 0 | 0 | 0 |
| 0 | 0 | 0 |
| 0 | 0 | 0 |
| 0 | 0 | 0 |
| 0 | 0 | 0 |
| 0 | 0 | 0 |
| 1 | 1 | 1 |
| 1 | 1 | 1 |
| 0 | 0 | 0 |
| 1 | 1 | 1 |
| 1 | 1 | 1 |
| 1 | 1 | 1 |
| 1 | 1 | 1 |
| 1 | 1 | 1 |
| 1 | 0 | 0 |
| 0 | 1 | 0 |
| 0 | 1 | 0 |
| 1 | 0 | 0 |

| | | |
|---|---|---|
| 0 | 0 | 0 |
| 1 | 1 | 1 |
| 0 | 0 | 0 |
| 0 | 1 | 0 |
| 1 | 0 | 0 |
| 0 | 0 | 0 |
| 0 | 0 | 0 |
| 0 | 0 | 0 |
| 1 | 0 | 0 |
| 1 | 0 | 0 |
| 0 | 0 | 0 |
| 1 | 1 | 1 |
| 1 | 0 | 0 |
| 1 | 0 | 0 |
| 0 | 0 | 0 |
| 0 | 0 | 0 |
| 0 | 0 | 0 |
| 0 | 0 | 0 |
| 1 | 1 | 1 |
| 0 | 0 | 0 |
| 0 | 0 | 0 |
| 0 | 0 | 0 |
| 0 | 0 | 0 |
| 0 | 0 | 0 |
| 1 | 1 | 1 |
| 0 | 0 | 0 |
| 1 | 1 | 1 |
| 0 | 1 | 0 |
| 1 | 1 | 1 |
| 0 | 0 | 0 |
| 1 | 1 | 1 |
| 1 | 1 | 1 |
| 0 | 0 | 0 |
| 0 | 0 | 0 |
| 1 | 1 | 1 |
| 1 | 1 | 1 |
| 1 | 1 | 1 |
| 0 | 1 | 0 |
| 1 | 0 | 0 |
| 0 | 0 | 0 |
| 0 | 0 | 0 |
| 0 | 0 | 0 |
| 0 | 0 | 0 |
| 0 | 0 | 0 |
| 0 | 0 | 0 |
| 0 | 1 | 0 |
| 0 | 1 | 0 |
| 0 | 0 | 0 |
| 0 | 0 | 0 |
| 1 | 1 | 1 |
| 0 | 0 | 0 |
| 0 | 0 | 0 |
| 0 | 1 | 0 |
| 1 | 1 | 1 |

| | | |
|---|---|---|
| 1 | 1 | 1 |
| 0 | 0 | 0 |
| 0 | 0 | 0 |
| 1 | 0 | 0 |
| 1 | 1 | 1 |
| 0 | 0 | 0 |
| 0 | 0 | 0 |
| 0 | 0 | 0 |
| 0 | 0 | 0 |
| 0 | 0 | 0 |
| 0 | 0 | 0 |
| 0 | 0 | 0 |
| 0 | 0 | 0 |
| 0 | 0 | 0 |
| 0 | 0 | 0 |
| 0 | 1 | 0 |
| 0 | 0 | 0 |
| 1 | 1 | 1 |
| 1 | 0 | 0 |
| 1 | 1 | 1 |
| 0 | 0 | 0 |
| 1 | 1 | 1 |
| 0 | 0 | 0 |
| 0 | 0 | 0 |
| 1 | 1 | 1 |
| 0 | 0 | 0 |
| 0 | 1 | 0 |
| 0 | 1 | 0 |
| 0 | 1 | 0 |
| 0 | 0 | 0 |
| 0 | 0 | 0 |
| 0 | 0 | 0 |
| 0 | 0 | 0 |
| 0 | 0 | 0 |
| 0 | 0 | 0 |
| 0 | 0 | 0 |
| 0 | 0 | 0 |
| 1 | 1 | 1 |
| 1 | 0 | 0 |
| 0 | 0 | 0 |
| 0 | 0 | 0 |
| 0 | 0 | 0 |
| 0 | 0 | 0 |
| 0 | 0 | 0 |
| 0 | 0 | 0 |
| 0 | 0 | 0 |
| 1 | 1 | 1 |
| 0 | 0 | 0 |
| 0 | 0 | 0 |
| 0 | 0 | 0 |
| 0 | 0 | 0 |
| 0 | 1 | 0 |
| 0 | 0 | 0 |
| 0 | 0 | 0 |

| | | |
|---|---|---|
| 0 | 0 | 0 |
| 0 | 0 | 0 |
| 0 | 0 | 0 |
| 0 | 0 | 0 |
| 0 | 0 | 0 |
| 0 | 0 | 0 |
| 0 | 0 | 0 |
| 0 | 0 | 0 |
| 0 | 0 | 0 |
| 0 | 0 | 0 |
| 1 | 1 | 1 |
| 0 | 0 | 0 |
| 1 | 0 | 0 |
| 0 | 0 | 0 |
| 0 | 1 | 0 |
| 1 | 1 | 1 |
| 1 | 1 | 1 |
| 1 | 1 | 1 |
| 0 | 0 | 0 |
| 0 | 0 | 0 |
| 1 | 1 | 1 |
| 0 | 0 | 0 |
| 0 | 0 | 0 |
| 1 | 1 | 1 |
| 0 | 0 | 0 |
| 0 | 0 | 0 |
| 0 | 0 | 0 |
| 1 | 0 | 0 |
| 0 | 0 | 0 |
| 0 | 0 | 0 |
| 1 | 0 | 0 |
| 0 | 0 | 0 |
| 1 | 0 | 0 |
| 1 | 0 | 0 |
| 0 | 0 | 0 |
| 0 | 0 | 0 |
| 0 | 0 | 0 |
| 1 | 0 | 0 |
| 1 | 0 | 0 |
| 0 | 0 | 0 |
| 0 | 0 | 0 |
| 0 | 0 | 0 |
| 0 | 0 | 0 |
| 0 | 0 | 0 |
| 0 | 0 | 0 |
| 0 | 0 | 0 |
| 0 | 0 | 0 |
| 0 | 0 | 0 |
| 0 | 0 | 0 |
| 0 | 0 | 0 |
| 0 | 0 | 0 |
| 0 | 0 | 0 |
| 0 | 0 | 0 |
| 0 | 0 | 0 |

| | | |
|---|---|---|
| 0 | 0 | 0 |
| 1 | 0 | 0 |
| 0 | 0 | 0 |
| 0 | 0 | 0 |
| 0 | 0 | 0 |
| 1 | 0 | 0 |
| 0 | 0 | 0 |
| 0 | 0 | 0 |
| 1 | 1 | 1 |
| 0 | 0 | 0 |
| 0 | 1 | 0 |
| 1 | 1 | 1 |
| 1 | 1 | 1 |
| 0 | 0 | 0 |
| 1 | 1 | 1 |
| 0 | 0 | 0 |
| 1 | 1 | 1 |
| 1 | 1 | 1 |
| 0 | 0 | 0 |
| 0 | 0 | 0 |
| 0 | 0 | 0 |
| 0 | 0 | 0 |
| 0 | 0 | 0 |
| 0 | 0 | 0 |
| 0 | 0 | 0 |
| 0 | 1 | 0 |
| 1 | 1 | 1 |
| 1 | 1 | 1 |
| 1 | 0 | 0 |
| 1 | 1 | 1 |
| 1 | 1 | 1 |
| 0 | 1 | 0 |
| 0 | 0 | 0 |
| 0 | 0 | 0 |
| 0 | 0 | 0 |
| 0 | 0 | 0 |
| 0 | 0 | 0 |
| 0 | 0 | 0 |
| 0 | 1 | 0 |
| 0 | 1 | 0 |
| 0 | 1 | 0 |
| 0 | 1 | 0 |
| 0 | 1 | 0 |
| 0 | 1 | 0 |
| 0 | 0 | 0 |
| 0 | 0 | 0 |
| 0 | 1 | 0 |
| 0 | 0 | 0 |
| 0 | 0 | 0 |
| 0 | 0 | 0 |
| 0 | 0 | 0 |
| 0 | 0 | 0 |
| 0 | 0 | 0 |
| 0 | 0 | 0 |

| | | |
|---|---|---|
| 1 | 1 | 1 |
| 1 | 1 | 1 |
| 0 | 0 | 0 |
| 0 | 0 | 0 |
| 1 | 1 | 1 |
| 0 | 0 | 0 |
| 0 | 0 | 0 |
| 1 | 1 | 1 |
| 1 | 1 | 1 |
| 0 | 0 | 0 |
| 1 | 1 | 1 |
| 1 | 1 | 1 |
| 0 | 0 | 0 |
| 0 | 0 | 0 |
| 0 | 0 | 0 |
| 0 | 0 | 0 |
| 0 | 1 | 0 |
| 0 | 1 | 0 |
| 1 | 1 | 1 |
| 1 | 1 | 1 |
| 1 | 1 | 1 |
| 0 | 1 | 0 |
| 1 | 1 | 1 |
| 1 | 1 | 1 |
| 1 | 1 | 1 |
| 0 | 0 | 0 |
| 1 | 1 | 1 |
| 1 | 1 | 1 |
| 1 | 1 | 1 |
| 0 | 1 | 0 |
| 0 | 1 | 0 |
| 1 | 1 | 1 |
| 1 | 1 | 1 |
| 1 | 1 | 1 |
| 1 | 1 | 1 |
| 1 | 1 | 1 |
| 1 | 1 | 1 |
| 0 | 0 | 0 |
| 0 | 0 | 0 |
| 1 | 0 | 0 |
| 0 | 0 | 0 |
| 1 | 1 | 1 |
| 1 | 1 | 1 |
| 0 | 0 | 0 |
| 0 | 0 | 0 |
| 0 | 1 | 0 |
| 0 | 1 | 0 |
| 0 | 1 | 0 |
| 1 | 1 | 1 |
| 1 | 1 | 1 |
| 1 | 1 | 1 |
| 0 | 0 | 0 |
| 0 | 0 | 0 |
| 0 | 0 | 0 |

| | | |
|---|---|---|
| 0 | 0 | 0 |
| 1 | 1 | 1 |
| 1 | 1 | 1 |
| 1 | 1 | 1 |
| 1 | 1 | 1 |
| 1 | 1 | 1 |
| 1 | 1 | 1 |
| 1 | 1 | 1 |
| 0 | 0 | 0 |
| 1 | 1 | 1 |
| 1 | 1 | 1 |
| 1 | 1 | 1 |
| 0 | 1 | 0 |
| 0 | 1 | 0 |
| 1 | 1 | 1 |
| 1 | 1 | 1 |
| 0 | 1 | 0 |
| 0 | 1 | 0 |
| 0 | 0 | 0 |
| 0 | 0 | 0 |
| 0 | 0 | 0 |
| 0 | 0 | 0 |
| 0 | 1 | 0 |
| 0 | 0 | 0 |
| 0 | 0 | 0 |
| 1 | 1 | 1 |
| 1 | 1 | 1 |
| 1 | 1 | 1 |
| 1 | 1 | 1 |
| 1 | 1 | 1 |
| 1 | 1 | 1 |
| 1 | 1 | 1 |
| 0 | 0 | 0 |
| 1 | 1 | 1 |
| 1 | 1 | 1 |
| 0 | 1 | 0 |
| 0 | 0 | 0 |
| 1 | 1 | 1 |
| 0 | 0 | 0 |
| 0 | 0 | 0 |
| 0 | 0 | 0 |
| 0 | 0 | 0 |
| 0 | 0 | 0 |
| 1 | 1 | 1 |
| 1 | 1 | 1 |
| 1 | 1 | 1 |
| 0 | 0 | 0 |
| 0 | 0 | 0 |
| 0 | 0 | 0 |
| 0 | 0 | 0 |
| 0 | 0 | 0 |
| 0 | 0 | 0 |
| 0 | 0 | 0 |
| 0 | 1 | 0 |

| | | |
|---|---|---|
| 0 | 0 | 0 |
| 0 | 0 | 0 |
| 0 | 1 | 0 |
| 0 | 0 | 0 |
| 0 | 0 | 0 |
| 1 | 1 | 1 |
| 1 | 1 | 1 |
| 1 | 0 | 0 |
| 1 | 1 | 1 |
| 1 | 1 | 1 |
| 1 | 1 | 1 |
| 0 | 0 | 0 |
| 1 | 1 | 1 |
| 1 | 1 | 1 |
| 0 | 0 | 0 |
| 1 | 1 | 1 |
| 0 | 0 | 0 |
| 1 | 1 | 1 |
| 0 | 0 | 0 |
| 0 | 0 | 0 |
| 1 | 1 | 1 |
| 1 | 1 | 1 |
| 0 | 0 | 0 |
| 1 | 1 | 1 |
| 0 | 0 | 0 |
| 1 | 1 | 1 |
| 1 | 1 | 1 |
| 0 | 0 | 0 |
| 1 | 1 | 1 |
| 0 | 0 | 0 |
| 0 | 1 | 0 |
| 0 | 0 | 0 |
| 0 | 0 | 0 |
| 0 | 0 | 0 |
| 1 | 1 | 1 |
| 0 | 0 | 0 |
| 1 | 1 | 1 |
| 1 | 1 | 1 |
| 0 | 0 | 0 |
| 1 | 1 | 1 |
| 1 | 1 | 1 |
| 1 | 1 | 1 |
| 1 | 1 | 1 |
| 0 | 0 | 0 |
| 0 | 0 | 0 |
| 1 | 1 | 1 |
| 1 | 1 | 1 |
| 1 | 0 | 0 |
| 0 | 0 | 0 |
| 1 | 0 | 0 |
| 0 | 0 | 0 |
| 1 | 1 | 1 |
| 1 | 1 | 1 |
| 0 | 0 | 0 |

| | | |
|---|---|---|
| 1 | 1 | 1 |
| 1 | 1 | 1 |
| 1 | 1 | 1 |
| 1 | 1 | 1 |
| 0 | 1 | 0 |
| 1 | 1 | 1 |
| 1 | 1 | 1 |
| 0 | 0 | 0 |
| 1 | 1 | 1 |
| 1 | 1 | 1 |
| 1 | 1 | 1 |
| 1 | 1 | 1 |
| 0 | 0 | 0 |
| 1 | 1 | 1 |
| 0 | 0 | 0 |
| 0 | 0 | 0 |
| 1 | 1 | 1 |
| 1 | 1 | 1 |
| 1 | 1 | 1 |
| 1 | 1 | 1 |
| 0 | 0 | 0 |
| 0 | 0 | 0 |
| 1 | 1 | 1 |
| 1 | 1 | 1 |
| 0 | 0 | 0 |
| 1 | 1 | 1 |
| 0 | 0 | 0 |
| 0 | 0 | 0 |
| 0 | 0 | 0 |
| 0 | 0 | 0 |
| 0 | 1 | 0 |
| 0 | 1 | 0 |
| 0 | 0 | 0 |
| 0 | 0 | 0 |
| 0 | 0 | 0 |
| 0 | 0 | 0 |
| 0 | 0 | 0 |
| 0 | 0 | 0 |
| 0 | 0 | 0 |
| 1 | 0 | 0 |
| 1 | 1 | 1 |
| 0 | 1 | 0 |
| 0 | 1 | 0 |
| 0 | 1 | 0 |
| 0 | 1 | 0 |
| 0 | 0 | 0 |
| 0 | 1 | 0 |
| 0 | 1 | 0 |
| 1 | 1 | 1 |
| 0 | 0 | 0 |
| 0 | 0 | 0 |
| 1 | 1 | 1 |
| 1 | 1 | 1 |
| 1 | 1 | 1 |

| | | |
|---|---|---|
| 1 | 1 | 1 |
| 1 | 1 | 1 |
| 1 | 1 | 1 |
| 1 | 1 | 1 |
| 0 | 0 | 0 |
| 1 | 1 | 1 |
| 0 | 0 | 0 |
| 0 | 0 | 0 |
| 0 | 0 | 0 |
| 0 | 0 | 0 |
| 0 | 0 | 0 |
| 1 | 1 | 1 |
| 0 | 0 | 0 |
| 0 | 0 | 0 |
| 1 | 1 | 1 |
| 1 | 1 | 1 |
| 1 | 1 | 1 |
| 1 | 1 | 1 |
| 1 | 1 | 1 |
| 1 | 1 | 1 |
| 1 | 1 | 1 |
| 1 | 1 | 1 |
| 1 | 1 | 1 |
| 0 | 0 | 0 |
| 1 | 1 | 1 |
| 1 | 1 | 1 |
| 1 | 1 | 1 |
| 0 | 0 | 0 |
| 1 | 0 | 0 |
| 1 | 0 | 0 |
| 0 | 0 | 0 |
| 0 | 0 | 0 |
| 1 | 1 | 1 |
| 0 | 0 | 0 |
| 0 | 1 | 0 |
| 1 | 1 | 1 |
| 1 | 1 | 1 |
| 1 | 1 | 1 |
| 1 | 1 | 1 |
| 1 | 1 | 1 |
| 1 | 1 | 1 |
| 1 | 1 | 1 |
| 1 | 1 | 1 |
| 1 | 1 | 1 |
| 1 | 1 | 1 |
| 1 | 1 | 1 |
| 1 | 1 | 1 |
| 1 | 1 | 1 |
| 0 | 0 | 0 |
| 0 | 1 | 0 |
| 1 | 1 | 1 |
| 1 | 0 | 0 |
| 1 | 0 | 0 |
| 1 | 1 | 1 |

| | | |
|---|---|---|
| 1 | 1 | 1 |
| 0 | 1 | 0 |
| 1 | 1 | 1 |
| 0 | 1 | 0 |
| 1 | 0 | 0 |
| 1 | 1 | 1 |
| 1 | 1 | 1 |
| 1 | 1 | 1 |
| 1 | 1 | 1 |
| 1 | 1 | 1 |
| 0 | 0 | 0 |
| 0 | 0 | 0 |
| 0 | 0 | 0 |
| 0 | 0 | 0 |
| 0 | 0 | 0 |
| 1 | 0 | 0 |
| 0 | 0 | 0 |
| 0 | 0 | 0 |
| 0 | 0 | 0 |
| 0 | 0 | 0 |
| 0 | 0 | 0 |
| 0 | 0 | 0 |
| 0 | 1 | 0 |
| 0 | 0 | 0 |
| 0 | 0 | 0 |
| 0 | 0 | 0 |
| 1 | 1 | 1 |
| 0 | 0 | 0 |
| 0 | 1 | 0 |
| 0 | 1 | 0 |
| 0 | 1 | 0 |
| 0 | 0 | 0 |
| 1 | 1 | 1 |
| 1 | 1 | 1 |
| 0 | 0 | 0 |
| 0 | 0 | 0 |
| 0 | 0 | 0 |
| 0 | 0 | 0 |
| 0 | 0 | 0 |
| 1 | 0 | 0 |
| 1 | 0 | 0 |
| 0 | 0 | 0 |
| 1 | 0 | 0 |
| 0 | 0 | 0 |
| 1 | 1 | 1 |
| 1 | 0 | 0 |
| 1 | 1 | 1 |
| 0 | 0 | 0 |
| 0 | 0 | 0 |
| 0 | 0 | 0 |
| 0 | 0 | 0 |
| 1 | 1 | 1 |
| 0 | 0 | 0 |
| 1 | 1 | 1 |

| | | |
|---|---|---|
| 1 | 1 | 1 |
| 1 | 1 | 1 |
| 1 | 1 | 1 |
| 1 | 1 | 1 |
| 1 | 1 | 1 |
| 0 | 0 | 0 |
| 0 | 0 | 0 |
| 0 | 0 | 0 |
| 0 | 0 | 0 |
| 0 | 1 | 0 |
| 0 | 0 | 0 |
| 0 | 0 | 0 |
| 0 | 0 | 0 |
| 0 | 0 | 0 |
| 0 | 0 | 0 |
| 0 | 0 | 0 |
| 1 | 1 | 1 |
| 0 | 1 | 0 |
| 1 | 1 | 1 |
| 1 | 1 | 1 |
| 1 | 1 | 1 |
| 1 | 1 | 1 |
| 1 | 1 | 1 |
| 1 | 1 | 1 |
| 1 | 1 | 1 |
| 1 | 1 | 1 |
| 1 | 1 | 1 |
| 1 | 1 | 1 |
| 0 | 0 | 0 |
| 0 | 1 | 0 |
| 0 | 1 | 0 |
| 1 | 0 | 0 |
| 1 | 0 | 0 |
| 0 | 0 | 0 |
| 1 | 1 | 1 |
| 0 | 0 | 0 |
| 1 | 0 | 0 |
| 0 | 0 | 0 |
| 1 | 1 | 1 |
| 0 | 0 | 0 |
| 1 | 1 | 1 |
| 1 | 1 | 1 |
| 0 | 1 | 0 |
| 0 | 0 | 0 |
| 1 | 1 | 1 |
| 1 | 1 | 1 |
| 1 | 1 | 1 |
| 1 | 1 | 1 |
| 1 | 1 | 1 |
| 1 | 1 | 1 |
| 1 | 1 | 1 |
| 1 | 0 | 0 |
| 0 | 0 | 0 |
| 0 | 0 | 0 |

| | | |
|---|---|---|
| 1 | 1 | 1 |
| 0 | 0 | 0 |
| 1 | 1 | 1 |
| 1 | 1 | 1 |
| 0 | 0 | 0 |
| 0 | 0 | 0 |
| 0 | 0 | 0 |
| 1 | 1 | 1 |
| 1 | 1 | 1 |
| 0 | 0 | 0 |
| 0 | 0 | 0 |
| 0 | 0 | 0 |
| 0 | 0 | 0 |
| 0 | 0 | 0 |
| 1 | 1 | 1 |
| 0 | 0 | 0 |
| 0 | 0 | 0 |
| 0 | 0 | 0 |
| 1 | 1 | 1 |
| 1 | 1 | 1 |
| 1 | 1 | 1 |
| 1 | 1 | 1 |
| 1 | 1 | 1 |
| 0 | 0 | 0 |
| 0 | 1 | 0 |
| 0 | 1 | 0 |
| 0 | 0 | 0 |
| 0 | 0 | 0 |
| 1 | 1 | 1 |
| 1 | 1 | 1 |
| 0 | 0 | 0 |
| 1 | 1 | 1 |
| 1 | 1 | 1 |
| 0 | 1 | 0 |
| 1 | 1 | 1 |
| 0 | 0 | 0 |
| 0 | 1 | 0 |
| 0 | 0 | 0 |
| 0 | 0 | 0 |
| 0 | 1 | 0 |
| 0 | 1 | 0 |
| 0 | 0 | 0 |
| 0 | 0 | 0 |
| 0 | 1 | 0 |
| 0 | 0 | 0 |
| 0 | 0 | 0 |
| 0 | 0 | 0 |
| 0 | 1 | 0 |
| 0 | 0 | 0 |
| 0 | 0 | 0 |
| 0 | 0 | 0 |
| 1 | 1 | 1 |
| 0 | 0 | 0 |
| 1 | 1 | 1 |

| | | |
|---|---|---|
| 1 | 1 | 1 |
| 1 | 1 | 1 |
| 1 | 1 | 1 |
| 1 | 1 | 1 |
| 0 | 0 | 0 |
| 0 | 0 | 0 |
| 1 | 1 | 1 |
| 1 | 0 | 0 |
| 1 | 1 | 1 |
| 0 | 0 | 0 |
| 1 | 0 | 0 |
| 1 | 0 | 0 |
| 0 | 0 | 0 |
| 1 | 1 | 1 |
| 1 | 1 | 1 |
| 1 | 1 | 1 |
| 1 | 1 | 1 |
| 1 | 1 | 1 |
| 1 | 1 | 1 |
| 1 | 1 | 1 |
| 1 | 1 | 1 |
| 0 | 1 | 0 |
| 1 | 1 | 1 |
| 0 | 0 | 0 |
| 1 | 1 | 1 |
| 0 | 0 | 0 |
| 1 | 1 | 1 |
| 0 | 0 | 0 |
| 1 | 1 | 1 |
| 0 | 0 | 0 |
| 0 | 0 | 0 |
| 0 | 0 | 0 |
| 0 | 0 | 0 |
| 0 | 0 | 0 |
| 0 | 1 | 0 |
| 0 | 0 | 0 |
| 1 | 1 | 1 |
| 0 | 0 | 0 |
| 1 | 1 | 1 |
| 1 | 1 | 1 |
| 0 | 0 | 0 |
| 0 | 0 | 0 |
| 0 | 0 | 0 |
| 0 | 0 | 0 |
| 0 | 0 | 0 |
| 0 | 0 | 0 |
| 0 | 0 | 0 |
| 0 | 0 | 0 |
| 1 | 1 | 1 |
| 0 | 0 | 0 |
| 1 | 0 | 0 |
| 1 | 0 | 0 |
| 1 | 1 | 1 |
| 1 | 1 | 1 |

| | | |
|---|---|---|
| 1 | 1 | 1 |
| 1 | 1 | 1 |
| 1 | 1 | 1 |
| 0 | 1 | 0 |
| 0 | 1 | 0 |
| 0 | 1 | 0 |
| 0 | 0 | 0 |
| 0 | 0 | 0 |
| 0 | 0 | 0 |
| 0 | 0 | 0 |
| 0 | 0 | 0 |
| 0 | 0 | 0 |
| 0 | 0 | 0 |
| 0 | 0 | 0 |
| 0 | 0 | 0 |
| 0 | 0 | 0 |
| 0 | 0 | 0 |
| 0 | 0 | 0 |
| 1 | 1 | 1 |
| 0 | 0 | 0 |
| 0 | 1 | 0 |
| 0 | 1 | 0 |
| 0 | 1 | 0 |
| 1 | 1 | 1 |
| 0 | 1 | 0 |
| 0 | 0 | 0 |
| 0 | 0 | 0 |
| 0 | 0 | 0 |
| 0 | 0 | 0 |
| 0 | 0 | 0 |
| 0 | 0 | 0 |
| 0 | 0 | 0 |
| 0 | 0 | 0 |
| 0 | 0 | 0 |
| 1 | 1 | 1 |
| 0 | 0 | 0 |
| 0 | 0 | 0 |
| 0 | 0 | 0 |
| 0 | 0 | 0 |
| 0 | 1 | 0 |
| 0 | 0 | 0 |
| 0 | 0 | 0 |
| 1 | 1 | 1 |
| 0 | 1 | 0 |
| 1 | 1 | 1 |
| 0 | 1 | 0 |
| 0 | 1 | 0 |
| 0 | 0 | 0 |
| 0 | 0 | 0 |
| 0 | 0 | 0 |
| 0 | 0 | 0 |
| 0 | 1 | 0 |
| 0 | 0 | 0 |
| 1 | 1 | 1 |

| | | |
|---|---|---|
| 0 | 1 | 0 |
| 1 | 1 | 1 |
| 1 | 1 | 1 |
| 1 | 1 | 1 |
| 1 | 1 | 1 |
| 0 | 0 | 0 |
| 0 | 1 | 0 |
| 1 | 1 | 1 |
| 1 | 1 | 1 |
| 1 | 1 | 1 |
| 0 | 0 | 0 |
| 0 | 0 | 0 |
| 0 | 0 | 0 |
| 0 | 0 | 0 |
| 0 | 0 | 0 |
| 0 | 1 | 0 |
| 1 | 0 | 0 |
| 0 | 0 | 0 |
| 1 | 1 | 1 |
| 1 | 1 | 1 |
| 1 | 1 | 1 |
| 1 | 1 | 1 |
| 0 | 0 | 0 |
| 1 | 1 | 1 |
| 0 | 0 | 0 |
| 0 | 0 | 0 |
| 0 | 0 | 0 |
| 0 | 0 | 0 |
| 1 | 1 | 1 |
| 0 | 0 | 0 |
| 1 | 1 | 1 |
| 0 | 1 | 0 |
| 0 | 1 | 0 |
| 0 | 1 | 0 |
| 1 | 1 | 1 |
| 1 | 1 | 1 |
| 0 | 1 | 0 |
| 0 | 0 | 0 |
| 0 | 0 | 0 |
| 0 | 0 | 0 |
| 0 | 0 | 0 |
| 0 | 0 | 0 |
| 1 | 1 | 1 |
| 0 | 0 | 0 |
| 0 | 0 | 0 |
| 1 | 1 | 1 |
| 0 | 0 | 0 |
| 0 | 0 | 0 |
| 0 | 0 | 0 |
| 0 | 0 | 0 |
| 1 | 1 | 1 |
| 1 | 1 | 1 |
| 1 | 1 | 1 |
| 0 | 0 | 0 |

| | | |
|---|---|---|
| 0 | 0 | 0 |
| 1 | 1 | 1 |
| 0 | 0 | 0 |
| 0 | 1 | 0 |
| 1 | 0 | 0 |
| 1 | 0 | 0 |
| 1 | 0 | 0 |
| 1 | 1 | 1 |
| 0 | 0 | 0 |
| 1 | 1 | 1 |
| 0 | 0 | 0 |
| 0 | 0 | 0 |
| 0 | 0 | 0 |
| 0 | 0 | 0 |
| 1 | 1 | 1 |
| 1 | 1 | 1 |
| 0 | 0 | 0 |
| 0 | 0 | 0 |
| 0 | 1 | 0 |
| 1 | 1 | 1 |
| 0 | 0 | 0 |
| 1 | 0 | 0 |
| 1 | 0 | 0 |
| 1 | 1 | 1 |
| 0 | 0 | 0 |
| 1 | 0 | 0 |
| 0 | 0 | 0 |
| 0 | 0 | 0 |
| 0 | 0 | 0 |
| 1 | 1 | 1 |
| 0 | 0 | 0 |
| 0 | 0 | 0 |
| 0 | 0 | 0 |
| 1 | 1 | 1 |
| 0 | 0 | 0 |
| 1 | 1 | 1 |
| 0 | 0 | 0 |
| 0 | 0 | 0 |
| 0 | 0 | 0 |
| 0 | 0 | 0 |
| 0 | 0 | 0 |
| 1 | 1 | 1 |
| 0 | 0 | 0 |
| 0 | 0 | 0 |
| 0 | 0 | 0 |
| 0 | 0 | 0 |
| 0 | 0 | 0 |
| 0 | 0 | 0 |
| 0 | 0 | 0 |
| 0 | 0 | 0 |
| 0 | 0 | 0 |
| 0 | 0 | 0 |
| 1 | 1 | 1 |
| 0 | 0 | 0 |

| | | |
|---|---|---|
| 0 | 1 | 0 |
| 1 | 1 | 1 |
| 1 | 1 | 1 |
| 1 | 1 | 1 |
| 1 | 1 | 1 |
| 1 | 1 | 1 |
| 1 | 1 | 1 |
| 1 | 1 | 1 |
| 1 | 1 | 1 |
| 0 | 0 | 0 |
| 0 | 0 | 0 |
| 1 | 1 | 1 |
| 1 | 1 | 1 |
| 1 | 1 | 1 |
| 0 | 0 | 0 |
| 1 | 1 | 1 |
| 1 | 1 | 1 |
| 0 | 0 | 0 |
| 0 | 1 | 0 |
| 0 | 1 | 0 |
| 0 | 1 | 0 |
| 0 | 0 | 0 |
| 0 | 0 | 0 |
| 0 | 0 | 0 |
| 0 | 0 | 0 |
| 0 | 0 | 0 |
| 0 | 0 | 0 |
| 0 | 0 | 0 |
| 1 | 1 | 1 |
| 0 | 0 | 0 |
| 0 | 0 | 0 |
| 1 | 1 | 1 |
| 1 | 1 | 1 |
| 1 | 1 | 1 |
| 1 | 1 | 1 |
| 1 | 1 | 1 |
| 1 | 1 | 1 |
| 1 | 1 | 1 |
| 1 | 1 | 1 |
| 1 | 1 | 1 |
| 1 | 1 | 1 |
| 1 | 1 | 1 |
| 1 | 1 | 1 |
| 0 | 0 | 0 |
| 1 | 1 | 1 |
| 0 | 0 | 0 |
| 1 | 1 | 1 |
| 1 | 1 | 1 |
| 1 | 1 | 1 |
| 1 | 1 | 1 |
| 0 | 0 | 0 |
| 1 | 1 | 1 |
| 1 | 1 | 1 |
| 0 | 0 | 0 |

| | | |
|---|---|---|
| 0 | 0 | 0 |
| 0 | 0 | 0 |
| 0 | 0 | 0 |
| 0 | 0 | 0 |
| 1 | 1 | 1 |
| 0 | 1 | 0 |
| 1 | 1 | 1 |
| 0 | 0 | 0 |
| 0 | 0 | 0 |
| 0 | 0 | 0 |
| 0 | 0 | 0 |
| 1 | 1 | 1 |
| 1 | 1 | 1 |
| 1 | 1 | 1 |
| 1 | 1 | 1 |
| 0 | 0 | 0 |
| 0 | 0 | 0 |
| 0 | 1 | 0 |
| 0 | 1 | 0 |
| 1 | 1 | 1 |
| 1 | 1 | 1 |
| 0 | 0 | 0 |
| 1 | 1 | 1 |
| 0 | 1 | 0 |
| 0 | 0 | 0 |
| 0 | 0 | 0 |
| 0 | 0 | 0 |
| 0 | 0 | 0 |
| 1 | 1 | 1 |
| 1 | 1 | 1 |
| 1 | 1 | 1 |
| 0 | 0 | 0 |
| 1 | 1 | 1 |
| 1 | 1 | 1 |
| 1 | 1 | 1 |
| 1 | 1 | 1 |
| 1 | 1 | 1 |
| 1 | 1 | 1 |
| 1 | 1 | 1 |
| 1 | 1 | 1 |
| 1 | 1 | 1 |
| 1 | 1 | 1 |
| 0 | 0 | 0 |
| 0 | 1 | 0 |
| 0 | 1 | 0 |
| 1 | 1 | 1 |
| 0 | 1 | 0 |
| 1 | 1 | 1 |
| 1 | 1 | 1 |
| 0 | 0 | 0 |
| 1 | 1 | 1 |
| 1 | 1 | 1 |
| 1 | 1 | 1 |
| 1 | 1 | 1 |

| | | |
|---|---|---|
| 0 | 0 | 0 |
| 1 | 1 | 1 |
| 0 | 0 | 0 |
| 0 | 1 | 0 |
| 1 | 1 | 1 |
| 1 | 1 | 1 |
| 1 | 1 | 1 |
| 0 | 0 | 0 |
| 1 | 1 | 1 |
| 1 | 1 | 1 |
| 0 | 0 | 0 |
| 0 | 0 | 0 |
| 0 | 0 | 0 |
| 1 | 1 | 1 |
| 0 | 0 | 0 |
| 0 | 0 | 0 |
| 0 | 0 | 0 |
| 0 | 0 | 0 |
| 0 | 1 | 0 |
| 0 | 0 | 0 |
| 0 | 0 | 0 |
| 0 | 0 | 0 |
| 0 | 0 | 0 |
| 0 | 0 | 0 |
| 0 | 0 | 0 |
| 0 | 0 | 0 |
| 0 | 0 | 0 |
| 0 | 0 | 0 |
| 0 | 0 | 0 |
| 0 | 0 | 0 |
| 0 | 0 | 0 |
| 0 | 0 | 0 |
| 1 | 1 | 1 |
| 0 | 0 | 0 |
| 1 | 1 | 1 |
| 0 | 0 | 0 |
| 0 | 0 | 0 |
| 1 | 1 | 1 |
| 1 | 1 | 1 |
| 0 | 1 | 0 |
| 0 | 1 | 0 |
| 0 | 0 | 0 |
| 0 | 0 | 0 |
| 0 | 0 | 0 |
| 1 | 1 | 1 |
| 0 | 0 | 0 |
| 1 | 1 | 1 |
| 0 | 0 | 0 |
| 0 | 1 | 0 |
| 0 | 1 | 0 |
| 0 | 1 | 0 |
| 0 | 1 | 0 |
| 0 | 1 | 0 |
| 1 | 1 | 1 |

| | | |
|---|---|---|
| 1 | 1 | 1 |
| 1 | 1 | 1 |
| 0 | 1 | 0 |
| 1 | 1 | 1 |
| 1 | 1 | 1 |
| 0 | 0 | 0 |
| 0 | 0 | 0 |
| 1 | 1 | 1 |
| 0 | 0 | 0 |
| 0 | 0 | 0 |
| 1 | 1 | 1 |
| 0 | 1 | 0 |
| 1 | 1 | 1 |
| 1 | 1 | 1 |
| 1 | 1 | 1 |
| 0 | 0 | 0 |
| 1 | 1 | 1 |
| 1 | 1 | 1 |
| 0 | 0 | 0 |
| 0 | 0 | 0 |
| 0 | 0 | 0 |
| 1 | 1 | 1 |
| 1 | 1 | 1 |
| 0 | 0 | 0 |
| 1 | 1 | 1 |
| 0 | 0 | 0 |
| 0 | 0 | 0 |
| 1 | 1 | 1 |
| 1 | 1 | 1 |
| 1 | 1 | 1 |
| 0 | 0 | 0 |
| 1 | 0 | 0 |
| 0 | 0 | 0 |
| 0 | 0 | 0 |
| 0 | 0 | 0 |
| 0 | 0 | 0 |
| 0 | 0 | 0 |
| 1 | 1 | 1 |
| 0 | 0 | 0 |
| 0 | 1 | 0 |
| 0 | 0 | 0 |
| 0 | 0 | 0 |
| 0 | 0 | 0 |
| 0 | 0 | 0 |
| 1 | 1 | 1 |
| 0 | 1 | 0 |
| 0 | 1 | 0 |
| 0 | 0 | 0 |
| 0 | 0 | 0 |
| 0 | 0 | 0 |
| 1 | 1 | 1 |
| 0 | 1 | 0 |
| 1 | 1 | 1 |
| 1 | 1 | 1 |

| | | |
|---|---|---|
| 1 | 1 | 1 |
| 0 | 1 | 0 |
| 0 | 1 | 0 |
| 0 | 0 | 0 |
| 0 | 0 | 0 |
| 1 | 1 | 1 |
| 0 | 0 | 0 |
| 1 | 1 | 1 |
| 1 | 1 | 1 |
| 1 | 1 | 1 |
| 1 | 1 | 1 |
| 0 | 0 | 0 |
| 1 | 0 | 0 |
| 0 | 0 | 0 |
| 1 | 1 | 1 |
| 0 | 0 | 0 |
| 0 | 0 | 0 |
| 0 | 0 | 0 |
| 1 | 1 | 1 |
| 0 | 0 | 0 |
| 0 | 0 | 0 |
| 0 | 0 | 0 |
| 0 | 0 | 0 |
| 0 | 0 | 0 |
| 0 | 0 | 0 |
| 1 | 1 | 1 |
| 0 | 0 | 0 |
| 0 | 1 | 0 |
| 0 | 1 | 0 |
| 0 | 1 | 0 |
| 0 | 1 | 0 |
| 0 | 1 | 0 |
| 0 | 0 | 0 |
| 0 | 0 | 0 |
| 1 | 1 | 1 |
| 1 | 1 | 1 |
| 0 | 0 | 0 |
| 0 | 0 | 0 |
| 0 | 0 | 0 |
| 1 | 0 | 0 |
| 0 | 0 | 0 |
| 0 | 0 | 0 |
| 1 | 1 | 1 |
| 0 | 0 | 0 |
| 1 | 1 | 1 |
| 1 | 1 | 1 |
| 1 | 1 | 1 |
| 1 | 1 | 1 |
| 1 | 1 | 1 |
| 0 | 0 | 0 |
| 1 | 1 | 1 |
| 1 | 1 | 1 |
| 1 | 1 | 1 |
| 0 | 1 | 0 |

| | | |
|---|---|---|
| 1 | 1 | 1 |
| 1 | 1 | 1 |
| 1 | 1 | 1 |
| 0 | 0 | 0 |
| 1 | 1 | 1 |
| 1 | 1 | 1 |
| 0 | 1 | 0 |
| 0 | 0 | 0 |
| 1 | 1 | 1 |
| 1 | 1 | 1 |
| 1 | 1 | 1 |
| 1 | 1 | 1 |
| 1 | 1 | 1 |
| 0 | 1 | 0 |
| 0 | 1 | 0 |
| 0 | 1 | 0 |
| 0 | 0 | 0 |
| 0 | 0 | 0 |
| 0 | 1 | 0 |
| 1 | 1 | 1 |
| 0 | 1 | 0 |
| 1 | 1 | 1 |
| 0 | 1 | 0 |
| 1 | 1 | 1 |
| 1 | 1 | 1 |
| 1 | 1 | 1 |
| 1 | 1 | 1 |
| 0 | 0 | 0 |
| 0 | 0 | 0 |
| 1 | 1 | 1 |
| 1 | 1 | 1 |
| 0 | 0 | 0 |
| 0 | 1 | 0 |
| 0 | 0 | 0 |
| 0 | 0 | 0 |
| 0 | 0 | 0 |
| 0 | 0 | 0 |
| 0 | 0 | 0 |
| 1 | 1 | 1 |
| 1 | 1 | 1 |
| 1 | 1 | 1 |
| 0 | 1 | 0 |
| 1 | 1 | 1 |
| 1 | 1 | 1 |
| 0 | 0 | 0 |
| 0 | 0 | 0 |
| 0 | 0 | 0 |
| 1 | 1 | 1 |
| 1 | 1 | 1 |
| 1 | 1 | 1 |
| 1 | 1 | 1 |
| 0 | 0 | 0 |
| 0 | 0 | 0 |
| 0 | 0 | 0 |

| | | |
|---|---|---|
| 0 | 0 | 0 |
| 0 | 0 | 0 |
| 1 | 1 | 1 |
| 0 | 0 | 0 |
| 0 | 0 | 0 |
| 0 | 0 | 0 |
| 0 | 0 | 0 |
| 1 | 1 | 1 |
| 1 | 1 | 1 |
| 1 | 1 | 1 |
| 1 | 1 | 1 |
| 1 | 1 | 1 |
| 0 | 1 | 0 |
| 0 | 1 | 0 |
| 1 | 1 | 1 |
| 1 | 1 | 1 |
| 1 | 1 | 1 |
| 1 | 1 | 1 |
| 1 | 1 | 1 |
| 1 | 1 | 1 |
| 0 | 0 | 0 |
| 0 | 0 | 0 |
| 0 | 0 | 0 |
| 0 | 0 | 0 |
| 0 | 0 | 0 |
| 1 | 1 | 1 |
| 1 | 1 | 1 |
| 0 | 0 | 0 |
| 0 | 1 | 0 |
| 0 | 1 | 0 |
| 0 | 0 | 0 |
| 0 | 0 | 0 |
| 0 | 0 | 0 |
| 1 | 1 | 1 |
| 0 | 0 | 0 |
| 1 | 1 | 1 |
| 0 | 1 | 0 |
| 0 | 0 | 0 |
| 1 | 1 | 1 |
| 1 | 1 | 1 |
| 0 | 1 | 0 |
| 0 | 1 | 0 |
| 1 | 1 | 1 |
| 1 | 1 | 1 |
| 1 | 1 | 1 |
| 1 | 1 | 1 |
| 1 | 1 | 1 |
| 1 | 1 | 1 |
| 0 | 0 | 0 |
| 1 | 1 | 1 |
| 0 | 0 | 0 |
| 0 | 0 | 0 |
| 1 | 1 | 1 |
| 0 | 1 | 0 |

| | | |
|---|---|---|
| 0 | 0 | 0 |
| 1 | 1 | 1 |
| 1 | 1 | 1 |
| 1 | 1 | 1 |
| 1 | 1 | 1 |
| 0 | 0 | 0 |
| 0 | 0 | 0 |
| 1 | 1 | 1 |
| 0 | 1 | 0 |
| 1 | 1 | 1 |
| 1 | 1 | 1 |
| 0 | 0 | 0 |
| 0 | 1 | 0 |
| 1 | 1 | 1 |
| 0 | 1 | 0 |
| 1 | 1 | 1 |
| 0 | 0 | 0 |
| 0 | 0 | 0 |
| 1 | 1 | 1 |
| 1 | 1 | 1 |
| 1 | 1 | 1 |
| 1 | 1 | 1 |
| 1 | 1 | 1 |
| 1 | 1 | 1 |
| 0 | 0 | 0 |
| 1 | 1 | 1 |
| 1 | 1 | 1 |
| 1 | 1 | 1 |
| 1 | 1 | 1 |
| 1 | 1 | 1 |
| 0 | 1 | 0 |
| 0 | 0 | 0 |
| 0 | 0 | 0 |
| 0 | 1 | 0 |
| 1 | 1 | 1 |
| 0 | 0 | 0 |
| 0 | 0 | 0 |
| 0 | 0 | 0 |
| 0 | 0 | 0 |
| 0 | 0 | 0 |
| 0 | 0 | 0 |
| 0 | 0 | 0 |
| 0 | 0 | 0 |
| 0 | 0 | 0 |
| 0 | 0 | 0 |
| 0 | 0 | 0 |
| 0 | 0 | 0 |
| 0 | 0 | 0 |
| 1 | 0 | 0 |
| 1 | 1 | 1 |
| 1 | 0 | 0 |
| 1 | 1 | 1 |
| 0 | 0 | 0 |
| 1 | 1 | 1 |

| | | |
|---|---|---|
| 1 | 1 | 1 |
| 1 | 1 | 1 |
| 1 | 1 | 1 |
| 1 | 1 | 1 |
| 1 | 1 | 1 |
| 1 | 1 | 1 |
| 1 | 1 | 1 |
| 0 | 0 | 0 |
| 0 | 0 | 0 |
| 1 | 0 | 0 |
| 1 | 1 | 1 |
| 1 | 1 | 1 |
| 1 | 1 | 1 |
| 0 | 0 | 0 |
| 1 | 1 | 1 |
| 1 | 1 | 1 |
| 0 | 0 | 0 |
| 1 | 1 | 1 |
| 0 | 0 | 0 |
| 0 | 1 | 0 |
| 1 | 1 | 1 |
| 1 | 1 | 1 |
| 1 | 1 | 1 |
| 0 | 0 | 0 |
| 1 | 1 | 1 |
| 1 | 1 | 1 |
| 0 | 1 | 0 |
| 0 | 1 | 0 |
| 0 | 1 | 0 |
| 0 | 1 | 0 |
| 1 | 1 | 1 |
| 0 | 0 | 0 |
| 1 | 1 | 1 |
| 0 | 1 | 0 |
| 1 | 1 | 1 |
| 1 | 1 | 1 |
| 1 | 1 | 1 |
| 1 | 1 | 1 |
| 0 | 0 | 0 |
| 1 | 1 | 1 |
| 1 | 1 | 1 |
| 1 | 1 | 1 |
| 0 | 1 | 0 |
| 1 | 1 | 1 |
| 1 | 1 | 1 |
| 0 | 0 | 0 |
| 0 | 1 | 0 |
| 0 | 1 | 0 |
| 1 | 0 | 0 |
| 1 | 0 | 0 |
| 0 | 1 | 0 |
| 1 | 0 | 0 |
| 0 | 0 | 0 |
| 1 | 0 | 0 |

| | | |
|---|---|---|
| 1 | 0 | 0 |
| 1 | 0 | 0 |
| 1 | 0 | 0 |
| 1 | 0 | 0 |
| 0 | 0 | 0 |
| 0 | 0 | 0 |
| 0 | 0 | 0 |
| 0 | 0 | 0 |
| 1 | 0 | 0 |
| 1 | 0 | 0 |
| 1 | 0 | 0 |
| 1 | 0 | 0 |
| 1 | 0 | 0 |
| 0 | 0 | 0 |
| 1 | 0 | 0 |
| 0 | 0 | 0 |
| 1 | 0 | 0 |
| 0 | 0 | 0 |
| 1 | 0 | 0 |
| 0 | 0 | 0 |
| 0 | 0 | 0 |
| 1 | 0 | 0 |
| 1 | 0 | 0 |
| 0 | 0 | 0 |
| 0 | 0 | 0 |
| 1 | 0 | 0 |
| 0 | 0 | 0 |
| 0 | 0 | 0 |
| 0 | 0 | 0 |
| 1 | 0 | 0 |
| 1 | 0 | 0 |
| 0 | 0 | 0 |
| 1 | 0 | 0 |
| 1 | 0 | 0 |
| 1 | 0 | 0 |
| 1 | 0 | 0 |
| 1 | 0 | 0 |
| 1 | 0 | 0 |
| 1 | 0 | 0 |
| 1 | 0 | 0 |
| 1 | 0 | 0 |
| 1 | 0 | 0 |
| 1 | 0 | 0 |
| 0 | 0 | 0 |
| 1 | 1 | 1 |
| 1 | 1 | 1 |
| 0 | 0 | 0 |
| 0 | 0 | 0 |
| 0 | 1 | 0 |
| 0 | 0 | 0 |
| 0 | 0 | 0 |
| 0 | 0 | 0 |
| 1 | 1 | 1 |
| 1 | 1 | 1 |

| | | |
|---|---|---|
| 1 | 1 | 1 |
| 1 | 1 | 1 |
| 1 | 1 | 1 |
| 1 | 1 | 1 |
| 1 | 1 | 1 |
| 1 | 1 | 1 |
| 1 | 1 | 1 |
| 0 | 0 | 0 |
| 1 | 1 | 1 |
| 1 | 1 | 1 |
| 0 | 1 | 0 |
| 0 | 1 | 0 |
| 1 | 0 | 0 |
| 1 | 1 | 1 |
| 0 | 0 | 0 |
| 0 | 0 | 0 |
| 0 | 1 | 0 |
| 0 | 0 | 0 |
| 1 | 1 | 1 |
| 1 | 1 | 1 |
| 0 | 0 | 0 |
| 0 | 1 | 0 |
| 0 | 1 | 0 |
| 1 | 1 | 1 |
| 0 | 0 | 0 |
| 1 | 1 | 1 |
| 0 | 1 | 0 |
| 0 | 0 | 0 |
| 0 | 0 | 0 |
| 0 | 0 | 0 |
| 0 | 0 | 0 |
| 1 | 1 | 1 |
| 1 | 1 | 1 |
| 0 | 1 | 0 |
| 1 | 1 | 1 |
| 1 | 1 | 1 |
| 0 | 0 | 0 |
| 0 | 0 | 0 |
| 1 | 1 | 1 |
| 1 | 1 | 1 |
| 1 | 1 | 1 |
| 1 | 1 | 1 |
| 0 | 0 | 0 |
| 0 | 1 | 0 |
| 1 | 1 | 1 |
| 1 | 1 | 1 |
| 1 | 1 | 1 |
| 1 | 1 | 1 |
| 0 | 0 | 0 |
| 0 | 0 | 0 |
| 0 | 0 | 0 |
| 0 | 0 | 0 |
| 0 | 0 | 0 |
| 1 | 1 | 1 |

| | | |
|---|---|---|
| 0 | 0 | 0 |
| 0 | 0 | 0 |
| 0 | 1 | 0 |
| 0 | 0 | 0 |
| 1 | 1 | 1 |
| 0 | 1 | 0 |
| 0 | 1 | 0 |
| 1 | 1 | 1 |
| 1 | 1 | 1 |
| 0 | 0 | 0 |
| 1 | 1 | 1 |
| 0 | 1 | 0 |
| 1 | 1 | 1 |
| 0 | 1 | 0 |
| 0 | 1 | 0 |
| 0 | 1 | 0 |
| 0 | 1 | 0 |
| 1 | 1 | 1 |
| 1 | 1 | 1 |
| 0 | 1 | 0 |
| 0 | 1 | 0 |
| 0 | 1 | 0 |
| 0 | 1 | 0 |
| 0 | 1 | 0 |
| 0 | 1 | 0 |
| 0 | 1 | 0 |
| 0 | 1 | 0 |
| 0 | 1 | 0 |
| 0 | 1 | 0 |
| 0 | 1 | 0 |
| 1 | 1 | 1 |
| 1 | 1 | 1 |
| 0 | 0 | 0 |
| 1 | 1 | 1 |
| 1 | 1 | 1 |
| 1 | 1 | 1 |
| 1 | 1 | 1 |
| 0 | 1 | 0 |
| 1 | 1 | 1 |
| 1 | 1 | 1 |
| 1 | 1 | 1 |
| 1 | 1 | 1 |
| 1 | 1 | 1 |
| 0 | 1 | 0 |
| 0 | 1 | 0 |
| 1 | 1 | 1 |
| 1 | 1 | 1 |
| 0 | 0 | 0 |
| 0 | 0 | 0 |
| 0 | 0 | 0 |
| 0 | 0 | 0 |
| 0 | 0 | 0 |
| 1 | 1 | 1 |
| 0 | 0 | 0 |

| | | |
|---|---|---|
| 1 | 1 | 1 |
| 0 | 0 | 0 |
| 1 | 1 | 1 |
| 0 | 1 | 0 |
| 0 | 1 | 0 |
| 0 | 0 | 0 |
| 0 | 0 | 0 |
| 0 | 0 | 0 |
| 0 | 0 | 0 |
| 0 | 0 | 0 |
| 0 | 1 | 0 |
| 0 | 0 | 0 |
| 0 | 1 | 0 |
| 1 | 1 | 1 |
| 0 | 1 | 0 |
| 0 | 0 | 0 |
| 1 | 1 | 1 |
| 1 | 1 | 1 |
| 0 | 0 | 0 |
| 0 | 0 | 0 |
| 0 | 0 | 0 |
| 0 | 0 | 0 |
| 0 | 0 | 0 |
| 0 | 0 | 0 |
| 0 | 1 | 0 |
| 0 | 0 | 0 |
| 0 | 0 | 0 |
| 0 | 0 | 0 |
| 0 | 0 | 0 |
| 0 | 0 | 0 |
| 0 | 0 | 0 |
| 1 | 1 | 1 |
| 1 | 1 | 1 |
| 0 | 0 | 0 |
| 0 | 0 | 0 |
| 1 | 0 | 0 |
| 0 | 0 | 0 |
| 0 | 0 | 0 |
| 0 | 0 | 0 |
| 0 | 0 | 0 |
| 0 | 0 | 0 |
| 1 | 0 | 0 |
| 1 | 1 | 1 |
| 1 | 1 | 1 |
| 0 | 0 | 0 |
| 0 | 0 | 0 |
| 0 | 0 | 0 |
| 1 | 1 | 1 |
| 1 | 1 | 1 |
| 1 | 1 | 1 |
| 1 | 1 | 1 |
| 1 | 1 | 1 |
| 1 | 1 | 1 |
| 0 | 0 | 0 |

| | | |
|---|---|---|
| 1 | 1 | 1 |
| 1 | 1 | 1 |
| 1 | 1 | 1 |
| 1 | 1 | 1 |
| 0 | 0 | 0 |
| 0 | 0 | 0 |
| 1 | 0 | 0 |
| 1 | 0 | 0 |
| 0 | 1 | 0 |
| 1 | 1 | 1 |
| 1 | 1 | 1 |
| 0 | 0 | 0 |
| 1 | 1 | 1 |
| 0 | 0 | 0 |
| 0 | 0 | 0 |
| 0 | 0 | 0 |
| 0 | 0 | 0 |
| 0 | 0 | 0 |
| 0 | 0 | 0 |
| 0 | 0 | 0 |
| 0 | 0 | 0 |
| 0 | 0 | 0 |
| 0 | 0 | 0 |
| 0 | 1 | 0 |
| 0 | 1 | 0 |
| 0 | 1 | 0 |
| 1 | 1 | 1 |
| 1 | 1 | 1 |
| 1 | 1 | 1 |
| 1 | 1 | 1 |
| 1 | 1 | 1 |
| 0 | 0 | 0 |
| 1 | 1 | 1 |
| 0 | 0 | 0 |
| 0 | 0 | 0 |
| 0 | 0 | 0 |
| 0 | 0 | 0 |
| 1 | 1 | 1 |
| 1 | 1 | 1 |
| 0 | 1 | 0 |
| 1 | 1 | 1 |
| 1 | 1 | 1 |
| 1 | 1 | 1 |
| 0 | 1 | 0 |
| 1 | 1 | 1 |
| 0 | 0 | 0 |
| 1 | 1 | 1 |
| 1 | 1 | 1 |
| 1 | 1 | 1 |
| 0 | 0 | 0 |
| 0 | 0 | 0 |
| 0 | 0 | 0 |
| 0 | 1 | 0 |
| 0 | 1 | 0 |

| | | |
|---|---|---|
| 0 | 1 | 0 |
| 0 | 1 | 0 |
| 0 | 1 | 0 |
| 0 | 0 | 0 |
| 0 | 1 | 0 |
| 1 | 1 | 1 |
| 0 | 0 | 0 |
| 0 | 1 | 0 |
| 1 | 1 | 1 |
| 1 | 1 | 1 |
| 0 | 0 | 0 |
| 0 | 0 | 0 |
| 1 | 1 | 1 |
| 1 | 1 | 1 |
| 0 | 1 | 0 |
| 0 | 1 | 0 |
| 0 | 0 | 0 |
| 0 | 0 | 0 |
| 1 | 1 | 1 |
| 0 | 0 | 0 |
| 0 | 0 | 0 |
| 0 | 0 | 0 |
| 0 | 1 | 0 |
| 0 | 0 | 0 |
| 0 | 0 | 0 |
| 1 | 1 | 1 |
| 0 | 0 | 0 |
| 0 | 1 | 0 |
| 1 | 1 | 1 |
| 0 | 0 | 0 |
| 0 | 1 | 0 |
| 1 | 1 | 1 |
| 0 | 0 | 0 |
| 0 | 1 | 0 |
| 0 | 1 | 0 |
| 0 | 0 | 0 |
| 1 | 1 | 1 |
| 0 | 0 | 0 |
| 1 | 1 | 1 |
| 0 | 0 | 0 |
| 0 | 0 | 0 |
| 1 | 0 | 0 |
| 0 | 0 | 0 |
| 0 | 0 | 0 |
| 1 | 1 | 1 |
| 1 | 0 | 0 |
| 0 | 1 | 0 |
| 1 | 1 | 1 |
| 1 | 1 | 1 |
| 1 | 1 | 1 |
| 1 | 1 | 1 |
| 0 | 1 | 0 |
| 0 | 1 | 0 |
| 0 | 1 | 0 |

| | | |
|---|---|---|
| 0 | 1 | 0 |
| 0 | 1 | 0 |
| 0 | 1 | 0 |
| 0 | 0 | 0 |
| 1 | 0 | 0 |
| 0 | 0 | 0 |
| 1 | 1 | 1 |
| 1 | 0 | 0 |
| 0 | 0 | 0 |
| 0 | 0 | 0 |
| 1 | 1 | 1 |
| 0 | 0 | 0 |
| 0 | 0 | 0 |
| 0 | 0 | 0 |
| 1 | 0 | 0 |
| 0 | 0 | 0 |
| 0 | 0 | 0 |
| 0 | 1 | 0 |
| 0 | 0 | 0 |
| 0 | 0 | 0 |
| 0 | 0 | 0 |
| 0 | 0 | 0 |
| 1 | 1 | 1 |
| 0 | 0 | 0 |
| 0 | 0 | 0 |
| 0 | 0 | 0 |
| 0 | 0 | 0 |
| 0 | 0 | 0 |
| 0 | 0 | 0 |
| 0 | 0 | 0 |
| 0 | 0 | 0 |
| 0 | 0 | 0 |
| 0 | 0 | 0 |
| 0 | 1 | 0 |
| 0 | 1 | 0 |
| 0 | 0 | 0 |
| 1 | 1 | 1 |
| 0 | 0 | 0 |
| 0 | 0 | 0 |
| 0 | 0 | 0 |
| 0 | 0 | 0 |
| 1 | 1 | 1 |
| 0 | 0 | 0 |
| 0 | 0 | 0 |
| 1 | 1 | 1 |
| 0 | 1 | 0 |
| 1 | 1 | 1 |
| 1 | 1 | 1 |
| 1 | 1 | 1 |
| 0 | 1 | 0 |
| 1 | 1 | 1 |
| 1 | 1 | 1 |
| 1 | 1 | 1 |
| 1 | 1 | 1 |

| | | |
|---|---|---|
| 1 | 1 | 1 |
| 0 | 0 | 0 |
| 1 | 1 | 1 |
| 1 | 1 | 1 |
| 1 | 1 | 1 |
| 1 | 1 | 1 |
| 1 | 1 | 1 |
| 1 | 1 | 1 |
| 1 | 1 | 1 |
| 0 | 1 | 0 |
| 1 | 1 | 1 |
| 0 | 0 | 0 |
| 1 | 0 | 0 |
| 0 | 1 | 0 |
| 0 | 0 | 0 |
| 0 | 1 | 0 |
| 0 | 1 | 0 |
| 1 | 1 | 1 |
| 1 | 1 | 1 |
| 1 | 1 | 1 |
| 0 | 0 | 0 |
| 0 | 1 | 0 |
| 0 | 0 | 0 |
| 0 | 0 | 0 |
| 1 | 1 | 1 |
| 0 | 0 | 0 |
| 1 | 1 | 1 |
| 1 | 1 | 1 |
| 1 | 1 | 1 |
| 1 | 1 | 1 |
| 1 | 1 | 1 |
| 0 | 0 | 0 |
| 1 | 1 | 1 |
| 0 | 0 | 0 |
| 1 | 1 | 1 |
| 0 | 1 | 0 |
| 0 | 1 | 0 |
| 0 | 1 | 0 |
| 1 | 1 | 1 |
| 0 | 0 | 0 |
| 1 | 1 | 1 |
| 1 | 1 | 1 |
| 0 | 1 | 0 |
| 0 | 0 | 0 |
| 1 | 1 | 1 |
| 1 | 1 | 1 |
| 1 | 1 | 1 |
| 0 | 0 | 0 |
| 0 | 0 | 0 |
| 1 | 1 | 1 |
| 1 | 1 | 1 |
| 0 | 0 | 0 |
| 0 | 0 | 0 |
| 0 | 0 | 0 |

| | | |
|---|---|---|
| 0 | 0 | 0 |
| 1 | 1 | 1 |
| 0 | 0 | 0 |
| 0 | 0 | 0 |
| 0 | 0 | 0 |
| 1 | 1 | 1 |
| 1 | 1 | 1 |
| 0 | 1 | 0 |
| 1 | 1 | 1 |
| 0 | 0 | 0 |
| 1 | 1 | 1 |
| 0 | 1 | 0 |
| 1 | 1 | 1 |
| 1 | 1 | 1 |
| 0 | 0 | 0 |
| 0 | 1 | 0 |
| 1 | 1 | 1 |
| 0 | 0 | 0 |
| 1 | 1 | 1 |
| 0 | 0 | 0 |
| 0 | 0 | 0 |
| 1 | 1 | 1 |
| 0 | 0 | 0 |
| 1 | 1 | 1 |
| 0 | 0 | 0 |
| 0 | 0 | 0 |
| 1 | 1 | 1 |
| 0 | 1 | 0 |
| 0 | 0 | 0 |
| 1 | 1 | 1 |
| 1 | 1 | 1 |
| 1 | 1 | 1 |
| 1 | 1 | 1 |
| 0 | 1 | 0 |
| 0 | 0 | 0 |
| 0 | 0 | 0 |
| 1 | 1 | 1 |
| 0 | 0 | 0 |
| 0 | 0 | 0 |
| 0 | 0 | 0 |
| 0 | 0 | 0 |
| 0 | 0 | 0 |
| 0 | 0 | 0 |
| 0 | 1 | 0 |
| 0 | 0 | 0 |
| 1 | 1 | 1 |
| 0 | 1 | 0 |
| 0 | 0 | 0 |
| 0 | 0 | 0 |
| 0 | 0 | 0 |
| 0 | 0 | 0 |
| 0 | 0 | 0 |
| 1 | 1 | 1 |
| 1 | 1 | 1 |

| | | |
|---|---|---|
| 1 | 1 | 1 |
| 0 | 0 | 0 |
| 0 | 0 | 0 |
| 0 | 0 | 0 |
| 0 | 0 | 0 |
| 0 | 0 | 0 |
| 0 | 0 | 0 |
| 1 | 0 | 0 |
| 1 | 0 | 0 |
| 1 | 1 | 1 |
| 0 | 0 | 0 |
| 0 | 0 | 0 |
| 1 | 1 | 1 |
| 0 | 0 | 0 |
| 1 | 1 | 1 |
| 0 | 0 | 0 |
| 1 | 1 | 1 |
| 1 | 1 | 1 |
| 0 | 1 | 0 |
| 0 | 1 | 0 |
| 0 | 1 | 0 |
| 1 | 1 | 1 |
| 0 | 1 | 0 |
| 0 | 0 | 0 |
| 0 | 0 | 0 |
| 1 | 1 | 1 |
| 1 | 1 | 1 |
| 1 | 1 | 1 |
| 1 | 1 | 1 |
| 0 | 0 | 0 |
| 0 | 0 | 0 |
| 0 | 0 | 0 |
| 0 | 0 | 0 |
| 1 | 1 | 1 |
| 0 | 0 | 0 |
| 0 | 0 | 0 |
| 0 | 0 | 0 |
| 0 | 0 | 0 |
| 0 | 0 | 0 |
| 1 | 1 | 1 |
| 1 | 1 | 1 |
| 0 | 0 | 0 |
| 1 | 1 | 1 |
| 1 | 1 | 1 |
| 0 | 0 | 0 |
| 1 | 0 | 0 |
| 1 | 1 | 1 |
| 0 | 1 | 0 |
| 1 | 0 | 0 |
| 0 | 1 | 0 |
| 0 | 0 | 0 |
| 1 | 1 | 1 |
| 0 | 0 | 0 |
| 0 | 0 | 0 |

| | | |
|---|---|---|
| 0 | 1 | 0 |
| 1 | 1 | 1 |
| 0 | 0 | 0 |
| 0 | 0 | 0 |
| 1 | 1 | 1 |
| 1 | 1 | 1 |
| 1 | 1 | 1 |
| 1 | 1 | 1 |
| 1 | 1 | 1 |
| 1 | 1 | 1 |
| 1 | 1 | 1 |
| 1 | 1 | 1 |
| 0 | 0 | 0 |
| 1 | 1 | 1 |
| 1 | 1 | 1 |
| 1 | 1 | 1 |
| 1 | 1 | 1 |
| 1 | 1 | 1 |
| 1 | 1 | 1 |
| 1 | 1 | 1 |
| 1 | 1 | 1 |
| 1 | 1 | 1 |
| 1 | 1 | 1 |
| 1 | 1 | 1 |
| 1 | 1 | 1 |
| 1 | 1 | 1 |
| 0 | 1 | 0 |
| 1 | 1 | 1 |
| 1 | 1 | 1 |
| 1 | 1 | 1 |
| 1 | 1 | 1 |
| 0 | 0 | 0 |
| 1 | 1 | 1 |
| 0 | 1 | 0 |
| 1 | 1 | 1 |
| 1 | 1 | 1 |
| 0 | 1 | 0 |
| 1 | 1 | 1 |
| 1 | 1 | 1 |
| 1 | 1 | 1 |
| 0 | 0 | 0 |
| 0 | 0 | 0 |
| 1 | 1 | 1 |
| 1 | 1 | 1 |
| 1 | 1 | 1 |
| 1 | 0 | 0 |
| 1 | 0 | 0 |
| 1 | 1 | 1 |
| 0 | 0 | 0 |
| 0 | 0 | 0 |
| 1 | 1 | 1 |
| 0 | 1 | 0 |
| 0 | 0 | 0 |
| 1 | 1 | 1 |

| | | |
|---|---|---|
| 0 | 0 | 0 |
| 0 | 0 | 0 |
| 0 | 0 | 0 |
| 1 | 1 | 1 |
| 1 | 1 | 1 |
| 1 | 1 | 1 |
| 1 | 1 | 1 |
| 0 | 0 | 0 |
| 0 | 0 | 0 |
| 0 | 0 | 0 |
| 0 | 0 | 0 |
| 1 | 1 | 1 |
| 0 | 0 | 0 |
| 1 | 1 | 1 |
| 1 | 1 | 1 |
| 1 | 1 | 1 |
| 1 | 1 | 1 |
| 0 | 0 | 0 |
| 1 | 1 | 1 |
| 1 | 1 | 1 |
| 1 | 1 | 1 |
| 1 | 1 | 1 |
| 0 | 1 | 0 |
| 1 | 1 | 1 |
| 1 | 1 | 1 |
| 0 | 1 | 0 |
| 1 | 1 | 1 |
| 1 | 1 | 1 |
| 1 | 1 | 1 |
| 1 | 1 | 1 |
| 0 | 0 | 0 |
| 0 | 0 | 0 |
| 1 | 1 | 1 |
| 1 | 1 | 1 |
| 1 | 1 | 1 |
| 1 | 1 | 1 |
| 1 | 1 | 1 |
| 1 | 1 | 1 |
| 1 | 1 | 1 |
| 1 | 1 | 1 |
| 1 | 1 | 1 |
| 0 | 0 | 0 |
| 1 | 1 | 1 |
| 1 | 1 | 1 |
| 0 | 0 | 0 |
| 0 | 0 | 0 |
| 1 | 1 | 1 |
| 1 | 1 | 1 |
| 0 | 0 | 0 |
| 0 | 0 | 0 |
| 0 | 0 | 0 |
| 0 | 0 | 0 |
| 0 | 0 | 0 |
| 0 | 0 | 0 |

| | | |
|---|---|---|
| 1 | 1 | 1 |
| 1 | 1 | 1 |
| 1 | 1 | 1 |
| 1 | 1 | 1 |
| 1 | 1 | 1 |
| 0 | 0 | 0 |
| 0 | 0 | 0 |
| 0 | 0 | 0 |
| 0 | 0 | 0 |
| 0 | 0 | 0 |
| 0 | 1 | 0 |
| 0 | 0 | 0 |
| 0 | 0 | 0 |
| 0 | 0 | 0 |
| 0 | 1 | 0 |
| 0 | 0 | 0 |
| 0 | 1 | 0 |
| 0 | 1 | 0 |
| 0 | 0 | 0 |
| 0 | 0 | 0 |
| 0 | 0 | 0 |
| 0 | 0 | 0 |
| 1 | 0 | 0 |
| 0 | 0 | 0 |
| 0 | 0 | 0 |
| 0 | 0 | 0 |
| 0 | 0 | 0 |
| 1 | 1 | 1 |
| 1 | 1 | 1 |
| 1 | 1 | 1 |
| 1 | 1 | 1 |
| 1 | 1 | 1 |
| 0 | 0 | 0 |
| 0 | 1 | 0 |
| 0 | 0 | 0 |
| 0 | 0 | 0 |
| 0 | 0 | 0 |
| 0 | 0 | 0 |
| 0 | 0 | 0 |
| 0 | 0 | 0 |
| 0 | 0 | 0 |
| 0 | 0 | 0 |
| 0 | 0 | 0 |
| 1 | 1 | 1 |
| 1 | 1 | 1 |
| 1 | 1 | 1 |
| 1 | 1 | 1 |
| 0 | 0 | 0 |
| 0 | 0 | 0 |
| 0 | 0 | 0 |
| 1 | 1 | 1 |
| 1 | 1 | 1 |
| 0 | 0 | 0 |
| 1 | 1 | 1 |

| | | |
|---|---|---|
| 1 | 1 | 1 |
| 0 | 0 | 0 |
| 0 | 1 | 0 |
| 0 | 1 | 0 |
| 1 | 1 | 1 |
| 1 | 1 | 1 |
| 0 | 1 | 0 |
| 1 | 1 | 1 |
| 0 | 1 | 0 |
| 1 | 1 | 1 |
| 1 | 1 | 1 |
| 1 | 1 | 1 |
| 0 | 0 | 0 |
| 0 | 0 | 0 |
| 0 | 0 | 0 |
| 0 | 0 | 0 |
| 0 | 0 | 0 |
| 0 | 0 | 0 |
| 0 | 0 | 0 |
| 0 | 1 | 0 |
| 0 | 0 | 0 |
| 0 | 0 | 0 |
| 0 | 0 | 0 |
| 0 | 0 | 0 |
| 0 | 0 | 0 |
| 1 | 1 | 1 |
| 1 | 1 | 1 |
| 0 | 0 | 0 |
| 0 | 1 | 0 |
| 0 | 0 | 0 |
| 0 | 0 | 0 |
| 1 | 1 | 1 |
| 0 | 0 | 0 |
| 1 | 1 | 1 |
| 1 | 1 | 1 |
| 1 | 1 | 1 |
| 1 | 0 | 0 |
| 1 | 1 | 1 |
| 1 | 1 | 1 |
| 1 | 1 | 1 |
| 0 | 0 | 0 |
| 1 | 1 | 1 |
| 1 | 1 | 1 |
| 0 | 0 | 0 |
| 0 | 0 | 0 |
| 1 | 1 | 1 |
| 0 | 0 | 0 |
| 0 | 0 | 0 |
| 1 | 1 | 1 |
| 1 | 1 | 1 |
| 0 | 0 | 0 |
| 1 | 1 | 1 |
| 1 | 1 | 1 |
| 1 | 1 | 1 |

| | | |
|---|---|---|
| 1 | 1 | 1 |
| 1 | 1 | 1 |
| 1 | 1 | 1 |
| 1 | 1 | 1 |
| 0 | 0 | 0 |
| 0 | 0 | 0 |
| 1 | 1 | 1 |
| 1 | 1 | 1 |
| 1 | 1 | 1 |
| 0 | 1 | 0 |
| 1 | 1 | 1 |
| 0 | 0 | 0 |
| 0 | 0 | 0 |
| 1 | 1 | 1 |
| 1 | 1 | 1 |
| 0 | 0 | 0 |
| 0 | 1 | 0 |
| 1 | 1 | 1 |
| 0 | 0 | 0 |
| 1 | 1 | 1 |
| 1 | 1 | 1 |
| 1 | 1 | 1 |
| 1 | 1 | 1 |
| 0 | 0 | 0 |
| 1 | 1 | 1 |
| 0 | 0 | 0 |
| 1 | 1 | 1 |
| 1 | 1 | 1 |
| 0 | 0 | 0 |
| 0 | 0 | 0 |
| 0 | 0 | 0 |
| 0 | 0 | 0 |
| 0 | 0 | 0 |
| 0 | 0 | 0 |
| 1 | 1 | 1 |
| 1 | 1 | 1 |
| 1 | 1 | 1 |
| 0 | 0 | 0 |
| 1 | 1 | 1 |
| 1 | 1 | 1 |
| 0 | 0 | 0 |
| 0 | 0 | 0 |
| 0 | 0 | 0 |
| 1 | 1 | 1 |
| 0 | 0 | 0 |
| 0 | 1 | 0 |
| 0 | 1 | 0 |
| 0 | 0 | 0 |
| 1 | 1 | 1 |
| 1 | 1 | 1 |
| 1 | 1 | 1 |
| 0 | 0 | 0 |
| 1 | 1 | 1 |
| 0 | 0 | 0 |

| | | |
|---|---|---|
| 1 | 1 | 1 |
| 0 | 0 | 0 |
| 0 | 0 | 0 |
| 0 | 1 | 0 |
| 1 | 1 | 1 |
| 1 | 1 | 1 |
| 0 | 0 | 0 |
| 1 | 1 | 1 |
| 0 | 1 | 0 |
| 0 | 1 | 0 |
| 1 | 1 | 1 |
| 1 | 1 | 1 |
| 1 | 1 | 1 |
| 1 | 1 | 1 |
| 1 | 1 | 1 |
| 1 | 1 | 1 |
| 1 | 1 | 1 |
| 1 | 1 | 1 |
| 1 | 1 | 1 |
| 1 | 1 | 1 |
| 1 | 1 | 1 |
| 0 | 0 | 0 |
| 0 | 1 | 0 |
| 0 | 1 | 0 |
| 1 | 1 | 1 |
| 1 | 1 | 1 |
| 0 | 0 | 0 |
| 1 | 1 | 1 |
| 0 | 0 | 0 |
| 0 | 1 | 0 |
| 1 | 1 | 1 |
| 0 | 0 | 0 |
| 0 | 0 | 0 |
| 1 | 0 | 0 |
| 1 | 1 | 1 |
| 1 | 1 | 1 |
| 1 | 1 | 1 |
| 1 | 1 | 1 |
| 0 | 0 | 0 |
| 0 | 0 | 0 |
| 0 | 0 | 0 |
| 1 | 1 | 1 |
| 1 | 1 | 1 |
| 0 | 1 | 0 |
| 0 | 1 | 0 |
| 0 | 0 | 0 |
| 1 | 1 | 1 |
| 0 | 0 | 0 |
| 1 | 1 | 1 |
| 0 | 0 | 0 |
| 0 | 0 | 0 |
| 0 | 0 | 0 |
| 1 | 1 | 1 |
| 1 | 1 | 1 |

| | | |
|---|---|---|
| 0 | 0 | 0 |
| 0 | 0 | 0 |
| 1 | 1 | 1 |
| 1 | 1 | 1 |
| 1 | 1 | 1 |
| 1 | 1 | 1 |
| 1 | 1 | 1 |
| 1 | 1 | 1 |
| 1 | 1 | 1 |
| 1 | 1 | 1 |
| 1 | 1 | 1 |
| 1 | 1 | 1 |
| 1 | 1 | 1 |
| 1 | 1 | 1 |
| 1 | 1 | 1 |
| 0 | 0 | 0 |
| 0 | 0 | 0 |
| 0 | 0 | 0 |
| 1 | 1 | 1 |
| 1 | 1 | 1 |
| 0 | 1 | 0 |
| 0 | 0 | 0 |
| 0 | 0 | 0 |
| 0 | 0 | 0 |
| 0 | 0 | 0 |
| 1 | 1 | 1 |
| 1 | 1 | 1 |
| 1 | 1 | 1 |
| 1 | 1 | 1 |
| 1 | 1 | 1 |
| 0 | 0 | 0 |
| 0 | 1 | 0 |
| 0 | 0 | 0 |
| 1 | 1 | 1 |
| 1 | 0 | 0 |
| 1 | 0 | 0 |
| 0 | 0 | 0 |
| 0 | 0 | 0 |
| 0 | 1 | 0 |
| 1 | 1 | 1 |
| 1 | 0 | 0 |
| 0 | 0 | 0 |
| 1 | 1 | 1 |
| 0 | 1 | 0 |
| 0 | 0 | 0 |
| 1 | 0 | 0 |
| 0 | 0 | 0 |
| 0 | 0 | 0 |
| 1 | 1 | 1 |
| 0 | 1 | 0 |
| 0 | 0 | 0 |
| 0 | 0 | 0 |
| 1 | 1 | 1 |
| 0 | 0 | 0 |

| | | |
|---|---|---|
| 0 | 1 | 0 |
| 0 | 0 | 0 |
| 1 | 0 | 0 |
| 0 | 0 | 0 |
| 1 | 1 | 1 |
| 1 | 1 | 1 |
| 1 | 1 | 1 |
| 0 | 1 | 0 |
| 0 | 0 | 0 |
| 1 | 1 | 1 |
| 0 | 1 | 0 |
| 0 | 1 | 0 |
| 0 | 0 | 0 |
| 0 | 0 | 0 |
| 0 | 0 | 0 |
| 1 | 1 | 1 |
| 0 | 0 | 0 |
| 0 | 1 | 0 |
| 0 | 0 | 0 |
| 0 | 0 | 0 |
| 0 | 0 | 0 |
| 0 | 0 | 0 |
| 0 | 0 | 0 |
| 0 | 0 | 0 |
| 0 | 1 | 0 |
| 0 | 1 | 0 |
| 0 | 0 | 0 |
| 0 | 0 | 0 |
| 0 | 0 | 0 |
| 0 | 0 | 0 |
| 1 | 1 | 1 |
| 1 | 1 | 1 |
| 1 | 1 | 1 |
| 1 | 1 | 1 |
| 1 | 1 | 1 |
| 0 | 0 | 0 |
| 0 | 0 | 0 |
| 0 | 1 | 0 |
| 1 | 1 | 1 |
| 1 | 1 | 1 |
| 0 | 1 | 0 |
| 1 | 1 | 1 |
| 1 | 1 | 1 |
| 0 | 0 | 0 |
| 0 | 0 | 0 |
| 0 | 0 | 0 |
| 1 | 1 | 1 |
| 0 | 0 | 0 |
| 0 | 0 | 0 |
| 0 | 0 | 0 |
| 1 | 1 | 1 |
| 1 | 1 | 1 |
| 0 | 1 | 0 |
| 0 | 1 | 0 |

| | | |
|---|---|---|
| 0 | 1 | 0 |
| 1 | 1 | 1 |
| 1 | 0 | 0 |
| 1 | 1 | 1 |
| 0 | 0 | 0 |
| 1 | 0 | 0 |
| 0 | 0 | 0 |
| 0 | 0 | 0 |
| 0 | 0 | 0 |
| 0 | 0 | 0 |
| 0 | 0 | 0 |
| 0 | 0 | 0 |
| 0 | 1 | 0 |
| 0 | 0 | 0 |
| 0 | 0 | 0 |
| 1 | 1 | 1 |
| 1 | 1 | 1 |
| 1 | 1 | 1 |
| 0 | 0 | 0 |
| 0 | 0 | 0 |
| 0 | 0 | 0 |
| 0 | 0 | 0 |
| 0 | 1 | 0 |
| 0 | 1 | 0 |
| 1 | 1 | 1 |
| 1 | 1 | 1 |
| 1 | 1 | 1 |
| 0 | 0 | 0 |
| 0 | 0 | 0 |
| 0 | 0 | 0 |
| 0 | 0 | 0 |
| 0 | 0 | 0 |
| 0 | 0 | 0 |
| 0 | 1 | 0 |
| 1 | 1 | 1 |
| 0 | 0 | 0 |
| 1 | 1 | 1 |
| 1 | 1 | 1 |
| 1 | 1 | 1 |
| 1 | 1 | 1 |
| 0 | 0 | 0 |
| 1 | 1 | 1 |
| 0 | 0 | 0 |
| 0 | 1 | 0 |
| 1 | 1 | 1 |
| 1 | 1 | 1 |
| 1 | 1 | 1 |
| 0 | 0 | 0 |
| 1 | 1 | 1 |
| 1 | 1 | 1 |
| 1 | 1 | 1 |
| 1 | 1 | 1 |
| 1 | 1 | 1 |
| 0 | 1 | 0 |

| | | |
|---|---|---|
| 0 | 1 | 0 |
| 1 | 1 | 1 |
| 1 | 1 | 1 |
| 0 | 1 | 0 |
| 1 | 1 | 1 |
| 0 | 1 | 0 |
| 0 | 1 | 0 |
| 0 | 1 | 0 |
| 0 | 1 | 0 |
| 0 | 1 | 0 |
| 0 | 1 | 0 |
| 0 | 0 | 0 |
| 0 | 1 | 0 |
| 0 | 1 | 0 |
| 0 | 1 | 0 |
| 0 | 1 | 0 |
| 0 | 1 | 0 |
| 0 | 0 | 0 |
| 0 | 0 | 0 |
| 0 | 0 | 0 |
| 0 | 0 | 0 |
| 0 | 1 | 0 |
| 0 | 0 | 0 |
| 1 | 1 | 1 |
| 1 | 1 | 1 |
| 1 | 1 | 1 |
| 1 | 1 | 1 |
| 0 | 1 | 0 |
| 0 | 0 | 0 |
| 0 | 0 | 0 |
| 1 | 1 | 1 |
| 1 | 1 | 1 |
| 0 | 0 | 0 |
| 1 | 1 | 1 |
| 0 | 0 | 0 |
| 0 | 0 | 0 |
| 0 | 0 | 0 |
| 1 | 1 | 1 |
| 0 | 0 | 0 |
| 1 | 1 | 1 |
| 0 | 0 | 0 |
| 1 | 1 | 1 |
| 0 | 0 | 0 |
| 1 | 0 | 0 |
| 1 | 0 | 0 |
| 1 | 0 | 0 |
| 1 | 0 | 0 |
| 0 | 1 | 0 |
| 0 | 1 | 0 |
| 0 | 0 | 0 |
| 1 | 0 | 0 |
| 0 | 0 | 0 |
| 1 | 0 | 0 |
| 1 | 1 | 1 |

| | | |
|---|---|---|
| 0 | 0 | 0 |
| 0 | 0 | 0 |
| 0 | 0 | 0 |
| 0 | 0 | 0 |
| 0 | 0 | 0 |
| 1 | 1 | 1 |
| 0 | 0 | 0 |
| 0 | 0 | 0 |
| 1 | 1 | 1 |
| 0 | 0 | 0 |
| 0 | 0 | 0 |
| 0 | 0 | 0 |
| 0 | 1 | 0 |
| 0 | 1 | 0 |
| 0 | 1 | 0 |
| 0 | 1 | 0 |
| 0 | 1 | 0 |
| 0 | 1 | 0 |
| 0 | 1 | 0 |
| 0 | 1 | 0 |
| 0 | 1 | 0 |
| 0 | 1 | 0 |
| 0 | 1 | 0 |
| 0 | 1 | 0 |
| 0 | 1 | 0 |
| 0 | 1 | 0 |
| 0 | 1 | 0 |
| 0 | 1 | 0 |
| 0 | 1 | 0 |
| 0 | 1 | 0 |
| 0 | 1 | 0 |
| 0 | 1 | 0 |
| 0 | 1 | 0 |
| 0 | 1 | 0 |
| 0 | 1 | 0 |
| 0 | 1 | 0 |
| 0 | 1 | 0 |
| 0 | 1 | 0 |
| 0 | 1 | 0 |
| 0 | 1 | 0 |
| 0 | 1 | 0 |
| 0 | 1 | 0 |
| 0 | 1 | 0 |
| 0 | 1 | 0 |
| 0 | 1 | 0 |
| 0 | 0 | 0 |
| 1 | 1 | 1 |
| 1 | 1 | 1 |
| 1 | 1 | 1 |
| 1 | 1 | 1 |
| 0 | 0 | 0 |
| 1 | 1 | 1 |
| 1 | 1 | 1 |
| 0 | 1 | 0 |

| | | |
|---|---|---|
| 1 | 1 | 1 |
| 1 | 1 | 1 |
| 1 | 1 | 1 |
| 1 | 1 | 1 |
| 0 | 1 | 0 |
| 0 | 1 | 0 |
| 0 | 1 | 0 |
| 0 | 0 | 0 |
| 0 | 0 | 0 |
| 1 | 1 | 1 |
| 0 | 1 | 0 |
| 1 | 1 | 1 |
| 1 | 1 | 1 |
| 1 | 1 | 1 |
| 0 | 0 | 0 |
| 0 | 0 | 0 |
| 0 | 0 | 0 |
| 1 | 1 | 1 |
| 1 | 0 | 0 |
| 1 | 1 | 1 |
| 0 | 0 | 0 |
| 0 | 0 | 0 |
| 0 | 0 | 0 |
| 0 | 0 | 0 |
| 0 | 0 | 0 |
| 0 | 0 | 0 |
| 0 | 1 | 0 |
| 1 | 1 | 1 |
| 0 | 0 | 0 |
| 1 | 1 | 1 |
| 1 | 1 | 1 |
| 1 | 1 | 1 |
| 0 | 0 | 0 |
| 1 | 1 | 1 |
| 1 | 1 | 1 |
| 1 | 1 | 1 |
| 0 | 0 | 0 |
| 0 | 0 | 0 |
| 0 | 0 | 0 |
| 1 | 1 | 1 |
| 1 | 1 | 1 |
| 1 | 1 | 1 |
| 1 | 1 | 1 |
| 1 | 1 | 1 |
| 0 | 1 | 0 |
| 0 | 0 | 0 |
| 1 | 1 | 1 |
| 0 | 1 | 0 |
| 1 | 1 | 1 |
| 0 | 1 | 0 |
| 1 | 1 | 1 |
| 1 | 1 | 1 |
| 0 | 1 | 0 |
| 1 | 1 | 1 |

| | | |
|---|---|---|
| 1 | 1 | 1 |
| 0 | 1 | 0 |
| 1 | 1 | 1 |
| 0 | 0 | 0 |
| 1 | 1 | 1 |
| 1 | 1 | 1 |
| 1 | 1 | 1 |
| 0 | 0 | 0 |
| 0 | 0 | 0 |
| 1 | 1 | 1 |
| 1 | 1 | 1 |
| 1 | 1 | 1 |
| 0 | 0 | 0 |
| 0 | 1 | 0 |
| 0 | 1 | 0 |
| 0 | 1 | 0 |
| 0 | 0 | 0 |
| 1 | 1 | 1 |
| 1 | 1 | 1 |
| 1 | 1 | 1 |
| 1 | 1 | 1 |
| 1 | 1 | 1 |
| 1 | 1 | 1 |
| 1 | 1 | 1 |
| 0 | 1 | 0 |
| 1 | 1 | 1 |
| 0 | 1 | 0 |
| 0 | 0 | 0 |
| 0 | 0 | 0 |
| 0 | 0 | 0 |
| 0 | 0 | 0 |
| 0 | 0 | 0 |
| 0 | 0 | 0 |
| 0 | 1 | 0 |
| 0 | 0 | 0 |
| 0 | 0 | 0 |
| 0 | 0 | 0 |
| 1 | 1 | 1 |
| 1 | 1 | 1 |
| 1 | 1 | 1 |
| 1 | 1 | 1 |
| 0 | 1 | 0 |
| 0 | 0 | 0 |
| 0 | 0 | 0 |
| 1 | 1 | 1 |
| 1 | 1 | 1 |
| 0 | 0 | 0 |
| 1 | 1 | 1 |
| 1 | 1 | 1 |
| 1 | 1 | 1 |
| 1 | 1 | 1 |
| 1 | 1 | 1 |
| 0 | 0 | 0 |
| 0 | 0 | 0 |

| | | |
|---|---|---|
| 0 | 0 | 0 |
| 0 | 0 | 0 |
| 0 | 0 | 0 |
| 0 | 0 | 0 |
| 0 | 0 | 0 |
| 0 | 0 | 0 |
| 0 | 0 | 0 |
| 0 | 0 | 0 |
| 1 | 1 | 1 |
| 0 | 0 | 0 |
| 1 | 1 | 1 |
| 1 | 1 | 1 |
| 1 | 1 | 1 |
| 1 | 1 | 1 |
| 0 | 0 | 0 |
| 1 | 1 | 1 |
| 1 | 1 | 1 |
| 1 | 1 | 1 |
| 1 | 1 | 1 |
| 0 | 0 | 0 |
| 0 | 0 | 0 |
| 1 | 1 | 1 |
| 0 | 0 | 0 |
| 0 | 0 | 0 |
| 0 | 0 | 0 |
| 0 | 0 | 0 |
| 0 | 0 | 0 |
| 0 | 1 | 0 |
| 0 | 1 | 0 |
| 0 | 1 | 0 |
| 0 | 0 | 0 |
| 1 | 1 | 1 |
| 1 | 1 | 1 |
| 1 | 1 | 1 |
| 1 | 1 | 1 |
| 1 | 1 | 1 |
| 1 | 1 | 1 |
| 1 | 1 | 1 |
| 1 | 1 | 1 |
| 1 | 1 | 1 |
| 0 | 1 | 0 |
| 0 | 0 | 0 |
| 1 | 1 | 1 |
| 1 | 1 | 1 |
| 1 | 1 | 1 |
| 0 | 1 | 0 |
| 1 | 1 | 1 |
| 1 | 1 | 1 |
| 0 | 0 | 0 |
| 0 | 1 | 0 |
| 1 | 1 | 1 |
| 1 | 1 | 1 |
| 0 | 0 | 0 |
| 0 | 0 | 0 |

| | | |
|---|---|---|
| 1 | 1 | 1 |
| 1 | 1 | 1 |
| 0 | 1 | 0 |
| 1 | 0 | 0 |
| 0 | 0 | 0 |
| 0 | 0 | 0 |
| 0 | 1 | 0 |
| 0 | 0 | 0 |
| 1 | 1 | 1 |
| 1 | 1 | 1 |
| 0 | 0 | 0 |
| 1 | 0 | 0 |
| 0 | 0 | 0 |
| 0 | 0 | 0 |
| 0 | 0 | 0 |
| 0 | 0 | 0 |
| 1 | 1 | 1 |
| 1 | 1 | 1 |
| 0 | 0 | 0 |
| 0 | 0 | 0 |
| 0 | 0 | 0 |
| 0 | 0 | 0 |
| 0 | 0 | 0 |
| 1 | 1 | 1 |
| 1 | 1 | 1 |
| 0 | 0 | 0 |
| 1 | 1 | 1 |
| 0 | 0 | 0 |
| 0 | 0 | 0 |
| 0 | 0 | 0 |
| 0 | 1 | 0 |
| 0 | 0 | 0 |
| 0 | 0 | 0 |
| 1 | 1 | 1 |
| 1 | 1 | 1 |
| 0 | 0 | 0 |
| 0 | 0 | 0 |
| 1 | 0 | 0 |
| 0 | 0 | 0 |
| 0 | 0 | 0 |
| 0 | 0 | 0 |
| 0 | 0 | 0 |
| 0 | 1 | 0 |
| 1 | 1 | 1 |
| 0 | 0 | 0 |
| 0 | 1 | 0 |
| 0 | 0 | 0 |
| 1 | 1 | 1 |
| 1 | 1 | 1 |
| 1 | 1 | 1 |
| 1 | 1 | 1 |
| 1 | 1 | 1 |
| 0 | 1 | 0 |
| 1 | 1 | 1 |

| | | |
|---|---|---|
| 1 | 1 | 1 |
| 1 | 1 | 1 |
| 1 | 1 | 1 |
| 0 | 1 | 0 |
| 1 | 1 | 1 |
| 1 | 1 | 1 |
| 0 | 1 | 0 |
| 0 | 0 | 0 |
| 1 | 1 | 1 |
| 0 | 0 | 0 |
| 0 | 0 | 0 |
| 0 | 0 | 0 |
| 0 | 1 | 0 |
| 1 | 1 | 1 |
| 0 | 0 | 0 |
| 0 | 1 | 0 |
| 0 | 1 | 0 |
| 0 | 1 | 0 |
| 0 | 1 | 0 |
| 0 | 1 | 0 |
| 1 | 1 | 1 |
| 1 | 1 | 1 |
| 1 | 1 | 1 |
| 0 | 1 | 0 |
| 1 | 1 | 1 |
| 1 | 1 | 1 |
| 1 | 1 | 1 |
| 1 | 0 | 0 |
| 1 | 1 | 1 |
| 0 | 0 | 0 |
| 0 | 0 | 0 |
| 1 | 1 | 1 |
| 1 | 1 | 1 |
| 0 | 1 | 0 |
| 0 | 1 | 0 |
| 0 | 0 | 0 |
| 1 | 1 | 1 |
| 1 | 1 | 1 |
| 1 | 1 | 1 |
| 1 | 1 | 1 |
| 1 | 1 | 1 |
| 0 | 0 | 0 |
| 0 | 0 | 0 |
| 0 | 0 | 0 |
| 1 | 1 | 1 |
| 1 | 1 | 1 |
| 1 | 1 | 1 |
| 1 | 1 | 1 |
| 1 | 1 | 1 |
| 1 | 1 | 1 |
| 1 | 1 | 1 |
| 1 | 1 | 1 |
| 1 | 1 | 1 |
| 0 | 0 | 0 |

| | | |
|---|---|---|
| 1 | 1 | 1 |
| 1 | 1 | 1 |
| 1 | 1 | 1 |
| 1 | 1 | 1 |
| 1 | 1 | 1 |
| 1 | 1 | 1 |
| 1 | 1 | 1 |
| 0 | 1 | 0 |
| 0 | 0 | 0 |
| 0 | 0 | 0 |
| 1 | 1 | 1 |
| 0 | 1 | 0 |
| 0 | 1 | 0 |
| 0 | 0 | 0 |
| 1 | 1 | 1 |
| 1 | 1 | 1 |
| 1 | 1 | 1 |
| 0 | 0 | 0 |
| 0 | 0 | 0 |
| 0 | 0 | 0 |
| 0 | 0 | 0 |
| 0 | 1 | 0 |
| 1 | 1 | 1 |
| 1 | 1 | 1 |
| 1 | 1 | 1 |
| 0 | 0 | 0 |
| 1 | 1 | 1 |
| 0 | 1 | 0 |
| 1 | 1 | 1 |
| 1 | 1 | 1 |
| 1 | 1 | 1 |
| 1 | 1 | 1 |
| 1 | 1 | 1 |
| 1 | 1 | 1 |
| 1 | 1 | 1 |
| 1 | 1 | 1 |
| 1 | 1 | 1 |
| 1 | 1 | 1 |
| 1 | 1 | 1 |
| 0 | 0 | 0 |
| 1 | 1 | 1 |
| 1 | 1 | 1 |
| 0 | 0 | 0 |
| 1 | 1 | 1 |
| 1 | 1 | 1 |
| 1 | 1 | 1 |
| 0 | 0 | 0 |
| 0 | 0 | 0 |
| 1 | 1 | 1 |
| 1 | 1 | 1 |
| 0 | 0 | 0 |
| 1 | 1 | 1 |
| 0 | 1 | 0 |
| 1 | 1 | 1 |

| | | |
|---|---|---|
| 1 | 1 | 1 |
| 1 | 1 | 1 |
| 1 | 1 | 1 |
| 1 | 1 | 1 |
| 1 | 1 | 1 |
| 1 | 1 | 1 |
| 1 | 1 | 1 |
| 0 | 1 | 0 |
| 1 | 1 | 1 |
| 1 | 1 | 1 |
| 1 | 1 | 1 |
| 1 | 1 | 1 |
| 1 | 1 | 1 |
| 0 | 0 | 0 |
| 0 | 0 | 0 |
| 0 | 0 | 0 |
| 0 | 0 | 0 |
| 0 | 1 | 0 |
| 1 | 1 | 1 |
| 0 | 0 | 0 |
| 1 | 1 | 1 |
| 1 | 0 | 0 |
| 1 | 1 | 1 |
| 1 | 1 | 1 |
| 1 | 1 | 1 |
| 0 | 0 | 0 |
| 0 | 1 | 0 |
| 1 | 1 | 1 |
| 1 | 1 | 1 |
| 0 | 0 | 0 |
| 0 | 0 | 0 |
| 1 | 1 | 1 |
| 1 | 1 | 1 |
| 1 | 1 | 1 |
| 1 | 1 | 1 |
| 1 | 1 | 1 |
| 1 | 1 | 1 |
| 1 | 1 | 1 |
| 0 | 0 | 0 |
| 1 | 1 | 1 |
| 1 | 1 | 1 |
| 0 | 0 | 0 |
| 0 | 0 | 0 |
| 0 | 0 | 0 |
| 0 | 0 | 0 |
| 1 | 1 | 1 |
| 0 | 0 | 0 |
| 1 | 1 | 1 |
| 1 | 1 | 1 |
| 1 | 1 | 1 |
| 0 | 1 | 0 |
| 1 | 0 | 0 |
| 1 | 0 | 0 |
| 1 | 0 | 0 |

| | | |
|---|---|---|
| 1 | 1 | 1 |
| 1 | 1 | 1 |
| 0 | 0 | 0 |
| 1 | 1 | 1 |
| 1 | 1 | 1 |
| 1 | 1 | 1 |
| 1 | 1 | 1 |
| 0 | 1 | 0 |
| 1 | 1 | 1 |
| 1 | 1 | 1 |
| 1 | 1 | 1 |
| 0 | 1 | 0 |
| 0 | 1 | 0 |
| 0 | 1 | 0 |
| 0 | 1 | 0 |
| 0 | 1 | 0 |
| 0 | 1 | 0 |
| 0 | 1 | 0 |
| 0 | 1 | 0 |
| 0 | 1 | 0 |
| 0 | 1 | 0 |
| 0 | 1 | 0 |
| 0 | 1 | 0 |
| 0 | 1 | 0 |
| 0 | 1 | 0 |
| 0 | 1 | 0 |
| 0 | 1 | 0 |
| 0 | 1 | 0 |
| 1 | 1 | 1 |
| 1 | 1 | 1 |
| 0 | 0 | 0 |
| 1 | 1 | 1 |
| 0 | 0 | 0 |
| 0 | 0 | 0 |
| 0 | 0 | 0 |
| 0 | 0 | 0 |
| 1 | 1 | 1 |
| 0 | 0 | 0 |
| 1 | 1 | 1 |
| 1 | 1 | 1 |
| 0 | 0 | 0 |
| 0 | 0 | 0 |
| 0 | 0 | 0 |
| 1 | 1 | 1 |
| 0 | 0 | 0 |
| 0 | 0 | 0 |
| 1 | 1 | 1 |
| 1 | 1 | 1 |
| 1 | 1 | 1 |
| 1 | 1 | 1 |
| 0 | 0 | 0 |
| 1 | 1 | 1 |
| 1 | 1 | 1 |
| 1 | 1 | 1 |

| | | |
|---|---|---|
| 1 | 1 | 1 |
| 1 | 1 | 1 |
| 1 | 1 | 1 |
| 1 | 1 | 1 |
| 1 | 1 | 1 |
| 0 | 0 | 0 |
| 1 | 1 | 1 |
| 0 | 0 | 0 |
| 0 | 0 | 0 |
| 0 | 0 | 0 |
| 1 | 1 | 1 |
| 0 | 0 | 0 |
| 1 | 1 | 1 |
| 1 | 1 | 1 |
| 1 | 1 | 1 |
| 1 | 0 | 0 |
| 1 | 1 | 1 |
| 0 | 0 | 0 |
| 1 | 1 | 1 |
| 0 | 0 | 0 |
| 1 | 1 | 1 |
| 0 | 0 | 0 |
| 0 | 0 | 0 |
| 0 | 0 | 0 |
| 1 | 1 | 1 |
| 0 | 1 | 0 |
| 1 | 1 | 1 |
| 1 | 1 | 1 |
| 0 | 0 | 0 |
| 1 | 1 | 1 |
| 1 | 1 | 1 |
| 1 | 1 | 1 |
| 1 | 1 | 1 |
| 1 | 1 | 1 |
| 0 | 0 | 0 |
| 0 | 0 | 0 |
| 1 | 1 | 1 |
| 0 | 0 | 0 |
| 0 | 0 | 0 |
| 0 | 1 | 0 |
| 0 | 0 | 0 |
| 0 | 0 | 0 |
| 0 | 1 | 0 |
| 0 | 1 | 0 |
| 1 | 1 | 1 |
| 0 | 1 | 0 |
| 1 | 1 | 1 |
| 1 | 1 | 1 |
| 1 | 1 | 1 |
| 1 | 1 | 1 |
| 1 | 1 | 1 |
| 1 | 1 | 1 |
| 1 | 1 | 1 |
| 1 | 1 | 1 |

| | | |
|---|---|---|
| 1 | 1 | 1 |
| 0 | 0 | 0 |
| 0 | 1 | 0 |
| 1 | 1 | 1 |
| 0 | 0 | 0 |
| 1 | 1 | 1 |
| 1 | 1 | 1 |
| 1 | 1 | 1 |
| 0 | 0 | 0 |
| 0 | 0 | 0 |
| 1 | 1 | 1 |
| 0 | 0 | 0 |
| 0 | 0 | 0 |
| 1 | 1 | 1 |
| 1 | 1 | 1 |
| 0 | 0 | 0 |
| 0 | 1 | 0 |
| 0 | 0 | 0 |
| 0 | 0 | 0 |
| 1 | 1 | 1 |
| 0 | 0 | 0 |
| 0 | 0 | 0 |
| 0 | 0 | 0 |
| 1 | 1 | 1 |
| 0 | 1 | 0 |
| 0 | 0 | 0 |
| 1 | 1 | 1 |
| 1 | 1 | 1 |
| 1 | 1 | 1 |
| 1 | 1 | 1 |
| 0 | 0 | 0 |
| 1 | 1 | 1 |
| 0 | 0 | 0 |
| 1 | 1 | 1 |
| 0 | 0 | 0 |
| 0 | 0 | 0 |
| 1 | 1 | 1 |
| 1 | 1 | 1 |
| 1 | 1 | 1 |
| 1 | 1 | 1 |
| 1 | 1 | 1 |
| 1 | 1 | 1 |
| 1 | 1 | 1 |
| 1 | 1 | 1 |
| 1 | 1 | 1 |
| 1 | 1 | 1 |
| 1 | 0 | 0 |
| 1 | 1 | 1 |
| 1 | 1 | 1 |
| 1 | 1 | 1 |
| 1 | 1 | 1 |
| 1 | 1 | 1 |
| 1 | 1 | 1 |
| 0 | 0 | 0 |

| | | |
|---|---|---|
| 1 | 1 | 1 |
| 1 | 1 | 1 |
| 1 | 1 | 1 |
| 1 | 1 | 1 |
| 0 | 0 | 0 |
| 1 | 1 | 1 |
| 0 | 1 | 0 |
| 0 | 1 | 0 |
| 0 | 1 | 0 |
| 1 | 1 | 1 |
| 0 | 0 | 0 |
| 1 | 1 | 1 |
| 1 | 1 | 1 |
| 1 | 1 | 1 |
| 0 | 0 | 0 |
| 0 | 0 | 0 |
| 0 | 0 | 0 |
| 0 | 0 | 0 |
| 0 | 1 | 0 |
| 0 | 0 | 0 |
| 0 | 0 | 0 |
| 0 | 0 | 0 |
| 1 | 1 | 1 |
| 0 | 0 | 0 |
| 0 | 0 | 0 |
| 1 | 1 | 1 |
| 0 | 0 | 0 |
| 0 | 0 | 0 |
| 0 | 0 | 0 |
| 0 | 0 | 0 |
| 0 | 0 | 0 |
| 1 | 1 | 1 |
| 1 | 1 | 1 |
| 0 | 1 | 0 |
| 1 | 1 | 1 |
| 1 | 1 | 1 |
| 1 | 1 | 1 |
| 0 | 0 | 0 |
| 0 | 0 | 0 |
| 0 | 1 | 0 |
| 1 | 1 | 1 |
| 0 | 0 | 0 |
| 0 | 0 | 0 |
| 0 | 1 | 0 |
| 0 | 0 | 0 |
| 1 | 1 | 1 |
| 1 | 1 | 1 |
| 1 | 1 | 1 |
| 0 | 0 | 0 |
| 0 | 0 | 0 |
| 0 | 0 | 0 |
| 1 | 1 | 1 |
| 0 | 1 | 0 |
| 0 | 0 | 0 |

| | | |
|---|---|---|
| 1 | 1 | 1 |
| 0 | 0 | 0 |
| 1 | 1 | 1 |
| 0 | 0 | 0 |
| 0 | 0 | 0 |
| 0 | 1 | 0 |
| 0 | 0 | 0 |
| 0 | 0 | 0 |
| 1 | 1 | 1 |
| 1 | 1 | 1 |
| 1 | 1 | 1 |
| 1 | 1 | 1 |
| 1 | 1 | 1 |
| 1 | 1 | 1 |
| 1 | 1 | 1 |
| 1 | 1 | 1 |
| 1 | 1 | 1 |
| 0 | 1 | 0 |
| 1 | 1 | 1 |
| 1 | 1 | 1 |
| 0 | 0 | 0 |
| 0 | 1 | 0 |
| 0 | 1 | 0 |
| 0 | 1 | 0 |
| 0 | 1 | 0 |
| 0 | 0 | 0 |
| 0 | 0 | 0 |
| 1 | 1 | 1 |
| 1 | 1 | 1 |
| 1 | 1 | 1 |
| 1 | 1 | 1 |
| 1 | 1 | 1 |
| 1 | 1 | 1 |
| 0 | 1 | 0 |
| 1 | 1 | 1 |
| 1 | 1 | 1 |
| 1 | 1 | 1 |
| 1 | 1 | 1 |
| 0 | 1 | 0 |
| 1 | 1 | 1 |
| 1 | 1 | 1 |
| 0 | 1 | 0 |
| 1 | 1 | 1 |
| 0 | 0 | 0 |
| 1 | 1 | 1 |
| 0 | 0 | 0 |
| 0 | 0 | 0 |
| 0 | 0 | 0 |
| 0 | 0 | 0 |
| 0 | 0 | 0 |
| 0 | 0 | 0 |
| 0 | 1 | 0 |
| 1 | 1 | 1 |
| 1 | 1 | 1 |

| | | |
|---|---|---|
| 1 | 1 | 1 |
| 1 | 1 | 1 |
| 1 | 1 | 1 |
| 0 | 0 | 0 |
| 1 | 1 | 1 |
| 0 | 0 | 0 |
| 1 | 1 | 1 |
| 1 | 1 | 1 |
| 0 | 0 | 0 |
| 1 | 1 | 1 |
| 1 | 1 | 1 |
| 0 | 1 | 0 |
| 1 | 1 | 1 |
| 1 | 1 | 1 |
| 1 | 1 | 1 |
| 0 | 0 | 0 |
| 0 | 0 | 0 |
| 0 | 0 | 0 |
| 0 | 0 | 0 |
| 0 | 0 | 0 |
| 1 | 1 | 1 |
| 1 | 1 | 1 |
| 1 | 1 | 1 |
| 1 | 1 | 1 |
| 0 | 0 | 0 |
| 1 | 1 | 1 |
| 1 | 1 | 1 |
| 1 | 1 | 1 |
| 0 | 0 | 0 |
| 1 | 1 | 1 |
| 0 | 0 | 0 |
| 0 | 0 | 0 |
| 0 | 0 | 0 |
| 0 | 0 | 0 |
| 0 | 0 | 0 |
| 1 | 1 | 1 |
| 0 | 0 | 0 |
| 0 | 0 | 0 |
| 0 | 0 | 0 |
| 0 | 0 | 0 |
| 0 | 0 | 0 |
| 0 | 0 | 0 |
| 0 | 0 | 0 |
| 0 | 0 | 0 |
| 0 | 0 | 0 |
| 0 | 0 | 0 |
| 0 | 0 | 0 |
| 0 | 0 | 0 |
| 1 | 1 | 1 |
| 0 | 0 | 0 |
| 1 | 1 | 1 |
| 1 | 1 | 1 |
| 0 | 1 | 0 |
| 0 | 0 | 0 |

| | | |
|---|---|---|
| 0 | 0 | 0 |
| 0 | 1 | 0 |
| 1 | 1 | 1 |
| 1 | 1 | 1 |
| 1 | 1 | 1 |
| 0 | 0 | 0 |
| 1 | 1 | 1 |
| 0 | 1 | 0 |
| 1 | 1 | 1 |
| 0 | 1 | 0 |
| 0 | 1 | 0 |
| 0 | 1 | 0 |
| 1 | 1 | 1 |
| 0 | 0 | 0 |
| 0 | 1 | 0 |
| 1 | 1 | 1 |
| 1 | 1 | 1 |
| 0 | 0 | 0 |
| 1 | 1 | 1 |
| 1 | 1 | 1 |
| 1 | 1 | 1 |
| 1 | 1 | 1 |
| 0 | 0 | 0 |
| 1 | 1 | 1 |
| 1 | 1 | 1 |
| 0 | 0 | 0 |
| 0 | 0 | 0 |
| 0 | 0 | 0 |
| 1 | 1 | 1 |
| 0 | 1 | 0 |
| 1 | 1 | 1 |
| 1 | 1 | 1 |
| 1 | 1 | 1 |
| 1 | 1 | 1 |
| 1 | 1 | 1 |
| 1 | 1 | 1 |
| 1 | 1 | 1 |
| 1 | 1 | 1 |
| 1 | 1 | 1 |
| 1 | 1 | 1 |
| 1 | 1 | 1 |
| 1 | 1 | 1 |
| 1 | 1 | 1 |
| 1 | 1 | 1 |
| 1 | 1 | 1 |
| 1 | 1 | 1 |
| 1 | 1 | 1 |
| 1 | 1 | 1 |
| 1 | 0 | 0 |
| 1 | 1 | 1 |
| 1 | 1 | 1 |
| 1 | 1 | 1 |
| 1 | 1 | 1 |
| 1 | 1 | 1 |

| | | |
|---|---|---|
| 1 | 1 | 1 |
| 0 | 0 | 0 |
| 1 | 1 | 1 |
| 1 | 1 | 1 |
| 1 | 1 | 1 |
| 0 | 0 | 0 |
| 0 | 0 | 0 |
| 0 | 0 | 0 |
| 1 | 1 | 1 |
| 1 | 1 | 1 |
| 0 | 1 | 0 |
| 1 | 1 | 1 |
| 1 | 1 | 1 |
| 1 | 1 | 1 |
| 0 | 0 | 0 |
| 0 | 0 | 0 |
| 0 | 0 | 0 |
| 1 | 1 | 1 |
| 0 | 0 | 0 |
| 0 | 0 | 0 |
| 0 | 0 | 0 |
| 0 | 0 | 0 |
| 0 | 0 | 0 |
| 0 | 0 | 0 |
| 0 | 1 | 0 |
| 1 | 1 | 1 |
| 1 | 1 | 1 |
| 0 | 1 | 0 |
| 0 | 1 | 0 |
| 1 | 1 | 1 |
| 1 | 1 | 1 |
| 1 | 1 | 1 |
| 0 | 0 | 0 |
| 1 | 1 | 1 |
| 0 | 1 | 0 |
| 0 | 0 | 0 |
| 0 | 0 | 0 |
| 0 | 0 | 0 |
| 0 | 0 | 0 |
| 1 | 1 | 1 |
| 0 | 1 | 0 |
| 1 | 1 | 1 |
| 1 | 1 | 1 |
| 0 | 0 | 0 |
| 0 | 0 | 0 |
| 0 | 0 | 0 |
| 0 | 1 | 0 |
| 1 | 1 | 1 |
| 0 | 0 | 0 |
| 1 | 1 | 1 |
| 1 | 1 | 1 |
| 0 | 0 | 0 |
| 1 | 1 | 1 |
| 0 | 1 | 0 |

| | | |
|---|---|---|
| 1 | 1 | 1 |
| 0 | 1 | 0 |
| 1 | 1 | 1 |
| 1 | 1 | 1 |
| 1 | 1 | 1 |
| 1 | 1 | 1 |
| 0 | 1 | 0 |
| 1 | 1 | 1 |
| 0 | 0 | 0 |
| 0 | 0 | 0 |
| 1 | 1 | 1 |
| 0 | 0 | 0 |
| 0 | 0 | 0 |
| 0 | 0 | 0 |
| 1 | 1 | 1 |
| 1 | 1 | 1 |
| 0 | 0 | 0 |
| 0 | 0 | 0 |
| 0 | 0 | 0 |
| 1 | 1 | 1 |
| 1 | 1 | 1 |
| 1 | 1 | 1 |
| 0 | 0 | 0 |
| 1 | 1 | 1 |
| 0 | 0 | 0 |
| 1 | 1 | 1 |
| 1 | 1 | 1 |
| 1 | 1 | 1 |
| 1 | 1 | 1 |
| 0 | 0 | 0 |
| 0 | 0 | 0 |
| 0 | 0 | 0 |
| 1 | 1 | 1 |
| 0 | 1 | 0 |
| 1 | 1 | 1 |
| 1 | 1 | 1 |
| 1 | 1 | 1 |
| 0 | 1 | 0 |
| 0 | 1 | 0 |
| 0 | 1 | 0 |
| 0 | 0 | 0 |
| 1 | 1 | 1 |
| 1 | 1 | 1 |
| 1 | 1 | 1 |
| 1 | 1 | 1 |
| 0 | 0 | 0 |
| 1 | 0 | 0 |
| 0 | 0 | 0 |
| 1 | 1 | 1 |
| 0 | 0 | 0 |
| 0 | 0 | 0 |
| 0 | 0 | 0 |
| 0 | 1 | 0 |
| 1 | 0 | 0 |

| | | |
|---|---|---|
| 1 | 0 | 0 |
| 0 | 0 | 0 |
| 0 | 1 | 0 |
| 1 | 1 | 1 |
| 0 | 1 | 0 |
| 0 | 1 | 0 |
| 0 | 1 | 0 |
| 0 | 0 | 0 |
| 0 | 0 | 0 |
| 0 | 1 | 0 |
| 0 | 0 | 0 |
| 1 | 1 | 1 |
| 1 | 1 | 1 |
| 1 | 1 | 1 |
| 0 | 1 | 0 |
| 1 | 1 | 1 |
| 1 | 1 | 1 |
| 1 | 1 | 1 |
| 1 | 1 | 1 |
| 0 | 1 | 0 |
| 1 | 1 | 1 |
| 1 | 1 | 1 |
| 1 | 1 | 1 |
| 1 | 1 | 1 |
| 0 | 1 | 0 |
| 0 | 1 | 0 |
| 0 | 1 | 0 |
| 0 | 1 | 0 |
| 0 | 0 | 0 |
| 0 | 1 | 0 |
| 0 | 0 | 0 |
| 1 | 1 | 1 |
| 0 | 0 | 0 |
| 0 | 0 | 0 |
| 0 | 1 | 0 |
| 0 | 0 | 0 |
| 0 | 0 | 0 |
| 1 | 1 | 1 |
| 0 | 1 | 0 |
| 0 | 1 | 0 |
| 1 | 1 | 1 |
| 0 | 0 | 0 |
| 0 | 0 | 0 |
| 1 | 1 | 1 |
| 1 | 1 | 1 |
| 0 | 0 | 0 |
| 0 | 0 | 0 |
| 0 | 0 | 0 |
| 0 | 0 | 0 |
| 0 | 1 | 0 |
| 0 | 0 | 0 |
| 0 | 0 | 0 |
| 1 | 1 | 1 |
| 1 | 1 | 1 |

| | | |
|---|---|---|
| 1 | 1 | 1 |
| 1 | 1 | 1 |
| 1 | 1 | 1 |
| 1 | 1 | 1 |
| 0 | 0 | 0 |
| 0 | 0 | 0 |
| 0 | 0 | 0 |
| 0 | 0 | 0 |
| 1 | 1 | 1 |
| 0 | 0 | 0 |
| 0 | 0 | 0 |
| 0 | 0 | 0 |
| 0 | 0 | 0 |
| 1 | 1 | 1 |
| 1 | 0 | 0 |
| 0 | 1 | 0 |
| 1 | 1 | 1 |
| 1 | 1 | 1 |
| 1 | 1 | 1 |
| 0 | 1 | 0 |
| 1 | 1 | 1 |
| 1 | 1 | 1 |
| 1 | 1 | 1 |
| 1 | 1 | 1 |
| 1 | 1 | 1 |
| 0 | 0 | 0 |
| 0 | 0 | 0 |
| 0 | 0 | 0 |
| 0 | 0 | 0 |
| 0 | 1 | 0 |
| 0 | 1 | 0 |
| 0 | 1 | 0 |
| 1 | 1 | 1 |
| 0 | 1 | 0 |
| 0 | 1 | 0 |
| 1 | 1 | 1 |
| 1 | 1 | 1 |
| 1 | 1 | 1 |
| 1 | 1 | 1 |
| 1 | 1 | 1 |
| 1 | 1 | 1 |
| 1 | 1 | 1 |
| 1 | 1 | 1 |
| 1 | 1 | 1 |
| 1 | 1 | 1 |
| 1 | 1 | 1 |
| 1 | 1 | 1 |
| 1 | 1 | 1 |
| 1 | 1 | 1 |
| 1 | 1 | 1 |
| 1 | 1 | 1 |
| 1 | 1 | 1 |
| 1 | 1 | 1 |
| 0 | 0 | 0 |

| | | |
|---|---|---|
| 1 | 1 | 1 |
| 1 | 1 | 1 |
| 1 | 1 | 1 |
| 0 | 1 | 0 |
| 0 | 1 | 0 |
| 0 | 1 | 0 |
| 0 | 1 | 0 |
| 0 | 1 | 0 |
| 1 | 1 | 1 |
| 1 | 1 | 1 |
| 1 | 1 | 1 |
| 1 | 1 | 1 |
| 1 | 1 | 1 |
| 1 | 1 | 1 |
| 1 | 1 | 1 |
| 1 | 1 | 1 |
| 1 | 1 | 1 |
| 1 | 1 | 1 |
| 1 | 1 | 1 |
| 1 | 1 | 1 |
| 1 | 1 | 1 |
| 1 | 1 | 1 |
| 1 | 1 | 1 |
| 1 | 1 | 1 |
| 1 | 1 | 1 |
| 1 | 1 | 1 |
| 1 | 1 | 1 |
| 1 | 1 | 1 |
| 1 | 1 | 1 |
| 1 | 1 | 1 |
| 1 | 1 | 1 |
| 1 | 1 | 1 |
| 1 | 1 | 1 |
| 0 | 1 | 0 |
| 1 | 1 | 1 |
| 1 | 1 | 1 |
| 1 | 1 | 1 |
| 1 | 1 | 1 |
| 1 | 1 | 1 |
| 1 | 1 | 1 |
| 0 | 1 | 0 |
| 1 | 1 | 1 |
| 1 | 1 | 1 |
| 1 | 1 | 1 |
| 0 | 1 | 0 |
| 1 | 0 | 0 |
| 0 | 0 | 0 |
| 1 | 1 | 1 |
| 0 | 1 | 0 |
| 1 | 1 | 1 |
| 1 | 1 | 1 |
| 0 | 1 | 0 |
| 0 | 1 | 0 |
| 0 | 1 | 0 |

| | | |
|---|---|---|
| 0 | 1 | 0 |
| 1 | 1 | 1 |
| 0 | 0 | 0 |
| 1 | 1 | 1 |
| 1 | 1 | 1 |
| 1 | 1 | 1 |
| 0 | 0 | 0 |
| 1 | 1 | 1 |
| 1 | 1 | 1 |
| 0 | 1 | 0 |
| 0 | 1 | 0 |
| 0 | 0 | 0 |
| 1 | 1 | 1 |
| 0 | 0 | 0 |
| 0 | 0 | 0 |
| 0 | 0 | 0 |
| 0 | 0 | 0 |
| 0 | 0 | 0 |
| 0 | 0 | 0 |
| 0 | 0 | 0 |
| 0 | 0 | 0 |

| organism_scientific | evo_cons | cluster_id | pdb_id | chain_id | resi | resn | change | uniprot_id |
|---|---|---|---|---|---|---|---|---|
| HOMO SAPIENS | 5 | 87 | 3uzw | A | 106 | LEU | p.Leu106Phe | P51857 |
| HOMO SAPIENS | 3 | 87 | 3uzw | A | 133 | PRO | p.Pro133Arg | P51857 |
| HOMO SAPIENS | 0 | 87 | 3uzw | A | 198 | PRO | p.Pro198Leu | P51857 |
| HOMO SAPIENS | 7 | 87 | 3uzw | A | 261 | ARG | p.Arg261Cys | P51857 |
| HOMO SAPIENS | 0 | 4693 | 3mk1 | A | 3 | PRO | p.Pro25Leu | P05187 |
| HOMO SAPIENS | 8 | 4830 | 2h64 | B | 101 | CYS | p.Cys124Arg | P36894 |
| HOMO SAPIENS | 2 | 4830 | 2h64 | B | 116 | PRO | p.Pro139Ser | P36894 |
| HOMO SAPIENS | 0 | 6937 | 3dar | A | 186 | MET | p.Met186Thr | P21802 |
| HOMO SAPIENS | 3 | 5 | 1aut | C | 30 | GLN | p.Gln226His | P04070 |
| HOMO SAPIENS | 1 | 5 | 1aut | C | 68 | LEU | p.Leu265Phe | P04070 |
| HOMO SAPIENS | 0 | 5 | 1aut | C | 75 | ARG | p.Arg272Cys | P04070 |
| HOMO SAPIENS | 6 | 5 | 1aut | C | 92 | PRO | p.Pro289Leu | P04070 |
| HOMO SAPIENS | 0 | 5 | 1aut | C | 104 | ALA | p.Ala301Val | P04070 |
| HOMO SAPIENS | 1 | 5 | 1aut | C | 112 | ALA | p.Ala309Thr | P04070 |
| HOMO SAPIENS | 2 | 5 | 1aut | C | 115 | SER | p.Ser312Leu | P04070 |
| HOMO SAPIENS | 1 | 5 | 1aut | C | 133 | GLY | p.Gly334Ser | P04070 |
| HOMO SAPIENS | 0 | 5 | 1aut | C | 147 | ARG | p.Arg348Ter | P04070 |
| HOMO SAPIENS | 8 | 5 | 1aut | C | 194 | ASP | p.Asp401Asn | P04070 |
| HOMO SAPIENS | 7 | 5 | 1aut | C | 237 | TRP | p.Trp444Cys | P04070 |
| HOMO SAPIENS | 0 | 5 | 1aut | C | 238 | ILE | p.Ile445Met | P04070 |
| HOMO SAPIENS | 8 | 43 | 1aiy | F | 10 | HIS | p.His34Asp | P01308 |
| HOMO SAPIENS | 2 | 43 | 1aiy | F | 22 | ARG | p.Arg46Gln | P01308 |
| HOMO SAPIENS | 4 | 43 | 1aiy | F | 23 | GLY | p.Gly47Val | P01308 |
| HOMO SAPIENS | 7 | 43 | 1aiy | F | 24 | PHE | p.Phe48Ser | P01308 |
| HOMO SAPIENS | 5 | 24 | 2fg8 | A | 33 | THR | p.Thr30Ile | P02792 |
| HOMO SAPIENS | 4 | 24 | 2fg8 | A | 99 | ALA | p.Ala96Thr | P02792 |
| HOMO SAPIENS | 5 | 24 | 2fg8 | A | 107 | GLU | p.Glu104Ter | P02792 |
| HOMO SAPIENS | 3 | 5409 | 5bnq | A | 199 | MET | p.Met199Lys | O14788 |
| HOMO SAPIENS | -1 | 839 | 3swc | A | 959 | GLN | p.Gln990His | Q9H9B1 |
| HOMO SAPIENS | -1 | 839 | 3swc | A | 968 | ALA | p.Ala999Val | Q9H9B1 |
| HOMO SAPIENS | -1 | 839 | 3swc | A | 973 | SER | p.Ser1004Ile | Q9H9B1 |
| HOMO SAPIENS | 8 | 17 | 1eou | A | 18 | LYS | p.Lys18Glu | P00918 |
| HOMO SAPIENS | 2 | 17 | 1eou | A | 107 | HIS | p.His107Tyr | P00918 |
| HOMO SAPIENS | 1 | 17 | 1eou | A | 236 | PRO | p.Pro236His | P00918 |
| HOMO SAPIENS | 6 | 17 | 1eou | A | 252 | ASN | p.Asn252Asp | P00918 |
| HOMO SAPIENS | 2 | 9140 | 5c4v | A | 445 | ARG | p.Arg445Ter | Q13485 |
| HOMO SAPIENS | 7 | 9140 | 5c4v | A | 496 | ARG | p.Arg496Cys | Q13485 |
| HOMO SAPIENS | -1 | 42 | 4pwi | A | 6 | GLY | p.Gly26Ser | P02766 |
| HOMO SAPIENS | 6 | 42 | 4pwi | A | 10 | CYS | p.Cys30Arg | P02766 |
| HOMO SAPIENS | 8 | 42 | 4pwi | A | 12 | LEU | p.Leu32Pro | P02766 |
| HOMO SAPIENS | 9 | 42 | 4pwi | A | 18 | ASP | p.Asp38Gly | P02766 |
| HOMO SAPIENS | 8 | 42 | 4pwi | A | 20 | VAL | p.Val40Ile | P02766 |
| HOMO SAPIENS | 4 | 42 | 4pwi | A | 30 | MET | p.Val50Ala | P02766 |
| HOMO SAPIENS | 4 | 42 | 4pwi | A | 30 | MET | p.Val50Leu | P02766 |
| HOMO SAPIENS | 6 | 42 | 4pwi | A | 33 | PHE | p.Phe53Ile | P02766 |
| HOMO SAPIENS | 7 | 42 | 4pwi | A | 36 | ALA | p.Ala56Pro | P02766 |
| HOMO SAPIENS | 7 | 42 | 4pwi | A | 42 | GLU | p.Glu62Gly | P02766 |
| HOMO SAPIENS | 8 | 42 | 4pwi | A | 44 | PHE | p.Phe64Ser | P02766 |
| HOMO SAPIENS | 4 | 42 | 4pwi | A | 47 | GLY | p.Gly67Ala | P02766 |
| HOMO SAPIENS | 6 | 42 | 4pwi | A | 49 | THR | p.Thr69Ala | P02766 |
| HOMO SAPIENS | 6 | 42 | 4pwi | A | 50 | SER | p.Ser70Ile | P02766 |
| HOMO SAPIENS | 6 | 42 | 4pwi | A | 50 | SER | p.Ser70Arg | P02766 |
| HOMO SAPIENS | 0 | 42 | 4pwi | A | 53 | GLY | p.Gly73Glu | P02766 |

| | | | | | | | | |
|---|---|---|---|---|---|---|---|---|
| HOMO SAPIENS | 7 | 42 | 4pwi | A | 55 | LEU | p.Leu75Pro | P02766 |
| HOMO SAPIENS | 6 | 42 | 4pwi | A | 58 | LEU | p.Leu78His | P02766 |
| HOMO SAPIENS | 7 | 42 | 4pwi | A | 61 | GLU | p.Glu81Lys | P02766 |
| HOMO SAPIENS | 8 | 42 | 4pwi | A | 64 | PHE | p.Phe84Leu | P02766 |
| HOMO SAPIENS | 8 | 42 | 4pwi | A | 68 | ILE | p.Ile88Leu | P02766 |
| HOMO SAPIENS | 8 | 42 | 4pwi | A | 69 | TYR | p.Tyr89His | P02766 |
| HOMO SAPIENS | 4 | 42 | 4pwi | A | 71 | VAL | p.Val91Ala | P02766 |
| HOMO SAPIENS | 6 | 42 | 4pwi | A | 77 | SER | p.Ser97Tyr | P02766 |
| HOMO SAPIENS | 4 | 42 | 4pwi | A | 84 | ILE | p.Ile104Ser | P02766 |
| HOMO SAPIENS | 8 | 42 | 4pwi | A | 89 | GLU | p.Glu109Gln | P02766 |
| HOMO SAPIENS | 8 | 42 | 4pwi | A | 97 | ALA | p.Ala117Gly | P02766 |
| HOMO SAPIENS | 7 | 42 | 4pwi | A | 99 | ASP | p.Asp119Asn | P02766 |
| HOMO SAPIENS | 9 | 42 | 4pwi | A | 107 | ILE | p.Ile127Val | P02766 |
| HOMO SAPIENS | 9 | 42 | 4pwi | A | 111 | LEU | p.Leu131Met | P02766 |
| HOMO SAPIENS | 7 | 42 | 4pwi | A | 114 | TYR | p.Tyr134Cys | P02766 |
| HOMO SAPIENS | 7 | 42 | 4pwi | A | 114 | TYR | p.Tyr134His | P02766 |
| HOMO SAPIENS | 7 | 42 | 4pwi | A | 122 | VAL | p.Val142Ile | P02766 |
| HOMO SAPIENS | -1 | 42 | 4pwi | A | 126 | LYS | p.Lys146Arg | P02766 |
| HOMO SAPIENS | -1 | 42 | 4pwi | A | 127 | GLU | p.Glu147Ter | P02766 |
| HOMO SAPIENS | -1 | 1312 | 4dce | A | 1275 | ARG | p.Arg1275Gln | Q9UM73 |
| HOMO SAPIENS | -1 | 1312 | 4dce | A | 1280 | ALA | p.Ala1280Val | Q9UM73 |
| HOMO SAPIENS | 1 | 20 | 1y0x | X | 234 | ALA | p.Ala234Thr | P10828 |
| HOMO SAPIENS | 0 | 20 | 1y0x | X | 243 | ARG | p.Arg243Trp | P10828 |
| HOMO SAPIENS | 3 | 20 | 1y0x | X | 316 | ARG | p.Arg316His | P10828 |
| HOMO SAPIENS | 1 | 20 | 1y0x | X | 317 | ALA | p.Ala317Thr | P10828 |
| HOMO SAPIENS | 2 | 20 | 1y0x | X | 320 | ARG | p.Arg320Cys | P10828 |
| HOMO SAPIENS | 2 | 20 | 1y0x | X | 320 | ARG | p.Arg320His | P10828 |
| HOMO SAPIENS | 0 | 20 | 1y0x | X | 332 | GLY | p.Gly332Arg | P10828 |
| HOMO SAPIENS | 1 | 20 | 1y0x | X | 338 | ARG | p.Arg338Trp | P10828 |
| HOMO SAPIENS | 0 | 20 | 1y0x | X | 340 | GLN | p.Gln340His | P10828 |
| HOMO SAPIENS | 0 | 20 | 1y0x | X | 345 | GLY | p.Gly345Val | P10828 |
| HOMO SAPIENS | 0 | 20 | 1y0x | X | 345 | GLY | p.Gly345Ser | P10828 |
| HOMO SAPIENS | 0 | 20 | 1y0x | X | 347 | GLY | p.Gly347Glu | P10828 |
| HOMO SAPIENS | 1 | 20 | 1y0x | X | 438 | ARG | p.Arg438His | P10828 |
| HOMO SAPIENS | 0 | 20 | 1y0x | X | 442 | MET | p.Met442Val | P10828 |
| HOMO SAPIENS | 0 | 20 | 1y0x | X | 443 | LYS | p.Lys443Glu | P10828 |
| HOMO SAPIENS | 0 | 20 | 1y0x | X | 446 | CYS | p.Cys446Arg | P10828 |
| HOMO SAPIENS | 2 | 20 | 1y0x | X | 453 | PRO | p.Pro453His | P10828 |
| HOMO SAPIENS | 2 | 20 | 1y0x | X | 453 | PRO | p.Pro453Thr | P10828 |
| HOMO SAPIENS | 3 | 20 | 1y0x | X | 459 | PHE | p.Phe459Cys | P10828 |
| HOMO SAPIENS | 0 | 4409 | 3qb7 | D | 87 | GLU | p.Glu109Lys | P31785 |
| HOMO SAPIENS | 1 | 4409 | 3qb7 | D | 92 | GLY | p.Gly114Asp | P31785 |
| HOMO SAPIENS | 1 | 4409 | 3qb7 | D | 93 | CYS | p.Cys115Arg | P31785 |
| HOMO SAPIENS | 0 | 4409 | 3qb7 | D | 131 | ILE | p.Ile153Asn | P31785 |
| HOMO SAPIENS | 9 | 4409 | 3qb7 | D | 200 | ARG | p.Arg222Cys | P31785 |
| HOMO SAPIENS | 0 | 3891 | 3ije | B | 33 | LEU | p.Leu59Pro | P05106 |
| HOMO SAPIENS | 0 | 3891 | 3ije | B | 117 | LEU | p.Leu143Trp | P05106 |
| HOMO SAPIENS | 3 | 3891 | 3ije | B | 119 | ASP | p.Asp145Tyr | P05106 |
| HOMO SAPIENS | 2 | 3891 | 3ije | B | 143 | ARG | p.Arg169Gln | P05106 |
| HOMO SAPIENS | 2 | 3891 | 3ije | B | 214 | ARG | p.Arg240Gln | P05106 |
| HOMO SAPIENS | 2 | 3891 | 3ije | B | 214 | ARG | p.Arg240Trp | P05106 |
| HOMO SAPIENS | 2 | 3891 | 3ije | B | 374 | CYS | p.Cys400Tyr | P05106 |
| HOMO SAPIENS | 1 | 3891 | 3ije | B | 407 | PRO | p.Pro433Ala | P05106 |
| HOMO SAPIENS | 0 | 3891 | 3ije | B | 489 | ARG | p.Arg515Gln | P05106 |

| | | | | | | | | |
|---|---|---|---|---|---|---|---|---|
| HOMO SAPIENS | -1 | 9613 | 5mke | B | 28 | ARG | p.Arg28Pro | Q13563 |
| HOMO SAPIENS | -1 | 9613 | 5mke | B | 190 | ALA | p.Ala190Thr | Q13563 |
| HOMO SAPIENS | 4 | 9613 | 5mke | B | 262 | VAL | p.Val262Met | Q13563 |
| HOMO SAPIENS | 0 | 9613 | 5mke | B | 482 | PHE | p.Phe482Cys | Q13563 |
| HOMO SAPIENS | 4 | 9613 | 5mke | B | 511 | ASP | p.Asp511Val | Q13563 |
| HOMO SAPIENS | 5 | 9613 | 5mke | B | 516 | VAL | p.Val516Leu | Q13563 |
| HOMO SAPIENS | -1 | 9613 | 5mke | B | 807 | ARG | p.Arg807Gln | Q13563 |
| HOMO SAPIENS | 4 | 2428 | 1olh | C | 337 | ARG | p.Arg337Cys | P04637 |
| HOMO SAPIENS | 4 | 2428 | 1olh | C | 337 | ARG | p.Arg337His | P04637 |
| HOMO SAPIENS | 8 | 2428 | 1olh | C | 338 | PHE | p.Phe338Leu | P04637 |
| HOMO SAPIENS | 8 | 2428 | 1olh | C | 339 | GLU | p.Glu339Lys | P04637 |
| HOMO SAPIENS | 7 | 2428 | 1olh | C | 344 | LEU | p.Leu344Pro | P04637 |
| HOMO SAPIENS | 0 | 2428 | 1olh | C | 360 | GLY | p.Gly360Ala | P04637 |
| HOMO SAPIENS | 7 | 4867 | 1mox | B | 404 | GLY | p.Gly428Asp | P00533 |
| HOMO SAPIENS | 5 | 4867 | 1mox | B | 497 | ARG | p.Arg521Lys | P00533 |
| HOMO SAPIENS | 1 | 4630 | 1fu1 | A | 43 | TRP | p.Trp43Arg | Q13426 |
| HOMO SAPIENS | 2 | 4630 | 1fu1 | A | 161 | ARG | p.Arg161Ter | Q13426 |
| HOMO SAPIENS | 2 | 15 | 1lfy | B | 2 | HIS | p.His3Gln | P68871 |
| HOMO SAPIENS | 2 | 15 | 1lfy | B | 6 | GLU | p.Glu7Ala | P68871 |
| HOMO SAPIENS | 2 | 15 | 1lfy | B | 6 | GLU | p.Glu7Lys | P68871 |
| HOMO SAPIENS | 4 | 15 | 1lfy | B | 9 | SER | p.Ser10Cys | P68871 |
| HOMO SAPIENS | 6 | 15 | 1lfy | B | 15 | TRP | p.Trp16Ter | P68871 |
| HOMO SAPIENS | 3 | 15 | 1lfy | B | 17 | LYS | p.Lys18Glu | P68871 |
| HOMO SAPIENS | 6 | 15 | 1lfy | B | 19 | ASN | p.Asn20Ser | P68871 |
| HOMO SAPIENS | 6 | 15 | 1lfy | B | 22 | GLU | p.Glu23Ala | P68871 |
| HOMO SAPIENS | 6 | 15 | 1lfy | B | 22 | GLU | p.Glu23Lys | P68871 |
| HOMO SAPIENS | 8 | 15 | 1lfy | B | 26 | GLU | p.Glu27Lys | P68871 |
| HOMO SAPIENS | 8 | 15 | 1lfy | B | 28 | LEU | p.Leu29Pro | P68871 |
| HOMO SAPIENS | 8 | 15 | 1lfy | B | 30 | ARG | p.Arg31Ser | P68871 |
| HOMO SAPIENS | 8 | 15 | 1lfy | B | 30 | ARG | p.Arg31Lys | P68871 |
| HOMO SAPIENS | 6 | 15 | 1lfy | B | 39 | GLN | p.Gln40Ter | P68871 |
| HOMO SAPIENS | 2 | 15 | 1lfy | B | 43 | GLU | p.Glu44Gln | P68871 |
| HOMO SAPIENS | 6 | 15 | 1lfy | B | 57 | ASN | p.Asn58Lys | P68871 |
| HOMO SAPIENS | 6 | 15 | 1lfy | B | 59 | LYS | p.Lys60Asn | P68871 |
| HOMO SAPIENS | 8 | 15 | 1lfy | B | 61 | LYS | p.Lys62Glu | P68871 |
| HOMO SAPIENS | 4 | 15 | 1lfy | B | 73 | ASP | p.Asp74Tyr | P68871 |
| HOMO SAPIENS | 4 | 15 | 1lfy | B | 87 | THR | p.Thr88Ile | P68871 |
| HOMO SAPIENS | 4 | 15 | 1lfy | B | 87 | THR | p.Thr88Pro | P68871 |
| HOMO SAPIENS | 5 | 15 | 1lfy | B | 90 | GLU | p.Glu91Lys | P68871 |
| HOMO SAPIENS | 6 | 15 | 1lfy | B | 91 | LEU | p.Leu92Pro | P68871 |
| HOMO SAPIENS | 5 | 15 | 1lfy | B | 97 | HIS | p.His98Gln | P68871 |
| HOMO SAPIENS | 6 | 15 | 1lfy | B | 110 | LEU | p.Leu111Pro | P68871 |
| HOMO SAPIENS | 6 | 15 | 1lfy | B | 113 | VAL | p.Val114Glu | P68871 |
| HOMO SAPIENS | 2 | 15 | 1lfy | B | 119 | GLY | p.Gly120Ala | P68871 |
| HOMO SAPIENS | 4 | 15 | 1lfy | B | 121 | GLU | p.Glu122Gln | P68871 |
| HOMO SAPIENS | 5 | 15 | 1lfy | B | 134 | VAL | p.Val135Glu | P68871 |
| HOMO SAPIENS | 3 | 15 | 1lfy | B | 143 | HIS | p.His144Gln | P68871 |
| HOMO SAPIENS | 4 | 91 | 5d9k | A | 75 | GLY | p.Gly75Val | P51812 |
| HOMO SAPIENS | 5 | 91 | 5d9k | A | 82 | VAL | p.Val82Phe | P51812 |
| HOMO SAPIENS | 0 | 91 | 5d9k | A | 114 | ARG | p.Arg114Trp | P51812 |
| HOMO SAPIENS | 4 | 91 | 5d9k | A | 189 | ILE | p.Ile189Lys | P51812 |
| HOMO SAPIENS | -1 | 91 | 5d9k | A | 227 | SER | p.Ser227Ala | P51812 |
| HOMO SAPIENS | 1 | 91 | 5d9k | A | 268 | PHE | p.Phe268Ser | P51812 |
| HOMO SAPIENS | 0 | 954 | 2vt0 | A | 15 | VAL | p.Val54Leu | P04062 |

| | | | | | | | | |
|---|---|---|---|---|---|---|---|---|
| HOMO SAPIENS | 5 | 954 | 2vt0 | A | 46 | GLY | p.Gly85Glu | P04062 |
| HOMO SAPIENS | 5 | 954 | 2vt0 | A | 48 | ARG | p.Arg87Trp | P04062 |
| HOMO SAPIENS | 6 | 954 | 2vt0 | A | 79 | LYS | p.Lys118Asn | P04062 |
| HOMO SAPIENS | 0 | 954 | 2vt0 | A | 120 | ARG | p.Arg159Gln | P04062 |
| HOMO SAPIENS | 0 | 954 | 2vt0 | A | 120 | ARG | p.Arg159Trp | P04062 |
| HOMO SAPIENS | 0 | 954 | 2vt0 | A | 122 | PRO | p.Pro161Ser | P04062 |
| HOMO SAPIENS | 5 | 954 | 2vt0 | A | 131 | ARG | p.Arg170Leu | P04062 |
| HOMO SAPIENS | 5 | 954 | 2vt0 | A | 135 | TYR | p.Tyr174Cys | P04062 |
| HOMO SAPIENS | 0 | 954 | 2vt0 | A | 157 | LYS | p.Lys196Gln | P04062 |
| HOMO SAPIENS | 5 | 954 | 2vt0 | A | 163 | ARG | p.Arg202Gln | P04062 |
| HOMO SAPIENS | 5 | 954 | 2vt0 | A | 170 | ARG | p.Arg209Cys | P04062 |
| HOMO SAPIENS | 5 | 954 | 2vt0 | A | 188 | ASN | p.Asn227Ser | P04062 |
| HOMO SAPIENS | 3 | 954 | 2vt0 | A | 196 | SER | p.Ser235Pro | P04062 |
| HOMO SAPIENS | 5 | 954 | 2vt0 | A | 202 | GLY | p.Gly241Arg | P04062 |
| HOMO SAPIENS | 0 | 954 | 2vt0 | A | 212 | TYR | p.Tyr251His | P04062 |
| HOMO SAPIENS | 0 | 954 | 2vt0 | A | 213 | PHE | p.Phe252Ile | P04062 |
| HOMO SAPIENS | 5 | 954 | 2vt0 | A | 216 | PHE | p.Phe255Tyr | P04062 |
| HOMO SAPIENS | 6 | 954 | 2vt0 | A | 251 | PHE | p.Phe290Leu | P04062 |
| HOMO SAPIENS | 5 | 954 | 2vt0 | A | 255 | HIS | p.His294Gln | P04062 |
| HOMO SAPIENS | 6 | 954 | 2vt0 | A | 257 | ARG | p.Arg296Gln | P04062 |
| HOMO SAPIENS | 5 | 954 | 2vt0 | A | 289 | PRO | p.Pro328Leu | P04062 |
| HOMO SAPIENS | 0 | 954 | 2vt0 | A | 309 | ALA | p.Ala348Val | P04062 |
| HOMO SAPIENS | 0 | 954 | 2vt0 | A | 311 | HIS | p.His350Arg | P04062 |
| HOMO SAPIENS | 6 | 954 | 2vt0 | A | 312 | TRP | p.Trp351Cys | P04062 |
| HOMO SAPIENS | 5 | 954 | 2vt0 | A | 323 | THR | p.Thr362Ile | P04062 |
| HOMO SAPIENS | 5 | 954 | 2vt0 | A | 325 | GLY | p.Gly364Arg | P04062 |
| HOMO SAPIENS | 6 | 954 | 2vt0 | A | 342 | CYS | p.Cys381Gly | P04062 |
| HOMO SAPIENS | 5 | 954 | 2vt0 | A | 353 | ARG | p.Arg392Gly | P04062 |
| HOMO SAPIENS | 3 | 954 | 2vt0 | A | 364 | SER | p.Ser403Thr | P04062 |
| HOMO SAPIENS | 5 | 954 | 2vt0 | A | 369 | THR | p.Thr408Met | P04062 |
| HOMO SAPIENS | 3 | 954 | 2vt0 | A | 370 | ASN | p.Asn409Ser | P04062 |
| HOMO SAPIENS | 0 | 954 | 2vt0 | A | 371 | LEU | p.Leu410Val | P04062 |
| HOMO SAPIENS | 0 | 954 | 2vt0 | A | 375 | VAL | p.Val414Leu | P04062 |
| HOMO SAPIENS | 0 | 954 | 2vt0 | A | 377 | GLY | p.Gly416Ser | P04062 |
| HOMO SAPIENS | 8 | 954 | 2vt0 | A | 398 | VAL | p.Val437Phe | P04062 |
| HOMO SAPIENS | 0 | 954 | 2vt0 | A | 401 | PRO | p.Pro440Leu | P04062 |
| HOMO SAPIENS | 3 | 954 | 2vt0 | A | 409 | ASP | p.Asp448His | P04062 |
| HOMO SAPIENS | 3 | 954 | 2vt0 | A | 409 | ASP | p.Asp448Val | P04062 |
| HOMO SAPIENS | 0 | 954 | 2vt0 | A | 415 | PRO | p.Pro454Arg | P04062 |
| HOMO SAPIENS | 5 | 954 | 2vt0 | A | 444 | LEU | p.Leu483Arg | P04062 |
| HOMO SAPIENS | 5 | 954 | 2vt0 | A | 456 | ALA | p.Ala495Pro | P04062 |
| HOMO SAPIENS | 5 | 954 | 2vt0 | A | 463 | ARG | p.Arg502Cys | P04062 |
| HOMO SAPIENS | 5 | 954 | 2vt0 | A | 463 | ARG | p.Arg502His | P04062 |
| HOMO SAPIENS | 5 | 954 | 2vt0 | A | 478 | GLY | p.Gly517Ser | P04062 |
| HOMO SAPIENS | 5 | 954 | 2vt0 | A | 496 | ARG | p.Arg535Cys | P04062 |
| HOMO SAPIENS | 5 | 954 | 2vt0 | A | 496 | ARG | p.Arg535His | P04062 |
| HOMO SAPIENS | 1 | 9037 | 2rqb | A | 946 | ALA | p.Ala946Thr | Q9BYX4 |
| HOMO SAPIENS | 0 | 253 | 5E+092 | A | 308 | LEU | p.Leu308Pro | P37173 |
| HOMO SAPIENS | 0 | 253 | 5E+092 | A | 315 | THR | p.Thr315Met | P37173 |
| HOMO SAPIENS | 0 | 253 | 5E+092 | A | 328 | HIS | p.His328Gln | P37173 |
| HOMO SAPIENS | 0 | 253 | 5E+092 | A | 336 | TYR | p.Tyr336Asn | P37173 |
| HOMO SAPIENS | 0 | 253 | 5E+092 | A | 348 | ARG | p.Arg348Cys | P37173 |
| HOMO SAPIENS | 0 | 253 | 5E+092 | A | 355 | ALA | p.Ala355Pro | P37173 |
| HOMO SAPIENS | 1 | 253 | 5E+092 | A | 356 | ARG | p.Arg356Gly | P37173 |

| | | | | | | | | |
|---|---|---|---|---|---|---|---|---|
| HOMO SAPIENS | 0 | 253 | 5E+092 | A | 357 | GLY | p.Gly357Trp | P37173 |
| HOMO SAPIENS | 1 | 253 | 5E+092 | A | 403 | ARG | p.Arg403His | P37173 |
| HOMO SAPIENS | 5 | 253 | 5E+092 | A | 449 | SER | p.Ser449Phe | P37173 |
| HOMO SAPIENS | 0 | 253 | 5E+092 | A | 460 | ARG | p.Arg460Cys | P37173 |
| HOMO SAPIENS | 0 | 253 | 5E+092 | A | 460 | ARG | p.Arg460His | P37173 |
| HOMO SAPIENS | 2 | 253 | 5E+092 | A | 526 | GLU | p.Glu526Gln | P37173 |
| HOMO SAPIENS | 5 | 253 | 5E+092 | A | 528 | ARG | p.Arg528Cys | P37173 |
| HOMO SAPIENS | 5 | 253 | 5E+092 | A | 528 | ARG | p.Arg528His | P37173 |
| HOMO SAPIENS | 6 | 158 | 5amc | A | 231 | ARG | p.Arg260Leu | P12821 |
| HOMO SAPIENS | 7 | 158 | 5amc | A | 237 | TYR | p.Tyr266Ter | P12821 |
| HOMO SAPIENS | 6 | 158 | 5amc | A | 325 | GLY | p.Gly354Arg | P12821 |
| HOMO SAPIENS | 0 | 158 | 5amc | A | 467 | ARG | p.Arg496Ter | P12821 |
| HOMO SAPIENS | 4 | 158 | 5amc | A | 479 | ARG | p.Arg508Ter | P12821 |
| HOMO SAPIENS | 0 | 158 | 5amc | A | 563 | ASP | p.Asp592Gly | P12821 |
| HOMO SAPIENS | -1 | 33062 | 1gr3 | A | 545 | GLY | p.Gly545Arg | Q03692 |
| HOMO SAPIENS | 5 | 33062 | 1gr3 | A | 591 | CYS | p.Cys591Arg | Q03692 |
| HOMO SAPIENS | 2 | 33062 | 1gr3 | A | 595 | GLY | p.Gly595Glu | Q03692 |
| HOMO SAPIENS | 0 | 33062 | 1gr3 | A | 597 | TYR | p.Tyr597Cys | Q03692 |
| HOMO SAPIENS | 5 | 33062 | 1gr3 | A | 598 | TYR | p.Tyr598Asp | Q03692 |
| HOMO SAPIENS | 7 | 33062 | 1gr3 | A | 600 | SER | p.Ser600Pro | Q03692 |
| HOMO SAPIENS | 1 | 33062 | 1gr3 | A | 614 | LEU | p.Leu614Pro | Q03692 |
| HOMO SAPIENS | 6 | 33062 | 1gr3 | A | 651 | TRP | p.Trp651Arg | Q03692 |
| HOMO SAPIENS | 7 | 33062 | 1gr3 | A | 671 | SER | p.Ser671Pro | Q03692 |
| HOMO SAPIENS | -1 | 38257 | 2l2i | B | 87 | PRO | p.Pro87Ala | Q13351 |
| HOMO SAPIENS | 2 | 2036 | 1ghq | A | 231 | GLY | p.Gly1224Asp | P01024 |
| HOMO SAPIENS | 5 | 3594 | 4cde | F | 388 | TYR | p.Tyr412Cys | P53634 |
| HOMO SAPIENS | 9 | 3594 | 4cde | F | 405 | TRP | p.Trp429Cys | P53634 |
| HOMO SAPIENS | 0 | 3594 | 4cde | F | 429 | ILE | p.Ile453Val | P53634 |
| HOMO SAPIENS | 7 | 18 | 1jkb | A | 34 | TRP | p.Trp52Ter | P61626 |
| HOMO SAPIENS | 9 | 18 | 1jkb | A | 70 | THR | p.Thr88Asn | P61626 |
| HOMO SAPIENS | 8 | 18 | 1jkb | A | 121 | VAL | p.Val139Ile | P61626 |
| HOMO SAPIENS | 2 | 414 | 3fj6 | A | 136 | ARG | p.Arg135Cys | Q02127 |
| HOMO SAPIENS | 1 | 414 | 3fj6 | A | 153 | GLY | p.Gly152Arg | Q02127 |
| HOMO SAPIENS | 0 | 414 | 3fj6 | A | 200 | ARG | p.Arg199Cys | Q02127 |
| HOMO SAPIENS | 0 | 414 | 3fj6 | A | 203 | GLY | p.Gly202Ala | Q02127 |
| HOMO SAPIENS | 0 | 414 | 3fj6 | A | 245 | ARG | p.Arg244Trp | Q02127 |
| HOMO SAPIENS | 0 | 414 | 3fj6 | A | 342 | ALA | p.Ala341Val | Q02127 |
| HOMO SAPIENS | 1 | 414 | 3fj6 | A | 347 | ARG | p.Arg346Trp | Q02127 |
| HOMO SAPIENS | 6 | 103 | 1u3w | A | 77 | GLY | p.Gly78Ter | P00326 |
| HOMO SAPIENS | -1 | 8369 | 4ux8 | F | 16 | ARG | p.Arg93Trp | P39905 |
| HOMO SAPIENS | 0 | 366 | 2zjc | A | 32 | GLY | p.Arg108Trp | P01375 |
| HOMO SAPIENS | 0 | 1457 | 3v9i | A | 352 | LEU | p.Ser352Leu | P30038 |
| HOMO SAPIENS | 0 | 1457 | 3v9i | A | 470 | VAL | p.Val470Ile | P30038 |
| HOMO SAPIENS | 0 | 1457 | 3v9i | A | 473 | THR | p.Thr473Ala | P30038 |
| HOMO SAPIENS | 1 | 2 | 4wo5 | A | 462 | ARG | p.Arg462Ile | P15056 |
| HOMO SAPIENS | 3 | 2 | 4wo5 | A | 463 | ILE | p.Ile463Ser | P15056 |
| HOMO SAPIENS | 1 | 2 | 4wo5 | A | 466 | GLY | p.Gly466Val | P15056 |
| HOMO SAPIENS | 1 | 2 | 4wo5 | A | 468 | PHE | p.Phe468Ser | P15056 |
| HOMO SAPIENS | 2 | 2 | 4wo5 | A | 469 | GLY | p.Gly469Ala | P15056 |
| HOMO SAPIENS | 2 | 2 | 4wo5 | A | 469 | GLY | p.Gly469Arg | P15056 |
| HOMO SAPIENS | 2 | 2 | 4wo5 | A | 485 | LEU | p.Leu485Phe | P15056 |
| HOMO SAPIENS | 0 | 2 | 4wo5 | A | 501 | GLU | p.Glu501Gly | P15056 |
| HOMO SAPIENS | 0 | 2 | 4wo5 | A | 501 | GLU | p.Glu501Lys | P15056 |
| HOMO SAPIENS | 7 | 2 | 4wo5 | A | 581 | ASN | p.Asn581Asp | P15056 |

| | | | | | | | | |
|---|---|---|---|---|---|---|---|---|
| HOMO SAPIENS | 8 | 2 | 4wo5 | A | 594 | ASP | p.Asp594Gly | P15056 |
| HOMO SAPIENS | 0 | 2 | 4wo5 | A | 597 | LEU | p.Leu597Arg | P15056 |
| HOMO SAPIENS | 0 | 2 | 4wo5 | A | 597 | LEU | p.Leu597Val | P15056 |
| HOMO SAPIENS | 0 | 2 | 4wo5 | A | 600 | VAL | p.Val600Glu | P15056 |
| HOMO SAPIENS | 2 | 2 | 4wo5 | A | 601 | LYS | p.Lys601Glu | P15056 |
| HOMO SAPIENS | 8 | 2 | 4wo5 | A | 638 | ASP | p.Asp638Glu | P15056 |
| HOMO SAPIENS | -1 | 30366 | 5bqd | B | 19 | ALA | p.Ala19Ser | Q99593 |
| HOMO SAPIENS | -1 | 30366 | 5bqd | B | 23 | PRO | p.Pro23His | Q99593 |
| HOMO SAPIENS | 1 | 30366 | 5bqd | B | 49 | GLN | p.Gln49Lys | Q99593 |
| HOMO SAPIENS | 2 | 30366 | 5bqd | B | 54 | ILE | p.Ile54Thr | Q99593 |
| HOMO SAPIENS | 8 | 30366 | 5bqd | B | 80 | GLY | p.Gly80Arg | Q99593 |
| HOMO SAPIENS | 6 | 30366 | 5bqd | B | 106 | ILE | p.Ile106Val | Q99593 |
| HOMO SAPIENS | 7 | 30366 | 5bqd | B | 111 | ASP | p.Asp111Tyr | Q99593 |
| HOMO SAPIENS | -1 | 30366 | 5bqd | B | 237 | ARG | p.Arg237Gln | Q99593 |
| HOMO SAPIENS | -1 | 30366 | 5bqd | B | 237 | ARG | p.Arg237Trp | Q99593 |
| HOMO SAPIENS | 5 | 4789 | 1n4m | A | 567 | SER | p.Ser567Leu | P06400 |
| HOMO SAPIENS | 0 | 4789 | 1n4m | A | 658 | ALA | p.Ala658Asp | P06400 |
| HOMO SAPIENS | 7 | 4789 | 1n4m | A | 661 | ARG | p.Arg661Trp | P06400 |
| HOMO SAPIENS | 6 | 4789 | 1n4m | A | 668 | ARG | p.Arg668His | P06400 |
| HOMO SAPIENS | 8 | 4789 | 1n4m | A | 697 | ASP | p.Asp697Glu | P06400 |
| HOMO SAPIENS | 6 | 4789 | 1n4m | A | 712 | CYS | p.Cys712Arg | P06400 |
| HOMO SAPIENS | 1 | 1798 | 2r5g | A | 101 | HIS | p.His101Gln | O15247 |
| HOMO SAPIENS | 0 | 1798 | 2r5g | A | 160 | PRO | p.Pro160Ala | O15247 |
| HOMO SAPIENS | 2 | 22 | 4x0t | A | 82 | ARG | p.Arg110Ter | P49419 |
| HOMO SAPIENS | 0 | 22 | 4x0t | A | 149 | ALA | p.Ala177Pro | P49419 |
| HOMO SAPIENS | 4 | 22 | 4x0t | A | 164 | THR | p.Thr192Met | P49419 |
| HOMO SAPIENS | 9 | 22 | 4x0t | A | 167 | ASN | p.Asn195Ser | P49419 |
| HOMO SAPIENS | 0 | 22 | 4x0t | A | 175 | TRP | p.Trp203Gly | P49419 |
| HOMO SAPIENS | 0 | 22 | 4x0t | A | 194 | THR | p.Thr222Ala | P49419 |
| HOMO SAPIENS | 0 | 22 | 4x0t | A | 273 | ASN | p.Asn301Ile | P49419 |
| HOMO SAPIENS | 0 | 22 | 4x0t | A | 307 | ARG | p.Arg335Gln | P49419 |
| HOMO SAPIENS | 3 | 22 | 4x0t | A | 313 | SER | p.Ser341Asn | P49419 |
| HOMO SAPIENS | 0 | 22 | 4x0t | A | 329 | ILE | p.Ile357Val | P49419 |
| HOMO SAPIENS | 1 | 22 | 4x0t | A | 380 | TYR | p.Tyr408Ter | P49419 |
| HOMO SAPIENS | 9 | 22 | 4x0t | A | 399 | GLU | p.Glu427Gln | P49419 |
| HOMO SAPIENS | 0 | 22 | 4x0t | A | 406 | TYR | p.Tyr434Cys | P49419 |
| HOMO SAPIENS | 0 | 22 | 4x0t | A | 431 | ILE | p.Ile459Phe | P49419 |
| HOMO SAPIENS | 4 | 22 | 4x0t | A | 477 | GLY | p.Gly505Arg | P49419 |
| HOMO SAPIENS | 0 | 22 | 4x0t | A | 495 | THR | p.Thr523Ala | P49419 |
| HOMO SAPIENS | 8 | 11042 | 1y6b | A | 1145 | PRO | p.Pro1147Ser | P35968 |
| HOMO SAPIENS | -1 | 3867 | 2xij | A | 18 | GLN | p.Gln18Ter | P22033 |
| HOMO SAPIENS | -1 | 3867 | 2xij | A | 31 | ARG | p.Arg31Ter | P22033 |
| HOMO SAPIENS | 2 | 3867 | 2xij | A | 93 | ARG | p.Arg93His | P22033 |
| HOMO SAPIENS | 0 | 3867 | 2xij | A | 94 | GLY | p.Gly94Arg | P22033 |
| HOMO SAPIENS | 2 | 3867 | 2xij | A | 95 | PRO | p.Pro95Leu | P22033 |
| HOMO SAPIENS | 2 | 3867 | 2xij | A | 105 | TRP | p.Trp105Arg | P22033 |
| HOMO SAPIENS | 2 | 3867 | 2xij | A | 108 | ARG | p.Arg108Cys | P22033 |
| HOMO SAPIENS | 3 | 3867 | 2xij | A | 154 | ARG | p.Arg154His | P22033 |
| HOMO SAPIENS | 3 | 3867 | 2xij | A | 156 | ASP | p.Asp156Val | P22033 |
| HOMO SAPIENS | 0 | 3867 | 2xij | A | 189 | ASN | p.Asn189Ile | P22033 |
| HOMO SAPIENS | 0 | 3867 | 2xij | A | 191 | ALA | p.Ala191Glu | P22033 |
| HOMO SAPIENS | 0 | 3867 | 2xij | A | 203 | GLY | p.Gly203Arg | P22033 |
| HOMO SAPIENS | 2 | 3867 | 2xij | A | 215 | GLY | p.Gly215Ser | P22033 |
| HOMO SAPIENS | 0 | 3867 | 2xij | A | 219 | ASN | p.Asn219Tyr | P22033 |

| | | | | | | | | |
|---|---|---|---|---|---|---|---|---|
| HOMO SAPIENS | 3 | 3867 | 2xij | A | 228 | ARG | p.Arg228Ter | P22033 |
| HOMO SAPIENS | 1 | 3867 | 2xij | A | 297 | THR | p.Thr297Ile | P22033 |
| HOMO SAPIENS | 0 | 3867 | 2xij | A | 326 | ARG | p.Arg326Lys | P22033 |
| HOMO SAPIENS | 3 | 3867 | 2xij | A | 369 | ARG | p.Arg369His | P22033 |
| HOMO SAPIENS | 3 | 3867 | 2xij | A | 369 | ARG | p.Arg369Cys | P22033 |
| HOMO SAPIENS | 0 | 3867 | 2xij | A | 370 | THR | p.Thr370Pro | P22033 |
| HOMO SAPIENS | 0 | 3867 | 2xij | A | 377 | ALA | p.Ala377Glu | P22033 |
| HOMO SAPIENS | 2 | 3867 | 2xij | A | 426 | GLY | p.Gly426Ala | P22033 |
| HOMO SAPIENS | 2 | 3867 | 2xij | A | 427 | GLY | p.Gly427Asp | P22033 |
| HOMO SAPIENS | 1 | 3867 | 2xij | A | 499 | THR | p.Ala499Thr | P22033 |
| HOMO SAPIENS | 2 | 3867 | 2xij | A | 532 | ARG | p.Arg532His | P22033 |
| HOMO SAPIENS | 3 | 3867 | 2xij | A | 555 | ALA | p.Ala555Thr | P22033 |
| HOMO SAPIENS | 0 | 3867 | 2xij | A | 573 | PHE | p.Phe573Ser | P22033 |
| HOMO SAPIENS | 3 | 3867 | 2xij | A | 623 | GLY | p.Gly623Arg | P22033 |
| HOMO SAPIENS | 5 | 3867 | 2xij | A | 671 | VAL | p.Ile671Val | P22033 |
| HOMO SAPIENS | 4 | 3867 | 2xij | A | 694 | ARG | p.Arg694Trp | P22033 |
| HOMO SAPIENS | 1 | 3867 | 2xij | A | 700 | MET | p.Met700Lys | P22033 |
| HOMO SAPIENS | 8 | 3867 | 2xij | A | 703 | GLY | p.Gly703Arg | P22033 |
| HOMO SAPIENS | 0 | 3867 | 2xij | A | 717 | GLY | p.Gly717Val | P22033 |
| HOMO SAPIENS | 1 | 3867 | 2xij | A | 727 | ARG | p.Arg727Ter | P22033 |
| HOMO SAPIENS | -1 | 52 | 3k91 | B | 72 | GLY | p.Gly73Ser | P00441 |
| HOMO SAPIENS | 6 | 52 | 3k91 | B | 96 | ASP | p.Asp97Asn | P00441 |
| HOMO SAPIENS | 5 | 52 | 3k91 | B | 112 | ILE | p.Ile113Thr | P00441 |
| HOMO SAPIENS | -1 | 52 | 3k91 | B | 134 | SER | p.Ser135Asn | P00441 |
| HOMO SAPIENS | 0 | 3950 | 4fom | A | 296 | ASN | p.Asn296His | Q9NQS3 |
| HOMO SAPIENS | 1 | 10 | 1de4 | A | 31 | VAL | p.Val53Met | Q30201 |
| HOMO SAPIENS | 0 | 10 | 1de4 | A | 37 | VAL | p.Val59Met | Q30201 |
| HOMO SAPIENS | 0 | 10 | 1de4 | A | 71 | GLY | p.Gly93Arg | Q30201 |
| HOMO SAPIENS | 0 | 10 | 1de4 | A | 83 | ILE | p.Ile105Thr | Q30201 |
| HOMO SAPIENS | 0 | 10 | 1de4 | A | 105 | GLN | p.Gln127His | Q30201 |
| HOMO SAPIENS | 1 | 10 | 1de4 | A | 146 | GLU | p.Glu168Ter | Q30201 |
| HOMO SAPIENS | 2 | 10 | 1de4 | A | 261 | GLN | p.Gln283Pro | Q30201 |
| HOMO SAPIENS | 0 | 8038 | 4x7n | A | 586 | ILE | p.Ile586Leu | Q9NZJ5 |
| HOMO SAPIENS | 0 | 8038 | 4x7n | A | 588 | ARG | p.Arg588Gln | Q9NZJ5 |
| HOMO SAPIENS | 0 | 35175 | 5ugw | A | 1062 | ALA | p.Ala1062Thr | O14746 |
| HOMO SAPIENS | 2 | 35175 | 5ugw | A | 1090 | VAL | p.Val1090Met | O14746 |
| HOMO SAPIENS | 0 | 1722 | 4z3k | C | 147 | ARG | p.Arg150Gly | P35270 |
| HOMO SAPIENS | 0 | 1722 | 4z3k | C | 160 | PRO | p.Pro163Leu | P35270 |
| HOMO SAPIENS | 0 | 1722 | 4z3k | C | 207 | ARG | p.Arg210Trp | P35270 |
| HOMO SAPIENS | 0 | 1722 | 4z3k | C | 216 | ARG | p.Arg219Ter | P35270 |
| HOMO SAPIENS | 0 | 1722 | 4z3k | C | 248 | LYS | p.Lys251Ter | P35270 |
| HOMO SAPIENS | 1 | 2832 | 4x8v | L | 115 | HIS | p.His175Gln | P08709 |
| HOMO SAPIENS | 0 | 825 | 1emh | A | 242 | PHE | p.Phe251Ser | P13051 |
| HOMO SAPIENS | 0 | 122 | 5uwk | A | 109 | ARG | p.Arg109Ter | P45452 |
| HOMO SAPIENS | 1 | 122 | 5uwk | A | 170 | LYS | p.Lys170Thr | P45452 |
| HOMO SAPIENS | 9 | 122 | 5uwk | A | 232 | HIS | p.His232Asn | P45452 |
| HOMO SAPIENS | 5 | 122 | 5uwk | A | 257 | ASP | p.Asp257Val | P45452 |
| HOMO SAPIENS | 9 | 1074 | 3p5i | A | 264 | TRP | p.Trp264Arg | Q9UJ71 |
| HOMO SAPIENS | 0 | 146 | 3r3g | A | 14C | GLU | p.Glu352Lys | P00734 |
| HOMO SAPIENS | 3 | 9522 | 1wyq | A | 83 | LEU | p.Leu253Pro | O15020 |
| HOMO SAPIENS | 8 | 67 | 2vgb | A | 130 | SER | p.Ser130Tyr | P30613 |
| HOMO SAPIENS | 8 | 67 | 2vgb | A | 163 | ARG | p.Arg163Cys | P30613 |
| HOMO SAPIENS | 4 | 67 | 2vgb | A | 241 | GLU | p.Glu241Ter | P30613 |
| HOMO SAPIENS | 9 | 67 | 2vgb | A | 384 | THR | p.Thr384Met | P30613 |

| | | | | | | | | |
|---|---|---|---|---|---|---|---|---|
| HOMO SAPIENS | 0 | 67 | 2vgb | A | 421 | GLN | p.Gln421Lys | P30613 |
| HOMO SAPIENS | 6 | 67 | 2vgb | A | 479 | ARG | p.Arg479His | P30613 |
| HOMO SAPIENS | 8 | 67 | 2vgb | A | 510 | ARG | p.Arg510Gln | P30613 |
| HOMO SAPIENS | 4 | 67 | 2vgb | A | 569 | ARG | p.Arg569Trp | P30613 |
| HOMO SAPIENS | 0 | 9088 | 5v44 | A | 232 | ASN | p.Asn232Lys | Q9NZJ4 |
| HOMO SAPIENS | 4 | 16 | 3lbn | A | 12 | GLY | p.Gly12Val | P01112 |
| HOMO SAPIENS | 4 | 16 | 3lbn | A | 12 | GLY | p.Gly12Ser | P01112 |
| HOMO SAPIENS | 4 | 16 | 3lbn | A | 13 | GLY | p.Gly13Cys | P01112 |
| HOMO SAPIENS | 4 | 16 | 3lbn | A | 13 | GLY | p.Gly13Asp | P01112 |
| HOMO SAPIENS | 5 | 16 | 3lbn | A | 22 | GLN | p.Gln22Lys | P01112 |
| HOMO SAPIENS | 5 | 16 | 3lbn | A | 58 | THR | p.Thr58Ile | P01112 |
| HOMO SAPIENS | 3 | 16 | 3lbn | A | 61 | GLN | p.Gln61Lys | P01112 |
| HOMO SAPIENS | 1 | 16 | 3lbn | A | 63 | GLU | p.Glu63Lys | P01112 |
| HOMO SAPIENS | 9 | 16 | 3lbn | A | 117 | LYS | p.Lys117Arg | P01112 |
| HOMO SAPIENS | 6 | 16 | 3lbn | A | 123 | ARG | p.Arg123Cys | P01112 |
| HOMO SAPIENS | 1 | 16 | 3lbn | A | 138 | GLY | p.Gly138Ser | P01112 |
| HOMO SAPIENS | 9 | 16 | 3lbn | A | 146 | ALA | p.Ala146Thr | P01112 |
| HOMO SAPIENS | 9 | 16 | 3lbn | A | 146 | ALA | p.Ala146Val | P01112 |
| HOMO SAPIENS | 1 | 359 | 5nvy | C | 63 | LEU | p.Leu63Pro | P40337 |
| HOMO SAPIENS | 1 | 359 | 5nvy | C | 64 | ARG | p.Arg64Pro | P40337 |
| HOMO SAPIENS | 1 | 359 | 5nvy | C | 70 | GLU | p.Glu70Lys | P40337 |
| HOMO SAPIENS | 0 | 359 | 5nvy | C | 84 | VAL | p.Val84Leu | P40337 |
| HOMO SAPIENS | 1 | 359 | 5nvy | C | 88 | TRP | p.Trp88Ser | P40337 |
| HOMO SAPIENS | 1 | 359 | 5nvy | C | 93 | GLY | p.Gly93Cys | P40337 |
| HOMO SAPIENS | 1 | 359 | 5nvy | C | 98 | TYR | p.Tyr98His | P40337 |
| HOMO SAPIENS | 1 | 359 | 5nvy | C | 111 | SER | p.Ser111Arg | P40337 |
| HOMO SAPIENS | 1 | 359 | 5nvy | C | 112 | TYR | p.Tyr112His | P40337 |
| HOMO SAPIENS | 1 | 359 | 5nvy | C | 126 | ASP | p.Asp126Tyr | P40337 |
| HOMO SAPIENS | 1 | 359 | 5nvy | C | 130 | VAL | p.Val130Leu | P40337 |
| HOMO SAPIENS | 1 | 359 | 5nvy | C | 135 | LEU | p.Leu135Phe | P40337 |
| HOMO SAPIENS | 9 | 359 | 5nvy | C | 161 | ARG | p.Arg161Gly | P40337 |
| HOMO SAPIENS | 7 | 359 | 5nvy | C | 162 | CYS | p.Cys162Phe | P40337 |
| HOMO SAPIENS | 4 | 359 | 5nvy | C | 163 | LEU | p.Leu163Pro | P40337 |
| HOMO SAPIENS | 3 | 359 | 5nvy | C | 164 | GLN | p.Gln164Arg | P40337 |
| HOMO SAPIENS | 9 | 359 | 5nvy | C | 166 | VAL | p.Val166Phe | P40337 |
| HOMO SAPIENS | 7 | 359 | 5nvy | C | 167 | ARG | p.Arg167Gly | P40337 |
| HOMO SAPIENS | 7 | 359 | 5nvy | C | 167 | ARG | p.Arg167Gln | P40337 |
| HOMO SAPIENS | 1 | 359 | 5nvy | C | 183 | SER | p.Ser183Ter | P40337 |
| HOMO SAPIENS | 2 | 359 | 5nvy | C | 188 | LEU | p.Leu188Val | P40337 |
| HOMO SAPIENS | 1 | 359 | 5nvy | C | 191 | HIS | p.His191Asp | P40337 |
| HOMO SAPIENS | 1 | 359 | 5nvy | C | 192 | PRO | p.Pro192Ser | P40337 |
| HOMO SAPIENS | 1 | 359 | 5nvy | C | 200 | ARG | p.Arg200Trp | P40337 |
| HOMO SAPIENS | -1 | 359 | 5nvy | C | 211 | MET | p.Met211Leu | P40337 |
| HOMO SAPIENS | 0 | 14099 | 4btm | A | 282 | ARG | p.Arg258Gln | Q5TCY1 |
| HOMO SAPIENS | 0 | 696 | 5h1b | A | 150 | ARG | p.Arg150Gln | Q06609 |
| HOMO SAPIENS | -1 | 208 | 2bxl | A | 1 | ASP | p.Asp25Val | P02768 |
| HOMO SAPIENS | 0 | 208 | 2bxl | A | 269 | ASP | p.Asp293Gly | P02768 |
| HOMO SAPIENS | 0 | 208 | 2bxl | A | 570 | GLU | p.Glu594Lys | P02768 |
| HOMO SAPIENS | 0 | 208 | 2bxl | A | 573 | LYS | p.Lys597Glu | P02768 |
| HOMO SAPIENS | -1 | 12813 | 2uzx | B | 35 | MET | p.Met35Leu | P08581 |
| HOMO SAPIENS | 1 | 12813 | 2uzx | B | 156 | SER | p.Ser156Leu | P08581 |
| HOMO SAPIENS | 0 | 12813 | 2uzx | B | 168 | GLU | p.Glu168Asp | P08581 |
| HOMO SAPIENS | 2 | 12813 | 2uzx | B | 211 | LEU | p.Leu211Trp | P08581 |
| HOMO SAPIENS | 2 | 12813 | 2uzx | B | 218 | ARG | p.Arg218Ser | P08581 |

| | | | | | | | | |
|---|---|---|---|---|---|---|---|---|
| HOMO SAPIENS | 1 | 12813 | 2uzx | B | 238 | LEU | p.Leu238Ser | P08581 |
| HOMO SAPIENS | 4 | 12813 | 2uzx | B | 320 | ALA | p.Ala320Val | P08581 |
| HOMO SAPIENS | 1 | 12813 | 2uzx | B | 375 | ASN | p.Asn375Ser | P08581 |
| HOMO SAPIENS | 0 | 12813 | 2uzx | B | 575 | LEU | p.Leu575Ile | P08581 |
| HOMO SAPIENS | 1 | 10780 | 4ejx | A | 348 | ARG | p.Arg367Cys | P00450 |
| HOMO SAPIENS | 2 | 10780 | 4ejx | A | 525 | ASP | p.Asp544Glu | P00450 |
| HOMO SAPIENS | 1 | 1407 | 5jn5 | A | 88 | ILE | p.Ile88Val | P36871 |
| HOMO SAPIENS | 7 | 1407 | 5jn5 | A | 115 | THR | p.Thr115Ala | P36871 |
| HOMO SAPIENS | 3 | 1407 | 5jn5 | A | 121 | GLY | p.Gly121Arg | P36871 |
| HOMO SAPIENS | 2 | 1407 | 5jn5 | A | 221 | ARG | p.Arg221Cys | P36871 |
| HOMO SAPIENS | 6 | 1407 | 5jn5 | A | 263 | TYR | p.Asp263Tyr | P36871 |
| HOMO SAPIENS | 3 | 1407 | 5jn5 | A | 420 | TYR | p.Tyr420His | P36871 |
| HOMO SAPIENS | 0 | 1407 | 5jn5 | A | 501 | VAL | p.Val501Ile | P36871 |
| HOMO SAPIENS | -1 | 1878 | 1yow | A | 255 | ARG | p.Arg255Leu | Q13285 |
| HOMO SAPIENS | 6 | 1878 | 1yow | A | 293 | ASP | p.Asp293Asn | Q13285 |
| HOMO SAPIENS | 1 | 1878 | 1yow | A | 437 | LEU | p.Leu437Gln | Q13285 |
| HOMO SAPIENS | 0 | 3071 | 2wpm | E | 145 | ARG | p.Arg191His | P00740 |
| HOMO SAPIENS | 0 | 3071 | 2wpm | E | 145 | ARG | p.Arg191Cys | P00740 |
| HOMO SAPIENS | 3 | 589 | 3ggc | B | 15 | GLY | p.Gly16Ser | P00492 |
| HOMO SAPIENS | 4 | 589 | 3ggc | B | 40 | LEU | p.Leu41Pro | P00492 |
| HOMO SAPIENS | 2 | 589 | 3ggc | B | 50 | ARG | p.Arg51Gly | P00492 |
| HOMO SAPIENS | 5 | 589 | 3ggc | B | 51 | ASP | p.Asp52Gly | P00492 |
| HOMO SAPIENS | 0 | 589 | 3ggc | B | 57 | GLY | p.Gly58Arg | P00492 |
| HOMO SAPIENS | 6 | 589 | 3ggc | B | 69 | GLY | p.Gly70Glu | P00492 |
| HOMO SAPIENS | 1 | 589 | 3ggc | B | 73 | PHE | p.Phe74Leu | P00492 |
| HOMO SAPIENS | 0 | 589 | 3ggc | B | 77 | LEU | p.Leu78Val | P00492 |
| HOMO SAPIENS | 3 | 589 | 3ggc | B | 79 | ASP | p.Asp80Val | P00492 |
| HOMO SAPIENS | 0 | 589 | 3ggc | B | 103 | SER | p.Ser104Arg | P00492 |
| HOMO SAPIENS | 0 | 589 | 3ggc | B | 109 | SER | p.Ser110Leu | P00492 |
| HOMO SAPIENS | 1 | 589 | 3ggc | B | 129 | VAL | p.Val130Asp | P00492 |
| HOMO SAPIENS | 4 | 589 | 3ggc | B | 131 | ILE | p.Ile132Met | P00492 |
| HOMO SAPIENS | 7 | 589 | 3ggc | B | 142 | MET | p.Met143Lys | P00492 |
| HOMO SAPIENS | 0 | 589 | 3ggc | B | 160 | ALA | p.Ala161Ser | P00492 |
| HOMO SAPIENS | 6 | 589 | 3ggc | B | 167 | THR | p.Thr168Ile | P00492 |
| HOMO SAPIENS | 0 | 589 | 3ggc | B | 169 | ARG | p.Arg170Ter | P00492 |
| HOMO SAPIENS | 5 | 589 | 3ggc | B | 175 | PRO | p.Pro176Leu | P00492 |
| HOMO SAPIENS | 8 | 589 | 3ggc | B | 193 | ASP | p.Asp194Glu | P00492 |
| HOMO SAPIENS | 8 | 589 | 3ggc | B | 193 | ASP | p.Asp194Asn | P00492 |
| HOMO SAPIENS | 8 | 589 | 3ggc | B | 198 | PHE | p.Phe199Val | P00492 |
| HOMO SAPIENS | 8 | 589 | 3ggc | B | 200 | ASP | p.Asp201Gly | P00492 |
| HOMO SAPIENS | 0 | 4269 | 2pla | B | 174 | GLU | p.Glu174Lys | Q8N335 |
| HOMO SAPIENS | 1 | 4269 | 2pla | B | 220 | ARG | p.Arg220His | Q8N335 |
| HOMO SAPIENS | 4 | 4269 | 2pla | B | 280 | ALA | p.Ala280Val | Q8N335 |
| HOMO SAPIENS | 0 | 2712 | 1ii4 | E | 252 | TRP | p.Ser252Leu | P21802 |
| HOMO SAPIENS | 0 | 2712 | 1ii4 | E | 253 | PRO | p.Pro253Arg | P21802 |
| HOMO SAPIENS | 5 | 2712 | 1ii4 | E | 267 | SER | p.Ser267Pro | P21802 |
| HOMO SAPIENS | 0 | 2712 | 1ii4 | E | 278 | CYS | p.Cys278Phe | P21802 |
| HOMO SAPIENS | 0 | 2712 | 1ii4 | E | 289 | GLN | p.Gln289Pro | P21802 |
| HOMO SAPIENS | 0 | 2712 | 1ii4 | E | 290 | TRP | p.Trp290Cys | P21802 |
| HOMO SAPIENS | 0 | 2712 | 1ii4 | E | 290 | TRP | p.Trp290Arg | P21802 |
| HOMO SAPIENS | 0 | 2712 | 1ii4 | E | 292 | LYS | p.Lys292Glu | P21802 |
| HOMO SAPIENS | 0 | 2712 | 1ii4 | E | 315 | ALA | p.Ala315Ser | P21802 |
| HOMO SAPIENS | 0 | 2712 | 1ii4 | E | 321 | ASP | p.Asp321Ala | P21802 |
| HOMO SAPIENS | 1 | 2712 | 1ii4 | E | 328 | TYR | p.Tyr328Cys | P21802 |

| | | | | | | | | |
|---|---|---|---|---|---|---|---|---|
| HOMO SAPIENS | 9 | 2712 | 1ii4 | E | 340 | TYR | p.Tyr340His | P21802 |
| HOMO SAPIENS | 1 | 2712 | 1ii4 | E | 341 | THR | p.Thr341Pro | P21802 |
| HOMO SAPIENS | 0 | 2712 | 1ii4 | E | 342 | CYS | p.Cys342Arg | P21802 |
| HOMO SAPIENS | 0 | 2712 | 1ii4 | E | 342 | CYS | p.Cys342Trp | P21802 |
| HOMO SAPIENS | 0 | 2712 | 1ii4 | E | 344 | ALA | p.Ala344Gly | P21802 |
| HOMO SAPIENS | 0 | 2712 | 1ii4 | E | 347 | SER | p.Ser347Cys | P21802 |
| HOMO SAPIENS | 0 | 2712 | 1ii4 | E | 351 | SER | p.Ser351Cys | P21802 |
| HOMO SAPIENS | 1 | 2712 | 1ii4 | E | 354 | SER | p.Ser354Cys | P21802 |
| HOMO SAPIENS | 2 | 53 | 4pj1 | H | 5 | ASP | p.Asp29Gly | P10809 |
| HOMO SAPIENS | 0 | 53 | 4pj1 | H | 74 | VAL | p.Val98Ile | P10809 |
| HOMO SAPIENS | 0 | 29968 | 2fdb | M | 71 | LYS | p.Lys89Glu | P55075 |
| HOMO SAPIENS | 2 | 29968 | 2fdb | M | 98 | ARG | p.Arg116Gly | P55075 |
| HOMO SAPIENS | 2 | 29968 | 2fdb | M | 122 | GLY | p.Gly140Ser | P55075 |
| HOMO SAPIENS | 8 | 5061 | 5f85 | B | 60 | GLY | p.Gly106Ser | P00740 |
| HOMO SAPIENS | 0 | 35944 | 1h6h | A | 85 | THR | p.Thr85Asn | Q15080 |
| HOMO SAPIENS | 8 | 35944 | 1h6h | A | 105 | ARG | p.Arg105Gln | Q15080 |
| HOMO SAPIENS | 7 | 12016 | 1hj7 | A | 295 | ASN | p.Asn316Ser | P01130 |
| HOMO SAPIENS | 5 | 12016 | 1hj7 | A | 302 | GLY | p.Gly323Ser | P01130 |
| HOMO SAPIENS | 6 | 12016 | 1hj7 | A | 319 | CYS | p.Cys340Trp | P01130 |
| HOMO SAPIENS | 7 | 12016 | 1hj7 | A | 329 | ARG | p.Arg350Ter | P01130 |
| HOMO SAPIENS | 5 | 12016 | 1hj7 | A | 331 | CYS | p.Cys352Ter | P01130 |
| HOMO SAPIENS | 2 | 12016 | 1hj7 | A | 345 | GLN | p.Gln366Arg | P01130 |
| HOMO SAPIENS | 2 | 1481 | 2i07 | A | 239 | TYR | p.Tyr261Ter | P01024 |
| HOMO SAPIENS | -1 | 1481 | 2i07 | A | 292 | LEU | p.Pro314Leu | P01024 |
| HOMO SAPIENS | 2 | 1481 | 2i07 | A | 326 | ILE | p.Ile348Val | P01024 |
| HOMO SAPIENS | 2 | 1481 | 2i07 | A | 413 | GLU | p.Glu435Lys | P01024 |
| HOMO SAPIENS | 4 | 1481 | 2i07 | A | 447 | GLU | p.Glu469Asp | P01024 |
| HOMO SAPIENS | 2 | 1481 | 2i07 | A | 597 | VAL | p.Val619Met | P01024 |
| HOMO SAPIENS | 5 | 1481 | 2i07 | A | 603 | ILE | p.Ile625Phe | P01024 |
| HOMO SAPIENS | 0 | 1481 | 2i07 | A | 611 | LYS | p.Lys633Arg | P01024 |
| HOMO SAPIENS | 0 | 11123 | 3c5n | A | 315 | GLY | p.Gly368Trp | O00294 |
| HOMO SAPIENS | 8 | 11123 | 3c5n | A | 329 | PHE | p.Phe382Ser | O00294 |
| HOMO SAPIENS | 5 | 11123 | 3c5n | A | 429 | ARG | p.Arg482Trp | O00294 |
| HOMO SAPIENS | 8 | 11123 | 3c5n | A | 438 | PHE | p.Phe491Leu | O00294 |
| HOMO SAPIENS | 3 | 4398 | 3fx0 | B | 1288 | ALA | p.Ala288Gly | Q9Y6K9 |
| HOMO SAPIENS | 7 | 4398 | 3fx0 | B | 1315 | GLU | p.Glu315Ala | Q9Y6K9 |
| HOMO SAPIENS | 8 | 4398 | 3fx0 | B | 1319 | ARG | p.Arg319Gln | Q9Y6K9 |
| HOMO SAPIENS | 6 | 3989 | 3qd6 | S | 83 | CYS | p.Cys83Arg | P25942 |
| HOMO SAPIENS | 1 | 3989 | 3qd6 | S | 124 | SER | p.Ser124Leu | P25942 |
| HOMO SAPIENS | 2 | 24003 | 2m0v | A | 225 | GLU | p.Glu225Lys | O14745 |
| HOMO SAPIENS | 2 | 46 | 4aqd | A | 70 | ASP | p.Asp98Gly | P06276 |
| HOMO SAPIENS | 4 | 46 | 4aqd | A | 115 | GLY | p.Gly143Asp | P06276 |
| HOMO SAPIENS | 3 | 46 | 4aqd | A | 184 | ALA | p.Ala212Val | P06276 |
| HOMO SAPIENS | 0 | 46 | 4aqd | A | 225 | GLY | p.Gly253Ter | P06276 |
| HOMO SAPIENS | 2 | 46 | 4aqd | A | 243 | THR | p.Thr271Met | P06276 |
| HOMO SAPIENS | 2 | 46 | 4aqd | A | 255 | GLU | p.Glu283Asp | P06276 |
| HOMO SAPIENS | 5 | 46 | 4aqd | A | 267 | LYS | p.Lys295Arg | P06276 |
| HOMO SAPIENS | 3 | 46 | 4aqd | A | 271 | GLU | p.Glu299Ter | P06276 |
| HOMO SAPIENS | 2 | 46 | 4aqd | A | 307 | LEU | p.Leu335Pro | P06276 |
| HOMO SAPIENS | 1 | 46 | 4aqd | A | 330 | LEU | p.Leu358Ile | P06276 |
| HOMO SAPIENS | 0 | 46 | 4aqd | A | 497 | GLU | p.Glu525Val | P06276 |
| HOMO SAPIENS | 3 | 786 | 1ols | B | 51 | GLY | p.Gly101Asp | P21953 |
| HOMO SAPIENS | 2 | 786 | 1ols | B | 87 | ALA | p.Ala137Val | P21953 |
| HOMO SAPIENS | 0 | 786 | 1ols | B | 110 | ILE | p.Ile160Ser | P21953 |

| | | | | | | | | |
|---|---|---|---|---|---|---|---|---|
| HOMO SAPIENS | 4 | 786 | 1ols | B | 120 | ARG | p.Arg170Cys | P21953 |
| HOMO SAPIENS | 2 | 786 | 1ols | B | 126 | ASN | p.Asn176Tyr | P21953 |
| HOMO SAPIENS | 0 | 786 | 1ols | B | 133 | ARG | p.Arg183Gly | P21953 |
| HOMO SAPIENS | 0 | 786 | 1ols | B | 133 | ARG | p.Arg183Pro | P21953 |
| HOMO SAPIENS | 0 | 786 | 1ols | B | 135 | PRO | p.Pro185Leu | P21953 |
| HOMO SAPIENS | 3 | 786 | 1ols | B | 156 | HIS | p.His206Tyr | P21953 |
| HOMO SAPIENS | 0 | 786 | 1ols | B | 228 | GLY | p.Gly278Ser | P21953 |
| HOMO SAPIENS | 1 | 786 | 1ols | B | 235 | ARG | p.Arg285Ter | P21953 |
| HOMO SAPIENS | 6 | 786 | 1ols | B | 274 | ARG | p.Arg324Ter | P21953 |
| HOMO SAPIENS | 1 | 831 | 4zt1 | B | 48 | VAL | p.Val202Ile | P12830 |
| HOMO SAPIENS | 6 | 831 | 4zt1 | B | 70 | ARG | p.Arg224Cys | P12830 |
| HOMO SAPIENS | 5 | 831 | 4zt1 | B | 79 | HIS | p.His233Gln | P12830 |
| HOMO SAPIENS | 4 | 831 | 4zt1 | B | 120 | GLY | p.Gly274Ser | P12830 |
| HOMO SAPIENS | 7 | 831 | 4zt1 | B | 187 | TYR | p.Tyr341Ter | P12830 |
| HOMO SAPIENS | 0 | 8497 | 3sla | A | 287 | ASN | p.Asn287Ser | P35222 |
| HOMO SAPIENS | 7 | 23986 | 2ke1 | A | 301 | VAL | p.Val301Met | O43918 |
| HOMO SAPIENS | 0 | 4967 | 1rj7 | B | 334 | ARG | p.Arg334His | Q92838 |
| HOMO SAPIENS | 0 | 4967 | 1rj7 | B | 338 | THR | p.Thr338Met | Q92838 |
| HOMO SAPIENS | 3 | 4967 | 1rj7 | B | 349 | ALA | p.Ala349Thr | Q92838 |
| HOMO SAPIENS | 2 | 4967 | 1rj7 | B | 358 | GLN | p.Gln358Glu | Q92838 |
| HOMO SAPIENS | 1 | 4967 | 1rj7 | B | 375 | LYS | p.Lys375Glu | Q92838 |
| HOMO SAPIENS | 9 | 7623 | 1d4t | A | 32 | ARG | p.Arg32Thr | O60880 |
| HOMO SAPIENS | 6 | 7623 | 1d4t | A | 55 | ARG | p.Arg55Leu | O60880 |
| HOMO SAPIENS | 0 | 7623 | 1d4t | A | 68 | THR | p.Thr68Ile | O60880 |
| HOMO SAPIENS | 6 | 705 | 3idh | A | 59 | PRO | p.Pro59Ser | P35557 |
| HOMO SAPIENS | 5 | 705 | 3idh | A | 124 | ASP | p.Asp124Asn | P35557 |
| HOMO SAPIENS | 4 | 705 | 3idh | A | 131 | SER | p.Ser131Pro | P35557 |
| HOMO SAPIENS | 0 | 705 | 3idh | A | 147 | GLY | p.Gly147Asp | P35557 |
| HOMO SAPIENS | 5 | 705 | 3idh | A | 186 | ARG | p.Arg186Ter | P35557 |
| HOMO SAPIENS | 0 | 705 | 3idh | A | 202 | MET | p.Met202Thr | P35557 |
| HOMO SAPIENS | 2 | 705 | 3idh | A | 210 | MET | p.Met210Lys | P35557 |
| HOMO SAPIENS | 1 | 705 | 3idh | A | 215 | TYR | p.Tyr215Ter | P35557 |
| HOMO SAPIENS | 0 | 705 | 3idh | A | 226 | VAL | p.Val226Met | P35557 |
| HOMO SAPIENS | 9 | 705 | 3idh | A | 228 | THR | p.Thr228Met | P35557 |
| HOMO SAPIENS | 9 | 705 | 3idh | A | 256 | GLU | p.Glu256Lys | P35557 |
| HOMO SAPIENS | 2 | 705 | 3idh | A | 261 | GLY | p.Gly261Arg | P35557 |
| HOMO SAPIENS | 6 | 705 | 3idh | A | 279 | GLU | p.Glu279Gln | P35557 |
| HOMO SAPIENS | 7 | 705 | 3idh | A | 299 | GLY | p.Gly299Arg | P35557 |
| HOMO SAPIENS | 2 | 705 | 3idh | A | 318 | GLY | p.Gly318Trp | P35557 |
| HOMO SAPIENS | 4 | 705 | 3idh | A | 378 | ALA | p.Ala378Thr | P35557 |
| HOMO SAPIENS | 0 | 705 | 3idh | A | 387 | ALA | p.Ala387Glu | P35557 |
| HOMO SAPIENS | 5 | 705 | 3idh | A | 414 | LYS | p.Lys414Glu | P35557 |
| HOMO SAPIENS | 0 | 705 | 3idh | A | 455 | VAL | p.Val455Met | P35557 |
| HOMO SAPIENS | -1 | 705 | 3idh | A | 462 | MET | p.Met462Ile | P35557 |
| HOMO SAPIENS | 1 | 509 | 4nef | A | 22 | LEU | p.Leu22Val | P41181 |
| HOMO SAPIENS | 4 | 509 | 4nef | A | 57 | GLN | p.Gln57Pro | P41181 |
| HOMO SAPIENS | 8 | 509 | 4nef | A | 64 | GLY | p.Gly64Arg | P41181 |
| HOMO SAPIENS | 7 | 509 | 4nef | A | 68 | ASN | p.Asn68Ser | P41181 |
| HOMO SAPIENS | 1 | 509 | 4nef | A | 100 | GLY | p.Gly100Val | P41181 |
| HOMO SAPIENS | 0 | 509 | 4nef | A | 125 | THR | p.Thr125Met | P41181 |
| HOMO SAPIENS | 0 | 509 | 4nef | A | 126 | THR | p.Thr126Met | P41181 |
| HOMO SAPIENS | 4 | 509 | 4nef | A | 146 | PHE | p.Phe146Leu | P41181 |
| HOMO SAPIENS | 0 | 509 | 4nef | A | 175 | GLY | p.Gly175Arg | P41181 |
| HOMO SAPIENS | 5 | 509 | 4nef | A | 181 | CYS | p.Cys181Trp | P41181 |

| | | | | | | | | | |
|---|---|---|---|---|---|---|---|---|---|
| HOMO SAPIENS | 9 | 509 | 4nef | A | | 187 | ARG | p.Arg187Cys | P41181 |
| HOMO SAPIENS | 0 | 509 | 4nef | A | | 190 | ALA | p.Ala190Thr | P41181 |
| HOMO SAPIENS | 0 | 509 | 4nef | A | | 216 | SER | p.Ser216Pro | P41181 |
| HOMO SAPIENS | 4 | 2242 | 1z70 | X | | 1155 | SER | p.Ser155Pro | Q8NBK3 |
| HOMO SAPIENS | 3 | 2242 | 1z70 | X | | 1218 | CYS | p.Cys218Tyr | Q8NBK3 |
| HOMO SAPIENS | 0 | 2242 | 1z70 | X | | 1279 | ALA | p.Ala279Val | Q8NBK3 |
| HOMO SAPIENS | -1 | 2242 | 1z70 | X | | 1336 | OCS | p.Cys336Arg | Q8NBK3 |
| HOMO SAPIENS | 4 | 2242 | 1z70 | X | | 1345 | ARG | p.Arg345Cys | Q8NBK3 |
| HOMO SAPIENS | 2 | 2242 | 1z70 | X | | 1348 | ALA | p.Ala348Pro | Q8NBK3 |
| HOMO SAPIENS | 5 | 2242 | 1z70 | X | | 1349 | ARG | p.Arg349Gln | Q8NBK3 |
| HOMO SAPIENS | 5 | 2242 | 1z70 | X | | 1349 | ARG | p.Arg349Trp | Q8NBK3 |
| HOMO SAPIENS | 3 | 2242 | 1z70 | X | | 1359 | SER | p.Ser359Ter | Q8NBK3 |
| HOMO SAPIENS | 5 | 8269 | 5wrr | B | | 136 | ARG | p.Arg136Ter | Q96MK3 |
| HOMO SAPIENS | 1 | 8269 | 5wrr | B | | 332 | LYS | p.Asn332Lys | Q96MK3 |
| HOMO SAPIENS | 3 | 8269 | 5wrr | B | | 478 | ARG | p.Arg478Ter | Q96MK3 |
| HOMO SAPIENS | 5 | 8269 | 5wrr | B | | 504 | THR | p.Thr504Ile | Q96MK3 |
| HOMO SAPIENS | 5 | 1411 | 4h1s | B | | 221 | SER | p.Ser221Ter | P21589 |
| HOMO SAPIENS | 1 | 1411 | 4h1s | B | | 358 | CYS | p.Cys358Tyr | P21589 |
| HOMO SAPIENS | 2 | 23 | 5vvd | B | | 298 | GLU | p.Glu298Asp | P29474 |
| HOMO SAPIENS | 0 | 19 | 5jlh | E | | 88 | THR | p.Thr89Ile | P63261 |
| HOMO SAPIENS | 6 | 19 | 5jlh | E | | 117 | LYS | p.Lys118Met | P63261 |
| HOMO SAPIENS | 5 | 19 | 5jlh | E | | 263 | PRO | p.Pro264Leu | P63261 |
| HOMO SAPIENS | 1 | 19 | 5jlh | E | | 277 | THR | p.Thr278Ile | P63261 |
| HOMO SAPIENS | 6 | 19 | 5jlh | E | | 331 | PRO | p.Pro332Ala | P63261 |
| HOMO SAPIENS | 5 | 19 | 5jlh | E | | 369 | VAL | p.Val370Ala | P63261 |
| HOMO SAPIENS | 0 | 14306 | 4w4o | C | | 92 | ARG | p.Arg92Ter | P12314 |
| HOMO SAPIENS | 3 | 603 | 1nuh | A | | 4 | THR | p.Thr5Ile | P06744 |
| HOMO SAPIENS | 2 | 603 | 1nuh | A | | 19 | HIS | p.His20Pro | P06744 |
| HOMO SAPIENS | 8 | 603 | 1nuh | A | | 158 | GLY | p.Gly159Ser | P06744 |
| HOMO SAPIENS | 6 | 603 | 1nuh | A | | 223 | THR | p.Thr224Met | P06744 |
| HOMO SAPIENS | 0 | 603 | 1nuh | A | | 338 | LEU | p.Leu339Pro | P06744 |
| HOMO SAPIENS | 6 | 603 | 1nuh | A | | 342 | GLN | p.Gln343Arg | P06744 |
| HOMO SAPIENS | 6 | 603 | 1nuh | A | | 374 | THR | p.Thr375Arg | P06744 |
| HOMO SAPIENS | 4 | 603 | 1nuh | A | | 524 | ILE | p.Ile525Thr | P06744 |
| HOMO SAPIENS | 4 | 603 | 1nuh | A | | 538 | ASP | p.Asp539Asn | P06744 |
| HOMO SAPIENS | 1 | 3107 | 3kzj | A | | 1197 | VAL | p.Val1197Ala | P08603 |
| HOMO SAPIENS | 9 | 1748 | 2noh | A | | 46 | ARG | p.Arg46Gln | O15527 |
| HOMO SAPIENS | 2 | 11750 | 3s4y | A | | 33 | ARG | p.Arg33His | Q9H3S4 |
| HOMO SAPIENS | 2 | 11750 | 3s4y | A | | 40 | LEU | p.Leu40Pro | Q9H3S4 |
| HOMO SAPIENS | 4 | 11750 | 3s4y | A | | 142 | LEU | p.Leu142Phe | Q9H3S4 |
| HOMO SAPIENS | 5 | 11750 | 3s4y | A | | 219 | ASN | p.Asn219Ser | Q9H3S4 |
| HOMO SAPIENS | 1 | 2591 | 5vnj | A | | 382 | PHE | p.Phe382Leu | Q15436 |
| HOMO SAPIENS | 2 | 2591 | 5vnj | A | | 702 | MET | p.Met702Val | Q15436 |
| HOMO SAPIENS | 1 | 2426 | 5m8t | A | | 87 | ARG | p.Arg87Trp | P17643 |
| HOMO SAPIENS | 1 | 2426 | 5m8t | A | | 166 | SER | p.Ser166Ter | P17643 |
| HOMO SAPIENS | 0 | 2426 | 5m8t | A | | 356 | ARG | p.Arg356Gln | P17643 |
| HOMO SAPIENS | 2 | 2426 | 5m8t | A | | 374 | SER | p.Arg374Ter | P17643 |
| HOMO SAPIENS | 6 | 2426 | 5m8t | A | | 378 | ASN | p.Asn378Ser | P17643 |
| HOMO SAPIENS | 6 | 2426 | 5m8t | A | | 382 | LEU | p.Leu382Pro | P17643 |
| HOMO SAPIENS | 7 | 10097 | 4n6v | | 1 | 46 | GLN | p.Gln46Ter | P04080 |
| HOMO SAPIENS | 3 | 10097 | 4n6v | | 1 | 65 | VAL | p.Val65Ile | P04080 |
| HOMO SAPIENS | 8 | 10097 | 4n6v | | 1 | 68 | ARG | p.Arg68Ter | P04080 |
| HOMO SAPIENS | 2 | 859 | 4z9a | A | | 105 | GLN | p.Gln106Arg | P24666 |
| HOMO SAPIENS | 8 | 1420 | 5nan | F | | 95 | SER | p.Ser95Leu | Q96PD4 |

| | | | | | | | | |
|---|---|---|---|---|---|---|---|---|
| HOMO SAPIENS | 6 | 1420 | 5nan | F | 126 | GLU | p.Glu126Gly | Q96PD4 |
| HOMO SAPIENS | 8 | 1420 | 5nan | F | 155 | VAL | p.Val155Ile | Q96PD4 |
| HOMO SAPIENS | 6 | 1420 | 5nan | F | 157 | PRO | p.Pro157Ser | Q96PD4 |
| HOMO SAPIENS | -1 | 1420 | 5nan | F | 161 | HIS | p.His161Arg | Q96PD4 |
| HOMO SAPIENS | 3 | 288 | 5i96 | B | 140 | GLN | p.Arg140Gly | P48735 |
| HOMO SAPIENS | 3 | 288 | 5i96 | B | 140 | GLN | p.Arg140Gln | P48735 |
| HOMO SAPIENS | 2 | 288 | 5i96 | B | 261 | ARG | p.Arg261His | P48735 |
| HOMO SAPIENS | 2 | 288 | 5i96 | B | 435 | THR | p.Thr435Met | P48735 |
| HOMO SAPIENS | 9 | 120 | 1tup | B | 107 | TYR | p.Tyr107His | P04637 |
| HOMO SAPIENS | 8 | 120 | 1tup | B | 110 | ARG | p.Arg110Leu | P04637 |
| HOMO SAPIENS | 5 | 120 | 1tup | B | 120 | LYS | p.Lys120Glu | P04637 |
| HOMO SAPIENS | 4 | 120 | 1tup | B | 133 | MET | p.Met133Thr | P04637 |
| HOMO SAPIENS | 3 | 120 | 1tup | B | 134 | PHE | p.Phe134Cys | P04637 |
| HOMO SAPIENS | 8 | 120 | 1tup | B | 136 | GLN | p.Gln136His | P04637 |
| HOMO SAPIENS | 9 | 120 | 1tup | B | 138 | ALA | p.Ala138Pro | P04637 |
| HOMO SAPIENS | 9 | 120 | 1tup | B | 151 | PRO | p.Pro151Ser | P04637 |
| HOMO SAPIENS | 9 | 120 | 1tup | B | 152 | PRO | p.Pro152Leu | P04637 |
| HOMO SAPIENS | 9 | 120 | 1tup | B | 158 | ARG | p.Arg158Cys | P04637 |
| HOMO SAPIENS | 9 | 120 | 1tup | B | 158 | ARG | p.Arg158His | P04637 |
| HOMO SAPIENS | 9 | 120 | 1tup | B | 175 | ARG | p.Arg175Cys | P04637 |
| HOMO SAPIENS | 9 | 120 | 1tup | B | 175 | ARG | p.Arg175Leu | P04637 |
| HOMO SAPIENS | 8 | 120 | 1tup | B | 181 | ARG | p.Arg181Leu | P04637 |
| HOMO SAPIENS | 9 | 120 | 1tup | B | 189 | ALA | p.Ala189Val | P04637 |
| HOMO SAPIENS | 9 | 120 | 1tup | B | 196 | ARG | p.Arg196Gly | P04637 |
| HOMO SAPIENS | 9 | 120 | 1tup | B | 213 | ARG | p.Arg213Gln | P04637 |
| HOMO SAPIENS | 9 | 120 | 1tup | B | 213 | ARG | p.Arg213Ter | P04637 |
| HOMO SAPIENS | 9 | 120 | 1tup | B | 214 | HIS | p.His214Gln | P04637 |
| HOMO SAPIENS | 9 | 120 | 1tup | B | 220 | TYR | p.Tyr220Cys | P04637 |
| HOMO SAPIENS | 0 | 120 | 1tup | B | 234 | TYR | p.Tyr234Cys | P04637 |
| HOMO SAPIENS | 9 | 120 | 1tup | B | 237 | MET | p.Met237Ile | P04637 |
| HOMO SAPIENS | 0 | 120 | 1tup | B | 238 | CYS | p.Cys238Tyr | P04637 |
| HOMO SAPIENS | 9 | 120 | 1tup | B | 241 | SER | p.Ser241Phe | P04637 |
| HOMO SAPIENS | 9 | 120 | 1tup | B | 242 | CYS | p.Cys242Ser | P04637 |
| HOMO SAPIENS | 9 | 120 | 1tup | B | 245 | GLY | p.Gly245Cys | P04637 |
| HOMO SAPIENS | 9 | 120 | 1tup | B | 245 | GLY | p.Gly245Asp | P04637 |
| HOMO SAPIENS | 9 | 120 | 1tup | B | 248 | ARG | p.Arg248Leu | P04637 |
| HOMO SAPIENS | 9 | 120 | 1tup | B | 248 | ARG | p.Arg248Trp | P04637 |
| HOMO SAPIENS | 9 | 120 | 1tup | B | 251 | ILE | p.Ile251Ser | P04637 |
| HOMO SAPIENS | 9 | 120 | 1tup | B | 252 | LEU | p.Leu252Pro | P04637 |
| HOMO SAPIENS | 9 | 120 | 1tup | B | 258 | GLU | p.Glu258Lys | P04637 |
| HOMO SAPIENS | 8 | 120 | 1tup | B | 272 | VAL | p.Val272Leu | P04637 |
| HOMO SAPIENS | 9 | 120 | 1tup | B | 273 | ARG | p.Arg273Cys | P04637 |
| HOMO SAPIENS | 9 | 120 | 1tup | B | 273 | ARG | p.Arg273His | P04637 |
| HOMO SAPIENS | 9 | 120 | 1tup | B | 277 | CYS | p.Cys277Phe | P04637 |
| HOMO SAPIENS | 9 | 120 | 1tup | B | 280 | ARG | p.Arg280Ile | P04637 |
| HOMO SAPIENS | 9 | 120 | 1tup | B | 282 | ARG | p.Arg282Trp | P04637 |
| HOMO SAPIENS | 9 | 120 | 1tup | B | 283 | ARG | p.Arg283His | P04637 |
| HOMO SAPIENS | 9 | 120 | 1tup | B | 285 | GLU | p.Glu285Val | P04637 |
| HOMO SAPIENS | -1 | 120 | 1tup | B | 292 | LYS | p.Lys292Ile | P04637 |
| HOMO SAPIENS | -1 | 120 | 1tup | B | 298 | GLU | p.Glu298Lys | P04637 |
| HOMO SAPIENS | -1 | 2791 | 4ygb | A | 39 | VAL | p.Val39Ala | P49257 |
| HOMO SAPIENS | 2 | 106 | 4x6j | A | 32 | GLY | p.Gly146Arg | P43235 |
| HOMO SAPIENS | 0 | 106 | 4x6j | A | 76 | GLN | p.Gln190Ter | P43235 |
| HOMO SAPIENS | 7 | 106 | 4x6j | A | 198 | ARG | p.Arg312Ter | P43235 |

| | | | | | | | | |
|---|---|---|---|---|---|---|---|---|
| HOMO SAPIENS | 5 | 490 | 2hlo | B | 172 | LEU | p.Leu202Gln | P02675 |
| HOMO SAPIENS | 0 | 490 | 2hlo | B | 353 | LEU | p.Leu383Arg | P02675 |
| HOMO SAPIENS | 4 | 490 | 2hlo | B | 400 | GLY | p.Gly430Asp | P02675 |
| HOMO SAPIENS | 7 | 490 | 2hlo | B | 448 | ARG | p.Arg478Lys | P02675 |
| HOMO SAPIENS | 1 | 289 | 4p13 | B | 18 | GLU | p.Glu43Lys | P11310 |
| HOMO SAPIENS | 2 | 289 | 4p13 | B | 20 | GLN | p.Gln45Arg | P11310 |
| HOMO SAPIENS | 5 | 289 | 4p13 | B | 28 | ARG | p.Arg53Cys | P11310 |
| HOMO SAPIENS | 1 | 289 | 4p13 | B | 42 | TYR | p.Tyr67His | P11310 |
| HOMO SAPIENS | 1 | 289 | 4p13 | B | 53 | ILE | p.Ile78Thr | P11310 |
| HOMO SAPIENS | 1 | 289 | 4p13 | B | 59 | LEU | p.Leu84Phe | P11310 |
| HOMO SAPIENS | 2 | 289 | 4p13 | B | 96 | THR | p.Thr121Ile | P11310 |
| HOMO SAPIENS | 0 | 289 | 4p13 | B | 124 | MET | p.Met149Ile | P11310 |
| HOMO SAPIENS | 5 | 289 | 4p13 | B | 143 | ASP | p.Asp168Ala | P11310 |
| HOMO SAPIENS | 7 | 289 | 4p13 | B | 168 | THR | p.Thr193Ala | P11310 |
| HOMO SAPIENS | 1 | 289 | 4p13 | B | 170 | GLY | p.Gly195Arg | P11310 |
| HOMO SAPIENS | 2 | 289 | 4p13 | B | 181 | ARG | p.Arg206Cys | P11310 |
| HOMO SAPIENS | 2 | 289 | 4p13 | B | 181 | ARG | p.Arg206His | P11310 |
| HOMO SAPIENS | 8 | 289 | 4p13 | B | 203 | THR | p.Thr228Ile | P11310 |
| HOMO SAPIENS | 2 | 289 | 4p13 | B | 208 | ILE | p.Ile233Asn | P11310 |
| HOMO SAPIENS | 1 | 289 | 4p13 | B | 219 | CYS | p.Cys244Arg | P11310 |
| HOMO SAPIENS | 6 | 289 | 4p13 | B | 220 | SER | p.Ser245Leu | P11310 |
| HOMO SAPIENS | 5 | 289 | 4p13 | B | 228 | GLU | p.Glu253Lys | P11310 |
| HOMO SAPIENS | 2 | 289 | 4p13 | B | 241 | ASP | p.Asp266Gly | P11310 |
| HOMO SAPIENS | 1 | 289 | 4p13 | B | 242 | GLY | p.Gly267Arg | P11310 |
| HOMO SAPIENS | 7 | 289 | 4p13 | B | 256 | ARG | p.Arg281Thr | P11310 |
| HOMO SAPIENS | 4 | 289 | 4p13 | B | 269 | ARG | p.Arg294Thr | P11310 |
| HOMO SAPIENS | 4 | 289 | 4p13 | B | 304 | GLU | p.Lys329Glu | P11310 |
| HOMO SAPIENS | 1 | 289 | 4p13 | B | 324 | ARG | p.Arg349Ter | P11310 |
| HOMO SAPIENS | 3 | 289 | 4p13 | B | 326 | THR | p.Thr351Ile | P11310 |
| HOMO SAPIENS | 7 | 289 | 4p13 | B | 350 | ILE | p.Ile375Thr | P11310 |
| HOMO SAPIENS | 3 | 289 | 4p13 | B | 359 | GLU | p.Glu384Ter | P11310 |
| HOMO SAPIENS | 1 | 4563 | 3i2b | A | 16 | ARG | p.Arg16Cys | Q03393 |
| HOMO SAPIENS | 5 | 4563 | 3i2b | A | 25 | ARG | p.Arg25Gln | Q03393 |
| HOMO SAPIENS | 1 | 4563 | 3i2b | A | 47 | ASN | p.Asn47Asp | Q03393 |
| HOMO SAPIENS | 8 | 4563 | 3i2b | A | 52 | ASN | p.Asn52Ser | Q03393 |
| HOMO SAPIENS | 1 | 4563 | 3i2b | A | 56 | VAL | p.Val56Met | Q03393 |
| HOMO SAPIENS | 1 | 4563 | 3i2b | A | 87 | PRO | p.Pro87Ser | Q03393 |
| HOMO SAPIENS | 1 | 4563 | 3i2b | A | 96 | ASP | p.Asp96Asn | Q03393 |
| HOMO SAPIENS | 0 | 4563 | 3i2b | A | 99 | TYR | p.Tyr99Ter | Q03393 |
| HOMO SAPIENS | 2 | 4563 | 3i2b | A | 116 | ASP | p.Asp116Gly | Q03393 |
| HOMO SAPIENS | 2 | 41 | 3exx | A | 1 | GLY | p.Gly90Cys | P01308 |
| HOMO SAPIENS | 5 | 41 | 3exx | A | 3 | VAL | p.Val92Leu | P01308 |
| HOMO SAPIENS | 4 | 41 | 3exx | A | 7 | CYS | p.Cys96Ser | P01308 |
| HOMO SAPIENS | 9 | 41 | 3exx | A | 19 | TYR | p.Tyr108Cys | P01308 |
| HOMO SAPIENS | 9 | 131 | 4mjo | D | 118 | ASP | p.Asp119Asn | P09467 |
| HOMO SAPIENS | 0 | 131 | 4mjo | D | 163 | GLY | p.Gly164Ser | P09467 |
| HOMO SAPIENS | 0 | 131 | 4mjo | D | 176 | ALA | p.Ala177Asp | P09467 |
| HOMO SAPIENS | 5 | 131 | 4mjo | D | 193 | PHE | p.Phe194Ser | P09467 |
| HOMO SAPIENS | 4 | 131 | 4mjo | D | 217 | ARG | p.Arg218Lys | P09467 |
| HOMO SAPIENS | 4 | 131 | 4mjo | D | 223 | VAL | p.Val224Ile | P09467 |
| HOMO SAPIENS | 4 | 131 | 4mjo | D | 232 | PHE | p.Phe233Ile | P09467 |
| HOMO SAPIENS | 5 | 131 | 4mjo | D | 243 | ARG | p.Arg244Trp | P09467 |
| HOMO SAPIENS | 0 | 131 | 4mjo | D | 283 | PRO | p.Pro284Arg | P09467 |
| HOMO SAPIENS | 0 | 131 | 4mjo | D | 288 | MET | p.Met289Leu | P09467 |

| | | | | | | | | |
|---|---|---|---|---|---|---|---|---|
| HOMO SAPIENS | 0 | 131 | 4mjo | D | 323 | ASP | p.Asp324Asn | P09467 |
| HOMO SAPIENS | -1 | 13122 | 3hfc | A | 583 | ARG | p.Arg583Cys | P51787 |
| HOMO SAPIENS | 1 | 13122 | 3hfc | A | 587 | THR | p.Thr587Met | P51787 |
| HOMO SAPIENS | 0 | 13122 | 3hfc | A | 589 | GLY | p.Gly589Asp | P51787 |
| HOMO SAPIENS | 4 | 3002 | 3nbq | B | 94 | ARG | p.Arg94Cys | Q16831 |
| HOMO SAPIENS | 1 | 3616 | 3dbp | A | 97 | ILE | p.Ile286Asn | P11172 |
| HOMO SAPIENS | 0 | 3616 | 3dbp | A | 240 | GLY | p.Gly429Arg | P11172 |
| HOMO SAPIENS | 0 | 3616 | 3dbp | A | 257 | ILE | p.Ile446Val | P11172 |
| HOMO SAPIENS | 1 | 584 | 1drf | A | 79 | LEU | p.Leu80Phe | P00374 |
| HOMO SAPIENS | 3 | 1609 | 3qa2 | B | 49 | VAL | p.Val49Ile | P50053 |
| HOMO SAPIENS | 0 | 20560 | 5mug | A | 101 | HIS | p.His101Gln | P49638 |
| HOMO SAPIENS | 2 | 20560 | 5mug | A | 120 | ALA | p.Ala120Thr | P49638 |
| HOMO SAPIENS | 3 | 20560 | 5mug | A | 192 | ARG | p.Arg192His | P49638 |
| HOMO SAPIENS | 3 | 20560 | 5mug | A | 221 | ARG | p.Arg221Trp | P49638 |
| HOMO SAPIENS | 4 | 256 | 3vtv | A | 65 | THR | p.Glu50Lys | Q9GZQ8 |
| HOMO SAPIENS | 4 | 256 | 3vtv | A | 113 | VAL | p.Met98Lys | Q9GZQ8 |
| HOMO SAPIENS | 4 | 5397 | 1m61 | A | 192 | ARG | p.Arg192Trp | P43403 |
| HOMO SAPIENS | 5 | 39881 | 1aut | L | 107 | HIS | p.His149Pro | P04070 |
| HOMO SAPIENS | 5 | 4010 | 4uf8 | I | 33 | ALA | p.Ala33Val | Q13501 |
| HOMO SAPIENS | 3 | 313 | 5w49 | A | 98 | PRO | p.Pro98Leu | P23526 |
| HOMO SAPIENS | 2 | 313 | 5w49 | A | 143 | TYR | p.Tyr143Cys | P23526 |
| HOMO SAPIENS | -1 | 1643 | 2m8t | A | 102 | PRO | p.Pro102Leu | P04156 |
| HOMO SAPIENS | -1 | 1643 | 2m8t | A | 105 | PRO | p.Pro105Leu | P04156 |
| HOMO SAPIENS | -1 | 1643 | 2m8t | A | 117 | ALA | p.Ala117Val | P04156 |
| HOMO SAPIENS | 5 | 1643 | 2m8t | A | 129 | MET | p.Met129Val | P04156 |
| HOMO SAPIENS | 5 | 1643 | 2m8t | A | 131 | GLY | p.Gly131Val | P04156 |
| HOMO SAPIENS | 2 | 1643 | 2m8t | A | 142 | GLY | p.Gly142Ser | P04156 |
| HOMO SAPIENS | 3 | 1643 | 2m8t | A | 171 | ASN | p.Asn171Ser | P04156 |
| HOMO SAPIENS | 9 | 1643 | 2m8t | A | 178 | ASP | p.Asp178Asn | P04156 |
| HOMO SAPIENS | 9 | 1643 | 2m8t | A | 180 | VAL | p.Val180Ile | P04156 |
| HOMO SAPIENS | 9 | 1643 | 2m8t | A | 183 | THR | p.Thr183Ala | P04156 |
| HOMO SAPIENS | 9 | 1643 | 2m8t | A | 187 | HIS | p.His187Arg | P04156 |
| HOMO SAPIENS | 8 | 1643 | 2m8t | A | 198 | PHE | p.Phe198Ser | P04156 |
| HOMO SAPIENS | 9 | 1643 | 2m8t | A | 200 | GLU | p.Glu200Lys | P04156 |
| HOMO SAPIENS | 9 | 1643 | 2m8t | A | 208 | ARG | p.Arg208His | P04156 |
| HOMO SAPIENS | 3 | 1643 | 2m8t | A | 210 | VAL | p.Val210Ile | P04156 |
| HOMO SAPIENS | 6 | 1643 | 2m8t | A | 217 | GLN | p.Gln217Arg | P04156 |
| HOMO SAPIENS | 1 | 29 | 5kos | A | 151 | GLY | p.Gly217Arg | P00797 |
| HOMO SAPIENS | 1 | 29 | 5kos | A | 164 | ARG | p.Arg230Lys | P00797 |
| HOMO SAPIENS | 3 | 504 | 4zkv | B | 37 | ARG | p.Arg37Pro | P49773 |
| HOMO SAPIENS | 2 | 504 | 4zkv | B | 62 | GLN | p.Gln62Ter | P49773 |
| HOMO SAPIENS | 2 | 504 | 4zkv | B | 84 | CYS | p.Cys84Arg | P49773 |
| HOMO SAPIENS | 4 | 504 | 4zkv | B | 93 | GLY | p.Gly93Asp | P49773 |
| HOMO SAPIENS | 9 | 504 | 4zkv | B | 112 | HIS | p.His112Asn | P49773 |
| HOMO SAPIENS | 0 | 74 | 5vwo | C | 54 | ASN | p.Asn54Lys | P04181 |
| HOMO SAPIENS | 0 | 74 | 5vwo | C | 55 | TYR | p.Tyr55His | P04181 |
| HOMO SAPIENS | 2 | 74 | 5vwo | C | 90 | GLN | p.Gln90Glu | P04181 |
| HOMO SAPIENS | 6 | 74 | 5vwo | C | 91 | GLY | p.Gly91Glu | P04181 |
| HOMO SAPIENS | 2 | 74 | 5vwo | C | 93 | CYS | p.Cys93Phe | P04181 |
| HOMO SAPIENS | 1 | 74 | 5vwo | C | 154 | ARG | p.Arg154Leu | P04181 |
| HOMO SAPIENS | 0 | 74 | 5vwo | C | 180 | ARG | p.Arg180Thr | P04181 |
| HOMO SAPIENS | 0 | 74 | 5vwo | C | 199 | PRO | p.Pro199Gln | P04181 |
| HOMO SAPIENS | 4 | 74 | 5vwo | C | 209 | TYR | p.Tyr209Ter | P04181 |
| HOMO SAPIENS | 0 | 74 | 5vwo | C | 226 | ALA | p.Ala226Val | P04181 |

| | | | | | | | | |
|---|---|---|---|---|---|---|---|---|
| HOMO SAPIENS | 5 | 74 | 5vwo | C | 233 | GLN | p.Gln233Arg | P04181 |
| HOMO SAPIENS | 5 | 74 | 5vwo | C | 241 | PRO | p.Pro241Leu | P04181 |
| HOMO SAPIENS | 2 | 74 | 5vwo | C | 245 | TYR | p.Tyr245Cys | P04181 |
| HOMO SAPIENS | 6 | 74 | 5vwo | C | 250 | ARG | p.Arg250Pro | P04181 |
| HOMO SAPIENS | 6 | 74 | 5vwo | C | 250 | ARG | p.Arg250Ter | P04181 |
| HOMO SAPIENS | 8 | 74 | 5vwo | C | 267 | THR | p.Thr267Ile | P04181 |
| HOMO SAPIENS | 0 | 74 | 5vwo | C | 270 | ALA | p.Ala270Pro | P04181 |
| HOMO SAPIENS | 0 | 74 | 5vwo | C | 271 | ARG | p.Arg271Lys | P04181 |
| HOMO SAPIENS | 1 | 74 | 5vwo | C | 299 | TYR | p.Tyr299Ter | P04181 |
| HOMO SAPIENS | 0 | 74 | 5vwo | C | 318 | GLU | p.Glu318Lys | P04181 |
| HOMO SAPIENS | 1 | 74 | 5vwo | C | 319 | HIS | p.His319Tyr | P04181 |
| HOMO SAPIENS | 1 | 74 | 5vwo | C | 331 | ARG | p.Arg331Ter | P04181 |
| HOMO SAPIENS | 0 | 74 | 5vwo | C | 332 | VAL | p.Val332Met | P04181 |
| HOMO SAPIENS | 1 | 74 | 5vwo | C | 353 | GLY | p.Gly353Asp | P04181 |
| HOMO SAPIENS | 2 | 74 | 5vwo | C | 375 | GLY | p.Gly375Ala | P04181 |
| HOMO SAPIENS | 0 | 74 | 5vwo | C | 394 | CYS | p.Cys394Arg | P04181 |
| HOMO SAPIENS | 0 | 74 | 5vwo | C | 402 | LEU | p.Leu402Pro | P04181 |
| HOMO SAPIENS | 0 | 74 | 5vwo | C | 436 | ILE | p.Ile436Asn | P04181 |
| HOMO SAPIENS | 7 | 24356 | 4az0 | A | 104 | VAL | p.Val132Met | P10619 |
| HOMO SAPIENS | 4 | 24356 | 4az0 | A | 221 | TYR | p.Tyr249Asn | P10619 |
| HOMO SAPIENS | 3 | 245 | 1rsz | A | 24 | ARG | p.Arg24Ter | P00491 |
| HOMO SAPIENS | 3 | 245 | 1rsz | A | 51 | SER | p.Gly51Ser | P00491 |
| HOMO SAPIENS | 1 | 245 | 1rsz | A | 58 | ARG | p.Arg58Ter | P00491 |
| HOMO SAPIENS | 4 | 245 | 1rsz | A | 89 | GLU | p.Glu89Lys | P00491 |
| HOMO SAPIENS | 6 | 245 | 1rsz | A | 128 | ASP | p.Asp128Gly | P00491 |
| HOMO SAPIENS | 0 | 245 | 1rsz | A | 174 | ALA | p.Ala174Pro | P00491 |
| HOMO SAPIENS | 8 | 245 | 1rsz | A | 192 | TYR | p.Tyr192Cys | P00491 |
| HOMO SAPIENS | 8 | 245 | 1rsz | A | 234 | ARG | p.Arg234Pro | P00491 |
| HOMO SAPIENS | 0 | 37 | 5tby | E | 22 | GLU | p.Glu22Lys | P10916 |
| HOMO SAPIENS | 1 | 37 | 5tby | E | 86 | PHE | p.Phe86Ser | P10916 |
| HOMO SAPIENS | 9 | 71 | 5m9j | A | 12 | GLN | p.Gln36Leu | P03950 |
| HOMO SAPIENS | 3 | 71 | 5m9j | A | 17 | LYS | p.Lys41Glu | P03950 |
| HOMO SAPIENS | 3 | 71 | 5m9j | A | 17 | LYS | p.Lys41Ile | P03950 |
| HOMO SAPIENS | 4 | 71 | 5m9j | A | 28 | SER | p.Ser52Asn | P03950 |
| HOMO SAPIENS | 7 | 71 | 5m9j | A | 31 | ARG | p.Arg55Lys | P03950 |
| HOMO SAPIENS | 9 | 71 | 5m9j | A | 39 | CYS | p.Cys63Trp | P03950 |
| HOMO SAPIENS | 9 | 71 | 5m9j | A | 40 | LYS | p.Lys64Ile | P03950 |
| HOMO SAPIENS | 2 | 71 | 5m9j | A | 60 | GLU | p.Lys84Glu | P03950 |
| HOMO SAPIENS | 0 | 71 | 5m9j | A | 112 | PRO | p.Pro136Leu | P03950 |
| HOMO SAPIENS | 9 | 71 | 5m9j | A | 113 | VAL | p.Val137Ile | P03950 |
| HOMO SAPIENS | 3 | 326 | 5dqi | A | 209 | GLY | p.Gly209Val | Q9Y253 |
| HOMO SAPIENS | 1 | 326 | 5dqi | A | 242 | SER | p.Ser242Ter | Q9Y253 |
| HOMO SAPIENS | 1 | 326 | 5dqi | A | 303 | ARG | p.Arg303Ter | Q9Y253 |
| HOMO SAPIENS | 1 | 8203 | 3iu3 | K | 132 | VAL | p.Val153Ile | P01589 |
| HOMO SAPIENS | 0 | 4995 | 5ffo | H | 234 | THR | p.Thr263Ile | P01137 |
| HOMO SAPIENS | 5 | 38609 | 2ruk | A | 49 | ASP | p.Asp49His | P04637 |
| HOMO SAPIENS | 5 | 38609 | 2ruk | A | 58 | PRO | p.Pro58Arg | P04637 |
| HOMO SAPIENS | 0 | 1607 | 6pah | A | 117 | THR | p.Thr117Ile | P00439 |
| HOMO SAPIENS | 3 | 1607 | 6pah | A | 145 | ASP | p.Asp145Val | P00439 |
| HOMO SAPIENS | 5 | 1607 | 6pah | A | 158 | ARG | p.Arg158Gln | P00439 |
| HOMO SAPIENS | 5 | 1607 | 6pah | A | 158 | ARG | p.Arg158Trp | P00439 |
| HOMO SAPIENS | 5 | 1607 | 6pah | A | 169 | ARG | p.Arg169His | P00439 |
| HOMO SAPIENS | 5 | 1607 | 6pah | A | 169 | ARG | p.Arg169Cys | P00439 |
| HOMO SAPIENS | 3 | 1607 | 6pah | A | 176 | ARG | p.Arg176Leu | P00439 |

| | | | | | | | | |
|---|---|---|---|---|---|---|---|---|
| HOMO SAPIENS | 3 | 1607 | 6pah | A | 177 | VAL | p.Val177Leu | P00439 |
| HOMO SAPIENS | 1 | 1607 | 6pah | A | 194 | LEU | p.Leu194Pro | P00439 |
| HOMO SAPIENS | 1 | 1607 | 6pah | A | 204 | TYR | p.Tyr204Cys | P00439 |
| HOMO SAPIENS | 1 | 1607 | 6pah | A | 204 | TYR | p.Tyr204Ter | P00439 |
| HOMO SAPIENS | 2 | 1607 | 6pah | A | 230 | VAL | p.Val230Ile | P00439 |
| HOMO SAPIENS | 4 | 1607 | 6pah | A | 232 | GLN | p.Gln232Ter | P00439 |
| HOMO SAPIENS | 4 | 1607 | 6pah | A | 241 | ARG | p.Arg241Cys | P00439 |
| HOMO SAPIENS | 4 | 1607 | 6pah | A | 241 | ARG | p.Arg241His | P00439 |
| HOMO SAPIENS | 5 | 1607 | 6pah | A | 243 | ARG | p.Arg243Gln | P00439 |
| HOMO SAPIENS | 5 | 1607 | 6pah | A | 243 | ARG | p.Arg243Ter | P00439 |
| HOMO SAPIENS | 7 | 1607 | 6pah | A | 249 | LEU | p.Leu249Phe | P00439 |
| HOMO SAPIENS | 5 | 1607 | 6pah | A | 252 | ARG | p.Arg252Gly | P00439 |
| HOMO SAPIENS | 5 | 1607 | 6pah | A | 261 | ARG | p.Arg261Ter | P00439 |
| HOMO SAPIENS | 4 | 1607 | 6pah | A | 267 | GLN | p.Gln267Leu | P00439 |
| HOMO SAPIENS | 1 | 1607 | 6pah | A | 272 | GLY | p.Gly272Ter | P00439 |
| HOMO SAPIENS | 2 | 1607 | 6pah | A | 277 | TYR | p.Tyr277Asp | P00439 |
| HOMO SAPIENS | 7 | 1607 | 6pah | A | 280 | GLU | p.Glu280Lys | P00439 |
| HOMO SAPIENS | 8 | 1607 | 6pah | A | 281 | PRO | p.Pro281Leu | P00439 |
| HOMO SAPIENS | 0 | 1607 | 6pah | A | 299 | PHE | p.Phe299Cys | P00439 |
| HOMO SAPIENS | 5 | 1607 | 6pah | A | 300 | ALA | p.Ala300Ser | P00439 |
| HOMO SAPIENS | 5 | 1607 | 6pah | A | 309 | ALA | p.Ala309Asp | P00439 |
| HOMO SAPIENS | 4 | 1607 | 6pah | A | 320 | LYS | p.Lys320Asn | P00439 |
| HOMO SAPIENS | 0 | 1607 | 6pah | A | 345 | ALA | p.Ala345Ser | P00439 |
| HOMO SAPIENS | 0 | 1607 | 6pah | A | 348 | LEU | p.Leu348Val | P00439 |
| HOMO SAPIENS | 6 | 1607 | 6pah | A | 356 | TYR | p.Tyr356Ter | P00439 |
| HOMO SAPIENS | 2 | 1607 | 6pah | A | 380 | THR | p.Thr380Met | P00439 |
| HOMO SAPIENS | 0 | 1607 | 6pah | A | 386 | TYR | p.Tyr386Cys | P00439 |
| HOMO SAPIENS | 5 | 1607 | 6pah | A | 390 | GLU | p.Glu390Gly | P00439 |
| HOMO SAPIENS | 5 | 1607 | 6pah | A | 403 | ALA | p.Ala403Val | P00439 |
| HOMO SAPIENS | 4 | 1607 | 6pah | A | 408 | ARG | p.Arg408Trp | P00439 |
| HOMO SAPIENS | 3 | 1607 | 6pah | A | 413 | ARG | p.Arg413Pro | P00439 |
| HOMO SAPIENS | 4 | 1607 | 6pah | A | 414 | TYR | p.Tyr414Cys | P00439 |
| HOMO SAPIENS | 4 | 1607 | 6pah | A | 414 | TYR | p.Tyr414His | P00439 |
| HOMO SAPIENS | 4 | 1607 | 6pah | A | 415 | ASP | p.Asp415Asn | P00439 |
| HOMO SAPIENS | -1 | 676 | 5dc5 | A | 7 | PRO | p.Pro7Leu | Q9BY41 |
| HOMO SAPIENS | 0 | 676 | 5dc5 | A | 311 | THR | p.Thr311Met | Q9BY41 |
| HOMO SAPIENS | 1 | 124 | 5ftk | E | 95 | ARG | p.Arg95Gly | P55072 |
| HOMO SAPIENS | 1 | 124 | 5ftk | E | 155 | ARG | p.Arg155Cys | P55072 |
| HOMO SAPIENS | 1 | 124 | 5ftk | E | 155 | ARG | p.Arg155His | P55072 |
| HOMO SAPIENS | 1 | 124 | 5ftk | E | 159 | ARG | p.Arg159His | P55072 |
| HOMO SAPIENS | 1 | 124 | 5ftk | E | 159 | ARG | p.Arg159Gly | P55072 |
| HOMO SAPIENS | 0 | 124 | 5ftk | E | 191 | ARG | p.Arg191Gln | P55072 |
| HOMO SAPIENS | 0 | 124 | 5ftk | E | 232 | ALA | p.Ala232Glu | P55072 |
| HOMO SAPIENS | 5 | 5442 | 1fgg | B | 223 | GLY | p.Gly223Ser | O94766 |
| HOMO SAPIENS | 0 | 5442 | 1fgg | B | 277 | ARG | p.Arg277Gln | O94766 |
| HOMO SAPIENS | 0 | 9072 | 2gys | B | 225 | GLU | p.Glu249Gln | P32927 |
| HOMO SAPIENS | 0 | 9072 | 2gys | B | 247 | SER | p.Ser271Ter | P32927 |
| HOMO SAPIENS | 0 | 9072 | 2gys | B | 294 | ASP | p.Asp312Asn | P32927 |
| HOMO SAPIENS | 0 | 1484 | 4zrj | A | 62 | PHE | p.Phe62Ser | P35240 |
| HOMO SAPIENS | 3 | 1484 | 4zrj | A | 205 | MET | p.Met205Val | P35240 |
| HOMO SAPIENS | 3 | 1484 | 4zrj | A | 301 | ILE | p.Ile301Met | P35240 |
| HOMO SAPIENS | 4 | 40923 | 2enq | A | 68 | ILE | p.Ile391Met | P42336 |
| HOMO SAPIENS | 1 | 6179 | 5ofv | A | 125 | THR | p.Thr125Met | O43175 |
| HOMO SAPIENS | 0 | 6179 | 5ofv | A | 135 | ARG | p.Arg135Trp | O43175 |

| | | | | | | | | |
|---|---|---|---|---|---|---|---|---|
| HOMO SAPIENS | 1 | 6179 | 5ofv | A | 163 | ARG | p.Arg163Gln | O43175 |
| HOMO SAPIENS | 9 | 6179 | 5ofv | A | 261 | VAL | p.Val261Met | O43175 |
| HOMO SAPIENS | 7 | 34118 | 2i5o | A | 24 | MET | p.Met647Leu | Q9Y253 |
| HOMO SAPIENS | 8 | 3234 | 5c84 | A | 311 | LYS | p.Lys311Asn | P98170 |
| HOMO SAPIENS | 3 | 3234 | 5c84 | A | 321 | ALA | p.Ala321Gly | P98170 |
| HOMO SAPIENS | -1 | 15129 | 1zy8 | K | 1 | GLY | p.Gly54Ser | O00330 |
| HOMO SAPIENS | -1 | 15129 | 1zy8 | K | 48 | THR | p.Thr101Ala | O00330 |
| HOMO SAPIENS | -1 | 15129 | 1zy8 | K | 105 | GLU | p.Glu158Asp | O00330 |
| HOMO SAPIENS | 8 | 15129 | 1zy8 | K | 136 | ARG | p.Arg189His | O00330 |
| HOMO SAPIENS | -1 | 15129 | 1zy8 | K | 195 | GLN | p.Gln248Ter | O00330 |
| HOMO SAPIENS | 8 | 202 | 4bzb | A | 123 | HIS | p.His123Pro | Q9Y3Z3 |
| HOMO SAPIENS | 4 | 202 | 4bzb | A | 145 | ARG | p.Arg145Ter | Q9Y3Z3 |
| HOMO SAPIENS | 6 | 202 | 4bzb | A | 164 | ARG | p.Arg164Ter | Q9Y3Z3 |
| HOMO SAPIENS | 3 | 202 | 4bzb | A | 201 | ILE | p.Ile201Asn | Q9Y3Z3 |
| HOMO SAPIENS | 0 | 202 | 4bzb | A | 209 | GLY | p.Gly209Ser | Q9Y3Z3 |
| HOMO SAPIENS | 0 | 202 | 4bzb | A | 254 | MET | p.Met254Val | Q9Y3Z3 |
| HOMO SAPIENS | 5 | 202 | 4bzb | A | 531 | ARG | p.Arg531Ser | Q9Y3Z3 |
| HOMO SAPIENS | 6 | 202 | 4bzb | A | 548 | GLN | p.Gln548Ter | Q9Y3Z3 |
| HOMO SAPIENS | 3 | 30947 | 5lgd | A | 90 | PRO | p.Pro90Ser | P16671 |
| HOMO SAPIENS | 6 | 30947 | 5lgd | A | 254 | PHE | p.Phe254Leu | P16671 |
| HOMO SAPIENS | 0 | 30947 | 5lgd | A | 413 | ILE | p.Ile413Leu | P16671 |
| HOMO SAPIENS | 2 | 3906 | 5jtc | A | 735 | ARG | p.Arg735Trp | Q12797 |
| HOMO SAPIENS | 1 | 45 | 5uyx | C | 146 | GLU | p.Glu146Val | P48643 |
| HOMO SAPIENS | 1 | 45 | 5uyx | C | 147 | HIS | p.His147Arg | P48643 |
| HOMO SAPIENS | 0 | 45 | 5uyx | C | 362 | ILE | p.Ile362Met | P48643 |
| HOMO SAPIENS | 8 | 39596 | 1mhn | A | 95 | GLY | p.Gly95Arg | Q16637 |
| HOMO SAPIENS | 4 | 39596 | 1mhn | A | 111 | ALA | p.Ala111Gly | Q16637 |
| HOMO SAPIENS | 9 | 39596 | 1mhn | A | 116 | ILE | p.Ile116Phe | Q16637 |
| HOMO SAPIENS | 1 | 39596 | 1mhn | A | 136 | GLN | p.Gln136Glu | Q16637 |
| HOMO SAPIENS | 1 | 6408 | 5xjy | A | 85 | PRO | p.Pro85Leu | O95477 |
| HOMO SAPIENS | 0 | 6408 | 5xjy | A | 219 | ARG | p.Arg219Lys | O95477 |
| HOMO SAPIENS | 0 | 6408 | 5xjy | A | 230 | ARG | p.Arg230Cys | O95477 |
| HOMO SAPIENS | 1 | 6408 | 5xjy | A | 281 | MET | p.Met281Thr | O95477 |
| HOMO SAPIENS | -1 | 6408 | 5xjy | A | 343 | ALA | p.Ala343Gly | O95477 |
| HOMO SAPIENS | 1 | 6408 | 5xjy | A | 399 | VAL | p.Val399Ala | O95477 |
| HOMO SAPIENS | 1 | 6408 | 5xjy | A | 496 | ARG | p.Arg496Trp | O95477 |
| HOMO SAPIENS | 1 | 6408 | 5xjy | A | 590 | TRP | p.Trp590Ser | O95477 |
| HOMO SAPIENS | 1 | 6408 | 5xjy | A | 638 | ARG | p.Arg638Gln | O95477 |
| HOMO SAPIENS | 4 | 6408 | 5xjy | A | 697 | ALA | p.Ala697Thr | O95477 |
| HOMO SAPIENS | 1 | 6408 | 5xjy | A | 771 | VAL | p.Val771Met | O95477 |
| HOMO SAPIENS | 1 | 6408 | 5xjy | A | 774 | THR | p.Thr774Pro | O95477 |
| HOMO SAPIENS | 1 | 6408 | 5xjy | A | 776 | LYS | p.Lys776Asn | O95477 |
| HOMO SAPIENS | 0 | 6408 | 5xjy | A | 825 | VAL | p.Val825Ile | O95477 |
| HOMO SAPIENS | -1 | 6408 | 5xjy | A | 868 | GLU | p.Glu868Lys | O95477 |
| HOMO SAPIENS | -1 | 6408 | 5xjy | A | 883 | ILE | p.Ile883Met | O95477 |
| HOMO SAPIENS | 0 | 6408 | 5xjy | A | 935 | ASN | p.Asn935Ser | O95477 |
| HOMO SAPIENS | 0 | 6408 | 5xjy | A | 935 | ASN | p.Asn935His | O95477 |
| HOMO SAPIENS | 0 | 6408 | 5xjy | A | 937 | ALA | p.Ala937Val | O95477 |
| HOMO SAPIENS | 0 | 6408 | 5xjy | A | 1018 | ASP | p.Asp1018Gly | O95477 |
| HOMO SAPIENS | 0 | 6408 | 5xjy | A | 1096 | VAL | p.Val1096Ile | O95477 |
| HOMO SAPIENS | 0 | 6408 | 5xjy | A | 1099 | ASP | p.Asp1099Tyr | O95477 |
| HOMO SAPIENS | 0 | 6408 | 5xjy | A | 1172 | GLU | p.Glu1172Asp | O95477 |
| HOMO SAPIENS | 0 | 6408 | 5xjy | A | 1181 | SER | p.Ser1181Phe | O95477 |
| HOMO SAPIENS | 0 | 6408 | 5xjy | A | 1182 | ALA | p.Ala1182Thr | O95477 |

| | | | | | | | | |
|---|---|---|---|---|---|---|---|---|
| HOMO SAPIENS | -1 | 6408 | 5xjy | A | 1289 | ASP | p.Asp1289Asn | O95477 |
| HOMO SAPIENS | 0 | 6408 | 5xjy | A | 1399 | THR | p.Thr1399Ser | O95477 |
| HOMO SAPIENS | 1 | 6408 | 5xjy | A | 1408 | LEU | p.Leu1408Phe | O95477 |
| HOMO SAPIENS | 0 | 6408 | 5xjy | A | 1587 | LYS | p.Lys1587Arg | O95477 |
| HOMO SAPIENS | 1 | 6408 | 5xjy | A | 1674 | VAL | p.Val1674Ile | O95477 |
| HOMO SAPIENS | 1 | 6408 | 5xjy | A | 1680 | ARG | p.Arg1680Trp | O95477 |
| HOMO SAPIENS | 1 | 6408 | 5xjy | A | 1680 | ARG | p.Arg1680Gln | O95477 |
| HOMO SAPIENS | 0 | 6408 | 5xjy | A | 1800 | ASN | p.Asn1800His | O95477 |
| HOMO SAPIENS | -1 | 6408 | 5xjy | A | 1925 | ARG | p.Arg1925Gln | O95477 |
| HOMO SAPIENS | 0 | 6408 | 5xjy | A | 2009 | PHE | p.Phe2009Ser | O95477 |
| HOMO SAPIENS | -1 | 6408 | 5xjy | A | 2243 | ASP | p.Asp2243Glu | O95477 |
| HOMO SAPIENS | 4 | 23760 | 2mut | A | 231 | LEU | p.Phe231Leu | P07992 |
| HOMO SAPIENS | 6 | 1460 | 2hre | B | 96 | ARG | p.Arg96Gln | P22830 |
| HOMO SAPIENS | 8 | 1460 | 2hre | B | 129 | GLY | p.Gly129Arg | P22830 |
| HOMO SAPIENS | 9 | 1460 | 2hre | B | 172 | GLU | p.Glu172Lys | P22830 |
| HOMO SAPIENS | 4 | 1460 | 2hre | B | 285 | GLN | p.Gln285Arg | P22830 |
| HOMO SAPIENS | 5 | 1460 | 2hre | B | 362 | VAL | p.Val362Gly | P22830 |
| HOMO SAPIENS | 6 | 1460 | 2hre | B | 411 | CYS | p.Cys411Gly | P22830 |
| HOMO SAPIENS | 6 | 1460 | 2hre | B | 417 | PHE | p.Phe417Ser | P22830 |
| HOMO SAPIENS | 0 | 2210 | 5bug | A | 19 | ASP | p.Asp19Asn | P60484 |
| HOMO SAPIENS | 0 | 2210 | 5bug | A | 35 | MET | p.Met35Arg | P60484 |
| HOMO SAPIENS | 1 | 2210 | 5bug | A | 61 | HIS | p.His61Asp | P60484 |
| HOMO SAPIENS | 0 | 2210 | 5bug | A | 68 | TYR | p.Tyr68His | P60484 |
| HOMO SAPIENS | 8 | 2210 | 5bug | A | 70 | LEU | p.Leu70Pro | P60484 |
| HOMO SAPIENS | 6 | 2210 | 5bug | A | 93 | HIS | p.His93Arg | P60484 |
| HOMO SAPIENS | 1 | 2210 | 5bug | A | 112 | LEU | p.Leu112Pro | P60484 |
| HOMO SAPIENS | 0 | 2210 | 5bug | A | 121 | ALA | p.Ala121Gly | P60484 |
| HOMO SAPIENS | 3 | 2210 | 5bug | A | 123 | HIS | p.His123Arg | P60484 |
| HOMO SAPIENS | 7 | 2210 | 5bug | A | 124 | CYS | p.Cys124Arg | P60484 |
| HOMO SAPIENS | 9 | 2210 | 5bug | A | 129 | GLY | p.Gly129Glu | P60484 |
| HOMO SAPIENS | 9 | 2210 | 5bug | A | 130 | ARG | p.Arg130Gln | P60484 |
| HOMO SAPIENS | 9 | 2210 | 5bug | A | 130 | ARG | p.Arg130Gly | P60484 |
| HOMO SAPIENS | 0 | 2210 | 5bug | A | 132 | GLY | p.Gly132Val | P60484 |
| HOMO SAPIENS | 0 | 2210 | 5bug | A | 170 | SER | p.Ser170Arg | P60484 |
| HOMO SAPIENS | 1 | 2210 | 5bug | A | 173 | ARG | p.Arg173His | P60484 |
| HOMO SAPIENS | 0 | 2210 | 5bug | A | 217 | VAL | p.Val217Ile | P60484 |
| HOMO SAPIENS | 0 | 2210 | 5bug | A | 241 | PHE | p.Phe241Ser | P60484 |
| HOMO SAPIENS | 0 | 2210 | 5bug | A | 250 | CYS | p.Cys250Ter | P60484 |
| HOMO SAPIENS | 1 | 2210 | 5bug | A | 252 | ASP | p.Asp252Gly | P60484 |
| HOMO SAPIENS | 3 | 123 | 2ibu | A | 93 | ASN | p.Asn93Ser | P24752 |
| HOMO SAPIENS | 0 | 123 | 2ibu | A | 145 | GLN | p.Gln145Glu | P24752 |
| HOMO SAPIENS | 0 | 123 | 2ibu | A | 152 | GLY | p.Gly152Ala | P24752 |
| HOMO SAPIENS | 2 | 123 | 2ibu | A | 158 | ASN | p.Asn158Asp | P24752 |
| HOMO SAPIENS | 2 | 123 | 2ibu | A | 158 | ASN | p.Asn158Ser | P24752 |
| HOMO SAPIENS | 5 | 123 | 2ibu | A | 183 | GLY | p.Gly183Arg | P24752 |
| HOMO SAPIENS | 3 | 123 | 2ibu | A | 272 | GLN | p.Gln272Ter | P24752 |
| HOMO SAPIENS | 3 | 123 | 2ibu | A | 312 | ILE | p.Ile312Thr | P24752 |
| HOMO SAPIENS | 1 | 123 | 2ibu | A | 333 | ALA | p.Ala333Pro | P24752 |
| HOMO SAPIENS | 5 | 123 | 2ibu | A | 379 | GLY | p.Gly379Val | P24752 |
| HOMO SAPIENS | 0 | 123 | 2ibu | A | 380 | ALA | p.Ala380Thr | P24752 |
| HOMO SAPIENS | 2 | 369 | 5nrw | A | 1175 | TYR | p.Tyr1175Cys | Q92793 |
| HOMO SAPIENS | 2 | 830 | 2vc2 | A | 116 | LEU | p.Leu147Val | P08514 |
| HOMO SAPIENS | 0 | 830 | 2vc2 | A | 183 | LEU | p.Leu214Pro | P08514 |
| HOMO SAPIENS | 4 | 830 | 2vc2 | A | 242 | GLY | p.Gly273Asp | P08514 |

| | | | | | | | | |
|---|---|---|---|---|---|---|---|---|
| HOMO SAPIENS | 6 | 830 | 2vc2 | A | 324 | GLU | p.Glu355Lys | P08514 |
| HOMO SAPIENS | -1 | 10353 | 5ejc | E | 989 | ALA | p.Ala989Ser | Q92574 |
| HOMO SAPIENS | 0 | 15448 | 3lgg | A | 21 | GLY | p.Gly47Arg | Q9NZK5 |
| HOMO SAPIENS | 0 | 15448 | 3lgg | A | 21 | GLY | p.Gly47Val | Q9NZK5 |
| HOMO SAPIENS | 9 | 15448 | 3lgg | A | 86 | HIS | p.His112Gln | Q9NZK5 |
| HOMO SAPIENS | 0 | 15448 | 3lgg | A | 143 | ARG | p.Arg169Gln | Q9NZK5 |
| HOMO SAPIENS | 3 | 15448 | 3lgg | A | 221 | ALA | p.Ala247Val | Q9NZK5 |
| HOMO SAPIENS | 0 | 15448 | 3lgg | A | 225 | PRO | p.Pro251Leu | Q9NZK5 |
| HOMO SAPIENS | 0 | 15448 | 3lgg | A | 283 | MET | p.Met309Ile | Q9NZK5 |
| HOMO SAPIENS | 2 | 15448 | 3lgg | A | 334 | THR | p.Thr360Ala | Q9NZK5 |
| HOMO SAPIENS | 7 | 15448 | 3lgg | A | 357 | GLY | p.Gly383Ser | Q9NZK5 |
| HOMO SAPIENS | 0 | 15448 | 3lgg | A | 427 | TYR | p.Tyr453Cys | Q9NZK5 |
| HOMO SAPIENS | 0 | 872 | 2y3i | A | 87 | LEU | p.Leu88Pro | P00558 |
| HOMO SAPIENS | 1 | 872 | 2y3i | A | 163 | ASP | p.Asp164Val | P00558 |
| HOMO SAPIENS | 4 | 872 | 2y3i | A | 205 | ARG | p.Arg206Pro | P00558 |
| HOMO SAPIENS | 4 | 872 | 2y3i | A | 267 | ASP | p.Asp268Asn | P00558 |
| HOMO SAPIENS | 8 | 872 | 2y3i | A | 284 | ASP | p.Asp285Val | P00558 |
| HOMO SAPIENS | 4 | 8 | 5jco | B | 188 | ILE | p.Ile188Leu | Q71U36 |
| HOMO SAPIENS | 7 | 8 | 5jco | B | 263 | PRO | p.Pro263Thr | Q71U36 |
| HOMO SAPIENS | 6 | 8 | 5jco | B | 264 | ARG | p.Arg264Cys | Q71U36 |
| HOMO SAPIENS | 7 | 8 | 5jco | B | 402 | ARG | p.Arg402His | Q71U36 |
| HOMO SAPIENS | 7 | 8 | 5jco | B | 419 | SER | p.Ser419Leu | Q71U36 |
| HOMO SAPIENS | -1 | 880 | 4nwf | A | 2 | THR | p.Thr2Ile | Q06124 |
| HOMO SAPIENS | 2 | 880 | 4nwf | A | 58 | ASN | p.Asn58Ser | Q06124 |
| HOMO SAPIENS | 4 | 880 | 4nwf | A | 61 | ASP | p.Asp61Gly | Q06124 |
| HOMO SAPIENS | 2 | 880 | 4nwf | A | 62 | TYR | p.Tyr62Asp | Q06124 |
| HOMO SAPIENS | 3 | 880 | 4nwf | A | 63 | TYR | p.Tyr63Cys | Q06124 |
| HOMO SAPIENS | 2 | 880 | 4nwf | A | 69 | GLU | p.Glu69Gln | Q06124 |
| HOMO SAPIENS | 2 | 880 | 4nwf | A | 72 | ALA | p.Ala72Gly | Q06124 |
| HOMO SAPIENS | 2 | 880 | 4nwf | A | 72 | ALA | p.Ala72Ser | Q06124 |
| HOMO SAPIENS | 3 | 880 | 4nwf | A | 73 | THR | p.Thr73Ile | Q06124 |
| HOMO SAPIENS | 5 | 880 | 4nwf | A | 76 | GLU | p.Glu76Ala | Q06124 |
| HOMO SAPIENS | 5 | 880 | 4nwf | A | 76 | GLU | p.Glu76Lys | Q06124 |
| HOMO SAPIENS | 3 | 880 | 4nwf | A | 79 | GLN | p.Gln79Arg | Q06124 |
| HOMO SAPIENS | 5 | 880 | 4nwf | A | 152 | ARG | p.Arg152His | Q06124 |
| HOMO SAPIENS | 2 | 880 | 4nwf | A | 268 | GLY | p.Gly268Cys | Q06124 |
| HOMO SAPIENS | 9 | 880 | 4nwf | A | 276 | LYS | p.Lys276Arg | Q06124 |
| HOMO SAPIENS | 9 | 880 | 4nwf | A | 279 | TYR | p.Tyr279Cys | Q06124 |
| HOMO SAPIENS | 5 | 880 | 4nwf | A | 285 | PHE | p.Phe285Ser | Q06124 |
| HOMO SAPIENS | 0 | 880 | 4nwf | A | 308 | ASP | p.Asn308Ser | Q06124 |
| HOMO SAPIENS | 0 | 880 | 4nwf | A | 308 | ASP | p.Asn308Asp | Q06124 |
| HOMO SAPIENS | 5 | 880 | 4nwf | A | 392 | ALA | p.Ala392Thr | Q06124 |
| HOMO SAPIENS | 1 | 880 | 4nwf | A | 411 | THR | p.Thr415Met | Q06124 |
| HOMO SAPIENS | 0 | 880 | 4nwf | A | 461 | ALA | p.Ala465Thr | Q06124 |
| HOMO SAPIENS | 0 | 880 | 4nwf | A | 464 | GLY | p.Gly468Ala | Q06124 |
| HOMO SAPIENS | 1 | 880 | 4nwf | A | 491 | PRO | p.Pro495Ser | Q06124 |
| HOMO SAPIENS | 1 | 880 | 4nwf | A | 502 | SER | p.Ser506Thr | Q06124 |
| HOMO SAPIENS | 8 | 880 | 4nwf | A | 510 | GLN | p.Gln514Pro | Q06124 |
| HOMO SAPIENS | 4 | 1872 | 3wf1 | D | 109 | ARG | p.Arg109Trp | P16278 |
| HOMO SAPIENS | 0 | 1872 | 3wf1 | D | 521 | ARG | p.Arg521Cys | P16278 |
| HOMO SAPIENS | 5 | 910 | 5xte | A | 45 | SER | p.Ser45Phe | O14949 |
| HOMO SAPIENS | 2 | 9017 | 5f0k | A | 382 | LYS | p.Lys382Arg | Q96QK1 |
| HOMO SAPIENS | 1 | 710 | 4lsj | A | 559 | ILE | p.Ile559Asn | P04150 |
| HOMO SAPIENS | 5 | 710 | 4lsj | A | 571 | VAL | p.Val571Ala | P04150 |

| | | | | | | | | |
|---|---|---|---|---|---|---|---|---|
| HOMO SAPIENS | 0 | 710 | 4lsj | A | 641 | ASP | p.Asp641Val | P04150 |
| HOMO SAPIENS | 1 | 710 | 4lsj | A | 678 | ASP | p.Asp678Glu | P04150 |
| HOMO SAPIENS | 2 | 710 | 4lsj | A | 679 | GLY | p.Gly679Ser | P04150 |
| HOMO SAPIENS | 1 | 710 | 4lsj | A | 737 | PHE | p.Phe737Leu | P04150 |
| HOMO SAPIENS | 1 | 710 | 4lsj | A | 747 | ILE | p.Ile747Met | P04150 |
| HOMO SAPIENS | 1 | 710 | 4lsj | A | 753 | LEU | p.Leu753Phe | P04150 |
| HOMO SAPIENS | 1 | 710 | 4lsj | A | 773 | LEU | p.Leu773Pro | P04150 |
| HOMO SAPIENS | 1 | 2145 | 1av1 | B | 71 | PHE | p.Phe95Tyr | P02647 |
| HOMO SAPIENS | 1 | 2145 | 1av1 | B | 128 | GLU | p.Glu152Lys | P02647 |
| HOMO SAPIENS | 1 | 2145 | 1av1 | B | 136 | GLU | p.Glu160Lys | P02647 |
| HOMO SAPIENS | 2 | 2145 | 1av1 | B | 143 | PRO | p.Pro167Arg | P02647 |
| HOMO SAPIENS | 6 | 2145 | 1av1 | B | 164 | ALA | p.Ala188Ser | P02647 |
| HOMO SAPIENS | 7 | 2145 | 1av1 | B | 173 | ARG | p.Arg197Cys | P02647 |
| HOMO SAPIENS | 2 | 2145 | 1av1 | B | 198 | GLU | p.Glu222Lys | P02647 |
| HOMO SAPIENS | -1 | 3432 | 2beu | A | 1 | SER | p.Ser46Ter | P12694 |
| HOMO SAPIENS | 5 | 3432 | 2beu | A | 79 | ARG | p.Arg124Trp | P12694 |
| HOMO SAPIENS | 7 | 3432 | 2beu | A | 114 | ARG | p.Arg159Gln | P12694 |
| HOMO SAPIENS | 6 | 3432 | 2beu | A | 166 | THR | p.Thr211Met | P12694 |
| HOMO SAPIENS | 4 | 3432 | 2beu | A | 175 | ALA | p.Ala220Val | P12694 |
| HOMO SAPIENS | 5 | 3432 | 2beu | A | 204 | GLY | p.Gly249Ser | P12694 |
| HOMO SAPIENS | 4 | 3432 | 2beu | A | 209 | ALA | p.Ala254Asp | P12694 |
| HOMO SAPIENS | 6 | 3432 | 2beu | A | 220 | ARG | p.Arg265Trp | P12694 |
| HOMO SAPIENS | 6 | 3432 | 2beu | A | 234 | TYR | p.Tyr279Cys | P12694 |
| HOMO SAPIENS | 5 | 3432 | 2beu | A | 240 | ALA | p.Ala285Pro | P12694 |
| HOMO SAPIENS | 4 | 3432 | 2beu | A | 245 | GLY | p.Gly290Arg | P12694 |
| HOMO SAPIENS | 5 | 3432 | 2beu | A | 269 | ARG | p.Arg314Ter | P12694 |
| HOMO SAPIENS | 3 | 3432 | 2beu | A | 301 | ARG | p.Arg346Cys | P12694 |
| HOMO SAPIENS | 5 | 3432 | 2beu | A | 359 | ASN | p.Asn404Ser | P12694 |
| HOMO SAPIENS | 4 | 3432 | 2beu | A | 393 | TYR | p.Tyr438Asn | P12694 |
| HOMO SAPIENS | 8 | 4541 | 1rr8 | C | 533 | ASP | p.Asp533Gly | P11387 |
| HOMO SAPIENS | 2 | 3247 | 1tzn | d | 1105 | GLY | p.Gly105Asp | P58335 |
| HOMO SAPIENS | 3 | 3247 | 1tzn | d | 1111 | ARG | p.Arg111Cys | P58335 |
| HOMO SAPIENS | 2 | 3247 | 1tzn | d | 1189 | ILE | p.Ile189Thr | P58335 |
| HOMO SAPIENS | 9 | 467 | 1suv | C | 58 | ASP | p.Asp77Asn | P02787 |
| HOMO SAPIENS | 2 | 467 | 1suv | C | 277 | ASP | p.Asp296Gly | P02787 |
| HOMO SAPIENS | 9 | 6485 | 1xwd | B | 51 | CYS | p.Cys69Gly | P01225 |
| HOMO SAPIENS | 1 | 694 | 1wuu | A | 28 | PRO | p.Pro28Thr | P51570 |
| HOMO SAPIENS | 0 | 694 | 1wuu | A | 32 | VAL | p.Val32Met | P51570 |
| HOMO SAPIENS | 0 | 694 | 1wuu | A | 198 | ALA | p.Ala198Val | P51570 |
| HOMO SAPIENS | 0 | 143 | 4nkx | A | 93 | PHE | p.Phe93Cys | P05093 |
| HOMO SAPIENS | 3 | 143 | 4nkx | A | 96 | ARG | p.Arg96Trp | P05093 |
| HOMO SAPIENS | 3 | 143 | 4nkx | A | 96 | ARG | p.Arg96Gln | P05093 |
| HOMO SAPIENS | 0 | 143 | 4nkx | A | 106 | SER | p.Ser106Pro | P05093 |
| HOMO SAPIENS | 1 | 143 | 4nkx | A | 114 | PHE | p.Phe114Val | P05093 |
| HOMO SAPIENS | 1 | 143 | 4nkx | A | 116 | ASP | p.Asp116Val | P05093 |
| HOMO SAPIENS | 3 | 143 | 4nkx | A | 125 | ARG | p.Arg125Gln | P05093 |
| HOMO SAPIENS | 0 | 143 | 4nkx | A | 201 | TYR | p.Tyr201Asp | P05093 |
| HOMO SAPIENS | 0 | 143 | 4nkx | A | 239 | ARG | p.Arg239Ter | P05093 |
| HOMO SAPIENS | 0 | 143 | 4nkx | A | 329 | TYR | p.Tyr329Asp | P05093 |
| HOMO SAPIENS | 1 | 143 | 4nkx | A | 342 | PRO | p.Pro342Thr | P05093 |
| HOMO SAPIENS | 2 | 143 | 4nkx | A | 347 | ARG | p.Arg347His | P05093 |
| HOMO SAPIENS | 2 | 143 | 4nkx | A | 347 | ARG | p.Arg347Cys | P05093 |
| HOMO SAPIENS | 1 | 143 | 4nkx | A | 358 | ARG | p.Arg358Gln | P05093 |
| HOMO SAPIENS | 0 | 143 | 4nkx | A | 362 | ARG | p.Arg362Cys | P05093 |

| | | | | | | | | |
|---|---|---|---|---|---|---|---|---|
| HOMO SAPIENS | 1 | 143 | 4nkx | A | 406 | TRP | p.Trp406Arg | P05093 |
| HOMO SAPIENS | 0 | 143 | 4nkx | A | 428 | PRO | p.Pro428Leu | P05093 |
| HOMO SAPIENS | 6 | 1441 | 1uuh | A | 46 | ARG | p.Arg46Gly | P16070 |
| HOMO SAPIENS | 3 | 82 | 4wlv | D | 37 | GLY | p.Gly37Arg | P40926 |
| HOMO SAPIENS | 0 | 82 | 4wlv | D | 133 | PRO | p.Pro133Leu | P40926 |
| HOMO SAPIENS | 2 | 392 | 5xtd | A | 56 | SER | p.Ser56Pro | P49821 |
| HOMO SAPIENS | 2 | 392 | 5xtd | A | 59 | ARG | p.Arg59Ter | P49821 |
| HOMO SAPIENS | 2 | 392 | 5xtd | A | 74 | ASP | p.Asp74Asn | P49821 |
| HOMO SAPIENS | 4 | 392 | 5xtd | A | 116 | ASN | p.Asn116Lys | P49821 |
| HOMO SAPIENS | 4 | 392 | 5xtd | A | 117 | ALA | p.Ala117Thr | P49821 |
| HOMO SAPIENS | 5 | 392 | 5xtd | A | 122 | PRO | p.Pro122Leu | P49821 |
| HOMO SAPIENS | 3 | 392 | 5xtd | A | 160 | GLY | p.Gly160Glu | P49821 |
| HOMO SAPIENS | 4 | 392 | 5xtd | A | 214 | GLU | p.Glu214Lys | P49821 |
| HOMO SAPIENS | 2 | 392 | 5xtd | A | 357 | MET | p.Met357Thr | P49821 |
| HOMO SAPIENS | 4 | 392 | 5xtd | A | 423 | THR | p.Thr423Met | P49821 |
| HOMO SAPIENS | 0 | 22593 | 3pdf | A | 8 | TYR | p.Tyr32Ter | P53634 |
| HOMO SAPIENS | 0 | 22593 | 3pdf | A | 15 | TRP | p.Trp39Ser | P53634 |
| HOMO SAPIENS | 0 | 22593 | 3pdf | A | 103 | HIS | p.His127Pro | P53634 |
| HOMO SAPIENS | 9 | 22593 | 3pdf | A | 228 | GLN | p.Gln252Leu | P53634 |
| HOMO SAPIENS | 3 | 22593 | 3pdf | A | 262 | GLN | p.Gln286Arg | P53634 |
| HOMO SAPIENS | 4 | 22593 | 3pdf | A | 277 | GLY | p.Gly301Ser | P53634 |
| HOMO SAPIENS | 0 | 22593 | 3pdf | A | 323 | TYR | p.Tyr347Cys | P53634 |
| HOMO SAPIENS | 5 | 3461 | 4c2a | A | 1306 | GLN | p.Arg1306Trp | P04275 |
| HOMO SAPIENS | 5 | 3461 | 4c2a | A | 1374 | ARG | p.Arg1374Cys | P04275 |
| HOMO SAPIENS | 5 | 3461 | 4c2a | A | 1379 | ARG | p.Arg1379Cys | P04275 |
| HOMO SAPIENS | 4 | 3461 | 4c2a | A | 1380 | ILE | p.Ile1380Val | P04275 |
| HOMO SAPIENS | 0 | 3461 | 4c2a | A | 1381 | ALA | p.Thr1381Ala | P04275 |
| HOMO SAPIENS | 5 | 3461 | 4c2a | A | 1399 | ARG | p.Arg1399His | P04275 |
| HOMO SAPIENS | 5 | 3461 | 4c2a | A | 1435 | ASN | p.Asn1435Ser | P04275 |
| HOMO SAPIENS | 0 | 4533 | 5uj7 | C | 174 | TYR | p.Tyr174Cys | O43929 |
| HOMO SAPIENS | 0 | 4533 | 5uj7 | C | 202 | LEU | p.Leu202Val | O43929 |
| HOMO SAPIENS | 0 | 1374 | 3j07 | O | 39 | PRO | p.Pro39Leu | P02511 |
| HOMO SAPIENS | 0 | 1374 | 3j07 | O | 107 | ARG | p.Arg107His | P02511 |
| HOMO SAPIENS | 0 | 1374 | 3j07 | O | 157 | ARG | p.Arg157His | P02511 |
| HOMO SAPIENS | -1 | 942 | 1s9i | B | 57 | PHE | p.Phe57Cys | P36507 |
| HOMO SAPIENS | -1 | 942 | 1s9i | B | 57 | PHE | p.Phe57Val | P36507 |
| HOMO SAPIENS | 2 | 942 | 1s9i | B | 128 | PRO | p.Pro128Gln | P36507 |
| HOMO SAPIENS | 1 | 942 | 1s9i | B | 134 | TYR | p.Tyr134His | P36507 |
| HOMO SAPIENS | 1 | 942 | 1s9i | B | 188 | HIS | p.His188Gln | P36507 |
| HOMO SAPIENS | 0 | 942 | 1s9i | B | 262 | VAL | p.Val262Phe | P36507 |
| HOMO SAPIENS | 0 | 942 | 1s9i | B | 263 | GLY | p.Gly263Arg | P36507 |
| HOMO SAPIENS | -1 | 942 | 1s9i | B | 298 | PRO | p.Pro298Leu | P36507 |
| HOMO SAPIENS | -1 | 942 | 1s9i | B | 305 | VAL | p.Val305Ile | P36507 |
| HOMO SAPIENS | 0 | 3245 | 1fbv | A | 367 | GLN | p.Gln367Pro | P22681 |
| HOMO SAPIENS | 0 | 3245 | 1fbv | A | 371 | TYR | p.Tyr371His | P22681 |
| HOMO SAPIENS | 1 | 3245 | 1fbv | A | 382 | LYS | p.Lys382Glu | P22681 |
| HOMO SAPIENS | 2 | 3245 | 1fbv | A | 390 | ASP | p.Asp390Tyr | P22681 |
| HOMO SAPIENS | 9 | 3245 | 1fbv | A | 396 | CYS | p.Cys396Arg | P22681 |
| HOMO SAPIENS | 7 | 3245 | 1fbv | A | 420 | ARG | p.Arg420Gln | P22681 |
| HOMO SAPIENS | 1 | 3245 | 1fbv | A | 429 | ILE | p.Ile429Met | P22681 |
| HOMO SAPIENS | 0 | 2604 | 2pbd | P | 111 | LEU | p.Leu112Met | P07737 |
| HOMO SAPIENS | 3 | 2604 | 2pbd | P | 116 | GLU | p.Glu117Gly | P07737 |
| HOMO SAPIENS | 4 | 2013 | 3b7w | A | 472 | ARG | p.Arg472Gly | Q08AH3 |
| HOMO SAPIENS | 0 | 24346 | 3sh5 | A | 85 | ARG | p.Arg4281His | P98160 |

| | | | | | | | | |
|---|---|---|---|---|---|---|---|---|
| HOMO SAPIENS | 0 | 24346 | 3sh5 | A | 122 | ARG | p.Arg4318Trp | P98160 |
| HOMO SAPIENS | 0 | 24346 | 3sh5 | A | 135 | SER | p.Ser4331Asn | P98160 |
| HOMO SAPIENS | 0 | 24346 | 3sh5 | A | 136 | VAL | p.Val4332Ile | P98160 |
| HOMO SAPIENS | 0 | 707 | 4p2g | A | 26 | MET | p.Met26Ile | Q99497 |
| HOMO SAPIENS | 5 | 707 | 4p2g | A | 39 | ALA | p.Ala39Ser | Q99497 |
| HOMO SAPIENS | 5 | 707 | 4p2g | A | 64 | GLU | p.Glu64Asp | Q99497 |
| HOMO SAPIENS | 4 | 707 | 4p2g | A | 149 | ASP | p.Asp149Ala | Q99497 |
| HOMO SAPIENS | 6 | 707 | 4p2g | A | 163 | GLU | p.Glu163Lys | Q99497 |
| HOMO SAPIENS | 6 | 707 | 4p2g | A | 166 | LEU | p.Leu166Pro | Q99497 |
| HOMO SAPIENS | 3 | 5121 | 4xss | E | 31 | GLY | p.Gly58Arg | P06213 |
| HOMO SAPIENS | 2 | 5121 | 4xss | E | 119 | ILE | p.Ile146Met | P06213 |
| HOMO SAPIENS | 8 | 5121 | 4xss | E | 209 | HIS | p.His236Arg | P06213 |
| HOMO SAPIENS | 2 | 5121 | 4xss | E | 234 | ASP | p.Asp261Glu | P06213 |
| HOMO SAPIENS | 1 | 5121 | 4xss | E | 276 | GLN | p.Gln303His | P06213 |
| HOMO SAPIENS | 1 | 27790 | 5erp | B | 140 | THR | p.Thr275Met | Q02487 |
| HOMO SAPIENS | 0 | 27790 | 5erp | B | 150 | ILE | p.Ile285Thr | Q02487 |
| HOMO SAPIENS | 1 | 27790 | 5erp | B | 205 | THR | p.Thr340Ala | Q02487 |
| HOMO SAPIENS | 1 | 27790 | 5erp | B | 223 | THR | p.Thr358Ile | Q02487 |
| HOMO SAPIENS | 0 | 27790 | 5erp | B | 301 | GLY | p.Gly436Ala | Q02487 |
| HOMO SAPIENS | -1 | 27790 | 5erp | B | 315 | ARG | p.Arg450Ser | Q02487 |
| HOMO SAPIENS | 0 | 27790 | 5erp | B | 369 | SER | p.Ser504Gly | Q02487 |
| HOMO SAPIENS | 0 | 27790 | 5erp | B | 385 | ILE | p.Ile520Thr | Q02487 |
| HOMO SAPIENS | 0 | 27790 | 5erp | B | 461 | ALA | p.Ala596Val | Q02487 |
| HOMO SAPIENS | 0 | 27790 | 5erp | B | 503 | GLN | p.Gln638His | Q02487 |
| HOMO SAPIENS | 5 | 6599 | 4nsc | A | 129 | ARG | p.Arg129Leu | Q9BPX6 |
| HOMO SAPIENS | -1 | 6599 | 4nsc | A | 183 | GLN | p.Gln183Ter | Q9BPX6 |
| HOMO SAPIENS | 0 | 20420 | 3ghm | A | 398 | ARG | p.Arg398His | Q76LX8 |
| HOMO SAPIENS | 0 | 20420 | 3ghm | A | 448 | GLN | p.Gln448Glu | Q76LX8 |
| HOMO SAPIENS | 1 | 20420 | 3ghm | A | 449 | GLN | p.Gln449Ter | Q76LX8 |
| HOMO SAPIENS | 0 | 20420 | 3ghm | A | 456 | GLN | p.Gln456His | Q76LX8 |
| HOMO SAPIENS | 1 | 20420 | 3ghm | A | 528 | ARG | p.Arg528Gly | Q76LX8 |
| HOMO SAPIENS | 0 | 20420 | 3ghm | A | 618 | PRO | p.Pro618Ala | Q76LX8 |
| HOMO SAPIENS | 0 | 20420 | 3ghm | A | 625 | ARG | p.Arg625His | Q76LX8 |
| HOMO SAPIENS | 0 | 275 | 4rmz | A | 293 | GLN | p.Gln293Ter | Q9NWZ3 |
| HOMO SAPIENS | 0 | 275 | 4rmz | A | 428 | ALA | p.Ala428Thr | Q9NWZ3 |
| HOMO SAPIENS | 2 | 10458 | 3puf | B | 138 | LEU | p.Leu138Phe | Q5TBB1 |
| HOMO SAPIENS | 6 | 10458 | 3puf | B | 177 | ALA | p.Ala177Thr | Q5TBB1 |
| HOMO SAPIENS | -1 | 10458 | 3puf | B | 185 | VAL | p.Val185Gly | Q5TBB1 |
| HOMO SAPIENS | 0 | 26648 | 2n90 | A | 36 | ARG | p.Arg444Gln | P04629 |
| HOMO SAPIENS | 2 | 977 | 1dnw | D | 166 | ALA | p.Ala332Val | P05164 |
| HOMO SAPIENS | 4 | 977 | 1dnw | D | 333 | ARG | p.Arg499Cys | P05164 |
| HOMO SAPIENS | 1 | 977 | 1dnw | D | 403 | ARG | p.Arg569Trp | P05164 |
| HOMO SAPIENS | 0 | 25704 | 3ajb | A | 82 | GLN | p.Gln82Arg | P56589 |
| HOMO SAPIENS | 3 | 255 | 3t1p | A | 38 | TYR | p.Tyr62Ter | P01009 |
| HOMO SAPIENS | 3 | 255 | 3t1p | A | 39 | ARG | p.Arg63Cys | P01009 |
| HOMO SAPIENS | 0 | 255 | 3t1p | A | 101 | ARG | p.Arg125His | P01009 |
| HOMO SAPIENS | 0 | 255 | 3t1p | A | 212 | GLN | p.Gln236Arg | P01009 |
| HOMO SAPIENS | 0 | 255 | 3t1p | A | 213 | VAL | p.Val237Ala | P01009 |
| HOMO SAPIENS | 1 | 255 | 3t1p | A | 269 | HIS | p.His293Gln | P01009 |
| HOMO SAPIENS | 8 | 255 | 3t1p | A | 341 | ASP | p.Asp365Asn | P01009 |
| HOMO SAPIENS | 9 | 255 | 3t1p | A | 342 | GLU | p.Glu366Lys | P01009 |
| HOMO SAPIENS | 0 | 255 | 3t1p | A | 369 | PRO | p.Pro393Ser | P01009 |
| HOMO SAPIENS | 0 | 255 | 3t1p | A | 376 | GLU | p.Glu400Asp | P01009 |
| HOMO SAPIENS | -1 | 5175 | 3zxi | A | 35 | ALA | p.Ala35Asp | Q9Y2Z4 |

| | | | | | | | | | |
|---|---|---|---|---|---|---|---|---|---|
| HOMO SAPIENS | 0 | 5175 | 3zxi | A | | 52 | PHE | p.Phe52Leu | Q9Y2Z4 |
| HOMO SAPIENS | -1 | 5175 | 3zxi | A | | 68 | GLY | p.Gly68Ser | Q9Y2Z4 |
| HOMO SAPIENS | 0 | 5175 | 3zxi | A | | 179 | LYS | p.Lys179Gln | Q9Y2Z4 |
| HOMO SAPIENS | 0 | 5175 | 3zxi | A | | 191 | GLY | p.Gly191Val | Q9Y2Z4 |
| HOMO SAPIENS | 0 | 5175 | 3zxi | A | | 251 | ILE | p.Ile251Val | Q9Y2Z4 |
| HOMO SAPIENS | 2 | 5175 | 3zxi | A | | 295 | ASP | p.Asp295Asn | Q9Y2Z4 |
| HOMO SAPIENS | 0 | 5175 | 3zxi | A | | 311 | ASP | p.Asp311Glu | Q9Y2Z4 |
| HOMO SAPIENS | 0 | 5175 | 3zxi | A | | 360 | ARG | p.Arg360Ter | Q9Y2Z4 |
| HOMO SAPIENS | 1 | 3016 | 3exg | | 3 | 4 | ASP | p.Asp33His | P08559 |
| HOMO SAPIENS | 1 | 3016 | 3exg | | 3 | 176 | PHE | p.Phe205Leu | P08559 |
| HOMO SAPIENS | 1 | 3016 | 3exg | | 3 | 214 | TYR | p.Tyr243Asn | P08559 |
| HOMO SAPIENS | 1 | 3016 | 3exg | | 3 | 229 | ASP | p.Asp258Ala | P08559 |
| HOMO SAPIENS | 0 | 3016 | 3exg | | 3 | 233 | VAL | p.Val262Ile | P08559 |
| HOMO SAPIENS | 1 | 3016 | 3exg | | 3 | 234 | ARG | p.Arg263Gly | P08559 |
| HOMO SAPIENS | 0 | 3016 | 3exg | | 3 | 253 | MET | p.Met282Leu | P08559 |
| HOMO SAPIENS | 7 | 3016 | 3exg | | 3 | 259 | ARG | p.Arg288His | P08559 |
| HOMO SAPIENS | -1 | 3016 | 3exg | | 3 | 273 | ARG | p.Arg302Cys | P08559 |
| HOMO SAPIENS | 2 | 3016 | 3exg | | 3 | 286 | ASP | p.Asp315Asn | P08559 |
| HOMO SAPIENS | 1 | 3016 | 3exg | | 3 | 349 | ARG | p.Arg378His | P08559 |
| HOMO SAPIENS | 5 | 6729 | 4a0l | E | | 572 | ARG | p.Arg572Cys | Q13620 |
| HOMO SAPIENS | 0 | 1067 | 4kyo | A | | 9 | THR | p.Thr9Asn | P21549 |
| HOMO SAPIENS | 0 | 1067 | 4kyo | A | | 11 | PRO | p.Pro11Leu | P21549 |
| HOMO SAPIENS | 0 | 1067 | 4kyo | A | | 22 | ASN | p.Asn22Ser | P21549 |
| HOMO SAPIENS | 4 | 1067 | 4kyo | A | | 25 | LEU | p.Leu25Arg | P21549 |
| HOMO SAPIENS | 7 | 1067 | 4kyo | A | | 36 | ARG | p.Arg36His | P21549 |
| HOMO SAPIENS | 3 | 1067 | 4kyo | A | | 41 | GLY | p.Gly41Arg | P21549 |
| HOMO SAPIENS | 0 | 1067 | 4kyo | A | | 42 | GLY | p.Gly42Glu | P21549 |
| HOMO SAPIENS | 3 | 1067 | 4kyo | A | | 56 | ILE | p.Ile56Asn | P21549 |
| HOMO SAPIENS | 3 | 1067 | 4kyo | A | | 59 | GLU | p.Glu59Lys | P21549 |
| HOMO SAPIENS | 3 | 1067 | 4kyo | A | | 66 | TYR | p.Tyr66Ter | P21549 |
| HOMO SAPIENS | 3 | 1067 | 4kyo | A | | 111 | ARG | p.Arg111Leu | P21549 |
| HOMO SAPIENS | 7 | 1067 | 4kyo | A | | 118 | ARG | p.Arg118His | P21549 |
| HOMO SAPIENS | 3 | 1067 | 4kyo | A | | 122 | ARG | p.Arg122Ter | P21549 |
| HOMO SAPIENS | 3 | 1067 | 4kyo | A | | 124 | HIS | p.His124Pro | P21549 |
| HOMO SAPIENS | 0 | 1067 | 4kyo | A | | 152 | PHE | p.Phe152Ile | P21549 |
| HOMO SAPIENS | 1 | 1067 | 4kyo | A | | 156 | GLY | p.Gly156Arg | P21549 |
| HOMO SAPIENS | 0 | 1067 | 4kyo | A | | 161 | GLY | p.Gly161Arg | P21549 |
| HOMO SAPIENS | 3 | 1067 | 4kyo | A | | 166 | LEU | p.Leu166Pro | P21549 |
| HOMO SAPIENS | 5 | 1067 | 4kyo | A | | 170 | GLY | p.Gly170Arg | P21549 |
| HOMO SAPIENS | 5 | 1067 | 4kyo | A | | 186 | ALA | p.Ala186Val | P21549 |
| HOMO SAPIENS | 3 | 1067 | 4kyo | A | | 187 | SER | p.Ser187Phe | P21549 |
| HOMO SAPIENS | 3 | 1067 | 4kyo | A | | 197 | ARG | p.Arg197Pro | P21549 |
| HOMO SAPIENS | 5 | 1067 | 4kyo | A | | 201 | ASP | p.Asp201Glu | P21549 |
| HOMO SAPIENS | 0 | 1067 | 4kyo | A | | 202 | ILE | p.Ile202Asn | P21549 |
| HOMO SAPIENS | 1 | 1067 | 4kyo | A | | 205 | SER | p.Ser205Pro | P21549 |
| HOMO SAPIENS | 1 | 1067 | 4kyo | A | | 205 | SER | p.Ser205Ter | P21549 |
| HOMO SAPIENS | 2 | 1067 | 4kyo | A | | 216 | GLY | p.Gly216Arg | P21549 |
| HOMO SAPIENS | 5 | 1067 | 4kyo | A | | 233 | ARG | p.Arg233Cys | P21549 |
| HOMO SAPIENS | 5 | 1067 | 4kyo | A | | 244 | ILE | p.Ile244Thr | P21549 |
| HOMO SAPIENS | 5 | 1067 | 4kyo | A | | 244 | ILE | p.Ile244Met | P21549 |
| HOMO SAPIENS | 4 | 1067 | 4kyo | A | | 253 | CYS | p.Cys253Arg | P21549 |
| HOMO SAPIENS | 7 | 1067 | 4kyo | A | | 274 | GLU | p.Glu274Asp | P21549 |
| HOMO SAPIENS | 7 | 1067 | 4kyo | A | | 279 | ILE | p.Ile279Thr | P21549 |
| HOMO SAPIENS | 6 | 1067 | 4kyo | A | | 295 | ALA | p.Ala295Thr | P21549 |

| | | | | | | | | |
|---|---|---|---|---|---|---|---|---|
| HOMO SAPIENS | 5 | 1067 | 4kyo | A | 340 | ILE | p.Ile340Met | P21549 |
| HOMO SAPIENS | 1 | 790 | 2ric | C | 270 | SER | p.Ser290Thr | P35247 |
| HOMO SAPIENS | 0 | 790 | 2ric | C | 289 | GLU | p.Glu309Lys | P35247 |
| HOMO SAPIENS | 1 | 212 | 5sxe | A | 545 | GLU | p.Glu545Lys | P42336 |
| HOMO SAPIENS | 3 | 212 | 5sxe | A | 546 | GLN | p.Gln546Glu | P42336 |
| HOMO SAPIENS | 2 | 212 | 5sxe | A | 1047 | HIS | p.His1047Leu | P42336 |
| HOMO SAPIENS | 7 | 152 | 2zqq | F | 197 | ARG | p.Arg197Ter | Q13825 |
| HOMO SAPIENS | 0 | 152 | 2zqq | F | 248 | ALA | p.Ala248Thr | Q13825 |
| HOMO SAPIENS | 0 | 152 | 2zqq | F | 275 | ALA | p.Ala275Val | Q13825 |
| HOMO SAPIENS | 5 | 152 | 2zqq | F | 331 | LYS | p.Lys331Ter | Q13825 |
| HOMO SAPIENS | 6 | 17801 | 2j8l | A | 283 | PHE | p.Phe301Leu | P03951 |
| HOMO SAPIENS | 3 | 17801 | 2j8l | A | 308 | ARG | p.Arg326Cys | P03951 |
| HOMO SAPIENS | 0 | 17801 | 2j8l | A | 321 | CYS | p.Cys339Phe | P03951 |
| HOMO SAPIENS | 3 | 17801 | 2j8l | A | 351 | TYR | p.Tyr369Ter | P03951 |
| HOMO SAPIENS | 7 | 1802 | 5db0 | A | 22 | LEU | p.Leu22Arg | O00255 |
| HOMO SAPIENS | 7 | 1802 | 5db0 | A | 26 | GLU | p.Glu26Lys | O00255 |
| HOMO SAPIENS | 7 | 1802 | 5db0 | A | 139 | HIS | p.His139Asp | O00255 |
| HOMO SAPIENS | 0 | 1802 | 5db0 | A | 171 | ARG | p.Arg176Gln | O00255 |
| HOMO SAPIENS | 0 | 1802 | 5db0 | A | 184 | VAL | p.Val189Glu | O00255 |
| HOMO SAPIENS | 7 | 1802 | 5db0 | A | 255 | GLU | p.Glu260Lys | O00255 |
| HOMO SAPIENS | 4 | 1802 | 5db0 | A | 418 | ASP | p.Asp423Asn | O00255 |
| HOMO SAPIENS | 0 | 1802 | 5db0 | A | 436 | TRP | p.Trp441Arg | O00255 |
| HOMO SAPIENS | -1 | 1802 | 5db0 | A | 541 | ALA | p.Thr546Ala | O00255 |
| HOMO SAPIENS | 4 | 1802 | 5db0 | A | 552 | THR | p.Thr557Ser | O00255 |
| HOMO SAPIENS | 0 | 3450 | 2yuz | A | 36 | PRO | p.Pro461Gln | Q00872 |
| HOMO SAPIENS | 1 | 3450 | 2yuz | A | 56 | GLN | p.His481Gln | Q00872 |
| HOMO SAPIENS | 2 | 3450 | 2yuz | A | 68 | ALA | p.Ala493Val | Q00872 |
| HOMO SAPIENS | 0 | 3211 | 3rqg | A | 72 | VAL | p.Val72Leu | Q9BUL8 |
| HOMO SAPIENS | 2 | 4168 | 4q4f | A | 121 | ARG | p.Arg121Ter | Q14108 |
| HOMO SAPIENS | 2 | 4168 | 4q4f | A | 121 | ARG | p.Arg121Gln | Q14108 |
| HOMO SAPIENS | 2 | 4168 | 4q4f | A | 159 | MET | p.Met159Val | Q14108 |
| HOMO SAPIENS | 4 | 4168 | 4q4f | A | 363 | HIS | p.His363Asn | Q14108 |
| HOMO SAPIENS | 2 | 4168 | 4q4f | A | 396 | VAL | p.Val396Ile | Q14108 |
| HOMO SAPIENS | 0 | 1448 | 1y97 | A | 87 | ARG | p.Arg130Ter | Q9BQ50 |
| HOMO SAPIENS | 0 | 3444 | 4iwp | A | 151 | SER | p.Ser151Phe | Q9UHD2 |
| HOMO SAPIENS | 0 | 3444 | 4iwp | A | 401 | LYS | p.Lys401Glu | Q9UHD2 |
| HOMO SAPIENS | 0 | 116 | 1zy8 | B | 194 | GLY | p.Gly229Cys | P09622 |
| HOMO SAPIENS | 0 | 116 | 1zy8 | B | 255 | ASP | p.Asp290Gly | P09622 |
| HOMO SAPIENS | 2 | 116 | 1zy8 | B | 326 | MET | p.Met361Val | P09622 |
| HOMO SAPIENS | 0 | 116 | 1zy8 | B | 340 | GLU | p.Glu375Lys | P09622 |
| HOMO SAPIENS | 2 | 116 | 1zy8 | B | 358 | ILE | p.Ile393Thr | P09622 |
| HOMO SAPIENS | 1 | 116 | 1zy8 | B | 453 | PRO | p.Pro488Leu | P09622 |
| HOMO SAPIENS | 7 | 127 | 4qo7 | B | 160 | SER | p.Ser161Arg | P00338 |
| HOMO SAPIENS | 0 | 2025 | 1wrl | A | 8 | ALA | p.Ala8Val | P63316 |
| HOMO SAPIENS | 8 | 2025 | 1wrl | A | 24 | PHE | p.Phe24Leu | P63316 |
| HOMO SAPIENS | 0 | 2025 | 1wrl | A | 84 | SER | p.Cys84Tyr | P63316 |
| HOMO SAPIENS | 8 | 3791 | 1mhd | B | 93 | ARG | p.Arg93Ter | P84022 |
| HOMO SAPIENS | 0 | 1575 | 4phc | A | 205 | GLY | p.Gly205Asp | P12081 |
| HOMO SAPIENS | 0 | 1575 | 4phc | A | 454 | TYR | p.Tyr454Ser | P12081 |
| HOMO SAPIENS | 0 | 39620 | 1ozs | B | 135 | ARG | p.Arg136Gln | P19429 |
| HOMO SAPIENS | 5 | 39620 | 1ozs | B | 140 | ARG | p.Arg141Trp | P19429 |
| HOMO SAPIENS | 5 | 39620 | 1ozs | B | 142 | THR | p.Thr143Asn | P19429 |
| HOMO SAPIENS | 5 | 39620 | 1ozs | B | 143 | LEU | p.Leu144Gln | P19429 |
| HOMO SAPIENS | 5 | 39620 | 1ozs | B | 144 | ARG | p.Arg145Gly | P19429 |

| | | | | | | | | |
|---|---|---|---|---|---|---|---|---|
| HOMO SAPIENS | 5 | 39620 | 1ozs | B | 144 | ARG | p.Arg145Gln | P19429 |
| HOMO SAPIENS | -1 | 20934 | 4nuv | D | 42 | GLY | p.Gly42Asp | Q16570 |
| HOMO SAPIENS | -1 | 40565 | 5lcw | S | 4 | VAL | p.Val4Met | O60566 |
| HOMO SAPIENS | 0 | 40565 | 5lcw | S | 40 | THR | p.Thr40Met | O60566 |
| HOMO SAPIENS | 4 | 40565 | 5lcw | S | 198 | ALA | p.Glu390Asp | O60566 |
| HOMO SAPIENS | 4 | 758 | 4oiu | A | 726 | ARG | p.Arg727Leu | P10275 |
| HOMO SAPIENS | 4 | 758 | 4oiu | A | 730 | VAL | p.Val731Met | P10275 |
| HOMO SAPIENS | 2 | 758 | 4oiu | A | 798 | GLN | p.Gln799Glu | P10275 |
| HOMO SAPIENS | 3 | 758 | 4oiu | A | 855 | ARG | p.Arg856His | P10275 |
| HOMO SAPIENS | 1 | 758 | 4oiu | A | 886 | MET | p.Met887Val | P10275 |
| HOMO SAPIENS | 0 | 3303 | 4ie0 | A | 134 | ALA | p.Ala134Thr | Q9C0B1 |
| HOMO SAPIENS | 9 | 3303 | 4ie0 | A | 316 | ARG | p.Arg316Gln | Q9C0B1 |
| HOMO SAPIENS | 1 | 41502 | 3fgh | A | 136 | PRO | p.Pro178Leu | Q00059 |
| HOMO SAPIENS | 5 | 175 | 3ceh | B | 58 | THR | p.Thr59Lys | P06737 |
| HOMO SAPIENS | 8 | 175 | 3ceh | B | 93 | ARG | p.Arg94Ter | P06737 |
| HOMO SAPIENS | 5 | 175 | 3ceh | B | 221 | VAL | p.Val222Ile | P06737 |
| HOMO SAPIENS | 0 | 175 | 3ceh | B | 338 | ASN | p.Asn339Ser | P06737 |
| HOMO SAPIENS | 1 | 175 | 3ceh | B | 372 | PHE | p.Phe373Leu | P06737 |
| HOMO SAPIENS | 9 | 175 | 3ceh | B | 376 | ASN | p.Asn377Lys | P06737 |
| HOMO SAPIENS | 8 | 175 | 3ceh | B | 398 | ARG | p.Arg399Ter | P06737 |
| HOMO SAPIENS | 5 | 175 | 3ceh | B | 424 | ARG | p.Arg425Pro | P06737 |
| HOMO SAPIENS | 9 | 175 | 3ceh | B | 455 | VAL | p.Val456Met | P06737 |
| HOMO SAPIENS | 8 | 175 | 3ceh | B | 490 | ARG | p.Arg491Cys | P06737 |
| HOMO SAPIENS | 8 | 175 | 3ceh | B | 631 | ASN | p.Asn632Ile | P06737 |
| HOMO SAPIENS | 9 | 175 | 3ceh | B | 648 | TYR | p.Tyr649Ter | P06737 |
| HOMO SAPIENS | 9 | 175 | 3ceh | B | 672 | GLU | p.Glu673Lys | P06737 |
| HOMO SAPIENS | 9 | 175 | 3ceh | B | 674 | SER | p.Ser675Leu | P06737 |
| HOMO SAPIENS | 0 | 175 | 3ceh | B | 805 | ILE | p.Ile806Leu | P06737 |
| HOMO SAPIENS | 0 | 175 | 3ceh | B | 820 | TYR | p.Tyr821His | P06737 |
| HOMO SAPIENS | 1 | 6750 | 4fdj | A | 47 | GLY | p.Gly47Arg | P34059 |
| HOMO SAPIENS | 9 | 6750 | 4fdj | A | 60 | ASP | p.Asp60Asn | P34059 |
| HOMO SAPIENS | 5 | 6750 | 4fdj | A | 67 | LEU | p.Leu67Met | P34059 |
| HOMO SAPIENS | 0 | 6750 | 4fdj | A | 69 | PHE | p.Phe69Val | P34059 |
| HOMO SAPIENS | 3 | 6750 | 4fdj | A | 94 | ARG | p.Arg94Gly | P34059 |
| HOMO SAPIENS | 0 | 6750 | 4fdj | A | 113 | ILE | p.Ile113Phe | P34059 |
| HOMO SAPIENS | 0 | 6750 | 4fdj | A | 138 | VAL | p.Val138Ala | P34059 |
| HOMO SAPIENS | 0 | 6750 | 4fdj | A | 139 | GLY | p.Gly139Ser | P34059 |
| HOMO SAPIENS | 0 | 6750 | 4fdj | A | 151 | PRO | p.Pro151Thr | P34059 |
| HOMO SAPIENS | 0 | 6750 | 4fdj | A | 155 | GLY | p.Gly155Arg | P34059 |
| HOMO SAPIENS | 0 | 6750 | 4fdj | A | 159 | TRP | p.Trp159Ter | P34059 |
| HOMO SAPIENS | 0 | 6750 | 4fdj | A | 162 | SER | p.Ser162Phe | P34059 |
| HOMO SAPIENS | 0 | 6750 | 4fdj | A | 200 | THR | p.Thr200Met | P34059 |
| HOMO SAPIENS | 0 | 6750 | 4fdj | A | 204 | ASN | p.Asn204Lys | P34059 |
| HOMO SAPIENS | 0 | 6750 | 4fdj | A | 231 | ALA | p.Ala231Gly | P34059 |
| HOMO SAPIENS | 2 | 6750 | 4fdj | A | 235 | THR | p.Thr235Lys | P34059 |
| HOMO SAPIENS | 1 | 6750 | 4fdj | A | 259 | ARG | p.Arg259Gln | P34059 |
| HOMO SAPIENS | 0 | 6750 | 4fdj | A | 287 | SER | p.Ser287Leu | P34059 |
| HOMO SAPIENS | 0 | 6750 | 4fdj | A | 291 | ALA | p.Ala291Thr | P34059 |
| HOMO SAPIENS | 3 | 6750 | 4fdj | A | 301 | GLY | p.Gly301Cys | P34059 |
| HOMO SAPIENS | 0 | 6750 | 4fdj | A | 312 | THR | p.Thr312Ser | P34059 |
| HOMO SAPIENS | 0 | 6750 | 4fdj | A | 318 | MET | p.Met318Arg | P34059 |
| HOMO SAPIENS | 0 | 6750 | 4fdj | A | 340 | GLY | p.Gly340Asp | P34059 |
| HOMO SAPIENS | 0 | 6750 | 4fdj | A | 386 | ARG | p.Arg386Cys | P34059 |
| HOMO SAPIENS | 0 | 6750 | 4fdj | A | 391 | MET | p.Met391Val | P34059 |

| | | | | | | | | |
|---|---|---|---|---|---|---|---|---|
| HOMO SAPIENS | 0 | 6750 | 4fdj | A | 392 | ALA | p.Ala392Val | P34059 |
| HOMO SAPIENS | 0 | 6750 | 4fdj | A | 393 | ALA | p.Ala393Ser | P34059 |
| HOMO SAPIENS | 0 | 6750 | 4fdj | A | 459 | ALA | p.Ala459Val | P34059 |
| HOMO SAPIENS | 0 | 6750 | 4fdj | A | 473 | GLN | p.Gln473Ter | P34059 |
| HOMO SAPIENS | 2 | 6750 | 4fdj | A | 480 | VAL | p.Val480Phe | P34059 |
| HOMO SAPIENS | 0 | 6750 | 4fdj | A | 487 | ASN | p.Asn487Ser | P34059 |
| HOMO SAPIENS | 0 | 6750 | 4fdj | A | 488 | VAL | p.Val488Met | P34059 |
| HOMO SAPIENS | 0 | 6750 | 4fdj | A | 520 | TRP | p.Trp520Ter | P34059 |
| HOMO SAPIENS | 4 | 1088 | 2p3u | A | 102 | GLU | p.Glu142Lys | P00742 |
| HOMO SAPIENS | 6 | 370 | 1nr1 | D | 73 | ASP | p.Asp126His | P00367 |
| HOMO SAPIENS | 3 | 370 | 1nr1 | D | 221 | ARG | p.Arg274Cys | P00367 |
| HOMO SAPIENS | 2 | 370 | 1nr1 | D | 265 | ARG | p.Arg318Lys | P00367 |
| HOMO SAPIENS | 1 | 370 | 1nr1 | D | 269 | ARG | p.Arg322His | P00367 |
| HOMO SAPIENS | 1 | 370 | 1nr1 | D | 296 | GLU | p.Glu349Ala | P00367 |
| HOMO SAPIENS | 1 | 370 | 1nr1 | D | 445 | SER | p.Ser498Leu | P00367 |
| HOMO SAPIENS | 1 | 370 | 1nr1 | D | 446 | GLY | p.Gly499Asp | P00367 |
| HOMO SAPIENS | 1 | 370 | 1nr1 | D | 446 | GLY | p.Gly499Ser | P00367 |
| HOMO SAPIENS | 1 | 370 | 1nr1 | D | 448 | SER | p.Ser501Pro | P00367 |
| HOMO SAPIENS | 2 | 370 | 1nr1 | D | 454 | HIS | p.His507Tyr | P00367 |
| HOMO SAPIENS | 1 | 7720 | 2he0 | A | 147 | HIS | p.His2018Gln | P46531 |
| HOMO SAPIENS | 2 | 11844 | 5l0v | A | 218 | ARG | p.Arg218Ter | Q8NBL1 |
| HOMO SAPIENS | 2 | 11844 | 5l0v | A | 279 | ARG | p.Arg279Trp | Q8NBL1 |
| HOMO SAPIENS | 9 | 1485 | 5va0 | A | 477 | ARG | p.Arg477His | P04150 |
| HOMO SAPIENS | 0 | 235 | 2fun | B | 2248 | ARG | p.Arg248Trp | Q14790 |
| HOMO SAPIENS | 4 | 235 | 2fun | B | 2285 | HIS | p.Asp285His | Q14790 |
| HOMO SAPIENS | 0 | 4822 | 5y3n | A | 307 | ARG | p.Arg307Gly | Q12931 |
| HOMO SAPIENS | 8 | 32928 | 4m6w | A | 1886 | HIS | p.His1886Tyr | Q8IYD8 |
| HOMO SAPIENS | 0 | 2711 | 4y5v | F | 22 | GLY | p.Gly46Glu | P19235 |
| HOMO SAPIENS | -1 | 2711 | 4y5v | F | 166 | ALA | p.Ala190Ser | P19235 |
| HOMO SAPIENS | 4 | 5161 | 5bpu | A | 42 | HIS | p.His42Arg | Q00604 |
| HOMO SAPIENS | 0 | 5161 | 5bpu | A | 44 | TYR | p.Tyr44Cys | Q00604 |
| HOMO SAPIENS | 0 | 5161 | 5bpu | A | 45 | VAL | p.Val45Glu | Q00604 |
| HOMO SAPIENS | 0 | 5161 | 5bpu | A | 60 | VAL | p.Val60Glu | Q00604 |
| HOMO SAPIENS | 0 | 5161 | 5bpu | A | 61 | LEU | p.Leu61Phe | Q00604 |
| HOMO SAPIENS | 7 | 5161 | 5bpu | A | 69 | CYS | p.Cys69Ser | Q00604 |
| HOMO SAPIENS | 0 | 5161 | 5bpu | A | 75 | SER | p.Ser75Cys | Q00604 |
| HOMO SAPIENS | 0 | 5161 | 5bpu | A | 90 | ARG | p.Arg90Pro | Q00604 |
| HOMO SAPIENS | 9 | 5161 | 5bpu | A | 96 | CYS | p.Cys96Tyr | Q00604 |
| HOMO SAPIENS | 9 | 5161 | 5bpu | A | 96 | CYS | p.Cys96Trp | Q00604 |
| HOMO SAPIENS | 0 | 5161 | 5bpu | A | 101 | SER | p.Ser101Phe | Q00604 |
| HOMO SAPIENS | 0 | 5161 | 5bpu | A | 105 | ALA | p.Ala105Thr | Q00604 |
| HOMO SAPIENS | 0 | 5161 | 5bpu | A | 110 | CYS | p.Cys110Gly | Q00604 |
| HOMO SAPIENS | 0 | 5161 | 5bpu | A | 121 | ARG | p.Arg121Trp | Q00604 |
| HOMO SAPIENS | 0 | 5161 | 5bpu | A | 121 | ARG | p.Arg121Leu | Q00604 |
| HOMO SAPIENS | 0 | 5161 | 5bpu | A | 124 | LEU | p.Leu124Phe | Q00604 |
| HOMO SAPIENS | 8 | 5161 | 5bpu | A | 128 | CYS | p.Cys128Ter | Q00604 |
| HOMO SAPIENS | 1 | 13944 | 5xqh | A | 238 | ARG | p.Arg238His | Q9GZN7 |
| HOMO SAPIENS | -1 | 7508 | 5jl6 | A | 39 | TRP | p.Trp39Arg | P11511 |
| HOMO SAPIENS | 0 | 7508 | 5jl6 | A | 201 | THR | p.Thr201Met | P11511 |
| HOMO SAPIENS | 0 | 7508 | 5jl6 | A | 210 | GLU | p.Glu210Lys | P11511 |
| HOMO SAPIENS | 1 | 7508 | 5jl6 | A | 264 | ARG | p.Arg264Cys | P11511 |
| HOMO SAPIENS | 2 | 7508 | 5jl6 | A | 365 | ARG | p.Arg365Gln | P11511 |
| HOMO SAPIENS | 7 | 7508 | 5jl6 | A | 375 | ARG | p.Arg375Cys | P11511 |
| HOMO SAPIENS | 8 | 7508 | 5jl6 | A | 435 | ARG | p.Arg435Cys | P11511 |

| | | | | | | | | |
|---|---|---|---|---|---|---|---|---|
| HOMO SAPIENS | 9 | 7508 | 5jl6 | A | 437 | CYS | p.Cys437Tyr | P11511 |
| HOMO SAPIENS | -1 | 30206 | 5l95 | B | 246 | ARG | p.Arg246Ter | Q9GZZ9 |
| HOMO SAPIENS | 4 | 875 | 3efh | B | 43 | GLU | p.Glu43Asp | P60891 |
| HOMO SAPIENS | 5 | 875 | 3efh | B | 65 | ASP | p.Asp65Asn | P60891 |
| HOMO SAPIENS | 0 | 875 | 3efh | B | 87 | ALA | p.Ala87Thr | P60891 |
| HOMO SAPIENS | 2 | 875 | 3efh | B | 115 | MET | p.Met115Thr | P60891 |
| HOMO SAPIENS | 8 | 875 | 3efh | B | 133 | GLN | p.Gln133Pro | P60891 |
| HOMO SAPIENS | 8 | 875 | 3efh | B | 152 | LEU | p.Leu152Pro | P60891 |
| HOMO SAPIENS | 7 | 875 | 3efh | B | 290 | ILE | p.Ile290Thr | P60891 |
| HOMO SAPIENS | 6 | 875 | 3efh | B | 306 | GLY | p.Gly306Arg | P60891 |
| HOMO SAPIENS | 0 | 7978 | 5edm | A | 122 | MET | p.Thr165Met | P00734 |
| HOMO SAPIENS | -1 | 7978 | 5edm | A | 257 | ARG | p.Arg314Cys | P00734 |
| HOMO SAPIENS | 0 | 1716 | 1b98 | M | 126 | ARG | p.Arg206Gln | P34130 |
| HOMO SAPIENS | 0 | 1716 | 1b98 | M | 126 | ARG | p.Arg206Trp | P34130 |
| HOMO SAPIENS | 9 | 3960 | 2qyp | B | 28 | ILE | p.Ile338Val | P07602 |
| HOMO SAPIENS | 1 | 3960 | 2qyp | B | 39 | LEU | p.Leu349Pro | P07602 |
| HOMO SAPIENS | 0 | 31414 | 5u1d | A | 333 | ILE | p.Ile393Val | Q03518 |
| HOMO SAPIENS | 0 | 31414 | 5u1d | A | 637 | ASP | p.Asp697Gly | Q03518 |
| HOMO SAPIENS | 0 | 31414 | 5u1d | A | 659 | ARG | p.Arg719Gln | Q03518 |
| HOMO SAPIENS | 3 | 119 | 2pho | A | 11 | ILE | p.Ile11Thr | P05089 |
| HOMO SAPIENS | 8 | 119 | 2pho | A | 21 | ARG | p.Arg21Ter | P05089 |
| HOMO SAPIENS | 0 | 119 | 2pho | A | 122 | TRP | p.Trp122Ter | P05089 |
| HOMO SAPIENS | 6 | 119 | 2pho | A | 138 | GLY | p.Gly138Val | P05089 |
| HOMO SAPIENS | 5 | 119 | 2pho | A | 290 | THR | p.Thr290Ser | P05089 |
| HOMO SAPIENS | 5 | 119 | 2pho | A | 291 | ARG | p.Arg291Ter | P05089 |
| HOMO SAPIENS | 0 | 3732 | 2i99 | A | 247 | TYR | p.Tyr247Ter | Q14894 |
| HOMO SAPIENS | -1 | 1382 | 4ffx | A | 2 | ALA | p.Ala2Val | P30566 |
| HOMO SAPIENS | 1 | 1382 | 4ffx | A | 72 | ILE | p.Ile72Met | P30566 |
| HOMO SAPIENS | 1 | 1382 | 4ffx | A | 100 | PRO | p.Pro100Ala | P30566 |
| HOMO SAPIENS | 0 | 1382 | 4ffx | A | 114 | TYR | p.Tyr114His | P30566 |
| HOMO SAPIENS | 1 | 1382 | 4ffx | A | 141 | ARG | p.Arg141Trp | P30566 |
| HOMO SAPIENS | 0 | 1382 | 4ffx | A | 147 | LYS | p.Lys147Met | P30566 |
| HOMO SAPIENS | 3 | 1382 | 4ffx | A | 190 | ARG | p.Arg190Gln | P30566 |
| HOMO SAPIENS | 3 | 1382 | 4ffx | A | 190 | ARG | p.Arg190Ter | P30566 |
| HOMO SAPIENS | 0 | 1382 | 4ffx | A | 217 | HIS | p.His217Asp | P30566 |
| HOMO SAPIENS | 2 | 1382 | 4ffx | A | 246 | LYS | p.Lys246Glu | P30566 |
| HOMO SAPIENS | 5 | 1382 | 4ffx | A | 303 | ARG | p.Arg303Cys | P30566 |
| HOMO SAPIENS | 1 | 1382 | 4ffx | A | 318 | PRO | p.Pro318Leu | P30566 |
| HOMO SAPIENS | 6 | 1382 | 4ffx | A | 332 | ASP | p.Asp332His | P30566 |
| HOMO SAPIENS | 6 | 1382 | 4ffx | A | 337 | ARG | p.Arg337Ter | P30566 |
| HOMO SAPIENS | 1 | 1382 | 4ffx | A | 396 | ARG | p.Arg396His | P30566 |
| HOMO SAPIENS | 1 | 1382 | 4ffx | A | 426 | ARG | p.Arg426His | P30566 |
| HOMO SAPIENS | 0 | 1382 | 4ffx | A | 438 | SER | p.Ser438Pro | P30566 |
| HOMO SAPIENS | 1 | 1382 | 4ffx | A | 447 | SER | p.Ser447Pro | P30566 |
| HOMO SAPIENS | 2 | 1382 | 4ffx | A | 448 | SER | p.Ser448Pro | P30566 |
| HOMO SAPIENS | 2 | 1382 | 4ffx | A | 450 | THR | p.Thr450Ser | P30566 |
| HOMO SAPIENS | 0 | 6212 | 1ao3 | A | 46 | SER | p.Ser1731Thr | P04275 |
| HOMO SAPIENS | 0 | 1711 | 3f9c | A | 279 | VAL | p.Val279Phe | Q13093 |
| HOMO SAPIENS | 0 | 6112 | 4fvl | A | 312 | HIS | p.His312Arg | P45452 |
| HOMO SAPIENS | 0 | 6112 | 4fvl | A | 323 | THR | p.Thr323Met | P45452 |
| HOMO SAPIENS | 0 | 6112 | 4fvl | A | 390 | ASP | p.Asp390Gly | P45452 |
| HOMO SAPIENS | 0 | 6112 | 4fvl | A | 458 | ARG | p.Arg458Cys | P45452 |
| HOMO SAPIENS | 2 | 6180 | 5ggn | A | 97 | ARG | p.Arg97Trp | Q8WZA1 |
| HOMO SAPIENS | 2 | 6180 | 5ggn | A | 215 | ARG | p.Arg215Ter | Q8WZA1 |

| | | | | | | | | |
|---|---|---|---|---|---|---|---|---|
| HOMO SAPIENS | -1 | 5526 | 4ak3 | A | 11 | ASP | p.Asp1232Asn | P02461 |
| HOMO SAPIENS | 6 | 5526 | 4ak3 | A | 38 | ALA | p.Ala1259Thr | P02461 |
| HOMO SAPIENS | 0 | 5526 | 4ak3 | A | 92 | LYS | p.Lys1313Arg | P02461 |
| HOMO SAPIENS | 0 | 5526 | 4ak3 | A | 132 | GLN | p.His1353Gln | P02461 |
| HOMO SAPIENS | 4 | 16616 | 1hci | B | 433 | SER | p.Ser433Leu | P35609 |
| HOMO SAPIENS | 1 | 16616 | 1hci | B | 475 | ASP | p.Asp475Asn | P35609 |
| HOMO SAPIENS | 1 | 16616 | 1hci | B | 476 | ALA | p.Ala476Ser | P35609 |
| HOMO SAPIENS | 2 | 16616 | 1hci | B | 507 | MET | p.Met507Val | P35609 |
| HOMO SAPIENS | 1 | 16616 | 1hci | B | 604 | MET | p.Met604Val | P35609 |
| HOMO SAPIENS | 2 | 16616 | 1hci | B | 640 | ARG | p.Arg640His | P35609 |
| HOMO SAPIENS | 1 | 16616 | 1hci | B | 721 | ARG | p.Arg721Cys | P35609 |
| HOMO SAPIENS | -1 | 3402 | 5iaa | B | 57 | MET | p.Met57Val | Q9GZZ9 |
| HOMO SAPIENS | 7 | 3250 | 3irq | B | 193 | PRO | p.Pro193Ala | P55265 |
| HOMO SAPIENS | 2 | 15200 | 2wns | A | 30 | SER | p.Ser30Gly | P11172 |
| HOMO SAPIENS | 0 | 15200 | 2wns | A | 96 | ARG | p.Arg96Gly | P11172 |
| HOMO SAPIENS | 0 | 15200 | 2wns | A | 109 | VAL | p.Val109Gly | P11172 |
| HOMO SAPIENS | 2 | 15759 | 4x4w | B | 140 | LEU | p.Leu166Ser | Q96Q11 |
| HOMO SAPIENS | 0 | 15759 | 4x4w | B | 197 | ILE | p.Ile223Thr | Q96Q11 |
| HOMO SAPIENS | 0 | 3156 | 3w03 | B | 14 | ALA | p.Ala14Thr | Q9H9Q4 |
| HOMO SAPIENS | 0 | 3156 | 3w03 | B | 57 | ARG | p.Arg57Gly | Q9H9Q4 |
| HOMO SAPIENS | 0 | 3156 | 3w03 | B | 123 | CYS | p.Cys123Arg | Q9H9Q4 |
| HOMO SAPIENS | 0 | 11693 | 3ojy | A | 63 | GLN | p.Gln93Lys | P07357 |
| HOMO SAPIENS | 0 | 1421 | 2qc8 | E | 275 | GLU | p.Glu275Asp | P15104 |
| HOMO SAPIENS | 9 | 1421 | 2qc8 | E | 324 | ARG | p.Arg324Cys | P15104 |
| HOMO SAPIENS | 6 | 1421 | 2qc8 | E | 341 | ARG | p.Arg341Cys | P15104 |
| HOMO SAPIENS | 6 | 2850 | 3qvb | A | 83 | THR | p.Thr83Met | P46976 |
| HOMO SAPIENS | 8 | 2850 | 3qvb | A | 102 | ASP | p.Asp102His | P46976 |
| HOMO SAPIENS | 1 | 24314 | 4m9p | A | 480 | PRO | p.Pro480Leu | P21333 |
| HOMO SAPIENS | 0 | 24314 | 4m9p | A | 528 | VAL | p.Val528Met | P21333 |
| HOMO SAPIENS | 0 | 24314 | 4m9p | A | 590 | VAL | p.Val590Ile | P21333 |
| HOMO SAPIENS | 0 | 24314 | 4m9p | A | 605 | ASP | p.Asp605Asn | P21333 |
| HOMO SAPIENS | 1 | 24314 | 4m9p | A | 637 | PRO | p.Pro637Gln | P21333 |
| HOMO SAPIENS | 0 | 24314 | 4m9p | A | 656 | LEU | p.Leu656Phe | P21333 |
| HOMO SAPIENS | 2 | 24314 | 4m9p | A | 711 | VAL | p.Val711Asp | P21333 |
| HOMO SAPIENS | 0 | 4951 | 5joe | A | 40 | THR | p.Thr9621Ser | Q8WZ42 |
| HOMO SAPIENS | 1 | 4951 | 5joe | A | 41 | GLU | p.Glu9622Gln | Q8WZ42 |
| HOMO SAPIENS | 0 | 4951 | 5joe | A | 82 | ALA | p.Ala9663Ser | Q8WZ42 |
| HOMO SAPIENS | 4 | 4951 | 5joe | A | 89 | ILE | p.Ile9670Asn | Q8WZ42 |
| HOMO SAPIENS | 0 | 9664 | 3c6m | A | 56 | GLY | p.Gly56Ser | P52788 |
| HOMO SAPIENS | 0 | 9664 | 3c6m | A | 132 | VAL | p.Val132Gly | P52788 |
| HOMO SAPIENS | 1 | 9664 | 3c6m | A | 190 | THR | p.Thr190Ile | P52788 |
| HOMO SAPIENS | -1 | 12118 | 5hcd | B | 19 | GLN | p.Gln19Ter | P01031 |
| HOMO SAPIENS | 0 | 1583 | 2b4x | L | 7 | ILE | p.Ile39Asn | P01008 |
| HOMO SAPIENS | 0 | 1583 | 2b4x | L | 24 | ARG | p.Arg56Cys | P01008 |
| HOMO SAPIENS | 0 | 1583 | 2b4x | L | 47 | ARG | p.Arg79Cys | P01008 |
| HOMO SAPIENS | 0 | 1583 | 2b4x | L | 47 | ARG | p.Arg79His | P01008 |
| HOMO SAPIENS | 1 | 1583 | 2b4x | L | 95 | CYS | p.Cys127Arg | P01008 |
| HOMO SAPIENS | 1 | 1583 | 2b4x | L | 99 | LEU | p.Leu131Phe | P01008 |
| HOMO SAPIENS | 0 | 1583 | 2b4x | L | 116 | SER | p.Ser148Pro | P01008 |
| HOMO SAPIENS | 0 | 1583 | 2b4x | L | 129 | ARG | p.Arg161Gln | P01008 |
| HOMO SAPIENS | 0 | 1583 | 2b4x | L | 135 | ASN | p.Asn167Thr | P01008 |
| HOMO SAPIENS | 2 | 1583 | 2b4x | L | 187 | ASN | p.Asn219Asp | P01008 |
| HOMO SAPIENS | 0 | 1583 | 2b4x | L | 191 | SER | p.Ser223Pro | P01008 |
| HOMO SAPIENS | 2 | 1583 | 2b4x | L | 349 | SER | p.Ser381Pro | P01008 |

| | | | | | | | | |
|---|---|---|---|---|---|---|---|---|
| HOMO SAPIENS | 0 | 1583 | 2b4x | L | 382 | ALA | p.Ala414Thr | P01008 |
| HOMO SAPIENS | 0 | 1583 | 2b4x | L | 387 | ALA | p.Ala419Val | P01008 |
| HOMO SAPIENS | 0 | 1583 | 2b4x | L | 392 | GLY | p.Gly424Asp | P01008 |
| HOMO SAPIENS | 0 | 1583 | 2b4x | L | 393 | ARG | p.Arg425His | P01008 |
| HOMO SAPIENS | 0 | 1583 | 2b4x | L | 393 | ARG | p.Arg425Cys | P01008 |
| HOMO SAPIENS | 0 | 1583 | 2b4x | L | 394 | SER | p.Ser426Leu | P01008 |
| HOMO SAPIENS | 0 | 1583 | 2b4x | L | 404 | ALA | p.Ala436Thr | P01008 |
| HOMO SAPIENS | 2 | 1583 | 2b4x | L | 407 | PRO | p.Pro439Leu | P01008 |
| HOMO SAPIENS | 2 | 1583 | 2b4x | L | 429 | PRO | p.Pro461Leu | P01008 |
| HOMO SAPIENS | 0 | 11647 | 2a1t | S | 78 | ALA | p.Ala78Ser | P38117 |
| HOMO SAPIENS | 0 | 11647 | 2a1t | S | 98 | ARG | p.Arg98Cys | P38117 |
| HOMO SAPIENS | 8 | 11647 | 2a1t | S | 128 | ASP | p.Asp128Asn | P38117 |
| HOMO SAPIENS | 5 | 11647 | 2a1t | S | 154 | THR | p.Thr154Met | P38117 |
| HOMO SAPIENS | 6 | 11647 | 2a1t | S | 164 | ARG | p.Arg164Gln | P38117 |
| HOMO SAPIENS | 5 | 11647 | 2a1t | S | 174 | ARG | p.Arg174His | P38117 |
| HOMO SAPIENS | -1 | 11200 | 4tx5 | A | 4 | ILE | p.Ile59Val | Q9NR28 |
| HOMO SAPIENS | 0 | 11200 | 4tx5 | A | 169 | GLY | p.Gly224Arg | Q9NR28 |
| HOMO SAPIENS | 3 | 417 | 2vr2 | A | 6 | ARG | p.Arg6Gln | Q14117 |
| HOMO SAPIENS | 1 | 417 | 2vr2 | A | 334 | GLN | p.Gln334Arg | Q14117 |
| HOMO SAPIENS | 2 | 417 | 2vr2 | A | 360 | TRP | p.Trp360Arg | Q14117 |
| HOMO SAPIENS | 0 | 417 | 2vr2 | A | 379 | SER | p.Ser379Arg | Q14117 |
| HOMO SAPIENS | 0 | 417 | 2vr2 | A | 435 | GLY | p.Gly435Arg | Q14117 |
| HOMO SAPIENS | 0 | 12498 | 3eo3 | B | 423 | ILE | p.Ile423Val | Q9Y223 |
| HOMO SAPIENS | 5 | 12498 | 3eo3 | B | 572 | VAL | p.Val572Leu | Q9Y223 |
| HOMO SAPIENS | 0 | 12498 | 3eo3 | B | 576 | GLY | p.Gly576Glu | Q9Y223 |
| HOMO SAPIENS | 0 | 12498 | 3eo3 | B | 587 | ILE | p.Ile587Thr | Q9Y223 |
| HOMO SAPIENS | 0 | 12498 | 3eo3 | B | 615 | SER | p.Ser615Ter | Q9Y223 |
| HOMO SAPIENS | 1 | 12498 | 3eo3 | B | 631 | ALA | p.Ala631Val | Q9Y223 |
| HOMO SAPIENS | 1 | 12498 | 3eo3 | B | 669 | GLY | p.Gly669Arg | Q9Y223 |
| HOMO SAPIENS | 1 | 12498 | 3eo3 | B | 696 | VAL | p.Val696Met | Q9Y223 |
| HOMO SAPIENS | 2 | 12498 | 3eo3 | B | 712 | MET | p.Met712Thr | Q9Y223 |
| HOMO SAPIENS | 0 | 1568 | 1r74 | A | 49 | LEU | p.Leu50Pro | Q14749 |
| HOMO SAPIENS | 0 | 1568 | 1r74 | A | 176 | HIS | p.His177Asn | Q14749 |
| HOMO SAPIENS | 3 | 147 | 5eig | A | 67 | THR | p.Thr67Ile | P32929 |
| HOMO SAPIENS | 3 | 147 | 5eig | A | 240 | GLN | p.Gln240Glu | P32929 |
| HOMO SAPIENS | -1 | 147 | 5eig | A | 403 | SER | p.Ser403Ile | P32929 |
| HOMO SAPIENS | 8 | 20036 | 2lsr | A | 31 | ARG | p.Arg31Ter | Q9Y6N9 |
| HOMO SAPIENS | -1 | 41382 | 4uis | B | 431 | UNK | p.Ala431Glu | P49768 |
| HOMO SAPIENS | -1 | 8536 | 3i6w | B | 85 | PRO | p.Pro85Leu | O96017 |
| HOMO SAPIENS | 0 | 8536 | 3i6w | B | 93 | TRP | p.Trp93Ter | O96017 |
| HOMO SAPIENS | 0 | 8536 | 3i6w | B | 95 | ARG | p.Arg95Ter | O96017 |
| HOMO SAPIENS | 0 | 8536 | 3i6w | B | 117 | ARG | p.Arg117Gly | O96017 |
| HOMO SAPIENS | 0 | 8536 | 3i6w | B | 119 | LYS | p.Lys119Arg | O96017 |
| HOMO SAPIENS | 0 | 8536 | 3i6w | B | 137 | ARG | p.Arg137Gln | O96017 |
| HOMO SAPIENS | 0 | 8536 | 3i6w | B | 137 | ARG | p.Arg137Ter | O96017 |
| HOMO SAPIENS | 0 | 8536 | 3i6w | B | 145 | ARG | p.Arg145Trp | O96017 |
| HOMO SAPIENS | 0 | 8536 | 3i6w | B | 160 | ILE | p.Ile160Met | O96017 |
| HOMO SAPIENS | 0 | 41322 | 4bxl | C | 46 | GLU | p.Glu46Lys | P37840 |
| HOMO SAPIENS | 5 | 41322 | 4bxl | C | 53 | ALA | p.Ala53Thr | P37840 |
| HOMO SAPIENS | 2 | 24966 | 3hve | A | 15 | ARG | p.Arg15Ser | Q9H2C0 |
| HOMO SAPIENS | 1 | 24966 | 3hve | A | 138 | ARG | p.Arg138His | Q9H2C0 |
| HOMO SAPIENS | 0 | 24966 | 3hve | A | 201 | ARG | p.Arg201Ter | Q9H2C0 |
| HOMO SAPIENS | 0 | 11680 | 5an9 | J | 8 | ASN | p.Asn8Lys | Q9Y3A5 |
| HOMO SAPIENS | 1 | 11680 | 5an9 | J | 33 | LYS | p.Lys33Arg | Q9Y3A5 |

| | | | | | | | | |
|---|---|---|---|---|---|---|---|---|
| HOMO SAPIENS | 1 | 11680 | 5an9 | J | 126 | ARG | p.Arg126Thr | Q9Y3A5 |
| HOMO SAPIENS | 1 | 11680 | 5an9 | J | 130 | VAL | p.Val130Met | Q9Y3A5 |
| HOMO SAPIENS | 1 | 11680 | 5an9 | J | 169 | ARG | p.Arg169Cys | Q9Y3A5 |
| HOMO SAPIENS | 1 | 11680 | 5an9 | J | 218 | ARG | p.Arg218Ter | Q9Y3A5 |
| HOMO SAPIENS | 4 | 978 | 3zs0 | B | 7 | TYR | p.Tyr173Cys | P05164 |
| HOMO SAPIENS | 1 | 978 | 3zs0 | B | 85 | MET | p.Met251Thr | P05164 |
| HOMO SAPIENS | 4 | 5140 | 2l4r | A | 52 | CYS | p.Cys52Ter | Q12809 |
| HOMO SAPIENS | 4 | 5140 | 2l4r | A | 63 | PRO | p.Pro63His | Q12809 |
| HOMO SAPIENS | 0 | 5140 | 2l4r | A | 100 | ARG | p.Arg100Gly | Q12809 |
| HOMO SAPIENS | 4 | 5140 | 2l4r | A | 129 | PHE | p.Phe129Leu | Q12809 |
| HOMO SAPIENS | 0 | 25836 | 2gy7 | B | 103 | GLU | p.Glu103Asp | Q02763 |
| HOMO SAPIENS | 0 | 25836 | 2gy7 | B | 148 | ILE | p.Ile148Thr | Q02763 |
| HOMO SAPIENS | 0 | 25836 | 2gy7 | B | 226 | ALA | p.Ala226Val | Q02763 |
| HOMO SAPIENS | 0 | 25836 | 2gy7 | B | 391 | THR | p.Thr391Ile | Q02763 |
| HOMO SAPIENS | 4 | 12563 | 5kwy | D | 20 | GLU | p.Glu20Ter | P61916 |
| HOMO SAPIENS | 2 | 12563 | 5kwy | D | 30 | VAL | p.Val30Met | P61916 |
| HOMO SAPIENS | 4 | 12563 | 5kwy | D | 39 | VAL | p.Val39Met | P61916 |
| HOMO SAPIENS | 8 | 12563 | 5kwy | D | 67 | SER | p.Ser67Pro | P61916 |
| HOMO SAPIENS | 8 | 12563 | 5kwy | D | 99 | CYS | p.Cys99Arg | P61916 |
| HOMO SAPIENS | 2 | 12563 | 5kwy | D | 118 | GLU | p.Glu118Ter | P61916 |
| HOMO SAPIENS | 2 | 12563 | 5kwy | D | 120 | PRO | p.Pro120Ser | P61916 |
| HOMO SAPIENS | -1 | 1679 | 1pmk | A | -2 | VAL | p.Val374Phe | P00747 |
| HOMO SAPIENS | -1 | 5315 | 2w2n | P | 53 | ALA | p.Ala53Val | Q8NBP7 |
| HOMO SAPIENS | 8 | 13496 | 2yad | A | 138 | THR | p.Asn138Thr | P11686 |
| HOMO SAPIENS | -1 | 13496 | 2yad | A | 173 | ARG | p.Arg167Gln | P11686 |
| HOMO SAPIENS | 2 | 13496 | 2yad | A | 186 | SER | p.Asn186Ser | P11686 |
| HOMO SAPIENS | 0 | 4426 | 3qd6 | C | 123 | ALA | p.Ala123Glu | P29965 |
| HOMO SAPIENS | 1 | 4426 | 3qd6 | C | 140 | TRP | p.Trp140Gly | P29965 |
| HOMO SAPIENS | 0 | 4426 | 3qd6 | C | 155 | LEU | p.Leu155Pro | P29965 |
| HOMO SAPIENS | 6 | 4426 | 3qd6 | C | 227 | GLY | p.Gly227Val | P29965 |
| HOMO SAPIENS | 0 | 4426 | 3qd6 | C | 235 | ALA | p.Ala235Pro | P29965 |
| HOMO SAPIENS | 0 | 10734 | 3dyd | B | 45 | PRO | p.Pro45Ser | P17735 |
| HOMO SAPIENS | -1 | 10734 | 3dyd | B | 57 | ARG | p.Arg57Ter | P17735 |
| HOMO SAPIENS | 8 | 10734 | 3dyd | B | 417 | ARG | p.Arg417Ter | P17735 |
| HOMO SAPIENS | 0 | 6956 | 4pzs | A | 1600 | LEU | p.Leu1600Phe | Q09472 |
| HOMO SAPIENS | 0 | 9370 | 3hfv | A | 21 | SER | p.Ser57Cys | O95363 |
| HOMO SAPIENS | 1 | 9370 | 3hfv | A | 25 | ASP | p.Asp61Glu | O95363 |
| HOMO SAPIENS | 1 | 9370 | 3hfv | A | 130 | ALA | p.Ala166Thr | O95363 |
| HOMO SAPIENS | 5 | 9370 | 3hfv | A | 148 | ASP | p.Asp184Asn | O95363 |
| HOMO SAPIENS | 0 | 9370 | 3hfv | A | 187 | ARG | p.Arg223Cys | O95363 |
| HOMO SAPIENS | 0 | 9370 | 3hfv | A | 210 | THR | p.Thr246Met | O95363 |
| HOMO SAPIENS | 1 | 9370 | 3hfv | A | 244 | ASN | p.Asn280Ser | O95363 |
| HOMO SAPIENS | 2 | 9370 | 3hfv | A | 288 | TYR | p.Tyr324Cys | O95363 |
| HOMO SAPIENS | 5 | 9370 | 3hfv | A | 293 | ILE | p.Ile329Thr | O95363 |
| HOMO SAPIENS | 1 | 9370 | 3hfv | A | 355 | ASP | p.Asp391Val | O95363 |
| HOMO SAPIENS | 1 | 9370 | 3hfv | A | 371 | THR | p.Thr407Met | O95363 |
| HOMO SAPIENS | 1 | 9370 | 3hfv | A | 383 | ARG | p.Arg419Cys | O95363 |
| HOMO SAPIENS | 1 | 9370 | 3hfv | A | 387 | ARG | p.Arg423Gln | O95363 |
| HOMO SAPIENS | 0 | 14401 | 1q3x | A | 371 | ASP | p.Asp371Tyr | O00187 |
| HOMO SAPIENS | 0 | 14401 | 1q3x | A | 377 | VAL | p.Val377Ala | O00187 |
| HOMO SAPIENS | 0 | 14401 | 1q3x | A | 439 | ARG | p.Arg439His | O00187 |
| HOMO SAPIENS | 0 | 14401 | 1q3x | A | 577 | GLN | p.Gln577His | O00187 |
| HOMO SAPIENS | 2 | 18270 | 3teq | A | 429 | ARG | p.Arg429Cys | Q13586 |
| HOMO SAPIENS | -1 | 16081 | 4mqt | A | 231 | VAL | p.Val231Ile | P08172 |

| | | | | | | | | |
|---|---|---|---|---|---|---|---|---|
| HOMO SAPIENS | 9 | 6607 | 3whu | B | 81 | ASP | p.Asp81His | Q8NI22 |
| HOMO SAPIENS | 9 | 6607 | 3whu | B | 129 | ASP | p.Asp129Glu | Q8NI22 |
| HOMO SAPIENS | 9 | 6607 | 3whu | B | 136 | ILE | p.Ile136Thr | Q8NI22 |
| HOMO SAPIENS | 0 | 7208 | 2e9w | D | 11 | ASN | p.Asn36Ser | P21583 |
| HOMO SAPIENS | 0 | 24566 | 4bdx | A | 10 | LEU | p.Leu140Val | P00748 |
| HOMO SAPIENS | 5 | 24566 | 4bdx | A | 77 | PRO | p.Ala207Pro | P00748 |
| HOMO SAPIENS | 7 | 606 | 3qnf | A | 319 | MET | p.Met319Val | Q9NZ08 |
| HOMO SAPIENS | 8 | 4097 | 1ydl | A | 21 | LEU | p.Leu21Pro | Q6ZYL4 |
| HOMO SAPIENS | 6 | 4097 | 1ydl | A | 56 | ARG | p.Arg56Ter | Q6ZYL4 |
| HOMO SAPIENS | 8 | 36691 | 1dqb | A | 42 | ARG | p.Arg403Lys | P07204 |
| HOMO SAPIENS | 0 | 5642 | 3j7y | X | 59 | ARG | p.Arg59Pro | Q13084 |
| HOMO SAPIENS | 1 | 5642 | 3j7y | X | 204 | VAL | p.Val204Met | Q13084 |
| HOMO SAPIENS | 1 | 1119 | 3nwu | B | 252 | ALA | p.Ala252Thr | Q92743 |
| HOMO SAPIENS | 3 | 1119 | 3nwu | B | 274 | ARG | p.Arg274Gln | Q92743 |
| HOMO SAPIENS | 9 | 1119 | 3nwu | B | 297 | VAL | p.Val297Met | Q92743 |
| HOMO SAPIENS | -1 | 1119 | 3nwu | B | 302 | ARG | p.Arg302Ter | Q92743 |
| HOMO SAPIENS | 9 | 1119 | 3nwu | B | 321 | ALA | p.Ala321Thr | Q92743 |
| HOMO SAPIENS | -1 | 1119 | 3nwu | B | 370 | ARG | p.Arg370Ter | Q92743 |
| HOMO SAPIENS | 0 | 416 | 4ycu | B | 105 | LEU | p.Leu105His | Q15046 |
| HOMO SAPIENS | 0 | 416 | 4ycu | B | 145 | TYR | p.Tyr145His | Q15046 |
| HOMO SAPIENS | 0 | 416 | 4ycu | B | 200 | PRO | p.Pro200Leu | Q15046 |
| HOMO SAPIENS | 0 | 416 | 4ycu | B | 201 | TYR | p.Tyr201His | Q15046 |
| HOMO SAPIENS | 0 | 416 | 4ycu | B | 420 | ARG | p.Arg420Cys | Q15046 |
| HOMO SAPIENS | 9 | 3297 | 4fpb | A | 680 | THR | p.Thr680Ser | P19367 |
| HOMO SAPIENS | 0 | 6771 | 2zt6 | A | 201 | ALA | p.Ala255Val | P41250 |
| HOMO SAPIENS | 0 | 6771 | 2zt6 | A | 202 | ASP | p.Asp256His | P41250 |
| HOMO SAPIENS | 0 | 6771 | 2zt6 | A | 334 | ARG | p.Arg388Gln | P41250 |
| HOMO SAPIENS | -1 | 6771 | 2zt6 | A | 439 | ASN | p.Asn493Ser | P41250 |
| HOMO SAPIENS | -1 | 6771 | 2zt6 | A | 500 | ASP | p.Asp554Asn | P41250 |
| HOMO SAPIENS | 1 | 6771 | 2zt6 | A | 581 | SER | p.Ser635Leu | P41250 |
| HOMO SAPIENS | -1 | 6771 | 2zt6 | A | 684 | GLU | p.Glu738Lys | P41250 |
| HOMO SAPIENS | 0 | 2687 | 4grw | B | 276 | VAL | p.Val298Phe | P29460 |
| HOMO SAPIENS | 6 | 913 | 1m75 | B | 45 | ASP | p.Asp57Glu | Q16836 |
| HOMO SAPIENS | 2 | 913 | 1m75 | B | 74 | PRO | p.Leu86Pro | Q16836 |
| HOMO SAPIENS | 3 | 913 | 1m75 | B | 214 | TYR | p.Tyr226His | Q16836 |
| HOMO SAPIENS | 3 | 913 | 1m75 | B | 224 | ARG | p.Arg236Ter | Q16836 |
| HOMO SAPIENS | 4 | 913 | 1m75 | B | 246 | PRO | p.Pro258Leu | Q16836 |
| HOMO SAPIENS | 2 | 4713 | 4djn | B | 40 | ARG | p.Arg40Cys | Q7LG56 |
| HOMO SAPIENS | 1 | 4713 | 4djn | B | 41 | ARG | p.Arg41Pro | Q7LG56 |
| HOMO SAPIENS | 2 | 4713 | 4djn | B | 110 | ARG | p.Arg110His | Q7LG56 |
| HOMO SAPIENS | 2 | 4713 | 4djn | B | 121 | ARG | p.Arg121His | Q7LG56 |
| HOMO SAPIENS | 9 | 4713 | 4djn | B | 131 | GLU | p.Glu131Lys | Q7LG56 |
| HOMO SAPIENS | 2 | 4713 | 4djn | B | 182 | THR | p.Thr182Ala | Q7LG56 |
| HOMO SAPIENS | 6 | 4713 | 4djn | B | 194 | GLU | p.Glu194Lys | Q7LG56 |
| HOMO SAPIENS | 2 | 4713 | 4djn | B | 211 | ARG | p.Arg211Lys | Q7LG56 |
| HOMO SAPIENS | 0 | 4713 | 4djn | B | 224 | ILE | p.Ile224Ser | Q7LG56 |
| HOMO SAPIENS | 3 | 4713 | 4djn | B | 229 | GLY | p.Gly229Val | Q7LG56 |
| HOMO SAPIENS | 2 | 4713 | 4djn | B | 236 | CYS | p.Cys236Phe | Q7LG56 |
| HOMO SAPIENS | 0 | 4713 | 4djn | B | 273 | GLY | p.Gly273Ser | Q7LG56 |
| HOMO SAPIENS | -1 | 4713 | 4djn | B | 282 | MSE | p.Met282Ile | Q7LG56 |
| HOMO SAPIENS | 3 | 4713 | 4djn | B | 284 | GLN | p.Gln284Ter | Q7LG56 |
| HOMO SAPIENS | 3 | 4713 | 4djn | B | 305 | ALA | p.Ala305Ser | Q7LG56 |
| HOMO SAPIENS | -1 | 8974 | 5n7i | A | 500 | ARG | p.Arg500Ter | Q8N4S9 |
| HOMO SAPIENS | 0 | 17072 | 3fxi | B | 299 | ASP | p.Asp299Gly | O00206 |

| | | | | | | | | |
|---|---|---|---|---|---|---|---|---|
| HOMO SAPIENS | 1 | 14355 | 2wwk | T | 55 | ILE | p.Ile34306Asn | Q8WZ42 |
| HOMO SAPIENS | 0 | 14355 | 2wwk | T | 64 | LEU | p.Leu34315Pro | Q8WZ42 |
| HOMO SAPIENS | 2 | 17797 | 1xu1 | S | 72 | ARG | p.Arg72His | O14836 |
| HOMO SAPIENS | -1 | 10836 | 4ztu | A | 39 | GLN | p.Gln39Arg | P54098 |
| HOMO SAPIENS | -1 | 10836 | 4ztu | A | 42 | ARG | p.Arg42Gln | P54098 |
| HOMO SAPIENS | -1 | 10836 | 4ztu | A | 45 | GLN | p.Gln45Arg | P54098 |
| HOMO SAPIENS | -1 | 10836 | 4ztu | A | 52 | GLN | p.Gln52Glu | P54098 |
| HOMO SAPIENS | 0 | 10836 | 4ztu | A | 88 | PHE | p.Phe88Leu | P54098 |
| HOMO SAPIENS | 1 | 10836 | 4ztu | A | 194 | ALA | p.Ala194Val | P54098 |
| HOMO SAPIENS | 1 | 10836 | 4ztu | A | 227 | ARG | p.Arg227Trp | P54098 |
| HOMO SAPIENS | 0 | 10836 | 4ztu | A | 232 | ARG | p.Arg232His | P54098 |
| HOMO SAPIENS | 1 | 10836 | 4ztu | A | 285 | GLN | p.Gln285His | P54098 |
| HOMO SAPIENS | 1 | 10836 | 4ztu | A | 434 | TYR | p.Tyr434His | P54098 |
| HOMO SAPIENS | -1 | 10836 | 4ztu | A | 511 | SER | p.Ser511Asn | P54098 |
| HOMO SAPIENS | 0 | 10836 | 4ztu | A | 533 | CYS | p.Cys533Trp | P54098 |
| HOMO SAPIENS | 1 | 10836 | 4ztu | A | 546 | ARG | p.Arg546Cys | P54098 |
| HOMO SAPIENS | 1 | 10836 | 4ztu | A | 587 | PRO | p.Pro587Leu | P54098 |
| HOMO SAPIENS | 0 | 10836 | 4ztu | A | 603 | MET | p.Met603Thr | P54098 |
| HOMO SAPIENS | 0 | 10836 | 4ztu | A | 623 | LEU | p.Leu623Trp | P54098 |
| HOMO SAPIENS | -1 | 10836 | 4ztu | A | 627 | ARG | p.Arg627Trp | P54098 |
| HOMO SAPIENS | -1 | 10836 | 4ztu | A | 628 | ARG | p.Arg628Pro | P54098 |
| HOMO SAPIENS | 0 | 10836 | 4ztu | A | 635 | PRO | p.Pro635Leu | P54098 |
| HOMO SAPIENS | 0 | 10836 | 4ztu | A | 662 | GLU | p.Glu662Lys | P54098 |
| HOMO SAPIENS | -1 | 10836 | 4ztu | A | 674 | GLY | p.Gly674Asp | P54098 |
| HOMO SAPIENS | -1 | 10836 | 4ztu | A | 722 | ARG | p.Arg722His | P54098 |
| HOMO SAPIENS | -1 | 10836 | 4ztu | A | 737 | GLY | p.Gly737Arg | P54098 |
| HOMO SAPIENS | 0 | 10836 | 4ztu | A | 748 | TRP | p.Trp748Ser | P54098 |
| HOMO SAPIENS | 0 | 10836 | 4ztu | A | 749 | PHE | p.Phe749Ser | P54098 |
| HOMO SAPIENS | 1 | 10836 | 4ztu | A | 807 | ARG | p.Arg807Cys | P54098 |
| HOMO SAPIENS | 1 | 10836 | 4ztu | A | 831 | TYR | p.Tyr831Cys | P54098 |
| HOMO SAPIENS | 8 | 10836 | 4ztu | A | 848 | GLY | p.Gly848Ser | P54098 |
| HOMO SAPIENS | 1 | 10836 | 4ztu | A | 852 | ARG | p.Arg852Cys | P54098 |
| HOMO SAPIENS | 9 | 10836 | 4ztu | A | 853 | ARG | p.Arg853Trp | P54098 |
| HOMO SAPIENS | 1 | 10836 | 4ztu | A | 864 | ASN | p.Asn864Ser | P54098 |
| HOMO SAPIENS | 1 | 10836 | 4ztu | A | 879 | GLN | p.Gln879Arg | P54098 |
| HOMO SAPIENS | 1 | 10836 | 4ztu | A | 914 | THR | p.Thr914Pro | P54098 |
| HOMO SAPIENS | 6 | 10836 | 4ztu | A | 932 | HIS | p.His932Tyr | P54098 |
| HOMO SAPIENS | 2 | 10836 | 4ztu | A | 944 | GLU | p.Glu944Lys | P54098 |
| HOMO SAPIENS | 6 | 10836 | 4ztu | A | 955 | TYR | p.Tyr955Cys | P54098 |
| HOMO SAPIENS | 1 | 10836 | 4ztu | A | 957 | ALA | p.Ala957Ser | P54098 |
| HOMO SAPIENS | 1 | 10836 | 4ztu | A | 957 | ALA | p.Ala957Val | P54098 |
| HOMO SAPIENS | 1 | 10836 | 4ztu | A | 964 | ARG | p.Arg964Cys | P54098 |
| HOMO SAPIENS | 1 | 10836 | 4ztu | A | 966 | LEU | p.Leu966Arg | P54098 |
| HOMO SAPIENS | 0 | 10836 | 4ztu | A | 1051 | GLY | p.Gly1051Arg | P54098 |
| HOMO SAPIENS | 1 | 10836 | 4ztu | A | 1073 | PRO | p.Pro1073Leu | P54098 |
| HOMO SAPIENS | 1 | 10836 | 4ztu | A | 1096 | ARG | p.Arg1096Cys | P54098 |
| HOMO SAPIENS | 1 | 10836 | 4ztu | A | 1096 | ARG | p.Arg1096His | P54098 |
| HOMO SAPIENS | 0 | 10836 | 4ztu | A | 1116 | MET | p.Met1116Thr | P54098 |
| HOMO SAPIENS | 1 | 10836 | 4ztu | A | 1142 | ARG | p.Arg1142Trp | P54098 |
| HOMO SAPIENS | 1 | 10836 | 4ztu | A | 1146 | ARG | p.Arg1146Cys | P54098 |
| HOMO SAPIENS | 1 | 10836 | 4ztu | A | 1157 | ASN | p.Asn1157Ser | P54098 |
| HOMO SAPIENS | -1 | 10836 | 4ztu | A | 1234 | ARG | p.Arg1234Gln | P54098 |
| HOMO SAPIENS | -1 | 10836 | 4ztu | A | 1236 | GLN | p.Gln1236His | P54098 |
| HOMO SAPIENS | 5 | 2666 | 1jph | A | 62 | PRO | p.Pro62Leu | P06132 |

| | | | | | | | | |
|---|---|---|---|---|---|---|---|---|
| HOMO SAPIENS | 5 | 2666 | 1jph | A | 116 | GLN | p.Gln116Ter | P06132 |
| HOMO SAPIENS | 0 | 2666 | 1jph | A | 165 | MET | p.Met165Arg | P06132 |
| HOMO SAPIENS | 6 | 2666 | 1jph | A | 167 | GLU | p.Glu167Lys | P06132 |
| HOMO SAPIENS | 0 | 2666 | 1jph | A | 195 | LEU | p.Leu195Phe | P06132 |
| HOMO SAPIENS | 7 | 2666 | 1jph | A | 253 | LEU | p.Leu253Gln | P06132 |
| HOMO SAPIENS | 0 | 2666 | 1jph | A | 281 | GLY | p.Gly281Glu | P06132 |
| HOMO SAPIENS | 9 | 2666 | 1jph | A | 304 | ASN | p.Asn304Lys | P06132 |
| HOMO SAPIENS | 8 | 2666 | 1jph | A | 311 | TYR | p.Tyr311Cys | P06132 |
| HOMO SAPIENS | 5 | 2666 | 1jph | A | 332 | ARG | p.Arg332His | P06132 |
| HOMO SAPIENS | 1 | 889 | 1m51 | A | 184 | LEU | p.Val184Leu | P35558 |
| HOMO SAPIENS | 0 | 889 | 1m51 | A | 267 | VAL | p.Ile267Val | P35558 |
| HOMO SAPIENS | 2 | 889 | 1m51 | A | 276 | GLU | p.Glu276Lys | P35558 |
| HOMO SAPIENS | 6 | 889 | 1m51 | A | 368 | VAL | p.Val368Ile | P35558 |
| HOMO SAPIENS | 0 | 889 | 1m51 | A | 579 | SER | p.Ser579Arg | P35558 |
| HOMO SAPIENS | 1 | 4084 | 2gjx | A | 28 | GLN | p.Gln28Ter | P06865 |
| HOMO SAPIENS | 0 | 4084 | 2gjx | A | 39 | LEU | p.Leu39Arg | P06865 |
| HOMO SAPIENS | 0 | 4084 | 2gjx | A | 127 | LEU | p.Leu127Arg | P06865 |
| HOMO SAPIENS | 6 | 4084 | 2gjx | A | 137 | ARG | p.Arg137Ter | P06865 |
| HOMO SAPIENS | 6 | 4084 | 2gjx | A | 170 | ARG | p.Arg170Gln | P06865 |
| HOMO SAPIENS | 9 | 4084 | 2gjx | A | 178 | ARG | p.Arg178Cys | P06865 |
| HOMO SAPIENS | 9 | 4084 | 2gjx | A | 178 | ARG | p.Arg178His | P06865 |
| HOMO SAPIENS | 5 | 4084 | 2gjx | A | 180 | TYR | p.Tyr180His | P06865 |
| HOMO SAPIENS | 5 | 4084 | 2gjx | A | 180 | TYR | p.Tyr180Ter | P06865 |
| HOMO SAPIENS | 1 | 4084 | 2gjx | A | 197 | LYS | p.Lys197Thr | P06865 |
| HOMO SAPIENS | 0 | 4084 | 2gjx | A | 200 | VAL | p.Val200Met | P06865 |
| HOMO SAPIENS | 9 | 4084 | 2gjx | A | 204 | HIS | p.His204Arg | P06865 |
| HOMO SAPIENS | 5 | 4084 | 2gjx | A | 210 | SER | p.Ser210Phe | P06865 |
| HOMO SAPIENS | 0 | 4084 | 2gjx | A | 211 | PHE | p.Phe211Ser | P06865 |
| HOMO SAPIENS | 2 | 4084 | 2gjx | A | 250 | GLY | p.Gly250Asp | P06865 |
| HOMO SAPIENS | 9 | 4084 | 2gjx | A | 258 | ASP | p.Asp258His | P06865 |
| HOMO SAPIENS | 0 | 4084 | 2gjx | A | 269 | GLY | p.Gly269Ser | P06865 |
| HOMO SAPIENS | 0 | 4084 | 2gjx | A | 301 | MET | p.Met301Arg | P06865 |
| HOMO SAPIENS | 3 | 4084 | 2gjx | A | 393 | ARG | p.Arg393Ter | P06865 |
| HOMO SAPIENS | 3 | 4084 | 2gjx | A | 393 | ARG | p.Arg393Pro | P06865 |
| HOMO SAPIENS | 0 | 4084 | 2gjx | A | 399 | ASN | p.Asn399Asp | P06865 |
| HOMO SAPIENS | 2 | 4084 | 2gjx | A | 420 | TRP | p.Trp420Cys | P06865 |
| HOMO SAPIENS | 1 | 4084 | 2gjx | A | 434 | PHE | p.Phe434Leu | P06865 |
| HOMO SAPIENS | 0 | 4084 | 2gjx | A | 454 | GLY | p.Gly454Ser | P06865 |
| HOMO SAPIENS | 1 | 4084 | 2gjx | A | 474 | TRP | p.Trp474Cys | P06865 |
| HOMO SAPIENS | 0 | 4084 | 2gjx | A | 479 | ALA | p.Ala479Thr | P06865 |
| HOMO SAPIENS | 5 | 4084 | 2gjx | A | 485 | TRP | p.Trp485Arg | P06865 |
| HOMO SAPIENS | 4 | 4084 | 2gjx | A | 499 | ARG | p.Arg499Cys | P06865 |
| HOMO SAPIENS | 4 | 4084 | 2gjx | A | 499 | ARG | p.Arg499His | P06865 |
| HOMO SAPIENS | 2 | 4084 | 2gjx | A | 504 | ARG | p.Arg504Cys | P06865 |
| HOMO SAPIENS | 2 | 4084 | 2gjx | A | 504 | ARG | p.Arg504His | P06865 |
| HOMO SAPIENS | -1 | 1943 | 1xmj | A | 417 | ASN | p.Asn417Lys | P13569 |
| HOMO SAPIENS | -1 | 1943 | 1xmj | A | 422 | SER | p.Ser422Phe | P13569 |
| HOMO SAPIENS | 1 | 1943 | 1xmj | A | 438 | THR | p.Thr438Ala | P13569 |
| HOMO SAPIENS | 0 | 1943 | 1xmj | A | 455 | ALA | p.Ala455Glu | P13569 |
| HOMO SAPIENS | 4 | 1943 | 1xmj | A | 456 | VAL | p.Val456Ala | P13569 |
| HOMO SAPIENS | 9 | 1943 | 1xmj | A | 458 | GLY | p.Gly458Val | P13569 |
| HOMO SAPIENS | 8 | 1943 | 1xmj | A | 466 | SER | p.Ser466Ter | P13569 |
| HOMO SAPIENS | 0 | 1943 | 1xmj | A | 467 | LEU | p.Leu467Pro | P13569 |
| HOMO SAPIENS | 1 | 1943 | 1xmj | A | 470 | MET | p.Val470Met | P13569 |

| | | | | | | | | |
|---|---|---|---|---|---|---|---|---|
| HOMO SAPIENS | 5 | 1943 | 1xmj | A | 480 | GLY | p.Gly480Cys | P13569 |
| HOMO SAPIENS | 3 | 1943 | 1xmj | A | 492 | SER | p.Ser492Phe | P13569 |
| HOMO SAPIENS | 8 | 1943 | 1xmj | A | 493 | GLN | p.Gln493Ter | P13569 |
| HOMO SAPIENS | 5 | 1943 | 1xmj | A | 506 | ILE | p.Ile506Met | P13569 |
| HOMO SAPIENS | 3 | 1943 | 1xmj | A | 520 | VAL | p.Val520Phe | P13569 |
| HOMO SAPIENS | 1 | 1943 | 1xmj | A | 528 | GLU | p.Glu528Asp | P13569 |
| HOMO SAPIENS | 1 | 1943 | 1xmj | A | 534 | ALA | p.Ala534Glu | P13569 |
| HOMO SAPIENS | 4 | 1943 | 1xmj | A | 542 | GLY | p.Gly542Ter | P13569 |
| HOMO SAPIENS | 6 | 1943 | 1xmj | A | 549 | SER | p.Ser549Asn | P13569 |
| HOMO SAPIENS | 6 | 1943 | 1xmj | A | 549 | SER | p.Ser549Arg | P13569 |
| HOMO SAPIENS | 6 | 1943 | 1xmj | A | 549 | SER | p.Ser549Arg | P13569 |
| HOMO SAPIENS | 6 | 1943 | 1xmj | A | 551 | GLY | p.Gly551Asp | P13569 |
| HOMO SAPIENS | 6 | 1943 | 1xmj | A | 551 | GLY | p.Gly551Ser | P13569 |
| HOMO SAPIENS | 6 | 1943 | 1xmj | A | 553 | GLN | p.Arg553Gln | P13569 |
| HOMO SAPIENS | 6 | 1943 | 1xmj | A | 553 | GLN | p.Arg553Ter | P13569 |
| HOMO SAPIENS | 0 | 1943 | 1xmj | A | 559 | ALA | p.Ala559Thr | P13569 |
| HOMO SAPIENS | 7 | 1943 | 1xmj | A | 560 | ARG | p.Arg560Lys | P13569 |
| HOMO SAPIENS | 4 | 1943 | 1xmj | A | 562 | VAL | p.Val562Ile | P13569 |
| HOMO SAPIENS | 3 | 1943 | 1xmj | A | 563 | TYR | p.Tyr563Asn | P13569 |
| HOMO SAPIENS | 3 | 1943 | 1xmj | A | 569 | TYR | p.Tyr569Asp | P13569 |
| HOMO SAPIENS | 3 | 1943 | 1xmj | A | 585 | GLU | p.Glu585Ter | P13569 |
| HOMO SAPIENS | 1 | 1943 | 1xmj | A | 618 | ILE | p.Ile618Thr | P13569 |
| HOMO SAPIENS | 9 | 1943 | 1xmj | A | 622 | GLY | p.Gly622Asp | P13569 |
| HOMO SAPIENS | -1 | 1943 | 1xmj | A | 648 | ASP | p.Asp648Val | P13569 |
| HOMO SAPIENS | 0 | 1943 | 1xmj | A | 673 | GLY | p.Gly673Ter | P13569 |
| HOMO SAPIENS | 5 | 5111 | 4rec | A | 749 | ARG | p.Arg749Ter | Q9Y2M0 |
| HOMO SAPIENS | -1 | 4580 | 4l0d | B | 2 | GLY | p.Pro2Leu | P35520 |
| HOMO SAPIENS | -1 | 4580 | 4l0d | B | 12 | PRO | p.Pro12Ser | P35520 |
| HOMO SAPIENS | -1 | 4580 | 4l0d | B | 18 | ARG | p.Arg18Cys | P35520 |
| HOMO SAPIENS | 1 | 4580 | 4l0d | B | 49 | PRO | p.Pro49Leu | P35520 |
| HOMO SAPIENS | 2 | 4580 | 4l0d | B | 70 | PRO | p.Pro70Leu | P35520 |
| HOMO SAPIENS | 1 | 4580 | 4l0d | B | 102 | LYS | p.Lys102Gln | P35520 |
| HOMO SAPIENS | 7 | 4580 | 4l0d | B | 114 | ALA | p.Ala114Val | P35520 |
| HOMO SAPIENS | 5 | 4580 | 4l0d | B | 116 | GLY | p.Gly116Arg | P35520 |
| HOMO SAPIENS | 8 | 4580 | 4l0d | B | 121 | ARG | p.Arg121Cys | P35520 |
| HOMO SAPIENS | 8 | 4580 | 4l0d | B | 121 | ARG | p.Arg121His | P35520 |
| HOMO SAPIENS | 5 | 4580 | 4l0d | B | 125 | ARG | p.Arg125Gln | P35520 |
| HOMO SAPIENS | 4 | 4580 | 4l0d | B | 134 | GLY | p.Gly134Arg | P35520 |
| HOMO SAPIENS | 4 | 4580 | 4l0d | B | 139 | GLY | p.Gly139Arg | P35520 |
| HOMO SAPIENS | 0 | 4580 | 4l0d | B | 145 | PRO | p.Pro145Leu | P35520 |
| HOMO SAPIENS | 0 | 4580 | 4l0d | B | 148 | GLY | p.Gly148Arg | P35520 |
| HOMO SAPIENS | 0 | 4580 | 4l0d | B | 168 | VAL | p.Val168Met | P35520 |
| HOMO SAPIENS | 2 | 4580 | 4l0d | B | 191 | THR | p.Thr191Met | P35520 |
| HOMO SAPIENS | 5 | 4580 | 4l0d | B | 234 | ASP | p.Asp234Asn | P35520 |
| HOMO SAPIENS | 0 | 4580 | 4l0d | B | 257 | THR | p.Thr257Met | P35520 |
| HOMO SAPIENS | 4 | 4580 | 4l0d | B | 262 | THR | p.Thr262Met | P35520 |
| HOMO SAPIENS | 0 | 4580 | 4l0d | B | 278 | ILE | p.Ile278Thr | P35520 |
| HOMO SAPIENS | 2 | 4580 | 4l0d | B | 307 | GLY | p.Gly307Ser | P35520 |
| HOMO SAPIENS | 4 | 4580 | 4l0d | B | 320 | VAL | p.Val320Ala | P35520 |
| HOMO SAPIENS | 2 | 4580 | 4l0d | B | 336 | ARG | p.Arg336Cys | P35520 |
| HOMO SAPIENS | 2 | 4580 | 4l0d | B | 336 | ARG | p.Arg336His | P35520 |
| HOMO SAPIENS | 0 | 4580 | 4l0d | B | 351 | GLY | p.Gly351Arg | P35520 |
| HOMO SAPIENS | 0 | 4580 | 4l0d | B | 353 | THR | p.Thr353Met | P35520 |
| HOMO SAPIENS | 0 | 4580 | 4l0d | B | 371 | VAL | p.Val371Met | P35520 |

| | | | | | | | | |
|---|---|---|---|---|---|---|---|---|
| HOMO SAPIENS | 2 | 4580 | 4l0d | B | 379 | ARG | p.Arg379Gln | P35520 |
| HOMO SAPIENS | 2 | 4580 | 4l0d | B | 379 | ARG | p.Arg379Trp | P35520 |
| HOMO SAPIENS | 2 | 4580 | 4l0d | B | 384 | LYS | p.Lys384Glu | P35520 |
| HOMO SAPIENS | 0 | 4580 | 4l0d | B | 422 | PRO | p.Pro422Leu | P35520 |
| HOMO SAPIENS | 0 | 4580 | 4l0d | B | 444 | ASP | p.Asp444Asn | P35520 |
| HOMO SAPIENS | 0 | 4580 | 4l0d | B | 466 | SER | p.Ser466Leu | P35520 |
| HOMO SAPIENS | 0 | 4580 | 4l0d | B | 539 | LEU | p.Leu539Ser | P35520 |
| HOMO SAPIENS | 0 | 4580 | 4l0d | B | 548 | ARG | p.Arg548Gln | P35520 |
| HOMO SAPIENS | 1 | 6536 | 3ulu | A | 412 | LEU | p.Leu412Phe | O15455 |
| HOMO SAPIENS | 1 | 28493 | 3ii6 | X | 814 | ARG | p.Arg814Ter | P49917 |
| HOMO SAPIENS | 1 | 28493 | 3ii6 | X | 857 | ALA | p.Ala857Thr | P49917 |
| HOMO SAPIENS | 9 | 3543 | 2q51 | A | 24 | GLU | p.Glu24Gly | P45381 |
| HOMO SAPIENS | 1 | 3543 | 2q51 | A | 27 | GLY | p.Gly27Arg | P45381 |
| HOMO SAPIENS | 9 | 3543 | 2q51 | A | 71 | ARG | p.Arg71His | P45381 |
| HOMO SAPIENS | -1 | 3543 | 2q51 | A | 82 | MSE | p.Met82Thr | P45381 |
| HOMO SAPIENS | 0 | 3543 | 2q51 | A | 143 | ILE | p.Ile143Phe | P45381 |
| HOMO SAPIENS | 1 | 3543 | 2q51 | A | 152 | CYS | p.Cys152Arg | P45381 |
| HOMO SAPIENS | 2 | 3543 | 2q51 | A | 218 | CYS | p.Cys218Ter | P45381 |
| HOMO SAPIENS | 1 | 3543 | 2q51 | A | 231 | TYR | p.Tyr231Cys | P45381 |
| HOMO SAPIENS | 1 | 3543 | 2q51 | A | 231 | TYR | p.Tyr231Ter | P45381 |
| HOMO SAPIENS | 2 | 3543 | 2q51 | A | 274 | GLY | p.Gly274Arg | P45381 |
| HOMO SAPIENS | 6 | 3543 | 2q51 | A | 285 | GLU | p.Glu285Ala | P45381 |
| HOMO SAPIENS | 5 | 3543 | 2q51 | A | 287 | ALA | p.Ala287Thr | P45381 |
| HOMO SAPIENS | 2 | 3543 | 2q51 | A | 305 | ALA | p.Ala305Glu | P45381 |
| HOMO SAPIENS | 2 | 24687 | 1hfi | A | 25 | SER | p.Ser890Ile | P08603 |
| HOMO SAPIENS | 0 | 1508 | 3wn5 | C | 45 | LEU | p.Leu66Arg | P08637 |
| HOMO SAPIENS | 0 | 1508 | 3wn5 | C | 155 | VAL | p.Phe176Val | P08637 |
| HOMO SAPIENS | 0 | 750 | 5tpa | A | 54 | VAL | p.Val452Met | Q12879 |
| HOMO SAPIENS | 9 | 750 | 5tpa | A | 120 | ARG | p.Arg518Cys | Q12879 |
| HOMO SAPIENS | 0 | 750 | 5tpa | A | 199 | ALA | p.Ala716Thr | Q12879 |
| HOMO SAPIENS | 4 | 1363 | 3hg2 | B | 40 | PRO | p.Pro40Ser | P06280 |
| HOMO SAPIENS | 3 | 1363 | 3hg2 | B | 65 | SER | p.Ser65Thr | P06280 |
| HOMO SAPIENS | 4 | 1363 | 3hg2 | B | 66 | GLU | p.Glu66Gln | P06280 |
| HOMO SAPIENS | 2 | 1363 | 3hg2 | B | 80 | GLY | p.Gly80Asp | P06280 |
| HOMO SAPIENS | 3 | 1363 | 3hg2 | B | 112 | ARG | p.Arg112Cys | P06280 |
| HOMO SAPIENS | 3 | 1363 | 3hg2 | B | 112 | ARG | p.Arg112His | P06280 |
| HOMO SAPIENS | 0 | 1363 | 3hg2 | B | 139 | ASN | p.Asn139Ser | P06280 |
| HOMO SAPIENS | 7 | 1363 | 3hg2 | B | 156 | ALA | p.Ala156Thr | P06280 |
| HOMO SAPIENS | 5 | 1363 | 3hg2 | B | 162 | TRP | p.Trp162Arg | P06280 |
| HOMO SAPIENS | 5 | 1363 | 3hg2 | B | 207 | TYR | p.Tyr207His | P06280 |
| HOMO SAPIENS | 0 | 1363 | 3hg2 | B | 215 | ASN | p.Asn215Ser | P06280 |
| HOMO SAPIENS | 0 | 1363 | 3hg2 | B | 264 | ASP | p.Asp264Val | P06280 |
| HOMO SAPIENS | 5 | 1363 | 3hg2 | B | 266 | ASP | p.Asp266Val | P06280 |
| HOMO SAPIENS | 2 | 1363 | 3hg2 | B | 269 | VAL | p.Val269Ala | P06280 |
| HOMO SAPIENS | 3 | 1363 | 3hg2 | B | 272 | ASN | p.Asn272Ser | P06280 |
| HOMO SAPIENS | 3 | 1363 | 3hg2 | B | 279 | GLN | p.Gln279Glu | P06280 |
| HOMO SAPIENS | 0 | 1363 | 3hg2 | B | 289 | ILE | p.Ile289Val | P06280 |
| HOMO SAPIENS | 0 | 1363 | 3hg2 | B | 296 | MET | p.Met296Val | P06280 |
| HOMO SAPIENS | 0 | 1363 | 3hg2 | B | 296 | MET | p.Met296Ile | P06280 |
| HOMO SAPIENS | 0 | 1363 | 3hg2 | B | 297 | SER | p.Ser297Phe | P06280 |
| HOMO SAPIENS | 2 | 1363 | 3hg2 | B | 301 | ARG | p.Arg301Gln | P06280 |
| HOMO SAPIENS | 3 | 1363 | 3hg2 | B | 327 | GLN | p.Gln327Lys | P06280 |
| HOMO SAPIENS | 0 | 1363 | 3hg2 | B | 328 | GLY | p.Gly328Arg | P06280 |
| HOMO SAPIENS | 0 | 1363 | 3hg2 | B | 328 | GLY | p.Gly328Ala | P06280 |

| | | | | | | | | |
|---|---|---|---|---|---|---|---|---|
| HOMO SAPIENS | 2 | 1363 | 3hg2 | B | 331 | LEU | p.Leu331Phe | P06280 |
| HOMO SAPIENS | 0 | 1363 | 3hg2 | B | 342 | ARG | p.Arg342Gln | P06280 |
| HOMO SAPIENS | 1 | 1363 | 3hg2 | B | 361 | GLY | p.Gly361Arg | P06280 |
| HOMO SAPIENS | 1 | 1363 | 3hg2 | B | 363 | ARG | p.Arg363His | P06280 |
| HOMO SAPIENS | 1 | 1363 | 3hg2 | B | 385 | THR | p.Thr385Ala | P06280 |
| HOMO SAPIENS | 1 | 1363 | 3hg2 | B | 399 | TRP | p.Trp399Ser | P06280 |
| HOMO SAPIENS | 1 | 1363 | 3hg2 | B | 410 | THR | p.Thr410Ala | P06280 |
| HOMO SAPIENS | 2 | 3904 | 3caa | B | 385 | MET | p.Met414Val | P01011 |
| HOMO SAPIENS | 0 | 558 | 5eub | A | 56 | VAL | p.Val56Ile | Q13126 |
| HOMO SAPIENS | 0 | 558 | 5eub | A | 189 | TRP | p.Trp189Leu | Q13126 |
| HOMO SAPIENS | 0 | 558 | 5eub | A | 218 | ALA | p.Ala218Gly | Q13126 |
| HOMO SAPIENS | 1 | 3865 | 1cb5 | B | 443 | ILE | p.Ile443Val | Q13867 |
| HOMO SAPIENS | 2 | 14329 | 4m92 | A | 24 | ILE | p.Ile65Val | Q13454 |
| HOMO SAPIENS | 1 | 7381 | 5kz5 | k | 50 | GLY | p.Gly50Glu | Q9H1K1 |
| HOMO SAPIENS | 0 | 1102 | 4fdh | E | 86 | PRO | p.Pro86Ala | P19099 |
| HOMO SAPIENS | 0 | 1102 | 4fdh | E | 118 | ALA | p.Ala118Thr | P19099 |
| HOMO SAPIENS | 0 | 1102 | 4fdh | E | 159 | PRO | p.Pro159Leu | P19099 |
| HOMO SAPIENS | 0 | 1102 | 4fdh | E | 173 | LYS | p.Lys173Arg | P19099 |
| HOMO SAPIENS | 0 | 1102 | 4fdh | E | 181 | ARG | p.Arg181Trp | P19099 |
| HOMO SAPIENS | 0 | 1102 | 4fdh | E | 185 | THR | p.Thr185Ile | P19099 |
| HOMO SAPIENS | 0 | 1102 | 4fdh | E | 198 | GLU | p.Glu198Asp | P19099 |
| HOMO SAPIENS | 0 | 1102 | 4fdh | E | 214 | HIS | p.His214Asp | P19099 |
| HOMO SAPIENS | 0 | 1102 | 4fdh | E | 251 | LYS | p.Lys251Arg | P19099 |
| HOMO SAPIENS | 0 | 1102 | 4fdh | E | 255 | GLU | p.Glu255Ter | P19099 |
| HOMO SAPIENS | 0 | 1102 | 4fdh | E | 281 | ASN | p.Asn281Ser | P19099 |
| HOMO SAPIENS | 0 | 1102 | 4fdh | E | 320 | ALA | p.Ala320Val | P19099 |
| HOMO SAPIENS | 0 | 1102 | 4fdh | E | 339 | ILE | p.Ile339Thr | P19099 |
| HOMO SAPIENS | 0 | 1102 | 4fdh | E | 341 | ARG | p.Arg341Cys | P19099 |
| HOMO SAPIENS | 2 | 1102 | 4fdh | E | 374 | ARG | p.Arg374Trp | P19099 |
| HOMO SAPIENS | 0 | 1102 | 4fdh | E | 386 | VAL | p.Val386Ala | P19099 |
| HOMO SAPIENS | 0 | 1102 | 4fdh | E | 390 | LEU | p.Leu390Phe | P19099 |
| HOMO SAPIENS | -1 | 1102 | 4fdh | E | 435 | GLY | p.Gly435Ser | P19099 |
| HOMO SAPIENS | 0 | 1102 | 4fdh | E | 461 | LEU | p.Leu461Pro | P19099 |
| HOMO SAPIENS | 0 | 1102 | 4fdh | E | 498 | THR | p.Thr498Ala | P19099 |
| HOMO SAPIENS | 6 | 10292 | 2of5 | F | 128 | GLY | p.Gly128Arg | P78560 |
| HOMO SAPIENS | 6 | 10292 | 2of5 | F | 170 | ARG | p.Arg170Cys | P78560 |
| HOMO SAPIENS | -1 | 18965 | 3ghg | A | 16 | ARG | p.Arg35Cys | P02671 |
| HOMO SAPIENS | -1 | 18965 | 3ghg | A | 16 | ARG | p.Arg35His | P02671 |
| HOMO SAPIENS | -1 | 18965 | 3ghg | A | 283 | PRO | p.Pro302Ala | P02671 |
| HOMO SAPIENS | -1 | 18965 | 3ghg | A | 312 | THR | p.Thr331Ala | P02671 |
| HOMO SAPIENS | -1 | 18965 | 3ghg | A | 381 | SER | p.Ser400Phe | P02671 |
| HOMO SAPIENS | -1 | 18965 | 3ghg | A | 463 | VAL | p.Val482Met | P02671 |
| HOMO SAPIENS | -1 | 18965 | 3ghg | A | 526 | GLU | p.Glu545Val | P02671 |
| HOMO SAPIENS | -1 | 18965 | 3ghg | A | 554 | ARG | p.Arg573Leu | P02671 |
| HOMO SAPIENS | 8 | 3023 | 4ttb | A | 101 | ARG | p.Arg101Trp | Q6PHW0 |
| HOMO SAPIENS | 2 | 3023 | 4ttb | A | 116 | ILE | p.Ile116Thr | Q6PHW0 |
| HOMO SAPIENS | 0 | 3023 | 4ttb | A | 220 | ALA | p.Ala220Thr | Q6PHW0 |
| HOMO SAPIENS | 2 | 3023 | 4ttb | A | 271 | GLU | p.Glu271Lys | Q6PHW0 |
| HOMO SAPIENS | 3 | 33226 | 1ya5 | T | 11 | SER | p.Ser11Ter | O15273 |
| HOMO SAPIENS | 6 | 33226 | 1ya5 | T | 64 | SER | p.Ser64Leu | O15273 |
| HOMO SAPIENS | 6 | 33226 | 1ya5 | T | 70 | ARG | p.Arg70Gln | O15273 |
| HOMO SAPIENS | 3 | 33226 | 1ya5 | T | 75 | GLY | p.Gly75Ser | O15273 |
| HOMO SAPIENS | 0 | 33226 | 1ya5 | T | 87 | ARG | p.Arg87Trp | O15273 |
| HOMO SAPIENS | 2 | 5667 | 3dax | A | 398 | PRO | p.Pro398Ala | P22680 |

| | | | | | | | | |
|---|---|---|---|---|---|---|---|---|
| HOMO SAPIENS | 0 | 12861 | 5mya | A | 464 | ASN | p.Asn464His | Q02763 |
| HOMO SAPIENS | 2 | 12861 | 5mya | A | 486 | VAL | p.Val486Ile | Q02763 |
| HOMO SAPIENS | 0 | 12861 | 5mya | A | 522 | ARG | p.Arg522His | Q02763 |
| HOMO SAPIENS | 2 | 12861 | 5mya | A | 600 | VAL | p.Val600Leu | Q02763 |
| HOMO SAPIENS | 0 | 12861 | 5mya | A | 634 | LEU | p.Leu634Phe | Q02763 |
| HOMO SAPIENS | 0 | 12861 | 5mya | A | 684 | HIS | p.His684Gln | Q02763 |
| HOMO SAPIENS | 0 | 12861 | 5mya | A | 724 | ALA | p.Ala724Thr | Q02763 |
| HOMO SAPIENS | 0 | 7743 | 3qzw | G | 90 | GLY | p.Gly111Ser | P01732 |
| HOMO SAPIENS | 2 | 9498 | 3kci | A | 4267 | ASP | p.Asp4267Glu | O95714 |
| HOMO SAPIENS | 5 | 201 | 3ips | B | 415 | ARG | p.Arg415Gln | Q13133 |
| HOMO SAPIENS | 3 | 229 | 2q50 | A | 68 | LEU | p.Leu68Pro | Q9UBQ7 |
| HOMO SAPIENS | 2 | 229 | 2q50 | A | 99 | ARG | p.Arg99Ter | Q9UBQ7 |
| HOMO SAPIENS | 3 | 229 | 2q50 | A | 160 | GLY | p.Gly160Arg | Q9UBQ7 |
| HOMO SAPIENS | 5 | 229 | 2q50 | A | 165 | GLY | p.Gly165Asp | Q9UBQ7 |
| HOMO SAPIENS | 4 | 229 | 2q50 | A | 302 | ARG | p.Arg302Cys | Q9UBQ7 |
| HOMO SAPIENS | 4 | 229 | 2q50 | A | 302 | ARG | p.Arg302His | Q9UBQ7 |
| HOMO SAPIENS | -1 | 229 | 2q50 | A | 322 | MSE | p.Met322Arg | Q9UBQ7 |
| HOMO SAPIENS | -1 | 4519 | 5clw | A | 40 | PRO | p.Pro40Thr | Q04446 |
| HOMO SAPIENS | 1 | 4519 | 5clw | A | 59 | ILE | p.Ile59Thr | Q04446 |
| HOMO SAPIENS | 0 | 4519 | 5clw | A | 190 | ARG | p.Arg190Gly | Q04446 |
| HOMO SAPIENS | 1 | 4519 | 5clw | A | 224 | LEU | p.Leu224Pro | Q04446 |
| HOMO SAPIENS | 0 | 4519 | 5clw | A | 236 | GLN | p.Gln236His | Q04446 |
| HOMO SAPIENS | 1 | 4519 | 5clw | A | 254 | THR | p.Thr254Ala | Q04446 |
| HOMO SAPIENS | 0 | 4519 | 5clw | A | 257 | PHE | p.Phe257Leu | Q04446 |
| HOMO SAPIENS | 1 | 4519 | 5clw | A | 262 | ARG | p.Arg262Cys | Q04446 |
| HOMO SAPIENS | 3 | 4519 | 5clw | A | 329 | TYR | p.Tyr329Ser | Q04446 |
| HOMO SAPIENS | 0 | 4519 | 5clw | A | 507 | THR | p.Thr507Ala | Q04446 |
| HOMO SAPIENS | 0 | 4519 | 5clw | A | 515 | ARG | p.Arg515Cys | Q04446 |
| HOMO SAPIENS | 0 | 4519 | 5clw | A | 515 | ARG | p.Arg515His | Q04446 |
| HOMO SAPIENS | 0 | 4519 | 5clw | A | 524 | ARG | p.Arg524Gln | Q04446 |
| HOMO SAPIENS | 0 | 4519 | 5clw | A | 524 | ARG | p.Arg524Ter | Q04446 |
| HOMO SAPIENS | 0 | 4519 | 5clw | A | 545 | HIS | p.His545Arg | Q04446 |
| HOMO SAPIENS | 2 | 4519 | 5clw | A | 548 | TRP | p.Trp548Ter | Q04446 |
| HOMO SAPIENS | 0 | 4519 | 5clw | A | 592 | GLU | p.Glu592Ter | Q04446 |
| HOMO SAPIENS | 0 | 4519 | 5clw | A | 628 | HIS | p.His628Arg | Q04446 |
| HOMO SAPIENS | 0 | 4519 | 5clw | A | 637 | ARG | p.Arg637Ter | Q04446 |
| HOMO SAPIENS | 0 | 4519 | 5clw | A | 673 | ALA | p.Ala673Thr | Q04446 |
| HOMO SAPIENS | 1 | 10405 | 3bg3 | D | 507 | VAL | p.Val507Ile | P11498 |
| HOMO SAPIENS | 0 | 10405 | 3bg3 | D | 583 | ARG | p.Arg583Leu | P11498 |
| HOMO SAPIENS | 1 | 10405 | 3bg3 | D | 589 | LYS | p.Lys589Asn | P11498 |
| HOMO SAPIENS | 6 | 10405 | 3bg3 | D | 610 | ALA | p.Ala610Thr | P11498 |
| HOMO SAPIENS | 2 | 10405 | 3bg3 | D | 631 | ARG | p.Arg631Gln | P11498 |
| HOMO SAPIENS | 1 | 10405 | 3bg3 | D | 732 | ARG | p.Arg732Gln | P11498 |
| HOMO SAPIENS | 6 | 10405 | 3bg3 | D | 743 | MET | p.Met743Ile | P11498 |
| HOMO SAPIENS | 0 | 10405 | 3bg3 | D | 750 | THR | p.Thr750Met | P11498 |
| HOMO SAPIENS | -1 | 13751 | 2o8d | B | 350 | ALA | p.Ala350Val | P52701 |
| HOMO SAPIENS | -1 | 13751 | 2o8d | B | 352 | VAL | p.Val352Ile | P52701 |
| HOMO SAPIENS | -1 | 13751 | 2o8d | B | 355 | GLY | p.Gly355Ser | P52701 |
| HOMO SAPIENS | 0 | 13751 | 2o8d | B | 385 | ARG | p.Arg385Trp | P52701 |
| HOMO SAPIENS | 0 | 13751 | 2o8d | B | 390 | ASP | p.Asp390Glu | P52701 |
| HOMO SAPIENS | 0 | 13751 | 2o8d | B | 396 | LEU | p.Leu396Val | P52701 |
| HOMO SAPIENS | 0 | 13751 | 2o8d | B | 449 | LEU | p.Leu449Pro | P52701 |
| HOMO SAPIENS | 0 | 13751 | 2o8d | B | 468 | ARG | p.Arg468His | P52701 |
| HOMO SAPIENS | 0 | 13751 | 2o8d | B | 503 | SER | p.Ser503Cys | P52701 |

| | | | | | | | | |
|---|---|---|---|---|---|---|---|---|
| HOMO SAPIENS | 1 | 13751 | 2o8d | B | 509 | VAL | p.Val509Ala | P52701 |
| HOMO SAPIENS | 0 | 13751 | 2o8d | B | 675 | GLU | p.Glu675Asp | P52701 |
| HOMO SAPIENS | 0 | 13751 | 2o8d | B | 795 | ILE | p.Ile795Thr | P52701 |
| HOMO SAPIENS | 0 | 13751 | 2o8d | B | 850 | TYR | p.Tyr850Ter | P52701 |
| HOMO SAPIENS | 0 | 13751 | 2o8d | B | 878 | VAL | p.Val878Ala | P52701 |
| HOMO SAPIENS | 0 | 13751 | 2o8d | B | 911 | ARG | p.Arg911Ter | P52701 |
| HOMO SAPIENS | 0 | 13751 | 2o8d | B | 922 | ARG | p.Arg922Ter | P52701 |
| HOMO SAPIENS | 0 | 13751 | 2o8d | B | 969 | TYR | p.Tyr969Cys | P52701 |
| HOMO SAPIENS | 0 | 13751 | 2o8d | B | 1005 | ARG | p.Arg1005Ter | P52701 |
| HOMO SAPIENS | 0 | 13751 | 2o8d | B | 1035 | ARG | p.Arg1035Ter | P52701 |
| HOMO SAPIENS | 1 | 13751 | 2o8d | B | 1048 | GLN | p.Gln1048Ter | P52701 |
| HOMO SAPIENS | 1 | 13751 | 2o8d | B | 1051 | VAL | p.Val1051Ile | P52701 |
| HOMO SAPIENS | 0 | 13751 | 2o8d | B | 1054 | ILE | p.Ile1054Phe | P52701 |
| HOMO SAPIENS | 0 | 13751 | 2o8d | B | 1054 | ILE | p.Ile1054Met | P52701 |
| HOMO SAPIENS | 0 | 13751 | 2o8d | B | 1163 | GLU | p.Glu1163Val | P52701 |
| HOMO SAPIENS | 0 | 13751 | 2o8d | B | 1237 | GLU | p.Glu1237Asp | P52701 |
| HOMO SAPIENS | 0 | 13751 | 2o8d | B | 1242 | ARG | p.Arg1242His | P52701 |
| HOMO SAPIENS | -1 | 13751 | 2o8d | B | 1278 | PRO | p.Pro1278Thr | P52701 |
| HOMO SAPIENS | 0 | 13751 | 2o8d | B | 1304 | ARG | p.Arg1304Lys | P52701 |
| HOMO SAPIENS | 0 | 13751 | 2o8d | B | 1317 | HIS | p.His1317Gln | P52701 |
| HOMO SAPIENS | 0 | 13751 | 2o8d | B | 1329 | SER | p.Ser1329Leu | P52701 |
| HOMO SAPIENS | 3 | 13273 | 4g8k | A | 265 | GLU | p.Glu265Ter | Q05823 |
| HOMO SAPIENS | -1 | 11088 | 5uj7 | A | 666 | ARG | p.Arg666Trp | Q13415 |
| HOMO SAPIENS | 4 | 11088 | 5uj7 | A | 720 | ARG | p.Arg720Gln | Q13415 |
| HOMO SAPIENS | 0 | 11088 | 5uj7 | A | 816 | MET | p.Met816Thr | Q13415 |
| HOMO SAPIENS | 1 | 4352 | 5va1 | A | 421 | THR | p.Thr421Met | Q12809 |
| HOMO SAPIENS | 0 | 4352 | 5va1 | A | 451 | PRO | p.Pro451Leu | Q12809 |
| HOMO SAPIENS | 0 | 4352 | 5va1 | A | 470 | ASN | p.Asn470Asp | Q12809 |
| HOMO SAPIENS | 0 | 4352 | 5va1 | A | 490 | ALA | p.Ala490Thr | Q12809 |
| HOMO SAPIENS | 0 | 4352 | 5va1 | A | 491 | VAL | p.Val491Ile | Q12809 |
| HOMO SAPIENS | 1 | 4352 | 5va1 | A | 533 | VAL | p.Val533Gly | Q12809 |
| HOMO SAPIENS | 0 | 4352 | 5va1 | A | 552 | LEU | p.Leu552Ser | Q12809 |
| HOMO SAPIENS | 0 | 4352 | 5va1 | A | 558 | ALA | p.Ala558Pro | Q12809 |
| HOMO SAPIENS | 1 | 4352 | 5va1 | A | 561 | ALA | p.Ala561Val | Q12809 |
| HOMO SAPIENS | 0 | 4352 | 5va1 | A | 572 | GLY | p.Gly572Arg | Q12809 |
| HOMO SAPIENS | -1 | 4352 | 5va1 | A | 582 | ARG | p.Arg582Cys | Q12809 |
| HOMO SAPIENS | 0 | 4352 | 5va1 | A | 584 | GLY | p.Gly584Ser | Q12809 |
| HOMO SAPIENS | 0 | 4352 | 5va1 | A | 588 | ASN | p.Asn588Lys | Q12809 |
| HOMO SAPIENS | 0 | 4352 | 5va1 | A | 593 | ILE | p.Ile593Arg | Q12809 |
| HOMO SAPIENS | 1 | 4352 | 5va1 | A | 627 | PHE | p.Phe627Leu | Q12809 |
| HOMO SAPIENS | 0 | 4352 | 5va1 | A | 630 | VAL | p.Val630Leu | Q12809 |
| HOMO SAPIENS | 0 | 4352 | 5va1 | A | 652 | TYR | p.Tyr652Ter | Q12809 |
| HOMO SAPIENS | 0 | 4352 | 5va1 | A | 711 | ILE | p.Ile711Val | Q12809 |
| HOMO SAPIENS | 0 | 4352 | 5va1 | A | 735 | SER | p.Ser735Leu | Q12809 |
| HOMO SAPIENS | 0 | 4352 | 5va1 | A | 744 | ARG | p.Arg744Ter | Q12809 |
| HOMO SAPIENS | 0 | 4352 | 5va1 | A | 784 | ARG | p.Arg784Trp | Q12809 |
| HOMO SAPIENS | 1 | 4352 | 5va1 | A | 820 | GLY | p.Gly820Arg | Q12809 |
| HOMO SAPIENS | 5 | 4352 | 5va1 | A | 822 | VAL | p.Val822Met | Q12809 |
| HOMO SAPIENS | 1 | 4352 | 5va1 | A | 863 | ARG | p.Arg863Ter | Q12809 |
| HOMO SAPIENS | 5 | 303 | 2obv | A | 55 | ALA | p.Ala55Asp | Q00266 |
| HOMO SAPIENS | 7 | 303 | 2obv | A | 259 | ALA | p.Ala259Gly | Q00266 |
| HOMO SAPIENS | 9 | 303 | 2obv | A | 264 | ARG | p.Arg264His | Q00266 |
| HOMO SAPIENS | 9 | 303 | 2obv | A | 264 | ARG | p.Arg264Cys | Q00266 |
| HOMO SAPIENS | 0 | 303 | 2obv | A | 305 | LEU | p.Leu305Pro | Q00266 |

| | | | | | | | | |
|---|---|---|---|---|---|---|---|---|
| HOMO SAPIENS | 7 | 303 | 2obv | A | 322 | ILE | p.Ile322Met | Q00266 |
| HOMO SAPIENS | 0 | 303 | 2obv | A | 336 | GLY | p.Gly336Arg | Q00266 |
| HOMO SAPIENS | 0 | 303 | 2obv | A | 357 | PRO | p.Pro357Leu | Q00266 |
| HOMO SAPIENS | 0 | 2077 | 4zxb | E | 462 | ASN | p.Asn489Ser | P06213 |
| HOMO SAPIENS | -1 | 2077 | 4zxb | E | 519 | ASP | p.Asp546Glu | P06213 |
| HOMO SAPIENS | 2 | 2077 | 4zxb | E | 819 | THR | p.Thr858Ala | P06213 |
| HOMO SAPIENS | 0 | 2077 | 4zxb | E | 907 | ASP | p.Asp946Glu | P06213 |
| HOMO SAPIENS | 7 | 3274 | 1wsv | B | 14 | HIS | p.His42Arg | P48728 |
| HOMO SAPIENS | 0 | 3274 | 1wsv | B | 19 | GLY | p.Gly47Arg | P48728 |
| HOMO SAPIENS | 4 | 3274 | 1wsv | B | 45 | ARG | p.Arg73Cys | P48728 |
| HOMO SAPIENS | 1 | 3274 | 1wsv | B | 49 | SER | p.Ser77Leu | P48728 |
| HOMO SAPIENS | 3 | 3274 | 1wsv | B | 241 | GLY | p.Gly269Asp | P48728 |
| HOMO SAPIENS | 7 | 3274 | 1wsv | B | 248 | ASP | p.Asp276His | P48728 |
| HOMO SAPIENS | 0 | 3274 | 1wsv | B | 272 | MET | p.Met300Val | P48728 |
| HOMO SAPIENS | 4 | 3274 | 1wsv | B | 292 | ARG | p.Arg320His | P48728 |
| HOMO SAPIENS | 0 | 3274 | 1wsv | B | 354 | ARG | p.Arg382Gln | P48728 |
| HOMO SAPIENS | -1 | 435 | 3uyt | A | 44 | THR | p.Thr44Ala | P48730 |
| HOMO SAPIENS | 1 | 4698 | 5ife | C | 666 | LYS | p.Lys666Asn | O75533 |
| HOMO SAPIENS | 1 | 16917 | 1apy | C | 37 | GLY | p.Gly60Asp | P20933 |
| HOMO SAPIENS | 0 | 16917 | 1apy | C | 49 | SER | p.Ser72Pro | P20933 |
| HOMO SAPIENS | 0 | 16917 | 1apy | C | 84 | ARG | p.Arg107Ter | P20933 |
| HOMO SAPIENS | 0 | 16917 | 1apy | C | 126 | SER | p.Thr149Ser | P20933 |
| HOMO SAPIENS | 0 | 16917 | 1apy | C | 135 | TRP | p.Trp158Ter | P20933 |
| HOMO SAPIENS | 0 | 16917 | 1apy | C | 140 | CYS | p.Cys163Ser | P20933 |
| HOMO SAPIENS | 0 | 16917 | 1apy | C | 145 | TRP | p.Trp168Ter | P20933 |
| HOMO SAPIENS | 0 | 10413 | 4frw | A | 185 | THR | p.Thr185Met | Q96NY8 |
| HOMO SAPIENS | 0 | 5217 | 5n6w | G | 96 | TRP | p.Trp96Arg | O15537 |
| HOMO SAPIENS | 1 | 5217 | 5n6w | G | 102 | ARG | p.Arg102Trp | O15537 |
| HOMO SAPIENS | 0 | 5217 | 5n6w | G | 109 | GLY | p.Gly109Arg | O15537 |
| HOMO SAPIENS | -1 | 5217 | 5n6w | G | 223 | CYS | p.Cys223Arg | O15537 |
| HOMO SAPIENS | 0 | 9485 | 4z6y | E | 138 | ALA | p.Ala138Asp | Q9P0N9 |
| HOMO SAPIENS | 3 | 4504 | 4mwt | B | 411 | GLY | p.Gly439Ser | P10619 |
| HOMO SAPIENS | 3 | 4504 | 4mwt | B | 412 | PHE | p.Phe440Val | P10619 |
| HOMO SAPIENS | 6 | 6391 | 4kxy | A | 318 | ARG | p.Arg318Cys | P29401 |
| HOMO SAPIENS | 6 | 19546 | 2bx6 | A | 87 | THR | p.Thr87Ile | O75695 |
| HOMO SAPIENS | 1 | 19546 | 2bx6 | A | 118 | ARG | p.Arg118His | O75695 |
| HOMO SAPIENS | 1 | 19546 | 2bx6 | A | 201 | ILE | p.Ile201Val | O75695 |
| HOMO SAPIENS | 1 | 19546 | 2bx6 | A | 282 | ARG | p.Arg282Trp | O75695 |
| HOMO SAPIENS | 1 | 5265 | 5usr | H | 68 | ARG | p.Arg68Leu | Q9HD34 |
| HOMO SAPIENS | 2 | 1012 | 4nyi | S | 569 | LEU | p.Leu569Val | Q07889 |
| HOMO SAPIENS | 2 | 1012 | 4nyi | S | 655 | PRO | p.Pro655Leu | Q07889 |
| HOMO SAPIENS | 4 | 1012 | 4nyi | S | 663 | ILE | p.Ile663Met | Q07889 |
| HOMO SAPIENS | 2 | 1012 | 4nyi | S | 702 | TYR | p.Tyr702Cys | Q07889 |
| HOMO SAPIENS | 2 | 1012 | 4nyi | S | 708 | ALA | p.Ala708Thr | Q07889 |
| HOMO SAPIENS | 4 | 1012 | 4nyi | S | 733 | ILE | p.Ile733Phe | Q07889 |
| HOMO SAPIENS | 1 | 1848 | 3g1n | A | 4013 | ARG | p.Arg4013Trp | Q7Z6Z7 |
| HOMO SAPIENS | 1 | 1848 | 3g1n | A | 4187 | ARG | p.Arg4187Cys | Q7Z6Z7 |
| HOMO SAPIENS | 1 | 77 | 4pod | A | 41 | CYS | p.Cys79Tyr | P60174 |
| HOMO SAPIENS | 9 | 77 | 4pod | A | 104 | GLU | p.Glu142Asp | P60174 |
| HOMO SAPIENS | 1 | 77 | 4pod | A | 122 | GLY | p.Gly160Arg | P60174 |
| HOMO SAPIENS | -1 | 8672 | 3thz | A | 8 | THR | p.Thr8Met | P43246 |
| HOMO SAPIENS | 0 | 8672 | 3thz | A | 19 | PHE | p.Phe19Leu | P43246 |
| HOMO SAPIENS | 0 | 8672 | 3thz | A | 23 | PHE | p.Phe23Leu | P43246 |
| HOMO SAPIENS | 0 | 8672 | 3thz | A | 96 | ARG | p.Arg96His | P43246 |

| | | | | | | | | |
|---|---|---|---|---|---|---|---|---|
| HOMO SAPIENS | 0 | 8672 | 3thz | A | 102 | VAL | p.Val102Ile | P43246 |
| HOMO SAPIENS | 0 | 8672 | 3thz | A | 106 | ARG | p.Arg106Lys | P43246 |
| HOMO SAPIENS | 0 | 8672 | 3thz | A | 127 | ASN | p.Asn127Ser | P43246 |
| HOMO SAPIENS | 0 | 8672 | 3thz | A | 128 | LEU | p.Leu128Val | P43246 |
| HOMO SAPIENS | 0 | 8672 | 3thz | A | 163 | VAL | p.Val163Gly | P43246 |
| HOMO SAPIENS | 0 | 8672 | 3thz | A | 169 | ILE | p.Ile169Val | P43246 |
| HOMO SAPIENS | 0 | 8672 | 3thz | A | 315 | GLY | p.Gly315Asp | P43246 |
| HOMO SAPIENS | -1 | 8672 | 3thz | A | 322 | GLY | p.Gly322Asp | P43246 |
| HOMO SAPIENS | 0 | 8672 | 3thz | A | 331 | ASN | p.Asn331Asp | P43246 |
| HOMO SAPIENS | 1 | 8672 | 3thz | A | 390 | LEU | p.Leu390Phe | P43246 |
| HOMO SAPIENS | 0 | 8672 | 3thz | A | 419 | GLN | p.Gln419Lys | P43246 |
| HOMO SAPIENS | 0 | 8672 | 3thz | A | 447 | PHE | p.Phe447Leu | P43246 |
| HOMO SAPIENS | 0 | 8672 | 3thz | A | 524 | ARG | p.Arg524Pro | P43246 |
| HOMO SAPIENS | 0 | 8672 | 3thz | A | 539 | PHE | p.Phe539Leu | P43246 |
| HOMO SAPIENS | -1 | 8672 | 3thz | A | 546 | LYS | p.Lys546Asn | P43246 |
| HOMO SAPIENS | 0 | 8672 | 3thz | A | 564 | THR | p.Thr564Ala | P43246 |
| HOMO SAPIENS | 0 | 8672 | 3thz | A | 577 | ILE | p.Ile577Thr | P43246 |
| HOMO SAPIENS | 0 | 8672 | 3thz | A | 588 | TYR | p.Tyr588Ter | P43246 |
| HOMO SAPIENS | 0 | 8672 | 3thz | A | 622 | PRO | p.Pro622Leu | P43246 |
| HOMO SAPIENS | 0 | 8672 | 3thz | A | 629 | GLN | p.Gln629Arg | P43246 |
| HOMO SAPIENS | 0 | 8672 | 3thz | A | 636 | ALA | p.Ala636Pro | P43246 |
| HOMO SAPIENS | 2 | 8672 | 3thz | A | 642 | VAL | p.Val642Ile | P43246 |
| HOMO SAPIENS | 1 | 8672 | 3thz | A | 654 | ASP | p.Asp654Glu | P43246 |
| HOMO SAPIENS | 0 | 8672 | 3thz | A | 662 | GLN | p.Gln662His | P43246 |
| HOMO SAPIENS | 2 | 8672 | 3thz | A | 681 | GLN | p.Gln681His | P43246 |
| HOMO SAPIENS | 0 | 8672 | 3thz | A | 707 | CYS | p.Cys707Ser | P43246 |
| HOMO SAPIENS | 0 | 8672 | 3thz | A | 809 | GLU | p.Glu809Lys | P43246 |
| HOMO SAPIENS | 0 | 8672 | 3thz | A | 813 | MET | p.Met813Val | P43246 |
| HOMO SAPIENS | 1 | 8672 | 3thz | A | 834 | ALA | p.Ala834Thr | P43246 |
| HOMO SAPIENS | 0 | 8672 | 3thz | A | 849 | LEU | p.Leu849Val | P43246 |
| HOMO SAPIENS | -1 | 8672 | 3thz | A | 860 | SER | p.Ser860Leu | P43246 |
| HOMO SAPIENS | -1 | 8672 | 3thz | A | 934 | THR | p.Thr934Lys | P43246 |
| HOMO SAPIENS | 2 | 292 | 1i3k | A | 34 | ASN | p.Asn34Ser | Q14376 |
| HOMO SAPIENS | 7 | 292 | 1i3k | A | 90 | GLY | p.Gly90Glu | Q14376 |
| HOMO SAPIENS | 2 | 292 | 1i3k | A | 103 | ASP | p.Asp103Gly | Q14376 |
| HOMO SAPIENS | 2 | 292 | 1i3k | A | 136 | TYR | p.Tyr136Ter | Q14376 |
| HOMO SAPIENS | 0 | 292 | 1i3k | A | 169 | ARG | p.Arg169Trp | Q14376 |
| HOMO SAPIENS | 0 | 292 | 1i3k | A | 183 | LEU | p.Leu183Pro | Q14376 |
| HOMO SAPIENS | 0 | 292 | 1i3k | A | 302 | GLY | p.Gly302Asp | Q14376 |
| HOMO SAPIENS | 0 | 292 | 1i3k | A | 313 | LEU | p.Leu313Met | Q14376 |
| HOMO SAPIENS | 1 | 2084 | 5vc8 | A | 295 | GLN | p.Gln295Leu | O96028 |
| HOMO SAPIENS | -1 | 10157 | 5wrw | A | 7 | GLY | p.Gly7Ala | O76094 |
| HOMO SAPIENS | -1 | 10157 | 5wrw | A | 7 | GLY | p.Gly7Arg | O76094 |
| HOMO SAPIENS | -1 | 10157 | 5wrw | A | 8 | GLY | p.Gly8Ala | O76094 |
| HOMO SAPIENS | 0 | 3113 | 3in5 | A | 29 | GLU | p.Glu29Lys | Q9UBT6 |
| HOMO SAPIENS | 5 | 4546 | 5in3 | B | 44 | VAL | p.Val44Leu | P07902 |
| HOMO SAPIENS | -1 | 4546 | 5in3 | B | 51 | ARG | p.Arg51Leu | P07902 |
| HOMO SAPIENS | -1 | 4546 | 5in3 | B | 62 | LEU | p.Leu62Met | P07902 |
| HOMO SAPIENS | 4 | 4546 | 5in3 | B | 66 | PRO | p.Pro66His | P07902 |
| HOMO SAPIENS | 2 | 4546 | 5in3 | B | 74 | LEU | p.Leu74Pro | P07902 |
| HOMO SAPIENS | 5 | 4546 | 5in3 | B | 97 | ASN | p.Asn97Ser | P07902 |
| HOMO SAPIENS | 1 | 4546 | 5in3 | B | 135 | SER | p.Ser135Leu | P07902 |
| HOMO SAPIENS | 5 | 4546 | 5in3 | B | 138 | THR | p.Thr138Met | P07902 |
| HOMO SAPIENS | 7 | 4546 | 5in3 | B | 139 | LEU | p.Leu139Pro | P07902 |

| | | | | | | | | |
|---|---|---|---|---|---|---|---|---|
| HOMO SAPIENS | 2 | 4546 | 5in3 | B | 142 | MET | p.Met142Lys | P07902 |
| HOMO SAPIENS | 1 | 4546 | 5in3 | B | 143 | SER | p.Ser143Leu | P07902 |
| HOMO SAPIENS | 2 | 4546 | 5in3 | B | 148 | ARG | p.Arg148Gly | P07902 |
| HOMO SAPIENS | 2 | 4546 | 5in3 | B | 148 | ARG | p.Arg148Gln | P07902 |
| HOMO SAPIENS | 5 | 4546 | 5in3 | B | 183 | PRO | p.Pro183Thr | P07902 |
| HOMO SAPIENS | 5 | 4546 | 5in3 | B | 185 | PRO | p.Pro185His | P07902 |
| HOMO SAPIENS | 5 | 4546 | 5in3 | B | 186 | HIS | p.His186Tyr | P07902 |
| HOMO SAPIENS | 7 | 4546 | 5in3 | B | 188 | GLN | p.Gln188Arg | P07902 |
| HOMO SAPIENS | 5 | 4546 | 5in3 | B | 195 | LEU | p.Leu195Pro | P07902 |
| HOMO SAPIENS | 4 | 4546 | 5in3 | B | 201 | ARG | p.Arg201His | P07902 |
| HOMO SAPIENS | 4 | 4546 | 5in3 | B | 201 | ARG | p.Arg201Cys | P07902 |
| HOMO SAPIENS | 4 | 4546 | 5in3 | B | 203 | GLU | p.Glu203Lys | P07902 |
| HOMO SAPIENS | 2 | 4546 | 5in3 | B | 209 | TYR | p.Tyr209Cys | P07902 |
| HOMO SAPIENS | 5 | 4546 | 5in3 | B | 223 | ARG | p.Arg223Ser | P07902 |
| HOMO SAPIENS | 5 | 4546 | 5in3 | B | 226 | LEU | p.Leu226Pro | P07902 |
| HOMO SAPIENS | 5 | 4546 | 5in3 | B | 226 | LEU | p.Leu226Val | P07902 |
| HOMO SAPIENS | 2 | 4546 | 5in3 | B | 227 | LEU | p.Leu227Pro | P07902 |
| HOMO SAPIENS | 8 | 4546 | 5in3 | B | 258 | ARG | p.Arg258Cys | P07902 |
| HOMO SAPIENS | 1 | 4546 | 5in3 | B | 265 | PRO | p.Pro265Ala | P07902 |
| HOMO SAPIENS | 8 | 4546 | 5in3 | B | 268 | THR | p.Thr268Asn | P07902 |
| HOMO SAPIENS | 7 | 4546 | 5in3 | B | 278 | ILE | p.Ile278Asn | P07902 |
| HOMO SAPIENS | 5 | 4546 | 5in3 | B | 285 | LYS | p.Lys285Asn | P07902 |
| HOMO SAPIENS | 1 | 4546 | 5in3 | B | 319 | HIS | p.His319Gln | P07902 |
| HOMO SAPIENS | 4 | 4546 | 5in3 | B | 321 | HIS | p.His321Tyr | P07902 |
| HOMO SAPIENS | 4 | 4546 | 5in3 | B | 325 | PRO | p.Pro325Leu | P07902 |
| HOMO SAPIENS | 2 | 4546 | 5in3 | B | 328 | ARG | p.Arg328His | P07902 |
| HOMO SAPIENS | 2 | 4546 | 5in3 | B | 329 | SER | p.Ser329Phe | P07902 |
| HOMO SAPIENS | 5 | 4546 | 5in3 | B | 334 | LYS | p.Lys334Arg | P07902 |
| HOMO SAPIENS | 6 | 1290 | 5uav | C | 119 | ARG | p.Arg119Gly | P32322 |
| HOMO SAPIENS | 6 | 1290 | 5uav | C | 119 | ARG | p.Arg119His | P32322 |
| HOMO SAPIENS | 3 | 1290 | 5uav | C | 251 | ARG | p.Arg251His | P32322 |
| HOMO SAPIENS | 7 | 1290 | 5uav | C | 266 | ARG | p.Arg266Gln | P32322 |
| HOMO SAPIENS | 0 | 9618 | 2r7e | A | 22 | GLY | p.Gly41Cys | P00451 |
| HOMO SAPIENS | 2 | 9618 | 2r7e | A | 113 | GLU | p.Glu132Asp | P00451 |
| HOMO SAPIENS | -1 | 9618 | 2r7e | A | 336 | ARG | p.Arg355Ter | P00451 |
| HOMO SAPIENS | 0 | 9618 | 2r7e | A | 372 | ARG | p.Arg391His | P00451 |
| HOMO SAPIENS | 0 | 9618 | 2r7e | A | 373 | SER | p.Ser392Leu | P00451 |
| HOMO SAPIENS | 0 | 9618 | 2r7e | A | 373 | SER | p.Ser392Pro | P00451 |
| HOMO SAPIENS | 3 | 9618 | 2r7e | A | 386 | ILE | p.Ile405Ser | P00451 |
| HOMO SAPIENS | 6 | 9618 | 2r7e | A | 390 | GLU | p.Glu409Gly | P00451 |
| HOMO SAPIENS | 2 | 9618 | 2r7e | A | 412 | LEU | p.Leu431Phe | P00451 |
| HOMO SAPIENS | 5 | 9618 | 2r7e | A | 425 | LYS | p.Lys444Arg | P00451 |
| HOMO SAPIENS | 0 | 9618 | 2r7e | A | 484 | ARG | p.Arg503His | P00451 |
| HOMO SAPIENS | 2 | 9618 | 2r7e | A | 531 | ARG | p.Arg550His | P00451 |
| HOMO SAPIENS | 3 | 9618 | 2r7e | A | 535 | SER | p.Ser554Gly | P00451 |
| HOMO SAPIENS | 2 | 9618 | 2r7e | A | 593 | ARG | p.Arg612Cys | P00451 |
| HOMO SAPIENS | 0 | 9618 | 2r7e | A | 704 | ALA | p.Ala723Thr | P00451 |
| HOMO SAPIENS | 0 | 9618 | 2r7e | A | 720 | GLU | p.Glu739Lys | P00451 |
| HOMO SAPIENS | 1 | 13085 | 3ud1 | A | 1075 | ILE | p.Thr1075Ile | P16157 |
| HOMO SAPIENS | -1 | 3289 | 2fy4 | A | 2 | ALA | p.Ala120Thr | P28329 |
| HOMO SAPIENS | 7 | 3289 | 2fy4 | A | 18 | VAL | p.Val136Met | P28329 |
| HOMO SAPIENS | 5 | 3289 | 2fy4 | A | 92 | LEU | p.Leu210Pro | P28329 |
| HOMO SAPIENS | 2 | 3289 | 2fy4 | A | 93 | PRO | p.Pro211Ala | P28329 |
| HOMO SAPIENS | 4 | 3289 | 2fy4 | A | 131 | LEU | p.Leu249Val | P28329 |

| | | | | | | | | |
|---|---|---|---|---|---|---|---|---|
| HOMO SAPIENS | 2 | 3289 | 2fy4 | A | 181 | PRO | p.Pro299Leu | P28329 |
| HOMO SAPIENS | 0 | 3289 | 2fy4 | A | 218 | ILE | p.Ile336Thr | P28329 |
| HOMO SAPIENS | 2 | 3289 | 2fy4 | A | 261 | ASP | p.Asp379His | P28329 |
| HOMO SAPIENS | 0 | 3289 | 2fy4 | A | 302 | ARG | p.Arg420Cys | P28329 |
| HOMO SAPIENS | 5 | 3289 | 2fy4 | A | 323 | GLU | p.Glu441Lys | P28329 |
| HOMO SAPIENS | 0 | 3289 | 2fy4 | A | 343 | MET | p.Val461Met | P28329 |
| HOMO SAPIENS | 2 | 3289 | 2fy4 | A | 364 | ARG | p.Arg482Gly | P28329 |
| HOMO SAPIENS | 3 | 3289 | 2fy4 | A | 380 | SER | p.Ser498Leu | P28329 |
| HOMO SAPIENS | 6 | 3289 | 2fy4 | A | 388 | VAL | p.Val506Leu | P28329 |
| HOMO SAPIENS | 7 | 3289 | 2fy4 | A | 429 | HIS | p.His547Gln | P28329 |
| HOMO SAPIENS | 8 | 3289 | 2fy4 | A | 442 | ARG | p.Arg560His | P28329 |
| HOMO SAPIENS | 3 | 3289 | 2fy4 | A | 443 | ARG | p.Arg561Gln | P28329 |
| HOMO SAPIENS | 0 | 4137 | 3hhr | B | 38 | CYS | p.Cys56Ter | P10912 |
| HOMO SAPIENS | 0 | 4137 | 3hhr | B | 43 | ARG | p.Arg61Ter | P10912 |
| HOMO SAPIENS | 0 | 4137 | 3hhr | B | 44 | GLU | p.Glu62Lys | P10912 |
| HOMO SAPIENS | 0 | 4137 | 3hhr | B | 96 | PHE | p.Phe114Ser | P10912 |
| HOMO SAPIENS | 5 | 4137 | 3hhr | B | 131 | PRO | p.Pro149Gln | P10912 |
| HOMO SAPIENS | 0 | 4137 | 3hhr | B | 144 | VAL | p.Val162Ile | P10912 |
| HOMO SAPIENS | 0 | 4137 | 3hhr | B | 150 | HIS | p.His168Gln | P10912 |
| HOMO SAPIENS | 0 | 4137 | 3hhr | B | 152 | ASP | p.Asp170His | P10912 |
| HOMO SAPIENS | 0 | 4137 | 3hhr | B | 153 | ILE | p.Ile171Thr | P10912 |
| HOMO SAPIENS | 0 | 4137 | 3hhr | B | 154 | GLN | p.Gln172Pro | P10912 |
| HOMO SAPIENS | 0 | 4137 | 3hhr | B | 155 | VAL | p.Val173Gly | P10912 |
| HOMO SAPIENS | 0 | 4137 | 3hhr | B | 217 | ARG | p.Arg235Ter | P10912 |
| HOMO SAPIENS | 5 | 4137 | 3hhr | B | 224 | GLU | p.Glu242Asp | P10912 |
| HOMO SAPIENS | 5 | 4137 | 3hhr | B | 224 | GLU | p.Glu242Ter | P10912 |
| HOMO SAPIENS | 6 | 2493 | 5bpb | A | 69 | HIS | p.His69Tyr | Q9ULV1 |
| HOMO SAPIENS | 6 | 2493 | 5bpb | A | 105 | MET | p.Met105Val | Q9ULV1 |
| HOMO SAPIENS | -1 | 2493 | 5bpb | A | 168 | PRO | p.Pro168Ser | Q9ULV1 |
| HOMO SAPIENS | 0 | 32123 | 4ccz | A | 314 | ASP | p.Asp314Asn | Q99707 |
| HOMO SAPIENS | 0 | 32123 | 4ccz | A | 345 | VAL | p.Val345Phe | Q99707 |
| HOMO SAPIENS | 1 | 32123 | 4ccz | A | 585 | ARG | p.Arg585Ter | Q99707 |
| HOMO SAPIENS | 4 | 4618 | 2ag4 | A | 26 | PRO | p.Pro55Leu | P17900 |
| HOMO SAPIENS | 4 | 4618 | 2ag4 | A | 30 | VAL | p.Ile59Val | P17900 |
| HOMO SAPIENS | 2 | 4618 | 2ag4 | A | 40 | VAL | p.Met69Val | P17900 |
| HOMO SAPIENS | 2 | 4618 | 2ag4 | A | 124 | VAL | p.Val153Ala | P17900 |
| HOMO SAPIENS | 8 | 4618 | 2ag4 | A | 140 | ARG | p.Arg169Pro | P17900 |
| HOMO SAPIENS | 5 | 1082 | 1hk0 | X | 14 | ARG | p.Arg15Cys | P07320 |
| HOMO SAPIENS | 7 | 1082 | 1hk0 | X | 23 | PRO | p.Pro24Thr | P07320 |
| HOMO SAPIENS | 6 | 1082 | 1hk0 | X | 36 | ARG | p.Arg37Ser | P07320 |
| HOMO SAPIENS | 5 | 1082 | 1hk0 | X | 55 | TYR | p.Tyr56Ter | P07320 |
| HOMO SAPIENS | 3 | 1082 | 1hk0 | X | 58 | ARG | p.Arg59His | P07320 |
| HOMO SAPIENS | 7 | 1082 | 1hk0 | X | 88 | HIS | p.His88Gln | P07320 |
| HOMO SAPIENS | 6 | 1082 | 1hk0 | X | 95 | ARG | p.Arg95Ser | P07320 |
| HOMO SAPIENS | 5 | 1082 | 1hk0 | X | 126 | VAL | p.Val126Met | P07320 |
| HOMO SAPIENS | 8 | 1082 | 1hk0 | X | 157 | TRP | p.Trp157Ter | P07320 |
| HOMO SAPIENS | 0 | 22879 | 3j0a | B | 644 | ILE | p.Ile644Phe | O60602 |
| HOMO SAPIENS | 8 | 6 | 2o23 | B | 122 | LEU | p.Leu122Val | Q99714 |
| HOMO SAPIENS | 1 | 6 | 2o23 | B | 130 | ARG | p.Arg130Cys | Q99714 |
| HOMO SAPIENS | 1 | 6 | 2o23 | B | 247 | ASN | p.Asn247Ser | Q99714 |
| HOMO SAPIENS | 0 | 6 | 2o23 | B | 249 | GLU | p.Glu249Gln | Q99714 |
| HOMO SAPIENS | 1 | 12910 | 4hl4 | A | 67 | ARG | p.Arg67Ter | Q96BZ9 |
| HOMO SAPIENS | 8 | 16105 | 1lmj | A | 7 | GLU | p.Glu1073Lys | P35555 |
| HOMO SAPIENS | 8 | 16105 | 1lmj | A | 8 | CYS | p.Cys1074Arg | P35555 |

| | | | | | | | | |
|---|---|---|---|---|---|---|---|---|
| HOMO SAPIENS | 9 | 16105 | 1lmj | A | 51 | CYS | p.Cys1117Tyr | P35555 |
| HOMO SAPIENS | 9 | 16105 | 1lmj | A | 63 | CYS | p.Cys1129Tyr | P35555 |
| HOMO SAPIENS | 9 | 16105 | 1lmj | A | 65 | ASN | p.Asn1131Tyr | P35555 |
| HOMO SAPIENS | 4 | 16105 | 1lmj | A | 71 | ARG | p.Arg1137Pro | P35555 |
| HOMO SAPIENS | 1 | 16105 | 1lmj | A | 82 | PRO | p.Pro1148Ala | P35555 |
| HOMO SAPIENS | 0 | 4795 | 3s5f | A | 106 | LEU | p.Leu106Ser | Q16595 |
| HOMO SAPIENS | 0 | 4795 | 3s5f | A | 130 | GLY | p.Gly130Val | Q16595 |
| HOMO SAPIENS | 0 | 4795 | 3s5f | A | 154 | ILE | p.Ile154Phe | Q16595 |
| HOMO SAPIENS | -1 | 4795 | 3s5f | A | 209 | ASP | p.Asp209Gly | Q16595 |
| HOMO SAPIENS | -1 | 2884 | 4lli | A | 1655 | ASP | p.Asp1655Glu | Q9Y4I1 |
| HOMO SAPIENS | 0 | 1786 | 7pck | C | 56P | TYR | p.Tyr71Ter | P43235 |
| HOMO SAPIENS | 0 | 1786 | 7pck | C | 64P | GLY | p.Gly79Glu | P43235 |
| HOMO SAPIENS | 7 | 38286 | 2l53 | B | 1914 | ARG | p.Arg1914Gly | Q14524 |
| HOMO SAPIENS | 1 | 38286 | 2l53 | B | 1924 | ALA | p.Ala1924Thr | Q14524 |
| HOMO SAPIENS | 0 | 31747 | 3ksy | A | 25 | PRO | p.Pro25Ser | Q07889 |
| HOMO SAPIENS | 3 | 31747 | 3ksy | A | 37 | THR | p.Thr37Ala | Q07889 |
| HOMO SAPIENS | 0 | 31747 | 3ksy | A | 54 | GLN | p.Gln54His | Q07889 |
| HOMO SAPIENS | -1 | 31747 | 3ksy | A | 190 | ASP | p.Asp190Glu | Q07889 |
| HOMO SAPIENS | 2 | 31747 | 3ksy | A | 266 | THR | p.Thr266Lys | Q07889 |
| HOMO SAPIENS | 1 | 31747 | 3ksy | A | 269 | MET | p.Met269Arg | Q07889 |
| HOMO SAPIENS | 1 | 31747 | 3ksy | A | 340 | PRO | p.Pro340Ser | Q07889 |
| HOMO SAPIENS | 1 | 31747 | 3ksy | A | 423 | ASN | p.Asn423Lys | Q07889 |
| HOMO SAPIENS | 1 | 31747 | 3ksy | A | 424 | GLU | p.Glu424Lys | Q07889 |
| HOMO SAPIENS | 1 | 31747 | 3ksy | A | 432 | TRP | p.Trp432Arg | Q07889 |
| HOMO SAPIENS | 1 | 31747 | 3ksy | A | 552 | ARG | p.Arg552Gly | Q07889 |
| HOMO SAPIENS | 1 | 31747 | 3ksy | A | 552 | ARG | p.Arg552Ser | Q07889 |
| HOMO SAPIENS | 3 | 7529 | 6jdw | A | 93 | ILE | p.Ile93Met | P50440 |
| HOMO SAPIENS | 2 | 7529 | 6jdw | A | 110 | GLN | p.Gln110His | P50440 |
| HOMO SAPIENS | 5 | 7529 | 6jdw | A | 149 | TRP | p.Trp149Ter | P50440 |
| HOMO SAPIENS | 0 | 7529 | 6jdw | A | 169 | ARG | p.Arg169Ter | P50440 |
| HOMO SAPIENS | 0 | 7529 | 6jdw | A | 260 | ARG | p.Arg260Ter | P50440 |
| HOMO SAPIENS | 3 | 7529 | 6jdw | A | 344 | ILE | p.Ile344Val | P50440 |
| HOMO SAPIENS | 5 | 7529 | 6jdw | A | 418 | LEU | p.Leu418Ile | P50440 |
| HOMO SAPIENS | 1 | 14788 | 1eak | A | 101 | ARG | p.Arg101His | P08253 |
| HOMO SAPIENS | 0 | 14788 | 1eak | A | 383 | ASP | p.Asp383Glu | P08253 |
| HOMO SAPIENS | 7 | 14788 | 1eak | A | 404 | GLN | p.Glu404Lys | P08253 |
| HOMO SAPIENS | 2 | 14788 | 1eak | A | 447 | ALA | p.Ala447Val | P08253 |
| HOMO SAPIENS | 0 | 1255 | 5xtc | p | 8 | PRO | p.Pro8His | Q9Y6M9 |
| HOMO SAPIENS | 0 | 1255 | 5xtc | p | 146 | PRO | p.Pro146Ser | Q9Y6M9 |
| HOMO SAPIENS | 3 | 6721 | 1a22 | A | 25 | PHE | p.Phe51Tyr | P01241 |
| HOMO SAPIENS | 2 | 6721 | 1a22 | A | 95 | SER | p.Ser121Arg | P01241 |
| HOMO SAPIENS | 2 | 6721 | 1a22 | A | 110 | VAL | p.Val136Ile | P01241 |
| HOMO SAPIENS | 0 | 1380 | 5i6z | A | 605 | LYS | p.Lys605Asn | P31645 |
| HOMO SAPIENS | 0 | 941 | 1qki | D | 48 | ILE | p.Ile48Thr | P11413 |
| HOMO SAPIENS | 3 | 941 | 1qki | D | 68 | VAL | p.Val68Met | P11413 |
| HOMO SAPIENS | 2 | 941 | 1qki | D | 126 | ASN | p.Asn126Asp | P11413 |
| HOMO SAPIENS | 8 | 941 | 1qki | D | 163 | GLY | p.Gly163Ser | P11413 |
| HOMO SAPIENS | 6 | 941 | 1qki | D | 165 | ASN | p.Asn165Asp | P11413 |
| HOMO SAPIENS | 4 | 941 | 1qki | D | 181 | ASP | p.Asp181Val | P11413 |
| HOMO SAPIENS | 4 | 941 | 1qki | D | 182 | ARG | p.Arg182Trp | P11413 |
| HOMO SAPIENS | 5 | 941 | 1qki | D | 188 | SER | p.Ser188Phe | P11413 |
| HOMO SAPIENS | 9 | 941 | 1qki | D | 198 | ARG | p.Arg198Cys | P11413 |
| HOMO SAPIENS | 8 | 941 | 1qki | D | 213 | VAL | p.Val213Leu | P11413 |
| HOMO SAPIENS | 5 | 941 | 1qki | D | 282 | ASP | p.Asp282His | P11413 |

| | | | | | | | | |
|---|---|---|---|---|---|---|---|---|
| HOMO SAPIENS | 7 | 941 | 1qki | D | 285 | ARG | p.Arg285His | P11413 |
| HOMO SAPIENS | 5 | 941 | 1qki | D | 291 | VAL | p.Val291Met | P11413 |
| HOMO SAPIENS | 8 | 941 | 1qki | D | 323 | LEU | p.Leu323Pro | P11413 |
| HOMO SAPIENS | 3 | 941 | 1qki | D | 335 | ALA | p.Ala335Thr | P11413 |
| HOMO SAPIENS | 6 | 941 | 1qki | D | 350 | ASP | p.Asp350His | P11413 |
| HOMO SAPIENS | 5 | 941 | 1qki | D | 393 | ARG | p.Arg393His | P11413 |
| HOMO SAPIENS | 5 | 941 | 1qki | D | 454 | ARG | p.Arg454Cys | P11413 |
| HOMO SAPIENS | 2 | 941 | 1qki | D | 459 | LEU | p.Arg459Leu | P11413 |
| HOMO SAPIENS | 4 | 941 | 1qki | D | 463 | ARG | p.Arg463His | P11413 |
| HOMO SAPIENS | 4 | 2483 | 1tij | A | 68 | LEU | p.Leu94Gln | P01034 |
| HOMO SAPIENS | 2 | 20632 | 3v5b | A | 331 | ARG | p.Arg331Pro | P02545 |
| HOMO SAPIENS | 2 | 20632 | 3v5b | A | 374 | HIS | p.His374Gln | P02545 |
| HOMO SAPIENS | 4 | 13929 | 5xtd | J | 75 | ARG | p.Arg75His | Q16795 |
| HOMO SAPIENS | 3 | 13929 | 5xtd | J | 243 | VAL | p.Val243Ala | Q16795 |
| HOMO SAPIENS | 1 | 13929 | 5xtd | J | 294 | PRO | p.Pro294Leu | Q16795 |
| HOMO SAPIENS | 2 | 13929 | 5xtd | J | 321 | ARG | p.Arg321Pro | Q16795 |
| HOMO SAPIENS | 0 | 9411 | 5thl | A | 41 | ARG | p.Gly41Arg | P54577 |
| HOMO SAPIENS | 0 | 9411 | 5thl | A | 196 | GLU | p.Glu196Lys | P54577 |
| HOMO SAPIENS | 0 | 9411 | 5thl | A | 316 | ALA | p.Ala316Ser | P54577 |
| HOMO SAPIENS | 0 | 9411 | 5thl | A | 339 | ALA | p.Ala339Thr | P54577 |
| HOMO SAPIENS | 4 | 16273 | 2e8n | A | 46 | THR | p.Thr940Ile | P29317 |
| HOMO SAPIENS | 7 | 16273 | 2e8n | A | 54 | GLY | p.Gly948Trp | P29317 |
| HOMO SAPIENS | 6 | 16273 | 2e8n | A | 65 | ALA | p.Ala959Thr | P29317 |
| HOMO SAPIENS | 7 | 30387 | 2k9y | A | 541 | VAL | p.Val541Met | P29317 |
| HOMO SAPIENS | 0 | 6961 | 4nha | A | 843 | ILE | p.Ile853Val | Q9UPY3 |
| HOMO SAPIENS | -1 | 6961 | 4nha | A | 862 | ALA | p.Ala872Thr | Q9UPY3 |
| HOMO SAPIENS | 3 | 15708 | 4pzg | A | 16 | ARG | p.Arg16Ter | Q9Y5X0 |
| HOMO SAPIENS | 4 | 15708 | 4pzg | A | 51 | ARG | p.Arg51Gln | Q9Y5X0 |
| HOMO SAPIENS | 3 | 30166 | 4o00 | A | 44 | ARG | p.Arg14257Ter | Q8WZ42 |
| HOMO SAPIENS | 2 | 30166 | 4o00 | A | 54 | ARG | p.Arg14267His | Q8WZ42 |
| HOMO SAPIENS | 8 | 11158 | 4hca | A | 275 | TRP | p.Trp274Arg | P23771 |
| HOMO SAPIENS | 2 | 4328 | 4e0s | A | 1476 | ARG | p.Arg1476Ter | P01031 |
| HOMO SAPIENS | 0 | 11462 | 1dx5 | I | 455 | ALA | p.Ala473Val | P07204 |
| HOMO SAPIENS | -1 | 11150 | 2fic | A | 45 | LYS | p.Lys35Asn | O00499 |
| HOMO SAPIENS | 6 | 11150 | 2fic | A | 161 | ASP | p.Asp151Asn | O00499 |
| HOMO SAPIENS | 8 | 27382 | 4mqv | B | 17 | ARG | p.Arg17Ter | Q9NZC9 |
| HOMO SAPIENS | -1 | 37991 | 1q0p | A | 231 | GLN | p.Gln256Ter | P00751 |
| HOMO SAPIENS | 1 | 443 | 5xte | U | 215 | LEU | p.Leu215Phe | P08574 |
| HOMO SAPIENS | 8 | 21471 | 5y4e | A | 1461 | ASN | p.Thr692Ala | Q01484 |
| HOMO SAPIENS | -1 | 35443 | 3um3 | B | 129 | ASP | p.Asp129Val | Q9H444 |
| HOMO SAPIENS | -1 | 35443 | 3um3 | B | 161 | GLU | p.Glu161Lys | Q9H444 |
| HOMO SAPIENS | 6 | 28781 | 2n2a | B | 111 | ILE | p.Ile654Val | P04626 |
| HOMO SAPIENS | 7 | 28781 | 2n2a | B | 112 | ILE | p.Ile655Val | P04626 |
| HOMO SAPIENS | -1 | 3346 | 4ztz | C | 61 | GLY | p.Gly61Arg | Q9UHN1 |
| HOMO SAPIENS | 4 | 3346 | 4ztz | C | 103 | GLY | p.Gly103Ser | Q9UHN1 |
| HOMO SAPIENS | 4 | 3346 | 4ztz | C | 133 | HIS | p.His133Tyr | Q9UHN1 |
| HOMO SAPIENS | -1 | 3346 | 4ztz | C | 153 | LEU | p.Leu153Val | Q9UHN1 |
| HOMO SAPIENS | -1 | 3346 | 4ztz | C | 169 | ALA | p.Ala169Thr | Q9UHN1 |
| HOMO SAPIENS | 7 | 3346 | 4ztz | C | 369 | ARG | p.Arg369Gly | Q9UHN1 |
| HOMO SAPIENS | 4 | 3346 | 4ztz | C | 386 | ASP | p.Asp386Glu | Q9UHN1 |
| HOMO SAPIENS | 4 | 3346 | 4ztz | C | 451 | GLY | p.Gly451Glu | Q9UHN1 |
| HOMO SAPIENS | 4 | 2131 | 5fo9 | E | 1549 | LEU | p.Leu1549Met | P01024 |
| HOMO SAPIENS | 2 | 2131 | 5fo9 | E | 1619 | SER | p.Ser1619Arg | P01024 |
| HOMO SAPIENS | 3 | 12662 | 4jhp | C | 75 | ILE | p.Ile75Val | Q92834 |

| | | | | | | | | |
|---|---|---|---|---|---|---|---|---|
| HOMO SAPIENS | 1 | 12662 | 4jhp | C | 173 | GLY | p.Gly173Arg | Q92834 |
| HOMO SAPIENS | 0 | 12662 | 4jhp | C | 184 | GLN | p.Gln184His | Q92834 |
| HOMO SAPIENS | 3 | 12662 | 4jhp | C | 262 | ALA | p.Ala262Gly | Q92834 |
| HOMO SAPIENS | 3 | 12662 | 4jhp | C | 302 | CYS | p.Cys302Tyr | Q92834 |
| HOMO SAPIENS | 5 | 1093 | 2jg9 | A | 186 | GLN | p.Gln208Ter | P02745 |
| HOMO SAPIENS | 0 | 13181 | 4r4l | A | 12 | MET | p.Met13Ile | Q13976 |
| HOMO SAPIENS | 8 | 25939 | 4iyp | A | 106 | ARG | p.Arg106Gln | P78318 |
| HOMO SAPIENS | 1 | 217 | 2xsx | A | 71 | SER | p.Asn71Ser | P13929 |
| HOMO SAPIENS | 0 | 217 | 2xsx | A | 76 | ALA | p.Ala76Pro | P13929 |
| HOMO SAPIENS | 1 | 217 | 2xsx | A | 85 | ALA | p.Val85Ala | P13929 |
| HOMO SAPIENS | 8 | 217 | 2xsx | A | 156 | GLY | p.Gly156Asp | P13929 |
| HOMO SAPIENS | 5 | 217 | 2xsx | A | 332 | VAL | p.Val332Ile | P13929 |
| HOMO SAPIENS | 7 | 217 | 2xsx | A | 374 | GLY | p.Gly374Glu | P13929 |
| HOMO SAPIENS | 1 | 5508 | 3szm | A | 682 | THR | p.Thr682Asn | Q8NEM0 |
| HOMO SAPIENS | 0 | 5508 | 3szm | A | 715 | TRP | p.Trp715Ter | Q8NEM0 |
| HOMO SAPIENS | 1 | 5508 | 3szm | A | 761 | VAL | p.Ala761Val | Q8NEM0 |
| HOMO SAPIENS | 1 | 5508 | 3szm | A | 828 | PRO | p.Pro828Ser | Q8NEM0 |
| HOMO SAPIENS | 4 | 5508 | 3szm | A | 829 | GLU | p.Glu829Asp | Q8NEM0 |
| HOMO SAPIENS | 2 | 16353 | 2q0z | X | 171 | GLY | p.Gly1978Ala | O75643 |
| HOMO SAPIENS | 2 | 16353 | 2q0z | X | 188 | ALA | p.Ala1995Thr | O75643 |
| HOMO SAPIENS | 0 | 4127 | 3swr | A | 860 | GLY | p.Gly860Arg | P26358 |
| HOMO SAPIENS | -1 | 4127 | 3swr | A | 1122 | SER | p.Ser1122Cys | P26358 |
| HOMO SAPIENS | 0 | 4127 | 3swr | A | 1216 | ASP | p.Asp1216Glu | P26358 |
| HOMO SAPIENS | 0 | 6382 | 3o96 | A | 17 | GLU | p.Glu17Lys | P31749 |
| HOMO SAPIENS | 6 | 14314 | 5xtc | Q | 71 | PRO | p.Pro71Leu | O75306 |
| HOMO SAPIENS | 2 | 5886 | 4fao | E | 22 | ARG | p.Arg40His | Q13705 |
| HOMO SAPIENS | 1 | 2534 | 3mp4 | A | 37 | GLU | p.Glu37Ter | P35914 |
| HOMO SAPIENS | 3 | 2534 | 3mp4 | A | 41 | MET | p.Arg41Gln | P35914 |
| HOMO SAPIENS | 1 | 2534 | 3mp4 | A | 70 | VAL | p.Val70Leu | P35914 |
| HOMO SAPIENS | 2 | 2196 | 3ifq | A | 142 | ARG | p.Arg142His | P14923 |
| HOMO SAPIENS | 2 | 2196 | 3ifq | A | 153 | ASP | p.Asp153Glu | P14923 |
| HOMO SAPIENS | 3 | 2196 | 3ifq | A | 176 | ARG | p.Arg176Gln | P14923 |
| HOMO SAPIENS | 3 | 2196 | 3ifq | A | 265 | ARG | p.Arg265His | P14923 |
| HOMO SAPIENS | 2 | 2196 | 3ifq | A | 301 | GLU | p.Glu301Gly | P14923 |
| HOMO SAPIENS | 0 | 2196 | 3ifq | A | 456 | VAL | p.Val456Ile | P14923 |
| HOMO SAPIENS | 3 | 2196 | 3ifq | A | 603 | VAL | p.Val603Leu | P14923 |
| HOMO SAPIENS | 0 | 2196 | 3ifq | A | 648 | VAL | p.Val648Ile | P14923 |
| HOMO SAPIENS | 1 | 8746 | 1cgj | I | 32 | PRO | p.Pro55Ser | P00995 |
| HOMO SAPIENS | 1 | 8746 | 1cgj | I | 44 | ARG | p.Arg67His | P00995 |
| HOMO SAPIENS | 1 | 8746 | 1cgj | I | 44 | ARG | p.Arg67Cys | P00995 |
| HOMO SAPIENS | 3 | 3212 | 1syq | A | 105 | ARG | p.Arg105Ter | P18206 |
| HOMO SAPIENS | 3 | 3212 | 1syq | A | 126 | PHE | p.Phe126Leu | P18206 |
| HOMO SAPIENS | 4 | 10028 | 3eto | A | 1609 | ARG | p.Arg1608His | P46531 |
| HOMO SAPIENS | 1 | 10028 | 3eto | A | 1672 | VAL | p.Val1671Ile | P46531 |
| HOMO SAPIENS | 5 | 10028 | 3eto | A | 1692 | GLN | p.Gln1691His | P46531 |
| HOMO SAPIENS | 3 | 3749 | 5xtc | Z | 70 | GLY | p.Gly70Ter | O43676 |
| HOMO SAPIENS | 2 | 17039 | 3brv | B | 57 | GLU | p.Glu57Lys | Q9Y6K9 |
| HOMO SAPIENS | 8 | 3589 | 2fh3 | A | 433 | VAL | p.Val460Leu | P06396 |
| HOMO SAPIENS | 1 | 3589 | 2fh3 | A | 454 | ARG | p.Arg481Cys | P06396 |
| HOMO SAPIENS | 4 | 3589 | 2fh3 | A | 498 | ARG | p.Arg525His | P06396 |
| HOMO SAPIENS | 2 | 3589 | 2fh3 | A | 520 | TYR | p.Tyr547Cys | P06396 |
| HOMO SAPIENS | 1 | 3589 | 2fh3 | A | 529 | GLY | p.Gly556Arg | P06396 |
| HOMO SAPIENS | 2 | 3589 | 2fh3 | A | 536 | THR | p.Thr563Ser | P06396 |
| HOMO SAPIENS | 2 | 3589 | 2fh3 | A | 579 | VAL | p.Val606Met | P06396 |

| | | | | | | | | |
|---|---|---|---|---|---|---|---|---|
| HOMO SAPIENS | 1 | 3589 | 2fh3 | A | 589 | THR | p.Thr616Met | P06396 |
| HOMO SAPIENS | 5 | 3589 | 2fh3 | A | 668 | THR | p.Thr695Met | P06396 |
| HOMO SAPIENS | 0 | 148 | 1lcl | A | 130 | ILE | p.Ile130Ser | Q05315 |
| HOMO SAPIENS | -1 | 24727 | 4fj3 | P | 237 | ALA | p.Ala237Thr | P04049 |
| HOMO SAPIENS | 0 | 24727 | 4fj3 | P | 257 | SER | p.Ser257Leu | P04049 |
| HOMO SAPIENS | 0 | 24727 | 4fj3 | P | 261 | PRO | p.Pro261Ala | P04049 |
| HOMO SAPIENS | 0 | 2949 | 5xtb | B | 77 | ARG | p.Arg77Gly | O00217 |
| HOMO SAPIENS | 0 | 2949 | 5xtb | B | 79 | PRO | p.Pro79Leu | O00217 |
| HOMO SAPIENS | 0 | 2949 | 5xtb | B | 100 | ALA | p.Ala100Val | O00217 |
| HOMO SAPIENS | 1 | 2949 | 5xtb | B | 115 | LYS | p.Lys115Glu | O00217 |
| HOMO SAPIENS | 0 | 2949 | 5xtb | B | 138 | ARG | p.Arg138His | O00217 |
| HOMO SAPIENS | 1 | 3959 | 5k5s | B | 25 | ARG | p.Arg25Ter | P41180 |
| HOMO SAPIENS | 4 | 3959 | 5k5s | B | 39 | PRO | p.Pro39Ala | P41180 |
| HOMO SAPIENS | 0 | 3959 | 5k5s | B | 47 | LYS | p.Lys47Asn | P41180 |
| HOMO SAPIENS | 2 | 3959 | 5k5s | B | 62 | ARG | p.Arg62Met | P41180 |
| HOMO SAPIENS | 2 | 3959 | 5k5s | B | 66 | ARG | p.Arg66Cys | P41180 |
| HOMO SAPIENS | 4 | 3959 | 5k5s | B | 90 | ASN | p.Asn90Thr | P41180 |
| HOMO SAPIENS | 1 | 3959 | 5k5s | B | 116 | ALA | p.Ala116Thr | P41180 |
| HOMO SAPIENS | 1 | 3959 | 5k5s | B | 118 | ASN | p.Asn118Lys | P41180 |
| HOMO SAPIENS | -1 | 3959 | 5k5s | B | 125 | LEU | p.Leu125Pro | P41180 |
| HOMO SAPIENS | -1 | 3959 | 5k5s | B | 127 | GLU | p.Glu127Ala | P41180 |
| HOMO SAPIENS | -1 | 3959 | 5k5s | B | 128 | PHE | p.Phe128Leu | P41180 |
| HOMO SAPIENS | -1 | 3959 | 5k5s | B | 131 | CYS | p.Cys131Trp | P41180 |
| HOMO SAPIENS | 1 | 3959 | 5k5s | B | 138 | THR | p.Thr138Met | P41180 |
| HOMO SAPIENS | 0 | 3959 | 5k5s | B | 143 | GLY | p.Gly143Glu | P41180 |
| HOMO SAPIENS | 5 | 3959 | 5k5s | B | 151 | THR | p.Thr151Met | P41180 |
| HOMO SAPIENS | 4 | 3959 | 5k5s | B | 180 | PHE | p.Phe180Cys | P41180 |
| HOMO SAPIENS | 0 | 3959 | 5k5s | B | 185 | ARG | p.Arg185Gln | P41180 |
| HOMO SAPIENS | 0 | 3959 | 5k5s | B | 185 | ARG | p.Arg185Ter | P41180 |
| HOMO SAPIENS | 2 | 3959 | 5k5s | B | 191 | GLU | p.Glu191Lys | P41180 |
| HOMO SAPIENS | 2 | 3959 | 5k5s | B | 227 | ARG | p.Arg227Leu | P41180 |
| HOMO SAPIENS | 2 | 3959 | 5k5s | B | 250 | GLU | p.Glu250Lys | P41180 |
| HOMO SAPIENS | 6 | 3959 | 5k5s | B | 297 | GLU | p.Glu297Lys | P41180 |
| HOMO SAPIENS | 1 | 3959 | 5k5s | B | 336 | LYS | p.Lys336Asn | P41180 |
| HOMO SAPIENS | 6 | 3959 | 5k5s | B | 429 | HIS | p.His429Tyr | P41180 |
| HOMO SAPIENS | 0 | 3959 | 5k5s | B | 445 | THR | p.Thr445Ala | P41180 |
| HOMO SAPIENS | 3 | 3959 | 5k5s | B | 509 | GLY | p.Gly509Arg | P41180 |
| HOMO SAPIENS | 1 | 3959 | 5k5s | B | 553 | GLY | p.Gly553Arg | P41180 |
| HOMO SAPIENS | 3 | 3959 | 5k5s | B | 592 | ASN | p.Asn592Ser | P41180 |
| HOMO SAPIENS | -1 | 3959 | 5k5s | B | 604 | GLU | p.Glu604Lys | P41180 |
| HOMO SAPIENS | 4 | 2564 | 5l0c | C | 975 | ARG | p.Arg975Trp | P18206 |
| HOMO SAPIENS | 4 | 2564 | 5l0c | C | 975 | ARG | p.Arg975Leu | P18206 |
| HOMO SAPIENS | 1 | 5695 | 5i8r | A | 294 | ARG | p.Arg294Gln | P17405 |
| HOMO SAPIENS | 0 | 5695 | 5i8r | A | 485 | ALA | p.Ala485Val | P17405 |
| HOMO SAPIENS | 1 | 5695 | 5i8r | A | 506 | GLY | p.Gly506Arg | P17405 |
| HOMO SAPIENS | -1 | 1978 | 3avs | A | 1060 | HIS | p.His1060Leu | O15550 |
| HOMO SAPIENS | 0 | 11173 | 4v0c | B | 362 | LYS | p.Lys362Arg | P51787 |
| HOMO SAPIENS | 3 | 11173 | 4v0c | B | 381 | CYS | p.Cys381Tyr | P51787 |
| HOMO SAPIENS | 7 | 11173 | 4v0c | B | 518 | ARG | p.Arg518Gly | P51787 |
| HOMO SAPIENS | 2 | 11173 | 4v0c | B | 525 | ALA | p.Ala525Thr | P51787 |
| HOMO SAPIENS | 0 | 11173 | 4v0c | B | 530 | GLN | p.Gln530Ter | P51787 |
| HOMO SAPIENS | -1 | 11173 | 4v0c | B | 536 | TYR | p.Tyr536Ter | P51787 |
| HOMO SAPIENS | 2 | 1113 | 5xtd | N | 60 | ARG | p.Arg60Ter | Q9UI09 |
| HOMO SAPIENS | 3 | 6061 | 2mmz | A | 148 | LEU | p.Leu148Ser | Q86SX6 |

| | | | | | | | | |
|---|---|---|---|---|---|---|---|---|
| HOMO SAPIENS | -1 | 26081 | 3eu7 | A | 837 | GLU | p.Glu837Ter | Q86YC2 |
| HOMO SAPIENS | 0 | 26081 | 3eu7 | A | 864 | PRO | p.Pro864Ser | Q86YC2 |
| HOMO SAPIENS | 5 | 26081 | 3eu7 | A | 891 | CYS | p.Cys891Trp | Q86YC2 |
| HOMO SAPIENS | 0 | 26081 | 3eu7 | A | 932 | VAL | p.Val932Met | Q86YC2 |
| HOMO SAPIENS | 5 | 26081 | 3eu7 | A | 961 | LEU | p.Leu961Met | Q86YC2 |
| HOMO SAPIENS | 5 | 26081 | 3eu7 | A | 1038 | TRP | p.Trp1038Ter | Q86YC2 |
| HOMO SAPIENS | -1 | 26081 | 3eu7 | A | 1086 | ARG | p.Arg1086Ter | Q86YC2 |
| HOMO SAPIENS | 0 | 26081 | 3eu7 | A | 1122 | ASP | p.Asp1122Glu | Q86YC2 |
| HOMO SAPIENS | 5 | 26081 | 3eu7 | A | 1158 | HIS | p.His1158Gln | Q86YC2 |
| HOMO SAPIENS | 5 | 26081 | 3eu7 | A | 1183 | TYR | p.Tyr1183Ter | Q86YC2 |
| HOMO SAPIENS | 4 | 199 | 1qo5 | C | 45 | ARG | p.Arg46Trp | P05062 |
| HOMO SAPIENS | 5 | 199 | 1qo5 | C | 147 | TRP | p.Trp148Arg | P05062 |
| HOMO SAPIENS | 0 | 199 | 1qo5 | C | 149 | ALA | p.Ala150Pro | P05062 |
| HOMO SAPIENS | 0 | 199 | 1qo5 | C | 174 | ALA | p.Ala175Asp | P05062 |
| HOMO SAPIENS | 0 | 199 | 1qo5 | C | 334 | ASN | p.Asn335Lys | P05062 |
| HOMO SAPIENS | 0 | 199 | 1qo5 | C | 337 | ALA | p.Ala338Val | P05062 |
| HOMO SAPIENS | 5 | 199 | 1qo5 | C | 342 | TYR | p.Tyr343His | P05062 |
| HOMO SAPIENS | 0 | 6910 | 4zht | B | 59 | ARG | p.Arg59Ter | Q9Y223 |
| HOMO SAPIENS | 4 | 6910 | 4zht | B | 129 | ARG | p.Arg129Ter | Q9Y223 |
| HOMO SAPIENS | 4 | 6910 | 4zht | B | 129 | ARG | p.Arg129Gln | Q9Y223 |
| HOMO SAPIENS | 1 | 6910 | 4zht | B | 162 | ARG | p.Arg162Cys | Q9Y223 |
| HOMO SAPIENS | 0 | 6910 | 4zht | B | 176 | ASP | p.Asp176Val | Q9Y223 |
| HOMO SAPIENS | 0 | 6910 | 4zht | B | 216 | VAL | p.Val216Ala | Q9Y223 |
| HOMO SAPIENS | 0 | 6910 | 4zht | B | 246 | ARG | p.Arg246Trp | Q9Y223 |
| HOMO SAPIENS | 0 | 6910 | 4zht | B | 263 | ARG | p.Arg263Leu | Q9Y223 |
| HOMO SAPIENS | 0 | 6910 | 4zht | B | 266 | ARG | p.Arg266Gln | Q9Y223 |
| HOMO SAPIENS | 0 | 6910 | 4zht | B | 266 | ARG | p.Arg266Trp | Q9Y223 |
| HOMO SAPIENS | 0 | 6910 | 4zht | B | 277 | ARG | p.Arg277Cys | Q9Y223 |
| HOMO SAPIENS | 0 | 6910 | 4zht | B | 378 | ASP | p.Asp378Tyr | Q9Y223 |
| HOMO SAPIENS | 3 | 211 | 5kxi | C | 262 | VAL | p.Val287Leu | P17787 |
| HOMO SAPIENS | 0 | 211 | 5kxi | C | 369 | PHE | p.Phe478Leu | P17787 |
| HOMO SAPIENS | 0 | 2454 | 5iy9 | V | null | PHE | p.Phe99Ser | P19447 |
| HOMO SAPIENS | 0 | 2454 | 5iy9 | V | null | ARG | p.Arg109Ter | P19447 |
| HOMO SAPIENS | 0 | 2454 | 5iy9 | V | null | THR | p.Thr119Pro | P19447 |
| HOMO SAPIENS | 0 | 2454 | 5iy9 | V | 425 | ARG | p.Arg425Ter | P19447 |
| HOMO SAPIENS | 1 | 2454 | 5iy9 | V | 545 | GLN | p.Gln545Ter | P19447 |
| HOMO SAPIENS | -1 | 3805 | 3rbq | A | 57 | LYS | p.Lys57Ter | Q13432 |
| HOMO SAPIENS | 2 | 2649 | 4ido | A | 43 | ASP | p.Asp43Glu | Q8WXF7 |
| HOMO SAPIENS | 2 | 2649 | 4ido | A | 157 | LEU | p.Leu157Trp | Q8WXF7 |
| HOMO SAPIENS | 2 | 2649 | 4ido | A | 193 | PHE | p.Phe193Cys | Q8WXF7 |
| HOMO SAPIENS | 7 | 2649 | 4ido | A | 217 | ARG | p.Arg217Gln | Q8WXF7 |
| HOMO SAPIENS | 4 | 2649 | 4ido | A | 239 | ARG | p.Arg239Cys | Q8WXF7 |
| HOMO SAPIENS | 3 | 2649 | 4ido | A | 258 | HIS | p.His258Arg | Q8WXF7 |
| HOMO SAPIENS | 4 | 2649 | 4ido | A | 259 | SER | p.Ser259Tyr | Q8WXF7 |
| HOMO SAPIENS | 4 | 2649 | 4ido | A | 408 | MET | p.Met408Val | Q8WXF7 |
| HOMO SAPIENS | 2 | 2649 | 4ido | A | 415 | ARG | p.Arg415Trp | Q8WXF7 |
| HOMO SAPIENS | 3 | 2649 | 4ido | A | 416 | ARG | p.Arg416Cys | Q8WXF7 |
| HOMO SAPIENS | 5 | 2203 | 3wff | A | 810 | LEU | p.Ser810Leu | P08235 |
| HOMO SAPIENS | 4 | 12648 | 5ep6 | B | 696 | GLU | p.Glu696Lys | Q9UHD2 |
| HOMO SAPIENS | 0 | 30622 | 2pe4 | A | 90 | GLU | p.Glu90Asp | Q12794 |
| HOMO SAPIENS | 0 | 30622 | 2pe4 | A | 256 | GLY | p.Gly256Arg | Q12794 |
| HOMO SAPIENS | 1 | 30622 | 2pe4 | A | 268 | GLU | p.Glu268Lys | Q12794 |
| HOMO SAPIENS | 0 | 30622 | 2pe4 | A | 392 | THR | p.Thr392Met | Q12794 |
| HOMO SAPIENS | 5 | 5411 | 1lya | A | 93 | SER | p.Ser157Leu | P07339 |

| | | | | | | | | |
|---|---|---|---|---|---|---|---|---|
| HOMO SAPIENS | 3 | 5743 | 2uzp | A | 189 | ASN | p.Asn189Lys | P05129 |
| HOMO SAPIENS | 1 | 12529 | 3kfd | J | 21 | ASN | p.Asn45Ser | P36897 |
| HOMO SAPIENS | 1 | 1063 | 2e1q | B | 149 | ARG | p.Arg149Cys | P47989 |
| HOMO SAPIENS | -1 | 1063 | 2e1q | B | 172 | GLY | p.Gly172Arg | P47989 |
| HOMO SAPIENS | 4 | 1063 | 2e1q | B | 228 | ARG | p.Arg228Ter | P47989 |
| HOMO SAPIENS | 5 | 1063 | 2e1q | B | 395 | LYS | p.Lys395Met | P47989 |
| HOMO SAPIENS | 2 | 1063 | 2e1q | B | 646 | ILE | p.Ile646Val | P47989 |
| HOMO SAPIENS | 4 | 1063 | 2e1q | B | 703 | ILE | p.Ile703Val | P47989 |
| HOMO SAPIENS | 8 | 2525 | 1kqn | F | 18 | ASN | p.Asn18Ser | Q9HAN9 |
| HOMO SAPIENS | 0 | 2525 | 1kqn | F | 151 | VAL | p.Val151Phe | Q9HAN9 |
| HOMO SAPIENS | 0 | 2525 | 1kqn | F | 153 | LEU | p.Leu153Val | Q9HAN9 |
| HOMO SAPIENS | 0 | 2525 | 1kqn | F | 169 | TRP | p.Trp169Ter | Q9HAN9 |
| HOMO SAPIENS | 0 | 2525 | 1kqn | F | 207 | ARG | p.Arg207Trp | Q9HAN9 |
| HOMO SAPIENS | 0 | 2525 | 1kqn | F | 273 | ASN | p.Asn273Asp | Q9HAN9 |
| HOMO SAPIENS | 0 | 12299 | 5duq | A | 432 | VAL | p.Val454Glu | P05155 |
| HOMO SAPIENS | 0 | 12299 | 5duq | A | 436 | ALA | p.Ala458Thr | P05155 |
| HOMO SAPIENS | 0 | 12299 | 5duq | A | 443 | ALA | p.Ala465Val | P05155 |
| HOMO SAPIENS | 0 | 12299 | 5duq | A | 444 | ARG | p.Arg466His | P05155 |
| HOMO SAPIENS | 0 | 12299 | 5duq | A | 458 | MET | p.Val480Met | P05155 |
| HOMO SAPIENS | 0 | 327 | 1uhp | A | 36 | ASP | p.Asp274Asn | Q9UPQ7 |
| HOMO SAPIENS | 0 | 64 | 2mo5 | A | 54 | ALA | p.Ala55Thr | P12104 |
| HOMO SAPIENS | 2 | 35969 | 1jrh | I | 60 | CYS | p.Cys77Tyr | P15260 |
| HOMO SAPIENS | 4 | 35969 | 1jrh | I | 70 | ILE | p.Ile87Thr | P15260 |
| HOMO SAPIENS | 2 | 1068 | 1mu7 | A | 304 | ARG | p.Arg304Gln | Q9NUW8 |
| HOMO SAPIENS | 8 | 1068 | 1mu7 | A | 493 | HIS | p.His493Arg | Q9NUW8 |
| HOMO SAPIENS | 0 | 32360 | 2kqp | A | 31 | ARG | p.Arg55Cys | P01308 |
| HOMO SAPIENS | 0 | 32360 | 2kqp | A | 65 | ARG | p.Arg89His | P01308 |
| HOMO SAPIENS | 0 | 32360 | 2kqp | A | 65 | ARG | p.Arg89Cys | P01308 |
| HOMO SAPIENS | 6 | 10805 | 3nks | A | 12 | ILE | p.Ile12Thr | P50336 |
| HOMO SAPIENS | 0 | 10805 | 3nks | A | 20 | HIS | p.His20Pro | P50336 |
| HOMO SAPIENS | 0 | 10805 | 3nks | A | 59 | ARG | p.Arg59Trp | P50336 |
| HOMO SAPIENS | 0 | 10805 | 3nks | A | 168 | ARG | p.Arg168Cys | P50336 |
| HOMO SAPIENS | 0 | 10805 | 3nks | A | 168 | ARG | p.Arg168His | P50336 |
| HOMO SAPIENS | 2 | 10805 | 3nks | A | 232 | GLY | p.Gly232Arg | P50336 |
| HOMO SAPIENS | 3 | 10805 | 3nks | A | 256 | PRO | p.Pro256Arg | P50336 |
| HOMO SAPIENS | 0 | 10805 | 3nks | A | 304 | ARG | p.Arg304His | P50336 |
| HOMO SAPIENS | 0 | 10805 | 3nks | A | 349 | ASP | p.Asp349Ala | P50336 |
| HOMO SAPIENS | 9 | 7008 | 3u2l | A | 125 | GLN | p.Gln125Ter | P55789 |
| HOMO SAPIENS | 4 | 7008 | 3u2l | A | 172 | HIS | p.His172Asn | P55789 |
| HOMO SAPIENS | 4 | 7008 | 3u2l | A | 194 | ARG | p.Arg194His | P55789 |
| HOMO SAPIENS | 3 | 7008 | 3u2l | A | 198 | GLY | p.Gly198Ser | P55789 |
| HOMO SAPIENS | 0 | 20821 | 3e0g | A | 65 | HIS | p.His116Tyr | P42702 |
| HOMO SAPIENS | 3 | 1151 | 5l9v | A | 278 | ASP | p.Asp278Asn | Q9GZT9 |
| HOMO SAPIENS | 4 | 1151 | 5l9v | A | 317 | CYS | p.Pro317Arg | Q9GZT9 |
| HOMO SAPIENS | 5 | 1151 | 5l9v | A | 371 | ARG | p.Arg371His | Q9GZT9 |
| HOMO SAPIENS | 2 | 3935 | 1gwb | A | 36 | ARG | p.Arg36Gly | P07359 |
| HOMO SAPIENS | 1 | 3935 | 1gwb | A | 73 | LEU | p.Leu73Phe | P07359 |
| HOMO SAPIENS | 1 | 3935 | 1gwb | A | 172 | ALA | p.Ala172Val | P07359 |
| HOMO SAPIENS | 0 | 3935 | 1gwb | A | 249 | GLY | p.Gly249Val | P07359 |
| HOMO SAPIENS | 0 | 3935 | 1gwb | A | 255 | MET | p.Met255Val | P07359 |
| HOMO SAPIENS | 2 | 14541 | 5cuz | A | 138 | GLU | p.Glu138Lys | Q9H3L0 |
| HOMO SAPIENS | 2 | 14541 | 5cuz | A | 143 | SER | p.Ser143Ile | Q9H3L0 |
| HOMO SAPIENS | 4 | 14541 | 5cuz | A | 182 | THR | p.Thr182Asn | Q9H3L0 |
| HOMO SAPIENS | 6 | 14541 | 5cuz | A | 206 | ASN | p.Asn206Ile | Q9H3L0 |

| | | | | | | | | |
|---|---|---|---|---|---|---|---|---|
| HOMO SAPIENS | 0 | 14541 | 5cuz | A | 249 | TYR | p.Tyr249Cys | Q9H3L0 |
| HOMO SAPIENS | 4 | 14541 | 5cuz | A | 250 | ARG | p.Arg250Ter | Q9H3L0 |
| HOMO SAPIENS | 2 | 14541 | 5cuz | A | 259 | LEU | p.Leu259Pro | Q9H3L0 |
| HOMO SAPIENS | 0 | 4206 | 4fao | D | 30 | PRO | p.Pro30Ser | P37023 |
| HOMO SAPIENS | 1 | 4206 | 4fao | D | 50 | TRP | p.Trp50Cys | P37023 |
| HOMO SAPIENS | 0 | 16553 | 3v30 | A | 251 | GLN | p.Gln251Glu | O14593 |
| HOMO SAPIENS | 0 | 5012 | 5hbf | A | 442 | ALA | p.Ala442Gly | Q13231 |
| HOMO SAPIENS | 5 | 4844 | 5tdc | A | 136 | HIS | p.His136Arg | Q8IWV7 |
| HOMO SAPIENS | 0 | 15435 | 4cgz | A | 643 | ARG | p.Arg643His | P54132 |
| HOMO SAPIENS | 1 | 15435 | 4cgz | A | 672 | GLN | p.Gln672Arg | P54132 |
| HOMO SAPIENS | 1 | 15435 | 4cgz | A | 700 | GLN | p.Gln700Ter | P54132 |
| HOMO SAPIENS | 0 | 15435 | 4cgz | A | 707 | PRO | p.Pro707Ser | P54132 |
| HOMO SAPIENS | 0 | 15435 | 4cgz | A | 755 | LYS | p.Lys755Glu | P54132 |
| HOMO SAPIENS | 0 | 15435 | 4cgz | A | 881 | TRP | p.Trp881Ter | P54132 |
| HOMO SAPIENS | 2 | 15435 | 4cgz | A | 899 | ARG | p.Arg899Ter | P54132 |
| HOMO SAPIENS | 0 | 15435 | 4cgz | A | 1036 | CYS | p.Cys1036Phe | P54132 |
| HOMO SAPIENS | 0 | 15435 | 4cgz | A | 1055 | CYS | p.Cys1055Ser | P54132 |
| HOMO SAPIENS | 0 | 15435 | 4cgz | A | 1139 | ARG | p.Arg1139Ter | P54132 |
| HOMO SAPIENS | -1 | 15435 | 4cgz | A | 1198 | VAL | p.Val1198Met | P54132 |
| HOMO SAPIENS | -1 | 15435 | 4cgz | A | 1205 | VAL | p.Val1205Ile | P54132 |
| HOMO SAPIENS | 0 | 15435 | 4cgz | A | 1251 | GLU | p.Glu1251Gln | P54132 |
| HOMO SAPIENS | 0 | 11185 | 1kb4 | A | 30 | ARG | p.Arg30Ter | P11473 |
| HOMO SAPIENS | 5 | 11185 | 1kb4 | A | 33 | GLY | p.Gly33Asp | P11473 |
| HOMO SAPIENS | 5 | 11185 | 1kb4 | A | 46 | GLY | p.Gly46Asp | P11473 |
| HOMO SAPIENS | 6 | 11185 | 1kb4 | A | 50 | ARG | p.Arg50Gln | P11473 |
| HOMO SAPIENS | 9 | 11185 | 1kb4 | A | 73 | ARG | p.Arg73Gln | P11473 |
| HOMO SAPIENS | 9 | 11185 | 1kb4 | A | 80 | ARG | p.Arg80Gln | P11473 |
| HOMO SAPIENS | 0 | 5382 | 3fl7 | A | 41 | LEU | p.Leu41Val | P29317 |
| HOMO SAPIENS | 2 | 5382 | 3fl7 | A | 175 | ARG | p.Arg175Cys | P29317 |
| HOMO SAPIENS | 1 | 5382 | 3fl7 | A | 195 | ARG | p.Arg195Cys | P29317 |
| HOMO SAPIENS | 0 | 5382 | 3fl7 | A | 232 | ASP | p.Asp232Gly | P29317 |
| HOMO SAPIENS | 0 | 5382 | 3fl7 | A | 277 | SER | p.Ser277Leu | P29317 |
| HOMO SAPIENS | 0 | 5382 | 3fl7 | A | 315 | ARG | p.Arg315Gln | P29317 |
| HOMO SAPIENS | 7 | 5382 | 3fl7 | A | 350 | PRO | p.Pro350Thr | P29317 |
| HOMO SAPIENS | 0 | 5382 | 3fl7 | A | 391 | GLY | p.Gly391Arg | P29317 |
| HOMO SAPIENS | 0 | 5382 | 3fl7 | A | 418 | VAL | p.Val418Ile | P29317 |
| HOMO SAPIENS | 0 | 5382 | 3fl7 | A | 511 | THR | p.Thr511Met | P29317 |
| HOMO SAPIENS | 2 | 29168 | 4p22 | B | 144 | VAL | p.Val144Ile | P22314 |
| HOMO SAPIENS | 5 | 4348 | 1bf5 | A | 241 | ARG | p.Arg241Gln | P42224 |
| HOMO SAPIENS | 2 | 4348 | 1bf5 | A | 266 | VAL | p.Val266Ile | P42224 |
| HOMO SAPIENS | 4 | 4348 | 1bf5 | A | 320 | GLU | p.Glu320Gln | P42224 |
| HOMO SAPIENS | 1 | 4348 | 1bf5 | A | 372 | ASN | p.Asn372Lys | P42224 |
| HOMO SAPIENS | 1 | 4348 | 1bf5 | A | 447 | ASP | p.Asp447Glu | P42224 |
| HOMO SAPIENS | 2 | 4348 | 1bf5 | A | 463 | GLN | p.Gln463His | P42224 |
| HOMO SAPIENS | 4 | 4348 | 1bf5 | A | 600 | LEU | p.Leu600Pro | P42224 |
| HOMO SAPIENS | 0 | 4348 | 1bf5 | A | 706 | LEU | p.Leu706Ser | P42224 |
| HOMO SAPIENS | 4 | 4991 | 4bej | B | 141 | ASN | p.Asn141Ser | O00429 |
| HOMO SAPIENS | 0 | 4991 | 4bej | B | 395 | ALA | p.Ala395Asp | O00429 |
| HOMO SAPIENS | 2 | 11792 | 1fvo | A | 40 | ARG | p.Arg40Cys | P00480 |
| HOMO SAPIENS | 2 | 11792 | 1fvo | A | 40 | ARG | p.Arg40His | P00480 |
| HOMO SAPIENS | 0 | 11792 | 1fvo | A | 45 | LEU | p.Leu45Pro | P00480 |
| HOMO SAPIENS | 1 | 11792 | 1fvo | A | 47 | ASN | p.Asn47Ile | P00480 |
| HOMO SAPIENS | 0 | 11792 | 1fvo | A | 50 | GLY | p.Gly50Arg | P00480 |
| HOMO SAPIENS | 5 | 11792 | 1fvo | A | 87 | GLU | p.Glu87Lys | P00480 |

| | | | | | | | | |
|---|---|---|---|---|---|---|---|---|
| HOMO SAPIENS | 3 | 11792 | 1fvo | A | 111 | LEU | p.Leu111Pro | P00480 |
| HOMO SAPIENS | 1 | 11792 | 1fvo | A | 125 | THR | p.Thr125Met | P00480 |
| HOMO SAPIENS | 3 | 11792 | 1fvo | A | 129 | ARG | p.Arg129His | P00480 |
| HOMO SAPIENS | 9 | 11792 | 1fvo | A | 141 | ARG | p.Arg141Pro | P00480 |
| HOMO SAPIENS | 0 | 11792 | 1fvo | A | 206 | MET | p.Met206Arg | P00480 |
| HOMO SAPIENS | 0 | 11792 | 1fvo | A | 207 | SER | p.Ser207Arg | P00480 |
| HOMO SAPIENS | 0 | 11792 | 1fvo | A | 216 | GLN | p.Gln216Glu | P00480 |
| HOMO SAPIENS | 1 | 11792 | 1fvo | A | 225 | PRO | p.Pro225Leu | P00480 |
| HOMO SAPIENS | 0 | 11792 | 1fvo | A | 270 | ARG | p.Gln270Arg | P00480 |
| HOMO SAPIENS | 0 | 11792 | 1fvo | A | 270 | ARG | p.Gln270Ter | P00480 |
| HOMO SAPIENS | 2 | 11792 | 1fvo | A | 277 | ARG | p.Arg277Trp | P00480 |
| HOMO SAPIENS | 0 | 3721 | 4r3z | B | 2 | ASP | p.Asp2Gly | P54136 |
| HOMO SAPIENS | 0 | 3721 | 4r3z | B | 456 | SER | p.Ser456Leu | P54136 |
| HOMO SAPIENS | 5 | 3721 | 4r3z | B | 512 | ARG | p.Arg512Gln | P54136 |
| HOMO SAPIENS | 3 | 5749 | 3hl2 | A | 239 | ALA | p.Ala239Thr | Q9HD40 |
| HOMO SAPIENS | 1 | 5749 | 3hl2 | A | 334 | TYR | p.Tyr334Cys | Q9HD40 |
| HOMO SAPIENS | 1 | 5749 | 3hl2 | A | 452 | LYS | p.Lys452Asn | Q9HD40 |
| HOMO SAPIENS | -1 | 5749 | 3hl2 | A | 465 | ARG | p.Arg465Leu | Q9HD40 |
| HOMO SAPIENS | -1 | 5749 | 3hl2 | A | 489 | ASP | p.Asp489Gly | Q9HD40 |
| HOMO SAPIENS | 5 | 1721 | 3k15 | A | 1652 | MET | p.Met1652Ile | P38398 |
| HOMO SAPIENS | 5 | 1721 | 3k15 | A | 1652 | MET | p.Met1652Lys | P38398 |
| HOMO SAPIENS | 5 | 1721 | 3k15 | A | 1662 | PHE | p.Phe1662Cys | P38398 |
| HOMO SAPIENS | 5 | 1721 | 3k15 | A | 1664 | LEU | p.Leu1664Pro | P38398 |
| HOMO SAPIENS | 0 | 1721 | 3k15 | A | 1665 | VAL | p.Val1665Met | P38398 |
| HOMO SAPIENS | 5 | 1721 | 3k15 | A | 1690 | LYS | p.Lys1690Gln | P38398 |
| HOMO SAPIENS | 5 | 1721 | 3k15 | A | 1692 | ASP | p.Asp1692Asn | P38398 |
| HOMO SAPIENS | 5 | 1721 | 3k15 | A | 1699 | ARG | p.Arg1699Gln | P38398 |
| HOMO SAPIENS | 5 | 1721 | 3k15 | A | 1699 | ARG | p.Arg1699Trp | P38398 |
| HOMO SAPIENS | 3 | 1721 | 3k15 | A | 1706 | GLY | p.Gly1706Glu | P38398 |
| HOMO SAPIENS | 5 | 1721 | 3k15 | A | 1720 | THR | p.Thr1720Ala | P38398 |
| HOMO SAPIENS | 5 | 1721 | 3k15 | A | 1751 | ARG | p.Arg1751Pro | P38398 |
| HOMO SAPIENS | 4 | 1721 | 3k15 | A | 1775 | MET | p.Met1775Arg | P38398 |
| HOMO SAPIENS | 2 | 1721 | 3k15 | A | 1788 | GLY | p.Gly1788Val | P38398 |
| HOMO SAPIENS | 5 | 1721 | 3k15 | A | 1794 | GLU | p.Glu1794Asp | P38398 |
| HOMO SAPIENS | 5 | 1721 | 3k15 | A | 1804 | VAL | p.Val1804Asp | P38398 |
| HOMO SAPIENS | 3 | 1721 | 3k15 | A | 1815 | TRP | p.Trp1815Ter | P38398 |
| HOMO SAPIENS | 1 | 1721 | 3k15 | A | 1819 | ASN | p.Asn1819Ser | P38398 |
| HOMO SAPIENS | 5 | 1721 | 3k15 | A | 1856 | PRO | p.Pro1856Ser | P38398 |
| HOMO SAPIENS | 5 | 1721 | 3k15 | A | 1858 | ILE | p.Ile1858Leu | P38398 |
| HOMO SAPIENS | 5 | 1721 | 3k15 | A | 1858 | ILE | p.Ile1858Thr | P38398 |
| HOMO SAPIENS | 5 | 1721 | 3k15 | A | 1859 | PRO | p.Pro1859Arg | P38398 |
| HOMO SAPIENS | 2 | 22086 | 3kq4 | B | 1297 | PRO | p.Pro1297Leu | O60494 |
| HOMO SAPIENS | 1 | 12135 | 2k9j | B | 724 | ARG | p.Arg750Ter | P05106 |
| HOMO SAPIENS | -1 | 24027 | 2qtl | A | 175 | LEU | p.Ser202Leu | Q9UBK8 |
| HOMO SAPIENS | 0 | 24027 | 2qtl | A | 257 | SER | p.Ser284Thr | Q9UBK8 |
| HOMO SAPIENS | 0 | 24027 | 2qtl | A | 333 | LEU | p.Leu360Val | Q9UBK8 |
| HOMO SAPIENS | 0 | 24027 | 2qtl | A | 350 | LYS | p.Lys377Arg | Q9UBK8 |
| HOMO SAPIENS | 0 | 24027 | 2qtl | A | 415 | ARG | p.Arg442Cys | Q9UBK8 |
| HOMO SAPIENS | 2 | 24027 | 2qtl | A | 450 | PRO | p.Pro477Arg | Q9UBK8 |
| HOMO SAPIENS | 4 | 24027 | 2qtl | A | 487 | GLY | p.Gly514Arg | Q9UBK8 |
| HOMO SAPIENS | 1 | 24027 | 2qtl | A | 595 | HIS | p.His622Tyr | Q9UBK8 |
| HOMO SAPIENS | 0 | 5758 | 3di2 | B | 46 | THR | p.Thr66Ile | P16871 |
| HOMO SAPIENS | 1 | 5758 | 3di2 | B | 118 | VAL | p.Ile138Val | P16871 |
| HOMO SAPIENS | 4 | 848 | 5oa3 | Y | 79 | LEU | p.Leu79Val | P62847 |

| | | | | | | | | |
|---|---|---|---|---|---|---|---|---|
| HOMO SAPIENS | 2 | 848 | 5oa3 | Y | 124 | ASN | p.Asn124Ser | P62847 |
| HOMO SAPIENS | 0 | 81 | 3s5o | A | 36 | ALA | p.Ala36Glu | Q86XE5 |
| HOMO SAPIENS | 0 | 81 | 3s5o | A | 39 | TYR | p.Tyr39Ter | Q86XE5 |
| HOMO SAPIENS | 0 | 81 | 3s5o | A | 45 | PRO | p.Pro45Leu | Q86XE5 |
| HOMO SAPIENS | 0 | 81 | 3s5o | A | 70 | ARG | p.Arg70Ter | Q86XE5 |
| HOMO SAPIENS | 0 | 81 | 3s5o | A | 70 | ARG | p.Arg70Pro | Q86XE5 |
| HOMO SAPIENS | 1 | 81 | 3s5o | A | 97 | ARG | p.Arg97Cys | Q86XE5 |
| HOMO SAPIENS | 4 | 81 | 3s5o | A | 113 | GLU | p.Glu113Lys | Q86XE5 |
| HOMO SAPIENS | 1 | 81 | 3s5o | A | 190 | PRO | p.Pro190Leu | Q86XE5 |
| HOMO SAPIENS | 0 | 81 | 3s5o | A | 245 | VAL | p.Val245Phe | Q86XE5 |
| HOMO SAPIENS | 0 | 81 | 3s5o | A | 257 | CYS | p.Cys257Gly | Q86XE5 |
| HOMO SAPIENS | 0 | 81 | 3s5o | A | 287 | GLY | p.Gly287Val | Q86XE5 |
| HOMO SAPIENS | 7 | 81 | 3s5o | A | 303 | ARG | p.Arg303Cys | Q86XE5 |
| HOMO SAPIENS | 0 | 81 | 3s5o | A | 325 | GLY | p.Gly325Ser | Q86XE5 |
| HOMO SAPIENS | -1 | 622 | 5era | A | 12 | GLY | p.Gly12Arg | P29033 |
| HOMO SAPIENS | -1 | 622 | 5era | A | 12 | GLY | p.Gly12Asp | P29033 |
| HOMO SAPIENS | -1 | 622 | 5era | A | 17 | SER | p.Ser17Phe | P29033 |
| HOMO SAPIENS | 5 | 622 | 5era | A | 24 | TRP | p.Trp24Ter | P29033 |
| HOMO SAPIENS | 0 | 622 | 5era | A | 32 | ARG | p.Arg32Cys | P29033 |
| HOMO SAPIENS | 5 | 622 | 5era | A | 34 | MET | p.Met34Thr | P29033 |
| HOMO SAPIENS | 5 | 622 | 5era | A | 36 | LEU | p.Leu36Pro | P29033 |
| HOMO SAPIENS | 5 | 622 | 5era | A | 37 | VAL | p.Val37Ile | P29033 |
| HOMO SAPIENS | 5 | 622 | 5era | A | 40 | ALA | p.Ala40Glu | P29033 |
| HOMO SAPIENS | 5 | 622 | 5era | A | 44 | TRP | p.Trp44Cys | P29033 |
| HOMO SAPIENS | 5 | 622 | 5era | A | 45 | GLY | p.Gly45Glu | P29033 |
| HOMO SAPIENS | 5 | 622 | 5era | A | 47 | GLU | p.Glu47Ter | P29033 |
| HOMO SAPIENS | 5 | 622 | 5era | A | 50 | ASP | p.Asp50Asn | P29033 |
| HOMO SAPIENS | 5 | 622 | 5era | A | 54 | ASN | p.Asn54Lys | P29033 |
| HOMO SAPIENS | 5 | 622 | 5era | A | 57 | GLN | p.Gln57Ter | P29033 |
| HOMO SAPIENS | 5 | 622 | 5era | A | 59 | GLY | p.Gly59Ala | P29033 |
| HOMO SAPIENS | 5 | 622 | 5era | A | 59 | GLY | p.Gly59Ser | P29033 |
| HOMO SAPIENS | 5 | 622 | 5era | A | 66 | ASP | p.Asp66His | P29033 |
| HOMO SAPIENS | 5 | 622 | 5era | A | 68 | TYR | p.Tyr68Cys | P29033 |
| HOMO SAPIENS | 5 | 622 | 5era | A | 70 | PRO | p.Pro70Ala | P29033 |
| HOMO SAPIENS | 5 | 622 | 5era | A | 73 | HIS | p.His73Arg | P29033 |
| HOMO SAPIENS | 5 | 622 | 5era | A | 75 | ARG | p.Arg75Trp | P29033 |
| HOMO SAPIENS | 5 | 622 | 5era | A | 75 | ARG | p.Arg75Gln | P29033 |
| HOMO SAPIENS | 2 | 622 | 5era | A | 77 | TRP | p.Trp77Arg | P29033 |
| HOMO SAPIENS | 2 | 622 | 5era | A | 77 | TRP | p.Trp77Ter | P29033 |
| HOMO SAPIENS | 0 | 622 | 5era | A | 80 | GLN | p.Gln80Ter | P29033 |
| HOMO SAPIENS | 0 | 622 | 5era | A | 80 | GLN | p.Gln80Pro | P29033 |
| HOMO SAPIENS | 5 | 622 | 5era | A | 81 | LEU | p.Leu81Val | P29033 |
| HOMO SAPIENS | 5 | 622 | 5era | A | 82 | ILE | p.Ile82Met | P29033 |
| HOMO SAPIENS | 5 | 622 | 5era | A | 83 | PHE | p.Phe83Leu | P29033 |
| HOMO SAPIENS | 5 | 622 | 5era | A | 84 | VAL | p.Val84Leu | P29033 |
| HOMO SAPIENS | 5 | 622 | 5era | A | 90 | LEU | p.Leu90Pro | P29033 |
| HOMO SAPIENS | 5 | 622 | 5era | A | 95 | VAL | p.Val95Met | P29033 |
| HOMO SAPIENS | -1 | 622 | 5era | A | 100 | HIS | p.His100Tyr | P29033 |
| HOMO SAPIENS | -1 | 622 | 5era | A | 113 | SER | p.Ser113Arg | P29033 |
| HOMO SAPIENS | -1 | 622 | 5era | A | 119 | GLU | p.Glu119Lys | P29033 |
| HOMO SAPIENS | -1 | 622 | 5era | A | 122 | LYS | p.Lys122Ile | P29033 |
| HOMO SAPIENS | -1 | 622 | 5era | A | 123 | THR | p.Thr123Asn | P29033 |
| HOMO SAPIENS | -1 | 622 | 5era | A | 124 | GLN | p.Gln124Ter | P29033 |
| HOMO SAPIENS | 5 | 622 | 5era | A | 139 | SER | p.Ser139Asn | P29033 |

| | | | | | | | | |
|---|---|---|---|---|---|---|---|---|
| HOMO SAPIENS | 5 | 622 | 5era | A | 143 | ARG | p.Arg143Trp | P29033 |
| HOMO SAPIENS | 5 | 622 | 5era | A | 143 | ARG | p.Arg143Gln | P29033 |
| HOMO SAPIENS | 0 | 622 | 5era | A | 147 | GLU | p.Glu147Lys | P29033 |
| HOMO SAPIENS | 7 | 622 | 5era | A | 155 | TYR | p.Tyr155Ter | P29033 |
| HOMO SAPIENS | 2 | 622 | 5era | A | 159 | ASP | p.Asp159Val | P29033 |
| HOMO SAPIENS | 5 | 622 | 5era | A | 163 | MET | p.Met163Val | P29033 |
| HOMO SAPIENS | 5 | 622 | 5era | A | 165 | ARG | p.Arg165Trp | P29033 |
| HOMO SAPIENS | 5 | 622 | 5era | A | 167 | VAL | p.Val167Met | P29033 |
| HOMO SAPIENS | 5 | 622 | 5era | A | 169 | CYS | p.Cys169Tyr | P29033 |
| HOMO SAPIENS | 5 | 622 | 5era | A | 179 | ASP | p.Asp179Asn | P29033 |
| HOMO SAPIENS | 0 | 622 | 5era | A | 199 | SER | p.Ser199Phe | P29033 |
| HOMO SAPIENS | 0 | 622 | 5era | A | 202 | CYS | p.Cys202Phe | P29033 |
| HOMO SAPIENS | 5 | 622 | 5era | A | 203 | ILE | p.Ile203Thr | P29033 |
| HOMO SAPIENS | 5 | 622 | 5era | A | 206 | ASN | p.Asn206Ser | P29033 |
| HOMO SAPIENS | 0 | 8103 | 5dot | A | 163 | GLY | p.Gly163Ter | P31327 |
| HOMO SAPIENS | 0 | 8103 | 5dot | A | 337 | HIS | p.His337Arg | P31327 |
| HOMO SAPIENS | 3 | 8103 | 5dot | A | 344 | THR | p.Thr344Ala | P31327 |
| HOMO SAPIENS | 1 | 8103 | 5dot | A | 405 | THR | p.Thr405Asn | P31327 |
| HOMO SAPIENS | 5 | 8103 | 5dot | A | 544 | THR | p.Thr544Met | P31327 |
| HOMO SAPIENS | 5 | 8103 | 5dot | A | 716 | ASN | p.Asn716Lys | P31327 |
| HOMO SAPIENS | -1 | 8103 | 5dot | A | 780 | ARG | p.Arg780His | P31327 |
| HOMO SAPIENS | 0 | 8103 | 5dot | A | 982 | GLY | p.Gly982Asp | P31327 |
| HOMO SAPIENS | 3 | 8103 | 5dot | A | 1119 | ALA | p.Ala1119Ser | P31327 |
| HOMO SAPIENS | 1 | 8103 | 5dot | A | 1376 | GLY | p.Gly1376Ser | P31327 |
| HOMO SAPIENS | 4 | 6833 | 1l8l | B | 27 | ARG | p.Arg27Ser | P78330 |
| HOMO SAPIENS | 3 | 6833 | 1l8l | B | 32 | ASP | p.Asp32Gly | P78330 |
| HOMO SAPIENS | 3 | 6833 | 1l8l | B | 32 | ASP | p.Asp32Asn | P78330 |
| HOMO SAPIENS | 1 | 6833 | 1l8l | B | 49 | ARG | p.Arg49Trp | P78330 |
| HOMO SAPIENS | 0 | 6833 | 1l8l | B | 52 | MET | p.Met52Thr | P78330 |
| HOMO SAPIENS | 0 | 6833 | 1l8l | B | 83 | GLN | p.Gln83His | P78330 |
| HOMO SAPIENS | 6 | 810 | 2iw2 | A | 185 | ARG | p.Arg184Gln | P12955 |
| HOMO SAPIENS | 0 | 810 | 2iw2 | A | 213 | ALA | p.Ala212Pro | P12955 |
| HOMO SAPIENS | 1 | 810 | 2iw2 | A | 266 | ARG | p.Arg265Ter | P12955 |
| HOMO SAPIENS | 9 | 810 | 2iw2 | A | 277 | ASP | p.Asp276Asn | P12955 |
| HOMO SAPIENS | 0 | 810 | 2iw2 | A | 279 | GLY | p.Gly278Asp | P12955 |
| HOMO SAPIENS | 9 | 810 | 2iw2 | A | 378 | HIS | p.His377Gln | P12955 |
| HOMO SAPIENS | 0 | 810 | 2iw2 | A | 389 | ARG | p.Arg388His | P12955 |
| HOMO SAPIENS | 0 | 810 | 2iw2 | A | 436 | PHE | p.Leu435Phe | P12955 |
| HOMO SAPIENS | 0 | 810 | 2iw2 | A | 449 | GLY | p.Gly448Arg | P12955 |
| HOMO SAPIENS | -1 | 1530 | 5fwj | A | 388 | ALA | p.Ala388Pro | P41229 |
| HOMO SAPIENS | 4 | 1530 | 5fwj | A | 451 | SER | p.Ser451Arg | P41229 |
| HOMO SAPIENS | 3 | 1530 | 5fwj | A | 731 | LEU | p.Leu731Phe | P41229 |
| HOMO SAPIENS | 3 | 578 | 5net | B | 123 | ALA | p.Ala143Thr | P18564 |
| HOMO SAPIENS | 6 | 578 | 5net | B | 176 | PRO | p.Pro196Thr | P18564 |
| HOMO SAPIENS | 3 | 578 | 5net | B | 280 | GLU | p.Glu300Lys | P18564 |
| HOMO SAPIENS | -1 | 5247 | 4j15 | A | 274 | ALA | p.Ala274Val | P14868 |
| HOMO SAPIENS | -1 | 5247 | 4j15 | A | 280 | HIS | p.His280Leu | P14868 |
| HOMO SAPIENS | 2 | 5247 | 4j15 | A | 460 | ARG | p.Arg460His | P14868 |
| HOMO SAPIENS | 5 | 5247 | 4j15 | A | 494 | ARG | p.Arg494Cys | P14868 |
| HOMO SAPIENS | 7 | 2566 | 1b55 | A | 28 | ARG | p.Arg28His | Q06187 |
| HOMO SAPIENS | 5 | 2566 | 1b55 | A | 33 | THR | p.Thr33Pro | Q06187 |
| HOMO SAPIENS | 0 | 2566 | 1b55 | A | 113 | VAL | p.Val113Asp | Q06187 |
| HOMO SAPIENS | 0 | 356 | 5a2s | A | 786 | ALA | p.Ala786Thr | P56524 |
| HOMO SAPIENS | 5 | 2960 | 4cak | A | 553 | ARG | p.Arg584Ter | P08514 |

| | | | | | | | | |
|---|---|---|---|---|---|---|---|---|
| HOMO SAPIENS | 5 | 2960 | 4cak | A | 565 | ILE | p.Ile596Thr | P08514 |
| HOMO SAPIENS | -1 | 2960 | 4cak | A | 841 | LEU | p.Leu872Met | P08514 |
| HOMO SAPIENS | -1 | 2960 | 4cak | A | 843 | ILE | p.Ile874Ser | P08514 |
| HOMO SAPIENS | 0 | 5054 | 4k8r | B | 216 | PHE | p.Phe216Leu | Q8NBP7 |
| HOMO SAPIENS | 1 | 5054 | 4k8r | B | 417 | HIS | p.His417Gln | Q8NBP7 |
| HOMO SAPIENS | 1 | 5054 | 4k8r | B | 425 | ASN | p.Asn425Ser | Q8NBP7 |
| HOMO SAPIENS | 1 | 5054 | 4k8r | B | 443 | ALA | p.Ala443Thr | Q8NBP7 |
| HOMO SAPIENS | 0 | 27283 | 3n5n | X | 71 | HIS | p.His82Arg | Q9UIF7 |
| HOMO SAPIENS | 1 | 27283 | 3n5n | X | 83 | ARG | p.Arg94Ter | Q9UIF7 |
| HOMO SAPIENS | 8 | 27283 | 3n5n | X | 89 | TRP | p.Trp100Ter | Q9UIF7 |
| HOMO SAPIENS | 8 | 27283 | 3n5n | X | 90 | TYR | p.Tyr101Ter | Q9UIF7 |
| HOMO SAPIENS | 7 | 27283 | 3n5n | X | 95 | ARG | p.Arg106Trp | Q9UIF7 |
| HOMO SAPIENS | 7 | 27283 | 3n5n | X | 117 | TRP | p.Trp128Arg | Q9UIF7 |
| HOMO SAPIENS | 0 | 27283 | 3n5n | X | 137 | GLY | p.Gly148Arg | Q9UIF7 |
| HOMO SAPIENS | 5 | 27283 | 3n5n | X | 142 | TRP | p.Trp153Ter | Q9UIF7 |
| HOMO SAPIENS | 0 | 27283 | 3n5n | X | 165 | TYR | p.Tyr176Cys | Q9UIF7 |
| HOMO SAPIENS | 0 | 27283 | 3n5n | X | 168 | ARG | p.Arg179His | Q9UIF7 |
| HOMO SAPIENS | 0 | 27283 | 3n5n | X | 168 | ARG | p.Arg179Cys | Q9UIF7 |
| HOMO SAPIENS | 0 | 27283 | 3n5n | X | 196 | GLN | p.Gln207Ter | Q9UIF7 |
| HOMO SAPIENS | 8 | 27283 | 3n5n | X | 201 | VAL | p.Val212Met | Q9UIF7 |
| HOMO SAPIENS | 9 | 27283 | 3n5n | X | 227 | ARG | p.Arg238Trp | Q9UIF7 |
| HOMO SAPIENS | 9 | 27283 | 3n5n | X | 231 | ARG | p.Arg242Cys | Q9UIF7 |
| HOMO SAPIENS | 9 | 27283 | 3n5n | X | 231 | ARG | p.Arg242His | Q9UIF7 |
| HOMO SAPIENS | 3 | 27283 | 3n5n | X | 232 | VAL | p.Val243Phe | Q9UIF7 |
| HOMO SAPIENS | 3 | 27283 | 3n5n | X | 246 | GLN | p.Gln257Ter | Q9UIF7 |
| HOMO SAPIENS | 9 | 27283 | 3n5n | X | 272 | GLY | p.Gly283Glu | Q9UIF7 |
| HOMO SAPIENS | 9 | 27283 | 3n5n | X | 281 | PRO | p.Pro292Leu | Q9UIF7 |
| HOMO SAPIENS | -1 | 27283 | 3n5n | X | 314 | ASP | p.Asp325Glu | Q9UIF7 |
| HOMO SAPIENS | 0 | 27283 | 3n5n | X | 324 | GLN | p.Gln335His | Q9UIF7 |
| HOMO SAPIENS | 0 | 27283 | 3n5n | X | 324 | GLN | p.Gln335Arg | Q9UIF7 |
| HOMO SAPIENS | 0 | 27283 | 3n5n | X | 324 | GLN | p.Gln335Ter | Q9UIF7 |
| HOMO SAPIENS | -1 | 20852 | 4p60 | B | 243 | ARG | p.Arg243Lys | O75746 |
| HOMO SAPIENS | -1 | 1875 | 5dd2 | G | 152 | VAL | p.Val179Leu | P06396 |
| HOMO SAPIENS | 9 | 2126 | 3n1p | B | 46 | PRO | p.Pro46Leu | Q14623 |
| HOMO SAPIENS | 8 | 2126 | 3n1p | B | 76 | GLU | p.Glu76Ala | Q14623 |
| HOMO SAPIENS | 9 | 2126 | 3n1p | B | 77 | ARG | p.Arg77Ser | Q14623 |
| HOMO SAPIENS | 6 | 2126 | 3n1p | B | 95 | GLU | p.Glu95Lys | Q14623 |
| HOMO SAPIENS | 9 | 2126 | 3n1p | B | 100 | ASP | p.Asp100Glu | Q14623 |
| HOMO SAPIENS | 9 | 2126 | 3n1p | B | 131 | GLU | p.Glu131Lys | Q14623 |
| HOMO SAPIENS | 0 | 2126 | 3n1p | B | 190 | VAL | p.Val190Ala | Q14623 |
| HOMO SAPIENS | 5 | 40410 | 3ioz | B | 70 | GLN | p.Gln70Leu | P20963 |
| HOMO SAPIENS | 1 | 5754 | 5lkz | A | 1108 | THR | p.Tyr1191Ter | Q09472 |
| HOMO SAPIENS | 3 | 8159 | 1tiu | A | 3 | GLU | p.Lys12679Glu | Q8WZ42 |
| HOMO SAPIENS | 4 | 3275 | 4ldq | A | 210 | ARG | p.Arg210Gln | Q9BV20 |
| HOMO SAPIENS | 0 | 8143 | 4miv | B | 66 | SER | p.Ser66Trp | P51688 |
| HOMO SAPIENS | 1 | 8143 | 4miv | B | 74 | ARG | p.Arg74Cys | P51688 |
| HOMO SAPIENS | 0 | 8143 | 4miv | B | 128 | PRO | p.Pro128Leu | P51688 |
| HOMO SAPIENS | 1 | 8143 | 4miv | B | 150 | ARG | p.Arg150Gln | P51688 |
| HOMO SAPIENS | 0 | 8143 | 4miv | B | 191 | GLY | p.Gly191Arg | P51688 |
| HOMO SAPIENS | 0 | 8143 | 4miv | B | 206 | ARG | p.Arg206Pro | P51688 |
| HOMO SAPIENS | 0 | 8143 | 4miv | B | 225 | PHE | p.Phe225Leu | P51688 |
| HOMO SAPIENS | 0 | 8143 | 4miv | B | 233 | ARG | p.Arg233Ter | P51688 |
| HOMO SAPIENS | 0 | 8143 | 4miv | B | 234 | ALA | p.Ala234Gly | P51688 |
| HOMO SAPIENS | 0 | 8143 | 4miv | B | 245 | ARG | p.Arg245His | P51688 |

| | | | | | | | | |
|---|---|---|---|---|---|---|---|---|
| HOMO SAPIENS | 0 | 8143 | 4miv | B | 293 | PRO | p.Pro293Ser | P51688 |
| HOMO SAPIENS | 1 | 8143 | 4miv | B | 298 | SER | p.Ser298Pro | P51688 |
| HOMO SAPIENS | 0 | 8143 | 4miv | B | 361 | VAL | p.Val361Ile | P51688 |
| HOMO SAPIENS | 0 | 8143 | 4miv | B | 369 | GLU | p.Glu369Lys | P51688 |
| HOMO SAPIENS | 0 | 8143 | 4miv | B | 372 | MET | p.Met372Ile | P51688 |
| HOMO SAPIENS | 1 | 8143 | 4miv | B | 380 | GLN | p.Gln380Arg | P51688 |
| HOMO SAPIENS | 0 | 8143 | 4miv | B | 387 | VAL | p.Val387Met | P51688 |
| HOMO SAPIENS | 0 | 8143 | 4miv | B | 394 | MET | p.Met394Ile | P51688 |
| HOMO SAPIENS | 0 | 8143 | 4miv | B | 433 | ARG | p.Arg433Gln | P51688 |
| HOMO SAPIENS | 1 | 8143 | 4miv | B | 446 | HIS | p.His446Arg | P51688 |
| HOMO SAPIENS | 5 | 8143 | 4miv | B | 447 | GLU | p.Glu447Lys | P51688 |
| HOMO SAPIENS | 2 | 8143 | 4miv | B | 456 | ARG | p.Arg456His | P51688 |
| HOMO SAPIENS | 1 | 21880 | 2oo9 | B | 857 | LEU | p.Leu857Phe | P22681 |
| HOMO SAPIENS | 0 | 9467 | 1s1d | A | 154 | LEU | p.Leu224Pro | Q8WVQ1 |
| HOMO SAPIENS | 0 | 9467 | 1s1d | A | 156 | VAL | p.Val226Met | Q8WVQ1 |
| HOMO SAPIENS | 0 | 9467 | 1s1d | A | 229 | PRO | p.Pro299Leu | Q8WVQ1 |
| HOMO SAPIENS | 4 | 9467 | 1s1d | A | 230 | ARG | p.Arg300Cys | Q8WVQ1 |
| HOMO SAPIENS | 4 | 9467 | 1s1d | A | 230 | ARG | p.Arg300His | Q8WVQ1 |
| HOMO SAPIENS | 0 | 9467 | 1s1d | A | 253 | ALA | p.Ala323Thr | Q8WVQ1 |
| HOMO SAPIENS | 2 | 9467 | 1s1d | A | 305 | MET | p.Met375Leu | Q8WVQ1 |
| HOMO SAPIENS | -1 | 2361 | 5llm | A | 5 | ASN | p.Asn5Lys | P43003 |
| HOMO SAPIENS | 0 | 2361 | 5llm | A | 71 | TYR | p.Tyr71Cys | P43003 |
| HOMO SAPIENS | -1 | 2361 | 5llm | A | 211 | LYS | p.Glu219Asp | Q15758 |
| HOMO SAPIENS | 8 | 2361 | 5llm | A | 270 | PRO | p.Pro290Arg | P43003 |
| HOMO SAPIENS | 1 | 2361 | 5llm | A | 321 | THR | p.Ala329Thr | Q15758 |
| HOMO SAPIENS | 1 | 2361 | 5llm | A | 365 | ARG | p.Arg385His | P43003 |
| HOMO SAPIENS | 1 | 2361 | 5llm | A | 479 | ARG | p.Arg499Gln | P43003 |
| HOMO SAPIENS | 3 | 5412 | 4obz | D | 165 | PHE | p.Phe229Ile | P07339 |
| HOMO SAPIENS | 2 | 5412 | 4obz | D | 187 | ASP | p.Asp251Asn | P07339 |
| HOMO SAPIENS | 0 | 5412 | 4obz | D | 218 | GLY | p.Gly282Arg | P07339 |
| HOMO SAPIENS | 0 | 5412 | 4obz | D | 245 | ARG | p.Arg309His | P07339 |
| HOMO SAPIENS | 2 | 5412 | 4obz | D | 319 | TRP | p.Trp383Cys | P07339 |
| HOMO SAPIENS | 0 | 3544 | 5h64 | A | 277 | MET | p.Met277Ile | P42345 |
| HOMO SAPIENS | 0 | 14230 | 2kbx | A | 145 | ARG | p.Arg145Ser | Q13418 |
| HOMO SAPIENS | 2 | 34970 | 4xhz | A | 834 | ARG | p.Arg855Trp | Q96QU1 |
| HOMO SAPIENS | 4 | 34970 | 4xhz | A | 854 | SER | p.Ser875Ter | Q96QU1 |
| HOMO SAPIENS | 1 | 34970 | 4xhz | A | 908 | ARG | p.Arg929Gln | Q96QU1 |
| HOMO SAPIENS | 1 | 34970 | 4xhz | A | 941 | ARG | p.Arg962Cys | Q96QU1 |
| HOMO SAPIENS | 0 | 34970 | 4xhz | A | 970 | ARG | p.Arg991Ter | Q96QU1 |
| HOMO SAPIENS | 0 | 34970 | 4xhz | A | 976 | GLU | p.Glu997Gly | Q96QU1 |
| HOMO SAPIENS | 0 | 25734 | 4asc | A | 345 | SER | p.Asn345Ser | Q2TBA0 |
| HOMO SAPIENS | 0 | 25734 | 4asc | A | 514 | VAL | p.Val514Met | Q2TBA0 |
| HOMO SAPIENS | 6 | 25734 | 4asc | A | 528 | GLU | p.Glu528Lys | Q2TBA0 |
| HOMO SAPIENS | 0 | 25734 | 4asc | A | 580 | LEU | p.Leu580Ile | Q2TBA0 |
| HOMO SAPIENS | 8 | 2356 | 3kse | D | 22 | LYS | p.Lys22Ter | P01040 |
| HOMO SAPIENS | 1 | 2356 | 3kse | D | 58 | ARG | p.Arg58Ter | P01040 |
| HOMO SAPIENS | 2 | 6000 | 1jqd | A | 60 | GLY | p.Gly60Asp | P50135 |
| HOMO SAPIENS | 9 | 6000 | 1jqd | A | 208 | LEU | p.Leu208Pro | P50135 |
| HOMO SAPIENS | 0 | 14741 | 5eh1 | A | 58 | THR | p.Thr58Arg | P38484 |
| HOMO SAPIENS | 0 | 14741 | 5eh1 | A | 168 | THR | p.Thr168Asn | P38484 |
| HOMO SAPIENS | 1 | 4814 | 4kxk | D | 427 | ARG | p.Arg427Ter | P50542 |
| HOMO SAPIENS | 0 | 188 | 2xt3 | A | 49 | ASP | p.Asp52Asn | Q2M1P5 |
| HOMO SAPIENS | 3 | 188 | 2xt3 | A | 142 | TYR | p.Tyr145Ser | Q2M1P5 |
| HOMO SAPIENS | 9 | 8681 | 5tby | D | 57 | ALA | p.Ala57Asp | P08590 |

| | | | | | | | | |
|---|---|---|---|---|---|---|---|---|
| HOMO SAPIENS | 3 | 8681 | 5tby | D | 62 | ASP | p.Asp62Asn | P08590 |
| HOMO SAPIENS | 0 | 8681 | 5tby | D | 65 | PRO | p.Pro65Arg | P08590 |
| HOMO SAPIENS | 7 | 8681 | 5tby | D | 79 | VAL | p.Val79Ile | P08590 |
| HOMO SAPIENS | 0 | 8681 | 5tby | D | 143 | GLU | p.Glu143Lys | P08590 |
| HOMO SAPIENS | 0 | 8681 | 5tby | D | 149 | MET | p.Met149Val | P08590 |
| HOMO SAPIENS | 3 | 8681 | 5tby | D | 154 | ARG | p.Arg154His | P08590 |
| HOMO SAPIENS | 6 | 30984 | 4xwh | A | 140 | TYR | p.Tyr140Cys | P54802 |
| HOMO SAPIENS | 4 | 30984 | 4xwh | A | 141 | SER | p.Ser141Thr | P54802 |
| HOMO SAPIENS | 2 | 30984 | 4xwh | A | 234 | ARG | p.Arg234Cys | P54802 |
| HOMO SAPIENS | 2 | 30984 | 4xwh | A | 336 | GLU | p.Glu336Ter | P54802 |
| HOMO SAPIENS | 6 | 30984 | 4xwh | A | 482 | ARG | p.Arg482Trp | P54802 |
| HOMO SAPIENS | 2 | 30984 | 4xwh | A | 521 | PRO | p.Pro521Leu | P54802 |
| HOMO SAPIENS | 0 | 30984 | 4xwh | A | 565 | ARG | p.Arg565Gln | P54802 |
| HOMO SAPIENS | 0 | 30984 | 4xwh | A | 565 | ARG | p.Arg565Trp | P54802 |
| HOMO SAPIENS | 6 | 30984 | 4xwh | A | 626 | ARG | p.Arg626Ter | P54802 |
| HOMO SAPIENS | 6 | 30984 | 4xwh | A | 643 | ARG | p.Arg643His | P54802 |
| HOMO SAPIENS | 6 | 30984 | 4xwh | A | 643 | ARG | p.Arg643Cys | P54802 |
| HOMO SAPIENS | 2 | 30984 | 4xwh | A | 674 | ARG | p.Arg674His | P54802 |
| HOMO SAPIENS | 2 | 30984 | 4xwh | A | 737 | ARG | p.Arg737Gly | P54802 |
| HOMO SAPIENS | 5 | 10545 | 1oqe | N | 6 | PRO | p.Pro21Arg | Q96RJ3 |
| HOMO SAPIENS | 1 | 16995 | 5d3a | B | 12 | MET | p.Met947Arg | Q7Z4S6 |
| HOMO SAPIENS | 1 | 16995 | 5d3a | B | 12 | MET | p.Met947Val | Q7Z4S6 |
| HOMO SAPIENS | 2 | 16995 | 5d3a | B | 19 | ARG | p.Arg954Gln | Q7Z4S6 |
| HOMO SAPIENS | 2 | 16995 | 5d3a | B | 19 | ARG | p.Arg954Trp | Q7Z4S6 |
| HOMO SAPIENS | 0 | 16995 | 5d3a | B | 75 | ILE | p.Ile1010Thr | Q7Z4S6 |
| HOMO SAPIENS | 0 | 17091 | 2yin | B | 1324 | GLN | p.Gln1324Ter | Q92608 |
| HOMO SAPIENS | 5 | 27606 | 1lm7 | B | 2366 | ARG | p.Arg2366Cys | P15924 |
| HOMO SAPIENS | 2 | 15731 | 1bi7 | B | 24 | ARG | p.Arg24Pro | P42771 |
| HOMO SAPIENS | 0 | 15731 | 1bi7 | B | 28 | VAL | p.Val28Gly | P42771 |
| HOMO SAPIENS | 0 | 15731 | 1bi7 | B | 49 | ILE | p.Ile49Thr | P42771 |
| HOMO SAPIENS | 0 | 15731 | 1bi7 | B | 53 | MET | p.Met53Ile | P42771 |
| HOMO SAPIENS | 2 | 15731 | 1bi7 | B | 56 | SER | p.Ser56Ile | P42771 |
| HOMO SAPIENS | 1 | 15731 | 1bi7 | B | 57 | ALA | p.Ala57Val | P42771 |
| HOMO SAPIENS | 4 | 15731 | 1bi7 | B | 59 | VAL | p.Val59Gly | P42771 |
| HOMO SAPIENS | 9 | 15731 | 1bi7 | B | 89 | GLY | p.Gly89Asp | P42771 |
| HOMO SAPIENS | 0 | 15731 | 1bi7 | B | 101 | GLY | p.Gly101Trp | P42771 |
| HOMO SAPIENS | 0 | 15731 | 1bi7 | B | 114 | PRO | p.Pro114Ser | P42771 |
| HOMO SAPIENS | 9 | 15731 | 1bi7 | B | 123 | HIS | p.His123Gln | P42771 |
| HOMO SAPIENS | 8 | 15731 | 1bi7 | B | 126 | VAL | p.Val126Asp | P42771 |
| HOMO SAPIENS | 7 | 15731 | 1bi7 | B | 127 | ALA | p.Ala127Ser | P42771 |
| HOMO SAPIENS | -1 | 15731 | 1bi7 | B | 148 | ALA | p.Ala148Thr | P42771 |
| HOMO SAPIENS | -1 | 19858 | 3jbk | M | 866 | ALA | p.Ala934Val | P18206 |
| HOMO SAPIENS | 0 | 19484 | 3ee6 | B | 62 | SER | p.Ser62Leu | O14773 |
| HOMO SAPIENS | 1 | 19484 | 3ee6 | B | 66 | GLN | p.Gln66Ter | O14773 |
| HOMO SAPIENS | 0 | 19484 | 3ee6 | B | 100 | GLN | p.Gln100Arg | O14773 |
| HOMO SAPIENS | 0 | 19484 | 3ee6 | B | 104 | LEU | p.Leu104Ter | O14773 |
| HOMO SAPIENS | 0 | 19484 | 3ee6 | B | 127 | ARG | p.Arg127Gln | O14773 |
| HOMO SAPIENS | 0 | 19484 | 3ee6 | B | 127 | ARG | p.Arg127Ter | O14773 |
| HOMO SAPIENS | -1 | 19484 | 3ee6 | B | 185 | ARG | p.Arg185Cys | O14773 |
| HOMO SAPIENS | 0 | 19484 | 3ee6 | B | 206 | ARG | p.Arg206Cys | O14773 |
| HOMO SAPIENS | 0 | 19484 | 3ee6 | B | 208 | ARG | p.Arg208Ter | O14773 |
| HOMO SAPIENS | 0 | 19484 | 3ee6 | B | 214 | GLN | p.Gln214Ter | O14773 |
| HOMO SAPIENS | 1 | 19484 | 3ee6 | B | 276 | ASP | p.Asp276Val | O14773 |
| HOMO SAPIENS | 0 | 19484 | 3ee6 | B | 284 | GLY | p.Gly284Val | O14773 |

| | | | | | | | | |
|---|---|---|---|---|---|---|---|---|
| HOMO SAPIENS | 0 | 19484 | 3ee6 | B | 286 | ASN | p.Asn286Ser | O14773 |
| HOMO SAPIENS | 0 | 19484 | 3ee6 | B | 339 | ARG | p.Arg339Gly | O14773 |
| HOMO SAPIENS | 2 | 19484 | 3ee6 | B | 345 | MET | p.Met345Leu | O14773 |
| HOMO SAPIENS | 0 | 19484 | 3ee6 | B | 365 | CYS | p.Cys365Arg | O14773 |
| HOMO SAPIENS | 0 | 19484 | 3ee6 | B | 365 | CYS | p.Cys365Tyr | O14773 |
| HOMO SAPIENS | 0 | 19484 | 3ee6 | B | 373 | GLN | p.Gln373Glu | O14773 |
| HOMO SAPIENS | 6 | 19484 | 3ee6 | B | 418 | ARG | p.Arg418Gln | O14773 |
| HOMO SAPIENS | 7 | 19484 | 3ee6 | B | 422 | GLN | p.Gln422His | O14773 |
| HOMO SAPIENS | 0 | 19484 | 3ee6 | B | 447 | ARG | p.Arg447His | O14773 |
| HOMO SAPIENS | 0 | 19484 | 3ee6 | B | 466 | VAL | p.Val466Met | O14773 |
| HOMO SAPIENS | 0 | 4654 | 5t5w | A | 47 | LYS | p.Lys47Glu | Q08334 |
| HOMO SAPIENS | 0 | 4654 | 5t5w | A | 141 | GLU | p.Glu141Ter | Q08334 |
| HOMO SAPIENS | -1 | 18815 | 4wj7 | A | 53 | VAL | p.Val53Ile | Q9BSQ5 |
| HOMO SAPIENS | 0 | 18815 | 4wj7 | A | 120 | VAL | p.Val120Ile | Q9BSQ5 |
| HOMO SAPIENS | 0 | 18815 | 4wj7 | A | 198 | LEU | p.Leu198Arg | Q9BSQ5 |
| HOMO SAPIENS | 0 | 4602 | 1mjd | A | 47 | SER | p.Ser47Arg | O43602 |
| HOMO SAPIENS | 6 | 4602 | 1mjd | A | 59 | ARG | p.Arg59Leu | O43602 |
| HOMO SAPIENS | 7 | 4602 | 1mjd | A | 62 | ASP | p.Asp62Asn | O43602 |
| HOMO SAPIENS | 7 | 4602 | 1mjd | A | 71 | ALA | p.Ala71Ser | O43602 |
| HOMO SAPIENS | 7 | 4602 | 1mjd | A | 78 | ARG | p.Arg78Leu | O43602 |
| HOMO SAPIENS | 7 | 4602 | 1mjd | A | 89 | ARG | p.Arg89Gly | O43602 |
| HOMO SAPIENS | 8 | 4602 | 1mjd | A | 104 | ILE | p.Ile104Phe | O43602 |
| HOMO SAPIENS | 7 | 4602 | 1mjd | A | 125 | TYR | p.Tyr125His | O43602 |
| HOMO SAPIENS | 5 | 34467 | 1n0w | B | 1529 | GLY | p.Gly1529Arg | P51587 |
| HOMO SAPIENS | 8 | 5738 | 1u31 | A | 130 | ASN | p.Asn1009Lys | Q13423 |
| HOMO SAPIENS | 0 | 5760 | 2i6s | A | 298 | GLU | p.Glu318Asp | P06681 |
| HOMO SAPIENS | 0 | 5760 | 2i6s | A | 573 | ARG | p.Arg593Gln | P06681 |
| HOMO SAPIENS | 0 | 5760 | 2i6s | A | 592 | SER | p.Ser612Asn | P06681 |
| HOMO SAPIENS | 0 | 5760 | 2i6s | A | 621 | VAL | p.Val641Ala | P06681 |
| HOMO SAPIENS | 1 | 10977 | 2fby | A | 114 | VAL | p.Val114Ile | Q14191 |
| HOMO SAPIENS | 0 | 10977 | 2fby | A | 135 | LYS | p.Lys135Glu | Q14191 |
| HOMO SAPIENS | -1 | 1175 | 1qrk | A | 34 | VAL | p.Val35Leu | P00488 |
| HOMO SAPIENS | 4 | 1175 | 1qrk | A | 564 | PRO | p.Pro565Leu | P00488 |
| HOMO SAPIENS | 0 | 1175 | 1qrk | A | 650 | VAL | p.Val651Ile | P00488 |
| HOMO SAPIENS | 0 | 1175 | 1qrk | A | 651 | GLU | p.Gln652Glu | P00488 |
| HOMO SAPIENS | 3 | 909 | 4h0w | A | 375 | GLU | p.Glu394Lys | P02787 |
| HOMO SAPIENS | 0 | 909 | 4h0w | A | 458 | ALA | p.Ala477Pro | P02787 |
| HOMO SAPIENS | 6 | 909 | 4h0w | A | 515 | TYR | p.Tyr534Ter | P02787 |
| HOMO SAPIENS | 7 | 909 | 4h0w | A | 570 | PRO | p.Pro589Ser | P02787 |
| HOMO SAPIENS | 3 | 909 | 4h0w | A | 652 | GLY | p.Gly671Glu | P02787 |
| HOMO SAPIENS | 5 | 1964 | 2car | A | 151 | TRP | p.Trp151Ter | Q9BY32 |
| HOMO SAPIENS | 7 | 1964 | 2car | A | 178 | ARG | p.Arg178Cys | Q9BY32 |
| HOMO SAPIENS | 0 | 144 | 2n0a | J | 30 | ALA | p.Ala30Pro | P37840 |
| HOMO SAPIENS | 1 | 12288 | 4q58 | D | 1225 | ARG | p.Arg1225His | P16144 |
| HOMO SAPIENS | 2 | 12288 | 4q58 | D | 1281 | ARG | p.Arg1281Trp | P16144 |
| HOMO SAPIENS | 2 | 12288 | 4q58 | D | 1287 | ASN | p.Asn1287Lys | P16144 |
| HOMO SAPIENS | 0 | 5085 | 3s94 | A | 360 | ARG | p.Arg360His | O75581 |
| HOMO SAPIENS | 0 | 5085 | 3s94 | A | 433 | ASN | p.Asn433Ser | O75581 |
| HOMO SAPIENS | 2 | 5085 | 3s94 | A | 611 | ARG | p.Arg611Cys | O75581 |
| HOMO SAPIENS | 0 | 15452 | 4dur | A | 19 | LYS | p.Lys38Glu | P00747 |
| HOMO SAPIENS | 2 | 15452 | 4dur | A | 216 | ARG | p.Arg235His | P00747 |
| HOMO SAPIENS | 7 | 8406 | 4wo1 | D | 28 | VAL | p.Val55Leu | O43914 |
| HOMO SAPIENS | 2 | 895 | 5xte | K | 183 | ARG | p.Arg183Trp | P22695 |
| HOMO SAPIENS | 2 | 4191 | 4wnd | A | 120 | ARG | p.Arg127Ter | P81274 |

| | | | | | | | | |
|---|---|---|---|---|---|---|---|---|
| HOMO SAPIENS | 2 | 4191 | 4wnd | A | 120 | ARG | p.Arg127Gln | P81274 |
| HOMO SAPIENS | 1 | 4191 | 4wnd | A | 349 | GLY | p.Gly356Arg | P81274 |
| HOMO SAPIENS | 0 | 41734 | 2kbi | A | 1777 | VAL | p.Val1777Met | Q14524 |
| HOMO SAPIENS | 1 | 41734 | 2kbi | A | 1784 | GLU | p.Glu1784Lys | Q14524 |
| HOMO SAPIENS | 5 | 41734 | 2kbi | A | 1795 | TYR | p.Tyr1795Cys | Q14524 |
| HOMO SAPIENS | 5 | 41734 | 2kbi | A | 1795 | TYR | p.Tyr1795His | Q14524 |
| HOMO SAPIENS | 7 | 41734 | 2kbi | A | 1826 | ARG | p.Arg1826Cys | Q14524 |
| HOMO SAPIENS | 3 | 2119 | 1pl0 | A | 426 | LYS | p.Lys426Arg | P31939 |
| HOMO SAPIENS | 5 | 3475 | 5u62 | C | 50 | LEU | p.Leu50Ser | Q15910 |
| HOMO SAPIENS | 3 | 379 | 4uez | A | 200 | TYR | p.Tyr200Ter | P15085 |
| HOMO SAPIENS | 0 | 8195 | 5xaw | B | 67 | ARG | p.Arg67Trp | Q8IWT1 |
| HOMO SAPIENS | 0 | 8195 | 5xaw | B | 94 | THR | p.Thr94Met | Q8IWT1 |
| HOMO SAPIENS | 1 | 5711 | 1fo3 | A | 265 | ARG | p.Arg265His | Q9UKM7 |
| HOMO SAPIENS | 4 | 5711 | 1fo3 | A | 334 | ARG | p.Arg334Cys | Q9UKM7 |
| HOMO SAPIENS | 1 | 5711 | 1fo3 | A | 478 | LYS | p.Lys478Arg | Q9UKM7 |
| HOMO SAPIENS | 1 | 5711 | 1fo3 | A | 682 | ASP | p.Asp682Glu | Q9UKM7 |
| HOMO SAPIENS | 0 | 170 | 2n3y | A | 41 | GLY | p.Gly42Ser | P99999 |
| HOMO SAPIENS | 0 | 4122 | 3w81 | A | 33 | HIS | p.His33Gln | P35475 |
| HOMO SAPIENS | 0 | 4122 | 3w81 | A | 64 | TYR | p.Tyr64Ter | P35475 |
| HOMO SAPIENS | 0 | 4122 | 3w81 | A | 70 | GLN | p.Gln70Ter | P35475 |
| HOMO SAPIENS | 0 | 4122 | 3w81 | A | 75 | ALA | p.Ala75Thr | P35475 |
| HOMO SAPIENS | 0 | 4122 | 3w81 | A | 79 | ALA | p.Ala79Val | P35475 |
| HOMO SAPIENS | 0 | 4122 | 3w81 | A | 79 | ALA | p.Ala79Thr | P35475 |
| HOMO SAPIENS | 0 | 4122 | 3w81 | A | 89 | ARG | p.Arg89Gln | P35475 |
| HOMO SAPIENS | 0 | 4122 | 3w81 | A | 99 | THR | p.Thr99Ile | P35475 |
| HOMO SAPIENS | 0 | 4122 | 3w81 | A | 105 | ARG | p.Arg105Gln | P35475 |
| HOMO SAPIENS | 0 | 4122 | 3w81 | A | 238 | LEU | p.Leu238Gln | P35475 |
| HOMO SAPIENS | 0 | 4122 | 3w81 | A | 300 | ALA | p.Ala300Thr | P35475 |
| HOMO SAPIENS | 0 | 4122 | 3w81 | A | 322 | VAL | p.Val322Glu | P35475 |
| HOMO SAPIENS | 0 | 4122 | 3w81 | A | 327 | ALA | p.Ala327Pro | P35475 |
| HOMO SAPIENS | 0 | 4122 | 3w81 | A | 346 | LEU | p.Leu346Arg | P35475 |
| HOMO SAPIENS | 0 | 4122 | 3w81 | A | 349 | ASP | p.Asp349Asn | P35475 |
| HOMO SAPIENS | 0 | 4122 | 3w81 | A | 361 | ALA | p.Ala361Thr | P35475 |
| HOMO SAPIENS | 0 | 4122 | 3w81 | A | 366 | THR | p.Thr366Pro | P35475 |
| HOMO SAPIENS | 0 | 4122 | 3w81 | A | 402 | TRP | p.Trp402Ter | P35475 |
| HOMO SAPIENS | 0 | 4122 | 3w81 | A | 409 | GLY | p.Gly409Arg | P35475 |
| HOMO SAPIENS | 0 | 4122 | 3w81 | A | 454 | VAL | p.Val454Ile | P35475 |
| HOMO SAPIENS | 1 | 4122 | 3w81 | A | 490 | LEU | p.Leu490Pro | P35475 |
| HOMO SAPIENS | 1 | 4122 | 3w81 | A | 492 | ARG | p.Arg492Pro | P35475 |
| HOMO SAPIENS | 0 | 4122 | 3w81 | A | 533 | PRO | p.Pro533Arg | P35475 |
| HOMO SAPIENS | 0 | 4122 | 3w81 | A | 591 | ALA | p.Ala591Thr | P35475 |
| HOMO SAPIENS | 7 | 4122 | 3w81 | A | 619 | ARG | p.Arg619Gly | P35475 |
| HOMO SAPIENS | 0 | 15325 | 3wvz | B | 54 | VAL | p.Val54Leu | Q53FT3 |
| HOMO SAPIENS | 3 | 3228 | 5iz8 | A | 421 | TRP | p.Trp421Ter | P25054 |
| HOMO SAPIENS | 2 | 3228 | 5iz8 | A | 437 | PRO | p.Pro437Arg | P25054 |
| HOMO SAPIENS | 3 | 3228 | 5iz8 | A | 499 | ARG | p.Arg499Ter | P25054 |
| HOMO SAPIENS | -1 | 29229 | 5buo | A | 539 | SER | p.Ser614Gly | P05067 |
| HOMO SAPIENS | -1 | 34931 | 4n7v | C | 54 | SER | p.Ser54Leu | O94986 |
| HOMO SAPIENS | 0 | 2450 | 5kzx | A | 123 | GLY | p.Gly123Glu | P10253 |
| HOMO SAPIENS | 1 | 2450 | 5kzx | A | 190 | ARG | p.Arg190His | P10253 |
| HOMO SAPIENS | -1 | 2450 | 5kzx | A | 199 | HIS | p.His199Arg | P10253 |
| HOMO SAPIENS | 0 | 2450 | 5kzx | A | 219 | GLY | p.Gly219Arg | P10253 |
| HOMO SAPIENS | 0 | 2450 | 5kzx | A | 220 | VAL | p.Val220Leu | P10253 |
| HOMO SAPIENS | 0 | 2450 | 5kzx | A | 222 | VAL | p.Val222Met | P10253 |

| | | | | | | | | |
|---|---|---|---|---|---|---|---|---|
| HOMO SAPIENS | 0 | 2450 | 5kzx | A | 237 | ALA | p.Ala237Val | P10253 |
| HOMO SAPIENS | 0 | 2450 | 5kzx | A | 251 | SER | p.Ser251Leu | P10253 |
| HOMO SAPIENS | 0 | 2450 | 5kzx | A | 254 | SER | p.Ser254Leu | P10253 |
| HOMO SAPIENS | 0 | 2450 | 5kzx | A | 262 | GLU | p.Glu262Lys | P10253 |
| HOMO SAPIENS | 0 | 2450 | 5kzx | A | 293 | GLY | p.Gly293Arg | P10253 |
| HOMO SAPIENS | 0 | 2450 | 5kzx | A | 299 | LEU | p.Leu299Arg | P10253 |
| HOMO SAPIENS | 0 | 2450 | 5kzx | A | 309 | GLY | p.Gly309Arg | P10253 |
| HOMO SAPIENS | 0 | 2450 | 5kzx | A | 318 | MET | p.Met318Thr | P10253 |
| HOMO SAPIENS | 0 | 2450 | 5kzx | A | 355 | LEU | p.Leu355Pro | P10253 |
| HOMO SAPIENS | 1 | 2450 | 5kzx | A | 361 | PRO | p.Pro361Leu | P10253 |
| HOMO SAPIENS | 4 | 2450 | 5kzx | A | 408 | MET | p.Met408Val | P10253 |
| HOMO SAPIENS | 1 | 2450 | 5kzx | A | 429 | GLN | p.Gln429Glu | P10253 |
| HOMO SAPIENS | 4 | 2450 | 5kzx | A | 437 | ARG | p.Arg437Cys | P10253 |
| HOMO SAPIENS | 0 | 2450 | 5kzx | A | 439 | MET | p.Met439Lys | P10253 |
| HOMO SAPIENS | 1 | 2450 | 5kzx | A | 448 | SER | p.Ser448Thr | P10253 |
| HOMO SAPIENS | 0 | 2450 | 5kzx | A | 451 | PRO | p.Pro451Arg | P10253 |
| HOMO SAPIENS | 1 | 2450 | 5kzx | A | 459 | ASP | p.Asp459His | P10253 |
| HOMO SAPIENS | 8 | 2450 | 5kzx | A | 481 | TRP | p.Trp481Arg | P10253 |
| HOMO SAPIENS | 1 | 2450 | 5kzx | A | 489 | ASP | p.Asp489Asn | P10253 |
| HOMO SAPIENS | 0 | 2450 | 5kzx | A | 521 | GLU | p.Glu521Lys | P10253 |
| HOMO SAPIENS | 0 | 2450 | 5kzx | A | 529 | SER | p.Ser529Val | P10253 |
| HOMO SAPIENS | 0 | 2450 | 5kzx | A | 545 | PRO | p.Pro545Leu | P10253 |
| HOMO SAPIENS | 0 | 2450 | 5kzx | A | 552 | LEU | p.Leu552Pro | P10253 |
| HOMO SAPIENS | 0 | 2450 | 5kzx | A | 576 | GLY | p.Gly576Ser | P10253 |
| HOMO SAPIENS | 9 | 2450 | 5kzx | A | 600 | ARG | p.Arg600His | P10253 |
| HOMO SAPIENS | 8 | 2450 | 5kzx | A | 615 | GLY | p.Gly615Arg | P10253 |
| HOMO SAPIENS | 3 | 2450 | 5kzx | A | 619 | SER | p.Ser619Asn | P10253 |
| HOMO SAPIENS | 0 | 2450 | 5kzx | A | 638 | GLY | p.Gly638Trp | P10253 |
| HOMO SAPIENS | 0 | 2450 | 5kzx | A | 643 | GLY | p.Gly643Arg | P10253 |
| HOMO SAPIENS | 9 | 2450 | 5kzx | A | 645 | ASP | p.Asp645Glu | P10253 |
| HOMO SAPIENS | 9 | 2450 | 5kzx | A | 645 | ASP | p.Asp645His | P10253 |
| HOMO SAPIENS | 0 | 2450 | 5kzx | A | 648 | GLY | p.Gly648Ser | P10253 |
| HOMO SAPIENS | 5 | 2450 | 5kzx | A | 660 | ARG | p.Arg660His | P10253 |
| HOMO SAPIENS | 8 | 2450 | 5kzx | A | 672 | ARG | p.Arg672Gln | P10253 |
| HOMO SAPIENS | 0 | 2450 | 5kzx | A | 702 | ARG | p.Arg702Leu | P10253 |
| HOMO SAPIENS | 3 | 2450 | 5kzx | A | 725 | ARG | p.Arg725Trp | P10253 |
| HOMO SAPIENS | 0 | 2450 | 5kzx | A | 746 | TRP | p.Trp746Cys | P10253 |
| HOMO SAPIENS | 0 | 2450 | 5kzx | A | 780 | ILE | p.Val780Ile | P10253 |
| HOMO SAPIENS | 1 | 2450 | 5kzx | A | 816 | VAL | p.Val816Ile | P10253 |
| HOMO SAPIENS | 0 | 2450 | 5kzx | A | 838 | GLN | p.Gln838Ter | P10253 |
| HOMO SAPIENS | 2 | 2450 | 5kzx | A | 854 | ARG | p.Arg854Ter | P10253 |
| HOMO SAPIENS | 3 | 2450 | 5kzx | A | 870 | ARG | p.Arg870Ter | P10253 |
| HOMO SAPIENS | 0 | 2450 | 5kzx | A | 890 | VAL | p.Val890Leu | P10253 |
| HOMO SAPIENS | 6 | 1882 | 4km6 | A | 175 | CYS | p.Cys175Ter | P15328 |
| HOMO SAPIENS | 3 | 14767 | 1lm5 | A | 2628 | LEU | p.Leu2628Pro | P15924 |
| HOMO SAPIENS | 5 | 14767 | 1lm5 | A | 2655 | ALA | p.Ala2655Gly | P15924 |
| HOMO SAPIENS | 3 | 14767 | 1lm5 | A | 2731 | TYR | p.Tyr2731His | P15924 |
| HOMO SAPIENS | 1 | 14767 | 1lm5 | A | 2767 | THR | p.Thr2767Asn | P15924 |
| HOMO SAPIENS | -1 | 14767 | 1lm5 | A | 2819 | MET | p.Met2819Leu | P15924 |
| HOMO SAPIENS | -1 | 11246 | 4y5o | A | 289 | SER | p.Ser289Asn | Q9BSQ5 |
| HOMO SAPIENS | 0 | 7380 | 5kz5 | I | 60 | ARG | p.Arg60His | Q16595 |
| HOMO SAPIENS | 0 | 7380 | 5kz5 | I | 76 | MET | p.Met76Val | Q16595 |
| HOMO SAPIENS | 2 | 22187 | 3fby | A | 328 | CYS | p.Cys328Arg | P49747 |
| HOMO SAPIENS | 2 | 22187 | 3fby | A | 342 | ASP | p.Asp342Tyr | P49747 |

| | | | | | | | | |
|---|---|---|---|---|---|---|---|---|
| HOMO SAPIENS | 2 | 22187 | 3fby | A | 348 | CYS | p.Cys348Arg | P49747 |
| HOMO SAPIENS | 2 | 22187 | 3fby | A | 386 | ASN | p.Asn386Asp | P49747 |
| HOMO SAPIENS | 2 | 22187 | 3fby | A | 453 | ASN | p.Asn453Ser | P49747 |
| HOMO SAPIENS | 2 | 22187 | 3fby | A | 468 | CYS | p.Cys468Tyr | P49747 |
| HOMO SAPIENS | 3 | 22187 | 3fby | A | 472 | ASP | p.Asp472Tyr | P49747 |
| HOMO SAPIENS | 3 | 22187 | 3fby | A | 473 | ASP | p.Asp473Gly | P49747 |
| HOMO SAPIENS | 2 | 22187 | 3fby | A | 523 | ASN | p.Asn523Lys | P49747 |
| HOMO SAPIENS | 1 | 22187 | 3fby | A | 718 | ARG | p.Arg718Trp | P49747 |
| HOMO SAPIENS | 1 | 22187 | 3fby | A | 719 | GLY | p.Gly719Asp | P49747 |
| HOMO SAPIENS | 0 | 22187 | 3fby | A | 756 | GLN | p.Gln756Arg | P49747 |
| HOMO SAPIENS | -1 | 2784 | 5kvh | A | 88 | MET | p.Met88Val | O95831 |
| HOMO SAPIENS | -1 | 2784 | 5kvh | A | 96 | ASN | p.Asn96Ser | O95831 |
| HOMO SAPIENS | 0 | 2784 | 5kvh | A | 282 | THR | p.Thr282Met | O95831 |
| HOMO SAPIENS | 1 | 2784 | 5kvh | A | 366 | ASN | p.Asn366Ser | O95831 |
| HOMO SAPIENS | 3 | 2784 | 5kvh | A | 463 | ARG | p.Arg463Ile | O95831 |
| HOMO SAPIENS | 0 | 2784 | 5kvh | A | 498 | VAL | p.Val498Met | O95831 |
| HOMO SAPIENS | -1 | 2784 | 5kvh | A | 549 | ALA | p.Ala549Val | O95831 |
| HOMO SAPIENS | 0 | 38316 | 2lnk | C | 1933 | ARG | p.Arg1933Ter | P35579 |
| HOMO SAPIENS | 0 | 33808 | 2miu | A | 29 | VAL | p.Val29Met | Q14192 |
| HOMO SAPIENS | 1 | 33808 | 2miu | A | 37 | ALA | p.Ala37Ser | Q14192 |
| HOMO SAPIENS | 2 | 20994 | 2hey | R | 65 | ARG | p.Arg65Cys | P43489 |
| HOMO SAPIENS | -1 | 5687 | 2xwj | I | 7 | ARG | p.Arg32Gln | P00751 |
| HOMO SAPIENS | -1 | 5687 | 2xwj | I | 7 | ARG | p.Arg32Trp | P00751 |
| HOMO SAPIENS | 0 | 5687 | 2xwj | I | 49 | ARG | p.Arg74His | P00751 |
| HOMO SAPIENS | 0 | 5687 | 2xwj | I | 177 | ARG | p.Arg202Gly | P00751 |
| HOMO SAPIENS | 0 | 5687 | 2xwj | I | 217 | ILE | p.Ile242Leu | P00751 |
| HOMO SAPIENS | -1 | 5687 | 2xwj | I | 227 | GLY | p.Gly252Ser | P00751 |
| HOMO SAPIENS | 0 | 5687 | 2xwj | I | 483 | HIS | p.His508Gln | P00751 |
| HOMO SAPIENS | 0 | 5687 | 2xwj | I | 508 | LYS | p.Lys533Arg | P00751 |
| HOMO SAPIENS | 0 | 5687 | 2xwj | I | 540 | LYS | p.Lys565Glu | P00751 |
| HOMO SAPIENS | 0 | 5687 | 2xwj | I | 541 | GLU | p.Glu566Ala | P00751 |
| HOMO SAPIENS | 2 | 953 | 3d4j | A | 249 | PHE | p.Phe249Ser | P53602 |
| HOMO SAPIENS | 1 | 953 | 3d4j | A | 292 | ASN | p.Asn292Ser | P53602 |
| HOMO SAPIENS | 0 | 3300 | 1s9c | A | 140 | ASN | p.Asn457Tyr | P51659 |
| HOMO SAPIENS | 2 | 3300 | 1s9c | A | 167 | ILE | p.Ile484Val | P51659 |
| HOMO SAPIENS | 0 | 3300 | 1s9c | A | 174 | ALA | p.Ala491Thr | P51659 |
| HOMO SAPIENS | 8 | 3300 | 1s9c | A | 193 | ASP | p.Asp510Asn | P51659 |
| HOMO SAPIENS | 3 | 3300 | 1s9c | A | 194 | TRP | p.Trp511Arg | P51659 |
| HOMO SAPIENS | 0 | 3300 | 1s9c | A | 242 | VAL | p.Ile559Val | P51659 |
| HOMO SAPIENS | 0 | 10810 | 5i25 | A | 5 | GLN | p.Gln23Ter | P03951 |
| HOMO SAPIENS | 0 | 10810 | 5i25 | A | 55 | TRP | p.Trp73Ter | P03951 |
| HOMO SAPIENS | 0 | 10810 | 5i25 | A | 91 | ALA | p.Ala109Thr | P03951 |
| HOMO SAPIENS | 0 | 10810 | 5i25 | A | 117 | GLU | p.Glu135Ter | P03951 |
| HOMO SAPIENS | 0 | 10810 | 5i25 | A | 118 | CYS | p.Cys136Ter | P03951 |
| HOMO SAPIENS | 0 | 10810 | 5i25 | A | 128 | CYS | p.Cys146Ter | P03951 |
| HOMO SAPIENS | 0 | 10810 | 5i25 | A | 210 | ARG | p.Arg228Ter | P03951 |
| HOMO SAPIENS | 0 | 10810 | 5i25 | A | 226 | GLN | p.Gln244Ter | P03951 |
| HOMO SAPIENS | 0 | 10810 | 5i25 | A | 233 | GLN | p.Gln251Ter | P03951 |
| HOMO SAPIENS | 0 | 16668 | 3lbx | A | 28 | ARG | p.Arg28Leu | P02549 |
| HOMO SAPIENS | 0 | 16668 | 3lbx | A | 28 | ARG | p.Arg28Cys | P02549 |
| HOMO SAPIENS | 1 | 16668 | 3lbx | A | 41 | ARG | p.Arg41Trp | P02549 |
| HOMO SAPIENS | 1 | 16668 | 3lbx | A | 45 | ARG | p.Arg45Ser | P02549 |
| HOMO SAPIENS | 2 | 16668 | 3lbx | A | 48 | LYS | p.Lys48Arg | P02549 |
| HOMO SAPIENS | 1 | 16668 | 3lbx | A | 152 | ASP | p.Asp152Asn | P02549 |

| | | | | | | | | |
|---|---|---|---|---|---|---|---|---|
| HOMO SAPIENS | 0 | 226 | 2ble | A | 256 | ILE | p.Phe256Ile | P36959 |
| HOMO SAPIENS | 2 | 36808 | 4c7n | A | 3 | SER | p.Ser357Pro | O75030 |
| HOMO SAPIENS | 0 | 4092 | 5jnx | A | 113 | CYS | p.Cys113Arg | O15118 |
| HOMO SAPIENS | 4 | 4092 | 5jnx | A | 215 | HIS | p.His215Arg | O15118 |
| HOMO SAPIENS | 6 | 4092 | 5jnx | A | 237 | PRO | p.Pro237Ser | O15118 |
| HOMO SAPIENS | 0 | 4092 | 5jnx | A | 363 | ALA | p.Ala363Val | O15118 |
| HOMO SAPIENS | 0 | 4092 | 5jnx | A | 378 | VAL | p.Val378Ala | O15118 |
| HOMO SAPIENS | 8 | 4092 | 5jnx | A | 404 | ARG | p.Arg404Gln | O15118 |
| HOMO SAPIENS | 8 | 4092 | 5jnx | A | 411 | ARG | p.Arg411Gln | O15118 |
| HOMO SAPIENS | 2 | 4092 | 5jnx | A | 474 | PRO | p.Pro474Leu | O15118 |
| HOMO SAPIENS | 8 | 4092 | 5jnx | A | 511 | THR | p.Thr511Met | O15118 |
| HOMO SAPIENS | 0 | 4092 | 5jnx | A | 642 | MET | p.Met642Ile | O15118 |
| HOMO SAPIENS | 2 | 4092 | 5jnx | A | 664 | VAL | p.Val664Met | O15118 |
| HOMO SAPIENS | 0 | 4092 | 5jnx | A | 734 | SER | p.Ser734Ile | O15118 |
| HOMO SAPIENS | -1 | 4092 | 5jnx | A | 775 | GLN | p.Gln775Pro | O15118 |
| HOMO SAPIENS | 0 | 4092 | 5jnx | A | 858 | ILE | p.Ile858Val | O15118 |
| HOMO SAPIENS | 0 | 4092 | 5jnx | A | 889 | VAL | p.Val889Met | O15118 |
| HOMO SAPIENS | 0 | 4092 | 5jnx | A | 910 | GLY | p.Gly910Ser | O15118 |
| HOMO SAPIENS | 0 | 4092 | 5jnx | A | 911 | GLY | p.Gly911Ser | O15118 |
| HOMO SAPIENS | 0 | 4092 | 5jnx | A | 928 | GLN | p.Gln928Pro | O15118 |
| HOMO SAPIENS | 0 | 4092 | 5jnx | A | 931 | ASN | p.Asn931Lys | O15118 |
| HOMO SAPIENS | 0 | 4092 | 5jnx | A | 940 | SER | p.Ser940Leu | O15118 |
| HOMO SAPIENS | 0 | 4092 | 5jnx | A | 950 | VAL | p.Val950Met | O15118 |
| HOMO SAPIENS | 0 | 4092 | 5jnx | A | 954 | SER | p.Ser954Leu | O15118 |
| HOMO SAPIENS | 0 | 4092 | 5jnx | A | 958 | ARG | p.Arg958Gln | O15118 |
| HOMO SAPIENS | 0 | 4092 | 5jnx | A | 958 | ARG | p.Arg958Ter | O15118 |
| HOMO SAPIENS | -1 | 4092 | 5jnx | A | 978 | ARG | p.Arg978Cys | O15118 |
| HOMO SAPIENS | 0 | 4092 | 5jnx | A | 992 | GLY | p.Gly992Trp | O15118 |
| HOMO SAPIENS | 0 | 4092 | 5jnx | A | 1007 | PRO | p.Pro1007Ala | O15118 |
| HOMO SAPIENS | 0 | 4092 | 5jnx | A | 1019 | TYR | p.Tyr1019Cys | O15118 |
| HOMO SAPIENS | 2 | 4092 | 5jnx | A | 1035 | ALA | p.Ala1035Val | O15118 |
| HOMO SAPIENS | 2 | 4092 | 5jnx | A | 1054 | ALA | p.Ala1054Thr | O15118 |
| HOMO SAPIENS | 2 | 4092 | 5jnx | A | 1061 | ILE | p.Ile1061Thr | O15118 |
| HOMO SAPIENS | 0 | 4092 | 5jnx | A | 1073 | GLY | p.Gly1073Ser | O15118 |
| HOMO SAPIENS | 2 | 4092 | 5jnx | A | 1088 | TYR | p.Tyr1088Cys | O15118 |
| HOMO SAPIENS | 2 | 4092 | 5jnx | A | 1142 | MET | p.Met1142Thr | O15118 |
| HOMO SAPIENS | 2 | 4092 | 5jnx | A | 1156 | ASN | p.Asn1156Ser | O15118 |
| HOMO SAPIENS | 0 | 4092 | 5jnx | A | 1179 | MET | p.Met1179Val | O15118 |
| HOMO SAPIENS | 0 | 4092 | 5jnx | A | 1186 | ARG | p.Arg1186His | O15118 |
| HOMO SAPIENS | 0 | 4092 | 5jnx | A | 1187 | ALA | p.Ala1187Gly | O15118 |
| HOMO SAPIENS | 2 | 4092 | 5jnx | A | 1189 | GLU | p.Glu1189Gly | O15118 |
| HOMO SAPIENS | 2 | 4092 | 5jnx | A | 1200 | SER | p.Ser1200Gly | O15118 |
| HOMO SAPIENS | 2 | 4092 | 5jnx | A | 1205 | THR | p.Thr1205Lys | O15118 |
| HOMO SAPIENS | 0 | 4092 | 5jnx | A | 1213 | LEU | p.Leu1213Phe | O15118 |
| HOMO SAPIENS | -1 | 4092 | 5jnx | A | 1266 | ARG | p.Arg1266Gln | O15118 |
| HOMO SAPIENS | 9 | 16498 | 4i6o | A | 64 | ARG | p.Arg735Trp | P01024 |
| HOMO SAPIENS | 1 | 21902 | 3un9 | C | 751 | ARG | p.Arg751His | Q86UT6 |
| HOMO SAPIENS | 7 | 20391 | 2h32 | B | 105 | PRO | p.Pro142Leu | P15814 |
| HOMO SAPIENS | 6 | 1635 | 4x45 | A | 65 | ASP | p.Asp65Val | P07741 |
| HOMO SAPIENS | 1 | 1635 | 4x45 | A | 98 | TRP | p.Trp98Ter | P07741 |
| HOMO SAPIENS | 0 | 1635 | 4x45 | A | 110 | LEU | p.Leu110Pro | P07741 |
| HOMO SAPIENS | 7 | 1635 | 4x45 | A | 136 | MET | p.Met136Thr | P07741 |
| HOMO SAPIENS | 1 | 1635 | 4x45 | A | 150 | VAL | p.Val150Phe | P07741 |
| HOMO SAPIENS | 4 | 34810 | 2rpc | A | 16 | CYS | p.Cys253Ser | O60481 |

| | | | | | | | | |
|---|---|---|---|---|---|---|---|---|
| HOMO SAPIENS | 1 | 34810 | 2rpc | A | 18 | TRP | p.Trp255Gly | O60481 |
| HOMO SAPIENS | 1 | 14359 | 4hxi | A | 78 | MET | p.Met78Val | Q9UH77 |
| HOMO SAPIENS | 0 | 5342 | 4d9t | A | 628 | GLU | p.Glu628Asp | P51812 |
| HOMO SAPIENS | -1 | 5342 | 4d9t | A | 723 | ARG | p.Arg723His | P51812 |
| HOMO SAPIENS | -1 | 5342 | 4d9t | A | 729 | ARG | p.Arg729Gln | P51812 |
| HOMO SAPIENS | 1 | 9619 | 3j2s | B | 1781 | ARG | p.Arg1800Cys | P00451 |
| HOMO SAPIENS | 0 | 9619 | 3j2s | B | 1823 | MET | p.Met1842Ile | P00451 |
| HOMO SAPIENS | 1 | 9619 | 3j2s | B | 1825 | PRO | p.Pro1844Ser | P00451 |
| HOMO SAPIENS | 0 | 9619 | 3j2s | B | 1826 | THR | p.Thr1845Pro | P00451 |
| HOMO SAPIENS | 0 | 9619 | 3j2s | B | 1834 | ALA | p.Ala1853Val | P00451 |
| HOMO SAPIENS | 0 | 9619 | 3j2s | B | 1846 | ASP | p.Asp1865Asn | P00451 |
| HOMO SAPIENS | 1 | 9619 | 3j2s | B | 1848 | HIS | p.His1867Arg | P00451 |
| HOMO SAPIENS | 0 | 9619 | 3j2s | B | 1854 | PRO | p.Pro1873Arg | P00451 |
| HOMO SAPIENS | 1 | 9619 | 3j2s | B | 1885 | GLU | p.Glu1904Lys | P00451 |
| HOMO SAPIENS | 0 | 9619 | 3j2s | B | 1922 | ASN | p.Asn1941Ser | P00451 |
| HOMO SAPIENS | 1 | 9619 | 3j2s | B | 1941 | ARG | p.Arg1960Gln | P00451 |
| HOMO SAPIENS | 4 | 9619 | 3j2s | B | 2105 | TYR | p.Tyr2124Cys | P00451 |
| HOMO SAPIENS | 6 | 9619 | 3j2s | B | 2150 | ARG | p.Arg2169His | P00451 |
| HOMO SAPIENS | 0 | 9619 | 3j2s | B | 2159 | ARG | p.Arg2178Cys | P00451 |
| HOMO SAPIENS | 0 | 9619 | 3j2s | B | 2159 | ARG | p.Arg2178His | P00451 |
| HOMO SAPIENS | 2 | 9619 | 3j2s | B | 2238 | MET | p.Met2257Val | P00451 |
| HOMO SAPIENS | 2 | 9619 | 3j2s | B | 2300 | PRO | p.Pro2319Ser | P00451 |
| HOMO SAPIENS | 1 | 667 | 1z2m | A | 127 | GLU | p.Glu127Ter | P05161 |
| HOMO SAPIENS | 0 | 15717 | 2nn6 | G | 30 | GLY | p.Gly31Ala | Q9NQT5 |
| HOMO SAPIENS | -1 | 15717 | 2nn6 | G | 55 | ASN | p.Asn56His | Q9NQT5 |
| HOMO SAPIENS | 7 | 15717 | 2nn6 | G | 131 | ASP | p.Asp132Ala | Q9NQT5 |
| HOMO SAPIENS | 7 | 784 | 3rkq | A | 161 | ARG | p.Arg161Pro | P52952 |
| HOMO SAPIENS | 6 | 784 | 3rkq | A | 178 | THR | p.Thr178Met | P52952 |
| HOMO SAPIENS | 7 | 784 | 3rkq | A | 183 | LYS | p.Lys183Glu | P52952 |
| HOMO SAPIENS | 8 | 784 | 3rkq | A | 190 | ARG | p.Arg190Cys | P52952 |
| HOMO SAPIENS | 6 | 385 | 30000 | C | 70 | ALA | p.Ala70Thr | P07954 |
| HOMO SAPIENS | 2 | 385 | 30000 | C | 101 | ARG | p.Arg101Pro | P07954 |
| HOMO SAPIENS | 2 | 385 | 30000 | C | 174 | PRO | p.Pro174Arg | P07954 |
| HOMO SAPIENS | 6 | 385 | 30000 | C | 187 | SER | p.Ser187Ter | P07954 |
| HOMO SAPIENS | 9 | 385 | 30000 | C | 233 | ARG | p.Arg233His | P07954 |
| HOMO SAPIENS | 3 | 385 | 30000 | C | 273 | ALA | p.Ala273Thr | P07954 |
| HOMO SAPIENS | 4 | 385 | 30000 | C | 334 | SER | p.Ser334Arg | P07954 |
| HOMO SAPIENS | 6 | 385 | 30000 | C | 340 | ASN | p.Asn340Lys | P07954 |
| HOMO SAPIENS | 8 | 385 | 30000 | C | 343 | ARG | p.Arg343Ter | P07954 |
| HOMO SAPIENS | 3 | 385 | 30000 | C | 350 | ARG | p.Arg350Trp | P07954 |
| HOMO SAPIENS | 4 | 385 | 30000 | C | 376 | GLN | p.Gln376Pro | P07954 |
| HOMO SAPIENS | 6 | 385 | 30000 | C | 434 | CYS | p.Cys434Ter | P07954 |
| HOMO SAPIENS | 0 | 3316 | 4f91 | B | 934 | THR | p.Thr934Ala | O75643 |
| HOMO SAPIENS | 0 | 3316 | 4f91 | B | 1002 | ASN | p.Asn1002Ser | O75643 |
| HOMO SAPIENS | 0 | 3316 | 4f91 | B | 1087 | SER | p.Ser1087Leu | O75643 |
| HOMO SAPIENS | 4 | 3316 | 4f91 | B | 1389 | VAL | p.Val1389Ile | O75643 |
| HOMO SAPIENS | 0 | 619 | 5hbt | B | 195 | ASP | p.Asp240His | P02708 |
| HOMO SAPIENS | 1 | 619 | 5hbt | B | 209 | ARG | p.Arg254Leu | P02708 |
| HOMO SAPIENS | 0 | 18171 | 2y25 | B | 1408 | ASP | p.Asp1408Asn | P52179 |
| HOMO SAPIENS | 0 | 18171 | 2y25 | B | 1453 | MET | p.Met1453Thr | P52179 |
| HOMO SAPIENS | 0 | 18171 | 2y25 | B | 1459 | ILE | p.Ile1459Lys | P52179 |
| HOMO SAPIENS | 0 | 18171 | 2y25 | B | 1558 | PHE | p.Phe1558Leu | P52179 |
| HOMO SAPIENS | 0 | 18171 | 2y25 | B | 1573 | ARG | p.Arg1573Gln | P52179 |
| HOMO SAPIENS | 7 | 18171 | 2y25 | B | 1663 | VAL | p.Val1663Met | P52179 |

| | | | | | | | | |
|---|---|---|---|---|---|---|---|---|
| HOMO SAPIENS | 8 | 14262 | 2fcw | B | 117 | SER | p.Ser138Ter | P01130 |
| HOMO SAPIENS | 8 | 14262 | 2fcw | B | 118 | ASP | p.Asp139Glu | P01130 |
| HOMO SAPIENS | 9 | 14262 | 2fcw | B | 119 | GLU | p.Glu140Lys | P01130 |
| HOMO SAPIENS | 7 | 14262 | 2fcw | B | 122 | CYS | p.Cys143Ter | P01130 |
| HOMO SAPIENS | 6 | 14262 | 2fcw | B | 134 | CYS | p.Cys155Ter | P01130 |
| HOMO SAPIENS | 7 | 14262 | 2fcw | B | 147 | ASP | p.Asp168Glu | P01130 |
| HOMO SAPIENS | 4 | 2624 | 3b7b | A | 855 | VAL | p.Val886Met | Q9H9B1 |
| HOMO SAPIENS | 7 | 2624 | 3b7b | A | 868 | ILE | p.Ile899Val | Q9H9B1 |
| HOMO SAPIENS | 0 | 2624 | 3b7b | A | 892 | GLU | p.Glu923Lys | Q9H9B1 |
| HOMO SAPIENS | 0 | 4791 | 5ctd | B | 68 | LEU | p.Pro549Ala | Q14055 |
| HOMO SAPIENS | 5 | 460 | 4ui1 | D | 80 | CYS | p.Cys80Arg | Q6ZVN8 |
| HOMO SAPIENS | 3 | 460 | 4ui1 | D | 101 | LEU | p.Leu101Pro | Q6ZVN8 |
| HOMO SAPIENS | 0 | 40782 | 1sy6 | A | 47 | LYS | p.Lys69Ter | P09693 |
| HOMO SAPIENS | -1 | 1092 | 2idx | A | 62 | THR | p.Thr62Met | Q96EY8 |
| HOMO SAPIENS | 6 | 1092 | 2idx | A | 183 | HIS | p.His183Leu | Q96EY8 |
| HOMO SAPIENS | 7 | 1092 | 2idx | A | 186 | ARG | p.Arg186Trp | Q96EY8 |
| HOMO SAPIENS | 6 | 1092 | 2idx | A | 190 | ARG | p.Arg190Cys | Q96EY8 |
| HOMO SAPIENS | 6 | 1092 | 2idx | A | 190 | ARG | p.Arg190His | Q96EY8 |
| HOMO SAPIENS | 7 | 1092 | 2idx | A | 191 | ARG | p.Arg191Trp | Q96EY8 |
| HOMO SAPIENS | 7 | 1092 | 2idx | A | 191 | ARG | p.Arg191Gln | Q96EY8 |
| HOMO SAPIENS | 6 | 1092 | 2idx | A | 195 | ARG | p.Arg195His | Q96EY8 |
| HOMO SAPIENS | 1 | 1092 | 2idx | A | 234 | GLN | p.Gln234Ter | Q96EY8 |
| HOMO SAPIENS | 0 | 1092 | 2idx | A | 239 | MET | p.Met239Lys | Q96EY8 |
| HOMO SAPIENS | -1 | 6418 | 5fur | G | 11 | THR | p.Thr11Ile | P21675 |
| HOMO SAPIENS | -1 | 6418 | 5fur | G | 290 | LEU | p.Leu269Val | P21675 |
| HOMO SAPIENS | 0 | 6100 | 1wiq | A | 240 | ARG | p.Arg265Trp | P01730 |
| HOMO SAPIENS | 0 | 13757 | 4ye6 | A | 45 | GLY | p.Gly45Val | P47897 |
| HOMO SAPIENS | 1 | 13757 | 4ye6 | A | 57 | TYR | p.Tyr57His | P47897 |
| HOMO SAPIENS | 8 | 13757 | 4ye6 | A | 403 | ARG | p.Arg403Trp | P47897 |
| HOMO SAPIENS | 7 | 13757 | 4ye6 | A | 524 | ARG | p.Arg524Trp | P47897 |
| HOMO SAPIENS | 6 | 13757 | 4ye6 | A | 690 | ARG | p.Arg690Cys | P47897 |
| HOMO SAPIENS | 0 | 26958 | 2lzl | A | 370 | GLY | p.Gly370Cys | P22607 |
| HOMO SAPIENS | 0 | 26958 | 2lzl | A | 371 | SER | p.Ser371Cys | P22607 |
| HOMO SAPIENS | 0 | 26958 | 2lzl | A | 373 | TYR | p.Tyr373Cys | P22607 |
| HOMO SAPIENS | 0 | 26958 | 2lzl | A | 380 | GLY | p.Gly380Arg | P22607 |
| HOMO SAPIENS | 0 | 26958 | 2lzl | A | 384 | PHE | p.Phe384Leu | P22607 |
| HOMO SAPIENS | 5 | 26958 | 2lzl | A | 391 | ALA | p.Ala391Glu | P22607 |
| HOMO SAPIENS | 6 | 11325 | 2wa6 | A | 121 | TYR | p.Tyr121Phe | O75369 |
| HOMO SAPIENS | 2 | 11325 | 2wa6 | A | 161 | PHE | p.Phe161Cys | O75369 |
| HOMO SAPIENS | 0 | 11325 | 2wa6 | A | 171 | LEU | p.Leu171Arg | O75369 |
| HOMO SAPIENS | 2 | 11325 | 2wa6 | A | 173 | ALA | p.Ala173Val | O75369 |
| HOMO SAPIENS | 0 | 11325 | 2wa6 | A | 202 | MET | p.Met202Val | O75369 |
| HOMO SAPIENS | 3 | 11325 | 2wa6 | A | 227 | GLU | p.Glu227Lys | O75369 |
| HOMO SAPIENS | 0 | 11325 | 2wa6 | A | 234 | LEU | p.Leu234Val | O75369 |
| HOMO SAPIENS | 2 | 11325 | 2wa6 | A | 235 | SER | p.Ser235Pro | O75369 |
| HOMO SAPIENS | 9 | 7240 | 3tgx | A | 137 | TYR | p.Tyr156Cys | Q9HBE5 |
| HOMO SAPIENS | 0 | 35479 | 4elj | A | 307 | THR | p.Thr307Ile | P06400 |
| HOMO SAPIENS | 0 | 35479 | 4elj | A | 310 | GLY | p.Gly310Glu | P06400 |
| HOMO SAPIENS | 0 | 35479 | 4elj | A | 377 | THR | p.Thr377Ser | P06400 |
| HOMO SAPIENS | 9 | 36706 | 1he7 | A | 359 | TYR | p.Tyr359Cys | P04629 |
| HOMO SAPIENS | 4 | 6538 | 4d0e | A | 504 | ARG | p.Arg504His | P46531 |
| HOMO SAPIENS | 1 | 26518 | 5fm5 | M | 600 | GLU | p.Glu600Val | P52179 |
| HOMO SAPIENS | -1 | 26518 | 5fm5 | M | 651 | ARG | p.Arg651Gln | P52179 |
| HOMO SAPIENS | -1 | 26518 | 5fm5 | M | 660 | PRO | p.Pro660Ala | P52179 |

| | | | | | | | | |
|---|---|---|---|---|---|---|---|---|
| HOMO SAPIENS | -1 | 26518 | 5fm5 | M | 688 | THR | p.Thr688Ser | P52179 |
| HOMO SAPIENS | -1 | 26518 | 5fm5 | M | 704 | GLU | p.Glu704Lys | P52179 |
| HOMO SAPIENS | -1 | 26518 | 5fm5 | M | 727 | THR | p.Thr727Ala | P52179 |
| HOMO SAPIENS | 2 | 27054 | 4rhp | B | 209 | LEU | p.Leu209Val | O75208 |
| HOMO SAPIENS | -1 | 27054 | 4rhp | B | 288 | LYS | p.Lys288Asn | O75208 |
| HOMO SAPIENS | 5 | 2384 | 5fqd | B | 419 | ARG | p.Arg419Ter | Q96SW2 |
| HOMO SAPIENS | 4 | 5154 | 2hl3 | A | 56 | LYS | p.Lys56Arg | Q14203 |
| HOMO SAPIENS | 0 | 5154 | 2hl3 | A | 59 | GLY | p.Gly59Ser | Q14203 |
| HOMO SAPIENS | 0 | 5154 | 2hl3 | A | 71 | GLY | p.Gly71Ala | Q14203 |
| HOMO SAPIENS | 0 | 5154 | 2hl3 | A | 71 | GLY | p.Gly71Arg | Q14203 |
| HOMO SAPIENS | 7 | 5154 | 2hl3 | A | 72 | THR | p.Thr72Pro | Q14203 |
| HOMO SAPIENS | 2 | 5154 | 2hl3 | A | 74 | GLN | p.Gln74Pro | Q14203 |
| HOMO SAPIENS | 2 | 20039 | 2l1g | A | 9 | GLY | p.Gly9Cys | Q9NVV9 |
| HOMO SAPIENS | 5 | 20039 | 2l1g | A | 81 | PHE | p.Phe81Leu | Q9NVV9 |
| HOMO SAPIENS | -1 | 97 | 5uz7 | A | 201 | ARG | p.Arg201His | P63092 |
| HOMO SAPIENS | -1 | 97 | 5uz7 | A | 201 | ARG | p.Arg201Cys | P63092 |
| HOMO SAPIENS | 0 | 3257 | 4b7w | B | 278 | GLU | p.Glu278Asp | P41235 |
| HOMO SAPIENS | 1 | 3257 | 4b7w | B | 331 | ARG | p.Arg331Cys | P41235 |
| HOMO SAPIENS | 0 | 2814 | 4qgk | B | 10 | GLN | p.Gln10Ter | P51648 |
| HOMO SAPIENS | 0 | 2814 | 4qgk | B | 177 | ARG | p.Arg177Ter | P51648 |
| HOMO SAPIENS | 9 | 2814 | 4qgk | B | 184 | THR | p.Thr184Met | P51648 |
| HOMO SAPIENS | 1 | 2814 | 4qgk | B | 188 | ALA | p.Ala188Val | P51648 |
| HOMO SAPIENS | 0 | 2814 | 4qgk | B | 214 | CYS | p.Cys214Tyr | P51648 |
| HOMO SAPIENS | 0 | 2814 | 4qgk | B | 245 | ASP | p.Asp245Asn | P51648 |
| HOMO SAPIENS | 0 | 2814 | 4qgk | B | 266 | LYS | p.Lys266Asn | P51648 |
| HOMO SAPIENS | 0 | 2814 | 4qgk | B | 315 | PRO | p.Pro315Ser | P51648 |
| HOMO SAPIENS | 0 | 2814 | 4qgk | B | 386 | ASN | p.Asn386Ser | P51648 |
| HOMO SAPIENS | -1 | 5486 | 5u6o | B | 47 | GLY | p.Gly47Val | O60741 |
| HOMO SAPIENS | 1 | 21475 | 3pdy | A | 679 | THR | p.Thr789Ile | Q15149 |
| HOMO SAPIENS | 0 | 32890 | 4djc | B | 1501 | LEU | p.Leu1501Val | Q14524 |
| HOMO SAPIENS | -1 | 32890 | 4djc | B | 1512 | ARG | p.Arg1512Trp | Q14524 |
| HOMO SAPIENS | 0 | 26683 | 2xze | B | 38 | ARG | p.Arg38Cys | O95630 |
| HOMO SAPIENS | 0 | 26683 | 2xze | B | 100 | PHE | p.Phe100Tyr | O95630 |
| HOMO SAPIENS | -1 | 23175 | 5dka | A | 163 | SER | p.Ser163Leu | Q96EQ8 |
| HOMO SAPIENS | -1 | 23175 | 5dka | A | 174 | ARG | p.Arg174Cys | Q96EQ8 |
| HOMO SAPIENS | 6 | 21177 | 3k1r | A | 155 | ARG | p.Arg155Ter | Q9Y6N9 |
| HOMO SAPIENS | 0 | 1237 | 2wwm | O | 40 | PRO | p.Pro40Arg | O75147 |
| HOMO SAPIENS | 1 | 2543 | 5ewl | B | 64 | VAL | p.Val64Leu | Q13224 |
| HOMO SAPIENS | 2 | 2543 | 5ewl | B | 271 | ALA | p.Ala271Glu | Q13224 |
| HOMO SAPIENS | 3 | 34262 | 2mng | A | 613 | PHE | p.Phe613Leu | Q9Y3Q4 |
| HOMO SAPIENS | 7 | 34262 | 2mng | A | 672 | SER | p.Ser672Arg | Q9Y3Q4 |
| HOMO SAPIENS | 4 | 1667 | 1f05 | A | 321 | LYS | p.Lys321Met | P37837 |
| HOMO SAPIENS | 0 | 25179 | 2knc | B | 752 | SER | p.Ser778Pro | P05106 |
| HOMO SAPIENS | 0 | 2185 | 2ig7 | A | 50 | TYR | p.Tyr50Cys | Q9Y259 |
| HOMO SAPIENS | 0 | 2185 | 2ig7 | A | 224 | ASN | p.Asn224His | Q9Y259 |
| HOMO SAPIENS | 0 | 2185 | 2ig7 | A | 270 | TYR | p.Tyr270Ter | Q9Y259 |
| HOMO SAPIENS | 0 | 2185 | 2ig7 | A | 328 | GLN | p.Gln328Arg | Q9Y259 |
| HOMO SAPIENS | 3 | 2201 | 2r0o | A | 179 | PRO | p.Pro179Leu | O43707 |
| HOMO SAPIENS | 3 | 2201 | 2r0o | A | 255 | GLU | p.Lys255Glu | O43707 |
| HOMO SAPIENS | 3 | 2201 | 2r0o | A | 259 | THR | p.Thr259Ile | O43707 |
| HOMO SAPIENS | 4 | 2201 | 2r0o | A | 262 | SER | p.Ser262Pro | O43707 |
| HOMO SAPIENS | -1 | 31221 | 2wvr | C | 44 | THR | p.Thr44Pro | Q9H211 |
| HOMO SAPIENS | -1 | 31221 | 2wvr | C | 135 | ALA | p.Ala135Val | Q9H211 |
| HOMO SAPIENS | -1 | 31221 | 2wvr | C | 456 | GLU | p.Glu456Ala | Q9H211 |

| | | | | | | | | |
|---|---|---|---|---|---|---|---|---|
| HOMO SAPIENS | -1 | 31221 | 2wvr | C | 462 | ARG | p.Arg462Gln | Q9H211 |
| HOMO SAPIENS | -1 | 31221 | 2wvr | C | 468 | GLU | p.Glu468Lys | Q9H211 |
| HOMO SAPIENS | -1 | 31221 | 2wvr | C | 537 | ALA | p.Ala537Val | Q9H211 |
| HOMO SAPIENS | 5 | 2576 | 5o3l | C | 363 | VAL | p.Val680Ile | P10636 |
| HOMO SAPIENS | 5 | 1345 | 1czt | A | 37 | ARG | p.Arg2102Cys | P12259 |
| HOMO SAPIENS | 1 | 1345 | 1czt | A | 83 | MET | p.Met2148Thr | P12259 |
| HOMO SAPIENS | 1 | 1345 | 1czt | A | 157 | ASP | p.Asp2222Gly | P12259 |
| HOMO SAPIENS | 8 | 35985 | 1lo1 | A | 110 | ALA | p.Ala110Val | O95718 |
| HOMO SAPIENS | 1 | 617 | 2ocp | B | 44 | GLU | p.Glu44Lys | Q16854 |
| HOMO SAPIENS | 5 | 617 | 2ocp | B | 46 | ASN | p.Asn46Ser | Q16854 |
| HOMO SAPIENS | 0 | 617 | 2ocp | B | 59 | THR | p.Thr59Met | Q16854 |
| HOMO SAPIENS | 7 | 617 | 2ocp | B | 142 | ARG | p.Arg142Lys | Q16854 |
| HOMO SAPIENS | 3 | 617 | 2ocp | B | 154 | ASN | p.Asn154Lys | Q16854 |
| HOMO SAPIENS | 3 | 617 | 2ocp | B | 170 | GLN | p.Gln170Arg | Q16854 |
| HOMO SAPIENS | 3 | 617 | 2ocp | B | 227 | GLU | p.Glu227Lys | Q16854 |
| HOMO SAPIENS | 3 | 617 | 2ocp | B | 233 | LYS | p.Lys233Glu | Q16854 |
| HOMO SAPIENS | 3 | 617 | 2ocp | B | 255 | ASP | p.Asp255Tyr | Q16854 |
| HOMO SAPIENS | -1 | 38975 | 5itz | D | 421 | GLN | p.Gln421His | Q9HC77 |
| HOMO SAPIENS | 2 | 6235 | 2f8v | A | 15 | VAL | p.Val15Ile | Q8WZ42 |
| HOMO SAPIENS | 0 | 6235 | 2f8v | A | 54 | VAL | p.Val54Met | Q8WZ42 |
| HOMO SAPIENS | 0 | 6235 | 2f8v | A | 59 | SER | p.Ser59Arg | Q8WZ42 |
| HOMO SAPIENS | 0 | 6235 | 2f8v | A | 60 | ASP | p.Asp60Tyr | Q8WZ42 |
| HOMO SAPIENS | 0 | 12630 | 1ext | B | 12 | ASP | p.Asp41Glu | P19438 |
| HOMO SAPIENS | 3 | 12630 | 1ext | B | 30 | CYS | p.Cys59Arg | P19438 |
| HOMO SAPIENS | 3 | 12630 | 1ext | B | 30 | CYS | p.Cys59Ser | P19438 |
| HOMO SAPIENS | 5 | 12630 | 1ext | B | 33 | CYS | p.Cys62Tyr | P19438 |
| HOMO SAPIENS | 5 | 12630 | 1ext | B | 33 | CYS | p.Cys62Gly | P19438 |
| HOMO SAPIENS | 1 | 12630 | 1ext | B | 46 | PRO | p.Pro75Leu | P19438 |
| HOMO SAPIENS | 3 | 12630 | 1ext | B | 50 | THR | p.Thr79Met | P19438 |
| HOMO SAPIENS | 6 | 12630 | 1ext | B | 52 | CYS | p.Cys81Phe | P19438 |
| HOMO SAPIENS | 6 | 12630 | 1ext | B | 70 | CYS | p.Cys99Ser | P19438 |
| HOMO SAPIENS | 5 | 12630 | 1ext | B | 88 | CYS | p.Cys117Arg | P19438 |
| HOMO SAPIENS | 5 | 12630 | 1ext | B | 88 | CYS | p.Cys117Tyr | P19438 |
| HOMO SAPIENS | -1 | 12630 | 1ext | B | 170 | ILE | p.Ile199Asn | P19438 |
| HOMO SAPIENS | 0 | 994 | 1aos | B | 95 | ARG | p.Arg95Cys | P04424 |
| HOMO SAPIENS | 0 | 994 | 1aos | B | 111 | ARG | p.Arg111Gln | P04424 |
| HOMO SAPIENS | 0 | 994 | 1aos | B | 131 | THR | p.Thr131Met | P04424 |
| HOMO SAPIENS | 6 | 994 | 1aos | B | 178 | VAL | p.Val178Met | P04424 |
| HOMO SAPIENS | 9 | 994 | 1aos | B | 182 | ARG | p.Arg182Gln | P04424 |
| HOMO SAPIENS | 9 | 994 | 1aos | B | 186 | ARG | p.Arg186Trp | P04424 |
| HOMO SAPIENS | 8 | 994 | 1aos | B | 193 | ARG | p.Arg193Gln | P04424 |
| HOMO SAPIENS | 5 | 994 | 1aos | B | 217 | ARG | p.Arg217Ter | P04424 |
| HOMO SAPIENS | 0 | 994 | 1aos | B | 286 | GLN | p.Gln286Arg | P04424 |
| HOMO SAPIENS | 5 | 994 | 1aos | B | 379 | ARG | p.Arg379Cys | P04424 |
| HOMO SAPIENS | 4 | 16066 | 5mv9 | A | 1719 | TYR | p.Tyr1719Cys | Q13402 |
| HOMO SAPIENS | -1 | 16066 | 5mv9 | A | 1739 | ARG | p.Arg1739Ter | Q13402 |
| HOMO SAPIENS | 4 | 16066 | 5mv9 | A | 1775 | ILE | p.Ile1775Thr | Q13402 |
| HOMO SAPIENS | 2 | 16066 | 5mv9 | A | 1786 | SER | p.Ser1786Thr | Q13402 |
| HOMO SAPIENS | 4 | 16066 | 5mv9 | A | 1798 | GLN | p.Gln1798Ter | Q13402 |
| HOMO SAPIENS | 0 | 16066 | 5mv9 | A | 1858 | LEU | p.Leu1858Pro | Q13402 |
| HOMO SAPIENS | 6 | 16066 | 5mv9 | A | 1873 | ARG | p.Arg1873Trp | Q13402 |
| HOMO SAPIENS | 6 | 16066 | 5mv9 | A | 1873 | ARG | p.Arg1873Leu | Q13402 |
| HOMO SAPIENS | 6 | 16066 | 5mv9 | A | 1883 | ARG | p.Arg1883Gln | Q13402 |
| HOMO SAPIENS | 4 | 16066 | 5mv9 | A | 1887 | PRO | p.Pro1887Leu | Q13402 |

| | | | | | | | | |
|---|---|---|---|---|---|---|---|---|
| HOMO SAPIENS | 5 | 16066 | 5mv9 | A | 1942 | GLY | p.Gly1942Ter | Q13402 |
| HOMO SAPIENS | 5 | 16066 | 5mv9 | A | 1954 | LEU | p.Leu1954Ile | Q13402 |
| HOMO SAPIENS | 5 | 16066 | 5mv9 | A | 1956 | VAL | p.Val1956Ile | Q13402 |
| HOMO SAPIENS | 5 | 16066 | 5mv9 | A | 1967 | ARG | p.Arg1967Ter | Q13402 |
| HOMO SAPIENS | 5 | 16066 | 5mv9 | A | 1990 | GLN | p.Gln1990Ter | Q13402 |
| HOMO SAPIENS | 2 | 16066 | 5mv9 | A | 2005 | LYS | p.Lys2005Glu | Q13402 |
| HOMO SAPIENS | 1 | 16066 | 5mv9 | A | 2072 | VAL | p.Val2072Ile | Q13402 |
| HOMO SAPIENS | 4 | 16066 | 5mv9 | A | 2109 | THR | p.Thr2109Ile | Q13402 |
| HOMO SAPIENS | 4 | 16066 | 5mv9 | A | 2142 | ASP | p.Asp2142Asn | Q13402 |
| HOMO SAPIENS | -1 | 16066 | 5mv9 | A | 2214 | GLY | p.Gly2214Ser | Q13402 |
| HOMO SAPIENS | 7 | 27942 | 1deb | A | 24 | ARG | p.Arg24Ter | P25054 |
| HOMO SAPIENS | 0 | 876 | 5hnr | B | 12 | PHE | p.Phe12Leu | P13716 |
| HOMO SAPIENS | 0 | 876 | 5hnr | B | 59 | LYS | p.Lys59Asn | P13716 |
| HOMO SAPIENS | -1 | 876 | 5hnr | B | 133 | GLY | p.Gly133Arg | P13716 |
| HOMO SAPIENS | 2 | 876 | 5hnr | B | 240 | ARG | p.Arg240Trp | P13716 |
| HOMO SAPIENS | 0 | 876 | 5hnr | B | 274 | ALA | p.Ala274Thr | P13716 |
| HOMO SAPIENS | 0 | 876 | 5hnr | B | 275 | VAL | p.Val275Met | P13716 |
| HOMO SAPIENS | 0 | 1142 | 3qe2 | A | 503 | VAL | p.Ala500Val | P16435 |
| HOMO SAPIENS | 8 | 14358 | 4dkd | C | 32 | VAL | p.Val32Gly | P07333 |
| HOMO SAPIENS | 0 | 14358 | 4dkd | C | 60 | LEU | p.Leu60Val | P07333 |
| HOMO SAPIENS | 7 | 14358 | 4dkd | C | 75 | THR | p.Thr75Ile | P07333 |
| HOMO SAPIENS | -1 | 14358 | 4dkd | C | 90 | PRO | p.Pro90Thr | P07333 |
| HOMO SAPIENS | 0 | 14358 | 4dkd | C | 106 | ARG | p.Arg106Trp | P07333 |
| HOMO SAPIENS | 0 | 14358 | 4dkd | C | 241 | ASN | p.Asn241Asp | P07333 |
| HOMO SAPIENS | 0 | 14358 | 4dkd | C | 245 | ALA | p.Ala245Ser | P07333 |
| HOMO SAPIENS | 0 | 14358 | 4dkd | C | 255 | ASN | p.Asn255Ile | P07333 |
| HOMO SAPIENS | 0 | 14358 | 4dkd | C | 268 | VAL | p.Val268Ile | P07333 |
| HOMO SAPIENS | 0 | 14358 | 4dkd | C | 279 | VAL | p.Val279Met | P07333 |
| HOMO SAPIENS | 0 | 14358 | 4dkd | C | 294 | ARG | p.Arg294Gln | P07333 |
| HOMO SAPIENS | 5 | 8968 | 3ppy | A | 1565 | VAL | p.Val1565Leu | P04275 |
| HOMO SAPIENS | 4 | 8968 | 3ppy | A | 1584 | TYR | p.Tyr1584Cys | P04275 |
| HOMO SAPIENS | 5 | 5341 | 3k3n | A | 279 | PHE | p.Phe315Ser | Q9UPP1 |
| HOMO SAPIENS | 0 | 29463 | 1q7l | A | 197 | ARG | p.Arg197Trp | Q03154 |
| HOMO SAPIENS | 0 | 33761 | 2leh | B | 30 | ASP | p.Asp30Asn | Q16637 |
| HOMO SAPIENS | 6 | 33761 | 2leh | B | 44 | ASP | p.Asp44Val | Q16637 |
| HOMO SAPIENS | -1 | 12249 | 2pnr | C | 224 | GLU | p.Glu310Lys | P10515 |
| HOMO SAPIENS | -1 | 12249 | 2pnr | C | 232 | VAL | p.Val318Ala | P10515 |
| HOMO SAPIENS | -1 | 7 | 5whk | B | -9 | ALA | p.Ala11Pro | P61769 |
| HOMO SAPIENS | 8 | 20298 | 4euw | A | 154 | PHE | p.Phe154Leu | P48436 |
| HOMO SAPIENS | 9 | 20298 | 4euw | A | 158 | ALA | p.Ala158Thr | P48436 |
| HOMO SAPIENS | 5 | 20298 | 4euw | A | 165 | HIS | p.His165Tyr | P48436 |
| HOMO SAPIENS | 3 | 20298 | 4euw | A | 169 | HIS | p.His169Gln | P48436 |
| HOMO SAPIENS | 0 | 20298 | 4euw | A | 173 | LYS | p.Lys173Glu | P48436 |
| HOMO SAPIENS | 1 | 16465 | 2jgx | A | 402 | TYR | p.Tyr402His | P08603 |
| HOMO SAPIENS | 0 | 8993 | 3v8o | A | 659 | LEU | p.Leu659Arg | Q14315 |
| HOMO SAPIENS | 6 | 8993 | 3v8o | A | 693 | ASP | p.Asp693Ala | Q14315 |
| HOMO SAPIENS | 0 | 8993 | 3v8o | A | 709 | ALA | p.Ala709Thr | Q14315 |
| HOMO SAPIENS | 2 | 3852 | 5es4 | B | 106 | ASP | p.Asp128Asn | P05107 |
| HOMO SAPIENS | 2 | 3852 | 5es4 | B | 116 | SER | p.Ser138Pro | P05107 |
| HOMO SAPIENS | 1 | 3852 | 5es4 | B | 127 | LEU | p.Leu149Pro | P05107 |
| HOMO SAPIENS | 0 | 3852 | 5es4 | B | 147 | GLY | p.Gly169Arg | P05107 |
| HOMO SAPIENS | 3 | 3852 | 5es4 | B | 156 | PRO | p.Pro178Leu | P05107 |
| HOMO SAPIENS | 0 | 3852 | 5es4 | B | 174 | LYS | p.Lys196Thr | P05107 |
| HOMO SAPIENS | 2 | 3852 | 5es4 | B | 251 | GLY | p.Gly273Arg | P05107 |

| | | | | | | | | |
|---|---|---|---|---|---|---|---|---|
| HOMO SAPIENS | 2 | 3852 | 5es4 | B | 329 | ASN | p.Asn351Ser | P05107 |
| HOMO SAPIENS | 2 | 3852 | 5es4 | B | 571 | ARG | p.Arg593Cys | P05107 |
| HOMO SAPIENS | 7 | 5962 | 5fn2 | B | 146 | MET | p.Met146Leu | P49768 |
| HOMO SAPIENS | 3 | 5962 | 5fn2 | B | 166 | LEU | p.Leu166Pro | P49768 |
| HOMO SAPIENS | 1 | 5962 | 5fn2 | B | 233 | MET | p.Met233Leu | P49768 |
| HOMO SAPIENS | 1 | 5962 | 5fn2 | B | 261 | VAL | p.Val261Gly | P49768 |
| HOMO SAPIENS | 1 | 5962 | 5fn2 | B | 266 | GLY | p.Gly266Ser | P49768 |
| HOMO SAPIENS | -1 | 5962 | 5fn2 | B | 333 | ASP | p.Asp333Gly | P49768 |
| HOMO SAPIENS | 6 | 1918 | 3dlx | A | 58 | THR | p.Thr58Met | P55809 |
| HOMO SAPIENS | 5 | 1918 | 3dlx | A | 133 | VAL | p.Val133Glu | P55809 |
| HOMO SAPIENS | 5 | 1918 | 3dlx | A | 219 | GLY | p.Gly219Glu | P55809 |
| HOMO SAPIENS | 0 | 1918 | 3dlx | A | 221 | VAL | p.Val221Met | P55809 |
| HOMO SAPIENS | 5 | 1918 | 3dlx | A | 283 | SER | p.Ser283Ter | P55809 |
| HOMO SAPIENS | 4 | 1918 | 3dlx | A | 324 | GLY | p.Gly324Glu | P55809 |
| HOMO SAPIENS | 0 | 1918 | 3dlx | A | 456 | CYS | p.Cys456Phe | P55809 |
| HOMO SAPIENS | 0 | 9010 | 2wfr | A | 162 | LEU | p.Leu162Pro | O43323 |
| HOMO SAPIENS | 5 | 3770 | 5n38 | A | 56 | VAL | p.Val56Glu | O60260 |
| HOMO SAPIENS | 1 | 3770 | 5n38 | A | 161 | LYS | p.Lys161Asn | O60260 |
| HOMO SAPIENS | 1 | 3770 | 5n38 | A | 167 | SER | p.Ser167Asn | O60260 |
| HOMO SAPIENS | 1 | 3770 | 5n38 | A | 211 | LYS | p.Lys211Asn | O60260 |
| HOMO SAPIENS | 1 | 3770 | 5n38 | A | 212 | CYS | p.Cys212Tyr | O60260 |
| HOMO SAPIENS | 1 | 3770 | 5n38 | A | 275 | ARG | p.Arg275Trp | O60260 |
| HOMO SAPIENS | 2 | 3770 | 5n38 | A | 334 | ARG | p.Arg334Cys | O60260 |
| HOMO SAPIENS | 1 | 3770 | 5n38 | A | 366 | ARG | p.Arg366Trp | O60260 |
| HOMO SAPIENS | -1 | 3770 | 5n38 | A | 380 | VAL | p.Val380Leu | O60260 |
| HOMO SAPIENS | 1 | 3770 | 5n38 | A | 394 | ASP | p.Asp394Asn | O60260 |
| HOMO SAPIENS | 1 | 3770 | 5n38 | A | 430 | GLY | p.Gly430Asp | O60260 |
| HOMO SAPIENS | -1 | 2467 | 4xx1 | A | 10 | PRO | p.Pro34Leu | P04180 |
| HOMO SAPIENS | 4 | 2467 | 4xx1 | A | 83 | TYR | p.Tyr107Ter | P04180 |
| HOMO SAPIENS | 4 | 2467 | 4xx1 | A | 93 | ALA | p.Ala117Thr | P04180 |
| HOMO SAPIENS | 4 | 2467 | 4xx1 | A | 123 | THR | p.Thr147Ile | P04180 |
| HOMO SAPIENS | 4 | 2467 | 4xx1 | A | 135 | ARG | p.Arg159Trp | P04180 |
| HOMO SAPIENS | 3 | 2467 | 4xx1 | A | 158 | ARG | p.Arg182Cys | P04180 |
| HOMO SAPIENS | 0 | 2467 | 4xx1 | A | 209 | LEU | p.Leu233Pro | P04180 |
| HOMO SAPIENS | 1 | 2467 | 4xx1 | A | 228 | ASN | p.Asn252Lys | P04180 |
| HOMO SAPIENS | 4 | 2467 | 4xx1 | A | 252 | MET | p.Met276Lys | P04180 |
| HOMO SAPIENS | 3 | 2467 | 4xx1 | A | 293 | MET | p.Met317Ile | P04180 |
| HOMO SAPIENS | 5 | 2467 | 4xx1 | A | 321 | THR | p.Thr345Met | P04180 |
| HOMO SAPIENS | 9 | 2467 | 4xx1 | A | 347 | THR | p.Thr371Met | P04180 |
| HOMO SAPIENS | 1 | 5834 | 3qrf | G | 371 | PHE | p.Phe371Cys | Q9BZS1 |
| HOMO SAPIENS | 1 | 5834 | 3qrf | G | 373 | PHE | p.Phe373Ala | Q9BZS1 |
| HOMO SAPIENS | 9 | 5834 | 3qrf | G | 384 | ALA | p.Ala384Thr | Q9BZS1 |
| HOMO SAPIENS | 6 | 5834 | 3qrf | G | 397 | ARG | p.Arg397Trp | Q9BZS1 |
| HOMO SAPIENS | -1 | 11869 | 4km8 | A | 15 | PRO | p.Pro15Leu | Q9UMX1 |
| HOMO SAPIENS | 5 | 262 | 4kbo | A | 126 | ARG | p.Arg126Trp | P38646 |
| HOMO SAPIENS | 3 | 262 | 4kbo | A | 128 | TYR | p.Tyr128Cys | P38646 |
| HOMO SAPIENS | -1 | 11347 | 4uf9 | D | 117 | ALA | p.Ala117Val | Q13501 |
| HOMO SAPIENS | 0 | 2173 | 4ah6 | A | 45 | SER | p.Ser45Gly | Q6PI48 |
| HOMO SAPIENS | 2 | 2173 | 4ah6 | A | 152 | CYS | p.Cys152Phe | Q6PI48 |
| HOMO SAPIENS | 4 | 2173 | 4ah6 | A | 179 | ARG | p.Arg179His | Q6PI48 |
| HOMO SAPIENS | 0 | 2173 | 4ah6 | A | 196 | LYS | p.Lys196Arg | Q6PI48 |
| HOMO SAPIENS | 8 | 2173 | 4ah6 | A | 263 | ARG | p.Arg263Gln | Q6PI48 |
| HOMO SAPIENS | 1 | 2173 | 4ah6 | A | 338 | GLY | p.Gly338Glu | Q6PI48 |
| HOMO SAPIENS | 0 | 2173 | 4ah6 | A | 425 | GLU | p.Glu425Ter | Q6PI48 |

| | | | | | | | | |
|---|---|---|---|---|---|---|---|---|
| HOMO SAPIENS | 1 | 2173 | 4ah6 | A | 548 | LEU | p.Leu548Met | Q6PI48 |
| HOMO SAPIENS | 0 | 2173 | 4ah6 | A | 609 | ARG | p.Arg609Trp | Q6PI48 |
| HOMO SAPIENS | 2 | 2173 | 4ah6 | A | 613 | LEU | p.Leu613Phe | Q6PI48 |
| HOMO SAPIENS | 0 | 2173 | 4ah6 | A | 626 | LEU | p.Leu626Gln | Q6PI48 |
| HOMO SAPIENS | 0 | 2173 | 4ah6 | A | 626 | LEU | p.Leu626Val | Q6PI48 |
| HOMO SAPIENS | 7 | 3203 | 2cn1 | A | 87 | ASP | p.Asp137Val | Q9H0P0 |
| HOMO SAPIENS | 8 | 3203 | 2cn1 | A | 166 | GLN | p.Gln216Ter | Q9H0P0 |
| HOMO SAPIENS | 8 | 3203 | 2cn1 | A | 179 | ASN | p.Asn229Ser | Q9H0P0 |
| HOMO SAPIENS | 7 | 3203 | 2cn1 | A | 230 | GLY | p.Gly280Arg | Q9H0P0 |
| HOMO SAPIENS | 2 | 3424 | 2a1t | R | 116 | GLY | p.Gly116Arg | P13804 |
| HOMO SAPIENS | 2 | 3424 | 2a1t | R | 157 | VAL | p.Val157Gly | P13804 |
| HOMO SAPIENS | 2 | 3424 | 2a1t | R | 171 | THR | p.Thr171Ile | P13804 |
| HOMO SAPIENS | 0 | 3424 | 2a1t | R | 178 | THR | p.Thr178Arg | P13804 |
| HOMO SAPIENS | 0 | 3424 | 2a1t | R | 194 | VAL | p.Val194Met | P13804 |
| HOMO SAPIENS | 1 | 3424 | 2a1t | R | 209 | ARG | p.Arg209Ter | P13804 |
| HOMO SAPIENS | 6 | 3424 | 2a1t | R | 266 | THR | p.Thr266Met | P13804 |
| HOMO SAPIENS | 0 | 27598 | 1g1c | A | 26 | ARG | p.Arg2098Gly | Q8WZ42 |
| HOMO SAPIENS | 1 | 27598 | 1g1c | A | 46 | ILE | p.Ile2118Thr | Q8WZ42 |
| HOMO SAPIENS | 0 | 32885 | 4bsk | A | 149 | ASN | p.Asn149Asp | P35916 |
| HOMO SAPIENS | -1 | 8074 | 3ghg | B | 17 | ARG | p.Arg47Ter | P02675 |
| HOMO SAPIENS | -1 | 447 | 5g4p | A | 190 | ALA | p.Ala190Thr | P84074 |
| HOMO SAPIENS | 0 | 10761 | 5u1d | B | 379 | VAL | p.Val379Ile | Q03519 |
| HOMO SAPIENS | 0 | 10761 | 5u1d | B | 665 | THR | p.Thr665Ala | Q03519 |
| HOMO SAPIENS | 5 | 29251 | 5fn7 | A | 79 | ILE | p.Ile294Leu | P08575 |
| HOMO SAPIENS | -1 | 27727 | 4ky9 | A | 56 | LYS | p.Lys56Glu | P02730 |
| HOMO SAPIENS | 2 | 27727 | 4ky9 | A | 72 | GLU | p.Glu72Asp | P02730 |
| HOMO SAPIENS | 2 | 27727 | 4ky9 | A | 90 | GLU | p.Glu90Lys | P02730 |
| HOMO SAPIENS | 1 | 27727 | 4ky9 | A | 130 | GLY | p.Gly130Arg | P02730 |
| HOMO SAPIENS | 2 | 27727 | 4ky9 | A | 150 | ARG | p.Arg150Ter | P02730 |
| HOMO SAPIENS | 0 | 27727 | 4ky9 | A | 327 | PRO | p.Pro327Arg | P02730 |
| HOMO SAPIENS | 2 | 13202 | 2fxo | B | 870 | ARG | p.Arg870His | P12883 |
| HOMO SAPIENS | 2 | 13202 | 2fxo | B | 870 | ARG | p.Arg870Cys | P12883 |
| HOMO SAPIENS | 4 | 13202 | 2fxo | B | 904 | ARG | p.Arg904Cys | P12883 |
| HOMO SAPIENS | 4 | 13202 | 2fxo | B | 906 | ASP | p.Asp906Gly | P12883 |
| HOMO SAPIENS | 2 | 13202 | 2fxo | B | 908 | LEU | p.Leu908Val | P12883 |
| HOMO SAPIENS | 2 | 13202 | 2fxo | B | 923 | ASN | p.Asn923Lys | P12883 |
| HOMO SAPIENS | 7 | 13202 | 2fxo | B | 935 | GLU | p.Glu935Lys | P12883 |
| HOMO SAPIENS | 7 | 13202 | 2fxo | B | 935 | GLU | p.Glu935Val | P12883 |
| HOMO SAPIENS | 8 | 13202 | 2fxo | B | 949 | GLU | p.Glu949Lys | P12883 |
| HOMO SAPIENS | 0 | 13200 | 2a1s | A | 349 | ARG | p.Arg349Trp | O95453 |
| HOMO SAPIENS | 0 | 13200 | 2a1s | A | 421 | LYS | p.Lys421Arg | O95453 |
| HOMO SAPIENS | 6 | 3196 | 5xtb | T | 115 | CYS | p.Cys115Tyr | O75380 |
| HOMO SAPIENS | 0 | 1108 | 5ukf | A | 89 | ARG | p.Arg89Gln | Q99986 |
| HOMO SAPIENS | 0 | 1108 | 5ukf | A | 119 | HIS | p.His119Arg | Q99986 |
| HOMO SAPIENS | 0 | 1108 | 5ukf | A | 133 | ARG | p.Arg133Cys | Q99986 |
| HOMO SAPIENS | 0 | 1108 | 5ukf | A | 236 | VAL | p.Val236Leu | Q99986 |
| HOMO SAPIENS | 0 | 1108 | 5ukf | A | 321 | ARG | p.Arg321Cys | Q99986 |
| HOMO SAPIENS | -1 | 1108 | 5ukf | A | 358 | ARG | p.Arg358Ter | Q99986 |
| HOMO SAPIENS | 0 | 5842 | 4yli | C | 204 | GLY | p.Gly204Ser | Q9BWP8 |
| HOMO SAPIENS | -1 | 31362 | 4wqo | A | 25 | PRO | p.Pro25Leu | P40337 |
| HOMO SAPIENS | 2 | 10107 | 2j8h | A | 34 | VAL | p.Val31886Ile | Q8WZ42 |
| HOMO SAPIENS | 8 | 20190 | 1wm5 | A | 77 | ARG | p.Arg77Gln | P19878 |
| HOMO SAPIENS | 1 | 20190 | 1wm5 | A | 100 | GLN | p.Gln100Ter | P19878 |
| HOMO SAPIENS | 0 | 20190 | 1wm5 | A | 102 | ARG | p.Arg102Ter | P19878 |

| | | | | | | | | | |
|---|---|---|---|---|---|---|---|---|---|
| HOMO SAPIENS | 0 | 20190 | 1wm5 | A | | 128 | ALA | p.Ala128Val | P19878 |
| HOMO SAPIENS | 0 | 20190 | 1wm5 | A | | 181 | LYS | p.Lys181Arg | P19878 |
| HOMO SAPIENS | 7 | 1278 | 5a2q | a | | 33 | ASP | p.Asp33Asn | P62854 |
| HOMO SAPIENS | 3 | 19515 | 2obd | A | | 11 | ILE | p.Ile28Met | P11597 |
| HOMO SAPIENS | 3 | 19515 | 2obd | A | | 89 | VAL | p.Val106Met | P11597 |
| HOMO SAPIENS | 3 | 19515 | 2obd | A | | 137 | ARG | p.Arg154Trp | P11597 |
| HOMO SAPIENS | 3 | 19515 | 2obd | A | | 139 | ASP | p.Asp156Asn | P11597 |
| HOMO SAPIENS | 3 | 19515 | 2obd | A | | 204 | SER | p.Ser221Arg | P11597 |
| HOMO SAPIENS | 0 | 19515 | 2obd | A | | 297 | GLU | p.Glu314Lys | P11597 |
| HOMO SAPIENS | 0 | 19515 | 2obd | A | | 314 | GLY | p.Gly331Ser | P11597 |
| HOMO SAPIENS | 0 | 19515 | 2obd | A | | 332 | SER | p.Ser349Tyr | P11597 |
| HOMO SAPIENS | 0 | 19515 | 2obd | A | | 368 | VAL | p.Val385Met | P11597 |
| HOMO SAPIENS | 0 | 19515 | 2obd | A | | 373 | ALA | p.Ala390Pro | P11597 |
| HOMO SAPIENS | 3 | 19515 | 2obd | A | | 405 | ILE | p.Val422Ile | P11597 |
| HOMO SAPIENS | 0 | 19515 | 2obd | A | | 451 | ARG | p.Arg468Gln | P11597 |
| HOMO SAPIENS | 0 | 19515 | 2obd | A | | 451 | ARG | p.Arg468Ter | P11597 |
| HOMO SAPIENS | 0 | 6626 | 3hop | A | | 39 | ALA | p.Ala39Gly | P21333 |
| HOMO SAPIENS | 1 | 6626 | 3hop | A | | 82 | GLU | p.Glu82Val | P21333 |
| HOMO SAPIENS | 1 | 6626 | 3hop | A | | 128 | ALA | p.Ala128Val | P21333 |
| HOMO SAPIENS | 2 | 6626 | 3hop | A | | 203 | ASP | p.Asp203Tyr | P21333 |
| HOMO SAPIENS | 1 | 6626 | 3hop | A | | 207 | PRO | p.Pro207Leu | P21333 |
| HOMO SAPIENS | 2 | 6626 | 3hop | A | | 210 | CYS | p.Cys210Phe | P21333 |
| HOMO SAPIENS | 7 | 6626 | 3hop | A | | 254 | GLU | p.Glu254Lys | P21333 |
| HOMO SAPIENS | 4 | 4210 | 2avg | A | | 8 | VAL | p.Val158Met | Q14896 |
| HOMO SAPIENS | 6 | 4210 | 2avg | A | | 15 | GLU | p.Glu165Asp | Q14896 |
| HOMO SAPIENS | 4 | 4210 | 2avg | A | | 27 | ARG | p.Arg177His | Q14896 |
| HOMO SAPIENS | 0 | 4210 | 2avg | A | | 28 | VAL | p.Val178Met | Q14896 |
| HOMO SAPIENS | 6 | 4210 | 2avg | A | | 39 | VAL | p.Val189Ile | Q14896 |
| HOMO SAPIENS | 2 | 4210 | 2avg | A | | 52 | LYS | p.Lys202Gln | Q14896 |
| HOMO SAPIENS | 4 | 4210 | 2avg | A | | 67 | SER | p.Ser217Gly | Q14896 |
| HOMO SAPIENS | 6 | 4210 | 2avg | A | | 69 | VAL | p.Val219Leu | Q14896 |
| HOMO SAPIENS | 2 | 4210 | 2avg | A | | 86 | GLY | p.Ser236Gly | Q14896 |
| HOMO SAPIENS | 4 | 4210 | 2avg | A | | 108 | GLU | p.Glu258Lys | Q14896 |
| HOMO SAPIENS | 0 | 22237 | 1w6k | A | | 588 | GLY | p.Gly588Ser | P48449 |
| HOMO SAPIENS | 0 | 10765 | 4a5w | B | | 98 | GLU | p.Glu98Ala | P13671 |
| HOMO SAPIENS | 0 | 10765 | 4a5w | B | | 596 | ARG | p.Arg596Ter | P13671 |
| HOMO SAPIENS | -1 | 15515 | 3hx3 | A | | 11 | MSE | p.Met10Lys | P12271 |
| HOMO SAPIENS | 8 | 15515 | 3hx3 | A | | 64 | ARG | p.Arg64Gln | P12271 |
| HOMO SAPIENS | 0 | 15515 | 3hx3 | A | | 182 | PHE | p.Phe182Cys | P12271 |
| HOMO SAPIENS | 0 | 15515 | 3hx3 | A | | 216 | ARG | p.Arg216Gln | P12271 |
| HOMO SAPIENS | -1 | 15515 | 3hx3 | A | | 226 | MSE | p.Met226Lys | P12271 |
| HOMO SAPIENS | 0 | 15515 | 3hx3 | A | | 234 | TRP | p.Arg234Trp | P12271 |
| HOMO SAPIENS | 0 | 15515 | 3hx3 | A | | 273 | LEU | p.Leu273Phe | P12271 |
| HOMO SAPIENS | 0 | 15515 | 3hx3 | A | | 292 | THR | p.Thr292Met | P12271 |
| HOMO SAPIENS | 0 | 5152 | 2www | D | | 95 | GLN | p.Gln95Ter | Q8IVH4 |
| HOMO SAPIENS | 0 | 5152 | 2www | D | | 133 | GLN | p.Gln133Ter | Q8IVH4 |
| HOMO SAPIENS | 0 | 5152 | 2www | D | | 145 | ARG | p.Arg145Ter | Q8IVH4 |
| HOMO SAPIENS | 0 | 5152 | 2www | D | | 207 | TYR | p.Tyr207Cys | Q8IVH4 |
| HOMO SAPIENS | 8 | 5152 | 2www | D | | 330 | ARG | p.Arg330Ter | Q8IVH4 |
| HOMO SAPIENS | 0 | 5152 | 2www | D | | 363 | GLN | p.Gln363His | Q8IVH4 |
| HOMO SAPIENS | 1 | 1626 | 1ilr | | 1 | 105 | SER | p.Ser130Arg | P18510 |
| HOMO SAPIENS | -1 | 929 | 5loh | A | | 12 | GLY | p.Gly12Arg | Q96GX5 |
| HOMO SAPIENS | 9 | 3206 | 1x63 | A | | 21 | CYS | p.Cys104Arg | Q13642 |
| HOMO SAPIENS | 8 | 3206 | 1x63 | A | | 39 | TRP | p.Trp122Ser | Q13642 |

| | | | | | | | | |
|---|---|---|---|---|---|---|---|---|
| HOMO SAPIENS | 9 | 3206 | 1x63 | A | 40 | HIS | p.His123Tyr | Q13642 |
| HOMO SAPIENS | 9 | 3206 | 1x63 | A | 40 | HIS | p.His123Leu | Q13642 |
| HOMO SAPIENS | 9 | 3206 | 1x63 | A | 40 | HIS | p.His123Gln | Q13642 |
| HOMO SAPIENS | 3 | 3206 | 1x63 | A | 49 | CYS | p.Cys132Phe | Q13642 |
| HOMO SAPIENS | 3 | 3206 | 1x63 | A | 67 | CYS | p.Cys150Tyr | Q13642 |
| HOMO SAPIENS | 3 | 3206 | 1x63 | A | 70 | CYS | p.Cys153Arg | Q13642 |
| HOMO SAPIENS | 3 | 3206 | 1x63 | A | 70 | CYS | p.Cys153Tyr | Q13642 |
| HOMO SAPIENS | 5 | 1052 | 3ecs | E | 208 | ASN | p.Asn208Tyr | Q14232 |
| HOMO SAPIENS | 2 | 1052 | 3ecs | E | 275 | TYR | p.Tyr275Cys | Q14232 |
| HOMO SAPIENS | 0 | 24253 | 5i35 | A | 271 | ARG | p.Arg271Cys | Q8NI60 |
| HOMO SAPIENS | 0 | 24253 | 5i35 | A | 272 | GLY | p.Gly272Asp | Q8NI60 |
| HOMO SAPIENS | 0 | 24253 | 5i35 | A | 299 | ARG | p.Arg299Trp | Q8NI60 |
| HOMO SAPIENS | 1 | 24253 | 5i35 | A | 339 | ALA | p.Ala339Thr | Q8NI60 |
| HOMO SAPIENS | 0 | 24253 | 5i35 | A | 348 | ARG | p.Arg348Ter | Q8NI60 |
| HOMO SAPIENS | 0 | 24253 | 5i35 | A | 395 | ILE | p.Ile395Met | Q8NI60 |
| HOMO SAPIENS | 0 | 24253 | 5i35 | A | 396 | ASP | p.Asp396Glu | Q8NI60 |
| HOMO SAPIENS | 0 | 24253 | 5i35 | A | 480 | PHE | p.Phe480Leu | Q8NI60 |
| HOMO SAPIENS | 2 | 24253 | 5i35 | A | 511 | THR | p.Thr511Met | Q8NI60 |
| HOMO SAPIENS | 0 | 24253 | 5i35 | A | 514 | TYR | p.Tyr514Cys | Q8NI60 |
| HOMO SAPIENS | 0 | 24253 | 5i35 | A | 538 | ARG | p.Arg538Gln | Q8NI60 |
| HOMO SAPIENS | 0 | 24253 | 5i35 | A | 549 | GLY | p.Gly549Ser | Q8NI60 |
| HOMO SAPIENS | 0 | 24253 | 5i35 | A | 551 | GLU | p.Glu551Lys | Q8NI60 |
| HOMO SAPIENS | -1 | 24253 | 5i35 | A | 555 | MSE | p.Met555Ile | Q8NI60 |
| HOMO SAPIENS | 0 | 24253 | 5i35 | A | 568 | GLU | p.Glu568Ter | Q8NI60 |
| HOMO SAPIENS | 0 | 24253 | 5i35 | A | 602 | PRO | p.Pro602Arg | Q8NI60 |
| HOMO SAPIENS | -1 | 15505 | 4goj | C | 22 | GLY | p.Gly22Val | Q13432 |
| HOMO SAPIENS | 5 | 23902 | 2lrr | A | 765 | HIS | p.His765Gln | P38935 |
| HOMO SAPIENS | 1 | 29650 | 2vj2 | A | 234 | CYS | p.Cys234Tyr | P78504 |
| HOMO SAPIENS | 0 | 29650 | 2vj2 | A | 255 | TYR | p.Tyr255Ter | P78504 |
| HOMO SAPIENS | 3 | 29650 | 2vj2 | A | 274 | GLY | p.Gly274Asp | P78504 |
| HOMO SAPIENS | 1 | 105 | 5he0 | B | 77 | GLY | p.Gly77Ser | P62873 |
| HOMO SAPIENS | 2 | 105 | 5he0 | B | 80 | ILE | p.Ile80Thr | P62873 |
| HOMO SAPIENS | 3 | 4587 | 3rgh | A | 159 | ASP | p.Asp1159Ala | P21333 |
| HOMO SAPIENS | 3 | 4587 | 3rgh | A | 184 | ASP | p.Asp1184Glu | P21333 |
| HOMO SAPIENS | 2 | 4587 | 3rgh | A | 186 | SER | p.Ser1186Leu | P21333 |
| HOMO SAPIENS | 4 | 4587 | 3rgh | A | 188 | ALA | p.Ala1188Thr | P21333 |
| HOMO SAPIENS | 2 | 4587 | 3rgh | A | 199 | SER | p.Ser1199Leu | P21333 |
| HOMO SAPIENS | 0 | 20283 | 4n3w | A | 1278 | GLU | p.Glu1278Lys | Q92793 |
| HOMO SAPIENS | 1 | 14294 | 2d9q | B | 208 | MET | p.Met231Thr | Q99062 |
| HOMO SAPIENS | 6 | 14294 | 2d9q | B | 297 | ASP | p.Asp320Asn | Q99062 |
| HOMO SAPIENS | 0 | 11642 | 3bpt | A | 66 | ARG | p.Arg66Trp | Q6NVY1 |
| HOMO SAPIENS | 0 | 11642 | 3bpt | A | 122 | TYR | p.Tyr122Cys | Q6NVY1 |
| HOMO SAPIENS | 0 | 11642 | 3bpt | A | 345 | GLY | p.Gly345Ser | Q6NVY1 |
| HOMO SAPIENS | 1 | 3942 | 3sby | A | 60 | ASP | p.Asp60His | Q9Y4U1 |
| HOMO SAPIENS | 0 | 3942 | 3sby | A | 111 | ARG | p.Arg111Ter | Q9Y4U1 |
| HOMO SAPIENS | 1 | 3942 | 3sby | A | 116 | LEU | p.Leu116Pro | Q9Y4U1 |
| HOMO SAPIENS | 1 | 3942 | 3sby | A | 130 | TYR | p.Tyr130Cys | Q9Y4U1 |
| HOMO SAPIENS | 1 | 3942 | 3sby | A | 132 | ARG | p.Arg132Ter | Q9Y4U1 |
| HOMO SAPIENS | 1 | 3942 | 3sby | A | 147 | GLY | p.Gly147Ala | Q9Y4U1 |
| HOMO SAPIENS | 1 | 3942 | 3sby | A | 161 | ARG | p.Arg161Gln | Q9Y4U1 |
| HOMO SAPIENS | 1 | 3942 | 3sby | A | 161 | ARG | p.Arg161Ter | Q9Y4U1 |
| HOMO SAPIENS | 1 | 3942 | 3sby | A | 203 | TRP | p.Trp203Ter | Q9Y4U1 |
| HOMO SAPIENS | 1 | 3942 | 3sby | A | 203 | TRP | p.Trp203Ter | Q9Y4U1 |
| HOMO SAPIENS | 1 | 3942 | 3sby | A | 205 | TYR | p.Tyr205Ter | Q9Y4U1 |

| | | | | | | | | | |
|---|---|---|---|---|---|---|---|---|---|
| HOMO SAPIENS | 0 | 38340 | 2mbh | B | | 39 | MET | p.Met39Leu | Q13351 |
| HOMO SAPIENS | 2 | 7360 | 4k30 | A | | 430 | LEU | p.Leu430Pro | Q8N159 |
| HOMO SAPIENS | 2 | 7360 | 4k30 | A | | 433 | GLU | p.Glu433Asp | Q8N159 |
| HOMO SAPIENS | 7 | 7360 | 4k30 | A | | 484 | TRP | p.Trp484Arg | Q8N159 |
| HOMO SAPIENS | 1 | 41903 | 2xs6 | A | | 151 | PRO | p.Pro151Ser | O00459 |
| HOMO SAPIENS | 0 | 172 | 5glh | A | | 183 | ALA | p.Ala183Gly | P24530 |
| HOMO SAPIENS | 0 | 172 | 5glh | A | | 201 | ARG | p.Arg201Ter | P24530 |
| HOMO SAPIENS | 0 | 172 | 5glh | A | | 244 | THR | p.Thr244Met | P24530 |
| HOMO SAPIENS | 0 | 172 | 5glh | A | | 253 | ARG | p.Arg253Ter | P24530 |
| HOMO SAPIENS | 9 | 172 | 5glh | A | | 1017 | ILE | p.Leu17Phe | P00720 |
| HOMO SAPIENS | -1 | 3342 | 1ljm | A | | 1058 | HIS | p.His58Asn | Q01196 |
| HOMO SAPIENS | 4 | 3342 | 1ljm | A | | 1174 | ARG | p.Arg174Gln | Q01196 |
| HOMO SAPIENS | 5 | 25607 | 1p22 | C | | 32 | ASP | p.Asp32Ala | P35222 |
| HOMO SAPIENS | 5 | 25607 | 1p22 | C | | 32 | ASP | p.Asp32Tyr | P35222 |
| HOMO SAPIENS | -1 | 25607 | 1p22 | C | | 33 | SEP | p.Ser33Phe | P35222 |
| HOMO SAPIENS | 5 | 25607 | 1p22 | C | | 34 | GLY | p.Gly34Glu | P35222 |
| HOMO SAPIENS | -1 | 25607 | 1p22 | C | | 37 | SEP | p.Ser37Cys | P35222 |
| HOMO SAPIENS | -1 | 25607 | 1p22 | C | | 41 | THR | p.Thr41Ile | P35222 |
| HOMO SAPIENS | 1 | 48 | 4uow | | 9 | 60 | GLN | p.Arg34111His | Q8WZ42 |
| HOMO SAPIENS | 0 | 48 | 4uow | | 9 | 64 | LEU | p.Val34115Ala | Q8WZ42 |
| HOMO SAPIENS | 0 | 1127 | 5c01 | A | | 638 | ARG | p.Arg638Ter | P29597 |
| HOMO SAPIENS | 0 | 1127 | 5c01 | A | | 684 | ILE | p.Ile684Ser | P29597 |
| HOMO SAPIENS | 0 | 1127 | 5c01 | A | | 703 | ARG | p.Arg703Trp | P29597 |
| HOMO SAPIENS | 5 | 16389 | 2aq0 | B | | 128 | ALA | p.Ala860Asp | Q92889 |
| HOMO SAPIENS | 2 | 16389 | 2aq0 | B | | 141 | ILE | p.Ile873Val | Q92889 |
| HOMO SAPIENS | 5 | 16389 | 2aq0 | B | | 143 | GLU | p.Glu875Gly | Q92889 |
| HOMO SAPIENS | 0 | 16389 | 2aq0 | B | | 180 | GLY | p.Gly912Arg | Q92889 |
| HOMO SAPIENS | 0 | 15816 | 2xrc | A | | 89 | THR | p.Thr107Ala | P05156 |
| HOMO SAPIENS | 0 | 15816 | 2xrc | A | | 133 | ASN | p.Asn151Ser | P05156 |
| HOMO SAPIENS | 0 | 15816 | 2xrc | A | | 185 | THR | p.Thr203Ile | P05156 |
| HOMO SAPIENS | 0 | 15816 | 2xrc | A | | 225 | GLY | p.Gly243Asp | P05156 |
| HOMO SAPIENS | 0 | 15816 | 2xrc | A | | 243 | GLY | p.Gly261Ala | P05156 |
| HOMO SAPIENS | 0 | 15816 | 2xrc | A | | 269 | GLY | p.Gly287Arg | P05156 |
| HOMO SAPIENS | -1 | 15816 | 2xrc | A | | 288 | ILE | p.Ile306Val | P05156 |
| HOMO SAPIENS | -1 | 15816 | 2xrc | A | | 388 | ARG | p.Arg406His | P05156 |
| HOMO SAPIENS | 7 | 15816 | 2xrc | A | | 398 | ILE | p.Ile416Leu | P05156 |
| HOMO SAPIENS | 9 | 15816 | 2xrc | A | | 400 | HIS | p.His418Leu | P05156 |
| HOMO SAPIENS | 0 | 15816 | 2xrc | A | | 423 | LYS | p.Lys441Arg | P05156 |
| HOMO SAPIENS | -1 | 15816 | 2xrc | A | | 530 | GLU | p.Glu548Gln | P05156 |
| HOMO SAPIENS | 0 | 15816 | 2xrc | A | | 535 | PRO | p.Pro553Ala | P05156 |
| HOMO SAPIENS | 0 | 15816 | 2xrc | A | | 536 | GLU | p.Glu554Gly | P05156 |
| HOMO SAPIENS | 1 | 12385 | 4zlp | B | | 1450 | ALA | p.Ala1450Pro | Q9UM47 |
| HOMO SAPIENS | 7 | 12385 | 4zlp | B | | 1482 | GLY | p.Gly1482Arg | Q9UM47 |
| HOMO SAPIENS | 2 | 12385 | 4zlp | B | | 1518 | LEU | p.Leu1518Met | Q9UM47 |
| HOMO SAPIENS | 0 | 12385 | 4zlp | B | | 1560 | ARG | p.Arg1560Pro | Q9UM47 |
| HOMO SAPIENS | 0 | 1227 | 4a04 | A | | 148 | PHE | p.Phe148Tyr | O43435 |
| HOMO SAPIENS | 4 | 1227 | 4a04 | A | | 194 | HIS | p.His194Gln | O43435 |
| HOMO SAPIENS | 2 | 8017 | 3ei4 | B | | 244 | LYS | p.Lys244Glu | Q92466 |
| HOMO SAPIENS | 1 | 8017 | 3ei4 | B | | 273 | ARG | p.Arg273His | Q92466 |
| HOMO SAPIENS | 3 | 36112 | 2k8f | B | | 9 | SER | p.Ser9Arg | P04637 |
| HOMO SAPIENS | 2 | 12880 | 5ggf | A | | 265 | ARG | p.Arg265His | Q8WZA1 |
| HOMO SAPIENS | 2 | 12880 | 5ggf | A | | 287 | ILE | p.Ile287Ser | Q8WZA1 |
| HOMO SAPIENS | 2 | 12880 | 5ggf | A | | 311 | ARG | p.Arg311Gln | Q8WZA1 |
| HOMO SAPIENS | 2 | 12880 | 5ggf | A | | 425 | TRP | p.Trp425Ser | Q8WZA1 |

| | | | | | | | | |
|---|---|---|---|---|---|---|---|---|
| HOMO SAPIENS | 2 | 12880 | 5ggf | A | 442 | ARG | p.Arg442Cys | Q8WZA1 |
| HOMO SAPIENS | 2 | 12880 | 5ggf | A | 448 | GLY | p.Gly448Arg | Q8WZA1 |
| HOMO SAPIENS | 0 | 12880 | 5ggf | A | 490 | CYS | p.Cys490Tyr | Q8WZA1 |
| HOMO SAPIENS | 0 | 12880 | 5ggf | A | 493 | PRO | p.Pro493Arg | Q8WZA1 |
| HOMO SAPIENS | 3 | 12880 | 5ggf | A | 550 | SER | p.Ser550Asn | Q8WZA1 |
| HOMO SAPIENS | 1 | 12880 | 5ggf | A | 556 | ASP | p.Asp556Asn | Q8WZA1 |
| HOMO SAPIENS | 0 | 12880 | 5ggf | A | 580 | ARG | p.Arg580Ter | Q8WZA1 |
| HOMO SAPIENS | 2 | 12880 | 5ggf | A | 605 | ARG | p.Arg605Pro | Q8WZA1 |
| HOMO SAPIENS | 0 | 12880 | 5ggf | A | 623 | VAL | p.Met623Val | Q8WZA1 |
| HOMO SAPIENS | 1 | 10187 | 4qw2 | A | 145 | ALA | p.Ala145Ser | Q06787 |
| HOMO SAPIENS | 1 | 10187 | 4qw2 | A | 174 | ASN | p.Asn174Lys | Q06787 |
| HOMO SAPIENS | 0 | 24002 | 2m0t | A | 110 | LEU | p.Leu110Val | O14745 |
| HOMO SAPIENS | 9 | 3126 | 4mjh | A | 99 | LEU | p.Leu99Met | P04792 |
| HOMO SAPIENS | 1 | 3126 | 4mjh | A | 127 | ARG | p.Arg127Trp | P04792 |
| HOMO SAPIENS | 1 | 3126 | 4mjh | A | 127 | ARG | p.Arg127Leu | P04792 |
| HOMO SAPIENS | 1 | 3126 | 4mjh | A | 135 | SER | p.Ser135Phe | P04792 |
| HOMO SAPIENS | 2 | 3126 | 4mjh | A | 136 | ARG | p.Arg136Trp | P04792 |
| HOMO SAPIENS | 1 | 3126 | 4mjh | A | 140 | ARG | p.Arg140Gly | P04792 |
| HOMO SAPIENS | 0 | 3126 | 4mjh | A | 151 | THR | p.Thr151Ile | P04792 |
| HOMO SAPIENS | -1 | 15780 | 2jlp | A | 213 | ARG | p.Arg231Gly | P08294 |
| HOMO SAPIENS | 8 | 3578 | 1apz | D | 203 | GLY | p.Gly226Asp | P20933 |
| HOMO SAPIENS | 1 | 3578 | 1apz | D | 279 | GLY | p.Gly302Arg | P20933 |
| HOMO SAPIENS | 0 | 3578 | 1apz | D | 283 | CYS | p.Cys306Arg | P20933 |
| HOMO SAPIENS | 5 | 1335 | 5xtd | C | 115 | ARG | p.Arg115Cys | O75251 |
| HOMO SAPIENS | 0 | 1335 | 5xtd | C | 122 | VAL | p.Val122Met | O75251 |
| HOMO SAPIENS | 2 | 5063 | 2le4 | A | 37 | ARG | p.Arg74Pro | P48431 |
| HOMO SAPIENS | 1 | 25870 | 2o10 | A | 44 | LEU | p.Leu44Pro | P50461 |
| HOMO SAPIENS | 4 | 25870 | 2o10 | A | 58 | CYS | p.Cys58Gly | P50461 |
| HOMO SAPIENS | 0 | 18748 | 5fii | C | 39 | PHE | p.Phe39Leu | P00439 |
| HOMO SAPIENS | 6 | 18748 | 5fii | C | 48 | LEU | p.Leu48Ser | P00439 |
| HOMO SAPIENS | 2 | 18748 | 5fii | C | 65 | ILE | p.Ile65Asn | P00439 |
| HOMO SAPIENS | 3 | 18748 | 5fii | C | 95 | ILE | p.Ile95Phe | P00439 |
| HOMO SAPIENS | 1 | 14261 | 2vay | B | 1539 | ARG | p.Arg1539Cys | Q13698 |
| HOMO SAPIENS | 0 | 2494 | 4dx1 | A | 183 | GLU | p.Glu183Asp | Q9HBA0 |
| HOMO SAPIENS | 0 | 2494 | 4dx1 | A | 217 | ALA | p.Ala217Ser | Q9HBA0 |
| HOMO SAPIENS | 0 | 2494 | 4dx1 | A | 217 | ALA | p.Ala217Val | Q9HBA0 |
| HOMO SAPIENS | 4 | 2494 | 4dx1 | A | 224 | ARG | p.Arg224Trp | Q9HBA0 |
| HOMO SAPIENS | 4 | 2494 | 4dx1 | A | 254 | VAL | p.Val254Leu | Q9HBA0 |
| HOMO SAPIENS | 0 | 2494 | 4dx1 | A | 269 | ARG | p.Arg269Cys | Q9HBA0 |
| HOMO SAPIENS | 0 | 2494 | 4dx1 | A | 269 | ARG | p.Arg269His | Q9HBA0 |
| HOMO SAPIENS | 0 | 2494 | 4dx1 | A | 278 | GLU | p.Glu278Lys | Q9HBA0 |
| HOMO SAPIENS | 0 | 2494 | 4dx1 | A | 315 | ARG | p.Arg315Trp | Q9HBA0 |
| HOMO SAPIENS | 0 | 2494 | 4dx1 | A | 316 | ARG | p.Arg316Cys | Q9HBA0 |
| HOMO SAPIENS | 0 | 2494 | 4dx1 | A | 331 | ILE | p.Ile331Phe | Q9HBA0 |
| HOMO SAPIENS | 0 | 2494 | 4dx1 | A | 333 | ASP | p.Asp333Gly | Q9HBA0 |
| HOMO SAPIENS | 1 | 2494 | 4dx1 | A | 365 | VAL | p.Val365Met | Q9HBA0 |
| HOMO SAPIENS | 0 | 23998 | 2l1r | B | 161 | ARG | p.Arg162Trp | P19429 |
| HOMO SAPIENS | 0 | 8354 | 4f0x | D | 187 | SER | p.Ser187Ter | O95822 |
| HOMO SAPIENS | 2 | 8354 | 4f0x | D | 259 | GLY | p.Gly259Ala | O95822 |
| HOMO SAPIENS | 2 | 8354 | 4f0x | D | 284 | THR | p.Thr284Ala | O95822 |
| HOMO SAPIENS | 3 | 8354 | 4f0x | D | 309 | LYS | p.Lys309Gln | O95822 |
| HOMO SAPIENS | 0 | 21182 | 4apo | A | 228 | GLN | p.Gln228Lys | O00170 |
| HOMO SAPIENS | 1 | 21182 | 4apo | A | 307 | GLN | p.Gln307Arg | O00170 |
| HOMO SAPIENS | 0 | 2287 | 5of4 | B | 112 | ARG | p.Arg112His | P18074 |

| | | | | | | | | |
|---|---|---|---|---|---|---|---|---|
| HOMO SAPIENS | 0 | 2287 | 5of4 | B | 312 | ASP | p.Asp312Asn | P18074 |
| HOMO SAPIENS | 3 | 2287 | 5of4 | B | 461 | LEU | p.Leu461Val | P18074 |
| HOMO SAPIENS | 0 | 2287 | 5of4 | B | 485 | LEU | p.Leu485Pro | P18074 |
| HOMO SAPIENS | 1 | 2287 | 5of4 | B | 541 | SER | p.Ser541Arg | P18074 |
| HOMO SAPIENS | 0 | 2287 | 5of4 | B | 616 | ARG | p.Arg616Pro | P18074 |
| HOMO SAPIENS | 0 | 2287 | 5of4 | B | 616 | ARG | p.Arg616Trp | P18074 |
| HOMO SAPIENS | 1 | 2287 | 5of4 | B | 658 | ARG | p.Arg658Cys | P18074 |
| HOMO SAPIENS | 1 | 2287 | 5of4 | B | 681 | ASP | p.Asp681Asn | P18074 |
| HOMO SAPIENS | 1 | 2287 | 5of4 | B | 683 | ARG | p.Arg683Trp | P18074 |
| HOMO SAPIENS | 0 | 2287 | 5of4 | B | 713 | GLY | p.Gly713Arg | P18074 |
| HOMO SAPIENS | 0 | 2287 | 5of4 | B | 717 | ALA | p.Ala717Gly | P18074 |
| HOMO SAPIENS | 0 | 2287 | 5of4 | B | 722 | ARG | p.Arg722Trp | P18074 |
| HOMO SAPIENS | 0 | 2287 | 5of4 | B | 725 | ALA | p.Ala725Pro | P18074 |
| HOMO SAPIENS | 4 | 12188 | 4its | A | 122 | ARG | p.Arg122Gly | Q9Y606 |
| HOMO SAPIENS | 4 | 12188 | 4its | A | 133 | ASP | p.Asp133Asn | Q9Y606 |
| HOMO SAPIENS | 4 | 12188 | 4its | A | 144 | ARG | p.Arg144Trp | Q9Y606 |
| HOMO SAPIENS | 0 | 31529 | 1kgd | A | 761 | ARG | p.Arg766Gln | O14936 |
| HOMO SAPIENS | 1 | 31529 | 1kgd | A | 824 | GLN | p.Gln829Arg | O14936 |
| HOMO SAPIENS | 0 | 10906 | 2h63 | C | 18 | ARG | p.Arg18Ter | P53004 |
| HOMO SAPIENS | 9 | 27767 | 4wyq | A | 280 | LEU | p.Leu278Phe | Q9UPY3 |
| HOMO SAPIENS | 0 | 27767 | 4wyq | A | 297 | SER | p.Ser295Cys | Q9UPY3 |
| HOMO SAPIENS | 1 | 27767 | 4wyq | A | 334 | TYR | p.Tyr332Ter | Q9UPY3 |
| HOMO SAPIENS | 7 | 7775 | 5sv7 | C | 136 | SER | p.Ser136Cys | Q9NZJ5 |
| HOMO SAPIENS | 2 | 7775 | 5sv7 | C | 166 | GLN | p.Gln166Arg | Q9NZJ5 |
| HOMO SAPIENS | 2 | 7775 | 5sv7 | C | 332 | GLU | p.Glu332Ter | Q9NZJ5 |
| HOMO SAPIENS | 0 | 21449 | 2e9w | B | 84 | THR | p.Thr84Met | P10721 |
| HOMO SAPIENS | 0 | 21449 | 2e9w | B | 168 | ALA | p.Ala168Glu | P10721 |
| HOMO SAPIENS | 0 | 21449 | 2e9w | B | 178 | ALA | p.Ala178Thr | P10721 |
| HOMO SAPIENS | 0 | 21449 | 2e9w | B | 293 | ASN | p.Asn293Ser | P10721 |
| HOMO SAPIENS | -1 | 21449 | 2e9w | B | 318 | MET | p.Met318Leu | P10721 |
| HOMO SAPIENS | 0 | 21449 | 2e9w | B | 374 | VAL | p.Val374Ile | P10721 |
| HOMO SAPIENS | 6 | 3010 | 3flg | A | 66 | SER | p.Ser270Pro | Q9UBC3 |
| HOMO SAPIENS | 1 | 33918 | 2xzz | A | 760 | ARG | p.Arg760Ter | P22735 |
| HOMO SAPIENS | 2 | 1254 | 5xtc | w | 65 | ASN | p.Asn65Ser | O95299 |
| HOMO SAPIENS | 1 | 1254 | 5xtc | w | 238 | GLU | p.Glu238Lys | O95299 |
| HOMO SAPIENS | 3 | 10245 | 3siv | B | 194 | ALA | p.Ala194Glu | Q8WWY3 |
| HOMO SAPIENS | 4 | 10245 | 3siv | B | 216 | ALA | p.Ala216Pro | Q8WWY3 |
| HOMO SAPIENS | 0 | 2670 | 3dwb | A | 341 | THR | p.Thr341Ile | P42892 |
| HOMO SAPIENS | 1 | 2670 | 3dwb | A | 462 | SER | p.Ser462Asn | P42892 |
| HOMO SAPIENS | 1 | 2670 | 3dwb | A | 754 | ARG | p.Arg754Cys | P42892 |
| HOMO SAPIENS | 1 | 2961 | 5xtb | O | 40 | VAL | p.Val40Met | P19404 |
| HOMO SAPIENS | 5 | 2961 | 5xtb | O | 172 | ILE | p.Ile172Thr | P19404 |
| HOMO SAPIENS | 5 | 2961 | 5xtb | O | 190 | ASP | p.Asp190Gly | P19404 |
| HOMO SAPIENS | 5 | 29655 | 2xdg | A | 57 | ALA | p.Ala57Thr | Q02643 |
| HOMO SAPIENS | 4 | 29655 | 2xdg | A | 72 | GLU | p.Glu72Ter | Q02643 |
| HOMO SAPIENS | 0 | 29655 | 2xdg | A | 121 | GLU | p.Glu121Asp | Q02643 |
| HOMO SAPIENS | -1 | 17935 | 1hyn | P | 38 | ASP | p.Asp38Ala | P02730 |
| HOMO SAPIENS | 7 | 12825 | 1g82 | D | 99 | SER | p.Ser99Asn | P31371 |
| HOMO SAPIENS | 6 | 24927 | 1jm7 | B | 85 | VAL | p.Val85Leu | Q99728 |
| HOMO SAPIENS | 1 | 24927 | 1jm7 | B | 112 | ARG | p.Arg112Ter | Q99728 |
| HOMO SAPIENS | 0 | 24927 | 1jm7 | B | 116 | HIS | p.His116Tyr | Q99728 |
| HOMO SAPIENS | 7 | 92 | 5hde | A | 61 | LYS | p.Lys61Arg | Q05209 |
| HOMO SAPIENS | 0 | 17246 | 1szb | A | 84 | ARG | p.Arg99Gln | O00187 |
| HOMO SAPIENS | 1 | 17246 | 1szb | A | 105 | ASP | p.Asp120Gly | O00187 |

| | | | | | | | | |
|---|---|---|---|---|---|---|---|---|
| HOMO SAPIENS | 0 | 17246 | 1szb | A | 111 | PRO | p.Pro126Leu | O00187 |
| HOMO SAPIENS | 9 | 17246 | 1szb | A | 141 | CYS | p.Cys156Tyr | O00187 |
| HOMO SAPIENS | 0 | 32473 | 1b72 | A | 207 | ARG | p.Arg207Cys | P14653 |
| HOMO SAPIENS | 0 | 1920 | 4yzf | B | 488 | VAL | p.Val488Met | P02730 |
| HOMO SAPIENS | 0 | 1920 | 4yzf | B | 589 | ARG | p.Arg589Cys | P02730 |
| HOMO SAPIENS | 0 | 1920 | 4yzf | B | 589 | ARG | p.Arg589His | P02730 |
| HOMO SAPIENS | 0 | 1920 | 4yzf | B | 602 | ARG | p.Arg602Pro | P02730 |
| HOMO SAPIENS | 0 | 1920 | 4yzf | B | 613 | SER | p.Ser613Phe | P02730 |
| HOMO SAPIENS | -1 | 1920 | 4yzf | B | 646 | ARG | p.Arg646Gln | P02730 |
| HOMO SAPIENS | -1 | 1920 | 4yzf | B | 646 | ARG | p.Arg646Trp | P02730 |
| HOMO SAPIENS | 0 | 1920 | 4yzf | B | 701 | GLY | p.Gly701Asp | P02730 |
| HOMO SAPIENS | 0 | 1920 | 4yzf | B | 760 | ARG | p.Arg760Gln | P02730 |
| HOMO SAPIENS | 0 | 1920 | 4yzf | B | 771 | GLY | p.Gly771Asp | P02730 |
| HOMO SAPIENS | 0 | 1920 | 4yzf | B | 773 | SER | p.Ser773Pro | P02730 |
| HOMO SAPIENS | 0 | 1920 | 4yzf | B | 837 | THR | p.Thr837Ala | P02730 |
| HOMO SAPIENS | 0 | 1920 | 4yzf | B | 854 | PRO | p.Pro854Leu | P02730 |
| HOMO SAPIENS | 0 | 1920 | 4yzf | B | 858 | ALA | p.Ala858Asp | P02730 |
| HOMO SAPIENS | 0 | 1920 | 4yzf | B | 862 | VAL | p.Val862Ile | P02730 |
| HOMO SAPIENS | 0 | 1920 | 4yzf | B | 868 | PRO | p.Pro868Leu | P02730 |
| HOMO SAPIENS | 0 | 1920 | 4yzf | B | 870 | ARG | p.Arg870Trp | P02730 |
| HOMO SAPIENS | 0 | 9863 | 2juj | A | 54 | VAL | p.Val904Ile | P22681 |
| HOMO SAPIENS | 0 | 21934 | 2na7 | A | 184 | ILE | p.Ile184Val | P25445 |
| HOMO SAPIENS | -1 | 6746 | 1lw3 | A | 3 | THR | p.Lys3Thr | Q13614 |
| HOMO SAPIENS | -1 | 6746 | 1lw3 | A | 44 | VAL | p.Val44Ile | Q13614 |
| HOMO SAPIENS | 3 | 6746 | 1lw3 | A | 183 | ALA | p.Ala183Ser | Q13614 |
| HOMO SAPIENS | 4 | 6746 | 1lw3 | A | 482 | GLN | p.Gln482Ter | Q13614 |
| HOMO SAPIENS | 4 | 6746 | 1lw3 | A | 502 | GLU | p.Glu502Gln | Q13614 |
| HOMO SAPIENS | 3 | 6746 | 1lw3 | A | 545 | ASN | p.Asn545Ser | Q13614 |
| HOMO SAPIENS | 0 | 6746 | 1lw3 | A | 586 | ARG | p.Arg586Trp | Q13614 |
| HOMO SAPIENS | -1 | 6746 | 1lw3 | A | 602 | ALA | p.Ala602Gly | Q13614 |
| HOMO SAPIENS | 1 | 319 | 4ch9 | B | 336 | ARG | p.Arg336Ile | Q9UH77 |
| HOMO SAPIENS | 3 | 319 | 4ch9 | B | 340 | ALA | p.Ala340Val | Q9UH77 |
| HOMO SAPIENS | 0 | 319 | 4ch9 | B | 384 | ARG | p.Arg384Gln | Q9UH77 |
| HOMO SAPIENS | 5 | 319 | 4ch9 | B | 426 | PRO | p.Pro426Leu | Q9UH77 |
| HOMO SAPIENS | 3 | 319 | 4ch9 | B | 431 | ARG | p.Arg431Gln | Q9UH77 |
| HOMO SAPIENS | 2 | 319 | 4ch9 | B | 575 | ARG | p.Arg575Trp | Q9UH77 |
| HOMO SAPIENS | 0 | 574 | 2qkk | A | 142 | VAL | p.Val142Ile | O60930 |
| HOMO SAPIENS | 0 | 24021 | 2n8y | A | 752 | ARG | p.Arg752Gln | P12814 |
| HOMO SAPIENS | 0 | 18693 | 2y7j | B | 44 | ARG | p.Arg44Ter | P15735 |
| HOMO SAPIENS | 0 | 18693 | 2y7j | B | 106 | VAL | p.Val106Glu | P15735 |
| HOMO SAPIENS | 3 | 18693 | 2y7j | B | 189 | GLY | p.Gly189Glu | P15735 |
| HOMO SAPIENS | 0 | 17087 | 2ifg | A | 237 | THR | p.Thr237Met | P04629 |
| HOMO SAPIENS | 0 | 7980 | 1qub | A | 247 | VAL | p.Val266Leu | P02749 |
| HOMO SAPIENS | -1 | 2205 | 1h7u | A | 18 | ILE | p.Ile18Val | P54278 |
| HOMO SAPIENS | -1 | 2205 | 1h7u | A | 20 | ARG | p.Arg20Gln | P54278 |
| HOMO SAPIENS | 9 | 2205 | 1h7u | A | 46 | SER | p.Ser46Ile | P54278 |
| HOMO SAPIENS | 0 | 2205 | 1h7u | A | 60 | ASP | p.Asp60Glu | P54278 |
| HOMO SAPIENS | 0 | 2205 | 1h7u | A | 80 | PHE | p.Phe80Leu | P54278 |
| HOMO SAPIENS | 0 | 2205 | 1h7u | A | 118 | SER | p.Ser118Arg | P54278 |
| HOMO SAPIENS | 0 | 2205 | 1h7u | A | 126 | HIS | p.His126Gln | P54278 |
| HOMO SAPIENS | 0 | 2205 | 1h7u | A | 128 | SER | p.Ser128Leu | P54278 |
| HOMO SAPIENS | 1 | 2205 | 1h7u | A | 134 | ARG | p.Arg134Ter | P54278 |
| HOMO SAPIENS | 0 | 2205 | 1h7u | A | 166 | LEU | p.Leu166Pro | P54278 |
| HOMO SAPIENS | 1 | 2205 | 1h7u | A | 191 | TYR | p.Tyr191Cys | P54278 |

| | | | | | | | | |
|---|---|---|---|---|---|---|---|---|
| HOMO SAPIENS | 0 | 2205 | 1h7u | A | 211 | ARG | p.Arg211Ter | P54278 |
| HOMO SAPIENS | 0 | 2205 | 1h7u | A | 233 | GLN | p.Gln233Ter | P54278 |
| HOMO SAPIENS | 0 | 2205 | 1h7u | A | 255 | TYR | p.Tyr255Ter | P54278 |
| HOMO SAPIENS | 0 | 2205 | 1h7u | A | 277 | THR | p.Thr277Lys | P54278 |
| HOMO SAPIENS | 0 | 2205 | 1h7u | A | 301 | LYS | p.Lys301Asn | P54278 |
| HOMO SAPIENS | 0 | 2205 | 1h7u | A | 312 | MET | p.Met312Ile | P54278 |
| HOMO SAPIENS | 0 | 2205 | 1h7u | A | 312 | MET | p.Met312Thr | P54278 |
| HOMO SAPIENS | 0 | 2205 | 1h7u | A | 315 | ARG | p.Arg315Ter | P54278 |
| HOMO SAPIENS | 0 | 2205 | 1h7u | A | 318 | TYR | p.Tyr318Cys | P54278 |
| HOMO SAPIENS | 0 | 2205 | 1h7u | A | 335 | ASN | p.Asn335Ile | P54278 |
| HOMO SAPIENS | 0 | 2205 | 1h7u | A | 360 | ILE | p.Ile360Met | P54278 |
| HOMO SAPIENS | 6 | 20754 | 5mpo | B | 7 | VAL | p.Val7Phe | O96033 |
| HOMO SAPIENS | 4 | 20754 | 5mpo | B | 64 | VAL | p.Val64Ile | O96033 |
| HOMO SAPIENS | 5 | 34944 | 4qam | B | 1114 | ASP | p.Asp1114Gly | Q96KN7 |
| HOMO SAPIENS | 5 | 34944 | 4qam | B | 1240 | GLY | p.Gly1240Glu | Q96KN7 |
| HOMO SAPIENS | 9 | 40496 | 3vnn | A | 278 | ARG | p.Arg278His | P49917 |
| HOMO SAPIENS | 6 | 30131 | 4dt7 | E | 40 | PRO | p.Pro210Leu | P04070 |
| HOMO SAPIENS | -1 | 30131 | 4dt7 | E | 50 | ARG | p.Arg220Trp | P04070 |
| HOMO SAPIENS | 7 | 15311 | 2pqt | A | 64 | ARG | p.Arg64Trp | P18440 |
| HOMO SAPIENS | 2 | 15311 | 2pqt | A | 149 | VAL | p.Val149Ile | P18440 |
| HOMO SAPIENS | -1 | 19581 | 3hy8 | A | 33 | ASP | p.Asp33Val | Q9NVS9 |
| HOMO SAPIENS | 5 | 19581 | 3hy8 | A | 116 | ARG | p.Arg116Gln | Q9NVS9 |
| HOMO SAPIENS | 4 | 19581 | 3hy8 | A | 225 | ARG | p.Arg225His | Q9NVS9 |
| HOMO SAPIENS | 2 | 19581 | 3hy8 | A | 229 | TRP | p.Arg229Trp | Q9NVS9 |
| HOMO SAPIENS | 2 | 19581 | 3hy8 | A | 229 | TRP | p.Arg229Gln | Q9NVS9 |
| HOMO SAPIENS | 0 | 10266 | 3t1i | A | 41 | ASP | p.Asp41His | P49959 |
| HOMO SAPIENS | 1 | 10266 | 3t1i | A | 305 | ARG | p.Arg305Trp | P49959 |
| HOMO SAPIENS | 0 | 10266 | 3t1i | A | 308 | PHE | p.Phe308Leu | P49959 |
| HOMO SAPIENS | 0 | 10266 | 3t1i | A | 334 | SER | p.Ser334Arg | P49959 |
| HOMO SAPIENS | 1 | 1600 | 3rbf | A | 61 | GLU | p.Glu61Asp | P20711 |
| HOMO SAPIENS | 0 | 1600 | 3rbf | A | 91 | ALA | p.Ala91Val | P20711 |
| HOMO SAPIENS | -1 | 1600 | 3rbf | A | 102 | GLY | p.Gly102Ser | P20711 |
| HOMO SAPIENS | 3 | 1600 | 3rbf | A | 147 | SER | p.Ser147Arg | P20711 |
| HOMO SAPIENS | 1 | 1600 | 3rbf | A | 210 | PRO | p.Pro210Leu | P20711 |
| HOMO SAPIENS | 0 | 1600 | 3rbf | A | 217 | MET | p.Met217Val | P20711 |
| HOMO SAPIENS | 4 | 1600 | 3rbf | A | 250 | SER | p.Ser250Phe | P20711 |
| HOMO SAPIENS | 2 | 1600 | 3rbf | A | 275 | ALA | p.Ala275Thr | P20711 |
| HOMO SAPIENS | 1 | 1600 | 3rbf | A | 309 | PHE | p.Phe309Leu | P20711 |
| HOMO SAPIENS | -1 | 1600 | 3rbf | A | 347 | ARG | p.Arg347Gln | P20711 |
| HOMO SAPIENS | 0 | 1600 | 3rbf | A | 462 | ARG | p.Arg462Gln | P20711 |
| HOMO SAPIENS | 6 | 20830 | 5y4f | A | 714 | THR | p.Thr715Ile | Q01484 |
| HOMO SAPIENS | 7 | 20830 | 5y4f | A | 719 | ASP | p.Asp720Glu | Q01484 |
| HOMO SAPIENS | 0 | 20830 | 5y4f | A | 814 | VAL | p.Val815Ile | Q01484 |
| HOMO SAPIENS | -1 | 302 | 2c9y | A | 9 | GLU | p.Glu9Ter | P54819 |
| HOMO SAPIENS | 0 | 302 | 2c9y | A | 103 | ARG | p.Arg103Trp | P54819 |
| HOMO SAPIENS | 3 | 302 | 2c9y | A | 165 | ASP | p.Asp165Gly | P54819 |
| HOMO SAPIENS | 0 | 302 | 2c9y | A | 182 | ALA | p.Ala182Asp | P54819 |
| HOMO SAPIENS | 2 | 302 | 2c9y | A | 186 | ARG | p.Arg186Cys | P54819 |
| HOMO SAPIENS | 0 | 302 | 2c9y | A | 209 | ALA | p.Ala209Thr | P54819 |
| HOMO SAPIENS | -1 | 16920 | 1jlj | B | 10 | ASN | p.Asn10Tyr | Q9NQX3 |
| HOMO SAPIENS | 9 | 8462 | 1zbq | B | 16 | GLY | p.Gly16Ser | P51659 |
| HOMO SAPIENS | 5 | 8462 | 1zbq | B | 19 | ALA | p.Ala19Gly | P51659 |
| HOMO SAPIENS | 1 | 8462 | 1zbq | B | 34 | ALA | p.Ala34Val | P51659 |
| HOMO SAPIENS | 1 | 8462 | 1zbq | B | 106 | ARG | p.Arg106His | P51659 |

| | | | | | | | | |
|---|---|---|---|---|---|---|---|---|
| HOMO SAPIENS | 0 | 8462 | 1zbq | B | 140 | LYS | p.Lys140Asn | P51659 |
| HOMO SAPIENS | 0 | 8462 | 1zbq | B | 292 | THR | p.Thr292Ser | P51659 |
| HOMO SAPIENS | 0 | 4375 | 4ndi | A | 199 | ARG | p.Arg213His | Q7Z2E3 |
| HOMO SAPIENS | 3 | 4375 | 4ndi | A | 201 | HIS | p.His215Arg | Q7Z2E3 |
| HOMO SAPIENS | 0 | 4375 | 4ndi | A | 206 | PRO | p.Pro220Leu | Q7Z2E3 |
| HOMO SAPIENS | 2 | 4375 | 4ndi | A | 223 | LEU | p.Leu237Phe | Q7Z2E3 |
| HOMO SAPIENS | 0 | 4375 | 4ndi | A | 245 | ARG | p.Arg259His | Q7Z2E3 |
| HOMO SAPIENS | 0 | 4375 | 4ndi | A | 263 | VAL | p.Val277Gly | Q7Z2E3 |
| HOMO SAPIENS | 0 | 4375 | 4ndi | A | 279 | TRP | p.Trp293Ter | Q7Z2E3 |
| HOMO SAPIENS | 0 | 27004 | 4ccg | Y | 321 | ASP | p.Asp321Glu | Q9NW38 |
| HOMO SAPIENS | -1 | 27004 | 4ccg | Y | 372 | GLY | p.Gly372Ala | Q9NW38 |
| HOMO SAPIENS | -1 | 6383 | 5fai | A | 34 | ALA | p.Ala34Gly | Q92979 |
| HOMO SAPIENS | 8 | 6383 | 5fai | A | 86 | ASP | p.Asp86Gly | Q92979 |
| HOMO SAPIENS | 2 | 24585 | 4mi5 | A | 684 | ARG | p.Arg679Cys | Q15910 |
| HOMO SAPIENS | 0 | 15967 | 3nhb | A | 723 | ARG | p.Arg723Gln | Q9NP58 |
| HOMO SAPIENS | 5 | 18152 | 1oey | A | 389 | HIS | p.His389Gln | P19878 |
| HOMO SAPIENS | 5 | 18152 | 1oey | A | 395 | ARG | p.Arg395Trp | P19878 |
| HOMO SAPIENS | -1 | 1044 | 1li1 | F | 2 | VAL | p.Val1486Ile | P08572 |
| HOMO SAPIENS | 6 | 1044 | 1li1 | F | 37 | GLU | p.Glu1521Lys | P08572 |
| HOMO SAPIENS | 5 | 1044 | 1li1 | F | 157 | SER | p.Ser1639Leu | P08572 |
| HOMO SAPIENS | 5 | 1044 | 1li1 | F | 179 | ARG | p.Arg1662His | P08572 |
| HOMO SAPIENS | 5 | 1044 | 1li1 | F | 180 | GLY | p.Gly1663Arg | P08572 |
| HOMO SAPIENS | 6 | 20076 | 4uyi | A | 671 | LEU | p.Leu671Ser | Q8IY92 |
| HOMO SAPIENS | 0 | 19588 | 1jmj | A | 189 | ARG | p.Arg208His | P05546 |
| HOMO SAPIENS | -1 | 38023 | 1ry7 | B | 65 | GLY | p.Gly65Arg | P22607 |
| HOMO SAPIENS | -1 | 38023 | 1ry7 | B | 139 | ASP | p.Asp139Glu | P22607 |
| HOMO SAPIENS | 0 | 38023 | 1ry7 | B | 248 | ARG | p.Arg248Cys | P22607 |
| HOMO SAPIENS | 0 | 38023 | 1ry7 | B | 249 | SER | p.Ser249Cys | P22607 |
| HOMO SAPIENS | 0 | 38023 | 1ry7 | B | 250 | PRO | p.Pro250Arg | P22607 |
| HOMO SAPIENS | 0 | 38023 | 1ry7 | B | 322 | GLU | p.Glu322Lys | P22607 |
| HOMO SAPIENS | 5 | 16884 | 5eli | A | 33 | GLN | p.Gln33Ter | Q9NZC2 |
| HOMO SAPIENS | 0 | 16884 | 5eli | A | 44 | TRP | p.Trp44Ter | Q9NZC2 |
| HOMO SAPIENS | 0 | 16884 | 5eli | A | 47 | ARG | p.Arg47His | Q9NZC2 |
| HOMO SAPIENS | 0 | 16884 | 5eli | A | 66 | THR | p.Thr66Met | Q9NZC2 |
| HOMO SAPIENS | 0 | 16884 | 5eli | A | 78 | TRP | p.Trp78Ter | Q9NZC2 |
| HOMO SAPIENS | 1 | 16884 | 5eli | A | 96 | THR | p.Thr96Lys | Q9NZC2 |
| HOMO SAPIENS | 9 | 16884 | 5eli | A | 126 | VAL | p.Val126Gly | Q9NZC2 |
| HOMO SAPIENS | -1 | 12267 | 3w5o | A | 3 | ALA | p.Ala3Val | P49917 |
| HOMO SAPIENS | 1 | 12267 | 3w5o | A | 9 | THR | p.Thr9Ile | P49917 |
| HOMO SAPIENS | 2 | 12267 | 3w5o | A | 188 | ARG | p.Arg188Gln | P49917 |
| HOMO SAPIENS | 0 | 12267 | 3w5o | A | 469 | GLY | p.Gly469Glu | P49917 |
| HOMO SAPIENS | 1 | 12267 | 3w5o | A | 580 | ARG | p.Arg580Ter | P49917 |
| HOMO SAPIENS | 0 | 12267 | 3w5o | A | 600 | GLU | p.Glu600Lys | P49917 |
| HOMO SAPIENS | 9 | 2839 | 2hw5 | B | 54 | ARG | p.Arg54Cys | P30084 |
| HOMO SAPIENS | 9 | 2839 | 2hw5 | B | 59 | ASN | p.Asn59Ser | P30084 |
| HOMO SAPIENS | 0 | 2839 | 2hw5 | B | 273 | LYS | p.Lys273Glu | P30084 |
| HOMO SAPIENS | 1 | 12266 | 2o2k | B | 939 | GLY | p.Gly939Cys | Q99707 |
| HOMO SAPIENS | 5 | 12266 | 2o2k | B | 1164 | ARG | p.Arg1164His | Q99707 |
| HOMO SAPIENS | 1 | 12266 | 2o2k | B | 1166 | LEU | p.Leu1166Met | Q99707 |
| HOMO SAPIENS | 5 | 12266 | 2o2k | B | 1173 | PRO | p.Pro1173Leu | Q99707 |
| HOMO SAPIENS | 5 | 12266 | 2o2k | B | 1222 | ASN | p.Asn1222Ser | Q99707 |
| HOMO SAPIENS | 0 | 30891 | 3ch4 | B | 138 | ARG | p.Arg138Ter | Q15126 |
| HOMO SAPIENS | -1 | 4433 | 1qlz | A | 54 | GLY | p.Gly54Ser | P04156 |
| HOMO SAPIENS | 0 | 22568 | 3cxl | A | 20 | LEU | p.Leu20Phe | P15882 |

| | | | | | | | | |
|---|---|---|---|---|---|---|---|---|
| HOMO SAPIENS | 0 | 22568 | 3cxl | A | 126 | ILE | p.Ile126Met | P15882 |
| HOMO SAPIENS | 0 | 22568 | 3cxl | A | 141 | PRO | p.Pro141Leu | P15882 |
| HOMO SAPIENS | 0 | 22568 | 3cxl | A | 143 | TYR | p.Tyr143His | P15882 |
| HOMO SAPIENS | 0 | 22568 | 3cxl | A | 223 | ALA | p.Ala223Val | P15882 |
| HOMO SAPIENS | 0 | 22568 | 3cxl | A | 228 | GLY | p.Gly228Ser | P15882 |
| HOMO SAPIENS | 0 | 22568 | 3cxl | A | 252 | PRO | p.Pro252Gln | P15882 |
| HOMO SAPIENS | 0 | 22568 | 3cxl | A | 313 | GLU | p.Glu313Lys | P15882 |
| HOMO SAPIENS | 3 | 2668 | 5eqg | A | 34 | ASN | p.Asn34Ile | P11166 |
| HOMO SAPIENS | 3 | 2668 | 5eqg | A | 77 | MET | p.Met77Val | P11166 |
| HOMO SAPIENS | 6 | 2668 | 5eqg | A | 91 | GLY | p.Gly91Asp | P11166 |
| HOMO SAPIENS | 6 | 2668 | 5eqg | A | 92 | ARG | p.Arg92Trp | P11166 |
| HOMO SAPIENS | 6 | 2668 | 5eqg | A | 93 | ARG | p.Arg93Trp | P11166 |
| HOMO SAPIENS | 0 | 2668 | 5eqg | A | 95 | SER | p.Ser95Ile | P11166 |
| HOMO SAPIENS | 3 | 2668 | 5eqg | A | 126 | ARG | p.Arg126His | P11166 |
| HOMO SAPIENS | 3 | 2668 | 5eqg | A | 126 | ARG | p.Arg126Cys | P11166 |
| HOMO SAPIENS | 4 | 2668 | 5eqg | A | 232 | ARG | p.Arg232Cys | P11166 |
| HOMO SAPIENS | 3 | 2668 | 5eqg | A | 249 | ARG | p.Arg249Gln | P11166 |
| HOMO SAPIENS | 3 | 2668 | 5eqg | A | 255 | LYS | p.Lys255Arg | P11166 |
| HOMO SAPIENS | 0 | 2668 | 5eqg | A | 275 | ALA | p.Ala275Thr | P11166 |
| HOMO SAPIENS | 4 | 2668 | 5eqg | A | 299 | GLU | p.Glu299Lys | P11166 |
| HOMO SAPIENS | 6 | 2668 | 5eqg | A | 314 | GLY | p.Gly314Ser | P11166 |
| HOMO SAPIENS | 7 | 2668 | 5eqg | A | 315 | ILE | p.Ile315Met | P11166 |
| HOMO SAPIENS | -1 | 2668 | 5eqg | A | 458 | ARG | p.Arg458Trp | P11166 |
| HOMO SAPIENS | -1 | 2668 | 5eqg | A | 468 | ARG | p.Arg468Trp | P11166 |
| HOMO SAPIENS | 3 | 2291 | 2amy | A | 31 | PHE | p.Phe31Leu | O15305 |
| HOMO SAPIENS | 7 | 2291 | 2amy | A | 32 | LEU | p.Leu32Arg | O15305 |
| HOMO SAPIENS | 7 | 2291 | 2amy | A | 65 | ASP | p.Asp65Tyr | O15305 |
| HOMO SAPIENS | 1 | 2291 | 2amy | A | 108 | ALA | p.Ala108Val | O15305 |
| HOMO SAPIENS | 3 | 2291 | 2amy | A | 113 | PRO | p.Pro113Leu | O15305 |
| HOMO SAPIENS | 3 | 2291 | 2amy | A | 113 | PRO | p.Pro113Ser | O15305 |
| HOMO SAPIENS | 6 | 2291 | 2amy | A | 117 | GLY | p.Gly117Arg | O15305 |
| HOMO SAPIENS | 6 | 2291 | 2amy | A | 119 | PHE | p.Phe119Leu | O15305 |
| HOMO SAPIENS | 6 | 2291 | 2amy | A | 123 | ARG | p.Arg123Gln | O15305 |
| HOMO SAPIENS | 0 | 2291 | 2amy | A | 129 | VAL | p.Val129Met | O15305 |
| HOMO SAPIENS | 7 | 2291 | 2amy | A | 139 | GLU | p.Glu139Lys | O15305 |
| HOMO SAPIENS | 5 | 2291 | 2amy | A | 141 | ARG | p.Arg141His | O15305 |
| HOMO SAPIENS | 7 | 2291 | 2amy | A | 144 | PHE | p.Phe144Leu | O15305 |
| HOMO SAPIENS | 6 | 2291 | 2amy | A | 148 | ASP | p.Asp148Asn | O15305 |
| HOMO SAPIENS | 6 | 2291 | 2amy | A | 153 | ILE | p.Ile153Thr | O15305 |
| HOMO SAPIENS | 2 | 2291 | 2amy | A | 157 | PHE | p.Phe157Ser | O15305 |
| HOMO SAPIENS | 1 | 2291 | 2amy | A | 162 | ARG | p.Arg162Trp | O15305 |
| HOMO SAPIENS | 6 | 2291 | 2amy | A | 188 | ASP | p.Asp188Gly | O15305 |
| HOMO SAPIENS | 5 | 2291 | 2amy | A | 194 | ARG | p.Arg194Ter | O15305 |
| HOMO SAPIENS | 0 | 2291 | 2amy | A | 207 | PHE | p.Phe207Ser | O15305 |
| HOMO SAPIENS | 5 | 2291 | 2amy | A | 208 | GLY | p.Gly208Ala | O15305 |
| HOMO SAPIENS | 1 | 2291 | 2amy | A | 216 | ASN | p.Asn216Ile | O15305 |
| HOMO SAPIENS | 5 | 2291 | 2amy | A | 218 | HIS | p.His218Leu | O15305 |
| HOMO SAPIENS | 6 | 2291 | 2amy | A | 223 | ASP | p.Asp223Glu | O15305 |
| HOMO SAPIENS | 6 | 2291 | 2amy | A | 226 | THR | p.Thr226Ser | O15305 |
| HOMO SAPIENS | 5 | 2291 | 2amy | A | 231 | VAL | p.Val231Met | O15305 |
| HOMO SAPIENS | 1 | 2291 | 2amy | A | 237 | THR | p.Thr237Met | O15305 |
| HOMO SAPIENS | 1 | 2291 | 2amy | A | 241 | CYS | p.Cys241Ser | O15305 |
| HOMO SAPIENS | 5 | 35907 | 1d6g | A | 21 | GLY | p.Gly21Arg | P32238 |
| HOMO SAPIENS | -1 | 33209 | 1x9d | A | 197 | PRO | p.Pro197Leu | Q9UKM7 |

| | | | | | | | | |
|---|---|---|---|---|---|---|---|---|
| HOMO SAPIENS | -1 | 33209 | 1x9d | A | 201 | PRO | p.Pro201Leu | Q9UKM7 |
| HOMO SAPIENS | 6 | 21305 | 4yep | B | 1326 | ARG | p.Arg1326Ter | P24043 |
| HOMO SAPIENS | 6 | 21305 | 4yep | B | 1350 | ARG | p.Arg1350Ter | P24043 |
| HOMO SAPIENS | 0 | 21553 | 3qyb | A | 1120 | VAL | p.Val1119Ala | O60343 |
| HOMO SAPIENS | 2 | 21553 | 3qyb | A | 1148 | THR | p.Thr1147Met | O60343 |
| HOMO SAPIENS | -1 | 30645 | 3m0c | C | -17 | TRP | p.Trp4Ter | P01130 |
| HOMO SAPIENS | -1 | 30645 | 3m0c | C | -17 | TRP | p.Trp4Ter | P01130 |
| HOMO SAPIENS | -1 | 30645 | 3m0c | C | 12 | GLN | p.Gln33Ter | P01130 |
| HOMO SAPIENS | -1 | 30645 | 3m0c | C | 15 | ASP | p.Asp36Glu | P01130 |
| HOMO SAPIENS | -1 | 30645 | 3m0c | C | 28 | SER | p.Ser49Arg | P01130 |
| HOMO SAPIENS | -1 | 30645 | 3m0c | C | 41 | THR | p.Thr62Met | P01130 |
| HOMO SAPIENS | -1 | 30645 | 3m0c | C | 69 | ASP | p.Asp90Asn | P01130 |
| HOMO SAPIENS | -1 | 30645 | 3m0c | C | 76 | ASN | p.Asn97Lys | P01130 |
| HOMO SAPIENS | -1 | 30645 | 3m0c | C | 78 | SER | p.Ser99Ter | P01130 |
| HOMO SAPIENS | -1 | 30645 | 3m0c | C | 81 | GLN | p.Gln102Ter | P01130 |
| HOMO SAPIENS | -1 | 30645 | 3m0c | C | 84 | PRO | p.Pro105Ser | P01130 |
| HOMO SAPIENS | -1 | 30645 | 3m0c | C | 194 | ARG | p.Arg215Cys | P01130 |
| HOMO SAPIENS | -1 | 30645 | 3m0c | C | 198 | GLY | p.Gly219Asp | P01130 |
| HOMO SAPIENS | -1 | 30645 | 3m0c | C | 201 | CYS | p.Cys222Tyr | P01130 |
| HOMO SAPIENS | -1 | 30645 | 3m0c | C | 201 | CYS | p.Cys222Ter | P01130 |
| HOMO SAPIENS | -1 | 30645 | 3m0c | C | 206 | ASP | p.Asp227Glu | P01130 |
| HOMO SAPIENS | -1 | 30645 | 3m0c | C | 249 | CYS | p.Cys270Ter | P01130 |
| HOMO SAPIENS | -1 | 30645 | 3m0c | C | 250 | VAL | p.Val271Ile | P01130 |
| HOMO SAPIENS | 0 | 30645 | 3m0c | C | 255 | CYS | p.Cys276Tyr | P01130 |
| HOMO SAPIENS | 0 | 30645 | 3m0c | C | 255 | CYS | p.Cys276Ter | P01130 |
| HOMO SAPIENS | 0 | 30645 | 3m0c | C | 256 | GLU | p.Glu277Lys | P01130 |
| HOMO SAPIENS | 0 | 30645 | 3m0c | C | 265 | SER | p.Ser286Arg | P01130 |
| HOMO SAPIENS | 0 | 30645 | 3m0c | C | 284 | TRP | p.Trp305Ter | P01130 |
| HOMO SAPIENS | 0 | 30645 | 3m0c | C | 292 | CYS | p.Cys313Ter | P01130 |
| HOMO SAPIENS | 1 | 30645 | 3m0c | C | 385 | ARG | p.Arg406Trp | P01130 |
| HOMO SAPIENS | 0 | 30645 | 3m0c | C | 395 | ARG | p.Arg416Gln | P01130 |
| HOMO SAPIENS | 0 | 30645 | 3m0c | C | 398 | TYR | p.Tyr419Ter | P01130 |
| HOMO SAPIENS | 1 | 30645 | 3m0c | C | 415 | VAL | p.Val436Ala | P01130 |
| HOMO SAPIENS | 1 | 30645 | 3m0c | C | 424 | ASP | p.Asp445Glu | P01130 |
| HOMO SAPIENS | 1 | 30645 | 3m0c | C | 425 | LEU | p.Leu446Val | P01130 |
| HOMO SAPIENS | -1 | 30645 | 3m0c | C | 440 | GLY | p.Gly461Cys | P01130 |
| HOMO SAPIENS | -1 | 30645 | 3m0c | C | 441 | VAL | p.Val462Ile | P01130 |
| HOMO SAPIENS | 1 | 30645 | 3m0c | C | 447 | VAL | p.Val468Ile | P01130 |
| HOMO SAPIENS | 1 | 30645 | 3m0c | C | 489 | THR | p.Thr510Met | P01130 |
| HOMO SAPIENS | 0 | 30645 | 3m0c | C | 524 | GLY | p.Gly545Glu | P01130 |
| HOMO SAPIENS | 1 | 30645 | 3m0c | C | 615 | GLY | p.Gly636Ser | P01130 |
| HOMO SAPIENS | 1 | 30645 | 3m0c | C | 625 | LEU | p.Leu646Ile | P01130 |
| HOMO SAPIENS | 1 | 30645 | 3m0c | C | 630 | ASP | p.Asp651Gly | P01130 |
| HOMO SAPIENS | 3 | 30645 | 3m0c | C | 646 | CYS | p.Cys667Tyr | P01130 |
| HOMO SAPIENS | 4 | 30645 | 3m0c | C | 654 | GLY | p.Gly675Ser | P01130 |
| HOMO SAPIENS | 6 | 30645 | 3m0c | C | 660 | CYS | p.Cys681Ter | P01130 |
| HOMO SAPIENS | 5 | 30645 | 3m0c | C | 676 | ALA | p.Ala697Pro | P01130 |
| HOMO SAPIENS | -1 | 30645 | 3m0c | C | 749 | GLN | p.Gln770Ter | P01130 |
| HOMO SAPIENS | -1 | 41323 | 4bxo | A | 1812 | PRO | p.Pro1812Ala | Q8IYD8 |
| HOMO SAPIENS | 5 | 8335 | 3h0t | C | 12 | GLY | p.Gly71Asp | P81172 |
| HOMO SAPIENS | 6 | 609 | 1bhg | B | 148 | PRO | p.Pro148Ser | P08236 |
| HOMO SAPIENS | 1 | 609 | 1bhg | B | 176 | LEU | p.Leu176Phe | P08236 |
| HOMO SAPIENS | 6 | 609 | 1bhg | B | 216 | ARG | p.Arg216Trp | P08236 |
| HOMO SAPIENS | 0 | 609 | 1bhg | B | 350 | LYS | p.Lys350Asn | P08236 |

| | | | | | | | | |
|---|---|---|---|---|---|---|---|---|
| HOMO SAPIENS | 1 | 609 | 1bhg | B | 354 | ALA | p.Ala354Val | P08236 |
| HOMO SAPIENS | 1 | 609 | 1bhg | B | 357 | ARG | p.Arg357Ter | P08236 |
| HOMO SAPIENS | 0 | 609 | 1bhg | B | 382 | ARG | p.Arg382Cys | P08236 |
| HOMO SAPIENS | 0 | 609 | 1bhg | B | 495 | TYR | p.Tyr495Cys | P08236 |
| HOMO SAPIENS | 2 | 609 | 1bhg | B | 507 | TRP | p.Trp507Ter | P08236 |
| HOMO SAPIENS | 0 | 609 | 1bhg | B | 577 | ARG | p.Arg577Leu | P08236 |
| HOMO SAPIENS | 1 | 609 | 1bhg | B | 611 | ARG | p.Arg611Trp | P08236 |
| HOMO SAPIENS | 0 | 609 | 1bhg | B | 619 | ALA | p.Ala619Val | P08236 |
| HOMO SAPIENS | 0 | 609 | 1bhg | B | 627 | TRP | p.Trp627Cys | P08236 |
| HOMO SAPIENS | 5 | 3422 | 1x9n | A | 566 | GLU | p.Glu566Lys | P18858 |
| HOMO SAPIENS | 2 | 3422 | 1x9n | A | 771 | ARG | p.Arg771Trp | P18858 |
| HOMO SAPIENS | 0 | 3279 | 5tby | B | 797 | ALA | p.Ala797Thr | P12883 |
| HOMO SAPIENS | -1 | 3279 | 5tby | B | 982 | MET | p.Met982Thr | P12883 |
| HOMO SAPIENS | -1 | 3279 | 5tby | B | 992 | LEU | p.Leu992Met | P12883 |
| HOMO SAPIENS | -1 | 3279 | 5tby | B | 1004 | GLN | p.Gln1004Glu | P12883 |
| HOMO SAPIENS | -1 | 3279 | 5tby | B | 1013 | GLU | p.Glu1013Lys | P12883 |
| HOMO SAPIENS | -1 | 3279 | 5tby | B | 1032 | ASP | p.Asp1032Asn | P12883 |
| HOMO SAPIENS | -1 | 3279 | 5tby | B | 1053 | ARG | p.Arg1053Gln | P12883 |
| HOMO SAPIENS | -1 | 3279 | 5tby | B | 1117 | GLU | p.Glu1117Ter | P12883 |
| HOMO SAPIENS | -1 | 3279 | 5tby | B | 1193 | ARG | p.Arg1193His | P12883 |
| HOMO SAPIENS | -1 | 3279 | 5tby | B | 1205 | GLU | p.Glu1205Lys | P12883 |
| HOMO SAPIENS | -1 | 3279 | 5tby | B | 1207 | ILE | p.Ile1207Met | P12883 |
| HOMO SAPIENS | -1 | 3279 | 5tby | B | 1250 | ARG | p.Arg1250Pro | P12883 |
| HOMO SAPIENS | -1 | 3279 | 5tby | B | 1259 | HIS | p.His1259Gln | P12883 |
| HOMO SAPIENS | -1 | 3279 | 5tby | B | 1272 | ASP | p.Asp1272Asn | P12883 |
| HOMO SAPIENS | -1 | 3279 | 5tby | B | 1295 | GLU | p.Glu1295Lys | P12883 |
| HOMO SAPIENS | -1 | 3279 | 5tby | B | 1335 | SER | p.Ser1335Leu | P12883 |
| HOMO SAPIENS | -1 | 3279 | 5tby | B | 1399 | GLU | p.Glu1399Asp | P12883 |
| HOMO SAPIENS | -1 | 3279 | 5tby | B | 1413 | SER | p.Ser1413Ter | P12883 |
| HOMO SAPIENS | -1 | 3279 | 5tby | B | 1459 | LYS | p.Lys1459Asn | P12883 |
| HOMO SAPIENS | -1 | 3279 | 5tby | B | 1467 | LEU | p.Leu1467Val | P12883 |
| HOMO SAPIENS | -1 | 3279 | 5tby | B | 1491 | SER | p.Ser1491Cys | P12883 |
| HOMO SAPIENS | -1 | 3279 | 5tby | B | 1496 | GLU | p.Glu1496Ala | P12883 |
| HOMO SAPIENS | -1 | 3279 | 5tby | B | 1500 | ARG | p.Arg1500Pro | P12883 |
| HOMO SAPIENS | -1 | 3279 | 5tby | B | 1606 | ARG | p.Arg1606His | P12883 |
| HOMO SAPIENS | -1 | 3279 | 5tby | B | 1664 | ASN | p.Asn1664Lys | P12883 |
| HOMO SAPIENS | -1 | 3279 | 5tby | B | 1712 | ARG | p.Arg1712Trp | P12883 |
| HOMO SAPIENS | -1 | 3279 | 5tby | B | 1769 | LEU | p.Leu1769Met | P12883 |
| HOMO SAPIENS | -1 | 3279 | 5tby | B | 1808 | GLY | p.Gly1808Ser | P12883 |
| HOMO SAPIENS | -1 | 3279 | 5tby | B | 1818 | ARG | p.Arg1818Trp | P12883 |
| HOMO SAPIENS | -1 | 3279 | 5tby | B | 1820 | ARG | p.Arg1820Trp | P12883 |
| HOMO SAPIENS | -1 | 3279 | 5tby | B | 1844 | GLU | p.Glu1844Lys | P12883 |
| HOMO SAPIENS | -1 | 3279 | 5tby | B | 1845 | ARG | p.Arg1845Trp | P12883 |
| HOMO SAPIENS | -1 | 3279 | 5tby | B | 1854 | THR | p.Thr1854Met | P12883 |
| HOMO SAPIENS | -1 | 3279 | 5tby | B | 1909 | ARG | p.Arg1909Trp | P12883 |
| HOMO SAPIENS | 2 | 2260 | 4kl1 | A | 553 | ASP | p.Asp553Asn | Q9Y3Q4 |
| HOMO SAPIENS | 0 | 40570 | 5lst | A | 522 | ARG | p.Arg522Cys | O94761 |
| HOMO SAPIENS | 0 | 40570 | 5lst | A | 522 | ARG | p.Arg522His | O94761 |
| HOMO SAPIENS | 0 | 40570 | 5lst | A | 591 | PRO | p.Pro591Leu | O94761 |
| HOMO SAPIENS | 0 | 40570 | 5lst | A | 793 | PRO | p.Pro793Ser | O94761 |
| HOMO SAPIENS | 4 | 40570 | 5lst | A | 799 | VAL | p.Val799Met | O94761 |
| HOMO SAPIENS | 0 | 40570 | 5lst | A | 1004 | ARG | p.Arg1004Trp | O94761 |
| HOMO SAPIENS | 0 | 40570 | 5lst | A | 1005 | ARG | p.Arg1005Gln | O94761 |
| HOMO SAPIENS | 0 | 40570 | 5lst | A | 1021 | ARG | p.Arg1021Gln | O94761 |

| | | | | | | | | |
|---|---|---|---|---|---|---|---|---|
| HOMO SAPIENS | 0 | 40570 | 5lst | A | 1043 | LEU | p.Leu1043Pro | O94761 |
| HOMO SAPIENS | 0 | 40570 | 5lst | A | 1045 | ALA | p.Ala1045Thr | O94761 |
| HOMO SAPIENS | 0 | 40570 | 5lst | A | 1105 | GLY | p.Gly1105Asp | O94761 |
| HOMO SAPIENS | 0 | 40570 | 5lst | A | 1105 | GLY | p.Gly1105Ser | O94761 |
| HOMO SAPIENS | -1 | 40570 | 5lst | A | 1113 | GLY | p.Gly1113Arg | O94761 |
| HOMO SAPIENS | -1 | 18865 | 1gxc | A | 64 | GLU | p.Glu64Lys | O96017 |
| HOMO SAPIENS | -1 | 11673 | 3dzu | D | 85 | PRO | p.Pro113Gln | P37231 |
| HOMO SAPIENS | 0 | 19383 | 3b96 | A | 63 | LYS | p.Lys103Arg | P49748 |
| HOMO SAPIENS | 4 | 19383 | 3b96 | A | 140 | ALA | p.Ala180Thr | P49748 |
| HOMO SAPIENS | 1 | 19383 | 3b96 | A | 145 | GLY | p.Gly185Ser | P49748 |
| HOMO SAPIENS | 0 | 19383 | 3b96 | A | 220 | THR | p.Thr260Met | P49748 |
| HOMO SAPIENS | 5 | 19383 | 3b96 | A | 243 | VAL | p.Val283Ala | P49748 |
| HOMO SAPIENS | 0 | 19383 | 3b96 | A | 249 | GLY | p.Gly289Arg | P49748 |
| HOMO SAPIENS | 0 | 19383 | 3b96 | A | 254 | GLY | p.Gly294Glu | P49748 |
| HOMO SAPIENS | 7 | 19383 | 3b96 | A | 326 | ARG | p.Arg366Cys | P49748 |
| HOMO SAPIENS | 7 | 19383 | 3b96 | A | 326 | ARG | p.Arg366His | P49748 |
| HOMO SAPIENS | 5 | 19383 | 3b96 | A | 342 | LYS | p.Lys382Gln | P49748 |
| HOMO SAPIENS | 0 | 19383 | 3b96 | A | 369 | THR | p.Thr409Met | P49748 |
| HOMO SAPIENS | 5 | 19383 | 3b96 | A | 376 | ALA | p.Ala416Ser | P49748 |
| HOMO SAPIENS | 0 | 19383 | 3b96 | A | 385 | ALA | p.Ala425Thr | P49748 |
| HOMO SAPIENS | 5 | 19383 | 3b96 | A | 399 | GLY | p.Gly439Asp | P49748 |
| HOMO SAPIENS | 7 | 19383 | 3b96 | A | 401 | GLY | p.Gly441Asp | P49748 |
| HOMO SAPIENS | 9 | 19383 | 3b96 | A | 413 | ARG | p.Arg453Gln | P49748 |
| HOMO SAPIENS | 2 | 19383 | 3b96 | A | 418 | PHE | p.Phe458Leu | P49748 |
| HOMO SAPIENS | 6 | 19383 | 3b96 | A | 419 | ARG | p.Arg459Trp | P49748 |
| HOMO SAPIENS | 6 | 19383 | 3b96 | A | 419 | ARG | p.Arg459Gln | P49748 |
| HOMO SAPIENS | -1 | 19383 | 3b96 | A | 471 | ARG | p.Arg511Gln | P49748 |
| HOMO SAPIENS | 0 | 19383 | 3b96 | A | 573 | ARG | p.Arg613Trp | P49748 |
| HOMO SAPIENS | 0 | 19383 | 3b96 | A | 575 | ARG | p.Arg615Gln | P49748 |
| HOMO SAPIENS | 0 | 22286 | 3k2s | A | 3 | PRO | p.Pro27His | P02647 |
| HOMO SAPIENS | 0 | 22286 | 3k2s | A | 4 | PRO | p.Pro28Arg | P02647 |
| HOMO SAPIENS | 0 | 22286 | 3k2s | A | 10 | ARG | p.Arg34Leu | P02647 |
| HOMO SAPIENS | 0 | 22286 | 3k2s | A | 26 | GLY | p.Gly50Arg | P02647 |
| HOMO SAPIENS | 0 | 22286 | 3k2s | A | 37 | ALA | p.Ala61Thr | P02647 |
| HOMO SAPIENS | 1 | 1622 | 2qrv | D | 631 | ARG | p.Arg635Trp | Q9Y6K1 |
| HOMO SAPIENS | 1 | 1622 | 2qrv | D | 900 | PRO | p.Pro904Leu | Q9Y6K1 |
| HOMO SAPIENS | 6 | 40796 | 1v18 | B | 1509 | LEU | p.Leu1509Val | P25054 |
| HOMO SAPIENS | 1 | 22375 | 4pki | G | 1214 | LYS | p.Asp214Asn | P28289 |
| HOMO SAPIENS | 1 | 16307 | 5mv8 | B | 483 | ALA | p.Ala483Ser | Q9Y6N9 |
| HOMO SAPIENS | 6 | 16307 | 5mv8 | B | 506 | ASN | p.Asn506Lys | Q9Y6N9 |
| HOMO SAPIENS | 4 | 16307 | 5mv8 | B | 519 | ASP | p.Glu519Asp | Q9Y6N9 |
| HOMO SAPIENS | 0 | 16307 | 5mv8 | B | 530 | GLY | p.Gly530Arg | Q9Y6N9 |
| HOMO SAPIENS | 4 | 36221 | 4cvo | A | 127 | VAL | p.Val127Ile | Q03468 |
| HOMO SAPIENS | 7 | 36221 | 4cvo | A | 134 | ARG | p.Arg134Trp | Q03468 |
| HOMO SAPIENS | 0 | 36221 | 4cvo | A | 156 | GLN | p.Gln156Ter | Q03468 |
| HOMO SAPIENS | 0 | 10829 | 4a11 | B | 13 | GLU | p.Glu13Ter | Q13216 |
| HOMO SAPIENS | 1 | 10829 | 4a11 | B | 100 | TYR | p.Tyr100Ter | Q13216 |
| HOMO SAPIENS | 0 | 10829 | 4a11 | B | 160 | ALA | p.Ala160Val | Q13216 |
| HOMO SAPIENS | 1 | 6759 | 5lev | A | 29 | ILE | p.Ile29Phe | Q9H3H5 |
| HOMO SAPIENS | 0 | 6759 | 5lev | A | 108 | MET | p.Met108Ile | Q9H3H5 |
| HOMO SAPIENS | 2 | 6759 | 5lev | A | 117 | VAL | p.Val117Ile | Q9H3H5 |
| HOMO SAPIENS | 1 | 6759 | 5lev | A | 170 | TYR | p.Tyr170Cys | Q9H3H5 |
| HOMO SAPIENS | 0 | 6759 | 5lev | A | 264 | GLY | p.Val264Gly | Q9H3H5 |
| HOMO SAPIENS | 0 | 6759 | 5lev | A | 332 | PHE | p.Phe332Val | Q9H3H5 |

| | | | | | | | | |
|---|---|---|---|---|---|---|---|---|
| HOMO SAPIENS | 7 | 6759 | 5lev | A | 393 | ILE | p.Ile393Val | Q9H3H5 |
| HOMO SAPIENS | -1 | 33473 | 5m5e | C | 9 | ASN | p.Asn31Ser | P31785 |
| HOMO SAPIENS | 0 | 26155 | 1ic8 | B | 98 | ALA | p.Ala98Val | P20823 |
| HOMO SAPIENS | 0 | 15329 | 4iqr | B | 127 | ARG | p.Arg136Trp | P41235 |
| HOMO SAPIENS | 0 | 20725 | 4bsj | A | 350 | GLU | p.Glu350Val | P35916 |
| HOMO SAPIENS | 0 | 20725 | 4bsj | A | 494 | THR | p.Thr494Ala | P35916 |
| HOMO SAPIENS | 0 | 20725 | 4bsj | A | 527 | ASN | p.Asn527Ser | P35916 |
| HOMO SAPIENS | 6 | 29959 | 1xiw | B | 47 | ARG | p.Arg68Ter | P04234 |
| HOMO SAPIENS | 2 | 29959 | 1xiw | B | 72 | CYS | p.Cys93Ter | P04234 |
| HOMO SAPIENS | 3 | 20242 | 5tjc | A | 102 | ARG | p.Arg102Ter | Q9GZU1 |
| HOMO SAPIENS | 3 | 20242 | 5tjc | A | 232 | THR | p.Thr232Pro | Q9GZU1 |
| HOMO SAPIENS | 7 | 9590 | 4ap8 | A | 50 | THR | p.Thr50Ala | O96007 |
| HOMO SAPIENS | 4 | 9590 | 4ap8 | A | 168 | GLU | p.Glu168Lys | O96007 |
| HOMO SAPIENS | -1 | 4297 | 5do7 | B | 19 | ASP | p.Asp19His | Q9H221 |
| HOMO SAPIENS | -1 | 4297 | 5do7 | B | 54 | TYR | p.Tyr54Cys | Q9H221 |
| HOMO SAPIENS | -1 | 4297 | 5do7 | B | 80 | CYS | p.Cys80Tyr | Q9H221 |
| HOMO SAPIENS | 5 | 4297 | 5do7 | B | 107 | SER | p.Ser107Ter | Q9H221 |
| HOMO SAPIENS | 1 | 4297 | 5do7 | B | 164 | ARG | p.Arg164Pro | Q9H221 |
| HOMO SAPIENS | 0 | 4297 | 5do7 | B | 210 | VAL | p.Val210Met | Q9H221 |
| HOMO SAPIENS | 0 | 4297 | 5do7 | B | 231 | PRO | p.Pro231Thr | Q9H221 |
| HOMO SAPIENS | 0 | 4297 | 5do7 | B | 263 | ARG | p.Arg263Gln | Q9H221 |
| HOMO SAPIENS | 0 | 4297 | 5do7 | B | 361 | TRP | p.Trp361Ter | Q9H221 |
| HOMO SAPIENS | 0 | 4297 | 5do7 | B | 400 | THR | p.Thr400Lys | Q9H221 |
| HOMO SAPIENS | 0 | 4297 | 5do7 | B | 549 | ALA | p.Ala549Pro | Q9H221 |
| HOMO SAPIENS | 0 | 4297 | 5do7 | B | 596 | LEU | p.Leu596Arg | Q9H221 |
| HOMO SAPIENS | 0 | 4297 | 5do7 | B | 632 | VAL | p.Val632Ala | Q9H221 |
| HOMO SAPIENS | 0 | 4297 | 5do7 | B | 658 | TYR | p.Tyr658Ter | Q9H221 |
| HOMO SAPIENS | 2 | 9356 | 4igg | B | 206 | GLN | p.Gln206His | P35221 |
| HOMO SAPIENS | 6 | 2179 | 2wk6 | A | 44 | ARG | p.Arg44Gln | P19971 |
| HOMO SAPIENS | 6 | 2179 | 2wk6 | A | 81 | ARG | p.Arg81Gln | P19971 |
| HOMO SAPIENS | 6 | 2179 | 2wk6 | A | 145 | GLY | p.Gly145Arg | P19971 |
| HOMO SAPIENS | 7 | 2179 | 2wk6 | A | 146 | ARG | p.Arg146His | P19971 |
| HOMO SAPIENS | 7 | 2179 | 2wk6 | A | 153 | GLY | p.Gly153Ser | P19971 |
| HOMO SAPIENS | 0 | 2179 | 2wk6 | A | 173 | MET | p.Met173Ile | P19971 |
| HOMO SAPIENS | 6 | 2179 | 2wk6 | A | 202 | ARG | p.Arg202Lys | P19971 |
| HOMO SAPIENS | 6 | 2179 | 2wk6 | A | 208 | VAL | p.Val208Met | P19971 |
| HOMO SAPIENS | 5 | 2179 | 2wk6 | A | 222 | LYS | p.Lys222Arg | P19971 |
| HOMO SAPIENS | 5 | 2179 | 2wk6 | A | 289 | GLU | p.Glu289Ala | P19971 |
| HOMO SAPIENS | 2 | 2179 | 2wk6 | A | 311 | GLY | p.Gly311Arg | P19971 |
| HOMO SAPIENS | 2 | 2179 | 2wk6 | A | 376 | ARG | p.Arg376Pro | P19971 |
| HOMO SAPIENS | 2 | 2179 | 2wk6 | A | 465 | ALA | p.Ala465Thr | P19971 |
| HOMO SAPIENS | -1 | 11856 | 3orh | A | 3 | ALA | p.Ala3Ser | Q14353 |
| HOMO SAPIENS | 1 | 11856 | 3orh | A | 20 | TRP | p.Trp20Ser | Q14353 |
| HOMO SAPIENS | 1 | 11856 | 3orh | A | 50 | MET | p.Met50Leu | Q14353 |
| HOMO SAPIENS | 1 | 11856 | 3orh | A | 110 | VAL | p.Val110Phe | Q14353 |
| HOMO SAPIENS | 8 | 11856 | 3orh | A | 164 | GLY | p.Gly164Asp | Q14353 |
| HOMO SAPIENS | 1 | 11856 | 3orh | A | 169 | CYS | p.Cys169Tyr | Q14353 |
| HOMO SAPIENS | 1 | 11856 | 3orh | A | 174 | TRP | p.Trp174Ter | Q14353 |
| HOMO SAPIENS | 2 | 11856 | 3orh | A | 196 | ALA | p.Ala196Val | Q14353 |
| HOMO SAPIENS | 1 | 11856 | 3orh | A | 209 | THR | p.Thr209Met | Q14353 |
| HOMO SAPIENS | 1 | 11856 | 3orh | A | 219 | ASP | p.Asp219Asn | Q14353 |
| HOMO SAPIENS | 1 | 11856 | 3orh | A | 224 | ALA | p.Ala224Thr | Q14353 |
| HOMO SAPIENS | 1 | 13244 | 2m6i | D | 218 | ARG | p.Arg246Leu | P23415 |
| HOMO SAPIENS | 1 | 13244 | 2m6i | D | 244 | ILE | p.Ile272Asn | P23415 |

| | | | | | | | | |
|---|---|---|---|---|---|---|---|---|
| HOMO SAPIENS | 5 | 13244 | 2m6i | D | 250 | PRO | p.Pro278Thr | P23415 |
| HOMO SAPIENS | 6 | 13244 | 2m6i | D | 266 | GLN | p.Gln294His | P23415 |
| HOMO SAPIENS | 1 | 13244 | 2m6i | D | 271 | ARG | p.Arg299Leu | P23415 |
| HOMO SAPIENS | 1 | 13244 | 2m6i | D | 276 | LYS | p.Lys304Glu | P23415 |
| HOMO SAPIENS | 1 | 13244 | 2m6i | D | 279 | TYR | p.Tyr307Cys | P23415 |
| HOMO SAPIENS | 0 | 13244 | 2m6i | D | 316 | GLY | p.Gly378Ser | P23415 |
| HOMO SAPIENS | 1 | 13244 | 2m6i | D | 400 | ARG | p.Arg428His | P23415 |
| HOMO SAPIENS | 3 | 1967 | 2i6b | A | 301 | ALA | p.Ala318Glu | P55263 |
| HOMO SAPIENS | 4 | 11096 | 1h8b | A | 61 | ALA | p.Ala882Thr | P35609 |
| HOMO SAPIENS | 0 | 15423 | 5ngz | A | 2 | GLN | p.Gln2Glu | Q9NPD8 |
| HOMO SAPIENS | 3 | 30475 | 3lpw | A | 21 | SER | p.Ser22894Tyr | Q8WZ42 |
| HOMO SAPIENS | 6 | 32687 | 2arf | A | 1041 | ARG | p.Arg1041Trp | P35670 |
| HOMO SAPIENS | 8 | 32687 | 2arf | A | 1061 | GLY | p.Gly1061Glu | P35670 |
| HOMO SAPIENS | 6 | 32687 | 2arf | A | 1064 | GLU | p.Glu1064Ala | P35670 |
| HOMO SAPIENS | 2 | 32687 | 2arf | A | 1069 | HIS | p.His1069Gln | P35670 |
| HOMO SAPIENS | 2 | 32687 | 2arf | A | 1088 | LEU | p.Leu1088Ter | P35670 |
| HOMO SAPIENS | 0 | 32687 | 2arf | A | 1099 | GLY | p.Gly1099Ser | P35670 |
| HOMO SAPIENS | 4 | 32687 | 2arf | A | 1106 | VAL | p.Val1106Asp | P35670 |
| HOMO SAPIENS | 2 | 32687 | 2arf | A | 1142 | GLN | p.Gln1142His | P35670 |
| HOMO SAPIENS | 8 | 32687 | 2arf | A | 1148 | ILE | p.Ile1148Thr | P35670 |
| HOMO SAPIENS | 6 | 32687 | 2arf | A | 1151 | ARG | p.Arg1151Cys | P35670 |
| HOMO SAPIENS | 4 | 32687 | 2arf | A | 1169 | MET | p.Met1169Val | P35670 |
| HOMO SAPIENS | 4 | 32687 | 2arf | A | 1173 | GLU | p.Glu1173Lys | P35670 |
| HOMO SAPIENS | 2 | 32687 | 2arf | A | 1176 | GLY | p.Gly1176Arg | P35670 |
| HOMO SAPIENS | 3 | 15544 | 5uq3 | C | 575 | PRO | p.Pro69Leu | P46527 |
| HOMO SAPIENS | 6 | 15544 | 5uq3 | C | 592 | GLU | p.Glu86Asp | P46527 |
| HOMO SAPIENS | 6 | 15544 | 5uq3 | C | 599 | ARG | p.Arg93Trp | P46527 |
| HOMO SAPIENS | -1 | 15544 | 5uq3 | C | 615 | VAL | p.Val109Gly | P46527 |
| HOMO SAPIENS | -1 | 15544 | 5uq3 | C | 631 | SER | p.Ser125Pro | P46527 |
| HOMO SAPIENS | -1 | 15544 | 5uq3 | C | 639 | PRO | p.Pro133Ser | P46527 |
| HOMO SAPIENS | 2 | 2703 | 4wxq | A | 245 | CYS | p.Cys245Tyr | Q99972 |
| HOMO SAPIENS | 0 | 2703 | 4wxq | A | 252 | GLY | p.Gly252Arg | Q99972 |
| HOMO SAPIENS | 1 | 2703 | 4wxq | A | 337 | GLN | p.Gln337Arg | Q99972 |
| HOMO SAPIENS | 5 | 2703 | 4wxq | A | 347 | TYR | p.Tyr347Ter | Q99972 |
| HOMO SAPIENS | 3 | 2703 | 4wxq | A | 364 | GLY | p.Gly364Val | Q99972 |
| HOMO SAPIENS | 2 | 2703 | 4wxq | A | 367 | GLY | p.Gly367Arg | Q99972 |
| HOMO SAPIENS | 2 | 2703 | 4wxq | A | 368 | GLN | p.Gln368Ter | Q99972 |
| HOMO SAPIENS | 6 | 2703 | 4wxq | A | 370 | PRO | p.Pro370Leu | Q99972 |
| HOMO SAPIENS | 1 | 2703 | 4wxq | A | 380 | ASP | p.Asp380His | Q99972 |
| HOMO SAPIENS | 8 | 2703 | 4wxq | A | 399 | GLY | p.Gly399Val | Q99972 |
| HOMO SAPIENS | 0 | 2703 | 4wxq | A | 423 | LYS | p.Lys423Glu | Q99972 |
| HOMO SAPIENS | 3 | 2703 | 4wxq | A | 433 | CYS | p.Cys433Arg | Q99972 |
| HOMO SAPIENS | 3 | 2703 | 4wxq | A | 437 | TYR | p.Tyr437His | Q99972 |
| HOMO SAPIENS | 2 | 2703 | 4wxq | A | 477 | ILE | p.Ile477Ser | Q99972 |
| HOMO SAPIENS | 3 | 2703 | 4wxq | A | 480 | ASN | p.Asn480Lys | Q99972 |
| HOMO SAPIENS | -1 | 34100 | 2fot | C | 1171 | GLN | p.Gln1173Pro | Q13813 |
| HOMO SAPIENS | 0 | 8773 | 1kw2 | A | 432 | ASP | p.Asp432Glu | P02774 |
| HOMO SAPIENS | 0 | 8773 | 1kw2 | A | 436 | LYS | p.Lys436Thr | P02774 |
| HOMO SAPIENS | 0 | 8402 | 4dzo | B | 649 | TYR | p.Tyr649Ter | Q9Y6D9 |
| HOMO SAPIENS | 0 | 39962 | 1nst | A | 611 | GLY | p.Gly611Ser | P52848 |
| HOMO SAPIENS | 2 | 22578 | 3dof | B | 45 | MET | p.Met45Arg | Q9Y2Y0 |
| HOMO SAPIENS | 1 | 23290 | 5j1s | A | 292 | ILE | p.Ile292Thr | O14656 |
| HOMO SAPIENS | 4 | 9505 | 4j1y | A | 534 | ARG | p.Arg534Trp | P09871 |
| HOMO SAPIENS | 9 | 260 | 2h57 | A | 31 | THR | p.Thr31Met | Q9H0F7 |

| | | | | | | | | |
|---|---|---|---|---|---|---|---|---|
| HOMO SAPIENS | 0 | 260 | 2h57 | A | 91 | ILE | p.Ile91Thr | Q9H0F7 |
| HOMO SAPIENS | 0 | 260 | 2h57 | A | 121 | ARG | p.Arg121His | Q9H0F7 |
| HOMO SAPIENS | 0 | 260 | 2h57 | A | 122 | ARG | p.Arg122Ter | Q9H0F7 |
| HOMO SAPIENS | 2 | 260 | 2h57 | A | 169 | GLY | p.Gly169Ala | Q9H0F7 |
| HOMO SAPIENS | 4 | 260 | 2h57 | A | 170 | LEU | p.Leu170Trp | Q9H0F7 |
| HOMO SAPIENS | 4 | 34023 | 2cr3 | A | 18 | GLY | p.Gly48Ser | P11362 |
| HOMO SAPIENS | 0 | 37667 | 3ii7 | A | 420 | ARG | p.Arg420Cys | Q8IXQ5 |
| HOMO SAPIENS | 0 | 41720 | 2k9u | A | 68 | ARG | p.Arg2364His | Q14315 |
| HOMO SAPIENS | 1 | 29250 | 5fmv | A | 568 | HIS | p.His568Gln | P08575 |
| HOMO SAPIENS | 1 | 1280 | 1fb1 | A | 69 | PRO | p.Pro69Leu | P30793 |
| HOMO SAPIENS | 3 | 1280 | 1fb1 | A | 88 | ARG | p.Arg88Trp | P30793 |
| HOMO SAPIENS | 2 | 1280 | 1fb1 | A | 108 | GLY | p.Gly108Asp | P30793 |
| HOMO SAPIENS | 4 | 1280 | 1fb1 | A | 134 | ASP | p.Asp134Val | P30793 |
| HOMO SAPIENS | 6 | 1280 | 1fb1 | A | 135 | ILE | p.Ile135Lys | P30793 |
| HOMO SAPIENS | 9 | 1280 | 1fb1 | A | 144 | HIS | p.His144Pro | P30793 |
| HOMO SAPIENS | 9 | 1280 | 1fb1 | A | 184 | ARG | p.Arg184His | P30793 |
| HOMO SAPIENS | 1 | 1280 | 1fb1 | A | 196 | ALA | p.Ala196Ser | P30793 |
| HOMO SAPIENS | 3 | 1280 | 1fb1 | A | 201 | GLY | p.Gly201Glu | P30793 |
| HOMO SAPIENS | 2 | 1280 | 1fb1 | A | 204 | VAL | p.Val204Ile | P30793 |
| HOMO SAPIENS | 2 | 1280 | 1fb1 | A | 211 | MET | p.Met211Ile | P30793 |
| HOMO SAPIENS | 0 | 1280 | 1fb1 | A | 221 | MET | p.Met221Thr | P30793 |
| HOMO SAPIENS | 3 | 1280 | 1fb1 | A | 224 | LYS | p.Lys224Arg | P30793 |
| HOMO SAPIENS | 2 | 1280 | 1fb1 | A | 249 | ARG | p.Arg249Ser | P30793 |
| HOMO SAPIENS | 8 | 41352 | 4k12 | A | 47 | ILE | p.Ile551Thr | P08603 |
| HOMO SAPIENS | 9 | 20703 | 5nir | A | 57 | CYS | p.Cys57Tyr | P02458 |
| HOMO SAPIENS | 9 | 20703 | 5nir | A | 64 | CYS | p.Cys64Ter | P02458 |
| HOMO SAPIENS | 0 | 20087 | 4cmn | A | 299 | MET | p.Met299Ile | Q01968 |
| HOMO SAPIENS | 1 | 20819 | 3aaf | A | 979 | ILE | p.Ile979Met | Q14191 |
| HOMO SAPIENS | 1 | 20819 | 3aaf | A | 984 | LEU | p.Leu984Ile | Q14191 |
| HOMO SAPIENS | 3 | 20819 | 3aaf | A | 987 | ARG | p.Arg987Ter | Q14191 |
| HOMO SAPIENS | -1 | 20819 | 3aaf | A | 1074 | PHE | p.Leu1074Phe | Q14191 |
| HOMO SAPIENS | 0 | 3619 | 3dem | A | 206 | ASN | p.Asn225Ser | P48740 |
| HOMO SAPIENS | 1 | 3619 | 3dem | A | 269 | ARG | p.Arg288Leu | P48740 |
| HOMO SAPIENS | 0 | 3105 | 3ooi | A | 164 | TYR | p.Tyr2015His | Q96L73 |
| HOMO SAPIENS | 6 | 15828 | 2lkx | A | 16 | LEU | p.Leu100Gln | Q99697 |
| HOMO SAPIENS | 8 | 15828 | 2lkx | A | 24 | ARG | p.Arg108His | Q99697 |
| HOMO SAPIENS | 2 | 15828 | 2lkx | A | 30 | THR | p.Thr114Pro | Q99697 |
| HOMO SAPIENS | 8 | 15828 | 2lkx | A | 45 | VAL | p.Val129Leu | Q99697 |
| HOMO SAPIENS | 8 | 15828 | 2lkx | A | 46 | ARG | p.Arg130Trp | Q99697 |
| HOMO SAPIENS | 9 | 15828 | 2lkx | A | 53 | ARG | p.Arg137Pro | Q99697 |
| HOMO SAPIENS | 9 | 13017 | 4irq | A | 141 | ARG | p.Arg141Trp | Q9UBV7 |
| HOMO SAPIENS | 0 | 13017 | 4irq | A | 186 | ALA | p.Ala186Asp | Q9UBV7 |
| HOMO SAPIENS | 0 | 13017 | 4irq | A | 206 | LEU | p.Leu206Pro | Q9UBV7 |
| HOMO SAPIENS | 1 | 13017 | 4irq | A | 214 | CYS | p.Cys214Tyr | Q9UBV7 |
| HOMO SAPIENS | 0 | 13017 | 4irq | A | 270 | ARG | p.Arg270Cys | Q9UBV7 |
| HOMO SAPIENS | -1 | 13053 | 4mqw | X | 307 | ALA | p.Ala307Thr | P23945 |
| HOMO SAPIENS | 0 | 19406 | 4b3g | A | 43 | ARG | p.Arg43Ter | P38935 |
| HOMO SAPIENS | 0 | 19406 | 4b3g | A | 46 | CYS | p.Cys46Ter | P38935 |
| HOMO SAPIENS | 0 | 19406 | 4b3g | A | 75 | ALA | p.Ala75Thr | P38935 |
| HOMO SAPIENS | 0 | 19406 | 4b3g | A | 201 | LEU | p.Leu201Ser | P38935 |
| HOMO SAPIENS | 3 | 19406 | 4b3g | A | 213 | HIS | p.His213Arg | P38935 |
| HOMO SAPIENS | 0 | 19406 | 4b3g | A | 275 | ILE | p.Ile275Val | P38935 |
| HOMO SAPIENS | -1 | 19406 | 4b3g | A | 320 | ARG | p.Arg320Ter | P38935 |
| HOMO SAPIENS | 0 | 19406 | 4b3g | A | 361 | LEU | p.Leu361Pro | P38935 |

| | | | | | | | | |
|---|---|---|---|---|---|---|---|---|
| HOMO SAPIENS | 0 | 19406 | 4b3g | A | 369 | PHE | p.Phe369Leu | P38935 |
| HOMO SAPIENS | 0 | 19406 | 4b3g | A | 493 | THR | p.Thr493Ile | P38935 |
| HOMO SAPIENS | 0 | 19406 | 4b3g | A | 496 | CYS | p.Cys496Ter | P38935 |
| HOMO SAPIENS | 0 | 19406 | 4b3g | A | 514 | GLU | p.Glu514Lys | P38935 |
| HOMO SAPIENS | 0 | 19406 | 4b3g | A | 528 | ALA | p.Ala528Thr | P38935 |
| HOMO SAPIENS | 0 | 19406 | 4b3g | A | 531 | PRO | p.Pro531Thr | P38935 |
| HOMO SAPIENS | 0 | 19406 | 4b3g | A | 580 | VAL | p.Val580Ile | P38935 |
| HOMO SAPIENS | 6 | 19406 | 4b3g | A | 603 | ARG | p.Arg603His | P38935 |
| HOMO SAPIENS | 0 | 21501 | 5jde | A | 40 | SER | p.Ser2831Asn | Q8WZ42 |
| HOMO SAPIENS | 5 | 21501 | 5jde | A | 189 | ALA | p.Ala2980Thr | Q8WZ42 |
| HOMO SAPIENS | 6 | 15008 | 3mop | H | 12 | ARG | p.Arg12Cys | Q9NWZ3 |
| HOMO SAPIENS | 8 | 17455 | 3jtc | C | 15 | ARG | p.Arg57Trp | P04070 |
| HOMO SAPIENS | -1 | 17455 | 3jtc | C | 20 | CGU | p.Glu62Ala | P04070 |
| HOMO SAPIENS | 0 | 27795 | 5gox | B | 625 | TYR | p.Tyr625Ter | Q92878 |
| HOMO SAPIENS | 0 | 27795 | 5gox | B | 653 | SER | p.Ser653Ter | Q92878 |
| HOMO SAPIENS | -1 | 6558 | 5nug | A | 137 | VAL | p.Val137Ile | Q14204 |
| HOMO SAPIENS | -1 | 6558 | 5nug | A | 567 | ARG | p.Arg567His | Q14204 |
| HOMO SAPIENS | -1 | 6558 | 5nug | A | 1140 | THR | p.Thr1140Met | Q14204 |
| HOMO SAPIENS | -1 | 6558 | 5nug | A | 1991 | ASP | p.Asp1991Asn | Q14204 |
| HOMO SAPIENS | -1 | 6558 | 5nug | A | 2401 | LYS | p.Lys2401Asn | Q14204 |
| HOMO SAPIENS | 0 | 6558 | 5nug | A | 2488 | ARG | p.Arg2488His | Q14204 |
| HOMO SAPIENS | 0 | 6558 | 5nug | A | 3907 | HIS | p.His3907Gln | Q14204 |
| HOMO SAPIENS | 0 | 6558 | 5nug | A | 3981 | THR | p.Thr3981Arg | Q14204 |
| HOMO SAPIENS | 0 | 6558 | 5nug | A | 4394 | THR | p.Thr4394Met | Q14204 |
| HOMO SAPIENS | 0 | 6558 | 5nug | A | 4573 | ASN | p.Asn4573Ser | Q14204 |
| HOMO SAPIENS | 0 | 6558 | 5nug | A | 4603 | SER | p.Ser4603Ile | Q14204 |
| HOMO SAPIENS | 1 | 4244 | 1umk | A | 57 | ARG | p.Arg58Gln | P00387 |
| HOMO SAPIENS | 1 | 4244 | 1umk | A | 72 | LEU | p.Leu73Pro | P00387 |
| HOMO SAPIENS | 1 | 4244 | 1umk | A | 105 | VAL | p.Val106Met | P00387 |
| HOMO SAPIENS | 0 | 4244 | 1umk | A | 116 | THR | p.Thr117Ser | P00387 |
| HOMO SAPIENS | 8 | 4244 | 1umk | A | 127 | SER | p.Ser128Pro | P00387 |
| HOMO SAPIENS | 0 | 4244 | 1umk | A | 148 | LEU | p.Leu149Pro | P00387 |
| HOMO SAPIENS | 0 | 4244 | 1umk | A | 159 | ARG | p.Arg160Ter | P00387 |
| HOMO SAPIENS | 0 | 4244 | 1umk | A | 203 | CYS | p.Cys204Arg | P00387 |
| HOMO SAPIENS | 0 | 4244 | 1umk | A | 203 | CYS | p.Cys204Tyr | P00387 |
| HOMO SAPIENS | 2 | 4244 | 1umk | A | 239 | ASP | p.Asp240Gly | P00387 |
| HOMO SAPIENS | 0 | 4244 | 1umk | A | 291 | GLY | p.Gly292Asp | P00387 |
| HOMO SAPIENS | 4 | 34495 | 1s35 | A | 1151 | ASP | p.Asn1151Asp | P11277 |
| HOMO SAPIENS | 3 | 30110 | 3zqs | A | 224 | THR | p.Thr224Ala | Q9NW38 |
| HOMO SAPIENS | 3 | 17588 | 5dse | B | 53 | LEU | p.Leu53Pro | Q9BYI3 |
| HOMO SAPIENS | 0 | 1564 | 4yqf | B | 72 | HIS | p.His347Gln | Q9UHD8 |
| HOMO SAPIENS | 0 | 17673 | 4jnw | A | 32 | ARG | p.Arg32186His | Q8WZ42 |
| HOMO SAPIENS | 3 | 17673 | 4jnw | A | 140 | ILE | p.Ile32294Val | Q8WZ42 |
| HOMO SAPIENS | 5 | 17673 | 4jnw | A | 269 | ALA | p.Ala32423Thr | Q8WZ42 |
| HOMO SAPIENS | 3 | 12858 | 3s97 | C | 302 | LEU | p.Leu302Pro | Q12860 |
| HOMO SAPIENS | 0 | 6147 | 4nn0 | A | 114 | ARG | p.Arg114Gln | Q9BXJ0 |
| HOMO SAPIENS | 0 | 7549 | 1tr2 | A | 277 | LEU | p.Leu277Met | P18206 |
| HOMO SAPIENS | 1 | 7549 | 1tr2 | A | 413 | ALA | p.Ala413Thr | P18206 |
| HOMO SAPIENS | 1 | 7549 | 1tr2 | A | 519 | ILE | p.Ile519Leu | P18206 |
| HOMO SAPIENS | 5 | 7549 | 1tr2 | A | 547 | ARG | p.Arg547Ter | P18206 |
| HOMO SAPIENS | 2 | 7549 | 1tr2 | A | 636 | HIS | p.His636Arg | P18206 |
| HOMO SAPIENS | 2 | 7549 | 1tr2 | A | 824 | ILE | p.Ile824Met | P18206 |
| HOMO SAPIENS | 2 | 13900 | 3gro | A | 45 | CYS | p.Cys45Tyr | P50897 |
| HOMO SAPIENS | 2 | 13900 | 3gro | A | 75 | THR | p.Thr75Pro | P50897 |

| | | | | | | | | |
|---|---|---|---|---|---|---|---|---|
| HOMO SAPIENS | 2 | 13900 | 3gro | A | 79 | ASP | p.Asp79Gly | P50897 |
| HOMO SAPIENS | 5 | 13900 | 3gro | A | 91 | GLN | p.Gln91Pro | P50897 |
| HOMO SAPIENS | 5 | 13900 | 3gro | A | 96 | CYS | p.Cys96Tyr | P50897 |
| HOMO SAPIENS | 0 | 13900 | 3gro | A | 104 | LYS | p.Lys104Ter | P50897 |
| HOMO SAPIENS | 2 | 13900 | 3gro | A | 108 | GLY | p.Gly108Arg | P50897 |
| HOMO SAPIENS | 2 | 13900 | 3gro | A | 109 | TYR | p.Tyr109Asp | P50897 |
| HOMO SAPIENS | 0 | 13900 | 3gro | A | 122 | ARG | p.Arg122Trp | P50897 |
| HOMO SAPIENS | 0 | 13900 | 3gro | A | 138 | SER | p.Ser138Leu | P50897 |
| HOMO SAPIENS | 5 | 13900 | 3gro | A | 151 | ARG | p.Arg151Ter | P50897 |
| HOMO SAPIENS | 7 | 13900 | 3gro | A | 164 | ARG | p.Arg164Ter | P50897 |
| HOMO SAPIENS | 0 | 13900 | 3gro | A | 177 | GLN | p.Gln177Glu | P50897 |
| HOMO SAPIENS | 0 | 13900 | 3gro | A | 181 | VAL | p.Val181Leu | P50897 |
| HOMO SAPIENS | 0 | 13900 | 3gro | A | 184 | GLU | p.Glu184Lys | P50897 |
| HOMO SAPIENS | 5 | 13900 | 3gro | A | 186 | TRP | p.Trp186Ter | P50897 |
| HOMO SAPIENS | 0 | 13900 | 3gro | A | 219 | LEU | p.Leu219Gln | P50897 |
| HOMO SAPIENS | 0 | 13900 | 3gro | A | 222 | LEU | p.Leu222Pro | P50897 |
| HOMO SAPIENS | 5 | 13900 | 3gro | A | 296 | TRP | p.Trp296Ter | P50897 |
| HOMO SAPIENS | 6 | 35028 | 5mwb | A | 444 | CYS | p.Cys444Tyr | Q04721 |
| HOMO SAPIENS | 0 | 35028 | 5mwb | A | 514 | ASN | p.Asn514Ser | Q04721 |
| HOMO SAPIENS | 5 | 1483 | 4gl2 | A | 822 | ARG | p.Arg822Gln | Q9BYX4 |
| HOMO SAPIENS | 0 | 1483 | 4gl2 | A | 843 | ARG | p.His843Arg | Q9BYX4 |
| HOMO SAPIENS | -1 | 28838 | 3cl3 | D | 153 | LEU | p.Leu153Arg | Q9Y6K9 |
| HOMO SAPIENS | 2 | 22563 | 4zel | B | 79 | ARG | p.Arg79Trp | P09172 |
| HOMO SAPIENS | 2 | 22563 | 4zel | B | 101 | VAL | p.Val101Met | P09172 |
| HOMO SAPIENS | 2 | 22563 | 4zel | B | 114 | ASP | p.Asp114Glu | P09172 |
| HOMO SAPIENS | 0 | 22563 | 4zel | B | 201 | ASN | p.Asn201Ser | P09172 |
| HOMO SAPIENS | 2 | 22563 | 4zel | B | 211 | ALA | p.Ala211Thr | P09172 |
| HOMO SAPIENS | -1 | 22563 | 4zel | B | 290 | ASP | p.Asp290Asn | P09172 |
| HOMO SAPIENS | 2 | 22563 | 4zel | B | 318 | ALA | p.Ala318Ser | P09172 |
| HOMO SAPIENS | 2 | 22563 | 4zel | B | 345 | ASP | p.Asp345Asn | P09172 |
| HOMO SAPIENS | 2 | 22563 | 4zel | B | 362 | ALA | p.Ala362Glu | P09172 |
| HOMO SAPIENS | 2 | 22563 | 4zel | B | 549 | ARG | p.Arg549Cys | P09172 |
| HOMO SAPIENS | 9 | 9780 | 1y6k | R | 36 | TYR | p.Tyr57Cys | Q13651 |
| HOMO SAPIENS | 1 | 9780 | 1y6k | R | 40 | LEU | p.Leu61Val | Q13651 |
| HOMO SAPIENS | 1 | 9780 | 1y6k | R | 63 | THR | p.Thr84Ile | Q13651 |
| HOMO SAPIENS | 9 | 9780 | 1y6k | R | 80 | ARG | p.Arg101Trp | Q13651 |
| HOMO SAPIENS | 0 | 9780 | 1y6k | R | 120 | GLY | p.Gly141Arg | Q13651 |
| HOMO SAPIENS | 0 | 9780 | 1y6k | R | 138 | SER | p.Ser159Gly | Q13651 |
| HOMO SAPIENS | 0 | 9780 | 1y6k | R | 203 | ILE | p.Ile224Val | Q13651 |
| HOMO SAPIENS | 0 | 16019 | 3rep | A | 211 | ARG | p.Arg211Cys | Q13418 |
| HOMO SAPIENS | 3 | 7879 | 5ec3 | A | 33 | ALA | p.Ala33Thr | P32754 |
| HOMO SAPIENS | 4 | 7879 | 5ec3 | A | 113 | ARG | p.Arg113Gln | P32754 |
| HOMO SAPIENS | 3 | 7879 | 5ec3 | A | 340 | VAL | p.Val340Leu | P32754 |
| HOMO SAPIENS | 0 | 8904 | 1qk9 | A | 6 | LYS | p.Lys82Arg | P51608 |
| HOMO SAPIENS | 4 | 8904 | 1qk9 | A | 24 | LEU | p.Leu100Val | P51608 |
| HOMO SAPIENS | 7 | 8904 | 1qk9 | A | 30 | ARG | p.Arg106Trp | P51608 |
| HOMO SAPIENS | 6 | 8904 | 1qk9 | A | 57 | ARG | p.Arg133Cys | P51608 |
| HOMO SAPIENS | 3 | 8904 | 1qk9 | A | 61 | GLU | p.Glu137Gly | P51608 |
| HOMO SAPIENS | 2 | 8904 | 1qk9 | A | 64 | ALA | p.Ala140Val | P51608 |
| HOMO SAPIENS | 5 | 8904 | 1qk9 | A | 79 | PHE | p.Phe155Ser | P51608 |
| HOMO SAPIENS | 3 | 8904 | 1qk9 | A | 82 | THR | p.Thr158Met | P51608 |
| HOMO SAPIENS | 1 | 23121 | 3e9k | A | 198 | THR | p.Thr198Ala | Q16719 |
| HOMO SAPIENS | -1 | 26090 | 3tje | F | 19 | LEU | p.Leu35Met | P25445 |
| HOMO SAPIENS | 8 | 26090 | 3tje | F | 106 | THR | p.Thr122Ile | P25445 |

| | | | | | | | | |
|---|---|---|---|---|---|---|---|---|
| HOMO SAPIENS | 2 | 36404 | 5cqr | A | 765 | GLY | p.Gly765Glu | O95163 |
| HOMO SAPIENS | 2 | 36404 | 5cqr | A | 816 | ILE | p.Ile816Leu | O95163 |
| HOMO SAPIENS | 2 | 36404 | 5cqr | A | 830 | ILE | p.Ile830Met | O95163 |
| HOMO SAPIENS | 5 | 36404 | 5cqr | A | 939 | TYR | p.Tyr939Ter | O95163 |
| HOMO SAPIENS | 2 | 36404 | 5cqr | A | 952 | LYS | p.Lys952Ile | O95163 |
| HOMO SAPIENS | 5 | 36404 | 5cqr | A | 1013 | GLY | p.Gly1013Ser | O95163 |
| HOMO SAPIENS | 5 | 36404 | 5cqr | A | 1072 | CYS | p.Cys1072Ser | O95163 |
| HOMO SAPIENS | -1 | 36404 | 5cqr | A | 1158 | PRO | p.Pro1158Leu | O95163 |
| HOMO SAPIENS | 1 | 34149 | 2k40 | A | 18 | ASN | p.Asn125Ser | Q9UBX0 |
| HOMO SAPIENS | 0 | 34149 | 2k40 | A | 42 | LEU | p.Glu149Lys | Q9UBX0 |
| HOMO SAPIENS | 9 | 34149 | 2k40 | A | 53 | ARG | p.Arg160Cys | Q9UBX0 |
| HOMO SAPIENS | 0 | 34149 | 2k40 | A | 63 | SER | p.Ser170Leu | Q9UBX0 |
| HOMO SAPIENS | 2 | 2468 | 3vp6 | A | 484 | LEU | p.Leu484Met | Q99259 |
| HOMO SAPIENS | 0 | 2468 | 3vp6 | A | 532 | ARG | p.Arg532Gln | Q99259 |
| HOMO SAPIENS | 5 | 1253 | 5xtd | S | 8 | GLY | p.Gly8Arg | O15239 |
| HOMO SAPIENS | 7 | 1253 | 5xtd | S | 32 | GLY | p.Gly32Arg | O15239 |
| HOMO SAPIENS | 7 | 1253 | 5xtd | S | 37 | ARG | p.Arg37Ser | O15239 |
| HOMO SAPIENS | -1 | 13292 | 3u1k | A | 121 | VAL | p.Ile121Val | Q8TCS8 |
| HOMO SAPIENS | 2 | 13292 | 3u1k | A | 165 | PRO | p.Pro165Ser | Q8TCS8 |
| HOMO SAPIENS | 2 | 13292 | 3u1k | A | 192 | ARG | p.Arg192Gln | Q8TCS8 |
| HOMO SAPIENS | 1 | 13292 | 3u1k | A | 315 | LYS | p.Lys315Arg | Q8TCS8 |
| HOMO SAPIENS | 4 | 13292 | 3u1k | A | 509 | VAL | p.Val509Ile | Q8TCS8 |
| HOMO SAPIENS | 7 | 13292 | 3u1k | A | 531 | THR | p.Thr531Arg | Q8TCS8 |
| HOMO SAPIENS | 0 | 13292 | 3u1k | A | 590 | ASN | p.Asn590Asp | Q8TCS8 |
| HOMO SAPIENS | 0 | 27048 | 4p5w | B | 184 | ARG | p.Arg184Ter | Q9UJS0 |
| HOMO SAPIENS | 0 | 27048 | 4p5w | B | 225 | SER | p.Ser225Ter | Q9UJS0 |
| HOMO SAPIENS | -1 | 27048 | 4p5w | B | 631 | ALA | p.Arg360Ter | Q9UJS0 |
| HOMO SAPIENS | 0 | 24240 | 4mdr | A | 318 | ARG | p.Arg318Ter | O00189 |
| HOMO SAPIENS | 0 | 24240 | 4mdr | A | 338 | ARG | p.Arg338Ter | O00189 |
| HOMO SAPIENS | 0 | 13429 | 4cg4 | A | 479 | PHE | p.Phe479Leu | O15553 |
| HOMO SAPIENS | 0 | 13429 | 4cg4 | A | 487 | VAL | p.Val487Leu | O15553 |
| HOMO SAPIENS | 3 | 13429 | 4cg4 | A | 653 | ARG | p.Arg653His | O15553 |
| HOMO SAPIENS | 1 | 13429 | 4cg4 | A | 680 | MET | p.Met680Ile | O15553 |
| HOMO SAPIENS | 1 | 13429 | 4cg4 | A | 688 | TYR | p.Tyr688Ter | O15553 |
| HOMO SAPIENS | 0 | 13429 | 4cg4 | A | 694 | MET | p.Met694Ile | O15553 |
| HOMO SAPIENS | 0 | 13429 | 4cg4 | A | 694 | MET | p.Met694Val | O15553 |
| HOMO SAPIENS | 1 | 13429 | 4cg4 | A | 726 | VAL | p.Val726Ala | O15553 |
| HOMO SAPIENS | 0 | 13429 | 4cg4 | A | 761 | ARG | p.Arg761His | O15553 |
| HOMO SAPIENS | 0 | 2079 | 4okh | A | 691 | GLY | p.Gly691Arg | P20807 |
| HOMO SAPIENS | 2 | 2079 | 4okh | A | 748 | ARG | p.Arg748Ter | P20807 |
| HOMO SAPIENS | 0 | 2079 | 4okh | A | 763 | TYR | p.Tyr763Cys | P20807 |
| HOMO SAPIENS | 1 | 2079 | 4okh | A | 769 | ARG | p.Arg769Gln | P20807 |
| HOMO SAPIENS | 9 | 2079 | 4okh | A | 777 | ILE | p.Ile777Val | P20807 |
| HOMO SAPIENS | 8 | 2079 | 4okh | A | 778 | ASP | p.Asp778His | P20807 |
| HOMO SAPIENS | 5 | 2079 | 4okh | A | 780 | ASP | p.Asp780His | P20807 |
| HOMO SAPIENS | 1 | 1200 | 2gi7 | A | 142 | ARG | p.Arg162Trp | Q9HCN6 |
| HOMO SAPIENS | 4 | 9199 | 3ezq | C | 260 | ASP | p.Asp260Val | P25445 |
| HOMO SAPIENS | 0 | 36522 | 5ms9 | A | 122 | ARG | p.Arg122Cys | P35555 |
| HOMO SAPIENS | 0 | 36522 | 5ms9 | A | 240 | ARG | p.Arg240Cys | P35555 |
| HOMO SAPIENS | 0 | 17448 | 3g2f | A | 225 | ARG | p.Arg225His | Q13873 |
| HOMO SAPIENS | 0 | 17448 | 3g2f | A | 347 | CYS | p.Cys347Tyr | Q13873 |
| HOMO SAPIENS | 3 | 17448 | 3g2f | A | 485 | ASP | p.Asp485Gly | Q13873 |
| HOMO SAPIENS | 7 | 17448 | 3g2f | A | 491 | ARG | p.Arg491Gln | Q13873 |
| HOMO SAPIENS | 7 | 17448 | 3g2f | A | 491 | ARG | p.Arg491Trp | Q13873 |

| | | | | | | | | |
|---|---|---|---|---|---|---|---|---|
| HOMO SAPIENS | 0 | 4861 | 4ux8 | A | 56 | LEU | p.Leu56Met | P07949 |
| HOMO SAPIENS | 1 | 4861 | 4ux8 | A | 88 | ILE | p.Ile88Leu | P07949 |
| HOMO SAPIENS | 0 | 4861 | 4ux8 | A | 114 | ARG | p.Arg114His | P07949 |
| HOMO SAPIENS | 5 | 4861 | 4ux8 | A | 180 | ARG | p.Arg180Gln | P07949 |
| HOMO SAPIENS | 0 | 4861 | 4ux8 | A | 208 | GLU | p.Glu208Asp | P07949 |
| HOMO SAPIENS | 0 | 4861 | 4ux8 | A | 373 | ALA | p.Ala373Val | P07949 |
| HOMO SAPIENS | 0 | 4861 | 4ux8 | A | 375 | LEU | p.Leu375Gln | P07949 |
| HOMO SAPIENS | -1 | 4861 | 4ux8 | A | 386 | ALA | p.Ala386Val | P07949 |
| HOMO SAPIENS | 0 | 4861 | 4ux8 | A | 446 | GLY | p.Gly446Arg | P07949 |
| HOMO SAPIENS | 0 | 4861 | 4ux8 | A | 480 | GLU | p.Glu480Lys | P07949 |
| HOMO SAPIENS | 0 | 4861 | 4ux8 | A | 489 | ASP | p.Asp489Asn | P07949 |
| HOMO SAPIENS | -1 | 4861 | 4ux8 | A | 510 | ALA | p.Ala510Val | P07949 |
| HOMO SAPIENS | -1 | 4861 | 4ux8 | A | 533 | GLY | p.Gly533Cys | P07949 |
| HOMO SAPIENS | -1 | 4861 | 4ux8 | A | 567 | ASP | p.Asp567Asn | P07949 |
| HOMO SAPIENS | -1 | 4861 | 4ux8 | A | 609 | CYS | p.Cys609Tyr | P07949 |
| HOMO SAPIENS | -1 | 4861 | 4ux8 | A | 609 | CYS | p.Cys609Gly | P07949 |
| HOMO SAPIENS | -1 | 4861 | 4ux8 | A | 611 | CYS | p.Cys611Trp | P07949 |
| HOMO SAPIENS | -1 | 4861 | 4ux8 | A | 618 | CYS | p.Cys618Gly | P07949 |
| HOMO SAPIENS | -1 | 4861 | 4ux8 | A | 618 | CYS | p.Cys618Phe | P07949 |
| HOMO SAPIENS | -1 | 4861 | 4ux8 | A | 620 | CYS | p.Cys620Gly | P07949 |
| HOMO SAPIENS | -1 | 4861 | 4ux8 | A | 620 | CYS | p.Cys620Ser | P07949 |
| HOMO SAPIENS | -1 | 4861 | 4ux8 | A | 620 | CYS | p.Cys620Trp | P07949 |
| HOMO SAPIENS | -1 | 4861 | 4ux8 | A | 623 | GLU | p.Glu623Lys | P07949 |
| HOMO SAPIENS | -1 | 4861 | 4ux8 | A | 631 | ASP | p.Asp631Gly | P07949 |
| HOMO SAPIENS | -1 | 4861 | 4ux8 | A | 631 | ASP | p.Asp631Glu | P07949 |
| HOMO SAPIENS | -1 | 4861 | 4ux8 | A | 634 | CYS | p.Cys634Gly | P07949 |
| HOMO SAPIENS | -1 | 4861 | 4ux8 | A | 634 | CYS | p.Cys634Phe | P07949 |
| HOMO SAPIENS | -1 | 4861 | 4ux8 | A | 634 | CYS | p.Cys634Trp | P07949 |
| HOMO SAPIENS | 0 | 36308 | 4nkq | B | 123 | PRO | p.Pro142Leu | P15509 |
| HOMO SAPIENS | 0 | 36308 | 4nkq | B | 145 | ARG | p.Arg164Gln | P15509 |
| HOMO SAPIENS | -1 | 36308 | 4nkq | B | 177 | GLY | p.Gly196Arg | P15509 |
| HOMO SAPIENS | 0 | 36006 | 1oqa | A | 107 | HIS | p.His1860Pro | P38398 |
| HOMO SAPIENS | 7 | 15394 | 2ozo | A | 465 | ARG | p.Arg465His | P43403 |
| HOMO SAPIENS | 7 | 15394 | 2ozo | A | 465 | ARG | p.Arg465Cys | P43403 |
| HOMO SAPIENS | 4 | 15394 | 2ozo | A | 518 | SER | p.Ser518Arg | P43403 |
| HOMO SAPIENS | 1 | 20813 | 3o5z | A | 19 | ARG | p.Arg19Trp | O00459 |
| HOMO SAPIENS | 2 | 4329 | 2bb5 | A | 5 | ILE | p.Ile23Val | P20062 |
| HOMO SAPIENS | 4 | 4329 | 2bb5 | A | 59 | LYS | p.Lys77Met | P20062 |
| HOMO SAPIENS | 4 | 4329 | 2bb5 | A | 197 | ARG | p.Arg215Trp | P20062 |
| HOMO SAPIENS | 1 | 4329 | 2bb5 | A | 241 | PRO | p.Arg259Pro | P20062 |
| HOMO SAPIENS | 2 | 4329 | 2bb5 | A | 315 | THR | p.Thr333Lys | P20062 |
| HOMO SAPIENS | 5 | 4329 | 2bb5 | A | 330 | SER | p.Ser348Phe | P20062 |
| HOMO SAPIENS | 4 | 4329 | 2bb5 | A | 358 | LEU | p.Leu376Ser | P20062 |
| HOMO SAPIENS | 5 | 4329 | 2bb5 | A | 381 | ARG | p.Arg399Gln | P20062 |
| HOMO SAPIENS | 4 | 18294 | 4r30 | A | 163 | ASN | p.Asn163Asp | O95278 |
| HOMO SAPIENS | 6 | 18294 | 4r30 | A | 227 | ALA | p.Ala227Val | O95278 |
| HOMO SAPIENS | 4 | 18294 | 4r30 | A | 241 | ARG | p.Arg241Ter | O95278 |
| HOMO SAPIENS | 4 | 18294 | 4r30 | A | 325 | ARG | p.Arg325His | O95278 |
| HOMO SAPIENS | 0 | 2632 | 4xem | A | 22 | GLU | p.Glu22Lys | P49588 |
| HOMO SAPIENS | 0 | 2632 | 4xem | A | 275 | GLY | p.Gly275Asp | P49588 |
| HOMO SAPIENS | 1 | 2632 | 4xem | A | 329 | ARG | p.Arg329His | P49588 |
| HOMO SAPIENS | 0 | 11431 | 5t4n | A | 284 | GLN | p.Gln305Glu | Q96QU1 |
| HOMO SAPIENS | 0 | 11431 | 5t4n | A | 315 | ARG | p.Arg336Ter | Q96QU1 |
| HOMO SAPIENS | 0 | 11431 | 5t4n | A | 322 | ARG | p.Arg343Lys | Q96QU1 |

| | | | | | | | | |
|---|---|---|---|---|---|---|---|---|
| HOMO SAPIENS | 1 | 11431 | 5t4n | A | 326 | LEU | p.Leu347Phe | Q96QU1 |
| HOMO SAPIENS | 0 | 11431 | 5t4n | A | 373 | TYR | p.Tyr394Cys | Q96QU1 |
| HOMO SAPIENS | 0 | 11431 | 5t4n | A | 398 | ASN | p.Asn419Ser | Q96QU1 |
| HOMO SAPIENS | 0 | 11431 | 5t4n | A | 414 | ALA | p.Asp435Ala | Q96QU1 |
| HOMO SAPIENS | 0 | 11431 | 5t4n | A | 433 | VAL | p.Val454Phe | Q96QU1 |
| HOMO SAPIENS | 1 | 11431 | 5t4n | A | 507 | VAL | p.Val528Asp | Q96QU1 |
| HOMO SAPIENS | 0 | 11431 | 5t4n | A | 547 | ALA | p.Ala568Ser | Q96QU1 |
| HOMO SAPIENS | 3 | 3194 | 5xtd | L | 84 | ARG | p.Arg84His | O43181 |
| HOMO SAPIENS | 1 | 3194 | 5xtd | L | 130 | THR | p.Thr130Ser | O43181 |
| HOMO SAPIENS | 8 | 3194 | 5xtd | L | 136 | SER | p.Ser136Ala | O43181 |
| HOMO SAPIENS | 0 | 40749 | 1nun | A | 106 | CYS | p.Cys106Phe | O15520 |
| HOMO SAPIENS | 9 | 40749 | 1nun | A | 156 | ILE | p.Ile156Arg | O15520 |
| HOMO SAPIENS | 1 | 40749 | 1nun | A | 193 | ARG | p.Arg193Ter | O15520 |
| HOMO SAPIENS | 0 | 40749 | 1nun | A | 207 | HIS | p.His207Pro | O15520 |
| HOMO SAPIENS | 0 | 7026 | 3n1f | D | 875 | SER | p.Ser875Gly | Q4KMG0 |
| HOMO SAPIENS | 6 | 23802 | 3rzv | A | 424 | ARG | p.Arg424Ter | O95630 |
| HOMO SAPIENS | 0 | 36720 | 1jm7 | A | 12 | GLN | p.Gln12Ter | P38398 |
| HOMO SAPIENS | 3 | 36720 | 1jm7 | A | 23 | GLU | p.Glu23Asp | P38398 |
| HOMO SAPIENS | 1 | 36720 | 1jm7 | A | 45 | LYS | p.Lys45Gln | P38398 |
| HOMO SAPIENS | 8 | 36720 | 1jm7 | A | 47 | CYS | p.Cys47Ter | P38398 |
| HOMO SAPIENS | 0 | 36720 | 1jm7 | A | 59 | SER | p.Ser59Ter | P38398 |
| HOMO SAPIENS | 8 | 36720 | 1jm7 | A | 61 | CYS | p.Cys61Gly | P38398 |
| HOMO SAPIENS | 8 | 36720 | 1jm7 | A | 63 | LEU | p.Leu63Ter | P38398 |
| HOMO SAPIENS | 7 | 36720 | 1jm7 | A | 64 | CYS | p.Cys64Gly | P38398 |
| HOMO SAPIENS | 0 | 36720 | 1jm7 | A | 101 | TYR | p.Tyr101Ter | P38398 |
| HOMO SAPIENS | 0 | 36720 | 1jm7 | A | null | TYR | p.Tyr105Cys | P38398 |
| HOMO SAPIENS | 0 | 28500 | 3kuz | A | 1217 | CYS | p.Cys1217Arg | O60486 |
| HOMO SAPIENS | 5 | 8097 | 3p56 | C | 69 | ARG | p.Arg69Trp | Q8TDP1 |
| HOMO SAPIENS | 5 | 8097 | 3p56 | C | 143 | LYS | p.Lys143Ile | Q8TDP1 |
| HOMO SAPIENS | 2 | 30073 | 3nmz | A | 380 | ARG | p.Arg380Gln | P25054 |
| HOMO SAPIENS | 1 | 22573 | 1cjy | A | 111 | SER | p.Ser111Pro | P47712 |
| HOMO SAPIENS | 0 | 22573 | 1cjy | A | 485 | ARG | p.Arg485His | P47712 |
| HOMO SAPIENS | 2 | 965 | 3E+077 | A | 99 | ALA | p.Ala99Val | Q9Y617 |
| HOMO SAPIENS | 2 | 965 | 3E+077 | A | 100 | ASP | p.Asp100Ala | Q9Y617 |
| HOMO SAPIENS | 4 | 1503 | 3kxc | A | 62 | ARG | p.Arg62Trp | O43617 |
| HOMO SAPIENS | 6 | 9418 | 1hdr | A | 22 | GLY | p.Gly23Asp | P09417 |
| HOMO SAPIENS | 2 | 9418 | 1hdr | A | 35 | TRP | p.Trp36Arg | P09417 |
| HOMO SAPIENS | 2 | 9418 | 1hdr | A | 107 | TRP | p.Trp108Gly | P09417 |
| HOMO SAPIENS | 6 | 9418 | 1hdr | A | 149 | TYR | p.Tyr150Cys | P09417 |
| HOMO SAPIENS | -1 | 12079 | 3tnu | B | 373 | GLY | p.Gly373Ala | P13647 |
| HOMO SAPIENS | 1 | 12079 | 3tnu | B | 463 | LEU | p.Leu463Pro | P13647 |
| HOMO SAPIENS | 1 | 12079 | 3tnu | B | 471 | ARG | p.Arg471Cys | P13647 |
| HOMO SAPIENS | 0 | 8135 | 3rbn | B | 89 | SER | p.Ser556Ile | P40692 |
| HOMO SAPIENS | 0 | 8135 | 3rbn | B | 111 | GLU | p.Glu578Gly | P40692 |
| HOMO SAPIENS | 2 | 8135 | 3rbn | B | 114 | PRO | p.Pro581Leu | P40692 |
| HOMO SAPIENS | 0 | 8135 | 3rbn | B | 133 | GLU | p.Glu600Gly | P40692 |
| HOMO SAPIENS | 2 | 8135 | 3rbn | B | 136 | PRO | p.Pro603Arg | P40692 |
| HOMO SAPIENS | 2 | 8135 | 3rbn | B | 136 | PRO | p.Pro603Leu | P40692 |
| HOMO SAPIENS | 7 | 8135 | 3rbn | B | 140 | LEU | p.Leu607His | P40692 |
| HOMO SAPIENS | 2 | 8135 | 3rbn | B | 151 | LYS | p.Lys618Ala | P40692 |
| HOMO SAPIENS | 2 | 8135 | 3rbn | B | 151 | LYS | p.Lys618Thr | P40692 |
| HOMO SAPIENS | 5 | 8135 | 3rbn | B | 181 | PRO | p.Pro648Ser | P40692 |
| HOMO SAPIENS | 2 | 8135 | 3rbn | B | 188 | ILE | p.Ile655Val | P40692 |
| HOMO SAPIENS | 2 | 8135 | 3rbn | B | 214 | ALA | p.Ala681Thr | P40692 |

| | | | | | | | | |
|---|---|---|---|---|---|---|---|---|
| HOMO SAPIENS | 2 | 8135 | 3rbn | B | 220 | ARG | p.Arg687Trp | P40692 |
| HOMO SAPIENS | 2 | 8135 | 3rbn | B | 222 | GLN | p.Gln689Arg | P40692 |
| HOMO SAPIENS | -1 | 8135 | 3rbn | B | 236 | GLN | p.Gln701Ter | P40692 |
| HOMO SAPIENS | 7 | 8135 | 3rbn | B | 251 | HIS | p.His718Tyr | P40692 |
| HOMO SAPIENS | 0 | 8135 | 3rbn | B | 284 | LYS | p.Lys751Arg | P40692 |
| HOMO SAPIENS | 1 | 21998 | 5d68 | A | 355 | LEU | p.Leu355Phe | O00522 |
| HOMO SAPIENS | 3 | 21998 | 5d68 | A | 380 | THR | p.Thr380Met | O00522 |
| HOMO SAPIENS | 0 | 21998 | 5d68 | A | 415 | ILE | p.Ile415Met | O00522 |
| HOMO SAPIENS | 0 | 33789 | 2m0c | A | 13 | ARG | p.Arg218Gln | Q9H161 |
| HOMO SAPIENS | 7 | 33789 | 2m0c | A | 52 | ARG | p.Arg257Thr | Q9H161 |
| HOMO SAPIENS | 0 | 33789 | 2m0c | A | 67 | ARG | p.Arg272Pro | Q9H161 |
| HOMO SAPIENS | 6 | 8263 | 2osg | A | 51 | GLY | p.Gly355Arg | Q9UDY2 |
| HOMO SAPIENS | 5 | 8263 | 2osg | A | 60 | THR | p.Thr364Met | Q9UDY2 |
| HOMO SAPIENS | 1 | 2475 | 5E+026 | B | 234 | THR | p.Thr234Ala | Q9BZ23 |
| HOMO SAPIENS | 2 | 2475 | 5E+026 | B | 264 | ARG | p.Arg264Trp | Q9BZ23 |
| HOMO SAPIENS | 4 | 2475 | 5E+026 | B | 286 | ARG | p.Arg286Cys | Q9BZ23 |
| HOMO SAPIENS | 0 | 2475 | 5E+026 | B | 357 | ARG | p.Arg357Trp | Q9BZ23 |
| HOMO SAPIENS | 2 | 2475 | 5E+026 | B | 481 | ARG | p.Arg481Ter | Q9BZ23 |
| HOMO SAPIENS | 8 | 2475 | 5E+026 | B | 521 | GLY | p.Gly521Arg | Q9BZ23 |
| HOMO SAPIENS | 2 | 2475 | 5E+026 | B | 528 | THR | p.Thr528Met | Q9BZ23 |
| HOMO SAPIENS | 1 | 3742 | 5xtd | E | 108 | HIS | p.His134Tyr | P56556 |
| HOMO SAPIENS | 1 | 16725 | 3g04 | C | 36 | ASP | p.Asp36His | P16473 |
| HOMO SAPIENS | 3 | 16725 | 3g04 | C | 52 | PRO | p.Pro52Thr | P16473 |
| HOMO SAPIENS | 1 | 16725 | 3g04 | C | 162 | PRO | p.Pro162Ala | P16473 |
| HOMO SAPIENS | -1 | 12366 | 5d2d | C | 750 | PRO | p.Pro750Leu | P13569 |
| HOMO SAPIENS | 0 | 12366 | 5d2d | C | 764 | ARG | p.Arg764Ter | P13569 |
| HOMO SAPIENS | 0 | 34845 | 2y23 | A | 1151 | ASP | p.Asp1151Glu | P52179 |
| HOMO SAPIENS | 0 | 34845 | 2y23 | A | 1180 | ASP | p.Asp1180Ala | P52179 |
| HOMO SAPIENS | 6 | 36740 | 3lbx | B | 2053 | ALA | p.Ala2053Pro | P11277 |
| HOMO SAPIENS | 0 | 19289 | 3ihj | A | 404 | ARG | p.Arg404Ter | Q8TD30 |
| HOMO SAPIENS | 5 | 1034 | 5xtd | Q | 228 | ARG | p.Arg228Gln | O75306 |
| HOMO SAPIENS | 5 | 1034 | 5xtd | Q | 229 | PRO | p.Pro229Gln | O75306 |
| HOMO SAPIENS | 4 | 1034 | 5xtd | Q | 323 | ARG | p.Arg323Gln | O75306 |
| HOMO SAPIENS | 4 | 1034 | 5xtd | Q | 352 | PRO | p.Pro352Ala | O75306 |
| HOMO SAPIENS | 5 | 1034 | 5xtd | Q | 413 | SER | p.Ser413Pro | O75306 |
| HOMO SAPIENS | 1 | 2772 | 2vaf | A | 1047 | THR | p.Thr66Ala | O14958 |
| HOMO SAPIENS | 5 | 2772 | 2vaf | A | 1057 | VAL | p.Val76Met | O14958 |
| HOMO SAPIENS | 0 | 2772 | 2vaf | A | 1148 | LEU | p.Leu167His | O14958 |
| HOMO SAPIENS | 1 | 2772 | 2vaf | A | 1149 | ILE | p.Ile168Thr | O14958 |
| HOMO SAPIENS | 8 | 2772 | 2vaf | A | 1225 | HIS | p.His244Arg | O14958 |
| HOMO SAPIENS | 8 | 2772 | 2vaf | A | 1225 | HIS | p.His244Tyr | O14958 |
| HOMO SAPIENS | 7 | 2772 | 2vaf | A | 1251 | ILE | p.Ile270Thr | O14958 |
| HOMO SAPIENS | 3 | 2772 | 2vaf | A | 1288 | ASP | p.Asp307His | O14958 |
| HOMO SAPIENS | 5 | 2772 | 2vaf | A | 1289 | PRO | p.Pro308Leu | O14958 |
| HOMO SAPIENS | 1 | 31885 | 2aex | A | 132 | ALA | p.Ala132Val | P36551 |
| HOMO SAPIENS | 1 | 31885 | 2aex | A | 272 | ASN | p.Asn272His | P36551 |
| HOMO SAPIENS | 8 | 31885 | 2aex | A | 279 | GLY | p.Gly279Arg | P36551 |
| HOMO SAPIENS | 5 | 31885 | 2aex | A | 294 | VAL | p.Val294Ile | P36551 |
| HOMO SAPIENS | 5 | 31885 | 2aex | A | 295 | HIS | p.His295Asp | P36551 |
| HOMO SAPIENS | 1 | 31885 | 2aex | A | 331 | ARG | p.Arg331Trp | P36551 |
| HOMO SAPIENS | 3 | 31885 | 2aex | A | 352 | ARG | p.Arg352Cys | P36551 |
| HOMO SAPIENS | 1 | 31885 | 2aex | A | 404 | LYS | p.Lys404Glu | P36551 |
| HOMO SAPIENS | 2 | 31885 | 2aex | A | 447 | ARG | p.Arg447Cys | P36551 |
| HOMO SAPIENS | 5 | 11721 | 2r3v | A | 20 | HIS | p.His20Pro | Q03426 |

| | | | | | | | | |
|---|---|---|---|---|---|---|---|---|
| HOMO SAPIENS | 1 | 11721 | 2r3v | A | 116 | TYR | p.Tyr116His | Q03426 |
| HOMO SAPIENS | 1 | 11721 | 2r3v | A | 148 | ALA | p.Ala148Thr | Q03426 |
| HOMO SAPIENS | 1 | 11721 | 2r3v | A | 167 | PRO | p.Pro167Leu | Q03426 |
| HOMO SAPIENS | 6 | 11721 | 2r3v | A | 203 | VAL | p.Val203Ala | Q03426 |
| HOMO SAPIENS | 2 | 11721 | 2r3v | A | 215 | ARG | p.Arg215Ter | Q03426 |
| HOMO SAPIENS | 2 | 11721 | 2r3v | A | 237 | THR | p.Thr237Ser | Q03426 |
| HOMO SAPIENS | 0 | 11721 | 2r3v | A | 268 | ILE | p.Ile268Thr | Q03426 |
| HOMO SAPIENS | 0 | 11721 | 2r3v | A | 268 | ILE | p.Ile268Val | Q03426 |
| HOMO SAPIENS | 0 | 11721 | 2r3v | A | 301 | ASN | p.Asn301Ser | Q03426 |
| HOMO SAPIENS | 6 | 11721 | 2r3v | A | 310 | VAL | p.Val310Met | Q03426 |
| HOMO SAPIENS | 5 | 11721 | 2r3v | A | 334 | ALA | p.Ala334Thr | Q03426 |
| HOMO SAPIENS | 2 | 11721 | 2r3v | A | 377 | VAL | p.Val377Ile | Q03426 |
| HOMO SAPIENS | 1 | 11721 | 2r3v | A | 386 | ASP | p.Asp386Asn | Q03426 |
| HOMO SAPIENS | 1 | 11721 | 2r3v | A | 388 | ARG | p.Arg388Ter | Q03426 |
| HOMO SAPIENS | 8 | 5010 | 2hgs | A | 125 | ARG | p.Arg125Cys | P48637 |
| HOMO SAPIENS | 1 | 5010 | 2hgs | A | 164 | ARG | p.Arg164Gln | P48637 |
| HOMO SAPIENS | 0 | 5010 | 2hgs | A | 219 | ASP | p.Asp219Gly | P48637 |
| HOMO SAPIENS | 7 | 5010 | 2hgs | A | 267 | ARG | p.Arg267Trp | P48637 |
| HOMO SAPIENS | 0 | 5010 | 2hgs | A | 283 | ARG | p.Arg283Cys | P48637 |
| HOMO SAPIENS | 0 | 6892 | 5erd | B | 7 | VAL | p.Val56Met | Q14126 |
| HOMO SAPIENS | 1 | 6892 | 5erd | B | 25 | HIS | p.His74Arg | Q14126 |
| HOMO SAPIENS | 0 | 6892 | 5erd | B | 40 | TYR | p.Tyr89Cys | Q14126 |
| HOMO SAPIENS | 0 | 6892 | 5erd | B | 97 | ARG | p.Arg146Leu | Q14126 |
| HOMO SAPIENS | 7 | 6892 | 5erd | B | 217 | ASN | p.Asn266Ser | Q14126 |
| HOMO SAPIENS | 0 | 6892 | 5erd | B | 243 | ARG | p.Arg292His | Q14126 |
| HOMO SAPIENS | 0 | 6892 | 5erd | B | 244 | ILE | p.Ile293Val | Q14126 |
| HOMO SAPIENS | 0 | 6892 | 5erd | B | 343 | VAL | p.Val392Ile | Q14126 |
| HOMO SAPIENS | 4 | 6892 | 5erd | B | 350 | GLU | p.Glu399Lys | Q14126 |
| HOMO SAPIENS | 0 | 6892 | 5erd | B | 410 | TYR | p.Tyr459Cys | Q14126 |
| HOMO SAPIENS | 0 | 6892 | 5erd | B | 466 | VAL | p.Val515Ile | Q14126 |
| HOMO SAPIENS | 0 | 6892 | 5erd | B | 482 | PHE | p.Phe531Cys | Q14126 |
| HOMO SAPIENS | 0 | 6892 | 5erd | B | 484 | VAL | p.Val533Ile | Q14126 |
| HOMO SAPIENS | 0 | 6892 | 5erd | B | 545 | LEU | p.Leu594Pro | Q14126 |
| HOMO SAPIENS | 0 | 1338 | 2ax4 | A | 58 | THR | p.Thr48Arg | O95340 |
| HOMO SAPIENS | 7 | 8675 | 2zkm | X | 385 | LYS | p.Lys385Arg | Q00722 |
| HOMO SAPIENS | 1 | 11769 | 2wya | B | 54 | VAL | p.Val54Met | P54868 |
| HOMO SAPIENS | 1 | 11769 | 2wya | B | 116 | ARG | p.Arg116Cys | P54868 |
| HOMO SAPIENS | 5 | 11769 | 2wya | B | 167 | TYR | p.Tyr167Cys | P54868 |
| HOMO SAPIENS | 1 | 11769 | 2wya | B | 174 | PHE | p.Phe174Leu | P54868 |
| HOMO SAPIENS | 0 | 11769 | 2wya | B | 212 | GLY | p.Gly212Arg | P54868 |
| HOMO SAPIENS | 1 | 11769 | 2wya | B | 288 | ARG | p.Arg288Ter | P54868 |
| HOMO SAPIENS | 0 | 11769 | 2wya | B | 375 | SER | p.Ser375Cys | P54868 |
| HOMO SAPIENS | 4 | 11769 | 2wya | B | 500 | ARG | p.Arg500His | P54868 |
| HOMO SAPIENS | 0 | 3102 | 2iwo | A | 1880 | ARG | p.Arg1908Lys | O75970 |
| HOMO SAPIENS | 6 | 14296 | 2m9w | A | 38 | GLY | p.Gly296Ser | P43694 |
| HOMO SAPIENS | 0 | 14296 | 2m9w | A | 58 | GLN | p.Gln316Glu | P43694 |
| HOMO SAPIENS | 2 | 704 | 5ekd | B | 239 | ARG | p.Arg239Ter | Q9UGM6 |
| HOMO SAPIENS | 2 | 704 | 5ekd | B | 264 | TYR | p.Tyr264Cys | Q9UGM6 |
| HOMO SAPIENS | 6 | 2228 | 1ey2 | A | 47 | SER | p.Ser47Ter | Q93099 |
| HOMO SAPIENS | 6 | 2228 | 1ey2 | A | 48 | ALA | p.Ala48Ser | Q93099 |
| HOMO SAPIENS | 6 | 2228 | 1ey2 | A | 53 | ARG | p.Arg53Gln | Q93099 |
| HOMO SAPIENS | 3 | 2228 | 1ey2 | A | 80 | HIS | p.Gln80His | Q93099 |
| HOMO SAPIENS | 0 | 2228 | 1ey2 | A | 120 | CYS | p.Cys120Trp | Q93099 |
| HOMO SAPIENS | 3 | 2228 | 1ey2 | A | 122 | ALA | p.Ala122Val | Q93099 |

| | | | | | | | | |
|---|---|---|---|---|---|---|---|---|
| HOMO SAPIENS | 3 | 2228 | 1ey2 | A | 161 | GLY | p.Gly161Arg | Q93099 |
| HOMO SAPIENS | 6 | 2228 | 1ey2 | A | 225 | ARG | p.Arg225His | Q93099 |
| HOMO SAPIENS | 0 | 2228 | 1ey2 | A | 230 | PRO | p.Pro230Ser | Q93099 |
| HOMO SAPIENS | 6 | 2228 | 1ey2 | A | 270 | GLY | p.Gly270Arg | Q93099 |
| HOMO SAPIENS | 6 | 2228 | 1ey2 | A | 330 | ARG | p.Arg330Ser | Q93099 |
| HOMO SAPIENS | -1 | 2228 | 1ey2 | A | 368 | MSE | p.Met368Val | Q93099 |
| HOMO SAPIENS | 6 | 2228 | 1ey2 | A | 371 | HIS | p.His371Arg | Q93099 |
| HOMO SAPIENS | 6 | 2228 | 1ey2 | A | 401 | GLU | p.Glu401Gln | Q93099 |
| HOMO SAPIENS | 5 | 34648 | 2k27 | A | 31 | ARG | p.Arg31His | Q06710 |
| HOMO SAPIENS | 7 | 34648 | 2k27 | A | 40 | GLN | p.Gln40Pro | Q06710 |
| HOMO SAPIENS | 5 | 34648 | 2k27 | A | 57 | CYS | p.Cys57Tyr | Q06710 |
| HOMO SAPIENS | 5 | 34648 | 2k27 | A | 62 | LEU | p.Leu62Arg | Q06710 |
| HOMO SAPIENS | 1 | 21399 | 3edu | A | 1819 | GLU | p.Glu1819Ter | P11277 |
| HOMO SAPIENS | 1 | 18153 | 1oey | J | 245 | TYR | p.Tyr245Ter | Q15080 |
| HOMO SAPIENS | 2 | 41567 | 3sw0 | X | 1050 | ASN | p.Asn1050Tyr | P08603 |
| HOMO SAPIENS | 5 | 41567 | 3sw0 | X | 1059 | ILE | p.Ile1059Thr | P08603 |
| HOMO SAPIENS | 0 | 36362 | 4wrm | A | 362 | HIS | p.His362Arg | P07333 |
| HOMO SAPIENS | -1 | 36362 | 4wrm | A | 413 | GLY | p.Gly413Ser | P07333 |
| HOMO SAPIENS | 0 | 31255 | 2w2j | A | 100 | SER | p.Ser100Pro | P35219 |
| HOMO SAPIENS | 0 | 31255 | 2w2j | A | 100 | SER | p.Ser100Leu | P35219 |
| HOMO SAPIENS | 5 | 31255 | 2w2j | A | 237 | ARG | p.Arg237Gln | P35219 |
| HOMO SAPIENS | -1 | 26450 | 1th1 | C | 1395 | SER | p.Ser1395Cys | P25054 |
| HOMO SAPIENS | -1 | 26450 | 1th1 | C | 1413 | MET | p.Met1413Val | P25054 |
| HOMO SAPIENS | -1 | 26450 | 1th1 | C | 1446 | ALA | p.Ala1446Thr | P25054 |
| HOMO SAPIENS | -1 | 26450 | 1th1 | C | 1454 | LYS | p.Lys1454Glu | P25054 |
| HOMO SAPIENS | 0 | 26450 | 1th1 | C | 1474 | ALA | p.Ala1474Thr | P25054 |
| HOMO SAPIENS | 1 | 981 | 5xtb | F | 14 | LEU | p.Leu14Val | O43678 |
| HOMO SAPIENS | 1 | 11069 | 3kq4 | A | 46 | SER | p.Ser46Leu | P27352 |
| HOMO SAPIENS | 1 | 11069 | 3kq4 | A | 97 | MET | p.Met97Thr | P27352 |
| HOMO SAPIENS | 1 | 11069 | 3kq4 | A | 255 | ASN | p.Asn255Ser | P27352 |
| HOMO SAPIENS | 1 | 11069 | 3kq4 | A | 330 | ASN | p.Asn330Lys | P27352 |
| HOMO SAPIENS | 0 | 41489 | 3erb | A | 53 | PRO | p.Pro73Leu | P06681 |
| HOMO SAPIENS | 6 | 8715 | 2bq8 | X | 23 | THR | p.Thr44Met | P13686 |
| HOMO SAPIENS | 0 | 8715 | 2bq8 | X | 88 | GLY | p.Gly109Arg | P13686 |
| HOMO SAPIENS | 6 | 8715 | 2bq8 | X | 127 | VAL | p.Val148Met | P13686 |
| HOMO SAPIENS | 2 | 8715 | 2bq8 | X | 179 | VAL | p.Val200Met | P13686 |
| HOMO SAPIENS | 6 | 8715 | 2bq8 | X | 251 | ARG | p.Arg272Cys | P13686 |
| HOMO SAPIENS | 7 | 8112 | 3p56 | A | 37 | GLY | p.Gly37Ser | O75792 |
| HOMO SAPIENS | -1 | 8112 | 3p56 | A | 202 | LEU | p.Leu202Ser | O75792 |
| HOMO SAPIENS | 1 | 8112 | 3p56 | A | 205 | ASP | p.Asp205Glu | O75792 |
| HOMO SAPIENS | 4 | 8112 | 3p56 | A | 212 | ASN | p.Asn212Ile | O75792 |
| HOMO SAPIENS | 0 | 8112 | 3p56 | A | 235 | ARG | p.Arg235Gln | O75792 |
| HOMO SAPIENS | 4 | 8112 | 3p56 | A | 240 | THR | p.Thr240Met | O75792 |
| HOMO SAPIENS | 0 | 8112 | 3p56 | A | 291 | ARG | p.Arg291His | O75792 |
| HOMO SAPIENS | 0 | 37603 | 2yqg | A | 10 | ARG | p.Arg49His | Q14126 |
| HOMO SAPIENS | 0 | 21178 | 3k1r | B | 458 | ASP | p.Asp458Val | Q495M9 |
| HOMO SAPIENS | -1 | 36828 | 4fq3 | B | 507 | GLY | p.Gly507Asp | P35637 |
| HOMO SAPIENS | 5 | 36828 | 4fq3 | B | 517 | HIS | p.His517Gln | P35637 |
| HOMO SAPIENS | 8 | 36828 | 4fq3 | B | 518 | ARG | p.Arg518Lys | P35637 |
| HOMO SAPIENS | 6 | 36828 | 4fq3 | B | 521 | ARG | p.Arg521Cys | P35637 |
| HOMO SAPIENS | 6 | 36828 | 4fq3 | B | 521 | ARG | p.Arg521His | P35637 |
| HOMO SAPIENS | 0 | 6363 | 3j7y | c | null | HIS | p.His13Arg | Q9H9J2 |
| HOMO SAPIENS | 2 | 6363 | 3j7y | c | 105 | ARG | p.Arg105Cys | Q9H9J2 |
| HOMO SAPIENS | 2 | 6363 | 3j7y | c | 138 | THR | p.Thr138Ile | Q9H9J2 |

| | | | | | | | | |
|---|---|---|---|---|---|---|---|---|
| HOMO SAPIENS | 1 | 6363 | 3j7y | c | 156 | LEU | p.Leu156Arg | Q9H9J2 |
| HOMO SAPIENS | 1 | 6782 | 5l46 | A | 109 | HIS | p.His109Arg | Q9UI17 |
| HOMO SAPIENS | 0 | 23951 | 5t76 | A | 175 | GLY | p.Gly931Ser | P49588 |
| HOMO SAPIENS | 8 | 12604 | 3p0l | A | 182 | ARG | p.Arg182Leu | P49675 |
| HOMO SAPIENS | 2 | 12604 | 3p0l | A | 187 | VAL | p.Val187Met | P49675 |
| HOMO SAPIENS | 2 | 12604 | 3p0l | A | 188 | ARG | p.Arg188Cys | P49675 |
| HOMO SAPIENS | 0 | 12604 | 3p0l | A | 193 | ARG | p.Arg193Ter | P49675 |
| HOMO SAPIENS | 9 | 12604 | 3p0l | A | 217 | ARG | p.Arg217Thr | P49675 |
| HOMO SAPIENS | 4 | 12604 | 3p0l | A | 218 | ALA | p.Ala218Val | P49675 |
| HOMO SAPIENS | 0 | 12604 | 3p0l | A | 250 | TRP | p.Trp250Ter | P49675 |
| HOMO SAPIENS | 1 | 12604 | 3p0l | A | 258 | GLN | p.Gln258Ter | P49675 |
| HOMO SAPIENS | -1 | 27765 | 4wkr | A | 4 | GLU | p.Glu4Lys | Q4G0J3 |
| HOMO SAPIENS | 2 | 3587 | 4kit | C | 2301 | PRO | p.Pro2301Thr | Q6P2Q9 |
| HOMO SAPIENS | 3 | 3587 | 4kit | C | 2304 | PHE | p.Phe2304Leu | Q6P2Q9 |
| HOMO SAPIENS | 3 | 3587 | 4kit | C | 2309 | HIS | p.His2309Pro | Q6P2Q9 |
| HOMO SAPIENS | 3 | 3587 | 4kit | C | 2310 | ARG | p.Arg2310Lys | Q6P2Q9 |
| HOMO SAPIENS | 0 | 9073 | 1n69 | B | 21 | ASN | p.Asn215His | P07602 |
| HOMO SAPIENS | 0 | 9073 | 1n69 | B | 23 | THR | p.Thr217Ile | P07602 |
| HOMO SAPIENS | 0 | 9073 | 1n69 | B | 47 | CYS | p.Cys241Ser | P07602 |
| HOMO SAPIENS | 0 | 475 | 2vd5 | A | 225 | THR | p.Thr225Met | Q09013 |
| HOMO SAPIENS | 1 | 9066 | 4mzv | A | 31 | ASN | p.Asn31Lys | P16422 |
| HOMO SAPIENS | 1 | 9066 | 4mzv | A | 47 | GLN | p.Gln47His | P16422 |
| HOMO SAPIENS | 3 | 9066 | 4mzv | A | 66 | CYS | p.Cys66Tyr | P16422 |
| HOMO SAPIENS | 1 | 9066 | 4mzv | A | 102 | SER | p.Ser102Gly | P16422 |
| HOMO SAPIENS | 5 | 9066 | 4mzv | A | 115 | THR | p.Met115Thr | P16422 |
| HOMO SAPIENS | 1 | 9066 | 4mzv | A | 138 | ARG | p.Arg138Ter | P16422 |
| HOMO SAPIENS | 1 | 9066 | 4mzv | A | 153 | ARG | p.Arg153Thr | P16422 |
| HOMO SAPIENS | 1 | 9066 | 4mzv | A | 165 | ALA | p.Ala165Thr | P16422 |
| HOMO SAPIENS | 8 | 1827 | 2nz2 | A | 14 | GLY | p.Gly14Ser | P00966 |
| HOMO SAPIENS | 5 | 1827 | 2nz2 | A | 18 | SER | p.Ser18Leu | P00966 |
| HOMO SAPIENS | 3 | 1827 | 2nz2 | A | 86 | ARG | p.Arg86Cys | P00966 |
| HOMO SAPIENS | 3 | 1827 | 2nz2 | A | 86 | ARG | p.Arg86His | P00966 |
| HOMO SAPIENS | 1 | 1827 | 2nz2 | A | 179 | TRP | p.Trp179Arg | P00966 |
| HOMO SAPIENS | 7 | 1827 | 2nz2 | A | 180 | SER | p.Ser180Asn | P00966 |
| HOMO SAPIENS | 7 | 1827 | 2nz2 | A | 191 | GLU | p.Glu191Lys | P00966 |
| HOMO SAPIENS | 4 | 1827 | 2nz2 | A | 256 | GLU | p.Glu256Lys | P00966 |
| HOMO SAPIENS | 4 | 1827 | 2nz2 | A | 263 | VAL | p.Val263Met | P00966 |
| HOMO SAPIENS | 5 | 1827 | 2nz2 | A | 265 | ARG | p.Arg265Cys | P00966 |
| HOMO SAPIENS | 5 | 1827 | 2nz2 | A | 268 | ILE | p.Ile268Met | P00966 |
| HOMO SAPIENS | 6 | 1827 | 2nz2 | A | 269 | VAL | p.Val269Met | P00966 |
| HOMO SAPIENS | 7 | 1827 | 2nz2 | A | 272 | ARG | p.Arg272Cys | P00966 |
| HOMO SAPIENS | 3 | 1827 | 2nz2 | A | 304 | ARG | p.Arg304Trp | P00966 |
| HOMO SAPIENS | 3 | 1827 | 2nz2 | A | 307 | ARG | p.Arg307Cys | P00966 |
| HOMO SAPIENS | 3 | 1827 | 2nz2 | A | 310 | LYS | p.Lys310Gln | P00966 |
| HOMO SAPIENS | 2 | 1827 | 2nz2 | A | 362 | GLY | p.Gly362Val | P00966 |
| HOMO SAPIENS | 2 | 1827 | 2nz2 | A | 363 | ARG | p.Arg363Trp | P00966 |
| HOMO SAPIENS | 2 | 1827 | 2nz2 | A | 363 | ARG | p.Arg363Gln | P00966 |
| HOMO SAPIENS | 1 | 1827 | 2nz2 | A | 390 | GLY | p.Gly390Arg | P00966 |
| HOMO SAPIENS | 0 | 29524 | 5vbn | B | 2165 | ARG | p.Arg2165His | Q07864 |
| HOMO SAPIENS | 2 | 29524 | 5vbn | B | 2255 | ILE | p.Ile2255Phe | Q07864 |
| HOMO SAPIENS | 2 | 29524 | 5vbn | B | 2274 | LEU | p.Leu2274Val | Q07864 |
| HOMO SAPIENS | 0 | 31897 | 3t0o | A | 184 | CYS | p.Cys184Arg | O00584 |
| HOMO SAPIENS | -1 | 31897 | 3t0o | A | 215 | GLU | p.Glu215Lys | O00584 |
| HOMO SAPIENS | 0 | 31897 | 3t0o | A | 236 | ARG | p.Arg236Trp | O00584 |

| | | | | | | | | |
|---|---|---|---|---|---|---|---|---|
| HOMO SAPIENS | 7 | 18490 | 4pcw | B | 31 | GLU | p.Glu32Ter | P20930 |
| HOMO SAPIENS | -1 | 39865 | 2h9e | L | 152 | GLY | p.Gly192Arg | P00742 |
| HOMO SAPIENS | 2 | 4309 | 2x0b | F | 174 | THR | p.Thr207Met | P01019 |
| HOMO SAPIENS | 3 | 4309 | 2x0b | F | 342 | ARG | p.Arg375Gln | P01019 |
| HOMO SAPIENS | 2 | 3094 | 1zll | B | 9 | ARG | p.Arg9Cys | P26678 |
| HOMO SAPIENS | 9 | 3094 | 1zll | B | 39 | LEU | p.Leu39Ter | P26678 |
| HOMO SAPIENS | 5 | 3094 | 1zll | B | 49 | VAL | p.Val49Met | P26678 |
| HOMO SAPIENS | -1 | 41498 | 3ffd | P | 8 | LEU | p.Leu44Pro | P12272 |
| HOMO SAPIENS | 1 | 20697 | 1igr | A | 407 | ARG | p.Arg437His | P08069 |
| HOMO SAPIENS | 0 | 20697 | 1igr | A | 416 | MET | p.Met446Val | P08069 |
| HOMO SAPIENS | -1 | 20697 | 1igr | A | 462 | VAL | p.Val492Ile | P08069 |
| HOMO SAPIENS | 0 | 3973 | 4cbz | A | 37 | LEU | p.Leu37Ser | P78504 |
| HOMO SAPIENS | 0 | 3973 | 4cbz | A | 45 | VAL | p.Val45Leu | P78504 |
| HOMO SAPIENS | 0 | 3973 | 4cbz | A | 134 | LEU | p.Leu134Met | P78504 |
| HOMO SAPIENS | -1 | 3973 | 4cbz | A | 146 | VAL | p.Val146Ile | P78504 |
| HOMO SAPIENS | 0 | 3973 | 4cbz | A | 176 | VAL | p.Val176Ile | P78504 |
| HOMO SAPIENS | 0 | 3973 | 4cbz | A | 184 | ARG | p.Arg184Cys | P78504 |
| HOMO SAPIENS | 6 | 1697 | 4hmy | S | 33 | ARG | p.Arg33Trp | Q96PC3 |
| HOMO SAPIENS | 2 | 23752 | 5j4o | A | 13 | ARG | p.Arg1611Ser | P02549 |
| HOMO SAPIENS | 0 | 23752 | 5j4o | A | 42 | ALA | p.Ala1640Gly | P02549 |
| HOMO SAPIENS | 0 | 23752 | 5j4o | A | 108 | HIS | p.His1706Gln | P02549 |
| HOMO SAPIENS | 6 | 23752 | 5j4o | A | 206 | LEU | p.Leu1804Phe | P02549 |
| HOMO SAPIENS | 8 | 531 | 2qz4 | A | 349 | GLY | p.Gly349Ser | Q9UQ90 |
| HOMO SAPIENS | 0 | 531 | 2qz4 | A | 486 | ARG | p.Arg486Gln | Q9UQ90 |
| HOMO SAPIENS | 1 | 531 | 2qz4 | A | 510 | ALA | p.Ala510Val | Q9UQ90 |
| HOMO SAPIENS | 0 | 531 | 2qz4 | A | 529 | ALA | p.Ala529Val | Q9UQ90 |
| HOMO SAPIENS | 0 | 531 | 2qz4 | A | 558 | LYS | p.Lys558Ter | Q9UQ90 |
| HOMO SAPIENS | 1 | 531 | 2qz4 | A | 559 | LYS | p.Lys559Ter | Q9UQ90 |
| HOMO SAPIENS | 2 | 7736 | 1fnh | A | 114 | TRP | p.Trp1834Arg | P02751 |
| HOMO SAPIENS | 0 | 7736 | 1fnh | A | 163 | LEU | p.Leu1883Arg | P02751 |
| HOMO SAPIENS | 3 | 20337 | 5wy5 | B | 264 | LEU | p.Leu264Phe | Q96MG7 |
| HOMO SAPIENS | 8 | 36096 | 2jku | A | 643 | GLY | p.Gly668Arg | P05165 |
| HOMO SAPIENS | 6 | 36096 | 2jku | A | 663 | CYS | p.Cys688Arg | P05165 |
| HOMO SAPIENS | 0 | 9850 | 3kbt | A | 1606 | TYR | p.Tyr1606Ter | P11277 |
| HOMO SAPIENS | 4 | 6876 | 4jkq | A | 217 | ILE | p.Ile217Val | Q92736 |
| HOMO SAPIENS | 5 | 6876 | 4jkq | A | 420 | ARG | p.Arg420Trp | Q92736 |
| HOMO SAPIENS | 2 | 27603 | 1itv | B | 62 | PRO | p.Arg574Pro | P14780 |
| HOMO SAPIENS | 4 | 27603 | 1itv | B | 156 | ARG | p.Arg668Gln | P14780 |
| HOMO SAPIENS | 2 | 22954 | 4f52 | E | 48 | ILE | p.Ile48Met | Q92990 |
| HOMO SAPIENS | 0 | 22954 | 4f52 | E | 91 | ASP | p.Asp91Asn | Q92990 |
| HOMO SAPIENS | 0 | 22954 | 4f52 | E | 146 | ALA | p.Ala146Thr | Q92990 |
| HOMO SAPIENS | 0 | 22954 | 4f52 | E | 181 | ILE | p.Ile181Lys | Q92990 |
| HOMO SAPIENS | 0 | 22954 | 4f52 | E | 194 | ASN | p.Asn194Ser | Q92990 |
| HOMO SAPIENS | 2 | 22954 | 4f52 | E | 254 | PRO | p.Pro254Arg | Q92990 |
| HOMO SAPIENS | 2 | 22954 | 4f52 | E | 300 | ILE | p.Ile300Thr | Q92990 |
| HOMO SAPIENS | 2 | 22954 | 4f52 | E | 336 | LEU | p.Leu336Ser | Q92990 |
| HOMO SAPIENS | 0 | 22954 | 4f52 | E | 353 | TYR | p.Tyr353His | Q92990 |
| HOMO SAPIENS | 0 | 22954 | 4f52 | E | 387 | MET | p.Met387Val | Q92990 |
| HOMO SAPIENS | 4 | 2946 | 5gl7 | A | 344 | TYR | p.Tyr344Cys | P56192 |
| HOMO SAPIENS | 0 | 2946 | 5gl7 | A | 370 | PHE | p.Phe370Leu | P56192 |
| HOMO SAPIENS | 0 | 2946 | 5gl7 | A | 393 | ALA | p.Ala393Thr | P56192 |
| HOMO SAPIENS | 0 | 2946 | 5gl7 | A | 523 | ILE | p.Ile523Thr | P56192 |
| HOMO SAPIENS | 0 | 2946 | 5gl7 | A | 567 | SER | p.Ser567Leu | P56192 |
| HOMO SAPIENS | 0 | 2946 | 5gl7 | A | 800 | PRO | p.Pro800Thr | P56192 |

| | | | | | | | | |
|---|---|---|---|---|---|---|---|---|
| HOMO SAPIENS | 3 | 34961 | 4wy4 | C | 44 | TYR | p.Tyr44His | O95721 |
| HOMO SAPIENS | 8 | 34961 | 4wy4 | C | 89 | GLU | p.Glu89Lys | O95721 |
| HOMO SAPIENS | 0 | 9427 | 4chl | B | 21 | ALA | p.Ala21Ser | O95571 |
| HOMO SAPIENS | 2 | 9427 | 4chl | B | 62 | ALA | p.Ala62Thr | O95571 |
| HOMO SAPIENS | 0 | 9427 | 4chl | B | 114 | ILE | p.Ile114Phe | O95571 |
| HOMO SAPIENS | 2 | 9427 | 4chl | B | 159 | ARG | p.Arg159His | O95571 |
| HOMO SAPIENS | 8 | 9427 | 4chl | B | 163 | ARG | p.Arg163Trp | O95571 |
| HOMO SAPIENS | 8 | 9427 | 4chl | B | 163 | ARG | p.Arg163Gln | O95571 |
| HOMO SAPIENS | 2 | 9427 | 4chl | B | 185 | LEU | p.Leu185Arg | O95571 |
| HOMO SAPIENS | 0 | 40917 | 2e8o | A | 11 | PRO | p.Pro26Leu | Q9Y3Z3 |
| HOMO SAPIENS | -1 | 39190 | 4gow | A | 556 | TYR | p.Tyr556Ter | P56696 |
| HOMO SAPIENS | 0 | 2399 | 2mkv | A | 8 | GLY | p.Gly10Ser | P54710 |
| HOMO SAPIENS | 1 | 2399 | 2mkv | A | 24 | VAL | p.Val26Ile | P54710 |
| HOMO SAPIENS | 4 | 2399 | 2mkv | A | 39 | GLY | p.Gly41Arg | P54710 |
| HOMO SAPIENS | 0 | 20801 | 2zej | B | 1441 | ARG | p.Arg1441Cys | Q5S007 |
| HOMO SAPIENS | 0 | 20801 | 2zej | B | 1441 | ARG | p.Arg1441His | Q5S007 |
| HOMO SAPIENS | 5 | 1019 | 5xtb | M | 62 | ARG | p.Arg62Gln | P28331 |
| HOMO SAPIENS | 1 | 1019 | 5xtb | M | 85 | ALA | p.Ala85Thr | P28331 |
| HOMO SAPIENS | 5 | 1019 | 5xtb | M | 231 | LEU | p.Leu231Val | P28331 |
| HOMO SAPIENS | 2 | 1019 | 5xtb | M | 241 | ARG | p.Arg241Trp | P28331 |
| HOMO SAPIENS | 1 | 1019 | 5xtb | M | 252 | ASP | p.Asp252Gly | P28331 |
| HOMO SAPIENS | 1 | 1019 | 5xtb | M | 408 | ARG | p.Arg408Cys | P28331 |
| HOMO SAPIENS | 2 | 1019 | 5xtb | M | 417 | ARG | p.Arg417Ser | P28331 |
| HOMO SAPIENS | 1 | 1019 | 5xtb | M | 557 | ARG | p.Arg557Ter | P28331 |
| HOMO SAPIENS | 0 | 1019 | 5xtb | M | 567 | ILE | p.Ile567Val | P28331 |
| HOMO SAPIENS | 1 | 1019 | 5xtb | M | 595 | THR | p.Thr595Ala | P28331 |
| HOMO SAPIENS | 1 | 1019 | 5xtb | M | 703 | ALA | p.Ala703Thr | P28331 |
| HOMO SAPIENS | -1 | 3437 | 5hyn | A | 185 | ASP | p.Asp185His | Q15910 |
| HOMO SAPIENS | -1 | 3437 | 5hyn | A | 482 | ALA | p.Ala482Thr | Q15910 |
| HOMO SAPIENS | 0 | 8201 | 2wfd | A | 373 | TYR | p.Tyr373Cys | Q9P2J5 |
| HOMO SAPIENS | 1 | 3590 | 3edi | A | 90 | VAL | p.Val238Ala | O43897 |
| HOMO SAPIENS | 6 | 4555 | 1ytq | A | 64 | ALA | p.Ala65Ser | P43320 |
| HOMO SAPIENS | 0 | 4555 | 1ytq | A | 194 | TRP | p.Trp195Gly | P43320 |
| HOMO SAPIENS | 0 | 1013 | 2nn6 | C | 272 | SER | p.Ser272Thr | Q96B26 |
| HOMO SAPIENS | -1 | 29026 | 5m73 | D | 530 | ILE | p.Ile530Asn | O76094 |
| HOMO SAPIENS | 8 | 2252 | 1d5v | A | 57 | SER | p.Ser125Leu | Q99958 |
| HOMO SAPIENS | 1 | 4337 | 3ecr | A | 29 | ARG | p.Arg26His | P08397 |
| HOMO SAPIENS | 2 | 4337 | 3ecr | A | 34 | ALA | p.Ala31Thr | P08397 |
| HOMO SAPIENS | 2 | 4337 | 3ecr | A | 37 | GLN | p.Gln34Lys | P08397 |
| HOMO SAPIENS | 1 | 4337 | 3ecr | A | 58 | ALA | p.Ala55Ser | P08397 |
| HOMO SAPIENS | 0 | 4337 | 3ecr | A | 84 | LEU | p.Leu81Pro | P08397 |
| HOMO SAPIENS | 0 | 4337 | 3ecr | A | 114 | GLY | p.Gly111Arg | P08397 |
| HOMO SAPIENS | 1 | 4337 | 3ecr | A | 119 | ARG | p.Arg116Trp | P08397 |
| HOMO SAPIENS | 1 | 4337 | 3ecr | A | 152 | ARG | p.Arg149Gln | P08397 |
| HOMO SAPIENS | 1 | 4337 | 3ecr | A | 170 | ARG | p.Arg167Trp | P08397 |
| HOMO SAPIENS | 1 | 4337 | 3ecr | A | 176 | ARG | p.Arg173Gln | P08397 |
| HOMO SAPIENS | 0 | 4337 | 3ecr | A | 180 | LEU | p.Leu177Arg | P08397 |
| HOMO SAPIENS | 0 | 4337 | 3ecr | A | 181 | ASP | p.Asp178Asn | P08397 |
| HOMO SAPIENS | 1 | 4337 | 3ecr | A | 198 | ARG | p.Arg195Cys | P08397 |
| HOMO SAPIENS | 0 | 4337 | 3ecr | A | 201 | TRP | p.Trp198Ter | P08397 |
| HOMO SAPIENS | 1 | 4337 | 3ecr | A | 204 | ARG | p.Arg201Trp | P08397 |
| HOMO SAPIENS | 1 | 4337 | 3ecr | A | 219 | GLY | p.Gly216Asp | P08397 |
| HOMO SAPIENS | 1 | 4337 | 3ecr | A | 226 | GLU | p.Glu223Lys | P08397 |
| HOMO SAPIENS | 0 | 4337 | 3ecr | A | 248 | LEU | p.Leu245Arg | P08397 |

| | | | | | | | | |
|---|---|---|---|---|---|---|---|---|
| HOMO SAPIENS | 1 | 4337 | 3ecr | A | 250 | CYS | p.Cys247Arg | P08397 |
| HOMO SAPIENS | 0 | 4337 | 3ecr | A | 253 | GLU | p.Glu250Lys | P08397 |
| HOMO SAPIENS | 0 | 4337 | 3ecr | A | 255 | ALA | p.Ala252Thr | P08397 |
| HOMO SAPIENS | 0 | 4337 | 3ecr | A | 255 | ALA | p.Ala252Val | P08397 |
| HOMO SAPIENS | 0 | 4337 | 3ecr | A | 259 | HIS | p.His256Asn | P08397 |
| HOMO SAPIENS | 0 | 4337 | 3ecr | A | 324 | ARG | p.Arg321His | P08397 |
| HOMO SAPIENS | -1 | 4337 | 3ecr | A | 362 | ASP | p.Asp359Asn | P08397 |
| HOMO SAPIENS | 1 | 1204 | 5lwv | A | 854 | ILE | p.Ala864Thr | O15294 |
| HOMO SAPIENS | 1 | 25016 | 1m4u | A | 35 | PRO | p.Pro35Arg | Q13253 |
| HOMO SAPIENS | 1 | 25016 | 1m4u | A | 35 | PRO | p.Pro35Ser | Q13253 |
| HOMO SAPIENS | 1 | 25016 | 1m4u | A | 167 | ARG | p.Arg167Gly | Q13253 |
| HOMO SAPIENS | 1 | 25016 | 1m4u | A | 184 | CYS | p.Cys184Tyr | Q13253 |
| HOMO SAPIENS | 0 | 25016 | 1m4u | A | 189 | GLY | p.Gly189Cys | Q13253 |
| HOMO SAPIENS | 1 | 25016 | 1m4u | A | 204 | ARG | p.Arg204Leu | Q13253 |
| HOMO SAPIENS | 1 | 25016 | 1m4u | A | 205 | TRP | p.Trp205Cys | Q13253 |
| HOMO SAPIENS | 1 | 25016 | 1m4u | A | 217 | TRP | p.Trp217Gly | Q13253 |
| HOMO SAPIENS | 0 | 25016 | 1m4u | A | 222 | TYR | p.Tyr222Cys | Q13253 |
| HOMO SAPIENS | 0 | 25016 | 1m4u | A | 222 | TYR | p.Tyr222Asp | Q13253 |
| HOMO SAPIENS | 1 | 25016 | 1m4u | A | 223 | PRO | p.Pro223Leu | Q13253 |
| HOMO SAPIENS | 0 | 1394 | 1um7 | A | 95 | ALA | p.Ala463Val | Q92796 |
| HOMO SAPIENS | 0 | 21294 | 4jpz | B | 1788 | GLU | p.Glu1788Asp | Q99250 |
| HOMO SAPIENS | 9 | 24068 | 2db6 | A | 47 | ARG | p.Arg119Cys | Q96MF2 |
| HOMO SAPIENS | 5 | 1869 | 4omt | A | 39 | ARG | p.Arg39Leu | P08237 |
| HOMO SAPIENS | 2 | 1869 | 4omt | A | 100 | ARG | p.Arg100Gln | P08237 |
| HOMO SAPIENS | 7 | 1869 | 4omt | A | 543 | ASP | p.Asp543Ala | P08237 |
| HOMO SAPIENS | 1 | 1869 | 4omt | A | 686 | TRP | p.Trp686Cys | P08237 |
| HOMO SAPIENS | 1 | 1869 | 4omt | A | 696 | ARG | p.Arg696His | P08237 |
| HOMO SAPIENS | 0 | 12163 | 4wxx | A | 495 | TYR | p.Tyr495Cys | P26358 |
| HOMO SAPIENS | 0 | 12163 | 4wxx | A | 528 | ILE | p.Ile528Met | P26358 |
| HOMO SAPIENS | 0 | 12391 | 1j1d | C | 36 | LYS | p.Lys36Gln | P19429 |
| HOMO SAPIENS | 0 | 12391 | 1j1d | C | 82 | PRO | p.Pro82Ser | P19429 |
| HOMO SAPIENS | 7 | 12391 | 1j1d | C | 102 | ALA | p.Ala102Thr | P19429 |
| HOMO SAPIENS | 1 | 5000 | 4cca | A | 17 | GLY | p.Gly17Arg | Q15833 |
| HOMO SAPIENS | 0 | 5000 | 4cca | A | 205 | VAL | p.Val205Ile | Q15833 |
| HOMO SAPIENS | 0 | 5000 | 4cca | A | 209 | LEU | p.Leu209Pro | Q15833 |
| HOMO SAPIENS | 1 | 5000 | 4cca | A | 459 | ARG | p.Arg459Trp | Q15833 |
| HOMO SAPIENS | 7 | 5000 | 4cca | A | 477 | PRO | p.Pro477Leu | Q15833 |
| HOMO SAPIENS | -1 | 5000 | 4cca | A | 526 | ILE | p.Ile526Val | Q15833 |
| HOMO SAPIENS | 0 | 5000 | 4cca | A | 541 | GLY | p.Gly541Ser | Q15833 |
| HOMO SAPIENS | 0 | 5000 | 4cca | A | 555 | ARG | p.Arg555Gly | Q15833 |
| HOMO SAPIENS | 5 | 28551 | 3vcm | P | 26 | ARG | p.Arg49Ter | P00797 |
| HOMO SAPIENS | 6 | 36453 | 5dzz | A | 2070 | ASP | p.Asp2070His | P15924 |
| HOMO SAPIENS | 0 | 36453 | 5dzz | A | 2166 | ARG | p.Arg2166Ter | P15924 |
| HOMO SAPIENS | 0 | 20268 | 2da6 | A | 96 | ASN | p.Asn321Ser | P35680 |
| HOMO SAPIENS | 1 | 1405 | 3iar | A | 8 | ASP | p.Asp8Asn | P00813 |
| HOMO SAPIENS | 8 | 1405 | 3iar | A | 15 | HIS | p.His15Asp | P00813 |
| HOMO SAPIENS | 6 | 1405 | 3iar | A | 74 | GLY | p.Gly74Val | P00813 |
| HOMO SAPIENS | 6 | 1405 | 3iar | A | 80 | LYS | p.Lys80Arg | P00813 |
| HOMO SAPIENS | 5 | 1405 | 3iar | A | 97 | TYR | p.Tyr97Cys | P00813 |
| HOMO SAPIENS | 5 | 1405 | 3iar | A | 101 | ARG | p.Arg101Leu | P00813 |
| HOMO SAPIENS | 5 | 1405 | 3iar | A | 101 | ARG | p.Arg101Trp | P00813 |
| HOMO SAPIENS | 5 | 1405 | 3iar | A | 106 | LEU | p.Leu106Val | P00813 |
| HOMO SAPIENS | 0 | 1405 | 3iar | A | 107 | LEU | p.Leu107Pro | P00813 |
| HOMO SAPIENS | 9 | 1405 | 3iar | A | 156 | ARG | p.Arg156Cys | P00813 |

| | | | | | | | | |
|---|---|---|---|---|---|---|---|---|
| HOMO SAPIENS | 6 | 1405 | 3iar | A | 211 | ARG | p.Arg211His | P00813 |
| HOMO SAPIENS | 0 | 1405 | 3iar | A | 216 | GLY | p.Gly216Arg | P00813 |
| HOMO SAPIENS | 0 | 1405 | 3iar | A | 291 | SER | p.Ser291Leu | P00813 |
| HOMO SAPIENS | 6 | 1405 | 3iar | A | 304 | LEU | p.Leu304Arg | P00813 |
| HOMO SAPIENS | 0 | 1405 | 3iar | A | 329 | ALA | p.Ala329Val | P00813 |
| HOMO SAPIENS | -1 | 31340 | 4atu | I | 186 | ARG | p.Arg186His | O43602 |
| HOMO SAPIENS | -1 | 31340 | 4atu | I | 192 | ARG | p.Arg192Trp | O43602 |
| HOMO SAPIENS | -1 | 31340 | 4atu | I | 196 | ARG | p.Arg196His | O43602 |
| HOMO SAPIENS | -1 | 31340 | 4atu | I | 203 | THR | p.Thr203Arg | O43602 |
| HOMO SAPIENS | -1 | 31340 | 4atu | I | 286 | ALA | p.Ala286Ser | O43602 |
| HOMO SAPIENS | 4 | 30734 | 1fsu | A | 95 | ARG | p.Arg95Gln | P15848 |
| HOMO SAPIENS | 0 | 30734 | 1fsu | A | 117 | CYS | p.Cys117Arg | P15848 |
| HOMO SAPIENS | 0 | 30734 | 1fsu | A | 137 | GLY | p.Gly137Val | P15848 |
| HOMO SAPIENS | 0 | 30734 | 1fsu | A | 210 | TYR | p.Tyr210Cys | P15848 |
| HOMO SAPIENS | 0 | 30734 | 1fsu | A | 236 | LEU | p.Leu236Pro | P15848 |
| HOMO SAPIENS | 1 | 30734 | 1fsu | A | 315 | ARG | p.Arg315Gln | P15848 |
| HOMO SAPIENS | 0 | 30734 | 1fsu | A | 358 | VAL | p.Val358Met | P15848 |
| HOMO SAPIENS | 2 | 30734 | 1fsu | A | 376 | VAL | p.Val376Met | P15848 |
| HOMO SAPIENS | 0 | 30734 | 1fsu | A | 384 | SER | p.Ser384Asn | P15848 |
| HOMO SAPIENS | 0 | 30734 | 1fsu | A | 393 | HIS | p.His393Pro | P15848 |
| HOMO SAPIENS | 0 | 30734 | 1fsu | A | 405 | CYS | p.Cys405Tyr | P15848 |
| HOMO SAPIENS | 0 | 30734 | 1fsu | A | 458 | ASN | p.Asn458Ser | P15848 |
| HOMO SAPIENS | 1 | 30734 | 1fsu | A | 484 | ARG | p.Arg484Gly | P15848 |
| HOMO SAPIENS | 0 | 38072 | 1wmv | A | 51 | PRO | p.Pro98Leu | Q9NZC7 |
| HOMO SAPIENS | -1 | 5674 | 1j1e | C | 185 | ASN | p.Asn185Lys | P19429 |
| HOMO SAPIENS | -1 | 5674 | 1j1e | C | 192 | ARG | p.Arg192His | P19429 |
| HOMO SAPIENS | -1 | 5674 | 1j1e | C | 196 | ASP | p.Asp196Asn | P19429 |
| HOMO SAPIENS | -1 | 5674 | 1j1e | C | 206 | LYS | p.Lys206Gln | P19429 |
| HOMO SAPIENS | 0 | 38067 | 1wlp | A | 13 | PRO | p.Pro156Gln | P13498 |
| HOMO SAPIENS | -1 | 20441 | 3vfd | A | 289 | HIS | p.His289Tyr | Q9UBP0 |
| HOMO SAPIENS | 2 | 20441 | 3vfd | A | 344 | ILE | p.Ile344Lys | Q9UBP0 |
| HOMO SAPIENS | 0 | 20441 | 3vfd | A | 362 | SER | p.Ser362Cys | Q9UBP0 |
| HOMO SAPIENS | 1 | 20441 | 3vfd | A | 386 | ASN | p.Asn386Ser | Q9UBP0 |
| HOMO SAPIENS | 9 | 20441 | 3vfd | A | 441 | ASP | p.Asp441Gly | Q9UBP0 |
| HOMO SAPIENS | -1 | 20441 | 3vfd | A | 448 | CYS | p.Cys448Tyr | Q9UBP0 |
| HOMO SAPIENS | 2 | 20441 | 3vfd | A | 500 | ARG | p.Arg499Cys | Q9UBP0 |
| HOMO SAPIENS | 1 | 20441 | 3vfd | A | 563 | ARG | p.Arg562Gly | Q9UBP0 |
| HOMO SAPIENS | 3 | 18471 | 4dx8 | J | 26 | ARG | p.Arg26Gln | O00522 |
| HOMO SAPIENS | -1 | 18471 | 4dx8 | J | 101 | GLY | p.Gly101Val | O00522 |
| HOMO SAPIENS | -1 | 34735 | 2ly4 | B | 83 | ALA | p.Ala83Glu | P04637 |
| HOMO SAPIENS | 0 | 31128 | 4cvh | A | 116 | ARG | p.Arg116Cys | A4D126 |
| HOMO SAPIENS | 8 | 31128 | 4cvh | A | 215 | GLN | p.Gln215Ter | A4D126 |
| HOMO SAPIENS | 0 | 31128 | 4cvh | A | 268 | ARG | p.Arg268Ter | A4D126 |
| HOMO SAPIENS | 0 | 31128 | 4cvh | A | 316 | THR | p.Thr316Lys | A4D126 |
| HOMO SAPIENS | 1 | 7164 | 3mt5 | A | 369 | ASP | p.Asp434Gly | Q12791 |
| HOMO SAPIENS | 2 | 26812 | 2wtk | F | 49 | TYR | p.Tyr49Asp | Q15831 |
| HOMO SAPIENS | 0 | 26812 | 2wtk | F | 67 | LEU | p.Leu67Pro | Q15831 |
| HOMO SAPIENS | 4 | 26812 | 2wtk | F | 104 | ARG | p.Arg104Ser | Q15831 |
| HOMO SAPIENS | 0 | 26812 | 2wtk | F | 135 | GLY | p.Gly135Arg | Q15831 |
| HOMO SAPIENS | 0 | 26812 | 2wtk | F | 163 | GLY | p.Gly163Asp | Q15831 |
| HOMO SAPIENS | 2 | 26812 | 2wtk | F | 194 | ALA | p.Asp194Asn | Q15831 |
| HOMO SAPIENS | 2 | 1357 | 5xtd | W | 57 | ARG | p.Arg57His | Q9P0J0 |
| HOMO SAPIENS | 0 | 28086 | 4nn2 | A | 229 | HIS | p.His229Arg | Q8IWS0 |
| HOMO SAPIENS | 0 | 28086 | 4nn2 | A | 234 | LYS | p.Lys234Glu | Q8IWS0 |

| | | | | | | | | |
|---|---|---|---|---|---|---|---|---|
| HOMO SAPIENS | 0 | 28086 | 4nn2 | A | 257 | ARG | p.Arg257Gly | Q8IWS0 |
| HOMO SAPIENS | 5 | 31182 | 2a1x | A | 176 | GLN | p.Gln176Lys | O14832 |
| HOMO SAPIENS | 1 | 31182 | 2a1x | A | 204 | GLY | p.Gly204Ser | O14832 |
| HOMO SAPIENS | 0 | 31182 | 2a1x | A | 269 | ASN | p.Asn269His | O14832 |
| HOMO SAPIENS | 3 | 31182 | 2a1x | A | 275 | ARG | p.Arg275Gln | O14832 |
| HOMO SAPIENS | 3 | 31182 | 2a1x | A | 275 | ARG | p.Arg275Trp | O14832 |
| HOMO SAPIENS | 4 | 2823 | 5xtb | P | 145 | THR | p.Thr145Ile | O75489 |
| HOMO SAPIENS | 5 | 2823 | 5xtb | P | 199 | ARG | p.Arg199Trp | O75489 |
| HOMO SAPIENS | 0 | 8645 | 2h5g | A | 372 | SER | p.Ser372Tyr | P54886 |
| HOMO SAPIENS | 3 | 8645 | 2h5g | A | 715 | ASP | p.Asp715His | P54886 |
| HOMO SAPIENS | 0 | 8645 | 2h5g | A | 765 | ARG | p.Arg765Gln | P54886 |
| HOMO SAPIENS | 0 | 8645 | 2h5g | A | 782 | TYR | p.Tyr782Cys | P54886 |
| HOMO SAPIENS | 0 | 8645 | 2h5g | A | 784 | HIS | p.His784Tyr | P54886 |
| HOMO SAPIENS | 1 | 13992 | 2cue | A | 39 | ARG | p.Arg242Thr | P26367 |
| HOMO SAPIENS | 9 | 13992 | 2cue | A | 55 | PHE | p.Phe258Ser | P26367 |
| HOMO SAPIENS | 0 | 30418 | 2xsw | A | 303 | VAL | p.Val303Met | Q9NRR6 |
| HOMO SAPIENS | 0 | 30418 | 2xsw | A | 315 | PRO | p.Pro315Leu | Q9NRR6 |
| HOMO SAPIENS | 0 | 30418 | 2xsw | A | 341 | GLY | p.Gly341Ser | Q9NRR6 |
| HOMO SAPIENS | 0 | 30418 | 2xsw | A | 378 | ARG | p.Arg378Cys | Q9NRR6 |
| HOMO SAPIENS | 0 | 30418 | 2xsw | A | 435 | ARG | p.Arg435Gln | Q9NRR6 |
| HOMO SAPIENS | 0 | 30418 | 2xsw | A | 515 | ARG | p.Arg515Trp | Q9NRR6 |
| HOMO SAPIENS | 1 | 30418 | 2xsw | A | 526 | PRO | p.Pro526Leu | Q9NRR6 |
| HOMO SAPIENS | 0 | 30418 | 2xsw | A | 563 | ARG | p.Arg563His | Q9NRR6 |
| HOMO SAPIENS | 2 | 30418 | 2xsw | A | 585 | ARG | p.Arg585His | Q9NRR6 |
| HOMO SAPIENS | 2 | 11167 | 4y99 | B | 205 | ARG | p.Arg215Leu | P45379 |
| HOMO SAPIENS | 5 | 11167 | 4y99 | B | 244 | GLU | p.Glu254Asp | P45379 |
| HOMO SAPIENS | 1 | 11167 | 4y99 | B | 253 | LYS | p.Lys263Arg | P45379 |
| HOMO SAPIENS | 8 | 11167 | 4y99 | B | 265 | ARG | p.Arg275Gln | P45379 |
| HOMO SAPIENS | 3 | 11167 | 4y99 | B | 270 | ASP | p.Asp280Asn | P45379 |
| HOMO SAPIENS | -1 | 11167 | 4y99 | B | 278 | ARG | p.Arg288Pro | P45379 |
| HOMO SAPIENS | -1 | 11167 | 4y99 | B | 286 | ARG | p.Arg296Cys | P45379 |
| HOMO SAPIENS | -1 | 11167 | 4y99 | B | 286 | ARG | p.Arg296His | P45379 |
| HOMO SAPIENS | 0 | 15993 | 4po6 | A | 197 | ARG | p.Arg197His | P29597 |
| HOMO SAPIENS | -1 | 15993 | 4po6 | A | 362 | VAL | p.Val362Phe | P29597 |
| HOMO SAPIENS | -1 | 15993 | 4po6 | A | 363 | GLY | p.Gly363Ser | P29597 |
| HOMO SAPIENS | 0 | 5646 | 5uak | A | 7 | GLU | p.Glu7Ter | P13569 |
| HOMO SAPIENS | 0 | 5646 | 5uak | A | 27 | GLY | p.Gly27Ter | P13569 |
| HOMO SAPIENS | 0 | 5646 | 5uak | A | 46 | ALA | p.Ala46Asp | P13569 |
| HOMO SAPIENS | 0 | 5646 | 5uak | A | 60 | GLU | p.Glu60Ter | P13569 |
| HOMO SAPIENS | 0 | 5646 | 5uak | A | 67 | PRO | p.Pro67Leu | P13569 |
| HOMO SAPIENS | 0 | 5646 | 5uak | A | 74 | ARG | p.Arg74Trp | P13569 |
| HOMO SAPIENS | 0 | 5646 | 5uak | A | 75 | ARG | p.Arg75Ter | P13569 |
| HOMO SAPIENS | 0 | 5646 | 5uak | A | 85 | GLY | p.Gly85Glu | P13569 |
| HOMO SAPIENS | 0 | 5646 | 5uak | A | 91 | GLY | p.Gly91Arg | P13569 |
| HOMO SAPIENS | 0 | 5646 | 5uak | A | 92 | GLU | p.Glu92Lys | P13569 |
| HOMO SAPIENS | 0 | 5646 | 5uak | A | 98 | GLN | p.Gln98Arg | P13569 |
| HOMO SAPIENS | 0 | 5646 | 5uak | A | 98 | GLN | p.Gln98Ter | P13569 |
| HOMO SAPIENS | 0 | 5646 | 5uak | A | 109 | TYR | p.Tyr109Cys | P13569 |
| HOMO SAPIENS | 0 | 5646 | 5uak | A | 110 | ASP | p.Asp110His | P13569 |
| HOMO SAPIENS | 0 | 5646 | 5uak | A | 117 | ARG | p.Arg117Cys | P13569 |
| HOMO SAPIENS | 0 | 5646 | 5uak | A | 117 | ARG | p.Arg117His | P13569 |
| HOMO SAPIENS | 0 | 5646 | 5uak | A | 125 | ILE | p.Ile125Thr | P13569 |
| HOMO SAPIENS | 0 | 5646 | 5uak | A | 148 | ILE | p.Ile148Thr | P13569 |
| HOMO SAPIENS | 0 | 5646 | 5uak | A | 170 | ARG | p.Arg170His | P13569 |

| | | | | | | | | |
|---|---|---|---|---|---|---|---|---|
| HOMO SAPIENS | 0 | 5646 | 5uak | A | 178 | GLY | p.Gly178Arg | P13569 |
| HOMO SAPIENS | 0 | 5646 | 5uak | A | 206 | LEU | p.Leu206Trp | P13569 |
| HOMO SAPIENS | 0 | 5646 | 5uak | A | 373 | ASP | p.Asp373Asn | P13569 |
| HOMO SAPIENS | -1 | 5646 | 5uak | A | 709 | ARG | p.Arg709Ter | P13569 |
| HOMO SAPIENS | -1 | 5646 | 5uak | A | 732 | LEU | p.Leu732Ter | P13569 |
| HOMO SAPIENS | -1 | 5646 | 5uak | A | 785 | ARG | p.Arg785Ter | P13569 |
| HOMO SAPIENS | -1 | 5646 | 5uak | A | 831 | GLU | p.Glu831Ter | P13569 |
| HOMO SAPIENS | -1 | 5646 | 5uak | A | 836 | ASP | p.Asp836Tyr | P13569 |
| HOMO SAPIENS | 0 | 5646 | 5uak | A | 851 | ARG | p.Arg851Ter | P13569 |
| HOMO SAPIENS | 0 | 5646 | 5uak | A | 913 | TYR | p.Tyr913Cys | P13569 |
| HOMO SAPIENS | 0 | 5646 | 5uak | A | 913 | TYR | p.Tyr913Ter | P13569 |
| HOMO SAPIENS | 0 | 5646 | 5uak | A | 945 | SER | p.Ser945Leu | P13569 |
| HOMO SAPIENS | 0 | 5646 | 5uak | A | 949 | HIS | p.His949Tyr | P13569 |
| HOMO SAPIENS | 0 | 5646 | 5uak | A | 970 | GLY | p.Gly970Asp | P13569 |
| HOMO SAPIENS | 0 | 5646 | 5uak | A | 1027 | ILE | p.Ile1027Thr | P13569 |
| HOMO SAPIENS | 0 | 5646 | 5uak | A | 1032 | TYR | p.Tyr1032Cys | P13569 |
| HOMO SAPIENS | 0 | 5646 | 5uak | A | 1061 | GLY | p.Gly1061Arg | P13569 |
| HOMO SAPIENS | 1 | 5646 | 5uak | A | 1065 | LEU | p.Leu1065Pro | P13569 |
| HOMO SAPIENS | 1 | 5646 | 5uak | A | 1066 | ARG | p.Arg1066Cys | P13569 |
| HOMO SAPIENS | 1 | 5646 | 5uak | A | 1066 | ARG | p.Arg1066His | P13569 |
| HOMO SAPIENS | 1 | 5646 | 5uak | A | 1067 | ALA | p.Ala1067Thr | P13569 |
| HOMO SAPIENS | 0 | 5646 | 5uak | A | 1069 | GLY | p.Gly1069Arg | P13569 |
| HOMO SAPIENS | 0 | 5646 | 5uak | A | 1070 | ARG | p.Arg1070Trp | P13569 |
| HOMO SAPIENS | 1 | 5646 | 5uak | A | 1071 | GLN | p.Gln1071Pro | P13569 |
| HOMO SAPIENS | 0 | 5646 | 5uak | A | 1074 | PHE | p.Phe1074Leu | P13569 |
| HOMO SAPIENS | 0 | 5646 | 5uak | A | 1077 | LEU | p.Leu1077Pro | P13569 |
| HOMO SAPIENS | 0 | 5646 | 5uak | A | 1089 | TRP | p.Trp1089Ter | P13569 |
| HOMO SAPIENS | 0 | 5646 | 5uak | A | 1092 | TYR | p.Tyr1092Ter | P13569 |
| HOMO SAPIENS | 0 | 5646 | 5uak | A | 1101 | MET | p.Met1101Lys | P13569 |
| HOMO SAPIENS | 0 | 5646 | 5uak | A | 1156 | LEU | p.Leu1156Phe | P13569 |
| HOMO SAPIENS | 0 | 5646 | 5uak | A | 1158 | ARG | p.Arg1158Ter | P13569 |
| HOMO SAPIENS | 0 | 5646 | 5uak | A | 1162 | ARG | p.Arg1162Leu | P13569 |
| HOMO SAPIENS | 0 | 5646 | 5uak | A | 1162 | ARG | p.Arg1162Ter | P13569 |
| HOMO SAPIENS | -1 | 5646 | 5uak | A | 1476 | GLN | p.Gln1476Ter | P13569 |
| HOMO SAPIENS | 0 | 22763 | 1imv | A | 52 | MET | p.Thr72Met | P36955 |
| HOMO SAPIENS | 0 | 22763 | 1imv | A | 301 | TYR | p.Tyr321Ter | P36955 |
| HOMO SAPIENS | 0 | 23179 | 5fql | A | 160 | PRO | p.Pro160Arg | P22304 |
| HOMO SAPIENS | 0 | 23179 | 5fql | A | 172 | ARG | p.Arg172Ter | P22304 |
| HOMO SAPIENS | 0 | 23179 | 5fql | A | 214 | THR | p.Thr214Met | P22304 |
| HOMO SAPIENS | 0 | 23179 | 5fql | A | 309 | THR | p.Thr309Ala | P22304 |
| HOMO SAPIENS | 0 | 23179 | 5fql | A | 333 | SER | p.Ser333Leu | P22304 |
| HOMO SAPIENS | 0 | 23179 | 5fql | A | 443 | ARG | p.Arg443Ter | P22304 |
| HOMO SAPIENS | 0 | 23179 | 5fql | A | 468 | ARG | p.Arg468Leu | P22304 |
| HOMO SAPIENS | 0 | 23179 | 5fql | A | 468 | ARG | p.Arg468Trp | P22304 |
| HOMO SAPIENS | 0 | 23179 | 5fql | A | 502 | TRP | p.Trp502Ser | P22304 |
| HOMO SAPIENS | 9 | 20989 | 1nfi | F | 185 | THR | p.Thr185Met | P25963 |
| HOMO SAPIENS | 4 | 20989 | 1nfi | F | 194 | GLY | p.Gly194Ala | P25963 |
| HOMO SAPIENS | 0 | 11538 | 3eab | A | 162 | VAL | p.Val162Ile | Q9UBP0 |
| HOMO SAPIENS | 0 | 33399 | 2p39 | A | 71 | SER | p.Ser71Gly | Q9GZV9 |
| HOMO SAPIENS | 3 | 33399 | 2p39 | A | 96 | MET | p.Met96Thr | Q9GZV9 |
| HOMO SAPIENS | 0 | 33399 | 2p39 | A | 129 | SER | p.Ser129Phe | Q9GZV9 |
| HOMO SAPIENS | -1 | 33399 | 2p39 | A | 172 | PRO | p.Pro172Leu | Q9GZV9 |
| HOMO SAPIENS | -1 | 33399 | 2p39 | A | 176 | ARG | p.Arg176Gln | Q9GZV9 |
| HOMO SAPIENS | -1 | 33399 | 2p39 | A | 179 | ARG | p.Arg179Trp | Q9GZV9 |

| | | | | | | | | |
|---|---|---|---|---|---|---|---|---|
| HOMO SAPIENS | -1 | 33399 | 2p39 | A | 179 | ARG | p.Arg179Gln | Q9GZV9 |
| HOMO SAPIENS | 0 | 16217 | 2dne | A | 79 | ILE | p.Ile163Leu | P10515 |
| HOMO SAPIENS | 8 | 7666 | 4kbq | A | 65 | ASN | p.Asn65Ser | Q9UNE7 |
| HOMO SAPIENS | 1 | 6407 | 1ikt | A | 71 | THR | p.Thr687Ile | P51659 |
| HOMO SAPIENS | 0 | 6407 | 1ikt | A | 112 | MET | p.Met728Val | P51659 |
| HOMO SAPIENS | 0 | 5849 | 4ihb | A | 67 | VAL | p.Val67Asp | O75923 |
| HOMO SAPIENS | 6 | 17703 | 5fcn | A | 20 | HIS | p.His99Gln | Q66GS9 |
| HOMO SAPIENS | 8 | 17703 | 5fcn | A | 33 | ARG | p.Arg112His | Q66GS9 |
| HOMO SAPIENS | 0 | 34095 | 2E+073 | A | 101 | HIS | p.His101Tyr | P05129 |
| HOMO SAPIENS | -1 | 41358 | 4liq | E | 507 | THR | p.Thr507Met | P07333 |
| HOMO SAPIENS | -1 | 21290 | 1fch | A | 235 | MET | p.Met272Arg | P50542 |
| HOMO SAPIENS | 5 | 23431 | 6pax | A | 23 | ARG | p.Arg26Gly | P26367 |
| HOMO SAPIENS | 1 | 23431 | 6pax | A | 61 | GLY | p.Gly64Val | P26367 |
| HOMO SAPIENS | 1 | 23431 | 6pax | A | 65 | PRO | p.Pro68Ser | P26367 |
| HOMO SAPIENS | 1 | 23431 | 6pax | A | 116 | SER | p.Ser119Arg | P26367 |
| HOMO SAPIENS | 9 | 23431 | 6pax | A | 123 | VAL | p.Val126Asp | P26367 |
| HOMO SAPIENS | 9 | 23431 | 6pax | A | 125 | ARG | p.Arg128Cys | P26367 |
| HOMO SAPIENS | 9 | 33003 | 4zrj | B | 535 | LEU | p.Leu535Pro | P35240 |
| HOMO SAPIENS | 7 | 33003 | 4zrj | B | 538 | GLN | p.Gln538Pro | P35240 |
| HOMO SAPIENS | 0 | 33003 | 4zrj | B | 585 | TRP | p.Ala585Thr | P35240 |
| HOMO SAPIENS | 0 | 3800 | 2nzj | A | 193 | VAL | p.Val193Met | O75628 |

| gene_name | max_cons | max_bs_id | max_pdb_id | max_chain_id |
|---|---|---|---|---|
| AKR1D1 | None | None | None | None |
| AKR1D1 | None | None | None | None |
| AKR1D1 | None | None | None | None |
| AKR1D1 | None | None | None | None |
| ALPP | None | None | None | None |
| BMPR1A | None | None | None | None |
| BMPR1A | None | None | None | None |
| FGFR2 | None | None | None | None |
| PROC | 0.587 | 3 | 1aut | C |
| PROC | None | None | None | None |
| PROC | None | None | None | None |
| PROC | None | None | None | None |
| PROC | None | None | None | None |
| PROC | None | None | None | None |
| PROC | None | None | None | None |
| PROC | None | None | None | None |
| PROC | None | None | None | None |
| PROC | 0.587 | 3 | 1aut | C |
| PROC | None | None | None | None |
| PROC | None | None | None | None |
| INS | 1 | 2 | 3zs2 | J |
| INS | 0.85 | 1 | 4ajx | D |
| INS | 1 | 1 | 3zs2 | J |
| INS | 1 | 1 | 3zs2 | J |
| FTL | 0.627 | 11 | 2fg8 | B |
| FTL | None | None | None | None |
| FTL | 0.544 | 24 | 2fg8 | D |
| TNFSF11 | None | None | None | None |
| EHMT1 | None | None | None | None |
| EHMT1 | None | None | None | None |
| EHMT1 | None | None | None | None |
| CA2 | 0.939 | 1 | 4idr | X |
| CA2 | 0.939 | 1 | 4idr | X |
| CA2 | 0.902 | 5 | 4idr | X |
| CA2 | 0.902 | 5 | 4idr | X |
| SMAD4 | None | None | None | None |
| SMAD4 | None | None | None | None |
| TTR | 0.691 | 12 | 1ttb | B |
| TTR | 0.961 | 7 | 3grb | A |
| TTR | 0.891 | 4 | 3dk2 | B |
| TTR | 0.947 | 3 | 1tta | A |
| TTR | 0.923 | 1 | 3grb | B |
| TTR | 0.891 | 1 | 4y9b | B |
| TTR | 0.891 | 1 | 4y9b | B |
| TTR | 0.882 | 15 | 3grb | A |
| TTR | 0.909 | 5 | 4tl4 | A |
| TTR | 0.836 | 7 | 1z7j | B |
| TTR | 0.882 | 15 | 3grb | A |
| TTR | 0.891 | 5 | 3i9i | B |
| TTR | 0.902 | 9 | 3m1o | B |
| TTR | 0.902 | 9 | 3m1o | B |
| TTR | 0.902 | 9 | 3m1o | B |
| TTR | 0.636 | 16 | 1thc | A |

| | | | | | |
|---|---|---|---|---|---|
| TTR | | 0.902 | 9 | 3m1o | B |
| TTR | | 0.836 | 7 | 1bze | A |
| TTR | | 0.846 | 16 | 3gps | A |
| TTR | | 0.846 | 16 | 3gps | A |
| TTR | | 0.909 | 5 | 4tl4 | A |
| TTR | | 0.909 | 5 | 4tl4 | A |
| TTR | | 0.673 | 15 | 3dk2 | B |
| TTR | | 0.945 | 4 | 4deu | A |
| TTR | | 0.941 | 3 | 3did | B |
| TTR | | 0.941 | 3 | 3did | B |
| TTR | | 0.868 | 6 | 4l1t | A |
| TTR | | 0.882 | 19 | 3gps | B |
| TTR | | 0.891 | 4 | 3dk2 | B |
| TTR | | 0.941 | 3 | 3did | B |
| TTR | | 0.923 | 1 | 4hiq | A |
| TTR | | 0.923 | 1 | 4hiq | A |
| TTR | | 0.961 | 7 | 3grb | A |
| TTR | | 0.839 | 7 | 1tsh | B |
| TTR | | 0.891 | 6 | 1ttb | B |
| ALK | | 0.682 | 1 | 4cd0 | A |
| ALK | None | None | | None | None |
| THRB | None | None | | None | None |
| THRB | None | None | | None | None |
| THRB | None | None | | None | None |
| THRB | None | None | | None | None |
| THRB | None | None | | None | None |
| THRB | None | None | | None | None |
| THRB | None | None | | None | None |
| THRB | None | None | | None | None |
| THRB | None | None | | None | None |
| THRB | None | None | | None | None |
| THRB | None | None | | None | None |
| THRB | None | None | | None | None |
| THRB | None | None | | None | None |
| THRB | None | None | | None | None |
| THRB | None | None | | None | None |
| THRB | None | None | | None | None |
| THRB | None | None | | None | None |
| THRB | None | None | | None | None |
| THRB | | 0.577 | 3 | 2j4a | A |
| IL2RG | None | None | | None | None |
| IL2RG | None | None | | None | None |
| IL2RG | None | None | | None | None |
| IL2RG | None | None | | None | None |
| IL2RG | None | None | | None | None |
| ITGB3 | None | None | | None | None |
| ITGB3 | None | None | | None | None |
| ITGB3 | None | None | | None | None |
| ITGB3 | None | None | | None | None |
| ITGB3 | None | None | | None | None |
| ITGB3 | None | None | | None | None |
| ITGB3 | None | None | | None | None |
| ITGB3 | None | None | | None | None |
| ITGB3 | None | None | | None | None |

| | | | | |
|---|---|---|---|---|
| PKD2 | None | None | None | None |
| PKD2 | None | None | None | None |
| PKD2 | None | None | None | None |
| PKD2 | None | None | None | None |
| PKD2 | None | None | None | None |
| PKD2 | None | None | None | None |
| PKD2 | None | None | None | None |
| TP53 | 0.571 | 1 | 1saf | B |
| TP53 | 0.571 | 1 | 1saf | B |
| TP53 | None | None | None | None |
| TP53 | 0.585 | 15 | 4mzr | B |
| TP53 | None | None | None | None |
| TP53 | None | None | None | None |
| EGFR | None | None | None | None |
| EGFR | None | None | None | None |
| XRCC4 | None | None | None | None |
| XRCC4 | None | None | None | None |
| HBB | 0.438 | 2 | 2w6v | D |
| HBB | 0.467 | 2 | 3qje | D |
| HBB | 0.467 | 2 | 3qje | D |
| HBB | None | None | None | None |
| HBB | None | None | None | None |
| HBB | None | None | None | None |
| HBB | 0.421 | 11 | 3ic2 | B |
| HBB | 0.459 | 2 | 5ee4 | D |
| HBB | 0.459 | 2 | 5ee4 | D |
| HBB | 0.459 | 2 | 5ee4 | D |
| HBB | 0.535 | 3 | 3r5i | B |
| HBB | None | None | None | None |
| HBB | None | None | None | None |
| HBB | 0.402 | 11 | 1m9p | D |
| HBB | None | None | None | None |
| HBB | 0.459 | 2 | 5ee4 | D |
| HBB | 0.566 | 1 | 1cbm | B |
| HBB | 0.51 | 3 | 1abw | B |
| HBB | None | None | None | None |
| HBB | 0.474 | 1 | 2h35 | B |
| HBB | 0.474 | 1 | 2h35 | B |
| HBB | None | None | None | None |
| HBB | 0.508 | 7 | 3ic2 | B |
| HBB | 0.545 | 3 | 1bij | D |
| HBB | 0.6 | 1 | 2w72 | D |
| HBB | None | None | None | None |
| HBB | None | None | None | None |
| HBB | None | None | None | None |
| HBB | 0.733 | 1 | 1yff | D |
| HBB | 0.692 | 2 | 1buw | B |
| RPS6KA3 | None | None | None | None |
| RPS6KA3 | 0.555 | 1 | 4nw5 | A |
| RPS6KA3 | None | None | None | None |
| RPS6KA3 | None | None | None | None |
| RPS6KA3 | None | None | None | None |
| RPS6KA3 | None | None | None | None |
| GBA | 1 | 15 | 3gxd | B |

| | | | | | |
|---|---|---|---|---|---|
| GBA | | 1 | 55 | 3gxd | B |
| GBA | | 1 | 15 | 3gxd | B |
| GBA | | 0.75 | 45 | 3ril | A |
| GBA | None | None | | None | None |
| GBA | None | None | | None | None |
| GBA | | 1 | 12 | 1ogs | A |
| GBA | | 0.75 | 122 | 3gxi | A |
| GBA | | 1 | 2 | 1ogs | A |
| GBA | | 1 | 11 | 1ogs | A |
| GBA | | 0.5 | 59 | 3gxf | B |
| GBA | | 0.75 | 45 | 3ril | A |
| GBA | | 1 | 18 | 1ogs | A |
| GBA | | 1 | 3 | 1ogs | A |
| GBA | None | None | | None | None |
| GBA | | 1 | 2 | 1ogs | A |
| GBA | None | None | | None | None |
| GBA | | 0.75 | 76 | 3gxm | C |
| GBA | | 1 | 3 | 1ogs | A |
| GBA | | 1 | 3 | 1ogs | A |
| GBA | | 0.75 | 62 | 3ril | A |
| GBA | | 1 | 28 | 1ogs | A |
| GBA | None | None | | None | None |
| GBA | | 0.5 | 78 | 3gxi | A |
| GBA | | 0.5 | 92 | 3rik | B |
| GBA | | 0.5 | 42 | 2xwe | A |
| GBA | None | None | | None | None |
| GBA | | 0.5 | 92 | 3rik | B |
| GBA | | 0.75 | 49 | 3ril | A |
| GBA | None | None | | None | None |
| GBA | | 0.667 | 60 | 1ogs | A |
| GBA | None | None | | None | None |
| GBA | | 0.5 | 11 | 2nt1 | A |
| GBA | | 0.75 | 21 | 3gxi | A |
| GBA | None | None | | None | None |
| GBA | | 0.5 | 35 | 2xwe | A |
| GBA | | 0.75 | 25 | 3keh | B |
| GBA | | 0.75 | 70 | 3ril | A |
| GBA | | 0.75 | 70 | 3ril | A |
| GBA | | 1 | 15 | 3gxd | B |
| GBA | | 1 | 44 | 3gxd | B |
| GBA | | 0.5 | 57 | 3gxf | D |
| GBA | | 1 | 44 | 3gxd | B |
| GBA | | 1 | 44 | 3gxd | B |
| GBA | None | None | | None | None |
| GBA | | 0.75 | 94 | 3gxi | A |
| GBA | | 0.75 | 94 | 3gxi | A |
| IFIH1 | None | None | | None | None |
| TGFBR2 | None | None | | None | None |
| TGFBR2 | None | None | | None | None |
| TGFBR2 | None | None | | None | None |
| TGFBR2 | None | None | | None | None |
| TGFBR2 | None | None | | None | None |
| TGFBR2 | None | None | | None | None |
| TGFBR2 | None | None | | None | None |

| | | | | | |
|---|---|---|---|---|---|
| TGFBR2 | None | None | None | | None |
| TGFBR2 | None | None | None | | None |
| TGFBR2 | 0.742 | | 1 | 5E+091 | A |
| TGFBR2 | None | None | None | | None |
| TGFBR2 | None | None | None | | None |
| TGFBR2 | None | None | None | | None |
| TGFBR2 | 0.676 | | 1 | 5e8v | A |
| TGFBR2 | 0.676 | | 1 | 5e8v | A |
| ACE | 0.684 | | 17 | 3nxq | A |
| ACE | 0.474 | | 25 | 4bxk | A |
| ACE | None | None | None | | None |
| ACE | None | None | None | | None |
| ACE | None | None | None | | None |
| ACE | None | None | None | | None |
| COL10A1 | None | None | None | | None |
| COL10A1 | None | None | None | | None |
| COL10A1 | None | None | None | | None |
| COL10A1 | 0.409 | | 1 | 1gr3 | A |
| COL10A1 | None | None | None | | None |
| COL10A1 | None | None | None | | None |
| COL10A1 | None | None | None | | None |
| COL10A1 | None | None | None | | None |
| COL10A1 | None | None | None | | None |
| KLF1 | None | None | None | | None |
| C3 | None | None | None | | None |
| CTSC | None | None | None | | None |
| CTSC | 0.934 | | 1 | 2djg | C |
| CTSC | None | None | None | | None |
| LYZ | 0.877 | | 3 | 1bb5 | A |
| LYZ | 0.923 | | 1 | 1lz4 | A |
| LYZ | 0.852 | | 2 | 1ubz | A |
| DHODH | None | None | None | | None |
| DHODH | None | None | None | | None |
| DHODH | None | None | None | | None |
| DHODH | None | None | None | | None |
| DHODH | None | None | None | | None |
| DHODH | None | None | None | | None |
| DHODH | None | None | None | | None |
| ADH1C | 0.527 | | 11 | 1ht0 | A |
| GDNF | None | None | None | | None |
| TNF | 0.5 | | 8 | 2e7a | B |
| ALDH4A1 | 0.636 | | 3 | 3v9h | A |
| ALDH4A1 | None | None | None | | None |
| ALDH4A1 | None | None | None | | None |
| BRAF | None | None | None | | None |
| BRAF | None | None | None | | None |
| BRAF | 0.752 | | 2 | 3psb | A |
| BRAF | 0.752 | | 2 | 3psb | A |
| BRAF | 0.737 | | 1 | 3pri | A |
| BRAF | 0.737 | | 1 | 3pri | A |
| BRAF | 0.737 | | 1 | 3pri | A |
| BRAF | 0.737 | | 1 | 3prf | A |
| BRAF | 0.737 | | 1 | 3prf | A |
| BRAF | 0.752 | | 2 | 3psb | A |

| | | | | |
|---|---|---|---|---|
| BRAF | 0.752 | 2 | 3psb | A |
| BRAF | 0.752 | 2 | 3psb | A |
| BRAF | 0.752 | 2 | 3psb | A |
| BRAF | 0.555 | 3 | 5vam | A |
| BRAF | 0.663 | 1 | 4ehe | B |
| BRAF | 0.734 | 1 | 5csx | A |
| TBX5 | None | None | None | None |
| TBX5 | None | None | None | None |
| TBX5 | None | None | None | None |
| TBX5 | None | None | None | None |
| TBX5 | None | None | None | None |
| TBX5 | None | None | None | None |
| TBX5 | None | None | None | None |
| TBX5 | None | None | None | None |
| TBX5 | None | None | None | None |
| RB1 | None | None | None | None |
| RB1 | None | None | None | None |
| RB1 | None | None | None | None |
| RB1 | None | None | None | None |
| RB1 | None | None | None | None |
| RB1 | None | None | None | None |
| CLIC2 | 0.5 | 3 | 2per | A |
| CLIC2 | None | None | None | None |
| ALDH7A1 | None | None | None | None |
| ALDH7A1 | None | None | None | None |
| ALDH7A1 | 0.405 | 4 | 2j6l | D |
| ALDH7A1 | None | None | None | None |
| ALDH7A1 | 0.409 | 3 | 4zvw | C |
| ALDH7A1 | None | None | None | None |
| ALDH7A1 | 0.534 | 3 | 2j6l | A |
| ALDH7A1 | None | None | None | None |
| ALDH7A1 | None | None | None | None |
| ALDH7A1 | None | None | None | None |
| ALDH7A1 | None | None | None | None |
| ALDH7A1 | 0.534 | 3 | 2j6l | A |
| ALDH7A1 | None | None | None | None |
| ALDH7A1 | None | None | None | None |
| ALDH7A1 | 0.632 | 1 | 4zuk | A |
| ALDH7A1 | None | None | None | None |
| KDR | 0.575 | 5 | 1y6a | A |
| MUT | None | None | None | None |
| MUT | None | None | None | None |
| MUT | None | None | None | None |
| MUT | None | None | None | None |
| MUT | None | None | None | None |
| MUT | None | None | None | None |
| MUT | None | None | None | None |
| MUT | None | None | None | None |
| MUT | None | None | None | None |
| MUT | None | None | None | None |
| MUT | None | None | None | None |
| MUT | None | None | None | None |
| MUT | None | None | None | None |
| MUT | None | None | None | None |

| | | | | |
|---|---|---|---|---|
| MUT | None | None | None | None |
| MUT | None | None | None | None |
| MUT | 0.6 | 1 | 3bic | A |
| MUT | 0.6 | 2 | 2xiq | A |
| MUT | 0.6 | 2 | 2xiq | A |
| MUT | None | None | None | None |
| MUT | None | None | None | None |
| MUT | 0.4 | 1 | 2xiq | A |
| MUT | None | None | None | None |
| MUT | None | None | None | None |
| MUT | None | None | None | None |
| MUT | None | None | None | None |
| MUT | None | None | None | None |
| MUT | None | None | None | None |
| MUT | None | None | None | None |
| MUT | None | None | None | None |
| MUT | None | None | None | None |
| MUT | None | None | None | None |
| MUT | None | None | None | None |
| MUT | None | None | None | None |
| SOD1 | 0.778 | 1 | 2gbu | A |
| SOD1 | 0.727 | 4 | 4xcr | A |
| SOD1 | None | None | None | None |
| SOD1 | 0.621 | 12 | 1hl5 | H |
| NECTIN3 | None | None | None | None |
| HFE | None | None | None | None |
| HFE | None | None | None | None |
| HFE | None | None | None | None |
| HFE | 0.5 | 1 | 1de4 | A |
| HFE | None | None | None | None |
| HFE | None | None | None | None |
| HFE | None | None | None | None |
| EIF2AK3 | None | None | None | None |
| EIF2AK3 | None | None | None | None |
| TERT | None | None | None | None |
| TERT | None | None | None | None |
| SPR | None | None | None | None |
| SPR | None | None | None | None |
| SPR | None | None | None | None |
| SPR | None | None | None | None |
| SPR | None | None | None | None |
| F7 | 0.625 | 2 | 1klj | L |
| UNG | None | None | None | None |
| MMP13 | None | None | None | None |
| MMP13 | 0.786 | 1 | 3tvc | A |
| MMP13 | 0.766 | 1 | 3zxh | B |
| MMP13 | 0.766 | 1 | 3zxh | B |
| CD207 | 0.621 | 1 | 3kqg | B |
| F2 | 1 | 2 | 1nrq | L |
| SPTBN2 | None | None | None | None |
| PKLR | 0.568 | 9 | 2vgb | A |
| PKLR | 0.486 | 17 | 2vgg | D |
| PKLR | None | None | None | None |
| PKLR | 0.486 | 8 | 2vgg | D |

| | | | | |
|---|---|---|---|---|
| PKLR | 0.568 | 3 | 4ip7 | D |
| PKLR | 0.432 | 13 | 2vgi | A |
| PKLR | None | None | None | None |
| PKLR | None | None | None | None |
| SACS | None | None | None | None |
| HRAS | 0.529 | 3 | 6q21 | A |
| HRAS | 0.529 | 3 | 6q21 | A |
| HRAS | 0.59 | 1 | 1gnq | A |
| HRAS | 0.59 | 1 | 1gnq | A |
| HRAS | 0.664 | 1 | 2lwi | A |
| HRAS | 0.742 | 1 | 1ctq | A |
| HRAS | 0.69 | 1 | 1xd2 | A |
| HRAS | 0.415 | 3 | 6q21 | D |
| HRAS | 0.529 | 3 | 6q21 | A |
| HRAS | 0.403 | 3 | 2n46 | A |
| HRAS | 0.521 | 4 | 4nyj | R |
| HRAS | 0.664 | 1 | 2lwi | A |
| HRAS | 0.664 | 1 | 2lwi | A |
| VHL | None | None | None | None |
| VHL | None | None | None | None |
| VHL | None | None | None | None |
| VHL | None | None | None | None |
| VHL | 0.5 | 22 | 4w9l | C |
| VHL | 0.5 | 22 | 5nvy | C |
| VHL | None | None | None | None |
| VHL | None | None | None | None |
| VHL | None | None | None | None |
| VHL | 0.5 | 2 | 5nvy | C |
| VHL | 0.5 | 1 | 5nvy | C |
| VHL | 0.5 | 1 | 5nvy | C |
| VHL | 0.5 | 4 | 5nvy | C |
| VHL | None | None | None | None |
| VHL | 0.5 | 6 | 5nvy | C |
| VHL | 0.5 | 2 | 5nvy | C |
| VHL | 0.5 | 8 | 3ztc | C |
| VHL | 0.5 | 2 | 5nvy | C |
| VHL | 0.5 | 2 | 5nvy | C |
| VHL | None | None | None | None |
| VHL | 0.5 | 7 | 4w9h | C |
| VHL | 0.5 | 2 | 5nvy | C |
| VHL | 0.5 | 2 | 5nvy | C |
| VHL | 0.5 | 19 | 4w9l | C |
| VHL | None | None | None | None |
| TTBK1 | None | None | None | None |
| RAD51 | None | None | None | None |
| ALB | None | None | None | None |
| ALB | 0.5 | 2 | 4n0u | D |
| ALB | None | None | None | None |
| ALB | None | None | None | None |
| MET | None | None | None | None |
| MET | None | None | None | None |
| MET | None | None | None | None |
| MET | None | None | None | None |
| MET | None | None | None | None |

| | | | | |
|---|---|---|---|---|
| MET | None | None | None | None |
| MET | None | None | None | None |
| MET | None | None | None | None |
| MET | None | None | None | None |
| CP | None | None | None | None |
| CP | None | None | None | None |
| PGM1 | None | None | None | None |
| PGM1 | 0.409 | 2 | 5f9c | A |
| PGM1 | 0.409 | 2 | 5f9c | A |
| PGM1 | None | None | None | None |
| PGM1 | None | None | None | None |
| PGM1 | None | None | None | None |
| PGM1 | None | None | None | None |
| NR5A1 | None | None | None | None |
| NR5A1 | None | None | None | None |
| NR5A1 | None | None | None | None |
| F9 | None | None | None | None |
| F9 | None | None | None | None |
| HPRT1 | None | None | None | None |
| HPRT1 | None | None | None | None |
| HPRT1 | None | None | None | None |
| HPRT1 | None | None | None | None |
| HPRT1 | None | None | None | None |
| HPRT1 | 0.871 | 3 | 4rac | B |
| HPRT1 | 0.806 | 2 | 4ran | D |
| HPRT1 | None | None | None | None |
| HPRT1 | None | None | None | None |
| HPRT1 | 0.44 | 3 | 1d6n | B |
| HPRT1 | None | None | None | None |
| HPRT1 | None | None | None | None |
| HPRT1 | None | None | None | None |
| HPRT1 | 0.703 | 4 | 5w8v | A |
| HPRT1 | None | None | None | None |
| HPRT1 | None | None | None | None |
| HPRT1 | None | None | None | None |
| HPRT1 | None | None | None | None |
| HPRT1 | 0.871 | 3 | 4rac | B |
| HPRT1 | 0.871 | 3 | 4rac | B |
| HPRT1 | 0.765 | 2 | 4rao | D |
| HPRT1 | 0.774 | 2 | 4rac | B |
| GPD1L | None | None | None | None |
| GPD1L | None | None | None | None |
| GPD1L | None | None | None | None |
| FGFR2 | None | None | None | None |
| FGFR2 | None | None | None | None |
| FGFR2 | None | None | None | None |
| FGFR2 | None | None | None | None |
| FGFR2 | None | None | None | None |
| FGFR2 | None | None | None | None |
| FGFR2 | None | None | None | None |
| FGFR2 | None | None | None | None |
| FGFR2 | None | None | None | None |
| FGFR2 | None | None | None | None |
| FGFR2 | None | None | None | None |

| | | | | |
|---|---|---|---|---|
| FGFR2 | None | None | None | None |
| FGFR2 | None | None | None | None |
| FGFR2 | None | None | None | None |
| FGFR2 | None | None | None | None |
| FGFR2 | None | None | None | None |
| FGFR2 | None | None | None | None |
| FGFR2 | None | None | None | None |
| FGFR2 | None | None | None | None |
| HSPD1 | None | None | None | None |
| HSPD1 | None | None | None | None |
| FGF8 | None | None | None | None |
| FGF8 | None | None | None | None |
| FGF8 | None | None | None | None |
| F9 | 0.438 | 1 | 5vyg | B |
| NCF4 | None | None | None | None |
| NCF4 | None | None | None | None |
| LDLR | None | None | None | None |
| LDLR | None | None | None | None |
| LDLR | 0.478 | 1 | 3gcx | E |
| LDLR | 0.478 | 1 | 3gcx | E |
| LDLR | 0.478 | 1 | 3gcx | E |
| LDLR | None | None | None | None |
| C3 | None | None | None | None |
| C3 | None | None | None | None |
| C3 | None | None | None | None |
| C3 | None | None | None | None |
| C3 | None | None | None | None |
| C3 | None | None | None | None |
| C3 | None | None | None | None |
| C3 | None | None | None | None |
| TULP1 | None | None | None | None |
| TULP1 | None | None | None | None |
| TULP1 | None | None | None | None |
| TULP1 | None | None | None | None |
| IKBKG | None | None | None | None |
| IKBKG | None | None | None | None |
| IKBKG | None | None | None | None |
| CD40 | None | None | None | None |
| CD40 | None | None | None | None |
| SLC9A3R1 | None | None | None | None |
| BCHE | None | None | None | None |
| BCHE | 0.439 | 5 | 2wil | B |
| BCHE | 0.544 | 11 | 1p0q | A |
| BCHE | None | None | None | None |
| BCHE | None | None | None | None |
| BCHE | None | None | None | None |
| BCHE | 0.63 | 4 | 2xqg | A |
| BCHE | None | None | None | None |
| BCHE | None | None | None | None |
| BCHE | None | None | None | None |
| BCHE | None | None | None | None |
| BCKDHB | 0.833 | 1 | 1v11 | B |
| BCKDHB | 0.5 | 18 | 2bfc | B |
| BCKDHB | None | None | None | None |

| | | | | | |
|---|---|---|---|---|---|
| BCKDHB | | 0.667 | 11 | 2bfd | B |
| BCKDHB | | 0.833 | 10 | 1olx | B |
| BCKDHB | | 0.833 | 1 | 1v11 | B |
| BCKDHB | | 0.833 | 1 | 1v11 | B |
| BCKDHB | | 0.625 | 2 | 1x7z | B |
| BCKDHB | | 0.429 | 16 | 1ols | B |
| BCKDHB | None | None | | None | None |
| BCKDHB | None | None | | None | None |
| BCKDHB | | 0.857 | 1 | 2bff | B |
| CDH1 | None | None | | None | None |
| CDH1 | | 0.575 | 1 | 3ff8 | A |
| CDH1 | | 0.543 | 1 | 3ff7 | A |
| CDH1 | None | None | | None | None |
| CDH1 | None | None | | None | None |
| CTNNB1 | None | None | | None | None |
| AIRE | None | None | | None | None |
| EDA | | 1 | 1 | 1rj8 | A |
| EDA | None | None | | None | None |
| EDA | None | None | | None | None |
| EDA | None | None | | None | None |
| EDA | None | None | | None | None |
| SH2D1A | | 0.444 | 5 | 1ka7 | A |
| SH2D1A | None | None | | None | None |
| SH2D1A | None | None | | None | None |
| GCK | | 0.556 | 37 | 3idh | A |
| GCK | None | None | | None | None |
| GCK | None | None | | None | None |
| GCK | None | None | | None | None |
| GCK | None | None | | None | None |
| GCK | | 0.611 | 13 | 3h1v | X |
| GCK | None | None | | None | None |
| GCK | None | None | | None | None |
| GCK | None | None | | None | None |
| GCK | | 0.778 | 5 | 3id8 | A |
| GCK | | 0.833 | 1 | 4dhy | A |
| GCK | None | None | | None | None |
| GCK | None | None | | None | None |
| GCK | | 0.611 | 6 | 3id8 | A |
| GCK | None | None | | None | None |
| GCK | None | None | | None | None |
| GCK | None | None | | None | None |
| GCK | | 0.5 | 5 | 3vey | A |
| GCK | None | None | | None | None |
| GCK | None | None | | None | None |
| AQP2 | None | None | | None | None |
| AQP2 | None | None | | None | None |
| AQP2 | None | None | | None | None |
| AQP2 | None | None | | None | None |
| AQP2 | None | None | | None | None |
| AQP2 | None | None | | None | None |
| AQP2 | None | None | | None | None |
| AQP2 | None | None | | None | None |
| AQP2 | None | None | | None | None |
| AQP2 | None | None | | None | None |

| | | | | |
|---|---|---|---|---|
| AQP2 | None | None | None | None |
| AQP2 | None | None | None | None |
| AQP2 | None | None | None | None |
| SUMF1 | 0.75 | 6 | 1z70 | X |
| SUMF1 | 0.556 | 30 | 1y1i | X |
| SUMF1 | 0.5 | 38 | 2aij | X |
| SUMF1 | 0.625 | 18 | 2aij | X |
| SUMF1 | 0.875 | 35 | 2afy | X |
| SUMF1 | 0.667 | 19 | 1y1i | X |
| SUMF1 | 0.75 | 17 | 1y1j | X |
| SUMF1 | 0.75 | 17 | 1y1j | X |
| SUMF1 | 0.625 | 27 | 1y1g | X |
| FAM20A | None | None | None | None |
| FAM20A | None | None | None | None |
| FAM20A | None | None | None | None |
| FAM20A | None | None | None | None |
| NT5E | None | None | None | None |
| NT5E | None | None | None | None |
| NOS3 | 0.564 | 54 | 1m9k | B |
| ACTG1 | None | None | None | None |
| ACTG1 | None | None | None | None |
| ACTG1 | None | None | None | None |
| ACTG1 | None | None | None | None |
| ACTG1 | 0.571 | 8 | 5jlh | E |
| ACTG1 | None | None | None | None |
| FCGR1A | None | None | None | None |
| GPI | None | None | None | None |
| GPI | None | None | None | None |
| GPI | 1 | 3 | 1jiq | A |
| GPI | None | None | None | None |
| GPI | 0.696 | 22 | 1iat | A |
| GPI | 0.522 | 22 | 1nuh | A |
| GPI | 0.696 | 27 | 1iat | A |
| GPI | None | None | None | None |
| GPI | None | None | None | None |
| CFH | None | None | None | None |
| OGG1 | 0.895 | 13 | 1m3h | A |
| TPK1 | None | None | None | None |
| TPK1 | None | None | None | None |
| TPK1 | None | None | None | None |
| TPK1 | None | None | None | None |
| SEC23A | None | None | None | None |
| SEC23A | None | None | None | None |
| TYRP1 | None | None | None | None |
| TYRP1 | None | None | None | None |
| TYRP1 | None | None | None | None |
| TYRP1 | None | None | None | None |
| TYRP1 | None | None | None | None |
| TYRP1 | None | None | None | None |
| CSTB | None | None | None | None |
| CSTB | None | None | None | None |
| CSTB | None | None | None | None |
| ACP1 | None | None | None | None |
| IL17F | None | None | None | None |

| | | | | |
|---|---|---|---|---|
| IL17F | None | None | None | None |
| IL17F | None | None | None | None |
| IL17F | None | None | None | None |
| IL17F | None | None | None | None |
| IDH2 | 0.5 | 2 | 5svn | B |
| IDH2 | 0.5 | 2 | 5svn | B |
| IDH2 | 0.48 | 13 | 5i96 | A |
| IDH2 | None | None | None | None |
| TP53 | 0.925 | 11 | 2ocj | A |
| TP53 | 0.8 | 16 | 2ahi | D |
| TP53 | 0.65 | 49 | 4ibq | A |
| TP53 | 0.725 | 27 | 5g4o | A |
| TP53 | 0.455 | 21 | 3q06 | A |
| TP53 | 0.9 | 1 | 4agp | A |
| TP53 | 0.9 | 1 | 4agp | A |
| TP53 | 0.925 | 10 | 4hje | A |
| TP53 | 0.7 | 29 | 3d05 | A |
| TP53 | 0.875 | 7 | 3kmd | B |
| TP53 | 0.875 | 7 | 3kmd | B |
| TP53 | 0.925 | 6 | 3kz8 | B |
| TP53 | 0.925 | 6 | 3kz8 | B |
| TP53 | 0.609 | 22 | 3q01 | B |
| TP53 | 0.925 | 5 | 3kmd | B |
| TP53 | 0.925 | 5 | 3kmd | B |
| TP53 | 0.925 | 8 | 3kz8 | B |
| TP53 | 0.925 | 8 | 3kz8 | B |
| TP53 | 0.925 | 1 | 3kmd | B |
| TP53 | 0.9 | 11 | 2pcx | A |
| TP53 | 0.9 | 1 | 5aol | A |
| TP53 | 0.85 | 5 | 5lgy | A |
| TP53 | 0.9 | 1 | 3q06 | B |
| TP53 | 0.725 | 19 | 5lgy | A |
| TP53 | 0.775 | 14 | 4mzi | A |
| TP53 | 0.825 | 8 | 1kzy | A |
| TP53 | 0.825 | 8 | 1kzy | A |
| TP53 | 0.8 | 11 | 4ibt | A |
| TP53 | 0.8 | 11 | 4ibt | A |
| TP53 | 0.725 | 15 | 5lgy | A |
| TP53 | 0.725 | 29 | 2h1l | O |
| TP53 | 0.875 | 7 | 3kmd | B |
| TP53 | 0.725 | 15 | 5lgy | A |
| TP53 | 0.8 | 10 | 4agp | A |
| TP53 | 0.8 | 10 | 4agp | A |
| TP53 | 0.8 | 10 | 4agp | A |
| TP53 | 0.8 | 10 | 4agp | A |
| TP53 | 0.675 | 35 | 4ibq | A |
| TP53 | 0.575 | 37 | 4agm | A |
| TP53 | 0.675 | 17 | 3kz8 | B |
| TP53 | None | None | None | None |
| TP53 | None | None | None | None |
| LMAN1 | None | None | None | None |
| CTSK | 0.75 | 2 | 4n79 | A |
| CTSK | None | None | None | None |
| CTSK | 0.413 | 12 | 4x6j | A |

| | | | | |
|---|---|---|---|---|
| FGB | None | None | None | None |
| FGB | None | None | None | None |
| FGB | 0.688 | 1 | 1fzf | E |
| FGB | None | None | None | None |
| ACADM | None | None | None | None |
| ACADM | None | None | None | None |
| ACADM | None | None | None | None |
| ACADM | 0.471 | 3 | 4p13 | D |
| ACADM | None | None | None | None |
| ACADM | None | None | None | None |
| ACADM | None | None | None | None |
| ACADM | None | None | None | None |
| ACADM | None | None | None | None |
| ACADM | None | None | None | None |
| ACADM | None | None | None | None |
| ACADM | None | None | None | None |
| ACADM | None | None | None | None |
| ACADM | None | None | None | None |
| ACADM | None | None | None | None |
| ACADM | 0.481 | 2 | 4p13 | C |
| ACADM | None | None | None | None |
| ACADM | None | None | None | None |
| ACADM | None | None | None | None |
| ACADM | None | None | None | None |
| ACADM | 0.564 | 1 | 1t9g | C |
| ACADM | None | None | None | None |
| ACADM | None | None | None | None |
| ACADM | None | None | None | None |
| ACADM | None | None | None | None |
| ACADM | None | None | None | None |
| ACADM | None | None | None | None |
| PTS | None | None | None | None |
| PTS | 0.417 | 2 | 3i2b | E |
| PTS | None | None | None | None |
| PTS | 0.778 | 1 | 3i2b | D |
| PTS | None | None | None | None |
| PTS | None | None | None | None |
| PTS | None | None | None | None |
| PTS | None | None | None | None |
| PTS | None | None | None | None |
| INS | 0.75 | 2 | 3zqr | E |
| INS | 1 | 5 | 2l1z | A |
| INS | 1 | 5 | 2l1z | A |
| INS | 1 | 1 | 1his | A |
| FBP1 | 0.71 | 26 | 2y5l | C |
| FBP1 | None | None | None | None |
| FBP1 | 0.5 | 18 | 1fta | A |
| FBP1 | 0.7 | 10 | 2y5k | A |
| FBP1 | 0.516 | 38 | 3kc1 | B |
| FBP1 | 0.733 | 10 | 3kc1 | A |
| FBP1 | 0.633 | 43 | 2jjk | A |
| FBP1 | 0.8 | 11 | 2y5k | A |
| FBP1 | 0.548 | 41 | 2vt5 | D |
| FBP1 | 0.438 | 25 | 2fie | A |

| | | | | |
|---|---|---|---|---|
| FBP1 | 0.643 | 7 | 4mjo | B |
| KCNQ1 | None | None | None | None |
| KCNQ1 | None | None | None | None |
| KCNQ1 | None | None | None | None |
| UPP1 | 0.706 | 4 | 3euf | A |
| UMPS | 1 | 3 | 2qcl | A |
| UMPS | 0.556 | 59 | 2qcl | B |
| UMPS | None | None | None | None |
| DHFR | None | None | None | None |
| KHK | 0.667 | 2 | 2hw1 | A |
| TTPA | None | None | None | None |
| TTPA | None | None | None | None |
| TTPA | 0.5 | 1 | 1r5l | A |
| TTPA | None | None | None | None |
| MAP1LC3B | None | None | None | None |
| MAP1LC3B | None | None | None | None |
| ZAP70 | None | None | None | None |
| PROC | 0.536 | 1 | 3f6u | L |
| SQSTM1 | None | None | None | None |
| AHCY | None | None | None | None |
| AHCY | None | None | None | None |
| PRNP | None | None | None | None |
| PRNP | None | None | None | None |
| PRNP | None | None | None | None |
| PRNP | None | None | None | None |
| PRNP | None | None | None | None |
| PRNP | None | None | None | None |
| PRNP | None | None | None | None |
| PRNP | None | None | None | None |
| PRNP | None | None | None | None |
| PRNP | None | None | None | None |
| PRNP | None | None | None | None |
| PRNP | None | None | None | None |
| PRNP | None | None | None | None |
| PRNP | None | None | None | None |
| PRNP | None | None | None | None |
| PRNP | None | None | None | None |
| REN | 0.431 | 2 | 2v13 | A |
| REN | 0.431 | 2 | 2v13 | A |
| HINT1 | 0.556 | 2 | 5ed6 | B |
| HINT1 | 0.897 | 1 | 5km4 | A |
| HINT1 | None | None | None | None |
| HINT1 | 0.667 | 11 | 5ed6 | B |
| HINT1 | 0.923 | 1 | 5ipc | A |
| OAT | None | None | None | None |
| OAT | None | None | None | None |
| OAT | None | None | None | None |
| OAT | None | None | None | None |
| OAT | None | None | None | None |
| OAT | None | None | None | None |
| OAT | None | None | None | None |
| OAT | None | None | None | None |
| OAT | None | None | None | None |
| OAT | None | None | None | None |

| | | | | |
|---|---|---|---|---|
| OAT | 0.675 | 2 | 2byl | B |
| OAT | None | None | None | None |
| OAT | None | None | None | None |
| OAT | None | None | None | None |
| OAT | None | None | None | None |
| OAT | 0.758 | 1 | 2byl | A |
| OAT | None | None | None | None |
| OAT | 0.525 | 4 | 2byl | B |
| OAT | 0.429 | 6 | 2byj | B |
| OAT | None | None | None | None |
| OAT | None | None | None | None |
| OAT | None | None | None | None |
| OAT | None | None | None | None |
| OAT | None | None | None | None |
| OAT | None | None | None | None |
| OAT | None | None | None | None |
| OAT | None | None | None | None |
| OAT | None | None | None | None |
| CTSA | None | None | None | None |
| CTSA | None | None | None | None |
| PNP | None | None | None | None |
| PNP | None | None | None | None |
| PNP | 0.406 | 43 | 3phb | E |
| PNP | 0.706 | 5 | 4ear | C |
| PNP | None | None | None | None |
| PNP | 0.5 | 17 | 1pf7 | E |
| PNP | 0.933 | 1 | 3ggs | C |
| PNP | 0.833 | 1 | 3d1v | A |
| MYL2 | None | None | None | None |
| MYL2 | None | None | None | None |
| ANG | 0.548 | 4 | 1k5b | A |
| ANG | 0.419 | 8 | 4ahi | A |
| ANG | 0.419 | 8 | 4ahi | A |
| ANG | None | None | None | None |
| ANG | 0.761 | 1 | 5eop | A |
| ANG | 0.576 | 4 | 5m9v | A |
| ANG | 0.579 | 4 | 5m9q | A |
| ANG | None | None | None | None |
| ANG | None | None | None | None |
| ANG | 0.52 | 5 | 5m9g | A |
| POLH | None | None | None | None |
| POLH | None | None | None | None |
| POLH | None | None | None | None |
| IL2RA | None | None | None | None |
| TGFB1 | None | None | None | None |
| TP53 | None | None | None | None |
| TP53 | None | None | None | None |
| PAH | None | None | None | None |
| PAH | None | None | None | None |
| PAH | 0.44 | 11 | 2pah | A |
| PAH | 0.44 | 11 | 2pah | A |
| PAH | None | None | None | None |
| PAH | None | None | None | None |
| PAH | 0.48 | 2 | 1pah | A |

| | | | | |
|---|---|---|---|---|
| PAH | 0.48 | 2 | 1pah | A |
| PAH | 0.4 | 11 | 3pah | A |
| PAH | None | None | None | None |
| PAH | None | None | None | None |
| PAH | None | None | None | None |
| PAH | 0.44 | 9 | 1dmw | A |
| PAH | 0.48 | 1 | 1dmw | A |
| PAH | 0.48 | 1 | 1dmw | A |
| PAH | 0.44 | 8 | 2pah | B |
| PAH | 0.44 | 8 | 2pah | B |
| PAH | 0.417 | 3 | 1j8t | A |
| PAH | 0.48 | 2 | 4anp | A |
| PAH | 0.48 | 1 | 1dmw | A |
| PAH | 0.48 | 4 | 2pah | B |
| PAH | 0.44 | 11 | 2pah | A |
| PAH | 0.48 | 4 | 1mmt | A |
| PAH | 0.68 | 6 | 2pah | A |
| PAH | 0.76 | 1 | 1tdw | A |
| PAH | None | None | None | None |
| PAH | None | None | None | None |
| PAH | 0.4 | 14 | 1j8u | A |
| PAH | None | None | None | None |
| PAH | None | None | None | None |
| PAH | None | None | None | None |
| PAH | None | None | None | None |
| PAH | 0.64 | 2 | 1lrm | A |
| PAH | 0.4 | 11 | 2pah | B |
| PAH | None | None | None | None |
| PAH | 0.4 | 14 | 1j8u | A |
| PAH | 0.4 | 14 | 1j8u | A |
| PAH | None | None | None | None |
| PAH | 0.48 | 1 | 1dmw | A |
| PAH | 0.48 | 1 | 1dmw | A |
| PAH | None | None | None | None |
| HDAC8 | None | None | None | None |
| HDAC8 | 0.732 | 3 | 3rqd | A |
| VCP | 0.438 | 4 | 3tiw | A |
| VCP | 0.5 | 2 | 3qq8 | A |
| VCP | 0.5 | 2 | 3qq8 | A |
| VCP | None | None | None | None |
| VCP | None | None | None | None |
| VCP | None | None | None | None |
| VCP | None | None | None | None |
| B3GAT3 | None | None | None | None |
| B3GAT3 | None | None | None | None |
| CSF2RB | None | None | None | None |
| CSF2RB | None | None | None | None |
| CSF2RB | None | None | None | None |
| NF2 | 0.5 | 6 | 1h4r | B |
| NF2 | None | None | None | None |
| NF2 | None | None | None | None |
| PIK3CA | None | None | None | None |
| PHGDH | None | None | None | None |
| PHGDH | None | None | None | None |

| | | | | |
|---|---|---|---|---|
| PHGDH | None | None | None | None |
| PHGDH | 0.66 | 1 | 5n53 | A |
| POLH | None | None | None | None |
| XIAP | 0.5 | 7 | 5c7b | A |
| XIAP | None | None | None | None |
| PDHX | None | None | None | None |
| PDHX | None | None | None | None |
| PDHX | None | None | None | None |
| PDHX | None | None | None | None |
| PDHX | None | None | None | None |
| SAMHD1 | 0.5 | 12 | 5ao0 | A |
| SAMHD1 | None | None | None | None |
| SAMHD1 | 0.429 | 7 | 4tnr | D |
| SAMHD1 | None | None | None | None |
| SAMHD1 | 0.571 | 3 | 4qfy | D |
| SAMHD1 | None | None | None | None |
| SAMHD1 | None | None | None | None |
| SAMHD1 | None | None | None | None |
| CD36 | None | None | None | None |
| CD36 | None | None | None | None |
| CD36 | None | None | None | None |
| ASPH | None | None | None | None |
| CCT5 | None | None | None | None |
| CCT5 | None | None | None | None |
| CCT5 | None | None | None | None |
| SMN1 | None | None | None | None |
| SMN1 | None | None | None | None |
| SMN1 | None | None | None | None |
| SMN1 | None | None | None | None |
| ABCA1 | None | None | None | None |
| ABCA1 | None | None | None | None |
| ABCA1 | None | None | None | None |
| ABCA1 | None | None | None | None |
| ABCA1 | None | None | None | None |
| ABCA1 | None | None | None | None |
| ABCA1 | None | None | None | None |
| ABCA1 | None | None | None | None |
| ABCA1 | None | None | None | None |
| ABCA1 | None | None | None | None |
| ABCA1 | None | None | None | None |
| ABCA1 | None | None | None | None |
| ABCA1 | None | None | None | None |
| ABCA1 | None | None | None | None |
| ABCA1 | None | None | None | None |
| ABCA1 | None | None | None | None |
| ABCA1 | None | None | None | None |
| ABCA1 | None | None | None | None |
| ABCA1 | None | None | None | None |
| ABCA1 | None | None | None | None |
| ABCA1 | None | None | None | None |
| ABCA1 | None | None | None | None |
| ABCA1 | None | None | None | None |
| ABCA1 | None | None | None | None |

| | | | | |
|---|---|---|---|---|
| ABCA1 | None | None | None | None |
| ABCA1 | None | None | None | None |
| ABCA1 | None | None | None | None |
| ABCA1 | None | None | None | None |
| ABCA1 | None | None | None | None |
| ABCA1 | None | None | None | None |
| ABCA1 | None | None | None | None |
| ABCA1 | None | None | None | None |
| ABCA1 | None | None | None | None |
| ABCA1 | None | None | None | None |
| ABCA1 | None | None | None | None |
| ERCC1 | None | None | None | None |
| FECH | 0.524 | 27 | 2po5 | A |
| FECH | 0.6 | 60 | 2hre | A |
| FECH | 0.636 | 20 | 2po7 | A |
| FECH | 0.476 | 61 | 4klr | A |
| FECH | 0.5 | 84 | 2hrc | A |
| FECH | 0.667 | 10 | 2po5 | A |
| FECH | 0.455 | 30 | 4kla | B |
| PTEN | None | None | None | None |
| PTEN | None | None | None | None |
| PTEN | None | None | None | None |
| PTEN | None | None | None | None |
| PTEN | None | None | None | None |
| PTEN | None | None | None | None |
| PTEN | None | None | None | None |
| PTEN | None | None | None | None |
| PTEN | None | None | None | None |
| PTEN | None | None | None | None |
| PTEN | None | None | None | None |
| PTEN | None | None | None | None |
| PTEN | None | None | None | None |
| PTEN | None | None | None | None |
| PTEN | None | None | None | None |
| PTEN | None | None | None | None |
| PTEN | None | None | None | None |
| PTEN | None | None | None | None |
| PTEN | None | None | None | None |
| PTEN | None | None | None | None |
| ACAT1 | 0.667 | 1 | 2ib9 | C |
| ACAT1 | None | None | None | None |
| ACAT1 | None | None | None | None |
| ACAT1 | 0.667 | 1 | 2ib9 | C |
| ACAT1 | 0.667 | 1 | 2ib9 | C |
| ACAT1 | 0.472 | 11 | 2f2s | A |
| ACAT1 | 0.833 | 6 | 2ib7 | C |
| ACAT1 | None | None | None | None |
| ACAT1 | None | None | None | None |
| ACAT1 | 0.528 | 9 | 2ibw | D |
| ACAT1 | 0.703 | 16 | 2ibu | A |
| CREBBP | None | None | None | None |
| ITGA2B | None | None | None | None |
| ITGA2B | None | None | None | None |
| ITGA2B | None | None | None | None |

| | | | | | |
|---|---|---|---|---|---|
| ITGA2B | | 0.538 | 1 | 3nig | A |
| TSC1 | None | None | | None | None |
| ADA2 | None | None | | None | None |
| ADA2 | None | None | | None | None |
| ADA2 | | 0.75 | 1 | 3lgd | A |
| ADA2 | None | None | | None | None |
| ADA2 | None | None | | None | None |
| ADA2 | None | None | | None | None |
| ADA2 | None | None | | None | None |
| ADA2 | None | None | | None | None |
| ADA2 | | 1 | 1 | 3lgd | B |
| ADA2 | None | None | | None | None |
| PGK1 | None | None | | None | None |
| PGK1 | | 0.536 | 8 | 2xe8 | A |
| PGK1 | None | None | | None | None |
| PGK1 | None | None | | None | None |
| PGK1 | | 0.571 | 3 | 2ybe | A |
| TUBA1A | None | None | | None | None |
| TUBA1A | None | None | | None | None |
| TUBA1A | None | None | | None | None |
| TUBA1A | None | None | | None | None |
| TUBA1A | None | None | | None | None |
| PTPN11 | None | None | | None | None |
| PTPN11 | None | None | | None | None |
| PTPN11 | None | None | | None | None |
| PTPN11 | None | None | | None | None |
| PTPN11 | None | None | | None | None |
| PTPN11 | None | None | | None | None |
| PTPN11 | None | None | | None | None |
| PTPN11 | None | None | | None | None |
| PTPN11 | None | None | | None | None |
| PTPN11 | | 0.404 | 5 | 3tl0 | A |
| PTPN11 | | 0.404 | 5 | 3tl0 | A |
| PTPN11 | | 0.404 | 5 | 3tl0 | A |
| PTPN11 | None | None | | None | None |
| PTPN11 | | 0.466 | 7 | 3zm1 | A |
| PTPN11 | | 0.621 | 2 | 3b7o | A |
| PTPN11 | | 0.776 | 1 | 3zm1 | A |
| PTPN11 | | 0.453 | 5 | 4ohd | A |
| PTPN11 | None | None | | None | None |
| PTPN11 | None | None | | None | None |
| PTPN11 | None | None | | None | None |
| PTPN11 | None | None | | None | None |
| PTPN11 | | 0.776 | 1 | 3zm1 | A |
| PTPN11 | | 0.621 | 6 | 3zm1 | A |
| PTPN11 | None | None | | None | None |
| PTPN11 | None | None | | None | None |
| PTPN11 | | 0.621 | 6 | 3zm1 | A |
| GLB1 | None | None | | None | None |
| GLB1 | None | None | | None | None |
| UQCRQ | None | None | | None | None |
| VPS35 | None | None | | None | None |
| NR3C1 | None | None | | None | None |
| NR3C1 | None | None | | None | None |

| | | | | |
|---|---|---|---|---|
| NR3C1 | None | None | None | None |
| NR3C1 | None | None | None | None |
| NR3C1 | 0.42 | 6 | 4mdd | A |
| NR3C1 | None | None | None | None |
| NR3C1 | None | None | None | None |
| NR3C1 | None | None | None | None |
| NR3C1 | 0.422 | 14 | 1nhz | A |
| APOA1 | None | None | None | None |
| APOA1 | None | None | None | None |
| APOA1 | None | None | None | None |
| APOA1 | None | None | None | None |
| APOA1 | None | None | None | None |
| APOA1 | None | None | None | None |
| APOA1 | None | None | None | None |
| BCKDHA | None | None | None | None |
| BCKDHA | 0.583 | 76 | 1x7x | A |
| BCKDHA | 0.963 | 1 | 1ols | A |
| BCKDHA | 0.696 | 3 | 1v1r | A |
| BCKDHA | 0.625 | 16 | 2beu | A |
| BCKDHA | None | None | None | None |
| BCKDHA | 0.652 | 11 | 1v1r | A |
| BCKDHA | 1 | 1 | 2beu | A |
| BCKDHA | 0.84 | 2 | 2j9f | A |
| BCKDHA | 0.556 | 42 | 1ols | A |
| BCKDHA | 0.609 | 90 | 1v1r | A |
| BCKDHA | 0.652 | 25 | 1v1r | A |
| BCKDHA | None | None | None | None |
| BCKDHA | 0.625 | 60 | 2beu | A |
| BCKDHA | 0.625 | 28 | 2beu | A |
| TOP1 | None | None | None | None |
| ANTXR2 | None | None | None | None |
| ANTXR2 | None | None | None | None |
| ANTXR2 | None | None | None | None |
| TF | 0.625 | 35 | 5dyh | A |
| TF | None | None | None | None |
| FSHB | None | None | None | None |
| GALK1 | None | None | None | None |
| GALK1 | None | None | None | None |
| GALK1 | None | None | None | None |
| CYP17A1 | None | None | None | None |
| CYP17A1 | None | None | None | None |
| CYP17A1 | None | None | None | None |
| CYP17A1 | None | None | None | None |
| CYP17A1 | None | None | None | None |
| CYP17A1 | None | None | None | None |
| CYP17A1 | None | None | None | None |
| CYP17A1 | None | None | None | None |
| CYP17A1 | None | None | None | None |
| CYP17A1 | None | None | None | None |
| CYP17A1 | None | None | None | None |
| CYP17A1 | None | None | None | None |
| CYP17A1 | None | None | None | None |
| CYP17A1 | None | None | None | None |
| CYP17A1 | None | None | None | None |

| | | | | |
|---|---|---|---|---|
| CYP17A1 | None | None | None | None |
| CYP17A1 | None | None | None | None |
| CD44 | 0.667 | 24 | 1uuh | A |
| MDH2 | 0.426 | 5 | 4wlo | B |
| MDH2 | None | None | None | None |
| NDUFV1 | None | None | None | None |
| NDUFV1 | None | None | None | None |
| NDUFV1 | None | None | None | None |
| NDUFV1 | None | None | None | None |
| NDUFV1 | None | None | None | None |
| NDUFV1 | None | None | None | None |
| NDUFV1 | None | None | None | None |
| NDUFV1 | None | None | None | None |
| NDUFV1 | None | None | None | None |
| NDUFV1 | None | None | None | None |
| CTSC | None | None | None | None |
| CTSC | None | None | None | None |
| CTSC | None | None | None | None |
| CTSC | None | None | None | None |
| CTSC | 0.643 | 1 | 4oel | A |
| CTSC | None | None | None | None |
| CTSC | None | None | None | None |
| VWF | None | None | None | None |
| VWF | None | None | None | None |
| VWF | None | None | None | None |
| VWF | None | None | None | None |
| VWF | None | None | None | None |
| VWF | None | None | None | None |
| VWF | None | None | None | None |
| ORC4 | None | None | None | None |
| ORC4 | None | None | None | None |
| CRYAB | None | None | None | None |
| CRYAB | None | None | None | None |
| CRYAB | None | None | None | None |
| MAP2K2 | None | None | None | None |
| MAP2K2 | None | None | None | None |
| MAP2K2 | None | None | None | None |
| MAP2K2 | None | None | None | None |
| MAP2K2 | None | None | None | None |
| MAP2K2 | None | None | None | None |
| MAP2K2 | None | None | None | None |
| MAP2K2 | None | None | None | None |
| MAP2K2 | None | None | None | None |
| CBL | None | None | None | None |
| CBL | None | None | None | None |
| CBL | None | None | None | None |
| CBL | None | None | None | None |
| CBL | None | None | None | None |
| CBL | None | None | None | None |
| CBL | None | None | None | None |
| PFN1 | None | None | None | None |
| PFN1 | 0.75 | 2 | 1fik | A |
| ACSM2A | None | None | None | None |
| HSPG2 | None | None | None | None |

| | | | | |
|---|---|---|---|---|
| HSPG2 | None | None | None | None |
| HSPG2 | None | None | None | None |
| HSPG2 | None | None | None | None |
| PARK7 | 0.444 | 48 | 3cyf | A |
| PARK7 | 0.769 | 4 | 1pe0 | A |
| PARK7 | 0.808 | 2 | 1pe0 | A |
| PARK7 | 0.667 | 8 | 4rky | A |
| PARK7 | 0.889 | 1 | 3cz9 | A |
| PARK7 | 0.75 | 2 | 1pdw | B |
| INSR | None | None | None | None |
| INSR | None | None | None | None |
| INSR | None | None | None | None |
| INSR | None | None | None | None |
| INSR | None | None | None | None |
| DSC2 | None | None | None | None |
| DSC2 | None | None | None | None |
| DSC2 | None | None | None | None |
| DSC2 | None | None | None | None |
| DSC2 | None | None | None | None |
| DSC2 | None | None | None | None |
| DSC2 | None | None | None | None |
| DSC2 | None | None | None | None |
| DSC2 | None | None | None | None |
| DSC2 | None | None | None | None |
| MICU1 | None | None | None | None |
| MICU1 | None | None | None | None |
| ADAMTS13 | None | None | None | None |
| ADAMTS13 | None | None | None | None |
| ADAMTS13 | None | None | None | None |
| ADAMTS13 | None | None | None | None |
| ADAMTS13 | None | None | None | None |
| ADAMTS13 | None | None | None | None |
| ADAMTS13 | None | None | None | None |
| IRAK4 | None | None | None | None |
| IRAK4 | None | None | None | None |
| RNASEH2B | None | None | None | None |
| RNASEH2B | None | None | None | None |
| RNASEH2B | None | None | None | None |
| NTRK1 | None | None | None | None |
| MPO | 0.75 | 3 | 1d5l | C |
| MPO | 0.6 | 25 | 1myp | D |
| MPO | None | None | None | None |
| PEX3 | None | None | None | None |
| SERPINA1 | None | None | None | None |
| SERPINA1 | None | None | None | None |
| SERPINA1 | None | None | None | None |
| SERPINA1 | None | None | None | None |
| SERPINA1 | None | None | None | None |
| SERPINA1 | None | None | None | None |
| SERPINA1 | None | None | None | None |
| SERPINA1 | 0.6 | 1 | 1ezx | A |
| SERPINA1 | None | None | None | None |
| SERPINA1 | None | None | None | None |
| YARS2 | None | None | None | None |

| | | | | |
|---|---|---|---|---|
| YARS2 | None | None | None | None |
| YARS2 | None | None | None | None |
| YARS2 | None | None | None | None |
| YARS2 | None | None | None | None |
| YARS2 | 0.781 | 1 | 2pid | B |
| YARS2 | None | None | None | None |
| YARS2 | None | None | None | None |
| YARS2 | None | None | None | None |
| PDHA1 | None | None | None | None |
| PDHA1 | None | None | None | None |
| PDHA1 | None | None | None | None |
| PDHA1 | None | None | None | None |
| PDHA1 | None | None | None | None |
| PDHA1 | None | None | None | None |
| PDHA1 | None | None | None | None |
| PDHA1 | 0.96 | 1 | 3exh | C |
| PDHA1 | None | None | None | None |
| PDHA1 | None | None | None | None |
| PDHA1 | None | None | None | None |
| CUL4B | None | None | None | None |
| AGXT | None | None | None | None |
| AGXT | None | None | None | None |
| AGXT | None | None | None | None |
| AGXT | 0.692 | 3 | 2yob | A |
| AGXT | 0.786 | 7 | 4kxk | C |
| AGXT | 0.538 | 21 | 3r9a | A |
| AGXT | None | None | None | None |
| AGXT | None | None | None | None |
| AGXT | None | None | None | None |
| AGXT | 0.611 | 4 | 5hhy | B |
| AGXT | None | None | None | None |
| AGXT | None | None | None | None |
| AGXT | None | None | None | None |
| AGXT | None | None | None | None |
| AGXT | 0.429 | 28 | 4kxk | C |
| AGXT | 0.684 | 1 | 1h0c | A |
| AGXT | 0.5 | 2 | 4cbr | A |
| AGXT | 0.538 | 19 | 4kyo | A |
| AGXT | 0.538 | 19 | 4kyo | A |
| AGXT | 0.75 | 2 | 5luc | B |
| AGXT | 0.722 | 3 | 5hhy | B |
| AGXT | 0.571 | 3 | 4kxk | C |
| AGXT | 0.571 | 5 | 4kxk | C |
| AGXT | None | None | None | None |
| AGXT | 0.778 | 1 | 5hhy | B |
| AGXT | 0.778 | 1 | 5hhy | B |
| AGXT | 0.421 | 12 | 1h0c | A |
| AGXT | 0.692 | 7 | 4kyo | A |
| AGXT | 0.5 | 18 | 4cbr | A |
| AGXT | 0.5 | 18 | 4cbr | A |
| AGXT | 0.615 | 13 | 3r9a | A |
| AGXT | 0.786 | 7 | 4kxk | C |
| AGXT | 0.846 | 1 | 4kxk | A |
| AGXT | 0.615 | 36 | 3r9a | A |

| | | | | | |
|---|---|---|---|---|---|
| AGXT | | 0.571 | 10 | 4kxk | C |
| SFTPD | | 0.509 | 2 | 3g83 | B |
| SFTPD | None | None | | None | None |
| PIK3CA | None | None | | None | None |
| PIK3CA | None | None | | None | None |
| PIK3CA | None | None | | None | None |
| AUH | None | None | | None | None |
| AUH | None | None | | None | None |
| AUH | None | None | | None | None |
| AUH | None | None | | None | None |
| F11 | None | None | | None | None |
| F11 | None | None | | None | None |
| F11 | None | None | | None | None |
| F11 | None | None | | None | None |
| MEN1 | | 0.6 | 81 | 5db0 | A |
| MEN1 | | 0.8 | 54 | 4i80 | A |
| MEN1 | | 0.8 | 84 | 5db0 | A |
| MEN1 | | 0.8 | 30 | 5ddb | A |
| MEN1 | | 1 | 2 | 4i80 | A |
| MEN1 | | 0.8 | 17 | 5db0 | A |
| MEN1 | | 0.6 | 17 | 5ddb | A |
| MEN1 | | 0.6 | 31 | 3u88 | A |
| MEN1 | None | None | | None | None |
| MEN1 | | 0.6 | 56 | 5ddb | A |
| MYBPC1 | None | None | | None | None |
| MYBPC1 | None | None | | None | None |
| MYBPC1 | None | None | | None | None |
| PDCD10 | None | None | | None | None |
| SCARB2 | None | None | | None | None |
| SCARB2 | None | None | | None | None |
| SCARB2 | None | None | | None | None |
| SCARB2 | None | None | | None | None |
| SCARB2 | None | None | | None | None |
| TREX2 | None | None | | None | None |
| TBK1 | None | None | | None | None |
| TBK1 | None | None | | None | None |
| DLD | None | None | | None | None |
| DLD | None | None | | None | None |
| DLD | | 0.505 | 3 | 1zy8 | B |
| DLD | None | None | | None | None |
| DLD | None | None | | None | None |
| DLD | None | None | | None | None |
| LDHA | | 0.802 | 1 | 4qt0 | A |
| TNNC1 | None | None | | None | None |
| TNNC1 | | 0.461 | 2 | 1wrl | A |
| TNNC1 | None | None | | None | None |
| SMAD3 | None | None | | None | None |
| HARS | None | None | | None | None |
| HARS | None | None | | None | None |
| TNNI3 | None | None | | None | None |
| TNNI3 | None | None | | None | None |
| TNNI3 | None | None | | None | None |
| TNNI3 | None | None | | None | None |
| TNNI3 | None | None | | None | None |

| | | | | |
|---|---|---|---|---|
| TNNI3 | None | None | None | None |
| ACKR1 | None | None | None | None |
| BUB1B | None | None | None | None |
| BUB1B | None | None | None | None |
| BUB1B | None | None | None | None |
| AR | 0.522 | 6 | 1xj7 | A |
| AR | None | None | None | None |
| AR | None | None | None | None |
| AR | None | None | None | None |
| AR | None | None | None | None |
| FTO | None | None | None | None |
| FTO | None | None | None | None |
| TFAM | None | None | None | None |
| PYGL | None | None | None | None |
| PYGL | 0.778 | 3 | 1fa9 | A |
| PYGL | None | None | None | None |
| PYGL | 0.722 | 2 | 1fa9 | A |
| PYGL | 0.5 | 19 | 3dds | A |
| PYGL | 0.611 | 25 | 3ceh | B |
| PYGL | 0.611 | 32 | 1l7x | B |
| PYGL | None | None | None | None |
| PYGL | 0.722 | 2 | 1fc0 | B |
| PYGL | 0.778 | 3 | 1fa9 | A |
| PYGL | None | None | None | None |
| PYGL | 0.722 | 12 | 2ati | A |
| PYGL | 0.722 | 1 | 3ceh | B |
| PYGL | 0.611 | 11 | 3ceh | B |
| PYGL | 0.611 | 18 | 1fc0 | B |
| PYGL | 0.722 | 2 | 1fc0 | B |
| GALNS | None | None | None | None |
| GALNS | None | None | None | None |
| GALNS | None | None | None | None |
| GALNS | None | None | None | None |
| GALNS | None | None | None | None |
| GALNS | None | None | None | None |
| GALNS | None | None | None | None |
| GALNS | None | None | None | None |
| GALNS | None | None | None | None |
| GALNS | None | None | None | None |
| GALNS | None | None | None | None |
| GALNS | None | None | None | None |
| GALNS | None | None | None | None |
| GALNS | None | None | None | None |
| GALNS | None | None | None | None |
| GALNS | None | None | None | None |
| GALNS | None | None | None | None |
| GALNS | None | None | None | None |
| GALNS | None | None | None | None |
| GALNS | None | None | None | None |
| GALNS | None | None | None | None |
| GALNS | None | None | None | None |
| GALNS | None | None | None | None |
| GALNS | None | None | None | None |

| | | | | |
|---|---|---|---|---|
| GALNS | None | None | None | None |
| GALNS | None | None | None | None |
| GALNS | None | None | None | None |
| GALNS | None | None | None | None |
| GALNS | None | None | None | None |
| GALNS | None | None | None | None |
| GALNS | None | None | None | None |
| GALNS | None | None | None | None |
| F10 | None | None | None | None |
| GLUD1 | None | None | None | None |
| GLUD1 | None | None | None | None |
| GLUD1 | None | None | None | None |
| GLUD1 | None | None | None | None |
| GLUD1 | None | None | None | None |
| GLUD1 | None | None | None | None |
| GLUD1 | None | None | None | None |
| GLUD1 | None | None | None | None |
| GLUD1 | None | None | None | None |
| GLUD1 | None | None | None | None |
| NOTCH1 | 0.528 | 3 | 3v79 | K |
| POGLUT1 | None | None | None | None |
| POGLUT1 | None | None | None | None |
| NR3C1 | 0.455 | 1 | 4hn5 | B |
| CASP8 | 0.535 | 6 | 1f9e | C |
| CASP8 | 0.592 | 8 | 1qdu | C |
| TRAP1 | None | None | None | None |
| FANCM | None | None | None | None |
| EPOR | None | None | None | None |
| EPOR | None | None | None | None |
| NDP | None | None | None | None |
| NDP | None | None | None | None |
| NDP | None | None | None | None |
| NDP | None | None | None | None |
| NDP | None | None | None | None |
| NDP | None | None | None | None |
| NDP | None | None | None | None |
| NDP | None | None | None | None |
| NDP | None | None | None | None |
| NDP | None | None | None | None |
| NDP | None | None | None | None |
| NDP | None | None | None | None |
| NDP | None | None | None | None |
| NDP | None | None | None | None |
| NDP | None | None | None | None |
| NDP | None | None | None | None |
| NDP | None | None | None | None |
| ROGDI | None | None | None | None |
| CYP19A1 | None | None | None | None |
| CYP19A1 | None | None | None | None |
| CYP19A1 | None | None | None | None |
| CYP19A1 | None | None | None | None |
| CYP19A1 | None | None | None | None |
| CYP19A1 | 0.487 | 1 | 3eqm | A |
| CYP19A1 | 0.487 | 1 | 3eqm | A |

| | | | | |
|---|---|---|---|---|
| CYP19A1 | None | None | None | None |
| UBA5 | None | None | None | None |
| PRPS1 | None | None | None | None |
| PRPS1 | None | None | None | None |
| PRPS1 | None | None | None | None |
| PRPS1 | None | None | None | None |
| PRPS1 | None | None | None | None |
| PRPS1 | None | None | None | None |
| PRPS1 | None | None | None | None |
| PRPS1 | None | None | None | None |
| F2 | None | None | None | None |
| F2 | None | None | None | None |
| NTF4 | None | None | None | None |
| NTF4 | None | None | None | None |
| PSAP | None | None | None | None |
| PSAP | None | None | None | None |
| TAP1 | None | None | None | None |
| TAP1 | None | None | None | None |
| TAP1 | None | None | None | None |
| ARG1 | None | None | None | None |
| ARG1 | 0.714 | 1 | 2pha | A |
| ARG1 | None | None | None | None |
| ARG1 | 0.471 | 10 | 3th7 | A |
| ARG1 | None | None | None | None |
| ARG1 | None | None | None | None |
| CRYM | None | None | None | None |
| ADSL | None | None | None | None |
| ADSL | None | None | None | None |
| ADSL | None | None | None | None |
| ADSL | None | None | None | None |
| ADSL | None | None | None | None |
| ADSL | None | None | None | None |
| ADSL | None | None | None | None |
| ADSL | None | None | None | None |
| ADSL | None | None | None | None |
| ADSL | None | None | None | None |
| ADSL | None | None | None | None |
| ADSL | None | None | None | None |
| ADSL | 0.647 | 1 | 2vd6 | D |
| ADSL | 0.435 | 1 | 4ffx | A |
| ADSL | None | None | None | None |
| ADSL | None | None | None | None |
| ADSL | None | None | None | None |
| ADSL | None | None | None | None |
| ADSL | None | None | None | None |
| ADSL | None | None | None | None |
| VWF | None | None | None | None |
| PLA2G7 | None | None | None | None |
| MMP13 | None | None | None | None |
| MMP13 | None | None | None | None |
| MMP13 | None | None | None | None |
| MMP13 | None | None | None | None |
| POMGNT1 | None | None | None | None |
| POMGNT1 | None | None | None | None |

| | | | | |
|---|---|---|---|---|
| COL3A1 | None | None | None | None |
| COL3A1 | None | None | None | None |
| COL3A1 | None | None | None | None |
| COL3A1 | None | None | None | None |
| ACTN2 | None | None | None | None |
| ACTN2 | None | None | None | None |
| ACTN2 | None | None | None | None |
| ACTN2 | None | None | None | None |
| ACTN2 | None | None | None | None |
| ACTN2 | None | None | None | None |
| ACTN2 | None | None | None | None |
| UBA5 | None | None | None | None |
| ADAR | 0.75 | 1 | 1qbj | B |
| UMPS | None | None | None | None |
| UMPS | None | None | None | None |
| UMPS | None | None | None | None |
| TRNT1 | None | None | None | None |
| TRNT1 | None | None | None | None |
| NHEJ1 | None | None | None | None |
| NHEJ1 | None | None | None | None |
| NHEJ1 | None | None | None | None |
| C8A | None | None | None | None |
| GLUL | None | None | None | None |
| GLUL | None | None | None | None |
| GLUL | 0.467 | 9 | 2qc8 | B |
| GYG1 | 0.615 | 45 | 3rmv | A |
| GYG1 | 0.923 | 1 | 3u2u | B |
| FLNA | None | None | None | None |
| FLNA | None | None | None | None |
| FLNA | None | None | None | None |
| FLNA | None | None | None | None |
| FLNA | None | None | None | None |
| FLNA | None | None | None | None |
| FLNA | None | None | None | None |
| TTN | None | None | None | None |
| TTN | None | None | None | None |
| TTN | None | None | None | None |
| TTN | None | None | None | None |
| SMS | None | None | None | None |
| SMS | None | None | None | None |
| SMS | None | None | None | None |
| C5 | None | None | None | None |
| SERPINC1 | None | None | None | None |
| SERPINC1 | None | None | None | None |
| SERPINC1 | None | None | None | None |
| SERPINC1 | None | None | None | None |
| SERPINC1 | None | None | None | None |
| SERPINC1 | None | None | None | None |
| SERPINC1 | None | None | None | None |
| SERPINC1 | None | None | None | None |
| SERPINC1 | None | None | None | None |
| SERPINC1 | None | None | None | None |
| SERPINC1 | None | None | None | None |
| SERPINC1 | None | None | None | None |

| | | | | |
|---|---|---|---|---|
| SERPINC1 | None | None | None | None |
| SERPINC1 | None | None | None | None |
| SERPINC1 | None | None | None | None |
| SERPINC1 | None | None | None | None |
| SERPINC1 | None | None | None | None |
| SERPINC1 | None | None | None | None |
| SERPINC1 | None | None | None | None |
| SERPINC1 | None | None | None | None |
| SERPINC1 | None | None | None | None |
| ETFB | None | None | None | None |
| ETFB | None | None | None | None |
| ETFB | None | None | None | None |
| ETFB | None | None | None | None |
| ETFB | None | None | None | None |
| ETFB | None | None | None | None |
| DIABLO | None | None | None | None |
| DIABLO | None | None | None | None |
| DPYS | None | None | None | None |
| DPYS | None | None | None | None |
| DPYS | None | None | None | None |
| DPYS | 0.533 | 7 | 2vr2 | A |
| DPYS | None | None | None | None |
| GNE | None | None | None | None |
| GNE | None | None | None | None |
| GNE | None | None | None | None |
| GNE | None | None | None | None |
| GNE | None | None | None | None |
| GNE | None | None | None | None |
| GNE | None | None | None | None |
| GNE | None | None | None | None |
| GNE | None | None | None | None |
| GNMT | None | None | None | None |
| GNMT | None | None | None | None |
| CTH | None | None | None | None |
| CTH | None | None | None | None |
| CTH | None | None | None | None |
| USH1C | None | None | None | None |
| PSEN1 | None | None | None | None |
| CHEK2 | None | None | None | None |
| CHEK2 | None | None | None | None |
| CHEK2 | None | None | None | None |
| CHEK2 | None | None | None | None |
| CHEK2 | None | None | None | None |
| CHEK2 | None | None | None | None |
| CHEK2 | None | None | None | None |
| CHEK2 | None | None | None | None |
| CHEK2 | None | None | None | None |
| SNCA | None | None | None | None |
| SNCA | None | None | None | None |
| GAN | None | None | None | None |
| GAN | None | None | None | None |
| GAN | None | None | None | None |
| SBDS | None | None | None | None |
| SBDS | None | None | None | None |

| | | | | | |
|---|---|---|---|---|---|
| SBDS | None | None | None | None |
| SBDS | None | None | None | None |
| SBDS | None | None | None | None |
| SBDS | None | None | None | None |
| MPO | | 1 | 3 | 3f9p | A |
| MPO | | 0.6 | 12 | 5mfa | A |
| KCNH2 | None | None | None | None |
| KCNH2 | None | None | None | None |
| KCNH2 | | 0.75 | 1 | 4hp9 | A |
| KCNH2 | None | None | None | None |
| TEK | None | None | None | None |
| TEK | None | None | None | None |
| TEK | None | None | None | None |
| TEK | None | None | None | None |
| NPC2 | None | None | None | None |
| NPC2 | None | None | None | None |
| NPC2 | None | None | None | None |
| NPC2 | None | None | None | None |
| NPC2 | None | None | None | None |
| NPC2 | None | None | None | None |
| NPC2 | None | None | None | None |
| PLG | None | None | None | None |
| PCSK9 | None | None | None | None |
| SFTPC | None | None | None | None |
| SFTPC | None | None | None | None |
| SFTPC | None | None | None | None |
| CD40LG | None | None | None | None |
| CD40LG | None | None | None | None |
| CD40LG | None | None | None | None |
| CD40LG | None | None | None | None |
| CD40LG | None | None | None | None |
| TAT | None | None | None | None |
| TAT | None | None | None | None |
| TAT | None | None | None | None |
| EP300 | None | None | None | None |
| FARS2 | None | None | None | None |
| FARS2 | None | None | None | None |
| FARS2 | None | None | None | None |
| FARS2 | None | None | None | None |
| FARS2 | None | None | None | None |
| FARS2 | None | None | None | None |
| FARS2 | None | None | None | None |
| FARS2 | None | None | None | None |
| FARS2 | | 0.5 | 1 | 5mgw | A |
| FARS2 | None | None | None | None |
| FARS2 | None | None | None | None |
| FARS2 | None | None | None | None |
| FARS2 | None | None | None | None |
| MASP2 | None | None | None | None |
| MASP2 | None | None | None | None |
| MASP2 | None | None | None | None |
| MASP2 | None | None | None | None |
| STIM1 | None | None | None | None |
| CHRM2 | None | None | None | None |

| | | | | | |
|---|---|---|---|---|---|
| MCFD2 | None | None | None | None |
| MCFD2 | 0.574 | | 1 | 4yge | B |
| MCFD2 | 0.623 | | 1 | 3wnx | B |
| KITLG | None | None | None | None |
| F12 | None | None | None | None |
| F12 | None | None | None | None |
| ERAP1 | None | None | None | None |
| GTF2H5 | None | None | None | None |
| GTF2H5 | None | None | None | None |
| THBD | None | None | None | None |
| MRPL28 | None | None | None | None |
| MRPL28 | None | None | None | None |
| HTRA1 | None | None | None | None |
| HTRA1 | None | None | None | None |
| HTRA1 | None | None | None | None |
| HTRA1 | None | None | None | None |
| HTRA1 | None | None | None | None |
| HTRA1 | None | None | None | None |
| KARS | None | None | None | None |
| KARS | None | None | None | None |
| KARS | None | None | None | None |
| KARS | None | None | None | None |
| KARS | None | None | None | None |
| HK1 | 0.722 | | 11 | 4fpb | B |
| GARS | None | None | None | None |
| GARS | None | None | None | None |
| GARS | None | None | None | None |
| GARS | None | None | None | None |
| GARS | None | None | None | None |
| GARS | None | None | None | None |
| GARS | None | None | None | None |
| IL12B | None | None | None | None |
| HADH | None | None | None | None |
| HADH | None | None | None | None |
| HADH | 0.52 | | 4 | 1lsj | A |
| HADH | 0.458 | | 3 | 1m75 | A |
| HADH | 0.52 | | 4 | 1lsj | A |
| RRM2B | None | None | None | None |
| RRM2B | None | None | None | None |
| RRM2B | None | None | None | None |
| RRM2B | None | None | None | None |
| RRM2B | None | None | None | None |
| RRM2B | None | None | None | None |
| RRM2B | None | None | None | None |
| RRM2B | None | None | None | None |
| RRM2B | None | None | None | None |
| RRM2B | None | None | None | None |
| RRM2B | None | None | None | None |
| RRM2B | None | None | None | None |
| RRM2B | None | None | None | None |
| RRM2B | None | None | None | None |
| RRM2B | None | None | None | None |
| MARVELD2 | None | None | None | None |
| TLR4 | None | None | None | None |

| | | | | |
|---|---|---|---|---|
| TTN | None | None | None | None |
| TTN | None | None | None | None |
| TNFRSF13B | None | None | None | None |
| POLG | None | None | None | None |
| POLG | None | None | None | None |
| POLG | None | None | None | None |
| POLG | None | None | None | None |
| POLG | None | None | None | None |
| POLG | None | None | None | None |
| POLG | None | None | None | None |
| POLG | None | None | None | None |
| POLG | None | None | None | None |
| POLG | None | None | None | None |
| POLG | None | None | None | None |
| POLG | None | None | None | None |
| POLG | None | None | None | None |
| POLG | None | None | None | None |
| POLG | None | None | None | None |
| POLG | None | None | None | None |
| POLG | None | None | None | None |
| POLG | None | None | None | None |
| POLG | None | None | None | None |
| POLG | None | None | None | None |
| POLG | None | None | None | None |
| POLG | None | None | None | None |
| POLG | None | None | None | None |
| POLG | None | None | None | None |
| POLG | None | None | None | None |
| POLG | None | None | None | None |
| POLG | None | None | None | None |
| POLG | 0.467 | 4 | 5c53 | A |
| POLG | 0.533 | 1 | 4ztu | A |
| POLG | None | None | None | None |
| POLG | None | None | None | None |
| POLG | None | None | None | None |
| POLG | None | None | None | None |
| POLG | None | None | None | None |
| POLG | 0.533 | 1 | 4ztu | A |
| POLG | None | None | None | None |
| POLG | None | None | None | None |
| POLG | None | None | None | None |
| POLG | None | None | None | None |
| POLG | None | None | None | None |
| POLG | None | None | None | None |
| POLG | None | None | None | None |
| POLG | None | None | None | None |
| POLG | None | None | None | None |
| POLG | None | None | None | None |
| POLG | None | None | None | None |
| POLG | None | None | None | None |
| POLG | None | None | None | None |
| POLG | None | None | None | None |
| UROD | 0.632 | 24 | 1jpk | A |

| | | | | |
|---|---|---|---|---|
| UROD | 0.579 | 33 | 1r3v | A |
| UROD | None | None | None | None |
| UROD | 0.526 | 49 | 1jph | A |
| UROD | None | None | None | None |
| UROD | 0.632 | 13 | 3gvw | A |
| UROD | 0.789 | 6 | 3gvw | A |
| UROD | 0.842 | 2 | 3gvv | A |
| UROD | 0.737 | 31 | 1r3q | A |
| UROD | 0.667 | 14 | 2q71 | A |
| PCK1 | 0.429 | 154 | 1khg | A |
| PCK1 | None | None | None | None |
| PCK1 | None | None | None | None |
| PCK1 | None | None | None | None |
| PCK1 | 0.524 | 60 | 1nhx | A |
| HEXA | None | None | None | None |
| HEXA | None | None | None | None |
| HEXA | None | None | None | None |
| HEXA | None | None | None | None |
| HEXA | None | None | None | None |
| HEXA | None | None | None | None |
| HEXA | None | None | None | None |
| HEXA | None | None | None | None |
| HEXA | None | None | None | None |
| HEXA | None | None | None | None |
| HEXA | None | None | None | None |
| HEXA | None | None | None | None |
| HEXA | None | None | None | None |
| HEXA | None | None | None | None |
| HEXA | None | None | None | None |
| HEXA | 0.455 | 1 | 2gjx | A |
| HEXA | None | None | None | None |
| HEXA | None | None | None | None |
| HEXA | None | None | None | None |
| HEXA | None | None | None | None |
| HEXA | None | None | None | None |
| HEXA | None | None | None | None |
| HEXA | None | None | None | None |
| HEXA | None | None | None | None |
| HEXA | None | None | None | None |
| HEXA | None | None | None | None |
| HEXA | None | None | None | None |
| HEXA | 0.455 | 3 | 2gjx | A |
| HEXA | 0.455 | 3 | 2gjx | A |
| HEXA | None | None | None | None |
| HEXA | None | None | None | None |
| CFTR | None | None | None | None |
| CFTR | None | None | None | None |
| CFTR | None | None | None | None |
| CFTR | None | None | None | None |
| CFTR | None | None | None | None |
| CFTR | None | None | None | None |
| CFTR | None | None | None | None |
| CFTR | None | None | None | None |
| CFTR | None | None | None | None |

| | | | | |
|---|---|---|---|---|
| CFTR | None | None | None | None |
| CFTR | 0.462 | 1 | 2pzg | A |
| CFTR | None | None | None | None |
| CFTR | 0.462 | 2 | 2pzg | A |
| CFTR | None | None | None | None |
| CFTR | None | None | None | None |
| CFTR | None | None | None | None |
| CFTR | None | None | None | None |
| CFTR | None | None | None | None |
| CFTR | None | None | None | None |
| CFTR | None | None | None | None |
| CFTR | None | None | None | None |
| CFTR | None | None | None | None |
| CFTR | None | None | None | None |
| CFTR | None | None | None | None |
| CFTR | None | None | None | None |
| CFTR | 0.462 | 1 | 2pzg | A |
| CFTR | None | None | None | None |
| CFTR | 0.462 | 2 | 2pzg | A |
| CFTR | None | None | None | None |
| CFTR | None | None | None | None |
| CFTR | None | None | None | None |
| CFTR | None | None | None | None |
| CFTR | None | None | None | None |
| CFTR | None | None | None | None |
| FAN1 | None | None | None | None |
| CBS | None | None | None | None |
| CBS | None | None | None | None |
| CBS | None | None | None | None |
| CBS | None | None | None | None |
| CBS | 0.406 | 6 | 4pcu | B |
| CBS | None | None | None | None |
| CBS | 0.469 | 6 | 4pcu | A |
| CBS | 0.597 | 3 | 5mms | C |
| CBS | 0.54 | 2 | 5mms | A |
| CBS | 0.54 | 2 | 5mms | A |
| CBS | None | None | None | None |
| CBS | None | None | None | None |
| CBS | None | None | None | None |
| CBS | None | None | None | None |
| CBS | 0.516 | 4 | 5mms | C |
| CBS | None | None | None | None |
| CBS | None | None | None | None |
| CBS | None | None | None | None |
| CBS | 0.698 | 2 | 1m54 | B |
| CBS | None | None | None | None |
| CBS | None | None | None | None |
| CBS | None | None | None | None |
| CBS | None | None | None | None |
| CBS | None | None | None | None |
| CBS | None | None | None | None |
| CBS | None | None | None | None |
| CBS | 0.424 | 6 | 4coo | B |
| CBS | None | None | None | None |

| | | | | |
|---|---|---|---|---|
| CBS | 0.603 | 3 | 4coo | A |
| CBS | 0.603 | 3 | 4coo | A |
| CBS | None | None | None | None |
| CBS | None | None | None | None |
| CBS | None | None | None | None |
| CBS | None | None | None | None |
| CBS | None | None | None | None |
| CBS | None | None | None | None |
| TLR3 | None | None | None | None |
| LIG4 | None | None | None | None |
| LIG4 | None | None | None | None |
| ASPA | 0.769 | 1 | 2i3c | B |
| ASPA | 0.769 | 1 | 2i3c | B |
| ASPA | None | None | None | None |
| ASPA | None | None | None | None |
| ASPA | None | None | None | None |
| ASPA | None | None | None | None |
| ASPA | None | None | None | None |
| ASPA | None | None | None | None |
| ASPA | None | None | None | None |
| ASPA | None | None | None | None |
| ASPA | 0.583 | 2 | 4tnu | A |
| ASPA | 0.583 | 2 | 4tnu | A |
| ASPA | None | None | None | None |
| CFH | None | None | None | None |
| FCGR3A | None | None | None | None |
| FCGR3A | None | None | None | None |
| GRIN2A | None | None | None | None |
| GRIN2A | 0.658 | 1 | 5i2n | A |
| GRIN2A | None | None | None | None |
| GLA | 0.467 | 10 | 1r46 | B |
| GLA | None | None | None | None |
| GLA | None | None | None | None |
| GLA | None | None | None | None |
| GLA | None | None | None | None |
| GLA | None | None | None | None |
| GLA | None | None | None | None |
| GLA | 0.647 | 2 | 3s5y | A |
| GLA | 0.562 | 20 | 3gxt | B |
| GLA | None | None | None | None |
| GLA | None | None | None | None |
| GLA | 0.75 | 3 | 3gxn | B |
| GLA | 0.75 | 3 | 3gxn | B |
| GLA | 0.438 | 7 | 1r47 | B |
| GLA | None | None | None | None |
| GLA | None | None | None | None |
| GLA | None | None | None | None |
| GLA | None | None | None | None |
| GLA | None | None | None | None |
| GLA | None | None | None | None |
| GLA | None | None | None | None |
| GLA | 0.4 | 5 | 3lxa | A |
| GLA | None | None | None | None |
| GLA | None | None | None | None |

| | | | | |
|---|---|---|---|---|
| GLA | None | None | None | None |
| GLA | 0.467 | 22 | 3gxn | A |
| GLA | None | None | None | None |
| GLA | None | None | None | None |
| GLA | None | None | None | None |
| GLA | None | None | None | None |
| GLA | None | None | None | None |
| SERPINA3 | None | None | None | None |
| MTAP | None | None | None | None |
| MTAP | 0.462 | 25 | 3oze | B |
| MTAP | None | None | None | None |
| BLMH | None | None | None | None |
| TUSC3 | None | None | None | None |
| ISCU | None | None | None | None |
| CYP11B2 | None | None | None | None |
| CYP11B2 | None | None | None | None |
| CYP11B2 | None | None | None | None |
| CYP11B2 | None | None | None | None |
| CYP11B2 | None | None | None | None |
| CYP11B2 | None | None | None | None |
| CYP11B2 | None | None | None | None |
| CYP11B2 | None | None | None | None |
| CYP11B2 | None | None | None | None |
| CYP11B2 | None | None | None | None |
| CYP11B2 | None | None | None | None |
| CYP11B2 | None | None | None | None |
| CYP11B2 | None | None | None | None |
| CYP11B2 | None | None | None | None |
| CYP11B2 | None | None | None | None |
| CYP11B2 | None | None | None | None |
| CYP11B2 | None | None | None | None |
| CYP11B2 | None | None | None | None |
| CYP11B2 | None | None | None | None |
| CYP11B2 | None | None | None | None |
| CRADD | None | None | None | None |
| CRADD | None | None | None | None |
| FGA | None | None | None | None |
| FGA | None | None | None | None |
| FGA | None | None | None | None |
| FGA | None | None | None | None |
| FGA | None | None | None | None |
| FGA | None | None | None | None |
| FGA | None | None | None | None |
| FGA | None | None | None | None |
| IYD | 1 | 1 | 4ttb | A |
| IYD | None | None | None | None |
| IYD | None | None | None | None |
| IYD | None | None | None | None |
| TCAP | None | None | None | None |
| TCAP | None | None | None | None |
| TCAP | None | None | None | None |
| TCAP | None | None | None | None |
| TCAP | None | None | None | None |
| CYP7A1 | None | None | None | None |

| | | | | |
|---|---|---|---|---|
| TEK | None | None | None | None |
| TEK | None | None | None | None |
| TEK | None | None | None | None |
| TEK | None | None | None | None |
| TEK | None | None | None | None |
| TEK | None | None | None | None |
| TEK | None | None | None | None |
| CD8A | None | None | None | None |
| HERC2 | None | None | None | None |
| NR1H3 | 0.537 | 3 | 3ipq | A |
| GRHPR | None | None | None | None |
| GRHPR | None | None | None | None |
| GRHPR | 0.458 | 2 | 2q50 | A |
| GRHPR | 0.458 | 2 | 2q50 | A |
| GRHPR | 0.697 | 1 | 2wwr | D |
| GRHPR | 0.697 | 1 | 2wwr | D |
| GRHPR | None | None | None | None |
| GBE1 | None | None | None | None |
| GBE1 | None | None | None | None |
| GBE1 | None | None | None | None |
| GBE1 | None | None | None | None |
| GBE1 | None | None | None | None |
| GBE1 | None | None | None | None |
| GBE1 | None | None | None | None |
| GBE1 | None | None | None | None |
| GBE1 | None | None | None | None |
| GBE1 | None | None | None | None |
| GBE1 | None | None | None | None |
| GBE1 | None | None | None | None |
| GBE1 | None | None | None | None |
| GBE1 | None | None | None | None |
| GBE1 | None | None | None | None |
| GBE1 | None | None | None | None |
| GBE1 | None | None | None | None |
| GBE1 | None | None | None | None |
| GBE1 | None | None | None | None |
| PC | None | None | None | None |
| PC | 0.429 | 1 | 3bg3 | A |
| PC | None | None | None | None |
| PC | 0.4 | 2 | 3bg9 | B |
| PC | None | None | None | None |
| PC | None | None | None | None |
| PC | 0.4 | 6 | 3bg3 | B |
| PC | None | None | None | None |
| MSH6 | None | None | None | None |
| MSH6 | None | None | None | None |
| MSH6 | None | None | None | None |
| MSH6 | None | None | None | None |
| MSH6 | None | None | None | None |
| MSH6 | None | None | None | None |
| MSH6 | None | None | None | None |
| MSH6 | None | None | None | None |
| MSH6 | None | None | None | None |

| | | | | |
|---|---|---|---|---|
| MSH6 | None | None | None | None |
| MSH6 | None | None | None | None |
| MSH6 | None | None | None | None |
| MSH6 | None | None | None | None |
| MSH6 | None | None | None | None |
| MSH6 | None | None | None | None |
| MSH6 | None | None | None | None |
| MSH6 | None | None | None | None |
| MSH6 | None | None | None | None |
| MSH6 | None | None | None | None |
| MSH6 | None | None | None | None |
| MSH6 | None | None | None | None |
| MSH6 | None | None | None | None |
| MSH6 | None | None | None | None |
| MSH6 | None | None | None | None |
| MSH6 | None | None | None | None |
| MSH6 | None | None | None | None |
| MSH6 | None | None | None | None |
| MSH6 | None | None | None | None |
| MSH6 | None | None | None | None |
| MSH6 | None | None | None | None |
| RNASEL | None | None | None | None |
| ORC1 | None | None | None | None |
| ORC1 | None | None | None | None |
| ORC1 | None | None | None | None |
| KCNH2 | None | None | None | None |
| KCNH2 | None | None | None | None |
| KCNH2 | None | None | None | None |
| KCNH2 | None | None | None | None |
| KCNH2 | None | None | None | None |
| KCNH2 | None | None | None | None |
| KCNH2 | None | None | None | None |
| KCNH2 | None | None | None | None |
| KCNH2 | None | None | None | None |
| KCNH2 | None | None | None | None |
| KCNH2 | None | None | None | None |
| KCNH2 | None | None | None | None |
| KCNH2 | None | None | None | None |
| KCNH2 | None | None | None | None |
| KCNH2 | None | None | None | None |
| KCNH2 | None | None | None | None |
| KCNH2 | None | None | None | None |
| KCNH2 | None | None | None | None |
| KCNH2 | None | None | None | None |
| KCNH2 | None | None | None | None |
| KCNH2 | None | None | None | None |
| KCNH2 | None | None | None | None |
| KCNH2 | None | None | None | None |
| KCNH2 | None | None | None | None |
| KCNH2 | None | None | None | None |
| MAT1A | 0.429 | 3 | 2obv | A |
| MAT1A | 0.429 | 14 | 2obv | A |
| MAT1A | 0.571 | 1 | 2obv | A |
| MAT1A | 0.571 | 1 | 2obv | A |
| MAT1A | None | None | None | None |

| | | | | |
|---|---|---|---|---|
| MAT1A | None | None | None | None |
| MAT1A | None | None | None | None |
| MAT1A | None | None | None | None |
| INSR | None | None | None | None |
| INSR | None | None | None | None |
| INSR | None | None | None | None |
| INSR | None | None | None | None |
| AMT | 0.545 | 5 | 1wsr | A |
| AMT | None | None | None | None |
| AMT | None | None | None | None |
| AMT | None | None | None | None |
| AMT | None | None | None | None |
| AMT | None | None | None | None |
| AMT | None | None | None | None |
| AMT | None | None | None | None |
| AMT | None | None | None | None |
| CSNK1D | None | None | None | None |
| SF3B1 | None | None | None | None |
| AGA | None | None | None | None |
| AGA | None | None | None | None |
| AGA | None | None | None | None |
| AGA | None | None | None | None |
| AGA | None | None | None | None |
| AGA | None | None | None | None |
| AGA | None | None | None | None |
| NECTIN4 | None | None | None | None |
| RS1 | None | None | None | None |
| RS1 | None | None | None | None |
| RS1 | None | None | None | None |
| RS1 | None | None | None | None |
| TBC1D7 | None | None | None | None |
| CTSA | None | None | None | None |
| CTSA | None | None | None | None |
| TKT | None | None | None | None |
| RP2 | None | None | None | None |
| RP2 | None | None | None | None |
| RP2 | None | None | None | None |
| RP2 | None | None | None | None |
| LYRM4 | None | None | None | None |
| SOS1 | None | None | None | None |
| SOS1 | None | None | None | None |
| SOS1 | None | None | None | None |
| SOS1 | None | None | None | None |
| SOS1 | None | None | None | None |
| SOS1 | None | None | None | None |
| HUWE1 | None | None | None | None |
| HUWE1 | None | None | None | None |
| TPI1 | None | None | None | None |
| TPI1 | 0.886 | 1 | 4unk | B |
| TPI1 | None | None | None | None |
| MSH2 | None | None | None | None |
| MSH2 | None | None | None | None |
| MSH2 | None | None | None | None |
| MSH2 | None | None | None | None |

| | | | | |
|---|---|---|---|---|
| MSH2 | None | None | None | None |
| MSH2 | None | None | None | None |
| MSH2 | None | None | None | None |
| MSH2 | None | None | None | None |
| MSH2 | None | None | None | None |
| MSH2 | None | None | None | None |
| MSH2 | None | None | None | None |
| MSH2 | None | None | None | None |
| MSH2 | None | None | None | None |
| MSH2 | None | None | None | None |
| MSH2 | None | None | None | None |
| MSH2 | None | None | None | None |
| MSH2 | None | None | None | None |
| MSH2 | None | None | None | None |
| MSH2 | None | None | None | None |
| MSH2 | None | None | None | None |
| MSH2 | None | None | None | None |
| MSH2 | None | None | None | None |
| MSH2 | None | None | None | None |
| MSH2 | None | None | None | None |
| MSH2 | None | None | None | None |
| MSH2 | None | None | None | None |
| MSH2 | None | None | None | None |
| MSH2 | None | None | None | None |
| MSH2 | None | None | None | None |
| MSH2 | None | None | None | None |
| MSH2 | None | None | None | None |
| MSH2 | None | None | None | None |
| MSH2 | None | None | None | None |
| MSH2 | None | None | None | None |
| MSH2 | None | None | None | None |
| MSH2 | None | None | None | None |
| GALE | None | None | None | None |
| GALE | 0.78 | 1 | 1i3n | A |
| GALE | None | None | None | None |
| GALE | None | None | None | None |
| GALE | None | None | None | None |
| GALE | None | None | None | None |
| GALE | None | None | None | None |
| GALE | None | None | None | None |
| NSD2 | None | None | None | None |
| SRP72 | None | None | None | None |
| SRP72 | None | None | None | None |
| SRP72 | None | None | None | None |
| POLK | None | None | None | None |
| GALT | None | None | None | None |
| GALT | None | None | None | None |
| GALT | None | None | None | None |
| GALT | None | None | None | None |
| GALT | None | None | None | None |
| GALT | None | None | None | None |
| GALT | None | None | None | None |
| GALT | None | None | None | None |
| GALT | None | None | None | None |

| | | | | |
|---|---|---|---|---|
| GALT | None | None | None | None |
| GALT | None | None | None | None |
| GALT | None | None | None | None |
| GALT | None | None | None | None |
| GALT | None | None | None | None |
| GALT | None | None | None | None |
| GALT | None | None | None | None |
| GALT | None | None | None | None |
| GALT | None | None | None | None |
| GALT | None | None | None | None |
| GALT | None | None | None | None |
| GALT | None | None | None | None |
| GALT | None | None | None | None |
| GALT | None | None | None | None |
| GALT | None | None | None | None |
| GALT | None | None | None | None |
| GALT | None | None | None | None |
| GALT | None | None | None | None |
| GALT | None | None | None | None |
| GALT | None | None | None | None |
| GALT | None | None | None | None |
| GALT | None | None | None | None |
| GALT | None | None | None | None |
| GALT | None | None | None | None |
| GALT | None | None | None | None |
| GALT | None | None | None | None |
| GALT | None | None | None | None |
| GALT | None | None | None | None |
| PYCR1 | 0.75 | 11 | 5uav | A |
| PYCR1 | 0.75 | 11 | 5uav | A |
| PYCR1 | 0.5 | 2 | 2izz | A |
| PYCR1 | 0.556 | 2 | 5uax | B |
| F8 | None | None | None | None |
| F8 | None | None | None | None |
| F8 | None | None | None | None |
| F8 | None | None | None | None |
| F8 | None | None | None | None |
| F8 | None | None | None | None |
| F8 | None | None | None | None |
| F8 | None | None | None | None |
| F8 | None | None | None | None |
| F8 | None | None | None | None |
| F8 | None | None | None | None |
| F8 | None | None | None | None |
| F8 | None | None | None | None |
| F8 | None | None | None | None |
| F8 | None | None | None | None |
| F8 | None | None | None | None |
| ANK1 | None | None | None | None |
| CHAT | None | None | None | None |
| CHAT | None | None | None | None |
| CHAT | 0.615 | 8 | 2fy5 | A |
| CHAT | 0.538 | 2 | 2fy5 | A |
| CHAT | None | None | None | None |

| | | | | |
|---|---|---|---|---|
| CHAT | None | None | None | None |
| CHAT | None | None | None | None |
| CHAT | None | None | None | None |
| CHAT | None | None | None | None |
| CHAT | 0.538 | 6 | 2fy5 | A |
| CHAT | None | None | None | None |
| CHAT | None | None | None | None |
| CHAT | None | None | None | None |
| CHAT | None | None | None | None |
| CHAT | None | None | None | None |
| CHAT | None | None | None | None |
| CHAT | None | None | None | None |
| GHR | None | None | None | None |
| GHR | None | None | None | None |
| GHR | None | None | None | None |
| GHR | None | None | None | None |
| GHR | None | None | None | None |
| GHR | None | None | None | None |
| GHR | None | None | None | None |
| GHR | None | None | None | None |
| GHR | None | None | None | None |
| GHR | None | None | None | None |
| GHR | None | None | None | None |
| GHR | None | None | None | None |
| GHR | None | None | None | None |
| GHR | None | None | None | None |
| FZD4 | None | None | None | None |
| FZD4 | None | None | None | None |
| FZD4 | None | None | None | None |
| MTR | None | None | None | None |
| MTR | None | None | None | None |
| MTR | None | None | None | None |
| GM2A | None | None | None | None |
| GM2A | None | None | None | None |
| GM2A | None | None | None | None |
| GM2A | None | None | None | None |
| GM2A | None | None | None | None |
| CRYGD | 0.444 | 3 | 2g98 | A |
| CRYGD | None | None | None | None |
| CRYGD | 0.522 | 1 | 2kfb | A |
| CRYGD | 0.4 | 3 | 4gr7 | X |
| CRYGD | None | None | None | None |
| CRYGD | None | None | None | None |
| CRYGD | None | None | None | None |
| CRYGD | None | None | None | None |
| CRYGD | 0.556 | 2 | 2g98 | A |
| TLR5 | None | None | None | None |
| HSD17B10 | 0.855 | 1 | 1u7t | B |
| HSD17B10 | None | None | None | None |
| HSD17B10 | 0.542 | 1 | 1so8 | A |
| HSD17B10 | None | None | None | None |
| TBC1D20 | None | None | None | None |
| FBN1 | None | None | None | None |
| FBN1 | None | None | None | None |

| | | | | |
|---|---|---|---|---|
| FBN1 | None | None | None | None |
| FBN1 | None | None | None | None |
| FBN1 | None | None | None | None |
| FBN1 | None | None | None | None |
| FBN1 | None | None | None | None |
| FXN | None | None | None | None |
| FXN | None | None | None | None |
| FXN | None | None | None | None |
| FXN | None | None | None | None |
| MYO5A | None | None | None | None |
| CTSK | None | None | None | None |
| CTSK | None | None | None | None |
| SCN5A | None | None | None | None |
| SCN5A | None | None | None | None |
| SOS1 | None | None | None | None |
| SOS1 | None | None | None | None |
| SOS1 | None | None | None | None |
| SOS1 | None | None | None | None |
| SOS1 | None | None | None | None |
| SOS1 | None | None | None | None |
| SOS1 | None | None | None | None |
| SOS1 | None | None | None | None |
| SOS1 | None | None | None | None |
| SOS1 | None | None | None | None |
| SOS1 | None | None | None | None |
| SOS1 | None | None | None | None |
| GATM | None | None | None | None |
| GATM | None | None | None | None |
| GATM | 0.75 | 4 | 4jdw | A |
| GATM | None | None | None | None |
| GATM | None | None | None | None |
| GATM | 0.75 | 6 | 6jdw | A |
| GATM | 0.75 | 11 | 6jdw | A |
| MMP2 | None | None | None | None |
| MMP2 | None | None | None | None |
| MMP2 | 0.407 | 7 | 1qib | A |
| MMP2 | None | None | None | None |
| NDUFB9 | None | None | None | None |
| NDUFB9 | None | None | None | None |
| GH1 | None | None | None | None |
| GH1 | None | None | None | None |
| GH1 | None | None | None | None |
| SLC6A4 | None | None | None | None |
| G6PD | None | None | None | None |
| G6PD | None | None | None | None |
| G6PD | None | None | None | None |
| G6PD | None | None | None | None |
| G6PD | None | None | None | None |
| G6PD | None | None | None | None |
| G6PD | None | None | None | None |
| G6PD | None | None | None | None |
| G6PD | 0.545 | 1 | 1qki | B |
| G6PD | None | None | None | None |
| G6PD | None | None | None | None |

| | | | | |
|---|---|---|---|---|
| G6PD | None | None | None | None |
| G6PD | None | None | None | None |
| G6PD | None | None | None | None |
| G6PD | None | None | None | None |
| G6PD | 0.455 | 1 | 1qki | D |
| G6PD | None | None | None | None |
| G6PD | None | None | None | None |
| G6PD | None | None | None | None |
| G6PD | None | None | None | None |
| CST3 | None | None | None | None |
| LMNA | None | None | None | None |
| LMNA | None | None | None | None |
| NDUFA9 | None | None | None | None |
| NDUFA9 | None | None | None | None |
| NDUFA9 | None | None | None | None |
| NDUFA9 | None | None | None | None |
| YARS | 0.581 | 1 | 5thl | A |
| YARS | None | None | None | None |
| YARS | None | None | None | None |
| YARS | None | None | None | None |
| EPHA2 | None | None | None | None |
| EPHA2 | None | None | None | None |
| EPHA2 | None | None | None | None |
| EPHA2 | None | None | None | None |
| DICER1 | None | None | None | None |
| DICER1 | None | None | None | None |
| SNX10 | None | None | None | None |
| SNX10 | None | None | None | None |
| TTN | None | None | None | None |
| TTN | None | None | None | None |
| GATA3 | None | None | None | None |
| C5 | None | None | None | None |
| THBD | None | None | None | None |
| BIN1 | None | None | None | None |
| BIN1 | None | None | None | None |
| SMARCAL1 | None | None | None | None |
| CFB | None | None | None | None |
| CYC1 | None | None | None | None |
| ANK2 | None | None | None | None |
| CHMP4B | None | None | None | None |
| CHMP4B | None | None | None | None |
| ERBB2 | None | None | None | None |
| ERBB2 | None | None | None | None |
| POLG2 | None | None | None | None |
| POLG2 | None | None | None | None |
| POLG2 | None | None | None | None |
| POLG2 | None | None | None | None |
| POLG2 | None | None | None | None |
| POLG2 | None | None | None | None |
| POLG2 | None | None | None | None |
| POLG2 | None | None | None | None |
| C3 | None | None | None | None |
| C3 | None | None | None | None |
| RPGR | None | None | None | None |

| | | | | |
|---|---|---|---|---|
| RPGR | None | None | None | None |
| RPGR | None | None | None | None |
| RPGR | None | None | None | None |
| RPGR | None | None | None | None |
| C1QA | None | None | None | None |
| PRKG1 | None | None | None | None |
| IGBP1 | None | None | None | None |
| ENO3 | None | None | None | None |
| ENO3 | None | None | None | None |
| ENO3 | 0.41 | 43 | 2xsx | A |
| ENO3 | 0.487 | 39 | 2xsx | A |
| ENO3 | None | None | None | None |
| ENO3 | 0.897 | 2 | 2xsx | B |
| MCPH1 | None | None | None | None |
| MCPH1 | None | None | None | None |
| MCPH1 | None | None | None | None |
| MCPH1 | None | None | None | None |
| MCPH1 | None | None | None | None |
| SNRNP200 | None | None | None | None |
| SNRNP200 | None | None | None | None |
| DNMT1 | None | None | None | None |
| DNMT1 | None | None | None | None |
| DNMT1 | None | None | None | None |
| AKT1 | None | None | None | None |
| NDUFS2 | None | None | None | None |
| ACVR2B | None | None | None | None |
| HMGCL | None | None | None | None |
| HMGCL | None | None | None | None |
| HMGCL | None | None | None | None |
| JUP | None | None | None | None |
| JUP | None | None | None | None |
| JUP | None | None | None | None |
| JUP | None | None | None | None |
| JUP | None | None | None | None |
| JUP | None | None | None | None |
| JUP | None | None | None | None |
| JUP | None | None | None | None |
| SPINK1 | None | None | None | None |
| SPINK1 | None | None | None | None |
| SPINK1 | None | None | None | None |
| VCL | None | None | None | None |
| VCL | None | None | None | None |
| NOTCH1 | None | None | None | None |
| NOTCH1 | None | None | None | None |
| NOTCH1 | None | None | None | None |
| NDUFB3 | None | None | None | None |
| IKBKG | None | None | None | None |
| GSN | 0.529 | 2 | 2fh1 | A |
| GSN | None | None | None | None |
| GSN | None | None | None | None |
| GSN | 0.5 | 3 | 1p8x | A |
| GSN | None | None | None | None |
| GSN | None | None | None | None |
| GSN | None | None | None | None |

| | | | | |
|---|---|---|---|---|
| GSN | None | None | None | None |
| GSN | None | None | None | None |
| CLC | None | None | None | None |
| RAF1 | None | None | None | None |
| RAF1 | None | None | None | None |
| RAF1 | None | None | None | None |
| NDUFS8 | None | None | None | None |
| NDUFS8 | None | None | None | None |
| NDUFS8 | None | None | None | None |
| NDUFS8 | None | None | None | None |
| NDUFS8 | None | None | None | None |
| CASR | None | None | None | None |
| CASR | None | None | None | None |
| CASR | None | None | None | None |
| CASR | None | None | None | None |
| CASR | None | None | None | None |
| CASR | None | None | None | None |
| CASR | None | None | None | None |
| CASR | None | None | None | None |
| CASR | None | None | None | None |
| CASR | None | None | None | None |
| CASR | None | None | None | None |
| CASR | None | None | None | None |
| CASR | None | None | None | None |
| CASR | None | None | None | None |
| CASR | None | None | None | None |
| CASR | None | None | None | None |
| CASR | None | None | None | None |
| CASR | None | None | None | None |
| CASR | None | None | None | None |
| CASR | None | None | None | None |
| CASR | None | None | None | None |
| CASR | None | None | None | None |
| CASR | None | None | None | None |
| CASR | None | None | None | None |
| CASR | None | None | None | None |
| CASR | None | None | None | None |
| CASR | None | None | None | None |
| CASR | None | None | None | None |
| CASR | None | None | None | None |
| VCL | None | None | None | None |
| VCL | None | None | None | None |
| SMPD1 | None | None | None | None |
| SMPD1 | None | None | None | None |
| SMPD1 | None | None | None | None |
| KDM6A | None | None | None | None |
| KCNQ1 | None | None | None | None |
| KCNQ1 | None | None | None | None |
| KCNQ1 | None | None | None | None |
| KCNQ1 | None | None | None | None |
| KCNQ1 | None | None | None | None |
| KCNQ1 | None | None | None | None |
| NDUFA12 | None | None | None | None |
| GLRX5 | None | None | None | None |

| | | | | |
|---|---|---|---|---|
| PALB2 | None | None | None | None |
| PALB2 | None | None | None | None |
| PALB2 | None | None | None | None |
| PALB2 | None | None | None | None |
| PALB2 | None | None | None | None |
| PALB2 | None | None | None | None |
| PALB2 | None | None | None | None |
| PALB2 | None | None | None | None |
| PALB2 | None | None | None | None |
| PALB2 | None | None | None | None |
| ALDOB | 0.462 | 89 | 1qo5 | C |
| ALDOB | 0.731 | 2 | 1xdm | A |
| ALDOB | 0.731 | 1 | 1qo5 | B |
| ALDOB | 0.538 | 6 | 1xdm | A |
| ALDOB | 0.692 | 7 | 1qo5 | C |
| ALDOB | None | None | None | None |
| ALDOB | 0.577 | 25 | 1qo5 | B |
| GNE | None | None | None | None |
| GNE | None | None | None | None |
| GNE | None | None | None | None |
| GNE | None | None | None | None |
| GNE | None | None | None | None |
| GNE | None | None | None | None |
| GNE | None | None | None | None |
| GNE | None | None | None | None |
| GNE | None | None | None | None |
| GNE | None | None | None | None |
| GNE | None | None | None | None |
| GNE | None | None | None | None |
| CHRNB2 | None | None | None | None |
| CHRNB2 | None | None | None | None |
| ERCC3 | None | None | None | None |
| ERCC3 | None | None | None | None |
| ERCC3 | None | None | None | None |
| ERCC3 | None | None | None | None |
| ERCC3 | None | None | None | None |
| UNC119 | None | None | None | None |
| ATL1 | None | None | None | None |
| ATL1 | None | None | None | None |
| ATL1 | None | None | None | None |
| ATL1 | 0.8 | 1 | 3q5d | A |
| ATL1 | None | None | None | None |
| ATL1 | None | None | None | None |
| ATL1 | None | None | None | None |
| ATL1 | None | None | None | None |
| ATL1 | None | None | None | None |
| ATL1 | None | None | None | None |
| NR3C2 | 0.701 | 1 | 2oax | A |
| TBK1 | None | None | None | None |
| HYAL1 | None | None | None | None |
| HYAL1 | None | None | None | None |
| HYAL1 | None | None | None | None |
| HYAL1 | None | None | None | None |
| CTSD | 0.633 | 1 | 4obz | A |

| | | | | |
|---|---|---|---|---|
| PRKCG | None | None | None | None |
| TGFBR1 | None | None | None | None |
| XDH | None | None | None | None |
| XDH | None | None | None | None |
| XDH | None | None | None | None |
| XDH | None | None | None | None |
| XDH | None | None | None | None |
| XDH | None | None | None | None |
| NMNAT1 | 0.833 | 1 | 1kqn | E |
| NMNAT1 | None | None | None | None |
| NMNAT1 | None | None | None | None |
| NMNAT1 | None | None | None | None |
| NMNAT1 | None | None | None | None |
| NMNAT1 | None | None | None | None |
| SERPING1 | None | None | None | None |
| SERPING1 | None | None | None | None |
| SERPING1 | None | None | None | None |
| SERPING1 | None | None | None | None |
| SERPING1 | None | None | None | None |
| PDZRN3 | None | None | None | None |
| FABP2 | None | None | None | None |
| IFNGR1 | None | None | None | None |
| IFNGR1 | None | None | None | None |
| TDP1 | None | None | None | None |
| TDP1 | None | None | None | None |
| INS | None | None | None | None |
| INS | 0.769 | 1 | 2r34 | C |
| INS | 0.769 | 1 | 2r34 | C |
| PPOX | 0.857 | 1 | 4ivm | B |
| PPOX | None | None | None | None |
| PPOX | None | None | None | None |
| PPOX | None | None | None | None |
| PPOX | None | None | None | None |
| PPOX | None | None | None | None |
| PPOX | None | None | None | None |
| PPOX | None | None | None | None |
| PPOX | None | None | None | None |
| GFER | None | None | None | None |
| GFER | 0.769 | 4 | 3tk0 | A |
| GFER | None | None | None | None |
| GFER | None | None | None | None |
| LIFR | None | None | None | None |
| EGLN1 | 0.556 | 21 | 5a3u | C |
| EGLN1 | 0.889 | 6 | 5v18 | A |
| EGLN1 | 0.944 | 2 | 5a3u | C |
| GP1BA | None | None | None | None |
| GP1BA | None | None | None | None |
| GP1BA | None | None | None | None |
| GP1BA | None | None | None | None |
| GP1BA | None | None | None | None |
| MMADHC | None | None | None | None |
| MMADHC | None | None | None | None |
| MMADHC | None | None | None | None |
| MMADHC | None | None | None | None |

| | | | | |
|---|---|---|---|---|
| MMADHC | None | None | None | None |
| MMADHC | None | None | None | None |
| MMADHC | None | None | None | None |
| ACVRL1 | None | None | None | None |
| ACVRL1 | None | None | None | None |
| RFXANK | None | None | None | None |
| CHIT1 | None | None | None | None |
| UBR1 | None | None | None | None |
| BLM | None | None | None | None |
| BLM | None | None | None | None |
| BLM | None | None | None | None |
| BLM | None | None | None | None |
| BLM | None | None | None | None |
| BLM | None | None | None | None |
| BLM | None | None | None | None |
| BLM | None | None | None | None |
| BLM | None | None | None | None |
| BLM | None | None | None | None |
| BLM | None | None | None | None |
| BLM | None | None | None | None |
| BLM | None | None | None | None |
| VDR | None | None | None | None |
| VDR | None | None | None | None |
| VDR | None | None | None | None |
| VDR | 0.578 | 1 | 1kb2 | B |
| VDR | 0.578 | 1 | 1kb2 | B |
| VDR | 0.578 | 1 | 1kb2 | B |
| EPHA2 | None | None | None | None |
| EPHA2 | None | None | None | None |
| EPHA2 | None | None | None | None |
| EPHA2 | None | None | None | None |
| EPHA2 | None | None | None | None |
| EPHA2 | None | None | None | None |
| EPHA2 | None | None | None | None |
| EPHA2 | None | None | None | None |
| EPHA2 | None | None | None | None |
| EPHA2 | None | None | None | None |
| UBA1 | None | None | None | None |
| STAT1 | None | None | None | None |
| STAT1 | None | None | None | None |
| STAT1 | None | None | None | None |
| STAT1 | None | None | None | None |
| STAT1 | None | None | None | None |
| STAT1 | None | None | None | None |
| STAT1 | None | None | None | None |
| STAT1 | None | None | None | None |
| DNM1L | None | None | None | None |
| DNM1L | None | None | None | None |
| OTC | None | None | None | None |
| OTC | None | None | None | None |
| OTC | None | None | None | None |
| OTC | None | None | None | None |
| OTC | None | None | None | None |
| OTC | None | None | None | None |

| | | | | |
|---|---|---|---|---|
| OTC | None | None | None | None |
| OTC | None | None | None | None |
| OTC | None | None | None | None |
| OTC | None | None | None | None |
| OTC | None | None | None | None |
| OTC | None | None | None | None |
| OTC | None | None | None | None |
| OTC | None | None | None | None |
| OTC | None | None | None | None |
| OTC | None | None | None | None |
| OTC | None | None | None | None |
| RARS | None | None | None | None |
| RARS | None | None | None | None |
| RARS | None | None | None | None |
| SEPSECS | None | None | None | None |
| SEPSECS | None | None | None | None |
| SEPSECS | None | None | None | None |
| SEPSECS | None | None | None | None |
| SEPSECS | None | None | None | None |
| BRCA1 | None | None | None | None |
| BRCA1 | None | None | None | None |
| BRCA1 | 0.538 | 1 | 3pxc | X |
| BRCA1 | None | None | None | None |
| BRCA1 | None | None | None | None |
| BRCA1 | 0.462 | 1 | 1t2u | A |
| BRCA1 | None | None | None | None |
| BRCA1 | None | None | None | None |
| BRCA1 | None | None | None | None |
| BRCA1 | None | None | None | None |
| BRCA1 | None | None | None | None |
| BRCA1 | None | None | None | None |
| BRCA1 | None | None | None | None |
| BRCA1 | None | None | None | None |
| BRCA1 | None | None | None | None |
| BRCA1 | None | None | None | None |
| BRCA1 | None | None | None | None |
| BRCA1 | None | None | None | None |
| BRCA1 | None | None | None | None |
| BRCA1 | None | None | None | None |
| BRCA1 | None | None | None | None |
| BRCA1 | None | None | None | None |
| CUBN | None | None | None | None |
| ITGB3 | None | None | None | None |
| MTRR | None | None | None | None |
| MTRR | None | None | None | None |
| MTRR | None | None | None | None |
| MTRR | None | None | None | None |
| MTRR | None | None | None | None |
| MTRR | None | None | None | None |
| MTRR | None | None | None | None |
| MTRR | None | None | None | None |
| IL7R | None | None | None | None |
| IL7R | None | None | None | None |
| RPS24 | None | None | None | None |

| | | | | |
|---|---|---|---|---|
| RPS24 | None | None | None | None |
| HOGA1 | None | None | None | None |
| HOGA1 | None | None | None | None |
| HOGA1 | None | None | None | None |
| HOGA1 | None | None | None | None |
| HOGA1 | None | None | None | None |
| HOGA1 | None | None | None | None |
| HOGA1 | 0.791 | 1 | 3s5n | A |
| HOGA1 | None | None | None | None |
| HOGA1 | None | None | None | None |
| HOGA1 | None | None | None | None |
| HOGA1 | None | None | None | None |
| HOGA1 | None | None | None | None |
| HOGA1 | None | None | None | None |
| GJB2 | None | None | None | None |
| GJB2 | None | None | None | None |
| GJB2 | None | None | None | None |
| GJB2 | None | None | None | None |
| GJB2 | None | None | None | None |
| GJB2 | None | None | None | None |
| GJB2 | None | None | None | None |
| GJB2 | None | None | None | None |
| GJB2 | None | None | None | None |
| GJB2 | None | None | None | None |
| GJB2 | None | None | None | None |
| GJB2 | None | None | None | None |
| GJB2 | None | None | None | None |
| GJB2 | None | None | None | None |
| GJB2 | None | None | None | None |
| GJB2 | None | None | None | None |
| GJB2 | None | None | None | None |
| GJB2 | None | None | None | None |
| GJB2 | None | None | None | None |
| GJB2 | None | None | None | None |
| GJB2 | None | None | None | None |
| GJB2 | None | None | None | None |
| GJB2 | None | None | None | None |
| GJB2 | None | None | None | None |
| GJB2 | None | None | None | None |
| GJB2 | None | None | None | None |
| GJB2 | None | None | None | None |
| GJB2 | None | None | None | None |
| GJB2 | None | None | None | None |
| GJB2 | None | None | None | None |
| GJB2 | None | None | None | None |
| GJB2 | None | None | None | None |
| GJB2 | None | None | None | None |
| GJB2 | None | None | None | None |
| GJB2 | None | None | None | None |
| GJB2 | None | None | None | None |
| GJB2 | None | None | None | None |
| GJB2 | None | None | None | None |
| GJB2 | None | None | None | None |
| GJB2 | None | None | None | None |

| | | | | |
|---|---|---|---|---|
| GJB2 | None | None | None | None |
| GJB2 | None | None | None | None |
| GJB2 | None | None | None | None |
| GJB2 | None | None | None | None |
| GJB2 | None | None | None | None |
| GJB2 | None | None | None | None |
| GJB2 | None | None | None | None |
| GJB2 | None | None | None | None |
| GJB2 | None | None | None | None |
| GJB2 | None | None | None | None |
| GJB2 | None | None | None | None |
| GJB2 | None | None | None | None |
| GJB2 | None | None | None | None |
| GJB2 | None | None | None | None |
| CPS1 | None | None | None | None |
| CPS1 | 0.545 | 127 | 5dou | A |
| CPS1 | None | None | None | None |
| CPS1 | None | None | None | None |
| CPS1 | None | None | None | None |
| CPS1 | None | None | None | None |
| CPS1 | None | None | None | None |
| CPS1 | None | None | None | None |
| CPS1 | None | None | None | None |
| CPS1 | None | None | None | None |
| PSPH | None | None | None | None |
| PSPH | None | None | None | None |
| PSPH | None | None | None | None |
| PSPH | None | None | None | None |
| PSPH | None | None | None | None |
| PSPH | None | None | None | None |
| PEPD | 0.512 | 2 | 5m4l | B |
| PEPD | None | None | None | None |
| PEPD | None | None | None | None |
| PEPD | 0.643 | 2 | 5m4j | B |
| PEPD | 0.643 | 2 | 5m4j | B |
| PEPD | 0.419 | 5 | 5m4l | B |
| PEPD | None | None | None | None |
| PEPD | None | None | None | None |
| PEPD | None | None | None | None |
| KDM5C | None | None | None | None |
| KDM5C | None | None | None | None |
| KDM5C | None | None | None | None |
| ITGB6 | None | None | None | None |
| ITGB6 | None | None | None | None |
| ITGB6 | None | None | None | None |
| DARS | None | None | None | None |
| DARS | None | None | None | None |
| DARS | None | None | None | None |
| DARS | None | None | None | None |
| BTK | None | None | None | None |
| BTK | None | None | None | None |
| BTK | None | None | None | None |
| HDAC4 | None | None | None | None |
| ITGA2B | None | None | None | None |

| | | | | |
|---|---|---|---|---|
| ITGA2B | None | None | None | None |
| ITGA2B | None | None | None | None |
| ITGA2B | None | None | None | None |
| PCSK9 | None | None | None | None |
| PCSK9 | None | None | None | None |
| PCSK9 | None | None | None | None |
| PCSK9 | None | None | None | None |
| MUTYH | None | None | None | None |
| MUTYH | None | None | None | None |
| MUTYH | None | None | None | None |
| MUTYH | None | None | None | None |
| MUTYH | None | None | None | None |
| MUTYH | None | None | None | None |
| MUTYH | None | None | None | None |
| MUTYH | None | None | None | None |
| MUTYH | None | None | None | None |
| MUTYH | None | None | None | None |
| MUTYH | None | None | None | None |
| MUTYH | None | None | None | None |
| MUTYH | None | None | None | None |
| MUTYH | None | None | None | None |
| MUTYH | None | None | None | None |
| MUTYH | None | None | None | None |
| MUTYH | None | None | None | None |
| MUTYH | None | None | None | None |
| MUTYH | None | None | None | None |
| MUTYH | None | None | None | None |
| MUTYH | None | None | None | None |
| MUTYH | None | None | None | None |
| MUTYH | None | None | None | None |
| MUTYH | None | None | None | None |
| SLC25A12 | None | None | None | None |
| GSN | None | None | None | None |
| IHH | 0.909 | 1 | 3n1p | B |
| IHH | None | None | None | None |
| IHH | None | None | None | None |
| IHH | 0.818 | 2 | 3n1p | B |
| IHH | 0.727 | 12 | 3n1p | B |
| IHH | 0.636 | 15 | 3n1p | B |
| IHH | 0.818 | 7 | 3n1p | B |
| CD247 | None | None | None | None |
| EP300 | None | None | None | None |
| TTN | None | None | None | None |
| MRI1 | None | None | None | None |
| SGSH | None | None | None | None |
| SGSH | None | None | None | None |
| SGSH | None | None | None | None |
| SGSH | None | None | None | None |
| SGSH | None | None | None | None |
| SGSH | None | None | None | None |
| SGSH | None | None | None | None |
| SGSH | None | None | None | None |
| SGSH | None | None | None | None |
| SGSH | None | None | None | None |

| | | | | |
|---|---|---|---|---|
| SGSH | None | None | None | None |
| SGSH | None | None | None | None |
| SGSH | None | None | None | None |
| SGSH | None | None | None | None |
| SGSH | None | None | None | None |
| SGSH | None | None | None | None |
| SGSH | None | None | None | None |
| SGSH | None | None | None | None |
| SGSH | None | None | None | None |
| SGSH | None | None | None | None |
| SGSH | None | None | None | None |
| SGSH | None | None | None | None |
| CBL | None | None | None | None |
| CANT1 | None | None | None | None |
| CANT1 | None | None | None | None |
| CANT1 | None | None | None | None |
| CANT1 | None | None | None | None |
| CANT1 | None | None | None | None |
| CANT1 | None | None | None | None |
| CANT1 | None | None | None | None |
| SLC1A3 | None | None | None | None |
| SLC1A3 | None | None | None | None |
| SLC1A5 | None | None | None | None |
| SLC1A3 | None | None | None | None |
| SLC1A5 | None | None | None | None |
| SLC1A3 | None | None | None | None |
| SLC1A3 | None | None | None | None |
| CTSD | 0.423 | 15 | 4od9 | D |
| CTSD | None | None | None | None |
| CTSD | None | None | None | None |
| CTSD | None | None | None | None |
| CTSD | None | None | None | None |
| MTOR | None | None | None | None |
| ILK | None | None | None | None |
| PCDH15 | None | None | None | None |
| PCDH15 | None | None | None | None |
| PCDH15 | None | None | None | None |
| PCDH15 | None | None | None | None |
| PCDH15 | None | None | None | None |
| PCDH15 | None | None | None | None |
| KLHL40 | None | None | None | None |
| KLHL40 | None | None | None | None |
| KLHL40 | None | None | None | None |
| KLHL40 | None | None | None | None |
| CSTA | None | None | None | None |
| CSTA | None | None | None | None |
| HNMT | None | None | None | None |
| HNMT | None | None | None | None |
| IFNGR2 | None | None | None | None |
| IFNGR2 | None | None | None | None |
| PEX5 | None | None | None | None |
| KIF7 | None | None | None | None |
| KIF7 | None | None | None | None |
| MYL3 | None | None | None | None |

| | | | | |
|---|---|---|---|---|
| MYL3 | None | None | None | None |
| MYL3 | None | None | None | None |
| MYL3 | None | None | None | None |
| MYL3 | None | None | None | None |
| MYL3 | None | None | None | None |
| MYL3 | None | None | None | None |
| NAGLU | None | None | None | None |
| NAGLU | None | None | None | None |
| NAGLU | None | None | None | None |
| NAGLU | None | None | None | None |
| NAGLU | None | None | None | None |
| NAGLU | None | None | None | None |
| NAGLU | None | None | None | None |
| NAGLU | None | None | None | None |
| NAGLU | None | None | None | None |
| NAGLU | None | None | None | None |
| NAGLU | None | None | None | None |
| NAGLU | None | None | None | None |
| NAGLU | None | None | None | None |
| TNFRSF13C | None | None | None | None |
| KIF21A | None | None | None | None |
| KIF21A | None | None | None | None |
| KIF21A | None | None | None | None |
| KIF21A | None | None | None | None |
| KIF21A | None | None | None | None |
| DOCK2 | None | None | None | None |
| DSP | None | None | None | None |
| CDKN2A | None | None | None | None |
| CDKN2A | None | None | None | None |
| CDKN2A | None | None | None | None |
| CDKN2A | None | None | None | None |
| CDKN2A | None | None | None | None |
| CDKN2A | None | None | None | None |
| CDKN2A | None | None | None | None |
| CDKN2A | None | None | None | None |
| CDKN2A | None | None | None | None |
| CDKN2A | None | None | None | None |
| CDKN2A | None | None | None | None |
| CDKN2A | None | None | None | None |
| CDKN2A | None | None | None | None |
| CDKN2A | None | None | None | None |
| VCL | None | None | None | None |
| TPP1 | None | None | None | None |
| TPP1 | None | None | None | None |
| TPP1 | None | None | None | None |
| TPP1 | None | None | None | None |
| TPP1 | None | None | None | None |
| TPP1 | None | None | None | None |
| TPP1 | None | None | None | None |
| TPP1 | None | None | None | None |
| TPP1 | None | None | None | None |
| TPP1 | None | None | None | None |
| TPP1 | None | None | None | None |
| TPP1 | None | None | None | None |

| | | | | |
|---|---|---|---|---|
| TPP1 | None | None | None | None |
| TPP1 | None | None | None | None |
| TPP1 | None | None | None | None |
| TPP1 | None | None | None | None |
| TPP1 | None | None | None | None |
| TPP1 | None | None | None | None |
| TPP1 | None | None | None | None |
| TPP1 | None | None | None | None |
| TPP1 | None | None | None | None |
| TPP1 | None | None | None | None |
| IL10RB | None | None | None | None |
| IL10RB | None | None | None | None |
| CCM2 | None | None | None | None |
| CCM2 | None | None | None | None |
| CCM2 | None | None | None | None |
| DCX | None | None | None | None |
| DCX | None | None | None | None |
| DCX | None | None | None | None |
| DCX | None | None | None | None |
| DCX | None | None | None | None |
| DCX | None | None | None | None |
| DCX | None | None | None | None |
| DCX | None | None | None | None |
| BRCA2 | None | None | None | None |
| NNT | None | None | None | None |
| C2 | None | None | None | None |
| C2 | None | None | None | None |
| C2 | None | None | None | None |
| C2 | None | None | None | None |
| WRN | None | None | None | None |
| WRN | None | None | None | None |
| F13A1 | None | None | None | None |
| F13A1 | None | None | None | None |
| F13A1 | None | None | None | None |
| F13A1 | None | None | None | None |
| TF | None | None | None | None |
| TF | None | None | None | None |
| TF | 0.516 | 2 | 3ve1 | B |
| TF | 0.562 | 39 | 3skp | A |
| TF | None | None | None | None |
| ITPA | None | None | None | None |
| ITPA | None | None | None | None |
| SNCA | None | None | None | None |
| ITGB4 | None | None | None | None |
| ITGB4 | None | None | None | None |
| ITGB4 | None | None | None | None |
| LRP6 | None | None | None | None |
| LRP6 | None | None | None | None |
| LRP6 | None | None | None | None |
| PLG | None | None | None | None |
| PLG | 0.61 | 1 | 1i5k | A |
| TYROBP | None | None | None | None |
| UQCRC2 | None | None | None | None |
| GPSM2 | None | None | None | None |

| | | | | |
|---|---|---|---|---|
| GPSM2 | None | None | None | None |
| GPSM2 | None | None | None | None |
| SCN5A | None | None | None | None |
| SCN5A | None | None | None | None |
| SCN5A | None | None | None | None |
| SCN5A | None | None | None | None |
| SCN5A | None | None | None | None |
| ATIC | 0.6 | 6 | 1p4r | A |
| EZH2 | None | None | None | None |
| CPA1 | None | None | None | None |
| SCN4B | None | None | None | None |
| SCN4B | None | None | None | None |
| MAN1B1 | None | None | None | None |
| MAN1B1 | 0.6 | 9 | 1fmi | A |
| MAN1B1 | None | None | None | None |
| MAN1B1 | None | None | None | None |
| CYCS | 0.74 | 1 | 2n9i | A |
| IDUA | None | None | None | None |
| IDUA | None | None | None | None |
| IDUA | None | None | None | None |
| IDUA | None | None | None | None |
| IDUA | None | None | None | None |
| IDUA | None | None | None | None |
| IDUA | None | None | None | None |
| IDUA | None | None | None | None |
| IDUA | None | None | None | None |
| IDUA | None | None | None | None |
| IDUA | None | None | None | None |
| IDUA | None | None | None | None |
| IDUA | None | None | None | None |
| IDUA | None | None | None | None |
| IDUA | None | None | None | None |
| IDUA | None | None | None | None |
| IDUA | None | None | None | None |
| IDUA | None | None | None | None |
| IDUA | None | None | None | None |
| IDUA | None | None | None | None |
| IDUA | None | None | None | None |
| IDUA | None | None | None | None |
| IDUA | None | None | None | None |
| IDUA | None | None | None | None |
| IDUA | None | None | None | None |
| HIKESHI | None | None | None | None |
| APC | None | None | None | None |
| APC | None | None | None | None |
| APC | None | None | None | None |
| APP | None | None | None | None |
| CEP152 | None | None | None | None |
| GAA | 0.609 | 1 | 5kzx | A |
| GAA | 0.609 | 1 | 5kzx | A |
| GAA | None | None | None | None |
| GAA | None | None | None | None |
| GAA | None | None | None | None |
| GAA | 0.609 | 1 | 5kzx | A |

| | | | | |
|---|---|---|---|---|
| GAA | 0.609 | 1 | 5kzx | A |
| GAA | 0.609 | 1 | 5kzx | A |
| GAA | 0.609 | 1 | 5kzx | A |
| GAA | 0.609 | 1 | 5kzx | A |
| GAA | 0.609 | 1 | 5kzx | A |
| GAA | 0.609 | 1 | 5kzx | A |
| GAA | 0.609 | 1 | 5kzx | A |
| GAA | 0.609 | 1 | 5kzx | A |
| GAA | 0.609 | 1 | 5kzx | A |
| GAA | 0.609 | 1 | 5kzx | A |
| GAA | 0.609 | 1 | 5kzx | A |
| GAA | 0.609 | 1 | 5kzx | A |
| GAA | 0.609 | 1 | 5kzx | A |
| GAA | 0.609 | 1 | 5kzx | A |
| GAA | 0.609 | 1 | 5kzx | A |
| GAA | 0.609 | 1 | 5kzx | A |
| GAA | 0.609 | 1 | 5kzx | A |
| GAA | 0.609 | 1 | 5kzx | A |
| GAA | 0.609 | 1 | 5kzx | A |
| GAA | 0.609 | 1 | 5kzx | A |
| GAA | 0.609 | 1 | 5kzx | A |
| GAA | 0.609 | 1 | 5kzx | A |
| GAA | 0.609 | 1 | 5kzx | A |
| GAA | 0.609 | 1 | 5kzx | A |
| GAA | 0.609 | 1 | 5kzx | A |
| GAA | 0.609 | 1 | 5kzx | A |
| GAA | 0.609 | 1 | 5kzx | A |
| GAA | 0.609 | 1 | 5kzx | A |
| GAA | 0.609 | 1 | 5kzx | A |
| GAA | 0.609 | 1 | 5kzx | A |
| GAA | 0.609 | 1 | 5kzx | A |
| GAA | 0.609 | 1 | 5kzx | A |
| GAA | 0.609 | 1 | 5kzx | A |
| GAA | 0.609 | 1 | 5kzx | A |
| GAA | 0.609 | 1 | 5kzx | A |
| GAA | 0.609 | 1 | 5kzx | A |
| GAA | 0.609 | 1 | 5kzx | A |
| GAA | 0.609 | 1 | 5kzx | A |
| GAA | 0.609 | 1 | 5kzx | A |
| GAA | 0.609 | 1 | 5kzx | A |
| GAA | 0.609 | 1 | 5kzx | A |
| GAA | 0.609 | 1 | 5kzx | A |
| GAA | None | None | None | None |
| FOLR1 | None | None | None | None |
| DSP | None | None | None | None |
| DSP | None | None | None | None |
| DSP | None | None | None | None |
| DSP | None | None | None | None |
| DSP | None | None | None | None |
| CCM2 | None | None | None | None |
| FXN | None | None | None | None |
| FXN | None | None | None | None |
| COMP | None | None | None | None |
| COMP | None | None | None | None |

| | | | | |
|---|---|---|---|---|
| COMP | None | None | None | None |
| COMP | None | None | None | None |
| COMP | None | None | None | None |
| COMP | None | None | None | None |
| COMP | None | None | None | None |
| COMP | None | None | None | None |
| COMP | None | None | None | None |
| COMP | None | None | None | None |
| COMP | None | None | None | None |
| COMP | None | None | None | None |
| AIFM1 | None | None | None | None |
| AIFM1 | None | None | None | None |
| AIFM1 | None | None | None | None |
| AIFM1 | None | None | None | None |
| AIFM1 | None | None | None | None |
| AIFM1 | None | None | None | None |
| AIFM1 | None | None | None | None |
| MYH9 | None | None | None | None |
| FHL2 | None | None | None | None |
| FHL2 | None | None | None | None |
| TNFRSF4 | None | None | None | None |
| CFB | None | None | None | None |
| CFB | None | None | None | None |
| CFB | None | None | None | None |
| CFB | None | None | None | None |
| CFB | None | None | None | None |
| CFB | None | None | None | None |
| CFB | None | None | None | None |
| CFB | None | None | None | None |
| CFB | None | None | None | None |
| CFB | None | None | None | None |
| MVD | None | None | None | None |
| MVD | None | None | None | None |
| HSD17B4 | None | None | None | None |
| HSD17B4 | None | None | None | None |
| HSD17B4 | None | None | None | None |
| HSD17B4 | 0.429 | 2 | 1s9c | A |
| HSD17B4 | None | None | None | None |
| HSD17B4 | None | None | None | None |
| F11 | None | None | None | None |
| F11 | None | None | None | None |
| F11 | None | None | None | None |
| F11 | None | None | None | None |
| F11 | None | None | None | None |
| F11 | None | None | None | None |
| F11 | None | None | None | None |
| F11 | None | None | None | None |
| F11 | None | None | None | None |
| SPTA1 | None | None | None | None |
| SPTA1 | None | None | None | None |
| SPTA1 | None | None | None | None |
| SPTA1 | None | None | None | None |
| SPTA1 | None | None | None | None |
| SPTA1 | None | None | None | None |

| | | | | |
|---|---|---|---|---|
| GMPR | None | None | None | None |
| MITF | None | None | None | None |
| NPC1 | None | None | None | None |
| NPC1 | None | None | None | None |
| NPC1 | None | None | None | None |
| NPC1 | None | None | None | None |
| NPC1 | None | None | None | None |
| NPC1 | None | None | None | None |
| NPC1 | None | None | None | None |
| NPC1 | None | None | None | None |
| NPC1 | None | None | None | None |
| NPC1 | None | None | None | None |
| NPC1 | None | None | None | None |
| NPC1 | None | None | None | None |
| NPC1 | None | None | None | None |
| NPC1 | None | None | None | None |
| NPC1 | None | None | None | None |
| NPC1 | None | None | None | None |
| NPC1 | None | None | None | None |
| NPC1 | None | None | None | None |
| NPC1 | None | None | None | None |
| NPC1 | None | None | None | None |
| NPC1 | None | None | None | None |
| NPC1 | None | None | None | None |
| NPC1 | None | None | None | None |
| NPC1 | None | None | None | None |
| NPC1 | None | None | None | None |
| NPC1 | None | None | None | None |
| NPC1 | None | None | None | None |
| NPC1 | None | None | None | None |
| NPC1 | None | None | None | None |
| NPC1 | None | None | None | None |
| NPC1 | None | None | None | None |
| NPC1 | None | None | None | None |
| NPC1 | None | None | None | None |
| NPC1 | None | None | None | None |
| NPC1 | None | None | None | None |
| NPC1 | None | None | None | None |
| NPC1 | None | None | None | None |
| NPC1 | None | None | None | None |
| NPC1 | None | None | None | None |
| NPC1 | None | None | None | None |
| NPC1 | None | None | None | None |
| NPC1 | None | None | None | None |
| NPC1 | None | None | None | None |
| C3 | None | None | None | None |
| NLRX1 | None | None | None | None |
| IGLL1 | None | None | None | None |
| APRT | 0.769 | 4 | 1zn9 | B |
| APRT | 0.417 | 5 | 4x44 | A |
| APRT | None | None | None | None |
| APRT | 0.923 | 1 | 1zn8 | A |
| APRT | None | None | None | None |
| ZIC3 | None | None | None | None |

| | | | | | |
|---|---|---|---|---|---|
| ZIC3 | None | None | None | | None |
| KLHL3 | None | None | None | | None |
| RPS6KA3 | None | None | None | | None |
| RPS6KA3 | None | None | None | | None |
| RPS6KA3 | None | None | None | | None |
| F8 | None | None | None | | None |
| F8 | None | None | None | | None |
| F8 | None | None | None | | None |
| F8 | None | None | None | | None |
| F8 | None | None | None | | None |
| F8 | None | None | None | | None |
| F8 | None | None | None | | None |
| F8 | None | None | None | | None |
| F8 | None | None | None | | None |
| F8 | None | None | None | | None |
| F8 | None | None | None | | None |
| F8 | None | None | None | | None |
| F8 | None | None | None | | None |
| F8 | None | None | None | | None |
| F8 | None | None | None | | None |
| F8 | None | None | None | | None |
| F8 | None | None | None | | None |
| ISG15 | None | None | None | | None |
| EXOSC3 | None | None | None | | None |
| EXOSC3 | None | None | None | | None |
| EXOSC3 | None | None | None | | None |
| NKX2-5 | 0.403 | 1 | 4s0h | | B |
| NKX2-5 | None | None | None | | None |
| NKX2-5 | 0.403 | 1 | 4s0h | | B |
| NKX2-5 | 0.403 | 1 | 4s0h | | B |
| FH | None | None | None | | None |
| FH | None | None | None | | None |
| FH | None | None | None | | None |
| FH | None | None | None | | None |
| FH | 0.621 | 4 | | 30000 | A |
| FH | 0.462 | 6 | | 30000 | C |
| FH | None | None | None | | None |
| FH | None | None | None | | None |
| FH | 0.64 | 6 | 5d6b | | A |
| FH | None | None | None | | None |
| FH | 0.64 | 1 | 5d6b | | A |
| FH | 0.64 | 1 | 5d6b | | A |
| SNRNP200 | None | None | None | | None |
| SNRNP200 | None | None | None | | None |
| SNRNP200 | None | None | None | | None |
| SNRNP200 | None | None | None | | None |
| CHRNA1 | None | None | None | | None |
| CHRNA1 | None | None | None | | None |
| MYOM1 | None | None | None | | None |
| MYOM1 | None | None | None | | None |
| MYOM1 | None | None | None | | None |
| MYOM1 | None | None | None | | None |
| MYOM1 | None | None | None | | None |
| MYOM1 | None | None | None | | None |

| | | | | |
|---|---|---|---|---|
| LDLR | None | None | None | None |
| LDLR | None | None | None | None |
| LDLR | None | None | None | None |
| LDLR | None | None | None | None |
| LDLR | None | None | None | None |
| LDLR | None | None | None | None |
| EHMT1 | None | None | None | None |
| EHMT1 | 0.545 | 3 | 3b7b | A |
| EHMT1 | None | None | None | None |
| COL9A2 | None | None | None | None |
| HFE2 | None | None | None | None |
| HFE2 | None | None | None | None |
| CD3G | None | None | None | None |
| MMAB | None | None | None | None |
| MMAB | 0.5 | 3 | 2idx | C |
| MMAB | 0.45 | 7 | 2idx | C |
| MMAB | 0.65 | 3 | 2idx | A |
| MMAB | 0.65 | 3 | 2idx | A |
| MMAB | 0.7 | 2 | 2idx | B |
| MMAB | 0.7 | 2 | 2idx | B |
| MMAB | None | None | None | None |
| MMAB | 0.4 | 11 | 2idx | C |
| MMAB | None | None | None | None |
| TAF1 | None | None | None | None |
| TAF1 | None | None | None | None |
| CD4 | None | None | None | None |
| QARS | None | None | None | None |
| QARS | None | None | None | None |
| QARS | None | None | None | None |
| QARS | None | None | None | None |
| QARS | None | None | None | None |
| FGFR3 | None | None | None | None |
| FGFR3 | None | None | None | None |
| FGFR3 | None | None | None | None |
| FGFR3 | None | None | None | None |
| FGFR3 | None | None | None | None |
| FGFR3 | None | None | None | None |
| FLNB | None | None | None | None |
| FLNB | None | None | None | None |
| FLNB | None | None | None | None |
| FLNB | None | None | None | None |
| FLNB | None | None | None | None |
| FLNB | None | None | None | None |
| FLNB | None | None | None | None |
| FLNB | None | None | None | None |
| IL21R | None | None | None | None |
| RB1 | None | None | None | None |
| RB1 | None | None | None | None |
| RB1 | None | None | None | None |
| NTRK1 | 0.533 | 1 | 1he7 | A |
| NOTCH1 | None | None | None | None |
| MYOM1 | None | None | None | None |
| MYOM1 | None | None | None | None |
| MYOM1 | None | None | None | None |

| | | | | |
|---|---|---|---|---|
| MYOM1 | None | None | None | None |
| MYOM1 | None | None | None | None |
| MYOM1 | None | None | None | None |
| COQ9 | None | None | None | None |
| COQ9 | None | None | None | None |
| CRBN | None | None | None | None |
| DCTN1 | None | None | None | None |
| DCTN1 | None | None | None | None |
| DCTN1 | None | None | None | None |
| DCTN1 | None | None | None | None |
| DCTN1 | None | None | None | None |
| DCTN1 | None | None | None | None |
| THAP1 | None | None | None | None |
| THAP1 | None | None | None | None |
| GNAS | None | None | None | None |
| GNAS | None | None | None | None |
| HNF4A | None | None | None | None |
| HNF4A | None | None | None | None |
| ALDH3A2 | None | None | None | None |
| ALDH3A2 | None | None | None | None |
| ALDH3A2 | 0.412 | 5 | 4qgk | B |
| ALDH3A2 | 0.557 | 2 | 4qgk | A |
| ALDH3A2 | None | None | None | None |
| ALDH3A2 | 0.569 | 3 | 4qgk | B |
| ALDH3A2 | None | None | None | None |
| ALDH3A2 | None | None | None | None |
| ALDH3A2 | None | None | None | None |
| HCN1 | None | None | None | None |
| PLEC | None | None | None | None |
| SCN5A | None | None | None | None |
| SCN5A | None | None | None | None |
| STAMBP | None | None | None | None |
| STAMBP | None | None | None | None |
| RNF125 | None | None | None | None |
| RNF125 | None | None | None | None |
| USH1C | None | None | None | None |
| OBSL1 | None | None | None | None |
| GRIN2B | None | None | None | None |
| GRIN2B | None | None | None | None |
| HCN4 | None | None | None | None |
| HCN4 | None | None | None | None |
| TALDO1 | None | None | None | None |
| ITGB3 | None | None | None | None |
| CHKB | None | None | None | None |
| CHKB | None | None | None | None |
| CHKB | None | None | None | None |
| CHKB | None | None | None | None |
| ACTN4 | None | None | None | None |
| ACTN4 | None | None | None | None |
| ACTN4 | None | None | None | None |
| ACTN4 | None | None | None | None |
| CDT1 | None | None | None | None |
| CDT1 | None | None | None | None |
| CDT1 | None | None | None | None |

| | | | | |
|---|---|---|---|---|
| CDT1 | None | None | None | None |
| CDT1 | None | None | None | None |
| CDT1 | None | None | None | None |
| MAPT | None | None | None | None |
| F5 | 0.517 | 4 | 1czs | A |
| F5 | None | None | None | None |
| F5 | None | None | None | None |
| ESRRB | None | None | None | None |
| DGUOK | None | None | None | None |
| DGUOK | 0.471 | 8 | 2ocp | A |
| DGUOK | None | None | None | None |
| DGUOK | None | None | None | None |
| DGUOK | 0.765 | 1 | 2ocp | A |
| DGUOK | 0.438 | 11 | 2ocp | B |
| DGUOK | None | None | None | None |
| DGUOK | None | None | None | None |
| DGUOK | None | None | None | None |
| CENPJ | None | None | None | None |
| TTN | None | None | None | None |
| TTN | None | None | None | None |
| TTN | None | None | None | None |
| TTN | None | None | None | None |
| TNFRSF1A | None | None | None | None |
| TNFRSF1A | None | None | None | None |
| TNFRSF1A | None | None | None | None |
| TNFRSF1A | None | None | None | None |
| TNFRSF1A | None | None | None | None |
| TNFRSF1A | None | None | None | None |
| TNFRSF1A | None | None | None | None |
| TNFRSF1A | None | None | None | None |
| TNFRSF1A | None | None | None | None |
| TNFRSF1A | None | None | None | None |
| TNFRSF1A | None | None | None | None |
| TNFRSF1A | None | None | None | None |
| ASL | None | None | None | None |
| ASL | 0.625 | 25 | 1aos | A |
| ASL | None | None | None | None |
| ASL | None | None | None | None |
| ASL | 0.438 | 4 | 1k62 | B |
| ASL | 0.562 | 30 | 1k62 | A |
| ASL | 0.625 | 1 | 1aos | A |
| ASL | None | None | None | None |
| ASL | None | None | None | None |
| ASL | None | None | None | None |
| MYO7A | None | None | None | None |
| MYO7A | None | None | None | None |
| MYO7A | None | None | None | None |
| MYO7A | None | None | None | None |
| MYO7A | None | None | None | None |
| MYO7A | None | None | None | None |
| MYO7A | None | None | None | None |
| MYO7A | None | None | None | None |
| MYO7A | None | None | None | None |
| MYO7A | None | None | None | None |

| | | | | |
|---|---|---|---|---|
| MYO7A | None | None | None | None |
| MYO7A | None | None | None | None |
| MYO7A | None | None | None | None |
| MYO7A | None | None | None | None |
| MYO7A | None | None | None | None |
| MYO7A | None | None | None | None |
| MYO7A | None | None | None | None |
| MYO7A | None | None | None | None |
| MYO7A | None | None | None | None |
| MYO7A | None | None | None | None |
| APC | None | None | None | None |
| ALAD | None | None | None | None |
| ALAD | None | None | None | None |
| ALAD | None | None | None | None |
| ALAD | 0.542 | 1 | 1E+051 | A |
| ALAD | None | None | None | None |
| ALAD | None | None | None | None |
| POR | None | None | None | None |
| CSF1R | None | None | None | None |
| CSF1R | None | None | None | None |
| CSF1R | None | None | None | None |
| CSF1R | None | None | None | None |
| CSF1R | None | None | None | None |
| CSF1R | None | None | None | None |
| CSF1R | None | None | None | None |
| CSF1R | None | None | None | None |
| CSF1R | None | None | None | None |
| CSF1R | None | None | None | None |
| CSF1R | None | None | None | None |
| VWF | None | None | None | None |
| VWF | None | None | None | None |
| PHF8 | None | None | None | None |
| ACY1 | None | None | None | None |
| SMN1 | None | None | None | None |
| SMN1 | None | None | None | None |
| DLAT | None | None | None | None |
| DLAT | None | None | None | None |
| B2M | None | None | None | None |
| SOX9 | None | None | None | None |
| SOX9 | None | None | None | None |
| SOX9 | None | None | None | None |
| SOX9 | None | None | None | None |
| SOX9 | None | None | None | None |
| CFH | None | None | None | None |
| FLNC | None | None | None | None |
| FLNC | None | None | None | None |
| FLNC | None | None | None | None |
| ITGB2 | None | None | None | None |
| ITGB2 | None | None | None | None |
| ITGB2 | None | None | None | None |
| ITGB2 | None | None | None | None |
| ITGB2 | None | None | None | None |
| ITGB2 | None | None | None | None |
| ITGB2 | None | None | None | None |

| | | | | |
|---|---|---|---|---|
| ITGB2 | None | None | None | None |
| ITGB2 | None | None | None | None |
| PSEN1 | None | None | None | None |
| PSEN1 | None | None | None | None |
| PSEN1 | None | None | None | None |
| PSEN1 | None | None | None | None |
| PSEN1 | None | None | None | None |
| PSEN1 | None | None | None | None |
| OXCT1 | None | None | None | None |
| OXCT1 | None | None | None | None |
| OXCT1 | None | None | None | None |
| OXCT1 | None | None | None | None |
| OXCT1 | None | None | None | None |
| OXCT1 | None | None | None | None |
| OXCT1 | None | None | None | None |
| DHH | None | None | None | None |
| PRKN | None | None | None | None |
| PRKN | None | None | None | None |
| PRKN | None | None | None | None |
| PRKN | None | None | None | None |
| PRKN | None | None | None | None |
| PRKN | None | None | None | None |
| PRKN | None | None | None | None |
| PRKN | None | None | None | None |
| PRKN | None | None | None | None |
| PRKN | None | None | None | None |
| PRKN | None | None | None | None |
| LCAT | None | None | None | None |
| LCAT | None | None | None | None |
| LCAT | None | None | None | None |
| LCAT | None | None | None | None |
| LCAT | None | None | None | None |
| LCAT | None | None | None | None |
| LCAT | None | None | None | None |
| LCAT | None | None | None | None |
| LCAT | None | None | None | None |
| LCAT | None | None | None | None |
| LCAT | None | None | None | None |
| LCAT | None | None | None | None |
| FOXP3 | None | None | None | None |
| FOXP3 | None | None | None | None |
| FOXP3 | None | None | None | None |
| FOXP3 | None | None | None | None |
| SUFU | None | None | None | None |
| HSPA9 | 0.571 | 14 | 4kbo | A |
| HSPA9 | None | None | None | None |
| SQSTM1 | None | None | None | None |
| DARS2 | None | None | None | None |
| DARS2 | None | None | None | None |
| DARS2 | None | None | None | None |
| DARS2 | None | None | None | None |
| DARS2 | 0.412 | 2 | 4ah6 | A |
| DARS2 | None | None | None | None |
| DARS2 | None | None | None | None |

| | | | | |
|---|---|---|---|---|
| DARS2 | None | None | None | None |
| DARS2 | None | None | None | None |
| DARS2 | None | None | None | None |
| DARS2 | None | None | None | None |
| DARS2 | None | None | None | None |
| NT5C3A | None | None | None | None |
| NT5C3A | | 0.4 | 18 | 2vkq | A |
| NT5C3A | | 0.4 | 7 | 2vkq | A |
| NT5C3A | | 0.6 | 14 | 2vkq | A |
| ETFA | None | None | None | None |
| ETFA | None | None | None | None |
| ETFA | None | None | None | None |
| ETFA | None | None | None | None |
| ETFA | None | None | None | None |
| ETFA | None | None | None | None |
| ETFA | None | None | None | None |
| TTN | None | None | None | None |
| TTN | None | None | None | None |
| FLT4 | None | None | None | None |
| FGB | None | None | None | None |
| HPCA | None | None | None | None |
| TAP2 | None | None | None | None |
| TAP2 | None | None | None | None |
| PTPRC | None | None | None | None |
| SLC4A1 | None | None | None | None |
| SLC4A1 | None | None | None | None |
| SLC4A1 | None | None | None | None |
| SLC4A1 | None | None | None | None |
| SLC4A1 | None | None | None | None |
| SLC4A1 | None | None | None | None |
| MYH7 | None | None | None | None |
| MYH7 | None | None | None | None |
| MYH7 | None | None | None | None |
| MYH7 | None | None | None | None |
| MYH7 | None | None | None | None |
| MYH7 | None | None | None | None |
| MYH7 | None | None | None | None |
| MYH7 | None | None | None | None |
| MYH7 | None | None | None | None |
| PARN | None | None | None | None |
| PARN | None | None | None | None |
| NDUFS6 | None | None | None | None |
| VRK1 | None | None | None | None |
| VRK1 | None | None | None | None |
| VRK1 | None | None | None | None |
| VRK1 | None | None | None | None |
| VRK1 | None | None | None | None |
| VRK1 | None | None | None | None |
| COLEC11 | None | None | None | None |
| VHL | None | None | None | None |
| TTN | None | None | None | None |
| NCF2 | None | None | None | None |
| NCF2 | None | None | None | None |
| NCF2 | None | None | None | None |

| | | | | |
|---|---|---|---|---|
| NCF2 | None | None | None | None |
| NCF2 | None | None | None | None |
| RPS26 | None | None | None | None |
| CETP | None | None | None | None |
| CETP | None | None | None | None |
| CETP | None | None | None | None |
| CETP | None | None | None | None |
| CETP | None | None | None | None |
| CETP | None | None | None | None |
| CETP | None | None | None | None |
| CETP | None | None | None | None |
| CETP | None | None | None | None |
| CETP | None | None | None | None |
| CETP | None | None | None | None |
| CETP | None | None | None | None |
| CETP | None | None | None | None |
| FLNA | None | None | None | None |
| FLNA | None | None | None | None |
| FLNA | None | None | None | None |
| FLNA | None | None | None | None |
| FLNA | None | None | None | None |
| FLNA | None | None | None | None |
| FLNA | None | None | None | None |
| MYBPC3 | None | None | None | None |
| MYBPC3 | None | None | None | None |
| MYBPC3 | None | None | None | None |
| MYBPC3 | None | None | None | None |
| MYBPC3 | None | None | None | None |
| MYBPC3 | None | None | None | None |
| MYBPC3 | None | None | None | None |
| MYBPC3 | None | None | None | None |
| MYBPC3 | None | None | None | None |
| MYBPC3 | None | None | None | None |
| LSS | None | None | None | None |
| C6 | None | None | None | None |
| C6 | None | None | None | None |
| RLBP1 | None | None | None | None |
| RLBP1 | None | None | None | None |
| RLBP1 | None | None | None | None |
| RLBP1 | None | None | None | None |
| RLBP1 | None | None | None | None |
| RLBP1 | None | None | None | None |
| RLBP1 | None | None | None | None |
| RLBP1 | None | None | None | None |
| MMAA | None | None | None | None |
| MMAA | None | None | None | None |
| MMAA | None | None | None | None |
| MMAA | None | None | None | None |
| MMAA | None | None | None | None |
| MMAA | None | None | None | None |
| IL1RN | None | None | None | None |
| MASTL | None | None | None | None |
| FHL1 | None | None | None | None |
| FHL1 | None | None | None | None |

| | | | | |
|---|---|---|---|---|
| FHL1 | None | None | None | None |
| FHL1 | None | None | None | None |
| FHL1 | None | None | None | None |
| FHL1 | None | None | None | None |
| FHL1 | None | None | None | None |
| FHL1 | None | None | None | None |
| FHL1 | None | None | None | None |
| EIF2B1 | 0.636 | 1 | 3ecs | H |
| EIF2B1 | 0.636 | 1 | 3ecs | H |
| COQ8A | None | None | None | None |
| COQ8A | None | None | None | None |
| COQ8A | None | None | None | None |
| COQ8A | None | None | None | None |
| COQ8A | None | None | None | None |
| COQ8A | None | None | None | None |
| COQ8A | None | None | None | None |
| COQ8A | None | None | None | None |
| COQ8A | None | None | None | None |
| COQ8A | None | None | None | None |
| COQ8A | None | None | None | None |
| COQ8A | None | None | None | None |
| COQ8A | None | None | None | None |
| COQ8A | None | None | None | None |
| COQ8A | None | None | None | None |
| COQ8A | None | None | None | None |
| UNC119 | None | None | None | None |
| IGHMBP2 | None | None | None | None |
| JAG1 | None | None | None | None |
| JAG1 | None | None | None | None |
| JAG1 | None | None | None | None |
| GNB1 | None | None | None | None |
| GNB1 | None | None | None | None |
| FLNA | None | None | None | None |
| FLNA | None | None | None | None |
| FLNA | None | None | None | None |
| FLNA | None | None | None | None |
| FLNA | None | None | None | None |
| CREBBP | None | None | None | None |
| CSF3R | None | None | None | None |
| CSF3R | None | None | None | None |
| HIBCH | None | None | None | None |
| HIBCH | None | None | None | None |
| HIBCH | None | None | None | None |
| MMACHC | None | None | None | None |
| MMACHC | None | None | None | None |
| MMACHC | None | None | None | None |
| MMACHC | None | None | None | None |
| MMACHC | None | None | None | None |
| MMACHC | None | None | None | None |
| MMACHC | None | None | None | None |
| MMACHC | None | None | None | None |
| MMACHC | None | None | None | None |
| MMACHC | None | None | None | None |
| MMACHC | None | None | None | None |

| | | | | |
|---|---|---|---|---|
| KLF1 | None | None | None | None |
| NAGS | None | None | None | None |
| NAGS | None | None | None | None |
| NAGS | None | None | None | None |
| PIK3R2 | None | None | None | None |
| EDNRB | None | None | None | None |
| EDNRB | None | None | None | None |
| EDNRB | None | None | None | None |
| EDNRB | None | None | None | None |
| E | None | None | None | None |
| RUNX1 | None | None | None | None |
| RUNX1 | None | None | None | None |
| CTNNB1 | None | None | None | None |
| CTNNB1 | None | None | None | None |
| CTNNB1 | None | None | None | None |
| CTNNB1 | None | None | None | None |
| CTNNB1 | None | None | None | None |
| CTNNB1 | None | None | None | None |
| TTN | None | None | None | None |
| TTN | None | None | None | None |
| TYK2 | None | None | None | None |
| TYK2 | None | None | None | None |
| TYK2 | None | None | None | None |
| ERCC4 | None | None | None | None |
| ERCC4 | None | None | None | None |
| ERCC4 | None | None | None | None |
| ERCC4 | None | None | None | None |
| CFI | None | None | None | None |
| CFI | None | None | None | None |
| CFI | None | None | None | None |
| CFI | None | None | None | None |
| CFI | None | None | None | None |
| CFI | None | None | None | None |
| CFI | None | None | None | None |
| CFI | None | None | None | None |
| CFI | None | None | None | None |
| CFI | None | None | None | None |
| CFI | None | None | None | None |
| CFI | 0.691 | 2 | 5o32 | J |
| CFI | 0.691 | 2 | 5o32 | J |
| CFI | 0.524 | 5 | 5o32 | I |
| NOTCH3 | None | None | None | None |
| NOTCH3 | None | None | None | None |
| NOTCH3 | None | None | None | None |
| NOTCH3 | None | None | None | None |
| TBX1 | None | None | None | None |
| TBX1 | None | None | None | None |
| DDB2 | None | None | None | None |
| DDB2 | None | None | None | None |
| TP53 | None | None | None | None |
| POMGNT1 | None | None | None | None |
| POMGNT1 | None | None | None | None |
| POMGNT1 | None | None | None | None |
| POMGNT1 | None | None | None | None |

| | | | | |
|---|---|---|---|---|
| POMGNT1 | None | None | None | None |
| POMGNT1 | None | None | None | None |
| POMGNT1 | None | None | None | None |
| POMGNT1 | None | None | None | None |
| POMGNT1 | None | None | None | None |
| POMGNT1 | None | None | None | None |
| POMGNT1 | None | None | None | None |
| POMGNT1 | None | None | None | None |
| POMGNT1 | None | None | None | None |
| FMR1 | None | None | None | None |
| FMR1 | None | None | None | None |
| SLC9A3R1 | None | None | None | None |
| HSPB1 | None | None | None | None |
| HSPB1 | None | None | None | None |
| HSPB1 | None | None | None | None |
| HSPB1 | None | None | None | None |
| HSPB1 | None | None | None | None |
| HSPB1 | None | None | None | None |
| HSPB1 | None | None | None | None |
| SOD3 | None | None | None | None |
| AGA | 1 | 2 | 1apz | D |
| AGA | None | None | None | None |
| AGA | None | None | None | None |
| NDUFS7 | None | None | None | None |
| NDUFS7 | None | None | None | None |
| SOX2 | None | None | None | None |
| CSRP3 | None | None | None | None |
| CSRP3 | None | None | None | None |
| PAH | None | None | None | None |
| PAH | None | None | None | None |
| PAH | None | None | None | None |
| PAH | None | None | None | None |
| CACNA1S | None | None | None | None |
| TRPV4 | None | None | None | None |
| TRPV4 | None | None | None | None |
| TRPV4 | None | None | None | None |
| TRPV4 | 0.58 | 3 | 4dx2 | A |
| TRPV4 | None | None | None | None |
| TRPV4 | None | None | None | None |
| TRPV4 | None | None | None | None |
| TRPV4 | None | None | None | None |
| TRPV4 | None | None | None | None |
| TRPV4 | None | None | None | None |
| TRPV4 | None | None | None | None |
| TRPV4 | None | None | None | None |
| TRPV4 | None | None | None | None |
| TNNI3 | None | None | None | None |
| MLYCD | None | None | None | None |
| MLYCD | None | None | None | None |
| MLYCD | None | None | None | None |
| MLYCD | None | None | None | None |
| AIP | None | None | None | None |
| AIP | None | None | None | None |
| ERCC2 | None | None | None | None |

| | | | | |
|---|---|---|---|---|
| ERCC2 | None | None | None | None |
| ERCC2 | None | None | None | None |
| ERCC2 | None | None | None | None |
| ERCC2 | None | None | None | None |
| ERCC2 | None | None | None | None |
| ERCC2 | None | None | None | None |
| ERCC2 | None | None | None | None |
| ERCC2 | None | None | None | None |
| ERCC2 | None | None | None | None |
| ERCC2 | None | None | None | None |
| ERCC2 | None | None | None | None |
| ERCC2 | None | None | None | None |
| ERCC2 | None | None | None | None |
| PUS1 | None | None | None | None |
| PUS1 | None | None | None | None |
| PUS1 | None | None | None | None |
| CASK | None | None | None | None |
| CASK | None | None | None | None |
| BLVRA | None | None | None | None |
| DICER1 | None | None | None | None |
| DICER1 | None | None | None | None |
| DICER1 | None | None | None | None |
| EIF2AK3 | None | None | None | None |
| EIF2AK3 | None | None | None | None |
| EIF2AK3 | None | None | None | None |
| KIT | None | None | None | None |
| KIT | None | None | None | None |
| KIT | None | None | None | None |
| KIT | None | None | None | None |
| KIT | None | None | None | None |
| KIT | None | None | None | None |
| DNMT3B | None | None | None | None |
| TGM1 | None | None | None | None |
| NDUFA10 | None | None | None | None |
| NDUFA10 | None | None | None | None |
| PRPF31 | None | None | None | None |
| PRPF31 | None | None | None | None |
| ECE1 | None | None | None | None |
| ECE1 | None | None | None | None |
| ECE1 | None | None | None | None |
| NDUFV2 | None | None | None | None |
| NDUFV2 | None | None | None | None |
| NDUFV2 | None | None | None | None |
| GHRHR | None | None | None | None |
| GHRHR | None | None | None | None |
| GHRHR | None | None | None | None |
| SLC4A1 | None | None | None | None |
| FGF9 | None | None | None | None |
| BARD1 | None | None | None | None |
| BARD1 | None | None | None | None |
| BARD1 | None | None | None | None |
| PTPN12 | 0.741 | 2 | 5j8r | D |
| MASP2 | None | None | None | None |
| MASP2 | None | None | None | None |

| | | | | |
|---|---|---|---|---|
| MASP2 | None | None | None | None |
| MASP2 | None | None | None | None |
| HOXB1 | None | None | None | None |
| SLC4A1 | None | None | None | None |
| SLC4A1 | None | None | None | None |
| SLC4A1 | None | None | None | None |
| SLC4A1 | None | None | None | None |
| SLC4A1 | None | None | None | None |
| SLC4A1 | None | None | None | None |
| SLC4A1 | None | None | None | None |
| SLC4A1 | None | None | None | None |
| SLC4A1 | None | None | None | None |
| SLC4A1 | None | None | None | None |
| SLC4A1 | None | None | None | None |
| SLC4A1 | None | None | None | None |
| SLC4A1 | None | None | None | None |
| SLC4A1 | None | None | None | None |
| SLC4A1 | None | None | None | None |
| SLC4A1 | None | None | None | None |
| SLC4A1 | None | None | None | None |
| CBL | None | None | None | None |
| FAS | None | None | None | None |
| MTMR2 | None | None | None | None |
| MTMR2 | None | None | None | None |
| MTMR2 | None | None | None | None |
| MTMR2 | None | None | None | None |
| MTMR2 | None | None | None | None |
| MTMR2 | None | None | None | None |
| MTMR2 | None | None | None | None |
| MTMR2 | None | None | None | None |
| KLHL3 | None | None | None | None |
| KLHL3 | None | None | None | None |
| KLHL3 | 0.667 | 1 | 4ch9 | B |
| KLHL3 | 0.733 | 1 | 5nkp | A |
| KLHL3 | None | None | None | None |
| KLHL3 | None | None | None | None |
| RNASEH1 | None | None | None | None |
| ACTN1 | None | None | None | None |
| PHKG2 | None | None | None | None |
| PHKG2 | None | None | None | None |
| PHKG2 | 0.472 | 2 | 2y7j | A |
| NTRK1 | None | None | None | None |
| APOH | None | None | None | None |
| PMS2 | None | None | None | None |
| PMS2 | None | None | None | None |
| PMS2 | 0.797 | 1 | 1ea6 | A |
| PMS2 | None | None | None | None |
| PMS2 | None | None | None | None |
| PMS2 | None | None | None | None |
| PMS2 | None | None | None | None |
| PMS2 | None | None | None | None |
| PMS2 | None | None | None | None |
| PMS2 | None | None | None | None |
| PMS2 | None | None | None | None |

| | | | | | |
|---|---|---|---|---|---|
| PMS2 | None | None | None | | None |
| PMS2 | None | None | None | | None |
| PMS2 | None | None | None | | None |
| PMS2 | None | None | None | | None |
| PMS2 | None | None | None | | None |
| PMS2 | None | None | None | | None |
| PMS2 | None | None | None | | None |
| PMS2 | None | None | None | | None |
| PMS2 | None | None | None | | None |
| PMS2 | None | None | None | | None |
| PMS2 | None | None | None | | None |
| MOCS2 | None | None | None | | None |
| MOCS2 | None | None | None | | None |
| RPGRIP1 | None | None | None | | None |
| RPGRIP1 | None | None | None | | None |
| LIG4 | None | None | None | | None |
| PROC | None | None | None | | None |
| PROC | None | None | None | | None |
| NAT1 | None | None | None | | None |
| NAT1 | None | None | None | | None |
| PNPO | None | None | None | | None |
| PNPO | 0.5 | | 2 | 1nrg | A |
| PNPO | None | None | None | | None |
| PNPO | None | None | None | | None |
| PNPO | None | None | None | | None |
| MRE11 | None | None | None | | None |
| MRE11 | None | None | None | | None |
| MRE11 | None | None | None | | None |
| MRE11 | None | None | None | | None |
| DDC | None | None | None | | None |
| DDC | None | None | None | | None |
| DDC | None | None | None | | None |
| DDC | None | None | None | | None |
| DDC | None | None | None | | None |
| DDC | 0.909 | | 7 | 3rch | B |
| DDC | 0.909 | | 7 | 3rch | B |
| DDC | 0.643 | | 4 | 3rbf | A |
| DDC | None | None | None | | None |
| DDC | None | None | None | | None |
| DDC | None | None | None | | None |
| ANK2 | None | None | None | | None |
| ANK2 | None | None | None | | None |
| ANK2 | None | None | None | | None |
| AK2 | None | None | None | | None |
| AK2 | None | None | None | | None |
| AK2 | None | None | None | | None |
| AK2 | None | None | None | | None |
| AK2 | None | None | None | | None |
| AK2 | None | None | None | | None |
| GPHN | None | None | None | | None |
| HSD17B4 | 0.66 | | 1 | 1zbq | A |
| HSD17B4 | 0.66 | | 1 | 1zbq | A |
| HSD17B4 | None | None | None | | None |
| HSD17B4 | None | None | None | | None |

| | | | | |
|---|---|---|---|---|
| HSD17B4 | None | None | None | None |
| HSD17B4 | None | None | None | None |
| APTX | 0.6 | 5 | 4ndg | A |
| APTX | 0.6 | 5 | 4ndg | A |
| APTX | None | None | None | None |
| APTX | None | None | None | None |
| APTX | None | None | None | None |
| APTX | None | None | None | None |
| APTX | None | None | None | None |
| FANCL | None | None | None | None |
| FANCL | None | None | None | None |
| EMG1 | None | None | None | None |
| EMG1 | None | None | None | None |
| EZH2 | None | None | None | None |
| ABCB6 | None | None | None | None |
| NCF2 | None | None | None | None |
| NCF2 | None | None | None | None |
| COL4A2 | None | None | None | None |
| COL4A2 | 1 | 1 | 1li1 | F |
| COL4A2 | None | None | None | None |
| COL4A2 | None | None | None | None |
| COL4A2 | 0.75 | 16 | 1li1 | C |
| SLX4 | None | None | None | None |
| SERPIND1 | None | None | None | None |
| FGFR3 | None | None | None | None |
| FGFR3 | None | None | None | None |
| FGFR3 | None | None | None | None |
| FGFR3 | None | None | None | None |
| FGFR3 | None | None | None | None |
| FGFR3 | None | None | None | None |
| TREM2 | None | None | None | None |
| TREM2 | None | None | None | None |
| TREM2 | None | None | None | None |
| TREM2 | None | None | None | None |
| TREM2 | None | None | None | None |
| TREM2 | None | None | None | None |
| TREM2 | None | None | None | None |
| LIG4 | None | None | None | None |
| LIG4 | None | None | None | None |
| LIG4 | None | None | None | None |
| LIG4 | None | None | None | None |
| LIG4 | None | None | None | None |
| LIG4 | None | None | None | None |
| ECHS1 | None | None | None | None |
| ECHS1 | None | None | None | None |
| ECHS1 | None | None | None | None |
| MTR | None | None | None | None |
| MTR | None | None | None | None |
| MTR | None | None | None | None |
| MTR | None | None | None | None |
| MTR | None | None | None | None |
| PMVK | None | None | None | None |
| PRNP | None | None | None | None |
| CHN1 | None | None | None | None |

| | | | | |
|---|---|---|---|---|
| CHN1 | None | None | None | None |
| CHN1 | None | None | None | None |
| CHN1 | None | None | None | None |
| CHN1 | None | None | None | None |
| CHN1 | None | None | None | None |
| CHN1 | None | None | None | None |
| CHN1 | None | None | None | None |
| SLC2A1 | None | None | None | None |
| SLC2A1 | None | None | None | None |
| SLC2A1 | None | None | None | None |
| SLC2A1 | None | None | None | None |
| SLC2A1 | None | None | None | None |
| SLC2A1 | None | None | None | None |
| SLC2A1 | None | None | None | None |
| SLC2A1 | None | None | None | None |
| SLC2A1 | None | None | None | None |
| SLC2A1 | None | None | None | None |
| SLC2A1 | None | None | None | None |
| SLC2A1 | None | None | None | None |
| SLC2A1 | None | None | None | None |
| SLC2A1 | None | None | None | None |
| SLC2A1 | None | None | None | None |
| SLC2A1 | None | None | None | None |
| SLC2A1 | None | None | None | None |
| PMM2 | None | None | None | None |
| PMM2 | None | None | None | None |
| PMM2 | None | None | None | None |
| PMM2 | None | None | None | None |
| PMM2 | None | None | None | None |
| PMM2 | None | None | None | None |
| PMM2 | None | None | None | None |
| PMM2 | None | None | None | None |
| PMM2 | None | None | None | None |
| PMM2 | None | None | None | None |
| PMM2 | None | None | None | None |
| PMM2 | None | None | None | None |
| PMM2 | None | None | None | None |
| PMM2 | None | None | None | None |
| PMM2 | None | None | None | None |
| PMM2 | None | None | None | None |
| PMM2 | None | None | None | None |
| PMM2 | None | None | None | None |
| PMM2 | None | None | None | None |
| PMM2 | None | None | None | None |
| PMM2 | None | None | None | None |
| PMM2 | None | None | None | None |
| PMM2 | None | None | None | None |
| PMM2 | None | None | None | None |
| PMM2 | None | None | None | None |
| PMM2 | None | None | None | None |
| PMM2 | None | None | None | None |
| PMM2 | None | None | None | None |
| CCKAR | None | None | None | None |
| MAN1B1 | None | None | None | None |

| | | | | |
|---|---|---|---|---|
| MAN1B1 | None | None | None | None |
| LAMA2 | None | None | None | None |
| LAMA2 | None | None | None | None |
| TBC1D4 | None | None | None | None |
| TBC1D4 | None | None | None | None |
| LDLR | None | None | None | None |
| LDLR | None | None | None | None |
| LDLR | None | None | None | None |
| LDLR | None | None | None | None |
| LDLR | None | None | None | None |
| LDLR | None | None | None | None |
| LDLR | None | None | None | None |
| LDLR | None | None | None | None |
| LDLR | None | None | None | None |
| LDLR | None | None | None | None |
| LDLR | None | None | None | None |
| LDLR | None | None | None | None |
| LDLR | None | None | None | None |
| LDLR | None | None | None | None |
| LDLR | None | None | None | None |
| LDLR | None | None | None | None |
| LDLR | None | None | None | None |
| LDLR | None | None | None | None |
| LDLR | None | None | None | None |
| LDLR | None | None | None | None |
| LDLR | None | None | None | None |
| LDLR | None | None | None | None |
| LDLR | None | None | None | None |
| LDLR | None | None | None | None |
| LDLR | None | None | None | None |
| LDLR | None | None | None | None |
| LDLR | None | None | None | None |
| LDLR | None | None | None | None |
| LDLR | None | None | None | None |
| LDLR | None | None | None | None |
| LDLR | None | None | None | None |
| LDLR | None | None | None | None |
| LDLR | None | None | None | None |
| LDLR | None | None | None | None |
| LDLR | None | None | None | None |
| LDLR | None | None | None | None |
| LDLR | None | None | None | None |
| LDLR | None | None | None | None |
| LDLR | None | None | None | None |
| LDLR | None | None | None | None |
| LDLR | None | None | None | None |
| LDLR | 0.5 | 1 | 1ijq | A |
| LDLR | None | None | None | None |
| FANCM | None | None | None | None |
| HAMP | None | None | None | None |
| GUSB | 0.462 | 8 | 3hn3 | A |
| GUSB | None | None | None | None |
| GUSB | 0.45 | 16 | 3hn3 | B |
| GUSB | None | None | None | None |

| | | | | |
|---|---|---|---|---|
| GUSB | None | None | None | None |
| GUSB | None | None | None | None |
| GUSB | None | None | None | None |
| GUSB | None | None | None | None |
| GUSB | None | None | None | None |
| GUSB | None | None | None | None |
| GUSB | None | None | None | None |
| GUSB | None | None | None | None |
| GUSB | None | None | None | None |
| LIG1 | None | None | None | None |
| LIG1 | None | None | None | None |
| MYH7 | None | None | None | None |
| MYH7 | None | None | None | None |
| MYH7 | None | None | None | None |
| MYH7 | None | None | None | None |
| MYH7 | None | None | None | None |
| MYH7 | None | None | None | None |
| MYH7 | None | None | None | None |
| MYH7 | None | None | None | None |
| MYH7 | None | None | None | None |
| MYH7 | None | None | None | None |
| MYH7 | None | None | None | None |
| MYH7 | None | None | None | None |
| MYH7 | None | None | None | None |
| MYH7 | None | None | None | None |
| MYH7 | None | None | None | None |
| MYH7 | None | None | None | None |
| MYH7 | None | None | None | None |
| MYH7 | None | None | None | None |
| MYH7 | None | None | None | None |
| MYH7 | None | None | None | None |
| MYH7 | None | None | None | None |
| MYH7 | None | None | None | None |
| MYH7 | None | None | None | None |
| MYH7 | None | None | None | None |
| MYH7 | None | None | None | None |
| MYH7 | None | None | None | None |
| MYH7 | None | None | None | None |
| MYH7 | None | None | None | None |
| MYH7 | None | None | None | None |
| MYH7 | None | None | None | None |
| MYH7 | None | None | None | None |
| MYH7 | None | None | None | None |
| MYH7 | None | None | None | None |
| MYH7 | None | None | None | None |
| HCN4 | None | None | None | None |
| RECQL4 | None | None | None | None |
| RECQL4 | None | None | None | None |
| RECQL4 | None | None | None | None |
| RECQL4 | None | None | None | None |
| RECQL4 | None | None | None | None |
| RECQL4 | None | None | None | None |
| RECQL4 | None | None | None | None |
| RECQL4 | None | None | None | None |

| | | | | |
|---|---|---|---|---|
| RECQL4 | None | None | None | None |
| RECQL4 | None | None | None | None |
| RECQL4 | None | None | None | None |
| RECQL4 | None | None | None | None |
| RECQL4 | None | None | None | None |
| CHEK2 | None | None | None | None |
| PPARG | None | None | None | None |
| ACADVL | None | None | None | None |
| ACADVL | None | None | None | None |
| ACADVL | None | None | None | None |
| ACADVL | None | None | None | None |
| ACADVL | None | None | None | None |
| ACADVL | None | None | None | None |
| ACADVL | None | None | None | None |
| ACADVL | None | None | None | None |
| ACADVL | None | None | None | None |
| ACADVL | None | None | None | None |
| ACADVL | None | None | None | None |
| ACADVL | None | None | None | None |
| ACADVL | None | None | None | None |
| ACADVL | None | None | None | None |
| ACADVL | None | None | None | None |
| ACADVL | 0.423 | 2 | 3b96 | A |
| ACADVL | None | None | None | None |
| ACADVL | None | None | None | None |
| ACADVL | None | None | None | None |
| ACADVL | None | None | None | None |
| ACADVL | None | None | None | None |
| ACADVL | None | None | None | None |
| APOA1 | None | None | None | None |
| APOA1 | None | None | None | None |
| APOA1 | None | None | None | None |
| APOA1 | None | None | None | None |
| APOA1 | None | None | None | None |
| DNMT3A | None | None | None | None |
| DNMT3A | None | None | None | None |
| APC | None | None | None | None |
| TMOD1 | None | None | None | None |
| USH1C | None | None | None | None |
| USH1C | None | None | None | None |
| USH1C | None | None | None | None |
| USH1C | None | None | None | None |
| ERCC6 | None | None | None | None |
| ERCC6 | None | None | None | None |
| ERCC6 | None | None | None | None |
| ERCC8 | None | None | None | None |
| ERCC8 | None | None | None | None |
| ERCC8 | None | None | None | None |
| DPAGT1 | None | None | None | None |
| DPAGT1 | None | None | None | None |
| DPAGT1 | None | None | None | None |
| DPAGT1 | None | None | None | None |
| DPAGT1 | None | None | None | None |
| DPAGT1 | None | None | None | None |

| | | | | |
|---|---|---|---|---|
| DPAGT1 | None | None | None | None |
| IL2RG | None | None | None | None |
| HNF1A | None | None | None | None |
| HNF4A | None | None | None | None |
| FLT4 | None | None | None | None |
| FLT4 | None | None | None | None |
| FLT4 | None | None | None | None |
| CD3D | None | None | None | None |
| CD3D | None | None | None | None |
| MCOLN1 | None | None | None | None |
| MCOLN1 | None | None | None | None |
| MOCS2 | None | None | None | None |
| MOCS2 | None | None | None | None |
| ABCG8 | None | None | None | None |
| ABCG8 | None | None | None | None |
| ABCG8 | None | None | None | None |
| ABCG8 | None | None | None | None |
| ABCG8 | None | None | None | None |
| ABCG8 | None | None | None | None |
| ABCG8 | None | None | None | None |
| ABCG8 | None | None | None | None |
| ABCG8 | None | None | None | None |
| ABCG8 | None | None | None | None |
| ABCG8 | None | None | None | None |
| ABCG8 | None | None | None | None |
| ABCG8 | None | None | None | None |
| ABCG8 | None | None | None | None |
| CTNNA1 | None | None | None | None |
| TYMP | None | None | None | None |
| TYMP | None | None | None | None |
| TYMP | None | None | None | None |
| TYMP | None | None | None | None |
| TYMP | None | None | None | None |
| TYMP | None | None | None | None |
| TYMP | None | None | None | None |
| TYMP | None | None | None | None |
| TYMP | None | None | None | None |
| TYMP | None | None | None | None |
| TYMP | None | None | None | None |
| TYMP | None | None | None | None |
| TYMP | None | None | None | None |
| GAMT | None | None | None | None |
| GAMT | None | None | None | None |
| GAMT | None | None | None | None |
| GAMT | None | None | None | None |
| GAMT | None | None | None | None |
| GAMT | None | None | None | None |
| GAMT | None | None | None | None |
| GAMT | None | None | None | None |
| GAMT | None | None | None | None |
| GAMT | None | None | None | None |
| GAMT | None | None | None | None |
| GLRA1 | None | None | None | None |
| GLRA1 | None | None | None | None |

| | | | | |
|---|---|---|---|---|
| GLRA1 | None | None | None | None |
| GLRA1 | None | None | None | None |
| GLRA1 | None | None | None | None |
| GLRA1 | None | None | None | None |
| GLRA1 | None | None | None | None |
| GLRA1 | None | None | None | None |
| GLRA1 | None | None | None | None |
| ADK | None | None | None | None |
| ACTN2 | None | None | None | None |
| UBE2T | None | None | None | None |
| TTN | None | None | None | None |
| ATP7B | None | None | None | None |
| ATP7B | None | None | None | None |
| ATP7B | None | None | None | None |
| ATP7B | None | None | None | None |
| ATP7B | None | None | None | None |
| ATP7B | None | None | None | None |
| ATP7B | None | None | None | None |
| ATP7B | None | None | None | None |
| ATP7B | None | None | None | None |
| ATP7B | None | None | None | None |
| ATP7B | None | None | None | None |
| ATP7B | None | None | None | None |
| ATP7B | None | None | None | None |
| CDKN1B | None | None | None | None |
| CDKN1B | None | None | None | None |
| CDKN1B | None | None | None | None |
| CDKN1B | None | None | None | None |
| CDKN1B | None | None | None | None |
| CDKN1B | None | None | None | None |
| MYOC | None | None | None | None |
| MYOC | None | None | None | None |
| MYOC | None | None | None | None |
| MYOC | None | None | None | None |
| MYOC | None | None | None | None |
| MYOC | None | None | None | None |
| MYOC | None | None | None | None |
| MYOC | None | None | None | None |
| MYOC | None | None | None | None |
| MYOC | None | None | None | None |
| MYOC | None | None | None | None |
| MYOC | None | None | None | None |
| MYOC | None | None | None | None |
| MYOC | None | None | None | None |
| MYOC | None | None | None | None |
| SPTAN1 | None | None | None | None |
| GC | None | None | None | None |
| GC | None | None | None | None |
| MAD1L1 | None | None | None | None |
| NDST1 | None | None | None | None |
| ARL2BP | None | None | None | None |
| TOR1A | None | None | None | None |
| C1S | None | None | None | None |
| ARL6 | 0.508 | 1 | 2h57 | A |

| | | | | |
|---|---|---|---|---|
| ARL6 | None | None | None | None |
| ARL6 | None | None | None | None |
| ARL6 | None | None | None | None |
| ARL6 | None | None | None | None |
| ARL6 | None | None | None | None |
| FGFR1 | None | None | None | None |
| KLHL7 | None | None | None | None |
| FLNC | None | None | None | None |
| PTPRC | None | None | None | None |
| GCH1 | None | None | None | None |
| GCH1 | None | None | None | None |
| GCH1 | None | None | None | None |
| GCH1 | None | None | None | None |
| GCH1 | None | None | None | None |
| GCH1 | None | None | None | None |
| GCH1 | None | None | None | None |
| GCH1 | None | None | None | None |
| GCH1 | None | None | None | None |
| GCH1 | None | None | None | None |
| GCH1 | None | None | None | None |
| GCH1 | None | None | None | None |
| GCH1 | None | None | None | None |
| GCH1 | None | None | None | None |
| CFH | None | None | None | None |
| COL2A1 | None | None | None | None |
| COL2A1 | None | None | None | None |
| OCRL | None | None | None | None |
| WRN | None | None | None | None |
| WRN | None | None | None | None |
| WRN | None | None | None | None |
| WRN | None | None | None | None |
| MASP1 | None | None | None | None |
| MASP1 | None | None | None | None |
| NSD1 | None | None | None | None |
| PITX2 | None | None | None | None |
| PITX2 | None | None | None | None |
| PITX2 | None | None | None | None |
| PITX2 | None | None | None | None |
| PITX2 | None | None | None | None |
| PITX2 | None | None | None | None |
| B4GALT7 | 0.917 | 1 | 4irq | A |
| B4GALT7 | None | None | None | None |
| B4GALT7 | None | None | None | None |
| B4GALT7 | None | None | None | None |
| B4GALT7 | None | None | None | None |
| FSHR | None | None | None | None |
| IGHMBP2 | None | None | None | None |
| IGHMBP2 | None | None | None | None |
| IGHMBP2 | None | None | None | None |
| IGHMBP2 | None | None | None | None |
| IGHMBP2 | None | None | None | None |
| IGHMBP2 | None | None | None | None |
| IGHMBP2 | None | None | None | None |
| IGHMBP2 | None | None | None | None |

| | | | | |
|---|---|---|---|---|
| IGHMBP2 | None | None | None | None |
| IGHMBP2 | None | None | None | None |
| IGHMBP2 | None | None | None | None |
| IGHMBP2 | None | None | None | None |
| IGHMBP2 | None | None | None | None |
| IGHMBP2 | None | None | None | None |
| IGHMBP2 | None | None | None | None |
| IGHMBP2 | None | None | None | None |
| TTN | None | None | None | None |
| TTN | None | None | None | None |
| IRAK4 | None | None | None | None |
| PROC | None | None | None | None |
| PROC | None | None | None | None |
| RAD50 | None | None | None | None |
| RAD50 | None | None | None | None |
| DYNC1H1 | None | None | None | None |
| DYNC1H1 | None | None | None | None |
| DYNC1H1 | None | None | None | None |
| DYNC1H1 | None | None | None | None |
| DYNC1H1 | None | None | None | None |
| DYNC1H1 | None | None | None | None |
| DYNC1H1 | None | None | None | None |
| DYNC1H1 | None | None | None | None |
| DYNC1H1 | None | None | None | None |
| DYNC1H1 | None | None | None | None |
| DYNC1H1 | None | None | None | None |
| CYB5R3 | None | None | None | None |
| CYB5R3 | None | None | None | None |
| CYB5R3 | None | None | None | None |
| CYB5R3 | None | None | None | None |
| CYB5R3 | None | None | None | None |
| CYB5R3 | None | None | None | None |
| CYB5R3 | None | None | None | None |
| CYB5R3 | None | None | None | None |
| CYB5R3 | None | None | None | None |
| CYB5R3 | None | None | None | None |
| CYB5R3 | None | None | None | None |
| SPTB | None | None | None | None |
| FANCL | None | None | None | None |
| FAM126A | None | None | None | None |
| SEPT9 | None | None | None | None |
| TTN | None | None | None | None |
| TTN | None | None | None | None |
| TTN | None | None | None | None |
| CNTN1 | None | None | None | None |
| C1QTNF5 | None | None | None | None |
| VCL | None | None | None | None |
| VCL | None | None | None | None |
| VCL | None | None | None | None |
| VCL | None | None | None | None |
| VCL | None | None | None | None |
| VCL | None | None | None | None |
| PPT1 | None | None | None | None |
| PPT1 | None | None | None | None |

| | | | | |
|---|---|---|---|---|
| PPT1 | None | None | None | None |
| PPT1 | None | None | None | None |
| PPT1 | None | None | None | None |
| PPT1 | None | None | None | None |
| PPT1 | None | None | None | None |
| PPT1 | None | None | None | None |
| PPT1 | None | None | None | None |
| PPT1 | None | None | None | None |
| PPT1 | None | None | None | None |
| PPT1 | None | None | None | None |
| PPT1 | None | None | None | None |
| PPT1 | None | None | None | None |
| PPT1 | None | None | None | None |
| PPT1 | None | None | None | None |
| PPT1 | None | None | None | None |
| PPT1 | None | None | None | None |
| PPT1 | None | None | None | None |
| NOTCH2 | None | None | None | None |
| NOTCH2 | None | None | None | None |
| IFIH1 | None | None | None | None |
| IFIH1 | None | None | None | None |
| IKBKG | None | None | None | None |
| DBH | None | None | None | None |
| DBH | None | None | None | None |
| DBH | None | None | None | None |
| DBH | None | None | None | None |
| DBH | None | None | None | None |
| DBH | None | None | None | None |
| DBH | None | None | None | None |
| DBH | None | None | None | None |
| DBH | None | None | None | None |
| DBH | None | None | None | None |
| IL10RA | None | None | None | None |
| IL10RA | None | None | None | None |
| IL10RA | None | None | None | None |
| IL10RA | None | None | None | None |
| IL10RA | None | None | None | None |
| IL10RA | None | None | None | None |
| IL10RA | None | None | None | None |
| ILK | None | None | None | None |
| HPD | None | None | None | None |
| HPD | None | None | None | None |
| HPD | None | None | None | None |
| MECP2 | None | None | None | None |
| MECP2 | None | None | None | None |
| MECP2 | None | None | None | None |
| MECP2 | None | None | None | None |
| MECP2 | None | None | None | None |
| MECP2 | None | None | None | None |
| MECP2 | None | None | None | None |
| MECP2 | None | None | None | None |
| KYNU | None | None | None | None |
| FAS | None | None | None | None |
| FAS | None | None | None | None |

| | | | | |
|---|---|---|---|---|
| ELP1 | None | None | None | None |
| ELP1 | None | None | None | None |
| ELP1 | None | None | None | None |
| ELP1 | None | None | None | None |
| ELP1 | None | None | None | None |
| ELP1 | None | None | None | None |
| ELP1 | None | None | None | None |
| ELP1 | None | None | None | None |
| HESX1 | None | None | None | None |
| HESX1 | None | None | None | None |
| HESX1 | None | None | None | None |
| HESX1 | None | None | None | None |
| GAD1 | None | None | None | None |
| GAD1 | None | None | None | None |
| NDUFA1 | None | None | None | None |
| NDUFA1 | None | None | None | None |
| NDUFA1 | None | None | None | None |
| PNPT1 | None | None | None | None |
| PNPT1 | None | None | None | None |
| PNPT1 | None | None | None | None |
| PNPT1 | None | None | None | None |
| PNPT1 | None | None | None | None |
| PNPT1 | None | None | None | None |
| PNPT1 | None | None | None | None |
| SLC25A13 | None | None | None | None |
| SLC25A13 | None | None | None | None |
| SLC25A13 | None | None | None | None |
| AP4M1 | None | None | None | None |
| AP4M1 | None | None | None | None |
| MEFV | None | None | None | None |
| MEFV | None | None | None | None |
| MEFV | None | None | None | None |
| MEFV | None | None | None | None |
| MEFV | None | None | None | None |
| MEFV | None | None | None | None |
| MEFV | None | None | None | None |
| MEFV | None | None | None | None |
| MEFV | None | None | None | None |
| CAPN3 | None | None | None | None |
| CAPN3 | None | None | None | None |
| CAPN3 | None | None | None | None |
| CAPN3 | None | None | None | None |
| CAPN3 | 0.474 | 2 | 4okh | A |
| CAPN3 | 0.474 | 2 | 4okh | A |
| CAPN3 | None | None | None | None |
| GP6 | None | None | None | None |
| FAS | None | None | None | None |
| FBN1 | None | None | None | None |
| FBN1 | None | None | None | None |
| BMPR2 | None | None | None | None |
| BMPR2 | None | None | None | None |
| BMPR2 | None | None | None | None |
| BMPR2 | None | None | None | None |
| BMPR2 | None | None | None | None |

| | | | | |
|---|---|---|---|---|
| RET | None | None | None | None |
| RET | None | None | None | None |
| RET | None | None | None | None |
| RET | None | None | None | None |
| RET | None | None | None | None |
| RET | None | None | None | None |
| RET | None | None | None | None |
| RET | None | None | None | None |
| RET | None | None | None | None |
| RET | None | None | None | None |
| RET | None | None | None | None |
| RET | None | None | None | None |
| RET | None | None | None | None |
| RET | None | None | None | None |
| RET | None | None | None | None |
| RET | None | None | None | None |
| RET | None | None | None | None |
| RET | None | None | None | None |
| RET | None | None | None | None |
| RET | None | None | None | None |
| RET | None | None | None | None |
| RET | None | None | None | None |
| RET | None | None | None | None |
| RET | None | None | None | None |
| RET | None | None | None | None |
| RET | None | None | None | None |
| RET | None | None | None | None |
| RET | None | None | None | None |
| CSF2RA | None | None | None | None |
| CSF2RA | None | None | None | None |
| CSF2RA | None | None | None | None |
| BRCA1 | None | None | None | None |
| ZAP70 | 0.636 | 1 | 2ozo | A |
| ZAP70 | 0.636 | 1 | 2ozo | A |
| ZAP70 | None | None | None | None |
| PIK3R2 | None | None | None | None |
| TCN2 | None | None | None | None |
| TCN2 | None | None | None | None |
| TCN2 | None | None | None | None |
| TCN2 | None | None | None | None |
| TCN2 | None | None | None | None |
| TCN2 | None | None | None | None |
| TCN2 | None | None | None | None |
| TCN2 | None | None | None | None |
| EPM2A | None | None | None | None |
| EPM2A | None | None | None | None |
| EPM2A | None | None | None | None |
| EPM2A | None | None | None | None |
| AARS | None | None | None | None |
| AARS | None | None | None | None |
| AARS | None | None | None | None |
| PCDH15 | None | None | None | None |
| PCDH15 | None | None | None | None |
| PCDH15 | None | None | None | None |

| | | | | |
|---|---|---|---|---|
| PCDH15 | None | None | None | None |
| PCDH15 | None | None | None | None |
| PCDH15 | None | None | None | None |
| PCDH15 | None | None | None | None |
| PCDH15 | None | None | None | None |
| PCDH15 | None | None | None | None |
| PCDH15 | None | None | None | None |
| NDUFS4 | None | None | None | None |
| NDUFS4 | None | None | None | None |
| NDUFS4 | None | None | None | None |
| FGF10 | None | None | None | None |
| FGF10 | 0.558 | 3 | 1nun | A |
| FGF10 | None | None | None | None |
| FGF10 | None | None | None | None |
| CDON | None | None | None | None |
| STAMBP | 0.909 | 2 | 3rzv | A |
| BRCA1 | None | None | None | None |
| BRCA1 | None | None | None | None |
| BRCA1 | None | None | None | None |
| BRCA1 | None | None | None | None |
| BRCA1 | None | None | None | None |
| BRCA1 | None | None | None | None |
| BRCA1 | None | None | None | None |
| BRCA1 | None | None | None | None |
| BRCA1 | None | None | None | None |
| BRCA1 | None | None | None | None |
| PLXNC1 | None | None | None | None |
| RNASEH2C | None | None | None | None |
| RNASEH2C | None | None | None | None |
| APC | None | None | None | None |
| PLA2G4A | None | None | None | None |
| PLA2G4A | None | None | None | None |
| PSAT1 | None | None | None | None |
| PSAT1 | None | None | None | None |
| TRAPPC3 | None | None | None | None |
| QDPR | None | None | None | None |
| QDPR | None | None | None | None |
| QDPR | None | None | None | None |
| QDPR | None | None | None | None |
| KRT5 | None | None | None | None |
| KRT5 | None | None | None | None |
| KRT5 | None | None | None | None |
| MLH1 | None | None | None | None |
| MLH1 | None | None | None | None |
| MLH1 | None | None | None | None |
| MLH1 | None | None | None | None |
| MLH1 | None | None | None | None |
| MLH1 | None | None | None | None |
| MLH1 | None | None | None | None |
| MLH1 | None | None | None | None |
| MLH1 | None | None | None | None |
| MLH1 | None | None | None | None |
| MLH1 | None | None | None | None |
| MLH1 | None | None | None | None |

| | | | | |
|---|---|---|---|---|
| MLH1 | None | None | None | None |
| MLH1 | None | None | None | None |
| MLH1 | None | None | None | None |
| MLH1 | None | None | None | None |
| MLH1 | None | None | None | None |
| KRIT1 | None | None | None | None |
| KRIT1 | None | None | None | None |
| KRIT1 | None | None | None | None |
| ALX4 | None | None | None | None |
| ALX4 | None | None | None | None |
| ALX4 | None | None | None | None |
| TJP2 | None | None | None | None |
| TJP2 | None | None | None | None |
| PANK2 | None | None | None | None |
| PANK2 | None | None | None | None |
| PANK2 | None | None | None | None |
| PANK2 | None | None | None | None |
| PANK2 | None | None | None | None |
| PANK2 | None | None | None | None |
| PANK2 | None | None | None | None |
| NDUFA6 | None | None | None | None |
| TSHR | None | None | None | None |
| TSHR | None | None | None | None |
| TSHR | None | None | None | None |
| CFTR | None | None | None | None |
| CFTR | None | None | None | None |
| MYOM1 | None | None | None | None |
| MYOM1 | None | None | None | None |
| SPTB | None | None | None | None |
| GPT2 | None | None | None | None |
| NDUFS2 | None | None | None | None |
| NDUFS2 | None | None | None | None |
| NDUFS2 | None | None | None | None |
| NDUFS2 | None | None | None | None |
| NDUFS2 | None | None | None | None |
| CASQ2 | None | None | None | None |
| CASQ2 | None | None | None | None |
| CASQ2 | None | None | None | None |
| CASQ2 | None | None | None | None |
| CASQ2 | None | None | None | None |
| CASQ2 | None | None | None | None |
| CASQ2 | None | None | None | None |
| CASQ2 | None | None | None | None |
| CASQ2 | None | None | None | None |
| CPOX | None | None | None | None |
| CPOX | None | None | None | None |
| CPOX | 0.857 | 2 | 2aex | A |
| CPOX | None | None | None | None |
| CPOX | None | None | None | None |
| CPOX | None | None | None | None |
| CPOX | None | None | None | None |
| CPOX | None | None | None | None |
| CPOX | None | None | None | None |
| MVK | None | None | None | None |

| | | | | |
|---|---|---|---|---|
| MVK | None | None | None | None |
| MVK | None | None | None | None |
| MVK | None | None | None | None |
| MVK | None | None | None | None |
| MVK | None | None | None | None |
| MVK | None | None | None | None |
| MVK | None | None | None | None |
| MVK | None | None | None | None |
| MVK | None | None | None | None |
| MVK | None | None | None | None |
| MVK | None | None | None | None |
| MVK | None | None | None | None |
| MVK | None | None | None | None |
| MVK | None | None | None | None |
| GSS | None | None | None | None |
| GSS | None | None | None | None |
| GSS | None | None | None | None |
| GSS | None | None | None | None |
| GSS | None | None | None | None |
| DSG2 | None | None | None | None |
| DSG2 | None | None | None | None |
| DSG2 | None | None | None | None |
| DSG2 | None | None | None | None |
| DSG2 | None | None | None | None |
| DSG2 | None | None | None | None |
| DSG2 | None | None | None | None |
| DSG2 | None | None | None | None |
| DSG2 | None | None | None | None |
| DSG2 | None | None | None | None |
| DSG2 | None | None | None | None |
| DSG2 | None | None | None | None |
| DSG2 | None | None | None | None |
| DSG2 | None | None | None | None |
| PAPSS2 | None | None | None | None |
| PLCB2 | None | None | None | None |
| HMGCS2 | None | None | None | None |
| HMGCS2 | None | None | None | None |
| HMGCS2 | 0.938 | 1 | 2wya | C |
| HMGCS2 | None | None | None | None |
| HMGCS2 | None | None | None | None |
| HMGCS2 | None | None | None | None |
| HMGCS2 | 0.562 | 4 | 2wya | C |
| HMGCS2 | None | None | None | None |
| MPDZ | None | None | None | None |
| GATA4 | None | None | None | None |
| GATA4 | None | None | None | None |
| WARS2 | None | None | None | None |
| WARS2 | None | None | None | None |
| HGD | 1 | 47 | 1eyb | A |
| HGD | 1 | 47 | 1eyb | A |
| HGD | 1 | 70 | 1eyb | A |
| HGD | None | None | None | None |
| HGD | None | None | None | None |
| HGD | None | None | None | None |

| | | | | |
|---|---|---|---|---|
| HGD | None | None | None | None |
| HGD | 0.5 | 51 | 1ey2 | A |
| HGD | None | None | None | None |
| HGD | 1 | 1 | 1ey2 | A |
| HGD | 1 | 9 | 1ey2 | A |
| HGD | None | None | None | None |
| HGD | 1 | 10 | 1eyb | A |
| HGD | 1 | 7 | 1ey2 | A |
| PAX8 | None | None | None | None |
| PAX8 | None | None | None | None |
| PAX8 | None | None | None | None |
| PAX8 | None | None | None | None |
| SPTB | None | None | None | None |
| NCF4 | None | None | None | None |
| CFH | None | None | None | None |
| CFH | None | None | None | None |
| CSF1R | None | None | None | None |
| CSF1R | None | None | None | None |
| CA8 | None | None | None | None |
| CA8 | None | None | None | None |
| CA8 | 0.748 | 7 | 2w2j | A |
| APC | None | None | None | None |
| APC | None | None | None | None |
| APC | None | None | None | None |
| APC | None | None | None | None |
| APC | None | None | None | None |
| NDUFA2 | None | None | None | None |
| GIF | None | None | None | None |
| GIF | None | None | None | None |
| GIF | None | None | None | None |
| GIF | None | None | None | None |
| C2 | None | None | None | None |
| ACP5 | None | None | None | None |
| ACP5 | None | None | None | None |
| ACP5 | None | None | None | None |
| ACP5 | None | None | None | None |
| ACP5 | None | None | None | None |
| RNASEH2A | None | None | None | None |
| RNASEH2A | None | None | None | None |
| RNASEH2A | None | None | None | None |
| RNASEH2A | None | None | None | None |
| RNASEH2A | None | None | None | None |
| RNASEH2A | None | None | None | None |
| RNASEH2A | None | None | None | None |
| DSG2 | None | None | None | None |
| USH1G | None | None | None | None |
| FUS | None | None | None | None |
| FUS | None | None | None | None |
| FUS | None | None | None | None |
| FUS | None | None | None | None |
| FUS | None | None | None | None |
| MRPL44 | None | None | None | None |
| MRPL44 | None | None | None | None |
| MRPL44 | None | None | None | None |

| | | | | |
|---|---|---|---|---|
| MRPL44 | None | None | None | None |
| DMGDH | None | None | None | None |
| AARS | None | None | None | None |
| STAR | None | None | None | None |
| STAR | None | None | None | None |
| STAR | None | None | None | None |
| STAR | None | None | None | None |
| STAR | None | None | None | None |
| STAR | None | None | None | None |
| STAR | None | None | None | None |
| STAR | None | None | None | None |
| LARP7 | None | None | None | None |
| PRPF8 | None | None | None | None |
| PRPF8 | None | None | None | None |
| PRPF8 | None | None | None | None |
| PRPF8 | None | None | None | None |
| PSAP | None | None | None | None |
| PSAP | None | None | None | None |
| PSAP | None | None | None | None |
| DMPK | None | None | None | None |
| EPCAM | None | None | None | None |
| EPCAM | None | None | None | None |
| EPCAM | None | None | None | None |
| EPCAM | None | None | None | None |
| EPCAM | None | None | None | None |
| EPCAM | None | None | None | None |
| EPCAM | None | None | None | None |
| EPCAM | None | None | None | None |
| ASS1 | None | None | None | None |
| ASS1 | None | None | None | None |
| ASS1 | 0.571 | 3 | 2nz2 | A |
| ASS1 | 0.571 | 3 | 2nz2 | A |
| ASS1 | None | None | None | None |
| ASS1 | 0.857 | 20 | 2nz2 | A |
| ASS1 | 0.857 | 9 | 2nz2 | A |
| ASS1 | None | None | None | None |
| ASS1 | 0.429 | 13 | 2nz2 | A |
| ASS1 | 0.571 | 4 | 2nz2 | A |
| ASS1 | 0.857 | 9 | 2nz2 | A |
| ASS1 | 0.857 | 9 | 2nz2 | A |
| ASS1 | 0.857 | 9 | 2nz2 | A |
| ASS1 | None | None | None | None |
| ASS1 | None | None | None | None |
| ASS1 | None | None | None | None |
| ASS1 | None | None | None | None |
| ASS1 | None | None | None | None |
| ASS1 | None | None | None | None |
| ASS1 | None | None | None | None |
| POLE | None | None | None | None |
| POLE | None | None | None | None |
| POLE | None | None | None | None |
| RNASET2 | None | None | None | None |
| RNASET2 | None | None | None | None |
| RNASET2 | None | None | None | None |

| | | | | |
|---|---|---|---|---|
| FLG | None | None | None | None |
| F10 | None | None | None | None |
| AGT | None | None | None | None |
| AGT | None | None | None | None |
| PLN | None | None | None | None |
| PLN | None | None | None | None |
| PLN | None | None | None | None |
| PTHLH | None | None | None | None |
| IGF1R | None | None | None | None |
| IGF1R | None | None | None | None |
| IGF1R | None | None | None | None |
| JAG1 | None | None | None | None |
| JAG1 | None | None | None | None |
| JAG1 | None | None | None | None |
| JAG1 | None | None | None | None |
| JAG1 | None | None | None | None |
| JAG1 | None | None | None | None |
| AP1S3 | None | None | None | None |
| SPTA1 | None | None | None | None |
| SPTA1 | None | None | None | None |
| SPTA1 | None | None | None | None |
| SPTA1 | None | None | None | None |
| SPG7 | None | None | None | None |
| SPG7 | None | None | None | None |
| SPG7 | None | None | None | None |
| SPG7 | None | None | None | None |
| SPG7 | None | None | None | None |
| SPG7 | None | None | None | None |
| FN1 | None | None | None | None |
| FN1 | None | None | None | None |
| NSMCE3 | None | None | None | None |
| PCCA | None | None | None | None |
| PCCA | None | None | None | None |
| SPTB | None | None | None | None |
| RYR2 | None | None | None | None |
| RYR2 | None | None | None | None |
| MMP9 | None | None | None | None |
| MMP9 | None | None | None | None |
| GLMN | None | None | None | None |
| GLMN | None | None | None | None |
| GLMN | None | None | None | None |
| GLMN | None | None | None | None |
| GLMN | None | None | None | None |
| GLMN | None | None | None | None |
| GLMN | None | None | None | None |
| GLMN | None | None | None | None |
| GLMN | None | None | None | None |
| GLMN | None | None | None | None |
| MARS | None | None | None | None |
| MARS | None | None | None | None |
| MARS | None | None | None | None |
| MARS | None | None | None | None |
| MARS | None | None | None | None |
| MARS | None | None | None | None |

| | | | | |
|---|---|---|---|---|
| SNAP29 | None | None | None | None |
| SNAP29 | None | None | None | None |
| ETHE1 | None | None | None | None |
| ETHE1 | None | None | None | None |
| ETHE1 | None | None | None | None |
| ETHE1 | None | None | None | None |
| ETHE1 | None | None | None | None |
| ETHE1 | None | None | None | None |
| ETHE1 | None | None | None | None |
| SAMHD1 | None | None | None | None |
| KCNQ4 | None | None | None | None |
| FXYD2 | None | None | None | None |
| FXYD2 | None | None | None | None |
| FXYD2 | None | None | None | None |
| LRRK2 | None | None | None | None |
| LRRK2 | None | None | None | None |
| NDUFS1 | None | None | None | None |
| NDUFS1 | None | None | None | None |
| NDUFS1 | None | None | None | None |
| NDUFS1 | None | None | None | None |
| NDUFS1 | None | None | None | None |
| NDUFS1 | None | None | None | None |
| NDUFS1 | None | None | None | None |
| NDUFS1 | None | None | None | None |
| NDUFS1 | None | None | None | None |
| NDUFS1 | None | None | None | None |
| NDUFS1 | None | None | None | None |
| EZH2 | None | None | None | None |
| EZH2 | None | None | None | None |
| LARS | None | None | None | None |
| TLL1 | None | None | None | None |
| CRYBB2 | None | None | None | None |
| CRYBB2 | None | None | None | None |
| EXOSC8 | None | None | None | None |
| SRP72 | None | None | None | None |
| FOXC2 | None | None | None | None |
| HMBS | None | None | None | None |
| HMBS | None | None | None | None |
| HMBS | None | None | None | None |
| HMBS | None | None | None | None |
| HMBS | None | None | None | None |
| HMBS | None | None | None | None |
| HMBS | None | None | None | None |
| HMBS | None | None | None | None |
| HMBS | None | None | None | None |
| HMBS | None | None | None | None |
| HMBS | None | None | None | None |
| HMBS | None | None | None | None |
| HMBS | None | None | None | None |
| HMBS | None | None | None | None |
| HMBS | None | None | None | None |
| HMBS | None | None | None | None |
| HMBS | None | None | None | None |
| HMBS | None | None | None | None |

| | | | | |
|---|---|---|---|---|
| HMBS | None | None | None | None |
| HMBS | None | None | None | None |
| HMBS | None | None | None | None |
| HMBS | None | None | None | None |
| HMBS | None | None | None | None |
| HMBS | None | None | None | None |
| HMBS | None | None | None | None |
| OGT | None | None | None | None |
| NOG | None | None | None | None |
| NOG | None | None | None | None |
| NOG | None | None | None | None |
| NOG | None | None | None | None |
| NOG | None | None | None | None |
| NOG | None | None | None | None |
| NOG | None | None | None | None |
| NOG | None | None | None | None |
| NOG | None | None | None | None |
| NOG | None | None | None | None |
| NOG | None | None | None | None |
| DLG3 | None | None | None | None |
| SCN2A | None | None | None | None |
| STAC3 | None | None | None | None |
| PFKM | None | None | None | None |
| PFKM | None | None | None | None |
| PFKM | None | None | None | None |
| PFKM | None | None | None | None |
| PFKM | None | None | None | None |
| DNMT1 | None | None | None | None |
| DNMT1 | None | None | None | None |
| TNNI3 | None | None | None | None |
| TNNI3 | None | None | None | None |
| TNNI3 | None | None | None | None |
| STXBP2 | None | None | None | None |
| STXBP2 | None | None | None | None |
| STXBP2 | None | None | None | None |
| STXBP2 | None | None | None | None |
| STXBP2 | None | None | None | None |
| STXBP2 | None | None | None | None |
| STXBP2 | None | None | None | None |
| STXBP2 | None | None | None | None |
| REN | None | None | None | None |
| DSP | None | None | None | None |
| DSP | None | None | None | None |
| HNF1B | None | None | None | None |
| ADA | None | None | None | None |
| ADA | None | None | None | None |
| ADA | 0.615 | 7 | 3iar | A |
| ADA | None | None | None | None |
| ADA | None | None | None | None |
| ADA | 0.538 | 5 | 3iar | A |
| ADA | 0.538 | 5 | 3iar | A |
| ADA | None | None | None | None |
| ADA | None | None | None | None |
| ADA | 0.615 | 1 | 3iar | A |

| | | | | |
|---|---|---|---|---|
| ADA | 0.462 | 6 | 3iar | A |
| ADA | 0.615 | 1 | 3iar | A |
| ADA | None | None | None | None |
| ADA | None | None | None | None |
| ADA | None | None | None | None |
| DCX | None | None | None | None |
| DCX | None | None | None | None |
| DCX | None | None | None | None |
| DCX | None | None | None | None |
| DCX | None | None | None | None |
| ARSB | None | None | None | None |
| ARSB | None | None | None | None |
| ARSB | None | None | None | None |
| ARSB | None | None | None | None |
| ARSB | None | None | None | None |
| ARSB | None | None | None | None |
| ARSB | None | None | None | None |
| ARSB | None | None | None | None |
| ARSB | None | None | None | None |
| ARSB | None | None | None | None |
| ARSB | None | None | None | None |
| ARSB | None | None | None | None |
| ARSB | None | None | None | None |
| WWOX | None | None | None | None |
| TNNI3 | None | None | None | None |
| TNNI3 | None | None | None | None |
| TNNI3 | None | None | None | None |
| TNNI3 | None | None | None | None |
| CYBA | None | None | None | None |
| SPAST | None | None | None | None |
| SPAST | None | None | None | None |
| SPAST | None | None | None | None |
| SPAST | None | None | None | None |
| SPAST | None | None | None | None |
| SPAST | None | None | None | None |
| SPAST | None | None | None | None |
| SPAST | None | None | None | None |
| KRIT1 | None | None | None | None |
| KRIT1 | None | None | None | None |
| TP53 | None | None | None | None |
| ISPD | None | None | None | None |
| ISPD | None | None | None | None |
| ISPD | None | None | None | None |
| ISPD | None | None | None | None |
| KCNMA1 | None | None | None | None |
| STK11 | None | None | None | None |
| STK11 | None | None | None | None |
| STK11 | None | None | None | None |
| STK11 | None | None | None | None |
| STK11 | None | None | None | None |
| STK11 | 0.412 | 2 | 2wtk | F |
| NDUFA13 | None | None | None | None |
| PHF6 | None | None | None | None |
| PHF6 | None | None | None | None |

| | | | | |
|---|---|---|---|---|
| PHF6 | None | None | None | None |
| PHYH | None | None | None | None |
| PHYH | None | None | None | None |
| PHYH | None | None | None | None |
| PHYH | None | None | None | None |
| PHYH | None | None | None | None |
| NDUFS3 | None | None | None | None |
| NDUFS3 | None | None | None | None |
| ALDH18A1 | None | None | None | None |
| ALDH18A1 | None | None | None | None |
| ALDH18A1 | None | None | None | None |
| ALDH18A1 | None | None | None | None |
| ALDH18A1 | None | None | None | None |
| PAX6 | None | None | None | None |
| PAX6 | None | None | None | None |
| INPP5E | None | None | None | None |
| INPP5E | None | None | None | None |
| INPP5E | None | None | None | None |
| INPP5E | None | None | None | None |
| INPP5E | None | None | None | None |
| INPP5E | None | None | None | None |
| INPP5E | None | None | None | None |
| INPP5E | None | None | None | None |
| INPP5E | None | None | None | None |
| TNNT2 | None | None | None | None |
| TNNT2 | None | None | None | None |
| TNNT2 | None | None | None | None |
| TNNT2 | None | None | None | None |
| TNNT2 | None | None | None | None |
| TNNT2 | None | None | None | None |
| TNNT2 | None | None | None | None |
| TNNT2 | None | None | None | None |
| TYK2 | None | None | None | None |
| TYK2 | None | None | None | None |
| TYK2 | None | None | None | None |
| CFTR | None | None | None | None |
| CFTR | None | None | None | None |
| CFTR | None | None | None | None |
| CFTR | None | None | None | None |
| CFTR | None | None | None | None |
| CFTR | None | None | None | None |
| CFTR | None | None | None | None |
| CFTR | None | None | None | None |
| CFTR | None | None | None | None |
| CFTR | None | None | None | None |
| CFTR | None | None | None | None |
| CFTR | None | None | None | None |
| CFTR | None | None | None | None |
| CFTR | None | None | None | None |
| CFTR | None | None | None | None |
| CFTR | None | None | None | None |
| CFTR | None | None | None | None |
| CFTR | None | None | None | None |
| CFTR | None | None | None | None |

| | | | | |
|---|---|---|---|---|
| CFTR | None | None | None | None |
| CFTR | None | None | None | None |
| CFTR | None | None | None | None |
| CFTR | None | None | None | None |
| CFTR | None | None | None | None |
| CFTR | None | None | None | None |
| CFTR | None | None | None | None |
| CFTR | None | None | None | None |
| CFTR | None | None | None | None |
| CFTR | None | None | None | None |
| CFTR | None | None | None | None |
| CFTR | None | None | None | None |
| CFTR | None | None | None | None |
| CFTR | None | None | None | None |
| CFTR | None | None | None | None |
| CFTR | None | None | None | None |
| CFTR | None | None | None | None |
| CFTR | None | None | None | None |
| CFTR | None | None | None | None |
| CFTR | None | None | None | None |
| CFTR | None | None | None | None |
| CFTR | None | None | None | None |
| CFTR | None | None | None | None |
| CFTR | None | None | None | None |
| CFTR | None | None | None | None |
| CFTR | None | None | None | None |
| CFTR | None | None | None | None |
| CFTR | None | None | None | None |
| CFTR | None | None | None | None |
| CFTR | None | None | None | None |
| CFTR | None | None | None | None |
| CFTR | None | None | None | None |
| CFTR | None | None | None | None |
| CFTR | None | None | None | None |
| SERPINF1 | None | None | None | None |
| SERPINF1 | None | None | None | None |
| IDS | None | None | None | None |
| IDS | None | None | None | None |
| IDS | None | None | None | None |
| IDS | None | None | None | None |
| IDS | None | None | None | None |
| IDS | None | None | None | None |
| IDS | None | None | None | None |
| IDS | None | None | None | None |
| IDS | None | None | None | None |
| NFKBIA | 0.7 | 1 | 1nfi | F |
| NFKBIA | None | None | None | None |
| SPAST | None | None | None | None |
| FGF23 | None | None | None | None |
| FGF23 | None | None | None | None |
| FGF23 | None | None | None | None |
| FGF23 | None | None | None | None |
| FGF23 | None | None | None | None |
| FGF23 | None | None | None | None |

| | | | | |
|---|---|---|---|---|
| FGF23 | None | None | None | None |
| DLAT | None | None | None | None |
| STUB1 | None | None | None | None |
| HSD17B4 | None | None | None | None |
| HSD17B4 | None | None | None | None |
| DYSF | None | None | None | None |
| CEP135 | None | None | None | None |
| CEP135 | None | None | None | None |
| PRKCG | None | None | None | None |
| CSF1R | None | None | None | None |
| PEX5 | None | None | None | None |
| PAX6 | None | None | None | None |
| PAX6 | None | None | None | None |
| PAX6 | None | None | None | None |
| PAX6 | None | None | None | None |
| PAX6 | None | None | None | None |
| PAX6 | None | None | None | None |
| NF2 | None | None | None | None |
| NF2 | None | None | None | None |
| NF2 | None | None | None | None |
| REM1 | None | None | None | None |

| source_id | classification | binding_site | binding_site_single_water | protein_binding_site |
|---|---|---|---|---|
| rs121918343 | pathogenic | 1 | 1 | 0 |
| rs267606649 | pathogenic | 1 | 1 | 1 |
| rs121918342 | pathogenic | 0 | 1 | 0 |
| rs267606650 | pathogenic | 1 | 1 | 1 |
| rs1130335 | benign | 1 | 1 | 1 |
| rs199476087 | pathogenic | 1 | 1 | 1 |
| rs772163112 | benign | 0 | 0 | 0 |
| rs755793 | benign | 0 | 0 | 0 |
| rs121918155 | pathogenic | 1 | 1 | 0 |
| rs121918156 | pathogenic | 1 | 1 | 0 |
| rs121918154 | pathogenic | 1 | 1 | 1 |
| rs121918151 | pathogenic | 1 | 1 | 1 |
| rs121918144 | pathogenic | 0 | 1 | 0 |
| rs121918146 | pathogenic | 1 | 1 | 1 |
| rs121918160 | pathogenic | 1 | 1 | 1 |
| rs121918150 | pathogenic | 1 | 1 | 1 |
| rs121918141 | pathogenic | 1 | 1 | 1 |
| rs142742242 | pathogenic | 1 | 1 | 1 |
| rs121918142 | pathogenic | 1 | 1 | 1 |
| rs121918157 | pathogenic | 1 | 1 | 1 |
| rs121918101 | pathogenic | 1 | 1 | 1 |
| rs121908260 | pathogenic | 1 | 1 | 1 |
| rs80356667 | pathogenic | 1 | 1 | 1 |
| rs80356668 | pathogenic | 1 | 1 | 1 |
| rs397514540 | pathogenic | 1 | 1 | 1 |
| rs104894685 | pathogenic | 0 | 1 | 0 |
| rs199869995 | pathogenic | 1 | 1 | 1 |
| rs121909072 | pathogenic | 0 | 0 | 0 |
| rs143669310 | benign | 1 | 1 | 1 |
| rs747347498 | benign | 0 | 0 | 0 |
| rs138283222 | benign | 0 | 0 | 0 |
| rs118203931 | pathogenic | 1 | 1 | 1 |
| rs118203933 | pathogenic | 1 | 1 | 0 |
| rs118203932 | pathogenic | 1 | 1 | 0 |
| rs2228063 | benign | 1 | 1 | 1 |
| rs377767360 | pathogenic | 0 | 0 | 0 |
| rs397518413 | pathogenic | 1 | 1 | 1 |
| rs1800458 | benign | 1 | 1 | 0 |
| rs121918083 | pathogenic | 1 | 1 | 1 |
| rs121918094 | pathogenic | 1 | 1 | 1 |
| rs121918098 | pathogenic | 1 | 1 | 1 |
| rs121918093 | pathogenic | 1 | 1 | 1 |
| rs79977247 | pathogenic | 1 | 1 | 1 |
| rs28933979 | pathogenic | 1 | 1 | 1 |
| rs121918068 | pathogenic | 1 | 1 | 1 |
| rs121918077 | pathogenic | 1 | 1 | 1 |
| rs11541796 | pathogenic | 1 | 1 | 1 |
| rs104894665 | pathogenic | 1 | 1 | 1 |
| rs121918090 | pathogenic | 1 | 1 | 1 |
| rs121918081 | pathogenic | 1 | 1 | 1 |
| rs121918080 | pathogenic | 1 | 1 | 1 |
| rs121918076 | pathogenic | 1 | 1 | 1 |
| rs121918097 | pathogenic | 1 | 1 | 1 |

| | | | | |
|---|---|---|---|---|
| rs121918079 | pathogenic | 1 | 1 | 1 |
| rs121918069 | pathogenic | 1 | 1 | 1 |
| rs121918086 | pathogenic | 1 | 1 | 1 |
| rs121918091 | pathogenic | 1 | 1 | 1 |
| rs121918085 | pathogenic | 1 | 1 | 1 |
| rs121918100 | pathogenic | 1 | 1 | 1 |
| rs121918084 | pathogenic | 1 | 1 | 1 |
| rs121918071 | pathogenic | 1 | 1 | 1 |
| rs121918072 | pathogenic | 1 | 1 | 1 |
| rs121918082 | pathogenic | 1 | 1 | 1 |
| rs121918087 | pathogenic | 1 | 1 | 1 |
| rs76410435 | benign | 1 | 1 | 1 |
| rs121918089 | pathogenic | 1 | 1 | 1 |
| rs121918073 | pathogenic | 1 | 1 | 1 |
| rs121918075 | pathogenic | 1 | 1 | 1 |
| rs121918088 | pathogenic | 1 | 1 | 1 |
| rs76992529 | pathogenic | 1 | 1 | 1 |
| rs536294863 | benign | 1 | 1 | 1 |
| rs730881162 | benign | 1 | 1 | 1 |
| rs113994087 | pathogenic | 1 | 1 | 1 |
| rs74716434 | benign | 1 | 1 | 1 |
| rs121918694 | pathogenic | 1 | 1 | 1 |
| rs121918707 | pathogenic | 1 | 1 | 1 |
| rs121918695 | pathogenic | 1 | 1 | 0 |
| rs121918690 | pathogenic | 1 | 1 | 0 |
| rs121918696 | pathogenic | 1 | 1 | 0 |
| rs121918693 | pathogenic | 1 | 1 | 0 |
| rs28999969 | pathogenic | 1 | 1 | 1 |
| rs121918697 | pathogenic | 1 | 1 | 1 |
| rs121918688 | pathogenic | 1 | 1 | 0 |
| rs28999970 | pathogenic | 1 | 1 | 0 |
| rs121918686 | pathogenic | 1 | 1 | 0 |
| rs28999971 | pathogenic | 1 | 1 | 1 |
| rs121918698 | pathogenic | 1 | 1 | 1 |
| rs121918691 | pathogenic | 1 | 1 | 0 |
| rs121918692 | pathogenic | 1 | 1 | 1 |
| rs121918703 | pathogenic | 1 | 1 | 1 |
| rs121918687 | pathogenic | 1 | 1 | 1 |
| rs28933408 | pathogenic | 1 | 1 | 1 |
| rs121918702 | pathogenic | 1 | 1 | 1 |
| rs17875899 | benign | 1 | 1 | 1 |
| rs111033620 | pathogenic | 1 | 1 | 1 |
| rs111033622 | pathogenic | 1 | 1 | 1 |
| rs111033621 | pathogenic | 1 | 1 | 1 |
| rs111033618 | pathogenic | 1 | 1 | 1 |
| rs5918 | benign | 0 | 0 | 0 |
| rs121918452 | pathogenic | 0 | 0 | 0 |
| rs121918445 | pathogenic | 1 | 1 | 1 |
| rs5917 | benign | 1 | 1 | 1 |
| rs121918444 | pathogenic | 1 | 1 | 1 |
| rs121918446 | pathogenic | 1 | 1 | 1 |
| rs121918449 | pathogenic | 0 | 0 | 0 |
| rs121918448 | benign | 0 | 0 | 0 |
| rs13306487 | benign | 1 | 1 | 1 |

| | | | | |
|---|---|---|---|---|
| rs1805044 | benign | 0 | 0 | 0 |
| rs117078377 | benign | 0 | 0 | 0 |
| rs138132026 | benign | 1 | 1 | 1 |
| rs75762896 | benign | 1 | 1 | 0 |
| rs121918043 | pathogenic | 1 | 1 | 0 |
| rs143581690 | benign | 1 | 1 | 0 |
| rs147654263 | benign | 0 | 0 | 0 |
| rs587782529 | pathogenic | 1 | 1 | 1 |
| rs121912664 | pathogenic | 1 | 1 | 1 |
| rs150293825 | benign | 1 | 1 | 1 |
| rs17882252 | benign | 1 | 1 | 1 |
| rs121912662 | pathogenic | 1 | 1 | 1 |
| rs35993958 | benign | 1 | 1 | 1 |
| rs606231253 | pathogenic | 1 | 1 | 1 |
| rs2227983 | benign | 1 | 1 | 1 |
| rs587779351 | pathogenic | 0 | 0 | 0 |
| rs779773463 | pathogenic | 1 | 1 | 1 |
| rs713040 | benign | 1 | 1 | 1 |
| rs334 | pathogenic | 1 | 1 | 1 |
| rs33930165 | pathogenic | 1 | 1 | 1 |
| rs33918131 | benign | 1 | 1 | 1 |
| rs63750783 | pathogenic | 1 | 1 | 1 |
| rs33986703 | pathogenic | 1 | 1 | 1 |
| rs33972047 | pathogenic | 1 | 1 | 1 |
| rs33936254 | pathogenic | 1 | 1 | 1 |
| rs33959855 | benign | 1 | 1 | 1 |
| rs33950507 | pathogenic | 1 | 1 | 1 |
| rs33916412 | pathogenic | 1 | 1 | 0 |
| rs1135071 | pathogenic | 1 | 1 | 1 |
| rs33960103 | pathogenic | 1 | 1 | 1 |
| rs11549407 | pathogenic | 1 | 1 | 1 |
| rs33922842 | pathogenic | 1 | 1 | 1 |
| rs35278874 | benign | 1 | 1 | 1 |
| rs34621955 | benign | 1 | 1 | 1 |
| rs33995148 | pathogenic | 1 | 1 | 1 |
| rs33945705 | benign | 1 | 1 | 1 |
| rs33993568 | benign | 1 | 1 | 1 |
| rs35553496 | benign | 1 | 1 | 1 |
| rs33913712 | pathogenic | 1 | 1 | 1 |
| rs33917785 | pathogenic | 1 | 1 | 1 |
| rs34515413 | benign | 1 | 1 | 1 |
| rs35256489 | pathogenic | 1 | 1 | 0 |
| rs34484056 | pathogenic | 1 | 1 | 1 |
| rs33947020 | benign | 1 | 1 | 1 |
| rs33946267 | pathogenic | 1 | 1 | 1 |
| rs33966761 | pathogenic | 1 | 1 | 1 |
| rs36020563 | benign | 1 | 1 | 1 |
| rs122454124 | pathogenic | 1 | 1 | 1 |
| rs122454126 | pathogenic | 1 | 1 | 0 |
| rs122454127 | pathogenic | 1 | 1 | 1 |
| rs122454130 | pathogenic | 1 | 1 | 1 |
| rs122454125 | pathogenic | 1 | 1 | 1 |
| rs122454131 | pathogenic | 1 | 1 | 0 |
| rs121908302 | pathogenic | 0 | 1 | 0 |

| | | | | |
|---|---|---|---|---|
| rs77829017 | pathogenic | 0 | 1 | 0 |
| rs1141814 | pathogenic | 0 | 1 | 0 |
| rs121908312 | pathogenic | 0 | 1 | 0 |
| rs79653797 | pathogenic | 1 | 1 | 0 |
| rs439898 | pathogenic | 1 | 1 | 0 |
| rs121908299 | pathogenic | 0 | 1 | 0 |
| rs80356763 | pathogenic | 1 | 1 | 1 |
| rs781152868 | pathogenic | 0 | 1 | 0 |
| rs121908297 | pathogenic | 0 | 1 | 0 |
| rs398123531 | benign | 0 | 1 | 0 |
| rs398123532 | pathogenic | 0 | 1 | 0 |
| rs364897 | pathogenic | 1 | 1 | 1 |
| rs1064644 | pathogenic | 0 | 1 | 0 |
| rs409652 | pathogenic | 1 | 1 | 1 |
| rs121908300 | pathogenic | 0 | 1 | 0 |
| rs381737 | pathogenic | 0 | 1 | 0 |
| rs74500255 | pathogenic | 0 | 1 | 0 |
| rs121908313 | pathogenic | 0 | 1 | 0 |
| rs367968666 | pathogenic | 0 | 1 | 0 |
| rs78973108 | pathogenic | 1 | 1 | 1 |
| rs121908298 | pathogenic | 1 | 1 | 1 |
| rs78396650 | pathogenic | 0 | 0 | 0 |
| rs78198234 | pathogenic | 1 | 1 | 0 |
| rs121908304 | pathogenic | 1 | 1 | 1 |
| rs76539814 | pathogenic | 1 | 1 | 1 |
| rs121908305 | pathogenic | 0 | 1 | 0 |
| rs121908306 | pathogenic | 1 | 1 | 0 |
| rs121908308 | pathogenic | 1 | 1 | 1 |
| rs121908307 | pathogenic | 0 | 1 | 0 |
| rs75548401 | benign | 0 | 1 | 0 |
| rs76763715 | pathogenic | 0 | 0 | 0 |
| rs121908314 | pathogenic | 0 | 1 | 0 |
| rs398123528 | pathogenic | 0 | 1 | 0 |
| rs121908311 | pathogenic | 0 | 0 | 0 |
| rs121908310 | pathogenic | 1 | 1 | 0 |
| rs74598136 | pathogenic | 0 | 1 | 0 |
| rs1064651 | pathogenic | 1 | 1 | 1 |
| rs77369218 | pathogenic | 1 | 1 | 1 |
| rs121908295 | pathogenic | 0 | 1 | 0 |
| rs421016 | pathogenic | 0 | 1 | 0 |
| rs368060 | pathogenic | 0 | 1 | 0 |
| rs80356771 | pathogenic | 1 | 1 | 1 |
| rs80356772 | pathogenic | 1 | 1 | 1 |
| rs121908301 | pathogenic | 0 | 0 | 0 |
| rs747506979 | pathogenic | 0 | 1 | 0 |
| rs75822236 | pathogenic | 0 | 1 | 0 |
| rs1990760 | benign | 1 | 1 | 0 |
| rs28934568 | pathogenic | 1 | 1 | 0 |
| rs34833812 | benign | 1 | 1 | 0 |
| rs193922666 | benign | 1 | 1 | 0 |
| rs104893812 | pathogenic | 1 | 1 | 0 |
| rs144701411 | benign | 0 | 0 | 0 |
| rs104893813 | pathogenic | 0 | 1 | 0 |
| rs199660234 | pathogenic | 1 | 1 | 0 |

| | | | | |
|---|---|---|---|---|
| rs104893814 | pathogenic | 1 | 1 | 0 |
| rs143095746 | benign | 1 | 1 | 1 |
| rs104893807 | pathogenic | 1 | 1 | 0 |
| rs104893811 | pathogenic | 0 | 0 | 0 |
| rs104893816 | pathogenic | 0 | 0 | 0 |
| rs121918714 | pathogenic | 0 | 1 | 0 |
| rs104893810 | pathogenic | 1 | 1 | 0 |
| rs104893815 | pathogenic | 1 | 1 | 0 |
| rs4303 | benign | 0 | 1 | 0 |
| rs121912704 | pathogenic | 0 | 1 | 0 |
| rs56394458 | benign | 1 | 1 | 0 |
| rs397514688 | pathogenic | 0 | 0 | 0 |
| rs367797185 | pathogenic | 1 | 1 | 0 |
| rs12709426 | benign | 0 | 1 | 0 |
| rs2228547 | benign | 0 | 0 | 0 |
| rs111033546 | pathogenic | 0 | 0 | 0 |
| rs111033553 | pathogenic | 0 | 0 | 0 |
| rs111033554 | pathogenic | 1 | 1 | 1 |
| rs111033544 | pathogenic | 1 | 1 | 1 |
| rs111033555 | pathogenic | 1 | 1 | 1 |
| rs111033545 | pathogenic | 1 | 1 | 1 |
| rs111033549 | pathogenic | 0 | 1 | 0 |
| rs111033552 | pathogenic | 1 | 1 | 1 |
| rs752204035 | benign | 0 | 0 | 0 |
| rs11569534 | benign | 0 | 0 | 0 |
| rs28937571 | pathogenic | 1 | 1 | 1 |
| rs104894215 | pathogenic | 1 | 1 | 1 |
| rs3888798 | benign | 1 | 1 | 1 |
| rs764263034 | benign | 1 | 1 | 1 |
| rs1800973 | benign | 1 | 1 | 1 |
| rs367623154 | benign | 1 | 1 | 1 |
| rs201230446 | pathogenic | 1 | 1 | 1 |
| rs267606766 | pathogenic | 0 | 1 | 0 |
| rs267606769 | pathogenic | 0 | 1 | 0 |
| rs267606767 | pathogenic | 1 | 1 | 0 |
| rs267606768 | pathogenic | 0 | 1 | 0 |
| rs61733129 | benign | 0 | 1 | 0 |
| rs201947120 | pathogenic | 1 | 1 | 1 |
| rs283413 | benign | 0 | 1 | 0 |
| rs36119840 | benign | 0 | 0 | 0 |
| rs281865419 | pathogenic | 1 | 1 | 1 |
| rs137852937 | pathogenic | 0 | 1 | 0 |
| rs2230709 | benign | 0 | 1 | 0 |
| rs6695033 | benign | 0 | 1 | 0 |
| rs180177032 | pathogenic | 1 | 1 | 1 |
| rs180177033 | pathogenic | 1 | 1 | 1 |
| rs121913351 | pathogenic | 1 | 1 | 1 |
| rs397507473 | pathogenic | 1 | 1 | 1 |
| rs121913355 | pathogenic | 1 | 1 | 1 |
| rs121913357 | pathogenic | 1 | 1 | 1 |
| rs180177036 | pathogenic | 1 | 1 | 1 |
| rs180177039 | pathogenic | 1 | 1 | 1 |
| rs180177038 | pathogenic | 1 | 1 | 1 |
| rs180177040 | pathogenic | 1 | 1 | 1 |

| | | | | |
|---|---|---|---|---|
| rs121913338 | pathogenic | 1 | 1 | 1 |
| rs121913366 | pathogenic | 1 | 1 | 1 |
| rs121913369 | pathogenic | 1 | 1 | 1 |
| rs113488022 | pathogenic | 1 | 1 | 1 |
| rs121913364 | pathogenic | 1 | 1 | 1 |
| rs180177042 | pathogenic | 1 | 1 | 1 |
| rs200461617 | benign | 0 | 0 | 0 |
| rs141609745 | benign | 0 | 0 | 0 |
| rs104894383 | pathogenic | 1 | 1 | 1 |
| rs104894384 | pathogenic | 1 | 1 | 1 |
| rs104894381 | pathogenic | 1 | 1 | 0 |
| rs147710408 | benign | 1 | 1 | 1 |
| rs77357563 | benign | 1 | 1 | 1 |
| rs104894378 | pathogenic | 0 | 0 | 0 |
| rs104894382 | pathogenic | 0 | 0 | 0 |
| rs137853292 | pathogenic | 1 | 1 | 1 |
| rs587778834 | pathogenic | 0 | 0 | 0 |
| rs137853294 | pathogenic | 1 | 1 | 1 |
| rs551747882 | benign | 0 | 0 | 0 |
| rs3092903 | benign | 1 | 1 | 1 |
| rs137853296 | pathogenic | 0 | 0 | 0 |
| rs398122917 | pathogenic | 0 | 1 | 0 |
| rs41304992 | benign | 0 | 0 | 0 |
| rs121912708 | pathogenic | 1 | 1 | 1 |
| rs140102105 | pathogenic | 1 | 1 | 1 |
| rs376917645 | pathogenic | 1 | 1 | 0 |
| rs372660425 | pathogenic | 1 | 1 | 0 |
| rs555896752 | pathogenic | 1 | 1 | 1 |
| rs777829351 | pathogenic | 1 | 1 | 0 |
| rs121912711 | pathogenic | 1 | 1 | 0 |
| rs754449549 | pathogenic | 0 | 1 | 0 |
| rs141701364 | benign | 0 | 1 | 0 |
| rs761011399 | benign | 1 | 1 | 1 |
| rs121912710 | benign | 1 | 1 | 0 |
| rs121912707 | pathogenic | 1 | 1 | 0 |
| rs747597620 | pathogenic | 0 | 1 | 0 |
| rs186558364 | pathogenic | 1 | 1 | 1 |
| rs556400964 | pathogenic | 1 | 1 | 1 |
| rs61757684 | benign | 1 | 1 | 1 |
| rs121917766 | pathogenic | 1 | 1 | 1 |
| rs121918248 | pathogenic | 0 | 0 | 0 |
| rs398123278 | pathogenic | 0 | 0 | 0 |
| rs121918251 | pathogenic | 1 | 1 | 1 |
| rs727504022 | pathogenic | 0 | 0 | 0 |
| rs190834116 | pathogenic | 1 | 1 | 1 |
| rs121918249 | pathogenic | 1 | 1 | 0 |
| rs121918257 | pathogenic | 1 | 1 | 0 |
| rs142068898 | benign | 0 | 0 | 0 |
| rs757000253 | pathogenic | 0 | 0 | 0 |
| rs200908035 | pathogenic | 1 | 1 | 0 |
| rs760782399 | pathogenic | 0 | 0 | 0 |
| rs778702777 | pathogenic | 0 | 0 | 0 |
| rs121918258 | pathogenic | 1 | 1 | 0 |
| rs121918256 | pathogenic | 0 | 0 | 0 |

| | | | | |
|---|---|---|---|---|
| rs200596762 | pathogenic | 1 | 1 | 0 |
| rs547709692 | pathogenic | 0 | 0 | 0 |
| rs758577372 | pathogenic | 0 | 1 | 0 |
| rs564069299 | pathogenic | 1 | 1 | 1 |
| rs772552898 | pathogenic | 1 | 1 | 1 |
| rs368790885 | pathogenic | 1 | 1 | 1 |
| rs121918250 | pathogenic | 0 | 0 | 0 |
| rs533755473 | pathogenic | 1 | 1 | 1 |
| rs753288303 | pathogenic | 1 | 1 | 1 |
| rs2229385 | benign | 0 | 0 | 0 |
| rs1141321 | benign | 0 | 0 | 0 |
| rs753564352 | pathogenic | 0 | 0 | 0 |
| rs775593146 | pathogenic | 0 | 0 | 0 |
| rs121918254 | pathogenic | 0 | 0 | 0 |
| rs8589 | benign | 1 | 1 | 0 |
| rs777758903 | pathogenic | 0 | 0 | 0 |
| rs140600746 | pathogenic | 1 | 1 | 0 |
| rs121918255 | pathogenic | 1 | 1 | 0 |
| rs121918252 | pathogenic | 0 | 0 | 0 |
| rs779990936 | pathogenic | 1 | 1 | 0 |
| rs121912455 | pathogenic | 1 | 1 | 1 |
| rs121912459 | pathogenic | 1 | 1 | 1 |
| rs74315452 | pathogenic | 1 | 1 | 1 |
| rs121912451 | pathogenic | 1 | 1 | 1 |
| rs79006549 | pathogenic | 0 | 0 | 0 |
| rs28934889 | benign | 0 | 1 | 0 |
| rs111033557 | benign | 1 | 1 | 1 |
| rs28934597 | pathogenic | 1 | 1 | 1 |
| rs28934596 | pathogenic | 1 | 1 | 1 |
| rs28934595 | pathogenic | 1 | 1 | 1 |
| rs146519482 | pathogenic | 1 | 1 | 1 |
| rs111033563 | pathogenic | 1 | 1 | 1 |
| rs75385605 | benign | 1 | 1 | 1 |
| rs121908569 | pathogenic | 1 | 1 | 1 |
| rs35719940 | benign | 0 | 0 | 0 |
| rs121918664 | pathogenic | 0 | 0 | 0 |
| rs104893665 | pathogenic | 1 | 1 | 1 |
| rs104893666 | pathogenic | 1 | 1 | 1 |
| rs200845457 | benign | 1 | 1 | 1 |
| rs779204655 | pathogenic | 1 | 1 | 1 |
| rs121917747 | pathogenic | 1 | 1 | 1 |
| rs6042 | benign | 1 | 1 | 1 |
| rs104894380 | pathogenic | 0 | 0 | 0 |
| rs369083541 | pathogenic | 1 | 1 | 1 |
| rs145243532 | benign | 1 | 1 | 1 |
| rs121909500 | pathogenic | 1 | 1 | 1 |
| rs140993310 | benign | 1 | 1 | 1 |
| rs200837270 | pathogenic | 1 | 1 | 1 |
| rs121918484 | pathogenic | 1 | 1 | 1 |
| rs121918306 | pathogenic | 0 | 0 | 0 |
| rs118204089 | pathogenic | 1 | 1 | 0 |
| rs118204083 | pathogenic | 1 | 1 | 0 |
| rs201953584 | pathogenic | 1 | 1 | 1 |
| rs74315362 | pathogenic | 1 | 1 | 1 |

| | | | | |
|---|---|---|---|---|
| rs118204084 | pathogenic | 1 | 1 | 0 |
| rs118204085 | pathogenic | 1 | 1 | 1 |
| rs113403872 | pathogenic | 1 | 1 | 0 |
| rs1052176 | benign | 1 | 1 | 1 |
| rs2031640 | benign | 0 | 0 | 0 |
| rs104894230 | pathogenic | 1 | 1 | 1 |
| rs104894229 | pathogenic | 1 | 1 | 1 |
| rs104894228 | pathogenic | 1 | 1 | 1 |
| rs104894226 | pathogenic | 1 | 1 | 1 |
| rs121917757 | pathogenic | 1 | 1 | 1 |
| rs121917758 | pathogenic | 1 | 1 | 1 |
| rs28933406 | pathogenic | 1 | 1 | 1 |
| rs121917756 | pathogenic | 1 | 1 | 1 |
| rs104894227 | pathogenic | 1 | 1 | 1 |
| rs369106578 | pathogenic | 1 | 1 | 1 |
| rs397517142 | benign | 1 | 1 | 1 |
| rs104894231 | pathogenic | 1 | 1 | 1 |
| rs121917759 | pathogenic | 1 | 1 | 1 |
| rs104893827 | pathogenic | 0 | 0 | 0 |
| rs104893826 | pathogenic | 1 | 1 | 1 |
| rs5030802 | pathogenic | 1 | 1 | 1 |
| rs5030827 | pathogenic | 1 | 1 | 0 |
| rs119103277 | pathogenic | 1 | 1 | 1 |
| rs5030808 | pathogenic | 0 | 1 | 0 |
| rs5030809 | pathogenic | 1 | 1 | 1 |
| rs765978945 | pathogenic | 1 | 1 | 1 |
| rs104893824 | pathogenic | 1 | 1 | 1 |
| rs104893831 | pathogenic | 0 | 1 | 0 |
| rs104893830 | pathogenic | 0 | 1 | 0 |
| rs119103278 | pathogenic | 0 | 1 | 0 |
| rs5030818 | pathogenic | 1 | 1 | 1 |
| rs397516444 | pathogenic | 1 | 1 | 1 |
| rs28940297 | pathogenic | 1 | 1 | 1 |
| rs267607170 | pathogenic | 1 | 1 | 1 |
| rs104893825 | pathogenic | 1 | 1 | 1 |
| rs5030820 | pathogenic | 0 | 1 | 0 |
| rs5030821 | pathogenic | 0 | 1 | 0 |
| rs5030823 | pathogenic | 1 | 1 | 1 |
| rs5030824 | pathogenic | 1 | 1 | 1 |
| rs28940301 | pathogenic | 0 | 1 | 0 |
| rs28940300 | pathogenic | 0 | 1 | 0 |
| rs28940298 | pathogenic | 0 | 1 | 0 |
| rs200019083 | benign | 0 | 0 | 0 |
| rs759748655 | pathogenic | 0 | 0 | 0 |
| rs121917739 | pathogenic | 1 | 1 | 1 |
| rs75353611 | pathogenic | 1 | 1 | 1 |
| rs79744198 | benign | 1 | 1 | 1 |
| rs79228041 | pathogenic | 1 | 1 | 1 |
| rs80106970 | benign | 1 | 1 | 1 |
| rs375353223 | benign | 0 | 0 | 0 |
| rs56311081 | benign | 0 | 0 | 0 |
| rs55985569 | benign | 1 | 1 | 1 |
| rs45483396 | benign | 0 | 0 | 0 |
| rs35284565 | benign | 1 | 1 | 1 |

| | | | | |
|---|---|---|---|---|
| rs34349517 | benign | 0 | 0 | 0 |
| rs35776110 | benign | 0 | 0 | 0 |
| rs33917957 | benign | 1 | 1 | 1 |
| rs587778445 | benign | 0 | 0 | 0 |
| rs34624984 | benign | 0 | 0 | 0 |
| rs701753 | benign | 0 | 0 | 0 |
| rs855314 | benign | 0 | 0 | 0 |
| rs121918371 | pathogenic | 1 | 1 | 0 |
| rs398122912 | pathogenic | 0 | 1 | 0 |
| rs1126728 | benign | 0 | 0 | 0 |
| rs587777404 | pathogenic | 1 | 1 | 0 |
| rs11208257 | benign | 0 | 0 | 0 |
| rs6676290 | benign | 0 | 0 | 0 |
| rs104894118 | pathogenic | 0 | 0 | 0 |
| rs121918655 | pathogenic | 1 | 1 | 1 |
| rs104894120 | pathogenic | 1 | 1 | 1 |
| rs137852238 | pathogenic | 0 | 0 | 0 |
| rs137852237 | pathogenic | 0 | 0 | 0 |
| rs137852499 | pathogenic | 1 | 1 | 1 |
| rs137852480 | pathogenic | 1 | 1 | 1 |
| rs137852494 | pathogenic | 1 | 1 | 1 |
| rs137852502 | pathogenic | 1 | 1 | 1 |
| rs137852500 | pathogenic | 1 | 1 | 1 |
| rs137852487 | pathogenic | 1 | 1 | 1 |
| rs137852481 | pathogenic | 1 | 1 | 1 |
| rs137852501 | pathogenic | 0 | 0 | 0 |
| rs137852478 | pathogenic | 1 | 1 | 1 |
| rs137852485 | pathogenic | 1 | 1 | 1 |
| rs137852482 | pathogenic | 1 | 1 | 0 |
| rs137852483 | pathogenic | 0 | 1 | 0 |
| rs137852477 | pathogenic | 1 | 1 | 0 |
| rs137852496 | pathogenic | 1 | 1 | 0 |
| rs137852484 | pathogenic | 0 | 1 | 0 |
| rs137852498 | pathogenic | 1 | 1 | 0 |
| rs137852497 | pathogenic | 1 | 1 | 0 |
| rs137852493 | pathogenic | 1 | 1 | 0 |
| rs137852504 | pathogenic | 1 | 1 | 1 |
| rs267606863 | pathogenic | 1 | 1 | 1 |
| rs137852486 | pathogenic | 1 | 1 | 1 |
| rs137852479 | pathogenic | 1 | 1 | 1 |
| rs112122950 | benign | 0 | 0 | 0 |
| rs72546648 | benign | 1 | 1 | 1 |
| rs72552291 | pathogenic | 0 | 0 | 0 |
| rs79184941 | pathogenic | 1 | 1 | 1 |
| rs77543610 | pathogenic | 1 | 1 | 1 |
| rs121918505 | pathogenic | 1 | 1 | 1 |
| rs776587763 | pathogenic | 1 | 1 | 1 |
| rs121918497 | pathogenic | 1 | 1 | 1 |
| rs121918499 | pathogenic | 1 | 1 | 1 |
| rs121918501 | pathogenic | 1 | 1 | 1 |
| rs121918500 | pathogenic | 0 | 0 | 0 |
| rs121918504 | pathogenic | 1 | 1 | 1 |
| rs121918510 | pathogenic | 1 | 1 | 1 |
| rs121918493 | pathogenic | 1 | 1 | 1 |

| | | | | |
|---|---|---|---|---|
| rs121918489 | pathogenic | 0 | 0 | 0 |
| rs121918495 | pathogenic | 1 | 1 | 1 |
| rs121918488 | pathogenic | 0 | 0 | 0 |
| rs121918496 | pathogenic | 0 | 0 | 0 |
| rs121918492 | pathogenic | 1 | 1 | 1 |
| rs121918494 | pathogenic | 1 | 1 | 1 |
| rs121918502 | pathogenic | 1 | 1 | 1 |
| rs121918490 | pathogenic | 1 | 1 | 1 |
| rs72466451 | pathogenic | 1 | 1 | 1 |
| rs66468541 | pathogenic | 0 | 0 | 0 |
| rs137852662 | pathogenic | 0 | 0 | 0 |
| rs137852663 | pathogenic | 0 | 0 | 0 |
| rs606231407 | pathogenic | 0 | 0 | 0 |
| rs137852233 | pathogenic | 1 | 1 | 1 |
| rs112306225 | benign | 0 | 0 | 0 |
| rs387906808 | pathogenic | 1 | 1 | 0 |
| rs730882094 | pathogenic | 1 | 1 | 1 |
| rs373869746 | pathogenic | 1 | 1 | 1 |
| rs777524402 | pathogenic | 1 | 1 | 1 |
| rs769737896 | pathogenic | 0 | 1 | 0 |
| rs13306515 | benign | 1 | 1 | 0 |
| rs746982741 | pathogenic | 0 | 0 | 0 |
| rs2230200 | benign | 1 | 1 | 1 |
| rs1047286 | benign | 0 | 0 | 0 |
| rs141737564 | benign | 0 | 0 | 0 |
| rs774826179 | benign | 0 | 0 | 0 |
| rs11569422 | benign | 1 | 1 | 1 |
| rs146613648 | benign | 0 | 0 | 0 |
| rs144432231 | benign | 0 | 0 | 0 |
| rs140655115 | benign | 0 | 0 | 0 |
| rs387906837 | pathogenic | 1 | 1 | 0 |
| rs121909076 | pathogenic | 1 | 1 | 0 |
| rs121909077 | pathogenic | 0 | 0 | 0 |
| rs121909074 | pathogenic | 0 | 0 | 0 |
| rs137853330 | pathogenic | 1 | 1 | 1 |
| rs137853331 | pathogenic | 1 | 1 | 1 |
| rs137853332 | pathogenic | 1 | 1 | 1 |
| rs28931586 | pathogenic | 1 | 1 | 0 |
| rs11569321 | benign | 1 | 1 | 1 |
| rs119486097 | benign | 0 | 0 | 0 |
| rs1799807 | pathogenic | 1 | 1 | 1 |
| rs201820739 | pathogenic | 1 | 1 | 0 |
| rs114706984 | pathogenic | 1 | 1 | 0 |
| rs140080572 | pathogenic | 1 | 1 | 0 |
| rs28933389 | pathogenic | 0 | 1 | 0 |
| rs16849700 | benign | 1 | 1 | 0 |
| rs115624085 | pathogenic | 1 | 1 | 1 |
| rs747196387 | pathogenic | 1 | 1 | 1 |
| rs104893684 | pathogenic | 1 | 1 | 0 |
| rs121918557 | pathogenic | 1 | 1 | 0 |
| rs121918556 | pathogenic | 1 | 1 | 1 |
| rs398124571 | pathogenic | 1 | 1 | 1 |
| rs776631396 | pathogenic | 0 | 1 | 0 |
| rs398124576 | pathogenic | 1 | 1 | 1 |

| | | | | |
|---|---|---|---|---|
| rs398124581 | pathogenic | 1 | 1 | 1 |
| rs398124582 | pathogenic | 1 | 1 | 1 |
| rs149766077 | pathogenic | 0 | 1 | 0 |
| rs79761867 | pathogenic | 0 | 1 | 0 |
| rs148905512 | pathogenic | 0 | 1 | 0 |
| rs121965004 | pathogenic | 1 | 1 | 1 |
| rs386834233 | pathogenic | 0 | 1 | 0 |
| rs398124598 | pathogenic | 0 | 1 | 0 |
| rs398124603 | pathogenic | 1 | 1 | 1 |
| rs546716073 | benign | 1 | 1 | 1 |
| rs200310662 | benign | 1 | 1 | 1 |
| rs115494727 | benign | 1 | 1 | 1 |
| rs781513008 | benign | 0 | 0 | 0 |
| rs587776398 | benign | 1 | 1 | 0 |
| rs35288908 | benign | 1 | 1 | 1 |
| rs150634562 | pathogenic | 1 | 1 | 0 |
| rs142948132 | pathogenic | 1 | 1 | 1 |
| rs132630321 | pathogenic | 1 | 1 | 1 |
| rs132630317 | pathogenic | 1 | 1 | 1 |
| rs132630320 | pathogenic | 0 | 0 | 0 |
| rs781394318 | pathogenic | 1 | 1 | 1 |
| rs111033624 | pathogenic | 1 | 1 | 1 |
| rs111033630 | pathogenic | 1 | 1 | 1 |
| rs111033627 | pathogenic | 1 | 1 | 1 |
| rs193922287 | pathogenic | 1 | 1 | 1 |
| rs759072800 | pathogenic | 0 | 1 | 0 |
| rs104894010 | pathogenic | 0 | 0 | 0 |
| rs193922296 | pathogenic | 0 | 0 | 0 |
| rs104894006 | pathogenic | 1 | 1 | 1 |
| rs193922311 | pathogenic | 1 | 1 | 0 |
| rs80356654 | pathogenic | 1 | 1 | 0 |
| rs144723656 | benign | 1 | 1 | 1 |
| rs148311934 | pathogenic | 1 | 1 | 0 |
| rs80356655 | pathogenic | 1 | 1 | 1 |
| rs769268803 | pathogenic | 1 | 1 | 0 |
| rs104894008 | pathogenic | 0 | 1 | 0 |
| rs104894005 | pathogenic | 1 | 1 | 1 |
| rs104894009 | pathogenic | 1 | 1 | 0 |
| rs193922340 | pathogenic | 0 | 0 | 0 |
| rs104894016 | pathogenic | 0 | 1 | 0 |
| rs193921338 | pathogenic | 0 | 0 | 0 |
| rs193922272 | pathogenic | 1 | 1 | 1 |
| rs104894012 | pathogenic | 1 | 1 | 0 |
| rs193922285 | pathogenic | 1 | 1 | 1 |
| rs104894336 | pathogenic | 1 | 1 | 1 |
| rs28931580 | pathogenic | 1 | 1 | 1 |
| rs104894326 | pathogenic | 1 | 1 | 1 |
| rs104894331 | pathogenic | 0 | 0 | 0 |
| rs104894338 | pathogenic | 0 | 0 | 0 |
| rs104894333 | pathogenic | 1 | 1 | 1 |
| rs104894330 | pathogenic | 1 | 1 | 1 |
| rs143886391 | benign | 1 | 1 | 1 |
| rs104894335 | pathogenic | 1 | 1 | 1 |
| rs104894337 | pathogenic | 0 | 0 | 0 |

| | | | | |
|---|---|---|---|---|
| rs104894328 | pathogenic | 0 | 0 | 0 |
| rs104894341 | pathogenic | 1 | 1 | 0 |
| rs104894329 | pathogenic | 0 | 0 | 0 |
| rs137852850 | pathogenic | 1 | 1 | 1 |
| rs137852854 | pathogenic | 0 | 1 | 0 |
| rs137852849 | pathogenic | 0 | 1 | 0 |
| rs137852848 | pathogenic | 1 | 1 | 1 |
| rs137852852 | pathogenic | 0 | 1 | 0 |
| rs137852853 | pathogenic | 0 | 1 | 0 |
| rs137852847 | pathogenic | 0 | 1 | 0 |
| rs137852846 | pathogenic | 0 | 1 | 0 |
| rs137852844 | pathogenic | 0 | 1 | 0 |
| rs144411158 | pathogenic | 0 | 0 | 0 |
| rs2302234 | benign | 0 | 0 | 0 |
| rs139620139 | pathogenic | 0 | 0 | 0 |
| rs146689929 | benign | 0 | 0 | 0 |
| rs373328681 | pathogenic | 1 | 1 | 0 |
| rs387906620 | pathogenic | 1 | 1 | 1 |
| rs1799983 | benign | 0 | 1 | 0 |
| rs28999111 | pathogenic | 1 | 1 | 1 |
| rs104894544 | pathogenic | 1 | 1 | 1 |
| rs104894546 | pathogenic | 1 | 1 | 1 |
| rs28999112 | pathogenic | 0 | 1 | 0 |
| rs104894545 | pathogenic | 1 | 1 | 1 |
| rs104894547 | pathogenic | 1 | 1 | 1 |
| rs74315310 | pathogenic | 0 | 0 | 0 |
| rs267606852 | pathogenic | 0 | 1 | 0 |
| rs137853586 | pathogenic | 0 | 1 | 0 |
| rs137853582 | pathogenic | 1 | 1 | 1 |
| rs61754634 | pathogenic | 1 | 1 | 1 |
| rs137853587 | pathogenic | 0 | 1 | 0 |
| rs267606851 | pathogenic | 1 | 1 | 1 |
| rs267606853 | pathogenic | 1 | 1 | 1 |
| rs137853584 | pathogenic | 1 | 1 | 1 |
| rs137853585 | pathogenic | 1 | 1 | 1 |
| rs460184 | pathogenic | 1 | 1 | 1 |
| rs104893751 | pathogenic | 0 | 1 | 0 |
| rs77358162 | benign | 0 | 0 | 0 |
| rs387906936 | pathogenic | 0 | 0 | 0 |
| rs769525399 | pathogenic | 0 | 0 | 0 |
| rs371271054 | pathogenic | 1 | 1 | 0 |
| rs118204000 | pathogenic | 0 | 0 | 0 |
| rs138568622 | pathogenic | 1 | 1 | 1 |
| rs34509359 | benign | 0 | 0 | 0 |
| rs104894130 | pathogenic | 1 | 1 | 0 |
| rs281865424 | pathogenic | 0 | 0 | 0 |
| rs121912778 | pathogenic | 1 | 1 | 0 |
| rs771055145 | pathogenic | 1 | 1 | 0 |
| rs776174514 | pathogenic | 1 | 1 | 0 |
| rs545986367 | pathogenic | 1 | 1 | 1 |
| rs570768038 | benign | 1 | 1 | 1 |
| rs74315442 | pathogenic | 1 | 1 | 1 |
| rs79716074 | benign | 1 | 1 | 1 |
| rs748486078 | pathogenic | 1 | 1 | 1 |

| | | | | |
|---|---|---|---|---|
| rs2397084 | benign | 1 | 1 | 1 |
| rs11465553 | benign | 1 | 1 | 1 |
| rs144576902 | benign | 1 | 1 | 1 |
| rs763780 | benign | 1 | 1 | 1 |
| rs267606870 | pathogenic | 1 | 1 | 0 |
| rs121913502 | pathogenic | 1 | 1 | 0 |
| rs118101777 | benign | 1 | 1 | 1 |
| rs118053940 | benign | 0 | 0 | 0 |
| rs368771578 | benign | 1 | 1 | 1 |
| rs11540654 | pathogenic | 1 | 1 | 1 |
| rs121912658 | pathogenic | 1 | 1 | 1 |
| rs28934873 | pathogenic | 0 | 1 | 0 |
| rs780442292 | pathogenic | 1 | 1 | 1 |
| rs758781593 | benign | 1 | 1 | 1 |
| rs28934875 | pathogenic | 1 | 1 | 1 |
| rs28934874 | pathogenic | 1 | 1 | 0 |
| rs587782705 | pathogenic | 1 | 1 | 1 |
| rs587780068 | pathogenic | 1 | 1 | 1 |
| rs587782144 | pathogenic | 1 | 1 | 1 |
| rs138729528 | pathogenic | 1 | 1 | 1 |
| rs28934578 | pathogenic | 1 | 1 | 1 |
| rs397514495 | pathogenic | 1 | 1 | 1 |
| rs121912665 | pathogenic | 1 | 1 | 1 |
| rs397516435 | pathogenic | 1 | 1 | 1 |
| rs587778720 | pathogenic | 1 | 1 | 1 |
| rs397516436 | pathogenic | 1 | 1 | 1 |
| rs587781386 | benign | 1 | 1 | 1 |
| rs121912666 | pathogenic | 1 | 1 | 0 |
| rs587780073 | pathogenic | 1 | 1 | 1 |
| rs587782664 | pathogenic | 1 | 1 | 1 |
| rs730882005 | pathogenic | 1 | 1 | 1 |
| rs28934573 | pathogenic | 1 | 1 | 1 |
| rs121912655 | pathogenic | 1 | 1 | 1 |
| rs28934575 | pathogenic | 1 | 1 | 1 |
| rs121912656 | pathogenic | 1 | 1 | 1 |
| rs11540652 | pathogenic | 1 | 1 | 1 |
| rs121912651 | pathogenic | 1 | 1 | 1 |
| rs730882027 | pathogenic | 1 | 1 | 1 |
| rs121912653 | pathogenic | 1 | 1 | 1 |
| rs121912652 | pathogenic | 1 | 1 | 1 |
| rs121912657 | pathogenic | 0 | 1 | 0 |
| rs121913343 | pathogenic | 1 | 1 | 1 |
| rs28934576 | pathogenic | 1 | 1 | 1 |
| rs763098116 | pathogenic | 1 | 1 | 1 |
| rs121912660 | pathogenic | 1 | 1 | 1 |
| rs28934574 | pathogenic | 1 | 1 | 1 |
| rs371409680 | pathogenic | 1 | 1 | 1 |
| rs121912667 | pathogenic | 1 | 1 | 1 |
| rs121912663 | pathogenic | 1 | 1 | 1 |
| rs201744589 | pathogenic | 0 | 0 | 0 |
| rs33926449 | benign | 0 | 0 | 0 |
| rs74315302 | pathogenic | 0 | 1 | 0 |
| rs202040269 | pathogenic | 1 | 1 | 1 |
| rs375958814 | pathogenic | 1 | 1 | 1 |

| | | | | |
|---|---|---|---|---|
| rs121909624 | pathogenic | 1 | 1 | 1 |
| rs121909621 | pathogenic | 1 | 1 | 1 |
| rs121909622 | pathogenic | 1 | 1 | 0 |
| rs4220 | benign | 1 | 1 | 1 |
| rs147559466 | pathogenic | 1 | 1 | 1 |
| rs757434857 | pathogenic | 1 | 1 | 1 |
| rs398123072 | pathogenic | 1 | 1 | 1 |
| rs121434280 | pathogenic | 0 | 1 | 0 |
| rs398123074 | pathogenic | 0 | 1 | 0 |
| rs762114560 | pathogenic | 1 | 1 | 1 |
| rs121434283 | pathogenic | 1 | 1 | 0 |
| rs121434277 | pathogenic | 0 | 1 | 0 |
| rs745844469 | pathogenic | 1 | 1 | 1 |
| rs121434279 | pathogenic | 1 | 1 | 0 |
| rs121434278 | pathogenic | 1 | 1 | 1 |
| rs373715782 | pathogenic | 1 | 1 | 1 |
| rs200724875 | pathogenic | 1 | 1 | 1 |
| rs149678400 | pathogenic | 0 | 1 | 0 |
| rs770273135 | pathogenic | 1 | 1 | 1 |
| rs121434276 | pathogenic | 0 | 1 | 0 |
| rs121434281 | pathogenic | 1 | 1 | 0 |
| rs768884003 | pathogenic | 1 | 1 | 1 |
| rs201375579 | pathogenic | 0 | 0 | 0 |
| rs121434274 | pathogenic | 1 | 1 | 0 |
| rs121434282 | pathogenic | 1 | 1 | 0 |
| rs779759347 | pathogenic | 0 | 1 | 0 |
| rs77931234 | pathogenic | 1 | 1 | 1 |
| rs148207467 | pathogenic | 1 | 1 | 1 |
| rs766140986 | pathogenic | 1 | 1 | 1 |
| rs121434275 | pathogenic | 1 | 1 | 1 |
| rs748110745 | pathogenic | 1 | 1 | 1 |
| rs104894274 | pathogenic | 1 | 1 | 1 |
| rs104894273 | pathogenic | 0 | 1 | 0 |
| rs104894278 | pathogenic | 1 | 1 | 1 |
| rs104894275 | pathogenic | 1 | 1 | 1 |
| rs104894277 | pathogenic | 1 | 1 | 1 |
| rs104894276 | pathogenic | 0 | 0 | 0 |
| rs104894280 | pathogenic | 1 | 1 | 1 |
| rs145882709 | pathogenic | 1 | 1 | 1 |
| rs104894279 | pathogenic | 1 | 1 | 1 |
| rs80356670 | pathogenic | 1 | 1 | 1 |
| rs121918102 | pathogenic | 1 | 1 | 1 |
| rs80356671 | pathogenic | 1 | 1 | 1 |
| rs80356672 | pathogenic | 1 | 1 | 1 |
| rs758609113 | pathogenic | 1 | 1 | 0 |
| rs121918188 | pathogenic | 0 | 1 | 0 |
| rs121918189 | pathogenic | 1 | 1 | 0 |
| rs121918191 | pathogenic | 1 | 1 | 1 |
| rs1769259 | benign | 1 | 1 | 1 |
| rs140222720 | benign | 1 | 1 | 1 |
| rs2297085 | benign | 1 | 1 | 1 |
| rs200679026 | pathogenic | 1 | 1 | 1 |
| rs121918192 | pathogenic | 0 | 1 | 0 |
| rs145771944 | benign | 0 | 1 | 0 |

| | | | | |
|---|---|---|---|---|
| rs201508880 | pathogenic | 0 | 1 | 0 |
| rs17221854 | pathogenic | 0 | 0 | 0 |
| rs120074189 | pathogenic | 1 | 1 | 1 |
| rs120074190 | pathogenic | 1 | 1 | 1 |
| rs758600283 | benign | 1 | 1 | 1 |
| rs200305064 | pathogenic | 1 | 1 | 1 |
| rs121917891 | pathogenic | 1 | 1 | 1 |
| rs3772809 | benign | 0 | 1 | 0 |
| rs387906619 | pathogenic | 1 | 1 | 0 |
| rs2304681 | benign | 0 | 1 | 0 |
| rs121917849 | pathogenic | 0 | 0 | 0 |
| rs143010236 | pathogenic | 0 | 0 | 0 |
| rs121917850 | pathogenic | 1 | 1 | 0 |
| rs35916840 | pathogenic | 1 | 1 | 0 |
| rs28939688 | pathogenic | 1 | 1 | 1 |
| rs11258194 | benign | 1 | 1 | 1 |
| rs199840952 | pathogenic | 1 | 1 | 1 |
| rs121918159 | pathogenic | 1 | 1 | 1 |
| rs200396166 | pathogenic | 0 | 0 | 0 |
| rs757966746 | pathogenic | 0 | 1 | 0 |
| rs121918608 | pathogenic | 0 | 1 | 0 |
| rs74315401 | pathogenic | 1 | 1 | 1 |
| rs11538758 | pathogenic | 0 | 0 | 0 |
| rs74315402 | pathogenic | 1 | 1 | 1 |
| rs1799990 | pathogenic | 1 | 1 | 1 |
| rs74315410 | pathogenic | 1 | 1 | 1 |
| rs150351644 | benign | 1 | 1 | 1 |
| rs16990018 | benign | 1 | 1 | 1 |
| rs74315403 | pathogenic | 1 | 1 | 1 |
| rs74315408 | pathogenic | 1 | 1 | 1 |
| rs74315411 | pathogenic | 1 | 1 | 1 |
| rs74315413 | pathogenic | 1 | 1 | 1 |
| rs74315405 | pathogenic | 1 | 1 | 1 |
| rs28933385 | pathogenic | 1 | 1 | 1 |
| rs74315412 | pathogenic | 1 | 1 | 1 |
| rs74315407 | pathogenic | 1 | 1 | 1 |
| rs74315406 | pathogenic | 1 | 1 | 1 |
| rs11571117 | benign | 1 | 1 | 1 |
| rs121917742 | pathogenic | 1 | 1 | 1 |
| rs149782619 | pathogenic | 1 | 1 | 1 |
| rs397514492 | pathogenic | 1 | 1 | 1 |
| rs397514489 | pathogenic | 0 | 1 | 0 |
| rs397514493 | pathogenic | 1 | 1 | 1 |
| rs373849532 | pathogenic | 1 | 1 | 1 |
| rs121965048 | pathogenic | 1 | 1 | 1 |
| rs121965037 | pathogenic | 1 | 1 | 1 |
| rs121965060 | pathogenic | 0 | 1 | 0 |
| rs386833603 | pathogenic | 1 | 1 | 1 |
| rs121965038 | pathogenic | 1 | 1 | 1 |
| rs121965039 | pathogenic | 1 | 1 | 1 |
| rs121965040 | pathogenic | 1 | 1 | 1 |
| rs267606925 | pathogenic | 1 | 1 | 1 |
| rs121965056 | pathogenic | 0 | 1 | 0 |
| rs121965059 | pathogenic | 0 | 0 | 0 |

| | | | | |
|---|---|---|---|---|
| rs386833615 | pathogenic | 1 | 1 | 0 |
| rs121965051 | pathogenic | 1 | 1 | 0 |
| rs121965046 | pathogenic | 0 | 1 | 0 |
| rs121965052 | pathogenic | 0 | 1 | 0 |
| rs386833617 | pathogenic | 0 | 1 | 0 |
| rs386833618 | pathogenic | 1 | 1 | 0 |
| rs121965041 | pathogenic | 0 | 1 | 0 |
| rs121965042 | pathogenic | 0 | 1 | 0 |
| rs121965057 | pathogenic | 1 | 1 | 1 |
| rs386833621 | pathogenic | 1 | 1 | 1 |
| rs121965049 | pathogenic | 1 | 1 | 1 |
| rs386833623 | pathogenic | 1 | 1 | 1 |
| rs121965047 | pathogenic | 1 | 1 | 1 |
| rs121965053 | pathogenic | 0 | 1 | 0 |
| rs121965045 | pathogenic | 0 | 1 | 0 |
| rs121965054 | pathogenic | 0 | 1 | 0 |
| rs121965043 | pathogenic | 0 | 1 | 0 |
| rs386833598 | pathogenic | 0 | 0 | 0 |
| rs137854545 | pathogenic | 1 | 1 | 1 |
| rs137854544 | pathogenic | 1 | 1 | 1 |
| rs104894455 | pathogenic | 0 | 1 | 0 |
| rs1049564 | benign | 0 | 1 | 0 |
| rs104894460 | pathogenic | 1 | 1 | 1 |
| rs104894453 | pathogenic | 1 | 1 | 1 |
| rs104894450 | pathogenic | 0 | 1 | 0 |
| rs104894454 | pathogenic | 0 | 1 | 0 |
| rs104894452 | pathogenic | 1 | 1 | 1 |
| rs104894451 | pathogenic | 0 | 1 | 0 |
| rs104894368 | pathogenic | 1 | 1 | 1 |
| rs730880950 | pathogenic | 1 | 1 | 1 |
| rs121909535 | pathogenic | 1 | 1 | 1 |
| rs121909537 | pathogenic | 1 | 1 | 1 |
| rs121909536 | pathogenic | 1 | 1 | 1 |
| rs121909542 | pathogenic | 1 | 1 | 1 |
| rs121909538 | pathogenic | 1 | 1 | 1 |
| rs121909539 | pathogenic | 1 | 1 | 1 |
| rs121909540 | pathogenic | 1 | 1 | 1 |
| rs17560 | benign | 1 | 1 | 1 |
| rs121909543 | pathogenic | 0 | 1 | 0 |
| rs121909544 | pathogenic | 1 | 1 | 1 |
| rs2307456 | benign | 0 | 1 | 0 |
| rs745778317 | pathogenic | 0 | 1 | 0 |
| rs759607901 | pathogenic | 0 | 1 | 0 |
| rs201188114 | benign | 0 | 0 | 0 |
| rs1800472 | benign | 0 | 0 | 0 |
| rs587780728 | benign | 1 | 1 | 1 |
| rs144386518 | benign | 1 | 1 | 1 |
| rs281865439 | pathogenic | 1 | 1 | 1 |
| rs140175796 | pathogenic | 1 | 1 | 1 |
| rs5030843 | pathogenic | 1 | 1 | 0 |
| rs75166491 | pathogenic | 1 | 1 | 0 |
| rs199475679 | pathogenic | 1 | 1 | 1 |
| rs281865440 | pathogenic | 1 | 1 | 1 |
| rs74486803 | pathogenic | 0 | 1 | 0 |

| | | | | |
|---|---|---|---|---|
| rs199475602 | pathogenic | 0 | 1 | 0 |
| rs5030844 | pathogenic | 1 | 1 | 0 |
| rs62514927 | pathogenic | 1 | 1 | 1 |
| rs62514928 | pathogenic | 1 | 1 | 1 |
| rs62516152 | pathogenic | 0 | 1 | 0 |
| rs62507348 | pathogenic | 1 | 1 | 1 |
| rs76687508 | pathogenic | 1 | 1 | 1 |
| rs62508730 | pathogenic | 1 | 1 | 1 |
| rs62508588 | pathogenic | 0 | 1 | 0 |
| rs5030846 | pathogenic | 0 | 1 | 0 |
| rs74503222 | pathogenic | 1 | 1 | 0 |
| rs5030847 | pathogenic | 1 | 1 | 0 |
| rs5030850 | pathogenic | 1 | 1 | 1 |
| rs778154939 | pathogenic | 0 | 1 | 0 |
| rs62514952 | pathogenic | 1 | 1 | 1 |
| rs78655458 | pathogenic | 1 | 1 | 1 |
| rs62508698 | pathogenic | 1 | 1 | 0 |
| rs5030851 | pathogenic | 1 | 1 | 0 |
| rs62642933 | pathogenic | 1 | 1 | 1 |
| rs5030853 | pathogenic | 1 | 1 | 1 |
| rs62642935 | pathogenic | 1 | 1 | 1 |
| rs199475615 | pathogenic | 0 | 0 | 0 |
| rs62516062 | pathogenic | 1 | 1 | 0 |
| rs62516092 | pathogenic | 0 | 0 | 0 |
| rs62516095 | pathogenic | 1 | 1 | 1 |
| rs62642937 | pathogenic | 1 | 1 | 1 |
| rs62516141 | pathogenic | 0 | 1 | 0 |
| rs5030856 | pathogenic | 1 | 1 | 1 |
| rs5030857 | pathogenic | 1 | 1 | 1 |
| rs5030858 | pathogenic | 1 | 1 | 1 |
| rs79931499 | pathogenic | 1 | 1 | 1 |
| rs5030860 | pathogenic | 1 | 1 | 1 |
| rs281865437 | pathogenic | 1 | 1 | 1 |
| rs62644499 | pathogenic | 1 | 1 | 1 |
| rs147689487 | benign | 0 | 0 | 0 |
| rs397515417 | pathogenic | 1 | 1 | 1 |
| rs121909332 | pathogenic | 1 | 1 | 1 |
| rs121909330 | pathogenic | 1 | 1 | 0 |
| rs121909329 | pathogenic | 1 | 1 | 0 |
| rs121909335 | pathogenic | 1 | 1 | 1 |
| rs387906789 | pathogenic | 1 | 1 | 1 |
| rs121909334 | pathogenic | 1 | 1 | 1 |
| rs121909331 | pathogenic | 1 | 1 | 1 |
| rs372487178 | pathogenic | 1 | 1 | 1 |
| rs387906937 | pathogenic | 1 | 1 | 0 |
| rs16845 | benign | 1 | 1 | 1 |
| rs672601313 | pathogenic | 1 | 1 | 1 |
| rs117805308 | benign | 0 | 0 | 0 |
| rs121434261 | pathogenic | 1 | 1 | 1 |
| rs141629512 | benign | 0 | 0 | 0 |
| rs199957176 | benign | 1 | 1 | 1 |
| rs2230461 | benign | 1 | 1 | 1 |
| rs764618040 | pathogenic | 1 | 1 | 1 |
| rs267606949 | pathogenic | 1 | 1 | 1 |

| | | | | |
|---|---|---|---|---|
| rs587777483 | pathogenic | 1 | 1 | 1 |
| rs267606947 | pathogenic | 1 | 1 | 1 |
| rs6941583 | benign | 0 | 0 | 0 |
| rs749597959 | benign | 1 | 1 | 1 |
| rs182340753 | benign | 1 | 1 | 1 |
| rs146678221 | benign | 0 | 0 | 0 |
| rs11539202 | benign | 0 | 0 | 0 |
| rs148723565 | benign | 0 | 0 | 0 |
| rs61752925 | benign | 1 | 1 | 1 |
| rs113309941 | pathogenic | 0 | 0 | 0 |
| rs121434520 | pathogenic | 1 | 1 | 1 |
| rs121434517 | pathogenic | 1 | 1 | 1 |
| rs267607027 | pathogenic | 1 | 1 | 1 |
| rs138603088 | pathogenic | 0 | 0 | 0 |
| rs121434516 | pathogenic | 1 | 1 | 0 |
| rs121434521 | pathogenic | 0 | 0 | 0 |
| rs145735112 | benign | 1 | 1 | 1 |
| rs121434519 | pathogenic | 1 | 1 | 1 |
| rs75326924 | pathogenic | 1 | 1 | 0 |
| rs142186404 | pathogenic | 0 | 0 | 0 |
| rs121918035 | pathogenic | 0 | 0 | 0 |
| rs374385878 | pathogenic | 1 | 1 | 0 |
| rs11557652 | benign | 0 | 0 | 0 |
| rs118203986 | pathogenic | 1 | 1 | 1 |
| rs141675330 | benign | 1 | 1 | 1 |
| rs104893927 | pathogenic | 0 | 0 | 0 |
| rs104893935 | pathogenic | 0 | 0 | 0 |
| rs104893933 | pathogenic | 1 | 1 | 0 |
| rs104893934 | pathogenic | 0 | 0 | 0 |
| rs145183203 | benign | 1 | 1 | 0 |
| rs2230806 | benign | 0 | 0 | 0 |
| rs9282541 | benign | 0 | 0 | 0 |
| rs559110055 | benign | 0 | 0 | 0 |
| rs200030513 | benign | 0 | 0 | 0 |
| rs9282543 | benign | 0 | 0 | 0 |
| rs147675550 | benign | 0 | 0 | 0 |
| rs137854496 | pathogenic | 0 | 0 | 0 |
| rs374190304 | benign | 0 | 0 | 0 |
| rs114620717 | benign | 0 | 0 | 0 |
| rs2066718 | benign | 0 | 0 | 0 |
| rs35819696 | benign | 0 | 0 | 0 |
| rs138880920 | benign | 0 | 0 | 0 |
| rs2066715 | benign | 0 | 0 | 0 |
| rs35207495 | benign | 0 | 0 | 0 |
| rs2066714 | benign | 0 | 0 | 0 |
| rs28937313 | pathogenic | 0 | 0 | 0 |
| rs28937314 | pathogenic | 0 | 0 | 0 |
| rs137854495 | pathogenic | 0 | 0 | 0 |
| rs140365800 | benign | 0 | 0 | 0 |
| rs13306073 | benign | 0 | 0 | 0 |
| rs28933692 | pathogenic | 0 | 0 | 0 |
| rs33918808 | benign | 0 | 0 | 0 |
| rs76881554 | benign | 0 | 0 | 0 |
| rs143180998 | benign | 0 | 0 | 0 |

| | | | | |
|---|---|---|---|---|
| rs137854500 | pathogenic | 0 | 0 | 0 |
| rs538614702 | benign | 0 | 0 | 0 |
| rs201879964 | benign | 0 | 0 | 0 |
| rs2230808 | benign | 0 | 0 | 0 |
| rs138422574 | benign | 0 | 0 | 0 |
| rs137854498 | pathogenic | 0 | 0 | 0 |
| rs150125857 | benign | 0 | 0 | 0 |
| rs146292819 | pathogenic | 0 | 0 | 0 |
| rs142688906 | benign | 0 | 0 | 0 |
| rs137854499 | pathogenic | 0 | 0 | 0 |
| rs34879708 | benign | 0 | 0 | 0 |
| rs121913028 | pathogenic | 1 | 1 | 1 |
| rs1041951 | benign | 1 | 1 | 0 |
| rs138714094 | benign | 1 | 1 | 0 |
| rs200650502 | benign | 0 | 1 | 0 |
| rs370708663 | pathogenic | 1 | 1 | 1 |
| rs118204040 | pathogenic | 0 | 1 | 0 |
| rs146899669 | pathogenic | 1 | 1 | 0 |
| rs118204039 | pathogenic | 0 | 1 | 0 |
| rs121909233 | pathogenic | 0 | 0 | 0 |
| rs121909225 | pathogenic | 1 | 1 | 0 |
| rs121909236 | pathogenic | 0 | 0 | 0 |
| rs398123317 | pathogenic | 0 | 0 | 0 |
| rs121909226 | pathogenic | 1 | 1 | 0 |
| rs121909238 | pathogenic | 1 | 1 | 0 |
| rs121909230 | pathogenic | 0 | 0 | 0 |
| rs121909237 | pathogenic | 0 | 0 | 0 |
| rs121909222 | pathogenic | 1 | 1 | 0 |
| rs121909223 | pathogenic | 1 | 1 | 0 |
| rs121909218 | pathogenic | 1 | 1 | 0 |
| rs121909229 | pathogenic | 1 | 1 | 0 |
| rs121909224 | pathogenic | 1 | 1 | 0 |
| rs121909241 | pathogenic | 0 | 0 | 0 |
| rs121909221 | pathogenic | 0 | 0 | 0 |
| rs121913294 | pathogenic | 0 | 0 | 0 |
| rs121909234 | pathogenic | 0 | 0 | 0 |
| rs121909240 | pathogenic | 0 | 0 | 0 |
| rs767623493 | benign | 0 | 0 | 0 |
| rs121909239 | pathogenic | 0 | 0 | 0 |
| rs120074145 | pathogenic | 1 | 1 | 1 |
| rs120074148 | pathogenic | 1 | 1 | 1 |
| rs762991875 | pathogenic | 0 | 0 | 0 |
| rs148639841 | pathogenic | 1 | 1 | 1 |
| rs199524907 | pathogenic | 1 | 1 | 1 |
| rs120074141 | pathogenic | 1 | 1 | 1 |
| rs120074144 | pathogenic | 0 | 1 | 0 |
| rs120074146 | pathogenic | 1 | 1 | 1 |
| rs120074147 | pathogenic | 0 | 1 | 0 |
| rs120074143 | pathogenic | 0 | 1 | 0 |
| rs120074140 | pathogenic | 1 | 1 | 0 |
| rs28937315 | pathogenic | 1 | 1 | 1 |
| rs76066357 | benign | 1 | 1 | 1 |
| rs137852911 | pathogenic | 0 | 1 | 0 |
| rs137852907 | pathogenic | 1 | 1 | 0 |

| | | | | |
|---|---|---|---|---|
| rs137852910 | pathogenic | 1 | 1 | 1 |
| rs537585211 | benign | 0 | 0 | 0 |
| rs202134424 | pathogenic | 0 | 0 | 0 |
| rs200930463 | pathogenic | 0 | 0 | 0 |
| rs587777241 | pathogenic | 1 | 1 | 0 |
| rs77563738 | pathogenic | 0 | 0 | 0 |
| rs750868279 | pathogenic | 1 | 1 | 0 |
| rs148936893 | pathogenic | 0 | 0 | 0 |
| rs146597836 | pathogenic | 0 | 0 | 0 |
| rs775440641 | pathogenic | 1 | 1 | 1 |
| rs770689762 | pathogenic | 0 | 1 | 0 |
| rs376785840 | pathogenic | 0 | 0 | 0 |
| rs137852531 | pathogenic | 0 | 0 | 0 |
| rs137852538 | pathogenic | 1 | 1 | 0 |
| rs137852529 | pathogenic | 0 | 1 | 0 |
| rs137852528 | pathogenic | 0 | 1 | 0 |
| rs137852535 | pathogenic | 1 | 1 | 0 |
| rs137853045 | pathogenic | 0 | 1 | 0 |
| rs137853046 | pathogenic | 1 | 1 | 1 |
| rs137853043 | pathogenic | 1 | 1 | 1 |
| rs137853044 | pathogenic | 1 | 1 | 1 |
| rs137853047 | pathogenic | 1 | 1 | 1 |
| rs267606990 | pathogenic | 1 | 1 | 1 |
| rs751437780 | pathogenic | 1 | 1 | 0 |
| rs121918461 | pathogenic | 1 | 1 | 1 |
| rs121918460 | pathogenic | 1 | 1 | 1 |
| rs121918459 | pathogenic | 1 | 1 | 0 |
| rs397507511 | pathogenic | 1 | 1 | 1 |
| rs121918454 | pathogenic | 1 | 1 | 1 |
| rs121918453 | pathogenic | 1 | 1 | 1 |
| rs121918462 | pathogenic | 1 | 1 | 0 |
| rs121918465 | pathogenic | 1 | 1 | 0 |
| rs121918464 | pathogenic | 1 | 1 | 0 |
| rs121918466 | pathogenic | 1 | 1 | 1 |
| rs397507521 | pathogenic | 1 | 1 | 1 |
| rs397507527 | pathogenic | 1 | 1 | 1 |
| rs777603059 | pathogenic | 0 | 1 | 0 |
| rs121918456 | pathogenic | 1 | 1 | 1 |
| rs121918463 | pathogenic | 1 | 1 | 1 |
| rs121918455 | pathogenic | 0 | 1 | 0 |
| rs28933386 | pathogenic | 0 | 1 | 0 |
| rs774356443 | benign | 1 | 1 | 1 |
| rs121918467 | pathogenic | 1 | 1 | 1 |
| rs121918468 | pathogenic | 1 | 1 | 1 |
| rs121918469 | pathogenic | 1 | 1 | 1 |
| rs397507539 | pathogenic | 1 | 1 | 0 |
| rs121918458 | pathogenic | 1 | 1 | 1 |
| rs121918470 | pathogenic | 1 | 1 | 1 |
| rs35289681 | benign | 0 | 1 | 0 |
| rs4302331 | benign | 0 | 1 | 0 |
| rs11544803 | pathogenic | 1 | 1 | 1 |
| rs116254156 | benign | 0 | 0 | 0 |
| rs104893909 | pathogenic | 1 | 1 | 1 |
| rs104893911 | pathogenic | 1 | 1 | 1 |

| | | | | |
|---|---|---|---|---|
| rs104893908 | pathogenic | 1 | 1 | 0 |
| rs258751 | benign | 1 | 1 | 1 |
| rs104893914 | pathogenic | 1 | 1 | 0 |
| rs121909727 | pathogenic | 1 | 1 | 1 |
| rs104893910 | pathogenic | 1 | 1 | 1 |
| rs121909726 | pathogenic | 1 | 1 | 1 |
| rs104893912 | pathogenic | 1 | 1 | 1 |
| rs138407155 | benign | 1 | 1 | 1 |
| rs574061789 | benign | 1 | 1 | 1 |
| rs121912718 | pathogenic | 1 | 1 | 1 |
| rs121912719 | pathogenic | 1 | 1 | 1 |
| rs140770089 | benign | 1 | 1 | 1 |
| rs28931573 | pathogenic | 1 | 1 | 1 |
| rs121912717 | pathogenic | 1 | 1 | 1 |
| rs376456598 | pathogenic | 0 | 0 | 0 |
| rs398123499 | pathogenic | 1 | 1 | 1 |
| rs773048903 | pathogenic | 1 | 1 | 0 |
| rs398123503 | pathogenic | 1 | 1 | 1 |
| rs375785084 | pathogenic | 0 | 1 | 0 |
| rs137852874 | pathogenic | 1 | 1 | 1 |
| rs373713279 | pathogenic | 1 | 1 | 1 |
| rs137852873 | pathogenic | 1 | 1 | 0 |
| rs367823977 | pathogenic | 1 | 1 | 1 |
| rs398123508 | pathogenic | 1 | 1 | 1 |
| rs137852871 | pathogenic | 1 | 1 | 1 |
| rs753698250 | pathogenic | 0 | 1 | 0 |
| rs182923857 | pathogenic | 1 | 1 | 1 |
| rs148090804 | benign | 1 | 1 | 1 |
| rs137852870 | pathogenic | 1 | 1 | 1 |
| rs267607131 | pathogenic | 1 | 1 | 0 |
| rs137852902 | pathogenic | 0 | 0 | 0 |
| rs61741646 | benign | 1 | 1 | 1 |
| rs137852905 | pathogenic | 0 | 0 | 0 |
| rs121918681 | pathogenic | 0 | 1 | 0 |
| rs8177238 | benign | 1 | 1 | 0 |
| rs5030776 | pathogenic | 1 | 1 | 1 |
| rs104894572 | pathogenic | 0 | 0 | 0 |
| rs104894576 | pathogenic | 0 | 0 | 0 |
| rs80084721 | pathogenic | 0 | 0 | 0 |
| rs104894146 | pathogenic | 0 | 1 | 0 |
| rs104894138 | pathogenic | 1 | 1 | 0 |
| rs104894153 | pathogenic | 1 | 1 | 0 |
| rs104894135 | pathogenic | 1 | 1 | 0 |
| rs104894147 | pathogenic | 1 | 1 | 0 |
| rs104894148 | pathogenic | 1 | 1 | 0 |
| rs104894154 | pathogenic | 1 | 1 | 0 |
| rs104894150 | pathogenic | 1 | 1 | 0 |
| rs104894136 | pathogenic | 1 | 1 | 0 |
| rs104894144 | pathogenic | 0 | 0 | 0 |
| rs104894137 | pathogenic | 0 | 0 | 0 |
| rs61754278 | pathogenic | 1 | 1 | 0 |
| rs104894149 | pathogenic | 1 | 1 | 0 |
| rs104894139 | pathogenic | 1 | 1 | 0 |
| rs104894142 | pathogenic | 0 | 0 | 0 |

| | | | | |
|---|---|---|---|---|
| rs104894143 | pathogenic | 0 | 0 | 0 |
| rs104894145 | pathogenic | 0 | 0 | 0 |
| rs121909545 | benign | 0 | 1 | 0 |
| rs782308462 | pathogenic | 1 | 1 | 1 |
| rs375002796 | pathogenic | 0 | 1 | 0 |
| rs201727685 | pathogenic | 0 | 0 | 0 |
| rs768050261 | pathogenic | 1 | 1 | 1 |
| rs780851340 | benign | 0 | 0 | 0 |
| rs139824241 | benign | 1 | 1 | 0 |
| rs757486575 | pathogenic | 1 | 1 | 0 |
| rs750831299 | pathogenic | 1 | 1 | 1 |
| rs767602961 | pathogenic | 1 | 1 | 1 |
| rs121913661 | pathogenic | 1 | 1 | 1 |
| rs777692271 | pathogenic | 1 | 1 | 1 |
| rs121913659 | pathogenic | 1 | 1 | 1 |
| rs587777533 | pathogenic | 1 | 1 | 1 |
| rs104894210 | pathogenic | 0 | 0 | 0 |
| rs104894216 | pathogenic | 1 | 1 | 1 |
| rs104894207 | pathogenic | 1 | 1 | 1 |
| rs104894208 | pathogenic | 0 | 1 | 0 |
| rs104894214 | pathogenic | 1 | 1 | 1 |
| rs104894211 | pathogenic | 1 | 1 | 1 |
| rs61749384 | pathogenic | 0 | 0 | 0 |
| rs61750071 | pathogenic | 1 | 1 | 1 |
| rs61750074 | pathogenic | 0 | 0 | 0 |
| rs11063988 | benign | 0 | 0 | 0 |
| rs216311 | benign | 0 | 0 | 0 |
| rs1800382 | benign | 1 | 1 | 1 |
| rs11063987 | benign | 1 | 1 | 1 |
| rs387906847 | pathogenic | 1 | 1 | 1 |
| rs73003466 | benign | 0 | 0 | 0 |
| rs149787233 | benign | 1 | 1 | 1 |
| rs144451841 | pathogenic | 1 | 1 | 1 |
| rs141638421 | pathogenic | 1 | 1 | 1 |
| rs121434497 | pathogenic | 0 | 0 | 0 |
| rs121434498 | pathogenic | 0 | 0 | 0 |
| rs267607230 | pathogenic | 1 | 1 | 0 |
| rs121434499 | pathogenic | 1 | 1 | 0 |
| rs546749336 | benign | 1 | 1 | 1 |
| rs138873805 | benign | 0 | 0 | 0 |
| rs730880522 | pathogenic | 0 | 0 | 0 |
| rs200371894 | benign | 0 | 0 | 0 |
| rs730880509 | benign | 0 | 0 | 0 |
| rs267606704 | pathogenic | 1 | 1 | 1 |
| rs267606706 | pathogenic | 0 | 0 | 0 |
| rs267606705 | pathogenic | 1 | 1 | 1 |
| rs267606707 | pathogenic | 0 | 0 | 0 |
| rs387906665 | pathogenic | 1 | 1 | 1 |
| rs267606708 | pathogenic | 1 | 1 | 1 |
| rs148368481 | benign | 0 | 0 | 0 |
| rs13204 | benign | 0 | 0 | 0 |
| rs140547520 | pathogenic | 1 | 1 | 1 |
| rs755234990 | benign | 1 | 1 | 0 |
| rs151184052 | benign | 0 | 0 | 0 |

| | | | | |
|---|---|---|---|---|
| rs3736358 | benign | 0 | 0 | 0 |
| rs3736360 | benign | 0 | 0 | 0 |
| rs1138469 | benign | 0 | 0 | 0 |
| rs74315351 | pathogenic | 0 | 1 | 0 |
| rs137853051 | pathogenic | 1 | 1 | 0 |
| rs74315353 | pathogenic | 0 | 1 | 0 |
| rs74315352 | pathogenic | 0 | 1 | 0 |
| rs74315354 | pathogenic | 1 | 1 | 1 |
| rs28938172 | pathogenic | 1 | 1 | 1 |
| rs52836744 | pathogenic | 0 | 0 | 0 |
| rs121913159 | pathogenic | 1 | 1 | 1 |
| rs121913145 | pathogenic | 1 | 1 | 1 |
| rs891087 | benign | 1 | 1 | 1 |
| rs9282757 | benign | 1 | 1 | 1 |
| rs397517404 | pathogenic | 1 | 1 | 1 |
| rs199918720 | benign | 1 | 1 | 0 |
| rs368299411 | benign | 1 | 1 | 0 |
| rs139399951 | benign | 1 | 1 | 0 |
| rs763981974 | benign | 1 | 1 | 0 |
| rs144242114 | benign | 0 | 0 | 0 |
| rs750965801 | benign | 0 | 0 | 0 |
| rs561310777 | benign | 0 | 0 | 0 |
| rs148185335 | benign | 0 | 0 | 0 |
| rs147742157 | benign | 0 | 0 | 0 |
| rs375664373 | pathogenic | 1 | 1 | 1 |
| rs777327250 | pathogenic | 0 | 0 | 0 |
| rs121908471 | pathogenic | 1 | 1 | 0 |
| rs2301612 | pathogenic | 1 | 1 | 0 |
| rs121908476 | pathogenic | 0 | 0 | 0 |
| rs36220239 | benign | 0 | 0 | 0 |
| rs121908473 | pathogenic | 0 | 0 | 0 |
| rs28647808 | benign | 0 | 0 | 0 |
| rs36090624 | benign | 0 | 0 | 0 |
| rs121908002 | pathogenic | 1 | 1 | 1 |
| rs4251545 | benign | 1 | 1 | 1 |
| rs78705382 | pathogenic | 0 | 0 | 0 |
| rs75184679 | pathogenic | 1 | 1 | 1 |
| rs74555752 | pathogenic | 0 | 0 | 0 |
| rs56320207 | benign | 1 | 1 | 1 |
| rs28730837 | pathogenic | 1 | 1 | 1 |
| rs119469014 | pathogenic | 1 | 1 | 1 |
| rs119468010 | pathogenic | 1 | 1 | 1 |
| rs35220041 | benign | 0 | 0 | 0 |
| rs762321137 | pathogenic | 1 | 1 | 1 |
| rs28931570 | pathogenic | 1 | 1 | 1 |
| rs709932 | benign | 1 | 1 | 1 |
| rs746531546 | benign | 1 | 1 | 1 |
| rs6647 | benign | 1 | 1 | 1 |
| rs141095970 | benign | 1 | 1 | 1 |
| rs143370956 | benign | 1 | 1 | 1 |
| rs28929474 | pathogenic | 1 | 1 | 1 |
| rs61761869 | pathogenic | 1 | 1 | 1 |
| rs1303 | benign | 1 | 1 | 1 |
| rs149447502 | benign | 0 | 0 | 0 |

| | | | | |
|---|---|---|---|---|
| rs267607180 | pathogenic | 1 | 1 | 1 |
| rs150304121 | benign | 0 | 0 | 0 |
| rs147630375 | benign | 1 | 1 | 1 |
| rs11539445 | benign | 1 | 1 | 1 |
| rs372098364 | pathogenic | 1 | 1 | 0 |
| rs760072286 | benign | 0 | 0 | 0 |
| rs745595833 | pathogenic | 1 | 1 | 1 |
| rs587777214 | pathogenic | 0 | 0 | 0 |
| rs150318528 | benign | 0 | 0 | 0 |
| rs137853254 | pathogenic | 1 | 1 | 1 |
| rs137853255 | pathogenic | 0 | 0 | 0 |
| rs137853253 | pathogenic | 0 | 0 | 0 |
| rs202166915 | benign | 0 | 0 | 0 |
| rs137853259 | pathogenic | 0 | 0 | 0 |
| rs2229137 | benign | 0 | 0 | 0 |
| rs137853258 | pathogenic | 1 | 1 | 0 |
| rs137853252 | pathogenic | 1 | 1 | 1 |
| rs137853256 | pathogenic | 0 | 0 | 0 |
| rs137853250 | pathogenic | 1 | 1 | 1 |
| rs121434615 | pathogenic | 0 | 0 | 0 |
| rs115014558 | benign | 1 | 1 | 1 |
| rs34116584 | benign | 1 | 1 | 1 |
| rs34885252 | benign | 1 | 1 | 1 |
| rs180177262 | pathogenic | 1 | 1 | 1 |
| rs180177162 | pathogenic | 1 | 1 | 1 |
| rs121908523 | pathogenic | 1 | 1 | 1 |
| rs180177170 | pathogenic | 1 | 1 | 1 |
| rs180177180 | pathogenic | 1 | 1 | 1 |
| rs767586362 | pathogenic | 1 | 1 | 1 |
| rs121908521 | pathogenic | 1 | 1 | 1 |
| rs180177203 | pathogenic | 1 | 1 | 1 |
| rs138025751 | pathogenic | 1 | 1 | 1 |
| rs180177210 | pathogenic | 0 | 1 | 0 |
| rs180177211 | pathogenic | 0 | 1 | 0 |
| rs121908524 | pathogenic | 1 | 1 | 0 |
| rs121908530 | pathogenic | 1 | 1 | 0 |
| rs180177227 | pathogenic | 0 | 1 | 0 |
| rs180177230 | pathogenic | 1 | 1 | 1 |
| rs121908529 | pathogenic | 0 | 1 | 0 |
| rs117195882 | benign | 1 | 1 | 0 |
| rs180177238 | pathogenic | 1 | 1 | 0 |
| rs34664134 | benign | 1 | 1 | 1 |
| rs180177246 | pathogenic | 0 | 1 | 0 |
| rs536352238 | pathogenic | 0 | 0 | 0 |
| rs121908520 | pathogenic | 1 | 1 | 0 |
| rs180177248 | pathogenic | 1 | 1 | 0 |
| rs180177252 | pathogenic | 1 | 1 | 1 |
| rs121908526 | pathogenic | 1 | 1 | 1 |
| rs121908525 | pathogenic | 1 | 1 | 1 |
| rs147106773 | benign | 1 | 1 | 1 |
| rs180177264 | pathogenic | 0 | 1 | 0 |
| rs146525143 | pathogenic | 1 | 1 | 1 |
| rs140992177 | benign | 1 | 1 | 1 |
| rs13408961 | benign | 0 | 1 | 0 |

| | | | | |
|---|---|---|---|---|
| rs4426527 | benign | 1 | 1 | 1 |
| rs3088308 | benign | 0 | 1 | 0 |
| rs4469829 | benign | 1 | 1 | 0 |
| rs104886003 | pathogenic | 1 | 1 | 1 |
| rs121913286 | pathogenic | 1 | 1 | 1 |
| rs121913279 | pathogenic | 1 | 1 | 1 |
| rs121434636 | pathogenic | 1 | 1 | 1 |
| rs755299132 | pathogenic | 1 | 1 | 1 |
| rs748318386 | pathogenic | 1 | 1 | 1 |
| rs387906757 | pathogenic | 1 | 1 | 1 |
| rs121965064 | pathogenic | 1 | 1 | 1 |
| rs28934608 | pathogenic | 0 | 0 | 0 |
| rs5967 | benign | 1 | 1 | 1 |
| rs773905328 | pathogenic | 0 | 0 | 0 |
| rs104894256 | pathogenic | 0 | 1 | 0 |
| rs28931612 | pathogenic | 0 | 1 | 0 |
| rs104894263 | pathogenic | 1 | 1 | 1 |
| rs607969 | benign | 1 | 1 | 0 |
| rs104894262 | pathogenic | 0 | 1 | 0 |
| rs104894268 | pathogenic | 1 | 1 | 1 |
| rs104894264 | pathogenic | 0 | 1 | 0 |
| rs104894259 | pathogenic | 1 | 1 | 0 |
| rs2959656 | benign | 0 | 0 | 0 |
| rs121913035 | pathogenic | 1 | 1 | 0 |
| rs139605474 | benign | 0 | 0 | 0 |
| rs3817552 | benign | 0 | 0 | 0 |
| rs150541115 | benign | 0 | 0 | 0 |
| rs562763010 | benign | 1 | 1 | 1 |
| rs200053119 | pathogenic | 0 | 0 | 0 |
| rs73826386 | benign | 0 | 0 | 0 |
| rs143655258 | benign | 0 | 0 | 0 |
| rs758857853 | pathogenic | 1 | 1 | 0 |
| rs2228380 | benign | 0 | 0 | 0 |
| rs141078733 | benign | 1 | 1 | 1 |
| rs55824172 | pathogenic | 1 | 1 | 1 |
| rs756751089 | pathogenic | 1 | 1 | 1 |
| rs121964990 | pathogenic | 0 | 0 | 0 |
| rs143555186 | benign | 0 | 0 | 0 |
| rs121964993 | pathogenic | 1 | 1 | 0 |
| rs121964992 | pathogenic | 1 | 1 | 1 |
| rs121964991 | pathogenic | 1 | 1 | 0 |
| rs121964988 | pathogenic | 1 | 1 | 1 |
| rs4687 | benign | 1 | 1 | 1 |
| rs267607125 | pathogenic | 1 | 1 | 1 |
| rs751682878 | benign | 1 | 1 | 1 |
| rs267607126 | pathogenic | 1 | 1 | 1 |
| rs768713596 | pathogenic | 0 | 0 | 0 |
| rs147288996 | benign | 0 | 0 | 0 |
| rs387906639 | pathogenic | 1 | 1 | 1 |
| rs730881069 | pathogenic | 1 | 1 | 1 |
| rs730881071 | pathogenic | 1 | 1 | 1 |
| rs397516348 | pathogenic | 0 | 0 | 0 |
| rs121917760 | pathogenic | 0 | 0 | 0 |
| rs104894724 | pathogenic | 1 | 1 | 1 |

| | | | | |
|---|---|---|---|---|
| rs397516349 | pathogenic | 1 | 1 | 1 |
| rs12075 | benign | 0 | 0 | 0 |
| rs147549987 | benign | 0 | 0 | 0 |
| rs56079734 | benign | 0 | 0 | 0 |
| rs1017842 | benign | 1 | 1 | 1 |
| rs137852593 | pathogenic | 1 | 1 | 1 |
| rs137852571 | pathogenic | 1 | 1 | 1 |
| rs137852591 | pathogenic | 1 | 1 | 1 |
| rs9332971 | pathogenic | 1 | 1 | 1 |
| rs755226547 | pathogenic | 1 | 1 | 1 |
| rs79206939 | benign | 0 | 0 | 0 |
| rs121918214 | pathogenic | 1 | 1 | 0 |
| rs757075712 | pathogenic | 1 | 1 | 0 |
| rs150483902 | benign | 1 | 1 | 1 |
| rs113993973 | pathogenic | 0 | 1 | 0 |
| rs946616 | benign | 0 | 0 | 0 |
| rs113993976 | pathogenic | 1 | 1 | 0 |
| rs151022264 | benign | 0 | 1 | 0 |
| rs113993977 | pathogenic | 1 | 1 | 0 |
| rs113993978 | pathogenic | 1 | 1 | 0 |
| rs2228499 | benign | 0 | 1 | 0 |
| rs113993979 | pathogenic | 1 | 1 | 0 |
| rs113993980 | pathogenic | 1 | 1 | 0 |
| rs113993983 | pathogenic | 0 | 1 | 0 |
| rs150547274 | pathogenic | 1 | 1 | 0 |
| rs113993984 | pathogenic | 1 | 1 | 0 |
| rs113993986 | pathogenic | 1 | 1 | 0 |
| rs34313873 | benign | 0 | 1 | 0 |
| rs113993988 | pathogenic | 1 | 1 | 0 |
| rs199638097 | pathogenic | 1 | 1 | 1 |
| rs118204447 | pathogenic | 1 | 1 | 1 |
| rs11862754 | benign | 0 | 0 | 0 |
| rs118204445 | pathogenic | 0 | 0 | 0 |
| rs118204441 | pathogenic | 0 | 0 | 0 |
| rs118204438 | pathogenic | 0 | 0 | 0 |
| rs118204436 | pathogenic | 0 | 0 | 0 |
| rs146093755 | pathogenic | 0 | 0 | 0 |
| rs781439830 | pathogenic | 0 | 0 | 0 |
| rs398123438 | pathogenic | 0 | 0 | 0 |
| rs398123439 | pathogenic | 0 | 0 | 0 |
| rs118204444 | pathogenic | 0 | 0 | 0 |
| rs7187889 | benign | 0 | 0 | 0 |
| rs118204435 | pathogenic | 1 | 1 | 0 |
| rs34745339 | benign | 0 | 0 | 0 |
| rs398123440 | pathogenic | 0 | 0 | 0 |
| rs118204442 | pathogenic | 0 | 0 | 0 |
| rs770053354 | pathogenic | 0 | 0 | 0 |
| rs118204448 | pathogenic | 0 | 0 | 0 |
| rs118204443 | pathogenic | 0 | 0 | 0 |
| rs118204446 | pathogenic | 0 | 0 | 0 |
| rs746756997 | pathogenic | 0 | 0 | 0 |
| rs267606838 | pathogenic | 0 | 0 | 0 |
| rs118204437 | pathogenic | 0 | 0 | 0 |
| rs398123429 | pathogenic | 0 | 0 | 0 |

| | | | | |
|---|---|---|---|---|
| rs398123430 | pathogenic | 0 | 0 | 0 |
| rs2303269 | benign | 0 | 0 | 0 |
| rs114703967 | benign | 0 | 0 | 0 |
| rs118204439 | pathogenic | 0 | 0 | 0 |
| rs151296605 | benign | 0 | 0 | 0 |
| rs118204440 | pathogenic | 0 | 0 | 0 |
| rs78127134 | benign | 0 | 0 | 0 |
| rs372893383 | pathogenic | 0 | 0 | 0 |
| rs61753266 | pathogenic | 1 | 1 | 1 |
| rs139579928 | benign | 1 | 1 | 1 |
| rs56275071 | pathogenic | 1 | 1 | 1 |
| rs121909736 | pathogenic | 1 | 1 | 0 |
| rs121909737 | pathogenic | 1 | 1 | 0 |
| rs121909735 | pathogenic | 1 | 1 | 0 |
| rs121909731 | pathogenic | 1 | 1 | 1 |
| rs121909734 | pathogenic | 1 | 1 | 1 |
| rs121909733 | pathogenic | 1 | 1 | 1 |
| rs121909732 | pathogenic | 1 | 1 | 1 |
| rs121909730 | pathogenic | 1 | 1 | 1 |
| rs202198360 | benign | 1 | 1 | 1 |
| rs587777294 | pathogenic | 1 | 1 | 0 |
| rs587777297 | pathogenic | 1 | 1 | 0 |
| rs104893913 | pathogenic | 1 | 1 | 0 |
| rs17860424 | pathogenic | 1 | 1 | 1 |
| rs1045485 | benign | 1 | 1 | 1 |
| rs13926 | benign | 0 | 0 | 0 |
| rs79343837 | benign | 0 | 0 | 0 |
| rs45516306 | benign | 1 | 1 | 1 |
| rs139849355 | benign | 1 | 1 | 1 |
| rs104894874 | pathogenic | 1 | 1 | 1 |
| rs104894870 | pathogenic | 1 | 1 | 1 |
| rs137852221 | pathogenic | 1 | 1 | 1 |
| rs104894869 | pathogenic | 1 | 1 | 1 |
| rs104894880 | pathogenic | 1 | 1 | 1 |
| rs104894872 | pathogenic | 1 | 1 | 1 |
| rs104894868 | pathogenic | 1 | 1 | 1 |
| rs104894867 | pathogenic | 1 | 1 | 1 |
| rs104894871 | pathogenic | 1 | 1 | 1 |
| rs104894877 | pathogenic | 1 | 1 | 1 |
| rs104894883 | pathogenic | 1 | 1 | 1 |
| rs104894875 | pathogenic | 1 | 1 | 1 |
| rs104894876 | pathogenic | 1 | 1 | 1 |
| rs104894878 | pathogenic | 1 | 1 | 1 |
| rs137852220 | pathogenic | 1 | 1 | 1 |
| rs28933684 | pathogenic | 1 | 1 | 1 |
| rs104894873 | pathogenic | 1 | 1 | 1 |
| rs138409264 | benign | 0 | 0 | 0 |
| rs2236722 | benign | 0 | 0 | 0 |
| rs28757184 | benign | 0 | 0 | 0 |
| rs121434538 | pathogenic | 0 | 0 | 0 |
| rs700519 | benign | 0 | 0 | 0 |
| rs80051519 | pathogenic | 0 | 0 | 0 |
| rs121434536 | pathogenic | 1 | 1 | 0 |
| rs121434534 | pathogenic | 1 | 1 | 0 |

| | | | | |
|---|---|---|---|---|
| rs78310315 | pathogenic | 1 | 1 | 0 |
| rs540839115 | pathogenic | 1 | 1 | 1 |
| rs80338731 | pathogenic | 1 | 1 | 1 |
| rs180177151 | pathogenic | 1 | 1 | 1 |
| rs180177152 | pathogenic | 0 | 0 | 0 |
| rs80338732 | pathogenic | 1 | 1 | 1 |
| rs80338675 | pathogenic | 1 | 1 | 1 |
| rs80338676 | pathogenic | 1 | 1 | 1 |
| rs180177153 | pathogenic | 0 | 0 | 0 |
| rs180177154 | pathogenic | 1 | 1 | 1 |
| rs5896 | benign | 1 | 1 | 1 |
| rs121918477 | pathogenic | 0 | 0 | 0 |
| rs121918428 | pathogenic | 1 | 1 | 1 |
| rs121918427 | pathogenic | 1 | 1 | 1 |
| rs544300820 | benign | 1 | 1 | 1 |
| rs121918110 | pathogenic | 1 | 1 | 1 |
| rs1057141 | benign | 0 | 0 | 0 |
| rs1135216 | benign | 1 | 1 | 1 |
| rs121917702 | pathogenic | 1 | 1 | 1 |
| rs28941474 | pathogenic | 0 | 0 | 0 |
| rs104893944 | pathogenic | 1 | 1 | 1 |
| rs104893947 | pathogenic | 1 | 1 | 0 |
| rs104893943 | pathogenic | 1 | 1 | 1 |
| rs104893942 | pathogenic | 1 | 1 | 1 |
| rs104893940 | pathogenic | 1 | 1 | 1 |
| rs34045013 | benign | 0 | 0 | 0 |
| rs143083947 | pathogenic | 0 | 0 | 0 |
| rs145786986 | benign | 0 | 0 | 0 |
| rs119450942 | pathogenic | 0 | 0 | 0 |
| rs374259530 | pathogenic | 0 | 0 | 0 |
| rs756210458 | pathogenic | 0 | 0 | 0 |
| rs11089991 | benign | 0 | 0 | 0 |
| rs28941471 | pathogenic | 0 | 1 | 0 |
| rs750614500 | pathogenic | 0 | 1 | 0 |
| rs199761158 | benign | 1 | 1 | 1 |
| rs119450944 | pathogenic | 1 | 1 | 1 |
| rs373458753 | pathogenic | 1 | 1 | 1 |
| rs202064195 | pathogenic | 1 | 1 | 1 |
| rs776496275 | pathogenic | 1 | 1 | 1 |
| rs761493155 | pathogenic | 1 | 1 | 1 |
| rs763542069 | pathogenic | 0 | 0 | 0 |
| rs119450941 | pathogenic | 0 | 0 | 0 |
| rs119450940 | pathogenic | 0 | 0 | 0 |
| rs777821034 | pathogenic | 1 | 1 | 1 |
| rs771121666 | pathogenic | 1 | 1 | 1 |
| rs372895468 | pathogenic | 1 | 1 | 1 |
| rs61750603 | pathogenic | 0 | 0 | 0 |
| rs76863441 | pathogenic | 0 | 0 | 0 |
| rs142601143 | benign | 1 | 1 | 1 |
| rs185832993 | benign | 1 | 1 | 0 |
| rs17860568 | benign | 0 | 0 | 0 |
| rs142064825 | benign | 0 | 0 | 0 |
| rs564076183 | benign | 0 | 0 | 0 |
| rs386834034 | pathogenic | 0 | 0 | 0 |

| | | | | |
|---|---|---|---|---|
| rs147663769 | benign | 0 | 0 | 0 |
| rs776478974 | benign | 0 | 0 | 0 |
| rs111840783 | benign | 0 | 0 | 0 |
| rs1516446 | benign | 1 | 1 | 1 |
| rs143749154 | benign | 1 | 1 | 1 |
| rs80257412 | benign | 0 | 0 | 0 |
| rs142943120 | benign | 0 | 0 | 0 |
| rs144553482 | benign | 0 | 0 | 0 |
| rs35997569 | benign | 0 | 0 | 0 |
| rs772909106 | pathogenic | 0 | 0 | 0 |
| rs149433837 | benign | 1 | 1 | 1 |
| rs532178791 | pathogenic | 0 | 0 | 0 |
| rs145588689 | pathogenic | 1 | 1 | 0 |
| rs17843776 | benign | 1 | 1 | 0 |
| rs121917890 | pathogenic | 1 | 1 | 1 |
| rs121917892 | pathogenic | 0 | 0 | 0 |
| rs606231289 | pathogenic | 0 | 0 | 0 |
| rs370011798 | pathogenic | 0 | 0 | 0 |
| rs34689457 | benign | 0 | 0 | 0 |
| rs118204451 | pathogenic | 0 | 0 | 0 |
| rs118204452 | pathogenic | 1 | 1 | 1 |
| rs652785 | benign | 0 | 0 | 0 |
| rs35378126 | benign | 0 | 0 | 0 |
| rs80358214 | pathogenic | 1 | 1 | 1 |
| rs80358215 | pathogenic | 0 | 1 | 0 |
| rs267606858 | pathogenic | 1 | 1 | 1 |
| rs143137713 | pathogenic | 1 | 1 | 1 |
| rs782168634 | benign | 0 | 0 | 0 |
| rs143873938 | benign | 0 | 0 | 0 |
| rs782776511 | benign | 0 | 0 | 0 |
| rs201093148 | benign | 0 | 0 | 0 |
| rs267606815 | pathogenic | 0 | 0 | 0 |
| rs137853311 | pathogenic | 0 | 0 | 0 |
| rs267606817 | pathogenic | 0 | 0 | 0 |
| rs72650006 | benign | 0 | 0 | 0 |
| rs727503638 | benign | 1 | 1 | 0 |
| rs189286381 | benign | 0 | 0 | 0 |
| rs376644553 | benign | 0 | 0 | 0 |
| rs121434610 | pathogenic | 1 | 1 | 1 |
| rs267607076 | pathogenic | 1 | 1 | 1 |
| rs201275730 | benign | 0 | 0 | 0 |
| rs121909587 | pathogenic | 0 | 0 | 0 |
| rs121909558 | pathogenic | 1 | 1 | 0 |
| rs28929469 | pathogenic | 1 | 1 | 0 |
| rs121909547 | pathogenic | 1 | 1 | 0 |
| rs121909552 | pathogenic | 1 | 1 | 0 |
| rs121909573 | pathogenic | 1 | 1 | 1 |
| rs121909567 | pathogenic | 1 | 1 | 0 |
| rs121909569 | pathogenic | 0 | 0 | 0 |
| rs121909563 | pathogenic | 1 | 1 | 0 |
| rs121909570 | pathogenic | 1 | 1 | 1 |
| rs121909571 | pathogenic | 1 | 1 | 0 |
| rs121909572 | pathogenic | 1 | 1 | 1 |
| rs121909565 | pathogenic | 1 | 1 | 1 |

| | | | | |
|---|---|---|---|---|
| rs121909557 | pathogenic | 1 | 1 | 1 |
| rs121909568 | pathogenic | 1 | 1 | 1 |
| rs121909566 | pathogenic | 1 | 1 | 1 |
| rs121909549 | pathogenic | 1 | 1 | 1 |
| rs121909554 | pathogenic | 1 | 1 | 1 |
| rs121909550 | pathogenic | 1 | 1 | 1 |
| rs121909546 | pathogenic | 1 | 1 | 0 |
| rs121909555 | pathogenic | 1 | 1 | 1 |
| rs121909564 | pathogenic | 1 | 1 | 1 |
| rs548046212 | pathogenic | 0 | 0 | 0 |
| rs147353781 | benign | 0 | 0 | 0 |
| rs104894678 | pathogenic | 1 | 1 | 1 |
| rs1130426 | benign | 0 | 0 | 0 |
| rs104894677 | pathogenic | 1 | 1 | 1 |
| rs141917423 | benign | 1 | 1 | 1 |
| rs574777883 | benign | 0 | 0 | 0 |
| rs150199226 | benign | 1 | 1 | 1 |
| rs199618701 | benign | 0 | 1 | 0 |
| rs121964923 | pathogenic | 0 | 0 | 0 |
| rs121964924 | pathogenic | 0 | 0 | 0 |
| rs201258823 | pathogenic | 0 | 1 | 0 |
| rs267606773 | pathogenic | 0 | 0 | 0 |
| rs35638832 | benign | 0 | 0 | 0 |
| rs121908632 | pathogenic | 1 | 1 | 1 |
| rs121908625 | pathogenic | 1 | 1 | 1 |
| rs748949603 | pathogenic | 0 | 0 | 0 |
| rs757523840 | pathogenic | 0 | 0 | 0 |
| rs62541771 | pathogenic | 0 | 0 | 0 |
| rs776384541 | pathogenic | 1 | 1 | 0 |
| rs121908627 | pathogenic | 1 | 1 | 0 |
| rs28937594 | pathogenic | 0 | 0 | 0 |
| rs121907888 | pathogenic | 0 | 0 | 0 |
| rs121907889 | pathogenic | 0 | 1 | 0 |
| rs28941785 | pathogenic | 1 | 1 | 1 |
| rs28941786 | pathogenic | 1 | 1 | 1 |
| rs1021737 | benign | 0 | 0 | 0 |
| rs121908370 | pathogenic | 1 | 1 | 1 |
| rs63750083 | pathogenic | 0 | 0 | 0 |
| rs17883862 | benign | 0 | 0 | 0 |
| rs587782070 | pathogenic | 1 | 1 | 1 |
| rs587781269 | pathogenic | 1 | 1 | 1 |
| rs28909982 | pathogenic | 1 | 1 | 1 |
| rs373648967 | benign | 1 | 1 | 1 |
| rs368570187 | benign | 1 | 1 | 1 |
| rs730881701 | pathogenic | 1 | 1 | 1 |
| rs137853007 | pathogenic | 0 | 0 | 0 |
| rs575910805 | benign | 0 | 0 | 0 |
| rs104893875 | pathogenic | 1 | 1 | 1 |
| rs104893877 | pathogenic | 1 | 1 | 1 |
| rs119485093 | pathogenic | 1 | 1 | 1 |
| rs119485092 | pathogenic | 0 | 0 | 0 |
| rs119485090 | pathogenic | 0 | 0 | 0 |
| rs28942099 | pathogenic | 1 | 1 | 1 |
| rs373730800 | pathogenic | 1 | 1 | 1 |

| | | | | |
|---|---|---|---|---|
| rs113993995 | pathogenic | 0 | 0 | 0 |
| rs201070132 | pathogenic | 0 | 0 | 0 |
| rs113993996 | pathogenic | 0 | 0 | 0 |
| rs113993998 | pathogenic | 0 | 0 | 0 |
| rs78950939 | pathogenic | 1 | 1 | 1 |
| rs56378716 | pathogenic | 1 | 1 | 1 |
| rs754921704 | benign | 0 | 1 | 0 |
| rs766379103 | pathogenic | 1 | 1 | 0 |
| rs121912515 | pathogenic | 0 | 1 | 0 |
| rs764831888 | benign | 1 | 1 | 0 |
| rs572527340 | benign | 0 | 0 | 0 |
| rs35969327 | benign | 0 | 0 | 0 |
| rs35814893 | benign | 0 | 0 | 0 |
| rs34032300 | benign | 0 | 0 | 0 |
| rs80358260 | pathogenic | 0 | 0 | 0 |
| rs151220873 | benign | 0 | 0 | 0 |
| rs80358261 | pathogenic | 1 | 1 | 0 |
| rs11694 | pathogenic | 0 | 0 | 0 |
| rs80358264 | pathogenic | 0 | 0 | 0 |
| rs80358266 | pathogenic | 1 | 1 | 0 |
| rs104894458 | pathogenic | 1 | 1 | 1 |
| rs121918028 | pathogenic | 0 | 0 | 0 |
| rs11583680 | pathogenic | 0 | 0 | 0 |
| rs4715 | benign | 1 | 1 | 1 |
| rs34957318 | benign | 0 | 0 | 0 |
| rs1124 | benign | 1 | 1 | 1 |
| rs104894778 | pathogenic | 1 | 1 | 1 |
| rs104894777 | pathogenic | 1 | 1 | 1 |
| rs104894769 | pathogenic | 1 | 1 | 1 |
| rs104894768 | pathogenic | 1 | 1 | 1 |
| rs104894771 | pathogenic | 1 | 1 | 1 |
| rs575372249 | benign | 1 | 1 | 1 |
| rs118203914 | pathogenic | 0 | 0 | 0 |
| rs118203916 | pathogenic | 1 | 1 | 0 |
| rs140154690 | benign | 0 | 0 | 0 |
| rs34382405 | benign | 0 | 0 | 0 |
| rs73718082 | benign | 0 | 0 | 0 |
| rs764932724 | benign | 0 | 0 | 0 |
| rs554931092 | pathogenic | 1 | 1 | 0 |
| rs202060864 | pathogenic | 0 | 0 | 0 |
| rs116567033 | benign | 0 | 0 | 0 |
| rs11243011 | benign | 0 | 0 | 0 |
| rs142073519 | pathogenic | 0 | 0 | 0 |
| rs397514611 | pathogenic | 1 | 1 | 0 |
| rs397514612 | pathogenic | 0 | 0 | 0 |
| rs372054960 | pathogenic | 0 | 0 | 0 |
| rs775690041 | pathogenic | 0 | 0 | 0 |
| rs148921184 | pathogenic | 0 | 0 | 0 |
| rs12711521 | benign | 0 | 0 | 0 |
| rs2273346 | benign | 0 | 0 | 0 |
| rs12085877 | benign | 1 | 1 | 1 |
| rs144471433 | benign | 1 | 1 | 1 |
| rs397514671 | pathogenic | 1 | 1 | 1 |
| rs76394680 | benign | 0 | 0 | 0 |

| | | | | |
|---|---|---|---|---|
| rs78289603 | pathogenic | 1 | 1 | 1 |
| rs137852913 | pathogenic | 1 | 1 | 1 |
| rs137852914 | pathogenic | 1 | 1 | 1 |
| rs121918653 | pathogenic | 1 | 1 | 1 |
| rs35515200 | benign | 1 | 1 | 0 |
| rs17876030 | benign | 1 | 1 | 1 |
| rs556623443 | benign | 1 | 1 | 1 |
| rs121434365 | pathogenic | 1 | 1 | 1 |
| rs121434364 | pathogenic | 0 | 0 | 0 |
| rs41400249 | benign | 0 | 0 | 0 |
| rs149440376 | benign | 1 | 1 | 1 |
| rs181590179 | benign | 0 | 0 | 0 |
| rs113993968 | pathogenic | 0 | 0 | 0 |
| rs587776445 | pathogenic | 1 | 1 | 1 |
| rs113993969 | pathogenic | 1 | 1 | 1 |
| rs113993970 | pathogenic | 1 | 1 | 1 |
| rs587776449 | pathogenic | 1 | 1 | 1 |
| rs113993971 | pathogenic | 1 | 1 | 1 |
| rs267607194 | pathogenic | 1 | 1 | 1 |
| rs397514745 | pathogenic | 0 | 0 | 0 |
| rs201650281 | pathogenic | 0 | 0 | 0 |
| rs150529876 | benign | 1 | 1 | 1 |
| rs77573084 | benign | 0 | 0 | 0 |
| rs398122379 | pathogenic | 1 | 1 | 0 |
| rs765478968 | benign | 0 | 0 | 0 |
| rs201399681 | benign | 0 | 0 | 0 |
| rs17159287 | benign | 0 | 0 | 0 |
| rs538571144 | benign | 0 | 0 | 0 |
| rs137852647 | pathogenic | 0 | 0 | 0 |
| rs201358272 | pathogenic | 1 | 1 | 1 |
| rs181251337 | benign | 0 | 0 | 0 |
| rs3213119 | benign | 0 | 0 | 0 |
| rs137853102 | pathogenic | 1 | 1 | 0 |
| rs4956145 | benign | 0 | 0 | 0 |
| rs146036912 | pathogenic | 1 | 1 | 1 |
| rs375717077 | pathogenic | 1 | 1 | 1 |
| rs137853103 | pathogenic | 1 | 1 | 1 |
| rs776184830 | pathogenic | 1 | 1 | 1 |
| rs200273673 | pathogenic | 1 | 1 | 1 |
| rs267607025 | pathogenic | 1 | 1 | 1 |
| rs267607024 | pathogenic | 1 | 1 | 1 |
| rs515726188 | pathogenic | 1 | 1 | 1 |
| rs147315735 | benign | 0 | 0 | 0 |
| rs121918308 | pathogenic | 1 | 1 | 0 |
| rs515726195 | pathogenic | 0 | 0 | 0 |
| rs515726196 | pathogenic | 1 | 1 | 0 |
| rs121918311 | pathogenic | 0 | 1 | 0 |
| rs121918309 | pathogenic | 0 | 0 | 0 |
| rs387906891 | pathogenic | 0 | 0 | 0 |
| rs182614164 | pathogenic | 1 | 1 | 0 |
| rs121918307 | pathogenic | 0 | 0 | 0 |
| rs201440849 | benign | 0 | 0 | 0 |
| rs118203957 | pathogenic | 0 | 0 | 0 |
| rs4986790 | benign | 0 | 0 | 0 |

| | | | | |
|---|---|---|---|---|
| rs281864928 | pathogenic | 1 | 1 | 0 |
| rs267607156 | pathogenic | 0 | 0 | 0 |
| rs55916807 | benign | 0 | 0 | 0 |
| rs749750052 | benign | 0 | 0 | 0 |
| rs74382477 | benign | 0 | 0 | 0 |
| rs201016638 | benign | 0 | 0 | 0 |
| rs376683989 | benign | 0 | 0 | 0 |
| rs144439703 | pathogenic | 0 | 0 | 0 |
| rs779122022 | benign | 0 | 0 | 0 |
| rs121918056 | pathogenic | 0 | 0 | 0 |
| rs113994093 | pathogenic | 1 | 1 | 1 |
| rs141367015 | pathogenic | 0 | 0 | 0 |
| rs775538075 | pathogenic | 0 | 0 | 0 |
| rs121918055 | pathogenic | 0 | 0 | 0 |
| rs754246294 | benign | 1 | 1 | 1 |
| rs2307447 | benign | 1 | 1 | 1 |
| rs113994096 | pathogenic | 1 | 1 | 1 |
| rs367610201 | pathogenic | 1 | 1 | 1 |
| rs758438414 | pathogenic | 1 | 1 | 0 |
| rs121918046 | pathogenic | 0 | 0 | 0 |
| rs201871736 | benign | 0 | 0 | 0 |
| rs773994204 | benign | 0 | 0 | 0 |
| rs2307450 | benign | 1 | 1 | 1 |
| rs200257554 | benign | 0 | 0 | 0 |
| rs185645212 | benign | 0 | 0 | 0 |
| rs121918054 | pathogenic | 0 | 0 | 0 |
| rs113994097 | pathogenic | 0 | 0 | 0 |
| rs202037973 | pathogenic | 0 | 0 | 0 |
| rs769827124 | pathogenic | 1 | 1 | 0 |
| rs41549716 | benign | 1 | 1 | 1 |
| rs113994098 | pathogenic | 1 | 1 | 0 |
| rs144500145 | pathogenic | 1 | 1 | 0 |
| rs121918053 | pathogenic | 1 | 1 | 0 |
| rs121918050 | pathogenic | 1 | 1 | 0 |
| rs368587966 | pathogenic | 0 | 0 | 0 |
| rs139590686 | pathogenic | 0 | 0 | 0 |
| rs121918048 | pathogenic | 1 | 1 | 0 |
| rs768653086 | pathogenic | 1 | 1 | 0 |
| rs113994099 | pathogenic | 1 | 1 | 0 |
| rs121918051 | pathogenic | 1 | 1 | 0 |
| rs753160398 | pathogenic | 1 | 1 | 0 |
| rs201477273 | pathogenic | 1 | 1 | 0 |
| rs142347031 | pathogenic | 0 | 0 | 0 |
| rs121918049 | pathogenic | 0 | 0 | 0 |
| rs267606959 | pathogenic | 1 | 1 | 0 |
| rs201732356 | pathogenic | 0 | 0 | 0 |
| rs368435864 | pathogenic | 0 | 0 | 0 |
| rs764036283 | pathogenic | 0 | 0 | 0 |
| rs2307442 | benign | 0 | 0 | 0 |
| rs2307440 | benign | 0 | 0 | 0 |
| rs548076633 | pathogenic | 0 | 1 | 0 |
| rs750792237 | benign | 0 | 0 | 0 |
| rs3087374 | benign | 0 | 0 | 0 |
| rs121918060 | pathogenic | 0 | 1 | 0 |

| | | | | |
|---|---|---|---|---|
| rs397514765 | pathogenic | 1 | 1 | 1 |
| rs121918063 | pathogenic | 0 | 0 | 0 |
| rs121918058 | pathogenic | 1 | 1 | 1 |
| rs121918064 | pathogenic | 0 | 0 | 0 |
| rs36033115 | benign | 0 | 1 | 0 |
| rs121918057 | pathogenic | 0 | 1 | 0 |
| rs121918065 | pathogenic | 1 | 1 | 1 |
| rs121918061 | pathogenic | 1 | 1 | 1 |
| rs121918066 | pathogenic | 0 | 1 | 0 |
| rs707555 | benign | 0 | 1 | 0 |
| rs8192708 | benign | 0 | 1 | 0 |
| rs11552145 | benign | 0 | 0 | 0 |
| rs1804160 | benign | 0 | 1 | 0 |
| rs34103538 | benign | 0 | 1 | 0 |
| rs751393950 | pathogenic | 1 | 1 | 1 |
| rs121907979 | pathogenic | 1 | 1 | 1 |
| rs121907975 | pathogenic | 1 | 1 | 1 |
| rs121907962 | pathogenic | 1 | 1 | 1 |
| rs121907957 | pathogenic | 0 | 0 | 0 |
| rs121907953 | pathogenic | 1 | 1 | 1 |
| rs28941770 | pathogenic | 1 | 1 | 1 |
| rs28941771 | pathogenic | 1 | 1 | 1 |
| rs121907969 | pathogenic | 1 | 1 | 1 |
| rs121907973 | pathogenic | 1 | 1 | 1 |
| rs1800429 | pathogenic | 0 | 0 | 0 |
| rs121907976 | pathogenic | 1 | 1 | 0 |
| rs121907961 | pathogenic | 0 | 0 | 0 |
| rs121907974 | pathogenic | 0 | 0 | 0 |
| rs121907959 | pathogenic | 0 | 0 | 0 |
| rs121907971 | pathogenic | 1 | 1 | 0 |
| rs121907954 | pathogenic | 0 | 0 | 0 |
| rs121907977 | pathogenic | 0 | 0 | 0 |
| rs121907963 | pathogenic | 1 | 1 | 0 |
| rs370266293 | pathogenic | 1 | 1 | 0 |
| rs1800430 | benign | 0 | 0 | 0 |
| rs121907958 | pathogenic | 0 | 0 | 0 |
| rs202173526 | pathogenic | 0 | 0 | 0 |
| rs121907978 | pathogenic | 0 | 0 | 0 |
| rs121907981 | pathogenic | 0 | 1 | 0 |
| rs145012038 | benign | 1 | 1 | 1 |
| rs121907968 | pathogenic | 1 | 1 | 1 |
| rs121907966 | pathogenic | 1 | 1 | 1 |
| rs121907956 | pathogenic | 1 | 1 | 1 |
| rs28942071 | pathogenic | 1 | 1 | 1 |
| rs121907955 | pathogenic | 1 | 1 | 1 |
| rs4727853 | benign | 0 | 0 | 0 |
| rs201880593 | benign | 0 | 0 | 0 |
| rs201434579 | benign | 1 | 1 | 1 |
| rs74551128 | pathogenic | 1 | 1 | 0 |
| rs193922500 | pathogenic | 0 | 0 | 0 |
| rs121909009 | pathogenic | 1 | 1 | 0 |
| rs121908805 | pathogenic | 1 | 1 | 0 |
| rs139573311 | pathogenic | 0 | 0 | 0 |
| rs213950 | benign | 0 | 0 | 0 |

| | | | | |
|---|---|---|---|---|
| rs79282516 | pathogenic | 0 | 0 | 0 |
| rs121909017 | pathogenic | 1 | 1 | 1 |
| rs77101217 | pathogenic | 1 | 1 | 1 |
| rs1800092 | pathogenic | 0 | 1 | 0 |
| rs77646904 | pathogenic | 0 | 0 | 0 |
| rs1800095 | benign | 1 | 1 | 1 |
| rs387906368 | pathogenic | 1 | 1 | 1 |
| rs113993959 | pathogenic | 1 | 1 | 1 |
| rs121908755 | pathogenic | 1 | 1 | 1 |
| rs121909005 | pathogenic | 1 | 1 | 1 |
| rs121908757 | pathogenic | 1 | 1 | 1 |
| rs75527207 | pathogenic | 1 | 1 | 0 |
| rs121909013 | pathogenic | 1 | 1 | 0 |
| rs121909044 | pathogenic | 1 | 1 | 0 |
| rs74597325 | pathogenic | 1 | 1 | 0 |
| rs75549581 | pathogenic | 0 | 0 | 0 |
| rs80055610 | pathogenic | 1 | 1 | 1 |
| rs1800097 | pathogenic | 0 | 0 | 0 |
| rs121909006 | pathogenic | 0 | 1 | 0 |
| rs397508276 | pathogenic | 0 | 0 | 0 |
| rs397508296 | pathogenic | 1 | 1 | 1 |
| rs139468767 | pathogenic | 0 | 0 | 0 |
| rs121908759 | pathogenic | 1 | 1 | 0 |
| rs121909033 | pathogenic | 1 | 1 | 0 |
| rs397508331 | pathogenic | 0 | 0 | 0 |
| rs387907279 | pathogenic | 1 | 1 | 1 |
| rs546530618 | benign | 0 | 0 | 0 |
| rs558259739 | benign | 0 | 0 | 0 |
| rs201827340 | benign | 0 | 0 | 0 |
| rs148865119 | pathogenic | 1 | 1 | 0 |
| rs2229413 | benign | 1 | 1 | 1 |
| rs34040148 | benign | 1 | 1 | 1 |
| rs121964964 | pathogenic | 1 | 1 | 1 |
| rs760214620 | pathogenic | 1 | 1 | 1 |
| rs775992753 | pathogenic | 1 | 1 | 1 |
| rs770095972 | pathogenic | 1 | 1 | 1 |
| rs781444670 | pathogenic | 1 | 1 | 1 |
| rs147474549 | pathogenic | 0 | 0 | 0 |
| rs121964965 | pathogenic | 0 | 1 | 0 |
| rs121964963 | pathogenic | 1 | 1 | 1 |
| rs755952006 | pathogenic | 1 | 1 | 1 |
| rs121964970 | pathogenic | 1 | 1 | 1 |
| rs121964973 | pathogenic | 1 | 1 | 1 |
| rs773734233 | pathogenic | 1 | 1 | 1 |
| rs758236584 | pathogenic | 1 | 1 | 1 |
| rs149119723 | pathogenic | 1 | 1 | 0 |
| rs5742905 | pathogenic | 0 | 0 | 0 |
| rs121964962 | pathogenic | 1 | 1 | 1 |
| rs781567152 | pathogenic | 1 | 1 | 0 |
| rs398123151 | pathogenic | 1 | 1 | 1 |
| rs760417941 | pathogenic | 1 | 1 | 1 |
| rs774926464 | pathogenic | 0 | 0 | 0 |
| rs121964972 | pathogenic | 1 | 1 | 0 |
| rs372010465 | pathogenic | 1 | 1 | 1 |

| | | | | |
|---|---|---|---|---|
| rs763036586 | pathogenic | 1 | 1 | 1 |
| rs769080151 | pathogenic | 1 | 1 | 1 |
| rs121964967 | pathogenic | 1 | 1 | 1 |
| rs28934892 | pathogenic | 1 | 1 | 0 |
| rs28934891 | pathogenic | 1 | 1 | 1 |
| rs121964971 | pathogenic | 1 | 1 | 1 |
| rs121964968 | pathogenic | 1 | 1 | 1 |
| rs150828989 | benign | 1 | 1 | 1 |
| rs3775291 | benign | 1 | 1 | 0 |
| rs104894419 | pathogenic | 1 | 1 | 1 |
| rs2232642 | benign | 0 | 0 | 0 |
| rs104894551 | pathogenic | 1 | 1 | 0 |
| rs766328537 | pathogenic | 0 | 1 | 0 |
| rs104894553 | pathogenic | 1 | 1 | 0 |
| rs80099330 | benign | 0 | 0 | 0 |
| rs199565861 | pathogenic | 0 | 0 | 0 |
| rs104894548 | pathogenic | 0 | 0 | 0 |
| rs104894549 | pathogenic | 0 | 0 | 0 |
| rs104894550 | pathogenic | 1 | 1 | 1 |
| rs12948217 | pathogenic | 1 | 1 | 1 |
| rs761064915 | pathogenic | 0 | 0 | 0 |
| rs28940279 | pathogenic | 1 | 1 | 1 |
| rs774323189 | pathogenic | 1 | 1 | 1 |
| rs28940574 | pathogenic | 0 | 0 | 0 |
| rs515299 | benign | 1 | 1 | 1 |
| rs10127939 | benign | 1 | 1 | 1 |
| rs396991 | benign | 1 | 1 | 1 |
| rs145956175 | benign | 0 | 1 | 0 |
| rs747838255 | pathogenic | 1 | 1 | 1 |
| rs762659685 | pathogenic | 1 | 1 | 0 |
| rs104894831 | pathogenic | 0 | 1 | 0 |
| rs104894848 | pathogenic | 1 | 1 | 0 |
| rs104894833 | pathogenic | 0 | 1 | 0 |
| rs781838005 | pathogenic | 0 | 1 | 0 |
| rs104894834 | pathogenic | 1 | 1 | 0 |
| rs372966991 | pathogenic | 1 | 1 | 0 |
| rs138886989 | benign | 1 | 1 | 1 |
| rs28935195 | pathogenic | 0 | 1 | 0 |
| rs28935196 | pathogenic | 0 | 1 | 0 |
| rs372416832 | pathogenic | 1 | 1 | 1 |
| rs28935197 | pathogenic | 1 | 1 | 0 |
| rs28935486 | pathogenic | 1 | 1 | 0 |
| rs28935487 | pathogenic | 1 | 1 | 0 |
| rs28935488 | pathogenic | 1 | 1 | 1 |
| rs28935495 | pathogenic | 1 | 1 | 1 |
| rs28935485 | pathogenic | 1 | 1 | 1 |
| rs140329381 | pathogenic | 0 | 1 | 0 |
| rs104894830 | pathogenic | 0 | 1 | 0 |
| rs104894846 | pathogenic | 0 | 1 | 0 |
| rs28935489 | pathogenic | 0 | 1 | 0 |
| rs104894828 | pathogenic | 1 | 1 | 0 |
| rs28935491 | pathogenic | 1 | 1 | 0 |
| rs104894832 | pathogenic | 1 | 1 | 0 |
| rs28935492 | pathogenic | 1 | 1 | 0 |

| | | | | |
|---|---|---|---|---|
| rs730880437 | benign | 1 | 1 | 0 |
| rs28935493 | pathogenic | 0 | 1 | 0 |
| rs28935494 | pathogenic | 1 | 1 | 1 |
| rs111422676 | pathogenic | 1 | 1 | 1 |
| rs397515869 | benign | 1 | 1 | 0 |
| rs782449839 | pathogenic | 0 | 1 | 0 |
| rs104894852 | pathogenic | 1 | 1 | 1 |
| rs116929575 | pathogenic | 1 | 1 | 1 |
| rs7023954 | benign | 0 | 0 | 0 |
| rs184520335 | benign | 1 | 1 | 1 |
| rs143080527 | benign | 0 | 0 | 0 |
| rs1050565 | benign | 0 | 0 | 0 |
| rs11545035 | benign | 0 | 0 | 0 |
| rs267607190 | pathogenic | 1 | 1 | 1 |
| rs200283987 | benign | 0 | 0 | 0 |
| rs372556807 | benign | 1 | 1 | 0 |
| rs563073392 | benign | 0 | 0 | 0 |
| rs4539 | benign | 0 | 0 | 0 |
| rs28931609 | pathogenic | 1 | 1 | 1 |
| rs121912978 | pathogenic | 1 | 1 | 1 |
| rs104894072 | pathogenic | 1 | 1 | 1 |
| rs551933154 | benign | 1 | 1 | 1 |
| rs752962897 | benign | 1 | 1 | 0 |
| rs121912977 | pathogenic | 1 | 1 | 0 |
| rs4537 | benign | 1 | 1 | 1 |
| rs201830462 | benign | 0 | 0 | 0 |
| rs4544 | benign | 0 | 0 | 0 |
| rs562670189 | benign | 0 | 0 | 0 |
| rs4538 | benign | 0 | 1 | 0 |
| rs61757294 | pathogenic | 0 | 0 | 0 |
| rs5313 | benign | 0 | 0 | 0 |
| rs4545 | benign | 0 | 0 | 0 |
| rs72554627 | pathogenic | 1 | 1 | 1 |
| rs72554626 | pathogenic | 1 | 1 | 1 |
| rs387906861 | pathogenic | 1 | 1 | 1 |
| rs749655461 | pathogenic | 1 | 1 | 1 |
| rs121909606 | pathogenic | 1 | 1 | 1 |
| rs121909607 | pathogenic | 1 | 1 | 1 |
| rs200203992 | benign | 0 | 0 | 0 |
| rs6050 | benign | 0 | 0 | 0 |
| rs184635235 | benign | 0 | 0 | 0 |
| rs139146037 | benign | 0 | 0 | 0 |
| rs121909612 | pathogenic | 0 | 0 | 0 |
| rs78506343 | pathogenic | 0 | 0 | 0 |
| rs121918138 | pathogenic | 1 | 1 | 1 |
| rs121918139 | pathogenic | 1 | 1 | 1 |
| rs121918140 | pathogenic | 0 | 0 | 0 |
| rs36063028 | benign | 1 | 1 | 1 |
| rs45495192 | benign | 1 | 1 | 1 |
| rs45458802 | benign | 1 | 1 | 1 |
| rs552865793 | pathogenic | 1 | 1 | 1 |
| rs753744791 | pathogenic | 1 | 1 | 1 |
| rs777518512 | benign | 1 | 1 | 1 |
| rs142708991 | benign | 1 | 1 | 0 |

| | | | | |
|---|---|---|---|---|
| rs746069957 | benign | 0 | 0 | 0 |
| rs1334811 | benign | 0 | 0 | 0 |
| rs138894008 | benign | 1 | 1 | 0 |
| rs35030851 | benign | 1 | 1 | 0 |
| rs35378598 | benign | 0 | 0 | 0 |
| rs35818764 | benign | 1 | 1 | 1 |
| rs4631561 | benign | 1 | 1 | 1 |
| rs121918660 | pathogenic | 0 | 0 | 0 |
| rs373175587 | benign | 0 | 0 | 0 |
| rs61731956 | pathogenic | 1 | 1 | 1 |
| rs180177305 | pathogenic | 0 | 0 | 0 |
| rs119490108 | pathogenic | 0 | 0 | 0 |
| rs180177312 | pathogenic | 1 | 1 | 0 |
| rs180177314 | pathogenic | 1 | 1 | 0 |
| rs180177322 | pathogenic | 1 | 1 | 1 |
| rs180177323 | pathogenic | 1 | 1 | 1 |
| rs180177325 | pathogenic | 0 | 0 | 0 |
| rs35196441 | benign | 0 | 0 | 0 |
| rs28763904 | benign | 0 | 0 | 0 |
| rs2229519 | benign | 0 | 0 | 0 |
| rs137852886 | pathogenic | 0 | 0 | 0 |
| rs137852892 | pathogenic | 0 | 0 | 0 |
| rs770427750 | pathogenic | 0 | 0 | 0 |
| rs137852887 | pathogenic | 0 | 0 | 0 |
| rs137852893 | pathogenic | 0 | 0 | 0 |
| rs80338671 | pathogenic | 1 | 1 | 0 |
| rs2228389 | benign | 0 | 0 | 0 |
| rs80338672 | pathogenic | 0 | 0 | 0 |
| rs201958741 | pathogenic | 0 | 0 | 0 |
| rs80338673 | pathogenic | 0 | 0 | 0 |
| rs137852888 | pathogenic | 0 | 0 | 0 |
| rs137852889 | pathogenic | 0 | 0 | 0 |
| rs137852894 | pathogenic | 0 | 0 | 0 |
| rs137852890 | pathogenic | 0 | 0 | 0 |
| rs137852891 | pathogenic | 0 | 0 | 0 |
| rs766935302 | pathogenic | 0 | 0 | 0 |
| rs193074572 | benign | 0 | 0 | 0 |
| rs555210692 | benign | 1 | 1 | 0 |
| rs119103242 | pathogenic | 0 | 1 | 0 |
| rs143725101 | benign | 0 | 1 | 0 |
| rs28940589 | pathogenic | 1 | 1 | 1 |
| rs113994145 | pathogenic | 0 | 0 | 0 |
| rs773872994 | benign | 0 | 0 | 0 |
| rs28940590 | pathogenic | 1 | 1 | 0 |
| rs145400751 | benign | 1 | 1 | 1 |
| rs587782331 | benign | 0 | 0 | 0 |
| rs730881787 | benign | 0 | 0 | 0 |
| rs587778531 | benign | 0 | 0 | 0 |
| rs781652274 | benign | 0 | 0 | 0 |
| rs55882234 | benign | 0 | 0 | 0 |
| rs2020908 | benign | 0 | 0 | 0 |
| rs63750741 | pathogenic | 1 | 1 | 1 |
| rs41295268 | benign | 1 | 1 | 1 |
| rs63750897 | benign | 1 | 1 | 0 |

| | | | | |
|---|---|---|---|---|
| rs63751005 | benign | 1 | 1 | 0 |
| rs587779223 | benign | 0 | 0 | 0 |
| rs202127474 | benign | 0 | 0 | 0 |
| rs374230313 | benign | 0 | 0 | 0 |
| rs2020912 | benign | 0 | 0 | 0 |
| rs63751017 | pathogenic | 0 | 0 | 0 |
| rs587779246 | pathogenic | 0 | 0 | 0 |
| rs63749919 | pathogenic | 0 | 0 | 0 |
| rs63750563 | pathogenic | 1 | 1 | 1 |
| rs63749999 | pathogenic | 0 | 0 | 0 |
| rs200492211 | pathogenic | 0 | 0 | 0 |
| rs576269342 | benign | 0 | 0 | 0 |
| rs267608075 | benign | 0 | 0 | 0 |
| rs149605979 | benign | 0 | 0 | 0 |
| rs63750252 | benign | 0 | 0 | 0 |
| rs754289472 | benign | 0 | 0 | 0 |
| rs63750119 | pathogenic | 0 | 0 | 0 |
| rs587782109 | benign | 0 | 0 | 0 |
| rs34625968 | benign | 1 | 1 | 1 |
| rs764786814 | benign | 1 | 1 | 1 |
| rs199594809 | benign | 1 | 1 | 1 |
| rs74315364 | pathogenic | 1 | 1 | 1 |
| rs201253919 | pathogenic | 0 | 0 | 0 |
| rs387906828 | pathogenic | 1 | 1 | 1 |
| rs34521609 | benign | 1 | 1 | 1 |
| rs199472894 | pathogenic | 0 | 0 | 0 |
| rs199472902 | pathogenic | 0 | 0 | 0 |
| rs121912505 | pathogenic | 0 | 0 | 0 |
| rs28928905 | pathogenic | 0 | 0 | 0 |
| rs374376640 | benign | 0 | 0 | 0 |
| rs202102276 | benign | 0 | 0 | 0 |
| rs199472918 | pathogenic | 0 | 0 | 0 |
| rs121912516 | pathogenic | 1 | 1 | 1 |
| rs121912504 | pathogenic | 0 | 0 | 0 |
| rs9333649 | pathogenic | 0 | 0 | 0 |
| rs121912508 | pathogenic | 0 | 0 | 0 |
| rs199473428 | pathogenic | 1 | 1 | 1 |
| rs104894021 | pathogenic | 0 | 0 | 0 |
| rs28928904 | pathogenic | 1 | 1 | 1 |
| rs199473039 | benign | 1 | 1 | 1 |
| rs199472958 | pathogenic | 1 | 1 | 1 |
| rs1137617 | pathogenic | 1 | 1 | 1 |
| rs199473532 | pathogenic | 1 | 1 | 1 |
| rs199472988 | pathogenic | 0 | 0 | 0 |
| rs189014161 | pathogenic | 0 | 0 | 0 |
| rs12720441 | pathogenic | 0 | 0 | 0 |
| rs199473001 | pathogenic | 1 | 1 | 0 |
| rs121912506 | pathogenic | 1 | 1 | 0 |
| rs773724817 | pathogenic | 1 | 1 | 1 |
| rs118204002 | pathogenic | 1 | 1 | 1 |
| rs138556525 | pathogenic | 1 | 1 | 1 |
| rs72558181 | pathogenic | 1 | 1 | 1 |
| rs118204005 | pathogenic | 1 | 1 | 1 |
| rs118204004 | pathogenic | 0 | 0 | 0 |

| | | | | |
|---|---|---|---|---|
| rs118204001 | pathogenic | 1 | 1 | 1 |
| rs118204006 | pathogenic | 1 | 1 | 1 |
| rs118204003 | pathogenic | 0 | 0 | 0 |
| rs121913147 | pathogenic | 1 | 1 | 0 |
| rs2229429 | benign | 1 | 1 | 0 |
| rs182552223 | benign | 0 | 0 | 0 |
| rs146588336 | benign | 0 | 0 | 0 |
| rs121964983 | pathogenic | 0 | 1 | 0 |
| rs121964982 | pathogenic | 0 | 0 | 0 |
| rs386833679 | pathogenic | 0 | 0 | 0 |
| rs386833680 | pathogenic | 0 | 0 | 0 |
| rs121964981 | pathogenic | 1 | 1 | 0 |
| rs121964984 | pathogenic | 0 | 0 | 0 |
| rs144971200 | benign | 0 | 0 | 0 |
| rs121964985 | pathogenic | 1 | 1 | 0 |
| rs141246107 | benign | 0 | 0 | 0 |
| rs104894561 | pathogenic | 1 | 1 | 0 |
| rs377023736 | pathogenic | 0 | 0 | 0 |
| rs121964907 | pathogenic | 1 | 1 | 1 |
| rs121964909 | pathogenic | 1 | 1 | 1 |
| rs765070743 | pathogenic | 0 | 0 | 0 |
| rs2228119 | benign | 0 | 0 | 0 |
| rs745976989 | pathogenic | 1 | 1 | 1 |
| rs121964904 | pathogenic | 0 | 0 | 0 |
| rs386833430 | pathogenic | 1 | 1 | 1 |
| rs267606992 | pathogenic | 0 | 0 | 0 |
| rs61752063 | pathogenic | 0 | 0 | 0 |
| rs61752067 | pathogenic | 1 | 1 | 1 |
| rs104894934 | pathogenic | 0 | 0 | 0 |
| rs104894929 | pathogenic | 1 | 1 | 1 |
| rs80189640 | benign | 0 | 0 | 0 |
| rs137854547 | pathogenic | 1 | 1 | 1 |
| rs137854540 | pathogenic | 1 | 1 | 1 |
| rs782092363 | pathogenic | 1 | 1 | 1 |
| rs147783915 | benign | 0 | 0 | 0 |
| rs28933687 | pathogenic | 1 | 1 | 1 |
| rs149503319 | benign | 0 | 0 | 0 |
| rs1805147 | benign | 0 | 0 | 0 |
| rs587777218 | pathogenic | 1 | 1 | 1 |
| rs200786705 | benign | 0 | 0 | 0 |
| rs56219475 | benign | 0 | 0 | 0 |
| rs587781172 | benign | 0 | 0 | 0 |
| rs757094189 | pathogenic | 0 | 0 | 0 |
| rs140811086 | benign | 1 | 1 | 1 |
| rs574088829 | pathogenic | 1 | 1 | 1 |
| rs121918525 | pathogenic | 0 | 0 | 0 |
| rs121918527 | pathogenic | 0 | 0 | 0 |
| rs121964848 | pathogenic | 0 | 1 | 0 |
| rs121964845 | pathogenic | 1 | 1 | 1 |
| rs121964846 | pathogenic | 0 | 1 | 0 |
| rs17217716 | benign | 1 | 1 | 1 |
| rs141711342 | benign | 0 | 0 | 0 |
| rs372619120 | benign | 0 | 0 | 0 |
| rs63750002 | benign | 1 | 1 | 1 |

| | | | | |
|---|---|---|---|---|
| rs193922373 | benign | 0 | 0 | 0 |
| rs41295286 | benign | 0 | 0 | 0 |
| rs17217772 | benign | 0 | 0 | 0 |
| rs145649774 | benign | 0 | 0 | 0 |
| rs63750214 | pathogenic | 0 | 0 | 0 |
| rs63750716 | benign | 0 | 0 | 0 |
| rs202026056 | benign | 0 | 0 | 0 |
| rs4987188 | benign | 1 | 1 | 0 |
| rs267607938 | benign | 0 | 0 | 0 |
| rs17224367 | benign | 0 | 0 | 0 |
| rs63750006 | benign | 0 | 0 | 0 |
| rs587781373 | benign | 0 | 0 | 0 |
| rs63751207 | pathogenic | 1 | 1 | 0 |
| rs730881759 | benign | 0 | 0 | 0 |
| rs372350768 | benign | 1 | 1 | 1 |
| rs55778204 | benign | 1 | 1 | 0 |
| rs63749910 | benign | 0 | 0 | 0 |
| rs63750844 | benign | 0 | 0 | 0 |
| rs28929483 | pathogenic | 0 | 0 | 0 |
| rs61756468 | benign | 0 | 0 | 0 |
| rs63750875 | pathogenic | 1 | 1 | 0 |
| rs776528054 | benign | 1 | 1 | 0 |
| rs751939698 | benign | 0 | 0 | 0 |
| rs587780685 | benign | 0 | 0 | 0 |
| rs730881763 | benign | 1 | 1 | 0 |
| rs373226409 | benign | 0 | 0 | 0 |
| rs202145681 | benign | 1 | 1 | 1 |
| rs63749841 | benign | 1 | 1 | 1 |
| rs63750757 | benign | 1 | 1 | 1 |
| rs587778527 | benign | 1 | 1 | 1 |
| rs63750849 | benign | 0 | 0 | 0 |
| rs587779969 | benign | 0 | 0 | 0 |
| rs121908046 | pathogenic | 1 | 1 | 1 |
| rs28940882 | pathogenic | 1 | 1 | 0 |
| rs28940883 | pathogenic | 1 | 1 | 1 |
| rs727503943 | pathogenic | 1 | 1 | 1 |
| rs137853859 | pathogenic | 1 | 1 | 1 |
| rs121908045 | pathogenic | 1 | 1 | 0 |
| rs137853861 | pathogenic | 1 | 1 | 0 |
| rs3180383 | pathogenic | 0 | 1 | 0 |
| rs144431814 | benign | 0 | 0 | 0 |
| rs139502866 | benign | 0 | 0 | 0 |
| rs17524437 | benign | 0 | 0 | 0 |
| rs760960779 | benign | 0 | 0 | 0 |
| rs148960463 | pathogenic | 0 | 0 | 0 |
| rs111033647 | pathogenic | 1 | 1 | 1 |
| rs111033648 | pathogenic | 0 | 0 | 0 |
| rs1800461 | benign | 0 | 0 | 0 |
| rs111033656 | pathogenic | 0 | 0 | 0 |
| rs111033663 | pathogenic | 1 | 1 | 0 |
| rs111033669 | pathogenic | 1 | 1 | 1 |
| rs111033690 | pathogenic | 0 | 0 | 0 |
| rs111033686 | pathogenic | 0 | 0 | 0 |
| rs111033687 | pathogenic | 0 | 0 | 0 |

| | | | | |
|---|---|---|---|---|
| rs111033695 | pathogenic | 0 | 0 | 0 |
| rs111033697 | pathogenic | 0 | 0 | 0 |
| rs111033693 | pathogenic | 0 | 0 | 0 |
| rs111033694 | pathogenic | 0 | 0 | 0 |
| rs111033721 | pathogenic | 1 | 1 | 1 |
| rs111033722 | pathogenic | 0 | 0 | 0 |
| rs111033725 | pathogenic | 1 | 1 | 0 |
| rs75391579 | pathogenic | 1 | 1 | 1 |
| rs111033728 | pathogenic | 1 | 1 | 1 |
| rs111033735 | pathogenic | 1 | 1 | 1 |
| rs111033739 | pathogenic | 1 | 1 | 1 |
| rs111033736 | pathogenic | 0 | 0 | 0 |
| rs111033744 | pathogenic | 1 | 1 | 1 |
| rs111033750 | pathogenic | 1 | 1 | 1 |
| rs111033752 | pathogenic | 0 | 0 | 0 |
| rs111033751 | pathogenic | 0 | 0 | 0 |
| rs111033846 | pathogenic | 0 | 0 | 0 |
| rs368166217 | pathogenic | 0 | 0 | 0 |
| rs111033764 | pathogenic | 0 | 0 | 0 |
| rs111033847 | benign | 0 | 0 | 0 |
| rs111033778 | pathogenic | 0 | 0 | 0 |
| rs111033773 | pathogenic | 0 | 0 | 0 |
| rs111033792 | pathogenic | 0 | 0 | 0 |
| rs367543266 | pathogenic | 1 | 1 | 1 |
| rs111033794 | pathogenic | 0 | 0 | 0 |
| rs111033802 | pathogenic | 0 | 0 | 0 |
| rs111033803 | pathogenic | 0 | 0 | 0 |
| rs111033809 | pathogenic | 1 | 1 | 0 |
| rs121918376 | pathogenic | 0 | 1 | 0 |
| rs121918377 | pathogenic | 0 | 1 | 0 |
| rs121918378 | pathogenic | 1 | 1 | 1 |
| rs121918374 | pathogenic | 1 | 1 | 1 |
| rs137852379 | pathogenic | 0 | 0 | 0 |
| rs137852388 | pathogenic | 1 | 1 | 1 |
| rs137852368 | pathogenic | 0 | 0 | 0 |
| rs28935499 | pathogenic | 0 | 0 | 0 |
| rs28933668 | pathogenic | 0 | 0 | 0 |
| rs28933669 | pathogenic | 0 | 0 | 0 |
| rs28933670 | pathogenic | 0 | 0 | 0 |
| rs28933671 | pathogenic | 0 | 0 | 0 |
| rs28933672 | pathogenic | 0 | 0 | 0 |
| rs28937272 | pathogenic | 0 | 0 | 0 |
| rs35383156 | benign | 0 | 0 | 0 |
| rs137852418 | pathogenic | 0 | 0 | 0 |
| rs137852419 | pathogenic | 0 | 0 | 0 |
| rs137852428 | pathogenic | 0 | 0 | 0 |
| rs137852436 | pathogenic | 0 | 0 | 0 |
| rs28937285 | pathogenic | 0 | 0 | 0 |
| rs35213384 | benign | 1 | 1 | 1 |
| rs3810950 | benign | 0 | 0 | 0 |
| rs201479289 | pathogenic | 0 | 0 | 0 |
| rs121912820 | pathogenic | 0 | 1 | 0 |
| rs121912815 | pathogenic | 0 | 1 | 0 |
| rs115510708 | benign | 0 | 0 | 0 |

| | | | | |
|---|---|---|---|---|
| rs868749 | benign | 0 | 0 | 0 |
| rs121912823 | pathogenic | 0 | 0 | 0 |
| rs115212829 | benign | 0 | 0 | 0 |
| rs121912822 | pathogenic | 0 | 0 | 0 |
| rs121912816 | pathogenic | 0 | 1 | 0 |
| rs4838544 | benign | 0 | 0 | 0 |
| rs121912818 | pathogenic | 0 | 0 | 0 |
| rs121912821 | pathogenic | 0 | 0 | 0 |
| rs121912817 | pathogenic | 0 | 0 | 0 |
| rs8178992 | benign | 0 | 0 | 0 |
| rs121912819 | pathogenic | 1 | 1 | 0 |
| rs80097077 | benign | 1 | 1 | 0 |
| rs121909359 | pathogenic | 1 | 1 | 0 |
| rs121909358 | pathogenic | 1 | 1 | 1 |
| rs121909361 | pathogenic | 1 | 1 | 1 |
| rs121909357 | pathogenic | 0 | 1 | 0 |
| rs121909365 | pathogenic | 1 | 1 | 1 |
| rs6413484 | pathogenic | 1 | 1 | 1 |
| rs121909373 | pathogenic | 1 | 1 | 1 |
| rs121909366 | pathogenic | 1 | 1 | 1 |
| rs121909367 | pathogenic | 1 | 1 | 1 |
| rs121909368 | pathogenic | 1 | 1 | 1 |
| rs121909369 | pathogenic | 0 | 0 | 0 |
| rs121909363 | pathogenic | 1 | 1 | 1 |
| rs45588036 | pathogenic | 1 | 1 | 1 |
| rs121909364 | pathogenic | 1 | 1 | 1 |
| rs80358282 | benign | 0 | 0 | 0 |
| rs80358284 | pathogenic | 1 | 1 | 1 |
| rs61735303 | benign | 0 | 0 | 0 |
| rs2229274 | benign | 0 | 0 | 0 |
| rs145006491 | benign | 0 | 0 | 0 |
| rs121913580 | pathogenic | 1 | 1 | 0 |
| rs730882196 | pathogenic | 0 | 0 | 0 |
| rs153477 | benign | 0 | 0 | 0 |
| rs153478 | benign | 1 | 1 | 0 |
| rs61740602 | benign | 1 | 1 | 0 |
| rs104893892 | pathogenic | 1 | 1 | 0 |
| rs121909595 | pathogenic | 1 | 1 | 1 |
| rs28931605 | pathogenic | 1 | 1 | 1 |
| rs121909597 | pathogenic | 0 | 1 | 0 |
| rs202233735 | pathogenic | 0 | 1 | 0 |
| rs121909596 | pathogenic | 1 | 1 | 1 |
| rs75156162 | benign | 0 | 1 | 0 |
| rs2305430 | benign | 1 | 1 | 1 |
| rs150318966 | benign | 0 | 0 | 0 |
| rs121909598 | pathogenic | 0 | 1 | 0 |
| rs5744175 | benign | 0 | 0 | 0 |
| rs28935476 | pathogenic | 1 | 1 | 1 |
| rs28935475 | pathogenic | 1 | 1 | 1 |
| rs122461163 | pathogenic | 1 | 1 | 1 |
| rs62626305 | pathogenic | 1 | 1 | 1 |
| rs587777157 | pathogenic | 0 | 0 | 0 |
| rs137854478 | pathogenic | 1 | 1 | 0 |
| rs137854465 | pathogenic | 0 | 0 | 0 |

| | | | | |
|---|---|---|---|---|
| rs137854470 | pathogenic | 0 | 0 | 0 |
| rs137854482 | pathogenic | 0 | 0 | 0 |
| rs137854473 | pathogenic | 0 | 0 | 0 |
| rs137854456 | pathogenic | 0 | 0 | 0 |
| rs140598 | benign | 0 | 0 | 0 |
| rs104894105 | pathogenic | 1 | 1 | 1 |
| rs104894107 | pathogenic | 1 | 1 | 1 |
| rs104894106 | pathogenic | 1 | 1 | 1 |
| rs74621026 | benign | 1 | 1 | 1 |
| rs2290332 | benign | 0 | 0 | 0 |
| rs780202604 | pathogenic | 1 | 1 | 1 |
| rs74315305 | pathogenic | 1 | 1 | 0 |
| rs766751878 | pathogenic | 1 | 1 | 1 |
| rs137854603 | pathogenic | 1 | 1 | 1 |
| rs139592595 | benign | 1 | 1 | 0 |
| rs150565592 | benign | 1 | 1 | 1 |
| rs763520126 | benign | 0 | 0 | 0 |
| rs55980502 | benign | 0 | 0 | 0 |
| rs137852812 | pathogenic | 0 | 0 | 0 |
| rs137852813 | pathogenic | 0 | 0 | 0 |
| rs190222208 | benign | 0 | 0 | 0 |
| rs138459502 | pathogenic | 0 | 0 | 0 |
| rs730881041 | pathogenic | 0 | 0 | 0 |
| rs267607080 | pathogenic | 0 | 0 | 0 |
| rs137852814 | pathogenic | 0 | 0 | 0 |
| rs267607079 | pathogenic | 0 | 0 | 0 |
| rs192378417 | benign | 0 | 0 | 0 |
| rs1288775 | benign | 0 | 0 | 0 |
| rs80338737 | pathogenic | 1 | 1 | 1 |
| rs397514708 | pathogenic | 0 | 0 | 0 |
| rs775933965 | pathogenic | 0 | 0 | 0 |
| rs747608698 | benign | 0 | 1 | 0 |
| rs1145086 | benign | 0 | 1 | 0 |
| rs121912953 | pathogenic | 1 | 1 | 0 |
| rs243849 | benign | 0 | 0 | 0 |
| rs121912955 | pathogenic | 1 | 1 | 1 |
| rs17859943 | benign | 0 | 1 | 0 |
| rs755557988 | benign | 1 | 1 | 1 |
| rs10195 | benign | 0 | 0 | 0 |
| rs61735357 | benign | 1 | 1 | 1 |
| rs137882374 | benign | 0 | 0 | 0 |
| rs5388 | benign | 0 | 0 | 0 |
| rs6352 | benign | 0 | 0 | 0 |
| rs76645461 | pathogenic | 1 | 1 | 0 |
| rs1050828 | pathogenic | 0 | 0 | 0 |
| rs1050829 | pathogenic | 0 | 0 | 0 |
| rs137852314 | pathogenic | 0 | 0 | 0 |
| rs137852331 | pathogenic | 0 | 0 | 0 |
| rs5030872 | pathogenic | 0 | 0 | 0 |
| rs267606836 | pathogenic | 0 | 0 | 0 |
| rs5030868 | pathogenic | 0 | 0 | 0 |
| rs137852330 | pathogenic | 0 | 1 | 0 |
| rs137852326 | pathogenic | 1 | 1 | 1 |
| rs137852318 | pathogenic | 1 | 1 | 1 |

| | | | | |
|---|---|---|---|---|
| rs74575103 | benign | 0 | 0 | 0 |
| rs137852327 | pathogenic | 1 | 1 | 1 |
| rs76723693 | pathogenic | 0 | 0 | 0 |
| rs5030869 | pathogenic | 0 | 0 | 0 |
| rs34193178 | pathogenic | 1 | 1 | 1 |
| rs137852316 | pathogenic | 1 | 1 | 0 |
| rs398123546 | pathogenic | 1 | 1 | 1 |
| rs72554665 | pathogenic | 0 | 0 | 0 |
| rs72554664 | pathogenic | 0 | 0 | 0 |
| rs28939068 | pathogenic | 1 | 1 | 1 |
| rs59301204 | pathogenic | 1 | 1 | 1 |
| rs143715750 | benign | 1 | 1 | 1 |
| rs35263902 | pathogenic | 1 | 1 | 1 |
| rs202214518 | benign | 1 | 1 | 1 |
| rs34076756 | benign | 0 | 0 | 0 |
| rs199592341 | pathogenic | 1 | 1 | 0 |
| rs121908833 | pathogenic | 1 | 1 | 1 |
| rs121908834 | pathogenic | 1 | 1 | 0 |
| rs138454151 | benign | 0 | 0 | 0 |
| rs141323599 | benign | 0 | 0 | 0 |
| rs137853200 | pathogenic | 0 | 0 | 0 |
| rs137853199 | pathogenic | 0 | 0 | 0 |
| rs139787163 | benign | 0 | 0 | 0 |
| rs61731097 | benign | 1 | 1 | 1 |
| rs61729795 | benign | 0 | 0 | 0 |
| rs149242330 | benign | 0 | 0 | 0 |
| rs587777490 | pathogenic | 1 | 1 | 1 |
| rs398123011 | pathogenic | 1 | 1 | 0 |
| rs775186117 | pathogenic | 0 | 0 | 0 |
| rs72677237 | benign | 0 | 0 | 0 |
| rs104894163 | pathogenic | 1 | 1 | 1 |
| rs121909588 | pathogenic | 0 | 0 | 0 |
| rs1042579 | benign | 0 | 0 | 0 |
| rs121909273 | pathogenic | 0 | 0 | 0 |
| rs121909274 | pathogenic | 0 | 0 | 0 |
| rs119473034 | pathogenic | 1 | 1 | 1 |
| rs398123065 | pathogenic | 0 | 0 | 0 |
| rs587777042 | pathogenic | 1 | 1 | 0 |
| rs761413864 | pathogenic | 1 | 1 | 1 |
| rs118203965 | pathogenic | 0 | 0 | 0 |
| rs118203966 | pathogenic | 0 | 0 | 0 |
| rs1801201 | benign | 1 | 1 | 0 |
| rs1136201 | benign | 1 | 1 | 1 |
| rs144148008 | benign | 0 | 0 | 0 |
| rs200118378 | benign | 1 | 1 | 1 |
| rs112984118 | benign | 1 | 1 | 1 |
| rs149446102 | benign | 1 | 1 | 1 |
| rs1427463 | benign | 0 | 0 | 0 |
| rs201936720 | benign | 1 | 1 | 1 |
| rs61751983 | benign | 1 | 1 | 1 |
| rs104894632 | pathogenic | 1 | 1 | 1 |
| rs149202905 | benign | 0 | 0 | 0 |
| rs2230210 | benign | 1 | 1 | 1 |
| rs111631988 | benign | 0 | 0 | 0 |

| | | | | |
|---|---|---|---|---|
| rs137852550 | pathogenic | 0 | 0 | 0 |
| rs5963403 | benign | 0 | 0 | 0 |
| rs138018739 | benign | 0 | 0 | 0 |
| rs62640590 | benign | 0 | 0 | 0 |
| rs121909581 | pathogenic | 1 | 1 | 1 |
| rs202017913 | benign | 1 | 1 | 1 |
| rs143163513 | benign | 0 | 0 | 0 |
| rs238238 | benign | 0 | 1 | 0 |
| rs143945974 | benign | 0 | 0 | 0 |
| rs238239 | benign | 1 | 1 | 1 |
| rs121918403 | pathogenic | 0 | 1 | 0 |
| rs61735456 | benign | 0 | 1 | 0 |
| rs121918404 | pathogenic | 1 | 1 | 0 |
| rs12674488 | benign | 0 | 0 | 0 |
| rs201599657 | pathogenic | 0 | 0 | 0 |
| rs1057090 | benign | 0 | 0 | 0 |
| rs1057091 | benign | 0 | 0 | 0 |
| rs587783740 | benign | 0 | 0 | 0 |
| rs75956769 | benign | 0 | 0 | 0 |
| rs201691299 | benign | 0 | 0 | 0 |
| rs62621087 | benign | 0 | 0 | 0 |
| rs201308454 | benign | 0 | 0 | 0 |
| rs145657360 | benign | 0 | 0 | 0 |
| rs121434592 | pathogenic | 1 | 1 | 0 |
| rs375650413 | benign | 0 | 0 | 0 |
| rs121434437 | pathogenic | 1 | 1 | 0 |
| rs763494292 | pathogenic | 0 | 0 | 0 |
| rs121964997 | pathogenic | 1 | 1 | 0 |
| rs121964996 | pathogenic | 0 | 0 | 0 |
| rs41283425 | benign | 1 | 1 | 1 |
| rs376289528 | benign | 1 | 1 | 1 |
| rs144171604 | benign | 1 | 1 | 1 |
| rs782440692 | pathogenic | 1 | 1 | 1 |
| rs782058451 | pathogenic | 1 | 1 | 1 |
| rs78437817 | benign | 0 | 0 | 0 |
| rs200327969 | benign | 1 | 1 | 1 |
| rs143043662 | benign | 0 | 0 | 0 |
| rs111966833 | benign | 1 | 1 | 1 |
| rs35523678 | benign | 0 | 0 | 0 |
| rs515726208 | pathogenic | 0 | 0 | 0 |
| rs397517239 | pathogenic | 1 | 1 | 1 |
| rs148966602 | benign | 1 | 1 | 1 |
| rs76371972 | benign | 1 | 1 | 1 |
| rs2229968 | benign | 1 | 1 | 1 |
| rs61751538 | benign | 1 | 1 | 1 |
| rs200800978 | pathogenic | 1 | 1 | 1 |
| rs148695964 | benign | 1 | 1 | 1 |
| rs140042418 | benign | 1 | 1 | 1 |
| rs116185403 | benign | 0 | 0 | 0 |
| rs142828669 | benign | 1 | 1 | 1 |
| rs139832048 | benign | 0 | 1 | 0 |
| rs58750568 | benign | 0 | 1 | 0 |
| rs77681311 | benign | 0 | 0 | 0 |
| rs151208452 | benign | 0 | 0 | 0 |

| | | | | |
|---|---|---|---|---|
| rs76463933 | benign | 0 | 0 | 0 |
| rs144434647 | benign | 0 | 0 | 0 |
| rs117361485 | benign | 0 | 0 | 0 |
| rs587777588 | pathogenic | 0 | 0 | 0 |
| rs80338796 | pathogenic | 0 | 0 | 0 |
| rs121434594 | pathogenic | 0 | 0 | 0 |
| rs146766138 | pathogenic | 1 | 1 | 1 |
| rs28939679 | pathogenic | 1 | 1 | 1 |
| rs748754134 | pathogenic | 1 | 1 | 0 |
| rs764276946 | pathogenic | 1 | 1 | 1 |
| rs111033588 | pathogenic | 1 | 1 | 1 |
| rs201633414 | pathogenic | 1 | 1 | 1 |
| rs121909262 | pathogenic | 1 | 1 | 1 |
| rs104893702 | pathogenic | 1 | 1 | 1 |
| rs121909265 | pathogenic | 1 | 1 | 1 |
| rs121909266 | pathogenic | 1 | 1 | 1 |
| rs193922439 | pathogenic | 1 | 1 | 0 |
| rs104893691 | pathogenic | 1 | 1 | 1 |
| rs104893695 | pathogenic | 0 | 0 | 0 |
| rs104893708 | pathogenic | 1 | 1 | 1 |
| rs121909260 | pathogenic | 0 | 0 | 0 |
| rs104893696 | pathogenic | 0 | 0 | 0 |
| rs121909267 | pathogenic | 0 | 0 | 0 |
| rs121909263 | pathogenic | 0 | 0 | 0 |
| rs121909264 | pathogenic | 0 | 0 | 0 |
| rs104893694 | pathogenic | 1 | 1 | 1 |
| rs121909268 | pathogenic | 1 | 1 | 1 |
| rs104893689 | pathogenic | 1 | 1 | 0 |
| rs104893707 | pathogenic | 1 | 1 | 0 |
| rs104893697 | pathogenic | 1 | 1 | 1 |
| rs28936684 | pathogenic | 1 | 1 | 1 |
| rs62269092 | benign | 1 | 1 | 0 |
| rs121909259 | pathogenic | 1 | 1 | 0 |
| rs548403340 | benign | 1 | 1 | 0 |
| rs142818334 | benign | 0 | 0 | 0 |
| rs12493789 | benign | 1 | 1 | 1 |
| rs193922423 | pathogenic | 1 | 1 | 0 |
| rs104893719 | pathogenic | 1 | 1 | 1 |
| rs117375173 | benign | 1 | 1 | 0 |
| rs104893712 | pathogenic | 0 | 0 | 0 |
| rs121917776 | pathogenic | 1 | 1 | 1 |
| rs767325003 | benign | 1 | 1 | 1 |
| rs35824453 | benign | 0 | 0 | 0 |
| rs141641266 | benign | 0 | 0 | 0 |
| rs1050239 | benign | 0 | 0 | 0 |
| rs141303384 | benign | 0 | 0 | 0 |
| rs12720458 | pathogenic | 0 | 0 | 0 |
| rs368507376 | pathogenic | 1 | 1 | 1 |
| rs17215500 | pathogenic | 1 | 1 | 1 |
| rs120074188 | pathogenic | 1 | 1 | 1 |
| rs397508097 | pathogenic | 1 | 1 | 1 |
| rs138551008 | pathogenic | 0 | 0 | 0 |
| rs387907139 | pathogenic | 1 | 1 | 1 |
| rs765487627 | pathogenic | 0 | 0 | 0 |

| | | | | |
|---|---|---|---|---|
| rs587778587 | benign | 0 | 0 | 0 |
| rs45568339 | benign | 0 | 0 | 0 |
| rs78179744 | benign | 0 | 0 | 0 |
| rs45624036 | benign | 0 | 0 | 0 |
| rs61755166 | benign | 0 | 0 | 0 |
| rs180177132 | pathogenic | 1 | 1 | 1 |
| rs587776527 | pathogenic | 0 | 0 | 0 |
| rs373783514 | benign | 0 | 0 | 0 |
| rs730881896 | benign | 0 | 0 | 0 |
| rs118203998 | pathogenic | 0 | 0 | 0 |
| rs41281039 | pathogenic | 1 | 1 | 1 |
| rs118204430 | pathogenic | 1 | 1 | 1 |
| rs1800546 | pathogenic | 1 | 1 | 0 |
| rs76917243 | pathogenic | 0 | 1 | 0 |
| rs78340951 | pathogenic | 0 | 1 | 0 |
| rs77718928 | pathogenic | 0 | 0 | 0 |
| rs369586696 | pathogenic | 0 | 1 | 0 |
| rs745517517 | pathogenic | 1 | 1 | 0 |
| rs372872777 | pathogenic | 0 | 0 | 0 |
| rs748704459 | pathogenic | 0 | 0 | 0 |
| rs769215411 | pathogenic | 0 | 0 | 0 |
| rs139425890 | pathogenic | 0 | 0 | 0 |
| rs779694939 | pathogenic | 1 | 1 | 0 |
| rs773729410 | pathogenic | 0 | 0 | 0 |
| rs121908623 | pathogenic | 1 | 1 | 1 |
| rs121908622 | pathogenic | 1 | 1 | 1 |
| rs121908621 | pathogenic | 1 | 1 | 1 |
| rs762106720 | pathogenic | 1 | 1 | 0 |
| rs199877522 | pathogenic | 0 | 0 | 0 |
| rs74315291 | pathogenic | 1 | 1 | 1 |
| rs79137415 | benign | 0 | 0 | 0 |
| rs121913045 | pathogenic | 0 | 0 | 0 |
| rs34295337 | pathogenic | 0 | 0 | 0 |
| rs121913046 | pathogenic | 0 | 0 | 0 |
| rs121913047 | pathogenic | 0 | 0 | 0 |
| rs121913048 | pathogenic | 0 | 0 | 0 |
| rs267607166 | pathogenic | 0 | 0 | 0 |
| rs17850684 | benign | 1 | 1 | 1 |
| rs119476051 | pathogenic | 1 | 1 | 1 |
| rs17850683 | benign | 1 | 1 | 1 |
| rs119476049 | pathogenic | 1 | 1 | 1 |
| rs119476046 | pathogenic | 1 | 1 | 1 |
| rs119476048 | pathogenic | 0 | 0 | 0 |
| rs119476047 | pathogenic | 1 | 1 | 1 |
| rs28939094 | pathogenic | 1 | 1 | 1 |
| rs119476050 | pathogenic | 0 | 0 | 0 |
| rs387906941 | pathogenic | 1 | 1 | 1 |
| rs41511344 | pathogenic | 1 | 1 | 0 |
| rs748112833 | pathogenic | 1 | 1 | 1 |
| rs74342080 | benign | 1 | 1 | 0 |
| rs116482870 | benign | 0 | 0 | 0 |
| rs104893743 | pathogenic | 0 | 0 | 0 |
| rs117179004 | benign | 0 | 0 | 0 |
| rs587779409 | pathogenic | 1 | 1 | 1 |

| | | | | |
|---|---|---|---|---|
| rs3745406 | benign | 1 | 1 | 0 |
| rs387906696 | pathogenic | 0 | 0 | 0 |
| rs72549369 | pathogenic | 1 | 1 | 0 |
| rs45523133 | benign | 0 | 0 | 0 |
| rs119460972 | pathogenic | 0 | 1 | 0 |
| rs34929837 | benign | 0 | 1 | 0 |
| rs17323225 | benign | 0 | 1 | 0 |
| rs17011368 | benign | 0 | 0 | 0 |
| rs748902766 | pathogenic | 1 | 1 | 1 |
| rs387907292 | pathogenic | 0 | 0 | 0 |
| rs387907293 | pathogenic | 1 | 1 | 0 |
| rs371526758 | pathogenic | 1 | 1 | 1 |
| rs142968179 | pathogenic | 1 | 1 | 1 |
| rs387907291 | pathogenic | 1 | 1 | 0 |
| rs121907949 | pathogenic | 0 | 0 | 0 |
| rs121907947 | pathogenic | 0 | 0 | 0 |
| rs121907950 | pathogenic | 1 | 1 | 1 |
| rs121907948 | pathogenic | 1 | 1 | 1 |
| rs4926 | benign | 0 | 0 | 0 |
| rs748809996 | pathogenic | 0 | 0 | 0 |
| rs1799883 | benign | 1 | 1 | 0 |
| rs104893974 | pathogenic | 0 | 1 | 0 |
| rs104893973 | pathogenic | 0 | 1 | 0 |
| rs34452707 | benign | 0 | 0 | 0 |
| rs119467003 | pathogenic | 1 | 1 | 1 |
| rs121908261 | pathogenic | 0 | 1 | 0 |
| rs28933985 | pathogenic | 1 | 1 | 1 |
| rs80356669 | pathogenic | 1 | 1 | 1 |
| rs28936677 | pathogenic | 1 | 1 | 0 |
| rs121918326 | pathogenic | 0 | 0 | 0 |
| rs121918324 | pathogenic | 1 | 1 | 0 |
| rs121918325 | pathogenic | 1 | 1 | 0 |
| rs41270025 | pathogenic | 1 | 1 | 0 |
| rs121918323 | pathogenic | 0 | 0 | 0 |
| rs12735723 | pathogenic | 1 | 1 | 0 |
| rs36013429 | benign | 0 | 0 | 0 |
| rs28936676 | pathogenic | 1 | 1 | 0 |
| rs771809901 | pathogenic | 1 | 1 | 1 |
| rs758350837 | benign | 1 | 1 | 0 |
| rs121908192 | pathogenic | 1 | 1 | 1 |
| rs147568500 | pathogenic | 1 | 1 | 1 |
| rs3729734 | benign | 0 | 0 | 0 |
| rs371536347 | benign | 1 | 1 | 1 |
| rs80358193 | pathogenic | 1 | 1 | 1 |
| rs119476044 | pathogenic | 1 | 1 | 1 |
| rs111292798 | benign | 1 | 1 | 1 |
| rs121908063 | pathogenic | 1 | 1 | 0 |
| rs121908065 | pathogenic | 1 | 1 | 1 |
| rs121908062 | pathogenic | 1 | 1 | 1 |
| rs121908064 | pathogenic | 1 | 1 | 1 |
| rs61746421 | benign | 1 | 1 | 1 |
| rs34886916 | benign | 1 | 1 | 1 |
| rs118204045 | pathogenic | 0 | 0 | 0 |
| rs138607412 | benign | 1 | 1 | 1 |

| | | | | |
|---|---|---|---|---|
| rs118204046 | pathogenic | 0 | 0 | 0 |
| rs118204048 | pathogenic | 0 | 0 | 0 |
| rs118204044 | pathogenic | 0 | 0 | 0 |
| rs149664056 | benign | 0 | 0 | 0 |
| rs121909285 | pathogenic | 0 | 0 | 0 |
| rs1802498 | benign | 0 | 0 | 0 |
| rs1065761 | benign | 0 | 0 | 0 |
| rs119477054 | pathogenic | 0 | 0 | 0 |
| rs12720097 | benign | 0 | 0 | 0 |
| rs747281324 | pathogenic | 1 | 1 | 0 |
| rs367543028 | pathogenic | 1 | 1 | 0 |
| rs146077918 | benign | 0 | 0 | 0 |
| rs142551229 | benign | 0 | 0 | 0 |
| rs367543039 | pathogenic | 0 | 0 | 0 |
| rs587779884 | pathogenic | 1 | 1 | 0 |
| rs137853153 | pathogenic | 0 | 0 | 0 |
| rs367543029 | pathogenic | 0 | 0 | 0 |
| rs587783037 | pathogenic | 1 | 1 | 0 |
| rs142928725 | benign | 0 | 0 | 0 |
| rs28385141 | benign | 0 | 0 | 0 |
| rs587779888 | pathogenic | 0 | 0 | 0 |
| rs121909801 | pathogenic | 1 | 1 | 1 |
| rs121909790 | pathogenic | 1 | 1 | 0 |
| rs121909797 | pathogenic | 1 | 1 | 0 |
| rs121909794 | pathogenic | 1 | 1 | 1 |
| rs121909791 | pathogenic | 1 | 1 | 1 |
| rs121909793 | pathogenic | 1 | 1 | 1 |
| rs147977279 | benign | 1 | 1 | 1 |
| rs147352564 | benign | 0 | 0 | 0 |
| rs368758045 | benign | 1 | 1 | 1 |
| rs114498261 | benign | 1 | 1 | 0 |
| rs35484156 | benign | 0 | 0 | 0 |
| rs139176878 | benign | 0 | 0 | 0 |
| rs11543934 | benign | 0 | 0 | 0 |
| rs34192549 | benign | 0 | 0 | 0 |
| rs568736295 | benign | 0 | 0 | 0 |
| rs55747232 | benign | 0 | 0 | 0 |
| rs781887272 | benign | 0 | 0 | 0 |
| rs146273341 | benign | 0 | 0 | 0 |
| rs41473544 | benign | 0 | 0 | 0 |
| rs137852680 | pathogenic | 0 | 0 | 0 |
| rs55891000 | benign | 0 | 0 | 0 |
| rs140351189 | benign | 0 | 0 | 0 |
| rs137852679 | pathogenic | 1 | 1 | 0 |
| rs137852678 | pathogenic | 0 | 0 | 0 |
| rs137852677 | pathogenic | 1 | 1 | 1 |
| rs138620818 | benign | 0 | 0 | 0 |
| rs121908531 | pathogenic | 0 | 0 | 0 |
| rs72554307 | pathogenic | 0 | 0 | 0 |
| rs72554308 | pathogenic | 0 | 0 | 0 |
| rs72554312 | pathogenic | 1 | 1 | 0 |
| rs67939655 | pathogenic | 0 | 0 | 0 |
| rs67486158 | pathogenic | 0 | 0 | 0 |
| rs72554338 | pathogenic | 1 | 1 | 1 |

| | | | | |
|---|---|---|---|---|
| rs1800324 | pathogenic | 1 | 1 | 1 |
| rs72554356 | pathogenic | 1 | 1 | 1 |
| rs66656800 | pathogenic | 1 | 1 | 1 |
| rs68026851 | pathogenic | 1 | 1 | 1 |
| rs72558412 | pathogenic | 0 | 0 | 0 |
| rs72558415 | pathogenic | 0 | 0 | 0 |
| rs72558423 | pathogenic | 0 | 0 | 0 |
| rs67120076 | pathogenic | 1 | 1 | 1 |
| rs1800328 | pathogenic | 1 | 1 | 1 |
| rs72558451 | pathogenic | 1 | 1 | 1 |
| rs72558454 | pathogenic | 1 | 1 | 1 |
| rs672601372 | pathogenic | 0 | 0 | 0 |
| rs139644798 | pathogenic | 0 | 0 | 0 |
| rs369398935 | pathogenic | 0 | 0 | 0 |
| rs267607035 | pathogenic | 0 | 0 | 0 |
| rs267607036 | pathogenic | 1 | 1 | 1 |
| rs2302566 | benign | 1 | 1 | 0 |
| rs115959591 | benign | 0 | 0 | 0 |
| rs773876739 | pathogenic | 0 | 0 | 0 |
| rs1799967 | benign | 0 | 0 | 0 |
| rs80356968 | benign | 0 | 0 | 0 |
| rs28897695 | benign | 1 | 1 | 1 |
| rs80357314 | benign | 0 | 0 | 0 |
| rs80357169 | benign | 0 | 0 | 0 |
| rs397507239 | benign | 1 | 1 | 1 |
| rs80187739 | pathogenic | 1 | 1 | 1 |
| rs41293459 | pathogenic | 1 | 1 | 1 |
| rs55770810 | pathogenic | 1 | 1 | 1 |
| rs80356860 | pathogenic | 0 | 0 | 0 |
| rs56195342 | benign | 0 | 0 | 0 |
| rs80357442 | benign | 0 | 0 | 0 |
| rs41293463 | pathogenic | 1 | 1 | 1 |
| rs80357069 | pathogenic | 0 | 0 | 0 |
| rs397509275 | benign | 0 | 0 | 0 |
| rs80356920 | benign | 0 | 0 | 0 |
| rs397509284 | pathogenic | 0 | 0 | 0 |
| rs80357286 | benign | 0 | 0 | 0 |
| rs80357274 | benign | 0 | 0 | 0 |
| rs765656957 | benign | 0 | 0 | 0 |
| rs755427809 | benign | 0 | 0 | 0 |
| rs80357322 | benign | 0 | 0 | 0 |
| rs121434430 | pathogenic | 0 | 0 | 0 |
| rs121918450 | pathogenic | 1 | 1 | 1 |
| rs1532268 | benign | 0 | 0 | 0 |
| rs2303080 | benign | 0 | 0 | 0 |
| rs10064631 | benign | 0 | 0 | 0 |
| rs162036 | benign | 0 | 0 | 0 |
| rs2287780 | benign | 0 | 0 | 0 |
| rs16879334 | benign | 0 | 0 | 0 |
| rs137853061 | pathogenic | 1 | 1 | 0 |
| rs10380 | benign | 0 | 0 | 0 |
| rs1494558 | benign | 0 | 0 | 0 |
| rs1494555 | benign | 0 | 0 | 0 |
| rs192994060 | benign | 0 | 0 | 0 |

| | | | | |
|---|---|---|---|---|
| rs147128920 | benign | 1 | 1 | 0 |
| rs201803986 | pathogenic | 0 | 0 | 0 |
| rs746419489 | pathogenic | 0 | 1 | 0 |
| rs764396564 | pathogenic | 0 | 0 | 0 |
| rs758304537 | pathogenic | 0 | 0 | 0 |
| rs267606763 | pathogenic | 0 | 0 | 0 |
| rs267606762 | pathogenic | 0 | 0 | 0 |
| rs150702945 | pathogenic | 1 | 1 | 1 |
| rs202047589 | pathogenic | 0 | 0 | 0 |
| rs755562733 | pathogenic | 0 | 0 | 0 |
| rs267606764 | pathogenic | 1 | 1 | 1 |
| rs138207257 | pathogenic | 1 | 1 | 0 |
| rs149150736 | pathogenic | 1 | 1 | 1 |
| rs777046879 | pathogenic | 0 | 0 | 0 |
| rs104894408 | pathogenic | 1 | 1 | 1 |
| rs1801002 | pathogenic | 1 | 1 | 1 |
| rs28929485 | pathogenic | 1 | 1 | 1 |
| rs104894396 | pathogenic | 1 | 1 | 1 |
| rs371024165 | pathogenic | 0 | 0 | 0 |
| rs35887622 | pathogenic | 1 | 1 | 1 |
| rs587783644 | pathogenic | 1 | 1 | 1 |
| rs72474224 | pathogenic | 1 | 1 | 1 |
| rs111033296 | pathogenic | 1 | 1 | 1 |
| rs104894407 | pathogenic | 1 | 1 | 1 |
| rs72561723 | pathogenic | 1 | 1 | 1 |
| rs104894398 | pathogenic | 1 | 1 | 1 |
| rs28931594 | pathogenic | 1 | 1 | 1 |
| rs104894412 | pathogenic | 1 | 1 | 1 |
| rs111033297 | pathogenic | 1 | 1 | 1 |
| rs104894404 | pathogenic | 1 | 1 | 1 |
| rs104894410 | pathogenic | 1 | 1 | 1 |
| rs104894403 | pathogenic | 1 | 1 | 1 |
| rs397516870 | benign | 0 | 0 | 0 |
| rs200023879 | pathogenic | 1 | 1 | 1 |
| rs121912968 | pathogenic | 0 | 0 | 0 |
| rs104894402 | pathogenic | 1 | 1 | 1 |
| rs28931593 | pathogenic | 1 | 1 | 1 |
| rs104894397 | pathogenic | 0 | 0 | 0 |
| rs80338944 | pathogenic | 0 | 0 | 0 |
| rs199883710 | pathogenic | 0 | 0 | 0 |
| rs727504302 | pathogenic | 0 | 0 | 0 |
| rs145216882 | benign | 1 | 1 | 1 |
| rs781534323 | pathogenic | 1 | 1 | 1 |
| rs111033218 | benign | 1 | 1 | 1 |
| rs104894409 | pathogenic | 0 | 0 | 0 |
| rs80338945 | pathogenic | 1 | 1 | 1 |
| rs111033299 | pathogenic | 0 | 0 | 0 |
| rs143343083 | pathogenic | 1 | 1 | 1 |
| rs80338946 | pathogenic | 0 | 0 | 0 |
| rs150529554 | pathogenic | 0 | 0 | 0 |
| rs111033295 | pathogenic | 0 | 0 | 0 |
| rs111033188 | benign | 0 | 0 | 0 |
| rs397516874 | pathogenic | 0 | 0 | 0 |
| rs76434661 | pathogenic | 0 | 0 | 0 |

| | | | | |
|---|---|---|---|---|
| rs80338948 | pathogenic | 0 | 0 | 0 |
| rs104894401 | pathogenic | 0 | 0 | 0 |
| rs767178508 | pathogenic | 0 | 0 | 0 |
| rs772264564 | pathogenic | 0 | 0 | 0 |
| rs28931592 | pathogenic | 0 | 0 | 0 |
| rs80338949 | pathogenic | 1 | 1 | 1 |
| rs376898963 | benign | 1 | 1 | 1 |
| rs111033360 | benign | 0 | 0 | 0 |
| rs774518779 | pathogenic | 1 | 1 | 1 |
| rs28931595 | pathogenic | 1 | 1 | 1 |
| rs771748289 | pathogenic | 0 | 0 | 0 |
| rs104894406 | pathogenic | 0 | 0 | 0 |
| rs76838169 | benign | 0 | 0 | 0 |
| rs111033294 | pathogenic | 0 | 0 | 0 |
| rs200214298 | pathogenic | 0 | 0 | 0 |
| rs28940283 | pathogenic | 0 | 1 | 0 |
| rs1047883 | benign | 0 | 0 | 0 |
| rs138254425 | benign | 0 | 0 | 0 |
| rs121912592 | pathogenic | 1 | 1 | 0 |
| rs369061090 | pathogenic | 1 | 1 | 0 |
| rs758724746 | pathogenic | 0 | 1 | 0 |
| rs121912595 | pathogenic | 0 | 0 | 0 |
| rs76340296 | benign | 0 | 0 | 0 |
| rs140578009 | benign | 1 | 1 | 1 |
| rs74445297 | benign | 0 | 0 | 0 |
| rs78599516 | benign | 1 | 1 | 0 |
| rs104894035 | pathogenic | 1 | 1 | 0 |
| rs79451216 | benign | 0 | 0 | 0 |
| rs104894036 | pathogenic | 0 | 0 | 0 |
| rs73343757 | benign | 0 | 0 | 0 |
| rs121917722 | pathogenic | 1 | 1 | 1 |
| rs747700126 | pathogenic | 0 | 1 | 0 |
| rs121917725 | pathogenic | 1 | 1 | 1 |
| rs121917721 | pathogenic | 1 | 1 | 1 |
| rs121917723 | pathogenic | 1 | 1 | 0 |
| rs17569 | benign | 1 | 1 | 1 |
| rs2230062 | benign | 1 | 1 | 1 |
| rs17570 | benign | 0 | 0 | 0 |
| rs121917724 | pathogenic | 0 | 1 | 0 |
| rs199422235 | pathogenic | 0 | 0 | 0 |
| rs199422237 | pathogenic | 1 | 1 | 1 |
| rs199422234 | pathogenic | 1 | 1 | 0 |
| rs140015315 | pathogenic | 1 | 1 | 1 |
| rs730880298 | pathogenic | 1 | 1 | 1 |
| rs779692470 | pathogenic | 0 | 0 | 0 |
| rs369152939 | pathogenic | 0 | 0 | 0 |
| rs527236040 | pathogenic | 0 | 0 | 0 |
| rs587776985 | pathogenic | 1 | 1 | 1 |
| rs147077598 | pathogenic | 1 | 1 | 1 |
| rs128620185 | pathogenic | 1 | 1 | 1 |
| rs128620189 | pathogenic | 0 | 0 | 0 |
| rs128621190 | pathogenic | 0 | 0 | 0 |
| rs61754648 | benign | 0 | 0 | 0 |
| rs137852906 | pathogenic | 0 | 0 | 0 |

| | | | | |
|---|---|---|---|---|
| rs76811038 | pathogenic | 0 | 0 | 0 |
| rs149468422 | benign | 0 | 0 | 0 |
| rs5911 | benign | 0 | 0 | 0 |
| rs28942112 | pathogenic | 1 | 1 | 1 |
| rs143275858 | pathogenic | 1 | 1 | 0 |
| rs28362261 | benign | 0 | 0 | 0 |
| rs28362263 | benign | 1 | 1 | 1 |
| rs558707786 | pathogenic | 0 | 0 | 0 |
| rs138775799 | pathogenic | 0 | 0 | 0 |
| rs748170941 | pathogenic | 0 | 0 | 0 |
| rs121908380 | pathogenic | 0 | 0 | 0 |
| rs765123255 | pathogenic | 1 | 1 | 1 |
| rs730881832 | pathogenic | 0 | 1 | 0 |
| rs587782165 | benign | 1 | 1 | 1 |
| rs762307622 | pathogenic | 0 | 0 | 0 |
| rs34612342 | pathogenic | 1 | 1 | 0 |
| rs143353451 | pathogenic | 0 | 1 | 0 |
| rs747993448 | pathogenic | 0 | 1 | 0 |
| rs376561094 | pathogenic | 0 | 0 | 0 |
| rs776487884 | pathogenic | 1 | 1 | 0 |
| rs34126013 | pathogenic | 1 | 1 | 0 |
| rs200495564 | pathogenic | 1 | 1 | 0 |
| rs140342925 | pathogenic | 1 | 1 | 0 |
| rs587780749 | pathogenic | 1 | 1 | 0 |
| rs773087549 | pathogenic | 1 | 1 | 0 |
| rs730881833 | pathogenic | 0 | 0 | 0 |
| rs374950566 | pathogenic | 1 | 1 | 0 |
| rs587780752 | benign | 0 | 0 | 0 |
| rs3219489 | benign | 0 | 0 | 0 |
| rs199742231 | benign | 0 | 0 | 0 |
| rs587780082 | pathogenic | 0 | 0 | 0 |
| rs35881803 | benign | 0 | 0 | 0 |
| rs41305623 | benign | 1 | 1 | 1 |
| rs121917856 | pathogenic | 1 | 1 | 1 |
| rs576284122 | benign | 0 | 0 | 0 |
| rs142036701 | benign | 0 | 0 | 0 |
| rs121917852 | pathogenic | 1 | 1 | 1 |
| rs121917854 | pathogenic | 1 | 1 | 1 |
| rs121917853 | pathogenic | 1 | 1 | 1 |
| rs121917857 | pathogenic | 0 | 1 | 0 |
| rs193922740 | pathogenic | 1 | 1 | 1 |
| rs565779970 | pathogenic | 1 | 1 | 1 |
| rs6723526 | benign | 0 | 0 | 0 |
| rs141094096 | pathogenic | 1 | 1 | 1 |
| rs104894637 | pathogenic | 0 | 0 | 0 |
| rs104894636 | pathogenic | 0 | 0 | 0 |
| rs104894642 | pathogenic | 0 | 0 | 0 |
| rs104894638 | pathogenic | 0 | 0 | 0 |
| rs753666460 | pathogenic | 0 | 0 | 0 |
| rs104894643 | pathogenic | 1 | 1 | 0 |
| rs34520362 | pathogenic | 0 | 0 | 0 |
| rs374621913 | pathogenic | 0 | 0 | 0 |
| rs113641837 | benign | 0 | 0 | 0 |
| rs104894635 | pathogenic | 0 | 0 | 0 |

| | | | | |
|---|---|---|---|---|
| rs143947056 | pathogenic | 0 | 0 | 0 |
| rs138504221 | pathogenic | 0 | 0 | 0 |
| rs9894254 | benign | 0 | 0 | 0 |
| rs104894640 | pathogenic | 1 | 1 | 1 |
| rs58786455 | benign | 0 | 0 | 0 |
| rs144143780 | pathogenic | 0 | 0 | 0 |
| rs62620232 | benign | 0 | 0 | 0 |
| rs34297805 | benign | 1 | 1 | 1 |
| rs104894641 | pathogenic | 0 | 0 | 0 |
| rs148876719 | benign | 0 | 0 | 0 |
| rs104894639 | pathogenic | 0 | 0 | 0 |
| rs7503034 | benign | 0 | 0 | 0 |
| rs201631570 | benign | 0 | 0 | 0 |
| rs150181226 | pathogenic | 0 | 0 | 0 |
| rs377546036 | pathogenic | 0 | 0 | 0 |
| rs267606700 | pathogenic | 0 | 0 | 0 |
| rs267606701 | pathogenic | 1 | 1 | 0 |
| rs267606699 | pathogenic | 1 | 1 | 0 |
| rs9903215 | benign | 0 | 0 | 0 |
| rs139486406 | benign | 0 | 0 | 0 |
| rs143791614 | benign | 0 | 0 | 0 |
| rs573483474 | benign | 0 | 0 | 0 |
| rs2032892 | benign | 0 | 0 | 0 |
| rs137852619 | pathogenic | 0 | 0 | 0 |
| rs200243548 | benign | 0 | 0 | 0 |
| rs115702388 | benign | 0 | 0 | 0 |
| rs138085358 | pathogenic | 0 | 0 | 0 |
| rs121912789 | pathogenic | 1 | 1 | 1 |
| rs763407972 | pathogenic | 1 | 1 | 1 |
| rs147278302 | benign | 0 | 0 | 0 |
| rs374540411 | benign | 0 | 0 | 0 |
| rs121912790 | pathogenic | 1 | 1 | 0 |
| rs752049917 | benign | 0 | 0 | 0 |
| rs139640491 | benign | 1 | 1 | 1 |
| rs138010738 | benign | 1 | 1 | 0 |
| rs201328768 | pathogenic | 1 | 1 | 0 |
| rs2135720 | benign | 1 | 1 | 0 |
| rs201816080 | benign | 1 | 1 | 0 |
| rs754391973 | pathogenic | 0 | 0 | 0 |
| rs139087859 | benign | 0 | 0 | 0 |
| rs6805421 | benign | 0 | 0 | 0 |
| rs146266900 | benign | 0 | 0 | 0 |
| rs397509419 | pathogenic | 0 | 0 | 0 |
| rs149980411 | benign | 0 | 0 | 0 |
| rs747711488 | pathogenic | 1 | 1 | 1 |
| rs149474339 | pathogenic | 1 | 1 | 1 |
| rs758252808 | pathogenic | 1 | 1 | 0 |
| rs745756308 | pathogenic | 0 | 0 | 0 |
| rs4986958 | benign | 0 | 0 | 0 |
| rs74315444 | pathogenic | 0 | 0 | 0 |
| rs61752137 | pathogenic | 0 | 0 | 0 |
| rs8179065 | benign | 1 | 1 | 1 |
| rs758361736 | pathogenic | 1 | 1 | 0 |
| rs139794067 | pathogenic | 1 | 1 | 1 |

| | | | | |
|---|---|---|---|---|
| rs730880954 | pathogenic | 1 | 1 | 1 |
| rs730880955 | pathogenic | 1 | 1 | 1 |
| rs150634297 | pathogenic | 0 | 0 | 0 |
| rs104893750 | pathogenic | 1 | 1 | 1 |
| rs104893748 | pathogenic | 1 | 1 | 1 |
| rs104893749 | pathogenic | 1 | 1 | 1 |
| rs753520553 | pathogenic | 1 | 1 | 0 |
| rs61737294 | benign | 0 | 0 | 0 |
| rs104894601 | pathogenic | 0 | 0 | 0 |
| rs376090795 | pathogenic | 0 | 0 | 0 |
| rs104894596 | pathogenic | 0 | 0 | 0 |
| rs104894595 | pathogenic | 0 | 0 | 0 |
| rs104894598 | pathogenic | 0 | 0 | 0 |
| rs104894597 | pathogenic | 0 | 0 | 0 |
| rs104894591 | pathogenic | 0 | 0 | 0 |
| rs104894593 | pathogenic | 0 | 0 | 0 |
| rs104894594 | pathogenic | 0 | 0 | 0 |
| rs104894590 | pathogenic | 0 | 0 | 0 |
| rs86312 | benign | 0 | 0 | 0 |
| rs77874543 | benign | 0 | 0 | 0 |
| rs121912590 | pathogenic | 1 | 1 | 1 |
| rs121912589 | pathogenic | 1 | 1 | 1 |
| rs121912586 | pathogenic | 1 | 1 | 1 |
| rs121912585 | pathogenic | 1 | 1 | 1 |
| rs121912587 | pathogenic | 1 | 1 | 1 |
| rs762909359 | pathogenic | 0 | 0 | 0 |
| rs28931610 | pathogenic | 0 | 0 | 0 |
| rs104894097 | pathogenic | 1 | 1 | 1 |
| rs775176191 | benign | 0 | 0 | 0 |
| rs199907548 | pathogenic | 0 | 0 | 0 |
| rs104894095 | pathogenic | 1 | 1 | 1 |
| rs104894109 | pathogenic | 1 | 1 | 1 |
| rs372266620 | benign | 1 | 1 | 1 |
| rs104894099 | pathogenic | 0 | 0 | 0 |
| rs137854599 | pathogenic | 1 | 1 | 1 |
| rs104894094 | pathogenic | 0 | 0 | 0 |
| rs104894104 | pathogenic | 0 | 0 | 0 |
| rs6413463 | benign | 1 | 1 | 1 |
| rs104894098 | pathogenic | 0 | 0 | 0 |
| rs6413464 | benign | 0 | 0 | 0 |
| rs3731249 | benign | 0 | 0 | 0 |
| rs16931179 | benign | 0 | 0 | 0 |
| rs2734715 | benign | 0 | 0 | 0 |
| rs759080581 | pathogenic | 0 | 0 | 0 |
| rs1800746 | benign | 0 | 0 | 0 |
| rs202189057 | pathogenic | 0 | 0 | 0 |
| rs121908204 | pathogenic | 0 | 0 | 0 |
| rs756564767 | pathogenic | 0 | 0 | 0 |
| rs34758634 | benign | 1 | 1 | 1 |
| rs28940573 | pathogenic | 0 | 0 | 0 |
| rs119455955 | pathogenic | 1 | 1 | 0 |
| rs752164603 | pathogenic | 1 | 1 | 0 |
| rs763162812 | pathogenic | 0 | 0 | 0 |
| rs119455957 | pathogenic | 0 | 0 | 0 |

| | | | | |
|---|---|---|---|---|
| rs119455958 | pathogenic | 1 | 1 | 0 |
| rs750428882 | pathogenic | 0 | 0 | 0 |
| rs141482368 | benign | 0 | 0 | 0 |
| rs119455953 | pathogenic | 0 | 0 | 0 |
| rs119455954 | pathogenic | 0 | 0 | 0 |
| rs112812685 | benign | 1 | 1 | 0 |
| rs138744051 | benign | 1 | 1 | 0 |
| rs121908200 | pathogenic | 0 | 0 | 0 |
| rs119455956 | pathogenic | 0 | 0 | 0 |
| rs78970155 | benign | 0 | 0 | 0 |
| rs2834167 | benign | 0 | 0 | 0 |
| rs387907326 | pathogenic | 1 | 1 | 1 |
| rs2107732 | benign | 0 | 0 | 0 |
| rs11552377 | benign | 0 | 0 | 0 |
| rs137852843 | pathogenic | 0 | 0 | 0 |
| rs104894783 | pathogenic | 0 | 0 | 0 |
| rs122457137 | pathogenic | 1 | 1 | 1 |
| rs104894779 | pathogenic | 1 | 1 | 1 |
| rs104894786 | pathogenic | 0 | 0 | 0 |
| rs104894784 | pathogenic | 0 | 0 | 0 |
| rs104894785 | pathogenic | 0 | 0 | 0 |
| rs201870761 | pathogenic | 0 | 0 | 0 |
| rs104894781 | pathogenic | 0 | 0 | 0 |
| rs28897728 | benign | 0 | 0 | 0 |
| rs370273690 | pathogenic | 1 | 1 | 0 |
| rs9332739 | benign | 0 | 0 | 0 |
| rs150878060 | benign | 0 | 0 | 0 |
| rs573509224 | benign | 0 | 0 | 0 |
| rs36221133 | benign | 0 | 0 | 0 |
| rs2230009 | benign | 0 | 0 | 0 |
| rs267607008 | pathogenic | 0 | 0 | 0 |
| rs5985 | benign | 1 | 1 | 1 |
| rs5982 | benign | 1 | 1 | 1 |
| rs5987 | benign | 0 | 0 | 0 |
| rs5988 | benign | 0 | 0 | 0 |
| rs121918680 | pathogenic | 1 | 1 | 0 |
| rs121918679 | pathogenic | 1 | 1 | 0 |
| rs7845 | benign | 1 | 1 | 0 |
| rs1049296 | benign | 1 | 1 | 1 |
| rs121918677 | pathogenic | 1 | 1 | 0 |
| rs200086262 | pathogenic | 1 | 1 | 0 |
| rs746930990 | pathogenic | 1 | 1 | 0 |
| rs104893878 | pathogenic | 1 | 1 | 1 |
| rs121912468 | pathogenic | 1 | 1 | 1 |
| rs121912467 | pathogenic | 1 | 1 | 1 |
| rs145351926 | benign | 1 | 1 | 1 |
| rs141212743 | pathogenic | 1 | 1 | 1 |
| rs397515473 | pathogenic | 1 | 1 | 0 |
| rs121918313 | pathogenic | 0 | 1 | 0 |
| rs73015965 | pathogenic | 0 | 0 | 0 |
| rs121918030 | pathogenic | 0 | 1 | 0 |
| rs77782321 | benign | 0 | 0 | 0 |
| rs374661051 | pathogenic | 1 | 1 | 1 |
| rs267606854 | pathogenic | 0 | 0 | 0 |

| | | | | |
|---|---|---|---|---|
| rs41279678 | benign | 0 | 0 | 0 |
| rs61754640 | benign | 0 | 0 | 0 |
| rs199473314 | pathogenic | 0 | 0 | 0 |
| rs137854601 | pathogenic | 0 | 0 | 0 |
| rs137854614 | pathogenic | 0 | 0 | 0 |
| rs137854615 | pathogenic | 0 | 0 | 0 |
| rs199473635 | pathogenic | 0 | 0 | 0 |
| rs121434478 | pathogenic | 1 | 1 | 1 |
| rs775407864 | pathogenic | 1 | 1 | 1 |
| rs12706927 | benign | 0 | 1 | 0 |
| rs775614261 | benign | 0 | 0 | 0 |
| rs768926261 | benign | 0 | 0 | 0 |
| rs114057640 | benign | 0 | 0 | 0 |
| rs387906886 | pathogenic | 1 | 1 | 0 |
| rs138658585 | benign | 0 | 0 | 0 |
| rs4880091 | benign | 0 | 0 | 0 |
| rs121918552 | pathogenic | 1 | 1 | 1 |
| rs10794537 | benign | 0 | 0 | 0 |
| rs121965022 | pathogenic | 1 | 1 | 0 |
| rs121965020 | pathogenic | 0 | 0 | 0 |
| rs758452450 | pathogenic | 0 | 0 | 0 |
| rs747981483 | benign | 0 | 0 | 0 |
| rs58037052 | benign | 0 | 0 | 0 |
| rs121965029 | pathogenic | 1 | 1 | 0 |
| rs147490060 | benign | 0 | 0 | 0 |
| rs3755955 | benign | 0 | 0 | 0 |
| rs148789453 | pathogenic | 0 | 0 | 0 |
| rs121965030 | pathogenic | 0 | 0 | 0 |
| rs76722191 | benign | 0 | 0 | 0 |
| rs199801029 | pathogenic | 0 | 0 | 0 |
| rs121965033 | pathogenic | 0 | 0 | 0 |
| rs368454909 | pathogenic | 1 | 1 | 0 |
| rs6831280 | benign | 0 | 0 | 0 |
| rs121965024 | pathogenic | 0 | 0 | 0 |
| rs121965019 | pathogenic | 0 | 0 | 0 |
| rs11934801 | benign | 0 | 0 | 0 |
| rs73066479 | benign | 0 | 0 | 0 |
| rs121965027 | pathogenic | 0 | 0 | 0 |
| rs121965026 | pathogenic | 1 | 1 | 0 |
| rs121965021 | pathogenic | 0 | 0 | 0 |
| rs398123257 | benign | 0 | 0 | 0 |
| rs121965031 | pathogenic | 0 | 0 | 0 |
| rs202003795 | pathogenic | 0 | 0 | 0 |
| rs559510809 | pathogenic | 0 | 0 | 0 |
| rs762936223 | benign | 0 | 0 | 0 |
| rs137854580 | pathogenic | 1 | 1 | 1 |
| rs112263157 | benign | 0 | 0 | 0 |
| rs2289181 | benign | 0 | 0 | 0 |
| rs138034915 | benign | 0 | 1 | 0 |
| rs528367092 | pathogenic | 0 | 1 | 0 |
| rs1042393 | benign | 0 | 0 | 0 |
| rs370950728 | pathogenic | 0 | 1 | 0 |
| rs530478036 | benign | 0 | 1 | 0 |
| rs374569672 | benign | 0 | 1 | 0 |

| | | | | |
|---|---|---|---|---|
| rs121907944 | pathogenic | 0 | 1 | 0 |
| rs200856561 | pathogenic | 0 | 1 | 0 |
| rs577915581 | pathogenic | 0 | 1 | 0 |
| rs201896815 | pathogenic | 0 | 1 | 0 |
| rs121907945 | pathogenic | 0 | 1 | 0 |
| rs121907940 | pathogenic | 0 | 1 | 0 |
| rs543300039 | pathogenic | 0 | 1 | 0 |
| rs121907936 | pathogenic | 0 | 1 | 0 |
| rs766074609 | pathogenic | 0 | 1 | 0 |
| rs755253527 | pathogenic | 0 | 1 | 0 |
| rs560575383 | pathogenic | 0 | 1 | 0 |
| rs528369909 | benign | 0 | 1 | 0 |
| rs770610356 | pathogenic | 0 | 1 | 0 |
| rs747610090 | pathogenic | 0 | 1 | 0 |
| rs145712232 | benign | 0 | 1 | 0 |
| rs7215458 | benign | 1 | 1 | 0 |
| rs535644999 | pathogenic | 0 | 1 | 0 |
| rs772883420 | pathogenic | 1 | 1 | 0 |
| rs398123169 | pathogenic | 0 | 1 | 0 |
| rs121907937 | pathogenic | 0 | 1 | 0 |
| rs121907941 | pathogenic | 1 | 1 | 0 |
| rs121907942 | pathogenic | 0 | 1 | 0 |
| rs779556619 | pathogenic | 0 | 1 | 0 |
| rs1800307 | benign | 0 | 1 | 0 |
| rs377544304 | pathogenic | 1 | 1 | 0 |
| rs549029029 | pathogenic | 0 | 1 | 0 |
| rs753269119 | pathogenic | 1 | 1 | 1 |
| rs757617999 | pathogenic | 0 | 1 | 0 |
| rs28937909 | pathogenic | 0 | 1 | 0 |
| rs28940868 | pathogenic | 0 | 1 | 0 |
| rs368438393 | pathogenic | 0 | 1 | 0 |
| rs536906561 | pathogenic | 0 | 1 | 0 |
| rs374143224 | pathogenic | 0 | 1 | 0 |
| rs778418246 | pathogenic | 0 | 1 | 0 |
| rs398123172 | pathogenic | 0 | 1 | 0 |
| rs121907938 | pathogenic | 0 | 1 | 0 |
| rs1800312 | pathogenic | 0 | 1 | 0 |
| rs1126690 | benign | 0 | 1 | 0 |
| rs1800314 | benign | 0 | 1 | 0 |
| rs369532274 | pathogenic | 0 | 1 | 0 |
| rs121907943 | pathogenic | 0 | 1 | 0 |
| rs780321415 | pathogenic | 0 | 1 | 0 |
| rs377286472 | benign | 0 | 1 | 0 |
| rs121918406 | pathogenic | 1 | 1 | 1 |
| rs147484870 | benign | 0 | 0 | 0 |
| rs193922671 | pathogenic | 0 | 0 | 0 |
| rs201397978 | benign | 1 | 1 | 1 |
| rs34884895 | benign | 1 | 1 | 1 |
| rs138329459 | benign | 0 | 0 | 0 |
| rs2289366 | benign | 1 | 1 | 1 |
| rs141858334 | benign | 1 | 1 | 1 |
| rs59907886 | benign | 1 | 1 | 1 |
| rs137852653 | pathogenic | 0 | 0 | 0 |
| rs137852652 | pathogenic | 0 | 0 | 0 |

| | | | | |
|---|---|---|---|---|
| rs137852656 | pathogenic | 0 | 0 | 0 |
| rs61739916 | benign | 0 | 0 | 0 |
| rs28936668 | pathogenic | 0 | 0 | 0 |
| rs137852651 | pathogenic | 0 | 0 | 0 |
| rs137852650 | pathogenic | 0 | 0 | 0 |
| rs28936669 | pathogenic | 0 | 0 | 0 |
| rs137852654 | pathogenic | 0 | 0 | 0 |
| rs28936368 | pathogenic | 1 | 1 | 0 |
| rs137852655 | pathogenic | 0 | 0 | 0 |
| rs61752496 | benign | 0 | 0 | 0 |
| rs750098055 | benign | 0 | 0 | 0 |
| rs764149793 | benign | 0 | 0 | 0 |
| rs724160017 | pathogenic | 0 | 1 | 0 |
| rs724160019 | pathogenic | 0 | 0 | 0 |
| rs202219398 | pathogenic | 1 | 1 | 0 |
| rs724160023 | pathogenic | 1 | 1 | 0 |
| rs761953453 | pathogenic | 0 | 0 | 0 |
| rs80338835 | pathogenic | 1 | 1 | 1 |
| rs56284940 | benign | 0 | 0 | 0 |
| rs74575749 | benign | 0 | 0 | 0 |
| rs587777075 | pathogenic | 1 | 1 | 1 |
| rs641153 | benign | 0 | 0 | 0 |
| rs12614 | benign | 0 | 0 | 0 |
| rs117314762 | benign | 0 | 0 | 0 |
| rs537478097 | benign | 1 | 1 | 1 |
| rs144812066 | benign | 0 | 0 | 0 |
| rs4151651 | benign | 0 | 0 | 0 |
| rs138207668 | benign | 1 | 1 | 1 |
| rs149101394 | benign | 1 | 1 | 1 |
| rs4151659 | benign | 1 | 1 | 1 |
| rs45484591 | benign | 1 | 1 | 1 |
| rs761991070 | pathogenic | 0 | 0 | 0 |
| rs755948940 | pathogenic | 0 | 0 | 0 |
| rs137853097 | pathogenic | 0 | 0 | 0 |
| rs775297352 | benign | 0 | 0 | 0 |
| rs28943591 | benign | 0 | 0 | 0 |
| rs191468413 | pathogenic | 1 | 1 | 0 |
| rs11539471 | benign | 1 | 1 | 1 |
| rs11205 | benign | 0 | 0 | 0 |
| rs768409400 | pathogenic | 0 | 0 | 0 |
| rs762013077 | pathogenic | 0 | 0 | 0 |
| rs768474112 | pathogenic | 0 | 0 | 0 |
| rs121965063 | pathogenic | 1 | 1 | 1 |
| rs143648758 | pathogenic | 0 | 0 | 0 |
| rs121965066 | pathogenic | 0 | 0 | 0 |
| rs752907087 | pathogenic | 0 | 0 | 0 |
| rs747702749 | pathogenic | 0 | 0 | 0 |
| rs754087775 | pathogenic | 0 | 0 | 0 |
| rs121918641 | pathogenic | 1 | 1 | 1 |
| rs121918642 | pathogenic | 1 | 1 | 1 |
| rs121918640 | pathogenic | 1 | 1 | 1 |
| rs121918637 | pathogenic | 1 | 1 | 1 |
| rs121918644 | pathogenic | 1 | 1 | 1 |
| rs16840544 | benign | 0 | 0 | 0 |

| | | | | |
|---|---|---|---|---|
| rs1042391 | benign | 1 | 1 | 1 |
| rs104893744 | pathogenic | 1 | 1 | 1 |
| rs120074136 | pathogenic | 0 | 0 | 0 |
| rs1805081 | benign | 1 | 1 | 0 |
| rs80358251 | benign | 0 | 0 | 0 |
| rs774588125 | benign | 0 | 0 | 0 |
| rs120074134 | pathogenic | 0 | 0 | 0 |
| rs139751448 | pathogenic | 0 | 0 | 0 |
| rs77080672 | benign | 0 | 0 | 0 |
| rs372445155 | pathogenic | 1 | 1 | 0 |
| rs13381670 | benign | 0 | 0 | 0 |
| rs1788799 | benign | 0 | 0 | 0 |
| rs376213990 | pathogenic | 0 | 0 | 0 |
| rs757475924 | pathogenic | 0 | 0 | 0 |
| rs80358253 | pathogenic | 0 | 0 | 0 |
| rs1805082 | benign | 0 | 0 | 0 |
| rs120074130 | pathogenic | 0 | 0 | 0 |
| rs768999208 | pathogenic | 0 | 0 | 0 |
| rs34302553 | benign | 0 | 0 | 0 |
| rs28940897 | pathogenic | 0 | 0 | 0 |
| rs1140458 | benign | 0 | 0 | 0 |
| rs143124972 | pathogenic | 0 | 0 | 0 |
| rs120074135 | pathogenic | 0 | 0 | 0 |
| rs543206298 | pathogenic | 0 | 0 | 0 |
| rs120074132 | pathogenic | 0 | 0 | 0 |
| rs759826138 | pathogenic | 0 | 0 | 0 |
| rs28942108 | pathogenic | 0 | 0 | 0 |
| rs80358254 | pathogenic | 0 | 0 | 0 |
| rs80358257 | pathogenic | 0 | 0 | 0 |
| rs781261962 | pathogenic | 0 | 0 | 0 |
| rs28942107 | pathogenic | 0 | 0 | 0 |
| rs80358258 | pathogenic | 0 | 0 | 0 |
| rs80358259 | pathogenic | 1 | 1 | 0 |
| rs141440861 | benign | 0 | 0 | 0 |
| rs28942106 | pathogenic | 0 | 0 | 0 |
| rs778878523 | pathogenic | 0 | 0 | 0 |
| rs28942105 | pathogenic | 0 | 0 | 0 |
| rs61731969 | benign | 0 | 0 | 0 |
| rs200444084 | pathogenic | 0 | 0 | 0 |
| rs113371321 | pathogenic | 0 | 0 | 0 |
| rs369098773 | pathogenic | 0 | 0 | 0 |
| rs35248744 | benign | 0 | 0 | 0 |
| rs758902805 | pathogenic | 0 | 0 | 0 |
| rs120074131 | pathogenic | 0 | 0 | 0 |
| rs1805084 | benign | 0 | 0 | 0 |
| rs117793540 | benign | 0 | 0 | 0 |
| rs530058063 | benign | 0 | 0 | 0 |
| rs1064422 | pathogenic | 1 | 1 | 1 |
| rs104894506 | pathogenic | 1 | 1 | 1 |
| rs104894507 | pathogenic | 1 | 1 | 1 |
| rs104894508 | pathogenic | 1 | 1 | 0 |
| rs28999113 | pathogenic | 1 | 1 | 0 |
| rs281860266 | pathogenic | 0 | 0 | 0 |
| rs104894961 | pathogenic | 1 | 1 | 0 |

| | | | | |
|---|---|---|---|---|
| rs122463168 | pathogenic | 0 | 0 | 0 |
| rs199469624 | pathogenic | 1 | 1 | 1 |
| rs150107747 | benign | 0 | 0 | 0 |
| rs144984628 | pathogenic | 0 | 0 | 0 |
| rs28935171 | pathogenic | 0 | 0 | 0 |
| rs137852443 | pathogenic | 1 | 1 | 0 |
| rs28933674 | pathogenic | 1 | 1 | 1 |
| rs28933675 | pathogenic | 1 | 1 | 1 |
| rs28933676 | pathogenic | 1 | 1 | 1 |
| rs28933677 | pathogenic | 1 | 1 | 1 |
| rs28933678 | pathogenic | 0 | 0 | 0 |
| rs28933679 | pathogenic | 1 | 1 | 1 |
| rs28933680 | pathogenic | 0 | 0 | 0 |
| rs28933681 | pathogenic | 0 | 0 | 0 |
| rs28933682 | pathogenic | 0 | 0 | 0 |
| rs28937294 | pathogenic | 1 | 1 | 0 |
| rs137852459 | pathogenic | 0 | 0 | 0 |
| rs137852461 | pathogenic | 0 | 0 | 0 |
| rs137852464 | pathogenic | 1 | 1 | 1 |
| rs137852465 | pathogenic | 1 | 1 | 1 |
| rs1800297 | benign | 1 | 1 | 0 |
| rs137852374 | pathogenic | 1 | 1 | 1 |
| rs672601312 | pathogenic | 1 | 1 | 1 |
| rs387907196 | pathogenic | 1 | 1 | 1 |
| rs148348866 | benign | 0 | 0 | 0 |
| rs141138948 | pathogenic | 1 | 1 | 1 |
| rs137852685 | pathogenic | 1 | 1 | 0 |
| rs104893900 | pathogenic | 1 | 1 | 0 |
| rs137852686 | pathogenic | 1 | 1 | 1 |
| rs104893906 | pathogenic | 1 | 1 | 0 |
| rs587782207 | pathogenic | 1 | 1 | 1 |
| rs75086406 | pathogenic | 0 | 0 | 0 |
| rs199822819 | pathogenic | 0 | 0 | 0 |
| rs398123166 | pathogenic | 1 | 1 | 1 |
| rs121913123 | pathogenic | 1 | 1 | 1 |
| rs772190176 | pathogenic | 0 | 1 | 0 |
| rs587782216 | pathogenic | 1 | 1 | 1 |
| rs398123159 | pathogenic | 1 | 1 | 1 |
| rs121913122 | pathogenic | 1 | 1 | 1 |
| rs755436052 | pathogenic | 1 | 1 | 1 |
| rs200796606 | pathogenic | 1 | 1 | 1 |
| rs2070080 | pathogenic | 0 | 1 | 0 |
| rs149616320 | benign | 0 | 0 | 0 |
| rs143529458 | benign | 0 | 0 | 0 |
| rs267607077 | pathogenic | 1 | 1 | 1 |
| rs143898031 | benign | 1 | 1 | 0 |
| rs148304857 | benign | 1 | 1 | 1 |
| rs137852809 | pathogenic | 0 | 0 | 0 |
| rs3765623 | benign | 0 | 0 | 0 |
| rs16944397 | benign | 0 | 0 | 0 |
| rs730880167 | benign | 0 | 0 | 0 |
| rs187108957 | benign | 0 | 0 | 0 |
| rs117342470 | benign | 0 | 0 | 0 |
| rs751200138 | pathogenic | 1 | 1 | 1 |

| | | | | |
|---|---|---|---|---|
| rs755799528 | pathogenic | 0 | 0 | 0 |
| rs537484504 | benign | 0 | 0 | 0 |
| rs748944640 | pathogenic | 0 | 0 | 0 |
| rs199774121 | pathogenic | 0 | 0 | 0 |
| rs766094434 | pathogenic | 1 | 1 | 0 |
| rs777321035 | pathogenic | 1 | 1 | 1 |
| rs146269774 | benign | 0 | 1 | 0 |
| rs144085805 | benign | 0 | 1 | 0 |
| rs367643028 | benign | 0 | 0 | 0 |
| rs42524 | benign | 0 | 0 | 0 |
| rs28940586 | pathogenic | 0 | 0 | 0 |
| rs74315327 | pathogenic | 1 | 1 | 1 |
| rs199676861 | pathogenic | 0 | 0 | 0 |
| rs115802744 | benign | 1 | 1 | 0 |
| rs752866643 | pathogenic | 1 | 1 | 1 |
| rs28941784 | pathogenic | 1 | 1 | 1 |
| rs398124434 | pathogenic | 1 | 1 | 1 |
| rs756414548 | pathogenic | 1 | 1 | 1 |
| rs376128990 | pathogenic | 1 | 1 | 1 |
| rs746219370 | pathogenic | 1 | 1 | 1 |
| rs756195708 | pathogenic | 1 | 1 | 1 |
| rs369296618 | pathogenic | 1 | 1 | 1 |
| rs9593 | benign | 1 | 1 | 1 |
| rs139510844 | benign | 0 | 0 | 0 |
| rs28382158 | benign | 0 | 0 | 0 |
| rs28919570 | pathogenic | 0 | 0 | 0 |
| rs587777331 | pathogenic | 0 | 0 | 0 |
| rs587777333 | pathogenic | 0 | 0 | 0 |
| rs587777332 | pathogenic | 0 | 0 | 0 |
| rs774980346 | pathogenic | 0 | 0 | 0 |
| rs370934093 | pathogenic | 0 | 0 | 0 |
| rs121913479 | pathogenic | 1 | 1 | 1 |
| rs121913484 | pathogenic | 0 | 0 | 0 |
| rs121913485 | pathogenic | 1 | 1 | 1 |
| rs28931614 | pathogenic | 1 | 1 | 1 |
| rs17881656 | benign | 1 | 1 | 1 |
| rs28931615 | pathogenic | 1 | 1 | 1 |
| rs147846832 | benign | 0 | 0 | 0 |
| rs80356506 | pathogenic | 0 | 0 | 0 |
| rs80356494 | pathogenic | 0 | 0 | 0 |
| rs121908894 | pathogenic | 0 | 0 | 0 |
| rs121908895 | pathogenic | 0 | 0 | 0 |
| rs80356508 | pathogenic | 0 | 0 | 0 |
| rs80356507 | pathogenic | 0 | 0 | 0 |
| rs121908896 | pathogenic | 0 | 0 | 0 |
| rs745372589 | pathogenic | 1 | 1 | 0 |
| rs183898408 | benign | 0 | 0 | 0 |
| rs200844292 | benign | 0 | 0 | 0 |
| rs146897002 | benign | 0 | 0 | 0 |
| rs121964869 | pathogenic | 1 | 1 | 1 |
| rs201768800 | benign | 1 | 1 | 1 |
| rs9807556 | benign | 1 | 1 | 1 |
| rs184721031 | benign | 0 | 0 | 0 |
| rs201104206 | benign | 0 | 0 | 0 |

| | | | | |
|---|---|---|---|---|
| rs188677538 | benign | 0 | 0 | 0 |
| rs149528866 | benign | 0 | 0 | 0 |
| rs115382168 | benign | 0 | 0 | 0 |
| rs78846023 | benign | 1 | 1 | 1 |
| rs61730662 | benign | 0 | 0 | 0 |
| rs121918368 | pathogenic | 0 | 0 | 0 |
| rs566433112 | pathogenic | 1 | 1 | 1 |
| rs121909342 | pathogenic | 0 | 0 | 0 |
| rs67586389 | pathogenic | 0 | 0 | 0 |
| rs72466485 | pathogenic | 0 | 0 | 0 |
| rs72466486 | pathogenic | 1 | 1 | 1 |
| rs72466487 | pathogenic | 1 | 1 | 1 |
| rs267607112 | pathogenic | 0 | 0 | 0 |
| rs118204013 | pathogenic | 0 | 0 | 0 |
| rs121913495 | pathogenic | 0 | 0 | 0 |
| rs11554273 | pathogenic | 0 | 0 | 0 |
| rs193922477 | pathogenic | 1 | 1 | 1 |
| rs193922479 | pathogenic | 1 | 1 | 1 |
| rs72547554 | pathogenic | 0 | 0 | 0 |
| rs72547561 | pathogenic | 1 | 1 | 1 |
| rs72547562 | pathogenic | 1 | 1 | 0 |
| rs115977487 | benign | 1 | 1 | 0 |
| rs72547564 | pathogenic | 0 | 0 | 0 |
| rs72547568 | pathogenic | 1 | 1 | 0 |
| rs72547569 | pathogenic | 1 | 1 | 1 |
| rs72547571 | pathogenic | 0 | 0 | 0 |
| rs72547575 | pathogenic | 1 | 1 | 1 |
| rs544994462 | benign | 0 | 0 | 0 |
| rs190470017 | benign | 0 | 0 | 0 |
| rs199473266 | pathogenic | 1 | 1 | 1 |
| rs137854602 | pathogenic | 0 | 0 | 0 |
| rs143739249 | pathogenic | 0 | 0 | 0 |
| rs397514697 | pathogenic | 1 | 1 | 1 |
| rs373764886 | pathogenic | 0 | 0 | 0 |
| rs370242930 | pathogenic | 0 | 0 | 0 |
| rs377145777 | pathogenic | 1 | 1 | 1 |
| rs191256186 | benign | 0 | 0 | 0 |
| rs150070901 | benign | 1 | 1 | 1 |
| rs138098032 | benign | 0 | 0 | 0 |
| rs117731813 | benign | 1 | 1 | 1 |
| rs104894488 | pathogenic | 1 | 1 | 1 |
| rs11302 | benign | 0 | 1 | 0 |
| rs121918447 | pathogenic | 1 | 1 | 1 |
| rs138205828 | benign | 1 | 1 | 0 |
| rs149858290 | benign | 0 | 0 | 0 |
| rs750764003 | pathogenic | 0 | 1 | 0 |
| rs141381896 | benign | 0 | 0 | 0 |
| rs149027682 | benign | 0 | 0 | 0 |
| rs121908415 | pathogenic | 0 | 0 | 0 |
| rs121908416 | pathogenic | 0 | 0 | 0 |
| rs121908417 | pathogenic | 0 | 0 | 0 |
| rs587780306 | benign | 0 | 0 | 0 |
| rs3218725 | benign | 0 | 0 | 0 |
| rs3218729 | benign | 0 | 0 | 0 |

| | | | | |
|---|---|---|---|---|
| rs387906917 | pathogenic | 0 | 0 | 0 |
| rs200652608 | pathogenic | 0 | 0 | 0 |
| rs3218721 | benign | 0 | 0 | 0 |
| rs63750869 | pathogenic | 1 | 1 | 1 |
| rs118203910 | pathogenic | 0 | 1 | 0 |
| rs9332701 | benign | 1 | 1 | 0 |
| rs6027 | benign | 0 | 0 | 0 |
| rs121909110 | pathogenic | 0 | 1 | 0 |
| rs762550967 | pathogenic | 1 | 1 | 0 |
| rs763615602 | pathogenic | 1 | 1 | 0 |
| rs144907403 | benign | 0 | 0 | 0 |
| rs104893631 | pathogenic | 1 | 1 | 0 |
| rs144181978 | pathogenic | 1 | 1 | 0 |
| rs74874677 | benign | 1 | 1 | 1 |
| rs104893632 | pathogenic | 0 | 1 | 0 |
| rs371342464 | benign | 1 | 1 | 1 |
| rs104893633 | pathogenic | 1 | 1 | 0 |
| rs201088712 | pathogenic | 0 | 0 | 0 |
| rs201857541 | benign | 1 | 1 | 1 |
| rs139517732 | pathogenic | 1 | 1 | 0 |
| rs191057824 | benign | 1 | 1 | 0 |
| rs35683768 | benign | 1 | 1 | 0 |
| rs104895271 | pathogenic | 0 | 0 | 0 |
| rs104895217 | pathogenic | 1 | 1 | 1 |
| rs104895223 | pathogenic | 1 | 1 | 1 |
| rs104895218 | pathogenic | 1 | 1 | 1 |
| rs104895225 | pathogenic | 1 | 1 | 1 |
| rs4149637 | benign | 1 | 1 | 1 |
| rs104895219 | pathogenic | 1 | 1 | 1 |
| rs104895220 | pathogenic | 1 | 1 | 1 |
| rs104895228 | pathogenic | 1 | 1 | 1 |
| rs104895221 | pathogenic | 1 | 1 | 1 |
| rs104895222 | pathogenic | 1 | 1 | 1 |
| rs104895247 | pathogenic | 0 | 0 | 0 |
| rs28940585 | pathogenic | 0 | 0 | 0 |
| rs561367199 | pathogenic | 1 | 1 | 1 |
| rs143793815 | benign | 0 | 0 | 0 |
| rs28941473 | pathogenic | 1 | 1 | 1 |
| rs751590073 | pathogenic | 1 | 1 | 1 |
| rs111407265 | benign | 1 | 1 | 1 |
| rs373697663 | pathogenic | 1 | 1 | 1 |
| rs369879957 | pathogenic | 0 | 0 | 0 |
| rs28941472 | pathogenic | 1 | 1 | 1 |
| rs28940287 | pathogenic | 1 | 1 | 1 |
| rs77625410 | benign | 0 | 0 | 0 |
| rs111033477 | benign | 0 | 0 | 0 |
| rs115123584 | benign | 0 | 0 | 0 |
| rs374576916 | benign | 0 | 0 | 0 |
| rs397516317 | pathogenic | 0 | 0 | 0 |
| rs368657015 | pathogenic | 0 | 0 | 0 |
| rs397516321 | pathogenic | 0 | 0 | 0 |
| rs397516322 | pathogenic | 0 | 0 | 0 |
| rs111033215 | pathogenic | 0 | 0 | 0 |
| rs199606180 | pathogenic | 0 | 0 | 0 |

| | | | | |
|---|---|---|---|---|
| rs111033192 | pathogenic | 0 | 0 | 0 |
| rs948962 | benign | 1 | 1 | 1 |
| rs142293185 | benign | 0 | 0 | 0 |
| rs376764423 | pathogenic | 0 | 0 | 0 |
| rs773844428 | pathogenic | 0 | 0 | 0 |
| rs186644871 | benign | 0 | 0 | 0 |
| rs200313391 | benign | 0 | 0 | 0 |
| rs377670513 | pathogenic | 0 | 0 | 0 |
| rs1132036 | benign | 0 | 0 | 0 |
| rs111033231 | benign | 0 | 0 | 0 |
| rs145945630 | pathogenic | 0 | 0 | 0 |
| rs121912984 | pathogenic | 1 | 1 | 1 |
| rs1800435 | benign | 0 | 1 | 0 |
| rs121912980 | pathogenic | 0 | 0 | 0 |
| rs121912982 | pathogenic | 1 | 1 | 1 |
| rs121912983 | pathogenic | 0 | 0 | 0 |
| rs121912981 | pathogenic | 0 | 0 | 0 |
| rs1057868 | benign | 0 | 0 | 0 |
| rs56048668 | benign | 0 | 0 | 0 |
| rs55865465 | benign | 0 | 0 | 0 |
| rs748096324 | benign | 1 | 1 | 0 |
| rs150475750 | benign | 0 | 0 | 0 |
| rs143025739 | benign | 1 | 1 | 0 |
| rs79702016 | benign | 1 | 1 | 0 |
| rs41338945 | benign | 0 | 0 | 0 |
| rs146406037 | benign | 1 | 1 | 1 |
| rs143274491 | benign | 0 | 0 | 0 |
| rs3829986 | benign | 0 | 0 | 0 |
| rs149168939 | benign | 0 | 0 | 0 |
| rs1800385 | benign | 0 | 0 | 0 |
| rs1800386 | pathogenic | 0 | 0 | 0 |
| rs121918524 | pathogenic | 1 | 1 | 1 |
| rs121912700 | pathogenic | 0 | 0 | 0 |
| rs104893930 | pathogenic | 1 | 1 | 1 |
| rs104893931 | pathogenic | 0 | 0 | 0 |
| rs116125936 | benign | 0 | 0 | 0 |
| rs627441 | benign | 0 | 0 | 0 |
| rs104894481 | pathogenic | 0 | 0 | 0 |
| rs137853129 | pathogenic | 0 | 0 | 0 |
| rs137853130 | pathogenic | 1 | 1 | 0 |
| rs28940282 | pathogenic | 0 | 0 | 0 |
| rs2229989 | benign | 0 | 0 | 0 |
| rs104894647 | pathogenic | 0 | 0 | 0 |
| rs1061170 | benign | 1 | 1 | 1 |
| rs576402053 | benign | 0 | 0 | 0 |
| rs34972246 | benign | 0 | 0 | 0 |
| rs192725607 | benign | 0 | 0 | 0 |
| rs137852615 | pathogenic | 1 | 1 | 0 |
| rs137852617 | pathogenic | 1 | 1 | 1 |
| rs137852611 | pathogenic | 0 | 0 | 0 |
| rs137852612 | pathogenic | 0 | 0 | 0 |
| rs137852614 | pathogenic | 1 | 1 | 1 |
| rs137852610 | pathogenic | 1 | 1 | 1 |
| rs137852618 | pathogenic | 1 | 1 | 1 |

| | | | | |
|---|---|---|---|---|
| rs137852613 | pathogenic | 1 | 1 | 0 |
| rs137852609 | pathogenic | 0 | 0 | 0 |
| rs63750306 | pathogenic | 1 | 1 | 1 |
| rs63750265 | pathogenic | 1 | 1 | 1 |
| rs63751287 | pathogenic | 1 | 1 | 1 |
| rs199723282 | benign | 0 | 0 | 0 |
| rs121917807 | pathogenic | 0 | 0 | 0 |
| rs121917809 | pathogenic | 0 | 0 | 0 |
| rs75134564 | pathogenic | 0 | 0 | 0 |
| rs267606930 | pathogenic | 1 | 1 | 0 |
| rs121909302 | pathogenic | 0 | 0 | 0 |
| rs121909303 | pathogenic | 0 | 0 | 0 |
| rs121909299 | pathogenic | 0 | 0 | 0 |
| rs121909301 | pathogenic | 0 | 0 | 0 |
| rs121909300 | pathogenic | 1 | 1 | 0 |
| rs111033589 | pathogenic | 0 | 0 | 0 |
| rs137853059 | pathogenic | 0 | 0 | 0 |
| rs137853057 | pathogenic | 1 | 1 | 1 |
| rs1801474 | benign | 0 | 0 | 0 |
| rs137853060 | pathogenic | 1 | 1 | 1 |
| rs137853058 | pathogenic | 1 | 1 | 1 |
| rs34424986 | pathogenic | 0 | 0 | 0 |
| rs199657839 | benign | 1 | 1 | 1 |
| rs56092260 | benign | 0 | 0 | 0 |
| rs1801582 | benign | 1 | 1 | 1 |
| rs1801334 | benign | 1 | 1 | 1 |
| rs191486604 | pathogenic | 0 | 0 | 0 |
| rs121908051 | pathogenic | 0 | 0 | 0 |
| rs121908055 | pathogenic | 1 | 1 | 0 |
| rs28940886 | pathogenic | 0 | 0 | 0 |
| rs121908050 | pathogenic | 0 | 0 | 0 |
| rs28940887 | pathogenic | 0 | 0 | 0 |
| rs387906300 | pathogenic | 0 | 0 | 0 |
| rs28942087 | pathogenic | 0 | 0 | 0 |
| rs121908049 | pathogenic | 1 | 1 | 0 |
| rs121908054 | pathogenic | 1 | 1 | 0 |
| rs121908048 | pathogenic | 0 | 0 | 0 |
| rs28940888 | pathogenic | 1 | 1 | 1 |
| rs121908053 | pathogenic | 1 | 1 | 0 |
| rs122467169 | pathogenic | 1 | 1 | 1 |
| rs122467172 | pathogenic | 1 | 1 | 1 |
| rs122467170 | pathogenic | 1 | 1 | 1 |
| rs28935477 | pathogenic | 1 | 1 | 0 |
| rs28942088 | pathogenic | 0 | 0 | 0 |
| rs751478142 | pathogenic | 1 | 1 | 0 |
| rs765368797 | pathogenic | 0 | 0 | 0 |
| rs147810437 | benign | 0 | 0 | 0 |
| rs121918209 | pathogenic | 1 | 1 | 1 |
| rs121918208 | pathogenic | 1 | 1 | 1 |
| rs121918210 | pathogenic | 1 | 1 | 1 |
| rs35515638 | benign | 1 | 1 | 1 |
| rs121918207 | pathogenic | 1 | 1 | 1 |
| rs141298312 | benign | 0 | 0 | 0 |
| rs121918211 | pathogenic | 0 | 0 | 0 |

| | | | | |
|---|---|---|---|---|
| rs149660059 | benign | 0 | 0 | 0 |
| rs200670286 | pathogenic | 1 | 1 | 1 |
| rs121918212 | pathogenic | 1 | 1 | 1 |
| rs121918213 | pathogenic | 1 | 1 | 1 |
| rs121918205 | pathogenic | 1 | 1 | 1 |
| rs104894025 | pathogenic | 1 | 1 | 1 |
| rs104894026 | pathogenic | 1 | 1 | 1 |
| rs104894028 | pathogenic | 0 | 1 | 0 |
| rs104894029 | pathogenic | 0 | 1 | 0 |
| rs119458971 | pathogenic | 0 | 0 | 0 |
| rs119458969 | pathogenic | 1 | 1 | 1 |
| rs1801591 | benign | 1 | 1 | 1 |
| rs140169311 | benign | 0 | 0 | 0 |
| rs753363278 | benign | 1 | 1 | 1 |
| rs199763682 | pathogenic | 0 | 0 | 0 |
| rs119458970 | pathogenic | 1 | 1 | 1 |
| rs763722868 | pathogenic | 0 | 0 | 0 |
| rs56404770 | benign | 0 | 0 | 0 |
| rs34221241 | benign | 0 | 0 | 0 |
| rs121909625 | pathogenic | 0 | 0 | 0 |
| rs550921485 | pathogenic | 0 | 0 | 0 |
| rs1800454 | benign | 0 | 0 | 0 |
| rs241447 | benign | 0 | 0 | 0 |
| rs2230606 | benign | 1 | 1 | 0 |
| rs5036 | pathogenic | 1 | 1 | 1 |
| rs13306788 | benign | 0 | 0 | 0 |
| rs28929480 | pathogenic | 0 | 0 | 0 |
| rs121912749 | pathogenic | 0 | 0 | 0 |
| rs56361140 | pathogenic | 0 | 0 | 0 |
| rs28931583 | pathogenic | 1 | 1 | 1 |
| rs36211715 | pathogenic | 1 | 1 | 1 |
| rs138049878 | pathogenic | 1 | 1 | 1 |
| rs727503253 | pathogenic | 1 | 1 | 1 |
| rs267606908 | pathogenic | 0 | 0 | 0 |
| rs121913631 | pathogenic | 1 | 1 | 1 |
| rs36211716 | benign | 1 | 1 | 1 |
| rs121913639 | pathogenic | 1 | 1 | 1 |
| rs730880761 | pathogenic | 1 | 1 | 1 |
| rs121913629 | pathogenic | 1 | 1 | 1 |
| rs754368658 | pathogenic | 0 | 0 | 0 |
| rs777090017 | pathogenic | 0 | 0 | 0 |
| rs267606913 | pathogenic | 1 | 1 | 1 |
| rs773138218 | pathogenic | 0 | 0 | 0 |
| rs371295780 | pathogenic | 0 | 0 | 0 |
| rs387906830 | pathogenic | 1 | 1 | 0 |
| rs771364038 | pathogenic | 0 | 1 | 0 |
| rs772731615 | pathogenic | 0 | 0 | 0 |
| rs137853063 | pathogenic | 0 | 0 | 0 |
| rs387907076 | pathogenic | 0 | 1 | 0 |
| rs35460768 | benign | 0 | 0 | 0 |
| rs2278196 | benign | 1 | 1 | 1 |
| rs119103275 | pathogenic | 0 | 0 | 0 |
| rs119103276 | pathogenic | 1 | 1 | 1 |
| rs374402066 | pathogenic | 1 | 1 | 1 |

| | | | | |
|---|---|---|---|---|
| rs119103274 | pathogenic | 0 | 0 | 0 |
| rs2274064 | benign | 0 | 0 | 0 |
| rs267607023 | pathogenic | 1 | 1 | 0 |
| rs142117489 | benign | 1 | 1 | 0 |
| rs371258270 | benign | 0 | 0 | 0 |
| rs34716057 | benign | 0 | 0 | 0 |
| rs201778606 | benign | 0 | 0 | 0 |
| rs201438792 | benign | 0 | 0 | 0 |
| rs140547417 | benign | 0 | 0 | 0 |
| rs5881 | benign | 0 | 0 | 0 |
| rs752298084 | benign | 0 | 0 | 0 |
| rs34855278 | benign | 0 | 0 | 0 |
| rs5880 | benign | 1 | 1 | 0 |
| rs5882 | benign | 0 | 0 | 0 |
| rs1800777 | benign | 0 | 0 | 0 |
| rs577301331 | benign | 0 | 0 | 0 |
| rs137853313 | pathogenic | 0 | 0 | 0 |
| rs28935169 | pathogenic | 0 | 0 | 0 |
| rs137853315 | pathogenic | 0 | 0 | 0 |
| rs137853314 | pathogenic | 0 | 0 | 0 |
| rs28935469 | pathogenic | 0 | 0 | 0 |
| rs137853318 | pathogenic | 0 | 0 | 0 |
| rs28935470 | pathogenic | 0 | 0 | 0 |
| rs3729986 | benign | 0 | 0 | 0 |
| rs730880619 | pathogenic | 0 | 0 | 0 |
| rs201012766 | benign | 0 | 0 | 0 |
| rs727503218 | pathogenic | 0 | 0 | 0 |
| rs11570052 | benign | 0 | 0 | 0 |
| rs730880623 | pathogenic | 0 | 0 | 0 |
| rs138753870 | benign | 0 | 0 | 0 |
| rs397516068 | pathogenic | 0 | 0 | 0 |
| rs3729989 | benign | 1 | 1 | 1 |
| rs397516074 | pathogenic | 0 | 0 | 0 |
| rs561449819 | pathogenic | 0 | 0 | 0 |
| rs121917779 | benign | 0 | 0 | 0 |
| rs142881576 | pathogenic | 0 | 0 | 0 |
| rs77384282 | benign | 0 | 0 | 0 |
| rs201865787 | benign | 0 | 0 | 0 |
| rs142244640 | benign | 0 | 0 | 0 |
| rs200488706 | benign | 0 | 0 | 0 |
| rs137853291 | pathogenic | 1 | 1 | 0 |
| rs28933990 | pathogenic | 1 | 1 | 0 |
| rs140570189 | benign | 0 | 0 | 0 |
| rs201588178 | benign | 0 | 0 | 0 |
| rs104893846 | pathogenic | 0 | 0 | 0 |
| rs754545360 | pathogenic | 0 | 1 | 0 |
| rs104893851 | pathogenic | 0 | 1 | 0 |
| rs104893849 | pathogenic | 0 | 0 | 0 |
| rs571038432 | pathogenic | 1 | 1 | 0 |
| rs2270655 | benign | 1 | 1 | 1 |
| rs315952 | benign | 0 | 0 | 0 |
| rs151176548 | benign | 0 | 0 | 0 |
| rs122459147 | pathogenic | 1 | 1 | 0 |
| rs122458140 | pathogenic | 0 | 0 | 0 |

| | | | | |
|---|---|---|---|---|
| rs122458142 | pathogenic | 1 | 1 | 0 |
| rs267606812 | pathogenic | 1 | 1 | 0 |
| rs267606813 | pathogenic | 1 | 1 | 0 |
| rs122458143 | pathogenic | 0 | 0 | 0 |
| rs122459146 | pathogenic | 0 | 0 | 0 |
| rs122458144 | pathogenic | 0 | 0 | 0 |
| rs122458145 | pathogenic | 0 | 0 | 0 |
| rs113994007 | pathogenic | 1 | 1 | 0 |
| rs758746181 | pathogenic | 1 | 1 | 1 |
| rs145034527 | pathogenic | 1 | 1 | 0 |
| rs119468006 | pathogenic | 1 | 1 | 0 |
| rs201908721 | pathogenic | 0 | 0 | 0 |
| rs755933881 | pathogenic | 1 | 1 | 0 |
| rs771578775 | pathogenic | 0 | 0 | 0 |
| rs17849927 | benign | 0 | 0 | 0 |
| rs139133094 | benign | 0 | 0 | 0 |
| rs12593 | benign | 0 | 0 | 0 |
| rs578189699 | pathogenic | 1 | 1 | 0 |
| rs119468008 | pathogenic | 0 | 0 | 0 |
| rs768524626 | benign | 0 | 0 | 0 |
| rs119468009 | pathogenic | 0 | 0 | 0 |
| rs119468004 | pathogenic | 0 | 0 | 0 |
| rs199874519 | pathogenic | 0 | 0 | 0 |
| rs376478331 | pathogenic | 0 | 0 | 0 |
| rs61995958 | pathogenic | 0 | 0 | 0 |
| rs199714731 | benign | 0 | 0 | 0 |
| rs149185954 | benign | 0 | 0 | 0 |
| rs121918353 | pathogenic | 0 | 0 | 0 |
| rs1131695 | benign | 0 | 0 | 0 |
| rs28939668 | pathogenic | 0 | 0 | 0 |
| rs758432471 | pathogenic | 1 | 1 | 1 |
| rs752746786 | pathogenic | 1 | 1 | 1 |
| rs28935471 | pathogenic | 0 | 0 | 0 |
| rs80338837 | pathogenic | 0 | 0 | 0 |
| rs137853312 | pathogenic | 0 | 0 | 0 |
| rs28935472 | pathogenic | 0 | 0 | 0 |
| rs28935473 | pathogenic | 0 | 0 | 0 |
| rs267606752 | pathogenic | 0 | 0 | 0 |
| rs3917973 | benign | 0 | 0 | 0 |
| rs3918018 | benign | 0 | 0 | 0 |
| rs757976755 | pathogenic | 0 | 0 | 0 |
| rs121918329 | pathogenic | 1 | 1 | 0 |
| rs770114459 | pathogenic | 0 | 0 | 0 |
| rs6662272 | benign | 0 | 0 | 0 |
| rs121918242 | pathogenic | 1 | 1 | 0 |
| rs121918240 | pathogenic | 1 | 1 | 0 |
| rs200094982 | pathogenic | 0 | 0 | 0 |
| rs121918241 | pathogenic | 0 | 0 | 0 |
| rs140522266 | pathogenic | 1 | 1 | 0 |
| rs121918243 | pathogenic | 1 | 1 | 0 |
| rs370596113 | pathogenic | 1 | 1 | 0 |
| rs398124295 | pathogenic | 1 | 1 | 0 |
| rs587776889 | pathogenic | 1 | 1 | 0 |
| rs747527726 | pathogenic | 1 | 1 | 0 |

| | | | | |
|---|---|---|---|---|
| rs112631212 | benign | 0 | 0 | 0 |
| rs104894605 | pathogenic | 0 | 0 | 0 |
| rs104894607 | pathogenic | 0 | 0 | 0 |
| rs104894606 | pathogenic | 0 | 0 | 0 |
| rs143873873 | benign | 0 | 0 | 0 |
| rs104894388 | pathogenic | 1 | 1 | 0 |
| rs104894391 | pathogenic | 1 | 1 | 0 |
| rs5350 | benign | 1 | 1 | 1 |
| rs104894390 | pathogenic | 1 | 1 | 1 |
| rs5346 | benign | 1 | 1 | 1 |
| rs121912500 | pathogenic | 0 | 0 | 0 |
| rs74315450 | pathogenic | 1 | 1 | 0 |
| rs121913396 | pathogenic | 1 | 1 | 1 |
| rs28931588 | pathogenic | 1 | 1 | 1 |
| rs121913400 | pathogenic | 0 | 0 | 0 |
| rs28931589 | pathogenic | 1 | 1 | 1 |
| rs121913403 | pathogenic | 0 | 0 | 0 |
| rs121913413 | pathogenic | 0 | 0 | 0 |
| rs760107623 | pathogenic | 1 | 1 | 0 |
| rs16866378 | benign | 1 | 1 | 0 |
| rs201025290 | pathogenic | 1 | 1 | 0 |
| rs12720356 | benign | 1 | 1 | 0 |
| rs55882956 | benign | 0 | 1 | 0 |
| rs4986933 | benign | 1 | 1 | 0 |
| rs2020957 | benign | 1 | 1 | 1 |
| rs1800124 | benign | 1 | 1 | 1 |
| rs150077735 | benign | 1 | 1 | 1 |
| rs201419000 | benign | 0 | 0 | 0 |
| rs772044176 | pathogenic | 0 | 0 | 0 |
| rs138346388 | benign | 0 | 0 | 0 |
| rs121964916 | pathogenic | 0 | 0 | 0 |
| rs112534524 | pathogenic | 0 | 0 | 0 |
| rs182078921 | benign | 1 | 1 | 1 |
| rs113273712 | benign | 0 | 0 | 0 |
| rs74817407 | benign | 1 | 1 | 1 |
| rs61733901 | benign | 1 | 1 | 1 |
| rs121964912 | pathogenic | 1 | 1 | 1 |
| rs41278047 | benign | 1 | 1 | 1 |
| rs7437875 | benign | 1 | 1 | 1 |
| rs113460688 | benign | 1 | 1 | 1 |
| rs754572081 | benign | 1 | 1 | 1 |
| rs201082692 | benign | 1 | 1 | 1 |
| rs202085803 | benign | 1 | 1 | 1 |
| rs141320511 | benign | 0 | 0 | 0 |
| rs78501403 | benign | 0 | 0 | 0 |
| rs28939675 | pathogenic | 0 | 0 | 0 |
| rs74315522 | pathogenic | 0 | 0 | 0 |
| rs121434639 | pathogenic | 1 | 1 | 0 |
| rs121434640 | pathogenic | 1 | 1 | 1 |
| rs757282628 | benign | 1 | 1 | 1 |
| rs386834010 | pathogenic | 0 | 0 | 0 |
| rs200863680 | pathogenic | 0 | 0 | 0 |
| rs193919336 | pathogenic | 1 | 1 | 0 |
| rs386834011 | pathogenic | 0 | 0 | 0 |

| | | | | |
|---|---|---|---|---|
| rs28940869 | pathogenic | 0 | 0 | 0 |
| rs386834014 | pathogenic | 1 | 1 | 0 |
| rs267606960 | pathogenic | 0 | 0 | 0 |
| rs28942068 | pathogenic | 0 | 0 | 0 |
| rs193919335 | pathogenic | 0 | 0 | 0 |
| rs74374973 | pathogenic | 0 | 0 | 0 |
| rs386834018 | pathogenic | 0 | 0 | 0 |
| rs267606962 | pathogenic | 0 | 0 | 0 |
| rs6659553 | benign | 0 | 0 | 0 |
| rs29281 | benign | 0 | 0 | 0 |
| rs782265090 | benign | 0 | 0 | 0 |
| rs35910969 | pathogenic | 0 | 0 | 0 |
| rs121909113 | pathogenic | 1 | 1 | 1 |
| rs29001571 | pathogenic | 1 | 1 | 1 |
| rs587781250 | pathogenic | 1 | 1 | 1 |
| rs28939680 | pathogenic | 1 | 1 | 1 |
| rs28939681 | pathogenic | 1 | 1 | 1 |
| rs121909112 | pathogenic | 1 | 1 | 1 |
| rs28937568 | pathogenic | 1 | 1 | 1 |
| rs1799895 | pathogenic | 0 | 0 | 0 |
| rs386833431 | pathogenic | 1 | 1 | 1 |
| rs121964905 | pathogenic | 1 | 1 | 1 |
| rs121964906 | pathogenic | 1 | 1 | 1 |
| rs11551664 | pathogenic | 1 | 1 | 1 |
| rs104894705 | pathogenic | 1 | 1 | 1 |
| rs104893805 | pathogenic | 0 | 0 | 0 |
| rs104894205 | pathogenic | 0 | 0 | 0 |
| rs104894204 | pathogenic | 1 | 1 | 0 |
| rs62642926 | pathogenic | 0 | 0 | 0 |
| rs5030841 | pathogenic | 1 | 1 | 1 |
| rs75193786 | pathogenic | 1 | 1 | 1 |
| rs62508682 | pathogenic | 0 | 0 | 0 |
| rs3850625 | benign | 1 | 1 | 1 |
| rs141908793 | benign | 0 | 0 | 0 |
| rs187864727 | benign | 0 | 0 | 0 |
| rs548909101 | benign | 0 | 0 | 0 |
| rs3825394 | benign | 0 | 1 | 0 |
| rs143548402 | benign | 0 | 0 | 0 |
| rs267607146 | pathogenic | 0 | 0 | 0 |
| rs267607144 | pathogenic | 0 | 0 | 0 |
| rs267607148 | pathogenic | 1 | 1 | 1 |
| rs267607143 | pathogenic | 0 | 0 | 0 |
| rs267607145 | pathogenic | 0 | 0 | 0 |
| rs121912636 | pathogenic | 1 | 1 | 1 |
| rs121912634 | pathogenic | 1 | 1 | 1 |
| rs570675468 | benign | 0 | 0 | 0 |
| rs368861241 | pathogenic | 1 | 1 | 1 |
| rs104894528 | pathogenic | 1 | 1 | 1 |
| rs77493891 | benign | 0 | 0 | 0 |
| rs62640903 | benign | 0 | 0 | 0 |
| rs149926091 | benign | 1 | 1 | 1 |
| rs641081 | benign | 0 | 0 | 0 |
| rs4930199 | benign | 0 | 0 | 0 |
| rs121913020 | pathogenic | 0 | 0 | 0 |

| | | | | |
|---|---|---|---|---|
| rs1799793 | benign | 0 | 0 | 0 |
| rs121913016 | pathogenic | 0 | 0 | 0 |
| rs121913025 | pathogenic | 0 | 0 | 0 |
| rs121913019 | pathogenic | 0 | 0 | 0 |
| rs376556895 | pathogenic | 1 | 1 | 1 |
| rs121913024 | pathogenic | 1 | 1 | 1 |
| rs121913021 | pathogenic | 0 | 0 | 0 |
| rs121913023 | pathogenic | 0 | 0 | 0 |
| rs41556519 | pathogenic | 0 | 0 | 0 |
| rs121913022 | pathogenic | 0 | 0 | 0 |
| rs144564120 | pathogenic | 0 | 0 | 0 |
| rs121913026 | pathogenic | 1 | 1 | 1 |
| rs121913018 | pathogenic | 1 | 1 | 1 |
| rs142954643 | benign | 0 | 0 | 0 |
| rs76655496 | benign | 0 | 0 | 0 |
| rs104894371 | pathogenic | 1 | 1 | 0 |
| rs137964936 | benign | 0 | 0 | 0 |
| rs759736132 | pathogenic | 0 | 0 | 0 |
| rs387906595 | pathogenic | 1 | 1 | 0 |
| rs768248216 | pathogenic | 0 | 0 | 0 |
| rs548231008 | benign | 0 | 0 | 0 |
| rs757094384 | benign | 0 | 0 | 0 |
| rs867529 | benign | 0 | 0 | 0 |
| rs13045 | benign | 1 | 1 | 1 |
| rs121908570 | pathogenic | 0 | 0 | 0 |
| rs201872586 | benign | 1 | 1 | 0 |
| rs149092990 | benign | 0 | 0 | 0 |
| rs115585711 | benign | 0 | 0 | 0 |
| rs137909416 | benign | 1 | 1 | 0 |
| rs143388949 | benign | 1 | 1 | 1 |
| rs73137716 | benign | 1 | 1 | 1 |
| rs121908947 | pathogenic | 1 | 1 | 1 |
| rs398122904 | pathogenic | 1 | 1 | 1 |
| rs35715497 | benign | 1 | 1 | 0 |
| rs35462421 | benign | 0 | 0 | 0 |
| rs119475043 | pathogenic | 0 | 0 | 0 |
| rs119475042 | pathogenic | 0 | 0 | 0 |
| rs1076669 | benign | 1 | 1 | 0 |
| rs141146885 | benign | 0 | 0 | 0 |
| rs3026906 | pathogenic | 0 | 0 | 0 |
| rs75362221 | benign | 1 | 1 | 1 |
| rs138472116 | benign | 1 | 1 | 1 |
| rs371040282 | pathogenic | 1 | 1 | 1 |
| rs4988496 | benign | 0 | 0 | 0 |
| rs121918117 | pathogenic | 1 | 1 | 1 |
| rs4988498 | benign | 0 | 0 | 0 |
| rs5035 | benign | 0 | 0 | 0 |
| rs121918322 | pathogenic | 1 | 1 | 1 |
| rs370359540 | benign | 0 | 0 | 0 |
| rs758972589 | pathogenic | 0 | 0 | 0 |
| rs144856889 | benign | 0 | 0 | 0 |
| rs121434623 | pathogenic | 1 | 1 | 1 |
| rs61735600 | benign | 0 | 0 | 0 |
| rs72550870 | benign | 1 | 1 | 1 |

| | | | | |
|---|---|---|---|---|
| rs56392418 | benign | 0 | 0 | 0 |
| rs41307788 | benign | 1 | 1 | 1 |
| rs387907239 | pathogenic | 1 | 1 | 0 |
| rs28931584 | pathogenic | 0 | 0 | 0 |
| rs121912745 | pathogenic | 0 | 0 | 0 |
| rs121912744 | pathogenic | 0 | 0 | 0 |
| rs121912754 | pathogenic | 1 | 1 | 1 |
| rs121912746 | pathogenic | 0 | 0 | 0 |
| rs121912757 | pathogenic | 0 | 0 | 0 |
| rs121912758 | pathogenic | 0 | 0 | 0 |
| rs121912748 | pathogenic | 0 | 0 | 0 |
| rs121912755 | pathogenic | 0 | 0 | 0 |
| rs121912741 | pathogenic | 0 | 0 | 0 |
| rs121912753 | pathogenic | 0 | 0 | 0 |
| rs121912750 | pathogenic | 0 | 0 | 0 |
| rs2285644 | pathogenic | 0 | 0 | 0 |
| rs121912751 | pathogenic | 0 | 0 | 0 |
| rs5026 | benign | 0 | 0 | 0 |
| rs121912759 | pathogenic | 0 | 0 | 0 |
| rs28931585 | pathogenic | 0 | 0 | 0 |
| rs17122769 | benign | 0 | 0 | 0 |
| rs28362322 | benign | 1 | 1 | 1 |
| rs3824874 | benign | 0 | 0 | 0 |
| rs146004831 | benign | 0 | 0 | 0 |
| rs142155860 | benign | 0 | 0 | 0 |
| rs121434404 | pathogenic | 0 | 0 | 0 |
| rs61735578 | benign | 1 | 1 | 1 |
| rs558018 | benign | 0 | 0 | 0 |
| rs61735577 | benign | 1 | 1 | 1 |
| rs76784113 | benign | 0 | 0 | 0 |
| rs199469640 | pathogenic | 0 | 0 | 0 |
| rs199469628 | pathogenic | 1 | 1 | 0 |
| rs199469629 | pathogenic | 0 | 1 | 0 |
| rs387907156 | pathogenic | 0 | 1 | 0 |
| rs199469643 | pathogenic | 0 | 0 | 0 |
| rs199469646 | pathogenic | 0 | 0 | 0 |
| rs766294940 | pathogenic | 0 | 0 | 0 |
| rs387907347 | pathogenic | 0 | 0 | 0 |
| rs137853590 | pathogenic | 1 | 1 | 1 |
| rs137853589 | pathogenic | 1 | 1 | 1 |
| rs137853588 | pathogenic | 1 | 1 | 1 |
| rs55909005 | benign | 0 | 0 | 0 |
| rs4581 | benign | 0 | 0 | 0 |
| rs63750123 | benign | 0 | 0 | 0 |
| rs10254120 | benign | 0 | 0 | 0 |
| rs121434629 | pathogenic | 1 | 1 | 0 |
| rs200313585 | benign | 0 | 0 | 0 |
| rs143162541 | benign | 1 | 1 | 0 |
| rs760615315 | benign | 0 | 0 | 0 |
| rs768488890 | benign | 1 | 1 | 1 |
| rs116373169 | benign | 1 | 1 | 0 |
| rs63750871 | pathogenic | 1 | 1 | 0 |
| rs116349687 | benign | 0 | 0 | 0 |
| rs375289386 | benign | 0 | 0 | 0 |

| | | | | |
|---|---|---|---|---|
| rs760228510 | pathogenic | 1 | 1 | 1 |
| rs587779343 | pathogenic | 1 | 1 | 1 |
| rs573125799 | benign | 0 | 0 | 0 |
| rs1805322 | benign | 0 | 0 | 0 |
| rs267608153 | pathogenic | 0 | 0 | 0 |
| rs139194813 | benign | 0 | 0 | 0 |
| rs530021751 | benign | 0 | 0 | 0 |
| rs200640585 | pathogenic | 0 | 0 | 0 |
| rs139438201 | benign | 0 | 0 | 0 |
| rs200513014 | benign | 0 | 0 | 0 |
| rs567102013 | benign | 0 | 0 | 0 |
| rs121908608 | pathogenic | 0 | 0 | 0 |
| rs121908606 | pathogenic | 0 | 0 | 0 |
| rs17103671 | benign | 0 | 0 | 0 |
| rs34725281 | benign | 0 | 0 | 0 |
| rs104894421 | pathogenic | 1 | 1 | 0 |
| rs121918145 | pathogenic | 1 | 1 | 1 |
| rs121918152 | pathogenic | 1 | 1 | 1 |
| rs56379106 | benign | 0 | 0 | 0 |
| rs4987076 | benign | 0 | 0 | 0 |
| rs370243877 | pathogenic | 0 | 0 | 0 |
| rs17679445 | benign | 1 | 1 | 1 |
| rs550423482 | pathogenic | 1 | 1 | 1 |
| rs104894629 | pathogenic | 1 | 1 | 1 |
| rs773450573 | pathogenic | 1 | 1 | 1 |
| rs116679717 | benign | 0 | 0 | 0 |
| rs372000848 | pathogenic | 0 | 0 | 0 |
| rs138595973 | benign | 0 | 0 | 0 |
| rs115244417 | benign | 0 | 0 | 0 |
| rs11575292 | benign | 0 | 0 | 0 |
| rs137853211 | pathogenic | 1 | 1 | 1 |
| rs137853207 | pathogenic | 0 | 0 | 0 |
| rs137853210 | pathogenic | 1 | 1 | 1 |
| rs6262 | benign | 0 | 0 | 0 |
| rs6263 | benign | 0 | 1 | 0 |
| rs137853208 | pathogenic | 0 | 1 | 0 |
| rs137853212 | pathogenic | 1 | 1 | 0 |
| rs137853209 | pathogenic | 1 | 1 | 1 |
| rs201951824 | pathogenic | 0 | 0 | 0 |
| rs11575542 | benign | 0 | 0 | 0 |
| rs199681788 | benign | 0 | 1 | 0 |
| rs748835902 | benign | 1 | 1 | 1 |
| rs753719949 | benign | 0 | 0 | 0 |
| rs267606647 | pathogenic | 0 | 0 | 0 |
| rs267606648 | pathogenic | 1 | 1 | 1 |
| rs267606643 | pathogenic | 0 | 1 | 0 |
| rs559947967 | pathogenic | 1 | 1 | 0 |
| rs267606645 | pathogenic | 1 | 1 | 1 |
| rs12116440 | benign | 0 | 0 | 0 |
| rs121908539 | pathogenic | 0 | 0 | 0 |
| rs137853096 | pathogenic | 1 | 1 | 0 |
| rs148363262 | benign | 1 | 1 | 0 |
| rs587777442 | pathogenic | 0 | 0 | 0 |
| rs25640 | benign | 1 | 1 | 1 |

| | | | | |
|---|---|---|---|---|
| rs28943589 | benign | 0 | 0 | 0 |
| rs1143650 | benign | 1 | 1 | 1 |
| rs150886026 | benign | 0 | 1 | 0 |
| rs121908133 | pathogenic | 1 | 1 | 0 |
| rs121908131 | pathogenic | 0 | 0 | 0 |
| rs754477154 | pathogenic | 0 | 0 | 0 |
| rs747334007 | benign | 0 | 0 | 0 |
| rs121908132 | pathogenic | 0 | 0 | 0 |
| rs104894103 | pathogenic | 1 | 1 | 0 |
| rs140088149 | benign | 0 | 0 | 0 |
| rs149803148 | benign | 0 | 0 | 0 |
| rs11064480 | benign | 0 | 0 | 0 |
| rs74435397 | pathogenic | 1 | 1 | 1 |
| rs587783626 | pathogenic | 1 | 1 | 1 |
| rs148211042 | pathogenic | 1 | 1 | 1 |
| rs17849502 | benign | 0 | 0 | 0 |
| rs13306575 | pathogenic | 1 | 1 | 1 |
| rs201426733 | benign | 0 | 0 | 0 |
| rs182278784 | benign | 1 | 1 | 1 |
| rs77786415 | benign | 1 | 1 | 1 |
| rs200192119 | benign | 1 | 1 | 1 |
| rs12877501 | benign | 1 | 1 | 1 |
| rs77985244 | benign | 1 | 1 | 1 |
| rs5907 | pathogenic | 1 | 1 | 0 |
| rs2305178 | benign | 0 | 0 | 0 |
| rs3135867 | benign | 0 | 0 | 0 |
| rs121913482 | pathogenic | 1 | 1 | 1 |
| rs121913483 | pathogenic | 1 | 1 | 1 |
| rs4647924 | pathogenic | 1 | 1 | 1 |
| rs121913111 | pathogenic | 1 | 1 | 1 |
| rs104894002 | pathogenic | 0 | 0 | 0 |
| rs104894001 | pathogenic | 0 | 0 | 0 |
| rs75932628 | benign | 0 | 0 | 0 |
| rs201258663 | pathogenic | 0 | 0 | 0 |
| rs104893998 | pathogenic | 0 | 0 | 0 |
| rs2234253 | benign | 1 | 1 | 0 |
| rs121908402 | pathogenic | 1 | 1 | 0 |
| rs1805389 | benign | 0 | 0 | 0 |
| rs1805388 | benign | 0 | 0 | 0 |
| rs748385144 | pathogenic | 0 | 0 | 0 |
| rs104894420 | pathogenic | 0 | 0 | 0 |
| rs104894418 | pathogenic | 0 | 0 | 0 |
| rs61731910 | benign | 0 | 0 | 0 |
| rs758723288 | pathogenic | 1 | 1 | 0 |
| rs201865375 | pathogenic | 1 | 1 | 0 |
| rs565090080 | pathogenic | 1 | 1 | 1 |
| rs113042166 | benign | 0 | 0 | 0 |
| rs61736326 | benign | 1 | 1 | 1 |
| rs12030699 | benign | 1 | 1 | 1 |
| rs121913578 | pathogenic | 1 | 1 | 1 |
| rs61739582 | benign | 0 | 0 | 0 |
| rs745983207 | pathogenic | 0 | 0 | 0 |
| rs763524380 | benign | 0 | 0 | 0 |
| rs121912792 | pathogenic | 0 | 0 | 0 |

| | | | | |
|---|---|---|---|---|
| rs121912793 | pathogenic | 0 | 0 | 0 |
| rs387906599 | pathogenic | 0 | 0 | 0 |
| rs121912794 | pathogenic | 0 | 1 | 0 |
| rs121912795 | pathogenic | 0 | 0 | 0 |
| rs121912796 | pathogenic | 1 | 1 | 0 |
| rs121912797 | pathogenic | 0 | 1 | 0 |
| rs121912798 | pathogenic | 1 | 1 | 1 |
| rs80359812 | pathogenic | 1 | 1 | 0 |
| rs776583130 | benign | 1 | 1 | 0 |
| rs80359814 | pathogenic | 1 | 1 | 0 |
| rs202060209 | pathogenic | 1 | 1 | 0 |
| rs267607061 | pathogenic | 1 | 1 | 0 |
| rs267607060 | pathogenic | 0 | 0 | 0 |
| rs80359816 | pathogenic | 1 | 1 | 0 |
| rs80359818 | pathogenic | 1 | 1 | 0 |
| rs387907313 | pathogenic | 0 | 0 | 0 |
| rs587784395 | benign | 0 | 0 | 0 |
| rs5811 | benign | 0 | 0 | 0 |
| rs121909740 | pathogenic | 1 | 1 | 0 |
| rs148518827 | benign | 1 | 1 | 0 |
| rs121909739 | pathogenic | 1 | 1 | 0 |
| rs780528770 | benign | 1 | 1 | 0 |
| rs13306758 | pathogenic | 0 | 0 | 0 |
| rs267607059 | pathogenic | 0 | 0 | 0 |
| rs61730638 | benign | 0 | 0 | 0 |
| rs104894533 | pathogenic | 0 | 0 | 0 |
| rs104894527 | pathogenic | 0 | 0 | 0 |
| rs200503569 | pathogenic | 1 | 1 | 1 |
| rs80338700 | pathogenic | 1 | 1 | 1 |
| rs765433263 | pathogenic | 1 | 1 | 1 |
| rs104894530 | pathogenic | 1 | 1 | 1 |
| rs80338701 | pathogenic | 1 | 1 | 1 |
| rs141498002 | pathogenic | 1 | 1 | 1 |
| rs104894525 | pathogenic | 0 | 0 | 0 |
| rs80338703 | pathogenic | 0 | 0 | 0 |
| rs28936415 | pathogenic | 1 | 1 | 0 |
| rs150719105 | pathogenic | 0 | 0 | 0 |
| rs148032587 | pathogenic | 0 | 0 | 0 |
| rs150577656 | pathogenic | 0 | 0 | 0 |
| rs190521996 | pathogenic | 0 | 0 | 0 |
| rs104894526 | pathogenic | 1 | 1 | 1 |
| rs80338704 | pathogenic | 0 | 0 | 0 |
| rs199562225 | pathogenic | 1 | 1 | 1 |
| rs532870929 | pathogenic | 0 | 0 | 0 |
| rs398123309 | pathogenic | 0 | 0 | 0 |
| rs78290141 | pathogenic | 0 | 0 | 0 |
| rs80338705 | pathogenic | 0 | 0 | 0 |
| rs104894531 | pathogenic | 1 | 1 | 1 |
| rs80338706 | pathogenic | 0 | 0 | 0 |
| rs80338707 | pathogenic | 0 | 0 | 0 |
| rs80338708 | pathogenic | 0 | 0 | 0 |
| rs80338709 | pathogenic | 0 | 0 | 0 |
| rs104893833 | benign | 0 | 0 | 0 |
| rs61744585 | benign | 0 | 0 | 0 |

| | | | | |
|---|---|---|---|---|
| rs73569515 | benign | 0 | 0 | 0 |
| rs398123373 | pathogenic | 0 | 0 | 0 |
| rs756854513 | pathogenic | 0 | 0 | 0 |
| rs58232698 | benign | 0 | 0 | 0 |
| rs9600455 | benign | 0 | 0 | 0 |
| rs201016593 | pathogenic | 0 | 0 | 0 |
| rs756039188 | pathogenic | 0 | 0 | 0 |
| rs121908024 | pathogenic | 0 | 0 | 0 |
| rs373144619 | benign | 0 | 0 | 0 |
| rs138078086 | benign | 0 | 0 | 0 |
| rs376207800 | pathogenic | 0 | 0 | 0 |
| rs749038326 | pathogenic | 1 | 1 | 0 |
| rs372845091 | benign | 0 | 0 | 0 |
| rs377271627 | pathogenic | 0 | 0 | 0 |
| rs563390335 | pathogenic | 0 | 0 | 0 |
| rs13306510 | benign | 0 | 0 | 0 |
| rs764042910 | pathogenic | 0 | 0 | 0 |
| rs201384282 | benign | 1 | 1 | 1 |
| rs730882086 | pathogenic | 1 | 1 | 1 |
| rs756613387 | pathogenic | 1 | 1 | 1 |
| rs121908028 | benign | 1 | 1 | 0 |
| rs773328511 | benign | 0 | 0 | 0 |
| rs749220643 | pathogenic | 1 | 1 | 0 |
| rs730882089 | pathogenic | 0 | 0 | 0 |
| rs146651743 | benign | 0 | 0 | 0 |
| rs148698650 | pathogenic | 0 | 0 | 0 |
| rs140241383 | benign | 0 | 0 | 0 |
| rs757252110 | pathogenic | 0 | 0 | 0 |
| rs13306512 | pathogenic | 1 | 1 | 1 |
| rs121908043 | pathogenic | 1 | 1 | 1 |
| rs773658037 | pathogenic | 0 | 0 | 0 |
| rs774439908 | pathogenic | 0 | 0 | 0 |
| rs779732323 | pathogenic | 1 | 1 | 0 |
| rs749780672 | pathogenic | 0 | 0 | 0 |
| rs375651668 | benign | 1 | 1 | 1 |
| rs193922568 | pathogenic | 1 | 1 | 0 |
| rs750363970 | benign | 1 | 1 | 0 |
| rs5932 | benign | 0 | 0 | 0 |
| rs755154048 | pathogenic | 1 | 1 | 0 |
| rs759876319 | pathogenic | 0 | 0 | 0 |
| rs764550980 | pathogenic | 1 | 1 | 0 |
| rs779940524 | pathogenic | 0 | 0 | 0 |
| rs563382937 | pathogenic | 1 | 1 | 1 |
| rs28942083 | pathogenic | 0 | 1 | 0 |
| rs770744861 | pathogenic | 0 | 1 | 0 |
| rs121908031 | pathogenic | 0 | 1 | 0 |
| rs776217028 | pathogenic | 0 | 1 | 0 |
| rs763625913 | pathogenic | 0 | 0 | 0 |
| rs3736772 | benign | 0 | 0 | 0 |
| rs104894696 | benign | 1 | 1 | 1 |
| rs121918177 | pathogenic | 0 | 1 | 0 |
| rs121918181 | pathogenic | 1 | 1 | 0 |
| rs121918174 | pathogenic | 0 | 1 | 0 |
| rs121918182 | pathogenic | 0 | 0 | 0 |

| | | | | |
|---|---|---|---|---|
| rs121918175 | pathogenic | 1 | 1 | 1 |
| rs121918185 | pathogenic | 1 | 1 | 1 |
| rs121918173 | pathogenic | 1 | 1 | 0 |
| rs121918178 | pathogenic | 1 | 1 | 1 |
| rs121918179 | pathogenic | 1 | 1 | 1 |
| rs121918183 | pathogenic | 0 | 0 | 0 |
| rs121918176 | pathogenic | 1 | 1 | 1 |
| rs121918172 | pathogenic | 0 | 1 | 0 |
| rs121918184 | pathogenic | 0 | 0 | 0 |
| rs121434560 | pathogenic | 1 | 1 | 0 |
| rs121434561 | pathogenic | 1 | 1 | 0 |
| rs3218716 | pathogenic | 1 | 1 | 1 |
| rs145532615 | benign | 0 | 0 | 0 |
| rs149840927 | pathogenic | 0 | 0 | 0 |
| rs730880762 | pathogenic | 0 | 0 | 0 |
| rs730880764 | pathogenic | 0 | 0 | 0 |
| rs730880765 | pathogenic | 0 | 0 | 0 |
| rs587782962 | pathogenic | 0 | 0 | 0 |
| rs141735183 | pathogenic | 0 | 0 | 0 |
| rs397516187 | pathogenic | 0 | 0 | 0 |
| rs727505026 | pathogenic | 0 | 0 | 0 |
| rs529700838 | pathogenic | 0 | 0 | 0 |
| rs540263945 | pathogenic | 0 | 0 | 0 |
| rs149103761 | benign | 0 | 0 | 0 |
| rs730880906 | pathogenic | 0 | 0 | 0 |
| rs730880785 | pathogenic | 0 | 0 | 0 |
| rs397516199 | pathogenic | 0 | 0 | 0 |
| rs730880725 | benign | 0 | 0 | 0 |
| rs730880796 | pathogenic | 0 | 0 | 0 |
| rs201307101 | benign | 0 | 0 | 0 |
| rs397516214 | pathogenic | 0 | 0 | 0 |
| rs3729823 | benign | 0 | 0 | 0 |
| rs397516218 | pathogenic | 0 | 0 | 0 |
| rs121913647 | pathogenic | 0 | 0 | 0 |
| rs373514686 | pathogenic | 0 | 0 | 0 |
| rs763538103 | pathogenic | 0 | 0 | 0 |
| rs121913650 | pathogenic | 0 | 0 | 0 |
| rs139222507 | pathogenic | 0 | 0 | 0 |
| rs369940645 | pathogenic | 0 | 0 | 0 |
| rs763073072 | pathogenic | 0 | 0 | 0 |
| rs145734640 | pathogenic | 0 | 0 | 0 |
| rs730880821 | pathogenic | 0 | 0 | 0 |
| rs28933098 | pathogenic | 0 | 0 | 0 |
| rs372381770 | pathogenic | 0 | 0 | 0 |
| rs146796870 | benign | 0 | 0 | 0 |
| rs104894485 | pathogenic | 1 | 1 | 1 |
| rs35407712 | benign | 0 | 0 | 0 |
| rs35842750 | benign | 0 | 0 | 0 |
| rs2721191 | benign | 0 | 0 | 0 |
| rs35098923 | benign | 0 | 0 | 0 |
| rs34293591 | benign | 1 | 1 | 0 |
| rs36023964 | benign | 0 | 0 | 0 |
| rs4251691 | benign | 0 | 0 | 0 |
| rs34666647 | benign | 0 | 0 | 0 |

| | | | | |
|---|---|---|---|---|
| rs4925828 | benign | 0 | 0 | 0 |
| rs35348691 | benign | 0 | 0 | 0 |
| rs36078464 | benign | 0 | 0 | 0 |
| rs34915097 | benign | 0 | 0 | 0 |
| rs35101495 | benign | 0 | 0 | 0 |
| rs141568342 | pathogenic | 0 | 0 | 0 |
| rs1800571 | pathogenic | 0 | 0 | 0 |
| rs140566084 | benign | 1 | 1 | 1 |
| rs727503791 | pathogenic | 1 | 1 | 0 |
| rs545215807 | pathogenic | 0 | 0 | 0 |
| rs113994168 | pathogenic | 0 | 0 | 0 |
| rs113994167 | pathogenic | 0 | 0 | 0 |
| rs200788251 | pathogenic | 0 | 0 | 0 |
| rs200573371 | pathogenic | 0 | 0 | 0 |
| rs771874163 | pathogenic | 1 | 1 | 1 |
| rs112406105 | pathogenic | 1 | 1 | 1 |
| rs118204015 | pathogenic | 1 | 1 | 1 |
| rs113994169 | pathogenic | 0 | 0 | 0 |
| rs118204018 | pathogenic | 0 | 1 | 0 |
| rs138834083 | pathogenic | 0 | 0 | 0 |
| rs533055438 | pathogenic | 1 | 1 | 1 |
| rs2309689 | pathogenic | 1 | 1 | 1 |
| rs138058572 | pathogenic | 1 | 1 | 1 |
| rs118204017 | pathogenic | 1 | 1 | 0 |
| rs766742117 | pathogenic | 1 | 1 | 1 |
| rs751995154 | pathogenic | 1 | 1 | 1 |
| rs200771970 | pathogenic | 0 | 0 | 0 |
| rs118204014 | pathogenic | 0 | 0 | 0 |
| rs148584617 | pathogenic | 1 | 1 | 1 |
| rs121912720 | pathogenic | 0 | 0 | 0 |
| rs121912721 | pathogenic | 0 | 0 | 0 |
| rs28929476 | pathogenic | 0 | 0 | 0 |
| rs28931574 | pathogenic | 1 | 1 | 1 |
| rs12718465 | benign | 0 | 0 | 0 |
| rs144689354 | pathogenic | 0 | 0 | 0 |
| rs149095705 | pathogenic | 0 | 0 | 0 |
| rs572839648 | benign | 1 | 1 | 1 |
| rs121909715 | pathogenic | 1 | 1 | 0 |
| rs34077456 | benign | 1 | 1 | 1 |
| rs397517876 | benign | 0 | 0 | 0 |
| rs1064074 | benign | 0 | 0 | 0 |
| rs142751309 | benign | 0 | 0 | 0 |
| rs116275562 | benign | 0 | 0 | 0 |
| rs148095899 | benign | 0 | 0 | 0 |
| rs751838040 | pathogenic | 0 | 0 | 0 |
| rs121434324 | pathogenic | 1 | 1 | 1 |
| rs143367518 | pathogenic | 0 | 0 | 0 |
| rs121434325 | pathogenic | 0 | 0 | 0 |
| rs397515328 | pathogenic | 0 | 0 | 0 |
| rs376039938 | pathogenic | 0 | 0 | 0 |
| rs387907243 | pathogenic | 0 | 0 | 0 |
| rs28934876 | pathogenic | 0 | 0 | 0 |
| rs387907245 | pathogenic | 0 | 0 | 0 |
| rs138544311 | benign | 0 | 0 | 0 |

| | | | | |
|---|---|---|---|---|
| rs643788 | benign | 0 | 0 | 0 |
| rs749977756 | benign | 0 | 0 | 0 |
| rs1800574 | benign | 0 | 0 | 0 |
| rs137853336 | pathogenic | 1 | 1 | 1 |
| rs111916034 | benign | 0 | 0 | 0 |
| rs307826 | benign | 0 | 0 | 0 |
| rs35874891 | benign | 0 | 0 | 0 |
| rs111033580 | pathogenic | 1 | 1 | 1 |
| rs111033581 | pathogenic | 1 | 1 | 1 |
| rs121908373 | pathogenic | 0 | 0 | 0 |
| rs767122713 | pathogenic | 0 | 0 | 0 |
| rs2233213 | benign | 0 | 0 | 0 |
| rs121908605 | pathogenic | 1 | 1 | 0 |
| rs11887534 | benign | 0 | 0 | 0 |
| rs4148211 | benign | 0 | 0 | 0 |
| rs80025980 | benign | 0 | 0 | 0 |
| rs137854891 | pathogenic | 1 | 1 | 0 |
| rs74358901 | benign | 0 | 0 | 0 |
| rs9282574 | benign | 0 | 0 | 0 |
| rs137852993 | pathogenic | 0 | 0 | 0 |
| rs137852990 | pathogenic | 0 | 0 | 0 |
| rs137852987 | pathogenic | 0 | 0 | 0 |
| rs4148217 | benign | 0 | 0 | 0 |
| rs140690030 | benign | 0 | 0 | 0 |
| rs137852992 | pathogenic | 0 | 0 | 0 |
| rs6544718 | benign | 0 | 0 | 0 |
| rs137852989 | pathogenic | 0 | 0 | 0 |
| rs150893072 | benign | 1 | 1 | 1 |
| rs28931613 | pathogenic | 1 | 1 | 1 |
| rs143789597 | benign | 1 | 1 | 1 |
| rs121913037 | pathogenic | 1 | 1 | 0 |
| rs188802138 | pathogenic | 1 | 1 | 0 |
| rs121913038 | pathogenic | 1 | 1 | 0 |
| rs146835206 | benign | 0 | 0 | 0 |
| rs121913041 | pathogenic | 1 | 1 | 1 |
| rs121913039 | pathogenic | 1 | 1 | 1 |
| rs149977726 | pathogenic | 1 | 1 | 0 |
| rs121913036 | pathogenic | 0 | 0 | 0 |
| rs121913040 | pathogenic | 0 | 0 | 0 |
| rs765007158 | benign | 0 | 0 | 0 |
| rs112723255 | benign | 0 | 0 | 0 |
| rs574164748 | benign | 0 | 0 | 0 |
| rs80338734 | pathogenic | 1 | 1 | 0 |
| rs104894694 | pathogenic | 1 | 1 | 0 |
| rs753198836 | pathogenic | 0 | 0 | 0 |
| rs760101382 | pathogenic | 0 | 0 | 0 |
| rs121909272 | pathogenic | 1 | 1 | 0 |
| rs370421531 | pathogenic | 0 | 0 | 0 |
| rs565109128 | benign | 0 | 0 | 0 |
| rs17851582 | benign | 0 | 0 | 0 |
| rs753228876 | benign | 0 | 0 | 0 |
| rs141471799 | benign | 0 | 0 | 0 |
| rs281864916 | pathogenic | 1 | 1 | 0 |
| rs121918409 | pathogenic | 1 | 1 | 1 |

| | | | | |
|---|---|---|---|---|
| rs121918413 | pathogenic | 1 | 1 | 1 |
| rs121918411 | pathogenic | 1 | 1 | 1 |
| rs121918408 | pathogenic | 1 | 1 | 1 |
| rs121918412 | pathogenic | 1 | 1 | 1 |
| rs121918410 | pathogenic | 1 | 1 | 1 |
| rs116474260 | benign | 1 | 1 | 1 |
| rs281864919 | pathogenic | 1 | 1 | 1 |
| rs397514452 | pathogenic | 1 | 1 | 0 |
| rs747843638 | benign | 0 | 0 | 0 |
| rs774357609 | pathogenic | 1 | 1 | 1 |
| rs201804005 | benign | 0 | 0 | 0 |
| rs746485916 | pathogenic | 0 | 0 | 0 |
| rs764131178 | pathogenic | 0 | 0 | 0 |
| rs374094065 | pathogenic | 0 | 0 | 0 |
| rs76151636 | pathogenic | 1 | 1 | 0 |
| rs753250853 | pathogenic | 0 | 0 | 0 |
| rs761632029 | pathogenic | 0 | 0 | 0 |
| rs775541743 | pathogenic | 0 | 0 | 0 |
| rs778749563 | pathogenic | 0 | 0 | 0 |
| rs60431989 | pathogenic | 1 | 1 | 0 |
| rs755554442 | pathogenic | 1 | 1 | 0 |
| rs749085322 | pathogenic | 0 | 0 | 0 |
| rs756029120 | pathogenic | 1 | 1 | 0 |
| rs137853279 | pathogenic | 1 | 1 | 0 |
| rs777354267 | pathogenic | 1 | 1 | 1 |
| rs142940486 | benign | 1 | 1 | 1 |
| rs140167393 | benign | 1 | 1 | 1 |
| rs2066827 | benign | 0 | 0 | 0 |
| rs765681672 | benign | 0 | 0 | 0 |
| rs137985549 | benign | 0 | 0 | 0 |
| rs74315340 | pathogenic | 1 | 1 | 1 |
| rs74315341 | pathogenic | 0 | 0 | 0 |
| rs74315335 | pathogenic | 0 | 0 | 0 |
| rs61730974 | benign | 0 | 0 | 0 |
| rs121909193 | pathogenic | 0 | 0 | 0 |
| rs74315334 | pathogenic | 0 | 0 | 0 |
| rs74315329 | pathogenic | 0 | 0 | 0 |
| rs74315330 | pathogenic | 0 | 0 | 0 |
| rs121909194 | pathogenic | 0 | 0 | 0 |
| rs28936694 | pathogenic | 0 | 0 | 0 |
| rs74315336 | pathogenic | 0 | 0 | 0 |
| rs74315338 | pathogenic | 1 | 1 | 1 |
| rs74315328 | pathogenic | 0 | 0 | 0 |
| rs74315331 | pathogenic | 0 | 0 | 0 |
| rs74315332 | pathogenic | 0 | 0 | 0 |
| rs141696651 | benign | 0 | 0 | 0 |
| rs7041 | benign | 0 | 0 | 0 |
| rs4588 | benign | 0 | 0 | 0 |
| rs121908981 | pathogenic | 1 | 1 | 1 |
| rs606231459 | pathogenic | 0 | 0 | 0 |
| rs398123053 | pathogenic | 0 | 0 | 0 |
| rs774259802 | benign | 0 | 0 | 0 |
| rs121909582 | pathogenic | 1 | 1 | 1 |
| rs104893680 | pathogenic | 1 | 1 | 1 |

| | | | | |
|---|---|---|---|---|
| rs137854907 | pathogenic | 0 | 0 | 0 |
| rs765715798 | pathogenic | 0 | 0 | 0 |
| rs104893678 | pathogenic | 0 | 0 | 0 |
| rs104893679 | pathogenic | 0 | 1 | 0 |
| rs104893681 | pathogenic | 1 | 1 | 0 |
| rs121909640 | pathogenic | 1 | 1 | 0 |
| rs780705654 | pathogenic | 0 | 0 | 0 |
| rs201672146 | benign | 1 | 1 | 0 |
| rs12136658 | benign | 0 | 0 | 0 |
| rs56127440 | benign | 0 | 0 | 0 |
| rs104894433 | pathogenic | 1 | 1 | 1 |
| rs104894435 | pathogenic | 0 | 1 | 0 |
| rs104894437 | pathogenic | 1 | 1 | 1 |
| rs104894441 | pathogenic | 1 | 1 | 1 |
| rs104894440 | pathogenic | 1 | 1 | 1 |
| rs104894445 | pathogenic | 1 | 1 | 1 |
| rs104894436 | pathogenic | 0 | 0 | 0 |
| rs104894438 | pathogenic | 0 | 0 | 0 |
| rs200891969 | pathogenic | 1 | 1 | 0 |
| rs104894443 | pathogenic | 1 | 1 | 1 |
| rs104894434 | pathogenic | 1 | 1 | 1 |
| rs41298442 | pathogenic | 1 | 1 | 1 |
| rs104894442 | pathogenic | 1 | 1 | 1 |
| rs35453854 | benign | 0 | 0 | 0 |
| rs121912898 | pathogenic | 0 | 0 | 0 |
| rs121912897 | pathogenic | 1 | 1 | 0 |
| rs138260625 | benign | 0 | 0 | 0 |
| rs147802438 | benign | 1 | 1 | 0 |
| rs146443310 | benign | 1 | 1 | 0 |
| rs747319628 | pathogenic | 1 | 1 | 0 |
| rs1801195 | benign | 0 | 0 | 0 |
| rs145045341 | benign | 1 | 1 | 0 |
| rs116001173 | benign | 1 | 1 | 0 |
| rs587784175 | pathogenic | 0 | 0 | 0 |
| rs104893857 | pathogenic | 0 | 0 | 0 |
| rs104893862 | pathogenic | 1 | 1 | 0 |
| rs104893858 | pathogenic | 1 | 1 | 1 |
| rs121909249 | pathogenic | 0 | 1 | 0 |
| rs121909248 | pathogenic | 1 | 1 | 1 |
| rs104893859 | pathogenic | 1 | 1 | 0 |
| rs187063864 | pathogenic | 1 | 1 | 0 |
| rs121917817 | pathogenic | 1 | 1 | 0 |
| rs121917818 | pathogenic | 0 | 0 | 0 |
| rs753594601 | pathogenic | 0 | 0 | 0 |
| rs28937869 | pathogenic | 1 | 1 | 0 |
| rs6165 | benign | 0 | 0 | 0 |
| rs200089714 | pathogenic | 1 | 1 | 0 |
| rs372000714 | pathogenic | 0 | 0 | 0 |
| rs2228206 | benign | 0 | 0 | 0 |
| rs560096 | benign | 0 | 0 | 0 |
| rs137852666 | pathogenic | 0 | 0 | 0 |
| rs10896380 | benign | 0 | 0 | 0 |
| rs773690764 | pathogenic | 0 | 0 | 0 |
| rs201060167 | pathogenic | 0 | 0 | 0 |

| | | | | |
|---|---|---|---|---|
| rs137852670 | pathogenic | 0 | 0 | 0 |
| rs780594709 | pathogenic | 0 | 0 | 0 |
| rs145226920 | pathogenic | 0 | 0 | 0 |
| rs137852665 | pathogenic | 0 | 0 | 0 |
| rs724159960 | pathogenic | 0 | 0 | 0 |
| rs756985703 | pathogenic | 0 | 0 | 0 |
| rs137852667 | pathogenic | 0 | 0 | 0 |
| rs151079750 | pathogenic | 1 | 1 | 0 |
| rs2306636 | benign | 0 | 0 | 0 |
| rs72647885 | benign | 1 | 1 | 0 |
| rs377584435 | pathogenic | 1 | 1 | 1 |
| rs757583846 | pathogenic | 1 | 1 | 0 |
| rs121918148 | pathogenic | 0 | 0 | 0 |
| rs149201802 | benign | 0 | 0 | 0 |
| rs587781904 | pathogenic | 1 | 1 | 1 |
| rs143421325 | benign | 0 | 0 | 0 |
| rs587781253 | pathogenic | 0 | 0 | 0 |
| rs587780331 | benign | 0 | 0 | 0 |
| rs151001016 | benign | 0 | 0 | 0 |
| rs150888094 | benign | 0 | 0 | 0 |
| rs149581331 | benign | 0 | 0 | 0 |
| rs774840535 | benign | 0 | 0 | 0 |
| rs138428684 | benign | 0 | 0 | 0 |
| rs149300055 | benign | 0 | 0 | 0 |
| rs571005578 | benign | 0 | 0 | 0 |
| rs200731839 | benign | 0 | 0 | 0 |
| rs121965007 | pathogenic | 0 | 0 | 0 |
| rs121965013 | pathogenic | 0 | 1 | 0 |
| rs121965009 | pathogenic | 0 | 0 | 0 |
| rs1800457 | benign | 1 | 1 | 0 |
| rs121965006 | pathogenic | 1 | 1 | 0 |
| rs121965008 | pathogenic | 0 | 0 | 0 |
| rs61732609 | pathogenic | 0 | 0 | 0 |
| rs121965011 | pathogenic | 0 | 0 | 0 |
| rs121965015 | pathogenic | 0 | 0 | 0 |
| rs121965018 | pathogenic | 1 | 1 | 0 |
| rs121965016 | pathogenic | 0 | 0 | 0 |
| rs77806 | benign | 0 | 0 | 0 |
| rs149731356 | benign | 0 | 0 | 0 |
| rs72549407 | pathogenic | 0 | 0 | 0 |
| rs200363929 | benign | 0 | 0 | 0 |
| rs376403708 | benign | 1 | 1 | 0 |
| rs56376197 | benign | 1 | 1 | 1 |
| rs200237973 | benign | 1 | 1 | 1 |
| rs548330075 | benign | 0 | 0 | 0 |
| rs148291350 | benign | 0 | 0 | 0 |
| rs71579353 | pathogenic | 0 | 0 | 0 |
| rs146278697 | benign | 0 | 0 | 0 |
| rs141033098 | benign | 0 | 0 | 0 |
| rs397517234 | pathogenic | 0 | 0 | 0 |
| rs71579374 | benign | 0 | 0 | 0 |
| rs140381835 | benign | 0 | 0 | 0 |
| rs137852702 | pathogenic | 0 | 0 | 0 |
| rs137852696 | pathogenic | 1 | 1 | 1 |

| | | | | |
|---|---|---|---|---|
| rs137852697 | pathogenic | 0 | 0 | 0 |
| rs386833639 | pathogenic | 0 | 0 | 0 |
| rs386833640 | pathogenic | 0 | 0 | 0 |
| rs386833641 | benign | 0 | 0 | 0 |
| rs137852701 | pathogenic | 0 | 0 | 0 |
| rs386833642 | pathogenic | 0 | 0 | 0 |
| rs137852695 | pathogenic | 0 | 0 | 0 |
| rs386833646 | pathogenic | 0 | 0 | 0 |
| rs137852700 | pathogenic | 1 | 1 | 0 |
| rs386833649 | pathogenic | 1 | 1 | 0 |
| rs386833650 | pathogenic | 0 | 0 | 0 |
| rs148412181 | pathogenic | 0 | 0 | 0 |
| rs386833655 | pathogenic | 0 | 0 | 0 |
| rs386833656 | pathogenic | 1 | 1 | 0 |
| rs137852698 | pathogenic | 0 | 0 | 0 |
| rs386833661 | pathogenic | 0 | 0 | 0 |
| rs386833670 | pathogenic | 0 | 0 | 0 |
| rs111033632 | pathogenic | 1 | 1 | 1 |
| rs34308573 | benign | 1 | 1 | 1 |
| rs376048533 | pathogenic | 1 | 1 | 0 |
| rs3747517 | benign | 1 | 1 | 0 |
| rs137853328 | pathogenic | 0 | 0 | 0 |
| rs77273740 | benign | 0 | 0 | 0 |
| rs267606760 | pathogenic | 0 | 0 | 0 |
| rs77576840 | pathogenic | 0 | 0 | 0 |
| rs45465204 | benign | 1 | 1 | 0 |
| rs5320 | benign | 0 | 0 | 0 |
| rs5324 | benign | 0 | 0 | 0 |
| rs4531 | benign | 0 | 0 | 0 |
| rs267606761 | pathogenic | 0 | 0 | 0 |
| rs75215331 | pathogenic | 0 | 0 | 0 |
| rs6271 | benign | 0 | 0 | 0 |
| rs201643277 | pathogenic | 0 | 0 | 0 |
| rs4252250 | benign | 0 | 0 | 0 |
| rs137853580 | pathogenic | 0 | 0 | 0 |
| rs368287711 | pathogenic | 0 | 0 | 0 |
| rs137853579 | pathogenic | 0 | 0 | 0 |
| rs3135932 | benign | 0 | 0 | 0 |
| rs2228055 | benign | 0 | 0 | 0 |
| rs140322345 | benign | 1 | 1 | 1 |
| rs1154510 | pathogenic | 0 | 0 | 0 |
| rs11833399 | benign | 0 | 0 | 0 |
| rs36023382 | benign | 1 | 1 | 1 |
| rs61754444 | benign | 0 | 0 | 0 |
| rs28935168 | pathogenic | 0 | 0 | 0 |
| rs28934907 | pathogenic | 0 | 0 | 0 |
| rs28934904 | pathogenic | 1 | 1 | 0 |
| rs61748392 | pathogenic | 1 | 1 | 0 |
| rs28934908 | pathogenic | 0 | 0 | 0 |
| rs28934905 | pathogenic | 0 | 0 | 0 |
| rs28934906 | pathogenic | 1 | 1 | 0 |
| rs606231307 | pathogenic | 0 | 0 | 0 |
| rs9333296 | benign | 0 | 0 | 0 |
| rs3218614 | benign | 0 | 0 | 0 |

| | | | | |
|---|---|---|---|---|
| rs2230792 | benign | 0 | 0 | 0 |
| rs2230793 | benign | 0 | 0 | 0 |
| rs2230794 | benign | 0 | 0 | 0 |
| rs749052963 | pathogenic | 0 | 0 | 0 |
| rs2230798 | benign | 0 | 0 | 0 |
| rs2230795 | benign | 0 | 0 | 0 |
| rs3204145 | benign | 0 | 0 | 0 |
| rs1538660 | benign | 0 | 0 | 0 |
| rs9878928 | benign | 0 | 0 | 0 |
| rs104893742 | pathogenic | 0 | 0 | 0 |
| rs28936702 | pathogenic | 0 | 0 | 0 |
| rs28936703 | pathogenic | 0 | 0 | 0 |
| rs769392 | benign | 0 | 0 | 0 |
| rs769402 | benign | 1 | 1 | 1 |
| rs104894884 | pathogenic | 1 | 1 | 1 |
| rs1801316 | benign | 1 | 1 | 1 |
| rs104894885 | pathogenic | 1 | 1 | 1 |
| rs782572 | benign | 0 | 0 | 0 |
| rs151166046 | pathogenic | 0 | 0 | 0 |
| rs772153760 | pathogenic | 1 | 1 | 0 |
| rs35405862 | benign | 0 | 0 | 0 |
| rs146571352 | pathogenic | 0 | 1 | 0 |
| rs374698153 | pathogenic | 0 | 0 | 0 |
| rs7594497 | benign | 0 | 0 | 0 |
| rs80338716 | pathogenic | 0 | 0 | 0 |
| rs80338719 | pathogenic | 1 | 1 | 1 |
| rs80338721 | pathogenic | 0 | 0 | 0 |
| rs730882249 | pathogenic | 0 | 0 | 0 |
| rs146262009 | pathogenic | 0 | 0 | 0 |
| rs104895083 | pathogenic | 1 | 1 | 1 |
| rs104895100 | benign | 1 | 1 | 1 |
| rs104895085 | pathogenic | 0 | 0 | 0 |
| rs28940580 | pathogenic | 0 | 0 | 0 |
| rs104895098 | pathogenic | 0 | 0 | 0 |
| rs28940578 | pathogenic | 0 | 0 | 0 |
| rs61752717 | pathogenic | 0 | 0 | 0 |
| rs28940579 | pathogenic | 0 | 0 | 0 |
| rs104895097 | pathogenic | 0 | 0 | 0 |
| rs140425651 | benign | 1 | 1 | 0 |
| rs768090444 | pathogenic | 0 | 0 | 0 |
| rs764459544 | pathogenic | 0 | 0 | 0 |
| rs80338802 | pathogenic | 1 | 1 | 1 |
| rs149969786 | pathogenic | 1 | 1 | 0 |
| rs115311625 | pathogenic | 1 | 1 | 1 |
| rs778768583 | pathogenic | 1 | 1 | 1 |
| rs892090 | benign | 0 | 0 | 0 |
| rs28929498 | pathogenic | 1 | 1 | 1 |
| rs137854467 | pathogenic | 0 | 0 | 0 |
| rs137854480 | pathogenic | 0 | 0 | 0 |
| rs148682262 | benign | 0 | 0 | 0 |
| rs137852744 | pathogenic | 0 | 0 | 0 |
| rs137852745 | pathogenic | 0 | 1 | 0 |
| rs137852749 | pathogenic | 0 | 1 | 0 |
| rs137852746 | pathogenic | 0 | 1 | 0 |

| | | | | |
|---|---|---|---|---|
| rs145633958 | benign | 0 | 0 | 0 |
| rs141679950 | benign | 0 | 0 | 0 |
| rs76397662 | benign | 1 | 1 | 1 |
| rs370736139 | benign | 0 | 0 | 0 |
| rs781750106 | benign | 0 | 0 | 0 |
| rs546866208 | benign | 0 | 0 | 0 |
| rs142338976 | benign | 0 | 0 | 0 |
| rs115272158 | benign | 0 | 0 | 0 |
| rs115423919 | benign | 0 | 0 | 0 |
| rs537874538 | benign | 0 | 0 | 0 |
| rs9282834 | benign | 0 | 0 | 0 |
| rs201745826 | benign | 0 | 0 | 0 |
| rs75873440 | pathogenic | 0 | 0 | 0 |
| rs147219360 | benign | 0 | 0 | 0 |
| rs77939446 | pathogenic | 0 | 0 | 0 |
| rs77558292 | pathogenic | 0 | 0 | 0 |
| rs80069458 | pathogenic | 0 | 0 | 0 |
| rs76262710 | pathogenic | 0 | 0 | 0 |
| rs79781594 | pathogenic | 0 | 0 | 0 |
| rs77316810 | pathogenic | 0 | 0 | 0 |
| rs77503355 | pathogenic | 0 | 0 | 0 |
| rs79890926 | pathogenic | 0 | 0 | 0 |
| rs377767402 | benign | 0 | 0 | 0 |
| rs121913308 | pathogenic | 0 | 0 | 0 |
| rs55846256 | benign | 0 | 0 | 0 |
| rs75076352 | pathogenic | 0 | 0 | 0 |
| rs75996173 | pathogenic | 0 | 0 | 0 |
| rs77709286 | pathogenic | 0 | 0 | 0 |
| rs151058706 | benign | 0 | 0 | 0 |
| rs138964358 | benign | 0 | 0 | 0 |
| rs137852353 | pathogenic | 1 | 1 | 0 |
| rs201196020 | benign | 0 | 0 | 0 |
| rs137853201 | pathogenic | 1 | 1 | 1 |
| rs113994174 | pathogenic | 1 | 1 | 1 |
| rs104893674 | pathogenic | 1 | 1 | 1 |
| rs779519249 | benign | 1 | 1 | 1 |
| rs9606756 | benign | 0 | 0 | 0 |
| rs75680863 | benign | 1 | 1 | 1 |
| rs35838082 | benign | 0 | 0 | 0 |
| rs1801198 | benign | 0 | 0 | 0 |
| rs117458738 | benign | 1 | 1 | 0 |
| rs9621049 | benign | 1 | 1 | 0 |
| rs1131603 | benign | 1 | 1 | 0 |
| rs4820889 | benign | 0 | 0 | 0 |
| rs777767978 | pathogenic | 1 | 1 | 0 |
| rs147399860 | benign | 1 | 1 | 1 |
| rs104893950 | pathogenic | 0 | 0 | 0 |
| rs777308612 | benign | 0 | 0 | 0 |
| rs151091410 | benign | 0 | 0 | 0 |
| rs11537667 | benign | 0 | 0 | 0 |
| rs267606621 | pathogenic | 1 | 1 | 0 |
| rs143058902 | benign | 0 | 0 | 0 |
| rs370261904 | pathogenic | 0 | 0 | 0 |
| rs141169746 | benign | 0 | 0 | 0 |

| | | | | |
|---|---|---|---|---|
| rs111033436 | benign | 0 | 0 | 0 |
| rs148533341 | benign | 0 | 0 | 0 |
| rs143827620 | benign | 0 | 0 | 0 |
| rs4935502 | benign | 0 | 0 | 0 |
| rs61735473 | benign | 0 | 0 | 0 |
| rs267606932 | pathogenic | 1 | 1 | 0 |
| rs61730754 | benign | 0 | 0 | 0 |
| rs374491359 | pathogenic | 1 | 1 | 1 |
| rs761800077 | benign | 1 | 1 | 1 |
| rs760391381 | benign | 1 | 1 | 1 |
| rs104893885 | pathogenic | 0 | 0 | 0 |
| rs104893886 | pathogenic | 1 | 1 | 1 |
| rs104893884 | pathogenic | 0 | 0 | 0 |
| rs147715509 | benign | 0 | 0 | 0 |
| rs115533243 | benign | 0 | 0 | 0 |
| rs397509390 | pathogenic | 0 | 1 | 0 |
| rs80357134 | pathogenic | 0 | 0 | 0 |
| rs766004110 | benign | 1 | 1 | 1 |
| rs769650474 | benign | 0 | 0 | 0 |
| rs398122635 | benign | 1 | 1 | 0 |
| rs199522616 | pathogenic | 0 | 0 | 0 |
| rs28897672 | pathogenic | 1 | 1 | 0 |
| rs80357086 | pathogenic | 1 | 1 | 1 |
| rs80357064 | pathogenic | 1 | 1 | 0 |
| rs80356936 | benign | 0 | 0 | 0 |
| rs28897673 | benign | 0 | 0 | 0 |
| rs146747324 | benign | 0 | 0 | 0 |
| rs78635798 | pathogenic | 0 | 0 | 0 |
| rs75146158 | pathogenic | 1 | 1 | 1 |
| rs587782886 | benign | 0 | 0 | 0 |
| rs121434634 | pathogenic | 0 | 0 | 0 |
| rs121434635 | pathogenic | 0 | 0 | 0 |
| rs587777778 | pathogenic | 0 | 0 | 0 |
| rs118203967 | pathogenic | 0 | 0 | 0 |
| rs751375244 | pathogenic | 1 | 1 | 1 |
| rs104893863 | pathogenic | 1 | 1 | 0 |
| rs104893865 | pathogenic | 0 | 0 | 0 |
| rs104893864 | pathogenic | 1 | 1 | 1 |
| rs104893866 | pathogenic | 1 | 1 | 0 |
| rs561738497 | benign | 0 | 0 | 0 |
| rs57599352 | pathogenic | 1 | 1 | 1 |
| rs267607448 | pathogenic | 1 | 1 | 1 |
| rs63751596 | pathogenic | 0 | 0 | 0 |
| rs63751612 | benign | 0 | 0 | 0 |
| rs63751684 | benign | 0 | 0 | 0 |
| rs267607861 | benign | 0 | 0 | 0 |
| rs35831931 | benign | 0 | 0 | 0 |
| rs63750876 | benign | 0 | 0 | 0 |
| rs41295284 | benign | 0 | 0 | 0 |
| rs35502531 | benign | 0 | 0 | 0 |
| rs63750449 | benign | 0 | 0 | 0 |
| rs63750899 | pathogenic | 0 | 0 | 0 |
| rs55907433 | benign | 0 | 0 | 0 |
| rs63750217 | pathogenic | 0 | 0 | 0 |

| | | | | |
|---|---|---|---|---|
| rs63751275 | pathogenic | 0 | 0 | 0 |
| rs63750702 | benign | 1 | 1 | 1 |
| rs63750114 | benign | 0 | 0 | 0 |
| rs2020873 | benign | 0 | 0 | 0 |
| rs140195825 | benign | 1 | 1 | 1 |
| rs182763805 | benign | 0 | 0 | 0 |
| rs533696377 | benign | 0 | 0 | 0 |
| rs41278788 | benign | 0 | 0 | 0 |
| rs104894193 | pathogenic | 1 | 1 | 0 |
| rs3824915 | benign | 1 | 1 | 1 |
| rs104894196 | pathogenic | 0 | 0 | 0 |
| rs78681604 | benign | 1 | 1 | 1 |
| rs77321498 | benign | 1 | 1 | 1 |
| rs137852965 | pathogenic | 0 | 0 | 0 |
| rs137852961 | pathogenic | 0 | 0 | 0 |
| rs137852962 | pathogenic | 1 | 1 | 0 |
| rs753376100 | pathogenic | 0 | 0 | 0 |
| rs137852968 | pathogenic | 1 | 1 | 1 |
| rs137852959 | pathogenic | 1 | 1 | 0 |
| rs137852967 | pathogenic | 0 | 0 | 0 |
| rs113437301 | benign | 1 | 1 | 1 |
| rs61747482 | benign | 1 | 1 | 1 |
| rs2234919 | benign | 1 | 1 | 0 |
| rs121908863 | pathogenic | 0 | 0 | 0 |
| rs140455771 | pathogenic | 0 | 0 | 0 |
| rs121908810 | pathogenic | 1 | 1 | 1 |
| rs143879853 | benign | 0 | 0 | 0 |
| rs188319622 | benign | 0 | 0 | 0 |
| rs121918645 | pathogenic | 0 | 0 | 0 |
| rs115352435 | pathogenic | 0 | 0 | 0 |
| rs121434427 | pathogenic | 1 | 1 | 1 |
| rs121434428 | pathogenic | 0 | 0 | 0 |
| rs35086265 | benign | 1 | 1 | 1 |
| rs11576415 | benign | 0 | 0 | 0 |
| rs121434429 | pathogenic | 1 | 1 | 1 |
| rs4074536 | benign | 1 | 1 | 1 |
| rs10801999 | benign | 0 | 0 | 0 |
| rs121434550 | pathogenic | 0 | 0 | 0 |
| rs148057999 | pathogenic | 0 | 0 | 0 |
| rs28730716 | benign | 0 | 0 | 0 |
| rs142036299 | benign | 0 | 0 | 0 |
| rs762381137 | pathogenic | 0 | 0 | 0 |
| rs121434549 | pathogenic | 0 | 0 | 0 |
| rs139228801 | pathogenic | 0 | 0 | 0 |
| rs147219463 | benign | 0 | 0 | 0 |
| rs1131857 | benign | 0 | 0 | 0 |
| rs121917874 | pathogenic | 0 | 1 | 0 |
| rs2228056 | benign | 0 | 0 | 0 |
| rs121917870 | pathogenic | 0 | 0 | 0 |
| rs121917866 | pathogenic | 0 | 0 | 0 |
| rs11921054 | benign | 0 | 0 | 0 |
| rs121917868 | pathogenic | 0 | 0 | 0 |
| rs28931603 | pathogenic | 0 | 0 | 0 |
| rs104895295 | pathogenic | 1 | 1 | 0 |

| | | | | |
|---|---|---|---|---|
| rs104895382 | pathogenic | 0 | 0 | 0 |
| rs104895298 | pathogenic | 0 | 0 | 0 |
| rs104895300 | pathogenic | 0 | 0 | 0 |
| rs104895332 | pathogenic | 0 | 0 | 0 |
| rs758026399 | pathogenic | 1 | 1 | 1 |
| rs104895366 | pathogenic | 0 | 0 | 0 |
| rs104895304 | pathogenic | 0 | 0 | 0 |
| rs759997079 | pathogenic | 0 | 0 | 0 |
| rs121917789 | pathogenic | 0 | 0 | 0 |
| rs104895319 | pathogenic | 0 | 0 | 0 |
| rs104895317 | pathogenic | 1 | 1 | 0 |
| rs28934897 | pathogenic | 0 | 0 | 0 |
| rs104895380 | benign | 0 | 0 | 0 |
| rs104895360 | pathogenic | 0 | 0 | 0 |
| rs28936396 | pathogenic | 1 | 1 | 0 |
| rs121909307 | pathogenic | 1 | 1 | 1 |
| rs28938472 | pathogenic | 1 | 1 | 0 |
| rs121909308 | pathogenic | 1 | 1 | 0 |
| rs121909309 | pathogenic | 0 | 0 | 0 |
| rs121913013 | benign | 1 | 1 | 1 |
| rs201855245 | benign | 1 | 1 | 1 |
| rs2230232 | benign | 1 | 1 | 0 |
| rs113451409 | benign | 0 | 0 | 0 |
| rs121913011 | pathogenic | 1 | 1 | 0 |
| rs185821167 | pathogenic | 1 | 1 | 0 |
| rs2230234 | benign | 0 | 0 | 0 |
| rs193922639 | benign | 1 | 1 | 0 |
| rs774863785 | pathogenic | 1 | 1 | 0 |
| rs576404380 | pathogenic | 1 | 1 | 0 |
| rs2230235 | benign | 1 | 1 | 0 |
| rs200484060 | pathogenic | 1 | 1 | 0 |
| rs199761749 | benign | 0 | 0 | 0 |
| rs199681901 | benign | 0 | 0 | 0 |
| rs121908951 | pathogenic | 0 | 0 | 0 |
| rs769251460 | benign | 0 | 0 | 0 |
| rs28937320 | pathogenic | 0 | 0 | 0 |
| rs200607527 | pathogenic | 0 | 0 | 0 |
| rs137852640 | pathogenic | 1 | 1 | 1 |
| rs137852636 | pathogenic | 1 | 1 | 1 |
| rs137852638 | pathogenic | 0 | 0 | 0 |
| rs142637231 | pathogenic | 0 | 0 | 0 |
| rs151328418 | benign | 0 | 1 | 0 |
| rs137852639 | pathogenic | 1 | 1 | 1 |
| rs34605667 | benign | 0 | 0 | 0 |
| rs104894073 | pathogenic | 1 | 1 | 0 |
| rs56298569 | pathogenic | 0 | 0 | 0 |
| rs757600616 | pathogenic | 0 | 0 | 0 |
| rs139194636 | pathogenic | 0 | 0 | 0 |
| rs369517993 | pathogenic | 1 | 1 | 1 |
| rs138846036 | benign | 1 | 1 | 1 |
| rs200808744 | pathogenic | 1 | 1 | 1 |
| rs2255543 | benign | 0 | 0 | 0 |
| rs149165166 | pathogenic | 0 | 0 | 0 |
| rs544956641 | pathogenic | 1 | 1 | 1 |

| | | | | |
|---|---|---|---|---|
| rs28941783 | pathogenic | 0 | 0 | 0 |
| rs562853291 | pathogenic | 1 | 1 | 1 |
| rs28942100 | pathogenic | 0 | 0 | 0 |
| rs120074174 | pathogenic | 1 | 1 | 1 |
| rs120074171 | pathogenic | 1 | 1 | 1 |
| rs120074173 | pathogenic | 0 | 0 | 0 |
| rs120074172 | pathogenic | 1 | 1 | 1 |
| rs767159114 | pathogenic | 1 | 1 | 1 |
| rs104893657 | pathogenic | 0 | 0 | 0 |
| rs104893656 | pathogenic | 0 | 0 | 0 |
| rs104893659 | pathogenic | 0 | 0 | 0 |
| rs104893658 | pathogenic | 0 | 0 | 0 |
| rs200386310 | pathogenic | 0 | 0 | 0 |
| rs2072712 | benign | 0 | 0 | 0 |
| rs35274867 | benign | 0 | 0 | 0 |
| rs35343172 | benign | 0 | 0 | 0 |
| rs10079250 | benign | 0 | 0 | 0 |
| rs34951517 | benign | 0 | 0 | 0 |
| rs267606695 | pathogenic | 0 | 0 | 0 |
| rs149391728 | benign | 0 | 0 | 0 |
| rs387906598 | pathogenic | 1 | 1 | 0 |
| rs137854578 | pathogenic | 0 | 0 | 0 |
| rs141519952 | benign | 0 | 0 | 0 |
| rs146572883 | benign | 0 | 0 | 0 |
| rs111866410 | benign | 0 | 0 | 0 |
| rs139387758 | benign | 1 | 1 | 1 |
| rs35672074 | benign | 1 | 1 | 1 |
| rs121434322 | pathogenic | 0 | 0 | 0 |
| rs150884181 | benign | 0 | 0 | 0 |
| rs35867471 | benign | 0 | 0 | 0 |
| rs2867802 | benign | 0 | 0 | 0 |
| rs137902889 | benign | 0 | 0 | 0 |
| rs369804864 | pathogenic | 0 | 0 | 0 |
| rs781050795 | pathogenic | 0 | 0 | 0 |
| rs2305799 | benign | 0 | 0 | 0 |
| rs2229531 | benign | 0 | 0 | 0 |
| rs147025508 | benign | 0 | 0 | 0 |
| rs76857106 | pathogenic | 0 | 0 | 0 |
| rs7247284 | benign | 0 | 0 | 0 |
| rs62619782 | benign | 0 | 0 | 0 |
| rs377244188 | pathogenic | 0 | 0 | 0 |
| rs75718910 | pathogenic | 0 | 0 | 0 |
| rs79843600 | pathogenic | 0 | 0 | 0 |
| rs75037667 | pathogenic | 1 | 1 | 1 |
| rs121913006 | pathogenic | 0 | 0 | 0 |
| rs397517925 | pathogenic | 1 | 1 | 1 |
| rs267606831 | pathogenic | 1 | 1 | 1 |
| rs121909667 | pathogenic | 1 | 1 | 1 |
| rs121909669 | pathogenic | 1 | 1 | 1 |
| rs121909668 | pathogenic | 1 | 1 | 1 |
| rs121909671 | pathogenic | 1 | 1 | 1 |
| rs534596036 | benign | 0 | 0 | 0 |
| rs144037629 | pathogenic | 0 | 0 | 0 |
| rs11546406 | benign | 0 | 0 | 0 |

| | | | | |
|---|---|---|---|---|
| rs143697995 | pathogenic | 0 | 0 | 0 |
| rs121908331 | pathogenic | 0 | 0 | 0 |
| rs149377346 | benign | 0 | 0 | 0 |
| rs104894086 | pathogenic | 1 | 1 | 0 |
| rs104894089 | pathogenic | 0 | 0 | 0 |
| rs104894090 | pathogenic | 1 | 1 | 0 |
| rs387907235 | pathogenic | 0 | 0 | 0 |
| rs137852689 | pathogenic | 0 | 0 | 0 |
| rs137852690 | pathogenic | 1 | 1 | 0 |
| rs104894087 | pathogenic | 0 | 0 | 0 |
| rs104894085 | pathogenic | 0 | 0 | 0 |
| rs79383654 | benign | 0 | 0 | 0 |
| rs121434239 | pathogenic | 0 | 0 | 0 |
| rs121434240 | pathogenic | 0 | 0 | 0 |
| rs121434236 | pathogenic | 0 | 0 | 0 |
| rs121434238 | pathogenic | 1 | 1 | 1 |
| rs121918107 | pathogenic | 1 | 1 | 1 |
| rs121918103 | pathogenic | 1 | 1 | 1 |
| rs121918104 | pathogenic | 1 | 1 | 1 |
| rs61747614 | benign | 1 | 1 | 1 |
| rs555329870 | benign | 0 | 0 | 0 |
| rs186094931 | benign | 0 | 0 | 0 |
| rs267606785 | pathogenic | 0 | 0 | 0 |
| rs34474955 | benign | 0 | 0 | 0 |
| rs1126497 | benign | 1 | 1 | 1 |
| rs397514661 | pathogenic | 0 | 0 | 0 |
| rs189732445 | benign | 0 | 0 | 0 |
| rs146685071 | benign | 0 | 0 | 0 |
| rs121908636 | pathogenic | 1 | 1 | 0 |
| rs121908643 | pathogenic | 0 | 0 | 0 |
| rs121908644 | pathogenic | 1 | 1 | 1 |
| rs575001023 | pathogenic | 1 | 1 | 1 |
| rs121908646 | pathogenic | 1 | 1 | 0 |
| rs121908638 | pathogenic | 1 | 1 | 1 |
| rs777828000 | pathogenic | 1 | 1 | 1 |
| rs74923032 | benign | 0 | 0 | 0 |
| rs192838388 | pathogenic | 1 | 1 | 1 |
| rs148918985 | pathogenic | 1 | 1 | 1 |
| rs773540764 | benign | 1 | 1 | 1 |
| rs370595480 | pathogenic | 1 | 1 | 1 |
| rs762387914 | pathogenic | 1 | 1 | 1 |
| rs121908642 | pathogenic | 1 | 1 | 1 |
| rs183276875 | pathogenic | 1 | 1 | 1 |
| rs121908648 | pathogenic | 1 | 1 | 1 |
| rs121908647 | pathogenic | 1 | 1 | 1 |
| rs121908640 | pathogenic | 1 | 1 | 1 |
| rs771937610 | pathogenic | 1 | 1 | 1 |
| rs121908641 | pathogenic | 1 | 1 | 1 |
| rs5745068 | benign | 0 | 0 | 0 |
| rs73155056 | benign | 1 | 1 | 1 |
| rs148788180 | benign | 0 | 0 | 0 |
| rs121918137 | pathogenic | 0 | 0 | 0 |
| rs41269593 | benign | 0 | 0 | 0 |
| rs11159 | benign | 1 | 1 | 0 |

| | | | | |
|---|---|---|---|---|
| rs114733570 | pathogenic | 1 | 1 | 0 |
| rs3211783 | benign | 0 | 0 | 0 |
| rs4762 | benign | 0 | 0 | 0 |
| rs74315283 | pathogenic | 0 | 0 | 0 |
| rs111033559 | pathogenic | 0 | 0 | 0 |
| rs111033560 | pathogenic | 1 | 1 | 1 |
| rs749962743 | benign | 0 | 0 | 0 |
| rs267606986 | pathogenic | 0 | 0 | 0 |
| rs34516635 | benign | 0 | 0 | 0 |
| rs117440569 | benign | 0 | 0 | 0 |
| rs61740877 | benign | 0 | 0 | 0 |
| rs121918352 | pathogenic | 0 | 0 | 0 |
| rs183974372 | benign | 0 | 0 | 0 |
| rs140511204 | benign | 1 | 1 | 0 |
| rs6040067 | benign | 0 | 0 | 0 |
| rs199674138 | benign | 0 | 0 | 0 |
| rs121918350 | pathogenic | 0 | 0 | 0 |
| rs138292988 | benign | 1 | 1 | 1 |
| rs145288947 | benign | 0 | 0 | 0 |
| rs369714529 | benign | 0 | 0 | 0 |
| rs200945419 | benign | 0 | 0 | 0 |
| rs116959874 | benign | 0 | 0 | 0 |
| rs141659620 | pathogenic | 1 | 1 | 0 |
| rs111475461 | benign | 0 | 0 | 0 |
| rs61755320 | pathogenic | 0 | 0 | 0 |
| rs748600162 | pathogenic | 1 | 1 | 1 |
| rs369227537 | pathogenic | 1 | 1 | 1 |
| rs372981030 | pathogenic | 1 | 1 | 1 |
| rs137854486 | pathogenic | 0 | 0 | 0 |
| rs137854487 | pathogenic | 0 | 0 | 0 |
| rs199905054 | pathogenic | 0 | 0 | 0 |
| rs771438170 | pathogenic | 0 | 0 | 0 |
| rs774949844 | pathogenic | 0 | 0 | 0 |
| rs4899145 | benign | 0 | 0 | 0 |
| rs200642525 | benign | 0 | 0 | 0 |
| rs190140598 | pathogenic | 0 | 0 | 0 |
| rs2250889 | benign | 0 | 0 | 0 |
| rs17577 | benign | 0 | 0 | 0 |
| rs142032681 | benign | 0 | 0 | 0 |
| rs144577963 | benign | 0 | 0 | 0 |
| rs61754623 | benign | 0 | 0 | 0 |
| rs139873200 | benign | 0 | 0 | 0 |
| rs532530906 | benign | 0 | 0 | 0 |
| rs145762716 | benign | 0 | 0 | 0 |
| rs34062020 | benign | 0 | 0 | 0 |
| rs35258161 | benign | 0 | 0 | 0 |
| rs149792649 | benign | 0 | 0 | 0 |
| rs78101774 | benign | 0 | 0 | 0 |
| rs766466297 | pathogenic | 0 | 1 | 0 |
| rs140467171 | pathogenic | 0 | 0 | 0 |
| rs141340466 | pathogenic | 0 | 0 | 0 |
| rs201555303 | pathogenic | 0 | 0 | 0 |
| rs143592405 | pathogenic | 0 | 0 | 0 |
| rs781249411 | pathogenic | 0 | 0 | 0 |

| | | | | |
|---|---|---|---|---|
| rs116644127 | benign | 0 | 0 | 0 |
| rs151146863 | benign | 1 | 1 | 1 |
| rs116440799 | benign | 0 | 0 | 0 |
| rs138958351 | benign | 0 | 0 | 0 |
| rs143319354 | benign | 0 | 0 | 0 |
| rs768669208 | pathogenic | 1 | 1 | 1 |
| rs28940289 | pathogenic | 1 | 1 | 0 |
| rs745656120 | pathogenic | 1 | 1 | 0 |
| rs387906987 | pathogenic | 0 | 0 | 0 |
| rs147240777 | benign | 0 | 0 | 0 |
| rs140945833 | benign | 0 | 0 | 0 |
| rs146614981 | benign | 0 | 0 | 0 |
| rs149878562 | benign | 0 | 0 | 0 |
| rs28938168 | pathogenic | 0 | 0 | 0 |
| rs33939927 | pathogenic | 1 | 1 | 1 |
| rs34995376 | pathogenic | 1 | 1 | 1 |
| rs201034481 | pathogenic | 1 | 1 | 1 |
| rs774069983 | benign | 0 | 0 | 0 |
| rs199422226 | pathogenic | 1 | 1 | 0 |
| rs199422225 | pathogenic | 1 | 1 | 1 |
| rs199422224 | pathogenic | 1 | 1 | 0 |
| rs149271416 | pathogenic | 1 | 1 | 0 |
| rs1801318 | benign | 1 | 1 | 1 |
| rs372691318 | pathogenic | 1 | 1 | 1 |
| rs147685849 | benign | 0 | 0 | 0 |
| rs387907199 | pathogenic | 0 | 0 | 0 |
| rs767122069 | pathogenic | 0 | 0 | 0 |
| rs2302427 | benign | 0 | 0 | 0 |
| rs201135441 | benign | 0 | 0 | 0 |
| rs201861847 | pathogenic | 0 | 0 | 0 |
| rs137852952 | pathogenic | 0 | 0 | 0 |
| rs16986560 | benign | 0 | 0 | 0 |
| rs147344332 | pathogenic | 1 | 1 | 1 |
| rs36027220 | pathogenic | 0 | 0 | 0 |
| rs192401229 | benign | 0 | 0 | 0 |
| rs121909106 | pathogenic | 1 | 1 | 0 |
| rs118204103 | pathogenic | 1 | 1 | 0 |
| rs118204104 | pathogenic | 1 | 1 | 0 |
| rs118204105 | pathogenic | 1 | 1 | 0 |
| rs118204106 | pathogenic | 0 | 0 | 0 |
| rs118204119 | pathogenic | 0 | 0 | 0 |
| rs118204107 | pathogenic | 0 | 0 | 0 |
| rs118204094 | pathogenic | 0 | 0 | 0 |
| rs118204098 | pathogenic | 1 | 1 | 0 |
| rs118204101 | pathogenic | 1 | 1 | 0 |
| rs118204096 | pathogenic | 1 | 1 | 0 |
| rs118204108 | pathogenic | 0 | 0 | 0 |
| rs536814318 | pathogenic | 0 | 0 | 0 |
| rs34413634 | pathogenic | 1 | 1 | 0 |
| rs118204100 | pathogenic | 0 | 0 | 0 |
| rs118204109 | pathogenic | 0 | 0 | 0 |
| rs118204116 | pathogenic | 1 | 1 | 0 |
| rs118204110 | pathogenic | 0 | 0 | 0 |
| rs118204099 | pathogenic | 0 | 0 | 0 |

| | | | | |
|---|---|---|---|---|
| rs118204111 | pathogenic | 0 | 0 | 0 |
| rs118204112 | pathogenic | 0 | 0 | 0 |
| rs118204113 | pathogenic | 0 | 0 | 0 |
| rs118204114 | pathogenic | 0 | 0 | 0 |
| rs118204115 | pathogenic | 0 | 0 | 0 |
| rs150428209 | benign | 0 | 0 | 0 |
| rs144949995 | benign | 0 | 0 | 0 |
| rs190023981 | benign | 0 | 0 | 0 |
| rs104894611 | pathogenic | 1 | 1 | 1 |
| rs28937580 | pathogenic | 1 | 1 | 1 |
| rs121908949 | pathogenic | 0 | 0 | 0 |
| rs104894612 | pathogenic | 1 | 1 | 1 |
| rs104894609 | pathogenic | 1 | 1 | 1 |
| rs104894610 | pathogenic | 1 | 1 | 1 |
| rs104894615 | pathogenic | 0 | 0 | 0 |
| rs104894603 | pathogenic | 0 | 0 | 0 |
| rs104894602 | pathogenic | 1 | 1 | 1 |
| rs121908948 | pathogenic | 1 | 1 | 1 |
| rs104894608 | pathogenic | 1 | 1 | 1 |
| rs371545673 | benign | 0 | 0 | 0 |
| rs199925238 | benign | 0 | 0 | 0 |
| rs146313451 | benign | 0 | 0 | 0 |
| rs121918193 | pathogenic | 1 | 1 | 0 |
| rs2228500 | benign | 0 | 0 | 0 |
| rs121918194 | pathogenic | 1 | 1 | 0 |
| rs121918196 | pathogenic | 0 | 0 | 0 |
| rs41291971 | benign | 0 | 0 | 0 |
| rs199473690 | pathogenic | 0 | 0 | 0 |
| rs2228613 | benign | 0 | 0 | 0 |
| rs267607130 | pathogenic | 1 | 1 | 1 |
| rs77615401 | benign | 1 | 1 | 1 |
| rs374618872 | benign | 1 | 1 | 1 |
| rs146165014 | benign | 0 | 0 | 0 |
| rs144586070 | benign | 0 | 0 | 0 |
| rs121918541 | pathogenic | 0 | 0 | 0 |
| rs142105943 | benign | 0 | 0 | 0 |
| rs121918540 | pathogenic | 0 | 0 | 0 |
| rs6791 | benign | 0 | 0 | 0 |
| rs61736587 | pathogenic | 0 | 0 | 0 |
| rs61736586 | benign | 0 | 0 | 0 |
| rs121917741 | pathogenic | 1 | 1 | 1 |
| rs41302885 | benign | 0 | 0 | 0 |
| rs141026028 | benign | 0 | 0 | 0 |
| rs193922493 | pathogenic | 0 | 0 | 0 |
| rs73598374 | benign | 0 | 0 | 0 |
| rs121908725 | pathogenic | 1 | 1 | 0 |
| rs199422328 | pathogenic | 0 | 1 | 0 |
| rs11555566 | benign | 1 | 1 | 1 |
| rs267606634 | pathogenic | 0 | 0 | 0 |
| rs121908714 | pathogenic | 1 | 1 | 0 |
| rs121908717 | pathogenic | 1 | 1 | 0 |
| rs267606635 | pathogenic | 1 | 1 | 0 |
| rs121908739 | pathogenic | 0 | 0 | 0 |
| rs121908735 | pathogenic | 0 | 1 | 0 |

| | | | | |
|---|---|---|---|---|
| rs121908716 | pathogenic | 0 | 1 | 0 |
| rs121908723 | pathogenic | 0 | 1 | 0 |
| rs121908721 | pathogenic | 0 | 0 | 0 |
| rs199422327 | pathogenic | 0 | 0 | 0 |
| rs121908715 | pathogenic | 0 | 0 | 0 |
| rs587783563 | pathogenic | 0 | 0 | 0 |
| rs104894780 | pathogenic | 0 | 0 | 0 |
| rs56030372 | pathogenic | 1 | 1 | 1 |
| rs104894782 | pathogenic | 1 | 1 | 1 |
| rs149495971 | benign | 0 | 0 | 0 |
| rs118203942 | pathogenic | 1 | 1 | 0 |
| rs118203939 | pathogenic | 0 | 0 | 0 |
| rs118203938 | pathogenic | 0 | 0 | 0 |
| rs118203943 | pathogenic | 0 | 0 | 0 |
| rs118203940 | pathogenic | 0 | 0 | 0 |
| rs727503809 | pathogenic | 0 | 0 | 0 |
| rs1065757 | benign | 0 | 0 | 0 |
| rs1071598 | benign | 0 | 0 | 0 |
| rs25414 | benign | 0 | 0 | 0 |
| rs118203944 | pathogenic | 0 | 0 | 0 |
| rs118203941 | pathogenic | 0 | 0 | 0 |
| rs144222167 | benign | 0 | 0 | 0 |
| rs201101343 | pathogenic | 0 | 0 | 0 |
| rs144601717 | benign | 0 | 0 | 0 |
| rs267607129 | pathogenic | 0 | 0 | 0 |
| rs104894729 | pathogenic | 0 | 0 | 0 |
| rs104894727 | pathogenic | 0 | 0 | 0 |
| rs104894725 | pathogenic | 0 | 0 | 0 |
| rs104894515 | pathogenic | 1 | 1 | 1 |
| rs74688377 | benign | 0 | 0 | 0 |
| rs121908513 | pathogenic | 1 | 1 | 0 |
| rs121908509 | pathogenic | 0 | 0 | 0 |
| rs121908514 | pathogenic | 1 | 1 | 0 |
| rs121908512 | pathogenic | 1 | 1 | 0 |
| rs121908510 | pathogenic | 0 | 0 | 0 |
| rs121908511 | pathogenic | 1 | 1 | 1 |
| rs121908518 | pathogenic | 1 | 1 | 1 |
| rs34358665 | benign | 0 | 0 | 0 |
| rs147568834 | benign | 0 | 0 | 0 |
| rs201717599 | benign | 0 | 0 | 0 |
| rs61744487 | benign | 0 | 0 | 0 |
| rs370627877 | pathogenic | 1 | 1 | 1 |
| rs368593151 | pathogenic | 1 | 1 | 1 |
| rs114363936 | benign | 0 | 0 | 0 |
| rs137853333 | pathogenic | 0 | 0 | 0 |
| rs137853080 | pathogenic | 1 | 1 | 1 |
| rs137853077 | pathogenic | 1 | 1 | 1 |
| rs780749732 | benign | 1 | 1 | 1 |
| rs137853081 | pathogenic | 1 | 1 | 0 |
| rs137853078 | pathogenic | 0 | 0 | 0 |
| rs121913315 | pathogenic | 1 | 1 | 1 |
| rs752513525 | pathogenic | 1 | 1 | 1 |
| rs104894918 | pathogenic | 0 | 0 | 0 |
| rs104894917 | pathogenic | 0 | 0 | 0 |

| | | | | |
|---|---|---|---|---|
| rs104894919 | pathogenic | 0 | 0 | 0 |
| rs28939672 | pathogenic | 0 | 0 | 0 |
| rs104894173 | pathogenic | 0 | 0 | 0 |
| rs104894179 | pathogenic | 0 | 0 | 0 |
| rs104894174 | pathogenic | 1 | 1 | 0 |
| rs104894178 | pathogenic | 1 | 1 | 0 |
| rs28939714 | pathogenic | 1 | 1 | 1 |
| rs104894270 | pathogenic | 1 | 1 | 1 |
| rs3765571 | benign | 0 | 0 | 0 |
| rs752669339 | pathogenic | 1 | 1 | 1 |
| rs537043237 | pathogenic | 1 | 1 | 1 |
| rs774047299 | pathogenic | 1 | 1 | 1 |
| rs121434583 | pathogenic | 1 | 1 | 1 |
| rs121907927 | pathogenic | 0 | 0 | 0 |
| rs121907925 | pathogenic | 0 | 1 | 0 |
| rs746212325 | pathogenic | 0 | 0 | 0 |
| rs754637179 | pathogenic | 0 | 0 | 0 |
| rs780882740 | pathogenic | 0 | 0 | 0 |
| rs121918130 | pathogenic | 0 | 0 | 0 |
| rs121918129 | pathogenic | 0 | 0 | 0 |
| rs13297509 | pathogenic | 0 | 0 | 0 |
| rs746867724 | pathogenic | 0 | 0 | 0 |
| rs121918128 | pathogenic | 0 | 0 | 0 |
| rs752300607 | pathogenic | 0 | 0 | 0 |
| rs121964860 | pathogenic | 0 | 0 | 0 |
| rs45466197 | pathogenic | 1 | 1 | 1 |
| rs3730238 | benign | 1 | 1 | 1 |
| rs730881125 | pathogenic | 1 | 1 | 1 |
| rs121964861 | pathogenic | 1 | 1 | 1 |
| rs397516484 | pathogenic | 0 | 0 | 0 |
| rs367785431 | pathogenic | 0 | 0 | 0 |
| rs141121678 | pathogenic | 0 | 0 | 0 |
| rs12720263 | benign | 0 | 0 | 0 |
| rs2304256 | benign | 0 | 0 | 0 |
| rs2304255 | benign | 0 | 0 | 0 |
| rs121909045 | pathogenic | 0 | 0 | 0 |
| rs397508796 | pathogenic | 0 | 0 | 0 |
| rs151020603 | pathogenic | 0 | 0 | 0 |
| rs77284892 | pathogenic | 0 | 0 | 0 |
| rs368505753 | pathogenic | 0 | 0 | 0 |
| rs115545701 | pathogenic | 0 | 0 | 0 |
| rs121908749 | pathogenic | 0 | 0 | 0 |
| rs75961395 | pathogenic | 0 | 0 | 0 |
| rs121908750 | pathogenic | 0 | 0 | 0 |
| rs121908751 | pathogenic | 0 | 0 | 0 |
| rs397508464 | pathogenic | 0 | 0 | 0 |
| rs397508461 | pathogenic | 0 | 0 | 0 |
| rs121909031 | pathogenic | 0 | 0 | 0 |
| rs113993958 | pathogenic | 0 | 0 | 0 |
| rs77834169 | pathogenic | 0 | 0 | 0 |
| rs78655421 | pathogenic | 0 | 0 | 0 |
| rs141723617 | benign | 0 | 0 | 0 |
| rs35516286 | benign | 0 | 0 | 0 |
| rs1800079 | pathogenic | 1 | 1 | 0 |

| | | | | |
|---|---|---|---|---|
| rs80282562 | pathogenic | 0 | 0 | 0 |
| rs121908752 | pathogenic | 0 | 0 | 0 |
| rs556880586 | pathogenic | 0 | 0 | 0 |
| rs121908760 | pathogenic | 0 | 0 | 0 |
| rs397508350 | pathogenic | 0 | 0 | 0 |
| rs374946172 | pathogenic | 0 | 0 | 0 |
| rs397508387 | pathogenic | 0 | 0 | 0 |
| rs201386642 | pathogenic | 0 | 0 | 0 |
| rs121909012 | pathogenic | 0 | 0 | 0 |
| rs121909008 | pathogenic | 0 | 0 | 0 |
| rs149790377 | pathogenic | 0 | 0 | 0 |
| rs397508442 | pathogenic | 0 | 0 | 0 |
| rs121909035 | pathogenic | 0 | 0 | 0 |
| rs386134230 | pathogenic | 0 | 0 | 0 |
| rs1800112 | benign | 0 | 0 | 0 |
| rs144055758 | pathogenic | 0 | 0 | 0 |
| rs142394380 | pathogenic | 0 | 0 | 0 |
| rs121909036 | pathogenic | 0 | 0 | 0 |
| rs78194216 | pathogenic | 0 | 0 | 0 |
| rs121909019 | pathogenic | 0 | 0 | 0 |
| rs121909020 | pathogenic | 0 | 0 | 0 |
| rs200321110 | pathogenic | 0 | 0 | 0 |
| rs202179988 | pathogenic | 0 | 0 | 0 |
| rs121909037 | pathogenic | 0 | 0 | 0 |
| rs186045772 | pathogenic | 0 | 0 | 0 |
| rs139304906 | pathogenic | 0 | 0 | 0 |
| rs78802634 | pathogenic | 0 | 0 | 0 |
| rs121908761 | pathogenic | 0 | 0 | 0 |
| rs36210737 | pathogenic | 0 | 0 | 0 |
| rs139729994 | pathogenic | 0 | 0 | 0 |
| rs79850223 | pathogenic | 0 | 0 | 0 |
| rs1800120 | benign | 0 | 0 | 0 |
| rs74767530 | pathogenic | 0 | 0 | 0 |
| rs374705585 | pathogenic | 0 | 0 | 0 |
| rs1136287 | benign | 0 | 0 | 0 |
| rs6828 | benign | 0 | 0 | 0 |
| rs104894856 | pathogenic | 0 | 0 | 0 |
| rs104894860 | pathogenic | 0 | 0 | 0 |
| rs61736892 | pathogenic | 0 | 0 | 0 |
| rs145807417 | benign | 0 | 0 | 0 |
| rs104894853 | pathogenic | 0 | 0 | 0 |
| rs199422227 | pathogenic | 0 | 0 | 0 |
| rs113993946 | pathogenic | 0 | 0 | 0 |
| rs199422231 | pathogenic | 0 | 0 | 0 |
| rs199422228 | pathogenic | 0 | 0 | 0 |
| rs142195196 | benign | 0 | 1 | 0 |
| rs148656104 | benign | 1 | 1 | 1 |
| rs141944844 | benign | 0 | 0 | 0 |
| rs104894342 | pathogenic | 0 | 0 | 0 |
| rs104894343 | pathogenic | 0 | 0 | 0 |
| rs104894344 | pathogenic | 0 | 0 | 0 |
| rs573322878 | benign | 0 | 0 | 0 |
| rs104894347 | pathogenic | 0 | 0 | 0 |
| rs28937882 | pathogenic | 0 | 0 | 0 |

| | | | | |
|---|---|---|---|---|
| rs193922702 | pathogenic | 0 | 0 | 0 |
| rs200786104 | benign | 0 | 0 | 0 |
| rs690016544 | pathogenic | 1 | 1 | 1 |
| rs28943592 | benign | 0 | 0 | 0 |
| rs28943594 | benign | 0 | 0 | 0 |
| rs121908957 | pathogenic | 0 | 0 | 0 |
| rs115408010 | benign | 1 | 1 | 1 |
| rs77591659 | benign | 0 | 0 | 0 |
| rs121918511 | pathogenic | 0 | 0 | 0 |
| rs538359948 | benign | 0 | 0 | 0 |
| rs76708142 | benign | 0 | 0 | 0 |
| rs121907913 | pathogenic | 1 | 1 | 0 |
| rs121907920 | pathogenic | 0 | 0 | 0 |
| rs121907923 | pathogenic | 1 | 1 | 0 |
| rs121907928 | pathogenic | 1 | 1 | 0 |
| rs121907919 | pathogenic | 1 | 1 | 0 |
| rs121907918 | pathogenic | 1 | 1 | 0 |
| rs74315493 | pathogenic | 1 | 1 | 1 |
| rs74315494 | pathogenic | 1 | 1 | 1 |
| rs145446060 | benign | 1 | 1 | 1 |
| rs550409143 | benign | 1 | 1 | 0 |

| compound_binding_site | cofactor_binding_site | nucleic_binding_site | ion_binding_site |
|---|---|---|---|
| 1 | 0 | 0 | 0 |
| 1 | 0 | 0 | 0 |
| 0 | 0 | 0 | 0 |
| 1 | 0 | 0 | 0 |
| 0 | 0 | 0 | 0 |
| 0 | 0 | 0 | 0 |
| 0 | 0 | 0 | 0 |
| 0 | 0 | 0 | 0 |
| 1 | 0 | 0 | 1 |
| 1 | 0 | 0 | 0 |
| 0 | 0 | 0 | 1 |
| 1 | 0 | 1 | 0 |
| 0 | 0 | 0 | 0 |
| 0 | 0 | 0 | 0 |
| 0 | 0 | 0 | 0 |
| 0 | 0 | 0 | 0 |
| 0 | 0 | 0 | 0 |
| 1 | 0 | 0 | 1 |
| 1 | 0 | 1 | 0 |
| 0 | 0 | 0 | 0 |
| 1 | 0 | 0 | 1 |
| 1 | 0 | 0 | 0 |
| 1 | 0 | 0 | 0 |
| 1 | 0 | 0 | 0 |
| 0 | 0 | 0 | 0 |
| 0 | 0 | 0 | 0 |
| 1 | 0 | 0 | 1 |
| 0 | 0 | 0 | 0 |
| 0 | 0 | 0 | 0 |
| 0 | 0 | 0 | 0 |
| 0 | 0 | 0 | 0 |
| 1 | 0 | 0 | 0 |
| 1 | 0 | 0 | 1 |
| 1 | 0 | 0 | 0 |
| 0 | 0 | 0 | 0 |
| 0 | 0 | 0 | 0 |
| 0 | 0 | 0 | 0 |
| 1 | 0 | 0 | 1 |
| 1 | 0 | 0 | 1 |
| 1 | 0 | 0 | 1 |
| 1 | 0 | 0 | 0 |
| 1 | 0 | 0 | 0 |
| 1 | 0 | 0 | 0 |
| 1 | 0 | 0 | 0 |
| 1 | 0 | 0 | 0 |
| 1 | 0 | 0 | 0 |
| 1 | 0 | 0 | 0 |
| 1 | 0 | 0 | 0 |
| 1 | 0 | 0 | 0 |
| 1 | 0 | 0 | 0 |
| 1 | 0 | 0 | 0 |
| 1 | 0 | 0 | 0 |
| 1 | 0 | 0 | 0 |

| | | | |
|---|---|---|---|
| 1 | 0 | 0 | 1 |
| 1 | 0 | 0 | 0 |
| 1 | 0 | 0 | 0 |
| 1 | 0 | 0 | 0 |
| 0 | 0 | 0 | 0 |
| 1 | 0 | 0 | 0 |
| 0 | 0 | 0 | 0 |
| 1 | 0 | 0 | 0 |
| 1 | 0 | 0 | 0 |
| 1 | 0 | 0 | 1 |
| 1 | 0 | 0 | 0 |
| 1 | 0 | 0 | 0 |
| 1 | 0 | 0 | 0 |
| 1 | 0 | 0 | 0 |
| 1 | 0 | 0 | 0 |
| 1 | 0 | 0 | 0 |
| 1 | 0 | 0 | 0 |
| 1 | 0 | 0 | 1 |
| 1 | 0 | 0 | 1 |
| 1 | 1 | 0 | 1 |
| 1 | 0 | 0 | 0 |
| 1 | 0 | 0 | 0 |
| 1 | 0 | 0 | 0 |
| 1 | 0 | 0 | 0 |
| 1 | 1 | 0 | 0 |
| 1 | 0 | 0 | 0 |
| 1 | 0 | 0 | 0 |
| 1 | 0 | 0 | 0 |
| 1 | 0 | 0 | 0 |
| 1 | 0 | 0 | 0 |
| 1 | 1 | 0 | 0 |
| 1 | 1 | 0 | 0 |
| 1 | 1 | 0 | 0 |
| 1 | 1 | 0 | 0 |
| 1 | 1 | 0 | 0 |
| 1 | 0 | 0 | 0 |
| 1 | 1 | 0 | 0 |
| 1 | 0 | 0 | 0 |
| 1 | 0 | 0 | 0 |
| 1 | 1 | 0 | 0 |
| 0 | 0 | 0 | 0 |
| 0 | 0 | 0 | 0 |
| 0 | 0 | 0 | 0 |
| 0 | 0 | 0 | 0 |
| 0 | 0 | 0 | 0 |
| 0 | 0 | 0 | 0 |
| 0 | 0 | 0 | 0 |
| 1 | 0 | 0 | 1 |
| 0 | 0 | 0 | 0 |
| 1 | 0 | 0 | 0 |
| 1 | 0 | 0 | 0 |
| 0 | 0 | 0 | 0 |
| 0 | 0 | 0 | 0 |
| 0 | 0 | 0 | 0 |

| | | | |
|---|---|---|---|
| 0 | 0 | 0 | 0 |
| 0 | 0 | 0 | 0 |
| 0 | 0 | 0 | 0 |
| 1 | 0 | 0 | 0 |
| 1 | 0 | 0 | 0 |
| 1 | 0 | 0 | 0 |
| 0 | 0 | 0 | 0 |
| 0 | 0 | 1 | 0 |
| 0 | 0 | 1 | 0 |
| 0 | 0 | 1 | 0 |
| 0 | 0 | 1 | 0 |
| 0 | 0 | 1 | 0 |
| 0 | 0 | 0 | 0 |
| 0 | 0 | 0 | 0 |
| 1 | 0 | 0 | 0 |
| 0 | 0 | 0 | 0 |
| 0 | 0 | 0 | 0 |
| 1 | 0 | 0 | 0 |
| 1 | 0 | 0 | 0 |
| 1 | 0 | 0 | 0 |
| 1 | 0 | 0 | 0 |
| 1 | 1 | 0 | 0 |
| 1 | 0 | 0 | 0 |
| 1 | 0 | 0 | 0 |
| 0 | 0 | 0 | 0 |
| 0 | 0 | 0 | 0 |
| 1 | 0 | 0 | 0 |
| 1 | 1 | 0 | 0 |
| 1 | 0 | 0 | 0 |
| 1 | 0 | 0 | 0 |
| 1 | 1 | 0 | 0 |
| 1 | 1 | 0 | 0 |
| 1 | 1 | 0 | 0 |
| 1 | 1 | 0 | 0 |
| 1 | 1 | 0 | 0 |
| 1 | 1 | 0 | 0 |
| 1 | 1 | 0 | 0 |
| 1 | 1 | 0 | 0 |
| 1 | 1 | 0 | 0 |
| 1 | 1 | 0 | 0 |
| 1 | 1 | 0 | 0 |
| 1 | 1 | 0 | 0 |
| 1 | 0 | 0 | 0 |
| 1 | 0 | 0 | 0 |
| 1 | 0 | 0 | 0 |
| 1 | 1 | 0 | 0 |
| 1 | 0 | 0 | 0 |
| 1 | 1 | 1 | 0 |
| 1 | 1 | 1 | 0 |
| 1 | 1 | 0 | 0 |
| 1 | 1 | 0 | 0 |
| 1 | 0 | 0 | 0 |
| 1 | 0 | 0 | 0 |
| 0 | 0 | 0 | 0 |

| | | | |
|---|---|---|---|
| 0 | 0 | 0 | 0 |
| 0 | 0 | 0 | 0 |
| 0 | 0 | 0 | 0 |
| 1 | 0 | 0 | 0 |
| 1 | 0 | 0 | 0 |
| 0 | 0 | 0 | 0 |
| 0 | 0 | 0 | 0 |
| 0 | 0 | 0 | 0 |
| 0 | 0 | 0 | 0 |
| 0 | 0 | 0 | 0 |
| 0 | 0 | 0 | 0 |
| 0 | 0 | 0 | 0 |
| 0 | 0 | 0 | 0 |
| 0 | 0 | 0 | 0 |
| 0 | 0 | 0 | 0 |
| 0 | 0 | 0 | 0 |
| 0 | 0 | 0 | 0 |
| 0 | 0 | 0 | 0 |
| 0 | 0 | 0 | 0 |
| 0 | 0 | 0 | 0 |
| 0 | 0 | 0 | 0 |
| 0 | 0 | 0 | 0 |
| 1 | 0 | 0 | 0 |
| 1 | 0 | 0 | 0 |
| 0 | 0 | 0 | 0 |
| 0 | 0 | 0 | 0 |
| 1 | 0 | 0 | 0 |
| 0 | 0 | 0 | 0 |
| 0 | 0 | 0 | 0 |
| 0 | 0 | 0 | 0 |
| 0 | 0 | 0 | 0 |
| 0 | 0 | 0 | 0 |
| 0 | 0 | 0 | 0 |
| 0 | 0 | 0 | 0 |
| 1 | 0 | 0 | 0 |
| 0 | 0 | 0 | 0 |
| 0 | 0 | 0 | 0 |
| 0 | 0 | 0 | 0 |
| 0 | 0 | 0 | 0 |
| 0 | 0 | 0 | 0 |
| 0 | 0 | 0 | 0 |
| 0 | 0 | 0 | 0 |
| 0 | 0 | 0 | 0 |
| 0 | 0 | 0 | 0 |
| 0 | 0 | 0 | 0 |
| 0 | 0 | 0 | 0 |
| 0 | 0 | 1 | 0 |
| 1 | 0 | 0 | 0 |
| 1 | 0 | 0 | 0 |
| 1 | 1 | 0 | 0 |
| 1 | 0 | 0 | 0 |
| 0 | 0 | 0 | 0 |
| 0 | 0 | 0 | 0 |
| 1 | 0 | 0 | 0 |

| | | | |
|---|---|---|---|
| 1 | 0 | 0 | 0 |
| 1 | 0 | 0 | 0 |
| 1 | 0 | 0 | 0 |
| 0 | 0 | 0 | 0 |
| 0 | 0 | 0 | 0 |
| 0 | 0 | 0 | 0 |
| 1 | 0 | 0 | 0 |
| 1 | 0 | 0 | 0 |
| 0 | 0 | 0 | 0 |
| 0 | 0 | 0 | 0 |
| 1 | 0 | 0 | 0 |
| 0 | 0 | 0 | 0 |
| 1 | 0 | 0 | 0 |
| 0 | 0 | 0 | 0 |
| 0 | 0 | 0 | 0 |
| 0 | 0 | 0 | 0 |
| 0 | 0 | 0 | 0 |
| 0 | 0 | 0 | 0 |
| 0 | 0 | 0 | 0 |
| 0 | 0 | 0 | 0 |
| 0 | 0 | 0 | 0 |
| 0 | 0 | 0 | 0 |
| 0 | 0 | 0 | 0 |
| 0 | 0 | 0 | 0 |
| 0 | 0 | 0 | 0 |
| 0 | 0 | 0 | 0 |
| 1 | 0 | 0 | 0 |
| 1 | 0 | 0 | 0 |
| 1 | 0 | 0 | 0 |
| 1 | 0 | 0 | 0 |
| 1 | 0 | 0 | 0 |
| 1 | 0 | 0 | 0 |
| 0 | 0 | 0 | 0 |
| 0 | 0 | 0 | 0 |
| 1 | 0 | 0 | 0 |
| 0 | 0 | 0 | 0 |
| 0 | 0 | 0 | 0 |
| 0 | 0 | 0 | 0 |
| 0 | 0 | 0 | 0 |
| 0 | 0 | 0 | 0 |
| 0 | 0 | 0 | 0 |
| 0 | 0 | 0 | 0 |
| 0 | 0 | 0 | 0 |
| 0 | 0 | 0 | 0 |
| 1 | 1 | 0 | 0 |
| 1 | 1 | 0 | 0 |
| 1 | 1 | 0 | 1 |
| 1 | 1 | 0 | 1 |
| 1 | 1 | 0 | 1 |
| 1 | 1 | 0 | 1 |
| 1 | 1 | 0 | 0 |
| 1 | 1 | 0 | 1 |
| 1 | 1 | 0 | 1 |
| 1 | 1 | 0 | 1 |

| | | | |
|---|---|---|---|
| 1 | 1 | 0 | 1 |
| 1 | 1 | 0 | 1 |
| 1 | 1 | 0 | 1 |
| 1 | 1 | 0 | 1 |
| 1 | 1 | 0 | 1 |
| 1 | 0 | 0 | 0 |
| 0 | 0 | 0 | 0 |
| 0 | 0 | 0 | 0 |
| 0 | 0 | 0 | 0 |
| 1 | 0 | 0 | 0 |
| 0 | 0 | 1 | 0 |
| 0 | 0 | 0 | 0 |
| 0 | 0 | 0 | 0 |
| 0 | 0 | 0 | 0 |
| 0 | 0 | 0 | 0 |
| 0 | 0 | 0 | 0 |
| 0 | 0 | 0 | 0 |
| 0 | 0 | 0 | 0 |
| 0 | 0 | 0 | 0 |
| 0 | 0 | 0 | 0 |
| 0 | 0 | 0 | 0 |
| 0 | 0 | 0 | 0 |
| 0 | 0 | 0 | 0 |
| 0 | 0 | 0 | 0 |
| 0 | 0 | 0 | 0 |
| 1 | 1 | 0 | 0 |
| 1 | 1 | 0 | 0 |
| 1 | 1 | 0 | 0 |
| 0 | 1 | 0 | 0 |
| 1 | 0 | 0 | 0 |
| 0 | 0 | 0 | 0 |
| 0 | 0 | 0 | 0 |
| 0 | 0 | 0 | 0 |
| 1 | 0 | 0 | 0 |
| 1 | 1 | 0 | 0 |
| 0 | 0 | 0 | 0 |
| 0 | 0 | 0 | 0 |
| 0 | 0 | 0 | 0 |
| 0 | 0 | 0 | 0 |
| 1 | 0 | 0 | 0 |
| 0 | 0 | 0 | 0 |
| 0 | 0 | 0 | 0 |
| 0 | 0 | 0 | 0 |
| 0 | 0 | 0 | 0 |
| 0 | 0 | 0 | 0 |
| 0 | 1 | 0 | 0 |
| 0 | 1 | 0 | 0 |
| 0 | 0 | 0 | 0 |
| 0 | 0 | 0 | 0 |
| 0 | 1 | 0 | 0 |
| 0 | 0 | 0 | 0 |
| 0 | 0 | 0 | 0 |
| 0 | 1 | 0 | 0 |
| 0 | 0 | 0 | 0 |

| | | | |
|---|---|---|---|
| 1 | 1 | 0 | 0 |
| 0 | 0 | 0 | 0 |
| 0 | 0 | 0 | 0 |
| 0 | 0 | 0 | 0 |
| 0 | 0 | 0 | 0 |
| 0 | 0 | 0 | 0 |
| 0 | 0 | 0 | 0 |
| 0 | 0 | 0 | 0 |
| 0 | 0 | 0 | 0 |
| 0 | 0 | 0 | 0 |
| 0 | 0 | 0 | 0 |
| 0 | 0 | 0 | 0 |
| 0 | 0 | 0 | 0 |
| 0 | 0 | 0 | 0 |
| 0 | 1 | 0 | 0 |
| 0 | 0 | 0 | 0 |
| 0 | 1 | 0 | 0 |
| 0 | 1 | 0 | 0 |
| 0 | 0 | 0 | 0 |
| 0 | 1 | 0 | 0 |
| 1 | 0 | 0 | 1 |
| 1 | 1 | 0 | 0 |
| 1 | 0 | 0 | 0 |
| 1 | 0 | 0 | 1 |
| 0 | 0 | 0 | 0 |
| 0 | 0 | 0 | 0 |
| 0 | 0 | 0 | 0 |
| 0 | 0 | 0 | 0 |
| 1 | 0 | 0 | 0 |
| 0 | 0 | 0 | 0 |
| 1 | 0 | 0 | 0 |
| 0 | 0 | 0 | 0 |
| 0 | 0 | 0 | 0 |
| 0 | 0 | 0 | 0 |
| 0 | 0 | 0 | 0 |
| 0 | 0 | 0 | 0 |
| 0 | 0 | 0 | 0 |
| 1 | 1 | 0 | 0 |
| 1 | 1 | 0 | 0 |
| 1 | 1 | 0 | 0 |
| 0 | 0 | 0 | 0 |
| 0 | 0 | 0 | 0 |
| 0 | 0 | 0 | 0 |
| 1 | 0 | 0 | 0 |
| 1 | 0 | 0 | 1 |
| 1 | 0 | 0 | 1 |
| 1 | 0 | 0 | 0 |
| 1 | 0 | 0 | 1 |
| 1 | 0 | 0 | 0 |
| 0 | 0 | 0 | 0 |
| 1 | 1 | 0 | 0 |
| 1 | 1 | 0 | 0 |
| 0 | 0 | 0 | 0 |
| 0 | 0 | 0 | 0 |

| | | | |
|---|---|---|---|
| 1 | 0 | 0 | 0 |
| 1 | 0 | 0 | 0 |
| 1 | 0 | 0 | 0 |
| 0 | 0 | 0 | 0 |
| 0 | 0 | 0 | 0 |
| 1 | 1 | 0 | 1 |
| 1 | 1 | 0 | 1 |
| 1 | 1 | 0 | 1 |
| 1 | 1 | 0 | 1 |
| 1 | 1 | 0 | 1 |
| 1 | 1 | 0 | 1 |
| 1 | 1 | 0 | 1 |
| 1 | 1 | 0 | 0 |
| 1 | 1 | 0 | 0 |
| 1 | 1 | 1 | 1 |
| 0 | 0 | 0 | 0 |
| 1 | 1 | 0 | 0 |
| 1 | 1 | 0 | 0 |
| 0 | 0 | 0 | 0 |
| 0 | 0 | 0 | 0 |
| 0 | 0 | 0 | 0 |
| 1 | 0 | 0 | 0 |
| 1 | 0 | 0 | 0 |
| 0 | 0 | 0 | 0 |
| 1 | 0 | 0 | 0 |
| 1 | 0 | 0 | 0 |
| 1 | 0 | 0 | 0 |
| 0 | 0 | 0 | 0 |
| 0 | 0 | 0 | 0 |
| 0 | 0 | 0 | 0 |
| 0 | 0 | 0 | 0 |
| 0 | 0 | 0 | 0 |
| 0 | 0 | 0 | 0 |
| 0 | 0 | 0 | 0 |
| 0 | 0 | 0 | 0 |
| 0 | 0 | 0 | 0 |
| 0 | 0 | 0 | 0 |
| 0 | 0 | 0 | 0 |
| 0 | 0 | 0 | 0 |
| 0 | 0 | 0 | 0 |
| 0 | 0 | 0 | 0 |
| 0 | 0 | 0 | 0 |
| 0 | 0 | 0 | 0 |
| 0 | 0 | 0 | 0 |
| 0 | 0 | 0 | 0 |
| 1 | 0 | 0 | 0 |
| 1 | 0 | 0 | 0 |
| 1 | 0 | 0 | 0 |
| 1 | 0 | 0 | 0 |
| 0 | 0 | 0 | 0 |
| 0 | 0 | 0 | 0 |
| 0 | 0 | 0 | 0 |
| 0 | 0 | 0 | 0 |
| 0 | 0 | 0 | 0 |

| | | | |
|---|---|---|---|
| 0 | 0 | 0 | 0 |
| 0 | 0 | 0 | 0 |
| 0 | 0 | 0 | 0 |
| 0 | 0 | 0 | 0 |
| 0 | 0 | 0 | 0 |
| 0 | 0 | 0 | 0 |
| 0 | 0 | 0 | 0 |
| 1 | 0 | 0 | 1 |
| 0 | 0 | 0 | 0 |
| 0 | 0 | 0 | 0 |
| 1 | 0 | 0 | 0 |
| 0 | 0 | 0 | 0 |
| 0 | 0 | 0 | 0 |
| 0 | 0 | 0 | 0 |
| 0 | 0 | 0 | 0 |
| 1 | 0 | 0 | 0 |
| 0 | 0 | 0 | 0 |
| 0 | 0 | 0 | 0 |
| 0 | 0 | 0 | 0 |
| 0 | 0 | 0 | 0 |
| 0 | 0 | 0 | 0 |
| 0 | 0 | 0 | 0 |
| 0 | 0 | 0 | 0 |
| 1 | 1 | 0 | 1 |
| 0 | 0 | 0 | 0 |
| 0 | 0 | 0 | 0 |
| 0 | 0 | 0 | 0 |
| 1 | 1 | 0 | 1 |
| 1 | 1 | 0 | 0 |
| 0 | 0 | 0 | 0 |
| 1 | 0 | 0 | 0 |
| 1 | 1 | 0 | 0 |
| 0 | 0 | 0 | 0 |
| 1 | 0 | 0 | 0 |
| 1 | 1 | 0 | 0 |
| 1 | 0 | 0 | 0 |
| 1 | 1 | 0 | 1 |
| 1 | 1 | 0 | 1 |
| 1 | 0 | 0 | 0 |
| 0 | 0 | 0 | 0 |
| 0 | 0 | 0 | 0 |
| 0 | 0 | 0 | 0 |
| 0 | 0 | 0 | 0 |
| 0 | 0 | 0 | 0 |
| 0 | 0 | 0 | 0 |
| 0 | 0 | 0 | 0 |
| 0 | 0 | 0 | 0 |
| 0 | 0 | 0 | 0 |
| 0 | 0 | 0 | 0 |
| 0 | 0 | 0 | 0 |
| 0 | 0 | 0 | 0 |
| 0 | 0 | 0 | 0 |
| 0 | 0 | 0 | 0 |
| 0 | 0 | 0 | 0 |

| | | | |
|---|---|---|---|
| 0 | 0 | 0 | 0 |
| 0 | 0 | 0 | 0 |
| 0 | 0 | 0 | 0 |
| 0 | 0 | 0 | 0 |
| 0 | 0 | 0 | 0 |
| 0 | 0 | 0 | 0 |
| 0 | 0 | 0 | 0 |
| 0 | 0 | 0 | 0 |
| 0 | 0 | 0 | 0 |
| 0 | 0 | 0 | 0 |
| 0 | 0 | 0 | 0 |
| 0 | 0 | 0 | 0 |
| 0 | 0 | 0 | 0 |
| 1 | 1 | 0 | 0 |
| 0 | 0 | 0 | 0 |
| 1 | 0 | 0 | 0 |
| 0 | 0 | 0 | 1 |
| 0 | 0 | 0 | 0 |
| 0 | 0 | 0 | 0 |
| 0 | 0 | 0 | 0 |
| 0 | 0 | 0 | 0 |
| 0 | 0 | 0 | 0 |
| 0 | 0 | 0 | 0 |
| 0 | 0 | 0 | 0 |
| 0 | 0 | 0 | 0 |
| 0 | 0 | 0 | 0 |
| 0 | 0 | 0 | 0 |
| 0 | 0 | 0 | 0 |
| 0 | 0 | 0 | 0 |
| 0 | 0 | 0 | 0 |
| 1 | 0 | 0 | 0 |
| 1 | 0 | 0 | 0 |
| 0 | 0 | 0 | 0 |
| 0 | 0 | 0 | 0 |
| 0 | 0 | 0 | 0 |
| 0 | 0 | 0 | 0 |
| 0 | 0 | 0 | 0 |
| 0 | 0 | 0 | 0 |
| 0 | 0 | 0 | 0 |
| 0 | 0 | 0 | 0 |
| 1 | 0 | 0 | 0 |
| 1 | 0 | 0 | 0 |
| 1 | 0 | 0 | 0 |
| 1 | 0 | 0 | 0 |
| 0 | 0 | 0 | 0 |
| 1 | 0 | 0 | 0 |
| 0 | 0 | 0 | 0 |
| 0 | 0 | 0 | 0 |
| 1 | 0 | 0 | 0 |
| 1 | 0 | 0 | 0 |
| 0 | 0 | 0 | 0 |
| 0 | 0 | 0 | 0 |
| 0 | 0 | 0 | 0 |
| 0 | 0 | 0 | 0 |

| | | | |
|---|---|---|---|
| 0 | 0 | 0 | 0 |
| 0 | 0 | 0 | 0 |
| 0 | 0 | 0 | 0 |
| 0 | 0 | 0 | 0 |
| 0 | 0 | 0 | 0 |
| 0 | 0 | 0 | 0 |
| 0 | 0 | 0 | 0 |
| 0 | 0 | 0 | 0 |
| 0 | 0 | 0 | 0 |
| 0 | 0 | 0 | 0 |
| 0 | 0 | 0 | 0 |
| 0 | 0 | 0 | 0 |
| 0 | 0 | 0 | 0 |
| 0 | 0 | 0 | 0 |
| 0 | 0 | 0 | 0 |
| 1 | 0 | 0 | 1 |
| 0 | 0 | 0 | 0 |
| 0 | 0 | 0 | 0 |
| 0 | 0 | 0 | 0 |
| 0 | 0 | 0 | 0 |
| 0 | 0 | 0 | 0 |
| 1 | 1 | 0 | 0 |
| 1 | 1 | 0 | 0 |
| 1 | 0 | 0 | 0 |
| 1 | 0 | 0 | 0 |
| 0 | 0 | 0 | 0 |
| 0 | 0 | 0 | 0 |
| 0 | 0 | 0 | 0 |
| 0 | 0 | 0 | 0 |
| 1 | 0 | 0 | 0 |
| 1 | 0 | 0 | 0 |
| 1 | 0 | 0 | 0 |
| 1 | 1 | 0 | 0 |
| 1 | 1 | 0 | 0 |
| 1 | 0 | 0 | 0 |
| 0 | 0 | 0 | 0 |
| 0 | 0 | 0 | 0 |
| 0 | 1 | 0 | 0 |
| 0 | 0 | 0 | 0 |
| 0 | 0 | 0 | 0 |
| 0 | 0 | 0 | 0 |
| 1 | 1 | 0 | 0 |
| 1 | 0 | 0 | 0 |
| 1 | 0 | 0 | 0 |
| 0 | 0 | 0 | 0 |
| 0 | 0 | 0 | 0 |
| 0 | 0 | 0 | 0 |
| 0 | 0 | 0 | 0 |
| 0 | 0 | 0 | 0 |
| 0 | 0 | 0 | 0 |
| 0 | 0 | 0 | 0 |
| 0 | 0 | 0 | 0 |
| 0 | 0 | 0 | 0 |
| 0 | 0 | 0 | 0 |

| | | | |
|---|---|---|---|
| 0 | 0 | 0 | 0 |
| 1 | 0 | 0 | 0 |
| 0 | 0 | 0 | 0 |
| 0 | 0 | 0 | 0 |
| 0 | 0 | 0 | 0 |
| 0 | 0 | 0 | 0 |
| 1 | 0 | 0 | 0 |
| 0 | 0 | 0 | 0 |
| 0 | 0 | 0 | 0 |
| 0 | 0 | 0 | 0 |
| 0 | 0 | 0 | 0 |
| 0 | 0 | 0 | 0 |
| 0 | 0 | 0 | 0 |
| 0 | 0 | 0 | 0 |
| 0 | 0 | 0 | 0 |
| 0 | 0 | 0 | 0 |
| 0 | 1 | 0 | 0 |
| 0 | 0 | 0 | 0 |
| 0 | 0 | 0 | 0 |
| 0 | 0 | 0 | 0 |
| 1 | 0 | 0 | 0 |
| 0 | 0 | 0 | 0 |
| 0 | 0 | 0 | 0 |
| 1 | 1 | 0 | 0 |
| 0 | 0 | 0 | 0 |
| 0 | 0 | 0 | 0 |
| 0 | 0 | 0 | 0 |
| 0 | 0 | 0 | 0 |
| 1 | 0 | 0 | 0 |
| 0 | 0 | 0 | 0 |
| 0 | 0 | 0 | 0 |
| 0 | 0 | 0 | 0 |
| 0 | 0 | 0 | 0 |
| 0 | 0 | 0 | 0 |
| 0 | 0 | 0 | 0 |
| 1 | 0 | 0 | 0 |
| 0 | 0 | 0 | 0 |
| 0 | 0 | 0 | 0 |
| 0 | 0 | 0 | 0 |
| 0 | 0 | 0 | 0 |
| 1 | 1 | 0 | 0 |
| 0 | 0 | 0 | 0 |
| 0 | 0 | 0 | 0 |
| 0 | 0 | 0 | 0 |
| 0 | 0 | 0 | 0 |
| 0 | 0 | 0 | 0 |
| 1 | 0 | 0 | 0 |
| 1 | 0 | 0 | 1 |
| 1 | 0 | 0 | 0 |
| 0 | 0 | 0 | 0 |
| 0 | 0 | 0 | 0 |
| 0 | 0 | 0 | 0 |
| 1 | 0 | 0 | 0 |
| 1 | 0 | 0 | 0 |

| | | | |
|---|---|---|---|
| 1 | 0 | 0 | 0 |
| 0 | 0 | 0 | 0 |
| 0 | 0 | 0 | 0 |
| 0 | 0 | 0 | 0 |
| 1 | 1 | 0 | 0 |
| 1 | 1 | 0 | 0 |
| 0 | 0 | 0 | 0 |
| 0 | 0 | 0 | 0 |
| 1 | 0 | 0 | 0 |
| 1 | 0 | 0 | 0 |
| 1 | 0 | 1 | 0 |
| 0 | 0 | 0 | 0 |
| 0 | 0 | 1 | 0 |
| 0 | 0 | 1 | 0 |
| 0 | 0 | 1 | 0 |
| 1 | 0 | 0 | 0 |
| 1 | 0 | 0 | 0 |
| 0 | 0 | 0 | 0 |
| 0 | 0 | 0 | 0 |
| 1 | 0 | 0 | 1 |
| 1 | 0 | 0 | 1 |
| 1 | 0 | 0 | 0 |
| 0 | 0 | 0 | 0 |
| 0 | 0 | 0 | 0 |
| 0 | 0 | 0 | 0 |
| 0 | 0 | 0 | 0 |
| 0 | 0 | 0 | 0 |
| 1 | 0 | 0 | 0 |
| 0 | 0 | 0 | 0 |
| 0 | 0 | 0 | 1 |
| 0 | 0 | 0 | 1 |
| 0 | 0 | 1 | 1 |
| 0 | 0 | 1 | 1 |
| 0 | 0 | 1 | 1 |
| 0 | 0 | 1 | 1 |
| 0 | 0 | 1 | 0 |
| 0 | 0 | 1 | 0 |
| 0 | 0 | 0 | 0 |
| 0 | 0 | 0 | 0 |
| 0 | 0 | 0 | 0 |
| 0 | 0 | 0 | 0 |
| 0 | 0 | 1 | 0 |
| 0 | 0 | 1 | 0 |
| 0 | 0 | 1 | 0 |
| 0 | 0 | 1 | 0 |
| 0 | 0 | 1 | 0 |
| 0 | 0 | 1 | 0 |
| 0 | 0 | 1 | 0 |
| 0 | 0 | 1 | 0 |
| 0 | 0 | 0 | 0 |
| 0 | 0 | 0 | 0 |
| 0 | 0 | 0 | 0 |
| 0 | 0 | 0 | 0 |
| 1 | 0 | 0 | 0 |

| | | | |
|---|---|---|---|
| 0 | 0 | 0 | 0 |
| 0 | 0 | 0 | 0 |
| 0 | 0 | 0 | 0 |
| 1 | 0 | 0 | 0 |
| 0 | 0 | 0 | 0 |
| 0 | 0 | 0 | 0 |
| 0 | 0 | 0 | 0 |
| 0 | 0 | 0 | 0 |
| 0 | 0 | 0 | 0 |
| 0 | 0 | 0 | 0 |
| 1 | 1 | 0 | 0 |
| 0 | 0 | 0 | 0 |
| 1 | 1 | 0 | 0 |
| 1 | 1 | 0 | 0 |
| 0 | 1 | 0 | 0 |
| 0 | 0 | 0 | 0 |
| 0 | 0 | 0 | 0 |
| 0 | 0 | 0 | 0 |
| 0 | 0 | 0 | 0 |
| 0 | 0 | 0 | 0 |
| 1 | 1 | 0 | 0 |
| 0 | 1 | 0 | 0 |
| 0 | 0 | 0 | 0 |
| 0 | 1 | 0 | 0 |
| 1 | 1 | 0 | 0 |
| 0 | 0 | 0 | 0 |
| 0 | 0 | 0 | 0 |
| 1 | 1 | 0 | 0 |
| 1 | 1 | 0 | 0 |
| 0 | 1 | 0 | 0 |
| 0 | 0 | 0 | 0 |
| 0 | 0 | 0 | 0 |
| 0 | 0 | 0 | 0 |
| 0 | 0 | 0 | 0 |
| 0 | 0 | 0 | 0 |
| 0 | 0 | 0 | 0 |
| 0 | 0 | 0 | 0 |
| 0 | 0 | 0 | 0 |
| 0 | 0 | 0 | 0 |
| 0 | 0 | 0 | 0 |
| 1 | 0 | 0 | 0 |
| 1 | 0 | 0 | 0 |
| 1 | 0 | 0 | 0 |
| 1 | 0 | 0 | 0 |
| 1 | 1 | 0 | 1 |
| 0 | 0 | 0 | 0 |
| 1 | 0 | 0 | 0 |
| 0 | 0 | 0 | 0 |
| 1 | 0 | 0 | 0 |
| 0 | 0 | 0 | 0 |
| 1 | 0 | 0 | 0 |
| 1 | 0 | 0 | 0 |
| 0 | 0 | 0 | 0 |
| 0 | 0 | 0 | 0 |

| | | | |
|---|---|---|---|
| 0 | 0 | 0 | 0 |
| 0 | 0 | 0 | 0 |
| 0 | 0 | 0 | 0 |
| 0 | 0 | 0 | 0 |
| 1 | 0 | 0 | 0 |
| 1 | 0 | 0 | 0 |
| 1 | 0 | 0 | 0 |
| 0 | 0 | 0 | 0 |
| 1 | 1 | 0 | 0 |
| 0 | 0 | 0 | 0 |
| 0 | 0 | 0 | 0 |
| 0 | 0 | 0 | 0 |
| 1 | 1 | 0 | 0 |
| 1 | 0 | 0 | 0 |
| 0 | 0 | 0 | 0 |
| 0 | 0 | 0 | 0 |
| 1 | 1 | 0 | 0 |
| 0 | 0 | 0 | 0 |
| 0 | 0 | 0 | 0 |
| 0 | 0 | 0 | 0 |
| 0 | 0 | 0 | 0 |
| 0 | 0 | 0 | 0 |
| 0 | 0 | 0 | 0 |
| 0 | 0 | 0 | 0 |
| 1 | 0 | 0 | 0 |
| 0 | 0 | 0 | 0 |
| 1 | 0 | 0 | 0 |
| 0 | 0 | 0 | 0 |
| 0 | 0 | 0 | 0 |
| 0 | 0 | 0 | 0 |
| 1 | 0 | 0 | 0 |
| 1 | 0 | 0 | 0 |
| 0 | 0 | 0 | 0 |
| 0 | 0 | 0 | 0 |
| 1 | 0 | 0 | 0 |
| 0 | 0 | 0 | 0 |
| 0 | 0 | 0 | 0 |
| 1 | 0 | 0 | 0 |
| 0 | 0 | 0 | 0 |
| 1 | 0 | 0 | 0 |
| 1 | 0 | 0 | 0 |
| 0 | 0 | 0 | 0 |
| 0 | 0 | 0 | 0 |
| 1 | 1 | 0 | 0 |
| 1 | 0 | 0 | 0 |
| 1 | 1 | 0 | 0 |
| 0 | 0 | 0 | 0 |
| 0 | 0 | 0 | 0 |
| 0 | 0 | 0 | 0 |
| 0 | 0 | 0 | 0 |
| 1 | 1 | 0 | 0 |
| 0 | 0 | 0 | 0 |
| 0 | 0 | 0 | 0 |
| 0 | 0 | 0 | 0 |

| | | | |
|---|---|---|---|
| 0 | 1 | 0 | 0 |
| 1 | 0 | 0 | 0 |
| 0 | 0 | 0 | 0 |
| 0 | 0 | 0 | 0 |
| 0 | 0 | 0 | 0 |
| 1 | 1 | 0 | 0 |
| 0 | 0 | 0 | 0 |
| 0 | 0 | 0 | 0 |
| 0 | 0 | 0 | 0 |
| 1 | 0 | 0 | 0 |
| 1 | 0 | 0 | 0 |
| 0 | 0 | 0 | 0 |
| 0 | 0 | 0 | 0 |
| 0 | 0 | 0 | 0 |
| 0 | 0 | 0 | 0 |
| 0 | 0 | 0 | 0 |
| 0 | 0 | 0 | 0 |
| 0 | 0 | 0 | 0 |
| 0 | 0 | 0 | 0 |
| 0 | 0 | 0 | 0 |
| 0 | 0 | 0 | 0 |
| 0 | 0 | 0 | 0 |
| 0 | 0 | 0 | 0 |
| 1 | 1 | 0 | 0 |
| 0 | 0 | 0 | 0 |
| 0 | 0 | 0 | 0 |
| 1 | 1 | 1 | 0 |
| 0 | 0 | 0 | 0 |
| 0 | 0 | 0 | 0 |
| 0 | 0 | 0 | 0 |
| 1 | 1 | 1 | 0 |
| 1 | 0 | 0 | 0 |
| 1 | 0 | 0 | 0 |
| 1 | 0 | 0 | 0 |
| 1 | 0 | 0 | 0 |
| 1 | 1 | 1 | 0 |
| 1 | 1 | 1 | 0 |
| 0 | 0 | 0 | 0 |
| 0 | 0 | 0 | 0 |
| 1 | 1 | 1 | 0 |
| 0 | 0 | 0 | 0 |
| 0 | 0 | 0 | 0 |
| 0 | 0 | 0 | 0 |
| 0 | 0 | 0 | 0 |
| 0 | 0 | 0 | 0 |
| 0 | 0 | 0 | 0 |
| 0 | 0 | 0 | 0 |
| 0 | 0 | 0 | 0 |
| 1 | 0 | 0 | 0 |
| 1 | 0 | 0 | 0 |
| 1 | 0 | 0 | 0 |
| 0 | 0 | 0 | 0 |
| 0 | 0 | 0 | 0 |
| 0 | 0 | 0 | 0 |

| | | | |
|---|---|---|---|
| 0 | 0 | 0 | 0 |
| 1 | 0 | 0 | 0 |
| 0 | 0 | 0 | 0 |
| 0 | 0 | 0 | 0 |
| 0 | 0 | 0 | 0 |
| 0 | 0 | 0 | 0 |
| 0 | 0 | 0 | 0 |
| 0 | 0 | 0 | 0 |
| 0 | 0 | 0 | 0 |
| 0 | 0 | 0 | 0 |
| 1 | 1 | 0 | 0 |
| 1 | 1 | 0 | 0 |
| 0 | 0 | 0 | 0 |
| 0 | 0 | 0 | 0 |
| 0 | 0 | 0 | 0 |
| 1 | 0 | 0 | 0 |
| 1 | 0 | 0 | 0 |
| 1 | 1 | 0 | 1 |
| 0 | 0 | 0 | 0 |
| 0 | 0 | 0 | 0 |
| 0 | 0 | 0 | 0 |
| 0 | 0 | 0 | 0 |
| 1 | 0 | 0 | 1 |
| 0 | 0 | 0 | 0 |
| 0 | 0 | 0 | 0 |
| 1 | 0 | 0 | 0 |
| 0 | 0 | 0 | 0 |
| 0 | 0 | 0 | 0 |
| 0 | 0 | 0 | 0 |
| 0 | 0 | 0 | 0 |
| 0 | 0 | 0 | 0 |
| 0 | 0 | 0 | 0 |
| 0 | 0 | 0 | 0 |
| 0 | 0 | 0 | 0 |
| 0 | 0 | 0 | 0 |
| 1 | 0 | 0 | 0 |
| 1 | 0 | 0 | 0 |
| 1 | 1 | 0 | 0 |
| 1 | 1 | 0 | 0 |
| 0 | 0 | 0 | 0 |
| 0 | 0 | 0 | 0 |
| 1 | 0 | 0 | 0 |
| 0 | 0 | 0 | 0 |
| 1 | 0 | 0 | 0 |
| 1 | 1 | 0 | 0 |
| 0 | 0 | 0 | 0 |
| 0 | 0 | 0 | 0 |
| 0 | 0 | 0 | 0 |
| 0 | 0 | 0 | 0 |
| 0 | 0 | 0 | 0 |
| 0 | 0 | 0 | 0 |
| 1 | 0 | 0 | 0 |
| 0 | 0 | 0 | 0 |
| 0 | 0 | 0 | 0 |

| | | | |
|---|---|---|---|
| 0 | 0 | 0 | 0 |
| 1 | 1 | 0 | 0 |
| 0 | 0 | 0 | 0 |
| 1 | 0 | 0 | 0 |
| 1 | 0 | 0 | 0 |
| 0 | 0 | 0 | 0 |
| 0 | 0 | 0 | 0 |
| 0 | 0 | 0 | 0 |
| 0 | 0 | 0 | 0 |
| 0 | 0 | 0 | 0 |
| 0 | 0 | 0 | 0 |
| 1 | 1 | 0 | 0 |
| 1 | 1 | 0 | 1 |
| 0 | 0 | 0 | 0 |
| 0 | 1 | 0 | 0 |
| 0 | 0 | 0 | 0 |
| 0 | 0 | 0 | 0 |
| 0 | 0 | 0 | 0 |
| 0 | 0 | 0 | 0 |
| 0 | 0 | 0 | 0 |
| 0 | 0 | 0 | 0 |
| 1 | 0 | 0 | 0 |
| 0 | 0 | 0 | 0 |
| 0 | 0 | 0 | 0 |
| 0 | 0 | 0 | 0 |
| 0 | 0 | 0 | 0 |
| 0 | 0 | 0 | 0 |
| 1 | 0 | 0 | 0 |
| 0 | 0 | 0 | 0 |
| 0 | 0 | 0 | 0 |
| 0 | 0 | 0 | 0 |
| 0 | 0 | 0 | 0 |
| 0 | 0 | 0 | 0 |
| 0 | 0 | 0 | 0 |
| 0 | 0 | 0 | 0 |
| 0 | 0 | 0 | 0 |
| 0 | 0 | 0 | 0 |
| 0 | 0 | 0 | 0 |
| 0 | 0 | 0 | 0 |
| 0 | 0 | 0 | 0 |
| 0 | 0 | 0 | 0 |
| 0 | 0 | 0 | 0 |
| 0 | 0 | 0 | 0 |
| 0 | 0 | 0 | 0 |
| 0 | 0 | 0 | 0 |
| 0 | 0 | 0 | 0 |
| 0 | 0 | 0 | 0 |
| 0 | 0 | 0 | 0 |
| 0 | 0 | 0 | 0 |
| 0 | 0 | 0 | 0 |
| 0 | 0 | 0 | 0 |
| 0 | 0 | 0 | 0 |
| 0 | 0 | 0 | 0 |
| 0 | 0 | 0 | 0 |

| | | | |
|---|---|---|---|
| 0 | 0 | 0 | 0 |
| 0 | 0 | 0 | 0 |
| 0 | 0 | 0 | 0 |
| 0 | 0 | 0 | 0 |
| 0 | 0 | 0 | 0 |
| 0 | 0 | 0 | 0 |
| 0 | 0 | 0 | 0 |
| 0 | 0 | 0 | 0 |
| 0 | 0 | 0 | 0 |
| 0 | 0 | 0 | 0 |
| 0 | 0 | 0 | 0 |
| 0 | 0 | 0 | 0 |
| 1 | 0 | 0 | 0 |
| 1 | 1 | 0 | 0 |
| 0 | 0 | 0 | 0 |
| 1 | 0 | 0 | 0 |
| 0 | 0 | 0 | 0 |
| 1 | 0 | 0 | 0 |
| 0 | 0 | 0 | 0 |
| 0 | 0 | 0 | 0 |
| 1 | 0 | 0 | 0 |
| 0 | 0 | 0 | 0 |
| 0 | 0 | 0 | 0 |
| 1 | 0 | 0 | 0 |
| 1 | 0 | 0 | 0 |
| 0 | 0 | 0 | 0 |
| 0 | 0 | 0 | 0 |
| 1 | 0 | 0 | 0 |
| 1 | 0 | 0 | 0 |
| 1 | 0 | 0 | 0 |
| 1 | 0 | 0 | 0 |
| 1 | 0 | 0 | 0 |
| 0 | 0 | 0 | 0 |
| 0 | 0 | 0 | 0 |
| 0 | 0 | 0 | 0 |
| 0 | 0 | 0 | 0 |
| 0 | 0 | 0 | 0 |
| 0 | 0 | 0 | 0 |
| 0 | 0 | 0 | 0 |
| 0 | 0 | 0 | 0 |
| 0 | 0 | 0 | 0 |
| 0 | 0 | 0 | 0 |
| 0 | 0 | 0 | 0 |
| 0 | 0 | 0 | 0 |
| 1 | 1 | 0 | 0 |
| 0 | 0 | 0 | 0 |
| 0 | 0 | 0 | 0 |
| 0 | 0 | 0 | 0 |
| 0 | 0 | 0 | 0 |
| 1 | 1 | 0 | 0 |
| 1 | 0 | 0 | 0 |
| 0 | 0 | 0 | 0 |
| 0 | 0 | 0 | 0 |
| 0 | 0 | 0 | 1 |

| | | | |
|---|---|---|---|
| 0 | 0 | 0 | 0 |
| 0 | 0 | 0 | 0 |
| 0 | 0 | 0 | 0 |
| 0 | 0 | 0 | 0 |
| 1 | 0 | 0 | 1 |
| 0 | 0 | 0 | 0 |
| 1 | 0 | 0 | 0 |
| 0 | 0 | 0 | 0 |
| 0 | 0 | 0 | 0 |
| 0 | 0 | 0 | 0 |
| 0 | 0 | 0 | 0 |
| 0 | 0 | 0 | 0 |
| 0 | 0 | 0 | 0 |
| 1 | 0 | 0 | 0 |
| 0 | 0 | 0 | 0 |
| 0 | 0 | 0 | 0 |
| 1 | 1 | 0 | 0 |
| 0 | 0 | 0 | 0 |
| 1 | 0 | 0 | 0 |
| 1 | 0 | 0 | 0 |
| 0 | 0 | 0 | 0 |
| 0 | 0 | 0 | 0 |
| 0 | 0 | 0 | 0 |
| 1 | 0 | 0 | 0 |
| 1 | 0 | 0 | 0 |
| 1 | 0 | 0 | 0 |
| 1 | 0 | 0 | 0 |
| 1 | 0 | 0 | 0 |
| 1 | 0 | 0 | 0 |
| 1 | 0 | 0 | 0 |
| 1 | 0 | 0 | 0 |
| 1 | 0 | 0 | 0 |
| 1 | 0 | 0 | 0 |
| 1 | 0 | 0 | 0 |
| 0 | 0 | 0 | 0 |
| 0 | 0 | 0 | 0 |
| 0 | 0 | 0 | 0 |
| 1 | 0 | 0 | 0 |
| 1 | 0 | 0 | 0 |
| 0 | 0 | 0 | 0 |
| 0 | 0 | 0 | 0 |
| 1 | 0 | 0 | 0 |
| 0 | 0 | 0 | 0 |
| 1 | 0 | 0 | 0 |
| 1 | 0 | 0 | 0 |
| 1 | 0 | 0 | 0 |
| 1 | 0 | 0 | 0 |
| 1 | 0 | 0 | 0 |
| 0 | 0 | 0 | 0 |
| 0 | 0 | 0 | 0 |
| 1 | 0 | 0 | 0 |
| 0 | 0 | 0 | 0 |
| 1 | 0 | 0 | 0 |
| 1 | 0 | 0 | 0 |

| | | | |
|---|---|---|---|
| 1 | 0 | 0 | 0 |
| 1 | 0 | 0 | 0 |
| 1 | 0 | 0 | 0 |
| 1 | 0 | 0 | 0 |
| 1 | 0 | 0 | 0 |
| 1 | 0 | 0 | 0 |
| 0 | 0 | 0 | 0 |
| 1 | 0 | 0 | 0 |
| 1 | 0 | 0 | 0 |
| 1 | 0 | 0 | 0 |
| 1 | 0 | 0 | 0 |
| 1 | 0 | 0 | 0 |
| 1 | 0 | 0 | 0 |
| 1 | 0 | 0 | 0 |
| 0 | 0 | 0 | 0 |
| 0 | 0 | 0 | 0 |
| 1 | 1 | 0 | 0 |
| 1 | 1 | 0 | 0 |
| 0 | 0 | 0 | 0 |
| 0 | 0 | 0 | 0 |
| 0 | 0 | 0 | 0 |
| 1 | 1 | 0 | 1 |
| 0 | 0 | 0 | 0 |
| 0 | 0 | 0 | 0 |
| 0 | 0 | 0 | 0 |
| 0 | 0 | 0 | 0 |
| 0 | 0 | 0 | 0 |
| 0 | 0 | 0 | 0 |
| 0 | 0 | 0 | 0 |
| 1 | 0 | 1 | 0 |
| 0 | 0 | 0 | 0 |
| 0 | 0 | 0 | 0 |
| 0 | 0 | 0 | 0 |
| 0 | 0 | 0 | 0 |
| 0 | 0 | 0 | 0 |
| 0 | 0 | 0 | 0 |
| 0 | 0 | 0 | 0 |
| 0 | 0 | 0 | 0 |
| 0 | 0 | 0 | 0 |
| 0 | 0 | 0 | 0 |
| 1 | 1 | 0 | 0 |
| 1 | 1 | 0 | 0 |
| 1 | 0 | 0 | 0 |
| 1 | 1 | 0 | 0 |
| 1 | 0 | 0 | 0 |
| 1 | 1 | 0 | 0 |
| 1 | 0 | 0 | 0 |
| 1 | 0 | 0 | 0 |
| 0 | 0 | 0 | 0 |
| 0 | 0 | 0 | 0 |
| 1 | 0 | 0 | 0 |
| 1 | 0 | 0 | 0 |
| 1 | 0 | 0 | 0 |
| 0 | 0 | 0 | 0 |

| | | | |
|---|---|---|---|
| 0 | 0 | 0 | 0 |
| 0 | 0 | 0 | 0 |
| 0 | 0 | 0 | 0 |
| 1 | 1 | 1 | 0 |
| 0 | 0 | 0 | 0 |
| 0 | 0 | 0 | 0 |
| 0 | 0 | 0 | 0 |
| 0 | 0 | 0 | 0 |
| 0 | 1 | 0 | 0 |
| 0 | 1 | 0 | 0 |
| 0 | 1 | 0 | 0 |
| 0 | 0 | 0 | 0 |
| 0 | 0 | 0 | 0 |
| 0 | 0 | 0 | 0 |
| 1 | 0 | 0 | 0 |
| 1 | 0 | 0 | 0 |
| 0 | 0 | 0 | 0 |
| 0 | 0 | 0 | 0 |
| 1 | 0 | 0 | 0 |
| 0 | 0 | 0 | 0 |
| 1 | 0 | 0 | 0 |
| 1 | 0 | 0 | 0 |
| 0 | 0 | 0 | 0 |
| 0 | 0 | 0 | 0 |
| 0 | 0 | 0 | 0 |
| 0 | 0 | 0 | 0 |
| 0 | 0 | 0 | 0 |
| 0 | 0 | 1 | 0 |
| 0 | 0 | 0 | 0 |
| 0 | 1 | 0 | 0 |
| 0 | 0 | 0 | 0 |
| 0 | 0 | 0 | 0 |
| 0 | 0 | 0 | 0 |
| 0 | 0 | 0 | 0 |
| 0 | 0 | 0 | 0 |
| 0 | 0 | 0 | 0 |
| 1 | 0 | 0 | 0 |
| 1 | 0 | 0 | 0 |
| 1 | 0 | 0 | 0 |
| 0 | 0 | 0 | 0 |
| 0 | 0 | 0 | 0 |
| 0 | 0 | 0 | 0 |
| 0 | 0 | 0 | 0 |
| 0 | 0 | 0 | 0 |
| 0 | 0 | 0 | 0 |
| 0 | 0 | 0 | 0 |
| 0 | 0 | 0 | 0 |
| 0 | 0 | 0 | 1 |
| 0 | 0 | 0 | 0 |
| 0 | 0 | 0 | 0 |
| 0 | 0 | 0 | 0 |
| 0 | 0 | 0 | 0 |
| 1 | 1 | 0 | 0 |
| 0 | 0 | 0 | 0 |

| | | | |
|---|---|---|---|
| 0 | 0 | 0 | 0 |
| 0 | 0 | 0 | 0 |
| 0 | 0 | 0 | 0 |
| 0 | 0 | 0 | 0 |
| 1 | 0 | 0 | 0 |
| 0 | 0 | 0 | 0 |
| 0 | 0 | 0 | 0 |
| 0 | 0 | 0 | 0 |
| 0 | 0 | 0 | 0 |
| 0 | 0 | 0 | 0 |
| 0 | 0 | 0 | 0 |
| 0 | 0 | 0 | 0 |
| 0 | 0 | 0 | 0 |
| 0 | 0 | 0 | 0 |
| 0 | 0 | 0 | 0 |
| 0 | 0 | 0 | 0 |
| 0 | 0 | 0 | 0 |
| 0 | 0 | 0 | 0 |
| 0 | 0 | 0 | 0 |
| 0 | 0 | 0 | 0 |
| 0 | 0 | 0 | 0 |
| 0 | 0 | 0 | 0 |
| 0 | 0 | 0 | 0 |
| 0 | 0 | 0 | 0 |
| 0 | 0 | 0 | 0 |
| 0 | 0 | 0 | 0 |
| 0 | 0 | 0 | 0 |
| 0 | 0 | 0 | 0 |
| 0 | 0 | 0 | 0 |
| 0 | 0 | 0 | 0 |
| 0 | 0 | 0 | 0 |
| 0 | 0 | 0 | 0 |
| 0 | 0 | 0 | 0 |
| 0 | 0 | 0 | 0 |
| 0 | 0 | 0 | 0 |
| 0 | 0 | 0 | 0 |
| 0 | 0 | 0 | 0 |
| 0 | 0 | 0 | 0 |
| 0 | 0 | 0 | 0 |
| 0 | 0 | 0 | 0 |
| 1 | 1 | 0 | 0 |
| 0 | 0 | 0 | 0 |
| 0 | 0 | 0 | 0 |
| 0 | 0 | 0 | 0 |
| 0 | 0 | 0 | 0 |
| 1 | 0 | 0 | 0 |
| 1 | 0 | 0 | 0 |
| 0 | 0 | 0 | 0 |
| 1 | 0 | 0 | 0 |
| 0 | 0 | 0 | 0 |
| 0 | 0 | 0 | 0 |
| 1 | 0 | 0 | 0 |
| 1 | 0 | 0 | 0 |
| 0 | 0 | 0 | 0 |

| | | | |
|---|---|---|---|
| 0 | 0 | 0 | 0 |
| 0 | 0 | 0 | 0 |
| 0 | 0 | 0 | 0 |
| 0 | 0 | 0 | 0 |
| 1 | 0 | 0 | 0 |
| 0 | 0 | 0 | 0 |
| 0 | 0 | 0 | 0 |
| 0 | 0 | 0 | 0 |
| 0 | 0 | 0 | 0 |
| 0 | 0 | 0 | 0 |
| 0 | 0 | 0 | 0 |
| 0 | 0 | 0 | 0 |
| 0 | 0 | 0 | 0 |
| 0 | 0 | 0 | 0 |
| 0 | 0 | 0 | 0 |
| 1 | 1 | 0 | 1 |
| 0 | 0 | 0 | 0 |
| 0 | 0 | 0 | 0 |
| 0 | 0 | 0 | 0 |
| 0 | 0 | 0 | 0 |
| 0 | 0 | 0 | 0 |
| 0 | 0 | 0 | 0 |
| 0 | 0 | 0 | 0 |
| 0 | 0 | 0 | 0 |
| 0 | 0 | 0 | 0 |
| 0 | 0 | 0 | 0 |
| 0 | 0 | 0 | 0 |
| 1 | 0 | 0 | 0 |
| 1 | 0 | 0 | 0 |
| 0 | 0 | 0 | 0 |
| 1 | 1 | 0 | 0 |
| 0 | 0 | 0 | 0 |
| 0 | 0 | 0 | 0 |
| 0 | 0 | 0 | 0 |
| 1 | 1 | 0 | 0 |
| 1 | 1 | 0 | 0 |
| 0 | 0 | 0 | 0 |
| 1 | 0 | 0 | 0 |
| 0 | 0 | 0 | 0 |
| 1 | 1 | 0 | 0 |
| 0 | 1 | 0 | 0 |
| 0 | 0 | 0 | 0 |
| 0 | 0 | 0 | 0 |
| 0 | 0 | 0 | 0 |
| 1 | 1 | 0 | 0 |
| 1 | 1 | 0 | 0 |
| 1 | 1 | 0 | 0 |
| 0 | 0 | 0 | 0 |
| 0 | 0 | 0 | 0 |
| 0 | 0 | 0 | 0 |
| 0 | 0 | 0 | 0 |
| 0 | 0 | 0 | 0 |
| 0 | 0 | 0 | 0 |
| 0 | 0 | 0 | 0 |

| | | | |
|---|---|---|---|
| 0 | 0 | 0 | 0 |
| 0 | 0 | 0 | 0 |
| 1 | 0 | 0 | 0 |
| 1 | 0 | 0 | 0 |
| 0 | 0 | 0 | 0 |
| 1 | 0 | 0 | 0 |
| 0 | 0 | 0 | 0 |
| 0 | 0 | 0 | 0 |
| 0 | 0 | 0 | 0 |
| 0 | 1 | 0 | 0 |
| 0 | 0 | 0 | 0 |
| 0 | 0 | 0 | 0 |
| 0 | 0 | 0 | 0 |
| 0 | 0 | 0 | 0 |
| 0 | 0 | 0 | 0 |
| 0 | 0 | 0 | 0 |
| 0 | 0 | 0 | 0 |
| 1 | 0 | 0 | 0 |
| 0 | 0 | 0 | 0 |
| 1 | 0 | 0 | 0 |
| 0 | 0 | 0 | 0 |
| 1 | 0 | 0 | 0 |
| 0 | 0 | 0 | 0 |
| 1 | 0 | 0 | 0 |
| 0 | 0 | 0 | 0 |
| 0 | 0 | 0 | 0 |
| 0 | 0 | 0 | 0 |
| 0 | 0 | 0 | 0 |
| 0 | 0 | 0 | 0 |
| 0 | 0 | 0 | 0 |
| 0 | 0 | 0 | 0 |
| 1 | 0 | 0 | 0 |
| 0 | 0 | 0 | 0 |
| 0 | 0 | 0 | 0 |
| 0 | 0 | 0 | 0 |
| 0 | 0 | 0 | 0 |
| 0 | 0 | 0 | 0 |
| 0 | 0 | 0 | 0 |
| 1 | 1 | 0 | 0 |
| 1 | 1 | 0 | 0 |
| 1 | 1 | 0 | 0 |
| 1 | 1 | 0 | 0 |
| 1 | 1 | 0 | 0 |
| 1 | 0 | 0 | 0 |
| 1 | 0 | 0 | 0 |
| 1 | 0 | 0 | 0 |
| 0 | 0 | 0 | 0 |
| 0 | 0 | 0 | 0 |
| 0 | 0 | 0 | 0 |
| 0 | 0 | 0 | 0 |
| 0 | 0 | 0 | 0 |
| 0 | 0 | 0 | 0 |
| 0 | 0 | 0 | 0 |
| 0 | 0 | 0 | 0 |

| | | | |
|---|---|---|---|
| 0 | 0 | 0 | 0 |
| 0 | 0 | 0 | 0 |
| 0 | 0 | 0 | 0 |
| 0 | 0 | 0 | 0 |
| 0 | 0 | 0 | 0 |
| 1 | 0 | 0 | 0 |
| 1 | 0 | 0 | 0 |
| 0 | 0 | 0 | 0 |
| 0 | 0 | 0 | 0 |
| 0 | 0 | 0 | 0 |
| 0 | 0 | 0 | 0 |
| 1 | 0 | 0 | 0 |
| 0 | 0 | 1 | 0 |
| 0 | 0 | 0 | 0 |
| 0 | 0 | 0 | 0 |
| 0 | 0 | 0 | 0 |
| 1 | 0 | 0 | 0 |
| 0 | 0 | 0 | 0 |
| 1 | 0 | 0 | 0 |
| 1 | 0 | 0 | 0 |
| 0 | 0 | 0 | 0 |
| 1 | 0 | 0 | 0 |
| 1 | 1 | 0 | 0 |
| 0 | 0 | 0 | 0 |
| 1 | 1 | 0 | 0 |
| 1 | 1 | 0 | 0 |
| 1 | 1 | 0 | 0 |
| 0 | 0 | 0 | 0 |
| 1 | 0 | 0 | 0 |
| 0 | 0 | 0 | 0 |
| 0 | 0 | 0 | 0 |
| 0 | 0 | 0 | 0 |
| 0 | 0 | 0 | 0 |
| 0 | 0 | 0 | 0 |
| 0 | 0 | 0 | 0 |
| 0 | 0 | 0 | 0 |
| 0 | 0 | 0 | 0 |
| 0 | 0 | 0 | 0 |
| 0 | 0 | 0 | 0 |
| 0 | 0 | 0 | 0 |
| 0 | 0 | 0 | 0 |
| 0 | 0 | 0 | 0 |
| 0 | 0 | 0 | 0 |
| 0 | 0 | 0 | 0 |
| 0 | 0 | 0 | 0 |
| 0 | 0 | 0 | 0 |
| 0 | 0 | 0 | 0 |
| 0 | 0 | 0 | 0 |
| 0 | 0 | 0 | 0 |
| 0 | 0 | 0 | 0 |
| 0 | 0 | 0 | 0 |
| 0 | 0 | 0 | 0 |
| 0 | 0 | 0 | 0 |
| 0 | 0 | 0 | 0 |

| | | | |
|---|---|---|---|
| 0 | 0 | 0 | 0 |
| 0 | 0 | 0 | 0 |
| 0 | 0 | 0 | 0 |
| 0 | 0 | 0 | 0 |
| 0 | 0 | 0 | 0 |
| 0 | 0 | 0 | 0 |
| 0 | 0 | 0 | 0 |
| 0 | 0 | 0 | 0 |
| 0 | 0 | 0 | 0 |
| 0 | 0 | 0 | 0 |
| 0 | 1 | 0 | 0 |
| 0 | 1 | 0 | 0 |
| 0 | 1 | 0 | 0 |
| 0 | 1 | 0 | 0 |
| 0 | 0 | 0 | 0 |
| 0 | 1 | 0 | 0 |
| 0 | 1 | 0 | 0 |
| 1 | 1 | 0 | 0 |
| 0 | 1 | 0 | 0 |
| 1 | 0 | 0 | 0 |
| 1 | 1 | 0 | 0 |
| 1 | 1 | 0 | 0 |
| 0 | 0 | 1 | 1 |
| 0 | 0 | 0 | 0 |
| 1 | 0 | 0 | 0 |
| 0 | 0 | 0 | 0 |
| 0 | 0 | 0 | 0 |
| 0 | 0 | 0 | 0 |
| 0 | 0 | 0 | 0 |
| 0 | 0 | 0 | 0 |
| 0 | 0 | 0 | 0 |
| 0 | 0 | 0 | 0 |
| 0 | 0 | 0 | 0 |
| 0 | 0 | 0 | 0 |
| 0 | 0 | 0 | 0 |
| 0 | 0 | 0 | 0 |
| 0 | 0 | 0 | 0 |
| 0 | 0 | 0 | 0 |
| 0 | 0 | 0 | 0 |
| 0 | 0 | 0 | 0 |
| 1 | 0 | 0 | 0 |
| 0 | 0 | 0 | 0 |
| 1 | 0 | 0 | 0 |
| 1 | 0 | 0 | 0 |
| 0 | 0 | 0 | 0 |
| 1 | 0 | 0 | 0 |
| 0 | 0 | 0 | 0 |
| 0 | 0 | 0 | 0 |
| 0 | 0 | 0 | 0 |
| 0 | 0 | 0 | 0 |
| 0 | 0 | 0 | 0 |
| 0 | 0 | 0 | 0 |
| 1 | 1 | 0 | 0 |
| 1 | 1 | 0 | 0 |

| | | | |
|---|---|---|---|
| 1 | 1 | 0 | 0 |
| 0 | 0 | 0 | 0 |
| 0 | 0 | 0 | 0 |
| 0 | 0 | 0 | 0 |
| 0 | 0 | 0 | 0 |
| 0 | 0 | 0 | 0 |
| 0 | 1 | 0 | 0 |
| 0 | 0 | 0 | 0 |
| 0 | 0 | 0 | 0 |
| 0 | 1 | 0 | 0 |
| 1 | 0 | 0 | 0 |
| 0 | 0 | 0 | 0 |
| 0 | 0 | 1 | 0 |
| 0 | 0 | 1 | 0 |
| 0 | 0 | 0 | 0 |
| 0 | 0 | 0 | 0 |
| 0 | 0 | 0 | 0 |
| 0 | 1 | 0 | 0 |
| 0 | 0 | 0 | 0 |
| 0 | 0 | 0 | 0 |
| 1 | 0 | 0 | 0 |
| 0 | 0 | 0 | 1 |
| 0 | 0 | 0 | 0 |
| 0 | 0 | 0 | 0 |
| 0 | 0 | 0 | 0 |
| 0 | 0 | 0 | 0 |
| 0 | 0 | 0 | 0 |
| 0 | 0 | 0 | 0 |
| 0 | 0 | 0 | 0 |
| 0 | 0 | 0 | 0 |
| 0 | 0 | 0 | 0 |
| 0 | 0 | 0 | 0 |
| 0 | 0 | 0 | 0 |
| 0 | 0 | 0 | 0 |
| 0 | 0 | 0 | 0 |
| 0 | 0 | 0 | 0 |
| 1 | 0 | 0 | 0 |
| 0 | 0 | 0 | 0 |
| 1 | 0 | 0 | 0 |
| 0 | 0 | 0 | 0 |
| 0 | 0 | 0 | 0 |
| 0 | 0 | 0 | 0 |
| 0 | 0 | 0 | 0 |
| 0 | 0 | 0 | 0 |
| 0 | 0 | 0 | 0 |
| 0 | 0 | 0 | 0 |
| 0 | 0 | 0 | 0 |
| 0 | 0 | 0 | 0 |
| 0 | 0 | 0 | 0 |
| 1 | 0 | 0 | 0 |
| 0 | 0 | 0 | 0 |
| 0 | 0 | 0 | 0 |
| 0 | 0 | 0 | 0 |
| 0 | 0 | 0 | 0 |

| | | | |
|---|---|---|---|
| 0 | 0 | 0 | 0 |
| 0 | 0 | 0 | 0 |
| 0 | 0 | 0 | 0 |
| 0 | 0 | 0 | 0 |
| 0 | 0 | 0 | 0 |
| 0 | 0 | 0 | 0 |
| 0 | 0 | 0 | 0 |
| 0 | 0 | 0 | 0 |
| 0 | 0 | 0 | 0 |
| 0 | 0 | 0 | 0 |
| 0 | 0 | 0 | 0 |
| 0 | 0 | 0 | 0 |
| 0 | 0 | 1 | 0 |
| 1 | 0 | 0 | 0 |
| 1 | 0 | 0 | 0 |
| 0 | 0 | 0 | 0 |
| 0 | 0 | 0 | 0 |
| 0 | 0 | 0 | 0 |
| 0 | 0 | 0 | 0 |
| 0 | 0 | 0 | 0 |
| 0 | 0 | 0 | 0 |
| 0 | 0 | 0 | 0 |
| 0 | 0 | 0 | 0 |
| 1 | 1 | 0 | 1 |
| 0 | 0 | 0 | 0 |
| 1 | 1 | 0 | 0 |
| 1 | 1 | 0 | 1 |
| 0 | 0 | 0 | 0 |
| 0 | 0 | 0 | 0 |
| 0 | 0 | 0 | 0 |
| 0 | 0 | 0 | 0 |
| 0 | 0 | 0 | 0 |
| 0 | 0 | 0 | 0 |
| 0 | 0 | 0 | 0 |
| 0 | 0 | 0 | 0 |
| 1 | 0 | 0 | 0 |
| 0 | 0 | 0 | 0 |
| 0 | 0 | 0 | 0 |
| 0 | 0 | 0 | 0 |
| 0 | 0 | 0 | 0 |
| 0 | 0 | 0 | 0 |
| 0 | 0 | 0 | 0 |
| 1 | 0 | 0 | 0 |
| 1 | 0 | 0 | 0 |
| 1 | 0 | 0 | 0 |
| 1 | 0 | 0 | 0 |
| 1 | 0 | 0 | 0 |
| 0 | 0 | 0 | 0 |
| 0 | 0 | 0 | 0 |
| 1 | 0 | 0 | 0 |
| 1 | 0 | 0 | 0 |
| 0 | 0 | 0 | 0 |
| 1 | 0 | 0 | 0 |
| 1 | 0 | 0 | 0 |

| | | | |
|---|---|---|---|
| 0 | 0 | 0 | 0 |
| 1 | 0 | 0 | 0 |
| 0 | 0 | 0 | 0 |
| 0 | 0 | 0 | 0 |
| 0 | 0 | 0 | 0 |
| 0 | 0 | 0 | 0 |
| 1 | 0 | 0 | 0 |
| 0 | 0 | 0 | 0 |
| 0 | 0 | 0 | 0 |
| 0 | 0 | 0 | 0 |
| 0 | 0 | 0 | 0 |
| 1 | 1 | 0 | 0 |
| 0 | 0 | 0 | 0 |
| 0 | 0 | 0 | 0 |
| 0 | 0 | 0 | 0 |
| 0 | 0 | 0 | 0 |
| 0 | 0 | 0 | 0 |
| 0 | 0 | 0 | 0 |
| 0 | 0 | 0 | 0 |
| 0 | 0 | 0 | 0 |
| 0 | 0 | 0 | 0 |
| 0 | 0 | 0 | 0 |
| 0 | 0 | 0 | 0 |
| 0 | 0 | 0 | 0 |
| 0 | 0 | 0 | 0 |
| 0 | 0 | 0 | 0 |
| 0 | 0 | 0 | 0 |
| 0 | 0 | 0 | 0 |
| 1 | 1 | 0 | 0 |
| 1 | 0 | 0 | 0 |
| 0 | 0 | 0 | 0 |
| 0 | 0 | 0 | 0 |
| 0 | 0 | 0 | 0 |
| 0 | 0 | 0 | 0 |
| 0 | 0 | 0 | 0 |
| 0 | 0 | 0 | 0 |
| 0 | 0 | 0 | 0 |
| 0 | 0 | 0 | 0 |
| 0 | 0 | 0 | 0 |
| 0 | 0 | 0 | 0 |
| 0 | 0 | 0 | 0 |
| 0 | 0 | 0 | 0 |
| 0 | 0 | 0 | 0 |
| 0 | 0 | 0 | 0 |
| 0 | 0 | 0 | 0 |
| 0 | 0 | 0 | 0 |
| 0 | 0 | 0 | 0 |
| 0 | 0 | 0 | 0 |
| 0 | 0 | 0 | 0 |
| 0 | 0 | 0 | 0 |
| 0 | 0 | 0 | 0 |
| 0 | 0 | 0 | 0 |
| 0 | 0 | 1 | 0 |
| 0 | 0 | 0 | 0 |

| | | | |
|---|---|---|---|
| 0 | 0 | 0 | 0 |
| 0 | 0 | 0 | 0 |
| 0 | 0 | 0 | 0 |
| 0 | 0 | 0 | 0 |
| 1 | 0 | 0 | 0 |
| 0 | 0 | 0 | 0 |
| 0 | 0 | 0 | 0 |
| 1 | 1 | 0 | 0 |
| 0 | 0 | 0 | 0 |
| 1 | 1 | 0 | 0 |
| 0 | 0 | 0 | 0 |
| 0 | 0 | 0 | 0 |
| 0 | 0 | 0 | 0 |
| 0 | 0 | 0 | 0 |
| 0 | 0 | 0 | 0 |
| 0 | 0 | 0 | 0 |
| 1 | 0 | 0 | 0 |
| 0 | 0 | 0 | 0 |
| 0 | 0 | 0 | 0 |
| 1 | 0 | 0 | 0 |
| 1 | 0 | 0 | 0 |
| 0 | 0 | 0 | 0 |
| 0 | 0 | 0 | 0 |
| 0 | 0 | 0 | 0 |
| 0 | 0 | 0 | 0 |
| 0 | 0 | 0 | 0 |
| 1 | 0 | 0 | 0 |
| 0 | 0 | 0 | 0 |
| 0 | 0 | 0 | 0 |
| 1 | 0 | 0 | 0 |
| 0 | 0 | 0 | 0 |
| 0 | 0 | 0 | 0 |
| 0 | 0 | 0 | 0 |
| 1 | 1 | 0 | 0 |
| 0 | 0 | 0 | 0 |
| 0 | 0 | 0 | 0 |
| 0 | 0 | 0 | 0 |
| 0 | 0 | 0 | 0 |
| 0 | 1 | 0 | 0 |
| 0 | 0 | 0 | 0 |
| 0 | 0 | 0 | 0 |
| 0 | 0 | 0 | 0 |
| 0 | 0 | 0 | 0 |
| 1 | 1 | 0 | 0 |
| 0 | 0 | 0 | 0 |
| 0 | 0 | 0 | 0 |
| 0 | 0 | 0 | 0 |
| 0 | 0 | 0 | 0 |
| 0 | 0 | 0 | 0 |
| 0 | 0 | 0 | 0 |
| 0 | 0 | 0 | 0 |
| 1 | 0 | 0 | 0 |
| 0 | 0 | 0 | 0 |
| 0 | 0 | 0 | 0 |

| | | | |
|---|---|---|---|
| 1 | 0 | 0 | 1 |
| 1 | 0 | 0 | 1 |
| 1 | 0 | 0 | 1 |
| 0 | 0 | 0 | 0 |
| 0 | 0 | 0 | 0 |
| 0 | 0 | 0 | 0 |
| 1 | 0 | 0 | 0 |
| 0 | 0 | 0 | 0 |
| 0 | 0 | 0 | 0 |
| 0 | 0 | 0 | 0 |
| 0 | 0 | 1 | 0 |
| 0 | 0 | 0 | 0 |
| 0 | 0 | 0 | 0 |
| 0 | 0 | 0 | 0 |
| 0 | 0 | 0 | 0 |
| 0 | 0 | 0 | 0 |
| 1 | 0 | 0 | 0 |
| 0 | 0 | 0 | 0 |
| 0 | 0 | 0 | 0 |
| 0 | 0 | 0 | 0 |
| 0 | 0 | 0 | 0 |
| 0 | 0 | 0 | 0 |
| 0 | 0 | 0 | 0 |
| 1 | 1 | 0 | 0 |
| 0 | 0 | 0 | 0 |
| 0 | 0 | 0 | 0 |
| 0 | 0 | 0 | 0 |
| 0 | 0 | 0 | 0 |
| 0 | 0 | 0 | 0 |
| 0 | 0 | 0 | 0 |
| 0 | 0 | 0 | 0 |
| 0 | 0 | 0 | 0 |
| 0 | 1 | 0 | 0 |
| 0 | 0 | 0 | 0 |
| 1 | 1 | 0 | 0 |
| 0 | 0 | 0 | 0 |
| 0 | 0 | 0 | 0 |
| 0 | 0 | 0 | 0 |
| 0 | 0 | 0 | 0 |
| 0 | 0 | 0 | 0 |
| 0 | 0 | 0 | 0 |
| 1 | 0 | 0 | 1 |
| 0 | 0 | 0 | 0 |
| 1 | 0 | 0 | 1 |
| 0 | 0 | 0 | 0 |
| 1 | 0 | 0 | 1 |
| 0 | 0 | 0 | 0 |
| 0 | 0 | 0 | 0 |
| 0 | 0 | 0 | 0 |
| 1 | 0 | 0 | 0 |
| 0 | 0 | 0 | 0 |
| 0 | 0 | 0 | 0 |
| 0 | 0 | 0 | 0 |
| 0 | 0 | 0 | 0 |

| | | | |
|---|---|---|---|
| 0 | 0 | 0 | 0 |
| 0 | 0 | 0 | 0 |
| 0 | 0 | 0 | 0 |
| 0 | 0 | 0 | 0 |
| 0 | 0 | 0 | 0 |
| 0 | 0 | 0 | 0 |
| 0 | 0 | 0 | 0 |
| 0 | 0 | 0 | 0 |
| 0 | 0 | 0 | 0 |
| 0 | 0 | 0 | 0 |
| 0 | 0 | 0 | 0 |
| 0 | 0 | 0 | 0 |
| 0 | 0 | 0 | 0 |
| 0 | 0 | 0 | 0 |
| 0 | 0 | 0 | 0 |
| 0 | 0 | 0 | 0 |
| 0 | 0 | 0 | 0 |
| 0 | 0 | 0 | 0 |
| 0 | 0 | 1 | 0 |
| 0 | 0 | 0 | 0 |
| 0 | 0 | 0 | 0 |
| 0 | 0 | 0 | 0 |
| 0 | 0 | 0 | 0 |
| 0 | 0 | 0 | 0 |
| 0 | 0 | 0 | 0 |
| 0 | 0 | 0 | 0 |
| 0 | 0 | 0 | 0 |
| 0 | 0 | 0 | 0 |
| 0 | 0 | 1 | 0 |
| 0 | 0 | 0 | 0 |
| 0 | 0 | 1 | 0 |
| 1 | 0 | 0 | 0 |
| 1 | 1 | 1 | 0 |
| 1 | 1 | 1 | 0 |
| 0 | 0 | 0 | 0 |
| 0 | 0 | 0 | 0 |
| 0 | 1 | 0 | 0 |
| 1 | 1 | 1 | 0 |
| 1 | 1 | 1 | 0 |
| 0 | 0 | 1 | 0 |
| 0 | 0 | 1 | 0 |
| 1 | 0 | 0 | 0 |
| 0 | 0 | 0 | 0 |
| 0 | 0 | 0 | 0 |
| 1 | 1 | 0 | 0 |
| 0 | 0 | 0 | 0 |
| 0 | 0 | 0 | 0 |
| 0 | 0 | 0 | 0 |
| 0 | 0 | 0 | 0 |
| 0 | 0 | 0 | 0 |
| 0 | 0 | 0 | 0 |
| 0 | 0 | 0 | 0 |
| 0 | 0 | 0 | 0 |
| 0 | 0 | 0 | 0 |

| | | | |
|---|---|---|---|
| 0 | 0 | 0 | 0 |
| 0 | 0 | 0 | 0 |
| 0 | 0 | 0 | 0 |
| 0 | 0 | 0 | 0 |
| 0 | 0 | 0 | 0 |
| 0 | 0 | 0 | 0 |
| 0 | 0 | 0 | 0 |
| 0 | 0 | 0 | 0 |
| 0 | 0 | 0 | 0 |
| 0 | 0 | 0 | 0 |
| 0 | 0 | 0 | 0 |
| 0 | 0 | 0 | 0 |
| 0 | 0 | 0 | 0 |
| 0 | 0 | 0 | 0 |
| 0 | 0 | 0 | 0 |
| 0 | 0 | 0 | 0 |
| 0 | 0 | 0 | 0 |
| 0 | 0 | 0 | 0 |
| 0 | 0 | 0 | 0 |
| 1 | 0 | 0 | 0 |
| 1 | 0 | 0 | 0 |
| 0 | 0 | 0 | 0 |
| 0 | 0 | 0 | 0 |
| 0 | 0 | 0 | 0 |
| 0 | 0 | 0 | 0 |
| 1 | 0 | 0 | 0 |
| 0 | 0 | 0 | 0 |
| 0 | 0 | 0 | 0 |
| 0 | 0 | 0 | 0 |
| 1 | 0 | 0 | 0 |
| 0 | 0 | 0 | 0 |
| 0 | 0 | 0 | 0 |
| 1 | 0 | 0 | 0 |
| 1 | 0 | 0 | 0 |
| 0 | 0 | 0 | 0 |
| 0 | 0 | 0 | 0 |
| 0 | 0 | 0 | 0 |
| 0 | 0 | 0 | 0 |
| 0 | 0 | 0 | 0 |
| 0 | 0 | 0 | 0 |
| 0 | 0 | 0 | 0 |
| 0 | 0 | 0 | 0 |
| 0 | 0 | 0 | 0 |
| 0 | 0 | 0 | 0 |
| 0 | 0 | 0 | 0 |
| 0 | 0 | 0 | 0 |
| 0 | 0 | 0 | 0 |
| 0 | 1 | 0 | 0 |
| 0 | 1 | 0 | 0 |
| 0 | 0 | 0 | 0 |
| 0 | 1 | 0 | 0 |
| 0 | 1 | 0 | 0 |
| 0 | 0 | 0 | 0 |
| 0 | 0 | 0 | 0 |

| | | | |
|---|---|---|---|
| 0 | 0 | 0 | 0 |
| 0 | 0 | 0 | 0 |
| 1 | 1 | 0 | 1 |
| 0 | 0 | 0 | 0 |
| 0 | 0 | 0 | 0 |
| 0 | 0 | 0 | 0 |
| 0 | 0 | 0 | 0 |
| 0 | 1 | 0 | 0 |
| 1 | 1 | 0 | 0 |
| 1 | 1 | 0 | 0 |
| 1 | 1 | 0 | 0 |
| 1 | 1 | 0 | 0 |
| 1 | 1 | 0 | 0 |
| 0 | 1 | 0 | 0 |
| 0 | 1 | 0 | 0 |
| 0 | 0 | 0 | 0 |
| 0 | 0 | 0 | 0 |
| 0 | 0 | 0 | 0 |
| 0 | 0 | 0 | 0 |
| 0 | 0 | 0 | 0 |
| 0 | 0 | 0 | 0 |
| 0 | 0 | 0 | 0 |
| 0 | 1 | 0 | 0 |
| 0 | 1 | 0 | 0 |
| 0 | 0 | 0 | 0 |
| 0 | 0 | 1 | 0 |
| 0 | 0 | 0 | 0 |
| 0 | 0 | 0 | 0 |
| 0 | 0 | 0 | 0 |
| 0 | 1 | 0 | 0 |
| 0 | 0 | 0 | 0 |
| 0 | 0 | 0 | 0 |
| 0 | 0 | 0 | 0 |
| 0 | 0 | 0 | 0 |
| 0 | 0 | 0 | 0 |
| 0 | 0 | 0 | 0 |
| 0 | 0 | 0 | 0 |
| 0 | 0 | 0 | 0 |
| 0 | 0 | 0 | 0 |
| 1 | 0 | 0 | 0 |
| 1 | 1 | 0 | 0 |
| 1 | 0 | 0 | 0 |
| 0 | 0 | 0 | 0 |
| 0 | 1 | 0 | 0 |
| 1 | 1 | 0 | 0 |
| 0 | 1 | 0 | 0 |
| 0 | 0 | 0 | 0 |
| 1 | 0 | 0 | 0 |
| 0 | 1 | 0 | 0 |
| 0 | 0 | 0 | 0 |
| 0 | 0 | 0 | 0 |
| 0 | 0 | 0 | 0 |
| 1 | 1 | 0 | 0 |
| 0 | 0 | 0 | 0 |

| | | | |
|---|---|---|---|
| 0 | 0 | 0 | 0 |
| 0 | 0 | 0 | 0 |
| 1 | 0 | 0 | 0 |
| 1 | 1 | 0 | 0 |
| 1 | 1 | 0 | 0 |
| 0 | 0 | 0 | 0 |
| 0 | 0 | 0 | 0 |
| 0 | 0 | 0 | 0 |
| 1 | 0 | 0 | 0 |
| 0 | 0 | 0 | 0 |
| 0 | 0 | 0 | 0 |
| 1 | 0 | 0 | 1 |
| 0 | 0 | 0 | 0 |
| 1 | 0 | 0 | 0 |
| 0 | 0 | 0 | 0 |
| 0 | 0 | 0 | 0 |
| 0 | 0 | 0 | 0 |
| 0 | 0 | 0 | 0 |
| 0 | 0 | 0 | 0 |
| 0 | 0 | 0 | 0 |
| 0 | 0 | 0 | 0 |
| 1 | 0 | 0 | 1 |
| 1 | 0 | 0 | 0 |
| 0 | 0 | 0 | 0 |
| 0 | 0 | 0 | 0 |
| 0 | 0 | 0 | 0 |
| 0 | 0 | 0 | 0 |
| 0 | 0 | 0 | 0 |
| 1 | 0 | 0 | 0 |
| 1 | 0 | 0 | 0 |
| 0 | 0 | 0 | 0 |
| 0 | 0 | 0 | 0 |
| 0 | 0 | 0 | 0 |
| 0 | 0 | 0 | 0 |
| 0 | 0 | 0 | 0 |
| 0 | 0 | 0 | 0 |
| 1 | 0 | 0 | 0 |
| 0 | 0 | 0 | 0 |
| 0 | 0 | 0 | 0 |
| 1 | 0 | 0 | 0 |
| 1 | 0 | 0 | 0 |
| 1 | 0 | 0 | 0 |
| 1 | 0 | 0 | 0 |
| 0 | 0 | 0 | 0 |
| 0 | 0 | 0 | 0 |
| 1 | 0 | 0 | 0 |
| 0 | 0 | 0 | 0 |
| 0 | 0 | 0 | 0 |
| 0 | 0 | 0 | 0 |
| 0 | 0 | 0 | 0 |
| 0 | 0 | 0 | 0 |
| 1 | 0 | 0 | 0 |
| 1 | 0 | 0 | 0 |
| 1 | 0 | 0 | 0 |

| | | | |
|---|---|---|---|
| 1 | 0 | 0 | 0 |
| 0 | 0 | 0 | 0 |
| 0 | 0 | 0 | 0 |
| 0 | 0 | 0 | 0 |
| 1 | 0 | 0 | 0 |
| 0 | 0 | 0 | 0 |
| 1 | 0 | 0 | 0 |
| 0 | 0 | 0 | 0 |
| 0 | 0 | 0 | 0 |
| 0 | 0 | 0 | 0 |
| 0 | 0 | 0 | 0 |
| 0 | 0 | 0 | 0 |
| 0 | 0 | 0 | 0 |
| 0 | 0 | 0 | 0 |
| 0 | 0 | 0 | 0 |
| 1 | 0 | 0 | 0 |
| 0 | 0 | 0 | 0 |
| 0 | 0 | 0 | 0 |
| 0 | 0 | 0 | 0 |
| 0 | 0 | 0 | 0 |
| 0 | 0 | 0 | 0 |
| 0 | 0 | 0 | 0 |
| 1 | 0 | 0 | 0 |
| 1 | 0 | 0 | 0 |
| 0 | 0 | 0 | 0 |
| 0 | 0 | 0 | 0 |
| 0 | 0 | 0 | 0 |
| 0 | 0 | 0 | 0 |
| 0 | 0 | 0 | 0 |
| 0 | 0 | 0 | 0 |
| 0 | 0 | 0 | 0 |
| 0 | 0 | 0 | 0 |
| 1 | 0 | 0 | 0 |
| 0 | 0 | 0 | 0 |
| 0 | 0 | 0 | 0 |
| 0 | 0 | 0 | 0 |
| 0 | 0 | 0 | 0 |
| 0 | 0 | 0 | 0 |
| 0 | 0 | 0 | 0 |
| 0 | 0 | 0 | 0 |
| 0 | 0 | 0 | 0 |
| 0 | 0 | 0 | 0 |
| 0 | 0 | 0 | 0 |
| 0 | 0 | 0 | 0 |
| 0 | 1 | 0 | 0 |
| 0 | 0 | 0 | 0 |
| 0 | 0 | 0 | 0 |
| 0 | 0 | 0 | 0 |
| 0 | 0 | 0 | 0 |
| 0 | 0 | 0 | 0 |
| 0 | 0 | 0 | 0 |
| 0 | 0 | 0 | 0 |
| 0 | 0 | 0 | 0 |
| 1 | 0 | 0 | 0 |

| | | | |
|---|---|---|---|
| 0 | 0 | 0 | 0 |
| 0 | 0 | 0 | 0 |
| 0 | 0 | 0 | 0 |
| 0 | 0 | 0 | 0 |
| 0 | 0 | 0 | 0 |
| 0 | 0 | 0 | 0 |
| 0 | 0 | 0 | 0 |
| 0 | 0 | 0 | 0 |
| 0 | 0 | 0 | 0 |
| 1 | 0 | 0 | 0 |
| 0 | 0 | 0 | 0 |
| 0 | 0 | 0 | 0 |
| 1 | 1 | 0 | 0 |
| 1 | 1 | 0 | 0 |
| 1 | 1 | 0 | 0 |
| 1 | 1 | 0 | 0 |
| 0 | 0 | 0 | 0 |
| 0 | 0 | 0 | 0 |
| 0 | 0 | 0 | 0 |
| 0 | 0 | 0 | 0 |
| 0 | 0 | 0 | 0 |
| 0 | 0 | 0 | 0 |
| 0 | 0 | 0 | 0 |
| 0 | 0 | 0 | 0 |
| 0 | 0 | 0 | 0 |
| 0 | 0 | 0 | 1 |
| 0 | 0 | 0 | 0 |
| 0 | 0 | 0 | 0 |
| 0 | 0 | 0 | 0 |
| 0 | 0 | 0 | 0 |
| 0 | 0 | 0 | 0 |
| 0 | 0 | 0 | 0 |
| 0 | 0 | 0 | 0 |
| 0 | 0 | 0 | 0 |
| 0 | 0 | 0 | 0 |
| 0 | 0 | 0 | 0 |
| 0 | 0 | 0 | 0 |
| 1 | 1 | 0 | 0 |
| 0 | 0 | 0 | 0 |
| 0 | 0 | 0 | 0 |
| 1 | 1 | 0 | 0 |
| 0 | 0 | 0 | 0 |
| 0 | 0 | 0 | 0 |
| 1 | 0 | 0 | 1 |
| 0 | 0 | 0 | 0 |
| 0 | 0 | 0 | 0 |
| 0 | 0 | 0 | 0 |
| 0 | 0 | 0 | 0 |
| 0 | 0 | 0 | 0 |
| 0 | 0 | 0 | 0 |
| 0 | 0 | 0 | 0 |
| 0 | 0 | 1 | 0 |
| 0 | 0 | 1 | 0 |
| 0 | 0 | 1 | 0 |

| | | | |
|---|---|---|---|
| 0 | 0 | 1 | 0 |
| 0 | 0 | 0 | 0 |
| 0 | 0 | 0 | 0 |
| 0 | 0 | 0 | 0 |
| 0 | 0 | 0 | 0 |
| 0 | 0 | 0 | 0 |
| 0 | 0 | 0 | 0 |
| 0 | 0 | 0 | 0 |
| 0 | 0 | 0 | 0 |
| 0 | 0 | 0 | 0 |
| 0 | 0 | 0 | 0 |
| 0 | 0 | 0 | 0 |
| 0 | 0 | 0 | 0 |
| 0 | 0 | 0 | 0 |
| 0 | 0 | 0 | 0 |
| 0 | 0 | 0 | 0 |
| 0 | 0 | 0 | 0 |
| 0 | 0 | 0 | 0 |
| 0 | 0 | 0 | 0 |
| 0 | 0 | 0 | 0 |
| 0 | 0 | 0 | 0 |
| 0 | 0 | 0 | 0 |
| 0 | 0 | 0 | 0 |
| 0 | 1 | 0 | 1 |
| 0 | 0 | 0 | 0 |
| 0 | 0 | 0 | 0 |
| 0 | 0 | 0 | 0 |
| 0 | 0 | 0 | 0 |
| 0 | 0 | 0 | 0 |
| 0 | 0 | 0 | 0 |
| 0 | 0 | 0 | 0 |
| 0 | 0 | 0 | 0 |
| 0 | 0 | 0 | 0 |
| 0 | 0 | 0 | 0 |
| 0 | 0 | 0 | 0 |
| 0 | 0 | 0 | 0 |
| 0 | 0 | 0 | 0 |
| 0 | 0 | 0 | 0 |
| 0 | 0 | 0 | 0 |
| 0 | 0 | 0 | 0 |
| 0 | 0 | 0 | 0 |
| 0 | 0 | 0 | 0 |
| 0 | 0 | 0 | 0 |
| 0 | 0 | 0 | 0 |
| 0 | 0 | 0 | 0 |
| 0 | 0 | 0 | 0 |
| 1 | 0 | 0 | 0 |
| 1 | 0 | 0 | 0 |
| 0 | 0 | 0 | 0 |
| 0 | 1 | 0 | 0 |
| 1 | 1 | 0 | 1 |
| 1 | 1 | 0 | 1 |
| 1 | 1 | 0 | 1 |
| 0 | 0 | 0 | 0 |

| | | | |
|---|---|---|---|
| 1 | 1 | 0 | 0 |
| 0 | 0 | 0 | 0 |
| 0 | 0 | 0 | 0 |
| 0 | 0 | 0 | 0 |
| 0 | 0 | 0 | 0 |
| 0 | 0 | 0 | 0 |
| 0 | 0 | 0 | 0 |
| 0 | 0 | 0 | 0 |
| 0 | 0 | 0 | 0 |
| 0 | 0 | 0 | 0 |
| 0 | 0 | 0 | 0 |
| 0 | 1 | 0 | 0 |
| 0 | 0 | 0 | 0 |
| 0 | 0 | 0 | 0 |
| 0 | 1 | 0 | 0 |
| 0 | 0 | 0 | 0 |
| 1 | 1 | 0 | 0 |
| 0 | 0 | 0 | 0 |
| 0 | 0 | 0 | 0 |
| 1 | 0 | 0 | 0 |
| 0 | 0 | 0 | 0 |
| 0 | 0 | 0 | 0 |
| 0 | 0 | 0 | 0 |
| 0 | 0 | 0 | 0 |
| 0 | 0 | 0 | 0 |
| 0 | 0 | 0 | 0 |
| 0 | 0 | 0 | 0 |
| 0 | 0 | 0 | 0 |
| 0 | 0 | 0 | 0 |
| 0 | 0 | 0 | 0 |
| 0 | 0 | 0 | 0 |
| 0 | 0 | 0 | 0 |
| 0 | 0 | 0 | 0 |
| 1 | 0 | 0 | 0 |
| 0 | 0 | 0 | 0 |
| 0 | 1 | 0 | 0 |
| 0 | 0 | 0 | 0 |
| 0 | 0 | 0 | 0 |
| 0 | 0 | 0 | 0 |
| 0 | 0 | 0 | 0 |
| 0 | 0 | 0 | 0 |
| 0 | 0 | 0 | 0 |
| 0 | 0 | 0 | 0 |
| 0 | 0 | 0 | 0 |
| 0 | 0 | 0 | 0 |
| 0 | 0 | 0 | 0 |
| 0 | 0 | 0 | 0 |
| 0 | 0 | 0 | 0 |
| 1 | 1 | 0 | 0 |
| 0 | 0 | 0 | 0 |
| 0 | 0 | 0 | 0 |
| 0 | 0 | 0 | 0 |
| 0 | 0 | 0 | 0 |
| 0 | 0 | 0 | 0 |

| | | | |
|---|---|---|---|
| 0 | 0 | 0 | 0 |
| 0 | 0 | 0 | 0 |
| 0 | 0 | 0 | 0 |
| 0 | 0 | 0 | 0 |
| 0 | 0 | 0 | 0 |
| 0 | 0 | 0 | 0 |
| 0 | 0 | 0 | 0 |
| 0 | 1 | 0 | 0 |
| 0 | 0 | 0 | 0 |
| 0 | 0 | 0 | 0 |
| 0 | 0 | 0 | 0 |
| 0 | 0 | 0 | 0 |
| 0 | 0 | 1 | 0 |
| 0 | 0 | 0 | 0 |
| 0 | 0 | 1 | 0 |
| 0 | 0 | 1 | 0 |
| 0 | 0 | 0 | 0 |
| 0 | 0 | 0 | 0 |
| 0 | 0 | 0 | 0 |
| 0 | 0 | 0 | 0 |
| 0 | 1 | 0 | 0 |
| 0 | 1 | 0 | 0 |
| 0 | 0 | 0 | 0 |
| 0 | 0 | 0 | 0 |
| 0 | 1 | 0 | 0 |
| 0 | 0 | 0 | 0 |
| 0 | 0 | 0 | 0 |
| 0 | 1 | 0 | 0 |
| 0 | 0 | 0 | 0 |
| 0 | 0 | 0 | 0 |
| 0 | 0 | 0 | 0 |
| 0 | 0 | 0 | 0 |
| 1 | 1 | 0 | 0 |
| 1 | 1 | 0 | 0 |
| 1 | 1 | 0 | 0 |
| 0 | 1 | 0 | 0 |
| 1 | 0 | 0 | 0 |
| 0 | 1 | 0 | 0 |
| 1 | 1 | 0 | 0 |
| 0 | 0 | 0 | 0 |
| 0 | 0 | 0 | 0 |
| 0 | 0 | 0 | 0 |
| 0 | 0 | 0 | 0 |
| 0 | 0 | 0 | 0 |
| 0 | 0 | 0 | 0 |
| 0 | 0 | 0 | 0 |
| 0 | 0 | 0 | 0 |
| 0 | 0 | 0 | 0 |
| 0 | 0 | 0 | 0 |
| 1 | 1 | 0 | 0 |
| 1 | 1 | 0 | 0 |
| 0 | 0 | 0 | 0 |
| 0 | 0 | 0 | 0 |
| 0 | 0 | 0 | 0 |

| | | | |
|---|---|---|---|
| 0 | 0 | 0 | 0 |
| 0 | 0 | 0 | 0 |
| 0 | 0 | 0 | 0 |
| 0 | 0 | 0 | 0 |
| 0 | 0 | 0 | 0 |
| 0 | 0 | 0 | 0 |
| 1 | 1 | 0 | 0 |
| 1 | 1 | 0 | 0 |
| 0 | 0 | 0 | 0 |
| 0 | 0 | 0 | 0 |
| 0 | 0 | 0 | 0 |
| 0 | 0 | 0 | 0 |
| 0 | 0 | 0 | 0 |
| 0 | 0 | 0 | 0 |
| 0 | 0 | 0 | 0 |
| 0 | 0 | 0 | 0 |
| 0 | 0 | 0 | 0 |
| 0 | 0 | 0 | 0 |
| 0 | 0 | 0 | 0 |
| 0 | 0 | 0 | 0 |
| 0 | 0 | 0 | 0 |
| 0 | 0 | 0 | 0 |
| 0 | 0 | 0 | 0 |
| 0 | 0 | 0 | 0 |
| 0 | 0 | 0 | 0 |
| 0 | 0 | 0 | 0 |
| 0 | 0 | 0 | 0 |
| 1 | 1 | 0 | 0 |
| 0 | 0 | 0 | 0 |
| 0 | 0 | 0 | 0 |
| 0 | 0 | 0 | 0 |
| 0 | 0 | 0 | 0 |
| 0 | 0 | 0 | 0 |
| 0 | 0 | 0 | 0 |
| 0 | 0 | 0 | 0 |
| 0 | 0 | 0 | 0 |
| 0 | 0 | 0 | 0 |
| 0 | 0 | 0 | 0 |
| 0 | 0 | 0 | 0 |
| 0 | 0 | 0 | 0 |
| 0 | 0 | 0 | 0 |
| 0 | 0 | 0 | 0 |
| 0 | 0 | 0 | 0 |
| 0 | 0 | 0 | 0 |
| 0 | 0 | 0 | 0 |
| 0 | 0 | 0 | 0 |
| 0 | 0 | 0 | 0 |
| 0 | 0 | 0 | 0 |
| 0 | 0 | 0 | 0 |
| 0 | 0 | 0 | 0 |
| 0 | 0 | 0 | 0 |
| 0 | 0 | 0 | 0 |
| 0 | 0 | 0 | 0 |
| 0 | 0 | 0 | 0 |

| | | | |
|---|---|---|---|
| 0 | 0 | 0 | 0 |
| 0 | 0 | 0 | 0 |
| 0 | 0 | 0 | 0 |
| 0 | 0 | 0 | 0 |
| 0 | 0 | 0 | 0 |
| 0 | 0 | 0 | 0 |
| 0 | 0 | 0 | 0 |
| 0 | 0 | 0 | 0 |
| 0 | 0 | 0 | 0 |
| 0 | 0 | 0 | 0 |
| 1 | 0 | 0 | 0 |
| 1 | 1 | 0 | 0 |
| 1 | 0 | 0 | 0 |
| 0 | 0 | 0 | 0 |
| 0 | 0 | 0 | 0 |
| 0 | 0 | 0 | 0 |
| 0 | 0 | 0 | 0 |
| 0 | 0 | 0 | 0 |
| 0 | 0 | 0 | 0 |
| 0 | 0 | 0 | 0 |
| 0 | 0 | 0 | 0 |
| 0 | 0 | 0 | 0 |
| 0 | 0 | 0 | 0 |
| 0 | 0 | 0 | 0 |
| 0 | 0 | 0 | 0 |
| 0 | 0 | 0 | 0 |
| 0 | 0 | 0 | 0 |
| 0 | 0 | 0 | 0 |
| 0 | 0 | 0 | 0 |
| 0 | 0 | 0 | 0 |
| 0 | 0 | 0 | 0 |
| 0 | 1 | 0 | 0 |
| 0 | 0 | 0 | 0 |
| 0 | 0 | 0 | 0 |
| 1 | 0 | 0 | 0 |
| 1 | 0 | 0 | 0 |
| 1 | 0 | 0 | 0 |
| 0 | 0 | 0 | 0 |
| 0 | 0 | 0 | 0 |
| 0 | 0 | 0 | 0 |
| 0 | 0 | 0 | 0 |
| 0 | 0 | 0 | 0 |
| 0 | 0 | 0 | 0 |
| 0 | 0 | 0 | 0 |
| 0 | 0 | 0 | 0 |
| 0 | 0 | 0 | 0 |
| 0 | 0 | 0 | 0 |
| 0 | 0 | 0 | 0 |
| 0 | 0 | 0 | 0 |
| 0 | 0 | 0 | 0 |
| 0 | 0 | 0 | 0 |
| 0 | 0 | 0 | 0 |
| 0 | 0 | 0 | 1 |
| 0 | 0 | 0 | 0 |

| | | | |
|---|---|---|---|
| 0 | 0 | 0 | 0 |
| 0 | 0 | 0 | 0 |
| 0 | 0 | 0 | 0 |
| 0 | 0 | 0 | 0 |
| 0 | 0 | 0 | 0 |
| 0 | 0 | 0 | 0 |
| 0 | 0 | 0 | 0 |
| 0 | 0 | 0 | 0 |
| 0 | 0 | 0 | 0 |
| 0 | 0 | 0 | 0 |
| 0 | 0 | 0 | 0 |
| 1 | 0 | 0 | 0 |
| 0 | 0 | 0 | 0 |
| 0 | 0 | 0 | 0 |
| 0 | 0 | 1 | 0 |
| 0 | 0 | 0 | 0 |
| 0 | 0 | 0 | 0 |
| 0 | 0 | 0 | 0 |
| 0 | 0 | 0 | 0 |
| 0 | 0 | 0 | 0 |
| 0 | 0 | 0 | 0 |
| 0 | 0 | 0 | 0 |
| 0 | 0 | 0 | 0 |
| 0 | 0 | 0 | 0 |
| 0 | 0 | 0 | 0 |
| 0 | 0 | 0 | 0 |
| 0 | 0 | 0 | 0 |
| 0 | 0 | 0 | 0 |
| 0 | 0 | 0 | 0 |
| 0 | 0 | 0 | 0 |
| 0 | 0 | 0 | 0 |
| 0 | 0 | 0 | 0 |
| 0 | 0 | 0 | 0 |
| 1 | 0 | 0 | 0 |
| 0 | 0 | 0 | 0 |
| 1 | 0 | 0 | 1 |
| 0 | 0 | 0 | 0 |
| 0 | 0 | 0 | 0 |
| 0 | 0 | 0 | 0 |
| 0 | 0 | 0 | 0 |
| 0 | 0 | 0 | 0 |
| 0 | 0 | 0 | 0 |
| 0 | 0 | 0 | 0 |
| 0 | 1 | 0 | 0 |
| 0 | 0 | 0 | 0 |
| 0 | 0 | 0 | 0 |
| 0 | 0 | 0 | 0 |
| 0 | 0 | 0 | 0 |
| 0 | 0 | 0 | 0 |
| 0 | 0 | 0 | 0 |
| 0 | 0 | 0 | 0 |
| 0 | 0 | 0 | 0 |
| 0 | 0 | 0 | 0 |
| 0 | 0 | 0 | 0 |

| | | | |
|---|---|---|---|
| 0 | 0 | 0 | 0 |
| 0 | 0 | 0 | 0 |
| 0 | 0 | 0 | 0 |
| 0 | 0 | 0 | 0 |
| 0 | 0 | 0 | 0 |
| 0 | 1 | 0 | 0 |
| 0 | 0 | 0 | 0 |
| 0 | 0 | 0 | 0 |
| 0 | 0 | 0 | 0 |
| 0 | 0 | 0 | 0 |
| 0 | 0 | 0 | 0 |
| 0 | 0 | 0 | 0 |
| 0 | 0 | 0 | 0 |
| 0 | 0 | 0 | 0 |
| 0 | 0 | 0 | 0 |
| 0 | 1 | 0 | 0 |
| 1 | 1 | 0 | 0 |
| 0 | 0 | 1 | 0 |
| 0 | 0 | 0 | 0 |
| 0 | 0 | 0 | 0 |
| 0 | 0 | 0 | 0 |
| 0 | 0 | 0 | 0 |
| 0 | 0 | 0 | 0 |
| 0 | 0 | 0 | 0 |
| 0 | 0 | 0 | 0 |
| 0 | 0 | 0 | 0 |
| 0 | 0 | 0 | 0 |
| 1 | 0 | 0 | 0 |
| 0 | 0 | 0 | 0 |
| 0 | 0 | 0 | 0 |
| 0 | 0 | 1 | 0 |
| 0 | 0 | 0 | 0 |
| 0 | 0 | 0 | 0 |
| 0 | 0 | 0 | 0 |
| 0 | 0 | 0 | 0 |
| 0 | 0 | 0 | 0 |
| 0 | 0 | 0 | 0 |
| 1 | 1 | 0 | 0 |
| 0 | 0 | 0 | 0 |
| 0 | 0 | 0 | 0 |
| 0 | 0 | 0 | 0 |
| 1 | 0 | 0 | 0 |
| 1 | 0 | 0 | 0 |
| 0 | 0 | 0 | 0 |
| 0 | 0 | 0 | 0 |
| 0 | 0 | 0 | 0 |
| 0 | 0 | 0 | 0 |
| 0 | 0 | 0 | 0 |
| 0 | 0 | 0 | 0 |
| 0 | 0 | 0 | 0 |
| 0 | 0 | 0 | 0 |
| 0 | 0 | 0 | 0 |
| 0 | 0 | 0 | 0 |
| 0 | 0 | 0 | 0 |

| | | | |
|---|---|---|---|
| 0 | 0 | 0 | 0 |
| 0 | 0 | 0 | 0 |
| 0 | 0 | 0 | 0 |
| 0 | 0 | 0 | 0 |
| 0 | 0 | 0 | 0 |
| 0 | 0 | 0 | 0 |
| 0 | 0 | 0 | 0 |
| 0 | 0 | 0 | 0 |
| 0 | 0 | 0 | 0 |
| 0 | 0 | 0 | 0 |
| 0 | 0 | 0 | 0 |
| 0 | 0 | 0 | 0 |
| 1 | 0 | 0 | 0 |
| 0 | 0 | 0 | 0 |
| 0 | 0 | 0 | 0 |
| 0 | 0 | 0 | 0 |
| 0 | 0 | 0 | 0 |
| 0 | 0 | 0 | 0 |
| 0 | 0 | 0 | 0 |
| 0 | 0 | 0 | 0 |
| 0 | 0 | 0 | 0 |
| 0 | 0 | 0 | 0 |
| 0 | 0 | 0 | 0 |
| 1 | 0 | 0 | 0 |
| 0 | 0 | 0 | 0 |
| 0 | 0 | 0 | 0 |
| 0 | 0 | 0 | 0 |
| 1 | 1 | 0 | 1 |
| 0 | 0 | 0 | 0 |
| 0 | 0 | 0 | 0 |
| 0 | 0 | 0 | 0 |
| 0 | 0 | 0 | 0 |
| 0 | 0 | 0 | 0 |
| 0 | 0 | 0 | 0 |
| 0 | 0 | 0 | 0 |
| 0 | 0 | 0 | 0 |
| 0 | 0 | 0 | 0 |
| 0 | 0 | 0 | 0 |
| 0 | 0 | 0 | 0 |
| 0 | 0 | 0 | 0 |
| 0 | 0 | 0 | 0 |
| 0 | 0 | 0 | 0 |
| 0 | 0 | 0 | 0 |
| 0 | 0 | 0 | 0 |
| 0 | 0 | 0 | 0 |
| 0 | 0 | 0 | 0 |
| 0 | 0 | 0 | 0 |
| 0 | 0 | 0 | 0 |
| 0 | 0 | 0 | 0 |
| 0 | 0 | 0 | 0 |
| 0 | 0 | 0 | 0 |
| 0 | 0 | 0 | 0 |
| 0 | 0 | 0 | 0 |
| 0 | 0 | 0 | 0 |

| | | | |
|---|---|---|---|
| 0 | 0 | 0 | 0 |
| 0 | 0 | 0 | 0 |
| 0 | 0 | 0 | 0 |
| 0 | 0 | 0 | 0 |
| 0 | 0 | 0 | 0 |
| 0 | 0 | 0 | 0 |
| 0 | 0 | 0 | 0 |
| 0 | 0 | 0 | 0 |
| 1 | 0 | 0 | 0 |
| 1 | 0 | 0 | 0 |
| 1 | 0 | 0 | 0 |
| 0 | 0 | 0 | 0 |
| 1 | 0 | 0 | 0 |
| 0 | 0 | 0 | 0 |
| 0 | 0 | 0 | 0 |
| 1 | 0 | 0 | 0 |
| 0 | 0 | 0 | 0 |
| 0 | 0 | 0 | 0 |
| 0 | 0 | 0 | 0 |
| 0 | 0 | 0 | 0 |
| 0 | 0 | 0 | 0 |
| 0 | 0 | 0 | 0 |
| 0 | 0 | 0 | 0 |
| 0 | 0 | 0 | 0 |
| 0 | 0 | 0 | 0 |
| 0 | 0 | 0 | 0 |
| 0 | 0 | 0 | 0 |
| 1 | 0 | 0 | 0 |
| 1 | 0 | 0 | 0 |
| 0 | 0 | 0 | 0 |
| 0 | 0 | 0 | 0 |
| 0 | 0 | 0 | 0 |
| 1 | 0 | 0 | 0 |
| 0 | 0 | 0 | 0 |
| 0 | 0 | 0 | 0 |
| 0 | 0 | 0 | 0 |
| 0 | 0 | 0 | 0 |
| 0 | 0 | 0 | 0 |
| 0 | 0 | 0 | 0 |
| 0 | 0 | 0 | 0 |
| 1 | 0 | 0 | 0 |
| 1 | 0 | 0 | 0 |
| 0 | 0 | 0 | 0 |
| 0 | 0 | 0 | 0 |
| 0 | 0 | 0 | 0 |
| 0 | 0 | 0 | 0 |
| 0 | 0 | 0 | 0 |
| 0 | 0 | 0 | 0 |
| 0 | 0 | 0 | 0 |
| 0 | 0 | 0 | 0 |
| 0 | 0 | 0 | 0 |
| 0 | 0 | 0 | 0 |
| 0 | 0 | 0 | 0 |
| 0 | 0 | 0 | 0 |

| | | | |
|---|---|---|---|
| 0 | 0 | 0 | 0 |
| 0 | 0 | 0 | 0 |
| 0 | 0 | 0 | 0 |
| 0 | 0 | 0 | 0 |
| 0 | 0 | 0 | 0 |
| 0 | 0 | 0 | 0 |
| 0 | 0 | 0 | 0 |
| 0 | 0 | 0 | 0 |
| 0 | 0 | 0 | 0 |
| 0 | 0 | 0 | 0 |
| 1 | 0 | 0 | 0 |
| 0 | 0 | 0 | 0 |
| 1 | 0 | 0 | 0 |
| 0 | 0 | 0 | 0 |
| 0 | 0 | 0 | 0 |
| 0 | 0 | 0 | 0 |
| 0 | 0 | 0 | 0 |
| 0 | 1 | 0 | 0 |
| 0 | 0 | 0 | 0 |
| 0 | 0 | 0 | 0 |
| 0 | 0 | 0 | 0 |
| 0 | 0 | 0 | 0 |
| 1 | 0 | 0 | 0 |
| 0 | 0 | 0 | 0 |
| 0 | 0 | 0 | 0 |
| 1 | 0 | 0 | 0 |
| 1 | 0 | 0 | 0 |
| 1 | 0 | 0 | 0 |
| 0 | 0 | 0 | 0 |
| 0 | 0 | 0 | 0 |
| 0 | 0 | 0 | 0 |
| 0 | 0 | 0 | 0 |
| 0 | 0 | 0 | 0 |
| 0 | 0 | 0 | 0 |
| 0 | 0 | 0 | 0 |
| 0 | 0 | 0 | 0 |
| 0 | 0 | 0 | 0 |
| 0 | 0 | 0 | 0 |
| 0 | 0 | 0 | 0 |
| 0 | 0 | 0 | 0 |
| 1 | 1 | 0 | 0 |
| 0 | 0 | 0 | 0 |
| 0 | 0 | 0 | 0 |
| 0 | 0 | 0 | 0 |
| 0 | 0 | 0 | 0 |
| 0 | 0 | 0 | 0 |
| 0 | 0 | 0 | 0 |
| 1 | 0 | 0 | 0 |
| 0 | 0 | 0 | 0 |
| 0 | 0 | 0 | 0 |
| 0 | 0 | 0 | 0 |
| 0 | 0 | 0 | 0 |
| 0 | 0 | 0 | 0 |
| 0 | 0 | 0 | 0 |

| | | | |
|---|---|---|---|
| 1 | 0 | 0 | 0 |
| 0 | 0 | 0 | 0 |
| 1 | 0 | 0 | 0 |
| 0 | 0 | 0 | 0 |
| 0 | 0 | 0 | 0 |
| 0 | 0 | 0 | 0 |
| 0 | 0 | 0 | 0 |
| 0 | 0 | 0 | 0 |
| 1 | 1 | 0 | 0 |
| 0 | 0 | 0 | 0 |
| 1 | 1 | 0 | 0 |
| 1 | 1 | 0 | 0 |
| 0 | 0 | 0 | 0 |
| 1 | 1 | 0 | 0 |
| 0 | 0 | 0 | 0 |
| 0 | 0 | 0 | 0 |
| 0 | 0 | 0 | 0 |
| 0 | 0 | 0 | 0 |
| 0 | 0 | 0 | 0 |
| 0 | 0 | 0 | 0 |
| 1 | 0 | 0 | 0 |
| 0 | 0 | 0 | 0 |
| 0 | 0 | 0 | 0 |
| 0 | 0 | 0 | 0 |
| 1 | 0 | 0 | 0 |
| 0 | 0 | 0 | 0 |
| 1 | 0 | 0 | 0 |
| 1 | 0 | 0 | 0 |
| 1 | 1 | 0 | 0 |
| 0 | 0 | 0 | 0 |
| 1 | 1 | 0 | 0 |
| 1 | 1 | 0 | 0 |
| 1 | 1 | 0 | 0 |
| 0 | 0 | 0 | 0 |
| 1 | 1 | 0 | 0 |
| 0 | 0 | 0 | 0 |
| 1 | 0 | 0 | 0 |
| 0 | 0 | 0 | 0 |
| 1 | 1 | 0 | 0 |
| 1 | 1 | 0 | 0 |
| 1 | 0 | 0 | 0 |
| 0 | 0 | 0 | 0 |
| 0 | 0 | 0 | 0 |
| 1 | 0 | 0 | 0 |
| 0 | 0 | 0 | 0 |
| 0 | 0 | 0 | 0 |
| 1 | 0 | 0 | 0 |
| 0 | 0 | 0 | 0 |
| 1 | 0 | 0 | 0 |
| 0 | 0 | 0 | 0 |
| 0 | 0 | 0 | 0 |
| 0 | 0 | 0 | 0 |
| 0 | 0 | 0 | 0 |
| 0 | 0 | 0 | 0 |

| | | | |
|---|---|---|---|
| 0 | 0 | 0 | 0 |
| 0 | 0 | 0 | 0 |
| 0 | 0 | 0 | 0 |
| 0 | 0 | 0 | 0 |
| 0 | 0 | 0 | 0 |
| 0 | 0 | 0 | 0 |
| 0 | 0 | 0 | 0 |
| 0 | 0 | 0 | 0 |
| 0 | 0 | 0 | 0 |
| 1 | 1 | 0 | 0 |
| 1 | 1 | 0 | 0 |
| 0 | 0 | 0 | 0 |
| 0 | 0 | 0 | 0 |
| 0 | 0 | 0 | 0 |
| 1 | 0 | 1 | 0 |
| 0 | 0 | 0 | 0 |
| 0 | 0 | 0 | 0 |
| 1 | 0 | 1 | 0 |
| 0 | 0 | 0 | 0 |
| 0 | 0 | 0 | 0 |
| 0 | 0 | 0 | 0 |
| 0 | 0 | 1 | 0 |
| 1 | 0 | 1 | 0 |
| 0 | 0 | 1 | 0 |
| 0 | 0 | 1 | 0 |
| 0 | 0 | 1 | 1 |
| 0 | 0 | 1 | 1 |
| 0 | 0 | 0 | 0 |
| 0 | 0 | 0 | 0 |
| 0 | 0 | 0 | 0 |
| 0 | 0 | 0 | 0 |
| 0 | 0 | 0 | 0 |
| 0 | 0 | 0 | 0 |
| 0 | 0 | 0 | 0 |
| 0 | 0 | 0 | 0 |
| 0 | 0 | 0 | 0 |
| 0 | 0 | 0 | 0 |
| 0 | 0 | 0 | 0 |
| 0 | 0 | 0 | 0 |
| 0 | 0 | 0 | 0 |
| 0 | 0 | 0 | 0 |
| 0 | 0 | 0 | 0 |
| 0 | 0 | 0 | 0 |
| 0 | 0 | 1 | 0 |
| 0 | 0 | 0 | 0 |
| 0 | 0 | 0 | 0 |
| 0 | 0 | 0 | 0 |
| 0 | 0 | 0 | 0 |
| 0 | 0 | 0 | 0 |
| 0 | 0 | 0 | 0 |
| 1 | 0 | 0 | 0 |
| 0 | 0 | 0 | 0 |
| 0 | 0 | 0 | 0 |
| 0 | 0 | 0 | 0 |

| | | | |
|---|---|---|---|
| 0 | 0 | 0 | 0 |
| 0 | 0 | 0 | 0 |
| 0 | 0 | 0 | 0 |
| 1 | 0 | 0 | 0 |
| 0 | 0 | 0 | 0 |
| 0 | 0 | 0 | 0 |
| 0 | 0 | 0 | 0 |
| 0 | 0 | 0 | 0 |
| 1 | 0 | 0 | 0 |
| 1 | 0 | 0 | 0 |
| 0 | 0 | 0 | 0 |
| 0 | 0 | 0 | 0 |
| 0 | 0 | 0 | 0 |
| 0 | 0 | 0 | 0 |
| 0 | 0 | 0 | 0 |
| 0 | 0 | 0 | 0 |
| 0 | 0 | 1 | 0 |
| 0 | 0 | 0 | 0 |
| 0 | 0 | 0 | 0 |
| 0 | 0 | 0 | 0 |
| 0 | 0 | 0 | 0 |
| 0 | 0 | 0 | 0 |
| 0 | 0 | 0 | 0 |
| 0 | 0 | 0 | 0 |
| 0 | 0 | 0 | 0 |
| 0 | 0 | 0 | 0 |
| 0 | 0 | 0 | 0 |
| 0 | 0 | 0 | 0 |
| 0 | 0 | 0 | 0 |
| 0 | 0 | 0 | 0 |
| 0 | 0 | 0 | 0 |
| 0 | 0 | 0 | 0 |
| 0 | 0 | 0 | 0 |
| 0 | 0 | 0 | 0 |
| 0 | 0 | 0 | 0 |
| 0 | 0 | 0 | 0 |
| 0 | 0 | 0 | 0 |
| 0 | 0 | 0 | 0 |
| 0 | 0 | 0 | 0 |
| 0 | 0 | 0 | 0 |
| 0 | 0 | 0 | 0 |
| 0 | 0 | 0 | 0 |
| 0 | 0 | 0 | 0 |
| 0 | 0 | 0 | 0 |
| 0 | 0 | 0 | 0 |
| 0 | 0 | 0 | 0 |
| 0 | 0 | 0 | 0 |
| 0 | 0 | 0 | 0 |
| 0 | 0 | 0 | 0 |
| 0 | 1 | 0 | 0 |
| 0 | 0 | 0 | 0 |
| 0 | 0 | 0 | 0 |
| 0 | 0 | 0 | 0 |
| 0 | 0 | 0 | 0 |

| | | | |
|---|---|---|---|
| 0 | 0 | 1 | 0 |
| 0 | 0 | 0 | 0 |
| 0 | 0 | 0 | 0 |
| 0 | 0 | 0 | 0 |
| 0 | 0 | 0 | 0 |
| 0 | 0 | 0 | 0 |
| 0 | 0 | 0 | 0 |
| 1 | 0 | 0 | 0 |
| 0 | 0 | 0 | 0 |
| 0 | 0 | 0 | 0 |
| 0 | 0 | 0 | 0 |
| 1 | 0 | 0 | 0 |
| 0 | 0 | 0 | 0 |
| 0 | 0 | 0 | 0 |
| 0 | 0 | 0 | 0 |
| 0 | 0 | 0 | 0 |
| 0 | 0 | 0 | 0 |
| 0 | 0 | 0 | 0 |
| 0 | 0 | 0 | 0 |
| 0 | 0 | 0 | 0 |
| 0 | 0 | 0 | 0 |
| 0 | 0 | 0 | 0 |
| 0 | 0 | 0 | 0 |
| 0 | 0 | 0 | 0 |
| 0 | 0 | 0 | 0 |
| 0 | 0 | 0 | 0 |
| 0 | 0 | 0 | 0 |
| 0 | 0 | 0 | 0 |
| 0 | 0 | 0 | 0 |
| 0 | 0 | 0 | 0 |
| 0 | 0 | 0 | 0 |
| 0 | 0 | 0 | 0 |
| 0 | 0 | 0 | 0 |
| 0 | 0 | 0 | 0 |
| 0 | 0 | 0 | 0 |
| 0 | 0 | 0 | 0 |
| 0 | 0 | 0 | 0 |
| 0 | 0 | 0 | 0 |
| 0 | 0 | 0 | 0 |
| 0 | 0 | 0 | 0 |
| 0 | 0 | 0 | 0 |
| 0 | 0 | 0 | 0 |
| 0 | 0 | 0 | 0 |
| 0 | 0 | 0 | 0 |
| 0 | 0 | 0 | 0 |
| 0 | 0 | 0 | 0 |
| 0 | 0 | 0 | 0 |
| 0 | 0 | 0 | 0 |
| 0 | 0 | 0 | 0 |
| 0 | 0 | 0 | 0 |
| 0 | 0 | 0 | 0 |
| 0 | 0 | 0 | 0 |
| 0 | 0 | 0 | 0 |
| 0 | 0 | 0 | 0 |

| | | | |
|---|---|---|---|
| 0 | 0 | 0 | 0 |
| 0 | 0 | 0 | 0 |
| 0 | 0 | 0 | 0 |
| 0 | 0 | 0 | 0 |
| 0 | 0 | 0 | 0 |
| 0 | 0 | 0 | 0 |
| 0 | 0 | 0 | 0 |
| 0 | 0 | 0 | 0 |
| 0 | 0 | 0 | 0 |
| 0 | 0 | 0 | 0 |
| 0 | 0 | 0 | 0 |
| 0 | 0 | 0 | 0 |
| 0 | 0 | 0 | 0 |
| 0 | 0 | 0 | 0 |
| 0 | 0 | 0 | 0 |
| 0 | 0 | 0 | 0 |
| 0 | 0 | 0 | 0 |
| 0 | 0 | 0 | 0 |
| 0 | 0 | 0 | 1 |
| 0 | 1 | 0 | 1 |
| 0 | 0 | 0 | 0 |
| 0 | 0 | 0 | 0 |
| 0 | 0 | 0 | 0 |
| 0 | 0 | 0 | 0 |
| 0 | 0 | 0 | 0 |
| 1 | 0 | 0 | 0 |
| 1 | 0 | 0 | 0 |
| 0 | 0 | 0 | 0 |
| 0 | 0 | 0 | 0 |
| 0 | 0 | 0 | 0 |
| 0 | 0 | 0 | 0 |
| 0 | 0 | 0 | 0 |
| 0 | 0 | 0 | 0 |
| 1 | 0 | 0 | 1 |
| 1 | 0 | 0 | 0 |
| 1 | 0 | 0 | 1 |
| 1 | 0 | 0 | 0 |
| 0 | 0 | 0 | 0 |
| 0 | 0 | 0 | 0 |
| 0 | 0 | 0 | 0 |
| 0 | 0 | 0 | 0 |
| 1 | 0 | 0 | 0 |
| 1 | 0 | 0 | 1 |
| 1 | 0 | 0 | 0 |
| 0 | 0 | 0 | 0 |
| 0 | 0 | 0 | 0 |
| 0 | 0 | 0 | 0 |
| 0 | 0 | 0 | 0 |
| 0 | 0 | 0 | 0 |
| 1 | 0 | 0 | 0 |
| 0 | 0 | 0 | 0 |
| 0 | 0 | 0 | 0 |
| 0 | 0 | 0 | 0 |
| 0 | 0 | 0 | 0 |

| | | | |
|---|---|---|---|
| 0 | 0 | 0 | 0 |
| 0 | 0 | 0 | 0 |
| 0 | 0 | 0 | 0 |
| 1 | 0 | 0 | 0 |
| 1 | 0 | 0 | 0 |
| 0 | 0 | 0 | 0 |
| 1 | 0 | 0 | 0 |
| 0 | 0 | 0 | 0 |
| 0 | 0 | 0 | 0 |
| 0 | 0 | 0 | 0 |
| 0 | 0 | 0 | 0 |
| 1 | 0 | 1 | 0 |
| 0 | 0 | 0 | 0 |
| 0 | 0 | 0 | 0 |
| 0 | 0 | 0 | 0 |
| 0 | 0 | 1 | 0 |
| 0 | 0 | 0 | 0 |
| 0 | 0 | 0 | 0 |
| 0 | 0 | 0 | 0 |
| 0 | 0 | 1 | 0 |
| 1 | 0 | 1 | 0 |
| 1 | 0 | 0 | 0 |
| 1 | 0 | 0 | 0 |
| 1 | 0 | 0 | 0 |
| 0 | 0 | 1 | 0 |
| 0 | 0 | 0 | 0 |
| 1 | 0 | 0 | 0 |
| 0 | 0 | 0 | 0 |
| 0 | 0 | 0 | 0 |
| 0 | 0 | 0 | 0 |
| 0 | 0 | 0 | 0 |
| 0 | 0 | 0 | 0 |
| 0 | 0 | 0 | 0 |
| 0 | 0 | 0 | 0 |
| 0 | 0 | 0 | 0 |
| 0 | 0 | 0 | 0 |
| 0 | 0 | 0 | 1 |
| 0 | 0 | 0 | 1 |
| 0 | 0 | 0 | 1 |
| 0 | 0 | 0 | 0 |
| 0 | 0 | 0 | 0 |
| 0 | 0 | 0 | 0 |
| 0 | 0 | 0 | 0 |
| 0 | 0 | 0 | 0 |
| 0 | 0 | 0 | 0 |
| 0 | 0 | 0 | 0 |
| 0 | 0 | 0 | 0 |
| 0 | 0 | 0 | 0 |
| 0 | 0 | 0 | 0 |
| 0 | 0 | 0 | 0 |
| 0 | 0 | 0 | 0 |
| 0 | 0 | 0 | 0 |
| 0 | 0 | 0 | 0 |
| 0 | 0 | 0 | 0 |

| | | | |
|---|---|---|---|
| 0 | 0 | 0 | 0 |
| 0 | 0 | 0 | 0 |
| 0 | 0 | 0 | 0 |
| 0 | 0 | 0 | 0 |
| 0 | 0 | 0 | 0 |
| 0 | 0 | 0 | 0 |
| 0 | 0 | 0 | 0 |
| 0 | 0 | 0 | 0 |
| 0 | 0 | 0 | 0 |
| 0 | 0 | 0 | 0 |
| 0 | 0 | 0 | 0 |
| 0 | 0 | 0 | 0 |
| 0 | 0 | 0 | 0 |
| 0 | 0 | 0 | 0 |
| 0 | 0 | 0 | 0 |
| 0 | 0 | 0 | 0 |
| 0 | 1 | 0 | 0 |
| 0 | 1 | 0 | 0 |
| 0 | 0 | 0 | 0 |
| 0 | 0 | 0 | 0 |
| 0 | 0 | 0 | 0 |
| 0 | 0 | 0 | 0 |
| 0 | 0 | 0 | 0 |
| 0 | 0 | 0 | 0 |
| 0 | 0 | 0 | 0 |
| 0 | 0 | 0 | 0 |
| 0 | 0 | 0 | 0 |
| 0 | 0 | 0 | 0 |
| 0 | 0 | 0 | 0 |
| 0 | 0 | 0 | 0 |
| 0 | 0 | 0 | 0 |
| 1 | 0 | 0 | 0 |
| 0 | 0 | 0 | 0 |
| 0 | 0 | 0 | 0 |
| 0 | 0 | 0 | 0 |
| 0 | 0 | 0 | 0 |
| 0 | 0 | 0 | 0 |
| 0 | 0 | 0 | 0 |
| 0 | 0 | 0 | 0 |
| 0 | 0 | 0 | 0 |
| 0 | 0 | 0 | 0 |
| 0 | 0 | 0 | 0 |
| 0 | 0 | 0 | 0 |
| 0 | 0 | 0 | 0 |
| 0 | 0 | 0 | 0 |
| 0 | 0 | 0 | 0 |
| 1 | 1 | 0 | 0 |
| 0 | 0 | 0 | 0 |
| 0 | 0 | 0 | 0 |
| 0 | 0 | 0 | 0 |
| 0 | 0 | 0 | 0 |
| 0 | 0 | 0 | 0 |
| 1 | 0 | 0 | 0 |
| 0 | 0 | 0 | 0 |

| | | | |
|---|---|---|---|
| 0 | 0 | 0 | 0 |
| 0 | 0 | 0 | 0 |
| 0 | 0 | 0 | 0 |
| 0 | 0 | 0 | 0 |
| 0 | 0 | 0 | 0 |
| 0 | 0 | 0 | 0 |
| 1 | 0 | 0 | 0 |
| 0 | 0 | 0 | 0 |
| 0 | 0 | 0 | 0 |
| 0 | 0 | 0 | 0 |
| 0 | 0 | 0 | 0 |
| 0 | 0 | 0 | 0 |
| 0 | 0 | 0 | 0 |
| 0 | 0 | 0 | 0 |
| 0 | 0 | 0 | 0 |
| 0 | 0 | 0 | 0 |
| 0 | 0 | 0 | 0 |
| 0 | 0 | 0 | 0 |
| 0 | 0 | 0 | 0 |
| 0 | 0 | 0 | 0 |
| 0 | 0 | 0 | 0 |
| 0 | 0 | 0 | 0 |
| 0 | 0 | 0 | 0 |
| 0 | 0 | 0 | 0 |
| 0 | 0 | 0 | 0 |
| 0 | 0 | 0 | 0 |
| 0 | 0 | 0 | 0 |
| 0 | 0 | 0 | 0 |
| 0 | 0 | 0 | 0 |
| 0 | 0 | 0 | 0 |
| 0 | 0 | 0 | 0 |
| 0 | 0 | 0 | 0 |
| 0 | 0 | 0 | 0 |
| 0 | 0 | 0 | 0 |
| 0 | 0 | 0 | 0 |
| 0 | 0 | 0 | 0 |
| 0 | 0 | 0 | 0 |
| 0 | 0 | 0 | 0 |
| 0 | 0 | 0 | 0 |
| 0 | 0 | 0 | 0 |
| 0 | 0 | 0 | 0 |
| 0 | 0 | 0 | 0 |
| 0 | 0 | 0 | 0 |
| 0 | 0 | 0 | 0 |
| 0 | 0 | 0 | 0 |
| 0 | 0 | 0 | 0 |
| 0 | 0 | 0 | 0 |
| 0 | 0 | 0 | 0 |
| 0 | 0 | 0 | 0 |
| 0 | 0 | 0 | 0 |
| 0 | 0 | 0 | 0 |
| 0 | 0 | 0 | 0 |
| 0 | 0 | 0 | 0 |
| 0 | 0 | 0 | 0 |

| | | | |
|---|---|---|---|
| 0 | 0 | 0 | 0 |
| 0 | 0 | 0 | 0 |
| 0 | 0 | 0 | 0 |
| 0 | 0 | 0 | 0 |
| 0 | 0 | 0 | 0 |
| 0 | 0 | 0 | 0 |
| 0 | 0 | 0 | 0 |
| 0 | 0 | 0 | 0 |
| 0 | 0 | 0 | 0 |
| 0 | 0 | 0 | 0 |
| 0 | 0 | 0 | 0 |
| 0 | 0 | 0 | 0 |
| 0 | 0 | 0 | 0 |
| 0 | 0 | 0 | 0 |
| 0 | 0 | 0 | 0 |
| 0 | 0 | 0 | 0 |
| 0 | 0 | 0 | 0 |
| 0 | 0 | 0 | 0 |
| 0 | 0 | 0 | 0 |
| 0 | 0 | 0 | 0 |
| 0 | 0 | 0 | 0 |
| 0 | 0 | 0 | 0 |
| 0 | 0 | 0 | 0 |
| 0 | 0 | 0 | 0 |
| 1 | 1 | 0 | 0 |
| 0 | 0 | 0 | 0 |
| 0 | 0 | 0 | 0 |
| 0 | 0 | 0 | 0 |
| 0 | 0 | 0 | 0 |
| 0 | 0 | 0 | 0 |
| 0 | 0 | 0 | 0 |
| 0 | 0 | 0 | 0 |
| 0 | 0 | 0 | 0 |
| 0 | 0 | 0 | 0 |
| 0 | 0 | 0 | 0 |
| 1 | 0 | 0 | 0 |
| 1 | 0 | 0 | 1 |
| 1 | 0 | 0 | 0 |
| 0 | 0 | 0 | 0 |
| 1 | 0 | 0 | 0 |
| 1 | 1 | 0 | 0 |
| 1 | 1 | 0 | 0 |
| 0 | 0 | 0 | 0 |
| 0 | 0 | 0 | 0 |
| 0 | 0 | 0 | 0 |
| 0 | 0 | 0 | 0 |
| 0 | 0 | 0 | 0 |
| 0 | 0 | 0 | 0 |
| 0 | 0 | 0 | 0 |
| 0 | 0 | 0 | 0 |
| 0 | 0 | 0 | 0 |
| 0 | 0 | 0 | 0 |
| 0 | 0 | 0 | 0 |
| 0 | 0 | 0 | 0 |

| | | | |
|---|---|---|---|
| 0 | 0 | 0 | 0 |
| 0 | 0 | 0 | 0 |
| 0 | 0 | 0 | 0 |
| 0 | 0 | 0 | 0 |
| 0 | 0 | 0 | 0 |
| 0 | 0 | 0 | 0 |
| 0 | 0 | 0 | 0 |
| 0 | 0 | 0 | 0 |
| 0 | 0 | 0 | 0 |
| 0 | 0 | 0 | 0 |
| 0 | 0 | 0 | 0 |
| 0 | 0 | 0 | 0 |
| 0 | 0 | 0 | 0 |
| 1 | 0 | 0 | 0 |
| 0 | 0 | 0 | 0 |
| 0 | 0 | 0 | 0 |
| 1 | 1 | 0 | 0 |
| 0 | 0 | 0 | 0 |
| 0 | 0 | 0 | 0 |
| 0 | 0 | 0 | 0 |
| 0 | 0 | 0 | 0 |
| 0 | 0 | 0 | 0 |
| 0 | 0 | 0 | 0 |
| 1 | 0 | 0 | 0 |
| 0 | 0 | 0 | 0 |
| 0 | 0 | 0 | 0 |
| 0 | 0 | 0 | 0 |
| 0 | 0 | 0 | 0 |
| 0 | 0 | 0 | 0 |
| 0 | 0 | 0 | 0 |
| 0 | 0 | 0 | 0 |
| 1 | 0 | 0 | 0 |
| 0 | 0 | 0 | 0 |
| 0 | 0 | 0 | 0 |
| 0 | 0 | 0 | 0 |
| 0 | 0 | 0 | 0 |
| 0 | 0 | 0 | 0 |
| 0 | 0 | 0 | 0 |
| 0 | 0 | 0 | 0 |
| 0 | 0 | 0 | 0 |
| 0 | 0 | 0 | 0 |
| 0 | 0 | 0 | 0 |
| 0 | 0 | 0 | 0 |
| 0 | 0 | 0 | 0 |
| 0 | 0 | 0 | 0 |
| 0 | 0 | 0 | 0 |
| 0 | 0 | 0 | 0 |
| 0 | 0 | 0 | 0 |
| 0 | 0 | 0 | 0 |
| 0 | 0 | 0 | 0 |
| 0 | 0 | 0 | 0 |
| 0 | 0 | 0 | 0 |
| 0 | 0 | 0 | 0 |
| 0 | 0 | 0 | 0 |

| | | | |
|---|---|---|---|
| 0 | 0 | 0 | 0 |
| 0 | 0 | 0 | 0 |
| 0 | 0 | 0 | 0 |
| 0 | 0 | 0 | 0 |
| 0 | 0 | 0 | 0 |
| 0 | 0 | 0 | 0 |
| 0 | 0 | 0 | 0 |
| 0 | 0 | 0 | 0 |
| 0 | 0 | 0 | 0 |
| 0 | 0 | 0 | 0 |
| 0 | 0 | 0 | 0 |
| 0 | 0 | 0 | 0 |
| 0 | 0 | 0 | 0 |
| 0 | 0 | 0 | 0 |
| 0 | 0 | 0 | 0 |
| 0 | 0 | 0 | 0 |
| 0 | 0 | 0 | 0 |
| 1 | 0 | 0 | 0 |
| 0 | 0 | 0 | 0 |
| 0 | 0 | 0 | 0 |
| 0 | 0 | 0 | 0 |
| 0 | 0 | 0 | 0 |
| 0 | 0 | 0 | 0 |
| 0 | 0 | 0 | 0 |
| 1 | 0 | 0 | 0 |
| 0 | 0 | 0 | 0 |
| 0 | 0 | 0 | 0 |
| 0 | 0 | 0 | 0 |
| 0 | 0 | 0 | 0 |
| 0 | 0 | 0 | 0 |
| 0 | 0 | 0 | 0 |
| 0 | 0 | 0 | 0 |
| 0 | 0 | 0 | 0 |
| 0 | 0 | 0 | 0 |
| 0 | 0 | 0 | 0 |
| 0 | 0 | 0 | 0 |
| 0 | 0 | 0 | 0 |
| 0 | 0 | 0 | 0 |
| 0 | 0 | 0 | 0 |
| 0 | 0 | 0 | 0 |
| 0 | 0 | 0 | 0 |
| 0 | 0 | 0 | 0 |
| 0 | 0 | 0 | 0 |
| 0 | 0 | 0 | 0 |
| 0 | 0 | 0 | 0 |
| 0 | 0 | 0 | 0 |
| 0 | 0 | 0 | 0 |
| 0 | 0 | 0 | 0 |
| 0 | 0 | 0 | 0 |
| 0 | 0 | 0 | 0 |
| 0 | 0 | 0 | 0 |
| 0 | 0 | 0 | 0 |
| 0 | 0 | 0 | 0 |
| 0 | 0 | 0 | 0 |

| | | | |
|---|---|---|---|
| 0 | 0 | 0 | 0 |
| 0 | 0 | 0 | 0 |
| 0 | 0 | 0 | 0 |
| 0 | 0 | 0 | 0 |
| 0 | 0 | 0 | 0 |
| 0 | 0 | 0 | 0 |
| 0 | 0 | 0 | 0 |
| 0 | 0 | 0 | 0 |
| 0 | 0 | 0 | 0 |
| 0 | 0 | 0 | 0 |
| 0 | 0 | 0 | 0 |
| 0 | 0 | 0 | 0 |
| 0 | 0 | 0 | 0 |
| 0 | 0 | 0 | 0 |
| 1 | 1 | 0 | 0 |
| 1 | 1 | 0 | 0 |
| 0 | 0 | 0 | 0 |
| 0 | 0 | 0 | 0 |
| 0 | 0 | 0 | 0 |
| 0 | 0 | 0 | 0 |
| 0 | 0 | 0 | 0 |
| 0 | 0 | 0 | 0 |
| 0 | 0 | 0 | 0 |
| 0 | 0 | 0 | 0 |
| 0 | 0 | 0 | 0 |
| 0 | 0 | 0 | 0 |
| 0 | 0 | 0 | 0 |
| 1 | 0 | 0 | 0 |
| 0 | 0 | 0 | 0 |
| 1 | 0 | 0 | 0 |
| 0 | 0 | 0 | 0 |
| 0 | 0 | 0 | 0 |
| 0 | 0 | 0 | 0 |
| 0 | 0 | 0 | 0 |
| 0 | 0 | 0 | 0 |
| 0 | 0 | 0 | 0 |
| 1 | 0 | 0 | 0 |
| 0 | 0 | 0 | 0 |
| 0 | 0 | 0 | 0 |
| 0 | 0 | 0 | 0 |
| 0 | 0 | 0 | 0 |
| 0 | 0 | 0 | 0 |
| 0 | 0 | 0 | 0 |
| 0 | 0 | 0 | 0 |
| 0 | 0 | 0 | 0 |
| 0 | 0 | 0 | 0 |
| 0 | 0 | 0 | 0 |
| 0 | 0 | 0 | 0 |
| 0 | 0 | 0 | 0 |
| 0 | 0 | 0 | 0 |
| 0 | 0 | 0 | 0 |
| 0 | 0 | 0 | 0 |
| 0 | 0 | 0 | 0 |
| 0 | 0 | 0 | 0 |

| | | | |
|---|---|---|---|
| 0 | 0 | 0 | 0 |
| 0 | 0 | 0 | 0 |
| 0 | 0 | 0 | 0 |
| 0 | 0 | 0 | 0 |
| 0 | 0 | 0 | 0 |
| 0 | 0 | 0 | 0 |
| 0 | 0 | 0 | 0 |
| 0 | 0 | 0 | 0 |
| 0 | 0 | 0 | 0 |
| 0 | 0 | 0 | 0 |
| 0 | 0 | 0 | 0 |
| 0 | 0 | 0 | 0 |
| 0 | 0 | 0 | 0 |
| 0 | 0 | 0 | 0 |
| 0 | 0 | 0 | 0 |
| 0 | 0 | 0 | 0 |
| 0 | 0 | 0 | 0 |
| 0 | 0 | 0 | 0 |
| 0 | 0 | 0 | 0 |
| 0 | 0 | 0 | 0 |
| 0 | 0 | 0 | 0 |
| 0 | 0 | 0 | 0 |
| 0 | 0 | 0 | 0 |
| 0 | 0 | 0 | 0 |
| 0 | 0 | 0 | 0 |
| 0 | 0 | 0 | 0 |
| 0 | 0 | 0 | 0 |
| 0 | 0 | 0 | 0 |
| 0 | 0 | 0 | 0 |
| 0 | 0 | 0 | 0 |
| 0 | 0 | 0 | 0 |
| 0 | 0 | 0 | 0 |
| 0 | 0 | 0 | 0 |
| 0 | 0 | 0 | 0 |
| 0 | 0 | 0 | 0 |
| 0 | 0 | 0 | 0 |
| 0 | 0 | 0 | 0 |
| 0 | 0 | 0 | 0 |
| 0 | 0 | 0 | 0 |
| 0 | 0 | 0 | 0 |
| 0 | 0 | 0 | 0 |
| 0 | 0 | 0 | 0 |
| 0 | 0 | 0 | 0 |
| 0 | 0 | 0 | 0 |
| 0 | 0 | 0 | 0 |
| 0 | 0 | 0 | 0 |
| 0 | 0 | 0 | 0 |
| 0 | 0 | 0 | 0 |
| 1 | 0 | 0 | 1 |
| 0 | 0 | 0 | 0 |
| 1 | 0 | 0 | 0 |
| 1 | 0 | 0 | 0 |
| 0 | 0 | 0 | 0 |
| 0 | 0 | 0 | 1 |

| | | | |
|---|---|---|---|
| 0 | 0 | 0 | 0 |
| 0 | 0 | 0 | 0 |
| 0 | 0 | 0 | 0 |
| 0 | 0 | 0 | 0 |
| 0 | 0 | 0 | 0 |
| 0 | 0 | 0 | 0 |
| 0 | 0 | 0 | 0 |
| 0 | 0 | 0 | 0 |
| 0 | 0 | 0 | 0 |
| 0 | 0 | 0 | 0 |
| 0 | 0 | 0 | 0 |
| 0 | 0 | 0 | 0 |
| 0 | 0 | 0 | 0 |
| 0 | 0 | 0 | 0 |
| 0 | 0 | 0 | 0 |
| 0 | 0 | 0 | 0 |
| 0 | 0 | 0 | 0 |
| 0 | 0 | 0 | 0 |
| 0 | 0 | 0 | 0 |
| 0 | 0 | 0 | 0 |
| 0 | 0 | 0 | 0 |
| 0 | 0 | 0 | 0 |
| 0 | 0 | 0 | 0 |
| 0 | 0 | 0 | 0 |
| 0 | 0 | 0 | 0 |
| 0 | 0 | 0 | 0 |
| 0 | 0 | 1 | 0 |
| 0 | 0 | 1 | 0 |
| 0 | 0 | 1 | 0 |
| 0 | 0 | 1 | 0 |
| 0 | 0 | 0 | 0 |
| 0 | 0 | 0 | 0 |
| 0 | 0 | 0 | 0 |
| 1 | 0 | 0 | 0 |
| 1 | 0 | 0 | 0 |
| 0 | 0 | 0 | 0 |
| 0 | 0 | 0 | 0 |
| 0 | 0 | 0 | 0 |
| 0 | 0 | 0 | 0 |
| 1 | 0 | 0 | 0 |
| 0 | 0 | 0 | 0 |
| 0 | 0 | 0 | 0 |
| 0 | 0 | 0 | 0 |
| 0 | 0 | 0 | 0 |
| 0 | 0 | 0 | 0 |
| 0 | 1 | 0 | 0 |
| 0 | 0 | 0 | 0 |
| 0 | 0 | 0 | 0 |
| 0 | 0 | 0 | 0 |
| 0 | 0 | 0 | 0 |
| 0 | 0 | 0 | 0 |
| 0 | 0 | 0 | 0 |
| 0 | 0 | 0 | 0 |
| 0 | 0 | 0 | 0 |

| | | | |
|---|---|---|---|
| 0 | 0 | 0 | 0 |
| 0 | 0 | 0 | 0 |
| 0 | 0 | 0 | 0 |
| 0 | 0 | 0 | 0 |
| 0 | 0 | 0 | 0 |
| 0 | 0 | 0 | 1 |
| 0 | 0 | 0 | 0 |
| 0 | 0 | 0 | 0 |
| 0 | 0 | 0 | 0 |
| 0 | 0 | 0 | 0 |
| 0 | 0 | 0 | 0 |
| 0 | 0 | 0 | 0 |
| 0 | 0 | 0 | 0 |
| 0 | 1 | 0 | 0 |
| 0 | 1 | 0 | 0 |
| 0 | 1 | 0 | 0 |
| 0 | 1 | 0 | 0 |
| 0 | 1 | 0 | 0 |
| 0 | 0 | 0 | 0 |
| 0 | 0 | 0 | 0 |
| 0 | 0 | 0 | 0 |
| 0 | 0 | 0 | 0 |
| 0 | 0 | 0 | 0 |
| 0 | 0 | 0 | 0 |
| 0 | 0 | 0 | 0 |
| 0 | 0 | 0 | 0 |
| 0 | 0 | 0 | 0 |
| 0 | 0 | 0 | 0 |
| 0 | 0 | 0 | 0 |
| 0 | 0 | 0 | 0 |
| 0 | 0 | 0 | 0 |
| 0 | 0 | 0 | 0 |
| 0 | 0 | 0 | 0 |
| 0 | 0 | 0 | 0 |
| 0 | 0 | 0 | 0 |
| 0 | 0 | 0 | 0 |
| 0 | 0 | 0 | 0 |
| 0 | 0 | 0 | 0 |
| 0 | 0 | 0 | 0 |
| 0 | 0 | 0 | 0 |
| 0 | 0 | 0 | 0 |
| 0 | 0 | 0 | 0 |
| 0 | 0 | 0 | 0 |
| 0 | 0 | 0 | 0 |
| 0 | 0 | 0 | 0 |
| 0 | 0 | 0 | 0 |
| 0 | 0 | 0 | 0 |
| 0 | 0 | 0 | 0 |
| 0 | 0 | 0 | 0 |
| 0 | 0 | 0 | 0 |
| 0 | 0 | 0 | 0 |
| 0 | 0 | 0 | 0 |
| 0 | 0 | 0 | 0 |
| 0 | 0 | 0 | 0 |

| | | | |
|---|---|---|---|
| 0 | 0 | 0 | 0 |
| 0 | 0 | 0 | 0 |
| 0 | 0 | 0 | 0 |
| 1 | 0 | 0 | 0 |
| 0 | 0 | 0 | 0 |
| 0 | 0 | 0 | 0 |
| 0 | 0 | 0 | 0 |
| 0 | 0 | 0 | 0 |
| 0 | 0 | 0 | 0 |
| 0 | 0 | 0 | 0 |
| 0 | 0 | 0 | 0 |
| 0 | 0 | 0 | 0 |
| 0 | 0 | 0 | 0 |
| 0 | 0 | 0 | 0 |
| 0 | 0 | 0 | 0 |
| 0 | 0 | 0 | 0 |
| 0 | 0 | 0 | 0 |
| 0 | 0 | 0 | 0 |
| 0 | 0 | 0 | 0 |
| 0 | 0 | 0 | 0 |
| 1 | 1 | 0 | 0 |
| 1 | 1 | 0 | 0 |
| 0 | 0 | 0 | 0 |
| 0 | 1 | 0 | 0 |
| 0 | 0 | 0 | 0 |
| 0 | 0 | 0 | 0 |
| 0 | 0 | 0 | 0 |
| 0 | 0 | 0 | 0 |
| 0 | 0 | 0 | 0 |
| 0 | 0 | 0 | 0 |
| 0 | 0 | 0 | 0 |
| 0 | 0 | 0 | 0 |
| 0 | 0 | 0 | 0 |
| 0 | 0 | 0 | 0 |
| 0 | 0 | 0 | 0 |
| 0 | 0 | 0 | 0 |
| 0 | 0 | 0 | 0 |
| 0 | 0 | 0 | 0 |
| 0 | 0 | 0 | 0 |
| 1 | 0 | 0 | 0 |
| 1 | 0 | 0 | 0 |
| 0 | 0 | 0 | 0 |
| 0 | 0 | 0 | 0 |
| 1 | 0 | 0 | 0 |
| 0 | 0 | 0 | 0 |
| 0 | 0 | 0 | 0 |
| 0 | 0 | 0 | 0 |
| 0 | 0 | 0 | 0 |
| 0 | 0 | 0 | 0 |
| 0 | 0 | 0 | 0 |
| 0 | 0 | 0 | 0 |
| 0 | 0 | 0 | 0 |
| 0 | 0 | 0 | 0 |
| 0 | 0 | 0 | 0 |

| | | | |
|---|---|---|---|
| 0 | 0 | 0 | 0 |
| 0 | 0 | 0 | 0 |
| 0 | 0 | 0 | 0 |
| 0 | 0 | 0 | 0 |
| 0 | 0 | 0 | 0 |
| 1 | 0 | 0 | 0 |
| 0 | 0 | 0 | 0 |
| 0 | 0 | 0 | 0 |
| 0 | 1 | 0 | 0 |
| 1 | 1 | 0 | 0 |
| 0 | 0 | 0 | 0 |
| 1 | 1 | 0 | 0 |
| 1 | 0 | 0 | 0 |
| 0 | 0 | 0 | 0 |
| 0 | 0 | 0 | 0 |
| 0 | 0 | 0 | 0 |
| 1 | 1 | 0 | 0 |
| 0 | 0 | 0 | 0 |
| 0 | 0 | 0 | 0 |
| 1 | 0 | 0 | 0 |
| 0 | 0 | 0 | 0 |
| 0 | 0 | 0 | 0 |
| 0 | 0 | 0 | 0 |
| 0 | 0 | 0 | 0 |
| 0 | 0 | 0 | 0 |
| 0 | 0 | 0 | 0 |
| 0 | 0 | 0 | 0 |
| 0 | 0 | 0 | 0 |
| 0 | 0 | 0 | 0 |
| 0 | 0 | 0 | 0 |
| 1 | 0 | 0 | 0 |
| 0 | 0 | 0 | 0 |
| 0 | 0 | 0 | 0 |
| 0 | 0 | 0 | 0 |
| 0 | 0 | 0 | 0 |
| 1 | 0 | 0 | 0 |
| 0 | 0 | 0 | 0 |
| 0 | 0 | 0 | 0 |
| 0 | 0 | 0 | 0 |
| 0 | 0 | 0 | 0 |
| 0 | 0 | 0 | 0 |
| 0 | 0 | 0 | 0 |
| 0 | 0 | 0 | 0 |
| 0 | 0 | 0 | 0 |
| 0 | 0 | 0 | 0 |
| 0 | 0 | 0 | 0 |
| 0 | 0 | 0 | 0 |
| 0 | 0 | 0 | 0 |
| 0 | 0 | 0 | 0 |
| 0 | 0 | 0 | 0 |
| 0 | 0 | 0 | 0 |
| 0 | 0 | 0 | 0 |
| 0 | 0 | 0 | 0 |
| 0 | 0 | 0 | 0 |

| | | | |
|---|---|---|---|
| 0 | 0 | 0 | 0 |
| 0 | 0 | 0 | 0 |
| 0 | 0 | 0 | 0 |
| 0 | 0 | 0 | 0 |
| 0 | 0 | 0 | 0 |
| 0 | 0 | 0 | 0 |
| 0 | 0 | 0 | 0 |
| 0 | 0 | 0 | 0 |
| 0 | 0 | 0 | 0 |
| 0 | 0 | 0 | 0 |
| 0 | 0 | 0 | 0 |
| 0 | 0 | 0 | 0 |
| 0 | 0 | 0 | 0 |
| 0 | 0 | 0 | 0 |
| 0 | 0 | 0 | 1 |
| 0 | 0 | 0 | 0 |
| 0 | 0 | 0 | 0 |
| 0 | 0 | 0 | 0 |
| 0 | 0 | 0 | 0 |
| 0 | 0 | 0 | 0 |
| 0 | 0 | 0 | 0 |
| 0 | 0 | 0 | 0 |
| 0 | 0 | 0 | 0 |
| 0 | 0 | 0 | 0 |
| 0 | 0 | 0 | 0 |
| 0 | 0 | 0 | 0 |
| 0 | 0 | 0 | 0 |
| 0 | 0 | 0 | 0 |
| 0 | 0 | 0 | 0 |
| 0 | 0 | 0 | 0 |
| 0 | 0 | 0 | 0 |
| 0 | 0 | 0 | 0 |
| 0 | 0 | 0 | 0 |
| 0 | 0 | 0 | 0 |
| 0 | 0 | 0 | 0 |
| 0 | 0 | 0 | 0 |
| 0 | 0 | 0 | 0 |
| 0 | 0 | 0 | 0 |
| 0 | 0 | 0 | 0 |
| 0 | 0 | 1 | 0 |
| 0 | 0 | 0 | 0 |
| 0 | 0 | 0 | 0 |
| 0 | 0 | 0 | 0 |
| 1 | 0 | 0 | 0 |
| 0 | 0 | 0 | 0 |
| 0 | 0 | 0 | 0 |
| 0 | 0 | 0 | 0 |
| 0 | 0 | 0 | 0 |
| 1 | 0 | 0 | 1 |
| 0 | 0 | 0 | 0 |
| 0 | 0 | 0 | 0 |
| 0 | 0 | 0 | 0 |
| 0 | 0 | 0 | 0 |
| 0 | 0 | 0 | 0 |

| | | | |
|---|---|---|---|
| 0 | 0 | 0 | 0 |
| 0 | 0 | 0 | 0 |
| 0 | 0 | 0 | 0 |
| 0 | 0 | 0 | 0 |
| 0 | 0 | 0 | 0 |
| 0 | 0 | 0 | 0 |
| 0 | 0 | 0 | 0 |
| 0 | 0 | 0 | 0 |
| 0 | 0 | 0 | 0 |
| 1 | 0 | 0 | 0 |
| 0 | 0 | 0 | 0 |
| 0 | 0 | 0 | 0 |
| 0 | 0 | 0 | 0 |
| 0 | 0 | 0 | 0 |
| 1 | 0 | 0 | 0 |
| 0 | 0 | 0 | 0 |
| 0 | 0 | 0 | 0 |
| 0 | 0 | 0 | 0 |
| 0 | 0 | 0 | 0 |
| 0 | 0 | 0 | 0 |
| 0 | 0 | 0 | 0 |
| 0 | 0 | 0 | 0 |
| 0 | 0 | 0 | 0 |
| 0 | 0 | 0 | 0 |
| 1 | 0 | 0 | 0 |
| 0 | 0 | 0 | 0 |
| 0 | 0 | 0 | 0 |
| 0 | 0 | 0 | 0 |
| 0 | 0 | 0 | 0 |
| 0 | 0 | 0 | 0 |
| 0 | 0 | 0 | 0 |
| 0 | 0 | 0 | 0 |
| 0 | 0 | 0 | 0 |
| 0 | 0 | 0 | 0 |
| 1 | 0 | 0 | 0 |
| 1 | 0 | 0 | 0 |
| 0 | 0 | 0 | 0 |
| 0 | 0 | 0 | 0 |
| 1 | 0 | 0 | 0 |
| 0 | 0 | 1 | 0 |
| 0 | 0 | 1 | 0 |
| 0 | 0 | 1 | 0 |
| 0 | 0 | 1 | 0 |
| 0 | 0 | 0 | 0 |
| 1 | 0 | 0 | 0 |
| 0 | 0 | 0 | 0 |
| 0 | 0 | 0 | 0 |
| 0 | 0 | 0 | 0 |
| 0 | 0 | 1 | 0 |
| 0 | 0 | 0 | 0 |
| 0 | 0 | 0 | 0 |
| 0 | 0 | 0 | 0 |
| 0 | 0 | 0 | 0 |
| 0 | 0 | 0 | 0 |

| | | | |
|---|---|---|---|
| 0 | 0 | 0 | 0 |
| 0 | 0 | 0 | 0 |
| 0 | 0 | 0 | 0 |
| 0 | 0 | 0 | 0 |
| 0 | 0 | 0 | 0 |
| 0 | 0 | 0 | 0 |
| 0 | 0 | 0 | 0 |
| 0 | 0 | 0 | 0 |
| 0 | 0 | 0 | 0 |
| 0 | 0 | 0 | 0 |
| 0 | 1 | 0 | 0 |
| 0 | 0 | 0 | 0 |
| 0 | 0 | 0 | 0 |
| 0 | 0 | 0 | 0 |
| 0 | 0 | 0 | 0 |
| 0 | 1 | 0 | 0 |
| 0 | 0 | 0 | 0 |
| 0 | 0 | 0 | 0 |
| 0 | 0 | 0 | 0 |
| 0 | 0 | 0 | 0 |
| 0 | 0 | 0 | 0 |
| 0 | 0 | 0 | 0 |
| 0 | 0 | 0 | 0 |
| 0 | 0 | 0 | 0 |
| 0 | 0 | 0 | 0 |
| 0 | 0 | 0 | 0 |
| 0 | 0 | 0 | 0 |
| 0 | 0 | 0 | 0 |
| 0 | 0 | 0 | 0 |
| 0 | 0 | 0 | 0 |
| 0 | 0 | 0 | 0 |
| 0 | 0 | 0 | 0 |
| 0 | 0 | 0 | 0 |
| 0 | 0 | 0 | 0 |
| 0 | 0 | 0 | 0 |
| 0 | 0 | 0 | 0 |
| 0 | 0 | 0 | 0 |
| 0 | 0 | 0 | 0 |
| 0 | 0 | 0 | 0 |
| 0 | 0 | 0 | 0 |
| 0 | 0 | 0 | 0 |
| 0 | 0 | 0 | 0 |
| 0 | 0 | 0 | 0 |
| 0 | 0 | 0 | 0 |
| 1 | 1 | 0 | 0 |
| 0 | 0 | 0 | 0 |
| 0 | 0 | 0 | 0 |
| 0 | 0 | 0 | 0 |
| 0 | 0 | 0 | 0 |
| 0 | 0 | 0 | 0 |
| 0 | 0 | 0 | 0 |
| 0 | 0 | 0 | 0 |
| 0 | 0 | 0 | 0 |
| 0 | 0 | 0 | 0 |

| | | | |
|---|---|---|---|
| 0 | 0 | 0 | 0 |
| 0 | 0 | 0 | 0 |
| 0 | 0 | 1 | 0 |
| 1 | 0 | 0 | 0 |
| 0 | 0 | 0 | 0 |
| 0 | 0 | 0 | 0 |
| 0 | 0 | 0 | 0 |
| 0 | 0 | 0 | 0 |
| 0 | 0 | 0 | 0 |
| 0 | 0 | 0 | 0 |
| 0 | 0 | 0 | 0 |
| 0 | 0 | 0 | 0 |
| 1 | 0 | 0 | 0 |
| 0 | 0 | 0 | 0 |
| 0 | 0 | 0 | 0 |
| 0 | 0 | 0 | 0 |
| 0 | 0 | 0 | 0 |
| 0 | 0 | 0 | 0 |
| 0 | 0 | 0 | 0 |
| 0 | 0 | 0 | 0 |
| 0 | 0 | 0 | 0 |
| 0 | 0 | 0 | 0 |
| 0 | 0 | 0 | 0 |
| 0 | 0 | 0 | 0 |
| 0 | 0 | 0 | 0 |
| 0 | 0 | 0 | 0 |
| 0 | 0 | 0 | 0 |
| 0 | 0 | 0 | 0 |
| 0 | 0 | 0 | 0 |
| 0 | 0 | 0 | 0 |
| 0 | 0 | 0 | 0 |
| 0 | 0 | 0 | 0 |
| 0 | 0 | 0 | 0 |
| 0 | 0 | 0 | 0 |
| 0 | 0 | 0 | 0 |
| 0 | 0 | 0 | 0 |
| 0 | 0 | 0 | 0 |
| 0 | 0 | 0 | 0 |
| 0 | 0 | 0 | 0 |
| 0 | 0 | 0 | 0 |
| 0 | 1 | 0 | 0 |
| 1 | 1 | 0 | 0 |
| 0 | 0 | 0 | 0 |
| 0 | 0 | 0 | 0 |
| 0 | 0 | 0 | 0 |
| 0 | 0 | 0 | 0 |
| 0 | 0 | 0 | 0 |
| 0 | 0 | 0 | 0 |
| 0 | 1 | 0 | 0 |
| 0 | 0 | 0 | 0 |
| 0 | 0 | 0 | 0 |
| 0 | 0 | 0 | 0 |
| 0 | 0 | 0 | 1 |
| 0 | 0 | 0 | 0 |

| | | | |
|---|---|---|---|
| 0 | 0 | 0 | 1 |
| 0 | 0 | 0 | 1 |
| 0 | 0 | 0 | 1 |
| 0 | 0 | 0 | 0 |
| 0 | 0 | 0 | 0 |
| 0 | 0 | 0 | 0 |
| 0 | 0 | 0 | 0 |
| 1 | 0 | 0 | 0 |
| 0 | 0 | 0 | 0 |
| 1 | 1 | 0 | 0 |
| 0 | 1 | 0 | 0 |
| 0 | 0 | 0 | 0 |
| 1 | 1 | 0 | 0 |
| 0 | 0 | 0 | 0 |
| 0 | 0 | 0 | 0 |
| 0 | 0 | 0 | 0 |
| 0 | 0 | 0 | 0 |
| 1 | 0 | 0 | 0 |
| 0 | 0 | 0 | 0 |
| 0 | 0 | 0 | 0 |
| 0 | 0 | 0 | 0 |
| 0 | 0 | 0 | 0 |
| 0 | 0 | 0 | 0 |
| 0 | 0 | 0 | 0 |
| 0 | 0 | 0 | 0 |
| 0 | 0 | 0 | 0 |
| 0 | 0 | 0 | 0 |
| 0 | 0 | 0 | 0 |
| 0 | 0 | 0 | 0 |
| 0 | 0 | 0 | 0 |
| 0 | 0 | 0 | 0 |
| 1 | 0 | 0 | 0 |
| 0 | 0 | 0 | 0 |
| 0 | 0 | 0 | 0 |
| 0 | 0 | 0 | 0 |
| 0 | 0 | 0 | 0 |
| 0 | 0 | 0 | 0 |
| 0 | 0 | 0 | 0 |
| 0 | 0 | 0 | 0 |
| 0 | 0 | 0 | 0 |
| 0 | 0 | 0 | 0 |
| 1 | 1 | 0 | 0 |
| 0 | 0 | 0 | 0 |
| 0 | 0 | 0 | 0 |
| 1 | 0 | 0 | 0 |
| 0 | 1 | 0 | 0 |
| 0 | 0 | 0 | 0 |
| 0 | 0 | 0 | 0 |
| 0 | 1 | 0 | 0 |
| 1 | 1 | 0 | 0 |
| 1 | 1 | 0 | 0 |
| 1 | 1 | 0 | 0 |
| 1 | 1 | 0 | 0 |
| 1 | 1 | 0 | 0 |

| | | | |
|---|---|---|---|
| 0 | 0 | 0 | 0 |
| 0 | 0 | 0 | 0 |
| 0 | 0 | 0 | 0 |
| 0 | 0 | 0 | 0 |
| 0 | 0 | 0 | 0 |
| 1 | 0 | 0 | 0 |
| 1 | 0 | 0 | 0 |
| 0 | 0 | 0 | 0 |
| 0 | 0 | 0 | 0 |
| 1 | 0 | 0 | 0 |
| 0 | 0 | 0 | 0 |
| 0 | 0 | 1 | 0 |
| 0 | 0 | 0 | 0 |
| 0 | 0 | 0 | 0 |
| 0 | 0 | 0 | 0 |
| 0 | 0 | 0 | 0 |
| 0 | 0 | 0 | 0 |
| 0 | 0 | 0 | 0 |
| 0 | 0 | 0 | 0 |
| 0 | 0 | 0 | 0 |
| 1 | 1 | 0 | 0 |
| 1 | 0 | 0 | 0 |
| 0 | 0 | 0 | 0 |
| 0 | 0 | 1 | 0 |
| 0 | 0 | 0 | 0 |
| 0 | 0 | 1 | 0 |
| 0 | 0 | 0 | 0 |
| 0 | 0 | 0 | 0 |
| 0 | 0 | 0 | 0 |
| 0 | 0 | 0 | 0 |
| 0 | 0 | 0 | 0 |
| 0 | 0 | 0 | 0 |
| 0 | 0 | 0 | 0 |
| 0 | 0 | 0 | 0 |
| 0 | 0 | 0 | 0 |
| 0 | 0 | 0 | 0 |
| 1 | 0 | 0 | 0 |
| 0 | 0 | 0 | 0 |
| 1 | 0 | 0 | 0 |
| 1 | 0 | 0 | 0 |
| 1 | 0 | 0 | 0 |
| 0 | 0 | 0 | 0 |
| 0 | 0 | 0 | 0 |
| 0 | 0 | 0 | 0 |
| 0 | 0 | 0 | 0 |
| 0 | 0 | 0 | 0 |
| 0 | 0 | 0 | 0 |
| 1 | 0 | 1 | 0 |
| 0 | 0 | 0 | 0 |
| 0 | 0 | 0 | 0 |
| 0 | 0 | 0 | 0 |
| 0 | 0 | 0 | 0 |
| 1 | 1 | 0 | 0 |
| 0 | 0 | 0 | 0 |

| | | | |
|---|---|---|---|
| 0 | 0 | 0 | 0 |
| 0 | 1 | 0 | 0 |
| 0 | 0 | 0 | 0 |
| 0 | 0 | 0 | 0 |
| 0 | 0 | 0 | 0 |
| 0 | 0 | 0 | 0 |
| 0 | 0 | 0 | 0 |
| 0 | 0 | 0 | 0 |
| 0 | 0 | 0 | 0 |
| 0 | 0 | 0 | 0 |
| 0 | 0 | 0 | 0 |
| 0 | 0 | 0 | 0 |
| 0 | 0 | 0 | 0 |
| 0 | 0 | 0 | 0 |
| 0 | 0 | 0 | 0 |
| 0 | 0 | 0 | 0 |
| 0 | 0 | 0 | 0 |
| 0 | 0 | 0 | 0 |
| 0 | 0 | 0 | 0 |
| 0 | 0 | 0 | 0 |
| 1 | 0 | 0 | 0 |
| 0 | 0 | 0 | 0 |
| 0 | 0 | 0 | 0 |
| 0 | 0 | 0 | 0 |
| 1 | 0 | 0 | 0 |
| 0 | 0 | 0 | 0 |
| 0 | 0 | 0 | 0 |
| 0 | 0 | 0 | 1 |
| 0 | 0 | 0 | 0 |
| 0 | 0 | 0 | 0 |
| 0 | 0 | 0 | 0 |
| 0 | 0 | 0 | 0 |
| 0 | 0 | 0 | 0 |
| 0 | 0 | 0 | 0 |
| 0 | 0 | 0 | 0 |
| 0 | 0 | 0 | 0 |
| 0 | 0 | 0 | 0 |
| 0 | 0 | 0 | 0 |
| 0 | 0 | 0 | 0 |
| 0 | 0 | 0 | 0 |
| 0 | 0 | 0 | 0 |
| 0 | 0 | 0 | 0 |
| 0 | 0 | 0 | 0 |
| 0 | 0 | 0 | 0 |
| 0 | 0 | 0 | 0 |
| 0 | 0 | 0 | 0 |
| 0 | 0 | 0 | 0 |
| 0 | 0 | 0 | 0 |
| 0 | 0 | 0 | 0 |
| 0 | 0 | 0 | 0 |
| 0 | 0 | 0 | 0 |
| 0 | 0 | 0 | 0 |
| 0 | 0 | 0 | 0 |
| 0 | 0 | 0 | 0 |

| | | | |
|---|---|---|---|
| 0 | 0 | 0 | 0 |
| 0 | 0 | 0 | 0 |
| 0 | 0 | 0 | 0 |
| 0 | 0 | 0 | 0 |
| 0 | 0 | 0 | 0 |
| 0 | 0 | 0 | 0 |
| 0 | 0 | 0 | 0 |
| 0 | 0 | 0 | 0 |
| 0 | 0 | 0 | 0 |
| 0 | 0 | 0 | 0 |
| 0 | 0 | 0 | 0 |
| 0 | 0 | 0 | 0 |
| 0 | 0 | 0 | 0 |
| 0 | 0 | 0 | 0 |
| 0 | 0 | 0 | 0 |
| 1 | 0 | 0 | 0 |
| 0 | 0 | 0 | 0 |
| 0 | 0 | 0 | 0 |
| 0 | 1 | 0 | 0 |
| 0 | 0 | 0 | 0 |
| 0 | 0 | 0 | 0 |
| 0 | 0 | 0 | 0 |
| 0 | 0 | 0 | 0 |
| 0 | 0 | 0 | 0 |
| 0 | 0 | 0 | 0 |
| 1 | 0 | 0 | 0 |
| 0 | 0 | 0 | 0 |
| 0 | 0 | 0 | 0 |
| 0 | 0 | 0 | 0 |
| 0 | 0 | 0 | 0 |
| 0 | 0 | 0 | 0 |
| 0 | 0 | 0 | 0 |
| 0 | 0 | 0 | 0 |
| 1 | 1 | 0 | 0 |
| 0 | 0 | 0 | 0 |
| 0 | 0 | 0 | 0 |
| 0 | 0 | 0 | 0 |
| 0 | 0 | 0 | 0 |
| 0 | 0 | 0 | 0 |
| 0 | 0 | 0 | 0 |
| 0 | 0 | 0 | 0 |
| 0 | 0 | 0 | 0 |
| 0 | 0 | 0 | 0 |
| 0 | 0 | 0 | 0 |
| 0 | 0 | 0 | 0 |
| 0 | 0 | 0 | 0 |
| 0 | 0 | 0 | 0 |
| 1 | 0 | 0 | 0 |
| 0 | 0 | 0 | 0 |
| 0 | 0 | 0 | 0 |
| 0 | 0 | 0 | 0 |
| 0 | 0 | 0 | 0 |
| 0 | 0 | 0 | 0 |
| 1 | 0 | 0 | 1 |

| | | | |
|---|---|---|---|
| 0 | 0 | 0 | 0 |
| 0 | 0 | 0 | 0 |
| 0 | 0 | 1 | 0 |
| 0 | 0 | 0 | 0 |
| 0 | 0 | 0 | 0 |
| 0 | 0 | 0 | 0 |
| 0 | 0 | 0 | 0 |
| 0 | 0 | 0 | 0 |
| 0 | 0 | 0 | 0 |
| 0 | 0 | 0 | 0 |
| 0 | 0 | 0 | 0 |
| 0 | 0 | 0 | 0 |
| 0 | 0 | 0 | 0 |
| 0 | 0 | 0 | 0 |
| 0 | 0 | 0 | 0 |
| 0 | 0 | 0 | 0 |
| 0 | 0 | 0 | 0 |
| 0 | 0 | 0 | 0 |
| 0 | 0 | 0 | 0 |
| 0 | 0 | 0 | 0 |
| 0 | 0 | 0 | 0 |
| 0 | 0 | 0 | 0 |
| 0 | 0 | 0 | 0 |
| 0 | 0 | 0 | 0 |
| 0 | 0 | 0 | 0 |
| 0 | 0 | 0 | 0 |
| 0 | 0 | 0 | 0 |
| 0 | 0 | 0 | 0 |
| 0 | 0 | 0 | 0 |
| 0 | 0 | 0 | 0 |
| 0 | 0 | 0 | 0 |
| 1 | 0 | 0 | 0 |
| 0 | 0 | 0 | 0 |
| 0 | 0 | 0 | 0 |
| 0 | 0 | 0 | 0 |
| 0 | 0 | 0 | 0 |
| 0 | 0 | 0 | 0 |
| 0 | 0 | 0 | 0 |
| 0 | 0 | 0 | 0 |
| 0 | 0 | 0 | 0 |
| 1 | 1 | 0 | 0 |
| 0 | 0 | 0 | 0 |
| 0 | 0 | 0 | 0 |
| 0 | 0 | 0 | 0 |
| 0 | 0 | 0 | 0 |
| 1 | 1 | 0 | 0 |
| 0 | 0 | 0 | 0 |
| 1 | 1 | 0 | 0 |
| 0 | 0 | 0 | 0 |
| 1 | 0 | 0 | 0 |
| 1 | 0 | 0 | 0 |
| 1 | 0 | 0 | 0 |
| 0 | 0 | 0 | 0 |
| 0 | 0 | 0 | 0 |

| | | | |
|---|---|---|---|
| 1 | 0 | 0 | 0 |
| 0 | 0 | 0 | 0 |
| 0 | 0 | 0 | 0 |
| 0 | 0 | 0 | 0 |
| 0 | 0 | 0 | 0 |
| 0 | 0 | 0 | 0 |
| 0 | 0 | 0 | 0 |
| 0 | 0 | 0 | 0 |
| 0 | 0 | 0 | 0 |
| 0 | 0 | 0 | 0 |
| 0 | 0 | 0 | 0 |
| 0 | 0 | 0 | 0 |
| 0 | 0 | 0 | 0 |
| 0 | 0 | 0 | 0 |
| 0 | 0 | 0 | 0 |
| 1 | 1 | 1 | 0 |
| 0 | 0 | 0 | 0 |
| 1 | 0 | 0 | 0 |
| 0 | 0 | 0 | 0 |
| 0 | 0 | 0 | 0 |
| 0 | 0 | 0 | 0 |
| 0 | 1 | 0 | 0 |
| 0 | 0 | 0 | 0 |
| 0 | 1 | 0 | 0 |
| 0 | 1 | 0 | 0 |
| 0 | 0 | 0 | 0 |
| 0 | 0 | 0 | 0 |
| 0 | 0 | 0 | 0 |
| 0 | 0 | 0 | 0 |
| 0 | 0 | 0 | 0 |
| 0 | 0 | 0 | 0 |
| 0 | 0 | 0 | 0 |
| 1 | 1 | 0 | 0 |
| 0 | 0 | 0 | 0 |
| 0 | 0 | 0 | 0 |
| 0 | 0 | 0 | 0 |
| 0 | 1 | 0 | 0 |
| 1 | 1 | 0 | 0 |
| 0 | 0 | 0 | 0 |
| 0 | 0 | 0 | 0 |
| 0 | 0 | 0 | 0 |
| 0 | 0 | 0 | 0 |
| 0 | 0 | 0 | 0 |
| 0 | 0 | 0 | 0 |
| 1 | 1 | 0 | 0 |
| 0 | 0 | 0 | 0 |
| 1 | 0 | 0 | 0 |
| 1 | 1 | 0 | 0 |
| 0 | 0 | 0 | 0 |
| 0 | 0 | 0 | 0 |
| 1 | 1 | 0 | 0 |
| 1 | 1 | 0 | 0 |
| 0 | 0 | 0 | 0 |
| 1 | 1 | 0 | 0 |

| | | | |
|---|---|---|---|
| 0 | 0 | 0 | 0 |
| 0 | 0 | 0 | 0 |
| 0 | 0 | 0 | 0 |
| 1 | 1 | 0 | 0 |
| 0 | 0 | 0 | 0 |
| 0 | 0 | 0 | 0 |
| 0 | 0 | 0 | 0 |
| 0 | 0 | 0 | 0 |
| 1 | 0 | 0 | 0 |
| 0 | 0 | 0 | 0 |
| 0 | 0 | 0 | 0 |
| 0 | 0 | 0 | 0 |
| 1 | 0 | 1 | 0 |
| 1 | 0 | 0 | 0 |
| 0 | 0 | 0 | 0 |
| 0 | 0 | 0 | 0 |
| 0 | 0 | 0 | 0 |
| 0 | 0 | 0 | 0 |
| 0 | 0 | 0 | 0 |
| 0 | 0 | 0 | 0 |
| 0 | 0 | 0 | 0 |
| 0 | 0 | 0 | 0 |
| 0 | 0 | 0 | 0 |
| 1 | 0 | 0 | 0 |
| 0 | 0 | 0 | 0 |
| 0 | 0 | 0 | 0 |
| 0 | 0 | 0 | 0 |
| 0 | 0 | 0 | 0 |
| 0 | 0 | 0 | 0 |
| 0 | 0 | 0 | 0 |
| 0 | 0 | 0 | 0 |
| 0 | 0 | 0 | 0 |
| 0 | 0 | 0 | 0 |
| 0 | 0 | 0 | 0 |
| 0 | 0 | 0 | 0 |
| 0 | 0 | 0 | 0 |
| 0 | 0 | 0 | 0 |
| 0 | 0 | 0 | 0 |
| 0 | 0 | 0 | 0 |
| 0 | 0 | 0 | 0 |
| 0 | 0 | 0 | 0 |
| 0 | 0 | 0 | 0 |
| 0 | 0 | 0 | 0 |
| 1 | 1 | 0 | 0 |
| 1 | 1 | 0 | 0 |
| 0 | 0 | 0 | 0 |
| 0 | 0 | 0 | 0 |
| 0 | 1 | 0 | 0 |
| 0 | 1 | 0 | 0 |
| 0 | 1 | 0 | 0 |
| 0 | 0 | 0 | 0 |
| 0 | 0 | 0 | 0 |
| 0 | 0 | 0 | 0 |
| 0 | 0 | 0 | 0 |

| | | | |
|---|---|---|---|
| 0 | 0 | 0 | 0 |
| 0 | 0 | 0 | 0 |
| 0 | 0 | 0 | 0 |
| 0 | 0 | 0 | 0 |
| 1 | 0 | 0 | 0 |
| 0 | 0 | 0 | 0 |
| 0 | 0 | 0 | 0 |
| 1 | 0 | 0 | 0 |
| 1 | 0 | 0 | 0 |
| 1 | 0 | 0 | 0 |
| 1 | 0 | 0 | 0 |
| 1 | 0 | 0 | 0 |
| 0 | 0 | 0 | 0 |
| 1 | 0 | 0 | 0 |
| 1 | 0 | 0 | 0 |
| 0 | 0 | 0 | 0 |
| 0 | 0 | 0 | 0 |
| 0 | 0 | 0 | 0 |
| 1 | 0 | 0 | 0 |
| 1 | 0 | 0 | 0 |
| 1 | 0 | 0 | 0 |
| 1 | 0 | 0 | 0 |
| 0 | 0 | 0 | 0 |
| 0 | 0 | 0 | 0 |
| 0 | 0 | 0 | 0 |
| 0 | 0 | 0 | 0 |
| 0 | 0 | 0 | 0 |
| 0 | 0 | 0 | 0 |
| 0 | 0 | 0 | 0 |
| 0 | 0 | 0 | 0 |
| 0 | 0 | 0 | 0 |
| 0 | 0 | 0 | 0 |
| 1 | 0 | 0 | 0 |
| 0 | 0 | 0 | 0 |
| 0 | 0 | 0 | 0 |
| 1 | 0 | 0 | 0 |
| 0 | 0 | 0 | 0 |
| 0 | 0 | 0 | 0 |
| 0 | 0 | 0 | 0 |
| 0 | 0 | 0 | 0 |
| 0 | 0 | 0 | 0 |
| 0 | 0 | 0 | 0 |
| 0 | 0 | 0 | 0 |
| 0 | 0 | 0 | 0 |
| 0 | 0 | 0 | 0 |
| 0 | 0 | 0 | 0 |
| 0 | 0 | 0 | 0 |
| 0 | 0 | 0 | 0 |
| 0 | 0 | 0 | 0 |
| 0 | 0 | 0 | 0 |
| 0 | 0 | 0 | 0 |
| 0 | 0 | 0 | 0 |
| 0 | 0 | 0 | 0 |
| 0 | 0 | 0 | 0 |

| | | | |
|---|---|---|---|
| 0 | 0 | 0 | 0 |
| 0 | 0 | 0 | 0 |
| 0 | 0 | 0 | 0 |
| 0 | 0 | 0 | 0 |
| 0 | 0 | 0 | 0 |
| 0 | 0 | 0 | 0 |
| 0 | 0 | 0 | 0 |
| 0 | 0 | 0 | 0 |
| 0 | 0 | 0 | 0 |
| 0 | 0 | 0 | 0 |
| 0 | 0 | 0 | 0 |
| 0 | 0 | 0 | 1 |
| 0 | 0 | 0 | 0 |
| 0 | 0 | 0 | 0 |
| 0 | 0 | 0 | 0 |
| 0 | 0 | 0 | 0 |
| 0 | 0 | 0 | 0 |
| 0 | 0 | 0 | 1 |
| 0 | 0 | 0 | 0 |
| 0 | 0 | 0 | 0 |
| 0 | 0 | 0 | 1 |
| 0 | 0 | 0 | 0 |
| 0 | 0 | 0 | 0 |
| 0 | 0 | 0 | 0 |
| 0 | 0 | 0 | 0 |
| 0 | 0 | 0 | 0 |
| 0 | 0 | 0 | 0 |
| 0 | 0 | 0 | 0 |
| 0 | 0 | 0 | 1 |
| 0 | 0 | 0 | 0 |
| 0 | 0 | 0 | 0 |
| 0 | 0 | 0 | 0 |
| 0 | 0 | 0 | 0 |
| 0 | 0 | 0 | 0 |
| 0 | 0 | 0 | 0 |
| 0 | 0 | 0 | 0 |
| 0 | 0 | 0 | 0 |
| 0 | 0 | 0 | 0 |
| 0 | 0 | 0 | 0 |
| 0 | 0 | 0 | 0 |
| 0 | 0 | 0 | 0 |
| 0 | 0 | 0 | 0 |
| 0 | 0 | 0 | 0 |
| 0 | 0 | 0 | 0 |
| 0 | 0 | 0 | 0 |
| 0 | 0 | 0 | 0 |
| 0 | 0 | 0 | 0 |
| 0 | 0 | 0 | 0 |
| 0 | 0 | 0 | 0 |
| 0 | 0 | 0 | 0 |
| 0 | 0 | 0 | 0 |
| 0 | 0 | 0 | 0 |
| 0 | 0 | 0 | 0 |
| 0 | 0 | 0 | 0 |

| | | | |
|---|---|---|---|
| 0 | 0 | 0 | 0 |
| 0 | 0 | 0 | 0 |
| 1 | 0 | 0 | 0 |
| 0 | 0 | 0 | 0 |
| 1 | 0 | 0 | 0 |
| 0 | 0 | 0 | 0 |
| 0 | 0 | 0 | 0 |
| 0 | 0 | 0 | 0 |
| 0 | 0 | 0 | 0 |
| 1 | 1 | 0 | 0 |
| 0 | 0 | 1 | 0 |
| 0 | 0 | 0 | 0 |
| 0 | 0 | 0 | 0 |
| 0 | 0 | 0 | 0 |
| 0 | 0 | 0 | 0 |
| 0 | 0 | 0 | 0 |
| 0 | 0 | 0 | 0 |
| 0 | 0 | 0 | 0 |
| 0 | 0 | 0 | 0 |
| 0 | 0 | 0 | 0 |
| 0 | 0 | 0 | 0 |
| 0 | 0 | 0 | 0 |
| 0 | 0 | 0 | 0 |
| 0 | 0 | 0 | 0 |
| 0 | 0 | 0 | 0 |
| 0 | 0 | 0 | 0 |
| 0 | 0 | 0 | 0 |
| 0 | 0 | 0 | 0 |
| 0 | 0 | 0 | 0 |
| 0 | 0 | 0 | 0 |
| 0 | 0 | 0 | 0 |
| 0 | 0 | 0 | 0 |
| 0 | 0 | 0 | 0 |
| 0 | 0 | 0 | 0 |
| 0 | 0 | 0 | 0 |
| 0 | 0 | 0 | 0 |
| 0 | 0 | 0 | 0 |
| 0 | 0 | 0 | 0 |
| 0 | 0 | 0 | 0 |
| 0 | 0 | 0 | 0 |
| 0 | 0 | 0 | 0 |
| 0 | 0 | 0 | 0 |
| 0 | 0 | 0 | 0 |
| 0 | 0 | 0 | 0 |
| 0 | 0 | 0 | 0 |
| 1 | 0 | 0 | 0 |
| 0 | 0 | 0 | 0 |
| 0 | 0 | 0 | 0 |
| 0 | 0 | 0 | 0 |
| 0 | 0 | 0 | 0 |
| 1 | 0 | 0 | 0 |
| 0 | 0 | 0 | 0 |
| 0 | 0 | 0 | 0 |
| 0 | 0 | 0 | 0 |

| | | | |
|---|---|---|---|
| 0 | 0 | 0 | 0 |
| 0 | 0 | 0 | 0 |
| 0 | 0 | 0 | 0 |
| 0 | 0 | 0 | 0 |
| 0 | 0 | 0 | 0 |
| 0 | 0 | 0 | 0 |
| 0 | 0 | 0 | 0 |
| 0 | 0 | 0 | 0 |
| 1 | 1 | 0 | 0 |
| 0 | 0 | 0 | 0 |
| 0 | 0 | 0 | 0 |
| 0 | 0 | 0 | 0 |
| 0 | 0 | 0 | 0 |
| 0 | 0 | 0 | 0 |
| 0 | 1 | 0 | 0 |
| 0 | 1 | 0 | 0 |
| 0 | 1 | 0 | 0 |
| 0 | 0 | 0 | 0 |
| 0 | 0 | 0 | 0 |
| 0 | 0 | 0 | 0 |
| 0 | 1 | 0 | 0 |
| 0 | 0 | 0 | 0 |
| 0 | 0 | 0 | 0 |
| 1 | 1 | 0 | 0 |
| 0 | 1 | 0 | 0 |
| 0 | 1 | 0 | 0 |
| 0 | 0 | 0 | 0 |
| 0 | 0 | 0 | 0 |
| 0 | 0 | 0 | 0 |
| 0 | 0 | 0 | 0 |
| 0 | 0 | 0 | 0 |
| 0 | 0 | 0 | 0 |
| 0 | 0 | 0 | 0 |
| 0 | 0 | 0 | 0 |
| 0 | 0 | 0 | 0 |
| 0 | 0 | 0 | 0 |
| 0 | 0 | 0 | 0 |
| 1 | 0 | 0 | 0 |
| 0 | 0 | 0 | 0 |
| 0 | 0 | 0 | 0 |
| 0 | 0 | 0 | 0 |
| 0 | 0 | 0 | 0 |
| 0 | 0 | 0 | 0 |
| 0 | 0 | 0 | 0 |
| 0 | 0 | 0 | 0 |
| 0 | 0 | 0 | 0 |
| 0 | 0 | 0 | 0 |
| 0 | 0 | 0 | 0 |
| 0 | 0 | 0 | 0 |
| 0 | 0 | 0 | 0 |
| 0 | 0 | 0 | 0 |
| 0 | 0 | 0 | 0 |
| 0 | 0 | 0 | 0 |
| 0 | 0 | 0 | 0 |

| | | | |
|---|---|---|---|
| 0 | 0 | 0 | 0 |
| 0 | 0 | 0 | 0 |
| 0 | 0 | 0 | 0 |
| 0 | 0 | 1 | 0 |
| 0 | 0 | 0 | 0 |
| 0 | 0 | 0 | 0 |
| 0 | 0 | 0 | 0 |
| 0 | 0 | 0 | 0 |
| 0 | 0 | 0 | 0 |
| 0 | 0 | 0 | 0 |
| 0 | 0 | 0 | 0 |
| 0 | 0 | 0 | 0 |
| 1 | 0 | 0 | 0 |
| 0 | 0 | 0 | 0 |
| 0 | 0 | 0 | 0 |
| 0 | 0 | 0 | 0 |
| 0 | 1 | 0 | 0 |
| 0 | 0 | 0 | 0 |
| 0 | 0 | 0 | 0 |
| 0 | 0 | 0 | 0 |
| 0 | 0 | 0 | 0 |
| 0 | 0 | 0 | 0 |
| 0 | 0 | 0 | 0 |
| 0 | 0 | 0 | 0 |
| 0 | 0 | 0 | 0 |
| 0 | 0 | 0 | 0 |
| 0 | 0 | 0 | 0 |
| 0 | 0 | 0 | 0 |
| 0 | 0 | 0 | 0 |
| 0 | 0 | 0 | 0 |
| 1 | 0 | 0 | 0 |
| 1 | 0 | 0 | 0 |
| 1 | 0 | 0 | 0 |
| 0 | 0 | 0 | 0 |
| 1 | 0 | 0 | 0 |
| 1 | 0 | 0 | 0 |
| 1 | 0 | 0 | 0 |
| 0 | 0 | 0 | 0 |
| 0 | 0 | 0 | 0 |
| 0 | 0 | 0 | 0 |
| 0 | 0 | 0 | 0 |
| 0 | 0 | 0 | 0 |
| 0 | 1 | 0 | 0 |
| 0 | 1 | 0 | 0 |
| 0 | 0 | 0 | 0 |
| 0 | 0 | 0 | 0 |
| 1 | 0 | 0 | 0 |
| 0 | 0 | 0 | 0 |
| 0 | 0 | 0 | 0 |
| 0 | 0 | 0 | 0 |
| 0 | 0 | 0 | 0 |
| 0 | 0 | 0 | 0 |
| 1 | 0 | 0 | 0 |
| 0 | 0 | 0 | 0 |

| | | | |
|---|---|---|---|
| 0 | 0 | 0 | 0 |
| 1 | 0 | 0 | 0 |
| 1 | 0 | 0 | 0 |
| 0 | 0 | 0 | 0 |
| 0 | 0 | 0 | 0 |
| 0 | 0 | 0 | 0 |
| 0 | 0 | 0 | 0 |
| 0 | 1 | 0 | 0 |
| 0 | 0 | 0 | 0 |
| 0 | 0 | 0 | 0 |
| 0 | 0 | 0 | 0 |
| 0 | 0 | 0 | 0 |
| 0 | 0 | 0 | 0 |
| 0 | 0 | 0 | 0 |
| 1 | 0 | 0 | 0 |
| 0 | 0 | 0 | 0 |
| 0 | 0 | 0 | 0 |
| 0 | 0 | 0 | 0 |
| 0 | 0 | 0 | 0 |
| 1 | 0 | 0 | 0 |
| 1 | 0 | 0 | 0 |
| 0 | 0 | 0 | 0 |
| 1 | 0 | 0 | 0 |
| 1 | 0 | 0 | 0 |
| 0 | 0 | 0 | 0 |
| 0 | 0 | 0 | 0 |
| 0 | 0 | 0 | 0 |
| 0 | 0 | 0 | 0 |
| 0 | 0 | 0 | 0 |
| 0 | 0 | 0 | 0 |
| 0 | 0 | 0 | 0 |
| 0 | 0 | 0 | 0 |
| 0 | 0 | 0 | 0 |
| 0 | 0 | 0 | 0 |
| 0 | 0 | 0 | 0 |
| 0 | 0 | 0 | 0 |
| 0 | 0 | 0 | 0 |
| 0 | 0 | 0 | 0 |
| 0 | 0 | 0 | 0 |
| 0 | 0 | 0 | 0 |
| 0 | 0 | 0 | 0 |
| 0 | 0 | 0 | 0 |
| 0 | 0 | 0 | 0 |
| 0 | 0 | 0 | 0 |
| 0 | 0 | 0 | 0 |
| 0 | 0 | 0 | 0 |
| 0 | 0 | 0 | 0 |
| 0 | 0 | 0 | 0 |
| 0 | 0 | 0 | 0 |
| 0 | 0 | 0 | 0 |
| 0 | 0 | 0 | 0 |
| 0 | 0 | 0 | 0 |
| 0 | 0 | 0 | 0 |
| 1 | 1 | 0 | 1 |

| | | | |
|---|---|---|---|
| 0 | 0 | 0 | 0 |
| 0 | 0 | 0 | 0 |
| 0 | 0 | 0 | 0 |
| 0 | 0 | 0 | 0 |
| 0 | 1 | 0 | 0 |
| 0 | 0 | 0 | 0 |
| 0 | 0 | 0 | 0 |
| 0 | 1 | 0 | 0 |
| 0 | 0 | 0 | 0 |
| 0 | 0 | 0 | 0 |
| 0 | 0 | 0 | 0 |
| 0 | 0 | 0 | 0 |
| 0 | 0 | 0 | 0 |
| 0 | 0 | 0 | 0 |
| 0 | 1 | 0 | 1 |
| 0 | 1 | 0 | 0 |
| 0 | 0 | 0 | 0 |
| 0 | 0 | 0 | 0 |
| 0 | 1 | 0 | 0 |
| 0 | 1 | 0 | 1 |
| 0 | 0 | 0 | 0 |
| 0 | 0 | 0 | 0 |
| 0 | 0 | 0 | 0 |
| 0 | 0 | 0 | 0 |
| 0 | 0 | 0 | 0 |
| 1 | 0 | 0 | 0 |
| 0 | 0 | 0 | 0 |
| 0 | 0 | 1 | 0 |
| 0 | 0 | 1 | 0 |
| 0 | 0 | 1 | 0 |
| 0 | 0 | 0 | 0 |
| 0 | 0 | 0 | 0 |
| 0 | 0 | 0 | 0 |
| 0 | 0 | 0 | 0 |
| 0 | 0 | 0 | 0 |
| 0 | 0 | 1 | 0 |
| 0 | 0 | 0 | 0 |
| 0 | 0 | 0 | 0 |
| 1 | 0 | 1 | 0 |
| 0 | 0 | 1 | 0 |
| 1 | 1 | 0 | 0 |
| 0 | 1 | 0 | 0 |
| 0 | 0 | 0 | 0 |
| 0 | 0 | 0 | 0 |
| 1 | 0 | 0 | 0 |
| 0 | 0 | 0 | 0 |
| 0 | 0 | 1 | 0 |
| 0 | 0 | 0 | 0 |
| 0 | 0 | 0 | 0 |
| 0 | 0 | 0 | 0 |
| 0 | 0 | 0 | 0 |
| 0 | 0 | 0 | 0 |
| 0 | 0 | 0 | 0 |
| 0 | 0 | 0 | 0 |

| | | | |
|---|---|---|---|
| 0 | 0 | 0 | 0 |
| 0 | 0 | 0 | 0 |
| 0 | 0 | 0 | 0 |
| 0 | 0 | 0 | 0 |
| 0 | 0 | 0 | 0 |
| 0 | 0 | 0 | 0 |
| 0 | 0 | 0 | 0 |
| 1 | 1 | 0 | 0 |
| 0 | 0 | 0 | 0 |
| 0 | 0 | 0 | 0 |
| 0 | 0 | 0 | 0 |
| 0 | 0 | 0 | 1 |
| 0 | 0 | 0 | 0 |
| 0 | 0 | 0 | 0 |
| 0 | 0 | 0 | 0 |
| 0 | 0 | 0 | 0 |
| 0 | 0 | 0 | 0 |
| 0 | 0 | 0 | 0 |
| 0 | 0 | 0 | 0 |
| 0 | 0 | 0 | 0 |
| 0 | 0 | 0 | 0 |
| 0 | 0 | 0 | 0 |
| 0 | 0 | 0 | 0 |
| 0 | 0 | 0 | 0 |
| 0 | 0 | 0 | 0 |
| 0 | 0 | 0 | 0 |
| 0 | 0 | 0 | 0 |
| 0 | 0 | 0 | 0 |
| 0 | 0 | 0 | 0 |
| 0 | 1 | 0 | 0 |
| 0 | 1 | 0 | 0 |
| 0 | 0 | 0 | 0 |
| 0 | 0 | 0 | 0 |
| 0 | 0 | 0 | 0 |
| 0 | 0 | 0 | 0 |
| 0 | 1 | 0 | 0 |
| 0 | 0 | 0 | 0 |
| 0 | 0 | 0 | 0 |
| 0 | 0 | 0 | 0 |
| 0 | 0 | 0 | 0 |
| 0 | 0 | 0 | 0 |
| 1 | 1 | 0 | 0 |
| 1 | 0 | 0 | 0 |
| 0 | 0 | 0 | 0 |
| 0 | 0 | 0 | 0 |
| 0 | 0 | 0 | 0 |
| 0 | 0 | 0 | 0 |
| 0 | 0 | 0 | 0 |
| 0 | 0 | 0 | 0 |
| 0 | 0 | 0 | 0 |
| 0 | 0 | 0 | 0 |
| 0 | 0 | 0 | 0 |
| 0 | 0 | 0 | 0 |
| 0 | 0 | 0 | 0 |

| | | | |
|---|---|---|---|
| 0 | 0 | 0 | 0 |
| 0 | 0 | 0 | 0 |
| 0 | 0 | 0 | 0 |
| 0 | 0 | 0 | 0 |
| 0 | 0 | 0 | 0 |
| 0 | 0 | 0 | 0 |
| 0 | 0 | 0 | 0 |
| 0 | 0 | 0 | 0 |
| 1 | 0 | 0 | 0 |
| 1 | 0 | 0 | 0 |
| 0 | 0 | 0 | 0 |
| 0 | 0 | 0 | 0 |
| 0 | 0 | 0 | 0 |
| 1 | 0 | 0 | 0 |
| 0 | 0 | 0 | 0 |
| 0 | 0 | 0 | 0 |
| 0 | 0 | 0 | 0 |
| 0 | 0 | 0 | 0 |
| 0 | 0 | 0 | 0 |
| 1 | 1 | 0 | 0 |
| 0 | 0 | 1 | 0 |
| 0 | 0 | 0 | 0 |
| 0 | 0 | 0 | 0 |
| 0 | 0 | 0 | 0 |
| 0 | 0 | 0 | 0 |
| 0 | 0 | 0 | 0 |
| 0 | 0 | 0 | 0 |
| 0 | 0 | 0 | 0 |
| 0 | 0 | 0 | 0 |
| 0 | 0 | 0 | 0 |
| 0 | 0 | 0 | 0 |
| 0 | 0 | 0 | 0 |
| 0 | 0 | 0 | 0 |
| 0 | 0 | 0 | 0 |
| 0 | 0 | 0 | 0 |
| 0 | 0 | 0 | 0 |
| 0 | 0 | 0 | 0 |
| 0 | 0 | 0 | 0 |
| 0 | 0 | 0 | 0 |
| 0 | 0 | 0 | 0 |
| 0 | 0 | 0 | 0 |
| 0 | 0 | 0 | 0 |
| 0 | 0 | 0 | 0 |
| 0 | 0 | 0 | 0 |
| 0 | 0 | 0 | 0 |
| 1 | 0 | 1 | 0 |
| 1 | 0 | 0 | 0 |
| 0 | 0 | 0 | 0 |
| 0 | 0 | 0 | 0 |
| 0 | 0 | 1 | 0 |
| 0 | 0 | 0 | 0 |
| 0 | 0 | 0 | 0 |
| 0 | 0 | 0 | 0 |

| | | | |
|---|---|---|---|
| 0 | 0 | 0 | 0 |
| 0 | 0 | 0 | 0 |
| 0 | 0 | 0 | 0 |
| 0 | 0 | 0 | 0 |
| 0 | 0 | 0 | 0 |
| 0 | 0 | 0 | 0 |
| 0 | 0 | 0 | 0 |
| 0 | 0 | 0 | 0 |
| 0 | 0 | 0 | 0 |
| 0 | 0 | 0 | 0 |
| 0 | 0 | 0 | 0 |
| 0 | 0 | 0 | 0 |
| 0 | 0 | 0 | 0 |
| 0 | 0 | 0 | 0 |
| 0 | 0 | 0 | 0 |
| 0 | 0 | 0 | 0 |
| 0 | 0 | 0 | 0 |
| 0 | 0 | 0 | 0 |
| 0 | 0 | 0 | 0 |
| 1 | 0 | 0 | 0 |
| 0 | 0 | 0 | 0 |
| 0 | 0 | 0 | 0 |
| 0 | 0 | 0 | 0 |
| 0 | 0 | 0 | 0 |
| 0 | 0 | 0 | 0 |
| 0 | 0 | 0 | 0 |
| 0 | 0 | 0 | 0 |
| 0 | 0 | 0 | 0 |
| 0 | 0 | 0 | 0 |
| 0 | 0 | 0 | 0 |
| 0 | 0 | 0 | 0 |
| 0 | 0 | 0 | 0 |
| 0 | 0 | 0 | 0 |
| 0 | 0 | 0 | 0 |
| 0 | 0 | 0 | 0 |
| 0 | 0 | 0 | 0 |
| 0 | 0 | 0 | 0 |
| 0 | 0 | 0 | 0 |
| 1 | 0 | 0 | 1 |
| 0 | 0 | 0 | 0 |
| 0 | 0 | 0 | 0 |
| 0 | 0 | 0 | 0 |
| 0 | 0 | 0 | 1 |
| 0 | 0 | 0 | 1 |
| 0 | 0 | 0 | 1 |
| 0 | 0 | 0 | 0 |
| 0 | 0 | 0 | 0 |
| 0 | 0 | 0 | 0 |
| 0 | 0 | 0 | 0 |
| 0 | 0 | 0 | 0 |
| 0 | 0 | 0 | 0 |
| 0 | 0 | 0 | 0 |
| 0 | 0 | 0 | 0 |
| 0 | 0 | 0 | 0 |

| | | | |
|---|---|---|---|
| 0 | 0 | 0 | 0 |
| 0 | 0 | 0 | 0 |
| 0 | 0 | 0 | 0 |
| 0 | 0 | 0 | 0 |
| 0 | 0 | 0 | 0 |
| 0 | 0 | 0 | 0 |
| 0 | 0 | 0 | 0 |
| 0 | 0 | 0 | 0 |
| 0 | 0 | 0 | 0 |
| 0 | 0 | 0 | 0 |
| 0 | 0 | 0 | 0 |
| 0 | 0 | 0 | 0 |
| 0 | 0 | 0 | 0 |
| 0 | 0 | 0 | 0 |
| 0 | 0 | 0 | 0 |
| 0 | 0 | 0 | 0 |
| 0 | 0 | 0 | 0 |
| 0 | 0 | 0 | 0 |
| 0 | 0 | 0 | 0 |
| 0 | 0 | 0 | 0 |
| 0 | 0 | 0 | 0 |
| 0 | 0 | 0 | 0 |
| 0 | 0 | 0 | 0 |
| 0 | 0 | 0 | 0 |
| 0 | 0 | 0 | 0 |
| 0 | 0 | 0 | 0 |
| 0 | 0 | 0 | 0 |
| 0 | 0 | 0 | 0 |
| 0 | 0 | 0 | 0 |
| 0 | 0 | 0 | 0 |
| 0 | 0 | 0 | 0 |
| 0 | 0 | 0 | 0 |
| 1 | 1 | 0 | 1 |
| 1 | 1 | 0 | 1 |
| 1 | 0 | 0 | 0 |
| 0 | 0 | 0 | 0 |
| 0 | 0 | 0 | 0 |
| 0 | 0 | 0 | 0 |
| 0 | 0 | 0 | 0 |
| 0 | 0 | 0 | 0 |
| 0 | 0 | 0 | 0 |
| 0 | 0 | 0 | 0 |
| 1 | 1 | 0 | 0 |
| 0 | 0 | 0 | 0 |
| 1 | 0 | 0 | 0 |
| 0 | 0 | 0 | 0 |
| 0 | 0 | 0 | 0 |
| 0 | 0 | 0 | 0 |
| 0 | 0 | 0 | 0 |
| 0 | 0 | 0 | 0 |
| 1 | 0 | 0 | 0 |
| 0 | 0 | 0 | 0 |
| 0 | 0 | 0 | 0 |
| 0 | 0 | 0 | 0 |

| | | | |
|---|---|---|---|
| 0 | 0 | 0 | 0 |
| 0 | 0 | 0 | 0 |
| 0 | 0 | 0 | 0 |
| 0 | 0 | 0 | 0 |
| 0 | 0 | 0 | 0 |
| 0 | 0 | 0 | 0 |
| 0 | 0 | 0 | 0 |
| 0 | 0 | 0 | 0 |
| 0 | 0 | 0 | 0 |
| 0 | 0 | 0 | 0 |
| 0 | 0 | 0 | 0 |
| 0 | 0 | 0 | 0 |
| 0 | 0 | 0 | 0 |
| 0 | 0 | 0 | 0 |
| 0 | 0 | 0 | 0 |
| 0 | 0 | 0 | 0 |
| 0 | 0 | 0 | 0 |
| 0 | 0 | 0 | 0 |
| 0 | 0 | 0 | 0 |
| 0 | 0 | 0 | 1 |
| 0 | 0 | 0 | 0 |
| 0 | 0 | 0 | 1 |
| 0 | 0 | 0 | 1 |
| 0 | 0 | 0 | 1 |
| 0 | 0 | 0 | 0 |
| 0 | 0 | 0 | 0 |
| 0 | 0 | 0 | 0 |
| 0 | 0 | 0 | 0 |
| 0 | 0 | 0 | 0 |
| 0 | 0 | 0 | 0 |
| 0 | 0 | 0 | 0 |
| 0 | 0 | 0 | 0 |
| 0 | 0 | 0 | 0 |
| 0 | 0 | 0 | 0 |
| 0 | 0 | 0 | 0 |
| 0 | 1 | 0 | 0 |
| 0 | 0 | 0 | 0 |
| 0 | 0 | 0 | 0 |
| 0 | 1 | 0 | 0 |
| 0 | 0 | 0 | 0 |
| 0 | 0 | 0 | 0 |
| 0 | 0 | 0 | 0 |
| 0 | 0 | 0 | 0 |
| 0 | 0 | 0 | 0 |
| 0 | 0 | 0 | 0 |
| 0 | 0 | 0 | 0 |
| 0 | 0 | 0 | 0 |
| 0 | 0 | 0 | 0 |
| 0 | 0 | 0 | 0 |
| 0 | 0 | 0 | 0 |
| 0 | 0 | 0 | 0 |
| 0 | 0 | 0 | 0 |
| 0 | 0 | 0 | 0 |
| 0 | 0 | 0 | 0 |

| | | | |
|---|---|---|---|
| 0 | 0 | 0 | 0 |
| 0 | 0 | 0 | 0 |
| 0 | 0 | 0 | 0 |
| 0 | 0 | 0 | 0 |
| 0 | 0 | 0 | 0 |
| 0 | 0 | 0 | 0 |
| 0 | 0 | 0 | 0 |
| 0 | 0 | 0 | 0 |
| 0 | 0 | 1 | 0 |
| 0 | 0 | 1 | 0 |
| 0 | 0 | 0 | 0 |
| 0 | 0 | 0 | 0 |
| 0 | 0 | 0 | 0 |
| 0 | 0 | 0 | 0 |
| 0 | 0 | 0 | 0 |
| 0 | 1 | 0 | 0 |
| 0 | 0 | 0 | 0 |
| 0 | 0 | 0 | 0 |
| 1 | 1 | 0 | 0 |
| 0 | 0 | 0 | 0 |
| 0 | 0 | 0 | 0 |
| 0 | 0 | 0 | 0 |
| 0 | 0 | 0 | 0 |
| 0 | 0 | 0 | 0 |
| 0 | 0 | 0 | 0 |
| 0 | 0 | 0 | 0 |
| 0 | 0 | 0 | 0 |
| 0 | 0 | 0 | 0 |
| 0 | 0 | 0 | 0 |
| 0 | 0 | 0 | 0 |
| 0 | 0 | 0 | 0 |
| 0 | 0 | 0 | 0 |
| 0 | 0 | 0 | 0 |
| 0 | 0 | 0 | 0 |
| 0 | 0 | 0 | 0 |
| 0 | 0 | 0 | 0 |
| 0 | 0 | 0 | 0 |
| 0 | 0 | 0 | 0 |
| 0 | 0 | 0 | 0 |
| 0 | 0 | 0 | 0 |
| 0 | 0 | 0 | 0 |
| 0 | 0 | 0 | 0 |
| 0 | 0 | 0 | 0 |
| 0 | 0 | 0 | 0 |
| 0 | 0 | 0 | 0 |
| 0 | 0 | 0 | 0 |
| 0 | 0 | 0 | 0 |
| 0 | 0 | 0 | 0 |
| 0 | 0 | 0 | 0 |
| 0 | 0 | 0 | 0 |
| 0 | 0 | 0 | 0 |
| 0 | 0 | 0 | 0 |
| 0 | 0 | 0 | 0 |
| 0 | 0 | 0 | 0 |
| 1 | 0 | 0 | 0 |

| | | | |
|---|---|---|---|
| 0 | 0 | 0 | 0 |
| 0 | 0 | 0 | 0 |
| 0 | 0 | 0 | 0 |
| 0 | 0 | 0 | 0 |
| 0 | 0 | 0 | 0 |
| 0 | 0 | 0 | 0 |
| 0 | 0 | 0 | 0 |
| 0 | 0 | 0 | 0 |
| 0 | 0 | 0 | 0 |
| 0 | 0 | 0 | 0 |
| 1 | 0 | 0 | 0 |
| 0 | 0 | 0 | 0 |
| 0 | 0 | 0 | 0 |
| 0 | 0 | 0 | 0 |
| 1 | 1 | 0 | 1 |
| 0 | 0 | 0 | 0 |
| 1 | 0 | 0 | 0 |
| 1 | 1 | 0 | 0 |
| 0 | 0 | 0 | 0 |
| 0 | 0 | 0 | 0 |
| 0 | 0 | 0 | 0 |
| 0 | 0 | 0 | 0 |
| 0 | 0 | 0 | 0 |
| 0 | 0 | 0 | 1 |
| 0 | 0 | 0 | 0 |
| 0 | 0 | 0 | 0 |
| 0 | 0 | 0 | 0 |
| 0 | 0 | 0 | 1 |
| 0 | 0 | 0 | 0 |
| 0 | 0 | 0 | 0 |
| 0 | 0 | 0 | 0 |
| 0 | 0 | 0 | 0 |
| 0 | 0 | 0 | 0 |
| 0 | 0 | 0 | 0 |
| 0 | 0 | 0 | 0 |
| 0 | 0 | 0 | 0 |
| 0 | 0 | 0 | 0 |
| 0 | 0 | 0 | 0 |
| 0 | 0 | 0 | 0 |
| 0 | 0 | 0 | 0 |
| 0 | 0 | 0 | 0 |
| 0 | 0 | 0 | 0 |
| 0 | 0 | 0 | 0 |
| 0 | 0 | 0 | 0 |
| 0 | 0 | 1 | 0 |
| 0 | 0 | 0 | 0 |
| 0 | 0 | 0 | 0 |
| 0 | 0 | 0 | 0 |
| 0 | 0 | 0 | 0 |
| 0 | 0 | 0 | 0 |
| 0 | 0 | 0 | 0 |
| 0 | 0 | 0 | 0 |
| 0 | 0 | 0 | 0 |
| 0 | 0 | 0 | 0 |

| | | | |
|---|---|---|---|
| 0 | 0 | 0 | 0 |
| 0 | 0 | 0 | 0 |
| 0 | 0 | 0 | 0 |
| 0 | 0 | 0 | 0 |
| 0 | 0 | 0 | 0 |
| 0 | 0 | 0 | 0 |
| 1 | 0 | 0 | 1 |
| 1 | 0 | 0 | 0 |
| 0 | 0 | 0 | 0 |
| 0 | 0 | 0 | 0 |
| 0 | 0 | 0 | 0 |
| 0 | 0 | 0 | 0 |
| 0 | 0 | 0 | 0 |
| 0 | 0 | 0 | 0 |
| 0 | 0 | 0 | 0 |
| 0 | 0 | 0 | 0 |
| 0 | 0 | 0 | 0 |
| 0 | 0 | 0 | 0 |
| 0 | 0 | 0 | 0 |
| 0 | 0 | 0 | 0 |
| 1 | 0 | 0 | 0 |
| 0 | 0 | 0 | 0 |
| 0 | 0 | 0 | 0 |
| 0 | 0 | 0 | 0 |
| 0 | 0 | 0 | 0 |
| 0 | 0 | 0 | 0 |
| 0 | 0 | 0 | 0 |
| 0 | 0 | 0 | 0 |
| 0 | 0 | 0 | 0 |
| 0 | 0 | 0 | 0 |
| 0 | 0 | 0 | 0 |
| 0 | 0 | 0 | 0 |
| 0 | 0 | 0 | 0 |
| 0 | 0 | 0 | 0 |
| 0 | 0 | 0 | 0 |
| 0 | 0 | 0 | 0 |
| 0 | 0 | 0 | 0 |
| 0 | 0 | 0 | 0 |
| 0 | 0 | 0 | 0 |
| 0 | 0 | 0 | 0 |
| 0 | 0 | 0 | 0 |
| 0 | 0 | 0 | 0 |
| 0 | 0 | 0 | 0 |
| 0 | 0 | 0 | 0 |
| 0 | 0 | 0 | 0 |
| 0 | 0 | 0 | 0 |
| 0 | 0 | 0 | 0 |
| 0 | 0 | 0 | 0 |
| 0 | 0 | 0 | 0 |
| 0 | 0 | 0 | 0 |
| 0 | 0 | 0 | 0 |
| 0 | 0 | 0 | 0 |
| 0 | 0 | 0 | 0 |
| 0 | 0 | 0 | 0 |

| | | | |
|---|---|---|---|
| 0 | 0 | 0 | 0 |
| 0 | 0 | 0 | 0 |
| 0 | 0 | 0 | 0 |
| 1 | 0 | 0 | 0 |
| 0 | 0 | 0 | 0 |
| 1 | 0 | 0 | 0 |
| 0 | 0 | 0 | 0 |
| 0 | 0 | 0 | 0 |
| 1 | 0 | 0 | 0 |
| 0 | 0 | 0 | 0 |
| 0 | 0 | 0 | 0 |
| 0 | 0 | 0 | 0 |
| 0 | 0 | 0 | 0 |
| 0 | 0 | 0 | 0 |
| 0 | 0 | 0 | 0 |
| 0 | 0 | 0 | 0 |
| 0 | 0 | 0 | 0 |
| 0 | 0 | 0 | 0 |
| 1 | 0 | 0 | 0 |
| 0 | 0 | 0 | 0 |
| 0 | 0 | 0 | 0 |
| 0 | 0 | 0 | 0 |
| 0 | 0 | 0 | 0 |
| 0 | 0 | 0 | 0 |
| 0 | 0 | 0 | 0 |
| 0 | 0 | 0 | 0 |
| 0 | 0 | 0 | 0 |
| 0 | 0 | 0 | 0 |
| 0 | 1 | 0 | 0 |
| 0 | 0 | 0 | 0 |
| 0 | 0 | 0 | 0 |
| 0 | 0 | 0 | 0 |
| 0 | 1 | 0 | 0 |
| 1 | 1 | 0 | 0 |
| 1 | 0 | 0 | 0 |
| 0 | 0 | 0 | 0 |
| 0 | 0 | 0 | 0 |
| 0 | 0 | 0 | 0 |
| 0 | 0 | 0 | 0 |
| 0 | 0 | 0 | 0 |
| 1 | 0 | 0 | 0 |
| 0 | 0 | 0 | 0 |
| 0 | 0 | 0 | 0 |
| 0 | 0 | 0 | 0 |
| 0 | 0 | 0 | 0 |
| 0 | 0 | 0 | 0 |
| 0 | 0 | 0 | 0 |
| 0 | 0 | 0 | 0 |
| 0 | 0 | 0 | 0 |
| 0 | 0 | 0 | 0 |
| 0 | 0 | 0 | 0 |
| 0 | 0 | 0 | 0 |
| 0 | 0 | 0 | 0 |
| 1 | 0 | 0 | 0 |

| | | | |
|---|---|---|---|
| 1 | 0 | 0 | 1 |
| 0 | 0 | 0 | 0 |
| 0 | 0 | 0 | 0 |
| 0 | 0 | 0 | 0 |
| 0 | 0 | 0 | 0 |
| 0 | 0 | 0 | 0 |
| 0 | 0 | 0 | 0 |
| 0 | 0 | 0 | 0 |
| 0 | 0 | 0 | 0 |
| 0 | 0 | 0 | 0 |
| 0 | 0 | 0 | 0 |
| 0 | 0 | 0 | 0 |
| 0 | 0 | 0 | 0 |
| 0 | 0 | 0 | 0 |
| 0 | 0 | 0 | 0 |
| 0 | 0 | 0 | 0 |
| 0 | 0 | 0 | 0 |
| 0 | 0 | 0 | 0 |
| 0 | 0 | 0 | 0 |
| 0 | 0 | 0 | 0 |
| 0 | 0 | 0 | 0 |
| 0 | 0 | 0 | 0 |
| 0 | 1 | 0 | 0 |
| 0 | 0 | 0 | 0 |
| 0 | 0 | 0 | 0 |
| 0 | 0 | 0 | 0 |
| 0 | 0 | 0 | 0 |
| 0 | 0 | 0 | 0 |
| 0 | 0 | 0 | 0 |
| 0 | 0 | 0 | 0 |
| 0 | 0 | 0 | 0 |
| 0 | 0 | 0 | 0 |
| 0 | 0 | 0 | 0 |
| 0 | 0 | 0 | 0 |
| 0 | 0 | 0 | 0 |
| 0 | 0 | 0 | 0 |
| 0 | 0 | 0 | 0 |
| 0 | 0 | 0 | 0 |
| 0 | 0 | 0 | 0 |
| 0 | 0 | 0 | 0 |
| 0 | 0 | 0 | 0 |
| 0 | 0 | 0 | 0 |
| 0 | 0 | 0 | 0 |
| 0 | 0 | 0 | 0 |
| 0 | 0 | 0 | 0 |
| 0 | 0 | 0 | 0 |
| 0 | 0 | 0 | 0 |
| 0 | 0 | 0 | 0 |
| 0 | 0 | 0 | 0 |
| 0 | 0 | 0 | 0 |
| 0 | 0 | 0 | 0 |
| 0 | 0 | 0 | 0 |
| 0 | 0 | 0 | 0 |
| 0 | 0 | 0 | 0 |

| | | | |
|---|---|---|---|
| 0 | 0 | 0 | 0 |
| 0 | 0 | 0 | 0 |
| 0 | 0 | 0 | 0 |
| 0 | 0 | 0 | 0 |
| 0 | 0 | 0 | 0 |
| 0 | 0 | 0 | 0 |
| 0 | 1 | 0 | 0 |
| 0 | 1 | 0 | 0 |
| 0 | 0 | 0 | 0 |
| 0 | 0 | 0 | 0 |
| 0 | 0 | 0 | 0 |
| 0 | 0 | 0 | 0 |
| 0 | 0 | 0 | 0 |
| 0 | 0 | 0 | 0 |
| 0 | 0 | 0 | 0 |
| 0 | 0 | 0 | 0 |
| 1 | 0 | 0 | 0 |
| 0 | 0 | 0 | 0 |
| 1 | 0 | 0 | 0 |
| 1 | 0 | 0 | 0 |
| 1 | 0 | 0 | 0 |
| 1 | 0 | 0 | 0 |
| 0 | 0 | 0 | 0 |
| 0 | 0 | 0 | 0 |
| 0 | 0 | 0 | 0 |
| 0 | 0 | 0 | 0 |
| 0 | 0 | 0 | 0 |
| 0 | 0 | 0 | 0 |
| 0 | 0 | 0 | 0 |
| 0 | 0 | 0 | 0 |
| 0 | 0 | 0 | 0 |
| 0 | 0 | 0 | 0 |
| 0 | 0 | 0 | 0 |
| 0 | 0 | 0 | 0 |
| 0 | 0 | 0 | 0 |
| 0 | 0 | 1 | 0 |
| 1 | 0 | 0 | 0 |
| 1 | 0 | 0 | 0 |
| 1 | 0 | 0 | 0 |
| 0 | 0 | 0 | 0 |
| 0 | 0 | 0 | 0 |
| 0 | 0 | 0 | 0 |
| 0 | 0 | 0 | 0 |
| 1 | 0 | 0 | 0 |
| 1 | 0 | 0 | 0 |
| 1 | 0 | 0 | 0 |
| 0 | 0 | 0 | 0 |
| 0 | 0 | 0 | 0 |
| 1 | 0 | 0 | 0 |
| 0 | 0 | 0 | 0 |
| 0 | 0 | 0 | 0 |
| 1 | 0 | 0 | 0 |
| 0 | 0 | 0 | 0 |
| 0 | 0 | 0 | 0 |

| | | | |
|---|---|---|---|
| 0 | 0 | 0 | 0 |
| 0 | 0 | 0 | 0 |
| 0 | 0 | 0 | 0 |
| 0 | 0 | 0 | 0 |
| 0 | 0 | 0 | 0 |
| 0 | 0 | 0 | 0 |
| 0 | 0 | 0 | 0 |
| 0 | 0 | 0 | 0 |
| 0 | 0 | 0 | 0 |
| 0 | 0 | 0 | 0 |
| 0 | 0 | 0 | 0 |
| 0 | 0 | 0 | 0 |
| 0 | 0 | 0 | 0 |
| 0 | 0 | 0 | 0 |
| 0 | 0 | 0 | 0 |
| 0 | 0 | 0 | 0 |
| 0 | 0 | 0 | 0 |
| 0 | 0 | 0 | 0 |
| 0 | 0 | 0 | 0 |
| 0 | 0 | 0 | 0 |
| 0 | 0 | 0 | 0 |
| 0 | 0 | 0 | 0 |
| 1 | 1 | 0 | 0 |
| 0 | 0 | 0 | 0 |
| 0 | 1 | 0 | 0 |
| 0 | 0 | 0 | 0 |
| 0 | 0 | 0 | 0 |
| 0 | 0 | 0 | 0 |
| 0 | 0 | 0 | 0 |
| 0 | 0 | 0 | 0 |
| 0 | 0 | 0 | 0 |
| 0 | 0 | 0 | 0 |
| 0 | 0 | 0 | 0 |
| 0 | 0 | 0 | 0 |
| 0 | 0 | 0 | 0 |
| 0 | 0 | 0 | 0 |
| 0 | 0 | 0 | 0 |
| 0 | 0 | 0 | 0 |
| 0 | 0 | 0 | 0 |
| 0 | 0 | 0 | 0 |
| 0 | 0 | 0 | 0 |
| 0 | 0 | 0 | 0 |
| 0 | 0 | 0 | 0 |
| 0 | 0 | 0 | 0 |
| 0 | 0 | 0 | 0 |
| 1 | 0 | 0 | 1 |
| 0 | 0 | 0 | 0 |
| 0 | 0 | 0 | 0 |
| 0 | 0 | 0 | 0 |
| 1 | 0 | 0 | 0 |
| 1 | 0 | 0 | 0 |
| 1 | 0 | 0 | 0 |
| 0 | 0 | 0 | 0 |
| 0 | 0 | 0 | 0 |

| | | | |
|---|---|---|---|
| 0 | 0 | 0 | 0 |
| 0 | 0 | 0 | 0 |
| 0 | 0 | 0 | 0 |
| 0 | 0 | 0 | 0 |
| 0 | 0 | 0 | 0 |
| 0 | 0 | 0 | 0 |
| 0 | 0 | 0 | 0 |
| 0 | 0 | 0 | 0 |
| 0 | 0 | 0 | 0 |
| 0 | 0 | 0 | 0 |
| 0 | 0 | 0 | 1 |
| 0 | 0 | 0 | 0 |
| 0 | 0 | 0 | 0 |
| 0 | 0 | 0 | 0 |
| 0 | 0 | 0 | 0 |
| 0 | 0 | 0 | 0 |
| 0 | 0 | 0 | 0 |
| 0 | 0 | 0 | 0 |
| 0 | 0 | 0 | 0 |
| 0 | 0 | 0 | 0 |
| 0 | 0 | 0 | 0 |
| 0 | 0 | 0 | 0 |
| 0 | 0 | 0 | 0 |
| 0 | 0 | 0 | 0 |
| 0 | 0 | 0 | 0 |
| 0 | 0 | 0 | 0 |
| 0 | 0 | 0 | 0 |
| 0 | 0 | 0 | 0 |
| 0 | 0 | 0 | 0 |
| 0 | 0 | 0 | 0 |
| 0 | 1 | 0 | 0 |
| 0 | 0 | 0 | 0 |
| 1 | 1 | 0 | 0 |
| 1 | 1 | 0 | 0 |
| 0 | 0 | 0 | 0 |
| 0 | 0 | 0 | 0 |
| 0 | 0 | 0 | 0 |
| 0 | 0 | 0 | 0 |
| 0 | 0 | 0 | 0 |
| 0 | 0 | 0 | 0 |
| 0 | 0 | 0 | 0 |
| 0 | 1 | 0 | 0 |
| 0 | 0 | 0 | 0 |
| 0 | 0 | 0 | 0 |
| 0 | 0 | 0 | 0 |
| 1 | 0 | 0 | 0 |
| 1 | 0 | 0 | 0 |
| 1 | 0 | 0 | 0 |
| 1 | 1 | 1 | 0 |
| 0 | 0 | 0 | 0 |
| 1 | 1 | 1 | 1 |
| 0 | 0 | 0 | 0 |
| 0 | 0 | 0 | 0 |
| 0 | 0 | 0 | 0 |

| | | | |
|---|---|---|---|
| 0 | 0 | 0 | 0 |
| 0 | 0 | 0 | 0 |
| 0 | 0 | 0 | 0 |
| 0 | 0 | 0 | 0 |
| 1 | 0 | 0 | 0 |
| 1 | 0 | 0 | 0 |
| 0 | 0 | 0 | 0 |
| 0 | 0 | 0 | 0 |
| 0 | 0 | 0 | 0 |
| 0 | 0 | 0 | 0 |
| 0 | 0 | 0 | 0 |
| 0 | 0 | 0 | 0 |
| 0 | 0 | 0 | 0 |
| 0 | 0 | 0 | 0 |
| 0 | 0 | 0 | 0 |
| 0 | 0 | 0 | 0 |
| 0 | 0 | 0 | 0 |
| 0 | 0 | 0 | 0 |
| 0 | 0 | 0 | 0 |
| 0 | 0 | 0 | 0 |
| 0 | 0 | 0 | 0 |
| 0 | 0 | 0 | 0 |
| 0 | 0 | 0 | 0 |
| 0 | 0 | 0 | 0 |
| 0 | 0 | 0 | 0 |
| 0 | 0 | 0 | 0 |
| 0 | 0 | 0 | 0 |
| 0 | 0 | 0 | 0 |
| 0 | 0 | 0 | 0 |
| 0 | 0 | 0 | 0 |
| 0 | 0 | 0 | 0 |
| 0 | 0 | 0 | 0 |
| 0 | 0 | 0 | 0 |
| 0 | 0 | 0 | 0 |
| 0 | 0 | 0 | 0 |
| 0 | 0 | 0 | 0 |
| 0 | 0 | 0 | 0 |
| 0 | 0 | 0 | 0 |
| 0 | 0 | 0 | 0 |
| 0 | 0 | 0 | 0 |
| 0 | 0 | 0 | 0 |
| 0 | 0 | 0 | 0 |
| 0 | 0 | 0 | 0 |
| 0 | 0 | 0 | 0 |
| 0 | 0 | 0 | 0 |
| 0 | 0 | 0 | 0 |
| 0 | 0 | 0 | 0 |
| 0 | 0 | 0 | 0 |
| 0 | 0 | 0 | 0 |
| 0 | 0 | 0 | 0 |
| 0 | 0 | 0 | 0 |
| 0 | 0 | 0 | 0 |
| 0 | 0 | 0 | 0 |
| 0 | 1 | 0 | 0 |

| | | | |
|---|---|---|---|
| 0 | 0 | 0 | 0 |
| 0 | 0 | 0 | 0 |
| 0 | 0 | 0 | 0 |
| 0 | 0 | 0 | 0 |
| 0 | 0 | 0 | 0 |
| 0 | 0 | 0 | 0 |
| 0 | 0 | 0 | 0 |
| 0 | 0 | 0 | 0 |
| 0 | 0 | 0 | 0 |
| 0 | 0 | 0 | 0 |
| 0 | 0 | 0 | 0 |
| 0 | 0 | 0 | 0 |
| 0 | 0 | 0 | 0 |
| 0 | 0 | 0 | 0 |
| 0 | 0 | 0 | 0 |
| 0 | 0 | 0 | 0 |
| 0 | 0 | 0 | 0 |
| 0 | 0 | 0 | 0 |
| 0 | 0 | 0 | 0 |
| 0 | 0 | 0 | 0 |
| 0 | 0 | 0 | 0 |
| 0 | 0 | 0 | 0 |
| 0 | 0 | 0 | 0 |
| 0 | 0 | 0 | 0 |
| 0 | 0 | 0 | 0 |
| 0 | 0 | 0 | 0 |
| 0 | 0 | 0 | 0 |
| 0 | 0 | 0 | 0 |
| 0 | 0 | 0 | 0 |
| 0 | 0 | 0 | 0 |
| 0 | 0 | 0 | 0 |
| 0 | 0 | 0 | 0 |
| 0 | 0 | 0 | 0 |
| 0 | 0 | 0 | 0 |
| 0 | 0 | 0 | 0 |
| 0 | 0 | 0 | 0 |
| 0 | 0 | 0 | 0 |
| 0 | 0 | 0 | 0 |
| 0 | 0 | 0 | 0 |
| 0 | 0 | 0 | 0 |
| 0 | 0 | 0 | 0 |
| 0 | 0 | 0 | 0 |
| 0 | 0 | 0 | 0 |
| 0 | 0 | 0 | 0 |
| 0 | 0 | 0 | 0 |
| 0 | 0 | 0 | 0 |
| 0 | 0 | 0 | 0 |
| 0 | 0 | 0 | 0 |
| 0 | 0 | 0 | 0 |
| 0 | 0 | 0 | 0 |
| 0 | 0 | 0 | 0 |
| 0 | 0 | 0 | 0 |
| 0 | 0 | 0 | 0 |
| 0 | 0 | 0 | 0 |

| | | | |
|---|---|---|---|
| 0 | 0 | 0 | 0 |
| 0 | 0 | 0 | 0 |
| 0 | 0 | 0 | 0 |
| 0 | 0 | 0 | 0 |
| 0 | 0 | 0 | 0 |
| 0 | 0 | 0 | 0 |
| 0 | 0 | 0 | 0 |
| 0 | 0 | 0 | 0 |
| 0 | 0 | 0 | 0 |
| 0 | 0 | 0 | 0 |
| 0 | 0 | 0 | 0 |
| 0 | 0 | 1 | 0 |
| 0 | 0 | 0 | 0 |
| 0 | 0 | 1 | 0 |
| 0 | 0 | 1 | 0 |
| 0 | 0 | 1 | 0 |
| 0 | 0 | 1 | 0 |
| 0 | 0 | 0 | 0 |
| 0 | 0 | 0 | 0 |
| 0 | 0 | 0 | 0 |
| 1 | 0 | 0 | 0 |

| glycan_binding_site | water_binding_site | water_only_binding_site | non_binding_site |
|---|---|---|---|
| 0 | 0 | 0 | 0 |
| 0 | 0 | 0 | 0 |
| 0 | 0 | 0 | 1 |
| 0 | 0 | 0 | 0 |
| 0 | 0 | 0 | 0 |
| 0 | 0 | 0 | 0 |
| 0 | 0 | 0 | 1 |
| 0 | 0 | 0 | 1 |
| 0 | 1 | 0 | 0 |
| 0 | 0 | 0 | 0 |
| 0 | 0 | 0 | 0 |
| 0 | 0 | 0 | 0 |
| 0 | 0 | 0 | 1 |
| 1 | 0 | 0 | 0 |
| 0 | 0 | 0 | 0 |
| 0 | 0 | 0 | 0 |
| 0 | 0 | 0 | 0 |
| 0 | 1 | 0 | 0 |
| 0 | 0 | 0 | 0 |
| 0 | 0 | 0 | 0 |
| 0 | 1 | 0 | 0 |
| 1 | 1 | 0 | 0 |
| 1 | 1 | 0 | 0 |
| 0 | 1 | 0 | 0 |
| 0 | 1 | 0 | 0 |
| 0 | 0 | 0 | 1 |
| 0 | 1 | 0 | 0 |
| 0 | 0 | 0 | 1 |
| 0 | 0 | 0 | 0 |
| 0 | 0 | 0 | 1 |
| 0 | 0 | 0 | 1 |
| 0 | 1 | 0 | 0 |
| 0 | 1 | 0 | 0 |
| 0 | 1 | 0 | 0 |
| 0 | 1 | 0 | 0 |
| 0 | 0 | 0 | 1 |
| 0 | 0 | 0 | 0 |
| 0 | 1 | 0 | 0 |
| 0 | 1 | 0 | 0 |
| 0 | 1 | 0 | 0 |
| 0 | 1 | 0 | 0 |
| 0 | 1 | 0 | 0 |
| 0 | 1 | 0 | 0 |
| 0 | 1 | 0 | 0 |
| 0 | 1 | 0 | 0 |
| 0 | 1 | 0 | 0 |
| 0 | 1 | 0 | 0 |
| 0 | 1 | 0 | 0 |
| 0 | 1 | 0 | 0 |
| 0 | 1 | 0 | 0 |
| 0 | 1 | 0 | 0 |
| 0 | 1 | 0 | 0 |
| 0 | 1 | 0 | 0 |

| | | | |
|---|---|---|---|
| 0 | 1 | 0 | 0 |
| 0 | 1 | 0 | 0 |
| 0 | 1 | 0 | 0 |
| 0 | 1 | 0 | 0 |
| 0 | 1 | 0 | 0 |
| 0 | 1 | 0 | 0 |
| 0 | 1 | 0 | 0 |
| 0 | 1 | 0 | 0 |
| 0 | 1 | 0 | 0 |
| 0 | 1 | 0 | 0 |
| 0 | 1 | 0 | 0 |
| 0 | 1 | 0 | 0 |
| 0 | 1 | 0 | 0 |
| 0 | 1 | 0 | 0 |
| 0 | 1 | 0 | 0 |
| 0 | 1 | 0 | 0 |
| 0 | 1 | 0 | 0 |
| 0 | 1 | 0 | 0 |
| 0 | 1 | 0 | 0 |
| 0 | 1 | 0 | 0 |
| 0 | 0 | 0 | 0 |
| 0 | 0 | 0 | 0 |
| 0 | 0 | 0 | 0 |
| 0 | 0 | 0 | 0 |
| 0 | 0 | 0 | 0 |
| 0 | 0 | 0 | 0 |
| 0 | 0 | 0 | 0 |
| 0 | 0 | 0 | 0 |
| 0 | 0 | 0 | 0 |
| 0 | 0 | 0 | 0 |
| 0 | 0 | 0 | 0 |
| 0 | 0 | 0 | 0 |
| 0 | 0 | 0 | 0 |
| 0 | 0 | 0 | 0 |
| 0 | 0 | 0 | 0 |
| 0 | 0 | 0 | 0 |
| 0 | 0 | 0 | 0 |
| 0 | 0 | 0 | 0 |
| 0 | 0 | 0 | 0 |
| 0 | 1 | 0 | 0 |
| 1 | 0 | 0 | 0 |
| 0 | 0 | 0 | 0 |
| 1 | 0 | 0 | 0 |
| 1 | 0 | 0 | 0 |
| 0 | 0 | 0 | 0 |
| 0 | 0 | 0 | 1 |
| 0 | 0 | 0 | 1 |
| 0 | 0 | 0 | 0 |
| 1 | 0 | 0 | 0 |
| 0 | 0 | 0 | 0 |
| 0 | 0 | 0 | 0 |
| 0 | 0 | 0 | 1 |
| 0 | 0 | 0 | 1 |
| 0 | 0 | 0 | 0 |

| | | | |
|---|---|---|---|
| 0 | 0 | 0 | 1 |
| 0 | 0 | 0 | 1 |
| 0 | 0 | 0 | 0 |
| 0 | 0 | 0 | 0 |
| 0 | 0 | 0 | 0 |
| 0 | 0 | 0 | 0 |
| 0 | 0 | 0 | 1 |
| 0 | 1 | 0 | 0 |
| 0 | 1 | 0 | 0 |
| 0 | 0 | 0 | 0 |
| 0 | 1 | 0 | 0 |
| 0 | 0 | 0 | 0 |
| 0 | 0 | 0 | 0 |
| 1 | 0 | 0 | 0 |
| 1 | 0 | 0 | 0 |
| 0 | 0 | 0 | 1 |
| 0 | 0 | 0 | 0 |
| 0 | 0 | 0 | 0 |
| 0 | 0 | 0 | 0 |
| 0 | 0 | 0 | 0 |
| 0 | 0 | 0 | 0 |
| 0 | 0 | 0 | 0 |
| 0 | 0 | 0 | 0 |
| 0 | 0 | 0 | 0 |
| 0 | 0 | 0 | 0 |
| 0 | 0 | 0 | 0 |
| 0 | 0 | 0 | 0 |
| 0 | 1 | 0 | 0 |
| 0 | 0 | 0 | 0 |
| 0 | 0 | 0 | 0 |
| 0 | 0 | 0 | 0 |
| 0 | 0 | 0 | 0 |
| 0 | 0 | 0 | 0 |
| 0 | 1 | 0 | 0 |
| 0 | 1 | 0 | 0 |
| 0 | 0 | 0 | 0 |
| 0 | 0 | 0 | 0 |
| 0 | 0 | 0 | 0 |
| 0 | 0 | 0 | 0 |
| 0 | 1 | 0 | 0 |
| 0 | 1 | 0 | 0 |
| 0 | 1 | 0 | 0 |
| 0 | 0 | 0 | 0 |
| 0 | 0 | 0 | 0 |
| 0 | 0 | 0 | 0 |
| 0 | 1 | 0 | 0 |
| 1 | 1 | 0 | 0 |
| 0 | 0 | 0 | 0 |
| 0 | 1 | 0 | 0 |
| 0 | 0 | 0 | 0 |
| 0 | 0 | 0 | 0 |
| 0 | 0 | 0 | 0 |
| 0 | 0 | 0 | 0 |
| 0 | 1 | 1 | 1 |

| | | | |
|---|---|---|---|
| 0 | 1 | 1 | 1 |
| 0 | 1 | 1 | 1 |
| 0 | 1 | 1 | 1 |
| 0 | 0 | 0 | 0 |
| 0 | 0 | 0 | 0 |
| 0 | 1 | 1 | 1 |
| 0 | 1 | 0 | 0 |
| 0 | 1 | 1 | 1 |
| 0 | 1 | 1 | 1 |
| 0 | 1 | 1 | 1 |
| 0 | 1 | 1 | 1 |
| 0 | 1 | 0 | 0 |
| 0 | 1 | 1 | 1 |
| 0 | 0 | 0 | 0 |
| 0 | 1 | 1 | 1 |
| 0 | 0 | 0 | 1 |
| 0 | 1 | 1 | 1 |
| 0 | 1 | 1 | 1 |
| 0 | 1 | 1 | 1 |
| 0 | 1 | 0 | 0 |
| 0 | 1 | 0 | 0 |
| 0 | 0 | 0 | 1 |
| 0 | 1 | 0 | 0 |
| 0 | 1 | 0 | 0 |
| 0 | 1 | 0 | 0 |
| 0 | 0 | 0 | 1 |
| 0 | 1 | 0 | 0 |
| 0 | 1 | 0 | 0 |
| 0 | 0 | 0 | 1 |
| 0 | 1 | 1 | 1 |
| 0 | 0 | 0 | 1 |
| 0 | 1 | 1 | 1 |
| 0 | 1 | 1 | 1 |
| 0 | 0 | 0 | 1 |
| 0 | 1 | 0 | 0 |
| 0 | 1 | 1 | 1 |
| 1 | 1 | 0 | 0 |
| 1 | 1 | 0 | 0 |
| 0 | 1 | 1 | 1 |
| 0 | 1 | 1 | 1 |
| 0 | 1 | 1 | 1 |
| 0 | 1 | 0 | 0 |
| 0 | 1 | 0 | 0 |
| 0 | 0 | 0 | 1 |
| 0 | 1 | 1 | 1 |
| 0 | 1 | 1 | 1 |
| 0 | 0 | 0 | 0 |
| 0 | 0 | 0 | 0 |
| 0 | 0 | 0 | 0 |
| 0 | 0 | 0 | 0 |
| 0 | 0 | 0 | 0 |
| 0 | 0 | 0 | 1 |
| 0 | 0 | 0 | 1 |
| 0 | 0 | 0 | 0 |

| | | | |
|---|---|---|---|
| 0 | 0 | 0 | 0 |
| 0 | 0 | 0 | 0 |
| 0 | 1 | 0 | 0 |
| 0 | 0 | 0 | 1 |
| 0 | 0 | 0 | 1 |
| 0 | 0 | 0 | 1 |
| 0 | 1 | 0 | 0 |
| 0 | 1 | 0 | 0 |
| 0 | 1 | 1 | 1 |
| 0 | 0 | 0 | 1 |
| 1 | 0 | 0 | 0 |
| 0 | 0 | 0 | 1 |
| 1 | 0 | 0 | 0 |
| 0 | 0 | 0 | 1 |
| 0 | 0 | 0 | 1 |
| 0 | 0 | 0 | 1 |
| 0 | 0 | 0 | 1 |
| 0 | 0 | 0 | 0 |
| 0 | 0 | 0 | 0 |
| 0 | 0 | 0 | 0 |
| 0 | 0 | 0 | 0 |
| 0 | 0 | 0 | 1 |
| 0 | 0 | 0 | 0 |
| 0 | 0 | 0 | 1 |
| 0 | 0 | 0 | 1 |
| 0 | 0 | 0 | 0 |
| 0 | 1 | 0 | 0 |
| 0 | 0 | 0 | 0 |
| 0 | 1 | 0 | 0 |
| 0 | 1 | 0 | 0 |
| 0 | 1 | 0 | 0 |
| 0 | 0 | 0 | 0 |
| 0 | 0 | 0 | 1 |
| 0 | 0 | 0 | 1 |
| 0 | 0 | 0 | 0 |
| 0 | 0 | 0 | 1 |
| 0 | 0 | 0 | 1 |
| 0 | 0 | 0 | 0 |
| 0 | 1 | 1 | 1 |
| 0 | 0 | 0 | 1 |
| 0 | 1 | 0 | 0 |
| 0 | 1 | 1 | 1 |
| 0 | 0 | 0 | 1 |
| 0 | 0 | 0 | 1 |
| 0 | 0 | 0 | 0 |
| 0 | 0 | 0 | 0 |
| 0 | 1 | 0 | 0 |
| 0 | 1 | 0 | 0 |
| 0 | 1 | 0 | 0 |
| 0 | 1 | 0 | 0 |
| 0 | 1 | 0 | 0 |
| 0 | 1 | 0 | 0 |
| 0 | 1 | 0 | 0 |
| 0 | 1 | 0 | 0 |

| | | | |
|---|---|---|---|
| 0 | 1 | 0 | 0 |
| 0 | 1 | 0 | 0 |
| 0 | 1 | 0 | 0 |
| 0 | 1 | 0 | 0 |
| 0 | 1 | 0 | 0 |
| 0 | 1 | 0 | 0 |
| 0 | 0 | 0 | 1 |
| 0 | 0 | 0 | 1 |
| 0 | 0 | 0 | 0 |
| 0 | 0 | 0 | 0 |
| 0 | 0 | 0 | 0 |
| 0 | 0 | 0 | 0 |
| 0 | 0 | 0 | 0 |
| 0 | 0 | 0 | 1 |
| 0 | 0 | 0 | 1 |
| 0 | 0 | 0 | 0 |
| 0 | 0 | 0 | 1 |
| 0 | 0 | 0 | 0 |
| 0 | 0 | 0 | 1 |
| 0 | 0 | 0 | 0 |
| 0 | 0 | 0 | 1 |
| 0 | 1 | 1 | 1 |
| 0 | 0 | 0 | 1 |
| 0 | 0 | 0 | 0 |
| 0 | 0 | 0 | 0 |
| 0 | 0 | 0 | 0 |
| 0 | 0 | 0 | 0 |
| 0 | 0 | 0 | 0 |
| 0 | 0 | 0 | 0 |
| 0 | 1 | 0 | 0 |
| 0 | 0 | 0 | 1 |
| 0 | 0 | 0 | 1 |
| 0 | 0 | 0 | 0 |
| 0 | 0 | 0 | 0 |
| 0 | 1 | 0 | 0 |
| 0 | 0 | 0 | 1 |
| 0 | 0 | 0 | 0 |
| 0 | 1 | 0 | 0 |
| 0 | 0 | 0 | 0 |
| 0 | 1 | 0 | 0 |
| 0 | 0 | 0 | 1 |
| 0 | 0 | 0 | 1 |
| 0 | 0 | 0 | 0 |
| 0 | 0 | 0 | 1 |
| 0 | 0 | 0 | 0 |
| 0 | 0 | 0 | 0 |
| 0 | 0 | 0 | 0 |
| 0 | 0 | 0 | 1 |
| 0 | 0 | 0 | 1 |
| 0 | 0 | 0 | 0 |
| 0 | 0 | 0 | 1 |
| 0 | 0 | 0 | 1 |
| 0 | 0 | 0 | 0 |
| 0 | 0 | 0 | 1 |

| | | | |
|---|---|---|---|
| 0 | 0 | 0 | 0 |
| 0 | 0 | 0 | 1 |
| 0 | 1 | 1 | 1 |
| 0 | 1 | 0 | 0 |
| 0 | 1 | 0 | 0 |
| 0 | 0 | 0 | 0 |
| 0 | 0 | 0 | 1 |
| 0 | 0 | 0 | 0 |
| 0 | 0 | 0 | 0 |
| 0 | 0 | 0 | 1 |
| 0 | 0 | 0 | 1 |
| 0 | 0 | 0 | 1 |
| 0 | 0 | 0 | 1 |
| 0 | 0 | 0 | 1 |
| 0 | 0 | 0 | 0 |
| 0 | 0 | 0 | 1 |
| 0 | 0 | 0 | 0 |
| 0 | 0 | 0 | 0 |
| 0 | 0 | 0 | 1 |
| 0 | 0 | 0 | 0 |
| 0 | 1 | 0 | 0 |
| 0 | 1 | 0 | 0 |
| 0 | 0 | 0 | 0 |
| 0 | 1 | 0 | 0 |
| 0 | 0 | 0 | 1 |
| 0 | 0 | 0 | 1 |
| 0 | 0 | 0 | 0 |
| 0 | 0 | 0 | 0 |
| 0 | 1 | 0 | 0 |
| 0 | 0 | 0 | 0 |
| 0 | 0 | 0 | 0 |
| 0 | 0 | 0 | 0 |
| 0 | 0 | 0 | 0 |
| 0 | 0 | 0 | 0 |
| 0 | 0 | 0 | 1 |
| 0 | 0 | 0 | 1 |
| 0 | 0 | 0 | 0 |
| 0 | 0 | 0 | 0 |
| 0 | 0 | 0 | 0 |
| 0 | 0 | 0 | 0 |
| 0 | 0 | 0 | 0 |
| 0 | 1 | 0 | 0 |
| 0 | 0 | 0 | 1 |
| 0 | 0 | 0 | 0 |
| 0 | 1 | 0 | 0 |
| 0 | 1 | 0 | 0 |
| 0 | 1 | 0 | 0 |
| 0 | 1 | 0 | 0 |
| 1 | 1 | 0 | 0 |
| 0 | 0 | 0 | 1 |
| 0 | 1 | 0 | 0 |
| 0 | 0 | 0 | 0 |
| 0 | 0 | 0 | 0 |
| 0 | 0 | 0 | 0 |

| | | | |
|---|---|---|---|
| 0 | 1 | 0 | 0 |
| 0 | 0 | 0 | 0 |
| 0 | 0 | 0 | 0 |
| 0 | 0 | 0 | 0 |
| 0 | 0 | 0 | 1 |
| 0 | 1 | 0 | 0 |
| 0 | 1 | 0 | 0 |
| 0 | 1 | 0 | 0 |
| 0 | 1 | 0 | 0 |
| 0 | 1 | 0 | 0 |
| 0 | 1 | 0 | 0 |
| 0 | 1 | 0 | 0 |
| 0 | 0 | 0 | 0 |
| 0 | 1 | 0 | 0 |
| 0 | 0 | 0 | 0 |
| 0 | 1 | 0 | 0 |
| 0 | 1 | 0 | 0 |
| 0 | 1 | 0 | 0 |
| 0 | 0 | 0 | 1 |
| 0 | 0 | 0 | 0 |
| 0 | 0 | 0 | 0 |
| 0 | 0 | 0 | 0 |
| 0 | 1 | 0 | 0 |
| 0 | 1 | 1 | 1 |
| 0 | 0 | 0 | 0 |
| 0 | 0 | 0 | 0 |
| 0 | 0 | 0 | 0 |
| 0 | 1 | 1 | 1 |
| 0 | 1 | 1 | 1 |
| 0 | 1 | 1 | 1 |
| 0 | 1 | 0 | 0 |
| 0 | 0 | 0 | 0 |
| 0 | 1 | 0 | 0 |
| 0 | 1 | 0 | 0 |
| 0 | 1 | 0 | 0 |
| 0 | 1 | 1 | 1 |
| 0 | 1 | 1 | 1 |
| 0 | 0 | 0 | 0 |
| 0 | 1 | 0 | 0 |
| 0 | 1 | 1 | 1 |
| 0 | 1 | 1 | 1 |
| 0 | 1 | 1 | 1 |
| 0 | 0 | 0 | 1 |
| 0 | 0 | 0 | 1 |
| 0 | 0 | 0 | 0 |
| 0 | 0 | 0 | 0 |
| 0 | 1 | 0 | 0 |
| 0 | 0 | 0 | 0 |
| 0 | 0 | 0 | 0 |
| 0 | 0 | 0 | 1 |
| 0 | 0 | 0 | 1 |
| 0 | 0 | 0 | 0 |
| 0 | 0 | 0 | 1 |
| 0 | 0 | 0 | 0 |

| | | | |
|---|---|---|---|
| 0 | 0 | 0 | 1 |
| 0 | 0 | 0 | 1 |
| 0 | 0 | 0 | 0 |
| 0 | 0 | 0 | 1 |
| 0 | 0 | 0 | 1 |
| 0 | 0 | 0 | 1 |
| 0 | 0 | 0 | 1 |
| 0 | 0 | 0 | 0 |
| 0 | 0 | 0 | 1 |
| 0 | 0 | 0 | 1 |
| 0 | 0 | 0 | 0 |
| 0 | 0 | 0 | 1 |
| 0 | 0 | 0 | 1 |
| 0 | 0 | 0 | 1 |
| 0 | 0 | 0 | 0 |
| 0 | 0 | 0 | 0 |
| 0 | 0 | 0 | 1 |
| 0 | 0 | 0 | 1 |
| 0 | 0 | 0 | 0 |
| 0 | 0 | 0 | 0 |
| 0 | 0 | 0 | 0 |
| 0 | 0 | 0 | 0 |
| 0 | 0 | 0 | 0 |
| 0 | 1 | 0 | 0 |
| 0 | 1 | 0 | 0 |
| 0 | 0 | 0 | 1 |
| 0 | 0 | 0 | 0 |
| 0 | 0 | 0 | 0 |
| 0 | 0 | 0 | 0 |
| 0 | 0 | 0 | 1 |
| 0 | 0 | 0 | 0 |
| 0 | 1 | 0 | 0 |
| 0 | 0 | 0 | 1 |
| 0 | 0 | 0 | 0 |
| 0 | 0 | 0 | 0 |
| 0 | 0 | 0 | 0 |
| 0 | 1 | 0 | 0 |
| 0 | 1 | 0 | 0 |
| 0 | 1 | 0 | 0 |
| 0 | 1 | 0 | 0 |
| 0 | 0 | 0 | 1 |
| 0 | 0 | 0 | 0 |
| 0 | 0 | 0 | 1 |
| 0 | 0 | 0 | 0 |
| 0 | 0 | 0 | 0 |
| 1 | 0 | 0 | 0 |
| 0 | 0 | 0 | 0 |
| 1 | 0 | 0 | 0 |
| 0 | 0 | 0 | 0 |
| 0 | 0 | 0 | 0 |
| 0 | 0 | 0 | 1 |
| 0 | 0 | 0 | 0 |
| 0 | 0 | 0 | 0 |
| 0 | 0 | 0 | 0 |

| | | | |
|---|---|---|---|
| 0 | 0 | 0 | 1 |
| 0 | 0 | 0 | 0 |
| 0 | 0 | 0 | 1 |
| 0 | 0 | 0 | 1 |
| 1 | 0 | 0 | 0 |
| 0 | 0 | 0 | 0 |
| 1 | 0 | 0 | 0 |
| 1 | 0 | 0 | 0 |
| 0 | 0 | 0 | 0 |
| 0 | 0 | 0 | 1 |
| 0 | 0 | 0 | 1 |
| 0 | 0 | 0 | 1 |
| 0 | 0 | 0 | 1 |
| 1 | 0 | 0 | 0 |
| 0 | 0 | 0 | 1 |
| 0 | 0 | 0 | 0 |
| 0 | 0 | 0 | 0 |
| 0 | 0 | 0 | 0 |
| 1 | 0 | 0 | 0 |
| 0 | 0 | 0 | 1 |
| 1 | 0 | 0 | 0 |
| 0 | 0 | 0 | 1 |
| 0 | 0 | 0 | 0 |
| 0 | 0 | 0 | 1 |
| 0 | 0 | 0 | 1 |
| 0 | 0 | 0 | 1 |
| 0 | 0 | 0 | 0 |
| 0 | 0 | 0 | 1 |
| 0 | 0 | 0 | 1 |
| 0 | 0 | 0 | 1 |
| 0 | 0 | 0 | 0 |
| 0 | 0 | 0 | 0 |
| 0 | 0 | 0 | 1 |
| 0 | 0 | 0 | 1 |
| 0 | 0 | 0 | 0 |
| 0 | 0 | 0 | 0 |
| 0 | 0 | 0 | 0 |
| 1 | 0 | 0 | 0 |
| 0 | 0 | 0 | 0 |
| 0 | 0 | 0 | 1 |
| 0 | 0 | 0 | 0 |
| 0 | 0 | 0 | 0 |
| 0 | 1 | 0 | 0 |
| 0 | 0 | 0 | 0 |
| 0 | 0 | 0 | 1 |
| 1 | 0 | 0 | 0 |
| 0 | 1 | 0 | 0 |
| 0 | 0 | 0 | 0 |
| 0 | 0 | 0 | 0 |
| 1 | 0 | 0 | 0 |
| 0 | 0 | 0 | 0 |
| 0 | 1 | 0 | 0 |
| 0 | 1 | 1 | 1 |
| 0 | 0 | 0 | 0 |

| | | | |
|---|---|---|---|
| 0 | 1 | 0 | 0 |
| 0 | 1 | 0 | 0 |
| 0 | 1 | 1 | 1 |
| 0 | 1 | 1 | 1 |
| 0 | 1 | 1 | 1 |
| 0 | 0 | 0 | 0 |
| 0 | 0 | 0 | 1 |
| 0 | 0 | 0 | 1 |
| 0 | 1 | 0 | 0 |
| 0 | 0 | 0 | 0 |
| 1 | 1 | 0 | 0 |
| 1 | 1 | 0 | 0 |
| 0 | 0 | 0 | 1 |
| 1 | 0 | 0 | 0 |
| 0 | 0 | 0 | 0 |
| 0 | 0 | 0 | 0 |
| 0 | 1 | 0 | 0 |
| 0 | 0 | 0 | 0 |
| 0 | 0 | 0 | 0 |
| 0 | 0 | 0 | 1 |
| 0 | 0 | 0 | 0 |
| 0 | 0 | 0 | 0 |
| 0 | 0 | 0 | 0 |
| 0 | 0 | 0 | 0 |
| 0 | 1 | 0 | 0 |
| 0 | 0 | 0 | 1 |
| 0 | 0 | 0 | 1 |
| 0 | 0 | 0 | 1 |
| 0 | 0 | 0 | 0 |
| 0 | 1 | 0 | 0 |
| 0 | 0 | 0 | 0 |
| 0 | 0 | 0 | 0 |
| 0 | 0 | 0 | 0 |
| 0 | 1 | 0 | 0 |
| 0 | 1 | 0 | 0 |
| 0 | 0 | 0 | 1 |
| 0 | 0 | 0 | 0 |
| 0 | 1 | 0 | 0 |
| 0 | 0 | 0 | 1 |
| 0 | 0 | 0 | 1 |
| 0 | 0 | 0 | 1 |
| 0 | 1 | 0 | 0 |
| 0 | 0 | 0 | 0 |
| 0 | 0 | 0 | 0 |
| 0 | 0 | 0 | 0 |
| 0 | 0 | 0 | 0 |
| 0 | 0 | 0 | 0 |
| 0 | 0 | 0 | 1 |
| 0 | 0 | 0 | 1 |
| 0 | 0 | 0 | 0 |
| 0 | 0 | 0 | 0 |
| 0 | 0 | 0 | 0 |
| 0 | 0 | 0 | 0 |
| 0 | 0 | 0 | 1 |

| | | | |
|---|---|---|---|
| 0 | 0 | 0 | 1 |
| 0 | 0 | 0 | 0 |
| 0 | 0 | 0 | 1 |
| 0 | 1 | 0 | 0 |
| 0 | 1 | 1 | 1 |
| 0 | 1 | 1 | 1 |
| 0 | 1 | 0 | 0 |
| 0 | 1 | 1 | 1 |
| 0 | 1 | 1 | 1 |
| 0 | 1 | 1 | 1 |
| 0 | 1 | 1 | 1 |
| 0 | 1 | 1 | 1 |
| 0 | 0 | 0 | 1 |
| 0 | 0 | 0 | 1 |
| 0 | 0 | 0 | 1 |
| 0 | 0 | 0 | 1 |
| 0 | 0 | 0 | 0 |
| 0 | 0 | 0 | 0 |
| 0 | 1 | 1 | 1 |
| 0 | 0 | 0 | 0 |
| 0 | 0 | 0 | 0 |
| 0 | 0 | 0 | 0 |
| 0 | 0 | 0 | 1 |
| 0 | 1 | 0 | 0 |
| 0 | 0 | 0 | 0 |
| 0 | 0 | 0 | 1 |
| 0 | 0 | 0 | 1 |
| 0 | 0 | 0 | 1 |
| 0 | 1 | 0 | 0 |
| 0 | 0 | 0 | 0 |
| 0 | 1 | 1 | 1 |
| 0 | 1 | 0 | 0 |
| 0 | 1 | 0 | 0 |
| 0 | 0 | 0 | 0 |
| 0 | 0 | 0 | 0 |
| 0 | 0 | 0 | 0 |
| 0 | 1 | 1 | 1 |
| 0 | 0 | 0 | 1 |
| 0 | 0 | 0 | 1 |
| 0 | 0 | 0 | 1 |
| 0 | 0 | 0 | 0 |
| 0 | 0 | 0 | 1 |
| 0 | 0 | 0 | 0 |
| 0 | 0 | 0 | 1 |
| 1 | 0 | 0 | 0 |
| 0 | 0 | 0 | 1 |
| 0 | 0 | 0 | 0 |
| 0 | 0 | 0 | 0 |
| 1 | 0 | 0 | 0 |
| 0 | 0 | 0 | 0 |
| 0 | 0 | 0 | 0 |
| 0 | 0 | 0 | 0 |
| 0 | 0 | 0 | 0 |
| 0 | 0 | 0 | 0 |

| | | | |
|---|---|---|---|
| 0 | 0 | 0 | 0 |
| 0 | 0 | 0 | 0 |
| 0 | 0 | 0 | 0 |
| 0 | 0 | 0 | 0 |
| 0 | 1 | 0 | 0 |
| 0 | 1 | 0 | 0 |
| 0 | 0 | 0 | 0 |
| 0 | 0 | 0 | 1 |
| 0 | 1 | 0 | 0 |
| 0 | 1 | 0 | 0 |
| 0 | 1 | 0 | 0 |
| 0 | 1 | 1 | 1 |
| 0 | 0 | 0 | 0 |
| 0 | 1 | 0 | 0 |
| 0 | 1 | 0 | 0 |
| 0 | 1 | 0 | 0 |
| 0 | 1 | 0 | 0 |
| 0 | 1 | 0 | 0 |
| 0 | 1 | 0 | 0 |
| 0 | 1 | 0 | 0 |
| 0 | 1 | 0 | 0 |
| 0 | 1 | 0 | 0 |
| 0 | 1 | 0 | 0 |
| 0 | 1 | 0 | 0 |
| 0 | 1 | 0 | 0 |
| 0 | 1 | 0 | 0 |
| 0 | 1 | 0 | 0 |
| 0 | 1 | 0 | 0 |
| 0 | 1 | 0 | 0 |
| 0 | 1 | 0 | 0 |
| 0 | 1 | 0 | 0 |
| 0 | 1 | 0 | 0 |
| 0 | 1 | 0 | 0 |
| 0 | 1 | 0 | 0 |
| 0 | 1 | 0 | 0 |
| 0 | 1 | 0 | 0 |
| 0 | 1 | 0 | 0 |
| 0 | 1 | 0 | 0 |
| 0 | 1 | 0 | 0 |
| 0 | 1 | 1 | 1 |
| 0 | 1 | 0 | 0 |
| 0 | 1 | 0 | 0 |
| 0 | 1 | 0 | 0 |
| 0 | 1 | 0 | 0 |
| 0 | 1 | 0 | 0 |
| 0 | 1 | 0 | 0 |
| 0 | 1 | 0 | 0 |
| 0 | 0 | 0 | 0 |
| 0 | 0 | 0 | 1 |
| 0 | 0 | 0 | 1 |
| 0 | 1 | 1 | 1 |
| 1 | 0 | 0 | 0 |
| 0 | 0 | 0 | 0 |

| | | | |
|---|---|---|---|
| 0 | 0 | 0 | 0 |
| 0 | 0 | 0 | 0 |
| 1 | 1 | 0 | 0 |
| 1 | 0 | 0 | 0 |
| 0 | 0 | 0 | 0 |
| 0 | 0 | 0 | 0 |
| 0 | 0 | 0 | 0 |
| 0 | 0 | 0 | 1 |
| 0 | 0 | 0 | 1 |
| 0 | 0 | 0 | 0 |
| 0 | 0 | 0 | 0 |
| 0 | 0 | 0 | 1 |
| 0 | 0 | 0 | 0 |
| 0 | 0 | 0 | 0 |
| 0 | 0 | 0 | 0 |
| 0 | 0 | 0 | 0 |
| 0 | 0 | 0 | 0 |
| 0 | 0 | 0 | 1 |
| 0 | 0 | 0 | 0 |
| 0 | 0 | 0 | 1 |
| 0 | 0 | 0 | 0 |
| 0 | 0 | 0 | 0 |
| 0 | 0 | 0 | 1 |
| 0 | 0 | 0 | 0 |
| 0 | 1 | 0 | 0 |
| 0 | 0 | 0 | 1 |
| 0 | 0 | 0 | 0 |
| 0 | 0 | 0 | 0 |
| 0 | 0 | 0 | 0 |
| 0 | 0 | 0 | 0 |
| 0 | 0 | 0 | 0 |
| 0 | 0 | 0 | 0 |
| 0 | 0 | 0 | 1 |
| 0 | 0 | 0 | 0 |
| 0 | 1 | 0 | 0 |
| 0 | 0 | 0 | 0 |
| 0 | 0 | 0 | 1 |
| 0 | 0 | 0 | 0 |
| 0 | 0 | 0 | 0 |
| 0 | 0 | 0 | 0 |
| 0 | 1 | 0 | 0 |
| 0 | 1 | 0 | 0 |
| 0 | 1 | 0 | 0 |
| 0 | 1 | 0 | 0 |
| 0 | 1 | 0 | 0 |
| 0 | 0 | 0 | 1 |
| 0 | 1 | 0 | 0 |
| 0 | 1 | 0 | 0 |
| 0 | 1 | 0 | 0 |
| 0 | 1 | 0 | 0 |
| 0 | 1 | 0 | 0 |
| 0 | 1 | 0 | 0 |
| 0 | 1 | 1 | 1 |
| 0 | 0 | 0 | 1 |

| | | | |
|---|---|---|---|
| 0 | 1 | 1 | 1 |
| 0 | 0 | 0 | 1 |
| 0 | 0 | 0 | 0 |
| 0 | 0 | 0 | 0 |
| 0 | 1 | 0 | 0 |
| 0 | 1 | 0 | 0 |
| 0 | 1 | 0 | 0 |
| 0 | 0 | 0 | 1 |
| 0 | 0 | 0 | 0 |
| 0 | 1 | 1 | 1 |
| 0 | 0 | 0 | 1 |
| 0 | 0 | 0 | 1 |
| 0 | 1 | 0 | 0 |
| 0 | 0 | 0 | 0 |
| 0 | 0 | 0 | 0 |
| 0 | 0 | 0 | 0 |
| 0 | 0 | 0 | 0 |
| 1 | 1 | 0 | 0 |
| 0 | 0 | 0 | 1 |
| 0 | 0 | 0 | 1 |
| 0 | 0 | 0 | 1 |
| 0 | 0 | 0 | 0 |
| 0 | 0 | 0 | 1 |
| 0 | 0 | 0 | 0 |
| 0 | 0 | 0 | 0 |
| 0 | 0 | 0 | 0 |
| 0 | 0 | 0 | 0 |
| 0 | 0 | 0 | 0 |
| 0 | 0 | 0 | 0 |
| 0 | 0 | 0 | 0 |
| 0 | 0 | 0 | 0 |
| 0 | 0 | 0 | 0 |
| 0 | 0 | 0 | 0 |
| 0 | 0 | 0 | 0 |
| 0 | 0 | 0 | 0 |
| 0 | 0 | 0 | 0 |
| 0 | 0 | 0 | 0 |
| 0 | 0 | 0 | 0 |
| 1 | 0 | 0 | 0 |
| 0 | 1 | 0 | 0 |
| 0 | 1 | 0 | 0 |
| 0 | 0 | 0 | 1 |
| 0 | 1 | 0 | 0 |
| 0 | 1 | 0 | 0 |
| 0 | 0 | 0 | 0 |
| 0 | 0 | 0 | 0 |
| 0 | 0 | 0 | 1 |
| 0 | 0 | 0 | 0 |
| 0 | 0 | 0 | 0 |
| 0 | 0 | 0 | 0 |
| 0 | 0 | 0 | 0 |
| 0 | 0 | 0 | 0 |
| 0 | 0 | 0 | 1 |
| 0 | 0 | 0 | 1 |

| | | | |
|---|---|---|---|
| 0 | 1 | 0 | 0 |
| 0 | 0 | 0 | 0 |
| 0 | 0 | 0 | 1 |
| 0 | 0 | 0 | 1 |
| 0 | 0 | 0 | 1 |
| 0 | 1 | 0 | 0 |
| 0 | 0 | 0 | 1 |
| 0 | 1 | 1 | 1 |
| 0 | 0 | 0 | 0 |
| 0 | 0 | 0 | 0 |
| 0 | 0 | 0 | 0 |
| 0 | 0 | 0 | 0 |
| 0 | 0 | 0 | 0 |
| 0 | 0 | 0 | 1 |
| 0 | 0 | 0 | 1 |
| 0 | 0 | 0 | 1 |
| 0 | 0 | 0 | 1 |
| 0 | 0 | 0 | 1 |
| 0 | 0 | 0 | 0 |
| 0 | 0 | 0 | 0 |
| 0 | 0 | 0 | 1 |
| 0 | 0 | 0 | 1 |
| 0 | 0 | 0 | 0 |
| 0 | 1 | 0 | 0 |
| 0 | 0 | 0 | 1 |
| 0 | 1 | 1 | 1 |
| 0 | 1 | 0 | 0 |
| 0 | 1 | 1 | 1 |
| 0 | 0 | 0 | 0 |
| 0 | 0 | 0 | 0 |
| 0 | 1 | 0 | 0 |
| 0 | 0 | 0 | 0 |
| 0 | 0 | 0 | 0 |
| 1 | 0 | 0 | 0 |
| 1 | 1 | 0 | 0 |
| 0 | 1 | 0 | 0 |
| 0 | 1 | 0 | 0 |
| 0 | 0 | 0 | 0 |
| 0 | 0 | 0 | 1 |
| 0 | 1 | 0 | 0 |
| 0 | 0 | 0 | 1 |
| 0 | 0 | 0 | 1 |
| 0 | 0 | 0 | 1 |
| 0 | 0 | 0 | 1 |
| 0 | 0 | 0 | 1 |
| 0 | 0 | 0 | 0 |
| 0 | 0 | 0 | 0 |
| 0 | 0 | 0 | 0 |
| 0 | 0 | 0 | 0 |
| 0 | 0 | 0 | 0 |
| 0 | 0 | 0 | 0 |
| 0 | 0 | 0 | 0 |
| 0 | 0 | 0 | 0 |
| 0 | 0 | 0 | 1 |

| | | | |
|---|---|---|---|
| 0 | 0 | 0 | 1 |
| 0 | 0 | 0 | 0 |
| 0 | 0 | 0 | 0 |
| 0 | 0 | 0 | 0 |
| 0 | 0 | 0 | 1 |
| 0 | 0 | 0 | 0 |
| 0 | 0 | 0 | 0 |
| 0 | 0 | 0 | 0 |
| 0 | 0 | 0 | 1 |
| 0 | 0 | 0 | 1 |
| 0 | 0 | 0 | 0 |
| 0 | 0 | 0 | 0 |
| 0 | 0 | 0 | 0 |
| 0 | 0 | 0 | 1 |
| 0 | 0 | 0 | 0 |
| 0 | 0 | 0 | 0 |
| 0 | 1 | 0 | 0 |
| 0 | 1 | 0 | 0 |
| 0 | 0 | 0 | 0 |
| 0 | 0 | 0 | 0 |
| 0 | 0 | 0 | 0 |
| 0 | 0 | 0 | 1 |
| 0 | 0 | 0 | 0 |
| 0 | 0 | 0 | 1 |
| 0 | 0 | 0 | 0 |
| 0 | 1 | 0 | 0 |
| 0 | 0 | 0 | 1 |
| 0 | 0 | 0 | 0 |
| 0 | 0 | 0 | 0 |
| 0 | 0 | 0 | 0 |
| 0 | 0 | 0 | 0 |
| 0 | 0 | 0 | 0 |
| 0 | 0 | 0 | 0 |
| 0 | 0 | 0 | 0 |
| 0 | 0 | 0 | 1 |
| 0 | 1 | 0 | 0 |
| 0 | 0 | 0 | 0 |
| 0 | 1 | 0 | 0 |
| 0 | 1 | 0 | 0 |
| 0 | 0 | 0 | 0 |
| 0 | 0 | 0 | 0 |
| 0 | 0 | 0 | 0 |
| 0 | 0 | 0 | 0 |
| 0 | 0 | 0 | 0 |
| 0 | 0 | 0 | 0 |
| 0 | 0 | 0 | 0 |
| 0 | 0 | 0 | 0 |
| 0 | 0 | 0 | 1 |
| 0 | 1 | 0 | 0 |
| 0 | 0 | 0 | 1 |
| 0 | 0 | 0 | 0 |
| 0 | 0 | 0 | 0 |
| 0 | 0 | 0 | 0 |
| 0 | 0 | 0 | 0 |

| | | | |
|---|---|---|---|
| 0 | 0 | 0 | 0 |
| 0 | 1 | 0 | 0 |
| 0 | 0 | 0 | 1 |
| 0 | 1 | 0 | 0 |
| 0 | 0 | 0 | 0 |
| 0 | 0 | 0 | 1 |
| 0 | 0 | 0 | 1 |
| 0 | 0 | 0 | 1 |
| 0 | 0 | 0 | 0 |
| 0 | 0 | 0 | 1 |
| 0 | 1 | 0 | 0 |
| 0 | 0 | 0 | 0 |
| 0 | 0 | 0 | 0 |
| 0 | 0 | 0 | 1 |
| 0 | 1 | 0 | 0 |
| 0 | 0 | 0 | 1 |
| 0 | 0 | 0 | 0 |
| 0 | 0 | 0 | 0 |
| 1 | 0 | 0 | 0 |
| 0 | 0 | 0 | 1 |
| 0 | 0 | 0 | 1 |
| 0 | 0 | 0 | 0 |
| 0 | 0 | 0 | 1 |
| 0 | 0 | 0 | 0 |
| 0 | 0 | 0 | 0 |
| 0 | 0 | 0 | 1 |
| 0 | 0 | 0 | 1 |
| 0 | 0 | 0 | 0 |
| 0 | 0 | 0 | 1 |
| 1 | 0 | 0 | 0 |
| 0 | 0 | 0 | 1 |
| 0 | 0 | 0 | 1 |
| 0 | 0 | 0 | 1 |
| 0 | 0 | 0 | 1 |
| 0 | 0 | 0 | 1 |
| 0 | 0 | 0 | 1 |
| 0 | 0 | 0 | 1 |
| 0 | 0 | 0 | 1 |
| 0 | 0 | 0 | 1 |
| 0 | 0 | 0 | 1 |
| 0 | 0 | 0 | 1 |
| 0 | 0 | 0 | 1 |
| 0 | 0 | 0 | 1 |
| 0 | 0 | 0 | 1 |
| 0 | 0 | 0 | 1 |
| 0 | 0 | 0 | 1 |
| 0 | 0 | 0 | 1 |
| 0 | 0 | 0 | 1 |
| 0 | 0 | 0 | 1 |
| 0 | 0 | 0 | 1 |
| 0 | 0 | 0 | 1 |
| 0 | 0 | 0 | 1 |
| 0 | 0 | 0 | 1 |
| 0 | 0 | 0 | 1 |

| | | | |
|---|---|---|---|
| 0 | 0 | 0 | 1 |
| 0 | 0 | 0 | 1 |
| 0 | 0 | 0 | 1 |
| 0 | 0 | 0 | 1 |
| 0 | 0 | 0 | 1 |
| 0 | 0 | 0 | 1 |
| 0 | 0 | 0 | 1 |
| 0 | 0 | 0 | 1 |
| 0 | 0 | 0 | 1 |
| 0 | 0 | 0 | 1 |
| 0 | 0 | 0 | 1 |
| 0 | 0 | 0 | 0 |
| 0 | 1 | 0 | 0 |
| 0 | 1 | 0 | 0 |
| 0 | 1 | 1 | 1 |
| 0 | 0 | 0 | 0 |
| 0 | 1 | 1 | 1 |
| 0 | 1 | 0 | 0 |
| 0 | 0 | 0 | 1 |
| 0 | 0 | 0 | 1 |
| 0 | 0 | 0 | 0 |
| 0 | 0 | 0 | 1 |
| 0 | 0 | 0 | 1 |
| 0 | 0 | 0 | 0 |
| 0 | 0 | 0 | 0 |
| 0 | 0 | 0 | 1 |
| 0 | 0 | 0 | 1 |
| 0 | 0 | 0 | 0 |
| 0 | 0 | 0 | 0 |
| 0 | 0 | 0 | 0 |
| 0 | 0 | 0 | 0 |
| 0 | 0 | 0 | 0 |
| 0 | 0 | 0 | 1 |
| 0 | 0 | 0 | 1 |
| 0 | 0 | 0 | 1 |
| 0 | 0 | 0 | 1 |
| 0 | 0 | 0 | 1 |
| 0 | 0 | 0 | 1 |
| 0 | 0 | 0 | 1 |
| 0 | 1 | 0 | 0 |
| 0 | 0 | 0 | 0 |
| 0 | 0 | 0 | 1 |
| 0 | 1 | 0 | 0 |
| 0 | 1 | 0 | 0 |
| 0 | 0 | 0 | 0 |
| 0 | 1 | 1 | 1 |
| 0 | 0 | 0 | 0 |
| 0 | 0 | 0 | 1 |
| 0 | 1 | 1 | 1 |
| 0 | 1 | 0 | 0 |
| 0 | 0 | 0 | 0 |
| 0 | 0 | 0 | 0 |
| 0 | 0 | 0 | 1 |
| 0 | 0 | 0 | 0 |

| | | | |
|---|---|---|---|
| 0 | 1 | 0 | 0 |
| 0 | 0 | 0 | 1 |
| 0 | 0 | 0 | 1 |
| 0 | 0 | 0 | 1 |
| 0 | 1 | 0 | 0 |
| 0 | 0 | 0 | 1 |
| 0 | 0 | 0 | 0 |
| 0 | 0 | 0 | 1 |
| 0 | 0 | 0 | 1 |
| 0 | 0 | 0 | 0 |
| 0 | 1 | 1 | 1 |
| 0 | 0 | 0 | 1 |
| 0 | 0 | 0 | 1 |
| 0 | 1 | 0 | 0 |
| 0 | 0 | 0 | 1 |
| 0 | 0 | 0 | 1 |
| 0 | 1 | 0 | 0 |
| 0 | 0 | 0 | 1 |
| 0 | 0 | 0 | 0 |
| 0 | 0 | 0 | 0 |
| 0 | 0 | 0 | 0 |
| 0 | 0 | 0 | 0 |
| 0 | 0 | 0 | 0 |
| 0 | 0 | 0 | 0 |
| 0 | 0 | 0 | 0 |
| 0 | 0 | 0 | 0 |
| 0 | 0 | 0 | 0 |
| 0 | 0 | 0 | 0 |
| 0 | 0 | 0 | 0 |
| 0 | 0 | 0 | 0 |
| 0 | 0 | 0 | 0 |
| 0 | 0 | 0 | 0 |
| 0 | 0 | 0 | 0 |
| 0 | 0 | 0 | 0 |
| 0 | 0 | 0 | 0 |
| 0 | 0 | 0 | 0 |
| 0 | 1 | 1 | 1 |
| 0 | 1 | 0 | 0 |
| 0 | 0 | 0 | 0 |
| 0 | 0 | 0 | 1 |
| 0 | 0 | 0 | 1 |
| 0 | 0 | 0 | 0 |
| 0 | 0 | 0 | 0 |
| 0 | 1 | 0 | 0 |
| 0 | 1 | 0 | 0 |
| 0 | 0 | 0 | 0 |
| 0 | 0 | 0 | 0 |
| 0 | 1 | 0 | 0 |
| 0 | 0 | 0 | 1 |
| 0 | 0 | 0 | 1 |
| 0 | 0 | 0 | 0 |
| 0 | 0 | 0 | 1 |
| 0 | 0 | 0 | 0 |
| 0 | 0 | 0 | 0 |

| | | | |
|---|---|---|---|
| 0 | 0 | 0 | 0 |
| 0 | 0 | 0 | 0 |
| 0 | 0 | 0 | 0 |
| 0 | 0 | 0 | 0 |
| 0 | 0 | 0 | 0 |
| 0 | 0 | 0 | 0 |
| 0 | 0 | 0 | 0 |
| 0 | 0 | 0 | 0 |
| 0 | 0 | 0 | 0 |
| 0 | 0 | 0 | 0 |
| 0 | 0 | 0 | 0 |
| 0 | 0 | 0 | 0 |
| 0 | 0 | 0 | 0 |
| 0 | 0 | 0 | 0 |
| 0 | 0 | 0 | 1 |
| 0 | 1 | 0 | 0 |
| 0 | 1 | 0 | 0 |
| 0 | 1 | 0 | 0 |
| 0 | 1 | 1 | 1 |
| 0 | 0 | 0 | 0 |
| 0 | 1 | 0 | 0 |
| 0 | 1 | 0 | 0 |
| 0 | 1 | 0 | 0 |
| 0 | 1 | 0 | 0 |
| 0 | 1 | 0 | 0 |
| 0 | 1 | 1 | 1 |
| 0 | 0 | 0 | 0 |
| 0 | 1 | 0 | 0 |
| 0 | 1 | 0 | 0 |
| 0 | 0 | 0 | 0 |
| 0 | 0 | 0 | 1 |
| 0 | 0 | 0 | 0 |
| 0 | 0 | 0 | 1 |
| 0 | 1 | 1 | 1 |
| 1 | 0 | 0 | 0 |
| 0 | 0 | 0 | 0 |
| 0 | 0 | 0 | 1 |
| 0 | 0 | 0 | 1 |
| 0 | 0 | 0 | 1 |
| 0 | 0 | 0 | 1 |
| 0 | 0 | 0 | 0 |
| 0 | 0 | 0 | 0 |
| 0 | 0 | 0 | 0 |
| 0 | 0 | 0 | 0 |
| 0 | 0 | 0 | 0 |
| 0 | 0 | 0 | 0 |
| 0 | 0 | 0 | 0 |
| 0 | 0 | 0 | 0 |
| 0 | 0 | 0 | 1 |
| 0 | 0 | 0 | 1 |
| 0 | 0 | 0 | 0 |
| 0 | 0 | 0 | 0 |
| 0 | 0 | 0 | 0 |
| 0 | 0 | 0 | 1 |

| | | | |
|---|---|---|---|
| 0 | 0 | 0 | 1 |
| 0 | 0 | 0 | 1 |
| 0 | 1 | 1 | 1 |
| 0 | 0 | 0 | 0 |
| 0 | 0 | 0 | 1 |
| 0 | 0 | 0 | 1 |
| 0 | 0 | 0 | 0 |
| 0 | 0 | 0 | 1 |
| 0 | 0 | 0 | 0 |
| 0 | 0 | 0 | 0 |
| 0 | 0 | 0 | 0 |
| 0 | 0 | 0 | 0 |
| 0 | 0 | 0 | 0 |
| 0 | 0 | 0 | 0 |
| 0 | 0 | 0 | 0 |
| 0 | 0 | 0 | 0 |
| 0 | 0 | 0 | 1 |
| 0 | 0 | 0 | 0 |
| 0 | 0 | 0 | 0 |
| 0 | 1 | 1 | 1 |
| 0 | 0 | 0 | 0 |
| 0 | 0 | 0 | 0 |
| 0 | 0 | 0 | 1 |
| 0 | 0 | 0 | 0 |
| 0 | 0 | 0 | 1 |
| 0 | 0 | 0 | 1 |
| 0 | 0 | 0 | 1 |
| 0 | 0 | 0 | 0 |
| 0 | 0 | 0 | 0 |
| 0 | 0 | 0 | 0 |
| 0 | 0 | 0 | 1 |
| 0 | 0 | 0 | 0 |
| 0 | 0 | 0 | 0 |
| 0 | 0 | 0 | 0 |
| 0 | 0 | 0 | 1 |
| 0 | 0 | 0 | 1 |
| 0 | 0 | 0 | 0 |
| 0 | 0 | 0 | 0 |
| 0 | 0 | 0 | 0 |
| 0 | 0 | 0 | 1 |
| 0 | 0 | 0 | 1 |
| 0 | 0 | 0 | 1 |
| 0 | 0 | 0 | 1 |
| 0 | 0 | 0 | 0 |
| 0 | 0 | 0 | 1 |
| 0 | 0 | 0 | 0 |
| 0 | 0 | 0 | 1 |
| 0 | 0 | 0 | 0 |
| 0 | 0 | 0 | 0 |
| 0 | 0 | 0 | 1 |
| 0 | 0 | 0 | 1 |
| 0 | 1 | 0 | 0 |
| 0 | 0 | 0 | 0 |
| 0 | 0 | 0 | 1 |

| | | | |
|---|---|---|---|
| 0 | 0 | 0 | 1 |
| 0 | 0 | 0 | 1 |
| 0 | 0 | 0 | 1 |
| 0 | 0 | 0 | 1 |
| 0 | 1 | 0 | 0 |
| 0 | 1 | 1 | 1 |
| 0 | 1 | 1 | 1 |
| 0 | 1 | 0 | 0 |
| 0 | 1 | 0 | 0 |
| 0 | 0 | 0 | 1 |
| 0 | 0 | 0 | 0 |
| 0 | 0 | 0 | 0 |
| 1 | 0 | 0 | 0 |
| 1 | 0 | 0 | 0 |
| 0 | 0 | 0 | 0 |
| 1 | 0 | 0 | 0 |
| 1 | 0 | 0 | 0 |
| 1 | 0 | 0 | 0 |
| 1 | 0 | 0 | 0 |
| 0 | 0 | 0 | 1 |
| 0 | 0 | 0 | 1 |
| 0 | 0 | 0 | 1 |
| 0 | 0 | 0 | 1 |
| 0 | 0 | 0 | 1 |
| 0 | 0 | 0 | 0 |
| 0 | 0 | 0 | 1 |
| 1 | 0 | 0 | 0 |
| 1 | 0 | 0 | 0 |
| 0 | 0 | 0 | 1 |
| 0 | 0 | 0 | 1 |
| 0 | 0 | 0 | 1 |
| 0 | 0 | 0 | 1 |
| 0 | 0 | 0 | 1 |
| 0 | 0 | 0 | 0 |
| 0 | 0 | 0 | 0 |
| 0 | 0 | 0 | 1 |
| 0 | 0 | 0 | 0 |
| 0 | 0 | 0 | 1 |
| 0 | 0 | 0 | 0 |
| 0 | 1 | 0 | 0 |
| 0 | 1 | 0 | 0 |
| 0 | 0 | 0 | 0 |
| 0 | 0 | 0 | 1 |
| 0 | 0 | 0 | 0 |
| 0 | 0 | 0 | 0 |
| 1 | 0 | 0 | 0 |
| 1 | 0 | 0 | 0 |
| 1 | 0 | 0 | 0 |
| 0 | 0 | 0 | 0 |
| 0 | 0 | 0 | 0 |
| 0 | 1 | 0 | 0 |
| 1 | 0 | 0 | 0 |
| 1 | 0 | 0 | 0 |
| 0 | 0 | 0 | 1 |

| | | | |
|---|---|---|---|
| 0 | 0 | 0 | 0 |
| 0 | 0 | 0 | 1 |
| 0 | 0 | 0 | 0 |
| 0 | 0 | 0 | 0 |
| 0 | 1 | 0 | 0 |
| 0 | 0 | 0 | 1 |
| 0 | 0 | 0 | 0 |
| 0 | 0 | 0 | 1 |
| 0 | 0 | 0 | 1 |
| 0 | 0 | 0 | 0 |
| 0 | 0 | 0 | 1 |
| 0 | 0 | 0 | 1 |
| 0 | 0 | 0 | 1 |
| 0 | 0 | 0 | 1 |
| 0 | 0 | 0 | 1 |
| 0 | 1 | 0 | 0 |
| 0 | 0 | 0 | 0 |
| 0 | 0 | 0 | 1 |
| 0 | 0 | 0 | 0 |
| 0 | 0 | 0 | 1 |
| 0 | 0 | 0 | 0 |
| 0 | 0 | 0 | 0 |
| 0 | 0 | 0 | 0 |
| 0 | 1 | 0 | 0 |
| 0 | 1 | 0 | 0 |
| 0 | 1 | 0 | 0 |
| 0 | 0 | 0 | 0 |
| 0 | 0 | 0 | 0 |
| 0 | 0 | 0 | 0 |
| 0 | 1 | 0 | 0 |
| 0 | 0 | 0 | 0 |
| 0 | 0 | 0 | 0 |
| 0 | 0 | 0 | 1 |
| 0 | 0 | 0 | 1 |
| 0 | 0 | 0 | 0 |
| 0 | 1 | 0 | 0 |
| 0 | 1 | 1 | 1 |
| 0 | 1 | 0 | 0 |
| 0 | 1 | 1 | 1 |
| 0 | 1 | 0 | 0 |
| 0 | 1 | 0 | 0 |
| 0 | 1 | 0 | 0 |
| 0 | 1 | 1 | 1 |
| 0 | 0 | 0 | 1 |
| 0 | 1 | 0 | 0 |
| 0 | 1 | 0 | 0 |
| 0 | 0 | 0 | 0 |
| 0 | 1 | 0 | 0 |
| 0 | 1 | 0 | 0 |
| 0 | 1 | 0 | 0 |
| 0 | 1 | 1 | 1 |
| 0 | 1 | 0 | 0 |
| 0 | 1 | 0 | 0 |
| 0 | 1 | 1 | 1 |

| | | | |
|---|---|---|---|
| 0 | 1 | 0 | 0 |
| 0 | 1 | 1 | 1 |
| 0 | 0 | 0 | 0 |
| 0 | 0 | 0 | 0 |
| 0 | 0 | 0 | 0 |
| 0 | 0 | 0 | 0 |
| 0 | 0 | 0 | 0 |
| 0 | 0 | 0 | 0 |
| 0 | 0 | 0 | 0 |
| 0 | 0 | 0 | 0 |
| 0 | 0 | 0 | 0 |
| 0 | 0 | 0 | 1 |
| 0 | 0 | 0 | 0 |
| 0 | 0 | 0 | 1 |
| 0 | 1 | 1 | 1 |
| 0 | 1 | 1 | 1 |
| 0 | 1 | 0 | 0 |
| 0 | 1 | 0 | 0 |
| 0 | 1 | 1 | 1 |
| 0 | 1 | 0 | 0 |
| 0 | 1 | 1 | 1 |
| 0 | 1 | 0 | 0 |
| 0 | 0 | 0 | 1 |
| 0 | 1 | 0 | 0 |
| 0 | 0 | 0 | 1 |
| 0 | 0 | 0 | 1 |
| 0 | 0 | 0 | 1 |
| 0 | 0 | 0 | 0 |
| 0 | 0 | 0 | 1 |
| 0 | 0 | 0 | 1 |
| 0 | 0 | 0 | 1 |
| 1 | 0 | 0 | 0 |
| 0 | 0 | 0 | 1 |
| 0 | 0 | 0 | 0 |
| 0 | 0 | 0 | 0 |
| 0 | 0 | 0 | 0 |
| 0 | 0 | 0 | 1 |
| 0 | 0 | 0 | 1 |
| 0 | 1 | 0 | 0 |
| 0 | 0 | 0 | 0 |
| 0 | 0 | 0 | 0 |
| 0 | 0 | 0 | 0 |
| 0 | 1 | 0 | 0 |
| 0 | 0 | 0 | 0 |
| 0 | 0 | 0 | 0 |
| 0 | 0 | 0 | 0 |
| 0 | 0 | 0 | 1 |
| 0 | 0 | 0 | 1 |
| 0 | 0 | 0 | 0 |
| 0 | 0 | 0 | 0 |
| 0 | 0 | 0 | 0 |
| 0 | 0 | 0 | 1 |
| 0 | 0 | 0 | 1 |
| 0 | 0 | 0 | 0 |

| | | | |
|---|---|---|---|
| 0 | 0 | 0 | 0 |
| 0 | 0 | 0 | 1 |
| 0 | 0 | 0 | 1 |
| 0 | 0 | 0 | 1 |
| 0 | 0 | 0 | 0 |
| 0 | 1 | 0 | 0 |
| 0 | 0 | 0 | 0 |
| 0 | 0 | 0 | 0 |
| 0 | 0 | 0 | 0 |
| 0 | 0 | 0 | 0 |
| 0 | 0 | 0 | 1 |
| 0 | 0 | 0 | 0 |
| 0 | 0 | 0 | 0 |
| 0 | 0 | 0 | 0 |
| 0 | 1 | 1 | 1 |
| 0 | 0 | 0 | 1 |
| 0 | 1 | 0 | 0 |
| 0 | 1 | 1 | 1 |
| 0 | 1 | 0 | 0 |
| 0 | 1 | 0 | 0 |
| 0 | 0 | 0 | 1 |
| 0 | 1 | 0 | 0 |
| 0 | 1 | 0 | 0 |
| 0 | 0 | 0 | 1 |
| 0 | 1 | 0 | 0 |
| 0 | 1 | 0 | 0 |
| 0 | 1 | 0 | 0 |
| 0 | 1 | 1 | 1 |
| 0 | 1 | 0 | 0 |
| 0 | 0 | 0 | 0 |
| 0 | 0 | 0 | 0 |
| 0 | 0 | 0 | 1 |
| 0 | 0 | 0 | 1 |
| 0 | 0 | 0 | 1 |
| 0 | 0 | 0 | 1 |
| 0 | 0 | 0 | 1 |
| 0 | 0 | 0 | 1 |
| 0 | 0 | 0 | 1 |
| 0 | 0 | 0 | 1 |
| 0 | 0 | 0 | 1 |
| 0 | 0 | 0 | 1 |
| 0 | 0 | 0 | 1 |
| 1 | 0 | 0 | 0 |
| 0 | 0 | 0 | 1 |
| 0 | 0 | 0 | 1 |
| 0 | 0 | 0 | 1 |
| 0 | 0 | 0 | 1 |
| 0 | 0 | 0 | 1 |
| 0 | 0 | 0 | 1 |
| 0 | 0 | 0 | 1 |
| 0 | 0 | 0 | 1 |
| 0 | 0 | 0 | 1 |
| 0 | 0 | 0 | 1 |
| 0 | 0 | 0 | 1 |

| | | | |
|---|---|---|---|
| 0 | 0 | 0 | 1 |
| 0 | 0 | 0 | 1 |
| 0 | 0 | 0 | 1 |
| 0 | 0 | 0 | 1 |
| 0 | 0 | 0 | 1 |
| 0 | 0 | 0 | 1 |
| 0 | 0 | 0 | 1 |
| 0 | 0 | 0 | 1 |
| 0 | 0 | 0 | 0 |
| 0 | 0 | 0 | 0 |
| 0 | 0 | 0 | 0 |
| 0 | 0 | 0 | 0 |
| 0 | 0 | 0 | 0 |
| 0 | 0 | 0 | 0 |
| 0 | 0 | 0 | 0 |
| 0 | 0 | 0 | 0 |
| 0 | 0 | 0 | 0 |
| 0 | 0 | 0 | 0 |
| 0 | 0 | 0 | 0 |
| 0 | 1 | 0 | 0 |
| 1 | 0 | 0 | 0 |
| 1 | 0 | 0 | 0 |
| 0 | 0 | 0 | 0 |
| 0 | 1 | 0 | 0 |
| 0 | 1 | 0 | 0 |
| 0 | 0 | 0 | 1 |
| 0 | 0 | 0 | 1 |
| 0 | 0 | 0 | 0 |
| 0 | 0 | 0 | 0 |
| 1 | 0 | 0 | 0 |
| 1 | 0 | 0 | 0 |
| 0 | 0 | 0 | 0 |
| 1 | 0 | 0 | 0 |
| 1 | 0 | 0 | 0 |
| 0 | 0 | 0 | 0 |
| 0 | 0 | 0 | 0 |
| 0 | 0 | 0 | 0 |
| 0 | 0 | 0 | 0 |
| 0 | 0 | 0 | 0 |
| 0 | 0 | 0 | 0 |
| 0 | 0 | 0 | 0 |
| 0 | 0 | 0 | 0 |
| 1 | 0 | 0 | 0 |
| 1 | 0 | 0 | 0 |
| 1 | 0 | 0 | 0 |
| 0 | 0 | 0 | 0 |
| 0 | 0 | 0 | 1 |
| 0 | 0 | 0 | 1 |
| 0 | 0 | 0 | 1 |
| 0 | 0 | 0 | 1 |
| 0 | 0 | 0 | 1 |
| 0 | 0 | 0 | 1 |
| 0 | 0 | 0 | 0 |
| 0 | 0 | 0 | 0 |

| | | | |
|---|---|---|---|
| 0 | 0 | 0 | 0 |
| 0 | 0 | 0 | 0 |
| 0 | 0 | 0 | 0 |
| 0 | 0 | 0 | 0 |
| 0 | 0 | 0 | 1 |
| 0 | 0 | 0 | 0 |
| 0 | 0 | 0 | 0 |
| 0 | 0 | 0 | 0 |
| 0 | 0 | 0 | 1 |
| 0 | 0 | 0 | 0 |
| 1 | 0 | 0 | 0 |
| 0 | 0 | 0 | 1 |
| 0 | 0 | 0 | 0 |
| 0 | 0 | 0 | 0 |
| 0 | 0 | 0 | 0 |
| 0 | 0 | 0 | 0 |
| 0 | 0 | 0 | 1 |
| 0 | 0 | 0 | 0 |
| 0 | 0 | 0 | 0 |
| 0 | 0 | 0 | 1 |
| 0 | 1 | 0 | 0 |
| 0 | 0 | 0 | 0 |
| 0 | 0 | 0 | 0 |
| 0 | 0 | 0 | 0 |
| 0 | 0 | 0 | 0 |
| 0 | 0 | 0 | 1 |
| 0 | 0 | 0 | 1 |
| 0 | 0 | 0 | 1 |
| 0 | 0 | 0 | 1 |
| 0 | 0 | 0 | 1 |
| 0 | 0 | 0 | 1 |
| 0 | 0 | 0 | 1 |
| 0 | 0 | 0 | 1 |
| 0 | 0 | 0 | 1 |
| 0 | 0 | 0 | 0 |
| 0 | 0 | 0 | 0 |
| 0 | 0 | 0 | 0 |
| 0 | 0 | 0 | 0 |
| 0 | 1 | 0 | 0 |
| 0 | 0 | 0 | 0 |
| 0 | 0 | 0 | 1 |
| 0 | 0 | 0 | 1 |
| 0 | 0 | 0 | 1 |
| 0 | 0 | 0 | 0 |
| 0 | 0 | 0 | 0 |
| 0 | 0 | 0 | 0 |
| 0 | 0 | 0 | 1 |
| 0 | 0 | 0 | 1 |
| 0 | 0 | 0 | 0 |
| 0 | 0 | 0 | 0 |
| 0 | 0 | 0 | 1 |
| 0 | 0 | 0 | 1 |
| 0 | 0 | 0 | 1 |
| 0 | 0 | 0 | 1 |

| | | | |
|---|---|---|---|
| 0 | 0 | 0 | 1 |
| 0 | 0 | 0 | 1 |
| 0 | 0 | 0 | 1 |
| 0 | 0 | 0 | 0 |
| 0 | 0 | 0 | 0 |
| 0 | 0 | 0 | 1 |
| 0 | 0 | 0 | 1 |
| 0 | 0 | 0 | 1 |
| 0 | 0 | 0 | 1 |
| 0 | 0 | 0 | 1 |
| 0 | 0 | 0 | 0 |
| 0 | 0 | 0 | 1 |
| 0 | 1 | 0 | 0 |
| 0 | 0 | 0 | 0 |
| 0 | 0 | 0 | 0 |
| 0 | 0 | 0 | 1 |
| 0 | 0 | 0 | 1 |
| 0 | 0 | 0 | 1 |
| 0 | 0 | 0 | 1 |
| 0 | 0 | 0 | 1 |
| 0 | 0 | 0 | 0 |
| 0 | 0 | 0 | 1 |
| 0 | 0 | 0 | 1 |
| 0 | 0 | 0 | 0 |
| 0 | 0 | 0 | 1 |
| 0 | 1 | 0 | 0 |
| 0 | 1 | 0 | 0 |
| 0 | 0 | 0 | 1 |
| 0 | 0 | 0 | 1 |
| 0 | 0 | 0 | 1 |
| 0 | 0 | 0 | 1 |
| 0 | 0 | 0 | 1 |
| 0 | 0 | 0 | 1 |
| 0 | 0 | 0 | 1 |
| 0 | 0 | 0 | 1 |
| 0 | 0 | 0 | 0 |
| 0 | 0 | 0 | 1 |
| 0 | 0 | 0 | 1 |
| 0 | 0 | 0 | 0 |
| 0 | 0 | 0 | 0 |
| 0 | 0 | 0 | 1 |
| 0 | 0 | 0 | 1 |
| 1 | 0 | 0 | 0 |
| 1 | 0 | 0 | 0 |
| 0 | 0 | 0 | 0 |
| 0 | 0 | 0 | 0 |
| 1 | 0 | 0 | 0 |
| 1 | 0 | 0 | 0 |
| 0 | 0 | 0 | 1 |
| 1 | 0 | 0 | 0 |
| 1 | 0 | 0 | 0 |
| 1 | 0 | 0 | 0 |
| 1 | 0 | 0 | 0 |
| 1 | 0 | 0 | 0 |

| | | | |
|---|---|---|---|
| 0 | 0 | 0 | 0 |
| 0 | 0 | 0 | 0 |
| 1 | 0 | 0 | 0 |
| 1 | 0 | 0 | 0 |
| 1 | 0 | 0 | 0 |
| 1 | 0 | 0 | 0 |
| 0 | 0 | 0 | 0 |
| 0 | 0 | 0 | 0 |
| 0 | 0 | 0 | 0 |
| 0 | 0 | 0 | 1 |
| 0 | 0 | 0 | 1 |
| 0 | 0 | 0 | 0 |
| 0 | 0 | 0 | 1 |
| 0 | 0 | 0 | 0 |
| 0 | 0 | 0 | 0 |
| 0 | 0 | 0 | 1 |
| 0 | 0 | 0 | 0 |
| 0 | 0 | 0 | 1 |
| 0 | 0 | 0 | 1 |
| 0 | 0 | 0 | 1 |
| 0 | 1 | 1 | 1 |
| 0 | 0 | 0 | 1 |
| 0 | 0 | 0 | 1 |
| 0 | 0 | 0 | 0 |
| 0 | 0 | 0 | 0 |
| 0 | 0 | 0 | 1 |
| 0 | 0 | 0 | 1 |
| 0 | 0 | 0 | 1 |
| 0 | 0 | 0 | 0 |
| 0 | 0 | 0 | 0 |
| 0 | 0 | 0 | 1 |
| 0 | 0 | 0 | 1 |
| 0 | 0 | 0 | 1 |
| 0 | 0 | 0 | 0 |
| 0 | 0 | 0 | 0 |
| 0 | 0 | 0 | 1 |
| 0 | 0 | 0 | 0 |
| 0 | 0 | 0 | 1 |
| 0 | 0 | 0 | 1 |
| 0 | 0 | 0 | 0 |
| 0 | 0 | 0 | 0 |
| 0 | 0 | 0 | 0 |
| 0 | 0 | 0 | 0 |
| 0 | 0 | 0 | 0 |
| 0 | 0 | 0 | 0 |
| 0 | 0 | 0 | 1 |
| 0 | 0 | 0 | 1 |
| 0 | 0 | 0 | 0 |
| 0 | 0 | 0 | 0 |
| 0 | 0 | 0 | 0 |
| 0 | 0 | 0 | 1 |
| 0 | 0 | 0 | 1 |
| 0 | 0 | 0 | 0 |
| 0 | 0 | 0 | 0 |

| | | | |
|---|---|---|---|
| 0 | 0 | 0 | 1 |
| 0 | 0 | 0 | 1 |
| 0 | 0 | 0 | 1 |
| 0 | 0 | 0 | 1 |
| 0 | 1 | 0 | 0 |
| 0 | 1 | 0 | 0 |
| 0 | 0 | 0 | 1 |
| 0 | 0 | 0 | 0 |
| 0 | 1 | 1 | 1 |
| 0 | 0 | 0 | 0 |
| 0 | 0 | 0 | 1 |
| 0 | 0 | 0 | 1 |
| 0 | 0 | 0 | 1 |
| 0 | 0 | 0 | 1 |
| 0 | 0 | 0 | 1 |
| 0 | 0 | 0 | 1 |
| 0 | 0 | 0 | 0 |
| 0 | 0 | 0 | 1 |
| 0 | 0 | 0 | 1 |
| 0 | 0 | 0 | 0 |
| 0 | 0 | 0 | 0 |
| 0 | 0 | 0 | 1 |
| 0 | 0 | 0 | 1 |
| 0 | 0 | 0 | 0 |
| 0 | 0 | 0 | 1 |
| 0 | 0 | 0 | 0 |
| 0 | 0 | 0 | 0 |
| 0 | 0 | 0 | 0 |
| 0 | 0 | 0 | 0 |
| 0 | 0 | 0 | 0 |
| 0 | 0 | 0 | 0 |
| 0 | 0 | 0 | 0 |
| 0 | 0 | 0 | 1 |
| 0 | 0 | 0 | 0 |
| 0 | 0 | 0 | 1 |
| 0 | 0 | 0 | 1 |
| 0 | 0 | 0 | 1 |
| 0 | 0 | 0 | 1 |
| 0 | 0 | 0 | 0 |
| 0 | 0 | 0 | 1 |
| 0 | 0 | 0 | 1 |
| 0 | 0 | 0 | 1 |
| 0 | 0 | 0 | 1 |
| 0 | 1 | 0 | 0 |
| 0 | 0 | 0 | 1 |
| 0 | 0 | 0 | 1 |
| 0 | 0 | 0 | 1 |
| 0 | 0 | 0 | 1 |
| 0 | 0 | 0 | 1 |
| 0 | 0 | 0 | 1 |
| 0 | 0 | 0 | 0 |
| 0 | 0 | 0 | 0 |
| 0 | 0 | 0 | 0 |
| 0 | 0 | 0 | 1 |

| | | | |
|---|---|---|---|
| 0 | 0 | 0 | 0 |
| 0 | 1 | 0 | 0 |
| 0 | 1 | 0 | 0 |
| 0 | 0 | 0 | 0 |
| 1 | 0 | 0 | 0 |
| 1 | 0 | 0 | 0 |
| 0 | 0 | 0 | 0 |
| 0 | 0 | 0 | 0 |
| 0 | 0 | 0 | 1 |
| 0 | 0 | 0 | 1 |
| 0 | 0 | 0 | 0 |
| 0 | 0 | 0 | 1 |
| 0 | 0 | 0 | 1 |
| 0 | 0 | 0 | 0 |
| 0 | 0 | 0 | 0 |
| 0 | 0 | 0 | 0 |
| 0 | 0 | 0 | 0 |
| 0 | 0 | 0 | 0 |
| 0 | 0 | 0 | 0 |
| 0 | 0 | 0 | 1 |
| 0 | 0 | 0 | 1 |
| 0 | 0 | 0 | 0 |
| 0 | 0 | 0 | 1 |
| 0 | 1 | 0 | 0 |
| 0 | 0 | 0 | 1 |
| 0 | 0 | 0 | 1 |
| 0 | 0 | 0 | 1 |
| 0 | 0 | 0 | 1 |
| 0 | 0 | 0 | 1 |
| 0 | 0 | 0 | 0 |
| 0 | 0 | 0 | 1 |
| 0 | 0 | 0 | 1 |
| 0 | 0 | 0 | 0 |
| 0 | 0 | 0 | 1 |
| 0 | 1 | 0 | 0 |
| 0 | 0 | 0 | 0 |
| 0 | 1 | 0 | 0 |
| 0 | 0 | 0 | 0 |
| 0 | 0 | 0 | 0 |
| 0 | 0 | 0 | 0 |
| 0 | 0 | 0 | 0 |
| 0 | 0 | 0 | 0 |
| 0 | 0 | 0 | 1 |
| 0 | 0 | 0 | 0 |
| 0 | 0 | 0 | 1 |
| 0 | 0 | 0 | 0 |
| 0 | 0 | 0 | 1 |
| 0 | 0 | 0 | 1 |
| 0 | 0 | 0 | 1 |
| 0 | 0 | 0 | 0 |
| 0 | 0 | 0 | 1 |
| 0 | 0 | 0 | 1 |
| 0 | 0 | 0 | 1 |
| 0 | 0 | 0 | 1 |

| | | | |
|---|---|---|---|
| 1 | 0 | 0 | 0 |
| 0 | 0 | 0 | 1 |
| 0 | 0 | 0 | 1 |
| 0 | 0 | 0 | 1 |
| 0 | 0 | 0 | 1 |
| 0 | 0 | 0 | 1 |
| 0 | 0 | 0 | 1 |
| 0 | 0 | 0 | 1 |
| 0 | 0 | 0 | 1 |
| 0 | 0 | 0 | 1 |
| 0 | 0 | 0 | 0 |
| 0 | 0 | 0 | 1 |
| 0 | 0 | 0 | 1 |
| 0 | 0 | 0 | 1 |
| 0 | 0 | 0 | 0 |
| 0 | 0 | 0 | 0 |
| 0 | 0 | 0 | 0 |
| 0 | 0 | 0 | 0 |
| 0 | 0 | 0 | 0 |
| 0 | 0 | 0 | 1 |
| 0 | 0 | 0 | 1 |
| 0 | 0 | 0 | 1 |
| 0 | 0 | 0 | 0 |
| 0 | 0 | 0 | 1 |
| 0 | 0 | 0 | 1 |
| 0 | 0 | 0 | 1 |
| 0 | 0 | 0 | 1 |
| 0 | 0 | 0 | 1 |
| 0 | 0 | 0 | 0 |
| 0 | 0 | 0 | 0 |
| 0 | 0 | 0 | 0 |
| 0 | 0 | 0 | 0 |
| 0 | 1 | 0 | 0 |
| 0 | 0 | 0 | 0 |
| 0 | 0 | 0 | 1 |
| 0 | 0 | 0 | 1 |
| 0 | 0 | 0 | 0 |
| 0 | 0 | 0 | 0 |
| 0 | 1 | 0 | 0 |
| 0 | 0 | 0 | 0 |
| 0 | 0 | 0 | 0 |
| 0 | 0 | 0 | 0 |
| 0 | 0 | 0 | 1 |
| 0 | 0 | 0 | 1 |
| 0 | 0 | 0 | 0 |
| 0 | 0 | 0 | 1 |
| 0 | 0 | 0 | 1 |
| 0 | 0 | 0 | 1 |
| 0 | 0 | 0 | 1 |
| 0 | 0 | 0 | 1 |
| 0 | 0 | 0 | 1 |
| 0 | 0 | 0 | 1 |
| 0 | 0 | 0 | 1 |
| 0 | 1 | 1 | 1 |

| | | | |
|---|---|---|---|
| 0 | 1 | 0 | 0 |
| 0 | 0 | 0 | 1 |
| 0 | 1 | 0 | 0 |
| 0 | 0 | 0 | 1 |
| 0 | 1 | 1 | 1 |
| 0 | 1 | 1 | 1 |
| 0 | 1 | 0 | 0 |
| 0 | 1 | 0 | 0 |
| 0 | 1 | 1 | 1 |
| 0 | 0 | 0 | 1 |
| 0 | 0 | 0 | 1 |
| 0 | 0 | 0 | 1 |
| 0 | 0 | 0 | 1 |
| 0 | 1 | 1 | 1 |
| 0 | 0 | 0 | 0 |
| 0 | 0 | 0 | 0 |
| 0 | 0 | 0 | 0 |
| 0 | 0 | 0 | 0 |
| 0 | 0 | 0 | 1 |
| 0 | 0 | 0 | 0 |
| 0 | 0 | 0 | 0 |
| 0 | 0 | 0 | 0 |
| 0 | 0 | 0 | 0 |
| 0 | 0 | 0 | 0 |
| 0 | 0 | 0 | 1 |
| 0 | 0 | 0 | 0 |
| 0 | 0 | 0 | 1 |
| 0 | 0 | 0 | 1 |
| 0 | 0 | 0 | 1 |
| 0 | 0 | 0 | 0 |
| 0 | 0 | 0 | 1 |
| 0 | 0 | 0 | 1 |
| 0 | 0 | 0 | 0 |
| 0 | 0 | 0 | 0 |
| 0 | 0 | 0 | 1 |
| 0 | 0 | 0 | 1 |
| 0 | 0 | 0 | 1 |
| 0 | 0 | 0 | 1 |
| 0 | 0 | 0 | 1 |
| 0 | 0 | 0 | 0 |
| 0 | 0 | 0 | 0 |
| 0 | 0 | 0 | 0 |
| 0 | 0 | 0 | 0 |
| 0 | 0 | 0 | 0 |
| 0 | 0 | 0 | 0 |
| 0 | 0 | 0 | 1 |
| 0 | 0 | 0 | 1 |
| 0 | 0 | 0 | 0 |
| 0 | 0 | 0 | 0 |
| 0 | 0 | 0 | 1 |
| 0 | 0 | 0 | 0 |
| 0 | 0 | 0 | 0 |
| 0 | 0 | 0 | 1 |
| 0 | 0 | 0 | 1 |

| | | | |
|---|---|---|---|
| 0 | 0 | 0 | 1 |
| 0 | 0 | 0 | 0 |
| 0 | 0 | 0 | 0 |
| 0 | 0 | 0 | 1 |
| 0 | 0 | 0 | 1 |
| 0 | 0 | 0 | 0 |
| 0 | 0 | 0 | 0 |
| 0 | 0 | 0 | 0 |
| 0 | 0 | 0 | 0 |
| 0 | 0 | 0 | 0 |
| 0 | 0 | 0 | 0 |
| 0 | 0 | 0 | 0 |
| 0 | 0 | 0 | 0 |
| 0 | 0 | 0 | 0 |
| 0 | 0 | 0 | 0 |
| 0 | 0 | 0 | 1 |
| 0 | 0 | 0 | 0 |
| 0 | 0 | 0 | 1 |
| 0 | 0 | 0 | 1 |
| 0 | 0 | 0 | 1 |
| 0 | 0 | 0 | 0 |
| 0 | 0 | 0 | 1 |
| 0 | 0 | 0 | 0 |
| 0 | 0 | 0 | 0 |
| 0 | 0 | 0 | 1 |
| 0 | 0 | 0 | 0 |
| 0 | 0 | 0 | 1 |
| 0 | 0 | 0 | 1 |
| 0 | 0 | 0 | 1 |
| 0 | 0 | 0 | 0 |
| 0 | 0 | 0 | 0 |
| 0 | 0 | 0 | 0 |
| 0 | 0 | 0 | 0 |
| 0 | 1 | 0 | 0 |
| 0 | 1 | 0 | 0 |
| 0 | 1 | 0 | 0 |
| 0 | 0 | 0 | 0 |
| 0 | 0 | 0 | 1 |
| 0 | 0 | 0 | 1 |
| 0 | 0 | 0 | 0 |
| 0 | 1 | 0 | 0 |
| 0 | 0 | 0 | 0 |
| 0 | 0 | 0 | 0 |
| 0 | 0 | 0 | 0 |
| 0 | 1 | 0 | 0 |
| 0 | 0 | 0 | 0 |
| 0 | 0 | 0 | 1 |
| 0 | 0 | 0 | 0 |
| 0 | 0 | 0 | 0 |
| 0 | 0 | 0 | 0 |
| 0 | 0 | 0 | 0 |
| 0 | 0 | 0 | 1 |
| 0 | 0 | 0 | 0 |
| 0 | 0 | 0 | 0 |

| | | | |
|---|---|---|---|
| 0 | 1 | 0 | 0 |
| 0 | 1 | 0 | 0 |
| 0 | 0 | 0 | 0 |
| 0 | 0 | 0 | 0 |
| 0 | 0 | 0 | 0 |
| 0 | 0 | 0 | 0 |
| 0 | 0 | 0 | 0 |
| 0 | 0 | 0 | 0 |
| 1 | 0 | 0 | 0 |
| 0 | 0 | 0 | 0 |
| 0 | 0 | 0 | 1 |
| 0 | 1 | 0 | 0 |
| 0 | 1 | 1 | 1 |
| 0 | 0 | 0 | 0 |
| 0 | 0 | 0 | 1 |
| 0 | 0 | 0 | 1 |
| 0 | 0 | 0 | 1 |
| 0 | 0 | 0 | 1 |
| 0 | 0 | 0 | 0 |
| 0 | 0 | 0 | 0 |
| 0 | 0 | 0 | 1 |
| 0 | 1 | 0 | 0 |
| 0 | 1 | 0 | 0 |
| 0 | 0 | 0 | 1 |
| 0 | 0 | 0 | 0 |
| 0 | 0 | 0 | 0 |
| 0 | 0 | 0 | 0 |
| 0 | 0 | 0 | 1 |
| 0 | 1 | 0 | 0 |
| 0 | 0 | 0 | 0 |
| 0 | 0 | 0 | 1 |
| 1 | 0 | 0 | 0 |
| 0 | 0 | 0 | 1 |
| 0 | 0 | 0 | 1 |
| 1 | 0 | 0 | 0 |
| 1 | 0 | 0 | 0 |
| 1 | 0 | 0 | 0 |
| 0 | 1 | 1 | 1 |
| 0 | 1 | 1 | 1 |
| 0 | 0 | 0 | 0 |
| 1 | 0 | 0 | 0 |
| 0 | 1 | 0 | 0 |
| 0 | 1 | 0 | 0 |
| 1 | 0 | 0 | 0 |
| 1 | 0 | 0 | 0 |
| 0 | 0 | 0 | 0 |
| 0 | 0 | 0 | 1 |
| 0 | 0 | 0 | 1 |
| 0 | 0 | 0 | 1 |
| 0 | 0 | 0 | 1 |
| 1 | 0 | 0 | 0 |
| 0 | 0 | 0 | 0 |
| 0 | 0 | 0 | 0 |
| 0 | 0 | 0 | 0 |

| | | | |
|---|---|---|---|
| 0 | 0 | 0 | 0 |
| 0 | 0 | 0 | 1 |
| 0 | 0 | 0 | 0 |
| 0 | 0 | 0 | 0 |
| 0 | 0 | 0 | 0 |
| 0 | 0 | 0 | 1 |
| 0 | 0 | 0 | 0 |
| 0 | 0 | 0 | 0 |
| 0 | 0 | 0 | 1 |
| 0 | 0 | 0 | 0 |
| 0 | 0 | 0 | 1 |
| 0 | 0 | 0 | 1 |
| 0 | 0 | 0 | 1 |
| 0 | 0 | 0 | 0 |
| 0 | 0 | 0 | 1 |
| 0 | 0 | 0 | 0 |
| 0 | 0 | 0 | 1 |
| 0 | 0 | 0 | 1 |
| 0 | 0 | 0 | 0 |
| 0 | 0 | 0 | 0 |
| 0 | 0 | 0 | 0 |
| 0 | 0 | 0 | 0 |
| 0 | 0 | 0 | 0 |
| 0 | 0 | 0 | 0 |
| 0 | 0 | 0 | 0 |
| 0 | 0 | 0 | 1 |
| 0 | 0 | 0 | 1 |
| 0 | 0 | 0 | 1 |
| 0 | 0 | 0 | 1 |
| 0 | 0 | 0 | 1 |
| 0 | 0 | 0 | 1 |
| 0 | 0 | 0 | 1 |
| 0 | 0 | 0 | 0 |
| 0 | 0 | 0 | 0 |
| 0 | 0 | 0 | 0 |
| 0 | 0 | 0 | 0 |
| 0 | 0 | 0 | 0 |
| 0 | 0 | 0 | 0 |
| 0 | 0 | 0 | 1 |
| 0 | 0 | 0 | 1 |
| 0 | 0 | 0 | 1 |
| 0 | 0 | 0 | 1 |
| 0 | 0 | 0 | 1 |
| 0 | 0 | 0 | 1 |
| 0 | 1 | 0 | 0 |
| 0 | 0 | 0 | 0 |
| 0 | 0 | 0 | 1 |
| 0 | 0 | 0 | 0 |
| 0 | 0 | 0 | 0 |
| 0 | 0 | 0 | 0 |
| 0 | 0 | 0 | 0 |
| 0 | 0 | 0 | 0 |
| 0 | 0 | 0 | 0 |
| 0 | 0 | 0 | 0 |

| | | | |
|---|---|---|---|
| 0 | 0 | 0 | 1 |
| 0 | 0 | 0 | 1 |
| 1 | 0 | 0 | 0 |
| 1 | 0 | 0 | 0 |
| 0 | 0 | 0 | 1 |
| 0 | 0 | 0 | 0 |
| 0 | 0 | 0 | 0 |
| 0 | 0 | 0 | 1 |
| 0 | 0 | 0 | 1 |
| 0 | 1 | 0 | 0 |
| 0 | 0 | 0 | 1 |
| 0 | 0 | 0 | 1 |
| 0 | 0 | 0 | 0 |
| 0 | 0 | 0 | 0 |
| 0 | 1 | 0 | 0 |
| 0 | 1 | 0 | 0 |
| 0 | 0 | 0 | 1 |
| 0 | 0 | 0 | 1 |
| 0 | 0 | 0 | 1 |
| 0 | 0 | 0 | 1 |
| 0 | 0 | 0 | 1 |
| 0 | 0 | 0 | 1 |
| 0 | 0 | 0 | 1 |
| 0 | 0 | 0 | 1 |
| 0 | 0 | 0 | 1 |
| 0 | 0 | 0 | 0 |
| 0 | 0 | 0 | 1 |
| 0 | 0 | 0 | 1 |
| 0 | 0 | 0 | 1 |
| 0 | 0 | 0 | 1 |
| 0 | 0 | 0 | 1 |
| 0 | 0 | 0 | 1 |
| 0 | 0 | 0 | 1 |
| 0 | 0 | 0 | 1 |
| 0 | 0 | 0 | 1 |
| 0 | 0 | 0 | 1 |
| 0 | 0 | 0 | 1 |
| 0 | 0 | 0 | 0 |
| 0 | 0 | 0 | 1 |
| 0 | 0 | 0 | 1 |
| 0 | 0 | 0 | 0 |
| 0 | 0 | 0 | 1 |
| 0 | 0 | 0 | 1 |
| 0 | 0 | 0 | 0 |
| 0 | 0 | 0 | 0 |
| 0 | 0 | 0 | 1 |
| 0 | 0 | 0 | 1 |
| 0 | 0 | 0 | 1 |
| 0 | 0 | 0 | 1 |
| 0 | 0 | 0 | 1 |
| 0 | 0 | 0 | 1 |
| 0 | 0 | 0 | 0 |
| 0 | 0 | 0 | 0 |
| 0 | 0 | 0 | 0 |

| | | | |
|---|---|---|---|
| 0 | 0 | 0 | 0 |
| 0 | 0 | 0 | 1 |
| 0 | 0 | 0 | 1 |
| 0 | 0 | 0 | 1 |
| 0 | 0 | 0 | 1 |
| 0 | 0 | 0 | 1 |
| 0 | 0 | 0 | 1 |
| 0 | 0 | 0 | 1 |
| 0 | 0 | 0 | 0 |
| 0 | 0 | 0 | 1 |
| 0 | 0 | 0 | 1 |
| 0 | 0 | 0 | 1 |
| 0 | 0 | 0 | 1 |
| 0 | 0 | 0 | 1 |
| 0 | 0 | 0 | 1 |
| 0 | 0 | 0 | 1 |
| 0 | 0 | 0 | 1 |
| 0 | 0 | 0 | 1 |
| 0 | 0 | 0 | 0 |
| 0 | 0 | 0 | 0 |
| 0 | 0 | 0 | 0 |
| 0 | 0 | 0 | 0 |
| 0 | 0 | 0 | 1 |
| 0 | 0 | 0 | 0 |
| 0 | 0 | 0 | 0 |
| 0 | 0 | 0 | 1 |
| 0 | 0 | 0 | 1 |
| 0 | 0 | 0 | 1 |
| 0 | 0 | 0 | 1 |
| 0 | 0 | 0 | 1 |
| 0 | 0 | 0 | 1 |
| 0 | 0 | 0 | 1 |
| 0 | 0 | 0 | 0 |
| 0 | 0 | 0 | 1 |
| 0 | 0 | 0 | 1 |
| 0 | 0 | 0 | 1 |
| 0 | 0 | 0 | 0 |
| 0 | 0 | 0 | 1 |
| 0 | 0 | 0 | 0 |
| 0 | 0 | 0 | 0 |
| 0 | 0 | 0 | 0 |
| 0 | 0 | 0 | 0 |
| 0 | 0 | 0 | 0 |
| 0 | 0 | 0 | 1 |
| 0 | 0 | 0 | 1 |
| 0 | 0 | 0 | 1 |
| 0 | 0 | 0 | 0 |
| 0 | 0 | 0 | 0 |
| 0 | 0 | 0 | 0 |
| 0 | 0 | 0 | 0 |
| 0 | 0 | 0 | 0 |
| 0 | 1 | 0 | 0 |
| 0 | 1 | 0 | 0 |
| 0 | 0 | 0 | 1 |

| | | | |
|---|---|---|---|
| 0 | 0 | 0 | 0 |
| 0 | 0 | 0 | 0 |
| 0 | 0 | 0 | 1 |
| 1 | 0 | 0 | 0 |
| 1 | 0 | 0 | 0 |
| 0 | 0 | 0 | 1 |
| 0 | 0 | 0 | 1 |
| 0 | 1 | 1 | 1 |
| 0 | 0 | 0 | 1 |
| 0 | 0 | 0 | 1 |
| 0 | 0 | 0 | 1 |
| 0 | 0 | 0 | 0 |
| 0 | 0 | 0 | 1 |
| 0 | 0 | 0 | 1 |
| 0 | 0 | 0 | 0 |
| 0 | 0 | 0 | 1 |
| 0 | 0 | 0 | 0 |
| 0 | 0 | 0 | 1 |
| 0 | 0 | 0 | 0 |
| 1 | 0 | 0 | 0 |
| 0 | 0 | 0 | 1 |
| 0 | 0 | 0 | 1 |
| 0 | 0 | 0 | 0 |
| 0 | 0 | 0 | 1 |
| 0 | 0 | 0 | 0 |
| 0 | 0 | 0 | 1 |
| 0 | 0 | 0 | 1 |
| 0 | 0 | 0 | 0 |
| 0 | 0 | 0 | 1 |
| 0 | 0 | 0 | 0 |
| 0 | 0 | 0 | 1 |
| 0 | 0 | 0 | 0 |
| 0 | 0 | 0 | 0 |
| 0 | 0 | 0 | 0 |
| 0 | 0 | 0 | 1 |
| 0 | 0 | 0 | 0 |
| 0 | 0 | 0 | 1 |
| 0 | 0 | 0 | 1 |
| 0 | 0 | 0 | 0 |
| 0 | 0 | 0 | 1 |
| 0 | 0 | 0 | 1 |
| 0 | 0 | 0 | 1 |
| 0 | 0 | 0 | 1 |
| 0 | 0 | 0 | 0 |
| 0 | 0 | 0 | 0 |
| 0 | 0 | 0 | 1 |
| 0 | 0 | 0 | 1 |
| 0 | 0 | 0 | 1 |
| 0 | 1 | 0 | 0 |
| 0 | 0 | 0 | 1 |
| 0 | 0 | 0 | 0 |
| 0 | 0 | 0 | 1 |
| 0 | 0 | 0 | 1 |
| 0 | 0 | 0 | 0 |

| | | | |
|---|---|---|---|
| 0 | 0 | 0 | 1 |
| 0 | 0 | 0 | 1 |
| 0 | 0 | 0 | 1 |
| 0 | 0 | 0 | 1 |
| 0 | 0 | 0 | 1 |
| 0 | 0 | 0 | 1 |
| 0 | 0 | 0 | 1 |
| 0 | 0 | 0 | 0 |
| 0 | 0 | 0 | 1 |
| 0 | 0 | 0 | 1 |
| 0 | 0 | 0 | 1 |
| 0 | 0 | 0 | 1 |
| 0 | 0 | 0 | 0 |
| 0 | 0 | 0 | 1 |
| 0 | 0 | 0 | 0 |
| 0 | 0 | 0 | 0 |
| 0 | 0 | 0 | 1 |
| 0 | 0 | 0 | 1 |
| 0 | 0 | 0 | 1 |
| 0 | 0 | 0 | 1 |
| 0 | 0 | 0 | 0 |
| 0 | 0 | 0 | 0 |
| 0 | 0 | 0 | 1 |
| 0 | 0 | 0 | 1 |
| 0 | 0 | 0 | 0 |
| 0 | 0 | 0 | 1 |
| 0 | 0 | 0 | 0 |
| 0 | 0 | 0 | 0 |
| 0 | 0 | 0 | 0 |
| 0 | 0 | 0 | 0 |
| 0 | 0 | 0 | 1 |
| 0 | 0 | 0 | 1 |
| 0 | 0 | 0 | 0 |
| 0 | 1 | 0 | 0 |
| 0 | 0 | 0 | 0 |
| 0 | 0 | 0 | 0 |
| 0 | 0 | 0 | 0 |
| 0 | 0 | 0 | 0 |
| 0 | 0 | 0 | 0 |
| 0 | 0 | 0 | 1 |
| 0 | 0 | 0 | 1 |
| 0 | 0 | 0 | 1 |
| 0 | 0 | 0 | 1 |
| 0 | 0 | 0 | 1 |
| 0 | 0 | 0 | 1 |
| 0 | 0 | 0 | 0 |
| 0 | 0 | 0 | 1 |
| 0 | 0 | 0 | 1 |
| 0 | 0 | 0 | 1 |
| 0 | 0 | 0 | 0 |
| 0 | 0 | 0 | 0 |
| 0 | 0 | 0 | 1 |
| 0 | 0 | 0 | 1 |
| 0 | 0 | 0 | 1 |

| | | | |
|---|---|---|---|
| 0 | 0 | 0 | 1 |
| 0 | 0 | 0 | 1 |
| 0 | 0 | 0 | 1 |
| 0 | 0 | 0 | 1 |
| 0 | 0 | 0 | 0 |
| 0 | 0 | 0 | 1 |
| 0 | 0 | 0 | 0 |
| 0 | 0 | 0 | 0 |
| 0 | 0 | 0 | 0 |
| 0 | 0 | 0 | 0 |
| 0 | 0 | 0 | 0 |
| 0 | 0 | 0 | 1 |
| 0 | 0 | 0 | 0 |
| 0 | 0 | 0 | 0 |
| 0 | 0 | 0 | 1 |
| 0 | 0 | 0 | 1 |
| 0 | 0 | 0 | 1 |
| 0 | 0 | 0 | 1 |
| 0 | 0 | 0 | 1 |
| 0 | 0 | 0 | 1 |
| 0 | 0 | 0 | 1 |
| 0 | 0 | 0 | 1 |
| 0 | 0 | 0 | 1 |
| 0 | 0 | 0 | 0 |
| 0 | 0 | 0 | 1 |
| 0 | 0 | 0 | 1 |
| 0 | 0 | 0 | 1 |
| 0 | 0 | 0 | 0 |
| 0 | 1 | 1 | 1 |
| 0 | 1 | 1 | 1 |
| 0 | 1 | 0 | 0 |
| 0 | 1 | 0 | 0 |
| 0 | 0 | 0 | 1 |
| 0 | 0 | 0 | 0 |
| 0 | 0 | 0 | 1 |
| 0 | 0 | 0 | 1 |
| 0 | 0 | 0 | 1 |
| 0 | 0 | 0 | 1 |
| 0 | 0 | 0 | 1 |
| 0 | 0 | 0 | 1 |
| 0 | 0 | 0 | 1 |
| 0 | 0 | 0 | 1 |
| 0 | 0 | 0 | 1 |
| 0 | 0 | 0 | 1 |
| 0 | 0 | 0 | 1 |
| 0 | 0 | 0 | 1 |
| 0 | 0 | 0 | 1 |
| 0 | 0 | 0 | 1 |
| 0 | 0 | 0 | 0 |
| 0 | 0 | 0 | 1 |
| 0 | 0 | 0 | 1 |
| 0 | 1 | 1 | 1 |
| 0 | 1 | 1 | 1 |
| 0 | 0 | 0 | 1 |

| | | | |
|---|---|---|---|
| 0 | 0 | 0 | 1 |
| 0 | 0 | 0 | 1 |
| 0 | 0 | 0 | 1 |
| 0 | 0 | 0 | 1 |
| 0 | 1 | 1 | 1 |
| 0 | 0 | 0 | 1 |
| 0 | 0 | 0 | 1 |
| 0 | 0 | 0 | 1 |
| 0 | 0 | 0 | 1 |
| 0 | 0 | 0 | 1 |
| 0 | 0 | 0 | 0 |
| 0 | 0 | 0 | 0 |
| 0 | 0 | 0 | 0 |
| 1 | 0 | 0 | 0 |
| 1 | 0 | 0 | 0 |
| 0 | 0 | 0 | 1 |
| 1 | 0 | 0 | 0 |
| 0 | 0 | 0 | 0 |
| 0 | 0 | 0 | 0 |
| 0 | 0 | 0 | 0 |
| 0 | 0 | 0 | 0 |
| 1 | 0 | 0 | 0 |
| 0 | 0 | 0 | 1 |
| 0 | 0 | 0 | 0 |
| 1 | 0 | 0 | 0 |
| 1 | 0 | 0 | 0 |
| 0 | 0 | 0 | 1 |
| 0 | 0 | 0 | 0 |
| 0 | 0 | 0 | 1 |
| 0 | 0 | 0 | 1 |
| 0 | 0 | 0 | 1 |
| 0 | 0 | 0 | 0 |
| 0 | 0 | 0 | 1 |
| 0 | 0 | 0 | 1 |
| 0 | 0 | 0 | 0 |
| 0 | 0 | 0 | 0 |
| 0 | 0 | 0 | 0 |
| 0 | 0 | 0 | 0 |
| 0 | 0 | 0 | 0 |
| 0 | 1 | 1 | 1 |
| 0 | 0 | 0 | 1 |
| 0 | 0 | 0 | 0 |
| 0 | 0 | 0 | 1 |
| 0 | 0 | 0 | 0 |
| 0 | 0 | 0 | 1 |
| 0 | 1 | 1 | 1 |
| 0 | 0 | 0 | 1 |
| 0 | 1 | 0 | 0 |
| 0 | 0 | 0 | 0 |
| 0 | 1 | 0 | 0 |
| 0 | 0 | 0 | 0 |
| 0 | 0 | 0 | 1 |
| 0 | 0 | 0 | 0 |
| 0 | 0 | 0 | 1 |

| | | | |
|---|---|---|---|
| 0 | 0 | 0 | 1 |
| 0 | 0 | 0 | 1 |
| 0 | 0 | 0 | 1 |
| 0 | 0 | 0 | 1 |
| 0 | 0 | 0 | 1 |
| 0 | 0 | 0 | 0 |
| 0 | 0 | 0 | 0 |
| 0 | 0 | 0 | 0 |
| 0 | 0 | 0 | 0 |
| 0 | 0 | 0 | 1 |
| 1 | 0 | 0 | 0 |
| 1 | 0 | 0 | 0 |
| 0 | 0 | 0 | 0 |
| 0 | 0 | 0 | 0 |
| 0 | 0 | 0 | 0 |
| 0 | 0 | 0 | 0 |
| 0 | 0 | 0 | 1 |
| 0 | 0 | 0 | 1 |
| 0 | 0 | 0 | 1 |
| 0 | 0 | 0 | 1 |
| 0 | 0 | 0 | 1 |
| 0 | 0 | 0 | 1 |
| 0 | 0 | 0 | 1 |
| 0 | 0 | 0 | 1 |
| 0 | 0 | 0 | 1 |
| 0 | 0 | 0 | 1 |
| 0 | 0 | 0 | 1 |
| 0 | 0 | 0 | 1 |
| 0 | 1 | 0 | 0 |
| 0 | 0 | 0 | 1 |
| 0 | 0 | 0 | 1 |
| 0 | 1 | 1 | 1 |
| 0 | 1 | 1 | 1 |
| 0 | 0 | 0 | 0 |
| 0 | 0 | 0 | 1 |
| 0 | 0 | 0 | 0 |
| 0 | 0 | 0 | 1 |
| 0 | 0 | 0 | 0 |
| 0 | 0 | 0 | 1 |
| 0 | 0 | 0 | 0 |
| 0 | 0 | 0 | 1 |
| 0 | 0 | 0 | 1 |
| 0 | 0 | 0 | 1 |
| 0 | 0 | 0 | 0 |
| 0 | 0 | 0 | 1 |
| 0 | 0 | 0 | 1 |
| 0 | 0 | 0 | 1 |
| 0 | 0 | 0 | 1 |
| 0 | 0 | 0 | 1 |
| 0 | 0 | 0 | 1 |
| 0 | 0 | 0 | 1 |
| 0 | 1 | 1 | 1 |
| 0 | 0 | 0 | 0 |
| 0 | 0 | 0 | 0 |

| | | | |
|---|---|---|---|
| 0 | 0 | 0 | 1 |
| 0 | 0 | 0 | 0 |
| 0 | 0 | 0 | 1 |
| 0 | 0 | 0 | 1 |
| 0 | 0 | 0 | 0 |
| 0 | 0 | 0 | 0 |
| 0 | 0 | 0 | 0 |
| 0 | 0 | 0 | 1 |
| 0 | 0 | 0 | 1 |
| 0 | 0 | 0 | 0 |
| 0 | 0 | 0 | 0 |
| 0 | 0 | 0 | 0 |
| 0 | 0 | 0 | 0 |
| 0 | 0 | 0 | 0 |
| 0 | 0 | 0 | 1 |
| 0 | 0 | 0 | 0 |
| 0 | 1 | 0 | 0 |
| 0 | 0 | 0 | 0 |
| 0 | 0 | 0 | 1 |
| 0 | 0 | 0 | 1 |
| 0 | 0 | 0 | 1 |
| 0 | 0 | 0 | 1 |
| 0 | 0 | 0 | 1 |
| 0 | 0 | 0 | 0 |
| 0 | 0 | 0 | 1 |
| 0 | 0 | 0 | 1 |
| 0 | 0 | 0 | 0 |
| 0 | 0 | 0 | 0 |
| 0 | 0 | 0 | 1 |
| 0 | 0 | 0 | 1 |
| 0 | 0 | 0 | 0 |
| 0 | 0 | 0 | 1 |
| 0 | 0 | 0 | 1 |
| 0 | 0 | 0 | 1 |
| 0 | 0 | 0 | 1 |
| 0 | 0 | 0 | 0 |
| 0 | 0 | 0 | 1 |
| 0 | 0 | 0 | 0 |
| 0 | 0 | 0 | 0 |
| 0 | 0 | 0 | 1 |
| 0 | 0 | 0 | 1 |
| 0 | 0 | 0 | 0 |
| 0 | 0 | 0 | 0 |
| 0 | 0 | 0 | 1 |
| 0 | 0 | 0 | 0 |
| 0 | 0 | 0 | 0 |
| 0 | 0 | 0 | 0 |
| 0 | 0 | 0 | 1 |
| 0 | 0 | 0 | 0 |
| 0 | 0 | 0 | 0 |
| 0 | 0 | 0 | 0 |
| 0 | 0 | 0 | 1 |
| 0 | 0 | 0 | 0 |
| 0 | 0 | 0 | 1 |

| | | | |
|---|---|---|---|
| 0 | 0 | 0 | 1 |
| 0 | 0 | 0 | 1 |
| 0 | 0 | 0 | 1 |
| 0 | 0 | 0 | 1 |
| 0 | 0 | 0 | 0 |
| 0 | 0 | 0 | 0 |
| 0 | 0 | 0 | 1 |
| 0 | 0 | 0 | 1 |
| 0 | 0 | 0 | 1 |
| 0 | 0 | 0 | 0 |
| 0 | 0 | 0 | 1 |
| 0 | 0 | 0 | 1 |
| 0 | 1 | 0 | 0 |
| 0 | 0 | 0 | 1 |
| 0 | 0 | 0 | 1 |
| 0 | 0 | 0 | 1 |
| 0 | 0 | 0 | 1 |
| 0 | 0 | 0 | 1 |
| 0 | 0 | 0 | 1 |
| 0 | 0 | 0 | 1 |
| 0 | 0 | 0 | 1 |
| 0 | 0 | 0 | 1 |
| 0 | 0 | 0 | 1 |
| 0 | 0 | 0 | 0 |
| 0 | 0 | 0 | 1 |
| 1 | 0 | 0 | 0 |
| 0 | 0 | 0 | 1 |
| 0 | 0 | 0 | 0 |
| 0 | 0 | 0 | 1 |
| 0 | 0 | 0 | 0 |
| 0 | 0 | 0 | 0 |
| 0 | 0 | 0 | 0 |
| 0 | 0 | 0 | 0 |
| 0 | 0 | 0 | 0 |
| 0 | 0 | 0 | 1 |
| 0 | 0 | 0 | 0 |
| 0 | 0 | 0 | 1 |
| 0 | 0 | 0 | 0 |
| 0 | 0 | 0 | 1 |
| 0 | 0 | 0 | 1 |
| 0 | 0 | 0 | 0 |
| 0 | 0 | 0 | 0 |
| 0 | 0 | 0 | 0 |
| 0 | 0 | 0 | 0 |
| 0 | 0 | 0 | 0 |
| 0 | 0 | 0 | 0 |
| 0 | 0 | 0 | 0 |
| 0 | 1 | 0 | 0 |
| 0 | 0 | 0 | 1 |
| 0 | 0 | 0 | 0 |
| 0 | 1 | 1 | 1 |
| 0 | 0 | 0 | 1 |
| 0 | 0 | 0 | 1 |
| 0 | 0 | 0 | 1 |

| | | | |
|---|---|---|---|
| 0 | 0 | 0 | 1 |
| 0 | 0 | 0 | 1 |
| 0 | 0 | 0 | 1 |
| 0 | 0 | 0 | 1 |
| 0 | 0 | 0 | 1 |
| 0 | 0 | 0 | 1 |
| 0 | 0 | 0 | 0 |
| 0 | 0 | 0 | 0 |
| 0 | 0 | 0 | 0 |
| 0 | 0 | 0 | 0 |
| 0 | 0 | 0 | 0 |
| 0 | 0 | 0 | 0 |
| 0 | 0 | 0 | 0 |
| 0 | 0 | 0 | 0 |
| 0 | 0 | 0 | 0 |
| 0 | 0 | 0 | 0 |
| 1 | 0 | 0 | 0 |
| 0 | 0 | 0 | 0 |
| 0 | 0 | 0 | 1 |
| 0 | 0 | 0 | 0 |
| 0 | 0 | 0 | 1 |
| 0 | 0 | 0 | 1 |
| 0 | 0 | 0 | 1 |
| 0 | 0 | 0 | 1 |
| 0 | 0 | 0 | 1 |
| 0 | 0 | 0 | 0 |
| 0 | 0 | 0 | 0 |
| 0 | 0 | 0 | 0 |
| 0 | 0 | 0 | 0 |
| 0 | 0 | 0 | 0 |
| 0 | 0 | 0 | 0 |
| 1 | 0 | 0 | 0 |
| 0 | 0 | 0 | 0 |
| 1 | 0 | 0 | 0 |
| 0 | 0 | 0 | 1 |
| 1 | 0 | 0 | 0 |
| 1 | 0 | 0 | 0 |
| 0 | 0 | 0 | 0 |
| 1 | 0 | 0 | 0 |
| 0 | 0 | 0 | 1 |
| 0 | 0 | 0 | 0 |
| 0 | 0 | 0 | 0 |
| 0 | 0 | 0 | 1 |
| 0 | 0 | 0 | 1 |
| 0 | 0 | 0 | 1 |
| 0 | 0 | 0 | 1 |
| 0 | 0 | 0 | 1 |
| 0 | 0 | 0 | 0 |
| 0 | 0 | 0 | 0 |
| 0 | 0 | 0 | 0 |
| 0 | 0 | 0 | 0 |
| 0 | 0 | 0 | 1 |
| 0 | 0 | 0 | 0 |
| 0 | 0 | 0 | 1 |

| | | | |
|---|---|---|---|
| 0 | 0 | 0 | 1 |
| 0 | 0 | 0 | 1 |
| 0 | 0 | 0 | 1 |
| 0 | 0 | 0 | 1 |
| 0 | 0 | 0 | 1 |
| 0 | 0 | 0 | 0 |
| 0 | 0 | 0 | 1 |
| 0 | 0 | 0 | 1 |
| 0 | 0 | 0 | 1 |
| 0 | 0 | 0 | 1 |
| 0 | 0 | 0 | 0 |
| 0 | 1 | 0 | 0 |
| 0 | 1 | 0 | 0 |
| 0 | 1 | 1 | 1 |
| 0 | 1 | 1 | 1 |
| 0 | 0 | 0 | 1 |
| 0 | 1 | 1 | 1 |
| 0 | 0 | 0 | 0 |
| 0 | 0 | 0 | 1 |
| 0 | 0 | 0 | 1 |
| 0 | 0 | 0 | 1 |
| 0 | 0 | 0 | 1 |
| 0 | 0 | 0 | 0 |
| 0 | 0 | 0 | 1 |
| 0 | 0 | 0 | 0 |
| 0 | 0 | 0 | 0 |
| 0 | 0 | 0 | 0 |
| 0 | 0 | 0 | 0 |
| 0 | 0 | 0 | 1 |
| 0 | 0 | 0 | 0 |
| 0 | 0 | 0 | 1 |
| 0 | 0 | 0 | 1 |
| 0 | 0 | 0 | 1 |
| 0 | 0 | 0 | 1 |
| 0 | 0 | 0 | 1 |
| 0 | 0 | 0 | 1 |
| 0 | 0 | 0 | 1 |
| 0 | 0 | 0 | 0 |
| 0 | 0 | 0 | 0 |
| 0 | 0 | 0 | 0 |
| 0 | 1 | 0 | 0 |
| 0 | 0 | 0 | 0 |
| 0 | 0 | 0 | 1 |
| 0 | 0 | 0 | 0 |
| 0 | 0 | 0 | 0 |
| 0 | 0 | 0 | 1 |
| 0 | 0 | 0 | 0 |
| 0 | 1 | 0 | 0 |
| 0 | 0 | 0 | 0 |
| 1 | 0 | 0 | 0 |
| 0 | 0 | 0 | 1 |
| 0 | 0 | 0 | 1 |
| 0 | 0 | 0 | 1 |
| 0 | 1 | 0 | 0 |

| | | | |
|---|---|---|---|
| 0 | 0 | 0 | 0 |
| 0 | 0 | 0 | 1 |
| 0 | 0 | 0 | 0 |
| 0 | 0 | 0 | 1 |
| 0 | 0 | 0 | 1 |
| 0 | 0 | 0 | 1 |
| 0 | 0 | 0 | 1 |
| 0 | 0 | 0 | 1 |
| 0 | 1 | 0 | 0 |
| 0 | 0 | 0 | 1 |
| 0 | 0 | 0 | 0 |
| 0 | 0 | 0 | 0 |
| 0 | 0 | 0 | 0 |
| 0 | 0 | 0 | 0 |
| 0 | 0 | 0 | 1 |
| 0 | 0 | 0 | 1 |
| 0 | 0 | 0 | 0 |
| 0 | 0 | 0 | 0 |
| 0 | 0 | 0 | 1 |
| 0 | 0 | 0 | 1 |
| 0 | 0 | 0 | 0 |
| 0 | 0 | 0 | 1 |
| 0 | 0 | 0 | 1 |
| 0 | 0 | 0 | 1 |
| 0 | 0 | 0 | 0 |
| 0 | 0 | 0 | 1 |
| 0 | 1 | 0 | 0 |
| 0 | 1 | 0 | 0 |
| 0 | 1 | 0 | 0 |
| 0 | 0 | 0 | 1 |
| 0 | 0 | 0 | 0 |
| 0 | 0 | 0 | 0 |
| 0 | 0 | 0 | 0 |
| 0 | 0 | 0 | 1 |
| 0 | 0 | 0 | 0 |
| 0 | 0 | 0 | 1 |
| 0 | 0 | 0 | 0 |
| 0 | 0 | 0 | 0 |
| 0 | 1 | 0 | 0 |
| 0 | 0 | 0 | 0 |
| 0 | 0 | 0 | 0 |
| 0 | 0 | 0 | 1 |
| 0 | 1 | 0 | 0 |
| 0 | 1 | 0 | 0 |
| 0 | 1 | 0 | 0 |
| 1 | 0 | 0 | 0 |
| 0 | 0 | 0 | 0 |
| 1 | 0 | 0 | 0 |
| 1 | 0 | 0 | 0 |
| 1 | 0 | 0 | 0 |
| 0 | 0 | 0 | 0 |
| 0 | 0 | 0 | 0 |
| 0 | 0 | 0 | 1 |
| 0 | 0 | 0 | 0 |

| | | | |
|---|---|---|---|
| 0 | 0 | 0 | 1 |
| 0 | 0 | 0 | 1 |
| 0 | 0 | 0 | 1 |
| 0 | 0 | 0 | 1 |
| 0 | 0 | 0 | 1 |
| 0 | 0 | 0 | 1 |
| 0 | 0 | 0 | 1 |
| 0 | 0 | 0 | 1 |
| 0 | 0 | 0 | 1 |
| 0 | 0 | 0 | 0 |
| 0 | 0 | 0 | 0 |
| 0 | 0 | 0 | 1 |
| 0 | 0 | 0 | 1 |
| 0 | 0 | 0 | 1 |
| 0 | 0 | 0 | 0 |
| 0 | 0 | 0 | 1 |
| 0 | 0 | 0 | 1 |
| 0 | 0 | 0 | 0 |
| 0 | 0 | 0 | 1 |
| 0 | 0 | 0 | 1 |
| 0 | 0 | 0 | 1 |
| 0 | 0 | 0 | 0 |
| 0 | 0 | 0 | 0 |
| 0 | 0 | 0 | 0 |
| 0 | 1 | 0 | 0 |
| 0 | 1 | 0 | 0 |
| 0 | 1 | 0 | 0 |
| 0 | 0 | 0 | 0 |
| 0 | 0 | 0 | 1 |
| 0 | 0 | 0 | 0 |
| 1 | 0 | 0 | 0 |
| 0 | 0 | 0 | 1 |
| 0 | 0 | 0 | 1 |
| 0 | 0 | 0 | 1 |
| 0 | 0 | 0 | 1 |
| 0 | 0 | 0 | 1 |
| 0 | 0 | 0 | 1 |
| 0 | 0 | 0 | 1 |
| 0 | 0 | 0 | 1 |
| 0 | 0 | 0 | 1 |
| 0 | 0 | 0 | 1 |
| 0 | 0 | 0 | 1 |
| 0 | 0 | 0 | 1 |
| 0 | 0 | 0 | 0 |
| 0 | 0 | 0 | 1 |
| 0 | 0 | 0 | 0 |
| 0 | 0 | 0 | 1 |
| 0 | 0 | 0 | 1 |
| 0 | 0 | 0 | 1 |
| 0 | 0 | 0 | 1 |
| 0 | 0 | 0 | 0 |
| 0 | 0 | 0 | 1 |
| 0 | 0 | 0 | 1 |
| 0 | 0 | 0 | 0 |

| | | | |
|---|---|---|---|
| 0 | 0 | 0 | 0 |
| 0 | 0 | 0 | 0 |
| 0 | 0 | 0 | 0 |
| 0 | 0 | 0 | 0 |
| 0 | 0 | 0 | 1 |
| 0 | 0 | 0 | 1 |
| 0 | 0 | 0 | 1 |
| 0 | 0 | 0 | 0 |
| 0 | 0 | 0 | 0 |
| 0 | 0 | 0 | 0 |
| 0 | 0 | 0 | 0 |
| 0 | 0 | 0 | 1 |
| 0 | 0 | 0 | 1 |
| 0 | 0 | 0 | 1 |
| 0 | 0 | 0 | 1 |
| 0 | 0 | 0 | 0 |
| 0 | 0 | 0 | 0 |
| 0 | 0 | 0 | 1 |
| 0 | 0 | 0 | 1 |
| 0 | 0 | 0 | 1 |
| 0 | 0 | 0 | 1 |
| 0 | 1 | 0 | 0 |
| 0 | 0 | 0 | 1 |
| 0 | 0 | 0 | 1 |
| 0 | 0 | 0 | 0 |
| 0 | 0 | 0 | 0 |
| 0 | 0 | 0 | 0 |
| 0 | 0 | 0 | 0 |
| 0 | 0 | 0 | 1 |
| 0 | 0 | 0 | 1 |
| 0 | 0 | 0 | 1 |
| 0 | 0 | 0 | 0 |
| 0 | 0 | 0 | 1 |
| 0 | 0 | 0 | 1 |
| 0 | 0 | 0 | 1 |
| 0 | 0 | 0 | 1 |
| 0 | 0 | 0 | 1 |
| 0 | 0 | 0 | 1 |
| 0 | 0 | 0 | 1 |
| 0 | 0 | 0 | 1 |
| 0 | 0 | 0 | 1 |
| 0 | 0 | 0 | 1 |
| 0 | 0 | 0 | 0 |
| 0 | 0 | 0 | 1 |
| 0 | 0 | 0 | 1 |
| 0 | 0 | 0 | 1 |
| 0 | 0 | 0 | 1 |
| 0 | 0 | 0 | 1 |
| 0 | 0 | 0 | 1 |
| 0 | 0 | 0 | 0 |
| 0 | 0 | 0 | 1 |
| 0 | 0 | 0 | 1 |
| 0 | 0 | 0 | 1 |
| 0 | 0 | 0 | 1 |

| | | | |
|---|---|---|---|
| 0 | 0 | 0 | 0 |
| 0 | 0 | 0 | 1 |
| 0 | 0 | 0 | 1 |
| 0 | 0 | 0 | 1 |
| 0 | 0 | 0 | 1 |
| 0 | 0 | 0 | 1 |
| 0 | 0 | 0 | 1 |
| 0 | 1 | 0 | 0 |
| 0 | 0 | 0 | 1 |
| 0 | 0 | 0 | 1 |
| 0 | 0 | 0 | 0 |
| 0 | 0 | 0 | 0 |
| 0 | 0 | 0 | 0 |
| 0 | 0 | 0 | 1 |
| 0 | 0 | 0 | 0 |
| 0 | 0 | 0 | 0 |
| 0 | 0 | 0 | 0 |
| 0 | 0 | 0 | 0 |
| 0 | 0 | 0 | 1 |
| 0 | 0 | 0 | 0 |
| 0 | 0 | 0 | 0 |
| 0 | 0 | 0 | 0 |
| 0 | 0 | 0 | 0 |
| 0 | 0 | 0 | 0 |
| 0 | 0 | 0 | 0 |
| 0 | 0 | 0 | 0 |
| 0 | 0 | 0 | 0 |
| 0 | 0 | 0 | 0 |
| 0 | 0 | 0 | 0 |
| 0 | 0 | 0 | 0 |
| 0 | 0 | 0 | 0 |
| 0 | 0 | 0 | 0 |
| 0 | 0 | 0 | 1 |
| 0 | 0 | 0 | 0 |
| 0 | 0 | 0 | 1 |
| 0 | 0 | 0 | 0 |
| 0 | 0 | 0 | 0 |
| 0 | 0 | 0 | 1 |
| 0 | 0 | 0 | 1 |
| 0 | 0 | 0 | 1 |
| 0 | 0 | 0 | 1 |
| 0 | 0 | 0 | 0 |
| 0 | 0 | 0 | 0 |
| 0 | 0 | 0 | 0 |
| 0 | 0 | 0 | 1 |
| 0 | 0 | 0 | 0 |
| 0 | 0 | 0 | 1 |
| 0 | 0 | 0 | 0 |
| 0 | 0 | 0 | 1 |
| 0 | 0 | 0 | 1 |
| 0 | 0 | 0 | 1 |
| 0 | 0 | 0 | 1 |
| 0 | 0 | 0 | 1 |
| 0 | 0 | 0 | 1 |

| | | | |
|---|---|---|---|
| 0 | 0 | 0 | 1 |
| 0 | 0 | 0 | 1 |
| 0 | 0 | 0 | 1 |
| 0 | 0 | 0 | 1 |
| 0 | 0 | 0 | 1 |
| 0 | 0 | 0 | 0 |
| 0 | 0 | 0 | 0 |
| 0 | 0 | 0 | 1 |
| 0 | 0 | 0 | 0 |
| 0 | 0 | 0 | 0 |
| 0 | 0 | 0 | 1 |
| 0 | 0 | 0 | 1 |
| 0 | 0 | 0 | 1 |
| 0 | 0 | 0 | 1 |
| 0 | 0 | 0 | 1 |
| 0 | 1 | 1 | 1 |
| 0 | 0 | 0 | 1 |
| 0 | 0 | 0 | 1 |
| 0 | 0 | 0 | 0 |
| 0 | 0 | 0 | 0 |
| 0 | 0 | 0 | 1 |
| 0 | 0 | 0 | 1 |
| 0 | 0 | 0 | 1 |
| 0 | 0 | 0 | 0 |
| 0 | 0 | 0 | 1 |
| 0 | 0 | 0 | 0 |
| 0 | 0 | 0 | 0 |
| 0 | 0 | 0 | 1 |
| 0 | 0 | 0 | 1 |
| 0 | 0 | 0 | 1 |
| 0 | 1 | 0 | 0 |
| 0 | 0 | 0 | 1 |
| 0 | 0 | 0 | 0 |
| 0 | 1 | 0 | 0 |
| 0 | 1 | 0 | 0 |
| 0 | 0 | 0 | 0 |
| 0 | 0 | 0 | 0 |
| 0 | 0 | 0 | 1 |
| 0 | 0 | 0 | 1 |
| 0 | 0 | 0 | 1 |
| 0 | 0 | 0 | 0 |
| 0 | 0 | 0 | 0 |
| 0 | 0 | 0 | 0 |
| 0 | 0 | 0 | 0 |
| 0 | 0 | 0 | 1 |
| 0 | 0 | 0 | 1 |
| 0 | 0 | 0 | 1 |
| 0 | 0 | 0 | 0 |
| 0 | 0 | 0 | 0 |
| 0 | 0 | 0 | 0 |
| 0 | 0 | 0 | 1 |
| 0 | 0 | 0 | 1 |
| 0 | 0 | 0 | 1 |
| 0 | 0 | 0 | 1 |

| | | | |
|---|---|---|---|
| 0 | 0 | 0 | 1 |
| 0 | 0 | 0 | 1 |
| 0 | 0 | 0 | 1 |
| 0 | 0 | 0 | 0 |
| 0 | 0 | 0 | 0 |
| 0 | 0 | 0 | 1 |
| 0 | 0 | 0 | 0 |
| 0 | 0 | 0 | 1 |
| 0 | 0 | 0 | 1 |
| 0 | 0 | 0 | 1 |
| 0 | 0 | 0 | 1 |
| 0 | 0 | 0 | 0 |
| 0 | 0 | 0 | 1 |
| 0 | 0 | 0 | 0 |
| 0 | 0 | 0 | 1 |
| 0 | 0 | 0 | 0 |
| 0 | 0 | 0 | 1 |
| 0 | 0 | 0 | 1 |
| 0 | 0 | 0 | 1 |
| 0 | 0 | 0 | 0 |
| 0 | 0 | 0 | 0 |
| 0 | 0 | 0 | 0 |
| 0 | 0 | 0 | 0 |
| 0 | 0 | 0 | 0 |
| 0 | 0 | 0 | 0 |
| 0 | 0 | 0 | 1 |
| 0 | 0 | 0 | 0 |
| 0 | 0 | 0 | 1 |
| 0 | 0 | 0 | 1 |
| 0 | 0 | 0 | 1 |
| 0 | 0 | 0 | 1 |
| 0 | 0 | 0 | 1 |
| 0 | 0 | 0 | 0 |
| 0 | 1 | 0 | 0 |
| 0 | 0 | 0 | 1 |
| 0 | 0 | 0 | 1 |
| 0 | 1 | 0 | 0 |
| 0 | 1 | 0 | 0 |
| 0 | 1 | 0 | 0 |
| 0 | 1 | 1 | 1 |
| 0 | 0 | 0 | 0 |
| 0 | 0 | 0 | 0 |
| 0 | 0 | 0 | 1 |
| 0 | 0 | 0 | 0 |
| 0 | 0 | 0 | 1 |
| 0 | 0 | 0 | 1 |
| 0 | 0 | 0 | 1 |
| 0 | 0 | 0 | 1 |
| 0 | 0 | 0 | 1 |
| 1 | 0 | 0 | 0 |
| 0 | 0 | 0 | 1 |
| 0 | 0 | 0 | 1 |
| 0 | 0 | 0 | 1 |
| 0 | 0 | 0 | 1 |

| | | | |
|---|---|---|---|
| 0 | 0 | 0 | 1 |
| 0 | 0 | 0 | 1 |
| 0 | 0 | 0 | 1 |
| 0 | 0 | 0 | 0 |
| 0 | 0 | 0 | 1 |
| 0 | 0 | 0 | 1 |
| 0 | 0 | 0 | 1 |
| 0 | 0 | 0 | 0 |
| 0 | 0 | 0 | 1 |
| 0 | 0 | 0 | 1 |
| 0 | 0 | 0 | 1 |
| 0 | 0 | 0 | 1 |
| 0 | 0 | 0 | 1 |
| 0 | 0 | 0 | 1 |
| 0 | 0 | 0 | 1 |
| 0 | 0 | 0 | 1 |
| 0 | 0 | 0 | 0 |
| 0 | 0 | 0 | 0 |
| 0 | 0 | 0 | 1 |
| 0 | 0 | 0 | 1 |
| 0 | 0 | 0 | 1 |
| 0 | 0 | 0 | 1 |
| 0 | 0 | 0 | 1 |
| 0 | 0 | 0 | 1 |
| 0 | 0 | 0 | 1 |
| 0 | 0 | 0 | 1 |
| 0 | 0 | 0 | 1 |
| 0 | 0 | 0 | 0 |
| 0 | 0 | 0 | 0 |
| 0 | 0 | 0 | 1 |
| 0 | 0 | 0 | 1 |
| 0 | 0 | 0 | 0 |
| 0 | 0 | 0 | 1 |
| 0 | 0 | 0 | 0 |
| 1 | 0 | 0 | 0 |
| 1 | 0 | 0 | 0 |
| 1 | 0 | 0 | 0 |
| 1 | 0 | 0 | 0 |
| 0 | 0 | 0 | 1 |
| 0 | 0 | 0 | 1 |
| 0 | 0 | 0 | 1 |
| 0 | 0 | 0 | 1 |
| 0 | 0 | 0 | 1 |
| 0 | 0 | 0 | 1 |
| 0 | 0 | 0 | 0 |
| 0 | 0 | 0 | 0 |
| 0 | 0 | 0 | 0 |
| 0 | 0 | 0 | 1 |
| 0 | 0 | 0 | 1 |
| 0 | 0 | 0 | 1 |
| 0 | 0 | 0 | 1 |
| 0 | 0 | 0 | 0 |
| 0 | 0 | 0 | 0 |
| 0 | 0 | 0 | 0 |

| | | | |
|---|---|---|---|
| 0 | 0 | 0 | 0 |
| 0 | 0 | 0 | 0 |
| 0 | 0 | 0 | 1 |
| 0 | 0 | 0 | 0 |
| 0 | 0 | 0 | 0 |
| 0 | 0 | 0 | 0 |
| 0 | 0 | 0 | 0 |
| 0 | 0 | 0 | 1 |
| 0 | 0 | 0 | 1 |
| 0 | 0 | 0 | 1 |
| 0 | 0 | 0 | 1 |
| 0 | 0 | 0 | 1 |
| 0 | 0 | 0 | 1 |
| 0 | 0 | 0 | 1 |
| 0 | 0 | 0 | 1 |
| 0 | 0 | 0 | 1 |
| 0 | 0 | 0 | 1 |
| 0 | 0 | 0 | 1 |
| 0 | 0 | 0 | 1 |
| 0 | 0 | 0 | 1 |
| 0 | 0 | 0 | 0 |
| 0 | 0 | 0 | 0 |
| 0 | 0 | 0 | 0 |
| 0 | 0 | 0 | 0 |
| 0 | 0 | 0 | 0 |
| 0 | 0 | 0 | 1 |
| 0 | 0 | 0 | 1 |
| 0 | 0 | 0 | 0 |
| 0 | 0 | 0 | 1 |
| 0 | 0 | 0 | 1 |
| 0 | 0 | 0 | 0 |
| 0 | 0 | 0 | 0 |
| 0 | 0 | 0 | 0 |
| 0 | 0 | 0 | 1 |
| 0 | 0 | 0 | 0 |
| 0 | 0 | 0 | 1 |
| 0 | 0 | 0 | 1 |
| 0 | 0 | 0 | 0 |
| 0 | 0 | 0 | 1 |
| 0 | 0 | 0 | 1 |
| 0 | 0 | 0 | 1 |
| 0 | 0 | 0 | 1 |
| 0 | 0 | 0 | 1 |
| 0 | 0 | 0 | 1 |
| 0 | 0 | 0 | 1 |
| 0 | 0 | 0 | 1 |
| 0 | 0 | 0 | 1 |
| 0 | 0 | 0 | 1 |
| 0 | 0 | 0 | 0 |
| 0 | 0 | 0 | 1 |
| 1 | 0 | 0 | 0 |
| 1 | 0 | 0 | 0 |
| 0 | 0 | 0 | 1 |
| 0 | 0 | 0 | 1 |

| | | | |
|---|---|---|---|
| 1 | 0 | 0 | 0 |
| 0 | 0 | 0 | 1 |
| 0 | 0 | 0 | 1 |
| 0 | 0 | 0 | 1 |
| 0 | 0 | 0 | 1 |
| 1 | 0 | 0 | 0 |
| 1 | 0 | 0 | 0 |
| 0 | 0 | 0 | 1 |
| 0 | 0 | 0 | 1 |
| 0 | 0 | 0 | 1 |
| 0 | 0 | 0 | 1 |
| 0 | 0 | 0 | 0 |
| 0 | 0 | 0 | 1 |
| 0 | 0 | 0 | 1 |
| 0 | 0 | 0 | 1 |
| 0 | 0 | 0 | 1 |
| 0 | 0 | 0 | 0 |
| 0 | 0 | 0 | 0 |
| 0 | 0 | 0 | 1 |
| 0 | 0 | 0 | 1 |
| 0 | 0 | 0 | 1 |
| 0 | 0 | 0 | 1 |
| 0 | 0 | 0 | 1 |
| 0 | 0 | 0 | 1 |
| 0 | 0 | 0 | 0 |
| 0 | 0 | 0 | 1 |
| 0 | 0 | 0 | 1 |
| 0 | 0 | 0 | 1 |
| 0 | 0 | 0 | 1 |
| 0 | 0 | 0 | 1 |
| 0 | 0 | 0 | 1 |
| 0 | 0 | 0 | 0 |
| 0 | 0 | 0 | 0 |
| 0 | 0 | 0 | 1 |
| 0 | 0 | 0 | 1 |
| 0 | 0 | 0 | 0 |
| 1 | 0 | 0 | 0 |
| 0 | 1 | 0 | 0 |
| 1 | 1 | 0 | 0 |
| 0 | 0 | 0 | 0 |
| 0 | 0 | 0 | 0 |
| 0 | 0 | 0 | 0 |
| 0 | 0 | 0 | 0 |
| 0 | 0 | 0 | 0 |
| 0 | 0 | 0 | 0 |
| 0 | 0 | 0 | 0 |
| 0 | 0 | 0 | 0 |
| 1 | 0 | 0 | 0 |
| 0 | 0 | 0 | 1 |
| 0 | 0 | 0 | 1 |
| 0 | 1 | 1 | 1 |
| 0 | 0 | 0 | 1 |
| 0 | 0 | 0 | 0 |
| 0 | 0 | 0 | 1 |

| | | | |
|---|---|---|---|
| 0 | 0 | 0 | 1 |
| 0 | 0 | 0 | 1 |
| 0 | 0 | 0 | 1 |
| 0 | 0 | 0 | 1 |
| 0 | 0 | 0 | 1 |
| 0 | 0 | 0 | 1 |
| 0 | 0 | 0 | 1 |
| 0 | 1 | 0 | 0 |
| 0 | 0 | 0 | 0 |
| 0 | 0 | 0 | 1 |
| 0 | 0 | 0 | 1 |
| 0 | 0 | 0 | 1 |
| 0 | 0 | 0 | 1 |
| 0 | 1 | 0 | 0 |
| 0 | 0 | 0 | 1 |
| 0 | 0 | 0 | 1 |
| 0 | 1 | 0 | 0 |
| 0 | 0 | 0 | 1 |
| 1 | 0 | 0 | 0 |
| 0 | 0 | 0 | 1 |
| 0 | 0 | 0 | 1 |
| 0 | 0 | 0 | 1 |
| 0 | 0 | 0 | 1 |
| 0 | 0 | 0 | 0 |
| 0 | 0 | 0 | 1 |
| 0 | 0 | 0 | 1 |
| 0 | 0 | 0 | 1 |
| 0 | 0 | 0 | 1 |
| 0 | 0 | 0 | 1 |
| 0 | 0 | 0 | 1 |
| 0 | 0 | 0 | 1 |
| 1 | 0 | 0 | 0 |
| 0 | 0 | 0 | 1 |
| 0 | 0 | 0 | 1 |
| 0 | 0 | 0 | 1 |
| 0 | 0 | 0 | 1 |
| 0 | 0 | 0 | 1 |
| 0 | 0 | 0 | 1 |
| 1 | 0 | 0 | 0 |
| 0 | 0 | 0 | 1 |
| 0 | 0 | 0 | 1 |
| 0 | 0 | 0 | 1 |
| 0 | 0 | 0 | 1 |
| 0 | 0 | 0 | 1 |
| 0 | 0 | 0 | 1 |
| 0 | 0 | 0 | 0 |
| 0 | 0 | 0 | 1 |
| 0 | 0 | 0 | 1 |
| 0 | 1 | 1 | 1 |
| 0 | 1 | 1 | 1 |
| 0 | 0 | 0 | 1 |
| 0 | 0 | 0 | 1 |
| 0 | 0 | 0 | 1 |
| 0 | 1 | 1 | 1 |

| | | | |
|---|---|---|---|
| 0 | 1 | 1 | 1 |
| 0 | 1 | 1 | 1 |
| 0 | 1 | 1 | 1 |
| 0 | 1 | 1 | 1 |
| 0 | 1 | 1 | 1 |
| 0 | 1 | 1 | 1 |
| 0 | 1 | 1 | 1 |
| 0 | 1 | 1 | 1 |
| 0 | 1 | 1 | 1 |
| 0 | 1 | 1 | 1 |
| 0 | 1 | 1 | 1 |
| 0 | 1 | 1 | 1 |
| 0 | 1 | 1 | 1 |
| 0 | 1 | 1 | 1 |
| 0 | 1 | 1 | 1 |
| 1 | 1 | 0 | 0 |
| 0 | 1 | 1 | 1 |
| 0 | 1 | 0 | 0 |
| 0 | 1 | 1 | 1 |
| 0 | 1 | 1 | 1 |
| 1 | 1 | 0 | 0 |
| 0 | 1 | 1 | 1 |
| 0 | 1 | 1 | 1 |
| 0 | 1 | 1 | 1 |
| 0 | 1 | 0 | 0 |
| 0 | 1 | 1 | 1 |
| 0 | 1 | 0 | 0 |
| 0 | 1 | 1 | 1 |
| 0 | 1 | 1 | 1 |
| 0 | 1 | 1 | 1 |
| 0 | 1 | 1 | 1 |
| 0 | 1 | 1 | 1 |
| 0 | 1 | 1 | 1 |
| 0 | 1 | 1 | 1 |
| 0 | 1 | 1 | 1 |
| 0 | 1 | 1 | 1 |
| 0 | 1 | 1 | 1 |
| 0 | 1 | 1 | 1 |
| 0 | 1 | 1 | 1 |
| 0 | 1 | 1 | 1 |
| 0 | 1 | 1 | 1 |
| 0 | 1 | 1 | 1 |
| 0 | 0 | 0 | 1 |
| 0 | 0 | 0 | 0 |
| 0 | 0 | 0 | 1 |
| 0 | 0 | 0 | 1 |
| 0 | 0 | 0 | 0 |
| 0 | 0 | 0 | 0 |
| 0 | 0 | 0 | 1 |
| 0 | 0 | 0 | 0 |
| 0 | 0 | 0 | 0 |
| 0 | 0 | 0 | 0 |
| 0 | 0 | 0 | 1 |
| 0 | 0 | 0 | 1 |

| | | | |
|---|---|---|---|
| 0 | 0 | 0 | 1 |
| 0 | 0 | 0 | 1 |
| 0 | 0 | 0 | 1 |
| 0 | 0 | 0 | 1 |
| 0 | 0 | 0 | 1 |
| 0 | 0 | 0 | 1 |
| 0 | 0 | 0 | 1 |
| 1 | 0 | 0 | 0 |
| 0 | 0 | 0 | 1 |
| 0 | 0 | 0 | 1 |
| 0 | 0 | 0 | 1 |
| 0 | 0 | 0 | 1 |
| 0 | 0 | 0 | 1 |
| 0 | 0 | 0 | 1 |
| 0 | 0 | 0 | 0 |
| 0 | 0 | 0 | 0 |
| 0 | 0 | 0 | 1 |
| 0 | 0 | 0 | 0 |
| 0 | 0 | 0 | 1 |
| 0 | 0 | 0 | 1 |
| 0 | 0 | 0 | 0 |
| 0 | 0 | 0 | 1 |
| 0 | 0 | 0 | 1 |
| 0 | 0 | 0 | 1 |
| 0 | 0 | 0 | 0 |
| 0 | 0 | 0 | 1 |
| 0 | 0 | 0 | 1 |
| 1 | 0 | 0 | 0 |
| 0 | 0 | 0 | 0 |
| 1 | 0 | 0 | 0 |
| 1 | 0 | 0 | 0 |
| 0 | 0 | 0 | 1 |
| 0 | 0 | 0 | 1 |
| 0 | 0 | 0 | 1 |
| 0 | 0 | 0 | 1 |
| 0 | 0 | 0 | 1 |
| 0 | 0 | 0 | 0 |
| 0 | 0 | 0 | 0 |
| 0 | 0 | 0 | 1 |
| 0 | 0 | 0 | 1 |
| 0 | 0 | 0 | 1 |
| 0 | 0 | 0 | 1 |
| 0 | 0 | 0 | 0 |
| 0 | 0 | 0 | 1 |
| 0 | 0 | 0 | 1 |
| 0 | 0 | 0 | 1 |
| 0 | 0 | 0 | 1 |
| 0 | 0 | 0 | 1 |
| 0 | 0 | 0 | 0 |
| 0 | 0 | 0 | 0 |
| 0 | 0 | 0 | 0 |
| 0 | 0 | 0 | 0 |
| 0 | 0 | 0 | 0 |
| 0 | 0 | 0 | 1 |

| | | | |
|---|---|---|---|
| 0 | 0 | 0 | 0 |
| 0 | 0 | 0 | 0 |
| 0 | 0 | 0 | 1 |
| 1 | 0 | 0 | 0 |
| 0 | 0 | 0 | 1 |
| 0 | 0 | 0 | 1 |
| 0 | 0 | 0 | 1 |
| 0 | 0 | 0 | 1 |
| 0 | 0 | 0 | 1 |
| 1 | 0 | 0 | 0 |
| 0 | 0 | 0 | 1 |
| 0 | 0 | 0 | 1 |
| 0 | 0 | 0 | 1 |
| 0 | 0 | 0 | 1 |
| 0 | 0 | 0 | 1 |
| 0 | 0 | 0 | 1 |
| 0 | 0 | 0 | 1 |
| 0 | 0 | 0 | 1 |
| 0 | 0 | 0 | 1 |
| 0 | 0 | 0 | 1 |
| 0 | 0 | 0 | 1 |
| 0 | 0 | 0 | 1 |
| 0 | 0 | 0 | 1 |
| 0 | 0 | 0 | 1 |
| 0 | 0 | 0 | 1 |
| 0 | 0 | 0 | 1 |
| 0 | 0 | 0 | 1 |
| 0 | 0 | 0 | 1 |
| 0 | 0 | 0 | 1 |
| 0 | 0 | 0 | 1 |
| 0 | 0 | 0 | 1 |
| 0 | 0 | 0 | 1 |
| 1 | 0 | 0 | 0 |
| 0 | 0 | 0 | 1 |
| 0 | 0 | 0 | 1 |
| 0 | 0 | 0 | 1 |
| 0 | 0 | 0 | 1 |
| 0 | 0 | 0 | 1 |
| 0 | 0 | 0 | 1 |
| 0 | 0 | 0 | 1 |
| 0 | 0 | 0 | 1 |
| 0 | 0 | 0 | 1 |
| 0 | 0 | 0 | 1 |
| 0 | 0 | 0 | 1 |
| 0 | 0 | 0 | 1 |
| 0 | 0 | 0 | 1 |
| 0 | 0 | 0 | 1 |
| 0 | 0 | 0 | 0 |
| 0 | 1 | 0 | 0 |
| 0 | 0 | 0 | 0 |
| 0 | 0 | 0 | 0 |
| 0 | 1 | 0 | 0 |
| 0 | 0 | 0 | 1 |
| 0 | 0 | 0 | 0 |

| | | | |
|---|---|---|---|
| 0 | 0 | 0 | 1 |
| 0 | 0 | 0 | 0 |
| 0 | 0 | 0 | 1 |
| 0 | 0 | 0 | 1 |
| 0 | 0 | 0 | 1 |
| 1 | 0 | 0 | 0 |
| 0 | 0 | 0 | 0 |
| 0 | 0 | 0 | 0 |
| 1 | 0 | 0 | 0 |
| 0 | 0 | 0 | 0 |
| 0 | 0 | 0 | 1 |
| 0 | 0 | 0 | 0 |
| 0 | 0 | 0 | 1 |
| 0 | 0 | 0 | 1 |
| 0 | 0 | 0 | 1 |
| 1 | 0 | 0 | 0 |
| 0 | 0 | 0 | 1 |
| 0 | 0 | 0 | 1 |
| 0 | 0 | 0 | 0 |
| 0 | 0 | 0 | 0 |
| 1 | 0 | 0 | 0 |
| 1 | 0 | 0 | 0 |
| 0 | 0 | 0 | 0 |
| 0 | 0 | 0 | 0 |
| 0 | 0 | 0 | 1 |
| 0 | 0 | 0 | 0 |
| 0 | 0 | 0 | 0 |
| 0 | 0 | 0 | 0 |
| 0 | 0 | 0 | 0 |
| 0 | 0 | 0 | 0 |
| 0 | 0 | 0 | 0 |
| 0 | 0 | 0 | 1 |
| 0 | 0 | 0 | 1 |
| 0 | 0 | 0 | 0 |
| 0 | 1 | 0 | 0 |
| 0 | 0 | 0 | 1 |
| 0 | 0 | 0 | 0 |
| 0 | 0 | 0 | 0 |
| 0 | 1 | 0 | 0 |
| 0 | 0 | 0 | 0 |
| 0 | 1 | 0 | 0 |
| 0 | 1 | 1 | 1 |
| 0 | 0 | 0 | 1 |
| 0 | 0 | 0 | 1 |
| 0 | 0 | 0 | 0 |
| 0 | 0 | 0 | 0 |
| 0 | 0 | 0 | 0 |
| 0 | 0 | 0 | 1 |
| 0 | 0 | 0 | 1 |
| 0 | 0 | 0 | 1 |
| 0 | 0 | 0 | 1 |
| 0 | 0 | 0 | 1 |
| 0 | 0 | 0 | 1 |
| 0 | 0 | 0 | 0 |

| | | | |
|---|---|---|---|
| 0 | 0 | 0 | 1 |
| 0 | 0 | 0 | 1 |
| 0 | 0 | 0 | 1 |
| 0 | 0 | 0 | 1 |
| 1 | 0 | 0 | 0 |
| 0 | 0 | 0 | 0 |
| 0 | 0 | 0 | 1 |
| 0 | 1 | 1 | 1 |
| 0 | 0 | 0 | 1 |
| 0 | 0 | 0 | 1 |
| 0 | 0 | 0 | 1 |
| 0 | 0 | 0 | 0 |
| 0 | 0 | 0 | 1 |
| 0 | 0 | 0 | 0 |
| 0 | 1 | 0 | 0 |
| 0 | 0 | 0 | 0 |
| 0 | 1 | 0 | 0 |
| 0 | 1 | 0 | 0 |
| 0 | 1 | 0 | 0 |
| 0 | 1 | 0 | 0 |
| 0 | 0 | 0 | 0 |
| 0 | 0 | 0 | 0 |
| 0 | 0 | 0 | 0 |
| 0 | 0 | 0 | 1 |
| 0 | 0 | 0 | 1 |
| 0 | 0 | 0 | 1 |
| 0 | 0 | 0 | 1 |
| 0 | 0 | 0 | 1 |
| 0 | 0 | 0 | 1 |
| 0 | 0 | 0 | 1 |
| 0 | 0 | 0 | 1 |
| 0 | 0 | 0 | 0 |
| 0 | 0 | 0 | 1 |
| 0 | 0 | 0 | 0 |
| 0 | 0 | 0 | 0 |
| 0 | 0 | 0 | 0 |
| 0 | 0 | 0 | 0 |
| 0 | 0 | 0 | 1 |
| 0 | 0 | 0 | 1 |
| 0 | 0 | 0 | 1 |
| 0 | 0 | 0 | 1 |
| 0 | 0 | 0 | 1 |
| 0 | 0 | 0 | 1 |
| 0 | 0 | 0 | 1 |
| 0 | 0 | 0 | 1 |
| 1 | 0 | 0 | 0 |
| 0 | 0 | 0 | 1 |
| 0 | 0 | 0 | 1 |
| 0 | 0 | 0 | 1 |
| 1 | 1 | 0 | 0 |
| 1 | 0 | 0 | 0 |
| 0 | 0 | 0 | 0 |
| 0 | 0 | 0 | 1 |
| 0 | 0 | 0 | 1 |

| | | | |
|---|---|---|---|
| 0 | 0 | 0 | 1 |
| 0 | 0 | 0 | 1 |
| 0 | 0 | 0 | 1 |
| 0 | 0 | 0 | 0 |
| 0 | 0 | 0 | 1 |
| 0 | 0 | 0 | 1 |
| 0 | 0 | 0 | 0 |
| 0 | 0 | 0 | 1 |
| 0 | 0 | 0 | 1 |
| 0 | 0 | 0 | 1 |
| 0 | 0 | 0 | 0 |
| 0 | 0 | 0 | 0 |
| 0 | 0 | 0 | 1 |
| 0 | 0 | 0 | 1 |
| 0 | 0 | 0 | 1 |
| 0 | 0 | 0 | 1 |
| 0 | 0 | 0 | 0 |
| 0 | 0 | 0 | 0 |
| 0 | 0 | 0 | 1 |
| 0 | 0 | 0 | 0 |
| 0 | 0 | 0 | 0 |
| 0 | 1 | 0 | 0 |
| 0 | 0 | 0 | 1 |
| 0 | 1 | 0 | 0 |
| 0 | 0 | 0 | 0 |
| 0 | 0 | 0 | 1 |
| 0 | 0 | 0 | 0 |
| 0 | 0 | 0 | 1 |
| 0 | 0 | 0 | 1 |
| 0 | 0 | 0 | 0 |
| 0 | 0 | 0 | 1 |
| 0 | 0 | 0 | 1 |
| 0 | 0 | 0 | 0 |
| 0 | 0 | 0 | 1 |
| 0 | 0 | 0 | 1 |
| 0 | 0 | 0 | 0 |
| 0 | 0 | 0 | 1 |
| 0 | 0 | 0 | 0 |
| 0 | 0 | 0 | 1 |
| 0 | 0 | 0 | 0 |
| 0 | 0 | 0 | 0 |
| 0 | 0 | 0 | 1 |
| 0 | 0 | 0 | 0 |
| 0 | 0 | 0 | 0 |
| 0 | 0 | 0 | 1 |
| 0 | 0 | 0 | 1 |
| 0 | 0 | 0 | 1 |
| 0 | 0 | 0 | 1 |
| 0 | 0 | 0 | 1 |
| 0 | 0 | 0 | 1 |
| 0 | 0 | 0 | 1 |
| 0 | 0 | 0 | 1 |
| 0 | 0 | 0 | 1 |
| 0 | 0 | 0 | 1 |

| | | | |
|---|---|---|---|
| 0 | 0 | 0 | 1 |
| 0 | 0 | 0 | 1 |
| 0 | 0 | 0 | 1 |
| 0 | 0 | 0 | 0 |
| 0 | 1 | 1 | 1 |
| 0 | 0 | 0 | 0 |
| 0 | 0 | 0 | 1 |
| 0 | 0 | 0 | 1 |
| 0 | 0 | 0 | 0 |
| 0 | 0 | 0 | 0 |
| 0 | 0 | 0 | 1 |
| 0 | 0 | 0 | 0 |
| 0 | 1 | 0 | 0 |
| 0 | 0 | 0 | 0 |
| 0 | 0 | 0 | 1 |
| 0 | 0 | 0 | 0 |
| 0 | 0 | 0 | 0 |
| 0 | 0 | 0 | 1 |
| 1 | 0 | 0 | 0 |
| 1 | 0 | 0 | 0 |
| 1 | 0 | 0 | 0 |
| 1 | 0 | 0 | 0 |
| 0 | 0 | 0 | 1 |
| 0 | 0 | 0 | 0 |
| 0 | 0 | 0 | 0 |
| 0 | 0 | 0 | 0 |
| 0 | 0 | 0 | 0 |
| 0 | 0 | 0 | 0 |
| 0 | 0 | 0 | 0 |
| 0 | 0 | 0 | 0 |
| 0 | 0 | 0 | 0 |
| 0 | 0 | 0 | 0 |
| 0 | 0 | 0 | 0 |
| 0 | 0 | 0 | 1 |
| 0 | 0 | 0 | 1 |
| 0 | 1 | 0 | 0 |
| 0 | 0 | 0 | 1 |
| 0 | 0 | 0 | 0 |
| 0 | 0 | 0 | 0 |
| 0 | 1 | 0 | 0 |
| 0 | 1 | 0 | 0 |
| 0 | 0 | 0 | 1 |
| 0 | 0 | 0 | 0 |
| 0 | 0 | 0 | 0 |
| 0 | 0 | 0 | 1 |
| 0 | 0 | 0 | 1 |
| 0 | 0 | 0 | 1 |
| 0 | 0 | 0 | 1 |
| 0 | 0 | 0 | 1 |
| 0 | 0 | 0 | 1 |
| 0 | 0 | 0 | 1 |
| 0 | 0 | 0 | 1 |
| 0 | 0 | 0 | 1 |
| 0 | 0 | 0 | 1 |

| | | | |
|---|---|---|---|
| 0 | 0 | 0 | 1 |
| 0 | 0 | 0 | 0 |
| 0 | 0 | 0 | 1 |
| 0 | 0 | 0 | 1 |
| 0 | 0 | 0 | 1 |
| 0 | 0 | 0 | 1 |
| 0 | 0 | 0 | 1 |
| 0 | 0 | 0 | 1 |
| 0 | 0 | 0 | 1 |
| 0 | 0 | 0 | 1 |
| 0 | 0 | 0 | 1 |
| 0 | 0 | 0 | 0 |
| 0 | 0 | 0 | 1 |
| 0 | 0 | 0 | 1 |
| 0 | 1 | 0 | 0 |
| 0 | 0 | 0 | 1 |
| 0 | 0 | 0 | 1 |
| 0 | 0 | 0 | 1 |
| 0 | 0 | 0 | 1 |
| 0 | 0 | 0 | 1 |
| 1 | 0 | 0 | 0 |
| 0 | 0 | 0 | 1 |
| 1 | 0 | 0 | 0 |
| 1 | 0 | 0 | 0 |
| 0 | 0 | 0 | 1 |
| 0 | 0 | 0 | 0 |
| 0 | 0 | 0 | 1 |
| 0 | 0 | 0 | 1 |
| 0 | 0 | 0 | 1 |
| 0 | 0 | 0 | 1 |
| 0 | 0 | 0 | 1 |
| 0 | 0 | 0 | 0 |
| 0 | 0 | 0 | 1 |
| 0 | 0 | 0 | 0 |
| 0 | 0 | 0 | 1 |
| 0 | 0 | 0 | 1 |
| 0 | 0 | 0 | 1 |
| 0 | 0 | 0 | 1 |
| 0 | 0 | 0 | 1 |
| 0 | 0 | 0 | 0 |
| 0 | 0 | 0 | 1 |
| 0 | 0 | 0 | 1 |
| 0 | 0 | 0 | 1 |
| 0 | 0 | 0 | 0 |
| 0 | 0 | 0 | 1 |
| 0 | 0 | 0 | 1 |
| 0 | 0 | 0 | 1 |
| 1 | 0 | 0 | 0 |
| 0 | 0 | 0 | 0 |
| 0 | 0 | 0 | 1 |
| 0 | 0 | 0 | 1 |
| 0 | 0 | 0 | 0 |
| 1 | 0 | 0 | 0 |
| 0 | 0 | 0 | 0 |

| | | | |
|---|---|---|---|
| 1 | 0 | 0 | 0 |
| 0 | 0 | 0 | 1 |
| 0 | 0 | 0 | 0 |
| 0 | 0 | 0 | 0 |
| 0 | 0 | 0 | 0 |
| 0 | 0 | 0 | 1 |
| 0 | 0 | 0 | 1 |
| 0 | 0 | 0 | 1 |
| 0 | 0 | 0 | 1 |
| 0 | 0 | 0 | 0 |
| 0 | 0 | 0 | 1 |
| 0 | 0 | 0 | 1 |
| 0 | 0 | 0 | 1 |
| 0 | 0 | 0 | 1 |
| 0 | 0 | 0 | 0 |
| 0 | 0 | 0 | 1 |
| 0 | 0 | 0 | 1 |
| 0 | 0 | 0 | 0 |
| 0 | 0 | 0 | 1 |
| 0 | 0 | 0 | 0 |
| 0 | 0 | 0 | 0 |
| 0 | 0 | 0 | 1 |
| 0 | 0 | 0 | 0 |
| 0 | 0 | 0 | 1 |
| 0 | 0 | 0 | 0 |
| 0 | 0 | 0 | 0 |
| 0 | 0 | 0 | 1 |
| 0 | 0 | 0 | 1 |
| 1 | 0 | 0 | 0 |
| 0 | 0 | 0 | 1 |
| 0 | 0 | 0 | 1 |
| 0 | 0 | 0 | 1 |
| 0 | 0 | 0 | 1 |
| 0 | 0 | 0 | 1 |
| 0 | 0 | 0 | 0 |
| 0 | 0 | 0 | 0 |
| 0 | 0 | 0 | 1 |
| 0 | 0 | 0 | 0 |
| 0 | 0 | 0 | 0 |
| 0 | 0 | 0 | 0 |
| 0 | 0 | 0 | 0 |
| 0 | 0 | 0 | 0 |
| 0 | 0 | 0 | 0 |
| 0 | 0 | 0 | 1 |
| 0 | 1 | 0 | 0 |
| 0 | 0 | 0 | 1 |
| 0 | 0 | 0 | 1 |
| 0 | 0 | 0 | 0 |
| 0 | 0 | 0 | 0 |
| 0 | 0 | 0 | 0 |
| 0 | 0 | 0 | 0 |
| 0 | 0 | 0 | 0 |
| 0 | 0 | 0 | 1 |
| 0 | 0 | 0 | 1 |

| | | | |
|---|---|---|---|
| 0 | 0 | 0 | 1 |
| 0 | 0 | 0 | 0 |
| 0 | 0 | 0 | 0 |
| 0 | 0 | 0 | 0 |
| 0 | 0 | 0 | 0 |
| 0 | 0 | 0 | 0 |
| 0 | 0 | 0 | 0 |
| 0 | 0 | 0 | 1 |
| 0 | 1 | 1 | 1 |
| 0 | 0 | 0 | 1 |
| 0 | 0 | 0 | 0 |
| 0 | 0 | 0 | 0 |
| 0 | 0 | 0 | 1 |
| 0 | 0 | 0 | 0 |
| 0 | 0 | 0 | 1 |
| 0 | 0 | 0 | 0 |
| 0 | 0 | 0 | 1 |
| 0 | 0 | 0 | 1 |
| 0 | 0 | 0 | 1 |
| 0 | 0 | 0 | 1 |
| 0 | 0 | 0 | 1 |
| 0 | 0 | 0 | 1 |
| 0 | 0 | 0 | 1 |
| 1 | 0 | 0 | 0 |
| 0 | 0 | 0 | 0 |
| 0 | 0 | 0 | 1 |
| 0 | 0 | 0 | 1 |
| 0 | 0 | 0 | 1 |
| 0 | 0 | 0 | 1 |
| 0 | 0 | 0 | 0 |
| 0 | 0 | 0 | 0 |
| 0 | 0 | 0 | 0 |
| 0 | 0 | 0 | 0 |
| 0 | 0 | 0 | 1 |
| 0 | 0 | 0 | 0 |
| 0 | 0 | 0 | 0 |
| 0 | 0 | 0 | 0 |
| 0 | 0 | 0 | 0 |
| 0 | 0 | 0 | 0 |
| 0 | 0 | 0 | 1 |
| 0 | 0 | 0 | 1 |
| 0 | 0 | 0 | 0 |
| 0 | 0 | 0 | 1 |
| 0 | 0 | 0 | 1 |
| 0 | 0 | 0 | 0 |
| 0 | 0 | 0 | 1 |
| 0 | 0 | 0 | 1 |
| 0 | 0 | 0 | 1 |
| 0 | 0 | 0 | 1 |
| 0 | 0 | 0 | 1 |
| 0 | 0 | 0 | 0 |
| 0 | 0 | 0 | 1 |
| 0 | 0 | 0 | 0 |
| 0 | 0 | 0 | 0 |

| | | | |
|---|---|---|---|
| 0 | 0 | 0 | 1 |
| 0 | 0 | 0 | 1 |
| 0 | 0 | 0 | 0 |
| 0 | 0 | 0 | 0 |
| 0 | 0 | 0 | 1 |
| 0 | 0 | 0 | 1 |
| 0 | 0 | 0 | 1 |
| 0 | 0 | 0 | 1 |
| 0 | 0 | 0 | 1 |
| 0 | 0 | 0 | 1 |
| 0 | 0 | 0 | 1 |
| 0 | 0 | 0 | 1 |
| 0 | 0 | 0 | 0 |
| 0 | 0 | 0 | 1 |
| 0 | 0 | 0 | 1 |
| 0 | 0 | 0 | 1 |
| 0 | 0 | 0 | 1 |
| 0 | 0 | 0 | 1 |
| 0 | 0 | 0 | 1 |
| 0 | 0 | 0 | 1 |
| 0 | 0 | 0 | 1 |
| 0 | 0 | 0 | 1 |
| 0 | 0 | 0 | 1 |
| 0 | 0 | 0 | 1 |
| 0 | 0 | 0 | 1 |
| 0 | 0 | 0 | 1 |
| 0 | 0 | 0 | 1 |
| 0 | 0 | 0 | 1 |
| 0 | 0 | 0 | 1 |
| 0 | 0 | 0 | 1 |
| 0 | 0 | 0 | 1 |
| 0 | 0 | 0 | 0 |
| 0 | 0 | 0 | 1 |
| 0 | 0 | 0 | 1 |
| 0 | 0 | 0 | 1 |
| 0 | 0 | 0 | 1 |
| 0 | 0 | 0 | 1 |
| 0 | 0 | 0 | 1 |
| 0 | 0 | 0 | 1 |
| 0 | 0 | 0 | 1 |
| 0 | 0 | 0 | 0 |
| 0 | 0 | 0 | 0 |
| 0 | 0 | 0 | 1 |
| 0 | 0 | 0 | 1 |
| 0 | 0 | 0 | 1 |
| 0 | 0 | 0 | 1 |
| 0 | 0 | 0 | 1 |
| 0 | 0 | 0 | 1 |
| 0 | 0 | 0 | 0 |
| 0 | 0 | 0 | 0 |
| 0 | 0 | 0 | 1 |
| 0 | 0 | 0 | 1 |
| 0 | 0 | 0 | 0 |
| 0 | 0 | 0 | 1 |

| | | | |
|---|---|---|---|
| 0 | 0 | 0 | 0 |
| 0 | 0 | 0 | 0 |
| 0 | 0 | 0 | 0 |
| 0 | 0 | 0 | 1 |
| 0 | 0 | 0 | 1 |
| 0 | 0 | 0 | 1 |
| 0 | 0 | 0 | 1 |
| 0 | 1 | 0 | 0 |
| 0 | 1 | 0 | 0 |
| 0 | 0 | 0 | 0 |
| 0 | 0 | 0 | 0 |
| 0 | 0 | 0 | 1 |
| 0 | 0 | 0 | 0 |
| 0 | 0 | 0 | 1 |
| 0 | 0 | 0 | 1 |
| 0 | 0 | 0 | 1 |
| 0 | 0 | 0 | 1 |
| 0 | 0 | 0 | 0 |
| 0 | 0 | 0 | 1 |
| 0 | 0 | 0 | 1 |
| 0 | 0 | 0 | 1 |
| 0 | 0 | 0 | 1 |
| 0 | 0 | 0 | 1 |
| 0 | 0 | 0 | 1 |
| 0 | 0 | 0 | 1 |
| 0 | 0 | 0 | 1 |
| 0 | 0 | 0 | 1 |
| 0 | 0 | 0 | 1 |
| 0 | 0 | 0 | 1 |
| 0 | 0 | 0 | 1 |
| 0 | 0 | 0 | 0 |
| 0 | 0 | 0 | 0 |
| 0 | 0 | 0 | 1 |
| 0 | 0 | 0 | 1 |
| 0 | 0 | 0 | 1 |
| 0 | 0 | 0 | 1 |
| 0 | 0 | 0 | 1 |
| 0 | 0 | 0 | 1 |
| 0 | 0 | 0 | 1 |
| 0 | 0 | 0 | 1 |
| 0 | 0 | 0 | 1 |
| 0 | 0 | 0 | 0 |
| 0 | 0 | 0 | 1 |
| 0 | 0 | 0 | 1 |
| 0 | 0 | 0 | 0 |
| 0 | 0 | 0 | 0 |
| 0 | 0 | 0 | 1 |
| 0 | 0 | 0 | 1 |
| 0 | 0 | 0 | 0 |
| 0 | 0 | 0 | 0 |
| 0 | 0 | 0 | 0 |
| 0 | 0 | 0 | 0 |
| 0 | 0 | 0 | 0 |
| 0 | 0 | 0 | 0 |

| | | | |
|---|---|---|---|
| 0 | 0 | 0 | 1 |
| 0 | 0 | 0 | 1 |
| 0 | 0 | 0 | 1 |
| 0 | 0 | 0 | 1 |
| 0 | 0 | 0 | 1 |
| 0 | 0 | 0 | 0 |
| 0 | 0 | 0 | 0 |
| 0 | 0 | 0 | 0 |
| 0 | 0 | 0 | 0 |
| 0 | 0 | 0 | 0 |
| 0 | 0 | 0 | 1 |
| 0 | 0 | 0 | 0 |
| 0 | 0 | 0 | 0 |
| 0 | 0 | 0 | 0 |
| 0 | 0 | 0 | 1 |
| 0 | 0 | 0 | 0 |
| 0 | 0 | 0 | 1 |
| 0 | 0 | 0 | 1 |
| 1 | 0 | 0 | 0 |
| 1 | 0 | 0 | 0 |
| 0 | 0 | 0 | 0 |
| 0 | 0 | 0 | 0 |
| 0 | 0 | 0 | 1 |
| 0 | 0 | 0 | 0 |
| 0 | 0 | 0 | 0 |
| 0 | 0 | 0 | 0 |
| 0 | 0 | 0 | 0 |
| 0 | 0 | 0 | 1 |
| 0 | 0 | 0 | 1 |
| 0 | 0 | 0 | 1 |
| 0 | 0 | 0 | 1 |
| 0 | 0 | 0 | 1 |
| 0 | 0 | 0 | 0 |
| 0 | 0 | 0 | 1 |
| 0 | 0 | 0 | 0 |
| 0 | 0 | 0 | 0 |
| 0 | 0 | 0 | 0 |
| 0 | 0 | 0 | 0 |
| 0 | 1 | 0 | 0 |
| 0 | 1 | 0 | 0 |
| 0 | 1 | 0 | 0 |
| 1 | 0 | 0 | 0 |
| 0 | 0 | 0 | 0 |
| 0 | 0 | 0 | 1 |
| 0 | 0 | 0 | 1 |
| 0 | 0 | 0 | 1 |
| 0 | 0 | 0 | 1 |
| 0 | 0 | 0 | 0 |
| 0 | 0 | 0 | 0 |
| 0 | 0 | 0 | 0 |
| 0 | 0 | 0 | 1 |
| 0 | 0 | 0 | 1 |
| 0 | 0 | 0 | 0 |
| 0 | 0 | 0 | 1 |

| | | | |
|---|---|---|---|
| 0 | 0 | 0 | 1 |
| 0 | 0 | 0 | 0 |
| 0 | 0 | 0 | 1 |
| 0 | 0 | 0 | 1 |
| 0 | 0 | 0 | 1 |
| 0 | 0 | 0 | 1 |
| 0 | 0 | 0 | 1 |
| 0 | 0 | 0 | 1 |
| 0 | 0 | 0 | 1 |
| 0 | 0 | 0 | 1 |
| 0 | 0 | 0 | 1 |
| 0 | 0 | 0 | 1 |
| 0 | 0 | 0 | 0 |
| 0 | 0 | 0 | 0 |
| 0 | 0 | 0 | 0 |
| 0 | 0 | 0 | 0 |
| 0 | 0 | 0 | 0 |
| 0 | 0 | 0 | 0 |
| 0 | 0 | 0 | 0 |
| 0 | 0 | 0 | 1 |
| 1 | 1 | 0 | 0 |
| 0 | 0 | 0 | 0 |
| 0 | 0 | 0 | 0 |
| 0 | 0 | 0 | 0 |
| 0 | 0 | 0 | 0 |
| 0 | 0 | 0 | 1 |
| 0 | 0 | 0 | 1 |
| 0 | 0 | 0 | 0 |
| 0 | 0 | 0 | 1 |
| 0 | 0 | 0 | 0 |
| 0 | 0 | 0 | 0 |
| 0 | 0 | 0 | 1 |
| 0 | 0 | 0 | 0 |
| 0 | 0 | 0 | 1 |
| 0 | 0 | 0 | 1 |
| 0 | 0 | 0 | 1 |
| 0 | 1 | 1 | 1 |
| 0 | 0 | 0 | 1 |
| 0 | 0 | 0 | 1 |
| 0 | 0 | 0 | 1 |
| 0 | 0 | 0 | 0 |
| 0 | 0 | 0 | 1 |
| 0 | 0 | 0 | 1 |
| 0 | 0 | 0 | 0 |
| 0 | 0 | 0 | 0 |
| 0 | 0 | 0 | 1 |
| 0 | 0 | 0 | 0 |
| 0 | 0 | 0 | 0 |
| 0 | 0 | 0 | 1 |
| 0 | 0 | 0 | 1 |
| 0 | 0 | 0 | 0 |
| 0 | 0 | 0 | 1 |
| 0 | 0 | 0 | 1 |
| 0 | 0 | 0 | 1 |

| | | | |
|---|---|---|---|
| 0 | 0 | 0 | 1 |
| 0 | 0 | 0 | 1 |
| 0 | 0 | 0 | 1 |
| 0 | 0 | 0 | 1 |
| 0 | 0 | 0 | 0 |
| 0 | 0 | 0 | 0 |
| 0 | 0 | 0 | 1 |
| 0 | 0 | 0 | 1 |
| 0 | 0 | 0 | 1 |
| 0 | 0 | 0 | 1 |
| 0 | 0 | 0 | 1 |
| 0 | 0 | 0 | 0 |
| 0 | 0 | 0 | 0 |
| 0 | 0 | 0 | 1 |
| 0 | 0 | 0 | 1 |
| 0 | 0 | 0 | 0 |
| 0 | 0 | 0 | 1 |
| 0 | 0 | 0 | 1 |
| 0 | 0 | 0 | 0 |
| 0 | 0 | 0 | 1 |
| 0 | 0 | 0 | 1 |
| 0 | 0 | 0 | 1 |
| 0 | 0 | 0 | 1 |
| 0 | 0 | 0 | 0 |
| 0 | 0 | 0 | 1 |
| 0 | 0 | 0 | 0 |
| 0 | 0 | 0 | 1 |
| 0 | 0 | 0 | 1 |
| 1 | 0 | 0 | 0 |
| 1 | 0 | 0 | 0 |
| 0 | 0 | 0 | 0 |
| 0 | 0 | 0 | 0 |
| 0 | 0 | 0 | 0 |
| 0 | 0 | 0 | 0 |
| 0 | 0 | 0 | 1 |
| 0 | 0 | 0 | 1 |
| 0 | 0 | 0 | 1 |
| 1 | 0 | 0 | 0 |
| 0 | 0 | 0 | 1 |
| 0 | 0 | 0 | 1 |
| 0 | 0 | 0 | 0 |
| 0 | 0 | 0 | 0 |
| 0 | 0 | 0 | 0 |
| 0 | 0 | 0 | 1 |
| 0 | 0 | 0 | 0 |
| 0 | 0 | 0 | 1 |
| 0 | 0 | 0 | 1 |
| 0 | 0 | 0 | 0 |
| 0 | 0 | 0 | 1 |
| 0 | 0 | 0 | 1 |
| 0 | 0 | 0 | 1 |
| 0 | 1 | 0 | 0 |
| 0 | 0 | 0 | 1 |
| 0 | 0 | 0 | 0 |

| | | | |
|---|---|---|---|
| 0 | 0 | 0 | 1 |
| 0 | 0 | 0 | 0 |
| 0 | 0 | 0 | 0 |
| 0 | 0 | 0 | 1 |
| 0 | 0 | 0 | 1 |
| 0 | 0 | 0 | 1 |
| 0 | 0 | 0 | 0 |
| 0 | 0 | 0 | 1 |
| 0 | 0 | 0 | 1 |
| 0 | 0 | 0 | 1 |
| 0 | 0 | 0 | 1 |
| 0 | 0 | 0 | 1 |
| 0 | 0 | 0 | 1 |
| 0 | 0 | 0 | 1 |
| 0 | 0 | 0 | 1 |
| 0 | 0 | 0 | 1 |
| 0 | 0 | 0 | 1 |
| 0 | 0 | 0 | 1 |
| 0 | 0 | 0 | 1 |
| 0 | 0 | 0 | 1 |
| 0 | 0 | 0 | 1 |
| 0 | 0 | 0 | 0 |
| 0 | 0 | 0 | 1 |
| 0 | 0 | 0 | 1 |
| 0 | 0 | 0 | 1 |
| 0 | 0 | 0 | 1 |
| 0 | 0 | 0 | 0 |
| 0 | 0 | 0 | 1 |
| 0 | 0 | 0 | 0 |
| 0 | 0 | 0 | 1 |
| 0 | 0 | 0 | 1 |
| 0 | 0 | 0 | 0 |
| 0 | 1 | 1 | 1 |
| 0 | 1 | 1 | 1 |
| 0 | 0 | 0 | 1 |
| 0 | 0 | 0 | 1 |
| 0 | 0 | 0 | 1 |
| 0 | 0 | 0 | 1 |
| 0 | 0 | 0 | 0 |
| 0 | 0 | 0 | 0 |
| 0 | 0 | 0 | 0 |
| 0 | 0 | 0 | 1 |
| 0 | 0 | 0 | 1 |
| 0 | 0 | 0 | 1 |
| 0 | 0 | 0 | 1 |
| 0 | 1 | 0 | 0 |
| 0 | 0 | 0 | 1 |
| 0 | 0 | 0 | 0 |
| 0 | 0 | 0 | 1 |
| 0 | 0 | 0 | 0 |
| 0 | 0 | 0 | 0 |
| 0 | 0 | 0 | 0 |
| 0 | 0 | 0 | 1 |
| 0 | 0 | 0 | 1 |

| | | | |
|---|---|---|---|
| 0 | 0 | 0 | 0 |
| 0 | 0 | 0 | 0 |
| 0 | 0 | 0 | 1 |
| 0 | 0 | 0 | 1 |
| 0 | 0 | 0 | 1 |
| 0 | 0 | 0 | 1 |
| 0 | 0 | 0 | 1 |
| 0 | 0 | 0 | 1 |
| 0 | 0 | 0 | 1 |
| 0 | 0 | 0 | 1 |
| 0 | 0 | 0 | 1 |
| 0 | 0 | 0 | 1 |
| 0 | 0 | 0 | 1 |
| 0 | 0 | 0 | 1 |
| 0 | 0 | 0 | 1 |
| 0 | 0 | 0 | 0 |
| 0 | 0 | 0 | 0 |
| 0 | 0 | 0 | 0 |
| 0 | 0 | 0 | 1 |
| 0 | 0 | 0 | 1 |
| 0 | 0 | 0 | 1 |
| 0 | 1 | 0 | 0 |
| 0 | 0 | 0 | 0 |
| 0 | 0 | 0 | 0 |
| 0 | 0 | 0 | 0 |
| 0 | 0 | 0 | 1 |
| 0 | 0 | 0 | 1 |
| 0 | 0 | 0 | 1 |
| 0 | 0 | 0 | 1 |
| 0 | 0 | 0 | 1 |
| 0 | 0 | 0 | 0 |
| 0 | 0 | 0 | 1 |
| 0 | 0 | 0 | 0 |
| 0 | 0 | 0 | 1 |
| 0 | 1 | 1 | 1 |
| 0 | 1 | 1 | 1 |
| 0 | 1 | 0 | 0 |
| 0 | 0 | 0 | 0 |
| 0 | 0 | 0 | 1 |
| 0 | 0 | 0 | 1 |
| 0 | 0 | 0 | 1 |
| 0 | 0 | 0 | 0 |
| 0 | 0 | 0 | 1 |
| 0 | 0 | 0 | 1 |
| 0 | 0 | 0 | 0 |
| 0 | 0 | 0 | 1 |
| 0 | 0 | 0 | 0 |
| 0 | 0 | 0 | 0 |
| 0 | 0 | 0 | 1 |
| 0 | 0 | 0 | 1 |
| 0 | 1 | 0 | 0 |
| 0 | 1 | 0 | 0 |
| 0 | 0 | 0 | 1 |
| 0 | 0 | 0 | 0 |

| | | | |
|---|---|---|---|
| 0 | 0 | 0 | 1 |
| 0 | 0 | 0 | 0 |
| 0 | 1 | 1 | 1 |
| 0 | 1 | 0 | 0 |
| 0 | 0 | 0 | 1 |
| 0 | 0 | 0 | 1 |
| 0 | 0 | 0 | 1 |
| 0 | 0 | 0 | 1 |
| 0 | 0 | 0 | 0 |
| 0 | 0 | 0 | 1 |
| 0 | 0 | 0 | 1 |
| 0 | 0 | 0 | 1 |
| 0 | 0 | 0 | 0 |
| 0 | 0 | 0 | 0 |
| 0 | 0 | 0 | 0 |
| 0 | 0 | 0 | 1 |
| 0 | 0 | 0 | 0 |
| 0 | 0 | 0 | 1 |
| 0 | 1 | 0 | 0 |
| 0 | 0 | 0 | 0 |
| 0 | 0 | 0 | 0 |
| 0 | 1 | 0 | 0 |
| 0 | 0 | 0 | 0 |
| 1 | 0 | 0 | 0 |
| 0 | 0 | 0 | 1 |
| 0 | 0 | 0 | 1 |
| 0 | 0 | 0 | 0 |
| 0 | 0 | 0 | 0 |
| 0 | 0 | 0 | 0 |
| 0 | 0 | 0 | 0 |
| 0 | 0 | 0 | 1 |
| 0 | 0 | 0 | 1 |
| 0 | 0 | 0 | 1 |
| 0 | 0 | 0 | 1 |
| 0 | 0 | 0 | 1 |
| 1 | 0 | 0 | 0 |
| 1 | 0 | 0 | 0 |
| 0 | 0 | 0 | 1 |
| 0 | 0 | 0 | 1 |
| 0 | 0 | 0 | 1 |
| 0 | 0 | 0 | 1 |
| 0 | 0 | 0 | 1 |
| 0 | 0 | 0 | 1 |
| 0 | 0 | 0 | 0 |
| 0 | 0 | 0 | 0 |
| 0 | 0 | 0 | 0 |
| 0 | 0 | 0 | 1 |
| 0 | 0 | 0 | 0 |
| 0 | 0 | 0 | 0 |
| 0 | 0 | 0 | 0 |
| 0 | 0 | 0 | 1 |
| 0 | 0 | 0 | 1 |
| 0 | 0 | 0 | 1 |
| 0 | 0 | 0 | 1 |

| | | | |
|---|---|---|---|
| 0 | 0 | 0 | 1 |
| 0 | 0 | 0 | 1 |
| 0 | 0 | 0 | 1 |
| 0 | 0 | 0 | 1 |
| 0 | 0 | 0 | 0 |
| 0 | 0 | 0 | 1 |
| 0 | 0 | 0 | 0 |
| 0 | 0 | 0 | 0 |
| 0 | 0 | 0 | 0 |
| 0 | 0 | 0 | 0 |
| 0 | 0 | 0 | 0 |
| 0 | 0 | 0 | 0 |
| 0 | 0 | 0 | 1 |
| 0 | 0 | 0 | 0 |
| 0 | 0 | 0 | 0 |
| 0 | 0 | 0 | 1 |
| 0 | 0 | 0 | 1 |
| 0 | 0 | 0 | 1 |
| 0 | 0 | 0 | 0 |
| 0 | 0 | 0 | 0 |
| 0 | 0 | 0 | 0 |
| 0 | 0 | 0 | 0 |
| 0 | 0 | 0 | 1 |
| 0 | 0 | 0 | 1 |
| 0 | 0 | 0 | 1 |
| 0 | 0 | 0 | 1 |
| 0 | 0 | 0 | 1 |
| 0 | 0 | 0 | 0 |
| 0 | 0 | 0 | 0 |
| 0 | 0 | 0 | 0 |
| 0 | 0 | 0 | 0 |
| 0 | 0 | 0 | 0 |
| 0 | 0 | 0 | 0 |
| 0 | 0 | 0 | 1 |
| 0 | 0 | 0 | 1 |
| 0 | 0 | 0 | 0 |
| 0 | 0 | 0 | 1 |
| 0 | 0 | 0 | 1 |
| 0 | 0 | 0 | 1 |
| 0 | 0 | 0 | 1 |
| 0 | 0 | 0 | 0 |
| 0 | 0 | 0 | 1 |
| 0 | 0 | 0 | 0 |
| 0 | 0 | 0 | 1 |
| 0 | 0 | 0 | 1 |
| 0 | 0 | 0 | 1 |
| 0 | 0 | 0 | 1 |
| 0 | 0 | 0 | 0 |
| 0 | 0 | 0 | 1 |
| 0 | 0 | 0 | 1 |
| 0 | 0 | 0 | 1 |
| 0 | 0 | 0 | 1 |
| 0 | 0 | 0 | 1 |
| 0 | 0 | 0 | 1 |

| | | | |
|---|---|---|---|
| 0 | 0 | 0 | 1 |
| 0 | 0 | 0 | 1 |
| 0 | 0 | 0 | 1 |
| 0 | 0 | 0 | 1 |
| 0 | 0 | 0 | 1 |
| 0 | 0 | 0 | 1 |
| 0 | 0 | 0 | 1 |
| 0 | 0 | 0 | 1 |
| 0 | 0 | 0 | 1 |
| 0 | 0 | 0 | 1 |
| 0 | 0 | 0 | 1 |
| 0 | 0 | 0 | 0 |
| 0 | 0 | 0 | 1 |
| 0 | 0 | 0 | 1 |
| 0 | 0 | 0 | 1 |
| 0 | 0 | 0 | 1 |
| 0 | 0 | 0 | 1 |
| 0 | 0 | 0 | 0 |
| 0 | 0 | 0 | 0 |
| 0 | 0 | 0 | 0 |
| 0 | 0 | 0 | 0 |
| 0 | 0 | 0 | 1 |
| 1 | 0 | 0 | 0 |
| 0 | 0 | 0 | 1 |
| 0 | 0 | 0 | 1 |
| 0 | 0 | 0 | 1 |
| 0 | 0 | 0 | 1 |
| 0 | 0 | 0 | 1 |
| 0 | 0 | 0 | 0 |
| 0 | 0 | 0 | 0 |
| 0 | 0 | 0 | 1 |
| 0 | 0 | 0 | 1 |
| 1 | 0 | 0 | 0 |
| 0 | 0 | 0 | 1 |
| 0 | 0 | 0 | 0 |
| 1 | 0 | 0 | 0 |
| 1 | 0 | 0 | 0 |
| 0 | 0 | 0 | 1 |
| 1 | 0 | 0 | 0 |
| 0 | 0 | 0 | 1 |
| 1 | 0 | 0 | 0 |
| 0 | 0 | 0 | 1 |
| 0 | 0 | 0 | 0 |
| 0 | 0 | 0 | 1 |
| 0 | 0 | 0 | 1 |
| 0 | 0 | 0 | 1 |
| 0 | 1 | 1 | 1 |
| 0 | 0 | 0 | 1 |
| 0 | 0 | 0 | 1 |
| 0 | 0 | 0 | 0 |
| 0 | 0 | 0 | 1 |
| 1 | 0 | 0 | 0 |
| 0 | 0 | 0 | 1 |
| 0 | 0 | 0 | 1 |

| | | | |
|---|---|---|---|
| 0 | 0 | 0 | 0 |
| 0 | 0 | 0 | 0 |
| 0 | 0 | 0 | 0 |
| 0 | 0 | 0 | 0 |
| 0 | 0 | 0 | 0 |
| 0 | 0 | 0 | 1 |
| 0 | 0 | 0 | 0 |
| 0 | 0 | 0 | 1 |
| 0 | 0 | 0 | 1 |
| 0 | 0 | 0 | 0 |
| 0 | 0 | 0 | 0 |
| 0 | 0 | 0 | 0 |
| 0 | 0 | 0 | 1 |
| 0 | 0 | 0 | 1 |
| 0 | 0 | 0 | 1 |
| 0 | 0 | 0 | 1 |
| 0 | 0 | 0 | 1 |
| 0 | 0 | 0 | 1 |
| 0 | 0 | 0 | 1 |
| 0 | 0 | 0 | 1 |
| 0 | 0 | 0 | 1 |
| 0 | 0 | 0 | 1 |
| 0 | 0 | 0 | 1 |
| 0 | 0 | 0 | 1 |
| 0 | 0 | 0 | 1 |
| 0 | 0 | 0 | 1 |
| 0 | 0 | 0 | 1 |
| 0 | 0 | 0 | 1 |
| 0 | 0 | 0 | 1 |
| 0 | 0 | 0 | 1 |
| 0 | 0 | 0 | 1 |
| 0 | 0 | 0 | 1 |
| 0 | 0 | 0 | 1 |
| 0 | 0 | 0 | 1 |
| 0 | 0 | 0 | 1 |
| 0 | 0 | 0 | 1 |
| 0 | 0 | 0 | 1 |
| 0 | 0 | 0 | 1 |
| 0 | 0 | 0 | 1 |
| 0 | 0 | 0 | 1 |
| 0 | 0 | 0 | 1 |
| 0 | 0 | 0 | 1 |
| 0 | 0 | 0 | 1 |
| 0 | 0 | 0 | 1 |
| 0 | 0 | 0 | 1 |
| 0 | 0 | 0 | 0 |
| 0 | 0 | 0 | 1 |
| 0 | 0 | 0 | 1 |
| 0 | 0 | 0 | 1 |
| 0 | 0 | 0 | 1 |
| 0 | 0 | 0 | 0 |
| 0 | 0 | 0 | 1 |
| 0 | 0 | 0 | 1 |
| 0 | 0 | 0 | 1 |

| | | | |
|---|---|---|---|
| 0 | 0 | 0 | 1 |
| 0 | 0 | 0 | 1 |
| 0 | 0 | 0 | 1 |
| 0 | 0 | 0 | 1 |
| 0 | 0 | 0 | 1 |
| 0 | 0 | 0 | 1 |
| 0 | 0 | 0 | 1 |
| 0 | 0 | 0 | 0 |
| 0 | 0 | 0 | 0 |
| 0 | 0 | 0 | 1 |
| 0 | 0 | 0 | 1 |
| 0 | 0 | 0 | 1 |
| 0 | 0 | 0 | 1 |
| 0 | 0 | 0 | 1 |
| 0 | 0 | 0 | 0 |
| 0 | 0 | 0 | 0 |
| 0 | 0 | 0 | 0 |
| 0 | 0 | 0 | 1 |
| 0 | 0 | 0 | 1 |
| 0 | 0 | 0 | 1 |
| 0 | 0 | 0 | 0 |
| 0 | 0 | 0 | 0 |
| 0 | 0 | 0 | 0 |
| 0 | 0 | 0 | 0 |
| 0 | 0 | 0 | 0 |
| 0 | 0 | 0 | 0 |
| 0 | 0 | 0 | 1 |
| 0 | 0 | 0 | 1 |
| 0 | 0 | 0 | 0 |
| 0 | 0 | 0 | 1 |
| 0 | 0 | 0 | 1 |
| 0 | 0 | 0 | 1 |
| 0 | 0 | 0 | 0 |
| 0 | 0 | 0 | 1 |
| 0 | 0 | 0 | 1 |
| 0 | 0 | 0 | 1 |
| 0 | 0 | 0 | 0 |
| 0 | 0 | 0 | 0 |
| 0 | 0 | 0 | 0 |
| 0 | 0 | 0 | 1 |
| 0 | 0 | 0 | 1 |
| 0 | 0 | 0 | 1 |
| 0 | 0 | 0 | 1 |
| 0 | 0 | 0 | 1 |
| 0 | 0 | 0 | 1 |
| 0 | 0 | 0 | 0 |
| 0 | 0 | 0 | 1 |
| 0 | 0 | 0 | 1 |
| 0 | 0 | 0 | 1 |
| 0 | 0 | 0 | 1 |
| 0 | 0 | 0 | 1 |
| 0 | 0 | 0 | 1 |
| 0 | 0 | 0 | 1 |
| 0 | 0 | 0 | 1 |

| | | | |
|---|---|---|---|
| 0 | 0 | 0 | 1 |
| 0 | 0 | 0 | 1 |
| 0 | 0 | 0 | 1 |
| 0 | 0 | 0 | 0 |
| 0 | 0 | 0 | 1 |
| 0 | 0 | 0 | 1 |
| 0 | 0 | 0 | 1 |
| 0 | 0 | 0 | 0 |
| 0 | 0 | 0 | 0 |
| 0 | 0 | 0 | 1 |
| 0 | 0 | 0 | 1 |
| 0 | 0 | 0 | 1 |
| 0 | 0 | 0 | 0 |
| 0 | 0 | 0 | 1 |
| 0 | 0 | 0 | 1 |
| 0 | 0 | 0 | 1 |
| 0 | 0 | 0 | 0 |
| 0 | 0 | 0 | 1 |
| 0 | 0 | 0 | 1 |
| 0 | 0 | 0 | 1 |
| 0 | 0 | 0 | 1 |
| 0 | 0 | 0 | 1 |
| 0 | 0 | 0 | 1 |
| 0 | 0 | 0 | 1 |
| 0 | 0 | 0 | 1 |
| 0 | 0 | 0 | 1 |
| 0 | 0 | 0 | 1 |
| 0 | 0 | 0 | 0 |
| 0 | 0 | 0 | 0 |
| 0 | 0 | 0 | 0 |
| 0 | 0 | 0 | 0 |
| 0 | 0 | 0 | 0 |
| 0 | 0 | 0 | 0 |
| 0 | 0 | 0 | 1 |
| 0 | 0 | 0 | 0 |
| 0 | 0 | 0 | 0 |
| 0 | 0 | 0 | 0 |
| 0 | 0 | 0 | 1 |
| 0 | 0 | 0 | 1 |
| 0 | 0 | 0 | 1 |
| 0 | 0 | 0 | 1 |
| 0 | 0 | 0 | 1 |
| 0 | 0 | 0 | 0 |
| 0 | 0 | 0 | 0 |
| 0 | 0 | 0 | 1 |
| 0 | 0 | 0 | 1 |
| 0 | 0 | 0 | 0 |
| 0 | 0 | 0 | 1 |
| 0 | 0 | 0 | 1 |
| 0 | 0 | 0 | 1 |
| 0 | 0 | 0 | 1 |
| 0 | 0 | 0 | 1 |
| 0 | 0 | 0 | 0 |
| 0 | 0 | 0 | 0 |

| | | | |
|---|---|---|---|
| 0 | 0 | 0 | 0 |
| 0 | 0 | 0 | 0 |
| 0 | 0 | 0 | 0 |
| 0 | 0 | 0 | 0 |
| 0 | 0 | 0 | 0 |
| 0 | 0 | 0 | 0 |
| 0 | 0 | 0 | 0 |
| 0 | 0 | 0 | 0 |
| 0 | 0 | 0 | 1 |
| 0 | 0 | 0 | 0 |
| 0 | 0 | 0 | 1 |
| 0 | 0 | 0 | 1 |
| 0 | 0 | 0 | 1 |
| 0 | 0 | 0 | 1 |
| 0 | 0 | 0 | 0 |
| 0 | 0 | 0 | 1 |
| 0 | 0 | 0 | 1 |
| 0 | 0 | 0 | 1 |
| 0 | 0 | 0 | 1 |
| 0 | 0 | 0 | 0 |
| 0 | 0 | 0 | 0 |
| 0 | 0 | 0 | 1 |
| 0 | 0 | 0 | 0 |
| 0 | 0 | 0 | 0 |
| 0 | 0 | 0 | 0 |
| 0 | 0 | 0 | 0 |
| 0 | 0 | 0 | 0 |
| 0 | 0 | 0 | 1 |
| 0 | 0 | 0 | 1 |
| 0 | 0 | 0 | 1 |
| 0 | 0 | 0 | 0 |
| 0 | 0 | 0 | 1 |
| 0 | 0 | 0 | 1 |
| 0 | 0 | 0 | 1 |
| 0 | 0 | 0 | 1 |
| 0 | 0 | 0 | 1 |
| 0 | 0 | 0 | 1 |
| 0 | 0 | 0 | 1 |
| 0 | 0 | 0 | 1 |
| 0 | 0 | 0 | 1 |
| 0 | 0 | 0 | 1 |
| 0 | 0 | 0 | 0 |
| 0 | 0 | 0 | 1 |
| 0 | 0 | 0 | 1 |
| 0 | 0 | 0 | 1 |
| 0 | 0 | 0 | 1 |
| 0 | 0 | 0 | 1 |
| 0 | 0 | 0 | 1 |
| 0 | 0 | 0 | 0 |
| 0 | 0 | 0 | 1 |
| 0 | 0 | 0 | 1 |
| 0 | 0 | 0 | 1 |
| 0 | 0 | 0 | 0 |
| 0 | 1 | 0 | 0 |

| | | | |
|---|---|---|---|
| 0 | 0 | 0 | 1 |
| 0 | 0 | 0 | 1 |
| 0 | 0 | 0 | 1 |
| 0 | 0 | 0 | 1 |
| 0 | 0 | 0 | 0 |
| 1 | 0 | 0 | 0 |
| 0 | 0 | 0 | 1 |
| 0 | 0 | 0 | 0 |
| 0 | 0 | 0 | 1 |
| 0 | 0 | 0 | 1 |
| 0 | 0 | 0 | 0 |
| 0 | 0 | 0 | 1 |
| 0 | 0 | 0 | 0 |
| 0 | 0 | 0 | 0 |
| 0 | 0 | 0 | 0 |
| 0 | 0 | 0 | 0 |
| 0 | 0 | 0 | 1 |
| 0 | 0 | 0 | 1 |
| 0 | 0 | 0 | 0 |
| 0 | 0 | 0 | 0 |
| 0 | 0 | 0 | 0 |
| 0 | 0 | 0 | 0 |
| 0 | 0 | 0 | 0 |
| 0 | 0 | 0 | 1 |
| 0 | 0 | 0 | 1 |
| 0 | 0 | 0 | 0 |
| 0 | 0 | 0 | 1 |
| 0 | 0 | 0 | 0 |
| 0 | 0 | 0 | 0 |
| 0 | 0 | 0 | 0 |
| 0 | 0 | 0 | 1 |
| 1 | 0 | 0 | 0 |
| 1 | 0 | 0 | 0 |
| 0 | 0 | 0 | 1 |
| 0 | 0 | 0 | 1 |
| 0 | 0 | 0 | 0 |
| 0 | 0 | 0 | 0 |
| 0 | 0 | 0 | 1 |
| 0 | 0 | 0 | 0 |
| 0 | 0 | 0 | 0 |
| 0 | 1 | 0 | 0 |
| 0 | 0 | 0 | 0 |
| 0 | 0 | 0 | 1 |
| 0 | 0 | 0 | 1 |
| 0 | 0 | 0 | 0 |
| 0 | 0 | 0 | 1 |
| 0 | 0 | 0 | 0 |
| 0 | 0 | 0 | 1 |
| 0 | 0 | 0 | 1 |
| 0 | 0 | 0 | 1 |
| 0 | 0 | 0 | 1 |
| 0 | 0 | 0 | 1 |
| 0 | 0 | 0 | 1 |
| 0 | 0 | 0 | 1 |

| | | | |
|---|---|---|---|
| 0 | 0 | 0 | 1 |
| 0 | 0 | 0 | 1 |
| 0 | 0 | 0 | 1 |
| 0 | 0 | 0 | 1 |
| 0 | 0 | 0 | 1 |
| 0 | 0 | 0 | 1 |
| 0 | 0 | 0 | 1 |
| 0 | 0 | 0 | 0 |
| 0 | 0 | 0 | 1 |
| 1 | 0 | 0 | 0 |
| 0 | 0 | 0 | 0 |
| 0 | 0 | 0 | 0 |
| 0 | 0 | 0 | 1 |
| 0 | 0 | 0 | 1 |
| 0 | 0 | 0 | 0 |
| 0 | 0 | 0 | 1 |
| 0 | 0 | 0 | 1 |
| 0 | 0 | 0 | 1 |
| 0 | 0 | 0 | 1 |
| 0 | 0 | 0 | 1 |
| 0 | 0 | 0 | 1 |
| 0 | 0 | 0 | 1 |
| 0 | 0 | 0 | 1 |
| 0 | 0 | 0 | 1 |
| 0 | 0 | 0 | 1 |
| 0 | 0 | 0 | 1 |
| 0 | 0 | 0 | 1 |
| 0 | 0 | 0 | 1 |
| 0 | 0 | 0 | 1 |
| 0 | 0 | 0 | 0 |
| 0 | 0 | 0 | 0 |
| 0 | 0 | 0 | 1 |
| 0 | 0 | 0 | 1 |
| 0 | 0 | 0 | 1 |
| 0 | 0 | 0 | 1 |
| 0 | 0 | 0 | 0 |
| 0 | 0 | 0 | 1 |
| 0 | 0 | 0 | 1 |
| 0 | 0 | 0 | 1 |
| 0 | 0 | 0 | 1 |
| 0 | 0 | 0 | 1 |
| 0 | 0 | 0 | 0 |
| 0 | 0 | 0 | 0 |
| 0 | 0 | 0 | 0 |
| 0 | 0 | 0 | 1 |
| 0 | 0 | 0 | 1 |
| 0 | 0 | 0 | 1 |
| 0 | 0 | 0 | 1 |
| 0 | 0 | 0 | 1 |
| 0 | 0 | 0 | 1 |
| 0 | 0 | 0 | 1 |
| 0 | 0 | 0 | 1 |
| 0 | 0 | 0 | 1 |
| 0 | 0 | 0 | 0 |

| | | | |
|---|---|---|---|
| 0 | 0 | 0 | 1 |
| 0 | 0 | 0 | 1 |
| 0 | 0 | 0 | 1 |
| 0 | 0 | 0 | 1 |
| 0 | 0 | 0 | 1 |
| 0 | 0 | 0 | 1 |
| 0 | 0 | 0 | 1 |
| 0 | 0 | 0 | 1 |
| 0 | 0 | 0 | 0 |
| 0 | 0 | 0 | 0 |
| 0 | 0 | 0 | 1 |
| 0 | 0 | 0 | 1 |
| 0 | 0 | 0 | 1 |
| 0 | 0 | 0 | 0 |
| 0 | 0 | 0 | 1 |
| 0 | 0 | 0 | 1 |
| 0 | 0 | 0 | 1 |
| 1 | 0 | 0 | 0 |
| 1 | 0 | 0 | 0 |
| 0 | 0 | 0 | 0 |
| 0 | 0 | 0 | 0 |
| 0 | 0 | 0 | 1 |
| 0 | 0 | 0 | 1 |
| 0 | 0 | 0 | 1 |
| 0 | 0 | 0 | 1 |
| 1 | 0 | 0 | 0 |
| 0 | 0 | 0 | 1 |
| 0 | 0 | 0 | 1 |
| 0 | 0 | 0 | 1 |
| 0 | 0 | 0 | 1 |
| 0 | 0 | 0 | 1 |
| 0 | 0 | 0 | 1 |
| 0 | 0 | 0 | 1 |
| 0 | 0 | 0 | 1 |
| 0 | 0 | 0 | 1 |
| 0 | 0 | 0 | 1 |
| 0 | 0 | 0 | 1 |
| 0 | 0 | 0 | 1 |
| 0 | 0 | 0 | 1 |
| 0 | 0 | 0 | 0 |
| 0 | 0 | 0 | 1 |
| 0 | 0 | 0 | 1 |
| 0 | 0 | 0 | 0 |
| 0 | 0 | 0 | 1 |
| 0 | 0 | 0 | 1 |
| 0 | 0 | 0 | 1 |
| 0 | 0 | 0 | 0 |
| 0 | 0 | 0 | 0 |
| 0 | 0 | 0 | 1 |
| 0 | 0 | 0 | 1 |
| 0 | 0 | 0 | 0 |
| 0 | 0 | 0 | 1 |
| 0 | 0 | 0 | 1 |
| 0 | 0 | 0 | 1 |

| | | | |
|---|---|---|---|
| 0 | 0 | 0 | 1 |
| 0 | 0 | 0 | 1 |
| 0 | 0 | 0 | 1 |
| 0 | 0 | 0 | 1 |
| 0 | 0 | 0 | 1 |
| 0 | 0 | 0 | 1 |
| 0 | 0 | 0 | 1 |
| 0 | 0 | 0 | 1 |
| 0 | 0 | 0 | 1 |
| 0 | 0 | 0 | 1 |
| 0 | 0 | 0 | 1 |
| 0 | 0 | 0 | 1 |
| 0 | 0 | 0 | 1 |
| 0 | 0 | 0 | 0 |
| 0 | 0 | 0 | 0 |
| 0 | 0 | 0 | 0 |
| 0 | 0 | 0 | 0 |
| 0 | 0 | 0 | 1 |
| 0 | 0 | 0 | 1 |
| 0 | 0 | 0 | 0 |
| 0 | 0 | 0 | 1 |
| 0 | 0 | 0 | 1 |
| 0 | 0 | 0 | 1 |
| 0 | 0 | 0 | 1 |
| 0 | 0 | 0 | 1 |
| 0 | 0 | 0 | 0 |
| 0 | 0 | 0 | 1 |
| 0 | 0 | 0 | 1 |
| 0 | 0 | 0 | 1 |
| 0 | 0 | 0 | 0 |
| 0 | 0 | 0 | 0 |
| 0 | 0 | 0 | 1 |
| 0 | 0 | 0 | 1 |
| 0 | 0 | 0 | 1 |
| 0 | 0 | 0 | 1 |
| 0 | 0 | 0 | 1 |
| 0 | 0 | 0 | 1 |
| 0 | 0 | 0 | 1 |
| 0 | 0 | 0 | 0 |
| 0 | 0 | 0 | 1 |
| 0 | 0 | 0 | 1 |
| 0 | 0 | 0 | 0 |
| 0 | 0 | 0 | 0 |
| 0 | 0 | 0 | 0 |
| 0 | 0 | 0 | 0 |
| 0 | 0 | 0 | 1 |
| 0 | 0 | 0 | 0 |
| 0 | 0 | 0 | 1 |
| 0 | 0 | 0 | 1 |
| 0 | 0 | 0 | 1 |
| 0 | 0 | 0 | 1 |
| 0 | 0 | 0 | 1 |
| 0 | 0 | 0 | 1 |
| 0 | 0 | 0 | 1 |

| | | | |
|---|---|---|---|
| 0 | 0 | 0 | 1 |
| 0 | 0 | 0 | 1 |
| 0 | 0 | 0 | 0 |
| 0 | 0 | 0 | 1 |
| 0 | 0 | 0 | 1 |
| 0 | 0 | 0 | 1 |
| 0 | 0 | 0 | 1 |
| 0 | 0 | 0 | 1 |
| 0 | 0 | 0 | 1 |
| 0 | 0 | 0 | 1 |
| 0 | 0 | 0 | 1 |
| 0 | 0 | 0 | 1 |
| 0 | 0 | 0 | 1 |
| 0 | 0 | 0 | 1 |
| 0 | 0 | 0 | 1 |
| 0 | 0 | 0 | 1 |
| 0 | 0 | 0 | 1 |
| 0 | 0 | 0 | 1 |
| 0 | 0 | 0 | 1 |
| 0 | 0 | 0 | 1 |
| 0 | 0 | 0 | 1 |
| 0 | 0 | 0 | 1 |
| 0 | 0 | 0 | 1 |
| 0 | 0 | 0 | 1 |
| 0 | 0 | 0 | 1 |
| 0 | 0 | 0 | 1 |
| 0 | 0 | 0 | 1 |
| 0 | 0 | 0 | 1 |
| 0 | 0 | 0 | 1 |
| 0 | 0 | 0 | 1 |
| 1 | 0 | 0 | 0 |
| 0 | 0 | 0 | 1 |
| 0 | 1 | 0 | 0 |
| 0 | 1 | 0 | 0 |
| 0 | 0 | 0 | 0 |
| 0 | 0 | 0 | 0 |
| 0 | 0 | 0 | 1 |
| 0 | 0 | 0 | 0 |
| 0 | 0 | 0 | 1 |
| 0 | 0 | 0 | 1 |
| 1 | 0 | 0 | 0 |
| 1 | 0 | 0 | 0 |
| 0 | 0 | 0 | 0 |
| 0 | 0 | 0 | 1 |
| 0 | 0 | 0 | 0 |
| 0 | 0 | 0 | 0 |
| 0 | 0 | 0 | 1 |
| 0 | 0 | 0 | 1 |
| 0 | 0 | 0 | 1 |
| 0 | 0 | 0 | 1 |
| 0 | 0 | 0 | 0 |
| 0 | 0 | 0 | 1 |
| 0 | 0 | 0 | 1 |
| 0 | 0 | 0 | 1 |

| | | | |
|---|---|---|---|
| 0 | 0 | 0 | 1 |
| 0 | 0 | 0 | 1 |
| 0 | 0 | 0 | 1 |
| 0 | 0 | 0 | 1 |
| 0 | 0 | 0 | 1 |
| 1 | 0 | 0 | 0 |
| 0 | 0 | 0 | 1 |
| 0 | 0 | 0 | 0 |
| 0 | 0 | 0 | 0 |
| 0 | 0 | 0 | 0 |
| 0 | 0 | 0 | 1 |
| 0 | 1 | 0 | 0 |
| 0 | 0 | 0 | 1 |
| 0 | 0 | 0 | 1 |
| 0 | 0 | 0 | 1 |
| 0 | 1 | 1 | 1 |
| 0 | 0 | 0 | 1 |
| 0 | 0 | 0 | 0 |
| 0 | 0 | 0 | 1 |
| 0 | 0 | 0 | 0 |
| 0 | 0 | 0 | 1 |
| 0 | 0 | 0 | 0 |
| 0 | 0 | 0 | 0 |
| 0 | 0 | 0 | 0 |
| 0 | 0 | 0 | 1 |
| 0 | 0 | 0 | 1 |
| 0 | 0 | 0 | 1 |
| 0 | 0 | 0 | 1 |
| 0 | 0 | 0 | 0 |
| 0 | 0 | 0 | 1 |
| 0 | 0 | 0 | 1 |
| 0 | 0 | 0 | 1 |
| 0 | 0 | 0 | 1 |
| 0 | 0 | 0 | 1 |
| 0 | 0 | 0 | 0 |
| 0 | 0 | 0 | 0 |
| 0 | 0 | 0 | 1 |
| 0 | 0 | 0 | 0 |
| 0 | 0 | 0 | 0 |
| 0 | 0 | 0 | 1 |
| 0 | 0 | 0 | 0 |
| 0 | 0 | 0 | 0 |
| 0 | 0 | 0 | 1 |
| 0 | 0 | 0 | 1 |
| 0 | 0 | 0 | 1 |
| 0 | 0 | 0 | 1 |
| 0 | 0 | 0 | 1 |
| 0 | 0 | 0 | 1 |
| 0 | 0 | 0 | 1 |
| 0 | 0 | 0 | 1 |
| 0 | 0 | 0 | 1 |
| 0 | 0 | 0 | 1 |
| 0 | 0 | 0 | 1 |
| 0 | 0 | 0 | 1 |

| | | | |
|---|---|---|---|
| 0 | 0 | 0 | 1 |
| 0 | 0 | 0 | 0 |
| 0 | 0 | 0 | 1 |
| 0 | 0 | 0 | 1 |
| 0 | 0 | 0 | 0 |
| 0 | 0 | 0 | 1 |
| 0 | 0 | 0 | 1 |
| 0 | 0 | 0 | 1 |
| 0 | 0 | 0 | 0 |
| 0 | 0 | 0 | 0 |
| 0 | 0 | 0 | 1 |
| 0 | 0 | 0 | 0 |
| 0 | 0 | 0 | 0 |
| 0 | 0 | 0 | 1 |
| 0 | 0 | 0 | 1 |
| 0 | 0 | 0 | 0 |
| 0 | 0 | 0 | 1 |
| 0 | 0 | 0 | 0 |
| 0 | 0 | 0 | 0 |
| 0 | 0 | 0 | 1 |
| 0 | 0 | 0 | 0 |
| 0 | 0 | 0 | 0 |
| 1 | 0 | 0 | 0 |
| 0 | 0 | 0 | 1 |
| 0 | 0 | 0 | 1 |
| 0 | 0 | 0 | 0 |
| 0 | 0 | 0 | 1 |
| 0 | 0 | 0 | 1 |
| 0 | 0 | 0 | 1 |
| 0 | 0 | 0 | 1 |
| 0 | 0 | 0 | 0 |
| 0 | 0 | 0 | 1 |
| 0 | 0 | 0 | 0 |
| 0 | 0 | 0 | 1 |
| 0 | 0 | 0 | 0 |
| 0 | 0 | 0 | 0 |
| 0 | 0 | 0 | 1 |
| 0 | 0 | 0 | 1 |
| 0 | 0 | 0 | 1 |
| 0 | 0 | 0 | 1 |
| 0 | 0 | 0 | 1 |
| 0 | 0 | 0 | 1 |
| 0 | 0 | 0 | 1 |
| 0 | 0 | 0 | 1 |
| 0 | 0 | 0 | 1 |
| 0 | 0 | 0 | 1 |
| 0 | 1 | 1 | 1 |
| 0 | 0 | 0 | 1 |
| 0 | 0 | 0 | 1 |
| 0 | 0 | 0 | 1 |
| 0 | 0 | 0 | 1 |
| 0 | 0 | 0 | 1 |
| 0 | 0 | 0 | 1 |
| 0 | 0 | 0 | 0 |

| | | | |
|---|---|---|---|
| 0 | 0 | 0 | 1 |
| 0 | 0 | 0 | 1 |
| 0 | 0 | 0 | 1 |
| 0 | 0 | 0 | 1 |
| 0 | 0 | 0 | 0 |
| 0 | 0 | 0 | 1 |
| 0 | 0 | 0 | 1 |
| 0 | 0 | 0 | 1 |
| 0 | 0 | 0 | 1 |
| 0 | 0 | 0 | 1 |
| 0 | 0 | 0 | 0 |
| 0 | 0 | 0 | 1 |
| 0 | 0 | 0 | 1 |
| 0 | 0 | 0 | 1 |
| 0 | 0 | 0 | 0 |
| 0 | 0 | 0 | 0 |
| 0 | 0 | 0 | 0 |
| 0 | 0 | 0 | 0 |
| 0 | 0 | 0 | 1 |
| 0 | 0 | 0 | 0 |
| 0 | 0 | 0 | 0 |
| 1 | 0 | 0 | 0 |
| 0 | 0 | 0 | 1 |
| 0 | 0 | 0 | 0 |
| 1 | 0 | 0 | 0 |
| 0 | 0 | 0 | 1 |
| 1 | 0 | 0 | 0 |
| 0 | 0 | 0 | 0 |
| 1 | 0 | 0 | 0 |
| 1 | 0 | 0 | 0 |
| 1 | 0 | 0 | 0 |
| 0 | 0 | 0 | 1 |
| 0 | 0 | 0 | 1 |
| 0 | 0 | 0 | 1 |
| 0 | 0 | 0 | 1 |
| 0 | 0 | 0 | 1 |
| 0 | 0 | 0 | 1 |
| 0 | 1 | 0 | 0 |
| 0 | 0 | 0 | 0 |
| 0 | 0 | 0 | 1 |
| 0 | 0 | 0 | 1 |
| 0 | 1 | 1 | 1 |
| 0 | 0 | 0 | 0 |
| 0 | 0 | 0 | 1 |
| 0 | 0 | 0 | 0 |
| 0 | 0 | 0 | 1 |
| 0 | 0 | 0 | 1 |
| 0 | 0 | 0 | 1 |
| 0 | 1 | 0 | 0 |
| 0 | 1 | 0 | 0 |
| 0 | 1 | 0 | 0 |
| 0 | 0 | 0 | 1 |
| 0 | 0 | 0 | 1 |
| 0 | 0 | 0 | 0 |

| | | | |
|---|---|---|---|
| 0 | 0 | 0 | 1 |
| 0 | 1 | 0 | 0 |
| 0 | 0 | 0 | 1 |
| 0 | 1 | 0 | 0 |
| 0 | 1 | 0 | 0 |
| 0 | 0 | 0 | 1 |
| 0 | 1 | 0 | 0 |
| 0 | 1 | 0 | 0 |
| 0 | 0 | 0 | 1 |
| 0 | 0 | 0 | 1 |
| 0 | 0 | 0 | 1 |
| 0 | 0 | 0 | 1 |
| 0 | 0 | 0 | 1 |
| 0 | 0 | 0 | 1 |
| 0 | 0 | 0 | 1 |
| 0 | 0 | 0 | 1 |
| 0 | 0 | 0 | 1 |
| 0 | 0 | 0 | 1 |
| 0 | 0 | 0 | 1 |
| 0 | 0 | 0 | 1 |
| 1 | 1 | 0 | 0 |
| 0 | 0 | 0 | 1 |
| 0 | 0 | 0 | 1 |
| 0 | 0 | 0 | 1 |
| 0 | 0 | 0 | 1 |
| 0 | 0 | 0 | 0 |
| 0 | 0 | 0 | 0 |
| 0 | 0 | 0 | 1 |
| 0 | 0 | 0 | 1 |
| 0 | 0 | 0 | 1 |
| 0 | 0 | 0 | 1 |
| 0 | 0 | 0 | 1 |
| 0 | 0 | 0 | 1 |
| 0 | 0 | 0 | 1 |
| 0 | 0 | 0 | 1 |
| 0 | 0 | 0 | 1 |
| 0 | 0 | 0 | 1 |
| 0 | 0 | 0 | 1 |
| 0 | 0 | 0 | 1 |
| 0 | 0 | 0 | 1 |
| 0 | 0 | 0 | 1 |
| 0 | 0 | 0 | 1 |
| 0 | 0 | 0 | 1 |
| 0 | 0 | 0 | 0 |
| 0 | 0 | 0 | 1 |
| 0 | 0 | 0 | 0 |
| 0 | 0 | 0 | 0 |
| 0 | 0 | 0 | 0 |
| 0 | 0 | 0 | 0 |
| 0 | 0 | 0 | 0 |
| 0 | 0 | 0 | 0 |
| 0 | 0 | 0 | 1 |
| 0 | 0 | 0 | 1 |
| 0 | 0 | 0 | 1 |

| | | | |
|---|---|---|---|
| 0 | 0 | 0 | 1 |
| 0 | 0 | 0 | 1 |
| 0 | 0 | 0 | 1 |
| 0 | 0 | 0 | 0 |
| 0 | 0 | 0 | 1 |
| 0 | 0 | 0 | 0 |
| 0 | 0 | 0 | 1 |
| 0 | 0 | 0 | 1 |
| 0 | 0 | 0 | 0 |
| 0 | 0 | 0 | 1 |
| 0 | 0 | 0 | 1 |
| 0 | 0 | 0 | 1 |
| 0 | 0 | 0 | 1 |
| 0 | 0 | 0 | 1 |
| 0 | 0 | 0 | 1 |
| 0 | 0 | 0 | 0 |
| 0 | 0 | 0 | 0 |
| 0 | 0 | 0 | 0 |
| 0 | 0 | 0 | 0 |
| 0 | 0 | 0 | 0 |
| 0 | 0 | 0 | 1 |
| 0 | 0 | 0 | 1 |
| 0 | 0 | 0 | 1 |
| 0 | 0 | 0 | 1 |
| 0 | 0 | 0 | 0 |
| 0 | 0 | 0 | 1 |
| 0 | 0 | 0 | 1 |
| 0 | 0 | 0 | 1 |
| 0 | 0 | 0 | 0 |
| 0 | 0 | 0 | 1 |
| 0 | 1 | 0 | 0 |
| 0 | 1 | 0 | 0 |
| 0 | 0 | 0 | 0 |
| 0 | 1 | 0 | 0 |
| 0 | 1 | 0 | 0 |
| 0 | 0 | 0 | 1 |
| 0 | 0 | 0 | 0 |
| 0 | 1 | 0 | 0 |
| 0 | 1 | 0 | 0 |
| 0 | 1 | 0 | 0 |
| 0 | 1 | 0 | 0 |
| 0 | 0 | 0 | 0 |
| 0 | 0 | 0 | 0 |
| 0 | 0 | 0 | 0 |
| 0 | 0 | 0 | 0 |
| 0 | 0 | 0 | 0 |
| 0 | 0 | 0 | 0 |
| 0 | 0 | 0 | 0 |
| 0 | 0 | 0 | 1 |
| 0 | 0 | 0 | 0 |
| 0 | 0 | 0 | 1 |
| 0 | 0 | 0 | 1 |
| 0 | 0 | 0 | 1 |
| 0 | 0 | 0 | 0 |

| | | | |
|---|---|---|---|
| 0 | 0 | 0 | 0 |
| 0 | 0 | 0 | 1 |
| 0 | 0 | 0 | 1 |
| 0 | 0 | 0 | 1 |
| 0 | 0 | 0 | 1 |
| 0 | 0 | 0 | 0 |
| 0 | 0 | 0 | 1 |
| 0 | 0 | 0 | 1 |
| 0 | 0 | 0 | 1 |
| 0 | 0 | 0 | 1 |
| 0 | 0 | 0 | 1 |
| 0 | 0 | 0 | 1 |
| 0 | 0 | 0 | 1 |
| 1 | 0 | 0 | 0 |
| 0 | 0 | 0 | 1 |
| 0 | 0 | 0 | 1 |
| 0 | 0 | 0 | 1 |
| 0 | 0 | 0 | 0 |
| 0 | 0 | 0 | 1 |
| 0 | 0 | 0 | 1 |
| 0 | 0 | 0 | 1 |
| 0 | 0 | 0 | 1 |
| 0 | 0 | 0 | 0 |
| 0 | 0 | 0 | 1 |
| 0 | 0 | 0 | 1 |
| 0 | 0 | 0 | 0 |
| 0 | 0 | 0 | 0 |
| 0 | 0 | 0 | 0 |
| 0 | 0 | 0 | 1 |
| 0 | 0 | 0 | 1 |
| 0 | 0 | 0 | 1 |
| 0 | 0 | 0 | 1 |
| 0 | 0 | 0 | 1 |
| 0 | 0 | 0 | 1 |
| 0 | 0 | 0 | 1 |
| 0 | 0 | 0 | 1 |
| 0 | 0 | 0 | 1 |
| 0 | 0 | 0 | 1 |
| 0 | 0 | 0 | 1 |
| 0 | 0 | 0 | 1 |
| 0 | 0 | 0 | 1 |
| 0 | 0 | 0 | 1 |
| 0 | 0 | 0 | 1 |
| 0 | 0 | 0 | 1 |
| 0 | 0 | 0 | 1 |
| 0 | 0 | 0 | 1 |
| 0 | 0 | 0 | 1 |
| 0 | 0 | 0 | 1 |
| 0 | 0 | 0 | 1 |
| 0 | 0 | 0 | 1 |
| 0 | 0 | 0 | 1 |
| 0 | 0 | 0 | 1 |
| 0 | 0 | 0 | 1 |

| | | | |
|---|---|---|---|
| 0 | 0 | 0 | 1 |
| 0 | 0 | 0 | 0 |
| 0 | 0 | 0 | 1 |
| 0 | 0 | 0 | 1 |
| 0 | 0 | 0 | 1 |
| 0 | 0 | 0 | 0 |
| 0 | 0 | 0 | 0 |
| 0 | 0 | 0 | 0 |
| 0 | 0 | 0 | 1 |
| 0 | 0 | 0 | 1 |
| 0 | 0 | 0 | 1 |
| 0 | 0 | 0 | 1 |
| 0 | 0 | 0 | 1 |
| 0 | 0 | 0 | 1 |
| 0 | 0 | 0 | 0 |
| 0 | 0 | 0 | 0 |
| 0 | 0 | 0 | 0 |
| 0 | 0 | 0 | 1 |
| 0 | 0 | 0 | 0 |
| 0 | 0 | 0 | 0 |
| 0 | 0 | 0 | 0 |
| 0 | 0 | 0 | 0 |
| 0 | 0 | 0 | 0 |
| 0 | 0 | 0 | 0 |
| 0 | 0 | 0 | 1 |
| 0 | 0 | 0 | 1 |
| 0 | 0 | 0 | 1 |
| 0 | 0 | 0 | 1 |
| 0 | 0 | 0 | 1 |
| 0 | 0 | 0 | 1 |
| 0 | 0 | 0 | 1 |
| 0 | 0 | 0 | 1 |
| 0 | 0 | 0 | 0 |
| 0 | 0 | 0 | 1 |
| 0 | 0 | 0 | 1 |
| 0 | 0 | 0 | 0 |
| 0 | 0 | 0 | 0 |
| 0 | 0 | 0 | 0 |
| 0 | 0 | 0 | 0 |
| 0 | 0 | 0 | 1 |
| 0 | 0 | 0 | 1 |
| 0 | 0 | 0 | 1 |
| 0 | 0 | 0 | 1 |
| 0 | 0 | 0 | 0 |
| 0 | 0 | 0 | 0 |
| 0 | 0 | 0 | 0 |
| 0 | 0 | 0 | 1 |
| 0 | 0 | 0 | 1 |
| 0 | 0 | 0 | 0 |
| 0 | 0 | 0 | 1 |
| 0 | 0 | 0 | 1 |
| 0 | 0 | 0 | 0 |
| 0 | 0 | 0 | 1 |
| 0 | 0 | 0 | 1 |

| | | | |
|---|---|---|---|
| 0 | 0 | 0 | 1 |
| 0 | 0 | 0 | 1 |
| 0 | 0 | 0 | 1 |
| 0 | 0 | 0 | 1 |
| 0 | 0 | 0 | 1 |
| 0 | 0 | 0 | 1 |
| 0 | 0 | 0 | 1 |
| 0 | 0 | 0 | 1 |
| 0 | 0 | 0 | 0 |
| 0 | 0 | 0 | 0 |
| 0 | 0 | 0 | 1 |
| 0 | 0 | 0 | 0 |
| 0 | 0 | 0 | 0 |
| 0 | 0 | 0 | 0 |
| 0 | 0 | 0 | 1 |
| 0 | 0 | 0 | 1 |
| 0 | 0 | 0 | 0 |
| 0 | 0 | 0 | 0 |
| 0 | 0 | 0 | 0 |
| 0 | 0 | 0 | 1 |
| 0 | 0 | 0 | 1 |
| 0 | 0 | 0 | 1 |
| 0 | 0 | 0 | 0 |
| 0 | 0 | 0 | 1 |
| 0 | 0 | 0 | 0 |
| 0 | 0 | 0 | 1 |
| 0 | 0 | 0 | 1 |
| 0 | 0 | 0 | 1 |
| 0 | 0 | 0 | 1 |
| 0 | 0 | 0 | 0 |
| 0 | 0 | 0 | 0 |
| 0 | 0 | 0 | 0 |
| 0 | 0 | 0 | 1 |
| 0 | 0 | 0 | 1 |
| 0 | 0 | 0 | 1 |
| 0 | 0 | 0 | 1 |
| 0 | 0 | 0 | 1 |
| 0 | 0 | 0 | 1 |
| 0 | 0 | 0 | 1 |
| 0 | 0 | 0 | 1 |
| 0 | 0 | 0 | 0 |
| 0 | 0 | 0 | 1 |
| 0 | 0 | 0 | 1 |
| 0 | 0 | 0 | 1 |
| 0 | 0 | 0 | 1 |
| 0 | 0 | 0 | 0 |
| 0 | 1 | 1 | 1 |
| 0 | 0 | 0 | 0 |
| 0 | 0 | 0 | 1 |
| 0 | 1 | 0 | 0 |
| 0 | 1 | 0 | 0 |
| 0 | 0 | 0 | 0 |
| 0 | 0 | 0 | 1 |
| 0 | 1 | 1 | 1 |

| | | | |
|---|---|---|---|
| 0 | 0 | 0 | 1 |
| 0 | 1 | 1 | 1 |
| 0 | 0 | 0 | 1 |
| 0 | 0 | 0 | 1 |
| 0 | 0 | 0 | 1 |
| 0 | 0 | 0 | 1 |
| 0 | 0 | 0 | 1 |
| 0 | 0 | 0 | 0 |
| 0 | 0 | 0 | 0 |
| 0 | 0 | 0 | 1 |
| 0 | 0 | 0 | 0 |
| 0 | 0 | 0 | 1 |
| 0 | 0 | 0 | 1 |
| 0 | 0 | 0 | 1 |
| 0 | 0 | 0 | 1 |
| 0 | 0 | 0 | 1 |
| 0 | 0 | 0 | 1 |
| 0 | 0 | 0 | 1 |
| 0 | 0 | 0 | 1 |
| 0 | 0 | 0 | 1 |
| 0 | 0 | 0 | 1 |
| 0 | 0 | 0 | 1 |
| 0 | 0 | 0 | 1 |
| 0 | 0 | 0 | 1 |
| 0 | 0 | 0 | 1 |
| 0 | 0 | 0 | 1 |
| 0 | 0 | 0 | 1 |
| 0 | 0 | 0 | 1 |
| 0 | 0 | 0 | 0 |
| 0 | 0 | 0 | 1 |
| 0 | 0 | 0 | 0 |
| 0 | 0 | 0 | 1 |
| 0 | 0 | 0 | 0 |
| 0 | 0 | 0 | 0 |
| 0 | 0 | 0 | 1 |
| 0 | 0 | 0 | 0 |
| 0 | 0 | 0 | 0 |
| 0 | 0 | 0 | 1 |
| 0 | 0 | 0 | 1 |
| 0 | 0 | 0 | 1 |
| 0 | 0 | 0 | 1 |
| 0 | 0 | 0 | 0 |
| 0 | 0 | 0 | 0 |
| 0 | 0 | 0 | 1 |
| 0 | 0 | 0 | 1 |
| 0 | 0 | 0 | 0 |
| 0 | 0 | 0 | 0 |
| 0 | 0 | 0 | 0 |
| 0 | 0 | 0 | 0 |
| 0 | 0 | 0 | 1 |
| 0 | 0 | 0 | 0 |
| 0 | 0 | 0 | 0 |
| 0 | 0 | 0 | 1 |
| 0 | 0 | 0 | 1 |

| | | | |
|---|---|---|---|
| 0 | 0 | 0 | 1 |
| 0 | 0 | 0 | 1 |
| 0 | 0 | 0 | 1 |
| 0 | 0 | 0 | 1 |
| 0 | 0 | 0 | 0 |
| 0 | 0 | 0 | 0 |
| 0 | 0 | 0 | 0 |
| 0 | 0 | 0 | 0 |
| 0 | 0 | 0 | 1 |
| 0 | 0 | 0 | 0 |
| 0 | 0 | 0 | 0 |
| 0 | 0 | 0 | 0 |
| 0 | 0 | 0 | 0 |
| 0 | 0 | 0 | 1 |
| 0 | 0 | 0 | 1 |
| 0 | 0 | 0 | 1 |
| 0 | 0 | 0 | 1 |
| 0 | 0 | 0 | 1 |
| 0 | 0 | 0 | 1 |
| 0 | 0 | 0 | 1 |
| 0 | 0 | 0 | 1 |
| 0 | 0 | 0 | 1 |
| 0 | 0 | 0 | 1 |
| 0 | 0 | 0 | 1 |
| 0 | 0 | 0 | 1 |
| 0 | 0 | 0 | 0 |
| 0 | 0 | 0 | 0 |
| 0 | 0 | 0 | 0 |
| 0 | 0 | 0 | 0 |
| 0 | 0 | 0 | 1 |
| 0 | 0 | 0 | 1 |
| 0 | 0 | 0 | 1 |
| 0 | 0 | 0 | 1 |
| 0 | 0 | 0 | 1 |
| 0 | 0 | 0 | 1 |
| 0 | 0 | 0 | 1 |
| 0 | 0 | 0 | 1 |
| 0 | 0 | 0 | 1 |
| 0 | 0 | 0 | 1 |
| 0 | 0 | 0 | 1 |
| 0 | 0 | 0 | 1 |
| 0 | 0 | 0 | 1 |
| 0 | 0 | 0 | 1 |
| 0 | 0 | 0 | 1 |
| 0 | 0 | 0 | 1 |
| 0 | 0 | 0 | 1 |
| 0 | 0 | 0 | 1 |
| 0 | 0 | 0 | 1 |
| 0 | 0 | 0 | 1 |
| 0 | 0 | 0 | 1 |
| 0 | 0 | 0 | 1 |
| 0 | 0 | 0 | 1 |
| 0 | 0 | 0 | 1 |
| 0 | 0 | 0 | 0 |

| | | | |
|---|---|---|---|
| 0 | 0 | 0 | 1 |
| 0 | 0 | 0 | 1 |
| 0 | 0 | 0 | 1 |
| 0 | 0 | 0 | 1 |
| 0 | 0 | 0 | 1 |
| 0 | 0 | 0 | 1 |
| 0 | 0 | 0 | 1 |
| 0 | 0 | 0 | 1 |
| 0 | 0 | 0 | 1 |
| 0 | 0 | 0 | 1 |
| 0 | 0 | 0 | 1 |
| 0 | 0 | 0 | 1 |
| 0 | 0 | 0 | 1 |
| 0 | 0 | 0 | 1 |
| 0 | 0 | 0 | 1 |
| 0 | 0 | 0 | 1 |
| 0 | 0 | 0 | 1 |
| 0 | 0 | 0 | 1 |
| 0 | 0 | 0 | 1 |
| 0 | 0 | 0 | 1 |
| 0 | 0 | 0 | 1 |
| 0 | 0 | 0 | 1 |
| 0 | 0 | 0 | 1 |
| 0 | 0 | 0 | 1 |
| 0 | 0 | 0 | 1 |
| 0 | 0 | 0 | 1 |
| 0 | 0 | 0 | 1 |
| 0 | 0 | 0 | 1 |
| 0 | 0 | 0 | 1 |
| 0 | 0 | 0 | 1 |
| 0 | 0 | 0 | 1 |
| 0 | 0 | 0 | 1 |
| 0 | 0 | 0 | 1 |
| 0 | 0 | 0 | 1 |
| 0 | 0 | 0 | 1 |
| 0 | 0 | 0 | 1 |
| 0 | 0 | 0 | 1 |
| 0 | 0 | 0 | 1 |
| 0 | 0 | 0 | 1 |
| 0 | 0 | 0 | 1 |
| 0 | 0 | 0 | 1 |
| 0 | 0 | 0 | 1 |
| 0 | 0 | 0 | 1 |
| 0 | 0 | 0 | 1 |
| 0 | 0 | 0 | 1 |
| 0 | 0 | 0 | 1 |
| 0 | 0 | 0 | 1 |
| 0 | 1 | 1 | 1 |
| 0 | 0 | 0 | 0 |
| 0 | 0 | 0 | 1 |
| 0 | 0 | 0 | 1 |
| 0 | 0 | 0 | 1 |
| 0 | 0 | 0 | 1 |
| 0 | 0 | 0 | 1 |
| 0 | 0 | 0 | 1 |
| 0 | 0 | 0 | 1 |

| | | | |
|---|---|---|---|
| 0 | 0 | 0 | 1 |
| 0 | 0 | 0 | 1 |
| 0 | 0 | 0 | 0 |
| 0 | 0 | 0 | 1 |
| 0 | 0 | 0 | 1 |
| 0 | 0 | 0 | 1 |
| 0 | 0 | 0 | 0 |
| 0 | 0 | 0 | 1 |
| 0 | 0 | 0 | 1 |
| 0 | 0 | 0 | 1 |
| 0 | 0 | 0 | 1 |
| 0 | 0 | 0 | 0 |
| 0 | 0 | 0 | 1 |
| 0 | 0 | 0 | 0 |
| 0 | 0 | 0 | 0 |
| 0 | 0 | 0 | 0 |
| 0 | 0 | 0 | 0 |
| 0 | 0 | 0 | 0 |
| 0 | 0 | 0 | 0 |
| 0 | 0 | 0 | 0 |
| 0 | 0 | 0 | 0 |

| surface_non_binding_site | non_binding_site_single_water | surface_non_binding_site_single_water |
|---|---|---|
| 0 | 0 | 0 |
| 0 | 0 | 0 |
| 0 | 0 | 0 |
| 0 | 0 | 0 |
| 0 | 0 | 0 |
| 0 | 0 | 0 |
| 1 | 1 | 1 |
| 1 | 1 | 1 |
| 0 | 0 | 0 |
| 0 | 0 | 0 |
| 0 | 0 | 0 |
| 0 | 0 | 0 |
| 0 | 0 | 0 |
| 0 | 0 | 0 |
| 0 | 0 | 0 |
| 0 | 0 | 0 |
| 0 | 0 | 0 |
| 0 | 0 | 0 |
| 0 | 0 | 0 |
| 0 | 0 | 0 |
| 0 | 0 | 0 |
| 0 | 0 | 0 |
| 0 | 0 | 0 |
| 0 | 0 | 0 |
| 0 | 0 | 0 |
| 1 | 0 | 0 |
| 0 | 0 | 0 |
| 1 | 1 | 1 |
| 0 | 0 | 0 |
| 0 | 1 | 0 |
| 0 | 1 | 0 |
| 0 | 0 | 0 |
| 0 | 0 | 0 |
| 0 | 0 | 0 |
| 0 | 0 | 0 |
| 1 | 1 | 1 |
| 0 | 0 | 0 |
| 0 | 0 | 0 |
| 0 | 0 | 0 |
| 0 | 0 | 0 |
| 0 | 0 | 0 |
| 0 | 0 | 0 |
| 0 | 0 | 0 |
| 0 | 0 | 0 |
| 0 | 0 | 0 |
| 0 | 0 | 0 |
| 0 | 0 | 0 |
| 0 | 0 | 0 |
| 0 | 0 | 0 |
| 0 | 0 | 0 |
| 0 | 0 | 0 |
| 0 | 0 | 0 |
| 0 | 0 | 0 |

| | | |
|---|---|---|
| 0 | 0 | 0 |
| 0 | 0 | 0 |
| 0 | 0 | 0 |
| 0 | 0 | 0 |
| 0 | 0 | 0 |
| 0 | 0 | 0 |
| 0 | 0 | 0 |
| 0 | 0 | 0 |
| 0 | 0 | 0 |
| 0 | 0 | 0 |
| 0 | 0 | 0 |
| 0 | 0 | 0 |
| 0 | 0 | 0 |
| 0 | 0 | 0 |
| 0 | 0 | 0 |
| 0 | 0 | 0 |
| 0 | 0 | 0 |
| 0 | 0 | 0 |
| 0 | 0 | 0 |
| 0 | 0 | 0 |
| 0 | 0 | 0 |
| 0 | 0 | 0 |
| 0 | 0 | 0 |
| 0 | 0 | 0 |
| 0 | 0 | 0 |
| 0 | 0 | 0 |
| 0 | 0 | 0 |
| 0 | 0 | 0 |
| 0 | 0 | 0 |
| 0 | 0 | 0 |
| 0 | 0 | 0 |
| 0 | 0 | 0 |
| 0 | 0 | 0 |
| 0 | 0 | 0 |
| 0 | 0 | 0 |
| 0 | 0 | 0 |
| 0 | 0 | 0 |
| 0 | 0 | 0 |
| 0 | 0 | 0 |
| 0 | 0 | 0 |
| 0 | 0 | 0 |
| 0 | 0 | 0 |
| 0 | 0 | 0 |
| 0 | 0 | 0 |
| 0 | 0 | 0 |
| 1 | 1 | 1 |
| 0 | 1 | 0 |
| 0 | 0 | 0 |
| 0 | 0 | 0 |
| 0 | 0 | 0 |
| 0 | 0 | 0 |
| 1 | 1 | 1 |
| 1 | 1 | 1 |
| 0 | 0 | 0 |

| | | |
|---|---|---|
| 0 | 1 | 0 |
| 0 | 1 | 0 |
| 0 | 0 | 0 |
| 0 | 0 | 0 |
| 0 | 0 | 0 |
| 0 | 0 | 0 |
| 0 | 1 | 0 |
| 0 | 0 | 0 |
| 0 | 0 | 0 |
| 0 | 0 | 0 |
| 0 | 0 | 0 |
| 0 | 0 | 0 |
| 0 | 0 | 0 |
| 0 | 0 | 0 |
| 0 | 0 | 0 |
| 1 | 1 | 1 |
| 0 | 0 | 0 |
| 0 | 0 | 0 |
| 0 | 0 | 0 |
| 0 | 0 | 0 |
| 0 | 0 | 0 |
| 0 | 0 | 0 |
| 0 | 0 | 0 |
| 0 | 0 | 0 |
| 0 | 0 | 0 |
| 0 | 0 | 0 |
| 0 | 0 | 0 |
| 0 | 0 | 0 |
| 0 | 0 | 0 |
| 0 | 0 | 0 |
| 0 | 0 | 0 |
| 0 | 0 | 0 |
| 0 | 0 | 0 |
| 0 | 0 | 0 |
| 0 | 0 | 0 |
| 0 | 0 | 0 |
| 0 | 0 | 0 |
| 0 | 0 | 0 |
| 0 | 0 | 0 |
| 0 | 0 | 0 |
| 0 | 0 | 0 |
| 0 | 0 | 0 |
| 0 | 0 | 0 |
| 0 | 0 | 0 |
| 0 | 0 | 0 |
| 0 | 0 | 0 |
| 0 | 0 | 0 |
| 0 | 0 | 0 |
| 0 | 0 | 0 |
| 0 | 0 | 0 |
| 0 | 0 | 0 |
| 0 | 0 | 0 |
| 0 | 0 | 0 |
| 0 | 0 | 0 |

| | | |
|---|---|---|
| 1 | 0 | 0 |
| 1 | 0 | 0 |
| 1 | 0 | 0 |
| 0 | 0 | 0 |
| 0 | 0 | 0 |
| 0 | 0 | 0 |
| 0 | 0 | 0 |
| 1 | 0 | 0 |
| 0 | 0 | 0 |
| 1 | 0 | 0 |
| 1 | 0 | 0 |
| 0 | 0 | 0 |
| 1 | 0 | 0 |
| 0 | 0 | 0 |
| 0 | 0 | 0 |
| 1 | 0 | 0 |
| 1 | 0 | 0 |
| 1 | 0 | 0 |
| 1 | 0 | 0 |
| 0 | 0 | 0 |
| 0 | 0 | 0 |
| 0 | 1 | 0 |
| 0 | 0 | 0 |
| 0 | 0 | 0 |
| 0 | 0 | 0 |
| 1 | 0 | 0 |
| 0 | 0 | 0 |
| 0 | 0 | 0 |
| 1 | 0 | 0 |
| 1 | 0 | 0 |
| 1 | 1 | 1 |
| 0 | 0 | 0 |
| 0 | 0 | 0 |
| 0 | 1 | 0 |
| 0 | 0 | 0 |
| 0 | 0 | 0 |
| 0 | 0 | 0 |
| 0 | 0 | 0 |
| 0 | 0 | 0 |
| 1 | 0 | 0 |
| 1 | 0 | 0 |
| 0 | 0 | 0 |
| 0 | 0 | 0 |
| 1 | 1 | 1 |
| 1 | 0 | 0 |
| 1 | 0 | 0 |
| 0 | 0 | 0 |
| 0 | 0 | 0 |
| 0 | 0 | 0 |
| 0 | 0 | 0 |
| 0 | 0 | 0 |
| 0 | 1 | 0 |
| 1 | 0 | 0 |
| 0 | 0 | 0 |

| | | |
|---|---|---|
| 0 | 0 | 0 |
| 0 | 0 | 0 |
| 0 | 0 | 0 |
| 1 | 1 | 1 |
| 1 | 1 | 1 |
| 1 | 0 | 0 |
| 0 | 0 | 0 |
| 0 | 0 | 0 |
| 1 | 0 | 0 |
| 1 | 0 | 0 |
| 0 | 0 | 0 |
| 0 | 1 | 0 |
| 0 | 0 | 0 |
| 1 | 0 | 0 |
| 0 | 1 | 0 |
| 1 | 1 | 1 |
| 1 | 1 | 1 |
| 0 | 0 | 0 |
| 0 | 0 | 0 |
| 0 | 0 | 0 |
| 0 | 0 | 0 |
| 1 | 0 | 0 |
| 0 | 0 | 0 |
| 0 | 1 | 0 |
| 1 | 1 | 1 |
| 0 | 0 | 0 |
| 0 | 0 | 0 |
| 0 | 0 | 0 |
| 0 | 0 | 0 |
| 0 | 0 | 0 |
| 0 | 0 | 0 |
| 0 | 0 | 0 |
| 1 | 0 | 0 |
| 1 | 0 | 0 |
| 0 | 0 | 0 |
| 1 | 0 | 0 |
| 0 | 0 | 0 |
| 0 | 0 | 0 |
| 1 | 0 | 0 |
| 0 | 1 | 0 |
| 0 | 0 | 0 |
| 1 | 0 | 0 |
| 1 | 0 | 0 |
| 1 | 0 | 0 |
| 0 | 0 | 0 |
| 0 | 0 | 0 |
| 0 | 0 | 0 |
| 0 | 0 | 0 |
| 0 | 0 | 0 |
| 0 | 0 | 0 |
| 0 | 0 | 0 |
| 0 | 0 | 0 |
| 0 | 0 | 0 |
| 0 | 0 | 0 |

| | | |
|---|---|---|
| 0 | 0 | 0 |
| 0 | 0 | 0 |
| 0 | 0 | 0 |
| 0 | 0 | 0 |
| 0 | 0 | 0 |
| 0 | 0 | 0 |
| 0 | 1 | 0 |
| 0 | 1 | 0 |
| 0 | 0 | 0 |
| 0 | 0 | 0 |
| 0 | 0 | 0 |
| 0 | 0 | 0 |
| 0 | 0 | 0 |
| 0 | 1 | 0 |
| 0 | 1 | 0 |
| 0 | 0 | 0 |
| 0 | 1 | 0 |
| 0 | 0 | 0 |
| 1 | 1 | 1 |
| 0 | 0 | 0 |
| 1 | 1 | 1 |
| 1 | 0 | 0 |
| 1 | 1 | 1 |
| 0 | 0 | 0 |
| 0 | 0 | 0 |
| 0 | 0 | 0 |
| 0 | 0 | 0 |
| 0 | 0 | 0 |
| 0 | 0 | 0 |
| 0 | 0 | 0 |
| 0 | 0 | 0 |
| 1 | 0 | 0 |
| 0 | 0 | 0 |
| 0 | 0 | 0 |
| 0 | 0 | 0 |
| 0 | 0 | 0 |
| 0 | 0 | 0 |
| 0 | 0 | 0 |
| 0 | 0 | 0 |
| 0 | 0 | 0 |
| 0 | 1 | 0 |
| 0 | 1 | 0 |
| 0 | 0 | 0 |
| 0 | 1 | 0 |
| 0 | 0 | 0 |
| 0 | 0 | 0 |
| 0 | 0 | 0 |
| 1 | 1 | 1 |
| 1 | 1 | 1 |
| 0 | 0 | 0 |
| 0 | 1 | 0 |
| 1 | 1 | 1 |
| 0 | 0 | 0 |
| 0 | 1 | 0 |

| | | |
|---|---|---|
| 0 | 0 | 0 |
| 1 | 1 | 1 |
| 0 | 0 | 0 |
| 0 | 0 | 0 |
| 0 | 0 | 0 |
| 0 | 0 | 0 |
| 0 | 1 | 0 |
| 0 | 0 | 0 |
| 0 | 0 | 0 |
| 1 | 1 | 1 |
| 1 | 1 | 1 |
| 1 | 1 | 1 |
| 1 | 1 | 1 |
| 1 | 1 | 1 |
| 0 | 0 | 0 |
| 1 | 1 | 1 |
| 0 | 0 | 0 |
| 0 | 0 | 0 |
| 1 | 1 | 1 |
| 0 | 0 | 0 |
| 0 | 0 | 0 |
| 0 | 0 | 0 |
| 0 | 0 | 0 |
| 0 | 0 | 0 |
| 1 | 1 | 1 |
| 1 | 0 | 0 |
| 0 | 0 | 0 |
| 0 | 0 | 0 |
| 0 | 0 | 0 |
| 0 | 0 | 0 |
| 0 | 0 | 0 |
| 0 | 0 | 0 |
| 0 | 0 | 0 |
| 0 | 0 | 0 |
| 1 | 1 | 1 |
| 1 | 1 | 1 |
| 0 | 0 | 0 |
| 0 | 0 | 0 |
| 0 | 0 | 0 |
| 0 | 0 | 0 |
| 0 | 0 | 0 |
| 0 | 0 | 0 |
| 0 | 1 | 0 |
| 0 | 0 | 0 |
| 0 | 0 | 0 |
| 0 | 0 | 0 |
| 0 | 0 | 0 |
| 0 | 0 | 0 |
| 0 | 0 | 0 |
| 1 | 1 | 1 |
| 0 | 0 | 0 |
| 0 | 0 | 0 |
| 0 | 0 | 0 |
| 0 | 0 | 0 |

| | | |
|---|---|---|
| 0 | 0 | 0 |
| 0 | 0 | 0 |
| 0 | 0 | 0 |
| 0 | 0 | 0 |
| 1 | 1 | 1 |
| 0 | 0 | 0 |
| 0 | 0 | 0 |
| 0 | 0 | 0 |
| 0 | 0 | 0 |
| 0 | 0 | 0 |
| 0 | 0 | 0 |
| 0 | 0 | 0 |
| 0 | 0 | 0 |
| 0 | 0 | 0 |
| 0 | 0 | 0 |
| 0 | 0 | 0 |
| 0 | 0 | 0 |
| 0 | 0 | 0 |
| 1 | 1 | 1 |
| 0 | 0 | 0 |
| 0 | 0 | 0 |
| 0 | 0 | 0 |
| 0 | 0 | 0 |
| 1 | 0 | 0 |
| 0 | 0 | 0 |
| 0 | 0 | 0 |
| 0 | 0 | 0 |
| 1 | 0 | 0 |
| 1 | 0 | 0 |
| 1 | 0 | 0 |
| 0 | 0 | 0 |
| 0 | 0 | 0 |
| 0 | 0 | 0 |
| 0 | 0 | 0 |
| 0 | 0 | 0 |
| 1 | 0 | 0 |
| 1 | 0 | 0 |
| 0 | 0 | 0 |
| 0 | 0 | 0 |
| 1 | 0 | 0 |
| 1 | 0 | 0 |
| 1 | 0 | 0 |
| 0 | 1 | 0 |
| 1 | 1 | 1 |
| 0 | 0 | 0 |
| 0 | 0 | 0 |
| 0 | 0 | 0 |
| 0 | 0 | 0 |
| 0 | 0 | 0 |
| 0 | 1 | 0 |
| 1 | 1 | 1 |
| 0 | 0 | 0 |
| 1 | 1 | 1 |
| 0 | 0 | 0 |

| | | |
|---|---|---|
| 1 | 1 | 1 |
| 1 | 1 | 1 |
| 0 | 0 | 0 |
| 1 | 1 | 1 |
| 1 | 1 | 1 |
| 1 | 1 | 1 |
| 1 | 1 | 1 |
| 0 | 0 | 0 |
| 1 | 0 | 0 |
| 1 | 1 | 1 |
| 0 | 0 | 0 |
| 1 | 1 | 1 |
| 1 | 1 | 1 |
| 0 | 1 | 0 |
| 0 | 0 | 0 |
| 0 | 0 | 0 |
| 1 | 1 | 1 |
| 1 | 1 | 1 |
| 0 | 0 | 0 |
| 0 | 0 | 0 |
| 0 | 0 | 0 |
| 0 | 0 | 0 |
| 0 | 0 | 0 |
| 0 | 0 | 0 |
| 0 | 0 | 0 |
| 1 | 1 | 1 |
| 0 | 0 | 0 |
| 0 | 0 | 0 |
| 0 | 0 | 0 |
| 1 | 0 | 0 |
| 0 | 0 | 0 |
| 0 | 0 | 0 |
| 0 | 0 | 0 |
| 0 | 0 | 0 |
| 0 | 0 | 0 |
| 0 | 0 | 0 |
| 0 | 0 | 0 |
| 0 | 0 | 0 |
| 0 | 0 | 0 |
| 0 | 0 | 0 |
| 0 | 1 | 0 |
| 0 | 0 | 0 |
| 1 | 1 | 1 |
| 0 | 0 | 0 |
| 0 | 0 | 0 |
| 0 | 0 | 0 |
| 0 | 0 | 0 |
| 0 | 0 | 0 |
| 0 | 0 | 0 |
| 0 | 0 | 0 |
| 0 | 1 | 0 |
| 0 | 0 | 0 |
| 0 | 0 | 0 |
| 0 | 0 | 0 |

| | | |
|---|---|---|
| 1 | 1 | 1 |
| 0 | 0 | 0 |
| 0 | 1 | 0 |
| 0 | 1 | 0 |
| 0 | 0 | 0 |
| 0 | 0 | 0 |
| 0 | 0 | 0 |
| 0 | 0 | 0 |
| 0 | 0 | 0 |
| 0 | 1 | 0 |
| 1 | 1 | 1 |
| 1 | 1 | 1 |
| 1 | 1 | 1 |
| 0 | 0 | 0 |
| 1 | 1 | 1 |
| 0 | 0 | 0 |
| 0 | 0 | 0 |
| 0 | 0 | 0 |
| 0 | 0 | 0 |
| 1 | 0 | 0 |
| 0 | 0 | 0 |
| 1 | 1 | 1 |
| 0 | 0 | 0 |
| 0 | 1 | 0 |
| 1 | 1 | 1 |
| 1 | 1 | 1 |
| 0 | 0 | 0 |
| 1 | 1 | 1 |
| 1 | 1 | 1 |
| 1 | 1 | 1 |
| 0 | 0 | 0 |
| 0 | 0 | 0 |
| 1 | 1 | 1 |
| 1 | 1 | 1 |
| 0 | 0 | 0 |
| 0 | 0 | 0 |
| 0 | 0 | 0 |
| 0 | 0 | 0 |
| 0 | 0 | 0 |
| 1 | 1 | 1 |
| 0 | 0 | 0 |
| 0 | 0 | 0 |
| 0 | 0 | 0 |
| 0 | 0 | 0 |
| 1 | 0 | 0 |
| 0 | 0 | 0 |
| 0 | 0 | 0 |
| 0 | 0 | 0 |
| 0 | 0 | 0 |
| 0 | 0 | 0 |
| 0 | 0 | 0 |
| 0 | 0 | 0 |
| 1 | 0 | 0 |
| 0 | 0 | 0 |

| | | |
|---|---|---|
| 0 | 0 | 0 |
| 0 | 0 | 0 |
| 0 | 0 | 0 |
| 0 | 0 | 0 |
| 0 | 0 | 0 |
| 0 | 0 | 0 |
| 0 | 0 | 0 |
| 1 | 0 | 0 |
| 0 | 0 | 0 |
| 0 | 0 | 0 |
| 0 | 0 | 0 |
| 0 | 0 | 0 |
| 1 | 1 | 1 |
| 0 | 0 | 0 |
| 0 | 0 | 0 |
| 0 | 0 | 0 |
| 0 | 0 | 0 |
| 0 | 0 | 0 |
| 0 | 0 | 0 |
| 1 | 1 | 1 |
| 0 | 0 | 0 |
| 0 | 0 | 0 |
| 0 | 0 | 0 |
| 0 | 0 | 0 |
| 0 | 0 | 0 |
| 1 | 0 | 0 |
| 1 | 1 | 1 |
| 0 | 1 | 0 |
| 0 | 0 | 0 |
| 0 | 0 | 0 |
| 0 | 0 | 0 |
| 0 | 0 | 0 |
| 0 | 0 | 0 |
| 0 | 0 | 0 |
| 0 | 0 | 0 |
| 1 | 0 | 0 |
| 0 | 0 | 0 |
| 0 | 0 | 0 |
| 1 | 1 | 1 |
| 1 | 0 | 0 |
| 0 | 1 | 0 |
| 0 | 0 | 0 |
| 0 | 0 | 0 |
| 0 | 0 | 0 |
| 0 | 0 | 0 |
| 0 | 0 | 0 |
| 0 | 0 | 0 |
| 1 | 1 | 1 |
| 1 | 1 | 1 |
| 0 | 0 | 0 |
| 0 | 0 | 0 |
| 0 | 0 | 0 |
| 0 | 0 | 0 |
| 1 | 1 | 1 |

| | | |
|---|---|---|
| 1 | 1 | 1 |
| 0 | 0 | 0 |
| 0 | 1 | 0 |
| 0 | 0 | 0 |
| 1 | 0 | 0 |
| 1 | 0 | 0 |
| 0 | 0 | 0 |
| 1 | 0 | 0 |
| 1 | 0 | 0 |
| 1 | 0 | 0 |
| 1 | 0 | 0 |
| 1 | 0 | 0 |
| 1 | 1 | 1 |
| 1 | 1 | 1 |
| 1 | 1 | 1 |
| 1 | 1 | 1 |
| 0 | 0 | 0 |
| 0 | 0 | 0 |
| 1 | 0 | 0 |
| 0 | 0 | 0 |
| 0 | 0 | 0 |
| 0 | 0 | 0 |
| 1 | 0 | 0 |
| 0 | 0 | 0 |
| 0 | 0 | 0 |
| 1 | 1 | 1 |
| 1 | 0 | 0 |
| 1 | 0 | 0 |
| 0 | 0 | 0 |
| 0 | 0 | 0 |
| 0 | 0 | 0 |
| 0 | 0 | 0 |
| 0 | 0 | 0 |
| 0 | 0 | 0 |
| 0 | 0 | 0 |
| 0 | 0 | 0 |
| 1 | 0 | 0 |
| 1 | 1 | 1 |
| 1 | 1 | 1 |
| 1 | 1 | 1 |
| 0 | 0 | 0 |
| 1 | 1 | 1 |
| 0 | 0 | 0 |
| 1 | 1 | 1 |
| 0 | 0 | 0 |
| 0 | 1 | 0 |
| 0 | 0 | 0 |
| 0 | 0 | 0 |
| 0 | 0 | 0 |
| 0 | 0 | 0 |
| 0 | 0 | 0 |
| 0 | 0 | 0 |
| 0 | 0 | 0 |
| 0 | 0 | 0 |

| | | |
|---|---|---|
| 0 | 0 | 0 |
| 0 | 0 | 0 |
| 0 | 0 | 0 |
| 0 | 0 | 0 |
| 0 | 0 | 0 |
| 0 | 0 | 0 |
| 0 | 0 | 0 |
| 1 | 1 | 1 |
| 0 | 0 | 0 |
| 0 | 0 | 0 |
| 0 | 0 | 0 |
| 1 | 0 | 0 |
| 0 | 0 | 0 |
| 0 | 0 | 0 |
| 0 | 0 | 0 |
| 0 | 0 | 0 |
| 0 | 0 | 0 |
| 0 | 0 | 0 |
| 0 | 0 | 0 |
| 0 | 0 | 0 |
| 0 | 0 | 0 |
| 0 | 0 | 0 |
| 0 | 0 | 0 |
| 0 | 0 | 0 |
| 0 | 0 | 0 |
| 0 | 0 | 0 |
| 0 | 0 | 0 |
| 0 | 0 | 0 |
| 0 | 0 | 0 |
| 0 | 0 | 0 |
| 0 | 0 | 0 |
| 0 | 0 | 0 |
| 0 | 0 | 0 |
| 0 | 0 | 0 |
| 0 | 0 | 0 |
| 0 | 0 | 0 |
| 0 | 0 | 0 |
| 0 | 0 | 0 |
| 0 | 0 | 0 |
| 0 | 0 | 0 |
| 1 | 0 | 0 |
| 0 | 0 | 0 |
| 0 | 0 | 0 |
| 0 | 0 | 0 |
| 0 | 0 | 0 |
| 0 | 0 | 0 |
| 0 | 0 | 0 |
| 0 | 0 | 0 |
| 0 | 0 | 0 |
| 0 | 1 | 0 |
| 0 | 1 | 0 |
| 1 | 0 | 0 |
| 0 | 0 | 0 |
| 0 | 0 | 0 |

| | | |
|---|---|---|
| 0 | 0 | 0 |
| 0 | 0 | 0 |
| 0 | 0 | 0 |
| 0 | 0 | 0 |
| 0 | 0 | 0 |
| 0 | 0 | 0 |
| 0 | 0 | 0 |
| 1 | 0 | 0 |
| 1 | 0 | 0 |
| 0 | 0 | 0 |
| 0 | 0 | 0 |
| 1 | 0 | 0 |
| 0 | 0 | 0 |
| 0 | 0 | 0 |
| 0 | 0 | 0 |
| 0 | 0 | 0 |
| 0 | 0 | 0 |
| 1 | 0 | 0 |
| 0 | 0 | 0 |
| 1 | 0 | 0 |
| 0 | 0 | 0 |
| 0 | 0 | 0 |
| 1 | 1 | 1 |
| 0 | 0 | 0 |
| 0 | 0 | 0 |
| 1 | 0 | 0 |
| 0 | 0 | 0 |
| 0 | 0 | 0 |
| 0 | 0 | 0 |
| 0 | 0 | 0 |
| 0 | 0 | 0 |
| 0 | 0 | 0 |
| 1 | 0 | 0 |
| 0 | 0 | 0 |
| 0 | 0 | 0 |
| 0 | 0 | 0 |
| 1 | 1 | 1 |
| 0 | 0 | 0 |
| 0 | 0 | 0 |
| 0 | 0 | 0 |
| 0 | 0 | 0 |
| 0 | 0 | 0 |
| 0 | 0 | 0 |
| 0 | 0 | 0 |
| 0 | 0 | 0 |
| 0 | 0 | 0 |
| 0 | 0 | 0 |
| 0 | 0 | 0 |
| 0 | 0 | 0 |
| 0 | 0 | 0 |
| 0 | 0 | 0 |
| 0 | 0 | 0 |
| 0 | 0 | 0 |
| 0 | 0 | 0 |

| | | |
|---|---|---|
| 1 | 0 | 0 |
| 0 | 1 | 0 |
| 0 | 0 | 0 |
| 0 | 0 | 0 |
| 0 | 0 | 0 |
| 0 | 0 | 0 |
| 0 | 0 | 0 |
| 0 | 0 | 0 |
| 0 | 0 | 0 |
| 1 | 0 | 0 |
| 0 | 1 | 0 |
| 1 | 1 | 1 |
| 0 | 0 | 0 |
| 0 | 0 | 0 |
| 0 | 0 | 0 |
| 0 | 0 | 0 |
| 0 | 0 | 0 |
| 0 | 0 | 0 |
| 1 | 1 | 1 |
| 1 | 0 | 0 |
| 1 | 0 | 0 |
| 0 | 0 | 0 |
| 0 | 1 | 0 |
| 0 | 0 | 0 |
| 0 | 0 | 0 |
| 0 | 0 | 0 |
| 0 | 0 | 0 |
| 0 | 0 | 0 |
| 0 | 0 | 0 |
| 0 | 0 | 0 |
| 0 | 0 | 0 |
| 0 | 0 | 0 |
| 0 | 0 | 0 |
| 0 | 0 | 0 |
| 0 | 0 | 0 |
| 0 | 0 | 0 |
| 0 | 0 | 0 |
| 0 | 0 | 0 |
| 0 | 0 | 0 |
| 0 | 0 | 0 |
| 0 | 0 | 0 |
| 1 | 0 | 0 |
| 0 | 0 | 0 |
| 0 | 0 | 0 |
| 0 | 0 | 0 |
| 0 | 0 | 0 |
| 1 | 0 | 0 |
| 0 | 0 | 0 |
| 0 | 0 | 0 |
| 0 | 0 | 0 |
| 0 | 0 | 0 |
| 0 | 0 | 0 |
| 1 | 0 | 0 |
| 0 | 1 | 0 |

| | | |
|---|---|---|
| 0 | 0 | 0 |
| 0 | 0 | 0 |
| 1 | 0 | 0 |
| 1 | 0 | 0 |
| 1 | 0 | 0 |
| 0 | 0 | 0 |
| 1 | 0 | 0 |
| 0 | 0 | 0 |
| 0 | 0 | 0 |
| 0 | 0 | 0 |
| 0 | 0 | 0 |
| 0 | 0 | 0 |
| 0 | 0 | 0 |
| 1 | 0 | 0 |
| 1 | 0 | 0 |
| 1 | 0 | 0 |
| 0 | 0 | 0 |
| 1 | 1 | 1 |
| 0 | 0 | 0 |
| 0 | 0 | 0 |
| 1 | 0 | 0 |
| 1 | 0 | 0 |
| 0 | 0 | 0 |
| 0 | 0 | 0 |
| 1 | 0 | 0 |
| 0 | 0 | 0 |
| 0 | 0 | 0 |
| 1 | 0 | 0 |
| 0 | 0 | 0 |
| 0 | 0 | 0 |
| 0 | 0 | 0 |
| 0 | 0 | 0 |
| 0 | 0 | 0 |
| 0 | 0 | 0 |
| 0 | 0 | 0 |
| 0 | 0 | 0 |
| 0 | 0 | 0 |
| 0 | 0 | 0 |
| 0 | 0 | 0 |
| 0 | 0 | 0 |
| 1 | 0 | 0 |
| 1 | 0 | 0 |
| 1 | 0 | 0 |
| 1 | 1 | 1 |
| 1 | 1 | 1 |
| 0 | 0 | 0 |
| 0 | 0 | 0 |
| 0 | 0 | 0 |
| 0 | 0 | 0 |
| 0 | 0 | 0 |
| 0 | 0 | 0 |
| 0 | 0 | 0 |
| 0 | 0 | 0 |
| 1 | 0 | 0 |

| | | |
|---|---|---|
| 1 | 0 | 0 |
| 0 | 0 | 0 |
| 0 | 0 | 0 |
| 0 | 0 | 0 |
| 1 | 0 | 0 |
| 0 | 0 | 0 |
| 0 | 0 | 0 |
| 0 | 0 | 0 |
| 1 | 0 | 0 |
| 1 | 0 | 0 |
| 0 | 0 | 0 |
| 0 | 0 | 0 |
| 0 | 0 | 0 |
| 1 | 0 | 0 |
| 0 | 0 | 0 |
| 0 | 0 | 0 |
| 0 | 0 | 0 |
| 0 | 0 | 0 |
| 0 | 0 | 0 |
| 0 | 0 | 0 |
| 0 | 0 | 0 |
| 1 | 1 | 1 |
| 0 | 0 | 0 |
| 0 | 1 | 0 |
| 0 | 0 | 0 |
| 0 | 0 | 0 |
| 0 | 0 | 0 |
| 0 | 0 | 0 |
| 0 | 0 | 0 |
| 0 | 0 | 0 |
| 0 | 0 | 0 |
| 0 | 0 | 0 |
| 0 | 0 | 0 |
| 0 | 0 | 0 |
| 0 | 1 | 0 |
| 0 | 0 | 0 |
| 0 | 0 | 0 |
| 0 | 0 | 0 |
| 0 | 0 | 0 |
| 0 | 0 | 0 |
| 0 | 0 | 0 |
| 0 | 0 | 0 |
| 0 | 0 | 0 |
| 0 | 0 | 0 |
| 0 | 0 | 0 |
| 0 | 0 | 0 |
| 0 | 0 | 0 |
| 1 | 1 | 1 |
| 0 | 0 | 0 |
| 1 | 1 | 1 |
| 0 | 0 | 0 |
| 0 | 0 | 0 |
| 0 | 0 | 0 |
| 0 | 0 | 0 |

| | | |
|---|---|---|
| 0 | 0 | 0 |
| 0 | 0 | 0 |
| 1 | 1 | 1 |
| 0 | 0 | 0 |
| 0 | 0 | 0 |
| 0 | 1 | 0 |
| 0 | 1 | 0 |
| 0 | 1 | 0 |
| 0 | 0 | 0 |
| 0 | 1 | 0 |
| 0 | 0 | 0 |
| 0 | 0 | 0 |
| 0 | 0 | 0 |
| 1 | 1 | 1 |
| 0 | 0 | 0 |
| 0 | 1 | 0 |
| 0 | 0 | 0 |
| 0 | 0 | 0 |
| 0 | 0 | 0 |
| 1 | 1 | 1 |
| 1 | 1 | 1 |
| 0 | 0 | 0 |
| 1 | 1 | 1 |
| 0 | 0 | 0 |
| 0 | 0 | 0 |
| 1 | 1 | 1 |
| 1 | 1 | 1 |
| 0 | 0 | 0 |
| 1 | 1 | 1 |
| 0 | 0 | 0 |
| 0 | 1 | 0 |
| 0 | 1 | 0 |
| 1 | 1 | 1 |
| 0 | 1 | 0 |
| 1 | 1 | 1 |
| 1 | 1 | 1 |
| 1 | 1 | 1 |
| 1 | 1 | 1 |
| 1 | 1 | 1 |
| 1 | 1 | 1 |
| 1 | 1 | 1 |
| 1 | 1 | 1 |
| 0 | 1 | 0 |
| 0 | 1 | 0 |
| 0 | 1 | 0 |
| 0 | 1 | 0 |
| 0 | 1 | 0 |
| 0 | 1 | 0 |
| 0 | 1 | 0 |
| 0 | 1 | 0 |
| 0 | 1 | 0 |
| 0 | 1 | 0 |
| 0 | 1 | 0 |
| 0 | 1 | 0 |

| | | |
|---|---|---|
| 0 | 1 | 0 |
| 1 | 1 | 1 |
| 1 | 1 | 1 |
| 1 | 1 | 1 |
| 1 | 1 | 1 |
| 1 | 1 | 1 |
| 1 | 1 | 1 |
| 0 | 1 | 0 |
| 0 | 1 | 0 |
| 0 | 1 | 0 |
| 0 | 1 | 0 |
| 0 | 0 | 0 |
| 0 | 0 | 0 |
| 0 | 0 | 0 |
| 1 | 0 | 0 |
| 0 | 0 | 0 |
| 1 | 0 | 0 |
| 0 | 0 | 0 |
| 1 | 0 | 0 |
| 1 | 1 | 1 |
| 0 | 0 | 0 |
| 1 | 1 | 1 |
| 1 | 1 | 1 |
| 0 | 0 | 0 |
| 0 | 0 | 0 |
| 1 | 1 | 1 |
| 1 | 1 | 1 |
| 0 | 0 | 0 |
| 0 | 0 | 0 |
| 0 | 0 | 0 |
| 0 | 0 | 0 |
| 0 | 0 | 0 |
| 0 | 1 | 0 |
| 0 | 1 | 0 |
| 1 | 1 | 1 |
| 1 | 1 | 1 |
| 1 | 1 | 1 |
| 1 | 1 | 1 |
| 1 | 1 | 1 |
| 0 | 0 | 0 |
| 0 | 0 | 0 |
| 0 | 1 | 0 |
| 0 | 0 | 0 |
| 0 | 0 | 0 |
| 0 | 0 | 0 |
| 1 | 0 | 0 |
| 0 | 0 | 0 |
| 1 | 0 | 0 |
| 1 | 0 | 0 |
| 0 | 0 | 0 |
| 0 | 0 | 0 |
| 0 | 0 | 0 |
| 0 | 0 | 0 |
| 0 | 0 | 0 |

| | | |
|---|---|---|
| 0 | 0 | 0 |
| 0 | 1 | 0 |
| 1 | 1 | 1 |
| 1 | 1 | 1 |
| 0 | 0 | 0 |
| 1 | 1 | 1 |
| 0 | 0 | 0 |
| 1 | 1 | 1 |
| 1 | 1 | 1 |
| 0 | 0 | 0 |
| 1 | 0 | 0 |
| 1 | 1 | 1 |
| 1 | 1 | 1 |
| 0 | 0 | 0 |
| 1 | 0 | 0 |
| 1 | 0 | 0 |
| 0 | 0 | 0 |
| 1 | 0 | 0 |
| 0 | 0 | 0 |
| 0 | 0 | 0 |
| 0 | 0 | 0 |
| 0 | 0 | 0 |
| 0 | 0 | 0 |
| 0 | 0 | 0 |
| 0 | 0 | 0 |
| 0 | 0 | 0 |
| 0 | 0 | 0 |
| 0 | 0 | 0 |
| 0 | 0 | 0 |
| 0 | 0 | 0 |
| 0 | 0 | 0 |
| 0 | 0 | 0 |
| 0 | 0 | 0 |
| 0 | 0 | 0 |
| 0 | 0 | 0 |
| 0 | 0 | 0 |
| 1 | 0 | 0 |
| 0 | 0 | 0 |
| 0 | 0 | 0 |
| 0 | 0 | 0 |
| 0 | 0 | 0 |
| 0 | 0 | 0 |
| 0 | 0 | 0 |
| 0 | 0 | 0 |
| 0 | 0 | 0 |
| 0 | 0 | 0 |
| 0 | 0 | 0 |
| 0 | 0 | 0 |
| 1 | 0 | 0 |
| 1 | 0 | 0 |
| 0 | 0 | 0 |
| 1 | 1 | 1 |
| 0 | 0 | 0 |
| 0 | 0 | 0 |

| | | |
|---|---|---|
| 0 | 0 | 0 |
| 0 | 0 | 0 |
| 0 | 0 | 0 |
| 0 | 0 | 0 |
| 0 | 0 | 0 |
| 0 | 0 | 0 |
| 0 | 0 | 0 |
| 0 | 0 | 0 |
| 0 | 0 | 0 |
| 0 | 0 | 0 |
| 0 | 0 | 0 |
| 0 | 0 | 0 |
| 0 | 0 | 0 |
| 0 | 0 | 0 |
| 0 | 1 | 0 |
| 0 | 0 | 0 |
| 0 | 0 | 0 |
| 0 | 0 | 0 |
| 1 | 0 | 0 |
| 0 | 0 | 0 |
| 0 | 0 | 0 |
| 0 | 0 | 0 |
| 0 | 0 | 0 |
| 0 | 0 | 0 |
| 0 | 0 | 0 |
| 1 | 0 | 0 |
| 0 | 0 | 0 |
| 0 | 0 | 0 |
| 0 | 0 | 0 |
| 0 | 0 | 0 |
| 1 | 1 | 1 |
| 0 | 0 | 0 |
| 1 | 1 | 1 |
| 1 | 0 | 0 |
| 0 | 0 | 0 |
| 0 | 0 | 0 |
| 1 | 1 | 1 |
| 1 | 1 | 1 |
| 1 | 1 | 1 |
| 1 | 0 | 0 |
| 0 | 0 | 0 |
| 0 | 0 | 0 |
| 0 | 0 | 0 |
| 0 | 0 | 0 |
| 0 | 0 | 0 |
| 0 | 0 | 0 |
| 0 | 0 | 0 |
| 0 | 0 | 0 |
| 1 | 1 | 1 |
| 1 | 1 | 1 |
| 0 | 0 | 0 |
| 0 | 0 | 0 |
| 0 | 0 | 0 |
| 0 | 1 | 0 |

| | | |
|---|---|---|
| 1 | 1 | 1 |
| 1 | 1 | 1 |
| 1 | 0 | 0 |
| 0 | 0 | 0 |
| 1 | 0 | 0 |
| 1 | 1 | 1 |
| 0 | 0 | 0 |
| 1 | 1 | 1 |
| 0 | 0 | 0 |
| 0 | 0 | 0 |
| 0 | 0 | 0 |
| 0 | 0 | 0 |
| 0 | 0 | 0 |
| 0 | 0 | 0 |
| 0 | 0 | 0 |
| 0 | 0 | 0 |
| 0 | 1 | 0 |
| 0 | 0 | 0 |
| 0 | 0 | 0 |
| 1 | 0 | 0 |
| 0 | 0 | 0 |
| 0 | 0 | 0 |
| 1 | 1 | 1 |
| 0 | 0 | 0 |
| 1 | 1 | 1 |
| 1 | 1 | 1 |
| 0 | 1 | 0 |
| 0 | 0 | 0 |
| 0 | 0 | 0 |
| 0 | 0 | 0 |
| 1 | 1 | 1 |
| 0 | 0 | 0 |
| 0 | 0 | 0 |
| 0 | 0 | 0 |
| 0 | 1 | 0 |
| 0 | 1 | 0 |
| 0 | 0 | 0 |
| 0 | 0 | 0 |
| 0 | 0 | 0 |
| 1 | 1 | 1 |
| 1 | 1 | 1 |
| 0 | 1 | 0 |
| 0 | 1 | 0 |
| 0 | 0 | 0 |
| 1 | 1 | 1 |
| 0 | 0 | 0 |
| 1 | 1 | 1 |
| 0 | 0 | 0 |
| 0 | 0 | 0 |
| 1 | 1 | 1 |
| 0 | 1 | 0 |
| 0 | 0 | 0 |
| 0 | 0 | 0 |
| 1 | 1 | 1 |

| | | |
|---|---|---|
| 1 | 1 | 1 |
| 1 | 1 | 1 |
| 0 | 1 | 0 |
| 0 | 0 | 0 |
| 0 | 0 | 0 |
| 1 | 0 | 0 |
| 1 | 0 | 0 |
| 0 | 0 | 0 |
| 0 | 0 | 0 |
| 1 | 1 | 1 |
| 0 | 0 | 0 |
| 0 | 0 | 0 |
| 0 | 0 | 0 |
| 0 | 0 | 0 |
| 0 | 0 | 0 |
| 0 | 0 | 0 |
| 0 | 0 | 0 |
| 0 | 0 | 0 |
| 0 | 0 | 0 |
| 0 | 1 | 0 |
| 1 | 1 | 1 |
| 1 | 1 | 1 |
| 1 | 1 | 1 |
| 1 | 1 | 1 |
| 0 | 0 | 0 |
| 0 | 1 | 0 |
| 0 | 0 | 0 |
| 0 | 0 | 0 |
| 1 | 1 | 1 |
| 1 | 1 | 1 |
| 1 | 1 | 1 |
| 1 | 1 | 1 |
| 1 | 1 | 1 |
| 0 | 0 | 0 |
| 0 | 0 | 0 |
| 1 | 1 | 1 |
| 0 | 0 | 0 |
| 0 | 1 | 0 |
| 0 | 0 | 0 |
| 0 | 0 | 0 |
| 0 | 0 | 0 |
| 0 | 0 | 0 |
| 0 | 1 | 0 |
| 0 | 0 | 0 |
| 0 | 0 | 0 |
| 0 | 0 | 0 |
| 0 | 0 | 0 |
| 0 | 0 | 0 |
| 0 | 0 | 0 |
| 0 | 0 | 0 |
| 0 | 0 | 0 |
| 0 | 0 | 0 |
| 0 | 0 | 0 |
| 0 | 1 | 0 |

| | | |
|---|---|---|
| 0 | 0 | 0 |
| 0 | 1 | 0 |
| 0 | 0 | 0 |
| 0 | 0 | 0 |
| 0 | 0 | 0 |
| 1 | 1 | 1 |
| 0 | 0 | 0 |
| 1 | 1 | 1 |
| 0 | 1 | 0 |
| 0 | 0 | 0 |
| 0 | 1 | 0 |
| 0 | 1 | 0 |
| 0 | 1 | 0 |
| 0 | 1 | 0 |
| 0 | 1 | 0 |
| 0 | 0 | 0 |
| 0 | 0 | 0 |
| 0 | 1 | 0 |
| 0 | 0 | 0 |
| 1 | 1 | 1 |
| 0 | 0 | 0 |
| 0 | 0 | 0 |
| 0 | 0 | 0 |
| 0 | 0 | 0 |
| 0 | 0 | 0 |
| 0 | 0 | 0 |
| 0 | 0 | 0 |
| 0 | 0 | 0 |
| 0 | 0 | 0 |
| 0 | 0 | 0 |
| 0 | 0 | 0 |
| 0 | 0 | 0 |
| 1 | 0 | 0 |
| 1 | 0 | 0 |
| 0 | 0 | 0 |
| 0 | 0 | 0 |
| 0 | 0 | 0 |
| 0 | 0 | 0 |
| 1 | 0 | 0 |
| 0 | 0 | 0 |
| 0 | 0 | 0 |
| 0 | 0 | 0 |
| 1 | 0 | 0 |
| 0 | 1 | 0 |
| 0 | 0 | 0 |
| 0 | 0 | 0 |
| 0 | 0 | 0 |
| 0 | 0 | 0 |
| 0 | 0 | 0 |
| 0 | 0 | 0 |
| 1 | 0 | 0 |
| 0 | 0 | 0 |
| 0 | 0 | 0 |
| 1 | 0 | 0 |

| | | |
|---|---|---|
| 0 | 0 | 0 |
| 1 | 0 | 0 |
| 0 | 0 | 0 |
| 0 | 0 | 0 |
| 0 | 0 | 0 |
| 0 | 0 | 0 |
| 0 | 0 | 0 |
| 0 | 0 | 0 |
| 0 | 0 | 0 |
| 0 | 0 | 0 |
| 0 | 0 | 0 |
| 1 | 1 | 1 |
| 0 | 0 | 0 |
| 1 | 1 | 1 |
| 1 | 0 | 0 |
| 1 | 0 | 0 |
| 0 | 0 | 0 |
| 0 | 0 | 0 |
| 0 | 0 | 0 |
| 0 | 0 | 0 |
| 1 | 0 | 0 |
| 0 | 0 | 0 |
| 0 | 1 | 0 |
| 0 | 0 | 0 |
| 1 | 1 | 1 |
| 1 | 1 | 1 |
| 1 | 1 | 1 |
| 0 | 0 | 0 |
| 1 | 1 | 1 |
| 1 | 1 | 1 |
| 1 | 1 | 1 |
| 0 | 0 | 0 |
| 1 | 1 | 1 |
| 0 | 0 | 0 |
| 0 | 0 | 0 |
| 0 | 0 | 0 |
| 0 | 1 | 0 |
| 1 | 1 | 1 |
| 0 | 0 | 0 |
| 0 | 0 | 0 |
| 0 | 0 | 0 |
| 0 | 0 | 0 |
| 0 | 0 | 0 |
| 0 | 0 | 0 |
| 0 | 0 | 0 |
| 0 | 0 | 0 |
| 1 | 1 | 1 |
| 1 | 1 | 1 |
| 0 | 0 | 0 |
| 0 | 0 | 0 |
| 0 | 0 | 0 |
| 1 | 1 | 1 |
| 1 | 1 | 1 |
| 0 | 0 | 0 |

| | | |
|---|---|---|
| 0 | 0 | 0 |
| 0 | 1 | 0 |
| 0 | 1 | 0 |
| 1 | 1 | 1 |
| 0 | 0 | 0 |
| 0 | 0 | 0 |
| 0 | 0 | 0 |
| 0 | 0 | 0 |
| 0 | 0 | 0 |
| 0 | 0 | 0 |
| 1 | 1 | 1 |
| 0 | 0 | 0 |
| 0 | 0 | 0 |
| 0 | 0 | 0 |
| 1 | 0 | 0 |
| 1 | 1 | 1 |
| 0 | 0 | 0 |
| 1 | 0 | 0 |
| 0 | 0 | 0 |
| 0 | 0 | 0 |
| 1 | 0 | 0 |
| 0 | 0 | 0 |
| 0 | 0 | 0 |
| 1 | 0 | 0 |
| 0 | 0 | 0 |
| 0 | 0 | 0 |
| 0 | 0 | 0 |
| 1 | 0 | 0 |
| 0 | 0 | 0 |
| 0 | 0 | 0 |
| 0 | 0 | 0 |
| 1 | 1 | 1 |
| 1 | 1 | 1 |
| 1 | 1 | 1 |
| 0 | 1 | 0 |
| 0 | 1 | 0 |
| 0 | 1 | 0 |
| 1 | 1 | 1 |
| 1 | 1 | 1 |
| 1 | 1 | 1 |
| 0 | 1 | 0 |
| 1 | 1 | 1 |
| 0 | 0 | 0 |
| 0 | 1 | 0 |
| 1 | 1 | 1 |
| 1 | 1 | 1 |
| 0 | 1 | 0 |
| 1 | 1 | 1 |
| 1 | 1 | 1 |
| 1 | 1 | 1 |
| 1 | 1 | 1 |
| 1 | 1 | 1 |
| 1 | 1 | 1 |
| 1 | 1 | 1 |

| | | |
|---|---|---|
| 0 | 1 | 0 |
| 0 | 1 | 0 |
| 1 | 1 | 1 |
| 1 | 1 | 1 |
| 1 | 1 | 1 |
| 1 | 1 | 1 |
| 1 | 1 | 1 |
| 1 | 1 | 1 |
| 0 | 0 | 0 |
| 0 | 0 | 0 |
| 0 | 0 | 0 |
| 0 | 0 | 0 |
| 0 | 0 | 0 |
| 0 | 0 | 0 |
| 0 | 0 | 0 |
| 0 | 0 | 0 |
| 0 | 0 | 0 |
| 0 | 0 | 0 |
| 0 | 0 | 0 |
| 0 | 0 | 0 |
| 0 | 0 | 0 |
| 0 | 0 | 0 |
| 0 | 0 | 0 |
| 0 | 0 | 0 |
| 0 | 0 | 0 |
| 1 | 1 | 1 |
| 1 | 1 | 1 |
| 0 | 0 | 0 |
| 0 | 0 | 0 |
| 0 | 0 | 0 |
| 0 | 0 | 0 |
| 0 | 0 | 0 |
| 0 | 0 | 0 |
| 0 | 0 | 0 |
| 0 | 0 | 0 |
| 0 | 0 | 0 |
| 0 | 0 | 0 |
| 0 | 0 | 0 |
| 0 | 0 | 0 |
| 0 | 0 | 0 |
| 0 | 0 | 0 |
| 0 | 0 | 0 |
| 0 | 0 | 0 |
| 0 | 0 | 0 |
| 0 | 0 | 0 |
| 0 | 0 | 0 |
| 1 | 1 | 1 |
| 0 | 1 | 0 |
| 1 | 1 | 1 |
| 1 | 1 | 1 |
| 1 | 1 | 1 |
| 1 | 1 | 1 |
| 0 | 0 | 0 |
| 0 | 0 | 0 |

| | | |
|---|---|---|
| 0 | 0 | 0 |
| 0 | 0 | 0 |
| 0 | 0 | 0 |
| 0 | 0 | 0 |
| 0 | 1 | 0 |
| 0 | 0 | 0 |
| 0 | 0 | 0 |
| 0 | 0 | 0 |
| 1 | 1 | 1 |
| 0 | 0 | 0 |
| 0 | 0 | 0 |
| 0 | 1 | 0 |
| 0 | 0 | 0 |
| 0 | 0 | 0 |
| 0 | 0 | 0 |
| 0 | 0 | 0 |
| 1 | 1 | 1 |
| 0 | 0 | 0 |
| 0 | 0 | 0 |
| 1 | 1 | 1 |
| 0 | 0 | 0 |
| 0 | 0 | 0 |
| 0 | 0 | 0 |
| 0 | 0 | 0 |
| 0 | 0 | 0 |
| 0 | 1 | 0 |
| 0 | 1 | 0 |
| 1 | 1 | 1 |
| 1 | 1 | 1 |
| 0 | 1 | 0 |
| 1 | 1 | 1 |
| 1 | 1 | 1 |
| 1 | 0 | 0 |
| 1 | 0 | 0 |
| 0 | 0 | 0 |
| 0 | 0 | 0 |
| 0 | 0 | 0 |
| 0 | 0 | 0 |
| 0 | 0 | 0 |
| 0 | 0 | 0 |
| 1 | 1 | 1 |
| 1 | 1 | 1 |
| 1 | 1 | 1 |
| 0 | 0 | 0 |
| 0 | 0 | 0 |
| 0 | 0 | 0 |
| 1 | 1 | 1 |
| 0 | 1 | 0 |
| 0 | 0 | 0 |
| 0 | 0 | 0 |
| 1 | 1 | 1 |
| 1 | 1 | 1 |
| 1 | 1 | 1 |
| 1 | 1 | 1 |

| | | |
|---|---|---|
| 0 | 1 | 0 |
| 1 | 1 | 1 |
| 0 | 1 | 0 |
| 0 | 0 | 0 |
| 0 | 0 | 0 |
| 1 | 1 | 1 |
| 1 | 1 | 1 |
| 1 | 1 | 1 |
| 1 | 1 | 1 |
| 1 | 1 | 1 |
| 0 | 0 | 0 |
| 0 | 1 | 0 |
| 0 | 0 | 0 |
| 0 | 0 | 0 |
| 0 | 0 | 0 |
| 1 | 1 | 1 |
| 1 | 1 | 1 |
| 0 | 1 | 0 |
| 1 | 1 | 1 |
| 1 | 1 | 1 |
| 0 | 0 | 0 |
| 1 | 1 | 1 |
| 1 | 1 | 1 |
| 0 | 0 | 0 |
| 1 | 0 | 0 |
| 0 | 0 | 0 |
| 0 | 0 | 0 |
| 1 | 1 | 1 |
| 1 | 1 | 1 |
| 1 | 1 | 1 |
| 1 | 1 | 1 |
| 1 | 1 | 1 |
| 1 | 1 | 1 |
| 1 | 1 | 1 |
| 1 | 1 | 1 |
| 0 | 0 | 0 |
| 1 | 1 | 1 |
| 1 | 1 | 1 |
| 0 | 0 | 0 |
| 0 | 0 | 0 |
| 1 | 1 | 1 |
| 0 | 1 | 0 |
| 0 | 0 | 0 |
| 0 | 0 | 0 |
| 0 | 0 | 0 |
| 0 | 0 | 0 |
| 0 | 0 | 0 |
| 0 | 0 | 0 |
| 1 | 1 | 1 |
| 0 | 0 | 0 |
| 0 | 0 | 0 |
| 0 | 0 | 0 |
| 0 | 0 | 0 |
| 0 | 0 | 0 |

| | | |
|---|---|---|
| 0 | 0 | 0 |
| 0 | 0 | 0 |
| 0 | 0 | 0 |
| 0 | 0 | 0 |
| 0 | 0 | 0 |
| 0 | 0 | 0 |
| 0 | 0 | 0 |
| 0 | 0 | 0 |
| 0 | 0 | 0 |
| 0 | 1 | 0 |
| 0 | 1 | 0 |
| 0 | 0 | 0 |
| 1 | 1 | 1 |
| 0 | 0 | 0 |
| 0 | 0 | 0 |
| 0 | 1 | 0 |
| 0 | 0 | 0 |
| 1 | 0 | 0 |
| 1 | 1 | 1 |
| 1 | 1 | 1 |
| 0 | 0 | 0 |
| 0 | 1 | 0 |
| 0 | 1 | 0 |
| 0 | 0 | 0 |
| 0 | 0 | 0 |
| 0 | 1 | 0 |
| 0 | 1 | 0 |
| 1 | 1 | 1 |
| 0 | 0 | 0 |
| 0 | 0 | 0 |
| 1 | 1 | 1 |
| 1 | 1 | 1 |
| 0 | 0 | 0 |
| 0 | 0 | 0 |
| 0 | 0 | 0 |
| 0 | 1 | 0 |
| 0 | 0 | 0 |
| 0 | 1 | 0 |
| 0 | 1 | 0 |
| 0 | 0 | 0 |
| 0 | 0 | 0 |
| 0 | 0 | 0 |
| 0 | 0 | 0 |
| 0 | 0 | 0 |
| 0 | 0 | 0 |
| 1 | 1 | 1 |
| 0 | 1 | 0 |
| 0 | 0 | 0 |
| 0 | 0 | 0 |
| 0 | 0 | 0 |
| 1 | 1 | 1 |
| 1 | 1 | 1 |
| 0 | 0 | 0 |
| 0 | 0 | 0 |

| | | |
|---|---|---|
| 1 | 1 | 1 |
| 1 | 1 | 1 |
| 1 | 1 | 1 |
| 1 | 1 | 1 |
| 0 | 0 | 0 |
| 0 | 0 | 0 |
| 1 | 0 | 0 |
| 0 | 0 | 0 |
| 1 | 0 | 0 |
| 0 | 0 | 0 |
| 1 | 1 | 1 |
| 1 | 1 | 1 |
| 1 | 1 | 1 |
| 1 | 1 | 1 |
| 1 | 1 | 1 |
| 1 | 1 | 1 |
| 0 | 0 | 0 |
| 1 | 1 | 1 |
| 1 | 1 | 1 |
| 0 | 0 | 0 |
| 0 | 0 | 0 |
| 0 | 1 | 0 |
| 0 | 1 | 0 |
| 0 | 0 | 0 |
| 0 | 1 | 0 |
| 0 | 0 | 0 |
| 0 | 0 | 0 |
| 0 | 0 | 0 |
| 0 | 0 | 0 |
| 0 | 0 | 0 |
| 0 | 0 | 0 |
| 0 | 0 | 0 |
| 0 | 1 | 0 |
| 0 | 0 | 0 |
| 1 | 1 | 1 |
| 1 | 1 | 1 |
| 1 | 1 | 1 |
| 1 | 1 | 1 |
| 0 | 0 | 0 |
| 1 | 1 | 1 |
| 1 | 1 | 1 |
| 1 | 1 | 1 |
| 1 | 1 | 1 |
| 0 | 0 | 0 |
| 1 | 1 | 1 |
| 1 | 1 | 1 |
| 1 | 1 | 1 |
| 1 | 1 | 1 |
| 1 | 1 | 1 |
| 1 | 1 | 1 |
| 0 | 0 | 0 |
| 0 | 0 | 0 |
| 0 | 0 | 0 |
| 0 | 1 | 0 |

| | | |
|---|---|---|
| 0 | 0 | 0 |
| 0 | 0 | 0 |
| 0 | 0 | 0 |
| 0 | 0 | 0 |
| 0 | 0 | 0 |
| 0 | 0 | 0 |
| 0 | 0 | 0 |
| 0 | 0 | 0 |
| 1 | 1 | 1 |
| 1 | 1 | 1 |
| 0 | 0 | 0 |
| 1 | 1 | 1 |
| 1 | 1 | 1 |
| 0 | 0 | 0 |
| 0 | 0 | 0 |
| 0 | 0 | 0 |
| 0 | 0 | 0 |
| 0 | 0 | 0 |
| 0 | 0 | 0 |
| 1 | 1 | 1 |
| 1 | 1 | 1 |
| 0 | 0 | 0 |
| 1 | 1 | 1 |
| 0 | 0 | 0 |
| 1 | 1 | 1 |
| 1 | 1 | 1 |
| 1 | 1 | 1 |
| 0 | 1 | 0 |
| 0 | 1 | 0 |
| 0 | 0 | 0 |
| 0 | 1 | 0 |
| 1 | 1 | 1 |
| 0 | 0 | 0 |
| 1 | 1 | 1 |
| 0 | 0 | 0 |
| 0 | 0 | 0 |
| 0 | 0 | 0 |
| 0 | 0 | 0 |
| 0 | 0 | 0 |
| 0 | 0 | 0 |
| 0 | 0 | 0 |
| 0 | 0 | 0 |
| 1 | 1 | 1 |
| 0 | 0 | 0 |
| 1 | 1 | 1 |
| 0 | 0 | 0 |
| 1 | 0 | 0 |
| 1 | 1 | 1 |
| 1 | 1 | 1 |
| 0 | 0 | 0 |
| 1 | 1 | 1 |
| 1 | 1 | 1 |
| 0 | 1 | 0 |
| 1 | 1 | 1 |

| | | |
|---|---|---|
| 0 | 0 | 0 |
| 0 | 1 | 0 |
| 1 | 1 | 1 |
| 0 | 1 | 0 |
| 0 | 1 | 0 |
| 0 | 1 | 0 |
| 0 | 1 | 0 |
| 1 | 1 | 1 |
| 1 | 1 | 1 |
| 1 | 1 | 1 |
| 0 | 0 | 0 |
| 1 | 1 | 1 |
| 1 | 1 | 1 |
| 0 | 1 | 0 |
| 0 | 0 | 0 |
| 0 | 0 | 0 |
| 0 | 0 | 0 |
| 0 | 0 | 0 |
| 0 | 0 | 0 |
| 0 | 1 | 0 |
| 0 | 1 | 0 |
| 1 | 1 | 1 |
| 0 | 0 | 0 |
| 0 | 1 | 0 |
| 0 | 1 | 0 |
| 0 | 1 | 0 |
| 1 | 1 | 1 |
| 1 | 1 | 1 |
| 0 | 0 | 0 |
| 0 | 0 | 0 |
| 0 | 0 | 0 |
| 0 | 0 | 0 |
| 0 | 0 | 0 |
| 0 | 0 | 0 |
| 1 | 1 | 1 |
| 1 | 1 | 1 |
| 0 | 0 | 0 |
| 0 | 0 | 0 |
| 0 | 0 | 0 |
| 0 | 0 | 0 |
| 0 | 0 | 0 |
| 0 | 0 | 0 |
| 1 | 1 | 1 |
| 1 | 1 | 1 |
| 0 | 0 | 0 |
| 1 | 1 | 1 |
| 1 | 1 | 1 |
| 1 | 1 | 1 |
| 1 | 1 | 1 |
| 1 | 1 | 1 |
| 1 | 0 | 0 |
| 0 | 1 | 0 |
| 0 | 1 | 0 |
| 1 | 0 | 0 |

| | | |
|---|---|---|
| 0 | 0 | 0 |
| 1 | 1 | 1 |
| 0 | 0 | 0 |
| 0 | 1 | 0 |
| 1 | 0 | 0 |
| 0 | 0 | 0 |
| 0 | 0 | 0 |
| 0 | 0 | 0 |
| 1 | 0 | 0 |
| 1 | 0 | 0 |
| 0 | 0 | 0 |
| 1 | 1 | 1 |
| 1 | 0 | 0 |
| 1 | 0 | 0 |
| 0 | 0 | 0 |
| 0 | 0 | 0 |
| 0 | 0 | 0 |
| 0 | 0 | 0 |
| 1 | 1 | 1 |
| 0 | 0 | 0 |
| 0 | 0 | 0 |
| 0 | 0 | 0 |
| 0 | 0 | 0 |
| 0 | 0 | 0 |
| 1 | 1 | 1 |
| 0 | 0 | 0 |
| 1 | 1 | 1 |
| 0 | 1 | 0 |
| 1 | 1 | 1 |
| 0 | 0 | 0 |
| 1 | 1 | 1 |
| 1 | 1 | 1 |
| 0 | 0 | 0 |
| 0 | 0 | 0 |
| 1 | 1 | 1 |
| 1 | 1 | 1 |
| 1 | 1 | 1 |
| 0 | 1 | 0 |
| 1 | 0 | 0 |
| 0 | 0 | 0 |
| 0 | 0 | 0 |
| 0 | 0 | 0 |
| 0 | 0 | 0 |
| 0 | 0 | 0 |
| 0 | 0 | 0 |
| 0 | 1 | 0 |
| 0 | 1 | 0 |
| 0 | 0 | 0 |
| 0 | 0 | 0 |
| 1 | 1 | 1 |
| 0 | 0 | 0 |
| 0 | 0 | 0 |
| 0 | 1 | 0 |
| 1 | 1 | 1 |

| | | |
|---|---|---|
| 1 | 1 | 1 |
| 0 | 0 | 0 |
| 0 | 0 | 0 |
| 1 | 0 | 0 |
| 1 | 1 | 1 |
| 0 | 0 | 0 |
| 0 | 0 | 0 |
| 0 | 0 | 0 |
| 0 | 0 | 0 |
| 0 | 0 | 0 |
| 0 | 0 | 0 |
| 0 | 0 | 0 |
| 0 | 0 | 0 |
| 0 | 0 | 0 |
| 0 | 0 | 0 |
| 0 | 1 | 0 |
| 0 | 0 | 0 |
| 1 | 1 | 1 |
| 1 | 0 | 0 |
| 1 | 1 | 1 |
| 0 | 0 | 0 |
| 1 | 1 | 1 |
| 0 | 0 | 0 |
| 0 | 0 | 0 |
| 1 | 1 | 1 |
| 0 | 0 | 0 |
| 0 | 1 | 0 |
| 0 | 1 | 0 |
| 0 | 1 | 0 |
| 0 | 0 | 0 |
| 0 | 0 | 0 |
| 0 | 0 | 0 |
| 0 | 0 | 0 |
| 0 | 0 | 0 |
| 0 | 0 | 0 |
| 0 | 0 | 0 |
| 0 | 0 | 0 |
| 1 | 1 | 1 |
| 1 | 0 | 0 |
| 0 | 0 | 0 |
| 0 | 0 | 0 |
| 0 | 0 | 0 |
| 0 | 0 | 0 |
| 0 | 0 | 0 |
| 0 | 0 | 0 |
| 0 | 0 | 0 |
| 1 | 1 | 1 |
| 0 | 0 | 0 |
| 0 | 0 | 0 |
| 0 | 0 | 0 |
| 0 | 0 | 0 |
| 0 | 1 | 0 |
| 0 | 0 | 0 |
| 0 | 0 | 0 |

| | | |
|---|---|---|
| 0 | 0 | 0 |
| 0 | 0 | 0 |
| 0 | 0 | 0 |
| 0 | 0 | 0 |
| 0 | 0 | 0 |
| 0 | 0 | 0 |
| 0 | 0 | 0 |
| 0 | 0 | 0 |
| 0 | 0 | 0 |
| 0 | 0 | 0 |
| 1 | 1 | 1 |
| 0 | 0 | 0 |
| 1 | 0 | 0 |
| 0 | 0 | 0 |
| 0 | 1 | 0 |
| 1 | 1 | 1 |
| 1 | 1 | 1 |
| 1 | 1 | 1 |
| 0 | 0 | 0 |
| 0 | 0 | 0 |
| 1 | 1 | 1 |
| 0 | 0 | 0 |
| 0 | 0 | 0 |
| 1 | 1 | 1 |
| 0 | 0 | 0 |
| 0 | 0 | 0 |
| 0 | 0 | 0 |
| 1 | 0 | 0 |
| 0 | 0 | 0 |
| 0 | 0 | 0 |
| 1 | 0 | 0 |
| 0 | 0 | 0 |
| 1 | 0 | 0 |
| 1 | 0 | 0 |
| 0 | 0 | 0 |
| 0 | 0 | 0 |
| 0 | 0 | 0 |
| 1 | 0 | 0 |
| 1 | 0 | 0 |
| 0 | 0 | 0 |
| 0 | 0 | 0 |
| 0 | 0 | 0 |
| 0 | 0 | 0 |
| 0 | 0 | 0 |
| 0 | 0 | 0 |
| 0 | 0 | 0 |
| 0 | 0 | 0 |
| 0 | 0 | 0 |
| 0 | 0 | 0 |
| 0 | 0 | 0 |
| 0 | 0 | 0 |
| 0 | 0 | 0 |
| 0 | 0 | 0 |
| 0 | 0 | 0 |

| | | |
|---|---|---|
| 0 | 0 | 0 |
| 1 | 0 | 0 |
| 0 | 0 | 0 |
| 0 | 0 | 0 |
| 0 | 0 | 0 |
| 1 | 0 | 0 |
| 0 | 0 | 0 |
| 0 | 0 | 0 |
| 1 | 1 | 1 |
| 0 | 0 | 0 |
| 0 | 1 | 0 |
| 1 | 1 | 1 |
| 1 | 1 | 1 |
| 0 | 0 | 0 |
| 1 | 1 | 1 |
| 0 | 0 | 0 |
| 1 | 1 | 1 |
| 1 | 1 | 1 |
| 0 | 0 | 0 |
| 0 | 0 | 0 |
| 0 | 0 | 0 |
| 0 | 0 | 0 |
| 0 | 0 | 0 |
| 0 | 0 | 0 |
| 0 | 0 | 0 |
| 0 | 1 | 0 |
| 1 | 1 | 1 |
| 1 | 1 | 1 |
| 1 | 0 | 0 |
| 1 | 1 | 1 |
| 1 | 1 | 1 |
| 0 | 1 | 0 |
| 0 | 0 | 0 |
| 0 | 0 | 0 |
| 0 | 0 | 0 |
| 0 | 0 | 0 |
| 0 | 0 | 0 |
| 0 | 0 | 0 |
| 0 | 1 | 0 |
| 0 | 1 | 0 |
| 0 | 1 | 0 |
| 0 | 1 | 0 |
| 0 | 1 | 0 |
| 0 | 1 | 0 |
| 0 | 0 | 0 |
| 0 | 0 | 0 |
| 0 | 1 | 0 |
| 0 | 0 | 0 |
| 0 | 0 | 0 |
| 0 | 0 | 0 |
| 0 | 0 | 0 |
| 0 | 0 | 0 |
| 0 | 0 | 0 |
| 0 | 0 | 0 |

| | | |
|---|---|---|
| 1 | 1 | 1 |
| 1 | 1 | 1 |
| 0 | 0 | 0 |
| 0 | 0 | 0 |
| 1 | 1 | 1 |
| 0 | 0 | 0 |
| 0 | 0 | 0 |
| 1 | 1 | 1 |
| 1 | 1 | 1 |
| 0 | 0 | 0 |
| 1 | 1 | 1 |
| 1 | 1 | 1 |
| 0 | 0 | 0 |
| 0 | 0 | 0 |
| 0 | 0 | 0 |
| 0 | 0 | 0 |
| 0 | 1 | 0 |
| 0 | 1 | 0 |
| 1 | 1 | 1 |
| 1 | 1 | 1 |
| 1 | 1 | 1 |
| 0 | 1 | 0 |
| 1 | 1 | 1 |
| 1 | 1 | 1 |
| 1 | 1 | 1 |
| 0 | 0 | 0 |
| 1 | 1 | 1 |
| 1 | 1 | 1 |
| 1 | 1 | 1 |
| 0 | 1 | 0 |
| 0 | 1 | 0 |
| 1 | 1 | 1 |
| 1 | 1 | 1 |
| 1 | 1 | 1 |
| 1 | 1 | 1 |
| 1 | 1 | 1 |
| 1 | 1 | 1 |
| 0 | 0 | 0 |
| 0 | 0 | 0 |
| 1 | 0 | 0 |
| 0 | 0 | 0 |
| 1 | 1 | 1 |
| 1 | 1 | 1 |
| 0 | 0 | 0 |
| 0 | 0 | 0 |
| 0 | 1 | 0 |
| 0 | 1 | 0 |
| 0 | 1 | 0 |
| 1 | 1 | 1 |
| 1 | 1 | 1 |
| 1 | 1 | 1 |
| 0 | 0 | 0 |
| 0 | 0 | 0 |
| 0 | 0 | 0 |

| | | |
|---|---|---|
| 0 | 0 | 0 |
| 1 | 1 | 1 |
| 1 | 1 | 1 |
| 1 | 1 | 1 |
| 1 | 1 | 1 |
| 1 | 1 | 1 |
| 1 | 1 | 1 |
| 1 | 1 | 1 |
| 0 | 0 | 0 |
| 1 | 1 | 1 |
| 1 | 1 | 1 |
| 1 | 1 | 1 |
| 0 | 1 | 0 |
| 0 | 1 | 0 |
| 1 | 1 | 1 |
| 1 | 1 | 1 |
| 0 | 1 | 0 |
| 0 | 1 | 0 |
| 0 | 0 | 0 |
| 0 | 0 | 0 |
| 0 | 0 | 0 |
| 0 | 0 | 0 |
| 0 | 1 | 0 |
| 0 | 0 | 0 |
| 0 | 0 | 0 |
| 1 | 1 | 1 |
| 1 | 1 | 1 |
| 1 | 1 | 1 |
| 1 | 1 | 1 |
| 1 | 1 | 1 |
| 1 | 1 | 1 |
| 1 | 1 | 1 |
| 0 | 0 | 0 |
| 1 | 1 | 1 |
| 1 | 1 | 1 |
| 0 | 1 | 0 |
| 0 | 0 | 0 |
| 1 | 1 | 1 |
| 0 | 0 | 0 |
| 0 | 0 | 0 |
| 0 | 0 | 0 |
| 0 | 0 | 0 |
| 0 | 0 | 0 |
| 1 | 1 | 1 |
| 1 | 1 | 1 |
| 1 | 1 | 1 |
| 0 | 0 | 0 |
| 0 | 0 | 0 |
| 0 | 0 | 0 |
| 0 | 0 | 0 |
| 0 | 0 | 0 |
| 0 | 0 | 0 |
| 0 | 0 | 0 |
| 0 | 1 | 0 |

| | | |
|---|---|---|
| 0 | 0 | 0 |
| 0 | 0 | 0 |
| 0 | 1 | 0 |
| 0 | 0 | 0 |
| 0 | 0 | 0 |
| 1 | 1 | 1 |
| 1 | 1 | 1 |
| 1 | 0 | 0 |
| 1 | 1 | 1 |
| 1 | 1 | 1 |
| 1 | 1 | 1 |
| 0 | 0 | 0 |
| 1 | 1 | 1 |
| 1 | 1 | 1 |
| 0 | 0 | 0 |
| 1 | 1 | 1 |
| 0 | 0 | 0 |
| 1 | 1 | 1 |
| 0 | 0 | 0 |
| 0 | 0 | 0 |
| 1 | 1 | 1 |
| 1 | 1 | 1 |
| 0 | 0 | 0 |
| 1 | 1 | 1 |
| 0 | 0 | 0 |
| 1 | 1 | 1 |
| 1 | 1 | 1 |
| 0 | 0 | 0 |
| 1 | 1 | 1 |
| 0 | 0 | 0 |
| 0 | 1 | 0 |
| 0 | 0 | 0 |
| 0 | 0 | 0 |
| 0 | 0 | 0 |
| 1 | 1 | 1 |
| 0 | 0 | 0 |
| 1 | 1 | 1 |
| 1 | 1 | 1 |
| 0 | 0 | 0 |
| 1 | 1 | 1 |
| 1 | 1 | 1 |
| 1 | 1 | 1 |
| 1 | 1 | 1 |
| 0 | 0 | 0 |
| 0 | 0 | 0 |
| 1 | 1 | 1 |
| 1 | 1 | 1 |
| 1 | 0 | 0 |
| 0 | 0 | 0 |
| 1 | 0 | 0 |
| 0 | 0 | 0 |
| 1 | 1 | 1 |
| 1 | 1 | 1 |
| 0 | 0 | 0 |

| | | |
|---|---|---|
| 1 | 1 | 1 |
| 1 | 1 | 1 |
| 1 | 1 | 1 |
| 1 | 1 | 1 |
| 0 | 1 | 0 |
| 1 | 1 | 1 |
| 1 | 1 | 1 |
| 0 | 0 | 0 |
| 1 | 1 | 1 |
| 1 | 1 | 1 |
| 1 | 1 | 1 |
| 1 | 1 | 1 |
| 0 | 0 | 0 |
| 1 | 1 | 1 |
| 0 | 0 | 0 |
| 0 | 0 | 0 |
| 1 | 1 | 1 |
| 1 | 1 | 1 |
| 1 | 1 | 1 |
| 1 | 1 | 1 |
| 0 | 0 | 0 |
| 0 | 0 | 0 |
| 1 | 1 | 1 |
| 1 | 1 | 1 |
| 0 | 0 | 0 |
| 1 | 1 | 1 |
| 0 | 0 | 0 |
| 0 | 0 | 0 |
| 0 | 0 | 0 |
| 0 | 0 | 0 |
| 0 | 1 | 0 |
| 0 | 1 | 0 |
| 0 | 0 | 0 |
| 0 | 0 | 0 |
| 0 | 0 | 0 |
| 0 | 0 | 0 |
| 0 | 0 | 0 |
| 0 | 0 | 0 |
| 0 | 0 | 0 |
| 1 | 0 | 0 |
| 1 | 1 | 1 |
| 0 | 1 | 0 |
| 0 | 1 | 0 |
| 0 | 1 | 0 |
| 0 | 1 | 0 |
| 0 | 0 | 0 |
| 0 | 1 | 0 |
| 0 | 1 | 0 |
| 1 | 1 | 1 |
| 0 | 0 | 0 |
| 0 | 0 | 0 |
| 1 | 1 | 1 |
| 1 | 1 | 1 |
| 1 | 1 | 1 |

| | | |
|---|---|---|
| 1 | 1 | 1 |
| 1 | 1 | 1 |
| 1 | 1 | 1 |
| 1 | 1 | 1 |
| 0 | 0 | 0 |
| 1 | 1 | 1 |
| 0 | 0 | 0 |
| 0 | 0 | 0 |
| 0 | 0 | 0 |
| 0 | 0 | 0 |
| 0 | 0 | 0 |
| 1 | 1 | 1 |
| 0 | 0 | 0 |
| 0 | 0 | 0 |
| 1 | 1 | 1 |
| 1 | 1 | 1 |
| 1 | 1 | 1 |
| 1 | 1 | 1 |
| 1 | 1 | 1 |
| 1 | 1 | 1 |
| 1 | 1 | 1 |
| 1 | 1 | 1 |
| 1 | 1 | 1 |
| 0 | 0 | 0 |
| 1 | 1 | 1 |
| 1 | 1 | 1 |
| 1 | 1 | 1 |
| 0 | 0 | 0 |
| 1 | 0 | 0 |
| 1 | 0 | 0 |
| 0 | 0 | 0 |
| 0 | 0 | 0 |
| 1 | 1 | 1 |
| 0 | 0 | 0 |
| 0 | 1 | 0 |
| 1 | 1 | 1 |
| 1 | 1 | 1 |
| 1 | 1 | 1 |
| 1 | 1 | 1 |
| 1 | 1 | 1 |
| 1 | 1 | 1 |
| 1 | 1 | 1 |
| 1 | 1 | 1 |
| 1 | 1 | 1 |
| 1 | 1 | 1 |
| 1 | 1 | 1 |
| 1 | 1 | 1 |
| 1 | 1 | 1 |
| 0 | 0 | 0 |
| 0 | 1 | 0 |
| 1 | 1 | 1 |
| 1 | 0 | 0 |
| 1 | 0 | 0 |
| 1 | 1 | 1 |

| | | |
|---|---|---|
| 1 | 1 | 1 |
| 0 | 1 | 0 |
| 1 | 1 | 1 |
| 0 | 1 | 0 |
| 1 | 0 | 0 |
| 1 | 1 | 1 |
| 1 | 1 | 1 |
| 1 | 1 | 1 |
| 1 | 1 | 1 |
| 1 | 1 | 1 |
| 0 | 0 | 0 |
| 0 | 0 | 0 |
| 0 | 0 | 0 |
| 0 | 0 | 0 |
| 0 | 0 | 0 |
| 1 | 0 | 0 |
| 0 | 0 | 0 |
| 0 | 0 | 0 |
| 0 | 0 | 0 |
| 0 | 0 | 0 |
| 0 | 0 | 0 |
| 0 | 0 | 0 |
| 0 | 1 | 0 |
| 0 | 0 | 0 |
| 0 | 0 | 0 |
| 0 | 0 | 0 |
| 1 | 1 | 1 |
| 0 | 0 | 0 |
| 0 | 1 | 0 |
| 0 | 1 | 0 |
| 0 | 1 | 0 |
| 0 | 0 | 0 |
| 1 | 1 | 1 |
| 1 | 1 | 1 |
| 0 | 0 | 0 |
| 0 | 0 | 0 |
| 0 | 0 | 0 |
| 0 | 0 | 0 |
| 0 | 0 | 0 |
| 1 | 0 | 0 |
| 1 | 0 | 0 |
| 0 | 0 | 0 |
| 1 | 0 | 0 |
| 0 | 0 | 0 |
| 1 | 1 | 1 |
| 1 | 0 | 0 |
| 1 | 1 | 1 |
| 0 | 0 | 0 |
| 0 | 0 | 0 |
| 0 | 0 | 0 |
| 0 | 0 | 0 |
| 1 | 1 | 1 |
| 0 | 0 | 0 |
| 1 | 1 | 1 |

| | | |
|---|---|---|
| 1 | 1 | 1 |
| 1 | 1 | 1 |
| 1 | 1 | 1 |
| 1 | 1 | 1 |
| 1 | 1 | 1 |
| 0 | 0 | 0 |
| 0 | 0 | 0 |
| 0 | 0 | 0 |
| 0 | 0 | 0 |
| 0 | 1 | 0 |
| 0 | 0 | 0 |
| 0 | 0 | 0 |
| 0 | 0 | 0 |
| 0 | 0 | 0 |
| 0 | 0 | 0 |
| 0 | 0 | 0 |
| 1 | 1 | 1 |
| 0 | 1 | 0 |
| 1 | 1 | 1 |
| 1 | 1 | 1 |
| 1 | 1 | 1 |
| 1 | 1 | 1 |
| 1 | 1 | 1 |
| 1 | 1 | 1 |
| 1 | 1 | 1 |
| 1 | 1 | 1 |
| 1 | 1 | 1 |
| 1 | 1 | 1 |
| 0 | 0 | 0 |
| 0 | 1 | 0 |
| 0 | 1 | 0 |
| 1 | 0 | 0 |
| 1 | 0 | 0 |
| 0 | 0 | 0 |
| 1 | 1 | 1 |
| 0 | 0 | 0 |
| 1 | 0 | 0 |
| 0 | 0 | 0 |
| 1 | 1 | 1 |
| 0 | 0 | 0 |
| 1 | 1 | 1 |
| 1 | 1 | 1 |
| 0 | 1 | 0 |
| 0 | 0 | 0 |
| 1 | 1 | 1 |
| 1 | 1 | 1 |
| 1 | 1 | 1 |
| 1 | 1 | 1 |
| 1 | 1 | 1 |
| 1 | 1 | 1 |
| 1 | 1 | 1 |
| 1 | 0 | 0 |
| 0 | 0 | 0 |
| 0 | 0 | 0 |

| | | |
|---|---|---|
| 1 | 1 | 1 |
| 0 | 0 | 0 |
| 1 | 1 | 1 |
| 1 | 1 | 1 |
| 0 | 0 | 0 |
| 0 | 0 | 0 |
| 0 | 0 | 0 |
| 1 | 1 | 1 |
| 1 | 1 | 1 |
| 0 | 0 | 0 |
| 0 | 0 | 0 |
| 0 | 0 | 0 |
| 0 | 0 | 0 |
| 0 | 0 | 0 |
| 1 | 1 | 1 |
| 0 | 0 | 0 |
| 0 | 0 | 0 |
| 0 | 0 | 0 |
| 1 | 1 | 1 |
| 1 | 1 | 1 |
| 1 | 1 | 1 |
| 1 | 1 | 1 |
| 1 | 1 | 1 |
| 0 | 0 | 0 |
| 0 | 1 | 0 |
| 0 | 1 | 0 |
| 0 | 0 | 0 |
| 0 | 0 | 0 |
| 1 | 1 | 1 |
| 1 | 1 | 1 |
| 0 | 0 | 0 |
| 1 | 1 | 1 |
| 1 | 1 | 1 |
| 0 | 1 | 0 |
| 1 | 1 | 1 |
| 0 | 0 | 0 |
| 0 | 1 | 0 |
| 0 | 0 | 0 |
| 0 | 0 | 0 |
| 0 | 1 | 0 |
| 0 | 1 | 0 |
| 0 | 0 | 0 |
| 0 | 0 | 0 |
| 0 | 1 | 0 |
| 0 | 0 | 0 |
| 0 | 0 | 0 |
| 0 | 0 | 0 |
| 0 | 1 | 0 |
| 0 | 0 | 0 |
| 0 | 0 | 0 |
| 0 | 0 | 0 |
| 1 | 1 | 1 |
| 0 | 0 | 0 |
| 1 | 1 | 1 |

| | | |
|---|---|---|
| 1 | 1 | 1 |
| 1 | 1 | 1 |
| 1 | 1 | 1 |
| 1 | 1 | 1 |
| 0 | 0 | 0 |
| 0 | 0 | 0 |
| 1 | 1 | 1 |
| 1 | 0 | 0 |
| 1 | 1 | 1 |
| 0 | 0 | 0 |
| 1 | 0 | 0 |
| 1 | 0 | 0 |
| 0 | 0 | 0 |
| 1 | 1 | 1 |
| 1 | 1 | 1 |
| 1 | 1 | 1 |
| 1 | 1 | 1 |
| 1 | 1 | 1 |
| 1 | 1 | 1 |
| 1 | 1 | 1 |
| 1 | 1 | 1 |
| 0 | 1 | 0 |
| 1 | 1 | 1 |
| 0 | 0 | 0 |
| 1 | 1 | 1 |
| 0 | 0 | 0 |
| 1 | 1 | 1 |
| 0 | 0 | 0 |
| 1 | 1 | 1 |
| 0 | 0 | 0 |
| 0 | 0 | 0 |
| 0 | 0 | 0 |
| 0 | 0 | 0 |
| 0 | 0 | 0 |
| 0 | 1 | 0 |
| 0 | 0 | 0 |
| 1 | 1 | 1 |
| 0 | 0 | 0 |
| 1 | 1 | 1 |
| 1 | 1 | 1 |
| 0 | 0 | 0 |
| 0 | 0 | 0 |
| 0 | 0 | 0 |
| 0 | 0 | 0 |
| 0 | 0 | 0 |
| 0 | 0 | 0 |
| 0 | 0 | 0 |
| 0 | 0 | 0 |
| 1 | 1 | 1 |
| 0 | 0 | 0 |
| 1 | 0 | 0 |
| 1 | 0 | 0 |
| 1 | 1 | 1 |
| 1 | 1 | 1 |

| | | |
|---|---|---|
| 1 | 1 | 1 |
| 1 | 1 | 1 |
| 1 | 1 | 1 |
| 0 | 1 | 0 |
| 0 | 1 | 0 |
| 0 | 1 | 0 |
| 0 | 0 | 0 |
| 0 | 0 | 0 |
| 0 | 0 | 0 |
| 0 | 0 | 0 |
| 0 | 0 | 0 |
| 0 | 0 | 0 |
| 0 | 0 | 0 |
| 0 | 0 | 0 |
| 0 | 0 | 0 |
| 0 | 0 | 0 |
| 0 | 0 | 0 |
| 0 | 0 | 0 |
| 1 | 1 | 1 |
| 0 | 0 | 0 |
| 0 | 1 | 0 |
| 0 | 1 | 0 |
| 0 | 1 | 0 |
| 1 | 1 | 1 |
| 0 | 1 | 0 |
| 0 | 0 | 0 |
| 0 | 0 | 0 |
| 0 | 0 | 0 |
| 0 | 0 | 0 |
| 0 | 0 | 0 |
| 0 | 0 | 0 |
| 0 | 0 | 0 |
| 0 | 0 | 0 |
| 0 | 0 | 0 |
| 1 | 1 | 1 |
| 0 | 0 | 0 |
| 0 | 0 | 0 |
| 0 | 0 | 0 |
| 0 | 0 | 0 |
| 0 | 1 | 0 |
| 0 | 0 | 0 |
| 0 | 0 | 0 |
| 1 | 1 | 1 |
| 0 | 1 | 0 |
| 1 | 1 | 1 |
| 0 | 1 | 0 |
| 0 | 1 | 0 |
| 0 | 0 | 0 |
| 0 | 0 | 0 |
| 0 | 0 | 0 |
| 0 | 0 | 0 |
| 0 | 1 | 0 |
| 0 | 0 | 0 |
| 1 | 1 | 1 |

| | | |
|---|---|---|
| 0 | 1 | 0 |
| 1 | 1 | 1 |
| 1 | 1 | 1 |
| 1 | 1 | 1 |
| 1 | 1 | 1 |
| 0 | 0 | 0 |
| 0 | 1 | 0 |
| 1 | 1 | 1 |
| 1 | 1 | 1 |
| 1 | 1 | 1 |
| 0 | 0 | 0 |
| 0 | 0 | 0 |
| 0 | 0 | 0 |
| 0 | 0 | 0 |
| 0 | 0 | 0 |
| 0 | 1 | 0 |
| 1 | 0 | 0 |
| 0 | 0 | 0 |
| 1 | 1 | 1 |
| 1 | 1 | 1 |
| 1 | 1 | 1 |
| 1 | 1 | 1 |
| 0 | 0 | 0 |
| 1 | 1 | 1 |
| 0 | 0 | 0 |
| 0 | 0 | 0 |
| 0 | 0 | 0 |
| 0 | 0 | 0 |
| 1 | 1 | 1 |
| 0 | 0 | 0 |
| 1 | 1 | 1 |
| 0 | 1 | 0 |
| 0 | 1 | 0 |
| 0 | 1 | 0 |
| 1 | 1 | 1 |
| 1 | 1 | 1 |
| 0 | 1 | 0 |
| 0 | 0 | 0 |
| 0 | 0 | 0 |
| 0 | 0 | 0 |
| 0 | 0 | 0 |
| 0 | 0 | 0 |
| 1 | 1 | 1 |
| 0 | 0 | 0 |
| 0 | 0 | 0 |
| 1 | 1 | 1 |
| 0 | 0 | 0 |
| 0 | 0 | 0 |
| 0 | 0 | 0 |
| 0 | 0 | 0 |
| 1 | 1 | 1 |
| 1 | 1 | 1 |
| 1 | 1 | 1 |
| 0 | 0 | 0 |

| | | |
|---|---|---|
| 0 | 0 | 0 |
| 1 | 1 | 1 |
| 0 | 0 | 0 |
| 0 | 1 | 0 |
| 1 | 0 | 0 |
| 1 | 0 | 0 |
| 1 | 0 | 0 |
| 1 | 1 | 1 |
| 0 | 0 | 0 |
| 1 | 1 | 1 |
| 0 | 0 | 0 |
| 0 | 0 | 0 |
| 0 | 0 | 0 |
| 0 | 0 | 0 |
| 1 | 1 | 1 |
| 1 | 1 | 1 |
| 0 | 0 | 0 |
| 0 | 0 | 0 |
| 0 | 1 | 0 |
| 1 | 1 | 1 |
| 0 | 0 | 0 |
| 1 | 0 | 0 |
| 1 | 0 | 0 |
| 1 | 1 | 1 |
| 0 | 0 | 0 |
| 1 | 0 | 0 |
| 0 | 0 | 0 |
| 0 | 0 | 0 |
| 0 | 0 | 0 |
| 1 | 1 | 1 |
| 0 | 0 | 0 |
| 0 | 0 | 0 |
| 0 | 0 | 0 |
| 1 | 1 | 1 |
| 0 | 0 | 0 |
| 1 | 1 | 1 |
| 0 | 0 | 0 |
| 0 | 0 | 0 |
| 0 | 0 | 0 |
| 0 | 0 | 0 |
| 0 | 0 | 0 |
| 1 | 1 | 1 |
| 0 | 0 | 0 |
| 0 | 0 | 0 |
| 0 | 0 | 0 |
| 0 | 0 | 0 |
| 0 | 0 | 0 |
| 0 | 0 | 0 |
| 0 | 0 | 0 |
| 0 | 0 | 0 |
| 0 | 0 | 0 |
| 0 | 0 | 0 |
| 1 | 1 | 1 |
| 0 | 0 | 0 |

| | | |
|---|---|---|
| 0 | 1 | 0 |
| 1 | 1 | 1 |
| 1 | 1 | 1 |
| 1 | 1 | 1 |
| 1 | 1 | 1 |
| 1 | 1 | 1 |
| 1 | 1 | 1 |
| 1 | 1 | 1 |
| 1 | 1 | 1 |
| 0 | 0 | 0 |
| 0 | 0 | 0 |
| 1 | 1 | 1 |
| 1 | 1 | 1 |
| 1 | 1 | 1 |
| 0 | 0 | 0 |
| 1 | 1 | 1 |
| 1 | 1 | 1 |
| 0 | 0 | 0 |
| 0 | 1 | 0 |
| 0 | 1 | 0 |
| 0 | 1 | 0 |
| 0 | 0 | 0 |
| 0 | 0 | 0 |
| 0 | 0 | 0 |
| 0 | 0 | 0 |
| 0 | 0 | 0 |
| 0 | 0 | 0 |
| 0 | 0 | 0 |
| 1 | 1 | 1 |
| 0 | 0 | 0 |
| 0 | 0 | 0 |
| 1 | 1 | 1 |
| 1 | 1 | 1 |
| 1 | 1 | 1 |
| 1 | 1 | 1 |
| 1 | 1 | 1 |
| 1 | 1 | 1 |
| 1 | 1 | 1 |
| 1 | 1 | 1 |
| 1 | 1 | 1 |
| 1 | 1 | 1 |
| 1 | 1 | 1 |
| 1 | 1 | 1 |
| 0 | 0 | 0 |
| 1 | 1 | 1 |
| 0 | 0 | 0 |
| 1 | 1 | 1 |
| 1 | 1 | 1 |
| 1 | 1 | 1 |
| 1 | 1 | 1 |
| 0 | 0 | 0 |
| 1 | 1 | 1 |
| 1 | 1 | 1 |
| 0 | 0 | 0 |

| | | |
|---|---|---|
| 0 | 0 | 0 |
| 0 | 0 | 0 |
| 0 | 0 | 0 |
| 0 | 0 | 0 |
| 1 | 1 | 1 |
| 0 | 1 | 0 |
| 1 | 1 | 1 |
| 0 | 0 | 0 |
| 0 | 0 | 0 |
| 0 | 0 | 0 |
| 0 | 0 | 0 |
| 1 | 1 | 1 |
| 1 | 1 | 1 |
| 1 | 1 | 1 |
| 1 | 1 | 1 |
| 0 | 0 | 0 |
| 0 | 0 | 0 |
| 0 | 1 | 0 |
| 0 | 1 | 0 |
| 1 | 1 | 1 |
| 1 | 1 | 1 |
| 0 | 0 | 0 |
| 1 | 1 | 1 |
| 0 | 1 | 0 |
| 0 | 0 | 0 |
| 0 | 0 | 0 |
| 0 | 0 | 0 |
| 0 | 0 | 0 |
| 1 | 1 | 1 |
| 1 | 1 | 1 |
| 1 | 1 | 1 |
| 0 | 0 | 0 |
| 1 | 1 | 1 |
| 1 | 1 | 1 |
| 1 | 1 | 1 |
| 1 | 1 | 1 |
| 1 | 1 | 1 |
| 1 | 1 | 1 |
| 1 | 1 | 1 |
| 1 | 1 | 1 |
| 1 | 1 | 1 |
| 1 | 1 | 1 |
| 0 | 0 | 0 |
| 0 | 1 | 0 |
| 0 | 1 | 0 |
| 1 | 1 | 1 |
| 0 | 1 | 0 |
| 1 | 1 | 1 |
| 1 | 1 | 1 |
| 0 | 0 | 0 |
| 1 | 1 | 1 |
| 1 | 1 | 1 |
| 1 | 1 | 1 |
| 1 | 1 | 1 |

| | | |
|---|---|---|
| 0 | 0 | 0 |
| 1 | 1 | 1 |
| 0 | 0 | 0 |
| 0 | 1 | 0 |
| 1 | 1 | 1 |
| 1 | 1 | 1 |
| 1 | 1 | 1 |
| 0 | 0 | 0 |
| 1 | 1 | 1 |
| 1 | 1 | 1 |
| 0 | 0 | 0 |
| 0 | 0 | 0 |
| 0 | 0 | 0 |
| 1 | 1 | 1 |
| 0 | 0 | 0 |
| 0 | 0 | 0 |
| 0 | 0 | 0 |
| 0 | 0 | 0 |
| 0 | 1 | 0 |
| 0 | 0 | 0 |
| 0 | 0 | 0 |
| 0 | 0 | 0 |
| 0 | 0 | 0 |
| 0 | 0 | 0 |
| 0 | 0 | 0 |
| 0 | 0 | 0 |
| 0 | 0 | 0 |
| 0 | 0 | 0 |
| 0 | 0 | 0 |
| 0 | 0 | 0 |
| 0 | 0 | 0 |
| 0 | 0 | 0 |
| 1 | 1 | 1 |
| 0 | 0 | 0 |
| 1 | 1 | 1 |
| 0 | 0 | 0 |
| 0 | 0 | 0 |
| 1 | 1 | 1 |
| 1 | 1 | 1 |
| 0 | 1 | 0 |
| 0 | 1 | 0 |
| 0 | 0 | 0 |
| 0 | 0 | 0 |
| 0 | 0 | 0 |
| 1 | 1 | 1 |
| 0 | 0 | 0 |
| 1 | 1 | 1 |
| 0 | 0 | 0 |
| 0 | 1 | 0 |
| 0 | 1 | 0 |
| 0 | 1 | 0 |
| 0 | 1 | 0 |
| 0 | 1 | 0 |
| 1 | 1 | 1 |

| | | |
|---|---|---|
| 1 | 1 | 1 |
| 1 | 1 | 1 |
| 0 | 1 | 0 |
| 1 | 1 | 1 |
| 1 | 1 | 1 |
| 0 | 0 | 0 |
| 0 | 0 | 0 |
| 1 | 1 | 1 |
| 0 | 0 | 0 |
| 0 | 0 | 0 |
| 1 | 1 | 1 |
| 0 | 1 | 0 |
| 1 | 1 | 1 |
| 1 | 1 | 1 |
| 1 | 1 | 1 |
| 0 | 0 | 0 |
| 1 | 1 | 1 |
| 1 | 1 | 1 |
| 0 | 0 | 0 |
| 0 | 0 | 0 |
| 0 | 0 | 0 |
| 1 | 1 | 1 |
| 1 | 1 | 1 |
| 0 | 0 | 0 |
| 1 | 1 | 1 |
| 0 | 0 | 0 |
| 0 | 0 | 0 |
| 1 | 1 | 1 |
| 1 | 1 | 1 |
| 1 | 1 | 1 |
| 0 | 0 | 0 |
| 1 | 0 | 0 |
| 0 | 0 | 0 |
| 0 | 0 | 0 |
| 0 | 0 | 0 |
| 0 | 0 | 0 |
| 0 | 0 | 0 |
| 1 | 1 | 1 |
| 0 | 0 | 0 |
| 0 | 1 | 0 |
| 0 | 0 | 0 |
| 0 | 0 | 0 |
| 0 | 0 | 0 |
| 0 | 0 | 0 |
| 1 | 1 | 1 |
| 0 | 1 | 0 |
| 0 | 1 | 0 |
| 0 | 0 | 0 |
| 0 | 0 | 0 |
| 0 | 0 | 0 |
| 1 | 1 | 1 |
| 0 | 1 | 0 |
| 1 | 1 | 1 |
| 1 | 1 | 1 |

| | | |
|---|---|---|
| 1 | 1 | 1 |
| 0 | 1 | 0 |
| 0 | 1 | 0 |
| 0 | 0 | 0 |
| 0 | 0 | 0 |
| 1 | 1 | 1 |
| 0 | 0 | 0 |
| 1 | 1 | 1 |
| 1 | 1 | 1 |
| 1 | 1 | 1 |
| 1 | 1 | 1 |
| 0 | 0 | 0 |
| 1 | 0 | 0 |
| 0 | 0 | 0 |
| 1 | 1 | 1 |
| 0 | 0 | 0 |
| 0 | 0 | 0 |
| 0 | 0 | 0 |
| 1 | 1 | 1 |
| 0 | 0 | 0 |
| 0 | 0 | 0 |
| 0 | 0 | 0 |
| 0 | 0 | 0 |
| 0 | 0 | 0 |
| 0 | 0 | 0 |
| 1 | 1 | 1 |
| 0 | 0 | 0 |
| 0 | 1 | 0 |
| 0 | 1 | 0 |
| 0 | 1 | 0 |
| 0 | 1 | 0 |
| 0 | 1 | 0 |
| 0 | 0 | 0 |
| 0 | 0 | 0 |
| 1 | 1 | 1 |
| 1 | 1 | 1 |
| 0 | 0 | 0 |
| 0 | 0 | 0 |
| 0 | 0 | 0 |
| 1 | 0 | 0 |
| 0 | 0 | 0 |
| 0 | 0 | 0 |
| 1 | 1 | 1 |
| 0 | 0 | 0 |
| 1 | 1 | 1 |
| 1 | 1 | 1 |
| 1 | 1 | 1 |
| 1 | 1 | 1 |
| 1 | 1 | 1 |
| 0 | 0 | 0 |
| 1 | 1 | 1 |
| 1 | 1 | 1 |
| 1 | 1 | 1 |
| 0 | 1 | 0 |

| | | |
|---|---|---|
| 1 | 1 | 1 |
| 1 | 1 | 1 |
| 1 | 1 | 1 |
| 0 | 0 | 0 |
| 1 | 1 | 1 |
| 1 | 1 | 1 |
| 0 | 1 | 0 |
| 0 | 0 | 0 |
| 1 | 1 | 1 |
| 1 | 1 | 1 |
| 1 | 1 | 1 |
| 1 | 1 | 1 |
| 1 | 1 | 1 |
| 0 | 1 | 0 |
| 0 | 1 | 0 |
| 0 | 1 | 0 |
| 0 | 0 | 0 |
| 0 | 0 | 0 |
| 0 | 1 | 0 |
| 1 | 1 | 1 |
| 0 | 1 | 0 |
| 1 | 1 | 1 |
| 0 | 1 | 0 |
| 1 | 1 | 1 |
| 1 | 1 | 1 |
| 1 | 1 | 1 |
| 1 | 1 | 1 |
| 0 | 0 | 0 |
| 0 | 0 | 0 |
| 1 | 1 | 1 |
| 1 | 1 | 1 |
| 0 | 0 | 0 |
| 0 | 1 | 0 |
| 0 | 0 | 0 |
| 0 | 0 | 0 |
| 0 | 0 | 0 |
| 0 | 0 | 0 |
| 0 | 0 | 0 |
| 1 | 1 | 1 |
| 1 | 1 | 1 |
| 1 | 1 | 1 |
| 0 | 1 | 0 |
| 1 | 1 | 1 |
| 1 | 1 | 1 |
| 0 | 0 | 0 |
| 0 | 0 | 0 |
| 0 | 0 | 0 |
| 1 | 1 | 1 |
| 1 | 1 | 1 |
| 1 | 1 | 1 |
| 1 | 1 | 1 |
| 0 | 0 | 0 |
| 0 | 0 | 0 |
| 0 | 0 | 0 |

| | | |
|---|---|---|
| 0 | 0 | 0 |
| 0 | 0 | 0 |
| 1 | 1 | 1 |
| 0 | 0 | 0 |
| 0 | 0 | 0 |
| 0 | 0 | 0 |
| 0 | 0 | 0 |
| 1 | 1 | 1 |
| 1 | 1 | 1 |
| 1 | 1 | 1 |
| 1 | 1 | 1 |
| 1 | 1 | 1 |
| 0 | 1 | 0 |
| 0 | 1 | 0 |
| 1 | 1 | 1 |
| 1 | 1 | 1 |
| 1 | 1 | 1 |
| 1 | 1 | 1 |
| 1 | 1 | 1 |
| 1 | 1 | 1 |
| 0 | 0 | 0 |
| 0 | 0 | 0 |
| 0 | 0 | 0 |
| 0 | 0 | 0 |
| 0 | 0 | 0 |
| 1 | 1 | 1 |
| 1 | 1 | 1 |
| 0 | 0 | 0 |
| 0 | 1 | 0 |
| 0 | 1 | 0 |
| 0 | 0 | 0 |
| 0 | 0 | 0 |
| 0 | 0 | 0 |
| 1 | 1 | 1 |
| 0 | 0 | 0 |
| 1 | 1 | 1 |
| 0 | 1 | 0 |
| 0 | 0 | 0 |
| 1 | 1 | 1 |
| 1 | 1 | 1 |
| 0 | 1 | 0 |
| 0 | 1 | 0 |
| 1 | 1 | 1 |
| 1 | 1 | 1 |
| 1 | 1 | 1 |
| 1 | 1 | 1 |
| 1 | 1 | 1 |
| 1 | 1 | 1 |
| 0 | 0 | 0 |
| 1 | 1 | 1 |
| 0 | 0 | 0 |
| 0 | 0 | 0 |
| 1 | 1 | 1 |
| 0 | 1 | 0 |

| | | |
|---|---|---|
| 0 | 0 | 0 |
| 1 | 1 | 1 |
| 1 | 1 | 1 |
| 1 | 1 | 1 |
| 1 | 1 | 1 |
| 0 | 0 | 0 |
| 0 | 0 | 0 |
| 1 | 1 | 1 |
| 0 | 1 | 0 |
| 1 | 1 | 1 |
| 1 | 1 | 1 |
| 0 | 0 | 0 |
| 0 | 1 | 0 |
| 1 | 1 | 1 |
| 0 | 1 | 0 |
| 1 | 1 | 1 |
| 0 | 0 | 0 |
| 0 | 0 | 0 |
| 1 | 1 | 1 |
| 1 | 1 | 1 |
| 1 | 1 | 1 |
| 1 | 1 | 1 |
| 1 | 1 | 1 |
| 1 | 1 | 1 |
| 0 | 0 | 0 |
| 1 | 1 | 1 |
| 1 | 1 | 1 |
| 1 | 1 | 1 |
| 1 | 1 | 1 |
| 1 | 1 | 1 |
| 0 | 1 | 0 |
| 0 | 0 | 0 |
| 0 | 0 | 0 |
| 0 | 1 | 0 |
| 1 | 1 | 1 |
| 0 | 0 | 0 |
| 0 | 0 | 0 |
| 0 | 0 | 0 |
| 0 | 0 | 0 |
| 0 | 0 | 0 |
| 0 | 0 | 0 |
| 0 | 0 | 0 |
| 0 | 0 | 0 |
| 0 | 0 | 0 |
| 0 | 0 | 0 |
| 0 | 0 | 0 |
| 0 | 0 | 0 |
| 0 | 0 | 0 |
| 1 | 0 | 0 |
| 1 | 1 | 1 |
| 1 | 0 | 0 |
| 1 | 1 | 1 |
| 0 | 0 | 0 |
| 1 | 1 | 1 |

| | | |
|---|---|---|
| 1 | 1 | 1 |
| 1 | 1 | 1 |
| 1 | 1 | 1 |
| 1 | 1 | 1 |
| 1 | 1 | 1 |
| 1 | 1 | 1 |
| 1 | 1 | 1 |
| 0 | 0 | 0 |
| 0 | 0 | 0 |
| 1 | 0 | 0 |
| 1 | 1 | 1 |
| 1 | 1 | 1 |
| 1 | 1 | 1 |
| 0 | 0 | 0 |
| 1 | 1 | 1 |
| 1 | 1 | 1 |
| 0 | 0 | 0 |
| 1 | 1 | 1 |
| 0 | 0 | 0 |
| 0 | 1 | 0 |
| 1 | 1 | 1 |
| 1 | 1 | 1 |
| 1 | 1 | 1 |
| 0 | 0 | 0 |
| 1 | 1 | 1 |
| 1 | 1 | 1 |
| 0 | 1 | 0 |
| 0 | 1 | 0 |
| 0 | 1 | 0 |
| 0 | 1 | 0 |
| 1 | 1 | 1 |
| 0 | 0 | 0 |
| 1 | 1 | 1 |
| 0 | 1 | 0 |
| 1 | 1 | 1 |
| 1 | 1 | 1 |
| 1 | 1 | 1 |
| 1 | 1 | 1 |
| 0 | 0 | 0 |
| 1 | 1 | 1 |
| 1 | 1 | 1 |
| 1 | 1 | 1 |
| 0 | 1 | 0 |
| 1 | 1 | 1 |
| 1 | 1 | 1 |
| 0 | 0 | 0 |
| 0 | 1 | 0 |
| 0 | 1 | 0 |
| 1 | 0 | 0 |
| 1 | 0 | 0 |
| 0 | 1 | 0 |
| 1 | 0 | 0 |
| 0 | 0 | 0 |
| 1 | 0 | 0 |

| | | |
|---|---|---|
| 1 | 0 | 0 |
| 1 | 0 | 0 |
| 1 | 0 | 0 |
| 1 | 0 | 0 |
| 0 | 0 | 0 |
| 0 | 0 | 0 |
| 0 | 0 | 0 |
| 0 | 0 | 0 |
| 1 | 0 | 0 |
| 1 | 0 | 0 |
| 1 | 0 | 0 |
| 1 | 0 | 0 |
| 1 | 0 | 0 |
| 0 | 0 | 0 |
| 1 | 0 | 0 |
| 0 | 0 | 0 |
| 1 | 0 | 0 |
| 0 | 0 | 0 |
| 1 | 0 | 0 |
| 0 | 0 | 0 |
| 0 | 0 | 0 |
| 1 | 0 | 0 |
| 1 | 0 | 0 |
| 0 | 0 | 0 |
| 0 | 0 | 0 |
| 1 | 0 | 0 |
| 0 | 0 | 0 |
| 0 | 0 | 0 |
| 0 | 0 | 0 |
| 1 | 0 | 0 |
| 1 | 0 | 0 |
| 0 | 0 | 0 |
| 1 | 0 | 0 |
| 1 | 0 | 0 |
| 1 | 0 | 0 |
| 1 | 0 | 0 |
| 1 | 0 | 0 |
| 1 | 0 | 0 |
| 1 | 0 | 0 |
| 1 | 0 | 0 |
| 1 | 0 | 0 |
| 1 | 0 | 0 |
| 1 | 0 | 0 |
| 0 | 0 | 0 |
| 1 | 1 | 1 |
| 1 | 1 | 1 |
| 0 | 0 | 0 |
| 0 | 0 | 0 |
| 0 | 1 | 0 |
| 0 | 0 | 0 |
| 0 | 0 | 0 |
| 0 | 0 | 0 |
| 1 | 1 | 1 |
| 1 | 1 | 1 |

| | | |
|---|---|---|
| 1 | 1 | 1 |
| 1 | 1 | 1 |
| 1 | 1 | 1 |
| 1 | 1 | 1 |
| 1 | 1 | 1 |
| 1 | 1 | 1 |
| 1 | 1 | 1 |
| 0 | 0 | 0 |
| 1 | 1 | 1 |
| 1 | 1 | 1 |
| 0 | 1 | 0 |
| 0 | 1 | 0 |
| 1 | 0 | 0 |
| 1 | 1 | 1 |
| 0 | 0 | 0 |
| 0 | 0 | 0 |
| 0 | 1 | 0 |
| 0 | 0 | 0 |
| 1 | 1 | 1 |
| 1 | 1 | 1 |
| 0 | 0 | 0 |
| 0 | 1 | 0 |
| 0 | 1 | 0 |
| 1 | 1 | 1 |
| 0 | 0 | 0 |
| 1 | 1 | 1 |
| 0 | 1 | 0 |
| 0 | 0 | 0 |
| 0 | 0 | 0 |
| 0 | 0 | 0 |
| 0 | 0 | 0 |
| 1 | 1 | 1 |
| 1 | 1 | 1 |
| 0 | 1 | 0 |
| 1 | 1 | 1 |
| 1 | 1 | 1 |
| 0 | 0 | 0 |
| 0 | 0 | 0 |
| 1 | 1 | 1 |
| 1 | 1 | 1 |
| 1 | 1 | 1 |
| 1 | 1 | 1 |
| 0 | 0 | 0 |
| 0 | 1 | 0 |
| 1 | 1 | 1 |
| 1 | 1 | 1 |
| 1 | 1 | 1 |
| 1 | 1 | 1 |
| 0 | 0 | 0 |
| 0 | 0 | 0 |
| 0 | 0 | 0 |
| 0 | 0 | 0 |
| 0 | 0 | 0 |
| 1 | 1 | 1 |

| | | |
|---|---|---|
| 0 | 0 | 0 |
| 0 | 0 | 0 |
| 0 | 1 | 0 |
| 0 | 0 | 0 |
| 1 | 1 | 1 |
| 0 | 1 | 0 |
| 0 | 1 | 0 |
| 1 | 1 | 1 |
| 1 | 1 | 1 |
| 0 | 0 | 0 |
| 1 | 1 | 1 |
| 0 | 1 | 0 |
| 1 | 1 | 1 |
| 0 | 1 | 0 |
| 0 | 1 | 0 |
| 0 | 1 | 0 |
| 0 | 1 | 0 |
| 1 | 1 | 1 |
| 1 | 1 | 1 |
| 0 | 1 | 0 |
| 0 | 1 | 0 |
| 0 | 1 | 0 |
| 0 | 1 | 0 |
| 0 | 1 | 0 |
| 0 | 1 | 0 |
| 0 | 1 | 0 |
| 0 | 1 | 0 |
| 0 | 1 | 0 |
| 0 | 1 | 0 |
| 0 | 1 | 0 |
| 1 | 1 | 1 |
| 1 | 1 | 1 |
| 0 | 0 | 0 |
| 1 | 1 | 1 |
| 1 | 1 | 1 |
| 1 | 1 | 1 |
| 1 | 1 | 1 |
| 0 | 1 | 0 |
| 1 | 1 | 1 |
| 1 | 1 | 1 |
| 1 | 1 | 1 |
| 1 | 1 | 1 |
| 1 | 1 | 1 |
| 0 | 1 | 0 |
| 0 | 1 | 0 |
| 1 | 1 | 1 |
| 1 | 1 | 1 |
| 0 | 0 | 0 |
| 0 | 0 | 0 |
| 0 | 0 | 0 |
| 0 | 0 | 0 |
| 0 | 0 | 0 |
| 1 | 1 | 1 |
| 0 | 0 | 0 |

| | | |
|---|---|---|
| 1 | 1 | 1 |
| 0 | 0 | 0 |
| 1 | 1 | 1 |
| 0 | 1 | 0 |
| 0 | 1 | 0 |
| 0 | 0 | 0 |
| 0 | 0 | 0 |
| 0 | 0 | 0 |
| 0 | 0 | 0 |
| 0 | 0 | 0 |
| 0 | 1 | 0 |
| 0 | 0 | 0 |
| 0 | 1 | 0 |
| 1 | 1 | 1 |
| 0 | 1 | 0 |
| 0 | 0 | 0 |
| 1 | 1 | 1 |
| 1 | 1 | 1 |
| 0 | 0 | 0 |
| 0 | 0 | 0 |
| 0 | 0 | 0 |
| 0 | 0 | 0 |
| 0 | 0 | 0 |
| 0 | 0 | 0 |
| 0 | 1 | 0 |
| 0 | 0 | 0 |
| 0 | 0 | 0 |
| 0 | 0 | 0 |
| 0 | 0 | 0 |
| 0 | 0 | 0 |
| 0 | 0 | 0 |
| 1 | 1 | 1 |
| 1 | 1 | 1 |
| 0 | 0 | 0 |
| 0 | 0 | 0 |
| 1 | 0 | 0 |
| 0 | 0 | 0 |
| 0 | 0 | 0 |
| 0 | 0 | 0 |
| 0 | 0 | 0 |
| 0 | 0 | 0 |
| 1 | 0 | 0 |
| 1 | 1 | 1 |
| 1 | 1 | 1 |
| 0 | 0 | 0 |
| 0 | 0 | 0 |
| 0 | 0 | 0 |
| 1 | 1 | 1 |
| 1 | 1 | 1 |
| 1 | 1 | 1 |
| 1 | 1 | 1 |
| 1 | 1 | 1 |
| 1 | 1 | 1 |
| 0 | 0 | 0 |

| | | |
|---|---|---|
| 1 | 1 | 1 |
| 1 | 1 | 1 |
| 1 | 1 | 1 |
| 1 | 1 | 1 |
| 0 | 0 | 0 |
| 0 | 0 | 0 |
| 1 | 0 | 0 |
| 1 | 0 | 0 |
| 0 | 1 | 0 |
| 1 | 1 | 1 |
| 1 | 1 | 1 |
| 0 | 0 | 0 |
| 1 | 1 | 1 |
| 0 | 0 | 0 |
| 0 | 0 | 0 |
| 0 | 0 | 0 |
| 0 | 0 | 0 |
| 0 | 0 | 0 |
| 0 | 0 | 0 |
| 0 | 0 | 0 |
| 0 | 0 | 0 |
| 0 | 0 | 0 |
| 0 | 0 | 0 |
| 0 | 1 | 0 |
| 0 | 1 | 0 |
| 0 | 1 | 0 |
| 1 | 1 | 1 |
| 1 | 1 | 1 |
| 1 | 1 | 1 |
| 1 | 1 | 1 |
| 1 | 1 | 1 |
| 0 | 0 | 0 |
| 1 | 1 | 1 |
| 0 | 0 | 0 |
| 0 | 0 | 0 |
| 0 | 0 | 0 |
| 0 | 0 | 0 |
| 1 | 1 | 1 |
| 1 | 1 | 1 |
| 0 | 1 | 0 |
| 1 | 1 | 1 |
| 1 | 1 | 1 |
| 1 | 1 | 1 |
| 0 | 1 | 0 |
| 1 | 1 | 1 |
| 0 | 0 | 0 |
| 1 | 1 | 1 |
| 1 | 1 | 1 |
| 1 | 1 | 1 |
| 0 | 0 | 0 |
| 0 | 0 | 0 |
| 0 | 0 | 0 |
| 0 | 1 | 0 |
| 0 | 1 | 0 |

| | | |
|---|---|---|
| 0 | 1 | 0 |
| 0 | 1 | 0 |
| 0 | 1 | 0 |
| 0 | 0 | 0 |
| 0 | 1 | 0 |
| 1 | 1 | 1 |
| 0 | 0 | 0 |
| 0 | 1 | 0 |
| 1 | 1 | 1 |
| 1 | 1 | 1 |
| 0 | 0 | 0 |
| 0 | 0 | 0 |
| 1 | 1 | 1 |
| 1 | 1 | 1 |
| 0 | 1 | 0 |
| 0 | 1 | 0 |
| 0 | 0 | 0 |
| 0 | 0 | 0 |
| 1 | 1 | 1 |
| 0 | 0 | 0 |
| 0 | 0 | 0 |
| 0 | 0 | 0 |
| 0 | 1 | 0 |
| 0 | 0 | 0 |
| 0 | 0 | 0 |
| 1 | 1 | 1 |
| 0 | 0 | 0 |
| 0 | 1 | 0 |
| 1 | 1 | 1 |
| 0 | 0 | 0 |
| 0 | 1 | 0 |
| 1 | 1 | 1 |
| 0 | 0 | 0 |
| 0 | 1 | 0 |
| 0 | 1 | 0 |
| 0 | 0 | 0 |
| 1 | 1 | 1 |
| 0 | 0 | 0 |
| 1 | 1 | 1 |
| 0 | 0 | 0 |
| 0 | 0 | 0 |
| 1 | 0 | 0 |
| 0 | 0 | 0 |
| 0 | 0 | 0 |
| 1 | 1 | 1 |
| 1 | 0 | 0 |
| 0 | 1 | 0 |
| 1 | 1 | 1 |
| 1 | 1 | 1 |
| 1 | 1 | 1 |
| 1 | 1 | 1 |
| 0 | 1 | 0 |
| 0 | 1 | 0 |
| 0 | 1 | 0 |

| | | |
|---|---|---|
| 0 | 1 | 0 |
| 0 | 1 | 0 |
| 0 | 1 | 0 |
| 0 | 0 | 0 |
| 1 | 0 | 0 |
| 0 | 0 | 0 |
| 1 | 1 | 1 |
| 1 | 0 | 0 |
| 0 | 0 | 0 |
| 0 | 0 | 0 |
| 1 | 1 | 1 |
| 0 | 0 | 0 |
| 0 | 0 | 0 |
| 0 | 0 | 0 |
| 1 | 0 | 0 |
| 0 | 0 | 0 |
| 0 | 0 | 0 |
| 0 | 1 | 0 |
| 0 | 0 | 0 |
| 0 | 0 | 0 |
| 0 | 0 | 0 |
| 0 | 0 | 0 |
| 1 | 1 | 1 |
| 0 | 0 | 0 |
| 0 | 0 | 0 |
| 0 | 0 | 0 |
| 0 | 0 | 0 |
| 0 | 0 | 0 |
| 0 | 0 | 0 |
| 0 | 0 | 0 |
| 0 | 0 | 0 |
| 0 | 0 | 0 |
| 0 | 0 | 0 |
| 0 | 1 | 0 |
| 0 | 1 | 0 |
| 0 | 0 | 0 |
| 1 | 1 | 1 |
| 0 | 0 | 0 |
| 0 | 0 | 0 |
| 0 | 0 | 0 |
| 0 | 0 | 0 |
| 1 | 1 | 1 |
| 0 | 0 | 0 |
| 0 | 0 | 0 |
| 1 | 1 | 1 |
| 0 | 1 | 0 |
| 1 | 1 | 1 |
| 1 | 1 | 1 |
| 1 | 1 | 1 |
| 0 | 1 | 0 |
| 1 | 1 | 1 |
| 1 | 1 | 1 |
| 1 | 1 | 1 |
| 1 | 1 | 1 |

| | | |
|---|---|---|
| 1 | 1 | 1 |
| 0 | 0 | 0 |
| 1 | 1 | 1 |
| 1 | 1 | 1 |
| 1 | 1 | 1 |
| 1 | 1 | 1 |
| 1 | 1 | 1 |
| 1 | 1 | 1 |
| 1 | 1 | 1 |
| 0 | 1 | 0 |
| 1 | 1 | 1 |
| 0 | 0 | 0 |
| 1 | 0 | 0 |
| 0 | 1 | 0 |
| 0 | 0 | 0 |
| 0 | 1 | 0 |
| 0 | 1 | 0 |
| 1 | 1 | 1 |
| 1 | 1 | 1 |
| 1 | 1 | 1 |
| 0 | 0 | 0 |
| 0 | 1 | 0 |
| 0 | 0 | 0 |
| 0 | 0 | 0 |
| 1 | 1 | 1 |
| 0 | 0 | 0 |
| 1 | 1 | 1 |
| 1 | 1 | 1 |
| 1 | 1 | 1 |
| 1 | 1 | 1 |
| 1 | 1 | 1 |
| 0 | 0 | 0 |
| 1 | 1 | 1 |
| 0 | 0 | 0 |
| 1 | 1 | 1 |
| 0 | 1 | 0 |
| 0 | 1 | 0 |
| 0 | 1 | 0 |
| 1 | 1 | 1 |
| 0 | 0 | 0 |
| 1 | 1 | 1 |
| 1 | 1 | 1 |
| 0 | 1 | 0 |
| 0 | 0 | 0 |
| 1 | 1 | 1 |
| 1 | 1 | 1 |
| 1 | 1 | 1 |
| 0 | 0 | 0 |
| 0 | 0 | 0 |
| 1 | 1 | 1 |
| 1 | 1 | 1 |
| 0 | 0 | 0 |
| 0 | 0 | 0 |
| 0 | 0 | 0 |

| | | |
|---|---|---|
| 0 | 0 | 0 |
| 1 | 1 | 1 |
| 0 | 0 | 0 |
| 0 | 0 | 0 |
| 0 | 0 | 0 |
| 1 | 1 | 1 |
| 1 | 1 | 1 |
| 0 | 1 | 0 |
| 1 | 1 | 1 |
| 0 | 0 | 0 |
| 1 | 1 | 1 |
| 0 | 1 | 0 |
| 1 | 1 | 1 |
| 1 | 1 | 1 |
| 0 | 0 | 0 |
| 0 | 1 | 0 |
| 1 | 1 | 1 |
| 0 | 0 | 0 |
| 1 | 1 | 1 |
| 0 | 0 | 0 |
| 0 | 0 | 0 |
| 1 | 1 | 1 |
| 0 | 0 | 0 |
| 1 | 1 | 1 |
| 0 | 0 | 0 |
| 0 | 0 | 0 |
| 1 | 1 | 1 |
| 0 | 1 | 0 |
| 0 | 0 | 0 |
| 1 | 1 | 1 |
| 1 | 1 | 1 |
| 1 | 1 | 1 |
| 1 | 1 | 1 |
| 0 | 1 | 0 |
| 0 | 0 | 0 |
| 0 | 0 | 0 |
| 1 | 1 | 1 |
| 0 | 0 | 0 |
| 0 | 0 | 0 |
| 0 | 0 | 0 |
| 0 | 0 | 0 |
| 0 | 0 | 0 |
| 0 | 0 | 0 |
| 0 | 1 | 0 |
| 0 | 0 | 0 |
| 1 | 1 | 1 |
| 0 | 1 | 0 |
| 0 | 0 | 0 |
| 0 | 0 | 0 |
| 0 | 0 | 0 |
| 0 | 0 | 0 |
| 0 | 0 | 0 |
| 1 | 1 | 1 |
| 1 | 1 | 1 |

| | | |
|---|---|---|
| 1 | 1 | 1 |
| 0 | 0 | 0 |
| 0 | 0 | 0 |
| 0 | 0 | 0 |
| 0 | 0 | 0 |
| 0 | 0 | 0 |
| 0 | 0 | 0 |
| 1 | 0 | 0 |
| 1 | 0 | 0 |
| 1 | 1 | 1 |
| 0 | 0 | 0 |
| 0 | 0 | 0 |
| 1 | 1 | 1 |
| 0 | 0 | 0 |
| 1 | 1 | 1 |
| 0 | 0 | 0 |
| 1 | 1 | 1 |
| 1 | 1 | 1 |
| 0 | 1 | 0 |
| 0 | 1 | 0 |
| 0 | 1 | 0 |
| 1 | 1 | 1 |
| 0 | 1 | 0 |
| 0 | 0 | 0 |
| 0 | 0 | 0 |
| 1 | 1 | 1 |
| 1 | 1 | 1 |
| 1 | 1 | 1 |
| 1 | 1 | 1 |
| 0 | 0 | 0 |
| 0 | 0 | 0 |
| 0 | 0 | 0 |
| 0 | 0 | 0 |
| 1 | 1 | 1 |
| 0 | 0 | 0 |
| 0 | 0 | 0 |
| 0 | 0 | 0 |
| 0 | 0 | 0 |
| 0 | 0 | 0 |
| 1 | 1 | 1 |
| 1 | 1 | 1 |
| 0 | 0 | 0 |
| 1 | 1 | 1 |
| 1 | 1 | 1 |
| 0 | 0 | 0 |
| 1 | 0 | 0 |
| 1 | 1 | 1 |
| 0 | 1 | 0 |
| 1 | 0 | 0 |
| 0 | 1 | 0 |
| 0 | 0 | 0 |
| 1 | 1 | 1 |
| 0 | 0 | 0 |
| 0 | 0 | 0 |

| | | |
|---|---|---|
| 0 | 1 | 0 |
| 1 | 1 | 1 |
| 0 | 0 | 0 |
| 0 | 0 | 0 |
| 1 | 1 | 1 |
| 1 | 1 | 1 |
| 1 | 1 | 1 |
| 1 | 1 | 1 |
| 1 | 1 | 1 |
| 1 | 1 | 1 |
| 1 | 1 | 1 |
| 1 | 1 | 1 |
| 0 | 0 | 0 |
| 1 | 1 | 1 |
| 1 | 1 | 1 |
| 1 | 1 | 1 |
| 1 | 1 | 1 |
| 1 | 1 | 1 |
| 1 | 1 | 1 |
| 1 | 1 | 1 |
| 1 | 1 | 1 |
| 1 | 1 | 1 |
| 1 | 1 | 1 |
| 1 | 1 | 1 |
| 1 | 1 | 1 |
| 1 | 1 | 1 |
| 0 | 1 | 0 |
| 1 | 1 | 1 |
| 1 | 1 | 1 |
| 1 | 1 | 1 |
| 1 | 1 | 1 |
| 0 | 0 | 0 |
| 1 | 1 | 1 |
| 0 | 1 | 0 |
| 1 | 1 | 1 |
| 1 | 1 | 1 |
| 0 | 1 | 0 |
| 1 | 1 | 1 |
| 1 | 1 | 1 |
| 1 | 1 | 1 |
| 0 | 0 | 0 |
| 0 | 0 | 0 |
| 1 | 1 | 1 |
| 1 | 1 | 1 |
| 1 | 1 | 1 |
| 1 | 0 | 0 |
| 1 | 0 | 0 |
| 1 | 1 | 1 |
| 0 | 0 | 0 |
| 0 | 0 | 0 |
| 1 | 1 | 1 |
| 0 | 1 | 0 |
| 0 | 0 | 0 |
| 1 | 1 | 1 |

| | | |
|---|---|---|
| 0 | 0 | 0 |
| 0 | 0 | 0 |
| 0 | 0 | 0 |
| 1 | 1 | 1 |
| 1 | 1 | 1 |
| 1 | 1 | 1 |
| 1 | 1 | 1 |
| 0 | 0 | 0 |
| 0 | 0 | 0 |
| 0 | 0 | 0 |
| 0 | 0 | 0 |
| 1 | 1 | 1 |
| 0 | 0 | 0 |
| 1 | 1 | 1 |
| 1 | 1 | 1 |
| 1 | 1 | 1 |
| 1 | 1 | 1 |
| 0 | 0 | 0 |
| 1 | 1 | 1 |
| 1 | 1 | 1 |
| 1 | 1 | 1 |
| 1 | 1 | 1 |
| 0 | 1 | 0 |
| 1 | 1 | 1 |
| 1 | 1 | 1 |
| 0 | 1 | 0 |
| 1 | 1 | 1 |
| 1 | 1 | 1 |
| 1 | 1 | 1 |
| 1 | 1 | 1 |
| 0 | 0 | 0 |
| 0 | 0 | 0 |
| 1 | 1 | 1 |
| 1 | 1 | 1 |
| 1 | 1 | 1 |
| 1 | 1 | 1 |
| 1 | 1 | 1 |
| 1 | 1 | 1 |
| 1 | 1 | 1 |
| 1 | 1 | 1 |
| 1 | 1 | 1 |
| 0 | 0 | 0 |
| 1 | 1 | 1 |
| 1 | 1 | 1 |
| 0 | 0 | 0 |
| 0 | 0 | 0 |
| 1 | 1 | 1 |
| 1 | 1 | 1 |
| 0 | 0 | 0 |
| 0 | 0 | 0 |
| 0 | 0 | 0 |
| 0 | 0 | 0 |
| 0 | 0 | 0 |
| 0 | 0 | 0 |

| | | |
|---|---|---|
| 1 | 1 | 1 |
| 1 | 1 | 1 |
| 1 | 1 | 1 |
| 1 | 1 | 1 |
| 1 | 1 | 1 |
| 0 | 0 | 0 |
| 0 | 0 | 0 |
| 0 | 0 | 0 |
| 0 | 0 | 0 |
| 0 | 0 | 0 |
| 0 | 1 | 0 |
| 0 | 0 | 0 |
| 0 | 0 | 0 |
| 0 | 0 | 0 |
| 0 | 1 | 0 |
| 0 | 0 | 0 |
| 0 | 1 | 0 |
| 0 | 1 | 0 |
| 0 | 0 | 0 |
| 0 | 0 | 0 |
| 0 | 0 | 0 |
| 0 | 0 | 0 |
| 1 | 0 | 0 |
| 0 | 0 | 0 |
| 0 | 0 | 0 |
| 0 | 0 | 0 |
| 0 | 0 | 0 |
| 1 | 1 | 1 |
| 1 | 1 | 1 |
| 1 | 1 | 1 |
| 1 | 1 | 1 |
| 1 | 1 | 1 |
| 0 | 0 | 0 |
| 0 | 1 | 0 |
| 0 | 0 | 0 |
| 0 | 0 | 0 |
| 0 | 0 | 0 |
| 0 | 0 | 0 |
| 0 | 0 | 0 |
| 0 | 0 | 0 |
| 0 | 0 | 0 |
| 0 | 0 | 0 |
| 0 | 0 | 0 |
| 1 | 1 | 1 |
| 1 | 1 | 1 |
| 1 | 1 | 1 |
| 1 | 1 | 1 |
| 0 | 0 | 0 |
| 0 | 0 | 0 |
| 0 | 0 | 0 |
| 1 | 1 | 1 |
| 1 | 1 | 1 |
| 0 | 0 | 0 |
| 1 | 1 | 1 |

| | | |
|---|---|---|
| 1 | 1 | 1 |
| 0 | 0 | 0 |
| 0 | 1 | 0 |
| 0 | 1 | 0 |
| 1 | 1 | 1 |
| 1 | 1 | 1 |
| 0 | 1 | 0 |
| 1 | 1 | 1 |
| 0 | 1 | 0 |
| 1 | 1 | 1 |
| 1 | 1 | 1 |
| 1 | 1 | 1 |
| 0 | 0 | 0 |
| 0 | 0 | 0 |
| 0 | 0 | 0 |
| 0 | 0 | 0 |
| 0 | 0 | 0 |
| 0 | 0 | 0 |
| 0 | 0 | 0 |
| 0 | 1 | 0 |
| 0 | 0 | 0 |
| 0 | 0 | 0 |
| 0 | 0 | 0 |
| 0 | 0 | 0 |
| 0 | 0 | 0 |
| 1 | 1 | 1 |
| 1 | 1 | 1 |
| 0 | 0 | 0 |
| 0 | 1 | 0 |
| 0 | 0 | 0 |
| 0 | 0 | 0 |
| 1 | 1 | 1 |
| 0 | 0 | 0 |
| 1 | 1 | 1 |
| 1 | 1 | 1 |
| 1 | 1 | 1 |
| 1 | 0 | 0 |
| 1 | 1 | 1 |
| 1 | 1 | 1 |
| 1 | 1 | 1 |
| 0 | 0 | 0 |
| 1 | 1 | 1 |
| 1 | 1 | 1 |
| 0 | 0 | 0 |
| 0 | 0 | 0 |
| 1 | 1 | 1 |
| 0 | 0 | 0 |
| 0 | 0 | 0 |
| 1 | 1 | 1 |
| 1 | 1 | 1 |
| 0 | 0 | 0 |
| 1 | 1 | 1 |
| 1 | 1 | 1 |
| 1 | 1 | 1 |

| | | |
|---|---|---|
| 1 | 1 | 1 |
| 1 | 1 | 1 |
| 1 | 1 | 1 |
| 1 | 1 | 1 |
| 0 | 0 | 0 |
| 0 | 0 | 0 |
| 1 | 1 | 1 |
| 1 | 1 | 1 |
| 1 | 1 | 1 |
| 0 | 1 | 0 |
| 1 | 1 | 1 |
| 0 | 0 | 0 |
| 0 | 0 | 0 |
| 1 | 1 | 1 |
| 1 | 1 | 1 |
| 0 | 0 | 0 |
| 0 | 1 | 0 |
| 1 | 1 | 1 |
| 0 | 0 | 0 |
| 1 | 1 | 1 |
| 1 | 1 | 1 |
| 1 | 1 | 1 |
| 1 | 1 | 1 |
| 0 | 0 | 0 |
| 1 | 1 | 1 |
| 0 | 0 | 0 |
| 1 | 1 | 1 |
| 1 | 1 | 1 |
| 0 | 0 | 0 |
| 0 | 0 | 0 |
| 0 | 0 | 0 |
| 0 | 0 | 0 |
| 0 | 0 | 0 |
| 0 | 0 | 0 |
| 1 | 1 | 1 |
| 1 | 1 | 1 |
| 1 | 1 | 1 |
| 0 | 0 | 0 |
| 1 | 1 | 1 |
| 1 | 1 | 1 |
| 0 | 0 | 0 |
| 0 | 0 | 0 |
| 0 | 0 | 0 |
| 1 | 1 | 1 |
| 0 | 0 | 0 |
| 0 | 1 | 0 |
| 0 | 1 | 0 |
| 0 | 0 | 0 |
| 1 | 1 | 1 |
| 1 | 1 | 1 |
| 1 | 1 | 1 |
| 0 | 0 | 0 |
| 1 | 1 | 1 |
| 0 | 0 | 0 |

| | | |
|---|---|---|
| 1 | 1 | 1 |
| 0 | 0 | 0 |
| 0 | 0 | 0 |
| 0 | 1 | 0 |
| 1 | 1 | 1 |
| 1 | 1 | 1 |
| 0 | 0 | 0 |
| 1 | 1 | 1 |
| 0 | 1 | 0 |
| 0 | 1 | 0 |
| 1 | 1 | 1 |
| 1 | 1 | 1 |
| 1 | 1 | 1 |
| 1 | 1 | 1 |
| 1 | 1 | 1 |
| 1 | 1 | 1 |
| 1 | 1 | 1 |
| 1 | 1 | 1 |
| 1 | 1 | 1 |
| 1 | 1 | 1 |
| 1 | 1 | 1 |
| 0 | 0 | 0 |
| 0 | 1 | 0 |
| 0 | 1 | 0 |
| 1 | 1 | 1 |
| 1 | 1 | 1 |
| 0 | 0 | 0 |
| 1 | 1 | 1 |
| 0 | 0 | 0 |
| 0 | 1 | 0 |
| 1 | 1 | 1 |
| 0 | 0 | 0 |
| 0 | 0 | 0 |
| 1 | 0 | 0 |
| 1 | 1 | 1 |
| 1 | 1 | 1 |
| 1 | 1 | 1 |
| 1 | 1 | 1 |
| 0 | 0 | 0 |
| 0 | 0 | 0 |
| 0 | 0 | 0 |
| 1 | 1 | 1 |
| 1 | 1 | 1 |
| 0 | 1 | 0 |
| 0 | 1 | 0 |
| 0 | 0 | 0 |
| 1 | 1 | 1 |
| 0 | 0 | 0 |
| 1 | 1 | 1 |
| 0 | 0 | 0 |
| 0 | 0 | 0 |
| 0 | 0 | 0 |
| 1 | 1 | 1 |
| 1 | 1 | 1 |

| | | |
|---|---|---|
| 0 | 0 | 0 |
| 0 | 0 | 0 |
| 1 | 1 | 1 |
| 1 | 1 | 1 |
| 1 | 1 | 1 |
| 1 | 1 | 1 |
| 1 | 1 | 1 |
| 1 | 1 | 1 |
| 1 | 1 | 1 |
| 1 | 1 | 1 |
| 1 | 1 | 1 |
| 1 | 1 | 1 |
| 1 | 1 | 1 |
| 1 | 1 | 1 |
| 1 | 1 | 1 |
| 0 | 0 | 0 |
| 0 | 0 | 0 |
| 0 | 0 | 0 |
| 1 | 1 | 1 |
| 1 | 1 | 1 |
| 0 | 1 | 0 |
| 0 | 0 | 0 |
| 0 | 0 | 0 |
| 0 | 0 | 0 |
| 0 | 0 | 0 |
| 1 | 1 | 1 |
| 1 | 1 | 1 |
| 1 | 1 | 1 |
| 1 | 1 | 1 |
| 1 | 1 | 1 |
| 0 | 0 | 0 |
| 0 | 1 | 0 |
| 0 | 0 | 0 |
| 1 | 1 | 1 |
| 1 | 0 | 0 |
| 1 | 0 | 0 |
| 0 | 0 | 0 |
| 0 | 0 | 0 |
| 0 | 1 | 0 |
| 1 | 1 | 1 |
| 1 | 0 | 0 |
| 0 | 0 | 0 |
| 1 | 1 | 1 |
| 0 | 1 | 0 |
| 0 | 0 | 0 |
| 1 | 0 | 0 |
| 0 | 0 | 0 |
| 0 | 0 | 0 |
| 1 | 1 | 1 |
| 0 | 1 | 0 |
| 0 | 0 | 0 |
| 0 | 0 | 0 |
| 1 | 1 | 1 |
| 0 | 0 | 0 |

| | | |
|---|---|---|
| 0 | 1 | 0 |
| 0 | 0 | 0 |
| 1 | 0 | 0 |
| 0 | 0 | 0 |
| 1 | 1 | 1 |
| 1 | 1 | 1 |
| 1 | 1 | 1 |
| 0 | 1 | 0 |
| 0 | 0 | 0 |
| 1 | 1 | 1 |
| 0 | 1 | 0 |
| 0 | 1 | 0 |
| 0 | 0 | 0 |
| 0 | 0 | 0 |
| 0 | 0 | 0 |
| 1 | 1 | 1 |
| 0 | 0 | 0 |
| 0 | 1 | 0 |
| 0 | 0 | 0 |
| 0 | 0 | 0 |
| 0 | 0 | 0 |
| 0 | 0 | 0 |
| 0 | 0 | 0 |
| 0 | 0 | 0 |
| 0 | 1 | 0 |
| 0 | 1 | 0 |
| 0 | 0 | 0 |
| 0 | 0 | 0 |
| 0 | 0 | 0 |
| 0 | 0 | 0 |
| 1 | 1 | 1 |
| 1 | 1 | 1 |
| 1 | 1 | 1 |
| 1 | 1 | 1 |
| 1 | 1 | 1 |
| 0 | 0 | 0 |
| 0 | 0 | 0 |
| 0 | 1 | 0 |
| 1 | 1 | 1 |
| 1 | 1 | 1 |
| 0 | 1 | 0 |
| 1 | 1 | 1 |
| 1 | 1 | 1 |
| 0 | 0 | 0 |
| 0 | 0 | 0 |
| 0 | 0 | 0 |
| 1 | 1 | 1 |
| 0 | 0 | 0 |
| 0 | 0 | 0 |
| 0 | 0 | 0 |
| 1 | 1 | 1 |
| 1 | 1 | 1 |
| 0 | 1 | 0 |
| 0 | 1 | 0 |

| | | |
|---|---|---|
| 0 | 1 | 0 |
| 1 | 1 | 1 |
| 1 | 0 | 0 |
| 1 | 1 | 1 |
| 0 | 0 | 0 |
| 1 | 0 | 0 |
| 0 | 0 | 0 |
| 0 | 0 | 0 |
| 0 | 0 | 0 |
| 0 | 0 | 0 |
| 0 | 0 | 0 |
| 0 | 0 | 0 |
| 0 | 1 | 0 |
| 0 | 0 | 0 |
| 0 | 0 | 0 |
| 1 | 1 | 1 |
| 1 | 1 | 1 |
| 1 | 1 | 1 |
| 0 | 0 | 0 |
| 0 | 0 | 0 |
| 0 | 0 | 0 |
| 0 | 0 | 0 |
| 0 | 1 | 0 |
| 0 | 1 | 0 |
| 1 | 1 | 1 |
| 1 | 1 | 1 |
| 1 | 1 | 1 |
| 0 | 0 | 0 |
| 0 | 0 | 0 |
| 0 | 0 | 0 |
| 0 | 0 | 0 |
| 0 | 0 | 0 |
| 0 | 0 | 0 |
| 0 | 1 | 0 |
| 1 | 1 | 1 |
| 0 | 0 | 0 |
| 1 | 1 | 1 |
| 1 | 1 | 1 |
| 1 | 1 | 1 |
| 1 | 1 | 1 |
| 0 | 0 | 0 |
| 1 | 1 | 1 |
| 0 | 0 | 0 |
| 0 | 1 | 0 |
| 1 | 1 | 1 |
| 1 | 1 | 1 |
| 1 | 1 | 1 |
| 0 | 0 | 0 |
| 1 | 1 | 1 |
| 1 | 1 | 1 |
| 1 | 1 | 1 |
| 1 | 1 | 1 |
| 1 | 1 | 1 |
| 0 | 1 | 0 |

| | | |
|---|---|---|
| 0 | 1 | 0 |
| 1 | 1 | 1 |
| 1 | 1 | 1 |
| 0 | 1 | 0 |
| 1 | 1 | 1 |
| 0 | 1 | 0 |
| 0 | 1 | 0 |
| 0 | 1 | 0 |
| 0 | 1 | 0 |
| 0 | 1 | 0 |
| 0 | 1 | 0 |
| 0 | 0 | 0 |
| 0 | 1 | 0 |
| 0 | 1 | 0 |
| 0 | 1 | 0 |
| 0 | 1 | 0 |
| 0 | 1 | 0 |
| 0 | 0 | 0 |
| 0 | 0 | 0 |
| 0 | 0 | 0 |
| 0 | 0 | 0 |
| 0 | 1 | 0 |
| 0 | 0 | 0 |
| 1 | 1 | 1 |
| 1 | 1 | 1 |
| 1 | 1 | 1 |
| 1 | 1 | 1 |
| 0 | 1 | 0 |
| 0 | 0 | 0 |
| 0 | 0 | 0 |
| 1 | 1 | 1 |
| 1 | 1 | 1 |
| 0 | 0 | 0 |
| 1 | 1 | 1 |
| 0 | 0 | 0 |
| 0 | 0 | 0 |
| 0 | 0 | 0 |
| 1 | 1 | 1 |
| 0 | 0 | 0 |
| 1 | 1 | 1 |
| 0 | 0 | 0 |
| 1 | 1 | 1 |
| 0 | 0 | 0 |
| 1 | 0 | 0 |
| 1 | 0 | 0 |
| 1 | 0 | 0 |
| 1 | 0 | 0 |
| 0 | 1 | 0 |
| 0 | 1 | 0 |
| 0 | 0 | 0 |
| 1 | 0 | 0 |
| 0 | 0 | 0 |
| 1 | 0 | 0 |
| 1 | 1 | 1 |

| | | |
|---|---|---|
| 0 | 0 | 0 |
| 0 | 0 | 0 |
| 0 | 0 | 0 |
| 0 | 0 | 0 |
| 0 | 0 | 0 |
| 1 | 1 | 1 |
| 0 | 0 | 0 |
| 0 | 0 | 0 |
| 1 | 1 | 1 |
| 0 | 0 | 0 |
| 0 | 0 | 0 |
| 0 | 0 | 0 |
| 0 | 1 | 0 |
| 0 | 1 | 0 |
| 0 | 1 | 0 |
| 0 | 1 | 0 |
| 0 | 1 | 0 |
| 0 | 1 | 0 |
| 0 | 1 | 0 |
| 0 | 1 | 0 |
| 0 | 1 | 0 |
| 0 | 1 | 0 |
| 0 | 1 | 0 |
| 0 | 1 | 0 |
| 0 | 1 | 0 |
| 0 | 1 | 0 |
| 0 | 1 | 0 |
| 0 | 1 | 0 |
| 0 | 1 | 0 |
| 0 | 1 | 0 |
| 0 | 1 | 0 |
| 0 | 1 | 0 |
| 0 | 1 | 0 |
| 0 | 1 | 0 |
| 0 | 1 | 0 |
| 0 | 1 | 0 |
| 0 | 1 | 0 |
| 0 | 1 | 0 |
| 0 | 1 | 0 |
| 0 | 1 | 0 |
| 0 | 1 | 0 |
| 0 | 1 | 0 |
| 0 | 1 | 0 |
| 0 | 1 | 0 |
| 0 | 1 | 0 |
| 0 | 0 | 0 |
| 1 | 1 | 1 |
| 1 | 1 | 1 |
| 1 | 1 | 1 |
| 1 | 1 | 1 |
| 0 | 0 | 0 |
| 1 | 1 | 1 |
| 1 | 1 | 1 |
| 0 | 1 | 0 |

| | | |
|---|---|---|
| 1 | 1 | 1 |
| 1 | 1 | 1 |
| 1 | 1 | 1 |
| 1 | 1 | 1 |
| 0 | 1 | 0 |
| 0 | 1 | 0 |
| 0 | 1 | 0 |
| 0 | 0 | 0 |
| 0 | 0 | 0 |
| 1 | 1 | 1 |
| 0 | 1 | 0 |
| 1 | 1 | 1 |
| 1 | 1 | 1 |
| 1 | 1 | 1 |
| 0 | 0 | 0 |
| 0 | 0 | 0 |
| 0 | 0 | 0 |
| 1 | 1 | 1 |
| 1 | 0 | 0 |
| 1 | 1 | 1 |
| 0 | 0 | 0 |
| 0 | 0 | 0 |
| 0 | 0 | 0 |
| 0 | 0 | 0 |
| 0 | 0 | 0 |
| 0 | 0 | 0 |
| 0 | 1 | 0 |
| 1 | 1 | 1 |
| 0 | 0 | 0 |
| 1 | 1 | 1 |
| 1 | 1 | 1 |
| 1 | 1 | 1 |
| 0 | 0 | 0 |
| 1 | 1 | 1 |
| 1 | 1 | 1 |
| 1 | 1 | 1 |
| 0 | 0 | 0 |
| 0 | 0 | 0 |
| 0 | 0 | 0 |
| 1 | 1 | 1 |
| 1 | 1 | 1 |
| 1 | 1 | 1 |
| 1 | 1 | 1 |
| 1 | 1 | 1 |
| 0 | 1 | 0 |
| 0 | 0 | 0 |
| 1 | 1 | 1 |
| 0 | 1 | 0 |
| 1 | 1 | 1 |
| 0 | 1 | 0 |
| 1 | 1 | 1 |
| 1 | 1 | 1 |
| 0 | 1 | 0 |
| 1 | 1 | 1 |

| | | |
|---|---|---|
| 1 | 1 | 1 |
| 0 | 1 | 0 |
| 1 | 1 | 1 |
| 0 | 0 | 0 |
| 1 | 1 | 1 |
| 1 | 1 | 1 |
| 1 | 1 | 1 |
| 0 | 0 | 0 |
| 0 | 0 | 0 |
| 1 | 1 | 1 |
| 1 | 1 | 1 |
| 1 | 1 | 1 |
| 0 | 0 | 0 |
| 0 | 1 | 0 |
| 0 | 1 | 0 |
| 0 | 1 | 0 |
| 0 | 0 | 0 |
| 1 | 1 | 1 |
| 1 | 1 | 1 |
| 1 | 1 | 1 |
| 1 | 1 | 1 |
| 1 | 1 | 1 |
| 1 | 1 | 1 |
| 1 | 1 | 1 |
| 0 | 1 | 0 |
| 1 | 1 | 1 |
| 0 | 1 | 0 |
| 0 | 0 | 0 |
| 0 | 0 | 0 |
| 0 | 0 | 0 |
| 0 | 0 | 0 |
| 0 | 0 | 0 |
| 0 | 0 | 0 |
| 0 | 1 | 0 |
| 0 | 0 | 0 |
| 0 | 0 | 0 |
| 0 | 0 | 0 |
| 1 | 1 | 1 |
| 1 | 1 | 1 |
| 1 | 1 | 1 |
| 1 | 1 | 1 |
| 0 | 1 | 0 |
| 0 | 0 | 0 |
| 0 | 0 | 0 |
| 1 | 1 | 1 |
| 1 | 1 | 1 |
| 0 | 0 | 0 |
| 1 | 1 | 1 |
| 1 | 1 | 1 |
| 1 | 1 | 1 |
| 1 | 1 | 1 |
| 1 | 1 | 1 |
| 0 | 0 | 0 |
| 0 | 0 | 0 |

| | | |
|---|---|---|
| 0 | 0 | 0 |
| 0 | 0 | 0 |
| 0 | 0 | 0 |
| 0 | 0 | 0 |
| 0 | 0 | 0 |
| 0 | 0 | 0 |
| 0 | 0 | 0 |
| 0 | 0 | 0 |
| 1 | 1 | 1 |
| 0 | 0 | 0 |
| 1 | 1 | 1 |
| 1 | 1 | 1 |
| 1 | 1 | 1 |
| 1 | 1 | 1 |
| 0 | 0 | 0 |
| 1 | 1 | 1 |
| 1 | 1 | 1 |
| 1 | 1 | 1 |
| 1 | 1 | 1 |
| 0 | 0 | 0 |
| 0 | 0 | 0 |
| 1 | 1 | 1 |
| 0 | 0 | 0 |
| 0 | 0 | 0 |
| 0 | 0 | 0 |
| 0 | 0 | 0 |
| 0 | 0 | 0 |
| 0 | 1 | 0 |
| 0 | 1 | 0 |
| 0 | 1 | 0 |
| 0 | 0 | 0 |
| 1 | 1 | 1 |
| 1 | 1 | 1 |
| 1 | 1 | 1 |
| 1 | 1 | 1 |
| 1 | 1 | 1 |
| 1 | 1 | 1 |
| 1 | 1 | 1 |
| 1 | 1 | 1 |
| 1 | 1 | 1 |
| 0 | 1 | 0 |
| 0 | 0 | 0 |
| 1 | 1 | 1 |
| 1 | 1 | 1 |
| 1 | 1 | 1 |
| 0 | 1 | 0 |
| 1 | 1 | 1 |
| 1 | 1 | 1 |
| 0 | 0 | 0 |
| 0 | 1 | 0 |
| 1 | 1 | 1 |
| 1 | 1 | 1 |
| 0 | 0 | 0 |
| 0 | 0 | 0 |

| | | |
|---|---|---|
| 1 | 1 | 1 |
| 1 | 1 | 1 |
| 0 | 1 | 0 |
| 1 | 0 | 0 |
| 0 | 0 | 0 |
| 0 | 0 | 0 |
| 0 | 1 | 0 |
| 0 | 0 | 0 |
| 1 | 1 | 1 |
| 1 | 1 | 1 |
| 0 | 0 | 0 |
| 1 | 0 | 0 |
| 0 | 0 | 0 |
| 0 | 0 | 0 |
| 0 | 0 | 0 |
| 0 | 0 | 0 |
| 1 | 1 | 1 |
| 1 | 1 | 1 |
| 0 | 0 | 0 |
| 0 | 0 | 0 |
| 0 | 0 | 0 |
| 0 | 0 | 0 |
| 0 | 0 | 0 |
| 1 | 1 | 1 |
| 1 | 1 | 1 |
| 0 | 0 | 0 |
| 1 | 1 | 1 |
| 0 | 0 | 0 |
| 0 | 0 | 0 |
| 0 | 0 | 0 |
| 0 | 1 | 0 |
| 0 | 0 | 0 |
| 0 | 0 | 0 |
| 1 | 1 | 1 |
| 1 | 1 | 1 |
| 0 | 0 | 0 |
| 0 | 0 | 0 |
| 1 | 0 | 0 |
| 0 | 0 | 0 |
| 0 | 0 | 0 |
| 0 | 0 | 0 |
| 0 | 0 | 0 |
| 0 | 1 | 0 |
| 1 | 1 | 1 |
| 0 | 0 | 0 |
| 0 | 1 | 0 |
| 0 | 0 | 0 |
| 1 | 1 | 1 |
| 1 | 1 | 1 |
| 1 | 1 | 1 |
| 1 | 1 | 1 |
| 1 | 1 | 1 |
| 0 | 1 | 0 |
| 1 | 1 | 1 |

| | | |
|---|---|---|
| 1 | 1 | 1 |
| 1 | 1 | 1 |
| 1 | 1 | 1 |
| 0 | 1 | 0 |
| 1 | 1 | 1 |
| 1 | 1 | 1 |
| 0 | 1 | 0 |
| 0 | 0 | 0 |
| 1 | 1 | 1 |
| 0 | 0 | 0 |
| 0 | 0 | 0 |
| 0 | 0 | 0 |
| 0 | 1 | 0 |
| 1 | 1 | 1 |
| 0 | 0 | 0 |
| 0 | 1 | 0 |
| 0 | 1 | 0 |
| 0 | 1 | 0 |
| 0 | 1 | 0 |
| 0 | 1 | 0 |
| 1 | 1 | 1 |
| 1 | 1 | 1 |
| 1 | 1 | 1 |
| 0 | 1 | 0 |
| 1 | 1 | 1 |
| 1 | 1 | 1 |
| 1 | 1 | 1 |
| 1 | 0 | 0 |
| 1 | 1 | 1 |
| 0 | 0 | 0 |
| 0 | 0 | 0 |
| 1 | 1 | 1 |
| 1 | 1 | 1 |
| 0 | 1 | 0 |
| 0 | 1 | 0 |
| 0 | 0 | 0 |
| 1 | 1 | 1 |
| 1 | 1 | 1 |
| 1 | 1 | 1 |
| 1 | 1 | 1 |
| 1 | 1 | 1 |
| 0 | 0 | 0 |
| 0 | 0 | 0 |
| 0 | 0 | 0 |
| 1 | 1 | 1 |
| 1 | 1 | 1 |
| 1 | 1 | 1 |
| 1 | 1 | 1 |
| 1 | 1 | 1 |
| 1 | 1 | 1 |
| 1 | 1 | 1 |
| 1 | 1 | 1 |
| 1 | 1 | 1 |
| 0 | 0 | 0 |

| | | |
|---|---|---|
| 1 | 1 | 1 |
| 1 | 1 | 1 |
| 1 | 1 | 1 |
| 1 | 1 | 1 |
| 1 | 1 | 1 |
| 1 | 1 | 1 |
| 1 | 1 | 1 |
| 0 | 1 | 0 |
| 0 | 0 | 0 |
| 0 | 0 | 0 |
| 1 | 1 | 1 |
| 0 | 1 | 0 |
| 0 | 1 | 0 |
| 0 | 0 | 0 |
| 1 | 1 | 1 |
| 1 | 1 | 1 |
| 1 | 1 | 1 |
| 0 | 0 | 0 |
| 0 | 0 | 0 |
| 0 | 0 | 0 |
| 0 | 0 | 0 |
| 0 | 1 | 0 |
| 1 | 1 | 1 |
| 1 | 1 | 1 |
| 1 | 1 | 1 |
| 0 | 0 | 0 |
| 1 | 1 | 1 |
| 0 | 1 | 0 |
| 1 | 1 | 1 |
| 1 | 1 | 1 |
| 1 | 1 | 1 |
| 1 | 1 | 1 |
| 1 | 1 | 1 |
| 1 | 1 | 1 |
| 1 | 1 | 1 |
| 1 | 1 | 1 |
| 1 | 1 | 1 |
| 1 | 1 | 1 |
| 1 | 1 | 1 |
| 0 | 0 | 0 |
| 1 | 1 | 1 |
| 1 | 1 | 1 |
| 0 | 0 | 0 |
| 1 | 1 | 1 |
| 1 | 1 | 1 |
| 1 | 1 | 1 |
| 0 | 0 | 0 |
| 0 | 0 | 0 |
| 1 | 1 | 1 |
| 1 | 1 | 1 |
| 0 | 0 | 0 |
| 1 | 1 | 1 |
| 0 | 1 | 0 |
| 1 | 1 | 1 |

| | | |
|---|---|---|
| 1 | 1 | 1 |
| 1 | 1 | 1 |
| 1 | 1 | 1 |
| 1 | 1 | 1 |
| 1 | 1 | 1 |
| 1 | 1 | 1 |
| 1 | 1 | 1 |
| 0 | 1 | 0 |
| 1 | 1 | 1 |
| 1 | 1 | 1 |
| 1 | 1 | 1 |
| 1 | 1 | 1 |
| 1 | 1 | 1 |
| 0 | 0 | 0 |
| 0 | 0 | 0 |
| 0 | 0 | 0 |
| 0 | 0 | 0 |
| 0 | 1 | 0 |
| 1 | 1 | 1 |
| 0 | 0 | 0 |
| 1 | 1 | 1 |
| 1 | 0 | 0 |
| 1 | 1 | 1 |
| 1 | 1 | 1 |
| 1 | 1 | 1 |
| 0 | 0 | 0 |
| 0 | 1 | 0 |
| 1 | 1 | 1 |
| 1 | 1 | 1 |
| 0 | 0 | 0 |
| 0 | 0 | 0 |
| 1 | 1 | 1 |
| 1 | 1 | 1 |
| 1 | 1 | 1 |
| 1 | 1 | 1 |
| 1 | 1 | 1 |
| 1 | 1 | 1 |
| 1 | 1 | 1 |
| 0 | 0 | 0 |
| 1 | 1 | 1 |
| 1 | 1 | 1 |
| 0 | 0 | 0 |
| 0 | 0 | 0 |
| 0 | 0 | 0 |
| 0 | 0 | 0 |
| 1 | 1 | 1 |
| 0 | 0 | 0 |
| 1 | 1 | 1 |
| 1 | 1 | 1 |
| 1 | 1 | 1 |
| 0 | 1 | 0 |
| 1 | 0 | 0 |
| 1 | 0 | 0 |
| 1 | 0 | 0 |

| | | |
|---|---|---|
| 1 | 1 | 1 |
| 1 | 1 | 1 |
| 0 | 0 | 0 |
| 1 | 1 | 1 |
| 1 | 1 | 1 |
| 1 | 1 | 1 |
| 1 | 1 | 1 |
| 0 | 1 | 0 |
| 1 | 1 | 1 |
| 1 | 1 | 1 |
| 1 | 1 | 1 |
| 0 | 1 | 0 |
| 0 | 1 | 0 |
| 0 | 1 | 0 |
| 0 | 1 | 0 |
| 0 | 1 | 0 |
| 0 | 1 | 0 |
| 0 | 1 | 0 |
| 0 | 1 | 0 |
| 0 | 1 | 0 |
| 0 | 1 | 0 |
| 0 | 1 | 0 |
| 0 | 1 | 0 |
| 0 | 1 | 0 |
| 0 | 1 | 0 |
| 0 | 1 | 0 |
| 0 | 1 | 0 |
| 0 | 1 | 0 |
| 1 | 1 | 1 |
| 1 | 1 | 1 |
| 0 | 0 | 0 |
| 1 | 1 | 1 |
| 0 | 0 | 0 |
| 0 | 0 | 0 |
| 0 | 0 | 0 |
| 0 | 0 | 0 |
| 1 | 1 | 1 |
| 0 | 0 | 0 |
| 1 | 1 | 1 |
| 1 | 1 | 1 |
| 0 | 0 | 0 |
| 0 | 0 | 0 |
| 0 | 0 | 0 |
| 1 | 1 | 1 |
| 0 | 0 | 0 |
| 0 | 0 | 0 |
| 1 | 1 | 1 |
| 1 | 1 | 1 |
| 1 | 1 | 1 |
| 1 | 1 | 1 |
| 0 | 0 | 0 |
| 1 | 1 | 1 |
| 1 | 1 | 1 |
| 1 | 1 | 1 |

| | | |
|---|---|---|
| 1 | 1 | 1 |
| 1 | 1 | 1 |
| 1 | 1 | 1 |
| 1 | 1 | 1 |
| 1 | 1 | 1 |
| 0 | 0 | 0 |
| 1 | 1 | 1 |
| 0 | 0 | 0 |
| 0 | 0 | 0 |
| 0 | 0 | 0 |
| 1 | 1 | 1 |
| 0 | 0 | 0 |
| 1 | 1 | 1 |
| 1 | 1 | 1 |
| 1 | 1 | 1 |
| 1 | 0 | 0 |
| 1 | 1 | 1 |
| 0 | 0 | 0 |
| 1 | 1 | 1 |
| 0 | 0 | 0 |
| 1 | 1 | 1 |
| 0 | 0 | 0 |
| 0 | 0 | 0 |
| 0 | 0 | 0 |
| 1 | 1 | 1 |
| 0 | 1 | 0 |
| 1 | 1 | 1 |
| 1 | 1 | 1 |
| 0 | 0 | 0 |
| 1 | 1 | 1 |
| 1 | 1 | 1 |
| 1 | 1 | 1 |
| 1 | 1 | 1 |
| 1 | 1 | 1 |
| 0 | 0 | 0 |
| 0 | 0 | 0 |
| 1 | 1 | 1 |
| 0 | 0 | 0 |
| 0 | 0 | 0 |
| 0 | 1 | 0 |
| 0 | 0 | 0 |
| 0 | 0 | 0 |
| 0 | 1 | 0 |
| 0 | 1 | 0 |
| 1 | 1 | 1 |
| 0 | 1 | 0 |
| 1 | 1 | 1 |
| 1 | 1 | 1 |
| 1 | 1 | 1 |
| 1 | 1 | 1 |
| 1 | 1 | 1 |
| 1 | 1 | 1 |
| 1 | 1 | 1 |
| 1 | 1 | 1 |

| | | |
|---|---|---|
| 1 | 1 | 1 |
| 0 | 0 | 0 |
| 0 | 1 | 0 |
| 1 | 1 | 1 |
| 0 | 0 | 0 |
| 1 | 1 | 1 |
| 1 | 1 | 1 |
| 1 | 1 | 1 |
| 0 | 0 | 0 |
| 0 | 0 | 0 |
| 1 | 1 | 1 |
| 0 | 0 | 0 |
| 0 | 0 | 0 |
| 1 | 1 | 1 |
| 1 | 1 | 1 |
| 0 | 0 | 0 |
| 0 | 1 | 0 |
| 0 | 0 | 0 |
| 0 | 0 | 0 |
| 1 | 1 | 1 |
| 0 | 0 | 0 |
| 0 | 0 | 0 |
| 0 | 0 | 0 |
| 1 | 1 | 1 |
| 0 | 1 | 0 |
| 0 | 0 | 0 |
| 1 | 1 | 1 |
| 1 | 1 | 1 |
| 1 | 1 | 1 |
| 1 | 1 | 1 |
| 0 | 0 | 0 |
| 1 | 1 | 1 |
| 0 | 0 | 0 |
| 1 | 1 | 1 |
| 0 | 0 | 0 |
| 0 | 0 | 0 |
| 1 | 1 | 1 |
| 1 | 1 | 1 |
| 1 | 1 | 1 |
| 1 | 1 | 1 |
| 1 | 1 | 1 |
| 1 | 1 | 1 |
| 1 | 1 | 1 |
| 1 | 1 | 1 |
| 1 | 1 | 1 |
| 1 | 1 | 1 |
| 1 | 0 | 0 |
| 1 | 1 | 1 |
| 1 | 1 | 1 |
| 1 | 1 | 1 |
| 1 | 1 | 1 |
| 1 | 1 | 1 |
| 1 | 1 | 1 |
| 0 | 0 | 0 |

| | | |
|---|---|---|
| 1 | 1 | 1 |
| 1 | 1 | 1 |
| 1 | 1 | 1 |
| 1 | 1 | 1 |
| 0 | 0 | 0 |
| 1 | 1 | 1 |
| 0 | 1 | 0 |
| 0 | 1 | 0 |
| 0 | 1 | 0 |
| 1 | 1 | 1 |
| 0 | 0 | 0 |
| 1 | 1 | 1 |
| 1 | 1 | 1 |
| 1 | 1 | 1 |
| 0 | 0 | 0 |
| 0 | 0 | 0 |
| 0 | 0 | 0 |
| 0 | 0 | 0 |
| 0 | 1 | 0 |
| 0 | 0 | 0 |
| 0 | 0 | 0 |
| 0 | 0 | 0 |
| 1 | 1 | 1 |
| 0 | 0 | 0 |
| 0 | 0 | 0 |
| 1 | 1 | 1 |
| 0 | 0 | 0 |
| 0 | 0 | 0 |
| 0 | 0 | 0 |
| 0 | 0 | 0 |
| 0 | 0 | 0 |
| 1 | 1 | 1 |
| 1 | 1 | 1 |
| 0 | 1 | 0 |
| 1 | 1 | 1 |
| 1 | 1 | 1 |
| 1 | 1 | 1 |
| 0 | 0 | 0 |
| 0 | 0 | 0 |
| 0 | 1 | 0 |
| 1 | 1 | 1 |
| 0 | 0 | 0 |
| 0 | 0 | 0 |
| 0 | 1 | 0 |
| 0 | 0 | 0 |
| 1 | 1 | 1 |
| 1 | 1 | 1 |
| 1 | 1 | 1 |
| 0 | 0 | 0 |
| 0 | 0 | 0 |
| 0 | 0 | 0 |
| 1 | 1 | 1 |
| 0 | 1 | 0 |
| 0 | 0 | 0 |

| | | |
|---|---|---|
| 1 | 1 | 1 |
| 0 | 0 | 0 |
| 1 | 1 | 1 |
| 0 | 0 | 0 |
| 0 | 0 | 0 |
| 0 | 1 | 0 |
| 0 | 0 | 0 |
| 0 | 0 | 0 |
| 1 | 1 | 1 |
| 1 | 1 | 1 |
| 1 | 1 | 1 |
| 1 | 1 | 1 |
| 1 | 1 | 1 |
| 1 | 1 | 1 |
| 1 | 1 | 1 |
| 1 | 1 | 1 |
| 1 | 1 | 1 |
| 0 | 1 | 0 |
| 1 | 1 | 1 |
| 1 | 1 | 1 |
| 0 | 0 | 0 |
| 0 | 1 | 0 |
| 0 | 1 | 0 |
| 0 | 1 | 0 |
| 0 | 1 | 0 |
| 0 | 0 | 0 |
| 0 | 0 | 0 |
| 1 | 1 | 1 |
| 1 | 1 | 1 |
| 1 | 1 | 1 |
| 1 | 1 | 1 |
| 1 | 1 | 1 |
| 1 | 1 | 1 |
| 0 | 1 | 0 |
| 1 | 1 | 1 |
| 1 | 1 | 1 |
| 1 | 1 | 1 |
| 1 | 1 | 1 |
| 0 | 1 | 0 |
| 1 | 1 | 1 |
| 1 | 1 | 1 |
| 0 | 1 | 0 |
| 1 | 1 | 1 |
| 0 | 0 | 0 |
| 1 | 1 | 1 |
| 0 | 0 | 0 |
| 0 | 0 | 0 |
| 0 | 0 | 0 |
| 0 | 0 | 0 |
| 0 | 0 | 0 |
| 0 | 0 | 0 |
| 0 | 1 | 0 |
| 1 | 1 | 1 |
| 1 | 1 | 1 |

| | | |
|---|---|---|
| 1 | 1 | 1 |
| 1 | 1 | 1 |
| 1 | 1 | 1 |
| 0 | 0 | 0 |
| 1 | 1 | 1 |
| 0 | 0 | 0 |
| 1 | 1 | 1 |
| 1 | 1 | 1 |
| 0 | 0 | 0 |
| 1 | 1 | 1 |
| 1 | 1 | 1 |
| 0 | 1 | 0 |
| 1 | 1 | 1 |
| 1 | 1 | 1 |
| 1 | 1 | 1 |
| 0 | 0 | 0 |
| 0 | 0 | 0 |
| 0 | 0 | 0 |
| 0 | 0 | 0 |
| 0 | 0 | 0 |
| 1 | 1 | 1 |
| 1 | 1 | 1 |
| 1 | 1 | 1 |
| 1 | 1 | 1 |
| 0 | 0 | 0 |
| 1 | 1 | 1 |
| 1 | 1 | 1 |
| 1 | 1 | 1 |
| 0 | 0 | 0 |
| 1 | 1 | 1 |
| 0 | 0 | 0 |
| 0 | 0 | 0 |
| 0 | 0 | 0 |
| 0 | 0 | 0 |
| 0 | 0 | 0 |
| 1 | 1 | 1 |
| 0 | 0 | 0 |
| 0 | 0 | 0 |
| 0 | 0 | 0 |
| 0 | 0 | 0 |
| 0 | 0 | 0 |
| 0 | 0 | 0 |
| 0 | 0 | 0 |
| 0 | 0 | 0 |
| 0 | 0 | 0 |
| 0 | 0 | 0 |
| 0 | 0 | 0 |
| 0 | 0 | 0 |
| 1 | 1 | 1 |
| 0 | 0 | 0 |
| 1 | 1 | 1 |
| 1 | 1 | 1 |
| 0 | 1 | 0 |
| 0 | 0 | 0 |

| | | |
|---|---|---|
| 0 | 0 | 0 |
| 0 | 1 | 0 |
| 1 | 1 | 1 |
| 1 | 1 | 1 |
| 1 | 1 | 1 |
| 0 | 0 | 0 |
| 1 | 1 | 1 |
| 0 | 1 | 0 |
| 1 | 1 | 1 |
| 0 | 1 | 0 |
| 0 | 1 | 0 |
| 0 | 1 | 0 |
| 1 | 1 | 1 |
| 0 | 0 | 0 |
| 0 | 1 | 0 |
| 1 | 1 | 1 |
| 1 | 1 | 1 |
| 0 | 0 | 0 |
| 1 | 1 | 1 |
| 1 | 1 | 1 |
| 1 | 1 | 1 |
| 1 | 1 | 1 |
| 0 | 0 | 0 |
| 1 | 1 | 1 |
| 1 | 1 | 1 |
| 0 | 0 | 0 |
| 0 | 0 | 0 |
| 0 | 0 | 0 |
| 1 | 1 | 1 |
| 0 | 1 | 0 |
| 1 | 1 | 1 |
| 1 | 1 | 1 |
| 1 | 1 | 1 |
| 1 | 1 | 1 |
| 1 | 1 | 1 |
| 1 | 1 | 1 |
| 1 | 1 | 1 |
| 1 | 1 | 1 |
| 1 | 1 | 1 |
| 1 | 1 | 1 |
| 1 | 1 | 1 |
| 1 | 1 | 1 |
| 1 | 1 | 1 |
| 1 | 1 | 1 |
| 1 | 1 | 1 |
| 1 | 1 | 1 |
| 1 | 1 | 1 |
| 1 | 1 | 1 |
| 1 | 0 | 0 |
| 1 | 1 | 1 |
| 1 | 1 | 1 |
| 1 | 1 | 1 |
| 1 | 1 | 1 |
| 1 | 1 | 1 |

| | | |
|---|---|---|
| 1 | 1 | 1 |
| 0 | 0 | 0 |
| 1 | 1 | 1 |
| 1 | 1 | 1 |
| 1 | 1 | 1 |
| 0 | 0 | 0 |
| 0 | 0 | 0 |
| 0 | 0 | 0 |
| 1 | 1 | 1 |
| 1 | 1 | 1 |
| 0 | 1 | 0 |
| 1 | 1 | 1 |
| 1 | 1 | 1 |
| 1 | 1 | 1 |
| 0 | 0 | 0 |
| 0 | 0 | 0 |
| 0 | 0 | 0 |
| 1 | 1 | 1 |
| 0 | 0 | 0 |
| 0 | 0 | 0 |
| 0 | 0 | 0 |
| 0 | 0 | 0 |
| 0 | 0 | 0 |
| 0 | 0 | 0 |
| 0 | 1 | 0 |
| 1 | 1 | 1 |
| 1 | 1 | 1 |
| 0 | 1 | 0 |
| 0 | 1 | 0 |
| 1 | 1 | 1 |
| 1 | 1 | 1 |
| 1 | 1 | 1 |
| 0 | 0 | 0 |
| 1 | 1 | 1 |
| 0 | 1 | 0 |
| 0 | 0 | 0 |
| 0 | 0 | 0 |
| 0 | 0 | 0 |
| 0 | 0 | 0 |
| 1 | 1 | 1 |
| 0 | 1 | 0 |
| 1 | 1 | 1 |
| 1 | 1 | 1 |
| 0 | 0 | 0 |
| 0 | 0 | 0 |
| 0 | 0 | 0 |
| 0 | 1 | 0 |
| 1 | 1 | 1 |
| 0 | 0 | 0 |
| 1 | 1 | 1 |
| 1 | 1 | 1 |
| 0 | 0 | 0 |
| 1 | 1 | 1 |
| 0 | 1 | 0 |

| | | |
|---|---|---|
| 1 | 1 | 1 |
| 0 | 1 | 0 |
| 1 | 1 | 1 |
| 1 | 1 | 1 |
| 1 | 1 | 1 |
| 1 | 1 | 1 |
| 0 | 1 | 0 |
| 1 | 1 | 1 |
| 0 | 0 | 0 |
| 0 | 0 | 0 |
| 1 | 1 | 1 |
| 0 | 0 | 0 |
| 0 | 0 | 0 |
| 0 | 0 | 0 |
| 1 | 1 | 1 |
| 1 | 1 | 1 |
| 0 | 0 | 0 |
| 0 | 0 | 0 |
| 0 | 0 | 0 |
| 1 | 1 | 1 |
| 1 | 1 | 1 |
| 1 | 1 | 1 |
| 0 | 0 | 0 |
| 1 | 1 | 1 |
| 0 | 0 | 0 |
| 1 | 1 | 1 |
| 1 | 1 | 1 |
| 1 | 1 | 1 |
| 1 | 1 | 1 |
| 0 | 0 | 0 |
| 0 | 0 | 0 |
| 0 | 0 | 0 |
| 1 | 1 | 1 |
| 0 | 1 | 0 |
| 1 | 1 | 1 |
| 1 | 1 | 1 |
| 1 | 1 | 1 |
| 0 | 1 | 0 |
| 0 | 1 | 0 |
| 0 | 1 | 0 |
| 0 | 0 | 0 |
| 1 | 1 | 1 |
| 1 | 1 | 1 |
| 1 | 1 | 1 |
| 1 | 1 | 1 |
| 0 | 0 | 0 |
| 1 | 0 | 0 |
| 0 | 0 | 0 |
| 1 | 1 | 1 |
| 0 | 0 | 0 |
| 0 | 0 | 0 |
| 0 | 0 | 0 |
| 0 | 1 | 0 |
| 1 | 0 | 0 |

| | | |
|---|---|---|
| 1 | 0 | 0 |
| 0 | 0 | 0 |
| 0 | 1 | 0 |
| 1 | 1 | 1 |
| 0 | 1 | 0 |
| 0 | 1 | 0 |
| 0 | 1 | 0 |
| 0 | 0 | 0 |
| 0 | 0 | 0 |
| 0 | 1 | 0 |
| 0 | 0 | 0 |
| 1 | 1 | 1 |
| 1 | 1 | 1 |
| 1 | 1 | 1 |
| 0 | 1 | 0 |
| 1 | 1 | 1 |
| 1 | 1 | 1 |
| 1 | 1 | 1 |
| 1 | 1 | 1 |
| 0 | 1 | 0 |
| 1 | 1 | 1 |
| 1 | 1 | 1 |
| 1 | 1 | 1 |
| 1 | 1 | 1 |
| 0 | 1 | 0 |
| 0 | 1 | 0 |
| 0 | 1 | 0 |
| 0 | 1 | 0 |
| 0 | 0 | 0 |
| 0 | 1 | 0 |
| 0 | 0 | 0 |
| 1 | 1 | 1 |
| 0 | 0 | 0 |
| 0 | 0 | 0 |
| 0 | 1 | 0 |
| 0 | 0 | 0 |
| 0 | 0 | 0 |
| 1 | 1 | 1 |
| 0 | 1 | 0 |
| 0 | 1 | 0 |
| 1 | 1 | 1 |
| 0 | 0 | 0 |
| 0 | 0 | 0 |
| 1 | 1 | 1 |
| 1 | 1 | 1 |
| 0 | 0 | 0 |
| 0 | 0 | 0 |
| 0 | 0 | 0 |
| 0 | 0 | 0 |
| 0 | 1 | 0 |
| 0 | 0 | 0 |
| 0 | 0 | 0 |
| 1 | 1 | 1 |
| 1 | 1 | 1 |

| | | |
|---|---|---|
| 1 | 1 | 1 |
| 1 | 1 | 1 |
| 1 | 1 | 1 |
| 1 | 1 | 1 |
| 0 | 0 | 0 |
| 0 | 0 | 0 |
| 0 | 0 | 0 |
| 0 | 0 | 0 |
| 1 | 1 | 1 |
| 0 | 0 | 0 |
| 0 | 0 | 0 |
| 0 | 0 | 0 |
| 0 | 0 | 0 |
| 1 | 1 | 1 |
| 1 | 0 | 0 |
| 0 | 1 | 0 |
| 1 | 1 | 1 |
| 1 | 1 | 1 |
| 1 | 1 | 1 |
| 0 | 1 | 0 |
| 1 | 1 | 1 |
| 1 | 1 | 1 |
| 1 | 1 | 1 |
| 1 | 1 | 1 |
| 1 | 1 | 1 |
| 0 | 0 | 0 |
| 0 | 0 | 0 |
| 0 | 0 | 0 |
| 0 | 0 | 0 |
| 0 | 1 | 0 |
| 0 | 1 | 0 |
| 0 | 1 | 0 |
| 1 | 1 | 1 |
| 0 | 1 | 0 |
| 0 | 1 | 0 |
| 1 | 1 | 1 |
| 1 | 1 | 1 |
| 1 | 1 | 1 |
| 1 | 1 | 1 |
| 1 | 1 | 1 |
| 1 | 1 | 1 |
| 1 | 1 | 1 |
| 1 | 1 | 1 |
| 1 | 1 | 1 |
| 1 | 1 | 1 |
| 1 | 1 | 1 |
| 1 | 1 | 1 |
| 1 | 1 | 1 |
| 1 | 1 | 1 |
| 1 | 1 | 1 |
| 1 | 1 | 1 |
| 1 | 1 | 1 |
| 1 | 1 | 1 |
| 0 | 0 | 0 |

| | | |
|---|---|---|
| 1 | 1 | 1 |
| 1 | 1 | 1 |
| 1 | 1 | 1 |
| 0 | 1 | 0 |
| 0 | 1 | 0 |
| 0 | 1 | 0 |
| 0 | 1 | 0 |
| 0 | 1 | 0 |
| 1 | 1 | 1 |
| 1 | 1 | 1 |
| 1 | 1 | 1 |
| 1 | 1 | 1 |
| 1 | 1 | 1 |
| 1 | 1 | 1 |
| 1 | 1 | 1 |
| 1 | 1 | 1 |
| 1 | 1 | 1 |
| 1 | 1 | 1 |
| 1 | 1 | 1 |
| 1 | 1 | 1 |
| 1 | 1 | 1 |
| 1 | 1 | 1 |
| 1 | 1 | 1 |
| 1 | 1 | 1 |
| 1 | 1 | 1 |
| 1 | 1 | 1 |
| 1 | 1 | 1 |
| 1 | 1 | 1 |
| 1 | 1 | 1 |
| 1 | 1 | 1 |
| 1 | 1 | 1 |
| 1 | 1 | 1 |
| 1 | 1 | 1 |
| 0 | 1 | 0 |
| 1 | 1 | 1 |
| 1 | 1 | 1 |
| 1 | 1 | 1 |
| 1 | 1 | 1 |
| 1 | 1 | 1 |
| 1 | 1 | 1 |
| 0 | 1 | 0 |
| 1 | 1 | 1 |
| 1 | 1 | 1 |
| 1 | 1 | 1 |
| 0 | 1 | 0 |
| 1 | 0 | 0 |
| 0 | 0 | 0 |
| 1 | 1 | 1 |
| 0 | 1 | 0 |
| 1 | 1 | 1 |
| 1 | 1 | 1 |
| 0 | 1 | 0 |
| 0 | 1 | 0 |
| 0 | 1 | 0 |

| | | |
|---|---|---|
| 0 | 1 | 0 |
| 1 | 1 | 1 |
| 0 | 0 | 0 |
| 1 | 1 | 1 |
| 1 | 1 | 1 |
| 1 | 1 | 1 |
| 0 | 0 | 0 |
| 1 | 1 | 1 |
| 1 | 1 | 1 |
| 0 | 1 | 0 |
| 0 | 1 | 0 |
| 0 | 0 | 0 |
| 1 | 1 | 1 |
| 0 | 0 | 0 |
| 0 | 0 | 0 |
| 0 | 0 | 0 |
| 0 | 0 | 0 |
| 0 | 0 | 0 |
| 0 | 0 | 0 |
| 0 | 0 | 0 |
| 0 | 0 | 0 |

| organism_scientific | evo_cons | cluster_id | pdb_id | chain_id | resi | resn | change | uniprot_id |
|---|---|---|---|---|---|---|---|---|
| HOMO SAPIENS | 5 | 87 | 3uzw | A | 106 | LEU | p.Leu106Phe | P51857 |
| HOMO SAPIENS | 3 | 87 | 3uzw | A | 133 | PRO | p.Pro133Arg | P51857 |
| HOMO SAPIENS | 0 | 87 | 3uzw | A | 198 | PRO | p.Pro198Leu | P51857 |
| HOMO SAPIENS | 7 | 87 | 3uzw | A | 261 | ARG | p.Arg261Cys | P51857 |
| HOMO SAPIENS | 0 | 4693 | 3mk1 | A | 3 | PRO | p.Pro25Leu | P05187 |
| HOMO SAPIENS | 8 | 4830 | 2h64 | B | 101 | CYS | p.Cys124Arg | P36894 |
| HOMO SAPIENS | 2 | 4830 | 2h64 | B | 116 | PRO | p.Pro139Ser | P36894 |
| HOMO SAPIENS | 0 | 6937 | 3dar | A | 186 | MET | p.Met186Thr | P21802 |
| HOMO SAPIENS | 3 | 5 | 1aut | C | 30 | GLN | p.Gln226His | P04070 |
| HOMO SAPIENS | 1 | 5 | 1aut | C | 68 | LEU | p.Leu265Phe | P04070 |
| HOMO SAPIENS | 0 | 5 | 1aut | C | 75 | ARG | p.Arg272Cys | P04070 |
| HOMO SAPIENS | 6 | 5 | 1aut | C | 92 | PRO | p.Pro289Leu | P04070 |
| HOMO SAPIENS | 0 | 5 | 1aut | C | 104 | ALA | p.Ala301Val | P04070 |
| HOMO SAPIENS | 1 | 5 | 1aut | C | 112 | ALA | p.Ala309Thr | P04070 |
| HOMO SAPIENS | 2 | 5 | 1aut | C | 115 | SER | p.Ser312Leu | P04070 |
| HOMO SAPIENS | 1 | 5 | 1aut | C | 133 | GLY | p.Gly334Ser | P04070 |
| HOMO SAPIENS | 0 | 5 | 1aut | C | 147 | ARG | p.Arg348Ter | P04070 |
| HOMO SAPIENS | 8 | 5 | 1aut | C | 194 | ASP | p.Asp401Asn | P04070 |
| HOMO SAPIENS | 7 | 5 | 1aut | C | 237 | TRP | p.Trp444Cys | P04070 |
| HOMO SAPIENS | 0 | 5 | 1aut | C | 238 | ILE | p.Ile445Met | P04070 |
| HOMO SAPIENS | 8 | 43 | 1aiy | F | 10 | HIS | p.His34Asp | P01308 |
| HOMO SAPIENS | 2 | 43 | 1aiy | F | 22 | ARG | p.Arg46Gln | P01308 |
| HOMO SAPIENS | 4 | 43 | 1aiy | F | 23 | GLY | p.Gly47Val | P01308 |
| HOMO SAPIENS | 7 | 43 | 1aiy | F | 24 | PHE | p.Phe48Ser | P01308 |
| HOMO SAPIENS | 5 | 24 | 2fg8 | A | 33 | THR | p.Thr30Ile | P02792 |
| HOMO SAPIENS | 4 | 24 | 2fg8 | A | 99 | ALA | p.Ala96Thr | P02792 |
| HOMO SAPIENS | 5 | 24 | 2fg8 | A | 107 | GLU | p.Glu104Ter | P02792 |
| HOMO SAPIENS | 3 | 5409 | 5bnq | A | 199 | MET | p.Met199Lys | O14788 |
| HOMO SAPIENS | -1 | 839 | 3swc | A | 959 | GLN | p.Gln990His | Q9H9B1 |
| HOMO SAPIENS | -1 | 839 | 3swc | A | 968 | ALA | p.Ala999Val | Q9H9B1 |
| HOMO SAPIENS | -1 | 839 | 3swc | A | 973 | SER | p.Ser1004Ile | Q9H9B1 |
| HOMO SAPIENS | 8 | 17 | 1eou | A | 18 | LYS | p.Lys18Glu | P00918 |
| HOMO SAPIENS | 2 | 17 | 1eou | A | 107 | HIS | p.His107Tyr | P00918 |
| HOMO SAPIENS | 1 | 17 | 1eou | A | 236 | PRO | p.Pro236His | P00918 |
| HOMO SAPIENS | 6 | 17 | 1eou | A | 252 | ASN | p.Asn252Asp | P00918 |
| HOMO SAPIENS | 2 | 9140 | 5c4v | A | 445 | ARG | p.Arg445Ter | Q13485 |
| HOMO SAPIENS | 7 | 9140 | 5c4v | A | 496 | ARG | p.Arg496Cys | Q13485 |
| HOMO SAPIENS | -1 | 42 | 4pwi | A | 6 | GLY | p.Gly26Ser | P02766 |
| HOMO SAPIENS | 6 | 42 | 4pwi | A | 10 | CYS | p.Cys30Arg | P02766 |
| HOMO SAPIENS | 8 | 42 | 4pwi | A | 12 | LEU | p.Leu32Pro | P02766 |
| HOMO SAPIENS | 9 | 42 | 4pwi | A | 18 | ASP | p.Asp38Gly | P02766 |
| HOMO SAPIENS | 8 | 42 | 4pwi | A | 20 | VAL | p.Val40Ile | P02766 |
| HOMO SAPIENS | 4 | 42 | 4pwi | A | 30 | MET | p.Val50Ala | P02766 |
| HOMO SAPIENS | 4 | 42 | 4pwi | A | 30 | MET | p.Val50Leu | P02766 |
| HOMO SAPIENS | 6 | 42 | 4pwi | A | 33 | PHE | p.Phe53Ile | P02766 |
| HOMO SAPIENS | 7 | 42 | 4pwi | A | 36 | ALA | p.Ala56Pro | P02766 |
| HOMO SAPIENS | 7 | 42 | 4pwi | A | 42 | GLU | p.Glu62Gly | P02766 |
| HOMO SAPIENS | 8 | 42 | 4pwi | A | 44 | PHE | p.Phe64Ser | P02766 |
| HOMO SAPIENS | 4 | 42 | 4pwi | A | 47 | GLY | p.Gly67Ala | P02766 |
| HOMO SAPIENS | 6 | 42 | 4pwi | A | 49 | THR | p.Thr69Ala | P02766 |
| HOMO SAPIENS | 6 | 42 | 4pwi | A | 50 | SER | p.Ser70Ile | P02766 |
| HOMO SAPIENS | 6 | 42 | 4pwi | A | 50 | SER | p.Ser70Arg | P02766 |
| HOMO SAPIENS | 0 | 42 | 4pwi | A | 53 | GLY | p.Gly73Glu | P02766 |

| | | | | | | | | |
|---|---|---|---|---|---|---|---|---|
| HOMO SAPIENS | 7 | 42 | 4pwi | A | 55 | LEU | p.Leu75Pro | P02766 |
| HOMO SAPIENS | 6 | 42 | 4pwi | A | 58 | LEU | p.Leu78His | P02766 |
| HOMO SAPIENS | 7 | 42 | 4pwi | A | 61 | GLU | p.Glu81Lys | P02766 |
| HOMO SAPIENS | 8 | 42 | 4pwi | A | 64 | PHE | p.Phe84Leu | P02766 |
| HOMO SAPIENS | 8 | 42 | 4pwi | A | 68 | ILE | p.Ile88Leu | P02766 |
| HOMO SAPIENS | 8 | 42 | 4pwi | A | 69 | TYR | p.Tyr89His | P02766 |
| HOMO SAPIENS | 4 | 42 | 4pwi | A | 71 | VAL | p.Val91Ala | P02766 |
| HOMO SAPIENS | 6 | 42 | 4pwi | A | 77 | SER | p.Ser97Tyr | P02766 |
| HOMO SAPIENS | 4 | 42 | 4pwi | A | 84 | ILE | p.Ile104Ser | P02766 |
| HOMO SAPIENS | 8 | 42 | 4pwi | A | 89 | GLU | p.Glu109Gln | P02766 |
| HOMO SAPIENS | 8 | 42 | 4pwi | A | 97 | ALA | p.Ala117Gly | P02766 |
| HOMO SAPIENS | 7 | 42 | 4pwi | A | 99 | ASP | p.Asp119Asn | P02766 |
| HOMO SAPIENS | 9 | 42 | 4pwi | A | 107 | ILE | p.Ile127Val | P02766 |
| HOMO SAPIENS | 9 | 42 | 4pwi | A | 111 | LEU | p.Leu131Met | P02766 |
| HOMO SAPIENS | 7 | 42 | 4pwi | A | 114 | TYR | p.Tyr134Cys | P02766 |
| HOMO SAPIENS | 7 | 42 | 4pwi | A | 114 | TYR | p.Tyr134His | P02766 |
| HOMO SAPIENS | 7 | 42 | 4pwi | A | 122 | VAL | p.Val142Ile | P02766 |
| HOMO SAPIENS | -1 | 42 | 4pwi | A | 126 | LYS | p.Lys146Arg | P02766 |
| HOMO SAPIENS | -1 | 42 | 4pwi | A | 127 | GLU | p.Glu147Ter | P02766 |
| HOMO SAPIENS | -1 | 1312 | 4dce | A | 1275 | ARG | p.Arg1275Gln | Q9UM73 |
| HOMO SAPIENS | -1 | 1312 | 4dce | A | 1280 | ALA | p.Ala1280Val | Q9UM73 |
| HOMO SAPIENS | 1 | 20 | 1y0x | X | 234 | ALA | p.Ala234Thr | P10828 |
| HOMO SAPIENS | 0 | 20 | 1y0x | X | 243 | ARG | p.Arg243Trp | P10828 |
| HOMO SAPIENS | 3 | 20 | 1y0x | X | 316 | ARG | p.Arg316His | P10828 |
| HOMO SAPIENS | 1 | 20 | 1y0x | X | 317 | ALA | p.Ala317Thr | P10828 |
| HOMO SAPIENS | 2 | 20 | 1y0x | X | 320 | ARG | p.Arg320Cys | P10828 |
| HOMO SAPIENS | 2 | 20 | 1y0x | X | 320 | ARG | p.Arg320His | P10828 |
| HOMO SAPIENS | 0 | 20 | 1y0x | X | 332 | GLY | p.Gly332Arg | P10828 |
| HOMO SAPIENS | 1 | 20 | 1y0x | X | 338 | ARG | p.Arg338Trp | P10828 |
| HOMO SAPIENS | 0 | 20 | 1y0x | X | 340 | GLN | p.Gln340His | P10828 |
| HOMO SAPIENS | 0 | 20 | 1y0x | X | 345 | GLY | p.Gly345Val | P10828 |
| HOMO SAPIENS | 0 | 20 | 1y0x | X | 345 | GLY | p.Gly345Ser | P10828 |
| HOMO SAPIENS | 0 | 20 | 1y0x | X | 347 | GLY | p.Gly347Glu | P10828 |
| HOMO SAPIENS | 1 | 20 | 1y0x | X | 438 | ARG | p.Arg438His | P10828 |
| HOMO SAPIENS | 0 | 20 | 1y0x | X | 442 | MET | p.Met442Val | P10828 |
| HOMO SAPIENS | 0 | 20 | 1y0x | X | 443 | LYS | p.Lys443Glu | P10828 |
| HOMO SAPIENS | 0 | 20 | 1y0x | X | 446 | CYS | p.Cys446Arg | P10828 |
| HOMO SAPIENS | 2 | 20 | 1y0x | X | 453 | PRO | p.Pro453His | P10828 |
| HOMO SAPIENS | 2 | 20 | 1y0x | X | 453 | PRO | p.Pro453Thr | P10828 |
| HOMO SAPIENS | 3 | 20 | 1y0x | X | 459 | PHE | p.Phe459Cys | P10828 |
| HOMO SAPIENS | 0 | 4409 | 3qb7 | D | 87 | GLU | p.Glu109Lys | P31785 |
| HOMO SAPIENS | 1 | 4409 | 3qb7 | D | 92 | GLY | p.Gly114Asp | P31785 |
| HOMO SAPIENS | 1 | 4409 | 3qb7 | D | 93 | CYS | p.Cys115Arg | P31785 |
| HOMO SAPIENS | 0 | 4409 | 3qb7 | D | 131 | ILE | p.Ile153Asn | P31785 |
| HOMO SAPIENS | 9 | 4409 | 3qb7 | D | 200 | ARG | p.Arg222Cys | P31785 |
| HOMO SAPIENS | 0 | 3891 | 3ije | B | 33 | LEU | p.Leu59Pro | P05106 |
| HOMO SAPIENS | 0 | 3891 | 3ije | B | 117 | LEU | p.Leu143Trp | P05106 |
| HOMO SAPIENS | 3 | 3891 | 3ije | B | 119 | ASP | p.Asp145Tyr | P05106 |
| HOMO SAPIENS | 2 | 3891 | 3ije | B | 143 | ARG | p.Arg169Gln | P05106 |
| HOMO SAPIENS | 2 | 3891 | 3ije | B | 214 | ARG | p.Arg240Gln | P05106 |
| HOMO SAPIENS | 2 | 3891 | 3ije | B | 214 | ARG | p.Arg240Trp | P05106 |
| HOMO SAPIENS | 2 | 3891 | 3ije | B | 374 | CYS | p.Cys400Tyr | P05106 |
| HOMO SAPIENS | 1 | 3891 | 3ije | B | 407 | PRO | p.Pro433Ala | P05106 |
| HOMO SAPIENS | 0 | 3891 | 3ije | B | 489 | ARG | p.Arg515Gln | P05106 |

| | | | | | | | | |
|---|---|---|---|---|---|---|---|---|
| HOMO SAPIENS | -1 | 9613 | 5mke | B | 28 | ARG | p.Arg28Pro | Q13563 |
| HOMO SAPIENS | -1 | 9613 | 5mke | B | 190 | ALA | p.Ala190Thr | Q13563 |
| HOMO SAPIENS | 4 | 9613 | 5mke | B | 262 | VAL | p.Val262Met | Q13563 |
| HOMO SAPIENS | 0 | 9613 | 5mke | B | 482 | PHE | p.Phe482Cys | Q13563 |
| HOMO SAPIENS | 4 | 9613 | 5mke | B | 511 | ASP | p.Asp511Val | Q13563 |
| HOMO SAPIENS | 5 | 9613 | 5mke | B | 516 | VAL | p.Val516Leu | Q13563 |
| HOMO SAPIENS | -1 | 9613 | 5mke | B | 807 | ARG | p.Arg807Gln | Q13563 |
| HOMO SAPIENS | 4 | 2428 | 1olh | C | 337 | ARG | p.Arg337Cys | P04637 |
| HOMO SAPIENS | 4 | 2428 | 1olh | C | 337 | ARG | p.Arg337His | P04637 |
| HOMO SAPIENS | 8 | 2428 | 1olh | C | 338 | PHE | p.Phe338Leu | P04637 |
| HOMO SAPIENS | 8 | 2428 | 1olh | C | 339 | GLU | p.Glu339Lys | P04637 |
| HOMO SAPIENS | 7 | 2428 | 1olh | C | 344 | LEU | p.Leu344Pro | P04637 |
| HOMO SAPIENS | 0 | 2428 | 1olh | C | 360 | GLY | p.Gly360Ala | P04637 |
| HOMO SAPIENS | 7 | 4867 | 1mox | B | 404 | GLY | p.Gly428Asp | P00533 |
| HOMO SAPIENS | 5 | 4867 | 1mox | B | 497 | ARG | p.Arg521Lys | P00533 |
| HOMO SAPIENS | 1 | 4630 | 1fu1 | A | 43 | TRP | p.Trp43Arg | Q13426 |
| HOMO SAPIENS | 2 | 4630 | 1fu1 | A | 161 | ARG | p.Arg161Ter | Q13426 |
| HOMO SAPIENS | 2 | 15 | 1lfy | B | 2 | HIS | p.His3Gln | P68871 |
| HOMO SAPIENS | 2 | 15 | 1lfy | B | 6 | GLU | p.Glu7Ala | P68871 |
| HOMO SAPIENS | 2 | 15 | 1lfy | B | 6 | GLU | p.Glu7Lys | P68871 |
| HOMO SAPIENS | 4 | 15 | 1lfy | B | 9 | SER | p.Ser10Cys | P68871 |
| HOMO SAPIENS | 6 | 15 | 1lfy | B | 15 | TRP | p.Trp16Ter | P68871 |
| HOMO SAPIENS | 3 | 15 | 1lfy | B | 17 | LYS | p.Lys18Glu | P68871 |
| HOMO SAPIENS | 6 | 15 | 1lfy | B | 19 | ASN | p.Asn20Ser | P68871 |
| HOMO SAPIENS | 6 | 15 | 1lfy | B | 22 | GLU | p.Glu23Ala | P68871 |
| HOMO SAPIENS | 6 | 15 | 1lfy | B | 22 | GLU | p.Glu23Lys | P68871 |
| HOMO SAPIENS | 8 | 15 | 1lfy | B | 26 | GLU | p.Glu27Lys | P68871 |
| HOMO SAPIENS | 8 | 15 | 1lfy | B | 28 | LEU | p.Leu29Pro | P68871 |
| HOMO SAPIENS | 8 | 15 | 1lfy | B | 30 | ARG | p.Arg31Ser | P68871 |
| HOMO SAPIENS | 8 | 15 | 1lfy | B | 30 | ARG | p.Arg31Lys | P68871 |
| HOMO SAPIENS | 6 | 15 | 1lfy | B | 39 | GLN | p.Gln40Ter | P68871 |
| HOMO SAPIENS | 2 | 15 | 1lfy | B | 43 | GLU | p.Glu44Gln | P68871 |
| HOMO SAPIENS | 6 | 15 | 1lfy | B | 57 | ASN | p.Asn58Lys | P68871 |
| HOMO SAPIENS | 6 | 15 | 1lfy | B | 59 | LYS | p.Lys60Asn | P68871 |
| HOMO SAPIENS | 8 | 15 | 1lfy | B | 61 | LYS | p.Lys62Glu | P68871 |
| HOMO SAPIENS | 4 | 15 | 1lfy | B | 73 | ASP | p.Asp74Tyr | P68871 |
| HOMO SAPIENS | 4 | 15 | 1lfy | B | 87 | THR | p.Thr88Ile | P68871 |
| HOMO SAPIENS | 4 | 15 | 1lfy | B | 87 | THR | p.Thr88Pro | P68871 |
| HOMO SAPIENS | 5 | 15 | 1lfy | B | 90 | GLU | p.Glu91Lys | P68871 |
| HOMO SAPIENS | 6 | 15 | 1lfy | B | 91 | LEU | p.Leu92Pro | P68871 |
| HOMO SAPIENS | 5 | 15 | 1lfy | B | 97 | HIS | p.His98Gln | P68871 |
| HOMO SAPIENS | 6 | 15 | 1lfy | B | 110 | LEU | p.Leu111Pro | P68871 |
| HOMO SAPIENS | 6 | 15 | 1lfy | B | 113 | VAL | p.Val114Glu | P68871 |
| HOMO SAPIENS | 2 | 15 | 1lfy | B | 119 | GLY | p.Gly120Ala | P68871 |
| HOMO SAPIENS | 4 | 15 | 1lfy | B | 121 | GLU | p.Glu122Gln | P68871 |
| HOMO SAPIENS | 5 | 15 | 1lfy | B | 134 | VAL | p.Val135Glu | P68871 |
| HOMO SAPIENS | 3 | 15 | 1lfy | B | 143 | HIS | p.His144Gln | P68871 |
| HOMO SAPIENS | 4 | 91 | 5d9k | A | 75 | GLY | p.Gly75Val | P51812 |
| HOMO SAPIENS | 5 | 91 | 5d9k | A | 82 | VAL | p.Val82Phe | P51812 |
| HOMO SAPIENS | 0 | 91 | 5d9k | A | 114 | ARG | p.Arg114Trp | P51812 |
| HOMO SAPIENS | 4 | 91 | 5d9k | A | 189 | ILE | p.Ile189Lys | P51812 |
| HOMO SAPIENS | -1 | 91 | 5d9k | A | 227 | SER | p.Ser227Ala | P51812 |
| HOMO SAPIENS | 1 | 91 | 5d9k | A | 268 | PHE | p.Phe268Ser | P51812 |
| HOMO SAPIENS | 0 | 954 | 2vt0 | A | 15 | VAL | p.Val54Leu | P04062 |

| | | | | | | | | |
|---|---|---|---|---|---|---|---|---|
| HOMO SAPIENS | 5 | 954 | 2vt0 | A | 46 | GLY | p.Gly85Glu | P04062 |
| HOMO SAPIENS | 5 | 954 | 2vt0 | A | 48 | ARG | p.Arg87Trp | P04062 |
| HOMO SAPIENS | 6 | 954 | 2vt0 | A | 79 | LYS | p.Lys118Asn | P04062 |
| HOMO SAPIENS | 0 | 954 | 2vt0 | A | 120 | ARG | p.Arg159Gln | P04062 |
| HOMO SAPIENS | 0 | 954 | 2vt0 | A | 120 | ARG | p.Arg159Trp | P04062 |
| HOMO SAPIENS | 0 | 954 | 2vt0 | A | 122 | PRO | p.Pro161Ser | P04062 |
| HOMO SAPIENS | 5 | 954 | 2vt0 | A | 131 | ARG | p.Arg170Leu | P04062 |
| HOMO SAPIENS | 5 | 954 | 2vt0 | A | 135 | TYR | p.Tyr174Cys | P04062 |
| HOMO SAPIENS | 0 | 954 | 2vt0 | A | 157 | LYS | p.Lys196Gln | P04062 |
| HOMO SAPIENS | 5 | 954 | 2vt0 | A | 163 | ARG | p.Arg202Gln | P04062 |
| HOMO SAPIENS | 5 | 954 | 2vt0 | A | 170 | ARG | p.Arg209Cys | P04062 |
| HOMO SAPIENS | 5 | 954 | 2vt0 | A | 188 | ASN | p.Asn227Ser | P04062 |
| HOMO SAPIENS | 3 | 954 | 2vt0 | A | 196 | SER | p.Ser235Pro | P04062 |
| HOMO SAPIENS | 5 | 954 | 2vt0 | A | 202 | GLY | p.Gly241Arg | P04062 |
| HOMO SAPIENS | 0 | 954 | 2vt0 | A | 212 | TYR | p.Tyr251His | P04062 |
| HOMO SAPIENS | 0 | 954 | 2vt0 | A | 213 | PHE | p.Phe252Ile | P04062 |
| HOMO SAPIENS | 5 | 954 | 2vt0 | A | 216 | PHE | p.Phe255Tyr | P04062 |
| HOMO SAPIENS | 6 | 954 | 2vt0 | A | 251 | PHE | p.Phe290Leu | P04062 |
| HOMO SAPIENS | 5 | 954 | 2vt0 | A | 255 | HIS | p.His294Gln | P04062 |
| HOMO SAPIENS | 6 | 954 | 2vt0 | A | 257 | ARG | p.Arg296Gln | P04062 |
| HOMO SAPIENS | 5 | 954 | 2vt0 | A | 289 | PRO | p.Pro328Leu | P04062 |
| HOMO SAPIENS | 0 | 954 | 2vt0 | A | 309 | ALA | p.Ala348Val | P04062 |
| HOMO SAPIENS | 0 | 954 | 2vt0 | A | 311 | HIS | p.His350Arg | P04062 |
| HOMO SAPIENS | 6 | 954 | 2vt0 | A | 312 | TRP | p.Trp351Cys | P04062 |
| HOMO SAPIENS | 5 | 954 | 2vt0 | A | 323 | THR | p.Thr362Ile | P04062 |
| HOMO SAPIENS | 5 | 954 | 2vt0 | A | 325 | GLY | p.Gly364Arg | P04062 |
| HOMO SAPIENS | 6 | 954 | 2vt0 | A | 342 | CYS | p.Cys381Gly | P04062 |
| HOMO SAPIENS | 5 | 954 | 2vt0 | A | 353 | ARG | p.Arg392Gly | P04062 |
| HOMO SAPIENS | 3 | 954 | 2vt0 | A | 364 | SER | p.Ser403Thr | P04062 |
| HOMO SAPIENS | 5 | 954 | 2vt0 | A | 369 | THR | p.Thr408Met | P04062 |
| HOMO SAPIENS | 3 | 954 | 2vt0 | A | 370 | ASN | p.Asn409Ser | P04062 |
| HOMO SAPIENS | 0 | 954 | 2vt0 | A | 371 | LEU | p.Leu410Val | P04062 |
| HOMO SAPIENS | 0 | 954 | 2vt0 | A | 375 | VAL | p.Val414Leu | P04062 |
| HOMO SAPIENS | 0 | 954 | 2vt0 | A | 377 | GLY | p.Gly416Ser | P04062 |
| HOMO SAPIENS | 8 | 954 | 2vt0 | A | 398 | VAL | p.Val437Phe | P04062 |
| HOMO SAPIENS | 0 | 954 | 2vt0 | A | 401 | PRO | p.Pro440Leu | P04062 |
| HOMO SAPIENS | 3 | 954 | 2vt0 | A | 409 | ASP | p.Asp448His | P04062 |
| HOMO SAPIENS | 3 | 954 | 2vt0 | A | 409 | ASP | p.Asp448Val | P04062 |
| HOMO SAPIENS | 0 | 954 | 2vt0 | A | 415 | PRO | p.Pro454Arg | P04062 |
| HOMO SAPIENS | 5 | 954 | 2vt0 | A | 444 | LEU | p.Leu483Arg | P04062 |
| HOMO SAPIENS | 5 | 954 | 2vt0 | A | 456 | ALA | p.Ala495Pro | P04062 |
| HOMO SAPIENS | 5 | 954 | 2vt0 | A | 463 | ARG | p.Arg502Cys | P04062 |
| HOMO SAPIENS | 5 | 954 | 2vt0 | A | 463 | ARG | p.Arg502His | P04062 |
| HOMO SAPIENS | 5 | 954 | 2vt0 | A | 478 | GLY | p.Gly517Ser | P04062 |
| HOMO SAPIENS | 5 | 954 | 2vt0 | A | 496 | ARG | p.Arg535Cys | P04062 |
| HOMO SAPIENS | 5 | 954 | 2vt0 | A | 496 | ARG | p.Arg535His | P04062 |
| HOMO SAPIENS | 1 | 9037 | 2rqb | A | 946 | ALA | p.Ala946Thr | Q9BYX4 |
| HOMO SAPIENS | 0 | 253 | 5E+092 | A | 308 | LEU | p.Leu308Pro | P37173 |
| HOMO SAPIENS | 0 | 253 | 5E+092 | A | 315 | THR | p.Thr315Met | P37173 |
| HOMO SAPIENS | 0 | 253 | 5E+092 | A | 328 | HIS | p.His328Gln | P37173 |
| HOMO SAPIENS | 0 | 253 | 5E+092 | A | 336 | TYR | p.Tyr336Asn | P37173 |
| HOMO SAPIENS | 0 | 253 | 5E+092 | A | 348 | ARG | p.Arg348Cys | P37173 |
| HOMO SAPIENS | 0 | 253 | 5E+092 | A | 355 | ALA | p.Ala355Pro | P37173 |
| HOMO SAPIENS | 1 | 253 | 5E+092 | A | 356 | ARG | p.Arg356Gly | P37173 |

| | | | | | | | | |
|---|---|---|---|---|---|---|---|---|
| HOMO SAPIENS | 0 | 253 | 5E+092 | A | 357 | GLY | p.Gly357Trp | P37173 |
| HOMO SAPIENS | 1 | 253 | 5E+092 | A | 403 | ARG | p.Arg403His | P37173 |
| HOMO SAPIENS | 5 | 253 | 5E+092 | A | 449 | SER | p.Ser449Phe | P37173 |
| HOMO SAPIENS | 0 | 253 | 5E+092 | A | 460 | ARG | p.Arg460Cys | P37173 |
| HOMO SAPIENS | 0 | 253 | 5E+092 | A | 460 | ARG | p.Arg460His | P37173 |
| HOMO SAPIENS | 2 | 253 | 5E+092 | A | 526 | GLU | p.Glu526Gln | P37173 |
| HOMO SAPIENS | 5 | 253 | 5E+092 | A | 528 | ARG | p.Arg528Cys | P37173 |
| HOMO SAPIENS | 5 | 253 | 5E+092 | A | 528 | ARG | p.Arg528His | P37173 |
| HOMO SAPIENS | 6 | 158 | 5amc | A | 231 | ARG | p.Arg260Leu | P12821 |
| HOMO SAPIENS | 7 | 158 | 5amc | A | 237 | TYR | p.Tyr266Ter | P12821 |
| HOMO SAPIENS | 6 | 158 | 5amc | A | 325 | GLY | p.Gly354Arg | P12821 |
| HOMO SAPIENS | 0 | 158 | 5amc | A | 467 | ARG | p.Arg496Ter | P12821 |
| HOMO SAPIENS | 4 | 158 | 5amc | A | 479 | ARG | p.Arg508Ter | P12821 |
| HOMO SAPIENS | 0 | 158 | 5amc | A | 563 | ASP | p.Asp592Gly | P12821 |
| HOMO SAPIENS | -1 | 33062 | 1gr3 | A | 545 | GLY | p.Gly545Arg | Q03692 |
| HOMO SAPIENS | 5 | 33062 | 1gr3 | A | 591 | CYS | p.Cys591Arg | Q03692 |
| HOMO SAPIENS | 2 | 33062 | 1gr3 | A | 595 | GLY | p.Gly595Glu | Q03692 |
| HOMO SAPIENS | 0 | 33062 | 1gr3 | A | 597 | TYR | p.Tyr597Cys | Q03692 |
| HOMO SAPIENS | 5 | 33062 | 1gr3 | A | 598 | TYR | p.Tyr598Asp | Q03692 |
| HOMO SAPIENS | 7 | 33062 | 1gr3 | A | 600 | SER | p.Ser600Pro | Q03692 |
| HOMO SAPIENS | 1 | 33062 | 1gr3 | A | 614 | LEU | p.Leu614Pro | Q03692 |
| HOMO SAPIENS | 6 | 33062 | 1gr3 | A | 651 | TRP | p.Trp651Arg | Q03692 |
| HOMO SAPIENS | 7 | 33062 | 1gr3 | A | 671 | SER | p.Ser671Pro | Q03692 |
| HOMO SAPIENS | -1 | 38257 | 2l2i | B | 87 | PRO | p.Pro87Ala | Q13351 |
| HOMO SAPIENS | 2 | 2036 | 1ghq | A | 231 | GLY | p.Gly1224Asp | P01024 |
| HOMO SAPIENS | 5 | 3594 | 4cde | F | 388 | TYR | p.Tyr412Cys | P53634 |
| HOMO SAPIENS | 9 | 3594 | 4cde | F | 405 | TRP | p.Trp429Cys | P53634 |
| HOMO SAPIENS | 0 | 3594 | 4cde | F | 429 | ILE | p.Ile453Val | P53634 |
| HOMO SAPIENS | 7 | 18 | 1jkb | A | 34 | TRP | p.Trp52Ter | P61626 |
| HOMO SAPIENS | 9 | 18 | 1jkb | A | 70 | THR | p.Thr88Asn | P61626 |
| HOMO SAPIENS | 8 | 18 | 1jkb | A | 121 | VAL | p.Val139Ile | P61626 |
| HOMO SAPIENS | 2 | 414 | 3fj6 | A | 136 | ARG | p.Arg135Cys | Q02127 |
| HOMO SAPIENS | 1 | 414 | 3fj6 | A | 153 | GLY | p.Gly152Arg | Q02127 |
| HOMO SAPIENS | 0 | 414 | 3fj6 | A | 200 | ARG | p.Arg199Cys | Q02127 |
| HOMO SAPIENS | 0 | 414 | 3fj6 | A | 203 | GLY | p.Gly202Ala | Q02127 |
| HOMO SAPIENS | 0 | 414 | 3fj6 | A | 245 | ARG | p.Arg244Trp | Q02127 |
| HOMO SAPIENS | 0 | 414 | 3fj6 | A | 342 | ALA | p.Ala341Val | Q02127 |
| HOMO SAPIENS | 1 | 414 | 3fj6 | A | 347 | ARG | p.Arg346Trp | Q02127 |
| HOMO SAPIENS | 6 | 103 | 1u3w | A | 77 | GLY | p.Gly78Ter | P00326 |
| HOMO SAPIENS | -1 | 8369 | 4ux8 | F | 16 | ARG | p.Arg93Trp | P39905 |
| HOMO SAPIENS | 0 | 366 | 2zjc | A | 32 | GLY | p.Arg108Trp | P01375 |
| HOMO SAPIENS | 0 | 1457 | 3v9i | A | 352 | LEU | p.Ser352Leu | P30038 |
| HOMO SAPIENS | 0 | 1457 | 3v9i | A | 470 | VAL | p.Val470Ile | P30038 |
| HOMO SAPIENS | 0 | 1457 | 3v9i | A | 473 | THR | p.Thr473Ala | P30038 |
| HOMO SAPIENS | 1 | 2 | 4wo5 | A | 462 | ARG | p.Arg462Ile | P15056 |
| HOMO SAPIENS | 3 | 2 | 4wo5 | A | 463 | ILE | p.Ile463Ser | P15056 |
| HOMO SAPIENS | 1 | 2 | 4wo5 | A | 466 | GLY | p.Gly466Val | P15056 |
| HOMO SAPIENS | 1 | 2 | 4wo5 | A | 468 | PHE | p.Phe468Ser | P15056 |
| HOMO SAPIENS | 2 | 2 | 4wo5 | A | 469 | GLY | p.Gly469Ala | P15056 |
| HOMO SAPIENS | 2 | 2 | 4wo5 | A | 469 | GLY | p.Gly469Arg | P15056 |
| HOMO SAPIENS | 2 | 2 | 4wo5 | A | 485 | LEU | p.Leu485Phe | P15056 |
| HOMO SAPIENS | 0 | 2 | 4wo5 | A | 501 | GLU | p.Glu501Gly | P15056 |
| HOMO SAPIENS | 0 | 2 | 4wo5 | A | 501 | GLU | p.Glu501Lys | P15056 |
| HOMO SAPIENS | 7 | 2 | 4wo5 | A | 581 | ASN | p.Asn581Asp | P15056 |

| | | | | | | | | |
|---|---|---|---|---|---|---|---|---|
| HOMO SAPIENS | 8 | 2 | 4wo5 | A | 594 | ASP | p.Asp594Gly | P15056 |
| HOMO SAPIENS | 0 | 2 | 4wo5 | A | 597 | LEU | p.Leu597Arg | P15056 |
| HOMO SAPIENS | 0 | 2 | 4wo5 | A | 597 | LEU | p.Leu597Val | P15056 |
| HOMO SAPIENS | 0 | 2 | 4wo5 | A | 600 | VAL | p.Val600Glu | P15056 |
| HOMO SAPIENS | 2 | 2 | 4wo5 | A | 601 | LYS | p.Lys601Glu | P15056 |
| HOMO SAPIENS | 8 | 2 | 4wo5 | A | 638 | ASP | p.Asp638Glu | P15056 |
| HOMO SAPIENS | -1 | 30366 | 5bqd | B | 19 | ALA | p.Ala19Ser | Q99593 |
| HOMO SAPIENS | -1 | 30366 | 5bqd | B | 23 | PRO | p.Pro23His | Q99593 |
| HOMO SAPIENS | 1 | 30366 | 5bqd | B | 49 | GLN | p.Gln49Lys | Q99593 |
| HOMO SAPIENS | 2 | 30366 | 5bqd | B | 54 | ILE | p.Ile54Thr | Q99593 |
| HOMO SAPIENS | 8 | 30366 | 5bqd | B | 80 | GLY | p.Gly80Arg | Q99593 |
| HOMO SAPIENS | 6 | 30366 | 5bqd | B | 106 | ILE | p.Ile106Val | Q99593 |
| HOMO SAPIENS | 7 | 30366 | 5bqd | B | 111 | ASP | p.Asp111Tyr | Q99593 |
| HOMO SAPIENS | -1 | 30366 | 5bqd | B | 237 | ARG | p.Arg237Gln | Q99593 |
| HOMO SAPIENS | -1 | 30366 | 5bqd | B | 237 | ARG | p.Arg237Trp | Q99593 |
| HOMO SAPIENS | 5 | 4789 | 1n4m | A | 567 | SER | p.Ser567Leu | P06400 |
| HOMO SAPIENS | 0 | 4789 | 1n4m | A | 658 | ALA | p.Ala658Asp | P06400 |
| HOMO SAPIENS | 7 | 4789 | 1n4m | A | 661 | ARG | p.Arg661Trp | P06400 |
| HOMO SAPIENS | 6 | 4789 | 1n4m | A | 668 | ARG | p.Arg668His | P06400 |
| HOMO SAPIENS | 8 | 4789 | 1n4m | A | 697 | ASP | p.Asp697Glu | P06400 |
| HOMO SAPIENS | 6 | 4789 | 1n4m | A | 712 | CYS | p.Cys712Arg | P06400 |
| HOMO SAPIENS | 1 | 1798 | 2r5g | A | 101 | HIS | p.His101Gln | O15247 |
| HOMO SAPIENS | 0 | 1798 | 2r5g | A | 160 | PRO | p.Pro160Ala | O15247 |
| HOMO SAPIENS | 2 | 22 | 4x0t | A | 82 | ARG | p.Arg110Ter | P49419 |
| HOMO SAPIENS | 0 | 22 | 4x0t | A | 149 | ALA | p.Ala177Pro | P49419 |
| HOMO SAPIENS | 4 | 22 | 4x0t | A | 164 | THR | p.Thr192Met | P49419 |
| HOMO SAPIENS | 9 | 22 | 4x0t | A | 167 | ASN | p.Asn195Ser | P49419 |
| HOMO SAPIENS | 0 | 22 | 4x0t | A | 175 | TRP | p.Trp203Gly | P49419 |
| HOMO SAPIENS | 0 | 22 | 4x0t | A | 194 | THR | p.Thr222Ala | P49419 |
| HOMO SAPIENS | 0 | 22 | 4x0t | A | 273 | ASN | p.Asn301Ile | P49419 |
| HOMO SAPIENS | 0 | 22 | 4x0t | A | 307 | ARG | p.Arg335Gln | P49419 |
| HOMO SAPIENS | 3 | 22 | 4x0t | A | 313 | SER | p.Ser341Asn | P49419 |
| HOMO SAPIENS | 0 | 22 | 4x0t | A | 329 | ILE | p.Ile357Val | P49419 |
| HOMO SAPIENS | 1 | 22 | 4x0t | A | 380 | TYR | p.Tyr408Ter | P49419 |
| HOMO SAPIENS | 9 | 22 | 4x0t | A | 399 | GLU | p.Glu427Gln | P49419 |
| HOMO SAPIENS | 0 | 22 | 4x0t | A | 406 | TYR | p.Tyr434Cys | P49419 |
| HOMO SAPIENS | 0 | 22 | 4x0t | A | 431 | ILE | p.Ile459Phe | P49419 |
| HOMO SAPIENS | 4 | 22 | 4x0t | A | 477 | GLY | p.Gly505Arg | P49419 |
| HOMO SAPIENS | 0 | 22 | 4x0t | A | 495 | THR | p.Thr523Ala | P49419 |
| HOMO SAPIENS | 8 | 11042 | 1y6b | A | 1145 | PRO | p.Pro1147Ser | P35968 |
| HOMO SAPIENS | -1 | 3867 | 2xij | A | 18 | GLN | p.Gln18Ter | P22033 |
| HOMO SAPIENS | -1 | 3867 | 2xij | A | 31 | ARG | p.Arg31Ter | P22033 |
| HOMO SAPIENS | 2 | 3867 | 2xij | A | 93 | ARG | p.Arg93His | P22033 |
| HOMO SAPIENS | 0 | 3867 | 2xij | A | 94 | GLY | p.Gly94Arg | P22033 |
| HOMO SAPIENS | 2 | 3867 | 2xij | A | 95 | PRO | p.Pro95Leu | P22033 |
| HOMO SAPIENS | 2 | 3867 | 2xij | A | 105 | TRP | p.Trp105Arg | P22033 |
| HOMO SAPIENS | 2 | 3867 | 2xij | A | 108 | ARG | p.Arg108Cys | P22033 |
| HOMO SAPIENS | 3 | 3867 | 2xij | A | 154 | ARG | p.Arg154His | P22033 |
| HOMO SAPIENS | 3 | 3867 | 2xij | A | 156 | ASP | p.Asp156Val | P22033 |
| HOMO SAPIENS | 0 | 3867 | 2xij | A | 189 | ASN | p.Asn189Ile | P22033 |
| HOMO SAPIENS | 0 | 3867 | 2xij | A | 191 | ALA | p.Ala191Glu | P22033 |
| HOMO SAPIENS | 0 | 3867 | 2xij | A | 203 | GLY | p.Gly203Arg | P22033 |
| HOMO SAPIENS | 2 | 3867 | 2xij | A | 215 | GLY | p.Gly215Ser | P22033 |
| HOMO SAPIENS | 0 | 3867 | 2xij | A | 219 | ASN | p.Asn219Tyr | P22033 |

| | | | | | | | | |
|---|---|---|---|---|---|---|---|---|
| HOMO SAPIENS | 3 | 3867 | 2xij | A | 228 | ARG | p.Arg228Ter | P22033 |
| HOMO SAPIENS | 1 | 3867 | 2xij | A | 297 | THR | p.Thr297Ile | P22033 |
| HOMO SAPIENS | 0 | 3867 | 2xij | A | 326 | ARG | p.Arg326Lys | P22033 |
| HOMO SAPIENS | 3 | 3867 | 2xij | A | 369 | ARG | p.Arg369His | P22033 |
| HOMO SAPIENS | 3 | 3867 | 2xij | A | 369 | ARG | p.Arg369Cys | P22033 |
| HOMO SAPIENS | 0 | 3867 | 2xij | A | 370 | THR | p.Thr370Pro | P22033 |
| HOMO SAPIENS | 0 | 3867 | 2xij | A | 377 | ALA | p.Ala377Glu | P22033 |
| HOMO SAPIENS | 2 | 3867 | 2xij | A | 426 | GLY | p.Gly426Ala | P22033 |
| HOMO SAPIENS | 2 | 3867 | 2xij | A | 427 | GLY | p.Gly427Asp | P22033 |
| HOMO SAPIENS | 1 | 3867 | 2xij | A | 499 | THR | p.Ala499Thr | P22033 |
| HOMO SAPIENS | 2 | 3867 | 2xij | A | 532 | ARG | p.Arg532His | P22033 |
| HOMO SAPIENS | 3 | 3867 | 2xij | A | 555 | ALA | p.Ala555Thr | P22033 |
| HOMO SAPIENS | 0 | 3867 | 2xij | A | 573 | PHE | p.Phe573Ser | P22033 |
| HOMO SAPIENS | 3 | 3867 | 2xij | A | 623 | GLY | p.Gly623Arg | P22033 |
| HOMO SAPIENS | 5 | 3867 | 2xij | A | 671 | VAL | p.Ile671Val | P22033 |
| HOMO SAPIENS | 4 | 3867 | 2xij | A | 694 | ARG | p.Arg694Trp | P22033 |
| HOMO SAPIENS | 1 | 3867 | 2xij | A | 700 | MET | p.Met700Lys | P22033 |
| HOMO SAPIENS | 8 | 3867 | 2xij | A | 703 | GLY | p.Gly703Arg | P22033 |
| HOMO SAPIENS | 0 | 3867 | 2xij | A | 717 | GLY | p.Gly717Val | P22033 |
| HOMO SAPIENS | 1 | 3867 | 2xij | A | 727 | ARG | p.Arg727Ter | P22033 |
| HOMO SAPIENS | -1 | 52 | 3k91 | B | 72 | GLY | p.Gly73Ser | P00441 |
| HOMO SAPIENS | 6 | 52 | 3k91 | B | 96 | ASP | p.Asp97Asn | P00441 |
| HOMO SAPIENS | 5 | 52 | 3k91 | B | 112 | ILE | p.Ile113Thr | P00441 |
| HOMO SAPIENS | -1 | 52 | 3k91 | B | 134 | SER | p.Ser135Asn | P00441 |
| HOMO SAPIENS | 0 | 3950 | 4fom | A | 296 | ASN | p.Asn296His | Q9NQS3 |
| HOMO SAPIENS | 1 | 10 | 1de4 | A | 31 | VAL | p.Val53Met | Q30201 |
| HOMO SAPIENS | 0 | 10 | 1de4 | A | 37 | VAL | p.Val59Met | Q30201 |
| HOMO SAPIENS | 0 | 10 | 1de4 | A | 71 | GLY | p.Gly93Arg | Q30201 |
| HOMO SAPIENS | 0 | 10 | 1de4 | A | 83 | ILE | p.Ile105Thr | Q30201 |
| HOMO SAPIENS | 0 | 10 | 1de4 | A | 105 | GLN | p.Gln127His | Q30201 |
| HOMO SAPIENS | 1 | 10 | 1de4 | A | 146 | GLU | p.Glu168Ter | Q30201 |
| HOMO SAPIENS | 2 | 10 | 1de4 | A | 261 | GLN | p.Gln283Pro | Q30201 |
| HOMO SAPIENS | 0 | 8038 | 4x7n | A | 586 | ILE | p.Ile586Leu | Q9NZJ5 |
| HOMO SAPIENS | 0 | 8038 | 4x7n | A | 588 | ARG | p.Arg588Gln | Q9NZJ5 |
| HOMO SAPIENS | 0 | 35175 | 5ugw | A | 1062 | ALA | p.Ala1062Thr | O14746 |
| HOMO SAPIENS | 2 | 35175 | 5ugw | A | 1090 | VAL | p.Val1090Met | O14746 |
| HOMO SAPIENS | 0 | 1722 | 4z3k | C | 147 | ARG | p.Arg150Gly | P35270 |
| HOMO SAPIENS | 0 | 1722 | 4z3k | C | 160 | PRO | p.Pro163Leu | P35270 |
| HOMO SAPIENS | 0 | 1722 | 4z3k | C | 207 | ARG | p.Arg210Trp | P35270 |
| HOMO SAPIENS | 0 | 1722 | 4z3k | C | 216 | ARG | p.Arg219Ter | P35270 |
| HOMO SAPIENS | 0 | 1722 | 4z3k | C | 248 | LYS | p.Lys251Ter | P35270 |
| HOMO SAPIENS | 1 | 2832 | 4x8v | L | 115 | HIS | p.His175Gln | P08709 |
| HOMO SAPIENS | 0 | 825 | 1emh | A | 242 | PHE | p.Phe251Ser | P13051 |
| HOMO SAPIENS | 0 | 122 | 5uwk | A | 109 | ARG | p.Arg109Ter | P45452 |
| HOMO SAPIENS | 1 | 122 | 5uwk | A | 170 | LYS | p.Lys170Thr | P45452 |
| HOMO SAPIENS | 9 | 122 | 5uwk | A | 232 | HIS | p.His232Asn | P45452 |
| HOMO SAPIENS | 5 | 122 | 5uwk | A | 257 | ASP | p.Asp257Val | P45452 |
| HOMO SAPIENS | 9 | 1074 | 3p5i | A | 264 | TRP | p.Trp264Arg | Q9UJ71 |
| HOMO SAPIENS | 0 | 146 | 3r3g | A | 14C | GLU | p.Glu352Lys | P00734 |
| HOMO SAPIENS | 3 | 9522 | 1wyq | A | 83 | LEU | p.Leu253Pro | O15020 |
| HOMO SAPIENS | 8 | 67 | 2vgb | A | 130 | SER | p.Ser130Tyr | P30613 |
| HOMO SAPIENS | 8 | 67 | 2vgb | A | 163 | ARG | p.Arg163Cys | P30613 |
| HOMO SAPIENS | 4 | 67 | 2vgb | A | 241 | GLU | p.Glu241Ter | P30613 |
| HOMO SAPIENS | 9 | 67 | 2vgb | A | 384 | THR | p.Thr384Met | P30613 |

| | | | | | | | | |
|---|---|---|---|---|---|---|---|---|
| HOMO SAPIENS | 0 | 67 | 2vgb | A | 421 | GLN | p.Gln421Lys | P30613 |
| HOMO SAPIENS | 6 | 67 | 2vgb | A | 479 | ARG | p.Arg479His | P30613 |
| HOMO SAPIENS | 8 | 67 | 2vgb | A | 510 | ARG | p.Arg510Gln | P30613 |
| HOMO SAPIENS | 4 | 67 | 2vgb | A | 569 | ARG | p.Arg569Trp | P30613 |
| HOMO SAPIENS | 0 | 9088 | 5v44 | A | 232 | ASN | p.Asn232Lys | Q9NZJ4 |
| HOMO SAPIENS | 4 | 16 | 3lbn | A | 12 | GLY | p.Gly12Val | P01112 |
| HOMO SAPIENS | 4 | 16 | 3lbn | A | 12 | GLY | p.Gly12Ser | P01112 |
| HOMO SAPIENS | 4 | 16 | 3lbn | A | 13 | GLY | p.Gly13Cys | P01112 |
| HOMO SAPIENS | 4 | 16 | 3lbn | A | 13 | GLY | p.Gly13Asp | P01112 |
| HOMO SAPIENS | 5 | 16 | 3lbn | A | 22 | GLN | p.Gln22Lys | P01112 |
| HOMO SAPIENS | 5 | 16 | 3lbn | A | 58 | THR | p.Thr58Ile | P01112 |
| HOMO SAPIENS | 3 | 16 | 3lbn | A | 61 | GLN | p.Gln61Lys | P01112 |
| HOMO SAPIENS | 1 | 16 | 3lbn | A | 63 | GLU | p.Glu63Lys | P01112 |
| HOMO SAPIENS | 9 | 16 | 3lbn | A | 117 | LYS | p.Lys117Arg | P01112 |
| HOMO SAPIENS | 6 | 16 | 3lbn | A | 123 | ARG | p.Arg123Cys | P01112 |
| HOMO SAPIENS | 1 | 16 | 3lbn | A | 138 | GLY | p.Gly138Ser | P01112 |
| HOMO SAPIENS | 9 | 16 | 3lbn | A | 146 | ALA | p.Ala146Thr | P01112 |
| HOMO SAPIENS | 9 | 16 | 3lbn | A | 146 | ALA | p.Ala146Val | P01112 |
| HOMO SAPIENS | 1 | 359 | 5nvy | C | 63 | LEU | p.Leu63Pro | P40337 |
| HOMO SAPIENS | 1 | 359 | 5nvy | C | 64 | ARG | p.Arg64Pro | P40337 |
| HOMO SAPIENS | 1 | 359 | 5nvy | C | 70 | GLU | p.Glu70Lys | P40337 |
| HOMO SAPIENS | 0 | 359 | 5nvy | C | 84 | VAL | p.Val84Leu | P40337 |
| HOMO SAPIENS | 1 | 359 | 5nvy | C | 88 | TRP | p.Trp88Ser | P40337 |
| HOMO SAPIENS | 1 | 359 | 5nvy | C | 93 | GLY | p.Gly93Cys | P40337 |
| HOMO SAPIENS | 1 | 359 | 5nvy | C | 98 | TYR | p.Tyr98His | P40337 |
| HOMO SAPIENS | 1 | 359 | 5nvy | C | 111 | SER | p.Ser111Arg | P40337 |
| HOMO SAPIENS | 1 | 359 | 5nvy | C | 112 | TYR | p.Tyr112His | P40337 |
| HOMO SAPIENS | 1 | 359 | 5nvy | C | 126 | ASP | p.Asp126Tyr | P40337 |
| HOMO SAPIENS | 1 | 359 | 5nvy | C | 130 | VAL | p.Val130Leu | P40337 |
| HOMO SAPIENS | 1 | 359 | 5nvy | C | 135 | LEU | p.Leu135Phe | P40337 |
| HOMO SAPIENS | 9 | 359 | 5nvy | C | 161 | ARG | p.Arg161Gly | P40337 |
| HOMO SAPIENS | 7 | 359 | 5nvy | C | 162 | CYS | p.Cys162Phe | P40337 |
| HOMO SAPIENS | 4 | 359 | 5nvy | C | 163 | LEU | p.Leu163Pro | P40337 |
| HOMO SAPIENS | 3 | 359 | 5nvy | C | 164 | GLN | p.Gln164Arg | P40337 |
| HOMO SAPIENS | 9 | 359 | 5nvy | C | 166 | VAL | p.Val166Phe | P40337 |
| HOMO SAPIENS | 7 | 359 | 5nvy | C | 167 | ARG | p.Arg167Gly | P40337 |
| HOMO SAPIENS | 7 | 359 | 5nvy | C | 167 | ARG | p.Arg167Gln | P40337 |
| HOMO SAPIENS | 1 | 359 | 5nvy | C | 183 | SER | p.Ser183Ter | P40337 |
| HOMO SAPIENS | 2 | 359 | 5nvy | C | 188 | LEU | p.Leu188Val | P40337 |
| HOMO SAPIENS | 1 | 359 | 5nvy | C | 191 | HIS | p.His191Asp | P40337 |
| HOMO SAPIENS | 1 | 359 | 5nvy | C | 192 | PRO | p.Pro192Ser | P40337 |
| HOMO SAPIENS | 1 | 359 | 5nvy | C | 200 | ARG | p.Arg200Trp | P40337 |
| HOMO SAPIENS | -1 | 359 | 5nvy | C | 211 | MET | p.Met211Leu | P40337 |
| HOMO SAPIENS | 0 | 14099 | 4btm | A | 282 | ARG | p.Arg258Gln | Q5TCY1 |
| HOMO SAPIENS | 0 | 696 | 5h1b | A | 150 | ARG | p.Arg150Gln | Q06609 |
| HOMO SAPIENS | -1 | 208 | 2bxl | A | 1 | ASP | p.Asp25Val | P02768 |
| HOMO SAPIENS | 0 | 208 | 2bxl | A | 269 | ASP | p.Asp293Gly | P02768 |
| HOMO SAPIENS | 0 | 208 | 2bxl | A | 570 | GLU | p.Glu594Lys | P02768 |
| HOMO SAPIENS | 0 | 208 | 2bxl | A | 573 | LYS | p.Lys597Glu | P02768 |
| HOMO SAPIENS | -1 | 12813 | 2uzx | B | 35 | MET | p.Met35Leu | P08581 |
| HOMO SAPIENS | 1 | 12813 | 2uzx | B | 156 | SER | p.Ser156Leu | P08581 |
| HOMO SAPIENS | 0 | 12813 | 2uzx | B | 168 | GLU | p.Glu168Asp | P08581 |
| HOMO SAPIENS | 2 | 12813 | 2uzx | B | 211 | LEU | p.Leu211Trp | P08581 |
| HOMO SAPIENS | 2 | 12813 | 2uzx | B | 218 | ARG | p.Arg218Ser | P08581 |

| | | | | | | | | |
|---|---|---|---|---|---|---|---|---|
| HOMO SAPIENS | 1 | 12813 | 2uzx | B | 238 | LEU | p.Leu238Ser | P08581 |
| HOMO SAPIENS | 4 | 12813 | 2uzx | B | 320 | ALA | p.Ala320Val | P08581 |
| HOMO SAPIENS | 1 | 12813 | 2uzx | B | 375 | ASN | p.Asn375Ser | P08581 |
| HOMO SAPIENS | 0 | 12813 | 2uzx | B | 575 | LEU | p.Leu575Ile | P08581 |
| HOMO SAPIENS | 1 | 10780 | 4ejx | A | 348 | ARG | p.Arg367Cys | P00450 |
| HOMO SAPIENS | 2 | 10780 | 4ejx | A | 525 | ASP | p.Asp544Glu | P00450 |
| HOMO SAPIENS | 1 | 1407 | 5jn5 | A | 88 | ILE | p.Ile88Val | P36871 |
| HOMO SAPIENS | 7 | 1407 | 5jn5 | A | 115 | THR | p.Thr115Ala | P36871 |
| HOMO SAPIENS | 3 | 1407 | 5jn5 | A | 121 | GLY | p.Gly121Arg | P36871 |
| HOMO SAPIENS | 2 | 1407 | 5jn5 | A | 221 | ARG | p.Arg221Cys | P36871 |
| HOMO SAPIENS | 6 | 1407 | 5jn5 | A | 263 | TYR | p.Asp263Tyr | P36871 |
| HOMO SAPIENS | 3 | 1407 | 5jn5 | A | 420 | TYR | p.Tyr420His | P36871 |
| HOMO SAPIENS | 0 | 1407 | 5jn5 | A | 501 | VAL | p.Val501Ile | P36871 |
| HOMO SAPIENS | -1 | 1878 | 1yow | A | 255 | ARG | p.Arg255Leu | Q13285 |
| HOMO SAPIENS | 6 | 1878 | 1yow | A | 293 | ASP | p.Asp293Asn | Q13285 |
| HOMO SAPIENS | 1 | 1878 | 1yow | A | 437 | LEU | p.Leu437Gln | Q13285 |
| HOMO SAPIENS | 0 | 3071 | 2wpm | E | 145 | ARG | p.Arg191His | P00740 |
| HOMO SAPIENS | 0 | 3071 | 2wpm | E | 145 | ARG | p.Arg191Cys | P00740 |
| HOMO SAPIENS | 3 | 589 | 3ggc | B | 15 | GLY | p.Gly16Ser | P00492 |
| HOMO SAPIENS | 4 | 589 | 3ggc | B | 40 | LEU | p.Leu41Pro | P00492 |
| HOMO SAPIENS | 2 | 589 | 3ggc | B | 50 | ARG | p.Arg51Gly | P00492 |
| HOMO SAPIENS | 5 | 589 | 3ggc | B | 51 | ASP | p.Asp52Gly | P00492 |
| HOMO SAPIENS | 0 | 589 | 3ggc | B | 57 | GLY | p.Gly58Arg | P00492 |
| HOMO SAPIENS | 6 | 589 | 3ggc | B | 69 | GLY | p.Gly70Glu | P00492 |
| HOMO SAPIENS | 1 | 589 | 3ggc | B | 73 | PHE | p.Phe74Leu | P00492 |
| HOMO SAPIENS | 0 | 589 | 3ggc | B | 77 | LEU | p.Leu78Val | P00492 |
| HOMO SAPIENS | 3 | 589 | 3ggc | B | 79 | ASP | p.Asp80Val | P00492 |
| HOMO SAPIENS | 0 | 589 | 3ggc | B | 103 | SER | p.Ser104Arg | P00492 |
| HOMO SAPIENS | 0 | 589 | 3ggc | B | 109 | SER | p.Ser110Leu | P00492 |
| HOMO SAPIENS | 1 | 589 | 3ggc | B | 129 | VAL | p.Val130Asp | P00492 |
| HOMO SAPIENS | 4 | 589 | 3ggc | B | 131 | ILE | p.Ile132Met | P00492 |
| HOMO SAPIENS | 7 | 589 | 3ggc | B | 142 | MET | p.Met143Lys | P00492 |
| HOMO SAPIENS | 0 | 589 | 3ggc | B | 160 | ALA | p.Ala161Ser | P00492 |
| HOMO SAPIENS | 6 | 589 | 3ggc | B | 167 | THR | p.Thr168Ile | P00492 |
| HOMO SAPIENS | 0 | 589 | 3ggc | B | 169 | ARG | p.Arg170Ter | P00492 |
| HOMO SAPIENS | 5 | 589 | 3ggc | B | 175 | PRO | p.Pro176Leu | P00492 |
| HOMO SAPIENS | 8 | 589 | 3ggc | B | 193 | ASP | p.Asp194Glu | P00492 |
| HOMO SAPIENS | 8 | 589 | 3ggc | B | 193 | ASP | p.Asp194Asn | P00492 |
| HOMO SAPIENS | 8 | 589 | 3ggc | B | 198 | PHE | p.Phe199Val | P00492 |
| HOMO SAPIENS | 8 | 589 | 3ggc | B | 200 | ASP | p.Asp201Gly | P00492 |
| HOMO SAPIENS | 0 | 4269 | 2pla | B | 174 | GLU | p.Glu174Lys | Q8N335 |
| HOMO SAPIENS | 1 | 4269 | 2pla | B | 220 | ARG | p.Arg220His | Q8N335 |
| HOMO SAPIENS | 4 | 4269 | 2pla | B | 280 | ALA | p.Ala280Val | Q8N335 |
| HOMO SAPIENS | 0 | 2712 | 1ii4 | E | 252 | TRP | p.Ser252Leu | P21802 |
| HOMO SAPIENS | 0 | 2712 | 1ii4 | E | 253 | PRO | p.Pro253Arg | P21802 |
| HOMO SAPIENS | 5 | 2712 | 1ii4 | E | 267 | SER | p.Ser267Pro | P21802 |
| HOMO SAPIENS | 0 | 2712 | 1ii4 | E | 278 | CYS | p.Cys278Phe | P21802 |
| HOMO SAPIENS | 0 | 2712 | 1ii4 | E | 289 | GLN | p.Gln289Pro | P21802 |
| HOMO SAPIENS | 0 | 2712 | 1ii4 | E | 290 | TRP | p.Trp290Cys | P21802 |
| HOMO SAPIENS | 0 | 2712 | 1ii4 | E | 290 | TRP | p.Trp290Arg | P21802 |
| HOMO SAPIENS | 0 | 2712 | 1ii4 | E | 292 | LYS | p.Lys292Glu | P21802 |
| HOMO SAPIENS | 0 | 2712 | 1ii4 | E | 315 | ALA | p.Ala315Ser | P21802 |
| HOMO SAPIENS | 0 | 2712 | 1ii4 | E | 321 | ASP | p.Asp321Ala | P21802 |
| HOMO SAPIENS | 1 | 2712 | 1ii4 | E | 328 | TYR | p.Tyr328Cys | P21802 |

| | | | | | | | | |
|---|---|---|---|---|---|---|---|---|
| HOMO SAPIENS | 9 | 2712 | 1ii4 | E | 340 | TYR | p.Tyr340His | P21802 |
| HOMO SAPIENS | 1 | 2712 | 1ii4 | E | 341 | THR | p.Thr341Pro | P21802 |
| HOMO SAPIENS | 0 | 2712 | 1ii4 | E | 342 | CYS | p.Cys342Arg | P21802 |
| HOMO SAPIENS | 0 | 2712 | 1ii4 | E | 342 | CYS | p.Cys342Trp | P21802 |
| HOMO SAPIENS | 0 | 2712 | 1ii4 | E | 344 | ALA | p.Ala344Gly | P21802 |
| HOMO SAPIENS | 0 | 2712 | 1ii4 | E | 347 | SER | p.Ser347Cys | P21802 |
| HOMO SAPIENS | 0 | 2712 | 1ii4 | E | 351 | SER | p.Ser351Cys | P21802 |
| HOMO SAPIENS | 1 | 2712 | 1ii4 | E | 354 | SER | p.Ser354Cys | P21802 |
| HOMO SAPIENS | 2 | 53 | 4pj1 | H | 5 | ASP | p.Asp29Gly | P10809 |
| HOMO SAPIENS | 0 | 53 | 4pj1 | H | 74 | VAL | p.Val98Ile | P10809 |
| HOMO SAPIENS | 0 | 29968 | 2fdb | M | 71 | LYS | p.Lys89Glu | P55075 |
| HOMO SAPIENS | 2 | 29968 | 2fdb | M | 98 | ARG | p.Arg116Gly | P55075 |
| HOMO SAPIENS | 2 | 29968 | 2fdb | M | 122 | GLY | p.Gly140Ser | P55075 |
| HOMO SAPIENS | 8 | 5061 | 5f85 | B | 60 | GLY | p.Gly106Ser | P00740 |
| HOMO SAPIENS | 0 | 35944 | 1h6h | A | 85 | THR | p.Thr85Asn | Q15080 |
| HOMO SAPIENS | 8 | 35944 | 1h6h | A | 105 | ARG | p.Arg105Gln | Q15080 |
| HOMO SAPIENS | 7 | 12016 | 1hj7 | A | 295 | ASN | p.Asn316Ser | P01130 |
| HOMO SAPIENS | 5 | 12016 | 1hj7 | A | 302 | GLY | p.Gly323Ser | P01130 |
| HOMO SAPIENS | 6 | 12016 | 1hj7 | A | 319 | CYS | p.Cys340Trp | P01130 |
| HOMO SAPIENS | 7 | 12016 | 1hj7 | A | 329 | ARG | p.Arg350Ter | P01130 |
| HOMO SAPIENS | 5 | 12016 | 1hj7 | A | 331 | CYS | p.Cys352Ter | P01130 |
| HOMO SAPIENS | 2 | 12016 | 1hj7 | A | 345 | GLN | p.Gln366Arg | P01130 |
| HOMO SAPIENS | 2 | 1481 | 2i07 | A | 239 | TYR | p.Tyr261Ter | P01024 |
| HOMO SAPIENS | -1 | 1481 | 2i07 | A | 292 | LEU | p.Pro314Leu | P01024 |
| HOMO SAPIENS | 2 | 1481 | 2i07 | A | 326 | ILE | p.Ile348Val | P01024 |
| HOMO SAPIENS | 2 | 1481 | 2i07 | A | 413 | GLU | p.Glu435Lys | P01024 |
| HOMO SAPIENS | 4 | 1481 | 2i07 | A | 447 | GLU | p.Glu469Asp | P01024 |
| HOMO SAPIENS | 2 | 1481 | 2i07 | A | 597 | VAL | p.Val619Met | P01024 |
| HOMO SAPIENS | 5 | 1481 | 2i07 | A | 603 | ILE | p.Ile625Phe | P01024 |
| HOMO SAPIENS | 0 | 1481 | 2i07 | A | 611 | LYS | p.Lys633Arg | P01024 |
| HOMO SAPIENS | 0 | 11123 | 3c5n | A | 315 | GLY | p.Gly368Trp | O00294 |
| HOMO SAPIENS | 8 | 11123 | 3c5n | A | 329 | PHE | p.Phe382Ser | O00294 |
| HOMO SAPIENS | 5 | 11123 | 3c5n | A | 429 | ARG | p.Arg482Trp | O00294 |
| HOMO SAPIENS | 8 | 11123 | 3c5n | A | 438 | PHE | p.Phe491Leu | O00294 |
| HOMO SAPIENS | 3 | 4398 | 3fx0 | B | 1288 | ALA | p.Ala288Gly | Q9Y6K9 |
| HOMO SAPIENS | 7 | 4398 | 3fx0 | B | 1315 | GLU | p.Glu315Ala | Q9Y6K9 |
| HOMO SAPIENS | 8 | 4398 | 3fx0 | B | 1319 | ARG | p.Arg319Gln | Q9Y6K9 |
| HOMO SAPIENS | 6 | 3989 | 3qd6 | S | 83 | CYS | p.Cys83Arg | P25942 |
| HOMO SAPIENS | 1 | 3989 | 3qd6 | S | 124 | SER | p.Ser124Leu | P25942 |
| HOMO SAPIENS | 2 | 24003 | 2m0v | A | 225 | GLU | p.Glu225Lys | O14745 |
| HOMO SAPIENS | 2 | 46 | 4aqd | A | 70 | ASP | p.Asp98Gly | P06276 |
| HOMO SAPIENS | 4 | 46 | 4aqd | A | 115 | GLY | p.Gly143Asp | P06276 |
| HOMO SAPIENS | 3 | 46 | 4aqd | A | 184 | ALA | p.Ala212Val | P06276 |
| HOMO SAPIENS | 0 | 46 | 4aqd | A | 225 | GLY | p.Gly253Ter | P06276 |
| HOMO SAPIENS | 2 | 46 | 4aqd | A | 243 | THR | p.Thr271Met | P06276 |
| HOMO SAPIENS | 2 | 46 | 4aqd | A | 255 | GLU | p.Glu283Asp | P06276 |
| HOMO SAPIENS | 5 | 46 | 4aqd | A | 267 | LYS | p.Lys295Arg | P06276 |
| HOMO SAPIENS | 3 | 46 | 4aqd | A | 271 | GLU | p.Glu299Ter | P06276 |
| HOMO SAPIENS | 2 | 46 | 4aqd | A | 307 | LEU | p.Leu335Pro | P06276 |
| HOMO SAPIENS | 1 | 46 | 4aqd | A | 330 | LEU | p.Leu358Ile | P06276 |
| HOMO SAPIENS | 0 | 46 | 4aqd | A | 497 | GLU | p.Glu525Val | P06276 |
| HOMO SAPIENS | 3 | 786 | 1ols | B | 51 | GLY | p.Gly101Asp | P21953 |
| HOMO SAPIENS | 2 | 786 | 1ols | B | 87 | ALA | p.Ala137Val | P21953 |
| HOMO SAPIENS | 0 | 786 | 1ols | B | 110 | ILE | p.Ile160Ser | P21953 |

| | | | | | | | | |
|---|---|---|---|---|---|---|---|---|
| HOMO SAPIENS | 4 | 786 | 1ols | B | 120 | ARG | p.Arg170Cys | P21953 |
| HOMO SAPIENS | 2 | 786 | 1ols | B | 126 | ASN | p.Asn176Tyr | P21953 |
| HOMO SAPIENS | 0 | 786 | 1ols | B | 133 | ARG | p.Arg183Gly | P21953 |
| HOMO SAPIENS | 0 | 786 | 1ols | B | 133 | ARG | p.Arg183Pro | P21953 |
| HOMO SAPIENS | 0 | 786 | 1ols | B | 135 | PRO | p.Pro185Leu | P21953 |
| HOMO SAPIENS | 3 | 786 | 1ols | B | 156 | HIS | p.His206Tyr | P21953 |
| HOMO SAPIENS | 0 | 786 | 1ols | B | 228 | GLY | p.Gly278Ser | P21953 |
| HOMO SAPIENS | 1 | 786 | 1ols | B | 235 | ARG | p.Arg285Ter | P21953 |
| HOMO SAPIENS | 6 | 786 | 1ols | B | 274 | ARG | p.Arg324Ter | P21953 |
| HOMO SAPIENS | 1 | 831 | 4zt1 | B | 48 | VAL | p.Val202Ile | P12830 |
| HOMO SAPIENS | 6 | 831 | 4zt1 | B | 70 | ARG | p.Arg224Cys | P12830 |
| HOMO SAPIENS | 5 | 831 | 4zt1 | B | 79 | HIS | p.His233Gln | P12830 |
| HOMO SAPIENS | 4 | 831 | 4zt1 | B | 120 | GLY | p.Gly274Ser | P12830 |
| HOMO SAPIENS | 7 | 831 | 4zt1 | B | 187 | TYR | p.Tyr341Ter | P12830 |
| HOMO SAPIENS | 0 | 8497 | 3sla | A | 287 | ASN | p.Asn287Ser | P35222 |
| HOMO SAPIENS | 7 | 23986 | 2ke1 | A | 301 | VAL | p.Val301Met | O43918 |
| HOMO SAPIENS | 0 | 4967 | 1rj7 | B | 334 | ARG | p.Arg334His | Q92838 |
| HOMO SAPIENS | 0 | 4967 | 1rj7 | B | 338 | THR | p.Thr338Met | Q92838 |
| HOMO SAPIENS | 3 | 4967 | 1rj7 | B | 349 | ALA | p.Ala349Thr | Q92838 |
| HOMO SAPIENS | 2 | 4967 | 1rj7 | B | 358 | GLN | p.Gln358Glu | Q92838 |
| HOMO SAPIENS | 1 | 4967 | 1rj7 | B | 375 | LYS | p.Lys375Glu | Q92838 |
| HOMO SAPIENS | 9 | 7623 | 1d4t | A | 32 | ARG | p.Arg32Thr | O60880 |
| HOMO SAPIENS | 6 | 7623 | 1d4t | A | 55 | ARG | p.Arg55Leu | O60880 |
| HOMO SAPIENS | 0 | 7623 | 1d4t | A | 68 | THR | p.Thr68Ile | O60880 |
| HOMO SAPIENS | 6 | 705 | 3idh | A | 59 | PRO | p.Pro59Ser | P35557 |
| HOMO SAPIENS | 5 | 705 | 3idh | A | 124 | ASP | p.Asp124Asn | P35557 |
| HOMO SAPIENS | 4 | 705 | 3idh | A | 131 | SER | p.Ser131Pro | P35557 |
| HOMO SAPIENS | 0 | 705 | 3idh | A | 147 | GLY | p.Gly147Asp | P35557 |
| HOMO SAPIENS | 5 | 705 | 3idh | A | 186 | ARG | p.Arg186Ter | P35557 |
| HOMO SAPIENS | 0 | 705 | 3idh | A | 202 | MET | p.Met202Thr | P35557 |
| HOMO SAPIENS | 2 | 705 | 3idh | A | 210 | MET | p.Met210Lys | P35557 |
| HOMO SAPIENS | 1 | 705 | 3idh | A | 215 | TYR | p.Tyr215Ter | P35557 |
| HOMO SAPIENS | 0 | 705 | 3idh | A | 226 | VAL | p.Val226Met | P35557 |
| HOMO SAPIENS | 9 | 705 | 3idh | A | 228 | THR | p.Thr228Met | P35557 |
| HOMO SAPIENS | 9 | 705 | 3idh | A | 256 | GLU | p.Glu256Lys | P35557 |
| HOMO SAPIENS | 2 | 705 | 3idh | A | 261 | GLY | p.Gly261Arg | P35557 |
| HOMO SAPIENS | 6 | 705 | 3idh | A | 279 | GLU | p.Glu279Gln | P35557 |
| HOMO SAPIENS | 7 | 705 | 3idh | A | 299 | GLY | p.Gly299Arg | P35557 |
| HOMO SAPIENS | 2 | 705 | 3idh | A | 318 | GLY | p.Gly318Trp | P35557 |
| HOMO SAPIENS | 4 | 705 | 3idh | A | 378 | ALA | p.Ala378Thr | P35557 |
| HOMO SAPIENS | 0 | 705 | 3idh | A | 387 | ALA | p.Ala387Glu | P35557 |
| HOMO SAPIENS | 5 | 705 | 3idh | A | 414 | LYS | p.Lys414Glu | P35557 |
| HOMO SAPIENS | 0 | 705 | 3idh | A | 455 | VAL | p.Val455Met | P35557 |
| HOMO SAPIENS | -1 | 705 | 3idh | A | 462 | MET | p.Met462Ile | P35557 |
| HOMO SAPIENS | 1 | 509 | 4nef | A | 22 | LEU | p.Leu22Val | P41181 |
| HOMO SAPIENS | 4 | 509 | 4nef | A | 57 | GLN | p.Gln57Pro | P41181 |
| HOMO SAPIENS | 8 | 509 | 4nef | A | 64 | GLY | p.Gly64Arg | P41181 |
| HOMO SAPIENS | 7 | 509 | 4nef | A | 68 | ASN | p.Asn68Ser | P41181 |
| HOMO SAPIENS | 1 | 509 | 4nef | A | 100 | GLY | p.Gly100Val | P41181 |
| HOMO SAPIENS | 0 | 509 | 4nef | A | 125 | THR | p.Thr125Met | P41181 |
| HOMO SAPIENS | 0 | 509 | 4nef | A | 126 | THR | p.Thr126Met | P41181 |
| HOMO SAPIENS | 4 | 509 | 4nef | A | 146 | PHE | p.Phe146Leu | P41181 |
| HOMO SAPIENS | 0 | 509 | 4nef | A | 175 | GLY | p.Gly175Arg | P41181 |
| HOMO SAPIENS | 5 | 509 | 4nef | A | 181 | CYS | p.Cys181Trp | P41181 |

| | | | | | | | | | |
|---|---|---|---|---|---|---|---|---|---|
| HOMO SAPIENS | 9 | 509 | 4nef | A | | 187 | ARG | p.Arg187Cys | P41181 |
| HOMO SAPIENS | 0 | 509 | 4nef | A | | 190 | ALA | p.Ala190Thr | P41181 |
| HOMO SAPIENS | 0 | 509 | 4nef | A | | 216 | SER | p.Ser216Pro | P41181 |
| HOMO SAPIENS | 4 | 2242 | 1z70 | X | | 1155 | SER | p.Ser155Pro | Q8NBK3 |
| HOMO SAPIENS | 3 | 2242 | 1z70 | X | | 1218 | CYS | p.Cys218Tyr | Q8NBK3 |
| HOMO SAPIENS | 0 | 2242 | 1z70 | X | | 1279 | ALA | p.Ala279Val | Q8NBK3 |
| HOMO SAPIENS | -1 | 2242 | 1z70 | X | | 1336 | OCS | p.Cys336Arg | Q8NBK3 |
| HOMO SAPIENS | 4 | 2242 | 1z70 | X | | 1345 | ARG | p.Arg345Cys | Q8NBK3 |
| HOMO SAPIENS | 2 | 2242 | 1z70 | X | | 1348 | ALA | p.Ala348Pro | Q8NBK3 |
| HOMO SAPIENS | 5 | 2242 | 1z70 | X | | 1349 | ARG | p.Arg349Gln | Q8NBK3 |
| HOMO SAPIENS | 5 | 2242 | 1z70 | X | | 1349 | ARG | p.Arg349Trp | Q8NBK3 |
| HOMO SAPIENS | 3 | 2242 | 1z70 | X | | 1359 | SER | p.Ser359Ter | Q8NBK3 |
| HOMO SAPIENS | 5 | 8269 | 5wrr | B | | 136 | ARG | p.Arg136Ter | Q96MK3 |
| HOMO SAPIENS | 1 | 8269 | 5wrr | B | | 332 | LYS | p.Asn332Lys | Q96MK3 |
| HOMO SAPIENS | 3 | 8269 | 5wrr | B | | 478 | ARG | p.Arg478Ter | Q96MK3 |
| HOMO SAPIENS | 5 | 8269 | 5wrr | B | | 504 | THR | p.Thr504Ile | Q96MK3 |
| HOMO SAPIENS | 5 | 1411 | 4h1s | B | | 221 | SER | p.Ser221Ter | P21589 |
| HOMO SAPIENS | 1 | 1411 | 4h1s | B | | 358 | CYS | p.Cys358Tyr | P21589 |
| HOMO SAPIENS | 2 | 23 | 5vvd | B | | 298 | GLU | p.Glu298Asp | P29474 |
| HOMO SAPIENS | 0 | 19 | 5jlh | E | | 88 | THR | p.Thr89Ile | P63261 |
| HOMO SAPIENS | 6 | 19 | 5jlh | E | | 117 | LYS | p.Lys118Met | P63261 |
| HOMO SAPIENS | 5 | 19 | 5jlh | E | | 263 | PRO | p.Pro264Leu | P63261 |
| HOMO SAPIENS | 1 | 19 | 5jlh | E | | 277 | THR | p.Thr278Ile | P63261 |
| HOMO SAPIENS | 6 | 19 | 5jlh | E | | 331 | PRO | p.Pro332Ala | P63261 |
| HOMO SAPIENS | 5 | 19 | 5jlh | E | | 369 | VAL | p.Val370Ala | P63261 |
| HOMO SAPIENS | 0 | 14306 | 4w4o | C | | 92 | ARG | p.Arg92Ter | P12314 |
| HOMO SAPIENS | 3 | 603 | 1nuh | A | | 4 | THR | p.Thr5Ile | P06744 |
| HOMO SAPIENS | 2 | 603 | 1nuh | A | | 19 | HIS | p.His20Pro | P06744 |
| HOMO SAPIENS | 8 | 603 | 1nuh | A | | 158 | GLY | p.Gly159Ser | P06744 |
| HOMO SAPIENS | 6 | 603 | 1nuh | A | | 223 | THR | p.Thr224Met | P06744 |
| HOMO SAPIENS | 0 | 603 | 1nuh | A | | 338 | LEU | p.Leu339Pro | P06744 |
| HOMO SAPIENS | 6 | 603 | 1nuh | A | | 342 | GLN | p.Gln343Arg | P06744 |
| HOMO SAPIENS | 6 | 603 | 1nuh | A | | 374 | THR | p.Thr375Arg | P06744 |
| HOMO SAPIENS | 4 | 603 | 1nuh | A | | 524 | ILE | p.Ile525Thr | P06744 |
| HOMO SAPIENS | 4 | 603 | 1nuh | A | | 538 | ASP | p.Asp539Asn | P06744 |
| HOMO SAPIENS | 1 | 3107 | 3kzj | A | | 1197 | VAL | p.Val1197Ala | P08603 |
| HOMO SAPIENS | 9 | 1748 | 2noh | A | | 46 | ARG | p.Arg46Gln | O15527 |
| HOMO SAPIENS | 2 | 11750 | 3s4y | A | | 33 | ARG | p.Arg33His | Q9H3S4 |
| HOMO SAPIENS | 2 | 11750 | 3s4y | A | | 40 | LEU | p.Leu40Pro | Q9H3S4 |
| HOMO SAPIENS | 4 | 11750 | 3s4y | A | | 142 | LEU | p.Leu142Phe | Q9H3S4 |
| HOMO SAPIENS | 5 | 11750 | 3s4y | A | | 219 | ASN | p.Asn219Ser | Q9H3S4 |
| HOMO SAPIENS | 1 | 2591 | 5vnj | A | | 382 | PHE | p.Phe382Leu | Q15436 |
| HOMO SAPIENS | 2 | 2591 | 5vnj | A | | 702 | MET | p.Met702Val | Q15436 |
| HOMO SAPIENS | 1 | 2426 | 5m8t | A | | 87 | ARG | p.Arg87Trp | P17643 |
| HOMO SAPIENS | 1 | 2426 | 5m8t | A | | 166 | SER | p.Ser166Ter | P17643 |
| HOMO SAPIENS | 0 | 2426 | 5m8t | A | | 356 | ARG | p.Arg356Gln | P17643 |
| HOMO SAPIENS | 2 | 2426 | 5m8t | A | | 374 | SER | p.Arg374Ter | P17643 |
| HOMO SAPIENS | 6 | 2426 | 5m8t | A | | 378 | ASN | p.Asn378Ser | P17643 |
| HOMO SAPIENS | 6 | 2426 | 5m8t | A | | 382 | LEU | p.Leu382Pro | P17643 |
| HOMO SAPIENS | 7 | 10097 | 4n6v | | 1 | 46 | GLN | p.Gln46Ter | P04080 |
| HOMO SAPIENS | 3 | 10097 | 4n6v | | 1 | 65 | VAL | p.Val65Ile | P04080 |
| HOMO SAPIENS | 8 | 10097 | 4n6v | | 1 | 68 | ARG | p.Arg68Ter | P04080 |
| HOMO SAPIENS | 2 | 859 | 4z9a | A | | 105 | GLN | p.Gln106Arg | P24666 |
| HOMO SAPIENS | 8 | 1420 | 5nan | F | | 95 | SER | p.Ser95Leu | Q96PD4 |

| | | | | | | | | |
|---|---|---|---|---|---|---|---|---|
| HOMO SAPIENS | 6 | 1420 | 5nan | F | 126 | GLU | p.Glu126Gly | Q96PD4 |
| HOMO SAPIENS | 8 | 1420 | 5nan | F | 155 | VAL | p.Val155Ile | Q96PD4 |
| HOMO SAPIENS | 6 | 1420 | 5nan | F | 157 | PRO | p.Pro157Ser | Q96PD4 |
| HOMO SAPIENS | -1 | 1420 | 5nan | F | 161 | HIS | p.His161Arg | Q96PD4 |
| HOMO SAPIENS | 3 | 288 | 5i96 | B | 140 | GLN | p.Arg140Gly | P48735 |
| HOMO SAPIENS | 3 | 288 | 5i96 | B | 140 | GLN | p.Arg140Gln | P48735 |
| HOMO SAPIENS | 2 | 288 | 5i96 | B | 261 | ARG | p.Arg261His | P48735 |
| HOMO SAPIENS | 2 | 288 | 5i96 | B | 435 | THR | p.Thr435Met | P48735 |
| HOMO SAPIENS | 9 | 120 | 1tup | B | 107 | TYR | p.Tyr107His | P04637 |
| HOMO SAPIENS | 8 | 120 | 1tup | B | 110 | ARG | p.Arg110Leu | P04637 |
| HOMO SAPIENS | 5 | 120 | 1tup | B | 120 | LYS | p.Lys120Glu | P04637 |
| HOMO SAPIENS | 4 | 120 | 1tup | B | 133 | MET | p.Met133Thr | P04637 |
| HOMO SAPIENS | 3 | 120 | 1tup | B | 134 | PHE | p.Phe134Cys | P04637 |
| HOMO SAPIENS | 8 | 120 | 1tup | B | 136 | GLN | p.Gln136His | P04637 |
| HOMO SAPIENS | 9 | 120 | 1tup | B | 138 | ALA | p.Ala138Pro | P04637 |
| HOMO SAPIENS | 9 | 120 | 1tup | B | 151 | PRO | p.Pro151Ser | P04637 |
| HOMO SAPIENS | 9 | 120 | 1tup | B | 152 | PRO | p.Pro152Leu | P04637 |
| HOMO SAPIENS | 9 | 120 | 1tup | B | 158 | ARG | p.Arg158Cys | P04637 |
| HOMO SAPIENS | 9 | 120 | 1tup | B | 158 | ARG | p.Arg158His | P04637 |
| HOMO SAPIENS | 9 | 120 | 1tup | B | 175 | ARG | p.Arg175Cys | P04637 |
| HOMO SAPIENS | 9 | 120 | 1tup | B | 175 | ARG | p.Arg175Leu | P04637 |
| HOMO SAPIENS | 8 | 120 | 1tup | B | 181 | ARG | p.Arg181Leu | P04637 |
| HOMO SAPIENS | 9 | 120 | 1tup | B | 189 | ALA | p.Ala189Val | P04637 |
| HOMO SAPIENS | 9 | 120 | 1tup | B | 196 | ARG | p.Arg196Gly | P04637 |
| HOMO SAPIENS | 9 | 120 | 1tup | B | 213 | ARG | p.Arg213Gln | P04637 |
| HOMO SAPIENS | 9 | 120 | 1tup | B | 213 | ARG | p.Arg213Ter | P04637 |
| HOMO SAPIENS | 9 | 120 | 1tup | B | 214 | HIS | p.His214Gln | P04637 |
| HOMO SAPIENS | 9 | 120 | 1tup | B | 220 | TYR | p.Tyr220Cys | P04637 |
| HOMO SAPIENS | 0 | 120 | 1tup | B | 234 | TYR | p.Tyr234Cys | P04637 |
| HOMO SAPIENS | 9 | 120 | 1tup | B | 237 | MET | p.Met237Ile | P04637 |
| HOMO SAPIENS | 0 | 120 | 1tup | B | 238 | CYS | p.Cys238Tyr | P04637 |
| HOMO SAPIENS | 9 | 120 | 1tup | B | 241 | SER | p.Ser241Phe | P04637 |
| HOMO SAPIENS | 9 | 120 | 1tup | B | 242 | CYS | p.Cys242Ser | P04637 |
| HOMO SAPIENS | 9 | 120 | 1tup | B | 245 | GLY | p.Gly245Cys | P04637 |
| HOMO SAPIENS | 9 | 120 | 1tup | B | 245 | GLY | p.Gly245Asp | P04637 |
| HOMO SAPIENS | 9 | 120 | 1tup | B | 248 | ARG | p.Arg248Leu | P04637 |
| HOMO SAPIENS | 9 | 120 | 1tup | B | 248 | ARG | p.Arg248Trp | P04637 |
| HOMO SAPIENS | 9 | 120 | 1tup | B | 251 | ILE | p.Ile251Ser | P04637 |
| HOMO SAPIENS | 9 | 120 | 1tup | B | 252 | LEU | p.Leu252Pro | P04637 |
| HOMO SAPIENS | 9 | 120 | 1tup | B | 258 | GLU | p.Glu258Lys | P04637 |
| HOMO SAPIENS | 8 | 120 | 1tup | B | 272 | VAL | p.Val272Leu | P04637 |
| HOMO SAPIENS | 9 | 120 | 1tup | B | 273 | ARG | p.Arg273Cys | P04637 |
| HOMO SAPIENS | 9 | 120 | 1tup | B | 273 | ARG | p.Arg273His | P04637 |
| HOMO SAPIENS | 9 | 120 | 1tup | B | 277 | CYS | p.Cys277Phe | P04637 |
| HOMO SAPIENS | 9 | 120 | 1tup | B | 280 | ARG | p.Arg280Ile | P04637 |
| HOMO SAPIENS | 9 | 120 | 1tup | B | 282 | ARG | p.Arg282Trp | P04637 |
| HOMO SAPIENS | 9 | 120 | 1tup | B | 283 | ARG | p.Arg283His | P04637 |
| HOMO SAPIENS | 9 | 120 | 1tup | B | 285 | GLU | p.Glu285Val | P04637 |
| HOMO SAPIENS | -1 | 120 | 1tup | B | 292 | LYS | p.Lys292Ile | P04637 |
| HOMO SAPIENS | -1 | 120 | 1tup | B | 298 | GLU | p.Glu298Lys | P04637 |
| HOMO SAPIENS | -1 | 2791 | 4ygb | A | 39 | VAL | p.Val39Ala | P49257 |
| HOMO SAPIENS | 2 | 106 | 4x6j | A | 32 | GLY | p.Gly146Arg | P43235 |
| HOMO SAPIENS | 0 | 106 | 4x6j | A | 76 | GLN | p.Gln190Ter | P43235 |
| HOMO SAPIENS | 7 | 106 | 4x6j | A | 198 | ARG | p.Arg312Ter | P43235 |

| | | | | | | | | |
|---|---|---|---|---|---|---|---|---|
| HOMO SAPIENS | 5 | 490 | 2hlo | B | 172 | LEU | p.Leu202Gln | P02675 |
| HOMO SAPIENS | 0 | 490 | 2hlo | B | 353 | LEU | p.Leu383Arg | P02675 |
| HOMO SAPIENS | 4 | 490 | 2hlo | B | 400 | GLY | p.Gly430Asp | P02675 |
| HOMO SAPIENS | 7 | 490 | 2hlo | B | 448 | ARG | p.Arg478Lys | P02675 |
| HOMO SAPIENS | 1 | 289 | 4p13 | B | 18 | GLU | p.Glu43Lys | P11310 |
| HOMO SAPIENS | 2 | 289 | 4p13 | B | 20 | GLN | p.Gln45Arg | P11310 |
| HOMO SAPIENS | 5 | 289 | 4p13 | B | 28 | ARG | p.Arg53Cys | P11310 |
| HOMO SAPIENS | 1 | 289 | 4p13 | B | 42 | TYR | p.Tyr67His | P11310 |
| HOMO SAPIENS | 1 | 289 | 4p13 | B | 53 | ILE | p.Ile78Thr | P11310 |
| HOMO SAPIENS | 1 | 289 | 4p13 | B | 59 | LEU | p.Leu84Phe | P11310 |
| HOMO SAPIENS | 2 | 289 | 4p13 | B | 96 | THR | p.Thr121Ile | P11310 |
| HOMO SAPIENS | 0 | 289 | 4p13 | B | 124 | MET | p.Met149Ile | P11310 |
| HOMO SAPIENS | 5 | 289 | 4p13 | B | 143 | ASP | p.Asp168Ala | P11310 |
| HOMO SAPIENS | 7 | 289 | 4p13 | B | 168 | THR | p.Thr193Ala | P11310 |
| HOMO SAPIENS | 1 | 289 | 4p13 | B | 170 | GLY | p.Gly195Arg | P11310 |
| HOMO SAPIENS | 2 | 289 | 4p13 | B | 181 | ARG | p.Arg206Cys | P11310 |
| HOMO SAPIENS | 2 | 289 | 4p13 | B | 181 | ARG | p.Arg206His | P11310 |
| HOMO SAPIENS | 8 | 289 | 4p13 | B | 203 | THR | p.Thr228Ile | P11310 |
| HOMO SAPIENS | 2 | 289 | 4p13 | B | 208 | ILE | p.Ile233Asn | P11310 |
| HOMO SAPIENS | 1 | 289 | 4p13 | B | 219 | CYS | p.Cys244Arg | P11310 |
| HOMO SAPIENS | 6 | 289 | 4p13 | B | 220 | SER | p.Ser245Leu | P11310 |
| HOMO SAPIENS | 5 | 289 | 4p13 | B | 228 | GLU | p.Glu253Lys | P11310 |
| HOMO SAPIENS | 2 | 289 | 4p13 | B | 241 | ASP | p.Asp266Gly | P11310 |
| HOMO SAPIENS | 1 | 289 | 4p13 | B | 242 | GLY | p.Gly267Arg | P11310 |
| HOMO SAPIENS | 7 | 289 | 4p13 | B | 256 | ARG | p.Arg281Thr | P11310 |
| HOMO SAPIENS | 4 | 289 | 4p13 | B | 269 | ARG | p.Arg294Thr | P11310 |
| HOMO SAPIENS | 4 | 289 | 4p13 | B | 304 | GLU | p.Lys329Glu | P11310 |
| HOMO SAPIENS | 1 | 289 | 4p13 | B | 324 | ARG | p.Arg349Ter | P11310 |
| HOMO SAPIENS | 3 | 289 | 4p13 | B | 326 | THR | p.Thr351Ile | P11310 |
| HOMO SAPIENS | 7 | 289 | 4p13 | B | 350 | ILE | p.Ile375Thr | P11310 |
| HOMO SAPIENS | 3 | 289 | 4p13 | B | 359 | GLU | p.Glu384Ter | P11310 |
| HOMO SAPIENS | 1 | 4563 | 3i2b | A | 16 | ARG | p.Arg16Cys | Q03393 |
| HOMO SAPIENS | 5 | 4563 | 3i2b | A | 25 | ARG | p.Arg25Gln | Q03393 |
| HOMO SAPIENS | 1 | 4563 | 3i2b | A | 47 | ASN | p.Asn47Asp | Q03393 |
| HOMO SAPIENS | 8 | 4563 | 3i2b | A | 52 | ASN | p.Asn52Ser | Q03393 |
| HOMO SAPIENS | 1 | 4563 | 3i2b | A | 56 | VAL | p.Val56Met | Q03393 |
| HOMO SAPIENS | 1 | 4563 | 3i2b | A | 87 | PRO | p.Pro87Ser | Q03393 |
| HOMO SAPIENS | 1 | 4563 | 3i2b | A | 96 | ASP | p.Asp96Asn | Q03393 |
| HOMO SAPIENS | 0 | 4563 | 3i2b | A | 99 | TYR | p.Tyr99Ter | Q03393 |
| HOMO SAPIENS | 2 | 4563 | 3i2b | A | 116 | ASP | p.Asp116Gly | Q03393 |
| HOMO SAPIENS | 2 | 41 | 3exx | A | 1 | GLY | p.Gly90Cys | P01308 |
| HOMO SAPIENS | 5 | 41 | 3exx | A | 3 | VAL | p.Val92Leu | P01308 |
| HOMO SAPIENS | 4 | 41 | 3exx | A | 7 | CYS | p.Cys96Ser | P01308 |
| HOMO SAPIENS | 9 | 41 | 3exx | A | 19 | TYR | p.Tyr108Cys | P01308 |
| HOMO SAPIENS | 9 | 131 | 4mjo | D | 118 | ASP | p.Asp119Asn | P09467 |
| HOMO SAPIENS | 0 | 131 | 4mjo | D | 163 | GLY | p.Gly164Ser | P09467 |
| HOMO SAPIENS | 0 | 131 | 4mjo | D | 176 | ALA | p.Ala177Asp | P09467 |
| HOMO SAPIENS | 5 | 131 | 4mjo | D | 193 | PHE | p.Phe194Ser | P09467 |
| HOMO SAPIENS | 4 | 131 | 4mjo | D | 217 | ARG | p.Arg218Lys | P09467 |
| HOMO SAPIENS | 4 | 131 | 4mjo | D | 223 | VAL | p.Val224Ile | P09467 |
| HOMO SAPIENS | 4 | 131 | 4mjo | D | 232 | PHE | p.Phe233Ile | P09467 |
| HOMO SAPIENS | 5 | 131 | 4mjo | D | 243 | ARG | p.Arg244Trp | P09467 |
| HOMO SAPIENS | 0 | 131 | 4mjo | D | 283 | PRO | p.Pro284Arg | P09467 |
| HOMO SAPIENS | 0 | 131 | 4mjo | D | 288 | MET | p.Met289Leu | P09467 |

| | | | | | | | | |
|---|---|---|---|---|---|---|---|---|
| HOMO SAPIENS | 0 | 131 | 4mjo | D | 323 | ASP | p.Asp324Asn | P09467 |
| HOMO SAPIENS | -1 | 13122 | 3hfc | A | 583 | ARG | p.Arg583Cys | P51787 |
| HOMO SAPIENS | 1 | 13122 | 3hfc | A | 587 | THR | p.Thr587Met | P51787 |
| HOMO SAPIENS | 0 | 13122 | 3hfc | A | 589 | GLY | p.Gly589Asp | P51787 |
| HOMO SAPIENS | 4 | 3002 | 3nbq | B | 94 | ARG | p.Arg94Cys | Q16831 |
| HOMO SAPIENS | 1 | 3616 | 3dbp | A | 97 | ILE | p.Ile286Asn | P11172 |
| HOMO SAPIENS | 0 | 3616 | 3dbp | A | 240 | GLY | p.Gly429Arg | P11172 |
| HOMO SAPIENS | 0 | 3616 | 3dbp | A | 257 | ILE | p.Ile446Val | P11172 |
| HOMO SAPIENS | 1 | 584 | 1drf | A | 79 | LEU | p.Leu80Phe | P00374 |
| HOMO SAPIENS | 3 | 1609 | 3qa2 | B | 49 | VAL | p.Val49Ile | P50053 |
| HOMO SAPIENS | 0 | 20560 | 5mug | A | 101 | HIS | p.His101Gln | P49638 |
| HOMO SAPIENS | 2 | 20560 | 5mug | A | 120 | ALA | p.Ala120Thr | P49638 |
| HOMO SAPIENS | 3 | 20560 | 5mug | A | 192 | ARG | p.Arg192His | P49638 |
| HOMO SAPIENS | 3 | 20560 | 5mug | A | 221 | ARG | p.Arg221Trp | P49638 |
| HOMO SAPIENS | 4 | 256 | 3vtv | A | 65 | THR | p.Glu50Lys | Q9GZQ8 |
| HOMO SAPIENS | 4 | 256 | 3vtv | A | 113 | VAL | p.Met98Lys | Q9GZQ8 |
| HOMO SAPIENS | 4 | 5397 | 1m61 | A | 192 | ARG | p.Arg192Trp | P43403 |
| HOMO SAPIENS | 5 | 39881 | 1aut | L | 107 | HIS | p.His149Pro | P04070 |
| HOMO SAPIENS | 5 | 4010 | 4uf8 | I | 33 | ALA | p.Ala33Val | Q13501 |
| HOMO SAPIENS | 3 | 313 | 5w49 | A | 98 | PRO | p.Pro98Leu | P23526 |
| HOMO SAPIENS | 2 | 313 | 5w49 | A | 143 | TYR | p.Tyr143Cys | P23526 |
| HOMO SAPIENS | -1 | 1643 | 2m8t | A | 102 | PRO | p.Pro102Leu | P04156 |
| HOMO SAPIENS | -1 | 1643 | 2m8t | A | 105 | PRO | p.Pro105Leu | P04156 |
| HOMO SAPIENS | -1 | 1643 | 2m8t | A | 117 | ALA | p.Ala117Val | P04156 |
| HOMO SAPIENS | 5 | 1643 | 2m8t | A | 129 | MET | p.Met129Val | P04156 |
| HOMO SAPIENS | 5 | 1643 | 2m8t | A | 131 | GLY | p.Gly131Val | P04156 |
| HOMO SAPIENS | 2 | 1643 | 2m8t | A | 142 | GLY | p.Gly142Ser | P04156 |
| HOMO SAPIENS | 3 | 1643 | 2m8t | A | 171 | ASN | p.Asn171Ser | P04156 |
| HOMO SAPIENS | 9 | 1643 | 2m8t | A | 178 | ASP | p.Asp178Asn | P04156 |
| HOMO SAPIENS | 9 | 1643 | 2m8t | A | 180 | VAL | p.Val180Ile | P04156 |
| HOMO SAPIENS | 9 | 1643 | 2m8t | A | 183 | THR | p.Thr183Ala | P04156 |
| HOMO SAPIENS | 9 | 1643 | 2m8t | A | 187 | HIS | p.His187Arg | P04156 |
| HOMO SAPIENS | 8 | 1643 | 2m8t | A | 198 | PHE | p.Phe198Ser | P04156 |
| HOMO SAPIENS | 9 | 1643 | 2m8t | A | 200 | GLU | p.Glu200Lys | P04156 |
| HOMO SAPIENS | 9 | 1643 | 2m8t | A | 208 | ARG | p.Arg208His | P04156 |
| HOMO SAPIENS | 3 | 1643 | 2m8t | A | 210 | VAL | p.Val210Ile | P04156 |
| HOMO SAPIENS | 6 | 1643 | 2m8t | A | 217 | GLN | p.Gln217Arg | P04156 |
| HOMO SAPIENS | 1 | 29 | 5kos | A | 151 | GLY | p.Gly217Arg | P00797 |
| HOMO SAPIENS | 1 | 29 | 5kos | A | 164 | ARG | p.Arg230Lys | P00797 |
| HOMO SAPIENS | 3 | 504 | 4zkv | B | 37 | ARG | p.Arg37Pro | P49773 |
| HOMO SAPIENS | 2 | 504 | 4zkv | B | 62 | GLN | p.Gln62Ter | P49773 |
| HOMO SAPIENS | 2 | 504 | 4zkv | B | 84 | CYS | p.Cys84Arg | P49773 |
| HOMO SAPIENS | 4 | 504 | 4zkv | B | 93 | GLY | p.Gly93Asp | P49773 |
| HOMO SAPIENS | 9 | 504 | 4zkv | B | 112 | HIS | p.His112Asn | P49773 |
| HOMO SAPIENS | 0 | 74 | 5vwo | C | 54 | ASN | p.Asn54Lys | P04181 |
| HOMO SAPIENS | 0 | 74 | 5vwo | C | 55 | TYR | p.Tyr55His | P04181 |
| HOMO SAPIENS | 2 | 74 | 5vwo | C | 90 | GLN | p.Gln90Glu | P04181 |
| HOMO SAPIENS | 6 | 74 | 5vwo | C | 91 | GLY | p.Gly91Glu | P04181 |
| HOMO SAPIENS | 2 | 74 | 5vwo | C | 93 | CYS | p.Cys93Phe | P04181 |
| HOMO SAPIENS | 1 | 74 | 5vwo | C | 154 | ARG | p.Arg154Leu | P04181 |
| HOMO SAPIENS | 0 | 74 | 5vwo | C | 180 | ARG | p.Arg180Thr | P04181 |
| HOMO SAPIENS | 0 | 74 | 5vwo | C | 199 | PRO | p.Pro199Gln | P04181 |
| HOMO SAPIENS | 4 | 74 | 5vwo | C | 209 | TYR | p.Tyr209Ter | P04181 |
| HOMO SAPIENS | 0 | 74 | 5vwo | C | 226 | ALA | p.Ala226Val | P04181 |

| | | | | | | | | |
|---|---|---|---|---|---|---|---|---|
| HOMO SAPIENS | 5 | 74 | 5vwo | C | 233 | GLN | p.Gln233Arg | P04181 |
| HOMO SAPIENS | 5 | 74 | 5vwo | C | 241 | PRO | p.Pro241Leu | P04181 |
| HOMO SAPIENS | 2 | 74 | 5vwo | C | 245 | TYR | p.Tyr245Cys | P04181 |
| HOMO SAPIENS | 6 | 74 | 5vwo | C | 250 | ARG | p.Arg250Pro | P04181 |
| HOMO SAPIENS | 6 | 74 | 5vwo | C | 250 | ARG | p.Arg250Ter | P04181 |
| HOMO SAPIENS | 8 | 74 | 5vwo | C | 267 | THR | p.Thr267Ile | P04181 |
| HOMO SAPIENS | 0 | 74 | 5vwo | C | 270 | ALA | p.Ala270Pro | P04181 |
| HOMO SAPIENS | 0 | 74 | 5vwo | C | 271 | ARG | p.Arg271Lys | P04181 |
| HOMO SAPIENS | 1 | 74 | 5vwo | C | 299 | TYR | p.Tyr299Ter | P04181 |
| HOMO SAPIENS | 0 | 74 | 5vwo | C | 318 | GLU | p.Glu318Lys | P04181 |
| HOMO SAPIENS | 1 | 74 | 5vwo | C | 319 | HIS | p.His319Tyr | P04181 |
| HOMO SAPIENS | 1 | 74 | 5vwo | C | 331 | ARG | p.Arg331Ter | P04181 |
| HOMO SAPIENS | 0 | 74 | 5vwo | C | 332 | VAL | p.Val332Met | P04181 |
| HOMO SAPIENS | 1 | 74 | 5vwo | C | 353 | GLY | p.Gly353Asp | P04181 |
| HOMO SAPIENS | 2 | 74 | 5vwo | C | 375 | GLY | p.Gly375Ala | P04181 |
| HOMO SAPIENS | 0 | 74 | 5vwo | C | 394 | CYS | p.Cys394Arg | P04181 |
| HOMO SAPIENS | 0 | 74 | 5vwo | C | 402 | LEU | p.Leu402Pro | P04181 |
| HOMO SAPIENS | 0 | 74 | 5vwo | C | 436 | ILE | p.Ile436Asn | P04181 |
| HOMO SAPIENS | 7 | 24356 | 4az0 | A | 104 | VAL | p.Val132Met | P10619 |
| HOMO SAPIENS | 4 | 24356 | 4az0 | A | 221 | TYR | p.Tyr249Asn | P10619 |
| HOMO SAPIENS | 3 | 245 | 1rsz | A | 24 | ARG | p.Arg24Ter | P00491 |
| HOMO SAPIENS | 3 | 245 | 1rsz | A | 51 | SER | p.Gly51Ser | P00491 |
| HOMO SAPIENS | 1 | 245 | 1rsz | A | 58 | ARG | p.Arg58Ter | P00491 |
| HOMO SAPIENS | 4 | 245 | 1rsz | A | 89 | GLU | p.Glu89Lys | P00491 |
| HOMO SAPIENS | 6 | 245 | 1rsz | A | 128 | ASP | p.Asp128Gly | P00491 |
| HOMO SAPIENS | 0 | 245 | 1rsz | A | 174 | ALA | p.Ala174Pro | P00491 |
| HOMO SAPIENS | 8 | 245 | 1rsz | A | 192 | TYR | p.Tyr192Cys | P00491 |
| HOMO SAPIENS | 8 | 245 | 1rsz | A | 234 | ARG | p.Arg234Pro | P00491 |
| HOMO SAPIENS | 0 | 37 | 5tby | E | 22 | GLU | p.Glu22Lys | P10916 |
| HOMO SAPIENS | 1 | 37 | 5tby | E | 86 | PHE | p.Phe86Ser | P10916 |
| HOMO SAPIENS | 9 | 71 | 5m9j | A | 12 | GLN | p.Gln36Leu | P03950 |
| HOMO SAPIENS | 3 | 71 | 5m9j | A | 17 | LYS | p.Lys41Glu | P03950 |
| HOMO SAPIENS | 3 | 71 | 5m9j | A | 17 | LYS | p.Lys41Ile | P03950 |
| HOMO SAPIENS | 4 | 71 | 5m9j | A | 28 | SER | p.Ser52Asn | P03950 |
| HOMO SAPIENS | 7 | 71 | 5m9j | A | 31 | ARG | p.Arg55Lys | P03950 |
| HOMO SAPIENS | 9 | 71 | 5m9j | A | 39 | CYS | p.Cys63Trp | P03950 |
| HOMO SAPIENS | 9 | 71 | 5m9j | A | 40 | LYS | p.Lys64Ile | P03950 |
| HOMO SAPIENS | 2 | 71 | 5m9j | A | 60 | GLU | p.Lys84Glu | P03950 |
| HOMO SAPIENS | 0 | 71 | 5m9j | A | 112 | PRO | p.Pro136Leu | P03950 |
| HOMO SAPIENS | 9 | 71 | 5m9j | A | 113 | VAL | p.Val137Ile | P03950 |
| HOMO SAPIENS | 3 | 326 | 5dqi | A | 209 | GLY | p.Gly209Val | Q9Y253 |
| HOMO SAPIENS | 1 | 326 | 5dqi | A | 242 | SER | p.Ser242Ter | Q9Y253 |
| HOMO SAPIENS | 1 | 326 | 5dqi | A | 303 | ARG | p.Arg303Ter | Q9Y253 |
| HOMO SAPIENS | 1 | 8203 | 3iu3 | K | 132 | VAL | p.Val153Ile | P01589 |
| HOMO SAPIENS | 0 | 4995 | 5ffo | H | 234 | THR | p.Thr263Ile | P01137 |
| HOMO SAPIENS | 5 | 38609 | 2ruk | A | 49 | ASP | p.Asp49His | P04637 |
| HOMO SAPIENS | 5 | 38609 | 2ruk | A | 58 | PRO | p.Pro58Arg | P04637 |
| HOMO SAPIENS | 0 | 1607 | 6pah | A | 117 | THR | p.Thr117Ile | P00439 |
| HOMO SAPIENS | 3 | 1607 | 6pah | A | 145 | ASP | p.Asp145Val | P00439 |
| HOMO SAPIENS | 5 | 1607 | 6pah | A | 158 | ARG | p.Arg158Gln | P00439 |
| HOMO SAPIENS | 5 | 1607 | 6pah | A | 158 | ARG | p.Arg158Trp | P00439 |
| HOMO SAPIENS | 5 | 1607 | 6pah | A | 169 | ARG | p.Arg169His | P00439 |
| HOMO SAPIENS | 5 | 1607 | 6pah | A | 169 | ARG | p.Arg169Cys | P00439 |
| HOMO SAPIENS | 3 | 1607 | 6pah | A | 176 | ARG | p.Arg176Leu | P00439 |

| | | | | | | | | |
|---|---|---|---|---|---|---|---|---|
| HOMO SAPIENS | 3 | 1607 | 6pah | A | 177 | VAL | p.Val177Leu | P00439 |
| HOMO SAPIENS | 1 | 1607 | 6pah | A | 194 | LEU | p.Leu194Pro | P00439 |
| HOMO SAPIENS | 1 | 1607 | 6pah | A | 204 | TYR | p.Tyr204Cys | P00439 |
| HOMO SAPIENS | 1 | 1607 | 6pah | A | 204 | TYR | p.Tyr204Ter | P00439 |
| HOMO SAPIENS | 2 | 1607 | 6pah | A | 230 | VAL | p.Val230Ile | P00439 |
| HOMO SAPIENS | 4 | 1607 | 6pah | A | 232 | GLN | p.Gln232Ter | P00439 |
| HOMO SAPIENS | 4 | 1607 | 6pah | A | 241 | ARG | p.Arg241Cys | P00439 |
| HOMO SAPIENS | 4 | 1607 | 6pah | A | 241 | ARG | p.Arg241His | P00439 |
| HOMO SAPIENS | 5 | 1607 | 6pah | A | 243 | ARG | p.Arg243Gln | P00439 |
| HOMO SAPIENS | 5 | 1607 | 6pah | A | 243 | ARG | p.Arg243Ter | P00439 |
| HOMO SAPIENS | 7 | 1607 | 6pah | A | 249 | LEU | p.Leu249Phe | P00439 |
| HOMO SAPIENS | 5 | 1607 | 6pah | A | 252 | ARG | p.Arg252Gly | P00439 |
| HOMO SAPIENS | 5 | 1607 | 6pah | A | 261 | ARG | p.Arg261Ter | P00439 |
| HOMO SAPIENS | 4 | 1607 | 6pah | A | 267 | GLN | p.Gln267Leu | P00439 |
| HOMO SAPIENS | 1 | 1607 | 6pah | A | 272 | GLY | p.Gly272Ter | P00439 |
| HOMO SAPIENS | 2 | 1607 | 6pah | A | 277 | TYR | p.Tyr277Asp | P00439 |
| HOMO SAPIENS | 7 | 1607 | 6pah | A | 280 | GLU | p.Glu280Lys | P00439 |
| HOMO SAPIENS | 8 | 1607 | 6pah | A | 281 | PRO | p.Pro281Leu | P00439 |
| HOMO SAPIENS | 0 | 1607 | 6pah | A | 299 | PHE | p.Phe299Cys | P00439 |
| HOMO SAPIENS | 5 | 1607 | 6pah | A | 300 | ALA | p.Ala300Ser | P00439 |
| HOMO SAPIENS | 5 | 1607 | 6pah | A | 309 | ALA | p.Ala309Asp | P00439 |
| HOMO SAPIENS | 4 | 1607 | 6pah | A | 320 | LYS | p.Lys320Asn | P00439 |
| HOMO SAPIENS | 0 | 1607 | 6pah | A | 345 | ALA | p.Ala345Ser | P00439 |
| HOMO SAPIENS | 0 | 1607 | 6pah | A | 348 | LEU | p.Leu348Val | P00439 |
| HOMO SAPIENS | 6 | 1607 | 6pah | A | 356 | TYR | p.Tyr356Ter | P00439 |
| HOMO SAPIENS | 2 | 1607 | 6pah | A | 380 | THR | p.Thr380Met | P00439 |
| HOMO SAPIENS | 0 | 1607 | 6pah | A | 386 | TYR | p.Tyr386Cys | P00439 |
| HOMO SAPIENS | 5 | 1607 | 6pah | A | 390 | GLU | p.Glu390Gly | P00439 |
| HOMO SAPIENS | 5 | 1607 | 6pah | A | 403 | ALA | p.Ala403Val | P00439 |
| HOMO SAPIENS | 4 | 1607 | 6pah | A | 408 | ARG | p.Arg408Trp | P00439 |
| HOMO SAPIENS | 3 | 1607 | 6pah | A | 413 | ARG | p.Arg413Pro | P00439 |
| HOMO SAPIENS | 4 | 1607 | 6pah | A | 414 | TYR | p.Tyr414Cys | P00439 |
| HOMO SAPIENS | 4 | 1607 | 6pah | A | 414 | TYR | p.Tyr414His | P00439 |
| HOMO SAPIENS | 4 | 1607 | 6pah | A | 415 | ASP | p.Asp415Asn | P00439 |
| HOMO SAPIENS | -1 | 676 | 5dc5 | A | 7 | PRO | p.Pro7Leu | Q9BY41 |
| HOMO SAPIENS | 0 | 676 | 5dc5 | A | 311 | THR | p.Thr311Met | Q9BY41 |
| HOMO SAPIENS | 1 | 124 | 5ftk | E | 95 | ARG | p.Arg95Gly | P55072 |
| HOMO SAPIENS | 1 | 124 | 5ftk | E | 155 | ARG | p.Arg155Cys | P55072 |
| HOMO SAPIENS | 1 | 124 | 5ftk | E | 155 | ARG | p.Arg155His | P55072 |
| HOMO SAPIENS | 1 | 124 | 5ftk | E | 159 | ARG | p.Arg159His | P55072 |
| HOMO SAPIENS | 1 | 124 | 5ftk | E | 159 | ARG | p.Arg159Gly | P55072 |
| HOMO SAPIENS | 0 | 124 | 5ftk | E | 191 | ARG | p.Arg191Gln | P55072 |
| HOMO SAPIENS | 0 | 124 | 5ftk | E | 232 | ALA | p.Ala232Glu | P55072 |
| HOMO SAPIENS | 5 | 5442 | 1fgg | B | 223 | GLY | p.Gly223Ser | O94766 |
| HOMO SAPIENS | 0 | 5442 | 1fgg | B | 277 | ARG | p.Arg277Gln | O94766 |
| HOMO SAPIENS | 0 | 9072 | 2gys | B | 225 | GLU | p.Glu249Gln | P32927 |
| HOMO SAPIENS | 0 | 9072 | 2gys | B | 247 | SER | p.Ser271Ter | P32927 |
| HOMO SAPIENS | 0 | 9072 | 2gys | B | 294 | ASP | p.Asp312Asn | P32927 |
| HOMO SAPIENS | 0 | 1484 | 4zrj | A | 62 | PHE | p.Phe62Ser | P35240 |
| HOMO SAPIENS | 3 | 1484 | 4zrj | A | 205 | MET | p.Met205Val | P35240 |
| HOMO SAPIENS | 3 | 1484 | 4zrj | A | 301 | ILE | p.Ile301Met | P35240 |
| HOMO SAPIENS | 4 | 40923 | 2enq | A | 68 | ILE | p.Ile391Met | P42336 |
| HOMO SAPIENS | 1 | 6179 | 5ofv | A | 125 | THR | p.Thr125Met | O43175 |
| HOMO SAPIENS | 0 | 6179 | 5ofv | A | 135 | ARG | p.Arg135Trp | O43175 |

| | | | | | | | | |
|---|---|---|---|---|---|---|---|---|
| HOMO SAPIENS | 1 | 6179 | 5ofv | A | 163 | ARG | p.Arg163Gln | O43175 |
| HOMO SAPIENS | 9 | 6179 | 5ofv | A | 261 | VAL | p.Val261Met | O43175 |
| HOMO SAPIENS | 7 | 34118 | 2i5o | A | 24 | MET | p.Met647Leu | Q9Y253 |
| HOMO SAPIENS | 8 | 3234 | 5c84 | A | 311 | LYS | p.Lys311Asn | P98170 |
| HOMO SAPIENS | 3 | 3234 | 5c84 | A | 321 | ALA | p.Ala321Gly | P98170 |
| HOMO SAPIENS | -1 | 15129 | 1zy8 | K | 1 | GLY | p.Gly54Ser | O00330 |
| HOMO SAPIENS | -1 | 15129 | 1zy8 | K | 48 | THR | p.Thr101Ala | O00330 |
| HOMO SAPIENS | -1 | 15129 | 1zy8 | K | 105 | GLU | p.Glu158Asp | O00330 |
| HOMO SAPIENS | 8 | 15129 | 1zy8 | K | 136 | ARG | p.Arg189His | O00330 |
| HOMO SAPIENS | -1 | 15129 | 1zy8 | K | 195 | GLN | p.Gln248Ter | O00330 |
| HOMO SAPIENS | 8 | 202 | 4bzb | A | 123 | HIS | p.His123Pro | Q9Y3Z3 |
| HOMO SAPIENS | 4 | 202 | 4bzb | A | 145 | ARG | p.Arg145Ter | Q9Y3Z3 |
| HOMO SAPIENS | 6 | 202 | 4bzb | A | 164 | ARG | p.Arg164Ter | Q9Y3Z3 |
| HOMO SAPIENS | 3 | 202 | 4bzb | A | 201 | ILE | p.Ile201Asn | Q9Y3Z3 |
| HOMO SAPIENS | 0 | 202 | 4bzb | A | 209 | GLY | p.Gly209Ser | Q9Y3Z3 |
| HOMO SAPIENS | 0 | 202 | 4bzb | A | 254 | MET | p.Met254Val | Q9Y3Z3 |
| HOMO SAPIENS | 5 | 202 | 4bzb | A | 531 | ARG | p.Arg531Ser | Q9Y3Z3 |
| HOMO SAPIENS | 6 | 202 | 4bzb | A | 548 | GLN | p.Gln548Ter | Q9Y3Z3 |
| HOMO SAPIENS | 3 | 30947 | 5lgd | A | 90 | PRO | p.Pro90Ser | P16671 |
| HOMO SAPIENS | 6 | 30947 | 5lgd | A | 254 | PHE | p.Phe254Leu | P16671 |
| HOMO SAPIENS | 0 | 30947 | 5lgd | A | 413 | ILE | p.Ile413Leu | P16671 |
| HOMO SAPIENS | 2 | 3906 | 5jtc | A | 735 | ARG | p.Arg735Trp | Q12797 |
| HOMO SAPIENS | 1 | 45 | 5uyx | C | 146 | GLU | p.Glu146Val | P48643 |
| HOMO SAPIENS | 1 | 45 | 5uyx | C | 147 | HIS | p.His147Arg | P48643 |
| HOMO SAPIENS | 0 | 45 | 5uyx | C | 362 | ILE | p.Ile362Met | P48643 |
| HOMO SAPIENS | 8 | 39596 | 1mhn | A | 95 | GLY | p.Gly95Arg | Q16637 |
| HOMO SAPIENS | 4 | 39596 | 1mhn | A | 111 | ALA | p.Ala111Gly | Q16637 |
| HOMO SAPIENS | 9 | 39596 | 1mhn | A | 116 | ILE | p.Ile116Phe | Q16637 |
| HOMO SAPIENS | 1 | 39596 | 1mhn | A | 136 | GLN | p.Gln136Glu | Q16637 |
| HOMO SAPIENS | 1 | 6408 | 5xjy | A | 85 | PRO | p.Pro85Leu | O95477 |
| HOMO SAPIENS | 0 | 6408 | 5xjy | A | 219 | ARG | p.Arg219Lys | O95477 |
| HOMO SAPIENS | 0 | 6408 | 5xjy | A | 230 | ARG | p.Arg230Cys | O95477 |
| HOMO SAPIENS | 1 | 6408 | 5xjy | A | 281 | MET | p.Met281Thr | O95477 |
| HOMO SAPIENS | -1 | 6408 | 5xjy | A | 343 | ALA | p.Ala343Gly | O95477 |
| HOMO SAPIENS | 1 | 6408 | 5xjy | A | 399 | VAL | p.Val399Ala | O95477 |
| HOMO SAPIENS | 1 | 6408 | 5xjy | A | 496 | ARG | p.Arg496Trp | O95477 |
| HOMO SAPIENS | 1 | 6408 | 5xjy | A | 590 | TRP | p.Trp590Ser | O95477 |
| HOMO SAPIENS | 1 | 6408 | 5xjy | A | 638 | ARG | p.Arg638Gln | O95477 |
| HOMO SAPIENS | 4 | 6408 | 5xjy | A | 697 | ALA | p.Ala697Thr | O95477 |
| HOMO SAPIENS | 1 | 6408 | 5xjy | A | 771 | VAL | p.Val771Met | O95477 |
| HOMO SAPIENS | 1 | 6408 | 5xjy | A | 774 | THR | p.Thr774Pro | O95477 |
| HOMO SAPIENS | 1 | 6408 | 5xjy | A | 776 | LYS | p.Lys776Asn | O95477 |
| HOMO SAPIENS | 0 | 6408 | 5xjy | A | 825 | VAL | p.Val825Ile | O95477 |
| HOMO SAPIENS | -1 | 6408 | 5xjy | A | 868 | GLU | p.Glu868Lys | O95477 |
| HOMO SAPIENS | -1 | 6408 | 5xjy | A | 883 | ILE | p.Ile883Met | O95477 |
| HOMO SAPIENS | 0 | 6408 | 5xjy | A | 935 | ASN | p.Asn935Ser | O95477 |
| HOMO SAPIENS | 0 | 6408 | 5xjy | A | 935 | ASN | p.Asn935His | O95477 |
| HOMO SAPIENS | 0 | 6408 | 5xjy | A | 937 | ALA | p.Ala937Val | O95477 |
| HOMO SAPIENS | 0 | 6408 | 5xjy | A | 1018 | ASP | p.Asp1018Gly | O95477 |
| HOMO SAPIENS | 0 | 6408 | 5xjy | A | 1096 | VAL | p.Val1096Ile | O95477 |
| HOMO SAPIENS | 0 | 6408 | 5xjy | A | 1099 | ASP | p.Asp1099Tyr | O95477 |
| HOMO SAPIENS | 0 | 6408 | 5xjy | A | 1172 | GLU | p.Glu1172Asp | O95477 |
| HOMO SAPIENS | 0 | 6408 | 5xjy | A | 1181 | SER | p.Ser1181Phe | O95477 |
| HOMO SAPIENS | 0 | 6408 | 5xjy | A | 1182 | ALA | p.Ala1182Thr | O95477 |

| | | | | | | | | |
|---|---|---|---|---|---|---|---|---|
| HOMO SAPIENS | -1 | 6408 | 5xjy | A | 1289 | ASP | p.Asp1289Asn | O95477 |
| HOMO SAPIENS | 0 | 6408 | 5xjy | A | 1399 | THR | p.Thr1399Ser | O95477 |
| HOMO SAPIENS | 1 | 6408 | 5xjy | A | 1408 | LEU | p.Leu1408Phe | O95477 |
| HOMO SAPIENS | 0 | 6408 | 5xjy | A | 1587 | LYS | p.Lys1587Arg | O95477 |
| HOMO SAPIENS | 1 | 6408 | 5xjy | A | 1674 | VAL | p.Val1674Ile | O95477 |
| HOMO SAPIENS | 1 | 6408 | 5xjy | A | 1680 | ARG | p.Arg1680Trp | O95477 |
| HOMO SAPIENS | 1 | 6408 | 5xjy | A | 1680 | ARG | p.Arg1680Gln | O95477 |
| HOMO SAPIENS | 0 | 6408 | 5xjy | A | 1800 | ASN | p.Asn1800His | O95477 |
| HOMO SAPIENS | -1 | 6408 | 5xjy | A | 1925 | ARG | p.Arg1925Gln | O95477 |
| HOMO SAPIENS | 0 | 6408 | 5xjy | A | 2009 | PHE | p.Phe2009Ser | O95477 |
| HOMO SAPIENS | -1 | 6408 | 5xjy | A | 2243 | ASP | p.Asp2243Glu | O95477 |
| HOMO SAPIENS | 4 | 23760 | 2mut | A | 231 | LEU | p.Phe231Leu | P07992 |
| HOMO SAPIENS | 6 | 1460 | 2hre | B | 96 | ARG | p.Arg96Gln | P22830 |
| HOMO SAPIENS | 8 | 1460 | 2hre | B | 129 | GLY | p.Gly129Arg | P22830 |
| HOMO SAPIENS | 9 | 1460 | 2hre | B | 172 | GLU | p.Glu172Lys | P22830 |
| HOMO SAPIENS | 4 | 1460 | 2hre | B | 285 | GLN | p.Gln285Arg | P22830 |
| HOMO SAPIENS | 5 | 1460 | 2hre | B | 362 | VAL | p.Val362Gly | P22830 |
| HOMO SAPIENS | 6 | 1460 | 2hre | B | 411 | CYS | p.Cys411Gly | P22830 |
| HOMO SAPIENS | 6 | 1460 | 2hre | B | 417 | PHE | p.Phe417Ser | P22830 |
| HOMO SAPIENS | 0 | 2210 | 5bug | A | 19 | ASP | p.Asp19Asn | P60484 |
| HOMO SAPIENS | 0 | 2210 | 5bug | A | 35 | MET | p.Met35Arg | P60484 |
| HOMO SAPIENS | 1 | 2210 | 5bug | A | 61 | HIS | p.His61Asp | P60484 |
| HOMO SAPIENS | 0 | 2210 | 5bug | A | 68 | TYR | p.Tyr68His | P60484 |
| HOMO SAPIENS | 8 | 2210 | 5bug | A | 70 | LEU | p.Leu70Pro | P60484 |
| HOMO SAPIENS | 6 | 2210 | 5bug | A | 93 | HIS | p.His93Arg | P60484 |
| HOMO SAPIENS | 1 | 2210 | 5bug | A | 112 | LEU | p.Leu112Pro | P60484 |
| HOMO SAPIENS | 0 | 2210 | 5bug | A | 121 | ALA | p.Ala121Gly | P60484 |
| HOMO SAPIENS | 3 | 2210 | 5bug | A | 123 | HIS | p.His123Arg | P60484 |
| HOMO SAPIENS | 7 | 2210 | 5bug | A | 124 | CYS | p.Cys124Arg | P60484 |
| HOMO SAPIENS | 9 | 2210 | 5bug | A | 129 | GLY | p.Gly129Glu | P60484 |
| HOMO SAPIENS | 9 | 2210 | 5bug | A | 130 | ARG | p.Arg130Gln | P60484 |
| HOMO SAPIENS | 9 | 2210 | 5bug | A | 130 | ARG | p.Arg130Gly | P60484 |
| HOMO SAPIENS | 0 | 2210 | 5bug | A | 132 | GLY | p.Gly132Val | P60484 |
| HOMO SAPIENS | 0 | 2210 | 5bug | A | 170 | SER | p.Ser170Arg | P60484 |
| HOMO SAPIENS | 1 | 2210 | 5bug | A | 173 | ARG | p.Arg173His | P60484 |
| HOMO SAPIENS | 0 | 2210 | 5bug | A | 217 | VAL | p.Val217Ile | P60484 |
| HOMO SAPIENS | 0 | 2210 | 5bug | A | 241 | PHE | p.Phe241Ser | P60484 |
| HOMO SAPIENS | 0 | 2210 | 5bug | A | 250 | CYS | p.Cys250Ter | P60484 |
| HOMO SAPIENS | 1 | 2210 | 5bug | A | 252 | ASP | p.Asp252Gly | P60484 |
| HOMO SAPIENS | 3 | 123 | 2ibu | A | 93 | ASN | p.Asn93Ser | P24752 |
| HOMO SAPIENS | 0 | 123 | 2ibu | A | 145 | GLN | p.Gln145Glu | P24752 |
| HOMO SAPIENS | 0 | 123 | 2ibu | A | 152 | GLY | p.Gly152Ala | P24752 |
| HOMO SAPIENS | 2 | 123 | 2ibu | A | 158 | ASN | p.Asn158Asp | P24752 |
| HOMO SAPIENS | 2 | 123 | 2ibu | A | 158 | ASN | p.Asn158Ser | P24752 |
| HOMO SAPIENS | 5 | 123 | 2ibu | A | 183 | GLY | p.Gly183Arg | P24752 |
| HOMO SAPIENS | 3 | 123 | 2ibu | A | 272 | GLN | p.Gln272Ter | P24752 |
| HOMO SAPIENS | 3 | 123 | 2ibu | A | 312 | ILE | p.Ile312Thr | P24752 |
| HOMO SAPIENS | 1 | 123 | 2ibu | A | 333 | ALA | p.Ala333Pro | P24752 |
| HOMO SAPIENS | 5 | 123 | 2ibu | A | 379 | GLY | p.Gly379Val | P24752 |
| HOMO SAPIENS | 0 | 123 | 2ibu | A | 380 | ALA | p.Ala380Thr | P24752 |
| HOMO SAPIENS | 2 | 369 | 5nrw | A | 1175 | TYR | p.Tyr1175Cys | Q92793 |
| HOMO SAPIENS | 2 | 830 | 2vc2 | A | 116 | LEU | p.Leu147Val | P08514 |
| HOMO SAPIENS | 0 | 830 | 2vc2 | A | 183 | LEU | p.Leu214Pro | P08514 |
| HOMO SAPIENS | 4 | 830 | 2vc2 | A | 242 | GLY | p.Gly273Asp | P08514 |

| | | | | | | | | |
|---|---|---|---|---|---|---|---|---|
| HOMO SAPIENS | 6 | 830 | 2vc2 | A | 324 | GLU | p.Glu355Lys | P08514 |
| HOMO SAPIENS | -1 | 10353 | 5ejc | E | 989 | ALA | p.Ala989Ser | Q92574 |
| HOMO SAPIENS | 0 | 15448 | 3lgg | A | 21 | GLY | p.Gly47Arg | Q9NZK5 |
| HOMO SAPIENS | 0 | 15448 | 3lgg | A | 21 | GLY | p.Gly47Val | Q9NZK5 |
| HOMO SAPIENS | 9 | 15448 | 3lgg | A | 86 | HIS | p.His112Gln | Q9NZK5 |
| HOMO SAPIENS | 0 | 15448 | 3lgg | A | 143 | ARG | p.Arg169Gln | Q9NZK5 |
| HOMO SAPIENS | 3 | 15448 | 3lgg | A | 221 | ALA | p.Ala247Val | Q9NZK5 |
| HOMO SAPIENS | 0 | 15448 | 3lgg | A | 225 | PRO | p.Pro251Leu | Q9NZK5 |
| HOMO SAPIENS | 0 | 15448 | 3lgg | A | 283 | MET | p.Met309Ile | Q9NZK5 |
| HOMO SAPIENS | 2 | 15448 | 3lgg | A | 334 | THR | p.Thr360Ala | Q9NZK5 |
| HOMO SAPIENS | 7 | 15448 | 3lgg | A | 357 | GLY | p.Gly383Ser | Q9NZK5 |
| HOMO SAPIENS | 0 | 15448 | 3lgg | A | 427 | TYR | p.Tyr453Cys | Q9NZK5 |
| HOMO SAPIENS | 0 | 872 | 2y3i | A | 87 | LEU | p.Leu88Pro | P00558 |
| HOMO SAPIENS | 1 | 872 | 2y3i | A | 163 | ASP | p.Asp164Val | P00558 |
| HOMO SAPIENS | 4 | 872 | 2y3i | A | 205 | ARG | p.Arg206Pro | P00558 |
| HOMO SAPIENS | 4 | 872 | 2y3i | A | 267 | ASP | p.Asp268Asn | P00558 |
| HOMO SAPIENS | 8 | 872 | 2y3i | A | 284 | ASP | p.Asp285Val | P00558 |
| HOMO SAPIENS | 4 | 8 | 5jco | B | 188 | ILE | p.Ile188Leu | Q71U36 |
| HOMO SAPIENS | 7 | 8 | 5jco | B | 263 | PRO | p.Pro263Thr | Q71U36 |
| HOMO SAPIENS | 6 | 8 | 5jco | B | 264 | ARG | p.Arg264Cys | Q71U36 |
| HOMO SAPIENS | 7 | 8 | 5jco | B | 402 | ARG | p.Arg402His | Q71U36 |
| HOMO SAPIENS | 7 | 8 | 5jco | B | 419 | SER | p.Ser419Leu | Q71U36 |
| HOMO SAPIENS | -1 | 880 | 4nwf | A | 2 | THR | p.Thr2Ile | Q06124 |
| HOMO SAPIENS | 2 | 880 | 4nwf | A | 58 | ASN | p.Asn58Ser | Q06124 |
| HOMO SAPIENS | 4 | 880 | 4nwf | A | 61 | ASP | p.Asp61Gly | Q06124 |
| HOMO SAPIENS | 2 | 880 | 4nwf | A | 62 | TYR | p.Tyr62Asp | Q06124 |
| HOMO SAPIENS | 3 | 880 | 4nwf | A | 63 | TYR | p.Tyr63Cys | Q06124 |
| HOMO SAPIENS | 2 | 880 | 4nwf | A | 69 | GLU | p.Glu69Gln | Q06124 |
| HOMO SAPIENS | 2 | 880 | 4nwf | A | 72 | ALA | p.Ala72Gly | Q06124 |
| HOMO SAPIENS | 2 | 880 | 4nwf | A | 72 | ALA | p.Ala72Ser | Q06124 |
| HOMO SAPIENS | 3 | 880 | 4nwf | A | 73 | THR | p.Thr73Ile | Q06124 |
| HOMO SAPIENS | 5 | 880 | 4nwf | A | 76 | GLU | p.Glu76Ala | Q06124 |
| HOMO SAPIENS | 5 | 880 | 4nwf | A | 76 | GLU | p.Glu76Lys | Q06124 |
| HOMO SAPIENS | 3 | 880 | 4nwf | A | 79 | GLN | p.Gln79Arg | Q06124 |
| HOMO SAPIENS | 5 | 880 | 4nwf | A | 152 | ARG | p.Arg152His | Q06124 |
| HOMO SAPIENS | 2 | 880 | 4nwf | A | 268 | GLY | p.Gly268Cys | Q06124 |
| HOMO SAPIENS | 9 | 880 | 4nwf | A | 276 | LYS | p.Lys276Arg | Q06124 |
| HOMO SAPIENS | 9 | 880 | 4nwf | A | 279 | TYR | p.Tyr279Cys | Q06124 |
| HOMO SAPIENS | 5 | 880 | 4nwf | A | 285 | PHE | p.Phe285Ser | Q06124 |
| HOMO SAPIENS | 0 | 880 | 4nwf | A | 308 | ASP | p.Asn308Ser | Q06124 |
| HOMO SAPIENS | 0 | 880 | 4nwf | A | 308 | ASP | p.Asn308Asp | Q06124 |
| HOMO SAPIENS | 5 | 880 | 4nwf | A | 392 | ALA | p.Ala392Thr | Q06124 |
| HOMO SAPIENS | 1 | 880 | 4nwf | A | 411 | THR | p.Thr415Met | Q06124 |
| HOMO SAPIENS | 0 | 880 | 4nwf | A | 461 | ALA | p.Ala465Thr | Q06124 |
| HOMO SAPIENS | 0 | 880 | 4nwf | A | 464 | GLY | p.Gly468Ala | Q06124 |
| HOMO SAPIENS | 1 | 880 | 4nwf | A | 491 | PRO | p.Pro495Ser | Q06124 |
| HOMO SAPIENS | 1 | 880 | 4nwf | A | 502 | SER | p.Ser506Thr | Q06124 |
| HOMO SAPIENS | 8 | 880 | 4nwf | A | 510 | GLN | p.Gln514Pro | Q06124 |
| HOMO SAPIENS | 4 | 1872 | 3wf1 | D | 109 | ARG | p.Arg109Trp | P16278 |
| HOMO SAPIENS | 0 | 1872 | 3wf1 | D | 521 | ARG | p.Arg521Cys | P16278 |
| HOMO SAPIENS | 5 | 910 | 5xte | A | 45 | SER | p.Ser45Phe | O14949 |
| HOMO SAPIENS | 2 | 9017 | 5f0k | A | 382 | LYS | p.Lys382Arg | Q96QK1 |
| HOMO SAPIENS | 1 | 710 | 4lsj | A | 559 | ILE | p.Ile559Asn | P04150 |
| HOMO SAPIENS | 5 | 710 | 4lsj | A | 571 | VAL | p.Val571Ala | P04150 |

| | | | | | | | | |
|---|---|---|---|---|---|---|---|---|
| HOMO SAPIENS | 0 | 710 | 4lsj | A | 641 | ASP | p.Asp641Val | P04150 |
| HOMO SAPIENS | 1 | 710 | 4lsj | A | 678 | ASP | p.Asp678Glu | P04150 |
| HOMO SAPIENS | 2 | 710 | 4lsj | A | 679 | GLY | p.Gly679Ser | P04150 |
| HOMO SAPIENS | 1 | 710 | 4lsj | A | 737 | PHE | p.Phe737Leu | P04150 |
| HOMO SAPIENS | 1 | 710 | 4lsj | A | 747 | ILE | p.Ile747Met | P04150 |
| HOMO SAPIENS | 1 | 710 | 4lsj | A | 753 | LEU | p.Leu753Phe | P04150 |
| HOMO SAPIENS | 1 | 710 | 4lsj | A | 773 | LEU | p.Leu773Pro | P04150 |
| HOMO SAPIENS | 1 | 2145 | 1av1 | B | 71 | PHE | p.Phe95Tyr | P02647 |
| HOMO SAPIENS | 1 | 2145 | 1av1 | B | 128 | GLU | p.Glu152Lys | P02647 |
| HOMO SAPIENS | 1 | 2145 | 1av1 | B | 136 | GLU | p.Glu160Lys | P02647 |
| HOMO SAPIENS | 2 | 2145 | 1av1 | B | 143 | PRO | p.Pro167Arg | P02647 |
| HOMO SAPIENS | 6 | 2145 | 1av1 | B | 164 | ALA | p.Ala188Ser | P02647 |
| HOMO SAPIENS | 7 | 2145 | 1av1 | B | 173 | ARG | p.Arg197Cys | P02647 |
| HOMO SAPIENS | 2 | 2145 | 1av1 | B | 198 | GLU | p.Glu222Lys | P02647 |
| HOMO SAPIENS | -1 | 3432 | 2beu | A | 1 | SER | p.Ser46Ter | P12694 |
| HOMO SAPIENS | 5 | 3432 | 2beu | A | 79 | ARG | p.Arg124Trp | P12694 |
| HOMO SAPIENS | 7 | 3432 | 2beu | A | 114 | ARG | p.Arg159Gln | P12694 |
| HOMO SAPIENS | 6 | 3432 | 2beu | A | 166 | THR | p.Thr211Met | P12694 |
| HOMO SAPIENS | 4 | 3432 | 2beu | A | 175 | ALA | p.Ala220Val | P12694 |
| HOMO SAPIENS | 5 | 3432 | 2beu | A | 204 | GLY | p.Gly249Ser | P12694 |
| HOMO SAPIENS | 4 | 3432 | 2beu | A | 209 | ALA | p.Ala254Asp | P12694 |
| HOMO SAPIENS | 6 | 3432 | 2beu | A | 220 | ARG | p.Arg265Trp | P12694 |
| HOMO SAPIENS | 6 | 3432 | 2beu | A | 234 | TYR | p.Tyr279Cys | P12694 |
| HOMO SAPIENS | 5 | 3432 | 2beu | A | 240 | ALA | p.Ala285Pro | P12694 |
| HOMO SAPIENS | 4 | 3432 | 2beu | A | 245 | GLY | p.Gly290Arg | P12694 |
| HOMO SAPIENS | 5 | 3432 | 2beu | A | 269 | ARG | p.Arg314Ter | P12694 |
| HOMO SAPIENS | 3 | 3432 | 2beu | A | 301 | ARG | p.Arg346Cys | P12694 |
| HOMO SAPIENS | 5 | 3432 | 2beu | A | 359 | ASN | p.Asn404Ser | P12694 |
| HOMO SAPIENS | 4 | 3432 | 2beu | A | 393 | TYR | p.Tyr438Asn | P12694 |
| HOMO SAPIENS | 8 | 4541 | 1rr8 | C | 533 | ASP | p.Asp533Gly | P11387 |
| HOMO SAPIENS | 2 | 3247 | 1tzn | d | 1105 | GLY | p.Gly105Asp | P58335 |
| HOMO SAPIENS | 3 | 3247 | 1tzn | d | 1111 | ARG | p.Arg111Cys | P58335 |
| HOMO SAPIENS | 2 | 3247 | 1tzn | d | 1189 | ILE | p.Ile189Thr | P58335 |
| HOMO SAPIENS | 9 | 467 | 1suv | C | 58 | ASP | p.Asp77Asn | P02787 |
| HOMO SAPIENS | 2 | 467 | 1suv | C | 277 | ASP | p.Asp296Gly | P02787 |
| HOMO SAPIENS | 9 | 6485 | 1xwd | B | 51 | CYS | p.Cys69Gly | P01225 |
| HOMO SAPIENS | 1 | 694 | 1wuu | A | 28 | PRO | p.Pro28Thr | P51570 |
| HOMO SAPIENS | 0 | 694 | 1wuu | A | 32 | VAL | p.Val32Met | P51570 |
| HOMO SAPIENS | 0 | 694 | 1wuu | A | 198 | ALA | p.Ala198Val | P51570 |
| HOMO SAPIENS | 0 | 143 | 4nkx | A | 93 | PHE | p.Phe93Cys | P05093 |
| HOMO SAPIENS | 3 | 143 | 4nkx | A | 96 | ARG | p.Arg96Trp | P05093 |
| HOMO SAPIENS | 3 | 143 | 4nkx | A | 96 | ARG | p.Arg96Gln | P05093 |
| HOMO SAPIENS | 0 | 143 | 4nkx | A | 106 | SER | p.Ser106Pro | P05093 |
| HOMO SAPIENS | 1 | 143 | 4nkx | A | 114 | PHE | p.Phe114Val | P05093 |
| HOMO SAPIENS | 1 | 143 | 4nkx | A | 116 | ASP | p.Asp116Val | P05093 |
| HOMO SAPIENS | 3 | 143 | 4nkx | A | 125 | ARG | p.Arg125Gln | P05093 |
| HOMO SAPIENS | 0 | 143 | 4nkx | A | 201 | TYR | p.Tyr201Asp | P05093 |
| HOMO SAPIENS | 0 | 143 | 4nkx | A | 239 | ARG | p.Arg239Ter | P05093 |
| HOMO SAPIENS | 0 | 143 | 4nkx | A | 329 | TYR | p.Tyr329Asp | P05093 |
| HOMO SAPIENS | 1 | 143 | 4nkx | A | 342 | PRO | p.Pro342Thr | P05093 |
| HOMO SAPIENS | 2 | 143 | 4nkx | A | 347 | ARG | p.Arg347His | P05093 |
| HOMO SAPIENS | 2 | 143 | 4nkx | A | 347 | ARG | p.Arg347Cys | P05093 |
| HOMO SAPIENS | 1 | 143 | 4nkx | A | 358 | ARG | p.Arg358Gln | P05093 |
| HOMO SAPIENS | 0 | 143 | 4nkx | A | 362 | ARG | p.Arg362Cys | P05093 |

| | | | | | | | | |
|---|---|---|---|---|---|---|---|---|
| HOMO SAPIENS | 1 | 143 | 4nkx | A | 406 | TRP | p.Trp406Arg | P05093 |
| HOMO SAPIENS | 0 | 143 | 4nkx | A | 428 | PRO | p.Pro428Leu | P05093 |
| HOMO SAPIENS | 6 | 1441 | 1uuh | A | 46 | ARG | p.Arg46Gly | P16070 |
| HOMO SAPIENS | 3 | 82 | 4wlv | D | 37 | GLY | p.Gly37Arg | P40926 |
| HOMO SAPIENS | 0 | 82 | 4wlv | D | 133 | PRO | p.Pro133Leu | P40926 |
| HOMO SAPIENS | 2 | 392 | 5xtd | A | 56 | SER | p.Ser56Pro | P49821 |
| HOMO SAPIENS | 2 | 392 | 5xtd | A | 59 | ARG | p.Arg59Ter | P49821 |
| HOMO SAPIENS | 2 | 392 | 5xtd | A | 74 | ASP | p.Asp74Asn | P49821 |
| HOMO SAPIENS | 4 | 392 | 5xtd | A | 116 | ASN | p.Asn116Lys | P49821 |
| HOMO SAPIENS | 4 | 392 | 5xtd | A | 117 | ALA | p.Ala117Thr | P49821 |
| HOMO SAPIENS | 5 | 392 | 5xtd | A | 122 | PRO | p.Pro122Leu | P49821 |
| HOMO SAPIENS | 3 | 392 | 5xtd | A | 160 | GLY | p.Gly160Glu | P49821 |
| HOMO SAPIENS | 4 | 392 | 5xtd | A | 214 | GLU | p.Glu214Lys | P49821 |
| HOMO SAPIENS | 2 | 392 | 5xtd | A | 357 | MET | p.Met357Thr | P49821 |
| HOMO SAPIENS | 4 | 392 | 5xtd | A | 423 | THR | p.Thr423Met | P49821 |
| HOMO SAPIENS | 0 | 22593 | 3pdf | A | 8 | TYR | p.Tyr32Ter | P53634 |
| HOMO SAPIENS | 0 | 22593 | 3pdf | A | 15 | TRP | p.Trp39Ser | P53634 |
| HOMO SAPIENS | 0 | 22593 | 3pdf | A | 103 | HIS | p.His127Pro | P53634 |
| HOMO SAPIENS | 9 | 22593 | 3pdf | A | 228 | GLN | p.Gln252Leu | P53634 |
| HOMO SAPIENS | 3 | 22593 | 3pdf | A | 262 | GLN | p.Gln286Arg | P53634 |
| HOMO SAPIENS | 4 | 22593 | 3pdf | A | 277 | GLY | p.Gly301Ser | P53634 |
| HOMO SAPIENS | 0 | 22593 | 3pdf | A | 323 | TYR | p.Tyr347Cys | P53634 |
| HOMO SAPIENS | 5 | 3461 | 4c2a | A | 1306 | GLN | p.Arg1306Trp | P04275 |
| HOMO SAPIENS | 5 | 3461 | 4c2a | A | 1374 | ARG | p.Arg1374Cys | P04275 |
| HOMO SAPIENS | 5 | 3461 | 4c2a | A | 1379 | ARG | p.Arg1379Cys | P04275 |
| HOMO SAPIENS | 4 | 3461 | 4c2a | A | 1380 | ILE | p.Ile1380Val | P04275 |
| HOMO SAPIENS | 0 | 3461 | 4c2a | A | 1381 | ALA | p.Thr1381Ala | P04275 |
| HOMO SAPIENS | 5 | 3461 | 4c2a | A | 1399 | ARG | p.Arg1399His | P04275 |
| HOMO SAPIENS | 5 | 3461 | 4c2a | A | 1435 | ASN | p.Asn1435Ser | P04275 |
| HOMO SAPIENS | 0 | 4533 | 5uj7 | C | 174 | TYR | p.Tyr174Cys | O43929 |
| HOMO SAPIENS | 0 | 4533 | 5uj7 | C | 202 | LEU | p.Leu202Val | O43929 |
| HOMO SAPIENS | 0 | 1374 | 3j07 | O | 39 | PRO | p.Pro39Leu | P02511 |
| HOMO SAPIENS | 0 | 1374 | 3j07 | O | 107 | ARG | p.Arg107His | P02511 |
| HOMO SAPIENS | 0 | 1374 | 3j07 | O | 157 | ARG | p.Arg157His | P02511 |
| HOMO SAPIENS | -1 | 942 | 1s9i | B | 57 | PHE | p.Phe57Cys | P36507 |
| HOMO SAPIENS | -1 | 942 | 1s9i | B | 57 | PHE | p.Phe57Val | P36507 |
| HOMO SAPIENS | 2 | 942 | 1s9i | B | 128 | PRO | p.Pro128Gln | P36507 |
| HOMO SAPIENS | 1 | 942 | 1s9i | B | 134 | TYR | p.Tyr134His | P36507 |
| HOMO SAPIENS | 1 | 942 | 1s9i | B | 188 | HIS | p.His188Gln | P36507 |
| HOMO SAPIENS | 0 | 942 | 1s9i | B | 262 | VAL | p.Val262Phe | P36507 |
| HOMO SAPIENS | 0 | 942 | 1s9i | B | 263 | GLY | p.Gly263Arg | P36507 |
| HOMO SAPIENS | -1 | 942 | 1s9i | B | 298 | PRO | p.Pro298Leu | P36507 |
| HOMO SAPIENS | -1 | 942 | 1s9i | B | 305 | VAL | p.Val305Ile | P36507 |
| HOMO SAPIENS | 0 | 3245 | 1fbv | A | 367 | GLN | p.Gln367Pro | P22681 |
| HOMO SAPIENS | 0 | 3245 | 1fbv | A | 371 | TYR | p.Tyr371His | P22681 |
| HOMO SAPIENS | 1 | 3245 | 1fbv | A | 382 | LYS | p.Lys382Glu | P22681 |
| HOMO SAPIENS | 2 | 3245 | 1fbv | A | 390 | ASP | p.Asp390Tyr | P22681 |
| HOMO SAPIENS | 9 | 3245 | 1fbv | A | 396 | CYS | p.Cys396Arg | P22681 |
| HOMO SAPIENS | 7 | 3245 | 1fbv | A | 420 | ARG | p.Arg420Gln | P22681 |
| HOMO SAPIENS | 1 | 3245 | 1fbv | A | 429 | ILE | p.Ile429Met | P22681 |
| HOMO SAPIENS | 0 | 2604 | 2pbd | P | 111 | LEU | p.Leu112Met | P07737 |
| HOMO SAPIENS | 3 | 2604 | 2pbd | P | 116 | GLU | p.Glu117Gly | P07737 |
| HOMO SAPIENS | 4 | 2013 | 3b7w | A | 472 | ARG | p.Arg472Gly | Q08AH3 |
| HOMO SAPIENS | 0 | 24346 | 3sh5 | A | 85 | ARG | p.Arg4281His | P98160 |

| | | | | | | | | |
|---|---|---|---|---|---|---|---|---|
| HOMO SAPIENS | 0 | 24346 | 3sh5 | A | 122 | ARG | p.Arg4318Trp | P98160 |
| HOMO SAPIENS | 0 | 24346 | 3sh5 | A | 135 | SER | p.Ser4331Asn | P98160 |
| HOMO SAPIENS | 0 | 24346 | 3sh5 | A | 136 | VAL | p.Val4332Ile | P98160 |
| HOMO SAPIENS | 0 | 707 | 4p2g | A | 26 | MET | p.Met26Ile | Q99497 |
| HOMO SAPIENS | 5 | 707 | 4p2g | A | 39 | ALA | p.Ala39Ser | Q99497 |
| HOMO SAPIENS | 5 | 707 | 4p2g | A | 64 | GLU | p.Glu64Asp | Q99497 |
| HOMO SAPIENS | 4 | 707 | 4p2g | A | 149 | ASP | p.Asp149Ala | Q99497 |
| HOMO SAPIENS | 6 | 707 | 4p2g | A | 163 | GLU | p.Glu163Lys | Q99497 |
| HOMO SAPIENS | 6 | 707 | 4p2g | A | 166 | LEU | p.Leu166Pro | Q99497 |
| HOMO SAPIENS | 3 | 5121 | 4xss | E | 31 | GLY | p.Gly58Arg | P06213 |
| HOMO SAPIENS | 2 | 5121 | 4xss | E | 119 | ILE | p.Ile146Met | P06213 |
| HOMO SAPIENS | 8 | 5121 | 4xss | E | 209 | HIS | p.His236Arg | P06213 |
| HOMO SAPIENS | 2 | 5121 | 4xss | E | 234 | ASP | p.Asp261Glu | P06213 |
| HOMO SAPIENS | 1 | 5121 | 4xss | E | 276 | GLN | p.Gln303His | P06213 |
| HOMO SAPIENS | 1 | 27790 | 5erp | B | 140 | THR | p.Thr275Met | Q02487 |
| HOMO SAPIENS | 0 | 27790 | 5erp | B | 150 | ILE | p.Ile285Thr | Q02487 |
| HOMO SAPIENS | 1 | 27790 | 5erp | B | 205 | THR | p.Thr340Ala | Q02487 |
| HOMO SAPIENS | 1 | 27790 | 5erp | B | 223 | THR | p.Thr358Ile | Q02487 |
| HOMO SAPIENS | 0 | 27790 | 5erp | B | 301 | GLY | p.Gly436Ala | Q02487 |
| HOMO SAPIENS | -1 | 27790 | 5erp | B | 315 | ARG | p.Arg450Ser | Q02487 |
| HOMO SAPIENS | 0 | 27790 | 5erp | B | 369 | SER | p.Ser504Gly | Q02487 |
| HOMO SAPIENS | 0 | 27790 | 5erp | B | 385 | ILE | p.Ile520Thr | Q02487 |
| HOMO SAPIENS | 0 | 27790 | 5erp | B | 461 | ALA | p.Ala596Val | Q02487 |
| HOMO SAPIENS | 0 | 27790 | 5erp | B | 503 | GLN | p.Gln638His | Q02487 |
| HOMO SAPIENS | 5 | 6599 | 4nsc | A | 129 | ARG | p.Arg129Leu | Q9BPX6 |
| HOMO SAPIENS | -1 | 6599 | 4nsc | A | 183 | GLN | p.Gln183Ter | Q9BPX6 |
| HOMO SAPIENS | 0 | 20420 | 3ghm | A | 398 | ARG | p.Arg398His | Q76LX8 |
| HOMO SAPIENS | 0 | 20420 | 3ghm | A | 448 | GLN | p.Gln448Glu | Q76LX8 |
| HOMO SAPIENS | 1 | 20420 | 3ghm | A | 449 | GLN | p.Gln449Ter | Q76LX8 |
| HOMO SAPIENS | 0 | 20420 | 3ghm | A | 456 | GLN | p.Gln456His | Q76LX8 |
| HOMO SAPIENS | 1 | 20420 | 3ghm | A | 528 | ARG | p.Arg528Gly | Q76LX8 |
| HOMO SAPIENS | 0 | 20420 | 3ghm | A | 618 | PRO | p.Pro618Ala | Q76LX8 |
| HOMO SAPIENS | 0 | 20420 | 3ghm | A | 625 | ARG | p.Arg625His | Q76LX8 |
| HOMO SAPIENS | 0 | 275 | 4rmz | A | 293 | GLN | p.Gln293Ter | Q9NWZ3 |
| HOMO SAPIENS | 0 | 275 | 4rmz | A | 428 | ALA | p.Ala428Thr | Q9NWZ3 |
| HOMO SAPIENS | 2 | 10458 | 3puf | B | 138 | LEU | p.Leu138Phe | Q5TBB1 |
| HOMO SAPIENS | 6 | 10458 | 3puf | B | 177 | ALA | p.Ala177Thr | Q5TBB1 |
| HOMO SAPIENS | -1 | 10458 | 3puf | B | 185 | VAL | p.Val185Gly | Q5TBB1 |
| HOMO SAPIENS | 0 | 26648 | 2n90 | A | 36 | ARG | p.Arg444Gln | P04629 |
| HOMO SAPIENS | 2 | 977 | 1dnw | D | 166 | ALA | p.Ala332Val | P05164 |
| HOMO SAPIENS | 4 | 977 | 1dnw | D | 333 | ARG | p.Arg499Cys | P05164 |
| HOMO SAPIENS | 1 | 977 | 1dnw | D | 403 | ARG | p.Arg569Trp | P05164 |
| HOMO SAPIENS | 0 | 25704 | 3ajb | A | 82 | GLN | p.Gln82Arg | P56589 |
| HOMO SAPIENS | 3 | 255 | 3t1p | A | 38 | TYR | p.Tyr62Ter | P01009 |
| HOMO SAPIENS | 3 | 255 | 3t1p | A | 39 | ARG | p.Arg63Cys | P01009 |
| HOMO SAPIENS | 0 | 255 | 3t1p | A | 101 | ARG | p.Arg125His | P01009 |
| HOMO SAPIENS | 0 | 255 | 3t1p | A | 212 | GLN | p.Gln236Arg | P01009 |
| HOMO SAPIENS | 0 | 255 | 3t1p | A | 213 | VAL | p.Val237Ala | P01009 |
| HOMO SAPIENS | 1 | 255 | 3t1p | A | 269 | HIS | p.His293Gln | P01009 |
| HOMO SAPIENS | 8 | 255 | 3t1p | A | 341 | ASP | p.Asp365Asn | P01009 |
| HOMO SAPIENS | 9 | 255 | 3t1p | A | 342 | GLU | p.Glu366Lys | P01009 |
| HOMO SAPIENS | 0 | 255 | 3t1p | A | 369 | PRO | p.Pro393Ser | P01009 |
| HOMO SAPIENS | 0 | 255 | 3t1p | A | 376 | GLU | p.Glu400Asp | P01009 |
| HOMO SAPIENS | -1 | 5175 | 3zxi | A | 35 | ALA | p.Ala35Asp | Q9Y2Z4 |

| | | | | | | | | | |
|---|---|---|---|---|---|---|---|---|---|
| HOMO SAPIENS | 0 | 5175 | 3zxi | A | | 52 | PHE | p.Phe52Leu | Q9Y2Z4 |
| HOMO SAPIENS | -1 | 5175 | 3zxi | A | | 68 | GLY | p.Gly68Ser | Q9Y2Z4 |
| HOMO SAPIENS | 0 | 5175 | 3zxi | A | | 179 | LYS | p.Lys179Gln | Q9Y2Z4 |
| HOMO SAPIENS | 0 | 5175 | 3zxi | A | | 191 | GLY | p.Gly191Val | Q9Y2Z4 |
| HOMO SAPIENS | 0 | 5175 | 3zxi | A | | 251 | ILE | p.Ile251Val | Q9Y2Z4 |
| HOMO SAPIENS | 2 | 5175 | 3zxi | A | | 295 | ASP | p.Asp295Asn | Q9Y2Z4 |
| HOMO SAPIENS | 0 | 5175 | 3zxi | A | | 311 | ASP | p.Asp311Glu | Q9Y2Z4 |
| HOMO SAPIENS | 0 | 5175 | 3zxi | A | | 360 | ARG | p.Arg360Ter | Q9Y2Z4 |
| HOMO SAPIENS | 1 | 3016 | 3exg | | 3 | 4 | ASP | p.Asp33His | P08559 |
| HOMO SAPIENS | 1 | 3016 | 3exg | | 3 | 176 | PHE | p.Phe205Leu | P08559 |
| HOMO SAPIENS | 1 | 3016 | 3exg | | 3 | 214 | TYR | p.Tyr243Asn | P08559 |
| HOMO SAPIENS | 1 | 3016 | 3exg | | 3 | 229 | ASP | p.Asp258Ala | P08559 |
| HOMO SAPIENS | 0 | 3016 | 3exg | | 3 | 233 | VAL | p.Val262Ile | P08559 |
| HOMO SAPIENS | 1 | 3016 | 3exg | | 3 | 234 | ARG | p.Arg263Gly | P08559 |
| HOMO SAPIENS | 0 | 3016 | 3exg | | 3 | 253 | MET | p.Met282Leu | P08559 |
| HOMO SAPIENS | 7 | 3016 | 3exg | | 3 | 259 | ARG | p.Arg288His | P08559 |
| HOMO SAPIENS | -1 | 3016 | 3exg | | 3 | 273 | ARG | p.Arg302Cys | P08559 |
| HOMO SAPIENS | 2 | 3016 | 3exg | | 3 | 286 | ASP | p.Asp315Asn | P08559 |
| HOMO SAPIENS | 1 | 3016 | 3exg | | 3 | 349 | ARG | p.Arg378His | P08559 |
| HOMO SAPIENS | 5 | 6729 | 4a0l | E | | 572 | ARG | p.Arg572Cys | Q13620 |
| HOMO SAPIENS | 0 | 1067 | 4kyo | A | | 9 | THR | p.Thr9Asn | P21549 |
| HOMO SAPIENS | 0 | 1067 | 4kyo | A | | 11 | PRO | p.Pro11Leu | P21549 |
| HOMO SAPIENS | 0 | 1067 | 4kyo | A | | 22 | ASN | p.Asn22Ser | P21549 |
| HOMO SAPIENS | 4 | 1067 | 4kyo | A | | 25 | LEU | p.Leu25Arg | P21549 |
| HOMO SAPIENS | 7 | 1067 | 4kyo | A | | 36 | ARG | p.Arg36His | P21549 |
| HOMO SAPIENS | 3 | 1067 | 4kyo | A | | 41 | GLY | p.Gly41Arg | P21549 |
| HOMO SAPIENS | 0 | 1067 | 4kyo | A | | 42 | GLY | p.Gly42Glu | P21549 |
| HOMO SAPIENS | 3 | 1067 | 4kyo | A | | 56 | ILE | p.Ile56Asn | P21549 |
| HOMO SAPIENS | 3 | 1067 | 4kyo | A | | 59 | GLU | p.Glu59Lys | P21549 |
| HOMO SAPIENS | 3 | 1067 | 4kyo | A | | 66 | TYR | p.Tyr66Ter | P21549 |
| HOMO SAPIENS | 3 | 1067 | 4kyo | A | | 111 | ARG | p.Arg111Leu | P21549 |
| HOMO SAPIENS | 7 | 1067 | 4kyo | A | | 118 | ARG | p.Arg118His | P21549 |
| HOMO SAPIENS | 3 | 1067 | 4kyo | A | | 122 | ARG | p.Arg122Ter | P21549 |
| HOMO SAPIENS | 3 | 1067 | 4kyo | A | | 124 | HIS | p.His124Pro | P21549 |
| HOMO SAPIENS | 0 | 1067 | 4kyo | A | | 152 | PHE | p.Phe152Ile | P21549 |
| HOMO SAPIENS | 1 | 1067 | 4kyo | A | | 156 | GLY | p.Gly156Arg | P21549 |
| HOMO SAPIENS | 0 | 1067 | 4kyo | A | | 161 | GLY | p.Gly161Arg | P21549 |
| HOMO SAPIENS | 3 | 1067 | 4kyo | A | | 166 | LEU | p.Leu166Pro | P21549 |
| HOMO SAPIENS | 5 | 1067 | 4kyo | A | | 170 | GLY | p.Gly170Arg | P21549 |
| HOMO SAPIENS | 5 | 1067 | 4kyo | A | | 186 | ALA | p.Ala186Val | P21549 |
| HOMO SAPIENS | 3 | 1067 | 4kyo | A | | 187 | SER | p.Ser187Phe | P21549 |
| HOMO SAPIENS | 3 | 1067 | 4kyo | A | | 197 | ARG | p.Arg197Pro | P21549 |
| HOMO SAPIENS | 5 | 1067 | 4kyo | A | | 201 | ASP | p.Asp201Glu | P21549 |
| HOMO SAPIENS | 0 | 1067 | 4kyo | A | | 202 | ILE | p.Ile202Asn | P21549 |
| HOMO SAPIENS | 1 | 1067 | 4kyo | A | | 205 | SER | p.Ser205Pro | P21549 |
| HOMO SAPIENS | 1 | 1067 | 4kyo | A | | 205 | SER | p.Ser205Ter | P21549 |
| HOMO SAPIENS | 2 | 1067 | 4kyo | A | | 216 | GLY | p.Gly216Arg | P21549 |
| HOMO SAPIENS | 5 | 1067 | 4kyo | A | | 233 | ARG | p.Arg233Cys | P21549 |
| HOMO SAPIENS | 5 | 1067 | 4kyo | A | | 244 | ILE | p.Ile244Thr | P21549 |
| HOMO SAPIENS | 5 | 1067 | 4kyo | A | | 244 | ILE | p.Ile244Met | P21549 |
| HOMO SAPIENS | 4 | 1067 | 4kyo | A | | 253 | CYS | p.Cys253Arg | P21549 |
| HOMO SAPIENS | 7 | 1067 | 4kyo | A | | 274 | GLU | p.Glu274Asp | P21549 |
| HOMO SAPIENS | 7 | 1067 | 4kyo | A | | 279 | ILE | p.Ile279Thr | P21549 |
| HOMO SAPIENS | 6 | 1067 | 4kyo | A | | 295 | ALA | p.Ala295Thr | P21549 |

| | | | | | | | | |
|---|---|---|---|---|---|---|---|---|
| HOMO SAPIENS | 5 | 1067 | 4kyo | A | 340 | ILE | p.Ile340Met | P21549 |
| HOMO SAPIENS | 1 | 790 | 2ric | C | 270 | SER | p.Ser290Thr | P35247 |
| HOMO SAPIENS | 0 | 790 | 2ric | C | 289 | GLU | p.Glu309Lys | P35247 |
| HOMO SAPIENS | 1 | 212 | 5sxe | A | 545 | GLU | p.Glu545Lys | P42336 |
| HOMO SAPIENS | 3 | 212 | 5sxe | A | 546 | GLN | p.Gln546Glu | P42336 |
| HOMO SAPIENS | 2 | 212 | 5sxe | A | 1047 | HIS | p.His1047Leu | P42336 |
| HOMO SAPIENS | 7 | 152 | 2zqq | F | 197 | ARG | p.Arg197Ter | Q13825 |
| HOMO SAPIENS | 0 | 152 | 2zqq | F | 248 | ALA | p.Ala248Thr | Q13825 |
| HOMO SAPIENS | 0 | 152 | 2zqq | F | 275 | ALA | p.Ala275Val | Q13825 |
| HOMO SAPIENS | 5 | 152 | 2zqq | F | 331 | LYS | p.Lys331Ter | Q13825 |
| HOMO SAPIENS | 6 | 17801 | 2j8l | A | 283 | PHE | p.Phe301Leu | P03951 |
| HOMO SAPIENS | 3 | 17801 | 2j8l | A | 308 | ARG | p.Arg326Cys | P03951 |
| HOMO SAPIENS | 0 | 17801 | 2j8l | A | 321 | CYS | p.Cys339Phe | P03951 |
| HOMO SAPIENS | 3 | 17801 | 2j8l | A | 351 | TYR | p.Tyr369Ter | P03951 |
| HOMO SAPIENS | 7 | 1802 | 5db0 | A | 22 | LEU | p.Leu22Arg | O00255 |
| HOMO SAPIENS | 7 | 1802 | 5db0 | A | 26 | GLU | p.Glu26Lys | O00255 |
| HOMO SAPIENS | 7 | 1802 | 5db0 | A | 139 | HIS | p.His139Asp | O00255 |
| HOMO SAPIENS | 0 | 1802 | 5db0 | A | 171 | ARG | p.Arg176Gln | O00255 |
| HOMO SAPIENS | 0 | 1802 | 5db0 | A | 184 | VAL | p.Val189Glu | O00255 |
| HOMO SAPIENS | 7 | 1802 | 5db0 | A | 255 | GLU | p.Glu260Lys | O00255 |
| HOMO SAPIENS | 4 | 1802 | 5db0 | A | 418 | ASP | p.Asp423Asn | O00255 |
| HOMO SAPIENS | 0 | 1802 | 5db0 | A | 436 | TRP | p.Trp441Arg | O00255 |
| HOMO SAPIENS | -1 | 1802 | 5db0 | A | 541 | ALA | p.Thr546Ala | O00255 |
| HOMO SAPIENS | 4 | 1802 | 5db0 | A | 552 | THR | p.Thr557Ser | O00255 |
| HOMO SAPIENS | 0 | 3450 | 2yuz | A | 36 | PRO | p.Pro461Gln | Q00872 |
| HOMO SAPIENS | 1 | 3450 | 2yuz | A | 56 | GLN | p.His481Gln | Q00872 |
| HOMO SAPIENS | 2 | 3450 | 2yuz | A | 68 | ALA | p.Ala493Val | Q00872 |
| HOMO SAPIENS | 0 | 3211 | 3rqg | A | 72 | VAL | p.Val72Leu | Q9BUL8 |
| HOMO SAPIENS | 2 | 4168 | 4q4f | A | 121 | ARG | p.Arg121Ter | Q14108 |
| HOMO SAPIENS | 2 | 4168 | 4q4f | A | 121 | ARG | p.Arg121Gln | Q14108 |
| HOMO SAPIENS | 2 | 4168 | 4q4f | A | 159 | MET | p.Met159Val | Q14108 |
| HOMO SAPIENS | 4 | 4168 | 4q4f | A | 363 | HIS | p.His363Asn | Q14108 |
| HOMO SAPIENS | 2 | 4168 | 4q4f | A | 396 | VAL | p.Val396Ile | Q14108 |
| HOMO SAPIENS | 0 | 1448 | 1y97 | A | 87 | ARG | p.Arg130Ter | Q9BQ50 |
| HOMO SAPIENS | 0 | 3444 | 4iwp | A | 151 | SER | p.Ser151Phe | Q9UHD2 |
| HOMO SAPIENS | 0 | 3444 | 4iwp | A | 401 | LYS | p.Lys401Glu | Q9UHD2 |
| HOMO SAPIENS | 0 | 116 | 1zy8 | B | 194 | GLY | p.Gly229Cys | P09622 |
| HOMO SAPIENS | 0 | 116 | 1zy8 | B | 255 | ASP | p.Asp290Gly | P09622 |
| HOMO SAPIENS | 2 | 116 | 1zy8 | B | 326 | MET | p.Met361Val | P09622 |
| HOMO SAPIENS | 0 | 116 | 1zy8 | B | 340 | GLU | p.Glu375Lys | P09622 |
| HOMO SAPIENS | 2 | 116 | 1zy8 | B | 358 | ILE | p.Ile393Thr | P09622 |
| HOMO SAPIENS | 1 | 116 | 1zy8 | B | 453 | PRO | p.Pro488Leu | P09622 |
| HOMO SAPIENS | 7 | 127 | 4qo7 | B | 160 | SER | p.Ser161Arg | P00338 |
| HOMO SAPIENS | 0 | 2025 | 1wrl | A | 8 | ALA | p.Ala8Val | P63316 |
| HOMO SAPIENS | 8 | 2025 | 1wrl | A | 24 | PHE | p.Phe24Leu | P63316 |
| HOMO SAPIENS | 0 | 2025 | 1wrl | A | 84 | SER | p.Cys84Tyr | P63316 |
| HOMO SAPIENS | 8 | 3791 | 1mhd | B | 93 | ARG | p.Arg93Ter | P84022 |
| HOMO SAPIENS | 0 | 1575 | 4phc | A | 205 | GLY | p.Gly205Asp | P12081 |
| HOMO SAPIENS | 0 | 1575 | 4phc | A | 454 | TYR | p.Tyr454Ser | P12081 |
| HOMO SAPIENS | 0 | 39620 | 1ozs | B | 135 | ARG | p.Arg136Gln | P19429 |
| HOMO SAPIENS | 5 | 39620 | 1ozs | B | 140 | ARG | p.Arg141Trp | P19429 |
| HOMO SAPIENS | 5 | 39620 | 1ozs | B | 142 | THR | p.Thr143Asn | P19429 |
| HOMO SAPIENS | 5 | 39620 | 1ozs | B | 143 | LEU | p.Leu144Gln | P19429 |
| HOMO SAPIENS | 5 | 39620 | 1ozs | B | 144 | ARG | p.Arg145Gly | P19429 |

| | | | | | | | | |
|---|---|---|---|---|---|---|---|---|
| HOMO SAPIENS | 5 | 39620 | 1ozs | B | 144 | ARG | p.Arg145Gln | P19429 |
| HOMO SAPIENS | -1 | 20934 | 4nuv | D | 42 | GLY | p.Gly42Asp | Q16570 |
| HOMO SAPIENS | -1 | 40565 | 5lcw | S | 4 | VAL | p.Val4Met | O60566 |
| HOMO SAPIENS | 0 | 40565 | 5lcw | S | 40 | THR | p.Thr40Met | O60566 |
| HOMO SAPIENS | 4 | 40565 | 5lcw | S | 198 | ALA | p.Glu390Asp | O60566 |
| HOMO SAPIENS | 4 | 758 | 4oiu | A | 726 | ARG | p.Arg727Leu | P10275 |
| HOMO SAPIENS | 4 | 758 | 4oiu | A | 730 | VAL | p.Val731Met | P10275 |
| HOMO SAPIENS | 2 | 758 | 4oiu | A | 798 | GLN | p.Gln799Glu | P10275 |
| HOMO SAPIENS | 3 | 758 | 4oiu | A | 855 | ARG | p.Arg856His | P10275 |
| HOMO SAPIENS | 1 | 758 | 4oiu | A | 886 | MET | p.Met887Val | P10275 |
| HOMO SAPIENS | 0 | 3303 | 4ie0 | A | 134 | ALA | p.Ala134Thr | Q9C0B1 |
| HOMO SAPIENS | 9 | 3303 | 4ie0 | A | 316 | ARG | p.Arg316Gln | Q9C0B1 |
| HOMO SAPIENS | 1 | 41502 | 3fgh | A | 136 | PRO | p.Pro178Leu | Q00059 |
| HOMO SAPIENS | 5 | 175 | 3ceh | B | 58 | THR | p.Thr59Lys | P06737 |
| HOMO SAPIENS | 8 | 175 | 3ceh | B | 93 | ARG | p.Arg94Ter | P06737 |
| HOMO SAPIENS | 5 | 175 | 3ceh | B | 221 | VAL | p.Val222Ile | P06737 |
| HOMO SAPIENS | 0 | 175 | 3ceh | B | 338 | ASN | p.Asn339Ser | P06737 |
| HOMO SAPIENS | 1 | 175 | 3ceh | B | 372 | PHE | p.Phe373Leu | P06737 |
| HOMO SAPIENS | 9 | 175 | 3ceh | B | 376 | ASN | p.Asn377Lys | P06737 |
| HOMO SAPIENS | 8 | 175 | 3ceh | B | 398 | ARG | p.Arg399Ter | P06737 |
| HOMO SAPIENS | 5 | 175 | 3ceh | B | 424 | ARG | p.Arg425Pro | P06737 |
| HOMO SAPIENS | 9 | 175 | 3ceh | B | 455 | VAL | p.Val456Met | P06737 |
| HOMO SAPIENS | 8 | 175 | 3ceh | B | 490 | ARG | p.Arg491Cys | P06737 |
| HOMO SAPIENS | 8 | 175 | 3ceh | B | 631 | ASN | p.Asn632Ile | P06737 |
| HOMO SAPIENS | 9 | 175 | 3ceh | B | 648 | TYR | p.Tyr649Ter | P06737 |
| HOMO SAPIENS | 9 | 175 | 3ceh | B | 672 | GLU | p.Glu673Lys | P06737 |
| HOMO SAPIENS | 9 | 175 | 3ceh | B | 674 | SER | p.Ser675Leu | P06737 |
| HOMO SAPIENS | 0 | 175 | 3ceh | B | 805 | ILE | p.Ile806Leu | P06737 |
| HOMO SAPIENS | 0 | 175 | 3ceh | B | 820 | TYR | p.Tyr821His | P06737 |
| HOMO SAPIENS | 1 | 6750 | 4fdj | A | 47 | GLY | p.Gly47Arg | P34059 |
| HOMO SAPIENS | 9 | 6750 | 4fdj | A | 60 | ASP | p.Asp60Asn | P34059 |
| HOMO SAPIENS | 5 | 6750 | 4fdj | A | 67 | LEU | p.Leu67Met | P34059 |
| HOMO SAPIENS | 0 | 6750 | 4fdj | A | 69 | PHE | p.Phe69Val | P34059 |
| HOMO SAPIENS | 3 | 6750 | 4fdj | A | 94 | ARG | p.Arg94Gly | P34059 |
| HOMO SAPIENS | 0 | 6750 | 4fdj | A | 113 | ILE | p.Ile113Phe | P34059 |
| HOMO SAPIENS | 0 | 6750 | 4fdj | A | 138 | VAL | p.Val138Ala | P34059 |
| HOMO SAPIENS | 0 | 6750 | 4fdj | A | 139 | GLY | p.Gly139Ser | P34059 |
| HOMO SAPIENS | 0 | 6750 | 4fdj | A | 151 | PRO | p.Pro151Thr | P34059 |
| HOMO SAPIENS | 0 | 6750 | 4fdj | A | 155 | GLY | p.Gly155Arg | P34059 |
| HOMO SAPIENS | 0 | 6750 | 4fdj | A | 159 | TRP | p.Trp159Ter | P34059 |
| HOMO SAPIENS | 0 | 6750 | 4fdj | A | 162 | SER | p.Ser162Phe | P34059 |
| HOMO SAPIENS | 0 | 6750 | 4fdj | A | 200 | THR | p.Thr200Met | P34059 |
| HOMO SAPIENS | 0 | 6750 | 4fdj | A | 204 | ASN | p.Asn204Lys | P34059 |
| HOMO SAPIENS | 0 | 6750 | 4fdj | A | 231 | ALA | p.Ala231Gly | P34059 |
| HOMO SAPIENS | 2 | 6750 | 4fdj | A | 235 | THR | p.Thr235Lys | P34059 |
| HOMO SAPIENS | 1 | 6750 | 4fdj | A | 259 | ARG | p.Arg259Gln | P34059 |
| HOMO SAPIENS | 0 | 6750 | 4fdj | A | 287 | SER | p.Ser287Leu | P34059 |
| HOMO SAPIENS | 0 | 6750 | 4fdj | A | 291 | ALA | p.Ala291Thr | P34059 |
| HOMO SAPIENS | 3 | 6750 | 4fdj | A | 301 | GLY | p.Gly301Cys | P34059 |
| HOMO SAPIENS | 0 | 6750 | 4fdj | A | 312 | THR | p.Thr312Ser | P34059 |
| HOMO SAPIENS | 0 | 6750 | 4fdj | A | 318 | MET | p.Met318Arg | P34059 |
| HOMO SAPIENS | 0 | 6750 | 4fdj | A | 340 | GLY | p.Gly340Asp | P34059 |
| HOMO SAPIENS | 0 | 6750 | 4fdj | A | 386 | ARG | p.Arg386Cys | P34059 |
| HOMO SAPIENS | 0 | 6750 | 4fdj | A | 391 | MET | p.Met391Val | P34059 |

| | | | | | | | | |
|---|---|---|---|---|---|---|---|---|
| HOMO SAPIENS | 0 | 6750 | 4fdj | A | 392 | ALA | p.Ala392Val | P34059 |
| HOMO SAPIENS | 0 | 6750 | 4fdj | A | 393 | ALA | p.Ala393Ser | P34059 |
| HOMO SAPIENS | 0 | 6750 | 4fdj | A | 459 | ALA | p.Ala459Val | P34059 |
| HOMO SAPIENS | 0 | 6750 | 4fdj | A | 473 | GLN | p.Gln473Ter | P34059 |
| HOMO SAPIENS | 2 | 6750 | 4fdj | A | 480 | VAL | p.Val480Phe | P34059 |
| HOMO SAPIENS | 0 | 6750 | 4fdj | A | 487 | ASN | p.Asn487Ser | P34059 |
| HOMO SAPIENS | 0 | 6750 | 4fdj | A | 488 | VAL | p.Val488Met | P34059 |
| HOMO SAPIENS | 0 | 6750 | 4fdj | A | 520 | TRP | p.Trp520Ter | P34059 |
| HOMO SAPIENS | 4 | 1088 | 2p3u | A | 102 | GLU | p.Glu142Lys | P00742 |
| HOMO SAPIENS | 6 | 370 | 1nr1 | D | 73 | ASP | p.Asp126His | P00367 |
| HOMO SAPIENS | 3 | 370 | 1nr1 | D | 221 | ARG | p.Arg274Cys | P00367 |
| HOMO SAPIENS | 2 | 370 | 1nr1 | D | 265 | ARG | p.Arg318Lys | P00367 |
| HOMO SAPIENS | 1 | 370 | 1nr1 | D | 269 | ARG | p.Arg322His | P00367 |
| HOMO SAPIENS | 1 | 370 | 1nr1 | D | 296 | GLU | p.Glu349Ala | P00367 |
| HOMO SAPIENS | 1 | 370 | 1nr1 | D | 445 | SER | p.Ser498Leu | P00367 |
| HOMO SAPIENS | 1 | 370 | 1nr1 | D | 446 | GLY | p.Gly499Asp | P00367 |
| HOMO SAPIENS | 1 | 370 | 1nr1 | D | 446 | GLY | p.Gly499Ser | P00367 |
| HOMO SAPIENS | 1 | 370 | 1nr1 | D | 448 | SER | p.Ser501Pro | P00367 |
| HOMO SAPIENS | 2 | 370 | 1nr1 | D | 454 | HIS | p.His507Tyr | P00367 |
| HOMO SAPIENS | 1 | 7720 | 2he0 | A | 147 | HIS | p.His2018Gln | P46531 |
| HOMO SAPIENS | 2 | 11844 | 5l0v | A | 218 | ARG | p.Arg218Ter | Q8NBL1 |
| HOMO SAPIENS | 2 | 11844 | 5l0v | A | 279 | ARG | p.Arg279Trp | Q8NBL1 |
| HOMO SAPIENS | 9 | 1485 | 5va0 | A | 477 | ARG | p.Arg477His | P04150 |
| HOMO SAPIENS | 0 | 235 | 2fun | B | 2248 | ARG | p.Arg248Trp | Q14790 |
| HOMO SAPIENS | 4 | 235 | 2fun | B | 2285 | HIS | p.Asp285His | Q14790 |
| HOMO SAPIENS | 0 | 4822 | 5y3n | A | 307 | ARG | p.Arg307Gly | Q12931 |
| HOMO SAPIENS | 8 | 32928 | 4m6w | A | 1886 | HIS | p.His1886Tyr | Q8IYD8 |
| HOMO SAPIENS | 0 | 2711 | 4y5v | F | 22 | GLY | p.Gly46Glu | P19235 |
| HOMO SAPIENS | -1 | 2711 | 4y5v | F | 166 | ALA | p.Ala190Ser | P19235 |
| HOMO SAPIENS | 4 | 5161 | 5bpu | A | 42 | HIS | p.His42Arg | Q00604 |
| HOMO SAPIENS | 0 | 5161 | 5bpu | A | 44 | TYR | p.Tyr44Cys | Q00604 |
| HOMO SAPIENS | 0 | 5161 | 5bpu | A | 45 | VAL | p.Val45Glu | Q00604 |
| HOMO SAPIENS | 0 | 5161 | 5bpu | A | 60 | VAL | p.Val60Glu | Q00604 |
| HOMO SAPIENS | 0 | 5161 | 5bpu | A | 61 | LEU | p.Leu61Phe | Q00604 |
| HOMO SAPIENS | 7 | 5161 | 5bpu | A | 69 | CYS | p.Cys69Ser | Q00604 |
| HOMO SAPIENS | 0 | 5161 | 5bpu | A | 75 | SER | p.Ser75Cys | Q00604 |
| HOMO SAPIENS | 0 | 5161 | 5bpu | A | 90 | ARG | p.Arg90Pro | Q00604 |
| HOMO SAPIENS | 9 | 5161 | 5bpu | A | 96 | CYS | p.Cys96Tyr | Q00604 |
| HOMO SAPIENS | 9 | 5161 | 5bpu | A | 96 | CYS | p.Cys96Trp | Q00604 |
| HOMO SAPIENS | 0 | 5161 | 5bpu | A | 101 | SER | p.Ser101Phe | Q00604 |
| HOMO SAPIENS | 0 | 5161 | 5bpu | A | 105 | ALA | p.Ala105Thr | Q00604 |
| HOMO SAPIENS | 0 | 5161 | 5bpu | A | 110 | CYS | p.Cys110Gly | Q00604 |
| HOMO SAPIENS | 0 | 5161 | 5bpu | A | 121 | ARG | p.Arg121Trp | Q00604 |
| HOMO SAPIENS | 0 | 5161 | 5bpu | A | 121 | ARG | p.Arg121Leu | Q00604 |
| HOMO SAPIENS | 0 | 5161 | 5bpu | A | 124 | LEU | p.Leu124Phe | Q00604 |
| HOMO SAPIENS | 8 | 5161 | 5bpu | A | 128 | CYS | p.Cys128Ter | Q00604 |
| HOMO SAPIENS | 1 | 13944 | 5xqh | A | 238 | ARG | p.Arg238His | Q9GZN7 |
| HOMO SAPIENS | -1 | 7508 | 5jl6 | A | 39 | TRP | p.Trp39Arg | P11511 |
| HOMO SAPIENS | 0 | 7508 | 5jl6 | A | 201 | THR | p.Thr201Met | P11511 |
| HOMO SAPIENS | 0 | 7508 | 5jl6 | A | 210 | GLU | p.Glu210Lys | P11511 |
| HOMO SAPIENS | 1 | 7508 | 5jl6 | A | 264 | ARG | p.Arg264Cys | P11511 |
| HOMO SAPIENS | 2 | 7508 | 5jl6 | A | 365 | ARG | p.Arg365Gln | P11511 |
| HOMO SAPIENS | 7 | 7508 | 5jl6 | A | 375 | ARG | p.Arg375Cys | P11511 |
| HOMO SAPIENS | 8 | 7508 | 5jl6 | A | 435 | ARG | p.Arg435Cys | P11511 |

| | | | | | | | | |
|---|---|---|---|---|---|---|---|---|
| HOMO SAPIENS | 9 | 7508 | 5jl6 | A | 437 | CYS | p.Cys437Tyr | P11511 |
| HOMO SAPIENS | -1 | 30206 | 5l95 | B | 246 | ARG | p.Arg246Ter | Q9GZZ9 |
| HOMO SAPIENS | 4 | 875 | 3efh | B | 43 | GLU | p.Glu43Asp | P60891 |
| HOMO SAPIENS | 5 | 875 | 3efh | B | 65 | ASP | p.Asp65Asn | P60891 |
| HOMO SAPIENS | 0 | 875 | 3efh | B | 87 | ALA | p.Ala87Thr | P60891 |
| HOMO SAPIENS | 2 | 875 | 3efh | B | 115 | MET | p.Met115Thr | P60891 |
| HOMO SAPIENS | 8 | 875 | 3efh | B | 133 | GLN | p.Gln133Pro | P60891 |
| HOMO SAPIENS | 8 | 875 | 3efh | B | 152 | LEU | p.Leu152Pro | P60891 |
| HOMO SAPIENS | 7 | 875 | 3efh | B | 290 | ILE | p.Ile290Thr | P60891 |
| HOMO SAPIENS | 6 | 875 | 3efh | B | 306 | GLY | p.Gly306Arg | P60891 |
| HOMO SAPIENS | 0 | 7978 | 5edm | A | 122 | MET | p.Thr165Met | P00734 |
| HOMO SAPIENS | -1 | 7978 | 5edm | A | 257 | ARG | p.Arg314Cys | P00734 |
| HOMO SAPIENS | 0 | 1716 | 1b98 | M | 126 | ARG | p.Arg206Gln | P34130 |
| HOMO SAPIENS | 0 | 1716 | 1b98 | M | 126 | ARG | p.Arg206Trp | P34130 |
| HOMO SAPIENS | 9 | 3960 | 2qyp | B | 28 | ILE | p.Ile338Val | P07602 |
| HOMO SAPIENS | 1 | 3960 | 2qyp | B | 39 | LEU | p.Leu349Pro | P07602 |
| HOMO SAPIENS | 0 | 31414 | 5u1d | A | 333 | ILE | p.Ile393Val | Q03518 |
| HOMO SAPIENS | 0 | 31414 | 5u1d | A | 637 | ASP | p.Asp697Gly | Q03518 |
| HOMO SAPIENS | 0 | 31414 | 5u1d | A | 659 | ARG | p.Arg719Gln | Q03518 |
| HOMO SAPIENS | 3 | 119 | 2pho | A | 11 | ILE | p.Ile11Thr | P05089 |
| HOMO SAPIENS | 8 | 119 | 2pho | A | 21 | ARG | p.Arg21Ter | P05089 |
| HOMO SAPIENS | 0 | 119 | 2pho | A | 122 | TRP | p.Trp122Ter | P05089 |
| HOMO SAPIENS | 6 | 119 | 2pho | A | 138 | GLY | p.Gly138Val | P05089 |
| HOMO SAPIENS | 5 | 119 | 2pho | A | 290 | THR | p.Thr290Ser | P05089 |
| HOMO SAPIENS | 5 | 119 | 2pho | A | 291 | ARG | p.Arg291Ter | P05089 |
| HOMO SAPIENS | 0 | 3732 | 2i99 | A | 247 | TYR | p.Tyr247Ter | Q14894 |
| HOMO SAPIENS | -1 | 1382 | 4ffx | A | 2 | ALA | p.Ala2Val | P30566 |
| HOMO SAPIENS | 1 | 1382 | 4ffx | A | 72 | ILE | p.Ile72Met | P30566 |
| HOMO SAPIENS | 1 | 1382 | 4ffx | A | 100 | PRO | p.Pro100Ala | P30566 |
| HOMO SAPIENS | 0 | 1382 | 4ffx | A | 114 | TYR | p.Tyr114His | P30566 |
| HOMO SAPIENS | 1 | 1382 | 4ffx | A | 141 | ARG | p.Arg141Trp | P30566 |
| HOMO SAPIENS | 0 | 1382 | 4ffx | A | 147 | LYS | p.Lys147Met | P30566 |
| HOMO SAPIENS | 3 | 1382 | 4ffx | A | 190 | ARG | p.Arg190Gln | P30566 |
| HOMO SAPIENS | 3 | 1382 | 4ffx | A | 190 | ARG | p.Arg190Ter | P30566 |
| HOMO SAPIENS | 0 | 1382 | 4ffx | A | 217 | HIS | p.His217Asp | P30566 |
| HOMO SAPIENS | 2 | 1382 | 4ffx | A | 246 | LYS | p.Lys246Glu | P30566 |
| HOMO SAPIENS | 5 | 1382 | 4ffx | A | 303 | ARG | p.Arg303Cys | P30566 |
| HOMO SAPIENS | 1 | 1382 | 4ffx | A | 318 | PRO | p.Pro318Leu | P30566 |
| HOMO SAPIENS | 6 | 1382 | 4ffx | A | 332 | ASP | p.Asp332His | P30566 |
| HOMO SAPIENS | 6 | 1382 | 4ffx | A | 337 | ARG | p.Arg337Ter | P30566 |
| HOMO SAPIENS | 1 | 1382 | 4ffx | A | 396 | ARG | p.Arg396His | P30566 |
| HOMO SAPIENS | 1 | 1382 | 4ffx | A | 426 | ARG | p.Arg426His | P30566 |
| HOMO SAPIENS | 0 | 1382 | 4ffx | A | 438 | SER | p.Ser438Pro | P30566 |
| HOMO SAPIENS | 1 | 1382 | 4ffx | A | 447 | SER | p.Ser447Pro | P30566 |
| HOMO SAPIENS | 2 | 1382 | 4ffx | A | 448 | SER | p.Ser448Pro | P30566 |
| HOMO SAPIENS | 2 | 1382 | 4ffx | A | 450 | THR | p.Thr450Ser | P30566 |
| HOMO SAPIENS | 0 | 6212 | 1ao3 | A | 46 | SER | p.Ser1731Thr | P04275 |
| HOMO SAPIENS | 0 | 1711 | 3f9c | A | 279 | VAL | p.Val279Phe | Q13093 |
| HOMO SAPIENS | 0 | 6112 | 4fvl | A | 312 | HIS | p.His312Arg | P45452 |
| HOMO SAPIENS | 0 | 6112 | 4fvl | A | 323 | THR | p.Thr323Met | P45452 |
| HOMO SAPIENS | 0 | 6112 | 4fvl | A | 390 | ASP | p.Asp390Gly | P45452 |
| HOMO SAPIENS | 0 | 6112 | 4fvl | A | 458 | ARG | p.Arg458Cys | P45452 |
| HOMO SAPIENS | 2 | 6180 | 5ggn | A | 97 | ARG | p.Arg97Trp | Q8WZA1 |
| HOMO SAPIENS | 2 | 6180 | 5ggn | A | 215 | ARG | p.Arg215Ter | Q8WZA1 |

| | | | | | | | | |
|---|---|---|---|---|---|---|---|---|
| HOMO SAPIENS | -1 | 5526 | 4ak3 | A | 11 | ASP | p.Asp1232Asn | P02461 |
| HOMO SAPIENS | 6 | 5526 | 4ak3 | A | 38 | ALA | p.Ala1259Thr | P02461 |
| HOMO SAPIENS | 0 | 5526 | 4ak3 | A | 92 | LYS | p.Lys1313Arg | P02461 |
| HOMO SAPIENS | 0 | 5526 | 4ak3 | A | 132 | GLN | p.His1353Gln | P02461 |
| HOMO SAPIENS | 4 | 16616 | 1hci | B | 433 | SER | p.Ser433Leu | P35609 |
| HOMO SAPIENS | 1 | 16616 | 1hci | B | 475 | ASP | p.Asp475Asn | P35609 |
| HOMO SAPIENS | 1 | 16616 | 1hci | B | 476 | ALA | p.Ala476Ser | P35609 |
| HOMO SAPIENS | 2 | 16616 | 1hci | B | 507 | MET | p.Met507Val | P35609 |
| HOMO SAPIENS | 1 | 16616 | 1hci | B | 604 | MET | p.Met604Val | P35609 |
| HOMO SAPIENS | 2 | 16616 | 1hci | B | 640 | ARG | p.Arg640His | P35609 |
| HOMO SAPIENS | 1 | 16616 | 1hci | B | 721 | ARG | p.Arg721Cys | P35609 |
| HOMO SAPIENS | -1 | 3402 | 5iaa | B | 57 | MET | p.Met57Val | Q9GZZ9 |
| HOMO SAPIENS | 7 | 3250 | 3irq | B | 193 | PRO | p.Pro193Ala | P55265 |
| HOMO SAPIENS | 2 | 15200 | 2wns | A | 30 | SER | p.Ser30Gly | P11172 |
| HOMO SAPIENS | 0 | 15200 | 2wns | A | 96 | ARG | p.Arg96Gly | P11172 |
| HOMO SAPIENS | 0 | 15200 | 2wns | A | 109 | VAL | p.Val109Gly | P11172 |
| HOMO SAPIENS | 2 | 15759 | 4x4w | B | 140 | LEU | p.Leu166Ser | Q96Q11 |
| HOMO SAPIENS | 0 | 15759 | 4x4w | B | 197 | ILE | p.Ile223Thr | Q96Q11 |
| HOMO SAPIENS | 0 | 3156 | 3w03 | B | 14 | ALA | p.Ala14Thr | Q9H9Q4 |
| HOMO SAPIENS | 0 | 3156 | 3w03 | B | 57 | ARG | p.Arg57Gly | Q9H9Q4 |
| HOMO SAPIENS | 0 | 3156 | 3w03 | B | 123 | CYS | p.Cys123Arg | Q9H9Q4 |
| HOMO SAPIENS | 0 | 11693 | 3ojy | A | 63 | GLN | p.Gln93Lys | P07357 |
| HOMO SAPIENS | 0 | 1421 | 2qc8 | E | 275 | GLU | p.Glu275Asp | P15104 |
| HOMO SAPIENS | 9 | 1421 | 2qc8 | E | 324 | ARG | p.Arg324Cys | P15104 |
| HOMO SAPIENS | 6 | 1421 | 2qc8 | E | 341 | ARG | p.Arg341Cys | P15104 |
| HOMO SAPIENS | 6 | 2850 | 3qvb | A | 83 | THR | p.Thr83Met | P46976 |
| HOMO SAPIENS | 8 | 2850 | 3qvb | A | 102 | ASP | p.Asp102His | P46976 |
| HOMO SAPIENS | 1 | 24314 | 4m9p | A | 480 | PRO | p.Pro480Leu | P21333 |
| HOMO SAPIENS | 0 | 24314 | 4m9p | A | 528 | VAL | p.Val528Met | P21333 |
| HOMO SAPIENS | 0 | 24314 | 4m9p | A | 590 | VAL | p.Val590Ile | P21333 |
| HOMO SAPIENS | 0 | 24314 | 4m9p | A | 605 | ASP | p.Asp605Asn | P21333 |
| HOMO SAPIENS | 1 | 24314 | 4m9p | A | 637 | PRO | p.Pro637Gln | P21333 |
| HOMO SAPIENS | 0 | 24314 | 4m9p | A | 656 | LEU | p.Leu656Phe | P21333 |
| HOMO SAPIENS | 2 | 24314 | 4m9p | A | 711 | VAL | p.Val711Asp | P21333 |
| HOMO SAPIENS | 0 | 4951 | 5joe | A | 40 | THR | p.Thr9621Ser | Q8WZ42 |
| HOMO SAPIENS | 1 | 4951 | 5joe | A | 41 | GLU | p.Glu9622Gln | Q8WZ42 |
| HOMO SAPIENS | 0 | 4951 | 5joe | A | 82 | ALA | p.Ala9663Ser | Q8WZ42 |
| HOMO SAPIENS | 4 | 4951 | 5joe | A | 89 | ILE | p.Ile9670Asn | Q8WZ42 |
| HOMO SAPIENS | 0 | 9664 | 3c6m | A | 56 | GLY | p.Gly56Ser | P52788 |
| HOMO SAPIENS | 0 | 9664 | 3c6m | A | 132 | VAL | p.Val132Gly | P52788 |
| HOMO SAPIENS | 1 | 9664 | 3c6m | A | 190 | THR | p.Thr190Ile | P52788 |
| HOMO SAPIENS | -1 | 12118 | 5hcd | B | 19 | GLN | p.Gln19Ter | P01031 |
| HOMO SAPIENS | 0 | 1583 | 2b4x | L | 7 | ILE | p.Ile39Asn | P01008 |
| HOMO SAPIENS | 0 | 1583 | 2b4x | L | 24 | ARG | p.Arg56Cys | P01008 |
| HOMO SAPIENS | 0 | 1583 | 2b4x | L | 47 | ARG | p.Arg79Cys | P01008 |
| HOMO SAPIENS | 0 | 1583 | 2b4x | L | 47 | ARG | p.Arg79His | P01008 |
| HOMO SAPIENS | 1 | 1583 | 2b4x | L | 95 | CYS | p.Cys127Arg | P01008 |
| HOMO SAPIENS | 1 | 1583 | 2b4x | L | 99 | LEU | p.Leu131Phe | P01008 |
| HOMO SAPIENS | 0 | 1583 | 2b4x | L | 116 | SER | p.Ser148Pro | P01008 |
| HOMO SAPIENS | 0 | 1583 | 2b4x | L | 129 | ARG | p.Arg161Gln | P01008 |
| HOMO SAPIENS | 0 | 1583 | 2b4x | L | 135 | ASN | p.Asn167Thr | P01008 |
| HOMO SAPIENS | 2 | 1583 | 2b4x | L | 187 | ASN | p.Asn219Asp | P01008 |
| HOMO SAPIENS | 0 | 1583 | 2b4x | L | 191 | SER | p.Ser223Pro | P01008 |
| HOMO SAPIENS | 2 | 1583 | 2b4x | L | 349 | SER | p.Ser381Pro | P01008 |

| | | | | | | | | |
|---|---|---|---|---|---|---|---|---|
| HOMO SAPIENS | 0 | 1583 | 2b4x | L | 382 | ALA | p.Ala414Thr | P01008 |
| HOMO SAPIENS | 0 | 1583 | 2b4x | L | 387 | ALA | p.Ala419Val | P01008 |
| HOMO SAPIENS | 0 | 1583 | 2b4x | L | 392 | GLY | p.Gly424Asp | P01008 |
| HOMO SAPIENS | 0 | 1583 | 2b4x | L | 393 | ARG | p.Arg425His | P01008 |
| HOMO SAPIENS | 0 | 1583 | 2b4x | L | 393 | ARG | p.Arg425Cys | P01008 |
| HOMO SAPIENS | 0 | 1583 | 2b4x | L | 394 | SER | p.Ser426Leu | P01008 |
| HOMO SAPIENS | 0 | 1583 | 2b4x | L | 404 | ALA | p.Ala436Thr | P01008 |
| HOMO SAPIENS | 2 | 1583 | 2b4x | L | 407 | PRO | p.Pro439Leu | P01008 |
| HOMO SAPIENS | 2 | 1583 | 2b4x | L | 429 | PRO | p.Pro461Leu | P01008 |
| HOMO SAPIENS | 0 | 11647 | 2a1t | S | 78 | ALA | p.Ala78Ser | P38117 |
| HOMO SAPIENS | 0 | 11647 | 2a1t | S | 98 | ARG | p.Arg98Cys | P38117 |
| HOMO SAPIENS | 8 | 11647 | 2a1t | S | 128 | ASP | p.Asp128Asn | P38117 |
| HOMO SAPIENS | 5 | 11647 | 2a1t | S | 154 | THR | p.Thr154Met | P38117 |
| HOMO SAPIENS | 6 | 11647 | 2a1t | S | 164 | ARG | p.Arg164Gln | P38117 |
| HOMO SAPIENS | 5 | 11647 | 2a1t | S | 174 | ARG | p.Arg174His | P38117 |
| HOMO SAPIENS | -1 | 11200 | 4tx5 | A | 4 | ILE | p.Ile59Val | Q9NR28 |
| HOMO SAPIENS | 0 | 11200 | 4tx5 | A | 169 | GLY | p.Gly224Arg | Q9NR28 |
| HOMO SAPIENS | 3 | 417 | 2vr2 | A | 6 | ARG | p.Arg6Gln | Q14117 |
| HOMO SAPIENS | 1 | 417 | 2vr2 | A | 334 | GLN | p.Gln334Arg | Q14117 |
| HOMO SAPIENS | 2 | 417 | 2vr2 | A | 360 | TRP | p.Trp360Arg | Q14117 |
| HOMO SAPIENS | 0 | 417 | 2vr2 | A | 379 | SER | p.Ser379Arg | Q14117 |
| HOMO SAPIENS | 0 | 417 | 2vr2 | A | 435 | GLY | p.Gly435Arg | Q14117 |
| HOMO SAPIENS | 0 | 12498 | 3eo3 | B | 423 | ILE | p.Ile423Val | Q9Y223 |
| HOMO SAPIENS | 5 | 12498 | 3eo3 | B | 572 | VAL | p.Val572Leu | Q9Y223 |
| HOMO SAPIENS | 0 | 12498 | 3eo3 | B | 576 | GLY | p.Gly576Glu | Q9Y223 |
| HOMO SAPIENS | 0 | 12498 | 3eo3 | B | 587 | ILE | p.Ile587Thr | Q9Y223 |
| HOMO SAPIENS | 0 | 12498 | 3eo3 | B | 615 | SER | p.Ser615Ter | Q9Y223 |
| HOMO SAPIENS | 1 | 12498 | 3eo3 | B | 631 | ALA | p.Ala631Val | Q9Y223 |
| HOMO SAPIENS | 1 | 12498 | 3eo3 | B | 669 | GLY | p.Gly669Arg | Q9Y223 |
| HOMO SAPIENS | 1 | 12498 | 3eo3 | B | 696 | VAL | p.Val696Met | Q9Y223 |
| HOMO SAPIENS | 2 | 12498 | 3eo3 | B | 712 | MET | p.Met712Thr | Q9Y223 |
| HOMO SAPIENS | 0 | 1568 | 1r74 | A | 49 | LEU | p.Leu50Pro | Q14749 |
| HOMO SAPIENS | 0 | 1568 | 1r74 | A | 176 | HIS | p.His177Asn | Q14749 |
| HOMO SAPIENS | 3 | 147 | 5eig | A | 67 | THR | p.Thr67Ile | P32929 |
| HOMO SAPIENS | 3 | 147 | 5eig | A | 240 | GLN | p.Gln240Glu | P32929 |
| HOMO SAPIENS | -1 | 147 | 5eig | A | 403 | SER | p.Ser403Ile | P32929 |
| HOMO SAPIENS | 8 | 20036 | 2lsr | A | 31 | ARG | p.Arg31Ter | Q9Y6N9 |
| HOMO SAPIENS | -1 | 41382 | 4uis | B | 431 | UNK | p.Ala431Glu | P49768 |
| HOMO SAPIENS | -1 | 8536 | 3i6w | B | 85 | PRO | p.Pro85Leu | O96017 |
| HOMO SAPIENS | 0 | 8536 | 3i6w | B | 93 | TRP | p.Trp93Ter | O96017 |
| HOMO SAPIENS | 0 | 8536 | 3i6w | B | 95 | ARG | p.Arg95Ter | O96017 |
| HOMO SAPIENS | 0 | 8536 | 3i6w | B | 117 | ARG | p.Arg117Gly | O96017 |
| HOMO SAPIENS | 0 | 8536 | 3i6w | B | 119 | LYS | p.Lys119Arg | O96017 |
| HOMO SAPIENS | 0 | 8536 | 3i6w | B | 137 | ARG | p.Arg137Gln | O96017 |
| HOMO SAPIENS | 0 | 8536 | 3i6w | B | 137 | ARG | p.Arg137Ter | O96017 |
| HOMO SAPIENS | 0 | 8536 | 3i6w | B | 145 | ARG | p.Arg145Trp | O96017 |
| HOMO SAPIENS | 0 | 8536 | 3i6w | B | 160 | ILE | p.Ile160Met | O96017 |
| HOMO SAPIENS | 0 | 41322 | 4bxl | C | 46 | GLU | p.Glu46Lys | P37840 |
| HOMO SAPIENS | 5 | 41322 | 4bxl | C | 53 | ALA | p.Ala53Thr | P37840 |
| HOMO SAPIENS | 2 | 24966 | 3hve | A | 15 | ARG | p.Arg15Ser | Q9H2C0 |
| HOMO SAPIENS | 1 | 24966 | 3hve | A | 138 | ARG | p.Arg138His | Q9H2C0 |
| HOMO SAPIENS | 0 | 24966 | 3hve | A | 201 | ARG | p.Arg201Ter | Q9H2C0 |
| HOMO SAPIENS | 0 | 11680 | 5an9 | J | 8 | ASN | p.Asn8Lys | Q9Y3A5 |
| HOMO SAPIENS | 1 | 11680 | 5an9 | J | 33 | LYS | p.Lys33Arg | Q9Y3A5 |

| | | | | | | | | |
|---|---|---|---|---|---|---|---|---|
| HOMO SAPIENS | 1 | 11680 | 5an9 | J | 126 | ARG | p.Arg126Thr | Q9Y3A5 |
| HOMO SAPIENS | 1 | 11680 | 5an9 | J | 130 | VAL | p.Val130Met | Q9Y3A5 |
| HOMO SAPIENS | 1 | 11680 | 5an9 | J | 169 | ARG | p.Arg169Cys | Q9Y3A5 |
| HOMO SAPIENS | 1 | 11680 | 5an9 | J | 218 | ARG | p.Arg218Ter | Q9Y3A5 |
| HOMO SAPIENS | 4 | 978 | 3zs0 | B | 7 | TYR | p.Tyr173Cys | P05164 |
| HOMO SAPIENS | 1 | 978 | 3zs0 | B | 85 | MET | p.Met251Thr | P05164 |
| HOMO SAPIENS | 4 | 5140 | 2l4r | A | 52 | CYS | p.Cys52Ter | Q12809 |
| HOMO SAPIENS | 4 | 5140 | 2l4r | A | 63 | PRO | p.Pro63His | Q12809 |
| HOMO SAPIENS | 0 | 5140 | 2l4r | A | 100 | ARG | p.Arg100Gly | Q12809 |
| HOMO SAPIENS | 4 | 5140 | 2l4r | A | 129 | PHE | p.Phe129Leu | Q12809 |
| HOMO SAPIENS | 0 | 25836 | 2gy7 | B | 103 | GLU | p.Glu103Asp | Q02763 |
| HOMO SAPIENS | 0 | 25836 | 2gy7 | B | 148 | ILE | p.Ile148Thr | Q02763 |
| HOMO SAPIENS | 0 | 25836 | 2gy7 | B | 226 | ALA | p.Ala226Val | Q02763 |
| HOMO SAPIENS | 0 | 25836 | 2gy7 | B | 391 | THR | p.Thr391Ile | Q02763 |
| HOMO SAPIENS | 4 | 12563 | 5kwy | D | 20 | GLU | p.Glu20Ter | P61916 |
| HOMO SAPIENS | 2 | 12563 | 5kwy | D | 30 | VAL | p.Val30Met | P61916 |
| HOMO SAPIENS | 4 | 12563 | 5kwy | D | 39 | VAL | p.Val39Met | P61916 |
| HOMO SAPIENS | 8 | 12563 | 5kwy | D | 67 | SER | p.Ser67Pro | P61916 |
| HOMO SAPIENS | 8 | 12563 | 5kwy | D | 99 | CYS | p.Cys99Arg | P61916 |
| HOMO SAPIENS | 2 | 12563 | 5kwy | D | 118 | GLU | p.Glu118Ter | P61916 |
| HOMO SAPIENS | 2 | 12563 | 5kwy | D | 120 | PRO | p.Pro120Ser | P61916 |
| HOMO SAPIENS | -1 | 1679 | 1pmk | A | -2 | VAL | p.Val374Phe | P00747 |
| HOMO SAPIENS | -1 | 5315 | 2w2n | P | 53 | ALA | p.Ala53Val | Q8NBP7 |
| HOMO SAPIENS | 8 | 13496 | 2yad | A | 138 | THR | p.Asn138Thr | P11686 |
| HOMO SAPIENS | -1 | 13496 | 2yad | A | 173 | ARG | p.Arg167Gln | P11686 |
| HOMO SAPIENS | 2 | 13496 | 2yad | A | 186 | SER | p.Asn186Ser | P11686 |
| HOMO SAPIENS | 0 | 4426 | 3qd6 | C | 123 | ALA | p.Ala123Glu | P29965 |
| HOMO SAPIENS | 1 | 4426 | 3qd6 | C | 140 | TRP | p.Trp140Gly | P29965 |
| HOMO SAPIENS | 0 | 4426 | 3qd6 | C | 155 | LEU | p.Leu155Pro | P29965 |
| HOMO SAPIENS | 6 | 4426 | 3qd6 | C | 227 | GLY | p.Gly227Val | P29965 |
| HOMO SAPIENS | 0 | 4426 | 3qd6 | C | 235 | ALA | p.Ala235Pro | P29965 |
| HOMO SAPIENS | 0 | 10734 | 3dyd | B | 45 | PRO | p.Pro45Ser | P17735 |
| HOMO SAPIENS | -1 | 10734 | 3dyd | B | 57 | ARG | p.Arg57Ter | P17735 |
| HOMO SAPIENS | 8 | 10734 | 3dyd | B | 417 | ARG | p.Arg417Ter | P17735 |
| HOMO SAPIENS | 0 | 6956 | 4pzs | A | 1600 | LEU | p.Leu1600Phe | Q09472 |
| HOMO SAPIENS | 0 | 9370 | 3hfv | A | 21 | SER | p.Ser57Cys | O95363 |
| HOMO SAPIENS | 1 | 9370 | 3hfv | A | 25 | ASP | p.Asp61Glu | O95363 |
| HOMO SAPIENS | 1 | 9370 | 3hfv | A | 130 | ALA | p.Ala166Thr | O95363 |
| HOMO SAPIENS | 5 | 9370 | 3hfv | A | 148 | ASP | p.Asp184Asn | O95363 |
| HOMO SAPIENS | 0 | 9370 | 3hfv | A | 187 | ARG | p.Arg223Cys | O95363 |
| HOMO SAPIENS | 0 | 9370 | 3hfv | A | 210 | THR | p.Thr246Met | O95363 |
| HOMO SAPIENS | 1 | 9370 | 3hfv | A | 244 | ASN | p.Asn280Ser | O95363 |
| HOMO SAPIENS | 2 | 9370 | 3hfv | A | 288 | TYR | p.Tyr324Cys | O95363 |
| HOMO SAPIENS | 5 | 9370 | 3hfv | A | 293 | ILE | p.Ile329Thr | O95363 |
| HOMO SAPIENS | 1 | 9370 | 3hfv | A | 355 | ASP | p.Asp391Val | O95363 |
| HOMO SAPIENS | 1 | 9370 | 3hfv | A | 371 | THR | p.Thr407Met | O95363 |
| HOMO SAPIENS | 1 | 9370 | 3hfv | A | 383 | ARG | p.Arg419Cys | O95363 |
| HOMO SAPIENS | 1 | 9370 | 3hfv | A | 387 | ARG | p.Arg423Gln | O95363 |
| HOMO SAPIENS | 0 | 14401 | 1q3x | A | 371 | ASP | p.Asp371Tyr | O00187 |
| HOMO SAPIENS | 0 | 14401 | 1q3x | A | 377 | VAL | p.Val377Ala | O00187 |
| HOMO SAPIENS | 0 | 14401 | 1q3x | A | 439 | ARG | p.Arg439His | O00187 |
| HOMO SAPIENS | 0 | 14401 | 1q3x | A | 577 | GLN | p.Gln577His | O00187 |
| HOMO SAPIENS | 2 | 18270 | 3teq | A | 429 | ARG | p.Arg429Cys | Q13586 |
| HOMO SAPIENS | -1 | 16081 | 4mqt | A | 231 | VAL | p.Val231Ile | P08172 |

| | | | | | | | | |
|---|---|---|---|---|---|---|---|---|
| HOMO SAPIENS | 9 | 6607 | 3whu | B | 81 | ASP | p.Asp81His | Q8NI22 |
| HOMO SAPIENS | 9 | 6607 | 3whu | B | 129 | ASP | p.Asp129Glu | Q8NI22 |
| HOMO SAPIENS | 9 | 6607 | 3whu | B | 136 | ILE | p.Ile136Thr | Q8NI22 |
| HOMO SAPIENS | 0 | 7208 | 2e9w | D | 11 | ASN | p.Asn36Ser | P21583 |
| HOMO SAPIENS | 0 | 24566 | 4bdx | A | 10 | LEU | p.Leu140Val | P00748 |
| HOMO SAPIENS | 5 | 24566 | 4bdx | A | 77 | PRO | p.Ala207Pro | P00748 |
| HOMO SAPIENS | 7 | 606 | 3qnf | A | 319 | MET | p.Met319Val | Q9NZ08 |
| HOMO SAPIENS | 8 | 4097 | 1ydl | A | 21 | LEU | p.Leu21Pro | Q6ZYL4 |
| HOMO SAPIENS | 6 | 4097 | 1ydl | A | 56 | ARG | p.Arg56Ter | Q6ZYL4 |
| HOMO SAPIENS | 8 | 36691 | 1dqb | A | 42 | ARG | p.Arg403Lys | P07204 |
| HOMO SAPIENS | 0 | 5642 | 3j7y | X | 59 | ARG | p.Arg59Pro | Q13084 |
| HOMO SAPIENS | 1 | 5642 | 3j7y | X | 204 | VAL | p.Val204Met | Q13084 |
| HOMO SAPIENS | 1 | 1119 | 3nwu | B | 252 | ALA | p.Ala252Thr | Q92743 |
| HOMO SAPIENS | 3 | 1119 | 3nwu | B | 274 | ARG | p.Arg274Gln | Q92743 |
| HOMO SAPIENS | 9 | 1119 | 3nwu | B | 297 | VAL | p.Val297Met | Q92743 |
| HOMO SAPIENS | -1 | 1119 | 3nwu | B | 302 | ARG | p.Arg302Ter | Q92743 |
| HOMO SAPIENS | 9 | 1119 | 3nwu | B | 321 | ALA | p.Ala321Thr | Q92743 |
| HOMO SAPIENS | -1 | 1119 | 3nwu | B | 370 | ARG | p.Arg370Ter | Q92743 |
| HOMO SAPIENS | 0 | 416 | 4ycu | B | 105 | LEU | p.Leu105His | Q15046 |
| HOMO SAPIENS | 0 | 416 | 4ycu | B | 145 | TYR | p.Tyr145His | Q15046 |
| HOMO SAPIENS | 0 | 416 | 4ycu | B | 200 | PRO | p.Pro200Leu | Q15046 |
| HOMO SAPIENS | 0 | 416 | 4ycu | B | 201 | TYR | p.Tyr201His | Q15046 |
| HOMO SAPIENS | 0 | 416 | 4ycu | B | 420 | ARG | p.Arg420Cys | Q15046 |
| HOMO SAPIENS | 9 | 3297 | 4fpb | A | 680 | THR | p.Thr680Ser | P19367 |
| HOMO SAPIENS | 0 | 6771 | 2zt6 | A | 201 | ALA | p.Ala255Val | P41250 |
| HOMO SAPIENS | 0 | 6771 | 2zt6 | A | 202 | ASP | p.Asp256His | P41250 |
| HOMO SAPIENS | 0 | 6771 | 2zt6 | A | 334 | ARG | p.Arg388Gln | P41250 |
| HOMO SAPIENS | -1 | 6771 | 2zt6 | A | 439 | ASN | p.Asn493Ser | P41250 |
| HOMO SAPIENS | -1 | 6771 | 2zt6 | A | 500 | ASP | p.Asp554Asn | P41250 |
| HOMO SAPIENS | 1 | 6771 | 2zt6 | A | 581 | SER | p.Ser635Leu | P41250 |
| HOMO SAPIENS | -1 | 6771 | 2zt6 | A | 684 | GLU | p.Glu738Lys | P41250 |
| HOMO SAPIENS | 0 | 2687 | 4grw | B | 276 | VAL | p.Val298Phe | P29460 |
| HOMO SAPIENS | 6 | 913 | 1m75 | B | 45 | ASP | p.Asp57Glu | Q16836 |
| HOMO SAPIENS | 2 | 913 | 1m75 | B | 74 | PRO | p.Leu86Pro | Q16836 |
| HOMO SAPIENS | 3 | 913 | 1m75 | B | 214 | TYR | p.Tyr226His | Q16836 |
| HOMO SAPIENS | 3 | 913 | 1m75 | B | 224 | ARG | p.Arg236Ter | Q16836 |
| HOMO SAPIENS | 4 | 913 | 1m75 | B | 246 | PRO | p.Pro258Leu | Q16836 |
| HOMO SAPIENS | 2 | 4713 | 4djn | B | 40 | ARG | p.Arg40Cys | Q7LG56 |
| HOMO SAPIENS | 1 | 4713 | 4djn | B | 41 | ARG | p.Arg41Pro | Q7LG56 |
| HOMO SAPIENS | 2 | 4713 | 4djn | B | 110 | ARG | p.Arg110His | Q7LG56 |
| HOMO SAPIENS | 2 | 4713 | 4djn | B | 121 | ARG | p.Arg121His | Q7LG56 |
| HOMO SAPIENS | 9 | 4713 | 4djn | B | 131 | GLU | p.Glu131Lys | Q7LG56 |
| HOMO SAPIENS | 2 | 4713 | 4djn | B | 182 | THR | p.Thr182Ala | Q7LG56 |
| HOMO SAPIENS | 6 | 4713 | 4djn | B | 194 | GLU | p.Glu194Lys | Q7LG56 |
| HOMO SAPIENS | 2 | 4713 | 4djn | B | 211 | ARG | p.Arg211Lys | Q7LG56 |
| HOMO SAPIENS | 0 | 4713 | 4djn | B | 224 | ILE | p.Ile224Ser | Q7LG56 |
| HOMO SAPIENS | 3 | 4713 | 4djn | B | 229 | GLY | p.Gly229Val | Q7LG56 |
| HOMO SAPIENS | 2 | 4713 | 4djn | B | 236 | CYS | p.Cys236Phe | Q7LG56 |
| HOMO SAPIENS | 0 | 4713 | 4djn | B | 273 | GLY | p.Gly273Ser | Q7LG56 |
| HOMO SAPIENS | -1 | 4713 | 4djn | B | 282 | MSE | p.Met282Ile | Q7LG56 |
| HOMO SAPIENS | 3 | 4713 | 4djn | B | 284 | GLN | p.Gln284Ter | Q7LG56 |
| HOMO SAPIENS | 3 | 4713 | 4djn | B | 305 | ALA | p.Ala305Ser | Q7LG56 |
| HOMO SAPIENS | -1 | 8974 | 5n7i | A | 500 | ARG | p.Arg500Ter | Q8N4S9 |
| HOMO SAPIENS | 0 | 17072 | 3fxi | B | 299 | ASP | p.Asp299Gly | O00206 |

| | | | | | | | | |
|---|---|---|---|---|---|---|---|---|
| HOMO SAPIENS | 1 | 14355 | 2wwk | T | 55 | ILE | p.Ile34306Asn | Q8WZ42 |
| HOMO SAPIENS | 0 | 14355 | 2wwk | T | 64 | LEU | p.Leu34315Pro | Q8WZ42 |
| HOMO SAPIENS | 2 | 17797 | 1xu1 | S | 72 | ARG | p.Arg72His | O14836 |
| HOMO SAPIENS | -1 | 10836 | 4ztu | A | 39 | GLN | p.Gln39Arg | P54098 |
| HOMO SAPIENS | -1 | 10836 | 4ztu | A | 42 | ARG | p.Arg42Gln | P54098 |
| HOMO SAPIENS | -1 | 10836 | 4ztu | A | 45 | GLN | p.Gln45Arg | P54098 |
| HOMO SAPIENS | -1 | 10836 | 4ztu | A | 52 | GLN | p.Gln52Glu | P54098 |
| HOMO SAPIENS | 0 | 10836 | 4ztu | A | 88 | PHE | p.Phe88Leu | P54098 |
| HOMO SAPIENS | 1 | 10836 | 4ztu | A | 194 | ALA | p.Ala194Val | P54098 |
| HOMO SAPIENS | 1 | 10836 | 4ztu | A | 227 | ARG | p.Arg227Trp | P54098 |
| HOMO SAPIENS | 0 | 10836 | 4ztu | A | 232 | ARG | p.Arg232His | P54098 |
| HOMO SAPIENS | 1 | 10836 | 4ztu | A | 285 | GLN | p.Gln285His | P54098 |
| HOMO SAPIENS | 1 | 10836 | 4ztu | A | 434 | TYR | p.Tyr434His | P54098 |
| HOMO SAPIENS | -1 | 10836 | 4ztu | A | 511 | SER | p.Ser511Asn | P54098 |
| HOMO SAPIENS | 0 | 10836 | 4ztu | A | 533 | CYS | p.Cys533Trp | P54098 |
| HOMO SAPIENS | 1 | 10836 | 4ztu | A | 546 | ARG | p.Arg546Cys | P54098 |
| HOMO SAPIENS | 1 | 10836 | 4ztu | A | 587 | PRO | p.Pro587Leu | P54098 |
| HOMO SAPIENS | 0 | 10836 | 4ztu | A | 603 | MET | p.Met603Thr | P54098 |
| HOMO SAPIENS | 0 | 10836 | 4ztu | A | 623 | LEU | p.Leu623Trp | P54098 |
| HOMO SAPIENS | -1 | 10836 | 4ztu | A | 627 | ARG | p.Arg627Trp | P54098 |
| HOMO SAPIENS | -1 | 10836 | 4ztu | A | 628 | ARG | p.Arg628Pro | P54098 |
| HOMO SAPIENS | 0 | 10836 | 4ztu | A | 635 | PRO | p.Pro635Leu | P54098 |
| HOMO SAPIENS | 0 | 10836 | 4ztu | A | 662 | GLU | p.Glu662Lys | P54098 |
| HOMO SAPIENS | -1 | 10836 | 4ztu | A | 674 | GLY | p.Gly674Asp | P54098 |
| HOMO SAPIENS | -1 | 10836 | 4ztu | A | 722 | ARG | p.Arg722His | P54098 |
| HOMO SAPIENS | -1 | 10836 | 4ztu | A | 737 | GLY | p.Gly737Arg | P54098 |
| HOMO SAPIENS | 0 | 10836 | 4ztu | A | 748 | TRP | p.Trp748Ser | P54098 |
| HOMO SAPIENS | 0 | 10836 | 4ztu | A | 749 | PHE | p.Phe749Ser | P54098 |
| HOMO SAPIENS | 1 | 10836 | 4ztu | A | 807 | ARG | p.Arg807Cys | P54098 |
| HOMO SAPIENS | 1 | 10836 | 4ztu | A | 831 | TYR | p.Tyr831Cys | P54098 |
| HOMO SAPIENS | 8 | 10836 | 4ztu | A | 848 | GLY | p.Gly848Ser | P54098 |
| HOMO SAPIENS | 1 | 10836 | 4ztu | A | 852 | ARG | p.Arg852Cys | P54098 |
| HOMO SAPIENS | 9 | 10836 | 4ztu | A | 853 | ARG | p.Arg853Trp | P54098 |
| HOMO SAPIENS | 1 | 10836 | 4ztu | A | 864 | ASN | p.Asn864Ser | P54098 |
| HOMO SAPIENS | 1 | 10836 | 4ztu | A | 879 | GLN | p.Gln879Arg | P54098 |
| HOMO SAPIENS | 1 | 10836 | 4ztu | A | 914 | THR | p.Thr914Pro | P54098 |
| HOMO SAPIENS | 6 | 10836 | 4ztu | A | 932 | HIS | p.His932Tyr | P54098 |
| HOMO SAPIENS | 2 | 10836 | 4ztu | A | 944 | GLU | p.Glu944Lys | P54098 |
| HOMO SAPIENS | 6 | 10836 | 4ztu | A | 955 | TYR | p.Tyr955Cys | P54098 |
| HOMO SAPIENS | 1 | 10836 | 4ztu | A | 957 | ALA | p.Ala957Ser | P54098 |
| HOMO SAPIENS | 1 | 10836 | 4ztu | A | 957 | ALA | p.Ala957Val | P54098 |
| HOMO SAPIENS | 1 | 10836 | 4ztu | A | 964 | ARG | p.Arg964Cys | P54098 |
| HOMO SAPIENS | 1 | 10836 | 4ztu | A | 966 | LEU | p.Leu966Arg | P54098 |
| HOMO SAPIENS | 0 | 10836 | 4ztu | A | 1051 | GLY | p.Gly1051Arg | P54098 |
| HOMO SAPIENS | 1 | 10836 | 4ztu | A | 1073 | PRO | p.Pro1073Leu | P54098 |
| HOMO SAPIENS | 1 | 10836 | 4ztu | A | 1096 | ARG | p.Arg1096Cys | P54098 |
| HOMO SAPIENS | 1 | 10836 | 4ztu | A | 1096 | ARG | p.Arg1096His | P54098 |
| HOMO SAPIENS | 0 | 10836 | 4ztu | A | 1116 | MET | p.Met1116Thr | P54098 |
| HOMO SAPIENS | 1 | 10836 | 4ztu | A | 1142 | ARG | p.Arg1142Trp | P54098 |
| HOMO SAPIENS | 1 | 10836 | 4ztu | A | 1146 | ARG | p.Arg1146Cys | P54098 |
| HOMO SAPIENS | 1 | 10836 | 4ztu | A | 1157 | ASN | p.Asn1157Ser | P54098 |
| HOMO SAPIENS | -1 | 10836 | 4ztu | A | 1234 | ARG | p.Arg1234Gln | P54098 |
| HOMO SAPIENS | -1 | 10836 | 4ztu | A | 1236 | GLN | p.Gln1236His | P54098 |
| HOMO SAPIENS | 5 | 2666 | 1jph | A | 62 | PRO | p.Pro62Leu | P06132 |

| | | | | | | | | |
|---|---|---|---|---|---|---|---|---|
| HOMO SAPIENS | 5 | 2666 | 1jph | A | 116 | GLN | p.Gln116Ter | P06132 |
| HOMO SAPIENS | 0 | 2666 | 1jph | A | 165 | MET | p.Met165Arg | P06132 |
| HOMO SAPIENS | 6 | 2666 | 1jph | A | 167 | GLU | p.Glu167Lys | P06132 |
| HOMO SAPIENS | 0 | 2666 | 1jph | A | 195 | LEU | p.Leu195Phe | P06132 |
| HOMO SAPIENS | 7 | 2666 | 1jph | A | 253 | LEU | p.Leu253Gln | P06132 |
| HOMO SAPIENS | 0 | 2666 | 1jph | A | 281 | GLY | p.Gly281Glu | P06132 |
| HOMO SAPIENS | 9 | 2666 | 1jph | A | 304 | ASN | p.Asn304Lys | P06132 |
| HOMO SAPIENS | 8 | 2666 | 1jph | A | 311 | TYR | p.Tyr311Cys | P06132 |
| HOMO SAPIENS | 5 | 2666 | 1jph | A | 332 | ARG | p.Arg332His | P06132 |
| HOMO SAPIENS | 1 | 889 | 1m51 | A | 184 | LEU | p.Val184Leu | P35558 |
| HOMO SAPIENS | 0 | 889 | 1m51 | A | 267 | VAL | p.Ile267Val | P35558 |
| HOMO SAPIENS | 2 | 889 | 1m51 | A | 276 | GLU | p.Glu276Lys | P35558 |
| HOMO SAPIENS | 6 | 889 | 1m51 | A | 368 | VAL | p.Val368Ile | P35558 |
| HOMO SAPIENS | 0 | 889 | 1m51 | A | 579 | SER | p.Ser579Arg | P35558 |
| HOMO SAPIENS | 1 | 4084 | 2gjx | A | 28 | GLN | p.Gln28Ter | P06865 |
| HOMO SAPIENS | 0 | 4084 | 2gjx | A | 39 | LEU | p.Leu39Arg | P06865 |
| HOMO SAPIENS | 0 | 4084 | 2gjx | A | 127 | LEU | p.Leu127Arg | P06865 |
| HOMO SAPIENS | 6 | 4084 | 2gjx | A | 137 | ARG | p.Arg137Ter | P06865 |
| HOMO SAPIENS | 6 | 4084 | 2gjx | A | 170 | ARG | p.Arg170Gln | P06865 |
| HOMO SAPIENS | 9 | 4084 | 2gjx | A | 178 | ARG | p.Arg178Cys | P06865 |
| HOMO SAPIENS | 9 | 4084 | 2gjx | A | 178 | ARG | p.Arg178His | P06865 |
| HOMO SAPIENS | 5 | 4084 | 2gjx | A | 180 | TYR | p.Tyr180His | P06865 |
| HOMO SAPIENS | 5 | 4084 | 2gjx | A | 180 | TYR | p.Tyr180Ter | P06865 |
| HOMO SAPIENS | 1 | 4084 | 2gjx | A | 197 | LYS | p.Lys197Thr | P06865 |
| HOMO SAPIENS | 0 | 4084 | 2gjx | A | 200 | VAL | p.Val200Met | P06865 |
| HOMO SAPIENS | 9 | 4084 | 2gjx | A | 204 | HIS | p.His204Arg | P06865 |
| HOMO SAPIENS | 5 | 4084 | 2gjx | A | 210 | SER | p.Ser210Phe | P06865 |
| HOMO SAPIENS | 0 | 4084 | 2gjx | A | 211 | PHE | p.Phe211Ser | P06865 |
| HOMO SAPIENS | 2 | 4084 | 2gjx | A | 250 | GLY | p.Gly250Asp | P06865 |
| HOMO SAPIENS | 9 | 4084 | 2gjx | A | 258 | ASP | p.Asp258His | P06865 |
| HOMO SAPIENS | 0 | 4084 | 2gjx | A | 269 | GLY | p.Gly269Ser | P06865 |
| HOMO SAPIENS | 0 | 4084 | 2gjx | A | 301 | MET | p.Met301Arg | P06865 |
| HOMO SAPIENS | 3 | 4084 | 2gjx | A | 393 | ARG | p.Arg393Ter | P06865 |
| HOMO SAPIENS | 3 | 4084 | 2gjx | A | 393 | ARG | p.Arg393Pro | P06865 |
| HOMO SAPIENS | 0 | 4084 | 2gjx | A | 399 | ASN | p.Asn399Asp | P06865 |
| HOMO SAPIENS | 2 | 4084 | 2gjx | A | 420 | TRP | p.Trp420Cys | P06865 |
| HOMO SAPIENS | 1 | 4084 | 2gjx | A | 434 | PHE | p.Phe434Leu | P06865 |
| HOMO SAPIENS | 0 | 4084 | 2gjx | A | 454 | GLY | p.Gly454Ser | P06865 |
| HOMO SAPIENS | 1 | 4084 | 2gjx | A | 474 | TRP | p.Trp474Cys | P06865 |
| HOMO SAPIENS | 0 | 4084 | 2gjx | A | 479 | ALA | p.Ala479Thr | P06865 |
| HOMO SAPIENS | 5 | 4084 | 2gjx | A | 485 | TRP | p.Trp485Arg | P06865 |
| HOMO SAPIENS | 4 | 4084 | 2gjx | A | 499 | ARG | p.Arg499Cys | P06865 |
| HOMO SAPIENS | 4 | 4084 | 2gjx | A | 499 | ARG | p.Arg499His | P06865 |
| HOMO SAPIENS | 2 | 4084 | 2gjx | A | 504 | ARG | p.Arg504Cys | P06865 |
| HOMO SAPIENS | 2 | 4084 | 2gjx | A | 504 | ARG | p.Arg504His | P06865 |
| HOMO SAPIENS | -1 | 1943 | 1xmj | A | 417 | ASN | p.Asn417Lys | P13569 |
| HOMO SAPIENS | -1 | 1943 | 1xmj | A | 422 | SER | p.Ser422Phe | P13569 |
| HOMO SAPIENS | 1 | 1943 | 1xmj | A | 438 | THR | p.Thr438Ala | P13569 |
| HOMO SAPIENS | 0 | 1943 | 1xmj | A | 455 | ALA | p.Ala455Glu | P13569 |
| HOMO SAPIENS | 4 | 1943 | 1xmj | A | 456 | VAL | p.Val456Ala | P13569 |
| HOMO SAPIENS | 9 | 1943 | 1xmj | A | 458 | GLY | p.Gly458Val | P13569 |
| HOMO SAPIENS | 8 | 1943 | 1xmj | A | 466 | SER | p.Ser466Ter | P13569 |
| HOMO SAPIENS | 0 | 1943 | 1xmj | A | 467 | LEU | p.Leu467Pro | P13569 |
| HOMO SAPIENS | 1 | 1943 | 1xmj | A | 470 | MET | p.Val470Met | P13569 |

| | | | | | | | | |
|---|---|---|---|---|---|---|---|---|
| HOMO SAPIENS | 5 | 1943 | 1xmj | A | 480 | GLY | p.Gly480Cys | P13569 |
| HOMO SAPIENS | 3 | 1943 | 1xmj | A | 492 | SER | p.Ser492Phe | P13569 |
| HOMO SAPIENS | 8 | 1943 | 1xmj | A | 493 | GLN | p.Gln493Ter | P13569 |
| HOMO SAPIENS | 5 | 1943 | 1xmj | A | 506 | ILE | p.Ile506Met | P13569 |
| HOMO SAPIENS | 3 | 1943 | 1xmj | A | 520 | VAL | p.Val520Phe | P13569 |
| HOMO SAPIENS | 1 | 1943 | 1xmj | A | 528 | GLU | p.Glu528Asp | P13569 |
| HOMO SAPIENS | 1 | 1943 | 1xmj | A | 534 | ALA | p.Ala534Glu | P13569 |
| HOMO SAPIENS | 4 | 1943 | 1xmj | A | 542 | GLY | p.Gly542Ter | P13569 |
| HOMO SAPIENS | 6 | 1943 | 1xmj | A | 549 | SER | p.Ser549Asn | P13569 |
| HOMO SAPIENS | 6 | 1943 | 1xmj | A | 549 | SER | p.Ser549Arg | P13569 |
| HOMO SAPIENS | 6 | 1943 | 1xmj | A | 549 | SER | p.Ser549Arg | P13569 |
| HOMO SAPIENS | 6 | 1943 | 1xmj | A | 551 | GLY | p.Gly551Asp | P13569 |
| HOMO SAPIENS | 6 | 1943 | 1xmj | A | 551 | GLY | p.Gly551Ser | P13569 |
| HOMO SAPIENS | 6 | 1943 | 1xmj | A | 553 | GLN | p.Arg553Gln | P13569 |
| HOMO SAPIENS | 6 | 1943 | 1xmj | A | 553 | GLN | p.Arg553Ter | P13569 |
| HOMO SAPIENS | 0 | 1943 | 1xmj | A | 559 | ALA | p.Ala559Thr | P13569 |
| HOMO SAPIENS | 7 | 1943 | 1xmj | A | 560 | ARG | p.Arg560Lys | P13569 |
| HOMO SAPIENS | 4 | 1943 | 1xmj | A | 562 | VAL | p.Val562Ile | P13569 |
| HOMO SAPIENS | 3 | 1943 | 1xmj | A | 563 | TYR | p.Tyr563Asn | P13569 |
| HOMO SAPIENS | 3 | 1943 | 1xmj | A | 569 | TYR | p.Tyr569Asp | P13569 |
| HOMO SAPIENS | 3 | 1943 | 1xmj | A | 585 | GLU | p.Glu585Ter | P13569 |
| HOMO SAPIENS | 1 | 1943 | 1xmj | A | 618 | ILE | p.Ile618Thr | P13569 |
| HOMO SAPIENS | 9 | 1943 | 1xmj | A | 622 | GLY | p.Gly622Asp | P13569 |
| HOMO SAPIENS | -1 | 1943 | 1xmj | A | 648 | ASP | p.Asp648Val | P13569 |
| HOMO SAPIENS | 0 | 1943 | 1xmj | A | 673 | GLY | p.Gly673Ter | P13569 |
| HOMO SAPIENS | 5 | 5111 | 4rec | A | 749 | ARG | p.Arg749Ter | Q9Y2M0 |
| HOMO SAPIENS | -1 | 4580 | 4l0d | B | 2 | GLY | p.Pro2Leu | P35520 |
| HOMO SAPIENS | -1 | 4580 | 4l0d | B | 12 | PRO | p.Pro12Ser | P35520 |
| HOMO SAPIENS | -1 | 4580 | 4l0d | B | 18 | ARG | p.Arg18Cys | P35520 |
| HOMO SAPIENS | 1 | 4580 | 4l0d | B | 49 | PRO | p.Pro49Leu | P35520 |
| HOMO SAPIENS | 2 | 4580 | 4l0d | B | 70 | PRO | p.Pro70Leu | P35520 |
| HOMO SAPIENS | 1 | 4580 | 4l0d | B | 102 | LYS | p.Lys102Gln | P35520 |
| HOMO SAPIENS | 7 | 4580 | 4l0d | B | 114 | ALA | p.Ala114Val | P35520 |
| HOMO SAPIENS | 5 | 4580 | 4l0d | B | 116 | GLY | p.Gly116Arg | P35520 |
| HOMO SAPIENS | 8 | 4580 | 4l0d | B | 121 | ARG | p.Arg121Cys | P35520 |
| HOMO SAPIENS | 8 | 4580 | 4l0d | B | 121 | ARG | p.Arg121His | P35520 |
| HOMO SAPIENS | 5 | 4580 | 4l0d | B | 125 | ARG | p.Arg125Gln | P35520 |
| HOMO SAPIENS | 4 | 4580 | 4l0d | B | 134 | GLY | p.Gly134Arg | P35520 |
| HOMO SAPIENS | 4 | 4580 | 4l0d | B | 139 | GLY | p.Gly139Arg | P35520 |
| HOMO SAPIENS | 0 | 4580 | 4l0d | B | 145 | PRO | p.Pro145Leu | P35520 |
| HOMO SAPIENS | 0 | 4580 | 4l0d | B | 148 | GLY | p.Gly148Arg | P35520 |
| HOMO SAPIENS | 0 | 4580 | 4l0d | B | 168 | VAL | p.Val168Met | P35520 |
| HOMO SAPIENS | 2 | 4580 | 4l0d | B | 191 | THR | p.Thr191Met | P35520 |
| HOMO SAPIENS | 5 | 4580 | 4l0d | B | 234 | ASP | p.Asp234Asn | P35520 |
| HOMO SAPIENS | 0 | 4580 | 4l0d | B | 257 | THR | p.Thr257Met | P35520 |
| HOMO SAPIENS | 4 | 4580 | 4l0d | B | 262 | THR | p.Thr262Met | P35520 |
| HOMO SAPIENS | 0 | 4580 | 4l0d | B | 278 | ILE | p.Ile278Thr | P35520 |
| HOMO SAPIENS | 2 | 4580 | 4l0d | B | 307 | GLY | p.Gly307Ser | P35520 |
| HOMO SAPIENS | 4 | 4580 | 4l0d | B | 320 | VAL | p.Val320Ala | P35520 |
| HOMO SAPIENS | 2 | 4580 | 4l0d | B | 336 | ARG | p.Arg336Cys | P35520 |
| HOMO SAPIENS | 2 | 4580 | 4l0d | B | 336 | ARG | p.Arg336His | P35520 |
| HOMO SAPIENS | 0 | 4580 | 4l0d | B | 351 | GLY | p.Gly351Arg | P35520 |
| HOMO SAPIENS | 0 | 4580 | 4l0d | B | 353 | THR | p.Thr353Met | P35520 |
| HOMO SAPIENS | 0 | 4580 | 4l0d | B | 371 | VAL | p.Val371Met | P35520 |

| | | | | | | | | |
|---|---|---|---|---|---|---|---|---|
| HOMO SAPIENS | 2 | 4580 | 4l0d | B | 379 | ARG | p.Arg379Gln | P35520 |
| HOMO SAPIENS | 2 | 4580 | 4l0d | B | 379 | ARG | p.Arg379Trp | P35520 |
| HOMO SAPIENS | 2 | 4580 | 4l0d | B | 384 | LYS | p.Lys384Glu | P35520 |
| HOMO SAPIENS | 0 | 4580 | 4l0d | B | 422 | PRO | p.Pro422Leu | P35520 |
| HOMO SAPIENS | 0 | 4580 | 4l0d | B | 444 | ASP | p.Asp444Asn | P35520 |
| HOMO SAPIENS | 0 | 4580 | 4l0d | B | 466 | SER | p.Ser466Leu | P35520 |
| HOMO SAPIENS | 0 | 4580 | 4l0d | B | 539 | LEU | p.Leu539Ser | P35520 |
| HOMO SAPIENS | 0 | 4580 | 4l0d | B | 548 | ARG | p.Arg548Gln | P35520 |
| HOMO SAPIENS | 1 | 6536 | 3ulu | A | 412 | LEU | p.Leu412Phe | O15455 |
| HOMO SAPIENS | 1 | 28493 | 3ii6 | X | 814 | ARG | p.Arg814Ter | P49917 |
| HOMO SAPIENS | 1 | 28493 | 3ii6 | X | 857 | ALA | p.Ala857Thr | P49917 |
| HOMO SAPIENS | 9 | 3543 | 2q51 | A | 24 | GLU | p.Glu24Gly | P45381 |
| HOMO SAPIENS | 1 | 3543 | 2q51 | A | 27 | GLY | p.Gly27Arg | P45381 |
| HOMO SAPIENS | 9 | 3543 | 2q51 | A | 71 | ARG | p.Arg71His | P45381 |
| HOMO SAPIENS | -1 | 3543 | 2q51 | A | 82 | MSE | p.Met82Thr | P45381 |
| HOMO SAPIENS | 0 | 3543 | 2q51 | A | 143 | ILE | p.Ile143Phe | P45381 |
| HOMO SAPIENS | 1 | 3543 | 2q51 | A | 152 | CYS | p.Cys152Arg | P45381 |
| HOMO SAPIENS | 2 | 3543 | 2q51 | A | 218 | CYS | p.Cys218Ter | P45381 |
| HOMO SAPIENS | 1 | 3543 | 2q51 | A | 231 | TYR | p.Tyr231Cys | P45381 |
| HOMO SAPIENS | 1 | 3543 | 2q51 | A | 231 | TYR | p.Tyr231Ter | P45381 |
| HOMO SAPIENS | 2 | 3543 | 2q51 | A | 274 | GLY | p.Gly274Arg | P45381 |
| HOMO SAPIENS | 6 | 3543 | 2q51 | A | 285 | GLU | p.Glu285Ala | P45381 |
| HOMO SAPIENS | 5 | 3543 | 2q51 | A | 287 | ALA | p.Ala287Thr | P45381 |
| HOMO SAPIENS | 2 | 3543 | 2q51 | A | 305 | ALA | p.Ala305Glu | P45381 |
| HOMO SAPIENS | 2 | 24687 | 1hfi | A | 25 | SER | p.Ser890Ile | P08603 |
| HOMO SAPIENS | 0 | 1508 | 3wn5 | C | 45 | LEU | p.Leu66Arg | P08637 |
| HOMO SAPIENS | 0 | 1508 | 3wn5 | C | 155 | VAL | p.Phe176Val | P08637 |
| HOMO SAPIENS | 0 | 750 | 5tpa | A | 54 | VAL | p.Val452Met | Q12879 |
| HOMO SAPIENS | 9 | 750 | 5tpa | A | 120 | ARG | p.Arg518Cys | Q12879 |
| HOMO SAPIENS | 0 | 750 | 5tpa | A | 199 | ALA | p.Ala716Thr | Q12879 |
| HOMO SAPIENS | 4 | 1363 | 3hg2 | B | 40 | PRO | p.Pro40Ser | P06280 |
| HOMO SAPIENS | 3 | 1363 | 3hg2 | B | 65 | SER | p.Ser65Thr | P06280 |
| HOMO SAPIENS | 4 | 1363 | 3hg2 | B | 66 | GLU | p.Glu66Gln | P06280 |
| HOMO SAPIENS | 2 | 1363 | 3hg2 | B | 80 | GLY | p.Gly80Asp | P06280 |
| HOMO SAPIENS | 3 | 1363 | 3hg2 | B | 112 | ARG | p.Arg112Cys | P06280 |
| HOMO SAPIENS | 3 | 1363 | 3hg2 | B | 112 | ARG | p.Arg112His | P06280 |
| HOMO SAPIENS | 0 | 1363 | 3hg2 | B | 139 | ASN | p.Asn139Ser | P06280 |
| HOMO SAPIENS | 7 | 1363 | 3hg2 | B | 156 | ALA | p.Ala156Thr | P06280 |
| HOMO SAPIENS | 5 | 1363 | 3hg2 | B | 162 | TRP | p.Trp162Arg | P06280 |
| HOMO SAPIENS | 5 | 1363 | 3hg2 | B | 207 | TYR | p.Tyr207His | P06280 |
| HOMO SAPIENS | 0 | 1363 | 3hg2 | B | 215 | ASN | p.Asn215Ser | P06280 |
| HOMO SAPIENS | 0 | 1363 | 3hg2 | B | 264 | ASP | p.Asp264Val | P06280 |
| HOMO SAPIENS | 5 | 1363 | 3hg2 | B | 266 | ASP | p.Asp266Val | P06280 |
| HOMO SAPIENS | 2 | 1363 | 3hg2 | B | 269 | VAL | p.Val269Ala | P06280 |
| HOMO SAPIENS | 3 | 1363 | 3hg2 | B | 272 | ASN | p.Asn272Ser | P06280 |
| HOMO SAPIENS | 3 | 1363 | 3hg2 | B | 279 | GLN | p.Gln279Glu | P06280 |
| HOMO SAPIENS | 0 | 1363 | 3hg2 | B | 289 | ILE | p.Ile289Val | P06280 |
| HOMO SAPIENS | 0 | 1363 | 3hg2 | B | 296 | MET | p.Met296Val | P06280 |
| HOMO SAPIENS | 0 | 1363 | 3hg2 | B | 296 | MET | p.Met296Ile | P06280 |
| HOMO SAPIENS | 0 | 1363 | 3hg2 | B | 297 | SER | p.Ser297Phe | P06280 |
| HOMO SAPIENS | 2 | 1363 | 3hg2 | B | 301 | ARG | p.Arg301Gln | P06280 |
| HOMO SAPIENS | 3 | 1363 | 3hg2 | B | 327 | GLN | p.Gln327Lys | P06280 |
| HOMO SAPIENS | 0 | 1363 | 3hg2 | B | 328 | GLY | p.Gly328Arg | P06280 |
| HOMO SAPIENS | 0 | 1363 | 3hg2 | B | 328 | GLY | p.Gly328Ala | P06280 |

| | | | | | | | | |
|---|---|---|---|---|---|---|---|---|
| HOMO SAPIENS | 2 | 1363 | 3hg2 | B | 331 | LEU | p.Leu331Phe | P06280 |
| HOMO SAPIENS | 0 | 1363 | 3hg2 | B | 342 | ARG | p.Arg342Gln | P06280 |
| HOMO SAPIENS | 1 | 1363 | 3hg2 | B | 361 | GLY | p.Gly361Arg | P06280 |
| HOMO SAPIENS | 1 | 1363 | 3hg2 | B | 363 | ARG | p.Arg363His | P06280 |
| HOMO SAPIENS | 1 | 1363 | 3hg2 | B | 385 | THR | p.Thr385Ala | P06280 |
| HOMO SAPIENS | 1 | 1363 | 3hg2 | B | 399 | TRP | p.Trp399Ser | P06280 |
| HOMO SAPIENS | 1 | 1363 | 3hg2 | B | 410 | THR | p.Thr410Ala | P06280 |
| HOMO SAPIENS | 2 | 3904 | 3caa | B | 385 | MET | p.Met414Val | P01011 |
| HOMO SAPIENS | 0 | 558 | 5eub | A | 56 | VAL | p.Val56Ile | Q13126 |
| HOMO SAPIENS | 0 | 558 | 5eub | A | 189 | TRP | p.Trp189Leu | Q13126 |
| HOMO SAPIENS | 0 | 558 | 5eub | A | 218 | ALA | p.Ala218Gly | Q13126 |
| HOMO SAPIENS | 1 | 3865 | 1cb5 | B | 443 | ILE | p.Ile443Val | Q13867 |
| HOMO SAPIENS | 2 | 14329 | 4m92 | A | 24 | ILE | p.Ile65Val | Q13454 |
| HOMO SAPIENS | 1 | 7381 | 5kz5 | k | 50 | GLY | p.Gly50Glu | Q9H1K1 |
| HOMO SAPIENS | 0 | 1102 | 4fdh | E | 86 | PRO | p.Pro86Ala | P19099 |
| HOMO SAPIENS | 0 | 1102 | 4fdh | E | 118 | ALA | p.Ala118Thr | P19099 |
| HOMO SAPIENS | 0 | 1102 | 4fdh | E | 159 | PRO | p.Pro159Leu | P19099 |
| HOMO SAPIENS | 0 | 1102 | 4fdh | E | 173 | LYS | p.Lys173Arg | P19099 |
| HOMO SAPIENS | 0 | 1102 | 4fdh | E | 181 | ARG | p.Arg181Trp | P19099 |
| HOMO SAPIENS | 0 | 1102 | 4fdh | E | 185 | THR | p.Thr185Ile | P19099 |
| HOMO SAPIENS | 0 | 1102 | 4fdh | E | 198 | GLU | p.Glu198Asp | P19099 |
| HOMO SAPIENS | 0 | 1102 | 4fdh | E | 214 | HIS | p.His214Asp | P19099 |
| HOMO SAPIENS | 0 | 1102 | 4fdh | E | 251 | LYS | p.Lys251Arg | P19099 |
| HOMO SAPIENS | 0 | 1102 | 4fdh | E | 255 | GLU | p.Glu255Ter | P19099 |
| HOMO SAPIENS | 0 | 1102 | 4fdh | E | 281 | ASN | p.Asn281Ser | P19099 |
| HOMO SAPIENS | 0 | 1102 | 4fdh | E | 320 | ALA | p.Ala320Val | P19099 |
| HOMO SAPIENS | 0 | 1102 | 4fdh | E | 339 | ILE | p.Ile339Thr | P19099 |
| HOMO SAPIENS | 0 | 1102 | 4fdh | E | 341 | ARG | p.Arg341Cys | P19099 |
| HOMO SAPIENS | 2 | 1102 | 4fdh | E | 374 | ARG | p.Arg374Trp | P19099 |
| HOMO SAPIENS | 0 | 1102 | 4fdh | E | 386 | VAL | p.Val386Ala | P19099 |
| HOMO SAPIENS | 0 | 1102 | 4fdh | E | 390 | LEU | p.Leu390Phe | P19099 |
| HOMO SAPIENS | -1 | 1102 | 4fdh | E | 435 | GLY | p.Gly435Ser | P19099 |
| HOMO SAPIENS | 0 | 1102 | 4fdh | E | 461 | LEU | p.Leu461Pro | P19099 |
| HOMO SAPIENS | 0 | 1102 | 4fdh | E | 498 | THR | p.Thr498Ala | P19099 |
| HOMO SAPIENS | 6 | 10292 | 2of5 | F | 128 | GLY | p.Gly128Arg | P78560 |
| HOMO SAPIENS | 6 | 10292 | 2of5 | F | 170 | ARG | p.Arg170Cys | P78560 |
| HOMO SAPIENS | -1 | 18965 | 3ghg | A | 16 | ARG | p.Arg35Cys | P02671 |
| HOMO SAPIENS | -1 | 18965 | 3ghg | A | 16 | ARG | p.Arg35His | P02671 |
| HOMO SAPIENS | -1 | 18965 | 3ghg | A | 283 | PRO | p.Pro302Ala | P02671 |
| HOMO SAPIENS | -1 | 18965 | 3ghg | A | 312 | THR | p.Thr331Ala | P02671 |
| HOMO SAPIENS | -1 | 18965 | 3ghg | A | 381 | SER | p.Ser400Phe | P02671 |
| HOMO SAPIENS | -1 | 18965 | 3ghg | A | 463 | VAL | p.Val482Met | P02671 |
| HOMO SAPIENS | -1 | 18965 | 3ghg | A | 526 | GLU | p.Glu545Val | P02671 |
| HOMO SAPIENS | -1 | 18965 | 3ghg | A | 554 | ARG | p.Arg573Leu | P02671 |
| HOMO SAPIENS | 8 | 3023 | 4ttb | A | 101 | ARG | p.Arg101Trp | Q6PHW0 |
| HOMO SAPIENS | 2 | 3023 | 4ttb | A | 116 | ILE | p.Ile116Thr | Q6PHW0 |
| HOMO SAPIENS | 0 | 3023 | 4ttb | A | 220 | ALA | p.Ala220Thr | Q6PHW0 |
| HOMO SAPIENS | 2 | 3023 | 4ttb | A | 271 | GLU | p.Glu271Lys | Q6PHW0 |
| HOMO SAPIENS | 3 | 33226 | 1ya5 | T | 11 | SER | p.Ser11Ter | O15273 |
| HOMO SAPIENS | 6 | 33226 | 1ya5 | T | 64 | SER | p.Ser64Leu | O15273 |
| HOMO SAPIENS | 6 | 33226 | 1ya5 | T | 70 | ARG | p.Arg70Gln | O15273 |
| HOMO SAPIENS | 3 | 33226 | 1ya5 | T | 75 | GLY | p.Gly75Ser | O15273 |
| HOMO SAPIENS | 0 | 33226 | 1ya5 | T | 87 | ARG | p.Arg87Trp | O15273 |
| HOMO SAPIENS | 2 | 5667 | 3dax | A | 398 | PRO | p.Pro398Ala | P22680 |

| | | | | | | | | |
|---|---|---|---|---|---|---|---|---|
| HOMO SAPIENS | 0 | 12861 | 5mya | A | 464 | ASN | p.Asn464His | Q02763 |
| HOMO SAPIENS | 2 | 12861 | 5mya | A | 486 | VAL | p.Val486Ile | Q02763 |
| HOMO SAPIENS | 0 | 12861 | 5mya | A | 522 | ARG | p.Arg522His | Q02763 |
| HOMO SAPIENS | 2 | 12861 | 5mya | A | 600 | VAL | p.Val600Leu | Q02763 |
| HOMO SAPIENS | 0 | 12861 | 5mya | A | 634 | LEU | p.Leu634Phe | Q02763 |
| HOMO SAPIENS | 0 | 12861 | 5mya | A | 684 | HIS | p.His684Gln | Q02763 |
| HOMO SAPIENS | 0 | 12861 | 5mya | A | 724 | ALA | p.Ala724Thr | Q02763 |
| HOMO SAPIENS | 0 | 7743 | 3qzw | G | 90 | GLY | p.Gly111Ser | P01732 |
| HOMO SAPIENS | 2 | 9498 | 3kci | A | 4267 | ASP | p.Asp4267Glu | O95714 |
| HOMO SAPIENS | 5 | 201 | 3ips | B | 415 | ARG | p.Arg415Gln | Q13133 |
| HOMO SAPIENS | 3 | 229 | 2q50 | A | 68 | LEU | p.Leu68Pro | Q9UBQ7 |
| HOMO SAPIENS | 2 | 229 | 2q50 | A | 99 | ARG | p.Arg99Ter | Q9UBQ7 |
| HOMO SAPIENS | 3 | 229 | 2q50 | A | 160 | GLY | p.Gly160Arg | Q9UBQ7 |
| HOMO SAPIENS | 5 | 229 | 2q50 | A | 165 | GLY | p.Gly165Asp | Q9UBQ7 |
| HOMO SAPIENS | 4 | 229 | 2q50 | A | 302 | ARG | p.Arg302Cys | Q9UBQ7 |
| HOMO SAPIENS | 4 | 229 | 2q50 | A | 302 | ARG | p.Arg302His | Q9UBQ7 |
| HOMO SAPIENS | -1 | 229 | 2q50 | A | 322 | MSE | p.Met322Arg | Q9UBQ7 |
| HOMO SAPIENS | -1 | 4519 | 5clw | A | 40 | PRO | p.Pro40Thr | Q04446 |
| HOMO SAPIENS | 1 | 4519 | 5clw | A | 59 | ILE | p.Ile59Thr | Q04446 |
| HOMO SAPIENS | 0 | 4519 | 5clw | A | 190 | ARG | p.Arg190Gly | Q04446 |
| HOMO SAPIENS | 1 | 4519 | 5clw | A | 224 | LEU | p.Leu224Pro | Q04446 |
| HOMO SAPIENS | 0 | 4519 | 5clw | A | 236 | GLN | p.Gln236His | Q04446 |
| HOMO SAPIENS | 1 | 4519 | 5clw | A | 254 | THR | p.Thr254Ala | Q04446 |
| HOMO SAPIENS | 0 | 4519 | 5clw | A | 257 | PHE | p.Phe257Leu | Q04446 |
| HOMO SAPIENS | 1 | 4519 | 5clw | A | 262 | ARG | p.Arg262Cys | Q04446 |
| HOMO SAPIENS | 3 | 4519 | 5clw | A | 329 | TYR | p.Tyr329Ser | Q04446 |
| HOMO SAPIENS | 0 | 4519 | 5clw | A | 507 | THR | p.Thr507Ala | Q04446 |
| HOMO SAPIENS | 0 | 4519 | 5clw | A | 515 | ARG | p.Arg515Cys | Q04446 |
| HOMO SAPIENS | 0 | 4519 | 5clw | A | 515 | ARG | p.Arg515His | Q04446 |
| HOMO SAPIENS | 0 | 4519 | 5clw | A | 524 | ARG | p.Arg524Gln | Q04446 |
| HOMO SAPIENS | 0 | 4519 | 5clw | A | 524 | ARG | p.Arg524Ter | Q04446 |
| HOMO SAPIENS | 0 | 4519 | 5clw | A | 545 | HIS | p.His545Arg | Q04446 |
| HOMO SAPIENS | 2 | 4519 | 5clw | A | 548 | TRP | p.Trp548Ter | Q04446 |
| HOMO SAPIENS | 0 | 4519 | 5clw | A | 592 | GLU | p.Glu592Ter | Q04446 |
| HOMO SAPIENS | 0 | 4519 | 5clw | A | 628 | HIS | p.His628Arg | Q04446 |
| HOMO SAPIENS | 0 | 4519 | 5clw | A | 637 | ARG | p.Arg637Ter | Q04446 |
| HOMO SAPIENS | 0 | 4519 | 5clw | A | 673 | ALA | p.Ala673Thr | Q04446 |
| HOMO SAPIENS | 1 | 10405 | 3bg3 | D | 507 | VAL | p.Val507Ile | P11498 |
| HOMO SAPIENS | 0 | 10405 | 3bg3 | D | 583 | ARG | p.Arg583Leu | P11498 |
| HOMO SAPIENS | 1 | 10405 | 3bg3 | D | 589 | LYS | p.Lys589Asn | P11498 |
| HOMO SAPIENS | 6 | 10405 | 3bg3 | D | 610 | ALA | p.Ala610Thr | P11498 |
| HOMO SAPIENS | 2 | 10405 | 3bg3 | D | 631 | ARG | p.Arg631Gln | P11498 |
| HOMO SAPIENS | 1 | 10405 | 3bg3 | D | 732 | ARG | p.Arg732Gln | P11498 |
| HOMO SAPIENS | 6 | 10405 | 3bg3 | D | 743 | MET | p.Met743Ile | P11498 |
| HOMO SAPIENS | 0 | 10405 | 3bg3 | D | 750 | THR | p.Thr750Met | P11498 |
| HOMO SAPIENS | -1 | 13751 | 2o8d | B | 350 | ALA | p.Ala350Val | P52701 |
| HOMO SAPIENS | -1 | 13751 | 2o8d | B | 352 | VAL | p.Val352Ile | P52701 |
| HOMO SAPIENS | -1 | 13751 | 2o8d | B | 355 | GLY | p.Gly355Ser | P52701 |
| HOMO SAPIENS | 0 | 13751 | 2o8d | B | 385 | ARG | p.Arg385Trp | P52701 |
| HOMO SAPIENS | 0 | 13751 | 2o8d | B | 390 | ASP | p.Asp390Glu | P52701 |
| HOMO SAPIENS | 0 | 13751 | 2o8d | B | 396 | LEU | p.Leu396Val | P52701 |
| HOMO SAPIENS | 0 | 13751 | 2o8d | B | 449 | LEU | p.Leu449Pro | P52701 |
| HOMO SAPIENS | 0 | 13751 | 2o8d | B | 468 | ARG | p.Arg468His | P52701 |
| HOMO SAPIENS | 0 | 13751 | 2o8d | B | 503 | SER | p.Ser503Cys | P52701 |

| | | | | | | | | |
|---|---|---|---|---|---|---|---|---|
| HOMO SAPIENS | 1 | 13751 | 2o8d | B | 509 | VAL | p.Val509Ala | P52701 |
| HOMO SAPIENS | 0 | 13751 | 2o8d | B | 675 | GLU | p.Glu675Asp | P52701 |
| HOMO SAPIENS | 0 | 13751 | 2o8d | B | 795 | ILE | p.Ile795Thr | P52701 |
| HOMO SAPIENS | 0 | 13751 | 2o8d | B | 850 | TYR | p.Tyr850Ter | P52701 |
| HOMO SAPIENS | 0 | 13751 | 2o8d | B | 878 | VAL | p.Val878Ala | P52701 |
| HOMO SAPIENS | 0 | 13751 | 2o8d | B | 911 | ARG | p.Arg911Ter | P52701 |
| HOMO SAPIENS | 0 | 13751 | 2o8d | B | 922 | ARG | p.Arg922Ter | P52701 |
| HOMO SAPIENS | 0 | 13751 | 2o8d | B | 969 | TYR | p.Tyr969Cys | P52701 |
| HOMO SAPIENS | 0 | 13751 | 2o8d | B | 1005 | ARG | p.Arg1005Ter | P52701 |
| HOMO SAPIENS | 0 | 13751 | 2o8d | B | 1035 | ARG | p.Arg1035Ter | P52701 |
| HOMO SAPIENS | 1 | 13751 | 2o8d | B | 1048 | GLN | p.Gln1048Ter | P52701 |
| HOMO SAPIENS | 1 | 13751 | 2o8d | B | 1051 | VAL | p.Val1051Ile | P52701 |
| HOMO SAPIENS | 0 | 13751 | 2o8d | B | 1054 | ILE | p.Ile1054Phe | P52701 |
| HOMO SAPIENS | 0 | 13751 | 2o8d | B | 1054 | ILE | p.Ile1054Met | P52701 |
| HOMO SAPIENS | 0 | 13751 | 2o8d | B | 1163 | GLU | p.Glu1163Val | P52701 |
| HOMO SAPIENS | 0 | 13751 | 2o8d | B | 1237 | GLU | p.Glu1237Asp | P52701 |
| HOMO SAPIENS | 0 | 13751 | 2o8d | B | 1242 | ARG | p.Arg1242His | P52701 |
| HOMO SAPIENS | -1 | 13751 | 2o8d | B | 1278 | PRO | p.Pro1278Thr | P52701 |
| HOMO SAPIENS | 0 | 13751 | 2o8d | B | 1304 | ARG | p.Arg1304Lys | P52701 |
| HOMO SAPIENS | 0 | 13751 | 2o8d | B | 1317 | HIS | p.His1317Gln | P52701 |
| HOMO SAPIENS | 0 | 13751 | 2o8d | B | 1329 | SER | p.Ser1329Leu | P52701 |
| HOMO SAPIENS | 3 | 13273 | 4g8k | A | 265 | GLU | p.Glu265Ter | Q05823 |
| HOMO SAPIENS | -1 | 11088 | 5uj7 | A | 666 | ARG | p.Arg666Trp | Q13415 |
| HOMO SAPIENS | 4 | 11088 | 5uj7 | A | 720 | ARG | p.Arg720Gln | Q13415 |
| HOMO SAPIENS | 0 | 11088 | 5uj7 | A | 816 | MET | p.Met816Thr | Q13415 |
| HOMO SAPIENS | 1 | 4352 | 5va1 | A | 421 | THR | p.Thr421Met | Q12809 |
| HOMO SAPIENS | 0 | 4352 | 5va1 | A | 451 | PRO | p.Pro451Leu | Q12809 |
| HOMO SAPIENS | 0 | 4352 | 5va1 | A | 470 | ASN | p.Asn470Asp | Q12809 |
| HOMO SAPIENS | 0 | 4352 | 5va1 | A | 490 | ALA | p.Ala490Thr | Q12809 |
| HOMO SAPIENS | 0 | 4352 | 5va1 | A | 491 | VAL | p.Val491Ile | Q12809 |
| HOMO SAPIENS | 1 | 4352 | 5va1 | A | 533 | VAL | p.Val533Gly | Q12809 |
| HOMO SAPIENS | 0 | 4352 | 5va1 | A | 552 | LEU | p.Leu552Ser | Q12809 |
| HOMO SAPIENS | 0 | 4352 | 5va1 | A | 558 | ALA | p.Ala558Pro | Q12809 |
| HOMO SAPIENS | 1 | 4352 | 5va1 | A | 561 | ALA | p.Ala561Val | Q12809 |
| HOMO SAPIENS | 0 | 4352 | 5va1 | A | 572 | GLY | p.Gly572Arg | Q12809 |
| HOMO SAPIENS | -1 | 4352 | 5va1 | A | 582 | ARG | p.Arg582Cys | Q12809 |
| HOMO SAPIENS | 0 | 4352 | 5va1 | A | 584 | GLY | p.Gly584Ser | Q12809 |
| HOMO SAPIENS | 0 | 4352 | 5va1 | A | 588 | ASN | p.Asn588Lys | Q12809 |
| HOMO SAPIENS | 0 | 4352 | 5va1 | A | 593 | ILE | p.Ile593Arg | Q12809 |
| HOMO SAPIENS | 1 | 4352 | 5va1 | A | 627 | PHE | p.Phe627Leu | Q12809 |
| HOMO SAPIENS | 0 | 4352 | 5va1 | A | 630 | VAL | p.Val630Leu | Q12809 |
| HOMO SAPIENS | 0 | 4352 | 5va1 | A | 652 | TYR | p.Tyr652Ter | Q12809 |
| HOMO SAPIENS | 0 | 4352 | 5va1 | A | 711 | ILE | p.Ile711Val | Q12809 |
| HOMO SAPIENS | 0 | 4352 | 5va1 | A | 735 | SER | p.Ser735Leu | Q12809 |
| HOMO SAPIENS | 0 | 4352 | 5va1 | A | 744 | ARG | p.Arg744Ter | Q12809 |
| HOMO SAPIENS | 0 | 4352 | 5va1 | A | 784 | ARG | p.Arg784Trp | Q12809 |
| HOMO SAPIENS | 1 | 4352 | 5va1 | A | 820 | GLY | p.Gly820Arg | Q12809 |
| HOMO SAPIENS | 5 | 4352 | 5va1 | A | 822 | VAL | p.Val822Met | Q12809 |
| HOMO SAPIENS | 1 | 4352 | 5va1 | A | 863 | ARG | p.Arg863Ter | Q12809 |
| HOMO SAPIENS | 5 | 303 | 2obv | A | 55 | ALA | p.Ala55Asp | Q00266 |
| HOMO SAPIENS | 7 | 303 | 2obv | A | 259 | ALA | p.Ala259Gly | Q00266 |
| HOMO SAPIENS | 9 | 303 | 2obv | A | 264 | ARG | p.Arg264His | Q00266 |
| HOMO SAPIENS | 9 | 303 | 2obv | A | 264 | ARG | p.Arg264Cys | Q00266 |
| HOMO SAPIENS | 0 | 303 | 2obv | A | 305 | LEU | p.Leu305Pro | Q00266 |

| | | | | | | | | |
|---|---|---|---|---|---|---|---|---|
| HOMO SAPIENS | 7 | 303 | 2obv | A | 322 | ILE | p.Ile322Met | Q00266 |
| HOMO SAPIENS | 0 | 303 | 2obv | A | 336 | GLY | p.Gly336Arg | Q00266 |
| HOMO SAPIENS | 0 | 303 | 2obv | A | 357 | PRO | p.Pro357Leu | Q00266 |
| HOMO SAPIENS | 0 | 2077 | 4zxb | E | 462 | ASN | p.Asn489Ser | P06213 |
| HOMO SAPIENS | -1 | 2077 | 4zxb | E | 519 | ASP | p.Asp546Glu | P06213 |
| HOMO SAPIENS | 2 | 2077 | 4zxb | E | 819 | THR | p.Thr858Ala | P06213 |
| HOMO SAPIENS | 0 | 2077 | 4zxb | E | 907 | ASP | p.Asp946Glu | P06213 |
| HOMO SAPIENS | 7 | 3274 | 1wsv | B | 14 | HIS | p.His42Arg | P48728 |
| HOMO SAPIENS | 0 | 3274 | 1wsv | B | 19 | GLY | p.Gly47Arg | P48728 |
| HOMO SAPIENS | 4 | 3274 | 1wsv | B | 45 | ARG | p.Arg73Cys | P48728 |
| HOMO SAPIENS | 1 | 3274 | 1wsv | B | 49 | SER | p.Ser77Leu | P48728 |
| HOMO SAPIENS | 3 | 3274 | 1wsv | B | 241 | GLY | p.Gly269Asp | P48728 |
| HOMO SAPIENS | 7 | 3274 | 1wsv | B | 248 | ASP | p.Asp276His | P48728 |
| HOMO SAPIENS | 0 | 3274 | 1wsv | B | 272 | MET | p.Met300Val | P48728 |
| HOMO SAPIENS | 4 | 3274 | 1wsv | B | 292 | ARG | p.Arg320His | P48728 |
| HOMO SAPIENS | 0 | 3274 | 1wsv | B | 354 | ARG | p.Arg382Gln | P48728 |
| HOMO SAPIENS | -1 | 435 | 3uyt | A | 44 | THR | p.Thr44Ala | P48730 |
| HOMO SAPIENS | 1 | 4698 | 5ife | C | 666 | LYS | p.Lys666Asn | O75533 |
| HOMO SAPIENS | 1 | 16917 | 1apy | C | 37 | GLY | p.Gly60Asp | P20933 |
| HOMO SAPIENS | 0 | 16917 | 1apy | C | 49 | SER | p.Ser72Pro | P20933 |
| HOMO SAPIENS | 0 | 16917 | 1apy | C | 84 | ARG | p.Arg107Ter | P20933 |
| HOMO SAPIENS | 0 | 16917 | 1apy | C | 126 | SER | p.Thr149Ser | P20933 |
| HOMO SAPIENS | 0 | 16917 | 1apy | C | 135 | TRP | p.Trp158Ter | P20933 |
| HOMO SAPIENS | 0 | 16917 | 1apy | C | 140 | CYS | p.Cys163Ser | P20933 |
| HOMO SAPIENS | 0 | 16917 | 1apy | C | 145 | TRP | p.Trp168Ter | P20933 |
| HOMO SAPIENS | 0 | 10413 | 4frw | A | 185 | THR | p.Thr185Met | Q96NY8 |
| HOMO SAPIENS | 0 | 5217 | 5n6w | G | 96 | TRP | p.Trp96Arg | O15537 |
| HOMO SAPIENS | 1 | 5217 | 5n6w | G | 102 | ARG | p.Arg102Trp | O15537 |
| HOMO SAPIENS | 0 | 5217 | 5n6w | G | 109 | GLY | p.Gly109Arg | O15537 |
| HOMO SAPIENS | -1 | 5217 | 5n6w | G | 223 | CYS | p.Cys223Arg | O15537 |
| HOMO SAPIENS | 0 | 9485 | 4z6y | E | 138 | ALA | p.Ala138Asp | Q9P0N9 |
| HOMO SAPIENS | 3 | 4504 | 4mwt | B | 411 | GLY | p.Gly439Ser | P10619 |
| HOMO SAPIENS | 3 | 4504 | 4mwt | B | 412 | PHE | p.Phe440Val | P10619 |
| HOMO SAPIENS | 6 | 6391 | 4kxy | A | 318 | ARG | p.Arg318Cys | P29401 |
| HOMO SAPIENS | 6 | 19546 | 2bx6 | A | 87 | THR | p.Thr87Ile | O75695 |
| HOMO SAPIENS | 1 | 19546 | 2bx6 | A | 118 | ARG | p.Arg118His | O75695 |
| HOMO SAPIENS | 1 | 19546 | 2bx6 | A | 201 | ILE | p.Ile201Val | O75695 |
| HOMO SAPIENS | 1 | 19546 | 2bx6 | A | 282 | ARG | p.Arg282Trp | O75695 |
| HOMO SAPIENS | 1 | 5265 | 5usr | H | 68 | ARG | p.Arg68Leu | Q9HD34 |
| HOMO SAPIENS | 2 | 1012 | 4nyi | S | 569 | LEU | p.Leu569Val | Q07889 |
| HOMO SAPIENS | 2 | 1012 | 4nyi | S | 655 | PRO | p.Pro655Leu | Q07889 |
| HOMO SAPIENS | 4 | 1012 | 4nyi | S | 663 | ILE | p.Ile663Met | Q07889 |
| HOMO SAPIENS | 2 | 1012 | 4nyi | S | 702 | TYR | p.Tyr702Cys | Q07889 |
| HOMO SAPIENS | 2 | 1012 | 4nyi | S | 708 | ALA | p.Ala708Thr | Q07889 |
| HOMO SAPIENS | 4 | 1012 | 4nyi | S | 733 | ILE | p.Ile733Phe | Q07889 |
| HOMO SAPIENS | 1 | 1848 | 3g1n | A | 4013 | ARG | p.Arg4013Trp | Q7Z6Z7 |
| HOMO SAPIENS | 1 | 1848 | 3g1n | A | 4187 | ARG | p.Arg4187Cys | Q7Z6Z7 |
| HOMO SAPIENS | 1 | 77 | 4pod | A | 41 | CYS | p.Cys79Tyr | P60174 |
| HOMO SAPIENS | 9 | 77 | 4pod | A | 104 | GLU | p.Glu142Asp | P60174 |
| HOMO SAPIENS | 1 | 77 | 4pod | A | 122 | GLY | p.Gly160Arg | P60174 |
| HOMO SAPIENS | -1 | 8672 | 3thz | A | 8 | THR | p.Thr8Met | P43246 |
| HOMO SAPIENS | 0 | 8672 | 3thz | A | 19 | PHE | p.Phe19Leu | P43246 |
| HOMO SAPIENS | 0 | 8672 | 3thz | A | 23 | PHE | p.Phe23Leu | P43246 |
| HOMO SAPIENS | 0 | 8672 | 3thz | A | 96 | ARG | p.Arg96His | P43246 |

| | | | | | | | | |
|---|---|---|---|---|---|---|---|---|
| HOMO SAPIENS | 0 | 8672 | 3thz | A | 102 | VAL | p.Val102Ile | P43246 |
| HOMO SAPIENS | 0 | 8672 | 3thz | A | 106 | ARG | p.Arg106Lys | P43246 |
| HOMO SAPIENS | 0 | 8672 | 3thz | A | 127 | ASN | p.Asn127Ser | P43246 |
| HOMO SAPIENS | 0 | 8672 | 3thz | A | 128 | LEU | p.Leu128Val | P43246 |
| HOMO SAPIENS | 0 | 8672 | 3thz | A | 163 | VAL | p.Val163Gly | P43246 |
| HOMO SAPIENS | 0 | 8672 | 3thz | A | 169 | ILE | p.Ile169Val | P43246 |
| HOMO SAPIENS | 0 | 8672 | 3thz | A | 315 | GLY | p.Gly315Asp | P43246 |
| HOMO SAPIENS | -1 | 8672 | 3thz | A | 322 | GLY | p.Gly322Asp | P43246 |
| HOMO SAPIENS | 0 | 8672 | 3thz | A | 331 | ASN | p.Asn331Asp | P43246 |
| HOMO SAPIENS | 1 | 8672 | 3thz | A | 390 | LEU | p.Leu390Phe | P43246 |
| HOMO SAPIENS | 0 | 8672 | 3thz | A | 419 | GLN | p.Gln419Lys | P43246 |
| HOMO SAPIENS | 0 | 8672 | 3thz | A | 447 | PHE | p.Phe447Leu | P43246 |
| HOMO SAPIENS | 0 | 8672 | 3thz | A | 524 | ARG | p.Arg524Pro | P43246 |
| HOMO SAPIENS | 0 | 8672 | 3thz | A | 539 | PHE | p.Phe539Leu | P43246 |
| HOMO SAPIENS | -1 | 8672 | 3thz | A | 546 | LYS | p.Lys546Asn | P43246 |
| HOMO SAPIENS | 0 | 8672 | 3thz | A | 564 | THR | p.Thr564Ala | P43246 |
| HOMO SAPIENS | 0 | 8672 | 3thz | A | 577 | ILE | p.Ile577Thr | P43246 |
| HOMO SAPIENS | 0 | 8672 | 3thz | A | 588 | TYR | p.Tyr588Ter | P43246 |
| HOMO SAPIENS | 0 | 8672 | 3thz | A | 622 | PRO | p.Pro622Leu | P43246 |
| HOMO SAPIENS | 0 | 8672 | 3thz | A | 629 | GLN | p.Gln629Arg | P43246 |
| HOMO SAPIENS | 0 | 8672 | 3thz | A | 636 | ALA | p.Ala636Pro | P43246 |
| HOMO SAPIENS | 2 | 8672 | 3thz | A | 642 | VAL | p.Val642Ile | P43246 |
| HOMO SAPIENS | 1 | 8672 | 3thz | A | 654 | ASP | p.Asp654Glu | P43246 |
| HOMO SAPIENS | 0 | 8672 | 3thz | A | 662 | GLN | p.Gln662His | P43246 |
| HOMO SAPIENS | 2 | 8672 | 3thz | A | 681 | GLN | p.Gln681His | P43246 |
| HOMO SAPIENS | 0 | 8672 | 3thz | A | 707 | CYS | p.Cys707Ser | P43246 |
| HOMO SAPIENS | 0 | 8672 | 3thz | A | 809 | GLU | p.Glu809Lys | P43246 |
| HOMO SAPIENS | 0 | 8672 | 3thz | A | 813 | MET | p.Met813Val | P43246 |
| HOMO SAPIENS | 1 | 8672 | 3thz | A | 834 | ALA | p.Ala834Thr | P43246 |
| HOMO SAPIENS | 0 | 8672 | 3thz | A | 849 | LEU | p.Leu849Val | P43246 |
| HOMO SAPIENS | -1 | 8672 | 3thz | A | 860 | SER | p.Ser860Leu | P43246 |
| HOMO SAPIENS | -1 | 8672 | 3thz | A | 934 | THR | p.Thr934Lys | P43246 |
| HOMO SAPIENS | 2 | 292 | 1i3k | A | 34 | ASN | p.Asn34Ser | Q14376 |
| HOMO SAPIENS | 7 | 292 | 1i3k | A | 90 | GLY | p.Gly90Glu | Q14376 |
| HOMO SAPIENS | 2 | 292 | 1i3k | A | 103 | ASP | p.Asp103Gly | Q14376 |
| HOMO SAPIENS | 2 | 292 | 1i3k | A | 136 | TYR | p.Tyr136Ter | Q14376 |
| HOMO SAPIENS | 0 | 292 | 1i3k | A | 169 | ARG | p.Arg169Trp | Q14376 |
| HOMO SAPIENS | 0 | 292 | 1i3k | A | 183 | LEU | p.Leu183Pro | Q14376 |
| HOMO SAPIENS | 0 | 292 | 1i3k | A | 302 | GLY | p.Gly302Asp | Q14376 |
| HOMO SAPIENS | 0 | 292 | 1i3k | A | 313 | LEU | p.Leu313Met | Q14376 |
| HOMO SAPIENS | 1 | 2084 | 5vc8 | A | 295 | GLN | p.Gln295Leu | O96028 |
| HOMO SAPIENS | -1 | 10157 | 5wrw | A | 7 | GLY | p.Gly7Ala | O76094 |
| HOMO SAPIENS | -1 | 10157 | 5wrw | A | 7 | GLY | p.Gly7Arg | O76094 |
| HOMO SAPIENS | -1 | 10157 | 5wrw | A | 8 | GLY | p.Gly8Ala | O76094 |
| HOMO SAPIENS | 0 | 3113 | 3in5 | A | 29 | GLU | p.Glu29Lys | Q9UBT6 |
| HOMO SAPIENS | 5 | 4546 | 5in3 | B | 44 | VAL | p.Val44Leu | P07902 |
| HOMO SAPIENS | -1 | 4546 | 5in3 | B | 51 | ARG | p.Arg51Leu | P07902 |
| HOMO SAPIENS | -1 | 4546 | 5in3 | B | 62 | LEU | p.Leu62Met | P07902 |
| HOMO SAPIENS | 4 | 4546 | 5in3 | B | 66 | PRO | p.Pro66His | P07902 |
| HOMO SAPIENS | 2 | 4546 | 5in3 | B | 74 | LEU | p.Leu74Pro | P07902 |
| HOMO SAPIENS | 5 | 4546 | 5in3 | B | 97 | ASN | p.Asn97Ser | P07902 |
| HOMO SAPIENS | 1 | 4546 | 5in3 | B | 135 | SER | p.Ser135Leu | P07902 |
| HOMO SAPIENS | 5 | 4546 | 5in3 | B | 138 | THR | p.Thr138Met | P07902 |
| HOMO SAPIENS | 7 | 4546 | 5in3 | B | 139 | LEU | p.Leu139Pro | P07902 |

| | | | | | | | | |
|---|---|---|---|---|---|---|---|---|
| HOMO SAPIENS | 2 | 4546 | 5in3 | B | 142 | MET | p.Met142Lys | P07902 |
| HOMO SAPIENS | 1 | 4546 | 5in3 | B | 143 | SER | p.Ser143Leu | P07902 |
| HOMO SAPIENS | 2 | 4546 | 5in3 | B | 148 | ARG | p.Arg148Gly | P07902 |
| HOMO SAPIENS | 2 | 4546 | 5in3 | B | 148 | ARG | p.Arg148Gln | P07902 |
| HOMO SAPIENS | 5 | 4546 | 5in3 | B | 183 | PRO | p.Pro183Thr | P07902 |
| HOMO SAPIENS | 5 | 4546 | 5in3 | B | 185 | PRO | p.Pro185His | P07902 |
| HOMO SAPIENS | 5 | 4546 | 5in3 | B | 186 | HIS | p.His186Tyr | P07902 |
| HOMO SAPIENS | 7 | 4546 | 5in3 | B | 188 | GLN | p.Gln188Arg | P07902 |
| HOMO SAPIENS | 5 | 4546 | 5in3 | B | 195 | LEU | p.Leu195Pro | P07902 |
| HOMO SAPIENS | 4 | 4546 | 5in3 | B | 201 | ARG | p.Arg201His | P07902 |
| HOMO SAPIENS | 4 | 4546 | 5in3 | B | 201 | ARG | p.Arg201Cys | P07902 |
| HOMO SAPIENS | 4 | 4546 | 5in3 | B | 203 | GLU | p.Glu203Lys | P07902 |
| HOMO SAPIENS | 2 | 4546 | 5in3 | B | 209 | TYR | p.Tyr209Cys | P07902 |
| HOMO SAPIENS | 5 | 4546 | 5in3 | B | 223 | ARG | p.Arg223Ser | P07902 |
| HOMO SAPIENS | 5 | 4546 | 5in3 | B | 226 | LEU | p.Leu226Pro | P07902 |
| HOMO SAPIENS | 5 | 4546 | 5in3 | B | 226 | LEU | p.Leu226Val | P07902 |
| HOMO SAPIENS | 2 | 4546 | 5in3 | B | 227 | LEU | p.Leu227Pro | P07902 |
| HOMO SAPIENS | 8 | 4546 | 5in3 | B | 258 | ARG | p.Arg258Cys | P07902 |
| HOMO SAPIENS | 1 | 4546 | 5in3 | B | 265 | PRO | p.Pro265Ala | P07902 |
| HOMO SAPIENS | 8 | 4546 | 5in3 | B | 268 | THR | p.Thr268Asn | P07902 |
| HOMO SAPIENS | 7 | 4546 | 5in3 | B | 278 | ILE | p.Ile278Asn | P07902 |
| HOMO SAPIENS | 5 | 4546 | 5in3 | B | 285 | LYS | p.Lys285Asn | P07902 |
| HOMO SAPIENS | 1 | 4546 | 5in3 | B | 319 | HIS | p.His319Gln | P07902 |
| HOMO SAPIENS | 4 | 4546 | 5in3 | B | 321 | HIS | p.His321Tyr | P07902 |
| HOMO SAPIENS | 4 | 4546 | 5in3 | B | 325 | PRO | p.Pro325Leu | P07902 |
| HOMO SAPIENS | 2 | 4546 | 5in3 | B | 328 | ARG | p.Arg328His | P07902 |
| HOMO SAPIENS | 2 | 4546 | 5in3 | B | 329 | SER | p.Ser329Phe | P07902 |
| HOMO SAPIENS | 5 | 4546 | 5in3 | B | 334 | LYS | p.Lys334Arg | P07902 |
| HOMO SAPIENS | 6 | 1290 | 5uav | C | 119 | ARG | p.Arg119Gly | P32322 |
| HOMO SAPIENS | 6 | 1290 | 5uav | C | 119 | ARG | p.Arg119His | P32322 |
| HOMO SAPIENS | 3 | 1290 | 5uav | C | 251 | ARG | p.Arg251His | P32322 |
| HOMO SAPIENS | 7 | 1290 | 5uav | C | 266 | ARG | p.Arg266Gln | P32322 |
| HOMO SAPIENS | 0 | 9618 | 2r7e | A | 22 | GLY | p.Gly41Cys | P00451 |
| HOMO SAPIENS | 2 | 9618 | 2r7e | A | 113 | GLU | p.Glu132Asp | P00451 |
| HOMO SAPIENS | -1 | 9618 | 2r7e | A | 336 | ARG | p.Arg355Ter | P00451 |
| HOMO SAPIENS | 0 | 9618 | 2r7e | A | 372 | ARG | p.Arg391His | P00451 |
| HOMO SAPIENS | 0 | 9618 | 2r7e | A | 373 | SER | p.Ser392Leu | P00451 |
| HOMO SAPIENS | 0 | 9618 | 2r7e | A | 373 | SER | p.Ser392Pro | P00451 |
| HOMO SAPIENS | 3 | 9618 | 2r7e | A | 386 | ILE | p.Ile405Ser | P00451 |
| HOMO SAPIENS | 6 | 9618 | 2r7e | A | 390 | GLU | p.Glu409Gly | P00451 |
| HOMO SAPIENS | 2 | 9618 | 2r7e | A | 412 | LEU | p.Leu431Phe | P00451 |
| HOMO SAPIENS | 5 | 9618 | 2r7e | A | 425 | LYS | p.Lys444Arg | P00451 |
| HOMO SAPIENS | 0 | 9618 | 2r7e | A | 484 | ARG | p.Arg503His | P00451 |
| HOMO SAPIENS | 2 | 9618 | 2r7e | A | 531 | ARG | p.Arg550His | P00451 |
| HOMO SAPIENS | 3 | 9618 | 2r7e | A | 535 | SER | p.Ser554Gly | P00451 |
| HOMO SAPIENS | 2 | 9618 | 2r7e | A | 593 | ARG | p.Arg612Cys | P00451 |
| HOMO SAPIENS | 0 | 9618 | 2r7e | A | 704 | ALA | p.Ala723Thr | P00451 |
| HOMO SAPIENS | 0 | 9618 | 2r7e | A | 720 | GLU | p.Glu739Lys | P00451 |
| HOMO SAPIENS | 1 | 13085 | 3ud1 | A | 1075 | ILE | p.Thr1075Ile | P16157 |
| HOMO SAPIENS | -1 | 3289 | 2fy4 | A | 2 | ALA | p.Ala120Thr | P28329 |
| HOMO SAPIENS | 7 | 3289 | 2fy4 | A | 18 | VAL | p.Val136Met | P28329 |
| HOMO SAPIENS | 5 | 3289 | 2fy4 | A | 92 | LEU | p.Leu210Pro | P28329 |
| HOMO SAPIENS | 2 | 3289 | 2fy4 | A | 93 | PRO | p.Pro211Ala | P28329 |
| HOMO SAPIENS | 4 | 3289 | 2fy4 | A | 131 | LEU | p.Leu249Val | P28329 |

| | | | | | | | | |
|---|---|---|---|---|---|---|---|---|
| HOMO SAPIENS | 2 | 3289 | 2fy4 | A | 181 | PRO | p.Pro299Leu | P28329 |
| HOMO SAPIENS | 0 | 3289 | 2fy4 | A | 218 | ILE | p.Ile336Thr | P28329 |
| HOMO SAPIENS | 2 | 3289 | 2fy4 | A | 261 | ASP | p.Asp379His | P28329 |
| HOMO SAPIENS | 0 | 3289 | 2fy4 | A | 302 | ARG | p.Arg420Cys | P28329 |
| HOMO SAPIENS | 5 | 3289 | 2fy4 | A | 323 | GLU | p.Glu441Lys | P28329 |
| HOMO SAPIENS | 0 | 3289 | 2fy4 | A | 343 | MET | p.Val461Met | P28329 |
| HOMO SAPIENS | 2 | 3289 | 2fy4 | A | 364 | ARG | p.Arg482Gly | P28329 |
| HOMO SAPIENS | 3 | 3289 | 2fy4 | A | 380 | SER | p.Ser498Leu | P28329 |
| HOMO SAPIENS | 6 | 3289 | 2fy4 | A | 388 | VAL | p.Val506Leu | P28329 |
| HOMO SAPIENS | 7 | 3289 | 2fy4 | A | 429 | HIS | p.His547Gln | P28329 |
| HOMO SAPIENS | 8 | 3289 | 2fy4 | A | 442 | ARG | p.Arg560His | P28329 |
| HOMO SAPIENS | 3 | 3289 | 2fy4 | A | 443 | ARG | p.Arg561Gln | P28329 |
| HOMO SAPIENS | 0 | 4137 | 3hhr | B | 38 | CYS | p.Cys56Ter | P10912 |
| HOMO SAPIENS | 0 | 4137 | 3hhr | B | 43 | ARG | p.Arg61Ter | P10912 |
| HOMO SAPIENS | 0 | 4137 | 3hhr | B | 44 | GLU | p.Glu62Lys | P10912 |
| HOMO SAPIENS | 0 | 4137 | 3hhr | B | 96 | PHE | p.Phe114Ser | P10912 |
| HOMO SAPIENS | 5 | 4137 | 3hhr | B | 131 | PRO | p.Pro149Gln | P10912 |
| HOMO SAPIENS | 0 | 4137 | 3hhr | B | 144 | VAL | p.Val162Ile | P10912 |
| HOMO SAPIENS | 0 | 4137 | 3hhr | B | 150 | HIS | p.His168Gln | P10912 |
| HOMO SAPIENS | 0 | 4137 | 3hhr | B | 152 | ASP | p.Asp170His | P10912 |
| HOMO SAPIENS | 0 | 4137 | 3hhr | B | 153 | ILE | p.Ile171Thr | P10912 |
| HOMO SAPIENS | 0 | 4137 | 3hhr | B | 154 | GLN | p.Gln172Pro | P10912 |
| HOMO SAPIENS | 0 | 4137 | 3hhr | B | 155 | VAL | p.Val173Gly | P10912 |
| HOMO SAPIENS | 0 | 4137 | 3hhr | B | 217 | ARG | p.Arg235Ter | P10912 |
| HOMO SAPIENS | 5 | 4137 | 3hhr | B | 224 | GLU | p.Glu242Asp | P10912 |
| HOMO SAPIENS | 5 | 4137 | 3hhr | B | 224 | GLU | p.Glu242Ter | P10912 |
| HOMO SAPIENS | 6 | 2493 | 5bpb | A | 69 | HIS | p.His69Tyr | Q9ULV1 |
| HOMO SAPIENS | 6 | 2493 | 5bpb | A | 105 | MET | p.Met105Val | Q9ULV1 |
| HOMO SAPIENS | -1 | 2493 | 5bpb | A | 168 | PRO | p.Pro168Ser | Q9ULV1 |
| HOMO SAPIENS | 0 | 32123 | 4ccz | A | 314 | ASP | p.Asp314Asn | Q99707 |
| HOMO SAPIENS | 0 | 32123 | 4ccz | A | 345 | VAL | p.Val345Phe | Q99707 |
| HOMO SAPIENS | 1 | 32123 | 4ccz | A | 585 | ARG | p.Arg585Ter | Q99707 |
| HOMO SAPIENS | 4 | 4618 | 2ag4 | A | 26 | PRO | p.Pro55Leu | P17900 |
| HOMO SAPIENS | 4 | 4618 | 2ag4 | A | 30 | VAL | p.Ile59Val | P17900 |
| HOMO SAPIENS | 2 | 4618 | 2ag4 | A | 40 | VAL | p.Met69Val | P17900 |
| HOMO SAPIENS | 2 | 4618 | 2ag4 | A | 124 | VAL | p.Val153Ala | P17900 |
| HOMO SAPIENS | 8 | 4618 | 2ag4 | A | 140 | ARG | p.Arg169Pro | P17900 |
| HOMO SAPIENS | 5 | 1082 | 1hk0 | X | 14 | ARG | p.Arg15Cys | P07320 |
| HOMO SAPIENS | 7 | 1082 | 1hk0 | X | 23 | PRO | p.Pro24Thr | P07320 |
| HOMO SAPIENS | 6 | 1082 | 1hk0 | X | 36 | ARG | p.Arg37Ser | P07320 |
| HOMO SAPIENS | 5 | 1082 | 1hk0 | X | 55 | TYR | p.Tyr56Ter | P07320 |
| HOMO SAPIENS | 3 | 1082 | 1hk0 | X | 58 | ARG | p.Arg59His | P07320 |
| HOMO SAPIENS | 7 | 1082 | 1hk0 | X | 88 | HIS | p.His88Gln | P07320 |
| HOMO SAPIENS | 6 | 1082 | 1hk0 | X | 95 | ARG | p.Arg95Ser | P07320 |
| HOMO SAPIENS | 5 | 1082 | 1hk0 | X | 126 | VAL | p.Val126Met | P07320 |
| HOMO SAPIENS | 8 | 1082 | 1hk0 | X | 157 | TRP | p.Trp157Ter | P07320 |
| HOMO SAPIENS | 0 | 22879 | 3j0a | B | 644 | ILE | p.Ile644Phe | O60602 |
| HOMO SAPIENS | 8 | 6 | 2o23 | B | 122 | LEU | p.Leu122Val | Q99714 |
| HOMO SAPIENS | 1 | 6 | 2o23 | B | 130 | ARG | p.Arg130Cys | Q99714 |
| HOMO SAPIENS | 1 | 6 | 2o23 | B | 247 | ASN | p.Asn247Ser | Q99714 |
| HOMO SAPIENS | 0 | 6 | 2o23 | B | 249 | GLU | p.Glu249Gln | Q99714 |
| HOMO SAPIENS | 1 | 12910 | 4hl4 | A | 67 | ARG | p.Arg67Ter | Q96BZ9 |
| HOMO SAPIENS | 8 | 16105 | 1lmj | A | 7 | GLU | p.Glu1073Lys | P35555 |
| HOMO SAPIENS | 8 | 16105 | 1lmj | A | 8 | CYS | p.Cys1074Arg | P35555 |

| | | | | | | | | |
|---|---|---|---|---|---|---|---|---|
| HOMO SAPIENS | 9 | 16105 | 1lmj | A | 51 | CYS | p.Cys1117Tyr | P35555 |
| HOMO SAPIENS | 9 | 16105 | 1lmj | A | 63 | CYS | p.Cys1129Tyr | P35555 |
| HOMO SAPIENS | 9 | 16105 | 1lmj | A | 65 | ASN | p.Asn1131Tyr | P35555 |
| HOMO SAPIENS | 4 | 16105 | 1lmj | A | 71 | ARG | p.Arg1137Pro | P35555 |
| HOMO SAPIENS | 1 | 16105 | 1lmj | A | 82 | PRO | p.Pro1148Ala | P35555 |
| HOMO SAPIENS | 0 | 4795 | 3s5f | A | 106 | LEU | p.Leu106Ser | Q16595 |
| HOMO SAPIENS | 0 | 4795 | 3s5f | A | 130 | GLY | p.Gly130Val | Q16595 |
| HOMO SAPIENS | 0 | 4795 | 3s5f | A | 154 | ILE | p.Ile154Phe | Q16595 |
| HOMO SAPIENS | -1 | 4795 | 3s5f | A | 209 | ASP | p.Asp209Gly | Q16595 |
| HOMO SAPIENS | -1 | 2884 | 4lli | A | 1655 | ASP | p.Asp1655Glu | Q9Y4I1 |
| HOMO SAPIENS | 0 | 1786 | 7pck | C | 56P | TYR | p.Tyr71Ter | P43235 |
| HOMO SAPIENS | 0 | 1786 | 7pck | C | 64P | GLY | p.Gly79Glu | P43235 |
| HOMO SAPIENS | 7 | 38286 | 2l53 | B | 1914 | ARG | p.Arg1914Gly | Q14524 |
| HOMO SAPIENS | 1 | 38286 | 2l53 | B | 1924 | ALA | p.Ala1924Thr | Q14524 |
| HOMO SAPIENS | 0 | 31747 | 3ksy | A | 25 | PRO | p.Pro25Ser | Q07889 |
| HOMO SAPIENS | 3 | 31747 | 3ksy | A | 37 | THR | p.Thr37Ala | Q07889 |
| HOMO SAPIENS | 0 | 31747 | 3ksy | A | 54 | GLN | p.Gln54His | Q07889 |
| HOMO SAPIENS | -1 | 31747 | 3ksy | A | 190 | ASP | p.Asp190Glu | Q07889 |
| HOMO SAPIENS | 2 | 31747 | 3ksy | A | 266 | THR | p.Thr266Lys | Q07889 |
| HOMO SAPIENS | 1 | 31747 | 3ksy | A | 269 | MET | p.Met269Arg | Q07889 |
| HOMO SAPIENS | 1 | 31747 | 3ksy | A | 340 | PRO | p.Pro340Ser | Q07889 |
| HOMO SAPIENS | 1 | 31747 | 3ksy | A | 423 | ASN | p.Asn423Lys | Q07889 |
| HOMO SAPIENS | 1 | 31747 | 3ksy | A | 424 | GLU | p.Glu424Lys | Q07889 |
| HOMO SAPIENS | 1 | 31747 | 3ksy | A | 432 | TRP | p.Trp432Arg | Q07889 |
| HOMO SAPIENS | 1 | 31747 | 3ksy | A | 552 | ARG | p.Arg552Gly | Q07889 |
| HOMO SAPIENS | 1 | 31747 | 3ksy | A | 552 | ARG | p.Arg552Ser | Q07889 |
| HOMO SAPIENS | 3 | 7529 | 6jdw | A | 93 | ILE | p.Ile93Met | P50440 |
| HOMO SAPIENS | 2 | 7529 | 6jdw | A | 110 | GLN | p.Gln110His | P50440 |
| HOMO SAPIENS | 5 | 7529 | 6jdw | A | 149 | TRP | p.Trp149Ter | P50440 |
| HOMO SAPIENS | 0 | 7529 | 6jdw | A | 169 | ARG | p.Arg169Ter | P50440 |
| HOMO SAPIENS | 0 | 7529 | 6jdw | A | 260 | ARG | p.Arg260Ter | P50440 |
| HOMO SAPIENS | 3 | 7529 | 6jdw | A | 344 | ILE | p.Ile344Val | P50440 |
| HOMO SAPIENS | 5 | 7529 | 6jdw | A | 418 | LEU | p.Leu418Ile | P50440 |
| HOMO SAPIENS | 1 | 14788 | 1eak | A | 101 | ARG | p.Arg101His | P08253 |
| HOMO SAPIENS | 0 | 14788 | 1eak | A | 383 | ASP | p.Asp383Glu | P08253 |
| HOMO SAPIENS | 7 | 14788 | 1eak | A | 404 | GLN | p.Glu404Lys | P08253 |
| HOMO SAPIENS | 2 | 14788 | 1eak | A | 447 | ALA | p.Ala447Val | P08253 |
| HOMO SAPIENS | 0 | 1255 | 5xtc | p | 8 | PRO | p.Pro8His | Q9Y6M9 |
| HOMO SAPIENS | 0 | 1255 | 5xtc | p | 146 | PRO | p.Pro146Ser | Q9Y6M9 |
| HOMO SAPIENS | 3 | 6721 | 1a22 | A | 25 | PHE | p.Phe51Tyr | P01241 |
| HOMO SAPIENS | 2 | 6721 | 1a22 | A | 95 | SER | p.Ser121Arg | P01241 |
| HOMO SAPIENS | 2 | 6721 | 1a22 | A | 110 | VAL | p.Val136Ile | P01241 |
| HOMO SAPIENS | 0 | 1380 | 5i6z | A | 605 | LYS | p.Lys605Asn | P31645 |
| HOMO SAPIENS | 0 | 941 | 1qki | D | 48 | ILE | p.Ile48Thr | P11413 |
| HOMO SAPIENS | 3 | 941 | 1qki | D | 68 | VAL | p.Val68Met | P11413 |
| HOMO SAPIENS | 2 | 941 | 1qki | D | 126 | ASN | p.Asn126Asp | P11413 |
| HOMO SAPIENS | 8 | 941 | 1qki | D | 163 | GLY | p.Gly163Ser | P11413 |
| HOMO SAPIENS | 6 | 941 | 1qki | D | 165 | ASN | p.Asn165Asp | P11413 |
| HOMO SAPIENS | 4 | 941 | 1qki | D | 181 | ASP | p.Asp181Val | P11413 |
| HOMO SAPIENS | 4 | 941 | 1qki | D | 182 | ARG | p.Arg182Trp | P11413 |
| HOMO SAPIENS | 5 | 941 | 1qki | D | 188 | SER | p.Ser188Phe | P11413 |
| HOMO SAPIENS | 9 | 941 | 1qki | D | 198 | ARG | p.Arg198Cys | P11413 |
| HOMO SAPIENS | 8 | 941 | 1qki | D | 213 | VAL | p.Val213Leu | P11413 |
| HOMO SAPIENS | 5 | 941 | 1qki | D | 282 | ASP | p.Asp282His | P11413 |

| | | | | | | | | |
|---|---|---|---|---|---|---|---|---|
| HOMO SAPIENS | 7 | 941 | 1qki | D | 285 | ARG | p.Arg285His | P11413 |
| HOMO SAPIENS | 5 | 941 | 1qki | D | 291 | VAL | p.Val291Met | P11413 |
| HOMO SAPIENS | 8 | 941 | 1qki | D | 323 | LEU | p.Leu323Pro | P11413 |
| HOMO SAPIENS | 3 | 941 | 1qki | D | 335 | ALA | p.Ala335Thr | P11413 |
| HOMO SAPIENS | 6 | 941 | 1qki | D | 350 | ASP | p.Asp350His | P11413 |
| HOMO SAPIENS | 5 | 941 | 1qki | D | 393 | ARG | p.Arg393His | P11413 |
| HOMO SAPIENS | 5 | 941 | 1qki | D | 454 | ARG | p.Arg454Cys | P11413 |
| HOMO SAPIENS | 2 | 941 | 1qki | D | 459 | LEU | p.Arg459Leu | P11413 |
| HOMO SAPIENS | 4 | 941 | 1qki | D | 463 | ARG | p.Arg463His | P11413 |
| HOMO SAPIENS | 4 | 2483 | 1tij | A | 68 | LEU | p.Leu94Gln | P01034 |
| HOMO SAPIENS | 2 | 20632 | 3v5b | A | 331 | ARG | p.Arg331Pro | P02545 |
| HOMO SAPIENS | 2 | 20632 | 3v5b | A | 374 | HIS | p.His374Gln | P02545 |
| HOMO SAPIENS | 4 | 13929 | 5xtd | J | 75 | ARG | p.Arg75His | Q16795 |
| HOMO SAPIENS | 3 | 13929 | 5xtd | J | 243 | VAL | p.Val243Ala | Q16795 |
| HOMO SAPIENS | 1 | 13929 | 5xtd | J | 294 | PRO | p.Pro294Leu | Q16795 |
| HOMO SAPIENS | 2 | 13929 | 5xtd | J | 321 | ARG | p.Arg321Pro | Q16795 |
| HOMO SAPIENS | 0 | 9411 | 5thl | A | 41 | ARG | p.Gly41Arg | P54577 |
| HOMO SAPIENS | 0 | 9411 | 5thl | A | 196 | GLU | p.Glu196Lys | P54577 |
| HOMO SAPIENS | 0 | 9411 | 5thl | A | 316 | ALA | p.Ala316Ser | P54577 |
| HOMO SAPIENS | 0 | 9411 | 5thl | A | 339 | ALA | p.Ala339Thr | P54577 |
| HOMO SAPIENS | 4 | 16273 | 2e8n | A | 46 | THR | p.Thr940Ile | P29317 |
| HOMO SAPIENS | 7 | 16273 | 2e8n | A | 54 | GLY | p.Gly948Trp | P29317 |
| HOMO SAPIENS | 6 | 16273 | 2e8n | A | 65 | ALA | p.Ala959Thr | P29317 |
| HOMO SAPIENS | 7 | 30387 | 2k9y | A | 541 | VAL | p.Val541Met | P29317 |
| HOMO SAPIENS | 0 | 6961 | 4nha | A | 843 | ILE | p.Ile853Val | Q9UPY3 |
| HOMO SAPIENS | -1 | 6961 | 4nha | A | 862 | ALA | p.Ala872Thr | Q9UPY3 |
| HOMO SAPIENS | 3 | 15708 | 4pzg | A | 16 | ARG | p.Arg16Ter | Q9Y5X0 |
| HOMO SAPIENS | 4 | 15708 | 4pzg | A | 51 | ARG | p.Arg51Gln | Q9Y5X0 |
| HOMO SAPIENS | 3 | 30166 | 4o00 | A | 44 | ARG | p.Arg14257Ter | Q8WZ42 |
| HOMO SAPIENS | 2 | 30166 | 4o00 | A | 54 | ARG | p.Arg14267His | Q8WZ42 |
| HOMO SAPIENS | 8 | 11158 | 4hca | A | 275 | TRP | p.Trp274Arg | P23771 |
| HOMO SAPIENS | 2 | 4328 | 4e0s | A | 1476 | ARG | p.Arg1476Ter | P01031 |
| HOMO SAPIENS | 0 | 11462 | 1dx5 | I | 455 | ALA | p.Ala473Val | P07204 |
| HOMO SAPIENS | -1 | 11150 | 2fic | A | 45 | LYS | p.Lys35Asn | O00499 |
| HOMO SAPIENS | 6 | 11150 | 2fic | A | 161 | ASP | p.Asp151Asn | O00499 |
| HOMO SAPIENS | 8 | 27382 | 4mqv | B | 17 | ARG | p.Arg17Ter | Q9NZC9 |
| HOMO SAPIENS | -1 | 37991 | 1q0p | A | 231 | GLN | p.Gln256Ter | P00751 |
| HOMO SAPIENS | 1 | 443 | 5xte | U | 215 | LEU | p.Leu215Phe | P08574 |
| HOMO SAPIENS | 8 | 21471 | 5y4e | A | 1461 | ASN | p.Thr692Ala | Q01484 |
| HOMO SAPIENS | -1 | 35443 | 3um3 | B | 129 | ASP | p.Asp129Val | Q9H444 |
| HOMO SAPIENS | -1 | 35443 | 3um3 | B | 161 | GLU | p.Glu161Lys | Q9H444 |
| HOMO SAPIENS | 6 | 28781 | 2n2a | B | 111 | ILE | p.Ile654Val | P04626 |
| HOMO SAPIENS | 7 | 28781 | 2n2a | B | 112 | ILE | p.Ile655Val | P04626 |
| HOMO SAPIENS | -1 | 3346 | 4ztz | C | 61 | GLY | p.Gly61Arg | Q9UHN1 |
| HOMO SAPIENS | 4 | 3346 | 4ztz | C | 103 | GLY | p.Gly103Ser | Q9UHN1 |
| HOMO SAPIENS | 4 | 3346 | 4ztz | C | 133 | HIS | p.His133Tyr | Q9UHN1 |
| HOMO SAPIENS | -1 | 3346 | 4ztz | C | 153 | LEU | p.Leu153Val | Q9UHN1 |
| HOMO SAPIENS | -1 | 3346 | 4ztz | C | 169 | ALA | p.Ala169Thr | Q9UHN1 |
| HOMO SAPIENS | 7 | 3346 | 4ztz | C | 369 | ARG | p.Arg369Gly | Q9UHN1 |
| HOMO SAPIENS | 4 | 3346 | 4ztz | C | 386 | ASP | p.Asp386Glu | Q9UHN1 |
| HOMO SAPIENS | 4 | 3346 | 4ztz | C | 451 | GLY | p.Gly451Glu | Q9UHN1 |
| HOMO SAPIENS | 4 | 2131 | 5fo9 | E | 1549 | LEU | p.Leu1549Met | P01024 |
| HOMO SAPIENS | 2 | 2131 | 5fo9 | E | 1619 | SER | p.Ser1619Arg | P01024 |
| HOMO SAPIENS | 3 | 12662 | 4jhp | C | 75 | ILE | p.Ile75Val | Q92834 |

| | | | | | | | | |
|---|---|---|---|---|---|---|---|---|
| HOMO SAPIENS | 1 | 12662 | 4jhp | C | 173 | GLY | p.Gly173Arg | Q92834 |
| HOMO SAPIENS | 0 | 12662 | 4jhp | C | 184 | GLN | p.Gln184His | Q92834 |
| HOMO SAPIENS | 3 | 12662 | 4jhp | C | 262 | ALA | p.Ala262Gly | Q92834 |
| HOMO SAPIENS | 3 | 12662 | 4jhp | C | 302 | CYS | p.Cys302Tyr | Q92834 |
| HOMO SAPIENS | 5 | 1093 | 2jg9 | A | 186 | GLN | p.Gln208Ter | P02745 |
| HOMO SAPIENS | 0 | 13181 | 4r4l | A | 12 | MET | p.Met13Ile | Q13976 |
| HOMO SAPIENS | 8 | 25939 | 4iyp | A | 106 | ARG | p.Arg106Gln | P78318 |
| HOMO SAPIENS | 1 | 217 | 2xsx | A | 71 | SER | p.Asn71Ser | P13929 |
| HOMO SAPIENS | 0 | 217 | 2xsx | A | 76 | ALA | p.Ala76Pro | P13929 |
| HOMO SAPIENS | 1 | 217 | 2xsx | A | 85 | ALA | p.Val85Ala | P13929 |
| HOMO SAPIENS | 8 | 217 | 2xsx | A | 156 | GLY | p.Gly156Asp | P13929 |
| HOMO SAPIENS | 5 | 217 | 2xsx | A | 332 | VAL | p.Val332Ile | P13929 |
| HOMO SAPIENS | 7 | 217 | 2xsx | A | 374 | GLY | p.Gly374Glu | P13929 |
| HOMO SAPIENS | 1 | 5508 | 3szm | A | 682 | THR | p.Thr682Asn | Q8NEM0 |
| HOMO SAPIENS | 0 | 5508 | 3szm | A | 715 | TRP | p.Trp715Ter | Q8NEM0 |
| HOMO SAPIENS | 1 | 5508 | 3szm | A | 761 | VAL | p.Ala761Val | Q8NEM0 |
| HOMO SAPIENS | 1 | 5508 | 3szm | A | 828 | PRO | p.Pro828Ser | Q8NEM0 |
| HOMO SAPIENS | 4 | 5508 | 3szm | A | 829 | GLU | p.Glu829Asp | Q8NEM0 |
| HOMO SAPIENS | 2 | 16353 | 2q0z | X | 171 | GLY | p.Gly1978Ala | O75643 |
| HOMO SAPIENS | 2 | 16353 | 2q0z | X | 188 | ALA | p.Ala1995Thr | O75643 |
| HOMO SAPIENS | 0 | 4127 | 3swr | A | 860 | GLY | p.Gly860Arg | P26358 |
| HOMO SAPIENS | -1 | 4127 | 3swr | A | 1122 | SER | p.Ser1122Cys | P26358 |
| HOMO SAPIENS | 0 | 4127 | 3swr | A | 1216 | ASP | p.Asp1216Glu | P26358 |
| HOMO SAPIENS | 0 | 6382 | 3o96 | A | 17 | GLU | p.Glu17Lys | P31749 |
| HOMO SAPIENS | 6 | 14314 | 5xtc | Q | 71 | PRO | p.Pro71Leu | O75306 |
| HOMO SAPIENS | 2 | 5886 | 4fao | E | 22 | ARG | p.Arg40His | Q13705 |
| HOMO SAPIENS | 1 | 2534 | 3mp4 | A | 37 | GLU | p.Glu37Ter | P35914 |
| HOMO SAPIENS | 3 | 2534 | 3mp4 | A | 41 | MET | p.Arg41Gln | P35914 |
| HOMO SAPIENS | 1 | 2534 | 3mp4 | A | 70 | VAL | p.Val70Leu | P35914 |
| HOMO SAPIENS | 2 | 2196 | 3ifq | A | 142 | ARG | p.Arg142His | P14923 |
| HOMO SAPIENS | 2 | 2196 | 3ifq | A | 153 | ASP | p.Asp153Glu | P14923 |
| HOMO SAPIENS | 3 | 2196 | 3ifq | A | 176 | ARG | p.Arg176Gln | P14923 |
| HOMO SAPIENS | 3 | 2196 | 3ifq | A | 265 | ARG | p.Arg265His | P14923 |
| HOMO SAPIENS | 2 | 2196 | 3ifq | A | 301 | GLU | p.Glu301Gly | P14923 |
| HOMO SAPIENS | 0 | 2196 | 3ifq | A | 456 | VAL | p.Val456Ile | P14923 |
| HOMO SAPIENS | 3 | 2196 | 3ifq | A | 603 | VAL | p.Val603Leu | P14923 |
| HOMO SAPIENS | 0 | 2196 | 3ifq | A | 648 | VAL | p.Val648Ile | P14923 |
| HOMO SAPIENS | 1 | 8746 | 1cgj | I | 32 | PRO | p.Pro55Ser | P00995 |
| HOMO SAPIENS | 1 | 8746 | 1cgj | I | 44 | ARG | p.Arg67His | P00995 |
| HOMO SAPIENS | 1 | 8746 | 1cgj | I | 44 | ARG | p.Arg67Cys | P00995 |
| HOMO SAPIENS | 3 | 3212 | 1syq | A | 105 | ARG | p.Arg105Ter | P18206 |
| HOMO SAPIENS | 3 | 3212 | 1syq | A | 126 | PHE | p.Phe126Leu | P18206 |
| HOMO SAPIENS | 4 | 10028 | 3eto | A | 1609 | ARG | p.Arg1608His | P46531 |
| HOMO SAPIENS | 1 | 10028 | 3eto | A | 1672 | VAL | p.Val1671Ile | P46531 |
| HOMO SAPIENS | 5 | 10028 | 3eto | A | 1692 | GLN | p.Gln1691His | P46531 |
| HOMO SAPIENS | 3 | 3749 | 5xtc | Z | 70 | GLY | p.Gly70Ter | O43676 |
| HOMO SAPIENS | 2 | 17039 | 3brv | B | 57 | GLU | p.Glu57Lys | Q9Y6K9 |
| HOMO SAPIENS | 8 | 3589 | 2fh3 | A | 433 | VAL | p.Val460Leu | P06396 |
| HOMO SAPIENS | 1 | 3589 | 2fh3 | A | 454 | ARG | p.Arg481Cys | P06396 |
| HOMO SAPIENS | 4 | 3589 | 2fh3 | A | 498 | ARG | p.Arg525His | P06396 |
| HOMO SAPIENS | 2 | 3589 | 2fh3 | A | 520 | TYR | p.Tyr547Cys | P06396 |
| HOMO SAPIENS | 1 | 3589 | 2fh3 | A | 529 | GLY | p.Gly556Arg | P06396 |
| HOMO SAPIENS | 2 | 3589 | 2fh3 | A | 536 | THR | p.Thr563Ser | P06396 |
| HOMO SAPIENS | 2 | 3589 | 2fh3 | A | 579 | VAL | p.Val606Met | P06396 |

| | | | | | | | | |
|---|---|---|---|---|---|---|---|---|
| HOMO SAPIENS | 1 | 3589 | 2fh3 | A | 589 | THR | p.Thr616Met | P06396 |
| HOMO SAPIENS | 5 | 3589 | 2fh3 | A | 668 | THR | p.Thr695Met | P06396 |
| HOMO SAPIENS | 0 | 148 | 1lcl | A | 130 | ILE | p.Ile130Ser | Q05315 |
| HOMO SAPIENS | -1 | 24727 | 4fj3 | P | 237 | ALA | p.Ala237Thr | P04049 |
| HOMO SAPIENS | 0 | 24727 | 4fj3 | P | 257 | SER | p.Ser257Leu | P04049 |
| HOMO SAPIENS | 0 | 24727 | 4fj3 | P | 261 | PRO | p.Pro261Ala | P04049 |
| HOMO SAPIENS | 0 | 2949 | 5xtb | B | 77 | ARG | p.Arg77Gly | O00217 |
| HOMO SAPIENS | 0 | 2949 | 5xtb | B | 79 | PRO | p.Pro79Leu | O00217 |
| HOMO SAPIENS | 0 | 2949 | 5xtb | B | 100 | ALA | p.Ala100Val | O00217 |
| HOMO SAPIENS | 1 | 2949 | 5xtb | B | 115 | LYS | p.Lys115Glu | O00217 |
| HOMO SAPIENS | 0 | 2949 | 5xtb | B | 138 | ARG | p.Arg138His | O00217 |
| HOMO SAPIENS | 1 | 3959 | 5k5s | B | 25 | ARG | p.Arg25Ter | P41180 |
| HOMO SAPIENS | 4 | 3959 | 5k5s | B | 39 | PRO | p.Pro39Ala | P41180 |
| HOMO SAPIENS | 0 | 3959 | 5k5s | B | 47 | LYS | p.Lys47Asn | P41180 |
| HOMO SAPIENS | 2 | 3959 | 5k5s | B | 62 | ARG | p.Arg62Met | P41180 |
| HOMO SAPIENS | 2 | 3959 | 5k5s | B | 66 | ARG | p.Arg66Cys | P41180 |
| HOMO SAPIENS | 4 | 3959 | 5k5s | B | 90 | ASN | p.Asn90Thr | P41180 |
| HOMO SAPIENS | 1 | 3959 | 5k5s | B | 116 | ALA | p.Ala116Thr | P41180 |
| HOMO SAPIENS | 1 | 3959 | 5k5s | B | 118 | ASN | p.Asn118Lys | P41180 |
| HOMO SAPIENS | -1 | 3959 | 5k5s | B | 125 | LEU | p.Leu125Pro | P41180 |
| HOMO SAPIENS | -1 | 3959 | 5k5s | B | 127 | GLU | p.Glu127Ala | P41180 |
| HOMO SAPIENS | -1 | 3959 | 5k5s | B | 128 | PHE | p.Phe128Leu | P41180 |
| HOMO SAPIENS | -1 | 3959 | 5k5s | B | 131 | CYS | p.Cys131Trp | P41180 |
| HOMO SAPIENS | 1 | 3959 | 5k5s | B | 138 | THR | p.Thr138Met | P41180 |
| HOMO SAPIENS | 0 | 3959 | 5k5s | B | 143 | GLY | p.Gly143Glu | P41180 |
| HOMO SAPIENS | 5 | 3959 | 5k5s | B | 151 | THR | p.Thr151Met | P41180 |
| HOMO SAPIENS | 4 | 3959 | 5k5s | B | 180 | PHE | p.Phe180Cys | P41180 |
| HOMO SAPIENS | 0 | 3959 | 5k5s | B | 185 | ARG | p.Arg185Gln | P41180 |
| HOMO SAPIENS | 0 | 3959 | 5k5s | B | 185 | ARG | p.Arg185Ter | P41180 |
| HOMO SAPIENS | 2 | 3959 | 5k5s | B | 191 | GLU | p.Glu191Lys | P41180 |
| HOMO SAPIENS | 2 | 3959 | 5k5s | B | 227 | ARG | p.Arg227Leu | P41180 |
| HOMO SAPIENS | 2 | 3959 | 5k5s | B | 250 | GLU | p.Glu250Lys | P41180 |
| HOMO SAPIENS | 6 | 3959 | 5k5s | B | 297 | GLU | p.Glu297Lys | P41180 |
| HOMO SAPIENS | 1 | 3959 | 5k5s | B | 336 | LYS | p.Lys336Asn | P41180 |
| HOMO SAPIENS | 6 | 3959 | 5k5s | B | 429 | HIS | p.His429Tyr | P41180 |
| HOMO SAPIENS | 0 | 3959 | 5k5s | B | 445 | THR | p.Thr445Ala | P41180 |
| HOMO SAPIENS | 3 | 3959 | 5k5s | B | 509 | GLY | p.Gly509Arg | P41180 |
| HOMO SAPIENS | 1 | 3959 | 5k5s | B | 553 | GLY | p.Gly553Arg | P41180 |
| HOMO SAPIENS | 3 | 3959 | 5k5s | B | 592 | ASN | p.Asn592Ser | P41180 |
| HOMO SAPIENS | -1 | 3959 | 5k5s | B | 604 | GLU | p.Glu604Lys | P41180 |
| HOMO SAPIENS | 4 | 2564 | 5l0c | C | 975 | ARG | p.Arg975Trp | P18206 |
| HOMO SAPIENS | 4 | 2564 | 5l0c | C | 975 | ARG | p.Arg975Leu | P18206 |
| HOMO SAPIENS | 1 | 5695 | 5i8r | A | 294 | ARG | p.Arg294Gln | P17405 |
| HOMO SAPIENS | 0 | 5695 | 5i8r | A | 485 | ALA | p.Ala485Val | P17405 |
| HOMO SAPIENS | 1 | 5695 | 5i8r | A | 506 | GLY | p.Gly506Arg | P17405 |
| HOMO SAPIENS | -1 | 1978 | 3avs | A | 1060 | HIS | p.His1060Leu | O15550 |
| HOMO SAPIENS | 0 | 11173 | 4v0c | B | 362 | LYS | p.Lys362Arg | P51787 |
| HOMO SAPIENS | 3 | 11173 | 4v0c | B | 381 | CYS | p.Cys381Tyr | P51787 |
| HOMO SAPIENS | 7 | 11173 | 4v0c | B | 518 | ARG | p.Arg518Gly | P51787 |
| HOMO SAPIENS | 2 | 11173 | 4v0c | B | 525 | ALA | p.Ala525Thr | P51787 |
| HOMO SAPIENS | 0 | 11173 | 4v0c | B | 530 | GLN | p.Gln530Ter | P51787 |
| HOMO SAPIENS | -1 | 11173 | 4v0c | B | 536 | TYR | p.Tyr536Ter | P51787 |
| HOMO SAPIENS | 2 | 1113 | 5xtd | N | 60 | ARG | p.Arg60Ter | Q9UI09 |
| HOMO SAPIENS | 3 | 6061 | 2mmz | A | 148 | LEU | p.Leu148Ser | Q86SX6 |

| | | | | | | | | |
|---|---|---|---|---|---|---|---|---|
| HOMO SAPIENS | -1 | 26081 | 3eu7 | A | 837 | GLU | p.Glu837Ter | Q86YC2 |
| HOMO SAPIENS | 0 | 26081 | 3eu7 | A | 864 | PRO | p.Pro864Ser | Q86YC2 |
| HOMO SAPIENS | 5 | 26081 | 3eu7 | A | 891 | CYS | p.Cys891Trp | Q86YC2 |
| HOMO SAPIENS | 0 | 26081 | 3eu7 | A | 932 | VAL | p.Val932Met | Q86YC2 |
| HOMO SAPIENS | 5 | 26081 | 3eu7 | A | 961 | LEU | p.Leu961Met | Q86YC2 |
| HOMO SAPIENS | 5 | 26081 | 3eu7 | A | 1038 | TRP | p.Trp1038Ter | Q86YC2 |
| HOMO SAPIENS | -1 | 26081 | 3eu7 | A | 1086 | ARG | p.Arg1086Ter | Q86YC2 |
| HOMO SAPIENS | 0 | 26081 | 3eu7 | A | 1122 | ASP | p.Asp1122Glu | Q86YC2 |
| HOMO SAPIENS | 5 | 26081 | 3eu7 | A | 1158 | HIS | p.His1158Gln | Q86YC2 |
| HOMO SAPIENS | 5 | 26081 | 3eu7 | A | 1183 | TYR | p.Tyr1183Ter | Q86YC2 |
| HOMO SAPIENS | 4 | 199 | 1qo5 | C | 45 | ARG | p.Arg46Trp | P05062 |
| HOMO SAPIENS | 5 | 199 | 1qo5 | C | 147 | TRP | p.Trp148Arg | P05062 |
| HOMO SAPIENS | 0 | 199 | 1qo5 | C | 149 | ALA | p.Ala150Pro | P05062 |
| HOMO SAPIENS | 0 | 199 | 1qo5 | C | 174 | ALA | p.Ala175Asp | P05062 |
| HOMO SAPIENS | 0 | 199 | 1qo5 | C | 334 | ASN | p.Asn335Lys | P05062 |
| HOMO SAPIENS | 0 | 199 | 1qo5 | C | 337 | ALA | p.Ala338Val | P05062 |
| HOMO SAPIENS | 5 | 199 | 1qo5 | C | 342 | TYR | p.Tyr343His | P05062 |
| HOMO SAPIENS | 0 | 6910 | 4zht | B | 59 | ARG | p.Arg59Ter | Q9Y223 |
| HOMO SAPIENS | 4 | 6910 | 4zht | B | 129 | ARG | p.Arg129Ter | Q9Y223 |
| HOMO SAPIENS | 4 | 6910 | 4zht | B | 129 | ARG | p.Arg129Gln | Q9Y223 |
| HOMO SAPIENS | 1 | 6910 | 4zht | B | 162 | ARG | p.Arg162Cys | Q9Y223 |
| HOMO SAPIENS | 0 | 6910 | 4zht | B | 176 | ASP | p.Asp176Val | Q9Y223 |
| HOMO SAPIENS | 0 | 6910 | 4zht | B | 216 | VAL | p.Val216Ala | Q9Y223 |
| HOMO SAPIENS | 0 | 6910 | 4zht | B | 246 | ARG | p.Arg246Trp | Q9Y223 |
| HOMO SAPIENS | 0 | 6910 | 4zht | B | 263 | ARG | p.Arg263Leu | Q9Y223 |
| HOMO SAPIENS | 0 | 6910 | 4zht | B | 266 | ARG | p.Arg266Gln | Q9Y223 |
| HOMO SAPIENS | 0 | 6910 | 4zht | B | 266 | ARG | p.Arg266Trp | Q9Y223 |
| HOMO SAPIENS | 0 | 6910 | 4zht | B | 277 | ARG | p.Arg277Cys | Q9Y223 |
| HOMO SAPIENS | 0 | 6910 | 4zht | B | 378 | ASP | p.Asp378Tyr | Q9Y223 |
| HOMO SAPIENS | 3 | 211 | 5kxi | C | 262 | VAL | p.Val287Leu | P17787 |
| HOMO SAPIENS | 0 | 211 | 5kxi | C | 369 | PHE | p.Phe478Leu | P17787 |
| HOMO SAPIENS | 0 | 2454 | 5iy9 | V | null | PHE | p.Phe99Ser | P19447 |
| HOMO SAPIENS | 0 | 2454 | 5iy9 | V | null | ARG | p.Arg109Ter | P19447 |
| HOMO SAPIENS | 0 | 2454 | 5iy9 | V | null | THR | p.Thr119Pro | P19447 |
| HOMO SAPIENS | 0 | 2454 | 5iy9 | V | 425 | ARG | p.Arg425Ter | P19447 |
| HOMO SAPIENS | 1 | 2454 | 5iy9 | V | 545 | GLN | p.Gln545Ter | P19447 |
| HOMO SAPIENS | -1 | 3805 | 3rbq | A | 57 | LYS | p.Lys57Ter | Q13432 |
| HOMO SAPIENS | 2 | 2649 | 4ido | A | 43 | ASP | p.Asp43Glu | Q8WXF7 |
| HOMO SAPIENS | 2 | 2649 | 4ido | A | 157 | LEU | p.Leu157Trp | Q8WXF7 |
| HOMO SAPIENS | 2 | 2649 | 4ido | A | 193 | PHE | p.Phe193Cys | Q8WXF7 |
| HOMO SAPIENS | 7 | 2649 | 4ido | A | 217 | ARG | p.Arg217Gln | Q8WXF7 |
| HOMO SAPIENS | 4 | 2649 | 4ido | A | 239 | ARG | p.Arg239Cys | Q8WXF7 |
| HOMO SAPIENS | 3 | 2649 | 4ido | A | 258 | HIS | p.His258Arg | Q8WXF7 |
| HOMO SAPIENS | 4 | 2649 | 4ido | A | 259 | SER | p.Ser259Tyr | Q8WXF7 |
| HOMO SAPIENS | 4 | 2649 | 4ido | A | 408 | MET | p.Met408Val | Q8WXF7 |
| HOMO SAPIENS | 2 | 2649 | 4ido | A | 415 | ARG | p.Arg415Trp | Q8WXF7 |
| HOMO SAPIENS | 3 | 2649 | 4ido | A | 416 | ARG | p.Arg416Cys | Q8WXF7 |
| HOMO SAPIENS | 5 | 2203 | 3wff | A | 810 | LEU | p.Ser810Leu | P08235 |
| HOMO SAPIENS | 4 | 12648 | 5ep6 | B | 696 | GLU | p.Glu696Lys | Q9UHD2 |
| HOMO SAPIENS | 0 | 30622 | 2pe4 | A | 90 | GLU | p.Glu90Asp | Q12794 |
| HOMO SAPIENS | 0 | 30622 | 2pe4 | A | 256 | GLY | p.Gly256Arg | Q12794 |
| HOMO SAPIENS | 1 | 30622 | 2pe4 | A | 268 | GLU | p.Glu268Lys | Q12794 |
| HOMO SAPIENS | 0 | 30622 | 2pe4 | A | 392 | THR | p.Thr392Met | Q12794 |
| HOMO SAPIENS | 5 | 5411 | 1lya | A | 93 | SER | p.Ser157Leu | P07339 |

| | | | | | | | | |
|---|---|---|---|---|---|---|---|---|
| HOMO SAPIENS | 3 | 5743 | 2uzp | A | 189 | ASN | p.Asn189Lys | P05129 |
| HOMO SAPIENS | 1 | 12529 | 3kfd | J | 21 | ASN | p.Asn45Ser | P36897 |
| HOMO SAPIENS | 1 | 1063 | 2e1q | B | 149 | ARG | p.Arg149Cys | P47989 |
| HOMO SAPIENS | -1 | 1063 | 2e1q | B | 172 | GLY | p.Gly172Arg | P47989 |
| HOMO SAPIENS | 4 | 1063 | 2e1q | B | 228 | ARG | p.Arg228Ter | P47989 |
| HOMO SAPIENS | 5 | 1063 | 2e1q | B | 395 | LYS | p.Lys395Met | P47989 |
| HOMO SAPIENS | 2 | 1063 | 2e1q | B | 646 | ILE | p.Ile646Val | P47989 |
| HOMO SAPIENS | 4 | 1063 | 2e1q | B | 703 | ILE | p.Ile703Val | P47989 |
| HOMO SAPIENS | 8 | 2525 | 1kqn | F | 18 | ASN | p.Asn18Ser | Q9HAN9 |
| HOMO SAPIENS | 0 | 2525 | 1kqn | F | 151 | VAL | p.Val151Phe | Q9HAN9 |
| HOMO SAPIENS | 0 | 2525 | 1kqn | F | 153 | LEU | p.Leu153Val | Q9HAN9 |
| HOMO SAPIENS | 0 | 2525 | 1kqn | F | 169 | TRP | p.Trp169Ter | Q9HAN9 |
| HOMO SAPIENS | 0 | 2525 | 1kqn | F | 207 | ARG | p.Arg207Trp | Q9HAN9 |
| HOMO SAPIENS | 0 | 2525 | 1kqn | F | 273 | ASN | p.Asn273Asp | Q9HAN9 |
| HOMO SAPIENS | 0 | 12299 | 5duq | A | 432 | VAL | p.Val454Glu | P05155 |
| HOMO SAPIENS | 0 | 12299 | 5duq | A | 436 | ALA | p.Ala458Thr | P05155 |
| HOMO SAPIENS | 0 | 12299 | 5duq | A | 443 | ALA | p.Ala465Val | P05155 |
| HOMO SAPIENS | 0 | 12299 | 5duq | A | 444 | ARG | p.Arg466His | P05155 |
| HOMO SAPIENS | 0 | 12299 | 5duq | A | 458 | MET | p.Val480Met | P05155 |
| HOMO SAPIENS | 0 | 327 | 1uhp | A | 36 | ASP | p.Asp274Asn | Q9UPQ7 |
| HOMO SAPIENS | 0 | 64 | 2mo5 | A | 54 | ALA | p.Ala55Thr | P12104 |
| HOMO SAPIENS | 2 | 35969 | 1jrh | I | 60 | CYS | p.Cys77Tyr | P15260 |
| HOMO SAPIENS | 4 | 35969 | 1jrh | I | 70 | ILE | p.Ile87Thr | P15260 |
| HOMO SAPIENS | 2 | 1068 | 1mu7 | A | 304 | ARG | p.Arg304Gln | Q9NUW8 |
| HOMO SAPIENS | 8 | 1068 | 1mu7 | A | 493 | HIS | p.His493Arg | Q9NUW8 |
| HOMO SAPIENS | 0 | 32360 | 2kqp | A | 31 | ARG | p.Arg55Cys | P01308 |
| HOMO SAPIENS | 0 | 32360 | 2kqp | A | 65 | ARG | p.Arg89His | P01308 |
| HOMO SAPIENS | 0 | 32360 | 2kqp | A | 65 | ARG | p.Arg89Cys | P01308 |
| HOMO SAPIENS | 6 | 10805 | 3nks | A | 12 | ILE | p.Ile12Thr | P50336 |
| HOMO SAPIENS | 0 | 10805 | 3nks | A | 20 | HIS | p.His20Pro | P50336 |
| HOMO SAPIENS | 0 | 10805 | 3nks | A | 59 | ARG | p.Arg59Trp | P50336 |
| HOMO SAPIENS | 0 | 10805 | 3nks | A | 168 | ARG | p.Arg168Cys | P50336 |
| HOMO SAPIENS | 0 | 10805 | 3nks | A | 168 | ARG | p.Arg168His | P50336 |
| HOMO SAPIENS | 2 | 10805 | 3nks | A | 232 | GLY | p.Gly232Arg | P50336 |
| HOMO SAPIENS | 3 | 10805 | 3nks | A | 256 | PRO | p.Pro256Arg | P50336 |
| HOMO SAPIENS | 0 | 10805 | 3nks | A | 304 | ARG | p.Arg304His | P50336 |
| HOMO SAPIENS | 0 | 10805 | 3nks | A | 349 | ASP | p.Asp349Ala | P50336 |
| HOMO SAPIENS | 9 | 7008 | 3u2l | A | 125 | GLN | p.Gln125Ter | P55789 |
| HOMO SAPIENS | 4 | 7008 | 3u2l | A | 172 | HIS | p.His172Asn | P55789 |
| HOMO SAPIENS | 4 | 7008 | 3u2l | A | 194 | ARG | p.Arg194His | P55789 |
| HOMO SAPIENS | 3 | 7008 | 3u2l | A | 198 | GLY | p.Gly198Ser | P55789 |
| HOMO SAPIENS | 0 | 20821 | 3e0g | A | 65 | HIS | p.His116Tyr | P42702 |
| HOMO SAPIENS | 3 | 1151 | 5l9v | A | 278 | ASP | p.Asp278Asn | Q9GZT9 |
| HOMO SAPIENS | 4 | 1151 | 5l9v | A | 317 | CYS | p.Pro317Arg | Q9GZT9 |
| HOMO SAPIENS | 5 | 1151 | 5l9v | A | 371 | ARG | p.Arg371His | Q9GZT9 |
| HOMO SAPIENS | 2 | 3935 | 1gwb | A | 36 | ARG | p.Arg36Gly | P07359 |
| HOMO SAPIENS | 1 | 3935 | 1gwb | A | 73 | LEU | p.Leu73Phe | P07359 |
| HOMO SAPIENS | 1 | 3935 | 1gwb | A | 172 | ALA | p.Ala172Val | P07359 |
| HOMO SAPIENS | 0 | 3935 | 1gwb | A | 249 | GLY | p.Gly249Val | P07359 |
| HOMO SAPIENS | 0 | 3935 | 1gwb | A | 255 | MET | p.Met255Val | P07359 |
| HOMO SAPIENS | 2 | 14541 | 5cuz | A | 138 | GLU | p.Glu138Lys | Q9H3L0 |
| HOMO SAPIENS | 2 | 14541 | 5cuz | A | 143 | SER | p.Ser143Ile | Q9H3L0 |
| HOMO SAPIENS | 4 | 14541 | 5cuz | A | 182 | THR | p.Thr182Asn | Q9H3L0 |
| HOMO SAPIENS | 6 | 14541 | 5cuz | A | 206 | ASN | p.Asn206Ile | Q9H3L0 |

| | | | | | | | | |
|---|---|---|---|---|---|---|---|---|
| HOMO SAPIENS | 0 | 14541 | 5cuz | A | 249 | TYR | p.Tyr249Cys | Q9H3L0 |
| HOMO SAPIENS | 4 | 14541 | 5cuz | A | 250 | ARG | p.Arg250Ter | Q9H3L0 |
| HOMO SAPIENS | 2 | 14541 | 5cuz | A | 259 | LEU | p.Leu259Pro | Q9H3L0 |
| HOMO SAPIENS | 0 | 4206 | 4fao | D | 30 | PRO | p.Pro30Ser | P37023 |
| HOMO SAPIENS | 1 | 4206 | 4fao | D | 50 | TRP | p.Trp50Cys | P37023 |
| HOMO SAPIENS | 0 | 16553 | 3v30 | A | 251 | GLN | p.Gln251Glu | O14593 |
| HOMO SAPIENS | 0 | 5012 | 5hbf | A | 442 | ALA | p.Ala442Gly | Q13231 |
| HOMO SAPIENS | 5 | 4844 | 5tdc | A | 136 | HIS | p.His136Arg | Q8IWV7 |
| HOMO SAPIENS | 0 | 15435 | 4cgz | A | 643 | ARG | p.Arg643His | P54132 |
| HOMO SAPIENS | 1 | 15435 | 4cgz | A | 672 | GLN | p.Gln672Arg | P54132 |
| HOMO SAPIENS | 1 | 15435 | 4cgz | A | 700 | GLN | p.Gln700Ter | P54132 |
| HOMO SAPIENS | 0 | 15435 | 4cgz | A | 707 | PRO | p.Pro707Ser | P54132 |
| HOMO SAPIENS | 0 | 15435 | 4cgz | A | 755 | LYS | p.Lys755Glu | P54132 |
| HOMO SAPIENS | 0 | 15435 | 4cgz | A | 881 | TRP | p.Trp881Ter | P54132 |
| HOMO SAPIENS | 2 | 15435 | 4cgz | A | 899 | ARG | p.Arg899Ter | P54132 |
| HOMO SAPIENS | 0 | 15435 | 4cgz | A | 1036 | CYS | p.Cys1036Phe | P54132 |
| HOMO SAPIENS | 0 | 15435 | 4cgz | A | 1055 | CYS | p.Cys1055Ser | P54132 |
| HOMO SAPIENS | 0 | 15435 | 4cgz | A | 1139 | ARG | p.Arg1139Ter | P54132 |
| HOMO SAPIENS | -1 | 15435 | 4cgz | A | 1198 | VAL | p.Val1198Met | P54132 |
| HOMO SAPIENS | -1 | 15435 | 4cgz | A | 1205 | VAL | p.Val1205Ile | P54132 |
| HOMO SAPIENS | 0 | 15435 | 4cgz | A | 1251 | GLU | p.Glu1251Gln | P54132 |
| HOMO SAPIENS | 0 | 11185 | 1kb4 | A | 30 | ARG | p.Arg30Ter | P11473 |
| HOMO SAPIENS | 5 | 11185 | 1kb4 | A | 33 | GLY | p.Gly33Asp | P11473 |
| HOMO SAPIENS | 5 | 11185 | 1kb4 | A | 46 | GLY | p.Gly46Asp | P11473 |
| HOMO SAPIENS | 6 | 11185 | 1kb4 | A | 50 | ARG | p.Arg50Gln | P11473 |
| HOMO SAPIENS | 9 | 11185 | 1kb4 | A | 73 | ARG | p.Arg73Gln | P11473 |
| HOMO SAPIENS | 9 | 11185 | 1kb4 | A | 80 | ARG | p.Arg80Gln | P11473 |
| HOMO SAPIENS | 0 | 5382 | 3fl7 | A | 41 | LEU | p.Leu41Val | P29317 |
| HOMO SAPIENS | 2 | 5382 | 3fl7 | A | 175 | ARG | p.Arg175Cys | P29317 |
| HOMO SAPIENS | 1 | 5382 | 3fl7 | A | 195 | ARG | p.Arg195Cys | P29317 |
| HOMO SAPIENS | 0 | 5382 | 3fl7 | A | 232 | ASP | p.Asp232Gly | P29317 |
| HOMO SAPIENS | 0 | 5382 | 3fl7 | A | 277 | SER | p.Ser277Leu | P29317 |
| HOMO SAPIENS | 0 | 5382 | 3fl7 | A | 315 | ARG | p.Arg315Gln | P29317 |
| HOMO SAPIENS | 7 | 5382 | 3fl7 | A | 350 | PRO | p.Pro350Thr | P29317 |
| HOMO SAPIENS | 0 | 5382 | 3fl7 | A | 391 | GLY | p.Gly391Arg | P29317 |
| HOMO SAPIENS | 0 | 5382 | 3fl7 | A | 418 | VAL | p.Val418Ile | P29317 |
| HOMO SAPIENS | 0 | 5382 | 3fl7 | A | 511 | THR | p.Thr511Met | P29317 |
| HOMO SAPIENS | 2 | 29168 | 4p22 | B | 144 | VAL | p.Val144Ile | P22314 |
| HOMO SAPIENS | 5 | 4348 | 1bf5 | A | 241 | ARG | p.Arg241Gln | P42224 |
| HOMO SAPIENS | 2 | 4348 | 1bf5 | A | 266 | VAL | p.Val266Ile | P42224 |
| HOMO SAPIENS | 4 | 4348 | 1bf5 | A | 320 | GLU | p.Glu320Gln | P42224 |
| HOMO SAPIENS | 1 | 4348 | 1bf5 | A | 372 | ASN | p.Asn372Lys | P42224 |
| HOMO SAPIENS | 1 | 4348 | 1bf5 | A | 447 | ASP | p.Asp447Glu | P42224 |
| HOMO SAPIENS | 2 | 4348 | 1bf5 | A | 463 | GLN | p.Gln463His | P42224 |
| HOMO SAPIENS | 4 | 4348 | 1bf5 | A | 600 | LEU | p.Leu600Pro | P42224 |
| HOMO SAPIENS | 0 | 4348 | 1bf5 | A | 706 | LEU | p.Leu706Ser | P42224 |
| HOMO SAPIENS | 4 | 4991 | 4bej | B | 141 | ASN | p.Asn141Ser | O00429 |
| HOMO SAPIENS | 0 | 4991 | 4bej | B | 395 | ALA | p.Ala395Asp | O00429 |
| HOMO SAPIENS | 2 | 11792 | 1fvo | A | 40 | ARG | p.Arg40Cys | P00480 |
| HOMO SAPIENS | 2 | 11792 | 1fvo | A | 40 | ARG | p.Arg40His | P00480 |
| HOMO SAPIENS | 0 | 11792 | 1fvo | A | 45 | LEU | p.Leu45Pro | P00480 |
| HOMO SAPIENS | 1 | 11792 | 1fvo | A | 47 | ASN | p.Asn47Ile | P00480 |
| HOMO SAPIENS | 0 | 11792 | 1fvo | A | 50 | GLY | p.Gly50Arg | P00480 |
| HOMO SAPIENS | 5 | 11792 | 1fvo | A | 87 | GLU | p.Glu87Lys | P00480 |

| | | | | | | | | |
|---|---|---|---|---|---|---|---|---|
| HOMO SAPIENS | 3 | 11792 | 1fvo | A | 111 | LEU | p.Leu111Pro | P00480 |
| HOMO SAPIENS | 1 | 11792 | 1fvo | A | 125 | THR | p.Thr125Met | P00480 |
| HOMO SAPIENS | 3 | 11792 | 1fvo | A | 129 | ARG | p.Arg129His | P00480 |
| HOMO SAPIENS | 9 | 11792 | 1fvo | A | 141 | ARG | p.Arg141Pro | P00480 |
| HOMO SAPIENS | 0 | 11792 | 1fvo | A | 206 | MET | p.Met206Arg | P00480 |
| HOMO SAPIENS | 0 | 11792 | 1fvo | A | 207 | SER | p.Ser207Arg | P00480 |
| HOMO SAPIENS | 0 | 11792 | 1fvo | A | 216 | GLN | p.Gln216Glu | P00480 |
| HOMO SAPIENS | 1 | 11792 | 1fvo | A | 225 | PRO | p.Pro225Leu | P00480 |
| HOMO SAPIENS | 0 | 11792 | 1fvo | A | 270 | ARG | p.Gln270Arg | P00480 |
| HOMO SAPIENS | 0 | 11792 | 1fvo | A | 270 | ARG | p.Gln270Ter | P00480 |
| HOMO SAPIENS | 2 | 11792 | 1fvo | A | 277 | ARG | p.Arg277Trp | P00480 |
| HOMO SAPIENS | 0 | 3721 | 4r3z | B | 2 | ASP | p.Asp2Gly | P54136 |
| HOMO SAPIENS | 0 | 3721 | 4r3z | B | 456 | SER | p.Ser456Leu | P54136 |
| HOMO SAPIENS | 5 | 3721 | 4r3z | B | 512 | ARG | p.Arg512Gln | P54136 |
| HOMO SAPIENS | 3 | 5749 | 3hl2 | A | 239 | ALA | p.Ala239Thr | Q9HD40 |
| HOMO SAPIENS | 1 | 5749 | 3hl2 | A | 334 | TYR | p.Tyr334Cys | Q9HD40 |
| HOMO SAPIENS | 1 | 5749 | 3hl2 | A | 452 | LYS | p.Lys452Asn | Q9HD40 |
| HOMO SAPIENS | -1 | 5749 | 3hl2 | A | 465 | ARG | p.Arg465Leu | Q9HD40 |
| HOMO SAPIENS | -1 | 5749 | 3hl2 | A | 489 | ASP | p.Asp489Gly | Q9HD40 |
| HOMO SAPIENS | 5 | 1721 | 3k15 | A | 1652 | MET | p.Met1652Ile | P38398 |
| HOMO SAPIENS | 5 | 1721 | 3k15 | A | 1652 | MET | p.Met1652Lys | P38398 |
| HOMO SAPIENS | 5 | 1721 | 3k15 | A | 1662 | PHE | p.Phe1662Cys | P38398 |
| HOMO SAPIENS | 5 | 1721 | 3k15 | A | 1664 | LEU | p.Leu1664Pro | P38398 |
| HOMO SAPIENS | 0 | 1721 | 3k15 | A | 1665 | VAL | p.Val1665Met | P38398 |
| HOMO SAPIENS | 5 | 1721 | 3k15 | A | 1690 | LYS | p.Lys1690Gln | P38398 |
| HOMO SAPIENS | 5 | 1721 | 3k15 | A | 1692 | ASP | p.Asp1692Asn | P38398 |
| HOMO SAPIENS | 5 | 1721 | 3k15 | A | 1699 | ARG | p.Arg1699Gln | P38398 |
| HOMO SAPIENS | 5 | 1721 | 3k15 | A | 1699 | ARG | p.Arg1699Trp | P38398 |
| HOMO SAPIENS | 3 | 1721 | 3k15 | A | 1706 | GLY | p.Gly1706Glu | P38398 |
| HOMO SAPIENS | 5 | 1721 | 3k15 | A | 1720 | THR | p.Thr1720Ala | P38398 |
| HOMO SAPIENS | 5 | 1721 | 3k15 | A | 1751 | ARG | p.Arg1751Pro | P38398 |
| HOMO SAPIENS | 4 | 1721 | 3k15 | A | 1775 | MET | p.Met1775Arg | P38398 |
| HOMO SAPIENS | 2 | 1721 | 3k15 | A | 1788 | GLY | p.Gly1788Val | P38398 |
| HOMO SAPIENS | 5 | 1721 | 3k15 | A | 1794 | GLU | p.Glu1794Asp | P38398 |
| HOMO SAPIENS | 5 | 1721 | 3k15 | A | 1804 | VAL | p.Val1804Asp | P38398 |
| HOMO SAPIENS | 3 | 1721 | 3k15 | A | 1815 | TRP | p.Trp1815Ter | P38398 |
| HOMO SAPIENS | 1 | 1721 | 3k15 | A | 1819 | ASN | p.Asn1819Ser | P38398 |
| HOMO SAPIENS | 5 | 1721 | 3k15 | A | 1856 | PRO | p.Pro1856Ser | P38398 |
| HOMO SAPIENS | 5 | 1721 | 3k15 | A | 1858 | ILE | p.Ile1858Leu | P38398 |
| HOMO SAPIENS | 5 | 1721 | 3k15 | A | 1858 | ILE | p.Ile1858Thr | P38398 |
| HOMO SAPIENS | 5 | 1721 | 3k15 | A | 1859 | PRO | p.Pro1859Arg | P38398 |
| HOMO SAPIENS | 2 | 22086 | 3kq4 | B | 1297 | PRO | p.Pro1297Leu | O60494 |
| HOMO SAPIENS | 1 | 12135 | 2k9j | B | 724 | ARG | p.Arg750Ter | P05106 |
| HOMO SAPIENS | -1 | 24027 | 2qtl | A | 175 | LEU | p.Ser202Leu | Q9UBK8 |
| HOMO SAPIENS | 0 | 24027 | 2qtl | A | 257 | SER | p.Ser284Thr | Q9UBK8 |
| HOMO SAPIENS | 0 | 24027 | 2qtl | A | 333 | LEU | p.Leu360Val | Q9UBK8 |
| HOMO SAPIENS | 0 | 24027 | 2qtl | A | 350 | LYS | p.Lys377Arg | Q9UBK8 |
| HOMO SAPIENS | 0 | 24027 | 2qtl | A | 415 | ARG | p.Arg442Cys | Q9UBK8 |
| HOMO SAPIENS | 2 | 24027 | 2qtl | A | 450 | PRO | p.Pro477Arg | Q9UBK8 |
| HOMO SAPIENS | 4 | 24027 | 2qtl | A | 487 | GLY | p.Gly514Arg | Q9UBK8 |
| HOMO SAPIENS | 1 | 24027 | 2qtl | A | 595 | HIS | p.His622Tyr | Q9UBK8 |
| HOMO SAPIENS | 0 | 5758 | 3di2 | B | 46 | THR | p.Thr66Ile | P16871 |
| HOMO SAPIENS | 1 | 5758 | 3di2 | B | 118 | VAL | p.Ile138Val | P16871 |
| HOMO SAPIENS | 4 | 848 | 5oa3 | Y | 79 | LEU | p.Leu79Val | P62847 |

| | | | | | | | | |
|---|---|---|---|---|---|---|---|---|
| HOMO SAPIENS | 2 | 848 | 5oa3 | Y | 124 | ASN | p.Asn124Ser | P62847 |
| HOMO SAPIENS | 0 | 81 | 3s5o | A | 36 | ALA | p.Ala36Glu | Q86XE5 |
| HOMO SAPIENS | 0 | 81 | 3s5o | A | 39 | TYR | p.Tyr39Ter | Q86XE5 |
| HOMO SAPIENS | 0 | 81 | 3s5o | A | 45 | PRO | p.Pro45Leu | Q86XE5 |
| HOMO SAPIENS | 0 | 81 | 3s5o | A | 70 | ARG | p.Arg70Ter | Q86XE5 |
| HOMO SAPIENS | 0 | 81 | 3s5o | A | 70 | ARG | p.Arg70Pro | Q86XE5 |
| HOMO SAPIENS | 1 | 81 | 3s5o | A | 97 | ARG | p.Arg97Cys | Q86XE5 |
| HOMO SAPIENS | 4 | 81 | 3s5o | A | 113 | GLU | p.Glu113Lys | Q86XE5 |
| HOMO SAPIENS | 1 | 81 | 3s5o | A | 190 | PRO | p.Pro190Leu | Q86XE5 |
| HOMO SAPIENS | 0 | 81 | 3s5o | A | 245 | VAL | p.Val245Phe | Q86XE5 |
| HOMO SAPIENS | 0 | 81 | 3s5o | A | 257 | CYS | p.Cys257Gly | Q86XE5 |
| HOMO SAPIENS | 0 | 81 | 3s5o | A | 287 | GLY | p.Gly287Val | Q86XE5 |
| HOMO SAPIENS | 7 | 81 | 3s5o | A | 303 | ARG | p.Arg303Cys | Q86XE5 |
| HOMO SAPIENS | 0 | 81 | 3s5o | A | 325 | GLY | p.Gly325Ser | Q86XE5 |
| HOMO SAPIENS | -1 | 622 | 5era | A | 12 | GLY | p.Gly12Arg | P29033 |
| HOMO SAPIENS | -1 | 622 | 5era | A | 12 | GLY | p.Gly12Asp | P29033 |
| HOMO SAPIENS | -1 | 622 | 5era | A | 17 | SER | p.Ser17Phe | P29033 |
| HOMO SAPIENS | 5 | 622 | 5era | A | 24 | TRP | p.Trp24Ter | P29033 |
| HOMO SAPIENS | 0 | 622 | 5era | A | 32 | ARG | p.Arg32Cys | P29033 |
| HOMO SAPIENS | 5 | 622 | 5era | A | 34 | MET | p.Met34Thr | P29033 |
| HOMO SAPIENS | 5 | 622 | 5era | A | 36 | LEU | p.Leu36Pro | P29033 |
| HOMO SAPIENS | 5 | 622 | 5era | A | 37 | VAL | p.Val37Ile | P29033 |
| HOMO SAPIENS | 5 | 622 | 5era | A | 40 | ALA | p.Ala40Glu | P29033 |
| HOMO SAPIENS | 5 | 622 | 5era | A | 44 | TRP | p.Trp44Cys | P29033 |
| HOMO SAPIENS | 5 | 622 | 5era | A | 45 | GLY | p.Gly45Glu | P29033 |
| HOMO SAPIENS | 5 | 622 | 5era | A | 47 | GLU | p.Glu47Ter | P29033 |
| HOMO SAPIENS | 5 | 622 | 5era | A | 50 | ASP | p.Asp50Asn | P29033 |
| HOMO SAPIENS | 5 | 622 | 5era | A | 54 | ASN | p.Asn54Lys | P29033 |
| HOMO SAPIENS | 5 | 622 | 5era | A | 57 | GLN | p.Gln57Ter | P29033 |
| HOMO SAPIENS | 5 | 622 | 5era | A | 59 | GLY | p.Gly59Ala | P29033 |
| HOMO SAPIENS | 5 | 622 | 5era | A | 59 | GLY | p.Gly59Ser | P29033 |
| HOMO SAPIENS | 5 | 622 | 5era | A | 66 | ASP | p.Asp66His | P29033 |
| HOMO SAPIENS | 5 | 622 | 5era | A | 68 | TYR | p.Tyr68Cys | P29033 |
| HOMO SAPIENS | 5 | 622 | 5era | A | 70 | PRO | p.Pro70Ala | P29033 |
| HOMO SAPIENS | 5 | 622 | 5era | A | 73 | HIS | p.His73Arg | P29033 |
| HOMO SAPIENS | 5 | 622 | 5era | A | 75 | ARG | p.Arg75Trp | P29033 |
| HOMO SAPIENS | 5 | 622 | 5era | A | 75 | ARG | p.Arg75Gln | P29033 |
| HOMO SAPIENS | 2 | 622 | 5era | A | 77 | TRP | p.Trp77Arg | P29033 |
| HOMO SAPIENS | 2 | 622 | 5era | A | 77 | TRP | p.Trp77Ter | P29033 |
| HOMO SAPIENS | 0 | 622 | 5era | A | 80 | GLN | p.Gln80Ter | P29033 |
| HOMO SAPIENS | 0 | 622 | 5era | A | 80 | GLN | p.Gln80Pro | P29033 |
| HOMO SAPIENS | 5 | 622 | 5era | A | 81 | LEU | p.Leu81Val | P29033 |
| HOMO SAPIENS | 5 | 622 | 5era | A | 82 | ILE | p.Ile82Met | P29033 |
| HOMO SAPIENS | 5 | 622 | 5era | A | 83 | PHE | p.Phe83Leu | P29033 |
| HOMO SAPIENS | 5 | 622 | 5era | A | 84 | VAL | p.Val84Leu | P29033 |
| HOMO SAPIENS | 5 | 622 | 5era | A | 90 | LEU | p.Leu90Pro | P29033 |
| HOMO SAPIENS | 5 | 622 | 5era | A | 95 | VAL | p.Val95Met | P29033 |
| HOMO SAPIENS | -1 | 622 | 5era | A | 100 | HIS | p.His100Tyr | P29033 |
| HOMO SAPIENS | -1 | 622 | 5era | A | 113 | SER | p.Ser113Arg | P29033 |
| HOMO SAPIENS | -1 | 622 | 5era | A | 119 | GLU | p.Glu119Lys | P29033 |
| HOMO SAPIENS | -1 | 622 | 5era | A | 122 | LYS | p.Lys122Ile | P29033 |
| HOMO SAPIENS | -1 | 622 | 5era | A | 123 | THR | p.Thr123Asn | P29033 |
| HOMO SAPIENS | -1 | 622 | 5era | A | 124 | GLN | p.Gln124Ter | P29033 |
| HOMO SAPIENS | 5 | 622 | 5era | A | 139 | SER | p.Ser139Asn | P29033 |

| | | | | | | | | |
|---|---|---|---|---|---|---|---|---|
| HOMO SAPIENS | 5 | 622 | 5era | A | 143 | ARG | p.Arg143Trp | P29033 |
| HOMO SAPIENS | 5 | 622 | 5era | A | 143 | ARG | p.Arg143Gln | P29033 |
| HOMO SAPIENS | 0 | 622 | 5era | A | 147 | GLU | p.Glu147Lys | P29033 |
| HOMO SAPIENS | 7 | 622 | 5era | A | 155 | TYR | p.Tyr155Ter | P29033 |
| HOMO SAPIENS | 2 | 622 | 5era | A | 159 | ASP | p.Asp159Val | P29033 |
| HOMO SAPIENS | 5 | 622 | 5era | A | 163 | MET | p.Met163Val | P29033 |
| HOMO SAPIENS | 5 | 622 | 5era | A | 165 | ARG | p.Arg165Trp | P29033 |
| HOMO SAPIENS | 5 | 622 | 5era | A | 167 | VAL | p.Val167Met | P29033 |
| HOMO SAPIENS | 5 | 622 | 5era | A | 169 | CYS | p.Cys169Tyr | P29033 |
| HOMO SAPIENS | 5 | 622 | 5era | A | 179 | ASP | p.Asp179Asn | P29033 |
| HOMO SAPIENS | 0 | 622 | 5era | A | 199 | SER | p.Ser199Phe | P29033 |
| HOMO SAPIENS | 0 | 622 | 5era | A | 202 | CYS | p.Cys202Phe | P29033 |
| HOMO SAPIENS | 5 | 622 | 5era | A | 203 | ILE | p.Ile203Thr | P29033 |
| HOMO SAPIENS | 5 | 622 | 5era | A | 206 | ASN | p.Asn206Ser | P29033 |
| HOMO SAPIENS | 0 | 8103 | 5dot | A | 163 | GLY | p.Gly163Ter | P31327 |
| HOMO SAPIENS | 0 | 8103 | 5dot | A | 337 | HIS | p.His337Arg | P31327 |
| HOMO SAPIENS | 3 | 8103 | 5dot | A | 344 | THR | p.Thr344Ala | P31327 |
| HOMO SAPIENS | 1 | 8103 | 5dot | A | 405 | THR | p.Thr405Asn | P31327 |
| HOMO SAPIENS | 5 | 8103 | 5dot | A | 544 | THR | p.Thr544Met | P31327 |
| HOMO SAPIENS | 5 | 8103 | 5dot | A | 716 | ASN | p.Asn716Lys | P31327 |
| HOMO SAPIENS | -1 | 8103 | 5dot | A | 780 | ARG | p.Arg780His | P31327 |
| HOMO SAPIENS | 0 | 8103 | 5dot | A | 982 | GLY | p.Gly982Asp | P31327 |
| HOMO SAPIENS | 3 | 8103 | 5dot | A | 1119 | ALA | p.Ala1119Ser | P31327 |
| HOMO SAPIENS | 1 | 8103 | 5dot | A | 1376 | GLY | p.Gly1376Ser | P31327 |
| HOMO SAPIENS | 4 | 6833 | 1l8l | B | 27 | ARG | p.Arg27Ser | P78330 |
| HOMO SAPIENS | 3 | 6833 | 1l8l | B | 32 | ASP | p.Asp32Gly | P78330 |
| HOMO SAPIENS | 3 | 6833 | 1l8l | B | 32 | ASP | p.Asp32Asn | P78330 |
| HOMO SAPIENS | 1 | 6833 | 1l8l | B | 49 | ARG | p.Arg49Trp | P78330 |
| HOMO SAPIENS | 0 | 6833 | 1l8l | B | 52 | MET | p.Met52Thr | P78330 |
| HOMO SAPIENS | 0 | 6833 | 1l8l | B | 83 | GLN | p.Gln83His | P78330 |
| HOMO SAPIENS | 6 | 810 | 2iw2 | A | 185 | ARG | p.Arg184Gln | P12955 |
| HOMO SAPIENS | 0 | 810 | 2iw2 | A | 213 | ALA | p.Ala212Pro | P12955 |
| HOMO SAPIENS | 1 | 810 | 2iw2 | A | 266 | ARG | p.Arg265Ter | P12955 |
| HOMO SAPIENS | 9 | 810 | 2iw2 | A | 277 | ASP | p.Asp276Asn | P12955 |
| HOMO SAPIENS | 0 | 810 | 2iw2 | A | 279 | GLY | p.Gly278Asp | P12955 |
| HOMO SAPIENS | 9 | 810 | 2iw2 | A | 378 | HIS | p.His377Gln | P12955 |
| HOMO SAPIENS | 0 | 810 | 2iw2 | A | 389 | ARG | p.Arg388His | P12955 |
| HOMO SAPIENS | 0 | 810 | 2iw2 | A | 436 | PHE | p.Leu435Phe | P12955 |
| HOMO SAPIENS | 0 | 810 | 2iw2 | A | 449 | GLY | p.Gly448Arg | P12955 |
| HOMO SAPIENS | -1 | 1530 | 5fwj | A | 388 | ALA | p.Ala388Pro | P41229 |
| HOMO SAPIENS | 4 | 1530 | 5fwj | A | 451 | SER | p.Ser451Arg | P41229 |
| HOMO SAPIENS | 3 | 1530 | 5fwj | A | 731 | LEU | p.Leu731Phe | P41229 |
| HOMO SAPIENS | 3 | 578 | 5net | B | 123 | ALA | p.Ala143Thr | P18564 |
| HOMO SAPIENS | 6 | 578 | 5net | B | 176 | PRO | p.Pro196Thr | P18564 |
| HOMO SAPIENS | 3 | 578 | 5net | B | 280 | GLU | p.Glu300Lys | P18564 |
| HOMO SAPIENS | -1 | 5247 | 4j15 | A | 274 | ALA | p.Ala274Val | P14868 |
| HOMO SAPIENS | -1 | 5247 | 4j15 | A | 280 | HIS | p.His280Leu | P14868 |
| HOMO SAPIENS | 2 | 5247 | 4j15 | A | 460 | ARG | p.Arg460His | P14868 |
| HOMO SAPIENS | 5 | 5247 | 4j15 | A | 494 | ARG | p.Arg494Cys | P14868 |
| HOMO SAPIENS | 7 | 2566 | 1b55 | A | 28 | ARG | p.Arg28His | Q06187 |
| HOMO SAPIENS | 5 | 2566 | 1b55 | A | 33 | THR | p.Thr33Pro | Q06187 |
| HOMO SAPIENS | 0 | 2566 | 1b55 | A | 113 | VAL | p.Val113Asp | Q06187 |
| HOMO SAPIENS | 0 | 356 | 5a2s | A | 786 | ALA | p.Ala786Thr | P56524 |
| HOMO SAPIENS | 5 | 2960 | 4cak | A | 553 | ARG | p.Arg584Ter | P08514 |

| | | | | | | | | |
|---|---|---|---|---|---|---|---|---|
| HOMO SAPIENS | 5 | 2960 | 4cak | A | 565 | ILE | p.Ile596Thr | P08514 |
| HOMO SAPIENS | -1 | 2960 | 4cak | A | 841 | LEU | p.Leu872Met | P08514 |
| HOMO SAPIENS | -1 | 2960 | 4cak | A | 843 | ILE | p.Ile874Ser | P08514 |
| HOMO SAPIENS | 0 | 5054 | 4k8r | B | 216 | PHE | p.Phe216Leu | Q8NBP7 |
| HOMO SAPIENS | 1 | 5054 | 4k8r | B | 417 | HIS | p.His417Gln | Q8NBP7 |
| HOMO SAPIENS | 1 | 5054 | 4k8r | B | 425 | ASN | p.Asn425Ser | Q8NBP7 |
| HOMO SAPIENS | 1 | 5054 | 4k8r | B | 443 | ALA | p.Ala443Thr | Q8NBP7 |
| HOMO SAPIENS | 0 | 27283 | 3n5n | X | 71 | HIS | p.His82Arg | Q9UIF7 |
| HOMO SAPIENS | 1 | 27283 | 3n5n | X | 83 | ARG | p.Arg94Ter | Q9UIF7 |
| HOMO SAPIENS | 8 | 27283 | 3n5n | X | 89 | TRP | p.Trp100Ter | Q9UIF7 |
| HOMO SAPIENS | 8 | 27283 | 3n5n | X | 90 | TYR | p.Tyr101Ter | Q9UIF7 |
| HOMO SAPIENS | 7 | 27283 | 3n5n | X | 95 | ARG | p.Arg106Trp | Q9UIF7 |
| HOMO SAPIENS | 7 | 27283 | 3n5n | X | 117 | TRP | p.Trp128Arg | Q9UIF7 |
| HOMO SAPIENS | 0 | 27283 | 3n5n | X | 137 | GLY | p.Gly148Arg | Q9UIF7 |
| HOMO SAPIENS | 5 | 27283 | 3n5n | X | 142 | TRP | p.Trp153Ter | Q9UIF7 |
| HOMO SAPIENS | 0 | 27283 | 3n5n | X | 165 | TYR | p.Tyr176Cys | Q9UIF7 |
| HOMO SAPIENS | 0 | 27283 | 3n5n | X | 168 | ARG | p.Arg179His | Q9UIF7 |
| HOMO SAPIENS | 0 | 27283 | 3n5n | X | 168 | ARG | p.Arg179Cys | Q9UIF7 |
| HOMO SAPIENS | 0 | 27283 | 3n5n | X | 196 | GLN | p.Gln207Ter | Q9UIF7 |
| HOMO SAPIENS | 8 | 27283 | 3n5n | X | 201 | VAL | p.Val212Met | Q9UIF7 |
| HOMO SAPIENS | 9 | 27283 | 3n5n | X | 227 | ARG | p.Arg238Trp | Q9UIF7 |
| HOMO SAPIENS | 9 | 27283 | 3n5n | X | 231 | ARG | p.Arg242Cys | Q9UIF7 |
| HOMO SAPIENS | 9 | 27283 | 3n5n | X | 231 | ARG | p.Arg242His | Q9UIF7 |
| HOMO SAPIENS | 3 | 27283 | 3n5n | X | 232 | VAL | p.Val243Phe | Q9UIF7 |
| HOMO SAPIENS | 3 | 27283 | 3n5n | X | 246 | GLN | p.Gln257Ter | Q9UIF7 |
| HOMO SAPIENS | 9 | 27283 | 3n5n | X | 272 | GLY | p.Gly283Glu | Q9UIF7 |
| HOMO SAPIENS | 9 | 27283 | 3n5n | X | 281 | PRO | p.Pro292Leu | Q9UIF7 |
| HOMO SAPIENS | -1 | 27283 | 3n5n | X | 314 | ASP | p.Asp325Glu | Q9UIF7 |
| HOMO SAPIENS | 0 | 27283 | 3n5n | X | 324 | GLN | p.Gln335His | Q9UIF7 |
| HOMO SAPIENS | 0 | 27283 | 3n5n | X | 324 | GLN | p.Gln335Arg | Q9UIF7 |
| HOMO SAPIENS | 0 | 27283 | 3n5n | X | 324 | GLN | p.Gln335Ter | Q9UIF7 |
| HOMO SAPIENS | -1 | 20852 | 4p60 | B | 243 | ARG | p.Arg243Lys | O75746 |
| HOMO SAPIENS | -1 | 1875 | 5dd2 | G | 152 | VAL | p.Val179Leu | P06396 |
| HOMO SAPIENS | 9 | 2126 | 3n1p | B | 46 | PRO | p.Pro46Leu | Q14623 |
| HOMO SAPIENS | 8 | 2126 | 3n1p | B | 76 | GLU | p.Glu76Ala | Q14623 |
| HOMO SAPIENS | 9 | 2126 | 3n1p | B | 77 | ARG | p.Arg77Ser | Q14623 |
| HOMO SAPIENS | 6 | 2126 | 3n1p | B | 95 | GLU | p.Glu95Lys | Q14623 |
| HOMO SAPIENS | 9 | 2126 | 3n1p | B | 100 | ASP | p.Asp100Glu | Q14623 |
| HOMO SAPIENS | 9 | 2126 | 3n1p | B | 131 | GLU | p.Glu131Lys | Q14623 |
| HOMO SAPIENS | 0 | 2126 | 3n1p | B | 190 | VAL | p.Val190Ala | Q14623 |
| HOMO SAPIENS | 5 | 40410 | 3ioz | B | 70 | GLN | p.Gln70Leu | P20963 |
| HOMO SAPIENS | 1 | 5754 | 5lkz | A | 1108 | THR | p.Tyr1191Ter | Q09472 |
| HOMO SAPIENS | 3 | 8159 | 1tiu | A | 3 | GLU | p.Lys12679Glu | Q8WZ42 |
| HOMO SAPIENS | 4 | 3275 | 4ldq | A | 210 | ARG | p.Arg210Gln | Q9BV20 |
| HOMO SAPIENS | 0 | 8143 | 4miv | B | 66 | SER | p.Ser66Trp | P51688 |
| HOMO SAPIENS | 1 | 8143 | 4miv | B | 74 | ARG | p.Arg74Cys | P51688 |
| HOMO SAPIENS | 0 | 8143 | 4miv | B | 128 | PRO | p.Pro128Leu | P51688 |
| HOMO SAPIENS | 1 | 8143 | 4miv | B | 150 | ARG | p.Arg150Gln | P51688 |
| HOMO SAPIENS | 0 | 8143 | 4miv | B | 191 | GLY | p.Gly191Arg | P51688 |
| HOMO SAPIENS | 0 | 8143 | 4miv | B | 206 | ARG | p.Arg206Pro | P51688 |
| HOMO SAPIENS | 0 | 8143 | 4miv | B | 225 | PHE | p.Phe225Leu | P51688 |
| HOMO SAPIENS | 0 | 8143 | 4miv | B | 233 | ARG | p.Arg233Ter | P51688 |
| HOMO SAPIENS | 0 | 8143 | 4miv | B | 234 | ALA | p.Ala234Gly | P51688 |
| HOMO SAPIENS | 0 | 8143 | 4miv | B | 245 | ARG | p.Arg245His | P51688 |

| | | | | | | | | |
|---|---|---|---|---|---|---|---|---|
| HOMO SAPIENS | 0 | 8143 | 4miv | B | 293 | PRO | p.Pro293Ser | P51688 |
| HOMO SAPIENS | 1 | 8143 | 4miv | B | 298 | SER | p.Ser298Pro | P51688 |
| HOMO SAPIENS | 0 | 8143 | 4miv | B | 361 | VAL | p.Val361Ile | P51688 |
| HOMO SAPIENS | 0 | 8143 | 4miv | B | 369 | GLU | p.Glu369Lys | P51688 |
| HOMO SAPIENS | 0 | 8143 | 4miv | B | 372 | MET | p.Met372Ile | P51688 |
| HOMO SAPIENS | 1 | 8143 | 4miv | B | 380 | GLN | p.Gln380Arg | P51688 |
| HOMO SAPIENS | 0 | 8143 | 4miv | B | 387 | VAL | p.Val387Met | P51688 |
| HOMO SAPIENS | 0 | 8143 | 4miv | B | 394 | MET | p.Met394Ile | P51688 |
| HOMO SAPIENS | 0 | 8143 | 4miv | B | 433 | ARG | p.Arg433Gln | P51688 |
| HOMO SAPIENS | 1 | 8143 | 4miv | B | 446 | HIS | p.His446Arg | P51688 |
| HOMO SAPIENS | 5 | 8143 | 4miv | B | 447 | GLU | p.Glu447Lys | P51688 |
| HOMO SAPIENS | 2 | 8143 | 4miv | B | 456 | ARG | p.Arg456His | P51688 |
| HOMO SAPIENS | 1 | 21880 | 2oo9 | B | 857 | LEU | p.Leu857Phe | P22681 |
| HOMO SAPIENS | 0 | 9467 | 1s1d | A | 154 | LEU | p.Leu224Pro | Q8WVQ1 |
| HOMO SAPIENS | 0 | 9467 | 1s1d | A | 156 | VAL | p.Val226Met | Q8WVQ1 |
| HOMO SAPIENS | 0 | 9467 | 1s1d | A | 229 | PRO | p.Pro299Leu | Q8WVQ1 |
| HOMO SAPIENS | 4 | 9467 | 1s1d | A | 230 | ARG | p.Arg300Cys | Q8WVQ1 |
| HOMO SAPIENS | 4 | 9467 | 1s1d | A | 230 | ARG | p.Arg300His | Q8WVQ1 |
| HOMO SAPIENS | 0 | 9467 | 1s1d | A | 253 | ALA | p.Ala323Thr | Q8WVQ1 |
| HOMO SAPIENS | 2 | 9467 | 1s1d | A | 305 | MET | p.Met375Leu | Q8WVQ1 |
| HOMO SAPIENS | -1 | 2361 | 5llm | A | 5 | ASN | p.Asn5Lys | P43003 |
| HOMO SAPIENS | 0 | 2361 | 5llm | A | 71 | TYR | p.Tyr71Cys | P43003 |
| HOMO SAPIENS | -1 | 2361 | 5llm | A | 211 | LYS | p.Glu219Asp | Q15758 |
| HOMO SAPIENS | 8 | 2361 | 5llm | A | 270 | PRO | p.Pro290Arg | P43003 |
| HOMO SAPIENS | 1 | 2361 | 5llm | A | 321 | THR | p.Ala329Thr | Q15758 |
| HOMO SAPIENS | 1 | 2361 | 5llm | A | 365 | ARG | p.Arg385His | P43003 |
| HOMO SAPIENS | 1 | 2361 | 5llm | A | 479 | ARG | p.Arg499Gln | P43003 |
| HOMO SAPIENS | 3 | 5412 | 4obz | D | 165 | PHE | p.Phe229Ile | P07339 |
| HOMO SAPIENS | 2 | 5412 | 4obz | D | 187 | ASP | p.Asp251Asn | P07339 |
| HOMO SAPIENS | 0 | 5412 | 4obz | D | 218 | GLY | p.Gly282Arg | P07339 |
| HOMO SAPIENS | 0 | 5412 | 4obz | D | 245 | ARG | p.Arg309His | P07339 |
| HOMO SAPIENS | 2 | 5412 | 4obz | D | 319 | TRP | p.Trp383Cys | P07339 |
| HOMO SAPIENS | 0 | 3544 | 5h64 | A | 277 | MET | p.Met277Ile | P42345 |
| HOMO SAPIENS | 0 | 14230 | 2kbx | A | 145 | ARG | p.Arg145Ser | Q13418 |
| HOMO SAPIENS | 2 | 34970 | 4xhz | A | 834 | ARG | p.Arg855Trp | Q96QU1 |
| HOMO SAPIENS | 4 | 34970 | 4xhz | A | 854 | SER | p.Ser875Ter | Q96QU1 |
| HOMO SAPIENS | 1 | 34970 | 4xhz | A | 908 | ARG | p.Arg929Gln | Q96QU1 |
| HOMO SAPIENS | 1 | 34970 | 4xhz | A | 941 | ARG | p.Arg962Cys | Q96QU1 |
| HOMO SAPIENS | 0 | 34970 | 4xhz | A | 970 | ARG | p.Arg991Ter | Q96QU1 |
| HOMO SAPIENS | 0 | 34970 | 4xhz | A | 976 | GLU | p.Glu997Gly | Q96QU1 |
| HOMO SAPIENS | 0 | 25734 | 4asc | A | 345 | SER | p.Asn345Ser | Q2TBA0 |
| HOMO SAPIENS | 0 | 25734 | 4asc | A | 514 | VAL | p.Val514Met | Q2TBA0 |
| HOMO SAPIENS | 6 | 25734 | 4asc | A | 528 | GLU | p.Glu528Lys | Q2TBA0 |
| HOMO SAPIENS | 0 | 25734 | 4asc | A | 580 | LEU | p.Leu580Ile | Q2TBA0 |
| HOMO SAPIENS | 8 | 2356 | 3kse | D | 22 | LYS | p.Lys22Ter | P01040 |
| HOMO SAPIENS | 1 | 2356 | 3kse | D | 58 | ARG | p.Arg58Ter | P01040 |
| HOMO SAPIENS | 2 | 6000 | 1jqd | A | 60 | GLY | p.Gly60Asp | P50135 |
| HOMO SAPIENS | 9 | 6000 | 1jqd | A | 208 | LEU | p.Leu208Pro | P50135 |
| HOMO SAPIENS | 0 | 14741 | 5eh1 | A | 58 | THR | p.Thr58Arg | P38484 |
| HOMO SAPIENS | 0 | 14741 | 5eh1 | A | 168 | THR | p.Thr168Asn | P38484 |
| HOMO SAPIENS | 1 | 4814 | 4kxk | D | 427 | ARG | p.Arg427Ter | P50542 |
| HOMO SAPIENS | 0 | 188 | 2xt3 | A | 49 | ASP | p.Asp52Asn | Q2M1P5 |
| HOMO SAPIENS | 3 | 188 | 2xt3 | A | 142 | TYR | p.Tyr145Ser | Q2M1P5 |
| HOMO SAPIENS | 9 | 8681 | 5tby | D | 57 | ALA | p.Ala57Asp | P08590 |

| | | | | | | | | |
|---|---|---|---|---|---|---|---|---|
| HOMO SAPIENS | 3 | 8681 | 5tby | D | 62 | ASP | p.Asp62Asn | P08590 |
| HOMO SAPIENS | 0 | 8681 | 5tby | D | 65 | PRO | p.Pro65Arg | P08590 |
| HOMO SAPIENS | 7 | 8681 | 5tby | D | 79 | VAL | p.Val79Ile | P08590 |
| HOMO SAPIENS | 0 | 8681 | 5tby | D | 143 | GLU | p.Glu143Lys | P08590 |
| HOMO SAPIENS | 0 | 8681 | 5tby | D | 149 | MET | p.Met149Val | P08590 |
| HOMO SAPIENS | 3 | 8681 | 5tby | D | 154 | ARG | p.Arg154His | P08590 |
| HOMO SAPIENS | 6 | 30984 | 4xwh | A | 140 | TYR | p.Tyr140Cys | P54802 |
| HOMO SAPIENS | 4 | 30984 | 4xwh | A | 141 | SER | p.Ser141Thr | P54802 |
| HOMO SAPIENS | 2 | 30984 | 4xwh | A | 234 | ARG | p.Arg234Cys | P54802 |
| HOMO SAPIENS | 2 | 30984 | 4xwh | A | 336 | GLU | p.Glu336Ter | P54802 |
| HOMO SAPIENS | 6 | 30984 | 4xwh | A | 482 | ARG | p.Arg482Trp | P54802 |
| HOMO SAPIENS | 2 | 30984 | 4xwh | A | 521 | PRO | p.Pro521Leu | P54802 |
| HOMO SAPIENS | 0 | 30984 | 4xwh | A | 565 | ARG | p.Arg565Gln | P54802 |
| HOMO SAPIENS | 0 | 30984 | 4xwh | A | 565 | ARG | p.Arg565Trp | P54802 |
| HOMO SAPIENS | 6 | 30984 | 4xwh | A | 626 | ARG | p.Arg626Ter | P54802 |
| HOMO SAPIENS | 6 | 30984 | 4xwh | A | 643 | ARG | p.Arg643His | P54802 |
| HOMO SAPIENS | 6 | 30984 | 4xwh | A | 643 | ARG | p.Arg643Cys | P54802 |
| HOMO SAPIENS | 2 | 30984 | 4xwh | A | 674 | ARG | p.Arg674His | P54802 |
| HOMO SAPIENS | 2 | 30984 | 4xwh | A | 737 | ARG | p.Arg737Gly | P54802 |
| HOMO SAPIENS | 5 | 10545 | 1oqe | N | 6 | PRO | p.Pro21Arg | Q96RJ3 |
| HOMO SAPIENS | 1 | 16995 | 5d3a | B | 12 | MET | p.Met947Arg | Q7Z4S6 |
| HOMO SAPIENS | 1 | 16995 | 5d3a | B | 12 | MET | p.Met947Val | Q7Z4S6 |
| HOMO SAPIENS | 2 | 16995 | 5d3a | B | 19 | ARG | p.Arg954Gln | Q7Z4S6 |
| HOMO SAPIENS | 2 | 16995 | 5d3a | B | 19 | ARG | p.Arg954Trp | Q7Z4S6 |
| HOMO SAPIENS | 0 | 16995 | 5d3a | B | 75 | ILE | p.Ile1010Thr | Q7Z4S6 |
| HOMO SAPIENS | 0 | 17091 | 2yin | B | 1324 | GLN | p.Gln1324Ter | Q92608 |
| HOMO SAPIENS | 5 | 27606 | 1lm7 | B | 2366 | ARG | p.Arg2366Cys | P15924 |
| HOMO SAPIENS | 2 | 15731 | 1bi7 | B | 24 | ARG | p.Arg24Pro | P42771 |
| HOMO SAPIENS | 0 | 15731 | 1bi7 | B | 28 | VAL | p.Val28Gly | P42771 |
| HOMO SAPIENS | 0 | 15731 | 1bi7 | B | 49 | ILE | p.Ile49Thr | P42771 |
| HOMO SAPIENS | 0 | 15731 | 1bi7 | B | 53 | MET | p.Met53Ile | P42771 |
| HOMO SAPIENS | 2 | 15731 | 1bi7 | B | 56 | SER | p.Ser56Ile | P42771 |
| HOMO SAPIENS | 1 | 15731 | 1bi7 | B | 57 | ALA | p.Ala57Val | P42771 |
| HOMO SAPIENS | 4 | 15731 | 1bi7 | B | 59 | VAL | p.Val59Gly | P42771 |
| HOMO SAPIENS | 9 | 15731 | 1bi7 | B | 89 | GLY | p.Gly89Asp | P42771 |
| HOMO SAPIENS | 0 | 15731 | 1bi7 | B | 101 | GLY | p.Gly101Trp | P42771 |
| HOMO SAPIENS | 0 | 15731 | 1bi7 | B | 114 | PRO | p.Pro114Ser | P42771 |
| HOMO SAPIENS | 9 | 15731 | 1bi7 | B | 123 | HIS | p.His123Gln | P42771 |
| HOMO SAPIENS | 8 | 15731 | 1bi7 | B | 126 | VAL | p.Val126Asp | P42771 |
| HOMO SAPIENS | 7 | 15731 | 1bi7 | B | 127 | ALA | p.Ala127Ser | P42771 |
| HOMO SAPIENS | -1 | 15731 | 1bi7 | B | 148 | ALA | p.Ala148Thr | P42771 |
| HOMO SAPIENS | -1 | 19858 | 3jbk | M | 866 | ALA | p.Ala934Val | P18206 |
| HOMO SAPIENS | 0 | 19484 | 3ee6 | B | 62 | SER | p.Ser62Leu | O14773 |
| HOMO SAPIENS | 1 | 19484 | 3ee6 | B | 66 | GLN | p.Gln66Ter | O14773 |
| HOMO SAPIENS | 0 | 19484 | 3ee6 | B | 100 | GLN | p.Gln100Arg | O14773 |
| HOMO SAPIENS | 0 | 19484 | 3ee6 | B | 104 | LEU | p.Leu104Ter | O14773 |
| HOMO SAPIENS | 0 | 19484 | 3ee6 | B | 127 | ARG | p.Arg127Gln | O14773 |
| HOMO SAPIENS | 0 | 19484 | 3ee6 | B | 127 | ARG | p.Arg127Ter | O14773 |
| HOMO SAPIENS | -1 | 19484 | 3ee6 | B | 185 | ARG | p.Arg185Cys | O14773 |
| HOMO SAPIENS | 0 | 19484 | 3ee6 | B | 206 | ARG | p.Arg206Cys | O14773 |
| HOMO SAPIENS | 0 | 19484 | 3ee6 | B | 208 | ARG | p.Arg208Ter | O14773 |
| HOMO SAPIENS | 0 | 19484 | 3ee6 | B | 214 | GLN | p.Gln214Ter | O14773 |
| HOMO SAPIENS | 1 | 19484 | 3ee6 | B | 276 | ASP | p.Asp276Val | O14773 |
| HOMO SAPIENS | 0 | 19484 | 3ee6 | B | 284 | GLY | p.Gly284Val | O14773 |

| | | | | | | | | |
|---|---|---|---|---|---|---|---|---|
| HOMO SAPIENS | 0 | 19484 | 3ee6 | B | 286 | ASN | p.Asn286Ser | O14773 |
| HOMO SAPIENS | 0 | 19484 | 3ee6 | B | 339 | ARG | p.Arg339Gly | O14773 |
| HOMO SAPIENS | 2 | 19484 | 3ee6 | B | 345 | MET | p.Met345Leu | O14773 |
| HOMO SAPIENS | 0 | 19484 | 3ee6 | B | 365 | CYS | p.Cys365Arg | O14773 |
| HOMO SAPIENS | 0 | 19484 | 3ee6 | B | 365 | CYS | p.Cys365Tyr | O14773 |
| HOMO SAPIENS | 0 | 19484 | 3ee6 | B | 373 | GLN | p.Gln373Glu | O14773 |
| HOMO SAPIENS | 6 | 19484 | 3ee6 | B | 418 | ARG | p.Arg418Gln | O14773 |
| HOMO SAPIENS | 7 | 19484 | 3ee6 | B | 422 | GLN | p.Gln422His | O14773 |
| HOMO SAPIENS | 0 | 19484 | 3ee6 | B | 447 | ARG | p.Arg447His | O14773 |
| HOMO SAPIENS | 0 | 19484 | 3ee6 | B | 466 | VAL | p.Val466Met | O14773 |
| HOMO SAPIENS | 0 | 4654 | 5t5w | A | 47 | LYS | p.Lys47Glu | Q08334 |
| HOMO SAPIENS | 0 | 4654 | 5t5w | A | 141 | GLU | p.Glu141Ter | Q08334 |
| HOMO SAPIENS | -1 | 18815 | 4wj7 | A | 53 | VAL | p.Val53Ile | Q9BSQ5 |
| HOMO SAPIENS | 0 | 18815 | 4wj7 | A | 120 | VAL | p.Val120Ile | Q9BSQ5 |
| HOMO SAPIENS | 0 | 18815 | 4wj7 | A | 198 | LEU | p.Leu198Arg | Q9BSQ5 |
| HOMO SAPIENS | 0 | 4602 | 1mjd | A | 47 | SER | p.Ser47Arg | O43602 |
| HOMO SAPIENS | 6 | 4602 | 1mjd | A | 59 | ARG | p.Arg59Leu | O43602 |
| HOMO SAPIENS | 7 | 4602 | 1mjd | A | 62 | ASP | p.Asp62Asn | O43602 |
| HOMO SAPIENS | 7 | 4602 | 1mjd | A | 71 | ALA | p.Ala71Ser | O43602 |
| HOMO SAPIENS | 7 | 4602 | 1mjd | A | 78 | ARG | p.Arg78Leu | O43602 |
| HOMO SAPIENS | 7 | 4602 | 1mjd | A | 89 | ARG | p.Arg89Gly | O43602 |
| HOMO SAPIENS | 8 | 4602 | 1mjd | A | 104 | ILE | p.Ile104Phe | O43602 |
| HOMO SAPIENS | 7 | 4602 | 1mjd | A | 125 | TYR | p.Tyr125His | O43602 |
| HOMO SAPIENS | 5 | 34467 | 1n0w | B | 1529 | GLY | p.Gly1529Arg | P51587 |
| HOMO SAPIENS | 8 | 5738 | 1u31 | A | 130 | ASN | p.Asn1009Lys | Q13423 |
| HOMO SAPIENS | 0 | 5760 | 2i6s | A | 298 | GLU | p.Glu318Asp | P06681 |
| HOMO SAPIENS | 0 | 5760 | 2i6s | A | 573 | ARG | p.Arg593Gln | P06681 |
| HOMO SAPIENS | 0 | 5760 | 2i6s | A | 592 | SER | p.Ser612Asn | P06681 |
| HOMO SAPIENS | 0 | 5760 | 2i6s | A | 621 | VAL | p.Val641Ala | P06681 |
| HOMO SAPIENS | 1 | 10977 | 2fby | A | 114 | VAL | p.Val114Ile | Q14191 |
| HOMO SAPIENS | 0 | 10977 | 2fby | A | 135 | LYS | p.Lys135Glu | Q14191 |
| HOMO SAPIENS | -1 | 1175 | 1qrk | A | 34 | VAL | p.Val35Leu | P00488 |
| HOMO SAPIENS | 4 | 1175 | 1qrk | A | 564 | PRO | p.Pro565Leu | P00488 |
| HOMO SAPIENS | 0 | 1175 | 1qrk | A | 650 | VAL | p.Val651Ile | P00488 |
| HOMO SAPIENS | 0 | 1175 | 1qrk | A | 651 | GLU | p.Gln652Glu | P00488 |
| HOMO SAPIENS | 3 | 909 | 4h0w | A | 375 | GLU | p.Glu394Lys | P02787 |
| HOMO SAPIENS | 0 | 909 | 4h0w | A | 458 | ALA | p.Ala477Pro | P02787 |
| HOMO SAPIENS | 6 | 909 | 4h0w | A | 515 | TYR | p.Tyr534Ter | P02787 |
| HOMO SAPIENS | 7 | 909 | 4h0w | A | 570 | PRO | p.Pro589Ser | P02787 |
| HOMO SAPIENS | 3 | 909 | 4h0w | A | 652 | GLY | p.Gly671Glu | P02787 |
| HOMO SAPIENS | 5 | 1964 | 2car | A | 151 | TRP | p.Trp151Ter | Q9BY32 |
| HOMO SAPIENS | 7 | 1964 | 2car | A | 178 | ARG | p.Arg178Cys | Q9BY32 |
| HOMO SAPIENS | 0 | 144 | 2n0a | J | 30 | ALA | p.Ala30Pro | P37840 |
| HOMO SAPIENS | 1 | 12288 | 4q58 | D | 1225 | ARG | p.Arg1225His | P16144 |
| HOMO SAPIENS | 2 | 12288 | 4q58 | D | 1281 | ARG | p.Arg1281Trp | P16144 |
| HOMO SAPIENS | 2 | 12288 | 4q58 | D | 1287 | ASN | p.Asn1287Lys | P16144 |
| HOMO SAPIENS | 0 | 5085 | 3s94 | A | 360 | ARG | p.Arg360His | O75581 |
| HOMO SAPIENS | 0 | 5085 | 3s94 | A | 433 | ASN | p.Asn433Ser | O75581 |
| HOMO SAPIENS | 2 | 5085 | 3s94 | A | 611 | ARG | p.Arg611Cys | O75581 |
| HOMO SAPIENS | 0 | 15452 | 4dur | A | 19 | LYS | p.Lys38Glu | P00747 |
| HOMO SAPIENS | 2 | 15452 | 4dur | A | 216 | ARG | p.Arg235His | P00747 |
| HOMO SAPIENS | 7 | 8406 | 4wo1 | D | 28 | VAL | p.Val55Leu | O43914 |
| HOMO SAPIENS | 2 | 895 | 5xte | K | 183 | ARG | p.Arg183Trp | P22695 |
| HOMO SAPIENS | 2 | 4191 | 4wnd | A | 120 | ARG | p.Arg127Ter | P81274 |

| | | | | | | | | |
|---|---|---|---|---|---|---|---|---|
| HOMO SAPIENS | 2 | 4191 | 4wnd | A | 120 | ARG | p.Arg127Gln | P81274 |
| HOMO SAPIENS | 1 | 4191 | 4wnd | A | 349 | GLY | p.Gly356Arg | P81274 |
| HOMO SAPIENS | 0 | 41734 | 2kbi | A | 1777 | VAL | p.Val1777Met | Q14524 |
| HOMO SAPIENS | 1 | 41734 | 2kbi | A | 1784 | GLU | p.Glu1784Lys | Q14524 |
| HOMO SAPIENS | 5 | 41734 | 2kbi | A | 1795 | TYR | p.Tyr1795Cys | Q14524 |
| HOMO SAPIENS | 5 | 41734 | 2kbi | A | 1795 | TYR | p.Tyr1795His | Q14524 |
| HOMO SAPIENS | 7 | 41734 | 2kbi | A | 1826 | ARG | p.Arg1826Cys | Q14524 |
| HOMO SAPIENS | 3 | 2119 | 1pl0 | A | 426 | LYS | p.Lys426Arg | P31939 |
| HOMO SAPIENS | 5 | 3475 | 5u62 | C | 50 | LEU | p.Leu50Ser | Q15910 |
| HOMO SAPIENS | 3 | 379 | 4uez | A | 200 | TYR | p.Tyr200Ter | P15085 |
| HOMO SAPIENS | 0 | 8195 | 5xaw | B | 67 | ARG | p.Arg67Trp | Q8IWT1 |
| HOMO SAPIENS | 0 | 8195 | 5xaw | B | 94 | THR | p.Thr94Met | Q8IWT1 |
| HOMO SAPIENS | 1 | 5711 | 1fo3 | A | 265 | ARG | p.Arg265His | Q9UKM7 |
| HOMO SAPIENS | 4 | 5711 | 1fo3 | A | 334 | ARG | p.Arg334Cys | Q9UKM7 |
| HOMO SAPIENS | 1 | 5711 | 1fo3 | A | 478 | LYS | p.Lys478Arg | Q9UKM7 |
| HOMO SAPIENS | 1 | 5711 | 1fo3 | A | 682 | ASP | p.Asp682Glu | Q9UKM7 |
| HOMO SAPIENS | 0 | 170 | 2n3y | A | 41 | GLY | p.Gly42Ser | P99999 |
| HOMO SAPIENS | 0 | 4122 | 3w81 | A | 33 | HIS | p.His33Gln | P35475 |
| HOMO SAPIENS | 0 | 4122 | 3w81 | A | 64 | TYR | p.Tyr64Ter | P35475 |
| HOMO SAPIENS | 0 | 4122 | 3w81 | A | 70 | GLN | p.Gln70Ter | P35475 |
| HOMO SAPIENS | 0 | 4122 | 3w81 | A | 75 | ALA | p.Ala75Thr | P35475 |
| HOMO SAPIENS | 0 | 4122 | 3w81 | A | 79 | ALA | p.Ala79Val | P35475 |
| HOMO SAPIENS | 0 | 4122 | 3w81 | A | 79 | ALA | p.Ala79Thr | P35475 |
| HOMO SAPIENS | 0 | 4122 | 3w81 | A | 89 | ARG | p.Arg89Gln | P35475 |
| HOMO SAPIENS | 0 | 4122 | 3w81 | A | 99 | THR | p.Thr99Ile | P35475 |
| HOMO SAPIENS | 0 | 4122 | 3w81 | A | 105 | ARG | p.Arg105Gln | P35475 |
| HOMO SAPIENS | 0 | 4122 | 3w81 | A | 238 | LEU | p.Leu238Gln | P35475 |
| HOMO SAPIENS | 0 | 4122 | 3w81 | A | 300 | ALA | p.Ala300Thr | P35475 |
| HOMO SAPIENS | 0 | 4122 | 3w81 | A | 322 | VAL | p.Val322Glu | P35475 |
| HOMO SAPIENS | 0 | 4122 | 3w81 | A | 327 | ALA | p.Ala327Pro | P35475 |
| HOMO SAPIENS | 0 | 4122 | 3w81 | A | 346 | LEU | p.Leu346Arg | P35475 |
| HOMO SAPIENS | 0 | 4122 | 3w81 | A | 349 | ASP | p.Asp349Asn | P35475 |
| HOMO SAPIENS | 0 | 4122 | 3w81 | A | 361 | ALA | p.Ala361Thr | P35475 |
| HOMO SAPIENS | 0 | 4122 | 3w81 | A | 366 | THR | p.Thr366Pro | P35475 |
| HOMO SAPIENS | 0 | 4122 | 3w81 | A | 402 | TRP | p.Trp402Ter | P35475 |
| HOMO SAPIENS | 0 | 4122 | 3w81 | A | 409 | GLY | p.Gly409Arg | P35475 |
| HOMO SAPIENS | 0 | 4122 | 3w81 | A | 454 | VAL | p.Val454Ile | P35475 |
| HOMO SAPIENS | 1 | 4122 | 3w81 | A | 490 | LEU | p.Leu490Pro | P35475 |
| HOMO SAPIENS | 1 | 4122 | 3w81 | A | 492 | ARG | p.Arg492Pro | P35475 |
| HOMO SAPIENS | 0 | 4122 | 3w81 | A | 533 | PRO | p.Pro533Arg | P35475 |
| HOMO SAPIENS | 0 | 4122 | 3w81 | A | 591 | ALA | p.Ala591Thr | P35475 |
| HOMO SAPIENS | 7 | 4122 | 3w81 | A | 619 | ARG | p.Arg619Gly | P35475 |
| HOMO SAPIENS | 0 | 15325 | 3wvz | B | 54 | VAL | p.Val54Leu | Q53FT3 |
| HOMO SAPIENS | 3 | 3228 | 5iz8 | A | 421 | TRP | p.Trp421Ter | P25054 |
| HOMO SAPIENS | 2 | 3228 | 5iz8 | A | 437 | PRO | p.Pro437Arg | P25054 |
| HOMO SAPIENS | 3 | 3228 | 5iz8 | A | 499 | ARG | p.Arg499Ter | P25054 |
| HOMO SAPIENS | -1 | 29229 | 5buo | A | 539 | SER | p.Ser614Gly | P05067 |
| HOMO SAPIENS | -1 | 34931 | 4n7v | C | 54 | SER | p.Ser54Leu | O94986 |
| HOMO SAPIENS | 0 | 2450 | 5kzx | A | 123 | GLY | p.Gly123Glu | P10253 |
| HOMO SAPIENS | 1 | 2450 | 5kzx | A | 190 | ARG | p.Arg190His | P10253 |
| HOMO SAPIENS | -1 | 2450 | 5kzx | A | 199 | HIS | p.His199Arg | P10253 |
| HOMO SAPIENS | 0 | 2450 | 5kzx | A | 219 | GLY | p.Gly219Arg | P10253 |
| HOMO SAPIENS | 0 | 2450 | 5kzx | A | 220 | VAL | p.Val220Leu | P10253 |
| HOMO SAPIENS | 0 | 2450 | 5kzx | A | 222 | VAL | p.Val222Met | P10253 |

| | | | | | | | | |
|---|---|---|---|---|---|---|---|---|
| HOMO SAPIENS | 0 | 2450 | 5kzx | A | 237 | ALA | p.Ala237Val | P10253 |
| HOMO SAPIENS | 0 | 2450 | 5kzx | A | 251 | SER | p.Ser251Leu | P10253 |
| HOMO SAPIENS | 0 | 2450 | 5kzx | A | 254 | SER | p.Ser254Leu | P10253 |
| HOMO SAPIENS | 0 | 2450 | 5kzx | A | 262 | GLU | p.Glu262Lys | P10253 |
| HOMO SAPIENS | 0 | 2450 | 5kzx | A | 293 | GLY | p.Gly293Arg | P10253 |
| HOMO SAPIENS | 0 | 2450 | 5kzx | A | 299 | LEU | p.Leu299Arg | P10253 |
| HOMO SAPIENS | 0 | 2450 | 5kzx | A | 309 | GLY | p.Gly309Arg | P10253 |
| HOMO SAPIENS | 0 | 2450 | 5kzx | A | 318 | MET | p.Met318Thr | P10253 |
| HOMO SAPIENS | 0 | 2450 | 5kzx | A | 355 | LEU | p.Leu355Pro | P10253 |
| HOMO SAPIENS | 1 | 2450 | 5kzx | A | 361 | PRO | p.Pro361Leu | P10253 |
| HOMO SAPIENS | 4 | 2450 | 5kzx | A | 408 | MET | p.Met408Val | P10253 |
| HOMO SAPIENS | 1 | 2450 | 5kzx | A | 429 | GLN | p.Gln429Glu | P10253 |
| HOMO SAPIENS | 4 | 2450 | 5kzx | A | 437 | ARG | p.Arg437Cys | P10253 |
| HOMO SAPIENS | 0 | 2450 | 5kzx | A | 439 | MET | p.Met439Lys | P10253 |
| HOMO SAPIENS | 1 | 2450 | 5kzx | A | 448 | SER | p.Ser448Thr | P10253 |
| HOMO SAPIENS | 0 | 2450 | 5kzx | A | 451 | PRO | p.Pro451Arg | P10253 |
| HOMO SAPIENS | 1 | 2450 | 5kzx | A | 459 | ASP | p.Asp459His | P10253 |
| HOMO SAPIENS | 8 | 2450 | 5kzx | A | 481 | TRP | p.Trp481Arg | P10253 |
| HOMO SAPIENS | 1 | 2450 | 5kzx | A | 489 | ASP | p.Asp489Asn | P10253 |
| HOMO SAPIENS | 0 | 2450 | 5kzx | A | 521 | GLU | p.Glu521Lys | P10253 |
| HOMO SAPIENS | 0 | 2450 | 5kzx | A | 529 | SER | p.Ser529Val | P10253 |
| HOMO SAPIENS | 0 | 2450 | 5kzx | A | 545 | PRO | p.Pro545Leu | P10253 |
| HOMO SAPIENS | 0 | 2450 | 5kzx | A | 552 | LEU | p.Leu552Pro | P10253 |
| HOMO SAPIENS | 0 | 2450 | 5kzx | A | 576 | GLY | p.Gly576Ser | P10253 |
| HOMO SAPIENS | 9 | 2450 | 5kzx | A | 600 | ARG | p.Arg600His | P10253 |
| HOMO SAPIENS | 8 | 2450 | 5kzx | A | 615 | GLY | p.Gly615Arg | P10253 |
| HOMO SAPIENS | 3 | 2450 | 5kzx | A | 619 | SER | p.Ser619Asn | P10253 |
| HOMO SAPIENS | 0 | 2450 | 5kzx | A | 638 | GLY | p.Gly638Trp | P10253 |
| HOMO SAPIENS | 0 | 2450 | 5kzx | A | 643 | GLY | p.Gly643Arg | P10253 |
| HOMO SAPIENS | 9 | 2450 | 5kzx | A | 645 | ASP | p.Asp645Glu | P10253 |
| HOMO SAPIENS | 9 | 2450 | 5kzx | A | 645 | ASP | p.Asp645His | P10253 |
| HOMO SAPIENS | 0 | 2450 | 5kzx | A | 648 | GLY | p.Gly648Ser | P10253 |
| HOMO SAPIENS | 5 | 2450 | 5kzx | A | 660 | ARG | p.Arg660His | P10253 |
| HOMO SAPIENS | 8 | 2450 | 5kzx | A | 672 | ARG | p.Arg672Gln | P10253 |
| HOMO SAPIENS | 0 | 2450 | 5kzx | A | 702 | ARG | p.Arg702Leu | P10253 |
| HOMO SAPIENS | 3 | 2450 | 5kzx | A | 725 | ARG | p.Arg725Trp | P10253 |
| HOMO SAPIENS | 0 | 2450 | 5kzx | A | 746 | TRP | p.Trp746Cys | P10253 |
| HOMO SAPIENS | 0 | 2450 | 5kzx | A | 780 | ILE | p.Val780Ile | P10253 |
| HOMO SAPIENS | 1 | 2450 | 5kzx | A | 816 | VAL | p.Val816Ile | P10253 |
| HOMO SAPIENS | 0 | 2450 | 5kzx | A | 838 | GLN | p.Gln838Ter | P10253 |
| HOMO SAPIENS | 2 | 2450 | 5kzx | A | 854 | ARG | p.Arg854Ter | P10253 |
| HOMO SAPIENS | 3 | 2450 | 5kzx | A | 870 | ARG | p.Arg870Ter | P10253 |
| HOMO SAPIENS | 0 | 2450 | 5kzx | A | 890 | VAL | p.Val890Leu | P10253 |
| HOMO SAPIENS | 6 | 1882 | 4km6 | A | 175 | CYS | p.Cys175Ter | P15328 |
| HOMO SAPIENS | 3 | 14767 | 1lm5 | A | 2628 | LEU | p.Leu2628Pro | P15924 |
| HOMO SAPIENS | 5 | 14767 | 1lm5 | A | 2655 | ALA | p.Ala2655Gly | P15924 |
| HOMO SAPIENS | 3 | 14767 | 1lm5 | A | 2731 | TYR | p.Tyr2731His | P15924 |
| HOMO SAPIENS | 1 | 14767 | 1lm5 | A | 2767 | THR | p.Thr2767Asn | P15924 |
| HOMO SAPIENS | -1 | 14767 | 1lm5 | A | 2819 | MET | p.Met2819Leu | P15924 |
| HOMO SAPIENS | -1 | 11246 | 4y5o | A | 289 | SER | p.Ser289Asn | Q9BSQ5 |
| HOMO SAPIENS | 0 | 7380 | 5kz5 | I | 60 | ARG | p.Arg60His | Q16595 |
| HOMO SAPIENS | 0 | 7380 | 5kz5 | I | 76 | MET | p.Met76Val | Q16595 |
| HOMO SAPIENS | 2 | 22187 | 3fby | A | 328 | CYS | p.Cys328Arg | P49747 |
| HOMO SAPIENS | 2 | 22187 | 3fby | A | 342 | ASP | p.Asp342Tyr | P49747 |

| | | | | | | | | |
|---|---|---|---|---|---|---|---|---|
| HOMO SAPIENS | 2 | 22187 | 3fby | A | 348 | CYS | p.Cys348Arg | P49747 |
| HOMO SAPIENS | 2 | 22187 | 3fby | A | 386 | ASN | p.Asn386Asp | P49747 |
| HOMO SAPIENS | 2 | 22187 | 3fby | A | 453 | ASN | p.Asn453Ser | P49747 |
| HOMO SAPIENS | 2 | 22187 | 3fby | A | 468 | CYS | p.Cys468Tyr | P49747 |
| HOMO SAPIENS | 3 | 22187 | 3fby | A | 472 | ASP | p.Asp472Tyr | P49747 |
| HOMO SAPIENS | 3 | 22187 | 3fby | A | 473 | ASP | p.Asp473Gly | P49747 |
| HOMO SAPIENS | 2 | 22187 | 3fby | A | 523 | ASN | p.Asn523Lys | P49747 |
| HOMO SAPIENS | 1 | 22187 | 3fby | A | 718 | ARG | p.Arg718Trp | P49747 |
| HOMO SAPIENS | 1 | 22187 | 3fby | A | 719 | GLY | p.Gly719Asp | P49747 |
| HOMO SAPIENS | 0 | 22187 | 3fby | A | 756 | GLN | p.Gln756Arg | P49747 |
| HOMO SAPIENS | -1 | 2784 | 5kvh | A | 88 | MET | p.Met88Val | O95831 |
| HOMO SAPIENS | -1 | 2784 | 5kvh | A | 96 | ASN | p.Asn96Ser | O95831 |
| HOMO SAPIENS | 0 | 2784 | 5kvh | A | 282 | THR | p.Thr282Met | O95831 |
| HOMO SAPIENS | 1 | 2784 | 5kvh | A | 366 | ASN | p.Asn366Ser | O95831 |
| HOMO SAPIENS | 3 | 2784 | 5kvh | A | 463 | ARG | p.Arg463Ile | O95831 |
| HOMO SAPIENS | 0 | 2784 | 5kvh | A | 498 | VAL | p.Val498Met | O95831 |
| HOMO SAPIENS | -1 | 2784 | 5kvh | A | 549 | ALA | p.Ala549Val | O95831 |
| HOMO SAPIENS | 0 | 38316 | 2lnk | C | 1933 | ARG | p.Arg1933Ter | P35579 |
| HOMO SAPIENS | 0 | 33808 | 2miu | A | 29 | VAL | p.Val29Met | Q14192 |
| HOMO SAPIENS | 1 | 33808 | 2miu | A | 37 | ALA | p.Ala37Ser | Q14192 |
| HOMO SAPIENS | 2 | 20994 | 2hey | R | 65 | ARG | p.Arg65Cys | P43489 |
| HOMO SAPIENS | -1 | 5687 | 2xwj | I | 7 | ARG | p.Arg32Gln | P00751 |
| HOMO SAPIENS | -1 | 5687 | 2xwj | I | 7 | ARG | p.Arg32Trp | P00751 |
| HOMO SAPIENS | 0 | 5687 | 2xwj | I | 49 | ARG | p.Arg74His | P00751 |
| HOMO SAPIENS | 0 | 5687 | 2xwj | I | 177 | ARG | p.Arg202Gly | P00751 |
| HOMO SAPIENS | 0 | 5687 | 2xwj | I | 217 | ILE | p.Ile242Leu | P00751 |
| HOMO SAPIENS | -1 | 5687 | 2xwj | I | 227 | GLY | p.Gly252Ser | P00751 |
| HOMO SAPIENS | 0 | 5687 | 2xwj | I | 483 | HIS | p.His508Gln | P00751 |
| HOMO SAPIENS | 0 | 5687 | 2xwj | I | 508 | LYS | p.Lys533Arg | P00751 |
| HOMO SAPIENS | 0 | 5687 | 2xwj | I | 540 | LYS | p.Lys565Glu | P00751 |
| HOMO SAPIENS | 0 | 5687 | 2xwj | I | 541 | GLU | p.Glu566Ala | P00751 |
| HOMO SAPIENS | 2 | 953 | 3d4j | A | 249 | PHE | p.Phe249Ser | P53602 |
| HOMO SAPIENS | 1 | 953 | 3d4j | A | 292 | ASN | p.Asn292Ser | P53602 |
| HOMO SAPIENS | 0 | 3300 | 1s9c | A | 140 | ASN | p.Asn457Tyr | P51659 |
| HOMO SAPIENS | 2 | 3300 | 1s9c | A | 167 | ILE | p.Ile484Val | P51659 |
| HOMO SAPIENS | 0 | 3300 | 1s9c | A | 174 | ALA | p.Ala491Thr | P51659 |
| HOMO SAPIENS | 8 | 3300 | 1s9c | A | 193 | ASP | p.Asp510Asn | P51659 |
| HOMO SAPIENS | 3 | 3300 | 1s9c | A | 194 | TRP | p.Trp511Arg | P51659 |
| HOMO SAPIENS | 0 | 3300 | 1s9c | A | 242 | VAL | p.Ile559Val | P51659 |
| HOMO SAPIENS | 0 | 10810 | 5i25 | A | 5 | GLN | p.Gln23Ter | P03951 |
| HOMO SAPIENS | 0 | 10810 | 5i25 | A | 55 | TRP | p.Trp73Ter | P03951 |
| HOMO SAPIENS | 0 | 10810 | 5i25 | A | 91 | ALA | p.Ala109Thr | P03951 |
| HOMO SAPIENS | 0 | 10810 | 5i25 | A | 117 | GLU | p.Glu135Ter | P03951 |
| HOMO SAPIENS | 0 | 10810 | 5i25 | A | 118 | CYS | p.Cys136Ter | P03951 |
| HOMO SAPIENS | 0 | 10810 | 5i25 | A | 128 | CYS | p.Cys146Ter | P03951 |
| HOMO SAPIENS | 0 | 10810 | 5i25 | A | 210 | ARG | p.Arg228Ter | P03951 |
| HOMO SAPIENS | 0 | 10810 | 5i25 | A | 226 | GLN | p.Gln244Ter | P03951 |
| HOMO SAPIENS | 0 | 10810 | 5i25 | A | 233 | GLN | p.Gln251Ter | P03951 |
| HOMO SAPIENS | 0 | 16668 | 3lbx | A | 28 | ARG | p.Arg28Leu | P02549 |
| HOMO SAPIENS | 0 | 16668 | 3lbx | A | 28 | ARG | p.Arg28Cys | P02549 |
| HOMO SAPIENS | 1 | 16668 | 3lbx | A | 41 | ARG | p.Arg41Trp | P02549 |
| HOMO SAPIENS | 1 | 16668 | 3lbx | A | 45 | ARG | p.Arg45Ser | P02549 |
| HOMO SAPIENS | 2 | 16668 | 3lbx | A | 48 | LYS | p.Lys48Arg | P02549 |
| HOMO SAPIENS | 1 | 16668 | 3lbx | A | 152 | ASP | p.Asp152Asn | P02549 |

| | | | | | | | | |
|---|---|---|---|---|---|---|---|---|
| HOMO SAPIENS | 0 | 226 | 2ble | A | 256 | ILE | p.Phe256Ile | P36959 |
| HOMO SAPIENS | 2 | 36808 | 4c7n | A | 3 | SER | p.Ser357Pro | O75030 |
| HOMO SAPIENS | 0 | 4092 | 5jnx | A | 113 | CYS | p.Cys113Arg | O15118 |
| HOMO SAPIENS | 4 | 4092 | 5jnx | A | 215 | HIS | p.His215Arg | O15118 |
| HOMO SAPIENS | 6 | 4092 | 5jnx | A | 237 | PRO | p.Pro237Ser | O15118 |
| HOMO SAPIENS | 0 | 4092 | 5jnx | A | 363 | ALA | p.Ala363Val | O15118 |
| HOMO SAPIENS | 0 | 4092 | 5jnx | A | 378 | VAL | p.Val378Ala | O15118 |
| HOMO SAPIENS | 8 | 4092 | 5jnx | A | 404 | ARG | p.Arg404Gln | O15118 |
| HOMO SAPIENS | 8 | 4092 | 5jnx | A | 411 | ARG | p.Arg411Gln | O15118 |
| HOMO SAPIENS | 2 | 4092 | 5jnx | A | 474 | PRO | p.Pro474Leu | O15118 |
| HOMO SAPIENS | 8 | 4092 | 5jnx | A | 511 | THR | p.Thr511Met | O15118 |
| HOMO SAPIENS | 0 | 4092 | 5jnx | A | 642 | MET | p.Met642Ile | O15118 |
| HOMO SAPIENS | 2 | 4092 | 5jnx | A | 664 | VAL | p.Val664Met | O15118 |
| HOMO SAPIENS | 0 | 4092 | 5jnx | A | 734 | SER | p.Ser734Ile | O15118 |
| HOMO SAPIENS | -1 | 4092 | 5jnx | A | 775 | GLN | p.Gln775Pro | O15118 |
| HOMO SAPIENS | 0 | 4092 | 5jnx | A | 858 | ILE | p.Ile858Val | O15118 |
| HOMO SAPIENS | 0 | 4092 | 5jnx | A | 889 | VAL | p.Val889Met | O15118 |
| HOMO SAPIENS | 0 | 4092 | 5jnx | A | 910 | GLY | p.Gly910Ser | O15118 |
| HOMO SAPIENS | 0 | 4092 | 5jnx | A | 911 | GLY | p.Gly911Ser | O15118 |
| HOMO SAPIENS | 0 | 4092 | 5jnx | A | 928 | GLN | p.Gln928Pro | O15118 |
| HOMO SAPIENS | 0 | 4092 | 5jnx | A | 931 | ASN | p.Asn931Lys | O15118 |
| HOMO SAPIENS | 0 | 4092 | 5jnx | A | 940 | SER | p.Ser940Leu | O15118 |
| HOMO SAPIENS | 0 | 4092 | 5jnx | A | 950 | VAL | p.Val950Met | O15118 |
| HOMO SAPIENS | 0 | 4092 | 5jnx | A | 954 | SER | p.Ser954Leu | O15118 |
| HOMO SAPIENS | 0 | 4092 | 5jnx | A | 958 | ARG | p.Arg958Gln | O15118 |
| HOMO SAPIENS | 0 | 4092 | 5jnx | A | 958 | ARG | p.Arg958Ter | O15118 |
| HOMO SAPIENS | -1 | 4092 | 5jnx | A | 978 | ARG | p.Arg978Cys | O15118 |
| HOMO SAPIENS | 0 | 4092 | 5jnx | A | 992 | GLY | p.Gly992Trp | O15118 |
| HOMO SAPIENS | 0 | 4092 | 5jnx | A | 1007 | PRO | p.Pro1007Ala | O15118 |
| HOMO SAPIENS | 0 | 4092 | 5jnx | A | 1019 | TYR | p.Tyr1019Cys | O15118 |
| HOMO SAPIENS | 2 | 4092 | 5jnx | A | 1035 | ALA | p.Ala1035Val | O15118 |
| HOMO SAPIENS | 2 | 4092 | 5jnx | A | 1054 | ALA | p.Ala1054Thr | O15118 |
| HOMO SAPIENS | 2 | 4092 | 5jnx | A | 1061 | ILE | p.Ile1061Thr | O15118 |
| HOMO SAPIENS | 0 | 4092 | 5jnx | A | 1073 | GLY | p.Gly1073Ser | O15118 |
| HOMO SAPIENS | 2 | 4092 | 5jnx | A | 1088 | TYR | p.Tyr1088Cys | O15118 |
| HOMO SAPIENS | 2 | 4092 | 5jnx | A | 1142 | MET | p.Met1142Thr | O15118 |
| HOMO SAPIENS | 2 | 4092 | 5jnx | A | 1156 | ASN | p.Asn1156Ser | O15118 |
| HOMO SAPIENS | 0 | 4092 | 5jnx | A | 1179 | MET | p.Met1179Val | O15118 |
| HOMO SAPIENS | 0 | 4092 | 5jnx | A | 1186 | ARG | p.Arg1186His | O15118 |
| HOMO SAPIENS | 0 | 4092 | 5jnx | A | 1187 | ALA | p.Ala1187Gly | O15118 |
| HOMO SAPIENS | 2 | 4092 | 5jnx | A | 1189 | GLU | p.Glu1189Gly | O15118 |
| HOMO SAPIENS | 2 | 4092 | 5jnx | A | 1200 | SER | p.Ser1200Gly | O15118 |
| HOMO SAPIENS | 2 | 4092 | 5jnx | A | 1205 | THR | p.Thr1205Lys | O15118 |
| HOMO SAPIENS | 0 | 4092 | 5jnx | A | 1213 | LEU | p.Leu1213Phe | O15118 |
| HOMO SAPIENS | -1 | 4092 | 5jnx | A | 1266 | ARG | p.Arg1266Gln | O15118 |
| HOMO SAPIENS | 9 | 16498 | 4i6o | A | 64 | ARG | p.Arg735Trp | P01024 |
| HOMO SAPIENS | 1 | 21902 | 3un9 | C | 751 | ARG | p.Arg751His | Q86UT6 |
| HOMO SAPIENS | 7 | 20391 | 2h32 | B | 105 | PRO | p.Pro142Leu | P15814 |
| HOMO SAPIENS | 6 | 1635 | 4x45 | A | 65 | ASP | p.Asp65Val | P07741 |
| HOMO SAPIENS | 1 | 1635 | 4x45 | A | 98 | TRP | p.Trp98Ter | P07741 |
| HOMO SAPIENS | 0 | 1635 | 4x45 | A | 110 | LEU | p.Leu110Pro | P07741 |
| HOMO SAPIENS | 7 | 1635 | 4x45 | A | 136 | MET | p.Met136Thr | P07741 |
| HOMO SAPIENS | 1 | 1635 | 4x45 | A | 150 | VAL | p.Val150Phe | P07741 |
| HOMO SAPIENS | 4 | 34810 | 2rpc | A | 16 | CYS | p.Cys253Ser | O60481 |

| | | | | | | | | |
|---|---|---|---|---|---|---|---|---|
| HOMO SAPIENS | 1 | 34810 | 2rpc | A | 18 | TRP | p.Trp255Gly | O60481 |
| HOMO SAPIENS | 1 | 14359 | 4hxi | A | 78 | MET | p.Met78Val | Q9UH77 |
| HOMO SAPIENS | 0 | 5342 | 4d9t | A | 628 | GLU | p.Glu628Asp | P51812 |
| HOMO SAPIENS | -1 | 5342 | 4d9t | A | 723 | ARG | p.Arg723His | P51812 |
| HOMO SAPIENS | -1 | 5342 | 4d9t | A | 729 | ARG | p.Arg729Gln | P51812 |
| HOMO SAPIENS | 1 | 9619 | 3j2s | B | 1781 | ARG | p.Arg1800Cys | P00451 |
| HOMO SAPIENS | 0 | 9619 | 3j2s | B | 1823 | MET | p.Met1842Ile | P00451 |
| HOMO SAPIENS | 1 | 9619 | 3j2s | B | 1825 | PRO | p.Pro1844Ser | P00451 |
| HOMO SAPIENS | 0 | 9619 | 3j2s | B | 1826 | THR | p.Thr1845Pro | P00451 |
| HOMO SAPIENS | 0 | 9619 | 3j2s | B | 1834 | ALA | p.Ala1853Val | P00451 |
| HOMO SAPIENS | 0 | 9619 | 3j2s | B | 1846 | ASP | p.Asp1865Asn | P00451 |
| HOMO SAPIENS | 1 | 9619 | 3j2s | B | 1848 | HIS | p.His1867Arg | P00451 |
| HOMO SAPIENS | 0 | 9619 | 3j2s | B | 1854 | PRO | p.Pro1873Arg | P00451 |
| HOMO SAPIENS | 1 | 9619 | 3j2s | B | 1885 | GLU | p.Glu1904Lys | P00451 |
| HOMO SAPIENS | 0 | 9619 | 3j2s | B | 1922 | ASN | p.Asn1941Ser | P00451 |
| HOMO SAPIENS | 1 | 9619 | 3j2s | B | 1941 | ARG | p.Arg1960Gln | P00451 |
| HOMO SAPIENS | 4 | 9619 | 3j2s | B | 2105 | TYR | p.Tyr2124Cys | P00451 |
| HOMO SAPIENS | 6 | 9619 | 3j2s | B | 2150 | ARG | p.Arg2169His | P00451 |
| HOMO SAPIENS | 0 | 9619 | 3j2s | B | 2159 | ARG | p.Arg2178Cys | P00451 |
| HOMO SAPIENS | 0 | 9619 | 3j2s | B | 2159 | ARG | p.Arg2178His | P00451 |
| HOMO SAPIENS | 2 | 9619 | 3j2s | B | 2238 | MET | p.Met2257Val | P00451 |
| HOMO SAPIENS | 2 | 9619 | 3j2s | B | 2300 | PRO | p.Pro2319Ser | P00451 |
| HOMO SAPIENS | 1 | 667 | 1z2m | A | 127 | GLU | p.Glu127Ter | P05161 |
| HOMO SAPIENS | 0 | 15717 | 2nn6 | G | 30 | GLY | p.Gly31Ala | Q9NQT5 |
| HOMO SAPIENS | -1 | 15717 | 2nn6 | G | 55 | ASN | p.Asn56His | Q9NQT5 |
| HOMO SAPIENS | 7 | 15717 | 2nn6 | G | 131 | ASP | p.Asp132Ala | Q9NQT5 |
| HOMO SAPIENS | 7 | 784 | 3rkq | A | 161 | ARG | p.Arg161Pro | P52952 |
| HOMO SAPIENS | 6 | 784 | 3rkq | A | 178 | THR | p.Thr178Met | P52952 |
| HOMO SAPIENS | 7 | 784 | 3rkq | A | 183 | LYS | p.Lys183Glu | P52952 |
| HOMO SAPIENS | 8 | 784 | 3rkq | A | 190 | ARG | p.Arg190Cys | P52952 |
| HOMO SAPIENS | 6 | 385 | 30000 | C | 70 | ALA | p.Ala70Thr | P07954 |
| HOMO SAPIENS | 2 | 385 | 30000 | C | 101 | ARG | p.Arg101Pro | P07954 |
| HOMO SAPIENS | 2 | 385 | 30000 | C | 174 | PRO | p.Pro174Arg | P07954 |
| HOMO SAPIENS | 6 | 385 | 30000 | C | 187 | SER | p.Ser187Ter | P07954 |
| HOMO SAPIENS | 9 | 385 | 30000 | C | 233 | ARG | p.Arg233His | P07954 |
| HOMO SAPIENS | 3 | 385 | 30000 | C | 273 | ALA | p.Ala273Thr | P07954 |
| HOMO SAPIENS | 4 | 385 | 30000 | C | 334 | SER | p.Ser334Arg | P07954 |
| HOMO SAPIENS | 6 | 385 | 30000 | C | 340 | ASN | p.Asn340Lys | P07954 |
| HOMO SAPIENS | 8 | 385 | 30000 | C | 343 | ARG | p.Arg343Ter | P07954 |
| HOMO SAPIENS | 3 | 385 | 30000 | C | 350 | ARG | p.Arg350Trp | P07954 |
| HOMO SAPIENS | 4 | 385 | 30000 | C | 376 | GLN | p.Gln376Pro | P07954 |
| HOMO SAPIENS | 6 | 385 | 30000 | C | 434 | CYS | p.Cys434Ter | P07954 |
| HOMO SAPIENS | 0 | 3316 | 4f91 | B | 934 | THR | p.Thr934Ala | O75643 |
| HOMO SAPIENS | 0 | 3316 | 4f91 | B | 1002 | ASN | p.Asn1002Ser | O75643 |
| HOMO SAPIENS | 0 | 3316 | 4f91 | B | 1087 | SER | p.Ser1087Leu | O75643 |
| HOMO SAPIENS | 4 | 3316 | 4f91 | B | 1389 | VAL | p.Val1389Ile | O75643 |
| HOMO SAPIENS | 0 | 619 | 5hbt | B | 195 | ASP | p.Asp240His | P02708 |
| HOMO SAPIENS | 1 | 619 | 5hbt | B | 209 | ARG | p.Arg254Leu | P02708 |
| HOMO SAPIENS | 0 | 18171 | 2y25 | B | 1408 | ASP | p.Asp1408Asn | P52179 |
| HOMO SAPIENS | 0 | 18171 | 2y25 | B | 1453 | MET | p.Met1453Thr | P52179 |
| HOMO SAPIENS | 0 | 18171 | 2y25 | B | 1459 | ILE | p.Ile1459Lys | P52179 |
| HOMO SAPIENS | 0 | 18171 | 2y25 | B | 1558 | PHE | p.Phe1558Leu | P52179 |
| HOMO SAPIENS | 0 | 18171 | 2y25 | B | 1573 | ARG | p.Arg1573Gln | P52179 |
| HOMO SAPIENS | 7 | 18171 | 2y25 | B | 1663 | VAL | p.Val1663Met | P52179 |

| | | | | | | | | |
|---|---|---|---|---|---|---|---|---|
| HOMO SAPIENS | 8 | 14262 | 2fcw | B | 117 | SER | p.Ser138Ter | P01130 |
| HOMO SAPIENS | 8 | 14262 | 2fcw | B | 118 | ASP | p.Asp139Glu | P01130 |
| HOMO SAPIENS | 9 | 14262 | 2fcw | B | 119 | GLU | p.Glu140Lys | P01130 |
| HOMO SAPIENS | 7 | 14262 | 2fcw | B | 122 | CYS | p.Cys143Ter | P01130 |
| HOMO SAPIENS | 6 | 14262 | 2fcw | B | 134 | CYS | p.Cys155Ter | P01130 |
| HOMO SAPIENS | 7 | 14262 | 2fcw | B | 147 | ASP | p.Asp168Glu | P01130 |
| HOMO SAPIENS | 4 | 2624 | 3b7b | A | 855 | VAL | p.Val886Met | Q9H9B1 |
| HOMO SAPIENS | 7 | 2624 | 3b7b | A | 868 | ILE | p.Ile899Val | Q9H9B1 |
| HOMO SAPIENS | 0 | 2624 | 3b7b | A | 892 | GLU | p.Glu923Lys | Q9H9B1 |
| HOMO SAPIENS | 0 | 4791 | 5ctd | B | 68 | LEU | p.Pro549Ala | Q14055 |
| HOMO SAPIENS | 5 | 460 | 4ui1 | D | 80 | CYS | p.Cys80Arg | Q6ZVN8 |
| HOMO SAPIENS | 3 | 460 | 4ui1 | D | 101 | LEU | p.Leu101Pro | Q6ZVN8 |
| HOMO SAPIENS | 0 | 40782 | 1sy6 | A | 47 | LYS | p.Lys69Ter | P09693 |
| HOMO SAPIENS | -1 | 1092 | 2idx | A | 62 | THR | p.Thr62Met | Q96EY8 |
| HOMO SAPIENS | 6 | 1092 | 2idx | A | 183 | HIS | p.His183Leu | Q96EY8 |
| HOMO SAPIENS | 7 | 1092 | 2idx | A | 186 | ARG | p.Arg186Trp | Q96EY8 |
| HOMO SAPIENS | 6 | 1092 | 2idx | A | 190 | ARG | p.Arg190Cys | Q96EY8 |
| HOMO SAPIENS | 6 | 1092 | 2idx | A | 190 | ARG | p.Arg190His | Q96EY8 |
| HOMO SAPIENS | 7 | 1092 | 2idx | A | 191 | ARG | p.Arg191Trp | Q96EY8 |
| HOMO SAPIENS | 7 | 1092 | 2idx | A | 191 | ARG | p.Arg191Gln | Q96EY8 |
| HOMO SAPIENS | 6 | 1092 | 2idx | A | 195 | ARG | p.Arg195His | Q96EY8 |
| HOMO SAPIENS | 1 | 1092 | 2idx | A | 234 | GLN | p.Gln234Ter | Q96EY8 |
| HOMO SAPIENS | 0 | 1092 | 2idx | A | 239 | MET | p.Met239Lys | Q96EY8 |
| HOMO SAPIENS | -1 | 6418 | 5fur | G | 11 | THR | p.Thr11Ile | P21675 |
| HOMO SAPIENS | -1 | 6418 | 5fur | G | 290 | LEU | p.Leu269Val | P21675 |
| HOMO SAPIENS | 0 | 6100 | 1wiq | A | 240 | ARG | p.Arg265Trp | P01730 |
| HOMO SAPIENS | 0 | 13757 | 4ye6 | A | 45 | GLY | p.Gly45Val | P47897 |
| HOMO SAPIENS | 1 | 13757 | 4ye6 | A | 57 | TYR | p.Tyr57His | P47897 |
| HOMO SAPIENS | 8 | 13757 | 4ye6 | A | 403 | ARG | p.Arg403Trp | P47897 |
| HOMO SAPIENS | 7 | 13757 | 4ye6 | A | 524 | ARG | p.Arg524Trp | P47897 |
| HOMO SAPIENS | 6 | 13757 | 4ye6 | A | 690 | ARG | p.Arg690Cys | P47897 |
| HOMO SAPIENS | 0 | 26958 | 2lzl | A | 370 | GLY | p.Gly370Cys | P22607 |
| HOMO SAPIENS | 0 | 26958 | 2lzl | A | 371 | SER | p.Ser371Cys | P22607 |
| HOMO SAPIENS | 0 | 26958 | 2lzl | A | 373 | TYR | p.Tyr373Cys | P22607 |
| HOMO SAPIENS | 0 | 26958 | 2lzl | A | 380 | GLY | p.Gly380Arg | P22607 |
| HOMO SAPIENS | 0 | 26958 | 2lzl | A | 384 | PHE | p.Phe384Leu | P22607 |
| HOMO SAPIENS | 5 | 26958 | 2lzl | A | 391 | ALA | p.Ala391Glu | P22607 |
| HOMO SAPIENS | 6 | 11325 | 2wa6 | A | 121 | TYR | p.Tyr121Phe | O75369 |
| HOMO SAPIENS | 2 | 11325 | 2wa6 | A | 161 | PHE | p.Phe161Cys | O75369 |
| HOMO SAPIENS | 0 | 11325 | 2wa6 | A | 171 | LEU | p.Leu171Arg | O75369 |
| HOMO SAPIENS | 2 | 11325 | 2wa6 | A | 173 | ALA | p.Ala173Val | O75369 |
| HOMO SAPIENS | 0 | 11325 | 2wa6 | A | 202 | MET | p.Met202Val | O75369 |
| HOMO SAPIENS | 3 | 11325 | 2wa6 | A | 227 | GLU | p.Glu227Lys | O75369 |
| HOMO SAPIENS | 0 | 11325 | 2wa6 | A | 234 | LEU | p.Leu234Val | O75369 |
| HOMO SAPIENS | 2 | 11325 | 2wa6 | A | 235 | SER | p.Ser235Pro | O75369 |
| HOMO SAPIENS | 9 | 7240 | 3tgx | A | 137 | TYR | p.Tyr156Cys | Q9HBE5 |
| HOMO SAPIENS | 0 | 35479 | 4elj | A | 307 | THR | p.Thr307Ile | P06400 |
| HOMO SAPIENS | 0 | 35479 | 4elj | A | 310 | GLY | p.Gly310Glu | P06400 |
| HOMO SAPIENS | 0 | 35479 | 4elj | A | 377 | THR | p.Thr377Ser | P06400 |
| HOMO SAPIENS | 9 | 36706 | 1he7 | A | 359 | TYR | p.Tyr359Cys | P04629 |
| HOMO SAPIENS | 4 | 6538 | 4d0e | A | 504 | ARG | p.Arg504His | P46531 |
| HOMO SAPIENS | 1 | 26518 | 5fm5 | M | 600 | GLU | p.Glu600Val | P52179 |
| HOMO SAPIENS | -1 | 26518 | 5fm5 | M | 651 | ARG | p.Arg651Gln | P52179 |
| HOMO SAPIENS | -1 | 26518 | 5fm5 | M | 660 | PRO | p.Pro660Ala | P52179 |

| | | | | | | | | |
|---|---|---|---|---|---|---|---|---|
| HOMO SAPIENS | -1 | 26518 | 5fm5 | M | 688 | THR | p.Thr688Ser | P52179 |
| HOMO SAPIENS | -1 | 26518 | 5fm5 | M | 704 | GLU | p.Glu704Lys | P52179 |
| HOMO SAPIENS | -1 | 26518 | 5fm5 | M | 727 | THR | p.Thr727Ala | P52179 |
| HOMO SAPIENS | 2 | 27054 | 4rhp | B | 209 | LEU | p.Leu209Val | O75208 |
| HOMO SAPIENS | -1 | 27054 | 4rhp | B | 288 | LYS | p.Lys288Asn | O75208 |
| HOMO SAPIENS | 5 | 2384 | 5fqd | B | 419 | ARG | p.Arg419Ter | Q96SW2 |
| HOMO SAPIENS | 4 | 5154 | 2hl3 | A | 56 | LYS | p.Lys56Arg | Q14203 |
| HOMO SAPIENS | 0 | 5154 | 2hl3 | A | 59 | GLY | p.Gly59Ser | Q14203 |
| HOMO SAPIENS | 0 | 5154 | 2hl3 | A | 71 | GLY | p.Gly71Ala | Q14203 |
| HOMO SAPIENS | 0 | 5154 | 2hl3 | A | 71 | GLY | p.Gly71Arg | Q14203 |
| HOMO SAPIENS | 7 | 5154 | 2hl3 | A | 72 | THR | p.Thr72Pro | Q14203 |
| HOMO SAPIENS | 2 | 5154 | 2hl3 | A | 74 | GLN | p.Gln74Pro | Q14203 |
| HOMO SAPIENS | 2 | 20039 | 2l1g | A | 9 | GLY | p.Gly9Cys | Q9NVV9 |
| HOMO SAPIENS | 5 | 20039 | 2l1g | A | 81 | PHE | p.Phe81Leu | Q9NVV9 |
| HOMO SAPIENS | -1 | 97 | 5uz7 | A | 201 | ARG | p.Arg201His | P63092 |
| HOMO SAPIENS | -1 | 97 | 5uz7 | A | 201 | ARG | p.Arg201Cys | P63092 |
| HOMO SAPIENS | 0 | 3257 | 4b7w | B | 278 | GLU | p.Glu278Asp | P41235 |
| HOMO SAPIENS | 1 | 3257 | 4b7w | B | 331 | ARG | p.Arg331Cys | P41235 |
| HOMO SAPIENS | 0 | 2814 | 4qgk | B | 10 | GLN | p.Gln10Ter | P51648 |
| HOMO SAPIENS | 0 | 2814 | 4qgk | B | 177 | ARG | p.Arg177Ter | P51648 |
| HOMO SAPIENS | 9 | 2814 | 4qgk | B | 184 | THR | p.Thr184Met | P51648 |
| HOMO SAPIENS | 1 | 2814 | 4qgk | B | 188 | ALA | p.Ala188Val | P51648 |
| HOMO SAPIENS | 0 | 2814 | 4qgk | B | 214 | CYS | p.Cys214Tyr | P51648 |
| HOMO SAPIENS | 0 | 2814 | 4qgk | B | 245 | ASP | p.Asp245Asn | P51648 |
| HOMO SAPIENS | 0 | 2814 | 4qgk | B | 266 | LYS | p.Lys266Asn | P51648 |
| HOMO SAPIENS | 0 | 2814 | 4qgk | B | 315 | PRO | p.Pro315Ser | P51648 |
| HOMO SAPIENS | 0 | 2814 | 4qgk | B | 386 | ASN | p.Asn386Ser | P51648 |
| HOMO SAPIENS | -1 | 5486 | 5u6o | B | 47 | GLY | p.Gly47Val | O60741 |
| HOMO SAPIENS | 1 | 21475 | 3pdy | A | 679 | THR | p.Thr789Ile | Q15149 |
| HOMO SAPIENS | 0 | 32890 | 4djc | B | 1501 | LEU | p.Leu1501Val | Q14524 |
| HOMO SAPIENS | -1 | 32890 | 4djc | B | 1512 | ARG | p.Arg1512Trp | Q14524 |
| HOMO SAPIENS | 0 | 26683 | 2xze | B | 38 | ARG | p.Arg38Cys | O95630 |
| HOMO SAPIENS | 0 | 26683 | 2xze | B | 100 | PHE | p.Phe100Tyr | O95630 |
| HOMO SAPIENS | -1 | 23175 | 5dka | A | 163 | SER | p.Ser163Leu | Q96EQ8 |
| HOMO SAPIENS | -1 | 23175 | 5dka | A | 174 | ARG | p.Arg174Cys | Q96EQ8 |
| HOMO SAPIENS | 6 | 21177 | 3k1r | A | 155 | ARG | p.Arg155Ter | Q9Y6N9 |
| HOMO SAPIENS | 0 | 1237 | 2wwm | O | 40 | PRO | p.Pro40Arg | O75147 |
| HOMO SAPIENS | 1 | 2543 | 5ewl | B | 64 | VAL | p.Val64Leu | Q13224 |
| HOMO SAPIENS | 2 | 2543 | 5ewl | B | 271 | ALA | p.Ala271Glu | Q13224 |
| HOMO SAPIENS | 3 | 34262 | 2mng | A | 613 | PHE | p.Phe613Leu | Q9Y3Q4 |
| HOMO SAPIENS | 7 | 34262 | 2mng | A | 672 | SER | p.Ser672Arg | Q9Y3Q4 |
| HOMO SAPIENS | 4 | 1667 | 1f05 | A | 321 | LYS | p.Lys321Met | P37837 |
| HOMO SAPIENS | 0 | 25179 | 2knc | B | 752 | SER | p.Ser778Pro | P05106 |
| HOMO SAPIENS | 0 | 2185 | 2ig7 | A | 50 | TYR | p.Tyr50Cys | Q9Y259 |
| HOMO SAPIENS | 0 | 2185 | 2ig7 | A | 224 | ASN | p.Asn224His | Q9Y259 |
| HOMO SAPIENS | 0 | 2185 | 2ig7 | A | 270 | TYR | p.Tyr270Ter | Q9Y259 |
| HOMO SAPIENS | 0 | 2185 | 2ig7 | A | 328 | GLN | p.Gln328Arg | Q9Y259 |
| HOMO SAPIENS | 3 | 2201 | 2r0o | A | 179 | PRO | p.Pro179Leu | O43707 |
| HOMO SAPIENS | 3 | 2201 | 2r0o | A | 255 | GLU | p.Lys255Glu | O43707 |
| HOMO SAPIENS | 3 | 2201 | 2r0o | A | 259 | THR | p.Thr259Ile | O43707 |
| HOMO SAPIENS | 4 | 2201 | 2r0o | A | 262 | SER | p.Ser262Pro | O43707 |
| HOMO SAPIENS | -1 | 31221 | 2wvr | C | 44 | THR | p.Thr44Pro | Q9H211 |
| HOMO SAPIENS | -1 | 31221 | 2wvr | C | 135 | ALA | p.Ala135Val | Q9H211 |
| HOMO SAPIENS | -1 | 31221 | 2wvr | C | 456 | GLU | p.Glu456Ala | Q9H211 |

| | | | | | | | | |
|---|---|---|---|---|---|---|---|---|
| HOMO SAPIENS | -1 | 31221 | 2wvr | C | 462 | ARG | p.Arg462Gln | Q9H211 |
| HOMO SAPIENS | -1 | 31221 | 2wvr | C | 468 | GLU | p.Glu468Lys | Q9H211 |
| HOMO SAPIENS | -1 | 31221 | 2wvr | C | 537 | ALA | p.Ala537Val | Q9H211 |
| HOMO SAPIENS | 5 | 2576 | 5o3l | C | 363 | VAL | p.Val680Ile | P10636 |
| HOMO SAPIENS | 5 | 1345 | 1czt | A | 37 | ARG | p.Arg2102Cys | P12259 |
| HOMO SAPIENS | 1 | 1345 | 1czt | A | 83 | MET | p.Met2148Thr | P12259 |
| HOMO SAPIENS | 1 | 1345 | 1czt | A | 157 | ASP | p.Asp2222Gly | P12259 |
| HOMO SAPIENS | 8 | 35985 | 1lo1 | A | 110 | ALA | p.Ala110Val | O95718 |
| HOMO SAPIENS | 1 | 617 | 2ocp | B | 44 | GLU | p.Glu44Lys | Q16854 |
| HOMO SAPIENS | 5 | 617 | 2ocp | B | 46 | ASN | p.Asn46Ser | Q16854 |
| HOMO SAPIENS | 0 | 617 | 2ocp | B | 59 | THR | p.Thr59Met | Q16854 |
| HOMO SAPIENS | 7 | 617 | 2ocp | B | 142 | ARG | p.Arg142Lys | Q16854 |
| HOMO SAPIENS | 3 | 617 | 2ocp | B | 154 | ASN | p.Asn154Lys | Q16854 |
| HOMO SAPIENS | 3 | 617 | 2ocp | B | 170 | GLN | p.Gln170Arg | Q16854 |
| HOMO SAPIENS | 3 | 617 | 2ocp | B | 227 | GLU | p.Glu227Lys | Q16854 |
| HOMO SAPIENS | 3 | 617 | 2ocp | B | 233 | LYS | p.Lys233Glu | Q16854 |
| HOMO SAPIENS | 3 | 617 | 2ocp | B | 255 | ASP | p.Asp255Tyr | Q16854 |
| HOMO SAPIENS | -1 | 38975 | 5itz | D | 421 | GLN | p.Gln421His | Q9HC77 |
| HOMO SAPIENS | 2 | 6235 | 2f8v | A | 15 | VAL | p.Val15Ile | Q8WZ42 |
| HOMO SAPIENS | 0 | 6235 | 2f8v | A | 54 | VAL | p.Val54Met | Q8WZ42 |
| HOMO SAPIENS | 0 | 6235 | 2f8v | A | 59 | SER | p.Ser59Arg | Q8WZ42 |
| HOMO SAPIENS | 0 | 6235 | 2f8v | A | 60 | ASP | p.Asp60Tyr | Q8WZ42 |
| HOMO SAPIENS | 0 | 12630 | 1ext | B | 12 | ASP | p.Asp41Glu | P19438 |
| HOMO SAPIENS | 3 | 12630 | 1ext | B | 30 | CYS | p.Cys59Arg | P19438 |
| HOMO SAPIENS | 3 | 12630 | 1ext | B | 30 | CYS | p.Cys59Ser | P19438 |
| HOMO SAPIENS | 5 | 12630 | 1ext | B | 33 | CYS | p.Cys62Tyr | P19438 |
| HOMO SAPIENS | 5 | 12630 | 1ext | B | 33 | CYS | p.Cys62Gly | P19438 |
| HOMO SAPIENS | 1 | 12630 | 1ext | B | 46 | PRO | p.Pro75Leu | P19438 |
| HOMO SAPIENS | 3 | 12630 | 1ext | B | 50 | THR | p.Thr79Met | P19438 |
| HOMO SAPIENS | 6 | 12630 | 1ext | B | 52 | CYS | p.Cys81Phe | P19438 |
| HOMO SAPIENS | 6 | 12630 | 1ext | B | 70 | CYS | p.Cys99Ser | P19438 |
| HOMO SAPIENS | 5 | 12630 | 1ext | B | 88 | CYS | p.Cys117Arg | P19438 |
| HOMO SAPIENS | 5 | 12630 | 1ext | B | 88 | CYS | p.Cys117Tyr | P19438 |
| HOMO SAPIENS | -1 | 12630 | 1ext | B | 170 | ILE | p.Ile199Asn | P19438 |
| HOMO SAPIENS | 0 | 994 | 1aos | B | 95 | ARG | p.Arg95Cys | P04424 |
| HOMO SAPIENS | 0 | 994 | 1aos | B | 111 | ARG | p.Arg111Gln | P04424 |
| HOMO SAPIENS | 0 | 994 | 1aos | B | 131 | THR | p.Thr131Met | P04424 |
| HOMO SAPIENS | 6 | 994 | 1aos | B | 178 | VAL | p.Val178Met | P04424 |
| HOMO SAPIENS | 9 | 994 | 1aos | B | 182 | ARG | p.Arg182Gln | P04424 |
| HOMO SAPIENS | 9 | 994 | 1aos | B | 186 | ARG | p.Arg186Trp | P04424 |
| HOMO SAPIENS | 8 | 994 | 1aos | B | 193 | ARG | p.Arg193Gln | P04424 |
| HOMO SAPIENS | 5 | 994 | 1aos | B | 217 | ARG | p.Arg217Ter | P04424 |
| HOMO SAPIENS | 0 | 994 | 1aos | B | 286 | GLN | p.Gln286Arg | P04424 |
| HOMO SAPIENS | 5 | 994 | 1aos | B | 379 | ARG | p.Arg379Cys | P04424 |
| HOMO SAPIENS | 4 | 16066 | 5mv9 | A | 1719 | TYR | p.Tyr1719Cys | Q13402 |
| HOMO SAPIENS | -1 | 16066 | 5mv9 | A | 1739 | ARG | p.Arg1739Ter | Q13402 |
| HOMO SAPIENS | 4 | 16066 | 5mv9 | A | 1775 | ILE | p.Ile1775Thr | Q13402 |
| HOMO SAPIENS | 2 | 16066 | 5mv9 | A | 1786 | SER | p.Ser1786Thr | Q13402 |
| HOMO SAPIENS | 4 | 16066 | 5mv9 | A | 1798 | GLN | p.Gln1798Ter | Q13402 |
| HOMO SAPIENS | 0 | 16066 | 5mv9 | A | 1858 | LEU | p.Leu1858Pro | Q13402 |
| HOMO SAPIENS | 6 | 16066 | 5mv9 | A | 1873 | ARG | p.Arg1873Trp | Q13402 |
| HOMO SAPIENS | 6 | 16066 | 5mv9 | A | 1873 | ARG | p.Arg1873Leu | Q13402 |
| HOMO SAPIENS | 6 | 16066 | 5mv9 | A | 1883 | ARG | p.Arg1883Gln | Q13402 |
| HOMO SAPIENS | 4 | 16066 | 5mv9 | A | 1887 | PRO | p.Pro1887Leu | Q13402 |

| | | | | | | | | |
|---|---|---|---|---|---|---|---|---|
| HOMO SAPIENS | 5 | 16066 | 5mv9 | A | 1942 | GLY | p.Gly1942Ter | Q13402 |
| HOMO SAPIENS | 5 | 16066 | 5mv9 | A | 1954 | LEU | p.Leu1954Ile | Q13402 |
| HOMO SAPIENS | 5 | 16066 | 5mv9 | A | 1956 | VAL | p.Val1956Ile | Q13402 |
| HOMO SAPIENS | 5 | 16066 | 5mv9 | A | 1967 | ARG | p.Arg1967Ter | Q13402 |
| HOMO SAPIENS | 5 | 16066 | 5mv9 | A | 1990 | GLN | p.Gln1990Ter | Q13402 |
| HOMO SAPIENS | 2 | 16066 | 5mv9 | A | 2005 | LYS | p.Lys2005Glu | Q13402 |
| HOMO SAPIENS | 1 | 16066 | 5mv9 | A | 2072 | VAL | p.Val2072Ile | Q13402 |
| HOMO SAPIENS | 4 | 16066 | 5mv9 | A | 2109 | THR | p.Thr2109Ile | Q13402 |
| HOMO SAPIENS | 4 | 16066 | 5mv9 | A | 2142 | ASP | p.Asp2142Asn | Q13402 |
| HOMO SAPIENS | -1 | 16066 | 5mv9 | A | 2214 | GLY | p.Gly2214Ser | Q13402 |
| HOMO SAPIENS | 7 | 27942 | 1deb | A | 24 | ARG | p.Arg24Ter | P25054 |
| HOMO SAPIENS | 0 | 876 | 5hnr | B | 12 | PHE | p.Phe12Leu | P13716 |
| HOMO SAPIENS | 0 | 876 | 5hnr | B | 59 | LYS | p.Lys59Asn | P13716 |
| HOMO SAPIENS | -1 | 876 | 5hnr | B | 133 | GLY | p.Gly133Arg | P13716 |
| HOMO SAPIENS | 2 | 876 | 5hnr | B | 240 | ARG | p.Arg240Trp | P13716 |
| HOMO SAPIENS | 0 | 876 | 5hnr | B | 274 | ALA | p.Ala274Thr | P13716 |
| HOMO SAPIENS | 0 | 876 | 5hnr | B | 275 | VAL | p.Val275Met | P13716 |
| HOMO SAPIENS | 0 | 1142 | 3qe2 | A | 503 | VAL | p.Ala500Val | P16435 |
| HOMO SAPIENS | 8 | 14358 | 4dkd | C | 32 | VAL | p.Val32Gly | P07333 |
| HOMO SAPIENS | 0 | 14358 | 4dkd | C | 60 | LEU | p.Leu60Val | P07333 |
| HOMO SAPIENS | 7 | 14358 | 4dkd | C | 75 | THR | p.Thr75Ile | P07333 |
| HOMO SAPIENS | -1 | 14358 | 4dkd | C | 90 | PRO | p.Pro90Thr | P07333 |
| HOMO SAPIENS | 0 | 14358 | 4dkd | C | 106 | ARG | p.Arg106Trp | P07333 |
| HOMO SAPIENS | 0 | 14358 | 4dkd | C | 241 | ASN | p.Asn241Asp | P07333 |
| HOMO SAPIENS | 0 | 14358 | 4dkd | C | 245 | ALA | p.Ala245Ser | P07333 |
| HOMO SAPIENS | 0 | 14358 | 4dkd | C | 255 | ASN | p.Asn255Ile | P07333 |
| HOMO SAPIENS | 0 | 14358 | 4dkd | C | 268 | VAL | p.Val268Ile | P07333 |
| HOMO SAPIENS | 0 | 14358 | 4dkd | C | 279 | VAL | p.Val279Met | P07333 |
| HOMO SAPIENS | 0 | 14358 | 4dkd | C | 294 | ARG | p.Arg294Gln | P07333 |
| HOMO SAPIENS | 5 | 8968 | 3ppy | A | 1565 | VAL | p.Val1565Leu | P04275 |
| HOMO SAPIENS | 4 | 8968 | 3ppy | A | 1584 | TYR | p.Tyr1584Cys | P04275 |
| HOMO SAPIENS | 5 | 5341 | 3k3n | A | 279 | PHE | p.Phe315Ser | Q9UPP1 |
| HOMO SAPIENS | 0 | 29463 | 1q7l | A | 197 | ARG | p.Arg197Trp | Q03154 |
| HOMO SAPIENS | 0 | 33761 | 2leh | B | 30 | ASP | p.Asp30Asn | Q16637 |
| HOMO SAPIENS | 6 | 33761 | 2leh | B | 44 | ASP | p.Asp44Val | Q16637 |
| HOMO SAPIENS | -1 | 12249 | 2pnr | C | 224 | GLU | p.Glu310Lys | P10515 |
| HOMO SAPIENS | -1 | 12249 | 2pnr | C | 232 | VAL | p.Val318Ala | P10515 |
| HOMO SAPIENS | -1 | 7 | 5whk | B | -9 | ALA | p.Ala11Pro | P61769 |
| HOMO SAPIENS | 8 | 20298 | 4euw | A | 154 | PHE | p.Phe154Leu | P48436 |
| HOMO SAPIENS | 9 | 20298 | 4euw | A | 158 | ALA | p.Ala158Thr | P48436 |
| HOMO SAPIENS | 5 | 20298 | 4euw | A | 165 | HIS | p.His165Tyr | P48436 |
| HOMO SAPIENS | 3 | 20298 | 4euw | A | 169 | HIS | p.His169Gln | P48436 |
| HOMO SAPIENS | 0 | 20298 | 4euw | A | 173 | LYS | p.Lys173Glu | P48436 |
| HOMO SAPIENS | 1 | 16465 | 2jgx | A | 402 | TYR | p.Tyr402His | P08603 |
| HOMO SAPIENS | 0 | 8993 | 3v8o | A | 659 | LEU | p.Leu659Arg | Q14315 |
| HOMO SAPIENS | 6 | 8993 | 3v8o | A | 693 | ASP | p.Asp693Ala | Q14315 |
| HOMO SAPIENS | 0 | 8993 | 3v8o | A | 709 | ALA | p.Ala709Thr | Q14315 |
| HOMO SAPIENS | 2 | 3852 | 5es4 | B | 106 | ASP | p.Asp128Asn | P05107 |
| HOMO SAPIENS | 2 | 3852 | 5es4 | B | 116 | SER | p.Ser138Pro | P05107 |
| HOMO SAPIENS | 1 | 3852 | 5es4 | B | 127 | LEU | p.Leu149Pro | P05107 |
| HOMO SAPIENS | 0 | 3852 | 5es4 | B | 147 | GLY | p.Gly169Arg | P05107 |
| HOMO SAPIENS | 3 | 3852 | 5es4 | B | 156 | PRO | p.Pro178Leu | P05107 |
| HOMO SAPIENS | 0 | 3852 | 5es4 | B | 174 | LYS | p.Lys196Thr | P05107 |
| HOMO SAPIENS | 2 | 3852 | 5es4 | B | 251 | GLY | p.Gly273Arg | P05107 |

| | | | | | | | | |
|---|---|---|---|---|---|---|---|---|
| HOMO SAPIENS | 2 | 3852 | 5es4 | B | 329 | ASN | p.Asn351Ser | P05107 |
| HOMO SAPIENS | 2 | 3852 | 5es4 | B | 571 | ARG | p.Arg593Cys | P05107 |
| HOMO SAPIENS | 7 | 5962 | 5fn2 | B | 146 | MET | p.Met146Leu | P49768 |
| HOMO SAPIENS | 3 | 5962 | 5fn2 | B | 166 | LEU | p.Leu166Pro | P49768 |
| HOMO SAPIENS | 1 | 5962 | 5fn2 | B | 233 | MET | p.Met233Leu | P49768 |
| HOMO SAPIENS | 1 | 5962 | 5fn2 | B | 261 | VAL | p.Val261Gly | P49768 |
| HOMO SAPIENS | 1 | 5962 | 5fn2 | B | 266 | GLY | p.Gly266Ser | P49768 |
| HOMO SAPIENS | -1 | 5962 | 5fn2 | B | 333 | ASP | p.Asp333Gly | P49768 |
| HOMO SAPIENS | 6 | 1918 | 3dlx | A | 58 | THR | p.Thr58Met | P55809 |
| HOMO SAPIENS | 5 | 1918 | 3dlx | A | 133 | VAL | p.Val133Glu | P55809 |
| HOMO SAPIENS | 5 | 1918 | 3dlx | A | 219 | GLY | p.Gly219Glu | P55809 |
| HOMO SAPIENS | 0 | 1918 | 3dlx | A | 221 | VAL | p.Val221Met | P55809 |
| HOMO SAPIENS | 5 | 1918 | 3dlx | A | 283 | SER | p.Ser283Ter | P55809 |
| HOMO SAPIENS | 4 | 1918 | 3dlx | A | 324 | GLY | p.Gly324Glu | P55809 |
| HOMO SAPIENS | 0 | 1918 | 3dlx | A | 456 | CYS | p.Cys456Phe | P55809 |
| HOMO SAPIENS | 0 | 9010 | 2wfr | A | 162 | LEU | p.Leu162Pro | O43323 |
| HOMO SAPIENS | 5 | 3770 | 5n38 | A | 56 | VAL | p.Val56Glu | O60260 |
| HOMO SAPIENS | 1 | 3770 | 5n38 | A | 161 | LYS | p.Lys161Asn | O60260 |
| HOMO SAPIENS | 1 | 3770 | 5n38 | A | 167 | SER | p.Ser167Asn | O60260 |
| HOMO SAPIENS | 1 | 3770 | 5n38 | A | 211 | LYS | p.Lys211Asn | O60260 |
| HOMO SAPIENS | 1 | 3770 | 5n38 | A | 212 | CYS | p.Cys212Tyr | O60260 |
| HOMO SAPIENS | 1 | 3770 | 5n38 | A | 275 | ARG | p.Arg275Trp | O60260 |
| HOMO SAPIENS | 2 | 3770 | 5n38 | A | 334 | ARG | p.Arg334Cys | O60260 |
| HOMO SAPIENS | 1 | 3770 | 5n38 | A | 366 | ARG | p.Arg366Trp | O60260 |
| HOMO SAPIENS | -1 | 3770 | 5n38 | A | 380 | VAL | p.Val380Leu | O60260 |
| HOMO SAPIENS | 1 | 3770 | 5n38 | A | 394 | ASP | p.Asp394Asn | O60260 |
| HOMO SAPIENS | 1 | 3770 | 5n38 | A | 430 | GLY | p.Gly430Asp | O60260 |
| HOMO SAPIENS | -1 | 2467 | 4xx1 | A | 10 | PRO | p.Pro34Leu | P04180 |
| HOMO SAPIENS | 4 | 2467 | 4xx1 | A | 83 | TYR | p.Tyr107Ter | P04180 |
| HOMO SAPIENS | 4 | 2467 | 4xx1 | A | 93 | ALA | p.Ala117Thr | P04180 |
| HOMO SAPIENS | 4 | 2467 | 4xx1 | A | 123 | THR | p.Thr147Ile | P04180 |
| HOMO SAPIENS | 4 | 2467 | 4xx1 | A | 135 | ARG | p.Arg159Trp | P04180 |
| HOMO SAPIENS | 3 | 2467 | 4xx1 | A | 158 | ARG | p.Arg182Cys | P04180 |
| HOMO SAPIENS | 0 | 2467 | 4xx1 | A | 209 | LEU | p.Leu233Pro | P04180 |
| HOMO SAPIENS | 1 | 2467 | 4xx1 | A | 228 | ASN | p.Asn252Lys | P04180 |
| HOMO SAPIENS | 4 | 2467 | 4xx1 | A | 252 | MET | p.Met276Lys | P04180 |
| HOMO SAPIENS | 3 | 2467 | 4xx1 | A | 293 | MET | p.Met317Ile | P04180 |
| HOMO SAPIENS | 5 | 2467 | 4xx1 | A | 321 | THR | p.Thr345Met | P04180 |
| HOMO SAPIENS | 9 | 2467 | 4xx1 | A | 347 | THR | p.Thr371Met | P04180 |
| HOMO SAPIENS | 1 | 5834 | 3qrf | G | 371 | PHE | p.Phe371Cys | Q9BZS1 |
| HOMO SAPIENS | 1 | 5834 | 3qrf | G | 373 | PHE | p.Phe373Ala | Q9BZS1 |
| HOMO SAPIENS | 9 | 5834 | 3qrf | G | 384 | ALA | p.Ala384Thr | Q9BZS1 |
| HOMO SAPIENS | 6 | 5834 | 3qrf | G | 397 | ARG | p.Arg397Trp | Q9BZS1 |
| HOMO SAPIENS | -1 | 11869 | 4km8 | A | 15 | PRO | p.Pro15Leu | Q9UMX1 |
| HOMO SAPIENS | 5 | 262 | 4kbo | A | 126 | ARG | p.Arg126Trp | P38646 |
| HOMO SAPIENS | 3 | 262 | 4kbo | A | 128 | TYR | p.Tyr128Cys | P38646 |
| HOMO SAPIENS | -1 | 11347 | 4uf9 | D | 117 | ALA | p.Ala117Val | Q13501 |
| HOMO SAPIENS | 0 | 2173 | 4ah6 | A | 45 | SER | p.Ser45Gly | Q6PI48 |
| HOMO SAPIENS | 2 | 2173 | 4ah6 | A | 152 | CYS | p.Cys152Phe | Q6PI48 |
| HOMO SAPIENS | 4 | 2173 | 4ah6 | A | 179 | ARG | p.Arg179His | Q6PI48 |
| HOMO SAPIENS | 0 | 2173 | 4ah6 | A | 196 | LYS | p.Lys196Arg | Q6PI48 |
| HOMO SAPIENS | 8 | 2173 | 4ah6 | A | 263 | ARG | p.Arg263Gln | Q6PI48 |
| HOMO SAPIENS | 1 | 2173 | 4ah6 | A | 338 | GLY | p.Gly338Glu | Q6PI48 |
| HOMO SAPIENS | 0 | 2173 | 4ah6 | A | 425 | GLU | p.Glu425Ter | Q6PI48 |

| | | | | | | | | |
|---|---|---|---|---|---|---|---|---|
| HOMO SAPIENS | 1 | 2173 | 4ah6 | A | 548 | LEU | p.Leu548Met | Q6PI48 |
| HOMO SAPIENS | 0 | 2173 | 4ah6 | A | 609 | ARG | p.Arg609Trp | Q6PI48 |
| HOMO SAPIENS | 2 | 2173 | 4ah6 | A | 613 | LEU | p.Leu613Phe | Q6PI48 |
| HOMO SAPIENS | 0 | 2173 | 4ah6 | A | 626 | LEU | p.Leu626Gln | Q6PI48 |
| HOMO SAPIENS | 0 | 2173 | 4ah6 | A | 626 | LEU | p.Leu626Val | Q6PI48 |
| HOMO SAPIENS | 7 | 3203 | 2cn1 | A | 87 | ASP | p.Asp137Val | Q9H0P0 |
| HOMO SAPIENS | 8 | 3203 | 2cn1 | A | 166 | GLN | p.Gln216Ter | Q9H0P0 |
| HOMO SAPIENS | 8 | 3203 | 2cn1 | A | 179 | ASN | p.Asn229Ser | Q9H0P0 |
| HOMO SAPIENS | 7 | 3203 | 2cn1 | A | 230 | GLY | p.Gly280Arg | Q9H0P0 |
| HOMO SAPIENS | 2 | 3424 | 2a1t | R | 116 | GLY | p.Gly116Arg | P13804 |
| HOMO SAPIENS | 2 | 3424 | 2a1t | R | 157 | VAL | p.Val157Gly | P13804 |
| HOMO SAPIENS | 2 | 3424 | 2a1t | R | 171 | THR | p.Thr171Ile | P13804 |
| HOMO SAPIENS | 0 | 3424 | 2a1t | R | 178 | THR | p.Thr178Arg | P13804 |
| HOMO SAPIENS | 0 | 3424 | 2a1t | R | 194 | VAL | p.Val194Met | P13804 |
| HOMO SAPIENS | 1 | 3424 | 2a1t | R | 209 | ARG | p.Arg209Ter | P13804 |
| HOMO SAPIENS | 6 | 3424 | 2a1t | R | 266 | THR | p.Thr266Met | P13804 |
| HOMO SAPIENS | 0 | 27598 | 1g1c | A | 26 | ARG | p.Arg2098Gly | Q8WZ42 |
| HOMO SAPIENS | 1 | 27598 | 1g1c | A | 46 | ILE | p.Ile2118Thr | Q8WZ42 |
| HOMO SAPIENS | 0 | 32885 | 4bsk | A | 149 | ASN | p.Asn149Asp | P35916 |
| HOMO SAPIENS | -1 | 8074 | 3ghg | B | 17 | ARG | p.Arg47Ter | P02675 |
| HOMO SAPIENS | -1 | 447 | 5g4p | A | 190 | ALA | p.Ala190Thr | P84074 |
| HOMO SAPIENS | 0 | 10761 | 5u1d | B | 379 | VAL | p.Val379Ile | Q03519 |
| HOMO SAPIENS | 0 | 10761 | 5u1d | B | 665 | THR | p.Thr665Ala | Q03519 |
| HOMO SAPIENS | 5 | 29251 | 5fn7 | A | 79 | ILE | p.Ile294Leu | P08575 |
| HOMO SAPIENS | -1 | 27727 | 4ky9 | A | 56 | LYS | p.Lys56Glu | P02730 |
| HOMO SAPIENS | 2 | 27727 | 4ky9 | A | 72 | GLU | p.Glu72Asp | P02730 |
| HOMO SAPIENS | 2 | 27727 | 4ky9 | A | 90 | GLU | p.Glu90Lys | P02730 |
| HOMO SAPIENS | 1 | 27727 | 4ky9 | A | 130 | GLY | p.Gly130Arg | P02730 |
| HOMO SAPIENS | 2 | 27727 | 4ky9 | A | 150 | ARG | p.Arg150Ter | P02730 |
| HOMO SAPIENS | 0 | 27727 | 4ky9 | A | 327 | PRO | p.Pro327Arg | P02730 |
| HOMO SAPIENS | 2 | 13202 | 2fxo | B | 870 | ARG | p.Arg870His | P12883 |
| HOMO SAPIENS | 2 | 13202 | 2fxo | B | 870 | ARG | p.Arg870Cys | P12883 |
| HOMO SAPIENS | 4 | 13202 | 2fxo | B | 904 | ARG | p.Arg904Cys | P12883 |
| HOMO SAPIENS | 4 | 13202 | 2fxo | B | 906 | ASP | p.Asp906Gly | P12883 |
| HOMO SAPIENS | 2 | 13202 | 2fxo | B | 908 | LEU | p.Leu908Val | P12883 |
| HOMO SAPIENS | 2 | 13202 | 2fxo | B | 923 | ASN | p.Asn923Lys | P12883 |
| HOMO SAPIENS | 7 | 13202 | 2fxo | B | 935 | GLU | p.Glu935Lys | P12883 |
| HOMO SAPIENS | 7 | 13202 | 2fxo | B | 935 | GLU | p.Glu935Val | P12883 |
| HOMO SAPIENS | 8 | 13202 | 2fxo | B | 949 | GLU | p.Glu949Lys | P12883 |
| HOMO SAPIENS | 0 | 13200 | 2a1s | A | 349 | ARG | p.Arg349Trp | O95453 |
| HOMO SAPIENS | 0 | 13200 | 2a1s | A | 421 | LYS | p.Lys421Arg | O95453 |
| HOMO SAPIENS | 6 | 3196 | 5xtb | T | 115 | CYS | p.Cys115Tyr | O75380 |
| HOMO SAPIENS | 0 | 1108 | 5ukf | A | 89 | ARG | p.Arg89Gln | Q99986 |
| HOMO SAPIENS | 0 | 1108 | 5ukf | A | 119 | HIS | p.His119Arg | Q99986 |
| HOMO SAPIENS | 0 | 1108 | 5ukf | A | 133 | ARG | p.Arg133Cys | Q99986 |
| HOMO SAPIENS | 0 | 1108 | 5ukf | A | 236 | VAL | p.Val236Leu | Q99986 |
| HOMO SAPIENS | 0 | 1108 | 5ukf | A | 321 | ARG | p.Arg321Cys | Q99986 |
| HOMO SAPIENS | -1 | 1108 | 5ukf | A | 358 | ARG | p.Arg358Ter | Q99986 |
| HOMO SAPIENS | 0 | 5842 | 4yli | C | 204 | GLY | p.Gly204Ser | Q9BWP8 |
| HOMO SAPIENS | -1 | 31362 | 4wqo | A | 25 | PRO | p.Pro25Leu | P40337 |
| HOMO SAPIENS | 2 | 10107 | 2j8h | A | 34 | VAL | p.Val31886Ile | Q8WZ42 |
| HOMO SAPIENS | 8 | 20190 | 1wm5 | A | 77 | ARG | p.Arg77Gln | P19878 |
| HOMO SAPIENS | 1 | 20190 | 1wm5 | A | 100 | GLN | p.Gln100Ter | P19878 |
| HOMO SAPIENS | 0 | 20190 | 1wm5 | A | 102 | ARG | p.Arg102Ter | P19878 |

| | | | | | | | | | |
|---|---|---|---|---|---|---|---|---|---|
| HOMO SAPIENS | 0 | 20190 | 1wm5 | A | | 128 | ALA | p.Ala128Val | P19878 |
| HOMO SAPIENS | 0 | 20190 | 1wm5 | A | | 181 | LYS | p.Lys181Arg | P19878 |
| HOMO SAPIENS | 7 | 1278 | 5a2q | a | | 33 | ASP | p.Asp33Asn | P62854 |
| HOMO SAPIENS | 3 | 19515 | 2obd | A | | 11 | ILE | p.Ile28Met | P11597 |
| HOMO SAPIENS | 3 | 19515 | 2obd | A | | 89 | VAL | p.Val106Met | P11597 |
| HOMO SAPIENS | 3 | 19515 | 2obd | A | | 137 | ARG | p.Arg154Trp | P11597 |
| HOMO SAPIENS | 3 | 19515 | 2obd | A | | 139 | ASP | p.Asp156Asn | P11597 |
| HOMO SAPIENS | 3 | 19515 | 2obd | A | | 204 | SER | p.Ser221Arg | P11597 |
| HOMO SAPIENS | 0 | 19515 | 2obd | A | | 297 | GLU | p.Glu314Lys | P11597 |
| HOMO SAPIENS | 0 | 19515 | 2obd | A | | 314 | GLY | p.Gly331Ser | P11597 |
| HOMO SAPIENS | 0 | 19515 | 2obd | A | | 332 | SER | p.Ser349Tyr | P11597 |
| HOMO SAPIENS | 0 | 19515 | 2obd | A | | 368 | VAL | p.Val385Met | P11597 |
| HOMO SAPIENS | 0 | 19515 | 2obd | A | | 373 | ALA | p.Ala390Pro | P11597 |
| HOMO SAPIENS | 3 | 19515 | 2obd | A | | 405 | ILE | p.Val422Ile | P11597 |
| HOMO SAPIENS | 0 | 19515 | 2obd | A | | 451 | ARG | p.Arg468Gln | P11597 |
| HOMO SAPIENS | 0 | 19515 | 2obd | A | | 451 | ARG | p.Arg468Ter | P11597 |
| HOMO SAPIENS | 0 | 6626 | 3hop | A | | 39 | ALA | p.Ala39Gly | P21333 |
| HOMO SAPIENS | 1 | 6626 | 3hop | A | | 82 | GLU | p.Glu82Val | P21333 |
| HOMO SAPIENS | 1 | 6626 | 3hop | A | | 128 | ALA | p.Ala128Val | P21333 |
| HOMO SAPIENS | 2 | 6626 | 3hop | A | | 203 | ASP | p.Asp203Tyr | P21333 |
| HOMO SAPIENS | 1 | 6626 | 3hop | A | | 207 | PRO | p.Pro207Leu | P21333 |
| HOMO SAPIENS | 2 | 6626 | 3hop | A | | 210 | CYS | p.Cys210Phe | P21333 |
| HOMO SAPIENS | 7 | 6626 | 3hop | A | | 254 | GLU | p.Glu254Lys | P21333 |
| HOMO SAPIENS | 4 | 4210 | 2avg | A | | 8 | VAL | p.Val158Met | Q14896 |
| HOMO SAPIENS | 6 | 4210 | 2avg | A | | 15 | GLU | p.Glu165Asp | Q14896 |
| HOMO SAPIENS | 4 | 4210 | 2avg | A | | 27 | ARG | p.Arg177His | Q14896 |
| HOMO SAPIENS | 0 | 4210 | 2avg | A | | 28 | VAL | p.Val178Met | Q14896 |
| HOMO SAPIENS | 6 | 4210 | 2avg | A | | 39 | VAL | p.Val189Ile | Q14896 |
| HOMO SAPIENS | 2 | 4210 | 2avg | A | | 52 | LYS | p.Lys202Gln | Q14896 |
| HOMO SAPIENS | 4 | 4210 | 2avg | A | | 67 | SER | p.Ser217Gly | Q14896 |
| HOMO SAPIENS | 6 | 4210 | 2avg | A | | 69 | VAL | p.Val219Leu | Q14896 |
| HOMO SAPIENS | 2 | 4210 | 2avg | A | | 86 | GLY | p.Ser236Gly | Q14896 |
| HOMO SAPIENS | 4 | 4210 | 2avg | A | | 108 | GLU | p.Glu258Lys | Q14896 |
| HOMO SAPIENS | 0 | 22237 | 1w6k | A | | 588 | GLY | p.Gly588Ser | P48449 |
| HOMO SAPIENS | 0 | 10765 | 4a5w | B | | 98 | GLU | p.Glu98Ala | P13671 |
| HOMO SAPIENS | 0 | 10765 | 4a5w | B | | 596 | ARG | p.Arg596Ter | P13671 |
| HOMO SAPIENS | -1 | 15515 | 3hx3 | A | | 11 | MSE | p.Met10Lys | P12271 |
| HOMO SAPIENS | 8 | 15515 | 3hx3 | A | | 64 | ARG | p.Arg64Gln | P12271 |
| HOMO SAPIENS | 0 | 15515 | 3hx3 | A | | 182 | PHE | p.Phe182Cys | P12271 |
| HOMO SAPIENS | 0 | 15515 | 3hx3 | A | | 216 | ARG | p.Arg216Gln | P12271 |
| HOMO SAPIENS | -1 | 15515 | 3hx3 | A | | 226 | MSE | p.Met226Lys | P12271 |
| HOMO SAPIENS | 0 | 15515 | 3hx3 | A | | 234 | TRP | p.Arg234Trp | P12271 |
| HOMO SAPIENS | 0 | 15515 | 3hx3 | A | | 273 | LEU | p.Leu273Phe | P12271 |
| HOMO SAPIENS | 0 | 15515 | 3hx3 | A | | 292 | THR | p.Thr292Met | P12271 |
| HOMO SAPIENS | 0 | 5152 | 2www | D | | 95 | GLN | p.Gln95Ter | Q8IVH4 |
| HOMO SAPIENS | 0 | 5152 | 2www | D | | 133 | GLN | p.Gln133Ter | Q8IVH4 |
| HOMO SAPIENS | 0 | 5152 | 2www | D | | 145 | ARG | p.Arg145Ter | Q8IVH4 |
| HOMO SAPIENS | 0 | 5152 | 2www | D | | 207 | TYR | p.Tyr207Cys | Q8IVH4 |
| HOMO SAPIENS | 8 | 5152 | 2www | D | | 330 | ARG | p.Arg330Ter | Q8IVH4 |
| HOMO SAPIENS | 0 | 5152 | 2www | D | | 363 | GLN | p.Gln363His | Q8IVH4 |
| HOMO SAPIENS | 1 | 1626 | 1ilr | | 1 | 105 | SER | p.Ser130Arg | P18510 |
| HOMO SAPIENS | -1 | 929 | 5loh | A | | 12 | GLY | p.Gly12Arg | Q96GX5 |
| HOMO SAPIENS | 9 | 3206 | 1x63 | A | | 21 | CYS | p.Cys104Arg | Q13642 |
| HOMO SAPIENS | 8 | 3206 | 1x63 | A | | 39 | TRP | p.Trp122Ser | Q13642 |

| | | | | | | | | |
|---|---|---|---|---|---|---|---|---|
| HOMO SAPIENS | 9 | 3206 | 1x63 | A | 40 | HIS | p.His123Tyr | Q13642 |
| HOMO SAPIENS | 9 | 3206 | 1x63 | A | 40 | HIS | p.His123Leu | Q13642 |
| HOMO SAPIENS | 9 | 3206 | 1x63 | A | 40 | HIS | p.His123Gln | Q13642 |
| HOMO SAPIENS | 3 | 3206 | 1x63 | A | 49 | CYS | p.Cys132Phe | Q13642 |
| HOMO SAPIENS | 3 | 3206 | 1x63 | A | 67 | CYS | p.Cys150Tyr | Q13642 |
| HOMO SAPIENS | 3 | 3206 | 1x63 | A | 70 | CYS | p.Cys153Arg | Q13642 |
| HOMO SAPIENS | 3 | 3206 | 1x63 | A | 70 | CYS | p.Cys153Tyr | Q13642 |
| HOMO SAPIENS | 5 | 1052 | 3ecs | E | 208 | ASN | p.Asn208Tyr | Q14232 |
| HOMO SAPIENS | 2 | 1052 | 3ecs | E | 275 | TYR | p.Tyr275Cys | Q14232 |
| HOMO SAPIENS | 0 | 24253 | 5i35 | A | 271 | ARG | p.Arg271Cys | Q8NI60 |
| HOMO SAPIENS | 0 | 24253 | 5i35 | A | 272 | GLY | p.Gly272Asp | Q8NI60 |
| HOMO SAPIENS | 0 | 24253 | 5i35 | A | 299 | ARG | p.Arg299Trp | Q8NI60 |
| HOMO SAPIENS | 1 | 24253 | 5i35 | A | 339 | ALA | p.Ala339Thr | Q8NI60 |
| HOMO SAPIENS | 0 | 24253 | 5i35 | A | 348 | ARG | p.Arg348Ter | Q8NI60 |
| HOMO SAPIENS | 0 | 24253 | 5i35 | A | 395 | ILE | p.Ile395Met | Q8NI60 |
| HOMO SAPIENS | 0 | 24253 | 5i35 | A | 396 | ASP | p.Asp396Glu | Q8NI60 |
| HOMO SAPIENS | 0 | 24253 | 5i35 | A | 480 | PHE | p.Phe480Leu | Q8NI60 |
| HOMO SAPIENS | 2 | 24253 | 5i35 | A | 511 | THR | p.Thr511Met | Q8NI60 |
| HOMO SAPIENS | 0 | 24253 | 5i35 | A | 514 | TYR | p.Tyr514Cys | Q8NI60 |
| HOMO SAPIENS | 0 | 24253 | 5i35 | A | 538 | ARG | p.Arg538Gln | Q8NI60 |
| HOMO SAPIENS | 0 | 24253 | 5i35 | A | 549 | GLY | p.Gly549Ser | Q8NI60 |
| HOMO SAPIENS | 0 | 24253 | 5i35 | A | 551 | GLU | p.Glu551Lys | Q8NI60 |
| HOMO SAPIENS | -1 | 24253 | 5i35 | A | 555 | MSE | p.Met555Ile | Q8NI60 |
| HOMO SAPIENS | 0 | 24253 | 5i35 | A | 568 | GLU | p.Glu568Ter | Q8NI60 |
| HOMO SAPIENS | 0 | 24253 | 5i35 | A | 602 | PRO | p.Pro602Arg | Q8NI60 |
| HOMO SAPIENS | -1 | 15505 | 4goj | C | 22 | GLY | p.Gly22Val | Q13432 |
| HOMO SAPIENS | 5 | 23902 | 2lrr | A | 765 | HIS | p.His765Gln | P38935 |
| HOMO SAPIENS | 1 | 29650 | 2vj2 | A | 234 | CYS | p.Cys234Tyr | P78504 |
| HOMO SAPIENS | 0 | 29650 | 2vj2 | A | 255 | TYR | p.Tyr255Ter | P78504 |
| HOMO SAPIENS | 3 | 29650 | 2vj2 | A | 274 | GLY | p.Gly274Asp | P78504 |
| HOMO SAPIENS | 1 | 105 | 5he0 | B | 77 | GLY | p.Gly77Ser | P62873 |
| HOMO SAPIENS | 2 | 105 | 5he0 | B | 80 | ILE | p.Ile80Thr | P62873 |
| HOMO SAPIENS | 3 | 4587 | 3rgh | A | 159 | ASP | p.Asp1159Ala | P21333 |
| HOMO SAPIENS | 3 | 4587 | 3rgh | A | 184 | ASP | p.Asp1184Glu | P21333 |
| HOMO SAPIENS | 2 | 4587 | 3rgh | A | 186 | SER | p.Ser1186Leu | P21333 |
| HOMO SAPIENS | 4 | 4587 | 3rgh | A | 188 | ALA | p.Ala1188Thr | P21333 |
| HOMO SAPIENS | 2 | 4587 | 3rgh | A | 199 | SER | p.Ser1199Leu | P21333 |
| HOMO SAPIENS | 0 | 20283 | 4n3w | A | 1278 | GLU | p.Glu1278Lys | Q92793 |
| HOMO SAPIENS | 1 | 14294 | 2d9q | B | 208 | MET | p.Met231Thr | Q99062 |
| HOMO SAPIENS | 6 | 14294 | 2d9q | B | 297 | ASP | p.Asp320Asn | Q99062 |
| HOMO SAPIENS | 0 | 11642 | 3bpt | A | 66 | ARG | p.Arg66Trp | Q6NVY1 |
| HOMO SAPIENS | 0 | 11642 | 3bpt | A | 122 | TYR | p.Tyr122Cys | Q6NVY1 |
| HOMO SAPIENS | 0 | 11642 | 3bpt | A | 345 | GLY | p.Gly345Ser | Q6NVY1 |
| HOMO SAPIENS | 1 | 3942 | 3sby | A | 60 | ASP | p.Asp60His | Q9Y4U1 |
| HOMO SAPIENS | 0 | 3942 | 3sby | A | 111 | ARG | p.Arg111Ter | Q9Y4U1 |
| HOMO SAPIENS | 1 | 3942 | 3sby | A | 116 | LEU | p.Leu116Pro | Q9Y4U1 |
| HOMO SAPIENS | 1 | 3942 | 3sby | A | 130 | TYR | p.Tyr130Cys | Q9Y4U1 |
| HOMO SAPIENS | 1 | 3942 | 3sby | A | 132 | ARG | p.Arg132Ter | Q9Y4U1 |
| HOMO SAPIENS | 1 | 3942 | 3sby | A | 147 | GLY | p.Gly147Ala | Q9Y4U1 |
| HOMO SAPIENS | 1 | 3942 | 3sby | A | 161 | ARG | p.Arg161Gln | Q9Y4U1 |
| HOMO SAPIENS | 1 | 3942 | 3sby | A | 161 | ARG | p.Arg161Ter | Q9Y4U1 |
| HOMO SAPIENS | 1 | 3942 | 3sby | A | 203 | TRP | p.Trp203Ter | Q9Y4U1 |
| HOMO SAPIENS | 1 | 3942 | 3sby | A | 203 | TRP | p.Trp203Ter | Q9Y4U1 |
| HOMO SAPIENS | 1 | 3942 | 3sby | A | 205 | TYR | p.Tyr205Ter | Q9Y4U1 |

| | | | | | | | | | |
|---|---|---|---|---|---|---|---|---|---|
| HOMO SAPIENS | 0 | 38340 | 2mbh | B | | 39 | MET | p.Met39Leu | Q13351 |
| HOMO SAPIENS | 2 | 7360 | 4k30 | A | | 430 | LEU | p.Leu430Pro | Q8N159 |
| HOMO SAPIENS | 2 | 7360 | 4k30 | A | | 433 | GLU | p.Glu433Asp | Q8N159 |
| HOMO SAPIENS | 7 | 7360 | 4k30 | A | | 484 | TRP | p.Trp484Arg | Q8N159 |
| HOMO SAPIENS | 1 | 41903 | 2xs6 | A | | 151 | PRO | p.Pro151Ser | O00459 |
| HOMO SAPIENS | 0 | 172 | 5glh | A | | 183 | ALA | p.Ala183Gly | P24530 |
| HOMO SAPIENS | 0 | 172 | 5glh | A | | 201 | ARG | p.Arg201Ter | P24530 |
| HOMO SAPIENS | 0 | 172 | 5glh | A | | 244 | THR | p.Thr244Met | P24530 |
| HOMO SAPIENS | 0 | 172 | 5glh | A | | 253 | ARG | p.Arg253Ter | P24530 |
| HOMO SAPIENS | 9 | 172 | 5glh | A | | 1017 | ILE | p.Leu17Phe | P00720 |
| HOMO SAPIENS | -1 | 3342 | 1ljm | A | | 1058 | HIS | p.His58Asn | Q01196 |
| HOMO SAPIENS | 4 | 3342 | 1ljm | A | | 1174 | ARG | p.Arg174Gln | Q01196 |
| HOMO SAPIENS | 5 | 25607 | 1p22 | C | | 32 | ASP | p.Asp32Ala | P35222 |
| HOMO SAPIENS | 5 | 25607 | 1p22 | C | | 32 | ASP | p.Asp32Tyr | P35222 |
| HOMO SAPIENS | -1 | 25607 | 1p22 | C | | 33 | SEP | p.Ser33Phe | P35222 |
| HOMO SAPIENS | 5 | 25607 | 1p22 | C | | 34 | GLY | p.Gly34Glu | P35222 |
| HOMO SAPIENS | -1 | 25607 | 1p22 | C | | 37 | SEP | p.Ser37Cys | P35222 |
| HOMO SAPIENS | -1 | 25607 | 1p22 | C | | 41 | THR | p.Thr41Ile | P35222 |
| HOMO SAPIENS | 1 | 48 | 4uow | | 9 | 60 | GLN | p.Arg34111His | Q8WZ42 |
| HOMO SAPIENS | 0 | 48 | 4uow | | 9 | 64 | LEU | p.Val34115Ala | Q8WZ42 |
| HOMO SAPIENS | 0 | 1127 | 5c01 | A | | 638 | ARG | p.Arg638Ter | P29597 |
| HOMO SAPIENS | 0 | 1127 | 5c01 | A | | 684 | ILE | p.Ile684Ser | P29597 |
| HOMO SAPIENS | 0 | 1127 | 5c01 | A | | 703 | ARG | p.Arg703Trp | P29597 |
| HOMO SAPIENS | 5 | 16389 | 2aq0 | B | | 128 | ALA | p.Ala860Asp | Q92889 |
| HOMO SAPIENS | 2 | 16389 | 2aq0 | B | | 141 | ILE | p.Ile873Val | Q92889 |
| HOMO SAPIENS | 5 | 16389 | 2aq0 | B | | 143 | GLU | p.Glu875Gly | Q92889 |
| HOMO SAPIENS | 0 | 16389 | 2aq0 | B | | 180 | GLY | p.Gly912Arg | Q92889 |
| HOMO SAPIENS | 0 | 15816 | 2xrc | A | | 89 | THR | p.Thr107Ala | P05156 |
| HOMO SAPIENS | 0 | 15816 | 2xrc | A | | 133 | ASN | p.Asn151Ser | P05156 |
| HOMO SAPIENS | 0 | 15816 | 2xrc | A | | 185 | THR | p.Thr203Ile | P05156 |
| HOMO SAPIENS | 0 | 15816 | 2xrc | A | | 225 | GLY | p.Gly243Asp | P05156 |
| HOMO SAPIENS | 0 | 15816 | 2xrc | A | | 243 | GLY | p.Gly261Ala | P05156 |
| HOMO SAPIENS | 0 | 15816 | 2xrc | A | | 269 | GLY | p.Gly287Arg | P05156 |
| HOMO SAPIENS | -1 | 15816 | 2xrc | A | | 288 | ILE | p.Ile306Val | P05156 |
| HOMO SAPIENS | -1 | 15816 | 2xrc | A | | 388 | ARG | p.Arg406His | P05156 |
| HOMO SAPIENS | 7 | 15816 | 2xrc | A | | 398 | ILE | p.Ile416Leu | P05156 |
| HOMO SAPIENS | 9 | 15816 | 2xrc | A | | 400 | HIS | p.His418Leu | P05156 |
| HOMO SAPIENS | 0 | 15816 | 2xrc | A | | 423 | LYS | p.Lys441Arg | P05156 |
| HOMO SAPIENS | -1 | 15816 | 2xrc | A | | 530 | GLU | p.Glu548Gln | P05156 |
| HOMO SAPIENS | 0 | 15816 | 2xrc | A | | 535 | PRO | p.Pro553Ala | P05156 |
| HOMO SAPIENS | 0 | 15816 | 2xrc | A | | 536 | GLU | p.Glu554Gly | P05156 |
| HOMO SAPIENS | 1 | 12385 | 4zlp | B | | 1450 | ALA | p.Ala1450Pro | Q9UM47 |
| HOMO SAPIENS | 7 | 12385 | 4zlp | B | | 1482 | GLY | p.Gly1482Arg | Q9UM47 |
| HOMO SAPIENS | 2 | 12385 | 4zlp | B | | 1518 | LEU | p.Leu1518Met | Q9UM47 |
| HOMO SAPIENS | 0 | 12385 | 4zlp | B | | 1560 | ARG | p.Arg1560Pro | Q9UM47 |
| HOMO SAPIENS | 0 | 1227 | 4a04 | A | | 148 | PHE | p.Phe148Tyr | O43435 |
| HOMO SAPIENS | 4 | 1227 | 4a04 | A | | 194 | HIS | p.His194Gln | O43435 |
| HOMO SAPIENS | 2 | 8017 | 3ei4 | B | | 244 | LYS | p.Lys244Glu | Q92466 |
| HOMO SAPIENS | 1 | 8017 | 3ei4 | B | | 273 | ARG | p.Arg273His | Q92466 |
| HOMO SAPIENS | 3 | 36112 | 2k8f | B | | 9 | SER | p.Ser9Arg | P04637 |
| HOMO SAPIENS | 2 | 12880 | 5ggf | A | | 265 | ARG | p.Arg265His | Q8WZA1 |
| HOMO SAPIENS | 2 | 12880 | 5ggf | A | | 287 | ILE | p.Ile287Ser | Q8WZA1 |
| HOMO SAPIENS | 2 | 12880 | 5ggf | A | | 311 | ARG | p.Arg311Gln | Q8WZA1 |
| HOMO SAPIENS | 2 | 12880 | 5ggf | A | | 425 | TRP | p.Trp425Ser | Q8WZA1 |

| | | | | | | | | |
|---|---|---|---|---|---|---|---|---|
| HOMO SAPIENS | 2 | 12880 | 5ggf | A | 442 | ARG | p.Arg442Cys | Q8WZA1 |
| HOMO SAPIENS | 2 | 12880 | 5ggf | A | 448 | GLY | p.Gly448Arg | Q8WZA1 |
| HOMO SAPIENS | 0 | 12880 | 5ggf | A | 490 | CYS | p.Cys490Tyr | Q8WZA1 |
| HOMO SAPIENS | 0 | 12880 | 5ggf | A | 493 | PRO | p.Pro493Arg | Q8WZA1 |
| HOMO SAPIENS | 3 | 12880 | 5ggf | A | 550 | SER | p.Ser550Asn | Q8WZA1 |
| HOMO SAPIENS | 1 | 12880 | 5ggf | A | 556 | ASP | p.Asp556Asn | Q8WZA1 |
| HOMO SAPIENS | 0 | 12880 | 5ggf | A | 580 | ARG | p.Arg580Ter | Q8WZA1 |
| HOMO SAPIENS | 2 | 12880 | 5ggf | A | 605 | ARG | p.Arg605Pro | Q8WZA1 |
| HOMO SAPIENS | 0 | 12880 | 5ggf | A | 623 | VAL | p.Met623Val | Q8WZA1 |
| HOMO SAPIENS | 1 | 10187 | 4qw2 | A | 145 | ALA | p.Ala145Ser | Q06787 |
| HOMO SAPIENS | 1 | 10187 | 4qw2 | A | 174 | ASN | p.Asn174Lys | Q06787 |
| HOMO SAPIENS | 0 | 24002 | 2m0t | A | 110 | LEU | p.Leu110Val | O14745 |
| HOMO SAPIENS | 9 | 3126 | 4mjh | A | 99 | LEU | p.Leu99Met | P04792 |
| HOMO SAPIENS | 1 | 3126 | 4mjh | A | 127 | ARG | p.Arg127Trp | P04792 |
| HOMO SAPIENS | 1 | 3126 | 4mjh | A | 127 | ARG | p.Arg127Leu | P04792 |
| HOMO SAPIENS | 1 | 3126 | 4mjh | A | 135 | SER | p.Ser135Phe | P04792 |
| HOMO SAPIENS | 2 | 3126 | 4mjh | A | 136 | ARG | p.Arg136Trp | P04792 |
| HOMO SAPIENS | 1 | 3126 | 4mjh | A | 140 | ARG | p.Arg140Gly | P04792 |
| HOMO SAPIENS | 0 | 3126 | 4mjh | A | 151 | THR | p.Thr151Ile | P04792 |
| HOMO SAPIENS | -1 | 15780 | 2jlp | A | 213 | ARG | p.Arg231Gly | P08294 |
| HOMO SAPIENS | 8 | 3578 | 1apz | D | 203 | GLY | p.Gly226Asp | P20933 |
| HOMO SAPIENS | 1 | 3578 | 1apz | D | 279 | GLY | p.Gly302Arg | P20933 |
| HOMO SAPIENS | 0 | 3578 | 1apz | D | 283 | CYS | p.Cys306Arg | P20933 |
| HOMO SAPIENS | 5 | 1335 | 5xtd | C | 115 | ARG | p.Arg115Cys | O75251 |
| HOMO SAPIENS | 0 | 1335 | 5xtd | C | 122 | VAL | p.Val122Met | O75251 |
| HOMO SAPIENS | 2 | 5063 | 2le4 | A | 37 | ARG | p.Arg74Pro | P48431 |
| HOMO SAPIENS | 1 | 25870 | 2o10 | A | 44 | LEU | p.Leu44Pro | P50461 |
| HOMO SAPIENS | 4 | 25870 | 2o10 | A | 58 | CYS | p.Cys58Gly | P50461 |
| HOMO SAPIENS | 0 | 18748 | 5fii | C | 39 | PHE | p.Phe39Leu | P00439 |
| HOMO SAPIENS | 6 | 18748 | 5fii | C | 48 | LEU | p.Leu48Ser | P00439 |
| HOMO SAPIENS | 2 | 18748 | 5fii | C | 65 | ILE | p.Ile65Asn | P00439 |
| HOMO SAPIENS | 3 | 18748 | 5fii | C | 95 | ILE | p.Ile95Phe | P00439 |
| HOMO SAPIENS | 1 | 14261 | 2vay | B | 1539 | ARG | p.Arg1539Cys | Q13698 |
| HOMO SAPIENS | 0 | 2494 | 4dx1 | A | 183 | GLU | p.Glu183Asp | Q9HBA0 |
| HOMO SAPIENS | 0 | 2494 | 4dx1 | A | 217 | ALA | p.Ala217Ser | Q9HBA0 |
| HOMO SAPIENS | 0 | 2494 | 4dx1 | A | 217 | ALA | p.Ala217Val | Q9HBA0 |
| HOMO SAPIENS | 4 | 2494 | 4dx1 | A | 224 | ARG | p.Arg224Trp | Q9HBA0 |
| HOMO SAPIENS | 4 | 2494 | 4dx1 | A | 254 | VAL | p.Val254Leu | Q9HBA0 |
| HOMO SAPIENS | 0 | 2494 | 4dx1 | A | 269 | ARG | p.Arg269Cys | Q9HBA0 |
| HOMO SAPIENS | 0 | 2494 | 4dx1 | A | 269 | ARG | p.Arg269His | Q9HBA0 |
| HOMO SAPIENS | 0 | 2494 | 4dx1 | A | 278 | GLU | p.Glu278Lys | Q9HBA0 |
| HOMO SAPIENS | 0 | 2494 | 4dx1 | A | 315 | ARG | p.Arg315Trp | Q9HBA0 |
| HOMO SAPIENS | 0 | 2494 | 4dx1 | A | 316 | ARG | p.Arg316Cys | Q9HBA0 |
| HOMO SAPIENS | 0 | 2494 | 4dx1 | A | 331 | ILE | p.Ile331Phe | Q9HBA0 |
| HOMO SAPIENS | 0 | 2494 | 4dx1 | A | 333 | ASP | p.Asp333Gly | Q9HBA0 |
| HOMO SAPIENS | 1 | 2494 | 4dx1 | A | 365 | VAL | p.Val365Met | Q9HBA0 |
| HOMO SAPIENS | 0 | 23998 | 2l1r | B | 161 | ARG | p.Arg162Trp | P19429 |
| HOMO SAPIENS | 0 | 8354 | 4f0x | D | 187 | SER | p.Ser187Ter | O95822 |
| HOMO SAPIENS | 2 | 8354 | 4f0x | D | 259 | GLY | p.Gly259Ala | O95822 |
| HOMO SAPIENS | 2 | 8354 | 4f0x | D | 284 | THR | p.Thr284Ala | O95822 |
| HOMO SAPIENS | 3 | 8354 | 4f0x | D | 309 | LYS | p.Lys309Gln | O95822 |
| HOMO SAPIENS | 0 | 21182 | 4apo | A | 228 | GLN | p.Gln228Lys | O00170 |
| HOMO SAPIENS | 1 | 21182 | 4apo | A | 307 | GLN | p.Gln307Arg | O00170 |
| HOMO SAPIENS | 0 | 2287 | 5of4 | B | 112 | ARG | p.Arg112His | P18074 |

| | | | | | | | | |
|---|---|---|---|---|---|---|---|---|
| HOMO SAPIENS | 0 | 2287 | 5of4 | B | 312 | ASP | p.Asp312Asn | P18074 |
| HOMO SAPIENS | 3 | 2287 | 5of4 | B | 461 | LEU | p.Leu461Val | P18074 |
| HOMO SAPIENS | 0 | 2287 | 5of4 | B | 485 | LEU | p.Leu485Pro | P18074 |
| HOMO SAPIENS | 1 | 2287 | 5of4 | B | 541 | SER | p.Ser541Arg | P18074 |
| HOMO SAPIENS | 0 | 2287 | 5of4 | B | 616 | ARG | p.Arg616Pro | P18074 |
| HOMO SAPIENS | 0 | 2287 | 5of4 | B | 616 | ARG | p.Arg616Trp | P18074 |
| HOMO SAPIENS | 1 | 2287 | 5of4 | B | 658 | ARG | p.Arg658Cys | P18074 |
| HOMO SAPIENS | 1 | 2287 | 5of4 | B | 681 | ASP | p.Asp681Asn | P18074 |
| HOMO SAPIENS | 1 | 2287 | 5of4 | B | 683 | ARG | p.Arg683Trp | P18074 |
| HOMO SAPIENS | 0 | 2287 | 5of4 | B | 713 | GLY | p.Gly713Arg | P18074 |
| HOMO SAPIENS | 0 | 2287 | 5of4 | B | 717 | ALA | p.Ala717Gly | P18074 |
| HOMO SAPIENS | 0 | 2287 | 5of4 | B | 722 | ARG | p.Arg722Trp | P18074 |
| HOMO SAPIENS | 0 | 2287 | 5of4 | B | 725 | ALA | p.Ala725Pro | P18074 |
| HOMO SAPIENS | 4 | 12188 | 4its | A | 122 | ARG | p.Arg122Gly | Q9Y606 |
| HOMO SAPIENS | 4 | 12188 | 4its | A | 133 | ASP | p.Asp133Asn | Q9Y606 |
| HOMO SAPIENS | 4 | 12188 | 4its | A | 144 | ARG | p.Arg144Trp | Q9Y606 |
| HOMO SAPIENS | 0 | 31529 | 1kgd | A | 761 | ARG | p.Arg766Gln | O14936 |
| HOMO SAPIENS | 1 | 31529 | 1kgd | A | 824 | GLN | p.Gln829Arg | O14936 |
| HOMO SAPIENS | 0 | 10906 | 2h63 | C | 18 | ARG | p.Arg18Ter | P53004 |
| HOMO SAPIENS | 9 | 27767 | 4wyq | A | 280 | LEU | p.Leu278Phe | Q9UPY3 |
| HOMO SAPIENS | 0 | 27767 | 4wyq | A | 297 | SER | p.Ser295Cys | Q9UPY3 |
| HOMO SAPIENS | 1 | 27767 | 4wyq | A | 334 | TYR | p.Tyr332Ter | Q9UPY3 |
| HOMO SAPIENS | 7 | 7775 | 5sv7 | C | 136 | SER | p.Ser136Cys | Q9NZJ5 |
| HOMO SAPIENS | 2 | 7775 | 5sv7 | C | 166 | GLN | p.Gln166Arg | Q9NZJ5 |
| HOMO SAPIENS | 2 | 7775 | 5sv7 | C | 332 | GLU | p.Glu332Ter | Q9NZJ5 |
| HOMO SAPIENS | 0 | 21449 | 2e9w | B | 84 | THR | p.Thr84Met | P10721 |
| HOMO SAPIENS | 0 | 21449 | 2e9w | B | 168 | ALA | p.Ala168Glu | P10721 |
| HOMO SAPIENS | 0 | 21449 | 2e9w | B | 178 | ALA | p.Ala178Thr | P10721 |
| HOMO SAPIENS | 0 | 21449 | 2e9w | B | 293 | ASN | p.Asn293Ser | P10721 |
| HOMO SAPIENS | -1 | 21449 | 2e9w | B | 318 | MET | p.Met318Leu | P10721 |
| HOMO SAPIENS | 0 | 21449 | 2e9w | B | 374 | VAL | p.Val374Ile | P10721 |
| HOMO SAPIENS | 6 | 3010 | 3flg | A | 66 | SER | p.Ser270Pro | Q9UBC3 |
| HOMO SAPIENS | 1 | 33918 | 2xzz | A | 760 | ARG | p.Arg760Ter | P22735 |
| HOMO SAPIENS | 2 | 1254 | 5xtc | w | 65 | ASN | p.Asn65Ser | O95299 |
| HOMO SAPIENS | 1 | 1254 | 5xtc | w | 238 | GLU | p.Glu238Lys | O95299 |
| HOMO SAPIENS | 3 | 10245 | 3siv | B | 194 | ALA | p.Ala194Glu | Q8WWY3 |
| HOMO SAPIENS | 4 | 10245 | 3siv | B | 216 | ALA | p.Ala216Pro | Q8WWY3 |
| HOMO SAPIENS | 0 | 2670 | 3dwb | A | 341 | THR | p.Thr341Ile | P42892 |
| HOMO SAPIENS | 1 | 2670 | 3dwb | A | 462 | SER | p.Ser462Asn | P42892 |
| HOMO SAPIENS | 1 | 2670 | 3dwb | A | 754 | ARG | p.Arg754Cys | P42892 |
| HOMO SAPIENS | 1 | 2961 | 5xtb | O | 40 | VAL | p.Val40Met | P19404 |
| HOMO SAPIENS | 5 | 2961 | 5xtb | O | 172 | ILE | p.Ile172Thr | P19404 |
| HOMO SAPIENS | 5 | 2961 | 5xtb | O | 190 | ASP | p.Asp190Gly | P19404 |
| HOMO SAPIENS | 5 | 29655 | 2xdg | A | 57 | ALA | p.Ala57Thr | Q02643 |
| HOMO SAPIENS | 4 | 29655 | 2xdg | A | 72 | GLU | p.Glu72Ter | Q02643 |
| HOMO SAPIENS | 0 | 29655 | 2xdg | A | 121 | GLU | p.Glu121Asp | Q02643 |
| HOMO SAPIENS | -1 | 17935 | 1hyn | P | 38 | ASP | p.Asp38Ala | P02730 |
| HOMO SAPIENS | 7 | 12825 | 1g82 | D | 99 | SER | p.Ser99Asn | P31371 |
| HOMO SAPIENS | 6 | 24927 | 1jm7 | B | 85 | VAL | p.Val85Leu | Q99728 |
| HOMO SAPIENS | 1 | 24927 | 1jm7 | B | 112 | ARG | p.Arg112Ter | Q99728 |
| HOMO SAPIENS | 0 | 24927 | 1jm7 | B | 116 | HIS | p.His116Tyr | Q99728 |
| HOMO SAPIENS | 7 | 92 | 5hde | A | 61 | LYS | p.Lys61Arg | Q05209 |
| HOMO SAPIENS | 0 | 17246 | 1szb | A | 84 | ARG | p.Arg99Gln | O00187 |
| HOMO SAPIENS | 1 | 17246 | 1szb | A | 105 | ASP | p.Asp120Gly | O00187 |

| | | | | | | | | |
|---|---|---|---|---|---|---|---|---|
| HOMO SAPIENS | 0 | 17246 | 1szb | A | 111 | PRO | p.Pro126Leu | O00187 |
| HOMO SAPIENS | 9 | 17246 | 1szb | A | 141 | CYS | p.Cys156Tyr | O00187 |
| HOMO SAPIENS | 0 | 32473 | 1b72 | A | 207 | ARG | p.Arg207Cys | P14653 |
| HOMO SAPIENS | 0 | 1920 | 4yzf | B | 488 | VAL | p.Val488Met | P02730 |
| HOMO SAPIENS | 0 | 1920 | 4yzf | B | 589 | ARG | p.Arg589Cys | P02730 |
| HOMO SAPIENS | 0 | 1920 | 4yzf | B | 589 | ARG | p.Arg589His | P02730 |
| HOMO SAPIENS | 0 | 1920 | 4yzf | B | 602 | ARG | p.Arg602Pro | P02730 |
| HOMO SAPIENS | 0 | 1920 | 4yzf | B | 613 | SER | p.Ser613Phe | P02730 |
| HOMO SAPIENS | -1 | 1920 | 4yzf | B | 646 | ARG | p.Arg646Gln | P02730 |
| HOMO SAPIENS | -1 | 1920 | 4yzf | B | 646 | ARG | p.Arg646Trp | P02730 |
| HOMO SAPIENS | 0 | 1920 | 4yzf | B | 701 | GLY | p.Gly701Asp | P02730 |
| HOMO SAPIENS | 0 | 1920 | 4yzf | B | 760 | ARG | p.Arg760Gln | P02730 |
| HOMO SAPIENS | 0 | 1920 | 4yzf | B | 771 | GLY | p.Gly771Asp | P02730 |
| HOMO SAPIENS | 0 | 1920 | 4yzf | B | 773 | SER | p.Ser773Pro | P02730 |
| HOMO SAPIENS | 0 | 1920 | 4yzf | B | 837 | THR | p.Thr837Ala | P02730 |
| HOMO SAPIENS | 0 | 1920 | 4yzf | B | 854 | PRO | p.Pro854Leu | P02730 |
| HOMO SAPIENS | 0 | 1920 | 4yzf | B | 858 | ALA | p.Ala858Asp | P02730 |
| HOMO SAPIENS | 0 | 1920 | 4yzf | B | 862 | VAL | p.Val862Ile | P02730 |
| HOMO SAPIENS | 0 | 1920 | 4yzf | B | 868 | PRO | p.Pro868Leu | P02730 |
| HOMO SAPIENS | 0 | 1920 | 4yzf | B | 870 | ARG | p.Arg870Trp | P02730 |
| HOMO SAPIENS | 0 | 9863 | 2juj | A | 54 | VAL | p.Val904Ile | P22681 |
| HOMO SAPIENS | 0 | 21934 | 2na7 | A | 184 | ILE | p.Ile184Val | P25445 |
| HOMO SAPIENS | -1 | 6746 | 1lw3 | A | 3 | THR | p.Lys3Thr | Q13614 |
| HOMO SAPIENS | -1 | 6746 | 1lw3 | A | 44 | VAL | p.Val44Ile | Q13614 |
| HOMO SAPIENS | 3 | 6746 | 1lw3 | A | 183 | ALA | p.Ala183Ser | Q13614 |
| HOMO SAPIENS | 4 | 6746 | 1lw3 | A | 482 | GLN | p.Gln482Ter | Q13614 |
| HOMO SAPIENS | 4 | 6746 | 1lw3 | A | 502 | GLU | p.Glu502Gln | Q13614 |
| HOMO SAPIENS | 3 | 6746 | 1lw3 | A | 545 | ASN | p.Asn545Ser | Q13614 |
| HOMO SAPIENS | 0 | 6746 | 1lw3 | A | 586 | ARG | p.Arg586Trp | Q13614 |
| HOMO SAPIENS | -1 | 6746 | 1lw3 | A | 602 | ALA | p.Ala602Gly | Q13614 |
| HOMO SAPIENS | 1 | 319 | 4ch9 | B | 336 | ARG | p.Arg336Ile | Q9UH77 |
| HOMO SAPIENS | 3 | 319 | 4ch9 | B | 340 | ALA | p.Ala340Val | Q9UH77 |
| HOMO SAPIENS | 0 | 319 | 4ch9 | B | 384 | ARG | p.Arg384Gln | Q9UH77 |
| HOMO SAPIENS | 5 | 319 | 4ch9 | B | 426 | PRO | p.Pro426Leu | Q9UH77 |
| HOMO SAPIENS | 3 | 319 | 4ch9 | B | 431 | ARG | p.Arg431Gln | Q9UH77 |
| HOMO SAPIENS | 2 | 319 | 4ch9 | B | 575 | ARG | p.Arg575Trp | Q9UH77 |
| HOMO SAPIENS | 0 | 574 | 2qkk | A | 142 | VAL | p.Val142Ile | O60930 |
| HOMO SAPIENS | 0 | 24021 | 2n8y | A | 752 | ARG | p.Arg752Gln | P12814 |
| HOMO SAPIENS | 0 | 18693 | 2y7j | B | 44 | ARG | p.Arg44Ter | P15735 |
| HOMO SAPIENS | 0 | 18693 | 2y7j | B | 106 | VAL | p.Val106Glu | P15735 |
| HOMO SAPIENS | 3 | 18693 | 2y7j | B | 189 | GLY | p.Gly189Glu | P15735 |
| HOMO SAPIENS | 0 | 17087 | 2ifg | A | 237 | THR | p.Thr237Met | P04629 |
| HOMO SAPIENS | 0 | 7980 | 1qub | A | 247 | VAL | p.Val266Leu | P02749 |
| HOMO SAPIENS | -1 | 2205 | 1h7u | A | 18 | ILE | p.Ile18Val | P54278 |
| HOMO SAPIENS | -1 | 2205 | 1h7u | A | 20 | ARG | p.Arg20Gln | P54278 |
| HOMO SAPIENS | 9 | 2205 | 1h7u | A | 46 | SER | p.Ser46Ile | P54278 |
| HOMO SAPIENS | 0 | 2205 | 1h7u | A | 60 | ASP | p.Asp60Glu | P54278 |
| HOMO SAPIENS | 0 | 2205 | 1h7u | A | 80 | PHE | p.Phe80Leu | P54278 |
| HOMO SAPIENS | 0 | 2205 | 1h7u | A | 118 | SER | p.Ser118Arg | P54278 |
| HOMO SAPIENS | 0 | 2205 | 1h7u | A | 126 | HIS | p.His126Gln | P54278 |
| HOMO SAPIENS | 0 | 2205 | 1h7u | A | 128 | SER | p.Ser128Leu | P54278 |
| HOMO SAPIENS | 1 | 2205 | 1h7u | A | 134 | ARG | p.Arg134Ter | P54278 |
| HOMO SAPIENS | 0 | 2205 | 1h7u | A | 166 | LEU | p.Leu166Pro | P54278 |
| HOMO SAPIENS | 1 | 2205 | 1h7u | A | 191 | TYR | p.Tyr191Cys | P54278 |

| | | | | | | | | |
|---|---|---|---|---|---|---|---|---|
| HOMO SAPIENS | 0 | 2205 | 1h7u | A | 211 | ARG | p.Arg211Ter | P54278 |
| HOMO SAPIENS | 0 | 2205 | 1h7u | A | 233 | GLN | p.Gln233Ter | P54278 |
| HOMO SAPIENS | 0 | 2205 | 1h7u | A | 255 | TYR | p.Tyr255Ter | P54278 |
| HOMO SAPIENS | 0 | 2205 | 1h7u | A | 277 | THR | p.Thr277Lys | P54278 |
| HOMO SAPIENS | 0 | 2205 | 1h7u | A | 301 | LYS | p.Lys301Asn | P54278 |
| HOMO SAPIENS | 0 | 2205 | 1h7u | A | 312 | MET | p.Met312Ile | P54278 |
| HOMO SAPIENS | 0 | 2205 | 1h7u | A | 312 | MET | p.Met312Thr | P54278 |
| HOMO SAPIENS | 0 | 2205 | 1h7u | A | 315 | ARG | p.Arg315Ter | P54278 |
| HOMO SAPIENS | 0 | 2205 | 1h7u | A | 318 | TYR | p.Tyr318Cys | P54278 |
| HOMO SAPIENS | 0 | 2205 | 1h7u | A | 335 | ASN | p.Asn335Ile | P54278 |
| HOMO SAPIENS | 0 | 2205 | 1h7u | A | 360 | ILE | p.Ile360Met | P54278 |
| HOMO SAPIENS | 6 | 20754 | 5mpo | B | 7 | VAL | p.Val7Phe | O96033 |
| HOMO SAPIENS | 4 | 20754 | 5mpo | B | 64 | VAL | p.Val64Ile | O96033 |
| HOMO SAPIENS | 5 | 34944 | 4qam | B | 1114 | ASP | p.Asp1114Gly | Q96KN7 |
| HOMO SAPIENS | 5 | 34944 | 4qam | B | 1240 | GLY | p.Gly1240Glu | Q96KN7 |
| HOMO SAPIENS | 9 | 40496 | 3vnn | A | 278 | ARG | p.Arg278His | P49917 |
| HOMO SAPIENS | 6 | 30131 | 4dt7 | E | 40 | PRO | p.Pro210Leu | P04070 |
| HOMO SAPIENS | -1 | 30131 | 4dt7 | E | 50 | ARG | p.Arg220Trp | P04070 |
| HOMO SAPIENS | 7 | 15311 | 2pqt | A | 64 | ARG | p.Arg64Trp | P18440 |
| HOMO SAPIENS | 2 | 15311 | 2pqt | A | 149 | VAL | p.Val149Ile | P18440 |
| HOMO SAPIENS | -1 | 19581 | 3hy8 | A | 33 | ASP | p.Asp33Val | Q9NVS9 |
| HOMO SAPIENS | 5 | 19581 | 3hy8 | A | 116 | ARG | p.Arg116Gln | Q9NVS9 |
| HOMO SAPIENS | 4 | 19581 | 3hy8 | A | 225 | ARG | p.Arg225His | Q9NVS9 |
| HOMO SAPIENS | 2 | 19581 | 3hy8 | A | 229 | TRP | p.Arg229Trp | Q9NVS9 |
| HOMO SAPIENS | 2 | 19581 | 3hy8 | A | 229 | TRP | p.Arg229Gln | Q9NVS9 |
| HOMO SAPIENS | 0 | 10266 | 3t1i | A | 41 | ASP | p.Asp41His | P49959 |
| HOMO SAPIENS | 1 | 10266 | 3t1i | A | 305 | ARG | p.Arg305Trp | P49959 |
| HOMO SAPIENS | 0 | 10266 | 3t1i | A | 308 | PHE | p.Phe308Leu | P49959 |
| HOMO SAPIENS | 0 | 10266 | 3t1i | A | 334 | SER | p.Ser334Arg | P49959 |
| HOMO SAPIENS | 1 | 1600 | 3rbf | A | 61 | GLU | p.Glu61Asp | P20711 |
| HOMO SAPIENS | 0 | 1600 | 3rbf | A | 91 | ALA | p.Ala91Val | P20711 |
| HOMO SAPIENS | -1 | 1600 | 3rbf | A | 102 | GLY | p.Gly102Ser | P20711 |
| HOMO SAPIENS | 3 | 1600 | 3rbf | A | 147 | SER | p.Ser147Arg | P20711 |
| HOMO SAPIENS | 1 | 1600 | 3rbf | A | 210 | PRO | p.Pro210Leu | P20711 |
| HOMO SAPIENS | 0 | 1600 | 3rbf | A | 217 | MET | p.Met217Val | P20711 |
| HOMO SAPIENS | 4 | 1600 | 3rbf | A | 250 | SER | p.Ser250Phe | P20711 |
| HOMO SAPIENS | 2 | 1600 | 3rbf | A | 275 | ALA | p.Ala275Thr | P20711 |
| HOMO SAPIENS | 1 | 1600 | 3rbf | A | 309 | PHE | p.Phe309Leu | P20711 |
| HOMO SAPIENS | -1 | 1600 | 3rbf | A | 347 | ARG | p.Arg347Gln | P20711 |
| HOMO SAPIENS | 0 | 1600 | 3rbf | A | 462 | ARG | p.Arg462Gln | P20711 |
| HOMO SAPIENS | 6 | 20830 | 5y4f | A | 714 | THR | p.Thr715Ile | Q01484 |
| HOMO SAPIENS | 7 | 20830 | 5y4f | A | 719 | ASP | p.Asp720Glu | Q01484 |
| HOMO SAPIENS | 0 | 20830 | 5y4f | A | 814 | VAL | p.Val815Ile | Q01484 |
| HOMO SAPIENS | -1 | 302 | 2c9y | A | 9 | GLU | p.Glu9Ter | P54819 |
| HOMO SAPIENS | 0 | 302 | 2c9y | A | 103 | ARG | p.Arg103Trp | P54819 |
| HOMO SAPIENS | 3 | 302 | 2c9y | A | 165 | ASP | p.Asp165Gly | P54819 |
| HOMO SAPIENS | 0 | 302 | 2c9y | A | 182 | ALA | p.Ala182Asp | P54819 |
| HOMO SAPIENS | 2 | 302 | 2c9y | A | 186 | ARG | p.Arg186Cys | P54819 |
| HOMO SAPIENS | 0 | 302 | 2c9y | A | 209 | ALA | p.Ala209Thr | P54819 |
| HOMO SAPIENS | -1 | 16920 | 1jlj | B | 10 | ASN | p.Asn10Tyr | Q9NQX3 |
| HOMO SAPIENS | 9 | 8462 | 1zbq | B | 16 | GLY | p.Gly16Ser | P51659 |
| HOMO SAPIENS | 5 | 8462 | 1zbq | B | 19 | ALA | p.Ala19Gly | P51659 |
| HOMO SAPIENS | 1 | 8462 | 1zbq | B | 34 | ALA | p.Ala34Val | P51659 |
| HOMO SAPIENS | 1 | 8462 | 1zbq | B | 106 | ARG | p.Arg106His | P51659 |

| | | | | | | | | |
|---|---|---|---|---|---|---|---|---|
| HOMO SAPIENS | 0 | 8462 | 1zbq | B | 140 | LYS | p.Lys140Asn | P51659 |
| HOMO SAPIENS | 0 | 8462 | 1zbq | B | 292 | THR | p.Thr292Ser | P51659 |
| HOMO SAPIENS | 0 | 4375 | 4ndi | A | 199 | ARG | p.Arg213His | Q7Z2E3 |
| HOMO SAPIENS | 3 | 4375 | 4ndi | A | 201 | HIS | p.His215Arg | Q7Z2E3 |
| HOMO SAPIENS | 0 | 4375 | 4ndi | A | 206 | PRO | p.Pro220Leu | Q7Z2E3 |
| HOMO SAPIENS | 2 | 4375 | 4ndi | A | 223 | LEU | p.Leu237Phe | Q7Z2E3 |
| HOMO SAPIENS | 0 | 4375 | 4ndi | A | 245 | ARG | p.Arg259His | Q7Z2E3 |
| HOMO SAPIENS | 0 | 4375 | 4ndi | A | 263 | VAL | p.Val277Gly | Q7Z2E3 |
| HOMO SAPIENS | 0 | 4375 | 4ndi | A | 279 | TRP | p.Trp293Ter | Q7Z2E3 |
| HOMO SAPIENS | 0 | 27004 | 4ccg | Y | 321 | ASP | p.Asp321Glu | Q9NW38 |
| HOMO SAPIENS | -1 | 27004 | 4ccg | Y | 372 | GLY | p.Gly372Ala | Q9NW38 |
| HOMO SAPIENS | -1 | 6383 | 5fai | A | 34 | ALA | p.Ala34Gly | Q92979 |
| HOMO SAPIENS | 8 | 6383 | 5fai | A | 86 | ASP | p.Asp86Gly | Q92979 |
| HOMO SAPIENS | 2 | 24585 | 4mi5 | A | 684 | ARG | p.Arg679Cys | Q15910 |
| HOMO SAPIENS | 0 | 15967 | 3nhb | A | 723 | ARG | p.Arg723Gln | Q9NP58 |
| HOMO SAPIENS | 5 | 18152 | 1oey | A | 389 | HIS | p.His389Gln | P19878 |
| HOMO SAPIENS | 5 | 18152 | 1oey | A | 395 | ARG | p.Arg395Trp | P19878 |
| HOMO SAPIENS | -1 | 1044 | 1li1 | F | 2 | VAL | p.Val1486Ile | P08572 |
| HOMO SAPIENS | 6 | 1044 | 1li1 | F | 37 | GLU | p.Glu1521Lys | P08572 |
| HOMO SAPIENS | 5 | 1044 | 1li1 | F | 157 | SER | p.Ser1639Leu | P08572 |
| HOMO SAPIENS | 5 | 1044 | 1li1 | F | 179 | ARG | p.Arg1662His | P08572 |
| HOMO SAPIENS | 5 | 1044 | 1li1 | F | 180 | GLY | p.Gly1663Arg | P08572 |
| HOMO SAPIENS | 6 | 20076 | 4uyi | A | 671 | LEU | p.Leu671Ser | Q8IY92 |
| HOMO SAPIENS | 0 | 19588 | 1jmj | A | 189 | ARG | p.Arg208His | P05546 |
| HOMO SAPIENS | -1 | 38023 | 1ry7 | B | 65 | GLY | p.Gly65Arg | P22607 |
| HOMO SAPIENS | -1 | 38023 | 1ry7 | B | 139 | ASP | p.Asp139Glu | P22607 |
| HOMO SAPIENS | 0 | 38023 | 1ry7 | B | 248 | ARG | p.Arg248Cys | P22607 |
| HOMO SAPIENS | 0 | 38023 | 1ry7 | B | 249 | SER | p.Ser249Cys | P22607 |
| HOMO SAPIENS | 0 | 38023 | 1ry7 | B | 250 | PRO | p.Pro250Arg | P22607 |
| HOMO SAPIENS | 0 | 38023 | 1ry7 | B | 322 | GLU | p.Glu322Lys | P22607 |
| HOMO SAPIENS | 5 | 16884 | 5eli | A | 33 | GLN | p.Gln33Ter | Q9NZC2 |
| HOMO SAPIENS | 0 | 16884 | 5eli | A | 44 | TRP | p.Trp44Ter | Q9NZC2 |
| HOMO SAPIENS | 0 | 16884 | 5eli | A | 47 | ARG | p.Arg47His | Q9NZC2 |
| HOMO SAPIENS | 0 | 16884 | 5eli | A | 66 | THR | p.Thr66Met | Q9NZC2 |
| HOMO SAPIENS | 0 | 16884 | 5eli | A | 78 | TRP | p.Trp78Ter | Q9NZC2 |
| HOMO SAPIENS | 1 | 16884 | 5eli | A | 96 | THR | p.Thr96Lys | Q9NZC2 |
| HOMO SAPIENS | 9 | 16884 | 5eli | A | 126 | VAL | p.Val126Gly | Q9NZC2 |
| HOMO SAPIENS | -1 | 12267 | 3w5o | A | 3 | ALA | p.Ala3Val | P49917 |
| HOMO SAPIENS | 1 | 12267 | 3w5o | A | 9 | THR | p.Thr9Ile | P49917 |
| HOMO SAPIENS | 2 | 12267 | 3w5o | A | 188 | ARG | p.Arg188Gln | P49917 |
| HOMO SAPIENS | 0 | 12267 | 3w5o | A | 469 | GLY | p.Gly469Glu | P49917 |
| HOMO SAPIENS | 1 | 12267 | 3w5o | A | 580 | ARG | p.Arg580Ter | P49917 |
| HOMO SAPIENS | 0 | 12267 | 3w5o | A | 600 | GLU | p.Glu600Lys | P49917 |
| HOMO SAPIENS | 9 | 2839 | 2hw5 | B | 54 | ARG | p.Arg54Cys | P30084 |
| HOMO SAPIENS | 9 | 2839 | 2hw5 | B | 59 | ASN | p.Asn59Ser | P30084 |
| HOMO SAPIENS | 0 | 2839 | 2hw5 | B | 273 | LYS | p.Lys273Glu | P30084 |
| HOMO SAPIENS | 1 | 12266 | 2o2k | B | 939 | GLY | p.Gly939Cys | Q99707 |
| HOMO SAPIENS | 5 | 12266 | 2o2k | B | 1164 | ARG | p.Arg1164His | Q99707 |
| HOMO SAPIENS | 1 | 12266 | 2o2k | B | 1166 | LEU | p.Leu1166Met | Q99707 |
| HOMO SAPIENS | 5 | 12266 | 2o2k | B | 1173 | PRO | p.Pro1173Leu | Q99707 |
| HOMO SAPIENS | 5 | 12266 | 2o2k | B | 1222 | ASN | p.Asn1222Ser | Q99707 |
| HOMO SAPIENS | 0 | 30891 | 3ch4 | B | 138 | ARG | p.Arg138Ter | Q15126 |
| HOMO SAPIENS | -1 | 4433 | 1qlz | A | 54 | GLY | p.Gly54Ser | P04156 |
| HOMO SAPIENS | 0 | 22568 | 3cxl | A | 20 | LEU | p.Leu20Phe | P15882 |

| | | | | | | | | |
|---|---|---|---|---|---|---|---|---|
| HOMO SAPIENS | 0 | 22568 | 3cxl | A | 126 | ILE | p.Ile126Met | P15882 |
| HOMO SAPIENS | 0 | 22568 | 3cxl | A | 141 | PRO | p.Pro141Leu | P15882 |
| HOMO SAPIENS | 0 | 22568 | 3cxl | A | 143 | TYR | p.Tyr143His | P15882 |
| HOMO SAPIENS | 0 | 22568 | 3cxl | A | 223 | ALA | p.Ala223Val | P15882 |
| HOMO SAPIENS | 0 | 22568 | 3cxl | A | 228 | GLY | p.Gly228Ser | P15882 |
| HOMO SAPIENS | 0 | 22568 | 3cxl | A | 252 | PRO | p.Pro252Gln | P15882 |
| HOMO SAPIENS | 0 | 22568 | 3cxl | A | 313 | GLU | p.Glu313Lys | P15882 |
| HOMO SAPIENS | 3 | 2668 | 5eqg | A | 34 | ASN | p.Asn34Ile | P11166 |
| HOMO SAPIENS | 3 | 2668 | 5eqg | A | 77 | MET | p.Met77Val | P11166 |
| HOMO SAPIENS | 6 | 2668 | 5eqg | A | 91 | GLY | p.Gly91Asp | P11166 |
| HOMO SAPIENS | 6 | 2668 | 5eqg | A | 92 | ARG | p.Arg92Trp | P11166 |
| HOMO SAPIENS | 6 | 2668 | 5eqg | A | 93 | ARG | p.Arg93Trp | P11166 |
| HOMO SAPIENS | 0 | 2668 | 5eqg | A | 95 | SER | p.Ser95Ile | P11166 |
| HOMO SAPIENS | 3 | 2668 | 5eqg | A | 126 | ARG | p.Arg126His | P11166 |
| HOMO SAPIENS | 3 | 2668 | 5eqg | A | 126 | ARG | p.Arg126Cys | P11166 |
| HOMO SAPIENS | 4 | 2668 | 5eqg | A | 232 | ARG | p.Arg232Cys | P11166 |
| HOMO SAPIENS | 3 | 2668 | 5eqg | A | 249 | ARG | p.Arg249Gln | P11166 |
| HOMO SAPIENS | 3 | 2668 | 5eqg | A | 255 | LYS | p.Lys255Arg | P11166 |
| HOMO SAPIENS | 0 | 2668 | 5eqg | A | 275 | ALA | p.Ala275Thr | P11166 |
| HOMO SAPIENS | 4 | 2668 | 5eqg | A | 299 | GLU | p.Glu299Lys | P11166 |
| HOMO SAPIENS | 6 | 2668 | 5eqg | A | 314 | GLY | p.Gly314Ser | P11166 |
| HOMO SAPIENS | 7 | 2668 | 5eqg | A | 315 | ILE | p.Ile315Met | P11166 |
| HOMO SAPIENS | -1 | 2668 | 5eqg | A | 458 | ARG | p.Arg458Trp | P11166 |
| HOMO SAPIENS | -1 | 2668 | 5eqg | A | 468 | ARG | p.Arg468Trp | P11166 |
| HOMO SAPIENS | 3 | 2291 | 2amy | A | 31 | PHE | p.Phe31Leu | O15305 |
| HOMO SAPIENS | 7 | 2291 | 2amy | A | 32 | LEU | p.Leu32Arg | O15305 |
| HOMO SAPIENS | 7 | 2291 | 2amy | A | 65 | ASP | p.Asp65Tyr | O15305 |
| HOMO SAPIENS | 1 | 2291 | 2amy | A | 108 | ALA | p.Ala108Val | O15305 |
| HOMO SAPIENS | 3 | 2291 | 2amy | A | 113 | PRO | p.Pro113Leu | O15305 |
| HOMO SAPIENS | 3 | 2291 | 2amy | A | 113 | PRO | p.Pro113Ser | O15305 |
| HOMO SAPIENS | 6 | 2291 | 2amy | A | 117 | GLY | p.Gly117Arg | O15305 |
| HOMO SAPIENS | 6 | 2291 | 2amy | A | 119 | PHE | p.Phe119Leu | O15305 |
| HOMO SAPIENS | 6 | 2291 | 2amy | A | 123 | ARG | p.Arg123Gln | O15305 |
| HOMO SAPIENS | 0 | 2291 | 2amy | A | 129 | VAL | p.Val129Met | O15305 |
| HOMO SAPIENS | 7 | 2291 | 2amy | A | 139 | GLU | p.Glu139Lys | O15305 |
| HOMO SAPIENS | 5 | 2291 | 2amy | A | 141 | ARG | p.Arg141His | O15305 |
| HOMO SAPIENS | 7 | 2291 | 2amy | A | 144 | PHE | p.Phe144Leu | O15305 |
| HOMO SAPIENS | 6 | 2291 | 2amy | A | 148 | ASP | p.Asp148Asn | O15305 |
| HOMO SAPIENS | 6 | 2291 | 2amy | A | 153 | ILE | p.Ile153Thr | O15305 |
| HOMO SAPIENS | 2 | 2291 | 2amy | A | 157 | PHE | p.Phe157Ser | O15305 |
| HOMO SAPIENS | 1 | 2291 | 2amy | A | 162 | ARG | p.Arg162Trp | O15305 |
| HOMO SAPIENS | 6 | 2291 | 2amy | A | 188 | ASP | p.Asp188Gly | O15305 |
| HOMO SAPIENS | 5 | 2291 | 2amy | A | 194 | ARG | p.Arg194Ter | O15305 |
| HOMO SAPIENS | 0 | 2291 | 2amy | A | 207 | PHE | p.Phe207Ser | O15305 |
| HOMO SAPIENS | 5 | 2291 | 2amy | A | 208 | GLY | p.Gly208Ala | O15305 |
| HOMO SAPIENS | 1 | 2291 | 2amy | A | 216 | ASN | p.Asn216Ile | O15305 |
| HOMO SAPIENS | 5 | 2291 | 2amy | A | 218 | HIS | p.His218Leu | O15305 |
| HOMO SAPIENS | 6 | 2291 | 2amy | A | 223 | ASP | p.Asp223Glu | O15305 |
| HOMO SAPIENS | 6 | 2291 | 2amy | A | 226 | THR | p.Thr226Ser | O15305 |
| HOMO SAPIENS | 5 | 2291 | 2amy | A | 231 | VAL | p.Val231Met | O15305 |
| HOMO SAPIENS | 1 | 2291 | 2amy | A | 237 | THR | p.Thr237Met | O15305 |
| HOMO SAPIENS | 1 | 2291 | 2amy | A | 241 | CYS | p.Cys241Ser | O15305 |
| HOMO SAPIENS | 5 | 35907 | 1d6g | A | 21 | GLY | p.Gly21Arg | P32238 |
| HOMO SAPIENS | -1 | 33209 | 1x9d | A | 197 | PRO | p.Pro197Leu | Q9UKM7 |

| | | | | | | | | |
|---|---|---|---|---|---|---|---|---|
| HOMO SAPIENS | -1 | 33209 | 1x9d | A | 201 | PRO | p.Pro201Leu | Q9UKM7 |
| HOMO SAPIENS | 6 | 21305 | 4yep | B | 1326 | ARG | p.Arg1326Ter | P24043 |
| HOMO SAPIENS | 6 | 21305 | 4yep | B | 1350 | ARG | p.Arg1350Ter | P24043 |
| HOMO SAPIENS | 0 | 21553 | 3qyb | A | 1120 | VAL | p.Val1119Ala | O60343 |
| HOMO SAPIENS | 2 | 21553 | 3qyb | A | 1148 | THR | p.Thr1147Met | O60343 |
| HOMO SAPIENS | -1 | 30645 | 3m0c | C | -17 | TRP | p.Trp4Ter | P01130 |
| HOMO SAPIENS | -1 | 30645 | 3m0c | C | -17 | TRP | p.Trp4Ter | P01130 |
| HOMO SAPIENS | -1 | 30645 | 3m0c | C | 12 | GLN | p.Gln33Ter | P01130 |
| HOMO SAPIENS | -1 | 30645 | 3m0c | C | 15 | ASP | p.Asp36Glu | P01130 |
| HOMO SAPIENS | -1 | 30645 | 3m0c | C | 28 | SER | p.Ser49Arg | P01130 |
| HOMO SAPIENS | -1 | 30645 | 3m0c | C | 41 | THR | p.Thr62Met | P01130 |
| HOMO SAPIENS | -1 | 30645 | 3m0c | C | 69 | ASP | p.Asp90Asn | P01130 |
| HOMO SAPIENS | -1 | 30645 | 3m0c | C | 76 | ASN | p.Asn97Lys | P01130 |
| HOMO SAPIENS | -1 | 30645 | 3m0c | C | 78 | SER | p.Ser99Ter | P01130 |
| HOMO SAPIENS | -1 | 30645 | 3m0c | C | 81 | GLN | p.Gln102Ter | P01130 |
| HOMO SAPIENS | -1 | 30645 | 3m0c | C | 84 | PRO | p.Pro105Ser | P01130 |
| HOMO SAPIENS | -1 | 30645 | 3m0c | C | 194 | ARG | p.Arg215Cys | P01130 |
| HOMO SAPIENS | -1 | 30645 | 3m0c | C | 198 | GLY | p.Gly219Asp | P01130 |
| HOMO SAPIENS | -1 | 30645 | 3m0c | C | 201 | CYS | p.Cys222Tyr | P01130 |
| HOMO SAPIENS | -1 | 30645 | 3m0c | C | 201 | CYS | p.Cys222Ter | P01130 |
| HOMO SAPIENS | -1 | 30645 | 3m0c | C | 206 | ASP | p.Asp227Glu | P01130 |
| HOMO SAPIENS | -1 | 30645 | 3m0c | C | 249 | CYS | p.Cys270Ter | P01130 |
| HOMO SAPIENS | -1 | 30645 | 3m0c | C | 250 | VAL | p.Val271Ile | P01130 |
| HOMO SAPIENS | 0 | 30645 | 3m0c | C | 255 | CYS | p.Cys276Tyr | P01130 |
| HOMO SAPIENS | 0 | 30645 | 3m0c | C | 255 | CYS | p.Cys276Ter | P01130 |
| HOMO SAPIENS | 0 | 30645 | 3m0c | C | 256 | GLU | p.Glu277Lys | P01130 |
| HOMO SAPIENS | 0 | 30645 | 3m0c | C | 265 | SER | p.Ser286Arg | P01130 |
| HOMO SAPIENS | 0 | 30645 | 3m0c | C | 284 | TRP | p.Trp305Ter | P01130 |
| HOMO SAPIENS | 0 | 30645 | 3m0c | C | 292 | CYS | p.Cys313Ter | P01130 |
| HOMO SAPIENS | 1 | 30645 | 3m0c | C | 385 | ARG | p.Arg406Trp | P01130 |
| HOMO SAPIENS | 0 | 30645 | 3m0c | C | 395 | ARG | p.Arg416Gln | P01130 |
| HOMO SAPIENS | 0 | 30645 | 3m0c | C | 398 | TYR | p.Tyr419Ter | P01130 |
| HOMO SAPIENS | 1 | 30645 | 3m0c | C | 415 | VAL | p.Val436Ala | P01130 |
| HOMO SAPIENS | 1 | 30645 | 3m0c | C | 424 | ASP | p.Asp445Glu | P01130 |
| HOMO SAPIENS | 1 | 30645 | 3m0c | C | 425 | LEU | p.Leu446Val | P01130 |
| HOMO SAPIENS | -1 | 30645 | 3m0c | C | 440 | GLY | p.Gly461Cys | P01130 |
| HOMO SAPIENS | -1 | 30645 | 3m0c | C | 441 | VAL | p.Val462Ile | P01130 |
| HOMO SAPIENS | 1 | 30645 | 3m0c | C | 447 | VAL | p.Val468Ile | P01130 |
| HOMO SAPIENS | 1 | 30645 | 3m0c | C | 489 | THR | p.Thr510Met | P01130 |
| HOMO SAPIENS | 0 | 30645 | 3m0c | C | 524 | GLY | p.Gly545Glu | P01130 |
| HOMO SAPIENS | 1 | 30645 | 3m0c | C | 615 | GLY | p.Gly636Ser | P01130 |
| HOMO SAPIENS | 1 | 30645 | 3m0c | C | 625 | LEU | p.Leu646Ile | P01130 |
| HOMO SAPIENS | 1 | 30645 | 3m0c | C | 630 | ASP | p.Asp651Gly | P01130 |
| HOMO SAPIENS | 3 | 30645 | 3m0c | C | 646 | CYS | p.Cys667Tyr | P01130 |
| HOMO SAPIENS | 4 | 30645 | 3m0c | C | 654 | GLY | p.Gly675Ser | P01130 |
| HOMO SAPIENS | 6 | 30645 | 3m0c | C | 660 | CYS | p.Cys681Ter | P01130 |
| HOMO SAPIENS | 5 | 30645 | 3m0c | C | 676 | ALA | p.Ala697Pro | P01130 |
| HOMO SAPIENS | -1 | 30645 | 3m0c | C | 749 | GLN | p.Gln770Ter | P01130 |
| HOMO SAPIENS | -1 | 41323 | 4bxo | A | 1812 | PRO | p.Pro1812Ala | Q8IYD8 |
| HOMO SAPIENS | 5 | 8335 | 3h0t | C | 12 | GLY | p.Gly71Asp | P81172 |
| HOMO SAPIENS | 6 | 609 | 1bhg | B | 148 | PRO | p.Pro148Ser | P08236 |
| HOMO SAPIENS | 1 | 609 | 1bhg | B | 176 | LEU | p.Leu176Phe | P08236 |
| HOMO SAPIENS | 6 | 609 | 1bhg | B | 216 | ARG | p.Arg216Trp | P08236 |
| HOMO SAPIENS | 0 | 609 | 1bhg | B | 350 | LYS | p.Lys350Asn | P08236 |

| | | | | | | | | |
|---|---|---|---|---|---|---|---|---|
| HOMO SAPIENS | 1 | 609 | 1bhg | B | 354 | ALA | p.Ala354Val | P08236 |
| HOMO SAPIENS | 1 | 609 | 1bhg | B | 357 | ARG | p.Arg357Ter | P08236 |
| HOMO SAPIENS | 0 | 609 | 1bhg | B | 382 | ARG | p.Arg382Cys | P08236 |
| HOMO SAPIENS | 0 | 609 | 1bhg | B | 495 | TYR | p.Tyr495Cys | P08236 |
| HOMO SAPIENS | 2 | 609 | 1bhg | B | 507 | TRP | p.Trp507Ter | P08236 |
| HOMO SAPIENS | 0 | 609 | 1bhg | B | 577 | ARG | p.Arg577Leu | P08236 |
| HOMO SAPIENS | 1 | 609 | 1bhg | B | 611 | ARG | p.Arg611Trp | P08236 |
| HOMO SAPIENS | 0 | 609 | 1bhg | B | 619 | ALA | p.Ala619Val | P08236 |
| HOMO SAPIENS | 0 | 609 | 1bhg | B | 627 | TRP | p.Trp627Cys | P08236 |
| HOMO SAPIENS | 5 | 3422 | 1x9n | A | 566 | GLU | p.Glu566Lys | P18858 |
| HOMO SAPIENS | 2 | 3422 | 1x9n | A | 771 | ARG | p.Arg771Trp | P18858 |
| HOMO SAPIENS | 0 | 3279 | 5tby | B | 797 | ALA | p.Ala797Thr | P12883 |
| HOMO SAPIENS | -1 | 3279 | 5tby | B | 982 | MET | p.Met982Thr | P12883 |
| HOMO SAPIENS | -1 | 3279 | 5tby | B | 992 | LEU | p.Leu992Met | P12883 |
| HOMO SAPIENS | -1 | 3279 | 5tby | B | 1004 | GLN | p.Gln1004Glu | P12883 |
| HOMO SAPIENS | -1 | 3279 | 5tby | B | 1013 | GLU | p.Glu1013Lys | P12883 |
| HOMO SAPIENS | -1 | 3279 | 5tby | B | 1032 | ASP | p.Asp1032Asn | P12883 |
| HOMO SAPIENS | -1 | 3279 | 5tby | B | 1053 | ARG | p.Arg1053Gln | P12883 |
| HOMO SAPIENS | -1 | 3279 | 5tby | B | 1117 | GLU | p.Glu1117Ter | P12883 |
| HOMO SAPIENS | -1 | 3279 | 5tby | B | 1193 | ARG | p.Arg1193His | P12883 |
| HOMO SAPIENS | -1 | 3279 | 5tby | B | 1205 | GLU | p.Glu1205Lys | P12883 |
| HOMO SAPIENS | -1 | 3279 | 5tby | B | 1207 | ILE | p.Ile1207Met | P12883 |
| HOMO SAPIENS | -1 | 3279 | 5tby | B | 1250 | ARG | p.Arg1250Pro | P12883 |
| HOMO SAPIENS | -1 | 3279 | 5tby | B | 1259 | HIS | p.His1259Gln | P12883 |
| HOMO SAPIENS | -1 | 3279 | 5tby | B | 1272 | ASP | p.Asp1272Asn | P12883 |
| HOMO SAPIENS | -1 | 3279 | 5tby | B | 1295 | GLU | p.Glu1295Lys | P12883 |
| HOMO SAPIENS | -1 | 3279 | 5tby | B | 1335 | SER | p.Ser1335Leu | P12883 |
| HOMO SAPIENS | -1 | 3279 | 5tby | B | 1399 | GLU | p.Glu1399Asp | P12883 |
| HOMO SAPIENS | -1 | 3279 | 5tby | B | 1413 | SER | p.Ser1413Ter | P12883 |
| HOMO SAPIENS | -1 | 3279 | 5tby | B | 1459 | LYS | p.Lys1459Asn | P12883 |
| HOMO SAPIENS | -1 | 3279 | 5tby | B | 1467 | LEU | p.Leu1467Val | P12883 |
| HOMO SAPIENS | -1 | 3279 | 5tby | B | 1491 | SER | p.Ser1491Cys | P12883 |
| HOMO SAPIENS | -1 | 3279 | 5tby | B | 1496 | GLU | p.Glu1496Ala | P12883 |
| HOMO SAPIENS | -1 | 3279 | 5tby | B | 1500 | ARG | p.Arg1500Pro | P12883 |
| HOMO SAPIENS | -1 | 3279 | 5tby | B | 1606 | ARG | p.Arg1606His | P12883 |
| HOMO SAPIENS | -1 | 3279 | 5tby | B | 1664 | ASN | p.Asn1664Lys | P12883 |
| HOMO SAPIENS | -1 | 3279 | 5tby | B | 1712 | ARG | p.Arg1712Trp | P12883 |
| HOMO SAPIENS | -1 | 3279 | 5tby | B | 1769 | LEU | p.Leu1769Met | P12883 |
| HOMO SAPIENS | -1 | 3279 | 5tby | B | 1808 | GLY | p.Gly1808Ser | P12883 |
| HOMO SAPIENS | -1 | 3279 | 5tby | B | 1818 | ARG | p.Arg1818Trp | P12883 |
| HOMO SAPIENS | -1 | 3279 | 5tby | B | 1820 | ARG | p.Arg1820Trp | P12883 |
| HOMO SAPIENS | -1 | 3279 | 5tby | B | 1844 | GLU | p.Glu1844Lys | P12883 |
| HOMO SAPIENS | -1 | 3279 | 5tby | B | 1845 | ARG | p.Arg1845Trp | P12883 |
| HOMO SAPIENS | -1 | 3279 | 5tby | B | 1854 | THR | p.Thr1854Met | P12883 |
| HOMO SAPIENS | -1 | 3279 | 5tby | B | 1909 | ARG | p.Arg1909Trp | P12883 |
| HOMO SAPIENS | 2 | 2260 | 4kl1 | A | 553 | ASP | p.Asp553Asn | Q9Y3Q4 |
| HOMO SAPIENS | 0 | 40570 | 5lst | A | 522 | ARG | p.Arg522Cys | O94761 |
| HOMO SAPIENS | 0 | 40570 | 5lst | A | 522 | ARG | p.Arg522His | O94761 |
| HOMO SAPIENS | 0 | 40570 | 5lst | A | 591 | PRO | p.Pro591Leu | O94761 |
| HOMO SAPIENS | 0 | 40570 | 5lst | A | 793 | PRO | p.Pro793Ser | O94761 |
| HOMO SAPIENS | 4 | 40570 | 5lst | A | 799 | VAL | p.Val799Met | O94761 |
| HOMO SAPIENS | 0 | 40570 | 5lst | A | 1004 | ARG | p.Arg1004Trp | O94761 |
| HOMO SAPIENS | 0 | 40570 | 5lst | A | 1005 | ARG | p.Arg1005Gln | O94761 |
| HOMO SAPIENS | 0 | 40570 | 5lst | A | 1021 | ARG | p.Arg1021Gln | O94761 |

| | | | | | | | | |
|---|---|---|---|---|---|---|---|---|
| HOMO SAPIENS | 0 | 40570 | 5lst | A | 1043 | LEU | p.Leu1043Pro | O94761 |
| HOMO SAPIENS | 0 | 40570 | 5lst | A | 1045 | ALA | p.Ala1045Thr | O94761 |
| HOMO SAPIENS | 0 | 40570 | 5lst | A | 1105 | GLY | p.Gly1105Asp | O94761 |
| HOMO SAPIENS | 0 | 40570 | 5lst | A | 1105 | GLY | p.Gly1105Ser | O94761 |
| HOMO SAPIENS | -1 | 40570 | 5lst | A | 1113 | GLY | p.Gly1113Arg | O94761 |
| HOMO SAPIENS | -1 | 18865 | 1gxc | A | 64 | GLU | p.Glu64Lys | O96017 |
| HOMO SAPIENS | -1 | 11673 | 3dzu | D | 85 | PRO | p.Pro113Gln | P37231 |
| HOMO SAPIENS | 0 | 19383 | 3b96 | A | 63 | LYS | p.Lys103Arg | P49748 |
| HOMO SAPIENS | 4 | 19383 | 3b96 | A | 140 | ALA | p.Ala180Thr | P49748 |
| HOMO SAPIENS | 1 | 19383 | 3b96 | A | 145 | GLY | p.Gly185Ser | P49748 |
| HOMO SAPIENS | 0 | 19383 | 3b96 | A | 220 | THR | p.Thr260Met | P49748 |
| HOMO SAPIENS | 5 | 19383 | 3b96 | A | 243 | VAL | p.Val283Ala | P49748 |
| HOMO SAPIENS | 0 | 19383 | 3b96 | A | 249 | GLY | p.Gly289Arg | P49748 |
| HOMO SAPIENS | 0 | 19383 | 3b96 | A | 254 | GLY | p.Gly294Glu | P49748 |
| HOMO SAPIENS | 7 | 19383 | 3b96 | A | 326 | ARG | p.Arg366Cys | P49748 |
| HOMO SAPIENS | 7 | 19383 | 3b96 | A | 326 | ARG | p.Arg366His | P49748 |
| HOMO SAPIENS | 5 | 19383 | 3b96 | A | 342 | LYS | p.Lys382Gln | P49748 |
| HOMO SAPIENS | 0 | 19383 | 3b96 | A | 369 | THR | p.Thr409Met | P49748 |
| HOMO SAPIENS | 5 | 19383 | 3b96 | A | 376 | ALA | p.Ala416Ser | P49748 |
| HOMO SAPIENS | 0 | 19383 | 3b96 | A | 385 | ALA | p.Ala425Thr | P49748 |
| HOMO SAPIENS | 5 | 19383 | 3b96 | A | 399 | GLY | p.Gly439Asp | P49748 |
| HOMO SAPIENS | 7 | 19383 | 3b96 | A | 401 | GLY | p.Gly441Asp | P49748 |
| HOMO SAPIENS | 9 | 19383 | 3b96 | A | 413 | ARG | p.Arg453Gln | P49748 |
| HOMO SAPIENS | 2 | 19383 | 3b96 | A | 418 | PHE | p.Phe458Leu | P49748 |
| HOMO SAPIENS | 6 | 19383 | 3b96 | A | 419 | ARG | p.Arg459Trp | P49748 |
| HOMO SAPIENS | 6 | 19383 | 3b96 | A | 419 | ARG | p.Arg459Gln | P49748 |
| HOMO SAPIENS | -1 | 19383 | 3b96 | A | 471 | ARG | p.Arg511Gln | P49748 |
| HOMO SAPIENS | 0 | 19383 | 3b96 | A | 573 | ARG | p.Arg613Trp | P49748 |
| HOMO SAPIENS | 0 | 19383 | 3b96 | A | 575 | ARG | p.Arg615Gln | P49748 |
| HOMO SAPIENS | 0 | 22286 | 3k2s | A | 3 | PRO | p.Pro27His | P02647 |
| HOMO SAPIENS | 0 | 22286 | 3k2s | A | 4 | PRO | p.Pro28Arg | P02647 |
| HOMO SAPIENS | 0 | 22286 | 3k2s | A | 10 | ARG | p.Arg34Leu | P02647 |
| HOMO SAPIENS | 0 | 22286 | 3k2s | A | 26 | GLY | p.Gly50Arg | P02647 |
| HOMO SAPIENS | 0 | 22286 | 3k2s | A | 37 | ALA | p.Ala61Thr | P02647 |
| HOMO SAPIENS | 1 | 1622 | 2qrv | D | 631 | ARG | p.Arg635Trp | Q9Y6K1 |
| HOMO SAPIENS | 1 | 1622 | 2qrv | D | 900 | PRO | p.Pro904Leu | Q9Y6K1 |
| HOMO SAPIENS | 6 | 40796 | 1v18 | B | 1509 | LEU | p.Leu1509Val | P25054 |
| HOMO SAPIENS | 1 | 22375 | 4pki | G | 1214 | LYS | p.Asp214Asn | P28289 |
| HOMO SAPIENS | 1 | 16307 | 5mv8 | B | 483 | ALA | p.Ala483Ser | Q9Y6N9 |
| HOMO SAPIENS | 6 | 16307 | 5mv8 | B | 506 | ASN | p.Asn506Lys | Q9Y6N9 |
| HOMO SAPIENS | 4 | 16307 | 5mv8 | B | 519 | ASP | p.Glu519Asp | Q9Y6N9 |
| HOMO SAPIENS | 0 | 16307 | 5mv8 | B | 530 | GLY | p.Gly530Arg | Q9Y6N9 |
| HOMO SAPIENS | 4 | 36221 | 4cvo | A | 127 | VAL | p.Val127Ile | Q03468 |
| HOMO SAPIENS | 7 | 36221 | 4cvo | A | 134 | ARG | p.Arg134Trp | Q03468 |
| HOMO SAPIENS | 0 | 36221 | 4cvo | A | 156 | GLN | p.Gln156Ter | Q03468 |
| HOMO SAPIENS | 0 | 10829 | 4a11 | B | 13 | GLU | p.Glu13Ter | Q13216 |
| HOMO SAPIENS | 1 | 10829 | 4a11 | B | 100 | TYR | p.Tyr100Ter | Q13216 |
| HOMO SAPIENS | 0 | 10829 | 4a11 | B | 160 | ALA | p.Ala160Val | Q13216 |
| HOMO SAPIENS | 1 | 6759 | 5lev | A | 29 | ILE | p.Ile29Phe | Q9H3H5 |
| HOMO SAPIENS | 0 | 6759 | 5lev | A | 108 | MET | p.Met108Ile | Q9H3H5 |
| HOMO SAPIENS | 2 | 6759 | 5lev | A | 117 | VAL | p.Val117Ile | Q9H3H5 |
| HOMO SAPIENS | 1 | 6759 | 5lev | A | 170 | TYR | p.Tyr170Cys | Q9H3H5 |
| HOMO SAPIENS | 0 | 6759 | 5lev | A | 264 | GLY | p.Val264Gly | Q9H3H5 |
| HOMO SAPIENS | 0 | 6759 | 5lev | A | 332 | PHE | p.Phe332Val | Q9H3H5 |

| | | | | | | | | |
|---|---|---|---|---|---|---|---|---|
| HOMO SAPIENS | 7 | 6759 | 5lev | A | 393 | ILE | p.Ile393Val | Q9H3H5 |
| HOMO SAPIENS | -1 | 33473 | 5m5e | C | 9 | ASN | p.Asn31Ser | P31785 |
| HOMO SAPIENS | 0 | 26155 | 1ic8 | B | 98 | ALA | p.Ala98Val | P20823 |
| HOMO SAPIENS | 0 | 15329 | 4iqr | B | 127 | ARG | p.Arg136Trp | P41235 |
| HOMO SAPIENS | 0 | 20725 | 4bsj | A | 350 | GLU | p.Glu350Val | P35916 |
| HOMO SAPIENS | 0 | 20725 | 4bsj | A | 494 | THR | p.Thr494Ala | P35916 |
| HOMO SAPIENS | 0 | 20725 | 4bsj | A | 527 | ASN | p.Asn527Ser | P35916 |
| HOMO SAPIENS | 6 | 29959 | 1xiw | B | 47 | ARG | p.Arg68Ter | P04234 |
| HOMO SAPIENS | 2 | 29959 | 1xiw | B | 72 | CYS | p.Cys93Ter | P04234 |
| HOMO SAPIENS | 3 | 20242 | 5tjc | A | 102 | ARG | p.Arg102Ter | Q9GZU1 |
| HOMO SAPIENS | 3 | 20242 | 5tjc | A | 232 | THR | p.Thr232Pro | Q9GZU1 |
| HOMO SAPIENS | 7 | 9590 | 4ap8 | A | 50 | THR | p.Thr50Ala | O96007 |
| HOMO SAPIENS | 4 | 9590 | 4ap8 | A | 168 | GLU | p.Glu168Lys | O96007 |
| HOMO SAPIENS | -1 | 4297 | 5do7 | B | 19 | ASP | p.Asp19His | Q9H221 |
| HOMO SAPIENS | -1 | 4297 | 5do7 | B | 54 | TYR | p.Tyr54Cys | Q9H221 |
| HOMO SAPIENS | -1 | 4297 | 5do7 | B | 80 | CYS | p.Cys80Tyr | Q9H221 |
| HOMO SAPIENS | 5 | 4297 | 5do7 | B | 107 | SER | p.Ser107Ter | Q9H221 |
| HOMO SAPIENS | 1 | 4297 | 5do7 | B | 164 | ARG | p.Arg164Pro | Q9H221 |
| HOMO SAPIENS | 0 | 4297 | 5do7 | B | 210 | VAL | p.Val210Met | Q9H221 |
| HOMO SAPIENS | 0 | 4297 | 5do7 | B | 231 | PRO | p.Pro231Thr | Q9H221 |
| HOMO SAPIENS | 0 | 4297 | 5do7 | B | 263 | ARG | p.Arg263Gln | Q9H221 |
| HOMO SAPIENS | 0 | 4297 | 5do7 | B | 361 | TRP | p.Trp361Ter | Q9H221 |
| HOMO SAPIENS | 0 | 4297 | 5do7 | B | 400 | THR | p.Thr400Lys | Q9H221 |
| HOMO SAPIENS | 0 | 4297 | 5do7 | B | 549 | ALA | p.Ala549Pro | Q9H221 |
| HOMO SAPIENS | 0 | 4297 | 5do7 | B | 596 | LEU | p.Leu596Arg | Q9H221 |
| HOMO SAPIENS | 0 | 4297 | 5do7 | B | 632 | VAL | p.Val632Ala | Q9H221 |
| HOMO SAPIENS | 0 | 4297 | 5do7 | B | 658 | TYR | p.Tyr658Ter | Q9H221 |
| HOMO SAPIENS | 2 | 9356 | 4igg | B | 206 | GLN | p.Gln206His | P35221 |
| HOMO SAPIENS | 6 | 2179 | 2wk6 | A | 44 | ARG | p.Arg44Gln | P19971 |
| HOMO SAPIENS | 6 | 2179 | 2wk6 | A | 81 | ARG | p.Arg81Gln | P19971 |
| HOMO SAPIENS | 6 | 2179 | 2wk6 | A | 145 | GLY | p.Gly145Arg | P19971 |
| HOMO SAPIENS | 7 | 2179 | 2wk6 | A | 146 | ARG | p.Arg146His | P19971 |
| HOMO SAPIENS | 7 | 2179 | 2wk6 | A | 153 | GLY | p.Gly153Ser | P19971 |
| HOMO SAPIENS | 0 | 2179 | 2wk6 | A | 173 | MET | p.Met173Ile | P19971 |
| HOMO SAPIENS | 6 | 2179 | 2wk6 | A | 202 | ARG | p.Arg202Lys | P19971 |
| HOMO SAPIENS | 6 | 2179 | 2wk6 | A | 208 | VAL | p.Val208Met | P19971 |
| HOMO SAPIENS | 5 | 2179 | 2wk6 | A | 222 | LYS | p.Lys222Arg | P19971 |
| HOMO SAPIENS | 5 | 2179 | 2wk6 | A | 289 | GLU | p.Glu289Ala | P19971 |
| HOMO SAPIENS | 2 | 2179 | 2wk6 | A | 311 | GLY | p.Gly311Arg | P19971 |
| HOMO SAPIENS | 2 | 2179 | 2wk6 | A | 376 | ARG | p.Arg376Pro | P19971 |
| HOMO SAPIENS | 2 | 2179 | 2wk6 | A | 465 | ALA | p.Ala465Thr | P19971 |
| HOMO SAPIENS | -1 | 11856 | 3orh | A | 3 | ALA | p.Ala3Ser | Q14353 |
| HOMO SAPIENS | 1 | 11856 | 3orh | A | 20 | TRP | p.Trp20Ser | Q14353 |
| HOMO SAPIENS | 1 | 11856 | 3orh | A | 50 | MET | p.Met50Leu | Q14353 |
| HOMO SAPIENS | 1 | 11856 | 3orh | A | 110 | VAL | p.Val110Phe | Q14353 |
| HOMO SAPIENS | 8 | 11856 | 3orh | A | 164 | GLY | p.Gly164Asp | Q14353 |
| HOMO SAPIENS | 1 | 11856 | 3orh | A | 169 | CYS | p.Cys169Tyr | Q14353 |
| HOMO SAPIENS | 1 | 11856 | 3orh | A | 174 | TRP | p.Trp174Ter | Q14353 |
| HOMO SAPIENS | 2 | 11856 | 3orh | A | 196 | ALA | p.Ala196Val | Q14353 |
| HOMO SAPIENS | 1 | 11856 | 3orh | A | 209 | THR | p.Thr209Met | Q14353 |
| HOMO SAPIENS | 1 | 11856 | 3orh | A | 219 | ASP | p.Asp219Asn | Q14353 |
| HOMO SAPIENS | 1 | 11856 | 3orh | A | 224 | ALA | p.Ala224Thr | Q14353 |
| HOMO SAPIENS | 1 | 13244 | 2m6i | D | 218 | ARG | p.Arg246Leu | P23415 |
| HOMO SAPIENS | 1 | 13244 | 2m6i | D | 244 | ILE | p.Ile272Asn | P23415 |

| | | | | | | | | |
|---|---|---|---|---|---|---|---|---|
| HOMO SAPIENS | 5 | 13244 | 2m6i | D | 250 | PRO | p.Pro278Thr | P23415 |
| HOMO SAPIENS | 6 | 13244 | 2m6i | D | 266 | GLN | p.Gln294His | P23415 |
| HOMO SAPIENS | 1 | 13244 | 2m6i | D | 271 | ARG | p.Arg299Leu | P23415 |
| HOMO SAPIENS | 1 | 13244 | 2m6i | D | 276 | LYS | p.Lys304Glu | P23415 |
| HOMO SAPIENS | 1 | 13244 | 2m6i | D | 279 | TYR | p.Tyr307Cys | P23415 |
| HOMO SAPIENS | 0 | 13244 | 2m6i | D | 316 | GLY | p.Gly378Ser | P23415 |
| HOMO SAPIENS | 1 | 13244 | 2m6i | D | 400 | ARG | p.Arg428His | P23415 |
| HOMO SAPIENS | 3 | 1967 | 2i6b | A | 301 | ALA | p.Ala318Glu | P55263 |
| HOMO SAPIENS | 4 | 11096 | 1h8b | A | 61 | ALA | p.Ala882Thr | P35609 |
| HOMO SAPIENS | 0 | 15423 | 5ngz | A | 2 | GLN | p.Gln2Glu | Q9NPD8 |
| HOMO SAPIENS | 3 | 30475 | 3lpw | A | 21 | SER | p.Ser22894Tyr | Q8WZ42 |
| HOMO SAPIENS | 6 | 32687 | 2arf | A | 1041 | ARG | p.Arg1041Trp | P35670 |
| HOMO SAPIENS | 8 | 32687 | 2arf | A | 1061 | GLY | p.Gly1061Glu | P35670 |
| HOMO SAPIENS | 6 | 32687 | 2arf | A | 1064 | GLU | p.Glu1064Ala | P35670 |
| HOMO SAPIENS | 2 | 32687 | 2arf | A | 1069 | HIS | p.His1069Gln | P35670 |
| HOMO SAPIENS | 2 | 32687 | 2arf | A | 1088 | LEU | p.Leu1088Ter | P35670 |
| HOMO SAPIENS | 0 | 32687 | 2arf | A | 1099 | GLY | p.Gly1099Ser | P35670 |
| HOMO SAPIENS | 4 | 32687 | 2arf | A | 1106 | VAL | p.Val1106Asp | P35670 |
| HOMO SAPIENS | 2 | 32687 | 2arf | A | 1142 | GLN | p.Gln1142His | P35670 |
| HOMO SAPIENS | 8 | 32687 | 2arf | A | 1148 | ILE | p.Ile1148Thr | P35670 |
| HOMO SAPIENS | 6 | 32687 | 2arf | A | 1151 | ARG | p.Arg1151Cys | P35670 |
| HOMO SAPIENS | 4 | 32687 | 2arf | A | 1169 | MET | p.Met1169Val | P35670 |
| HOMO SAPIENS | 4 | 32687 | 2arf | A | 1173 | GLU | p.Glu1173Lys | P35670 |
| HOMO SAPIENS | 2 | 32687 | 2arf | A | 1176 | GLY | p.Gly1176Arg | P35670 |
| HOMO SAPIENS | 3 | 15544 | 5uq3 | C | 575 | PRO | p.Pro69Leu | P46527 |
| HOMO SAPIENS | 6 | 15544 | 5uq3 | C | 592 | GLU | p.Glu86Asp | P46527 |
| HOMO SAPIENS | 6 | 15544 | 5uq3 | C | 599 | ARG | p.Arg93Trp | P46527 |
| HOMO SAPIENS | -1 | 15544 | 5uq3 | C | 615 | VAL | p.Val109Gly | P46527 |
| HOMO SAPIENS | -1 | 15544 | 5uq3 | C | 631 | SER | p.Ser125Pro | P46527 |
| HOMO SAPIENS | -1 | 15544 | 5uq3 | C | 639 | PRO | p.Pro133Ser | P46527 |
| HOMO SAPIENS | 2 | 2703 | 4wxq | A | 245 | CYS | p.Cys245Tyr | Q99972 |
| HOMO SAPIENS | 0 | 2703 | 4wxq | A | 252 | GLY | p.Gly252Arg | Q99972 |
| HOMO SAPIENS | 1 | 2703 | 4wxq | A | 337 | GLN | p.Gln337Arg | Q99972 |
| HOMO SAPIENS | 5 | 2703 | 4wxq | A | 347 | TYR | p.Tyr347Ter | Q99972 |
| HOMO SAPIENS | 3 | 2703 | 4wxq | A | 364 | GLY | p.Gly364Val | Q99972 |
| HOMO SAPIENS | 2 | 2703 | 4wxq | A | 367 | GLY | p.Gly367Arg | Q99972 |
| HOMO SAPIENS | 2 | 2703 | 4wxq | A | 368 | GLN | p.Gln368Ter | Q99972 |
| HOMO SAPIENS | 6 | 2703 | 4wxq | A | 370 | PRO | p.Pro370Leu | Q99972 |
| HOMO SAPIENS | 1 | 2703 | 4wxq | A | 380 | ASP | p.Asp380His | Q99972 |
| HOMO SAPIENS | 8 | 2703 | 4wxq | A | 399 | GLY | p.Gly399Val | Q99972 |
| HOMO SAPIENS | 0 | 2703 | 4wxq | A | 423 | LYS | p.Lys423Glu | Q99972 |
| HOMO SAPIENS | 3 | 2703 | 4wxq | A | 433 | CYS | p.Cys433Arg | Q99972 |
| HOMO SAPIENS | 3 | 2703 | 4wxq | A | 437 | TYR | p.Tyr437His | Q99972 |
| HOMO SAPIENS | 2 | 2703 | 4wxq | A | 477 | ILE | p.Ile477Ser | Q99972 |
| HOMO SAPIENS | 3 | 2703 | 4wxq | A | 480 | ASN | p.Asn480Lys | Q99972 |
| HOMO SAPIENS | -1 | 34100 | 2fot | C | 1171 | GLN | p.Gln1173Pro | Q13813 |
| HOMO SAPIENS | 0 | 8773 | 1kw2 | A | 432 | ASP | p.Asp432Glu | P02774 |
| HOMO SAPIENS | 0 | 8773 | 1kw2 | A | 436 | LYS | p.Lys436Thr | P02774 |
| HOMO SAPIENS | 0 | 8402 | 4dzo | B | 649 | TYR | p.Tyr649Ter | Q9Y6D9 |
| HOMO SAPIENS | 0 | 39962 | 1nst | A | 611 | GLY | p.Gly611Ser | P52848 |
| HOMO SAPIENS | 2 | 22578 | 3dof | B | 45 | MET | p.Met45Arg | Q9Y2Y0 |
| HOMO SAPIENS | 1 | 23290 | 5j1s | A | 292 | ILE | p.Ile292Thr | O14656 |
| HOMO SAPIENS | 4 | 9505 | 4j1y | A | 534 | ARG | p.Arg534Trp | P09871 |
| HOMO SAPIENS | 9 | 260 | 2h57 | A | 31 | THR | p.Thr31Met | Q9H0F7 |

| | | | | | | | | |
|---|---|---|---|---|---|---|---|---|
| HOMO SAPIENS | 0 | 260 | 2h57 | A | 91 | ILE | p.Ile91Thr | Q9H0F7 |
| HOMO SAPIENS | 0 | 260 | 2h57 | A | 121 | ARG | p.Arg121His | Q9H0F7 |
| HOMO SAPIENS | 0 | 260 | 2h57 | A | 122 | ARG | p.Arg122Ter | Q9H0F7 |
| HOMO SAPIENS | 2 | 260 | 2h57 | A | 169 | GLY | p.Gly169Ala | Q9H0F7 |
| HOMO SAPIENS | 4 | 260 | 2h57 | A | 170 | LEU | p.Leu170Trp | Q9H0F7 |
| HOMO SAPIENS | 4 | 34023 | 2cr3 | A | 18 | GLY | p.Gly48Ser | P11362 |
| HOMO SAPIENS | 0 | 37667 | 3ii7 | A | 420 | ARG | p.Arg420Cys | Q8IXQ5 |
| HOMO SAPIENS | 0 | 41720 | 2k9u | A | 68 | ARG | p.Arg2364His | Q14315 |
| HOMO SAPIENS | 1 | 29250 | 5fmv | A | 568 | HIS | p.His568Gln | P08575 |
| HOMO SAPIENS | 1 | 1280 | 1fb1 | A | 69 | PRO | p.Pro69Leu | P30793 |
| HOMO SAPIENS | 3 | 1280 | 1fb1 | A | 88 | ARG | p.Arg88Trp | P30793 |
| HOMO SAPIENS | 2 | 1280 | 1fb1 | A | 108 | GLY | p.Gly108Asp | P30793 |
| HOMO SAPIENS | 4 | 1280 | 1fb1 | A | 134 | ASP | p.Asp134Val | P30793 |
| HOMO SAPIENS | 6 | 1280 | 1fb1 | A | 135 | ILE | p.Ile135Lys | P30793 |
| HOMO SAPIENS | 9 | 1280 | 1fb1 | A | 144 | HIS | p.His144Pro | P30793 |
| HOMO SAPIENS | 9 | 1280 | 1fb1 | A | 184 | ARG | p.Arg184His | P30793 |
| HOMO SAPIENS | 1 | 1280 | 1fb1 | A | 196 | ALA | p.Ala196Ser | P30793 |
| HOMO SAPIENS | 3 | 1280 | 1fb1 | A | 201 | GLY | p.Gly201Glu | P30793 |
| HOMO SAPIENS | 2 | 1280 | 1fb1 | A | 204 | VAL | p.Val204Ile | P30793 |
| HOMO SAPIENS | 2 | 1280 | 1fb1 | A | 211 | MET | p.Met211Ile | P30793 |
| HOMO SAPIENS | 0 | 1280 | 1fb1 | A | 221 | MET | p.Met221Thr | P30793 |
| HOMO SAPIENS | 3 | 1280 | 1fb1 | A | 224 | LYS | p.Lys224Arg | P30793 |
| HOMO SAPIENS | 2 | 1280 | 1fb1 | A | 249 | ARG | p.Arg249Ser | P30793 |
| HOMO SAPIENS | 8 | 41352 | 4k12 | A | 47 | ILE | p.Ile551Thr | P08603 |
| HOMO SAPIENS | 9 | 20703 | 5nir | A | 57 | CYS | p.Cys57Tyr | P02458 |
| HOMO SAPIENS | 9 | 20703 | 5nir | A | 64 | CYS | p.Cys64Ter | P02458 |
| HOMO SAPIENS | 0 | 20087 | 4cmn | A | 299 | MET | p.Met299Ile | Q01968 |
| HOMO SAPIENS | 1 | 20819 | 3aaf | A | 979 | ILE | p.Ile979Met | Q14191 |
| HOMO SAPIENS | 1 | 20819 | 3aaf | A | 984 | LEU | p.Leu984Ile | Q14191 |
| HOMO SAPIENS | 3 | 20819 | 3aaf | A | 987 | ARG | p.Arg987Ter | Q14191 |
| HOMO SAPIENS | -1 | 20819 | 3aaf | A | 1074 | PHE | p.Leu1074Phe | Q14191 |
| HOMO SAPIENS | 0 | 3619 | 3dem | A | 206 | ASN | p.Asn225Ser | P48740 |
| HOMO SAPIENS | 1 | 3619 | 3dem | A | 269 | ARG | p.Arg288Leu | P48740 |
| HOMO SAPIENS | 0 | 3105 | 3ooi | A | 164 | TYR | p.Tyr2015His | Q96L73 |
| HOMO SAPIENS | 6 | 15828 | 2lkx | A | 16 | LEU | p.Leu100Gln | Q99697 |
| HOMO SAPIENS | 8 | 15828 | 2lkx | A | 24 | ARG | p.Arg108His | Q99697 |
| HOMO SAPIENS | 2 | 15828 | 2lkx | A | 30 | THR | p.Thr114Pro | Q99697 |
| HOMO SAPIENS | 8 | 15828 | 2lkx | A | 45 | VAL | p.Val129Leu | Q99697 |
| HOMO SAPIENS | 8 | 15828 | 2lkx | A | 46 | ARG | p.Arg130Trp | Q99697 |
| HOMO SAPIENS | 9 | 15828 | 2lkx | A | 53 | ARG | p.Arg137Pro | Q99697 |
| HOMO SAPIENS | 9 | 13017 | 4irq | A | 141 | ARG | p.Arg141Trp | Q9UBV7 |
| HOMO SAPIENS | 0 | 13017 | 4irq | A | 186 | ALA | p.Ala186Asp | Q9UBV7 |
| HOMO SAPIENS | 0 | 13017 | 4irq | A | 206 | LEU | p.Leu206Pro | Q9UBV7 |
| HOMO SAPIENS | 1 | 13017 | 4irq | A | 214 | CYS | p.Cys214Tyr | Q9UBV7 |
| HOMO SAPIENS | 0 | 13017 | 4irq | A | 270 | ARG | p.Arg270Cys | Q9UBV7 |
| HOMO SAPIENS | -1 | 13053 | 4mqw | X | 307 | ALA | p.Ala307Thr | P23945 |
| HOMO SAPIENS | 0 | 19406 | 4b3g | A | 43 | ARG | p.Arg43Ter | P38935 |
| HOMO SAPIENS | 0 | 19406 | 4b3g | A | 46 | CYS | p.Cys46Ter | P38935 |
| HOMO SAPIENS | 0 | 19406 | 4b3g | A | 75 | ALA | p.Ala75Thr | P38935 |
| HOMO SAPIENS | 0 | 19406 | 4b3g | A | 201 | LEU | p.Leu201Ser | P38935 |
| HOMO SAPIENS | 3 | 19406 | 4b3g | A | 213 | HIS | p.His213Arg | P38935 |
| HOMO SAPIENS | 0 | 19406 | 4b3g | A | 275 | ILE | p.Ile275Val | P38935 |
| HOMO SAPIENS | -1 | 19406 | 4b3g | A | 320 | ARG | p.Arg320Ter | P38935 |
| HOMO SAPIENS | 0 | 19406 | 4b3g | A | 361 | LEU | p.Leu361Pro | P38935 |

| | | | | | | | | |
|---|---|---|---|---|---|---|---|---|
| HOMO SAPIENS | 0 | 19406 | 4b3g | A | 369 | PHE | p.Phe369Leu | P38935 |
| HOMO SAPIENS | 0 | 19406 | 4b3g | A | 493 | THR | p.Thr493Ile | P38935 |
| HOMO SAPIENS | 0 | 19406 | 4b3g | A | 496 | CYS | p.Cys496Ter | P38935 |
| HOMO SAPIENS | 0 | 19406 | 4b3g | A | 514 | GLU | p.Glu514Lys | P38935 |
| HOMO SAPIENS | 0 | 19406 | 4b3g | A | 528 | ALA | p.Ala528Thr | P38935 |
| HOMO SAPIENS | 0 | 19406 | 4b3g | A | 531 | PRO | p.Pro531Thr | P38935 |
| HOMO SAPIENS | 0 | 19406 | 4b3g | A | 580 | VAL | p.Val580Ile | P38935 |
| HOMO SAPIENS | 6 | 19406 | 4b3g | A | 603 | ARG | p.Arg603His | P38935 |
| HOMO SAPIENS | 0 | 21501 | 5jde | A | 40 | SER | p.Ser2831Asn | Q8WZ42 |
| HOMO SAPIENS | 5 | 21501 | 5jde | A | 189 | ALA | p.Ala2980Thr | Q8WZ42 |
| HOMO SAPIENS | 6 | 15008 | 3mop | H | 12 | ARG | p.Arg12Cys | Q9NWZ3 |
| HOMO SAPIENS | 8 | 17455 | 3jtc | C | 15 | ARG | p.Arg57Trp | P04070 |
| HOMO SAPIENS | -1 | 17455 | 3jtc | C | 20 | CGU | p.Glu62Ala | P04070 |
| HOMO SAPIENS | 0 | 27795 | 5gox | B | 625 | TYR | p.Tyr625Ter | Q92878 |
| HOMO SAPIENS | 0 | 27795 | 5gox | B | 653 | SER | p.Ser653Ter | Q92878 |
| HOMO SAPIENS | -1 | 6558 | 5nug | A | 137 | VAL | p.Val137Ile | Q14204 |
| HOMO SAPIENS | -1 | 6558 | 5nug | A | 567 | ARG | p.Arg567His | Q14204 |
| HOMO SAPIENS | -1 | 6558 | 5nug | A | 1140 | THR | p.Thr1140Met | Q14204 |
| HOMO SAPIENS | -1 | 6558 | 5nug | A | 1991 | ASP | p.Asp1991Asn | Q14204 |
| HOMO SAPIENS | -1 | 6558 | 5nug | A | 2401 | LYS | p.Lys2401Asn | Q14204 |
| HOMO SAPIENS | 0 | 6558 | 5nug | A | 2488 | ARG | p.Arg2488His | Q14204 |
| HOMO SAPIENS | 0 | 6558 | 5nug | A | 3907 | HIS | p.His3907Gln | Q14204 |
| HOMO SAPIENS | 0 | 6558 | 5nug | A | 3981 | THR | p.Thr3981Arg | Q14204 |
| HOMO SAPIENS | 0 | 6558 | 5nug | A | 4394 | THR | p.Thr4394Met | Q14204 |
| HOMO SAPIENS | 0 | 6558 | 5nug | A | 4573 | ASN | p.Asn4573Ser | Q14204 |
| HOMO SAPIENS | 0 | 6558 | 5nug | A | 4603 | SER | p.Ser4603Ile | Q14204 |
| HOMO SAPIENS | 1 | 4244 | 1umk | A | 57 | ARG | p.Arg58Gln | P00387 |
| HOMO SAPIENS | 1 | 4244 | 1umk | A | 72 | LEU | p.Leu73Pro | P00387 |
| HOMO SAPIENS | 1 | 4244 | 1umk | A | 105 | VAL | p.Val106Met | P00387 |
| HOMO SAPIENS | 0 | 4244 | 1umk | A | 116 | THR | p.Thr117Ser | P00387 |
| HOMO SAPIENS | 8 | 4244 | 1umk | A | 127 | SER | p.Ser128Pro | P00387 |
| HOMO SAPIENS | 0 | 4244 | 1umk | A | 148 | LEU | p.Leu149Pro | P00387 |
| HOMO SAPIENS | 0 | 4244 | 1umk | A | 159 | ARG | p.Arg160Ter | P00387 |
| HOMO SAPIENS | 0 | 4244 | 1umk | A | 203 | CYS | p.Cys204Arg | P00387 |
| HOMO SAPIENS | 0 | 4244 | 1umk | A | 203 | CYS | p.Cys204Tyr | P00387 |
| HOMO SAPIENS | 2 | 4244 | 1umk | A | 239 | ASP | p.Asp240Gly | P00387 |
| HOMO SAPIENS | 0 | 4244 | 1umk | A | 291 | GLY | p.Gly292Asp | P00387 |
| HOMO SAPIENS | 4 | 34495 | 1s35 | A | 1151 | ASP | p.Asn1151Asp | P11277 |
| HOMO SAPIENS | 3 | 30110 | 3zqs | A | 224 | THR | p.Thr224Ala | Q9NW38 |
| HOMO SAPIENS | 3 | 17588 | 5dse | B | 53 | LEU | p.Leu53Pro | Q9BYI3 |
| HOMO SAPIENS | 0 | 1564 | 4yqf | B | 72 | HIS | p.His347Gln | Q9UHD8 |
| HOMO SAPIENS | 0 | 17673 | 4jnw | A | 32 | ARG | p.Arg32186His | Q8WZ42 |
| HOMO SAPIENS | 3 | 17673 | 4jnw | A | 140 | ILE | p.Ile32294Val | Q8WZ42 |
| HOMO SAPIENS | 5 | 17673 | 4jnw | A | 269 | ALA | p.Ala32423Thr | Q8WZ42 |
| HOMO SAPIENS | 3 | 12858 | 3s97 | C | 302 | LEU | p.Leu302Pro | Q12860 |
| HOMO SAPIENS | 0 | 6147 | 4nn0 | A | 114 | ARG | p.Arg114Gln | Q9BXJ0 |
| HOMO SAPIENS | 0 | 7549 | 1tr2 | A | 277 | LEU | p.Leu277Met | P18206 |
| HOMO SAPIENS | 1 | 7549 | 1tr2 | A | 413 | ALA | p.Ala413Thr | P18206 |
| HOMO SAPIENS | 1 | 7549 | 1tr2 | A | 519 | ILE | p.Ile519Leu | P18206 |
| HOMO SAPIENS | 5 | 7549 | 1tr2 | A | 547 | ARG | p.Arg547Ter | P18206 |
| HOMO SAPIENS | 2 | 7549 | 1tr2 | A | 636 | HIS | p.His636Arg | P18206 |
| HOMO SAPIENS | 2 | 7549 | 1tr2 | A | 824 | ILE | p.Ile824Met | P18206 |
| HOMO SAPIENS | 2 | 13900 | 3gro | A | 45 | CYS | p.Cys45Tyr | P50897 |
| HOMO SAPIENS | 2 | 13900 | 3gro | A | 75 | THR | p.Thr75Pro | P50897 |

| | | | | | | | | |
|---|---|---|---|---|---|---|---|---|
| HOMO SAPIENS | 2 | 13900 | 3gro | A | 79 | ASP | p.Asp79Gly | P50897 |
| HOMO SAPIENS | 5 | 13900 | 3gro | A | 91 | GLN | p.Gln91Pro | P50897 |
| HOMO SAPIENS | 5 | 13900 | 3gro | A | 96 | CYS | p.Cys96Tyr | P50897 |
| HOMO SAPIENS | 0 | 13900 | 3gro | A | 104 | LYS | p.Lys104Ter | P50897 |
| HOMO SAPIENS | 2 | 13900 | 3gro | A | 108 | GLY | p.Gly108Arg | P50897 |
| HOMO SAPIENS | 2 | 13900 | 3gro | A | 109 | TYR | p.Tyr109Asp | P50897 |
| HOMO SAPIENS | 0 | 13900 | 3gro | A | 122 | ARG | p.Arg122Trp | P50897 |
| HOMO SAPIENS | 0 | 13900 | 3gro | A | 138 | SER | p.Ser138Leu | P50897 |
| HOMO SAPIENS | 5 | 13900 | 3gro | A | 151 | ARG | p.Arg151Ter | P50897 |
| HOMO SAPIENS | 7 | 13900 | 3gro | A | 164 | ARG | p.Arg164Ter | P50897 |
| HOMO SAPIENS | 0 | 13900 | 3gro | A | 177 | GLN | p.Gln177Glu | P50897 |
| HOMO SAPIENS | 0 | 13900 | 3gro | A | 181 | VAL | p.Val181Leu | P50897 |
| HOMO SAPIENS | 0 | 13900 | 3gro | A | 184 | GLU | p.Glu184Lys | P50897 |
| HOMO SAPIENS | 5 | 13900 | 3gro | A | 186 | TRP | p.Trp186Ter | P50897 |
| HOMO SAPIENS | 0 | 13900 | 3gro | A | 219 | LEU | p.Leu219Gln | P50897 |
| HOMO SAPIENS | 0 | 13900 | 3gro | A | 222 | LEU | p.Leu222Pro | P50897 |
| HOMO SAPIENS | 5 | 13900 | 3gro | A | 296 | TRP | p.Trp296Ter | P50897 |
| HOMO SAPIENS | 6 | 35028 | 5mwb | A | 444 | CYS | p.Cys444Tyr | Q04721 |
| HOMO SAPIENS | 0 | 35028 | 5mwb | A | 514 | ASN | p.Asn514Ser | Q04721 |
| HOMO SAPIENS | 5 | 1483 | 4gl2 | A | 822 | ARG | p.Arg822Gln | Q9BYX4 |
| HOMO SAPIENS | 0 | 1483 | 4gl2 | A | 843 | ARG | p.His843Arg | Q9BYX4 |
| HOMO SAPIENS | -1 | 28838 | 3cl3 | D | 153 | LEU | p.Leu153Arg | Q9Y6K9 |
| HOMO SAPIENS | 2 | 22563 | 4zel | B | 79 | ARG | p.Arg79Trp | P09172 |
| HOMO SAPIENS | 2 | 22563 | 4zel | B | 101 | VAL | p.Val101Met | P09172 |
| HOMO SAPIENS | 2 | 22563 | 4zel | B | 114 | ASP | p.Asp114Glu | P09172 |
| HOMO SAPIENS | 0 | 22563 | 4zel | B | 201 | ASN | p.Asn201Ser | P09172 |
| HOMO SAPIENS | 2 | 22563 | 4zel | B | 211 | ALA | p.Ala211Thr | P09172 |
| HOMO SAPIENS | -1 | 22563 | 4zel | B | 290 | ASP | p.Asp290Asn | P09172 |
| HOMO SAPIENS | 2 | 22563 | 4zel | B | 318 | ALA | p.Ala318Ser | P09172 |
| HOMO SAPIENS | 2 | 22563 | 4zel | B | 345 | ASP | p.Asp345Asn | P09172 |
| HOMO SAPIENS | 2 | 22563 | 4zel | B | 362 | ALA | p.Ala362Glu | P09172 |
| HOMO SAPIENS | 2 | 22563 | 4zel | B | 549 | ARG | p.Arg549Cys | P09172 |
| HOMO SAPIENS | 9 | 9780 | 1y6k | R | 36 | TYR | p.Tyr57Cys | Q13651 |
| HOMO SAPIENS | 1 | 9780 | 1y6k | R | 40 | LEU | p.Leu61Val | Q13651 |
| HOMO SAPIENS | 1 | 9780 | 1y6k | R | 63 | THR | p.Thr84Ile | Q13651 |
| HOMO SAPIENS | 9 | 9780 | 1y6k | R | 80 | ARG | p.Arg101Trp | Q13651 |
| HOMO SAPIENS | 0 | 9780 | 1y6k | R | 120 | GLY | p.Gly141Arg | Q13651 |
| HOMO SAPIENS | 0 | 9780 | 1y6k | R | 138 | SER | p.Ser159Gly | Q13651 |
| HOMO SAPIENS | 0 | 9780 | 1y6k | R | 203 | ILE | p.Ile224Val | Q13651 |
| HOMO SAPIENS | 0 | 16019 | 3rep | A | 211 | ARG | p.Arg211Cys | Q13418 |
| HOMO SAPIENS | 3 | 7879 | 5ec3 | A | 33 | ALA | p.Ala33Thr | P32754 |
| HOMO SAPIENS | 4 | 7879 | 5ec3 | A | 113 | ARG | p.Arg113Gln | P32754 |
| HOMO SAPIENS | 3 | 7879 | 5ec3 | A | 340 | VAL | p.Val340Leu | P32754 |
| HOMO SAPIENS | 0 | 8904 | 1qk9 | A | 6 | LYS | p.Lys82Arg | P51608 |
| HOMO SAPIENS | 4 | 8904 | 1qk9 | A | 24 | LEU | p.Leu100Val | P51608 |
| HOMO SAPIENS | 7 | 8904 | 1qk9 | A | 30 | ARG | p.Arg106Trp | P51608 |
| HOMO SAPIENS | 6 | 8904 | 1qk9 | A | 57 | ARG | p.Arg133Cys | P51608 |
| HOMO SAPIENS | 3 | 8904 | 1qk9 | A | 61 | GLU | p.Glu137Gly | P51608 |
| HOMO SAPIENS | 2 | 8904 | 1qk9 | A | 64 | ALA | p.Ala140Val | P51608 |
| HOMO SAPIENS | 5 | 8904 | 1qk9 | A | 79 | PHE | p.Phe155Ser | P51608 |
| HOMO SAPIENS | 3 | 8904 | 1qk9 | A | 82 | THR | p.Thr158Met | P51608 |
| HOMO SAPIENS | 1 | 23121 | 3e9k | A | 198 | THR | p.Thr198Ala | Q16719 |
| HOMO SAPIENS | -1 | 26090 | 3tje | F | 19 | LEU | p.Leu35Met | P25445 |
| HOMO SAPIENS | 8 | 26090 | 3tje | F | 106 | THR | p.Thr122Ile | P25445 |

| | | | | | | | | |
|---|---|---|---|---|---|---|---|---|
| HOMO SAPIENS | 2 | 36404 | 5cqr | A | 765 | GLY | p.Gly765Glu | O95163 |
| HOMO SAPIENS | 2 | 36404 | 5cqr | A | 816 | ILE | p.Ile816Leu | O95163 |
| HOMO SAPIENS | 2 | 36404 | 5cqr | A | 830 | ILE | p.Ile830Met | O95163 |
| HOMO SAPIENS | 5 | 36404 | 5cqr | A | 939 | TYR | p.Tyr939Ter | O95163 |
| HOMO SAPIENS | 2 | 36404 | 5cqr | A | 952 | LYS | p.Lys952Ile | O95163 |
| HOMO SAPIENS | 5 | 36404 | 5cqr | A | 1013 | GLY | p.Gly1013Ser | O95163 |
| HOMO SAPIENS | 5 | 36404 | 5cqr | A | 1072 | CYS | p.Cys1072Ser | O95163 |
| HOMO SAPIENS | -1 | 36404 | 5cqr | A | 1158 | PRO | p.Pro1158Leu | O95163 |
| HOMO SAPIENS | 1 | 34149 | 2k40 | A | 18 | ASN | p.Asn125Ser | Q9UBX0 |
| HOMO SAPIENS | 0 | 34149 | 2k40 | A | 42 | LEU | p.Glu149Lys | Q9UBX0 |
| HOMO SAPIENS | 9 | 34149 | 2k40 | A | 53 | ARG | p.Arg160Cys | Q9UBX0 |
| HOMO SAPIENS | 0 | 34149 | 2k40 | A | 63 | SER | p.Ser170Leu | Q9UBX0 |
| HOMO SAPIENS | 2 | 2468 | 3vp6 | A | 484 | LEU | p.Leu484Met | Q99259 |
| HOMO SAPIENS | 0 | 2468 | 3vp6 | A | 532 | ARG | p.Arg532Gln | Q99259 |
| HOMO SAPIENS | 5 | 1253 | 5xtd | S | 8 | GLY | p.Gly8Arg | O15239 |
| HOMO SAPIENS | 7 | 1253 | 5xtd | S | 32 | GLY | p.Gly32Arg | O15239 |
| HOMO SAPIENS | 7 | 1253 | 5xtd | S | 37 | ARG | p.Arg37Ser | O15239 |
| HOMO SAPIENS | -1 | 13292 | 3u1k | A | 121 | VAL | p.Ile121Val | Q8TCS8 |
| HOMO SAPIENS | 2 | 13292 | 3u1k | A | 165 | PRO | p.Pro165Ser | Q8TCS8 |
| HOMO SAPIENS | 2 | 13292 | 3u1k | A | 192 | ARG | p.Arg192Gln | Q8TCS8 |
| HOMO SAPIENS | 1 | 13292 | 3u1k | A | 315 | LYS | p.Lys315Arg | Q8TCS8 |
| HOMO SAPIENS | 4 | 13292 | 3u1k | A | 509 | VAL | p.Val509Ile | Q8TCS8 |
| HOMO SAPIENS | 7 | 13292 | 3u1k | A | 531 | THR | p.Thr531Arg | Q8TCS8 |
| HOMO SAPIENS | 0 | 13292 | 3u1k | A | 590 | ASN | p.Asn590Asp | Q8TCS8 |
| HOMO SAPIENS | 0 | 27048 | 4p5w | B | 184 | ARG | p.Arg184Ter | Q9UJS0 |
| HOMO SAPIENS | 0 | 27048 | 4p5w | B | 225 | SER | p.Ser225Ter | Q9UJS0 |
| HOMO SAPIENS | -1 | 27048 | 4p5w | B | 631 | ALA | p.Arg360Ter | Q9UJS0 |
| HOMO SAPIENS | 0 | 24240 | 4mdr | A | 318 | ARG | p.Arg318Ter | O00189 |
| HOMO SAPIENS | 0 | 24240 | 4mdr | A | 338 | ARG | p.Arg338Ter | O00189 |
| HOMO SAPIENS | 0 | 13429 | 4cg4 | A | 479 | PHE | p.Phe479Leu | O15553 |
| HOMO SAPIENS | 0 | 13429 | 4cg4 | A | 487 | VAL | p.Val487Leu | O15553 |
| HOMO SAPIENS | 3 | 13429 | 4cg4 | A | 653 | ARG | p.Arg653His | O15553 |
| HOMO SAPIENS | 1 | 13429 | 4cg4 | A | 680 | MET | p.Met680Ile | O15553 |
| HOMO SAPIENS | 1 | 13429 | 4cg4 | A | 688 | TYR | p.Tyr688Ter | O15553 |
| HOMO SAPIENS | 0 | 13429 | 4cg4 | A | 694 | MET | p.Met694Ile | O15553 |
| HOMO SAPIENS | 0 | 13429 | 4cg4 | A | 694 | MET | p.Met694Val | O15553 |
| HOMO SAPIENS | 1 | 13429 | 4cg4 | A | 726 | VAL | p.Val726Ala | O15553 |
| HOMO SAPIENS | 0 | 13429 | 4cg4 | A | 761 | ARG | p.Arg761His | O15553 |
| HOMO SAPIENS | 0 | 2079 | 4okh | A | 691 | GLY | p.Gly691Arg | P20807 |
| HOMO SAPIENS | 2 | 2079 | 4okh | A | 748 | ARG | p.Arg748Ter | P20807 |
| HOMO SAPIENS | 0 | 2079 | 4okh | A | 763 | TYR | p.Tyr763Cys | P20807 |
| HOMO SAPIENS | 1 | 2079 | 4okh | A | 769 | ARG | p.Arg769Gln | P20807 |
| HOMO SAPIENS | 9 | 2079 | 4okh | A | 777 | ILE | p.Ile777Val | P20807 |
| HOMO SAPIENS | 8 | 2079 | 4okh | A | 778 | ASP | p.Asp778His | P20807 |
| HOMO SAPIENS | 5 | 2079 | 4okh | A | 780 | ASP | p.Asp780His | P20807 |
| HOMO SAPIENS | 1 | 1200 | 2gi7 | A | 142 | ARG | p.Arg162Trp | Q9HCN6 |
| HOMO SAPIENS | 4 | 9199 | 3ezq | C | 260 | ASP | p.Asp260Val | P25445 |
| HOMO SAPIENS | 0 | 36522 | 5ms9 | A | 122 | ARG | p.Arg122Cys | P35555 |
| HOMO SAPIENS | 0 | 36522 | 5ms9 | A | 240 | ARG | p.Arg240Cys | P35555 |
| HOMO SAPIENS | 0 | 17448 | 3g2f | A | 225 | ARG | p.Arg225His | Q13873 |
| HOMO SAPIENS | 0 | 17448 | 3g2f | A | 347 | CYS | p.Cys347Tyr | Q13873 |
| HOMO SAPIENS | 3 | 17448 | 3g2f | A | 485 | ASP | p.Asp485Gly | Q13873 |
| HOMO SAPIENS | 7 | 17448 | 3g2f | A | 491 | ARG | p.Arg491Gln | Q13873 |
| HOMO SAPIENS | 7 | 17448 | 3g2f | A | 491 | ARG | p.Arg491Trp | Q13873 |

| | | | | | | | | |
|---|---|---|---|---|---|---|---|---|
| HOMO SAPIENS | 0 | 4861 | 4ux8 | A | 56 | LEU | p.Leu56Met | P07949 |
| HOMO SAPIENS | 1 | 4861 | 4ux8 | A | 88 | ILE | p.Ile88Leu | P07949 |
| HOMO SAPIENS | 0 | 4861 | 4ux8 | A | 114 | ARG | p.Arg114His | P07949 |
| HOMO SAPIENS | 5 | 4861 | 4ux8 | A | 180 | ARG | p.Arg180Gln | P07949 |
| HOMO SAPIENS | 0 | 4861 | 4ux8 | A | 208 | GLU | p.Glu208Asp | P07949 |
| HOMO SAPIENS | 0 | 4861 | 4ux8 | A | 373 | ALA | p.Ala373Val | P07949 |
| HOMO SAPIENS | 0 | 4861 | 4ux8 | A | 375 | LEU | p.Leu375Gln | P07949 |
| HOMO SAPIENS | -1 | 4861 | 4ux8 | A | 386 | ALA | p.Ala386Val | P07949 |
| HOMO SAPIENS | 0 | 4861 | 4ux8 | A | 446 | GLY | p.Gly446Arg | P07949 |
| HOMO SAPIENS | 0 | 4861 | 4ux8 | A | 480 | GLU | p.Glu480Lys | P07949 |
| HOMO SAPIENS | 0 | 4861 | 4ux8 | A | 489 | ASP | p.Asp489Asn | P07949 |
| HOMO SAPIENS | -1 | 4861 | 4ux8 | A | 510 | ALA | p.Ala510Val | P07949 |
| HOMO SAPIENS | -1 | 4861 | 4ux8 | A | 533 | GLY | p.Gly533Cys | P07949 |
| HOMO SAPIENS | -1 | 4861 | 4ux8 | A | 567 | ASP | p.Asp567Asn | P07949 |
| HOMO SAPIENS | -1 | 4861 | 4ux8 | A | 609 | CYS | p.Cys609Tyr | P07949 |
| HOMO SAPIENS | -1 | 4861 | 4ux8 | A | 609 | CYS | p.Cys609Gly | P07949 |
| HOMO SAPIENS | -1 | 4861 | 4ux8 | A | 611 | CYS | p.Cys611Trp | P07949 |
| HOMO SAPIENS | -1 | 4861 | 4ux8 | A | 618 | CYS | p.Cys618Gly | P07949 |
| HOMO SAPIENS | -1 | 4861 | 4ux8 | A | 618 | CYS | p.Cys618Phe | P07949 |
| HOMO SAPIENS | -1 | 4861 | 4ux8 | A | 620 | CYS | p.Cys620Gly | P07949 |
| HOMO SAPIENS | -1 | 4861 | 4ux8 | A | 620 | CYS | p.Cys620Ser | P07949 |
| HOMO SAPIENS | -1 | 4861 | 4ux8 | A | 620 | CYS | p.Cys620Trp | P07949 |
| HOMO SAPIENS | -1 | 4861 | 4ux8 | A | 623 | GLU | p.Glu623Lys | P07949 |
| HOMO SAPIENS | -1 | 4861 | 4ux8 | A | 631 | ASP | p.Asp631Gly | P07949 |
| HOMO SAPIENS | -1 | 4861 | 4ux8 | A | 631 | ASP | p.Asp631Glu | P07949 |
| HOMO SAPIENS | -1 | 4861 | 4ux8 | A | 634 | CYS | p.Cys634Gly | P07949 |
| HOMO SAPIENS | -1 | 4861 | 4ux8 | A | 634 | CYS | p.Cys634Phe | P07949 |
| HOMO SAPIENS | -1 | 4861 | 4ux8 | A | 634 | CYS | p.Cys634Trp | P07949 |
| HOMO SAPIENS | 0 | 36308 | 4nkq | B | 123 | PRO | p.Pro142Leu | P15509 |
| HOMO SAPIENS | 0 | 36308 | 4nkq | B | 145 | ARG | p.Arg164Gln | P15509 |
| HOMO SAPIENS | -1 | 36308 | 4nkq | B | 177 | GLY | p.Gly196Arg | P15509 |
| HOMO SAPIENS | 0 | 36006 | 1oqa | A | 107 | HIS | p.His1860Pro | P38398 |
| HOMO SAPIENS | 7 | 15394 | 2ozo | A | 465 | ARG | p.Arg465His | P43403 |
| HOMO SAPIENS | 7 | 15394 | 2ozo | A | 465 | ARG | p.Arg465Cys | P43403 |
| HOMO SAPIENS | 4 | 15394 | 2ozo | A | 518 | SER | p.Ser518Arg | P43403 |
| HOMO SAPIENS | 1 | 20813 | 3o5z | A | 19 | ARG | p.Arg19Trp | O00459 |
| HOMO SAPIENS | 2 | 4329 | 2bb5 | A | 5 | ILE | p.Ile23Val | P20062 |
| HOMO SAPIENS | 4 | 4329 | 2bb5 | A | 59 | LYS | p.Lys77Met | P20062 |
| HOMO SAPIENS | 4 | 4329 | 2bb5 | A | 197 | ARG | p.Arg215Trp | P20062 |
| HOMO SAPIENS | 1 | 4329 | 2bb5 | A | 241 | PRO | p.Arg259Pro | P20062 |
| HOMO SAPIENS | 2 | 4329 | 2bb5 | A | 315 | THR | p.Thr333Lys | P20062 |
| HOMO SAPIENS | 5 | 4329 | 2bb5 | A | 330 | SER | p.Ser348Phe | P20062 |
| HOMO SAPIENS | 4 | 4329 | 2bb5 | A | 358 | LEU | p.Leu376Ser | P20062 |
| HOMO SAPIENS | 5 | 4329 | 2bb5 | A | 381 | ARG | p.Arg399Gln | P20062 |
| HOMO SAPIENS | 4 | 18294 | 4r30 | A | 163 | ASN | p.Asn163Asp | O95278 |
| HOMO SAPIENS | 6 | 18294 | 4r30 | A | 227 | ALA | p.Ala227Val | O95278 |
| HOMO SAPIENS | 4 | 18294 | 4r30 | A | 241 | ARG | p.Arg241Ter | O95278 |
| HOMO SAPIENS | 4 | 18294 | 4r30 | A | 325 | ARG | p.Arg325His | O95278 |
| HOMO SAPIENS | 0 | 2632 | 4xem | A | 22 | GLU | p.Glu22Lys | P49588 |
| HOMO SAPIENS | 0 | 2632 | 4xem | A | 275 | GLY | p.Gly275Asp | P49588 |
| HOMO SAPIENS | 1 | 2632 | 4xem | A | 329 | ARG | p.Arg329His | P49588 |
| HOMO SAPIENS | 0 | 11431 | 5t4n | A | 284 | GLN | p.Gln305Glu | Q96QU1 |
| HOMO SAPIENS | 0 | 11431 | 5t4n | A | 315 | ARG | p.Arg336Ter | Q96QU1 |
| HOMO SAPIENS | 0 | 11431 | 5t4n | A | 322 | ARG | p.Arg343Lys | Q96QU1 |

| | | | | | | | | |
|---|---|---|---|---|---|---|---|---|
| HOMO SAPIENS | 1 | 11431 | 5t4n | A | 326 | LEU | p.Leu347Phe | Q96QU1 |
| HOMO SAPIENS | 0 | 11431 | 5t4n | A | 373 | TYR | p.Tyr394Cys | Q96QU1 |
| HOMO SAPIENS | 0 | 11431 | 5t4n | A | 398 | ASN | p.Asn419Ser | Q96QU1 |
| HOMO SAPIENS | 0 | 11431 | 5t4n | A | 414 | ALA | p.Asp435Ala | Q96QU1 |
| HOMO SAPIENS | 0 | 11431 | 5t4n | A | 433 | VAL | p.Val454Phe | Q96QU1 |
| HOMO SAPIENS | 1 | 11431 | 5t4n | A | 507 | VAL | p.Val528Asp | Q96QU1 |
| HOMO SAPIENS | 0 | 11431 | 5t4n | A | 547 | ALA | p.Ala568Ser | Q96QU1 |
| HOMO SAPIENS | 3 | 3194 | 5xtd | L | 84 | ARG | p.Arg84His | O43181 |
| HOMO SAPIENS | 1 | 3194 | 5xtd | L | 130 | THR | p.Thr130Ser | O43181 |
| HOMO SAPIENS | 8 | 3194 | 5xtd | L | 136 | SER | p.Ser136Ala | O43181 |
| HOMO SAPIENS | 0 | 40749 | 1nun | A | 106 | CYS | p.Cys106Phe | O15520 |
| HOMO SAPIENS | 9 | 40749 | 1nun | A | 156 | ILE | p.Ile156Arg | O15520 |
| HOMO SAPIENS | 1 | 40749 | 1nun | A | 193 | ARG | p.Arg193Ter | O15520 |
| HOMO SAPIENS | 0 | 40749 | 1nun | A | 207 | HIS | p.His207Pro | O15520 |
| HOMO SAPIENS | 0 | 7026 | 3n1f | D | 875 | SER | p.Ser875Gly | Q4KMG0 |
| HOMO SAPIENS | 6 | 23802 | 3rzv | A | 424 | ARG | p.Arg424Ter | O95630 |
| HOMO SAPIENS | 0 | 36720 | 1jm7 | A | 12 | GLN | p.Gln12Ter | P38398 |
| HOMO SAPIENS | 3 | 36720 | 1jm7 | A | 23 | GLU | p.Glu23Asp | P38398 |
| HOMO SAPIENS | 1 | 36720 | 1jm7 | A | 45 | LYS | p.Lys45Gln | P38398 |
| HOMO SAPIENS | 8 | 36720 | 1jm7 | A | 47 | CYS | p.Cys47Ter | P38398 |
| HOMO SAPIENS | 0 | 36720 | 1jm7 | A | 59 | SER | p.Ser59Ter | P38398 |
| HOMO SAPIENS | 8 | 36720 | 1jm7 | A | 61 | CYS | p.Cys61Gly | P38398 |
| HOMO SAPIENS | 8 | 36720 | 1jm7 | A | 63 | LEU | p.Leu63Ter | P38398 |
| HOMO SAPIENS | 7 | 36720 | 1jm7 | A | 64 | CYS | p.Cys64Gly | P38398 |
| HOMO SAPIENS | 0 | 36720 | 1jm7 | A | 101 | TYR | p.Tyr101Ter | P38398 |
| HOMO SAPIENS | 0 | 36720 | 1jm7 | A | null | TYR | p.Tyr105Cys | P38398 |
| HOMO SAPIENS | 0 | 28500 | 3kuz | A | 1217 | CYS | p.Cys1217Arg | O60486 |
| HOMO SAPIENS | 5 | 8097 | 3p56 | C | 69 | ARG | p.Arg69Trp | Q8TDP1 |
| HOMO SAPIENS | 5 | 8097 | 3p56 | C | 143 | LYS | p.Lys143Ile | Q8TDP1 |
| HOMO SAPIENS | 2 | 30073 | 3nmz | A | 380 | ARG | p.Arg380Gln | P25054 |
| HOMO SAPIENS | 1 | 22573 | 1cjy | A | 111 | SER | p.Ser111Pro | P47712 |
| HOMO SAPIENS | 0 | 22573 | 1cjy | A | 485 | ARG | p.Arg485His | P47712 |
| HOMO SAPIENS | 2 | 965 | 3E+077 | A | 99 | ALA | p.Ala99Val | Q9Y617 |
| HOMO SAPIENS | 2 | 965 | 3E+077 | A | 100 | ASP | p.Asp100Ala | Q9Y617 |
| HOMO SAPIENS | 4 | 1503 | 3kxc | A | 62 | ARG | p.Arg62Trp | O43617 |
| HOMO SAPIENS | 6 | 9418 | 1hdr | A | 22 | GLY | p.Gly23Asp | P09417 |
| HOMO SAPIENS | 2 | 9418 | 1hdr | A | 35 | TRP | p.Trp36Arg | P09417 |
| HOMO SAPIENS | 2 | 9418 | 1hdr | A | 107 | TRP | p.Trp108Gly | P09417 |
| HOMO SAPIENS | 6 | 9418 | 1hdr | A | 149 | TYR | p.Tyr150Cys | P09417 |
| HOMO SAPIENS | -1 | 12079 | 3tnu | B | 373 | GLY | p.Gly373Ala | P13647 |
| HOMO SAPIENS | 1 | 12079 | 3tnu | B | 463 | LEU | p.Leu463Pro | P13647 |
| HOMO SAPIENS | 1 | 12079 | 3tnu | B | 471 | ARG | p.Arg471Cys | P13647 |
| HOMO SAPIENS | 0 | 8135 | 3rbn | B | 89 | SER | p.Ser556Ile | P40692 |
| HOMO SAPIENS | 0 | 8135 | 3rbn | B | 111 | GLU | p.Glu578Gly | P40692 |
| HOMO SAPIENS | 2 | 8135 | 3rbn | B | 114 | PRO | p.Pro581Leu | P40692 |
| HOMO SAPIENS | 0 | 8135 | 3rbn | B | 133 | GLU | p.Glu600Gly | P40692 |
| HOMO SAPIENS | 2 | 8135 | 3rbn | B | 136 | PRO | p.Pro603Arg | P40692 |
| HOMO SAPIENS | 2 | 8135 | 3rbn | B | 136 | PRO | p.Pro603Leu | P40692 |
| HOMO SAPIENS | 7 | 8135 | 3rbn | B | 140 | LEU | p.Leu607His | P40692 |
| HOMO SAPIENS | 2 | 8135 | 3rbn | B | 151 | LYS | p.Lys618Ala | P40692 |
| HOMO SAPIENS | 2 | 8135 | 3rbn | B | 151 | LYS | p.Lys618Thr | P40692 |
| HOMO SAPIENS | 5 | 8135 | 3rbn | B | 181 | PRO | p.Pro648Ser | P40692 |
| HOMO SAPIENS | 2 | 8135 | 3rbn | B | 188 | ILE | p.Ile655Val | P40692 |
| HOMO SAPIENS | 2 | 8135 | 3rbn | B | 214 | ALA | p.Ala681Thr | P40692 |

| | | | | | | | | |
|---|---|---|---|---|---|---|---|---|
| HOMO SAPIENS | 2 | 8135 | 3rbn | B | 220 | ARG | p.Arg687Trp | P40692 |
| HOMO SAPIENS | 2 | 8135 | 3rbn | B | 222 | GLN | p.Gln689Arg | P40692 |
| HOMO SAPIENS | -1 | 8135 | 3rbn | B | 236 | GLN | p.Gln701Ter | P40692 |
| HOMO SAPIENS | 7 | 8135 | 3rbn | B | 251 | HIS | p.His718Tyr | P40692 |
| HOMO SAPIENS | 0 | 8135 | 3rbn | B | 284 | LYS | p.Lys751Arg | P40692 |
| HOMO SAPIENS | 1 | 21998 | 5d68 | A | 355 | LEU | p.Leu355Phe | O00522 |
| HOMO SAPIENS | 3 | 21998 | 5d68 | A | 380 | THR | p.Thr380Met | O00522 |
| HOMO SAPIENS | 0 | 21998 | 5d68 | A | 415 | ILE | p.Ile415Met | O00522 |
| HOMO SAPIENS | 0 | 33789 | 2m0c | A | 13 | ARG | p.Arg218Gln | Q9H161 |
| HOMO SAPIENS | 7 | 33789 | 2m0c | A | 52 | ARG | p.Arg257Thr | Q9H161 |
| HOMO SAPIENS | 0 | 33789 | 2m0c | A | 67 | ARG | p.Arg272Pro | Q9H161 |
| HOMO SAPIENS | 6 | 8263 | 2osg | A | 51 | GLY | p.Gly355Arg | Q9UDY2 |
| HOMO SAPIENS | 5 | 8263 | 2osg | A | 60 | THR | p.Thr364Met | Q9UDY2 |
| HOMO SAPIENS | 1 | 2475 | 5E+026 | B | 234 | THR | p.Thr234Ala | Q9BZ23 |
| HOMO SAPIENS | 2 | 2475 | 5E+026 | B | 264 | ARG | p.Arg264Trp | Q9BZ23 |
| HOMO SAPIENS | 4 | 2475 | 5E+026 | B | 286 | ARG | p.Arg286Cys | Q9BZ23 |
| HOMO SAPIENS | 0 | 2475 | 5E+026 | B | 357 | ARG | p.Arg357Trp | Q9BZ23 |
| HOMO SAPIENS | 2 | 2475 | 5E+026 | B | 481 | ARG | p.Arg481Ter | Q9BZ23 |
| HOMO SAPIENS | 8 | 2475 | 5E+026 | B | 521 | GLY | p.Gly521Arg | Q9BZ23 |
| HOMO SAPIENS | 2 | 2475 | 5E+026 | B | 528 | THR | p.Thr528Met | Q9BZ23 |
| HOMO SAPIENS | 1 | 3742 | 5xtd | E | 108 | HIS | p.His134Tyr | P56556 |
| HOMO SAPIENS | 1 | 16725 | 3g04 | C | 36 | ASP | p.Asp36His | P16473 |
| HOMO SAPIENS | 3 | 16725 | 3g04 | C | 52 | PRO | p.Pro52Thr | P16473 |
| HOMO SAPIENS | 1 | 16725 | 3g04 | C | 162 | PRO | p.Pro162Ala | P16473 |
| HOMO SAPIENS | -1 | 12366 | 5d2d | C | 750 | PRO | p.Pro750Leu | P13569 |
| HOMO SAPIENS | 0 | 12366 | 5d2d | C | 764 | ARG | p.Arg764Ter | P13569 |
| HOMO SAPIENS | 0 | 34845 | 2y23 | A | 1151 | ASP | p.Asp1151Glu | P52179 |
| HOMO SAPIENS | 0 | 34845 | 2y23 | A | 1180 | ASP | p.Asp1180Ala | P52179 |
| HOMO SAPIENS | 6 | 36740 | 3lbx | B | 2053 | ALA | p.Ala2053Pro | P11277 |
| HOMO SAPIENS | 0 | 19289 | 3ihj | A | 404 | ARG | p.Arg404Ter | Q8TD30 |
| HOMO SAPIENS | 5 | 1034 | 5xtd | Q | 228 | ARG | p.Arg228Gln | O75306 |
| HOMO SAPIENS | 5 | 1034 | 5xtd | Q | 229 | PRO | p.Pro229Gln | O75306 |
| HOMO SAPIENS | 4 | 1034 | 5xtd | Q | 323 | ARG | p.Arg323Gln | O75306 |
| HOMO SAPIENS | 4 | 1034 | 5xtd | Q | 352 | PRO | p.Pro352Ala | O75306 |
| HOMO SAPIENS | 5 | 1034 | 5xtd | Q | 413 | SER | p.Ser413Pro | O75306 |
| HOMO SAPIENS | 1 | 2772 | 2vaf | A | 1047 | THR | p.Thr66Ala | O14958 |
| HOMO SAPIENS | 5 | 2772 | 2vaf | A | 1057 | VAL | p.Val76Met | O14958 |
| HOMO SAPIENS | 0 | 2772 | 2vaf | A | 1148 | LEU | p.Leu167His | O14958 |
| HOMO SAPIENS | 1 | 2772 | 2vaf | A | 1149 | ILE | p.Ile168Thr | O14958 |
| HOMO SAPIENS | 8 | 2772 | 2vaf | A | 1225 | HIS | p.His244Arg | O14958 |
| HOMO SAPIENS | 8 | 2772 | 2vaf | A | 1225 | HIS | p.His244Tyr | O14958 |
| HOMO SAPIENS | 7 | 2772 | 2vaf | A | 1251 | ILE | p.Ile270Thr | O14958 |
| HOMO SAPIENS | 3 | 2772 | 2vaf | A | 1288 | ASP | p.Asp307His | O14958 |
| HOMO SAPIENS | 5 | 2772 | 2vaf | A | 1289 | PRO | p.Pro308Leu | O14958 |
| HOMO SAPIENS | 1 | 31885 | 2aex | A | 132 | ALA | p.Ala132Val | P36551 |
| HOMO SAPIENS | 1 | 31885 | 2aex | A | 272 | ASN | p.Asn272His | P36551 |
| HOMO SAPIENS | 8 | 31885 | 2aex | A | 279 | GLY | p.Gly279Arg | P36551 |
| HOMO SAPIENS | 5 | 31885 | 2aex | A | 294 | VAL | p.Val294Ile | P36551 |
| HOMO SAPIENS | 5 | 31885 | 2aex | A | 295 | HIS | p.His295Asp | P36551 |
| HOMO SAPIENS | 1 | 31885 | 2aex | A | 331 | ARG | p.Arg331Trp | P36551 |
| HOMO SAPIENS | 3 | 31885 | 2aex | A | 352 | ARG | p.Arg352Cys | P36551 |
| HOMO SAPIENS | 1 | 31885 | 2aex | A | 404 | LYS | p.Lys404Glu | P36551 |
| HOMO SAPIENS | 2 | 31885 | 2aex | A | 447 | ARG | p.Arg447Cys | P36551 |
| HOMO SAPIENS | 5 | 11721 | 2r3v | A | 20 | HIS | p.His20Pro | Q03426 |

| | | | | | | | | |
|---|---|---|---|---|---|---|---|---|
| HOMO SAPIENS | 1 | 11721 | 2r3v | A | 116 | TYR | p.Tyr116His | Q03426 |
| HOMO SAPIENS | 1 | 11721 | 2r3v | A | 148 | ALA | p.Ala148Thr | Q03426 |
| HOMO SAPIENS | 1 | 11721 | 2r3v | A | 167 | PRO | p.Pro167Leu | Q03426 |
| HOMO SAPIENS | 6 | 11721 | 2r3v | A | 203 | VAL | p.Val203Ala | Q03426 |
| HOMO SAPIENS | 2 | 11721 | 2r3v | A | 215 | ARG | p.Arg215Ter | Q03426 |
| HOMO SAPIENS | 2 | 11721 | 2r3v | A | 237 | THR | p.Thr237Ser | Q03426 |
| HOMO SAPIENS | 0 | 11721 | 2r3v | A | 268 | ILE | p.Ile268Thr | Q03426 |
| HOMO SAPIENS | 0 | 11721 | 2r3v | A | 268 | ILE | p.Ile268Val | Q03426 |
| HOMO SAPIENS | 0 | 11721 | 2r3v | A | 301 | ASN | p.Asn301Ser | Q03426 |
| HOMO SAPIENS | 6 | 11721 | 2r3v | A | 310 | VAL | p.Val310Met | Q03426 |
| HOMO SAPIENS | 5 | 11721 | 2r3v | A | 334 | ALA | p.Ala334Thr | Q03426 |
| HOMO SAPIENS | 2 | 11721 | 2r3v | A | 377 | VAL | p.Val377Ile | Q03426 |
| HOMO SAPIENS | 1 | 11721 | 2r3v | A | 386 | ASP | p.Asp386Asn | Q03426 |
| HOMO SAPIENS | 1 | 11721 | 2r3v | A | 388 | ARG | p.Arg388Ter | Q03426 |
| HOMO SAPIENS | 8 | 5010 | 2hgs | A | 125 | ARG | p.Arg125Cys | P48637 |
| HOMO SAPIENS | 1 | 5010 | 2hgs | A | 164 | ARG | p.Arg164Gln | P48637 |
| HOMO SAPIENS | 0 | 5010 | 2hgs | A | 219 | ASP | p.Asp219Gly | P48637 |
| HOMO SAPIENS | 7 | 5010 | 2hgs | A | 267 | ARG | p.Arg267Trp | P48637 |
| HOMO SAPIENS | 0 | 5010 | 2hgs | A | 283 | ARG | p.Arg283Cys | P48637 |
| HOMO SAPIENS | 0 | 6892 | 5erd | B | 7 | VAL | p.Val56Met | Q14126 |
| HOMO SAPIENS | 1 | 6892 | 5erd | B | 25 | HIS | p.His74Arg | Q14126 |
| HOMO SAPIENS | 0 | 6892 | 5erd | B | 40 | TYR | p.Tyr89Cys | Q14126 |
| HOMO SAPIENS | 0 | 6892 | 5erd | B | 97 | ARG | p.Arg146Leu | Q14126 |
| HOMO SAPIENS | 7 | 6892 | 5erd | B | 217 | ASN | p.Asn266Ser | Q14126 |
| HOMO SAPIENS | 0 | 6892 | 5erd | B | 243 | ARG | p.Arg292His | Q14126 |
| HOMO SAPIENS | 0 | 6892 | 5erd | B | 244 | ILE | p.Ile293Val | Q14126 |
| HOMO SAPIENS | 0 | 6892 | 5erd | B | 343 | VAL | p.Val392Ile | Q14126 |
| HOMO SAPIENS | 4 | 6892 | 5erd | B | 350 | GLU | p.Glu399Lys | Q14126 |
| HOMO SAPIENS | 0 | 6892 | 5erd | B | 410 | TYR | p.Tyr459Cys | Q14126 |
| HOMO SAPIENS | 0 | 6892 | 5erd | B | 466 | VAL | p.Val515Ile | Q14126 |
| HOMO SAPIENS | 0 | 6892 | 5erd | B | 482 | PHE | p.Phe531Cys | Q14126 |
| HOMO SAPIENS | 0 | 6892 | 5erd | B | 484 | VAL | p.Val533Ile | Q14126 |
| HOMO SAPIENS | 0 | 6892 | 5erd | B | 545 | LEU | p.Leu594Pro | Q14126 |
| HOMO SAPIENS | 0 | 1338 | 2ax4 | A | 58 | THR | p.Thr48Arg | O95340 |
| HOMO SAPIENS | 7 | 8675 | 2zkm | X | 385 | LYS | p.Lys385Arg | Q00722 |
| HOMO SAPIENS | 1 | 11769 | 2wya | B | 54 | VAL | p.Val54Met | P54868 |
| HOMO SAPIENS | 1 | 11769 | 2wya | B | 116 | ARG | p.Arg116Cys | P54868 |
| HOMO SAPIENS | 5 | 11769 | 2wya | B | 167 | TYR | p.Tyr167Cys | P54868 |
| HOMO SAPIENS | 1 | 11769 | 2wya | B | 174 | PHE | p.Phe174Leu | P54868 |
| HOMO SAPIENS | 0 | 11769 | 2wya | B | 212 | GLY | p.Gly212Arg | P54868 |
| HOMO SAPIENS | 1 | 11769 | 2wya | B | 288 | ARG | p.Arg288Ter | P54868 |
| HOMO SAPIENS | 0 | 11769 | 2wya | B | 375 | SER | p.Ser375Cys | P54868 |
| HOMO SAPIENS | 4 | 11769 | 2wya | B | 500 | ARG | p.Arg500His | P54868 |
| HOMO SAPIENS | 0 | 3102 | 2iwo | A | 1880 | ARG | p.Arg1908Lys | O75970 |
| HOMO SAPIENS | 6 | 14296 | 2m9w | A | 38 | GLY | p.Gly296Ser | P43694 |
| HOMO SAPIENS | 0 | 14296 | 2m9w | A | 58 | GLN | p.Gln316Glu | P43694 |
| HOMO SAPIENS | 2 | 704 | 5ekd | B | 239 | ARG | p.Arg239Ter | Q9UGM6 |
| HOMO SAPIENS | 2 | 704 | 5ekd | B | 264 | TYR | p.Tyr264Cys | Q9UGM6 |
| HOMO SAPIENS | 6 | 2228 | 1ey2 | A | 47 | SER | p.Ser47Ter | Q93099 |
| HOMO SAPIENS | 6 | 2228 | 1ey2 | A | 48 | ALA | p.Ala48Ser | Q93099 |
| HOMO SAPIENS | 6 | 2228 | 1ey2 | A | 53 | ARG | p.Arg53Gln | Q93099 |
| HOMO SAPIENS | 3 | 2228 | 1ey2 | A | 80 | HIS | p.Gln80His | Q93099 |
| HOMO SAPIENS | 0 | 2228 | 1ey2 | A | 120 | CYS | p.Cys120Trp | Q93099 |
| HOMO SAPIENS | 3 | 2228 | 1ey2 | A | 122 | ALA | p.Ala122Val | Q93099 |

| | | | | | | | | |
|---|---|---|---|---|---|---|---|---|
| HOMO SAPIENS | 3 | 2228 | 1ey2 | A | 161 | GLY | p.Gly161Arg | Q93099 |
| HOMO SAPIENS | 6 | 2228 | 1ey2 | A | 225 | ARG | p.Arg225His | Q93099 |
| HOMO SAPIENS | 0 | 2228 | 1ey2 | A | 230 | PRO | p.Pro230Ser | Q93099 |
| HOMO SAPIENS | 6 | 2228 | 1ey2 | A | 270 | GLY | p.Gly270Arg | Q93099 |
| HOMO SAPIENS | 6 | 2228 | 1ey2 | A | 330 | ARG | p.Arg330Ser | Q93099 |
| HOMO SAPIENS | -1 | 2228 | 1ey2 | A | 368 | MSE | p.Met368Val | Q93099 |
| HOMO SAPIENS | 6 | 2228 | 1ey2 | A | 371 | HIS | p.His371Arg | Q93099 |
| HOMO SAPIENS | 6 | 2228 | 1ey2 | A | 401 | GLU | p.Glu401Gln | Q93099 |
| HOMO SAPIENS | 5 | 34648 | 2k27 | A | 31 | ARG | p.Arg31His | Q06710 |
| HOMO SAPIENS | 7 | 34648 | 2k27 | A | 40 | GLN | p.Gln40Pro | Q06710 |
| HOMO SAPIENS | 5 | 34648 | 2k27 | A | 57 | CYS | p.Cys57Tyr | Q06710 |
| HOMO SAPIENS | 5 | 34648 | 2k27 | A | 62 | LEU | p.Leu62Arg | Q06710 |
| HOMO SAPIENS | 1 | 21399 | 3edu | A | 1819 | GLU | p.Glu1819Ter | P11277 |
| HOMO SAPIENS | 1 | 18153 | 1oey | J | 245 | TYR | p.Tyr245Ter | Q15080 |
| HOMO SAPIENS | 2 | 41567 | 3sw0 | X | 1050 | ASN | p.Asn1050Tyr | P08603 |
| HOMO SAPIENS | 5 | 41567 | 3sw0 | X | 1059 | ILE | p.Ile1059Thr | P08603 |
| HOMO SAPIENS | 0 | 36362 | 4wrm | A | 362 | HIS | p.His362Arg | P07333 |
| HOMO SAPIENS | -1 | 36362 | 4wrm | A | 413 | GLY | p.Gly413Ser | P07333 |
| HOMO SAPIENS | 0 | 31255 | 2w2j | A | 100 | SER | p.Ser100Pro | P35219 |
| HOMO SAPIENS | 0 | 31255 | 2w2j | A | 100 | SER | p.Ser100Leu | P35219 |
| HOMO SAPIENS | 5 | 31255 | 2w2j | A | 237 | ARG | p.Arg237Gln | P35219 |
| HOMO SAPIENS | -1 | 26450 | 1th1 | C | 1395 | SER | p.Ser1395Cys | P25054 |
| HOMO SAPIENS | -1 | 26450 | 1th1 | C | 1413 | MET | p.Met1413Val | P25054 |
| HOMO SAPIENS | -1 | 26450 | 1th1 | C | 1446 | ALA | p.Ala1446Thr | P25054 |
| HOMO SAPIENS | -1 | 26450 | 1th1 | C | 1454 | LYS | p.Lys1454Glu | P25054 |
| HOMO SAPIENS | 0 | 26450 | 1th1 | C | 1474 | ALA | p.Ala1474Thr | P25054 |
| HOMO SAPIENS | 1 | 981 | 5xtb | F | 14 | LEU | p.Leu14Val | O43678 |
| HOMO SAPIENS | 1 | 11069 | 3kq4 | A | 46 | SER | p.Ser46Leu | P27352 |
| HOMO SAPIENS | 1 | 11069 | 3kq4 | A | 97 | MET | p.Met97Thr | P27352 |
| HOMO SAPIENS | 1 | 11069 | 3kq4 | A | 255 | ASN | p.Asn255Ser | P27352 |
| HOMO SAPIENS | 1 | 11069 | 3kq4 | A | 330 | ASN | p.Asn330Lys | P27352 |
| HOMO SAPIENS | 0 | 41489 | 3erb | A | 53 | PRO | p.Pro73Leu | P06681 |
| HOMO SAPIENS | 6 | 8715 | 2bq8 | X | 23 | THR | p.Thr44Met | P13686 |
| HOMO SAPIENS | 0 | 8715 | 2bq8 | X | 88 | GLY | p.Gly109Arg | P13686 |
| HOMO SAPIENS | 6 | 8715 | 2bq8 | X | 127 | VAL | p.Val148Met | P13686 |
| HOMO SAPIENS | 2 | 8715 | 2bq8 | X | 179 | VAL | p.Val200Met | P13686 |
| HOMO SAPIENS | 6 | 8715 | 2bq8 | X | 251 | ARG | p.Arg272Cys | P13686 |
| HOMO SAPIENS | 7 | 8112 | 3p56 | A | 37 | GLY | p.Gly37Ser | O75792 |
| HOMO SAPIENS | -1 | 8112 | 3p56 | A | 202 | LEU | p.Leu202Ser | O75792 |
| HOMO SAPIENS | 1 | 8112 | 3p56 | A | 205 | ASP | p.Asp205Glu | O75792 |
| HOMO SAPIENS | 4 | 8112 | 3p56 | A | 212 | ASN | p.Asn212Ile | O75792 |
| HOMO SAPIENS | 0 | 8112 | 3p56 | A | 235 | ARG | p.Arg235Gln | O75792 |
| HOMO SAPIENS | 4 | 8112 | 3p56 | A | 240 | THR | p.Thr240Met | O75792 |
| HOMO SAPIENS | 0 | 8112 | 3p56 | A | 291 | ARG | p.Arg291His | O75792 |
| HOMO SAPIENS | 0 | 37603 | 2yqg | A | 10 | ARG | p.Arg49His | Q14126 |
| HOMO SAPIENS | 0 | 21178 | 3k1r | B | 458 | ASP | p.Asp458Val | Q495M9 |
| HOMO SAPIENS | -1 | 36828 | 4fq3 | B | 507 | GLY | p.Gly507Asp | P35637 |
| HOMO SAPIENS | 5 | 36828 | 4fq3 | B | 517 | HIS | p.His517Gln | P35637 |
| HOMO SAPIENS | 8 | 36828 | 4fq3 | B | 518 | ARG | p.Arg518Lys | P35637 |
| HOMO SAPIENS | 6 | 36828 | 4fq3 | B | 521 | ARG | p.Arg521Cys | P35637 |
| HOMO SAPIENS | 6 | 36828 | 4fq3 | B | 521 | ARG | p.Arg521His | P35637 |
| HOMO SAPIENS | 0 | 6363 | 3j7y | c | null | HIS | p.His13Arg | Q9H9J2 |
| HOMO SAPIENS | 2 | 6363 | 3j7y | c | 105 | ARG | p.Arg105Cys | Q9H9J2 |
| HOMO SAPIENS | 2 | 6363 | 3j7y | c | 138 | THR | p.Thr138Ile | Q9H9J2 |

| | | | | | | | | |
|---|---|---|---|---|---|---|---|---|
| HOMO SAPIENS | 1 | 6363 | 3j7y | c | 156 | LEU | p.Leu156Arg | Q9H9J2 |
| HOMO SAPIENS | 1 | 6782 | 5l46 | A | 109 | HIS | p.His109Arg | Q9UI17 |
| HOMO SAPIENS | 0 | 23951 | 5t76 | A | 175 | GLY | p.Gly931Ser | P49588 |
| HOMO SAPIENS | 8 | 12604 | 3p0l | A | 182 | ARG | p.Arg182Leu | P49675 |
| HOMO SAPIENS | 2 | 12604 | 3p0l | A | 187 | VAL | p.Val187Met | P49675 |
| HOMO SAPIENS | 2 | 12604 | 3p0l | A | 188 | ARG | p.Arg188Cys | P49675 |
| HOMO SAPIENS | 0 | 12604 | 3p0l | A | 193 | ARG | p.Arg193Ter | P49675 |
| HOMO SAPIENS | 9 | 12604 | 3p0l | A | 217 | ARG | p.Arg217Thr | P49675 |
| HOMO SAPIENS | 4 | 12604 | 3p0l | A | 218 | ALA | p.Ala218Val | P49675 |
| HOMO SAPIENS | 0 | 12604 | 3p0l | A | 250 | TRP | p.Trp250Ter | P49675 |
| HOMO SAPIENS | 1 | 12604 | 3p0l | A | 258 | GLN | p.Gln258Ter | P49675 |
| HOMO SAPIENS | -1 | 27765 | 4wkr | A | 4 | GLU | p.Glu4Lys | Q4G0J3 |
| HOMO SAPIENS | 2 | 3587 | 4kit | C | 2301 | PRO | p.Pro2301Thr | Q6P2Q9 |
| HOMO SAPIENS | 3 | 3587 | 4kit | C | 2304 | PHE | p.Phe2304Leu | Q6P2Q9 |
| HOMO SAPIENS | 3 | 3587 | 4kit | C | 2309 | HIS | p.His2309Pro | Q6P2Q9 |
| HOMO SAPIENS | 3 | 3587 | 4kit | C | 2310 | ARG | p.Arg2310Lys | Q6P2Q9 |
| HOMO SAPIENS | 0 | 9073 | 1n69 | B | 21 | ASN | p.Asn215His | P07602 |
| HOMO SAPIENS | 0 | 9073 | 1n69 | B | 23 | THR | p.Thr217Ile | P07602 |
| HOMO SAPIENS | 0 | 9073 | 1n69 | B | 47 | CYS | p.Cys241Ser | P07602 |
| HOMO SAPIENS | 0 | 475 | 2vd5 | A | 225 | THR | p.Thr225Met | Q09013 |
| HOMO SAPIENS | 1 | 9066 | 4mzv | A | 31 | ASN | p.Asn31Lys | P16422 |
| HOMO SAPIENS | 1 | 9066 | 4mzv | A | 47 | GLN | p.Gln47His | P16422 |
| HOMO SAPIENS | 3 | 9066 | 4mzv | A | 66 | CYS | p.Cys66Tyr | P16422 |
| HOMO SAPIENS | 1 | 9066 | 4mzv | A | 102 | SER | p.Ser102Gly | P16422 |
| HOMO SAPIENS | 5 | 9066 | 4mzv | A | 115 | THR | p.Met115Thr | P16422 |
| HOMO SAPIENS | 1 | 9066 | 4mzv | A | 138 | ARG | p.Arg138Ter | P16422 |
| HOMO SAPIENS | 1 | 9066 | 4mzv | A | 153 | ARG | p.Arg153Thr | P16422 |
| HOMO SAPIENS | 1 | 9066 | 4mzv | A | 165 | ALA | p.Ala165Thr | P16422 |
| HOMO SAPIENS | 8 | 1827 | 2nz2 | A | 14 | GLY | p.Gly14Ser | P00966 |
| HOMO SAPIENS | 5 | 1827 | 2nz2 | A | 18 | SER | p.Ser18Leu | P00966 |
| HOMO SAPIENS | 3 | 1827 | 2nz2 | A | 86 | ARG | p.Arg86Cys | P00966 |
| HOMO SAPIENS | 3 | 1827 | 2nz2 | A | 86 | ARG | p.Arg86His | P00966 |
| HOMO SAPIENS | 1 | 1827 | 2nz2 | A | 179 | TRP | p.Trp179Arg | P00966 |
| HOMO SAPIENS | 7 | 1827 | 2nz2 | A | 180 | SER | p.Ser180Asn | P00966 |
| HOMO SAPIENS | 7 | 1827 | 2nz2 | A | 191 | GLU | p.Glu191Lys | P00966 |
| HOMO SAPIENS | 4 | 1827 | 2nz2 | A | 256 | GLU | p.Glu256Lys | P00966 |
| HOMO SAPIENS | 4 | 1827 | 2nz2 | A | 263 | VAL | p.Val263Met | P00966 |
| HOMO SAPIENS | 5 | 1827 | 2nz2 | A | 265 | ARG | p.Arg265Cys | P00966 |
| HOMO SAPIENS | 5 | 1827 | 2nz2 | A | 268 | ILE | p.Ile268Met | P00966 |
| HOMO SAPIENS | 6 | 1827 | 2nz2 | A | 269 | VAL | p.Val269Met | P00966 |
| HOMO SAPIENS | 7 | 1827 | 2nz2 | A | 272 | ARG | p.Arg272Cys | P00966 |
| HOMO SAPIENS | 3 | 1827 | 2nz2 | A | 304 | ARG | p.Arg304Trp | P00966 |
| HOMO SAPIENS | 3 | 1827 | 2nz2 | A | 307 | ARG | p.Arg307Cys | P00966 |
| HOMO SAPIENS | 3 | 1827 | 2nz2 | A | 310 | LYS | p.Lys310Gln | P00966 |
| HOMO SAPIENS | 2 | 1827 | 2nz2 | A | 362 | GLY | p.Gly362Val | P00966 |
| HOMO SAPIENS | 2 | 1827 | 2nz2 | A | 363 | ARG | p.Arg363Trp | P00966 |
| HOMO SAPIENS | 2 | 1827 | 2nz2 | A | 363 | ARG | p.Arg363Gln | P00966 |
| HOMO SAPIENS | 1 | 1827 | 2nz2 | A | 390 | GLY | p.Gly390Arg | P00966 |
| HOMO SAPIENS | 0 | 29524 | 5vbn | B | 2165 | ARG | p.Arg2165His | Q07864 |
| HOMO SAPIENS | 2 | 29524 | 5vbn | B | 2255 | ILE | p.Ile2255Phe | Q07864 |
| HOMO SAPIENS | 2 | 29524 | 5vbn | B | 2274 | LEU | p.Leu2274Val | Q07864 |
| HOMO SAPIENS | 0 | 31897 | 3t0o | A | 184 | CYS | p.Cys184Arg | O00584 |
| HOMO SAPIENS | -1 | 31897 | 3t0o | A | 215 | GLU | p.Glu215Lys | O00584 |
| HOMO SAPIENS | 0 | 31897 | 3t0o | A | 236 | ARG | p.Arg236Trp | O00584 |

| | | | | | | | | |
|---|---|---|---|---|---|---|---|---|
| HOMO SAPIENS | 7 | 18490 | 4pcw | B | 31 | GLU | p.Glu32Ter | P20930 |
| HOMO SAPIENS | -1 | 39865 | 2h9e | L | 152 | GLY | p.Gly192Arg | P00742 |
| HOMO SAPIENS | 2 | 4309 | 2x0b | F | 174 | THR | p.Thr207Met | P01019 |
| HOMO SAPIENS | 3 | 4309 | 2x0b | F | 342 | ARG | p.Arg375Gln | P01019 |
| HOMO SAPIENS | 2 | 3094 | 1zll | B | 9 | ARG | p.Arg9Cys | P26678 |
| HOMO SAPIENS | 9 | 3094 | 1zll | B | 39 | LEU | p.Leu39Ter | P26678 |
| HOMO SAPIENS | 5 | 3094 | 1zll | B | 49 | VAL | p.Val49Met | P26678 |
| HOMO SAPIENS | -1 | 41498 | 3ffd | P | 8 | LEU | p.Leu44Pro | P12272 |
| HOMO SAPIENS | 1 | 20697 | 1igr | A | 407 | ARG | p.Arg437His | P08069 |
| HOMO SAPIENS | 0 | 20697 | 1igr | A | 416 | MET | p.Met446Val | P08069 |
| HOMO SAPIENS | -1 | 20697 | 1igr | A | 462 | VAL | p.Val492Ile | P08069 |
| HOMO SAPIENS | 0 | 3973 | 4cbz | A | 37 | LEU | p.Leu37Ser | P78504 |
| HOMO SAPIENS | 0 | 3973 | 4cbz | A | 45 | VAL | p.Val45Leu | P78504 |
| HOMO SAPIENS | 0 | 3973 | 4cbz | A | 134 | LEU | p.Leu134Met | P78504 |
| HOMO SAPIENS | -1 | 3973 | 4cbz | A | 146 | VAL | p.Val146Ile | P78504 |
| HOMO SAPIENS | 0 | 3973 | 4cbz | A | 176 | VAL | p.Val176Ile | P78504 |
| HOMO SAPIENS | 0 | 3973 | 4cbz | A | 184 | ARG | p.Arg184Cys | P78504 |
| HOMO SAPIENS | 6 | 1697 | 4hmy | S | 33 | ARG | p.Arg33Trp | Q96PC3 |
| HOMO SAPIENS | 2 | 23752 | 5j4o | A | 13 | ARG | p.Arg1611Ser | P02549 |
| HOMO SAPIENS | 0 | 23752 | 5j4o | A | 42 | ALA | p.Ala1640Gly | P02549 |
| HOMO SAPIENS | 0 | 23752 | 5j4o | A | 108 | HIS | p.His1706Gln | P02549 |
| HOMO SAPIENS | 6 | 23752 | 5j4o | A | 206 | LEU | p.Leu1804Phe | P02549 |
| HOMO SAPIENS | 8 | 531 | 2qz4 | A | 349 | GLY | p.Gly349Ser | Q9UQ90 |
| HOMO SAPIENS | 0 | 531 | 2qz4 | A | 486 | ARG | p.Arg486Gln | Q9UQ90 |
| HOMO SAPIENS | 1 | 531 | 2qz4 | A | 510 | ALA | p.Ala510Val | Q9UQ90 |
| HOMO SAPIENS | 0 | 531 | 2qz4 | A | 529 | ALA | p.Ala529Val | Q9UQ90 |
| HOMO SAPIENS | 0 | 531 | 2qz4 | A | 558 | LYS | p.Lys558Ter | Q9UQ90 |
| HOMO SAPIENS | 1 | 531 | 2qz4 | A | 559 | LYS | p.Lys559Ter | Q9UQ90 |
| HOMO SAPIENS | 2 | 7736 | 1fnh | A | 114 | TRP | p.Trp1834Arg | P02751 |
| HOMO SAPIENS | 0 | 7736 | 1fnh | A | 163 | LEU | p.Leu1883Arg | P02751 |
| HOMO SAPIENS | 3 | 20337 | 5wy5 | B | 264 | LEU | p.Leu264Phe | Q96MG7 |
| HOMO SAPIENS | 8 | 36096 | 2jku | A | 643 | GLY | p.Gly668Arg | P05165 |
| HOMO SAPIENS | 6 | 36096 | 2jku | A | 663 | CYS | p.Cys688Arg | P05165 |
| HOMO SAPIENS | 0 | 9850 | 3kbt | A | 1606 | TYR | p.Tyr1606Ter | P11277 |
| HOMO SAPIENS | 4 | 6876 | 4jkq | A | 217 | ILE | p.Ile217Val | Q92736 |
| HOMO SAPIENS | 5 | 6876 | 4jkq | A | 420 | ARG | p.Arg420Trp | Q92736 |
| HOMO SAPIENS | 2 | 27603 | 1itv | B | 62 | PRO | p.Arg574Pro | P14780 |
| HOMO SAPIENS | 4 | 27603 | 1itv | B | 156 | ARG | p.Arg668Gln | P14780 |
| HOMO SAPIENS | 2 | 22954 | 4f52 | E | 48 | ILE | p.Ile48Met | Q92990 |
| HOMO SAPIENS | 0 | 22954 | 4f52 | E | 91 | ASP | p.Asp91Asn | Q92990 |
| HOMO SAPIENS | 0 | 22954 | 4f52 | E | 146 | ALA | p.Ala146Thr | Q92990 |
| HOMO SAPIENS | 0 | 22954 | 4f52 | E | 181 | ILE | p.Ile181Lys | Q92990 |
| HOMO SAPIENS | 0 | 22954 | 4f52 | E | 194 | ASN | p.Asn194Ser | Q92990 |
| HOMO SAPIENS | 2 | 22954 | 4f52 | E | 254 | PRO | p.Pro254Arg | Q92990 |
| HOMO SAPIENS | 2 | 22954 | 4f52 | E | 300 | ILE | p.Ile300Thr | Q92990 |
| HOMO SAPIENS | 2 | 22954 | 4f52 | E | 336 | LEU | p.Leu336Ser | Q92990 |
| HOMO SAPIENS | 0 | 22954 | 4f52 | E | 353 | TYR | p.Tyr353His | Q92990 |
| HOMO SAPIENS | 0 | 22954 | 4f52 | E | 387 | MET | p.Met387Val | Q92990 |
| HOMO SAPIENS | 4 | 2946 | 5gl7 | A | 344 | TYR | p.Tyr344Cys | P56192 |
| HOMO SAPIENS | 0 | 2946 | 5gl7 | A | 370 | PHE | p.Phe370Leu | P56192 |
| HOMO SAPIENS | 0 | 2946 | 5gl7 | A | 393 | ALA | p.Ala393Thr | P56192 |
| HOMO SAPIENS | 0 | 2946 | 5gl7 | A | 523 | ILE | p.Ile523Thr | P56192 |
| HOMO SAPIENS | 0 | 2946 | 5gl7 | A | 567 | SER | p.Ser567Leu | P56192 |
| HOMO SAPIENS | 0 | 2946 | 5gl7 | A | 800 | PRO | p.Pro800Thr | P56192 |

| | | | | | | | | |
|---|---|---|---|---|---|---|---|---|
| HOMO SAPIENS | 3 | 34961 | 4wy4 | C | 44 | TYR | p.Tyr44His | O95721 |
| HOMO SAPIENS | 8 | 34961 | 4wy4 | C | 89 | GLU | p.Glu89Lys | O95721 |
| HOMO SAPIENS | 0 | 9427 | 4chl | B | 21 | ALA | p.Ala21Ser | O95571 |
| HOMO SAPIENS | 2 | 9427 | 4chl | B | 62 | ALA | p.Ala62Thr | O95571 |
| HOMO SAPIENS | 0 | 9427 | 4chl | B | 114 | ILE | p.Ile114Phe | O95571 |
| HOMO SAPIENS | 2 | 9427 | 4chl | B | 159 | ARG | p.Arg159His | O95571 |
| HOMO SAPIENS | 8 | 9427 | 4chl | B | 163 | ARG | p.Arg163Trp | O95571 |
| HOMO SAPIENS | 8 | 9427 | 4chl | B | 163 | ARG | p.Arg163Gln | O95571 |
| HOMO SAPIENS | 2 | 9427 | 4chl | B | 185 | LEU | p.Leu185Arg | O95571 |
| HOMO SAPIENS | 0 | 40917 | 2e8o | A | 11 | PRO | p.Pro26Leu | Q9Y3Z3 |
| HOMO SAPIENS | -1 | 39190 | 4gow | A | 556 | TYR | p.Tyr556Ter | P56696 |
| HOMO SAPIENS | 0 | 2399 | 2mkv | A | 8 | GLY | p.Gly10Ser | P54710 |
| HOMO SAPIENS | 1 | 2399 | 2mkv | A | 24 | VAL | p.Val26Ile | P54710 |
| HOMO SAPIENS | 4 | 2399 | 2mkv | A | 39 | GLY | p.Gly41Arg | P54710 |
| HOMO SAPIENS | 0 | 20801 | 2zej | B | 1441 | ARG | p.Arg1441Cys | Q5S007 |
| HOMO SAPIENS | 0 | 20801 | 2zej | B | 1441 | ARG | p.Arg1441His | Q5S007 |
| HOMO SAPIENS | 5 | 1019 | 5xtb | M | 62 | ARG | p.Arg62Gln | P28331 |
| HOMO SAPIENS | 1 | 1019 | 5xtb | M | 85 | ALA | p.Ala85Thr | P28331 |
| HOMO SAPIENS | 5 | 1019 | 5xtb | M | 231 | LEU | p.Leu231Val | P28331 |
| HOMO SAPIENS | 2 | 1019 | 5xtb | M | 241 | ARG | p.Arg241Trp | P28331 |
| HOMO SAPIENS | 1 | 1019 | 5xtb | M | 252 | ASP | p.Asp252Gly | P28331 |
| HOMO SAPIENS | 1 | 1019 | 5xtb | M | 408 | ARG | p.Arg408Cys | P28331 |
| HOMO SAPIENS | 2 | 1019 | 5xtb | M | 417 | ARG | p.Arg417Ser | P28331 |
| HOMO SAPIENS | 1 | 1019 | 5xtb | M | 557 | ARG | p.Arg557Ter | P28331 |
| HOMO SAPIENS | 0 | 1019 | 5xtb | M | 567 | ILE | p.Ile567Val | P28331 |
| HOMO SAPIENS | 1 | 1019 | 5xtb | M | 595 | THR | p.Thr595Ala | P28331 |
| HOMO SAPIENS | 1 | 1019 | 5xtb | M | 703 | ALA | p.Ala703Thr | P28331 |
| HOMO SAPIENS | -1 | 3437 | 5hyn | A | 185 | ASP | p.Asp185His | Q15910 |
| HOMO SAPIENS | -1 | 3437 | 5hyn | A | 482 | ALA | p.Ala482Thr | Q15910 |
| HOMO SAPIENS | 0 | 8201 | 2wfd | A | 373 | TYR | p.Tyr373Cys | Q9P2J5 |
| HOMO SAPIENS | 1 | 3590 | 3edi | A | 90 | VAL | p.Val238Ala | O43897 |
| HOMO SAPIENS | 6 | 4555 | 1ytq | A | 64 | ALA | p.Ala65Ser | P43320 |
| HOMO SAPIENS | 0 | 4555 | 1ytq | A | 194 | TRP | p.Trp195Gly | P43320 |
| HOMO SAPIENS | 0 | 1013 | 2nn6 | C | 272 | SER | p.Ser272Thr | Q96B26 |
| HOMO SAPIENS | -1 | 29026 | 5m73 | D | 530 | ILE | p.Ile530Asn | O76094 |
| HOMO SAPIENS | 8 | 2252 | 1d5v | A | 57 | SER | p.Ser125Leu | Q99958 |
| HOMO SAPIENS | 1 | 4337 | 3ecr | A | 29 | ARG | p.Arg26His | P08397 |
| HOMO SAPIENS | 2 | 4337 | 3ecr | A | 34 | ALA | p.Ala31Thr | P08397 |
| HOMO SAPIENS | 2 | 4337 | 3ecr | A | 37 | GLN | p.Gln34Lys | P08397 |
| HOMO SAPIENS | 1 | 4337 | 3ecr | A | 58 | ALA | p.Ala55Ser | P08397 |
| HOMO SAPIENS | 0 | 4337 | 3ecr | A | 84 | LEU | p.Leu81Pro | P08397 |
| HOMO SAPIENS | 0 | 4337 | 3ecr | A | 114 | GLY | p.Gly111Arg | P08397 |
| HOMO SAPIENS | 1 | 4337 | 3ecr | A | 119 | ARG | p.Arg116Trp | P08397 |
| HOMO SAPIENS | 1 | 4337 | 3ecr | A | 152 | ARG | p.Arg149Gln | P08397 |
| HOMO SAPIENS | 1 | 4337 | 3ecr | A | 170 | ARG | p.Arg167Trp | P08397 |
| HOMO SAPIENS | 1 | 4337 | 3ecr | A | 176 | ARG | p.Arg173Gln | P08397 |
| HOMO SAPIENS | 0 | 4337 | 3ecr | A | 180 | LEU | p.Leu177Arg | P08397 |
| HOMO SAPIENS | 0 | 4337 | 3ecr | A | 181 | ASP | p.Asp178Asn | P08397 |
| HOMO SAPIENS | 1 | 4337 | 3ecr | A | 198 | ARG | p.Arg195Cys | P08397 |
| HOMO SAPIENS | 0 | 4337 | 3ecr | A | 201 | TRP | p.Trp198Ter | P08397 |
| HOMO SAPIENS | 1 | 4337 | 3ecr | A | 204 | ARG | p.Arg201Trp | P08397 |
| HOMO SAPIENS | 1 | 4337 | 3ecr | A | 219 | GLY | p.Gly216Asp | P08397 |
| HOMO SAPIENS | 1 | 4337 | 3ecr | A | 226 | GLU | p.Glu223Lys | P08397 |
| HOMO SAPIENS | 0 | 4337 | 3ecr | A | 248 | LEU | p.Leu245Arg | P08397 |

| | | | | | | | | |
|---|---|---|---|---|---|---|---|---|
| HOMO SAPIENS | 1 | 4337 | 3ecr | A | 250 | CYS | p.Cys247Arg | P08397 |
| HOMO SAPIENS | 0 | 4337 | 3ecr | A | 253 | GLU | p.Glu250Lys | P08397 |
| HOMO SAPIENS | 0 | 4337 | 3ecr | A | 255 | ALA | p.Ala252Thr | P08397 |
| HOMO SAPIENS | 0 | 4337 | 3ecr | A | 255 | ALA | p.Ala252Val | P08397 |
| HOMO SAPIENS | 0 | 4337 | 3ecr | A | 259 | HIS | p.His256Asn | P08397 |
| HOMO SAPIENS | 0 | 4337 | 3ecr | A | 324 | ARG | p.Arg321His | P08397 |
| HOMO SAPIENS | -1 | 4337 | 3ecr | A | 362 | ASP | p.Asp359Asn | P08397 |
| HOMO SAPIENS | 1 | 1204 | 5lwv | A | 854 | ILE | p.Ala864Thr | O15294 |
| HOMO SAPIENS | 1 | 25016 | 1m4u | A | 35 | PRO | p.Pro35Arg | Q13253 |
| HOMO SAPIENS | 1 | 25016 | 1m4u | A | 35 | PRO | p.Pro35Ser | Q13253 |
| HOMO SAPIENS | 1 | 25016 | 1m4u | A | 167 | ARG | p.Arg167Gly | Q13253 |
| HOMO SAPIENS | 1 | 25016 | 1m4u | A | 184 | CYS | p.Cys184Tyr | Q13253 |
| HOMO SAPIENS | 0 | 25016 | 1m4u | A | 189 | GLY | p.Gly189Cys | Q13253 |
| HOMO SAPIENS | 1 | 25016 | 1m4u | A | 204 | ARG | p.Arg204Leu | Q13253 |
| HOMO SAPIENS | 1 | 25016 | 1m4u | A | 205 | TRP | p.Trp205Cys | Q13253 |
| HOMO SAPIENS | 1 | 25016 | 1m4u | A | 217 | TRP | p.Trp217Gly | Q13253 |
| HOMO SAPIENS | 0 | 25016 | 1m4u | A | 222 | TYR | p.Tyr222Cys | Q13253 |
| HOMO SAPIENS | 0 | 25016 | 1m4u | A | 222 | TYR | p.Tyr222Asp | Q13253 |
| HOMO SAPIENS | 1 | 25016 | 1m4u | A | 223 | PRO | p.Pro223Leu | Q13253 |
| HOMO SAPIENS | 0 | 1394 | 1um7 | A | 95 | ALA | p.Ala463Val | Q92796 |
| HOMO SAPIENS | 0 | 21294 | 4jpz | B | 1788 | GLU | p.Glu1788Asp | Q99250 |
| HOMO SAPIENS | 9 | 24068 | 2db6 | A | 47 | ARG | p.Arg119Cys | Q96MF2 |
| HOMO SAPIENS | 5 | 1869 | 4omt | A | 39 | ARG | p.Arg39Leu | P08237 |
| HOMO SAPIENS | 2 | 1869 | 4omt | A | 100 | ARG | p.Arg100Gln | P08237 |
| HOMO SAPIENS | 7 | 1869 | 4omt | A | 543 | ASP | p.Asp543Ala | P08237 |
| HOMO SAPIENS | 1 | 1869 | 4omt | A | 686 | TRP | p.Trp686Cys | P08237 |
| HOMO SAPIENS | 1 | 1869 | 4omt | A | 696 | ARG | p.Arg696His | P08237 |
| HOMO SAPIENS | 0 | 12163 | 4wxx | A | 495 | TYR | p.Tyr495Cys | P26358 |
| HOMO SAPIENS | 0 | 12163 | 4wxx | A | 528 | ILE | p.Ile528Met | P26358 |
| HOMO SAPIENS | 0 | 12391 | 1j1d | C | 36 | LYS | p.Lys36Gln | P19429 |
| HOMO SAPIENS | 0 | 12391 | 1j1d | C | 82 | PRO | p.Pro82Ser | P19429 |
| HOMO SAPIENS | 7 | 12391 | 1j1d | C | 102 | ALA | p.Ala102Thr | P19429 |
| HOMO SAPIENS | 1 | 5000 | 4cca | A | 17 | GLY | p.Gly17Arg | Q15833 |
| HOMO SAPIENS | 0 | 5000 | 4cca | A | 205 | VAL | p.Val205Ile | Q15833 |
| HOMO SAPIENS | 0 | 5000 | 4cca | A | 209 | LEU | p.Leu209Pro | Q15833 |
| HOMO SAPIENS | 1 | 5000 | 4cca | A | 459 | ARG | p.Arg459Trp | Q15833 |
| HOMO SAPIENS | 7 | 5000 | 4cca | A | 477 | PRO | p.Pro477Leu | Q15833 |
| HOMO SAPIENS | -1 | 5000 | 4cca | A | 526 | ILE | p.Ile526Val | Q15833 |
| HOMO SAPIENS | 0 | 5000 | 4cca | A | 541 | GLY | p.Gly541Ser | Q15833 |
| HOMO SAPIENS | 0 | 5000 | 4cca | A | 555 | ARG | p.Arg555Gly | Q15833 |
| HOMO SAPIENS | 5 | 28551 | 3vcm | P | 26 | ARG | p.Arg49Ter | P00797 |
| HOMO SAPIENS | 6 | 36453 | 5dzz | A | 2070 | ASP | p.Asp2070His | P15924 |
| HOMO SAPIENS | 0 | 36453 | 5dzz | A | 2166 | ARG | p.Arg2166Ter | P15924 |
| HOMO SAPIENS | 0 | 20268 | 2da6 | A | 96 | ASN | p.Asn321Ser | P35680 |
| HOMO SAPIENS | 1 | 1405 | 3iar | A | 8 | ASP | p.Asp8Asn | P00813 |
| HOMO SAPIENS | 8 | 1405 | 3iar | A | 15 | HIS | p.His15Asp | P00813 |
| HOMO SAPIENS | 6 | 1405 | 3iar | A | 74 | GLY | p.Gly74Val | P00813 |
| HOMO SAPIENS | 6 | 1405 | 3iar | A | 80 | LYS | p.Lys80Arg | P00813 |
| HOMO SAPIENS | 5 | 1405 | 3iar | A | 97 | TYR | p.Tyr97Cys | P00813 |
| HOMO SAPIENS | 5 | 1405 | 3iar | A | 101 | ARG | p.Arg101Leu | P00813 |
| HOMO SAPIENS | 5 | 1405 | 3iar | A | 101 | ARG | p.Arg101Trp | P00813 |
| HOMO SAPIENS | 5 | 1405 | 3iar | A | 106 | LEU | p.Leu106Val | P00813 |
| HOMO SAPIENS | 0 | 1405 | 3iar | A | 107 | LEU | p.Leu107Pro | P00813 |
| HOMO SAPIENS | 9 | 1405 | 3iar | A | 156 | ARG | p.Arg156Cys | P00813 |

| | | | | | | | | |
|---|---|---|---|---|---|---|---|---|
| HOMO SAPIENS | 6 | 1405 | 3iar | A | 211 | ARG | p.Arg211His | P00813 |
| HOMO SAPIENS | 0 | 1405 | 3iar | A | 216 | GLY | p.Gly216Arg | P00813 |
| HOMO SAPIENS | 0 | 1405 | 3iar | A | 291 | SER | p.Ser291Leu | P00813 |
| HOMO SAPIENS | 6 | 1405 | 3iar | A | 304 | LEU | p.Leu304Arg | P00813 |
| HOMO SAPIENS | 0 | 1405 | 3iar | A | 329 | ALA | p.Ala329Val | P00813 |
| HOMO SAPIENS | -1 | 31340 | 4atu | I | 186 | ARG | p.Arg186His | O43602 |
| HOMO SAPIENS | -1 | 31340 | 4atu | I | 192 | ARG | p.Arg192Trp | O43602 |
| HOMO SAPIENS | -1 | 31340 | 4atu | I | 196 | ARG | p.Arg196His | O43602 |
| HOMO SAPIENS | -1 | 31340 | 4atu | I | 203 | THR | p.Thr203Arg | O43602 |
| HOMO SAPIENS | -1 | 31340 | 4atu | I | 286 | ALA | p.Ala286Ser | O43602 |
| HOMO SAPIENS | 4 | 30734 | 1fsu | A | 95 | ARG | p.Arg95Gln | P15848 |
| HOMO SAPIENS | 0 | 30734 | 1fsu | A | 117 | CYS | p.Cys117Arg | P15848 |
| HOMO SAPIENS | 0 | 30734 | 1fsu | A | 137 | GLY | p.Gly137Val | P15848 |
| HOMO SAPIENS | 0 | 30734 | 1fsu | A | 210 | TYR | p.Tyr210Cys | P15848 |
| HOMO SAPIENS | 0 | 30734 | 1fsu | A | 236 | LEU | p.Leu236Pro | P15848 |
| HOMO SAPIENS | 1 | 30734 | 1fsu | A | 315 | ARG | p.Arg315Gln | P15848 |
| HOMO SAPIENS | 0 | 30734 | 1fsu | A | 358 | VAL | p.Val358Met | P15848 |
| HOMO SAPIENS | 2 | 30734 | 1fsu | A | 376 | VAL | p.Val376Met | P15848 |
| HOMO SAPIENS | 0 | 30734 | 1fsu | A | 384 | SER | p.Ser384Asn | P15848 |
| HOMO SAPIENS | 0 | 30734 | 1fsu | A | 393 | HIS | p.His393Pro | P15848 |
| HOMO SAPIENS | 0 | 30734 | 1fsu | A | 405 | CYS | p.Cys405Tyr | P15848 |
| HOMO SAPIENS | 0 | 30734 | 1fsu | A | 458 | ASN | p.Asn458Ser | P15848 |
| HOMO SAPIENS | 1 | 30734 | 1fsu | A | 484 | ARG | p.Arg484Gly | P15848 |
| HOMO SAPIENS | 0 | 38072 | 1wmv | A | 51 | PRO | p.Pro98Leu | Q9NZC7 |
| HOMO SAPIENS | -1 | 5674 | 1j1e | C | 185 | ASN | p.Asn185Lys | P19429 |
| HOMO SAPIENS | -1 | 5674 | 1j1e | C | 192 | ARG | p.Arg192His | P19429 |
| HOMO SAPIENS | -1 | 5674 | 1j1e | C | 196 | ASP | p.Asp196Asn | P19429 |
| HOMO SAPIENS | -1 | 5674 | 1j1e | C | 206 | LYS | p.Lys206Gln | P19429 |
| HOMO SAPIENS | 0 | 38067 | 1wlp | A | 13 | PRO | p.Pro156Gln | P13498 |
| HOMO SAPIENS | -1 | 20441 | 3vfd | A | 289 | HIS | p.His289Tyr | Q9UBP0 |
| HOMO SAPIENS | 2 | 20441 | 3vfd | A | 344 | ILE | p.Ile344Lys | Q9UBP0 |
| HOMO SAPIENS | 0 | 20441 | 3vfd | A | 362 | SER | p.Ser362Cys | Q9UBP0 |
| HOMO SAPIENS | 1 | 20441 | 3vfd | A | 386 | ASN | p.Asn386Ser | Q9UBP0 |
| HOMO SAPIENS | 9 | 20441 | 3vfd | A | 441 | ASP | p.Asp441Gly | Q9UBP0 |
| HOMO SAPIENS | -1 | 20441 | 3vfd | A | 448 | CYS | p.Cys448Tyr | Q9UBP0 |
| HOMO SAPIENS | 2 | 20441 | 3vfd | A | 500 | ARG | p.Arg499Cys | Q9UBP0 |
| HOMO SAPIENS | 1 | 20441 | 3vfd | A | 563 | ARG | p.Arg562Gly | Q9UBP0 |
| HOMO SAPIENS | 3 | 18471 | 4dx8 | J | 26 | ARG | p.Arg26Gln | O00522 |
| HOMO SAPIENS | -1 | 18471 | 4dx8 | J | 101 | GLY | p.Gly101Val | O00522 |
| HOMO SAPIENS | -1 | 34735 | 2ly4 | B | 83 | ALA | p.Ala83Glu | P04637 |
| HOMO SAPIENS | 0 | 31128 | 4cvh | A | 116 | ARG | p.Arg116Cys | A4D126 |
| HOMO SAPIENS | 8 | 31128 | 4cvh | A | 215 | GLN | p.Gln215Ter | A4D126 |
| HOMO SAPIENS | 0 | 31128 | 4cvh | A | 268 | ARG | p.Arg268Ter | A4D126 |
| HOMO SAPIENS | 0 | 31128 | 4cvh | A | 316 | THR | p.Thr316Lys | A4D126 |
| HOMO SAPIENS | 1 | 7164 | 3mt5 | A | 369 | ASP | p.Asp434Gly | Q12791 |
| HOMO SAPIENS | 2 | 26812 | 2wtk | F | 49 | TYR | p.Tyr49Asp | Q15831 |
| HOMO SAPIENS | 0 | 26812 | 2wtk | F | 67 | LEU | p.Leu67Pro | Q15831 |
| HOMO SAPIENS | 4 | 26812 | 2wtk | F | 104 | ARG | p.Arg104Ser | Q15831 |
| HOMO SAPIENS | 0 | 26812 | 2wtk | F | 135 | GLY | p.Gly135Arg | Q15831 |
| HOMO SAPIENS | 0 | 26812 | 2wtk | F | 163 | GLY | p.Gly163Asp | Q15831 |
| HOMO SAPIENS | 2 | 26812 | 2wtk | F | 194 | ALA | p.Asp194Asn | Q15831 |
| HOMO SAPIENS | 2 | 1357 | 5xtd | W | 57 | ARG | p.Arg57His | Q9P0J0 |
| HOMO SAPIENS | 0 | 28086 | 4nn2 | A | 229 | HIS | p.His229Arg | Q8IWS0 |
| HOMO SAPIENS | 0 | 28086 | 4nn2 | A | 234 | LYS | p.Lys234Glu | Q8IWS0 |

| | | | | | | | | |
|---|---|---|---|---|---|---|---|---|
| HOMO SAPIENS | 0 | 28086 | 4nn2 | A | 257 | ARG | p.Arg257Gly | Q8IWS0 |
| HOMO SAPIENS | 5 | 31182 | 2a1x | A | 176 | GLN | p.Gln176Lys | O14832 |
| HOMO SAPIENS | 1 | 31182 | 2a1x | A | 204 | GLY | p.Gly204Ser | O14832 |
| HOMO SAPIENS | 0 | 31182 | 2a1x | A | 269 | ASN | p.Asn269His | O14832 |
| HOMO SAPIENS | 3 | 31182 | 2a1x | A | 275 | ARG | p.Arg275Gln | O14832 |
| HOMO SAPIENS | 3 | 31182 | 2a1x | A | 275 | ARG | p.Arg275Trp | O14832 |
| HOMO SAPIENS | 4 | 2823 | 5xtb | P | 145 | THR | p.Thr145Ile | O75489 |
| HOMO SAPIENS | 5 | 2823 | 5xtb | P | 199 | ARG | p.Arg199Trp | O75489 |
| HOMO SAPIENS | 0 | 8645 | 2h5g | A | 372 | SER | p.Ser372Tyr | P54886 |
| HOMO SAPIENS | 3 | 8645 | 2h5g | A | 715 | ASP | p.Asp715His | P54886 |
| HOMO SAPIENS | 0 | 8645 | 2h5g | A | 765 | ARG | p.Arg765Gln | P54886 |
| HOMO SAPIENS | 0 | 8645 | 2h5g | A | 782 | TYR | p.Tyr782Cys | P54886 |
| HOMO SAPIENS | 0 | 8645 | 2h5g | A | 784 | HIS | p.His784Tyr | P54886 |
| HOMO SAPIENS | 1 | 13992 | 2cue | A | 39 | ARG | p.Arg242Thr | P26367 |
| HOMO SAPIENS | 9 | 13992 | 2cue | A | 55 | PHE | p.Phe258Ser | P26367 |
| HOMO SAPIENS | 0 | 30418 | 2xsw | A | 303 | VAL | p.Val303Met | Q9NRR6 |
| HOMO SAPIENS | 0 | 30418 | 2xsw | A | 315 | PRO | p.Pro315Leu | Q9NRR6 |
| HOMO SAPIENS | 0 | 30418 | 2xsw | A | 341 | GLY | p.Gly341Ser | Q9NRR6 |
| HOMO SAPIENS | 0 | 30418 | 2xsw | A | 378 | ARG | p.Arg378Cys | Q9NRR6 |
| HOMO SAPIENS | 0 | 30418 | 2xsw | A | 435 | ARG | p.Arg435Gln | Q9NRR6 |
| HOMO SAPIENS | 0 | 30418 | 2xsw | A | 515 | ARG | p.Arg515Trp | Q9NRR6 |
| HOMO SAPIENS | 1 | 30418 | 2xsw | A | 526 | PRO | p.Pro526Leu | Q9NRR6 |
| HOMO SAPIENS | 0 | 30418 | 2xsw | A | 563 | ARG | p.Arg563His | Q9NRR6 |
| HOMO SAPIENS | 2 | 30418 | 2xsw | A | 585 | ARG | p.Arg585His | Q9NRR6 |
| HOMO SAPIENS | 2 | 11167 | 4y99 | B | 205 | ARG | p.Arg215Leu | P45379 |
| HOMO SAPIENS | 5 | 11167 | 4y99 | B | 244 | GLU | p.Glu254Asp | P45379 |
| HOMO SAPIENS | 1 | 11167 | 4y99 | B | 253 | LYS | p.Lys263Arg | P45379 |
| HOMO SAPIENS | 8 | 11167 | 4y99 | B | 265 | ARG | p.Arg275Gln | P45379 |
| HOMO SAPIENS | 3 | 11167 | 4y99 | B | 270 | ASP | p.Asp280Asn | P45379 |
| HOMO SAPIENS | -1 | 11167 | 4y99 | B | 278 | ARG | p.Arg288Pro | P45379 |
| HOMO SAPIENS | -1 | 11167 | 4y99 | B | 286 | ARG | p.Arg296Cys | P45379 |
| HOMO SAPIENS | -1 | 11167 | 4y99 | B | 286 | ARG | p.Arg296His | P45379 |
| HOMO SAPIENS | 0 | 15993 | 4po6 | A | 197 | ARG | p.Arg197His | P29597 |
| HOMO SAPIENS | -1 | 15993 | 4po6 | A | 362 | VAL | p.Val362Phe | P29597 |
| HOMO SAPIENS | -1 | 15993 | 4po6 | A | 363 | GLY | p.Gly363Ser | P29597 |
| HOMO SAPIENS | 0 | 5646 | 5uak | A | 7 | GLU | p.Glu7Ter | P13569 |
| HOMO SAPIENS | 0 | 5646 | 5uak | A | 27 | GLY | p.Gly27Ter | P13569 |
| HOMO SAPIENS | 0 | 5646 | 5uak | A | 46 | ALA | p.Ala46Asp | P13569 |
| HOMO SAPIENS | 0 | 5646 | 5uak | A | 60 | GLU | p.Glu60Ter | P13569 |
| HOMO SAPIENS | 0 | 5646 | 5uak | A | 67 | PRO | p.Pro67Leu | P13569 |
| HOMO SAPIENS | 0 | 5646 | 5uak | A | 74 | ARG | p.Arg74Trp | P13569 |
| HOMO SAPIENS | 0 | 5646 | 5uak | A | 75 | ARG | p.Arg75Ter | P13569 |
| HOMO SAPIENS | 0 | 5646 | 5uak | A | 85 | GLY | p.Gly85Glu | P13569 |
| HOMO SAPIENS | 0 | 5646 | 5uak | A | 91 | GLY | p.Gly91Arg | P13569 |
| HOMO SAPIENS | 0 | 5646 | 5uak | A | 92 | GLU | p.Glu92Lys | P13569 |
| HOMO SAPIENS | 0 | 5646 | 5uak | A | 98 | GLN | p.Gln98Arg | P13569 |
| HOMO SAPIENS | 0 | 5646 | 5uak | A | 98 | GLN | p.Gln98Ter | P13569 |
| HOMO SAPIENS | 0 | 5646 | 5uak | A | 109 | TYR | p.Tyr109Cys | P13569 |
| HOMO SAPIENS | 0 | 5646 | 5uak | A | 110 | ASP | p.Asp110His | P13569 |
| HOMO SAPIENS | 0 | 5646 | 5uak | A | 117 | ARG | p.Arg117Cys | P13569 |
| HOMO SAPIENS | 0 | 5646 | 5uak | A | 117 | ARG | p.Arg117His | P13569 |
| HOMO SAPIENS | 0 | 5646 | 5uak | A | 125 | ILE | p.Ile125Thr | P13569 |
| HOMO SAPIENS | 0 | 5646 | 5uak | A | 148 | ILE | p.Ile148Thr | P13569 |
| HOMO SAPIENS | 0 | 5646 | 5uak | A | 170 | ARG | p.Arg170His | P13569 |

| | | | | | | | | |
|---|---|---|---|---|---|---|---|---|
| HOMO SAPIENS | 0 | 5646 | 5uak | A | 178 | GLY | p.Gly178Arg | P13569 |
| HOMO SAPIENS | 0 | 5646 | 5uak | A | 206 | LEU | p.Leu206Trp | P13569 |
| HOMO SAPIENS | 0 | 5646 | 5uak | A | 373 | ASP | p.Asp373Asn | P13569 |
| HOMO SAPIENS | -1 | 5646 | 5uak | A | 709 | ARG | p.Arg709Ter | P13569 |
| HOMO SAPIENS | -1 | 5646 | 5uak | A | 732 | LEU | p.Leu732Ter | P13569 |
| HOMO SAPIENS | -1 | 5646 | 5uak | A | 785 | ARG | p.Arg785Ter | P13569 |
| HOMO SAPIENS | -1 | 5646 | 5uak | A | 831 | GLU | p.Glu831Ter | P13569 |
| HOMO SAPIENS | -1 | 5646 | 5uak | A | 836 | ASP | p.Asp836Tyr | P13569 |
| HOMO SAPIENS | 0 | 5646 | 5uak | A | 851 | ARG | p.Arg851Ter | P13569 |
| HOMO SAPIENS | 0 | 5646 | 5uak | A | 913 | TYR | p.Tyr913Cys | P13569 |
| HOMO SAPIENS | 0 | 5646 | 5uak | A | 913 | TYR | p.Tyr913Ter | P13569 |
| HOMO SAPIENS | 0 | 5646 | 5uak | A | 945 | SER | p.Ser945Leu | P13569 |
| HOMO SAPIENS | 0 | 5646 | 5uak | A | 949 | HIS | p.His949Tyr | P13569 |
| HOMO SAPIENS | 0 | 5646 | 5uak | A | 970 | GLY | p.Gly970Asp | P13569 |
| HOMO SAPIENS | 0 | 5646 | 5uak | A | 1027 | ILE | p.Ile1027Thr | P13569 |
| HOMO SAPIENS | 0 | 5646 | 5uak | A | 1032 | TYR | p.Tyr1032Cys | P13569 |
| HOMO SAPIENS | 0 | 5646 | 5uak | A | 1061 | GLY | p.Gly1061Arg | P13569 |
| HOMO SAPIENS | 1 | 5646 | 5uak | A | 1065 | LEU | p.Leu1065Pro | P13569 |
| HOMO SAPIENS | 1 | 5646 | 5uak | A | 1066 | ARG | p.Arg1066Cys | P13569 |
| HOMO SAPIENS | 1 | 5646 | 5uak | A | 1066 | ARG | p.Arg1066His | P13569 |
| HOMO SAPIENS | 1 | 5646 | 5uak | A | 1067 | ALA | p.Ala1067Thr | P13569 |
| HOMO SAPIENS | 0 | 5646 | 5uak | A | 1069 | GLY | p.Gly1069Arg | P13569 |
| HOMO SAPIENS | 0 | 5646 | 5uak | A | 1070 | ARG | p.Arg1070Trp | P13569 |
| HOMO SAPIENS | 1 | 5646 | 5uak | A | 1071 | GLN | p.Gln1071Pro | P13569 |
| HOMO SAPIENS | 0 | 5646 | 5uak | A | 1074 | PHE | p.Phe1074Leu | P13569 |
| HOMO SAPIENS | 0 | 5646 | 5uak | A | 1077 | LEU | p.Leu1077Pro | P13569 |
| HOMO SAPIENS | 0 | 5646 | 5uak | A | 1089 | TRP | p.Trp1089Ter | P13569 |
| HOMO SAPIENS | 0 | 5646 | 5uak | A | 1092 | TYR | p.Tyr1092Ter | P13569 |
| HOMO SAPIENS | 0 | 5646 | 5uak | A | 1101 | MET | p.Met1101Lys | P13569 |
| HOMO SAPIENS | 0 | 5646 | 5uak | A | 1156 | LEU | p.Leu1156Phe | P13569 |
| HOMO SAPIENS | 0 | 5646 | 5uak | A | 1158 | ARG | p.Arg1158Ter | P13569 |
| HOMO SAPIENS | 0 | 5646 | 5uak | A | 1162 | ARG | p.Arg1162Leu | P13569 |
| HOMO SAPIENS | 0 | 5646 | 5uak | A | 1162 | ARG | p.Arg1162Ter | P13569 |
| HOMO SAPIENS | -1 | 5646 | 5uak | A | 1476 | GLN | p.Gln1476Ter | P13569 |
| HOMO SAPIENS | 0 | 22763 | 1imv | A | 52 | MET | p.Thr72Met | P36955 |
| HOMO SAPIENS | 0 | 22763 | 1imv | A | 301 | TYR | p.Tyr321Ter | P36955 |
| HOMO SAPIENS | 0 | 23179 | 5fql | A | 160 | PRO | p.Pro160Arg | P22304 |
| HOMO SAPIENS | 0 | 23179 | 5fql | A | 172 | ARG | p.Arg172Ter | P22304 |
| HOMO SAPIENS | 0 | 23179 | 5fql | A | 214 | THR | p.Thr214Met | P22304 |
| HOMO SAPIENS | 0 | 23179 | 5fql | A | 309 | THR | p.Thr309Ala | P22304 |
| HOMO SAPIENS | 0 | 23179 | 5fql | A | 333 | SER | p.Ser333Leu | P22304 |
| HOMO SAPIENS | 0 | 23179 | 5fql | A | 443 | ARG | p.Arg443Ter | P22304 |
| HOMO SAPIENS | 0 | 23179 | 5fql | A | 468 | ARG | p.Arg468Leu | P22304 |
| HOMO SAPIENS | 0 | 23179 | 5fql | A | 468 | ARG | p.Arg468Trp | P22304 |
| HOMO SAPIENS | 0 | 23179 | 5fql | A | 502 | TRP | p.Trp502Ser | P22304 |
| HOMO SAPIENS | 9 | 20989 | 1nfi | F | 185 | THR | p.Thr185Met | P25963 |
| HOMO SAPIENS | 4 | 20989 | 1nfi | F | 194 | GLY | p.Gly194Ala | P25963 |
| HOMO SAPIENS | 0 | 11538 | 3eab | A | 162 | VAL | p.Val162Ile | Q9UBP0 |
| HOMO SAPIENS | 0 | 33399 | 2p39 | A | 71 | SER | p.Ser71Gly | Q9GZV9 |
| HOMO SAPIENS | 3 | 33399 | 2p39 | A | 96 | MET | p.Met96Thr | Q9GZV9 |
| HOMO SAPIENS | 0 | 33399 | 2p39 | A | 129 | SER | p.Ser129Phe | Q9GZV9 |
| HOMO SAPIENS | -1 | 33399 | 2p39 | A | 172 | PRO | p.Pro172Leu | Q9GZV9 |
| HOMO SAPIENS | -1 | 33399 | 2p39 | A | 176 | ARG | p.Arg176Gln | Q9GZV9 |
| HOMO SAPIENS | -1 | 33399 | 2p39 | A | 179 | ARG | p.Arg179Trp | Q9GZV9 |

| | | | | | | | | |
|---|---|---|---|---|---|---|---|---|
| HOMO SAPIENS | -1 | 33399 | 2p39 | A | 179 | ARG | p.Arg179Gln | Q9GZV9 |
| HOMO SAPIENS | 0 | 16217 | 2dne | A | 79 | ILE | p.Ile163Leu | P10515 |
| HOMO SAPIENS | 8 | 7666 | 4kbq | A | 65 | ASN | p.Asn65Ser | Q9UNE7 |
| HOMO SAPIENS | 1 | 6407 | 1ikt | A | 71 | THR | p.Thr687Ile | P51659 |
| HOMO SAPIENS | 0 | 6407 | 1ikt | A | 112 | MET | p.Met728Val | P51659 |
| HOMO SAPIENS | 0 | 5849 | 4ihb | A | 67 | VAL | p.Val67Asp | O75923 |
| HOMO SAPIENS | 6 | 17703 | 5fcn | A | 20 | HIS | p.His99Gln | Q66GS9 |
| HOMO SAPIENS | 8 | 17703 | 5fcn | A | 33 | ARG | p.Arg112His | Q66GS9 |
| HOMO SAPIENS | 0 | 34095 | 2E+073 | A | 101 | HIS | p.His101Tyr | P05129 |
| HOMO SAPIENS | -1 | 41358 | 4liq | E | 507 | THR | p.Thr507Met | P07333 |
| HOMO SAPIENS | -1 | 21290 | 1fch | A | 235 | MET | p.Met272Arg | P50542 |
| HOMO SAPIENS | 5 | 23431 | 6pax | A | 23 | ARG | p.Arg26Gly | P26367 |
| HOMO SAPIENS | 1 | 23431 | 6pax | A | 61 | GLY | p.Gly64Val | P26367 |
| HOMO SAPIENS | 1 | 23431 | 6pax | A | 65 | PRO | p.Pro68Ser | P26367 |
| HOMO SAPIENS | 1 | 23431 | 6pax | A | 116 | SER | p.Ser119Arg | P26367 |
| HOMO SAPIENS | 9 | 23431 | 6pax | A | 123 | VAL | p.Val126Asp | P26367 |
| HOMO SAPIENS | 9 | 23431 | 6pax | A | 125 | ARG | p.Arg128Cys | P26367 |
| HOMO SAPIENS | 9 | 33003 | 4zrj | B | 535 | LEU | p.Leu535Pro | P35240 |
| HOMO SAPIENS | 7 | 33003 | 4zrj | B | 538 | GLN | p.Gln538Pro | P35240 |
| HOMO SAPIENS | 0 | 33003 | 4zrj | B | 585 | TRP | p.Ala585Thr | P35240 |
| HOMO SAPIENS | 0 | 3800 | 2nzj | A | 193 | VAL | p.Val193Met | O75628 |

| gene_name | max_cons | max_bs_id | max_pdb_id | max_chain_id |
|---|---|---|---|---|
| AKR1D1 | None | None | None | None |
| AKR1D1 | None | None | None | None |
| AKR1D1 | None | None | None | None |
| AKR1D1 | None | None | None | None |
| ALPP | None | None | None | None |
| BMPR1A | None | None | None | None |
| BMPR1A | None | None | None | None |
| FGFR2 | None | None | None | None |
| PROC | 0.587 | 3 | 1aut | C |
| PROC | None | None | None | None |
| PROC | None | None | None | None |
| PROC | None | None | None | None |
| PROC | None | None | None | None |
| PROC | None | None | None | None |
| PROC | None | None | None | None |
| PROC | None | None | None | None |
| PROC | None | None | None | None |
| PROC | 0.587 | 3 | 1aut | C |
| PROC | None | None | None | None |
| PROC | None | None | None | None |
| INS | 1 | 2 | 3zs2 | J |
| INS | 0.85 | 1 | 4ajx | D |
| INS | 1 | 1 | 3zs2 | J |
| INS | 1 | 1 | 3zs2 | J |
| FTL | 0.627 | 11 | 2fg8 | B |
| FTL | None | None | None | None |
| FTL | 0.544 | 24 | 2fg8 | D |
| TNFSF11 | None | None | None | None |
| EHMT1 | None | None | None | None |
| EHMT1 | None | None | None | None |
| EHMT1 | None | None | None | None |
| CA2 | 0.939 | 1 | 4idr | X |
| CA2 | 0.939 | 1 | 4idr | X |
| CA2 | 0.902 | 5 | 4idr | X |
| CA2 | 0.902 | 5 | 4idr | X |
| SMAD4 | None | None | None | None |
| SMAD4 | None | None | None | None |
| TTR | 0.691 | 12 | 1ttb | B |
| TTR | 0.961 | 7 | 3grb | A |
| TTR | 0.891 | 4 | 3dk2 | B |
| TTR | 0.947 | 3 | 1tta | A |
| TTR | 0.923 | 1 | 3grb | B |
| TTR | 0.891 | 1 | 4y9b | B |
| TTR | 0.891 | 1 | 4y9b | B |
| TTR | 0.882 | 15 | 3grb | A |
| TTR | 0.909 | 5 | 4tl4 | A |
| TTR | 0.836 | 7 | 1z7j | B |
| TTR | 0.882 | 15 | 3grb | A |
| TTR | 0.891 | 5 | 3i9i | B |
| TTR | 0.902 | 9 | 3m1o | B |
| TTR | 0.902 | 9 | 3m1o | B |
| TTR | 0.902 | 9 | 3m1o | B |
| TTR | 0.636 | 16 | 1thc | A |

| | | | | |
|---|---|---|---|---|
| TTR | 0.902 | 9 | 3m1o | B |
| TTR | 0.836 | 7 | 1bze | A |
| TTR | 0.846 | 16 | 3gps | A |
| TTR | 0.846 | 16 | 3gps | A |
| TTR | 0.909 | 5 | 4tl4 | A |
| TTR | 0.909 | 5 | 4tl4 | A |
| TTR | 0.673 | 15 | 3dk2 | B |
| TTR | 0.945 | 4 | 4deu | A |
| TTR | 0.941 | 3 | 3did | B |
| TTR | 0.941 | 3 | 3did | B |
| TTR | 0.868 | 6 | 4l1t | A |
| TTR | 0.882 | 19 | 3gps | B |
| TTR | 0.891 | 4 | 3dk2 | B |
| TTR | 0.941 | 3 | 3did | B |
| TTR | 0.923 | 1 | 4hiq | A |
| TTR | 0.923 | 1 | 4hiq | A |
| TTR | 0.961 | 7 | 3grb | A |
| TTR | 0.839 | 7 | 1tsh | B |
| TTR | 0.891 | 6 | 1ttb | B |
| ALK | 0.682 | 1 | 4cd0 | A |
| ALK | None | None | None | None |
| THRB | None | None | None | None |
| THRB | None | None | None | None |
| THRB | None | None | None | None |
| THRB | None | None | None | None |
| THRB | None | None | None | None |
| THRB | None | None | None | None |
| THRB | None | None | None | None |
| THRB | None | None | None | None |
| THRB | None | None | None | None |
| THRB | None | None | None | None |
| THRB | None | None | None | None |
| THRB | None | None | None | None |
| THRB | None | None | None | None |
| THRB | None | None | None | None |
| THRB | None | None | None | None |
| THRB | None | None | None | None |
| THRB | None | None | None | None |
| THRB | None | None | None | None |
| THRB | 0.577 | 3 | 2j4a | A |
| IL2RG | None | None | None | None |
| IL2RG | None | None | None | None |
| IL2RG | None | None | None | None |
| IL2RG | None | None | None | None |
| IL2RG | None | None | None | None |
| ITGB3 | None | None | None | None |
| ITGB3 | None | None | None | None |
| ITGB3 | None | None | None | None |
| ITGB3 | None | None | None | None |
| ITGB3 | None | None | None | None |
| ITGB3 | None | None | None | None |
| ITGB3 | None | None | None | None |
| ITGB3 | None | None | None | None |
| ITGB3 | None | None | None | None |

| | | | | | |
|---|---|---|---|---|---|
| PKD2 | None | None | | None | None |
| PKD2 | None | None | | None | None |
| PKD2 | None | None | | None | None |
| PKD2 | None | None | | None | None |
| PKD2 | None | None | | None | None |
| PKD2 | None | None | | None | None |
| PKD2 | None | None | | None | None |
| TP53 | 0.571 | | 1 | 1saf | B |
| TP53 | 0.571 | | 1 | 1saf | B |
| TP53 | None | None | | None | None |
| TP53 | 0.585 | | 15 | 4mzr | B |
| TP53 | None | None | | None | None |
| TP53 | None | None | | None | None |
| EGFR | None | None | | None | None |
| EGFR | None | None | | None | None |
| XRCC4 | None | None | | None | None |
| XRCC4 | None | None | | None | None |
| HBB | None | None | | None | None |
| HBB | None | None | | None | None |
| HBB | None | None | | None | None |
| HBB | None | None | | None | None |
| HBB | None | None | | None | None |
| HBB | None | None | | None | None |
| HBB | None | None | | None | None |
| HBB | None | None | | None | None |
| HBB | None | None | | None | None |
| HBB | None | None | | None | None |
| HBB | 0.535 | | 3 | 3r5i | B |
| HBB | None | None | | None | None |
| HBB | None | None | | None | None |
| HBB | None | None | | None | None |
| HBB | None | None | | None | None |
| HBB | None | None | | None | None |
| HBB | 0.566 | | 1 | 1cbm | B |
| HBB | 0.51 | | 3 | 1abw | B |
| HBB | None | None | | None | None |
| HBB | None | None | | None | None |
| HBB | None | None | | None | None |
| HBB | None | None | | None | None |
| HBB | 0.508 | | 7 | 3ic2 | B |
| HBB | 0.545 | | 3 | 1bij | D |
| HBB | 0.6 | | 1 | 2w72 | D |
| HBB | None | None | | None | None |
| HBB | None | None | | None | None |
| HBB | None | None | | None | None |
| HBB | 0.733 | | 1 | 1yff | D |
| HBB | 0.692 | | 2 | 1buw | B |
| RPS6KA3 | None | None | | None | None |
| RPS6KA3 | 0.555 | | 1 | 4nw5 | A |
| RPS6KA3 | None | None | | None | None |
| RPS6KA3 | None | None | | None | None |
| RPS6KA3 | None | None | | None | None |
| RPS6KA3 | None | None | | None | None |
| GBA | 1 | | 15 | 3gxd | B |

| | | | | | |
|---|---|---|---|---|---|
| GBA | | 1 | 55 | 3gxd | B |
| GBA | | 1 | 15 | 3gxd | B |
| GBA | | 0.75 | 45 | 3ril | A |
| GBA | None | None | None | | None |
| GBA | None | None | None | | None |
| GBA | | 1 | 12 | 1ogs | A |
| GBA | | 0.75 | 122 | 3gxi | A |
| GBA | | 1 | 2 | 1ogs | A |
| GBA | | 1 | 11 | 1ogs | A |
| GBA | | 0.5 | 59 | 3gxf | B |
| GBA | | 0.75 | 45 | 3ril | A |
| GBA | | 1 | 18 | 1ogs | A |
| GBA | | 1 | 3 | 1ogs | A |
| GBA | None | None | None | | None |
| GBA | | 1 | 2 | 1ogs | A |
| GBA | None | None | None | | None |
| GBA | | 0.75 | 76 | 3gxm | C |
| GBA | | 1 | 3 | 1ogs | A |
| GBA | | 1 | 3 | 1ogs | A |
| GBA | | 0.75 | 62 | 3ril | A |
| GBA | | 1 | 28 | 1ogs | A |
| GBA | None | None | None | | None |
| GBA | | 0.5 | 78 | 3gxi | A |
| GBA | | 0.5 | 92 | 3rik | B |
| GBA | | 0.5 | 42 | 2xwe | A |
| GBA | None | None | None | | None |
| GBA | | 0.5 | 92 | 3rik | B |
| GBA | | 0.75 | 49 | 3ril | A |
| GBA | None | None | None | | None |
| GBA | | 0.667 | 60 | 1ogs | A |
| GBA | None | None | None | | None |
| GBA | | 0.5 | 11 | 2nt1 | A |
| GBA | | 0.75 | 21 | 3gxi | A |
| GBA | None | None | None | | None |
| GBA | | 0.5 | 35 | 2xwe | A |
| GBA | | 0.75 | 25 | 3keh | B |
| GBA | | 0.75 | 70 | 3ril | A |
| GBA | | 0.75 | 70 | 3ril | A |
| GBA | | 1 | 15 | 3gxd | B |
| GBA | | 1 | 44 | 3gxd | B |
| GBA | | 0.5 | 57 | 3gxf | D |
| GBA | | 1 | 44 | 3gxd | B |
| GBA | | 1 | 44 | 3gxd | B |
| GBA | None | None | None | | None |
| GBA | | 0.75 | 94 | 3gxi | A |
| GBA | | 0.75 | 94 | 3gxi | A |
| IFIH1 | None | None | None | | None |
| TGFBR2 | None | None | None | | None |
| TGFBR2 | None | None | None | | None |
| TGFBR2 | None | None | None | | None |
| TGFBR2 | None | None | None | | None |
| TGFBR2 | None | None | None | | None |
| TGFBR2 | None | None | None | | None |
| TGFBR2 | None | None | None | | None |

| | | | | |
|---|---|---|---|---|
| TGFBR2 | None | None | None | None |
| TGFBR2 | None | None | None | None |
| TGFBR2 | 0.742 | 1 | 5E+091 | A |
| TGFBR2 | None | None | None | None |
| TGFBR2 | None | None | None | None |
| TGFBR2 | None | None | None | None |
| TGFBR2 | 0.676 | 1 | 5e8v | A |
| TGFBR2 | 0.676 | 1 | 5e8v | A |
| ACE | 0.684 | 17 | 3nxq | A |
| ACE | None | None | None | None |
| ACE | None | None | None | None |
| ACE | None | None | None | None |
| ACE | None | None | None | None |
| ACE | None | None | None | None |
| COL10A1 | None | None | None | None |
| COL10A1 | None | None | None | None |
| COL10A1 | None | None | None | None |
| COL10A1 | None | None | None | None |
| COL10A1 | None | None | None | None |
| COL10A1 | None | None | None | None |
| COL10A1 | None | None | None | None |
| COL10A1 | None | None | None | None |
| COL10A1 | None | None | None | None |
| KLF1 | None | None | None | None |
| C3 | None | None | None | None |
| CTSC | None | None | None | None |
| CTSC | 0.934 | 1 | 2djg | C |
| CTSC | None | None | None | None |
| LYZ | 0.877 | 3 | 1bb5 | A |
| LYZ | 0.923 | 1 | 1lz4 | A |
| LYZ | 0.852 | 2 | 1ubz | A |
| DHODH | None | None | None | None |
| DHODH | None | None | None | None |
| DHODH | None | None | None | None |
| DHODH | None | None | None | None |
| DHODH | None | None | None | None |
| DHODH | None | None | None | None |
| DHODH | None | None | None | None |
| ADH1C | 0.527 | 11 | 1ht0 | A |
| GDNF | None | None | None | None |
| TNF | 0.5 | 8 | 2e7a | B |
| ALDH4A1 | 0.636 | 3 | 3v9h | A |
| ALDH4A1 | None | None | None | None |
| ALDH4A1 | None | None | None | None |
| BRAF | None | None | None | None |
| BRAF | None | None | None | None |
| BRAF | 0.752 | 2 | 3psb | A |
| BRAF | 0.752 | 2 | 3psb | A |
| BRAF | 0.737 | 1 | 3pri | A |
| BRAF | 0.737 | 1 | 3pri | A |
| BRAF | 0.737 | 1 | 3pri | A |
| BRAF | 0.737 | 1 | 3prf | A |
| BRAF | 0.737 | 1 | 3prf | A |
| BRAF | 0.752 | 2 | 3psb | A |

| | | | | |
|---|---|---|---|---|
| BRAF | 0.752 | 2 | 3psb | A |
| BRAF | 0.752 | 2 | 3psb | A |
| BRAF | 0.752 | 2 | 3psb | A |
| BRAF | 0.555 | 3 | 5vam | A |
| BRAF | 0.663 | 1 | 4ehe | B |
| BRAF | 0.734 | 1 | 5csx | A |
| TBX5 | None | None | None | None |
| TBX5 | None | None | None | None |
| TBX5 | None | None | None | None |
| TBX5 | None | None | None | None |
| TBX5 | None | None | None | None |
| TBX5 | None | None | None | None |
| TBX5 | None | None | None | None |
| TBX5 | None | None | None | None |
| TBX5 | None | None | None | None |
| RB1 | None | None | None | None |
| RB1 | None | None | None | None |
| RB1 | None | None | None | None |
| RB1 | None | None | None | None |
| RB1 | None | None | None | None |
| RB1 | None | None | None | None |
| CLIC2 | 0.5 | 3 | 2per | A |
| CLIC2 | None | None | None | None |
| ALDH7A1 | None | None | None | None |
| ALDH7A1 | None | None | None | None |
| ALDH7A1 | None | None | None | None |
| ALDH7A1 | None | None | None | None |
| ALDH7A1 | None | None | None | None |
| ALDH7A1 | None | None | None | None |
| ALDH7A1 | 0.534 | 3 | 2j6l | A |
| ALDH7A1 | None | None | None | None |
| ALDH7A1 | None | None | None | None |
| ALDH7A1 | None | None | None | None |
| ALDH7A1 | None | None | None | None |
| ALDH7A1 | 0.534 | 3 | 2j6l | A |
| ALDH7A1 | None | None | None | None |
| ALDH7A1 | None | None | None | None |
| ALDH7A1 | 0.632 | 1 | 4zuk | A |
| ALDH7A1 | None | None | None | None |
| KDR | 0.575 | 5 | 1y6a | A |
| MUT | None | None | None | None |
| MUT | None | None | None | None |
| MUT | None | None | None | None |
| MUT | None | None | None | None |
| MUT | None | None | None | None |
| MUT | None | None | None | None |
| MUT | None | None | None | None |
| MUT | None | None | None | None |
| MUT | None | None | None | None |
| MUT | None | None | None | None |
| MUT | None | None | None | None |
| MUT | None | None | None | None |
| MUT | None | None | None | None |
| MUT | None | None | None | None |

| | | | | |
|---|---|---|---|---|
| MUT | None | None | None | None |
| MUT | None | None | None | None |
| MUT | 0.6 | 1 | 3bic | A |
| MUT | 0.6 | 2 | 2xiq | A |
| MUT | 0.6 | 2 | 2xiq | A |
| MUT | None | None | None | None |
| MUT | None | None | None | None |
| MUT | None | None | None | None |
| MUT | None | None | None | None |
| MUT | None | None | None | None |
| MUT | None | None | None | None |
| MUT | None | None | None | None |
| MUT | None | None | None | None |
| MUT | None | None | None | None |
| MUT | None | None | None | None |
| MUT | None | None | None | None |
| MUT | None | None | None | None |
| MUT | None | None | None | None |
| MUT | None | None | None | None |
| MUT | None | None | None | None |
| SOD1 | 0.778 | 1 | 2gbu | A |
| SOD1 | 0.727 | 4 | 4xcr | A |
| SOD1 | None | None | None | None |
| SOD1 | 0.621 | 12 | 1hl5 | H |
| NECTIN3 | None | None | None | None |
| HFE | None | None | None | None |
| HFE | None | None | None | None |
| HFE | None | None | None | None |
| HFE | 0.5 | 1 | 1de4 | A |
| HFE | None | None | None | None |
| HFE | None | None | None | None |
| HFE | None | None | None | None |
| EIF2AK3 | None | None | None | None |
| EIF2AK3 | None | None | None | None |
| TERT | None | None | None | None |
| TERT | None | None | None | None |
| SPR | None | None | None | None |
| SPR | None | None | None | None |
| SPR | None | None | None | None |
| SPR | None | None | None | None |
| SPR | None | None | None | None |
| F7 | 0.625 | 2 | 1klj | L |
| UNG | None | None | None | None |
| MMP13 | None | None | None | None |
| MMP13 | 0.786 | 1 | 3tvc | A |
| MMP13 | 0.766 | 1 | 3zxh | B |
| MMP13 | 0.766 | 1 | 3zxh | B |
| CD207 | 0.621 | 1 | 3kqg | B |
| F2 | 1 | 2 | 1nrq | L |
| SPTBN2 | None | None | None | None |
| PKLR | 0.568 | 9 | 2vgb | A |
| PKLR | None | None | None | None |
| PKLR | None | None | None | None |
| PKLR | None | None | None | None |

| | | | | |
|---|---|---|---|---|
| PKLR | 0.568 | 3 | 4ip7 | D |
| PKLR | None | None | None | None |
| PKLR | None | None | None | None |
| PKLR | None | None | None | None |
| SACS | None | None | None | None |
| HRAS | 0.529 | 3 | 6q21 | A |
| HRAS | 0.529 | 3 | 6q21 | A |
| HRAS | 0.59 | 1 | 1gnq | A |
| HRAS | 0.59 | 1 | 1gnq | A |
| HRAS | 0.664 | 1 | 2lwi | A |
| HRAS | 0.742 | 1 | 1ctq | A |
| HRAS | 0.69 | 1 | 1xd2 | A |
| HRAS | None | None | None | None |
| HRAS | 0.529 | 3 | 6q21 | A |
| HRAS | None | None | None | None |
| HRAS | 0.521 | 4 | 4nyj | R |
| HRAS | 0.664 | 1 | 2lwi | A |
| HRAS | 0.664 | 1 | 2lwi | A |
| VHL | None | None | None | None |
| VHL | None | None | None | None |
| VHL | None | None | None | None |
| VHL | None | None | None | None |
| VHL | 0.5 | 22 | 4w9l | C |
| VHL | 0.5 | 22 | 5nvy | C |
| VHL | None | None | None | None |
| VHL | None | None | None | None |
| VHL | None | None | None | None |
| VHL | 0.5 | 2 | 5nvy | C |
| VHL | 0.5 | 1 | 5nvy | C |
| VHL | 0.5 | 1 | 5nvy | C |
| VHL | 0.5 | 4 | 5nvy | C |
| VHL | None | None | None | None |
| VHL | 0.5 | 6 | 5nvy | C |
| VHL | 0.5 | 2 | 5nvy | C |
| VHL | 0.5 | 8 | 3ztc | C |
| VHL | 0.5 | 2 | 5nvy | C |
| VHL | 0.5 | 2 | 5nvy | C |
| VHL | None | None | None | None |
| VHL | 0.5 | 7 | 4w9h | C |
| VHL | 0.5 | 2 | 5nvy | C |
| VHL | 0.5 | 2 | 5nvy | C |
| VHL | 0.5 | 19 | 4w9l | C |
| VHL | None | None | None | None |
| TTBK1 | None | None | None | None |
| RAD51 | None | None | None | None |
| ALB | None | None | None | None |
| ALB | 0.5 | 2 | 4n0u | D |
| ALB | None | None | None | None |
| ALB | None | None | None | None |
| MET | None | None | None | None |
| MET | None | None | None | None |
| MET | None | None | None | None |
| MET | None | None | None | None |
| MET | None | None | None | None |

| | | | | |
|---|---|---|---|---|
| MET | None | None | None | None |
| MET | None | None | None | None |
| MET | None | None | None | None |
| MET | None | None | None | None |
| CP | None | None | None | None |
| CP | None | None | None | None |
| PGM1 | None | None | None | None |
| PGM1 | None | None | None | None |
| PGM1 | None | None | None | None |
| PGM1 | None | None | None | None |
| PGM1 | None | None | None | None |
| PGM1 | None | None | None | None |
| PGM1 | None | None | None | None |
| NR5A1 | None | None | None | None |
| NR5A1 | None | None | None | None |
| NR5A1 | None | None | None | None |
| F9 | None | None | None | None |
| F9 | None | None | None | None |
| HPRT1 | None | None | None | None |
| HPRT1 | None | None | None | None |
| HPRT1 | None | None | None | None |
| HPRT1 | None | None | None | None |
| HPRT1 | None | None | None | None |
| HPRT1 | 0.871 | 3 | 4rac | B |
| HPRT1 | 0.806 | 2 | 4ran | D |
| HPRT1 | None | None | None | None |
| HPRT1 | None | None | None | None |
| HPRT1 | None | None | None | None |
| HPRT1 | None | None | None | None |
| HPRT1 | None | None | None | None |
| HPRT1 | None | None | None | None |
| HPRT1 | 0.703 | 4 | 5w8v | A |
| HPRT1 | None | None | None | None |
| HPRT1 | None | None | None | None |
| HPRT1 | None | None | None | None |
| HPRT1 | None | None | None | None |
| HPRT1 | 0.871 | 3 | 4rac | B |
| HPRT1 | 0.871 | 3 | 4rac | B |
| HPRT1 | 0.765 | 2 | 4rao | D |
| HPRT1 | 0.774 | 2 | 4rac | B |
| GPD1L | None | None | None | None |
| GPD1L | None | None | None | None |
| GPD1L | None | None | None | None |
| FGFR2 | None | None | None | None |
| FGFR2 | None | None | None | None |
| FGFR2 | None | None | None | None |
| FGFR2 | None | None | None | None |
| FGFR2 | None | None | None | None |
| FGFR2 | None | None | None | None |
| FGFR2 | None | None | None | None |
| FGFR2 | None | None | None | None |
| FGFR2 | None | None | None | None |
| FGFR2 | None | None | None | None |
| FGFR2 | None | None | None | None |

| | | | | |
|---|---|---|---|---|
| FGFR2 | None | None | None | None |
| FGFR2 | None | None | None | None |
| FGFR2 | None | None | None | None |
| FGFR2 | None | None | None | None |
| FGFR2 | None | None | None | None |
| FGFR2 | None | None | None | None |
| FGFR2 | None | None | None | None |
| FGFR2 | None | None | None | None |
| HSPD1 | None | None | None | None |
| HSPD1 | None | None | None | None |
| FGF8 | None | None | None | None |
| FGF8 | None | None | None | None |
| FGF8 | None | None | None | None |
| F9 | None | None | None | None |
| NCF4 | None | None | None | None |
| NCF4 | None | None | None | None |
| LDLR | None | None | None | None |
| LDLR | None | None | None | None |
| LDLR | None | None | None | None |
| LDLR | None | None | None | None |
| LDLR | None | None | None | None |
| LDLR | None | None | None | None |
| C3 | None | None | None | None |
| C3 | None | None | None | None |
| C3 | None | None | None | None |
| C3 | None | None | None | None |
| C3 | None | None | None | None |
| C3 | None | None | None | None |
| C3 | None | None | None | None |
| C3 | None | None | None | None |
| TULP1 | None | None | None | None |
| TULP1 | None | None | None | None |
| TULP1 | None | None | None | None |
| TULP1 | None | None | None | None |
| IKBKG | None | None | None | None |
| IKBKG | None | None | None | None |
| IKBKG | None | None | None | None |
| CD40 | None | None | None | None |
| CD40 | None | None | None | None |
| SLC9A3R1 | None | None | None | None |
| BCHE | None | None | None | None |
| BCHE | None | None | None | None |
| BCHE | 0.544 | 11 | 1p0q | A |
| BCHE | None | None | None | None |
| BCHE | None | None | None | None |
| BCHE | None | None | None | None |
| BCHE | 0.63 | 4 | 2xqg | A |
| BCHE | None | None | None | None |
| BCHE | None | None | None | None |
| BCHE | None | None | None | None |
| BCHE | None | None | None | None |
| BCKDHB | 0.833 | 1 | 1v11 | B |
| BCKDHB | 0.5 | 18 | 2bfc | B |
| BCKDHB | None | None | None | None |

| | | | | |
|---|---|---|---|---|
| BCKDHB | 0.667 | 11 | 2bfd | B |
| BCKDHB | 0.833 | 10 | 1olx | B |
| BCKDHB | 0.833 | 1 | 1v11 | B |
| BCKDHB | 0.833 | 1 | 1v11 | B |
| BCKDHB | 0.625 | 2 | 1x7z | B |
| BCKDHB | None | None | None | None |
| BCKDHB | None | None | None | None |
| BCKDHB | None | None | None | None |
| BCKDHB | 0.857 | 1 | 2bff | B |
| CDH1 | None | None | None | None |
| CDH1 | 0.575 | 1 | 3ff8 | A |
| CDH1 | 0.543 | 1 | 3ff7 | A |
| CDH1 | None | None | None | None |
| CDH1 | None | None | None | None |
| CTNNB1 | None | None | None | None |
| AIRE | None | None | None | None |
| EDA | 1 | 1 | 1rj8 | A |
| EDA | None | None | None | None |
| EDA | None | None | None | None |
| EDA | None | None | None | None |
| EDA | None | None | None | None |
| SH2D1A | None | None | None | None |
| SH2D1A | None | None | None | None |
| SH2D1A | None | None | None | None |
| GCK | 0.556 | 37 | 3idh | A |
| GCK | None | None | None | None |
| GCK | None | None | None | None |
| GCK | None | None | None | None |
| GCK | None | None | None | None |
| GCK | 0.611 | 13 | 3h1v | X |
| GCK | None | None | None | None |
| GCK | None | None | None | None |
| GCK | None | None | None | None |
| GCK | 0.778 | 5 | 3id8 | A |
| GCK | 0.833 | 1 | 4dhy | A |
| GCK | None | None | None | None |
| GCK | None | None | None | None |
| GCK | 0.611 | 6 | 3id8 | A |
| GCK | None | None | None | None |
| GCK | None | None | None | None |
| GCK | None | None | None | None |
| GCK | 0.5 | 5 | 3vey | A |
| GCK | None | None | None | None |
| GCK | None | None | None | None |
| AQP2 | None | None | None | None |
| AQP2 | None | None | None | None |
| AQP2 | None | None | None | None |
| AQP2 | None | None | None | None |
| AQP2 | None | None | None | None |
| AQP2 | None | None | None | None |
| AQP2 | None | None | None | None |
| AQP2 | None | None | None | None |
| AQP2 | None | None | None | None |
| AQP2 | None | None | None | None |

| | | | | |
|---|---|---|---|---|
| AQP2 | None | None | None | None |
| AQP2 | None | None | None | None |
| AQP2 | None | None | None | None |
| SUMF1 | 0.75 | 6 | 1z70 | X |
| SUMF1 | 0.556 | 30 | 1y1i | X |
| SUMF1 | 0.5 | 38 | 2aij | X |
| SUMF1 | 0.625 | 18 | 2aij | X |
| SUMF1 | 0.875 | 35 | 2afy | X |
| SUMF1 | 0.667 | 19 | 1y1i | X |
| SUMF1 | 0.75 | 17 | 1y1j | X |
| SUMF1 | 0.75 | 17 | 1y1j | X |
| SUMF1 | 0.625 | 27 | 1y1g | X |
| FAM20A | None | None | None | None |
| FAM20A | None | None | None | None |
| FAM20A | None | None | None | None |
| FAM20A | None | None | None | None |
| NT5E | None | None | None | None |
| NT5E | None | None | None | None |
| NOS3 | 0.564 | 54 | 1m9k | B |
| ACTG1 | None | None | None | None |
| ACTG1 | None | None | None | None |
| ACTG1 | None | None | None | None |
| ACTG1 | None | None | None | None |
| ACTG1 | 0.571 | 8 | 5jlh | E |
| ACTG1 | None | None | None | None |
| FCGR1A | None | None | None | None |
| GPI | None | None | None | None |
| GPI | None | None | None | None |
| GPI | 1 | 3 | 1jiq | A |
| GPI | None | None | None | None |
| GPI | 0.696 | 22 | 1iat | A |
| GPI | 0.522 | 22 | 1nuh | A |
| GPI | 0.696 | 27 | 1iat | A |
| GPI | None | None | None | None |
| GPI | None | None | None | None |
| CFH | None | None | None | None |
| OGG1 | 0.895 | 13 | 1m3h | A |
| TPK1 | None | None | None | None |
| TPK1 | None | None | None | None |
| TPK1 | None | None | None | None |
| TPK1 | None | None | None | None |
| SEC23A | None | None | None | None |
| SEC23A | None | None | None | None |
| TYRP1 | None | None | None | None |
| TYRP1 | None | None | None | None |
| TYRP1 | None | None | None | None |
| TYRP1 | None | None | None | None |
| TYRP1 | None | None | None | None |
| TYRP1 | None | None | None | None |
| CSTB | None | None | None | None |
| CSTB | None | None | None | None |
| CSTB | None | None | None | None |
| ACP1 | None | None | None | None |
| IL17F | None | None | None | None |

| | | | | |
|---|---|---|---|---|
| IL17F | None | None | None | None |
| IL17F | None | None | None | None |
| IL17F | None | None | None | None |
| IL17F | None | None | None | None |
| IDH2 | 0.5 | 2 | 5svn | B |
| IDH2 | 0.5 | 2 | 5svn | B |
| IDH2 | None | None | None | None |
| IDH2 | None | None | None | None |
| TP53 | 0.925 | 11 | 2ocj | A |
| TP53 | 0.8 | 16 | 2ahi | D |
| TP53 | 0.65 | 49 | 4ibq | A |
| TP53 | 0.725 | 27 | 5g4o | A |
| TP53 | None | None | None | None |
| TP53 | 0.9 | 1 | 4agp | A |
| TP53 | 0.9 | 1 | 4agp | A |
| TP53 | 0.925 | 10 | 4hje | A |
| TP53 | 0.7 | 29 | 3d05 | A |
| TP53 | 0.875 | 7 | 3kmd | B |
| TP53 | 0.875 | 7 | 3kmd | B |
| TP53 | 0.925 | 6 | 3kz8 | B |
| TP53 | 0.925 | 6 | 3kz8 | B |
| TP53 | 0.609 | 22 | 3q01 | B |
| TP53 | 0.925 | 5 | 3kmd | B |
| TP53 | 0.925 | 5 | 3kmd | B |
| TP53 | 0.925 | 8 | 3kz8 | B |
| TP53 | 0.925 | 8 | 3kz8 | B |
| TP53 | 0.925 | 1 | 3kmd | B |
| TP53 | 0.9 | 11 | 2pcx | A |
| TP53 | 0.9 | 1 | 5aol | A |
| TP53 | 0.85 | 5 | 5lgy | A |
| TP53 | 0.9 | 1 | 3q06 | B |
| TP53 | 0.725 | 19 | 5lgy | A |
| TP53 | 0.775 | 14 | 4mzi | A |
| TP53 | 0.825 | 8 | 1kzy | A |
| TP53 | 0.825 | 8 | 1kzy | A |
| TP53 | 0.8 | 11 | 4ibt | A |
| TP53 | 0.8 | 11 | 4ibt | A |
| TP53 | 0.725 | 15 | 5lgy | A |
| TP53 | 0.725 | 29 | 2h1l | O |
| TP53 | 0.875 | 7 | 3kmd | B |
| TP53 | 0.725 | 15 | 5lgy | A |
| TP53 | 0.8 | 10 | 4agp | A |
| TP53 | 0.8 | 10 | 4agp | A |
| TP53 | 0.8 | 10 | 4agp | A |
| TP53 | 0.8 | 10 | 4agp | A |
| TP53 | 0.675 | 35 | 4ibq | A |
| TP53 | 0.575 | 37 | 4agm | A |
| TP53 | 0.675 | 17 | 3kz8 | B |
| TP53 | None | None | None | None |
| TP53 | None | None | None | None |
| LMAN1 | None | None | None | None |
| CTSK | 0.75 | 2 | 4n79 | A |
| CTSK | None | None | None | None |
| CTSK | None | None | None | None |

| | | | | |
|---|---|---|---|---|
| FGB | None | None | None | None |
| FGB | None | None | None | None |
| FGB | 0.688 | 1 | 1fzf | E |
| FGB | None | None | None | None |
| ACADM | None | None | None | None |
| ACADM | None | None | None | None |
| ACADM | None | None | None | None |
| ACADM | None | None | None | None |
| ACADM | None | None | None | None |
| ACADM | None | None | None | None |
| ACADM | None | None | None | None |
| ACADM | None | None | None | None |
| ACADM | None | None | None | None |
| ACADM | None | None | None | None |
| ACADM | None | None | None | None |
| ACADM | None | None | None | None |
| ACADM | None | None | None | None |
| ACADM | None | None | None | None |
| ACADM | None | None | None | None |
| ACADM | None | None | None | None |
| ACADM | None | None | None | None |
| ACADM | None | None | None | None |
| ACADM | None | None | None | None |
| ACADM | None | None | None | None |
| ACADM | 0.564 | 1 | 1t9g | C |
| ACADM | None | None | None | None |
| ACADM | None | None | None | None |
| ACADM | None | None | None | None |
| ACADM | None | None | None | None |
| ACADM | None | None | None | None |
| ACADM | None | None | None | None |
| PTS | None | None | None | None |
| PTS | None | None | None | None |
| PTS | None | None | None | None |
| PTS | 0.778 | 1 | 3i2b | D |
| PTS | None | None | None | None |
| PTS | None | None | None | None |
| PTS | None | None | None | None |
| PTS | None | None | None | None |
| PTS | None | None | None | None |
| INS | 0.75 | 2 | 3zqr | E |
| INS | 1 | 5 | 2l1z | A |
| INS | 1 | 5 | 2l1z | A |
| INS | 1 | 1 | 1his | A |
| FBP1 | 0.71 | 26 | 2y5l | C |
| FBP1 | None | None | None | None |
| FBP1 | 0.5 | 18 | 1fta | A |
| FBP1 | 0.7 | 10 | 2y5k | A |
| FBP1 | 0.516 | 38 | 3kc1 | B |
| FBP1 | 0.733 | 10 | 3kc1 | A |
| FBP1 | 0.633 | 43 | 2jjk | A |
| FBP1 | 0.8 | 11 | 2y5k | A |
| FBP1 | 0.548 | 41 | 2vt5 | D |
| FBP1 | None | None | None | None |

| | | | | |
|---|---|---|---|---|
| FBP1 | 0.643 | 7 | 4mjo | B |
| KCNQ1 | None | None | None | None |
| KCNQ1 | None | None | None | None |
| KCNQ1 | None | None | None | None |
| UPP1 | 0.706 | 4 | 3euf | A |
| UMPS | 1 | 3 | 2qcl | A |
| UMPS | 0.556 | 59 | 2qcl | B |
| UMPS | None | None | None | None |
| DHFR | None | None | None | None |
| KHK | 0.667 | 2 | 2hw1 | A |
| TTPA | None | None | None | None |
| TTPA | None | None | None | None |
| TTPA | 0.5 | 1 | 1r5l | A |
| TTPA | None | None | None | None |
| MAP1LC3B | None | None | None | None |
| MAP1LC3B | None | None | None | None |
| ZAP70 | None | None | None | None |
| PROC | 0.536 | 1 | 3f6u | L |
| SQSTM1 | None | None | None | None |
| AHCY | None | None | None | None |
| AHCY | None | None | None | None |
| PRNP | None | None | None | None |
| PRNP | None | None | None | None |
| PRNP | None | None | None | None |
| PRNP | None | None | None | None |
| PRNP | None | None | None | None |
| PRNP | None | None | None | None |
| PRNP | None | None | None | None |
| PRNP | None | None | None | None |
| PRNP | None | None | None | None |
| PRNP | None | None | None | None |
| PRNP | None | None | None | None |
| PRNP | None | None | None | None |
| PRNP | None | None | None | None |
| PRNP | None | None | None | None |
| PRNP | None | None | None | None |
| PRNP | None | None | None | None |
| REN | None | None | None | None |
| REN | None | None | None | None |
| HINT1 | 0.556 | 2 | 5ed6 | B |
| HINT1 | 0.897 | 1 | 5km4 | A |
| HINT1 | None | None | None | None |
| HINT1 | 0.667 | 11 | 5ed6 | B |
| HINT1 | 0.923 | 1 | 5ipc | A |
| OAT | None | None | None | None |
| OAT | None | None | None | None |
| OAT | None | None | None | None |
| OAT | None | None | None | None |
| OAT | None | None | None | None |
| OAT | None | None | None | None |
| OAT | None | None | None | None |
| OAT | None | None | None | None |
| OAT | None | None | None | None |
| OAT | None | None | None | None |

| | | | | | |
|---|---|---|---|---|---|
| OAT | 0.675 | | 2 | 2byl | B |
| OAT | None | None | | None | None |
| OAT | None | None | | None | None |
| OAT | None | None | | None | None |
| OAT | None | None | | None | None |
| OAT | 0.758 | | 1 | 2byl | A |
| OAT | None | None | | None | None |
| OAT | 0.525 | | 4 | 2byl | B |
| OAT | None | None | | None | None |
| OAT | None | None | | None | None |
| OAT | None | None | | None | None |
| OAT | None | None | | None | None |
| OAT | None | None | | None | None |
| OAT | None | None | | None | None |
| OAT | None | None | | None | None |
| OAT | None | None | | None | None |
| OAT | None | None | | None | None |
| OAT | None | None | | None | None |
| CTSA | None | None | | None | None |
| CTSA | None | None | | None | None |
| PNP | None | None | | None | None |
| PNP | None | None | | None | None |
| PNP | None | None | | None | None |
| PNP | 0.706 | | 5 | 4ear | C |
| PNP | None | None | | None | None |
| PNP | 0.5 | | 17 | 1pf7 | E |
| PNP | 0.933 | | 1 | 3ggs | C |
| PNP | 0.833 | | 1 | 3d1v | A |
| MYL2 | None | None | | None | None |
| MYL2 | None | None | | None | None |
| ANG | 0.548 | | 4 | 1k5b | A |
| ANG | None | None | | None | None |
| ANG | None | None | | None | None |
| ANG | None | None | | None | None |
| ANG | 0.761 | | 1 | 5eop | A |
| ANG | 0.576 | | 4 | 5m9v | A |
| ANG | 0.579 | | 4 | 5m9q | A |
| ANG | None | None | | None | None |
| ANG | None | None | | None | None |
| ANG | 0.52 | | 5 | 5m9g | A |
| POLH | None | None | | None | None |
| POLH | None | None | | None | None |
| POLH | None | None | | None | None |
| IL2RA | None | None | | None | None |
| TGFB1 | None | None | | None | None |
| TP53 | None | None | | None | None |
| TP53 | None | None | | None | None |
| PAH | None | None | | None | None |
| PAH | None | None | | None | None |
| PAH | None | None | | None | None |
| PAH | None | None | | None | None |
| PAH | None | None | | None | None |
| PAH | None | None | | None | None |
| PAH | None | None | | None | None |

| | | | | |
|---|---|---|---|---|
| PAH | None | None | None | None |
| PAH | None | None | None | None |
| PAH | None | None | None | None |
| PAH | None | None | None | None |
| PAH | None | None | None | None |
| PAH | None | None | None | None |
| PAH | None | None | None | None |
| PAH | None | None | None | None |
| PAH | None | None | None | None |
| PAH | None | None | None | None |
| PAH | None | None | None | None |
| PAH | None | None | None | None |
| PAH | None | None | None | None |
| PAH | None | None | None | None |
| PAH | None | None | None | None |
| PAH | None | None | None | None |
| PAH | 0.68 | 6 | 2pah | A |
| PAH | 0.76 | 1 | 1tdw | A |
| PAH | None | None | None | None |
| PAH | None | None | None | None |
| PAH | None | None | None | None |
| PAH | None | None | None | None |
| PAH | None | None | None | None |
| PAH | None | None | None | None |
| PAH | None | None | None | None |
| PAH | 0.64 | 2 | 1lrm | A |
| PAH | None | None | None | None |
| PAH | None | None | None | None |
| PAH | None | None | None | None |
| PAH | None | None | None | None |
| PAH | None | None | None | None |
| PAH | None | None | None | None |
| PAH | None | None | None | None |
| PAH | None | None | None | None |
| HDAC8 | None | None | None | None |
| HDAC8 | 0.732 | 3 | 3rqd | A |
| VCP | None | None | None | None |
| VCP | 0.5 | 2 | 3qq8 | A |
| VCP | 0.5 | 2 | 3qq8 | A |
| VCP | None | None | None | None |
| VCP | None | None | None | None |
| VCP | None | None | None | None |
| VCP | None | None | None | None |
| B3GAT3 | None | None | None | None |
| B3GAT3 | None | None | None | None |
| CSF2RB | None | None | None | None |
| CSF2RB | None | None | None | None |
| CSF2RB | None | None | None | None |
| NF2 | 0.5 | 6 | 1h4r | B |
| NF2 | None | None | None | None |
| NF2 | None | None | None | None |
| PIK3CA | None | None | None | None |
| PHGDH | None | None | None | None |
| PHGDH | None | None | None | None |

| | | | | |
|---|---|---|---|---|
| PHGDH | None | None | None | None |
| PHGDH | 0.66 | 1 | 5n53 | A |
| POLH | None | None | None | None |
| XIAP | 0.5 | 7 | 5c7b | A |
| XIAP | None | None | None | None |
| PDHX | None | None | None | None |
| PDHX | None | None | None | None |
| PDHX | None | None | None | None |
| PDHX | None | None | None | None |
| PDHX | None | None | None | None |
| SAMHD1 | 0.5 | 12 | 5ao0 | A |
| SAMHD1 | None | None | None | None |
| SAMHD1 | None | None | None | None |
| SAMHD1 | None | None | None | None |
| SAMHD1 | 0.571 | 3 | 4qfy | D |
| SAMHD1 | None | None | None | None |
| SAMHD1 | None | None | None | None |
| SAMHD1 | None | None | None | None |
| CD36 | None | None | None | None |
| CD36 | None | None | None | None |
| CD36 | None | None | None | None |
| ASPH | None | None | None | None |
| CCT5 | None | None | None | None |
| CCT5 | None | None | None | None |
| CCT5 | None | None | None | None |
| SMN1 | None | None | None | None |
| SMN1 | None | None | None | None |
| SMN1 | None | None | None | None |
| SMN1 | None | None | None | None |
| ABCA1 | None | None | None | None |
| ABCA1 | None | None | None | None |
| ABCA1 | None | None | None | None |
| ABCA1 | None | None | None | None |
| ABCA1 | None | None | None | None |
| ABCA1 | None | None | None | None |
| ABCA1 | None | None | None | None |
| ABCA1 | None | None | None | None |
| ABCA1 | None | None | None | None |
| ABCA1 | None | None | None | None |
| ABCA1 | None | None | None | None |
| ABCA1 | None | None | None | None |
| ABCA1 | None | None | None | None |
| ABCA1 | None | None | None | None |
| ABCA1 | None | None | None | None |
| ABCA1 | None | None | None | None |
| ABCA1 | None | None | None | None |
| ABCA1 | None | None | None | None |
| ABCA1 | None | None | None | None |
| ABCA1 | None | None | None | None |
| ABCA1 | None | None | None | None |
| ABCA1 | None | None | None | None |
| ABCA1 | None | None | None | None |
| ABCA1 | None | None | None | None |
| ABCA1 | None | None | None | None |

| | | | | |
|---|---|---|---|---|
| ABCA1 | None | None | None | None |
| ABCA1 | None | None | None | None |
| ABCA1 | None | None | None | None |
| ABCA1 | None | None | None | None |
| ABCA1 | None | None | None | None |
| ABCA1 | None | None | None | None |
| ABCA1 | None | None | None | None |
| ABCA1 | None | None | None | None |
| ABCA1 | None | None | None | None |
| ABCA1 | None | None | None | None |
| ABCA1 | None | None | None | None |
| ERCC1 | None | None | None | None |
| FECH | 0.524 | 27 | 2po5 | A |
| FECH | 0.6 | 60 | 2hre | A |
| FECH | 0.636 | 20 | 2po7 | A |
| FECH | None | None | None | None |
| FECH | 0.5 | 84 | 2hrc | A |
| FECH | 0.667 | 10 | 2po5 | A |
| FECH | None | None | None | None |
| PTEN | None | None | None | None |
| PTEN | None | None | None | None |
| PTEN | None | None | None | None |
| PTEN | None | None | None | None |
| PTEN | None | None | None | None |
| PTEN | None | None | None | None |
| PTEN | None | None | None | None |
| PTEN | None | None | None | None |
| PTEN | None | None | None | None |
| PTEN | None | None | None | None |
| PTEN | None | None | None | None |
| PTEN | None | None | None | None |
| PTEN | None | None | None | None |
| PTEN | None | None | None | None |
| PTEN | None | None | None | None |
| PTEN | None | None | None | None |
| PTEN | None | None | None | None |
| PTEN | None | None | None | None |
| PTEN | None | None | None | None |
| PTEN | None | None | None | None |
| ACAT1 | 0.667 | 1 | 2ib9 | C |
| ACAT1 | None | None | None | None |
| ACAT1 | None | None | None | None |
| ACAT1 | 0.667 | 1 | 2ib9 | C |
| ACAT1 | 0.667 | 1 | 2ib9 | C |
| ACAT1 | None | None | None | None |
| ACAT1 | 0.833 | 6 | 2ib7 | C |
| ACAT1 | None | None | None | None |
| ACAT1 | None | None | None | None |
| ACAT1 | 0.528 | 9 | 2ibw | D |
| ACAT1 | 0.703 | 16 | 2ibu | A |
| CREBBP | None | None | None | None |
| ITGA2B | None | None | None | None |
| ITGA2B | None | None | None | None |
| ITGA2B | None | None | None | None |

| | | | | |
|---|---|---|---|---|
| ITGA2B | 0.538 | 1 | 3nig | A |
| TSC1 | None | None | None | None |
| ADA2 | None | None | None | None |
| ADA2 | None | None | None | None |
| ADA2 | 0.75 | 1 | 3lgd | A |
| ADA2 | None | None | None | None |
| ADA2 | None | None | None | None |
| ADA2 | None | None | None | None |
| ADA2 | None | None | None | None |
| ADA2 | None | None | None | None |
| ADA2 | 1 | 1 | 3lgd | B |
| ADA2 | None | None | None | None |
| PGK1 | None | None | None | None |
| PGK1 | 0.536 | 8 | 2xe8 | A |
| PGK1 | None | None | None | None |
| PGK1 | None | None | None | None |
| PGK1 | 0.571 | 3 | 2ybe | A |
| TUBA1A | None | None | None | None |
| TUBA1A | None | None | None | None |
| TUBA1A | None | None | None | None |
| TUBA1A | None | None | None | None |
| TUBA1A | None | None | None | None |
| PTPN11 | None | None | None | None |
| PTPN11 | None | None | None | None |
| PTPN11 | None | None | None | None |
| PTPN11 | None | None | None | None |
| PTPN11 | None | None | None | None |
| PTPN11 | None | None | None | None |
| PTPN11 | None | None | None | None |
| PTPN11 | None | None | None | None |
| PTPN11 | None | None | None | None |
| PTPN11 | None | None | None | None |
| PTPN11 | None | None | None | None |
| PTPN11 | None | None | None | None |
| PTPN11 | None | None | None | None |
| PTPN11 | None | None | None | None |
| PTPN11 | 0.621 | 2 | 3b7o | A |
| PTPN11 | 0.776 | 1 | 3zm1 | A |
| PTPN11 | None | None | None | None |
| PTPN11 | None | None | None | None |
| PTPN11 | None | None | None | None |
| PTPN11 | None | None | None | None |
| PTPN11 | None | None | None | None |
| PTPN11 | 0.776 | 1 | 3zm1 | A |
| PTPN11 | 0.621 | 6 | 3zm1 | A |
| PTPN11 | None | None | None | None |
| PTPN11 | None | None | None | None |
| PTPN11 | 0.621 | 6 | 3zm1 | A |
| GLB1 | None | None | None | None |
| GLB1 | None | None | None | None |
| UQCRQ | None | None | None | None |
| VPS35 | None | None | None | None |
| NR3C1 | None | None | None | None |
| NR3C1 | None | None | None | None |

| | | | | |
|---|---|---|---|---|
| NR3C1 | None | None | None | None |
| NR3C1 | None | None | None | None |
| NR3C1 | None | None | None | None |
| NR3C1 | None | None | None | None |
| NR3C1 | None | None | None | None |
| NR3C1 | None | None | None | None |
| NR3C1 | None | None | None | None |
| APOA1 | None | None | None | None |
| APOA1 | None | None | None | None |
| APOA1 | None | None | None | None |
| APOA1 | None | None | None | None |
| APOA1 | None | None | None | None |
| APOA1 | None | None | None | None |
| APOA1 | None | None | None | None |
| BCKDHA | None | None | None | None |
| BCKDHA | 0.583 | 76 | 1x7x | A |
| BCKDHA | 0.963 | 1 | 1ols | A |
| BCKDHA | 0.696 | 3 | 1v1r | A |
| BCKDHA | 0.625 | 16 | 2beu | A |
| BCKDHA | None | None | None | None |
| BCKDHA | 0.652 | 11 | 1v1r | A |
| BCKDHA | 1 | 1 | 2beu | A |
| BCKDHA | 0.84 | 2 | 2j9f | A |
| BCKDHA | 0.556 | 42 | 1ols | A |
| BCKDHA | 0.609 | 90 | 1v1r | A |
| BCKDHA | 0.652 | 25 | 1v1r | A |
| BCKDHA | None | None | None | None |
| BCKDHA | 0.625 | 60 | 2beu | A |
| BCKDHA | 0.625 | 28 | 2beu | A |
| TOP1 | None | None | None | None |
| ANTXR2 | None | None | None | None |
| ANTXR2 | None | None | None | None |
| ANTXR2 | None | None | None | None |
| TF | 0.625 | 35 | 5dyh | A |
| TF | None | None | None | None |
| FSHB | None | None | None | None |
| GALK1 | None | None | None | None |
| GALK1 | None | None | None | None |
| GALK1 | None | None | None | None |
| CYP17A1 | None | None | None | None |
| CYP17A1 | None | None | None | None |
| CYP17A1 | None | None | None | None |
| CYP17A1 | None | None | None | None |
| CYP17A1 | None | None | None | None |
| CYP17A1 | None | None | None | None |
| CYP17A1 | None | None | None | None |
| CYP17A1 | None | None | None | None |
| CYP17A1 | None | None | None | None |
| CYP17A1 | None | None | None | None |
| CYP17A1 | None | None | None | None |
| CYP17A1 | None | None | None | None |
| CYP17A1 | None | None | None | None |
| CYP17A1 | None | None | None | None |
| CYP17A1 | None | None | None | None |

| | | | | |
|---|---|---|---|---|
| CYP17A1 | None | None | None | None |
| CYP17A1 | None | None | None | None |
| CD44 | 0.667 | 24 | 1uuh | A |
| MDH2 | None | None | None | None |
| MDH2 | None | None | None | None |
| NDUFV1 | None | None | None | None |
| NDUFV1 | None | None | None | None |
| NDUFV1 | None | None | None | None |
| NDUFV1 | None | None | None | None |
| NDUFV1 | None | None | None | None |
| NDUFV1 | None | None | None | None |
| NDUFV1 | None | None | None | None |
| NDUFV1 | None | None | None | None |
| NDUFV1 | None | None | None | None |
| NDUFV1 | None | None | None | None |
| CTSC | None | None | None | None |
| CTSC | None | None | None | None |
| CTSC | None | None | None | None |
| CTSC | None | None | None | None |
| CTSC | 0.643 | 1 | 4oel | A |
| CTSC | None | None | None | None |
| CTSC | None | None | None | None |
| VWF | None | None | None | None |
| VWF | None | None | None | None |
| VWF | None | None | None | None |
| VWF | None | None | None | None |
| VWF | None | None | None | None |
| VWF | None | None | None | None |
| VWF | None | None | None | None |
| ORC4 | None | None | None | None |
| ORC4 | None | None | None | None |
| CRYAB | None | None | None | None |
| CRYAB | None | None | None | None |
| CRYAB | None | None | None | None |
| MAP2K2 | None | None | None | None |
| MAP2K2 | None | None | None | None |
| MAP2K2 | None | None | None | None |
| MAP2K2 | None | None | None | None |
| MAP2K2 | None | None | None | None |
| MAP2K2 | None | None | None | None |
| MAP2K2 | None | None | None | None |
| MAP2K2 | None | None | None | None |
| MAP2K2 | None | None | None | None |
| CBL | None | None | None | None |
| CBL | None | None | None | None |
| CBL | None | None | None | None |
| CBL | None | None | None | None |
| CBL | None | None | None | None |
| CBL | None | None | None | None |
| CBL | None | None | None | None |
| PFN1 | None | None | None | None |
| PFN1 | 0.75 | 2 | 1fik | A |
| ACSM2A | None | None | None | None |
| HSPG2 | None | None | None | None |

| | | | | |
|---|---|---|---|---|
| HSPG2 | None | None | None | None |
| HSPG2 | None | None | None | None |
| HSPG2 | None | None | None | None |
| PARK7 | None | None | None | None |
| PARK7 | 0.769 | 4 | 1pe0 | A |
| PARK7 | 0.808 | 2 | 1pe0 | A |
| PARK7 | 0.667 | 8 | 4rky | A |
| PARK7 | 0.889 | 1 | 3cz9 | A |
| PARK7 | 0.75 | 2 | 1pdw | B |
| INSR | None | None | None | None |
| INSR | None | None | None | None |
| INSR | None | None | None | None |
| INSR | None | None | None | None |
| INSR | None | None | None | None |
| DSC2 | None | None | None | None |
| DSC2 | None | None | None | None |
| DSC2 | None | None | None | None |
| DSC2 | None | None | None | None |
| DSC2 | None | None | None | None |
| DSC2 | None | None | None | None |
| DSC2 | None | None | None | None |
| DSC2 | None | None | None | None |
| DSC2 | None | None | None | None |
| DSC2 | None | None | None | None |
| MICU1 | None | None | None | None |
| MICU1 | None | None | None | None |
| ADAMTS13 | None | None | None | None |
| ADAMTS13 | None | None | None | None |
| ADAMTS13 | None | None | None | None |
| ADAMTS13 | None | None | None | None |
| ADAMTS13 | None | None | None | None |
| ADAMTS13 | None | None | None | None |
| ADAMTS13 | None | None | None | None |
| IRAK4 | None | None | None | None |
| IRAK4 | None | None | None | None |
| RNASEH2B | None | None | None | None |
| RNASEH2B | None | None | None | None |
| RNASEH2B | None | None | None | None |
| NTRK1 | None | None | None | None |
| MPO | 0.75 | 3 | 1d5l | C |
| MPO | 0.6 | 25 | 1myp | D |
| MPO | None | None | None | None |
| PEX3 | None | None | None | None |
| SERPINA1 | None | None | None | None |
| SERPINA1 | None | None | None | None |
| SERPINA1 | None | None | None | None |
| SERPINA1 | None | None | None | None |
| SERPINA1 | None | None | None | None |
| SERPINA1 | None | None | None | None |
| SERPINA1 | None | None | None | None |
| SERPINA1 | 0.6 | 1 | 1ezx | A |
| SERPINA1 | None | None | None | None |
| SERPINA1 | None | None | None | None |
| YARS2 | None | None | None | None |

| | | | | |
|---|---|---|---|---|
| YARS2 | None | None | None | None |
| YARS2 | None | None | None | None |
| YARS2 | None | None | None | None |
| YARS2 | None | None | None | None |
| YARS2 | 0.781 | 1 | 2pid | B |
| YARS2 | None | None | None | None |
| YARS2 | None | None | None | None |
| YARS2 | None | None | None | None |
| PDHA1 | None | None | None | None |
| PDHA1 | None | None | None | None |
| PDHA1 | None | None | None | None |
| PDHA1 | None | None | None | None |
| PDHA1 | None | None | None | None |
| PDHA1 | None | None | None | None |
| PDHA1 | None | None | None | None |
| PDHA1 | 0.96 | 1 | 3exh | C |
| PDHA1 | None | None | None | None |
| PDHA1 | None | None | None | None |
| PDHA1 | None | None | None | None |
| CUL4B | None | None | None | None |
| AGXT | None | None | None | None |
| AGXT | None | None | None | None |
| AGXT | None | None | None | None |
| AGXT | 0.692 | 3 | 2yob | A |
| AGXT | 0.786 | 7 | 4kxk | C |
| AGXT | 0.538 | 21 | 3r9a | A |
| AGXT | None | None | None | None |
| AGXT | None | None | None | None |
| AGXT | None | None | None | None |
| AGXT | 0.611 | 4 | 5hhy | B |
| AGXT | None | None | None | None |
| AGXT | None | None | None | None |
| AGXT | None | None | None | None |
| AGXT | None | None | None | None |
| AGXT | None | None | None | None |
| AGXT | 0.684 | 1 | 1h0c | A |
| AGXT | 0.5 | 2 | 4cbr | A |
| AGXT | 0.538 | 19 | 4kyo | A |
| AGXT | 0.538 | 19 | 4kyo | A |
| AGXT | 0.75 | 2 | 5luc | B |
| AGXT | 0.722 | 3 | 5hhy | B |
| AGXT | 0.571 | 3 | 4kxk | C |
| AGXT | 0.571 | 5 | 4kxk | C |
| AGXT | None | None | None | None |
| AGXT | 0.778 | 1 | 5hhy | B |
| AGXT | 0.778 | 1 | 5hhy | B |
| AGXT | None | None | None | None |
| AGXT | 0.692 | 7 | 4kyo | A |
| AGXT | 0.5 | 18 | 4cbr | A |
| AGXT | 0.5 | 18 | 4cbr | A |
| AGXT | 0.615 | 13 | 3r9a | A |
| AGXT | 0.786 | 7 | 4kxk | C |
| AGXT | 0.846 | 1 | 4kxk | A |
| AGXT | 0.615 | 36 | 3r9a | A |

| | | | | |
|---|---|---|---|---|
| AGXT | 0.571 | 10 | 4kxk | C |
| SFTPD | 0.509 | 2 | 3g83 | B |
| SFTPD | None | None | None | None |
| PIK3CA | None | None | None | None |
| PIK3CA | None | None | None | None |
| PIK3CA | None | None | None | None |
| AUH | None | None | None | None |
| AUH | None | None | None | None |
| AUH | None | None | None | None |
| AUH | None | None | None | None |
| F11 | None | None | None | None |
| F11 | None | None | None | None |
| F11 | None | None | None | None |
| F11 | None | None | None | None |
| MEN1 | 0.6 | 81 | 5db0 | A |
| MEN1 | 0.8 | 54 | 4i80 | A |
| MEN1 | 0.8 | 84 | 5db0 | A |
| MEN1 | 0.8 | 30 | 5ddb | A |
| MEN1 | 1 | 2 | 4i80 | A |
| MEN1 | 0.8 | 17 | 5db0 | A |
| MEN1 | 0.6 | 17 | 5ddb | A |
| MEN1 | 0.6 | 31 | 3u88 | A |
| MEN1 | None | None | None | None |
| MEN1 | 0.6 | 56 | 5ddb | A |
| MYBPC1 | None | None | None | None |
| MYBPC1 | None | None | None | None |
| MYBPC1 | None | None | None | None |
| PDCD10 | None | None | None | None |
| SCARB2 | None | None | None | None |
| SCARB2 | None | None | None | None |
| SCARB2 | None | None | None | None |
| SCARB2 | None | None | None | None |
| SCARB2 | None | None | None | None |
| TREX2 | None | None | None | None |
| TBK1 | None | None | None | None |
| TBK1 | None | None | None | None |
| DLD | None | None | None | None |
| DLD | None | None | None | None |
| DLD | 0.505 | 3 | 1zy8 | B |
| DLD | None | None | None | None |
| DLD | None | None | None | None |
| DLD | None | None | None | None |
| LDHA | 0.802 | 1 | 4qt0 | A |
| TNNC1 | None | None | None | None |
| TNNC1 | None | None | None | None |
| TNNC1 | None | None | None | None |
| SMAD3 | None | None | None | None |
| HARS | None | None | None | None |
| HARS | None | None | None | None |
| TNNI3 | None | None | None | None |
| TNNI3 | None | None | None | None |
| TNNI3 | None | None | None | None |
| TNNI3 | None | None | None | None |
| TNNI3 | None | None | None | None |

| | | | | |
|---|---|---|---|---|
| TNNI3 | None | None | None | None |
| ACKR1 | None | None | None | None |
| BUB1B | None | None | None | None |
| BUB1B | None | None | None | None |
| BUB1B | None | None | None | None |
| AR | 0.522 | 6 | 1xj7 | A |
| AR | None | None | None | None |
| AR | None | None | None | None |
| AR | None | None | None | None |
| AR | None | None | None | None |
| FTO | None | None | None | None |
| FTO | None | None | None | None |
| TFAM | None | None | None | None |
| PYGL | None | None | None | None |
| PYGL | 0.778 | 3 | 1fa9 | A |
| PYGL | None | None | None | None |
| PYGL | 0.722 | 2 | 1fa9 | A |
| PYGL | 0.5 | 19 | 3dds | A |
| PYGL | 0.611 | 25 | 3ceh | B |
| PYGL | 0.611 | 32 | 1l7x | B |
| PYGL | None | None | None | None |
| PYGL | 0.722 | 2 | 1fc0 | B |
| PYGL | 0.778 | 3 | 1fa9 | A |
| PYGL | None | None | None | None |
| PYGL | 0.722 | 12 | 2ati | A |
| PYGL | 0.722 | 1 | 3ceh | B |
| PYGL | 0.611 | 11 | 3ceh | B |
| PYGL | 0.611 | 18 | 1fc0 | B |
| PYGL | 0.722 | 2 | 1fc0 | B |
| GALNS | None | None | None | None |
| GALNS | None | None | None | None |
| GALNS | None | None | None | None |
| GALNS | None | None | None | None |
| GALNS | None | None | None | None |
| GALNS | None | None | None | None |
| GALNS | None | None | None | None |
| GALNS | None | None | None | None |
| GALNS | None | None | None | None |
| GALNS | None | None | None | None |
| GALNS | None | None | None | None |
| GALNS | None | None | None | None |
| GALNS | None | None | None | None |
| GALNS | None | None | None | None |
| GALNS | None | None | None | None |
| GALNS | None | None | None | None |
| GALNS | None | None | None | None |
| GALNS | None | None | None | None |
| GALNS | None | None | None | None |
| GALNS | None | None | None | None |
| GALNS | None | None | None | None |
| GALNS | None | None | None | None |
| GALNS | None | None | None | None |
| GALNS | None | None | None | None |
| GALNS | None | None | None | None |

| | | | | |
|---|---|---|---|---|
| GALNS | None | None | None | None |
| GALNS | None | None | None | None |
| GALNS | None | None | None | None |
| GALNS | None | None | None | None |
| GALNS | None | None | None | None |
| GALNS | None | None | None | None |
| GALNS | None | None | None | None |
| GALNS | None | None | None | None |
| F10 | None | None | None | None |
| GLUD1 | None | None | None | None |
| GLUD1 | None | None | None | None |
| GLUD1 | None | None | None | None |
| GLUD1 | None | None | None | None |
| GLUD1 | None | None | None | None |
| GLUD1 | None | None | None | None |
| GLUD1 | None | None | None | None |
| GLUD1 | None | None | None | None |
| GLUD1 | None | None | None | None |
| GLUD1 | None | None | None | None |
| NOTCH1 | 0.528 | 3 | 3v79 | K |
| POGLUT1 | None | None | None | None |
| POGLUT1 | None | None | None | None |
| NR3C1 | None | None | None | None |
| CASP8 | 0.535 | 6 | 1f9e | C |
| CASP8 | 0.592 | 8 | 1qdu | C |
| TRAP1 | None | None | None | None |
| FANCM | None | None | None | None |
| EPOR | None | None | None | None |
| EPOR | None | None | None | None |
| NDP | None | None | None | None |
| NDP | None | None | None | None |
| NDP | None | None | None | None |
| NDP | None | None | None | None |
| NDP | None | None | None | None |
| NDP | None | None | None | None |
| NDP | None | None | None | None |
| NDP | None | None | None | None |
| NDP | None | None | None | None |
| NDP | None | None | None | None |
| NDP | None | None | None | None |
| NDP | None | None | None | None |
| NDP | None | None | None | None |
| NDP | None | None | None | None |
| NDP | None | None | None | None |
| NDP | None | None | None | None |
| NDP | None | None | None | None |
| ROGDI | None | None | None | None |
| CYP19A1 | None | None | None | None |
| CYP19A1 | None | None | None | None |
| CYP19A1 | None | None | None | None |
| CYP19A1 | None | None | None | None |
| CYP19A1 | None | None | None | None |
| CYP19A1 | None | None | None | None |
| CYP19A1 | None | None | None | None |

| | | | | |
|---|---|---|---|---|
| CYP19A1 | None | None | None | None |
| UBA5 | None | None | None | None |
| PRPS1 | None | None | None | None |
| PRPS1 | None | None | None | None |
| PRPS1 | None | None | None | None |
| PRPS1 | None | None | None | None |
| PRPS1 | None | None | None | None |
| PRPS1 | None | None | None | None |
| PRPS1 | None | None | None | None |
| PRPS1 | None | None | None | None |
| F2 | None | None | None | None |
| F2 | None | None | None | None |
| NTF4 | None | None | None | None |
| NTF4 | None | None | None | None |
| PSAP | None | None | None | None |
| PSAP | None | None | None | None |
| TAP1 | None | None | None | None |
| TAP1 | None | None | None | None |
| TAP1 | None | None | None | None |
| ARG1 | None | None | None | None |
| ARG1 | 0.714 | 1 | 2pha | A |
| ARG1 | None | None | None | None |
| ARG1 | None | None | None | None |
| ARG1 | None | None | None | None |
| ARG1 | None | None | None | None |
| CRYM | None | None | None | None |
| ADSL | None | None | None | None |
| ADSL | None | None | None | None |
| ADSL | None | None | None | None |
| ADSL | None | None | None | None |
| ADSL | None | None | None | None |
| ADSL | None | None | None | None |
| ADSL | None | None | None | None |
| ADSL | None | None | None | None |
| ADSL | None | None | None | None |
| ADSL | None | None | None | None |
| ADSL | None | None | None | None |
| ADSL | None | None | None | None |
| ADSL | 0.647 | 1 | 2vd6 | D |
| ADSL | None | None | None | None |
| ADSL | None | None | None | None |
| ADSL | None | None | None | None |
| ADSL | None | None | None | None |
| ADSL | None | None | None | None |
| ADSL | None | None | None | None |
| ADSL | None | None | None | None |
| VWF | None | None | None | None |
| PLA2G7 | None | None | None | None |
| MMP13 | None | None | None | None |
| MMP13 | None | None | None | None |
| MMP13 | None | None | None | None |
| MMP13 | None | None | None | None |
| POMGNT1 | None | None | None | None |
| POMGNT1 | None | None | None | None |

| | | | | |
|---|---|---|---|---|
| COL3A1 | None | None | None | None |
| COL3A1 | None | None | None | None |
| COL3A1 | None | None | None | None |
| COL3A1 | None | None | None | None |
| ACTN2 | None | None | None | None |
| ACTN2 | None | None | None | None |
| ACTN2 | None | None | None | None |
| ACTN2 | None | None | None | None |
| ACTN2 | None | None | None | None |
| ACTN2 | None | None | None | None |
| ACTN2 | None | None | None | None |
| UBA5 | None | None | None | None |
| ADAR | 0.75 | 1 | 1qbj | B |
| UMPS | None | None | None | None |
| UMPS | None | None | None | None |
| UMPS | None | None | None | None |
| TRNT1 | None | None | None | None |
| TRNT1 | None | None | None | None |
| NHEJ1 | None | None | None | None |
| NHEJ1 | None | None | None | None |
| NHEJ1 | None | None | None | None |
| C8A | None | None | None | None |
| GLUL | None | None | None | None |
| GLUL | None | None | None | None |
| GLUL | None | None | None | None |
| GYG1 | 0.615 | 45 | 3rmv | A |
| GYG1 | 0.923 | 1 | 3u2u | B |
| FLNA | None | None | None | None |
| FLNA | None | None | None | None |
| FLNA | None | None | None | None |
| FLNA | None | None | None | None |
| FLNA | None | None | None | None |
| FLNA | None | None | None | None |
| FLNA | None | None | None | None |
| TTN | None | None | None | None |
| TTN | None | None | None | None |
| TTN | None | None | None | None |
| TTN | None | None | None | None |
| SMS | None | None | None | None |
| SMS | None | None | None | None |
| SMS | None | None | None | None |
| C5 | None | None | None | None |
| SERPINC1 | None | None | None | None |
| SERPINC1 | None | None | None | None |
| SERPINC1 | None | None | None | None |
| SERPINC1 | None | None | None | None |
| SERPINC1 | None | None | None | None |
| SERPINC1 | None | None | None | None |
| SERPINC1 | None | None | None | None |
| SERPINC1 | None | None | None | None |
| SERPINC1 | None | None | None | None |
| SERPINC1 | None | None | None | None |
| SERPINC1 | None | None | None | None |
| SERPINC1 | None | None | None | None |

| | | | | |
|---|---|---|---|---|
| SERPINC1 | None | None | None | None |
| SERPINC1 | None | None | None | None |
| SERPINC1 | None | None | None | None |
| SERPINC1 | None | None | None | None |
| SERPINC1 | None | None | None | None |
| SERPINC1 | None | None | None | None |
| SERPINC1 | None | None | None | None |
| SERPINC1 | None | None | None | None |
| SERPINC1 | None | None | None | None |
| ETFB | None | None | None | None |
| ETFB | None | None | None | None |
| ETFB | None | None | None | None |
| ETFB | None | None | None | None |
| ETFB | None | None | None | None |
| ETFB | None | None | None | None |
| DIABLO | None | None | None | None |
| DIABLO | None | None | None | None |
| DPYS | None | None | None | None |
| DPYS | None | None | None | None |
| DPYS | None | None | None | None |
| DPYS | 0.533 | 7 | 2vr2 | A |
| DPYS | None | None | None | None |
| GNE | None | None | None | None |
| GNE | None | None | None | None |
| GNE | None | None | None | None |
| GNE | None | None | None | None |
| GNE | None | None | None | None |
| GNE | None | None | None | None |
| GNE | None | None | None | None |
| GNE | None | None | None | None |
| GNE | None | None | None | None |
| GNMT | None | None | None | None |
| GNMT | None | None | None | None |
| CTH | None | None | None | None |
| CTH | None | None | None | None |
| CTH | None | None | None | None |
| USH1C | None | None | None | None |
| PSEN1 | None | None | None | None |
| CHEK2 | None | None | None | None |
| CHEK2 | None | None | None | None |
| CHEK2 | None | None | None | None |
| CHEK2 | None | None | None | None |
| CHEK2 | None | None | None | None |
| CHEK2 | None | None | None | None |
| CHEK2 | None | None | None | None |
| CHEK2 | None | None | None | None |
| CHEK2 | None | None | None | None |
| SNCA | None | None | None | None |
| SNCA | None | None | None | None |
| GAN | None | None | None | None |
| GAN | None | None | None | None |
| GAN | None | None | None | None |
| SBDS | None | None | None | None |
| SBDS | None | None | None | None |

| | | | | |
|---|---|---|---|---|
| SBDS | None | None | None | None |
| SBDS | None | None | None | None |
| SBDS | None | None | None | None |
| SBDS | None | None | None | None |
| MPO | 1 | 3 | 3f9p | A |
| MPO | 0.6 | 12 | 5mfa | A |
| KCNH2 | None | None | None | None |
| KCNH2 | None | None | None | None |
| KCNH2 | 0.75 | 1 | 4hp9 | A |
| KCNH2 | None | None | None | None |
| TEK | None | None | None | None |
| TEK | None | None | None | None |
| TEK | None | None | None | None |
| TEK | None | None | None | None |
| NPC2 | None | None | None | None |
| NPC2 | None | None | None | None |
| NPC2 | None | None | None | None |
| NPC2 | None | None | None | None |
| NPC2 | None | None | None | None |
| NPC2 | None | None | None | None |
| NPC2 | None | None | None | None |
| PLG | None | None | None | None |
| PCSK9 | None | None | None | None |
| SFTPC | None | None | None | None |
| SFTPC | None | None | None | None |
| SFTPC | None | None | None | None |
| CD40LG | None | None | None | None |
| CD40LG | None | None | None | None |
| CD40LG | None | None | None | None |
| CD40LG | None | None | None | None |
| CD40LG | None | None | None | None |
| TAT | None | None | None | None |
| TAT | None | None | None | None |
| TAT | None | None | None | None |
| EP300 | None | None | None | None |
| FARS2 | None | None | None | None |
| FARS2 | None | None | None | None |
| FARS2 | None | None | None | None |
| FARS2 | None | None | None | None |
| FARS2 | None | None | None | None |
| FARS2 | None | None | None | None |
| FARS2 | None | None | None | None |
| FARS2 | None | None | None | None |
| FARS2 | 0.5 | 1 | 5mgw | A |
| FARS2 | None | None | None | None |
| FARS2 | None | None | None | None |
| FARS2 | None | None | None | None |
| FARS2 | None | None | None | None |
| MASP2 | None | None | None | None |
| MASP2 | None | None | None | None |
| MASP2 | None | None | None | None |
| MASP2 | None | None | None | None |
| STIM1 | None | None | None | None |
| CHRM2 | None | None | None | None |

| | | | | |
|---|---|---|---|---|
| MCFD2 | None | None | None | None |
| MCFD2 | 0.574 | 1 | 4yge | B |
| MCFD2 | 0.623 | 1 | 3wnx | B |
| KITLG | None | None | None | None |
| F12 | None | None | None | None |
| F12 | None | None | None | None |
| ERAP1 | None | None | None | None |
| GTF2H5 | None | None | None | None |
| GTF2H5 | None | None | None | None |
| THBD | None | None | None | None |
| MRPL28 | None | None | None | None |
| MRPL28 | None | None | None | None |
| HTRA1 | None | None | None | None |
| HTRA1 | None | None | None | None |
| HTRA1 | None | None | None | None |
| HTRA1 | None | None | None | None |
| HTRA1 | None | None | None | None |
| HTRA1 | None | None | None | None |
| KARS | None | None | None | None |
| KARS | None | None | None | None |
| KARS | None | None | None | None |
| KARS | None | None | None | None |
| KARS | None | None | None | None |
| HK1 | 0.722 | 11 | 4fpb | B |
| GARS | None | None | None | None |
| GARS | None | None | None | None |
| GARS | None | None | None | None |
| GARS | None | None | None | None |
| GARS | None | None | None | None |
| GARS | None | None | None | None |
| GARS | None | None | None | None |
| IL12B | None | None | None | None |
| HADH | None | None | None | None |
| HADH | None | None | None | None |
| HADH | 0.52 | 4 | 1lsj | A |
| HADH | None | None | None | None |
| HADH | 0.52 | 4 | 1lsj | A |
| RRM2B | None | None | None | None |
| RRM2B | None | None | None | None |
| RRM2B | None | None | None | None |
| RRM2B | None | None | None | None |
| RRM2B | None | None | None | None |
| RRM2B | None | None | None | None |
| RRM2B | None | None | None | None |
| RRM2B | None | None | None | None |
| RRM2B | None | None | None | None |
| RRM2B | None | None | None | None |
| RRM2B | None | None | None | None |
| RRM2B | None | None | None | None |
| RRM2B | None | None | None | None |
| RRM2B | None | None | None | None |
| RRM2B | None | None | None | None |
| MARVELD2 | None | None | None | None |
| TLR4 | None | None | None | None |

| | | | | |
|---|---|---|---|---|
| TTN | None | None | None | None |
| TTN | None | None | None | None |
| TNFRSF13B | None | None | None | None |
| POLG | None | None | None | None |
| POLG | None | None | None | None |
| POLG | None | None | None | None |
| POLG | None | None | None | None |
| POLG | None | None | None | None |
| POLG | None | None | None | None |
| POLG | None | None | None | None |
| POLG | None | None | None | None |
| POLG | None | None | None | None |
| POLG | None | None | None | None |
| POLG | None | None | None | None |
| POLG | None | None | None | None |
| POLG | None | None | None | None |
| POLG | None | None | None | None |
| POLG | None | None | None | None |
| POLG | None | None | None | None |
| POLG | None | None | None | None |
| POLG | None | None | None | None |
| POLG | None | None | None | None |
| POLG | None | None | None | None |
| POLG | None | None | None | None |
| POLG | None | None | None | None |
| POLG | None | None | None | None |
| POLG | None | None | None | None |
| POLG | None | None | None | None |
| POLG | None | None | None | None |
| POLG | None | None | None | None |
| POLG | None | None | None | None |
| POLG | 0.533 | 1 | 4ztu | A |
| POLG | None | None | None | None |
| POLG | None | None | None | None |
| POLG | None | None | None | None |
| POLG | None | None | None | None |
| POLG | None | None | None | None |
| POLG | 0.533 | 1 | 4ztu | A |
| POLG | None | None | None | None |
| POLG | None | None | None | None |
| POLG | None | None | None | None |
| POLG | None | None | None | None |
| POLG | None | None | None | None |
| POLG | None | None | None | None |
| POLG | None | None | None | None |
| POLG | None | None | None | None |
| POLG | None | None | None | None |
| POLG | None | None | None | None |
| POLG | None | None | None | None |
| POLG | None | None | None | None |
| POLG | None | None | None | None |
| POLG | None | None | None | None |
| UROD | 0.632 | 24 | 1jpk | A |

| | | | | | |
|---|---|---|---|---|---|
| UROD | | 0.579 | 33 | 1r3v | A |
| UROD | None | None | | None | None |
| UROD | | 0.526 | 49 | 1jph | A |
| UROD | None | None | | None | None |
| UROD | | 0.632 | 13 | 3gvw | A |
| UROD | | 0.789 | 6 | 3gvw | A |
| UROD | | 0.842 | 2 | 3gvv | A |
| UROD | | 0.737 | 31 | 1r3q | A |
| UROD | | 0.667 | 14 | 2q71 | A |
| PCK1 | None | None | | None | None |
| PCK1 | None | None | | None | None |
| PCK1 | None | None | | None | None |
| PCK1 | None | None | | None | None |
| PCK1 | | 0.524 | 60 | 1nhx | A |
| HEXA | None | None | | None | None |
| HEXA | None | None | | None | None |
| HEXA | None | None | | None | None |
| HEXA | None | None | | None | None |
| HEXA | None | None | | None | None |
| HEXA | None | None | | None | None |
| HEXA | None | None | | None | None |
| HEXA | None | None | | None | None |
| HEXA | None | None | | None | None |
| HEXA | None | None | | None | None |
| HEXA | None | None | | None | None |
| HEXA | None | None | | None | None |
| HEXA | None | None | | None | None |
| HEXA | None | None | | None | None |
| HEXA | None | None | | None | None |
| HEXA | None | None | | None | None |
| HEXA | None | None | | None | None |
| HEXA | None | None | | None | None |
| HEXA | None | None | | None | None |
| HEXA | None | None | | None | None |
| HEXA | None | None | | None | None |
| HEXA | None | None | | None | None |
| HEXA | None | None | | None | None |
| HEXA | None | None | | None | None |
| HEXA | None | None | | None | None |
| HEXA | None | None | | None | None |
| HEXA | None | None | | None | None |
| HEXA | None | None | | None | None |
| HEXA | None | None | | None | None |
| HEXA | None | None | | None | None |
| CFTR | None | None | | None | None |
| CFTR | None | None | | None | None |
| CFTR | None | None | | None | None |
| CFTR | None | None | | None | None |
| CFTR | None | None | | None | None |
| CFTR | None | None | | None | None |
| CFTR | None | None | | None | None |
| CFTR | None | None | | None | None |
| CFTR | None | None | | None | None |

| | | | | |
|---|---|---|---|---|
| CFTR | None | None | None | None |
| CFTR | None | None | None | None |
| CFTR | None | None | None | None |
| CFTR | None | None | None | None |
| CFTR | None | None | None | None |
| CFTR | None | None | None | None |
| CFTR | None | None | None | None |
| CFTR | None | None | None | None |
| CFTR | None | None | None | None |
| CFTR | None | None | None | None |
| CFTR | None | None | None | None |
| CFTR | None | None | None | None |
| CFTR | None | None | None | None |
| CFTR | None | None | None | None |
| CFTR | None | None | None | None |
| CFTR | None | None | None | None |
| CFTR | None | None | None | None |
| CFTR | None | None | None | None |
| CFTR | None | None | None | None |
| CFTR | None | None | None | None |
| CFTR | None | None | None | None |
| CFTR | None | None | None | None |
| CFTR | None | None | None | None |
| CFTR | None | None | None | None |
| CFTR | None | None | None | None |
| FAN1 | None | None | None | None |
| CBS | None | None | None | None |
| CBS | None | None | None | None |
| CBS | None | None | None | None |
| CBS | None | None | None | None |
| CBS | None | None | None | None |
| CBS | None | None | None | None |
| CBS | None | None | None | None |
| CBS | 0.597 | 3 | 5mms | C |
| CBS | 0.54 | 2 | 5mms | A |
| CBS | 0.54 | 2 | 5mms | A |
| CBS | None | None | None | None |
| CBS | None | None | None | None |
| CBS | None | None | None | None |
| CBS | None | None | None | None |
| CBS | 0.516 | 4 | 5mms | C |
| CBS | None | None | None | None |
| CBS | None | None | None | None |
| CBS | None | None | None | None |
| CBS | 0.698 | 2 | 1m54 | B |
| CBS | None | None | None | None |
| CBS | None | None | None | None |
| CBS | None | None | None | None |
| CBS | None | None | None | None |
| CBS | None | None | None | None |
| CBS | None | None | None | None |
| CBS | None | None | None | None |
| CBS | None | None | None | None |
| CBS | None | None | None | None |

| | | | | | |
|---|---|---|---|---|---|
| CBS | | 0.603 | 3 | 4coo | A |
| CBS | | 0.603 | 3 | 4coo | A |
| CBS | None | None | | None | None |
| CBS | None | None | | None | None |
| CBS | None | None | | None | None |
| CBS | None | None | | None | None |
| CBS | None | None | | None | None |
| CBS | None | None | | None | None |
| TLR3 | None | None | | None | None |
| LIG4 | None | None | | None | None |
| LIG4 | None | None | | None | None |
| ASPA | | 0.769 | 1 | 2i3c | B |
| ASPA | | 0.769 | 1 | 2i3c | B |
| ASPA | None | None | | None | None |
| ASPA | None | None | | None | None |
| ASPA | None | None | | None | None |
| ASPA | None | None | | None | None |
| ASPA | None | None | | None | None |
| ASPA | None | None | | None | None |
| ASPA | None | None | | None | None |
| ASPA | None | None | | None | None |
| ASPA | | 0.583 | 2 | 4tnu | A |
| ASPA | | 0.583 | 2 | 4tnu | A |
| ASPA | None | None | | None | None |
| CFH | None | None | | None | None |
| FCGR3A | None | None | | None | None |
| FCGR3A | None | None | | None | None |
| GRIN2A | None | None | | None | None |
| GRIN2A | | 0.658 | 1 | 5i2n | A |
| GRIN2A | None | None | | None | None |
| GLA | None | None | | None | None |
| GLA | None | None | | None | None |
| GLA | None | None | | None | None |
| GLA | None | None | | None | None |
| GLA | None | None | | None | None |
| GLA | None | None | | None | None |
| GLA | None | None | | None | None |
| GLA | | 0.647 | 2 | 3s5y | A |
| GLA | | 0.562 | 20 | 3gxt | B |
| GLA | None | None | | None | None |
| GLA | None | None | | None | None |
| GLA | | 0.75 | 3 | 3gxn | B |
| GLA | | 0.75 | 3 | 3gxn | B |
| GLA | None | None | | None | None |
| GLA | None | None | | None | None |
| GLA | None | None | | None | None |
| GLA | None | None | | None | None |
| GLA | None | None | | None | None |
| GLA | None | None | | None | None |
| GLA | None | None | | None | None |
| GLA | None | None | | None | None |
| GLA | None | None | | None | None |
| GLA | None | None | | None | None |
| GLA | None | None | | None | None |

| | | | | |
|---|---|---|---|---|
| GLA | None | None | None | None |
| GLA | None | None | None | None |
| GLA | None | None | None | None |
| GLA | None | None | None | None |
| GLA | None | None | None | None |
| GLA | None | None | None | None |
| GLA | None | None | None | None |
| SERPINA3 | None | None | None | None |
| MTAP | None | None | None | None |
| MTAP | None | None | None | None |
| MTAP | None | None | None | None |
| BLMH | None | None | None | None |
| TUSC3 | None | None | None | None |
| ISCU | None | None | None | None |
| CYP11B2 | None | None | None | None |
| CYP11B2 | None | None | None | None |
| CYP11B2 | None | None | None | None |
| CYP11B2 | None | None | None | None |
| CYP11B2 | None | None | None | None |
| CYP11B2 | None | None | None | None |
| CYP11B2 | None | None | None | None |
| CYP11B2 | None | None | None | None |
| CYP11B2 | None | None | None | None |
| CYP11B2 | None | None | None | None |
| CYP11B2 | None | None | None | None |
| CYP11B2 | None | None | None | None |
| CYP11B2 | None | None | None | None |
| CYP11B2 | None | None | None | None |
| CYP11B2 | None | None | None | None |
| CYP11B2 | None | None | None | None |
| CYP11B2 | None | None | None | None |
| CYP11B2 | None | None | None | None |
| CYP11B2 | None | None | None | None |
| CYP11B2 | None | None | None | None |
| CRADD | None | None | None | None |
| CRADD | None | None | None | None |
| FGA | None | None | None | None |
| FGA | None | None | None | None |
| FGA | None | None | None | None |
| FGA | None | None | None | None |
| FGA | None | None | None | None |
| FGA | None | None | None | None |
| FGA | None | None | None | None |
| FGA | None | None | None | None |
| IYD | 1 | 1 | 4ttb | A |
| IYD | None | None | None | None |
| IYD | None | None | None | None |
| IYD | None | None | None | None |
| TCAP | None | None | None | None |
| TCAP | None | None | None | None |
| TCAP | None | None | None | None |
| TCAP | None | None | None | None |
| TCAP | None | None | None | None |
| CYP7A1 | None | None | None | None |

| | | | | |
|---|---|---|---|---|
| TEK | None | None | None | None |
| TEK | None | None | None | None |
| TEK | None | None | None | None |
| TEK | None | None | None | None |
| TEK | None | None | None | None |
| TEK | None | None | None | None |
| TEK | None | None | None | None |
| CD8A | None | None | None | None |
| HERC2 | None | None | None | None |
| NR1H3 | 0.537 | 3 | 3ipq | A |
| GRHPR | None | None | None | None |
| GRHPR | None | None | None | None |
| GRHPR | None | None | None | None |
| GRHPR | None | None | None | None |
| GRHPR | 0.697 | 1 | 2wwr | D |
| GRHPR | 0.697 | 1 | 2wwr | D |
| GRHPR | None | None | None | None |
| GBE1 | None | None | None | None |
| GBE1 | None | None | None | None |
| GBE1 | None | None | None | None |
| GBE1 | None | None | None | None |
| GBE1 | None | None | None | None |
| GBE1 | None | None | None | None |
| GBE1 | None | None | None | None |
| GBE1 | None | None | None | None |
| GBE1 | None | None | None | None |
| GBE1 | None | None | None | None |
| GBE1 | None | None | None | None |
| GBE1 | None | None | None | None |
| GBE1 | None | None | None | None |
| GBE1 | None | None | None | None |
| GBE1 | None | None | None | None |
| GBE1 | None | None | None | None |
| GBE1 | None | None | None | None |
| GBE1 | None | None | None | None |
| GBE1 | None | None | None | None |
| GBE1 | None | None | None | None |
| PC | None | None | None | None |
| PC | None | None | None | None |
| PC | None | None | None | None |
| PC | None | None | None | None |
| PC | None | None | None | None |
| PC | None | None | None | None |
| PC | None | None | None | None |
| PC | None | None | None | None |
| MSH6 | None | None | None | None |
| MSH6 | None | None | None | None |
| MSH6 | None | None | None | None |
| MSH6 | None | None | None | None |
| MSH6 | None | None | None | None |
| MSH6 | None | None | None | None |
| MSH6 | None | None | None | None |
| MSH6 | None | None | None | None |
| MSH6 | None | None | None | None |

| | | | | |
|---|---|---|---|---|
| MSH6 | None | None | None | None |
| MSH6 | None | None | None | None |
| MSH6 | None | None | None | None |
| MSH6 | None | None | None | None |
| MSH6 | None | None | None | None |
| MSH6 | None | None | None | None |
| MSH6 | None | None | None | None |
| MSH6 | None | None | None | None |
| MSH6 | None | None | None | None |
| MSH6 | None | None | None | None |
| MSH6 | None | None | None | None |
| MSH6 | None | None | None | None |
| MSH6 | None | None | None | None |
| MSH6 | None | None | None | None |
| MSH6 | None | None | None | None |
| MSH6 | None | None | None | None |
| MSH6 | None | None | None | None |
| MSH6 | None | None | None | None |
| MSH6 | None | None | None | None |
| MSH6 | None | None | None | None |
| MSH6 | None | None | None | None |
| RNASEL | None | None | None | None |
| ORC1 | None | None | None | None |
| ORC1 | None | None | None | None |
| ORC1 | None | None | None | None |
| KCNH2 | None | None | None | None |
| KCNH2 | None | None | None | None |
| KCNH2 | None | None | None | None |
| KCNH2 | None | None | None | None |
| KCNH2 | None | None | None | None |
| KCNH2 | None | None | None | None |
| KCNH2 | None | None | None | None |
| KCNH2 | None | None | None | None |
| KCNH2 | None | None | None | None |
| KCNH2 | None | None | None | None |
| KCNH2 | None | None | None | None |
| KCNH2 | None | None | None | None |
| KCNH2 | None | None | None | None |
| KCNH2 | None | None | None | None |
| KCNH2 | None | None | None | None |
| KCNH2 | None | None | None | None |
| KCNH2 | None | None | None | None |
| KCNH2 | None | None | None | None |
| KCNH2 | None | None | None | None |
| KCNH2 | None | None | None | None |
| KCNH2 | None | None | None | None |
| KCNH2 | None | None | None | None |
| KCNH2 | None | None | None | None |
| KCNH2 | None | None | None | None |
| MAT1A | None | None | None | None |
| MAT1A | None | None | None | None |
| MAT1A | 0.571 | 1 | 2obv | A |
| MAT1A | 0.571 | 1 | 2obv | A |
| MAT1A | None | None | None | None |

| | | | | |
|---|---|---|---|---|
| MAT1A | None | None | None | None |
| MAT1A | None | None | None | None |
| MAT1A | None | None | None | None |
| INSR | None | None | None | None |
| INSR | None | None | None | None |
| INSR | None | None | None | None |
| INSR | None | None | None | None |
| AMT | 0.545 | 5 | 1wsr | A |
| AMT | None | None | None | None |
| AMT | None | None | None | None |
| AMT | None | None | None | None |
| AMT | None | None | None | None |
| AMT | None | None | None | None |
| AMT | None | None | None | None |
| AMT | None | None | None | None |
| AMT | None | None | None | None |
| CSNK1D | None | None | None | None |
| SF3B1 | None | None | None | None |
| AGA | None | None | None | None |
| AGA | None | None | None | None |
| AGA | None | None | None | None |
| AGA | None | None | None | None |
| AGA | None | None | None | None |
| AGA | None | None | None | None |
| AGA | None | None | None | None |
| NECTIN4 | None | None | None | None |
| RS1 | None | None | None | None |
| RS1 | None | None | None | None |
| RS1 | None | None | None | None |
| RS1 | None | None | None | None |
| TBC1D7 | None | None | None | None |
| CTSA | None | None | None | None |
| CTSA | None | None | None | None |
| TKT | None | None | None | None |
| RP2 | None | None | None | None |
| RP2 | None | None | None | None |
| RP2 | None | None | None | None |
| RP2 | None | None | None | None |
| LYRM4 | None | None | None | None |
| SOS1 | None | None | None | None |
| SOS1 | None | None | None | None |
| SOS1 | None | None | None | None |
| SOS1 | None | None | None | None |
| SOS1 | None | None | None | None |
| SOS1 | None | None | None | None |
| HUWE1 | None | None | None | None |
| HUWE1 | None | None | None | None |
| TPI1 | None | None | None | None |
| TPI1 | 0.886 | 1 | 4unk | B |
| TPI1 | None | None | None | None |
| MSH2 | None | None | None | None |
| MSH2 | None | None | None | None |
| MSH2 | None | None | None | None |
| MSH2 | None | None | None | None |

| | | | | |
|---|---|---|---|---|
| MSH2 | None | None | None | None |
| MSH2 | None | None | None | None |
| MSH2 | None | None | None | None |
| MSH2 | None | None | None | None |
| MSH2 | None | None | None | None |
| MSH2 | None | None | None | None |
| MSH2 | None | None | None | None |
| MSH2 | None | None | None | None |
| MSH2 | None | None | None | None |
| MSH2 | None | None | None | None |
| MSH2 | None | None | None | None |
| MSH2 | None | None | None | None |
| MSH2 | None | None | None | None |
| MSH2 | None | None | None | None |
| MSH2 | None | None | None | None |
| MSH2 | None | None | None | None |
| MSH2 | None | None | None | None |
| MSH2 | None | None | None | None |
| MSH2 | None | None | None | None |
| MSH2 | None | None | None | None |
| MSH2 | None | None | None | None |
| MSH2 | None | None | None | None |
| MSH2 | None | None | None | None |
| MSH2 | None | None | None | None |
| MSH2 | None | None | None | None |
| MSH2 | None | None | None | None |
| MSH2 | None | None | None | None |
| MSH2 | None | None | None | None |
| MSH2 | None | None | None | None |
| MSH2 | None | None | None | None |
| MSH2 | None | None | None | None |
| MSH2 | None | None | None | None |
| GALE | None | None | None | None |
| GALE | 0.78 | 1 | 1i3n | A |
| GALE | None | None | None | None |
| GALE | None | None | None | None |
| GALE | None | None | None | None |
| GALE | None | None | None | None |
| GALE | None | None | None | None |
| GALE | None | None | None | None |
| NSD2 | None | None | None | None |
| SRP72 | None | None | None | None |
| SRP72 | None | None | None | None |
| SRP72 | None | None | None | None |
| POLK | None | None | None | None |
| GALT | None | None | None | None |
| GALT | None | None | None | None |
| GALT | None | None | None | None |
| GALT | None | None | None | None |
| GALT | None | None | None | None |
| GALT | None | None | None | None |
| GALT | None | None | None | None |
| GALT | None | None | None | None |
| GALT | None | None | None | None |

| | | | | |
|---|---|---|---|---|
| GALT | None | None | None | None |
| GALT | None | None | None | None |
| GALT | None | None | None | None |
| GALT | None | None | None | None |
| GALT | None | None | None | None |
| GALT | None | None | None | None |
| GALT | None | None | None | None |
| GALT | None | None | None | None |
| GALT | None | None | None | None |
| GALT | None | None | None | None |
| GALT | None | None | None | None |
| GALT | None | None | None | None |
| GALT | None | None | None | None |
| GALT | None | None | None | None |
| GALT | None | None | None | None |
| GALT | None | None | None | None |
| GALT | None | None | None | None |
| GALT | None | None | None | None |
| GALT | None | None | None | None |
| GALT | None | None | None | None |
| GALT | None | None | None | None |
| GALT | None | None | None | None |
| GALT | None | None | None | None |
| GALT | None | None | None | None |
| GALT | None | None | None | None |
| GALT | None | None | None | None |
| GALT | None | None | None | None |
| GALT | None | None | None | None |
| PYCR1 | 0.75 | 11 | 5uav | A |
| PYCR1 | 0.75 | 11 | 5uav | A |
| PYCR1 | 0.5 | 2 | 2izz | A |
| PYCR1 | 0.556 | 2 | 5uax | B |
| F8 | None | None | None | None |
| F8 | None | None | None | None |
| F8 | None | None | None | None |
| F8 | None | None | None | None |
| F8 | None | None | None | None |
| F8 | None | None | None | None |
| F8 | None | None | None | None |
| F8 | None | None | None | None |
| F8 | None | None | None | None |
| F8 | None | None | None | None |
| F8 | None | None | None | None |
| F8 | None | None | None | None |
| F8 | None | None | None | None |
| F8 | None | None | None | None |
| F8 | None | None | None | None |
| F8 | None | None | None | None |
| ANK1 | None | None | None | None |
| CHAT | None | None | None | None |
| CHAT | None | None | None | None |
| CHAT | 0.615 | 8 | 2fy5 | A |
| CHAT | 0.538 | 2 | 2fy5 | A |
| CHAT | None | None | None | None |

| | | | | |
|---|---|---|---|---|
| CHAT | None | None | None | None |
| CHAT | None | None | None | None |
| CHAT | None | None | None | None |
| CHAT | None | None | None | None |
| CHAT | 0.538 | 6 | 2fy5 | A |
| CHAT | None | None | None | None |
| CHAT | None | None | None | None |
| CHAT | None | None | None | None |
| CHAT | None | None | None | None |
| CHAT | None | None | None | None |
| CHAT | None | None | None | None |
| CHAT | None | None | None | None |
| GHR | None | None | None | None |
| GHR | None | None | None | None |
| GHR | None | None | None | None |
| GHR | None | None | None | None |
| GHR | None | None | None | None |
| GHR | None | None | None | None |
| GHR | None | None | None | None |
| GHR | None | None | None | None |
| GHR | None | None | None | None |
| GHR | None | None | None | None |
| GHR | None | None | None | None |
| GHR | None | None | None | None |
| GHR | None | None | None | None |
| GHR | None | None | None | None |
| FZD4 | None | None | None | None |
| FZD4 | None | None | None | None |
| FZD4 | None | None | None | None |
| MTR | None | None | None | None |
| MTR | None | None | None | None |
| MTR | None | None | None | None |
| GM2A | None | None | None | None |
| GM2A | None | None | None | None |
| GM2A | None | None | None | None |
| GM2A | None | None | None | None |
| GM2A | None | None | None | None |
| CRYGD | None | None | None | None |
| CRYGD | None | None | None | None |
| CRYGD | 0.522 | 1 | 2kfb | A |
| CRYGD | None | None | None | None |
| CRYGD | None | None | None | None |
| CRYGD | None | None | None | None |
| CRYGD | None | None | None | None |
| CRYGD | None | None | None | None |
| CRYGD | 0.556 | 2 | 2g98 | A |
| TLR5 | None | None | None | None |
| HSD17B10 | 0.855 | 1 | 1u7t | B |
| HSD17B10 | None | None | None | None |
| HSD17B10 | 0.542 | 1 | 1so8 | A |
| HSD17B10 | None | None | None | None |
| TBC1D20 | None | None | None | None |
| FBN1 | None | None | None | None |
| FBN1 | None | None | None | None |

| | | | | |
|---|---|---|---|---|
| FBN1 | None | None | None | None |
| FBN1 | None | None | None | None |
| FBN1 | None | None | None | None |
| FBN1 | None | None | None | None |
| FBN1 | None | None | None | None |
| FXN | None | None | None | None |
| FXN | None | None | None | None |
| FXN | None | None | None | None |
| FXN | None | None | None | None |
| MYO5A | None | None | None | None |
| CTSK | None | None | None | None |
| CTSK | None | None | None | None |
| SCN5A | None | None | None | None |
| SCN5A | None | None | None | None |
| SOS1 | None | None | None | None |
| SOS1 | None | None | None | None |
| SOS1 | None | None | None | None |
| SOS1 | None | None | None | None |
| SOS1 | None | None | None | None |
| SOS1 | None | None | None | None |
| SOS1 | None | None | None | None |
| SOS1 | None | None | None | None |
| SOS1 | None | None | None | None |
| SOS1 | None | None | None | None |
| SOS1 | None | None | None | None |
| SOS1 | None | None | None | None |
| GATM | None | None | None | None |
| GATM | None | None | None | None |
| GATM | 0.75 | 4 | 4jdw | A |
| GATM | None | None | None | None |
| GATM | None | None | None | None |
| GATM | 0.75 | 6 | 6jdw | A |
| GATM | 0.75 | 11 | 6jdw | A |
| MMP2 | None | None | None | None |
| MMP2 | None | None | None | None |
| MMP2 | None | None | None | None |
| MMP2 | None | None | None | None |
| NDUFB9 | None | None | None | None |
| NDUFB9 | None | None | None | None |
| GH1 | None | None | None | None |
| GH1 | None | None | None | None |
| GH1 | None | None | None | None |
| SLC6A4 | None | None | None | None |
| G6PD | None | None | None | None |
| G6PD | None | None | None | None |
| G6PD | None | None | None | None |
| G6PD | None | None | None | None |
| G6PD | None | None | None | None |
| G6PD | None | None | None | None |
| G6PD | None | None | None | None |
| G6PD | None | None | None | None |
| G6PD | 0.545 | 1 | 1qki | B |
| G6PD | None | None | None | None |
| G6PD | None | None | None | None |

| | | | | |
|---|---|---|---|---|
| G6PD | None | None | None | None |
| G6PD | None | None | None | None |
| G6PD | None | None | None | None |
| G6PD | None | None | None | None |
| G6PD | None | None | None | None |
| G6PD | None | None | None | None |
| G6PD | None | None | None | None |
| G6PD | None | None | None | None |
| G6PD | None | None | None | None |
| CST3 | None | None | None | None |
| LMNA | None | None | None | None |
| LMNA | None | None | None | None |
| NDUFA9 | None | None | None | None |
| NDUFA9 | None | None | None | None |
| NDUFA9 | None | None | None | None |
| NDUFA9 | None | None | None | None |
| YARS | 0.581 | 1 | 5thl | A |
| YARS | None | None | None | None |
| YARS | None | None | None | None |
| YARS | None | None | None | None |
| EPHA2 | None | None | None | None |
| EPHA2 | None | None | None | None |
| EPHA2 | None | None | None | None |
| EPHA2 | None | None | None | None |
| DICER1 | None | None | None | None |
| DICER1 | None | None | None | None |
| SNX10 | None | None | None | None |
| SNX10 | None | None | None | None |
| TTN | None | None | None | None |
| TTN | None | None | None | None |
| GATA3 | None | None | None | None |
| C5 | None | None | None | None |
| THBD | None | None | None | None |
| BIN1 | None | None | None | None |
| BIN1 | None | None | None | None |
| SMARCAL1 | None | None | None | None |
| CFB | None | None | None | None |
| CYC1 | None | None | None | None |
| ANK2 | None | None | None | None |
| CHMP4B | None | None | None | None |
| CHMP4B | None | None | None | None |
| ERBB2 | None | None | None | None |
| ERBB2 | None | None | None | None |
| POLG2 | None | None | None | None |
| POLG2 | None | None | None | None |
| POLG2 | None | None | None | None |
| POLG2 | None | None | None | None |
| POLG2 | None | None | None | None |
| POLG2 | None | None | None | None |
| POLG2 | None | None | None | None |
| POLG2 | None | None | None | None |
| C3 | None | None | None | None |
| C3 | None | None | None | None |
| RPGR | None | None | None | None |

| | | | | |
|---|---|---|---|---|
| RPGR | None | None | None | None |
| RPGR | None | None | None | None |
| RPGR | None | None | None | None |
| RPGR | None | None | None | None |
| C1QA | None | None | None | None |
| PRKG1 | None | None | None | None |
| IGBP1 | None | None | None | None |
| ENO3 | None | None | None | None |
| ENO3 | None | None | None | None |
| ENO3 | None | None | None | None |
| ENO3 | None | None | None | None |
| ENO3 | None | None | None | None |
| ENO3 | 0.897 | 2 | 2xsx | B |
| MCPH1 | None | None | None | None |
| MCPH1 | None | None | None | None |
| MCPH1 | None | None | None | None |
| MCPH1 | None | None | None | None |
| MCPH1 | None | None | None | None |
| SNRNP200 | None | None | None | None |
| SNRNP200 | None | None | None | None |
| DNMT1 | None | None | None | None |
| DNMT1 | None | None | None | None |
| DNMT1 | None | None | None | None |
| AKT1 | None | None | None | None |
| NDUFS2 | None | None | None | None |
| ACVR2B | None | None | None | None |
| HMGCL | None | None | None | None |
| HMGCL | None | None | None | None |
| HMGCL | None | None | None | None |
| JUP | None | None | None | None |
| JUP | None | None | None | None |
| JUP | None | None | None | None |
| JUP | None | None | None | None |
| JUP | None | None | None | None |
| JUP | None | None | None | None |
| JUP | None | None | None | None |
| JUP | None | None | None | None |
| SPINK1 | None | None | None | None |
| SPINK1 | None | None | None | None |
| SPINK1 | None | None | None | None |
| VCL | None | None | None | None |
| VCL | None | None | None | None |
| NOTCH1 | None | None | None | None |
| NOTCH1 | None | None | None | None |
| NOTCH1 | None | None | None | None |
| NDUFB3 | None | None | None | None |
| IKBKG | None | None | None | None |
| GSN | 0.529 | 2 | 2fh1 | A |
| GSN | None | None | None | None |
| GSN | None | None | None | None |
| GSN | 0.5 | 3 | 1p8x | A |
| GSN | None | None | None | None |
| GSN | None | None | None | None |
| GSN | None | None | None | None |

| | | | | |
|---|---|---|---|---|
| GSN | None | None | None | None |
| GSN | None | None | None | None |
| CLC | None | None | None | None |
| RAF1 | None | None | None | None |
| RAF1 | None | None | None | None |
| RAF1 | None | None | None | None |
| NDUFS8 | None | None | None | None |
| NDUFS8 | None | None | None | None |
| NDUFS8 | None | None | None | None |
| NDUFS8 | None | None | None | None |
| NDUFS8 | None | None | None | None |
| CASR | None | None | None | None |
| CASR | None | None | None | None |
| CASR | None | None | None | None |
| CASR | None | None | None | None |
| CASR | None | None | None | None |
| CASR | None | None | None | None |
| CASR | None | None | None | None |
| CASR | None | None | None | None |
| CASR | None | None | None | None |
| CASR | None | None | None | None |
| CASR | None | None | None | None |
| CASR | None | None | None | None |
| CASR | None | None | None | None |
| CASR | None | None | None | None |
| CASR | None | None | None | None |
| CASR | None | None | None | None |
| CASR | None | None | None | None |
| CASR | None | None | None | None |
| CASR | None | None | None | None |
| CASR | None | None | None | None |
| CASR | None | None | None | None |
| CASR | None | None | None | None |
| CASR | None | None | None | None |
| CASR | None | None | None | None |
| CASR | None | None | None | None |
| CASR | None | None | None | None |
| CASR | None | None | None | None |
| CASR | None | None | None | None |
| CASR | None | None | None | None |
| CASR | None | None | None | None |
| VCL | None | None | None | None |
| VCL | None | None | None | None |
| SMPD1 | None | None | None | None |
| SMPD1 | None | None | None | None |
| SMPD1 | None | None | None | None |
| KDM6A | None | None | None | None |
| KCNQ1 | None | None | None | None |
| KCNQ1 | None | None | None | None |
| KCNQ1 | None | None | None | None |
| KCNQ1 | None | None | None | None |
| KCNQ1 | None | None | None | None |
| KCNQ1 | None | None | None | None |
| NDUFA12 | None | None | None | None |
| GLRX5 | None | None | None | None |

| | | | | |
|---|---|---|---|---|
| PALB2 | None | None | None | None |
| PALB2 | None | None | None | None |
| PALB2 | None | None | None | None |
| PALB2 | None | None | None | None |
| PALB2 | None | None | None | None |
| PALB2 | None | None | None | None |
| PALB2 | None | None | None | None |
| PALB2 | None | None | None | None |
| PALB2 | None | None | None | None |
| PALB2 | None | None | None | None |
| ALDOB | None | None | None | None |
| ALDOB | 0.731 | 2 | 1xdm | A |
| ALDOB | 0.731 | 1 | 1qo5 | B |
| ALDOB | 0.538 | 6 | 1xdm | A |
| ALDOB | 0.692 | 7 | 1qo5 | C |
| ALDOB | None | None | None | None |
| ALDOB | 0.577 | 25 | 1qo5 | B |
| GNE | None | None | None | None |
| GNE | None | None | None | None |
| GNE | None | None | None | None |
| GNE | None | None | None | None |
| GNE | None | None | None | None |
| GNE | None | None | None | None |
| GNE | None | None | None | None |
| GNE | None | None | None | None |
| GNE | None | None | None | None |
| GNE | None | None | None | None |
| GNE | None | None | None | None |
| GNE | None | None | None | None |
| CHRNB2 | None | None | None | None |
| CHRNB2 | None | None | None | None |
| ERCC3 | None | None | None | None |
| ERCC3 | None | None | None | None |
| ERCC3 | None | None | None | None |
| ERCC3 | None | None | None | None |
| ERCC3 | None | None | None | None |
| UNC119 | None | None | None | None |
| ATL1 | None | None | None | None |
| ATL1 | None | None | None | None |
| ATL1 | None | None | None | None |
| ATL1 | 0.8 | 1 | 3q5d | A |
| ATL1 | None | None | None | None |
| ATL1 | None | None | None | None |
| ATL1 | None | None | None | None |
| ATL1 | None | None | None | None |
| ATL1 | None | None | None | None |
| ATL1 | None | None | None | None |
| NR3C2 | 0.701 | 1 | 2oax | A |
| TBK1 | None | None | None | None |
| HYAL1 | None | None | None | None |
| HYAL1 | None | None | None | None |
| HYAL1 | None | None | None | None |
| HYAL1 | None | None | None | None |
| CTSD | 0.633 | 1 | 4obz | A |

| | | | | |
|---|---|---|---|---|
| PRKCG | None | None | None | None |
| TGFBR1 | None | None | None | None |
| XDH | None | None | None | None |
| XDH | None | None | None | None |
| XDH | None | None | None | None |
| XDH | None | None | None | None |
| XDH | None | None | None | None |
| XDH | None | None | None | None |
| NMNAT1 | 0.833 | 1 | 1kqn | E |
| NMNAT1 | None | None | None | None |
| NMNAT1 | None | None | None | None |
| NMNAT1 | None | None | None | None |
| NMNAT1 | None | None | None | None |
| NMNAT1 | None | None | None | None |
| SERPING1 | None | None | None | None |
| SERPING1 | None | None | None | None |
| SERPING1 | None | None | None | None |
| SERPING1 | None | None | None | None |
| SERPING1 | None | None | None | None |
| PDZRN3 | None | None | None | None |
| FABP2 | None | None | None | None |
| IFNGR1 | None | None | None | None |
| IFNGR1 | None | None | None | None |
| TDP1 | None | None | None | None |
| TDP1 | None | None | None | None |
| INS | None | None | None | None |
| INS | 0.769 | 1 | 2r34 | C |
| INS | 0.769 | 1 | 2r34 | C |
| PPOX | 0.857 | 1 | 4ivm | B |
| PPOX | None | None | None | None |
| PPOX | None | None | None | None |
| PPOX | None | None | None | None |
| PPOX | None | None | None | None |
| PPOX | None | None | None | None |
| PPOX | None | None | None | None |
| PPOX | None | None | None | None |
| PPOX | None | None | None | None |
| GFER | None | None | None | None |
| GFER | 0.769 | 4 | 3tk0 | A |
| GFER | None | None | None | None |
| GFER | None | None | None | None |
| LIFR | None | None | None | None |
| EGLN1 | 0.556 | 21 | 5a3u | C |
| EGLN1 | 0.889 | 6 | 5v18 | A |
| EGLN1 | 0.944 | 2 | 5a3u | C |
| GP1BA | None | None | None | None |
| GP1BA | None | None | None | None |
| GP1BA | None | None | None | None |
| GP1BA | None | None | None | None |
| GP1BA | None | None | None | None |
| MMADHC | None | None | None | None |
| MMADHC | None | None | None | None |
| MMADHC | None | None | None | None |
| MMADHC | None | None | None | None |

| | | | | |
|---|---|---|---|---|
| MMADHC | None | None | None | None |
| MMADHC | None | None | None | None |
| MMADHC | None | None | None | None |
| ACVRL1 | None | None | None | None |
| ACVRL1 | None | None | None | None |
| RFXANK | None | None | None | None |
| CHIT1 | None | None | None | None |
| UBR1 | None | None | None | None |
| BLM | None | None | None | None |
| BLM | None | None | None | None |
| BLM | None | None | None | None |
| BLM | None | None | None | None |
| BLM | None | None | None | None |
| BLM | None | None | None | None |
| BLM | None | None | None | None |
| BLM | None | None | None | None |
| BLM | None | None | None | None |
| BLM | None | None | None | None |
| BLM | None | None | None | None |
| BLM | None | None | None | None |
| BLM | None | None | None | None |
| VDR | None | None | None | None |
| VDR | None | None | None | None |
| VDR | None | None | None | None |
| VDR | 0.578 | 1 | 1kb2 | B |
| VDR | 0.578 | 1 | 1kb2 | B |
| VDR | 0.578 | 1 | 1kb2 | B |
| EPHA2 | None | None | None | None |
| EPHA2 | None | None | None | None |
| EPHA2 | None | None | None | None |
| EPHA2 | None | None | None | None |
| EPHA2 | None | None | None | None |
| EPHA2 | None | None | None | None |
| EPHA2 | None | None | None | None |
| EPHA2 | None | None | None | None |
| EPHA2 | None | None | None | None |
| EPHA2 | None | None | None | None |
| UBA1 | None | None | None | None |
| STAT1 | None | None | None | None |
| STAT1 | None | None | None | None |
| STAT1 | None | None | None | None |
| STAT1 | None | None | None | None |
| STAT1 | None | None | None | None |
| STAT1 | None | None | None | None |
| STAT1 | None | None | None | None |
| STAT1 | None | None | None | None |
| DNM1L | None | None | None | None |
| DNM1L | None | None | None | None |
| OTC | None | None | None | None |
| OTC | None | None | None | None |
| OTC | None | None | None | None |
| OTC | None | None | None | None |
| OTC | None | None | None | None |
| OTC | None | None | None | None |

| | | | | |
|---|---|---|---|---|
| OTC | None | None | None | None |
| OTC | None | None | None | None |
| OTC | None | None | None | None |
| OTC | None | None | None | None |
| OTC | None | None | None | None |
| OTC | None | None | None | None |
| OTC | None | None | None | None |
| OTC | None | None | None | None |
| OTC | None | None | None | None |
| OTC | None | None | None | None |
| OTC | None | None | None | None |
| RARS | None | None | None | None |
| RARS | None | None | None | None |
| RARS | None | None | None | None |
| SEPSECS | None | None | None | None |
| SEPSECS | None | None | None | None |
| SEPSECS | None | None | None | None |
| SEPSECS | None | None | None | None |
| SEPSECS | None | None | None | None |
| BRCA1 | None | None | None | None |
| BRCA1 | None | None | None | None |
| BRCA1 | 0.538 | 1 | 3pxc | X |
| BRCA1 | None | None | None | None |
| BRCA1 | None | None | None | None |
| BRCA1 | None | None | None | None |
| BRCA1 | None | None | None | None |
| BRCA1 | None | None | None | None |
| BRCA1 | None | None | None | None |
| BRCA1 | None | None | None | None |
| BRCA1 | None | None | None | None |
| BRCA1 | None | None | None | None |
| BRCA1 | None | None | None | None |
| BRCA1 | None | None | None | None |
| BRCA1 | None | None | None | None |
| BRCA1 | None | None | None | None |
| BRCA1 | None | None | None | None |
| BRCA1 | None | None | None | None |
| BRCA1 | None | None | None | None |
| BRCA1 | None | None | None | None |
| BRCA1 | None | None | None | None |
| BRCA1 | None | None | None | None |
| CUBN | None | None | None | None |
| ITGB3 | None | None | None | None |
| MTRR | None | None | None | None |
| MTRR | None | None | None | None |
| MTRR | None | None | None | None |
| MTRR | None | None | None | None |
| MTRR | None | None | None | None |
| MTRR | None | None | None | None |
| MTRR | None | None | None | None |
| MTRR | None | None | None | None |
| IL7R | None | None | None | None |
| IL7R | None | None | None | None |
| RPS24 | None | None | None | None |

| | | | | |
|---|---|---|---|---|
| RPS24 | None | None | None | None |
| HOGA1 | None | None | None | None |
| HOGA1 | None | None | None | None |
| HOGA1 | None | None | None | None |
| HOGA1 | None | None | None | None |
| HOGA1 | None | None | None | None |
| HOGA1 | None | None | None | None |
| HOGA1 | 0.791 | 1 | 3s5n | A |
| HOGA1 | None | None | None | None |
| HOGA1 | None | None | None | None |
| HOGA1 | None | None | None | None |
| HOGA1 | None | None | None | None |
| HOGA1 | None | None | None | None |
| HOGA1 | None | None | None | None |
| GJB2 | None | None | None | None |
| GJB2 | None | None | None | None |
| GJB2 | None | None | None | None |
| GJB2 | None | None | None | None |
| GJB2 | None | None | None | None |
| GJB2 | None | None | None | None |
| GJB2 | None | None | None | None |
| GJB2 | None | None | None | None |
| GJB2 | None | None | None | None |
| GJB2 | None | None | None | None |
| GJB2 | None | None | None | None |
| GJB2 | None | None | None | None |
| GJB2 | None | None | None | None |
| GJB2 | None | None | None | None |
| GJB2 | None | None | None | None |
| GJB2 | None | None | None | None |
| GJB2 | None | None | None | None |
| GJB2 | None | None | None | None |
| GJB2 | None | None | None | None |
| GJB2 | None | None | None | None |
| GJB2 | None | None | None | None |
| GJB2 | None | None | None | None |
| GJB2 | None | None | None | None |
| GJB2 | None | None | None | None |
| GJB2 | None | None | None | None |
| GJB2 | None | None | None | None |
| GJB2 | None | None | None | None |
| GJB2 | None | None | None | None |
| GJB2 | None | None | None | None |
| GJB2 | None | None | None | None |
| GJB2 | None | None | None | None |
| GJB2 | None | None | None | None |
| GJB2 | None | None | None | None |
| GJB2 | None | None | None | None |
| GJB2 | None | None | None | None |
| GJB2 | None | None | None | None |
| GJB2 | None | None | None | None |
| GJB2 | None | None | None | None |
| GJB2 | None | None | None | None |
| GJB2 | None | None | None | None |

| | | | | |
|---|---|---|---|---|
| GJB2 | None | None | None | None |
| GJB2 | None | None | None | None |
| GJB2 | None | None | None | None |
| GJB2 | None | None | None | None |
| GJB2 | None | None | None | None |
| GJB2 | None | None | None | None |
| GJB2 | None | None | None | None |
| GJB2 | None | None | None | None |
| GJB2 | None | None | None | None |
| GJB2 | None | None | None | None |
| GJB2 | None | None | None | None |
| GJB2 | None | None | None | None |
| GJB2 | None | None | None | None |
| GJB2 | None | None | None | None |
| CPS1 | None | None | None | None |
| CPS1 | 0.545 | 127 | 5dou | A |
| CPS1 | None | None | None | None |
| CPS1 | None | None | None | None |
| CPS1 | None | None | None | None |
| CPS1 | None | None | None | None |
| CPS1 | None | None | None | None |
| CPS1 | None | None | None | None |
| CPS1 | None | None | None | None |
| CPS1 | None | None | None | None |
| PSPH | None | None | None | None |
| PSPH | None | None | None | None |
| PSPH | None | None | None | None |
| PSPH | None | None | None | None |
| PSPH | None | None | None | None |
| PSPH | None | None | None | None |
| PEPD | 0.512 | 2 | 5m4l | B |
| PEPD | None | None | None | None |
| PEPD | None | None | None | None |
| PEPD | 0.643 | 2 | 5m4j | B |
| PEPD | 0.643 | 2 | 5m4j | B |
| PEPD | None | None | None | None |
| PEPD | None | None | None | None |
| PEPD | None | None | None | None |
| PEPD | None | None | None | None |
| KDM5C | None | None | None | None |
| KDM5C | None | None | None | None |
| KDM5C | None | None | None | None |
| ITGB6 | None | None | None | None |
| ITGB6 | None | None | None | None |
| ITGB6 | None | None | None | None |
| DARS | None | None | None | None |
| DARS | None | None | None | None |
| DARS | None | None | None | None |
| DARS | None | None | None | None |
| BTK | None | None | None | None |
| BTK | None | None | None | None |
| BTK | None | None | None | None |
| HDAC4 | None | None | None | None |
| ITGA2B | None | None | None | None |

| | | | | |
|---|---|---|---|---|
| ITGA2B | None | None | None | None |
| ITGA2B | None | None | None | None |
| ITGA2B | None | None | None | None |
| PCSK9 | None | None | None | None |
| PCSK9 | None | None | None | None |
| PCSK9 | None | None | None | None |
| PCSK9 | None | None | None | None |
| MUTYH | None | None | None | None |
| MUTYH | None | None | None | None |
| MUTYH | None | None | None | None |
| MUTYH | None | None | None | None |
| MUTYH | None | None | None | None |
| MUTYH | None | None | None | None |
| MUTYH | None | None | None | None |
| MUTYH | None | None | None | None |
| MUTYH | None | None | None | None |
| MUTYH | None | None | None | None |
| MUTYH | None | None | None | None |
| MUTYH | None | None | None | None |
| MUTYH | None | None | None | None |
| MUTYH | None | None | None | None |
| MUTYH | None | None | None | None |
| MUTYH | None | None | None | None |
| MUTYH | None | None | None | None |
| MUTYH | None | None | None | None |
| MUTYH | None | None | None | None |
| MUTYH | None | None | None | None |
| MUTYH | None | None | None | None |
| MUTYH | None | None | None | None |
| MUTYH | None | None | None | None |
| MUTYH | None | None | None | None |
| SLC25A12 | None | None | None | None |
| GSN | None | None | None | None |
| IHH | 0.909 | 1 | 3n1p | B |
| IHH | None | None | None | None |
| IHH | None | None | None | None |
| IHH | 0.818 | 2 | 3n1p | B |
| IHH | 0.727 | 12 | 3n1p | B |
| IHH | 0.636 | 15 | 3n1p | B |
| IHH | 0.818 | 7 | 3n1p | B |
| CD247 | None | None | None | None |
| EP300 | None | None | None | None |
| TTN | None | None | None | None |
| MRI1 | None | None | None | None |
| SGSH | None | None | None | None |
| SGSH | None | None | None | None |
| SGSH | None | None | None | None |
| SGSH | None | None | None | None |
| SGSH | None | None | None | None |
| SGSH | None | None | None | None |
| SGSH | None | None | None | None |
| SGSH | None | None | None | None |
| SGSH | None | None | None | None |
| SGSH | None | None | None | None |

| | | | | |
|---|---|---|---|---|
| SGSH | None | None | None | None |
| SGSH | None | None | None | None |
| SGSH | None | None | None | None |
| SGSH | None | None | None | None |
| SGSH | None | None | None | None |
| SGSH | None | None | None | None |
| SGSH | None | None | None | None |
| SGSH | None | None | None | None |
| SGSH | None | None | None | None |
| SGSH | None | None | None | None |
| SGSH | None | None | None | None |
| SGSH | None | None | None | None |
| CBL | None | None | None | None |
| CANT1 | None | None | None | None |
| CANT1 | None | None | None | None |
| CANT1 | None | None | None | None |
| CANT1 | None | None | None | None |
| CANT1 | None | None | None | None |
| CANT1 | None | None | None | None |
| CANT1 | None | None | None | None |
| SLC1A3 | None | None | None | None |
| SLC1A3 | None | None | None | None |
| SLC1A5 | None | None | None | None |
| SLC1A3 | None | None | None | None |
| SLC1A5 | None | None | None | None |
| SLC1A3 | None | None | None | None |
| SLC1A3 | None | None | None | None |
| CTSD | None | None | None | None |
| CTSD | None | None | None | None |
| CTSD | None | None | None | None |
| CTSD | None | None | None | None |
| CTSD | None | None | None | None |
| MTOR | None | None | None | None |
| ILK | None | None | None | None |
| PCDH15 | None | None | None | None |
| PCDH15 | None | None | None | None |
| PCDH15 | None | None | None | None |
| PCDH15 | None | None | None | None |
| PCDH15 | None | None | None | None |
| PCDH15 | None | None | None | None |
| KLHL40 | None | None | None | None |
| KLHL40 | None | None | None | None |
| KLHL40 | None | None | None | None |
| KLHL40 | None | None | None | None |
| CSTA | None | None | None | None |
| CSTA | None | None | None | None |
| HNMT | None | None | None | None |
| HNMT | None | None | None | None |
| IFNGR2 | None | None | None | None |
| IFNGR2 | None | None | None | None |
| PEX5 | None | None | None | None |
| KIF7 | None | None | None | None |
| KIF7 | None | None | None | None |
| MYL3 | None | None | None | None |

| | | | | |
|---|---|---|---|---|
| MYL3 | None | None | None | None |
| MYL3 | None | None | None | None |
| MYL3 | None | None | None | None |
| MYL3 | None | None | None | None |
| MYL3 | None | None | None | None |
| MYL3 | None | None | None | None |
| NAGLU | None | None | None | None |
| NAGLU | None | None | None | None |
| NAGLU | None | None | None | None |
| NAGLU | None | None | None | None |
| NAGLU | None | None | None | None |
| NAGLU | None | None | None | None |
| NAGLU | None | None | None | None |
| NAGLU | None | None | None | None |
| NAGLU | None | None | None | None |
| NAGLU | None | None | None | None |
| NAGLU | None | None | None | None |
| NAGLU | None | None | None | None |
| NAGLU | None | None | None | None |
| TNFRSF13C | None | None | None | None |
| KIF21A | None | None | None | None |
| KIF21A | None | None | None | None |
| KIF21A | None | None | None | None |
| KIF21A | None | None | None | None |
| KIF21A | None | None | None | None |
| DOCK2 | None | None | None | None |
| DSP | None | None | None | None |
| CDKN2A | None | None | None | None |
| CDKN2A | None | None | None | None |
| CDKN2A | None | None | None | None |
| CDKN2A | None | None | None | None |
| CDKN2A | None | None | None | None |
| CDKN2A | None | None | None | None |
| CDKN2A | None | None | None | None |
| CDKN2A | None | None | None | None |
| CDKN2A | None | None | None | None |
| CDKN2A | None | None | None | None |
| CDKN2A | None | None | None | None |
| CDKN2A | None | None | None | None |
| CDKN2A | None | None | None | None |
| CDKN2A | None | None | None | None |
| VCL | None | None | None | None |
| TPP1 | None | None | None | None |
| TPP1 | None | None | None | None |
| TPP1 | None | None | None | None |
| TPP1 | None | None | None | None |
| TPP1 | None | None | None | None |
| TPP1 | None | None | None | None |
| TPP1 | None | None | None | None |
| TPP1 | None | None | None | None |
| TPP1 | None | None | None | None |
| TPP1 | None | None | None | None |
| TPP1 | None | None | None | None |
| TPP1 | None | None | None | None |

| | | | | |
|---|---|---|---|---|
| TPP1 | None | None | None | None |
| TPP1 | None | None | None | None |
| TPP1 | None | None | None | None |
| TPP1 | None | None | None | None |
| TPP1 | None | None | None | None |
| TPP1 | None | None | None | None |
| TPP1 | None | None | None | None |
| TPP1 | None | None | None | None |
| TPP1 | None | None | None | None |
| TPP1 | None | None | None | None |
| IL10RB | None | None | None | None |
| IL10RB | None | None | None | None |
| CCM2 | None | None | None | None |
| CCM2 | None | None | None | None |
| CCM2 | None | None | None | None |
| DCX | None | None | None | None |
| DCX | None | None | None | None |
| DCX | None | None | None | None |
| DCX | None | None | None | None |
| DCX | None | None | None | None |
| DCX | None | None | None | None |
| DCX | None | None | None | None |
| DCX | None | None | None | None |
| BRCA2 | None | None | None | None |
| NNT | None | None | None | None |
| C2 | None | None | None | None |
| C2 | None | None | None | None |
| C2 | None | None | None | None |
| C2 | None | None | None | None |
| WRN | None | None | None | None |
| WRN | None | None | None | None |
| F13A1 | None | None | None | None |
| F13A1 | None | None | None | None |
| F13A1 | None | None | None | None |
| F13A1 | None | None | None | None |
| TF | None | None | None | None |
| TF | None | None | None | None |
| TF | 0.516 | 2 | 3ve1 | B |
| TF | 0.562 | 39 | 3skp | A |
| TF | None | None | None | None |
| ITPA | None | None | None | None |
| ITPA | None | None | None | None |
| SNCA | None | None | None | None |
| ITGB4 | None | None | None | None |
| ITGB4 | None | None | None | None |
| ITGB4 | None | None | None | None |
| LRP6 | None | None | None | None |
| LRP6 | None | None | None | None |
| LRP6 | None | None | None | None |
| PLG | None | None | None | None |
| PLG | 0.61 | 1 | 1i5k | A |
| TYROBP | None | None | None | None |
| UQCRC2 | None | None | None | None |
| GPSM2 | None | None | None | None |

| | | | | |
|---|---|---|---|---|
| GPSM2 | None | None | None | None |
| GPSM2 | None | None | None | None |
| SCN5A | None | None | None | None |
| SCN5A | None | None | None | None |
| SCN5A | None | None | None | None |
| SCN5A | None | None | None | None |
| SCN5A | None | None | None | None |
| ATIC | 0.6 | 6 | 1p4r | A |
| EZH2 | None | None | None | None |
| CPA1 | None | None | None | None |
| SCN4B | None | None | None | None |
| SCN4B | None | None | None | None |
| MAN1B1 | None | None | None | None |
| MAN1B1 | 0.6 | 9 | 1fmi | A |
| MAN1B1 | None | None | None | None |
| MAN1B1 | None | None | None | None |
| CYCS | 0.74 | 1 | 2n9i | A |
| IDUA | None | None | None | None |
| IDUA | None | None | None | None |
| IDUA | None | None | None | None |
| IDUA | None | None | None | None |
| IDUA | None | None | None | None |
| IDUA | None | None | None | None |
| IDUA | None | None | None | None |
| IDUA | None | None | None | None |
| IDUA | None | None | None | None |
| IDUA | None | None | None | None |
| IDUA | None | None | None | None |
| IDUA | None | None | None | None |
| IDUA | None | None | None | None |
| IDUA | None | None | None | None |
| IDUA | None | None | None | None |
| IDUA | None | None | None | None |
| IDUA | None | None | None | None |
| IDUA | None | None | None | None |
| IDUA | None | None | None | None |
| IDUA | None | None | None | None |
| IDUA | None | None | None | None |
| IDUA | None | None | None | None |
| IDUA | None | None | None | None |
| IDUA | None | None | None | None |
| IDUA | None | None | None | None |
| HIKESHI | None | None | None | None |
| APC | None | None | None | None |
| APC | None | None | None | None |
| APC | None | None | None | None |
| APP | None | None | None | None |
| CEP152 | None | None | None | None |
| GAA | 0.609 | 1 | 5kzx | A |
| GAA | 0.609 | 1 | 5kzx | A |
| GAA | None | None | None | None |
| GAA | None | None | None | None |
| GAA | None | None | None | None |
| GAA | 0.609 | 1 | 5kzx | A |

| | | | | |
|---|---|---|---|---|
| GAA | 0.609 | 1 | 5kzx | A |
| GAA | 0.609 | 1 | 5kzx | A |
| GAA | 0.609 | 1 | 5kzx | A |
| GAA | 0.609 | 1 | 5kzx | A |
| GAA | 0.609 | 1 | 5kzx | A |
| GAA | 0.609 | 1 | 5kzx | A |
| GAA | 0.609 | 1 | 5kzx | A |
| GAA | 0.609 | 1 | 5kzx | A |
| GAA | 0.609 | 1 | 5kzx | A |
| GAA | 0.609 | 1 | 5kzx | A |
| GAA | 0.609 | 1 | 5kzx | A |
| GAA | 0.609 | 1 | 5kzx | A |
| GAA | 0.609 | 1 | 5kzx | A |
| GAA | 0.609 | 1 | 5kzx | A |
| GAA | 0.609 | 1 | 5kzx | A |
| GAA | 0.609 | 1 | 5kzx | A |
| GAA | 0.609 | 1 | 5kzx | A |
| GAA | 0.609 | 1 | 5kzx | A |
| GAA | 0.609 | 1 | 5kzx | A |
| GAA | 0.609 | 1 | 5kzx | A |
| GAA | 0.609 | 1 | 5kzx | A |
| GAA | 0.609 | 1 | 5kzx | A |
| GAA | 0.609 | 1 | 5kzx | A |
| GAA | 0.609 | 1 | 5kzx | A |
| GAA | 0.609 | 1 | 5kzx | A |
| GAA | 0.609 | 1 | 5kzx | A |
| GAA | 0.609 | 1 | 5kzx | A |
| GAA | 0.609 | 1 | 5kzx | A |
| GAA | 0.609 | 1 | 5kzx | A |
| GAA | 0.609 | 1 | 5kzx | A |
| GAA | 0.609 | 1 | 5kzx | A |
| GAA | 0.609 | 1 | 5kzx | A |
| GAA | 0.609 | 1 | 5kzx | A |
| GAA | 0.609 | 1 | 5kzx | A |
| GAA | 0.609 | 1 | 5kzx | A |
| GAA | 0.609 | 1 | 5kzx | A |
| GAA | 0.609 | 1 | 5kzx | A |
| GAA | 0.609 | 1 | 5kzx | A |
| GAA | 0.609 | 1 | 5kzx | A |
| GAA | 0.609 | 1 | 5kzx | A |
| GAA | 0.609 | 1 | 5kzx | A |
| GAA | 0.609 | 1 | 5kzx | A |
| GAA | None | None | None | None |
| FOLR1 | None | None | None | None |
| DSP | None | None | None | None |
| DSP | None | None | None | None |
| DSP | None | None | None | None |
| DSP | None | None | None | None |
| DSP | None | None | None | None |
| CCM2 | None | None | None | None |
| FXN | None | None | None | None |
| FXN | None | None | None | None |
| COMP | None | None | None | None |
| COMP | None | None | None | None |

| | | | | |
|---|---|---|---|---|
| COMP | None | None | None | None |
| COMP | None | None | None | None |
| COMP | None | None | None | None |
| COMP | None | None | None | None |
| COMP | None | None | None | None |
| COMP | None | None | None | None |
| COMP | None | None | None | None |
| COMP | None | None | None | None |
| COMP | None | None | None | None |
| COMP | None | None | None | None |
| AIFM1 | None | None | None | None |
| AIFM1 | None | None | None | None |
| AIFM1 | None | None | None | None |
| AIFM1 | None | None | None | None |
| AIFM1 | None | None | None | None |
| AIFM1 | None | None | None | None |
| AIFM1 | None | None | None | None |
| MYH9 | None | None | None | None |
| FHL2 | None | None | None | None |
| FHL2 | None | None | None | None |
| TNFRSF4 | None | None | None | None |
| CFB | None | None | None | None |
| CFB | None | None | None | None |
| CFB | None | None | None | None |
| CFB | None | None | None | None |
| CFB | None | None | None | None |
| CFB | None | None | None | None |
| CFB | None | None | None | None |
| CFB | None | None | None | None |
| CFB | None | None | None | None |
| CFB | None | None | None | None |
| MVD | None | None | None | None |
| MVD | None | None | None | None |
| HSD17B4 | None | None | None | None |
| HSD17B4 | None | None | None | None |
| HSD17B4 | None | None | None | None |
| HSD17B4 | None | None | None | None |
| HSD17B4 | None | None | None | None |
| HSD17B4 | None | None | None | None |
| F11 | None | None | None | None |
| F11 | None | None | None | None |
| F11 | None | None | None | None |
| F11 | None | None | None | None |
| F11 | None | None | None | None |
| F11 | None | None | None | None |
| F11 | None | None | None | None |
| F11 | None | None | None | None |
| F11 | None | None | None | None |
| SPTA1 | None | None | None | None |
| SPTA1 | None | None | None | None |
| SPTA1 | None | None | None | None |
| SPTA1 | None | None | None | None |
| SPTA1 | None | None | None | None |
| SPTA1 | None | None | None | None |

| | | | | |
|---|---|---|---|---|
| GMPR | None | None | None | None |
| MITF | None | None | None | None |
| NPC1 | None | None | None | None |
| NPC1 | None | None | None | None |
| NPC1 | None | None | None | None |
| NPC1 | None | None | None | None |
| NPC1 | None | None | None | None |
| NPC1 | None | None | None | None |
| NPC1 | None | None | None | None |
| NPC1 | None | None | None | None |
| NPC1 | None | None | None | None |
| NPC1 | None | None | None | None |
| NPC1 | None | None | None | None |
| NPC1 | None | None | None | None |
| NPC1 | None | None | None | None |
| NPC1 | None | None | None | None |
| NPC1 | None | None | None | None |
| NPC1 | None | None | None | None |
| NPC1 | None | None | None | None |
| NPC1 | None | None | None | None |
| NPC1 | None | None | None | None |
| NPC1 | None | None | None | None |
| NPC1 | None | None | None | None |
| NPC1 | None | None | None | None |
| NPC1 | None | None | None | None |
| NPC1 | None | None | None | None |
| NPC1 | None | None | None | None |
| NPC1 | None | None | None | None |
| NPC1 | None | None | None | None |
| NPC1 | None | None | None | None |
| NPC1 | None | None | None | None |
| NPC1 | None | None | None | None |
| NPC1 | None | None | None | None |
| NPC1 | None | None | None | None |
| NPC1 | None | None | None | None |
| NPC1 | None | None | None | None |
| NPC1 | None | None | None | None |
| NPC1 | None | None | None | None |
| NPC1 | None | None | None | None |
| NPC1 | None | None | None | None |
| NPC1 | None | None | None | None |
| NPC1 | None | None | None | None |
| NPC1 | None | None | None | None |
| NPC1 | None | None | None | None |
| NPC1 | None | None | None | None |
| C3 | None | None | None | None |
| NLRX1 | None | None | None | None |
| IGLL1 | None | None | None | None |
| APRT | 0.769 | 4 | 1zn9 | B |
| APRT | None | None | None | None |
| APRT | None | None | None | None |
| APRT | 0.923 | 1 | 1zn8 | A |
| APRT | None | None | None | None |
| ZIC3 | None | None | None | None |

| | | | | |
|---|---|---|---|---|
| ZIC3 | None | None | None | None |
| KLHL3 | None | None | None | None |
| RPS6KA3 | None | None | None | None |
| RPS6KA3 | None | None | None | None |
| RPS6KA3 | None | None | None | None |
| F8 | None | None | None | None |
| F8 | None | None | None | None |
| F8 | None | None | None | None |
| F8 | None | None | None | None |
| F8 | None | None | None | None |
| F8 | None | None | None | None |
| F8 | None | None | None | None |
| F8 | None | None | None | None |
| F8 | None | None | None | None |
| F8 | None | None | None | None |
| F8 | None | None | None | None |
| F8 | None | None | None | None |
| F8 | None | None | None | None |
| F8 | None | None | None | None |
| F8 | None | None | None | None |
| F8 | None | None | None | None |
| F8 | None | None | None | None |
| ISG15 | None | None | None | None |
| EXOSC3 | None | None | None | None |
| EXOSC3 | None | None | None | None |
| EXOSC3 | None | None | None | None |
| NKX2-5 | None | None | None | None |
| NKX2-5 | None | None | None | None |
| NKX2-5 | None | None | None | None |
| NKX2-5 | None | None | None | None |
| FH | None | None | None | None |
| FH | None | None | None | None |
| FH | None | None | None | None |
| FH | None | None | None | None |
| FH | 0.621 | 4 | 30000 | A |
| FH | None | None | None | None |
| FH | None | None | None | None |
| FH | None | None | None | None |
| FH | 0.64 | 6 | 5d6b | A |
| FH | None | None | None | None |
| FH | 0.64 | 1 | 5d6b | A |
| FH | 0.64 | 1 | 5d6b | A |
| SNRNP200 | None | None | None | None |
| SNRNP200 | None | None | None | None |
| SNRNP200 | None | None | None | None |
| SNRNP200 | None | None | None | None |
| CHRNA1 | None | None | None | None |
| CHRNA1 | None | None | None | None |
| MYOM1 | None | None | None | None |
| MYOM1 | None | None | None | None |
| MYOM1 | None | None | None | None |
| MYOM1 | None | None | None | None |
| MYOM1 | None | None | None | None |
| MYOM1 | None | None | None | None |

| | | | | |
|---|---|---|---|---|
| LDLR | None | None | None | None |
| LDLR | None | None | None | None |
| LDLR | None | None | None | None |
| LDLR | None | None | None | None |
| LDLR | None | None | None | None |
| LDLR | None | None | None | None |
| EHMT1 | None | None | None | None |
| EHMT1 | 0.545 | 3 | 3b7b | A |
| EHMT1 | None | None | None | None |
| COL9A2 | None | None | None | None |
| HFE2 | None | None | None | None |
| HFE2 | None | None | None | None |
| CD3G | None | None | None | None |
| MMAB | None | None | None | None |
| MMAB | 0.5 | 3 | 2idx | C |
| MMAB | None | None | None | None |
| MMAB | 0.65 | 3 | 2idx | A |
| MMAB | 0.65 | 3 | 2idx | A |
| MMAB | 0.7 | 2 | 2idx | B |
| MMAB | 0.7 | 2 | 2idx | B |
| MMAB | None | None | None | None |
| MMAB | None | None | None | None |
| MMAB | None | None | None | None |
| TAF1 | None | None | None | None |
| TAF1 | None | None | None | None |
| CD4 | None | None | None | None |
| QARS | None | None | None | None |
| QARS | None | None | None | None |
| QARS | None | None | None | None |
| QARS | None | None | None | None |
| QARS | None | None | None | None |
| FGFR3 | None | None | None | None |
| FGFR3 | None | None | None | None |
| FGFR3 | None | None | None | None |
| FGFR3 | None | None | None | None |
| FGFR3 | None | None | None | None |
| FGFR3 | None | None | None | None |
| FLNB | None | None | None | None |
| FLNB | None | None | None | None |
| FLNB | None | None | None | None |
| FLNB | None | None | None | None |
| FLNB | None | None | None | None |
| FLNB | None | None | None | None |
| FLNB | None | None | None | None |
| FLNB | None | None | None | None |
| IL21R | None | None | None | None |
| RB1 | None | None | None | None |
| RB1 | None | None | None | None |
| RB1 | None | None | None | None |
| NTRK1 | 0.533 | 1 | 1he7 | A |
| NOTCH1 | None | None | None | None |
| MYOM1 | None | None | None | None |
| MYOM1 | None | None | None | None |
| MYOM1 | None | None | None | None |

| | | | | |
|---|---|---|---|---|
| MYOM1 | None | None | None | None |
| MYOM1 | None | None | None | None |
| MYOM1 | None | None | None | None |
| COQ9 | None | None | None | None |
| COQ9 | None | None | None | None |
| CRBN | None | None | None | None |
| DCTN1 | None | None | None | None |
| DCTN1 | None | None | None | None |
| DCTN1 | None | None | None | None |
| DCTN1 | None | None | None | None |
| DCTN1 | None | None | None | None |
| DCTN1 | None | None | None | None |
| THAP1 | None | None | None | None |
| THAP1 | None | None | None | None |
| GNAS | None | None | None | None |
| GNAS | None | None | None | None |
| HNF4A | None | None | None | None |
| HNF4A | None | None | None | None |
| ALDH3A2 | None | None | None | None |
| ALDH3A2 | None | None | None | None |
| ALDH3A2 | None | None | None | None |
| ALDH3A2 | 0.557 | 2 | 4qgk | A |
| ALDH3A2 | None | None | None | None |
| ALDH3A2 | 0.569 | 3 | 4qgk | B |
| ALDH3A2 | None | None | None | None |
| ALDH3A2 | None | None | None | None |
| ALDH3A2 | None | None | None | None |
| HCN1 | None | None | None | None |
| PLEC | None | None | None | None |
| SCN5A | None | None | None | None |
| SCN5A | None | None | None | None |
| STAMBP | None | None | None | None |
| STAMBP | None | None | None | None |
| RNF125 | None | None | None | None |
| RNF125 | None | None | None | None |
| USH1C | None | None | None | None |
| OBSL1 | None | None | None | None |
| GRIN2B | None | None | None | None |
| GRIN2B | None | None | None | None |
| HCN4 | None | None | None | None |
| HCN4 | None | None | None | None |
| TALDO1 | None | None | None | None |
| ITGB3 | None | None | None | None |
| CHKB | None | None | None | None |
| CHKB | None | None | None | None |
| CHKB | None | None | None | None |
| CHKB | None | None | None | None |
| ACTN4 | None | None | None | None |
| ACTN4 | None | None | None | None |
| ACTN4 | None | None | None | None |
| ACTN4 | None | None | None | None |
| CDT1 | None | None | None | None |
| CDT1 | None | None | None | None |
| CDT1 | None | None | None | None |

| | | | | |
|---|---|---|---|---|
| CDT1 | None | None | None | None |
| CDT1 | None | None | None | None |
| CDT1 | None | None | None | None |
| MAPT | None | None | None | None |
| F5 | 0.517 | 4 | 1czs | A |
| F5 | None | None | None | None |
| F5 | None | None | None | None |
| ESRRB | None | None | None | None |
| DGUOK | None | None | None | None |
| DGUOK | None | None | None | None |
| DGUOK | None | None | None | None |
| DGUOK | None | None | None | None |
| DGUOK | 0.765 | 1 | 2ocp | A |
| DGUOK | None | None | None | None |
| DGUOK | None | None | None | None |
| DGUOK | None | None | None | None |
| DGUOK | None | None | None | None |
| CENPJ | None | None | None | None |
| TTN | None | None | None | None |
| TTN | None | None | None | None |
| TTN | None | None | None | None |
| TTN | None | None | None | None |
| TNFRSF1A | None | None | None | None |
| TNFRSF1A | None | None | None | None |
| TNFRSF1A | None | None | None | None |
| TNFRSF1A | None | None | None | None |
| TNFRSF1A | None | None | None | None |
| TNFRSF1A | None | None | None | None |
| TNFRSF1A | None | None | None | None |
| TNFRSF1A | None | None | None | None |
| TNFRSF1A | None | None | None | None |
| TNFRSF1A | None | None | None | None |
| TNFRSF1A | None | None | None | None |
| TNFRSF1A | None | None | None | None |
| ASL | None | None | None | None |
| ASL | 0.625 | 25 | 1aos | A |
| ASL | None | None | None | None |
| ASL | None | None | None | None |
| ASL | None | None | None | None |
| ASL | 0.562 | 30 | 1k62 | A |
| ASL | 0.625 | 1 | 1aos | A |
| ASL | None | None | None | None |
| ASL | None | None | None | None |
| ASL | None | None | None | None |
| MYO7A | None | None | None | None |
| MYO7A | None | None | None | None |
| MYO7A | None | None | None | None |
| MYO7A | None | None | None | None |
| MYO7A | None | None | None | None |
| MYO7A | None | None | None | None |
| MYO7A | None | None | None | None |
| MYO7A | None | None | None | None |
| MYO7A | None | None | None | None |
| MYO7A | None | None | None | None |

| | | | | |
|---|---|---|---|---|
| MYO7A | None | None | None | None |
| MYO7A | None | None | None | None |
| MYO7A | None | None | None | None |
| MYO7A | None | None | None | None |
| MYO7A | None | None | None | None |
| MYO7A | None | None | None | None |
| MYO7A | None | None | None | None |
| MYO7A | None | None | None | None |
| MYO7A | None | None | None | None |
| MYO7A | None | None | None | None |
| APC | None | None | None | None |
| ALAD | None | None | None | None |
| ALAD | None | None | None | None |
| ALAD | None | None | None | None |
| ALAD | 0.542 | 1 | 1E+051 | A |
| ALAD | None | None | None | None |
| ALAD | None | None | None | None |
| POR | None | None | None | None |
| CSF1R | None | None | None | None |
| CSF1R | None | None | None | None |
| CSF1R | None | None | None | None |
| CSF1R | None | None | None | None |
| CSF1R | None | None | None | None |
| CSF1R | None | None | None | None |
| CSF1R | None | None | None | None |
| CSF1R | None | None | None | None |
| CSF1R | None | None | None | None |
| CSF1R | None | None | None | None |
| CSF1R | None | None | None | None |
| VWF | None | None | None | None |
| VWF | None | None | None | None |
| PHF8 | None | None | None | None |
| ACY1 | None | None | None | None |
| SMN1 | None | None | None | None |
| SMN1 | None | None | None | None |
| DLAT | None | None | None | None |
| DLAT | None | None | None | None |
| B2M | None | None | None | None |
| SOX9 | None | None | None | None |
| SOX9 | None | None | None | None |
| SOX9 | None | None | None | None |
| SOX9 | None | None | None | None |
| SOX9 | None | None | None | None |
| CFH | None | None | None | None |
| FLNC | None | None | None | None |
| FLNC | None | None | None | None |
| FLNC | None | None | None | None |
| ITGB2 | None | None | None | None |
| ITGB2 | None | None | None | None |
| ITGB2 | None | None | None | None |
| ITGB2 | None | None | None | None |
| ITGB2 | None | None | None | None |
| ITGB2 | None | None | None | None |
| ITGB2 | None | None | None | None |

| | | | | |
|---|---|---|---|---|
| ITGB2 | None | None | None | None |
| ITGB2 | None | None | None | None |
| PSEN1 | None | None | None | None |
| PSEN1 | None | None | None | None |
| PSEN1 | None | None | None | None |
| PSEN1 | None | None | None | None |
| PSEN1 | None | None | None | None |
| PSEN1 | None | None | None | None |
| OXCT1 | None | None | None | None |
| OXCT1 | None | None | None | None |
| OXCT1 | None | None | None | None |
| OXCT1 | None | None | None | None |
| OXCT1 | None | None | None | None |
| OXCT1 | None | None | None | None |
| OXCT1 | None | None | None | None |
| DHH | None | None | None | None |
| PRKN | None | None | None | None |
| PRKN | None | None | None | None |
| PRKN | None | None | None | None |
| PRKN | None | None | None | None |
| PRKN | None | None | None | None |
| PRKN | None | None | None | None |
| PRKN | None | None | None | None |
| PRKN | None | None | None | None |
| PRKN | None | None | None | None |
| PRKN | None | None | None | None |
| PRKN | None | None | None | None |
| LCAT | None | None | None | None |
| LCAT | None | None | None | None |
| LCAT | None | None | None | None |
| LCAT | None | None | None | None |
| LCAT | None | None | None | None |
| LCAT | None | None | None | None |
| LCAT | None | None | None | None |
| LCAT | None | None | None | None |
| LCAT | None | None | None | None |
| LCAT | None | None | None | None |
| LCAT | None | None | None | None |
| LCAT | None | None | None | None |
| FOXP3 | None | None | None | None |
| FOXP3 | None | None | None | None |
| FOXP3 | None | None | None | None |
| FOXP3 | None | None | None | None |
| SUFU | None | None | None | None |
| HSPA9 | 0.571 | 14 | 4kbo | A |
| HSPA9 | None | None | None | None |
| SQSTM1 | None | None | None | None |
| DARS2 | None | None | None | None |
| DARS2 | None | None | None | None |
| DARS2 | None | None | None | None |
| DARS2 | None | None | None | None |
| DARS2 | None | None | None | None |
| DARS2 | None | None | None | None |
| DARS2 | None | None | None | None |

| | | | | |
|---|---|---|---|---|
| DARS2 | None | None | None | None |
| DARS2 | None | None | None | None |
| DARS2 | None | None | None | None |
| DARS2 | None | None | None | None |
| DARS2 | None | None | None | None |
| NT5C3A | None | None | None | None |
| NT5C3A | None | None | None | None |
| NT5C3A | None | None | None | None |
| NT5C3A | 0.6 | 14 | 2vkq | A |
| ETFA | None | None | None | None |
| ETFA | None | None | None | None |
| ETFA | None | None | None | None |
| ETFA | None | None | None | None |
| ETFA | None | None | None | None |
| ETFA | None | None | None | None |
| ETFA | None | None | None | None |
| TTN | None | None | None | None |
| TTN | None | None | None | None |
| FLT4 | None | None | None | None |
| FGB | None | None | None | None |
| HPCA | None | None | None | None |
| TAP2 | None | None | None | None |
| TAP2 | None | None | None | None |
| PTPRC | None | None | None | None |
| SLC4A1 | None | None | None | None |
| SLC4A1 | None | None | None | None |
| SLC4A1 | None | None | None | None |
| SLC4A1 | None | None | None | None |
| SLC4A1 | None | None | None | None |
| SLC4A1 | None | None | None | None |
| MYH7 | None | None | None | None |
| MYH7 | None | None | None | None |
| MYH7 | None | None | None | None |
| MYH7 | None | None | None | None |
| MYH7 | None | None | None | None |
| MYH7 | None | None | None | None |
| MYH7 | None | None | None | None |
| MYH7 | None | None | None | None |
| MYH7 | None | None | None | None |
| PARN | None | None | None | None |
| PARN | None | None | None | None |
| NDUFS6 | None | None | None | None |
| VRK1 | None | None | None | None |
| VRK1 | None | None | None | None |
| VRK1 | None | None | None | None |
| VRK1 | None | None | None | None |
| VRK1 | None | None | None | None |
| VRK1 | None | None | None | None |
| COLEC11 | None | None | None | None |
| VHL | None | None | None | None |
| TTN | None | None | None | None |
| NCF2 | None | None | None | None |
| NCF2 | None | None | None | None |
| NCF2 | None | None | None | None |

| | | | | |
|---|---|---|---|---|
| NCF2 | None | None | None | None |
| NCF2 | None | None | None | None |
| RPS26 | None | None | None | None |
| CETP | None | None | None | None |
| CETP | None | None | None | None |
| CETP | None | None | None | None |
| CETP | None | None | None | None |
| CETP | None | None | None | None |
| CETP | None | None | None | None |
| CETP | None | None | None | None |
| CETP | None | None | None | None |
| CETP | None | None | None | None |
| CETP | None | None | None | None |
| CETP | None | None | None | None |
| CETP | None | None | None | None |
| CETP | None | None | None | None |
| FLNA | None | None | None | None |
| FLNA | None | None | None | None |
| FLNA | None | None | None | None |
| FLNA | None | None | None | None |
| FLNA | None | None | None | None |
| FLNA | None | None | None | None |
| FLNA | None | None | None | None |
| MYBPC3 | None | None | None | None |
| MYBPC3 | None | None | None | None |
| MYBPC3 | None | None | None | None |
| MYBPC3 | None | None | None | None |
| MYBPC3 | None | None | None | None |
| MYBPC3 | None | None | None | None |
| MYBPC3 | None | None | None | None |
| MYBPC3 | None | None | None | None |
| MYBPC3 | None | None | None | None |
| MYBPC3 | None | None | None | None |
| LSS | None | None | None | None |
| C6 | None | None | None | None |
| C6 | None | None | None | None |
| RLBP1 | None | None | None | None |
| RLBP1 | None | None | None | None |
| RLBP1 | None | None | None | None |
| RLBP1 | None | None | None | None |
| RLBP1 | None | None | None | None |
| RLBP1 | None | None | None | None |
| RLBP1 | None | None | None | None |
| RLBP1 | None | None | None | None |
| MMAA | None | None | None | None |
| MMAA | None | None | None | None |
| MMAA | None | None | None | None |
| MMAA | None | None | None | None |
| MMAA | None | None | None | None |
| MMAA | None | None | None | None |
| IL1RN | None | None | None | None |
| MASTL | None | None | None | None |
| FHL1 | None | None | None | None |
| FHL1 | None | None | None | None |

| | | | | |
|---|---|---|---|---|
| FHL1 | None | None | None | None |
| FHL1 | None | None | None | None |
| FHL1 | None | None | None | None |
| FHL1 | None | None | None | None |
| FHL1 | None | None | None | None |
| FHL1 | None | None | None | None |
| FHL1 | None | None | None | None |
| EIF2B1 | 0.636 | 1 | 3ecs | H |
| EIF2B1 | 0.636 | 1 | 3ecs | H |
| COQ8A | None | None | None | None |
| COQ8A | None | None | None | None |
| COQ8A | None | None | None | None |
| COQ8A | None | None | None | None |
| COQ8A | None | None | None | None |
| COQ8A | None | None | None | None |
| COQ8A | None | None | None | None |
| COQ8A | None | None | None | None |
| COQ8A | None | None | None | None |
| COQ8A | None | None | None | None |
| COQ8A | None | None | None | None |
| COQ8A | None | None | None | None |
| COQ8A | None | None | None | None |
| COQ8A | None | None | None | None |
| COQ8A | None | None | None | None |
| COQ8A | None | None | None | None |
| UNC119 | None | None | None | None |
| IGHMBP2 | None | None | None | None |
| JAG1 | None | None | None | None |
| JAG1 | None | None | None | None |
| JAG1 | None | None | None | None |
| GNB1 | None | None | None | None |
| GNB1 | None | None | None | None |
| FLNA | None | None | None | None |
| FLNA | None | None | None | None |
| FLNA | None | None | None | None |
| FLNA | None | None | None | None |
| FLNA | None | None | None | None |
| CREBBP | None | None | None | None |
| CSF3R | None | None | None | None |
| CSF3R | None | None | None | None |
| HIBCH | None | None | None | None |
| HIBCH | None | None | None | None |
| HIBCH | None | None | None | None |
| MMACHC | None | None | None | None |
| MMACHC | None | None | None | None |
| MMACHC | None | None | None | None |
| MMACHC | None | None | None | None |
| MMACHC | None | None | None | None |
| MMACHC | None | None | None | None |
| MMACHC | None | None | None | None |
| MMACHC | None | None | None | None |
| MMACHC | None | None | None | None |
| MMACHC | None | None | None | None |
| MMACHC | None | None | None | None |

| | | | | |
|---|---|---|---|---|
| KLF1 | None | None | None | None |
| NAGS | None | None | None | None |
| NAGS | None | None | None | None |
| NAGS | None | None | None | None |
| PIK3R2 | None | None | None | None |
| EDNRB | None | None | None | None |
| EDNRB | None | None | None | None |
| EDNRB | None | None | None | None |
| EDNRB | None | None | None | None |
| E | None | None | None | None |
| RUNX1 | None | None | None | None |
| RUNX1 | None | None | None | None |
| CTNNB1 | None | None | None | None |
| CTNNB1 | None | None | None | None |
| CTNNB1 | None | None | None | None |
| CTNNB1 | None | None | None | None |
| CTNNB1 | None | None | None | None |
| CTNNB1 | None | None | None | None |
| TTN | None | None | None | None |
| TTN | None | None | None | None |
| TYK2 | None | None | None | None |
| TYK2 | None | None | None | None |
| TYK2 | None | None | None | None |
| ERCC4 | None | None | None | None |
| ERCC4 | None | None | None | None |
| ERCC4 | None | None | None | None |
| ERCC4 | None | None | None | None |
| CFI | None | None | None | None |
| CFI | None | None | None | None |
| CFI | None | None | None | None |
| CFI | None | None | None | None |
| CFI | None | None | None | None |
| CFI | None | None | None | None |
| CFI | None | None | None | None |
| CFI | None | None | None | None |
| CFI | None | None | None | None |
| CFI | None | None | None | None |
| CFI | None | None | None | None |
| CFI | 0.691 | 2 | 5o32 | J |
| CFI | 0.691 | 2 | 5o32 | J |
| CFI | 0.524 | 5 | 5o32 | I |
| NOTCH3 | None | None | None | None |
| NOTCH3 | None | None | None | None |
| NOTCH3 | None | None | None | None |
| NOTCH3 | None | None | None | None |
| TBX1 | None | None | None | None |
| TBX1 | None | None | None | None |
| DDB2 | None | None | None | None |
| DDB2 | None | None | None | None |
| TP53 | None | None | None | None |
| POMGNT1 | None | None | None | None |
| POMGNT1 | None | None | None | None |
| POMGNT1 | None | None | None | None |
| POMGNT1 | None | None | None | None |

| | | | | |
|---|---|---|---|---|
| POMGNT1 | None | None | None | None |
| POMGNT1 | None | None | None | None |
| POMGNT1 | None | None | None | None |
| POMGNT1 | None | None | None | None |
| POMGNT1 | None | None | None | None |
| POMGNT1 | None | None | None | None |
| POMGNT1 | None | None | None | None |
| POMGNT1 | None | None | None | None |
| POMGNT1 | None | None | None | None |
| FMR1 | None | None | None | None |
| FMR1 | None | None | None | None |
| SLC9A3R1 | None | None | None | None |
| HSPB1 | None | None | None | None |
| HSPB1 | None | None | None | None |
| HSPB1 | None | None | None | None |
| HSPB1 | None | None | None | None |
| HSPB1 | None | None | None | None |
| HSPB1 | None | None | None | None |
| HSPB1 | None | None | None | None |
| SOD3 | None | None | None | None |
| AGA | 1 | 2 | 1apz | D |
| AGA | None | None | None | None |
| AGA | None | None | None | None |
| NDUFS7 | None | None | None | None |
| NDUFS7 | None | None | None | None |
| SOX2 | None | None | None | None |
| CSRP3 | None | None | None | None |
| CSRP3 | None | None | None | None |
| PAH | None | None | None | None |
| PAH | None | None | None | None |
| PAH | None | None | None | None |
| PAH | None | None | None | None |
| CACNA1S | None | None | None | None |
| TRPV4 | None | None | None | None |
| TRPV4 | None | None | None | None |
| TRPV4 | None | None | None | None |
| TRPV4 | 0.58 | 3 | 4dx2 | A |
| TRPV4 | None | None | None | None |
| TRPV4 | None | None | None | None |
| TRPV4 | None | None | None | None |
| TRPV4 | None | None | None | None |
| TRPV4 | None | None | None | None |
| TRPV4 | None | None | None | None |
| TRPV4 | None | None | None | None |
| TRPV4 | None | None | None | None |
| TRPV4 | None | None | None | None |
| TNNI3 | None | None | None | None |
| MLYCD | None | None | None | None |
| MLYCD | None | None | None | None |
| MLYCD | None | None | None | None |
| MLYCD | None | None | None | None |
| AIP | None | None | None | None |
| AIP | None | None | None | None |
| ERCC2 | None | None | None | None |

| | | | | |
|---|---|---|---|---|
| ERCC2 | None | None | None | None |
| ERCC2 | None | None | None | None |
| ERCC2 | None | None | None | None |
| ERCC2 | None | None | None | None |
| ERCC2 | None | None | None | None |
| ERCC2 | None | None | None | None |
| ERCC2 | None | None | None | None |
| ERCC2 | None | None | None | None |
| ERCC2 | None | None | None | None |
| ERCC2 | None | None | None | None |
| ERCC2 | None | None | None | None |
| ERCC2 | None | None | None | None |
| ERCC2 | None | None | None | None |
| PUS1 | None | None | None | None |
| PUS1 | None | None | None | None |
| PUS1 | None | None | None | None |
| CASK | None | None | None | None |
| CASK | None | None | None | None |
| BLVRA | None | None | None | None |
| DICER1 | None | None | None | None |
| DICER1 | None | None | None | None |
| DICER1 | None | None | None | None |
| EIF2AK3 | None | None | None | None |
| EIF2AK3 | None | None | None | None |
| EIF2AK3 | None | None | None | None |
| KIT | None | None | None | None |
| KIT | None | None | None | None |
| KIT | None | None | None | None |
| KIT | None | None | None | None |
| KIT | None | None | None | None |
| KIT | None | None | None | None |
| DNMT3B | None | None | None | None |
| TGM1 | None | None | None | None |
| NDUFA10 | None | None | None | None |
| NDUFA10 | None | None | None | None |
| PRPF31 | None | None | None | None |
| PRPF31 | None | None | None | None |
| ECE1 | None | None | None | None |
| ECE1 | None | None | None | None |
| ECE1 | None | None | None | None |
| NDUFV2 | None | None | None | None |
| NDUFV2 | None | None | None | None |
| NDUFV2 | None | None | None | None |
| GHRHR | None | None | None | None |
| GHRHR | None | None | None | None |
| GHRHR | None | None | None | None |
| SLC4A1 | None | None | None | None |
| FGF9 | None | None | None | None |
| BARD1 | None | None | None | None |
| BARD1 | None | None | None | None |
| BARD1 | None | None | None | None |
| PTPN12 | 0.741 | 2 | 5j8r | D |
| MASP2 | None | None | None | None |
| MASP2 | None | None | None | None |

| | | | | |
|---|---|---|---|---|
| MASP2 | None | None | None | None |
| MASP2 | None | None | None | None |
| HOXB1 | None | None | None | None |
| SLC4A1 | None | None | None | None |
| SLC4A1 | None | None | None | None |
| SLC4A1 | None | None | None | None |
| SLC4A1 | None | None | None | None |
| SLC4A1 | None | None | None | None |
| SLC4A1 | None | None | None | None |
| SLC4A1 | None | None | None | None |
| SLC4A1 | None | None | None | None |
| SLC4A1 | None | None | None | None |
| SLC4A1 | None | None | None | None |
| SLC4A1 | None | None | None | None |
| SLC4A1 | None | None | None | None |
| SLC4A1 | None | None | None | None |
| SLC4A1 | None | None | None | None |
| SLC4A1 | None | None | None | None |
| SLC4A1 | None | None | None | None |
| SLC4A1 | None | None | None | None |
| CBL | None | None | None | None |
| FAS | None | None | None | None |
| MTMR2 | None | None | None | None |
| MTMR2 | None | None | None | None |
| MTMR2 | None | None | None | None |
| MTMR2 | None | None | None | None |
| MTMR2 | None | None | None | None |
| MTMR2 | None | None | None | None |
| MTMR2 | None | None | None | None |
| MTMR2 | None | None | None | None |
| KLHL3 | None | None | None | None |
| KLHL3 | None | None | None | None |
| KLHL3 | 0.667 | 1 | 4ch9 | B |
| KLHL3 | 0.733 | 1 | 5nkp | A |
| KLHL3 | None | None | None | None |
| KLHL3 | None | None | None | None |
| RNASEH1 | None | None | None | None |
| ACTN1 | None | None | None | None |
| PHKG2 | None | None | None | None |
| PHKG2 | None | None | None | None |
| PHKG2 | None | None | None | None |
| NTRK1 | None | None | None | None |
| APOH | None | None | None | None |
| PMS2 | None | None | None | None |
| PMS2 | None | None | None | None |
| PMS2 | 0.797 | 1 | 1ea6 | A |
| PMS2 | None | None | None | None |
| PMS2 | None | None | None | None |
| PMS2 | None | None | None | None |
| PMS2 | None | None | None | None |
| PMS2 | None | None | None | None |
| PMS2 | None | None | None | None |
| PMS2 | None | None | None | None |
| PMS2 | None | None | None | None |

| | | | | |
|---|---|---|---|---|
| PMS2 | None | None | None | None |
| PMS2 | None | None | None | None |
| PMS2 | None | None | None | None |
| PMS2 | None | None | None | None |
| PMS2 | None | None | None | None |
| PMS2 | None | None | None | None |
| PMS2 | None | None | None | None |
| PMS2 | None | None | None | None |
| PMS2 | None | None | None | None |
| PMS2 | None | None | None | None |
| PMS2 | None | None | None | None |
| MOCS2 | None | None | None | None |
| MOCS2 | None | None | None | None |
| RPGRIP1 | None | None | None | None |
| RPGRIP1 | None | None | None | None |
| LIG4 | None | None | None | None |
| PROC | None | None | None | None |
| PROC | None | None | None | None |
| NAT1 | None | None | None | None |
| NAT1 | None | None | None | None |
| PNPO | None | None | None | None |
| PNPO | 0.5 | 2 | 1nrg | A |
| PNPO | None | None | None | None |
| PNPO | None | None | None | None |
| PNPO | None | None | None | None |
| MRE11 | None | None | None | None |
| MRE11 | None | None | None | None |
| MRE11 | None | None | None | None |
| MRE11 | None | None | None | None |
| DDC | None | None | None | None |
| DDC | None | None | None | None |
| DDC | None | None | None | None |
| DDC | None | None | None | None |
| DDC | None | None | None | None |
| DDC | 0.909 | 7 | 3rch | B |
| DDC | 0.909 | 7 | 3rch | B |
| DDC | 0.643 | 4 | 3rbf | A |
| DDC | None | None | None | None |
| DDC | None | None | None | None |
| DDC | None | None | None | None |
| ANK2 | None | None | None | None |
| ANK2 | None | None | None | None |
| ANK2 | None | None | None | None |
| AK2 | None | None | None | None |
| AK2 | None | None | None | None |
| AK2 | None | None | None | None |
| AK2 | None | None | None | None |
| AK2 | None | None | None | None |
| AK2 | None | None | None | None |
| GPHN | None | None | None | None |
| HSD17B4 | 0.66 | 1 | 1zbq | A |
| HSD17B4 | 0.66 | 1 | 1zbq | A |
| HSD17B4 | None | None | None | None |
| HSD17B4 | None | None | None | None |

| | | | | |
|---|---|---|---|---|
| HSD17B4 | None | None | None | None |
| HSD17B4 | None | None | None | None |
| APTX | 0.6 | 5 | 4ndg | A |
| APTX | 0.6 | 5 | 4ndg | A |
| APTX | None | None | None | None |
| APTX | None | None | None | None |
| APTX | None | None | None | None |
| APTX | None | None | None | None |
| APTX | None | None | None | None |
| FANCL | None | None | None | None |
| FANCL | None | None | None | None |
| EMG1 | None | None | None | None |
| EMG1 | None | None | None | None |
| EZH2 | None | None | None | None |
| ABCB6 | None | None | None | None |
| NCF2 | None | None | None | None |
| NCF2 | None | None | None | None |
| COL4A2 | None | None | None | None |
| COL4A2 | 1 | 1 | 1li1 | F |
| COL4A2 | None | None | None | None |
| COL4A2 | None | None | None | None |
| COL4A2 | 0.75 | 16 | 1li1 | C |
| SLX4 | None | None | None | None |
| SERPIND1 | None | None | None | None |
| FGFR3 | None | None | None | None |
| FGFR3 | None | None | None | None |
| FGFR3 | None | None | None | None |
| FGFR3 | None | None | None | None |
| FGFR3 | None | None | None | None |
| FGFR3 | None | None | None | None |
| TREM2 | None | None | None | None |
| TREM2 | None | None | None | None |
| TREM2 | None | None | None | None |
| TREM2 | None | None | None | None |
| TREM2 | None | None | None | None |
| TREM2 | None | None | None | None |
| TREM2 | None | None | None | None |
| LIG4 | None | None | None | None |
| LIG4 | None | None | None | None |
| LIG4 | None | None | None | None |
| LIG4 | None | None | None | None |
| LIG4 | None | None | None | None |
| LIG4 | None | None | None | None |
| ECHS1 | None | None | None | None |
| ECHS1 | None | None | None | None |
| ECHS1 | None | None | None | None |
| MTR | None | None | None | None |
| MTR | None | None | None | None |
| MTR | None | None | None | None |
| MTR | None | None | None | None |
| MTR | None | None | None | None |
| PMVK | None | None | None | None |
| PRNP | None | None | None | None |
| CHN1 | None | None | None | None |

| | | | | |
|---|---|---|---|---|
| CHN1 | None | None | None | None |
| CHN1 | None | None | None | None |
| CHN1 | None | None | None | None |
| CHN1 | None | None | None | None |
| CHN1 | None | None | None | None |
| CHN1 | None | None | None | None |
| CHN1 | None | None | None | None |
| SLC2A1 | None | None | None | None |
| SLC2A1 | None | None | None | None |
| SLC2A1 | None | None | None | None |
| SLC2A1 | None | None | None | None |
| SLC2A1 | None | None | None | None |
| SLC2A1 | None | None | None | None |
| SLC2A1 | None | None | None | None |
| SLC2A1 | None | None | None | None |
| SLC2A1 | None | None | None | None |
| SLC2A1 | None | None | None | None |
| SLC2A1 | None | None | None | None |
| SLC2A1 | None | None | None | None |
| SLC2A1 | None | None | None | None |
| SLC2A1 | None | None | None | None |
| SLC2A1 | None | None | None | None |
| SLC2A1 | None | None | None | None |
| SLC2A1 | None | None | None | None |
| PMM2 | None | None | None | None |
| PMM2 | None | None | None | None |
| PMM2 | None | None | None | None |
| PMM2 | None | None | None | None |
| PMM2 | None | None | None | None |
| PMM2 | None | None | None | None |
| PMM2 | None | None | None | None |
| PMM2 | None | None | None | None |
| PMM2 | None | None | None | None |
| PMM2 | None | None | None | None |
| PMM2 | None | None | None | None |
| PMM2 | None | None | None | None |
| PMM2 | None | None | None | None |
| PMM2 | None | None | None | None |
| PMM2 | None | None | None | None |
| PMM2 | None | None | None | None |
| PMM2 | None | None | None | None |
| PMM2 | None | None | None | None |
| PMM2 | None | None | None | None |
| PMM2 | None | None | None | None |
| PMM2 | None | None | None | None |
| PMM2 | None | None | None | None |
| PMM2 | None | None | None | None |
| PMM2 | None | None | None | None |
| PMM2 | None | None | None | None |
| PMM2 | None | None | None | None |
| PMM2 | None | None | None | None |
| PMM2 | None | None | None | None |
| CCKAR | None | None | None | None |
| MAN1B1 | None | None | None | None |

| | | | | |
|---|---|---|---|---|
| MAN1B1 | None | None | None | None |
| LAMA2 | None | None | None | None |
| LAMA2 | None | None | None | None |
| TBC1D4 | None | None | None | None |
| TBC1D4 | None | None | None | None |
| LDLR | None | None | None | None |
| LDLR | None | None | None | None |
| LDLR | None | None | None | None |
| LDLR | None | None | None | None |
| LDLR | None | None | None | None |
| LDLR | None | None | None | None |
| LDLR | None | None | None | None |
| LDLR | None | None | None | None |
| LDLR | None | None | None | None |
| LDLR | None | None | None | None |
| LDLR | None | None | None | None |
| LDLR | None | None | None | None |
| LDLR | None | None | None | None |
| LDLR | None | None | None | None |
| LDLR | None | None | None | None |
| LDLR | None | None | None | None |
| LDLR | None | None | None | None |
| LDLR | None | None | None | None |
| LDLR | None | None | None | None |
| LDLR | None | None | None | None |
| LDLR | None | None | None | None |
| LDLR | None | None | None | None |
| LDLR | None | None | None | None |
| LDLR | None | None | None | None |
| LDLR | None | None | None | None |
| LDLR | None | None | None | None |
| LDLR | None | None | None | None |
| LDLR | None | None | None | None |
| LDLR | None | None | None | None |
| LDLR | None | None | None | None |
| LDLR | None | None | None | None |
| LDLR | None | None | None | None |
| LDLR | None | None | None | None |
| LDLR | None | None | None | None |
| LDLR | None | None | None | None |
| LDLR | None | None | None | None |
| LDLR | None | None | None | None |
| LDLR | None | None | None | None |
| LDLR | None | None | None | None |
| LDLR | None | None | None | None |
| LDLR | None | None | None | None |
| LDLR | 0.5 | 1 | 1ijq | A |
| LDLR | None | None | None | None |
| FANCM | None | None | None | None |
| HAMP | None | None | None | None |
| GUSB | None | None | None | None |
| GUSB | None | None | None | None |
| GUSB | None | None | None | None |
| GUSB | None | None | None | None |

| | | | | |
|---|---|---|---|---|
| GUSB | None | None | None | None |
| GUSB | None | None | None | None |
| GUSB | None | None | None | None |
| GUSB | None | None | None | None |
| GUSB | None | None | None | None |
| GUSB | None | None | None | None |
| GUSB | None | None | None | None |
| GUSB | None | None | None | None |
| GUSB | None | None | None | None |
| LIG1 | None | None | None | None |
| LIG1 | None | None | None | None |
| MYH7 | None | None | None | None |
| MYH7 | None | None | None | None |
| MYH7 | None | None | None | None |
| MYH7 | None | None | None | None |
| MYH7 | None | None | None | None |
| MYH7 | None | None | None | None |
| MYH7 | None | None | None | None |
| MYH7 | None | None | None | None |
| MYH7 | None | None | None | None |
| MYH7 | None | None | None | None |
| MYH7 | None | None | None | None |
| MYH7 | None | None | None | None |
| MYH7 | None | None | None | None |
| MYH7 | None | None | None | None |
| MYH7 | None | None | None | None |
| MYH7 | None | None | None | None |
| MYH7 | None | None | None | None |
| MYH7 | None | None | None | None |
| MYH7 | None | None | None | None |
| MYH7 | None | None | None | None |
| MYH7 | None | None | None | None |
| MYH7 | None | None | None | None |
| MYH7 | None | None | None | None |
| MYH7 | None | None | None | None |
| MYH7 | None | None | None | None |
| MYH7 | None | None | None | None |
| MYH7 | None | None | None | None |
| MYH7 | None | None | None | None |
| MYH7 | None | None | None | None |
| MYH7 | None | None | None | None |
| MYH7 | None | None | None | None |
| MYH7 | None | None | None | None |
| MYH7 | None | None | None | None |
| MYH7 | None | None | None | None |
| HCN4 | None | None | None | None |
| RECQL4 | None | None | None | None |
| RECQL4 | None | None | None | None |
| RECQL4 | None | None | None | None |
| RECQL4 | None | None | None | None |
| RECQL4 | None | None | None | None |
| RECQL4 | None | None | None | None |
| RECQL4 | None | None | None | None |
| RECQL4 | None | None | None | None |

| | | | | |
|---|---|---|---|---|
| RECQL4 | None | None | None | None |
| RECQL4 | None | None | None | None |
| RECQL4 | None | None | None | None |
| RECQL4 | None | None | None | None |
| RECQL4 | None | None | None | None |
| CHEK2 | None | None | None | None |
| PPARG | None | None | None | None |
| ACADVL | None | None | None | None |
| ACADVL | None | None | None | None |
| ACADVL | None | None | None | None |
| ACADVL | None | None | None | None |
| ACADVL | None | None | None | None |
| ACADVL | None | None | None | None |
| ACADVL | None | None | None | None |
| ACADVL | None | None | None | None |
| ACADVL | None | None | None | None |
| ACADVL | None | None | None | None |
| ACADVL | None | None | None | None |
| ACADVL | None | None | None | None |
| ACADVL | None | None | None | None |
| ACADVL | None | None | None | None |
| ACADVL | None | None | None | None |
| ACADVL | None | None | None | None |
| ACADVL | None | None | None | None |
| ACADVL | None | None | None | None |
| ACADVL | None | None | None | None |
| ACADVL | None | None | None | None |
| ACADVL | None | None | None | None |
| ACADVL | None | None | None | None |
| APOA1 | None | None | None | None |
| APOA1 | None | None | None | None |
| APOA1 | None | None | None | None |
| APOA1 | None | None | None | None |
| APOA1 | None | None | None | None |
| DNMT3A | None | None | None | None |
| DNMT3A | None | None | None | None |
| APC | None | None | None | None |
| TMOD1 | None | None | None | None |
| USH1C | None | None | None | None |
| USH1C | None | None | None | None |
| USH1C | None | None | None | None |
| USH1C | None | None | None | None |
| ERCC6 | None | None | None | None |
| ERCC6 | None | None | None | None |
| ERCC6 | None | None | None | None |
| ERCC8 | None | None | None | None |
| ERCC8 | None | None | None | None |
| ERCC8 | None | None | None | None |
| DPAGT1 | None | None | None | None |
| DPAGT1 | None | None | None | None |
| DPAGT1 | None | None | None | None |
| DPAGT1 | None | None | None | None |
| DPAGT1 | None | None | None | None |
| DPAGT1 | None | None | None | None |

| | | | | |
|---|---|---|---|---|
| DPAGT1 | None | None | None | None |
| IL2RG | None | None | None | None |
| HNF1A | None | None | None | None |
| HNF4A | None | None | None | None |
| FLT4 | None | None | None | None |
| FLT4 | None | None | None | None |
| FLT4 | None | None | None | None |
| CD3D | None | None | None | None |
| CD3D | None | None | None | None |
| MCOLN1 | None | None | None | None |
| MCOLN1 | None | None | None | None |
| MOCS2 | None | None | None | None |
| MOCS2 | None | None | None | None |
| ABCG8 | None | None | None | None |
| ABCG8 | None | None | None | None |
| ABCG8 | None | None | None | None |
| ABCG8 | None | None | None | None |
| ABCG8 | None | None | None | None |
| ABCG8 | None | None | None | None |
| ABCG8 | None | None | None | None |
| ABCG8 | None | None | None | None |
| ABCG8 | None | None | None | None |
| ABCG8 | None | None | None | None |
| ABCG8 | None | None | None | None |
| ABCG8 | None | None | None | None |
| ABCG8 | None | None | None | None |
| ABCG8 | None | None | None | None |
| CTNNA1 | None | None | None | None |
| TYMP | None | None | None | None |
| TYMP | None | None | None | None |
| TYMP | None | None | None | None |
| TYMP | None | None | None | None |
| TYMP | None | None | None | None |
| TYMP | None | None | None | None |
| TYMP | None | None | None | None |
| TYMP | None | None | None | None |
| TYMP | None | None | None | None |
| TYMP | None | None | None | None |
| TYMP | None | None | None | None |
| TYMP | None | None | None | None |
| TYMP | None | None | None | None |
| GAMT | None | None | None | None |
| GAMT | None | None | None | None |
| GAMT | None | None | None | None |
| GAMT | None | None | None | None |
| GAMT | None | None | None | None |
| GAMT | None | None | None | None |
| GAMT | None | None | None | None |
| GAMT | None | None | None | None |
| GAMT | None | None | None | None |
| GAMT | None | None | None | None |
| GAMT | None | None | None | None |
| GLRA1 | None | None | None | None |
| GLRA1 | None | None | None | None |

| | | | | |
|---|---|---|---|---|
| GLRA1 | None | None | None | None |
| GLRA1 | None | None | None | None |
| GLRA1 | None | None | None | None |
| GLRA1 | None | None | None | None |
| GLRA1 | None | None | None | None |
| GLRA1 | None | None | None | None |
| GLRA1 | None | None | None | None |
| ADK | None | None | None | None |
| ACTN2 | None | None | None | None |
| UBE2T | None | None | None | None |
| TTN | None | None | None | None |
| ATP7B | None | None | None | None |
| ATP7B | None | None | None | None |
| ATP7B | None | None | None | None |
| ATP7B | None | None | None | None |
| ATP7B | None | None | None | None |
| ATP7B | None | None | None | None |
| ATP7B | None | None | None | None |
| ATP7B | None | None | None | None |
| ATP7B | None | None | None | None |
| ATP7B | None | None | None | None |
| ATP7B | None | None | None | None |
| ATP7B | None | None | None | None |
| ATP7B | None | None | None | None |
| CDKN1B | None | None | None | None |
| CDKN1B | None | None | None | None |
| CDKN1B | None | None | None | None |
| CDKN1B | None | None | None | None |
| CDKN1B | None | None | None | None |
| CDKN1B | None | None | None | None |
| MYOC | None | None | None | None |
| MYOC | None | None | None | None |
| MYOC | None | None | None | None |
| MYOC | None | None | None | None |
| MYOC | None | None | None | None |
| MYOC | None | None | None | None |
| MYOC | None | None | None | None |
| MYOC | None | None | None | None |
| MYOC | None | None | None | None |
| MYOC | None | None | None | None |
| MYOC | None | None | None | None |
| MYOC | None | None | None | None |
| MYOC | None | None | None | None |
| MYOC | None | None | None | None |
| MYOC | None | None | None | None |
| SPTAN1 | None | None | None | None |
| GC | None | None | None | None |
| GC | None | None | None | None |
| MAD1L1 | None | None | None | None |
| NDST1 | None | None | None | None |
| ARL2BP | None | None | None | None |
| TOR1A | None | None | None | None |
| C1S | None | None | None | None |
| ARL6 | 0.508 | 1 | 2h57 | A |

| | | | | |
|---|---|---|---|---|
| ARL6 | None | None | None | None |
| ARL6 | None | None | None | None |
| ARL6 | None | None | None | None |
| ARL6 | None | None | None | None |
| ARL6 | None | None | None | None |
| FGFR1 | None | None | None | None |
| KLHL7 | None | None | None | None |
| FLNC | None | None | None | None |
| PTPRC | None | None | None | None |
| GCH1 | None | None | None | None |
| GCH1 | None | None | None | None |
| GCH1 | None | None | None | None |
| GCH1 | None | None | None | None |
| GCH1 | None | None | None | None |
| GCH1 | None | None | None | None |
| GCH1 | None | None | None | None |
| GCH1 | None | None | None | None |
| GCH1 | None | None | None | None |
| GCH1 | None | None | None | None |
| GCH1 | None | None | None | None |
| GCH1 | None | None | None | None |
| GCH1 | None | None | None | None |
| GCH1 | None | None | None | None |
| CFH | None | None | None | None |
| COL2A1 | None | None | None | None |
| COL2A1 | None | None | None | None |
| OCRL | None | None | None | None |
| WRN | None | None | None | None |
| WRN | None | None | None | None |
| WRN | None | None | None | None |
| WRN | None | None | None | None |
| MASP1 | None | None | None | None |
| MASP1 | None | None | None | None |
| NSD1 | None | None | None | None |
| PITX2 | None | None | None | None |
| PITX2 | None | None | None | None |
| PITX2 | None | None | None | None |
| PITX2 | None | None | None | None |
| PITX2 | None | None | None | None |
| PITX2 | None | None | None | None |
| B4GALT7 | 0.917 | 1 | 4irq | A |
| B4GALT7 | None | None | None | None |
| B4GALT7 | None | None | None | None |
| B4GALT7 | None | None | None | None |
| B4GALT7 | None | None | None | None |
| FSHR | None | None | None | None |
| IGHMBP2 | None | None | None | None |
| IGHMBP2 | None | None | None | None |
| IGHMBP2 | None | None | None | None |
| IGHMBP2 | None | None | None | None |
| IGHMBP2 | None | None | None | None |
| IGHMBP2 | None | None | None | None |
| IGHMBP2 | None | None | None | None |
| IGHMBP2 | None | None | None | None |

| | | | | |
|---|---|---|---|---|
| IGHMBP2 | None | None | None | None |
| IGHMBP2 | None | None | None | None |
| IGHMBP2 | None | None | None | None |
| IGHMBP2 | None | None | None | None |
| IGHMBP2 | None | None | None | None |
| IGHMBP2 | None | None | None | None |
| IGHMBP2 | None | None | None | None |
| IGHMBP2 | None | None | None | None |
| TTN | None | None | None | None |
| TTN | None | None | None | None |
| IRAK4 | None | None | None | None |
| PROC | None | None | None | None |
| PROC | None | None | None | None |
| RAD50 | None | None | None | None |
| RAD50 | None | None | None | None |
| DYNC1H1 | None | None | None | None |
| DYNC1H1 | None | None | None | None |
| DYNC1H1 | None | None | None | None |
| DYNC1H1 | None | None | None | None |
| DYNC1H1 | None | None | None | None |
| DYNC1H1 | None | None | None | None |
| DYNC1H1 | None | None | None | None |
| DYNC1H1 | None | None | None | None |
| DYNC1H1 | None | None | None | None |
| DYNC1H1 | None | None | None | None |
| DYNC1H1 | None | None | None | None |
| CYB5R3 | None | None | None | None |
| CYB5R3 | None | None | None | None |
| CYB5R3 | None | None | None | None |
| CYB5R3 | None | None | None | None |
| CYB5R3 | None | None | None | None |
| CYB5R3 | None | None | None | None |
| CYB5R3 | None | None | None | None |
| CYB5R3 | None | None | None | None |
| CYB5R3 | None | None | None | None |
| CYB5R3 | None | None | None | None |
| CYB5R3 | None | None | None | None |
| SPTB | None | None | None | None |
| FANCL | None | None | None | None |
| FAM126A | None | None | None | None |
| SEPT9 | None | None | None | None |
| TTN | None | None | None | None |
| TTN | None | None | None | None |
| TTN | None | None | None | None |
| CNTN1 | None | None | None | None |
| C1QTNF5 | None | None | None | None |
| VCL | None | None | None | None |
| VCL | None | None | None | None |
| VCL | None | None | None | None |
| VCL | None | None | None | None |
| VCL | None | None | None | None |
| VCL | None | None | None | None |
| PPT1 | None | None | None | None |
| PPT1 | None | None | None | None |

| | | | | |
|---|---|---|---|---|
| PPT1 | None | None | None | None |
| PPT1 | None | None | None | None |
| PPT1 | None | None | None | None |
| PPT1 | None | None | None | None |
| PPT1 | None | None | None | None |
| PPT1 | None | None | None | None |
| PPT1 | None | None | None | None |
| PPT1 | None | None | None | None |
| PPT1 | None | None | None | None |
| PPT1 | None | None | None | None |
| PPT1 | None | None | None | None |
| PPT1 | None | None | None | None |
| PPT1 | None | None | None | None |
| PPT1 | None | None | None | None |
| PPT1 | None | None | None | None |
| PPT1 | None | None | None | None |
| PPT1 | None | None | None | None |
| NOTCH2 | None | None | None | None |
| NOTCH2 | None | None | None | None |
| IFIH1 | None | None | None | None |
| IFIH1 | None | None | None | None |
| IKBKG | None | None | None | None |
| DBH | None | None | None | None |
| DBH | None | None | None | None |
| DBH | None | None | None | None |
| DBH | None | None | None | None |
| DBH | None | None | None | None |
| DBH | None | None | None | None |
| DBH | None | None | None | None |
| DBH | None | None | None | None |
| DBH | None | None | None | None |
| DBH | None | None | None | None |
| IL10RA | None | None | None | None |
| IL10RA | None | None | None | None |
| IL10RA | None | None | None | None |
| IL10RA | None | None | None | None |
| IL10RA | None | None | None | None |
| IL10RA | None | None | None | None |
| IL10RA | None | None | None | None |
| ILK | None | None | None | None |
| HPD | None | None | None | None |
| HPD | None | None | None | None |
| HPD | None | None | None | None |
| MECP2 | None | None | None | None |
| MECP2 | None | None | None | None |
| MECP2 | None | None | None | None |
| MECP2 | None | None | None | None |
| MECP2 | None | None | None | None |
| MECP2 | None | None | None | None |
| MECP2 | None | None | None | None |
| MECP2 | None | None | None | None |
| KYNU | None | None | None | None |
| FAS | None | None | None | None |
| FAS | None | None | None | None |

| | | | | |
|---|---|---|---|---|
| ELP1 | None | None | None | None |
| ELP1 | None | None | None | None |
| ELP1 | None | None | None | None |
| ELP1 | None | None | None | None |
| ELP1 | None | None | None | None |
| ELP1 | None | None | None | None |
| ELP1 | None | None | None | None |
| ELP1 | None | None | None | None |
| HESX1 | None | None | None | None |
| HESX1 | None | None | None | None |
| HESX1 | None | None | None | None |
| HESX1 | None | None | None | None |
| GAD1 | None | None | None | None |
| GAD1 | None | None | None | None |
| NDUFA1 | None | None | None | None |
| NDUFA1 | None | None | None | None |
| NDUFA1 | None | None | None | None |
| PNPT1 | None | None | None | None |
| PNPT1 | None | None | None | None |
| PNPT1 | None | None | None | None |
| PNPT1 | None | None | None | None |
| PNPT1 | None | None | None | None |
| PNPT1 | None | None | None | None |
| PNPT1 | None | None | None | None |
| SLC25A13 | None | None | None | None |
| SLC25A13 | None | None | None | None |
| SLC25A13 | None | None | None | None |
| AP4M1 | None | None | None | None |
| AP4M1 | None | None | None | None |
| MEFV | None | None | None | None |
| MEFV | None | None | None | None |
| MEFV | None | None | None | None |
| MEFV | None | None | None | None |
| MEFV | None | None | None | None |
| MEFV | None | None | None | None |
| MEFV | None | None | None | None |
| MEFV | None | None | None | None |
| MEFV | None | None | None | None |
| CAPN3 | None | None | None | None |
| CAPN3 | None | None | None | None |
| CAPN3 | None | None | None | None |
| CAPN3 | None | None | None | None |
| CAPN3 | None | None | None | None |
| CAPN3 | None | None | None | None |
| CAPN3 | None | None | None | None |
| GP6 | None | None | None | None |
| FAS | None | None | None | None |
| FBN1 | None | None | None | None |
| FBN1 | None | None | None | None |
| BMPR2 | None | None | None | None |
| BMPR2 | None | None | None | None |
| BMPR2 | None | None | None | None |
| BMPR2 | None | None | None | None |
| BMPR2 | None | None | None | None |

| | | | | |
|---|---|---|---|---|
| RET | None | None | None | None |
| RET | None | None | None | None |
| RET | None | None | None | None |
| RET | None | None | None | None |
| RET | None | None | None | None |
| RET | None | None | None | None |
| RET | None | None | None | None |
| RET | None | None | None | None |
| RET | None | None | None | None |
| RET | None | None | None | None |
| RET | None | None | None | None |
| RET | None | None | None | None |
| RET | None | None | None | None |
| RET | None | None | None | None |
| RET | None | None | None | None |
| RET | None | None | None | None |
| RET | None | None | None | None |
| RET | None | None | None | None |
| RET | None | None | None | None |
| RET | None | None | None | None |
| RET | None | None | None | None |
| RET | None | None | None | None |
| RET | None | None | None | None |
| RET | None | None | None | None |
| RET | None | None | None | None |
| RET | None | None | None | None |
| RET | None | None | None | None |
| RET | None | None | None | None |
| CSF2RA | None | None | None | None |
| CSF2RA | None | None | None | None |
| CSF2RA | None | None | None | None |
| BRCA1 | None | None | None | None |
| ZAP70 | 0.636 | 1 | 2ozo | A |
| ZAP70 | 0.636 | 1 | 2ozo | A |
| ZAP70 | None | None | None | None |
| PIK3R2 | None | None | None | None |
| TCN2 | None | None | None | None |
| TCN2 | None | None | None | None |
| TCN2 | None | None | None | None |
| TCN2 | None | None | None | None |
| TCN2 | None | None | None | None |
| TCN2 | None | None | None | None |
| TCN2 | None | None | None | None |
| TCN2 | None | None | None | None |
| EPM2A | None | None | None | None |
| EPM2A | None | None | None | None |
| EPM2A | None | None | None | None |
| EPM2A | None | None | None | None |
| AARS | None | None | None | None |
| AARS | None | None | None | None |
| AARS | None | None | None | None |
| PCDH15 | None | None | None | None |
| PCDH15 | None | None | None | None |
| PCDH15 | None | None | None | None |

| | | | | |
|---|---|---|---|---|
| PCDH15 | None | None | None | None |
| PCDH15 | None | None | None | None |
| PCDH15 | None | None | None | None |
| PCDH15 | None | None | None | None |
| PCDH15 | None | None | None | None |
| PCDH15 | None | None | None | None |
| PCDH15 | None | None | None | None |
| NDUFS4 | None | None | None | None |
| NDUFS4 | None | None | None | None |
| NDUFS4 | None | None | None | None |
| FGF10 | None | None | None | None |
| FGF10 | 0.558 | 3 | 1nun | A |
| FGF10 | None | None | None | None |
| FGF10 | None | None | None | None |
| CDON | None | None | None | None |
| STAMBP | 0.909 | 2 | 3rzv | A |
| BRCA1 | None | None | None | None |
| BRCA1 | None | None | None | None |
| BRCA1 | None | None | None | None |
| BRCA1 | None | None | None | None |
| BRCA1 | None | None | None | None |
| BRCA1 | None | None | None | None |
| BRCA1 | None | None | None | None |
| BRCA1 | None | None | None | None |
| BRCA1 | None | None | None | None |
| BRCA1 | None | None | None | None |
| PLXNC1 | None | None | None | None |
| RNASEH2C | None | None | None | None |
| RNASEH2C | None | None | None | None |
| APC | None | None | None | None |
| PLA2G4A | None | None | None | None |
| PLA2G4A | None | None | None | None |
| PSAT1 | None | None | None | None |
| PSAT1 | None | None | None | None |
| TRAPPC3 | None | None | None | None |
| QDPR | None | None | None | None |
| QDPR | None | None | None | None |
| QDPR | None | None | None | None |
| QDPR | None | None | None | None |
| KRT5 | None | None | None | None |
| KRT5 | None | None | None | None |
| KRT5 | None | None | None | None |
| MLH1 | None | None | None | None |
| MLH1 | None | None | None | None |
| MLH1 | None | None | None | None |
| MLH1 | None | None | None | None |
| MLH1 | None | None | None | None |
| MLH1 | None | None | None | None |
| MLH1 | None | None | None | None |
| MLH1 | None | None | None | None |
| MLH1 | None | None | None | None |
| MLH1 | None | None | None | None |
| MLH1 | None | None | None | None |
| MLH1 | None | None | None | None |

| | | | | |
|---|---|---|---|---|
| MLH1 | None | None | None | None |
| MLH1 | None | None | None | None |
| MLH1 | None | None | None | None |
| MLH1 | None | None | None | None |
| MLH1 | None | None | None | None |
| KRIT1 | None | None | None | None |
| KRIT1 | None | None | None | None |
| KRIT1 | None | None | None | None |
| ALX4 | None | None | None | None |
| ALX4 | None | None | None | None |
| ALX4 | None | None | None | None |
| TJP2 | None | None | None | None |
| TJP2 | None | None | None | None |
| PANK2 | None | None | None | None |
| PANK2 | None | None | None | None |
| PANK2 | None | None | None | None |
| PANK2 | None | None | None | None |
| PANK2 | None | None | None | None |
| PANK2 | None | None | None | None |
| PANK2 | None | None | None | None |
| NDUFA6 | None | None | None | None |
| TSHR | None | None | None | None |
| TSHR | None | None | None | None |
| TSHR | None | None | None | None |
| CFTR | None | None | None | None |
| CFTR | None | None | None | None |
| MYOM1 | None | None | None | None |
| MYOM1 | None | None | None | None |
| SPTB | None | None | None | None |
| GPT2 | None | None | None | None |
| NDUFS2 | None | None | None | None |
| NDUFS2 | None | None | None | None |
| NDUFS2 | None | None | None | None |
| NDUFS2 | None | None | None | None |
| NDUFS2 | None | None | None | None |
| CASQ2 | None | None | None | None |
| CASQ2 | None | None | None | None |
| CASQ2 | None | None | None | None |
| CASQ2 | None | None | None | None |
| CASQ2 | None | None | None | None |
| CASQ2 | None | None | None | None |
| CASQ2 | None | None | None | None |
| CASQ2 | None | None | None | None |
| CASQ2 | None | None | None | None |
| CPOX | None | None | None | None |
| CPOX | None | None | None | None |
| CPOX | 0.857 | 2 | 2aex | A |
| CPOX | None | None | None | None |
| CPOX | None | None | None | None |
| CPOX | None | None | None | None |
| CPOX | None | None | None | None |
| CPOX | None | None | None | None |
| CPOX | None | None | None | None |
| MVK | None | None | None | None |

| | | | | |
|---|---|---|---|---|
| MVK | None | None | None | None |
| MVK | None | None | None | None |
| MVK | None | None | None | None |
| MVK | None | None | None | None |
| MVK | None | None | None | None |
| MVK | None | None | None | None |
| MVK | None | None | None | None |
| MVK | None | None | None | None |
| MVK | None | None | None | None |
| MVK | None | None | None | None |
| MVK | None | None | None | None |
| MVK | None | None | None | None |
| MVK | None | None | None | None |
| MVK | None | None | None | None |
| GSS | None | None | None | None |
| GSS | None | None | None | None |
| GSS | None | None | None | None |
| GSS | None | None | None | None |
| GSS | None | None | None | None |
| DSG2 | None | None | None | None |
| DSG2 | None | None | None | None |
| DSG2 | None | None | None | None |
| DSG2 | None | None | None | None |
| DSG2 | None | None | None | None |
| DSG2 | None | None | None | None |
| DSG2 | None | None | None | None |
| DSG2 | None | None | None | None |
| DSG2 | None | None | None | None |
| DSG2 | None | None | None | None |
| DSG2 | None | None | None | None |
| DSG2 | None | None | None | None |
| DSG2 | None | None | None | None |
| DSG2 | None | None | None | None |
| PAPSS2 | None | None | None | None |
| PLCB2 | None | None | None | None |
| HMGCS2 | None | None | None | None |
| HMGCS2 | None | None | None | None |
| HMGCS2 | 0.938 | 1 | 2wya | C |
| HMGCS2 | None | None | None | None |
| HMGCS2 | None | None | None | None |
| HMGCS2 | None | None | None | None |
| HMGCS2 | 0.562 | 4 | 2wya | C |
| HMGCS2 | None | None | None | None |
| MPDZ | None | None | None | None |
| GATA4 | None | None | None | None |
| GATA4 | None | None | None | None |
| WARS2 | None | None | None | None |
| WARS2 | None | None | None | None |
| HGD | 1 | 47 | 1eyb | A |
| HGD | 1 | 47 | 1eyb | A |
| HGD | 1 | 70 | 1eyb | A |
| HGD | None | None | None | None |
| HGD | None | None | None | None |
| HGD | None | None | None | None |

| | | | | |
|---|---|---|---|---|
| HGD | None | None | None | None |
| HGD | 0.5 | 51 | 1ey2 | A |
| HGD | None | None | None | None |
| HGD | 1 | 1 | 1ey2 | A |
| HGD | 1 | 9 | 1ey2 | A |
| HGD | None | None | None | None |
| HGD | 1 | 10 | 1eyb | A |
| HGD | 1 | 7 | 1ey2 | A |
| PAX8 | None | None | None | None |
| PAX8 | None | None | None | None |
| PAX8 | None | None | None | None |
| PAX8 | None | None | None | None |
| SPTB | None | None | None | None |
| NCF4 | None | None | None | None |
| CFH | None | None | None | None |
| CFH | None | None | None | None |
| CSF1R | None | None | None | None |
| CSF1R | None | None | None | None |
| CA8 | None | None | None | None |
| CA8 | None | None | None | None |
| CA8 | 0.748 | 7 | 2w2j | A |
| APC | None | None | None | None |
| APC | None | None | None | None |
| APC | None | None | None | None |
| APC | None | None | None | None |
| APC | None | None | None | None |
| NDUFA2 | None | None | None | None |
| GIF | None | None | None | None |
| GIF | None | None | None | None |
| GIF | None | None | None | None |
| GIF | None | None | None | None |
| C2 | None | None | None | None |
| ACP5 | None | None | None | None |
| ACP5 | None | None | None | None |
| ACP5 | None | None | None | None |
| ACP5 | None | None | None | None |
| ACP5 | None | None | None | None |
| RNASEH2A | None | None | None | None |
| RNASEH2A | None | None | None | None |
| RNASEH2A | None | None | None | None |
| RNASEH2A | None | None | None | None |
| RNASEH2A | None | None | None | None |
| RNASEH2A | None | None | None | None |
| RNASEH2A | None | None | None | None |
| DSG2 | None | None | None | None |
| USH1G | None | None | None | None |
| FUS | None | None | None | None |
| FUS | None | None | None | None |
| FUS | None | None | None | None |
| FUS | None | None | None | None |
| FUS | None | None | None | None |
| MRPL44 | None | None | None | None |
| MRPL44 | None | None | None | None |
| MRPL44 | None | None | None | None |

| | | | | |
|---|---|---|---|---|
| MRPL44 | None | None | None | None |
| DMGDH | None | None | None | None |
| AARS | None | None | None | None |
| STAR | None | None | None | None |
| STAR | None | None | None | None |
| STAR | None | None | None | None |
| STAR | None | None | None | None |
| STAR | None | None | None | None |
| STAR | None | None | None | None |
| STAR | None | None | None | None |
| STAR | None | None | None | None |
| LARP7 | None | None | None | None |
| PRPF8 | None | None | None | None |
| PRPF8 | None | None | None | None |
| PRPF8 | None | None | None | None |
| PRPF8 | None | None | None | None |
| PSAP | None | None | None | None |
| PSAP | None | None | None | None |
| PSAP | None | None | None | None |
| DMPK | None | None | None | None |
| EPCAM | None | None | None | None |
| EPCAM | None | None | None | None |
| EPCAM | None | None | None | None |
| EPCAM | None | None | None | None |
| EPCAM | None | None | None | None |
| EPCAM | None | None | None | None |
| EPCAM | None | None | None | None |
| EPCAM | None | None | None | None |
| ASS1 | None | None | None | None |
| ASS1 | None | None | None | None |
| ASS1 | 0.571 | 3 | 2nz2 | A |
| ASS1 | 0.571 | 3 | 2nz2 | A |
| ASS1 | None | None | None | None |
| ASS1 | 0.857 | 20 | 2nz2 | A |
| ASS1 | 0.857 | 9 | 2nz2 | A |
| ASS1 | None | None | None | None |
| ASS1 | None | None | None | None |
| ASS1 | 0.571 | 4 | 2nz2 | A |
| ASS1 | 0.857 | 9 | 2nz2 | A |
| ASS1 | 0.857 | 9 | 2nz2 | A |
| ASS1 | 0.857 | 9 | 2nz2 | A |
| ASS1 | None | None | None | None |
| ASS1 | None | None | None | None |
| ASS1 | None | None | None | None |
| ASS1 | None | None | None | None |
| ASS1 | None | None | None | None |
| ASS1 | None | None | None | None |
| ASS1 | None | None | None | None |
| POLE | None | None | None | None |
| POLE | None | None | None | None |
| POLE | None | None | None | None |
| RNASET2 | None | None | None | None |
| RNASET2 | None | None | None | None |
| RNASET2 | None | None | None | None |

| | | | | |
|---|---|---|---|---|
| FLG | None | None | None | None |
| F10 | None | None | None | None |
| AGT | None | None | None | None |
| AGT | None | None | None | None |
| PLN | None | None | None | None |
| PLN | None | None | None | None |
| PLN | None | None | None | None |
| PTHLH | None | None | None | None |
| IGF1R | None | None | None | None |
| IGF1R | None | None | None | None |
| IGF1R | None | None | None | None |
| JAG1 | None | None | None | None |
| JAG1 | None | None | None | None |
| JAG1 | None | None | None | None |
| JAG1 | None | None | None | None |
| JAG1 | None | None | None | None |
| JAG1 | None | None | None | None |
| AP1S3 | None | None | None | None |
| SPTA1 | None | None | None | None |
| SPTA1 | None | None | None | None |
| SPTA1 | None | None | None | None |
| SPTA1 | None | None | None | None |
| SPG7 | None | None | None | None |
| SPG7 | None | None | None | None |
| SPG7 | None | None | None | None |
| SPG7 | None | None | None | None |
| SPG7 | None | None | None | None |
| SPG7 | None | None | None | None |
| FN1 | None | None | None | None |
| FN1 | None | None | None | None |
| NSMCE3 | None | None | None | None |
| PCCA | None | None | None | None |
| PCCA | None | None | None | None |
| SPTB | None | None | None | None |
| RYR2 | None | None | None | None |
| RYR2 | None | None | None | None |
| MMP9 | None | None | None | None |
| MMP9 | None | None | None | None |
| GLMN | None | None | None | None |
| GLMN | None | None | None | None |
| GLMN | None | None | None | None |
| GLMN | None | None | None | None |
| GLMN | None | None | None | None |
| GLMN | None | None | None | None |
| GLMN | None | None | None | None |
| GLMN | None | None | None | None |
| GLMN | None | None | None | None |
| GLMN | None | None | None | None |
| MARS | None | None | None | None |
| MARS | None | None | None | None |
| MARS | None | None | None | None |
| MARS | None | None | None | None |
| MARS | None | None | None | None |
| MARS | None | None | None | None |

| | | | | |
|---|---|---|---|---|
| SNAP29 | None | None | None | None |
| SNAP29 | None | None | None | None |
| ETHE1 | None | None | None | None |
| ETHE1 | None | None | None | None |
| ETHE1 | None | None | None | None |
| ETHE1 | None | None | None | None |
| ETHE1 | None | None | None | None |
| ETHE1 | None | None | None | None |
| ETHE1 | None | None | None | None |
| SAMHD1 | None | None | None | None |
| KCNQ4 | None | None | None | None |
| FXYD2 | None | None | None | None |
| FXYD2 | None | None | None | None |
| FXYD2 | None | None | None | None |
| LRRK2 | None | None | None | None |
| LRRK2 | None | None | None | None |
| NDUFS1 | None | None | None | None |
| NDUFS1 | None | None | None | None |
| NDUFS1 | None | None | None | None |
| NDUFS1 | None | None | None | None |
| NDUFS1 | None | None | None | None |
| NDUFS1 | None | None | None | None |
| NDUFS1 | None | None | None | None |
| NDUFS1 | None | None | None | None |
| NDUFS1 | None | None | None | None |
| NDUFS1 | None | None | None | None |
| NDUFS1 | None | None | None | None |
| EZH2 | None | None | None | None |
| EZH2 | None | None | None | None |
| LARS | None | None | None | None |
| TLL1 | None | None | None | None |
| CRYBB2 | None | None | None | None |
| CRYBB2 | None | None | None | None |
| EXOSC8 | None | None | None | None |
| SRP72 | None | None | None | None |
| FOXC2 | None | None | None | None |
| HMBS | None | None | None | None |
| HMBS | None | None | None | None |
| HMBS | None | None | None | None |
| HMBS | None | None | None | None |
| HMBS | None | None | None | None |
| HMBS | None | None | None | None |
| HMBS | None | None | None | None |
| HMBS | None | None | None | None |
| HMBS | None | None | None | None |
| HMBS | None | None | None | None |
| HMBS | None | None | None | None |
| HMBS | None | None | None | None |
| HMBS | None | None | None | None |
| HMBS | None | None | None | None |
| HMBS | None | None | None | None |
| HMBS | None | None | None | None |
| HMBS | None | None | None | None |
| HMBS | None | None | None | None |

| | | | | |
|---|---|---|---|---|
| HMBS | None | None | None | None |
| HMBS | None | None | None | None |
| HMBS | None | None | None | None |
| HMBS | None | None | None | None |
| HMBS | None | None | None | None |
| HMBS | None | None | None | None |
| HMBS | None | None | None | None |
| OGT | None | None | None | None |
| NOG | None | None | None | None |
| NOG | None | None | None | None |
| NOG | None | None | None | None |
| NOG | None | None | None | None |
| NOG | None | None | None | None |
| NOG | None | None | None | None |
| NOG | None | None | None | None |
| NOG | None | None | None | None |
| NOG | None | None | None | None |
| NOG | None | None | None | None |
| NOG | None | None | None | None |
| DLG3 | None | None | None | None |
| SCN2A | None | None | None | None |
| STAC3 | None | None | None | None |
| PFKM | None | None | None | None |
| PFKM | None | None | None | None |
| PFKM | None | None | None | None |
| PFKM | None | None | None | None |
| PFKM | None | None | None | None |
| DNMT1 | None | None | None | None |
| DNMT1 | None | None | None | None |
| TNNI3 | None | None | None | None |
| TNNI3 | None | None | None | None |
| TNNI3 | None | None | None | None |
| STXBP2 | None | None | None | None |
| STXBP2 | None | None | None | None |
| STXBP2 | None | None | None | None |
| STXBP2 | None | None | None | None |
| STXBP2 | None | None | None | None |
| STXBP2 | None | None | None | None |
| STXBP2 | None | None | None | None |
| STXBP2 | None | None | None | None |
| REN | None | None | None | None |
| DSP | None | None | None | None |
| DSP | None | None | None | None |
| HNF1B | None | None | None | None |
| ADA | None | None | None | None |
| ADA | None | None | None | None |
| ADA | 0.615 | 7 | 3iar | A |
| ADA | None | None | None | None |
| ADA | None | None | None | None |
| ADA | 0.538 | 5 | 3iar | A |
| ADA | 0.538 | 5 | 3iar | A |
| ADA | None | None | None | None |
| ADA | None | None | None | None |
| ADA | 0.615 | 1 | 3iar | A |

| | | | | |
|---|---|---|---|---|
| ADA | None | None | None | None |
| ADA | 0.615 | | 1 3iar | A |
| ADA | None | None | None | None |
| ADA | None | None | None | None |
| ADA | None | None | None | None |
| DCX | None | None | None | None |
| DCX | None | None | None | None |
| DCX | None | None | None | None |
| DCX | None | None | None | None |
| DCX | None | None | None | None |
| ARSB | None | None | None | None |
| ARSB | None | None | None | None |
| ARSB | None | None | None | None |
| ARSB | None | None | None | None |
| ARSB | None | None | None | None |
| ARSB | None | None | None | None |
| ARSB | None | None | None | None |
| ARSB | None | None | None | None |
| ARSB | None | None | None | None |
| ARSB | None | None | None | None |
| ARSB | None | None | None | None |
| ARSB | None | None | None | None |
| ARSB | None | None | None | None |
| WWOX | None | None | None | None |
| TNNI3 | None | None | None | None |
| TNNI3 | None | None | None | None |
| TNNI3 | None | None | None | None |
| TNNI3 | None | None | None | None |
| CYBA | None | None | None | None |
| SPAST | None | None | None | None |
| SPAST | None | None | None | None |
| SPAST | None | None | None | None |
| SPAST | None | None | None | None |
| SPAST | None | None | None | None |
| SPAST | None | None | None | None |
| SPAST | None | None | None | None |
| SPAST | None | None | None | None |
| KRIT1 | None | None | None | None |
| KRIT1 | None | None | None | None |
| TP53 | None | None | None | None |
| ISPD | None | None | None | None |
| ISPD | None | None | None | None |
| ISPD | None | None | None | None |
| ISPD | None | None | None | None |
| KCNMA1 | None | None | None | None |
| STK11 | None | None | None | None |
| STK11 | None | None | None | None |
| STK11 | None | None | None | None |
| STK11 | None | None | None | None |
| STK11 | None | None | None | None |
| STK11 | None | None | None | None |
| NDUFA13 | None | None | None | None |
| PHF6 | None | None | None | None |
| PHF6 | None | None | None | None |

| | | | | |
|---|---|---|---|---|
| PHF6 | None | None | None | None |
| PHYH | None | None | None | None |
| PHYH | None | None | None | None |
| PHYH | None | None | None | None |
| PHYH | None | None | None | None |
| PHYH | None | None | None | None |
| NDUFS3 | None | None | None | None |
| NDUFS3 | None | None | None | None |
| ALDH18A1 | None | None | None | None |
| ALDH18A1 | None | None | None | None |
| ALDH18A1 | None | None | None | None |
| ALDH18A1 | None | None | None | None |
| ALDH18A1 | None | None | None | None |
| PAX6 | None | None | None | None |
| PAX6 | None | None | None | None |
| INPP5E | None | None | None | None |
| INPP5E | None | None | None | None |
| INPP5E | None | None | None | None |
| INPP5E | None | None | None | None |
| INPP5E | None | None | None | None |
| INPP5E | None | None | None | None |
| INPP5E | None | None | None | None |
| INPP5E | None | None | None | None |
| INPP5E | None | None | None | None |
| TNNT2 | None | None | None | None |
| TNNT2 | None | None | None | None |
| TNNT2 | None | None | None | None |
| TNNT2 | None | None | None | None |
| TNNT2 | None | None | None | None |
| TNNT2 | None | None | None | None |
| TNNT2 | None | None | None | None |
| TNNT2 | None | None | None | None |
| TYK2 | None | None | None | None |
| TYK2 | None | None | None | None |
| TYK2 | None | None | None | None |
| CFTR | None | None | None | None |
| CFTR | None | None | None | None |
| CFTR | None | None | None | None |
| CFTR | None | None | None | None |
| CFTR | None | None | None | None |
| CFTR | None | None | None | None |
| CFTR | None | None | None | None |
| CFTR | None | None | None | None |
| CFTR | None | None | None | None |
| CFTR | None | None | None | None |
| CFTR | None | None | None | None |
| CFTR | None | None | None | None |
| CFTR | None | None | None | None |
| CFTR | None | None | None | None |
| CFTR | None | None | None | None |
| CFTR | None | None | None | None |
| CFTR | None | None | None | None |
| CFTR | None | None | None | None |
| CFTR | None | None | None | None |

| | | | | |
|---|---|---|---|---|
| CFTR | None | None | None | None |
| CFTR | None | None | None | None |
| CFTR | None | None | None | None |
| CFTR | None | None | None | None |
| CFTR | None | None | None | None |
| CFTR | None | None | None | None |
| CFTR | None | None | None | None |
| CFTR | None | None | None | None |
| CFTR | None | None | None | None |
| CFTR | None | None | None | None |
| CFTR | None | None | None | None |
| CFTR | None | None | None | None |
| CFTR | None | None | None | None |
| CFTR | None | None | None | None |
| CFTR | None | None | None | None |
| CFTR | None | None | None | None |
| CFTR | None | None | None | None |
| CFTR | None | None | None | None |
| CFTR | None | None | None | None |
| CFTR | None | None | None | None |
| CFTR | None | None | None | None |
| CFTR | None | None | None | None |
| CFTR | None | None | None | None |
| CFTR | None | None | None | None |
| CFTR | None | None | None | None |
| CFTR | None | None | None | None |
| CFTR | None | None | None | None |
| CFTR | None | None | None | None |
| CFTR | None | None | None | None |
| CFTR | None | None | None | None |
| CFTR | None | None | None | None |
| CFTR | None | None | None | None |
| CFTR | None | None | None | None |
| CFTR | None | None | None | None |
| SERPINF1 | None | None | None | None |
| SERPINF1 | None | None | None | None |
| IDS | None | None | None | None |
| IDS | None | None | None | None |
| IDS | None | None | None | None |
| IDS | None | None | None | None |
| IDS | None | None | None | None |
| IDS | None | None | None | None |
| IDS | None | None | None | None |
| IDS | None | None | None | None |
| IDS | None | None | None | None |
| NFKBIA | 0.7 | 1 | 1nfi | F |
| NFKBIA | None | None | None | None |
| SPAST | None | None | None | None |
| FGF23 | None | None | None | None |
| FGF23 | None | None | None | None |
| FGF23 | None | None | None | None |
| FGF23 | None | None | None | None |
| FGF23 | None | None | None | None |
| FGF23 | None | None | None | None |

| | | | | |
|---|---|---|---|---|
| FGF23 | None | None | None | None |
| DLAT | None | None | None | None |
| STUB1 | None | None | None | None |
| HSD17B4 | None | None | None | None |
| HSD17B4 | None | None | None | None |
| DYSF | None | None | None | None |
| CEP135 | None | None | None | None |
| CEP135 | None | None | None | None |
| PRKCG | None | None | None | None |
| CSF1R | None | None | None | None |
| PEX5 | None | None | None | None |
| PAX6 | None | None | None | None |
| PAX6 | None | None | None | None |
| PAX6 | None | None | None | None |
| PAX6 | None | None | None | None |
| PAX6 | None | None | None | None |
| PAX6 | None | None | None | None |
| NF2 | None | None | None | None |
| NF2 | None | None | None | None |
| NF2 | None | None | None | None |
| REM1 | None | None | None | None |

| source_id | classification | binding_site | binding_site_single_water | protein_binding_site |
|---|---|---|---|---|
| rs121918343 | pathogenic | 1 | 1 | 0 |
| rs267606649 | pathogenic | 1 | 1 | 1 |
| rs121918342 | pathogenic | 0 | 1 | 0 |
| rs267606650 | pathogenic | 1 | 1 | 1 |
| rs1130335 | benign | 1 | 1 | 1 |
| rs199476087 | pathogenic | 1 | 1 | 1 |
| rs772163112 | benign | 0 | 0 | 0 |
| rs755793 | benign | 0 | 0 | 0 |
| rs121918155 | pathogenic | 1 | 1 | 0 |
| rs121918156 | pathogenic | 1 | 1 | 0 |
| rs121918154 | pathogenic | 1 | 1 | 1 |
| rs121918151 | pathogenic | 1 | 1 | 1 |
| rs121918144 | pathogenic | 0 | 1 | 0 |
| rs121918146 | pathogenic | 1 | 1 | 1 |
| rs121918160 | pathogenic | 1 | 1 | 1 |
| rs121918150 | pathogenic | 1 | 1 | 1 |
| rs121918141 | pathogenic | 1 | 1 | 1 |
| rs142742242 | pathogenic | 1 | 1 | 1 |
| rs121918142 | pathogenic | 1 | 1 | 1 |
| rs121918157 | pathogenic | 1 | 1 | 1 |
| rs121918101 | pathogenic | 1 | 1 | 1 |
| rs121908260 | pathogenic | 1 | 1 | 1 |
| rs80356667 | pathogenic | 1 | 1 | 1 |
| rs80356668 | pathogenic | 1 | 1 | 1 |
| rs397514540 | pathogenic | 1 | 1 | 1 |
| rs104894685 | pathogenic | 0 | 1 | 0 |
| rs199869995 | pathogenic | 1 | 1 | 1 |
| rs121909072 | pathogenic | 0 | 0 | 0 |
| rs143669310 | benign | 1 | 1 | 1 |
| rs747347498 | benign | 0 | 0 | 0 |
| rs138283222 | benign | 0 | 0 | 0 |
| rs118203931 | pathogenic | 1 | 1 | 1 |
| rs118203933 | pathogenic | 1 | 1 | 0 |
| rs118203932 | pathogenic | 1 | 1 | 0 |
| rs2228063 | benign | 1 | 1 | 1 |
| rs377767360 | pathogenic | 0 | 0 | 0 |
| rs397518413 | pathogenic | 1 | 1 | 1 |
| rs1800458 | benign | 1 | 1 | 0 |
| rs121918083 | pathogenic | 1 | 1 | 1 |
| rs121918094 | pathogenic | 1 | 1 | 1 |
| rs121918098 | pathogenic | 1 | 1 | 1 |
| rs121918093 | pathogenic | 1 | 1 | 1 |
| rs79977247 | pathogenic | 1 | 1 | 1 |
| rs28933979 | pathogenic | 1 | 1 | 1 |
| rs121918068 | pathogenic | 1 | 1 | 1 |
| rs121918077 | pathogenic | 1 | 1 | 1 |
| rs11541796 | pathogenic | 1 | 1 | 1 |
| rs104894665 | pathogenic | 1 | 1 | 1 |
| rs121918090 | pathogenic | 1 | 1 | 1 |
| rs121918081 | pathogenic | 1 | 1 | 1 |
| rs121918080 | pathogenic | 1 | 1 | 1 |
| rs121918076 | pathogenic | 1 | 1 | 1 |
| rs121918097 | pathogenic | 1 | 1 | 1 |

| | | | | |
|---|---|---|---|---|
| rs121918079 | pathogenic | 1 | 1 | 1 |
| rs121918069 | pathogenic | 1 | 1 | 1 |
| rs121918086 | pathogenic | 1 | 1 | 1 |
| rs121918091 | pathogenic | 1 | 1 | 1 |
| rs121918085 | pathogenic | 1 | 1 | 1 |
| rs121918100 | pathogenic | 1 | 1 | 1 |
| rs121918084 | pathogenic | 1 | 1 | 1 |
| rs121918071 | pathogenic | 1 | 1 | 1 |
| rs121918072 | pathogenic | 1 | 1 | 1 |
| rs121918082 | pathogenic | 1 | 1 | 1 |
| rs121918087 | pathogenic | 1 | 1 | 1 |
| rs76410435 | benign | 1 | 1 | 1 |
| rs121918089 | pathogenic | 1 | 1 | 1 |
| rs121918073 | pathogenic | 1 | 1 | 1 |
| rs121918075 | pathogenic | 1 | 1 | 1 |
| rs121918088 | pathogenic | 1 | 1 | 1 |
| rs76992529 | pathogenic | 1 | 1 | 1 |
| rs536294863 | benign | 1 | 1 | 1 |
| rs730881162 | benign | 1 | 1 | 1 |
| rs113994087 | pathogenic | 1 | 1 | 1 |
| rs74716434 | benign | 1 | 1 | 1 |
| rs121918694 | pathogenic | 1 | 1 | 1 |
| rs121918707 | pathogenic | 1 | 1 | 1 |
| rs121918695 | pathogenic | 1 | 1 | 0 |
| rs121918690 | pathogenic | 1 | 1 | 0 |
| rs121918696 | pathogenic | 1 | 1 | 0 |
| rs121918693 | pathogenic | 1 | 1 | 0 |
| rs28999969 | pathogenic | 1 | 1 | 1 |
| rs121918697 | pathogenic | 1 | 1 | 1 |
| rs121918688 | pathogenic | 1 | 1 | 0 |
| rs28999970 | pathogenic | 1 | 1 | 0 |
| rs121918686 | pathogenic | 1 | 1 | 0 |
| rs28999971 | pathogenic | 1 | 1 | 1 |
| rs121918698 | pathogenic | 1 | 1 | 1 |
| rs121918691 | pathogenic | 1 | 1 | 0 |
| rs121918692 | pathogenic | 1 | 1 | 1 |
| rs121918703 | pathogenic | 1 | 1 | 1 |
| rs121918687 | pathogenic | 1 | 1 | 1 |
| rs28933408 | pathogenic | 1 | 1 | 1 |
| rs121918702 | pathogenic | 1 | 1 | 1 |
| rs17875899 | benign | 1 | 1 | 1 |
| rs111033620 | pathogenic | 1 | 1 | 1 |
| rs111033622 | pathogenic | 1 | 1 | 1 |
| rs111033621 | pathogenic | 1 | 1 | 1 |
| rs111033618 | pathogenic | 1 | 1 | 1 |
| rs5918 | benign | 0 | 0 | 0 |
| rs121918452 | pathogenic | 0 | 0 | 0 |
| rs121918445 | pathogenic | 1 | 1 | 1 |
| rs5917 | benign | 1 | 1 | 1 |
| rs121918444 | pathogenic | 1 | 1 | 1 |
| rs121918446 | pathogenic | 1 | 1 | 1 |
| rs121918449 | pathogenic | 0 | 0 | 0 |
| rs121918448 | benign | 0 | 0 | 0 |
| rs13306487 | benign | 1 | 1 | 1 |

| | | | | |
|---|---|---|---|---|
| rs1805044 | benign | 0 | 0 | 0 |
| rs117078377 | benign | 0 | 0 | 0 |
| rs138132026 | benign | 1 | 1 | 1 |
| rs75762896 | benign | 1 | 1 | 0 |
| rs121918043 | pathogenic | 1 | 1 | 0 |
| rs143581690 | benign | 1 | 1 | 0 |
| rs147654263 | benign | 0 | 0 | 0 |
| rs587782529 | pathogenic | 1 | 1 | 1 |
| rs121912664 | pathogenic | 1 | 1 | 1 |
| rs150293825 | benign | 1 | 1 | 1 |
| rs17882252 | benign | 1 | 1 | 1 |
| rs121912662 | pathogenic | 1 | 1 | 1 |
| rs35993958 | benign | 1 | 1 | 1 |
| rs606231253 | pathogenic | 1 | 1 | 1 |
| rs2227983 | benign | 1 | 1 | 1 |
| rs587779351 | pathogenic | 0 | 0 | 0 |
| rs779773463 | pathogenic | 1 | 1 | 1 |
| rs713040 | benign | 1 | 1 | 1 |
| rs334 | pathogenic | 1 | 1 | 1 |
| rs33930165 | pathogenic | 1 | 1 | 1 |
| rs33918131 | benign | 1 | 1 | 1 |
| rs63750783 | pathogenic | 1 | 1 | 1 |
| rs33986703 | pathogenic | 1 | 1 | 1 |
| rs33972047 | pathogenic | 1 | 1 | 1 |
| rs33936254 | pathogenic | 1 | 1 | 1 |
| rs33959855 | benign | 1 | 1 | 1 |
| rs33950507 | pathogenic | 1 | 1 | 1 |
| rs33916412 | pathogenic | 1 | 1 | 0 |
| rs1135071 | pathogenic | 1 | 1 | 1 |
| rs33960103 | pathogenic | 1 | 1 | 1 |
| rs11549407 | pathogenic | 1 | 1 | 1 |
| rs33922842 | pathogenic | 1 | 1 | 1 |
| rs35278874 | benign | 1 | 1 | 1 |
| rs34621955 | benign | 1 | 1 | 1 |
| rs33995148 | pathogenic | 1 | 1 | 1 |
| rs33945705 | benign | 1 | 1 | 1 |
| rs33993568 | benign | 1 | 1 | 1 |
| rs35553496 | benign | 1 | 1 | 1 |
| rs33913712 | pathogenic | 1 | 1 | 1 |
| rs33917785 | pathogenic | 1 | 1 | 1 |
| rs34515413 | benign | 1 | 1 | 1 |
| rs35256489 | pathogenic | 1 | 1 | 0 |
| rs34484056 | pathogenic | 1 | 1 | 1 |
| rs33947020 | benign | 1 | 1 | 1 |
| rs33946267 | pathogenic | 1 | 1 | 1 |
| rs33966761 | pathogenic | 1 | 1 | 1 |
| rs36020563 | benign | 1 | 1 | 1 |
| rs122454124 | pathogenic | 1 | 1 | 1 |
| rs122454126 | pathogenic | 1 | 1 | 0 |
| rs122454127 | pathogenic | 1 | 1 | 1 |
| rs122454130 | pathogenic | 1 | 1 | 1 |
| rs122454125 | pathogenic | 1 | 1 | 1 |
| rs122454131 | pathogenic | 1 | 1 | 0 |
| rs121908302 | pathogenic | 0 | 1 | 0 |

| | | | | |
|---|---|---|---|---|
| rs77829017 | pathogenic | 0 | 1 | 0 |
| rs1141814 | pathogenic | 0 | 1 | 0 |
| rs121908312 | pathogenic | 0 | 1 | 0 |
| rs79653797 | pathogenic | 1 | 1 | 0 |
| rs439898 | pathogenic | 1 | 1 | 0 |
| rs121908299 | pathogenic | 0 | 1 | 0 |
| rs80356763 | pathogenic | 1 | 1 | 1 |
| rs781152868 | pathogenic | 0 | 1 | 0 |
| rs121908297 | pathogenic | 0 | 1 | 0 |
| rs398123531 | benign | 0 | 1 | 0 |
| rs398123532 | pathogenic | 0 | 1 | 0 |
| rs364897 | pathogenic | 1 | 1 | 1 |
| rs1064644 | pathogenic | 0 | 1 | 0 |
| rs409652 | pathogenic | 1 | 1 | 1 |
| rs121908300 | pathogenic | 0 | 1 | 0 |
| rs381737 | pathogenic | 0 | 1 | 0 |
| rs74500255 | pathogenic | 0 | 1 | 0 |
| rs121908313 | pathogenic | 0 | 1 | 0 |
| rs367968666 | pathogenic | 0 | 1 | 0 |
| rs78973108 | pathogenic | 1 | 1 | 1 |
| rs121908298 | pathogenic | 1 | 1 | 1 |
| rs78396650 | pathogenic | 0 | 0 | 0 |
| rs78198234 | pathogenic | 1 | 1 | 0 |
| rs121908304 | pathogenic | 1 | 1 | 1 |
| rs76539814 | pathogenic | 1 | 1 | 1 |
| rs121908305 | pathogenic | 0 | 1 | 0 |
| rs121908306 | pathogenic | 1 | 1 | 0 |
| rs121908308 | pathogenic | 1 | 1 | 1 |
| rs121908307 | pathogenic | 0 | 1 | 0 |
| rs75548401 | benign | 0 | 1 | 0 |
| rs76763715 | pathogenic | 0 | 0 | 0 |
| rs121908314 | pathogenic | 0 | 1 | 0 |
| rs398123528 | pathogenic | 0 | 1 | 0 |
| rs121908311 | pathogenic | 0 | 0 | 0 |
| rs121908310 | pathogenic | 1 | 1 | 0 |
| rs74598136 | pathogenic | 0 | 1 | 0 |
| rs1064651 | pathogenic | 1 | 1 | 1 |
| rs77369218 | pathogenic | 1 | 1 | 1 |
| rs121908295 | pathogenic | 0 | 1 | 0 |
| rs421016 | pathogenic | 0 | 1 | 0 |
| rs368060 | pathogenic | 0 | 1 | 0 |
| rs80356771 | pathogenic | 1 | 1 | 1 |
| rs80356772 | pathogenic | 1 | 1 | 1 |
| rs121908301 | pathogenic | 0 | 0 | 0 |
| rs747506979 | pathogenic | 0 | 1 | 0 |
| rs75822236 | pathogenic | 0 | 1 | 0 |
| rs1990760 | benign | 1 | 1 | 0 |
| rs28934568 | pathogenic | 1 | 1 | 0 |
| rs34833812 | benign | 1 | 1 | 0 |
| rs193922666 | benign | 1 | 1 | 0 |
| rs104893812 | pathogenic | 1 | 1 | 0 |
| rs144701411 | benign | 0 | 0 | 0 |
| rs104893813 | pathogenic | 0 | 1 | 0 |
| rs199660234 | pathogenic | 1 | 1 | 0 |

| | | | | |
|---|---|---|---|---|
| rs104893814 | pathogenic | 1 | 1 | 0 |
| rs143095746 | benign | 1 | 1 | 1 |
| rs104893807 | pathogenic | 1 | 1 | 0 |
| rs104893811 | pathogenic | 0 | 0 | 0 |
| rs104893816 | pathogenic | 0 | 0 | 0 |
| rs121918714 | pathogenic | 0 | 1 | 0 |
| rs104893810 | pathogenic | 1 | 1 | 0 |
| rs104893815 | pathogenic | 1 | 1 | 0 |
| rs4303 | benign | 0 | 1 | 0 |
| rs121912704 | pathogenic | 0 | 1 | 0 |
| rs56394458 | benign | 1 | 1 | 0 |
| rs397514688 | pathogenic | 0 | 0 | 0 |
| rs367797185 | pathogenic | 1 | 1 | 0 |
| rs12709426 | benign | 0 | 1 | 0 |
| rs2228547 | benign | 0 | 0 | 0 |
| rs111033546 | pathogenic | 0 | 0 | 0 |
| rs111033553 | pathogenic | 0 | 0 | 0 |
| rs111033554 | pathogenic | 1 | 1 | 1 |
| rs111033544 | pathogenic | 1 | 1 | 1 |
| rs111033555 | pathogenic | 1 | 1 | 1 |
| rs111033545 | pathogenic | 1 | 1 | 1 |
| rs111033549 | pathogenic | 0 | 1 | 0 |
| rs111033552 | pathogenic | 1 | 1 | 1 |
| rs752204035 | benign | 0 | 0 | 0 |
| rs11569534 | benign | 0 | 0 | 0 |
| rs28937571 | pathogenic | 1 | 1 | 1 |
| rs104894215 | pathogenic | 1 | 1 | 1 |
| rs3888798 | benign | 1 | 1 | 1 |
| rs764263034 | benign | 1 | 1 | 1 |
| rs1800973 | benign | 1 | 1 | 1 |
| rs367623154 | benign | 1 | 1 | 1 |
| rs201230446 | pathogenic | 1 | 1 | 1 |
| rs267606766 | pathogenic | 0 | 1 | 0 |
| rs267606769 | pathogenic | 0 | 1 | 0 |
| rs267606767 | pathogenic | 1 | 1 | 0 |
| rs267606768 | pathogenic | 0 | 1 | 0 |
| rs61733129 | benign | 0 | 1 | 0 |
| rs201947120 | pathogenic | 1 | 1 | 1 |
| rs283413 | benign | 0 | 1 | 0 |
| rs36119840 | benign | 0 | 0 | 0 |
| rs281865419 | pathogenic | 1 | 1 | 1 |
| rs137852937 | pathogenic | 0 | 1 | 0 |
| rs2230709 | benign | 0 | 1 | 0 |
| rs6695033 | benign | 0 | 1 | 0 |
| rs180177032 | pathogenic | 1 | 1 | 1 |
| rs180177033 | pathogenic | 1 | 1 | 1 |
| rs121913351 | pathogenic | 1 | 1 | 1 |
| rs397507473 | pathogenic | 1 | 1 | 1 |
| rs121913355 | pathogenic | 1 | 1 | 1 |
| rs121913357 | pathogenic | 1 | 1 | 1 |
| rs180177036 | pathogenic | 1 | 1 | 1 |
| rs180177039 | pathogenic | 1 | 1 | 1 |
| rs180177038 | pathogenic | 1 | 1 | 1 |
| rs180177040 | pathogenic | 1 | 1 | 1 |

| | | | | |
|---|---|---|---|---|
| rs121913338 | pathogenic | 1 | 1 | 1 |
| rs121913366 | pathogenic | 1 | 1 | 1 |
| rs121913369 | pathogenic | 1 | 1 | 1 |
| rs113488022 | pathogenic | 1 | 1 | 1 |
| rs121913364 | pathogenic | 1 | 1 | 1 |
| rs180177042 | pathogenic | 1 | 1 | 1 |
| rs200461617 | benign | 0 | 0 | 0 |
| rs141609745 | benign | 0 | 0 | 0 |
| rs104894383 | pathogenic | 1 | 1 | 1 |
| rs104894384 | pathogenic | 1 | 1 | 1 |
| rs104894381 | pathogenic | 1 | 1 | 0 |
| rs147710408 | benign | 1 | 1 | 1 |
| rs77357563 | benign | 1 | 1 | 1 |
| rs104894378 | pathogenic | 0 | 0 | 0 |
| rs104894382 | pathogenic | 0 | 0 | 0 |
| rs137853292 | pathogenic | 1 | 1 | 1 |
| rs587778834 | pathogenic | 0 | 0 | 0 |
| rs137853294 | pathogenic | 1 | 1 | 1 |
| rs551747882 | benign | 0 | 0 | 0 |
| rs3092903 | benign | 1 | 1 | 1 |
| rs137853296 | pathogenic | 0 | 0 | 0 |
| rs398122917 | pathogenic | 0 | 1 | 0 |
| rs41304992 | benign | 0 | 0 | 0 |
| rs121912708 | pathogenic | 1 | 1 | 1 |
| rs140102105 | pathogenic | 1 | 1 | 1 |
| rs376917645 | pathogenic | 1 | 1 | 0 |
| rs372660425 | pathogenic | 1 | 1 | 0 |
| rs555896752 | pathogenic | 1 | 1 | 1 |
| rs777829351 | pathogenic | 1 | 1 | 0 |
| rs121912711 | pathogenic | 1 | 1 | 0 |
| rs754449549 | pathogenic | 0 | 1 | 0 |
| rs141701364 | benign | 0 | 1 | 0 |
| rs761011399 | benign | 1 | 1 | 1 |
| rs121912710 | benign | 1 | 1 | 0 |
| rs121912707 | pathogenic | 1 | 1 | 0 |
| rs747597620 | pathogenic | 0 | 1 | 0 |
| rs186558364 | pathogenic | 1 | 1 | 1 |
| rs556400964 | pathogenic | 1 | 1 | 1 |
| rs61757684 | benign | 1 | 1 | 1 |
| rs121917766 | pathogenic | 1 | 1 | 1 |
| rs121918248 | pathogenic | 0 | 0 | 0 |
| rs398123278 | pathogenic | 0 | 0 | 0 |
| rs121918251 | pathogenic | 1 | 1 | 1 |
| rs727504022 | pathogenic | 0 | 0 | 0 |
| rs190834116 | pathogenic | 1 | 1 | 1 |
| rs121918249 | pathogenic | 1 | 1 | 0 |
| rs121918257 | pathogenic | 1 | 1 | 0 |
| rs142068898 | benign | 0 | 0 | 0 |
| rs757000253 | pathogenic | 0 | 0 | 0 |
| rs200908035 | pathogenic | 1 | 1 | 0 |
| rs760782399 | pathogenic | 0 | 0 | 0 |
| rs778702777 | pathogenic | 0 | 0 | 0 |
| rs121918258 | pathogenic | 1 | 1 | 0 |
| rs121918256 | pathogenic | 0 | 0 | 0 |

| | | | | |
|---|---|---|---|---|
| rs200596762 | pathogenic | 1 | 1 | 0 |
| rs547709692 | pathogenic | 0 | 0 | 0 |
| rs758577372 | pathogenic | 0 | 1 | 0 |
| rs564069299 | pathogenic | 1 | 1 | 1 |
| rs772552898 | pathogenic | 1 | 1 | 1 |
| rs368790885 | pathogenic | 1 | 1 | 1 |
| rs121918250 | pathogenic | 0 | 0 | 0 |
| rs533755473 | pathogenic | 1 | 1 | 1 |
| rs753288303 | pathogenic | 1 | 1 | 1 |
| rs2229385 | benign | 0 | 0 | 0 |
| rs1141321 | benign | 0 | 0 | 0 |
| rs753564352 | pathogenic | 0 | 0 | 0 |
| rs775593146 | pathogenic | 0 | 0 | 0 |
| rs121918254 | pathogenic | 0 | 0 | 0 |
| rs8589 | benign | 1 | 1 | 0 |
| rs777758903 | pathogenic | 0 | 0 | 0 |
| rs140600746 | pathogenic | 1 | 1 | 0 |
| rs121918255 | pathogenic | 1 | 1 | 0 |
| rs121918252 | pathogenic | 0 | 0 | 0 |
| rs779990936 | pathogenic | 1 | 1 | 0 |
| rs121912455 | pathogenic | 1 | 1 | 1 |
| rs121912459 | pathogenic | 1 | 1 | 1 |
| rs74315452 | pathogenic | 1 | 1 | 1 |
| rs121912451 | pathogenic | 1 | 1 | 1 |
| rs79006549 | pathogenic | 0 | 0 | 0 |
| rs28934889 | benign | 0 | 1 | 0 |
| rs111033557 | benign | 1 | 1 | 1 |
| rs28934597 | pathogenic | 1 | 1 | 1 |
| rs28934596 | pathogenic | 1 | 1 | 1 |
| rs28934595 | pathogenic | 1 | 1 | 1 |
| rs146519482 | pathogenic | 1 | 1 | 1 |
| rs111033563 | pathogenic | 1 | 1 | 1 |
| rs75385605 | benign | 1 | 1 | 1 |
| rs121908569 | pathogenic | 1 | 1 | 1 |
| rs35719940 | benign | 0 | 0 | 0 |
| rs121918664 | pathogenic | 0 | 0 | 0 |
| rs104893665 | pathogenic | 1 | 1 | 1 |
| rs104893666 | pathogenic | 1 | 1 | 1 |
| rs200845457 | benign | 1 | 1 | 1 |
| rs779204655 | pathogenic | 1 | 1 | 1 |
| rs121917747 | pathogenic | 1 | 1 | 1 |
| rs6042 | benign | 1 | 1 | 1 |
| rs104894380 | pathogenic | 0 | 0 | 0 |
| rs369083541 | pathogenic | 1 | 1 | 1 |
| rs145243532 | benign | 1 | 1 | 1 |
| rs121909500 | pathogenic | 1 | 1 | 1 |
| rs140993310 | benign | 1 | 1 | 1 |
| rs200837270 | pathogenic | 1 | 1 | 1 |
| rs121918484 | pathogenic | 1 | 1 | 1 |
| rs121918306 | pathogenic | 0 | 0 | 0 |
| rs118204089 | pathogenic | 1 | 1 | 0 |
| rs118204083 | pathogenic | 1 | 1 | 0 |
| rs201953584 | pathogenic | 1 | 1 | 1 |
| rs74315362 | pathogenic | 1 | 1 | 1 |

| | | | | |
|---|---|---|---|---|
| rs118204084 | pathogenic | 1 | 1 | 0 |
| rs118204085 | pathogenic | 1 | 1 | 1 |
| rs113403872 | pathogenic | 1 | 1 | 0 |
| rs1052176 | benign | 1 | 1 | 1 |
| rs2031640 | benign | 0 | 0 | 0 |
| rs104894230 | pathogenic | 1 | 1 | 1 |
| rs104894229 | pathogenic | 1 | 1 | 1 |
| rs104894228 | pathogenic | 1 | 1 | 1 |
| rs104894226 | pathogenic | 1 | 1 | 1 |
| rs121917757 | pathogenic | 1 | 1 | 1 |
| rs121917758 | pathogenic | 1 | 1 | 1 |
| rs28933406 | pathogenic | 1 | 1 | 1 |
| rs121917756 | pathogenic | 1 | 1 | 1 |
| rs104894227 | pathogenic | 1 | 1 | 1 |
| rs369106578 | pathogenic | 1 | 1 | 1 |
| rs397517142 | benign | 1 | 1 | 1 |
| rs104894231 | pathogenic | 1 | 1 | 1 |
| rs121917759 | pathogenic | 1 | 1 | 1 |
| rs104893827 | pathogenic | 0 | 0 | 0 |
| rs104893826 | pathogenic | 1 | 1 | 1 |
| rs5030802 | pathogenic | 1 | 1 | 1 |
| rs5030827 | pathogenic | 1 | 1 | 0 |
| rs119103277 | pathogenic | 1 | 1 | 1 |
| rs5030808 | pathogenic | 0 | 1 | 0 |
| rs5030809 | pathogenic | 1 | 1 | 1 |
| rs765978945 | pathogenic | 1 | 1 | 1 |
| rs104893824 | pathogenic | 1 | 1 | 1 |
| rs104893831 | pathogenic | 0 | 1 | 0 |
| rs104893830 | pathogenic | 0 | 1 | 0 |
| rs119103278 | pathogenic | 0 | 1 | 0 |
| rs5030818 | pathogenic | 1 | 1 | 1 |
| rs397516444 | pathogenic | 1 | 1 | 1 |
| rs28940297 | pathogenic | 1 | 1 | 1 |
| rs267607170 | pathogenic | 1 | 1 | 1 |
| rs104893825 | pathogenic | 1 | 1 | 1 |
| rs5030820 | pathogenic | 0 | 1 | 0 |
| rs5030821 | pathogenic | 0 | 1 | 0 |
| rs5030823 | pathogenic | 1 | 1 | 1 |
| rs5030824 | pathogenic | 1 | 1 | 1 |
| rs28940301 | pathogenic | 0 | 1 | 0 |
| rs28940300 | pathogenic | 0 | 1 | 0 |
| rs28940298 | pathogenic | 0 | 1 | 0 |
| rs200019083 | benign | 0 | 0 | 0 |
| rs759748655 | pathogenic | 0 | 0 | 0 |
| rs121917739 | pathogenic | 1 | 1 | 1 |
| rs75353611 | pathogenic | 1 | 1 | 1 |
| rs79744198 | benign | 1 | 1 | 1 |
| rs79228041 | pathogenic | 1 | 1 | 1 |
| rs80106970 | benign | 1 | 1 | 1 |
| rs375353223 | benign | 0 | 0 | 0 |
| rs56311081 | benign | 0 | 0 | 0 |
| rs55985569 | benign | 1 | 1 | 1 |
| rs45483396 | benign | 0 | 0 | 0 |
| rs35284565 | benign | 1 | 1 | 1 |

| | | | | |
|---|---|---|---|---|
| rs34349517 | benign | 0 | 0 | 0 |
| rs35776110 | benign | 0 | 0 | 0 |
| rs33917957 | benign | 1 | 1 | 1 |
| rs587778445 | benign | 0 | 0 | 0 |
| rs34624984 | benign | 0 | 0 | 0 |
| rs701753 | benign | 0 | 0 | 0 |
| rs855314 | benign | 0 | 0 | 0 |
| rs121918371 | pathogenic | 1 | 1 | 0 |
| rs398122912 | pathogenic | 0 | 1 | 0 |
| rs1126728 | benign | 0 | 0 | 0 |
| rs587777404 | pathogenic | 1 | 1 | 0 |
| rs11208257 | benign | 0 | 0 | 0 |
| rs6676290 | benign | 0 | 0 | 0 |
| rs104894118 | pathogenic | 0 | 0 | 0 |
| rs121918655 | pathogenic | 1 | 1 | 1 |
| rs104894120 | pathogenic | 1 | 1 | 1 |
| rs137852238 | pathogenic | 0 | 0 | 0 |
| rs137852237 | pathogenic | 0 | 0 | 0 |
| rs137852499 | pathogenic | 1 | 1 | 1 |
| rs137852480 | pathogenic | 1 | 1 | 1 |
| rs137852494 | pathogenic | 1 | 1 | 1 |
| rs137852502 | pathogenic | 1 | 1 | 1 |
| rs137852500 | pathogenic | 1 | 1 | 1 |
| rs137852487 | pathogenic | 1 | 1 | 1 |
| rs137852481 | pathogenic | 1 | 1 | 1 |
| rs137852501 | pathogenic | 0 | 0 | 0 |
| rs137852478 | pathogenic | 1 | 1 | 1 |
| rs137852485 | pathogenic | 1 | 1 | 1 |
| rs137852482 | pathogenic | 1 | 1 | 0 |
| rs137852483 | pathogenic | 0 | 1 | 0 |
| rs137852477 | pathogenic | 1 | 1 | 0 |
| rs137852496 | pathogenic | 1 | 1 | 0 |
| rs137852484 | pathogenic | 0 | 1 | 0 |
| rs137852498 | pathogenic | 1 | 1 | 0 |
| rs137852497 | pathogenic | 1 | 1 | 0 |
| rs137852493 | pathogenic | 1 | 1 | 0 |
| rs137852504 | pathogenic | 1 | 1 | 1 |
| rs267606863 | pathogenic | 1 | 1 | 1 |
| rs137852486 | pathogenic | 1 | 1 | 1 |
| rs137852479 | pathogenic | 1 | 1 | 1 |
| rs112122950 | benign | 0 | 0 | 0 |
| rs72546648 | benign | 1 | 1 | 1 |
| rs72552291 | pathogenic | 0 | 0 | 0 |
| rs79184941 | pathogenic | 1 | 1 | 1 |
| rs77543610 | pathogenic | 1 | 1 | 1 |
| rs121918505 | pathogenic | 1 | 1 | 1 |
| rs776587763 | pathogenic | 1 | 1 | 1 |
| rs121918497 | pathogenic | 1 | 1 | 1 |
| rs121918499 | pathogenic | 1 | 1 | 1 |
| rs121918501 | pathogenic | 1 | 1 | 1 |
| rs121918500 | pathogenic | 0 | 0 | 0 |
| rs121918504 | pathogenic | 1 | 1 | 1 |
| rs121918510 | pathogenic | 1 | 1 | 1 |
| rs121918493 | pathogenic | 1 | 1 | 1 |

| | | | | |
|---|---|---|---|---|
| rs121918489 | pathogenic | 0 | 0 | 0 |
| rs121918495 | pathogenic | 1 | 1 | 1 |
| rs121918488 | pathogenic | 0 | 0 | 0 |
| rs121918496 | pathogenic | 0 | 0 | 0 |
| rs121918492 | pathogenic | 1 | 1 | 1 |
| rs121918494 | pathogenic | 1 | 1 | 1 |
| rs121918502 | pathogenic | 1 | 1 | 1 |
| rs121918490 | pathogenic | 1 | 1 | 1 |
| rs72466451 | pathogenic | 1 | 1 | 1 |
| rs66468541 | pathogenic | 0 | 0 | 0 |
| rs137852662 | pathogenic | 0 | 0 | 0 |
| rs137852663 | pathogenic | 0 | 0 | 0 |
| rs606231407 | pathogenic | 0 | 0 | 0 |
| rs137852233 | pathogenic | 1 | 1 | 1 |
| rs112306225 | benign | 0 | 0 | 0 |
| rs387906808 | pathogenic | 1 | 1 | 0 |
| rs730882094 | pathogenic | 1 | 1 | 1 |
| rs373869746 | pathogenic | 1 | 1 | 1 |
| rs777524402 | pathogenic | 1 | 1 | 1 |
| rs769737896 | pathogenic | 0 | 1 | 0 |
| rs13306515 | benign | 1 | 1 | 0 |
| rs746982741 | pathogenic | 0 | 0 | 0 |
| rs2230200 | benign | 1 | 1 | 1 |
| rs1047286 | benign | 0 | 0 | 0 |
| rs141737564 | benign | 0 | 0 | 0 |
| rs774826179 | benign | 0 | 0 | 0 |
| rs11569422 | benign | 1 | 1 | 1 |
| rs146613648 | benign | 0 | 0 | 0 |
| rs144432231 | benign | 0 | 0 | 0 |
| rs140655115 | benign | 0 | 0 | 0 |
| rs387906837 | pathogenic | 1 | 1 | 0 |
| rs121909076 | pathogenic | 1 | 1 | 0 |
| rs121909077 | pathogenic | 0 | 0 | 0 |
| rs121909074 | pathogenic | 0 | 0 | 0 |
| rs137853330 | pathogenic | 1 | 1 | 1 |
| rs137853331 | pathogenic | 1 | 1 | 1 |
| rs137853332 | pathogenic | 1 | 1 | 1 |
| rs28931586 | pathogenic | 1 | 1 | 0 |
| rs11569321 | benign | 1 | 1 | 1 |
| rs119486097 | benign | 0 | 0 | 0 |
| rs1799807 | pathogenic | 1 | 1 | 1 |
| rs201820739 | pathogenic | 1 | 1 | 0 |
| rs114706984 | pathogenic | 1 | 1 | 0 |
| rs140080572 | pathogenic | 1 | 1 | 0 |
| rs28933389 | pathogenic | 0 | 1 | 0 |
| rs16849700 | benign | 1 | 1 | 0 |
| rs115624085 | pathogenic | 1 | 1 | 1 |
| rs747196387 | pathogenic | 1 | 1 | 1 |
| rs104893684 | pathogenic | 1 | 1 | 0 |
| rs121918557 | pathogenic | 1 | 1 | 0 |
| rs121918556 | pathogenic | 1 | 1 | 1 |
| rs398124571 | pathogenic | 1 | 1 | 1 |
| rs776631396 | pathogenic | 0 | 1 | 0 |
| rs398124576 | pathogenic | 1 | 1 | 1 |

| | | | | |
|---|---|---|---|---|
| rs398124581 | pathogenic | 1 | 1 | 1 |
| rs398124582 | pathogenic | 1 | 1 | 1 |
| rs149766077 | pathogenic | 0 | 1 | 0 |
| rs79761867 | pathogenic | 0 | 1 | 0 |
| rs148905512 | pathogenic | 0 | 1 | 0 |
| rs121965004 | pathogenic | 1 | 1 | 1 |
| rs386834233 | pathogenic | 0 | 1 | 0 |
| rs398124598 | pathogenic | 0 | 1 | 0 |
| rs398124603 | pathogenic | 1 | 1 | 1 |
| rs546716073 | benign | 1 | 1 | 1 |
| rs200310662 | benign | 1 | 1 | 1 |
| rs115494727 | benign | 1 | 1 | 1 |
| rs781513008 | benign | 0 | 0 | 0 |
| rs587776398 | benign | 1 | 1 | 0 |
| rs35288908 | benign | 1 | 1 | 1 |
| rs150634562 | pathogenic | 1 | 1 | 0 |
| rs142948132 | pathogenic | 1 | 1 | 1 |
| rs132630321 | pathogenic | 1 | 1 | 1 |
| rs132630317 | pathogenic | 1 | 1 | 1 |
| rs132630320 | pathogenic | 0 | 0 | 0 |
| rs781394318 | pathogenic | 1 | 1 | 1 |
| rs111033624 | pathogenic | 1 | 1 | 1 |
| rs111033630 | pathogenic | 1 | 1 | 1 |
| rs111033627 | pathogenic | 1 | 1 | 1 |
| rs193922287 | pathogenic | 1 | 1 | 1 |
| rs759072800 | pathogenic | 0 | 1 | 0 |
| rs104894010 | pathogenic | 0 | 0 | 0 |
| rs193922296 | pathogenic | 0 | 0 | 0 |
| rs104894006 | pathogenic | 1 | 1 | 1 |
| rs193922311 | pathogenic | 1 | 1 | 0 |
| rs80356654 | pathogenic | 1 | 1 | 0 |
| rs144723656 | benign | 1 | 1 | 1 |
| rs148311934 | pathogenic | 1 | 1 | 0 |
| rs80356655 | pathogenic | 1 | 1 | 1 |
| rs769268803 | pathogenic | 1 | 1 | 0 |
| rs104894008 | pathogenic | 0 | 1 | 0 |
| rs104894005 | pathogenic | 1 | 1 | 1 |
| rs104894009 | pathogenic | 1 | 1 | 0 |
| rs193922340 | pathogenic | 0 | 0 | 0 |
| rs104894016 | pathogenic | 0 | 1 | 0 |
| rs193921338 | pathogenic | 0 | 0 | 0 |
| rs193922272 | pathogenic | 1 | 1 | 1 |
| rs104894012 | pathogenic | 1 | 1 | 0 |
| rs193922285 | pathogenic | 1 | 1 | 1 |
| rs104894336 | pathogenic | 1 | 1 | 1 |
| rs28931580 | pathogenic | 1 | 1 | 1 |
| rs104894326 | pathogenic | 1 | 1 | 1 |
| rs104894331 | pathogenic | 0 | 0 | 0 |
| rs104894338 | pathogenic | 0 | 0 | 0 |
| rs104894333 | pathogenic | 1 | 1 | 1 |
| rs104894330 | pathogenic | 1 | 1 | 1 |
| rs143886391 | benign | 1 | 1 | 1 |
| rs104894335 | pathogenic | 1 | 1 | 1 |
| rs104894337 | pathogenic | 0 | 0 | 0 |

| | | | | |
|---|---|---|---|---|
| rs104894328 | pathogenic | 0 | 0 | 0 |
| rs104894341 | pathogenic | 1 | 1 | 0 |
| rs104894329 | pathogenic | 0 | 0 | 0 |
| rs137852850 | pathogenic | 1 | 1 | 1 |
| rs137852854 | pathogenic | 0 | 1 | 0 |
| rs137852849 | pathogenic | 0 | 1 | 0 |
| rs137852848 | pathogenic | 1 | 1 | 1 |
| rs137852852 | pathogenic | 0 | 1 | 0 |
| rs137852853 | pathogenic | 0 | 1 | 0 |
| rs137852847 | pathogenic | 0 | 1 | 0 |
| rs137852846 | pathogenic | 0 | 1 | 0 |
| rs137852844 | pathogenic | 0 | 1 | 0 |
| rs144411158 | pathogenic | 0 | 0 | 0 |
| rs2302234 | benign | 0 | 0 | 0 |
| rs139620139 | pathogenic | 0 | 0 | 0 |
| rs146689929 | benign | 0 | 0 | 0 |
| rs373328681 | pathogenic | 1 | 1 | 0 |
| rs387906620 | pathogenic | 1 | 1 | 1 |
| rs1799983 | benign | 0 | 1 | 0 |
| rs28999111 | pathogenic | 1 | 1 | 1 |
| rs104894544 | pathogenic | 1 | 1 | 1 |
| rs104894546 | pathogenic | 1 | 1 | 1 |
| rs28999112 | pathogenic | 0 | 1 | 0 |
| rs104894545 | pathogenic | 1 | 1 | 1 |
| rs104894547 | pathogenic | 1 | 1 | 1 |
| rs74315310 | pathogenic | 0 | 0 | 0 |
| rs267606852 | pathogenic | 0 | 1 | 0 |
| rs137853586 | pathogenic | 0 | 1 | 0 |
| rs137853582 | pathogenic | 1 | 1 | 1 |
| rs61754634 | pathogenic | 1 | 1 | 1 |
| rs137853587 | pathogenic | 0 | 1 | 0 |
| rs267606851 | pathogenic | 1 | 1 | 1 |
| rs267606853 | pathogenic | 1 | 1 | 1 |
| rs137853584 | pathogenic | 1 | 1 | 1 |
| rs137853585 | pathogenic | 1 | 1 | 1 |
| rs460184 | pathogenic | 1 | 1 | 1 |
| rs104893751 | pathogenic | 0 | 1 | 0 |
| rs77358162 | benign | 0 | 0 | 0 |
| rs387906936 | pathogenic | 0 | 0 | 0 |
| rs769525399 | pathogenic | 0 | 0 | 0 |
| rs371271054 | pathogenic | 1 | 1 | 0 |
| rs118204000 | pathogenic | 0 | 0 | 0 |
| rs138568622 | pathogenic | 1 | 1 | 1 |
| rs34509359 | benign | 0 | 0 | 0 |
| rs104894130 | pathogenic | 1 | 1 | 0 |
| rs281865424 | pathogenic | 0 | 0 | 0 |
| rs121912778 | pathogenic | 1 | 1 | 0 |
| rs771055145 | pathogenic | 1 | 1 | 0 |
| rs776174514 | pathogenic | 1 | 1 | 0 |
| rs545986367 | pathogenic | 1 | 1 | 1 |
| rs570768038 | benign | 1 | 1 | 1 |
| rs74315442 | pathogenic | 1 | 1 | 1 |
| rs79716074 | benign | 1 | 1 | 1 |
| rs748486078 | pathogenic | 1 | 1 | 1 |

| | | | | |
|---|---|---|---|---|
| rs2397084 | benign | 1 | 1 | 1 |
| rs11465553 | benign | 1 | 1 | 1 |
| rs144576902 | benign | 1 | 1 | 1 |
| rs763780 | benign | 1 | 1 | 1 |
| rs267606870 | pathogenic | 1 | 1 | 0 |
| rs121913502 | pathogenic | 1 | 1 | 0 |
| rs118101777 | benign | 1 | 1 | 1 |
| rs118053940 | benign | 0 | 0 | 0 |
| rs368771578 | benign | 1 | 1 | 1 |
| rs11540654 | pathogenic | 1 | 1 | 1 |
| rs121912658 | pathogenic | 1 | 1 | 1 |
| rs28934873 | pathogenic | 0 | 1 | 0 |
| rs780442292 | pathogenic | 1 | 1 | 1 |
| rs758781593 | benign | 1 | 1 | 1 |
| rs28934875 | pathogenic | 1 | 1 | 1 |
| rs28934874 | pathogenic | 1 | 1 | 0 |
| rs587782705 | pathogenic | 1 | 1 | 1 |
| rs587780068 | pathogenic | 1 | 1 | 1 |
| rs587782144 | pathogenic | 1 | 1 | 1 |
| rs138729528 | pathogenic | 1 | 1 | 1 |
| rs28934578 | pathogenic | 1 | 1 | 1 |
| rs397514495 | pathogenic | 1 | 1 | 1 |
| rs121912665 | pathogenic | 1 | 1 | 1 |
| rs397516435 | pathogenic | 1 | 1 | 1 |
| rs587778720 | pathogenic | 1 | 1 | 1 |
| rs397516436 | pathogenic | 1 | 1 | 1 |
| rs587781386 | benign | 1 | 1 | 1 |
| rs121912666 | pathogenic | 1 | 1 | 0 |
| rs587780073 | pathogenic | 1 | 1 | 1 |
| rs587782664 | pathogenic | 1 | 1 | 1 |
| rs730882005 | pathogenic | 1 | 1 | 1 |
| rs28934573 | pathogenic | 1 | 1 | 1 |
| rs121912655 | pathogenic | 1 | 1 | 1 |
| rs28934575 | pathogenic | 1 | 1 | 1 |
| rs121912656 | pathogenic | 1 | 1 | 1 |
| rs11540652 | pathogenic | 1 | 1 | 1 |
| rs121912651 | pathogenic | 1 | 1 | 1 |
| rs730882027 | pathogenic | 1 | 1 | 1 |
| rs121912653 | pathogenic | 1 | 1 | 1 |
| rs121912652 | pathogenic | 1 | 1 | 1 |
| rs121912657 | pathogenic | 0 | 1 | 0 |
| rs121913343 | pathogenic | 1 | 1 | 1 |
| rs28934576 | pathogenic | 1 | 1 | 1 |
| rs763098116 | pathogenic | 1 | 1 | 1 |
| rs121912660 | pathogenic | 1 | 1 | 1 |
| rs28934574 | pathogenic | 1 | 1 | 1 |
| rs371409680 | pathogenic | 1 | 1 | 1 |
| rs121912667 | pathogenic | 1 | 1 | 1 |
| rs121912663 | pathogenic | 1 | 1 | 1 |
| rs201744589 | pathogenic | 0 | 0 | 0 |
| rs33926449 | benign | 0 | 0 | 0 |
| rs74315302 | pathogenic | 0 | 1 | 0 |
| rs202040269 | pathogenic | 1 | 1 | 1 |
| rs375958814 | pathogenic | 1 | 1 | 1 |

| | | | | |
|---|---|---|---|---|
| rs121909624 | pathogenic | 1 | 1 | 1 |
| rs121909621 | pathogenic | 1 | 1 | 1 |
| rs121909622 | pathogenic | 1 | 1 | 0 |
| rs4220 | benign | 1 | 1 | 1 |
| rs147559466 | pathogenic | 1 | 1 | 1 |
| rs757434857 | pathogenic | 1 | 1 | 1 |
| rs398123072 | pathogenic | 1 | 1 | 1 |
| rs121434280 | pathogenic | 0 | 1 | 0 |
| rs398123074 | pathogenic | 0 | 1 | 0 |
| rs762114560 | pathogenic | 1 | 1 | 1 |
| rs121434283 | pathogenic | 1 | 1 | 0 |
| rs121434277 | pathogenic | 0 | 1 | 0 |
| rs745844469 | pathogenic | 1 | 1 | 1 |
| rs121434279 | pathogenic | 1 | 1 | 0 |
| rs121434278 | pathogenic | 1 | 1 | 1 |
| rs373715782 | pathogenic | 1 | 1 | 1 |
| rs200724875 | pathogenic | 1 | 1 | 1 |
| rs149678400 | pathogenic | 0 | 1 | 0 |
| rs770273135 | pathogenic | 1 | 1 | 1 |
| rs121434276 | pathogenic | 0 | 1 | 0 |
| rs121434281 | pathogenic | 1 | 1 | 0 |
| rs768884003 | pathogenic | 1 | 1 | 1 |
| rs201375579 | pathogenic | 0 | 0 | 0 |
| rs121434274 | pathogenic | 1 | 1 | 0 |
| rs121434282 | pathogenic | 1 | 1 | 0 |
| rs779759347 | pathogenic | 0 | 1 | 0 |
| rs77931234 | pathogenic | 1 | 1 | 1 |
| rs148207467 | pathogenic | 1 | 1 | 1 |
| rs766140986 | pathogenic | 1 | 1 | 1 |
| rs121434275 | pathogenic | 1 | 1 | 1 |
| rs748110745 | pathogenic | 1 | 1 | 1 |
| rs104894274 | pathogenic | 1 | 1 | 1 |
| rs104894273 | pathogenic | 0 | 1 | 0 |
| rs104894278 | pathogenic | 1 | 1 | 1 |
| rs104894275 | pathogenic | 1 | 1 | 1 |
| rs104894277 | pathogenic | 1 | 1 | 1 |
| rs104894276 | pathogenic | 0 | 0 | 0 |
| rs104894280 | pathogenic | 1 | 1 | 1 |
| rs145882709 | pathogenic | 1 | 1 | 1 |
| rs104894279 | pathogenic | 1 | 1 | 1 |
| rs80356670 | pathogenic | 1 | 1 | 1 |
| rs121918102 | pathogenic | 1 | 1 | 1 |
| rs80356671 | pathogenic | 1 | 1 | 1 |
| rs80356672 | pathogenic | 1 | 1 | 1 |
| rs758609113 | pathogenic | 1 | 1 | 0 |
| rs121918188 | pathogenic | 0 | 1 | 0 |
| rs121918189 | pathogenic | 1 | 1 | 0 |
| rs121918191 | pathogenic | 1 | 1 | 1 |
| rs1769259 | benign | 1 | 1 | 1 |
| rs140222720 | benign | 1 | 1 | 1 |
| rs2297085 | benign | 1 | 1 | 1 |
| rs200679026 | pathogenic | 1 | 1 | 1 |
| rs121918192 | pathogenic | 0 | 1 | 0 |
| rs145771944 | benign | 0 | 1 | 0 |

| | | | | |
|---|---|---|---|---|
| rs201508880 | pathogenic | 0 | 1 | 0 |
| rs17221854 | pathogenic | 0 | 0 | 0 |
| rs120074189 | pathogenic | 1 | 1 | 1 |
| rs120074190 | pathogenic | 1 | 1 | 1 |
| rs758600283 | benign | 1 | 1 | 1 |
| rs200305064 | pathogenic | 1 | 1 | 1 |
| rs121917891 | pathogenic | 1 | 1 | 1 |
| rs3772809 | benign | 0 | 1 | 0 |
| rs387906619 | pathogenic | 1 | 1 | 0 |
| rs2304681 | benign | 0 | 1 | 0 |
| rs121917849 | pathogenic | 0 | 0 | 0 |
| rs143010236 | pathogenic | 0 | 0 | 0 |
| rs121917850 | pathogenic | 1 | 1 | 0 |
| rs35916840 | pathogenic | 1 | 1 | 0 |
| rs28939688 | pathogenic | 1 | 1 | 1 |
| rs11258194 | benign | 1 | 1 | 1 |
| rs199840952 | pathogenic | 1 | 1 | 1 |
| rs121918159 | pathogenic | 1 | 1 | 1 |
| rs200396166 | pathogenic | 0 | 0 | 0 |
| rs757966746 | pathogenic | 0 | 1 | 0 |
| rs121918608 | pathogenic | 0 | 1 | 0 |
| rs74315401 | pathogenic | 1 | 1 | 1 |
| rs11538758 | pathogenic | 0 | 0 | 0 |
| rs74315402 | pathogenic | 1 | 1 | 1 |
| rs1799990 | pathogenic | 1 | 1 | 1 |
| rs74315410 | pathogenic | 1 | 1 | 1 |
| rs150351644 | benign | 1 | 1 | 1 |
| rs16990018 | benign | 1 | 1 | 1 |
| rs74315403 | pathogenic | 1 | 1 | 1 |
| rs74315408 | pathogenic | 1 | 1 | 1 |
| rs74315411 | pathogenic | 1 | 1 | 1 |
| rs74315413 | pathogenic | 1 | 1 | 1 |
| rs74315405 | pathogenic | 1 | 1 | 1 |
| rs28933385 | pathogenic | 1 | 1 | 1 |
| rs74315412 | pathogenic | 1 | 1 | 1 |
| rs74315407 | pathogenic | 1 | 1 | 1 |
| rs74315406 | pathogenic | 1 | 1 | 1 |
| rs11571117 | benign | 1 | 1 | 1 |
| rs121917742 | pathogenic | 1 | 1 | 1 |
| rs149782619 | pathogenic | 1 | 1 | 1 |
| rs397514492 | pathogenic | 1 | 1 | 1 |
| rs397514489 | pathogenic | 0 | 1 | 0 |
| rs397514493 | pathogenic | 1 | 1 | 1 |
| rs373849532 | pathogenic | 1 | 1 | 1 |
| rs121965048 | pathogenic | 1 | 1 | 1 |
| rs121965037 | pathogenic | 1 | 1 | 1 |
| rs121965060 | pathogenic | 0 | 1 | 0 |
| rs386833603 | pathogenic | 1 | 1 | 1 |
| rs121965038 | pathogenic | 1 | 1 | 1 |
| rs121965039 | pathogenic | 1 | 1 | 1 |
| rs121965040 | pathogenic | 1 | 1 | 1 |
| rs267606925 | pathogenic | 1 | 1 | 1 |
| rs121965056 | pathogenic | 0 | 1 | 0 |
| rs121965059 | pathogenic | 0 | 0 | 0 |

| | | | | |
|---|---|---|---|---|
| rs386833615 | pathogenic | 1 | 1 | 0 |
| rs121965051 | pathogenic | 1 | 1 | 0 |
| rs121965046 | pathogenic | 0 | 1 | 0 |
| rs121965052 | pathogenic | 0 | 1 | 0 |
| rs386833617 | pathogenic | 0 | 1 | 0 |
| rs386833618 | pathogenic | 1 | 1 | 0 |
| rs121965041 | pathogenic | 0 | 1 | 0 |
| rs121965042 | pathogenic | 0 | 1 | 0 |
| rs121965057 | pathogenic | 1 | 1 | 1 |
| rs386833621 | pathogenic | 1 | 1 | 1 |
| rs121965049 | pathogenic | 1 | 1 | 1 |
| rs386833623 | pathogenic | 1 | 1 | 1 |
| rs121965047 | pathogenic | 1 | 1 | 1 |
| rs121965053 | pathogenic | 0 | 1 | 0 |
| rs121965045 | pathogenic | 0 | 1 | 0 |
| rs121965054 | pathogenic | 0 | 1 | 0 |
| rs121965043 | pathogenic | 0 | 1 | 0 |
| rs386833598 | pathogenic | 0 | 0 | 0 |
| rs137854545 | pathogenic | 1 | 1 | 1 |
| rs137854544 | pathogenic | 1 | 1 | 1 |
| rs104894455 | pathogenic | 0 | 1 | 0 |
| rs1049564 | benign | 0 | 1 | 0 |
| rs104894460 | pathogenic | 1 | 1 | 1 |
| rs104894453 | pathogenic | 1 | 1 | 1 |
| rs104894450 | pathogenic | 0 | 1 | 0 |
| rs104894454 | pathogenic | 0 | 1 | 0 |
| rs104894452 | pathogenic | 1 | 1 | 1 |
| rs104894451 | pathogenic | 0 | 1 | 0 |
| rs104894368 | pathogenic | 1 | 1 | 1 |
| rs730880950 | pathogenic | 1 | 1 | 1 |
| rs121909535 | pathogenic | 1 | 1 | 1 |
| rs121909537 | pathogenic | 1 | 1 | 1 |
| rs121909536 | pathogenic | 1 | 1 | 1 |
| rs121909542 | pathogenic | 1 | 1 | 1 |
| rs121909538 | pathogenic | 1 | 1 | 1 |
| rs121909539 | pathogenic | 1 | 1 | 1 |
| rs121909540 | pathogenic | 1 | 1 | 1 |
| rs17560 | benign | 1 | 1 | 1 |
| rs121909543 | pathogenic | 0 | 1 | 0 |
| rs121909544 | pathogenic | 1 | 1 | 1 |
| rs2307456 | benign | 0 | 1 | 0 |
| rs745778317 | pathogenic | 0 | 1 | 0 |
| rs759607901 | pathogenic | 0 | 1 | 0 |
| rs201188114 | benign | 0 | 0 | 0 |
| rs1800472 | benign | 0 | 0 | 0 |
| rs587780728 | benign | 1 | 1 | 1 |
| rs144386518 | benign | 1 | 1 | 1 |
| rs281865439 | pathogenic | 1 | 1 | 1 |
| rs140175796 | pathogenic | 1 | 1 | 1 |
| rs5030843 | pathogenic | 1 | 1 | 0 |
| rs75166491 | pathogenic | 1 | 1 | 0 |
| rs199475679 | pathogenic | 1 | 1 | 1 |
| rs281865440 | pathogenic | 1 | 1 | 1 |
| rs74486803 | pathogenic | 0 | 1 | 0 |

| | | | | |
|---|---|---|---|---|
| rs199475602 | pathogenic | 0 | 1 | 0 |
| rs5030844 | pathogenic | 1 | 1 | 0 |
| rs62514927 | pathogenic | 1 | 1 | 1 |
| rs62514928 | pathogenic | 1 | 1 | 1 |
| rs62516152 | pathogenic | 0 | 1 | 0 |
| rs62507348 | pathogenic | 1 | 1 | 1 |
| rs76687508 | pathogenic | 1 | 1 | 1 |
| rs62508730 | pathogenic | 1 | 1 | 1 |
| rs62508588 | pathogenic | 0 | 1 | 0 |
| rs5030846 | pathogenic | 0 | 1 | 0 |
| rs74503222 | pathogenic | 1 | 1 | 0 |
| rs5030847 | pathogenic | 1 | 1 | 0 |
| rs5030850 | pathogenic | 1 | 1 | 1 |
| rs778154939 | pathogenic | 0 | 1 | 0 |
| rs62514952 | pathogenic | 1 | 1 | 1 |
| rs78655458 | pathogenic | 1 | 1 | 1 |
| rs62508698 | pathogenic | 1 | 1 | 0 |
| rs5030851 | pathogenic | 1 | 1 | 0 |
| rs62642933 | pathogenic | 1 | 1 | 1 |
| rs5030853 | pathogenic | 1 | 1 | 1 |
| rs62642935 | pathogenic | 1 | 1 | 1 |
| rs199475615 | pathogenic | 0 | 0 | 0 |
| rs62516062 | pathogenic | 1 | 1 | 0 |
| rs62516092 | pathogenic | 0 | 0 | 0 |
| rs62516095 | pathogenic | 1 | 1 | 1 |
| rs62642937 | pathogenic | 1 | 1 | 1 |
| rs62516141 | pathogenic | 0 | 1 | 0 |
| rs5030856 | pathogenic | 1 | 1 | 1 |
| rs5030857 | pathogenic | 1 | 1 | 1 |
| rs5030858 | pathogenic | 1 | 1 | 1 |
| rs79931499 | pathogenic | 1 | 1 | 1 |
| rs5030860 | pathogenic | 1 | 1 | 1 |
| rs281865437 | pathogenic | 1 | 1 | 1 |
| rs62644499 | pathogenic | 1 | 1 | 1 |
| rs147689487 | benign | 0 | 0 | 0 |
| rs397515417 | pathogenic | 1 | 1 | 1 |
| rs121909332 | pathogenic | 1 | 1 | 1 |
| rs121909330 | pathogenic | 1 | 1 | 0 |
| rs121909329 | pathogenic | 1 | 1 | 0 |
| rs121909335 | pathogenic | 1 | 1 | 1 |
| rs387906789 | pathogenic | 1 | 1 | 1 |
| rs121909334 | pathogenic | 1 | 1 | 1 |
| rs121909331 | pathogenic | 1 | 1 | 1 |
| rs372487178 | pathogenic | 1 | 1 | 1 |
| rs387906937 | pathogenic | 1 | 1 | 0 |
| rs16845 | benign | 1 | 1 | 1 |
| rs672601313 | pathogenic | 1 | 1 | 1 |
| rs117805308 | benign | 0 | 0 | 0 |
| rs121434261 | pathogenic | 1 | 1 | 1 |
| rs141629512 | benign | 0 | 0 | 0 |
| rs199957176 | benign | 1 | 1 | 1 |
| rs2230461 | benign | 1 | 1 | 1 |
| rs764618040 | pathogenic | 1 | 1 | 1 |
| rs267606949 | pathogenic | 1 | 1 | 1 |

| | | | | |
|---|---|---|---|---|
| rs587777483 | pathogenic | 1 | 1 | 1 |
| rs267606947 | pathogenic | 1 | 1 | 1 |
| rs6941583 | benign | 0 | 0 | 0 |
| rs749597959 | benign | 1 | 1 | 1 |
| rs182340753 | benign | 1 | 1 | 1 |
| rs146678221 | benign | 0 | 0 | 0 |
| rs11539202 | benign | 0 | 0 | 0 |
| rs148723565 | benign | 0 | 0 | 0 |
| rs61752925 | benign | 1 | 1 | 1 |
| rs113309941 | pathogenic | 0 | 0 | 0 |
| rs121434520 | pathogenic | 1 | 1 | 1 |
| rs121434517 | pathogenic | 1 | 1 | 1 |
| rs267607027 | pathogenic | 1 | 1 | 1 |
| rs138603088 | pathogenic | 0 | 0 | 0 |
| rs121434516 | pathogenic | 1 | 1 | 0 |
| rs121434521 | pathogenic | 0 | 0 | 0 |
| rs145735112 | benign | 1 | 1 | 1 |
| rs121434519 | pathogenic | 1 | 1 | 1 |
| rs75326924 | pathogenic | 1 | 1 | 0 |
| rs142186404 | pathogenic | 0 | 0 | 0 |
| rs121918035 | pathogenic | 0 | 0 | 0 |
| rs374385878 | pathogenic | 1 | 1 | 0 |
| rs11557652 | benign | 0 | 0 | 0 |
| rs118203986 | pathogenic | 1 | 1 | 1 |
| rs141675330 | benign | 1 | 1 | 1 |
| rs104893927 | pathogenic | 0 | 0 | 0 |
| rs104893935 | pathogenic | 0 | 0 | 0 |
| rs104893933 | pathogenic | 1 | 1 | 0 |
| rs104893934 | pathogenic | 0 | 0 | 0 |
| rs145183203 | benign | 1 | 1 | 0 |
| rs2230806 | benign | 0 | 0 | 0 |
| rs9282541 | benign | 0 | 0 | 0 |
| rs559110055 | benign | 0 | 0 | 0 |
| rs200030513 | benign | 0 | 0 | 0 |
| rs9282543 | benign | 0 | 0 | 0 |
| rs147675550 | benign | 0 | 0 | 0 |
| rs137854496 | pathogenic | 0 | 0 | 0 |
| rs374190304 | benign | 0 | 0 | 0 |
| rs114620717 | benign | 0 | 0 | 0 |
| rs2066718 | benign | 0 | 0 | 0 |
| rs35819696 | benign | 0 | 0 | 0 |
| rs138880920 | benign | 0 | 0 | 0 |
| rs2066715 | benign | 0 | 0 | 0 |
| rs35207495 | benign | 0 | 0 | 0 |
| rs2066714 | benign | 0 | 0 | 0 |
| rs28937313 | pathogenic | 0 | 0 | 0 |
| rs28937314 | pathogenic | 0 | 0 | 0 |
| rs137854495 | pathogenic | 0 | 0 | 0 |
| rs140365800 | benign | 0 | 0 | 0 |
| rs13306073 | benign | 0 | 0 | 0 |
| rs28933692 | pathogenic | 0 | 0 | 0 |
| rs33918808 | benign | 0 | 0 | 0 |
| rs76881554 | benign | 0 | 0 | 0 |
| rs143180998 | benign | 0 | 0 | 0 |

| | | | | |
|---|---|---|---|---|
| rs137854500 | pathogenic | 0 | 0 | 0 |
| rs538614702 | benign | 0 | 0 | 0 |
| rs201879964 | benign | 0 | 0 | 0 |
| rs2230808 | benign | 0 | 0 | 0 |
| rs138422574 | benign | 0 | 0 | 0 |
| rs137854498 | pathogenic | 0 | 0 | 0 |
| rs150125857 | benign | 0 | 0 | 0 |
| rs146292819 | pathogenic | 0 | 0 | 0 |
| rs142688906 | benign | 0 | 0 | 0 |
| rs137854499 | pathogenic | 0 | 0 | 0 |
| rs34879708 | benign | 0 | 0 | 0 |
| rs121913028 | pathogenic | 1 | 1 | 1 |
| rs1041951 | benign | 1 | 1 | 0 |
| rs138714094 | benign | 1 | 1 | 0 |
| rs200650502 | benign | 0 | 1 | 0 |
| rs370708663 | pathogenic | 1 | 1 | 1 |
| rs118204040 | pathogenic | 0 | 1 | 0 |
| rs146899669 | pathogenic | 1 | 1 | 0 |
| rs118204039 | pathogenic | 0 | 1 | 0 |
| rs121909233 | pathogenic | 0 | 0 | 0 |
| rs121909225 | pathogenic | 1 | 1 | 0 |
| rs121909236 | pathogenic | 0 | 0 | 0 |
| rs398123317 | pathogenic | 0 | 0 | 0 |
| rs121909226 | pathogenic | 1 | 1 | 0 |
| rs121909238 | pathogenic | 1 | 1 | 0 |
| rs121909230 | pathogenic | 0 | 0 | 0 |
| rs121909237 | pathogenic | 0 | 0 | 0 |
| rs121909222 | pathogenic | 1 | 1 | 0 |
| rs121909223 | pathogenic | 1 | 1 | 0 |
| rs121909218 | pathogenic | 1 | 1 | 0 |
| rs121909229 | pathogenic | 1 | 1 | 0 |
| rs121909224 | pathogenic | 1 | 1 | 0 |
| rs121909241 | pathogenic | 0 | 0 | 0 |
| rs121909221 | pathogenic | 0 | 0 | 0 |
| rs121913294 | pathogenic | 0 | 0 | 0 |
| rs121909234 | pathogenic | 0 | 0 | 0 |
| rs121909240 | pathogenic | 0 | 0 | 0 |
| rs767623493 | benign | 0 | 0 | 0 |
| rs121909239 | pathogenic | 0 | 0 | 0 |
| rs120074145 | pathogenic | 1 | 1 | 1 |
| rs120074148 | pathogenic | 1 | 1 | 1 |
| rs762991875 | pathogenic | 0 | 0 | 0 |
| rs148639841 | pathogenic | 1 | 1 | 1 |
| rs199524907 | pathogenic | 1 | 1 | 1 |
| rs120074141 | pathogenic | 1 | 1 | 1 |
| rs120074144 | pathogenic | 0 | 1 | 0 |
| rs120074146 | pathogenic | 1 | 1 | 1 |
| rs120074147 | pathogenic | 0 | 1 | 0 |
| rs120074143 | pathogenic | 0 | 1 | 0 |
| rs120074140 | pathogenic | 1 | 1 | 0 |
| rs28937315 | pathogenic | 1 | 1 | 1 |
| rs76066357 | benign | 1 | 1 | 1 |
| rs137852911 | pathogenic | 0 | 1 | 0 |
| rs137852907 | pathogenic | 1 | 1 | 0 |

| | | | | |
|---|---|---|---|---|
| rs137852910 | pathogenic | 1 | 1 | 1 |
| rs537585211 | benign | 0 | 0 | 0 |
| rs202134424 | pathogenic | 0 | 0 | 0 |
| rs200930463 | pathogenic | 0 | 0 | 0 |
| rs587777241 | pathogenic | 1 | 1 | 0 |
| rs77563738 | pathogenic | 0 | 0 | 0 |
| rs750868279 | pathogenic | 1 | 1 | 0 |
| rs148936893 | pathogenic | 0 | 0 | 0 |
| rs146597836 | pathogenic | 0 | 0 | 0 |
| rs775440641 | pathogenic | 1 | 1 | 1 |
| rs770689762 | pathogenic | 0 | 1 | 0 |
| rs376785840 | pathogenic | 0 | 0 | 0 |
| rs137852531 | pathogenic | 0 | 0 | 0 |
| rs137852538 | pathogenic | 1 | 1 | 0 |
| rs137852529 | pathogenic | 0 | 1 | 0 |
| rs137852528 | pathogenic | 0 | 1 | 0 |
| rs137852535 | pathogenic | 1 | 1 | 0 |
| rs137853045 | pathogenic | 0 | 1 | 0 |
| rs137853046 | pathogenic | 1 | 1 | 1 |
| rs137853043 | pathogenic | 1 | 1 | 1 |
| rs137853044 | pathogenic | 1 | 1 | 1 |
| rs137853047 | pathogenic | 1 | 1 | 1 |
| rs267606990 | pathogenic | 1 | 1 | 1 |
| rs751437780 | pathogenic | 1 | 1 | 0 |
| rs121918461 | pathogenic | 1 | 1 | 1 |
| rs121918460 | pathogenic | 1 | 1 | 1 |
| rs121918459 | pathogenic | 1 | 1 | 0 |
| rs397507511 | pathogenic | 1 | 1 | 1 |
| rs121918454 | pathogenic | 1 | 1 | 1 |
| rs121918453 | pathogenic | 1 | 1 | 1 |
| rs121918462 | pathogenic | 1 | 1 | 0 |
| rs121918465 | pathogenic | 1 | 1 | 0 |
| rs121918464 | pathogenic | 1 | 1 | 0 |
| rs121918466 | pathogenic | 1 | 1 | 1 |
| rs397507521 | pathogenic | 1 | 1 | 1 |
| rs397507527 | pathogenic | 1 | 1 | 1 |
| rs777603059 | pathogenic | 0 | 1 | 0 |
| rs121918456 | pathogenic | 1 | 1 | 1 |
| rs121918463 | pathogenic | 1 | 1 | 1 |
| rs121918455 | pathogenic | 0 | 1 | 0 |
| rs28933386 | pathogenic | 0 | 1 | 0 |
| rs774356443 | benign | 1 | 1 | 1 |
| rs121918467 | pathogenic | 1 | 1 | 1 |
| rs121918468 | pathogenic | 1 | 1 | 1 |
| rs121918469 | pathogenic | 1 | 1 | 1 |
| rs397507539 | pathogenic | 1 | 1 | 0 |
| rs121918458 | pathogenic | 1 | 1 | 1 |
| rs121918470 | pathogenic | 1 | 1 | 1 |
| rs35289681 | benign | 0 | 1 | 0 |
| rs4302331 | benign | 0 | 1 | 0 |
| rs11544803 | pathogenic | 1 | 1 | 1 |
| rs116254156 | benign | 0 | 0 | 0 |
| rs104893909 | pathogenic | 1 | 1 | 1 |
| rs104893911 | pathogenic | 1 | 1 | 1 |

| | | | | |
|---|---|---|---|---|
| rs104893908 | pathogenic | 1 | 1 | 0 |
| rs258751 | benign | 1 | 1 | 1 |
| rs104893914 | pathogenic | 1 | 1 | 0 |
| rs121909727 | pathogenic | 1 | 1 | 1 |
| rs104893910 | pathogenic | 1 | 1 | 1 |
| rs121909726 | pathogenic | 1 | 1 | 1 |
| rs104893912 | pathogenic | 1 | 1 | 1 |
| rs138407155 | benign | 1 | 1 | 1 |
| rs574061789 | benign | 1 | 1 | 1 |
| rs121912718 | pathogenic | 1 | 1 | 1 |
| rs121912719 | pathogenic | 1 | 1 | 1 |
| rs140770089 | benign | 1 | 1 | 1 |
| rs28931573 | pathogenic | 1 | 1 | 1 |
| rs121912717 | pathogenic | 1 | 1 | 1 |
| rs376456598 | pathogenic | 0 | 0 | 0 |
| rs398123499 | pathogenic | 1 | 1 | 1 |
| rs773048903 | pathogenic | 1 | 1 | 0 |
| rs398123503 | pathogenic | 1 | 1 | 1 |
| rs375785084 | pathogenic | 0 | 1 | 0 |
| rs137852874 | pathogenic | 1 | 1 | 1 |
| rs373713279 | pathogenic | 1 | 1 | 1 |
| rs137852873 | pathogenic | 1 | 1 | 0 |
| rs367823977 | pathogenic | 1 | 1 | 1 |
| rs398123508 | pathogenic | 1 | 1 | 1 |
| rs137852871 | pathogenic | 1 | 1 | 1 |
| rs753698250 | pathogenic | 0 | 1 | 0 |
| rs182923857 | pathogenic | 1 | 1 | 1 |
| rs148090804 | benign | 1 | 1 | 1 |
| rs137852870 | pathogenic | 1 | 1 | 1 |
| rs267607131 | pathogenic | 1 | 1 | 0 |
| rs137852902 | pathogenic | 0 | 0 | 0 |
| rs61741646 | benign | 1 | 1 | 1 |
| rs137852905 | pathogenic | 0 | 0 | 0 |
| rs121918681 | pathogenic | 0 | 1 | 0 |
| rs8177238 | benign | 1 | 1 | 0 |
| rs5030776 | pathogenic | 1 | 1 | 1 |
| rs104894572 | pathogenic | 0 | 0 | 0 |
| rs104894576 | pathogenic | 0 | 0 | 0 |
| rs80084721 | pathogenic | 0 | 0 | 0 |
| rs104894146 | pathogenic | 0 | 1 | 0 |
| rs104894138 | pathogenic | 1 | 1 | 0 |
| rs104894153 | pathogenic | 1 | 1 | 0 |
| rs104894135 | pathogenic | 1 | 1 | 0 |
| rs104894147 | pathogenic | 1 | 1 | 0 |
| rs104894148 | pathogenic | 1 | 1 | 0 |
| rs104894154 | pathogenic | 1 | 1 | 0 |
| rs104894150 | pathogenic | 1 | 1 | 0 |
| rs104894136 | pathogenic | 1 | 1 | 0 |
| rs104894144 | pathogenic | 0 | 0 | 0 |
| rs104894137 | pathogenic | 0 | 0 | 0 |
| rs61754278 | pathogenic | 1 | 1 | 0 |
| rs104894149 | pathogenic | 1 | 1 | 0 |
| rs104894139 | pathogenic | 1 | 1 | 0 |
| rs104894142 | pathogenic | 0 | 0 | 0 |

| | | | | |
|---|---|---|---|---|
| rs104894143 | pathogenic | 0 | 0 | 0 |
| rs104894145 | pathogenic | 0 | 0 | 0 |
| rs121909545 | benign | 0 | 1 | 0 |
| rs782308462 | pathogenic | 1 | 1 | 1 |
| rs375002796 | pathogenic | 0 | 1 | 0 |
| rs201727685 | pathogenic | 0 | 0 | 0 |
| rs768050261 | pathogenic | 1 | 1 | 1 |
| rs780851340 | benign | 0 | 0 | 0 |
| rs139824241 | benign | 1 | 1 | 0 |
| rs757486575 | pathogenic | 1 | 1 | 0 |
| rs750831299 | pathogenic | 1 | 1 | 1 |
| rs767602961 | pathogenic | 1 | 1 | 1 |
| rs121913661 | pathogenic | 1 | 1 | 1 |
| rs777692271 | pathogenic | 1 | 1 | 1 |
| rs121913659 | pathogenic | 1 | 1 | 1 |
| rs587777533 | pathogenic | 1 | 1 | 1 |
| rs104894210 | pathogenic | 0 | 0 | 0 |
| rs104894216 | pathogenic | 1 | 1 | 1 |
| rs104894207 | pathogenic | 1 | 1 | 1 |
| rs104894208 | pathogenic | 0 | 1 | 0 |
| rs104894214 | pathogenic | 1 | 1 | 1 |
| rs104894211 | pathogenic | 1 | 1 | 1 |
| rs61749384 | pathogenic | 0 | 0 | 0 |
| rs61750071 | pathogenic | 1 | 1 | 1 |
| rs61750074 | pathogenic | 0 | 0 | 0 |
| rs11063988 | benign | 0 | 0 | 0 |
| rs216311 | benign | 0 | 0 | 0 |
| rs1800382 | benign | 1 | 1 | 1 |
| rs11063987 | benign | 1 | 1 | 1 |
| rs387906847 | pathogenic | 1 | 1 | 1 |
| rs73003466 | benign | 0 | 0 | 0 |
| rs149787233 | benign | 1 | 1 | 1 |
| rs144451841 | pathogenic | 1 | 1 | 1 |
| rs141638421 | pathogenic | 1 | 1 | 1 |
| rs121434497 | pathogenic | 0 | 0 | 0 |
| rs121434498 | pathogenic | 0 | 0 | 0 |
| rs267607230 | pathogenic | 1 | 1 | 0 |
| rs121434499 | pathogenic | 1 | 1 | 0 |
| rs546749336 | benign | 1 | 1 | 1 |
| rs138873805 | benign | 0 | 0 | 0 |
| rs730880522 | pathogenic | 0 | 0 | 0 |
| rs200371894 | benign | 0 | 0 | 0 |
| rs730880509 | benign | 0 | 0 | 0 |
| rs267606704 | pathogenic | 1 | 1 | 1 |
| rs267606706 | pathogenic | 0 | 0 | 0 |
| rs267606705 | pathogenic | 1 | 1 | 1 |
| rs267606707 | pathogenic | 0 | 0 | 0 |
| rs387906665 | pathogenic | 1 | 1 | 1 |
| rs267606708 | pathogenic | 1 | 1 | 1 |
| rs148368481 | benign | 0 | 0 | 0 |
| rs13204 | benign | 0 | 0 | 0 |
| rs140547520 | pathogenic | 1 | 1 | 1 |
| rs755234990 | benign | 1 | 1 | 0 |
| rs151184052 | benign | 0 | 0 | 0 |

| | | | | |
|---|---|---|---|---|
| rs3736358 | benign | 0 | 0 | 0 |
| rs3736360 | benign | 0 | 0 | 0 |
| rs1138469 | benign | 0 | 0 | 0 |
| rs74315351 | pathogenic | 0 | 1 | 0 |
| rs137853051 | pathogenic | 1 | 1 | 0 |
| rs74315353 | pathogenic | 0 | 1 | 0 |
| rs74315352 | pathogenic | 0 | 1 | 0 |
| rs74315354 | pathogenic | 1 | 1 | 1 |
| rs28938172 | pathogenic | 1 | 1 | 1 |
| rs52836744 | pathogenic | 0 | 0 | 0 |
| rs121913159 | pathogenic | 1 | 1 | 1 |
| rs121913145 | pathogenic | 1 | 1 | 1 |
| rs891087 | benign | 1 | 1 | 1 |
| rs9282757 | benign | 1 | 1 | 1 |
| rs397517404 | pathogenic | 1 | 1 | 1 |
| rs199918720 | benign | 1 | 1 | 0 |
| rs368299411 | benign | 1 | 1 | 0 |
| rs139399951 | benign | 1 | 1 | 0 |
| rs763981974 | benign | 1 | 1 | 0 |
| rs144242114 | benign | 0 | 0 | 0 |
| rs750965801 | benign | 0 | 0 | 0 |
| rs561310777 | benign | 0 | 0 | 0 |
| rs148185335 | benign | 0 | 0 | 0 |
| rs147742157 | benign | 0 | 0 | 0 |
| rs375664373 | pathogenic | 1 | 1 | 1 |
| rs777327250 | pathogenic | 0 | 0 | 0 |
| rs121908471 | pathogenic | 1 | 1 | 0 |
| rs2301612 | pathogenic | 1 | 1 | 0 |
| rs121908476 | pathogenic | 0 | 0 | 0 |
| rs36220239 | benign | 0 | 0 | 0 |
| rs121908473 | pathogenic | 0 | 0 | 0 |
| rs28647808 | benign | 0 | 0 | 0 |
| rs36090624 | benign | 0 | 0 | 0 |
| rs121908002 | pathogenic | 1 | 1 | 1 |
| rs4251545 | benign | 1 | 1 | 1 |
| rs78705382 | pathogenic | 0 | 0 | 0 |
| rs75184679 | pathogenic | 1 | 1 | 1 |
| rs74555752 | pathogenic | 0 | 0 | 0 |
| rs56320207 | benign | 1 | 1 | 1 |
| rs28730837 | pathogenic | 1 | 1 | 1 |
| rs119469014 | pathogenic | 1 | 1 | 1 |
| rs119468010 | pathogenic | 1 | 1 | 1 |
| rs35220041 | benign | 0 | 0 | 0 |
| rs762321137 | pathogenic | 1 | 1 | 1 |
| rs28931570 | pathogenic | 1 | 1 | 1 |
| rs709932 | benign | 1 | 1 | 1 |
| rs746531546 | benign | 1 | 1 | 1 |
| rs6647 | benign | 1 | 1 | 1 |
| rs141095970 | benign | 1 | 1 | 1 |
| rs143370956 | benign | 1 | 1 | 1 |
| rs28929474 | pathogenic | 1 | 1 | 1 |
| rs61761869 | pathogenic | 1 | 1 | 1 |
| rs1303 | benign | 1 | 1 | 1 |
| rs149447502 | benign | 0 | 0 | 0 |

| | | | | |
|---|---|---|---|---|
| rs267607180 | pathogenic | 1 | 1 | 1 |
| rs150304121 | benign | 0 | 0 | 0 |
| rs147630375 | benign | 1 | 1 | 1 |
| rs11539445 | benign | 1 | 1 | 1 |
| rs372098364 | pathogenic | 1 | 1 | 0 |
| rs760072286 | benign | 0 | 0 | 0 |
| rs745595833 | pathogenic | 1 | 1 | 1 |
| rs587777214 | pathogenic | 0 | 0 | 0 |
| rs150318528 | benign | 0 | 0 | 0 |
| rs137853254 | pathogenic | 1 | 1 | 1 |
| rs137853255 | pathogenic | 0 | 0 | 0 |
| rs137853253 | pathogenic | 0 | 0 | 0 |
| rs202166915 | benign | 0 | 0 | 0 |
| rs137853259 | pathogenic | 0 | 0 | 0 |
| rs2229137 | benign | 0 | 0 | 0 |
| rs137853258 | pathogenic | 1 | 1 | 0 |
| rs137853252 | pathogenic | 1 | 1 | 1 |
| rs137853256 | pathogenic | 0 | 0 | 0 |
| rs137853250 | pathogenic | 1 | 1 | 1 |
| rs121434615 | pathogenic | 0 | 0 | 0 |
| rs115014558 | benign | 1 | 1 | 1 |
| rs34116584 | benign | 1 | 1 | 1 |
| rs34885252 | benign | 1 | 1 | 1 |
| rs180177262 | pathogenic | 1 | 1 | 1 |
| rs180177162 | pathogenic | 1 | 1 | 1 |
| rs121908523 | pathogenic | 1 | 1 | 1 |
| rs180177170 | pathogenic | 1 | 1 | 1 |
| rs180177180 | pathogenic | 1 | 1 | 1 |
| rs767586362 | pathogenic | 1 | 1 | 1 |
| rs121908521 | pathogenic | 1 | 1 | 1 |
| rs180177203 | pathogenic | 1 | 1 | 1 |
| rs138025751 | pathogenic | 1 | 1 | 1 |
| rs180177210 | pathogenic | 0 | 1 | 0 |
| rs180177211 | pathogenic | 0 | 1 | 0 |
| rs121908524 | pathogenic | 1 | 1 | 0 |
| rs121908530 | pathogenic | 1 | 1 | 0 |
| rs180177227 | pathogenic | 0 | 1 | 0 |
| rs180177230 | pathogenic | 1 | 1 | 1 |
| rs121908529 | pathogenic | 0 | 1 | 0 |
| rs117195882 | benign | 1 | 1 | 0 |
| rs180177238 | pathogenic | 1 | 1 | 0 |
| rs34664134 | benign | 1 | 1 | 1 |
| rs180177246 | pathogenic | 0 | 1 | 0 |
| rs536352238 | pathogenic | 0 | 0 | 0 |
| rs121908520 | pathogenic | 1 | 1 | 0 |
| rs180177248 | pathogenic | 1 | 1 | 0 |
| rs180177252 | pathogenic | 1 | 1 | 1 |
| rs121908526 | pathogenic | 1 | 1 | 1 |
| rs121908525 | pathogenic | 1 | 1 | 1 |
| rs147106773 | benign | 1 | 1 | 1 |
| rs180177264 | pathogenic | 0 | 1 | 0 |
| rs146525143 | pathogenic | 1 | 1 | 1 |
| rs140992177 | benign | 1 | 1 | 1 |
| rs13408961 | benign | 0 | 1 | 0 |

| | | | | |
|---|---|---|---|---|
| rs4426527 | benign | 1 | 1 | 1 |
| rs3088308 | benign | 0 | 1 | 0 |
| rs4469829 | benign | 1 | 1 | 0 |
| rs104886003 | pathogenic | 1 | 1 | 1 |
| rs121913286 | pathogenic | 1 | 1 | 1 |
| rs121913279 | pathogenic | 1 | 1 | 1 |
| rs121434636 | pathogenic | 1 | 1 | 1 |
| rs755299132 | pathogenic | 1 | 1 | 1 |
| rs748318386 | pathogenic | 1 | 1 | 1 |
| rs387906757 | pathogenic | 1 | 1 | 1 |
| rs121965064 | pathogenic | 1 | 1 | 1 |
| rs28934608 | pathogenic | 0 | 0 | 0 |
| rs5967 | benign | 1 | 1 | 1 |
| rs773905328 | pathogenic | 0 | 0 | 0 |
| rs104894256 | pathogenic | 0 | 1 | 0 |
| rs28931612 | pathogenic | 0 | 1 | 0 |
| rs104894263 | pathogenic | 1 | 1 | 1 |
| rs607969 | benign | 1 | 1 | 0 |
| rs104894262 | pathogenic | 0 | 1 | 0 |
| rs104894268 | pathogenic | 1 | 1 | 1 |
| rs104894264 | pathogenic | 0 | 1 | 0 |
| rs104894259 | pathogenic | 1 | 1 | 0 |
| rs2959656 | benign | 0 | 0 | 0 |
| rs121913035 | pathogenic | 1 | 1 | 0 |
| rs139605474 | benign | 0 | 0 | 0 |
| rs3817552 | benign | 0 | 0 | 0 |
| rs150541115 | benign | 0 | 0 | 0 |
| rs562763010 | benign | 1 | 1 | 1 |
| rs200053119 | pathogenic | 0 | 0 | 0 |
| rs73826386 | benign | 0 | 0 | 0 |
| rs143655258 | benign | 0 | 0 | 0 |
| rs758857853 | pathogenic | 1 | 1 | 0 |
| rs2228380 | benign | 0 | 0 | 0 |
| rs141078733 | benign | 1 | 1 | 1 |
| rs55824172 | pathogenic | 1 | 1 | 1 |
| rs756751089 | pathogenic | 1 | 1 | 1 |
| rs121964990 | pathogenic | 0 | 0 | 0 |
| rs143555186 | benign | 0 | 0 | 0 |
| rs121964993 | pathogenic | 1 | 1 | 0 |
| rs121964992 | pathogenic | 1 | 1 | 1 |
| rs121964991 | pathogenic | 1 | 1 | 0 |
| rs121964988 | pathogenic | 1 | 1 | 1 |
| rs4687 | benign | 1 | 1 | 1 |
| rs267607125 | pathogenic | 1 | 1 | 1 |
| rs751682878 | benign | 1 | 1 | 1 |
| rs267607126 | pathogenic | 1 | 1 | 1 |
| rs768713596 | pathogenic | 0 | 0 | 0 |
| rs147288996 | benign | 0 | 0 | 0 |
| rs387906639 | pathogenic | 1 | 1 | 1 |
| rs730881069 | pathogenic | 1 | 1 | 1 |
| rs730881071 | pathogenic | 1 | 1 | 1 |
| rs397516348 | pathogenic | 0 | 0 | 0 |
| rs121917760 | pathogenic | 0 | 0 | 0 |
| rs104894724 | pathogenic | 1 | 1 | 1 |

| | | | | |
|---|---|---|---|---|
| rs397516349 | pathogenic | 1 | 1 | 1 |
| rs12075 | benign | 0 | 0 | 0 |
| rs147549987 | benign | 0 | 0 | 0 |
| rs56079734 | benign | 0 | 0 | 0 |
| rs1017842 | benign | 1 | 1 | 1 |
| rs137852593 | pathogenic | 1 | 1 | 1 |
| rs137852571 | pathogenic | 1 | 1 | 1 |
| rs137852591 | pathogenic | 1 | 1 | 1 |
| rs9332971 | pathogenic | 1 | 1 | 1 |
| rs755226547 | pathogenic | 1 | 1 | 1 |
| rs79206939 | benign | 0 | 0 | 0 |
| rs121918214 | pathogenic | 1 | 1 | 0 |
| rs757075712 | pathogenic | 1 | 1 | 0 |
| rs150483902 | benign | 1 | 1 | 1 |
| rs113993973 | pathogenic | 0 | 1 | 0 |
| rs946616 | benign | 0 | 0 | 0 |
| rs113993976 | pathogenic | 1 | 1 | 0 |
| rs151022264 | benign | 0 | 1 | 0 |
| rs113993977 | pathogenic | 1 | 1 | 0 |
| rs113993978 | pathogenic | 1 | 1 | 0 |
| rs2228499 | benign | 0 | 1 | 0 |
| rs113993979 | pathogenic | 1 | 1 | 0 |
| rs113993980 | pathogenic | 1 | 1 | 0 |
| rs113993983 | pathogenic | 0 | 1 | 0 |
| rs150547274 | pathogenic | 1 | 1 | 0 |
| rs113993984 | pathogenic | 1 | 1 | 0 |
| rs113993986 | pathogenic | 1 | 1 | 0 |
| rs34313873 | benign | 0 | 1 | 0 |
| rs113993988 | pathogenic | 1 | 1 | 0 |
| rs199638097 | pathogenic | 1 | 1 | 1 |
| rs118204447 | pathogenic | 1 | 1 | 1 |
| rs11862754 | benign | 0 | 0 | 0 |
| rs118204445 | pathogenic | 0 | 0 | 0 |
| rs118204441 | pathogenic | 0 | 0 | 0 |
| rs118204438 | pathogenic | 0 | 0 | 0 |
| rs118204436 | pathogenic | 0 | 0 | 0 |
| rs146093755 | pathogenic | 0 | 0 | 0 |
| rs781439830 | pathogenic | 0 | 0 | 0 |
| rs398123438 | pathogenic | 0 | 0 | 0 |
| rs398123439 | pathogenic | 0 | 0 | 0 |
| rs118204444 | pathogenic | 0 | 0 | 0 |
| rs7187889 | benign | 0 | 0 | 0 |
| rs118204435 | pathogenic | 1 | 1 | 0 |
| rs34745339 | benign | 0 | 0 | 0 |
| rs398123440 | pathogenic | 0 | 0 | 0 |
| rs118204442 | pathogenic | 0 | 0 | 0 |
| rs770053354 | pathogenic | 0 | 0 | 0 |
| rs118204448 | pathogenic | 0 | 0 | 0 |
| rs118204443 | pathogenic | 0 | 0 | 0 |
| rs118204446 | pathogenic | 0 | 0 | 0 |
| rs746756997 | pathogenic | 0 | 0 | 0 |
| rs267606838 | pathogenic | 0 | 0 | 0 |
| rs118204437 | pathogenic | 0 | 0 | 0 |
| rs398123429 | pathogenic | 0 | 0 | 0 |

| | | | | |
|---|---|---|---|---|
| rs398123430 | pathogenic | 0 | 0 | 0 |
| rs2303269 | benign | 0 | 0 | 0 |
| rs114703967 | benign | 0 | 0 | 0 |
| rs118204439 | pathogenic | 0 | 0 | 0 |
| rs151296605 | benign | 0 | 0 | 0 |
| rs118204440 | pathogenic | 0 | 0 | 0 |
| rs78127134 | benign | 0 | 0 | 0 |
| rs372893383 | pathogenic | 0 | 0 | 0 |
| rs61753266 | pathogenic | 1 | 1 | 1 |
| rs139579928 | benign | 1 | 1 | 1 |
| rs56275071 | pathogenic | 1 | 1 | 1 |
| rs121909736 | pathogenic | 1 | 1 | 0 |
| rs121909737 | pathogenic | 1 | 1 | 0 |
| rs121909735 | pathogenic | 1 | 1 | 0 |
| rs121909731 | pathogenic | 1 | 1 | 1 |
| rs121909734 | pathogenic | 1 | 1 | 1 |
| rs121909733 | pathogenic | 1 | 1 | 1 |
| rs121909732 | pathogenic | 1 | 1 | 1 |
| rs121909730 | pathogenic | 1 | 1 | 1 |
| rs202198360 | benign | 1 | 1 | 1 |
| rs587777294 | pathogenic | 1 | 1 | 0 |
| rs587777297 | pathogenic | 1 | 1 | 0 |
| rs104893913 | pathogenic | 1 | 1 | 0 |
| rs17860424 | pathogenic | 1 | 1 | 1 |
| rs1045485 | benign | 1 | 1 | 1 |
| rs13926 | benign | 0 | 0 | 0 |
| rs79343837 | benign | 0 | 0 | 0 |
| rs45516306 | benign | 1 | 1 | 1 |
| rs139849355 | benign | 1 | 1 | 1 |
| rs104894874 | pathogenic | 1 | 1 | 1 |
| rs104894870 | pathogenic | 1 | 1 | 1 |
| rs137852221 | pathogenic | 1 | 1 | 1 |
| rs104894869 | pathogenic | 1 | 1 | 1 |
| rs104894880 | pathogenic | 1 | 1 | 1 |
| rs104894872 | pathogenic | 1 | 1 | 1 |
| rs104894868 | pathogenic | 1 | 1 | 1 |
| rs104894867 | pathogenic | 1 | 1 | 1 |
| rs104894871 | pathogenic | 1 | 1 | 1 |
| rs104894877 | pathogenic | 1 | 1 | 1 |
| rs104894883 | pathogenic | 1 | 1 | 1 |
| rs104894875 | pathogenic | 1 | 1 | 1 |
| rs104894876 | pathogenic | 1 | 1 | 1 |
| rs104894878 | pathogenic | 1 | 1 | 1 |
| rs137852220 | pathogenic | 1 | 1 | 1 |
| rs28933684 | pathogenic | 1 | 1 | 1 |
| rs104894873 | pathogenic | 1 | 1 | 1 |
| rs138409264 | benign | 0 | 0 | 0 |
| rs2236722 | benign | 0 | 0 | 0 |
| rs28757184 | benign | 0 | 0 | 0 |
| rs121434538 | pathogenic | 0 | 0 | 0 |
| rs700519 | benign | 0 | 0 | 0 |
| rs80051519 | pathogenic | 0 | 0 | 0 |
| rs121434536 | pathogenic | 1 | 1 | 0 |
| rs121434534 | pathogenic | 1 | 1 | 0 |

| | | | | |
|---|---|---|---|---|
| rs78310315 | pathogenic | 1 | 1 | 0 |
| rs540839115 | pathogenic | 1 | 1 | 1 |
| rs80338731 | pathogenic | 1 | 1 | 1 |
| rs180177151 | pathogenic | 1 | 1 | 1 |
| rs180177152 | pathogenic | 0 | 0 | 0 |
| rs80338732 | pathogenic | 1 | 1 | 1 |
| rs80338675 | pathogenic | 1 | 1 | 1 |
| rs80338676 | pathogenic | 1 | 1 | 1 |
| rs180177153 | pathogenic | 0 | 0 | 0 |
| rs180177154 | pathogenic | 1 | 1 | 1 |
| rs5896 | benign | 1 | 1 | 1 |
| rs121918477 | pathogenic | 0 | 0 | 0 |
| rs121918428 | pathogenic | 1 | 1 | 1 |
| rs121918427 | pathogenic | 1 | 1 | 1 |
| rs544300820 | benign | 1 | 1 | 1 |
| rs121918110 | pathogenic | 1 | 1 | 1 |
| rs1057141 | benign | 0 | 0 | 0 |
| rs1135216 | benign | 1 | 1 | 1 |
| rs121917702 | pathogenic | 1 | 1 | 1 |
| rs28941474 | pathogenic | 0 | 0 | 0 |
| rs104893944 | pathogenic | 1 | 1 | 1 |
| rs104893947 | pathogenic | 1 | 1 | 0 |
| rs104893943 | pathogenic | 1 | 1 | 1 |
| rs104893942 | pathogenic | 1 | 1 | 1 |
| rs104893940 | pathogenic | 1 | 1 | 1 |
| rs34045013 | benign | 0 | 0 | 0 |
| rs143083947 | pathogenic | 0 | 0 | 0 |
| rs145786986 | benign | 0 | 0 | 0 |
| rs119450942 | pathogenic | 0 | 0 | 0 |
| rs374259530 | pathogenic | 0 | 0 | 0 |
| rs756210458 | pathogenic | 0 | 0 | 0 |
| rs11089991 | benign | 0 | 0 | 0 |
| rs28941471 | pathogenic | 0 | 1 | 0 |
| rs750614500 | pathogenic | 0 | 1 | 0 |
| rs199761158 | benign | 1 | 1 | 1 |
| rs119450944 | pathogenic | 1 | 1 | 1 |
| rs373458753 | pathogenic | 1 | 1 | 1 |
| rs202064195 | pathogenic | 1 | 1 | 1 |
| rs776496275 | pathogenic | 1 | 1 | 1 |
| rs761493155 | pathogenic | 1 | 1 | 1 |
| rs763542069 | pathogenic | 0 | 0 | 0 |
| rs119450941 | pathogenic | 0 | 0 | 0 |
| rs119450940 | pathogenic | 0 | 0 | 0 |
| rs777821034 | pathogenic | 1 | 1 | 1 |
| rs771121666 | pathogenic | 1 | 1 | 1 |
| rs372895468 | pathogenic | 1 | 1 | 1 |
| rs61750603 | pathogenic | 0 | 0 | 0 |
| rs76863441 | pathogenic | 0 | 0 | 0 |
| rs142601143 | benign | 1 | 1 | 1 |
| rs185832993 | benign | 1 | 1 | 0 |
| rs17860568 | benign | 0 | 0 | 0 |
| rs142064825 | benign | 0 | 0 | 0 |
| rs564076183 | benign | 0 | 0 | 0 |
| rs386834034 | pathogenic | 0 | 0 | 0 |

| | | | | |
|---|---|---|---|---|
| rs147663769 | benign | 0 | 0 | 0 |
| rs776478974 | benign | 0 | 0 | 0 |
| rs111840783 | benign | 0 | 0 | 0 |
| rs1516446 | benign | 1 | 1 | 1 |
| rs143749154 | benign | 1 | 1 | 1 |
| rs80257412 | benign | 0 | 0 | 0 |
| rs142943120 | benign | 0 | 0 | 0 |
| rs144553482 | benign | 0 | 0 | 0 |
| rs35997569 | benign | 0 | 0 | 0 |
| rs772909106 | pathogenic | 0 | 0 | 0 |
| rs149433837 | benign | 1 | 1 | 1 |
| rs532178791 | pathogenic | 0 | 0 | 0 |
| rs145588689 | pathogenic | 1 | 1 | 0 |
| rs17843776 | benign | 1 | 1 | 0 |
| rs121917890 | pathogenic | 1 | 1 | 1 |
| rs121917892 | pathogenic | 0 | 0 | 0 |
| rs606231289 | pathogenic | 0 | 0 | 0 |
| rs370011798 | pathogenic | 0 | 0 | 0 |
| rs34689457 | benign | 0 | 0 | 0 |
| rs118204451 | pathogenic | 0 | 0 | 0 |
| rs118204452 | pathogenic | 1 | 1 | 1 |
| rs652785 | benign | 0 | 0 | 0 |
| rs35378126 | benign | 0 | 0 | 0 |
| rs80358214 | pathogenic | 1 | 1 | 1 |
| rs80358215 | pathogenic | 0 | 1 | 0 |
| rs267606858 | pathogenic | 1 | 1 | 1 |
| rs143137713 | pathogenic | 1 | 1 | 1 |
| rs782168634 | benign | 0 | 0 | 0 |
| rs143873938 | benign | 0 | 0 | 0 |
| rs782776511 | benign | 0 | 0 | 0 |
| rs201093148 | benign | 0 | 0 | 0 |
| rs267606815 | pathogenic | 0 | 0 | 0 |
| rs137853311 | pathogenic | 0 | 0 | 0 |
| rs267606817 | pathogenic | 0 | 0 | 0 |
| rs72650006 | benign | 0 | 0 | 0 |
| rs727503638 | benign | 1 | 1 | 0 |
| rs189286381 | benign | 0 | 0 | 0 |
| rs376644553 | benign | 0 | 0 | 0 |
| rs121434610 | pathogenic | 1 | 1 | 1 |
| rs267607076 | pathogenic | 1 | 1 | 1 |
| rs201275730 | benign | 0 | 0 | 0 |
| rs121909587 | pathogenic | 0 | 0 | 0 |
| rs121909558 | pathogenic | 1 | 1 | 0 |
| rs28929469 | pathogenic | 1 | 1 | 0 |
| rs121909547 | pathogenic | 1 | 1 | 0 |
| rs121909552 | pathogenic | 1 | 1 | 0 |
| rs121909573 | pathogenic | 1 | 1 | 1 |
| rs121909567 | pathogenic | 1 | 1 | 0 |
| rs121909569 | pathogenic | 0 | 0 | 0 |
| rs121909563 | pathogenic | 1 | 1 | 0 |
| rs121909570 | pathogenic | 1 | 1 | 1 |
| rs121909571 | pathogenic | 1 | 1 | 0 |
| rs121909572 | pathogenic | 1 | 1 | 1 |
| rs121909565 | pathogenic | 1 | 1 | 1 |

| | | | | |
|---|---|---|---|---|
| rs121909557 | pathogenic | 1 | 1 | 1 |
| rs121909568 | pathogenic | 1 | 1 | 1 |
| rs121909566 | pathogenic | 1 | 1 | 1 |
| rs121909549 | pathogenic | 1 | 1 | 1 |
| rs121909554 | pathogenic | 1 | 1 | 1 |
| rs121909550 | pathogenic | 1 | 1 | 1 |
| rs121909546 | pathogenic | 1 | 1 | 0 |
| rs121909555 | pathogenic | 1 | 1 | 1 |
| rs121909564 | pathogenic | 1 | 1 | 1 |
| rs548046212 | pathogenic | 0 | 0 | 0 |
| rs147353781 | benign | 0 | 0 | 0 |
| rs104894678 | pathogenic | 1 | 1 | 1 |
| rs1130426 | benign | 0 | 0 | 0 |
| rs104894677 | pathogenic | 1 | 1 | 1 |
| rs141917423 | benign | 1 | 1 | 1 |
| rs574777883 | benign | 0 | 0 | 0 |
| rs150199226 | benign | 1 | 1 | 1 |
| rs199618701 | benign | 0 | 1 | 0 |
| rs121964923 | pathogenic | 0 | 0 | 0 |
| rs121964924 | pathogenic | 0 | 0 | 0 |
| rs201258823 | pathogenic | 0 | 1 | 0 |
| rs267606773 | pathogenic | 0 | 0 | 0 |
| rs35638832 | benign | 0 | 0 | 0 |
| rs121908632 | pathogenic | 1 | 1 | 1 |
| rs121908625 | pathogenic | 1 | 1 | 1 |
| rs748949603 | pathogenic | 0 | 0 | 0 |
| rs757523840 | pathogenic | 0 | 0 | 0 |
| rs62541771 | pathogenic | 0 | 0 | 0 |
| rs776384541 | pathogenic | 1 | 1 | 0 |
| rs121908627 | pathogenic | 1 | 1 | 0 |
| rs28937594 | pathogenic | 0 | 0 | 0 |
| rs121907888 | pathogenic | 0 | 0 | 0 |
| rs121907889 | pathogenic | 0 | 1 | 0 |
| rs28941785 | pathogenic | 1 | 1 | 1 |
| rs28941786 | pathogenic | 1 | 1 | 1 |
| rs1021737 | benign | 0 | 0 | 0 |
| rs121908370 | pathogenic | 1 | 1 | 1 |
| rs63750083 | pathogenic | 0 | 0 | 0 |
| rs17883862 | benign | 0 | 0 | 0 |
| rs587782070 | pathogenic | 1 | 1 | 1 |
| rs587781269 | pathogenic | 1 | 1 | 1 |
| rs28909982 | pathogenic | 1 | 1 | 1 |
| rs373648967 | benign | 1 | 1 | 1 |
| rs368570187 | benign | 1 | 1 | 1 |
| rs730881701 | pathogenic | 1 | 1 | 1 |
| rs137853007 | pathogenic | 0 | 0 | 0 |
| rs575910805 | benign | 0 | 0 | 0 |
| rs104893875 | pathogenic | 1 | 1 | 1 |
| rs104893877 | pathogenic | 1 | 1 | 1 |
| rs119485093 | pathogenic | 1 | 1 | 1 |
| rs119485092 | pathogenic | 0 | 0 | 0 |
| rs119485090 | pathogenic | 0 | 0 | 0 |
| rs28942099 | pathogenic | 1 | 1 | 1 |
| rs373730800 | pathogenic | 1 | 1 | 1 |

| | | | | |
|---|---|---|---|---|
| rs113993995 | pathogenic | 0 | 0 | 0 |
| rs201070132 | pathogenic | 0 | 0 | 0 |
| rs113993996 | pathogenic | 0 | 0 | 0 |
| rs113993998 | pathogenic | 0 | 0 | 0 |
| rs78950939 | pathogenic | 1 | 1 | 1 |
| rs56378716 | pathogenic | 1 | 1 | 1 |
| rs754921704 | benign | 0 | 1 | 0 |
| rs766379103 | pathogenic | 1 | 1 | 0 |
| rs121912515 | pathogenic | 0 | 1 | 0 |
| rs764831888 | benign | 1 | 1 | 0 |
| rs572527340 | benign | 0 | 0 | 0 |
| rs35969327 | benign | 0 | 0 | 0 |
| rs35814893 | benign | 0 | 0 | 0 |
| rs34032300 | benign | 0 | 0 | 0 |
| rs80358260 | pathogenic | 0 | 0 | 0 |
| rs151220873 | benign | 0 | 0 | 0 |
| rs80358261 | pathogenic | 1 | 1 | 0 |
| rs11694 | pathogenic | 0 | 0 | 0 |
| rs80358264 | pathogenic | 0 | 0 | 0 |
| rs80358266 | pathogenic | 1 | 1 | 0 |
| rs104894458 | pathogenic | 1 | 1 | 1 |
| rs121918028 | pathogenic | 0 | 0 | 0 |
| rs11583680 | pathogenic | 0 | 0 | 0 |
| rs4715 | benign | 1 | 1 | 1 |
| rs34957318 | benign | 0 | 0 | 0 |
| rs1124 | benign | 1 | 1 | 1 |
| rs104894778 | pathogenic | 1 | 1 | 1 |
| rs104894777 | pathogenic | 1 | 1 | 1 |
| rs104894769 | pathogenic | 1 | 1 | 1 |
| rs104894768 | pathogenic | 1 | 1 | 1 |
| rs104894771 | pathogenic | 1 | 1 | 1 |
| rs575372249 | benign | 1 | 1 | 1 |
| rs118203914 | pathogenic | 0 | 0 | 0 |
| rs118203916 | pathogenic | 1 | 1 | 0 |
| rs140154690 | benign | 0 | 0 | 0 |
| rs34382405 | benign | 0 | 0 | 0 |
| rs73718082 | benign | 0 | 0 | 0 |
| rs764932724 | benign | 0 | 0 | 0 |
| rs554931092 | pathogenic | 1 | 1 | 0 |
| rs202060864 | pathogenic | 0 | 0 | 0 |
| rs116567033 | benign | 0 | 0 | 0 |
| rs11243011 | benign | 0 | 0 | 0 |
| rs142073519 | pathogenic | 0 | 0 | 0 |
| rs397514611 | pathogenic | 1 | 1 | 0 |
| rs397514612 | pathogenic | 0 | 0 | 0 |
| rs372054960 | pathogenic | 0 | 0 | 0 |
| rs775690041 | pathogenic | 0 | 0 | 0 |
| rs148921184 | pathogenic | 0 | 0 | 0 |
| rs12711521 | benign | 0 | 0 | 0 |
| rs2273346 | benign | 0 | 0 | 0 |
| rs12085877 | benign | 1 | 1 | 1 |
| rs144471433 | benign | 1 | 1 | 1 |
| rs397514671 | pathogenic | 1 | 1 | 1 |
| rs76394680 | benign | 0 | 0 | 0 |

| | | | | |
|---|---|---|---|---|
| rs78289603 | pathogenic | 1 | 1 | 1 |
| rs137852913 | pathogenic | 1 | 1 | 1 |
| rs137852914 | pathogenic | 1 | 1 | 1 |
| rs121918653 | pathogenic | 1 | 1 | 1 |
| rs35515200 | benign | 1 | 1 | 0 |
| rs17876030 | benign | 1 | 1 | 1 |
| rs556623443 | benign | 1 | 1 | 1 |
| rs121434365 | pathogenic | 1 | 1 | 1 |
| rs121434364 | pathogenic | 0 | 0 | 0 |
| rs41400249 | benign | 0 | 0 | 0 |
| rs149440376 | benign | 1 | 1 | 1 |
| rs181590179 | benign | 0 | 0 | 0 |
| rs113993968 | pathogenic | 0 | 0 | 0 |
| rs587776445 | pathogenic | 1 | 1 | 1 |
| rs113993969 | pathogenic | 1 | 1 | 1 |
| rs113993970 | pathogenic | 1 | 1 | 1 |
| rs587776449 | pathogenic | 1 | 1 | 1 |
| rs113993971 | pathogenic | 1 | 1 | 1 |
| rs267607194 | pathogenic | 1 | 1 | 1 |
| rs397514745 | pathogenic | 0 | 0 | 0 |
| rs201650281 | pathogenic | 0 | 0 | 0 |
| rs150529876 | benign | 1 | 1 | 1 |
| rs77573084 | benign | 0 | 0 | 0 |
| rs398122379 | pathogenic | 1 | 1 | 0 |
| rs765478968 | benign | 0 | 0 | 0 |
| rs201399681 | benign | 0 | 0 | 0 |
| rs17159287 | benign | 0 | 0 | 0 |
| rs538571144 | benign | 0 | 0 | 0 |
| rs137852647 | pathogenic | 0 | 0 | 0 |
| rs201358272 | pathogenic | 1 | 1 | 1 |
| rs181251337 | benign | 0 | 0 | 0 |
| rs3213119 | benign | 0 | 0 | 0 |
| rs137853102 | pathogenic | 1 | 1 | 0 |
| rs4956145 | benign | 0 | 0 | 0 |
| rs146036912 | pathogenic | 1 | 1 | 1 |
| rs375717077 | pathogenic | 1 | 1 | 1 |
| rs137853103 | pathogenic | 1 | 1 | 1 |
| rs776184830 | pathogenic | 1 | 1 | 1 |
| rs200273673 | pathogenic | 1 | 1 | 1 |
| rs267607025 | pathogenic | 1 | 1 | 1 |
| rs267607024 | pathogenic | 1 | 1 | 1 |
| rs515726188 | pathogenic | 1 | 1 | 1 |
| rs147315735 | benign | 0 | 0 | 0 |
| rs121918308 | pathogenic | 1 | 1 | 0 |
| rs515726195 | pathogenic | 0 | 0 | 0 |
| rs515726196 | pathogenic | 1 | 1 | 0 |
| rs121918311 | pathogenic | 0 | 1 | 0 |
| rs121918309 | pathogenic | 0 | 0 | 0 |
| rs387906891 | pathogenic | 0 | 0 | 0 |
| rs182614164 | pathogenic | 1 | 1 | 0 |
| rs121918307 | pathogenic | 0 | 0 | 0 |
| rs201440849 | benign | 0 | 0 | 0 |
| rs118203957 | pathogenic | 0 | 0 | 0 |
| rs4986790 | benign | 0 | 0 | 0 |

| | | | | |
|---|---|---|---|---|
| rs281864928 | pathogenic | 1 | 1 | 0 |
| rs267607156 | pathogenic | 0 | 0 | 0 |
| rs55916807 | benign | 0 | 0 | 0 |
| rs749750052 | benign | 0 | 0 | 0 |
| rs74382477 | benign | 0 | 0 | 0 |
| rs201016638 | benign | 0 | 0 | 0 |
| rs376683989 | benign | 0 | 0 | 0 |
| rs144439703 | pathogenic | 0 | 0 | 0 |
| rs779122022 | benign | 0 | 0 | 0 |
| rs121918056 | pathogenic | 0 | 0 | 0 |
| rs113994093 | pathogenic | 1 | 1 | 1 |
| rs141367015 | pathogenic | 0 | 0 | 0 |
| rs775538075 | pathogenic | 0 | 0 | 0 |
| rs121918055 | pathogenic | 0 | 0 | 0 |
| rs754246294 | benign | 1 | 1 | 1 |
| rs2307447 | benign | 1 | 1 | 1 |
| rs113994096 | pathogenic | 1 | 1 | 1 |
| rs367610201 | pathogenic | 1 | 1 | 1 |
| rs758438414 | pathogenic | 1 | 1 | 0 |
| rs121918046 | pathogenic | 0 | 0 | 0 |
| rs201871736 | benign | 0 | 0 | 0 |
| rs773994204 | benign | 0 | 0 | 0 |
| rs2307450 | benign | 1 | 1 | 1 |
| rs200257554 | benign | 0 | 0 | 0 |
| rs185645212 | benign | 0 | 0 | 0 |
| rs121918054 | pathogenic | 0 | 0 | 0 |
| rs113994097 | pathogenic | 0 | 0 | 0 |
| rs202037973 | pathogenic | 0 | 0 | 0 |
| rs769827124 | pathogenic | 1 | 1 | 0 |
| rs41549716 | benign | 1 | 1 | 1 |
| rs113994098 | pathogenic | 1 | 1 | 0 |
| rs144500145 | pathogenic | 1 | 1 | 0 |
| rs121918053 | pathogenic | 1 | 1 | 0 |
| rs121918050 | pathogenic | 1 | 1 | 0 |
| rs368587966 | pathogenic | 0 | 0 | 0 |
| rs139590686 | pathogenic | 0 | 0 | 0 |
| rs121918048 | pathogenic | 1 | 1 | 0 |
| rs768653086 | pathogenic | 1 | 1 | 0 |
| rs113994099 | pathogenic | 1 | 1 | 0 |
| rs121918051 | pathogenic | 1 | 1 | 0 |
| rs753160398 | pathogenic | 1 | 1 | 0 |
| rs201477273 | pathogenic | 1 | 1 | 0 |
| rs142347031 | pathogenic | 0 | 0 | 0 |
| rs121918049 | pathogenic | 0 | 0 | 0 |
| rs267606959 | pathogenic | 1 | 1 | 0 |
| rs201732356 | pathogenic | 0 | 0 | 0 |
| rs368435864 | pathogenic | 0 | 0 | 0 |
| rs764036283 | pathogenic | 0 | 0 | 0 |
| rs2307442 | benign | 0 | 0 | 0 |
| rs2307440 | benign | 0 | 0 | 0 |
| rs548076633 | pathogenic | 0 | 1 | 0 |
| rs750792237 | benign | 0 | 0 | 0 |
| rs3087374 | benign | 0 | 0 | 0 |
| rs121918060 | pathogenic | 0 | 1 | 0 |

| | | | | |
|---|---|---|---|---|
| rs397514765 | pathogenic | 1 | 1 | 1 |
| rs121918063 | pathogenic | 0 | 0 | 0 |
| rs121918058 | pathogenic | 1 | 1 | 1 |
| rs121918064 | pathogenic | 0 | 0 | 0 |
| rs36033115 | benign | 0 | 1 | 0 |
| rs121918057 | pathogenic | 0 | 1 | 0 |
| rs121918065 | pathogenic | 1 | 1 | 1 |
| rs121918061 | pathogenic | 1 | 1 | 1 |
| rs121918066 | pathogenic | 0 | 1 | 0 |
| rs707555 | benign | 0 | 1 | 0 |
| rs8192708 | benign | 0 | 1 | 0 |
| rs11552145 | benign | 0 | 0 | 0 |
| rs1804160 | benign | 0 | 1 | 0 |
| rs34103538 | benign | 0 | 1 | 0 |
| rs751393950 | pathogenic | 1 | 1 | 1 |
| rs121907979 | pathogenic | 1 | 1 | 1 |
| rs121907975 | pathogenic | 1 | 1 | 1 |
| rs121907962 | pathogenic | 1 | 1 | 1 |
| rs121907957 | pathogenic | 0 | 0 | 0 |
| rs121907953 | pathogenic | 1 | 1 | 1 |
| rs28941770 | pathogenic | 1 | 1 | 1 |
| rs28941771 | pathogenic | 1 | 1 | 1 |
| rs121907969 | pathogenic | 1 | 1 | 1 |
| rs121907973 | pathogenic | 1 | 1 | 1 |
| rs1800429 | pathogenic | 0 | 0 | 0 |
| rs121907976 | pathogenic | 1 | 1 | 0 |
| rs121907961 | pathogenic | 0 | 0 | 0 |
| rs121907974 | pathogenic | 0 | 0 | 0 |
| rs121907959 | pathogenic | 0 | 0 | 0 |
| rs121907971 | pathogenic | 1 | 1 | 0 |
| rs121907954 | pathogenic | 0 | 0 | 0 |
| rs121907977 | pathogenic | 0 | 0 | 0 |
| rs121907963 | pathogenic | 1 | 1 | 0 |
| rs370266293 | pathogenic | 1 | 1 | 0 |
| rs1800430 | benign | 0 | 0 | 0 |
| rs121907958 | pathogenic | 0 | 0 | 0 |
| rs202173526 | pathogenic | 0 | 0 | 0 |
| rs121907978 | pathogenic | 0 | 0 | 0 |
| rs121907981 | pathogenic | 0 | 1 | 0 |
| rs145012038 | benign | 1 | 1 | 1 |
| rs121907968 | pathogenic | 1 | 1 | 1 |
| rs121907966 | pathogenic | 1 | 1 | 1 |
| rs121907956 | pathogenic | 1 | 1 | 1 |
| rs28942071 | pathogenic | 1 | 1 | 1 |
| rs121907955 | pathogenic | 1 | 1 | 1 |
| rs4727853 | benign | 0 | 0 | 0 |
| rs201880593 | benign | 0 | 0 | 0 |
| rs201434579 | benign | 1 | 1 | 1 |
| rs74551128 | pathogenic | 1 | 1 | 0 |
| rs193922500 | pathogenic | 0 | 0 | 0 |
| rs121909009 | pathogenic | 1 | 1 | 0 |
| rs121908805 | pathogenic | 1 | 1 | 0 |
| rs139573311 | pathogenic | 0 | 0 | 0 |
| rs213950 | benign | 0 | 0 | 0 |

| | | | | |
|---|---|---|---|---|
| rs79282516 | pathogenic | 0 | 0 | 0 |
| rs121909017 | pathogenic | 1 | 1 | 1 |
| rs77101217 | pathogenic | 1 | 1 | 1 |
| rs1800092 | pathogenic | 0 | 1 | 0 |
| rs77646904 | pathogenic | 0 | 0 | 0 |
| rs1800095 | benign | 1 | 1 | 1 |
| rs387906368 | pathogenic | 1 | 1 | 1 |
| rs113993959 | pathogenic | 1 | 1 | 1 |
| rs121908755 | pathogenic | 1 | 1 | 1 |
| rs121909005 | pathogenic | 1 | 1 | 1 |
| rs121908757 | pathogenic | 1 | 1 | 1 |
| rs75527207 | pathogenic | 1 | 1 | 0 |
| rs121909013 | pathogenic | 1 | 1 | 0 |
| rs121909044 | pathogenic | 1 | 1 | 0 |
| rs74597325 | pathogenic | 1 | 1 | 0 |
| rs75549581 | pathogenic | 0 | 0 | 0 |
| rs80055610 | pathogenic | 1 | 1 | 1 |
| rs1800097 | pathogenic | 0 | 0 | 0 |
| rs121909006 | pathogenic | 0 | 1 | 0 |
| rs397508276 | pathogenic | 0 | 0 | 0 |
| rs397508296 | pathogenic | 1 | 1 | 1 |
| rs139468767 | pathogenic | 0 | 0 | 0 |
| rs121908759 | pathogenic | 1 | 1 | 0 |
| rs121909033 | pathogenic | 1 | 1 | 0 |
| rs397508331 | pathogenic | 0 | 0 | 0 |
| rs387907279 | pathogenic | 1 | 1 | 1 |
| rs546530618 | benign | 0 | 0 | 0 |
| rs558259739 | benign | 0 | 0 | 0 |
| rs201827340 | benign | 0 | 0 | 0 |
| rs148865119 | pathogenic | 1 | 1 | 0 |
| rs2229413 | benign | 1 | 1 | 1 |
| rs34040148 | benign | 1 | 1 | 1 |
| rs121964964 | pathogenic | 1 | 1 | 1 |
| rs760214620 | pathogenic | 1 | 1 | 1 |
| rs775992753 | pathogenic | 1 | 1 | 1 |
| rs770095972 | pathogenic | 1 | 1 | 1 |
| rs781444670 | pathogenic | 1 | 1 | 1 |
| rs147474549 | pathogenic | 0 | 0 | 0 |
| rs121964965 | pathogenic | 0 | 1 | 0 |
| rs121964963 | pathogenic | 1 | 1 | 1 |
| rs755952006 | pathogenic | 1 | 1 | 1 |
| rs121964970 | pathogenic | 1 | 1 | 1 |
| rs121964973 | pathogenic | 1 | 1 | 1 |
| rs773734233 | pathogenic | 1 | 1 | 1 |
| rs758236584 | pathogenic | 1 | 1 | 1 |
| rs149119723 | pathogenic | 1 | 1 | 0 |
| rs5742905 | pathogenic | 0 | 0 | 0 |
| rs121964962 | pathogenic | 1 | 1 | 1 |
| rs781567152 | pathogenic | 1 | 1 | 0 |
| rs398123151 | pathogenic | 1 | 1 | 1 |
| rs760417941 | pathogenic | 1 | 1 | 1 |
| rs774926464 | pathogenic | 0 | 0 | 0 |
| rs121964972 | pathogenic | 1 | 1 | 0 |
| rs372010465 | pathogenic | 1 | 1 | 1 |

| | | | | |
|---|---|---|---|---|
| rs763036586 | pathogenic | 1 | 1 | 1 |
| rs769080151 | pathogenic | 1 | 1 | 1 |
| rs121964967 | pathogenic | 1 | 1 | 1 |
| rs28934892 | pathogenic | 1 | 1 | 0 |
| rs28934891 | pathogenic | 1 | 1 | 1 |
| rs121964971 | pathogenic | 1 | 1 | 1 |
| rs121964968 | pathogenic | 1 | 1 | 1 |
| rs150828989 | benign | 1 | 1 | 1 |
| rs3775291 | benign | 1 | 1 | 0 |
| rs104894419 | pathogenic | 1 | 1 | 1 |
| rs2232642 | benign | 0 | 0 | 0 |
| rs104894551 | pathogenic | 1 | 1 | 0 |
| rs766328537 | pathogenic | 0 | 1 | 0 |
| rs104894553 | pathogenic | 1 | 1 | 0 |
| rs80099330 | benign | 0 | 0 | 0 |
| rs199565861 | pathogenic | 0 | 0 | 0 |
| rs104894548 | pathogenic | 0 | 0 | 0 |
| rs104894549 | pathogenic | 0 | 0 | 0 |
| rs104894550 | pathogenic | 1 | 1 | 1 |
| rs12948217 | pathogenic | 1 | 1 | 1 |
| rs761064915 | pathogenic | 0 | 0 | 0 |
| rs28940279 | pathogenic | 1 | 1 | 1 |
| rs774323189 | pathogenic | 1 | 1 | 1 |
| rs28940574 | pathogenic | 0 | 0 | 0 |
| rs515299 | benign | 1 | 1 | 1 |
| rs10127939 | benign | 1 | 1 | 1 |
| rs396991 | benign | 1 | 1 | 1 |
| rs145956175 | benign | 0 | 1 | 0 |
| rs747838255 | pathogenic | 1 | 1 | 1 |
| rs762659685 | pathogenic | 1 | 1 | 0 |
| rs104894831 | pathogenic | 0 | 1 | 0 |
| rs104894848 | pathogenic | 1 | 1 | 0 |
| rs104894833 | pathogenic | 0 | 1 | 0 |
| rs781838005 | pathogenic | 0 | 1 | 0 |
| rs104894834 | pathogenic | 1 | 1 | 0 |
| rs372966991 | pathogenic | 1 | 1 | 0 |
| rs138886989 | benign | 1 | 1 | 1 |
| rs28935195 | pathogenic | 0 | 1 | 0 |
| rs28935196 | pathogenic | 0 | 1 | 0 |
| rs372416832 | pathogenic | 1 | 1 | 1 |
| rs28935197 | pathogenic | 1 | 1 | 0 |
| rs28935486 | pathogenic | 1 | 1 | 0 |
| rs28935487 | pathogenic | 1 | 1 | 0 |
| rs28935488 | pathogenic | 1 | 1 | 1 |
| rs28935495 | pathogenic | 1 | 1 | 1 |
| rs28935485 | pathogenic | 1 | 1 | 1 |
| rs140329381 | pathogenic | 0 | 1 | 0 |
| rs104894830 | pathogenic | 0 | 1 | 0 |
| rs104894846 | pathogenic | 0 | 1 | 0 |
| rs28935489 | pathogenic | 0 | 1 | 0 |
| rs104894828 | pathogenic | 1 | 1 | 0 |
| rs28935491 | pathogenic | 1 | 1 | 0 |
| rs104894832 | pathogenic | 1 | 1 | 0 |
| rs28935492 | pathogenic | 1 | 1 | 0 |

| | | | | |
|---|---|---|---|---|
| rs730880437 | benign | 1 | 1 | 0 |
| rs28935493 | pathogenic | 0 | 1 | 0 |
| rs28935494 | pathogenic | 1 | 1 | 1 |
| rs111422676 | pathogenic | 1 | 1 | 1 |
| rs397515869 | benign | 1 | 1 | 0 |
| rs782449839 | pathogenic | 0 | 1 | 0 |
| rs104894852 | pathogenic | 1 | 1 | 1 |
| rs116929575 | pathogenic | 1 | 1 | 1 |
| rs7023954 | benign | 0 | 0 | 0 |
| rs184520335 | benign | 1 | 1 | 1 |
| rs143080527 | benign | 0 | 0 | 0 |
| rs1050565 | benign | 0 | 0 | 0 |
| rs11545035 | benign | 0 | 0 | 0 |
| rs267607190 | pathogenic | 1 | 1 | 1 |
| rs200283987 | benign | 0 | 0 | 0 |
| rs372556807 | benign | 1 | 1 | 0 |
| rs563073392 | benign | 0 | 0 | 0 |
| rs4539 | benign | 0 | 0 | 0 |
| rs28931609 | pathogenic | 1 | 1 | 1 |
| rs121912978 | pathogenic | 1 | 1 | 1 |
| rs104894072 | pathogenic | 1 | 1 | 1 |
| rs551933154 | benign | 1 | 1 | 1 |
| rs752962897 | benign | 1 | 1 | 0 |
| rs121912977 | pathogenic | 1 | 1 | 0 |
| rs4537 | benign | 1 | 1 | 1 |
| rs201830462 | benign | 0 | 0 | 0 |
| rs4544 | benign | 0 | 0 | 0 |
| rs562670189 | benign | 0 | 0 | 0 |
| rs4538 | benign | 0 | 1 | 0 |
| rs61757294 | pathogenic | 0 | 0 | 0 |
| rs5313 | benign | 0 | 0 | 0 |
| rs4545 | benign | 0 | 0 | 0 |
| rs72554627 | pathogenic | 1 | 1 | 1 |
| rs72554626 | pathogenic | 1 | 1 | 1 |
| rs387906861 | pathogenic | 1 | 1 | 1 |
| rs749655461 | pathogenic | 1 | 1 | 1 |
| rs121909606 | pathogenic | 1 | 1 | 1 |
| rs121909607 | pathogenic | 1 | 1 | 1 |
| rs200203992 | benign | 0 | 0 | 0 |
| rs6050 | benign | 0 | 0 | 0 |
| rs184635235 | benign | 0 | 0 | 0 |
| rs139146037 | benign | 0 | 0 | 0 |
| rs121909612 | pathogenic | 0 | 0 | 0 |
| rs78506343 | pathogenic | 0 | 0 | 0 |
| rs121918138 | pathogenic | 1 | 1 | 1 |
| rs121918139 | pathogenic | 1 | 1 | 1 |
| rs121918140 | pathogenic | 0 | 0 | 0 |
| rs36063028 | benign | 1 | 1 | 1 |
| rs45495192 | benign | 1 | 1 | 1 |
| rs45458802 | benign | 1 | 1 | 1 |
| rs552865793 | pathogenic | 1 | 1 | 1 |
| rs753744791 | pathogenic | 1 | 1 | 1 |
| rs777518512 | benign | 1 | 1 | 1 |
| rs142708991 | benign | 1 | 1 | 0 |

| | | | | |
|---|---|---|---|---|
| rs746069957 | benign | 0 | 0 | 0 |
| rs1334811 | benign | 0 | 0 | 0 |
| rs138894008 | benign | 1 | 1 | 0 |
| rs35030851 | benign | 1 | 1 | 0 |
| rs35378598 | benign | 0 | 0 | 0 |
| rs35818764 | benign | 1 | 1 | 1 |
| rs4631561 | benign | 1 | 1 | 1 |
| rs121918660 | pathogenic | 0 | 0 | 0 |
| rs373175587 | benign | 0 | 0 | 0 |
| rs61731956 | pathogenic | 1 | 1 | 1 |
| rs180177305 | pathogenic | 0 | 0 | 0 |
| rs119490108 | pathogenic | 0 | 0 | 0 |
| rs180177312 | pathogenic | 1 | 1 | 0 |
| rs180177314 | pathogenic | 1 | 1 | 0 |
| rs180177322 | pathogenic | 1 | 1 | 1 |
| rs180177323 | pathogenic | 1 | 1 | 1 |
| rs180177325 | pathogenic | 0 | 0 | 0 |
| rs35196441 | benign | 0 | 0 | 0 |
| rs28763904 | benign | 0 | 0 | 0 |
| rs2229519 | benign | 0 | 0 | 0 |
| rs137852886 | pathogenic | 0 | 0 | 0 |
| rs137852892 | pathogenic | 0 | 0 | 0 |
| rs770427750 | pathogenic | 0 | 0 | 0 |
| rs137852887 | pathogenic | 0 | 0 | 0 |
| rs137852893 | pathogenic | 0 | 0 | 0 |
| rs80338671 | pathogenic | 1 | 1 | 0 |
| rs2228389 | benign | 0 | 0 | 0 |
| rs80338672 | pathogenic | 0 | 0 | 0 |
| rs201958741 | pathogenic | 0 | 0 | 0 |
| rs80338673 | pathogenic | 0 | 0 | 0 |
| rs137852888 | pathogenic | 0 | 0 | 0 |
| rs137852889 | pathogenic | 0 | 0 | 0 |
| rs137852894 | pathogenic | 0 | 0 | 0 |
| rs137852890 | pathogenic | 0 | 0 | 0 |
| rs137852891 | pathogenic | 0 | 0 | 0 |
| rs766935302 | pathogenic | 0 | 0 | 0 |
| rs193074572 | benign | 0 | 0 | 0 |
| rs555210692 | benign | 1 | 1 | 0 |
| rs119103242 | pathogenic | 0 | 1 | 0 |
| rs143725101 | benign | 0 | 1 | 0 |
| rs28940589 | pathogenic | 1 | 1 | 1 |
| rs113994145 | pathogenic | 0 | 0 | 0 |
| rs773872994 | benign | 0 | 0 | 0 |
| rs28940590 | pathogenic | 1 | 1 | 0 |
| rs145400751 | benign | 1 | 1 | 1 |
| rs587782331 | benign | 0 | 0 | 0 |
| rs730881787 | benign | 0 | 0 | 0 |
| rs587778531 | benign | 0 | 0 | 0 |
| rs781652274 | benign | 0 | 0 | 0 |
| rs55882234 | benign | 0 | 0 | 0 |
| rs2020908 | benign | 0 | 0 | 0 |
| rs63750741 | pathogenic | 1 | 1 | 1 |
| rs41295268 | benign | 1 | 1 | 1 |
| rs63750897 | benign | 1 | 1 | 0 |

| | | | | |
|---|---|---|---|---|
| rs63751005 | benign | 1 | 1 | 0 |
| rs587779223 | benign | 0 | 0 | 0 |
| rs202127474 | benign | 0 | 0 | 0 |
| rs374230313 | benign | 0 | 0 | 0 |
| rs2020912 | benign | 0 | 0 | 0 |
| rs63751017 | pathogenic | 0 | 0 | 0 |
| rs587779246 | pathogenic | 0 | 0 | 0 |
| rs63749919 | pathogenic | 0 | 0 | 0 |
| rs63750563 | pathogenic | 1 | 1 | 1 |
| rs63749999 | pathogenic | 0 | 0 | 0 |
| rs200492211 | pathogenic | 0 | 0 | 0 |
| rs576269342 | benign | 0 | 0 | 0 |
| rs267608075 | benign | 0 | 0 | 0 |
| rs149605979 | benign | 0 | 0 | 0 |
| rs63750252 | benign | 0 | 0 | 0 |
| rs754289472 | benign | 0 | 0 | 0 |
| rs63750119 | pathogenic | 0 | 0 | 0 |
| rs587782109 | benign | 0 | 0 | 0 |
| rs34625968 | benign | 1 | 1 | 1 |
| rs764786814 | benign | 1 | 1 | 1 |
| rs199594809 | benign | 1 | 1 | 1 |
| rs74315364 | pathogenic | 1 | 1 | 1 |
| rs201253919 | pathogenic | 0 | 0 | 0 |
| rs387906828 | pathogenic | 1 | 1 | 1 |
| rs34521609 | benign | 1 | 1 | 1 |
| rs199472894 | pathogenic | 0 | 0 | 0 |
| rs199472902 | pathogenic | 0 | 0 | 0 |
| rs121912505 | pathogenic | 0 | 0 | 0 |
| rs28928905 | pathogenic | 0 | 0 | 0 |
| rs374376640 | benign | 0 | 0 | 0 |
| rs202102276 | benign | 0 | 0 | 0 |
| rs199472918 | pathogenic | 0 | 0 | 0 |
| rs121912516 | pathogenic | 1 | 1 | 1 |
| rs121912504 | pathogenic | 0 | 0 | 0 |
| rs9333649 | pathogenic | 0 | 0 | 0 |
| rs121912508 | pathogenic | 0 | 0 | 0 |
| rs199473428 | pathogenic | 1 | 1 | 1 |
| rs104894021 | pathogenic | 0 | 0 | 0 |
| rs28928904 | pathogenic | 1 | 1 | 1 |
| rs199473039 | benign | 1 | 1 | 1 |
| rs199472958 | pathogenic | 1 | 1 | 1 |
| rs1137617 | pathogenic | 1 | 1 | 1 |
| rs199473532 | pathogenic | 1 | 1 | 1 |
| rs199472988 | pathogenic | 0 | 0 | 0 |
| rs189014161 | pathogenic | 0 | 0 | 0 |
| rs12720441 | pathogenic | 0 | 0 | 0 |
| rs199473001 | pathogenic | 1 | 1 | 0 |
| rs121912506 | pathogenic | 1 | 1 | 0 |
| rs773724817 | pathogenic | 1 | 1 | 1 |
| rs118204002 | pathogenic | 1 | 1 | 1 |
| rs138556525 | pathogenic | 1 | 1 | 1 |
| rs72558181 | pathogenic | 1 | 1 | 1 |
| rs118204005 | pathogenic | 1 | 1 | 1 |
| rs118204004 | pathogenic | 0 | 0 | 0 |

| | | | | |
|---|---|---|---|---|
| rs118204001 | pathogenic | 1 | 1 | 1 |
| rs118204006 | pathogenic | 1 | 1 | 1 |
| rs118204003 | pathogenic | 0 | 0 | 0 |
| rs121913147 | pathogenic | 1 | 1 | 0 |
| rs2229429 | benign | 1 | 1 | 0 |
| rs182552223 | benign | 0 | 0 | 0 |
| rs146588336 | benign | 0 | 0 | 0 |
| rs121964983 | pathogenic | 0 | 1 | 0 |
| rs121964982 | pathogenic | 0 | 0 | 0 |
| rs386833679 | pathogenic | 0 | 0 | 0 |
| rs386833680 | pathogenic | 0 | 0 | 0 |
| rs121964981 | pathogenic | 1 | 1 | 0 |
| rs121964984 | pathogenic | 0 | 0 | 0 |
| rs144971200 | benign | 0 | 0 | 0 |
| rs121964985 | pathogenic | 1 | 1 | 0 |
| rs141246107 | benign | 0 | 0 | 0 |
| rs104894561 | pathogenic | 1 | 1 | 0 |
| rs377023736 | pathogenic | 0 | 0 | 0 |
| rs121964907 | pathogenic | 1 | 1 | 1 |
| rs121964909 | pathogenic | 1 | 1 | 1 |
| rs765070743 | pathogenic | 0 | 0 | 0 |
| rs2228119 | benign | 0 | 0 | 0 |
| rs745976989 | pathogenic | 1 | 1 | 1 |
| rs121964904 | pathogenic | 0 | 0 | 0 |
| rs386833430 | pathogenic | 1 | 1 | 1 |
| rs267606992 | pathogenic | 0 | 0 | 0 |
| rs61752063 | pathogenic | 0 | 0 | 0 |
| rs61752067 | pathogenic | 1 | 1 | 1 |
| rs104894934 | pathogenic | 0 | 0 | 0 |
| rs104894929 | pathogenic | 1 | 1 | 1 |
| rs80189640 | benign | 0 | 0 | 0 |
| rs137854547 | pathogenic | 1 | 1 | 1 |
| rs137854540 | pathogenic | 1 | 1 | 1 |
| rs782092363 | pathogenic | 1 | 1 | 1 |
| rs147783915 | benign | 0 | 0 | 0 |
| rs28933687 | pathogenic | 1 | 1 | 1 |
| rs149503319 | benign | 0 | 0 | 0 |
| rs1805147 | benign | 0 | 0 | 0 |
| rs587777218 | pathogenic | 1 | 1 | 1 |
| rs200786705 | benign | 0 | 0 | 0 |
| rs56219475 | benign | 0 | 0 | 0 |
| rs587781172 | benign | 0 | 0 | 0 |
| rs757094189 | pathogenic | 0 | 0 | 0 |
| rs140811086 | benign | 1 | 1 | 1 |
| rs574088829 | pathogenic | 1 | 1 | 1 |
| rs121918525 | pathogenic | 0 | 0 | 0 |
| rs121918527 | pathogenic | 0 | 0 | 0 |
| rs121964848 | pathogenic | 0 | 1 | 0 |
| rs121964845 | pathogenic | 1 | 1 | 1 |
| rs121964846 | pathogenic | 0 | 1 | 0 |
| rs17217716 | benign | 1 | 1 | 1 |
| rs141711342 | benign | 0 | 0 | 0 |
| rs372619120 | benign | 0 | 0 | 0 |
| rs63750002 | benign | 1 | 1 | 1 |

| | | | | |
|---|---|---|---|---|
| rs193922373 | benign | 0 | 0 | 0 |
| rs41295286 | benign | 0 | 0 | 0 |
| rs17217772 | benign | 0 | 0 | 0 |
| rs145649774 | benign | 0 | 0 | 0 |
| rs63750214 | pathogenic | 0 | 0 | 0 |
| rs63750716 | benign | 0 | 0 | 0 |
| rs202026056 | benign | 0 | 0 | 0 |
| rs4987188 | benign | 1 | 1 | 0 |
| rs267607938 | benign | 0 | 0 | 0 |
| rs17224367 | benign | 0 | 0 | 0 |
| rs63750006 | benign | 0 | 0 | 0 |
| rs587781373 | benign | 0 | 0 | 0 |
| rs63751207 | pathogenic | 1 | 1 | 0 |
| rs730881759 | benign | 0 | 0 | 0 |
| rs372350768 | benign | 1 | 1 | 1 |
| rs55778204 | benign | 1 | 1 | 0 |
| rs63749910 | benign | 0 | 0 | 0 |
| rs63750844 | benign | 0 | 0 | 0 |
| rs28929483 | pathogenic | 0 | 0 | 0 |
| rs61756468 | benign | 0 | 0 | 0 |
| rs63750875 | pathogenic | 1 | 1 | 0 |
| rs776528054 | benign | 1 | 1 | 0 |
| rs751939698 | benign | 0 | 0 | 0 |
| rs587780685 | benign | 0 | 0 | 0 |
| rs730881763 | benign | 1 | 1 | 0 |
| rs373226409 | benign | 0 | 0 | 0 |
| rs202145681 | benign | 1 | 1 | 1 |
| rs63749841 | benign | 1 | 1 | 1 |
| rs63750757 | benign | 1 | 1 | 1 |
| rs587778527 | benign | 1 | 1 | 1 |
| rs63750849 | benign | 0 | 0 | 0 |
| rs587779969 | benign | 0 | 0 | 0 |
| rs121908046 | pathogenic | 1 | 1 | 1 |
| rs28940882 | pathogenic | 1 | 1 | 0 |
| rs28940883 | pathogenic | 1 | 1 | 1 |
| rs727503943 | pathogenic | 1 | 1 | 1 |
| rs137853859 | pathogenic | 1 | 1 | 1 |
| rs121908045 | pathogenic | 1 | 1 | 0 |
| rs137853861 | pathogenic | 1 | 1 | 0 |
| rs3180383 | pathogenic | 0 | 1 | 0 |
| rs144431814 | benign | 0 | 0 | 0 |
| rs139502866 | benign | 0 | 0 | 0 |
| rs17524437 | benign | 0 | 0 | 0 |
| rs760960779 | benign | 0 | 0 | 0 |
| rs148960463 | pathogenic | 0 | 0 | 0 |
| rs111033647 | pathogenic | 1 | 1 | 1 |
| rs111033648 | pathogenic | 0 | 0 | 0 |
| rs1800461 | benign | 0 | 0 | 0 |
| rs111033656 | pathogenic | 0 | 0 | 0 |
| rs111033663 | pathogenic | 1 | 1 | 0 |
| rs111033669 | pathogenic | 1 | 1 | 1 |
| rs111033690 | pathogenic | 0 | 0 | 0 |
| rs111033686 | pathogenic | 0 | 0 | 0 |
| rs111033687 | pathogenic | 0 | 0 | 0 |

| | | | | |
|---|---|---|---|---|
| rs111033695 | pathogenic | 0 | 0 | 0 |
| rs111033697 | pathogenic | 0 | 0 | 0 |
| rs111033693 | pathogenic | 0 | 0 | 0 |
| rs111033694 | pathogenic | 0 | 0 | 0 |
| rs111033721 | pathogenic | 1 | 1 | 1 |
| rs111033722 | pathogenic | 0 | 0 | 0 |
| rs111033725 | pathogenic | 1 | 1 | 0 |
| rs75391579 | pathogenic | 1 | 1 | 1 |
| rs111033728 | pathogenic | 1 | 1 | 1 |
| rs111033735 | pathogenic | 1 | 1 | 1 |
| rs111033739 | pathogenic | 1 | 1 | 1 |
| rs111033736 | pathogenic | 0 | 0 | 0 |
| rs111033744 | pathogenic | 1 | 1 | 1 |
| rs111033750 | pathogenic | 1 | 1 | 1 |
| rs111033752 | pathogenic | 0 | 0 | 0 |
| rs111033751 | pathogenic | 0 | 0 | 0 |
| rs111033846 | pathogenic | 0 | 0 | 0 |
| rs368166217 | pathogenic | 0 | 0 | 0 |
| rs111033764 | pathogenic | 0 | 0 | 0 |
| rs111033847 | benign | 0 | 0 | 0 |
| rs111033778 | pathogenic | 0 | 0 | 0 |
| rs111033773 | pathogenic | 0 | 0 | 0 |
| rs111033792 | pathogenic | 0 | 0 | 0 |
| rs367543266 | pathogenic | 1 | 1 | 1 |
| rs111033794 | pathogenic | 0 | 0 | 0 |
| rs111033802 | pathogenic | 0 | 0 | 0 |
| rs111033803 | pathogenic | 0 | 0 | 0 |
| rs111033809 | pathogenic | 1 | 1 | 0 |
| rs121918376 | pathogenic | 0 | 1 | 0 |
| rs121918377 | pathogenic | 0 | 1 | 0 |
| rs121918378 | pathogenic | 1 | 1 | 1 |
| rs121918374 | pathogenic | 1 | 1 | 1 |
| rs137852379 | pathogenic | 0 | 0 | 0 |
| rs137852388 | pathogenic | 1 | 1 | 1 |
| rs137852368 | pathogenic | 0 | 0 | 0 |
| rs28935499 | pathogenic | 0 | 0 | 0 |
| rs28933668 | pathogenic | 0 | 0 | 0 |
| rs28933669 | pathogenic | 0 | 0 | 0 |
| rs28933670 | pathogenic | 0 | 0 | 0 |
| rs28933671 | pathogenic | 0 | 0 | 0 |
| rs28933672 | pathogenic | 0 | 0 | 0 |
| rs28937272 | pathogenic | 0 | 0 | 0 |
| rs35383156 | benign | 0 | 0 | 0 |
| rs137852418 | pathogenic | 0 | 0 | 0 |
| rs137852419 | pathogenic | 0 | 0 | 0 |
| rs137852428 | pathogenic | 0 | 0 | 0 |
| rs137852436 | pathogenic | 0 | 0 | 0 |
| rs28937285 | pathogenic | 0 | 0 | 0 |
| rs35213384 | benign | 1 | 1 | 1 |
| rs3810950 | benign | 0 | 0 | 0 |
| rs201479289 | pathogenic | 0 | 0 | 0 |
| rs121912820 | pathogenic | 0 | 1 | 0 |
| rs121912815 | pathogenic | 0 | 1 | 0 |
| rs115510708 | benign | 0 | 0 | 0 |

| | | | | |
|---|---|---|---|---|
| rs868749 | benign | 0 | 0 | 0 |
| rs121912823 | pathogenic | 0 | 0 | 0 |
| rs115212829 | benign | 0 | 0 | 0 |
| rs121912822 | pathogenic | 0 | 0 | 0 |
| rs121912816 | pathogenic | 0 | 1 | 0 |
| rs4838544 | benign | 0 | 0 | 0 |
| rs121912818 | pathogenic | 0 | 0 | 0 |
| rs121912821 | pathogenic | 0 | 0 | 0 |
| rs121912817 | pathogenic | 0 | 0 | 0 |
| rs8178992 | benign | 0 | 0 | 0 |
| rs121912819 | pathogenic | 1 | 1 | 0 |
| rs80097077 | benign | 1 | 1 | 0 |
| rs121909359 | pathogenic | 1 | 1 | 0 |
| rs121909358 | pathogenic | 1 | 1 | 1 |
| rs121909361 | pathogenic | 1 | 1 | 1 |
| rs121909357 | pathogenic | 0 | 1 | 0 |
| rs121909365 | pathogenic | 1 | 1 | 1 |
| rs6413484 | pathogenic | 1 | 1 | 1 |
| rs121909373 | pathogenic | 1 | 1 | 1 |
| rs121909366 | pathogenic | 1 | 1 | 1 |
| rs121909367 | pathogenic | 1 | 1 | 1 |
| rs121909368 | pathogenic | 1 | 1 | 1 |
| rs121909369 | pathogenic | 0 | 0 | 0 |
| rs121909363 | pathogenic | 1 | 1 | 1 |
| rs45588036 | pathogenic | 1 | 1 | 1 |
| rs121909364 | pathogenic | 1 | 1 | 1 |
| rs80358282 | benign | 0 | 0 | 0 |
| rs80358284 | pathogenic | 1 | 1 | 1 |
| rs61735303 | benign | 0 | 0 | 0 |
| rs2229274 | benign | 0 | 0 | 0 |
| rs145006491 | benign | 0 | 0 | 0 |
| rs121913580 | pathogenic | 1 | 1 | 0 |
| rs730882196 | pathogenic | 0 | 0 | 0 |
| rs153477 | benign | 0 | 0 | 0 |
| rs153478 | benign | 1 | 1 | 0 |
| rs61740602 | benign | 1 | 1 | 0 |
| rs104893892 | pathogenic | 1 | 1 | 0 |
| rs121909595 | pathogenic | 1 | 1 | 1 |
| rs28931605 | pathogenic | 1 | 1 | 1 |
| rs121909597 | pathogenic | 0 | 1 | 0 |
| rs202233735 | pathogenic | 0 | 1 | 0 |
| rs121909596 | pathogenic | 1 | 1 | 1 |
| rs75156162 | benign | 0 | 1 | 0 |
| rs2305430 | benign | 1 | 1 | 1 |
| rs150318966 | benign | 0 | 0 | 0 |
| rs121909598 | pathogenic | 0 | 1 | 0 |
| rs5744175 | benign | 0 | 0 | 0 |
| rs28935476 | pathogenic | 1 | 1 | 1 |
| rs28935475 | pathogenic | 1 | 1 | 1 |
| rs122461163 | pathogenic | 1 | 1 | 1 |
| rs62626305 | pathogenic | 1 | 1 | 1 |
| rs587777157 | pathogenic | 0 | 0 | 0 |
| rs137854478 | pathogenic | 1 | 1 | 0 |
| rs137854465 | pathogenic | 0 | 0 | 0 |

| | | | | |
|---|---|---|---|---|
| rs137854470 | pathogenic | 0 | 0 | 0 |
| rs137854482 | pathogenic | 0 | 0 | 0 |
| rs137854473 | pathogenic | 0 | 0 | 0 |
| rs137854456 | pathogenic | 0 | 0 | 0 |
| rs140598 | benign | 0 | 0 | 0 |
| rs104894105 | pathogenic | 1 | 1 | 1 |
| rs104894107 | pathogenic | 1 | 1 | 1 |
| rs104894106 | pathogenic | 1 | 1 | 1 |
| rs74621026 | benign | 1 | 1 | 1 |
| rs2290332 | benign | 0 | 0 | 0 |
| rs780202604 | pathogenic | 1 | 1 | 1 |
| rs74315305 | pathogenic | 1 | 1 | 0 |
| rs766751878 | pathogenic | 1 | 1 | 1 |
| rs137854603 | pathogenic | 1 | 1 | 1 |
| rs139592595 | benign | 1 | 1 | 0 |
| rs150565592 | benign | 1 | 1 | 1 |
| rs763520126 | benign | 0 | 0 | 0 |
| rs55980502 | benign | 0 | 0 | 0 |
| rs137852812 | pathogenic | 0 | 0 | 0 |
| rs137852813 | pathogenic | 0 | 0 | 0 |
| rs190222208 | benign | 0 | 0 | 0 |
| rs138459502 | pathogenic | 0 | 0 | 0 |
| rs730881041 | pathogenic | 0 | 0 | 0 |
| rs267607080 | pathogenic | 0 | 0 | 0 |
| rs137852814 | pathogenic | 0 | 0 | 0 |
| rs267607079 | pathogenic | 0 | 0 | 0 |
| rs192378417 | benign | 0 | 0 | 0 |
| rs1288775 | benign | 0 | 0 | 0 |
| rs80338737 | pathogenic | 1 | 1 | 1 |
| rs397514708 | pathogenic | 0 | 0 | 0 |
| rs775933965 | pathogenic | 0 | 0 | 0 |
| rs747608698 | benign | 0 | 1 | 0 |
| rs1145086 | benign | 0 | 1 | 0 |
| rs121912953 | pathogenic | 1 | 1 | 0 |
| rs243849 | benign | 0 | 0 | 0 |
| rs121912955 | pathogenic | 1 | 1 | 1 |
| rs17859943 | benign | 0 | 1 | 0 |
| rs755557988 | benign | 1 | 1 | 1 |
| rs10195 | benign | 0 | 0 | 0 |
| rs61735357 | benign | 1 | 1 | 1 |
| rs137882374 | benign | 0 | 0 | 0 |
| rs5388 | benign | 0 | 0 | 0 |
| rs6352 | benign | 0 | 0 | 0 |
| rs76645461 | pathogenic | 1 | 1 | 0 |
| rs1050828 | pathogenic | 0 | 0 | 0 |
| rs1050829 | pathogenic | 0 | 0 | 0 |
| rs137852314 | pathogenic | 0 | 0 | 0 |
| rs137852331 | pathogenic | 0 | 0 | 0 |
| rs5030872 | pathogenic | 0 | 0 | 0 |
| rs267606836 | pathogenic | 0 | 0 | 0 |
| rs5030868 | pathogenic | 0 | 0 | 0 |
| rs137852330 | pathogenic | 0 | 1 | 0 |
| rs137852326 | pathogenic | 1 | 1 | 1 |
| rs137852318 | pathogenic | 1 | 1 | 1 |

| | | | | |
|---|---|---|---|---|
| rs74575103 | benign | 0 | 0 | 0 |
| rs137852327 | pathogenic | 1 | 1 | 1 |
| rs76723693 | pathogenic | 0 | 0 | 0 |
| rs5030869 | pathogenic | 0 | 0 | 0 |
| rs34193178 | pathogenic | 1 | 1 | 1 |
| rs137852316 | pathogenic | 1 | 1 | 0 |
| rs398123546 | pathogenic | 1 | 1 | 1 |
| rs72554665 | pathogenic | 0 | 0 | 0 |
| rs72554664 | pathogenic | 0 | 0 | 0 |
| rs28939068 | pathogenic | 1 | 1 | 1 |
| rs59301204 | pathogenic | 1 | 1 | 1 |
| rs143715750 | benign | 1 | 1 | 1 |
| rs35263902 | pathogenic | 1 | 1 | 1 |
| rs202214518 | benign | 1 | 1 | 1 |
| rs34076756 | benign | 0 | 0 | 0 |
| rs199592341 | pathogenic | 1 | 1 | 0 |
| rs121908833 | pathogenic | 1 | 1 | 1 |
| rs121908834 | pathogenic | 1 | 1 | 0 |
| rs138454151 | benign | 0 | 0 | 0 |
| rs141323599 | benign | 0 | 0 | 0 |
| rs137853200 | pathogenic | 0 | 0 | 0 |
| rs137853199 | pathogenic | 0 | 0 | 0 |
| rs139787163 | benign | 0 | 0 | 0 |
| rs61731097 | benign | 1 | 1 | 1 |
| rs61729795 | benign | 0 | 0 | 0 |
| rs149242330 | benign | 0 | 0 | 0 |
| rs587777490 | pathogenic | 1 | 1 | 1 |
| rs398123011 | pathogenic | 1 | 1 | 0 |
| rs775186117 | pathogenic | 0 | 0 | 0 |
| rs72677237 | benign | 0 | 0 | 0 |
| rs104894163 | pathogenic | 1 | 1 | 1 |
| rs121909588 | pathogenic | 0 | 0 | 0 |
| rs1042579 | benign | 0 | 0 | 0 |
| rs121909273 | pathogenic | 0 | 0 | 0 |
| rs121909274 | pathogenic | 0 | 0 | 0 |
| rs119473034 | pathogenic | 1 | 1 | 1 |
| rs398123065 | pathogenic | 0 | 0 | 0 |
| rs587777042 | pathogenic | 1 | 1 | 0 |
| rs761413864 | pathogenic | 1 | 1 | 1 |
| rs118203965 | pathogenic | 0 | 0 | 0 |
| rs118203966 | pathogenic | 0 | 0 | 0 |
| rs1801201 | benign | 1 | 1 | 0 |
| rs1136201 | benign | 1 | 1 | 1 |
| rs144148008 | benign | 0 | 0 | 0 |
| rs200118378 | benign | 1 | 1 | 1 |
| rs112984118 | benign | 1 | 1 | 1 |
| rs149446102 | benign | 1 | 1 | 1 |
| rs1427463 | benign | 0 | 0 | 0 |
| rs201936720 | benign | 1 | 1 | 1 |
| rs61751983 | benign | 1 | 1 | 1 |
| rs104894632 | pathogenic | 1 | 1 | 1 |
| rs149202905 | benign | 0 | 0 | 0 |
| rs2230210 | benign | 1 | 1 | 1 |
| rs111631988 | benign | 0 | 0 | 0 |

| | | | | |
|---|---|---|---|---|
| rs137852550 | pathogenic | 0 | 0 | 0 |
| rs5963403 | benign | 0 | 0 | 0 |
| rs138018739 | benign | 0 | 0 | 0 |
| rs62640590 | benign | 0 | 0 | 0 |
| rs121909581 | pathogenic | 1 | 1 | 1 |
| rs202017913 | benign | 1 | 1 | 1 |
| rs143163513 | benign | 0 | 0 | 0 |
| rs238238 | benign | 0 | 1 | 0 |
| rs143945974 | benign | 0 | 0 | 0 |
| rs238239 | benign | 1 | 1 | 1 |
| rs121918403 | pathogenic | 0 | 1 | 0 |
| rs61735456 | benign | 0 | 1 | 0 |
| rs121918404 | pathogenic | 1 | 1 | 0 |
| rs12674488 | benign | 0 | 0 | 0 |
| rs201599657 | pathogenic | 0 | 0 | 0 |
| rs1057090 | benign | 0 | 0 | 0 |
| rs1057091 | benign | 0 | 0 | 0 |
| rs587783740 | benign | 0 | 0 | 0 |
| rs75956769 | benign | 0 | 0 | 0 |
| rs201691299 | benign | 0 | 0 | 0 |
| rs62621087 | benign | 0 | 0 | 0 |
| rs201308454 | benign | 0 | 0 | 0 |
| rs145657360 | benign | 0 | 0 | 0 |
| rs121434592 | pathogenic | 1 | 1 | 0 |
| rs375650413 | benign | 0 | 0 | 0 |
| rs121434437 | pathogenic | 1 | 1 | 0 |
| rs763494292 | pathogenic | 0 | 0 | 0 |
| rs121964997 | pathogenic | 1 | 1 | 0 |
| rs121964996 | pathogenic | 0 | 0 | 0 |
| rs41283425 | benign | 1 | 1 | 1 |
| rs376289528 | benign | 1 | 1 | 1 |
| rs144171604 | benign | 1 | 1 | 1 |
| rs782440692 | pathogenic | 1 | 1 | 1 |
| rs782058451 | pathogenic | 1 | 1 | 1 |
| rs78437817 | benign | 0 | 0 | 0 |
| rs200327969 | benign | 1 | 1 | 1 |
| rs143043662 | benign | 0 | 0 | 0 |
| rs111966833 | benign | 1 | 1 | 1 |
| rs35523678 | benign | 0 | 0 | 0 |
| rs515726208 | pathogenic | 0 | 0 | 0 |
| rs397517239 | pathogenic | 1 | 1 | 1 |
| rs148966602 | benign | 1 | 1 | 1 |
| rs76371972 | benign | 1 | 1 | 1 |
| rs2229968 | benign | 1 | 1 | 1 |
| rs61751538 | benign | 1 | 1 | 1 |
| rs200800978 | pathogenic | 1 | 1 | 1 |
| rs148695964 | benign | 1 | 1 | 1 |
| rs140042418 | benign | 1 | 1 | 1 |
| rs116185403 | benign | 0 | 0 | 0 |
| rs142828669 | benign | 1 | 1 | 1 |
| rs139832048 | benign | 0 | 1 | 0 |
| rs58750568 | benign | 0 | 1 | 0 |
| rs77681311 | benign | 0 | 0 | 0 |
| rs151208452 | benign | 0 | 0 | 0 |

| | | | | |
|---|---|---|---|---|
| rs76463933 | benign | 0 | 0 | 0 |
| rs144434647 | benign | 0 | 0 | 0 |
| rs117361485 | benign | 0 | 0 | 0 |
| rs587777588 | pathogenic | 0 | 0 | 0 |
| rs80338796 | pathogenic | 0 | 0 | 0 |
| rs121434594 | pathogenic | 0 | 0 | 0 |
| rs146766138 | pathogenic | 1 | 1 | 1 |
| rs28939679 | pathogenic | 1 | 1 | 1 |
| rs748754134 | pathogenic | 1 | 1 | 0 |
| rs764276946 | pathogenic | 1 | 1 | 1 |
| rs111033588 | pathogenic | 1 | 1 | 1 |
| rs201633414 | pathogenic | 1 | 1 | 1 |
| rs121909262 | pathogenic | 1 | 1 | 1 |
| rs104893702 | pathogenic | 1 | 1 | 1 |
| rs121909265 | pathogenic | 1 | 1 | 1 |
| rs121909266 | pathogenic | 1 | 1 | 1 |
| rs193922439 | pathogenic | 1 | 1 | 0 |
| rs104893691 | pathogenic | 1 | 1 | 1 |
| rs104893695 | pathogenic | 0 | 0 | 0 |
| rs104893708 | pathogenic | 1 | 1 | 1 |
| rs121909260 | pathogenic | 0 | 0 | 0 |
| rs104893696 | pathogenic | 0 | 0 | 0 |
| rs121909267 | pathogenic | 0 | 0 | 0 |
| rs121909263 | pathogenic | 0 | 0 | 0 |
| rs121909264 | pathogenic | 0 | 0 | 0 |
| rs104893694 | pathogenic | 1 | 1 | 1 |
| rs121909268 | pathogenic | 1 | 1 | 1 |
| rs104893689 | pathogenic | 1 | 1 | 0 |
| rs104893707 | pathogenic | 1 | 1 | 0 |
| rs104893697 | pathogenic | 1 | 1 | 1 |
| rs28936684 | pathogenic | 1 | 1 | 1 |
| rs62269092 | benign | 1 | 1 | 0 |
| rs121909259 | pathogenic | 1 | 1 | 0 |
| rs548403340 | benign | 1 | 1 | 0 |
| rs142818334 | benign | 0 | 0 | 0 |
| rs12493789 | benign | 1 | 1 | 1 |
| rs193922423 | pathogenic | 1 | 1 | 0 |
| rs104893719 | pathogenic | 1 | 1 | 1 |
| rs117375173 | benign | 1 | 1 | 0 |
| rs104893712 | pathogenic | 0 | 0 | 0 |
| rs121917776 | pathogenic | 1 | 1 | 1 |
| rs767325003 | benign | 1 | 1 | 1 |
| rs35824453 | benign | 0 | 0 | 0 |
| rs141641266 | benign | 0 | 0 | 0 |
| rs1050239 | benign | 0 | 0 | 0 |
| rs141303384 | benign | 0 | 0 | 0 |
| rs12720458 | pathogenic | 0 | 0 | 0 |
| rs368507376 | pathogenic | 1 | 1 | 1 |
| rs17215500 | pathogenic | 1 | 1 | 1 |
| rs120074188 | pathogenic | 1 | 1 | 1 |
| rs397508097 | pathogenic | 1 | 1 | 1 |
| rs138551008 | pathogenic | 0 | 0 | 0 |
| rs387907139 | pathogenic | 1 | 1 | 1 |
| rs765487627 | pathogenic | 0 | 0 | 0 |

| | | | | |
|---|---|---|---|---|
| rs587778587 | benign | 0 | 0 | 0 |
| rs45568339 | benign | 0 | 0 | 0 |
| rs78179744 | benign | 0 | 0 | 0 |
| rs45624036 | benign | 0 | 0 | 0 |
| rs61755166 | benign | 0 | 0 | 0 |
| rs180177132 | pathogenic | 1 | 1 | 1 |
| rs587776527 | pathogenic | 0 | 0 | 0 |
| rs373783514 | benign | 0 | 0 | 0 |
| rs730881896 | benign | 0 | 0 | 0 |
| rs118203998 | pathogenic | 0 | 0 | 0 |
| rs41281039 | pathogenic | 1 | 1 | 1 |
| rs118204430 | pathogenic | 1 | 1 | 1 |
| rs1800546 | pathogenic | 1 | 1 | 0 |
| rs76917243 | pathogenic | 0 | 1 | 0 |
| rs78340951 | pathogenic | 0 | 1 | 0 |
| rs77718928 | pathogenic | 0 | 0 | 0 |
| rs369586696 | pathogenic | 0 | 1 | 0 |
| rs745517517 | pathogenic | 1 | 1 | 0 |
| rs372872777 | pathogenic | 0 | 0 | 0 |
| rs748704459 | pathogenic | 0 | 0 | 0 |
| rs769215411 | pathogenic | 0 | 0 | 0 |
| rs139425890 | pathogenic | 0 | 0 | 0 |
| rs779694939 | pathogenic | 1 | 1 | 0 |
| rs773729410 | pathogenic | 0 | 0 | 0 |
| rs121908623 | pathogenic | 1 | 1 | 1 |
| rs121908622 | pathogenic | 1 | 1 | 1 |
| rs121908621 | pathogenic | 1 | 1 | 1 |
| rs762106720 | pathogenic | 1 | 1 | 0 |
| rs199877522 | pathogenic | 0 | 0 | 0 |
| rs74315291 | pathogenic | 1 | 1 | 1 |
| rs79137415 | benign | 0 | 0 | 0 |
| rs121913045 | pathogenic | 0 | 0 | 0 |
| rs34295337 | pathogenic | 0 | 0 | 0 |
| rs121913046 | pathogenic | 0 | 0 | 0 |
| rs121913047 | pathogenic | 0 | 0 | 0 |
| rs121913048 | pathogenic | 0 | 0 | 0 |
| rs267607166 | pathogenic | 0 | 0 | 0 |
| rs17850684 | benign | 1 | 1 | 1 |
| rs119476051 | pathogenic | 1 | 1 | 1 |
| rs17850683 | benign | 1 | 1 | 1 |
| rs119476049 | pathogenic | 1 | 1 | 1 |
| rs119476046 | pathogenic | 1 | 1 | 1 |
| rs119476048 | pathogenic | 0 | 0 | 0 |
| rs119476047 | pathogenic | 1 | 1 | 1 |
| rs28939094 | pathogenic | 1 | 1 | 1 |
| rs119476050 | pathogenic | 0 | 0 | 0 |
| rs387906941 | pathogenic | 1 | 1 | 1 |
| rs41511344 | pathogenic | 1 | 1 | 0 |
| rs748112833 | pathogenic | 1 | 1 | 1 |
| rs74342080 | benign | 1 | 1 | 0 |
| rs116482870 | benign | 0 | 0 | 0 |
| rs104893743 | pathogenic | 0 | 0 | 0 |
| rs117179004 | benign | 0 | 0 | 0 |
| rs587779409 | pathogenic | 1 | 1 | 1 |

| | | | | |
|---|---|---|---|---|
| rs3745406 | benign | 1 | 1 | 0 |
| rs387906696 | pathogenic | 0 | 0 | 0 |
| rs72549369 | pathogenic | 1 | 1 | 0 |
| rs45523133 | benign | 0 | 0 | 0 |
| rs119460972 | pathogenic | 0 | 1 | 0 |
| rs34929837 | benign | 0 | 1 | 0 |
| rs17323225 | benign | 0 | 1 | 0 |
| rs17011368 | benign | 0 | 0 | 0 |
| rs748902766 | pathogenic | 1 | 1 | 1 |
| rs387907292 | pathogenic | 0 | 0 | 0 |
| rs387907293 | pathogenic | 1 | 1 | 0 |
| rs371526758 | pathogenic | 1 | 1 | 1 |
| rs142968179 | pathogenic | 1 | 1 | 1 |
| rs387907291 | pathogenic | 1 | 1 | 0 |
| rs121907949 | pathogenic | 0 | 0 | 0 |
| rs121907947 | pathogenic | 0 | 0 | 0 |
| rs121907950 | pathogenic | 1 | 1 | 1 |
| rs121907948 | pathogenic | 1 | 1 | 1 |
| rs4926 | benign | 0 | 0 | 0 |
| rs748809996 | pathogenic | 0 | 0 | 0 |
| rs1799883 | benign | 1 | 1 | 0 |
| rs104893974 | pathogenic | 0 | 1 | 0 |
| rs104893973 | pathogenic | 0 | 1 | 0 |
| rs34452707 | benign | 0 | 0 | 0 |
| rs119467003 | pathogenic | 1 | 1 | 1 |
| rs121908261 | pathogenic | 0 | 1 | 0 |
| rs28933985 | pathogenic | 1 | 1 | 1 |
| rs80356669 | pathogenic | 1 | 1 | 1 |
| rs28936677 | pathogenic | 1 | 1 | 0 |
| rs121918326 | pathogenic | 0 | 0 | 0 |
| rs121918324 | pathogenic | 1 | 1 | 0 |
| rs121918325 | pathogenic | 1 | 1 | 0 |
| rs41270025 | pathogenic | 1 | 1 | 0 |
| rs121918323 | pathogenic | 0 | 0 | 0 |
| rs12735723 | pathogenic | 1 | 1 | 0 |
| rs36013429 | benign | 0 | 0 | 0 |
| rs28936676 | pathogenic | 1 | 1 | 0 |
| rs771809901 | pathogenic | 1 | 1 | 1 |
| rs758350837 | benign | 1 | 1 | 0 |
| rs121908192 | pathogenic | 1 | 1 | 1 |
| rs147568500 | pathogenic | 1 | 1 | 1 |
| rs3729734 | benign | 0 | 0 | 0 |
| rs371536347 | benign | 1 | 1 | 1 |
| rs80358193 | pathogenic | 1 | 1 | 1 |
| rs119476044 | pathogenic | 1 | 1 | 1 |
| rs111292798 | benign | 1 | 1 | 1 |
| rs121908063 | pathogenic | 1 | 1 | 0 |
| rs121908065 | pathogenic | 1 | 1 | 1 |
| rs121908062 | pathogenic | 1 | 1 | 1 |
| rs121908064 | pathogenic | 1 | 1 | 1 |
| rs61746421 | benign | 1 | 1 | 1 |
| rs34886916 | benign | 1 | 1 | 1 |
| rs118204045 | pathogenic | 0 | 0 | 0 |
| rs138607412 | benign | 1 | 1 | 1 |

| | | | | |
|---|---|---|---|---|
| rs118204046 | pathogenic | 0 | 0 | 0 |
| rs118204048 | pathogenic | 0 | 0 | 0 |
| rs118204044 | pathogenic | 0 | 0 | 0 |
| rs149664056 | benign | 0 | 0 | 0 |
| rs121909285 | pathogenic | 0 | 0 | 0 |
| rs1802498 | benign | 0 | 0 | 0 |
| rs1065761 | benign | 0 | 0 | 0 |
| rs119477054 | pathogenic | 0 | 0 | 0 |
| rs12720097 | benign | 0 | 0 | 0 |
| rs747281324 | pathogenic | 1 | 1 | 0 |
| rs367543028 | pathogenic | 1 | 1 | 0 |
| rs146077918 | benign | 0 | 0 | 0 |
| rs142551229 | benign | 0 | 0 | 0 |
| rs367543039 | pathogenic | 0 | 0 | 0 |
| rs587779884 | pathogenic | 1 | 1 | 0 |
| rs137853153 | pathogenic | 0 | 0 | 0 |
| rs367543029 | pathogenic | 0 | 0 | 0 |
| rs587783037 | pathogenic | 1 | 1 | 0 |
| rs142928725 | benign | 0 | 0 | 0 |
| rs28385141 | benign | 0 | 0 | 0 |
| rs587779888 | pathogenic | 0 | 0 | 0 |
| rs121909801 | pathogenic | 1 | 1 | 1 |
| rs121909790 | pathogenic | 1 | 1 | 0 |
| rs121909797 | pathogenic | 1 | 1 | 0 |
| rs121909794 | pathogenic | 1 | 1 | 1 |
| rs121909791 | pathogenic | 1 | 1 | 1 |
| rs121909793 | pathogenic | 1 | 1 | 1 |
| rs147977279 | benign | 1 | 1 | 1 |
| rs147352564 | benign | 0 | 0 | 0 |
| rs368758045 | benign | 1 | 1 | 1 |
| rs114498261 | benign | 1 | 1 | 0 |
| rs35484156 | benign | 0 | 0 | 0 |
| rs139176878 | benign | 0 | 0 | 0 |
| rs11543934 | benign | 0 | 0 | 0 |
| rs34192549 | benign | 0 | 0 | 0 |
| rs568736295 | benign | 0 | 0 | 0 |
| rs55747232 | benign | 0 | 0 | 0 |
| rs781887272 | benign | 0 | 0 | 0 |
| rs146273341 | benign | 0 | 0 | 0 |
| rs41473544 | benign | 0 | 0 | 0 |
| rs137852680 | pathogenic | 0 | 0 | 0 |
| rs55891000 | benign | 0 | 0 | 0 |
| rs140351189 | benign | 0 | 0 | 0 |
| rs137852679 | pathogenic | 1 | 1 | 0 |
| rs137852678 | pathogenic | 0 | 0 | 0 |
| rs137852677 | pathogenic | 1 | 1 | 1 |
| rs138620818 | benign | 0 | 0 | 0 |
| rs121908531 | pathogenic | 0 | 0 | 0 |
| rs72554307 | pathogenic | 0 | 0 | 0 |
| rs72554308 | pathogenic | 0 | 0 | 0 |
| rs72554312 | pathogenic | 1 | 1 | 0 |
| rs67939655 | pathogenic | 0 | 0 | 0 |
| rs67486158 | pathogenic | 0 | 0 | 0 |
| rs72554338 | pathogenic | 1 | 1 | 1 |

| | | | | |
|---|---|---|---|---|
| rs1800324 | pathogenic | 1 | 1 | 1 |
| rs72554356 | pathogenic | 1 | 1 | 1 |
| rs66656800 | pathogenic | 1 | 1 | 1 |
| rs68026851 | pathogenic | 1 | 1 | 1 |
| rs72558412 | pathogenic | 0 | 0 | 0 |
| rs72558415 | pathogenic | 0 | 0 | 0 |
| rs72558423 | pathogenic | 0 | 0 | 0 |
| rs67120076 | pathogenic | 1 | 1 | 1 |
| rs1800328 | pathogenic | 1 | 1 | 1 |
| rs72558451 | pathogenic | 1 | 1 | 1 |
| rs72558454 | pathogenic | 1 | 1 | 1 |
| rs672601372 | pathogenic | 0 | 0 | 0 |
| rs139644798 | pathogenic | 0 | 0 | 0 |
| rs369398935 | pathogenic | 0 | 0 | 0 |
| rs267607035 | pathogenic | 0 | 0 | 0 |
| rs267607036 | pathogenic | 1 | 1 | 1 |
| rs2302566 | benign | 1 | 1 | 0 |
| rs115959591 | benign | 0 | 0 | 0 |
| rs773876739 | pathogenic | 0 | 0 | 0 |
| rs1799967 | benign | 0 | 0 | 0 |
| rs80356968 | benign | 0 | 0 | 0 |
| rs28897695 | benign | 1 | 1 | 1 |
| rs80357314 | benign | 0 | 0 | 0 |
| rs80357169 | benign | 0 | 0 | 0 |
| rs397507239 | benign | 1 | 1 | 1 |
| rs80187739 | pathogenic | 1 | 1 | 1 |
| rs41293459 | pathogenic | 1 | 1 | 1 |
| rs55770810 | pathogenic | 1 | 1 | 1 |
| rs80356860 | pathogenic | 0 | 0 | 0 |
| rs56195342 | benign | 0 | 0 | 0 |
| rs80357442 | benign | 0 | 0 | 0 |
| rs41293463 | pathogenic | 1 | 1 | 1 |
| rs80357069 | pathogenic | 0 | 0 | 0 |
| rs397509275 | benign | 0 | 0 | 0 |
| rs80356920 | benign | 0 | 0 | 0 |
| rs397509284 | pathogenic | 0 | 0 | 0 |
| rs80357286 | benign | 0 | 0 | 0 |
| rs80357274 | benign | 0 | 0 | 0 |
| rs765656957 | benign | 0 | 0 | 0 |
| rs755427809 | benign | 0 | 0 | 0 |
| rs80357322 | benign | 0 | 0 | 0 |
| rs121434430 | pathogenic | 0 | 0 | 0 |
| rs121918450 | pathogenic | 1 | 1 | 1 |
| rs1532268 | benign | 0 | 0 | 0 |
| rs2303080 | benign | 0 | 0 | 0 |
| rs10064631 | benign | 0 | 0 | 0 |
| rs162036 | benign | 0 | 0 | 0 |
| rs2287780 | benign | 0 | 0 | 0 |
| rs16879334 | benign | 0 | 0 | 0 |
| rs137853061 | pathogenic | 1 | 1 | 0 |
| rs10380 | benign | 0 | 0 | 0 |
| rs1494558 | benign | 0 | 0 | 0 |
| rs1494555 | benign | 0 | 0 | 0 |
| rs192994060 | benign | 0 | 0 | 0 |

| | | | | |
|---|---|---|---|---|
| rs147128920 | benign | 1 | 1 | 0 |
| rs201803986 | pathogenic | 0 | 0 | 0 |
| rs746419489 | pathogenic | 0 | 1 | 0 |
| rs764396564 | pathogenic | 0 | 0 | 0 |
| rs758304537 | pathogenic | 0 | 0 | 0 |
| rs267606763 | pathogenic | 0 | 0 | 0 |
| rs267606762 | pathogenic | 0 | 0 | 0 |
| rs150702945 | pathogenic | 1 | 1 | 1 |
| rs202047589 | pathogenic | 0 | 0 | 0 |
| rs755562733 | pathogenic | 0 | 0 | 0 |
| rs267606764 | pathogenic | 1 | 1 | 1 |
| rs138207257 | pathogenic | 1 | 1 | 0 |
| rs149150736 | pathogenic | 1 | 1 | 1 |
| rs777046879 | pathogenic | 0 | 0 | 0 |
| rs104894408 | pathogenic | 1 | 1 | 1 |
| rs1801002 | pathogenic | 1 | 1 | 1 |
| rs28929485 | pathogenic | 1 | 1 | 1 |
| rs104894396 | pathogenic | 1 | 1 | 1 |
| rs371024165 | pathogenic | 0 | 0 | 0 |
| rs35887622 | pathogenic | 1 | 1 | 1 |
| rs587783644 | pathogenic | 1 | 1 | 1 |
| rs72474224 | pathogenic | 1 | 1 | 1 |
| rs111033296 | pathogenic | 1 | 1 | 1 |
| rs104894407 | pathogenic | 1 | 1 | 1 |
| rs72561723 | pathogenic | 1 | 1 | 1 |
| rs104894398 | pathogenic | 1 | 1 | 1 |
| rs28931594 | pathogenic | 1 | 1 | 1 |
| rs104894412 | pathogenic | 1 | 1 | 1 |
| rs111033297 | pathogenic | 1 | 1 | 1 |
| rs104894404 | pathogenic | 1 | 1 | 1 |
| rs104894410 | pathogenic | 1 | 1 | 1 |
| rs104894403 | pathogenic | 1 | 1 | 1 |
| rs397516870 | benign | 0 | 0 | 0 |
| rs200023879 | pathogenic | 1 | 1 | 1 |
| rs121912968 | pathogenic | 0 | 0 | 0 |
| rs104894402 | pathogenic | 1 | 1 | 1 |
| rs28931593 | pathogenic | 1 | 1 | 1 |
| rs104894397 | pathogenic | 0 | 0 | 0 |
| rs80338944 | pathogenic | 0 | 0 | 0 |
| rs199883710 | pathogenic | 0 | 0 | 0 |
| rs727504302 | pathogenic | 0 | 0 | 0 |
| rs145216882 | benign | 1 | 1 | 1 |
| rs781534323 | pathogenic | 1 | 1 | 1 |
| rs111033218 | benign | 1 | 1 | 1 |
| rs104894409 | pathogenic | 0 | 0 | 0 |
| rs80338945 | pathogenic | 1 | 1 | 1 |
| rs111033299 | pathogenic | 0 | 0 | 0 |
| rs143343083 | pathogenic | 1 | 1 | 1 |
| rs80338946 | pathogenic | 0 | 0 | 0 |
| rs150529554 | pathogenic | 0 | 0 | 0 |
| rs111033295 | pathogenic | 0 | 0 | 0 |
| rs111033188 | benign | 0 | 0 | 0 |
| rs397516874 | pathogenic | 0 | 0 | 0 |
| rs76434661 | pathogenic | 0 | 0 | 0 |

| | | | | |
|---|---|---|---|---|
| rs80338948 | pathogenic | 0 | 0 | 0 |
| rs104894401 | pathogenic | 0 | 0 | 0 |
| rs767178508 | pathogenic | 0 | 0 | 0 |
| rs772264564 | pathogenic | 0 | 0 | 0 |
| rs28931592 | pathogenic | 0 | 0 | 0 |
| rs80338949 | pathogenic | 1 | 1 | 1 |
| rs376898963 | benign | 1 | 1 | 1 |
| rs111033360 | benign | 0 | 0 | 0 |
| rs774518779 | pathogenic | 1 | 1 | 1 |
| rs28931595 | pathogenic | 1 | 1 | 1 |
| rs771748289 | pathogenic | 0 | 0 | 0 |
| rs104894406 | pathogenic | 0 | 0 | 0 |
| rs76838169 | benign | 0 | 0 | 0 |
| rs111033294 | pathogenic | 0 | 0 | 0 |
| rs200214298 | pathogenic | 0 | 0 | 0 |
| rs28940283 | pathogenic | 0 | 1 | 0 |
| rs1047883 | benign | 0 | 0 | 0 |
| rs138254425 | benign | 0 | 0 | 0 |
| rs121912592 | pathogenic | 1 | 1 | 0 |
| rs369061090 | pathogenic | 1 | 1 | 0 |
| rs758724746 | pathogenic | 0 | 1 | 0 |
| rs121912595 | pathogenic | 0 | 0 | 0 |
| rs76340296 | benign | 0 | 0 | 0 |
| rs140578009 | benign | 1 | 1 | 1 |
| rs74445297 | benign | 0 | 0 | 0 |
| rs78599516 | benign | 1 | 1 | 0 |
| rs104894035 | pathogenic | 1 | 1 | 0 |
| rs79451216 | benign | 0 | 0 | 0 |
| rs104894036 | pathogenic | 0 | 0 | 0 |
| rs73343757 | benign | 0 | 0 | 0 |
| rs121917722 | pathogenic | 1 | 1 | 1 |
| rs747700126 | pathogenic | 0 | 1 | 0 |
| rs121917725 | pathogenic | 1 | 1 | 1 |
| rs121917721 | pathogenic | 1 | 1 | 1 |
| rs121917723 | pathogenic | 1 | 1 | 0 |
| rs17569 | benign | 1 | 1 | 1 |
| rs2230062 | benign | 1 | 1 | 1 |
| rs17570 | benign | 0 | 0 | 0 |
| rs121917724 | pathogenic | 0 | 1 | 0 |
| rs199422235 | pathogenic | 0 | 0 | 0 |
| rs199422237 | pathogenic | 1 | 1 | 1 |
| rs199422234 | pathogenic | 1 | 1 | 0 |
| rs140015315 | pathogenic | 1 | 1 | 1 |
| rs730880298 | pathogenic | 1 | 1 | 1 |
| rs779692470 | pathogenic | 0 | 0 | 0 |
| rs369152939 | pathogenic | 0 | 0 | 0 |
| rs527236040 | pathogenic | 0 | 0 | 0 |
| rs587776985 | pathogenic | 1 | 1 | 1 |
| rs147077598 | pathogenic | 1 | 1 | 1 |
| rs128620185 | pathogenic | 1 | 1 | 1 |
| rs128620189 | pathogenic | 0 | 0 | 0 |
| rs128621190 | pathogenic | 0 | 0 | 0 |
| rs61754648 | benign | 0 | 0 | 0 |
| rs137852906 | pathogenic | 0 | 0 | 0 |

| | | | | |
|---|---|---|---|---|
| rs76811038 | pathogenic | 0 | 0 | 0 |
| rs149468422 | benign | 0 | 0 | 0 |
| rs5911 | benign | 0 | 0 | 0 |
| rs28942112 | pathogenic | 1 | 1 | 1 |
| rs143275858 | pathogenic | 1 | 1 | 0 |
| rs28362261 | benign | 0 | 0 | 0 |
| rs28362263 | benign | 1 | 1 | 1 |
| rs558707786 | pathogenic | 0 | 0 | 0 |
| rs138775799 | pathogenic | 0 | 0 | 0 |
| rs748170941 | pathogenic | 0 | 0 | 0 |
| rs121908380 | pathogenic | 0 | 0 | 0 |
| rs765123255 | pathogenic | 1 | 1 | 1 |
| rs730881832 | pathogenic | 0 | 1 | 0 |
| rs587782165 | benign | 1 | 1 | 1 |
| rs762307622 | pathogenic | 0 | 0 | 0 |
| rs34612342 | pathogenic | 1 | 1 | 0 |
| rs143353451 | pathogenic | 0 | 1 | 0 |
| rs747993448 | pathogenic | 0 | 1 | 0 |
| rs376561094 | pathogenic | 0 | 0 | 0 |
| rs776487884 | pathogenic | 1 | 1 | 0 |
| rs34126013 | pathogenic | 1 | 1 | 0 |
| rs200495564 | pathogenic | 1 | 1 | 0 |
| rs140342925 | pathogenic | 1 | 1 | 0 |
| rs587780749 | pathogenic | 1 | 1 | 0 |
| rs773087549 | pathogenic | 1 | 1 | 0 |
| rs730881833 | pathogenic | 0 | 0 | 0 |
| rs374950566 | pathogenic | 1 | 1 | 0 |
| rs587780752 | benign | 0 | 0 | 0 |
| rs3219489 | benign | 0 | 0 | 0 |
| rs199742231 | benign | 0 | 0 | 0 |
| rs587780082 | pathogenic | 0 | 0 | 0 |
| rs35881803 | benign | 0 | 0 | 0 |
| rs41305623 | benign | 1 | 1 | 1 |
| rs121917856 | pathogenic | 1 | 1 | 1 |
| rs576284122 | benign | 0 | 0 | 0 |
| rs142036701 | benign | 0 | 0 | 0 |
| rs121917852 | pathogenic | 1 | 1 | 1 |
| rs121917854 | pathogenic | 1 | 1 | 1 |
| rs121917853 | pathogenic | 1 | 1 | 1 |
| rs121917857 | pathogenic | 0 | 1 | 0 |
| rs193922740 | pathogenic | 1 | 1 | 1 |
| rs565779970 | pathogenic | 1 | 1 | 1 |
| rs6723526 | benign | 0 | 0 | 0 |
| rs141094096 | pathogenic | 1 | 1 | 1 |
| rs104894637 | pathogenic | 0 | 0 | 0 |
| rs104894636 | pathogenic | 0 | 0 | 0 |
| rs104894642 | pathogenic | 0 | 0 | 0 |
| rs104894638 | pathogenic | 0 | 0 | 0 |
| rs753666460 | pathogenic | 0 | 0 | 0 |
| rs104894643 | pathogenic | 1 | 1 | 0 |
| rs34520362 | pathogenic | 0 | 0 | 0 |
| rs374621913 | pathogenic | 0 | 0 | 0 |
| rs113641837 | benign | 0 | 0 | 0 |
| rs104894635 | pathogenic | 0 | 0 | 0 |

| | | | | |
|---|---|---|---|---|
| rs143947056 | pathogenic | 0 | 0 | 0 |
| rs138504221 | pathogenic | 0 | 0 | 0 |
| rs9894254 | benign | 0 | 0 | 0 |
| rs104894640 | pathogenic | 1 | 1 | 1 |
| rs58786455 | benign | 0 | 0 | 0 |
| rs144143780 | pathogenic | 0 | 0 | 0 |
| rs62620232 | benign | 0 | 0 | 0 |
| rs34297805 | benign | 1 | 1 | 1 |
| rs104894641 | pathogenic | 0 | 0 | 0 |
| rs148876719 | benign | 0 | 0 | 0 |
| rs104894639 | pathogenic | 0 | 0 | 0 |
| rs7503034 | benign | 0 | 0 | 0 |
| rs201631570 | benign | 0 | 0 | 0 |
| rs150181226 | pathogenic | 0 | 0 | 0 |
| rs377546036 | pathogenic | 0 | 0 | 0 |
| rs267606700 | pathogenic | 0 | 0 | 0 |
| rs267606701 | pathogenic | 1 | 1 | 0 |
| rs267606699 | pathogenic | 1 | 1 | 0 |
| rs9903215 | benign | 0 | 0 | 0 |
| rs139486406 | benign | 0 | 0 | 0 |
| rs143791614 | benign | 0 | 0 | 0 |
| rs573483474 | benign | 0 | 0 | 0 |
| rs2032892 | benign | 0 | 0 | 0 |
| rs137852619 | pathogenic | 0 | 0 | 0 |
| rs200243548 | benign | 0 | 0 | 0 |
| rs115702388 | benign | 0 | 0 | 0 |
| rs138085358 | pathogenic | 0 | 0 | 0 |
| rs121912789 | pathogenic | 1 | 1 | 1 |
| rs763407972 | pathogenic | 1 | 1 | 1 |
| rs147278302 | benign | 0 | 0 | 0 |
| rs374540411 | benign | 0 | 0 | 0 |
| rs121912790 | pathogenic | 1 | 1 | 0 |
| rs752049917 | benign | 0 | 0 | 0 |
| rs139640491 | benign | 1 | 1 | 1 |
| rs138010738 | benign | 1 | 1 | 0 |
| rs201328768 | pathogenic | 1 | 1 | 0 |
| rs2135720 | benign | 1 | 1 | 0 |
| rs201816080 | benign | 1 | 1 | 0 |
| rs754391973 | pathogenic | 0 | 0 | 0 |
| rs139087859 | benign | 0 | 0 | 0 |
| rs6805421 | benign | 0 | 0 | 0 |
| rs146266900 | benign | 0 | 0 | 0 |
| rs397509419 | pathogenic | 0 | 0 | 0 |
| rs149980411 | benign | 0 | 0 | 0 |
| rs747711488 | pathogenic | 1 | 1 | 1 |
| rs149474339 | pathogenic | 1 | 1 | 1 |
| rs758252808 | pathogenic | 1 | 1 | 0 |
| rs745756308 | pathogenic | 0 | 0 | 0 |
| rs4986958 | benign | 0 | 0 | 0 |
| rs74315444 | pathogenic | 0 | 0 | 0 |
| rs61752137 | pathogenic | 0 | 0 | 0 |
| rs8179065 | benign | 1 | 1 | 1 |
| rs758361736 | pathogenic | 1 | 1 | 0 |
| rs139794067 | pathogenic | 1 | 1 | 1 |

| | | | | |
|---|---|---|---|---|
| rs730880954 | pathogenic | 1 | 1 | 1 |
| rs730880955 | pathogenic | 1 | 1 | 1 |
| rs150634297 | pathogenic | 0 | 0 | 0 |
| rs104893750 | pathogenic | 1 | 1 | 1 |
| rs104893748 | pathogenic | 1 | 1 | 1 |
| rs104893749 | pathogenic | 1 | 1 | 1 |
| rs753520553 | pathogenic | 1 | 1 | 0 |
| rs61737294 | benign | 0 | 0 | 0 |
| rs104894601 | pathogenic | 0 | 0 | 0 |
| rs376090795 | pathogenic | 0 | 0 | 0 |
| rs104894596 | pathogenic | 0 | 0 | 0 |
| rs104894595 | pathogenic | 0 | 0 | 0 |
| rs104894598 | pathogenic | 0 | 0 | 0 |
| rs104894597 | pathogenic | 0 | 0 | 0 |
| rs104894591 | pathogenic | 0 | 0 | 0 |
| rs104894593 | pathogenic | 0 | 0 | 0 |
| rs104894594 | pathogenic | 0 | 0 | 0 |
| rs104894590 | pathogenic | 0 | 0 | 0 |
| rs86312 | benign | 0 | 0 | 0 |
| rs77874543 | benign | 0 | 0 | 0 |
| rs121912590 | pathogenic | 1 | 1 | 1 |
| rs121912589 | pathogenic | 1 | 1 | 1 |
| rs121912586 | pathogenic | 1 | 1 | 1 |
| rs121912585 | pathogenic | 1 | 1 | 1 |
| rs121912587 | pathogenic | 1 | 1 | 1 |
| rs762909359 | pathogenic | 0 | 0 | 0 |
| rs28931610 | pathogenic | 0 | 0 | 0 |
| rs104894097 | pathogenic | 1 | 1 | 1 |
| rs775176191 | benign | 0 | 0 | 0 |
| rs199907548 | pathogenic | 0 | 0 | 0 |
| rs104894095 | pathogenic | 1 | 1 | 1 |
| rs104894109 | pathogenic | 1 | 1 | 1 |
| rs372266620 | benign | 1 | 1 | 1 |
| rs104894099 | pathogenic | 0 | 0 | 0 |
| rs137854599 | pathogenic | 1 | 1 | 1 |
| rs104894094 | pathogenic | 0 | 0 | 0 |
| rs104894104 | pathogenic | 0 | 0 | 0 |
| rs6413463 | benign | 1 | 1 | 1 |
| rs104894098 | pathogenic | 0 | 0 | 0 |
| rs6413464 | benign | 0 | 0 | 0 |
| rs3731249 | benign | 0 | 0 | 0 |
| rs16931179 | benign | 0 | 0 | 0 |
| rs2734715 | benign | 0 | 0 | 0 |
| rs759080581 | pathogenic | 0 | 0 | 0 |
| rs1800746 | benign | 0 | 0 | 0 |
| rs202189057 | pathogenic | 0 | 0 | 0 |
| rs121908204 | pathogenic | 0 | 0 | 0 |
| rs756564767 | pathogenic | 0 | 0 | 0 |
| rs34758634 | benign | 1 | 1 | 1 |
| rs28940573 | pathogenic | 0 | 0 | 0 |
| rs119455955 | pathogenic | 1 | 1 | 0 |
| rs752164603 | pathogenic | 1 | 1 | 0 |
| rs763162812 | pathogenic | 0 | 0 | 0 |
| rs119455957 | pathogenic | 0 | 0 | 0 |

| | | | | |
|---|---|---|---|---|
| rs119455958 | pathogenic | 1 | 1 | 0 |
| rs750428882 | pathogenic | 0 | 0 | 0 |
| rs141482368 | benign | 0 | 0 | 0 |
| rs119455953 | pathogenic | 0 | 0 | 0 |
| rs119455954 | pathogenic | 0 | 0 | 0 |
| rs112812685 | benign | 1 | 1 | 0 |
| rs138744051 | benign | 1 | 1 | 0 |
| rs121908200 | pathogenic | 0 | 0 | 0 |
| rs119455956 | pathogenic | 0 | 0 | 0 |
| rs78970155 | benign | 0 | 0 | 0 |
| rs2834167 | benign | 0 | 0 | 0 |
| rs387907326 | pathogenic | 1 | 1 | 1 |
| rs2107732 | benign | 0 | 0 | 0 |
| rs11552377 | benign | 0 | 0 | 0 |
| rs137852843 | pathogenic | 0 | 0 | 0 |
| rs104894783 | pathogenic | 0 | 0 | 0 |
| rs122457137 | pathogenic | 1 | 1 | 1 |
| rs104894779 | pathogenic | 1 | 1 | 1 |
| rs104894786 | pathogenic | 0 | 0 | 0 |
| rs104894784 | pathogenic | 0 | 0 | 0 |
| rs104894785 | pathogenic | 0 | 0 | 0 |
| rs201870761 | pathogenic | 0 | 0 | 0 |
| rs104894781 | pathogenic | 0 | 0 | 0 |
| rs28897728 | benign | 0 | 0 | 0 |
| rs370273690 | pathogenic | 1 | 1 | 0 |
| rs9332739 | benign | 0 | 0 | 0 |
| rs150878060 | benign | 0 | 0 | 0 |
| rs573509224 | benign | 0 | 0 | 0 |
| rs36221133 | benign | 0 | 0 | 0 |
| rs2230009 | benign | 0 | 0 | 0 |
| rs267607008 | pathogenic | 0 | 0 | 0 |
| rs5985 | benign | 1 | 1 | 1 |
| rs5982 | benign | 1 | 1 | 1 |
| rs5987 | benign | 0 | 0 | 0 |
| rs5988 | benign | 0 | 0 | 0 |
| rs121918680 | pathogenic | 1 | 1 | 0 |
| rs121918679 | pathogenic | 1 | 1 | 0 |
| rs7845 | benign | 1 | 1 | 0 |
| rs1049296 | benign | 1 | 1 | 1 |
| rs121918677 | pathogenic | 1 | 1 | 0 |
| rs200086262 | pathogenic | 1 | 1 | 0 |
| rs746930990 | pathogenic | 1 | 1 | 0 |
| rs104893878 | pathogenic | 1 | 1 | 1 |
| rs121912468 | pathogenic | 1 | 1 | 1 |
| rs121912467 | pathogenic | 1 | 1 | 1 |
| rs145351926 | benign | 1 | 1 | 1 |
| rs141212743 | pathogenic | 1 | 1 | 1 |
| rs397515473 | pathogenic | 1 | 1 | 0 |
| rs121918313 | pathogenic | 0 | 1 | 0 |
| rs73015965 | pathogenic | 0 | 0 | 0 |
| rs121918030 | pathogenic | 0 | 1 | 0 |
| rs77782321 | benign | 0 | 0 | 0 |
| rs374661051 | pathogenic | 1 | 1 | 1 |
| rs267606854 | pathogenic | 0 | 0 | 0 |

| | | | | |
|---|---|---|---|---|
| rs41279678 | benign | 0 | 0 | 0 |
| rs61754640 | benign | 0 | 0 | 0 |
| rs199473314 | pathogenic | 0 | 0 | 0 |
| rs137854601 | pathogenic | 0 | 0 | 0 |
| rs137854614 | pathogenic | 0 | 0 | 0 |
| rs137854615 | pathogenic | 0 | 0 | 0 |
| rs199473635 | pathogenic | 0 | 0 | 0 |
| rs121434478 | pathogenic | 1 | 1 | 1 |
| rs775407864 | pathogenic | 1 | 1 | 1 |
| rs12706927 | benign | 0 | 1 | 0 |
| rs775614261 | benign | 0 | 0 | 0 |
| rs768926261 | benign | 0 | 0 | 0 |
| rs114057640 | benign | 0 | 0 | 0 |
| rs387906886 | pathogenic | 1 | 1 | 0 |
| rs138658585 | benign | 0 | 0 | 0 |
| rs4880091 | benign | 0 | 0 | 0 |
| rs121918552 | pathogenic | 1 | 1 | 1 |
| rs10794537 | benign | 0 | 0 | 0 |
| rs121965022 | pathogenic | 1 | 1 | 0 |
| rs121965020 | pathogenic | 0 | 0 | 0 |
| rs758452450 | pathogenic | 0 | 0 | 0 |
| rs747981483 | benign | 0 | 0 | 0 |
| rs58037052 | benign | 0 | 0 | 0 |
| rs121965029 | pathogenic | 1 | 1 | 0 |
| rs147490060 | benign | 0 | 0 | 0 |
| rs3755955 | benign | 0 | 0 | 0 |
| rs148789453 | pathogenic | 0 | 0 | 0 |
| rs121965030 | pathogenic | 0 | 0 | 0 |
| rs76722191 | benign | 0 | 0 | 0 |
| rs199801029 | pathogenic | 0 | 0 | 0 |
| rs121965033 | pathogenic | 0 | 0 | 0 |
| rs368454909 | pathogenic | 1 | 1 | 0 |
| rs6831280 | benign | 0 | 0 | 0 |
| rs121965024 | pathogenic | 0 | 0 | 0 |
| rs121965019 | pathogenic | 0 | 0 | 0 |
| rs11934801 | benign | 0 | 0 | 0 |
| rs73066479 | benign | 0 | 0 | 0 |
| rs121965027 | pathogenic | 0 | 0 | 0 |
| rs121965026 | pathogenic | 1 | 1 | 0 |
| rs121965021 | pathogenic | 0 | 0 | 0 |
| rs398123257 | benign | 0 | 0 | 0 |
| rs121965031 | pathogenic | 0 | 0 | 0 |
| rs202003795 | pathogenic | 0 | 0 | 0 |
| rs559510809 | pathogenic | 0 | 0 | 0 |
| rs762936223 | benign | 0 | 0 | 0 |
| rs137854580 | pathogenic | 1 | 1 | 1 |
| rs112263157 | benign | 0 | 0 | 0 |
| rs2289181 | benign | 0 | 0 | 0 |
| rs138034915 | benign | 0 | 1 | 0 |
| rs528367092 | pathogenic | 0 | 1 | 0 |
| rs1042393 | benign | 0 | 0 | 0 |
| rs370950728 | pathogenic | 0 | 1 | 0 |
| rs530478036 | benign | 0 | 1 | 0 |
| rs374569672 | benign | 0 | 1 | 0 |

| | | | | |
|---|---|---|---|---|
| rs121907944 | pathogenic | 0 | 1 | 0 |
| rs200856561 | pathogenic | 0 | 1 | 0 |
| rs577915581 | pathogenic | 0 | 1 | 0 |
| rs201896815 | pathogenic | 0 | 1 | 0 |
| rs121907945 | pathogenic | 0 | 1 | 0 |
| rs121907940 | pathogenic | 0 | 1 | 0 |
| rs543300039 | pathogenic | 0 | 1 | 0 |
| rs121907936 | pathogenic | 0 | 1 | 0 |
| rs766074609 | pathogenic | 0 | 1 | 0 |
| rs755253527 | pathogenic | 0 | 1 | 0 |
| rs560575383 | pathogenic | 0 | 1 | 0 |
| rs528369909 | benign | 0 | 1 | 0 |
| rs770610356 | pathogenic | 0 | 1 | 0 |
| rs747610090 | pathogenic | 0 | 1 | 0 |
| rs145712232 | benign | 0 | 1 | 0 |
| rs7215458 | benign | 1 | 1 | 0 |
| rs535644999 | pathogenic | 0 | 1 | 0 |
| rs772883420 | pathogenic | 1 | 1 | 0 |
| rs398123169 | pathogenic | 0 | 1 | 0 |
| rs121907937 | pathogenic | 0 | 1 | 0 |
| rs121907941 | pathogenic | 1 | 1 | 0 |
| rs121907942 | pathogenic | 0 | 1 | 0 |
| rs779556619 | pathogenic | 0 | 1 | 0 |
| rs1800307 | benign | 0 | 1 | 0 |
| rs377544304 | pathogenic | 1 | 1 | 0 |
| rs549029029 | pathogenic | 0 | 1 | 0 |
| rs753269119 | pathogenic | 1 | 1 | 1 |
| rs757617999 | pathogenic | 0 | 1 | 0 |
| rs28937909 | pathogenic | 0 | 1 | 0 |
| rs28940868 | pathogenic | 0 | 1 | 0 |
| rs368438393 | pathogenic | 0 | 1 | 0 |
| rs536906561 | pathogenic | 0 | 1 | 0 |
| rs374143224 | pathogenic | 0 | 1 | 0 |
| rs778418246 | pathogenic | 0 | 1 | 0 |
| rs398123172 | pathogenic | 0 | 1 | 0 |
| rs121907938 | pathogenic | 0 | 1 | 0 |
| rs1800312 | pathogenic | 0 | 1 | 0 |
| rs1126690 | benign | 0 | 1 | 0 |
| rs1800314 | benign | 0 | 1 | 0 |
| rs369532274 | pathogenic | 0 | 1 | 0 |
| rs121907943 | pathogenic | 0 | 1 | 0 |
| rs780321415 | pathogenic | 0 | 1 | 0 |
| rs377286472 | benign | 0 | 1 | 0 |
| rs121918406 | pathogenic | 1 | 1 | 1 |
| rs147484870 | benign | 0 | 0 | 0 |
| rs193922671 | pathogenic | 0 | 0 | 0 |
| rs201397978 | benign | 1 | 1 | 1 |
| rs34884895 | benign | 1 | 1 | 1 |
| rs138329459 | benign | 0 | 0 | 0 |
| rs2289366 | benign | 1 | 1 | 1 |
| rs141858334 | benign | 1 | 1 | 1 |
| rs59907886 | benign | 1 | 1 | 1 |
| rs137852653 | pathogenic | 0 | 0 | 0 |
| rs137852652 | pathogenic | 0 | 0 | 0 |

| | | | | |
|---|---|---|---|---|
| rs137852656 | pathogenic | 0 | 0 | 0 |
| rs61739916 | benign | 0 | 0 | 0 |
| rs28936668 | pathogenic | 0 | 0 | 0 |
| rs137852651 | pathogenic | 0 | 0 | 0 |
| rs137852650 | pathogenic | 0 | 0 | 0 |
| rs28936669 | pathogenic | 0 | 0 | 0 |
| rs137852654 | pathogenic | 0 | 0 | 0 |
| rs28936368 | pathogenic | 1 | 1 | 0 |
| rs137852655 | pathogenic | 0 | 0 | 0 |
| rs61752496 | benign | 0 | 0 | 0 |
| rs750098055 | benign | 0 | 0 | 0 |
| rs764149793 | benign | 0 | 0 | 0 |
| rs724160017 | pathogenic | 0 | 1 | 0 |
| rs724160019 | pathogenic | 0 | 0 | 0 |
| rs202219398 | pathogenic | 1 | 1 | 0 |
| rs724160023 | pathogenic | 1 | 1 | 0 |
| rs761953453 | pathogenic | 0 | 0 | 0 |
| rs80338835 | pathogenic | 1 | 1 | 1 |
| rs56284940 | benign | 0 | 0 | 0 |
| rs74575749 | benign | 0 | 0 | 0 |
| rs587777075 | pathogenic | 1 | 1 | 1 |
| rs641153 | benign | 0 | 0 | 0 |
| rs12614 | benign | 0 | 0 | 0 |
| rs117314762 | benign | 0 | 0 | 0 |
| rs537478097 | benign | 1 | 1 | 1 |
| rs144812066 | benign | 0 | 0 | 0 |
| rs4151651 | benign | 0 | 0 | 0 |
| rs138207668 | benign | 1 | 1 | 1 |
| rs149101394 | benign | 1 | 1 | 1 |
| rs4151659 | benign | 1 | 1 | 1 |
| rs45484591 | benign | 1 | 1 | 1 |
| rs761991070 | pathogenic | 0 | 0 | 0 |
| rs755948940 | pathogenic | 0 | 0 | 0 |
| rs137853097 | pathogenic | 0 | 0 | 0 |
| rs775297352 | benign | 0 | 0 | 0 |
| rs28943591 | benign | 0 | 0 | 0 |
| rs191468413 | pathogenic | 1 | 1 | 0 |
| rs11539471 | benign | 1 | 1 | 1 |
| rs11205 | benign | 0 | 0 | 0 |
| rs768409400 | pathogenic | 0 | 0 | 0 |
| rs762013077 | pathogenic | 0 | 0 | 0 |
| rs768474112 | pathogenic | 0 | 0 | 0 |
| rs121965063 | pathogenic | 1 | 1 | 1 |
| rs143648758 | pathogenic | 0 | 0 | 0 |
| rs121965066 | pathogenic | 0 | 0 | 0 |
| rs752907087 | pathogenic | 0 | 0 | 0 |
| rs747702749 | pathogenic | 0 | 0 | 0 |
| rs754087775 | pathogenic | 0 | 0 | 0 |
| rs121918641 | pathogenic | 1 | 1 | 1 |
| rs121918642 | pathogenic | 1 | 1 | 1 |
| rs121918640 | pathogenic | 1 | 1 | 1 |
| rs121918637 | pathogenic | 1 | 1 | 1 |
| rs121918644 | pathogenic | 1 | 1 | 1 |
| rs16840544 | benign | 0 | 0 | 0 |

| | | | | |
|---|---|---|---|---|
| rs1042391 | benign | 1 | 1 | 1 |
| rs104893744 | pathogenic | 1 | 1 | 1 |
| rs120074136 | pathogenic | 0 | 0 | 0 |
| rs1805081 | benign | 1 | 1 | 0 |
| rs80358251 | benign | 0 | 0 | 0 |
| rs774588125 | benign | 0 | 0 | 0 |
| rs120074134 | pathogenic | 0 | 0 | 0 |
| rs139751448 | pathogenic | 0 | 0 | 0 |
| rs77080672 | benign | 0 | 0 | 0 |
| rs372445155 | pathogenic | 1 | 1 | 0 |
| rs13381670 | benign | 0 | 0 | 0 |
| rs1788799 | benign | 0 | 0 | 0 |
| rs376213990 | pathogenic | 0 | 0 | 0 |
| rs757475924 | pathogenic | 0 | 0 | 0 |
| rs80358253 | pathogenic | 0 | 0 | 0 |
| rs1805082 | benign | 0 | 0 | 0 |
| rs120074130 | pathogenic | 0 | 0 | 0 |
| rs768999208 | pathogenic | 0 | 0 | 0 |
| rs34302553 | benign | 0 | 0 | 0 |
| rs28940897 | pathogenic | 0 | 0 | 0 |
| rs1140458 | benign | 0 | 0 | 0 |
| rs143124972 | pathogenic | 0 | 0 | 0 |
| rs120074135 | pathogenic | 0 | 0 | 0 |
| rs543206298 | pathogenic | 0 | 0 | 0 |
| rs120074132 | pathogenic | 0 | 0 | 0 |
| rs759826138 | pathogenic | 0 | 0 | 0 |
| rs28942108 | pathogenic | 0 | 0 | 0 |
| rs80358254 | pathogenic | 0 | 0 | 0 |
| rs80358257 | pathogenic | 0 | 0 | 0 |
| rs781261962 | pathogenic | 0 | 0 | 0 |
| rs28942107 | pathogenic | 0 | 0 | 0 |
| rs80358258 | pathogenic | 0 | 0 | 0 |
| rs80358259 | pathogenic | 1 | 1 | 0 |
| rs141440861 | benign | 0 | 0 | 0 |
| rs28942106 | pathogenic | 0 | 0 | 0 |
| rs778878523 | pathogenic | 0 | 0 | 0 |
| rs28942105 | pathogenic | 0 | 0 | 0 |
| rs61731969 | benign | 0 | 0 | 0 |
| rs200444084 | pathogenic | 0 | 0 | 0 |
| rs113371321 | pathogenic | 0 | 0 | 0 |
| rs369098773 | pathogenic | 0 | 0 | 0 |
| rs35248744 | benign | 0 | 0 | 0 |
| rs758902805 | pathogenic | 0 | 0 | 0 |
| rs120074131 | pathogenic | 0 | 0 | 0 |
| rs1805084 | benign | 0 | 0 | 0 |
| rs117793540 | benign | 0 | 0 | 0 |
| rs530058063 | benign | 0 | 0 | 0 |
| rs1064422 | pathogenic | 1 | 1 | 1 |
| rs104894506 | pathogenic | 1 | 1 | 1 |
| rs104894507 | pathogenic | 1 | 1 | 1 |
| rs104894508 | pathogenic | 1 | 1 | 0 |
| rs28999113 | pathogenic | 1 | 1 | 0 |
| rs281860266 | pathogenic | 0 | 0 | 0 |
| rs104894961 | pathogenic | 1 | 1 | 0 |

| | | | | |
|---|---|---|---|---|
| rs122463168 | pathogenic | 0 | 0 | 0 |
| rs199469624 | pathogenic | 1 | 1 | 1 |
| rs150107747 | benign | 0 | 0 | 0 |
| rs144984628 | pathogenic | 0 | 0 | 0 |
| rs28935171 | pathogenic | 0 | 0 | 0 |
| rs137852443 | pathogenic | 1 | 1 | 0 |
| rs28933674 | pathogenic | 1 | 1 | 1 |
| rs28933675 | pathogenic | 1 | 1 | 1 |
| rs28933676 | pathogenic | 1 | 1 | 1 |
| rs28933677 | pathogenic | 1 | 1 | 1 |
| rs28933678 | pathogenic | 0 | 0 | 0 |
| rs28933679 | pathogenic | 1 | 1 | 1 |
| rs28933680 | pathogenic | 0 | 0 | 0 |
| rs28933681 | pathogenic | 0 | 0 | 0 |
| rs28933682 | pathogenic | 0 | 0 | 0 |
| rs28937294 | pathogenic | 1 | 1 | 0 |
| rs137852459 | pathogenic | 0 | 0 | 0 |
| rs137852461 | pathogenic | 0 | 0 | 0 |
| rs137852464 | pathogenic | 1 | 1 | 1 |
| rs137852465 | pathogenic | 1 | 1 | 1 |
| rs1800297 | benign | 1 | 1 | 0 |
| rs137852374 | pathogenic | 1 | 1 | 1 |
| rs672601312 | pathogenic | 1 | 1 | 1 |
| rs387907196 | pathogenic | 1 | 1 | 1 |
| rs148348866 | benign | 0 | 0 | 0 |
| rs141138948 | pathogenic | 1 | 1 | 1 |
| rs137852685 | pathogenic | 1 | 1 | 0 |
| rs104893900 | pathogenic | 1 | 1 | 0 |
| rs137852686 | pathogenic | 1 | 1 | 1 |
| rs104893906 | pathogenic | 1 | 1 | 0 |
| rs587782207 | pathogenic | 1 | 1 | 1 |
| rs75086406 | pathogenic | 0 | 0 | 0 |
| rs199822819 | pathogenic | 0 | 0 | 0 |
| rs398123166 | pathogenic | 1 | 1 | 1 |
| rs121913123 | pathogenic | 1 | 1 | 1 |
| rs772190176 | pathogenic | 0 | 1 | 0 |
| rs587782216 | pathogenic | 1 | 1 | 1 |
| rs398123159 | pathogenic | 1 | 1 | 1 |
| rs121913122 | pathogenic | 1 | 1 | 1 |
| rs755436052 | pathogenic | 1 | 1 | 1 |
| rs200796606 | pathogenic | 1 | 1 | 1 |
| rs2070080 | pathogenic | 0 | 1 | 0 |
| rs149616320 | benign | 0 | 0 | 0 |
| rs143529458 | benign | 0 | 0 | 0 |
| rs267607077 | pathogenic | 1 | 1 | 1 |
| rs143898031 | benign | 1 | 1 | 0 |
| rs148304857 | benign | 1 | 1 | 1 |
| rs137852809 | pathogenic | 0 | 0 | 0 |
| rs3765623 | benign | 0 | 0 | 0 |
| rs16944397 | benign | 0 | 0 | 0 |
| rs730880167 | benign | 0 | 0 | 0 |
| rs187108957 | benign | 0 | 0 | 0 |
| rs117342470 | benign | 0 | 0 | 0 |
| rs751200138 | pathogenic | 1 | 1 | 1 |

| | | | | |
|---|---|---|---|---|
| rs755799528 | pathogenic | 0 | 0 | 0 |
| rs537484504 | benign | 0 | 0 | 0 |
| rs748944640 | pathogenic | 0 | 0 | 0 |
| rs199774121 | pathogenic | 0 | 0 | 0 |
| rs766094434 | pathogenic | 1 | 1 | 0 |
| rs777321035 | pathogenic | 1 | 1 | 1 |
| rs146269774 | benign | 0 | 1 | 0 |
| rs144085805 | benign | 0 | 1 | 0 |
| rs367643028 | benign | 0 | 0 | 0 |
| rs42524 | benign | 0 | 0 | 0 |
| rs28940586 | pathogenic | 0 | 0 | 0 |
| rs74315327 | pathogenic | 1 | 1 | 1 |
| rs199676861 | pathogenic | 0 | 0 | 0 |
| rs115802744 | benign | 1 | 1 | 0 |
| rs752866643 | pathogenic | 1 | 1 | 1 |
| rs28941784 | pathogenic | 1 | 1 | 1 |
| rs398124434 | pathogenic | 1 | 1 | 1 |
| rs756414548 | pathogenic | 1 | 1 | 1 |
| rs376128990 | pathogenic | 1 | 1 | 1 |
| rs746219370 | pathogenic | 1 | 1 | 1 |
| rs756195708 | pathogenic | 1 | 1 | 1 |
| rs369296618 | pathogenic | 1 | 1 | 1 |
| rs9593 | benign | 1 | 1 | 1 |
| rs139510844 | benign | 0 | 0 | 0 |
| rs28382158 | benign | 0 | 0 | 0 |
| rs28919570 | pathogenic | 0 | 0 | 0 |
| rs587777331 | pathogenic | 0 | 0 | 0 |
| rs587777333 | pathogenic | 0 | 0 | 0 |
| rs587777332 | pathogenic | 0 | 0 | 0 |
| rs774980346 | pathogenic | 0 | 0 | 0 |
| rs370934093 | pathogenic | 0 | 0 | 0 |
| rs121913479 | pathogenic | 1 | 1 | 1 |
| rs121913484 | pathogenic | 0 | 0 | 0 |
| rs121913485 | pathogenic | 1 | 1 | 1 |
| rs28931614 | pathogenic | 1 | 1 | 1 |
| rs17881656 | benign | 1 | 1 | 1 |
| rs28931615 | pathogenic | 1 | 1 | 1 |
| rs147846832 | benign | 0 | 0 | 0 |
| rs80356506 | pathogenic | 0 | 0 | 0 |
| rs80356494 | pathogenic | 0 | 0 | 0 |
| rs121908894 | pathogenic | 0 | 0 | 0 |
| rs121908895 | pathogenic | 0 | 0 | 0 |
| rs80356508 | pathogenic | 0 | 0 | 0 |
| rs80356507 | pathogenic | 0 | 0 | 0 |
| rs121908896 | pathogenic | 0 | 0 | 0 |
| rs745372589 | pathogenic | 1 | 1 | 0 |
| rs183898408 | benign | 0 | 0 | 0 |
| rs200844292 | benign | 0 | 0 | 0 |
| rs146897002 | benign | 0 | 0 | 0 |
| rs121964869 | pathogenic | 1 | 1 | 1 |
| rs201768800 | benign | 1 | 1 | 1 |
| rs9807556 | benign | 1 | 1 | 1 |
| rs184721031 | benign | 0 | 0 | 0 |
| rs201104206 | benign | 0 | 0 | 0 |

| | | | | |
|---|---|---|---|---|
| rs188677538 | benign | 0 | 0 | 0 |
| rs149528866 | benign | 0 | 0 | 0 |
| rs115382168 | benign | 0 | 0 | 0 |
| rs78846023 | benign | 1 | 1 | 1 |
| rs61730662 | benign | 0 | 0 | 0 |
| rs121918368 | pathogenic | 0 | 0 | 0 |
| rs566433112 | pathogenic | 1 | 1 | 1 |
| rs121909342 | pathogenic | 0 | 0 | 0 |
| rs67586389 | pathogenic | 0 | 0 | 0 |
| rs72466485 | pathogenic | 0 | 0 | 0 |
| rs72466486 | pathogenic | 1 | 1 | 1 |
| rs72466487 | pathogenic | 1 | 1 | 1 |
| rs267607112 | pathogenic | 0 | 0 | 0 |
| rs118204013 | pathogenic | 0 | 0 | 0 |
| rs121913495 | pathogenic | 0 | 0 | 0 |
| rs11554273 | pathogenic | 0 | 0 | 0 |
| rs193922477 | pathogenic | 1 | 1 | 1 |
| rs193922479 | pathogenic | 1 | 1 | 1 |
| rs72547554 | pathogenic | 0 | 0 | 0 |
| rs72547561 | pathogenic | 1 | 1 | 1 |
| rs72547562 | pathogenic | 1 | 1 | 0 |
| rs115977487 | benign | 1 | 1 | 0 |
| rs72547564 | pathogenic | 0 | 0 | 0 |
| rs72547568 | pathogenic | 1 | 1 | 0 |
| rs72547569 | pathogenic | 1 | 1 | 1 |
| rs72547571 | pathogenic | 0 | 0 | 0 |
| rs72547575 | pathogenic | 1 | 1 | 1 |
| rs544994462 | benign | 0 | 0 | 0 |
| rs190470017 | benign | 0 | 0 | 0 |
| rs199473266 | pathogenic | 1 | 1 | 1 |
| rs137854602 | pathogenic | 0 | 0 | 0 |
| rs143739249 | pathogenic | 0 | 0 | 0 |
| rs397514697 | pathogenic | 1 | 1 | 1 |
| rs373764886 | pathogenic | 0 | 0 | 0 |
| rs370242930 | pathogenic | 0 | 0 | 0 |
| rs377145777 | pathogenic | 1 | 1 | 1 |
| rs191256186 | benign | 0 | 0 | 0 |
| rs150070901 | benign | 1 | 1 | 1 |
| rs138098032 | benign | 0 | 0 | 0 |
| rs117731813 | benign | 1 | 1 | 1 |
| rs104894488 | pathogenic | 1 | 1 | 1 |
| rs11302 | benign | 0 | 1 | 0 |
| rs121918447 | pathogenic | 1 | 1 | 1 |
| rs138205828 | benign | 1 | 1 | 0 |
| rs149858290 | benign | 0 | 0 | 0 |
| rs750764003 | pathogenic | 0 | 1 | 0 |
| rs141381896 | benign | 0 | 0 | 0 |
| rs149027682 | benign | 0 | 0 | 0 |
| rs121908415 | pathogenic | 0 | 0 | 0 |
| rs121908416 | pathogenic | 0 | 0 | 0 |
| rs121908417 | pathogenic | 0 | 0 | 0 |
| rs587780306 | benign | 0 | 0 | 0 |
| rs3218725 | benign | 0 | 0 | 0 |
| rs3218729 | benign | 0 | 0 | 0 |

| | | | | |
|---|---|---|---|---|
| rs387906917 | pathogenic | 0 | 0 | 0 |
| rs200652608 | pathogenic | 0 | 0 | 0 |
| rs3218721 | benign | 0 | 0 | 0 |
| rs63750869 | pathogenic | 1 | 1 | 1 |
| rs118203910 | pathogenic | 0 | 1 | 0 |
| rs9332701 | benign | 1 | 1 | 0 |
| rs6027 | benign | 0 | 0 | 0 |
| rs121909110 | pathogenic | 0 | 1 | 0 |
| rs762550967 | pathogenic | 1 | 1 | 0 |
| rs763615602 | pathogenic | 1 | 1 | 0 |
| rs144907403 | benign | 0 | 0 | 0 |
| rs104893631 | pathogenic | 1 | 1 | 0 |
| rs144181978 | pathogenic | 1 | 1 | 0 |
| rs74874677 | benign | 1 | 1 | 1 |
| rs104893632 | pathogenic | 0 | 1 | 0 |
| rs371342464 | benign | 1 | 1 | 1 |
| rs104893633 | pathogenic | 1 | 1 | 0 |
| rs201088712 | pathogenic | 0 | 0 | 0 |
| rs201857541 | benign | 1 | 1 | 1 |
| rs139517732 | pathogenic | 1 | 1 | 0 |
| rs191057824 | benign | 1 | 1 | 0 |
| rs35683768 | benign | 1 | 1 | 0 |
| rs104895271 | pathogenic | 0 | 0 | 0 |
| rs104895217 | pathogenic | 1 | 1 | 1 |
| rs104895223 | pathogenic | 1 | 1 | 1 |
| rs104895218 | pathogenic | 1 | 1 | 1 |
| rs104895225 | pathogenic | 1 | 1 | 1 |
| rs4149637 | benign | 1 | 1 | 1 |
| rs104895219 | pathogenic | 1 | 1 | 1 |
| rs104895220 | pathogenic | 1 | 1 | 1 |
| rs104895228 | pathogenic | 1 | 1 | 1 |
| rs104895221 | pathogenic | 1 | 1 | 1 |
| rs104895222 | pathogenic | 1 | 1 | 1 |
| rs104895247 | pathogenic | 0 | 0 | 0 |
| rs28940585 | pathogenic | 0 | 0 | 0 |
| rs561367199 | pathogenic | 1 | 1 | 1 |
| rs143793815 | benign | 0 | 0 | 0 |
| rs28941473 | pathogenic | 1 | 1 | 1 |
| rs751590073 | pathogenic | 1 | 1 | 1 |
| rs111407265 | benign | 1 | 1 | 1 |
| rs373697663 | pathogenic | 1 | 1 | 1 |
| rs369879957 | pathogenic | 0 | 0 | 0 |
| rs28941472 | pathogenic | 1 | 1 | 1 |
| rs28940287 | pathogenic | 1 | 1 | 1 |
| rs77625410 | benign | 0 | 0 | 0 |
| rs111033477 | benign | 0 | 0 | 0 |
| rs115123584 | benign | 0 | 0 | 0 |
| rs374576916 | benign | 0 | 0 | 0 |
| rs397516317 | pathogenic | 0 | 0 | 0 |
| rs368657015 | pathogenic | 0 | 0 | 0 |
| rs397516321 | pathogenic | 0 | 0 | 0 |
| rs397516322 | pathogenic | 0 | 0 | 0 |
| rs111033215 | pathogenic | 0 | 0 | 0 |
| rs199606180 | pathogenic | 0 | 0 | 0 |

| | | | | |
|---|---|---|---|---|
| rs111033192 | pathogenic | 0 | 0 | 0 |
| rs948962 | benign | 1 | 1 | 1 |
| rs142293185 | benign | 0 | 0 | 0 |
| rs376764423 | pathogenic | 0 | 0 | 0 |
| rs773844428 | pathogenic | 0 | 0 | 0 |
| rs186644871 | benign | 0 | 0 | 0 |
| rs200313391 | benign | 0 | 0 | 0 |
| rs377670513 | pathogenic | 0 | 0 | 0 |
| rs1132036 | benign | 0 | 0 | 0 |
| rs111033231 | benign | 0 | 0 | 0 |
| rs145945630 | pathogenic | 0 | 0 | 0 |
| rs121912984 | pathogenic | 1 | 1 | 1 |
| rs1800435 | benign | 0 | 1 | 0 |
| rs121912980 | pathogenic | 0 | 0 | 0 |
| rs121912982 | pathogenic | 1 | 1 | 1 |
| rs121912983 | pathogenic | 0 | 0 | 0 |
| rs121912981 | pathogenic | 0 | 0 | 0 |
| rs1057868 | benign | 0 | 0 | 0 |
| rs56048668 | benign | 0 | 0 | 0 |
| rs55865465 | benign | 0 | 0 | 0 |
| rs748096324 | benign | 1 | 1 | 0 |
| rs150475750 | benign | 0 | 0 | 0 |
| rs143025739 | benign | 1 | 1 | 0 |
| rs79702016 | benign | 1 | 1 | 0 |
| rs41338945 | benign | 0 | 0 | 0 |
| rs146406037 | benign | 1 | 1 | 1 |
| rs143274491 | benign | 0 | 0 | 0 |
| rs3829986 | benign | 0 | 0 | 0 |
| rs149168939 | benign | 0 | 0 | 0 |
| rs1800385 | benign | 0 | 0 | 0 |
| rs1800386 | pathogenic | 0 | 0 | 0 |
| rs121918524 | pathogenic | 1 | 1 | 1 |
| rs121912700 | pathogenic | 0 | 0 | 0 |
| rs104893930 | pathogenic | 1 | 1 | 1 |
| rs104893931 | pathogenic | 0 | 0 | 0 |
| rs116125936 | benign | 0 | 0 | 0 |
| rs627441 | benign | 0 | 0 | 0 |
| rs104894481 | pathogenic | 0 | 0 | 0 |
| rs137853129 | pathogenic | 0 | 0 | 0 |
| rs137853130 | pathogenic | 1 | 1 | 0 |
| rs28940282 | pathogenic | 0 | 0 | 0 |
| rs2229989 | benign | 0 | 0 | 0 |
| rs104894647 | pathogenic | 0 | 0 | 0 |
| rs1061170 | benign | 1 | 1 | 1 |
| rs576402053 | benign | 0 | 0 | 0 |
| rs34972246 | benign | 0 | 0 | 0 |
| rs192725607 | benign | 0 | 0 | 0 |
| rs137852615 | pathogenic | 1 | 1 | 0 |
| rs137852617 | pathogenic | 1 | 1 | 1 |
| rs137852611 | pathogenic | 0 | 0 | 0 |
| rs137852612 | pathogenic | 0 | 0 | 0 |
| rs137852614 | pathogenic | 1 | 1 | 1 |
| rs137852610 | pathogenic | 1 | 1 | 1 |
| rs137852618 | pathogenic | 1 | 1 | 1 |

| | | | | |
|---|---|---|---|---|
| rs137852613 | pathogenic | 1 | 1 | 0 |
| rs137852609 | pathogenic | 0 | 0 | 0 |
| rs63750306 | pathogenic | 1 | 1 | 1 |
| rs63750265 | pathogenic | 1 | 1 | 1 |
| rs63751287 | pathogenic | 1 | 1 | 1 |
| rs199723282 | benign | 0 | 0 | 0 |
| rs121917807 | pathogenic | 0 | 0 | 0 |
| rs121917809 | pathogenic | 0 | 0 | 0 |
| rs75134564 | pathogenic | 0 | 0 | 0 |
| rs267606930 | pathogenic | 1 | 1 | 0 |
| rs121909302 | pathogenic | 0 | 0 | 0 |
| rs121909303 | pathogenic | 0 | 0 | 0 |
| rs121909299 | pathogenic | 0 | 0 | 0 |
| rs121909301 | pathogenic | 0 | 0 | 0 |
| rs121909300 | pathogenic | 1 | 1 | 0 |
| rs111033589 | pathogenic | 0 | 0 | 0 |
| rs137853059 | pathogenic | 0 | 0 | 0 |
| rs137853057 | pathogenic | 1 | 1 | 1 |
| rs1801474 | benign | 0 | 0 | 0 |
| rs137853060 | pathogenic | 1 | 1 | 1 |
| rs137853058 | pathogenic | 1 | 1 | 1 |
| rs34424986 | pathogenic | 0 | 0 | 0 |
| rs199657839 | benign | 1 | 1 | 1 |
| rs56092260 | benign | 0 | 0 | 0 |
| rs1801582 | benign | 1 | 1 | 1 |
| rs1801334 | benign | 1 | 1 | 1 |
| rs191486604 | pathogenic | 0 | 0 | 0 |
| rs121908051 | pathogenic | 0 | 0 | 0 |
| rs121908055 | pathogenic | 1 | 1 | 0 |
| rs28940886 | pathogenic | 0 | 0 | 0 |
| rs121908050 | pathogenic | 0 | 0 | 0 |
| rs28940887 | pathogenic | 0 | 0 | 0 |
| rs387906300 | pathogenic | 0 | 0 | 0 |
| rs28942087 | pathogenic | 0 | 0 | 0 |
| rs121908049 | pathogenic | 1 | 1 | 0 |
| rs121908054 | pathogenic | 1 | 1 | 0 |
| rs121908048 | pathogenic | 0 | 0 | 0 |
| rs28940888 | pathogenic | 1 | 1 | 1 |
| rs121908053 | pathogenic | 1 | 1 | 0 |
| rs122467169 | pathogenic | 1 | 1 | 1 |
| rs122467172 | pathogenic | 1 | 1 | 1 |
| rs122467170 | pathogenic | 1 | 1 | 1 |
| rs28935477 | pathogenic | 1 | 1 | 0 |
| rs28942088 | pathogenic | 0 | 0 | 0 |
| rs751478142 | pathogenic | 1 | 1 | 0 |
| rs765368797 | pathogenic | 0 | 0 | 0 |
| rs147810437 | benign | 0 | 0 | 0 |
| rs121918209 | pathogenic | 1 | 1 | 1 |
| rs121918208 | pathogenic | 1 | 1 | 1 |
| rs121918210 | pathogenic | 1 | 1 | 1 |
| rs35515638 | benign | 1 | 1 | 1 |
| rs121918207 | pathogenic | 1 | 1 | 1 |
| rs141298312 | benign | 0 | 0 | 0 |
| rs121918211 | pathogenic | 0 | 0 | 0 |

| | | | | |
|---|---|---|---|---|
| rs149660059 | benign | 0 | 0 | 0 |
| rs200670286 | pathogenic | 1 | 1 | 1 |
| rs121918212 | pathogenic | 1 | 1 | 1 |
| rs121918213 | pathogenic | 1 | 1 | 1 |
| rs121918205 | pathogenic | 1 | 1 | 1 |
| rs104894025 | pathogenic | 1 | 1 | 1 |
| rs104894026 | pathogenic | 1 | 1 | 1 |
| rs104894028 | pathogenic | 0 | 1 | 0 |
| rs104894029 | pathogenic | 0 | 1 | 0 |
| rs119458971 | pathogenic | 0 | 0 | 0 |
| rs119458969 | pathogenic | 1 | 1 | 1 |
| rs1801591 | benign | 1 | 1 | 1 |
| rs140169311 | benign | 0 | 0 | 0 |
| rs753363278 | benign | 1 | 1 | 1 |
| rs199763682 | pathogenic | 0 | 0 | 0 |
| rs119458970 | pathogenic | 1 | 1 | 1 |
| rs763722868 | pathogenic | 0 | 0 | 0 |
| rs56404770 | benign | 0 | 0 | 0 |
| rs34221241 | benign | 0 | 0 | 0 |
| rs121909625 | pathogenic | 0 | 0 | 0 |
| rs550921485 | pathogenic | 0 | 0 | 0 |
| rs1800454 | benign | 0 | 0 | 0 |
| rs241447 | benign | 0 | 0 | 0 |
| rs2230606 | benign | 1 | 1 | 0 |
| rs5036 | pathogenic | 1 | 1 | 1 |
| rs13306788 | benign | 0 | 0 | 0 |
| rs28929480 | pathogenic | 0 | 0 | 0 |
| rs121912749 | pathogenic | 0 | 0 | 0 |
| rs56361140 | pathogenic | 0 | 0 | 0 |
| rs28931583 | pathogenic | 1 | 1 | 1 |
| rs36211715 | pathogenic | 1 | 1 | 1 |
| rs138049878 | pathogenic | 1 | 1 | 1 |
| rs727503253 | pathogenic | 1 | 1 | 1 |
| rs267606908 | pathogenic | 0 | 0 | 0 |
| rs121913631 | pathogenic | 1 | 1 | 1 |
| rs36211716 | benign | 1 | 1 | 1 |
| rs121913639 | pathogenic | 1 | 1 | 1 |
| rs730880761 | pathogenic | 1 | 1 | 1 |
| rs121913629 | pathogenic | 1 | 1 | 1 |
| rs754368658 | pathogenic | 0 | 0 | 0 |
| rs777090017 | pathogenic | 0 | 0 | 0 |
| rs267606913 | pathogenic | 1 | 1 | 1 |
| rs773138218 | pathogenic | 0 | 0 | 0 |
| rs371295780 | pathogenic | 0 | 0 | 0 |
| rs387906830 | pathogenic | 1 | 1 | 0 |
| rs771364038 | pathogenic | 0 | 1 | 0 |
| rs772731615 | pathogenic | 0 | 0 | 0 |
| rs137853063 | pathogenic | 0 | 0 | 0 |
| rs387907076 | pathogenic | 0 | 1 | 0 |
| rs35460768 | benign | 0 | 0 | 0 |
| rs2278196 | benign | 1 | 1 | 1 |
| rs119103275 | pathogenic | 0 | 0 | 0 |
| rs119103276 | pathogenic | 1 | 1 | 1 |
| rs374402066 | pathogenic | 1 | 1 | 1 |

| | | | | |
|---|---|---|---|---|
| rs119103274 | pathogenic | 0 | 0 | 0 |
| rs2274064 | benign | 0 | 0 | 0 |
| rs267607023 | pathogenic | 1 | 1 | 0 |
| rs142117489 | benign | 1 | 1 | 0 |
| rs371258270 | benign | 0 | 0 | 0 |
| rs34716057 | benign | 0 | 0 | 0 |
| rs201778606 | benign | 0 | 0 | 0 |
| rs201438792 | benign | 0 | 0 | 0 |
| rs140547417 | benign | 0 | 0 | 0 |
| rs5881 | benign | 0 | 0 | 0 |
| rs752298084 | benign | 0 | 0 | 0 |
| rs34855278 | benign | 0 | 0 | 0 |
| rs5880 | benign | 1 | 1 | 0 |
| rs5882 | benign | 0 | 0 | 0 |
| rs1800777 | benign | 0 | 0 | 0 |
| rs577301331 | benign | 0 | 0 | 0 |
| rs137853313 | pathogenic | 0 | 0 | 0 |
| rs28935169 | pathogenic | 0 | 0 | 0 |
| rs137853315 | pathogenic | 0 | 0 | 0 |
| rs137853314 | pathogenic | 0 | 0 | 0 |
| rs28935469 | pathogenic | 0 | 0 | 0 |
| rs137853318 | pathogenic | 0 | 0 | 0 |
| rs28935470 | pathogenic | 0 | 0 | 0 |
| rs3729986 | benign | 0 | 0 | 0 |
| rs730880619 | pathogenic | 0 | 0 | 0 |
| rs201012766 | benign | 0 | 0 | 0 |
| rs727503218 | pathogenic | 0 | 0 | 0 |
| rs11570052 | benign | 0 | 0 | 0 |
| rs730880623 | pathogenic | 0 | 0 | 0 |
| rs138753870 | benign | 0 | 0 | 0 |
| rs397516068 | pathogenic | 0 | 0 | 0 |
| rs3729989 | benign | 1 | 1 | 1 |
| rs397516074 | pathogenic | 0 | 0 | 0 |
| rs561449819 | pathogenic | 0 | 0 | 0 |
| rs121917779 | benign | 0 | 0 | 0 |
| rs142881576 | pathogenic | 0 | 0 | 0 |
| rs77384282 | benign | 0 | 0 | 0 |
| rs201865787 | benign | 0 | 0 | 0 |
| rs142244640 | benign | 0 | 0 | 0 |
| rs200488706 | benign | 0 | 0 | 0 |
| rs137853291 | pathogenic | 1 | 1 | 0 |
| rs28933990 | pathogenic | 1 | 1 | 0 |
| rs140570189 | benign | 0 | 0 | 0 |
| rs201588178 | benign | 0 | 0 | 0 |
| rs104893846 | pathogenic | 0 | 0 | 0 |
| rs754545360 | pathogenic | 0 | 1 | 0 |
| rs104893851 | pathogenic | 0 | 1 | 0 |
| rs104893849 | pathogenic | 0 | 0 | 0 |
| rs571038432 | pathogenic | 1 | 1 | 0 |
| rs2270655 | benign | 1 | 1 | 1 |
| rs315952 | benign | 0 | 0 | 0 |
| rs151176548 | benign | 0 | 0 | 0 |
| rs122459147 | pathogenic | 1 | 1 | 0 |
| rs122458140 | pathogenic | 0 | 0 | 0 |

| | | | | |
|---|---|---|---|---|
| rs122458142 | pathogenic | 1 | 1 | 0 |
| rs267606812 | pathogenic | 1 | 1 | 0 |
| rs267606813 | pathogenic | 1 | 1 | 0 |
| rs122458143 | pathogenic | 0 | 0 | 0 |
| rs122459146 | pathogenic | 0 | 0 | 0 |
| rs122458144 | pathogenic | 0 | 0 | 0 |
| rs122458145 | pathogenic | 0 | 0 | 0 |
| rs113994007 | pathogenic | 1 | 1 | 0 |
| rs758746181 | pathogenic | 1 | 1 | 1 |
| rs145034527 | pathogenic | 1 | 1 | 0 |
| rs119468006 | pathogenic | 1 | 1 | 0 |
| rs201908721 | pathogenic | 0 | 0 | 0 |
| rs755933881 | pathogenic | 1 | 1 | 0 |
| rs771578775 | pathogenic | 0 | 0 | 0 |
| rs17849927 | benign | 0 | 0 | 0 |
| rs139133094 | benign | 0 | 0 | 0 |
| rs12593 | benign | 0 | 0 | 0 |
| rs578189699 | pathogenic | 1 | 1 | 0 |
| rs119468008 | pathogenic | 0 | 0 | 0 |
| rs768524626 | benign | 0 | 0 | 0 |
| rs119468009 | pathogenic | 0 | 0 | 0 |
| rs119468004 | pathogenic | 0 | 0 | 0 |
| rs199874519 | pathogenic | 0 | 0 | 0 |
| rs376478331 | pathogenic | 0 | 0 | 0 |
| rs61995958 | pathogenic | 0 | 0 | 0 |
| rs199714731 | benign | 0 | 0 | 0 |
| rs149185954 | benign | 0 | 0 | 0 |
| rs121918353 | pathogenic | 0 | 0 | 0 |
| rs1131695 | benign | 0 | 0 | 0 |
| rs28939668 | pathogenic | 0 | 0 | 0 |
| rs758432471 | pathogenic | 1 | 1 | 1 |
| rs752746786 | pathogenic | 1 | 1 | 1 |
| rs28935471 | pathogenic | 0 | 0 | 0 |
| rs80338837 | pathogenic | 0 | 0 | 0 |
| rs137853312 | pathogenic | 0 | 0 | 0 |
| rs28935472 | pathogenic | 0 | 0 | 0 |
| rs28935473 | pathogenic | 0 | 0 | 0 |
| rs267606752 | pathogenic | 0 | 0 | 0 |
| rs3917973 | benign | 0 | 0 | 0 |
| rs3918018 | benign | 0 | 0 | 0 |
| rs757976755 | pathogenic | 0 | 0 | 0 |
| rs121918329 | pathogenic | 1 | 1 | 0 |
| rs770114459 | pathogenic | 0 | 0 | 0 |
| rs6662272 | benign | 0 | 0 | 0 |
| rs121918242 | pathogenic | 1 | 1 | 0 |
| rs121918240 | pathogenic | 1 | 1 | 0 |
| rs200094982 | pathogenic | 0 | 0 | 0 |
| rs121918241 | pathogenic | 0 | 0 | 0 |
| rs140522266 | pathogenic | 1 | 1 | 0 |
| rs121918243 | pathogenic | 1 | 1 | 0 |
| rs370596113 | pathogenic | 1 | 1 | 0 |
| rs398124295 | pathogenic | 1 | 1 | 0 |
| rs587776889 | pathogenic | 1 | 1 | 0 |
| rs747527726 | pathogenic | 1 | 1 | 0 |

| | | | | |
|---|---|---|---|---|
| rs112631212 | benign | 0 | 0 | 0 |
| rs104894605 | pathogenic | 0 | 0 | 0 |
| rs104894607 | pathogenic | 0 | 0 | 0 |
| rs104894606 | pathogenic | 0 | 0 | 0 |
| rs143873873 | benign | 0 | 0 | 0 |
| rs104894388 | pathogenic | 1 | 1 | 0 |
| rs104894391 | pathogenic | 1 | 1 | 0 |
| rs5350 | benign | 1 | 1 | 1 |
| rs104894390 | pathogenic | 1 | 1 | 1 |
| rs5346 | benign | 1 | 1 | 1 |
| rs121912500 | pathogenic | 0 | 0 | 0 |
| rs74315450 | pathogenic | 1 | 1 | 0 |
| rs121913396 | pathogenic | 1 | 1 | 1 |
| rs28931588 | pathogenic | 1 | 1 | 1 |
| rs121913400 | pathogenic | 0 | 0 | 0 |
| rs28931589 | pathogenic | 1 | 1 | 1 |
| rs121913403 | pathogenic | 0 | 0 | 0 |
| rs121913413 | pathogenic | 0 | 0 | 0 |
| rs760107623 | pathogenic | 1 | 1 | 0 |
| rs16866378 | benign | 1 | 1 | 0 |
| rs201025290 | pathogenic | 1 | 1 | 0 |
| rs12720356 | benign | 1 | 1 | 0 |
| rs55882956 | benign | 0 | 1 | 0 |
| rs4986933 | benign | 1 | 1 | 0 |
| rs2020957 | benign | 1 | 1 | 1 |
| rs1800124 | benign | 1 | 1 | 1 |
| rs150077735 | benign | 1 | 1 | 1 |
| rs201419000 | benign | 0 | 0 | 0 |
| rs772044176 | pathogenic | 0 | 0 | 0 |
| rs138346388 | benign | 0 | 0 | 0 |
| rs121964916 | pathogenic | 0 | 0 | 0 |
| rs112534524 | pathogenic | 0 | 0 | 0 |
| rs182078921 | benign | 1 | 1 | 1 |
| rs113273712 | benign | 0 | 0 | 0 |
| rs74817407 | benign | 1 | 1 | 1 |
| rs61733901 | benign | 1 | 1 | 1 |
| rs121964912 | pathogenic | 1 | 1 | 1 |
| rs41278047 | benign | 1 | 1 | 1 |
| rs7437875 | benign | 1 | 1 | 1 |
| rs113460688 | benign | 1 | 1 | 1 |
| rs754572081 | benign | 1 | 1 | 1 |
| rs201082692 | benign | 1 | 1 | 1 |
| rs202085803 | benign | 1 | 1 | 1 |
| rs141320511 | benign | 0 | 0 | 0 |
| rs78501403 | benign | 0 | 0 | 0 |
| rs28939675 | pathogenic | 0 | 0 | 0 |
| rs74315522 | pathogenic | 0 | 0 | 0 |
| rs121434639 | pathogenic | 1 | 1 | 0 |
| rs121434640 | pathogenic | 1 | 1 | 1 |
| rs757282628 | benign | 1 | 1 | 1 |
| rs386834010 | pathogenic | 0 | 0 | 0 |
| rs200863680 | pathogenic | 0 | 0 | 0 |
| rs193919336 | pathogenic | 1 | 1 | 0 |
| rs386834011 | pathogenic | 0 | 0 | 0 |

| | | | | |
|---|---|---|---|---|
| rs28940869 | pathogenic | 0 | 0 | 0 |
| rs386834014 | pathogenic | 1 | 1 | 0 |
| rs267606960 | pathogenic | 0 | 0 | 0 |
| rs28942068 | pathogenic | 0 | 0 | 0 |
| rs193919335 | pathogenic | 0 | 0 | 0 |
| rs74374973 | pathogenic | 0 | 0 | 0 |
| rs386834018 | pathogenic | 0 | 0 | 0 |
| rs267606962 | pathogenic | 0 | 0 | 0 |
| rs6659553 | benign | 0 | 0 | 0 |
| rs29281 | benign | 0 | 0 | 0 |
| rs782265090 | benign | 0 | 0 | 0 |
| rs35910969 | pathogenic | 0 | 0 | 0 |
| rs121909113 | pathogenic | 1 | 1 | 1 |
| rs29001571 | pathogenic | 1 | 1 | 1 |
| rs587781250 | pathogenic | 1 | 1 | 1 |
| rs28939680 | pathogenic | 1 | 1 | 1 |
| rs28939681 | pathogenic | 1 | 1 | 1 |
| rs121909112 | pathogenic | 1 | 1 | 1 |
| rs28937568 | pathogenic | 1 | 1 | 1 |
| rs1799895 | pathogenic | 0 | 0 | 0 |
| rs386833431 | pathogenic | 1 | 1 | 1 |
| rs121964905 | pathogenic | 1 | 1 | 1 |
| rs121964906 | pathogenic | 1 | 1 | 1 |
| rs11551664 | pathogenic | 1 | 1 | 1 |
| rs104894705 | pathogenic | 1 | 1 | 1 |
| rs104893805 | pathogenic | 0 | 0 | 0 |
| rs104894205 | pathogenic | 0 | 0 | 0 |
| rs104894204 | pathogenic | 1 | 1 | 0 |
| rs62642926 | pathogenic | 0 | 0 | 0 |
| rs5030841 | pathogenic | 1 | 1 | 1 |
| rs75193786 | pathogenic | 1 | 1 | 1 |
| rs62508682 | pathogenic | 0 | 0 | 0 |
| rs3850625 | benign | 1 | 1 | 1 |
| rs141908793 | benign | 0 | 0 | 0 |
| rs187864727 | benign | 0 | 0 | 0 |
| rs548909101 | benign | 0 | 0 | 0 |
| rs3825394 | benign | 0 | 1 | 0 |
| rs143548402 | benign | 0 | 0 | 0 |
| rs267607146 | pathogenic | 0 | 0 | 0 |
| rs267607144 | pathogenic | 0 | 0 | 0 |
| rs267607148 | pathogenic | 1 | 1 | 1 |
| rs267607143 | pathogenic | 0 | 0 | 0 |
| rs267607145 | pathogenic | 0 | 0 | 0 |
| rs121912636 | pathogenic | 1 | 1 | 1 |
| rs121912634 | pathogenic | 1 | 1 | 1 |
| rs570675468 | benign | 0 | 0 | 0 |
| rs368861241 | pathogenic | 1 | 1 | 1 |
| rs104894528 | pathogenic | 1 | 1 | 1 |
| rs77493891 | benign | 0 | 0 | 0 |
| rs62640903 | benign | 0 | 0 | 0 |
| rs149926091 | benign | 1 | 1 | 1 |
| rs641081 | benign | 0 | 0 | 0 |
| rs4930199 | benign | 0 | 0 | 0 |
| rs121913020 | pathogenic | 0 | 0 | 0 |

| | | | | |
|---|---|---|---|---|
| rs1799793 | benign | 0 | 0 | 0 |
| rs121913016 | pathogenic | 0 | 0 | 0 |
| rs121913025 | pathogenic | 0 | 0 | 0 |
| rs121913019 | pathogenic | 0 | 0 | 0 |
| rs376556895 | pathogenic | 1 | 1 | 1 |
| rs121913024 | pathogenic | 1 | 1 | 1 |
| rs121913021 | pathogenic | 0 | 0 | 0 |
| rs121913023 | pathogenic | 0 | 0 | 0 |
| rs41556519 | pathogenic | 0 | 0 | 0 |
| rs121913022 | pathogenic | 0 | 0 | 0 |
| rs144564120 | pathogenic | 0 | 0 | 0 |
| rs121913026 | pathogenic | 1 | 1 | 1 |
| rs121913018 | pathogenic | 1 | 1 | 1 |
| rs142954643 | benign | 0 | 0 | 0 |
| rs76655496 | benign | 0 | 0 | 0 |
| rs104894371 | pathogenic | 1 | 1 | 0 |
| rs137964936 | benign | 0 | 0 | 0 |
| rs759736132 | pathogenic | 0 | 0 | 0 |
| rs387906595 | pathogenic | 1 | 1 | 0 |
| rs768248216 | pathogenic | 0 | 0 | 0 |
| rs548231008 | benign | 0 | 0 | 0 |
| rs757094384 | benign | 0 | 0 | 0 |
| rs867529 | benign | 0 | 0 | 0 |
| rs13045 | benign | 1 | 1 | 1 |
| rs121908570 | pathogenic | 0 | 0 | 0 |
| rs201872586 | benign | 1 | 1 | 0 |
| rs149092990 | benign | 0 | 0 | 0 |
| rs115585711 | benign | 0 | 0 | 0 |
| rs137909416 | benign | 1 | 1 | 0 |
| rs143388949 | benign | 1 | 1 | 1 |
| rs73137716 | benign | 1 | 1 | 1 |
| rs121908947 | pathogenic | 1 | 1 | 1 |
| rs398122904 | pathogenic | 1 | 1 | 1 |
| rs35715497 | benign | 1 | 1 | 0 |
| rs35462421 | benign | 0 | 0 | 0 |
| rs119475043 | pathogenic | 0 | 0 | 0 |
| rs119475042 | pathogenic | 0 | 0 | 0 |
| rs1076669 | benign | 1 | 1 | 0 |
| rs141146885 | benign | 0 | 0 | 0 |
| rs3026906 | pathogenic | 0 | 0 | 0 |
| rs75362221 | benign | 1 | 1 | 1 |
| rs138472116 | benign | 1 | 1 | 1 |
| rs371040282 | pathogenic | 1 | 1 | 1 |
| rs4988496 | benign | 0 | 0 | 0 |
| rs121918117 | pathogenic | 1 | 1 | 1 |
| rs4988498 | benign | 0 | 0 | 0 |
| rs5035 | benign | 0 | 0 | 0 |
| rs121918322 | pathogenic | 1 | 1 | 1 |
| rs370359540 | benign | 0 | 0 | 0 |
| rs758972589 | pathogenic | 0 | 0 | 0 |
| rs144856889 | benign | 0 | 0 | 0 |
| rs121434623 | pathogenic | 1 | 1 | 1 |
| rs61735600 | benign | 0 | 0 | 0 |
| rs72550870 | benign | 1 | 1 | 1 |

| | | | | |
|---|---|---|---|---|
| rs56392418 | benign | 0 | 0 | 0 |
| rs41307788 | benign | 1 | 1 | 1 |
| rs387907239 | pathogenic | 1 | 1 | 0 |
| rs28931584 | pathogenic | 0 | 0 | 0 |
| rs121912745 | pathogenic | 0 | 0 | 0 |
| rs121912744 | pathogenic | 0 | 0 | 0 |
| rs121912754 | pathogenic | 1 | 1 | 1 |
| rs121912746 | pathogenic | 0 | 0 | 0 |
| rs121912757 | pathogenic | 0 | 0 | 0 |
| rs121912758 | pathogenic | 0 | 0 | 0 |
| rs121912748 | pathogenic | 0 | 0 | 0 |
| rs121912755 | pathogenic | 0 | 0 | 0 |
| rs121912741 | pathogenic | 0 | 0 | 0 |
| rs121912753 | pathogenic | 0 | 0 | 0 |
| rs121912750 | pathogenic | 0 | 0 | 0 |
| rs2285644 | pathogenic | 0 | 0 | 0 |
| rs121912751 | pathogenic | 0 | 0 | 0 |
| rs5026 | benign | 0 | 0 | 0 |
| rs121912759 | pathogenic | 0 | 0 | 0 |
| rs28931585 | pathogenic | 0 | 0 | 0 |
| rs17122769 | benign | 0 | 0 | 0 |
| rs28362322 | benign | 1 | 1 | 1 |
| rs3824874 | benign | 0 | 0 | 0 |
| rs146004831 | benign | 0 | 0 | 0 |
| rs142155860 | benign | 0 | 0 | 0 |
| rs121434404 | pathogenic | 0 | 0 | 0 |
| rs61735578 | benign | 1 | 1 | 1 |
| rs558018 | benign | 0 | 0 | 0 |
| rs61735577 | benign | 1 | 1 | 1 |
| rs76784113 | benign | 0 | 0 | 0 |
| rs199469640 | pathogenic | 0 | 0 | 0 |
| rs199469628 | pathogenic | 1 | 1 | 0 |
| rs199469629 | pathogenic | 0 | 1 | 0 |
| rs387907156 | pathogenic | 0 | 1 | 0 |
| rs199469643 | pathogenic | 0 | 0 | 0 |
| rs199469646 | pathogenic | 0 | 0 | 0 |
| rs766294940 | pathogenic | 0 | 0 | 0 |
| rs387907347 | pathogenic | 0 | 0 | 0 |
| rs137853590 | pathogenic | 1 | 1 | 1 |
| rs137853589 | pathogenic | 1 | 1 | 1 |
| rs137853588 | pathogenic | 1 | 1 | 1 |
| rs55909005 | benign | 0 | 0 | 0 |
| rs4581 | benign | 0 | 0 | 0 |
| rs63750123 | benign | 0 | 0 | 0 |
| rs10254120 | benign | 0 | 0 | 0 |
| rs121434629 | pathogenic | 1 | 1 | 0 |
| rs200313585 | benign | 0 | 0 | 0 |
| rs143162541 | benign | 1 | 1 | 0 |
| rs760615315 | benign | 0 | 0 | 0 |
| rs768488890 | benign | 1 | 1 | 1 |
| rs116373169 | benign | 1 | 1 | 0 |
| rs63750871 | pathogenic | 1 | 1 | 0 |
| rs116349687 | benign | 0 | 0 | 0 |
| rs375289386 | benign | 0 | 0 | 0 |

| | | | | |
|---|---|---|---|---|
| rs760228510 | pathogenic | 1 | 1 | 1 |
| rs587779343 | pathogenic | 1 | 1 | 1 |
| rs573125799 | benign | 0 | 0 | 0 |
| rs1805322 | benign | 0 | 0 | 0 |
| rs267608153 | pathogenic | 0 | 0 | 0 |
| rs139194813 | benign | 0 | 0 | 0 |
| rs530021751 | benign | 0 | 0 | 0 |
| rs200640585 | pathogenic | 0 | 0 | 0 |
| rs139438201 | benign | 0 | 0 | 0 |
| rs200513014 | benign | 0 | 0 | 0 |
| rs567102013 | benign | 0 | 0 | 0 |
| rs121908608 | pathogenic | 0 | 0 | 0 |
| rs121908606 | pathogenic | 0 | 0 | 0 |
| rs17103671 | benign | 0 | 0 | 0 |
| rs34725281 | benign | 0 | 0 | 0 |
| rs104894421 | pathogenic | 1 | 1 | 0 |
| rs121918145 | pathogenic | 1 | 1 | 1 |
| rs121918152 | pathogenic | 1 | 1 | 1 |
| rs56379106 | benign | 0 | 0 | 0 |
| rs4987076 | benign | 0 | 0 | 0 |
| rs370243877 | pathogenic | 0 | 0 | 0 |
| rs17679445 | benign | 1 | 1 | 1 |
| rs550423482 | pathogenic | 1 | 1 | 1 |
| rs104894629 | pathogenic | 1 | 1 | 1 |
| rs773450573 | pathogenic | 1 | 1 | 1 |
| rs116679717 | benign | 0 | 0 | 0 |
| rs372000848 | pathogenic | 0 | 0 | 0 |
| rs138595973 | benign | 0 | 0 | 0 |
| rs115244417 | benign | 0 | 0 | 0 |
| rs11575292 | benign | 0 | 0 | 0 |
| rs137853211 | pathogenic | 1 | 1 | 1 |
| rs137853207 | pathogenic | 0 | 0 | 0 |
| rs137853210 | pathogenic | 1 | 1 | 1 |
| rs6262 | benign | 0 | 0 | 0 |
| rs6263 | benign | 0 | 1 | 0 |
| rs137853208 | pathogenic | 0 | 1 | 0 |
| rs137853212 | pathogenic | 1 | 1 | 0 |
| rs137853209 | pathogenic | 1 | 1 | 1 |
| rs201951824 | pathogenic | 0 | 0 | 0 |
| rs11575542 | benign | 0 | 0 | 0 |
| rs199681788 | benign | 0 | 1 | 0 |
| rs748835902 | benign | 1 | 1 | 1 |
| rs753719949 | benign | 0 | 0 | 0 |
| rs267606647 | pathogenic | 0 | 0 | 0 |
| rs267606648 | pathogenic | 1 | 1 | 1 |
| rs267606643 | pathogenic | 0 | 1 | 0 |
| rs559947967 | pathogenic | 1 | 1 | 0 |
| rs267606645 | pathogenic | 1 | 1 | 1 |
| rs12116440 | benign | 0 | 0 | 0 |
| rs121908539 | pathogenic | 0 | 0 | 0 |
| rs137853096 | pathogenic | 1 | 1 | 0 |
| rs148363262 | benign | 1 | 1 | 0 |
| rs587777442 | pathogenic | 0 | 0 | 0 |
| rs25640 | benign | 1 | 1 | 1 |

| | | | | |
|---|---|---|---|---|
| rs28943589 | benign | 0 | 0 | 0 |
| rs1143650 | benign | 1 | 1 | 1 |
| rs150886026 | benign | 0 | 1 | 0 |
| rs121908133 | pathogenic | 1 | 1 | 0 |
| rs121908131 | pathogenic | 0 | 0 | 0 |
| rs754477154 | pathogenic | 0 | 0 | 0 |
| rs747334007 | benign | 0 | 0 | 0 |
| rs121908132 | pathogenic | 0 | 0 | 0 |
| rs104894103 | pathogenic | 1 | 1 | 0 |
| rs140088149 | benign | 0 | 0 | 0 |
| rs149803148 | benign | 0 | 0 | 0 |
| rs11064480 | benign | 0 | 0 | 0 |
| rs74435397 | pathogenic | 1 | 1 | 1 |
| rs587783626 | pathogenic | 1 | 1 | 1 |
| rs148211042 | pathogenic | 1 | 1 | 1 |
| rs17849502 | benign | 0 | 0 | 0 |
| rs13306575 | pathogenic | 1 | 1 | 1 |
| rs201426733 | benign | 0 | 0 | 0 |
| rs182278784 | benign | 1 | 1 | 1 |
| rs77786415 | benign | 1 | 1 | 1 |
| rs200192119 | benign | 1 | 1 | 1 |
| rs12877501 | benign | 1 | 1 | 1 |
| rs77985244 | benign | 1 | 1 | 1 |
| rs5907 | pathogenic | 1 | 1 | 0 |
| rs2305178 | benign | 0 | 0 | 0 |
| rs3135867 | benign | 0 | 0 | 0 |
| rs121913482 | pathogenic | 1 | 1 | 1 |
| rs121913483 | pathogenic | 1 | 1 | 1 |
| rs4647924 | pathogenic | 1 | 1 | 1 |
| rs121913111 | pathogenic | 1 | 1 | 1 |
| rs104894002 | pathogenic | 0 | 0 | 0 |
| rs104894001 | pathogenic | 0 | 0 | 0 |
| rs75932628 | benign | 0 | 0 | 0 |
| rs201258663 | pathogenic | 0 | 0 | 0 |
| rs104893998 | pathogenic | 0 | 0 | 0 |
| rs2234253 | benign | 1 | 1 | 0 |
| rs121908402 | pathogenic | 1 | 1 | 0 |
| rs1805389 | benign | 0 | 0 | 0 |
| rs1805388 | benign | 0 | 0 | 0 |
| rs748385144 | pathogenic | 0 | 0 | 0 |
| rs104894420 | pathogenic | 0 | 0 | 0 |
| rs104894418 | pathogenic | 0 | 0 | 0 |
| rs61731910 | benign | 0 | 0 | 0 |
| rs758723288 | pathogenic | 1 | 1 | 0 |
| rs201865375 | pathogenic | 1 | 1 | 0 |
| rs565090080 | pathogenic | 1 | 1 | 1 |
| rs113042166 | benign | 0 | 0 | 0 |
| rs61736326 | benign | 1 | 1 | 1 |
| rs12030699 | benign | 1 | 1 | 1 |
| rs121913578 | pathogenic | 1 | 1 | 1 |
| rs61739582 | benign | 0 | 0 | 0 |
| rs745983207 | pathogenic | 0 | 0 | 0 |
| rs763524380 | benign | 0 | 0 | 0 |
| rs121912792 | pathogenic | 0 | 0 | 0 |

| | | | | |
|---|---|---|---|---|
| rs121912793 | pathogenic | 0 | 0 | 0 |
| rs387906599 | pathogenic | 0 | 0 | 0 |
| rs121912794 | pathogenic | 0 | 1 | 0 |
| rs121912795 | pathogenic | 0 | 0 | 0 |
| rs121912796 | pathogenic | 1 | 1 | 0 |
| rs121912797 | pathogenic | 0 | 1 | 0 |
| rs121912798 | pathogenic | 1 | 1 | 1 |
| rs80359812 | pathogenic | 1 | 1 | 0 |
| rs776583130 | benign | 1 | 1 | 0 |
| rs80359814 | pathogenic | 1 | 1 | 0 |
| rs202060209 | pathogenic | 1 | 1 | 0 |
| rs267607061 | pathogenic | 1 | 1 | 0 |
| rs267607060 | pathogenic | 0 | 0 | 0 |
| rs80359816 | pathogenic | 1 | 1 | 0 |
| rs80359818 | pathogenic | 1 | 1 | 0 |
| rs387907313 | pathogenic | 0 | 0 | 0 |
| rs587784395 | benign | 0 | 0 | 0 |
| rs5811 | benign | 0 | 0 | 0 |
| rs121909740 | pathogenic | 1 | 1 | 0 |
| rs148518827 | benign | 1 | 1 | 0 |
| rs121909739 | pathogenic | 1 | 1 | 0 |
| rs780528770 | benign | 1 | 1 | 0 |
| rs13306758 | pathogenic | 0 | 0 | 0 |
| rs267607059 | pathogenic | 0 | 0 | 0 |
| rs61730638 | benign | 0 | 0 | 0 |
| rs104894533 | pathogenic | 0 | 0 | 0 |
| rs104894527 | pathogenic | 0 | 0 | 0 |
| rs200503569 | pathogenic | 1 | 1 | 1 |
| rs80338700 | pathogenic | 1 | 1 | 1 |
| rs765433263 | pathogenic | 1 | 1 | 1 |
| rs104894530 | pathogenic | 1 | 1 | 1 |
| rs80338701 | pathogenic | 1 | 1 | 1 |
| rs141498002 | pathogenic | 1 | 1 | 1 |
| rs104894525 | pathogenic | 0 | 0 | 0 |
| rs80338703 | pathogenic | 0 | 0 | 0 |
| rs28936415 | pathogenic | 1 | 1 | 0 |
| rs150719105 | pathogenic | 0 | 0 | 0 |
| rs148032587 | pathogenic | 0 | 0 | 0 |
| rs150577656 | pathogenic | 0 | 0 | 0 |
| rs190521996 | pathogenic | 0 | 0 | 0 |
| rs104894526 | pathogenic | 1 | 1 | 1 |
| rs80338704 | pathogenic | 0 | 0 | 0 |
| rs199562225 | pathogenic | 1 | 1 | 1 |
| rs532870929 | pathogenic | 0 | 0 | 0 |
| rs398123309 | pathogenic | 0 | 0 | 0 |
| rs78290141 | pathogenic | 0 | 0 | 0 |
| rs80338705 | pathogenic | 0 | 0 | 0 |
| rs104894531 | pathogenic | 1 | 1 | 1 |
| rs80338706 | pathogenic | 0 | 0 | 0 |
| rs80338707 | pathogenic | 0 | 0 | 0 |
| rs80338708 | pathogenic | 0 | 0 | 0 |
| rs80338709 | pathogenic | 0 | 0 | 0 |
| rs104893833 | benign | 0 | 0 | 0 |
| rs61744585 | benign | 0 | 0 | 0 |

| | | | | |
|---|---|---|---|---|
| rs73569515 | benign | 0 | 0 | 0 |
| rs398123373 | pathogenic | 0 | 0 | 0 |
| rs756854513 | pathogenic | 0 | 0 | 0 |
| rs58232698 | benign | 0 | 0 | 0 |
| rs9600455 | benign | 0 | 0 | 0 |
| rs201016593 | pathogenic | 0 | 0 | 0 |
| rs756039188 | pathogenic | 0 | 0 | 0 |
| rs121908024 | pathogenic | 0 | 0 | 0 |
| rs373144619 | benign | 0 | 0 | 0 |
| rs138078086 | benign | 0 | 0 | 0 |
| rs376207800 | pathogenic | 0 | 0 | 0 |
| rs749038326 | pathogenic | 1 | 1 | 0 |
| rs372845091 | benign | 0 | 0 | 0 |
| rs377271627 | pathogenic | 0 | 0 | 0 |
| rs563390335 | pathogenic | 0 | 0 | 0 |
| rs13306510 | benign | 0 | 0 | 0 |
| rs764042910 | pathogenic | 0 | 0 | 0 |
| rs201384282 | benign | 1 | 1 | 1 |
| rs730882086 | pathogenic | 1 | 1 | 1 |
| rs756613387 | pathogenic | 1 | 1 | 1 |
| rs121908028 | benign | 1 | 1 | 0 |
| rs773328511 | benign | 0 | 0 | 0 |
| rs749220643 | pathogenic | 1 | 1 | 0 |
| rs730882089 | pathogenic | 0 | 0 | 0 |
| rs146651743 | benign | 0 | 0 | 0 |
| rs148698650 | pathogenic | 0 | 0 | 0 |
| rs140241383 | benign | 0 | 0 | 0 |
| rs757252110 | pathogenic | 0 | 0 | 0 |
| rs13306512 | pathogenic | 1 | 1 | 1 |
| rs121908043 | pathogenic | 1 | 1 | 1 |
| rs773658037 | pathogenic | 0 | 0 | 0 |
| rs774439908 | pathogenic | 0 | 0 | 0 |
| rs779732323 | pathogenic | 1 | 1 | 0 |
| rs749780672 | pathogenic | 0 | 0 | 0 |
| rs375651668 | benign | 1 | 1 | 1 |
| rs193922568 | pathogenic | 1 | 1 | 0 |
| rs750363970 | benign | 1 | 1 | 0 |
| rs5932 | benign | 0 | 0 | 0 |
| rs755154048 | pathogenic | 1 | 1 | 0 |
| rs759876319 | pathogenic | 0 | 0 | 0 |
| rs764550980 | pathogenic | 1 | 1 | 0 |
| rs779940524 | pathogenic | 0 | 0 | 0 |
| rs563382937 | pathogenic | 1 | 1 | 1 |
| rs28942083 | pathogenic | 0 | 1 | 0 |
| rs770744861 | pathogenic | 0 | 1 | 0 |
| rs121908031 | pathogenic | 0 | 1 | 0 |
| rs776217028 | pathogenic | 0 | 1 | 0 |
| rs763625913 | pathogenic | 0 | 0 | 0 |
| rs3736772 | benign | 0 | 0 | 0 |
| rs104894696 | benign | 1 | 1 | 1 |
| rs121918177 | pathogenic | 0 | 1 | 0 |
| rs121918181 | pathogenic | 1 | 1 | 0 |
| rs121918174 | pathogenic | 0 | 1 | 0 |
| rs121918182 | pathogenic | 0 | 0 | 0 |

| | | | | |
|---|---|---|---|---|
| rs121918175 | pathogenic | 1 | 1 | 1 |
| rs121918185 | pathogenic | 1 | 1 | 1 |
| rs121918173 | pathogenic | 1 | 1 | 0 |
| rs121918178 | pathogenic | 1 | 1 | 1 |
| rs121918179 | pathogenic | 1 | 1 | 1 |
| rs121918183 | pathogenic | 0 | 0 | 0 |
| rs121918176 | pathogenic | 1 | 1 | 1 |
| rs121918172 | pathogenic | 0 | 1 | 0 |
| rs121918184 | pathogenic | 0 | 0 | 0 |
| rs121434560 | pathogenic | 1 | 1 | 0 |
| rs121434561 | pathogenic | 1 | 1 | 0 |
| rs3218716 | pathogenic | 1 | 1 | 1 |
| rs145532615 | benign | 0 | 0 | 0 |
| rs149840927 | pathogenic | 0 | 0 | 0 |
| rs730880762 | pathogenic | 0 | 0 | 0 |
| rs730880764 | pathogenic | 0 | 0 | 0 |
| rs730880765 | pathogenic | 0 | 0 | 0 |
| rs587782962 | pathogenic | 0 | 0 | 0 |
| rs141735183 | pathogenic | 0 | 0 | 0 |
| rs397516187 | pathogenic | 0 | 0 | 0 |
| rs727505026 | pathogenic | 0 | 0 | 0 |
| rs529700838 | pathogenic | 0 | 0 | 0 |
| rs540263945 | pathogenic | 0 | 0 | 0 |
| rs149103761 | benign | 0 | 0 | 0 |
| rs730880906 | pathogenic | 0 | 0 | 0 |
| rs730880785 | pathogenic | 0 | 0 | 0 |
| rs397516199 | pathogenic | 0 | 0 | 0 |
| rs730880725 | benign | 0 | 0 | 0 |
| rs730880796 | pathogenic | 0 | 0 | 0 |
| rs201307101 | benign | 0 | 0 | 0 |
| rs397516214 | pathogenic | 0 | 0 | 0 |
| rs3729823 | benign | 0 | 0 | 0 |
| rs397516218 | pathogenic | 0 | 0 | 0 |
| rs121913647 | pathogenic | 0 | 0 | 0 |
| rs373514686 | pathogenic | 0 | 0 | 0 |
| rs763538103 | pathogenic | 0 | 0 | 0 |
| rs121913650 | pathogenic | 0 | 0 | 0 |
| rs139222507 | pathogenic | 0 | 0 | 0 |
| rs369940645 | pathogenic | 0 | 0 | 0 |
| rs763073072 | pathogenic | 0 | 0 | 0 |
| rs145734640 | pathogenic | 0 | 0 | 0 |
| rs730880821 | pathogenic | 0 | 0 | 0 |
| rs28933098 | pathogenic | 0 | 0 | 0 |
| rs372381770 | pathogenic | 0 | 0 | 0 |
| rs146796870 | benign | 0 | 0 | 0 |
| rs104894485 | pathogenic | 1 | 1 | 1 |
| rs35407712 | benign | 0 | 0 | 0 |
| rs35842750 | benign | 0 | 0 | 0 |
| rs2721191 | benign | 0 | 0 | 0 |
| rs35098923 | benign | 0 | 0 | 0 |
| rs34293591 | benign | 1 | 1 | 0 |
| rs36023964 | benign | 0 | 0 | 0 |
| rs4251691 | benign | 0 | 0 | 0 |
| rs34666647 | benign | 0 | 0 | 0 |

| | | | | |
|---|---|---|---|---|
| rs4925828 | benign | 0 | 0 | 0 |
| rs35348691 | benign | 0 | 0 | 0 |
| rs36078464 | benign | 0 | 0 | 0 |
| rs34915097 | benign | 0 | 0 | 0 |
| rs35101495 | benign | 0 | 0 | 0 |
| rs141568342 | pathogenic | 0 | 0 | 0 |
| rs1800571 | pathogenic | 0 | 0 | 0 |
| rs140566084 | benign | 1 | 1 | 1 |
| rs727503791 | pathogenic | 1 | 1 | 0 |
| rs545215807 | pathogenic | 0 | 0 | 0 |
| rs113994168 | pathogenic | 0 | 0 | 0 |
| rs113994167 | pathogenic | 0 | 0 | 0 |
| rs200788251 | pathogenic | 0 | 0 | 0 |
| rs200573371 | pathogenic | 0 | 0 | 0 |
| rs771874163 | pathogenic | 1 | 1 | 1 |
| rs112406105 | pathogenic | 1 | 1 | 1 |
| rs118204015 | pathogenic | 1 | 1 | 1 |
| rs113994169 | pathogenic | 0 | 0 | 0 |
| rs118204018 | pathogenic | 0 | 1 | 0 |
| rs138834083 | pathogenic | 0 | 0 | 0 |
| rs533055438 | pathogenic | 1 | 1 | 1 |
| rs2309689 | pathogenic | 1 | 1 | 1 |
| rs138058572 | pathogenic | 1 | 1 | 1 |
| rs118204017 | pathogenic | 1 | 1 | 0 |
| rs766742117 | pathogenic | 1 | 1 | 1 |
| rs751995154 | pathogenic | 1 | 1 | 1 |
| rs200771970 | pathogenic | 0 | 0 | 0 |
| rs118204014 | pathogenic | 0 | 0 | 0 |
| rs148584617 | pathogenic | 1 | 1 | 1 |
| rs121912720 | pathogenic | 0 | 0 | 0 |
| rs121912721 | pathogenic | 0 | 0 | 0 |
| rs28929476 | pathogenic | 0 | 0 | 0 |
| rs28931574 | pathogenic | 1 | 1 | 1 |
| rs12718465 | benign | 0 | 0 | 0 |
| rs144689354 | pathogenic | 0 | 0 | 0 |
| rs149095705 | pathogenic | 0 | 0 | 0 |
| rs572839648 | benign | 1 | 1 | 1 |
| rs121909715 | pathogenic | 1 | 1 | 0 |
| rs34077456 | benign | 1 | 1 | 1 |
| rs397517876 | benign | 0 | 0 | 0 |
| rs1064074 | benign | 0 | 0 | 0 |
| rs142751309 | benign | 0 | 0 | 0 |
| rs116275562 | benign | 0 | 0 | 0 |
| rs148095899 | benign | 0 | 0 | 0 |
| rs751838040 | pathogenic | 0 | 0 | 0 |
| rs121434324 | pathogenic | 1 | 1 | 1 |
| rs143367518 | pathogenic | 0 | 0 | 0 |
| rs121434325 | pathogenic | 0 | 0 | 0 |
| rs397515328 | pathogenic | 0 | 0 | 0 |
| rs376039938 | pathogenic | 0 | 0 | 0 |
| rs387907243 | pathogenic | 0 | 0 | 0 |
| rs28934876 | pathogenic | 0 | 0 | 0 |
| rs387907245 | pathogenic | 0 | 0 | 0 |
| rs138544311 | benign | 0 | 0 | 0 |

| | | | | |
|---|---|---|---|---|
| rs643788 | benign | 0 | 0 | 0 |
| rs749977756 | benign | 0 | 0 | 0 |
| rs1800574 | benign | 0 | 0 | 0 |
| rs137853336 | pathogenic | 1 | 1 | 1 |
| rs111916034 | benign | 0 | 0 | 0 |
| rs307826 | benign | 0 | 0 | 0 |
| rs35874891 | benign | 0 | 0 | 0 |
| rs111033580 | pathogenic | 1 | 1 | 1 |
| rs111033581 | pathogenic | 1 | 1 | 1 |
| rs121908373 | pathogenic | 0 | 0 | 0 |
| rs767122713 | pathogenic | 0 | 0 | 0 |
| rs2233213 | benign | 0 | 0 | 0 |
| rs121908605 | pathogenic | 1 | 1 | 0 |
| rs11887534 | benign | 0 | 0 | 0 |
| rs4148211 | benign | 0 | 0 | 0 |
| rs80025980 | benign | 0 | 0 | 0 |
| rs137854891 | pathogenic | 1 | 1 | 0 |
| rs74358901 | benign | 0 | 0 | 0 |
| rs9282574 | benign | 0 | 0 | 0 |
| rs137852993 | pathogenic | 0 | 0 | 0 |
| rs137852990 | pathogenic | 0 | 0 | 0 |
| rs137852987 | pathogenic | 0 | 0 | 0 |
| rs4148217 | benign | 0 | 0 | 0 |
| rs140690030 | benign | 0 | 0 | 0 |
| rs137852992 | pathogenic | 0 | 0 | 0 |
| rs6544718 | benign | 0 | 0 | 0 |
| rs137852989 | pathogenic | 0 | 0 | 0 |
| rs150893072 | benign | 1 | 1 | 1 |
| rs28931613 | pathogenic | 1 | 1 | 1 |
| rs143789597 | benign | 1 | 1 | 1 |
| rs121913037 | pathogenic | 1 | 1 | 0 |
| rs188802138 | pathogenic | 1 | 1 | 0 |
| rs121913038 | pathogenic | 1 | 1 | 0 |
| rs146835206 | benign | 0 | 0 | 0 |
| rs121913041 | pathogenic | 1 | 1 | 1 |
| rs121913039 | pathogenic | 1 | 1 | 1 |
| rs149977726 | pathogenic | 1 | 1 | 0 |
| rs121913036 | pathogenic | 0 | 0 | 0 |
| rs121913040 | pathogenic | 0 | 0 | 0 |
| rs765007158 | benign | 0 | 0 | 0 |
| rs112723255 | benign | 0 | 0 | 0 |
| rs574164748 | benign | 0 | 0 | 0 |
| rs80338734 | pathogenic | 1 | 1 | 0 |
| rs104894694 | pathogenic | 1 | 1 | 0 |
| rs753198836 | pathogenic | 0 | 0 | 0 |
| rs760101382 | pathogenic | 0 | 0 | 0 |
| rs121909272 | pathogenic | 1 | 1 | 0 |
| rs370421531 | pathogenic | 0 | 0 | 0 |
| rs565109128 | benign | 0 | 0 | 0 |
| rs17851582 | benign | 0 | 0 | 0 |
| rs753228876 | benign | 0 | 0 | 0 |
| rs141471799 | benign | 0 | 0 | 0 |
| rs281864916 | pathogenic | 1 | 1 | 0 |
| rs121918409 | pathogenic | 1 | 1 | 1 |

| | | | | |
|---|---|---|---|---|
| rs121918413 | pathogenic | 1 | 1 | 1 |
| rs121918411 | pathogenic | 1 | 1 | 1 |
| rs121918408 | pathogenic | 1 | 1 | 1 |
| rs121918412 | pathogenic | 1 | 1 | 1 |
| rs121918410 | pathogenic | 1 | 1 | 1 |
| rs116474260 | benign | 1 | 1 | 1 |
| rs281864919 | pathogenic | 1 | 1 | 1 |
| rs397514452 | pathogenic | 1 | 1 | 0 |
| rs747843638 | benign | 0 | 0 | 0 |
| rs774357609 | pathogenic | 1 | 1 | 1 |
| rs201804005 | benign | 0 | 0 | 0 |
| rs746485916 | pathogenic | 0 | 0 | 0 |
| rs764131178 | pathogenic | 0 | 0 | 0 |
| rs374094065 | pathogenic | 0 | 0 | 0 |
| rs76151636 | pathogenic | 1 | 1 | 0 |
| rs753250853 | pathogenic | 0 | 0 | 0 |
| rs761632029 | pathogenic | 0 | 0 | 0 |
| rs775541743 | pathogenic | 0 | 0 | 0 |
| rs778749563 | pathogenic | 0 | 0 | 0 |
| rs60431989 | pathogenic | 1 | 1 | 0 |
| rs755554442 | pathogenic | 1 | 1 | 0 |
| rs749085322 | pathogenic | 0 | 0 | 0 |
| rs756029120 | pathogenic | 1 | 1 | 0 |
| rs137853279 | pathogenic | 1 | 1 | 0 |
| rs777354267 | pathogenic | 1 | 1 | 1 |
| rs142940486 | benign | 1 | 1 | 1 |
| rs140167393 | benign | 1 | 1 | 1 |
| rs2066827 | benign | 0 | 0 | 0 |
| rs765681672 | benign | 0 | 0 | 0 |
| rs137985549 | benign | 0 | 0 | 0 |
| rs74315340 | pathogenic | 1 | 1 | 1 |
| rs74315341 | pathogenic | 0 | 0 | 0 |
| rs74315335 | pathogenic | 0 | 0 | 0 |
| rs61730974 | benign | 0 | 0 | 0 |
| rs121909193 | pathogenic | 0 | 0 | 0 |
| rs74315334 | pathogenic | 0 | 0 | 0 |
| rs74315329 | pathogenic | 0 | 0 | 0 |
| rs74315330 | pathogenic | 0 | 0 | 0 |
| rs121909194 | pathogenic | 0 | 0 | 0 |
| rs28936694 | pathogenic | 0 | 0 | 0 |
| rs74315336 | pathogenic | 0 | 0 | 0 |
| rs74315338 | pathogenic | 1 | 1 | 1 |
| rs74315328 | pathogenic | 0 | 0 | 0 |
| rs74315331 | pathogenic | 0 | 0 | 0 |
| rs74315332 | pathogenic | 0 | 0 | 0 |
| rs141696651 | benign | 0 | 0 | 0 |
| rs7041 | benign | 0 | 0 | 0 |
| rs4588 | benign | 0 | 0 | 0 |
| rs121908981 | pathogenic | 1 | 1 | 1 |
| rs606231459 | pathogenic | 0 | 0 | 0 |
| rs398123053 | pathogenic | 0 | 0 | 0 |
| rs774259802 | benign | 0 | 0 | 0 |
| rs121909582 | pathogenic | 1 | 1 | 1 |
| rs104893680 | pathogenic | 1 | 1 | 1 |

| | | | | |
|---|---|---|---|---|
| rs137854907 | pathogenic | 0 | 0 | 0 |
| rs765715798 | pathogenic | 0 | 0 | 0 |
| rs104893678 | pathogenic | 0 | 0 | 0 |
| rs104893679 | pathogenic | 0 | 1 | 0 |
| rs104893681 | pathogenic | 1 | 1 | 0 |
| rs121909640 | pathogenic | 1 | 1 | 0 |
| rs780705654 | pathogenic | 0 | 0 | 0 |
| rs201672146 | benign | 1 | 1 | 0 |
| rs12136658 | benign | 0 | 0 | 0 |
| rs56127440 | benign | 0 | 0 | 0 |
| rs104894433 | pathogenic | 1 | 1 | 1 |
| rs104894435 | pathogenic | 0 | 1 | 0 |
| rs104894437 | pathogenic | 1 | 1 | 1 |
| rs104894441 | pathogenic | 1 | 1 | 1 |
| rs104894440 | pathogenic | 1 | 1 | 1 |
| rs104894445 | pathogenic | 1 | 1 | 1 |
| rs104894436 | pathogenic | 0 | 0 | 0 |
| rs104894438 | pathogenic | 0 | 0 | 0 |
| rs200891969 | pathogenic | 1 | 1 | 0 |
| rs104894443 | pathogenic | 1 | 1 | 1 |
| rs104894434 | pathogenic | 1 | 1 | 1 |
| rs41298442 | pathogenic | 1 | 1 | 1 |
| rs104894442 | pathogenic | 1 | 1 | 1 |
| rs35453854 | benign | 0 | 0 | 0 |
| rs121912898 | pathogenic | 0 | 0 | 0 |
| rs121912897 | pathogenic | 1 | 1 | 0 |
| rs138260625 | benign | 0 | 0 | 0 |
| rs147802438 | benign | 1 | 1 | 0 |
| rs146443310 | benign | 1 | 1 | 0 |
| rs747319628 | pathogenic | 1 | 1 | 0 |
| rs1801195 | benign | 0 | 0 | 0 |
| rs145045341 | benign | 1 | 1 | 0 |
| rs116001173 | benign | 1 | 1 | 0 |
| rs587784175 | pathogenic | 0 | 0 | 0 |
| rs104893857 | pathogenic | 0 | 0 | 0 |
| rs104893862 | pathogenic | 1 | 1 | 0 |
| rs104893858 | pathogenic | 1 | 1 | 1 |
| rs121909249 | pathogenic | 0 | 1 | 0 |
| rs121909248 | pathogenic | 1 | 1 | 1 |
| rs104893859 | pathogenic | 1 | 1 | 0 |
| rs187063864 | pathogenic | 1 | 1 | 0 |
| rs121917817 | pathogenic | 1 | 1 | 0 |
| rs121917818 | pathogenic | 0 | 0 | 0 |
| rs753594601 | pathogenic | 0 | 0 | 0 |
| rs28937869 | pathogenic | 1 | 1 | 0 |
| rs6165 | benign | 0 | 0 | 0 |
| rs200089714 | pathogenic | 1 | 1 | 0 |
| rs372000714 | pathogenic | 0 | 0 | 0 |
| rs2228206 | benign | 0 | 0 | 0 |
| rs560096 | benign | 0 | 0 | 0 |
| rs137852666 | pathogenic | 0 | 0 | 0 |
| rs10896380 | benign | 0 | 0 | 0 |
| rs773690764 | pathogenic | 0 | 0 | 0 |
| rs201060167 | pathogenic | 0 | 0 | 0 |

| | | | | |
|---|---|---|---|---|
| rs137852670 | pathogenic | 0 | 0 | 0 |
| rs780594709 | pathogenic | 0 | 0 | 0 |
| rs145226920 | pathogenic | 0 | 0 | 0 |
| rs137852665 | pathogenic | 0 | 0 | 0 |
| rs724159960 | pathogenic | 0 | 0 | 0 |
| rs756985703 | pathogenic | 0 | 0 | 0 |
| rs137852667 | pathogenic | 0 | 0 | 0 |
| rs151079750 | pathogenic | 1 | 1 | 0 |
| rs2306636 | benign | 0 | 0 | 0 |
| rs72647885 | benign | 1 | 1 | 0 |
| rs377584435 | pathogenic | 1 | 1 | 1 |
| rs757583846 | pathogenic | 1 | 1 | 0 |
| rs121918148 | pathogenic | 0 | 0 | 0 |
| rs149201802 | benign | 0 | 0 | 0 |
| rs587781904 | pathogenic | 1 | 1 | 1 |
| rs143421325 | benign | 0 | 0 | 0 |
| rs587781253 | pathogenic | 0 | 0 | 0 |
| rs587780331 | benign | 0 | 0 | 0 |
| rs151001016 | benign | 0 | 0 | 0 |
| rs150888094 | benign | 0 | 0 | 0 |
| rs149581331 | benign | 0 | 0 | 0 |
| rs774840535 | benign | 0 | 0 | 0 |
| rs138428684 | benign | 0 | 0 | 0 |
| rs149300055 | benign | 0 | 0 | 0 |
| rs571005578 | benign | 0 | 0 | 0 |
| rs200731839 | benign | 0 | 0 | 0 |
| rs121965007 | pathogenic | 0 | 0 | 0 |
| rs121965013 | pathogenic | 0 | 1 | 0 |
| rs121965009 | pathogenic | 0 | 0 | 0 |
| rs1800457 | benign | 1 | 1 | 0 |
| rs121965006 | pathogenic | 1 | 1 | 0 |
| rs121965008 | pathogenic | 0 | 0 | 0 |
| rs61732609 | pathogenic | 0 | 0 | 0 |
| rs121965011 | pathogenic | 0 | 0 | 0 |
| rs121965015 | pathogenic | 0 | 0 | 0 |
| rs121965018 | pathogenic | 1 | 1 | 0 |
| rs121965016 | pathogenic | 0 | 0 | 0 |
| rs77806 | benign | 0 | 0 | 0 |
| rs149731356 | benign | 0 | 0 | 0 |
| rs72549407 | pathogenic | 0 | 0 | 0 |
| rs200363929 | benign | 0 | 0 | 0 |
| rs376403708 | benign | 1 | 1 | 0 |
| rs56376197 | benign | 1 | 1 | 1 |
| rs200237973 | benign | 1 | 1 | 1 |
| rs548330075 | benign | 0 | 0 | 0 |
| rs148291350 | benign | 0 | 0 | 0 |
| rs71579353 | pathogenic | 0 | 0 | 0 |
| rs146278697 | benign | 0 | 0 | 0 |
| rs141033098 | benign | 0 | 0 | 0 |
| rs397517234 | pathogenic | 0 | 0 | 0 |
| rs71579374 | benign | 0 | 0 | 0 |
| rs140381835 | benign | 0 | 0 | 0 |
| rs137852702 | pathogenic | 0 | 0 | 0 |
| rs137852696 | pathogenic | 1 | 1 | 1 |

| | | | | |
|---|---|---|---|---|
| rs137852697 | pathogenic | 0 | 0 | 0 |
| rs386833639 | pathogenic | 0 | 0 | 0 |
| rs386833640 | pathogenic | 0 | 0 | 0 |
| rs386833641 | benign | 0 | 0 | 0 |
| rs137852701 | pathogenic | 0 | 0 | 0 |
| rs386833642 | pathogenic | 0 | 0 | 0 |
| rs137852695 | pathogenic | 0 | 0 | 0 |
| rs386833646 | pathogenic | 0 | 0 | 0 |
| rs137852700 | pathogenic | 1 | 1 | 0 |
| rs386833649 | pathogenic | 1 | 1 | 0 |
| rs386833650 | pathogenic | 0 | 0 | 0 |
| rs148412181 | pathogenic | 0 | 0 | 0 |
| rs386833655 | pathogenic | 0 | 0 | 0 |
| rs386833656 | pathogenic | 1 | 1 | 0 |
| rs137852698 | pathogenic | 0 | 0 | 0 |
| rs386833661 | pathogenic | 0 | 0 | 0 |
| rs386833670 | pathogenic | 0 | 0 | 0 |
| rs111033632 | pathogenic | 1 | 1 | 1 |
| rs34308573 | benign | 1 | 1 | 1 |
| rs376048533 | pathogenic | 1 | 1 | 0 |
| rs3747517 | benign | 1 | 1 | 0 |
| rs137853328 | pathogenic | 0 | 0 | 0 |
| rs77273740 | benign | 0 | 0 | 0 |
| rs267606760 | pathogenic | 0 | 0 | 0 |
| rs77576840 | pathogenic | 0 | 0 | 0 |
| rs45465204 | benign | 1 | 1 | 0 |
| rs5320 | benign | 0 | 0 | 0 |
| rs5324 | benign | 0 | 0 | 0 |
| rs4531 | benign | 0 | 0 | 0 |
| rs267606761 | pathogenic | 0 | 0 | 0 |
| rs75215331 | pathogenic | 0 | 0 | 0 |
| rs6271 | benign | 0 | 0 | 0 |
| rs201643277 | pathogenic | 0 | 0 | 0 |
| rs4252250 | benign | 0 | 0 | 0 |
| rs137853580 | pathogenic | 0 | 0 | 0 |
| rs368287711 | pathogenic | 0 | 0 | 0 |
| rs137853579 | pathogenic | 0 | 0 | 0 |
| rs3135932 | benign | 0 | 0 | 0 |
| rs2228055 | benign | 0 | 0 | 0 |
| rs140322345 | benign | 1 | 1 | 1 |
| rs1154510 | pathogenic | 0 | 0 | 0 |
| rs11833399 | benign | 0 | 0 | 0 |
| rs36023382 | benign | 1 | 1 | 1 |
| rs61754444 | benign | 0 | 0 | 0 |
| rs28935168 | pathogenic | 0 | 0 | 0 |
| rs28934907 | pathogenic | 0 | 0 | 0 |
| rs28934904 | pathogenic | 1 | 1 | 0 |
| rs61748392 | pathogenic | 1 | 1 | 0 |
| rs28934908 | pathogenic | 0 | 0 | 0 |
| rs28934905 | pathogenic | 0 | 0 | 0 |
| rs28934906 | pathogenic | 1 | 1 | 0 |
| rs606231307 | pathogenic | 0 | 0 | 0 |
| rs9333296 | benign | 0 | 0 | 0 |
| rs3218614 | benign | 0 | 0 | 0 |

| | | | | |
|---|---|---|---|---|
| rs2230792 | benign | 0 | 0 | 0 |
| rs2230793 | benign | 0 | 0 | 0 |
| rs2230794 | benign | 0 | 0 | 0 |
| rs749052963 | pathogenic | 0 | 0 | 0 |
| rs2230798 | benign | 0 | 0 | 0 |
| rs2230795 | benign | 0 | 0 | 0 |
| rs3204145 | benign | 0 | 0 | 0 |
| rs1538660 | benign | 0 | 0 | 0 |
| rs9878928 | benign | 0 | 0 | 0 |
| rs104893742 | pathogenic | 0 | 0 | 0 |
| rs28936702 | pathogenic | 0 | 0 | 0 |
| rs28936703 | pathogenic | 0 | 0 | 0 |
| rs769392 | benign | 0 | 0 | 0 |
| rs769402 | benign | 1 | 1 | 1 |
| rs104894884 | pathogenic | 1 | 1 | 1 |
| rs1801316 | benign | 1 | 1 | 1 |
| rs104894885 | pathogenic | 1 | 1 | 1 |
| rs782572 | benign | 0 | 0 | 0 |
| rs151166046 | pathogenic | 0 | 0 | 0 |
| rs772153760 | pathogenic | 1 | 1 | 0 |
| rs35405862 | benign | 0 | 0 | 0 |
| rs146571352 | pathogenic | 0 | 1 | 0 |
| rs374698153 | pathogenic | 0 | 0 | 0 |
| rs7594497 | benign | 0 | 0 | 0 |
| rs80338716 | pathogenic | 0 | 0 | 0 |
| rs80338719 | pathogenic | 1 | 1 | 1 |
| rs80338721 | pathogenic | 0 | 0 | 0 |
| rs730882249 | pathogenic | 0 | 0 | 0 |
| rs146262009 | pathogenic | 0 | 0 | 0 |
| rs104895083 | pathogenic | 1 | 1 | 1 |
| rs104895100 | benign | 1 | 1 | 1 |
| rs104895085 | pathogenic | 0 | 0 | 0 |
| rs28940580 | pathogenic | 0 | 0 | 0 |
| rs104895098 | pathogenic | 0 | 0 | 0 |
| rs28940578 | pathogenic | 0 | 0 | 0 |
| rs61752717 | pathogenic | 0 | 0 | 0 |
| rs28940579 | pathogenic | 0 | 0 | 0 |
| rs104895097 | pathogenic | 0 | 0 | 0 |
| rs140425651 | benign | 1 | 1 | 0 |
| rs768090444 | pathogenic | 0 | 0 | 0 |
| rs764459544 | pathogenic | 0 | 0 | 0 |
| rs80338802 | pathogenic | 1 | 1 | 1 |
| rs149969786 | pathogenic | 1 | 1 | 0 |
| rs115311625 | pathogenic | 1 | 1 | 1 |
| rs778768583 | pathogenic | 1 | 1 | 1 |
| rs892090 | benign | 0 | 0 | 0 |
| rs28929498 | pathogenic | 1 | 1 | 1 |
| rs137854467 | pathogenic | 0 | 0 | 0 |
| rs137854480 | pathogenic | 0 | 0 | 0 |
| rs148682262 | benign | 0 | 0 | 0 |
| rs137852744 | pathogenic | 0 | 0 | 0 |
| rs137852745 | pathogenic | 0 | 1 | 0 |
| rs137852749 | pathogenic | 0 | 1 | 0 |
| rs137852746 | pathogenic | 0 | 1 | 0 |

| | | | | |
|---|---|---|---|---|
| rs145633958 | benign | 0 | 0 | 0 |
| rs141679950 | benign | 0 | 0 | 0 |
| rs76397662 | benign | 1 | 1 | 1 |
| rs370736139 | benign | 0 | 0 | 0 |
| rs781750106 | benign | 0 | 0 | 0 |
| rs546866208 | benign | 0 | 0 | 0 |
| rs142338976 | benign | 0 | 0 | 0 |
| rs115272158 | benign | 0 | 0 | 0 |
| rs115423919 | benign | 0 | 0 | 0 |
| rs537874538 | benign | 0 | 0 | 0 |
| rs9282834 | benign | 0 | 0 | 0 |
| rs201745826 | benign | 0 | 0 | 0 |
| rs75873440 | pathogenic | 0 | 0 | 0 |
| rs147219360 | benign | 0 | 0 | 0 |
| rs77939446 | pathogenic | 0 | 0 | 0 |
| rs77558292 | pathogenic | 0 | 0 | 0 |
| rs80069458 | pathogenic | 0 | 0 | 0 |
| rs76262710 | pathogenic | 0 | 0 | 0 |
| rs79781594 | pathogenic | 0 | 0 | 0 |
| rs77316810 | pathogenic | 0 | 0 | 0 |
| rs77503355 | pathogenic | 0 | 0 | 0 |
| rs79890926 | pathogenic | 0 | 0 | 0 |
| rs377767402 | benign | 0 | 0 | 0 |
| rs121913308 | pathogenic | 0 | 0 | 0 |
| rs55846256 | benign | 0 | 0 | 0 |
| rs75076352 | pathogenic | 0 | 0 | 0 |
| rs75996173 | pathogenic | 0 | 0 | 0 |
| rs77709286 | pathogenic | 0 | 0 | 0 |
| rs151058706 | benign | 0 | 0 | 0 |
| rs138964358 | benign | 0 | 0 | 0 |
| rs137852353 | pathogenic | 1 | 1 | 0 |
| rs201196020 | benign | 0 | 0 | 0 |
| rs137853201 | pathogenic | 1 | 1 | 1 |
| rs113994174 | pathogenic | 1 | 1 | 1 |
| rs104893674 | pathogenic | 1 | 1 | 1 |
| rs779519249 | benign | 1 | 1 | 1 |
| rs9606756 | benign | 0 | 0 | 0 |
| rs75680863 | benign | 1 | 1 | 1 |
| rs35838082 | benign | 0 | 0 | 0 |
| rs1801198 | benign | 0 | 0 | 0 |
| rs117458738 | benign | 1 | 1 | 0 |
| rs9621049 | benign | 1 | 1 | 0 |
| rs1131603 | benign | 1 | 1 | 0 |
| rs4820889 | benign | 0 | 0 | 0 |
| rs777767978 | pathogenic | 1 | 1 | 0 |
| rs147399860 | benign | 1 | 1 | 1 |
| rs104893950 | pathogenic | 0 | 0 | 0 |
| rs777308612 | benign | 0 | 0 | 0 |
| rs151091410 | benign | 0 | 0 | 0 |
| rs11537667 | benign | 0 | 0 | 0 |
| rs267606621 | pathogenic | 1 | 1 | 0 |
| rs143058902 | benign | 0 | 0 | 0 |
| rs370261904 | pathogenic | 0 | 0 | 0 |
| rs141169746 | benign | 0 | 0 | 0 |

| | | | | |
|---|---|---|---|---|
| rs111033436 | benign | 0 | 0 | 0 |
| rs148533341 | benign | 0 | 0 | 0 |
| rs143827620 | benign | 0 | 0 | 0 |
| rs4935502 | benign | 0 | 0 | 0 |
| rs61735473 | benign | 0 | 0 | 0 |
| rs267606932 | pathogenic | 1 | 1 | 0 |
| rs61730754 | benign | 0 | 0 | 0 |
| rs374491359 | pathogenic | 1 | 1 | 1 |
| rs761800077 | benign | 1 | 1 | 1 |
| rs760391381 | benign | 1 | 1 | 1 |
| rs104893885 | pathogenic | 0 | 0 | 0 |
| rs104893886 | pathogenic | 1 | 1 | 1 |
| rs104893884 | pathogenic | 0 | 0 | 0 |
| rs147715509 | benign | 0 | 0 | 0 |
| rs115533243 | benign | 0 | 0 | 0 |
| rs397509390 | pathogenic | 0 | 1 | 0 |
| rs80357134 | pathogenic | 0 | 0 | 0 |
| rs766004110 | benign | 1 | 1 | 1 |
| rs769650474 | benign | 0 | 0 | 0 |
| rs398122635 | benign | 1 | 1 | 0 |
| rs199522616 | pathogenic | 0 | 0 | 0 |
| rs28897672 | pathogenic | 1 | 1 | 0 |
| rs80357086 | pathogenic | 1 | 1 | 1 |
| rs80357064 | pathogenic | 1 | 1 | 0 |
| rs80356936 | benign | 0 | 0 | 0 |
| rs28897673 | benign | 0 | 0 | 0 |
| rs146747324 | benign | 0 | 0 | 0 |
| rs78635798 | pathogenic | 0 | 0 | 0 |
| rs75146158 | pathogenic | 1 | 1 | 1 |
| rs587782886 | benign | 0 | 0 | 0 |
| rs121434634 | pathogenic | 0 | 0 | 0 |
| rs121434635 | pathogenic | 0 | 0 | 0 |
| rs587777778 | pathogenic | 0 | 0 | 0 |
| rs118203967 | pathogenic | 0 | 0 | 0 |
| rs751375244 | pathogenic | 1 | 1 | 1 |
| rs104893863 | pathogenic | 1 | 1 | 0 |
| rs104893865 | pathogenic | 0 | 0 | 0 |
| rs104893864 | pathogenic | 1 | 1 | 1 |
| rs104893866 | pathogenic | 1 | 1 | 0 |
| rs561738497 | benign | 0 | 0 | 0 |
| rs57599352 | pathogenic | 1 | 1 | 1 |
| rs267607448 | pathogenic | 1 | 1 | 1 |
| rs63751596 | pathogenic | 0 | 0 | 0 |
| rs63751612 | benign | 0 | 0 | 0 |
| rs63751684 | benign | 0 | 0 | 0 |
| rs267607861 | benign | 0 | 0 | 0 |
| rs35831931 | benign | 0 | 0 | 0 |
| rs63750876 | benign | 0 | 0 | 0 |
| rs41295284 | benign | 0 | 0 | 0 |
| rs35502531 | benign | 0 | 0 | 0 |
| rs63750449 | benign | 0 | 0 | 0 |
| rs63750899 | pathogenic | 0 | 0 | 0 |
| rs55907433 | benign | 0 | 0 | 0 |
| rs63750217 | pathogenic | 0 | 0 | 0 |

| | | | | |
|---|---|---|---|---|
| rs63751275 | pathogenic | 0 | 0 | 0 |
| rs63750702 | benign | 1 | 1 | 1 |
| rs63750114 | benign | 0 | 0 | 0 |
| rs2020873 | benign | 0 | 0 | 0 |
| rs140195825 | benign | 1 | 1 | 1 |
| rs182763805 | benign | 0 | 0 | 0 |
| rs533696377 | benign | 0 | 0 | 0 |
| rs41278788 | benign | 0 | 0 | 0 |
| rs104894193 | pathogenic | 1 | 1 | 0 |
| rs3824915 | benign | 1 | 1 | 1 |
| rs104894196 | pathogenic | 0 | 0 | 0 |
| rs78681604 | benign | 1 | 1 | 1 |
| rs77321498 | benign | 1 | 1 | 1 |
| rs137852965 | pathogenic | 0 | 0 | 0 |
| rs137852961 | pathogenic | 0 | 0 | 0 |
| rs137852962 | pathogenic | 1 | 1 | 0 |
| rs753376100 | pathogenic | 0 | 0 | 0 |
| rs137852968 | pathogenic | 1 | 1 | 1 |
| rs137852959 | pathogenic | 1 | 1 | 0 |
| rs137852967 | pathogenic | 0 | 0 | 0 |
| rs113437301 | benign | 1 | 1 | 1 |
| rs61747482 | benign | 1 | 1 | 1 |
| rs2234919 | benign | 1 | 1 | 0 |
| rs121908863 | pathogenic | 0 | 0 | 0 |
| rs140455771 | pathogenic | 0 | 0 | 0 |
| rs121908810 | pathogenic | 1 | 1 | 1 |
| rs143879853 | benign | 0 | 0 | 0 |
| rs188319622 | benign | 0 | 0 | 0 |
| rs121918645 | pathogenic | 0 | 0 | 0 |
| rs115352435 | pathogenic | 0 | 0 | 0 |
| rs121434427 | pathogenic | 1 | 1 | 1 |
| rs121434428 | pathogenic | 0 | 0 | 0 |
| rs35086265 | benign | 1 | 1 | 1 |
| rs11576415 | benign | 0 | 0 | 0 |
| rs121434429 | pathogenic | 1 | 1 | 1 |
| rs4074536 | benign | 1 | 1 | 1 |
| rs10801999 | benign | 0 | 0 | 0 |
| rs121434550 | pathogenic | 0 | 0 | 0 |
| rs148057999 | pathogenic | 0 | 0 | 0 |
| rs28730716 | benign | 0 | 0 | 0 |
| rs142036299 | benign | 0 | 0 | 0 |
| rs762381137 | pathogenic | 0 | 0 | 0 |
| rs121434549 | pathogenic | 0 | 0 | 0 |
| rs139228801 | pathogenic | 0 | 0 | 0 |
| rs147219463 | benign | 0 | 0 | 0 |
| rs1131857 | benign | 0 | 0 | 0 |
| rs121917874 | pathogenic | 0 | 1 | 0 |
| rs2228056 | benign | 0 | 0 | 0 |
| rs121917870 | pathogenic | 0 | 0 | 0 |
| rs121917866 | pathogenic | 0 | 0 | 0 |
| rs11921054 | benign | 0 | 0 | 0 |
| rs121917868 | pathogenic | 0 | 0 | 0 |
| rs28931603 | pathogenic | 0 | 0 | 0 |
| rs104895295 | pathogenic | 1 | 1 | 0 |

| | | | | |
|---|---|---|---|---|
| rs104895382 | pathogenic | 0 | 0 | 0 |
| rs104895298 | pathogenic | 0 | 0 | 0 |
| rs104895300 | pathogenic | 0 | 0 | 0 |
| rs104895332 | pathogenic | 0 | 0 | 0 |
| rs758026399 | pathogenic | 1 | 1 | 1 |
| rs104895366 | pathogenic | 0 | 0 | 0 |
| rs104895304 | pathogenic | 0 | 0 | 0 |
| rs759997079 | pathogenic | 0 | 0 | 0 |
| rs121917789 | pathogenic | 0 | 0 | 0 |
| rs104895319 | pathogenic | 0 | 0 | 0 |
| rs104895317 | pathogenic | 1 | 1 | 0 |
| rs28934897 | pathogenic | 0 | 0 | 0 |
| rs104895380 | benign | 0 | 0 | 0 |
| rs104895360 | pathogenic | 0 | 0 | 0 |
| rs28936396 | pathogenic | 1 | 1 | 0 |
| rs121909307 | pathogenic | 1 | 1 | 1 |
| rs28938472 | pathogenic | 1 | 1 | 0 |
| rs121909308 | pathogenic | 1 | 1 | 0 |
| rs121909309 | pathogenic | 0 | 0 | 0 |
| rs121913013 | benign | 1 | 1 | 1 |
| rs201855245 | benign | 1 | 1 | 1 |
| rs2230232 | benign | 1 | 1 | 0 |
| rs113451409 | benign | 0 | 0 | 0 |
| rs121913011 | pathogenic | 1 | 1 | 0 |
| rs185821167 | pathogenic | 1 | 1 | 0 |
| rs2230234 | benign | 0 | 0 | 0 |
| rs193922639 | benign | 1 | 1 | 0 |
| rs774863785 | pathogenic | 1 | 1 | 0 |
| rs576404380 | pathogenic | 1 | 1 | 0 |
| rs2230235 | benign | 1 | 1 | 0 |
| rs200484060 | pathogenic | 1 | 1 | 0 |
| rs199761749 | benign | 0 | 0 | 0 |
| rs199681901 | benign | 0 | 0 | 0 |
| rs121908951 | pathogenic | 0 | 0 | 0 |
| rs769251460 | benign | 0 | 0 | 0 |
| rs28937320 | pathogenic | 0 | 0 | 0 |
| rs200607527 | pathogenic | 0 | 0 | 0 |
| rs137852640 | pathogenic | 1 | 1 | 1 |
| rs137852636 | pathogenic | 1 | 1 | 1 |
| rs137852638 | pathogenic | 0 | 0 | 0 |
| rs142637231 | pathogenic | 0 | 0 | 0 |
| rs151328418 | benign | 0 | 1 | 0 |
| rs137852639 | pathogenic | 1 | 1 | 1 |
| rs34605667 | benign | 0 | 0 | 0 |
| rs104894073 | pathogenic | 1 | 1 | 0 |
| rs56298569 | pathogenic | 0 | 0 | 0 |
| rs757600616 | pathogenic | 0 | 0 | 0 |
| rs139194636 | pathogenic | 0 | 0 | 0 |
| rs369517993 | pathogenic | 1 | 1 | 1 |
| rs138846036 | benign | 1 | 1 | 1 |
| rs200808744 | pathogenic | 1 | 1 | 1 |
| rs2255543 | benign | 0 | 0 | 0 |
| rs149165166 | pathogenic | 0 | 0 | 0 |
| rs544956641 | pathogenic | 1 | 1 | 1 |

| | | | | |
|---|---|---|---|---|
| rs28941783 | pathogenic | 0 | 0 | 0 |
| rs562853291 | pathogenic | 1 | 1 | 1 |
| rs28942100 | pathogenic | 0 | 0 | 0 |
| rs120074174 | pathogenic | 1 | 1 | 1 |
| rs120074171 | pathogenic | 1 | 1 | 1 |
| rs120074173 | pathogenic | 0 | 0 | 0 |
| rs120074172 | pathogenic | 1 | 1 | 1 |
| rs767159114 | pathogenic | 1 | 1 | 1 |
| rs104893657 | pathogenic | 0 | 0 | 0 |
| rs104893656 | pathogenic | 0 | 0 | 0 |
| rs104893659 | pathogenic | 0 | 0 | 0 |
| rs104893658 | pathogenic | 0 | 0 | 0 |
| rs200386310 | pathogenic | 0 | 0 | 0 |
| rs2072712 | benign | 0 | 0 | 0 |
| rs35274867 | benign | 0 | 0 | 0 |
| rs35343172 | benign | 0 | 0 | 0 |
| rs10079250 | benign | 0 | 0 | 0 |
| rs34951517 | benign | 0 | 0 | 0 |
| rs267606695 | pathogenic | 0 | 0 | 0 |
| rs149391728 | benign | 0 | 0 | 0 |
| rs387906598 | pathogenic | 1 | 1 | 0 |
| rs137854578 | pathogenic | 0 | 0 | 0 |
| rs141519952 | benign | 0 | 0 | 0 |
| rs146572883 | benign | 0 | 0 | 0 |
| rs111866410 | benign | 0 | 0 | 0 |
| rs139387758 | benign | 1 | 1 | 1 |
| rs35672074 | benign | 1 | 1 | 1 |
| rs121434322 | pathogenic | 0 | 0 | 0 |
| rs150884181 | benign | 0 | 0 | 0 |
| rs35867471 | benign | 0 | 0 | 0 |
| rs2867802 | benign | 0 | 0 | 0 |
| rs137902889 | benign | 0 | 0 | 0 |
| rs369804864 | pathogenic | 0 | 0 | 0 |
| rs781050795 | pathogenic | 0 | 0 | 0 |
| rs2305799 | benign | 0 | 0 | 0 |
| rs2229531 | benign | 0 | 0 | 0 |
| rs147025508 | benign | 0 | 0 | 0 |
| rs76857106 | pathogenic | 0 | 0 | 0 |
| rs7247284 | benign | 0 | 0 | 0 |
| rs62619782 | benign | 0 | 0 | 0 |
| rs377244188 | pathogenic | 0 | 0 | 0 |
| rs75718910 | pathogenic | 0 | 0 | 0 |
| rs79843600 | pathogenic | 0 | 0 | 0 |
| rs75037667 | pathogenic | 1 | 1 | 1 |
| rs121913006 | pathogenic | 0 | 0 | 0 |
| rs397517925 | pathogenic | 1 | 1 | 1 |
| rs267606831 | pathogenic | 1 | 1 | 1 |
| rs121909667 | pathogenic | 1 | 1 | 1 |
| rs121909669 | pathogenic | 1 | 1 | 1 |
| rs121909668 | pathogenic | 1 | 1 | 1 |
| rs121909671 | pathogenic | 1 | 1 | 1 |
| rs534596036 | benign | 0 | 0 | 0 |
| rs144037629 | pathogenic | 0 | 0 | 0 |
| rs11546406 | benign | 0 | 0 | 0 |

| | | | | |
|---|---|---|---|---|
| rs143697995 | pathogenic | 0 | 0 | 0 |
| rs121908331 | pathogenic | 0 | 0 | 0 |
| rs149377346 | benign | 0 | 0 | 0 |
| rs104894086 | pathogenic | 1 | 1 | 0 |
| rs104894089 | pathogenic | 0 | 0 | 0 |
| rs104894090 | pathogenic | 1 | 1 | 0 |
| rs387907235 | pathogenic | 0 | 0 | 0 |
| rs137852689 | pathogenic | 0 | 0 | 0 |
| rs137852690 | pathogenic | 1 | 1 | 0 |
| rs104894087 | pathogenic | 0 | 0 | 0 |
| rs104894085 | pathogenic | 0 | 0 | 0 |
| rs79383654 | benign | 0 | 0 | 0 |
| rs121434239 | pathogenic | 0 | 0 | 0 |
| rs121434240 | pathogenic | 0 | 0 | 0 |
| rs121434236 | pathogenic | 0 | 0 | 0 |
| rs121434238 | pathogenic | 1 | 1 | 1 |
| rs121918107 | pathogenic | 1 | 1 | 1 |
| rs121918103 | pathogenic | 1 | 1 | 1 |
| rs121918104 | pathogenic | 1 | 1 | 1 |
| rs61747614 | benign | 1 | 1 | 1 |
| rs555329870 | benign | 0 | 0 | 0 |
| rs186094931 | benign | 0 | 0 | 0 |
| rs267606785 | pathogenic | 0 | 0 | 0 |
| rs34474955 | benign | 0 | 0 | 0 |
| rs1126497 | benign | 1 | 1 | 1 |
| rs397514661 | pathogenic | 0 | 0 | 0 |
| rs189732445 | benign | 0 | 0 | 0 |
| rs146685071 | benign | 0 | 0 | 0 |
| rs121908636 | pathogenic | 1 | 1 | 0 |
| rs121908643 | pathogenic | 0 | 0 | 0 |
| rs121908644 | pathogenic | 1 | 1 | 1 |
| rs575001023 | pathogenic | 1 | 1 | 1 |
| rs121908646 | pathogenic | 1 | 1 | 0 |
| rs121908638 | pathogenic | 1 | 1 | 1 |
| rs777828000 | pathogenic | 1 | 1 | 1 |
| rs74923032 | benign | 0 | 0 | 0 |
| rs192838388 | pathogenic | 1 | 1 | 1 |
| rs148918985 | pathogenic | 1 | 1 | 1 |
| rs773540764 | benign | 1 | 1 | 1 |
| rs370595480 | pathogenic | 1 | 1 | 1 |
| rs762387914 | pathogenic | 1 | 1 | 1 |
| rs121908642 | pathogenic | 1 | 1 | 1 |
| rs183276875 | pathogenic | 1 | 1 | 1 |
| rs121908648 | pathogenic | 1 | 1 | 1 |
| rs121908647 | pathogenic | 1 | 1 | 1 |
| rs121908640 | pathogenic | 1 | 1 | 1 |
| rs771937610 | pathogenic | 1 | 1 | 1 |
| rs121908641 | pathogenic | 1 | 1 | 1 |
| rs5745068 | benign | 0 | 0 | 0 |
| rs73155056 | benign | 1 | 1 | 1 |
| rs148788180 | benign | 0 | 0 | 0 |
| rs121918137 | pathogenic | 0 | 0 | 0 |
| rs41269593 | benign | 0 | 0 | 0 |
| rs11159 | benign | 1 | 1 | 0 |

| | | | | |
|---|---|---|---|---|
| rs114733570 | pathogenic | 1 | 1 | 0 |
| rs3211783 | benign | 0 | 0 | 0 |
| rs4762 | benign | 0 | 0 | 0 |
| rs74315283 | pathogenic | 0 | 0 | 0 |
| rs111033559 | pathogenic | 0 | 0 | 0 |
| rs111033560 | pathogenic | 1 | 1 | 1 |
| rs749962743 | benign | 0 | 0 | 0 |
| rs267606986 | pathogenic | 0 | 0 | 0 |
| rs34516635 | benign | 0 | 0 | 0 |
| rs117440569 | benign | 0 | 0 | 0 |
| rs61740877 | benign | 0 | 0 | 0 |
| rs121918352 | pathogenic | 0 | 0 | 0 |
| rs183974372 | benign | 0 | 0 | 0 |
| rs140511204 | benign | 1 | 1 | 0 |
| rs6040067 | benign | 0 | 0 | 0 |
| rs199674138 | benign | 0 | 0 | 0 |
| rs121918350 | pathogenic | 0 | 0 | 0 |
| rs138292988 | benign | 1 | 1 | 1 |
| rs145288947 | benign | 0 | 0 | 0 |
| rs369714529 | benign | 0 | 0 | 0 |
| rs200945419 | benign | 0 | 0 | 0 |
| rs116959874 | benign | 0 | 0 | 0 |
| rs141659620 | pathogenic | 1 | 1 | 0 |
| rs111475461 | benign | 0 | 0 | 0 |
| rs61755320 | pathogenic | 0 | 0 | 0 |
| rs748600162 | pathogenic | 1 | 1 | 1 |
| rs369227537 | pathogenic | 1 | 1 | 1 |
| rs372981030 | pathogenic | 1 | 1 | 1 |
| rs137854486 | pathogenic | 0 | 0 | 0 |
| rs137854487 | pathogenic | 0 | 0 | 0 |
| rs199905054 | pathogenic | 0 | 0 | 0 |
| rs771438170 | pathogenic | 0 | 0 | 0 |
| rs774949844 | pathogenic | 0 | 0 | 0 |
| rs4899145 | benign | 0 | 0 | 0 |
| rs200642525 | benign | 0 | 0 | 0 |
| rs190140598 | pathogenic | 0 | 0 | 0 |
| rs2250889 | benign | 0 | 0 | 0 |
| rs17577 | benign | 0 | 0 | 0 |
| rs142032681 | benign | 0 | 0 | 0 |
| rs144577963 | benign | 0 | 0 | 0 |
| rs61754623 | benign | 0 | 0 | 0 |
| rs139873200 | benign | 0 | 0 | 0 |
| rs532530906 | benign | 0 | 0 | 0 |
| rs145762716 | benign | 0 | 0 | 0 |
| rs34062020 | benign | 0 | 0 | 0 |
| rs35258161 | benign | 0 | 0 | 0 |
| rs149792649 | benign | 0 | 0 | 0 |
| rs78101774 | benign | 0 | 0 | 0 |
| rs766466297 | pathogenic | 0 | 1 | 0 |
| rs140467171 | pathogenic | 0 | 0 | 0 |
| rs141340466 | pathogenic | 0 | 0 | 0 |
| rs201555303 | pathogenic | 0 | 0 | 0 |
| rs143592405 | pathogenic | 0 | 0 | 0 |
| rs781249411 | pathogenic | 0 | 0 | 0 |

| | | | | |
|---|---|---|---|---|
| rs116644127 | benign | 0 | 0 | 0 |
| rs151146863 | benign | 1 | 1 | 1 |
| rs116440799 | benign | 0 | 0 | 0 |
| rs138958351 | benign | 0 | 0 | 0 |
| rs143319354 | benign | 0 | 0 | 0 |
| rs768669208 | pathogenic | 1 | 1 | 1 |
| rs28940289 | pathogenic | 1 | 1 | 0 |
| rs745656120 | pathogenic | 1 | 1 | 0 |
| rs387906987 | pathogenic | 0 | 0 | 0 |
| rs147240777 | benign | 0 | 0 | 0 |
| rs140945833 | benign | 0 | 0 | 0 |
| rs146614981 | benign | 0 | 0 | 0 |
| rs149878562 | benign | 0 | 0 | 0 |
| rs28938168 | pathogenic | 0 | 0 | 0 |
| rs33939927 | pathogenic | 1 | 1 | 1 |
| rs34995376 | pathogenic | 1 | 1 | 1 |
| rs201034481 | pathogenic | 1 | 1 | 1 |
| rs774069983 | benign | 0 | 0 | 0 |
| rs199422226 | pathogenic | 1 | 1 | 0 |
| rs199422225 | pathogenic | 1 | 1 | 1 |
| rs199422224 | pathogenic | 1 | 1 | 0 |
| rs149271416 | pathogenic | 1 | 1 | 0 |
| rs1801318 | benign | 1 | 1 | 1 |
| rs372691318 | pathogenic | 1 | 1 | 1 |
| rs147685849 | benign | 0 | 0 | 0 |
| rs387907199 | pathogenic | 0 | 0 | 0 |
| rs767122069 | pathogenic | 0 | 0 | 0 |
| rs2302427 | benign | 0 | 0 | 0 |
| rs201135441 | benign | 0 | 0 | 0 |
| rs201861847 | pathogenic | 0 | 0 | 0 |
| rs137852952 | pathogenic | 0 | 0 | 0 |
| rs16986560 | benign | 0 | 0 | 0 |
| rs147344332 | pathogenic | 1 | 1 | 1 |
| rs36027220 | pathogenic | 0 | 0 | 0 |
| rs192401229 | benign | 0 | 0 | 0 |
| rs121909106 | pathogenic | 1 | 1 | 0 |
| rs118204103 | pathogenic | 1 | 1 | 0 |
| rs118204104 | pathogenic | 1 | 1 | 0 |
| rs118204105 | pathogenic | 1 | 1 | 0 |
| rs118204106 | pathogenic | 0 | 0 | 0 |
| rs118204119 | pathogenic | 0 | 0 | 0 |
| rs118204107 | pathogenic | 0 | 0 | 0 |
| rs118204094 | pathogenic | 0 | 0 | 0 |
| rs118204098 | pathogenic | 1 | 1 | 0 |
| rs118204101 | pathogenic | 1 | 1 | 0 |
| rs118204096 | pathogenic | 1 | 1 | 0 |
| rs118204108 | pathogenic | 0 | 0 | 0 |
| rs536814318 | pathogenic | 0 | 0 | 0 |
| rs34413634 | pathogenic | 1 | 1 | 0 |
| rs118204100 | pathogenic | 0 | 0 | 0 |
| rs118204109 | pathogenic | 0 | 0 | 0 |
| rs118204116 | pathogenic | 1 | 1 | 0 |
| rs118204110 | pathogenic | 0 | 0 | 0 |
| rs118204099 | pathogenic | 0 | 0 | 0 |

| | | | | |
|---|---|---|---|---|
| rs118204111 | pathogenic | 0 | 0 | 0 |
| rs118204112 | pathogenic | 0 | 0 | 0 |
| rs118204113 | pathogenic | 0 | 0 | 0 |
| rs118204114 | pathogenic | 0 | 0 | 0 |
| rs118204115 | pathogenic | 0 | 0 | 0 |
| rs150428209 | benign | 0 | 0 | 0 |
| rs144949995 | benign | 0 | 0 | 0 |
| rs190023981 | benign | 0 | 0 | 0 |
| rs104894611 | pathogenic | 1 | 1 | 1 |
| rs28937580 | pathogenic | 1 | 1 | 1 |
| rs121908949 | pathogenic | 0 | 0 | 0 |
| rs104894612 | pathogenic | 1 | 1 | 1 |
| rs104894609 | pathogenic | 1 | 1 | 1 |
| rs104894610 | pathogenic | 1 | 1 | 1 |
| rs104894615 | pathogenic | 0 | 0 | 0 |
| rs104894603 | pathogenic | 0 | 0 | 0 |
| rs104894602 | pathogenic | 1 | 1 | 1 |
| rs121908948 | pathogenic | 1 | 1 | 1 |
| rs104894608 | pathogenic | 1 | 1 | 1 |
| rs371545673 | benign | 0 | 0 | 0 |
| rs199925238 | benign | 0 | 0 | 0 |
| rs146313451 | benign | 0 | 0 | 0 |
| rs121918193 | pathogenic | 1 | 1 | 0 |
| rs2228500 | benign | 0 | 0 | 0 |
| rs121918194 | pathogenic | 1 | 1 | 0 |
| rs121918196 | pathogenic | 0 | 0 | 0 |
| rs41291971 | benign | 0 | 0 | 0 |
| rs199473690 | pathogenic | 0 | 0 | 0 |
| rs2228613 | benign | 0 | 0 | 0 |
| rs267607130 | pathogenic | 1 | 1 | 1 |
| rs77615401 | benign | 1 | 1 | 1 |
| rs374618872 | benign | 1 | 1 | 1 |
| rs146165014 | benign | 0 | 0 | 0 |
| rs144586070 | benign | 0 | 0 | 0 |
| rs121918541 | pathogenic | 0 | 0 | 0 |
| rs142105943 | benign | 0 | 0 | 0 |
| rs121918540 | pathogenic | 0 | 0 | 0 |
| rs6791 | benign | 0 | 0 | 0 |
| rs61736587 | pathogenic | 0 | 0 | 0 |
| rs61736586 | benign | 0 | 0 | 0 |
| rs121917741 | pathogenic | 1 | 1 | 1 |
| rs41302885 | benign | 0 | 0 | 0 |
| rs141026028 | benign | 0 | 0 | 0 |
| rs193922493 | pathogenic | 0 | 0 | 0 |
| rs73598374 | benign | 0 | 0 | 0 |
| rs121908725 | pathogenic | 1 | 1 | 0 |
| rs199422328 | pathogenic | 0 | 1 | 0 |
| rs11555566 | benign | 1 | 1 | 1 |
| rs267606634 | pathogenic | 0 | 0 | 0 |
| rs121908714 | pathogenic | 1 | 1 | 0 |
| rs121908717 | pathogenic | 1 | 1 | 0 |
| rs267606635 | pathogenic | 1 | 1 | 0 |
| rs121908739 | pathogenic | 0 | 0 | 0 |
| rs121908735 | pathogenic | 0 | 1 | 0 |

| | | | | |
|---|---|---|---|---|
| rs121908716 | pathogenic | 0 | 1 | 0 |
| rs121908723 | pathogenic | 0 | 1 | 0 |
| rs121908721 | pathogenic | 0 | 0 | 0 |
| rs199422327 | pathogenic | 0 | 0 | 0 |
| rs121908715 | pathogenic | 0 | 0 | 0 |
| rs587783563 | pathogenic | 0 | 0 | 0 |
| rs104894780 | pathogenic | 0 | 0 | 0 |
| rs56030372 | pathogenic | 1 | 1 | 1 |
| rs104894782 | pathogenic | 1 | 1 | 1 |
| rs149495971 | benign | 0 | 0 | 0 |
| rs118203942 | pathogenic | 1 | 1 | 0 |
| rs118203939 | pathogenic | 0 | 0 | 0 |
| rs118203938 | pathogenic | 0 | 0 | 0 |
| rs118203943 | pathogenic | 0 | 0 | 0 |
| rs118203940 | pathogenic | 0 | 0 | 0 |
| rs727503809 | pathogenic | 0 | 0 | 0 |
| rs1065757 | benign | 0 | 0 | 0 |
| rs1071598 | benign | 0 | 0 | 0 |
| rs25414 | benign | 0 | 0 | 0 |
| rs118203944 | pathogenic | 0 | 0 | 0 |
| rs118203941 | pathogenic | 0 | 0 | 0 |
| rs144222167 | benign | 0 | 0 | 0 |
| rs201101343 | pathogenic | 0 | 0 | 0 |
| rs144601717 | benign | 0 | 0 | 0 |
| rs267607129 | pathogenic | 0 | 0 | 0 |
| rs104894729 | pathogenic | 0 | 0 | 0 |
| rs104894727 | pathogenic | 0 | 0 | 0 |
| rs104894725 | pathogenic | 0 | 0 | 0 |
| rs104894515 | pathogenic | 1 | 1 | 1 |
| rs74688377 | benign | 0 | 0 | 0 |
| rs121908513 | pathogenic | 1 | 1 | 0 |
| rs121908509 | pathogenic | 0 | 0 | 0 |
| rs121908514 | pathogenic | 1 | 1 | 0 |
| rs121908512 | pathogenic | 1 | 1 | 0 |
| rs121908510 | pathogenic | 0 | 0 | 0 |
| rs121908511 | pathogenic | 1 | 1 | 1 |
| rs121908518 | pathogenic | 1 | 1 | 1 |
| rs34358665 | benign | 0 | 0 | 0 |
| rs147568834 | benign | 0 | 0 | 0 |
| rs201717599 | benign | 0 | 0 | 0 |
| rs61744487 | benign | 0 | 0 | 0 |
| rs370627877 | pathogenic | 1 | 1 | 1 |
| rs368593151 | pathogenic | 1 | 1 | 1 |
| rs114363936 | benign | 0 | 0 | 0 |
| rs137853333 | pathogenic | 0 | 0 | 0 |
| rs137853080 | pathogenic | 1 | 1 | 1 |
| rs137853077 | pathogenic | 1 | 1 | 1 |
| rs780749732 | benign | 1 | 1 | 1 |
| rs137853081 | pathogenic | 1 | 1 | 0 |
| rs137853078 | pathogenic | 0 | 0 | 0 |
| rs121913315 | pathogenic | 1 | 1 | 1 |
| rs752513525 | pathogenic | 1 | 1 | 1 |
| rs104894918 | pathogenic | 0 | 0 | 0 |
| rs104894917 | pathogenic | 0 | 0 | 0 |

| | | | | |
|---|---|---|---|---|
| rs104894919 | pathogenic | 0 | 0 | 0 |
| rs28939672 | pathogenic | 0 | 0 | 0 |
| rs104894173 | pathogenic | 0 | 0 | 0 |
| rs104894179 | pathogenic | 0 | 0 | 0 |
| rs104894174 | pathogenic | 1 | 1 | 0 |
| rs104894178 | pathogenic | 1 | 1 | 0 |
| rs28939714 | pathogenic | 1 | 1 | 1 |
| rs104894270 | pathogenic | 1 | 1 | 1 |
| rs3765571 | benign | 0 | 0 | 0 |
| rs752669339 | pathogenic | 1 | 1 | 1 |
| rs537043237 | pathogenic | 1 | 1 | 1 |
| rs774047299 | pathogenic | 1 | 1 | 1 |
| rs121434583 | pathogenic | 1 | 1 | 1 |
| rs121907927 | pathogenic | 0 | 0 | 0 |
| rs121907925 | pathogenic | 0 | 1 | 0 |
| rs746212325 | pathogenic | 0 | 0 | 0 |
| rs754637179 | pathogenic | 0 | 0 | 0 |
| rs780882740 | pathogenic | 0 | 0 | 0 |
| rs121918130 | pathogenic | 0 | 0 | 0 |
| rs121918129 | pathogenic | 0 | 0 | 0 |
| rs13297509 | pathogenic | 0 | 0 | 0 |
| rs746867724 | pathogenic | 0 | 0 | 0 |
| rs121918128 | pathogenic | 0 | 0 | 0 |
| rs752300607 | pathogenic | 0 | 0 | 0 |
| rs121964860 | pathogenic | 0 | 0 | 0 |
| rs45466197 | pathogenic | 1 | 1 | 1 |
| rs3730238 | benign | 1 | 1 | 1 |
| rs730881125 | pathogenic | 1 | 1 | 1 |
| rs121964861 | pathogenic | 1 | 1 | 1 |
| rs397516484 | pathogenic | 0 | 0 | 0 |
| rs367785431 | pathogenic | 0 | 0 | 0 |
| rs141121678 | pathogenic | 0 | 0 | 0 |
| rs12720263 | benign | 0 | 0 | 0 |
| rs2304256 | benign | 0 | 0 | 0 |
| rs2304255 | benign | 0 | 0 | 0 |
| rs121909045 | pathogenic | 0 | 0 | 0 |
| rs397508796 | pathogenic | 0 | 0 | 0 |
| rs151020603 | pathogenic | 0 | 0 | 0 |
| rs77284892 | pathogenic | 0 | 0 | 0 |
| rs368505753 | pathogenic | 0 | 0 | 0 |
| rs115545701 | pathogenic | 0 | 0 | 0 |
| rs121908749 | pathogenic | 0 | 0 | 0 |
| rs75961395 | pathogenic | 0 | 0 | 0 |
| rs121908750 | pathogenic | 0 | 0 | 0 |
| rs121908751 | pathogenic | 0 | 0 | 0 |
| rs397508464 | pathogenic | 0 | 0 | 0 |
| rs397508461 | pathogenic | 0 | 0 | 0 |
| rs121909031 | pathogenic | 0 | 0 | 0 |
| rs113993958 | pathogenic | 0 | 0 | 0 |
| rs77834169 | pathogenic | 0 | 0 | 0 |
| rs78655421 | pathogenic | 0 | 0 | 0 |
| rs141723617 | benign | 0 | 0 | 0 |
| rs35516286 | benign | 0 | 0 | 0 |
| rs1800079 | pathogenic | 1 | 1 | 0 |

| | | | | |
|---|---|---|---|---|
| rs80282562 | pathogenic | 0 | 0 | 0 |
| rs121908752 | pathogenic | 0 | 0 | 0 |
| rs556880586 | pathogenic | 0 | 0 | 0 |
| rs121908760 | pathogenic | 0 | 0 | 0 |
| rs397508350 | pathogenic | 0 | 0 | 0 |
| rs374946172 | pathogenic | 0 | 0 | 0 |
| rs397508387 | pathogenic | 0 | 0 | 0 |
| rs201386642 | pathogenic | 0 | 0 | 0 |
| rs121909012 | pathogenic | 0 | 0 | 0 |
| rs121909008 | pathogenic | 0 | 0 | 0 |
| rs149790377 | pathogenic | 0 | 0 | 0 |
| rs397508442 | pathogenic | 0 | 0 | 0 |
| rs121909035 | pathogenic | 0 | 0 | 0 |
| rs386134230 | pathogenic | 0 | 0 | 0 |
| rs1800112 | benign | 0 | 0 | 0 |
| rs144055758 | pathogenic | 0 | 0 | 0 |
| rs142394380 | pathogenic | 0 | 0 | 0 |
| rs121909036 | pathogenic | 0 | 0 | 0 |
| rs78194216 | pathogenic | 0 | 0 | 0 |
| rs121909019 | pathogenic | 0 | 0 | 0 |
| rs121909020 | pathogenic | 0 | 0 | 0 |
| rs200321110 | pathogenic | 0 | 0 | 0 |
| rs202179988 | pathogenic | 0 | 0 | 0 |
| rs121909037 | pathogenic | 0 | 0 | 0 |
| rs186045772 | pathogenic | 0 | 0 | 0 |
| rs139304906 | pathogenic | 0 | 0 | 0 |
| rs78802634 | pathogenic | 0 | 0 | 0 |
| rs121908761 | pathogenic | 0 | 0 | 0 |
| rs36210737 | pathogenic | 0 | 0 | 0 |
| rs139729994 | pathogenic | 0 | 0 | 0 |
| rs79850223 | pathogenic | 0 | 0 | 0 |
| rs1800120 | benign | 0 | 0 | 0 |
| rs74767530 | pathogenic | 0 | 0 | 0 |
| rs374705585 | pathogenic | 0 | 0 | 0 |
| rs1136287 | benign | 0 | 0 | 0 |
| rs6828 | benign | 0 | 0 | 0 |
| rs104894856 | pathogenic | 0 | 0 | 0 |
| rs104894860 | pathogenic | 0 | 0 | 0 |
| rs61736892 | pathogenic | 0 | 0 | 0 |
| rs145807417 | benign | 0 | 0 | 0 |
| rs104894853 | pathogenic | 0 | 0 | 0 |
| rs199422227 | pathogenic | 0 | 0 | 0 |
| rs113993946 | pathogenic | 0 | 0 | 0 |
| rs199422231 | pathogenic | 0 | 0 | 0 |
| rs199422228 | pathogenic | 0 | 0 | 0 |
| rs142195196 | benign | 0 | 1 | 0 |
| rs148656104 | benign | 1 | 1 | 1 |
| rs141944844 | benign | 0 | 0 | 0 |
| rs104894342 | pathogenic | 0 | 0 | 0 |
| rs104894343 | pathogenic | 0 | 0 | 0 |
| rs104894344 | pathogenic | 0 | 0 | 0 |
| rs573322878 | benign | 0 | 0 | 0 |
| rs104894347 | pathogenic | 0 | 0 | 0 |
| rs28937882 | pathogenic | 0 | 0 | 0 |

| | | | | |
|---|---|---|---|---|
| rs193922702 | pathogenic | 0 | 0 | 0 |
| rs200786104 | benign | 0 | 0 | 0 |
| rs690016544 | pathogenic | 1 | 1 | 1 |
| rs28943592 | benign | 0 | 0 | 0 |
| rs28943594 | benign | 0 | 0 | 0 |
| rs121908957 | pathogenic | 0 | 0 | 0 |
| rs115408010 | benign | 1 | 1 | 1 |
| rs77591659 | benign | 0 | 0 | 0 |
| rs121918511 | pathogenic | 0 | 0 | 0 |
| rs538359948 | benign | 0 | 0 | 0 |
| rs76708142 | benign | 0 | 0 | 0 |
| rs121907913 | pathogenic | 1 | 1 | 0 |
| rs121907920 | pathogenic | 0 | 0 | 0 |
| rs121907923 | pathogenic | 1 | 1 | 0 |
| rs121907928 | pathogenic | 1 | 1 | 0 |
| rs121907919 | pathogenic | 1 | 1 | 0 |
| rs121907918 | pathogenic | 1 | 1 | 0 |
| rs74315493 | pathogenic | 1 | 1 | 1 |
| rs74315494 | pathogenic | 1 | 1 | 1 |
| rs145446060 | benign | 1 | 1 | 1 |
| rs550409143 | benign | 1 | 1 | 0 |

| compound_binding_site | cofactor_binding_site | nucleic_binding_site | ion_binding_site |
|---|---|---|---|
| 1 | 0 | 0 | 0 |
| 1 | 0 | 0 | 0 |
| 0 | 0 | 0 | 0 |
| 1 | 0 | 0 | 0 |
| 0 | 0 | 0 | 0 |
| 0 | 0 | 0 | 0 |
| 0 | 0 | 0 | 0 |
| 0 | 0 | 0 | 0 |
| 1 | 0 | 0 | 1 |
| 1 | 0 | 0 | 0 |
| 0 | 0 | 0 | 1 |
| 1 | 0 | 1 | 0 |
| 0 | 0 | 0 | 0 |
| 0 | 0 | 0 | 0 |
| 0 | 0 | 0 | 0 |
| 0 | 0 | 0 | 0 |
| 0 | 0 | 0 | 0 |
| 1 | 0 | 0 | 1 |
| 1 | 0 | 1 | 0 |
| 0 | 0 | 0 | 0 |
| 1 | 0 | 0 | 1 |
| 1 | 0 | 0 | 0 |
| 1 | 0 | 0 | 0 |
| 1 | 0 | 0 | 0 |
| 0 | 0 | 0 | 0 |
| 0 | 0 | 0 | 0 |
| 1 | 0 | 0 | 1 |
| 0 | 0 | 0 | 0 |
| 0 | 0 | 0 | 0 |
| 0 | 0 | 0 | 0 |
| 0 | 0 | 0 | 0 |
| 1 | 0 | 0 | 0 |
| 1 | 0 | 0 | 1 |
| 1 | 0 | 0 | 0 |
| 0 | 0 | 0 | 0 |
| 0 | 0 | 0 | 0 |
| 0 | 0 | 0 | 0 |
| 1 | 0 | 0 | 1 |
| 1 | 0 | 0 | 1 |
| 1 | 0 | 0 | 1 |
| 1 | 0 | 0 | 0 |
| 1 | 0 | 0 | 0 |
| 1 | 0 | 0 | 0 |
| 1 | 0 | 0 | 0 |
| 1 | 0 | 0 | 0 |
| 1 | 0 | 0 | 0 |
| 1 | 0 | 0 | 0 |
| 1 | 0 | 0 | 0 |
| 1 | 0 | 0 | 0 |
| 1 | 0 | 0 | 0 |
| 1 | 0 | 0 | 0 |
| 1 | 0 | 0 | 0 |
| 1 | 0 | 0 | 0 |

| | | | |
|---|---|---|---|
| 1 | 0 | 0 | 1 |
| 1 | 0 | 0 | 0 |
| 1 | 0 | 0 | 0 |
| 1 | 0 | 0 | 0 |
| 0 | 0 | 0 | 0 |
| 1 | 0 | 0 | 0 |
| 0 | 0 | 0 | 0 |
| 1 | 0 | 0 | 0 |
| 1 | 0 | 0 | 0 |
| 1 | 0 | 0 | 1 |
| 1 | 0 | 0 | 0 |
| 1 | 0 | 0 | 0 |
| 1 | 0 | 0 | 0 |
| 1 | 0 | 0 | 0 |
| 1 | 0 | 0 | 0 |
| 1 | 0 | 0 | 0 |
| 1 | 0 | 0 | 0 |
| 1 | 0 | 0 | 1 |
| 1 | 0 | 0 | 1 |
| 1 | 1 | 0 | 1 |
| 1 | 0 | 0 | 0 |
| 1 | 0 | 0 | 0 |
| 1 | 0 | 0 | 0 |
| 1 | 0 | 0 | 0 |
| 1 | 1 | 0 | 0 |
| 1 | 0 | 0 | 0 |
| 1 | 0 | 0 | 0 |
| 1 | 0 | 0 | 0 |
| 1 | 0 | 0 | 0 |
| 1 | 0 | 0 | 0 |
| 1 | 1 | 0 | 0 |
| 1 | 1 | 0 | 0 |
| 1 | 1 | 0 | 0 |
| 1 | 1 | 0 | 0 |
| 1 | 1 | 0 | 0 |
| 1 | 0 | 0 | 0 |
| 1 | 1 | 0 | 0 |
| 1 | 0 | 0 | 0 |
| 1 | 0 | 0 | 0 |
| 1 | 1 | 0 | 0 |
| 0 | 0 | 0 | 0 |
| 0 | 0 | 0 | 0 |
| 0 | 0 | 0 | 0 |
| 0 | 0 | 0 | 0 |
| 0 | 0 | 0 | 0 |
| 0 | 0 | 0 | 0 |
| 0 | 0 | 0 | 0 |
| 1 | 0 | 0 | 1 |
| 0 | 0 | 0 | 0 |
| 1 | 0 | 0 | 0 |
| 1 | 0 | 0 | 0 |
| 0 | 0 | 0 | 0 |
| 0 | 0 | 0 | 0 |
| 0 | 0 | 0 | 0 |

| | | | |
|---|---|---|---|
| 0 | 0 | 0 | 0 |
| 0 | 0 | 0 | 0 |
| 0 | 0 | 0 | 0 |
| 1 | 0 | 0 | 0 |
| 1 | 0 | 0 | 0 |
| 1 | 0 | 0 | 0 |
| 0 | 0 | 0 | 0 |
| 0 | 0 | 1 | 0 |
| 0 | 0 | 1 | 0 |
| 0 | 0 | 1 | 0 |
| 0 | 0 | 1 | 0 |
| 0 | 0 | 1 | 0 |
| 0 | 0 | 0 | 0 |
| 0 | 0 | 0 | 0 |
| 1 | 0 | 0 | 0 |
| 0 | 0 | 0 | 0 |
| 0 | 0 | 0 | 0 |
| 1 | 0 | 0 | 0 |
| 1 | 0 | 0 | 0 |
| 1 | 0 | 0 | 0 |
| 1 | 0 | 0 | 0 |
| 1 | 1 | 0 | 0 |
| 1 | 0 | 0 | 0 |
| 1 | 0 | 0 | 0 |
| 0 | 0 | 0 | 0 |
| 0 | 0 | 0 | 0 |
| 1 | 0 | 0 | 0 |
| 1 | 1 | 0 | 0 |
| 1 | 0 | 0 | 0 |
| 1 | 0 | 0 | 0 |
| 1 | 1 | 0 | 0 |
| 1 | 1 | 0 | 0 |
| 1 | 1 | 0 | 0 |
| 1 | 1 | 0 | 0 |
| 1 | 1 | 0 | 0 |
| 1 | 1 | 0 | 0 |
| 1 | 1 | 0 | 0 |
| 1 | 1 | 0 | 0 |
| 1 | 1 | 0 | 0 |
| 1 | 1 | 0 | 0 |
| 1 | 1 | 0 | 0 |
| 1 | 1 | 0 | 0 |
| 1 | 0 | 0 | 0 |
| 1 | 0 | 0 | 0 |
| 1 | 0 | 0 | 0 |
| 1 | 1 | 0 | 0 |
| 1 | 0 | 0 | 0 |
| 1 | 1 | 1 | 0 |
| 1 | 1 | 1 | 0 |
| 1 | 1 | 0 | 0 |
| 1 | 1 | 0 | 0 |
| 1 | 0 | 0 | 0 |
| 1 | 0 | 0 | 0 |
| 0 | 0 | 0 | 0 |

| | | | |
|---|---|---|---|
| 0 | 0 | 0 | 0 |
| 0 | 0 | 0 | 0 |
| 0 | 0 | 0 | 0 |
| 1 | 0 | 0 | 0 |
| 1 | 0 | 0 | 0 |
| 0 | 0 | 0 | 0 |
| 0 | 0 | 0 | 0 |
| 0 | 0 | 0 | 0 |
| 0 | 0 | 0 | 0 |
| 0 | 0 | 0 | 0 |
| 0 | 0 | 0 | 0 |
| 0 | 0 | 0 | 0 |
| 0 | 0 | 0 | 0 |
| 0 | 0 | 0 | 0 |
| 0 | 0 | 0 | 0 |
| 0 | 0 | 0 | 0 |
| 0 | 0 | 0 | 0 |
| 0 | 0 | 0 | 0 |
| 0 | 0 | 0 | 0 |
| 0 | 0 | 0 | 0 |
| 0 | 0 | 0 | 0 |
| 0 | 0 | 0 | 0 |
| 1 | 0 | 0 | 0 |
| 1 | 0 | 0 | 0 |
| 0 | 0 | 0 | 0 |
| 0 | 0 | 0 | 0 |
| 1 | 0 | 0 | 0 |
| 0 | 0 | 0 | 0 |
| 0 | 0 | 0 | 0 |
| 0 | 0 | 0 | 0 |
| 0 | 0 | 0 | 0 |
| 0 | 0 | 0 | 0 |
| 0 | 0 | 0 | 0 |
| 0 | 0 | 0 | 0 |
| 1 | 0 | 0 | 0 |
| 0 | 0 | 0 | 0 |
| 0 | 0 | 0 | 0 |
| 0 | 0 | 0 | 0 |
| 0 | 0 | 0 | 0 |
| 0 | 0 | 0 | 0 |
| 0 | 0 | 0 | 0 |
| 0 | 0 | 0 | 0 |
| 0 | 0 | 0 | 0 |
| 0 | 0 | 0 | 0 |
| 0 | 0 | 0 | 0 |
| 0 | 0 | 0 | 0 |
| 0 | 0 | 1 | 0 |
| 1 | 0 | 0 | 0 |
| 1 | 0 | 0 | 0 |
| 1 | 1 | 0 | 0 |
| 1 | 0 | 0 | 0 |
| 0 | 0 | 0 | 0 |
| 0 | 0 | 0 | 0 |
| 1 | 0 | 0 | 0 |

| | | | |
|---|---|---|---|
| 1 | 0 | 0 | 0 |
| 1 | 0 | 0 | 0 |
| 1 | 0 | 0 | 0 |
| 0 | 0 | 0 | 0 |
| 0 | 0 | 0 | 0 |
| 0 | 0 | 0 | 0 |
| 1 | 0 | 0 | 0 |
| 1 | 0 | 0 | 0 |
| 0 | 0 | 0 | 0 |
| 0 | 0 | 0 | 0 |
| 1 | 0 | 0 | 0 |
| 0 | 0 | 0 | 0 |
| 1 | 0 | 0 | 0 |
| 0 | 0 | 0 | 0 |
| 0 | 0 | 0 | 0 |
| 0 | 0 | 0 | 0 |
| 0 | 0 | 0 | 0 |
| 0 | 0 | 0 | 0 |
| 0 | 0 | 0 | 0 |
| 0 | 0 | 0 | 0 |
| 0 | 0 | 0 | 0 |
| 0 | 0 | 0 | 0 |
| 0 | 0 | 0 | 0 |
| 0 | 0 | 0 | 0 |
| 0 | 0 | 0 | 0 |
| 0 | 0 | 0 | 0 |
| 1 | 0 | 0 | 0 |
| 1 | 0 | 0 | 0 |
| 1 | 0 | 0 | 0 |
| 1 | 0 | 0 | 0 |
| 1 | 0 | 0 | 0 |
| 1 | 0 | 0 | 0 |
| 0 | 0 | 0 | 0 |
| 0 | 0 | 0 | 0 |
| 1 | 0 | 0 | 0 |
| 0 | 0 | 0 | 0 |
| 0 | 0 | 0 | 0 |
| 0 | 0 | 0 | 0 |
| 0 | 0 | 0 | 0 |
| 0 | 0 | 0 | 0 |
| 0 | 0 | 0 | 0 |
| 0 | 0 | 0 | 0 |
| 0 | 0 | 0 | 0 |
| 0 | 0 | 0 | 0 |
| 1 | 1 | 0 | 0 |
| 1 | 1 | 0 | 0 |
| 1 | 1 | 0 | 1 |
| 1 | 1 | 0 | 1 |
| 1 | 1 | 0 | 1 |
| 1 | 1 | 0 | 1 |
| 1 | 1 | 0 | 0 |
| 1 | 1 | 0 | 1 |
| 1 | 1 | 0 | 1 |
| 1 | 1 | 0 | 1 |

| | | | |
|---|---|---|---|
| 1 | 1 | 0 | 1 |
| 1 | 1 | 0 | 1 |
| 1 | 1 | 0 | 1 |
| 1 | 1 | 0 | 1 |
| 1 | 1 | 0 | 1 |
| 1 | 0 | 0 | 0 |
| 0 | 0 | 0 | 0 |
| 0 | 0 | 0 | 0 |
| 0 | 0 | 0 | 0 |
| 1 | 0 | 0 | 0 |
| 0 | 0 | 1 | 0 |
| 0 | 0 | 0 | 0 |
| 0 | 0 | 0 | 0 |
| 0 | 0 | 0 | 0 |
| 0 | 0 | 0 | 0 |
| 0 | 0 | 0 | 0 |
| 0 | 0 | 0 | 0 |
| 0 | 0 | 0 | 0 |
| 0 | 0 | 0 | 0 |
| 0 | 0 | 0 | 0 |
| 0 | 0 | 0 | 0 |
| 0 | 0 | 0 | 0 |
| 0 | 0 | 0 | 0 |
| 0 | 0 | 0 | 0 |
| 0 | 0 | 0 | 0 |
| 1 | 1 | 0 | 0 |
| 1 | 1 | 0 | 0 |
| 1 | 1 | 0 | 0 |
| 0 | 1 | 0 | 0 |
| 1 | 0 | 0 | 0 |
| 0 | 0 | 0 | 0 |
| 0 | 0 | 0 | 0 |
| 0 | 0 | 0 | 0 |
| 1 | 0 | 0 | 0 |
| 1 | 1 | 0 | 0 |
| 0 | 0 | 0 | 0 |
| 0 | 0 | 0 | 0 |
| 0 | 0 | 0 | 0 |
| 0 | 0 | 0 | 0 |
| 1 | 0 | 0 | 0 |
| 0 | 0 | 0 | 0 |
| 0 | 0 | 0 | 0 |
| 0 | 0 | 0 | 0 |
| 0 | 0 | 0 | 0 |
| 0 | 0 | 0 | 0 |
| 0 | 1 | 0 | 0 |
| 0 | 1 | 0 | 0 |
| 0 | 0 | 0 | 0 |
| 0 | 0 | 0 | 0 |
| 0 | 1 | 0 | 0 |
| 0 | 0 | 0 | 0 |
| 0 | 0 | 0 | 0 |
| 0 | 1 | 0 | 0 |
| 0 | 0 | 0 | 0 |

| | | | |
|---|---|---|---|
| 1 | 1 | 0 | 0 |
| 0 | 0 | 0 | 0 |
| 0 | 0 | 0 | 0 |
| 0 | 0 | 0 | 0 |
| 0 | 0 | 0 | 0 |
| 0 | 0 | 0 | 0 |
| 0 | 0 | 0 | 0 |
| 0 | 0 | 0 | 0 |
| 0 | 0 | 0 | 0 |
| 0 | 0 | 0 | 0 |
| 0 | 0 | 0 | 0 |
| 0 | 0 | 0 | 0 |
| 0 | 0 | 0 | 0 |
| 0 | 0 | 0 | 0 |
| 0 | 1 | 0 | 0 |
| 0 | 0 | 0 | 0 |
| 0 | 1 | 0 | 0 |
| 0 | 1 | 0 | 0 |
| 0 | 0 | 0 | 0 |
| 0 | 1 | 0 | 0 |
| 1 | 0 | 0 | 1 |
| 1 | 1 | 0 | 0 |
| 1 | 0 | 0 | 0 |
| 1 | 0 | 0 | 1 |
| 0 | 0 | 0 | 0 |
| 0 | 0 | 0 | 0 |
| 0 | 0 | 0 | 0 |
| 0 | 0 | 0 | 0 |
| 1 | 0 | 0 | 0 |
| 0 | 0 | 0 | 0 |
| 1 | 0 | 0 | 0 |
| 0 | 0 | 0 | 0 |
| 0 | 0 | 0 | 0 |
| 0 | 0 | 0 | 0 |
| 0 | 0 | 0 | 0 |
| 0 | 0 | 0 | 0 |
| 0 | 0 | 0 | 0 |
| 1 | 1 | 0 | 0 |
| 1 | 1 | 0 | 0 |
| 1 | 1 | 0 | 0 |
| 0 | 0 | 0 | 0 |
| 0 | 0 | 0 | 0 |
| 0 | 0 | 0 | 0 |
| 1 | 0 | 0 | 0 |
| 1 | 0 | 0 | 1 |
| 1 | 0 | 0 | 1 |
| 1 | 0 | 0 | 0 |
| 1 | 0 | 0 | 1 |
| 1 | 0 | 0 | 0 |
| 0 | 0 | 0 | 0 |
| 1 | 1 | 0 | 0 |
| 1 | 1 | 0 | 0 |
| 0 | 0 | 0 | 0 |
| 0 | 0 | 0 | 0 |

| | | | |
|---|---|---|---|
| 1 | 0 | 0 | 0 |
| 1 | 0 | 0 | 0 |
| 1 | 0 | 0 | 0 |
| 0 | 0 | 0 | 0 |
| 0 | 0 | 0 | 0 |
| 1 | 1 | 0 | 1 |
| 1 | 1 | 0 | 1 |
| 1 | 1 | 0 | 1 |
| 1 | 1 | 0 | 1 |
| 1 | 1 | 0 | 1 |
| 1 | 1 | 0 | 1 |
| 1 | 1 | 0 | 1 |
| 1 | 1 | 0 | 0 |
| 1 | 1 | 0 | 0 |
| 1 | 1 | 1 | 1 |
| 0 | 0 | 0 | 0 |
| 1 | 1 | 0 | 0 |
| 1 | 1 | 0 | 0 |
| 0 | 0 | 0 | 0 |
| 0 | 0 | 0 | 0 |
| 0 | 0 | 0 | 0 |
| 1 | 0 | 0 | 0 |
| 1 | 0 | 0 | 0 |
| 0 | 0 | 0 | 0 |
| 1 | 0 | 0 | 0 |
| 1 | 0 | 0 | 0 |
| 1 | 0 | 0 | 0 |
| 0 | 0 | 0 | 0 |
| 0 | 0 | 0 | 0 |
| 0 | 0 | 0 | 0 |
| 0 | 0 | 0 | 0 |
| 0 | 0 | 0 | 0 |
| 0 | 0 | 0 | 0 |
| 0 | 0 | 0 | 0 |
| 0 | 0 | 0 | 0 |
| 0 | 0 | 0 | 0 |
| 0 | 0 | 0 | 0 |
| 0 | 0 | 0 | 0 |
| 0 | 0 | 0 | 0 |
| 0 | 0 | 0 | 0 |
| 0 | 0 | 0 | 0 |
| 0 | 0 | 0 | 0 |
| 0 | 0 | 0 | 0 |
| 0 | 0 | 0 | 0 |
| 0 | 0 | 0 | 0 |
| 1 | 0 | 0 | 0 |
| 1 | 0 | 0 | 0 |
| 1 | 0 | 0 | 0 |
| 1 | 0 | 0 | 0 |
| 0 | 0 | 0 | 0 |
| 0 | 0 | 0 | 0 |
| 0 | 0 | 0 | 0 |
| 0 | 0 | 0 | 0 |
| 0 | 0 | 0 | 0 |

| | | | |
|---|---|---|---|
| 0 | 0 | 0 | 0 |
| 0 | 0 | 0 | 0 |
| 0 | 0 | 0 | 0 |
| 0 | 0 | 0 | 0 |
| 0 | 0 | 0 | 0 |
| 0 | 0 | 0 | 0 |
| 0 | 0 | 0 | 0 |
| 1 | 0 | 0 | 1 |
| 0 | 0 | 0 | 0 |
| 0 | 0 | 0 | 0 |
| 1 | 0 | 0 | 0 |
| 0 | 0 | 0 | 0 |
| 0 | 0 | 0 | 0 |
| 0 | 0 | 0 | 0 |
| 0 | 0 | 0 | 0 |
| 1 | 0 | 0 | 0 |
| 0 | 0 | 0 | 0 |
| 0 | 0 | 0 | 0 |
| 0 | 0 | 0 | 0 |
| 0 | 0 | 0 | 0 |
| 0 | 0 | 0 | 0 |
| 0 | 0 | 0 | 0 |
| 0 | 0 | 0 | 0 |
| 1 | 1 | 0 | 1 |
| 0 | 0 | 0 | 0 |
| 0 | 0 | 0 | 0 |
| 0 | 0 | 0 | 0 |
| 1 | 1 | 0 | 1 |
| 1 | 1 | 0 | 0 |
| 0 | 0 | 0 | 0 |
| 1 | 0 | 0 | 0 |
| 1 | 1 | 0 | 0 |
| 0 | 0 | 0 | 0 |
| 1 | 0 | 0 | 0 |
| 1 | 1 | 0 | 0 |
| 1 | 0 | 0 | 0 |
| 1 | 1 | 0 | 1 |
| 1 | 1 | 0 | 1 |
| 1 | 0 | 0 | 0 |
| 0 | 0 | 0 | 0 |
| 0 | 0 | 0 | 0 |
| 0 | 0 | 0 | 0 |
| 0 | 0 | 0 | 0 |
| 0 | 0 | 0 | 0 |
| 0 | 0 | 0 | 0 |
| 0 | 0 | 0 | 0 |
| 0 | 0 | 0 | 0 |
| 0 | 0 | 0 | 0 |
| 0 | 0 | 0 | 0 |
| 0 | 0 | 0 | 0 |
| 0 | 0 | 0 | 0 |
| 0 | 0 | 0 | 0 |
| 0 | 0 | 0 | 0 |
| 0 | 0 | 0 | 0 |

| | | | |
|---|---|---|---|
| 0 | 0 | 0 | 0 |
| 0 | 0 | 0 | 0 |
| 0 | 0 | 0 | 0 |
| 0 | 0 | 0 | 0 |
| 0 | 0 | 0 | 0 |
| 0 | 0 | 0 | 0 |
| 0 | 0 | 0 | 0 |
| 0 | 0 | 0 | 0 |
| 0 | 0 | 0 | 0 |
| 0 | 0 | 0 | 0 |
| 0 | 0 | 0 | 0 |
| 0 | 0 | 0 | 0 |
| 0 | 0 | 0 | 0 |
| 1 | 1 | 0 | 0 |
| 0 | 0 | 0 | 0 |
| 1 | 0 | 0 | 0 |
| 0 | 0 | 0 | 1 |
| 0 | 0 | 0 | 0 |
| 0 | 0 | 0 | 0 |
| 0 | 0 | 0 | 0 |
| 0 | 0 | 0 | 0 |
| 0 | 0 | 0 | 0 |
| 0 | 0 | 0 | 0 |
| 0 | 0 | 0 | 0 |
| 0 | 0 | 0 | 0 |
| 0 | 0 | 0 | 0 |
| 0 | 0 | 0 | 0 |
| 0 | 0 | 0 | 0 |
| 0 | 0 | 0 | 0 |
| 0 | 0 | 0 | 0 |
| 1 | 0 | 0 | 0 |
| 1 | 0 | 0 | 0 |
| 0 | 0 | 0 | 0 |
| 0 | 0 | 0 | 0 |
| 0 | 0 | 0 | 0 |
| 0 | 0 | 0 | 0 |
| 0 | 0 | 0 | 0 |
| 0 | 0 | 0 | 0 |
| 0 | 0 | 0 | 0 |
| 0 | 0 | 0 | 0 |
| 1 | 0 | 0 | 0 |
| 1 | 0 | 0 | 0 |
| 1 | 0 | 0 | 0 |
| 1 | 0 | 0 | 0 |
| 0 | 0 | 0 | 0 |
| 1 | 0 | 0 | 0 |
| 0 | 0 | 0 | 0 |
| 0 | 0 | 0 | 0 |
| 1 | 0 | 0 | 0 |
| 1 | 0 | 0 | 0 |
| 0 | 0 | 0 | 0 |
| 0 | 0 | 0 | 0 |
| 0 | 0 | 0 | 0 |
| 0 | 0 | 0 | 0 |

| | | | |
|---|---|---|---|
| 0 | 0 | 0 | 0 |
| 0 | 0 | 0 | 0 |
| 0 | 0 | 0 | 0 |
| 0 | 0 | 0 | 0 |
| 0 | 0 | 0 | 0 |
| 0 | 0 | 0 | 0 |
| 0 | 0 | 0 | 0 |
| 0 | 0 | 0 | 0 |
| 0 | 0 | 0 | 0 |
| 0 | 0 | 0 | 0 |
| 0 | 0 | 0 | 0 |
| 0 | 0 | 0 | 0 |
| 0 | 0 | 0 | 0 |
| 0 | 0 | 0 | 0 |
| 0 | 0 | 0 | 0 |
| 1 | 0 | 0 | 1 |
| 0 | 0 | 0 | 0 |
| 0 | 0 | 0 | 0 |
| 0 | 0 | 0 | 0 |
| 0 | 0 | 0 | 0 |
| 0 | 0 | 0 | 0 |
| 1 | 1 | 0 | 0 |
| 1 | 1 | 0 | 0 |
| 1 | 0 | 0 | 0 |
| 1 | 0 | 0 | 0 |
| 0 | 0 | 0 | 0 |
| 0 | 0 | 0 | 0 |
| 0 | 0 | 0 | 0 |
| 0 | 0 | 0 | 0 |
| 1 | 0 | 0 | 0 |
| 1 | 0 | 0 | 0 |
| 1 | 0 | 0 | 0 |
| 1 | 1 | 0 | 0 |
| 1 | 1 | 0 | 0 |
| 1 | 0 | 0 | 0 |
| 0 | 0 | 0 | 0 |
| 0 | 0 | 0 | 0 |
| 0 | 1 | 0 | 0 |
| 0 | 0 | 0 | 0 |
| 0 | 0 | 0 | 0 |
| 0 | 0 | 0 | 0 |
| 1 | 1 | 0 | 0 |
| 1 | 0 | 0 | 0 |
| 1 | 0 | 0 | 0 |
| 0 | 0 | 0 | 0 |
| 0 | 0 | 0 | 0 |
| 0 | 0 | 0 | 0 |
| 0 | 0 | 0 | 0 |
| 0 | 0 | 0 | 0 |
| 0 | 0 | 0 | 0 |
| 0 | 0 | 0 | 0 |
| 0 | 0 | 0 | 0 |
| 0 | 0 | 0 | 0 |
| 0 | 0 | 0 | 0 |

| | | | |
|---|---|---|---|
| 0 | 0 | 0 | 0 |
| 1 | 0 | 0 | 0 |
| 0 | 0 | 0 | 0 |
| 0 | 0 | 0 | 0 |
| 0 | 0 | 0 | 0 |
| 0 | 0 | 0 | 0 |
| 1 | 0 | 0 | 0 |
| 0 | 0 | 0 | 0 |
| 0 | 0 | 0 | 0 |
| 0 | 0 | 0 | 0 |
| 0 | 0 | 0 | 0 |
| 0 | 0 | 0 | 0 |
| 0 | 0 | 0 | 0 |
| 0 | 0 | 0 | 0 |
| 0 | 0 | 0 | 0 |
| 0 | 0 | 0 | 0 |
| 0 | 1 | 0 | 0 |
| 0 | 0 | 0 | 0 |
| 0 | 0 | 0 | 0 |
| 0 | 0 | 0 | 0 |
| 1 | 0 | 0 | 0 |
| 0 | 0 | 0 | 0 |
| 0 | 0 | 0 | 0 |
| 1 | 1 | 0 | 0 |
| 0 | 0 | 0 | 0 |
| 0 | 0 | 0 | 0 |
| 0 | 0 | 0 | 0 |
| 0 | 0 | 0 | 0 |
| 1 | 0 | 0 | 0 |
| 0 | 0 | 0 | 0 |
| 0 | 0 | 0 | 0 |
| 0 | 0 | 0 | 0 |
| 0 | 0 | 0 | 0 |
| 0 | 0 | 0 | 0 |
| 0 | 0 | 0 | 0 |
| 1 | 0 | 0 | 0 |
| 0 | 0 | 0 | 0 |
| 0 | 0 | 0 | 0 |
| 0 | 0 | 0 | 0 |
| 0 | 0 | 0 | 0 |
| 1 | 1 | 0 | 0 |
| 0 | 0 | 0 | 0 |
| 0 | 0 | 0 | 0 |
| 0 | 0 | 0 | 0 |
| 0 | 0 | 0 | 0 |
| 0 | 0 | 0 | 0 |
| 1 | 0 | 0 | 0 |
| 1 | 0 | 0 | 1 |
| 1 | 0 | 0 | 0 |
| 0 | 0 | 0 | 0 |
| 0 | 0 | 0 | 0 |
| 0 | 0 | 0 | 0 |
| 1 | 0 | 0 | 0 |
| 1 | 0 | 0 | 0 |

| | | | |
|---|---|---|---|
| 1 | 0 | 0 | 0 |
| 0 | 0 | 0 | 0 |
| 0 | 0 | 0 | 0 |
| 0 | 0 | 0 | 0 |
| 1 | 1 | 0 | 0 |
| 1 | 1 | 0 | 0 |
| 0 | 0 | 0 | 0 |
| 0 | 0 | 0 | 0 |
| 1 | 0 | 0 | 0 |
| 1 | 0 | 0 | 0 |
| 1 | 0 | 1 | 0 |
| 0 | 0 | 0 | 0 |
| 0 | 0 | 1 | 0 |
| 0 | 0 | 1 | 0 |
| 0 | 0 | 1 | 0 |
| 1 | 0 | 0 | 0 |
| 1 | 0 | 0 | 0 |
| 0 | 0 | 0 | 0 |
| 0 | 0 | 0 | 0 |
| 1 | 0 | 0 | 1 |
| 1 | 0 | 0 | 1 |
| 1 | 0 | 0 | 0 |
| 0 | 0 | 0 | 0 |
| 0 | 0 | 0 | 0 |
| 0 | 0 | 0 | 0 |
| 0 | 0 | 0 | 0 |
| 0 | 0 | 0 | 0 |
| 1 | 0 | 0 | 0 |
| 0 | 0 | 0 | 0 |
| 0 | 0 | 0 | 1 |
| 0 | 0 | 0 | 1 |
| 0 | 0 | 1 | 1 |
| 0 | 0 | 1 | 1 |
| 0 | 0 | 1 | 1 |
| 0 | 0 | 1 | 1 |
| 0 | 0 | 1 | 0 |
| 0 | 0 | 1 | 0 |
| 0 | 0 | 0 | 0 |
| 0 | 0 | 0 | 0 |
| 0 | 0 | 0 | 0 |
| 0 | 0 | 0 | 0 |
| 0 | 0 | 1 | 0 |
| 0 | 0 | 1 | 0 |
| 0 | 0 | 1 | 0 |
| 0 | 0 | 1 | 0 |
| 0 | 0 | 1 | 0 |
| 0 | 0 | 1 | 0 |
| 0 | 0 | 1 | 0 |
| 0 | 0 | 1 | 0 |
| 0 | 0 | 0 | 0 |
| 0 | 0 | 0 | 0 |
| 0 | 0 | 0 | 0 |
| 0 | 0 | 0 | 0 |
| 1 | 0 | 0 | 0 |

| | | | |
|---|---|---|---|
| 0 | 0 | 0 | 0 |
| 0 | 0 | 0 | 0 |
| 0 | 0 | 0 | 0 |
| 1 | 0 | 0 | 0 |
| 0 | 0 | 0 | 0 |
| 0 | 0 | 0 | 0 |
| 0 | 0 | 0 | 0 |
| 0 | 0 | 0 | 0 |
| 0 | 0 | 0 | 0 |
| 0 | 0 | 0 | 0 |
| 1 | 1 | 0 | 0 |
| 0 | 0 | 0 | 0 |
| 1 | 1 | 0 | 0 |
| 1 | 1 | 0 | 0 |
| 0 | 1 | 0 | 0 |
| 0 | 0 | 0 | 0 |
| 0 | 0 | 0 | 0 |
| 0 | 0 | 0 | 0 |
| 0 | 0 | 0 | 0 |
| 0 | 0 | 0 | 0 |
| 1 | 1 | 0 | 0 |
| 0 | 1 | 0 | 0 |
| 0 | 0 | 0 | 0 |
| 0 | 1 | 0 | 0 |
| 1 | 1 | 0 | 0 |
| 0 | 0 | 0 | 0 |
| 0 | 0 | 0 | 0 |
| 1 | 1 | 0 | 0 |
| 1 | 1 | 0 | 0 |
| 0 | 1 | 0 | 0 |
| 0 | 0 | 0 | 0 |
| 0 | 0 | 0 | 0 |
| 0 | 0 | 0 | 0 |
| 0 | 0 | 0 | 0 |
| 0 | 0 | 0 | 0 |
| 0 | 0 | 0 | 0 |
| 0 | 0 | 0 | 0 |
| 0 | 0 | 0 | 0 |
| 0 | 0 | 0 | 0 |
| 0 | 0 | 0 | 0 |
| 1 | 0 | 0 | 0 |
| 1 | 0 | 0 | 0 |
| 1 | 0 | 0 | 0 |
| 1 | 0 | 0 | 0 |
| 1 | 1 | 0 | 1 |
| 0 | 0 | 0 | 0 |
| 1 | 0 | 0 | 0 |
| 0 | 0 | 0 | 0 |
| 1 | 0 | 0 | 0 |
| 0 | 0 | 0 | 0 |
| 1 | 0 | 0 | 0 |
| 1 | 0 | 0 | 0 |
| 0 | 0 | 0 | 0 |
| 0 | 0 | 0 | 0 |

| | | | |
|---|---|---|---|
| 0 | 0 | 0 | 0 |
| 0 | 0 | 0 | 0 |
| 0 | 0 | 0 | 0 |
| 0 | 0 | 0 | 0 |
| 1 | 0 | 0 | 0 |
| 1 | 0 | 0 | 0 |
| 1 | 0 | 0 | 0 |
| 0 | 0 | 0 | 0 |
| 1 | 1 | 0 | 0 |
| 0 | 0 | 0 | 0 |
| 0 | 0 | 0 | 0 |
| 0 | 0 | 0 | 0 |
| 1 | 1 | 0 | 0 |
| 1 | 0 | 0 | 0 |
| 0 | 0 | 0 | 0 |
| 0 | 0 | 0 | 0 |
| 1 | 1 | 0 | 0 |
| 0 | 0 | 0 | 0 |
| 0 | 0 | 0 | 0 |
| 0 | 0 | 0 | 0 |
| 0 | 0 | 0 | 0 |
| 0 | 0 | 0 | 0 |
| 0 | 0 | 0 | 0 |
| 0 | 0 | 0 | 0 |
| 1 | 0 | 0 | 0 |
| 0 | 0 | 0 | 0 |
| 1 | 0 | 0 | 0 |
| 0 | 0 | 0 | 0 |
| 0 | 0 | 0 | 0 |
| 0 | 0 | 0 | 0 |
| 1 | 0 | 0 | 0 |
| 1 | 0 | 0 | 0 |
| 0 | 0 | 0 | 0 |
| 0 | 0 | 0 | 0 |
| 1 | 0 | 0 | 0 |
| 0 | 0 | 0 | 0 |
| 0 | 0 | 0 | 0 |
| 1 | 0 | 0 | 0 |
| 0 | 0 | 0 | 0 |
| 1 | 0 | 0 | 0 |
| 1 | 0 | 0 | 0 |
| 0 | 0 | 0 | 0 |
| 0 | 0 | 0 | 0 |
| 1 | 1 | 0 | 0 |
| 1 | 0 | 0 | 0 |
| 1 | 1 | 0 | 0 |
| 0 | 0 | 0 | 0 |
| 0 | 0 | 0 | 0 |
| 0 | 0 | 0 | 0 |
| 0 | 0 | 0 | 0 |
| 1 | 1 | 0 | 0 |
| 0 | 0 | 0 | 0 |
| 0 | 0 | 0 | 0 |
| 0 | 0 | 0 | 0 |

| | | | |
|---|---|---|---|
| 0 | 1 | 0 | 0 |
| 1 | 0 | 0 | 0 |
| 0 | 0 | 0 | 0 |
| 0 | 0 | 0 | 0 |
| 0 | 0 | 0 | 0 |
| 1 | 1 | 0 | 0 |
| 0 | 0 | 0 | 0 |
| 0 | 0 | 0 | 0 |
| 0 | 0 | 0 | 0 |
| 1 | 0 | 0 | 0 |
| 1 | 0 | 0 | 0 |
| 0 | 0 | 0 | 0 |
| 0 | 0 | 0 | 0 |
| 0 | 0 | 0 | 0 |
| 0 | 0 | 0 | 0 |
| 0 | 0 | 0 | 0 |
| 0 | 0 | 0 | 0 |
| 0 | 0 | 0 | 0 |
| 0 | 0 | 0 | 0 |
| 0 | 0 | 0 | 0 |
| 0 | 0 | 0 | 0 |
| 0 | 0 | 0 | 0 |
| 0 | 0 | 0 | 0 |
| 1 | 1 | 0 | 0 |
| 0 | 0 | 0 | 0 |
| 0 | 0 | 0 | 0 |
| 1 | 1 | 1 | 0 |
| 0 | 0 | 0 | 0 |
| 0 | 0 | 0 | 0 |
| 0 | 0 | 0 | 0 |
| 1 | 1 | 1 | 0 |
| 1 | 0 | 0 | 0 |
| 1 | 0 | 0 | 0 |
| 1 | 0 | 0 | 0 |
| 1 | 0 | 0 | 0 |
| 1 | 1 | 1 | 0 |
| 1 | 1 | 1 | 0 |
| 0 | 0 | 0 | 0 |
| 0 | 0 | 0 | 0 |
| 1 | 1 | 1 | 0 |
| 0 | 0 | 0 | 0 |
| 0 | 0 | 0 | 0 |
| 0 | 0 | 0 | 0 |
| 0 | 0 | 0 | 0 |
| 0 | 0 | 0 | 0 |
| 0 | 0 | 0 | 0 |
| 0 | 0 | 0 | 0 |
| 0 | 0 | 0 | 0 |
| 1 | 0 | 0 | 0 |
| 1 | 0 | 0 | 0 |
| 1 | 0 | 0 | 0 |
| 0 | 0 | 0 | 0 |
| 0 | 0 | 0 | 0 |
| 0 | 0 | 0 | 0 |

| | | | |
|---|---|---|---|
| 0 | 0 | 0 | 0 |
| 1 | 0 | 0 | 0 |
| 0 | 0 | 0 | 0 |
| 0 | 0 | 0 | 0 |
| 0 | 0 | 0 | 0 |
| 0 | 0 | 0 | 0 |
| 0 | 0 | 0 | 0 |
| 0 | 0 | 0 | 0 |
| 0 | 0 | 0 | 0 |
| 0 | 0 | 0 | 0 |
| 1 | 1 | 0 | 0 |
| 1 | 1 | 0 | 0 |
| 0 | 0 | 0 | 0 |
| 0 | 0 | 0 | 0 |
| 0 | 0 | 0 | 0 |
| 1 | 0 | 0 | 0 |
| 1 | 0 | 0 | 0 |
| 1 | 1 | 0 | 1 |
| 0 | 0 | 0 | 0 |
| 0 | 0 | 0 | 0 |
| 0 | 0 | 0 | 0 |
| 0 | 0 | 0 | 0 |
| 1 | 0 | 0 | 1 |
| 0 | 0 | 0 | 0 |
| 0 | 0 | 0 | 0 |
| 1 | 0 | 0 | 0 |
| 0 | 0 | 0 | 0 |
| 0 | 0 | 0 | 0 |
| 0 | 0 | 0 | 0 |
| 0 | 0 | 0 | 0 |
| 0 | 0 | 0 | 0 |
| 0 | 0 | 0 | 0 |
| 0 | 0 | 0 | 0 |
| 0 | 0 | 0 | 0 |
| 0 | 0 | 0 | 0 |
| 1 | 0 | 0 | 0 |
| 1 | 0 | 0 | 0 |
| 1 | 1 | 0 | 0 |
| 1 | 1 | 0 | 0 |
| 0 | 0 | 0 | 0 |
| 0 | 0 | 0 | 0 |
| 1 | 0 | 0 | 0 |
| 0 | 0 | 0 | 0 |
| 1 | 0 | 0 | 0 |
| 1 | 1 | 0 | 0 |
| 0 | 0 | 0 | 0 |
| 0 | 0 | 0 | 0 |
| 0 | 0 | 0 | 0 |
| 0 | 0 | 0 | 0 |
| 0 | 0 | 0 | 0 |
| 0 | 0 | 0 | 0 |
| 1 | 0 | 0 | 0 |
| 0 | 0 | 0 | 0 |
| 0 | 0 | 0 | 0 |

| | | | |
|---|---|---|---|
| 0 | 0 | 0 | 0 |
| 1 | 1 | 0 | 0 |
| 0 | 0 | 0 | 0 |
| 1 | 0 | 0 | 0 |
| 1 | 0 | 0 | 0 |
| 0 | 0 | 0 | 0 |
| 0 | 0 | 0 | 0 |
| 0 | 0 | 0 | 0 |
| 0 | 0 | 0 | 0 |
| 0 | 0 | 0 | 0 |
| 0 | 0 | 0 | 0 |
| 1 | 1 | 0 | 0 |
| 1 | 1 | 0 | 1 |
| 0 | 0 | 0 | 0 |
| 0 | 1 | 0 | 0 |
| 0 | 0 | 0 | 0 |
| 0 | 0 | 0 | 0 |
| 0 | 0 | 0 | 0 |
| 0 | 0 | 0 | 0 |
| 0 | 0 | 0 | 0 |
| 0 | 0 | 0 | 0 |
| 1 | 0 | 0 | 0 |
| 0 | 0 | 0 | 0 |
| 0 | 0 | 0 | 0 |
| 0 | 0 | 0 | 0 |
| 0 | 0 | 0 | 0 |
| 0 | 0 | 0 | 0 |
| 1 | 0 | 0 | 0 |
| 0 | 0 | 0 | 0 |
| 0 | 0 | 0 | 0 |
| 0 | 0 | 0 | 0 |
| 0 | 0 | 0 | 0 |
| 0 | 0 | 0 | 0 |
| 0 | 0 | 0 | 0 |
| 0 | 0 | 0 | 0 |
| 0 | 0 | 0 | 0 |
| 0 | 0 | 0 | 0 |
| 0 | 0 | 0 | 0 |
| 0 | 0 | 0 | 0 |
| 0 | 0 | 0 | 0 |
| 0 | 0 | 0 | 0 |
| 0 | 0 | 0 | 0 |
| 0 | 0 | 0 | 0 |
| 0 | 0 | 0 | 0 |
| 0 | 0 | 0 | 0 |
| 0 | 0 | 0 | 0 |
| 0 | 0 | 0 | 0 |
| 0 | 0 | 0 | 0 |
| 0 | 0 | 0 | 0 |
| 0 | 0 | 0 | 0 |
| 0 | 0 | 0 | 0 |
| 0 | 0 | 0 | 0 |
| 0 | 0 | 0 | 0 |
| 0 | 0 | 0 | 0 |

| | | | |
|---|---|---|---|
| 0 | 0 | 0 | 0 |
| 0 | 0 | 0 | 0 |
| 0 | 0 | 0 | 0 |
| 0 | 0 | 0 | 0 |
| 0 | 0 | 0 | 0 |
| 0 | 0 | 0 | 0 |
| 0 | 0 | 0 | 0 |
| 0 | 0 | 0 | 0 |
| 0 | 0 | 0 | 0 |
| 0 | 0 | 0 | 0 |
| 0 | 0 | 0 | 0 |
| 0 | 0 | 0 | 0 |
| 1 | 0 | 0 | 0 |
| 1 | 1 | 0 | 0 |
| 0 | 0 | 0 | 0 |
| 1 | 0 | 0 | 0 |
| 0 | 0 | 0 | 0 |
| 1 | 0 | 0 | 0 |
| 0 | 0 | 0 | 0 |
| 0 | 0 | 0 | 0 |
| 1 | 0 | 0 | 0 |
| 0 | 0 | 0 | 0 |
| 0 | 0 | 0 | 0 |
| 1 | 0 | 0 | 0 |
| 1 | 0 | 0 | 0 |
| 0 | 0 | 0 | 0 |
| 0 | 0 | 0 | 0 |
| 1 | 0 | 0 | 0 |
| 1 | 0 | 0 | 0 |
| 1 | 0 | 0 | 0 |
| 1 | 0 | 0 | 0 |
| 1 | 0 | 0 | 0 |
| 0 | 0 | 0 | 0 |
| 0 | 0 | 0 | 0 |
| 0 | 0 | 0 | 0 |
| 0 | 0 | 0 | 0 |
| 0 | 0 | 0 | 0 |
| 0 | 0 | 0 | 0 |
| 0 | 0 | 0 | 0 |
| 0 | 0 | 0 | 0 |
| 0 | 0 | 0 | 0 |
| 0 | 0 | 0 | 0 |
| 0 | 0 | 0 | 0 |
| 0 | 0 | 0 | 0 |
| 1 | 1 | 0 | 0 |
| 0 | 0 | 0 | 0 |
| 0 | 0 | 0 | 0 |
| 0 | 0 | 0 | 0 |
| 0 | 0 | 0 | 0 |
| 1 | 1 | 0 | 0 |
| 1 | 0 | 0 | 0 |
| 0 | 0 | 0 | 0 |
| 0 | 0 | 0 | 0 |
| 0 | 0 | 0 | 1 |

| | | | |
|---|---|---|---|
| 0 | 0 | 0 | 0 |
| 0 | 0 | 0 | 0 |
| 0 | 0 | 0 | 0 |
| 0 | 0 | 0 | 0 |
| 1 | 0 | 0 | 1 |
| 0 | 0 | 0 | 0 |
| 1 | 0 | 0 | 0 |
| 0 | 0 | 0 | 0 |
| 0 | 0 | 0 | 0 |
| 0 | 0 | 0 | 0 |
| 0 | 0 | 0 | 0 |
| 0 | 0 | 0 | 0 |
| 0 | 0 | 0 | 0 |
| 1 | 0 | 0 | 0 |
| 0 | 0 | 0 | 0 |
| 0 | 0 | 0 | 0 |
| 1 | 1 | 0 | 0 |
| 0 | 0 | 0 | 0 |
| 1 | 0 | 0 | 0 |
| 1 | 0 | 0 | 0 |
| 0 | 0 | 0 | 0 |
| 0 | 0 | 0 | 0 |
| 0 | 0 | 0 | 0 |
| 1 | 0 | 0 | 0 |
| 1 | 0 | 0 | 0 |
| 1 | 0 | 0 | 0 |
| 1 | 0 | 0 | 0 |
| 1 | 0 | 0 | 0 |
| 1 | 0 | 0 | 0 |
| 1 | 0 | 0 | 0 |
| 1 | 0 | 0 | 0 |
| 1 | 0 | 0 | 0 |
| 1 | 0 | 0 | 0 |
| 1 | 0 | 0 | 0 |
| 0 | 0 | 0 | 0 |
| 0 | 0 | 0 | 0 |
| 0 | 0 | 0 | 0 |
| 1 | 0 | 0 | 0 |
| 1 | 0 | 0 | 0 |
| 0 | 0 | 0 | 0 |
| 0 | 0 | 0 | 0 |
| 1 | 0 | 0 | 0 |
| 0 | 0 | 0 | 0 |
| 1 | 0 | 0 | 0 |
| 1 | 0 | 0 | 0 |
| 1 | 0 | 0 | 0 |
| 1 | 0 | 0 | 0 |
| 1 | 0 | 0 | 0 |
| 0 | 0 | 0 | 0 |
| 0 | 0 | 0 | 0 |
| 1 | 0 | 0 | 0 |
| 0 | 0 | 0 | 0 |
| 1 | 0 | 0 | 0 |
| 1 | 0 | 0 | 0 |

| | | | |
|---|---|---|---|
| 1 | 0 | 0 | 0 |
| 1 | 0 | 0 | 0 |
| 1 | 0 | 0 | 0 |
| 1 | 0 | 0 | 0 |
| 1 | 0 | 0 | 0 |
| 1 | 0 | 0 | 0 |
| 0 | 0 | 0 | 0 |
| 1 | 0 | 0 | 0 |
| 1 | 0 | 0 | 0 |
| 1 | 0 | 0 | 0 |
| 1 | 0 | 0 | 0 |
| 1 | 0 | 0 | 0 |
| 1 | 0 | 0 | 0 |
| 1 | 0 | 0 | 0 |
| 0 | 0 | 0 | 0 |
| 0 | 0 | 0 | 0 |
| 1 | 1 | 0 | 0 |
| 1 | 1 | 0 | 0 |
| 0 | 0 | 0 | 0 |
| 0 | 0 | 0 | 0 |
| 0 | 0 | 0 | 0 |
| 1 | 1 | 0 | 1 |
| 0 | 0 | 0 | 0 |
| 0 | 0 | 0 | 0 |
| 0 | 0 | 0 | 0 |
| 0 | 0 | 0 | 0 |
| 0 | 0 | 0 | 0 |
| 0 | 0 | 0 | 0 |
| 0 | 0 | 0 | 0 |
| 1 | 0 | 1 | 0 |
| 0 | 0 | 0 | 0 |
| 0 | 0 | 0 | 0 |
| 0 | 0 | 0 | 0 |
| 0 | 0 | 0 | 0 |
| 0 | 0 | 0 | 0 |
| 0 | 0 | 0 | 0 |
| 0 | 0 | 0 | 0 |
| 0 | 0 | 0 | 0 |
| 0 | 0 | 0 | 0 |
| 0 | 0 | 0 | 0 |
| 1 | 1 | 0 | 0 |
| 1 | 1 | 0 | 0 |
| 1 | 0 | 0 | 0 |
| 1 | 1 | 0 | 0 |
| 1 | 0 | 0 | 0 |
| 1 | 1 | 0 | 0 |
| 1 | 0 | 0 | 0 |
| 1 | 0 | 0 | 0 |
| 0 | 0 | 0 | 0 |
| 0 | 0 | 0 | 0 |
| 1 | 0 | 0 | 0 |
| 1 | 0 | 0 | 0 |
| 1 | 0 | 0 | 0 |
| 0 | 0 | 0 | 0 |

| | | | |
|---|---|---|---|
| 0 | 0 | 0 | 0 |
| 0 | 0 | 0 | 0 |
| 0 | 0 | 0 | 0 |
| 1 | 1 | 1 | 0 |
| 0 | 0 | 0 | 0 |
| 0 | 0 | 0 | 0 |
| 0 | 0 | 0 | 0 |
| 0 | 0 | 0 | 0 |
| 0 | 1 | 0 | 0 |
| 0 | 1 | 0 | 0 |
| 0 | 1 | 0 | 0 |
| 0 | 0 | 0 | 0 |
| 0 | 0 | 0 | 0 |
| 0 | 0 | 0 | 0 |
| 1 | 0 | 0 | 0 |
| 1 | 0 | 0 | 0 |
| 0 | 0 | 0 | 0 |
| 0 | 0 | 0 | 0 |
| 1 | 0 | 0 | 0 |
| 0 | 0 | 0 | 0 |
| 1 | 0 | 0 | 0 |
| 1 | 0 | 0 | 0 |
| 0 | 0 | 0 | 0 |
| 0 | 0 | 0 | 0 |
| 0 | 0 | 0 | 0 |
| 0 | 0 | 0 | 0 |
| 0 | 0 | 0 | 0 |
| 0 | 0 | 1 | 0 |
| 0 | 0 | 0 | 0 |
| 0 | 1 | 0 | 0 |
| 0 | 0 | 0 | 0 |
| 0 | 0 | 0 | 0 |
| 0 | 0 | 0 | 0 |
| 0 | 0 | 0 | 0 |
| 0 | 0 | 0 | 0 |
| 0 | 0 | 0 | 0 |
| 1 | 0 | 0 | 0 |
| 1 | 0 | 0 | 0 |
| 1 | 0 | 0 | 0 |
| 0 | 0 | 0 | 0 |
| 0 | 0 | 0 | 0 |
| 0 | 0 | 0 | 0 |
| 0 | 0 | 0 | 0 |
| 0 | 0 | 0 | 0 |
| 0 | 0 | 0 | 0 |
| 0 | 0 | 0 | 0 |
| 0 | 0 | 0 | 0 |
| 0 | 0 | 0 | 1 |
| 0 | 0 | 0 | 0 |
| 0 | 0 | 0 | 0 |
| 0 | 0 | 0 | 0 |
| 0 | 0 | 0 | 0 |
| 1 | 1 | 0 | 0 |
| 0 | 0 | 0 | 0 |

| | | | |
|---|---|---|---|
| 0 | 0 | 0 | 0 |
| 0 | 0 | 0 | 0 |
| 0 | 0 | 0 | 0 |
| 0 | 0 | 0 | 0 |
| 1 | 0 | 0 | 0 |
| 0 | 0 | 0 | 0 |
| 0 | 0 | 0 | 0 |
| 0 | 0 | 0 | 0 |
| 0 | 0 | 0 | 0 |
| 0 | 0 | 0 | 0 |
| 0 | 0 | 0 | 0 |
| 0 | 0 | 0 | 0 |
| 0 | 0 | 0 | 0 |
| 0 | 0 | 0 | 0 |
| 0 | 0 | 0 | 0 |
| 0 | 0 | 0 | 0 |
| 0 | 0 | 0 | 0 |
| 0 | 0 | 0 | 0 |
| 0 | 0 | 0 | 0 |
| 0 | 0 | 0 | 0 |
| 0 | 0 | 0 | 0 |
| 0 | 0 | 0 | 0 |
| 0 | 0 | 0 | 0 |
| 0 | 0 | 0 | 0 |
| 0 | 0 | 0 | 0 |
| 0 | 0 | 0 | 0 |
| 0 | 0 | 0 | 0 |
| 0 | 0 | 0 | 0 |
| 0 | 0 | 0 | 0 |
| 0 | 0 | 0 | 0 |
| 0 | 0 | 0 | 0 |
| 0 | 0 | 0 | 0 |
| 0 | 0 | 0 | 0 |
| 0 | 0 | 0 | 0 |
| 0 | 0 | 0 | 0 |
| 0 | 0 | 0 | 0 |
| 0 | 0 | 0 | 0 |
| 0 | 0 | 0 | 0 |
| 0 | 0 | 0 | 0 |
| 0 | 0 | 0 | 0 |
| 1 | 1 | 0 | 0 |
| 0 | 0 | 0 | 0 |
| 0 | 0 | 0 | 0 |
| 0 | 0 | 0 | 0 |
| 0 | 0 | 0 | 0 |
| 1 | 0 | 0 | 0 |
| 1 | 0 | 0 | 0 |
| 0 | 0 | 0 | 0 |
| 1 | 0 | 0 | 0 |
| 0 | 0 | 0 | 0 |
| 0 | 0 | 0 | 0 |
| 1 | 0 | 0 | 0 |
| 1 | 0 | 0 | 0 |
| 0 | 0 | 0 | 0 |

| | | | |
|---|---|---|---|
| 0 | 0 | 0 | 0 |
| 0 | 0 | 0 | 0 |
| 0 | 0 | 0 | 0 |
| 0 | 0 | 0 | 0 |
| 1 | 0 | 0 | 0 |
| 0 | 0 | 0 | 0 |
| 0 | 0 | 0 | 0 |
| 0 | 0 | 0 | 0 |
| 0 | 0 | 0 | 0 |
| 0 | 0 | 0 | 0 |
| 0 | 0 | 0 | 0 |
| 0 | 0 | 0 | 0 |
| 0 | 0 | 0 | 0 |
| 0 | 0 | 0 | 0 |
| 0 | 0 | 0 | 0 |
| 1 | 1 | 0 | 1 |
| 0 | 0 | 0 | 0 |
| 0 | 0 | 0 | 0 |
| 0 | 0 | 0 | 0 |
| 0 | 0 | 0 | 0 |
| 0 | 0 | 0 | 0 |
| 0 | 0 | 0 | 0 |
| 0 | 0 | 0 | 0 |
| 0 | 0 | 0 | 0 |
| 0 | 0 | 0 | 0 |
| 0 | 0 | 0 | 0 |
| 0 | 0 | 0 | 0 |
| 1 | 0 | 0 | 0 |
| 1 | 0 | 0 | 0 |
| 0 | 0 | 0 | 0 |
| 1 | 1 | 0 | 0 |
| 0 | 0 | 0 | 0 |
| 0 | 0 | 0 | 0 |
| 0 | 0 | 0 | 0 |
| 1 | 1 | 0 | 0 |
| 1 | 1 | 0 | 0 |
| 0 | 0 | 0 | 0 |
| 1 | 0 | 0 | 0 |
| 0 | 0 | 0 | 0 |
| 1 | 1 | 0 | 0 |
| 0 | 1 | 0 | 0 |
| 0 | 0 | 0 | 0 |
| 0 | 0 | 0 | 0 |
| 0 | 0 | 0 | 0 |
| 1 | 1 | 0 | 0 |
| 1 | 1 | 0 | 0 |
| 1 | 1 | 0 | 0 |
| 0 | 0 | 0 | 0 |
| 0 | 0 | 0 | 0 |
| 0 | 0 | 0 | 0 |
| 0 | 0 | 0 | 0 |
| 0 | 0 | 0 | 0 |
| 0 | 0 | 0 | 0 |
| 0 | 0 | 0 | 0 |

| | | | |
|---|---|---|---|
| 0 | 0 | 0 | 0 |
| 0 | 0 | 0 | 0 |
| 1 | 0 | 0 | 0 |
| 1 | 0 | 0 | 0 |
| 0 | 0 | 0 | 0 |
| 1 | 0 | 0 | 0 |
| 0 | 0 | 0 | 0 |
| 0 | 0 | 0 | 0 |
| 0 | 0 | 0 | 0 |
| 0 | 1 | 0 | 0 |
| 0 | 0 | 0 | 0 |
| 0 | 0 | 0 | 0 |
| 0 | 0 | 0 | 0 |
| 0 | 0 | 0 | 0 |
| 0 | 0 | 0 | 0 |
| 0 | 0 | 0 | 0 |
| 0 | 0 | 0 | 0 |
| 1 | 0 | 0 | 0 |
| 0 | 0 | 0 | 0 |
| 1 | 0 | 0 | 0 |
| 0 | 0 | 0 | 0 |
| 1 | 0 | 0 | 0 |
| 0 | 0 | 0 | 0 |
| 1 | 0 | 0 | 0 |
| 0 | 0 | 0 | 0 |
| 0 | 0 | 0 | 0 |
| 0 | 0 | 0 | 0 |
| 0 | 0 | 0 | 0 |
| 0 | 0 | 0 | 0 |
| 0 | 0 | 0 | 0 |
| 0 | 0 | 0 | 0 |
| 1 | 0 | 0 | 0 |
| 0 | 0 | 0 | 0 |
| 0 | 0 | 0 | 0 |
| 0 | 0 | 0 | 0 |
| 0 | 0 | 0 | 0 |
| 0 | 0 | 0 | 0 |
| 0 | 0 | 0 | 0 |
| 1 | 1 | 0 | 0 |
| 1 | 1 | 0 | 0 |
| 1 | 1 | 0 | 0 |
| 1 | 1 | 0 | 0 |
| 1 | 1 | 0 | 0 |
| 1 | 0 | 0 | 0 |
| 1 | 0 | 0 | 0 |
| 1 | 0 | 0 | 0 |
| 0 | 0 | 0 | 0 |
| 0 | 0 | 0 | 0 |
| 0 | 0 | 0 | 0 |
| 0 | 0 | 0 | 0 |
| 0 | 0 | 0 | 0 |
| 0 | 0 | 0 | 0 |
| 0 | 0 | 0 | 0 |
| 0 | 0 | 0 | 0 |

| | | | |
|---|---|---|---|
| 0 | 0 | 0 | 0 |
| 0 | 0 | 0 | 0 |
| 0 | 0 | 0 | 0 |
| 0 | 0 | 0 | 0 |
| 0 | 0 | 0 | 0 |
| 1 | 0 | 0 | 0 |
| 1 | 0 | 0 | 0 |
| 0 | 0 | 0 | 0 |
| 0 | 0 | 0 | 0 |
| 0 | 0 | 0 | 0 |
| 0 | 0 | 0 | 0 |
| 1 | 0 | 0 | 0 |
| 0 | 0 | 1 | 0 |
| 0 | 0 | 0 | 0 |
| 0 | 0 | 0 | 0 |
| 0 | 0 | 0 | 0 |
| 1 | 0 | 0 | 0 |
| 0 | 0 | 0 | 0 |
| 1 | 0 | 0 | 0 |
| 1 | 0 | 0 | 0 |
| 0 | 0 | 0 | 0 |
| 1 | 0 | 0 | 0 |
| 1 | 1 | 0 | 0 |
| 0 | 0 | 0 | 0 |
| 1 | 1 | 0 | 0 |
| 1 | 1 | 0 | 0 |
| 1 | 1 | 0 | 0 |
| 0 | 0 | 0 | 0 |
| 1 | 0 | 0 | 0 |
| 0 | 0 | 0 | 0 |
| 0 | 0 | 0 | 0 |
| 0 | 0 | 0 | 0 |
| 0 | 0 | 0 | 0 |
| 0 | 0 | 0 | 0 |
| 0 | 0 | 0 | 0 |
| 0 | 0 | 0 | 0 |
| 0 | 0 | 0 | 0 |
| 0 | 0 | 0 | 0 |
| 0 | 0 | 0 | 0 |
| 0 | 0 | 0 | 0 |
| 0 | 0 | 0 | 0 |
| 0 | 0 | 0 | 0 |
| 0 | 0 | 0 | 0 |
| 0 | 0 | 0 | 0 |
| 0 | 0 | 0 | 0 |
| 0 | 0 | 0 | 0 |
| 0 | 0 | 0 | 0 |
| 0 | 0 | 0 | 0 |
| 0 | 0 | 0 | 0 |
| 0 | 0 | 0 | 0 |
| 0 | 0 | 0 | 0 |
| 0 | 0 | 0 | 0 |
| 0 | 0 | 0 | 0 |
| 0 | 0 | 0 | 0 |

| | | | |
|---|---|---|---|
| 0 | 0 | 0 | 0 |
| 0 | 0 | 0 | 0 |
| 0 | 0 | 0 | 0 |
| 0 | 0 | 0 | 0 |
| 0 | 0 | 0 | 0 |
| 0 | 0 | 0 | 0 |
| 0 | 0 | 0 | 0 |
| 0 | 0 | 0 | 0 |
| 0 | 0 | 0 | 0 |
| 0 | 0 | 0 | 0 |
| 0 | 1 | 0 | 0 |
| 0 | 1 | 0 | 0 |
| 0 | 1 | 0 | 0 |
| 0 | 1 | 0 | 0 |
| 0 | 0 | 0 | 0 |
| 0 | 1 | 0 | 0 |
| 0 | 1 | 0 | 0 |
| 1 | 1 | 0 | 0 |
| 0 | 1 | 0 | 0 |
| 1 | 0 | 0 | 0 |
| 1 | 1 | 0 | 0 |
| 1 | 1 | 0 | 0 |
| 0 | 0 | 1 | 1 |
| 0 | 0 | 0 | 0 |
| 1 | 0 | 0 | 0 |
| 0 | 0 | 0 | 0 |
| 0 | 0 | 0 | 0 |
| 0 | 0 | 0 | 0 |
| 0 | 0 | 0 | 0 |
| 0 | 0 | 0 | 0 |
| 0 | 0 | 0 | 0 |
| 0 | 0 | 0 | 0 |
| 0 | 0 | 0 | 0 |
| 0 | 0 | 0 | 0 |
| 0 | 0 | 0 | 0 |
| 0 | 0 | 0 | 0 |
| 0 | 0 | 0 | 0 |
| 0 | 0 | 0 | 0 |
| 0 | 0 | 0 | 0 |
| 0 | 0 | 0 | 0 |
| 1 | 0 | 0 | 0 |
| 0 | 0 | 0 | 0 |
| 1 | 0 | 0 | 0 |
| 1 | 0 | 0 | 0 |
| 0 | 0 | 0 | 0 |
| 1 | 0 | 0 | 0 |
| 0 | 0 | 0 | 0 |
| 0 | 0 | 0 | 0 |
| 0 | 0 | 0 | 0 |
| 0 | 0 | 0 | 0 |
| 0 | 0 | 0 | 0 |
| 0 | 0 | 0 | 0 |
| 1 | 1 | 0 | 0 |
| 1 | 1 | 0 | 0 |

| | | | |
|---|---|---|---|
| 1 | 1 | 0 | 0 |
| 0 | 0 | 0 | 0 |
| 0 | 0 | 0 | 0 |
| 0 | 0 | 0 | 0 |
| 0 | 0 | 0 | 0 |
| 0 | 0 | 0 | 0 |
| 0 | 1 | 0 | 0 |
| 0 | 0 | 0 | 0 |
| 0 | 0 | 0 | 0 |
| 0 | 1 | 0 | 0 |
| 1 | 0 | 0 | 0 |
| 0 | 0 | 0 | 0 |
| 0 | 0 | 1 | 0 |
| 0 | 0 | 1 | 0 |
| 0 | 0 | 0 | 0 |
| 0 | 0 | 0 | 0 |
| 0 | 0 | 0 | 0 |
| 0 | 1 | 0 | 0 |
| 0 | 0 | 0 | 0 |
| 0 | 0 | 0 | 0 |
| 1 | 0 | 0 | 0 |
| 0 | 0 | 0 | 1 |
| 0 | 0 | 0 | 0 |
| 0 | 0 | 0 | 0 |
| 0 | 0 | 0 | 0 |
| 0 | 0 | 0 | 0 |
| 0 | 0 | 0 | 0 |
| 0 | 0 | 0 | 0 |
| 0 | 0 | 0 | 0 |
| 0 | 0 | 0 | 0 |
| 0 | 0 | 0 | 0 |
| 0 | 0 | 0 | 0 |
| 0 | 0 | 0 | 0 |
| 0 | 0 | 0 | 0 |
| 0 | 0 | 0 | 0 |
| 0 | 0 | 0 | 0 |
| 1 | 0 | 0 | 0 |
| 0 | 0 | 0 | 0 |
| 1 | 0 | 0 | 0 |
| 0 | 0 | 0 | 0 |
| 0 | 0 | 0 | 0 |
| 0 | 0 | 0 | 0 |
| 0 | 0 | 0 | 0 |
| 0 | 0 | 0 | 0 |
| 0 | 0 | 0 | 0 |
| 0 | 0 | 0 | 0 |
| 0 | 0 | 0 | 0 |
| 0 | 0 | 0 | 0 |
| 0 | 0 | 0 | 0 |
| 1 | 0 | 0 | 0 |
| 0 | 0 | 0 | 0 |
| 0 | 0 | 0 | 0 |
| 0 | 0 | 0 | 0 |
| 0 | 0 | 0 | 0 |

| | | | |
|---|---|---|---|
| 0 | 0 | 0 | 0 |
| 0 | 0 | 0 | 0 |
| 0 | 0 | 0 | 0 |
| 0 | 0 | 0 | 0 |
| 0 | 0 | 0 | 0 |
| 0 | 0 | 0 | 0 |
| 0 | 0 | 0 | 0 |
| 0 | 0 | 0 | 0 |
| 0 | 0 | 0 | 0 |
| 0 | 0 | 0 | 0 |
| 0 | 0 | 0 | 0 |
| 0 | 0 | 0 | 0 |
| 0 | 0 | 1 | 0 |
| 1 | 0 | 0 | 0 |
| 1 | 0 | 0 | 0 |
| 0 | 0 | 0 | 0 |
| 0 | 0 | 0 | 0 |
| 0 | 0 | 0 | 0 |
| 0 | 0 | 0 | 0 |
| 0 | 0 | 0 | 0 |
| 0 | 0 | 0 | 0 |
| 0 | 0 | 0 | 0 |
| 0 | 0 | 0 | 0 |
| 1 | 1 | 0 | 1 |
| 0 | 0 | 0 | 0 |
| 1 | 1 | 0 | 0 |
| 1 | 1 | 0 | 1 |
| 0 | 0 | 0 | 0 |
| 0 | 0 | 0 | 0 |
| 0 | 0 | 0 | 0 |
| 0 | 0 | 0 | 0 |
| 0 | 0 | 0 | 0 |
| 0 | 0 | 0 | 0 |
| 0 | 0 | 0 | 0 |
| 0 | 0 | 0 | 0 |
| 1 | 0 | 0 | 0 |
| 0 | 0 | 0 | 0 |
| 0 | 0 | 0 | 0 |
| 0 | 0 | 0 | 0 |
| 0 | 0 | 0 | 0 |
| 0 | 0 | 0 | 0 |
| 0 | 0 | 0 | 0 |
| 1 | 0 | 0 | 0 |
| 1 | 0 | 0 | 0 |
| 1 | 0 | 0 | 0 |
| 1 | 0 | 0 | 0 |
| 1 | 0 | 0 | 0 |
| 0 | 0 | 0 | 0 |
| 0 | 0 | 0 | 0 |
| 1 | 0 | 0 | 0 |
| 1 | 0 | 0 | 0 |
| 0 | 0 | 0 | 0 |
| 1 | 0 | 0 | 0 |
| 1 | 0 | 0 | 0 |

| | | | |
|---|---|---|---|
| 0 | 0 | 0 | 0 |
| 1 | 0 | 0 | 0 |
| 0 | 0 | 0 | 0 |
| 0 | 0 | 0 | 0 |
| 0 | 0 | 0 | 0 |
| 0 | 0 | 0 | 0 |
| 1 | 0 | 0 | 0 |
| 0 | 0 | 0 | 0 |
| 0 | 0 | 0 | 0 |
| 0 | 0 | 0 | 0 |
| 0 | 0 | 0 | 0 |
| 1 | 1 | 0 | 0 |
| 0 | 0 | 0 | 0 |
| 0 | 0 | 0 | 0 |
| 0 | 0 | 0 | 0 |
| 0 | 0 | 0 | 0 |
| 0 | 0 | 0 | 0 |
| 0 | 0 | 0 | 0 |
| 0 | 0 | 0 | 0 |
| 0 | 0 | 0 | 0 |
| 0 | 0 | 0 | 0 |
| 0 | 0 | 0 | 0 |
| 0 | 0 | 0 | 0 |
| 0 | 0 | 0 | 0 |
| 0 | 0 | 0 | 0 |
| 0 | 0 | 0 | 0 |
| 0 | 0 | 0 | 0 |
| 0 | 0 | 0 | 0 |
| 1 | 1 | 0 | 0 |
| 1 | 0 | 0 | 0 |
| 0 | 0 | 0 | 0 |
| 0 | 0 | 0 | 0 |
| 0 | 0 | 0 | 0 |
| 0 | 0 | 0 | 0 |
| 0 | 0 | 0 | 0 |
| 0 | 0 | 0 | 0 |
| 0 | 0 | 0 | 0 |
| 0 | 0 | 0 | 0 |
| 0 | 0 | 0 | 0 |
| 0 | 0 | 0 | 0 |
| 0 | 0 | 0 | 0 |
| 0 | 0 | 0 | 0 |
| 0 | 0 | 0 | 0 |
| 0 | 0 | 0 | 0 |
| 0 | 0 | 0 | 0 |
| 0 | 0 | 0 | 0 |
| 0 | 0 | 0 | 0 |
| 0 | 0 | 0 | 0 |
| 0 | 0 | 0 | 0 |
| 0 | 0 | 0 | 0 |
| 0 | 0 | 0 | 0 |
| 0 | 0 | 0 | 0 |
| 0 | 0 | 1 | 0 |
| 0 | 0 | 0 | 0 |

| | | | |
|---|---|---|---|
| 0 | 0 | 0 | 0 |
| 0 | 0 | 0 | 0 |
| 0 | 0 | 0 | 0 |
| 0 | 0 | 0 | 0 |
| 1 | 0 | 0 | 0 |
| 0 | 0 | 0 | 0 |
| 0 | 0 | 0 | 0 |
| 1 | 1 | 0 | 0 |
| 0 | 0 | 0 | 0 |
| 1 | 1 | 0 | 0 |
| 0 | 0 | 0 | 0 |
| 0 | 0 | 0 | 0 |
| 0 | 0 | 0 | 0 |
| 0 | 0 | 0 | 0 |
| 0 | 0 | 0 | 0 |
| 0 | 0 | 0 | 0 |
| 1 | 0 | 0 | 0 |
| 0 | 0 | 0 | 0 |
| 0 | 0 | 0 | 0 |
| 1 | 0 | 0 | 0 |
| 1 | 0 | 0 | 0 |
| 0 | 0 | 0 | 0 |
| 0 | 0 | 0 | 0 |
| 0 | 0 | 0 | 0 |
| 0 | 0 | 0 | 0 |
| 0 | 0 | 0 | 0 |
| 1 | 0 | 0 | 0 |
| 0 | 0 | 0 | 0 |
| 0 | 0 | 0 | 0 |
| 1 | 0 | 0 | 0 |
| 0 | 0 | 0 | 0 |
| 0 | 0 | 0 | 0 |
| 0 | 0 | 0 | 0 |
| 1 | 1 | 0 | 0 |
| 0 | 0 | 0 | 0 |
| 0 | 0 | 0 | 0 |
| 0 | 0 | 0 | 0 |
| 0 | 0 | 0 | 0 |
| 0 | 1 | 0 | 0 |
| 0 | 0 | 0 | 0 |
| 0 | 0 | 0 | 0 |
| 0 | 0 | 0 | 0 |
| 0 | 0 | 0 | 0 |
| 1 | 1 | 0 | 0 |
| 0 | 0 | 0 | 0 |
| 0 | 0 | 0 | 0 |
| 0 | 0 | 0 | 0 |
| 0 | 0 | 0 | 0 |
| 0 | 0 | 0 | 0 |
| 0 | 0 | 0 | 0 |
| 0 | 0 | 0 | 0 |
| 1 | 0 | 0 | 0 |
| 0 | 0 | 0 | 0 |
| 0 | 0 | 0 | 0 |

| | | | |
|---|---|---|---|
| 1 | 0 | 0 | 1 |
| 1 | 0 | 0 | 1 |
| 1 | 0 | 0 | 1 |
| 0 | 0 | 0 | 0 |
| 0 | 0 | 0 | 0 |
| 0 | 0 | 0 | 0 |
| 1 | 0 | 0 | 0 |
| 0 | 0 | 0 | 0 |
| 0 | 0 | 0 | 0 |
| 0 | 0 | 0 | 0 |
| 0 | 0 | 1 | 0 |
| 0 | 0 | 0 | 0 |
| 0 | 0 | 0 | 0 |
| 0 | 0 | 0 | 0 |
| 0 | 0 | 0 | 0 |
| 0 | 0 | 0 | 0 |
| 1 | 0 | 0 | 0 |
| 0 | 0 | 0 | 0 |
| 0 | 0 | 0 | 0 |
| 0 | 0 | 0 | 0 |
| 0 | 0 | 0 | 0 |
| 0 | 0 | 0 | 0 |
| 0 | 0 | 0 | 0 |
| 1 | 1 | 0 | 0 |
| 0 | 0 | 0 | 0 |
| 0 | 0 | 0 | 0 |
| 0 | 0 | 0 | 0 |
| 0 | 0 | 0 | 0 |
| 0 | 0 | 0 | 0 |
| 0 | 0 | 0 | 0 |
| 0 | 0 | 0 | 0 |
| 0 | 0 | 0 | 0 |
| 0 | 1 | 0 | 0 |
| 0 | 0 | 0 | 0 |
| 1 | 1 | 0 | 0 |
| 0 | 0 | 0 | 0 |
| 0 | 0 | 0 | 0 |
| 0 | 0 | 0 | 0 |
| 0 | 0 | 0 | 0 |
| 0 | 0 | 0 | 0 |
| 0 | 0 | 0 | 0 |
| 1 | 0 | 0 | 1 |
| 0 | 0 | 0 | 0 |
| 1 | 0 | 0 | 1 |
| 0 | 0 | 0 | 0 |
| 1 | 0 | 0 | 1 |
| 0 | 0 | 0 | 0 |
| 0 | 0 | 0 | 0 |
| 0 | 0 | 0 | 0 |
| 1 | 0 | 0 | 0 |
| 0 | 0 | 0 | 0 |
| 0 | 0 | 0 | 0 |
| 0 | 0 | 0 | 0 |
| 0 | 0 | 0 | 0 |

| | | | |
|---|---|---|---|
| 0 | 0 | 0 | 0 |
| 0 | 0 | 0 | 0 |
| 0 | 0 | 0 | 0 |
| 0 | 0 | 0 | 0 |
| 0 | 0 | 0 | 0 |
| 0 | 0 | 0 | 0 |
| 0 | 0 | 0 | 0 |
| 0 | 0 | 0 | 0 |
| 0 | 0 | 0 | 0 |
| 0 | 0 | 0 | 0 |
| 0 | 0 | 0 | 0 |
| 0 | 0 | 0 | 0 |
| 0 | 0 | 0 | 0 |
| 0 | 0 | 0 | 0 |
| 0 | 0 | 0 | 0 |
| 0 | 0 | 0 | 0 |
| 0 | 0 | 0 | 0 |
| 0 | 0 | 0 | 0 |
| 0 | 0 | 1 | 0 |
| 0 | 0 | 0 | 0 |
| 0 | 0 | 0 | 0 |
| 0 | 0 | 0 | 0 |
| 0 | 0 | 0 | 0 |
| 0 | 0 | 0 | 0 |
| 0 | 0 | 0 | 0 |
| 0 | 0 | 0 | 0 |
| 0 | 0 | 0 | 0 |
| 0 | 0 | 0 | 0 |
| 0 | 0 | 1 | 0 |
| 0 | 0 | 0 | 0 |
| 0 | 0 | 1 | 0 |
| 1 | 0 | 0 | 0 |
| 1 | 1 | 1 | 0 |
| 1 | 1 | 1 | 0 |
| 0 | 0 | 0 | 0 |
| 0 | 0 | 0 | 0 |
| 0 | 1 | 0 | 0 |
| 1 | 1 | 1 | 0 |
| 1 | 1 | 1 | 0 |
| 0 | 0 | 1 | 0 |
| 0 | 0 | 1 | 0 |
| 1 | 0 | 0 | 0 |
| 0 | 0 | 0 | 0 |
| 0 | 0 | 0 | 0 |
| 1 | 1 | 0 | 0 |
| 0 | 0 | 0 | 0 |
| 0 | 0 | 0 | 0 |
| 0 | 0 | 0 | 0 |
| 0 | 0 | 0 | 0 |
| 0 | 0 | 0 | 0 |
| 0 | 0 | 0 | 0 |
| 0 | 0 | 0 | 0 |
| 0 | 0 | 0 | 0 |
| 0 | 0 | 0 | 0 |

| | | | |
|---|---|---|---|
| 0 | 0 | 0 | 0 |
| 0 | 0 | 0 | 0 |
| 0 | 0 | 0 | 0 |
| 0 | 0 | 0 | 0 |
| 0 | 0 | 0 | 0 |
| 0 | 0 | 0 | 0 |
| 0 | 0 | 0 | 0 |
| 0 | 0 | 0 | 0 |
| 0 | 0 | 0 | 0 |
| 0 | 0 | 0 | 0 |
| 0 | 0 | 0 | 0 |
| 0 | 0 | 0 | 0 |
| 0 | 0 | 0 | 0 |
| 0 | 0 | 0 | 0 |
| 0 | 0 | 0 | 0 |
| 0 | 0 | 0 | 0 |
| 0 | 0 | 0 | 0 |
| 0 | 0 | 0 | 0 |
| 0 | 0 | 0 | 0 |
| 1 | 0 | 0 | 0 |
| 1 | 0 | 0 | 0 |
| 0 | 0 | 0 | 0 |
| 0 | 0 | 0 | 0 |
| 0 | 0 | 0 | 0 |
| 0 | 0 | 0 | 0 |
| 1 | 0 | 0 | 0 |
| 0 | 0 | 0 | 0 |
| 0 | 0 | 0 | 0 |
| 0 | 0 | 0 | 0 |
| 1 | 0 | 0 | 0 |
| 0 | 0 | 0 | 0 |
| 0 | 0 | 0 | 0 |
| 1 | 0 | 0 | 0 |
| 1 | 0 | 0 | 0 |
| 0 | 0 | 0 | 0 |
| 0 | 0 | 0 | 0 |
| 0 | 0 | 0 | 0 |
| 0 | 0 | 0 | 0 |
| 0 | 0 | 0 | 0 |
| 0 | 0 | 0 | 0 |
| 0 | 0 | 0 | 0 |
| 0 | 0 | 0 | 0 |
| 0 | 0 | 0 | 0 |
| 0 | 0 | 0 | 0 |
| 0 | 0 | 0 | 0 |
| 0 | 0 | 0 | 0 |
| 0 | 0 | 0 | 0 |
| 0 | 1 | 0 | 0 |
| 0 | 1 | 0 | 0 |
| 0 | 0 | 0 | 0 |
| 0 | 1 | 0 | 0 |
| 0 | 1 | 0 | 0 |
| 0 | 0 | 0 | 0 |
| 0 | 0 | 0 | 0 |

| | | | |
|---|---|---|---|
| 0 | 0 | 0 | 0 |
| 0 | 0 | 0 | 0 |
| 1 | 1 | 0 | 1 |
| 0 | 0 | 0 | 0 |
| 0 | 0 | 0 | 0 |
| 0 | 0 | 0 | 0 |
| 0 | 0 | 0 | 0 |
| 0 | 1 | 0 | 0 |
| 1 | 1 | 0 | 0 |
| 1 | 1 | 0 | 0 |
| 1 | 1 | 0 | 0 |
| 1 | 1 | 0 | 0 |
| 1 | 1 | 0 | 0 |
| 0 | 1 | 0 | 0 |
| 0 | 1 | 0 | 0 |
| 0 | 0 | 0 | 0 |
| 0 | 0 | 0 | 0 |
| 0 | 0 | 0 | 0 |
| 0 | 0 | 0 | 0 |
| 0 | 0 | 0 | 0 |
| 0 | 0 | 0 | 0 |
| 0 | 0 | 0 | 0 |
| 0 | 1 | 0 | 0 |
| 0 | 1 | 0 | 0 |
| 0 | 0 | 0 | 0 |
| 0 | 0 | 1 | 0 |
| 0 | 0 | 0 | 0 |
| 0 | 0 | 0 | 0 |
| 0 | 0 | 0 | 0 |
| 0 | 1 | 0 | 0 |
| 0 | 0 | 0 | 0 |
| 0 | 0 | 0 | 0 |
| 0 | 0 | 0 | 0 |
| 0 | 0 | 0 | 0 |
| 0 | 0 | 0 | 0 |
| 0 | 0 | 0 | 0 |
| 0 | 0 | 0 | 0 |
| 0 | 0 | 0 | 0 |
| 0 | 0 | 0 | 0 |
| 1 | 0 | 0 | 0 |
| 1 | 1 | 0 | 0 |
| 1 | 0 | 0 | 0 |
| 0 | 0 | 0 | 0 |
| 0 | 1 | 0 | 0 |
| 1 | 1 | 0 | 0 |
| 0 | 1 | 0 | 0 |
| 0 | 0 | 0 | 0 |
| 1 | 0 | 0 | 0 |
| 0 | 1 | 0 | 0 |
| 0 | 0 | 0 | 0 |
| 0 | 0 | 0 | 0 |
| 0 | 0 | 0 | 0 |
| 1 | 1 | 0 | 0 |
| 0 | 0 | 0 | 0 |

| | | | |
|---|---|---|---|
| 0 | 0 | 0 | 0 |
| 0 | 0 | 0 | 0 |
| 1 | 0 | 0 | 0 |
| 1 | 1 | 0 | 0 |
| 1 | 1 | 0 | 0 |
| 0 | 0 | 0 | 0 |
| 0 | 0 | 0 | 0 |
| 0 | 0 | 0 | 0 |
| 1 | 0 | 0 | 0 |
| 0 | 0 | 0 | 0 |
| 0 | 0 | 0 | 0 |
| 1 | 0 | 0 | 1 |
| 0 | 0 | 0 | 0 |
| 1 | 0 | 0 | 0 |
| 0 | 0 | 0 | 0 |
| 0 | 0 | 0 | 0 |
| 0 | 0 | 0 | 0 |
| 0 | 0 | 0 | 0 |
| 0 | 0 | 0 | 0 |
| 0 | 0 | 0 | 0 |
| 0 | 0 | 0 | 0 |
| 1 | 0 | 0 | 1 |
| 1 | 0 | 0 | 0 |
| 0 | 0 | 0 | 0 |
| 0 | 0 | 0 | 0 |
| 0 | 0 | 0 | 0 |
| 0 | 0 | 0 | 0 |
| 0 | 0 | 0 | 0 |
| 1 | 0 | 0 | 0 |
| 1 | 0 | 0 | 0 |
| 0 | 0 | 0 | 0 |
| 0 | 0 | 0 | 0 |
| 0 | 0 | 0 | 0 |
| 0 | 0 | 0 | 0 |
| 0 | 0 | 0 | 0 |
| 0 | 0 | 0 | 0 |
| 1 | 0 | 0 | 0 |
| 0 | 0 | 0 | 0 |
| 0 | 0 | 0 | 0 |
| 1 | 0 | 0 | 0 |
| 1 | 0 | 0 | 0 |
| 1 | 0 | 0 | 0 |
| 1 | 0 | 0 | 0 |
| 0 | 0 | 0 | 0 |
| 0 | 0 | 0 | 0 |
| 1 | 0 | 0 | 0 |
| 0 | 0 | 0 | 0 |
| 0 | 0 | 0 | 0 |
| 0 | 0 | 0 | 0 |
| 0 | 0 | 0 | 0 |
| 0 | 0 | 0 | 0 |
| 1 | 0 | 0 | 0 |
| 1 | 0 | 0 | 0 |
| 1 | 0 | 0 | 0 |

| | | | |
|---|---|---|---|
| 1 | 0 | 0 | 0 |
| 0 | 0 | 0 | 0 |
| 0 | 0 | 0 | 0 |
| 0 | 0 | 0 | 0 |
| 1 | 0 | 0 | 0 |
| 0 | 0 | 0 | 0 |
| 1 | 0 | 0 | 0 |
| 0 | 0 | 0 | 0 |
| 0 | 0 | 0 | 0 |
| 0 | 0 | 0 | 0 |
| 0 | 0 | 0 | 0 |
| 0 | 0 | 0 | 0 |
| 0 | 0 | 0 | 0 |
| 0 | 0 | 0 | 0 |
| 0 | 0 | 0 | 0 |
| 1 | 0 | 0 | 0 |
| 0 | 0 | 0 | 0 |
| 0 | 0 | 0 | 0 |
| 0 | 0 | 0 | 0 |
| 0 | 0 | 0 | 0 |
| 0 | 0 | 0 | 0 |
| 0 | 0 | 0 | 0 |
| 1 | 0 | 0 | 0 |
| 1 | 0 | 0 | 0 |
| 0 | 0 | 0 | 0 |
| 0 | 0 | 0 | 0 |
| 0 | 0 | 0 | 0 |
| 0 | 0 | 0 | 0 |
| 0 | 0 | 0 | 0 |
| 0 | 0 | 0 | 0 |
| 0 | 0 | 0 | 0 |
| 0 | 0 | 0 | 0 |
| 1 | 0 | 0 | 0 |
| 0 | 0 | 0 | 0 |
| 0 | 0 | 0 | 0 |
| 0 | 0 | 0 | 0 |
| 0 | 0 | 0 | 0 |
| 0 | 0 | 0 | 0 |
| 0 | 0 | 0 | 0 |
| 0 | 0 | 0 | 0 |
| 0 | 0 | 0 | 0 |
| 0 | 0 | 0 | 0 |
| 0 | 0 | 0 | 0 |
| 0 | 0 | 0 | 0 |
| 0 | 1 | 0 | 0 |
| 0 | 0 | 0 | 0 |
| 0 | 0 | 0 | 0 |
| 0 | 0 | 0 | 0 |
| 0 | 0 | 0 | 0 |
| 0 | 0 | 0 | 0 |
| 0 | 0 | 0 | 0 |
| 0 | 0 | 0 | 0 |
| 0 | 0 | 0 | 0 |
| 1 | 0 | 0 | 0 |

| | | | |
|---|---|---|---|
| 0 | 0 | 0 | 0 |
| 0 | 0 | 0 | 0 |
| 0 | 0 | 0 | 0 |
| 0 | 0 | 0 | 0 |
| 0 | 0 | 0 | 0 |
| 0 | 0 | 0 | 0 |
| 0 | 0 | 0 | 0 |
| 0 | 0 | 0 | 0 |
| 0 | 0 | 0 | 0 |
| 1 | 0 | 0 | 0 |
| 0 | 0 | 0 | 0 |
| 0 | 0 | 0 | 0 |
| 1 | 1 | 0 | 0 |
| 1 | 1 | 0 | 0 |
| 1 | 1 | 0 | 0 |
| 1 | 1 | 0 | 0 |
| 0 | 0 | 0 | 0 |
| 0 | 0 | 0 | 0 |
| 0 | 0 | 0 | 0 |
| 0 | 0 | 0 | 0 |
| 0 | 0 | 0 | 0 |
| 0 | 0 | 0 | 0 |
| 0 | 0 | 0 | 0 |
| 0 | 0 | 0 | 0 |
| 0 | 0 | 0 | 0 |
| 0 | 0 | 0 | 1 |
| 0 | 0 | 0 | 0 |
| 0 | 0 | 0 | 0 |
| 0 | 0 | 0 | 0 |
| 0 | 0 | 0 | 0 |
| 0 | 0 | 0 | 0 |
| 0 | 0 | 0 | 0 |
| 0 | 0 | 0 | 0 |
| 0 | 0 | 0 | 0 |
| 0 | 0 | 0 | 0 |
| 0 | 0 | 0 | 0 |
| 0 | 0 | 0 | 0 |
| 1 | 1 | 0 | 0 |
| 0 | 0 | 0 | 0 |
| 0 | 0 | 0 | 0 |
| 1 | 1 | 0 | 0 |
| 0 | 0 | 0 | 0 |
| 0 | 0 | 0 | 0 |
| 1 | 0 | 0 | 1 |
| 0 | 0 | 0 | 0 |
| 0 | 0 | 0 | 0 |
| 0 | 0 | 0 | 0 |
| 0 | 0 | 0 | 0 |
| 0 | 0 | 0 | 0 |
| 0 | 0 | 0 | 0 |
| 0 | 0 | 0 | 0 |
| 0 | 0 | 1 | 0 |
| 0 | 0 | 1 | 0 |
| 0 | 0 | 1 | 0 |

| | | | |
|---|---|---|---|
| 0 | 0 | 1 | 0 |
| 0 | 0 | 0 | 0 |
| 0 | 0 | 0 | 0 |
| 0 | 0 | 0 | 0 |
| 0 | 0 | 0 | 0 |
| 0 | 0 | 0 | 0 |
| 0 | 0 | 0 | 0 |
| 0 | 0 | 0 | 0 |
| 0 | 0 | 0 | 0 |
| 0 | 0 | 0 | 0 |
| 0 | 0 | 0 | 0 |
| 0 | 0 | 0 | 0 |
| 0 | 0 | 0 | 0 |
| 0 | 0 | 0 | 0 |
| 0 | 0 | 0 | 0 |
| 0 | 0 | 0 | 0 |
| 0 | 0 | 0 | 0 |
| 0 | 0 | 0 | 0 |
| 0 | 0 | 0 | 0 |
| 0 | 0 | 0 | 0 |
| 0 | 0 | 0 | 0 |
| 0 | 0 | 0 | 0 |
| 0 | 0 | 0 | 0 |
| 0 | 1 | 0 | 1 |
| 0 | 0 | 0 | 0 |
| 0 | 0 | 0 | 0 |
| 0 | 0 | 0 | 0 |
| 0 | 0 | 0 | 0 |
| 0 | 0 | 0 | 0 |
| 0 | 0 | 0 | 0 |
| 0 | 0 | 0 | 0 |
| 0 | 0 | 0 | 0 |
| 0 | 0 | 0 | 0 |
| 0 | 0 | 0 | 0 |
| 0 | 0 | 0 | 0 |
| 0 | 0 | 0 | 0 |
| 0 | 0 | 0 | 0 |
| 0 | 0 | 0 | 0 |
| 0 | 0 | 0 | 0 |
| 0 | 0 | 0 | 0 |
| 0 | 0 | 0 | 0 |
| 0 | 0 | 0 | 0 |
| 0 | 0 | 0 | 0 |
| 0 | 0 | 0 | 0 |
| 0 | 0 | 0 | 0 |
| 0 | 0 | 0 | 0 |
| 1 | 0 | 0 | 0 |
| 1 | 0 | 0 | 0 |
| 0 | 0 | 0 | 0 |
| 0 | 1 | 0 | 0 |
| 1 | 1 | 0 | 1 |
| 1 | 1 | 0 | 1 |
| 1 | 1 | 0 | 1 |
| 0 | 0 | 0 | 0 |

| | | | |
|---|---|---|---|
| 1 | 1 | 0 | 0 |
| 0 | 0 | 0 | 0 |
| 0 | 0 | 0 | 0 |
| 0 | 0 | 0 | 0 |
| 0 | 0 | 0 | 0 |
| 0 | 0 | 0 | 0 |
| 0 | 0 | 0 | 0 |
| 0 | 0 | 0 | 0 |
| 0 | 0 | 0 | 0 |
| 0 | 0 | 0 | 0 |
| 0 | 0 | 0 | 0 |
| 0 | 1 | 0 | 0 |
| 0 | 0 | 0 | 0 |
| 0 | 0 | 0 | 0 |
| 0 | 1 | 0 | 0 |
| 0 | 0 | 0 | 0 |
| 1 | 1 | 0 | 0 |
| 0 | 0 | 0 | 0 |
| 0 | 0 | 0 | 0 |
| 1 | 0 | 0 | 0 |
| 0 | 0 | 0 | 0 |
| 0 | 0 | 0 | 0 |
| 0 | 0 | 0 | 0 |
| 0 | 0 | 0 | 0 |
| 0 | 0 | 0 | 0 |
| 0 | 0 | 0 | 0 |
| 0 | 0 | 0 | 0 |
| 0 | 0 | 0 | 0 |
| 0 | 0 | 0 | 0 |
| 0 | 0 | 0 | 0 |
| 0 | 0 | 0 | 0 |
| 0 | 0 | 0 | 0 |
| 0 | 0 | 0 | 0 |
| 1 | 0 | 0 | 0 |
| 0 | 0 | 0 | 0 |
| 0 | 1 | 0 | 0 |
| 0 | 0 | 0 | 0 |
| 0 | 0 | 0 | 0 |
| 0 | 0 | 0 | 0 |
| 0 | 0 | 0 | 0 |
| 0 | 0 | 0 | 0 |
| 0 | 0 | 0 | 0 |
| 0 | 0 | 0 | 0 |
| 0 | 0 | 0 | 0 |
| 0 | 0 | 0 | 0 |
| 0 | 0 | 0 | 0 |
| 0 | 0 | 0 | 0 |
| 0 | 0 | 0 | 0 |
| 1 | 1 | 0 | 0 |
| 0 | 0 | 0 | 0 |
| 0 | 0 | 0 | 0 |
| 0 | 0 | 0 | 0 |
| 0 | 0 | 0 | 0 |
| 0 | 0 | 0 | 0 |

| | | | |
|---|---|---|---|
| 0 | 0 | 0 | 0 |
| 0 | 0 | 0 | 0 |
| 0 | 0 | 0 | 0 |
| 0 | 0 | 0 | 0 |
| 0 | 0 | 0 | 0 |
| 0 | 0 | 0 | 0 |
| 0 | 0 | 0 | 0 |
| 0 | 1 | 0 | 0 |
| 0 | 0 | 0 | 0 |
| 0 | 0 | 0 | 0 |
| 0 | 0 | 0 | 0 |
| 0 | 0 | 0 | 0 |
| 0 | 0 | 1 | 0 |
| 0 | 0 | 0 | 0 |
| 0 | 0 | 1 | 0 |
| 0 | 0 | 1 | 0 |
| 0 | 0 | 0 | 0 |
| 0 | 0 | 0 | 0 |
| 0 | 0 | 0 | 0 |
| 0 | 0 | 0 | 0 |
| 0 | 1 | 0 | 0 |
| 0 | 1 | 0 | 0 |
| 0 | 0 | 0 | 0 |
| 0 | 0 | 0 | 0 |
| 0 | 1 | 0 | 0 |
| 0 | 0 | 0 | 0 |
| 0 | 0 | 0 | 0 |
| 0 | 1 | 0 | 0 |
| 0 | 0 | 0 | 0 |
| 0 | 0 | 0 | 0 |
| 0 | 0 | 0 | 0 |
| 0 | 0 | 0 | 0 |
| 1 | 1 | 0 | 0 |
| 1 | 1 | 0 | 0 |
| 1 | 1 | 0 | 0 |
| 0 | 1 | 0 | 0 |
| 1 | 0 | 0 | 0 |
| 0 | 1 | 0 | 0 |
| 1 | 1 | 0 | 0 |
| 0 | 0 | 0 | 0 |
| 0 | 0 | 0 | 0 |
| 0 | 0 | 0 | 0 |
| 0 | 0 | 0 | 0 |
| 0 | 0 | 0 | 0 |
| 0 | 0 | 0 | 0 |
| 0 | 0 | 0 | 0 |
| 0 | 0 | 0 | 0 |
| 0 | 0 | 0 | 0 |
| 0 | 0 | 0 | 0 |
| 1 | 1 | 0 | 0 |
| 1 | 1 | 0 | 0 |
| 0 | 0 | 0 | 0 |
| 0 | 0 | 0 | 0 |
| 0 | 0 | 0 | 0 |

| | | | |
|---|---|---|---|
| 0 | 0 | 0 | 0 |
| 0 | 0 | 0 | 0 |
| 0 | 0 | 0 | 0 |
| 0 | 0 | 0 | 0 |
| 0 | 0 | 0 | 0 |
| 0 | 0 | 0 | 0 |
| 1 | 1 | 0 | 0 |
| 1 | 1 | 0 | 0 |
| 0 | 0 | 0 | 0 |
| 0 | 0 | 0 | 0 |
| 0 | 0 | 0 | 0 |
| 0 | 0 | 0 | 0 |
| 0 | 0 | 0 | 0 |
| 0 | 0 | 0 | 0 |
| 0 | 0 | 0 | 0 |
| 0 | 0 | 0 | 0 |
| 0 | 0 | 0 | 0 |
| 0 | 0 | 0 | 0 |
| 0 | 0 | 0 | 0 |
| 0 | 0 | 0 | 0 |
| 0 | 0 | 0 | 0 |
| 0 | 0 | 0 | 0 |
| 0 | 0 | 0 | 0 |
| 0 | 0 | 0 | 0 |
| 0 | 0 | 0 | 0 |
| 0 | 0 | 0 | 0 |
| 0 | 0 | 0 | 0 |
| 1 | 1 | 0 | 0 |
| 0 | 0 | 0 | 0 |
| 0 | 0 | 0 | 0 |
| 0 | 0 | 0 | 0 |
| 0 | 0 | 0 | 0 |
| 0 | 0 | 0 | 0 |
| 0 | 0 | 0 | 0 |
| 0 | 0 | 0 | 0 |
| 0 | 0 | 0 | 0 |
| 0 | 0 | 0 | 0 |
| 0 | 0 | 0 | 0 |
| 0 | 0 | 0 | 0 |
| 0 | 0 | 0 | 0 |
| 0 | 0 | 0 | 0 |
| 0 | 0 | 0 | 0 |
| 0 | 0 | 0 | 0 |
| 0 | 0 | 0 | 0 |
| 0 | 0 | 0 | 0 |
| 0 | 0 | 0 | 0 |
| 0 | 0 | 0 | 0 |
| 0 | 0 | 0 | 0 |
| 0 | 0 | 0 | 0 |
| 0 | 0 | 0 | 0 |
| 0 | 0 | 0 | 0 |
| 0 | 0 | 0 | 0 |
| 0 | 0 | 0 | 0 |
| 0 | 0 | 0 | 0 |

| | | | |
|---|---|---|---|
| 0 | 0 | 0 | 0 |
| 0 | 0 | 0 | 0 |
| 0 | 0 | 0 | 0 |
| 0 | 0 | 0 | 0 |
| 0 | 0 | 0 | 0 |
| 0 | 0 | 0 | 0 |
| 0 | 0 | 0 | 0 |
| 0 | 0 | 0 | 0 |
| 0 | 0 | 0 | 0 |
| 0 | 0 | 0 | 0 |
| 1 | 0 | 0 | 0 |
| 1 | 1 | 0 | 0 |
| 1 | 0 | 0 | 0 |
| 0 | 0 | 0 | 0 |
| 0 | 0 | 0 | 0 |
| 0 | 0 | 0 | 0 |
| 0 | 0 | 0 | 0 |
| 0 | 0 | 0 | 0 |
| 0 | 0 | 0 | 0 |
| 0 | 0 | 0 | 0 |
| 0 | 0 | 0 | 0 |
| 0 | 0 | 0 | 0 |
| 0 | 0 | 0 | 0 |
| 0 | 0 | 0 | 0 |
| 0 | 0 | 0 | 0 |
| 0 | 0 | 0 | 0 |
| 0 | 0 | 0 | 0 |
| 0 | 0 | 0 | 0 |
| 0 | 0 | 0 | 0 |
| 0 | 0 | 0 | 0 |
| 0 | 0 | 0 | 0 |
| 0 | 1 | 0 | 0 |
| 0 | 0 | 0 | 0 |
| 0 | 0 | 0 | 0 |
| 1 | 0 | 0 | 0 |
| 1 | 0 | 0 | 0 |
| 1 | 0 | 0 | 0 |
| 0 | 0 | 0 | 0 |
| 0 | 0 | 0 | 0 |
| 0 | 0 | 0 | 0 |
| 0 | 0 | 0 | 0 |
| 0 | 0 | 0 | 0 |
| 0 | 0 | 0 | 0 |
| 0 | 0 | 0 | 0 |
| 0 | 0 | 0 | 0 |
| 0 | 0 | 0 | 0 |
| 0 | 0 | 0 | 0 |
| 0 | 0 | 0 | 0 |
| 0 | 0 | 0 | 0 |
| 0 | 0 | 0 | 0 |
| 0 | 0 | 0 | 0 |
| 0 | 0 | 0 | 0 |
| 0 | 0 | 0 | 1 |
| 0 | 0 | 0 | 0 |

| | | | |
|---|---|---|---|
| 0 | 0 | 0 | 0 |
| 0 | 0 | 0 | 0 |
| 0 | 0 | 0 | 0 |
| 0 | 0 | 0 | 0 |
| 0 | 0 | 0 | 0 |
| 0 | 0 | 0 | 0 |
| 0 | 0 | 0 | 0 |
| 0 | 0 | 0 | 0 |
| 0 | 0 | 0 | 0 |
| 0 | 0 | 0 | 0 |
| 0 | 0 | 0 | 0 |
| 1 | 0 | 0 | 0 |
| 0 | 0 | 0 | 0 |
| 0 | 0 | 0 | 0 |
| 0 | 0 | 1 | 0 |
| 0 | 0 | 0 | 0 |
| 0 | 0 | 0 | 0 |
| 0 | 0 | 0 | 0 |
| 0 | 0 | 0 | 0 |
| 0 | 0 | 0 | 0 |
| 0 | 0 | 0 | 0 |
| 0 | 0 | 0 | 0 |
| 0 | 0 | 0 | 0 |
| 0 | 0 | 0 | 0 |
| 0 | 0 | 0 | 0 |
| 0 | 0 | 0 | 0 |
| 0 | 0 | 0 | 0 |
| 0 | 0 | 0 | 0 |
| 0 | 0 | 0 | 0 |
| 0 | 0 | 0 | 0 |
| 0 | 0 | 0 | 0 |
| 0 | 0 | 0 | 0 |
| 0 | 0 | 0 | 0 |
| 1 | 0 | 0 | 0 |
| 0 | 0 | 0 | 0 |
| 1 | 0 | 0 | 1 |
| 0 | 0 | 0 | 0 |
| 0 | 0 | 0 | 0 |
| 0 | 0 | 0 | 0 |
| 0 | 0 | 0 | 0 |
| 0 | 0 | 0 | 0 |
| 0 | 0 | 0 | 0 |
| 0 | 0 | 0 | 0 |
| 0 | 1 | 0 | 0 |
| 0 | 0 | 0 | 0 |
| 0 | 0 | 0 | 0 |
| 0 | 0 | 0 | 0 |
| 0 | 0 | 0 | 0 |
| 0 | 0 | 0 | 0 |
| 0 | 0 | 0 | 0 |
| 0 | 0 | 0 | 0 |
| 0 | 0 | 0 | 0 |
| 0 | 0 | 0 | 0 |
| 0 | 0 | 0 | 0 |

| | | | |
|---|---|---|---|
| 0 | 0 | 0 | 0 |
| 0 | 0 | 0 | 0 |
| 0 | 0 | 0 | 0 |
| 0 | 0 | 0 | 0 |
| 0 | 0 | 0 | 0 |
| 0 | 1 | 0 | 0 |
| 0 | 0 | 0 | 0 |
| 0 | 0 | 0 | 0 |
| 0 | 0 | 0 | 0 |
| 0 | 0 | 0 | 0 |
| 0 | 0 | 0 | 0 |
| 0 | 0 | 0 | 0 |
| 0 | 0 | 0 | 0 |
| 0 | 0 | 0 | 0 |
| 0 | 0 | 0 | 0 |
| 0 | 1 | 0 | 0 |
| 1 | 1 | 0 | 0 |
| 0 | 0 | 1 | 0 |
| 0 | 0 | 0 | 0 |
| 0 | 0 | 0 | 0 |
| 0 | 0 | 0 | 0 |
| 0 | 0 | 0 | 0 |
| 0 | 0 | 0 | 0 |
| 0 | 0 | 0 | 0 |
| 0 | 0 | 0 | 0 |
| 0 | 0 | 0 | 0 |
| 0 | 0 | 0 | 0 |
| 1 | 0 | 0 | 0 |
| 0 | 0 | 0 | 0 |
| 0 | 0 | 0 | 0 |
| 0 | 0 | 1 | 0 |
| 0 | 0 | 0 | 0 |
| 0 | 0 | 0 | 0 |
| 0 | 0 | 0 | 0 |
| 0 | 0 | 0 | 0 |
| 0 | 0 | 0 | 0 |
| 0 | 0 | 0 | 0 |
| 1 | 1 | 0 | 0 |
| 0 | 0 | 0 | 0 |
| 0 | 0 | 0 | 0 |
| 0 | 0 | 0 | 0 |
| 1 | 0 | 0 | 0 |
| 1 | 0 | 0 | 0 |
| 0 | 0 | 0 | 0 |
| 0 | 0 | 0 | 0 |
| 0 | 0 | 0 | 0 |
| 0 | 0 | 0 | 0 |
| 0 | 0 | 0 | 0 |
| 0 | 0 | 0 | 0 |
| 0 | 0 | 0 | 0 |
| 0 | 0 | 0 | 0 |
| 0 | 0 | 0 | 0 |
| 0 | 0 | 0 | 0 |
| 0 | 0 | 0 | 0 |

| | | | |
|---|---|---|---|
| 0 | 0 | 0 | 0 |
| 0 | 0 | 0 | 0 |
| 0 | 0 | 0 | 0 |
| 0 | 0 | 0 | 0 |
| 0 | 0 | 0 | 0 |
| 0 | 0 | 0 | 0 |
| 0 | 0 | 0 | 0 |
| 0 | 0 | 0 | 0 |
| 0 | 0 | 0 | 0 |
| 0 | 0 | 0 | 0 |
| 0 | 0 | 0 | 0 |
| 0 | 0 | 0 | 0 |
| 1 | 0 | 0 | 0 |
| 0 | 0 | 0 | 0 |
| 0 | 0 | 0 | 0 |
| 0 | 0 | 0 | 0 |
| 0 | 0 | 0 | 0 |
| 0 | 0 | 0 | 0 |
| 0 | 0 | 0 | 0 |
| 0 | 0 | 0 | 0 |
| 0 | 0 | 0 | 0 |
| 0 | 0 | 0 | 0 |
| 0 | 0 | 0 | 0 |
| 1 | 0 | 0 | 0 |
| 0 | 0 | 0 | 0 |
| 0 | 0 | 0 | 0 |
| 0 | 0 | 0 | 0 |
| 1 | 1 | 0 | 1 |
| 0 | 0 | 0 | 0 |
| 0 | 0 | 0 | 0 |
| 0 | 0 | 0 | 0 |
| 0 | 0 | 0 | 0 |
| 0 | 0 | 0 | 0 |
| 0 | 0 | 0 | 0 |
| 0 | 0 | 0 | 0 |
| 0 | 0 | 0 | 0 |
| 0 | 0 | 0 | 0 |
| 0 | 0 | 0 | 0 |
| 0 | 0 | 0 | 0 |
| 0 | 0 | 0 | 0 |
| 0 | 0 | 0 | 0 |
| 0 | 0 | 0 | 0 |
| 0 | 0 | 0 | 0 |
| 0 | 0 | 0 | 0 |
| 0 | 0 | 0 | 0 |
| 0 | 0 | 0 | 0 |
| 0 | 0 | 0 | 0 |
| 0 | 0 | 0 | 0 |
| 0 | 0 | 0 | 0 |
| 0 | 0 | 0 | 0 |
| 0 | 0 | 0 | 0 |
| 0 | 0 | 0 | 0 |
| 0 | 0 | 0 | 0 |
| 0 | 0 | 0 | 0 |

| | | | |
|---|---|---|---|
| 0 | 0 | 0 | 0 |
| 0 | 0 | 0 | 0 |
| 0 | 0 | 0 | 0 |
| 0 | 0 | 0 | 0 |
| 0 | 0 | 0 | 0 |
| 0 | 0 | 0 | 0 |
| 0 | 0 | 0 | 0 |
| 0 | 0 | 0 | 0 |
| 1 | 0 | 0 | 0 |
| 1 | 0 | 0 | 0 |
| 1 | 0 | 0 | 0 |
| 0 | 0 | 0 | 0 |
| 1 | 0 | 0 | 0 |
| 0 | 0 | 0 | 0 |
| 0 | 0 | 0 | 0 |
| 1 | 0 | 0 | 0 |
| 0 | 0 | 0 | 0 |
| 0 | 0 | 0 | 0 |
| 0 | 0 | 0 | 0 |
| 0 | 0 | 0 | 0 |
| 0 | 0 | 0 | 0 |
| 0 | 0 | 0 | 0 |
| 0 | 0 | 0 | 0 |
| 0 | 0 | 0 | 0 |
| 0 | 0 | 0 | 0 |
| 0 | 0 | 0 | 0 |
| 0 | 0 | 0 | 0 |
| 1 | 0 | 0 | 0 |
| 1 | 0 | 0 | 0 |
| 0 | 0 | 0 | 0 |
| 0 | 0 | 0 | 0 |
| 0 | 0 | 0 | 0 |
| 1 | 0 | 0 | 0 |
| 0 | 0 | 0 | 0 |
| 0 | 0 | 0 | 0 |
| 0 | 0 | 0 | 0 |
| 0 | 0 | 0 | 0 |
| 0 | 0 | 0 | 0 |
| 0 | 0 | 0 | 0 |
| 0 | 0 | 0 | 0 |
| 1 | 0 | 0 | 0 |
| 1 | 0 | 0 | 0 |
| 0 | 0 | 0 | 0 |
| 0 | 0 | 0 | 0 |
| 0 | 0 | 0 | 0 |
| 0 | 0 | 0 | 0 |
| 0 | 0 | 0 | 0 |
| 0 | 0 | 0 | 0 |
| 0 | 0 | 0 | 0 |
| 0 | 0 | 0 | 0 |
| 0 | 0 | 0 | 0 |
| 0 | 0 | 0 | 0 |
| 0 | 0 | 0 | 0 |
| 0 | 0 | 0 | 0 |

| | | | |
|---|---|---|---|
| 0 | 0 | 0 | 0 |
| 0 | 0 | 0 | 0 |
| 0 | 0 | 0 | 0 |
| 0 | 0 | 0 | 0 |
| 0 | 0 | 0 | 0 |
| 0 | 0 | 0 | 0 |
| 0 | 0 | 0 | 0 |
| 0 | 0 | 0 | 0 |
| 0 | 0 | 0 | 0 |
| 0 | 0 | 0 | 0 |
| 1 | 0 | 0 | 0 |
| 0 | 0 | 0 | 0 |
| 1 | 0 | 0 | 0 |
| 0 | 0 | 0 | 0 |
| 0 | 0 | 0 | 0 |
| 0 | 0 | 0 | 0 |
| 0 | 0 | 0 | 0 |
| 0 | 1 | 0 | 0 |
| 0 | 0 | 0 | 0 |
| 0 | 0 | 0 | 0 |
| 0 | 0 | 0 | 0 |
| 0 | 0 | 0 | 0 |
| 1 | 0 | 0 | 0 |
| 0 | 0 | 0 | 0 |
| 0 | 0 | 0 | 0 |
| 1 | 0 | 0 | 0 |
| 1 | 0 | 0 | 0 |
| 1 | 0 | 0 | 0 |
| 0 | 0 | 0 | 0 |
| 0 | 0 | 0 | 0 |
| 0 | 0 | 0 | 0 |
| 0 | 0 | 0 | 0 |
| 0 | 0 | 0 | 0 |
| 0 | 0 | 0 | 0 |
| 0 | 0 | 0 | 0 |
| 0 | 0 | 0 | 0 |
| 0 | 0 | 0 | 0 |
| 0 | 0 | 0 | 0 |
| 0 | 0 | 0 | 0 |
| 0 | 0 | 0 | 0 |
| 1 | 1 | 0 | 0 |
| 0 | 0 | 0 | 0 |
| 0 | 0 | 0 | 0 |
| 0 | 0 | 0 | 0 |
| 0 | 0 | 0 | 0 |
| 0 | 0 | 0 | 0 |
| 0 | 0 | 0 | 0 |
| 1 | 0 | 0 | 0 |
| 0 | 0 | 0 | 0 |
| 0 | 0 | 0 | 0 |
| 0 | 0 | 0 | 0 |
| 0 | 0 | 0 | 0 |
| 0 | 0 | 0 | 0 |
| 0 | 0 | 0 | 0 |

| | | | |
|---|---|---|---|
| 1 | 0 | 0 | 0 |
| 0 | 0 | 0 | 0 |
| 1 | 0 | 0 | 0 |
| 0 | 0 | 0 | 0 |
| 0 | 0 | 0 | 0 |
| 0 | 0 | 0 | 0 |
| 0 | 0 | 0 | 0 |
| 0 | 0 | 0 | 0 |
| 1 | 1 | 0 | 0 |
| 0 | 0 | 0 | 0 |
| 1 | 1 | 0 | 0 |
| 1 | 1 | 0 | 0 |
| 0 | 0 | 0 | 0 |
| 1 | 1 | 0 | 0 |
| 0 | 0 | 0 | 0 |
| 0 | 0 | 0 | 0 |
| 0 | 0 | 0 | 0 |
| 0 | 0 | 0 | 0 |
| 0 | 0 | 0 | 0 |
| 0 | 0 | 0 | 0 |
| 1 | 0 | 0 | 0 |
| 0 | 0 | 0 | 0 |
| 0 | 0 | 0 | 0 |
| 0 | 0 | 0 | 0 |
| 1 | 0 | 0 | 0 |
| 0 | 0 | 0 | 0 |
| 1 | 0 | 0 | 0 |
| 1 | 0 | 0 | 0 |
| 1 | 1 | 0 | 0 |
| 0 | 0 | 0 | 0 |
| 1 | 1 | 0 | 0 |
| 1 | 1 | 0 | 0 |
| 1 | 1 | 0 | 0 |
| 0 | 0 | 0 | 0 |
| 1 | 1 | 0 | 0 |
| 0 | 0 | 0 | 0 |
| 1 | 0 | 0 | 0 |
| 0 | 0 | 0 | 0 |
| 1 | 1 | 0 | 0 |
| 1 | 1 | 0 | 0 |
| 1 | 0 | 0 | 0 |
| 0 | 0 | 0 | 0 |
| 0 | 0 | 0 | 0 |
| 1 | 0 | 0 | 0 |
| 0 | 0 | 0 | 0 |
| 0 | 0 | 0 | 0 |
| 1 | 0 | 0 | 0 |
| 0 | 0 | 0 | 0 |
| 1 | 0 | 0 | 0 |
| 0 | 0 | 0 | 0 |
| 0 | 0 | 0 | 0 |
| 0 | 0 | 0 | 0 |
| 0 | 0 | 0 | 0 |
| 0 | 0 | 0 | 0 |

| | | | |
|---|---|---|---|
| 0 | 0 | 0 | 0 |
| 0 | 0 | 0 | 0 |
| 0 | 0 | 0 | 0 |
| 0 | 0 | 0 | 0 |
| 0 | 0 | 0 | 0 |
| 0 | 0 | 0 | 0 |
| 0 | 0 | 0 | 0 |
| 0 | 0 | 0 | 0 |
| 0 | 0 | 0 | 0 |
| 1 | 1 | 0 | 0 |
| 1 | 1 | 0 | 0 |
| 0 | 0 | 0 | 0 |
| 0 | 0 | 0 | 0 |
| 0 | 0 | 0 | 0 |
| 1 | 0 | 1 | 0 |
| 0 | 0 | 0 | 0 |
| 0 | 0 | 0 | 0 |
| 1 | 0 | 1 | 0 |
| 0 | 0 | 0 | 0 |
| 0 | 0 | 0 | 0 |
| 0 | 0 | 0 | 0 |
| 0 | 0 | 1 | 0 |
| 1 | 0 | 1 | 0 |
| 0 | 0 | 1 | 0 |
| 0 | 0 | 1 | 0 |
| 0 | 0 | 1 | 1 |
| 0 | 0 | 1 | 1 |
| 0 | 0 | 0 | 0 |
| 0 | 0 | 0 | 0 |
| 0 | 0 | 0 | 0 |
| 0 | 0 | 0 | 0 |
| 0 | 0 | 0 | 0 |
| 0 | 0 | 0 | 0 |
| 0 | 0 | 0 | 0 |
| 0 | 0 | 0 | 0 |
| 0 | 0 | 0 | 0 |
| 0 | 0 | 0 | 0 |
| 0 | 0 | 0 | 0 |
| 0 | 0 | 0 | 0 |
| 0 | 0 | 0 | 0 |
| 0 | 0 | 0 | 0 |
| 0 | 0 | 0 | 0 |
| 0 | 0 | 0 | 0 |
| 0 | 0 | 1 | 0 |
| 0 | 0 | 0 | 0 |
| 0 | 0 | 0 | 0 |
| 0 | 0 | 0 | 0 |
| 0 | 0 | 0 | 0 |
| 0 | 0 | 0 | 0 |
| 0 | 0 | 0 | 0 |
| 1 | 0 | 0 | 0 |
| 0 | 0 | 0 | 0 |
| 0 | 0 | 0 | 0 |
| 0 | 0 | 0 | 0 |

| | | | |
|---|---|---|---|
| 0 | 0 | 0 | 0 |
| 0 | 0 | 0 | 0 |
| 0 | 0 | 0 | 0 |
| 1 | 0 | 0 | 0 |
| 0 | 0 | 0 | 0 |
| 0 | 0 | 0 | 0 |
| 0 | 0 | 0 | 0 |
| 0 | 0 | 0 | 0 |
| 1 | 0 | 0 | 0 |
| 1 | 0 | 0 | 0 |
| 0 | 0 | 0 | 0 |
| 0 | 0 | 0 | 0 |
| 0 | 0 | 0 | 0 |
| 0 | 0 | 0 | 0 |
| 0 | 0 | 0 | 0 |
| 0 | 0 | 0 | 0 |
| 0 | 0 | 1 | 0 |
| 0 | 0 | 0 | 0 |
| 0 | 0 | 0 | 0 |
| 0 | 0 | 0 | 0 |
| 0 | 0 | 0 | 0 |
| 0 | 0 | 0 | 0 |
| 0 | 0 | 0 | 0 |
| 0 | 0 | 0 | 0 |
| 0 | 0 | 0 | 0 |
| 0 | 0 | 0 | 0 |
| 0 | 0 | 0 | 0 |
| 0 | 0 | 0 | 0 |
| 0 | 0 | 0 | 0 |
| 0 | 0 | 0 | 0 |
| 0 | 0 | 0 | 0 |
| 0 | 0 | 0 | 0 |
| 0 | 0 | 0 | 0 |
| 0 | 0 | 0 | 0 |
| 0 | 0 | 0 | 0 |
| 0 | 0 | 0 | 0 |
| 0 | 0 | 0 | 0 |
| 0 | 0 | 0 | 0 |
| 0 | 0 | 0 | 0 |
| 0 | 0 | 0 | 0 |
| 0 | 0 | 0 | 0 |
| 0 | 0 | 0 | 0 |
| 0 | 0 | 0 | 0 |
| 0 | 0 | 0 | 0 |
| 0 | 0 | 0 | 0 |
| 0 | 0 | 0 | 0 |
| 0 | 0 | 0 | 0 |
| 0 | 0 | 0 | 0 |
| 0 | 0 | 0 | 0 |
| 0 | 1 | 0 | 0 |
| 0 | 0 | 0 | 0 |
| 0 | 0 | 0 | 0 |
| 0 | 0 | 0 | 0 |
| 0 | 0 | 0 | 0 |

| | | | |
|---|---|---|---|
| 0 | 0 | 1 | 0 |
| 0 | 0 | 0 | 0 |
| 0 | 0 | 0 | 0 |
| 0 | 0 | 0 | 0 |
| 0 | 0 | 0 | 0 |
| 0 | 0 | 0 | 0 |
| 0 | 0 | 0 | 0 |
| 1 | 0 | 0 | 0 |
| 0 | 0 | 0 | 0 |
| 0 | 0 | 0 | 0 |
| 0 | 0 | 0 | 0 |
| 1 | 0 | 0 | 0 |
| 0 | 0 | 0 | 0 |
| 0 | 0 | 0 | 0 |
| 0 | 0 | 0 | 0 |
| 0 | 0 | 0 | 0 |
| 0 | 0 | 0 | 0 |
| 0 | 0 | 0 | 0 |
| 0 | 0 | 0 | 0 |
| 0 | 0 | 0 | 0 |
| 0 | 0 | 0 | 0 |
| 0 | 0 | 0 | 0 |
| 0 | 0 | 0 | 0 |
| 0 | 0 | 0 | 0 |
| 0 | 0 | 0 | 0 |
| 0 | 0 | 0 | 0 |
| 0 | 0 | 0 | 0 |
| 0 | 0 | 0 | 0 |
| 0 | 0 | 0 | 0 |
| 0 | 0 | 0 | 0 |
| 0 | 0 | 0 | 0 |
| 0 | 0 | 0 | 0 |
| 0 | 0 | 0 | 0 |
| 0 | 0 | 0 | 0 |
| 0 | 0 | 0 | 0 |
| 0 | 0 | 0 | 0 |
| 0 | 0 | 0 | 0 |
| 0 | 0 | 0 | 0 |
| 0 | 0 | 0 | 0 |
| 0 | 0 | 0 | 0 |
| 0 | 0 | 0 | 0 |
| 0 | 0 | 0 | 0 |
| 0 | 0 | 0 | 0 |
| 0 | 0 | 0 | 0 |
| 0 | 0 | 0 | 0 |
| 0 | 0 | 0 | 0 |
| 0 | 0 | 0 | 0 |
| 0 | 0 | 0 | 0 |
| 0 | 0 | 0 | 0 |
| 0 | 0 | 0 | 0 |
| 0 | 0 | 0 | 0 |
| 0 | 0 | 0 | 0 |
| 0 | 0 | 0 | 0 |
| 0 | 0 | 0 | 0 |

| | | | |
|---|---|---|---|
| 0 | 0 | 0 | 0 |
| 0 | 0 | 0 | 0 |
| 0 | 0 | 0 | 0 |
| 0 | 0 | 0 | 0 |
| 0 | 0 | 0 | 0 |
| 0 | 0 | 0 | 0 |
| 0 | 0 | 0 | 0 |
| 0 | 0 | 0 | 0 |
| 0 | 0 | 0 | 0 |
| 0 | 0 | 0 | 0 |
| 0 | 0 | 0 | 0 |
| 0 | 0 | 0 | 0 |
| 0 | 0 | 0 | 0 |
| 0 | 0 | 0 | 0 |
| 0 | 0 | 0 | 0 |
| 0 | 0 | 0 | 0 |
| 0 | 0 | 0 | 0 |
| 0 | 0 | 0 | 0 |
| 0 | 0 | 0 | 1 |
| 0 | 1 | 0 | 1 |
| 0 | 0 | 0 | 0 |
| 0 | 0 | 0 | 0 |
| 0 | 0 | 0 | 0 |
| 0 | 0 | 0 | 0 |
| 0 | 0 | 0 | 0 |
| 1 | 0 | 0 | 0 |
| 1 | 0 | 0 | 0 |
| 0 | 0 | 0 | 0 |
| 0 | 0 | 0 | 0 |
| 0 | 0 | 0 | 0 |
| 0 | 0 | 0 | 0 |
| 0 | 0 | 0 | 0 |
| 0 | 0 | 0 | 0 |
| 1 | 0 | 0 | 1 |
| 1 | 0 | 0 | 0 |
| 1 | 0 | 0 | 1 |
| 1 | 0 | 0 | 0 |
| 0 | 0 | 0 | 0 |
| 0 | 0 | 0 | 0 |
| 0 | 0 | 0 | 0 |
| 0 | 0 | 0 | 0 |
| 1 | 0 | 0 | 0 |
| 1 | 0 | 0 | 1 |
| 1 | 0 | 0 | 0 |
| 0 | 0 | 0 | 0 |
| 0 | 0 | 0 | 0 |
| 0 | 0 | 0 | 0 |
| 0 | 0 | 0 | 0 |
| 0 | 0 | 0 | 0 |
| 1 | 0 | 0 | 0 |
| 0 | 0 | 0 | 0 |
| 0 | 0 | 0 | 0 |
| 0 | 0 | 0 | 0 |
| 0 | 0 | 0 | 0 |

| | | | |
|---|---|---|---|
| 0 | 0 | 0 | 0 |
| 0 | 0 | 0 | 0 |
| 0 | 0 | 0 | 0 |
| 1 | 0 | 0 | 0 |
| 1 | 0 | 0 | 0 |
| 0 | 0 | 0 | 0 |
| 1 | 0 | 0 | 0 |
| 0 | 0 | 0 | 0 |
| 0 | 0 | 0 | 0 |
| 0 | 0 | 0 | 0 |
| 0 | 0 | 0 | 0 |
| 1 | 0 | 1 | 0 |
| 0 | 0 | 0 | 0 |
| 0 | 0 | 0 | 0 |
| 0 | 0 | 0 | 0 |
| 0 | 0 | 1 | 0 |
| 0 | 0 | 0 | 0 |
| 0 | 0 | 0 | 0 |
| 0 | 0 | 0 | 0 |
| 0 | 0 | 1 | 0 |
| 1 | 0 | 1 | 0 |
| 1 | 0 | 0 | 0 |
| 1 | 0 | 0 | 0 |
| 1 | 0 | 0 | 0 |
| 0 | 0 | 1 | 0 |
| 0 | 0 | 0 | 0 |
| 1 | 0 | 0 | 0 |
| 0 | 0 | 0 | 0 |
| 0 | 0 | 0 | 0 |
| 0 | 0 | 0 | 0 |
| 0 | 0 | 0 | 0 |
| 0 | 0 | 0 | 0 |
| 0 | 0 | 0 | 0 |
| 0 | 0 | 0 | 0 |
| 0 | 0 | 0 | 0 |
| 0 | 0 | 0 | 0 |
| 0 | 0 | 0 | 1 |
| 0 | 0 | 0 | 1 |
| 0 | 0 | 0 | 1 |
| 0 | 0 | 0 | 0 |
| 0 | 0 | 0 | 0 |
| 0 | 0 | 0 | 0 |
| 0 | 0 | 0 | 0 |
| 0 | 0 | 0 | 0 |
| 0 | 0 | 0 | 0 |
| 0 | 0 | 0 | 0 |
| 0 | 0 | 0 | 0 |
| 0 | 0 | 0 | 0 |
| 0 | 0 | 0 | 0 |
| 0 | 0 | 0 | 0 |
| 0 | 0 | 0 | 0 |
| 0 | 0 | 0 | 0 |
| 0 | 0 | 0 | 0 |
| 0 | 0 | 0 | 0 |

| | | | |
|---|---|---|---|
| 0 | 0 | 0 | 0 |
| 0 | 0 | 0 | 0 |
| 0 | 0 | 0 | 0 |
| 0 | 0 | 0 | 0 |
| 0 | 0 | 0 | 0 |
| 0 | 0 | 0 | 0 |
| 0 | 0 | 0 | 0 |
| 0 | 0 | 0 | 0 |
| 0 | 0 | 0 | 0 |
| 0 | 0 | 0 | 0 |
| 0 | 0 | 0 | 0 |
| 0 | 0 | 0 | 0 |
| 0 | 0 | 0 | 0 |
| 0 | 0 | 0 | 0 |
| 0 | 0 | 0 | 0 |
| 0 | 0 | 0 | 0 |
| 0 | 1 | 0 | 0 |
| 0 | 1 | 0 | 0 |
| 0 | 0 | 0 | 0 |
| 0 | 0 | 0 | 0 |
| 0 | 0 | 0 | 0 |
| 0 | 0 | 0 | 0 |
| 0 | 0 | 0 | 0 |
| 0 | 0 | 0 | 0 |
| 0 | 0 | 0 | 0 |
| 0 | 0 | 0 | 0 |
| 0 | 0 | 0 | 0 |
| 0 | 0 | 0 | 0 |
| 0 | 0 | 0 | 0 |
| 0 | 0 | 0 | 0 |
| 0 | 0 | 0 | 0 |
| 1 | 0 | 0 | 0 |
| 0 | 0 | 0 | 0 |
| 0 | 0 | 0 | 0 |
| 0 | 0 | 0 | 0 |
| 0 | 0 | 0 | 0 |
| 0 | 0 | 0 | 0 |
| 0 | 0 | 0 | 0 |
| 0 | 0 | 0 | 0 |
| 0 | 0 | 0 | 0 |
| 0 | 0 | 0 | 0 |
| 0 | 0 | 0 | 0 |
| 0 | 0 | 0 | 0 |
| 0 | 0 | 0 | 0 |
| 0 | 0 | 0 | 0 |
| 0 | 0 | 0 | 0 |
| 1 | 1 | 0 | 0 |
| 0 | 0 | 0 | 0 |
| 0 | 0 | 0 | 0 |
| 0 | 0 | 0 | 0 |
| 0 | 0 | 0 | 0 |
| 0 | 0 | 0 | 0 |
| 1 | 0 | 0 | 0 |
| 0 | 0 | 0 | 0 |

| | | | |
|---|---|---|---|
| 0 | 0 | 0 | 0 |
| 0 | 0 | 0 | 0 |
| 0 | 0 | 0 | 0 |
| 0 | 0 | 0 | 0 |
| 0 | 0 | 0 | 0 |
| 0 | 0 | 0 | 0 |
| 1 | 0 | 0 | 0 |
| 0 | 0 | 0 | 0 |
| 0 | 0 | 0 | 0 |
| 0 | 0 | 0 | 0 |
| 0 | 0 | 0 | 0 |
| 0 | 0 | 0 | 0 |
| 0 | 0 | 0 | 0 |
| 0 | 0 | 0 | 0 |
| 0 | 0 | 0 | 0 |
| 0 | 0 | 0 | 0 |
| 0 | 0 | 0 | 0 |
| 0 | 0 | 0 | 0 |
| 0 | 0 | 0 | 0 |
| 0 | 0 | 0 | 0 |
| 0 | 0 | 0 | 0 |
| 0 | 0 | 0 | 0 |
| 0 | 0 | 0 | 0 |
| 0 | 0 | 0 | 0 |
| 0 | 0 | 0 | 0 |
| 0 | 0 | 0 | 0 |
| 0 | 0 | 0 | 0 |
| 0 | 0 | 0 | 0 |
| 0 | 0 | 0 | 0 |
| 0 | 0 | 0 | 0 |
| 0 | 0 | 0 | 0 |
| 0 | 0 | 0 | 0 |
| 0 | 0 | 0 | 0 |
| 0 | 0 | 0 | 0 |
| 0 | 0 | 0 | 0 |
| 0 | 0 | 0 | 0 |
| 0 | 0 | 0 | 0 |
| 0 | 0 | 0 | 0 |
| 0 | 0 | 0 | 0 |
| 0 | 0 | 0 | 0 |
| 0 | 0 | 0 | 0 |
| 0 | 0 | 0 | 0 |
| 0 | 0 | 0 | 0 |
| 0 | 0 | 0 | 0 |
| 0 | 0 | 0 | 0 |
| 0 | 0 | 0 | 0 |
| 0 | 0 | 0 | 0 |
| 0 | 0 | 0 | 0 |
| 0 | 0 | 0 | 0 |
| 0 | 0 | 0 | 0 |
| 0 | 0 | 0 | 0 |
| 0 | 0 | 0 | 0 |
| 0 | 0 | 0 | 0 |
| 0 | 0 | 0 | 0 |

| | | | |
|---|---|---|---|
| 0 | 0 | 0 | 0 |
| 0 | 0 | 0 | 0 |
| 0 | 0 | 0 | 0 |
| 0 | 0 | 0 | 0 |
| 0 | 0 | 0 | 0 |
| 0 | 0 | 0 | 0 |
| 0 | 0 | 0 | 0 |
| 0 | 0 | 0 | 0 |
| 0 | 0 | 0 | 0 |
| 0 | 0 | 0 | 0 |
| 0 | 0 | 0 | 0 |
| 0 | 0 | 0 | 0 |
| 0 | 0 | 0 | 0 |
| 0 | 0 | 0 | 0 |
| 0 | 0 | 0 | 0 |
| 0 | 0 | 0 | 0 |
| 0 | 0 | 0 | 0 |
| 0 | 0 | 0 | 0 |
| 0 | 0 | 0 | 0 |
| 0 | 0 | 0 | 0 |
| 0 | 0 | 0 | 0 |
| 0 | 0 | 0 | 0 |
| 0 | 0 | 0 | 0 |
| 0 | 0 | 0 | 0 |
| 1 | 1 | 0 | 0 |
| 0 | 0 | 0 | 0 |
| 0 | 0 | 0 | 0 |
| 0 | 0 | 0 | 0 |
| 0 | 0 | 0 | 0 |
| 0 | 0 | 0 | 0 |
| 0 | 0 | 0 | 0 |
| 0 | 0 | 0 | 0 |
| 0 | 0 | 0 | 0 |
| 0 | 0 | 0 | 0 |
| 0 | 0 | 0 | 0 |
| 1 | 0 | 0 | 0 |
| 1 | 0 | 0 | 1 |
| 1 | 0 | 0 | 0 |
| 0 | 0 | 0 | 0 |
| 1 | 0 | 0 | 0 |
| 1 | 1 | 0 | 0 |
| 1 | 1 | 0 | 0 |
| 0 | 0 | 0 | 0 |
| 0 | 0 | 0 | 0 |
| 0 | 0 | 0 | 0 |
| 0 | 0 | 0 | 0 |
| 0 | 0 | 0 | 0 |
| 0 | 0 | 0 | 0 |
| 0 | 0 | 0 | 0 |
| 0 | 0 | 0 | 0 |
| 0 | 0 | 0 | 0 |
| 0 | 0 | 0 | 0 |
| 0 | 0 | 0 | 0 |
| 0 | 0 | 0 | 0 |

| | | | |
|---|---|---|---|
| 0 | 0 | 0 | 0 |
| 0 | 0 | 0 | 0 |
| 0 | 0 | 0 | 0 |
| 0 | 0 | 0 | 0 |
| 0 | 0 | 0 | 0 |
| 0 | 0 | 0 | 0 |
| 0 | 0 | 0 | 0 |
| 0 | 0 | 0 | 0 |
| 0 | 0 | 0 | 0 |
| 0 | 0 | 0 | 0 |
| 0 | 0 | 0 | 0 |
| 0 | 0 | 0 | 0 |
| 0 | 0 | 0 | 0 |
| 1 | 0 | 0 | 0 |
| 0 | 0 | 0 | 0 |
| 0 | 0 | 0 | 0 |
| 1 | 1 | 0 | 0 |
| 0 | 0 | 0 | 0 |
| 0 | 0 | 0 | 0 |
| 0 | 0 | 0 | 0 |
| 0 | 0 | 0 | 0 |
| 0 | 0 | 0 | 0 |
| 0 | 0 | 0 | 0 |
| 1 | 0 | 0 | 0 |
| 0 | 0 | 0 | 0 |
| 0 | 0 | 0 | 0 |
| 0 | 0 | 0 | 0 |
| 0 | 0 | 0 | 0 |
| 0 | 0 | 0 | 0 |
| 0 | 0 | 0 | 0 |
| 0 | 0 | 0 | 0 |
| 1 | 0 | 0 | 0 |
| 0 | 0 | 0 | 0 |
| 0 | 0 | 0 | 0 |
| 0 | 0 | 0 | 0 |
| 0 | 0 | 0 | 0 |
| 0 | 0 | 0 | 0 |
| 0 | 0 | 0 | 0 |
| 0 | 0 | 0 | 0 |
| 0 | 0 | 0 | 0 |
| 0 | 0 | 0 | 0 |
| 0 | 0 | 0 | 0 |
| 0 | 0 | 0 | 0 |
| 0 | 0 | 0 | 0 |
| 0 | 0 | 0 | 0 |
| 0 | 0 | 0 | 0 |
| 0 | 0 | 0 | 0 |
| 0 | 0 | 0 | 0 |
| 0 | 0 | 0 | 0 |
| 0 | 0 | 0 | 0 |
| 0 | 0 | 0 | 0 |
| 0 | 0 | 0 | 0 |
| 0 | 0 | 0 | 0 |
| 0 | 0 | 0 | 0 |

| | | | |
|---|---|---|---|
| 0 | 0 | 0 | 0 |
| 0 | 0 | 0 | 0 |
| 0 | 0 | 0 | 0 |
| 0 | 0 | 0 | 0 |
| 0 | 0 | 0 | 0 |
| 0 | 0 | 0 | 0 |
| 0 | 0 | 0 | 0 |
| 0 | 0 | 0 | 0 |
| 0 | 0 | 0 | 0 |
| 0 | 0 | 0 | 0 |
| 0 | 0 | 0 | 0 |
| 0 | 0 | 0 | 0 |
| 0 | 0 | 0 | 0 |
| 0 | 0 | 0 | 0 |
| 0 | 0 | 0 | 0 |
| 0 | 0 | 0 | 0 |
| 0 | 0 | 0 | 0 |
| 1 | 0 | 0 | 0 |
| 0 | 0 | 0 | 0 |
| 0 | 0 | 0 | 0 |
| 0 | 0 | 0 | 0 |
| 0 | 0 | 0 | 0 |
| 0 | 0 | 0 | 0 |
| 0 | 0 | 0 | 0 |
| 1 | 0 | 0 | 0 |
| 0 | 0 | 0 | 0 |
| 0 | 0 | 0 | 0 |
| 0 | 0 | 0 | 0 |
| 0 | 0 | 0 | 0 |
| 0 | 0 | 0 | 0 |
| 0 | 0 | 0 | 0 |
| 0 | 0 | 0 | 0 |
| 0 | 0 | 0 | 0 |
| 0 | 0 | 0 | 0 |
| 0 | 0 | 0 | 0 |
| 0 | 0 | 0 | 0 |
| 0 | 0 | 0 | 0 |
| 0 | 0 | 0 | 0 |
| 0 | 0 | 0 | 0 |
| 0 | 0 | 0 | 0 |
| 0 | 0 | 0 | 0 |
| 0 | 0 | 0 | 0 |
| 0 | 0 | 0 | 0 |
| 0 | 0 | 0 | 0 |
| 0 | 0 | 0 | 0 |
| 0 | 0 | 0 | 0 |
| 0 | 0 | 0 | 0 |
| 0 | 0 | 0 | 0 |
| 0 | 0 | 0 | 0 |
| 0 | 0 | 0 | 0 |
| 0 | 0 | 0 | 0 |
| 0 | 0 | 0 | 0 |
| 0 | 0 | 0 | 0 |
| 0 | 0 | 0 | 0 |

| | | | |
|---|---|---|---|
| 0 | 0 | 0 | 0 |
| 0 | 0 | 0 | 0 |
| 0 | 0 | 0 | 0 |
| 0 | 0 | 0 | 0 |
| 0 | 0 | 0 | 0 |
| 0 | 0 | 0 | 0 |
| 0 | 0 | 0 | 0 |
| 0 | 0 | 0 | 0 |
| 0 | 0 | 0 | 0 |
| 0 | 0 | 0 | 0 |
| 0 | 0 | 0 | 0 |
| 0 | 0 | 0 | 0 |
| 0 | 0 | 0 | 0 |
| 0 | 0 | 0 | 0 |
| 1 | 1 | 0 | 0 |
| 1 | 1 | 0 | 0 |
| 0 | 0 | 0 | 0 |
| 0 | 0 | 0 | 0 |
| 0 | 0 | 0 | 0 |
| 0 | 0 | 0 | 0 |
| 0 | 0 | 0 | 0 |
| 0 | 0 | 0 | 0 |
| 0 | 0 | 0 | 0 |
| 0 | 0 | 0 | 0 |
| 0 | 0 | 0 | 0 |
| 0 | 0 | 0 | 0 |
| 0 | 0 | 0 | 0 |
| 1 | 0 | 0 | 0 |
| 0 | 0 | 0 | 0 |
| 1 | 0 | 0 | 0 |
| 0 | 0 | 0 | 0 |
| 0 | 0 | 0 | 0 |
| 0 | 0 | 0 | 0 |
| 0 | 0 | 0 | 0 |
| 0 | 0 | 0 | 0 |
| 0 | 0 | 0 | 0 |
| 1 | 0 | 0 | 0 |
| 0 | 0 | 0 | 0 |
| 0 | 0 | 0 | 0 |
| 0 | 0 | 0 | 0 |
| 0 | 0 | 0 | 0 |
| 0 | 0 | 0 | 0 |
| 0 | 0 | 0 | 0 |
| 0 | 0 | 0 | 0 |
| 0 | 0 | 0 | 0 |
| 0 | 0 | 0 | 0 |
| 0 | 0 | 0 | 0 |
| 0 | 0 | 0 | 0 |
| 0 | 0 | 0 | 0 |
| 0 | 0 | 0 | 0 |
| 0 | 0 | 0 | 0 |
| 0 | 0 | 0 | 0 |
| 0 | 0 | 0 | 0 |
| 0 | 0 | 0 | 0 |

| | | | |
|---|---|---|---|
| 0 | 0 | 0 | 0 |
| 0 | 0 | 0 | 0 |
| 0 | 0 | 0 | 0 |
| 0 | 0 | 0 | 0 |
| 0 | 0 | 0 | 0 |
| 0 | 0 | 0 | 0 |
| 0 | 0 | 0 | 0 |
| 0 | 0 | 0 | 0 |
| 0 | 0 | 0 | 0 |
| 0 | 0 | 0 | 0 |
| 0 | 0 | 0 | 0 |
| 0 | 0 | 0 | 0 |
| 0 | 0 | 0 | 0 |
| 0 | 0 | 0 | 0 |
| 0 | 0 | 0 | 0 |
| 0 | 0 | 0 | 0 |
| 0 | 0 | 0 | 0 |
| 0 | 0 | 0 | 0 |
| 0 | 0 | 0 | 0 |
| 0 | 0 | 0 | 0 |
| 0 | 0 | 0 | 0 |
| 0 | 0 | 0 | 0 |
| 0 | 0 | 0 | 0 |
| 0 | 0 | 0 | 0 |
| 0 | 0 | 0 | 0 |
| 0 | 0 | 0 | 0 |
| 0 | 0 | 0 | 0 |
| 0 | 0 | 0 | 0 |
| 0 | 0 | 0 | 0 |
| 0 | 0 | 0 | 0 |
| 0 | 0 | 0 | 0 |
| 0 | 0 | 0 | 0 |
| 0 | 0 | 0 | 0 |
| 0 | 0 | 0 | 0 |
| 0 | 0 | 0 | 0 |
| 0 | 0 | 0 | 0 |
| 0 | 0 | 0 | 0 |
| 0 | 0 | 0 | 0 |
| 0 | 0 | 0 | 0 |
| 0 | 0 | 0 | 0 |
| 0 | 0 | 0 | 0 |
| 0 | 0 | 0 | 0 |
| 0 | 0 | 0 | 0 |
| 0 | 0 | 0 | 0 |
| 0 | 0 | 0 | 0 |
| 0 | 0 | 0 | 0 |
| 0 | 0 | 0 | 0 |
| 0 | 0 | 0 | 0 |
| 1 | 0 | 0 | 1 |
| 0 | 0 | 0 | 0 |
| 1 | 0 | 0 | 0 |
| 1 | 0 | 0 | 0 |
| 0 | 0 | 0 | 0 |
| 0 | 0 | 0 | 1 |

| | | | |
|---|---|---|---|
| 0 | 0 | 0 | 0 |
| 0 | 0 | 0 | 0 |
| 0 | 0 | 0 | 0 |
| 0 | 0 | 0 | 0 |
| 0 | 0 | 0 | 0 |
| 0 | 0 | 0 | 0 |
| 0 | 0 | 0 | 0 |
| 0 | 0 | 0 | 0 |
| 0 | 0 | 0 | 0 |
| 0 | 0 | 0 | 0 |
| 0 | 0 | 0 | 0 |
| 0 | 0 | 0 | 0 |
| 0 | 0 | 0 | 0 |
| 0 | 0 | 0 | 0 |
| 0 | 0 | 0 | 0 |
| 0 | 0 | 0 | 0 |
| 0 | 0 | 0 | 0 |
| 0 | 0 | 0 | 0 |
| 0 | 0 | 0 | 0 |
| 0 | 0 | 0 | 0 |
| 0 | 0 | 0 | 0 |
| 0 | 0 | 0 | 0 |
| 0 | 0 | 0 | 0 |
| 0 | 0 | 0 | 0 |
| 0 | 0 | 0 | 0 |
| 0 | 0 | 0 | 0 |
| 0 | 0 | 1 | 0 |
| 0 | 0 | 1 | 0 |
| 0 | 0 | 1 | 0 |
| 0 | 0 | 1 | 0 |
| 0 | 0 | 0 | 0 |
| 0 | 0 | 0 | 0 |
| 0 | 0 | 0 | 0 |
| 1 | 0 | 0 | 0 |
| 1 | 0 | 0 | 0 |
| 0 | 0 | 0 | 0 |
| 0 | 0 | 0 | 0 |
| 0 | 0 | 0 | 0 |
| 0 | 0 | 0 | 0 |
| 1 | 0 | 0 | 0 |
| 0 | 0 | 0 | 0 |
| 0 | 0 | 0 | 0 |
| 0 | 0 | 0 | 0 |
| 0 | 0 | 0 | 0 |
| 0 | 0 | 0 | 0 |
| 0 | 1 | 0 | 0 |
| 0 | 0 | 0 | 0 |
| 0 | 0 | 0 | 0 |
| 0 | 0 | 0 | 0 |
| 0 | 0 | 0 | 0 |
| 0 | 0 | 0 | 0 |
| 0 | 0 | 0 | 0 |
| 0 | 0 | 0 | 0 |
| 0 | 0 | 0 | 0 |

| | | | |
|---|---|---|---|
| 0 | 0 | 0 | 0 |
| 0 | 0 | 0 | 0 |
| 0 | 0 | 0 | 0 |
| 0 | 0 | 0 | 0 |
| 0 | 0 | 0 | 0 |
| 0 | 0 | 0 | 1 |
| 0 | 0 | 0 | 0 |
| 0 | 0 | 0 | 0 |
| 0 | 0 | 0 | 0 |
| 0 | 0 | 0 | 0 |
| 0 | 0 | 0 | 0 |
| 0 | 0 | 0 | 0 |
| 0 | 0 | 0 | 0 |
| 0 | 1 | 0 | 0 |
| 0 | 1 | 0 | 0 |
| 0 | 1 | 0 | 0 |
| 0 | 1 | 0 | 0 |
| 0 | 1 | 0 | 0 |
| 0 | 0 | 0 | 0 |
| 0 | 0 | 0 | 0 |
| 0 | 0 | 0 | 0 |
| 0 | 0 | 0 | 0 |
| 0 | 0 | 0 | 0 |
| 0 | 0 | 0 | 0 |
| 0 | 0 | 0 | 0 |
| 0 | 0 | 0 | 0 |
| 0 | 0 | 0 | 0 |
| 0 | 0 | 0 | 0 |
| 0 | 0 | 0 | 0 |
| 0 | 0 | 0 | 0 |
| 0 | 0 | 0 | 0 |
| 0 | 0 | 0 | 0 |
| 0 | 0 | 0 | 0 |
| 0 | 0 | 0 | 0 |
| 0 | 0 | 0 | 0 |
| 0 | 0 | 0 | 0 |
| 0 | 0 | 0 | 0 |
| 0 | 0 | 0 | 0 |
| 0 | 0 | 0 | 0 |
| 0 | 0 | 0 | 0 |
| 0 | 0 | 0 | 0 |
| 0 | 0 | 0 | 0 |
| 0 | 0 | 0 | 0 |
| 0 | 0 | 0 | 0 |
| 0 | 0 | 0 | 0 |
| 0 | 0 | 0 | 0 |
| 0 | 0 | 0 | 0 |
| 0 | 0 | 0 | 0 |
| 0 | 0 | 0 | 0 |
| 0 | 0 | 0 | 0 |
| 0 | 0 | 0 | 0 |
| 0 | 0 | 0 | 0 |
| 0 | 0 | 0 | 0 |
| 0 | 0 | 0 | 0 |

| | | | |
|---|---|---|---|
| 0 | 0 | 0 | 0 |
| 0 | 0 | 0 | 0 |
| 0 | 0 | 0 | 0 |
| 1 | 0 | 0 | 0 |
| 0 | 0 | 0 | 0 |
| 0 | 0 | 0 | 0 |
| 0 | 0 | 0 | 0 |
| 0 | 0 | 0 | 0 |
| 0 | 0 | 0 | 0 |
| 0 | 0 | 0 | 0 |
| 0 | 0 | 0 | 0 |
| 0 | 0 | 0 | 0 |
| 0 | 0 | 0 | 0 |
| 0 | 0 | 0 | 0 |
| 0 | 0 | 0 | 0 |
| 0 | 0 | 0 | 0 |
| 0 | 0 | 0 | 0 |
| 0 | 0 | 0 | 0 |
| 0 | 0 | 0 | 0 |
| 0 | 0 | 0 | 0 |
| 1 | 1 | 0 | 0 |
| 1 | 1 | 0 | 0 |
| 0 | 0 | 0 | 0 |
| 0 | 1 | 0 | 0 |
| 0 | 0 | 0 | 0 |
| 0 | 0 | 0 | 0 |
| 0 | 0 | 0 | 0 |
| 0 | 0 | 0 | 0 |
| 0 | 0 | 0 | 0 |
| 0 | 0 | 0 | 0 |
| 0 | 0 | 0 | 0 |
| 0 | 0 | 0 | 0 |
| 0 | 0 | 0 | 0 |
| 0 | 0 | 0 | 0 |
| 0 | 0 | 0 | 0 |
| 0 | 0 | 0 | 0 |
| 0 | 0 | 0 | 0 |
| 0 | 0 | 0 | 0 |
| 0 | 0 | 0 | 0 |
| 1 | 0 | 0 | 0 |
| 1 | 0 | 0 | 0 |
| 0 | 0 | 0 | 0 |
| 0 | 0 | 0 | 0 |
| 1 | 0 | 0 | 0 |
| 0 | 0 | 0 | 0 |
| 0 | 0 | 0 | 0 |
| 0 | 0 | 0 | 0 |
| 0 | 0 | 0 | 0 |
| 0 | 0 | 0 | 0 |
| 0 | 0 | 0 | 0 |
| 0 | 0 | 0 | 0 |
| 0 | 0 | 0 | 0 |
| 0 | 0 | 0 | 0 |
| 0 | 0 | 0 | 0 |

| | | | |
|---|---|---|---|
| 0 | 0 | 0 | 0 |
| 0 | 0 | 0 | 0 |
| 0 | 0 | 0 | 0 |
| 0 | 0 | 0 | 0 |
| 0 | 0 | 0 | 0 |
| 1 | 0 | 0 | 0 |
| 0 | 0 | 0 | 0 |
| 0 | 0 | 0 | 0 |
| 0 | 1 | 0 | 0 |
| 1 | 1 | 0 | 0 |
| 0 | 0 | 0 | 0 |
| 1 | 1 | 0 | 0 |
| 1 | 0 | 0 | 0 |
| 0 | 0 | 0 | 0 |
| 0 | 0 | 0 | 0 |
| 0 | 0 | 0 | 0 |
| 1 | 1 | 0 | 0 |
| 0 | 0 | 0 | 0 |
| 0 | 0 | 0 | 0 |
| 1 | 0 | 0 | 0 |
| 0 | 0 | 0 | 0 |
| 0 | 0 | 0 | 0 |
| 0 | 0 | 0 | 0 |
| 0 | 0 | 0 | 0 |
| 0 | 0 | 0 | 0 |
| 0 | 0 | 0 | 0 |
| 0 | 0 | 0 | 0 |
| 0 | 0 | 0 | 0 |
| 0 | 0 | 0 | 0 |
| 0 | 0 | 0 | 0 |
| 1 | 0 | 0 | 0 |
| 0 | 0 | 0 | 0 |
| 0 | 0 | 0 | 0 |
| 0 | 0 | 0 | 0 |
| 0 | 0 | 0 | 0 |
| 1 | 0 | 0 | 0 |
| 0 | 0 | 0 | 0 |
| 0 | 0 | 0 | 0 |
| 0 | 0 | 0 | 0 |
| 0 | 0 | 0 | 0 |
| 0 | 0 | 0 | 0 |
| 0 | 0 | 0 | 0 |
| 0 | 0 | 0 | 0 |
| 0 | 0 | 0 | 0 |
| 0 | 0 | 0 | 0 |
| 0 | 0 | 0 | 0 |
| 0 | 0 | 0 | 0 |
| 0 | 0 | 0 | 0 |
| 0 | 0 | 0 | 0 |
| 0 | 0 | 0 | 0 |
| 0 | 0 | 0 | 0 |
| 0 | 0 | 0 | 0 |
| 0 | 0 | 0 | 0 |
| 0 | 0 | 0 | 0 |

| | | | |
|---|---|---|---|
| 0 | 0 | 0 | 0 |
| 0 | 0 | 0 | 0 |
| 0 | 0 | 0 | 0 |
| 0 | 0 | 0 | 0 |
| 0 | 0 | 0 | 0 |
| 0 | 0 | 0 | 0 |
| 0 | 0 | 0 | 0 |
| 0 | 0 | 0 | 0 |
| 0 | 0 | 0 | 0 |
| 0 | 0 | 0 | 0 |
| 0 | 0 | 0 | 0 |
| 0 | 0 | 0 | 0 |
| 0 | 0 | 0 | 0 |
| 0 | 0 | 0 | 0 |
| 0 | 0 | 0 | 1 |
| 0 | 0 | 0 | 0 |
| 0 | 0 | 0 | 0 |
| 0 | 0 | 0 | 0 |
| 0 | 0 | 0 | 0 |
| 0 | 0 | 0 | 0 |
| 0 | 0 | 0 | 0 |
| 0 | 0 | 0 | 0 |
| 0 | 0 | 0 | 0 |
| 0 | 0 | 0 | 0 |
| 0 | 0 | 0 | 0 |
| 0 | 0 | 0 | 0 |
| 0 | 0 | 0 | 0 |
| 0 | 0 | 0 | 0 |
| 0 | 0 | 0 | 0 |
| 0 | 0 | 0 | 0 |
| 0 | 0 | 0 | 0 |
| 0 | 0 | 0 | 0 |
| 0 | 0 | 0 | 0 |
| 0 | 0 | 0 | 0 |
| 0 | 0 | 0 | 0 |
| 0 | 0 | 0 | 0 |
| 0 | 0 | 0 | 0 |
| 0 | 0 | 0 | 0 |
| 0 | 0 | 0 | 0 |
| 0 | 0 | 1 | 0 |
| 0 | 0 | 0 | 0 |
| 0 | 0 | 0 | 0 |
| 0 | 0 | 0 | 0 |
| 1 | 0 | 0 | 0 |
| 0 | 0 | 0 | 0 |
| 0 | 0 | 0 | 0 |
| 0 | 0 | 0 | 0 |
| 0 | 0 | 0 | 0 |
| 1 | 0 | 0 | 1 |
| 0 | 0 | 0 | 0 |
| 0 | 0 | 0 | 0 |
| 0 | 0 | 0 | 0 |
| 0 | 0 | 0 | 0 |
| 0 | 0 | 0 | 0 |

| | | | |
|---|---|---|---|
| 0 | 0 | 0 | 0 |
| 0 | 0 | 0 | 0 |
| 0 | 0 | 0 | 0 |
| 0 | 0 | 0 | 0 |
| 0 | 0 | 0 | 0 |
| 0 | 0 | 0 | 0 |
| 0 | 0 | 0 | 0 |
| 0 | 0 | 0 | 0 |
| 0 | 0 | 0 | 0 |
| 1 | 0 | 0 | 0 |
| 0 | 0 | 0 | 0 |
| 0 | 0 | 0 | 0 |
| 0 | 0 | 0 | 0 |
| 0 | 0 | 0 | 0 |
| 1 | 0 | 0 | 0 |
| 0 | 0 | 0 | 0 |
| 0 | 0 | 0 | 0 |
| 0 | 0 | 0 | 0 |
| 0 | 0 | 0 | 0 |
| 0 | 0 | 0 | 0 |
| 0 | 0 | 0 | 0 |
| 0 | 0 | 0 | 0 |
| 0 | 0 | 0 | 0 |
| 0 | 0 | 0 | 0 |
| 1 | 0 | 0 | 0 |
| 0 | 0 | 0 | 0 |
| 0 | 0 | 0 | 0 |
| 0 | 0 | 0 | 0 |
| 0 | 0 | 0 | 0 |
| 0 | 0 | 0 | 0 |
| 0 | 0 | 0 | 0 |
| 0 | 0 | 0 | 0 |
| 0 | 0 | 0 | 0 |
| 0 | 0 | 0 | 0 |
| 1 | 0 | 0 | 0 |
| 1 | 0 | 0 | 0 |
| 0 | 0 | 0 | 0 |
| 0 | 0 | 0 | 0 |
| 1 | 0 | 0 | 0 |
| 0 | 0 | 1 | 0 |
| 0 | 0 | 1 | 0 |
| 0 | 0 | 1 | 0 |
| 0 | 0 | 1 | 0 |
| 0 | 0 | 0 | 0 |
| 1 | 0 | 0 | 0 |
| 0 | 0 | 0 | 0 |
| 0 | 0 | 0 | 0 |
| 0 | 0 | 0 | 0 |
| 0 | 0 | 1 | 0 |
| 0 | 0 | 0 | 0 |
| 0 | 0 | 0 | 0 |
| 0 | 0 | 0 | 0 |
| 0 | 0 | 0 | 0 |
| 0 | 0 | 0 | 0 |

| | | | |
|---|---|---|---|
| 0 | 0 | 0 | 0 |
| 0 | 0 | 0 | 0 |
| 0 | 0 | 0 | 0 |
| 0 | 0 | 0 | 0 |
| 0 | 0 | 0 | 0 |
| 0 | 0 | 0 | 0 |
| 0 | 0 | 0 | 0 |
| 0 | 0 | 0 | 0 |
| 0 | 0 | 0 | 0 |
| 0 | 0 | 0 | 0 |
| 0 | 1 | 0 | 0 |
| 0 | 0 | 0 | 0 |
| 0 | 0 | 0 | 0 |
| 0 | 0 | 0 | 0 |
| 0 | 0 | 0 | 0 |
| 0 | 1 | 0 | 0 |
| 0 | 0 | 0 | 0 |
| 0 | 0 | 0 | 0 |
| 0 | 0 | 0 | 0 |
| 0 | 0 | 0 | 0 |
| 0 | 0 | 0 | 0 |
| 0 | 0 | 0 | 0 |
| 0 | 0 | 0 | 0 |
| 0 | 0 | 0 | 0 |
| 0 | 0 | 0 | 0 |
| 0 | 0 | 0 | 0 |
| 0 | 0 | 0 | 0 |
| 0 | 0 | 0 | 0 |
| 0 | 0 | 0 | 0 |
| 0 | 0 | 0 | 0 |
| 0 | 0 | 0 | 0 |
| 0 | 0 | 0 | 0 |
| 0 | 0 | 0 | 0 |
| 0 | 0 | 0 | 0 |
| 0 | 0 | 0 | 0 |
| 0 | 0 | 0 | 0 |
| 0 | 0 | 0 | 0 |
| 0 | 0 | 0 | 0 |
| 0 | 0 | 0 | 0 |
| 0 | 0 | 0 | 0 |
| 0 | 0 | 0 | 0 |
| 0 | 0 | 0 | 0 |
| 0 | 0 | 0 | 0 |
| 0 | 0 | 0 | 0 |
| 0 | 0 | 0 | 0 |
| 1 | 1 | 0 | 0 |
| 0 | 0 | 0 | 0 |
| 0 | 0 | 0 | 0 |
| 0 | 0 | 0 | 0 |
| 0 | 0 | 0 | 0 |
| 0 | 0 | 0 | 0 |
| 0 | 0 | 0 | 0 |
| 0 | 0 | 0 | 0 |
| 0 | 0 | 0 | 0 |
| 0 | 0 | 0 | 0 |

| | | | |
|---|---|---|---|
| 0 | 0 | 0 | 0 |
| 0 | 0 | 0 | 0 |
| 0 | 0 | 1 | 0 |
| 1 | 0 | 0 | 0 |
| 0 | 0 | 0 | 0 |
| 0 | 0 | 0 | 0 |
| 0 | 0 | 0 | 0 |
| 0 | 0 | 0 | 0 |
| 0 | 0 | 0 | 0 |
| 0 | 0 | 0 | 0 |
| 0 | 0 | 0 | 0 |
| 0 | 0 | 0 | 0 |
| 1 | 0 | 0 | 0 |
| 0 | 0 | 0 | 0 |
| 0 | 0 | 0 | 0 |
| 0 | 0 | 0 | 0 |
| 0 | 0 | 0 | 0 |
| 0 | 0 | 0 | 0 |
| 0 | 0 | 0 | 0 |
| 0 | 0 | 0 | 0 |
| 0 | 0 | 0 | 0 |
| 0 | 0 | 0 | 0 |
| 0 | 0 | 0 | 0 |
| 0 | 0 | 0 | 0 |
| 0 | 0 | 0 | 0 |
| 0 | 0 | 0 | 0 |
| 0 | 0 | 0 | 0 |
| 0 | 0 | 0 | 0 |
| 0 | 0 | 0 | 0 |
| 0 | 0 | 0 | 0 |
| 0 | 0 | 0 | 0 |
| 0 | 0 | 0 | 0 |
| 0 | 0 | 0 | 0 |
| 0 | 0 | 0 | 0 |
| 0 | 0 | 0 | 0 |
| 0 | 0 | 0 | 0 |
| 0 | 0 | 0 | 0 |
| 0 | 0 | 0 | 0 |
| 0 | 0 | 0 | 0 |
| 0 | 0 | 0 | 0 |
| 0 | 1 | 0 | 0 |
| 1 | 1 | 0 | 0 |
| 0 | 0 | 0 | 0 |
| 0 | 0 | 0 | 0 |
| 0 | 0 | 0 | 0 |
| 0 | 0 | 0 | 0 |
| 0 | 0 | 0 | 0 |
| 0 | 0 | 0 | 0 |
| 0 | 1 | 0 | 0 |
| 0 | 0 | 0 | 0 |
| 0 | 0 | 0 | 0 |
| 0 | 0 | 0 | 0 |
| 0 | 0 | 0 | 1 |
| 0 | 0 | 0 | 0 |

| | | | |
|---|---|---|---|
| 0 | 0 | 0 | 1 |
| 0 | 0 | 0 | 1 |
| 0 | 0 | 0 | 1 |
| 0 | 0 | 0 | 0 |
| 0 | 0 | 0 | 0 |
| 0 | 0 | 0 | 0 |
| 0 | 0 | 0 | 0 |
| 1 | 0 | 0 | 0 |
| 0 | 0 | 0 | 0 |
| 1 | 1 | 0 | 0 |
| 0 | 1 | 0 | 0 |
| 0 | 0 | 0 | 0 |
| 1 | 1 | 0 | 0 |
| 0 | 0 | 0 | 0 |
| 0 | 0 | 0 | 0 |
| 0 | 0 | 0 | 0 |
| 0 | 0 | 0 | 0 |
| 1 | 0 | 0 | 0 |
| 0 | 0 | 0 | 0 |
| 0 | 0 | 0 | 0 |
| 0 | 0 | 0 | 0 |
| 0 | 0 | 0 | 0 |
| 0 | 0 | 0 | 0 |
| 0 | 0 | 0 | 0 |
| 0 | 0 | 0 | 0 |
| 0 | 0 | 0 | 0 |
| 0 | 0 | 0 | 0 |
| 0 | 0 | 0 | 0 |
| 0 | 0 | 0 | 0 |
| 0 | 0 | 0 | 0 |
| 0 | 0 | 0 | 0 |
| 1 | 0 | 0 | 0 |
| 0 | 0 | 0 | 0 |
| 0 | 0 | 0 | 0 |
| 0 | 0 | 0 | 0 |
| 0 | 0 | 0 | 0 |
| 0 | 0 | 0 | 0 |
| 0 | 0 | 0 | 0 |
| 0 | 0 | 0 | 0 |
| 0 | 0 | 0 | 0 |
| 0 | 0 | 0 | 0 |
| 1 | 1 | 0 | 0 |
| 0 | 0 | 0 | 0 |
| 0 | 0 | 0 | 0 |
| 1 | 0 | 0 | 0 |
| 0 | 1 | 0 | 0 |
| 0 | 0 | 0 | 0 |
| 0 | 0 | 0 | 0 |
| 0 | 1 | 0 | 0 |
| 1 | 1 | 0 | 0 |
| 1 | 1 | 0 | 0 |
| 1 | 1 | 0 | 0 |
| 1 | 1 | 0 | 0 |
| 1 | 1 | 0 | 0 |

| | | | |
|---|---|---|---|
| 0 | 0 | 0 | 0 |
| 0 | 0 | 0 | 0 |
| 0 | 0 | 0 | 0 |
| 0 | 0 | 0 | 0 |
| 0 | 0 | 0 | 0 |
| 1 | 0 | 0 | 0 |
| 1 | 0 | 0 | 0 |
| 0 | 0 | 0 | 0 |
| 0 | 0 | 0 | 0 |
| 1 | 0 | 0 | 0 |
| 0 | 0 | 0 | 0 |
| 0 | 0 | 1 | 0 |
| 0 | 0 | 0 | 0 |
| 0 | 0 | 0 | 0 |
| 0 | 0 | 0 | 0 |
| 0 | 0 | 0 | 0 |
| 0 | 0 | 0 | 0 |
| 0 | 0 | 0 | 0 |
| 0 | 0 | 0 | 0 |
| 0 | 0 | 0 | 0 |
| 1 | 1 | 0 | 0 |
| 1 | 0 | 0 | 0 |
| 0 | 0 | 0 | 0 |
| 0 | 0 | 1 | 0 |
| 0 | 0 | 0 | 0 |
| 0 | 0 | 1 | 0 |
| 0 | 0 | 0 | 0 |
| 0 | 0 | 0 | 0 |
| 0 | 0 | 0 | 0 |
| 0 | 0 | 0 | 0 |
| 0 | 0 | 0 | 0 |
| 0 | 0 | 0 | 0 |
| 0 | 0 | 0 | 0 |
| 0 | 0 | 0 | 0 |
| 0 | 0 | 0 | 0 |
| 0 | 0 | 0 | 0 |
| 1 | 0 | 0 | 0 |
| 0 | 0 | 0 | 0 |
| 1 | 0 | 0 | 0 |
| 1 | 0 | 0 | 0 |
| 1 | 0 | 0 | 0 |
| 0 | 0 | 0 | 0 |
| 0 | 0 | 0 | 0 |
| 0 | 0 | 0 | 0 |
| 0 | 0 | 0 | 0 |
| 0 | 0 | 0 | 0 |
| 0 | 0 | 0 | 0 |
| 1 | 0 | 1 | 0 |
| 0 | 0 | 0 | 0 |
| 0 | 0 | 0 | 0 |
| 0 | 0 | 0 | 0 |
| 0 | 0 | 0 | 0 |
| 1 | 1 | 0 | 0 |
| 0 | 0 | 0 | 0 |

| | | | |
|---|---|---|---|
| 0 | 0 | 0 | 0 |
| 0 | 1 | 0 | 0 |
| 0 | 0 | 0 | 0 |
| 0 | 0 | 0 | 0 |
| 0 | 0 | 0 | 0 |
| 0 | 0 | 0 | 0 |
| 0 | 0 | 0 | 0 |
| 0 | 0 | 0 | 0 |
| 0 | 0 | 0 | 0 |
| 0 | 0 | 0 | 0 |
| 0 | 0 | 0 | 0 |
| 0 | 0 | 0 | 0 |
| 0 | 0 | 0 | 0 |
| 0 | 0 | 0 | 0 |
| 0 | 0 | 0 | 0 |
| 0 | 0 | 0 | 0 |
| 0 | 0 | 0 | 0 |
| 0 | 0 | 0 | 0 |
| 0 | 0 | 0 | 0 |
| 0 | 0 | 0 | 0 |
| 1 | 0 | 0 | 0 |
| 0 | 0 | 0 | 0 |
| 0 | 0 | 0 | 0 |
| 0 | 0 | 0 | 0 |
| 1 | 0 | 0 | 0 |
| 0 | 0 | 0 | 0 |
| 0 | 0 | 0 | 0 |
| 0 | 0 | 0 | 1 |
| 0 | 0 | 0 | 0 |
| 0 | 0 | 0 | 0 |
| 0 | 0 | 0 | 0 |
| 0 | 0 | 0 | 0 |
| 0 | 0 | 0 | 0 |
| 0 | 0 | 0 | 0 |
| 0 | 0 | 0 | 0 |
| 0 | 0 | 0 | 0 |
| 0 | 0 | 0 | 0 |
| 0 | 0 | 0 | 0 |
| 0 | 0 | 0 | 0 |
| 0 | 0 | 0 | 0 |
| 0 | 0 | 0 | 0 |
| 0 | 0 | 0 | 0 |
| 0 | 0 | 0 | 0 |
| 0 | 0 | 0 | 0 |
| 0 | 0 | 0 | 0 |
| 0 | 0 | 0 | 0 |
| 0 | 0 | 0 | 0 |
| 0 | 0 | 0 | 0 |
| 0 | 0 | 0 | 0 |
| 0 | 0 | 0 | 0 |
| 0 | 0 | 0 | 0 |
| 0 | 0 | 0 | 0 |
| 0 | 0 | 0 | 0 |
| 0 | 0 | 0 | 0 |

| | | | |
|---|---|---|---|
| 0 | 0 | 0 | 0 |
| 0 | 0 | 0 | 0 |
| 0 | 0 | 0 | 0 |
| 0 | 0 | 0 | 0 |
| 0 | 0 | 0 | 0 |
| 0 | 0 | 0 | 0 |
| 0 | 0 | 0 | 0 |
| 0 | 0 | 0 | 0 |
| 0 | 0 | 0 | 0 |
| 0 | 0 | 0 | 0 |
| 0 | 0 | 0 | 0 |
| 0 | 0 | 0 | 0 |
| 0 | 0 | 0 | 0 |
| 0 | 0 | 0 | 0 |
| 0 | 0 | 0 | 0 |
| 1 | 0 | 0 | 0 |
| 0 | 0 | 0 | 0 |
| 0 | 0 | 0 | 0 |
| 0 | 1 | 0 | 0 |
| 0 | 0 | 0 | 0 |
| 0 | 0 | 0 | 0 |
| 0 | 0 | 0 | 0 |
| 0 | 0 | 0 | 0 |
| 0 | 0 | 0 | 0 |
| 0 | 0 | 0 | 0 |
| 1 | 0 | 0 | 0 |
| 0 | 0 | 0 | 0 |
| 0 | 0 | 0 | 0 |
| 0 | 0 | 0 | 0 |
| 0 | 0 | 0 | 0 |
| 0 | 0 | 0 | 0 |
| 0 | 0 | 0 | 0 |
| 0 | 0 | 0 | 0 |
| 1 | 1 | 0 | 0 |
| 0 | 0 | 0 | 0 |
| 0 | 0 | 0 | 0 |
| 0 | 0 | 0 | 0 |
| 0 | 0 | 0 | 0 |
| 0 | 0 | 0 | 0 |
| 0 | 0 | 0 | 0 |
| 0 | 0 | 0 | 0 |
| 0 | 0 | 0 | 0 |
| 0 | 0 | 0 | 0 |
| 0 | 0 | 0 | 0 |
| 0 | 0 | 0 | 0 |
| 0 | 0 | 0 | 0 |
| 0 | 0 | 0 | 0 |
| 1 | 0 | 0 | 0 |
| 0 | 0 | 0 | 0 |
| 0 | 0 | 0 | 0 |
| 0 | 0 | 0 | 0 |
| 0 | 0 | 0 | 0 |
| 0 | 0 | 0 | 0 |
| 1 | 0 | 0 | 1 |

| | | | |
|---|---|---|---|
| 0 | 0 | 0 | 0 |
| 0 | 0 | 0 | 0 |
| 0 | 0 | 1 | 0 |
| 0 | 0 | 0 | 0 |
| 0 | 0 | 0 | 0 |
| 0 | 0 | 0 | 0 |
| 0 | 0 | 0 | 0 |
| 0 | 0 | 0 | 0 |
| 0 | 0 | 0 | 0 |
| 0 | 0 | 0 | 0 |
| 0 | 0 | 0 | 0 |
| 0 | 0 | 0 | 0 |
| 0 | 0 | 0 | 0 |
| 0 | 0 | 0 | 0 |
| 0 | 0 | 0 | 0 |
| 0 | 0 | 0 | 0 |
| 0 | 0 | 0 | 0 |
| 0 | 0 | 0 | 0 |
| 0 | 0 | 0 | 0 |
| 0 | 0 | 0 | 0 |
| 0 | 0 | 0 | 0 |
| 0 | 0 | 0 | 0 |
| 0 | 0 | 0 | 0 |
| 0 | 0 | 0 | 0 |
| 0 | 0 | 0 | 0 |
| 0 | 0 | 0 | 0 |
| 0 | 0 | 0 | 0 |
| 0 | 0 | 0 | 0 |
| 0 | 0 | 0 | 0 |
| 0 | 0 | 0 | 0 |
| 0 | 0 | 0 | 0 |
| 1 | 0 | 0 | 0 |
| 0 | 0 | 0 | 0 |
| 0 | 0 | 0 | 0 |
| 0 | 0 | 0 | 0 |
| 0 | 0 | 0 | 0 |
| 0 | 0 | 0 | 0 |
| 0 | 0 | 0 | 0 |
| 0 | 0 | 0 | 0 |
| 0 | 0 | 0 | 0 |
| 1 | 1 | 0 | 0 |
| 0 | 0 | 0 | 0 |
| 0 | 0 | 0 | 0 |
| 0 | 0 | 0 | 0 |
| 0 | 0 | 0 | 0 |
| 1 | 1 | 0 | 0 |
| 0 | 0 | 0 | 0 |
| 1 | 1 | 0 | 0 |
| 0 | 0 | 0 | 0 |
| 1 | 0 | 0 | 0 |
| 1 | 0 | 0 | 0 |
| 1 | 0 | 0 | 0 |
| 0 | 0 | 0 | 0 |
| 0 | 0 | 0 | 0 |

| | | | |
|---|---|---|---|
| 1 | 0 | 0 | 0 |
| 0 | 0 | 0 | 0 |
| 0 | 0 | 0 | 0 |
| 0 | 0 | 0 | 0 |
| 0 | 0 | 0 | 0 |
| 0 | 0 | 0 | 0 |
| 0 | 0 | 0 | 0 |
| 0 | 0 | 0 | 0 |
| 0 | 0 | 0 | 0 |
| 0 | 0 | 0 | 0 |
| 0 | 0 | 0 | 0 |
| 0 | 0 | 0 | 0 |
| 0 | 0 | 0 | 0 |
| 0 | 0 | 0 | 0 |
| 0 | 0 | 0 | 0 |
| 1 | 1 | 1 | 0 |
| 0 | 0 | 0 | 0 |
| 1 | 0 | 0 | 0 |
| 0 | 0 | 0 | 0 |
| 0 | 0 | 0 | 0 |
| 0 | 0 | 0 | 0 |
| 0 | 1 | 0 | 0 |
| 0 | 0 | 0 | 0 |
| 0 | 1 | 0 | 0 |
| 0 | 1 | 0 | 0 |
| 0 | 0 | 0 | 0 |
| 0 | 0 | 0 | 0 |
| 0 | 0 | 0 | 0 |
| 0 | 0 | 0 | 0 |
| 0 | 0 | 0 | 0 |
| 0 | 0 | 0 | 0 |
| 0 | 0 | 0 | 0 |
| 1 | 1 | 0 | 0 |
| 0 | 0 | 0 | 0 |
| 0 | 0 | 0 | 0 |
| 0 | 0 | 0 | 0 |
| 0 | 1 | 0 | 0 |
| 1 | 1 | 0 | 0 |
| 0 | 0 | 0 | 0 |
| 0 | 0 | 0 | 0 |
| 0 | 0 | 0 | 0 |
| 0 | 0 | 0 | 0 |
| 0 | 0 | 0 | 0 |
| 0 | 0 | 0 | 0 |
| 1 | 1 | 0 | 0 |
| 0 | 0 | 0 | 0 |
| 1 | 0 | 0 | 0 |
| 1 | 1 | 0 | 0 |
| 0 | 0 | 0 | 0 |
| 0 | 0 | 0 | 0 |
| 1 | 1 | 0 | 0 |
| 1 | 1 | 0 | 0 |
| 0 | 0 | 0 | 0 |
| 1 | 1 | 0 | 0 |

| | | | |
|---|---|---|---|
| 0 | 0 | 0 | 0 |
| 0 | 0 | 0 | 0 |
| 0 | 0 | 0 | 0 |
| 1 | 1 | 0 | 0 |
| 0 | 0 | 0 | 0 |
| 0 | 0 | 0 | 0 |
| 0 | 0 | 0 | 0 |
| 0 | 0 | 0 | 0 |
| 1 | 0 | 0 | 0 |
| 0 | 0 | 0 | 0 |
| 0 | 0 | 0 | 0 |
| 0 | 0 | 0 | 0 |
| 1 | 0 | 1 | 0 |
| 1 | 0 | 0 | 0 |
| 0 | 0 | 0 | 0 |
| 0 | 0 | 0 | 0 |
| 0 | 0 | 0 | 0 |
| 0 | 0 | 0 | 0 |
| 0 | 0 | 0 | 0 |
| 0 | 0 | 0 | 0 |
| 0 | 0 | 0 | 0 |
| 0 | 0 | 0 | 0 |
| 0 | 0 | 0 | 0 |
| 1 | 0 | 0 | 0 |
| 0 | 0 | 0 | 0 |
| 0 | 0 | 0 | 0 |
| 0 | 0 | 0 | 0 |
| 0 | 0 | 0 | 0 |
| 0 | 0 | 0 | 0 |
| 0 | 0 | 0 | 0 |
| 0 | 0 | 0 | 0 |
| 0 | 0 | 0 | 0 |
| 0 | 0 | 0 | 0 |
| 0 | 0 | 0 | 0 |
| 0 | 0 | 0 | 0 |
| 0 | 0 | 0 | 0 |
| 0 | 0 | 0 | 0 |
| 0 | 0 | 0 | 0 |
| 0 | 0 | 0 | 0 |
| 0 | 0 | 0 | 0 |
| 0 | 0 | 0 | 0 |
| 0 | 0 | 0 | 0 |
| 0 | 0 | 0 | 0 |
| 1 | 1 | 0 | 0 |
| 1 | 1 | 0 | 0 |
| 0 | 0 | 0 | 0 |
| 0 | 0 | 0 | 0 |
| 0 | 1 | 0 | 0 |
| 0 | 1 | 0 | 0 |
| 0 | 1 | 0 | 0 |
| 0 | 0 | 0 | 0 |
| 0 | 0 | 0 | 0 |
| 0 | 0 | 0 | 0 |
| 0 | 0 | 0 | 0 |

| | | | |
|---|---|---|---|
| 0 | 0 | 0 | 0 |
| 0 | 0 | 0 | 0 |
| 0 | 0 | 0 | 0 |
| 0 | 0 | 0 | 0 |
| 1 | 0 | 0 | 0 |
| 0 | 0 | 0 | 0 |
| 0 | 0 | 0 | 0 |
| 1 | 0 | 0 | 0 |
| 1 | 0 | 0 | 0 |
| 1 | 0 | 0 | 0 |
| 1 | 0 | 0 | 0 |
| 1 | 0 | 0 | 0 |
| 0 | 0 | 0 | 0 |
| 1 | 0 | 0 | 0 |
| 1 | 0 | 0 | 0 |
| 0 | 0 | 0 | 0 |
| 0 | 0 | 0 | 0 |
| 0 | 0 | 0 | 0 |
| 1 | 0 | 0 | 0 |
| 1 | 0 | 0 | 0 |
| 1 | 0 | 0 | 0 |
| 1 | 0 | 0 | 0 |
| 0 | 0 | 0 | 0 |
| 0 | 0 | 0 | 0 |
| 0 | 0 | 0 | 0 |
| 0 | 0 | 0 | 0 |
| 0 | 0 | 0 | 0 |
| 0 | 0 | 0 | 0 |
| 0 | 0 | 0 | 0 |
| 0 | 0 | 0 | 0 |
| 0 | 0 | 0 | 0 |
| 0 | 0 | 0 | 0 |
| 1 | 0 | 0 | 0 |
| 0 | 0 | 0 | 0 |
| 0 | 0 | 0 | 0 |
| 1 | 0 | 0 | 0 |
| 0 | 0 | 0 | 0 |
| 0 | 0 | 0 | 0 |
| 0 | 0 | 0 | 0 |
| 0 | 0 | 0 | 0 |
| 0 | 0 | 0 | 0 |
| 0 | 0 | 0 | 0 |
| 0 | 0 | 0 | 0 |
| 0 | 0 | 0 | 0 |
| 0 | 0 | 0 | 0 |
| 0 | 0 | 0 | 0 |
| 0 | 0 | 0 | 0 |
| 0 | 0 | 0 | 0 |
| 0 | 0 | 0 | 0 |
| 0 | 0 | 0 | 0 |
| 0 | 0 | 0 | 0 |
| 0 | 0 | 0 | 0 |
| 0 | 0 | 0 | 0 |
| 0 | 0 | 0 | 0 |
| 0 | 0 | 0 | 0 |

| | | | |
|---|---|---|---|
| 0 | 0 | 0 | 0 |
| 0 | 0 | 0 | 0 |
| 0 | 0 | 0 | 0 |
| 0 | 0 | 0 | 0 |
| 0 | 0 | 0 | 0 |
| 0 | 0 | 0 | 0 |
| 0 | 0 | 0 | 0 |
| 0 | 0 | 0 | 0 |
| 0 | 0 | 0 | 0 |
| 0 | 0 | 0 | 0 |
| 0 | 0 | 0 | 0 |
| 0 | 0 | 0 | 1 |
| 0 | 0 | 0 | 0 |
| 0 | 0 | 0 | 0 |
| 0 | 0 | 0 | 0 |
| 0 | 0 | 0 | 0 |
| 0 | 0 | 0 | 0 |
| 0 | 0 | 0 | 1 |
| 0 | 0 | 0 | 0 |
| 0 | 0 | 0 | 0 |
| 0 | 0 | 0 | 1 |
| 0 | 0 | 0 | 0 |
| 0 | 0 | 0 | 0 |
| 0 | 0 | 0 | 0 |
| 0 | 0 | 0 | 0 |
| 0 | 0 | 0 | 0 |
| 0 | 0 | 0 | 0 |
| 0 | 0 | 0 | 0 |
| 0 | 0 | 0 | 1 |
| 0 | 0 | 0 | 0 |
| 0 | 0 | 0 | 0 |
| 0 | 0 | 0 | 0 |
| 0 | 0 | 0 | 0 |
| 0 | 0 | 0 | 0 |
| 0 | 0 | 0 | 0 |
| 0 | 0 | 0 | 0 |
| 0 | 0 | 0 | 0 |
| 0 | 0 | 0 | 0 |
| 0 | 0 | 0 | 0 |
| 0 | 0 | 0 | 0 |
| 0 | 0 | 0 | 0 |
| 0 | 0 | 0 | 0 |
| 0 | 0 | 0 | 0 |
| 0 | 0 | 0 | 0 |
| 0 | 0 | 0 | 0 |
| 0 | 0 | 0 | 0 |
| 0 | 0 | 0 | 0 |
| 0 | 0 | 0 | 0 |
| 0 | 0 | 0 | 0 |
| 0 | 0 | 0 | 0 |
| 0 | 0 | 0 | 0 |
| 0 | 0 | 0 | 0 |
| 0 | 0 | 0 | 0 |
| 0 | 0 | 0 | 0 |

| | | | |
|---|---|---|---|
| 0 | 0 | 0 | 0 |
| 0 | 0 | 0 | 0 |
| 1 | 0 | 0 | 0 |
| 0 | 0 | 0 | 0 |
| 1 | 0 | 0 | 0 |
| 0 | 0 | 0 | 0 |
| 0 | 0 | 0 | 0 |
| 0 | 0 | 0 | 0 |
| 0 | 0 | 0 | 0 |
| 1 | 1 | 0 | 0 |
| 0 | 0 | 1 | 0 |
| 0 | 0 | 0 | 0 |
| 0 | 0 | 0 | 0 |
| 0 | 0 | 0 | 0 |
| 0 | 0 | 0 | 0 |
| 0 | 0 | 0 | 0 |
| 0 | 0 | 0 | 0 |
| 0 | 0 | 0 | 0 |
| 0 | 0 | 0 | 0 |
| 0 | 0 | 0 | 0 |
| 0 | 0 | 0 | 0 |
| 0 | 0 | 0 | 0 |
| 0 | 0 | 0 | 0 |
| 0 | 0 | 0 | 0 |
| 0 | 0 | 0 | 0 |
| 0 | 0 | 0 | 0 |
| 0 | 0 | 0 | 0 |
| 0 | 0 | 0 | 0 |
| 0 | 0 | 0 | 0 |
| 0 | 0 | 0 | 0 |
| 0 | 0 | 0 | 0 |
| 0 | 0 | 0 | 0 |
| 0 | 0 | 0 | 0 |
| 0 | 0 | 0 | 0 |
| 0 | 0 | 0 | 0 |
| 0 | 0 | 0 | 0 |
| 0 | 0 | 0 | 0 |
| 0 | 0 | 0 | 0 |
| 0 | 0 | 0 | 0 |
| 0 | 0 | 0 | 0 |
| 0 | 0 | 0 | 0 |
| 0 | 0 | 0 | 0 |
| 0 | 0 | 0 | 0 |
| 0 | 0 | 0 | 0 |
| 0 | 0 | 0 | 0 |
| 1 | 0 | 0 | 0 |
| 0 | 0 | 0 | 0 |
| 0 | 0 | 0 | 0 |
| 0 | 0 | 0 | 0 |
| 0 | 0 | 0 | 0 |
| 1 | 0 | 0 | 0 |
| 0 | 0 | 0 | 0 |
| 0 | 0 | 0 | 0 |
| 0 | 0 | 0 | 0 |

| | | | |
|---|---|---|---|
| 0 | 0 | 0 | 0 |
| 0 | 0 | 0 | 0 |
| 0 | 0 | 0 | 0 |
| 0 | 0 | 0 | 0 |
| 0 | 0 | 0 | 0 |
| 0 | 0 | 0 | 0 |
| 0 | 0 | 0 | 0 |
| 0 | 0 | 0 | 0 |
| 1 | 1 | 0 | 0 |
| 0 | 0 | 0 | 0 |
| 0 | 0 | 0 | 0 |
| 0 | 0 | 0 | 0 |
| 0 | 0 | 0 | 0 |
| 0 | 0 | 0 | 0 |
| 0 | 1 | 0 | 0 |
| 0 | 1 | 0 | 0 |
| 0 | 1 | 0 | 0 |
| 0 | 0 | 0 | 0 |
| 0 | 0 | 0 | 0 |
| 0 | 0 | 0 | 0 |
| 0 | 1 | 0 | 0 |
| 0 | 0 | 0 | 0 |
| 0 | 0 | 0 | 0 |
| 1 | 1 | 0 | 0 |
| 0 | 1 | 0 | 0 |
| 0 | 1 | 0 | 0 |
| 0 | 0 | 0 | 0 |
| 0 | 0 | 0 | 0 |
| 0 | 0 | 0 | 0 |
| 0 | 0 | 0 | 0 |
| 0 | 0 | 0 | 0 |
| 0 | 0 | 0 | 0 |
| 0 | 0 | 0 | 0 |
| 0 | 0 | 0 | 0 |
| 0 | 0 | 0 | 0 |
| 0 | 0 | 0 | 0 |
| 0 | 0 | 0 | 0 |
| 1 | 0 | 0 | 0 |
| 0 | 0 | 0 | 0 |
| 0 | 0 | 0 | 0 |
| 0 | 0 | 0 | 0 |
| 0 | 0 | 0 | 0 |
| 0 | 0 | 0 | 0 |
| 0 | 0 | 0 | 0 |
| 0 | 0 | 0 | 0 |
| 0 | 0 | 0 | 0 |
| 0 | 0 | 0 | 0 |
| 0 | 0 | 0 | 0 |
| 0 | 0 | 0 | 0 |
| 0 | 0 | 0 | 0 |
| 0 | 0 | 0 | 0 |
| 0 | 0 | 0 | 0 |
| 0 | 0 | 0 | 0 |

| | | | |
|---|---|---|---|
| 0 | 0 | 0 | 0 |
| 0 | 0 | 0 | 0 |
| 0 | 0 | 0 | 0 |
| 0 | 0 | 1 | 0 |
| 0 | 0 | 0 | 0 |
| 0 | 0 | 0 | 0 |
| 0 | 0 | 0 | 0 |
| 0 | 0 | 0 | 0 |
| 0 | 0 | 0 | 0 |
| 0 | 0 | 0 | 0 |
| 0 | 0 | 0 | 0 |
| 0 | 0 | 0 | 0 |
| 1 | 0 | 0 | 0 |
| 0 | 0 | 0 | 0 |
| 0 | 0 | 0 | 0 |
| 0 | 0 | 0 | 0 |
| 0 | 1 | 0 | 0 |
| 0 | 0 | 0 | 0 |
| 0 | 0 | 0 | 0 |
| 0 | 0 | 0 | 0 |
| 0 | 0 | 0 | 0 |
| 0 | 0 | 0 | 0 |
| 0 | 0 | 0 | 0 |
| 0 | 0 | 0 | 0 |
| 0 | 0 | 0 | 0 |
| 0 | 0 | 0 | 0 |
| 0 | 0 | 0 | 0 |
| 0 | 0 | 0 | 0 |
| 0 | 0 | 0 | 0 |
| 0 | 0 | 0 | 0 |
| 1 | 0 | 0 | 0 |
| 1 | 0 | 0 | 0 |
| 1 | 0 | 0 | 0 |
| 0 | 0 | 0 | 0 |
| 1 | 0 | 0 | 0 |
| 1 | 0 | 0 | 0 |
| 1 | 0 | 0 | 0 |
| 0 | 0 | 0 | 0 |
| 0 | 0 | 0 | 0 |
| 0 | 0 | 0 | 0 |
| 0 | 0 | 0 | 0 |
| 0 | 0 | 0 | 0 |
| 0 | 1 | 0 | 0 |
| 0 | 1 | 0 | 0 |
| 0 | 0 | 0 | 0 |
| 0 | 0 | 0 | 0 |
| 1 | 0 | 0 | 0 |
| 0 | 0 | 0 | 0 |
| 0 | 0 | 0 | 0 |
| 0 | 0 | 0 | 0 |
| 0 | 0 | 0 | 0 |
| 0 | 0 | 0 | 0 |
| 1 | 0 | 0 | 0 |
| 0 | 0 | 0 | 0 |

| | | | |
|---|---|---|---|
| 0 | 0 | 0 | 0 |
| 1 | 0 | 0 | 0 |
| 1 | 0 | 0 | 0 |
| 0 | 0 | 0 | 0 |
| 0 | 0 | 0 | 0 |
| 0 | 0 | 0 | 0 |
| 0 | 0 | 0 | 0 |
| 0 | 1 | 0 | 0 |
| 0 | 0 | 0 | 0 |
| 0 | 0 | 0 | 0 |
| 0 | 0 | 0 | 0 |
| 0 | 0 | 0 | 0 |
| 0 | 0 | 0 | 0 |
| 0 | 0 | 0 | 0 |
| 1 | 0 | 0 | 0 |
| 0 | 0 | 0 | 0 |
| 0 | 0 | 0 | 0 |
| 0 | 0 | 0 | 0 |
| 0 | 0 | 0 | 0 |
| 1 | 0 | 0 | 0 |
| 1 | 0 | 0 | 0 |
| 0 | 0 | 0 | 0 |
| 1 | 0 | 0 | 0 |
| 1 | 0 | 0 | 0 |
| 0 | 0 | 0 | 0 |
| 0 | 0 | 0 | 0 |
| 0 | 0 | 0 | 0 |
| 0 | 0 | 0 | 0 |
| 0 | 0 | 0 | 0 |
| 0 | 0 | 0 | 0 |
| 0 | 0 | 0 | 0 |
| 0 | 0 | 0 | 0 |
| 0 | 0 | 0 | 0 |
| 0 | 0 | 0 | 0 |
| 0 | 0 | 0 | 0 |
| 0 | 0 | 0 | 0 |
| 0 | 0 | 0 | 0 |
| 0 | 0 | 0 | 0 |
| 0 | 0 | 0 | 0 |
| 0 | 0 | 0 | 0 |
| 0 | 0 | 0 | 0 |
| 0 | 0 | 0 | 0 |
| 0 | 0 | 0 | 0 |
| 0 | 0 | 0 | 0 |
| 0 | 0 | 0 | 0 |
| 0 | 0 | 0 | 0 |
| 0 | 0 | 0 | 0 |
| 0 | 0 | 0 | 0 |
| 0 | 0 | 0 | 0 |
| 0 | 0 | 0 | 0 |
| 0 | 0 | 0 | 0 |
| 0 | 0 | 0 | 0 |
| 0 | 0 | 0 | 0 |
| 0 | 0 | 0 | 0 |
| 1 | 1 | 0 | 1 |

| | | | |
|---|---|---|---|
| 0 | 0 | 0 | 0 |
| 0 | 0 | 0 | 0 |
| 0 | 0 | 0 | 0 |
| 0 | 0 | 0 | 0 |
| 0 | 1 | 0 | 0 |
| 0 | 0 | 0 | 0 |
| 0 | 0 | 0 | 0 |
| 0 | 1 | 0 | 0 |
| 0 | 0 | 0 | 0 |
| 0 | 0 | 0 | 0 |
| 0 | 0 | 0 | 0 |
| 0 | 0 | 0 | 0 |
| 0 | 0 | 0 | 0 |
| 0 | 0 | 0 | 0 |
| 0 | 1 | 0 | 1 |
| 0 | 1 | 0 | 0 |
| 0 | 0 | 0 | 0 |
| 0 | 0 | 0 | 0 |
| 0 | 1 | 0 | 0 |
| 0 | 1 | 0 | 1 |
| 0 | 0 | 0 | 0 |
| 0 | 0 | 0 | 0 |
| 0 | 0 | 0 | 0 |
| 0 | 0 | 0 | 0 |
| 0 | 0 | 0 | 0 |
| 1 | 0 | 0 | 0 |
| 0 | 0 | 0 | 0 |
| 0 | 0 | 1 | 0 |
| 0 | 0 | 1 | 0 |
| 0 | 0 | 1 | 0 |
| 0 | 0 | 0 | 0 |
| 0 | 0 | 0 | 0 |
| 0 | 0 | 0 | 0 |
| 0 | 0 | 0 | 0 |
| 0 | 0 | 0 | 0 |
| 0 | 0 | 1 | 0 |
| 0 | 0 | 0 | 0 |
| 0 | 0 | 0 | 0 |
| 1 | 0 | 1 | 0 |
| 0 | 0 | 1 | 0 |
| 1 | 1 | 0 | 0 |
| 0 | 1 | 0 | 0 |
| 0 | 0 | 0 | 0 |
| 0 | 0 | 0 | 0 |
| 1 | 0 | 0 | 0 |
| 0 | 0 | 0 | 0 |
| 0 | 0 | 1 | 0 |
| 0 | 0 | 0 | 0 |
| 0 | 0 | 0 | 0 |
| 0 | 0 | 0 | 0 |
| 0 | 0 | 0 | 0 |
| 0 | 0 | 0 | 0 |
| 0 | 0 | 0 | 0 |
| 0 | 0 | 0 | 0 |

| | | | |
|---|---|---|---|
| 0 | 0 | 0 | 0 |
| 0 | 0 | 0 | 0 |
| 0 | 0 | 0 | 0 |
| 0 | 0 | 0 | 0 |
| 0 | 0 | 0 | 0 |
| 0 | 0 | 0 | 0 |
| 0 | 0 | 0 | 0 |
| 1 | 1 | 0 | 0 |
| 0 | 0 | 0 | 0 |
| 0 | 0 | 0 | 0 |
| 0 | 0 | 0 | 0 |
| 0 | 0 | 0 | 1 |
| 0 | 0 | 0 | 0 |
| 0 | 0 | 0 | 0 |
| 0 | 0 | 0 | 0 |
| 0 | 0 | 0 | 0 |
| 0 | 0 | 0 | 0 |
| 0 | 0 | 0 | 0 |
| 0 | 0 | 0 | 0 |
| 0 | 0 | 0 | 0 |
| 0 | 0 | 0 | 0 |
| 0 | 0 | 0 | 0 |
| 0 | 0 | 0 | 0 |
| 0 | 0 | 0 | 0 |
| 0 | 0 | 0 | 0 |
| 0 | 0 | 0 | 0 |
| 0 | 0 | 0 | 0 |
| 0 | 0 | 0 | 0 |
| 0 | 0 | 0 | 0 |
| 0 | 1 | 0 | 0 |
| 0 | 1 | 0 | 0 |
| 0 | 0 | 0 | 0 |
| 0 | 0 | 0 | 0 |
| 0 | 0 | 0 | 0 |
| 0 | 0 | 0 | 0 |
| 0 | 1 | 0 | 0 |
| 0 | 0 | 0 | 0 |
| 0 | 0 | 0 | 0 |
| 0 | 0 | 0 | 0 |
| 0 | 0 | 0 | 0 |
| 0 | 0 | 0 | 0 |
| 1 | 1 | 0 | 0 |
| 1 | 0 | 0 | 0 |
| 0 | 0 | 0 | 0 |
| 0 | 0 | 0 | 0 |
| 0 | 0 | 0 | 0 |
| 0 | 0 | 0 | 0 |
| 0 | 0 | 0 | 0 |
| 0 | 0 | 0 | 0 |
| 0 | 0 | 0 | 0 |
| 0 | 0 | 0 | 0 |
| 0 | 0 | 0 | 0 |
| 0 | 0 | 0 | 0 |
| 0 | 0 | 0 | 0 |

| | | | |
|---|---|---|---|
| 0 | 0 | 0 | 0 |
| 0 | 0 | 0 | 0 |
| 0 | 0 | 0 | 0 |
| 0 | 0 | 0 | 0 |
| 0 | 0 | 0 | 0 |
| 0 | 0 | 0 | 0 |
| 0 | 0 | 0 | 0 |
| 0 | 0 | 0 | 0 |
| 1 | 0 | 0 | 0 |
| 1 | 0 | 0 | 0 |
| 0 | 0 | 0 | 0 |
| 0 | 0 | 0 | 0 |
| 0 | 0 | 0 | 0 |
| 1 | 0 | 0 | 0 |
| 0 | 0 | 0 | 0 |
| 0 | 0 | 0 | 0 |
| 0 | 0 | 0 | 0 |
| 0 | 0 | 0 | 0 |
| 0 | 0 | 0 | 0 |
| 1 | 1 | 0 | 0 |
| 0 | 0 | 1 | 0 |
| 0 | 0 | 0 | 0 |
| 0 | 0 | 0 | 0 |
| 0 | 0 | 0 | 0 |
| 0 | 0 | 0 | 0 |
| 0 | 0 | 0 | 0 |
| 0 | 0 | 0 | 0 |
| 0 | 0 | 0 | 0 |
| 0 | 0 | 0 | 0 |
| 0 | 0 | 0 | 0 |
| 0 | 0 | 0 | 0 |
| 0 | 0 | 0 | 0 |
| 0 | 0 | 0 | 0 |
| 0 | 0 | 0 | 0 |
| 0 | 0 | 0 | 0 |
| 0 | 0 | 0 | 0 |
| 0 | 0 | 0 | 0 |
| 0 | 0 | 0 | 0 |
| 0 | 0 | 0 | 0 |
| 0 | 0 | 0 | 0 |
| 0 | 0 | 0 | 0 |
| 0 | 0 | 0 | 0 |
| 0 | 0 | 0 | 0 |
| 0 | 0 | 0 | 0 |
| 0 | 0 | 0 | 0 |
| 0 | 0 | 0 | 0 |
| 1 | 0 | 1 | 0 |
| 1 | 0 | 0 | 0 |
| 0 | 0 | 0 | 0 |
| 0 | 0 | 0 | 0 |
| 0 | 0 | 1 | 0 |
| 0 | 0 | 0 | 0 |
| 0 | 0 | 0 | 0 |
| 0 | 0 | 0 | 0 |

| | | | |
|---|---|---|---|
| 0 | 0 | 0 | 0 |
| 0 | 0 | 0 | 0 |
| 0 | 0 | 0 | 0 |
| 0 | 0 | 0 | 0 |
| 0 | 0 | 0 | 0 |
| 0 | 0 | 0 | 0 |
| 0 | 0 | 0 | 0 |
| 0 | 0 | 0 | 0 |
| 0 | 0 | 0 | 0 |
| 0 | 0 | 0 | 0 |
| 0 | 0 | 0 | 0 |
| 0 | 0 | 0 | 0 |
| 0 | 0 | 0 | 0 |
| 0 | 0 | 0 | 0 |
| 0 | 0 | 0 | 0 |
| 0 | 0 | 0 | 0 |
| 0 | 0 | 0 | 0 |
| 0 | 0 | 0 | 0 |
| 0 | 0 | 0 | 0 |
| 1 | 0 | 0 | 0 |
| 0 | 0 | 0 | 0 |
| 0 | 0 | 0 | 0 |
| 0 | 0 | 0 | 0 |
| 0 | 0 | 0 | 0 |
| 0 | 0 | 0 | 0 |
| 0 | 0 | 0 | 0 |
| 0 | 0 | 0 | 0 |
| 0 | 0 | 0 | 0 |
| 0 | 0 | 0 | 0 |
| 0 | 0 | 0 | 0 |
| 0 | 0 | 0 | 0 |
| 0 | 0 | 0 | 0 |
| 0 | 0 | 0 | 0 |
| 0 | 0 | 0 | 0 |
| 0 | 0 | 0 | 0 |
| 0 | 0 | 0 | 0 |
| 0 | 0 | 0 | 0 |
| 0 | 0 | 0 | 0 |
| 1 | 0 | 0 | 1 |
| 0 | 0 | 0 | 0 |
| 0 | 0 | 0 | 0 |
| 0 | 0 | 0 | 0 |
| 0 | 0 | 0 | 1 |
| 0 | 0 | 0 | 1 |
| 0 | 0 | 0 | 1 |
| 0 | 0 | 0 | 0 |
| 0 | 0 | 0 | 0 |
| 0 | 0 | 0 | 0 |
| 0 | 0 | 0 | 0 |
| 0 | 0 | 0 | 0 |
| 0 | 0 | 0 | 0 |
| 0 | 0 | 0 | 0 |
| 0 | 0 | 0 | 0 |
| 0 | 0 | 0 | 0 |

| | | | |
|---|---|---|---|
| 0 | 0 | 0 | 0 |
| 0 | 0 | 0 | 0 |
| 0 | 0 | 0 | 0 |
| 0 | 0 | 0 | 0 |
| 0 | 0 | 0 | 0 |
| 0 | 0 | 0 | 0 |
| 0 | 0 | 0 | 0 |
| 0 | 0 | 0 | 0 |
| 0 | 0 | 0 | 0 |
| 0 | 0 | 0 | 0 |
| 0 | 0 | 0 | 0 |
| 0 | 0 | 0 | 0 |
| 0 | 0 | 0 | 0 |
| 0 | 0 | 0 | 0 |
| 0 | 0 | 0 | 0 |
| 0 | 0 | 0 | 0 |
| 0 | 0 | 0 | 0 |
| 0 | 0 | 0 | 0 |
| 0 | 0 | 0 | 0 |
| 0 | 0 | 0 | 0 |
| 0 | 0 | 0 | 0 |
| 0 | 0 | 0 | 0 |
| 0 | 0 | 0 | 0 |
| 0 | 0 | 0 | 0 |
| 0 | 0 | 0 | 0 |
| 0 | 0 | 0 | 0 |
| 0 | 0 | 0 | 0 |
| 0 | 0 | 0 | 0 |
| 0 | 0 | 0 | 0 |
| 0 | 0 | 0 | 0 |
| 0 | 0 | 0 | 0 |
| 0 | 0 | 0 | 0 |
| 1 | 1 | 0 | 1 |
| 1 | 1 | 0 | 1 |
| 1 | 0 | 0 | 0 |
| 0 | 0 | 0 | 0 |
| 0 | 0 | 0 | 0 |
| 0 | 0 | 0 | 0 |
| 0 | 0 | 0 | 0 |
| 0 | 0 | 0 | 0 |
| 0 | 0 | 0 | 0 |
| 0 | 0 | 0 | 0 |
| 1 | 1 | 0 | 0 |
| 0 | 0 | 0 | 0 |
| 1 | 0 | 0 | 0 |
| 0 | 0 | 0 | 0 |
| 0 | 0 | 0 | 0 |
| 0 | 0 | 0 | 0 |
| 0 | 0 | 0 | 0 |
| 0 | 0 | 0 | 0 |
| 1 | 0 | 0 | 0 |
| 0 | 0 | 0 | 0 |
| 0 | 0 | 0 | 0 |
| 0 | 0 | 0 | 0 |

| | | | |
|---|---|---|---|
| 0 | 0 | 0 | 0 |
| 0 | 0 | 0 | 0 |
| 0 | 0 | 0 | 0 |
| 0 | 0 | 0 | 0 |
| 0 | 0 | 0 | 0 |
| 0 | 0 | 0 | 0 |
| 0 | 0 | 0 | 0 |
| 0 | 0 | 0 | 0 |
| 0 | 0 | 0 | 0 |
| 0 | 0 | 0 | 0 |
| 0 | 0 | 0 | 0 |
| 0 | 0 | 0 | 0 |
| 0 | 0 | 0 | 0 |
| 0 | 0 | 0 | 0 |
| 0 | 0 | 0 | 0 |
| 0 | 0 | 0 | 0 |
| 0 | 0 | 0 | 0 |
| 0 | 0 | 0 | 0 |
| 0 | 0 | 0 | 0 |
| 0 | 0 | 0 | 1 |
| 0 | 0 | 0 | 0 |
| 0 | 0 | 0 | 1 |
| 0 | 0 | 0 | 1 |
| 0 | 0 | 0 | 1 |
| 0 | 0 | 0 | 0 |
| 0 | 0 | 0 | 0 |
| 0 | 0 | 0 | 0 |
| 0 | 0 | 0 | 0 |
| 0 | 0 | 0 | 0 |
| 0 | 0 | 0 | 0 |
| 0 | 0 | 0 | 0 |
| 0 | 0 | 0 | 0 |
| 0 | 0 | 0 | 0 |
| 0 | 0 | 0 | 0 |
| 0 | 0 | 0 | 0 |
| 0 | 1 | 0 | 0 |
| 0 | 0 | 0 | 0 |
| 0 | 0 | 0 | 0 |
| 0 | 1 | 0 | 0 |
| 0 | 0 | 0 | 0 |
| 0 | 0 | 0 | 0 |
| 0 | 0 | 0 | 0 |
| 0 | 0 | 0 | 0 |
| 0 | 0 | 0 | 0 |
| 0 | 0 | 0 | 0 |
| 0 | 0 | 0 | 0 |
| 0 | 0 | 0 | 0 |
| 0 | 0 | 0 | 0 |
| 0 | 0 | 0 | 0 |
| 0 | 0 | 0 | 0 |
| 0 | 0 | 0 | 0 |
| 0 | 0 | 0 | 0 |
| 0 | 0 | 0 | 0 |
| 0 | 0 | 0 | 0 |

| | | | |
|---|---|---|---|
| 0 | 0 | 0 | 0 |
| 0 | 0 | 0 | 0 |
| 0 | 0 | 0 | 0 |
| 0 | 0 | 0 | 0 |
| 0 | 0 | 0 | 0 |
| 0 | 0 | 0 | 0 |
| 0 | 0 | 0 | 0 |
| 0 | 0 | 0 | 0 |
| 0 | 0 | 1 | 0 |
| 0 | 0 | 1 | 0 |
| 0 | 0 | 0 | 0 |
| 0 | 0 | 0 | 0 |
| 0 | 0 | 0 | 0 |
| 0 | 0 | 0 | 0 |
| 0 | 0 | 0 | 0 |
| 0 | 1 | 0 | 0 |
| 0 | 0 | 0 | 0 |
| 0 | 0 | 0 | 0 |
| 1 | 1 | 0 | 0 |
| 0 | 0 | 0 | 0 |
| 0 | 0 | 0 | 0 |
| 0 | 0 | 0 | 0 |
| 0 | 0 | 0 | 0 |
| 0 | 0 | 0 | 0 |
| 0 | 0 | 0 | 0 |
| 0 | 0 | 0 | 0 |
| 0 | 0 | 0 | 0 |
| 0 | 0 | 0 | 0 |
| 0 | 0 | 0 | 0 |
| 0 | 0 | 0 | 0 |
| 0 | 0 | 0 | 0 |
| 0 | 0 | 0 | 0 |
| 0 | 0 | 0 | 0 |
| 0 | 0 | 0 | 0 |
| 0 | 0 | 0 | 0 |
| 0 | 0 | 0 | 0 |
| 0 | 0 | 0 | 0 |
| 0 | 0 | 0 | 0 |
| 0 | 0 | 0 | 0 |
| 0 | 0 | 0 | 0 |
| 0 | 0 | 0 | 0 |
| 0 | 0 | 0 | 0 |
| 0 | 0 | 0 | 0 |
| 0 | 0 | 0 | 0 |
| 0 | 0 | 0 | 0 |
| 0 | 0 | 0 | 0 |
| 0 | 0 | 0 | 0 |
| 0 | 0 | 0 | 0 |
| 0 | 0 | 0 | 0 |
| 0 | 0 | 0 | 0 |
| 0 | 0 | 0 | 0 |
| 0 | 0 | 0 | 0 |
| 0 | 0 | 0 | 0 |
| 1 | 0 | 0 | 0 |

| | | | |
|---|---|---|---|
| 0 | 0 | 0 | 0 |
| 0 | 0 | 0 | 0 |
| 0 | 0 | 0 | 0 |
| 0 | 0 | 0 | 0 |
| 0 | 0 | 0 | 0 |
| 0 | 0 | 0 | 0 |
| 0 | 0 | 0 | 0 |
| 0 | 0 | 0 | 0 |
| 0 | 0 | 0 | 0 |
| 0 | 0 | 0 | 0 |
| 1 | 0 | 0 | 0 |
| 0 | 0 | 0 | 0 |
| 0 | 0 | 0 | 0 |
| 0 | 0 | 0 | 0 |
| 1 | 1 | 0 | 1 |
| 0 | 0 | 0 | 0 |
| 1 | 0 | 0 | 0 |
| 1 | 1 | 0 | 0 |
| 0 | 0 | 0 | 0 |
| 0 | 0 | 0 | 0 |
| 0 | 0 | 0 | 0 |
| 0 | 0 | 0 | 0 |
| 0 | 0 | 0 | 0 |
| 0 | 0 | 0 | 1 |
| 0 | 0 | 0 | 0 |
| 0 | 0 | 0 | 0 |
| 0 | 0 | 0 | 0 |
| 0 | 0 | 0 | 1 |
| 0 | 0 | 0 | 0 |
| 0 | 0 | 0 | 0 |
| 0 | 0 | 0 | 0 |
| 0 | 0 | 0 | 0 |
| 0 | 0 | 0 | 0 |
| 0 | 0 | 0 | 0 |
| 0 | 0 | 0 | 0 |
| 0 | 0 | 0 | 0 |
| 0 | 0 | 0 | 0 |
| 0 | 0 | 0 | 0 |
| 0 | 0 | 0 | 0 |
| 0 | 0 | 0 | 0 |
| 0 | 0 | 0 | 0 |
| 0 | 0 | 0 | 0 |
| 0 | 0 | 0 | 0 |
| 0 | 0 | 0 | 0 |
| 0 | 0 | 1 | 0 |
| 0 | 0 | 0 | 0 |
| 0 | 0 | 0 | 0 |
| 0 | 0 | 0 | 0 |
| 0 | 0 | 0 | 0 |
| 0 | 0 | 0 | 0 |
| 0 | 0 | 0 | 0 |
| 0 | 0 | 0 | 0 |
| 0 | 0 | 0 | 0 |
| 0 | 0 | 0 | 0 |

| | | | |
|---|---|---|---|
| 0 | 0 | 0 | 0 |
| 0 | 0 | 0 | 0 |
| 0 | 0 | 0 | 0 |
| 0 | 0 | 0 | 0 |
| 0 | 0 | 0 | 0 |
| 0 | 0 | 0 | 0 |
| 1 | 0 | 0 | 1 |
| 1 | 0 | 0 | 0 |
| 0 | 0 | 0 | 0 |
| 0 | 0 | 0 | 0 |
| 0 | 0 | 0 | 0 |
| 0 | 0 | 0 | 0 |
| 0 | 0 | 0 | 0 |
| 0 | 0 | 0 | 0 |
| 0 | 0 | 0 | 0 |
| 0 | 0 | 0 | 0 |
| 0 | 0 | 0 | 0 |
| 0 | 0 | 0 | 0 |
| 0 | 0 | 0 | 0 |
| 0 | 0 | 0 | 0 |
| 1 | 0 | 0 | 0 |
| 0 | 0 | 0 | 0 |
| 0 | 0 | 0 | 0 |
| 0 | 0 | 0 | 0 |
| 0 | 0 | 0 | 0 |
| 0 | 0 | 0 | 0 |
| 0 | 0 | 0 | 0 |
| 0 | 0 | 0 | 0 |
| 0 | 0 | 0 | 0 |
| 0 | 0 | 0 | 0 |
| 0 | 0 | 0 | 0 |
| 0 | 0 | 0 | 0 |
| 0 | 0 | 0 | 0 |
| 0 | 0 | 0 | 0 |
| 0 | 0 | 0 | 0 |
| 0 | 0 | 0 | 0 |
| 0 | 0 | 0 | 0 |
| 0 | 0 | 0 | 0 |
| 0 | 0 | 0 | 0 |
| 0 | 0 | 0 | 0 |
| 0 | 0 | 0 | 0 |
| 0 | 0 | 0 | 0 |
| 0 | 0 | 0 | 0 |
| 0 | 0 | 0 | 0 |
| 0 | 0 | 0 | 0 |
| 0 | 0 | 0 | 0 |
| 0 | 0 | 0 | 0 |
| 0 | 0 | 0 | 0 |
| 0 | 0 | 0 | 0 |
| 0 | 0 | 0 | 0 |
| 0 | 0 | 0 | 0 |
| 0 | 0 | 0 | 0 |
| 0 | 0 | 0 | 0 |
| 0 | 0 | 0 | 0 |

| | | | |
|---|---|---|---|
| 0 | 0 | 0 | 0 |
| 0 | 0 | 0 | 0 |
| 0 | 0 | 0 | 0 |
| 1 | 0 | 0 | 0 |
| 0 | 0 | 0 | 0 |
| 1 | 0 | 0 | 0 |
| 0 | 0 | 0 | 0 |
| 0 | 0 | 0 | 0 |
| 1 | 0 | 0 | 0 |
| 0 | 0 | 0 | 0 |
| 0 | 0 | 0 | 0 |
| 0 | 0 | 0 | 0 |
| 0 | 0 | 0 | 0 |
| 0 | 0 | 0 | 0 |
| 0 | 0 | 0 | 0 |
| 0 | 0 | 0 | 0 |
| 0 | 0 | 0 | 0 |
| 0 | 0 | 0 | 0 |
| 1 | 0 | 0 | 0 |
| 0 | 0 | 0 | 0 |
| 0 | 0 | 0 | 0 |
| 0 | 0 | 0 | 0 |
| 0 | 0 | 0 | 0 |
| 0 | 0 | 0 | 0 |
| 0 | 0 | 0 | 0 |
| 0 | 0 | 0 | 0 |
| 0 | 0 | 0 | 0 |
| 0 | 0 | 0 | 0 |
| 0 | 1 | 0 | 0 |
| 0 | 0 | 0 | 0 |
| 0 | 0 | 0 | 0 |
| 0 | 0 | 0 | 0 |
| 0 | 1 | 0 | 0 |
| 1 | 1 | 0 | 0 |
| 1 | 0 | 0 | 0 |
| 0 | 0 | 0 | 0 |
| 0 | 0 | 0 | 0 |
| 0 | 0 | 0 | 0 |
| 0 | 0 | 0 | 0 |
| 0 | 0 | 0 | 0 |
| 1 | 0 | 0 | 0 |
| 0 | 0 | 0 | 0 |
| 0 | 0 | 0 | 0 |
| 0 | 0 | 0 | 0 |
| 0 | 0 | 0 | 0 |
| 0 | 0 | 0 | 0 |
| 0 | 0 | 0 | 0 |
| 0 | 0 | 0 | 0 |
| 0 | 0 | 0 | 0 |
| 0 | 0 | 0 | 0 |
| 0 | 0 | 0 | 0 |
| 0 | 0 | 0 | 0 |
| 0 | 0 | 0 | 0 |
| 1 | 0 | 0 | 0 |

| | | | |
|---|---|---|---|
| 1 | 0 | 0 | 1 |
| 0 | 0 | 0 | 0 |
| 0 | 0 | 0 | 0 |
| 0 | 0 | 0 | 0 |
| 0 | 0 | 0 | 0 |
| 0 | 0 | 0 | 0 |
| 0 | 0 | 0 | 0 |
| 0 | 0 | 0 | 0 |
| 0 | 0 | 0 | 0 |
| 0 | 0 | 0 | 0 |
| 0 | 0 | 0 | 0 |
| 0 | 0 | 0 | 0 |
| 0 | 0 | 0 | 0 |
| 0 | 0 | 0 | 0 |
| 0 | 0 | 0 | 0 |
| 0 | 0 | 0 | 0 |
| 0 | 0 | 0 | 0 |
| 0 | 0 | 0 | 0 |
| 0 | 0 | 0 | 0 |
| 0 | 0 | 0 | 0 |
| 0 | 0 | 0 | 0 |
| 0 | 0 | 0 | 0 |
| 0 | 1 | 0 | 0 |
| 0 | 0 | 0 | 0 |
| 0 | 0 | 0 | 0 |
| 0 | 0 | 0 | 0 |
| 0 | 0 | 0 | 0 |
| 0 | 0 | 0 | 0 |
| 0 | 0 | 0 | 0 |
| 0 | 0 | 0 | 0 |
| 0 | 0 | 0 | 0 |
| 0 | 0 | 0 | 0 |
| 0 | 0 | 0 | 0 |
| 0 | 0 | 0 | 0 |
| 0 | 0 | 0 | 0 |
| 0 | 0 | 0 | 0 |
| 0 | 0 | 0 | 0 |
| 0 | 0 | 0 | 0 |
| 0 | 0 | 0 | 0 |
| 0 | 0 | 0 | 0 |
| 0 | 0 | 0 | 0 |
| 0 | 0 | 0 | 0 |
| 0 | 0 | 0 | 0 |
| 0 | 0 | 0 | 0 |
| 0 | 0 | 0 | 0 |
| 0 | 0 | 0 | 0 |
| 0 | 0 | 0 | 0 |
| 0 | 0 | 0 | 0 |
| 0 | 0 | 0 | 0 |
| 0 | 0 | 0 | 0 |
| 0 | 0 | 0 | 0 |
| 0 | 0 | 0 | 0 |
| 0 | 0 | 0 | 0 |
| 0 | 0 | 0 | 0 |

| | | | |
|---|---|---|---|
| 0 | 0 | 0 | 0 |
| 0 | 0 | 0 | 0 |
| 0 | 0 | 0 | 0 |
| 0 | 0 | 0 | 0 |
| 0 | 0 | 0 | 0 |
| 0 | 0 | 0 | 0 |
| 0 | 1 | 0 | 0 |
| 0 | 1 | 0 | 0 |
| 0 | 0 | 0 | 0 |
| 0 | 0 | 0 | 0 |
| 0 | 0 | 0 | 0 |
| 0 | 0 | 0 | 0 |
| 0 | 0 | 0 | 0 |
| 0 | 0 | 0 | 0 |
| 0 | 0 | 0 | 0 |
| 0 | 0 | 0 | 0 |
| 1 | 0 | 0 | 0 |
| 0 | 0 | 0 | 0 |
| 1 | 0 | 0 | 0 |
| 1 | 0 | 0 | 0 |
| 1 | 0 | 0 | 0 |
| 1 | 0 | 0 | 0 |
| 0 | 0 | 0 | 0 |
| 0 | 0 | 0 | 0 |
| 0 | 0 | 0 | 0 |
| 0 | 0 | 0 | 0 |
| 0 | 0 | 0 | 0 |
| 0 | 0 | 0 | 0 |
| 0 | 0 | 0 | 0 |
| 0 | 0 | 0 | 0 |
| 0 | 0 | 0 | 0 |
| 0 | 0 | 0 | 0 |
| 0 | 0 | 0 | 0 |
| 0 | 0 | 0 | 0 |
| 0 | 0 | 0 | 0 |
| 0 | 0 | 1 | 0 |
| 1 | 0 | 0 | 0 |
| 1 | 0 | 0 | 0 |
| 1 | 0 | 0 | 0 |
| 0 | 0 | 0 | 0 |
| 0 | 0 | 0 | 0 |
| 0 | 0 | 0 | 0 |
| 0 | 0 | 0 | 0 |
| 1 | 0 | 0 | 0 |
| 1 | 0 | 0 | 0 |
| 1 | 0 | 0 | 0 |
| 0 | 0 | 0 | 0 |
| 0 | 0 | 0 | 0 |
| 1 | 0 | 0 | 0 |
| 0 | 0 | 0 | 0 |
| 0 | 0 | 0 | 0 |
| 1 | 0 | 0 | 0 |
| 0 | 0 | 0 | 0 |
| 0 | 0 | 0 | 0 |

| | | | |
|---|---|---|---|
| 0 | 0 | 0 | 0 |
| 0 | 0 | 0 | 0 |
| 0 | 0 | 0 | 0 |
| 0 | 0 | 0 | 0 |
| 0 | 0 | 0 | 0 |
| 0 | 0 | 0 | 0 |
| 0 | 0 | 0 | 0 |
| 0 | 0 | 0 | 0 |
| 0 | 0 | 0 | 0 |
| 0 | 0 | 0 | 0 |
| 0 | 0 | 0 | 0 |
| 0 | 0 | 0 | 0 |
| 0 | 0 | 0 | 0 |
| 0 | 0 | 0 | 0 |
| 0 | 0 | 0 | 0 |
| 0 | 0 | 0 | 0 |
| 0 | 0 | 0 | 0 |
| 0 | 0 | 0 | 0 |
| 0 | 0 | 0 | 0 |
| 0 | 0 | 0 | 0 |
| 0 | 0 | 0 | 0 |
| 0 | 0 | 0 | 0 |
| 1 | 1 | 0 | 0 |
| 0 | 0 | 0 | 0 |
| 0 | 1 | 0 | 0 |
| 0 | 0 | 0 | 0 |
| 0 | 0 | 0 | 0 |
| 0 | 0 | 0 | 0 |
| 0 | 0 | 0 | 0 |
| 0 | 0 | 0 | 0 |
| 0 | 0 | 0 | 0 |
| 0 | 0 | 0 | 0 |
| 0 | 0 | 0 | 0 |
| 0 | 0 | 0 | 0 |
| 0 | 0 | 0 | 0 |
| 0 | 0 | 0 | 0 |
| 0 | 0 | 0 | 0 |
| 0 | 0 | 0 | 0 |
| 0 | 0 | 0 | 0 |
| 0 | 0 | 0 | 0 |
| 0 | 0 | 0 | 0 |
| 0 | 0 | 0 | 0 |
| 0 | 0 | 0 | 0 |
| 0 | 0 | 0 | 0 |
| 0 | 0 | 0 | 0 |
| 1 | 0 | 0 | 1 |
| 0 | 0 | 0 | 0 |
| 0 | 0 | 0 | 0 |
| 0 | 0 | 0 | 0 |
| 1 | 0 | 0 | 0 |
| 1 | 0 | 0 | 0 |
| 1 | 0 | 0 | 0 |
| 0 | 0 | 0 | 0 |
| 0 | 0 | 0 | 0 |

| | | | |
|---|---|---|---|
| 0 | 0 | 0 | 0 |
| 0 | 0 | 0 | 0 |
| 0 | 0 | 0 | 0 |
| 0 | 0 | 0 | 0 |
| 0 | 0 | 0 | 0 |
| 0 | 0 | 0 | 0 |
| 0 | 0 | 0 | 0 |
| 0 | 0 | 0 | 0 |
| 0 | 0 | 0 | 0 |
| 0 | 0 | 0 | 0 |
| 0 | 0 | 0 | 1 |
| 0 | 0 | 0 | 0 |
| 0 | 0 | 0 | 0 |
| 0 | 0 | 0 | 0 |
| 0 | 0 | 0 | 0 |
| 0 | 0 | 0 | 0 |
| 0 | 0 | 0 | 0 |
| 0 | 0 | 0 | 0 |
| 0 | 0 | 0 | 0 |
| 0 | 0 | 0 | 0 |
| 0 | 0 | 0 | 0 |
| 0 | 0 | 0 | 0 |
| 0 | 0 | 0 | 0 |
| 0 | 0 | 0 | 0 |
| 0 | 0 | 0 | 0 |
| 0 | 0 | 0 | 0 |
| 0 | 0 | 0 | 0 |
| 0 | 0 | 0 | 0 |
| 0 | 0 | 0 | 0 |
| 0 | 0 | 0 | 0 |
| 0 | 1 | 0 | 0 |
| 0 | 0 | 0 | 0 |
| 1 | 1 | 0 | 0 |
| 1 | 1 | 0 | 0 |
| 0 | 0 | 0 | 0 |
| 0 | 0 | 0 | 0 |
| 0 | 0 | 0 | 0 |
| 0 | 0 | 0 | 0 |
| 0 | 0 | 0 | 0 |
| 0 | 0 | 0 | 0 |
| 0 | 0 | 0 | 0 |
| 0 | 1 | 0 | 0 |
| 0 | 0 | 0 | 0 |
| 0 | 0 | 0 | 0 |
| 0 | 0 | 0 | 0 |
| 1 | 0 | 0 | 0 |
| 1 | 0 | 0 | 0 |
| 1 | 0 | 0 | 0 |
| 1 | 1 | 1 | 0 |
| 0 | 0 | 0 | 0 |
| 1 | 1 | 1 | 1 |
| 0 | 0 | 0 | 0 |
| 0 | 0 | 0 | 0 |
| 0 | 0 | 0 | 0 |

| | | | |
|---|---|---|---|
| 0 | 0 | 0 | 0 |
| 0 | 0 | 0 | 0 |
| 0 | 0 | 0 | 0 |
| 0 | 0 | 0 | 0 |
| 1 | 0 | 0 | 0 |
| 1 | 0 | 0 | 0 |
| 0 | 0 | 0 | 0 |
| 0 | 0 | 0 | 0 |
| 0 | 0 | 0 | 0 |
| 0 | 0 | 0 | 0 |
| 0 | 0 | 0 | 0 |
| 0 | 0 | 0 | 0 |
| 0 | 0 | 0 | 0 |
| 0 | 0 | 0 | 0 |
| 0 | 0 | 0 | 0 |
| 0 | 0 | 0 | 0 |
| 0 | 0 | 0 | 0 |
| 0 | 0 | 0 | 0 |
| 0 | 0 | 0 | 0 |
| 0 | 0 | 0 | 0 |
| 0 | 0 | 0 | 0 |
| 0 | 0 | 0 | 0 |
| 0 | 0 | 0 | 0 |
| 0 | 0 | 0 | 0 |
| 0 | 0 | 0 | 0 |
| 0 | 0 | 0 | 0 |
| 0 | 0 | 0 | 0 |
| 0 | 0 | 0 | 0 |
| 0 | 0 | 0 | 0 |
| 0 | 0 | 0 | 0 |
| 0 | 0 | 0 | 0 |
| 0 | 0 | 0 | 0 |
| 0 | 0 | 0 | 0 |
| 0 | 0 | 0 | 0 |
| 0 | 0 | 0 | 0 |
| 0 | 0 | 0 | 0 |
| 0 | 0 | 0 | 0 |
| 0 | 0 | 0 | 0 |
| 0 | 0 | 0 | 0 |
| 0 | 0 | 0 | 0 |
| 0 | 0 | 0 | 0 |
| 0 | 0 | 0 | 0 |
| 0 | 0 | 0 | 0 |
| 0 | 0 | 0 | 0 |
| 0 | 0 | 0 | 0 |
| 0 | 0 | 0 | 0 |
| 0 | 0 | 0 | 0 |
| 0 | 0 | 0 | 0 |
| 0 | 0 | 0 | 0 |
| 0 | 0 | 0 | 0 |
| 0 | 0 | 0 | 0 |
| 0 | 0 | 0 | 0 |
| 0 | 0 | 0 | 0 |
| 0 | 1 | 0 | 0 |

| | | | |
|---|---|---|---|
| 0 | 0 | 0 | 0 |
| 0 | 0 | 0 | 0 |
| 0 | 0 | 0 | 0 |
| 0 | 0 | 0 | 0 |
| 0 | 0 | 0 | 0 |
| 0 | 0 | 0 | 0 |
| 0 | 0 | 0 | 0 |
| 0 | 0 | 0 | 0 |
| 0 | 0 | 0 | 0 |
| 0 | 0 | 0 | 0 |
| 0 | 0 | 0 | 0 |
| 0 | 0 | 0 | 0 |
| 0 | 0 | 0 | 0 |
| 0 | 0 | 0 | 0 |
| 0 | 0 | 0 | 0 |
| 0 | 0 | 0 | 0 |
| 0 | 0 | 0 | 0 |
| 0 | 0 | 0 | 0 |
| 0 | 0 | 0 | 0 |
| 0 | 0 | 0 | 0 |
| 0 | 0 | 0 | 0 |
| 0 | 0 | 0 | 0 |
| 0 | 0 | 0 | 0 |
| 0 | 0 | 0 | 0 |
| 0 | 0 | 0 | 0 |
| 0 | 0 | 0 | 0 |
| 0 | 0 | 0 | 0 |
| 0 | 0 | 0 | 0 |
| 0 | 0 | 0 | 0 |
| 0 | 0 | 0 | 0 |
| 0 | 0 | 0 | 0 |
| 0 | 0 | 0 | 0 |
| 0 | 0 | 0 | 0 |
| 0 | 0 | 0 | 0 |
| 0 | 0 | 0 | 0 |
| 0 | 0 | 0 | 0 |
| 0 | 0 | 0 | 0 |
| 0 | 0 | 0 | 0 |
| 0 | 0 | 0 | 0 |
| 0 | 0 | 0 | 0 |
| 0 | 0 | 0 | 0 |
| 0 | 0 | 0 | 0 |
| 0 | 0 | 0 | 0 |
| 0 | 0 | 0 | 0 |
| 0 | 0 | 0 | 0 |
| 0 | 0 | 0 | 0 |
| 0 | 0 | 0 | 0 |
| 0 | 0 | 0 | 0 |
| 0 | 0 | 0 | 0 |
| 0 | 0 | 0 | 0 |
| 0 | 0 | 0 | 0 |
| 0 | 0 | 0 | 0 |
| 0 | 0 | 0 | 0 |
| 0 | 0 | 0 | 0 |

| | | | |
|---|---|---|---|
| 0 | 0 | 0 | 0 |
| 0 | 0 | 0 | 0 |
| 0 | 0 | 0 | 0 |
| 0 | 0 | 0 | 0 |
| 0 | 0 | 0 | 0 |
| 0 | 0 | 0 | 0 |
| 0 | 0 | 0 | 0 |
| 0 | 0 | 0 | 0 |
| 0 | 0 | 0 | 0 |
| 0 | 0 | 0 | 0 |
| 0 | 0 | 0 | 0 |
| 0 | 0 | 1 | 0 |
| 0 | 0 | 0 | 0 |
| 0 | 0 | 1 | 0 |
| 0 | 0 | 1 | 0 |
| 0 | 0 | 1 | 0 |
| 0 | 0 | 1 | 0 |
| 0 | 0 | 0 | 0 |
| 0 | 0 | 0 | 0 |
| 0 | 0 | 0 | 0 |
| 1 | 0 | 0 | 0 |

| glycan_binding_site | water_binding_site | water_only_binding_site | non_binding_site |
|---|---|---|---|
| 0 | 0 | 0 | 0 |
| 0 | 0 | 0 | 0 |
| 0 | 0 | 0 | 1 |
| 0 | 0 | 0 | 0 |
| 0 | 0 | 0 | 0 |
| 0 | 0 | 0 | 0 |
| 0 | 0 | 0 | 1 |
| 0 | 0 | 0 | 1 |
| 0 | 0 | 0 | 0 |
| 0 | 0 | 0 | 0 |
| 0 | 0 | 0 | 0 |
| 0 | 0 | 0 | 0 |
| 0 | 0 | 0 | 1 |
| 1 | 0 | 0 | 0 |
| 0 | 0 | 0 | 0 |
| 0 | 0 | 0 | 0 |
| 0 | 0 | 0 | 0 |
| 0 | 0 | 0 | 0 |
| 0 | 0 | 0 | 0 |
| 0 | 0 | 0 | 0 |
| 0 | 1 | 0 | 0 |
| 1 | 1 | 0 | 0 |
| 1 | 1 | 0 | 0 |
| 0 | 1 | 0 | 0 |
| 0 | 1 | 0 | 0 |
| 0 | 0 | 0 | 1 |
| 0 | 0 | 0 | 0 |
| 0 | 0 | 0 | 1 |
| 0 | 0 | 0 | 0 |
| 0 | 0 | 0 | 1 |
| 0 | 0 | 0 | 1 |
| 0 | 1 | 0 | 0 |
| 0 | 1 | 0 | 0 |
| 0 | 1 | 0 | 0 |
| 0 | 1 | 0 | 0 |
| 0 | 0 | 0 | 1 |
| 0 | 0 | 0 | 0 |
| 0 | 1 | 0 | 0 |
| 0 | 1 | 0 | 0 |
| 0 | 1 | 0 | 0 |
| 0 | 1 | 0 | 0 |
| 0 | 1 | 0 | 0 |
| 0 | 1 | 0 | 0 |
| 0 | 1 | 0 | 0 |
| 0 | 1 | 0 | 0 |
| 0 | 1 | 0 | 0 |
| 0 | 1 | 0 | 0 |
| 0 | 1 | 0 | 0 |
| 0 | 1 | 0 | 0 |
| 0 | 1 | 0 | 0 |
| 0 | 1 | 0 | 0 |
| 0 | 1 | 0 | 0 |
| 0 | 1 | 0 | 0 |

| | | | |
|---|---|---|---|
| 0 | 1 | 0 | 0 |
| 0 | 1 | 0 | 0 |
| 0 | 1 | 0 | 0 |
| 0 | 1 | 0 | 0 |
| 0 | 1 | 0 | 0 |
| 0 | 1 | 0 | 0 |
| 0 | 1 | 0 | 0 |
| 0 | 1 | 0 | 0 |
| 0 | 1 | 0 | 0 |
| 0 | 1 | 0 | 0 |
| 0 | 1 | 0 | 0 |
| 0 | 1 | 0 | 0 |
| 0 | 1 | 0 | 0 |
| 0 | 1 | 0 | 0 |
| 0 | 1 | 0 | 0 |
| 0 | 1 | 0 | 0 |
| 0 | 1 | 0 | 0 |
| 0 | 1 | 0 | 0 |
| 0 | 1 | 0 | 0 |
| 0 | 1 | 0 | 0 |
| 0 | 0 | 0 | 0 |
| 0 | 0 | 0 | 0 |
| 0 | 0 | 0 | 0 |
| 0 | 0 | 0 | 0 |
| 0 | 0 | 0 | 0 |
| 0 | 0 | 0 | 0 |
| 0 | 0 | 0 | 0 |
| 0 | 0 | 0 | 0 |
| 0 | 0 | 0 | 0 |
| 0 | 0 | 0 | 0 |
| 0 | 0 | 0 | 0 |
| 0 | 0 | 0 | 0 |
| 0 | 0 | 0 | 0 |
| 0 | 0 | 0 | 0 |
| 0 | 0 | 0 | 0 |
| 0 | 0 | 0 | 0 |
| 0 | 0 | 0 | 0 |
| 0 | 0 | 0 | 0 |
| 0 | 0 | 0 | 0 |
| 0 | 0 | 0 | 0 |
| 1 | 0 | 0 | 0 |
| 0 | 0 | 0 | 0 |
| 1 | 0 | 0 | 0 |
| 1 | 0 | 0 | 0 |
| 0 | 0 | 0 | 0 |
| 0 | 0 | 0 | 1 |
| 0 | 0 | 0 | 1 |
| 0 | 0 | 0 | 0 |
| 1 | 0 | 0 | 0 |
| 0 | 0 | 0 | 0 |
| 0 | 0 | 0 | 0 |
| 0 | 0 | 0 | 1 |
| 0 | 0 | 0 | 1 |
| 0 | 0 | 0 | 0 |

| | | | |
|---|---|---|---|
| 0 | 0 | 0 | 1 |
| 0 | 0 | 0 | 1 |
| 0 | 0 | 0 | 0 |
| 0 | 0 | 0 | 0 |
| 0 | 0 | 0 | 0 |
| 0 | 0 | 0 | 0 |
| 0 | 0 | 0 | 1 |
| 0 | 0 | 0 | 0 |
| 0 | 0 | 0 | 0 |
| 0 | 0 | 0 | 0 |
| 0 | 0 | 0 | 0 |
| 0 | 0 | 0 | 0 |
| 0 | 0 | 0 | 0 |
| 1 | 0 | 0 | 0 |
| 1 | 0 | 0 | 0 |
| 0 | 0 | 0 | 1 |
| 0 | 0 | 0 | 0 |
| 0 | 0 | 0 | 0 |
| 0 | 0 | 0 | 0 |
| 0 | 0 | 0 | 0 |
| 0 | 0 | 0 | 0 |
| 0 | 0 | 0 | 0 |
| 0 | 0 | 0 | 0 |
| 0 | 0 | 0 | 0 |
| 0 | 0 | 0 | 0 |
| 0 | 0 | 0 | 0 |
| 0 | 0 | 0 | 0 |
| 0 | 0 | 0 | 0 |
| 0 | 0 | 0 | 0 |
| 0 | 0 | 0 | 0 |
| 0 | 0 | 0 | 0 |
| 0 | 0 | 0 | 0 |
| 0 | 0 | 0 | 0 |
| 0 | 0 | 0 | 0 |
| 0 | 0 | 0 | 0 |
| 0 | 0 | 0 | 0 |
| 0 | 0 | 0 | 0 |
| 0 | 0 | 0 | 0 |
| 0 | 0 | 0 | 0 |
| 0 | 0 | 0 | 0 |
| 0 | 1 | 0 | 0 |
| 0 | 0 | 0 | 0 |
| 0 | 0 | 0 | 0 |
| 0 | 0 | 0 | 0 |
| 0 | 1 | 0 | 0 |
| 1 | 1 | 0 | 0 |
| 0 | 0 | 0 | 0 |
| 0 | 0 | 0 | 0 |
| 0 | 0 | 0 | 0 |
| 0 | 0 | 0 | 0 |
| 0 | 0 | 0 | 0 |
| 0 | 0 | 0 | 0 |
| 0 | 1 | 1 | 1 |

| | | | |
|---|---|---|---|
| 0 | 1 | 1 | 1 |
| 0 | 1 | 1 | 1 |
| 0 | 1 | 1 | 1 |
| 0 | 0 | 0 | 0 |
| 0 | 0 | 0 | 0 |
| 0 | 1 | 1 | 1 |
| 0 | 1 | 0 | 0 |
| 0 | 1 | 1 | 1 |
| 0 | 1 | 1 | 1 |
| 0 | 0 | 0 | 1 |
| 0 | 1 | 1 | 1 |
| 0 | 1 | 0 | 0 |
| 0 | 1 | 1 | 1 |
| 0 | 0 | 0 | 0 |
| 0 | 1 | 1 | 1 |
| 0 | 0 | 0 | 1 |
| 0 | 1 | 1 | 1 |
| 0 | 1 | 1 | 1 |
| 0 | 1 | 1 | 1 |
| 0 | 1 | 0 | 0 |
| 0 | 1 | 0 | 0 |
| 0 | 0 | 0 | 1 |
| 0 | 0 | 0 | 0 |
| 0 | 0 | 0 | 0 |
| 0 | 0 | 0 | 0 |
| 0 | 0 | 0 | 1 |
| 0 | 0 | 0 | 0 |
| 0 | 1 | 0 | 0 |
| 0 | 0 | 0 | 1 |
| 0 | 1 | 1 | 1 |
| 0 | 0 | 0 | 1 |
| 0 | 0 | 0 | 1 |
| 0 | 1 | 1 | 1 |
| 0 | 0 | 0 | 1 |
| 0 | 0 | 0 | 0 |
| 0 | 1 | 1 | 1 |
| 1 | 1 | 0 | 0 |
| 1 | 1 | 0 | 0 |
| 0 | 1 | 1 | 1 |
| 0 | 1 | 1 | 1 |
| 0 | 0 | 0 | 1 |
| 0 | 1 | 0 | 0 |
| 0 | 1 | 0 | 0 |
| 0 | 0 | 0 | 1 |
| 0 | 1 | 1 | 1 |
| 0 | 1 | 1 | 1 |
| 0 | 0 | 0 | 0 |
| 0 | 0 | 0 | 0 |
| 0 | 0 | 0 | 0 |
| 0 | 0 | 0 | 0 |
| 0 | 0 | 0 | 0 |
| 0 | 0 | 0 | 1 |
| 0 | 0 | 0 | 1 |
| 0 | 0 | 0 | 0 |

| | | | |
|---|---|---|---|
| 0 | 0 | 0 | 0 |
| 0 | 0 | 0 | 0 |
| 0 | 1 | 0 | 0 |
| 0 | 0 | 0 | 1 |
| 0 | 0 | 0 | 1 |
| 0 | 0 | 0 | 1 |
| 0 | 1 | 0 | 0 |
| 0 | 1 | 0 | 0 |
| 0 | 1 | 1 | 1 |
| 0 | 0 | 0 | 1 |
| 1 | 0 | 0 | 0 |
| 0 | 0 | 0 | 1 |
| 1 | 0 | 0 | 0 |
| 0 | 0 | 0 | 1 |
| 0 | 0 | 0 | 1 |
| 0 | 0 | 0 | 1 |
| 0 | 0 | 0 | 1 |
| 0 | 0 | 0 | 0 |
| 0 | 0 | 0 | 0 |
| 0 | 0 | 0 | 0 |
| 0 | 0 | 0 | 0 |
| 0 | 0 | 0 | 1 |
| 0 | 0 | 0 | 0 |
| 0 | 0 | 0 | 1 |
| 0 | 0 | 0 | 1 |
| 0 | 0 | 0 | 0 |
| 0 | 1 | 0 | 0 |
| 0 | 0 | 0 | 0 |
| 0 | 1 | 0 | 0 |
| 0 | 1 | 0 | 0 |
| 0 | 1 | 0 | 0 |
| 0 | 0 | 0 | 0 |
| 0 | 0 | 0 | 1 |
| 0 | 0 | 0 | 1 |
| 0 | 0 | 0 | 0 |
| 0 | 0 | 0 | 1 |
| 0 | 0 | 0 | 1 |
| 0 | 0 | 0 | 0 |
| 0 | 0 | 0 | 1 |
| 0 | 0 | 0 | 1 |
| 0 | 0 | 0 | 0 |
| 0 | 1 | 1 | 1 |
| 0 | 0 | 0 | 1 |
| 0 | 0 | 0 | 1 |
| 0 | 0 | 0 | 0 |
| 0 | 0 | 0 | 0 |
| 0 | 1 | 0 | 0 |
| 0 | 1 | 0 | 0 |
| 0 | 1 | 0 | 0 |
| 0 | 1 | 0 | 0 |
| 0 | 1 | 0 | 0 |
| 0 | 1 | 0 | 0 |
| 0 | 1 | 0 | 0 |
| 0 | 1 | 0 | 0 |

| | | | |
|---|---|---|---|
| 0 | 1 | 0 | 0 |
| 0 | 1 | 0 | 0 |
| 0 | 1 | 0 | 0 |
| 0 | 0 | 0 | 0 |
| 0 | 1 | 0 | 0 |
| 0 | 1 | 0 | 0 |
| 0 | 0 | 0 | 1 |
| 0 | 0 | 0 | 1 |
| 0 | 0 | 0 | 0 |
| 0 | 0 | 0 | 0 |
| 0 | 0 | 0 | 0 |
| 0 | 0 | 0 | 0 |
| 0 | 0 | 0 | 0 |
| 0 | 0 | 0 | 1 |
| 0 | 0 | 0 | 1 |
| 0 | 0 | 0 | 0 |
| 0 | 0 | 0 | 1 |
| 0 | 0 | 0 | 0 |
| 0 | 0 | 0 | 1 |
| 0 | 0 | 0 | 0 |
| 0 | 0 | 0 | 1 |
| 0 | 0 | 0 | 1 |
| 0 | 0 | 0 | 1 |
| 0 | 0 | 0 | 0 |
| 0 | 0 | 0 | 0 |
| 0 | 0 | 0 | 0 |
| 0 | 0 | 0 | 0 |
| 0 | 0 | 0 | 0 |
| 0 | 0 | 0 | 0 |
| 0 | 0 | 0 | 0 |
| 0 | 0 | 0 | 1 |
| 0 | 0 | 0 | 1 |
| 0 | 0 | 0 | 0 |
| 0 | 0 | 0 | 0 |
| 0 | 0 | 0 | 0 |
| 0 | 0 | 0 | 1 |
| 0 | 0 | 0 | 0 |
| 0 | 1 | 0 | 0 |
| 0 | 0 | 0 | 0 |
| 0 | 0 | 0 | 0 |
| 0 | 0 | 0 | 1 |
| 0 | 0 | 0 | 1 |
| 0 | 0 | 0 | 0 |
| 0 | 0 | 0 | 1 |
| 0 | 0 | 0 | 0 |
| 0 | 0 | 0 | 0 |
| 0 | 0 | 0 | 0 |
| 0 | 0 | 0 | 1 |
| 0 | 0 | 0 | 1 |
| 0 | 0 | 0 | 0 |
| 0 | 0 | 0 | 1 |
| 0 | 0 | 0 | 1 |
| 0 | 0 | 0 | 0 |
| 0 | 0 | 0 | 1 |

| | | | |
|---|---|---|---|
| 0 | 0 | 0 | 0 |
| 0 | 0 | 0 | 1 |
| 0 | 1 | 1 | 1 |
| 0 | 1 | 0 | 0 |
| 0 | 1 | 0 | 0 |
| 0 | 0 | 0 | 0 |
| 0 | 0 | 0 | 1 |
| 0 | 0 | 0 | 0 |
| 0 | 0 | 0 | 0 |
| 0 | 0 | 0 | 1 |
| 0 | 0 | 0 | 1 |
| 0 | 0 | 0 | 1 |
| 0 | 0 | 0 | 1 |
| 0 | 0 | 0 | 1 |
| 0 | 0 | 0 | 0 |
| 0 | 0 | 0 | 1 |
| 0 | 0 | 0 | 0 |
| 0 | 0 | 0 | 0 |
| 0 | 0 | 0 | 1 |
| 0 | 0 | 0 | 0 |
| 0 | 1 | 0 | 0 |
| 0 | 1 | 0 | 0 |
| 0 | 0 | 0 | 0 |
| 0 | 1 | 0 | 0 |
| 0 | 0 | 0 | 1 |
| 0 | 0 | 0 | 1 |
| 0 | 0 | 0 | 0 |
| 0 | 0 | 0 | 0 |
| 0 | 0 | 0 | 0 |
| 0 | 0 | 0 | 0 |
| 0 | 0 | 0 | 0 |
| 0 | 0 | 0 | 0 |
| 0 | 0 | 0 | 0 |
| 0 | 0 | 0 | 0 |
| 0 | 0 | 0 | 1 |
| 0 | 0 | 0 | 1 |
| 0 | 0 | 0 | 0 |
| 0 | 0 | 0 | 0 |
| 0 | 0 | 0 | 0 |
| 0 | 0 | 0 | 0 |
| 0 | 0 | 0 | 0 |
| 0 | 1 | 0 | 0 |
| 0 | 0 | 0 | 1 |
| 0 | 0 | 0 | 0 |
| 0 | 1 | 0 | 0 |
| 0 | 1 | 0 | 0 |
| 0 | 1 | 0 | 0 |
| 0 | 1 | 0 | 0 |
| 1 | 1 | 0 | 0 |
| 0 | 0 | 0 | 1 |
| 0 | 0 | 0 | 0 |
| 0 | 0 | 0 | 0 |
| 0 | 0 | 0 | 0 |
| 0 | 0 | 0 | 0 |

| | | | |
|---|---|---|---|
| 0 | 0 | 0 | 0 |
| 0 | 0 | 0 | 0 |
| 0 | 0 | 0 | 0 |
| 0 | 0 | 0 | 0 |
| 0 | 0 | 0 | 1 |
| 0 | 0 | 0 | 0 |
| 0 | 0 | 0 | 0 |
| 0 | 0 | 0 | 0 |
| 0 | 0 | 0 | 0 |
| 0 | 1 | 0 | 0 |
| 0 | 1 | 0 | 0 |
| 0 | 1 | 0 | 0 |
| 0 | 0 | 0 | 0 |
| 0 | 0 | 0 | 0 |
| 0 | 0 | 0 | 0 |
| 0 | 0 | 0 | 0 |
| 0 | 1 | 0 | 0 |
| 0 | 1 | 0 | 0 |
| 0 | 0 | 0 | 1 |
| 0 | 0 | 0 | 0 |
| 0 | 0 | 0 | 0 |
| 0 | 0 | 0 | 0 |
| 0 | 0 | 0 | 0 |
| 0 | 0 | 0 | 1 |
| 0 | 0 | 0 | 0 |
| 0 | 0 | 0 | 0 |
| 0 | 0 | 0 | 0 |
| 0 | 0 | 0 | 1 |
| 0 | 0 | 0 | 1 |
| 0 | 0 | 0 | 1 |
| 0 | 0 | 0 | 0 |
| 0 | 0 | 0 | 0 |
| 0 | 0 | 0 | 0 |
| 0 | 0 | 0 | 0 |
| 0 | 0 | 0 | 0 |
| 0 | 0 | 0 | 1 |
| 0 | 0 | 0 | 1 |
| 0 | 0 | 0 | 0 |
| 0 | 0 | 0 | 0 |
| 0 | 0 | 0 | 1 |
| 0 | 0 | 0 | 1 |
| 0 | 0 | 0 | 1 |
| 0 | 0 | 0 | 1 |
| 0 | 0 | 0 | 1 |
| 0 | 0 | 0 | 0 |
| 0 | 0 | 0 | 0 |
| 0 | 0 | 0 | 0 |
| 0 | 0 | 0 | 0 |
| 0 | 0 | 0 | 0 |
| 0 | 0 | 0 | 1 |
| 0 | 0 | 0 | 1 |
| 0 | 0 | 0 | 0 |
| 0 | 0 | 0 | 1 |
| 0 | 0 | 0 | 0 |

| | | | |
|---|---|---|---|
| 0 | 0 | 0 | 1 |
| 0 | 0 | 0 | 1 |
| 0 | 0 | 0 | 0 |
| 0 | 0 | 0 | 1 |
| 0 | 0 | 0 | 1 |
| 0 | 0 | 0 | 1 |
| 0 | 0 | 0 | 1 |
| 0 | 0 | 0 | 0 |
| 0 | 0 | 0 | 1 |
| 0 | 0 | 0 | 1 |
| 0 | 0 | 0 | 0 |
| 0 | 0 | 0 | 1 |
| 0 | 0 | 0 | 1 |
| 0 | 0 | 0 | 1 |
| 0 | 0 | 0 | 0 |
| 0 | 0 | 0 | 0 |
| 0 | 0 | 0 | 1 |
| 0 | 0 | 0 | 1 |
| 0 | 0 | 0 | 0 |
| 0 | 0 | 0 | 0 |
| 0 | 0 | 0 | 0 |
| 0 | 0 | 0 | 0 |
| 0 | 0 | 0 | 0 |
| 0 | 1 | 0 | 0 |
| 0 | 1 | 0 | 0 |
| 0 | 0 | 0 | 1 |
| 0 | 0 | 0 | 0 |
| 0 | 0 | 0 | 0 |
| 0 | 0 | 0 | 0 |
| 0 | 0 | 0 | 1 |
| 0 | 0 | 0 | 0 |
| 0 | 1 | 0 | 0 |
| 0 | 0 | 0 | 1 |
| 0 | 0 | 0 | 0 |
| 0 | 0 | 0 | 0 |
| 0 | 0 | 0 | 0 |
| 0 | 1 | 0 | 0 |
| 0 | 1 | 0 | 0 |
| 0 | 1 | 0 | 0 |
| 0 | 1 | 0 | 0 |
| 0 | 0 | 0 | 1 |
| 0 | 0 | 0 | 0 |
| 0 | 0 | 0 | 1 |
| 0 | 0 | 0 | 0 |
| 0 | 0 | 0 | 0 |
| 1 | 0 | 0 | 0 |
| 0 | 0 | 0 | 0 |
| 1 | 0 | 0 | 0 |
| 0 | 0 | 0 | 0 |
| 0 | 0 | 0 | 0 |
| 0 | 0 | 0 | 1 |
| 0 | 0 | 0 | 0 |
| 0 | 0 | 0 | 0 |
| 0 | 0 | 0 | 0 |

| | | | |
|---|---|---|---|
| 0 | 0 | 0 | 1 |
| 0 | 0 | 0 | 0 |
| 0 | 0 | 0 | 1 |
| 0 | 0 | 0 | 1 |
| 1 | 0 | 0 | 0 |
| 0 | 0 | 0 | 0 |
| 1 | 0 | 0 | 0 |
| 1 | 0 | 0 | 0 |
| 0 | 0 | 0 | 0 |
| 0 | 0 | 0 | 1 |
| 0 | 0 | 0 | 1 |
| 0 | 0 | 0 | 1 |
| 0 | 0 | 0 | 1 |
| 1 | 0 | 0 | 0 |
| 0 | 0 | 0 | 1 |
| 0 | 0 | 0 | 0 |
| 0 | 0 | 0 | 0 |
| 0 | 0 | 0 | 0 |
| 1 | 0 | 0 | 0 |
| 0 | 0 | 0 | 1 |
| 1 | 0 | 0 | 0 |
| 0 | 0 | 0 | 1 |
| 0 | 0 | 0 | 0 |
| 0 | 0 | 0 | 1 |
| 0 | 0 | 0 | 1 |
| 0 | 0 | 0 | 1 |
| 0 | 0 | 0 | 0 |
| 0 | 0 | 0 | 1 |
| 0 | 0 | 0 | 1 |
| 0 | 0 | 0 | 1 |
| 0 | 0 | 0 | 0 |
| 0 | 0 | 0 | 0 |
| 0 | 0 | 0 | 1 |
| 0 | 0 | 0 | 1 |
| 0 | 0 | 0 | 0 |
| 0 | 0 | 0 | 0 |
| 0 | 0 | 0 | 0 |
| 1 | 0 | 0 | 0 |
| 0 | 0 | 0 | 0 |
| 0 | 0 | 0 | 1 |
| 0 | 0 | 0 | 0 |
| 0 | 0 | 0 | 0 |
| 0 | 0 | 0 | 0 |
| 0 | 0 | 0 | 0 |
| 0 | 0 | 0 | 1 |
| 1 | 0 | 0 | 0 |
| 0 | 1 | 0 | 0 |
| 0 | 0 | 0 | 0 |
| 0 | 0 | 0 | 0 |
| 1 | 0 | 0 | 0 |
| 0 | 0 | 0 | 0 |
| 0 | 1 | 0 | 0 |
| 0 | 0 | 0 | 1 |
| 0 | 0 | 0 | 0 |

| | | | |
|---|---|---|---|
| 0 | 1 | 0 | 0 |
| 0 | 1 | 0 | 0 |
| 0 | 1 | 1 | 1 |
| 0 | 1 | 1 | 1 |
| 0 | 1 | 1 | 1 |
| 0 | 0 | 0 | 0 |
| 0 | 0 | 0 | 1 |
| 0 | 0 | 0 | 1 |
| 0 | 1 | 0 | 0 |
| 0 | 0 | 0 | 0 |
| 1 | 0 | 0 | 0 |
| 1 | 0 | 0 | 0 |
| 0 | 0 | 0 | 1 |
| 1 | 0 | 0 | 0 |
| 0 | 0 | 0 | 0 |
| 0 | 0 | 0 | 0 |
| 0 | 1 | 0 | 0 |
| 0 | 0 | 0 | 0 |
| 0 | 0 | 0 | 0 |
| 0 | 0 | 0 | 1 |
| 0 | 0 | 0 | 0 |
| 0 | 0 | 0 | 0 |
| 0 | 0 | 0 | 0 |
| 0 | 0 | 0 | 0 |
| 0 | 0 | 0 | 0 |
| 0 | 0 | 0 | 1 |
| 0 | 0 | 0 | 1 |
| 0 | 0 | 0 | 1 |
| 0 | 0 | 0 | 0 |
| 0 | 1 | 0 | 0 |
| 0 | 0 | 0 | 0 |
| 0 | 0 | 0 | 0 |
| 0 | 0 | 0 | 0 |
| 0 | 1 | 0 | 0 |
| 0 | 1 | 0 | 0 |
| 0 | 0 | 0 | 1 |
| 0 | 0 | 0 | 0 |
| 0 | 1 | 0 | 0 |
| 0 | 0 | 0 | 1 |
| 0 | 0 | 0 | 1 |
| 0 | 0 | 0 | 1 |
| 0 | 0 | 0 | 0 |
| 0 | 0 | 0 | 0 |
| 0 | 0 | 0 | 0 |
| 0 | 0 | 0 | 0 |
| 0 | 0 | 0 | 0 |
| 0 | 0 | 0 | 0 |
| 0 | 0 | 0 | 1 |
| 0 | 0 | 0 | 1 |
| 0 | 0 | 0 | 0 |
| 0 | 0 | 0 | 0 |
| 0 | 0 | 0 | 0 |
| 0 | 0 | 0 | 0 |
| 0 | 0 | 0 | 1 |

| | | | |
|---|---|---|---|
| 0 | 0 | 0 | 1 |
| 0 | 0 | 0 | 0 |
| 0 | 0 | 0 | 1 |
| 0 | 1 | 0 | 0 |
| 0 | 0 | 0 | 1 |
| 0 | 0 | 0 | 1 |
| 0 | 1 | 0 | 0 |
| 0 | 1 | 1 | 1 |
| 0 | 1 | 1 | 1 |
| 0 | 1 | 1 | 1 |
| 0 | 1 | 1 | 1 |
| 0 | 1 | 1 | 1 |
| 0 | 0 | 0 | 1 |
| 0 | 0 | 0 | 1 |
| 0 | 0 | 0 | 1 |
| 0 | 0 | 0 | 1 |
| 0 | 0 | 0 | 0 |
| 0 | 0 | 0 | 0 |
| 0 | 0 | 0 | 1 |
| 0 | 0 | 0 | 0 |
| 0 | 0 | 0 | 0 |
| 0 | 0 | 0 | 0 |
| 0 | 0 | 0 | 1 |
| 0 | 0 | 0 | 0 |
| 0 | 0 | 0 | 0 |
| 0 | 0 | 0 | 1 |
| 0 | 0 | 0 | 1 |
| 0 | 0 | 0 | 1 |
| 0 | 1 | 0 | 0 |
| 0 | 0 | 0 | 0 |
| 0 | 1 | 1 | 1 |
| 0 | 0 | 0 | 0 |
| 0 | 1 | 0 | 0 |
| 0 | 0 | 0 | 0 |
| 0 | 0 | 0 | 0 |
| 0 | 0 | 0 | 0 |
| 0 | 1 | 1 | 1 |
| 0 | 0 | 0 | 1 |
| 0 | 0 | 0 | 1 |
| 0 | 0 | 0 | 1 |
| 0 | 0 | 0 | 0 |
| 0 | 0 | 0 | 1 |
| 0 | 0 | 0 | 0 |
| 0 | 0 | 0 | 1 |
| 1 | 0 | 0 | 0 |
| 0 | 0 | 0 | 1 |
| 0 | 0 | 0 | 0 |
| 0 | 0 | 0 | 0 |
| 1 | 0 | 0 | 0 |
| 0 | 0 | 0 | 0 |
| 0 | 0 | 0 | 0 |
| 0 | 0 | 0 | 0 |
| 0 | 0 | 0 | 0 |
| 0 | 0 | 0 | 0 |

| | | | |
|---|---|---|---|
| 0 | 0 | 0 | 0 |
| 0 | 0 | 0 | 0 |
| 0 | 0 | 0 | 0 |
| 0 | 0 | 0 | 0 |
| 0 | 0 | 0 | 0 |
| 0 | 0 | 0 | 0 |
| 0 | 0 | 0 | 0 |
| 0 | 0 | 0 | 1 |
| 0 | 1 | 0 | 0 |
| 0 | 1 | 0 | 0 |
| 0 | 1 | 0 | 0 |
| 0 | 1 | 1 | 1 |
| 0 | 0 | 0 | 0 |
| 0 | 1 | 0 | 0 |
| 0 | 1 | 0 | 0 |
| 0 | 1 | 0 | 0 |
| 0 | 1 | 0 | 0 |
| 0 | 1 | 0 | 0 |
| 0 | 1 | 0 | 0 |
| 0 | 1 | 0 | 0 |
| 0 | 1 | 0 | 0 |
| 0 | 1 | 0 | 0 |
| 0 | 1 | 0 | 0 |
| 0 | 1 | 0 | 0 |
| 0 | 1 | 0 | 0 |
| 0 | 1 | 0 | 0 |
| 0 | 1 | 0 | 0 |
| 0 | 1 | 0 | 0 |
| 0 | 1 | 0 | 0 |
| 0 | 1 | 0 | 0 |
| 0 | 1 | 0 | 0 |
| 0 | 1 | 0 | 0 |
| 0 | 1 | 0 | 0 |
| 0 | 1 | 0 | 0 |
| 0 | 1 | 0 | 0 |
| 0 | 1 | 0 | 0 |
| 0 | 1 | 0 | 0 |
| 0 | 1 | 0 | 0 |
| 0 | 1 | 0 | 0 |
| 0 | 1 | 0 | 0 |
| 0 | 1 | 1 | 1 |
| 0 | 1 | 0 | 0 |
| 0 | 1 | 0 | 0 |
| 0 | 1 | 0 | 0 |
| 0 | 1 | 0 | 0 |
| 0 | 1 | 0 | 0 |
| 0 | 0 | 0 | 0 |
| 0 | 1 | 0 | 0 |
| 0 | 0 | 0 | 0 |
| 0 | 0 | 0 | 1 |
| 0 | 0 | 0 | 1 |
| 0 | 1 | 1 | 1 |
| 1 | 0 | 0 | 0 |
| 0 | 0 | 0 | 0 |

| | | | |
|---|---|---|---|
| 0 | 0 | 0 | 0 |
| 0 | 0 | 0 | 0 |
| 1 | 1 | 0 | 0 |
| 1 | 0 | 0 | 0 |
| 0 | 0 | 0 | 0 |
| 0 | 0 | 0 | 0 |
| 0 | 0 | 0 | 0 |
| 0 | 0 | 0 | 1 |
| 0 | 0 | 0 | 1 |
| 0 | 0 | 0 | 0 |
| 0 | 0 | 0 | 0 |
| 0 | 0 | 0 | 1 |
| 0 | 0 | 0 | 0 |
| 0 | 0 | 0 | 0 |
| 0 | 0 | 0 | 0 |
| 0 | 0 | 0 | 0 |
| 0 | 0 | 0 | 0 |
| 0 | 0 | 0 | 1 |
| 0 | 0 | 0 | 0 |
| 0 | 0 | 0 | 1 |
| 0 | 0 | 0 | 0 |
| 0 | 0 | 0 | 0 |
| 0 | 0 | 0 | 1 |
| 0 | 0 | 0 | 0 |
| 0 | 0 | 0 | 0 |
| 0 | 0 | 0 | 1 |
| 0 | 0 | 0 | 0 |
| 0 | 0 | 0 | 0 |
| 0 | 0 | 0 | 0 |
| 0 | 0 | 0 | 0 |
| 0 | 0 | 0 | 0 |
| 0 | 0 | 0 | 0 |
| 0 | 0 | 0 | 1 |
| 0 | 0 | 0 | 0 |
| 0 | 1 | 0 | 0 |
| 0 | 0 | 0 | 0 |
| 0 | 0 | 0 | 1 |
| 0 | 0 | 0 | 0 |
| 0 | 0 | 0 | 0 |
| 0 | 0 | 0 | 0 |
| 0 | 1 | 0 | 0 |
| 0 | 1 | 0 | 0 |
| 0 | 1 | 0 | 0 |
| 0 | 1 | 0 | 0 |
| 0 | 1 | 0 | 0 |
| 0 | 0 | 0 | 1 |
| 0 | 0 | 0 | 0 |
| 0 | 1 | 0 | 0 |
| 0 | 0 | 0 | 0 |
| 0 | 1 | 0 | 0 |
| 0 | 1 | 0 | 0 |
| 0 | 1 | 0 | 0 |
| 0 | 0 | 0 | 1 |
| 0 | 0 | 0 | 1 |

| | | | |
|---|---|---|---|
| 0 | 1 | 1 | 1 |
| 0 | 0 | 0 | 1 |
| 0 | 0 | 0 | 0 |
| 0 | 0 | 0 | 0 |
| 0 | 1 | 0 | 0 |
| 0 | 1 | 0 | 0 |
| 0 | 0 | 0 | 0 |
| 0 | 0 | 0 | 1 |
| 0 | 0 | 0 | 0 |
| 0 | 1 | 1 | 1 |
| 0 | 0 | 0 | 1 |
| 0 | 0 | 0 | 1 |
| 0 | 0 | 0 | 0 |
| 0 | 0 | 0 | 0 |
| 0 | 0 | 0 | 0 |
| 0 | 0 | 0 | 0 |
| 0 | 0 | 0 | 0 |
| 1 | 0 | 0 | 0 |
| 0 | 0 | 0 | 1 |
| 0 | 0 | 0 | 1 |
| 0 | 0 | 0 | 1 |
| 0 | 0 | 0 | 0 |
| 0 | 0 | 0 | 1 |
| 0 | 0 | 0 | 0 |
| 0 | 0 | 0 | 0 |
| 0 | 0 | 0 | 0 |
| 0 | 0 | 0 | 0 |
| 0 | 0 | 0 | 0 |
| 0 | 0 | 0 | 0 |
| 0 | 0 | 0 | 0 |
| 0 | 0 | 0 | 0 |
| 0 | 0 | 0 | 0 |
| 0 | 0 | 0 | 0 |
| 0 | 0 | 0 | 0 |
| 0 | 0 | 0 | 0 |
| 0 | 0 | 0 | 0 |
| 0 | 0 | 0 | 0 |
| 0 | 0 | 0 | 0 |
| 1 | 0 | 0 | 0 |
| 0 | 0 | 0 | 0 |
| 0 | 1 | 0 | 0 |
| 0 | 0 | 0 | 1 |
| 0 | 1 | 0 | 0 |
| 0 | 1 | 0 | 0 |
| 0 | 0 | 0 | 0 |
| 0 | 0 | 0 | 0 |
| 0 | 0 | 0 | 1 |
| 0 | 0 | 0 | 0 |
| 0 | 0 | 0 | 0 |
| 0 | 0 | 0 | 0 |
| 0 | 0 | 0 | 0 |
| 0 | 0 | 0 | 0 |
| 0 | 0 | 0 | 1 |
| 0 | 0 | 0 | 1 |

| | | | |
|---|---|---|---|
| 0 | 1 | 0 | 0 |
| 0 | 0 | 0 | 0 |
| 0 | 0 | 0 | 1 |
| 0 | 0 | 0 | 1 |
| 0 | 0 | 0 | 1 |
| 0 | 1 | 0 | 0 |
| 0 | 0 | 0 | 1 |
| 0 | 0 | 0 | 1 |
| 0 | 0 | 0 | 0 |
| 0 | 0 | 0 | 0 |
| 0 | 0 | 0 | 0 |
| 0 | 0 | 0 | 0 |
| 0 | 0 | 0 | 0 |
| 0 | 0 | 0 | 1 |
| 0 | 0 | 0 | 1 |
| 0 | 0 | 0 | 1 |
| 0 | 0 | 0 | 1 |
| 0 | 0 | 0 | 1 |
| 0 | 0 | 0 | 0 |
| 0 | 0 | 0 | 0 |
| 0 | 0 | 0 | 1 |
| 0 | 0 | 0 | 1 |
| 0 | 0 | 0 | 0 |
| 0 | 1 | 0 | 0 |
| 0 | 0 | 0 | 1 |
| 0 | 0 | 0 | 1 |
| 0 | 1 | 0 | 0 |
| 0 | 1 | 1 | 1 |
| 0 | 0 | 0 | 0 |
| 0 | 0 | 0 | 0 |
| 0 | 0 | 0 | 0 |
| 0 | 0 | 0 | 0 |
| 0 | 0 | 0 | 0 |
| 1 | 0 | 0 | 0 |
| 1 | 1 | 0 | 0 |
| 0 | 0 | 0 | 0 |
| 0 | 0 | 0 | 0 |
| 0 | 0 | 0 | 0 |
| 0 | 0 | 0 | 1 |
| 0 | 0 | 0 | 0 |
| 0 | 0 | 0 | 1 |
| 0 | 0 | 0 | 1 |
| 0 | 0 | 0 | 1 |
| 0 | 0 | 0 | 1 |
| 0 | 0 | 0 | 1 |
| 0 | 0 | 0 | 0 |
| 0 | 0 | 0 | 0 |
| 0 | 0 | 0 | 0 |
| 0 | 0 | 0 | 0 |
| 0 | 0 | 0 | 0 |
| 0 | 0 | 0 | 0 |
| 0 | 0 | 0 | 0 |
| 0 | 0 | 0 | 0 |
| 0 | 0 | 0 | 1 |

| | | | |
|---|---|---|---|
| 0 | 0 | 0 | 1 |
| 0 | 0 | 0 | 0 |
| 0 | 0 | 0 | 0 |
| 0 | 0 | 0 | 0 |
| 0 | 0 | 0 | 1 |
| 0 | 0 | 0 | 0 |
| 0 | 0 | 0 | 0 |
| 0 | 0 | 0 | 0 |
| 0 | 0 | 0 | 1 |
| 0 | 0 | 0 | 1 |
| 0 | 0 | 0 | 0 |
| 0 | 0 | 0 | 0 |
| 0 | 0 | 0 | 0 |
| 0 | 0 | 0 | 1 |
| 0 | 0 | 0 | 0 |
| 0 | 0 | 0 | 0 |
| 0 | 1 | 0 | 0 |
| 0 | 1 | 0 | 0 |
| 0 | 0 | 0 | 0 |
| 0 | 0 | 0 | 0 |
| 0 | 0 | 0 | 0 |
| 0 | 0 | 0 | 1 |
| 0 | 0 | 0 | 0 |
| 0 | 0 | 0 | 1 |
| 0 | 0 | 0 | 0 |
| 0 | 1 | 0 | 0 |
| 0 | 0 | 0 | 1 |
| 0 | 0 | 0 | 0 |
| 0 | 0 | 0 | 0 |
| 0 | 0 | 0 | 0 |
| 0 | 0 | 0 | 0 |
| 0 | 0 | 0 | 0 |
| 0 | 0 | 0 | 0 |
| 0 | 0 | 0 | 0 |
| 0 | 0 | 0 | 1 |
| 0 | 1 | 0 | 0 |
| 0 | 0 | 0 | 0 |
| 0 | 0 | 0 | 0 |
| 0 | 0 | 0 | 0 |
| 0 | 0 | 0 | 0 |
| 0 | 0 | 0 | 0 |
| 0 | 0 | 0 | 0 |
| 0 | 0 | 0 | 0 |
| 0 | 0 | 0 | 0 |
| 0 | 0 | 0 | 0 |
| 0 | 0 | 0 | 0 |
| 0 | 0 | 0 | 0 |
| 0 | 0 | 0 | 1 |
| 0 | 0 | 0 | 0 |
| 0 | 0 | 0 | 1 |
| 0 | 0 | 0 | 0 |
| 0 | 0 | 0 | 0 |
| 0 | 0 | 0 | 0 |
| 0 | 0 | 0 | 0 |

| | | | |
|---|---|---|---|
| 0 | 0 | 0 | 0 |
| 0 | 1 | 0 | 0 |
| 0 | 0 | 0 | 1 |
| 0 | 0 | 0 | 0 |
| 0 | 0 | 0 | 0 |
| 0 | 0 | 0 | 1 |
| 0 | 0 | 0 | 1 |
| 0 | 0 | 0 | 1 |
| 0 | 0 | 0 | 0 |
| 0 | 0 | 0 | 1 |
| 0 | 0 | 0 | 0 |
| 0 | 0 | 0 | 0 |
| 0 | 0 | 0 | 0 |
| 0 | 0 | 0 | 1 |
| 0 | 0 | 0 | 0 |
| 0 | 0 | 0 | 1 |
| 0 | 0 | 0 | 0 |
| 0 | 0 | 0 | 0 |
| 1 | 0 | 0 | 0 |
| 0 | 0 | 0 | 1 |
| 0 | 0 | 0 | 1 |
| 0 | 0 | 0 | 0 |
| 0 | 0 | 0 | 1 |
| 0 | 0 | 0 | 0 |
| 0 | 0 | 0 | 0 |
| 0 | 0 | 0 | 1 |
| 0 | 0 | 0 | 1 |
| 0 | 0 | 0 | 0 |
| 0 | 0 | 0 | 1 |
| 1 | 0 | 0 | 0 |
| 0 | 0 | 0 | 1 |
| 0 | 0 | 0 | 1 |
| 0 | 0 | 0 | 1 |
| 0 | 0 | 0 | 1 |
| 0 | 0 | 0 | 1 |
| 0 | 0 | 0 | 1 |
| 0 | 0 | 0 | 1 |
| 0 | 0 | 0 | 1 |
| 0 | 0 | 0 | 1 |
| 0 | 0 | 0 | 1 |
| 0 | 0 | 0 | 1 |
| 0 | 0 | 0 | 1 |
| 0 | 0 | 0 | 1 |
| 0 | 0 | 0 | 1 |
| 0 | 0 | 0 | 1 |
| 0 | 0 | 0 | 1 |
| 0 | 0 | 0 | 1 |
| 0 | 0 | 0 | 1 |
| 0 | 0 | 0 | 1 |
| 0 | 0 | 0 | 1 |
| 0 | 0 | 0 | 1 |
| 0 | 0 | 0 | 1 |
| 0 | 0 | 0 | 1 |
| 0 | 0 | 0 | 1 |

| | | | |
|---|---|---|---|
| 0 | 0 | 0 | 1 |
| 0 | 0 | 0 | 1 |
| 0 | 0 | 0 | 1 |
| 0 | 0 | 0 | 1 |
| 0 | 0 | 0 | 1 |
| 0 | 0 | 0 | 1 |
| 0 | 0 | 0 | 1 |
| 0 | 0 | 0 | 1 |
| 0 | 0 | 0 | 1 |
| 0 | 0 | 0 | 1 |
| 0 | 0 | 0 | 1 |
| 0 | 0 | 0 | 0 |
| 0 | 0 | 0 | 0 |
| 0 | 1 | 0 | 0 |
| 0 | 1 | 1 | 1 |
| 0 | 0 | 0 | 0 |
| 0 | 0 | 0 | 1 |
| 0 | 1 | 0 | 0 |
| 0 | 0 | 0 | 1 |
| 0 | 0 | 0 | 1 |
| 0 | 0 | 0 | 0 |
| 0 | 0 | 0 | 1 |
| 0 | 0 | 0 | 1 |
| 0 | 0 | 0 | 0 |
| 0 | 0 | 0 | 0 |
| 0 | 0 | 0 | 1 |
| 0 | 0 | 0 | 1 |
| 0 | 0 | 0 | 0 |
| 0 | 0 | 0 | 0 |
| 0 | 0 | 0 | 0 |
| 0 | 0 | 0 | 0 |
| 0 | 0 | 0 | 0 |
| 0 | 0 | 0 | 1 |
| 0 | 0 | 0 | 1 |
| 0 | 0 | 0 | 1 |
| 0 | 0 | 0 | 1 |
| 0 | 0 | 0 | 1 |
| 0 | 0 | 0 | 1 |
| 0 | 0 | 0 | 1 |
| 0 | 1 | 0 | 0 |
| 0 | 0 | 0 | 0 |
| 0 | 0 | 0 | 1 |
| 0 | 1 | 0 | 0 |
| 0 | 1 | 0 | 0 |
| 0 | 0 | 0 | 0 |
| 0 | 1 | 1 | 1 |
| 0 | 0 | 0 | 0 |
| 0 | 0 | 0 | 1 |
| 0 | 0 | 0 | 1 |
| 0 | 1 | 0 | 0 |
| 0 | 0 | 0 | 0 |
| 0 | 0 | 0 | 0 |
| 0 | 0 | 0 | 1 |
| 0 | 0 | 0 | 0 |

| | | | |
|---|---|---|---|
| 0 | 0 | 0 | 0 |
| 0 | 0 | 0 | 1 |
| 0 | 0 | 0 | 1 |
| 0 | 0 | 0 | 1 |
| 0 | 1 | 0 | 0 |
| 0 | 0 | 0 | 1 |
| 0 | 0 | 0 | 0 |
| 0 | 0 | 0 | 1 |
| 0 | 0 | 0 | 1 |
| 0 | 0 | 0 | 0 |
| 0 | 1 | 1 | 1 |
| 0 | 0 | 0 | 1 |
| 0 | 0 | 0 | 1 |
| 0 | 0 | 0 | 0 |
| 0 | 0 | 0 | 1 |
| 0 | 0 | 0 | 1 |
| 0 | 0 | 0 | 0 |
| 0 | 0 | 0 | 1 |
| 0 | 0 | 0 | 0 |
| 0 | 0 | 0 | 0 |
| 0 | 0 | 0 | 0 |
| 0 | 0 | 0 | 0 |
| 0 | 0 | 0 | 0 |
| 0 | 0 | 0 | 0 |
| 0 | 0 | 0 | 0 |
| 0 | 0 | 0 | 0 |
| 0 | 0 | 0 | 0 |
| 0 | 0 | 0 | 0 |
| 0 | 0 | 0 | 0 |
| 0 | 0 | 0 | 0 |
| 0 | 0 | 0 | 0 |
| 0 | 0 | 0 | 0 |
| 0 | 0 | 0 | 0 |
| 0 | 0 | 0 | 0 |
| 0 | 0 | 0 | 0 |
| 0 | 0 | 0 | 0 |
| 0 | 1 | 1 | 1 |
| 0 | 1 | 0 | 0 |
| 0 | 0 | 0 | 0 |
| 0 | 0 | 0 | 1 |
| 0 | 0 | 0 | 1 |
| 0 | 0 | 0 | 0 |
| 0 | 0 | 0 | 0 |
| 0 | 1 | 0 | 0 |
| 0 | 1 | 0 | 0 |
| 0 | 0 | 0 | 0 |
| 0 | 0 | 0 | 0 |
| 0 | 1 | 0 | 0 |
| 0 | 0 | 0 | 1 |
| 0 | 0 | 0 | 1 |
| 0 | 0 | 0 | 0 |
| 0 | 0 | 0 | 1 |
| 0 | 0 | 0 | 0 |
| 0 | 0 | 0 | 0 |

| | | | |
|---|---|---|---|
| 0 | 0 | 0 | 0 |
| 0 | 0 | 0 | 0 |
| 0 | 0 | 0 | 0 |
| 0 | 0 | 0 | 0 |
| 0 | 0 | 0 | 0 |
| 0 | 0 | 0 | 0 |
| 0 | 0 | 0 | 0 |
| 0 | 0 | 0 | 0 |
| 0 | 0 | 0 | 0 |
| 0 | 0 | 0 | 0 |
| 0 | 0 | 0 | 0 |
| 0 | 0 | 0 | 0 |
| 0 | 0 | 0 | 0 |
| 0 | 0 | 0 | 0 |
| 0 | 0 | 0 | 1 |
| 0 | 0 | 0 | 0 |
| 0 | 1 | 0 | 0 |
| 0 | 1 | 0 | 0 |
| 0 | 1 | 1 | 1 |
| 0 | 0 | 0 | 0 |
| 0 | 1 | 0 | 0 |
| 0 | 1 | 0 | 0 |
| 0 | 1 | 0 | 0 |
| 0 | 0 | 0 | 0 |
| 0 | 1 | 0 | 0 |
| 0 | 1 | 1 | 1 |
| 0 | 0 | 0 | 0 |
| 0 | 1 | 0 | 0 |
| 0 | 1 | 0 | 0 |
| 0 | 0 | 0 | 0 |
| 0 | 0 | 0 | 1 |
| 0 | 0 | 0 | 0 |
| 0 | 0 | 0 | 1 |
| 0 | 1 | 1 | 1 |
| 1 | 0 | 0 | 0 |
| 0 | 0 | 0 | 0 |
| 0 | 0 | 0 | 1 |
| 0 | 0 | 0 | 1 |
| 0 | 0 | 0 | 1 |
| 0 | 0 | 0 | 1 |
| 0 | 0 | 0 | 0 |
| 0 | 0 | 0 | 0 |
| 0 | 0 | 0 | 0 |
| 0 | 0 | 0 | 0 |
| 0 | 0 | 0 | 0 |
| 0 | 0 | 0 | 0 |
| 0 | 0 | 0 | 0 |
| 0 | 0 | 0 | 0 |
| 0 | 0 | 0 | 1 |
| 0 | 0 | 0 | 1 |
| 0 | 0 | 0 | 0 |
| 0 | 0 | 0 | 0 |
| 0 | 0 | 0 | 0 |
| 0 | 0 | 0 | 1 |

| | | | |
|---|---|---|---|
| 0 | 0 | 0 | 1 |
| 0 | 0 | 0 | 1 |
| 0 | 1 | 1 | 1 |
| 0 | 0 | 0 | 0 |
| 0 | 0 | 0 | 1 |
| 0 | 0 | 0 | 1 |
| 0 | 0 | 0 | 0 |
| 0 | 0 | 0 | 1 |
| 0 | 0 | 0 | 0 |
| 0 | 0 | 0 | 0 |
| 0 | 0 | 0 | 0 |
| 0 | 0 | 0 | 0 |
| 0 | 0 | 0 | 0 |
| 0 | 0 | 0 | 0 |
| 0 | 0 | 0 | 0 |
| 0 | 0 | 0 | 0 |
| 0 | 0 | 0 | 1 |
| 0 | 0 | 0 | 0 |
| 0 | 0 | 0 | 0 |
| 0 | 1 | 1 | 1 |
| 0 | 0 | 0 | 0 |
| 0 | 0 | 0 | 0 |
| 0 | 0 | 0 | 1 |
| 0 | 0 | 0 | 0 |
| 0 | 0 | 0 | 1 |
| 0 | 0 | 0 | 1 |
| 0 | 0 | 0 | 1 |
| 0 | 0 | 0 | 0 |
| 0 | 0 | 0 | 0 |
| 0 | 0 | 0 | 0 |
| 0 | 0 | 0 | 1 |
| 0 | 0 | 0 | 0 |
| 0 | 0 | 0 | 0 |
| 0 | 0 | 0 | 0 |
| 0 | 0 | 0 | 1 |
| 0 | 0 | 0 | 1 |
| 0 | 0 | 0 | 0 |
| 0 | 0 | 0 | 0 |
| 0 | 0 | 0 | 0 |
| 0 | 0 | 0 | 1 |
| 0 | 0 | 0 | 1 |
| 0 | 0 | 0 | 1 |
| 0 | 0 | 0 | 1 |
| 0 | 0 | 0 | 0 |
| 0 | 0 | 0 | 1 |
| 0 | 0 | 0 | 0 |
| 0 | 0 | 0 | 1 |
| 0 | 0 | 0 | 0 |
| 0 | 0 | 0 | 0 |
| 0 | 0 | 0 | 1 |
| 0 | 0 | 0 | 1 |
| 0 | 1 | 0 | 0 |
| 0 | 0 | 0 | 0 |
| 0 | 0 | 0 | 1 |

| | | | |
|---|---|---|---|
| 0 | 0 | 0 | 1 |
| 0 | 0 | 0 | 1 |
| 0 | 0 | 0 | 1 |
| 0 | 0 | 0 | 1 |
| 0 | 1 | 0 | 0 |
| 0 | 1 | 1 | 1 |
| 0 | 1 | 1 | 1 |
| 0 | 1 | 0 | 0 |
| 0 | 1 | 0 | 0 |
| 0 | 0 | 0 | 1 |
| 0 | 0 | 0 | 0 |
| 0 | 0 | 0 | 0 |
| 1 | 0 | 0 | 0 |
| 1 | 0 | 0 | 0 |
| 0 | 0 | 0 | 0 |
| 1 | 0 | 0 | 0 |
| 1 | 0 | 0 | 0 |
| 1 | 0 | 0 | 0 |
| 1 | 0 | 0 | 0 |
| 0 | 0 | 0 | 1 |
| 0 | 0 | 0 | 1 |
| 0 | 0 | 0 | 1 |
| 0 | 0 | 0 | 1 |
| 0 | 0 | 0 | 1 |
| 0 | 0 | 0 | 0 |
| 0 | 0 | 0 | 1 |
| 1 | 0 | 0 | 0 |
| 1 | 0 | 0 | 0 |
| 0 | 0 | 0 | 1 |
| 0 | 0 | 0 | 1 |
| 0 | 0 | 0 | 1 |
| 0 | 0 | 0 | 1 |
| 0 | 0 | 0 | 1 |
| 0 | 0 | 0 | 0 |
| 0 | 0 | 0 | 0 |
| 0 | 0 | 0 | 1 |
| 0 | 0 | 0 | 0 |
| 0 | 0 | 0 | 1 |
| 0 | 0 | 0 | 0 |
| 0 | 1 | 0 | 0 |
| 0 | 1 | 0 | 0 |
| 0 | 0 | 0 | 0 |
| 0 | 0 | 0 | 1 |
| 0 | 0 | 0 | 0 |
| 0 | 0 | 0 | 0 |
| 1 | 0 | 0 | 0 |
| 1 | 0 | 0 | 0 |
| 1 | 0 | 0 | 0 |
| 0 | 0 | 0 | 0 |
| 0 | 0 | 0 | 0 |
| 0 | 1 | 0 | 0 |
| 1 | 0 | 0 | 0 |
| 1 | 0 | 0 | 0 |
| 0 | 0 | 0 | 1 |

| | | | |
|---|---|---|---|
| 0 | 0 | 0 | 0 |
| 0 | 0 | 0 | 1 |
| 0 | 0 | 0 | 0 |
| 0 | 0 | 0 | 0 |
| 0 | 1 | 0 | 0 |
| 0 | 0 | 0 | 1 |
| 0 | 0 | 0 | 0 |
| 0 | 0 | 0 | 1 |
| 0 | 0 | 0 | 1 |
| 0 | 0 | 0 | 0 |
| 0 | 0 | 0 | 1 |
| 0 | 0 | 0 | 1 |
| 0 | 0 | 0 | 1 |
| 0 | 0 | 0 | 1 |
| 0 | 0 | 0 | 1 |
| 0 | 1 | 0 | 0 |
| 0 | 0 | 0 | 0 |
| 0 | 0 | 0 | 1 |
| 0 | 0 | 0 | 0 |
| 0 | 0 | 0 | 1 |
| 0 | 0 | 0 | 0 |
| 0 | 0 | 0 | 0 |
| 0 | 0 | 0 | 0 |
| 0 | 1 | 0 | 0 |
| 0 | 1 | 0 | 0 |
| 0 | 0 | 0 | 0 |
| 0 | 0 | 0 | 0 |
| 0 | 0 | 0 | 0 |
| 0 | 0 | 0 | 0 |
| 0 | 1 | 0 | 0 |
| 0 | 0 | 0 | 0 |
| 0 | 0 | 0 | 0 |
| 0 | 0 | 0 | 1 |
| 0 | 0 | 0 | 1 |
| 0 | 0 | 0 | 0 |
| 0 | 1 | 0 | 0 |
| 0 | 0 | 0 | 1 |
| 0 | 0 | 0 | 0 |
| 0 | 0 | 0 | 1 |
| 0 | 1 | 0 | 0 |
| 0 | 1 | 0 | 0 |
| 0 | 0 | 0 | 0 |
| 0 | 0 | 0 | 1 |
| 0 | 0 | 0 | 1 |
| 0 | 1 | 0 | 0 |
| 0 | 1 | 0 | 0 |
| 0 | 0 | 0 | 0 |
| 0 | 1 | 0 | 0 |
| 0 | 0 | 0 | 0 |
| 0 | 0 | 0 | 0 |
| 0 | 1 | 1 | 1 |
| 0 | 1 | 0 | 0 |
| 0 | 1 | 0 | 0 |
| 0 | 1 | 1 | 1 |

| | | | |
|---|---|---|---|
| 0 | 0 | 0 | 0 |
| 0 | 0 | 0 | 1 |
| 0 | 0 | 0 | 0 |
| 0 | 0 | 0 | 0 |
| 0 | 0 | 0 | 0 |
| 0 | 0 | 0 | 0 |
| 0 | 0 | 0 | 0 |
| 0 | 0 | 0 | 0 |
| 0 | 0 | 0 | 0 |
| 0 | 0 | 0 | 0 |
| 0 | 0 | 0 | 0 |
| 0 | 0 | 0 | 1 |
| 0 | 0 | 0 | 0 |
| 0 | 0 | 0 | 1 |
| 0 | 1 | 1 | 1 |
| 0 | 1 | 1 | 1 |
| 0 | 1 | 0 | 0 |
| 0 | 1 | 0 | 0 |
| 0 | 1 | 1 | 1 |
| 0 | 1 | 0 | 0 |
| 0 | 1 | 1 | 1 |
| 0 | 1 | 0 | 0 |
| 0 | 0 | 0 | 1 |
| 0 | 1 | 0 | 0 |
| 0 | 0 | 0 | 1 |
| 0 | 0 | 0 | 1 |
| 0 | 0 | 0 | 1 |
| 0 | 0 | 0 | 0 |
| 0 | 0 | 0 | 1 |
| 0 | 0 | 0 | 1 |
| 0 | 0 | 0 | 1 |
| 1 | 0 | 0 | 0 |
| 0 | 0 | 0 | 1 |
| 0 | 0 | 0 | 0 |
| 0 | 0 | 0 | 0 |
| 0 | 0 | 0 | 0 |
| 0 | 0 | 0 | 1 |
| 0 | 0 | 0 | 1 |
| 0 | 0 | 0 | 0 |
| 0 | 0 | 0 | 0 |
| 0 | 0 | 0 | 0 |
| 0 | 0 | 0 | 0 |
| 0 | 1 | 0 | 0 |
| 0 | 0 | 0 | 0 |
| 0 | 0 | 0 | 0 |
| 0 | 0 | 0 | 0 |
| 0 | 0 | 0 | 1 |
| 0 | 0 | 0 | 1 |
| 0 | 0 | 0 | 0 |
| 0 | 0 | 0 | 0 |
| 0 | 0 | 0 | 0 |
| 0 | 0 | 0 | 1 |
| 0 | 0 | 0 | 1 |
| 0 | 0 | 0 | 0 |

| | | | |
|---|---|---|---|
| 0 | 0 | 0 | 0 |
| 0 | 0 | 0 | 1 |
| 0 | 0 | 0 | 1 |
| 0 | 0 | 0 | 1 |
| 0 | 0 | 0 | 0 |
| 0 | 0 | 0 | 0 |
| 0 | 0 | 0 | 0 |
| 0 | 0 | 0 | 0 |
| 0 | 0 | 0 | 0 |
| 0 | 0 | 0 | 0 |
| 0 | 0 | 0 | 1 |
| 0 | 0 | 0 | 0 |
| 0 | 0 | 0 | 0 |
| 0 | 0 | 0 | 0 |
| 0 | 1 | 1 | 1 |
| 0 | 0 | 0 | 1 |
| 0 | 1 | 0 | 0 |
| 0 | 0 | 0 | 1 |
| 0 | 1 | 0 | 0 |
| 0 | 1 | 0 | 0 |
| 0 | 0 | 0 | 1 |
| 0 | 1 | 0 | 0 |
| 0 | 1 | 0 | 0 |
| 0 | 0 | 0 | 1 |
| 0 | 1 | 0 | 0 |
| 0 | 1 | 0 | 0 |
| 0 | 1 | 0 | 0 |
| 0 | 1 | 1 | 1 |
| 0 | 1 | 0 | 0 |
| 0 | 0 | 0 | 0 |
| 0 | 0 | 0 | 0 |
| 0 | 0 | 0 | 1 |
| 0 | 0 | 0 | 1 |
| 0 | 0 | 0 | 1 |
| 0 | 0 | 0 | 1 |
| 0 | 0 | 0 | 1 |
| 0 | 0 | 0 | 1 |
| 0 | 0 | 0 | 1 |
| 0 | 0 | 0 | 1 |
| 0 | 0 | 0 | 1 |
| 0 | 0 | 0 | 1 |
| 0 | 0 | 0 | 1 |
| 1 | 0 | 0 | 0 |
| 0 | 0 | 0 | 1 |
| 0 | 0 | 0 | 1 |
| 0 | 0 | 0 | 1 |
| 0 | 0 | 0 | 1 |
| 0 | 0 | 0 | 1 |
| 0 | 0 | 0 | 1 |
| 0 | 0 | 0 | 1 |
| 0 | 0 | 0 | 1 |
| 0 | 0 | 0 | 1 |
| 0 | 0 | 0 | 1 |
| 0 | 0 | 0 | 1 |

| | | | |
|---|---|---|---|
| 0 | 0 | 0 | 1 |
| 0 | 0 | 0 | 1 |
| 0 | 0 | 0 | 1 |
| 0 | 0 | 0 | 1 |
| 0 | 0 | 0 | 1 |
| 0 | 0 | 0 | 1 |
| 0 | 0 | 0 | 1 |
| 0 | 0 | 0 | 1 |
| 0 | 0 | 0 | 0 |
| 0 | 0 | 0 | 0 |
| 0 | 0 | 0 | 0 |
| 0 | 0 | 0 | 0 |
| 0 | 0 | 0 | 0 |
| 0 | 0 | 0 | 0 |
| 0 | 0 | 0 | 0 |
| 0 | 0 | 0 | 0 |
| 0 | 0 | 0 | 0 |
| 0 | 0 | 0 | 0 |
| 0 | 0 | 0 | 0 |
| 0 | 0 | 0 | 0 |
| 1 | 0 | 0 | 0 |
| 1 | 0 | 0 | 0 |
| 0 | 0 | 0 | 0 |
| 0 | 0 | 0 | 0 |
| 0 | 0 | 0 | 0 |
| 0 | 0 | 0 | 1 |
| 0 | 0 | 0 | 1 |
| 0 | 0 | 0 | 0 |
| 0 | 0 | 0 | 0 |
| 1 | 0 | 0 | 0 |
| 1 | 0 | 0 | 0 |
| 0 | 0 | 0 | 0 |
| 1 | 0 | 0 | 0 |
| 1 | 0 | 0 | 0 |
| 0 | 0 | 0 | 0 |
| 0 | 0 | 0 | 0 |
| 0 | 0 | 0 | 0 |
| 0 | 0 | 0 | 0 |
| 0 | 0 | 0 | 0 |
| 0 | 0 | 0 | 0 |
| 0 | 0 | 0 | 0 |
| 0 | 0 | 0 | 0 |
| 1 | 0 | 0 | 0 |
| 1 | 0 | 0 | 0 |
| 1 | 0 | 0 | 0 |
| 0 | 0 | 0 | 0 |
| 0 | 0 | 0 | 1 |
| 0 | 0 | 0 | 1 |
| 0 | 0 | 0 | 1 |
| 0 | 0 | 0 | 1 |
| 0 | 0 | 0 | 1 |
| 0 | 0 | 0 | 1 |
| 0 | 0 | 0 | 0 |
| 0 | 0 | 0 | 0 |

| | | | |
|---|---|---|---|
| 0 | 0 | 0 | 0 |
| 0 | 0 | 0 | 0 |
| 0 | 0 | 0 | 0 |
| 0 | 0 | 0 | 0 |
| 0 | 0 | 0 | 1 |
| 0 | 0 | 0 | 0 |
| 0 | 0 | 0 | 0 |
| 0 | 0 | 0 | 0 |
| 0 | 0 | 0 | 1 |
| 0 | 0 | 0 | 0 |
| 1 | 0 | 0 | 0 |
| 0 | 0 | 0 | 1 |
| 0 | 0 | 0 | 0 |
| 0 | 0 | 0 | 0 |
| 0 | 0 | 0 | 0 |
| 0 | 0 | 0 | 0 |
| 0 | 0 | 0 | 1 |
| 0 | 0 | 0 | 0 |
| 0 | 0 | 0 | 0 |
| 0 | 0 | 0 | 1 |
| 0 | 1 | 0 | 0 |
| 0 | 0 | 0 | 0 |
| 0 | 0 | 0 | 0 |
| 0 | 0 | 0 | 0 |
| 0 | 0 | 0 | 0 |
| 0 | 0 | 0 | 1 |
| 0 | 0 | 0 | 1 |
| 0 | 0 | 0 | 1 |
| 0 | 0 | 0 | 1 |
| 0 | 0 | 0 | 1 |
| 0 | 0 | 0 | 1 |
| 0 | 0 | 0 | 1 |
| 0 | 0 | 0 | 1 |
| 0 | 0 | 0 | 1 |
| 0 | 0 | 0 | 0 |
| 0 | 0 | 0 | 0 |
| 0 | 0 | 0 | 0 |
| 0 | 0 | 0 | 0 |
| 0 | 1 | 0 | 0 |
| 0 | 0 | 0 | 0 |
| 0 | 0 | 0 | 1 |
| 0 | 0 | 0 | 1 |
| 0 | 0 | 0 | 1 |
| 0 | 0 | 0 | 0 |
| 0 | 0 | 0 | 0 |
| 0 | 0 | 0 | 0 |
| 0 | 0 | 0 | 1 |
| 0 | 0 | 0 | 1 |
| 0 | 0 | 0 | 0 |
| 0 | 0 | 0 | 0 |
| 0 | 0 | 0 | 1 |
| 0 | 0 | 0 | 1 |
| 0 | 0 | 0 | 1 |
| 0 | 0 | 0 | 1 |

| | | | |
|---|---|---|---|
| 0 | 0 | 0 | 1 |
| 0 | 0 | 0 | 1 |
| 0 | 0 | 0 | 1 |
| 0 | 0 | 0 | 0 |
| 0 | 0 | 0 | 0 |
| 0 | 0 | 0 | 1 |
| 0 | 0 | 0 | 1 |
| 0 | 0 | 0 | 1 |
| 0 | 0 | 0 | 1 |
| 0 | 0 | 0 | 1 |
| 0 | 0 | 0 | 0 |
| 0 | 0 | 0 | 1 |
| 0 | 1 | 0 | 0 |
| 0 | 0 | 0 | 0 |
| 0 | 0 | 0 | 0 |
| 0 | 0 | 0 | 1 |
| 0 | 0 | 0 | 1 |
| 0 | 0 | 0 | 1 |
| 0 | 0 | 0 | 1 |
| 0 | 0 | 0 | 1 |
| 0 | 0 | 0 | 0 |
| 0 | 0 | 0 | 1 |
| 0 | 0 | 0 | 1 |
| 0 | 0 | 0 | 0 |
| 0 | 0 | 0 | 1 |
| 0 | 1 | 0 | 0 |
| 0 | 1 | 0 | 0 |
| 0 | 0 | 0 | 1 |
| 0 | 0 | 0 | 1 |
| 0 | 0 | 0 | 1 |
| 0 | 0 | 0 | 1 |
| 0 | 0 | 0 | 1 |
| 0 | 0 | 0 | 1 |
| 0 | 0 | 0 | 1 |
| 0 | 0 | 0 | 1 |
| 0 | 0 | 0 | 0 |
| 0 | 0 | 0 | 1 |
| 0 | 0 | 0 | 1 |
| 0 | 0 | 0 | 0 |
| 0 | 0 | 0 | 0 |
| 0 | 0 | 0 | 1 |
| 0 | 0 | 0 | 1 |
| 1 | 0 | 0 | 0 |
| 1 | 0 | 0 | 0 |
| 0 | 0 | 0 | 0 |
| 0 | 0 | 0 | 0 |
| 1 | 0 | 0 | 0 |
| 1 | 0 | 0 | 0 |
| 0 | 0 | 0 | 1 |
| 1 | 0 | 0 | 0 |
| 1 | 0 | 0 | 0 |
| 1 | 0 | 0 | 0 |
| 1 | 0 | 0 | 0 |
| 1 | 0 | 0 | 0 |

| | | | |
|---|---|---|---|
| 0 | 0 | 0 | 0 |
| 0 | 0 | 0 | 0 |
| 1 | 0 | 0 | 0 |
| 1 | 0 | 0 | 0 |
| 1 | 0 | 0 | 0 |
| 1 | 0 | 0 | 0 |
| 0 | 0 | 0 | 0 |
| 0 | 0 | 0 | 0 |
| 0 | 0 | 0 | 0 |
| 0 | 0 | 0 | 1 |
| 0 | 0 | 0 | 1 |
| 0 | 0 | 0 | 0 |
| 0 | 0 | 0 | 1 |
| 0 | 0 | 0 | 0 |
| 0 | 0 | 0 | 0 |
| 0 | 0 | 0 | 1 |
| 0 | 0 | 0 | 0 |
| 0 | 0 | 0 | 1 |
| 0 | 0 | 0 | 1 |
| 0 | 0 | 0 | 1 |
| 0 | 0 | 0 | 1 |
| 0 | 0 | 0 | 1 |
| 0 | 0 | 0 | 1 |
| 0 | 0 | 0 | 0 |
| 0 | 0 | 0 | 0 |
| 0 | 0 | 0 | 1 |
| 0 | 0 | 0 | 1 |
| 0 | 0 | 0 | 1 |
| 0 | 0 | 0 | 0 |
| 0 | 0 | 0 | 0 |
| 0 | 0 | 0 | 1 |
| 0 | 0 | 0 | 1 |
| 0 | 0 | 0 | 1 |
| 0 | 0 | 0 | 0 |
| 0 | 0 | 0 | 0 |
| 0 | 0 | 0 | 1 |
| 0 | 0 | 0 | 0 |
| 0 | 0 | 0 | 1 |
| 0 | 0 | 0 | 1 |
| 0 | 0 | 0 | 0 |
| 0 | 0 | 0 | 0 |
| 0 | 0 | 0 | 0 |
| 0 | 0 | 0 | 0 |
| 0 | 0 | 0 | 0 |
| 0 | 0 | 0 | 0 |
| 0 | 0 | 0 | 1 |
| 0 | 0 | 0 | 1 |
| 0 | 0 | 0 | 0 |
| 0 | 0 | 0 | 0 |
| 0 | 0 | 0 | 0 |
| 0 | 0 | 0 | 1 |
| 0 | 0 | 0 | 1 |
| 0 | 0 | 0 | 0 |
| 0 | 0 | 0 | 0 |

| | | | |
|---|---|---|---|
| 0 | 0 | 0 | 1 |
| 0 | 0 | 0 | 1 |
| 0 | 0 | 0 | 1 |
| 0 | 0 | 0 | 1 |
| 0 | 1 | 0 | 0 |
| 0 | 1 | 0 | 0 |
| 0 | 0 | 0 | 1 |
| 0 | 0 | 0 | 0 |
| 0 | 1 | 1 | 1 |
| 0 | 0 | 0 | 0 |
| 0 | 0 | 0 | 1 |
| 0 | 0 | 0 | 1 |
| 0 | 0 | 0 | 1 |
| 0 | 0 | 0 | 1 |
| 0 | 0 | 0 | 1 |
| 0 | 0 | 0 | 1 |
| 0 | 0 | 0 | 0 |
| 0 | 0 | 0 | 1 |
| 0 | 0 | 0 | 1 |
| 0 | 0 | 0 | 0 |
| 0 | 0 | 0 | 0 |
| 0 | 0 | 0 | 1 |
| 0 | 0 | 0 | 1 |
| 0 | 0 | 0 | 0 |
| 0 | 0 | 0 | 1 |
| 0 | 0 | 0 | 0 |
| 0 | 0 | 0 | 0 |
| 0 | 0 | 0 | 0 |
| 0 | 0 | 0 | 0 |
| 0 | 0 | 0 | 0 |
| 0 | 0 | 0 | 0 |
| 0 | 0 | 0 | 0 |
| 0 | 0 | 0 | 1 |
| 0 | 0 | 0 | 0 |
| 0 | 0 | 0 | 1 |
| 0 | 0 | 0 | 1 |
| 0 | 0 | 0 | 1 |
| 0 | 0 | 0 | 1 |
| 0 | 0 | 0 | 0 |
| 0 | 0 | 0 | 1 |
| 0 | 0 | 0 | 1 |
| 0 | 0 | 0 | 1 |
| 0 | 0 | 0 | 1 |
| 0 | 0 | 0 | 0 |
| 0 | 0 | 0 | 1 |
| 0 | 0 | 0 | 1 |
| 0 | 0 | 0 | 1 |
| 0 | 0 | 0 | 1 |
| 0 | 0 | 0 | 1 |
| 0 | 0 | 0 | 1 |
| 0 | 0 | 0 | 0 |
| 0 | 0 | 0 | 0 |
| 0 | 0 | 0 | 0 |
| 0 | 0 | 0 | 1 |

| | | | |
|---|---|---|---|
| 0 | 0 | 0 | 0 |
| 0 | 0 | 0 | 0 |
| 0 | 1 | 0 | 0 |
| 0 | 0 | 0 | 0 |
| 1 | 0 | 0 | 0 |
| 1 | 0 | 0 | 0 |
| 0 | 0 | 0 | 0 |
| 0 | 0 | 0 | 0 |
| 0 | 0 | 0 | 1 |
| 0 | 0 | 0 | 1 |
| 0 | 0 | 0 | 0 |
| 0 | 0 | 0 | 1 |
| 0 | 0 | 0 | 1 |
| 0 | 0 | 0 | 0 |
| 0 | 0 | 0 | 0 |
| 0 | 0 | 0 | 0 |
| 0 | 0 | 0 | 0 |
| 0 | 0 | 0 | 0 |
| 0 | 0 | 0 | 0 |
| 0 | 0 | 0 | 1 |
| 0 | 0 | 0 | 1 |
| 0 | 0 | 0 | 0 |
| 0 | 0 | 0 | 1 |
| 0 | 1 | 0 | 0 |
| 0 | 0 | 0 | 1 |
| 0 | 0 | 0 | 1 |
| 0 | 0 | 0 | 1 |
| 0 | 0 | 0 | 1 |
| 0 | 0 | 0 | 1 |
| 0 | 0 | 0 | 0 |
| 0 | 0 | 0 | 1 |
| 0 | 0 | 0 | 1 |
| 0 | 0 | 0 | 0 |
| 0 | 0 | 0 | 1 |
| 0 | 0 | 0 | 0 |
| 0 | 0 | 0 | 0 |
| 0 | 0 | 0 | 0 |
| 0 | 0 | 0 | 0 |
| 0 | 0 | 0 | 0 |
| 0 | 0 | 0 | 0 |
| 0 | 0 | 0 | 0 |
| 0 | 0 | 0 | 0 |
| 0 | 0 | 0 | 1 |
| 0 | 0 | 0 | 0 |
| 0 | 0 | 0 | 1 |
| 0 | 0 | 0 | 0 |
| 0 | 0 | 0 | 1 |
| 0 | 0 | 0 | 1 |
| 0 | 0 | 0 | 1 |
| 0 | 0 | 0 | 0 |
| 0 | 0 | 0 | 1 |
| 0 | 0 | 0 | 1 |
| 0 | 0 | 0 | 1 |
| 0 | 0 | 0 | 1 |

| | | | |
|---|---|---|---|
| 1 | 0 | 0 | 0 |
| 0 | 0 | 0 | 1 |
| 0 | 0 | 0 | 1 |
| 0 | 0 | 0 | 1 |
| 0 | 0 | 0 | 1 |
| 0 | 0 | 0 | 1 |
| 0 | 0 | 0 | 1 |
| 0 | 0 | 0 | 1 |
| 0 | 0 | 0 | 1 |
| 0 | 0 | 0 | 1 |
| 0 | 0 | 0 | 0 |
| 0 | 0 | 0 | 1 |
| 0 | 0 | 0 | 1 |
| 0 | 0 | 0 | 1 |
| 0 | 0 | 0 | 0 |
| 0 | 0 | 0 | 0 |
| 0 | 0 | 0 | 0 |
| 0 | 0 | 0 | 0 |
| 0 | 0 | 0 | 0 |
| 0 | 0 | 0 | 1 |
| 0 | 0 | 0 | 1 |
| 0 | 0 | 0 | 1 |
| 0 | 0 | 0 | 0 |
| 0 | 0 | 0 | 1 |
| 0 | 0 | 0 | 1 |
| 0 | 0 | 0 | 1 |
| 0 | 0 | 0 | 1 |
| 0 | 0 | 0 | 1 |
| 0 | 0 | 0 | 0 |
| 0 | 0 | 0 | 0 |
| 0 | 0 | 0 | 0 |
| 0 | 0 | 0 | 0 |
| 0 | 0 | 0 | 0 |
| 0 | 0 | 0 | 0 |
| 0 | 0 | 0 | 1 |
| 0 | 0 | 0 | 1 |
| 0 | 0 | 0 | 0 |
| 0 | 0 | 0 | 0 |
| 0 | 0 | 0 | 0 |
| 0 | 0 | 0 | 0 |
| 0 | 0 | 0 | 0 |
| 0 | 0 | 0 | 0 |
| 0 | 0 | 0 | 1 |
| 0 | 0 | 0 | 1 |
| 0 | 0 | 0 | 0 |
| 0 | 0 | 0 | 1 |
| 0 | 0 | 0 | 1 |
| 0 | 0 | 0 | 1 |
| 0 | 0 | 0 | 1 |
| 0 | 0 | 0 | 1 |
| 0 | 0 | 0 | 1 |
| 0 | 0 | 0 | 1 |
| 0 | 0 | 0 | 1 |
| 0 | 1 | 1 | 1 |

| | | | |
|---|---|---|---|
| 0 | 0 | 0 | 0 |
| 0 | 0 | 0 | 1 |
| 0 | 0 | 0 | 0 |
| 0 | 0 | 0 | 1 |
| 0 | 1 | 1 | 1 |
| 0 | 1 | 1 | 1 |
| 0 | 1 | 0 | 0 |
| 0 | 1 | 0 | 0 |
| 0 | 1 | 1 | 1 |
| 0 | 0 | 0 | 1 |
| 0 | 0 | 0 | 1 |
| 0 | 0 | 0 | 1 |
| 0 | 0 | 0 | 1 |
| 0 | 0 | 0 | 1 |
| 0 | 0 | 0 | 0 |
| 0 | 0 | 0 | 0 |
| 0 | 0 | 0 | 0 |
| 0 | 0 | 0 | 0 |
| 0 | 0 | 0 | 1 |
| 0 | 0 | 0 | 0 |
| 0 | 0 | 0 | 0 |
| 0 | 0 | 0 | 0 |
| 0 | 0 | 0 | 0 |
| 0 | 0 | 0 | 0 |
| 0 | 0 | 0 | 1 |
| 0 | 0 | 0 | 0 |
| 0 | 0 | 0 | 1 |
| 0 | 0 | 0 | 1 |
| 0 | 0 | 0 | 1 |
| 0 | 0 | 0 | 0 |
| 0 | 0 | 0 | 1 |
| 0 | 0 | 0 | 1 |
| 0 | 0 | 0 | 0 |
| 0 | 0 | 0 | 0 |
| 0 | 0 | 0 | 1 |
| 0 | 0 | 0 | 1 |
| 0 | 0 | 0 | 1 |
| 0 | 0 | 0 | 1 |
| 0 | 0 | 0 | 1 |
| 0 | 0 | 0 | 0 |
| 0 | 0 | 0 | 0 |
| 0 | 0 | 0 | 0 |
| 0 | 0 | 0 | 0 |
| 0 | 0 | 0 | 0 |
| 0 | 0 | 0 | 0 |
| 0 | 0 | 0 | 1 |
| 0 | 0 | 0 | 1 |
| 0 | 0 | 0 | 0 |
| 0 | 0 | 0 | 0 |
| 0 | 0 | 0 | 1 |
| 0 | 0 | 0 | 0 |
| 0 | 0 | 0 | 0 |
| 0 | 0 | 0 | 1 |
| 0 | 0 | 0 | 1 |

| | | | |
|---|---|---|---|
| 0 | 0 | 0 | 1 |
| 0 | 0 | 0 | 0 |
| 0 | 0 | 0 | 0 |
| 0 | 0 | 0 | 1 |
| 0 | 0 | 0 | 1 |
| 0 | 0 | 0 | 0 |
| 0 | 0 | 0 | 0 |
| 0 | 0 | 0 | 0 |
| 0 | 0 | 0 | 0 |
| 0 | 0 | 0 | 0 |
| 0 | 0 | 0 | 0 |
| 0 | 0 | 0 | 0 |
| 0 | 0 | 0 | 0 |
| 0 | 0 | 0 | 0 |
| 0 | 0 | 0 | 0 |
| 0 | 0 | 0 | 1 |
| 0 | 0 | 0 | 0 |
| 0 | 0 | 0 | 1 |
| 0 | 0 | 0 | 1 |
| 0 | 0 | 0 | 1 |
| 0 | 0 | 0 | 0 |
| 0 | 0 | 0 | 1 |
| 0 | 0 | 0 | 0 |
| 0 | 0 | 0 | 0 |
| 0 | 0 | 0 | 1 |
| 0 | 0 | 0 | 0 |
| 0 | 0 | 0 | 1 |
| 0 | 0 | 0 | 1 |
| 0 | 0 | 0 | 1 |
| 0 | 0 | 0 | 0 |
| 0 | 0 | 0 | 0 |
| 0 | 0 | 0 | 0 |
| 0 | 0 | 0 | 0 |
| 0 | 0 | 0 | 0 |
| 0 | 0 | 0 | 0 |
| 0 | 0 | 0 | 0 |
| 0 | 0 | 0 | 0 |
| 0 | 0 | 0 | 1 |
| 0 | 0 | 0 | 1 |
| 0 | 0 | 0 | 0 |
| 0 | 0 | 0 | 0 |
| 0 | 0 | 0 | 0 |
| 0 | 0 | 0 | 0 |
| 0 | 0 | 0 | 0 |
| 0 | 1 | 0 | 0 |
| 0 | 0 | 0 | 0 |
| 0 | 0 | 0 | 1 |
| 0 | 0 | 0 | 0 |
| 0 | 0 | 0 | 0 |
| 0 | 0 | 0 | 0 |
| 0 | 0 | 0 | 0 |
| 0 | 0 | 0 | 1 |
| 0 | 0 | 0 | 0 |
| 0 | 0 | 0 | 0 |

| | | | |
|---|---|---|---|
| 0 | 1 | 0 | 0 |
| 0 | 1 | 0 | 0 |
| 0 | 0 | 0 | 0 |
| 0 | 0 | 0 | 0 |
| 0 | 0 | 0 | 0 |
| 0 | 0 | 0 | 0 |
| 0 | 0 | 0 | 0 |
| 0 | 0 | 0 | 0 |
| 1 | 0 | 0 | 0 |
| 0 | 0 | 0 | 0 |
| 0 | 0 | 0 | 1 |
| 0 | 1 | 0 | 0 |
| 0 | 1 | 1 | 1 |
| 0 | 0 | 0 | 0 |
| 0 | 0 | 0 | 1 |
| 0 | 0 | 0 | 1 |
| 0 | 0 | 0 | 1 |
| 0 | 0 | 0 | 1 |
| 0 | 0 | 0 | 0 |
| 0 | 0 | 0 | 0 |
| 0 | 0 | 0 | 1 |
| 0 | 0 | 0 | 0 |
| 0 | 0 | 0 | 0 |
| 0 | 0 | 0 | 1 |
| 0 | 0 | 0 | 0 |
| 0 | 0 | 0 | 0 |
| 0 | 0 | 0 | 0 |
| 0 | 0 | 0 | 1 |
| 0 | 1 | 0 | 0 |
| 0 | 0 | 0 | 0 |
| 0 | 0 | 0 | 1 |
| 1 | 0 | 0 | 0 |
| 0 | 0 | 0 | 1 |
| 0 | 0 | 0 | 1 |
| 1 | 0 | 0 | 0 |
| 1 | 0 | 0 | 0 |
| 1 | 0 | 0 | 0 |
| 0 | 1 | 1 | 1 |
| 0 | 0 | 0 | 1 |
| 0 | 0 | 0 | 0 |
| 1 | 0 | 0 | 0 |
| 0 | 1 | 0 | 0 |
| 0 | 1 | 0 | 0 |
| 1 | 0 | 0 | 0 |
| 1 | 0 | 0 | 0 |
| 0 | 0 | 0 | 0 |
| 0 | 0 | 0 | 1 |
| 0 | 0 | 0 | 1 |
| 0 | 0 | 0 | 1 |
| 0 | 0 | 0 | 1 |
| 1 | 0 | 0 | 0 |
| 0 | 0 | 0 | 0 |
| 0 | 0 | 0 | 0 |
| 0 | 0 | 0 | 0 |

| | | | |
|---|---|---|---|
| 0 | 0 | 0 | 0 |
| 0 | 0 | 0 | 1 |
| 0 | 0 | 0 | 0 |
| 0 | 0 | 0 | 0 |
| 0 | 0 | 0 | 0 |
| 0 | 0 | 0 | 1 |
| 0 | 0 | 0 | 0 |
| 0 | 0 | 0 | 0 |
| 0 | 0 | 0 | 1 |
| 0 | 0 | 0 | 0 |
| 0 | 0 | 0 | 1 |
| 0 | 0 | 0 | 1 |
| 0 | 0 | 0 | 1 |
| 0 | 0 | 0 | 0 |
| 0 | 0 | 0 | 1 |
| 0 | 0 | 0 | 0 |
| 0 | 0 | 0 | 1 |
| 0 | 0 | 0 | 1 |
| 0 | 0 | 0 | 0 |
| 0 | 0 | 0 | 0 |
| 0 | 0 | 0 | 0 |
| 0 | 0 | 0 | 0 |
| 0 | 0 | 0 | 0 |
| 0 | 0 | 0 | 0 |
| 0 | 0 | 0 | 0 |
| 0 | 0 | 0 | 1 |
| 0 | 0 | 0 | 1 |
| 0 | 0 | 0 | 1 |
| 0 | 0 | 0 | 1 |
| 0 | 0 | 0 | 1 |
| 0 | 0 | 0 | 1 |
| 0 | 0 | 0 | 1 |
| 0 | 0 | 0 | 0 |
| 0 | 0 | 0 | 0 |
| 0 | 0 | 0 | 0 |
| 0 | 0 | 0 | 0 |
| 0 | 0 | 0 | 0 |
| 0 | 0 | 0 | 0 |
| 0 | 0 | 0 | 1 |
| 0 | 0 | 0 | 1 |
| 0 | 0 | 0 | 1 |
| 0 | 0 | 0 | 1 |
| 0 | 0 | 0 | 1 |
| 0 | 0 | 0 | 1 |
| 0 | 1 | 0 | 0 |
| 0 | 0 | 0 | 0 |
| 0 | 0 | 0 | 1 |
| 0 | 0 | 0 | 0 |
| 0 | 0 | 0 | 0 |
| 0 | 0 | 0 | 0 |
| 0 | 0 | 0 | 0 |
| 0 | 0 | 0 | 0 |
| 0 | 0 | 0 | 0 |
| 0 | 0 | 0 | 0 |

| | | | |
|---|---|---|---|
| 0 | 0 | 0 | 1 |
| 0 | 0 | 0 | 1 |
| 1 | 0 | 0 | 0 |
| 1 | 0 | 0 | 0 |
| 0 | 0 | 0 | 1 |
| 0 | 0 | 0 | 0 |
| 0 | 0 | 0 | 0 |
| 0 | 0 | 0 | 1 |
| 0 | 0 | 0 | 1 |
| 0 | 0 | 0 | 0 |
| 0 | 0 | 0 | 1 |
| 0 | 0 | 0 | 1 |
| 0 | 0 | 0 | 0 |
| 0 | 0 | 0 | 0 |
| 0 | 1 | 0 | 0 |
| 0 | 1 | 0 | 0 |
| 0 | 0 | 0 | 1 |
| 0 | 0 | 0 | 1 |
| 0 | 0 | 0 | 1 |
| 0 | 0 | 0 | 1 |
| 0 | 0 | 0 | 1 |
| 0 | 0 | 0 | 1 |
| 0 | 0 | 0 | 1 |
| 0 | 0 | 0 | 1 |
| 0 | 0 | 0 | 1 |
| 0 | 0 | 0 | 0 |
| 0 | 0 | 0 | 1 |
| 0 | 0 | 0 | 1 |
| 0 | 0 | 0 | 1 |
| 0 | 0 | 0 | 1 |
| 0 | 0 | 0 | 1 |
| 0 | 0 | 0 | 1 |
| 0 | 0 | 0 | 1 |
| 0 | 0 | 0 | 1 |
| 0 | 0 | 0 | 1 |
| 0 | 0 | 0 | 1 |
| 0 | 0 | 0 | 1 |
| 0 | 0 | 0 | 0 |
| 0 | 0 | 0 | 1 |
| 0 | 0 | 0 | 1 |
| 0 | 0 | 0 | 0 |
| 0 | 0 | 0 | 1 |
| 0 | 0 | 0 | 1 |
| 0 | 0 | 0 | 0 |
| 0 | 0 | 0 | 0 |
| 0 | 0 | 0 | 1 |
| 0 | 0 | 0 | 1 |
| 0 | 0 | 0 | 1 |
| 0 | 0 | 0 | 1 |
| 0 | 0 | 0 | 1 |
| 0 | 0 | 0 | 1 |
| 0 | 0 | 0 | 0 |
| 0 | 0 | 0 | 0 |
| 0 | 0 | 0 | 0 |

| | | | |
|---|---|---|---|
| 0 | 0 | 0 | 0 |
| 0 | 0 | 0 | 1 |
| 0 | 0 | 0 | 1 |
| 0 | 0 | 0 | 1 |
| 0 | 0 | 0 | 1 |
| 0 | 0 | 0 | 1 |
| 0 | 0 | 0 | 1 |
| 0 | 0 | 0 | 1 |
| 0 | 0 | 0 | 0 |
| 0 | 0 | 0 | 1 |
| 0 | 0 | 0 | 1 |
| 0 | 0 | 0 | 1 |
| 0 | 0 | 0 | 1 |
| 0 | 0 | 0 | 1 |
| 0 | 0 | 0 | 1 |
| 0 | 0 | 0 | 1 |
| 0 | 0 | 0 | 1 |
| 0 | 0 | 0 | 1 |
| 0 | 0 | 0 | 0 |
| 0 | 0 | 0 | 0 |
| 0 | 0 | 0 | 0 |
| 0 | 0 | 0 | 0 |
| 0 | 0 | 0 | 1 |
| 0 | 0 | 0 | 0 |
| 0 | 0 | 0 | 0 |
| 0 | 0 | 0 | 1 |
| 0 | 0 | 0 | 1 |
| 0 | 0 | 0 | 1 |
| 0 | 0 | 0 | 1 |
| 0 | 0 | 0 | 1 |
| 0 | 0 | 0 | 1 |
| 0 | 0 | 0 | 1 |
| 0 | 0 | 0 | 0 |
| 0 | 0 | 0 | 1 |
| 0 | 0 | 0 | 1 |
| 0 | 0 | 0 | 1 |
| 0 | 0 | 0 | 0 |
| 0 | 0 | 0 | 1 |
| 0 | 0 | 0 | 0 |
| 0 | 0 | 0 | 0 |
| 0 | 0 | 0 | 0 |
| 0 | 0 | 0 | 0 |
| 0 | 0 | 0 | 0 |
| 0 | 0 | 0 | 1 |
| 0 | 0 | 0 | 1 |
| 0 | 0 | 0 | 1 |
| 0 | 0 | 0 | 0 |
| 0 | 0 | 0 | 0 |
| 0 | 0 | 0 | 0 |
| 0 | 0 | 0 | 0 |
| 0 | 0 | 0 | 0 |
| 0 | 0 | 0 | 0 |
| 0 | 0 | 0 | 0 |
| 0 | 0 | 0 | 1 |

| | | | |
|---|---|---|---|
| 0 | 0 | 0 | 0 |
| 0 | 0 | 0 | 0 |
| 0 | 0 | 0 | 1 |
| 1 | 0 | 0 | 0 |
| 1 | 0 | 0 | 0 |
| 0 | 0 | 0 | 1 |
| 0 | 0 | 0 | 1 |
| 0 | 0 | 0 | 1 |
| 0 | 0 | 0 | 1 |
| 0 | 0 | 0 | 1 |
| 0 | 0 | 0 | 1 |
| 0 | 0 | 0 | 0 |
| 0 | 0 | 0 | 1 |
| 0 | 0 | 0 | 1 |
| 0 | 0 | 0 | 0 |
| 0 | 0 | 0 | 1 |
| 0 | 0 | 0 | 0 |
| 0 | 0 | 0 | 1 |
| 0 | 0 | 0 | 0 |
| 1 | 0 | 0 | 0 |
| 0 | 0 | 0 | 1 |
| 0 | 0 | 0 | 1 |
| 0 | 0 | 0 | 0 |
| 0 | 0 | 0 | 1 |
| 0 | 0 | 0 | 0 |
| 0 | 0 | 0 | 1 |
| 0 | 0 | 0 | 1 |
| 0 | 0 | 0 | 0 |
| 0 | 0 | 0 | 1 |
| 0 | 0 | 0 | 0 |
| 0 | 0 | 0 | 1 |
| 0 | 0 | 0 | 0 |
| 0 | 0 | 0 | 0 |
| 0 | 0 | 0 | 0 |
| 0 | 0 | 0 | 1 |
| 0 | 0 | 0 | 0 |
| 0 | 0 | 0 | 1 |
| 0 | 0 | 0 | 1 |
| 0 | 0 | 0 | 0 |
| 0 | 0 | 0 | 1 |
| 0 | 0 | 0 | 1 |
| 0 | 0 | 0 | 1 |
| 0 | 0 | 0 | 1 |
| 0 | 0 | 0 | 0 |
| 0 | 0 | 0 | 0 |
| 0 | 0 | 0 | 1 |
| 0 | 0 | 0 | 1 |
| 0 | 0 | 0 | 1 |
| 0 | 1 | 0 | 0 |
| 0 | 0 | 0 | 1 |
| 0 | 0 | 0 | 0 |
| 0 | 0 | 0 | 1 |
| 0 | 0 | 0 | 1 |
| 0 | 0 | 0 | 0 |

| | | | |
|---|---|---|---|
| 0 | 0 | 0 | 1 |
| 0 | 0 | 0 | 1 |
| 0 | 0 | 0 | 1 |
| 0 | 0 | 0 | 1 |
| 0 | 0 | 0 | 1 |
| 0 | 0 | 0 | 1 |
| 0 | 0 | 0 | 1 |
| 0 | 0 | 0 | 0 |
| 0 | 0 | 0 | 1 |
| 0 | 0 | 0 | 1 |
| 0 | 0 | 0 | 1 |
| 0 | 0 | 0 | 1 |
| 0 | 0 | 0 | 0 |
| 0 | 0 | 0 | 1 |
| 0 | 0 | 0 | 0 |
| 0 | 0 | 0 | 0 |
| 0 | 0 | 0 | 1 |
| 0 | 0 | 0 | 1 |
| 0 | 0 | 0 | 1 |
| 0 | 0 | 0 | 1 |
| 0 | 0 | 0 | 0 |
| 0 | 0 | 0 | 0 |
| 0 | 0 | 0 | 1 |
| 0 | 0 | 0 | 1 |
| 0 | 0 | 0 | 0 |
| 0 | 0 | 0 | 1 |
| 0 | 0 | 0 | 0 |
| 0 | 0 | 0 | 0 |
| 0 | 0 | 0 | 0 |
| 0 | 0 | 0 | 0 |
| 0 | 0 | 0 | 1 |
| 0 | 0 | 0 | 1 |
| 0 | 0 | 0 | 0 |
| 0 | 1 | 0 | 0 |
| 0 | 0 | 0 | 0 |
| 0 | 0 | 0 | 0 |
| 0 | 0 | 0 | 0 |
| 0 | 0 | 0 | 0 |
| 0 | 0 | 0 | 0 |
| 0 | 0 | 0 | 1 |
| 0 | 0 | 0 | 1 |
| 0 | 0 | 0 | 1 |
| 0 | 0 | 0 | 1 |
| 0 | 0 | 0 | 1 |
| 0 | 0 | 0 | 1 |
| 0 | 0 | 0 | 0 |
| 0 | 0 | 0 | 1 |
| 0 | 0 | 0 | 1 |
| 0 | 0 | 0 | 1 |
| 0 | 0 | 0 | 0 |
| 0 | 0 | 0 | 0 |
| 0 | 0 | 0 | 1 |
| 0 | 0 | 0 | 1 |
| 0 | 0 | 0 | 1 |

| | | | |
|---|---|---|---|
| 0 | 0 | 0 | 1 |
| 0 | 0 | 0 | 1 |
| 0 | 0 | 0 | 1 |
| 0 | 0 | 0 | 1 |
| 0 | 0 | 0 | 0 |
| 0 | 0 | 0 | 1 |
| 0 | 0 | 0 | 0 |
| 0 | 0 | 0 | 0 |
| 0 | 0 | 0 | 0 |
| 0 | 0 | 0 | 0 |
| 0 | 0 | 0 | 0 |
| 0 | 0 | 0 | 1 |
| 0 | 0 | 0 | 0 |
| 0 | 0 | 0 | 0 |
| 0 | 0 | 0 | 1 |
| 0 | 0 | 0 | 1 |
| 0 | 0 | 0 | 1 |
| 0 | 0 | 0 | 1 |
| 0 | 0 | 0 | 1 |
| 0 | 0 | 0 | 1 |
| 0 | 0 | 0 | 1 |
| 0 | 0 | 0 | 1 |
| 0 | 0 | 0 | 1 |
| 0 | 0 | 0 | 0 |
| 0 | 0 | 0 | 1 |
| 0 | 0 | 0 | 1 |
| 0 | 0 | 0 | 1 |
| 0 | 0 | 0 | 0 |
| 0 | 1 | 1 | 1 |
| 0 | 1 | 1 | 1 |
| 0 | 0 | 0 | 0 |
| 0 | 0 | 0 | 0 |
| 0 | 0 | 0 | 1 |
| 0 | 0 | 0 | 0 |
| 0 | 0 | 0 | 1 |
| 0 | 0 | 0 | 1 |
| 0 | 0 | 0 | 1 |
| 0 | 0 | 0 | 1 |
| 0 | 0 | 0 | 1 |
| 0 | 0 | 0 | 1 |
| 0 | 0 | 0 | 1 |
| 0 | 0 | 0 | 1 |
| 0 | 0 | 0 | 1 |
| 0 | 0 | 0 | 1 |
| 0 | 0 | 0 | 1 |
| 0 | 0 | 0 | 1 |
| 0 | 0 | 0 | 1 |
| 0 | 0 | 0 | 1 |
| 0 | 0 | 0 | 0 |
| 0 | 0 | 0 | 1 |
| 0 | 0 | 0 | 1 |
| 0 | 1 | 1 | 1 |
| 0 | 0 | 0 | 1 |
| 0 | 0 | 0 | 1 |

| | | | |
|---|---|---|---|
| 0 | 0 | 0 | 1 |
| 0 | 0 | 0 | 1 |
| 0 | 0 | 0 | 1 |
| 0 | 0 | 0 | 1 |
| 0 | 0 | 0 | 1 |
| 0 | 0 | 0 | 1 |
| 0 | 0 | 0 | 1 |
| 0 | 0 | 0 | 1 |
| 0 | 0 | 0 | 1 |
| 0 | 0 | 0 | 1 |
| 0 | 0 | 0 | 0 |
| 0 | 0 | 0 | 0 |
| 0 | 0 | 0 | 0 |
| 1 | 0 | 0 | 0 |
| 1 | 0 | 0 | 0 |
| 0 | 0 | 0 | 1 |
| 1 | 0 | 0 | 0 |
| 0 | 0 | 0 | 0 |
| 0 | 0 | 0 | 0 |
| 0 | 0 | 0 | 0 |
| 0 | 0 | 0 | 0 |
| 1 | 0 | 0 | 0 |
| 0 | 0 | 0 | 1 |
| 0 | 0 | 0 | 0 |
| 1 | 0 | 0 | 0 |
| 1 | 0 | 0 | 0 |
| 0 | 0 | 0 | 1 |
| 0 | 0 | 0 | 0 |
| 0 | 0 | 0 | 1 |
| 0 | 0 | 0 | 1 |
| 0 | 0 | 0 | 1 |
| 0 | 0 | 0 | 0 |
| 0 | 0 | 0 | 1 |
| 0 | 0 | 0 | 1 |
| 0 | 0 | 0 | 0 |
| 0 | 0 | 0 | 0 |
| 0 | 0 | 0 | 0 |
| 0 | 0 | 0 | 0 |
| 0 | 0 | 0 | 0 |
| 0 | 0 | 0 | 1 |
| 0 | 0 | 0 | 1 |
| 0 | 0 | 0 | 0 |
| 0 | 0 | 0 | 1 |
| 0 | 0 | 0 | 0 |
| 0 | 0 | 0 | 1 |
| 0 | 0 | 0 | 1 |
| 0 | 0 | 0 | 1 |
| 0 | 1 | 0 | 0 |
| 0 | 0 | 0 | 0 |
| 0 | 0 | 0 | 0 |
| 0 | 0 | 0 | 0 |
| 0 | 0 | 0 | 1 |
| 0 | 0 | 0 | 0 |
| 0 | 0 | 0 | 1 |

| | | | |
|---|---|---|---|
| 0 | 0 | 0 | 1 |
| 0 | 0 | 0 | 1 |
| 0 | 0 | 0 | 1 |
| 0 | 0 | 0 | 1 |
| 0 | 0 | 0 | 1 |
| 0 | 0 | 0 | 0 |
| 0 | 0 | 0 | 0 |
| 0 | 0 | 0 | 0 |
| 0 | 0 | 0 | 0 |
| 0 | 0 | 0 | 1 |
| 1 | 0 | 0 | 0 |
| 1 | 0 | 0 | 0 |
| 0 | 0 | 0 | 0 |
| 0 | 0 | 0 | 0 |
| 0 | 0 | 0 | 0 |
| 0 | 0 | 0 | 0 |
| 0 | 0 | 0 | 1 |
| 0 | 0 | 0 | 1 |
| 0 | 0 | 0 | 1 |
| 0 | 0 | 0 | 1 |
| 0 | 0 | 0 | 1 |
| 0 | 0 | 0 | 1 |
| 0 | 0 | 0 | 1 |
| 0 | 0 | 0 | 1 |
| 0 | 0 | 0 | 1 |
| 0 | 0 | 0 | 1 |
| 0 | 0 | 0 | 1 |
| 0 | 0 | 0 | 1 |
| 0 | 1 | 0 | 0 |
| 0 | 0 | 0 | 1 |
| 0 | 0 | 0 | 1 |
| 0 | 1 | 1 | 1 |
| 0 | 1 | 1 | 1 |
| 0 | 0 | 0 | 0 |
| 0 | 0 | 0 | 1 |
| 0 | 0 | 0 | 0 |
| 0 | 0 | 0 | 1 |
| 0 | 0 | 0 | 0 |
| 0 | 0 | 0 | 1 |
| 0 | 0 | 0 | 0 |
| 0 | 0 | 0 | 1 |
| 0 | 0 | 0 | 1 |
| 0 | 0 | 0 | 1 |
| 0 | 0 | 0 | 0 |
| 0 | 0 | 0 | 1 |
| 0 | 0 | 0 | 1 |
| 0 | 0 | 0 | 1 |
| 0 | 0 | 0 | 1 |
| 0 | 0 | 0 | 1 |
| 0 | 0 | 0 | 1 |
| 0 | 0 | 0 | 1 |
| 0 | 0 | 0 | 1 |
| 0 | 0 | 0 | 0 |
| 0 | 0 | 0 | 0 |

| | | | |
|---|---|---|---|
| 0 | 0 | 0 | 1 |
| 0 | 0 | 0 | 0 |
| 0 | 0 | 0 | 1 |
| 0 | 0 | 0 | 1 |
| 0 | 0 | 0 | 0 |
| 0 | 0 | 0 | 0 |
| 0 | 0 | 0 | 0 |
| 0 | 0 | 0 | 1 |
| 0 | 0 | 0 | 1 |
| 0 | 0 | 0 | 0 |
| 0 | 0 | 0 | 0 |
| 0 | 0 | 0 | 0 |
| 0 | 0 | 0 | 0 |
| 0 | 0 | 0 | 0 |
| 0 | 0 | 0 | 1 |
| 0 | 0 | 0 | 0 |
| 0 | 0 | 0 | 0 |
| 0 | 0 | 0 | 0 |
| 0 | 0 | 0 | 1 |
| 0 | 0 | 0 | 1 |
| 0 | 0 | 0 | 1 |
| 0 | 0 | 0 | 1 |
| 0 | 0 | 0 | 1 |
| 0 | 0 | 0 | 0 |
| 0 | 0 | 0 | 1 |
| 0 | 0 | 0 | 1 |
| 0 | 0 | 0 | 0 |
| 0 | 0 | 0 | 0 |
| 0 | 0 | 0 | 1 |
| 0 | 0 | 0 | 1 |
| 0 | 0 | 0 | 0 |
| 0 | 0 | 0 | 1 |
| 0 | 0 | 0 | 1 |
| 0 | 0 | 0 | 1 |
| 0 | 0 | 0 | 1 |
| 0 | 0 | 0 | 0 |
| 0 | 0 | 0 | 1 |
| 0 | 0 | 0 | 0 |
| 0 | 0 | 0 | 0 |
| 0 | 0 | 0 | 1 |
| 0 | 0 | 0 | 1 |
| 0 | 0 | 0 | 0 |
| 0 | 0 | 0 | 0 |
| 0 | 0 | 0 | 1 |
| 0 | 0 | 0 | 0 |
| 0 | 0 | 0 | 0 |
| 0 | 0 | 0 | 0 |
| 0 | 0 | 0 | 1 |
| 0 | 0 | 0 | 0 |
| 0 | 0 | 0 | 0 |
| 0 | 0 | 0 | 0 |
| 0 | 0 | 0 | 1 |
| 0 | 0 | 0 | 0 |
| 0 | 0 | 0 | 1 |

| | | | |
|---|---|---|---|
| 0 | 0 | 0 | 1 |
| 0 | 0 | 0 | 1 |
| 0 | 0 | 0 | 1 |
| 0 | 0 | 0 | 1 |
| 0 | 0 | 0 | 0 |
| 0 | 0 | 0 | 0 |
| 0 | 0 | 0 | 1 |
| 0 | 0 | 0 | 1 |
| 0 | 0 | 0 | 1 |
| 0 | 0 | 0 | 0 |
| 0 | 0 | 0 | 1 |
| 0 | 0 | 0 | 1 |
| 0 | 1 | 0 | 0 |
| 0 | 0 | 0 | 1 |
| 0 | 0 | 0 | 1 |
| 0 | 0 | 0 | 1 |
| 0 | 0 | 0 | 1 |
| 0 | 0 | 0 | 1 |
| 0 | 0 | 0 | 1 |
| 0 | 0 | 0 | 1 |
| 0 | 0 | 0 | 1 |
| 0 | 0 | 0 | 1 |
| 0 | 0 | 0 | 1 |
| 0 | 0 | 0 | 0 |
| 0 | 0 | 0 | 1 |
| 1 | 0 | 0 | 0 |
| 0 | 0 | 0 | 1 |
| 0 | 0 | 0 | 0 |
| 0 | 0 | 0 | 1 |
| 0 | 0 | 0 | 0 |
| 0 | 0 | 0 | 0 |
| 0 | 0 | 0 | 0 |
| 0 | 0 | 0 | 0 |
| 0 | 0 | 0 | 0 |
| 0 | 0 | 0 | 1 |
| 0 | 0 | 0 | 0 |
| 0 | 0 | 0 | 1 |
| 0 | 0 | 0 | 0 |
| 0 | 0 | 0 | 1 |
| 0 | 0 | 0 | 1 |
| 0 | 0 | 0 | 0 |
| 0 | 0 | 0 | 0 |
| 0 | 0 | 0 | 0 |
| 0 | 0 | 0 | 0 |
| 0 | 0 | 0 | 0 |
| 0 | 0 | 0 | 0 |
| 0 | 0 | 0 | 0 |
| 0 | 0 | 0 | 0 |
| 0 | 0 | 0 | 1 |
| 0 | 0 | 0 | 0 |
| 0 | 0 | 0 | 1 |
| 0 | 0 | 0 | 1 |
| 0 | 0 | 0 | 1 |
| 0 | 0 | 0 | 1 |

| | | | |
|---|---|---|---|
| 0 | 0 | 0 | 1 |
| 0 | 0 | 0 | 1 |
| 0 | 0 | 0 | 1 |
| 0 | 0 | 0 | 1 |
| 0 | 0 | 0 | 1 |
| 0 | 0 | 0 | 1 |
| 0 | 0 | 0 | 0 |
| 0 | 0 | 0 | 0 |
| 0 | 0 | 0 | 0 |
| 0 | 0 | 0 | 0 |
| 0 | 0 | 0 | 0 |
| 0 | 0 | 0 | 0 |
| 0 | 0 | 0 | 0 |
| 0 | 0 | 0 | 0 |
| 0 | 0 | 0 | 0 |
| 0 | 0 | 0 | 0 |
| 1 | 0 | 0 | 0 |
| 0 | 0 | 0 | 0 |
| 0 | 0 | 0 | 1 |
| 0 | 0 | 0 | 0 |
| 0 | 0 | 0 | 1 |
| 0 | 0 | 0 | 1 |
| 0 | 0 | 0 | 1 |
| 0 | 0 | 0 | 1 |
| 0 | 0 | 0 | 1 |
| 0 | 0 | 0 | 0 |
| 0 | 0 | 0 | 0 |
| 0 | 0 | 0 | 0 |
| 0 | 0 | 0 | 0 |
| 0 | 0 | 0 | 0 |
| 0 | 0 | 0 | 0 |
| 1 | 0 | 0 | 0 |
| 0 | 0 | 0 | 0 |
| 1 | 0 | 0 | 0 |
| 0 | 0 | 0 | 1 |
| 1 | 0 | 0 | 0 |
| 1 | 0 | 0 | 0 |
| 0 | 0 | 0 | 0 |
| 1 | 0 | 0 | 0 |
| 0 | 0 | 0 | 1 |
| 0 | 0 | 0 | 0 |
| 0 | 0 | 0 | 0 |
| 0 | 0 | 0 | 1 |
| 0 | 0 | 0 | 1 |
| 0 | 0 | 0 | 1 |
| 0 | 0 | 0 | 1 |
| 0 | 0 | 0 | 1 |
| 0 | 0 | 0 | 0 |
| 0 | 0 | 0 | 0 |
| 0 | 0 | 0 | 0 |
| 0 | 0 | 0 | 0 |
| 0 | 0 | 0 | 1 |
| 0 | 0 | 0 | 0 |
| 0 | 0 | 0 | 1 |

| | | | |
|---|---|---|---|
| 0 | 0 | 0 | 1 |
| 0 | 0 | 0 | 1 |
| 0 | 0 | 0 | 1 |
| 0 | 0 | 0 | 1 |
| 0 | 0 | 0 | 1 |
| 0 | 0 | 0 | 0 |
| 0 | 0 | 0 | 1 |
| 0 | 0 | 0 | 1 |
| 0 | 0 | 0 | 1 |
| 0 | 0 | 0 | 1 |
| 0 | 0 | 0 | 0 |
| 0 | 1 | 0 | 0 |
| 0 | 1 | 0 | 0 |
| 0 | 0 | 0 | 1 |
| 0 | 1 | 1 | 1 |
| 0 | 0 | 0 | 1 |
| 0 | 0 | 0 | 1 |
| 0 | 0 | 0 | 0 |
| 0 | 0 | 0 | 1 |
| 0 | 0 | 0 | 1 |
| 0 | 0 | 0 | 1 |
| 0 | 0 | 0 | 1 |
| 0 | 0 | 0 | 0 |
| 0 | 0 | 0 | 1 |
| 0 | 0 | 0 | 0 |
| 0 | 0 | 0 | 0 |
| 0 | 0 | 0 | 0 |
| 0 | 0 | 0 | 0 |
| 0 | 0 | 0 | 1 |
| 0 | 0 | 0 | 0 |
| 0 | 0 | 0 | 1 |
| 0 | 0 | 0 | 1 |
| 0 | 0 | 0 | 1 |
| 0 | 0 | 0 | 1 |
| 0 | 0 | 0 | 1 |
| 0 | 0 | 0 | 1 |
| 0 | 0 | 0 | 1 |
| 0 | 0 | 0 | 0 |
| 0 | 0 | 0 | 0 |
| 0 | 0 | 0 | 0 |
| 0 | 1 | 0 | 0 |
| 0 | 0 | 0 | 0 |
| 0 | 0 | 0 | 1 |
| 0 | 0 | 0 | 0 |
| 0 | 0 | 0 | 0 |
| 0 | 0 | 0 | 1 |
| 0 | 0 | 0 | 0 |
| 0 | 1 | 0 | 0 |
| 0 | 0 | 0 | 0 |
| 1 | 0 | 0 | 0 |
| 0 | 0 | 0 | 1 |
| 0 | 0 | 0 | 1 |
| 0 | 0 | 0 | 1 |
| 0 | 1 | 0 | 0 |

| | | | |
|---|---|---|---|
| 0 | 0 | 0 | 0 |
| 0 | 0 | 0 | 1 |
| 0 | 0 | 0 | 0 |
| 0 | 0 | 0 | 1 |
| 0 | 0 | 0 | 1 |
| 0 | 0 | 0 | 1 |
| 0 | 0 | 0 | 1 |
| 0 | 0 | 0 | 1 |
| 0 | 1 | 0 | 0 |
| 0 | 0 | 0 | 1 |
| 0 | 0 | 0 | 0 |
| 0 | 0 | 0 | 0 |
| 0 | 0 | 0 | 0 |
| 0 | 0 | 0 | 0 |
| 0 | 0 | 0 | 1 |
| 0 | 0 | 0 | 1 |
| 0 | 0 | 0 | 0 |
| 0 | 0 | 0 | 0 |
| 0 | 0 | 0 | 1 |
| 0 | 0 | 0 | 1 |
| 0 | 0 | 0 | 0 |
| 0 | 0 | 0 | 1 |
| 0 | 0 | 0 | 1 |
| 0 | 0 | 0 | 1 |
| 0 | 0 | 0 | 0 |
| 0 | 0 | 0 | 1 |
| 0 | 1 | 0 | 0 |
| 0 | 1 | 0 | 0 |
| 0 | 1 | 0 | 0 |
| 0 | 0 | 0 | 1 |
| 0 | 0 | 0 | 0 |
| 0 | 0 | 0 | 0 |
| 0 | 0 | 0 | 0 |
| 0 | 0 | 0 | 1 |
| 0 | 0 | 0 | 0 |
| 0 | 0 | 0 | 1 |
| 0 | 0 | 0 | 0 |
| 0 | 0 | 0 | 0 |
| 0 | 1 | 0 | 0 |
| 0 | 0 | 0 | 0 |
| 0 | 0 | 0 | 0 |
| 0 | 0 | 0 | 1 |
| 0 | 0 | 0 | 0 |
| 0 | 1 | 0 | 0 |
| 0 | 1 | 0 | 0 |
| 1 | 0 | 0 | 0 |
| 0 | 0 | 0 | 0 |
| 1 | 0 | 0 | 0 |
| 1 | 0 | 0 | 0 |
| 1 | 0 | 0 | 0 |
| 0 | 0 | 0 | 0 |
| 0 | 0 | 0 | 0 |
| 0 | 0 | 0 | 1 |
| 0 | 0 | 0 | 0 |

| | | | |
|---|---|---|---|
| 0 | 0 | 0 | 1 |
| 0 | 0 | 0 | 1 |
| 0 | 0 | 0 | 1 |
| 0 | 0 | 0 | 1 |
| 0 | 0 | 0 | 1 |
| 0 | 0 | 0 | 1 |
| 0 | 0 | 0 | 1 |
| 0 | 0 | 0 | 1 |
| 0 | 0 | 0 | 1 |
| 0 | 0 | 0 | 0 |
| 0 | 0 | 0 | 0 |
| 0 | 0 | 0 | 1 |
| 0 | 0 | 0 | 1 |
| 0 | 0 | 0 | 1 |
| 0 | 0 | 0 | 0 |
| 0 | 0 | 0 | 1 |
| 0 | 0 | 0 | 1 |
| 0 | 0 | 0 | 0 |
| 0 | 0 | 0 | 1 |
| 0 | 0 | 0 | 1 |
| 0 | 0 | 0 | 1 |
| 0 | 0 | 0 | 0 |
| 0 | 0 | 0 | 0 |
| 0 | 0 | 0 | 0 |
| 0 | 0 | 0 | 0 |
| 0 | 0 | 0 | 0 |
| 0 | 0 | 0 | 0 |
| 0 | 0 | 0 | 0 |
| 0 | 0 | 0 | 1 |
| 0 | 0 | 0 | 0 |
| 1 | 0 | 0 | 0 |
| 0 | 0 | 0 | 1 |
| 0 | 0 | 0 | 1 |
| 0 | 0 | 0 | 1 |
| 0 | 0 | 0 | 1 |
| 0 | 0 | 0 | 1 |
| 0 | 0 | 0 | 1 |
| 0 | 0 | 0 | 1 |
| 0 | 0 | 0 | 1 |
| 0 | 0 | 0 | 1 |
| 0 | 0 | 0 | 1 |
| 0 | 0 | 0 | 1 |
| 0 | 0 | 0 | 1 |
| 0 | 0 | 0 | 0 |
| 0 | 0 | 0 | 1 |
| 0 | 0 | 0 | 0 |
| 0 | 0 | 0 | 1 |
| 0 | 0 | 0 | 1 |
| 0 | 0 | 0 | 1 |
| 0 | 0 | 0 | 1 |
| 0 | 0 | 0 | 0 |
| 0 | 0 | 0 | 1 |
| 0 | 0 | 0 | 1 |
| 0 | 0 | 0 | 0 |

| | | | |
|---|---|---|---|
| 0 | 0 | 0 | 0 |
| 0 | 0 | 0 | 0 |
| 0 | 0 | 0 | 0 |
| 0 | 0 | 0 | 0 |
| 0 | 0 | 0 | 1 |
| 0 | 0 | 0 | 1 |
| 0 | 0 | 0 | 1 |
| 0 | 0 | 0 | 0 |
| 0 | 0 | 0 | 0 |
| 0 | 0 | 0 | 0 |
| 0 | 0 | 0 | 0 |
| 0 | 0 | 0 | 1 |
| 0 | 0 | 0 | 1 |
| 0 | 0 | 0 | 1 |
| 0 | 0 | 0 | 1 |
| 0 | 0 | 0 | 0 |
| 0 | 0 | 0 | 0 |
| 0 | 0 | 0 | 1 |
| 0 | 0 | 0 | 1 |
| 0 | 0 | 0 | 1 |
| 0 | 0 | 0 | 1 |
| 0 | 0 | 0 | 0 |
| 0 | 0 | 0 | 1 |
| 0 | 0 | 0 | 1 |
| 0 | 0 | 0 | 0 |
| 0 | 0 | 0 | 0 |
| 0 | 0 | 0 | 0 |
| 0 | 0 | 0 | 0 |
| 0 | 0 | 0 | 1 |
| 0 | 0 | 0 | 1 |
| 0 | 0 | 0 | 1 |
| 0 | 0 | 0 | 0 |
| 0 | 0 | 0 | 1 |
| 0 | 0 | 0 | 1 |
| 0 | 0 | 0 | 1 |
| 0 | 0 | 0 | 1 |
| 0 | 0 | 0 | 1 |
| 0 | 0 | 0 | 1 |
| 0 | 0 | 0 | 1 |
| 0 | 0 | 0 | 1 |
| 0 | 0 | 0 | 1 |
| 0 | 0 | 0 | 1 |
| 0 | 0 | 0 | 0 |
| 0 | 0 | 0 | 1 |
| 0 | 0 | 0 | 1 |
| 0 | 0 | 0 | 1 |
| 0 | 0 | 0 | 1 |
| 0 | 0 | 0 | 1 |
| 0 | 0 | 0 | 1 |
| 0 | 0 | 0 | 0 |
| 0 | 0 | 0 | 1 |
| 0 | 0 | 0 | 1 |
| 0 | 0 | 0 | 1 |
| 0 | 0 | 0 | 1 |

| | | | |
|---|---|---|---|
| 0 | 0 | 0 | 0 |
| 0 | 0 | 0 | 1 |
| 0 | 0 | 0 | 1 |
| 0 | 0 | 0 | 1 |
| 0 | 0 | 0 | 1 |
| 0 | 0 | 0 | 1 |
| 0 | 0 | 0 | 1 |
| 0 | 1 | 0 | 0 |
| 0 | 0 | 0 | 1 |
| 0 | 0 | 0 | 1 |
| 0 | 0 | 0 | 0 |
| 0 | 0 | 0 | 0 |
| 0 | 0 | 0 | 0 |
| 0 | 0 | 0 | 1 |
| 0 | 0 | 0 | 0 |
| 0 | 0 | 0 | 0 |
| 0 | 0 | 0 | 0 |
| 0 | 0 | 0 | 0 |
| 0 | 0 | 0 | 1 |
| 0 | 0 | 0 | 0 |
| 0 | 0 | 0 | 0 |
| 0 | 0 | 0 | 0 |
| 0 | 0 | 0 | 0 |
| 0 | 0 | 0 | 0 |
| 0 | 0 | 0 | 0 |
| 0 | 0 | 0 | 0 |
| 0 | 0 | 0 | 0 |
| 0 | 0 | 0 | 0 |
| 0 | 0 | 0 | 0 |
| 0 | 0 | 0 | 0 |
| 0 | 0 | 0 | 0 |
| 0 | 0 | 0 | 0 |
| 0 | 0 | 0 | 1 |
| 0 | 0 | 0 | 0 |
| 0 | 0 | 0 | 1 |
| 0 | 0 | 0 | 0 |
| 0 | 0 | 0 | 0 |
| 0 | 0 | 0 | 1 |
| 0 | 0 | 0 | 1 |
| 0 | 0 | 0 | 1 |
| 0 | 0 | 0 | 1 |
| 0 | 0 | 0 | 0 |
| 0 | 0 | 0 | 0 |
| 0 | 0 | 0 | 0 |
| 0 | 0 | 0 | 1 |
| 0 | 0 | 0 | 0 |
| 0 | 0 | 0 | 1 |
| 0 | 0 | 0 | 0 |
| 0 | 0 | 0 | 1 |
| 0 | 0 | 0 | 1 |
| 0 | 0 | 0 | 1 |
| 0 | 0 | 0 | 1 |
| 0 | 0 | 0 | 1 |
| 0 | 0 | 0 | 1 |

| | | | |
|---|---|---|---|
| 0 | 0 | 0 | 1 |
| 0 | 0 | 0 | 1 |
| 0 | 0 | 0 | 1 |
| 0 | 0 | 0 | 1 |
| 0 | 0 | 0 | 1 |
| 0 | 0 | 0 | 0 |
| 0 | 0 | 0 | 0 |
| 0 | 0 | 0 | 1 |
| 0 | 0 | 0 | 0 |
| 0 | 0 | 0 | 0 |
| 0 | 0 | 0 | 1 |
| 0 | 0 | 0 | 1 |
| 0 | 0 | 0 | 1 |
| 0 | 0 | 0 | 1 |
| 0 | 0 | 0 | 1 |
| 0 | 0 | 0 | 1 |
| 0 | 0 | 0 | 1 |
| 0 | 0 | 0 | 1 |
| 0 | 0 | 0 | 0 |
| 0 | 0 | 0 | 0 |
| 0 | 0 | 0 | 1 |
| 0 | 0 | 0 | 1 |
| 0 | 0 | 0 | 1 |
| 0 | 0 | 0 | 0 |
| 0 | 0 | 0 | 1 |
| 0 | 0 | 0 | 0 |
| 0 | 0 | 0 | 0 |
| 0 | 0 | 0 | 1 |
| 0 | 0 | 0 | 1 |
| 0 | 0 | 0 | 1 |
| 0 | 0 | 0 | 0 |
| 0 | 0 | 0 | 1 |
| 0 | 0 | 0 | 0 |
| 0 | 1 | 0 | 0 |
| 0 | 1 | 0 | 0 |
| 0 | 0 | 0 | 0 |
| 0 | 0 | 0 | 0 |
| 0 | 0 | 0 | 1 |
| 0 | 0 | 0 | 1 |
| 0 | 0 | 0 | 1 |
| 0 | 0 | 0 | 0 |
| 0 | 0 | 0 | 0 |
| 0 | 0 | 0 | 0 |
| 0 | 0 | 0 | 0 |
| 0 | 0 | 0 | 1 |
| 0 | 0 | 0 | 1 |
| 0 | 0 | 0 | 1 |
| 0 | 0 | 0 | 0 |
| 0 | 0 | 0 | 0 |
| 0 | 0 | 0 | 0 |
| 0 | 0 | 0 | 1 |
| 0 | 0 | 0 | 1 |
| 0 | 0 | 0 | 1 |
| 0 | 0 | 0 | 1 |

| | | | |
|---|---|---|---|
| 0 | 0 | 0 | 1 |
| 0 | 0 | 0 | 1 |
| 0 | 0 | 0 | 1 |
| 0 | 0 | 0 | 0 |
| 0 | 0 | 0 | 0 |
| 0 | 0 | 0 | 1 |
| 0 | 0 | 0 | 0 |
| 0 | 0 | 0 | 1 |
| 0 | 0 | 0 | 1 |
| 0 | 0 | 0 | 1 |
| 0 | 0 | 0 | 1 |
| 0 | 0 | 0 | 0 |
| 0 | 0 | 0 | 1 |
| 0 | 0 | 0 | 0 |
| 0 | 0 | 0 | 1 |
| 0 | 0 | 0 | 0 |
| 0 | 0 | 0 | 1 |
| 0 | 0 | 0 | 1 |
| 0 | 0 | 0 | 1 |
| 0 | 0 | 0 | 0 |
| 0 | 0 | 0 | 0 |
| 0 | 0 | 0 | 0 |
| 0 | 0 | 0 | 0 |
| 0 | 0 | 0 | 0 |
| 0 | 0 | 0 | 0 |
| 0 | 0 | 0 | 1 |
| 0 | 0 | 0 | 0 |
| 0 | 0 | 0 | 1 |
| 0 | 0 | 0 | 1 |
| 0 | 0 | 0 | 1 |
| 0 | 0 | 0 | 1 |
| 0 | 0 | 0 | 1 |
| 0 | 0 | 0 | 0 |
| 0 | 1 | 0 | 0 |
| 0 | 0 | 0 | 1 |
| 0 | 0 | 0 | 1 |
| 0 | 1 | 0 | 0 |
| 0 | 1 | 0 | 0 |
| 0 | 1 | 0 | 0 |
| 0 | 1 | 1 | 1 |
| 0 | 0 | 0 | 0 |
| 0 | 0 | 0 | 0 |
| 0 | 0 | 0 | 1 |
| 0 | 0 | 0 | 0 |
| 0 | 0 | 0 | 1 |
| 0 | 0 | 0 | 1 |
| 0 | 0 | 0 | 1 |
| 0 | 0 | 0 | 1 |
| 0 | 0 | 0 | 1 |
| 1 | 0 | 0 | 0 |
| 0 | 0 | 0 | 1 |
| 0 | 0 | 0 | 1 |
| 0 | 0 | 0 | 1 |
| 0 | 0 | 0 | 1 |

| | | | |
|---|---|---|---|
| 0 | 0 | 0 | 1 |
| 0 | 0 | 0 | 1 |
| 0 | 0 | 0 | 1 |
| 0 | 0 | 0 | 0 |
| 0 | 0 | 0 | 1 |
| 0 | 0 | 0 | 1 |
| 0 | 0 | 0 | 1 |
| 0 | 0 | 0 | 0 |
| 0 | 0 | 0 | 1 |
| 0 | 0 | 0 | 1 |
| 0 | 0 | 0 | 1 |
| 0 | 0 | 0 | 1 |
| 0 | 0 | 0 | 1 |
| 0 | 0 | 0 | 1 |
| 0 | 0 | 0 | 1 |
| 0 | 0 | 0 | 1 |
| 0 | 0 | 0 | 0 |
| 0 | 0 | 0 | 0 |
| 0 | 0 | 0 | 1 |
| 0 | 0 | 0 | 1 |
| 0 | 0 | 0 | 1 |
| 0 | 0 | 0 | 1 |
| 0 | 0 | 0 | 1 |
| 0 | 0 | 0 | 1 |
| 0 | 0 | 0 | 1 |
| 0 | 0 | 0 | 1 |
| 0 | 0 | 0 | 1 |
| 0 | 0 | 0 | 0 |
| 0 | 0 | 0 | 0 |
| 0 | 0 | 0 | 1 |
| 0 | 0 | 0 | 1 |
| 0 | 0 | 0 | 0 |
| 0 | 0 | 0 | 1 |
| 0 | 0 | 0 | 0 |
| 1 | 0 | 0 | 0 |
| 1 | 0 | 0 | 0 |
| 1 | 0 | 0 | 0 |
| 1 | 0 | 0 | 0 |
| 0 | 0 | 0 | 1 |
| 0 | 0 | 0 | 1 |
| 0 | 0 | 0 | 1 |
| 0 | 0 | 0 | 1 |
| 0 | 0 | 0 | 1 |
| 0 | 0 | 0 | 1 |
| 0 | 0 | 0 | 0 |
| 0 | 0 | 0 | 0 |
| 0 | 0 | 0 | 0 |
| 0 | 0 | 0 | 1 |
| 0 | 0 | 0 | 1 |
| 0 | 0 | 0 | 1 |
| 0 | 0 | 0 | 1 |
| 0 | 0 | 0 | 0 |
| 0 | 0 | 0 | 0 |
| 0 | 0 | 0 | 0 |

| | | | |
|---|---|---|---|
| 0 | 0 | 0 | 0 |
| 0 | 0 | 0 | 0 |
| 0 | 0 | 0 | 1 |
| 0 | 0 | 0 | 0 |
| 0 | 0 | 0 | 0 |
| 0 | 0 | 0 | 0 |
| 0 | 0 | 0 | 0 |
| 0 | 0 | 0 | 1 |
| 0 | 0 | 0 | 1 |
| 0 | 0 | 0 | 1 |
| 0 | 0 | 0 | 1 |
| 0 | 0 | 0 | 1 |
| 0 | 0 | 0 | 1 |
| 0 | 0 | 0 | 1 |
| 0 | 0 | 0 | 1 |
| 0 | 0 | 0 | 1 |
| 0 | 0 | 0 | 1 |
| 0 | 0 | 0 | 1 |
| 0 | 0 | 0 | 1 |
| 0 | 0 | 0 | 1 |
| 0 | 0 | 0 | 0 |
| 0 | 0 | 0 | 0 |
| 0 | 0 | 0 | 0 |
| 0 | 0 | 0 | 0 |
| 0 | 0 | 0 | 0 |
| 0 | 0 | 0 | 1 |
| 0 | 0 | 0 | 1 |
| 0 | 0 | 0 | 0 |
| 0 | 0 | 0 | 1 |
| 0 | 0 | 0 | 1 |
| 0 | 0 | 0 | 0 |
| 0 | 0 | 0 | 0 |
| 0 | 0 | 0 | 0 |
| 0 | 0 | 0 | 1 |
| 0 | 0 | 0 | 0 |
| 0 | 0 | 0 | 1 |
| 0 | 0 | 0 | 1 |
| 0 | 0 | 0 | 0 |
| 0 | 0 | 0 | 1 |
| 0 | 0 | 0 | 1 |
| 0 | 0 | 0 | 1 |
| 0 | 0 | 0 | 1 |
| 0 | 0 | 0 | 1 |
| 0 | 0 | 0 | 1 |
| 0 | 0 | 0 | 1 |
| 0 | 0 | 0 | 1 |
| 0 | 0 | 0 | 1 |
| 0 | 0 | 0 | 1 |
| 0 | 0 | 0 | 0 |
| 0 | 0 | 0 | 1 |
| 1 | 0 | 0 | 0 |
| 1 | 0 | 0 | 0 |
| 0 | 0 | 0 | 1 |
| 0 | 0 | 0 | 1 |

| | | | |
|---|---|---|---|
| 1 | 0 | 0 | 0 |
| 0 | 0 | 0 | 1 |
| 0 | 0 | 0 | 1 |
| 0 | 0 | 0 | 1 |
| 0 | 0 | 0 | 1 |
| 1 | 0 | 0 | 0 |
| 1 | 0 | 0 | 0 |
| 0 | 0 | 0 | 1 |
| 0 | 0 | 0 | 1 |
| 0 | 0 | 0 | 1 |
| 0 | 0 | 0 | 1 |
| 0 | 0 | 0 | 0 |
| 0 | 0 | 0 | 1 |
| 0 | 0 | 0 | 1 |
| 0 | 0 | 0 | 1 |
| 0 | 0 | 0 | 1 |
| 0 | 0 | 0 | 0 |
| 0 | 0 | 0 | 0 |
| 0 | 0 | 0 | 1 |
| 0 | 0 | 0 | 1 |
| 0 | 0 | 0 | 1 |
| 0 | 0 | 0 | 1 |
| 0 | 0 | 0 | 1 |
| 0 | 0 | 0 | 1 |
| 0 | 0 | 0 | 0 |
| 0 | 0 | 0 | 1 |
| 0 | 0 | 0 | 1 |
| 0 | 0 | 0 | 1 |
| 0 | 0 | 0 | 1 |
| 0 | 0 | 0 | 1 |
| 0 | 0 | 0 | 1 |
| 0 | 0 | 0 | 0 |
| 0 | 0 | 0 | 0 |
| 0 | 0 | 0 | 1 |
| 0 | 0 | 0 | 1 |
| 0 | 0 | 0 | 0 |
| 1 | 0 | 0 | 0 |
| 0 | 0 | 0 | 0 |
| 1 | 0 | 0 | 0 |
| 0 | 0 | 0 | 0 |
| 0 | 0 | 0 | 0 |
| 0 | 0 | 0 | 0 |
| 0 | 0 | 0 | 0 |
| 0 | 0 | 0 | 0 |
| 0 | 0 | 0 | 0 |
| 0 | 0 | 0 | 0 |
| 0 | 0 | 0 | 0 |
| 1 | 0 | 0 | 0 |
| 0 | 0 | 0 | 1 |
| 0 | 0 | 0 | 1 |
| 0 | 1 | 1 | 1 |
| 0 | 0 | 0 | 1 |
| 0 | 0 | 0 | 0 |
| 0 | 0 | 0 | 1 |

| | | | |
|---|---|---|---|
| 0 | 0 | 0 | 1 |
| 0 | 0 | 0 | 1 |
| 0 | 0 | 0 | 1 |
| 0 | 0 | 0 | 1 |
| 0 | 0 | 0 | 1 |
| 0 | 0 | 0 | 1 |
| 0 | 0 | 0 | 1 |
| 0 | 1 | 0 | 0 |
| 0 | 0 | 0 | 0 |
| 0 | 0 | 0 | 1 |
| 0 | 0 | 0 | 1 |
| 0 | 0 | 0 | 1 |
| 0 | 0 | 0 | 1 |
| 0 | 1 | 0 | 0 |
| 0 | 0 | 0 | 1 |
| 0 | 0 | 0 | 1 |
| 0 | 1 | 0 | 0 |
| 0 | 0 | 0 | 1 |
| 1 | 0 | 0 | 0 |
| 0 | 0 | 0 | 1 |
| 0 | 0 | 0 | 1 |
| 0 | 0 | 0 | 1 |
| 0 | 0 | 0 | 1 |
| 0 | 0 | 0 | 0 |
| 0 | 0 | 0 | 1 |
| 0 | 0 | 0 | 1 |
| 0 | 0 | 0 | 1 |
| 0 | 0 | 0 | 1 |
| 0 | 0 | 0 | 1 |
| 0 | 0 | 0 | 1 |
| 0 | 0 | 0 | 1 |
| 1 | 0 | 0 | 0 |
| 0 | 0 | 0 | 1 |
| 0 | 0 | 0 | 1 |
| 0 | 0 | 0 | 1 |
| 0 | 0 | 0 | 1 |
| 0 | 0 | 0 | 1 |
| 0 | 0 | 0 | 1 |
| 1 | 0 | 0 | 0 |
| 0 | 0 | 0 | 1 |
| 0 | 0 | 0 | 1 |
| 0 | 0 | 0 | 1 |
| 0 | 0 | 0 | 1 |
| 0 | 0 | 0 | 1 |
| 0 | 0 | 0 | 1 |
| 0 | 0 | 0 | 0 |
| 0 | 0 | 0 | 1 |
| 0 | 0 | 0 | 1 |
| 0 | 1 | 1 | 1 |
| 0 | 1 | 1 | 1 |
| 0 | 0 | 0 | 1 |
| 0 | 0 | 0 | 1 |
| 0 | 0 | 0 | 1 |
| 0 | 1 | 1 | 1 |

| | | | |
|---|---|---|---|
| 0 | 1 | 1 | 1 |
| 0 | 1 | 1 | 1 |
| 0 | 1 | 1 | 1 |
| 0 | 1 | 1 | 1 |
| 0 | 1 | 1 | 1 |
| 0 | 1 | 1 | 1 |
| 0 | 1 | 1 | 1 |
| 0 | 1 | 1 | 1 |
| 0 | 1 | 1 | 1 |
| 0 | 1 | 1 | 1 |
| 0 | 1 | 1 | 1 |
| 0 | 1 | 1 | 1 |
| 0 | 1 | 1 | 1 |
| 0 | 1 | 1 | 1 |
| 0 | 1 | 1 | 1 |
| 1 | 1 | 0 | 0 |
| 0 | 1 | 1 | 1 |
| 0 | 1 | 0 | 0 |
| 0 | 1 | 1 | 1 |
| 0 | 1 | 1 | 1 |
| 1 | 1 | 0 | 0 |
| 0 | 1 | 1 | 1 |
| 0 | 1 | 1 | 1 |
| 0 | 1 | 1 | 1 |
| 0 | 1 | 0 | 0 |
| 0 | 1 | 1 | 1 |
| 0 | 1 | 0 | 0 |
| 0 | 1 | 1 | 1 |
| 0 | 1 | 1 | 1 |
| 0 | 1 | 1 | 1 |
| 0 | 1 | 1 | 1 |
| 0 | 1 | 1 | 1 |
| 0 | 1 | 1 | 1 |
| 0 | 1 | 1 | 1 |
| 0 | 1 | 1 | 1 |
| 0 | 1 | 1 | 1 |
| 0 | 1 | 1 | 1 |
| 0 | 1 | 1 | 1 |
| 0 | 1 | 1 | 1 |
| 0 | 1 | 1 | 1 |
| 0 | 1 | 1 | 1 |
| 0 | 1 | 1 | 1 |
| 0 | 0 | 0 | 1 |
| 0 | 0 | 0 | 0 |
| 0 | 0 | 0 | 1 |
| 0 | 0 | 0 | 1 |
| 0 | 0 | 0 | 0 |
| 0 | 0 | 0 | 0 |
| 0 | 0 | 0 | 1 |
| 0 | 0 | 0 | 0 |
| 0 | 0 | 0 | 0 |
| 0 | 0 | 0 | 0 |
| 0 | 0 | 0 | 1 |
| 0 | 0 | 0 | 1 |

| | | | |
|---|---|---|---|
| 0 | 0 | 0 | 1 |
| 0 | 0 | 0 | 1 |
| 0 | 0 | 0 | 1 |
| 0 | 0 | 0 | 1 |
| 0 | 0 | 0 | 1 |
| 0 | 0 | 0 | 1 |
| 0 | 0 | 0 | 1 |
| 1 | 0 | 0 | 0 |
| 0 | 0 | 0 | 1 |
| 0 | 0 | 0 | 1 |
| 0 | 0 | 0 | 1 |
| 0 | 0 | 0 | 1 |
| 0 | 0 | 0 | 1 |
| 0 | 0 | 0 | 1 |
| 0 | 0 | 0 | 0 |
| 0 | 0 | 0 | 0 |
| 0 | 0 | 0 | 1 |
| 0 | 0 | 0 | 0 |
| 0 | 0 | 0 | 1 |
| 0 | 0 | 0 | 1 |
| 0 | 0 | 0 | 0 |
| 0 | 0 | 0 | 1 |
| 0 | 0 | 0 | 1 |
| 0 | 0 | 0 | 1 |
| 0 | 0 | 0 | 0 |
| 0 | 0 | 0 | 1 |
| 0 | 0 | 0 | 1 |
| 1 | 0 | 0 | 0 |
| 0 | 0 | 0 | 0 |
| 1 | 0 | 0 | 0 |
| 1 | 0 | 0 | 0 |
| 0 | 0 | 0 | 1 |
| 0 | 0 | 0 | 1 |
| 0 | 0 | 0 | 1 |
| 0 | 0 | 0 | 1 |
| 0 | 0 | 0 | 1 |
| 0 | 0 | 0 | 0 |
| 0 | 0 | 0 | 0 |
| 0 | 0 | 0 | 1 |
| 0 | 0 | 0 | 1 |
| 0 | 0 | 0 | 1 |
| 0 | 0 | 0 | 1 |
| 0 | 0 | 0 | 0 |
| 0 | 0 | 0 | 1 |
| 0 | 0 | 0 | 1 |
| 0 | 0 | 0 | 1 |
| 0 | 0 | 0 | 1 |
| 0 | 0 | 0 | 1 |
| 0 | 0 | 0 | 0 |
| 0 | 0 | 0 | 0 |
| 0 | 0 | 0 | 0 |
| 0 | 0 | 0 | 0 |
| 0 | 0 | 0 | 0 |
| 0 | 0 | 0 | 1 |

| | | | |
|---|---|---|---|
| 0 | 0 | 0 | 0 |
| 0 | 0 | 0 | 0 |
| 0 | 0 | 0 | 1 |
| 1 | 0 | 0 | 0 |
| 0 | 0 | 0 | 1 |
| 0 | 0 | 0 | 1 |
| 0 | 0 | 0 | 1 |
| 0 | 0 | 0 | 1 |
| 0 | 0 | 0 | 1 |
| 1 | 0 | 0 | 0 |
| 0 | 0 | 0 | 1 |
| 0 | 0 | 0 | 1 |
| 0 | 0 | 0 | 1 |
| 0 | 0 | 0 | 1 |
| 0 | 0 | 0 | 1 |
| 0 | 0 | 0 | 1 |
| 0 | 0 | 0 | 1 |
| 0 | 0 | 0 | 1 |
| 0 | 0 | 0 | 1 |
| 0 | 0 | 0 | 1 |
| 0 | 0 | 0 | 1 |
| 0 | 0 | 0 | 1 |
| 0 | 0 | 0 | 1 |
| 0 | 0 | 0 | 1 |
| 0 | 0 | 0 | 1 |
| 0 | 0 | 0 | 1 |
| 0 | 0 | 0 | 1 |
| 0 | 0 | 0 | 1 |
| 0 | 0 | 0 | 1 |
| 0 | 0 | 0 | 1 |
| 0 | 0 | 0 | 1 |
| 0 | 0 | 0 | 1 |
| 1 | 0 | 0 | 0 |
| 0 | 0 | 0 | 1 |
| 0 | 0 | 0 | 1 |
| 0 | 0 | 0 | 1 |
| 0 | 0 | 0 | 1 |
| 0 | 0 | 0 | 1 |
| 0 | 0 | 0 | 1 |
| 0 | 0 | 0 | 1 |
| 0 | 0 | 0 | 1 |
| 0 | 0 | 0 | 1 |
| 0 | 0 | 0 | 1 |
| 0 | 0 | 0 | 1 |
| 0 | 0 | 0 | 1 |
| 0 | 0 | 0 | 1 |
| 0 | 0 | 0 | 1 |
| 0 | 0 | 0 | 0 |
| 0 | 1 | 0 | 0 |
| 0 | 0 | 0 | 0 |
| 0 | 0 | 0 | 0 |
| 0 | 1 | 0 | 0 |
| 0 | 0 | 0 | 1 |
| 0 | 0 | 0 | 0 |

| | | | |
|---|---|---|---|
| 0 | 0 | 0 | 1 |
| 0 | 0 | 0 | 0 |
| 0 | 0 | 0 | 1 |
| 0 | 0 | 0 | 1 |
| 0 | 0 | 0 | 1 |
| 1 | 0 | 0 | 0 |
| 0 | 0 | 0 | 0 |
| 0 | 0 | 0 | 0 |
| 1 | 0 | 0 | 0 |
| 0 | 0 | 0 | 0 |
| 0 | 0 | 0 | 1 |
| 0 | 0 | 0 | 0 |
| 0 | 0 | 0 | 1 |
| 0 | 0 | 0 | 1 |
| 0 | 0 | 0 | 1 |
| 1 | 0 | 0 | 0 |
| 0 | 0 | 0 | 1 |
| 0 | 0 | 0 | 1 |
| 0 | 0 | 0 | 0 |
| 0 | 0 | 0 | 0 |
| 1 | 0 | 0 | 0 |
| 1 | 0 | 0 | 0 |
| 0 | 0 | 0 | 0 |
| 0 | 0 | 0 | 0 |
| 0 | 0 | 0 | 1 |
| 0 | 0 | 0 | 0 |
| 0 | 0 | 0 | 0 |
| 0 | 0 | 0 | 0 |
| 0 | 0 | 0 | 0 |
| 0 | 0 | 0 | 0 |
| 0 | 0 | 0 | 0 |
| 0 | 0 | 0 | 1 |
| 0 | 0 | 0 | 1 |
| 0 | 0 | 0 | 0 |
| 0 | 1 | 0 | 0 |
| 0 | 0 | 0 | 1 |
| 0 | 0 | 0 | 0 |
| 0 | 0 | 0 | 0 |
| 0 | 1 | 0 | 0 |
| 0 | 0 | 0 | 0 |
| 0 | 1 | 0 | 0 |
| 0 | 1 | 1 | 1 |
| 0 | 0 | 0 | 1 |
| 0 | 0 | 0 | 1 |
| 0 | 0 | 0 | 0 |
| 0 | 0 | 0 | 0 |
| 0 | 0 | 0 | 0 |
| 0 | 0 | 0 | 1 |
| 0 | 0 | 0 | 1 |
| 0 | 0 | 0 | 1 |
| 0 | 0 | 0 | 1 |
| 0 | 0 | 0 | 1 |
| 0 | 0 | 0 | 1 |
| 0 | 0 | 0 | 0 |

| | | | |
|---|---|---|---|
| 0 | 0 | 0 | 1 |
| 0 | 0 | 0 | 1 |
| 0 | 0 | 0 | 1 |
| 0 | 0 | 0 | 1 |
| 1 | 0 | 0 | 0 |
| 0 | 0 | 0 | 0 |
| 0 | 0 | 0 | 1 |
| 0 | 0 | 0 | 1 |
| 0 | 0 | 0 | 1 |
| 0 | 0 | 0 | 1 |
| 0 | 0 | 0 | 1 |
| 0 | 0 | 0 | 0 |
| 0 | 0 | 0 | 1 |
| 0 | 0 | 0 | 0 |
| 0 | 0 | 0 | 0 |
| 0 | 0 | 0 | 0 |
| 0 | 1 | 0 | 0 |
| 0 | 1 | 0 | 0 |
| 0 | 1 | 0 | 0 |
| 0 | 1 | 0 | 0 |
| 0 | 0 | 0 | 0 |
| 0 | 0 | 0 | 0 |
| 0 | 0 | 0 | 0 |
| 0 | 0 | 0 | 1 |
| 0 | 0 | 0 | 1 |
| 0 | 0 | 0 | 1 |
| 0 | 0 | 0 | 1 |
| 0 | 0 | 0 | 1 |
| 0 | 0 | 0 | 1 |
| 0 | 0 | 0 | 1 |
| 0 | 0 | 0 | 1 |
| 0 | 0 | 0 | 0 |
| 0 | 0 | 0 | 1 |
| 0 | 0 | 0 | 0 |
| 0 | 0 | 0 | 0 |
| 0 | 0 | 0 | 0 |
| 0 | 0 | 0 | 0 |
| 0 | 0 | 0 | 1 |
| 0 | 0 | 0 | 1 |
| 0 | 0 | 0 | 1 |
| 0 | 0 | 0 | 1 |
| 0 | 0 | 0 | 1 |
| 0 | 0 | 0 | 1 |
| 0 | 0 | 0 | 1 |
| 0 | 0 | 0 | 1 |
| 1 | 0 | 0 | 0 |
| 0 | 0 | 0 | 1 |
| 0 | 0 | 0 | 1 |
| 0 | 0 | 0 | 1 |
| 1 | 0 | 0 | 0 |
| 1 | 0 | 0 | 0 |
| 0 | 0 | 0 | 0 |
| 0 | 0 | 0 | 1 |
| 0 | 0 | 0 | 1 |

| | | | |
|---|---|---|---|
| 0 | 0 | 0 | 1 |
| 0 | 0 | 0 | 1 |
| 0 | 0 | 0 | 1 |
| 0 | 0 | 0 | 0 |
| 0 | 0 | 0 | 1 |
| 0 | 0 | 0 | 1 |
| 0 | 0 | 0 | 0 |
| 0 | 0 | 0 | 1 |
| 0 | 0 | 0 | 1 |
| 0 | 0 | 0 | 1 |
| 0 | 0 | 0 | 0 |
| 0 | 0 | 0 | 0 |
| 0 | 0 | 0 | 1 |
| 0 | 0 | 0 | 1 |
| 0 | 0 | 0 | 1 |
| 0 | 0 | 0 | 1 |
| 0 | 0 | 0 | 0 |
| 0 | 0 | 0 | 0 |
| 0 | 0 | 0 | 1 |
| 0 | 0 | 0 | 0 |
| 0 | 0 | 0 | 0 |
| 0 | 0 | 0 | 0 |
| 0 | 0 | 0 | 1 |
| 0 | 0 | 0 | 0 |
| 0 | 0 | 0 | 0 |
| 0 | 0 | 0 | 1 |
| 0 | 0 | 0 | 0 |
| 0 | 0 | 0 | 1 |
| 0 | 0 | 0 | 1 |
| 0 | 0 | 0 | 0 |
| 0 | 0 | 0 | 1 |
| 0 | 0 | 0 | 1 |
| 0 | 0 | 0 | 0 |
| 0 | 0 | 0 | 1 |
| 0 | 0 | 0 | 1 |
| 0 | 0 | 0 | 0 |
| 0 | 0 | 0 | 1 |
| 0 | 0 | 0 | 0 |
| 0 | 0 | 0 | 1 |
| 0 | 0 | 0 | 0 |
| 0 | 0 | 0 | 0 |
| 0 | 0 | 0 | 1 |
| 0 | 0 | 0 | 0 |
| 0 | 0 | 0 | 0 |
| 0 | 0 | 0 | 1 |
| 0 | 0 | 0 | 1 |
| 0 | 0 | 0 | 1 |
| 0 | 0 | 0 | 1 |
| 0 | 0 | 0 | 1 |
| 0 | 0 | 0 | 1 |
| 0 | 0 | 0 | 1 |
| 0 | 0 | 0 | 1 |
| 0 | 0 | 0 | 1 |
| 0 | 0 | 0 | 1 |

| | | | |
|---|---|---|---|
| 0 | 0 | 0 | 1 |
| 0 | 0 | 0 | 1 |
| 0 | 0 | 0 | 1 |
| 0 | 0 | 0 | 0 |
| 0 | 0 | 0 | 1 |
| 0 | 0 | 0 | 0 |
| 0 | 0 | 0 | 1 |
| 0 | 0 | 0 | 1 |
| 0 | 0 | 0 | 0 |
| 0 | 0 | 0 | 0 |
| 0 | 0 | 0 | 1 |
| 0 | 0 | 0 | 0 |
| 0 | 1 | 0 | 0 |
| 0 | 0 | 0 | 0 |
| 0 | 0 | 0 | 1 |
| 0 | 0 | 0 | 0 |
| 0 | 0 | 0 | 0 |
| 0 | 0 | 0 | 1 |
| 1 | 0 | 0 | 0 |
| 1 | 0 | 0 | 0 |
| 1 | 0 | 0 | 0 |
| 1 | 0 | 0 | 0 |
| 0 | 0 | 0 | 1 |
| 0 | 0 | 0 | 0 |
| 0 | 0 | 0 | 0 |
| 0 | 0 | 0 | 0 |
| 0 | 0 | 0 | 0 |
| 0 | 0 | 0 | 0 |
| 0 | 0 | 0 | 0 |
| 0 | 0 | 0 | 0 |
| 0 | 0 | 0 | 0 |
| 0 | 0 | 0 | 0 |
| 0 | 0 | 0 | 0 |
| 0 | 0 | 0 | 1 |
| 0 | 0 | 0 | 1 |
| 0 | 1 | 0 | 0 |
| 0 | 0 | 0 | 1 |
| 0 | 0 | 0 | 0 |
| 0 | 0 | 0 | 0 |
| 0 | 0 | 0 | 0 |
| 0 | 1 | 0 | 0 |
| 0 | 0 | 0 | 1 |
| 0 | 0 | 0 | 0 |
| 0 | 0 | 0 | 0 |
| 0 | 0 | 0 | 1 |
| 0 | 0 | 0 | 1 |
| 0 | 0 | 0 | 1 |
| 0 | 0 | 0 | 1 |
| 0 | 0 | 0 | 1 |
| 0 | 0 | 0 | 1 |
| 0 | 0 | 0 | 1 |
| 0 | 0 | 0 | 1 |
| 0 | 0 | 0 | 1 |
| 0 | 0 | 0 | 1 |

| | | | |
|---|---|---|---|
| 0 | 0 | 0 | 1 |
| 0 | 0 | 0 | 0 |
| 0 | 0 | 0 | 1 |
| 0 | 0 | 0 | 1 |
| 0 | 0 | 0 | 1 |
| 0 | 0 | 0 | 1 |
| 0 | 0 | 0 | 1 |
| 0 | 0 | 0 | 1 |
| 0 | 0 | 0 | 1 |
| 0 | 0 | 0 | 1 |
| 0 | 0 | 0 | 1 |
| 0 | 0 | 0 | 0 |
| 0 | 0 | 0 | 1 |
| 0 | 0 | 0 | 1 |
| 0 | 0 | 0 | 0 |
| 0 | 0 | 0 | 1 |
| 0 | 0 | 0 | 1 |
| 0 | 0 | 0 | 1 |
| 0 | 0 | 0 | 1 |
| 0 | 0 | 0 | 1 |
| 1 | 0 | 0 | 0 |
| 0 | 0 | 0 | 1 |
| 1 | 0 | 0 | 0 |
| 1 | 0 | 0 | 0 |
| 0 | 0 | 0 | 1 |
| 0 | 0 | 0 | 0 |
| 0 | 0 | 0 | 1 |
| 0 | 0 | 0 | 1 |
| 0 | 0 | 0 | 1 |
| 0 | 0 | 0 | 1 |
| 0 | 0 | 0 | 1 |
| 0 | 0 | 0 | 0 |
| 0 | 0 | 0 | 1 |
| 0 | 0 | 0 | 0 |
| 0 | 0 | 0 | 1 |
| 0 | 0 | 0 | 1 |
| 0 | 0 | 0 | 1 |
| 0 | 0 | 0 | 1 |
| 0 | 0 | 0 | 1 |
| 0 | 0 | 0 | 0 |
| 0 | 0 | 0 | 1 |
| 0 | 0 | 0 | 1 |
| 0 | 0 | 0 | 1 |
| 0 | 0 | 0 | 0 |
| 0 | 0 | 0 | 1 |
| 0 | 0 | 0 | 1 |
| 0 | 0 | 0 | 1 |
| 1 | 0 | 0 | 0 |
| 0 | 0 | 0 | 0 |
| 0 | 0 | 0 | 1 |
| 0 | 0 | 0 | 1 |
| 0 | 0 | 0 | 0 |
| 1 | 0 | 0 | 0 |
| 0 | 0 | 0 | 0 |

| | | | |
|---|---|---|---|
| 1 | 0 | 0 | 0 |
| 0 | 0 | 0 | 1 |
| 0 | 0 | 0 | 0 |
| 0 | 0 | 0 | 0 |
| 0 | 0 | 0 | 0 |
| 0 | 0 | 0 | 1 |
| 0 | 0 | 0 | 1 |
| 0 | 0 | 0 | 1 |
| 0 | 0 | 0 | 1 |
| 0 | 0 | 0 | 0 |
| 0 | 0 | 0 | 1 |
| 0 | 0 | 0 | 1 |
| 0 | 0 | 0 | 1 |
| 0 | 0 | 0 | 1 |
| 0 | 0 | 0 | 0 |
| 0 | 0 | 0 | 1 |
| 0 | 0 | 0 | 1 |
| 0 | 0 | 0 | 0 |
| 0 | 0 | 0 | 1 |
| 0 | 0 | 0 | 0 |
| 0 | 0 | 0 | 0 |
| 0 | 0 | 0 | 1 |
| 0 | 0 | 0 | 0 |
| 0 | 0 | 0 | 1 |
| 0 | 0 | 0 | 0 |
| 0 | 0 | 0 | 0 |
| 0 | 0 | 0 | 1 |
| 0 | 0 | 0 | 1 |
| 1 | 0 | 0 | 0 |
| 0 | 0 | 0 | 1 |
| 0 | 0 | 0 | 1 |
| 0 | 0 | 0 | 1 |
| 0 | 0 | 0 | 1 |
| 0 | 0 | 0 | 1 |
| 0 | 0 | 0 | 0 |
| 0 | 0 | 0 | 0 |
| 0 | 0 | 0 | 1 |
| 0 | 0 | 0 | 0 |
| 0 | 0 | 0 | 0 |
| 0 | 0 | 0 | 0 |
| 0 | 0 | 0 | 0 |
| 0 | 0 | 0 | 0 |
| 0 | 0 | 0 | 0 |
| 0 | 0 | 0 | 1 |
| 0 | 0 | 0 | 0 |
| 0 | 0 | 0 | 1 |
| 0 | 0 | 0 | 1 |
| 0 | 0 | 0 | 0 |
| 0 | 0 | 0 | 0 |
| 0 | 0 | 0 | 0 |
| 0 | 0 | 0 | 0 |
| 0 | 0 | 0 | 0 |
| 0 | 0 | 0 | 1 |
| 0 | 0 | 0 | 1 |

| | | | |
|---|---|---|---|
| 0 | 0 | 0 | 1 |
| 0 | 0 | 0 | 0 |
| 0 | 0 | 0 | 0 |
| 0 | 0 | 0 | 0 |
| 0 | 0 | 0 | 0 |
| 0 | 0 | 0 | 0 |
| 0 | 0 | 0 | 0 |
| 0 | 0 | 0 | 1 |
| 0 | 1 | 1 | 1 |
| 0 | 0 | 0 | 1 |
| 0 | 0 | 0 | 0 |
| 0 | 0 | 0 | 0 |
| 0 | 0 | 0 | 1 |
| 0 | 0 | 0 | 0 |
| 0 | 0 | 0 | 1 |
| 0 | 0 | 0 | 0 |
| 0 | 0 | 0 | 1 |
| 0 | 0 | 0 | 1 |
| 0 | 0 | 0 | 1 |
| 0 | 0 | 0 | 1 |
| 0 | 0 | 0 | 1 |
| 0 | 0 | 0 | 1 |
| 0 | 0 | 0 | 1 |
| 1 | 0 | 0 | 0 |
| 0 | 0 | 0 | 0 |
| 0 | 0 | 0 | 1 |
| 0 | 0 | 0 | 1 |
| 0 | 0 | 0 | 1 |
| 0 | 0 | 0 | 1 |
| 0 | 0 | 0 | 0 |
| 0 | 0 | 0 | 0 |
| 0 | 0 | 0 | 0 |
| 0 | 0 | 0 | 0 |
| 0 | 0 | 0 | 1 |
| 0 | 0 | 0 | 0 |
| 0 | 0 | 0 | 0 |
| 0 | 0 | 0 | 0 |
| 0 | 0 | 0 | 0 |
| 0 | 0 | 0 | 0 |
| 0 | 0 | 0 | 1 |
| 0 | 0 | 0 | 1 |
| 0 | 0 | 0 | 0 |
| 0 | 0 | 0 | 1 |
| 0 | 0 | 0 | 1 |
| 0 | 0 | 0 | 0 |
| 0 | 0 | 0 | 1 |
| 0 | 0 | 0 | 1 |
| 0 | 0 | 0 | 1 |
| 0 | 0 | 0 | 1 |
| 0 | 0 | 0 | 1 |
| 0 | 0 | 0 | 0 |
| 0 | 0 | 0 | 1 |
| 0 | 0 | 0 | 0 |
| 0 | 0 | 0 | 0 |

| | | | |
|---|---|---|---|
| 0 | 0 | 0 | 1 |
| 0 | 0 | 0 | 1 |
| 0 | 0 | 0 | 0 |
| 0 | 0 | 0 | 0 |
| 0 | 0 | 0 | 1 |
| 0 | 0 | 0 | 1 |
| 0 | 0 | 0 | 1 |
| 0 | 0 | 0 | 1 |
| 0 | 0 | 0 | 1 |
| 0 | 0 | 0 | 1 |
| 0 | 0 | 0 | 1 |
| 0 | 0 | 0 | 1 |
| 0 | 0 | 0 | 0 |
| 0 | 0 | 0 | 1 |
| 0 | 0 | 0 | 1 |
| 0 | 0 | 0 | 1 |
| 0 | 0 | 0 | 1 |
| 0 | 0 | 0 | 1 |
| 0 | 0 | 0 | 1 |
| 0 | 0 | 0 | 1 |
| 0 | 0 | 0 | 1 |
| 0 | 0 | 0 | 1 |
| 0 | 0 | 0 | 1 |
| 0 | 0 | 0 | 1 |
| 0 | 0 | 0 | 1 |
| 0 | 0 | 0 | 1 |
| 0 | 0 | 0 | 1 |
| 0 | 0 | 0 | 1 |
| 0 | 0 | 0 | 1 |
| 0 | 0 | 0 | 1 |
| 0 | 0 | 0 | 1 |
| 0 | 0 | 0 | 0 |
| 0 | 0 | 0 | 1 |
| 0 | 0 | 0 | 1 |
| 0 | 0 | 0 | 1 |
| 0 | 0 | 0 | 1 |
| 0 | 0 | 0 | 1 |
| 0 | 0 | 0 | 1 |
| 0 | 0 | 0 | 1 |
| 0 | 0 | 0 | 1 |
| 0 | 0 | 0 | 0 |
| 0 | 0 | 0 | 0 |
| 0 | 0 | 0 | 1 |
| 0 | 0 | 0 | 1 |
| 0 | 0 | 0 | 1 |
| 0 | 0 | 0 | 1 |
| 0 | 0 | 0 | 1 |
| 0 | 0 | 0 | 1 |
| 0 | 0 | 0 | 0 |
| 0 | 0 | 0 | 0 |
| 0 | 0 | 0 | 1 |
| 0 | 0 | 0 | 1 |
| 0 | 0 | 0 | 0 |
| 0 | 0 | 0 | 1 |

| | | | |
|---|---|---|---|
| 0 | 0 | 0 | 0 |
| 0 | 0 | 0 | 0 |
| 0 | 0 | 0 | 0 |
| 0 | 0 | 0 | 1 |
| 0 | 0 | 0 | 1 |
| 0 | 0 | 0 | 1 |
| 0 | 0 | 0 | 1 |
| 0 | 1 | 0 | 0 |
| 0 | 1 | 0 | 0 |
| 0 | 0 | 0 | 0 |
| 0 | 0 | 0 | 0 |
| 0 | 0 | 0 | 1 |
| 0 | 0 | 0 | 0 |
| 0 | 0 | 0 | 1 |
| 0 | 0 | 0 | 1 |
| 0 | 0 | 0 | 1 |
| 0 | 0 | 0 | 1 |
| 0 | 0 | 0 | 0 |
| 0 | 0 | 0 | 1 |
| 0 | 0 | 0 | 1 |
| 0 | 0 | 0 | 1 |
| 0 | 0 | 0 | 1 |
| 0 | 0 | 0 | 1 |
| 0 | 0 | 0 | 1 |
| 0 | 0 | 0 | 1 |
| 0 | 0 | 0 | 1 |
| 0 | 0 | 0 | 1 |
| 0 | 0 | 0 | 1 |
| 0 | 0 | 0 | 1 |
| 0 | 0 | 0 | 1 |
| 0 | 0 | 0 | 0 |
| 0 | 0 | 0 | 0 |
| 0 | 0 | 0 | 1 |
| 0 | 0 | 0 | 1 |
| 0 | 0 | 0 | 1 |
| 0 | 0 | 0 | 1 |
| 0 | 0 | 0 | 1 |
| 0 | 0 | 0 | 1 |
| 0 | 0 | 0 | 1 |
| 0 | 0 | 0 | 1 |
| 0 | 0 | 0 | 1 |
| 0 | 0 | 0 | 0 |
| 0 | 0 | 0 | 1 |
| 0 | 0 | 0 | 1 |
| 0 | 0 | 0 | 0 |
| 0 | 0 | 0 | 0 |
| 0 | 0 | 0 | 1 |
| 0 | 0 | 0 | 1 |
| 0 | 0 | 0 | 0 |
| 0 | 0 | 0 | 0 |
| 0 | 0 | 0 | 0 |
| 0 | 0 | 0 | 0 |
| 0 | 0 | 0 | 0 |
| 0 | 0 | 0 | 0 |

| | | | |
|---|---|---|---|
| 0 | 0 | 0 | 1 |
| 0 | 0 | 0 | 1 |
| 0 | 0 | 0 | 1 |
| 0 | 0 | 0 | 1 |
| 0 | 0 | 0 | 1 |
| 0 | 0 | 0 | 0 |
| 0 | 0 | 0 | 0 |
| 0 | 0 | 0 | 0 |
| 0 | 0 | 0 | 0 |
| 0 | 0 | 0 | 0 |
| 0 | 0 | 0 | 1 |
| 0 | 0 | 0 | 0 |
| 0 | 0 | 0 | 0 |
| 0 | 0 | 0 | 0 |
| 0 | 0 | 0 | 1 |
| 0 | 0 | 0 | 0 |
| 0 | 0 | 0 | 1 |
| 0 | 0 | 0 | 1 |
| 1 | 0 | 0 | 0 |
| 1 | 0 | 0 | 0 |
| 0 | 0 | 0 | 0 |
| 0 | 0 | 0 | 0 |
| 0 | 0 | 0 | 1 |
| 0 | 0 | 0 | 0 |
| 0 | 0 | 0 | 0 |
| 0 | 0 | 0 | 0 |
| 0 | 0 | 0 | 0 |
| 0 | 0 | 0 | 1 |
| 0 | 0 | 0 | 1 |
| 0 | 0 | 0 | 1 |
| 0 | 0 | 0 | 1 |
| 0 | 0 | 0 | 1 |
| 0 | 0 | 0 | 0 |
| 0 | 0 | 0 | 1 |
| 0 | 0 | 0 | 0 |
| 0 | 0 | 0 | 0 |
| 0 | 0 | 0 | 0 |
| 0 | 0 | 0 | 0 |
| 0 | 1 | 0 | 0 |
| 0 | 1 | 0 | 0 |
| 0 | 0 | 0 | 0 |
| 1 | 0 | 0 | 0 |
| 0 | 0 | 0 | 0 |
| 0 | 0 | 0 | 1 |
| 0 | 0 | 0 | 1 |
| 0 | 0 | 0 | 1 |
| 0 | 0 | 0 | 1 |
| 0 | 0 | 0 | 0 |
| 0 | 0 | 0 | 0 |
| 0 | 0 | 0 | 0 |
| 0 | 0 | 0 | 1 |
| 0 | 0 | 0 | 1 |
| 0 | 0 | 0 | 0 |
| 0 | 0 | 0 | 1 |

| | | | |
|---|---|---|---|
| 0 | 0 | 0 | 1 |
| 0 | 0 | 0 | 0 |
| 0 | 0 | 0 | 1 |
| 0 | 0 | 0 | 1 |
| 0 | 0 | 0 | 1 |
| 0 | 0 | 0 | 1 |
| 0 | 0 | 0 | 1 |
| 0 | 0 | 0 | 1 |
| 0 | 0 | 0 | 1 |
| 0 | 0 | 0 | 1 |
| 0 | 0 | 0 | 1 |
| 0 | 0 | 0 | 1 |
| 0 | 0 | 0 | 0 |
| 0 | 0 | 0 | 0 |
| 0 | 0 | 0 | 0 |
| 0 | 0 | 0 | 0 |
| 0 | 0 | 0 | 0 |
| 0 | 0 | 0 | 0 |
| 0 | 0 | 0 | 0 |
| 0 | 0 | 0 | 1 |
| 1 | 1 | 0 | 0 |
| 0 | 0 | 0 | 0 |
| 0 | 0 | 0 | 0 |
| 0 | 0 | 0 | 0 |
| 0 | 0 | 0 | 0 |
| 0 | 0 | 0 | 1 |
| 0 | 0 | 0 | 1 |
| 0 | 0 | 0 | 0 |
| 0 | 0 | 0 | 1 |
| 0 | 0 | 0 | 0 |
| 0 | 0 | 0 | 0 |
| 0 | 0 | 0 | 1 |
| 0 | 0 | 0 | 0 |
| 0 | 0 | 0 | 1 |
| 0 | 0 | 0 | 1 |
| 0 | 0 | 0 | 1 |
| 0 | 0 | 0 | 1 |
| 0 | 0 | 0 | 1 |
| 0 | 0 | 0 | 1 |
| 0 | 0 | 0 | 1 |
| 0 | 0 | 0 | 0 |
| 0 | 0 | 0 | 1 |
| 0 | 0 | 0 | 1 |
| 0 | 0 | 0 | 0 |
| 0 | 0 | 0 | 0 |
| 0 | 0 | 0 | 1 |
| 0 | 0 | 0 | 0 |
| 0 | 0 | 0 | 0 |
| 0 | 0 | 0 | 1 |
| 0 | 0 | 0 | 1 |
| 0 | 0 | 0 | 0 |
| 0 | 0 | 0 | 1 |
| 0 | 0 | 0 | 1 |
| 0 | 0 | 0 | 1 |

| | | | |
|---|---|---|---|
| 0 | 0 | 0 | 1 |
| 0 | 0 | 0 | 1 |
| 0 | 0 | 0 | 1 |
| 0 | 0 | 0 | 1 |
| 0 | 0 | 0 | 0 |
| 0 | 0 | 0 | 0 |
| 0 | 0 | 0 | 1 |
| 0 | 0 | 0 | 1 |
| 0 | 0 | 0 | 1 |
| 0 | 0 | 0 | 1 |
| 0 | 0 | 0 | 1 |
| 0 | 0 | 0 | 0 |
| 0 | 0 | 0 | 0 |
| 0 | 0 | 0 | 1 |
| 0 | 0 | 0 | 1 |
| 0 | 0 | 0 | 0 |
| 0 | 0 | 0 | 1 |
| 0 | 0 | 0 | 1 |
| 0 | 0 | 0 | 0 |
| 0 | 0 | 0 | 1 |
| 0 | 0 | 0 | 1 |
| 0 | 0 | 0 | 1 |
| 0 | 0 | 0 | 1 |
| 0 | 0 | 0 | 0 |
| 0 | 0 | 0 | 1 |
| 0 | 0 | 0 | 0 |
| 0 | 0 | 0 | 1 |
| 0 | 0 | 0 | 1 |
| 1 | 0 | 0 | 0 |
| 1 | 0 | 0 | 0 |
| 0 | 0 | 0 | 0 |
| 0 | 0 | 0 | 0 |
| 0 | 0 | 0 | 0 |
| 0 | 0 | 0 | 0 |
| 0 | 0 | 0 | 1 |
| 0 | 0 | 0 | 1 |
| 0 | 0 | 0 | 1 |
| 1 | 0 | 0 | 0 |
| 0 | 0 | 0 | 1 |
| 0 | 0 | 0 | 1 |
| 0 | 0 | 0 | 0 |
| 0 | 0 | 0 | 0 |
| 0 | 0 | 0 | 0 |
| 0 | 0 | 0 | 1 |
| 0 | 0 | 0 | 0 |
| 0 | 0 | 0 | 1 |
| 0 | 0 | 0 | 1 |
| 0 | 0 | 0 | 0 |
| 0 | 0 | 0 | 1 |
| 0 | 0 | 0 | 1 |
| 0 | 0 | 0 | 1 |
| 0 | 1 | 0 | 0 |
| 0 | 0 | 0 | 1 |
| 0 | 0 | 0 | 0 |

| | | | |
|---|---|---|---|
| 0 | 0 | 0 | 1 |
| 0 | 0 | 0 | 0 |
| 0 | 0 | 0 | 0 |
| 0 | 0 | 0 | 1 |
| 0 | 0 | 0 | 1 |
| 0 | 0 | 0 | 1 |
| 0 | 0 | 0 | 0 |
| 0 | 0 | 0 | 1 |
| 0 | 0 | 0 | 1 |
| 0 | 0 | 0 | 1 |
| 0 | 0 | 0 | 1 |
| 0 | 0 | 0 | 1 |
| 0 | 0 | 0 | 1 |
| 0 | 0 | 0 | 1 |
| 0 | 0 | 0 | 1 |
| 0 | 0 | 0 | 1 |
| 0 | 0 | 0 | 1 |
| 0 | 0 | 0 | 1 |
| 0 | 0 | 0 | 1 |
| 0 | 0 | 0 | 1 |
| 0 | 0 | 0 | 1 |
| 0 | 0 | 0 | 0 |
| 0 | 0 | 0 | 1 |
| 0 | 0 | 0 | 1 |
| 0 | 0 | 0 | 1 |
| 0 | 0 | 0 | 1 |
| 0 | 0 | 0 | 0 |
| 0 | 0 | 0 | 1 |
| 0 | 0 | 0 | 0 |
| 0 | 0 | 0 | 1 |
| 0 | 0 | 0 | 1 |
| 0 | 0 | 0 | 0 |
| 0 | 1 | 1 | 1 |
| 0 | 1 | 1 | 1 |
| 0 | 0 | 0 | 1 |
| 0 | 0 | 0 | 1 |
| 0 | 0 | 0 | 1 |
| 0 | 0 | 0 | 1 |
| 0 | 0 | 0 | 0 |
| 0 | 0 | 0 | 0 |
| 0 | 0 | 0 | 0 |
| 0 | 0 | 0 | 1 |
| 0 | 0 | 0 | 1 |
| 0 | 0 | 0 | 1 |
| 0 | 0 | 0 | 1 |
| 0 | 1 | 0 | 0 |
| 0 | 0 | 0 | 1 |
| 0 | 0 | 0 | 0 |
| 0 | 0 | 0 | 1 |
| 0 | 0 | 0 | 0 |
| 0 | 0 | 0 | 0 |
| 0 | 0 | 0 | 0 |
| 0 | 0 | 0 | 1 |
| 0 | 0 | 0 | 1 |

| | | | |
|---|---|---|---|
| 0 | 0 | 0 | 0 |
| 0 | 0 | 0 | 0 |
| 0 | 0 | 0 | 1 |
| 0 | 0 | 0 | 1 |
| 0 | 0 | 0 | 1 |
| 0 | 0 | 0 | 1 |
| 0 | 0 | 0 | 1 |
| 0 | 0 | 0 | 1 |
| 0 | 0 | 0 | 1 |
| 0 | 0 | 0 | 1 |
| 0 | 0 | 0 | 1 |
| 0 | 0 | 0 | 1 |
| 0 | 0 | 0 | 1 |
| 0 | 0 | 0 | 1 |
| 0 | 0 | 0 | 1 |
| 0 | 0 | 0 | 0 |
| 0 | 0 | 0 | 0 |
| 0 | 0 | 0 | 0 |
| 0 | 0 | 0 | 1 |
| 0 | 0 | 0 | 1 |
| 0 | 0 | 0 | 1 |
| 0 | 0 | 0 | 0 |
| 0 | 0 | 0 | 0 |
| 0 | 0 | 0 | 0 |
| 0 | 0 | 0 | 0 |
| 0 | 0 | 0 | 1 |
| 0 | 0 | 0 | 1 |
| 0 | 0 | 0 | 1 |
| 0 | 0 | 0 | 1 |
| 0 | 0 | 0 | 1 |
| 0 | 0 | 0 | 0 |
| 0 | 0 | 0 | 1 |
| 0 | 0 | 0 | 0 |
| 0 | 0 | 0 | 1 |
| 0 | 1 | 1 | 1 |
| 0 | 1 | 1 | 1 |
| 0 | 1 | 0 | 0 |
| 0 | 0 | 0 | 0 |
| 0 | 0 | 0 | 1 |
| 0 | 0 | 0 | 1 |
| 0 | 0 | 0 | 1 |
| 0 | 0 | 0 | 0 |
| 0 | 0 | 0 | 1 |
| 0 | 0 | 0 | 1 |
| 0 | 0 | 0 | 0 |
| 0 | 0 | 0 | 1 |
| 0 | 0 | 0 | 0 |
| 0 | 0 | 0 | 0 |
| 0 | 0 | 0 | 1 |
| 0 | 0 | 0 | 1 |
| 0 | 1 | 0 | 0 |
| 0 | 1 | 0 | 0 |
| 0 | 0 | 0 | 1 |
| 0 | 0 | 0 | 0 |

| | | | |
|---|---|---|---|
| 0 | 0 | 0 | 1 |
| 0 | 0 | 0 | 0 |
| 0 | 1 | 1 | 1 |
| 0 | 1 | 0 | 0 |
| 0 | 0 | 0 | 1 |
| 0 | 0 | 0 | 1 |
| 0 | 0 | 0 | 1 |
| 0 | 0 | 0 | 1 |
| 0 | 0 | 0 | 0 |
| 0 | 0 | 0 | 1 |
| 0 | 0 | 0 | 1 |
| 0 | 0 | 0 | 1 |
| 0 | 0 | 0 | 0 |
| 0 | 0 | 0 | 0 |
| 0 | 0 | 0 | 0 |
| 0 | 0 | 0 | 1 |
| 0 | 0 | 0 | 0 |
| 0 | 0 | 0 | 1 |
| 0 | 1 | 0 | 0 |
| 0 | 0 | 0 | 0 |
| 0 | 0 | 0 | 0 |
| 0 | 1 | 0 | 0 |
| 0 | 0 | 0 | 0 |
| 1 | 0 | 0 | 0 |
| 0 | 0 | 0 | 1 |
| 0 | 0 | 0 | 1 |
| 0 | 0 | 0 | 0 |
| 0 | 0 | 0 | 0 |
| 0 | 0 | 0 | 0 |
| 0 | 0 | 0 | 0 |
| 0 | 0 | 0 | 1 |
| 0 | 0 | 0 | 1 |
| 0 | 0 | 0 | 1 |
| 0 | 0 | 0 | 1 |
| 0 | 0 | 0 | 1 |
| 1 | 0 | 0 | 0 |
| 1 | 0 | 0 | 0 |
| 0 | 0 | 0 | 1 |
| 0 | 0 | 0 | 1 |
| 0 | 0 | 0 | 1 |
| 0 | 0 | 0 | 1 |
| 0 | 0 | 0 | 1 |
| 0 | 0 | 0 | 1 |
| 0 | 0 | 0 | 0 |
| 0 | 0 | 0 | 0 |
| 0 | 0 | 0 | 0 |
| 0 | 0 | 0 | 1 |
| 0 | 0 | 0 | 0 |
| 0 | 0 | 0 | 0 |
| 0 | 0 | 0 | 0 |
| 0 | 0 | 0 | 1 |
| 0 | 0 | 0 | 1 |
| 0 | 0 | 0 | 1 |
| 0 | 0 | 0 | 1 |

| | | | |
|---|---|---|---|
| 0 | 0 | 0 | 1 |
| 0 | 0 | 0 | 1 |
| 0 | 0 | 0 | 1 |
| 0 | 0 | 0 | 1 |
| 0 | 0 | 0 | 0 |
| 0 | 0 | 0 | 1 |
| 0 | 0 | 0 | 0 |
| 0 | 0 | 0 | 0 |
| 0 | 0 | 0 | 0 |
| 0 | 0 | 0 | 0 |
| 0 | 0 | 0 | 0 |
| 0 | 0 | 0 | 0 |
| 0 | 0 | 0 | 1 |
| 0 | 0 | 0 | 0 |
| 0 | 0 | 0 | 0 |
| 0 | 0 | 0 | 1 |
| 0 | 0 | 0 | 1 |
| 0 | 0 | 0 | 1 |
| 0 | 0 | 0 | 0 |
| 0 | 0 | 0 | 0 |
| 0 | 0 | 0 | 0 |
| 0 | 0 | 0 | 0 |
| 0 | 0 | 0 | 1 |
| 0 | 0 | 0 | 1 |
| 0 | 0 | 0 | 1 |
| 0 | 0 | 0 | 1 |
| 0 | 0 | 0 | 1 |
| 0 | 0 | 0 | 0 |
| 0 | 0 | 0 | 0 |
| 0 | 0 | 0 | 0 |
| 0 | 0 | 0 | 0 |
| 0 | 0 | 0 | 0 |
| 0 | 0 | 0 | 0 |
| 0 | 0 | 0 | 1 |
| 0 | 0 | 0 | 1 |
| 0 | 0 | 0 | 0 |
| 0 | 0 | 0 | 1 |
| 0 | 0 | 0 | 1 |
| 0 | 0 | 0 | 1 |
| 0 | 0 | 0 | 1 |
| 0 | 0 | 0 | 0 |
| 0 | 0 | 0 | 1 |
| 0 | 0 | 0 | 0 |
| 0 | 0 | 0 | 1 |
| 0 | 0 | 0 | 1 |
| 0 | 0 | 0 | 1 |
| 0 | 0 | 0 | 1 |
| 0 | 0 | 0 | 0 |
| 0 | 0 | 0 | 1 |
| 0 | 0 | 0 | 1 |
| 0 | 0 | 0 | 1 |
| 0 | 0 | 0 | 1 |
| 0 | 0 | 0 | 1 |
| 0 | 0 | 0 | 1 |

| | | | |
|---|---|---|---|
| 0 | 0 | 0 | 1 |
| 0 | 0 | 0 | 1 |
| 0 | 0 | 0 | 1 |
| 0 | 0 | 0 | 1 |
| 0 | 0 | 0 | 1 |
| 0 | 0 | 0 | 1 |
| 0 | 0 | 0 | 1 |
| 0 | 0 | 0 | 1 |
| 0 | 0 | 0 | 1 |
| 0 | 0 | 0 | 1 |
| 0 | 0 | 0 | 1 |
| 0 | 0 | 0 | 0 |
| 0 | 0 | 0 | 1 |
| 0 | 0 | 0 | 1 |
| 0 | 0 | 0 | 1 |
| 0 | 0 | 0 | 1 |
| 0 | 0 | 0 | 1 |
| 0 | 0 | 0 | 0 |
| 0 | 0 | 0 | 0 |
| 0 | 0 | 0 | 0 |
| 0 | 0 | 0 | 0 |
| 0 | 0 | 0 | 1 |
| 1 | 0 | 0 | 0 |
| 0 | 0 | 0 | 1 |
| 0 | 0 | 0 | 1 |
| 0 | 0 | 0 | 1 |
| 0 | 0 | 0 | 1 |
| 0 | 0 | 0 | 1 |
| 0 | 0 | 0 | 0 |
| 0 | 0 | 0 | 0 |
| 0 | 0 | 0 | 1 |
| 0 | 0 | 0 | 1 |
| 1 | 0 | 0 | 0 |
| 0 | 0 | 0 | 1 |
| 0 | 0 | 0 | 0 |
| 1 | 0 | 0 | 0 |
| 1 | 0 | 0 | 0 |
| 0 | 0 | 0 | 1 |
| 1 | 0 | 0 | 0 |
| 0 | 0 | 0 | 1 |
| 1 | 0 | 0 | 0 |
| 0 | 0 | 0 | 1 |
| 0 | 0 | 0 | 0 |
| 0 | 0 | 0 | 1 |
| 0 | 0 | 0 | 1 |
| 0 | 0 | 0 | 1 |
| 0 | 0 | 0 | 1 |
| 0 | 0 | 0 | 1 |
| 0 | 0 | 0 | 1 |
| 0 | 0 | 0 | 0 |
| 0 | 0 | 0 | 1 |
| 1 | 0 | 0 | 0 |
| 0 | 0 | 0 | 1 |
| 0 | 0 | 0 | 1 |

| | | | |
|---|---|---|---|
| 0 | 0 | 0 | 0 |
| 0 | 0 | 0 | 0 |
| 0 | 0 | 0 | 0 |
| 0 | 0 | 0 | 0 |
| 0 | 0 | 0 | 0 |
| 0 | 0 | 0 | 1 |
| 0 | 0 | 0 | 0 |
| 0 | 0 | 0 | 1 |
| 0 | 0 | 0 | 1 |
| 0 | 0 | 0 | 0 |
| 0 | 0 | 0 | 0 |
| 0 | 0 | 0 | 0 |
| 0 | 0 | 0 | 1 |
| 0 | 0 | 0 | 1 |
| 0 | 0 | 0 | 1 |
| 0 | 0 | 0 | 1 |
| 0 | 0 | 0 | 1 |
| 0 | 0 | 0 | 1 |
| 0 | 0 | 0 | 1 |
| 0 | 0 | 0 | 1 |
| 0 | 0 | 0 | 1 |
| 0 | 0 | 0 | 1 |
| 0 | 0 | 0 | 1 |
| 0 | 0 | 0 | 1 |
| 0 | 0 | 0 | 1 |
| 0 | 0 | 0 | 1 |
| 0 | 0 | 0 | 1 |
| 0 | 0 | 0 | 1 |
| 0 | 0 | 0 | 1 |
| 0 | 0 | 0 | 1 |
| 0 | 0 | 0 | 1 |
| 0 | 0 | 0 | 1 |
| 0 | 0 | 0 | 1 |
| 0 | 0 | 0 | 1 |
| 0 | 0 | 0 | 1 |
| 0 | 0 | 0 | 1 |
| 0 | 0 | 0 | 1 |
| 0 | 0 | 0 | 1 |
| 0 | 0 | 0 | 1 |
| 0 | 0 | 0 | 1 |
| 0 | 0 | 0 | 1 |
| 0 | 0 | 0 | 1 |
| 0 | 0 | 0 | 1 |
| 0 | 0 | 0 | 1 |
| 0 | 0 | 0 | 1 |
| 0 | 0 | 0 | 0 |
| 0 | 0 | 0 | 1 |
| 0 | 0 | 0 | 1 |
| 0 | 0 | 0 | 1 |
| 0 | 0 | 0 | 1 |
| 0 | 0 | 0 | 0 |
| 0 | 0 | 0 | 1 |
| 0 | 0 | 0 | 1 |
| 0 | 0 | 0 | 1 |

| | | | |
|---|---|---|---|
| 0 | 0 | 0 | 1 |
| 0 | 0 | 0 | 1 |
| 0 | 0 | 0 | 1 |
| 0 | 0 | 0 | 1 |
| 0 | 0 | 0 | 1 |
| 0 | 0 | 0 | 1 |
| 0 | 0 | 0 | 1 |
| 0 | 0 | 0 | 0 |
| 0 | 0 | 0 | 0 |
| 0 | 0 | 0 | 1 |
| 0 | 0 | 0 | 1 |
| 0 | 0 | 0 | 1 |
| 0 | 0 | 0 | 1 |
| 0 | 0 | 0 | 1 |
| 0 | 0 | 0 | 0 |
| 0 | 0 | 0 | 0 |
| 0 | 0 | 0 | 0 |
| 0 | 0 | 0 | 1 |
| 0 | 0 | 0 | 1 |
| 0 | 0 | 0 | 1 |
| 0 | 0 | 0 | 0 |
| 0 | 0 | 0 | 0 |
| 0 | 0 | 0 | 0 |
| 0 | 0 | 0 | 0 |
| 0 | 0 | 0 | 0 |
| 0 | 0 | 0 | 0 |
| 0 | 0 | 0 | 1 |
| 0 | 0 | 0 | 1 |
| 0 | 0 | 0 | 0 |
| 0 | 0 | 0 | 1 |
| 0 | 0 | 0 | 1 |
| 0 | 0 | 0 | 1 |
| 0 | 0 | 0 | 0 |
| 0 | 0 | 0 | 1 |
| 0 | 0 | 0 | 1 |
| 0 | 0 | 0 | 1 |
| 0 | 0 | 0 | 0 |
| 0 | 0 | 0 | 0 |
| 0 | 0 | 0 | 0 |
| 0 | 0 | 0 | 1 |
| 0 | 0 | 0 | 1 |
| 0 | 0 | 0 | 1 |
| 0 | 0 | 0 | 1 |
| 0 | 0 | 0 | 1 |
| 0 | 0 | 0 | 1 |
| 0 | 0 | 0 | 0 |
| 0 | 0 | 0 | 1 |
| 0 | 0 | 0 | 1 |
| 0 | 0 | 0 | 1 |
| 0 | 0 | 0 | 1 |
| 0 | 0 | 0 | 1 |
| 0 | 0 | 0 | 1 |
| 0 | 0 | 0 | 1 |
| 0 | 0 | 0 | 1 |

| | | | |
|---|---|---|---|
| 0 | 0 | 0 | 1 |
| 0 | 0 | 0 | 1 |
| 0 | 0 | 0 | 1 |
| 0 | 0 | 0 | 0 |
| 0 | 0 | 0 | 1 |
| 0 | 0 | 0 | 1 |
| 0 | 0 | 0 | 1 |
| 0 | 0 | 0 | 0 |
| 0 | 0 | 0 | 0 |
| 0 | 0 | 0 | 1 |
| 0 | 0 | 0 | 1 |
| 0 | 0 | 0 | 1 |
| 0 | 0 | 0 | 0 |
| 0 | 0 | 0 | 1 |
| 0 | 0 | 0 | 1 |
| 0 | 0 | 0 | 1 |
| 0 | 0 | 0 | 0 |
| 0 | 0 | 0 | 1 |
| 0 | 0 | 0 | 1 |
| 0 | 0 | 0 | 1 |
| 0 | 0 | 0 | 1 |
| 0 | 0 | 0 | 1 |
| 0 | 0 | 0 | 1 |
| 0 | 0 | 0 | 1 |
| 0 | 0 | 0 | 1 |
| 0 | 0 | 0 | 1 |
| 0 | 0 | 0 | 1 |
| 0 | 0 | 0 | 0 |
| 0 | 0 | 0 | 0 |
| 0 | 0 | 0 | 0 |
| 0 | 0 | 0 | 0 |
| 0 | 0 | 0 | 0 |
| 0 | 0 | 0 | 0 |
| 0 | 0 | 0 | 1 |
| 0 | 0 | 0 | 0 |
| 0 | 0 | 0 | 0 |
| 0 | 0 | 0 | 0 |
| 0 | 0 | 0 | 1 |
| 0 | 0 | 0 | 1 |
| 0 | 0 | 0 | 1 |
| 0 | 0 | 0 | 1 |
| 0 | 0 | 0 | 1 |
| 0 | 0 | 0 | 0 |
| 0 | 0 | 0 | 0 |
| 0 | 0 | 0 | 1 |
| 0 | 0 | 0 | 1 |
| 0 | 0 | 0 | 0 |
| 0 | 0 | 0 | 1 |
| 0 | 0 | 0 | 1 |
| 0 | 0 | 0 | 1 |
| 0 | 0 | 0 | 1 |
| 0 | 0 | 0 | 1 |
| 0 | 0 | 0 | 0 |
| 0 | 0 | 0 | 0 |

| | | | |
|---|---|---|---|
| 0 | 0 | 0 | 0 |
| 0 | 0 | 0 | 0 |
| 0 | 0 | 0 | 0 |
| 0 | 0 | 0 | 0 |
| 0 | 0 | 0 | 0 |
| 0 | 0 | 0 | 0 |
| 0 | 0 | 0 | 0 |
| 0 | 0 | 0 | 0 |
| 0 | 0 | 0 | 1 |
| 0 | 0 | 0 | 0 |
| 0 | 0 | 0 | 1 |
| 0 | 0 | 0 | 1 |
| 0 | 0 | 0 | 1 |
| 0 | 0 | 0 | 1 |
| 0 | 0 | 0 | 0 |
| 0 | 0 | 0 | 1 |
| 0 | 0 | 0 | 1 |
| 0 | 0 | 0 | 1 |
| 0 | 0 | 0 | 1 |
| 0 | 0 | 0 | 0 |
| 0 | 0 | 0 | 0 |
| 0 | 0 | 0 | 1 |
| 0 | 0 | 0 | 0 |
| 0 | 0 | 0 | 0 |
| 0 | 0 | 0 | 0 |
| 0 | 0 | 0 | 0 |
| 0 | 0 | 0 | 0 |
| 0 | 0 | 0 | 1 |
| 0 | 0 | 0 | 1 |
| 0 | 0 | 0 | 1 |
| 0 | 0 | 0 | 0 |
| 0 | 0 | 0 | 1 |
| 0 | 0 | 0 | 1 |
| 0 | 0 | 0 | 1 |
| 0 | 0 | 0 | 1 |
| 0 | 0 | 0 | 1 |
| 0 | 0 | 0 | 1 |
| 0 | 0 | 0 | 1 |
| 0 | 0 | 0 | 1 |
| 0 | 0 | 0 | 1 |
| 0 | 0 | 0 | 1 |
| 0 | 0 | 0 | 0 |
| 0 | 0 | 0 | 1 |
| 0 | 0 | 0 | 1 |
| 0 | 0 | 0 | 1 |
| 0 | 0 | 0 | 1 |
| 0 | 0 | 0 | 1 |
| 0 | 0 | 0 | 1 |
| 0 | 0 | 0 | 0 |
| 0 | 0 | 0 | 1 |
| 0 | 0 | 0 | 1 |
| 0 | 0 | 0 | 1 |
| 0 | 0 | 0 | 0 |
| 0 | 0 | 0 | 0 |

| | | | |
|---|---|---|---|
| 0 | 0 | 0 | 1 |
| 0 | 0 | 0 | 1 |
| 0 | 0 | 0 | 1 |
| 0 | 0 | 0 | 1 |
| 0 | 0 | 0 | 0 |
| 1 | 0 | 0 | 0 |
| 0 | 0 | 0 | 1 |
| 0 | 0 | 0 | 0 |
| 0 | 0 | 0 | 1 |
| 0 | 0 | 0 | 1 |
| 0 | 0 | 0 | 0 |
| 0 | 0 | 0 | 1 |
| 0 | 0 | 0 | 0 |
| 0 | 0 | 0 | 0 |
| 0 | 0 | 0 | 0 |
| 0 | 0 | 0 | 0 |
| 0 | 0 | 0 | 1 |
| 0 | 0 | 0 | 1 |
| 0 | 0 | 0 | 0 |
| 0 | 0 | 0 | 0 |
| 0 | 0 | 0 | 0 |
| 0 | 0 | 0 | 0 |
| 0 | 0 | 0 | 0 |
| 0 | 0 | 0 | 1 |
| 0 | 0 | 0 | 1 |
| 0 | 0 | 0 | 0 |
| 0 | 0 | 0 | 1 |
| 0 | 0 | 0 | 0 |
| 0 | 0 | 0 | 0 |
| 0 | 0 | 0 | 0 |
| 0 | 0 | 0 | 1 |
| 1 | 0 | 0 | 0 |
| 1 | 0 | 0 | 0 |
| 0 | 0 | 0 | 1 |
| 0 | 0 | 0 | 1 |
| 0 | 0 | 0 | 0 |
| 0 | 0 | 0 | 0 |
| 0 | 0 | 0 | 1 |
| 0 | 0 | 0 | 0 |
| 0 | 0 | 0 | 0 |
| 0 | 1 | 0 | 0 |
| 0 | 0 | 0 | 0 |
| 0 | 0 | 0 | 1 |
| 0 | 0 | 0 | 1 |
| 0 | 0 | 0 | 0 |
| 0 | 0 | 0 | 1 |
| 0 | 0 | 0 | 0 |
| 0 | 0 | 0 | 1 |
| 0 | 0 | 0 | 1 |
| 0 | 0 | 0 | 1 |
| 0 | 0 | 0 | 1 |
| 0 | 0 | 0 | 1 |
| 0 | 0 | 0 | 1 |
| 0 | 0 | 0 | 1 |

| | | | |
|---|---|---|---|
| 0 | 0 | 0 | 1 |
| 0 | 0 | 0 | 1 |
| 0 | 0 | 0 | 1 |
| 0 | 0 | 0 | 1 |
| 0 | 0 | 0 | 1 |
| 0 | 0 | 0 | 1 |
| 0 | 0 | 0 | 1 |
| 0 | 0 | 0 | 0 |
| 0 | 0 | 0 | 1 |
| 1 | 0 | 0 | 0 |
| 0 | 0 | 0 | 0 |
| 0 | 0 | 0 | 0 |
| 0 | 0 | 0 | 1 |
| 0 | 0 | 0 | 1 |
| 0 | 0 | 0 | 0 |
| 0 | 0 | 0 | 1 |
| 0 | 0 | 0 | 1 |
| 0 | 0 | 0 | 1 |
| 0 | 0 | 0 | 1 |
| 0 | 0 | 0 | 1 |
| 0 | 0 | 0 | 1 |
| 0 | 0 | 0 | 1 |
| 0 | 0 | 0 | 1 |
| 0 | 0 | 0 | 1 |
| 0 | 0 | 0 | 1 |
| 0 | 0 | 0 | 1 |
| 0 | 0 | 0 | 1 |
| 0 | 0 | 0 | 1 |
| 0 | 0 | 0 | 1 |
| 0 | 0 | 0 | 0 |
| 0 | 0 | 0 | 0 |
| 0 | 0 | 0 | 1 |
| 0 | 0 | 0 | 1 |
| 0 | 0 | 0 | 1 |
| 0 | 0 | 0 | 1 |
| 0 | 0 | 0 | 0 |
| 0 | 0 | 0 | 1 |
| 0 | 0 | 0 | 1 |
| 0 | 0 | 0 | 1 |
| 0 | 0 | 0 | 1 |
| 0 | 0 | 0 | 1 |
| 0 | 0 | 0 | 0 |
| 0 | 0 | 0 | 0 |
| 0 | 0 | 0 | 0 |
| 0 | 0 | 0 | 1 |
| 0 | 0 | 0 | 1 |
| 0 | 0 | 0 | 1 |
| 0 | 0 | 0 | 1 |
| 0 | 0 | 0 | 1 |
| 0 | 0 | 0 | 1 |
| 0 | 0 | 0 | 1 |
| 0 | 0 | 0 | 1 |
| 0 | 0 | 0 | 1 |
| 0 | 0 | 0 | 0 |

| | | | |
|---|---|---|---|
| 0 | 0 | 0 | 1 |
| 0 | 0 | 0 | 1 |
| 0 | 0 | 0 | 1 |
| 0 | 0 | 0 | 1 |
| 0 | 0 | 0 | 1 |
| 0 | 0 | 0 | 1 |
| 0 | 0 | 0 | 1 |
| 0 | 0 | 0 | 1 |
| 0 | 0 | 0 | 0 |
| 0 | 0 | 0 | 0 |
| 0 | 0 | 0 | 1 |
| 0 | 0 | 0 | 1 |
| 0 | 0 | 0 | 1 |
| 0 | 0 | 0 | 0 |
| 0 | 0 | 0 | 1 |
| 0 | 0 | 0 | 1 |
| 0 | 0 | 0 | 1 |
| 1 | 0 | 0 | 0 |
| 1 | 0 | 0 | 0 |
| 0 | 0 | 0 | 0 |
| 0 | 0 | 0 | 0 |
| 0 | 0 | 0 | 1 |
| 0 | 0 | 0 | 1 |
| 0 | 0 | 0 | 1 |
| 0 | 0 | 0 | 1 |
| 1 | 0 | 0 | 0 |
| 0 | 0 | 0 | 1 |
| 0 | 0 | 0 | 1 |
| 0 | 0 | 0 | 1 |
| 0 | 0 | 0 | 1 |
| 0 | 0 | 0 | 1 |
| 0 | 0 | 0 | 1 |
| 0 | 0 | 0 | 1 |
| 0 | 0 | 0 | 1 |
| 0 | 0 | 0 | 1 |
| 0 | 0 | 0 | 1 |
| 0 | 0 | 0 | 1 |
| 0 | 0 | 0 | 1 |
| 0 | 0 | 0 | 1 |
| 0 | 0 | 0 | 0 |
| 0 | 0 | 0 | 1 |
| 0 | 0 | 0 | 1 |
| 0 | 0 | 0 | 0 |
| 0 | 0 | 0 | 1 |
| 0 | 0 | 0 | 1 |
| 0 | 0 | 0 | 1 |
| 0 | 0 | 0 | 0 |
| 0 | 0 | 0 | 0 |
| 0 | 0 | 0 | 1 |
| 0 | 0 | 0 | 1 |
| 0 | 0 | 0 | 0 |
| 0 | 0 | 0 | 1 |
| 0 | 0 | 0 | 1 |
| 0 | 0 | 0 | 1 |

| | | | |
|---|---|---|---|
| 0 | 0 | 0 | 1 |
| 0 | 0 | 0 | 1 |
| 0 | 0 | 0 | 1 |
| 0 | 0 | 0 | 1 |
| 0 | 0 | 0 | 1 |
| 0 | 0 | 0 | 1 |
| 0 | 0 | 0 | 1 |
| 0 | 0 | 0 | 1 |
| 0 | 0 | 0 | 1 |
| 0 | 0 | 0 | 1 |
| 0 | 0 | 0 | 1 |
| 0 | 0 | 0 | 1 |
| 0 | 0 | 0 | 1 |
| 0 | 0 | 0 | 0 |
| 0 | 0 | 0 | 0 |
| 0 | 0 | 0 | 0 |
| 0 | 0 | 0 | 0 |
| 0 | 0 | 0 | 1 |
| 0 | 0 | 0 | 1 |
| 0 | 0 | 0 | 0 |
| 0 | 0 | 0 | 1 |
| 0 | 0 | 0 | 1 |
| 0 | 0 | 0 | 1 |
| 0 | 0 | 0 | 1 |
| 0 | 0 | 0 | 1 |
| 0 | 0 | 0 | 0 |
| 0 | 0 | 0 | 1 |
| 0 | 0 | 0 | 1 |
| 0 | 0 | 0 | 1 |
| 0 | 0 | 0 | 0 |
| 0 | 0 | 0 | 0 |
| 0 | 0 | 0 | 1 |
| 0 | 0 | 0 | 1 |
| 0 | 0 | 0 | 1 |
| 0 | 0 | 0 | 1 |
| 0 | 0 | 0 | 1 |
| 0 | 0 | 0 | 1 |
| 0 | 0 | 0 | 1 |
| 0 | 0 | 0 | 0 |
| 0 | 0 | 0 | 1 |
| 0 | 0 | 0 | 1 |
| 0 | 0 | 0 | 0 |
| 0 | 0 | 0 | 0 |
| 0 | 0 | 0 | 0 |
| 0 | 0 | 0 | 0 |
| 0 | 0 | 0 | 1 |
| 0 | 0 | 0 | 0 |
| 0 | 0 | 0 | 1 |
| 0 | 0 | 0 | 1 |
| 0 | 0 | 0 | 1 |
| 0 | 0 | 0 | 1 |
| 0 | 0 | 0 | 1 |
| 0 | 0 | 0 | 1 |
| 0 | 0 | 0 | 1 |

| | | | |
|---|---|---|---|
| 0 | 0 | 0 | 1 |
| 0 | 0 | 0 | 1 |
| 0 | 0 | 0 | 0 |
| 0 | 0 | 0 | 1 |
| 0 | 0 | 0 | 1 |
| 0 | 0 | 0 | 1 |
| 0 | 0 | 0 | 1 |
| 0 | 0 | 0 | 1 |
| 0 | 0 | 0 | 1 |
| 0 | 0 | 0 | 1 |
| 0 | 0 | 0 | 1 |
| 0 | 0 | 0 | 1 |
| 0 | 0 | 0 | 1 |
| 0 | 0 | 0 | 1 |
| 0 | 0 | 0 | 1 |
| 0 | 0 | 0 | 1 |
| 0 | 0 | 0 | 1 |
| 0 | 0 | 0 | 1 |
| 0 | 0 | 0 | 1 |
| 0 | 0 | 0 | 1 |
| 0 | 0 | 0 | 1 |
| 0 | 0 | 0 | 1 |
| 0 | 0 | 0 | 1 |
| 0 | 0 | 0 | 1 |
| 0 | 0 | 0 | 1 |
| 0 | 0 | 0 | 1 |
| 0 | 0 | 0 | 1 |
| 0 | 0 | 0 | 1 |
| 0 | 0 | 0 | 1 |
| 0 | 0 | 0 | 1 |
| 1 | 0 | 0 | 0 |
| 0 | 0 | 0 | 1 |
| 0 | 1 | 0 | 0 |
| 0 | 1 | 0 | 0 |
| 0 | 0 | 0 | 0 |
| 0 | 0 | 0 | 0 |
| 0 | 0 | 0 | 1 |
| 0 | 0 | 0 | 0 |
| 0 | 0 | 0 | 1 |
| 0 | 0 | 0 | 1 |
| 1 | 0 | 0 | 0 |
| 1 | 0 | 0 | 0 |
| 0 | 0 | 0 | 0 |
| 0 | 0 | 0 | 1 |
| 0 | 0 | 0 | 0 |
| 0 | 0 | 0 | 0 |
| 0 | 0 | 0 | 1 |
| 0 | 0 | 0 | 1 |
| 0 | 0 | 0 | 1 |
| 0 | 0 | 0 | 1 |
| 0 | 0 | 0 | 0 |
| 0 | 0 | 0 | 1 |
| 0 | 0 | 0 | 1 |
| 0 | 0 | 0 | 1 |

| | | | |
|---|---|---|---|
| 0 | 0 | 0 | 1 |
| 0 | 0 | 0 | 1 |
| 0 | 0 | 0 | 1 |
| 0 | 0 | 0 | 1 |
| 0 | 0 | 0 | 1 |
| 1 | 0 | 0 | 0 |
| 0 | 0 | 0 | 1 |
| 0 | 0 | 0 | 0 |
| 0 | 0 | 0 | 0 |
| 0 | 0 | 0 | 0 |
| 0 | 0 | 0 | 1 |
| 0 | 0 | 0 | 0 |
| 0 | 0 | 0 | 1 |
| 0 | 0 | 0 | 1 |
| 0 | 0 | 0 | 1 |
| 0 | 1 | 1 | 1 |
| 0 | 0 | 0 | 1 |
| 0 | 0 | 0 | 0 |
| 0 | 0 | 0 | 1 |
| 0 | 0 | 0 | 0 |
| 0 | 0 | 0 | 1 |
| 0 | 0 | 0 | 0 |
| 0 | 0 | 0 | 0 |
| 0 | 0 | 0 | 0 |
| 0 | 0 | 0 | 1 |
| 0 | 0 | 0 | 1 |
| 0 | 0 | 0 | 1 |
| 0 | 0 | 0 | 1 |
| 0 | 0 | 0 | 0 |
| 0 | 0 | 0 | 1 |
| 0 | 0 | 0 | 1 |
| 0 | 0 | 0 | 1 |
| 0 | 0 | 0 | 1 |
| 0 | 0 | 0 | 1 |
| 0 | 0 | 0 | 0 |
| 0 | 0 | 0 | 0 |
| 0 | 0 | 0 | 1 |
| 0 | 0 | 0 | 0 |
| 0 | 0 | 0 | 0 |
| 0 | 0 | 0 | 1 |
| 0 | 0 | 0 | 0 |
| 0 | 0 | 0 | 0 |
| 0 | 0 | 0 | 1 |
| 0 | 0 | 0 | 1 |
| 0 | 0 | 0 | 1 |
| 0 | 0 | 0 | 1 |
| 0 | 0 | 0 | 1 |
| 0 | 0 | 0 | 1 |
| 0 | 0 | 0 | 1 |
| 0 | 0 | 0 | 1 |
| 0 | 0 | 0 | 1 |
| 0 | 0 | 0 | 1 |
| 0 | 0 | 0 | 1 |
| 0 | 0 | 0 | 1 |

| | | | |
|---|---|---|---|
| 0 | 0 | 0 | 1 |
| 0 | 0 | 0 | 0 |
| 0 | 0 | 0 | 1 |
| 0 | 0 | 0 | 1 |
| 0 | 0 | 0 | 0 |
| 0 | 0 | 0 | 1 |
| 0 | 0 | 0 | 1 |
| 0 | 0 | 0 | 1 |
| 0 | 0 | 0 | 0 |
| 0 | 0 | 0 | 0 |
| 0 | 0 | 0 | 1 |
| 0 | 0 | 0 | 0 |
| 0 | 0 | 0 | 0 |
| 0 | 0 | 0 | 1 |
| 0 | 0 | 0 | 1 |
| 0 | 0 | 0 | 0 |
| 0 | 0 | 0 | 1 |
| 0 | 0 | 0 | 0 |
| 0 | 0 | 0 | 0 |
| 0 | 0 | 0 | 1 |
| 0 | 0 | 0 | 0 |
| 0 | 0 | 0 | 0 |
| 1 | 0 | 0 | 0 |
| 0 | 0 | 0 | 1 |
| 0 | 0 | 0 | 1 |
| 0 | 0 | 0 | 0 |
| 0 | 0 | 0 | 1 |
| 0 | 0 | 0 | 1 |
| 0 | 0 | 0 | 1 |
| 0 | 0 | 0 | 1 |
| 0 | 0 | 0 | 0 |
| 0 | 0 | 0 | 1 |
| 0 | 0 | 0 | 0 |
| 0 | 0 | 0 | 1 |
| 0 | 0 | 0 | 0 |
| 0 | 0 | 0 | 0 |
| 0 | 0 | 0 | 1 |
| 0 | 0 | 0 | 1 |
| 0 | 0 | 0 | 1 |
| 0 | 0 | 0 | 1 |
| 0 | 0 | 0 | 1 |
| 0 | 0 | 0 | 1 |
| 0 | 0 | 0 | 1 |
| 0 | 0 | 0 | 1 |
| 0 | 0 | 0 | 1 |
| 0 | 0 | 0 | 1 |
| 0 | 1 | 1 | 1 |
| 0 | 0 | 0 | 1 |
| 0 | 0 | 0 | 1 |
| 0 | 0 | 0 | 1 |
| 0 | 0 | 0 | 1 |
| 0 | 0 | 0 | 1 |
| 0 | 0 | 0 | 1 |
| 0 | 0 | 0 | 0 |

| | | | |
|---|---|---|---|
| 0 | 0 | 0 | 1 |
| 0 | 0 | 0 | 1 |
| 0 | 0 | 0 | 1 |
| 0 | 0 | 0 | 1 |
| 0 | 0 | 0 | 0 |
| 0 | 0 | 0 | 1 |
| 0 | 0 | 0 | 1 |
| 0 | 0 | 0 | 1 |
| 0 | 0 | 0 | 1 |
| 0 | 0 | 0 | 1 |
| 0 | 0 | 0 | 0 |
| 0 | 0 | 0 | 1 |
| 0 | 0 | 0 | 1 |
| 0 | 0 | 0 | 1 |
| 0 | 0 | 0 | 0 |
| 0 | 0 | 0 | 0 |
| 0 | 0 | 0 | 0 |
| 0 | 0 | 0 | 0 |
| 0 | 0 | 0 | 1 |
| 0 | 0 | 0 | 0 |
| 0 | 0 | 0 | 0 |
| 1 | 0 | 0 | 0 |
| 0 | 0 | 0 | 1 |
| 0 | 0 | 0 | 0 |
| 1 | 0 | 0 | 0 |
| 0 | 0 | 0 | 1 |
| 1 | 0 | 0 | 0 |
| 0 | 0 | 0 | 0 |
| 1 | 0 | 0 | 0 |
| 1 | 0 | 0 | 0 |
| 1 | 0 | 0 | 0 |
| 0 | 0 | 0 | 1 |
| 0 | 0 | 0 | 1 |
| 0 | 0 | 0 | 1 |
| 0 | 0 | 0 | 1 |
| 0 | 0 | 0 | 1 |
| 0 | 0 | 0 | 1 |
| 0 | 1 | 0 | 0 |
| 0 | 0 | 0 | 0 |
| 0 | 0 | 0 | 1 |
| 0 | 0 | 0 | 1 |
| 0 | 0 | 0 | 1 |
| 0 | 0 | 0 | 0 |
| 0 | 0 | 0 | 1 |
| 0 | 0 | 0 | 0 |
| 0 | 0 | 0 | 1 |
| 0 | 0 | 0 | 1 |
| 0 | 0 | 0 | 1 |
| 0 | 1 | 0 | 0 |
| 0 | 1 | 0 | 0 |
| 0 | 1 | 0 | 0 |
| 0 | 0 | 0 | 1 |
| 0 | 0 | 0 | 1 |
| 0 | 0 | 0 | 0 |

| | | | |
|---|---|---|---|
| 0 | 0 | 0 | 1 |
| 0 | 0 | 0 | 0 |
| 0 | 0 | 0 | 1 |
| 0 | 1 | 0 | 0 |
| 0 | 1 | 0 | 0 |
| 0 | 0 | 0 | 1 |
| 0 | 1 | 0 | 0 |
| 0 | 1 | 0 | 0 |
| 0 | 0 | 0 | 1 |
| 0 | 0 | 0 | 1 |
| 0 | 0 | 0 | 1 |
| 0 | 0 | 0 | 1 |
| 0 | 0 | 0 | 1 |
| 0 | 0 | 0 | 1 |
| 0 | 0 | 0 | 1 |
| 0 | 0 | 0 | 1 |
| 0 | 0 | 0 | 1 |
| 0 | 0 | 0 | 1 |
| 0 | 0 | 0 | 1 |
| 0 | 0 | 0 | 1 |
| 1 | 1 | 0 | 0 |
| 0 | 0 | 0 | 1 |
| 0 | 0 | 0 | 1 |
| 0 | 0 | 0 | 1 |
| 0 | 0 | 0 | 1 |
| 0 | 0 | 0 | 0 |
| 0 | 0 | 0 | 0 |
| 0 | 0 | 0 | 1 |
| 0 | 0 | 0 | 1 |
| 0 | 0 | 0 | 1 |
| 0 | 0 | 0 | 1 |
| 0 | 0 | 0 | 1 |
| 0 | 0 | 0 | 1 |
| 0 | 0 | 0 | 1 |
| 0 | 0 | 0 | 1 |
| 0 | 0 | 0 | 1 |
| 0 | 0 | 0 | 1 |
| 0 | 0 | 0 | 1 |
| 0 | 0 | 0 | 1 |
| 0 | 0 | 0 | 1 |
| 0 | 0 | 0 | 1 |
| 0 | 0 | 0 | 1 |
| 0 | 0 | 0 | 1 |
| 0 | 0 | 0 | 0 |
| 0 | 0 | 0 | 1 |
| 0 | 0 | 0 | 0 |
| 0 | 0 | 0 | 0 |
| 0 | 0 | 0 | 0 |
| 0 | 0 | 0 | 0 |
| 0 | 0 | 0 | 0 |
| 0 | 0 | 0 | 0 |
| 0 | 0 | 0 | 1 |
| 0 | 0 | 0 | 1 |
| 0 | 0 | 0 | 1 |

| | | | |
|---|---|---|---|
| 0 | 0 | 0 | 1 |
| 0 | 0 | 0 | 1 |
| 0 | 0 | 0 | 1 |
| 0 | 0 | 0 | 0 |
| 0 | 0 | 0 | 1 |
| 0 | 0 | 0 | 0 |
| 0 | 0 | 0 | 1 |
| 0 | 0 | 0 | 1 |
| 0 | 0 | 0 | 0 |
| 0 | 0 | 0 | 1 |
| 0 | 0 | 0 | 1 |
| 0 | 0 | 0 | 1 |
| 0 | 0 | 0 | 1 |
| 0 | 0 | 0 | 1 |
| 0 | 0 | 0 | 1 |
| 0 | 0 | 0 | 0 |
| 0 | 0 | 0 | 0 |
| 0 | 0 | 0 | 0 |
| 0 | 0 | 0 | 0 |
| 0 | 0 | 0 | 0 |
| 0 | 0 | 0 | 1 |
| 0 | 0 | 0 | 1 |
| 0 | 0 | 0 | 1 |
| 0 | 0 | 0 | 1 |
| 0 | 0 | 0 | 0 |
| 0 | 0 | 0 | 1 |
| 0 | 0 | 0 | 1 |
| 0 | 0 | 0 | 1 |
| 0 | 0 | 0 | 0 |
| 0 | 0 | 0 | 1 |
| 0 | 0 | 0 | 0 |
| 0 | 0 | 0 | 0 |
| 0 | 0 | 0 | 0 |
| 0 | 1 | 0 | 0 |
| 0 | 1 | 0 | 0 |
| 0 | 0 | 0 | 1 |
| 0 | 0 | 0 | 0 |
| 0 | 0 | 0 | 0 |
| 0 | 1 | 0 | 0 |
| 0 | 1 | 0 | 0 |
| 0 | 1 | 0 | 0 |
| 0 | 0 | 0 | 0 |
| 0 | 0 | 0 | 0 |
| 0 | 0 | 0 | 0 |
| 0 | 0 | 0 | 0 |
| 0 | 0 | 0 | 0 |
| 0 | 0 | 0 | 0 |
| 0 | 0 | 0 | 0 |
| 0 | 0 | 0 | 1 |
| 0 | 0 | 0 | 0 |
| 0 | 0 | 0 | 1 |
| 0 | 0 | 0 | 1 |
| 0 | 0 | 0 | 1 |
| 0 | 0 | 0 | 0 |

| | | | |
|---|---|---|---|
| 0 | 0 | 0 | 0 |
| 0 | 0 | 0 | 1 |
| 0 | 0 | 0 | 1 |
| 0 | 0 | 0 | 1 |
| 0 | 0 | 0 | 1 |
| 0 | 0 | 0 | 0 |
| 0 | 0 | 0 | 1 |
| 0 | 0 | 0 | 1 |
| 0 | 0 | 0 | 1 |
| 0 | 0 | 0 | 1 |
| 0 | 0 | 0 | 1 |
| 0 | 0 | 0 | 1 |
| 0 | 0 | 0 | 1 |
| 1 | 0 | 0 | 0 |
| 0 | 0 | 0 | 1 |
| 0 | 0 | 0 | 1 |
| 0 | 0 | 0 | 1 |
| 0 | 0 | 0 | 0 |
| 0 | 0 | 0 | 1 |
| 0 | 0 | 0 | 1 |
| 0 | 0 | 0 | 1 |
| 0 | 0 | 0 | 1 |
| 0 | 0 | 0 | 0 |
| 0 | 0 | 0 | 1 |
| 0 | 0 | 0 | 1 |
| 0 | 0 | 0 | 0 |
| 0 | 0 | 0 | 0 |
| 0 | 0 | 0 | 0 |
| 0 | 0 | 0 | 1 |
| 0 | 0 | 0 | 1 |
| 0 | 0 | 0 | 1 |
| 0 | 0 | 0 | 1 |
| 0 | 0 | 0 | 1 |
| 0 | 0 | 0 | 1 |
| 0 | 0 | 0 | 1 |
| 0 | 0 | 0 | 1 |
| 0 | 0 | 0 | 1 |
| 0 | 0 | 0 | 1 |
| 0 | 0 | 0 | 1 |
| 0 | 0 | 0 | 1 |
| 0 | 0 | 0 | 1 |
| 0 | 0 | 0 | 1 |
| 0 | 0 | 0 | 1 |
| 0 | 0 | 0 | 1 |
| 0 | 0 | 0 | 1 |
| 0 | 0 | 0 | 1 |
| 0 | 0 | 0 | 1 |
| 0 | 0 | 0 | 1 |
| 0 | 0 | 0 | 1 |
| 0 | 0 | 0 | 1 |
| 0 | 0 | 0 | 1 |
| 0 | 0 | 0 | 1 |
| 0 | 0 | 0 | 1 |

| | | | |
|---|---|---|---|
| 0 | 0 | 0 | 1 |
| 0 | 0 | 0 | 0 |
| 0 | 0 | 0 | 1 |
| 0 | 0 | 0 | 1 |
| 0 | 0 | 0 | 1 |
| 0 | 0 | 0 | 0 |
| 0 | 0 | 0 | 0 |
| 0 | 0 | 0 | 0 |
| 0 | 0 | 0 | 1 |
| 0 | 0 | 0 | 1 |
| 0 | 0 | 0 | 1 |
| 0 | 0 | 0 | 1 |
| 0 | 0 | 0 | 1 |
| 0 | 0 | 0 | 1 |
| 0 | 0 | 0 | 0 |
| 0 | 0 | 0 | 0 |
| 0 | 0 | 0 | 0 |
| 0 | 0 | 0 | 1 |
| 0 | 0 | 0 | 0 |
| 0 | 0 | 0 | 0 |
| 0 | 0 | 0 | 0 |
| 0 | 0 | 0 | 0 |
| 0 | 0 | 0 | 0 |
| 0 | 0 | 0 | 0 |
| 0 | 0 | 0 | 1 |
| 0 | 0 | 0 | 1 |
| 0 | 0 | 0 | 1 |
| 0 | 0 | 0 | 1 |
| 0 | 0 | 0 | 1 |
| 0 | 0 | 0 | 1 |
| 0 | 0 | 0 | 1 |
| 0 | 0 | 0 | 1 |
| 0 | 0 | 0 | 0 |
| 0 | 0 | 0 | 1 |
| 0 | 0 | 0 | 1 |
| 0 | 0 | 0 | 0 |
| 0 | 0 | 0 | 0 |
| 0 | 0 | 0 | 0 |
| 0 | 0 | 0 | 0 |
| 0 | 0 | 0 | 1 |
| 0 | 0 | 0 | 1 |
| 0 | 0 | 0 | 1 |
| 0 | 0 | 0 | 1 |
| 0 | 0 | 0 | 0 |
| 0 | 0 | 0 | 0 |
| 0 | 0 | 0 | 0 |
| 0 | 0 | 0 | 1 |
| 0 | 0 | 0 | 1 |
| 0 | 0 | 0 | 0 |
| 0 | 0 | 0 | 1 |
| 0 | 0 | 0 | 1 |
| 0 | 0 | 0 | 0 |
| 0 | 0 | 0 | 1 |
| 0 | 0 | 0 | 1 |

| | | | |
|---|---|---|---|
| 0 | 0 | 0 | 1 |
| 0 | 0 | 0 | 1 |
| 0 | 0 | 0 | 1 |
| 0 | 0 | 0 | 1 |
| 0 | 0 | 0 | 1 |
| 0 | 0 | 0 | 1 |
| 0 | 0 | 0 | 1 |
| 0 | 0 | 0 | 1 |
| 0 | 0 | 0 | 0 |
| 0 | 0 | 0 | 0 |
| 0 | 0 | 0 | 1 |
| 0 | 0 | 0 | 0 |
| 0 | 0 | 0 | 0 |
| 0 | 0 | 0 | 0 |
| 0 | 0 | 0 | 1 |
| 0 | 0 | 0 | 1 |
| 0 | 0 | 0 | 0 |
| 0 | 0 | 0 | 0 |
| 0 | 0 | 0 | 0 |
| 0 | 0 | 0 | 1 |
| 0 | 0 | 0 | 1 |
| 0 | 0 | 0 | 1 |
| 0 | 0 | 0 | 0 |
| 0 | 0 | 0 | 1 |
| 0 | 0 | 0 | 0 |
| 0 | 0 | 0 | 1 |
| 0 | 0 | 0 | 1 |
| 0 | 0 | 0 | 1 |
| 0 | 0 | 0 | 1 |
| 0 | 0 | 0 | 0 |
| 0 | 0 | 0 | 0 |
| 0 | 0 | 0 | 0 |
| 0 | 0 | 0 | 1 |
| 0 | 0 | 0 | 1 |
| 0 | 0 | 0 | 1 |
| 0 | 0 | 0 | 1 |
| 0 | 0 | 0 | 1 |
| 0 | 0 | 0 | 1 |
| 0 | 0 | 0 | 1 |
| 0 | 0 | 0 | 1 |
| 0 | 0 | 0 | 0 |
| 0 | 0 | 0 | 1 |
| 0 | 0 | 0 | 1 |
| 0 | 0 | 0 | 1 |
| 0 | 0 | 0 | 1 |
| 0 | 0 | 0 | 0 |
| 0 | 1 | 1 | 1 |
| 0 | 0 | 0 | 0 |
| 0 | 0 | 0 | 1 |
| 0 | 0 | 0 | 0 |
| 0 | 0 | 0 | 0 |
| 0 | 0 | 0 | 0 |
| 0 | 0 | 0 | 1 |
| 0 | 1 | 1 | 1 |

| | | | |
|---|---|---|---|
| 0 | 0 | 0 | 1 |
| 0 | 1 | 1 | 1 |
| 0 | 0 | 0 | 1 |
| 0 | 0 | 0 | 1 |
| 0 | 0 | 0 | 1 |
| 0 | 0 | 0 | 1 |
| 0 | 0 | 0 | 1 |
| 0 | 0 | 0 | 0 |
| 0 | 0 | 0 | 0 |
| 0 | 0 | 0 | 1 |
| 0 | 0 | 0 | 0 |
| 0 | 0 | 0 | 1 |
| 0 | 0 | 0 | 1 |
| 0 | 0 | 0 | 1 |
| 0 | 0 | 0 | 1 |
| 0 | 0 | 0 | 1 |
| 0 | 0 | 0 | 1 |
| 0 | 0 | 0 | 1 |
| 0 | 0 | 0 | 1 |
| 0 | 0 | 0 | 1 |
| 0 | 0 | 0 | 1 |
| 0 | 0 | 0 | 1 |
| 0 | 0 | 0 | 1 |
| 0 | 0 | 0 | 1 |
| 0 | 0 | 0 | 1 |
| 0 | 0 | 0 | 1 |
| 0 | 0 | 0 | 1 |
| 0 | 0 | 0 | 1 |
| 0 | 0 | 0 | 0 |
| 0 | 0 | 0 | 1 |
| 0 | 0 | 0 | 0 |
| 0 | 0 | 0 | 1 |
| 0 | 0 | 0 | 0 |
| 0 | 0 | 0 | 0 |
| 0 | 0 | 0 | 1 |
| 0 | 0 | 0 | 0 |
| 0 | 0 | 0 | 0 |
| 0 | 0 | 0 | 1 |
| 0 | 0 | 0 | 1 |
| 0 | 0 | 0 | 1 |
| 0 | 0 | 0 | 1 |
| 0 | 0 | 0 | 0 |
| 0 | 0 | 0 | 0 |
| 0 | 0 | 0 | 1 |
| 0 | 0 | 0 | 1 |
| 0 | 0 | 0 | 0 |
| 0 | 0 | 0 | 0 |
| 0 | 0 | 0 | 0 |
| 0 | 0 | 0 | 0 |
| 0 | 0 | 0 | 1 |
| 0 | 0 | 0 | 0 |
| 0 | 0 | 0 | 0 |
| 0 | 0 | 0 | 1 |
| 0 | 0 | 0 | 1 |

| | | | |
|---|---|---|---|
| 0 | 0 | 0 | 1 |
| 0 | 0 | 0 | 1 |
| 0 | 0 | 0 | 1 |
| 0 | 0 | 0 | 1 |
| 0 | 0 | 0 | 0 |
| 0 | 0 | 0 | 0 |
| 0 | 0 | 0 | 0 |
| 0 | 0 | 0 | 0 |
| 0 | 0 | 0 | 1 |
| 0 | 0 | 0 | 0 |
| 0 | 0 | 0 | 0 |
| 0 | 0 | 0 | 0 |
| 0 | 0 | 0 | 0 |
| 0 | 0 | 0 | 1 |
| 0 | 0 | 0 | 1 |
| 0 | 0 | 0 | 1 |
| 0 | 0 | 0 | 1 |
| 0 | 0 | 0 | 1 |
| 0 | 0 | 0 | 1 |
| 0 | 0 | 0 | 1 |
| 0 | 0 | 0 | 1 |
| 0 | 0 | 0 | 1 |
| 0 | 0 | 0 | 1 |
| 0 | 0 | 0 | 1 |
| 0 | 0 | 0 | 1 |
| 0 | 0 | 0 | 0 |
| 0 | 0 | 0 | 0 |
| 0 | 0 | 0 | 0 |
| 0 | 0 | 0 | 0 |
| 0 | 0 | 0 | 1 |
| 0 | 0 | 0 | 1 |
| 0 | 0 | 0 | 1 |
| 0 | 0 | 0 | 1 |
| 0 | 0 | 0 | 1 |
| 0 | 0 | 0 | 1 |
| 0 | 0 | 0 | 1 |
| 0 | 0 | 0 | 1 |
| 0 | 0 | 0 | 1 |
| 0 | 0 | 0 | 1 |
| 0 | 0 | 0 | 1 |
| 0 | 0 | 0 | 1 |
| 0 | 0 | 0 | 1 |
| 0 | 0 | 0 | 1 |
| 0 | 0 | 0 | 1 |
| 0 | 0 | 0 | 1 |
| 0 | 0 | 0 | 1 |
| 0 | 0 | 0 | 1 |
| 0 | 0 | 0 | 1 |
| 0 | 0 | 0 | 1 |
| 0 | 0 | 0 | 1 |
| 0 | 0 | 0 | 1 |
| 0 | 0 | 0 | 1 |
| 0 | 0 | 0 | 1 |
| 0 | 0 | 0 | 0 |

| | | | |
|---|---|---|---|
| 0 | 0 | 0 | 1 |
| 0 | 0 | 0 | 1 |
| 0 | 0 | 0 | 1 |
| 0 | 0 | 0 | 1 |
| 0 | 0 | 0 | 1 |
| 0 | 0 | 0 | 1 |
| 0 | 0 | 0 | 1 |
| 0 | 0 | 0 | 1 |
| 0 | 0 | 0 | 1 |
| 0 | 0 | 0 | 1 |
| 0 | 0 | 0 | 1 |
| 0 | 0 | 0 | 1 |
| 0 | 0 | 0 | 1 |
| 0 | 0 | 0 | 1 |
| 0 | 0 | 0 | 1 |
| 0 | 0 | 0 | 1 |
| 0 | 0 | 0 | 1 |
| 0 | 0 | 0 | 1 |
| 0 | 0 | 0 | 1 |
| 0 | 0 | 0 | 1 |
| 0 | 0 | 0 | 1 |
| 0 | 0 | 0 | 1 |
| 0 | 0 | 0 | 1 |
| 0 | 0 | 0 | 1 |
| 0 | 0 | 0 | 1 |
| 0 | 0 | 0 | 1 |
| 0 | 0 | 0 | 1 |
| 0 | 0 | 0 | 1 |
| 0 | 0 | 0 | 1 |
| 0 | 0 | 0 | 1 |
| 0 | 0 | 0 | 1 |
| 0 | 0 | 0 | 1 |
| 0 | 0 | 0 | 1 |
| 0 | 0 | 0 | 1 |
| 0 | 0 | 0 | 1 |
| 0 | 0 | 0 | 1 |
| 0 | 0 | 0 | 1 |
| 0 | 0 | 0 | 1 |
| 0 | 0 | 0 | 1 |
| 0 | 0 | 0 | 1 |
| 0 | 0 | 0 | 1 |
| 0 | 0 | 0 | 1 |
| 0 | 0 | 0 | 1 |
| 0 | 0 | 0 | 1 |
| 0 | 0 | 0 | 1 |
| 0 | 0 | 0 | 1 |
| 0 | 1 | 1 | 1 |
| 0 | 0 | 0 | 0 |
| 0 | 0 | 0 | 1 |
| 0 | 0 | 0 | 1 |
| 0 | 0 | 0 | 1 |
| 0 | 0 | 0 | 1 |
| 0 | 0 | 0 | 1 |
| 0 | 0 | 0 | 1 |
| 0 | 0 | 0 | 1 |

| | | | |
|---|---|---|---|
| 0 | 0 | 0 | 1 |
| 0 | 0 | 0 | 1 |
| 0 | 0 | 0 | 0 |
| 0 | 0 | 0 | 1 |
| 0 | 0 | 0 | 1 |
| 0 | 0 | 0 | 1 |
| 0 | 0 | 0 | 0 |
| 0 | 0 | 0 | 1 |
| 0 | 0 | 0 | 1 |
| 0 | 0 | 0 | 1 |
| 0 | 0 | 0 | 1 |
| 0 | 0 | 0 | 0 |
| 0 | 0 | 0 | 1 |
| 0 | 0 | 0 | 0 |
| 0 | 0 | 0 | 0 |
| 0 | 0 | 0 | 0 |
| 0 | 0 | 0 | 0 |
| 0 | 0 | 0 | 0 |
| 0 | 0 | 0 | 0 |
| 0 | 0 | 0 | 0 |
| 0 | 0 | 0 | 0 |

| surface_non_binding_site | non_binding_site_single_water | surface_non_binding_site_single_water |
|---|---|---|
| 0 | 0 | 0 |
| 0 | 0 | 0 |
| 0 | 0 | 0 |
| 0 | 0 | 0 |
| 0 | 0 | 0 |
| 0 | 0 | 0 |
| 1 | 1 | 1 |
| 1 | 1 | 1 |
| 0 | 0 | 0 |
| 0 | 0 | 0 |
| 0 | 0 | 0 |
| 0 | 0 | 0 |
| 0 | 0 | 0 |
| 0 | 0 | 0 |
| 0 | 0 | 0 |
| 0 | 0 | 0 |
| 0 | 0 | 0 |
| 0 | 0 | 0 |
| 0 | 0 | 0 |
| 0 | 0 | 0 |
| 0 | 0 | 0 |
| 0 | 0 | 0 |
| 0 | 0 | 0 |
| 0 | 0 | 0 |
| 0 | 0 | 0 |
| 1 | 0 | 0 |
| 0 | 0 | 0 |
| 1 | 1 | 1 |
| 0 | 0 | 0 |
| 0 | 1 | 0 |
| 0 | 1 | 0 |
| 0 | 0 | 0 |
| 0 | 0 | 0 |
| 0 | 0 | 0 |
| 0 | 0 | 0 |
| 1 | 1 | 1 |
| 0 | 0 | 0 |
| 0 | 0 | 0 |
| 0 | 0 | 0 |
| 0 | 0 | 0 |
| 0 | 0 | 0 |
| 0 | 0 | 0 |
| 0 | 0 | 0 |
| 0 | 0 | 0 |
| 0 | 0 | 0 |
| 0 | 0 | 0 |
| 0 | 0 | 0 |
| 0 | 0 | 0 |
| 0 | 0 | 0 |
| 0 | 0 | 0 |
| 0 | 0 | 0 |
| 0 | 0 | 0 |
| 0 | 0 | 0 |

| | | |
|---|---|---|
| 0 | 0 | 0 |
| 0 | 0 | 0 |
| 0 | 0 | 0 |
| 0 | 0 | 0 |
| 0 | 0 | 0 |
| 0 | 0 | 0 |
| 0 | 0 | 0 |
| 0 | 0 | 0 |
| 0 | 0 | 0 |
| 0 | 0 | 0 |
| 0 | 0 | 0 |
| 0 | 0 | 0 |
| 0 | 0 | 0 |
| 0 | 0 | 0 |
| 0 | 0 | 0 |
| 0 | 0 | 0 |
| 0 | 0 | 0 |
| 0 | 0 | 0 |
| 0 | 0 | 0 |
| 0 | 0 | 0 |
| 0 | 0 | 0 |
| 0 | 0 | 0 |
| 0 | 0 | 0 |
| 0 | 0 | 0 |
| 0 | 0 | 0 |
| 0 | 0 | 0 |
| 0 | 0 | 0 |
| 0 | 0 | 0 |
| 0 | 0 | 0 |
| 0 | 0 | 0 |
| 0 | 0 | 0 |
| 0 | 0 | 0 |
| 0 | 0 | 0 |
| 0 | 0 | 0 |
| 0 | 0 | 0 |
| 0 | 0 | 0 |
| 0 | 0 | 0 |
| 0 | 0 | 0 |
| 0 | 0 | 0 |
| 0 | 0 | 0 |
| 0 | 0 | 0 |
| 0 | 0 | 0 |
| 0 | 0 | 0 |
| 0 | 0 | 0 |
| 0 | 0 | 0 |
| 1 | 1 | 1 |
| 0 | 1 | 0 |
| 0 | 0 | 0 |
| 0 | 0 | 0 |
| 0 | 0 | 0 |
| 0 | 0 | 0 |
| 1 | 1 | 1 |
| 1 | 1 | 1 |
| 0 | 0 | 0 |

| | | |
|---|---|---|
| 0 | 1 | 0 |
| 0 | 1 | 0 |
| 0 | 0 | 0 |
| 0 | 0 | 0 |
| 0 | 0 | 0 |
| 0 | 0 | 0 |
| 0 | 1 | 0 |
| 0 | 0 | 0 |
| 0 | 0 | 0 |
| 0 | 0 | 0 |
| 0 | 0 | 0 |
| 0 | 0 | 0 |
| 0 | 0 | 0 |
| 0 | 0 | 0 |
| 0 | 0 | 0 |
| 1 | 1 | 1 |
| 0 | 0 | 0 |
| 0 | 0 | 0 |
| 0 | 0 | 0 |
| 0 | 0 | 0 |
| 0 | 0 | 0 |
| 0 | 0 | 0 |
| 0 | 0 | 0 |
| 0 | 0 | 0 |
| 0 | 0 | 0 |
| 0 | 0 | 0 |
| 0 | 0 | 0 |
| 0 | 0 | 0 |
| 0 | 0 | 0 |
| 0 | 0 | 0 |
| 0 | 0 | 0 |
| 0 | 0 | 0 |
| 0 | 0 | 0 |
| 0 | 0 | 0 |
| 0 | 0 | 0 |
| 0 | 0 | 0 |
| 0 | 0 | 0 |
| 0 | 0 | 0 |
| 0 | 0 | 0 |
| 0 | 0 | 0 |
| 0 | 0 | 0 |
| 0 | 0 | 0 |
| 0 | 0 | 0 |
| 0 | 0 | 0 |
| 0 | 0 | 0 |
| 0 | 0 | 0 |
| 0 | 0 | 0 |
| 0 | 0 | 0 |
| 0 | 0 | 0 |
| 0 | 0 | 0 |
| 0 | 0 | 0 |
| 0 | 0 | 0 |
| 0 | 0 | 0 |
| 0 | 0 | 0 |

| | | |
|---|---|---|
| 1 | 0 | 0 |
| 1 | 0 | 0 |
| 1 | 0 | 0 |
| 0 | 0 | 0 |
| 0 | 0 | 0 |
| 0 | 0 | 0 |
| 0 | 0 | 0 |
| 1 | 0 | 0 |
| 0 | 0 | 0 |
| 1 | 0 | 0 |
| 1 | 0 | 0 |
| 0 | 0 | 0 |
| 1 | 0 | 0 |
| 0 | 0 | 0 |
| 0 | 0 | 0 |
| 1 | 0 | 0 |
| 1 | 0 | 0 |
| 1 | 0 | 0 |
| 1 | 0 | 0 |
| 0 | 0 | 0 |
| 0 | 0 | 0 |
| 0 | 1 | 0 |
| 0 | 0 | 0 |
| 0 | 0 | 0 |
| 0 | 0 | 0 |
| 1 | 0 | 0 |
| 0 | 0 | 0 |
| 0 | 0 | 0 |
| 1 | 0 | 0 |
| 1 | 0 | 0 |
| 1 | 1 | 1 |
| 0 | 0 | 0 |
| 0 | 0 | 0 |
| 0 | 1 | 0 |
| 0 | 0 | 0 |
| 0 | 0 | 0 |
| 0 | 0 | 0 |
| 0 | 0 | 0 |
| 0 | 0 | 0 |
| 1 | 0 | 0 |
| 1 | 0 | 0 |
| 0 | 0 | 0 |
| 0 | 0 | 0 |
| 1 | 1 | 1 |
| 1 | 0 | 0 |
| 1 | 0 | 0 |
| 0 | 0 | 0 |
| 0 | 0 | 0 |
| 0 | 0 | 0 |
| 0 | 0 | 0 |
| 0 | 0 | 0 |
| 0 | 1 | 0 |
| 1 | 0 | 0 |
| 0 | 0 | 0 |

| | | |
|---|---|---|
| 0 | 0 | 0 |
| 0 | 0 | 0 |
| 0 | 0 | 0 |
| 1 | 1 | 1 |
| 1 | 1 | 1 |
| 1 | 0 | 0 |
| 0 | 0 | 0 |
| 0 | 0 | 0 |
| 1 | 0 | 0 |
| 1 | 0 | 0 |
| 0 | 0 | 0 |
| 0 | 1 | 0 |
| 0 | 0 | 0 |
| 1 | 0 | 0 |
| 0 | 1 | 0 |
| 1 | 1 | 1 |
| 1 | 1 | 1 |
| 0 | 0 | 0 |
| 0 | 0 | 0 |
| 0 | 0 | 0 |
| 0 | 0 | 0 |
| 1 | 0 | 0 |
| 0 | 0 | 0 |
| 0 | 1 | 0 |
| 1 | 1 | 1 |
| 0 | 0 | 0 |
| 0 | 0 | 0 |
| 0 | 0 | 0 |
| 0 | 0 | 0 |
| 0 | 0 | 0 |
| 0 | 0 | 0 |
| 0 | 0 | 0 |
| 1 | 0 | 0 |
| 1 | 0 | 0 |
| 0 | 0 | 0 |
| 1 | 0 | 0 |
| 0 | 0 | 0 |
| 0 | 0 | 0 |
| 1 | 0 | 0 |
| 0 | 1 | 0 |
| 0 | 0 | 0 |
| 1 | 0 | 0 |
| 1 | 0 | 0 |
| 1 | 0 | 0 |
| 0 | 0 | 0 |
| 0 | 0 | 0 |
| 0 | 0 | 0 |
| 0 | 0 | 0 |
| 0 | 0 | 0 |
| 0 | 0 | 0 |
| 0 | 0 | 0 |
| 0 | 0 | 0 |
| 0 | 0 | 0 |
| 0 | 0 | 0 |

| | | |
|---|---|---|
| 0 | 0 | 0 |
| 0 | 0 | 0 |
| 0 | 0 | 0 |
| 0 | 0 | 0 |
| 0 | 0 | 0 |
| 0 | 0 | 0 |
| 0 | 1 | 0 |
| 0 | 1 | 0 |
| 0 | 0 | 0 |
| 0 | 0 | 0 |
| 0 | 0 | 0 |
| 0 | 0 | 0 |
| 0 | 0 | 0 |
| 0 | 1 | 0 |
| 0 | 1 | 0 |
| 0 | 0 | 0 |
| 0 | 1 | 0 |
| 0 | 0 | 0 |
| 1 | 1 | 1 |
| 0 | 0 | 0 |
| 1 | 1 | 1 |
| 1 | 0 | 0 |
| 1 | 1 | 1 |
| 0 | 0 | 0 |
| 0 | 0 | 0 |
| 0 | 0 | 0 |
| 0 | 0 | 0 |
| 0 | 0 | 0 |
| 0 | 0 | 0 |
| 0 | 0 | 0 |
| 0 | 0 | 0 |
| 1 | 0 | 0 |
| 0 | 0 | 0 |
| 0 | 0 | 0 |
| 0 | 0 | 0 |
| 0 | 0 | 0 |
| 0 | 0 | 0 |
| 0 | 0 | 0 |
| 0 | 0 | 0 |
| 0 | 0 | 0 |
| 0 | 1 | 0 |
| 0 | 1 | 0 |
| 0 | 0 | 0 |
| 0 | 1 | 0 |
| 0 | 0 | 0 |
| 0 | 0 | 0 |
| 0 | 0 | 0 |
| 1 | 1 | 1 |
| 1 | 1 | 1 |
| 0 | 0 | 0 |
| 0 | 1 | 0 |
| 1 | 1 | 1 |
| 0 | 0 | 0 |
| 0 | 1 | 0 |

| | | |
|---|---|---|
| 0 | 0 | 0 |
| 1 | 1 | 1 |
| 0 | 0 | 0 |
| 0 | 0 | 0 |
| 0 | 0 | 0 |
| 0 | 0 | 0 |
| 0 | 1 | 0 |
| 0 | 0 | 0 |
| 0 | 0 | 0 |
| 1 | 1 | 1 |
| 1 | 1 | 1 |
| 1 | 1 | 1 |
| 1 | 1 | 1 |
| 1 | 1 | 1 |
| 0 | 0 | 0 |
| 1 | 1 | 1 |
| 0 | 0 | 0 |
| 0 | 0 | 0 |
| 1 | 1 | 1 |
| 0 | 0 | 0 |
| 0 | 0 | 0 |
| 0 | 0 | 0 |
| 0 | 0 | 0 |
| 0 | 0 | 0 |
| 1 | 1 | 1 |
| 1 | 0 | 0 |
| 0 | 0 | 0 |
| 0 | 0 | 0 |
| 0 | 0 | 0 |
| 0 | 0 | 0 |
| 0 | 0 | 0 |
| 0 | 0 | 0 |
| 0 | 0 | 0 |
| 0 | 0 | 0 |
| 1 | 1 | 1 |
| 1 | 1 | 1 |
| 0 | 0 | 0 |
| 0 | 0 | 0 |
| 0 | 0 | 0 |
| 0 | 0 | 0 |
| 0 | 0 | 0 |
| 0 | 0 | 0 |
| 0 | 1 | 0 |
| 0 | 0 | 0 |
| 0 | 0 | 0 |
| 0 | 0 | 0 |
| 0 | 0 | 0 |
| 0 | 0 | 0 |
| 0 | 0 | 0 |
| 1 | 1 | 1 |
| 0 | 0 | 0 |
| 0 | 0 | 0 |
| 0 | 0 | 0 |
| 0 | 0 | 0 |

| | | |
|---|---|---|
| 0 | 0 | 0 |
| 0 | 0 | 0 |
| 0 | 0 | 0 |
| 0 | 0 | 0 |
| 1 | 1 | 1 |
| 0 | 0 | 0 |
| 0 | 0 | 0 |
| 0 | 0 | 0 |
| 0 | 0 | 0 |
| 0 | 0 | 0 |
| 0 | 0 | 0 |
| 0 | 0 | 0 |
| 0 | 0 | 0 |
| 0 | 0 | 0 |
| 0 | 0 | 0 |
| 0 | 0 | 0 |
| 0 | 0 | 0 |
| 0 | 0 | 0 |
| 1 | 1 | 1 |
| 0 | 0 | 0 |
| 0 | 0 | 0 |
| 0 | 0 | 0 |
| 0 | 0 | 0 |
| 1 | 0 | 0 |
| 0 | 0 | 0 |
| 0 | 0 | 0 |
| 0 | 0 | 0 |
| 1 | 0 | 0 |
| 1 | 0 | 0 |
| 1 | 0 | 0 |
| 0 | 0 | 0 |
| 0 | 0 | 0 |
| 0 | 0 | 0 |
| 0 | 0 | 0 |
| 0 | 0 | 0 |
| 1 | 0 | 0 |
| 1 | 0 | 0 |
| 0 | 0 | 0 |
| 0 | 0 | 0 |
| 1 | 0 | 0 |
| 1 | 0 | 0 |
| 1 | 0 | 0 |
| 0 | 1 | 0 |
| 1 | 1 | 1 |
| 0 | 0 | 0 |
| 0 | 0 | 0 |
| 0 | 0 | 0 |
| 0 | 0 | 0 |
| 0 | 0 | 0 |
| 0 | 1 | 0 |
| 1 | 1 | 1 |
| 0 | 0 | 0 |
| 1 | 1 | 1 |
| 0 | 0 | 0 |

| | | |
|---|---|---|
| 1 | 1 | 1 |
| 1 | 1 | 1 |
| 0 | 0 | 0 |
| 1 | 1 | 1 |
| 1 | 1 | 1 |
| 1 | 1 | 1 |
| 1 | 1 | 1 |
| 0 | 0 | 0 |
| 1 | 0 | 0 |
| 1 | 1 | 1 |
| 0 | 0 | 0 |
| 1 | 1 | 1 |
| 1 | 1 | 1 |
| 0 | 1 | 0 |
| 0 | 0 | 0 |
| 0 | 0 | 0 |
| 1 | 1 | 1 |
| 1 | 1 | 1 |
| 0 | 0 | 0 |
| 0 | 0 | 0 |
| 0 | 0 | 0 |
| 0 | 0 | 0 |
| 0 | 0 | 0 |
| 0 | 0 | 0 |
| 0 | 0 | 0 |
| 1 | 1 | 1 |
| 0 | 0 | 0 |
| 0 | 0 | 0 |
| 0 | 0 | 0 |
| 1 | 0 | 0 |
| 0 | 0 | 0 |
| 0 | 0 | 0 |
| 0 | 0 | 0 |
| 0 | 0 | 0 |
| 0 | 0 | 0 |
| 0 | 0 | 0 |
| 0 | 0 | 0 |
| 0 | 0 | 0 |
| 0 | 0 | 0 |
| 0 | 0 | 0 |
| 0 | 1 | 0 |
| 0 | 0 | 0 |
| 1 | 1 | 1 |
| 0 | 0 | 0 |
| 0 | 0 | 0 |
| 0 | 0 | 0 |
| 0 | 0 | 0 |
| 0 | 0 | 0 |
| 0 | 0 | 0 |
| 0 | 0 | 0 |
| 0 | 1 | 0 |
| 0 | 0 | 0 |
| 0 | 0 | 0 |
| 0 | 0 | 0 |

| | | |
|---|---|---|
| 1 | 1 | 1 |
| 0 | 0 | 0 |
| 0 | 1 | 0 |
| 0 | 1 | 0 |
| 0 | 0 | 0 |
| 0 | 0 | 0 |
| 0 | 0 | 0 |
| 0 | 0 | 0 |
| 0 | 0 | 0 |
| 0 | 1 | 0 |
| 1 | 1 | 1 |
| 1 | 1 | 1 |
| 1 | 1 | 1 |
| 0 | 0 | 0 |
| 1 | 1 | 1 |
| 0 | 0 | 0 |
| 0 | 0 | 0 |
| 0 | 0 | 0 |
| 0 | 0 | 0 |
| 1 | 0 | 0 |
| 0 | 0 | 0 |
| 1 | 1 | 1 |
| 0 | 0 | 0 |
| 0 | 1 | 0 |
| 1 | 1 | 1 |
| 1 | 1 | 1 |
| 0 | 0 | 0 |
| 1 | 1 | 1 |
| 1 | 1 | 1 |
| 1 | 1 | 1 |
| 0 | 0 | 0 |
| 0 | 0 | 0 |
| 1 | 1 | 1 |
| 1 | 1 | 1 |
| 0 | 0 | 0 |
| 0 | 0 | 0 |
| 0 | 0 | 0 |
| 0 | 0 | 0 |
| 0 | 0 | 0 |
| 1 | 1 | 1 |
| 0 | 0 | 0 |
| 0 | 0 | 0 |
| 0 | 0 | 0 |
| 0 | 0 | 0 |
| 1 | 0 | 0 |
| 0 | 0 | 0 |
| 0 | 0 | 0 |
| 0 | 0 | 0 |
| 0 | 0 | 0 |
| 0 | 0 | 0 |
| 0 | 0 | 0 |
| 0 | 0 | 0 |
| 1 | 0 | 0 |
| 0 | 0 | 0 |

| | | |
|---|---|---|
| 0 | 0 | 0 |
| 0 | 0 | 0 |
| 0 | 0 | 0 |
| 0 | 0 | 0 |
| 0 | 0 | 0 |
| 0 | 0 | 0 |
| 0 | 0 | 0 |
| 1 | 0 | 0 |
| 0 | 0 | 0 |
| 0 | 0 | 0 |
| 0 | 0 | 0 |
| 0 | 0 | 0 |
| 1 | 1 | 1 |
| 0 | 0 | 0 |
| 0 | 0 | 0 |
| 0 | 0 | 0 |
| 0 | 0 | 0 |
| 0 | 0 | 0 |
| 0 | 0 | 0 |
| 1 | 1 | 1 |
| 0 | 0 | 0 |
| 0 | 0 | 0 |
| 0 | 0 | 0 |
| 0 | 0 | 0 |
| 0 | 0 | 0 |
| 1 | 0 | 0 |
| 1 | 1 | 1 |
| 0 | 1 | 0 |
| 0 | 0 | 0 |
| 0 | 0 | 0 |
| 0 | 0 | 0 |
| 0 | 0 | 0 |
| 0 | 0 | 0 |
| 0 | 0 | 0 |
| 0 | 0 | 0 |
| 1 | 0 | 0 |
| 0 | 0 | 0 |
| 0 | 0 | 0 |
| 1 | 1 | 1 |
| 1 | 0 | 0 |
| 0 | 1 | 0 |
| 0 | 0 | 0 |
| 0 | 0 | 0 |
| 0 | 0 | 0 |
| 0 | 0 | 0 |
| 0 | 0 | 0 |
| 0 | 0 | 0 |
| 1 | 1 | 1 |
| 1 | 1 | 1 |
| 0 | 0 | 0 |
| 0 | 0 | 0 |
| 0 | 0 | 0 |
| 0 | 0 | 0 |
| 1 | 1 | 1 |

| | | |
|---|---|---|
| 1 | 1 | 1 |
| 0 | 0 | 0 |
| 0 | 1 | 0 |
| 0 | 0 | 0 |
| 1 | 0 | 0 |
| 1 | 0 | 0 |
| 0 | 0 | 0 |
| 1 | 0 | 0 |
| 1 | 0 | 0 |
| 1 | 0 | 0 |
| 1 | 0 | 0 |
| 1 | 0 | 0 |
| 1 | 1 | 1 |
| 1 | 1 | 1 |
| 1 | 1 | 1 |
| 1 | 1 | 1 |
| 0 | 0 | 0 |
| 0 | 0 | 0 |
| 1 | 0 | 0 |
| 0 | 0 | 0 |
| 0 | 0 | 0 |
| 0 | 0 | 0 |
| 1 | 0 | 0 |
| 0 | 0 | 0 |
| 0 | 0 | 0 |
| 1 | 1 | 1 |
| 1 | 0 | 0 |
| 1 | 0 | 0 |
| 0 | 0 | 0 |
| 0 | 0 | 0 |
| 0 | 0 | 0 |
| 0 | 0 | 0 |
| 0 | 0 | 0 |
| 0 | 0 | 0 |
| 0 | 0 | 0 |
| 0 | 0 | 0 |
| 1 | 0 | 0 |
| 1 | 1 | 1 |
| 1 | 1 | 1 |
| 1 | 1 | 1 |
| 0 | 0 | 0 |
| 1 | 1 | 1 |
| 0 | 0 | 0 |
| 1 | 1 | 1 |
| 0 | 0 | 0 |
| 0 | 1 | 0 |
| 0 | 0 | 0 |
| 0 | 0 | 0 |
| 0 | 0 | 0 |
| 0 | 0 | 0 |
| 0 | 0 | 0 |
| 0 | 0 | 0 |
| 0 | 0 | 0 |
| 0 | 0 | 0 |

| | | |
|---|---|---|
| 0 | 0 | 0 |
| 0 | 0 | 0 |
| 0 | 0 | 0 |
| 0 | 0 | 0 |
| 0 | 0 | 0 |
| 0 | 0 | 0 |
| 0 | 0 | 0 |
| 1 | 1 | 1 |
| 0 | 0 | 0 |
| 0 | 0 | 0 |
| 0 | 0 | 0 |
| 1 | 0 | 0 |
| 0 | 0 | 0 |
| 0 | 0 | 0 |
| 0 | 0 | 0 |
| 0 | 0 | 0 |
| 0 | 0 | 0 |
| 0 | 0 | 0 |
| 0 | 0 | 0 |
| 0 | 0 | 0 |
| 0 | 0 | 0 |
| 0 | 0 | 0 |
| 0 | 0 | 0 |
| 0 | 0 | 0 |
| 0 | 0 | 0 |
| 0 | 0 | 0 |
| 0 | 0 | 0 |
| 0 | 0 | 0 |
| 0 | 0 | 0 |
| 0 | 0 | 0 |
| 0 | 0 | 0 |
| 0 | 0 | 0 |
| 0 | 0 | 0 |
| 0 | 0 | 0 |
| 0 | 0 | 0 |
| 0 | 0 | 0 |
| 0 | 0 | 0 |
| 0 | 0 | 0 |
| 0 | 0 | 0 |
| 0 | 0 | 0 |
| 1 | 0 | 0 |
| 0 | 0 | 0 |
| 0 | 0 | 0 |
| 0 | 0 | 0 |
| 0 | 0 | 0 |
| 0 | 0 | 0 |
| 0 | 0 | 0 |
| 0 | 0 | 0 |
| 0 | 0 | 0 |
| 0 | 1 | 0 |
| 0 | 1 | 0 |
| 1 | 0 | 0 |
| 0 | 0 | 0 |
| 0 | 0 | 0 |

| | | |
|---|---|---|
| 0 | 0 | 0 |
| 0 | 0 | 0 |
| 0 | 0 | 0 |
| 0 | 0 | 0 |
| 0 | 0 | 0 |
| 0 | 0 | 0 |
| 0 | 0 | 0 |
| 1 | 0 | 0 |
| 1 | 0 | 0 |
| 0 | 0 | 0 |
| 0 | 0 | 0 |
| 1 | 0 | 0 |
| 0 | 0 | 0 |
| 0 | 0 | 0 |
| 0 | 0 | 0 |
| 0 | 0 | 0 |
| 0 | 0 | 0 |
| 1 | 0 | 0 |
| 0 | 0 | 0 |
| 1 | 0 | 0 |
| 0 | 0 | 0 |
| 0 | 0 | 0 |
| 1 | 1 | 1 |
| 0 | 0 | 0 |
| 0 | 0 | 0 |
| 1 | 0 | 0 |
| 0 | 0 | 0 |
| 0 | 0 | 0 |
| 0 | 0 | 0 |
| 0 | 0 | 0 |
| 0 | 0 | 0 |
| 0 | 0 | 0 |
| 1 | 0 | 0 |
| 0 | 0 | 0 |
| 0 | 0 | 0 |
| 0 | 0 | 0 |
| 1 | 1 | 1 |
| 0 | 0 | 0 |
| 0 | 0 | 0 |
| 0 | 0 | 0 |
| 0 | 0 | 0 |
| 0 | 0 | 0 |
| 0 | 0 | 0 |
| 0 | 0 | 0 |
| 0 | 0 | 0 |
| 0 | 0 | 0 |
| 0 | 0 | 0 |
| 0 | 0 | 0 |
| 0 | 0 | 0 |
| 0 | 0 | 0 |
| 0 | 0 | 0 |
| 0 | 0 | 0 |
| 0 | 0 | 0 |
| 0 | 0 | 0 |

| | | |
|---|---|---|
| 1 | 0 | 0 |
| 0 | 1 | 0 |
| 0 | 0 | 0 |
| 0 | 0 | 0 |
| 0 | 0 | 0 |
| 0 | 0 | 0 |
| 0 | 0 | 0 |
| 0 | 0 | 0 |
| 0 | 0 | 0 |
| 1 | 0 | 0 |
| 0 | 1 | 0 |
| 1 | 1 | 1 |
| 0 | 0 | 0 |
| 0 | 0 | 0 |
| 0 | 0 | 0 |
| 0 | 0 | 0 |
| 0 | 0 | 0 |
| 0 | 0 | 0 |
| 1 | 1 | 1 |
| 1 | 0 | 0 |
| 1 | 0 | 0 |
| 0 | 0 | 0 |
| 0 | 1 | 0 |
| 0 | 0 | 0 |
| 0 | 0 | 0 |
| 0 | 0 | 0 |
| 0 | 0 | 0 |
| 0 | 0 | 0 |
| 0 | 0 | 0 |
| 0 | 0 | 0 |
| 0 | 0 | 0 |
| 0 | 0 | 0 |
| 0 | 0 | 0 |
| 0 | 0 | 0 |
| 0 | 0 | 0 |
| 0 | 0 | 0 |
| 0 | 0 | 0 |
| 0 | 0 | 0 |
| 0 | 0 | 0 |
| 0 | 0 | 0 |
| 0 | 0 | 0 |
| 1 | 0 | 0 |
| 0 | 0 | 0 |
| 0 | 0 | 0 |
| 0 | 0 | 0 |
| 0 | 0 | 0 |
| 1 | 0 | 0 |
| 0 | 0 | 0 |
| 0 | 0 | 0 |
| 0 | 0 | 0 |
| 0 | 0 | 0 |
| 0 | 0 | 0 |
| 1 | 0 | 0 |
| 0 | 1 | 0 |

| | | |
|---|---|---|
| 0 | 0 | 0 |
| 0 | 0 | 0 |
| 1 | 0 | 0 |
| 1 | 0 | 0 |
| 1 | 0 | 0 |
| 0 | 0 | 0 |
| 1 | 0 | 0 |
| 0 | 0 | 0 |
| 0 | 0 | 0 |
| 0 | 0 | 0 |
| 0 | 0 | 0 |
| 0 | 0 | 0 |
| 0 | 0 | 0 |
| 1 | 0 | 0 |
| 1 | 0 | 0 |
| 1 | 0 | 0 |
| 0 | 0 | 0 |
| 1 | 1 | 1 |
| 0 | 0 | 0 |
| 0 | 0 | 0 |
| 1 | 0 | 0 |
| 1 | 0 | 0 |
| 0 | 0 | 0 |
| 0 | 0 | 0 |
| 1 | 0 | 0 |
| 0 | 0 | 0 |
| 0 | 0 | 0 |
| 1 | 0 | 0 |
| 0 | 0 | 0 |
| 0 | 0 | 0 |
| 0 | 0 | 0 |
| 0 | 0 | 0 |
| 0 | 0 | 0 |
| 0 | 0 | 0 |
| 0 | 0 | 0 |
| 0 | 0 | 0 |
| 0 | 0 | 0 |
| 0 | 0 | 0 |
| 0 | 0 | 0 |
| 0 | 0 | 0 |
| 1 | 0 | 0 |
| 1 | 0 | 0 |
| 1 | 0 | 0 |
| 1 | 1 | 1 |
| 1 | 1 | 1 |
| 0 | 0 | 0 |
| 0 | 0 | 0 |
| 0 | 0 | 0 |
| 0 | 0 | 0 |
| 0 | 0 | 0 |
| 0 | 0 | 0 |
| 0 | 0 | 0 |
| 0 | 0 | 0 |
| 1 | 0 | 0 |

| | | |
|---|---|---|
| 1 | 0 | 0 |
| 0 | 0 | 0 |
| 0 | 0 | 0 |
| 0 | 0 | 0 |
| 1 | 0 | 0 |
| 0 | 0 | 0 |
| 0 | 0 | 0 |
| 0 | 0 | 0 |
| 1 | 0 | 0 |
| 1 | 0 | 0 |
| 0 | 0 | 0 |
| 0 | 0 | 0 |
| 0 | 0 | 0 |
| 1 | 0 | 0 |
| 0 | 0 | 0 |
| 0 | 0 | 0 |
| 0 | 0 | 0 |
| 0 | 0 | 0 |
| 0 | 0 | 0 |
| 0 | 0 | 0 |
| 0 | 0 | 0 |
| 1 | 1 | 1 |
| 0 | 0 | 0 |
| 0 | 1 | 0 |
| 0 | 0 | 0 |
| 0 | 0 | 0 |
| 0 | 0 | 0 |
| 0 | 0 | 0 |
| 0 | 0 | 0 |
| 0 | 0 | 0 |
| 0 | 0 | 0 |
| 0 | 0 | 0 |
| 0 | 0 | 0 |
| 0 | 0 | 0 |
| 0 | 1 | 0 |
| 0 | 0 | 0 |
| 0 | 0 | 0 |
| 0 | 0 | 0 |
| 0 | 0 | 0 |
| 0 | 0 | 0 |
| 0 | 0 | 0 |
| 0 | 0 | 0 |
| 0 | 0 | 0 |
| 0 | 0 | 0 |
| 0 | 0 | 0 |
| 0 | 0 | 0 |
| 0 | 0 | 0 |
| 1 | 1 | 1 |
| 0 | 0 | 0 |
| 1 | 1 | 1 |
| 0 | 0 | 0 |
| 0 | 0 | 0 |
| 0 | 0 | 0 |
| 0 | 0 | 0 |

| | | |
|---|---|---|
| 0 | 0 | 0 |
| 0 | 0 | 0 |
| 1 | 1 | 1 |
| 0 | 0 | 0 |
| 0 | 0 | 0 |
| 0 | 1 | 0 |
| 0 | 1 | 0 |
| 0 | 1 | 0 |
| 0 | 0 | 0 |
| 0 | 1 | 0 |
| 0 | 0 | 0 |
| 0 | 0 | 0 |
| 0 | 0 | 0 |
| 1 | 1 | 1 |
| 0 | 0 | 0 |
| 0 | 1 | 0 |
| 0 | 0 | 0 |
| 0 | 0 | 0 |
| 0 | 0 | 0 |
| 1 | 1 | 1 |
| 1 | 1 | 1 |
| 0 | 0 | 0 |
| 1 | 1 | 1 |
| 0 | 0 | 0 |
| 0 | 0 | 0 |
| 1 | 1 | 1 |
| 1 | 1 | 1 |
| 0 | 0 | 0 |
| 1 | 1 | 1 |
| 0 | 0 | 0 |
| 0 | 1 | 0 |
| 0 | 1 | 0 |
| 1 | 1 | 1 |
| 0 | 1 | 0 |
| 1 | 1 | 1 |
| 1 | 1 | 1 |
| 1 | 1 | 1 |
| 1 | 1 | 1 |
| 1 | 1 | 1 |
| 1 | 1 | 1 |
| 1 | 1 | 1 |
| 1 | 1 | 1 |
| 0 | 1 | 0 |
| 0 | 1 | 0 |
| 0 | 1 | 0 |
| 0 | 1 | 0 |
| 0 | 1 | 0 |
| 0 | 1 | 0 |
| 0 | 1 | 0 |
| 0 | 1 | 0 |
| 0 | 1 | 0 |
| 0 | 1 | 0 |
| 0 | 1 | 0 |
| 0 | 1 | 0 |

| | | |
|---|---|---|
| 0 | 1 | 0 |
| 1 | 1 | 1 |
| 1 | 1 | 1 |
| 1 | 1 | 1 |
| 1 | 1 | 1 |
| 1 | 1 | 1 |
| 1 | 1 | 1 |
| 0 | 1 | 0 |
| 0 | 1 | 0 |
| 0 | 1 | 0 |
| 0 | 1 | 0 |
| 0 | 0 | 0 |
| 0 | 0 | 0 |
| 0 | 0 | 0 |
| 1 | 0 | 0 |
| 0 | 0 | 0 |
| 1 | 0 | 0 |
| 0 | 0 | 0 |
| 1 | 0 | 0 |
| 1 | 1 | 1 |
| 0 | 0 | 0 |
| 1 | 1 | 1 |
| 1 | 1 | 1 |
| 0 | 0 | 0 |
| 0 | 0 | 0 |
| 1 | 1 | 1 |
| 1 | 1 | 1 |
| 0 | 0 | 0 |
| 0 | 0 | 0 |
| 0 | 0 | 0 |
| 0 | 0 | 0 |
| 0 | 0 | 0 |
| 0 | 1 | 0 |
| 0 | 1 | 0 |
| 1 | 1 | 1 |
| 1 | 1 | 1 |
| 1 | 1 | 1 |
| 1 | 1 | 1 |
| 1 | 1 | 1 |
| 0 | 0 | 0 |
| 0 | 0 | 0 |
| 0 | 1 | 0 |
| 0 | 0 | 0 |
| 0 | 0 | 0 |
| 0 | 0 | 0 |
| 1 | 0 | 0 |
| 0 | 0 | 0 |
| 1 | 0 | 0 |
| 1 | 0 | 0 |
| 0 | 0 | 0 |
| 0 | 0 | 0 |
| 0 | 0 | 0 |
| 0 | 0 | 0 |
| 0 | 0 | 0 |

| | | |
|---|---|---|
| 0 | 0 | 0 |
| 0 | 1 | 0 |
| 1 | 1 | 1 |
| 1 | 1 | 1 |
| 0 | 0 | 0 |
| 1 | 1 | 1 |
| 0 | 0 | 0 |
| 1 | 1 | 1 |
| 1 | 1 | 1 |
| 0 | 0 | 0 |
| 1 | 0 | 0 |
| 1 | 1 | 1 |
| 1 | 1 | 1 |
| 0 | 0 | 0 |
| 1 | 0 | 0 |
| 1 | 0 | 0 |
| 0 | 0 | 0 |
| 1 | 0 | 0 |
| 0 | 0 | 0 |
| 0 | 0 | 0 |
| 0 | 0 | 0 |
| 0 | 0 | 0 |
| 0 | 0 | 0 |
| 0 | 0 | 0 |
| 0 | 0 | 0 |
| 0 | 0 | 0 |
| 0 | 0 | 0 |
| 0 | 0 | 0 |
| 0 | 0 | 0 |
| 0 | 0 | 0 |
| 0 | 0 | 0 |
| 0 | 0 | 0 |
| 0 | 0 | 0 |
| 0 | 0 | 0 |
| 0 | 0 | 0 |
| 0 | 0 | 0 |
| 1 | 0 | 0 |
| 0 | 0 | 0 |
| 0 | 0 | 0 |
| 0 | 0 | 0 |
| 0 | 0 | 0 |
| 0 | 0 | 0 |
| 0 | 0 | 0 |
| 0 | 0 | 0 |
| 0 | 0 | 0 |
| 0 | 0 | 0 |
| 0 | 0 | 0 |
| 0 | 0 | 0 |
| 1 | 0 | 0 |
| 1 | 0 | 0 |
| 0 | 0 | 0 |
| 1 | 1 | 1 |
| 0 | 0 | 0 |
| 0 | 0 | 0 |

| | | |
|---|---|---|
| 0 | 0 | 0 |
| 0 | 0 | 0 |
| 0 | 0 | 0 |
| 0 | 0 | 0 |
| 0 | 0 | 0 |
| 0 | 0 | 0 |
| 0 | 0 | 0 |
| 0 | 0 | 0 |
| 0 | 0 | 0 |
| 0 | 0 | 0 |
| 0 | 0 | 0 |
| 0 | 0 | 0 |
| 0 | 0 | 0 |
| 0 | 0 | 0 |
| 0 | 1 | 0 |
| 0 | 0 | 0 |
| 0 | 0 | 0 |
| 0 | 0 | 0 |
| 1 | 0 | 0 |
| 0 | 0 | 0 |
| 0 | 0 | 0 |
| 0 | 0 | 0 |
| 0 | 0 | 0 |
| 0 | 0 | 0 |
| 0 | 0 | 0 |
| 1 | 0 | 0 |
| 0 | 0 | 0 |
| 0 | 0 | 0 |
| 0 | 0 | 0 |
| 0 | 0 | 0 |
| 1 | 1 | 1 |
| 0 | 0 | 0 |
| 1 | 1 | 1 |
| 1 | 0 | 0 |
| 0 | 0 | 0 |
| 0 | 0 | 0 |
| 1 | 1 | 1 |
| 1 | 1 | 1 |
| 1 | 1 | 1 |
| 1 | 0 | 0 |
| 0 | 0 | 0 |
| 0 | 0 | 0 |
| 0 | 0 | 0 |
| 0 | 0 | 0 |
| 0 | 0 | 0 |
| 0 | 0 | 0 |
| 0 | 0 | 0 |
| 0 | 0 | 0 |
| 1 | 1 | 1 |
| 1 | 1 | 1 |
| 0 | 0 | 0 |
| 0 | 0 | 0 |
| 0 | 0 | 0 |
| 0 | 1 | 0 |

| | | |
|---|---|---|
| 1 | 1 | 1 |
| 1 | 1 | 1 |
| 1 | 0 | 0 |
| 0 | 0 | 0 |
| 1 | 0 | 0 |
| 1 | 1 | 1 |
| 0 | 0 | 0 |
| 1 | 1 | 1 |
| 0 | 0 | 0 |
| 0 | 0 | 0 |
| 0 | 0 | 0 |
| 0 | 0 | 0 |
| 0 | 0 | 0 |
| 0 | 0 | 0 |
| 0 | 0 | 0 |
| 0 | 0 | 0 |
| 0 | 1 | 0 |
| 0 | 0 | 0 |
| 0 | 0 | 0 |
| 1 | 0 | 0 |
| 0 | 0 | 0 |
| 0 | 0 | 0 |
| 1 | 1 | 1 |
| 0 | 0 | 0 |
| 1 | 1 | 1 |
| 1 | 1 | 1 |
| 0 | 1 | 0 |
| 0 | 0 | 0 |
| 0 | 0 | 0 |
| 0 | 0 | 0 |
| 1 | 1 | 1 |
| 0 | 0 | 0 |
| 0 | 0 | 0 |
| 0 | 0 | 0 |
| 0 | 1 | 0 |
| 0 | 1 | 0 |
| 0 | 0 | 0 |
| 0 | 0 | 0 |
| 0 | 0 | 0 |
| 1 | 1 | 1 |
| 1 | 1 | 1 |
| 0 | 1 | 0 |
| 0 | 1 | 0 |
| 0 | 0 | 0 |
| 1 | 1 | 1 |
| 0 | 0 | 0 |
| 1 | 1 | 1 |
| 0 | 0 | 0 |
| 0 | 0 | 0 |
| 1 | 1 | 1 |
| 0 | 1 | 0 |
| 0 | 0 | 0 |
| 0 | 0 | 0 |
| 1 | 1 | 1 |

| | | |
|---|---|---|
| 1 | 1 | 1 |
| 1 | 1 | 1 |
| 0 | 1 | 0 |
| 0 | 0 | 0 |
| 0 | 0 | 0 |
| 1 | 0 | 0 |
| 1 | 0 | 0 |
| 0 | 0 | 0 |
| 0 | 0 | 0 |
| 1 | 1 | 1 |
| 0 | 0 | 0 |
| 0 | 0 | 0 |
| 0 | 0 | 0 |
| 0 | 0 | 0 |
| 0 | 0 | 0 |
| 0 | 0 | 0 |
| 0 | 0 | 0 |
| 0 | 0 | 0 |
| 0 | 0 | 0 |
| 0 | 1 | 0 |
| 1 | 1 | 1 |
| 1 | 1 | 1 |
| 1 | 1 | 1 |
| 1 | 1 | 1 |
| 0 | 0 | 0 |
| 0 | 1 | 0 |
| 0 | 0 | 0 |
| 0 | 0 | 0 |
| 1 | 1 | 1 |
| 1 | 1 | 1 |
| 1 | 1 | 1 |
| 1 | 1 | 1 |
| 1 | 1 | 1 |
| 0 | 0 | 0 |
| 0 | 0 | 0 |
| 1 | 1 | 1 |
| 0 | 0 | 0 |
| 0 | 1 | 0 |
| 0 | 0 | 0 |
| 0 | 0 | 0 |
| 0 | 0 | 0 |
| 0 | 0 | 0 |
| 0 | 1 | 0 |
| 0 | 0 | 0 |
| 0 | 0 | 0 |
| 0 | 0 | 0 |
| 0 | 0 | 0 |
| 0 | 0 | 0 |
| 0 | 0 | 0 |
| 0 | 0 | 0 |
| 0 | 0 | 0 |
| 0 | 0 | 0 |
| 0 | 0 | 0 |
| 0 | 1 | 0 |

| | | |
|---|---|---|
| 0 | 0 | 0 |
| 0 | 1 | 0 |
| 0 | 0 | 0 |
| 0 | 0 | 0 |
| 0 | 0 | 0 |
| 1 | 1 | 1 |
| 0 | 0 | 0 |
| 1 | 1 | 1 |
| 0 | 1 | 0 |
| 0 | 0 | 0 |
| 0 | 1 | 0 |
| 0 | 1 | 0 |
| 0 | 1 | 0 |
| 0 | 1 | 0 |
| 0 | 1 | 0 |
| 0 | 0 | 0 |
| 0 | 0 | 0 |
| 0 | 1 | 0 |
| 0 | 0 | 0 |
| 1 | 1 | 1 |
| 0 | 0 | 0 |
| 0 | 0 | 0 |
| 0 | 0 | 0 |
| 0 | 0 | 0 |
| 0 | 0 | 0 |
| 0 | 0 | 0 |
| 0 | 0 | 0 |
| 0 | 0 | 0 |
| 0 | 0 | 0 |
| 0 | 0 | 0 |
| 0 | 0 | 0 |
| 0 | 0 | 0 |
| 1 | 0 | 0 |
| 1 | 0 | 0 |
| 0 | 0 | 0 |
| 0 | 0 | 0 |
| 0 | 0 | 0 |
| 0 | 0 | 0 |
| 1 | 0 | 0 |
| 0 | 0 | 0 |
| 0 | 0 | 0 |
| 0 | 0 | 0 |
| 1 | 0 | 0 |
| 0 | 1 | 0 |
| 0 | 0 | 0 |
| 0 | 0 | 0 |
| 0 | 0 | 0 |
| 0 | 0 | 0 |
| 0 | 0 | 0 |
| 0 | 0 | 0 |
| 1 | 0 | 0 |
| 0 | 0 | 0 |
| 0 | 0 | 0 |
| 1 | 0 | 0 |

| | | |
|---|---|---|
| 0 | 0 | 0 |
| 1 | 0 | 0 |
| 0 | 0 | 0 |
| 0 | 0 | 0 |
| 0 | 0 | 0 |
| 0 | 0 | 0 |
| 0 | 0 | 0 |
| 0 | 0 | 0 |
| 0 | 0 | 0 |
| 0 | 0 | 0 |
| 0 | 0 | 0 |
| 1 | 1 | 1 |
| 0 | 0 | 0 |
| 1 | 1 | 1 |
| 1 | 0 | 0 |
| 1 | 0 | 0 |
| 0 | 0 | 0 |
| 0 | 0 | 0 |
| 0 | 0 | 0 |
| 0 | 0 | 0 |
| 1 | 0 | 0 |
| 0 | 0 | 0 |
| 0 | 1 | 0 |
| 0 | 0 | 0 |
| 1 | 1 | 1 |
| 1 | 1 | 1 |
| 1 | 1 | 1 |
| 0 | 0 | 0 |
| 1 | 1 | 1 |
| 1 | 1 | 1 |
| 1 | 1 | 1 |
| 0 | 0 | 0 |
| 1 | 1 | 1 |
| 0 | 0 | 0 |
| 0 | 0 | 0 |
| 0 | 0 | 0 |
| 0 | 1 | 0 |
| 1 | 1 | 1 |
| 0 | 0 | 0 |
| 0 | 0 | 0 |
| 0 | 0 | 0 |
| 0 | 0 | 0 |
| 0 | 0 | 0 |
| 0 | 0 | 0 |
| 0 | 0 | 0 |
| 0 | 0 | 0 |
| 1 | 1 | 1 |
| 1 | 1 | 1 |
| 0 | 0 | 0 |
| 0 | 0 | 0 |
| 0 | 0 | 0 |
| 1 | 1 | 1 |
| 1 | 1 | 1 |
| 0 | 0 | 0 |

| | | |
|---|---|---|
| 0 | 0 | 0 |
| 0 | 1 | 0 |
| 0 | 1 | 0 |
| 1 | 1 | 1 |
| 0 | 0 | 0 |
| 0 | 0 | 0 |
| 0 | 0 | 0 |
| 0 | 0 | 0 |
| 0 | 0 | 0 |
| 0 | 0 | 0 |
| 1 | 1 | 1 |
| 0 | 0 | 0 |
| 0 | 0 | 0 |
| 0 | 0 | 0 |
| 1 | 0 | 0 |
| 1 | 1 | 1 |
| 0 | 0 | 0 |
| 1 | 0 | 0 |
| 0 | 0 | 0 |
| 0 | 0 | 0 |
| 1 | 0 | 0 |
| 0 | 0 | 0 |
| 0 | 0 | 0 |
| 1 | 0 | 0 |
| 0 | 0 | 0 |
| 0 | 0 | 0 |
| 0 | 0 | 0 |
| 1 | 0 | 0 |
| 0 | 0 | 0 |
| 0 | 0 | 0 |
| 0 | 0 | 0 |
| 1 | 1 | 1 |
| 1 | 1 | 1 |
| 1 | 1 | 1 |
| 0 | 1 | 0 |
| 0 | 1 | 0 |
| 0 | 1 | 0 |
| 1 | 1 | 1 |
| 1 | 1 | 1 |
| 1 | 1 | 1 |
| 0 | 1 | 0 |
| 1 | 1 | 1 |
| 0 | 0 | 0 |
| 0 | 1 | 0 |
| 1 | 1 | 1 |
| 1 | 1 | 1 |
| 0 | 1 | 0 |
| 1 | 1 | 1 |
| 1 | 1 | 1 |
| 1 | 1 | 1 |
| 1 | 1 | 1 |
| 1 | 1 | 1 |
| 1 | 1 | 1 |
| 1 | 1 | 1 |

| | | |
|---|---|---|
| 0 | 1 | 0 |
| 0 | 1 | 0 |
| 1 | 1 | 1 |
| 1 | 1 | 1 |
| 1 | 1 | 1 |
| 1 | 1 | 1 |
| 1 | 1 | 1 |
| 1 | 1 | 1 |
| 0 | 0 | 0 |
| 0 | 0 | 0 |
| 0 | 0 | 0 |
| 0 | 0 | 0 |
| 0 | 0 | 0 |
| 0 | 0 | 0 |
| 0 | 0 | 0 |
| 0 | 0 | 0 |
| 0 | 0 | 0 |
| 0 | 0 | 0 |
| 0 | 0 | 0 |
| 0 | 0 | 0 |
| 0 | 0 | 0 |
| 0 | 0 | 0 |
| 0 | 0 | 0 |
| 0 | 0 | 0 |
| 0 | 0 | 0 |
| 1 | 1 | 1 |
| 1 | 1 | 1 |
| 0 | 0 | 0 |
| 0 | 0 | 0 |
| 0 | 0 | 0 |
| 0 | 0 | 0 |
| 0 | 0 | 0 |
| 0 | 0 | 0 |
| 0 | 0 | 0 |
| 0 | 0 | 0 |
| 0 | 0 | 0 |
| 0 | 0 | 0 |
| 0 | 0 | 0 |
| 0 | 0 | 0 |
| 0 | 0 | 0 |
| 0 | 0 | 0 |
| 0 | 0 | 0 |
| 0 | 0 | 0 |
| 0 | 0 | 0 |
| 0 | 0 | 0 |
| 0 | 0 | 0 |
| 1 | 1 | 1 |
| 0 | 1 | 0 |
| 1 | 1 | 1 |
| 1 | 1 | 1 |
| 1 | 1 | 1 |
| 1 | 1 | 1 |
| 0 | 0 | 0 |
| 0 | 0 | 0 |

| | | |
|---|---|---|
| 0 | 0 | 0 |
| 0 | 0 | 0 |
| 0 | 0 | 0 |
| 0 | 0 | 0 |
| 0 | 1 | 0 |
| 0 | 0 | 0 |
| 0 | 0 | 0 |
| 0 | 0 | 0 |
| 1 | 1 | 1 |
| 0 | 0 | 0 |
| 0 | 0 | 0 |
| 0 | 1 | 0 |
| 0 | 0 | 0 |
| 0 | 0 | 0 |
| 0 | 0 | 0 |
| 0 | 0 | 0 |
| 1 | 1 | 1 |
| 0 | 0 | 0 |
| 0 | 0 | 0 |
| 1 | 1 | 1 |
| 0 | 0 | 0 |
| 0 | 0 | 0 |
| 0 | 0 | 0 |
| 0 | 0 | 0 |
| 0 | 0 | 0 |
| 0 | 1 | 0 |
| 0 | 1 | 0 |
| 1 | 1 | 1 |
| 1 | 1 | 1 |
| 0 | 1 | 0 |
| 1 | 1 | 1 |
| 1 | 1 | 1 |
| 1 | 0 | 0 |
| 1 | 0 | 0 |
| 0 | 0 | 0 |
| 0 | 0 | 0 |
| 0 | 0 | 0 |
| 0 | 0 | 0 |
| 0 | 0 | 0 |
| 0 | 0 | 0 |
| 1 | 1 | 1 |
| 1 | 1 | 1 |
| 1 | 1 | 1 |
| 0 | 0 | 0 |
| 0 | 0 | 0 |
| 0 | 0 | 0 |
| 1 | 1 | 1 |
| 0 | 1 | 0 |
| 0 | 0 | 0 |
| 0 | 0 | 0 |
| 1 | 1 | 1 |
| 1 | 1 | 1 |
| 1 | 1 | 1 |
| 1 | 1 | 1 |

| | | |
|---|---|---|
| 0 | 1 | 0 |
| 1 | 1 | 1 |
| 0 | 1 | 0 |
| 0 | 0 | 0 |
| 0 | 0 | 0 |
| 1 | 1 | 1 |
| 1 | 1 | 1 |
| 1 | 1 | 1 |
| 1 | 1 | 1 |
| 1 | 1 | 1 |
| 0 | 0 | 0 |
| 0 | 1 | 0 |
| 0 | 0 | 0 |
| 0 | 0 | 0 |
| 0 | 0 | 0 |
| 1 | 1 | 1 |
| 1 | 1 | 1 |
| 0 | 1 | 0 |
| 1 | 1 | 1 |
| 1 | 1 | 1 |
| 0 | 0 | 0 |
| 1 | 1 | 1 |
| 1 | 1 | 1 |
| 0 | 0 | 0 |
| 1 | 0 | 0 |
| 0 | 0 | 0 |
| 0 | 0 | 0 |
| 1 | 1 | 1 |
| 1 | 1 | 1 |
| 1 | 1 | 1 |
| 1 | 1 | 1 |
| 1 | 1 | 1 |
| 1 | 1 | 1 |
| 1 | 1 | 1 |
| 1 | 1 | 1 |
| 0 | 0 | 0 |
| 1 | 1 | 1 |
| 1 | 1 | 1 |
| 0 | 0 | 0 |
| 0 | 0 | 0 |
| 1 | 1 | 1 |
| 0 | 1 | 0 |
| 0 | 0 | 0 |
| 0 | 0 | 0 |
| 0 | 0 | 0 |
| 0 | 0 | 0 |
| 0 | 0 | 0 |
| 0 | 0 | 0 |
| 1 | 1 | 1 |
| 0 | 0 | 0 |
| 0 | 0 | 0 |
| 0 | 0 | 0 |
| 0 | 0 | 0 |
| 0 | 0 | 0 |

| | | |
|---|---|---|
| 0 | 0 | 0 |
| 0 | 0 | 0 |
| 0 | 0 | 0 |
| 0 | 0 | 0 |
| 0 | 0 | 0 |
| 0 | 0 | 0 |
| 0 | 0 | 0 |
| 0 | 0 | 0 |
| 0 | 0 | 0 |
| 0 | 1 | 0 |
| 0 | 1 | 0 |
| 0 | 0 | 0 |
| 1 | 1 | 1 |
| 0 | 0 | 0 |
| 0 | 0 | 0 |
| 0 | 1 | 0 |
| 0 | 0 | 0 |
| 1 | 0 | 0 |
| 1 | 1 | 1 |
| 1 | 1 | 1 |
| 0 | 0 | 0 |
| 0 | 1 | 0 |
| 0 | 1 | 0 |
| 0 | 0 | 0 |
| 0 | 0 | 0 |
| 0 | 1 | 0 |
| 0 | 1 | 0 |
| 1 | 1 | 1 |
| 0 | 0 | 0 |
| 0 | 0 | 0 |
| 1 | 1 | 1 |
| 1 | 1 | 1 |
| 0 | 0 | 0 |
| 0 | 0 | 0 |
| 0 | 0 | 0 |
| 0 | 1 | 0 |
| 0 | 0 | 0 |
| 0 | 1 | 0 |
| 0 | 1 | 0 |
| 0 | 0 | 0 |
| 0 | 0 | 0 |
| 0 | 0 | 0 |
| 0 | 0 | 0 |
| 0 | 0 | 0 |
| 0 | 0 | 0 |
| 1 | 1 | 1 |
| 0 | 1 | 0 |
| 0 | 0 | 0 |
| 0 | 0 | 0 |
| 0 | 0 | 0 |
| 1 | 1 | 1 |
| 1 | 1 | 1 |
| 0 | 0 | 0 |
| 0 | 0 | 0 |

| | | |
|---|---|---|
| 1 | 1 | 1 |
| 1 | 1 | 1 |
| 1 | 1 | 1 |
| 1 | 1 | 1 |
| 0 | 0 | 0 |
| 0 | 0 | 0 |
| 1 | 0 | 0 |
| 0 | 0 | 0 |
| 1 | 0 | 0 |
| 0 | 0 | 0 |
| 1 | 1 | 1 |
| 1 | 1 | 1 |
| 1 | 1 | 1 |
| 1 | 1 | 1 |
| 1 | 1 | 1 |
| 1 | 1 | 1 |
| 0 | 0 | 0 |
| 1 | 1 | 1 |
| 1 | 1 | 1 |
| 0 | 0 | 0 |
| 0 | 0 | 0 |
| 0 | 1 | 0 |
| 0 | 1 | 0 |
| 0 | 0 | 0 |
| 0 | 1 | 0 |
| 0 | 0 | 0 |
| 0 | 0 | 0 |
| 0 | 0 | 0 |
| 0 | 0 | 0 |
| 0 | 0 | 0 |
| 0 | 0 | 0 |
| 0 | 0 | 0 |
| 0 | 1 | 0 |
| 0 | 0 | 0 |
| 1 | 1 | 1 |
| 1 | 1 | 1 |
| 1 | 1 | 1 |
| 1 | 1 | 1 |
| 0 | 0 | 0 |
| 1 | 1 | 1 |
| 1 | 1 | 1 |
| 1 | 1 | 1 |
| 1 | 1 | 1 |
| 0 | 0 | 0 |
| 1 | 1 | 1 |
| 1 | 1 | 1 |
| 1 | 1 | 1 |
| 1 | 1 | 1 |
| 1 | 1 | 1 |
| 1 | 1 | 1 |
| 0 | 0 | 0 |
| 0 | 0 | 0 |
| 0 | 0 | 0 |
| 0 | 1 | 0 |

| | | |
|---|---|---|
| 0 | 0 | 0 |
| 0 | 0 | 0 |
| 0 | 0 | 0 |
| 0 | 0 | 0 |
| 0 | 0 | 0 |
| 0 | 0 | 0 |
| 0 | 0 | 0 |
| 0 | 0 | 0 |
| 1 | 1 | 1 |
| 1 | 1 | 1 |
| 0 | 0 | 0 |
| 1 | 1 | 1 |
| 1 | 1 | 1 |
| 0 | 0 | 0 |
| 0 | 0 | 0 |
| 0 | 0 | 0 |
| 0 | 0 | 0 |
| 0 | 0 | 0 |
| 0 | 0 | 0 |
| 1 | 1 | 1 |
| 1 | 1 | 1 |
| 0 | 0 | 0 |
| 1 | 1 | 1 |
| 0 | 0 | 0 |
| 1 | 1 | 1 |
| 1 | 1 | 1 |
| 1 | 1 | 1 |
| 0 | 1 | 0 |
| 0 | 1 | 0 |
| 0 | 0 | 0 |
| 0 | 1 | 0 |
| 1 | 1 | 1 |
| 0 | 0 | 0 |
| 1 | 1 | 1 |
| 0 | 0 | 0 |
| 0 | 0 | 0 |
| 0 | 0 | 0 |
| 0 | 0 | 0 |
| 0 | 0 | 0 |
| 0 | 0 | 0 |
| 0 | 0 | 0 |
| 0 | 0 | 0 |
| 1 | 1 | 1 |
| 0 | 0 | 0 |
| 1 | 1 | 1 |
| 0 | 0 | 0 |
| 1 | 0 | 0 |
| 1 | 1 | 1 |
| 1 | 1 | 1 |
| 0 | 0 | 0 |
| 1 | 1 | 1 |
| 1 | 1 | 1 |
| 0 | 1 | 0 |
| 1 | 1 | 1 |

| | | |
|---|---|---|
| 0 | 0 | 0 |
| 0 | 1 | 0 |
| 1 | 1 | 1 |
| 0 | 1 | 0 |
| 0 | 1 | 0 |
| 0 | 1 | 0 |
| 0 | 1 | 0 |
| 1 | 1 | 1 |
| 1 | 1 | 1 |
| 1 | 1 | 1 |
| 0 | 0 | 0 |
| 1 | 1 | 1 |
| 1 | 1 | 1 |
| 0 | 1 | 0 |
| 0 | 0 | 0 |
| 0 | 0 | 0 |
| 0 | 0 | 0 |
| 0 | 0 | 0 |
| 0 | 0 | 0 |
| 0 | 1 | 0 |
| 0 | 1 | 0 |
| 1 | 1 | 1 |
| 0 | 0 | 0 |
| 0 | 1 | 0 |
| 0 | 1 | 0 |
| 0 | 1 | 0 |
| 1 | 1 | 1 |
| 1 | 1 | 1 |
| 0 | 0 | 0 |
| 0 | 0 | 0 |
| 0 | 0 | 0 |
| 0 | 0 | 0 |
| 0 | 0 | 0 |
| 0 | 0 | 0 |
| 1 | 1 | 1 |
| 1 | 1 | 1 |
| 0 | 0 | 0 |
| 0 | 0 | 0 |
| 0 | 0 | 0 |
| 0 | 0 | 0 |
| 0 | 0 | 0 |
| 0 | 0 | 0 |
| 1 | 1 | 1 |
| 1 | 1 | 1 |
| 0 | 0 | 0 |
| 1 | 1 | 1 |
| 1 | 1 | 1 |
| 1 | 1 | 1 |
| 1 | 1 | 1 |
| 1 | 1 | 1 |
| 1 | 0 | 0 |
| 0 | 1 | 0 |
| 0 | 1 | 0 |
| 1 | 0 | 0 |

| | | |
|---|---|---|
| 0 | 0 | 0 |
| 1 | 1 | 1 |
| 0 | 0 | 0 |
| 0 | 1 | 0 |
| 1 | 0 | 0 |
| 0 | 0 | 0 |
| 0 | 0 | 0 |
| 0 | 0 | 0 |
| 1 | 0 | 0 |
| 1 | 0 | 0 |
| 0 | 0 | 0 |
| 1 | 1 | 1 |
| 1 | 0 | 0 |
| 1 | 0 | 0 |
| 0 | 0 | 0 |
| 0 | 0 | 0 |
| 0 | 0 | 0 |
| 0 | 0 | 0 |
| 1 | 1 | 1 |
| 0 | 0 | 0 |
| 0 | 0 | 0 |
| 0 | 0 | 0 |
| 0 | 0 | 0 |
| 0 | 0 | 0 |
| 1 | 1 | 1 |
| 0 | 0 | 0 |
| 1 | 1 | 1 |
| 0 | 1 | 0 |
| 1 | 1 | 1 |
| 0 | 0 | 0 |
| 1 | 1 | 1 |
| 1 | 1 | 1 |
| 0 | 0 | 0 |
| 0 | 0 | 0 |
| 1 | 1 | 1 |
| 1 | 1 | 1 |
| 1 | 1 | 1 |
| 0 | 1 | 0 |
| 1 | 0 | 0 |
| 0 | 0 | 0 |
| 0 | 0 | 0 |
| 0 | 0 | 0 |
| 0 | 0 | 0 |
| 0 | 0 | 0 |
| 0 | 0 | 0 |
| 0 | 1 | 0 |
| 0 | 1 | 0 |
| 0 | 0 | 0 |
| 0 | 0 | 0 |
| 1 | 1 | 1 |
| 0 | 0 | 0 |
| 0 | 0 | 0 |
| 0 | 1 | 0 |
| 1 | 1 | 1 |

| | | |
|---|---|---|
| 1 | 1 | 1 |
| 0 | 0 | 0 |
| 0 | 0 | 0 |
| 1 | 0 | 0 |
| 1 | 1 | 1 |
| 0 | 0 | 0 |
| 0 | 0 | 0 |
| 0 | 0 | 0 |
| 0 | 0 | 0 |
| 0 | 0 | 0 |
| 0 | 0 | 0 |
| 0 | 0 | 0 |
| 0 | 0 | 0 |
| 0 | 0 | 0 |
| 0 | 0 | 0 |
| 0 | 1 | 0 |
| 0 | 0 | 0 |
| 1 | 1 | 1 |
| 1 | 0 | 0 |
| 1 | 1 | 1 |
| 0 | 0 | 0 |
| 1 | 1 | 1 |
| 0 | 0 | 0 |
| 0 | 0 | 0 |
| 1 | 1 | 1 |
| 0 | 0 | 0 |
| 0 | 1 | 0 |
| 0 | 1 | 0 |
| 0 | 1 | 0 |
| 0 | 0 | 0 |
| 0 | 0 | 0 |
| 0 | 0 | 0 |
| 0 | 0 | 0 |
| 0 | 0 | 0 |
| 0 | 0 | 0 |
| 0 | 0 | 0 |
| 0 | 0 | 0 |
| 1 | 1 | 1 |
| 1 | 0 | 0 |
| 0 | 0 | 0 |
| 0 | 0 | 0 |
| 0 | 0 | 0 |
| 0 | 0 | 0 |
| 0 | 0 | 0 |
| 0 | 0 | 0 |
| 0 | 0 | 0 |
| 1 | 1 | 1 |
| 0 | 0 | 0 |
| 0 | 0 | 0 |
| 0 | 0 | 0 |
| 0 | 0 | 0 |
| 0 | 1 | 0 |
| 0 | 0 | 0 |
| 0 | 0 | 0 |

| | | |
|---|---|---|
| 0 | 0 | 0 |
| 0 | 0 | 0 |
| 0 | 0 | 0 |
| 0 | 0 | 0 |
| 0 | 0 | 0 |
| 0 | 0 | 0 |
| 0 | 0 | 0 |
| 0 | 0 | 0 |
| 0 | 0 | 0 |
| 0 | 0 | 0 |
| 1 | 1 | 1 |
| 0 | 0 | 0 |
| 1 | 0 | 0 |
| 0 | 0 | 0 |
| 0 | 1 | 0 |
| 1 | 1 | 1 |
| 1 | 1 | 1 |
| 1 | 1 | 1 |
| 0 | 0 | 0 |
| 0 | 0 | 0 |
| 1 | 1 | 1 |
| 0 | 0 | 0 |
| 0 | 0 | 0 |
| 1 | 1 | 1 |
| 0 | 0 | 0 |
| 0 | 0 | 0 |
| 0 | 0 | 0 |
| 1 | 0 | 0 |
| 0 | 0 | 0 |
| 0 | 0 | 0 |
| 1 | 0 | 0 |
| 0 | 0 | 0 |
| 1 | 0 | 0 |
| 1 | 0 | 0 |
| 0 | 0 | 0 |
| 0 | 0 | 0 |
| 0 | 0 | 0 |
| 1 | 0 | 0 |
| 1 | 0 | 0 |
| 0 | 0 | 0 |
| 0 | 0 | 0 |
| 0 | 0 | 0 |
| 0 | 0 | 0 |
| 0 | 0 | 0 |
| 0 | 0 | 0 |
| 0 | 0 | 0 |
| 0 | 0 | 0 |
| 0 | 0 | 0 |
| 0 | 0 | 0 |
| 0 | 0 | 0 |
| 0 | 0 | 0 |
| 0 | 0 | 0 |
| 0 | 0 | 0 |
| 0 | 0 | 0 |

| | | |
|---|---|---|
| 0 | 0 | 0 |
| 1 | 0 | 0 |
| 0 | 0 | 0 |
| 0 | 0 | 0 |
| 0 | 0 | 0 |
| 1 | 0 | 0 |
| 0 | 0 | 0 |
| 0 | 0 | 0 |
| 1 | 1 | 1 |
| 0 | 0 | 0 |
| 0 | 1 | 0 |
| 1 | 1 | 1 |
| 1 | 1 | 1 |
| 0 | 0 | 0 |
| 1 | 1 | 1 |
| 0 | 0 | 0 |
| 1 | 1 | 1 |
| 1 | 1 | 1 |
| 0 | 0 | 0 |
| 0 | 0 | 0 |
| 0 | 0 | 0 |
| 0 | 0 | 0 |
| 0 | 0 | 0 |
| 0 | 0 | 0 |
| 0 | 0 | 0 |
| 0 | 1 | 0 |
| 1 | 1 | 1 |
| 1 | 1 | 1 |
| 1 | 0 | 0 |
| 1 | 1 | 1 |
| 1 | 1 | 1 |
| 0 | 1 | 0 |
| 0 | 0 | 0 |
| 0 | 0 | 0 |
| 0 | 0 | 0 |
| 0 | 0 | 0 |
| 0 | 0 | 0 |
| 0 | 0 | 0 |
| 0 | 1 | 0 |
| 0 | 1 | 0 |
| 0 | 1 | 0 |
| 0 | 1 | 0 |
| 0 | 1 | 0 |
| 0 | 1 | 0 |
| 0 | 0 | 0 |
| 0 | 0 | 0 |
| 0 | 1 | 0 |
| 0 | 0 | 0 |
| 0 | 0 | 0 |
| 0 | 0 | 0 |
| 0 | 0 | 0 |
| 0 | 0 | 0 |
| 0 | 0 | 0 |
| 0 | 0 | 0 |

| | | |
|---|---|---|
| 1 | 1 | 1 |
| 1 | 1 | 1 |
| 0 | 0 | 0 |
| 0 | 0 | 0 |
| 1 | 1 | 1 |
| 0 | 0 | 0 |
| 0 | 0 | 0 |
| 1 | 1 | 1 |
| 1 | 1 | 1 |
| 0 | 0 | 0 |
| 1 | 1 | 1 |
| 1 | 1 | 1 |
| 0 | 0 | 0 |
| 0 | 0 | 0 |
| 0 | 0 | 0 |
| 0 | 0 | 0 |
| 0 | 1 | 0 |
| 0 | 1 | 0 |
| 1 | 1 | 1 |
| 1 | 1 | 1 |
| 1 | 1 | 1 |
| 0 | 1 | 0 |
| 1 | 1 | 1 |
| 1 | 1 | 1 |
| 1 | 1 | 1 |
| 0 | 0 | 0 |
| 1 | 1 | 1 |
| 1 | 1 | 1 |
| 1 | 1 | 1 |
| 0 | 1 | 0 |
| 0 | 1 | 0 |
| 1 | 1 | 1 |
| 1 | 1 | 1 |
| 1 | 1 | 1 |
| 1 | 1 | 1 |
| 1 | 1 | 1 |
| 1 | 1 | 1 |
| 0 | 0 | 0 |
| 0 | 0 | 0 |
| 1 | 0 | 0 |
| 0 | 0 | 0 |
| 1 | 1 | 1 |
| 1 | 1 | 1 |
| 0 | 0 | 0 |
| 0 | 0 | 0 |
| 0 | 1 | 0 |
| 0 | 1 | 0 |
| 0 | 1 | 0 |
| 1 | 1 | 1 |
| 1 | 1 | 1 |
| 1 | 1 | 1 |
| 0 | 0 | 0 |
| 0 | 0 | 0 |
| 0 | 0 | 0 |

| | | |
|---|---|---|
| 0 | 0 | 0 |
| 1 | 1 | 1 |
| 1 | 1 | 1 |
| 1 | 1 | 1 |
| 1 | 1 | 1 |
| 1 | 1 | 1 |
| 1 | 1 | 1 |
| 1 | 1 | 1 |
| 0 | 0 | 0 |
| 1 | 1 | 1 |
| 1 | 1 | 1 |
| 1 | 1 | 1 |
| 0 | 1 | 0 |
| 0 | 1 | 0 |
| 1 | 1 | 1 |
| 1 | 1 | 1 |
| 0 | 1 | 0 |
| 0 | 1 | 0 |
| 0 | 0 | 0 |
| 0 | 0 | 0 |
| 0 | 0 | 0 |
| 0 | 0 | 0 |
| 0 | 1 | 0 |
| 0 | 0 | 0 |
| 0 | 0 | 0 |
| 1 | 1 | 1 |
| 1 | 1 | 1 |
| 1 | 1 | 1 |
| 1 | 1 | 1 |
| 1 | 1 | 1 |
| 1 | 1 | 1 |
| 1 | 1 | 1 |
| 0 | 0 | 0 |
| 1 | 1 | 1 |
| 1 | 1 | 1 |
| 0 | 1 | 0 |
| 0 | 0 | 0 |
| 1 | 1 | 1 |
| 0 | 0 | 0 |
| 0 | 0 | 0 |
| 0 | 0 | 0 |
| 0 | 0 | 0 |
| 0 | 0 | 0 |
| 1 | 1 | 1 |
| 1 | 1 | 1 |
| 1 | 1 | 1 |
| 0 | 0 | 0 |
| 0 | 0 | 0 |
| 0 | 0 | 0 |
| 0 | 0 | 0 |
| 0 | 0 | 0 |
| 0 | 0 | 0 |
| 0 | 0 | 0 |
| 0 | 1 | 0 |

| | | |
|---|---|---|
| 0 | 0 | 0 |
| 0 | 0 | 0 |
| 0 | 1 | 0 |
| 0 | 0 | 0 |
| 0 | 0 | 0 |
| 1 | 1 | 1 |
| 1 | 1 | 1 |
| 1 | 0 | 0 |
| 1 | 1 | 1 |
| 1 | 1 | 1 |
| 1 | 1 | 1 |
| 0 | 0 | 0 |
| 1 | 1 | 1 |
| 1 | 1 | 1 |
| 0 | 0 | 0 |
| 1 | 1 | 1 |
| 0 | 0 | 0 |
| 1 | 1 | 1 |
| 0 | 0 | 0 |
| 0 | 0 | 0 |
| 1 | 1 | 1 |
| 1 | 1 | 1 |
| 0 | 0 | 0 |
| 1 | 1 | 1 |
| 0 | 0 | 0 |
| 1 | 1 | 1 |
| 1 | 1 | 1 |
| 0 | 0 | 0 |
| 1 | 1 | 1 |
| 0 | 0 | 0 |
| 0 | 1 | 0 |
| 0 | 0 | 0 |
| 0 | 0 | 0 |
| 0 | 0 | 0 |
| 1 | 1 | 1 |
| 0 | 0 | 0 |
| 1 | 1 | 1 |
| 1 | 1 | 1 |
| 0 | 0 | 0 |
| 1 | 1 | 1 |
| 1 | 1 | 1 |
| 1 | 1 | 1 |
| 1 | 1 | 1 |
| 0 | 0 | 0 |
| 0 | 0 | 0 |
| 1 | 1 | 1 |
| 1 | 1 | 1 |
| 1 | 0 | 0 |
| 0 | 0 | 0 |
| 1 | 0 | 0 |
| 0 | 0 | 0 |
| 1 | 1 | 1 |
| 1 | 1 | 1 |
| 0 | 0 | 0 |

| | | |
|---|---|---|
| 1 | 1 | 1 |
| 1 | 1 | 1 |
| 1 | 1 | 1 |
| 1 | 1 | 1 |
| 0 | 1 | 0 |
| 1 | 1 | 1 |
| 1 | 1 | 1 |
| 0 | 0 | 0 |
| 1 | 1 | 1 |
| 1 | 1 | 1 |
| 1 | 1 | 1 |
| 1 | 1 | 1 |
| 0 | 0 | 0 |
| 1 | 1 | 1 |
| 0 | 0 | 0 |
| 0 | 0 | 0 |
| 1 | 1 | 1 |
| 1 | 1 | 1 |
| 1 | 1 | 1 |
| 1 | 1 | 1 |
| 0 | 0 | 0 |
| 0 | 0 | 0 |
| 1 | 1 | 1 |
| 1 | 1 | 1 |
| 0 | 0 | 0 |
| 1 | 1 | 1 |
| 0 | 0 | 0 |
| 0 | 0 | 0 |
| 0 | 0 | 0 |
| 0 | 0 | 0 |
| 0 | 1 | 0 |
| 0 | 1 | 0 |
| 0 | 0 | 0 |
| 0 | 0 | 0 |
| 0 | 0 | 0 |
| 0 | 0 | 0 |
| 0 | 0 | 0 |
| 0 | 0 | 0 |
| 0 | 0 | 0 |
| 1 | 0 | 0 |
| 1 | 1 | 1 |
| 0 | 1 | 0 |
| 0 | 1 | 0 |
| 0 | 1 | 0 |
| 0 | 1 | 0 |
| 0 | 0 | 0 |
| 0 | 1 | 0 |
| 0 | 1 | 0 |
| 1 | 1 | 1 |
| 0 | 0 | 0 |
| 0 | 0 | 0 |
| 1 | 1 | 1 |
| 1 | 1 | 1 |
| 1 | 1 | 1 |

| | | |
|---|---|---|
| 1 | 1 | 1 |
| 1 | 1 | 1 |
| 1 | 1 | 1 |
| 1 | 1 | 1 |
| 0 | 0 | 0 |
| 1 | 1 | 1 |
| 0 | 0 | 0 |
| 0 | 0 | 0 |
| 0 | 0 | 0 |
| 0 | 0 | 0 |
| 0 | 0 | 0 |
| 1 | 1 | 1 |
| 0 | 0 | 0 |
| 0 | 0 | 0 |
| 1 | 1 | 1 |
| 1 | 1 | 1 |
| 1 | 1 | 1 |
| 1 | 1 | 1 |
| 1 | 1 | 1 |
| 1 | 1 | 1 |
| 1 | 1 | 1 |
| 1 | 1 | 1 |
| 1 | 1 | 1 |
| 0 | 0 | 0 |
| 1 | 1 | 1 |
| 1 | 1 | 1 |
| 1 | 1 | 1 |
| 0 | 0 | 0 |
| 1 | 0 | 0 |
| 1 | 0 | 0 |
| 0 | 0 | 0 |
| 0 | 0 | 0 |
| 1 | 1 | 1 |
| 0 | 0 | 0 |
| 0 | 1 | 0 |
| 1 | 1 | 1 |
| 1 | 1 | 1 |
| 1 | 1 | 1 |
| 1 | 1 | 1 |
| 1 | 1 | 1 |
| 1 | 1 | 1 |
| 1 | 1 | 1 |
| 1 | 1 | 1 |
| 1 | 1 | 1 |
| 1 | 1 | 1 |
| 1 | 1 | 1 |
| 1 | 1 | 1 |
| 1 | 1 | 1 |
| 0 | 0 | 0 |
| 0 | 1 | 0 |
| 1 | 1 | 1 |
| 1 | 0 | 0 |
| 1 | 0 | 0 |
| 1 | 1 | 1 |

| | | |
|---|---|---|
| 1 | 1 | 1 |
| 0 | 1 | 0 |
| 1 | 1 | 1 |
| 0 | 1 | 0 |
| 1 | 0 | 0 |
| 1 | 1 | 1 |
| 1 | 1 | 1 |
| 1 | 1 | 1 |
| 1 | 1 | 1 |
| 1 | 1 | 1 |
| 0 | 0 | 0 |
| 0 | 0 | 0 |
| 0 | 0 | 0 |
| 0 | 0 | 0 |
| 0 | 0 | 0 |
| 1 | 0 | 0 |
| 0 | 0 | 0 |
| 0 | 0 | 0 |
| 0 | 0 | 0 |
| 0 | 0 | 0 |
| 0 | 0 | 0 |
| 0 | 0 | 0 |
| 0 | 1 | 0 |
| 0 | 0 | 0 |
| 0 | 0 | 0 |
| 0 | 0 | 0 |
| 1 | 1 | 1 |
| 0 | 0 | 0 |
| 0 | 1 | 0 |
| 0 | 1 | 0 |
| 0 | 1 | 0 |
| 0 | 0 | 0 |
| 1 | 1 | 1 |
| 1 | 1 | 1 |
| 0 | 0 | 0 |
| 0 | 0 | 0 |
| 0 | 0 | 0 |
| 0 | 0 | 0 |
| 0 | 0 | 0 |
| 1 | 0 | 0 |
| 1 | 0 | 0 |
| 0 | 0 | 0 |
| 1 | 0 | 0 |
| 0 | 0 | 0 |
| 1 | 1 | 1 |
| 1 | 0 | 0 |
| 1 | 1 | 1 |
| 0 | 0 | 0 |
| 0 | 0 | 0 |
| 0 | 0 | 0 |
| 0 | 0 | 0 |
| 1 | 1 | 1 |
| 0 | 0 | 0 |
| 1 | 1 | 1 |

| | | |
|---|---|---|
| 1 | 1 | 1 |
| 1 | 1 | 1 |
| 1 | 1 | 1 |
| 1 | 1 | 1 |
| 1 | 1 | 1 |
| 0 | 0 | 0 |
| 0 | 0 | 0 |
| 0 | 0 | 0 |
| 0 | 0 | 0 |
| 0 | 1 | 0 |
| 0 | 0 | 0 |
| 0 | 0 | 0 |
| 0 | 0 | 0 |
| 0 | 0 | 0 |
| 0 | 0 | 0 |
| 0 | 0 | 0 |
| 1 | 1 | 1 |
| 0 | 1 | 0 |
| 1 | 1 | 1 |
| 1 | 1 | 1 |
| 1 | 1 | 1 |
| 1 | 1 | 1 |
| 1 | 1 | 1 |
| 1 | 1 | 1 |
| 1 | 1 | 1 |
| 1 | 1 | 1 |
| 1 | 1 | 1 |
| 1 | 1 | 1 |
| 0 | 0 | 0 |
| 0 | 1 | 0 |
| 0 | 1 | 0 |
| 1 | 0 | 0 |
| 1 | 0 | 0 |
| 0 | 0 | 0 |
| 1 | 1 | 1 |
| 0 | 0 | 0 |
| 1 | 0 | 0 |
| 0 | 0 | 0 |
| 1 | 1 | 1 |
| 0 | 0 | 0 |
| 1 | 1 | 1 |
| 1 | 1 | 1 |
| 0 | 1 | 0 |
| 0 | 0 | 0 |
| 1 | 1 | 1 |
| 1 | 1 | 1 |
| 1 | 1 | 1 |
| 1 | 1 | 1 |
| 1 | 1 | 1 |
| 1 | 1 | 1 |
| 1 | 1 | 1 |
| 1 | 0 | 0 |
| 0 | 0 | 0 |
| 0 | 0 | 0 |

| | | |
|---|---|---|
| 1 | 1 | 1 |
| 0 | 0 | 0 |
| 1 | 1 | 1 |
| 1 | 1 | 1 |
| 0 | 0 | 0 |
| 0 | 0 | 0 |
| 0 | 0 | 0 |
| 1 | 1 | 1 |
| 1 | 1 | 1 |
| 0 | 0 | 0 |
| 0 | 0 | 0 |
| 0 | 0 | 0 |
| 0 | 0 | 0 |
| 0 | 0 | 0 |
| 1 | 1 | 1 |
| 0 | 0 | 0 |
| 0 | 0 | 0 |
| 0 | 0 | 0 |
| 1 | 1 | 1 |
| 1 | 1 | 1 |
| 1 | 1 | 1 |
| 1 | 1 | 1 |
| 1 | 1 | 1 |
| 0 | 0 | 0 |
| 0 | 1 | 0 |
| 0 | 1 | 0 |
| 0 | 0 | 0 |
| 0 | 0 | 0 |
| 1 | 1 | 1 |
| 1 | 1 | 1 |
| 0 | 0 | 0 |
| 1 | 1 | 1 |
| 1 | 1 | 1 |
| 0 | 1 | 0 |
| 1 | 1 | 1 |
| 0 | 0 | 0 |
| 0 | 1 | 0 |
| 0 | 0 | 0 |
| 0 | 0 | 0 |
| 0 | 1 | 0 |
| 0 | 1 | 0 |
| 0 | 0 | 0 |
| 0 | 0 | 0 |
| 0 | 1 | 0 |
| 0 | 0 | 0 |
| 0 | 0 | 0 |
| 0 | 0 | 0 |
| 0 | 1 | 0 |
| 0 | 0 | 0 |
| 0 | 0 | 0 |
| 0 | 0 | 0 |
| 1 | 1 | 1 |
| 0 | 0 | 0 |
| 1 | 1 | 1 |

| | | |
|---|---|---|
| 1 | 1 | 1 |
| 1 | 1 | 1 |
| 1 | 1 | 1 |
| 1 | 1 | 1 |
| 0 | 0 | 0 |
| 0 | 0 | 0 |
| 1 | 1 | 1 |
| 1 | 0 | 0 |
| 1 | 1 | 1 |
| 0 | 0 | 0 |
| 1 | 0 | 0 |
| 1 | 0 | 0 |
| 0 | 0 | 0 |
| 1 | 1 | 1 |
| 1 | 1 | 1 |
| 1 | 1 | 1 |
| 1 | 1 | 1 |
| 1 | 1 | 1 |
| 1 | 1 | 1 |
| 1 | 1 | 1 |
| 1 | 1 | 1 |
| 0 | 1 | 0 |
| 1 | 1 | 1 |
| 0 | 0 | 0 |
| 1 | 1 | 1 |
| 0 | 0 | 0 |
| 1 | 1 | 1 |
| 0 | 0 | 0 |
| 1 | 1 | 1 |
| 0 | 0 | 0 |
| 0 | 0 | 0 |
| 0 | 0 | 0 |
| 0 | 0 | 0 |
| 0 | 0 | 0 |
| 0 | 1 | 0 |
| 0 | 0 | 0 |
| 1 | 1 | 1 |
| 0 | 0 | 0 |
| 1 | 1 | 1 |
| 1 | 1 | 1 |
| 0 | 0 | 0 |
| 0 | 0 | 0 |
| 0 | 0 | 0 |
| 0 | 0 | 0 |
| 0 | 0 | 0 |
| 0 | 0 | 0 |
| 0 | 0 | 0 |
| 0 | 0 | 0 |
| 1 | 1 | 1 |
| 0 | 0 | 0 |
| 1 | 0 | 0 |
| 1 | 0 | 0 |
| 1 | 1 | 1 |
| 1 | 1 | 1 |

| | | |
|---|---|---|
| 1 | 1 | 1 |
| 1 | 1 | 1 |
| 1 | 1 | 1 |
| 0 | 1 | 0 |
| 0 | 1 | 0 |
| 0 | 1 | 0 |
| 0 | 0 | 0 |
| 0 | 0 | 0 |
| 0 | 0 | 0 |
| 0 | 0 | 0 |
| 0 | 0 | 0 |
| 0 | 0 | 0 |
| 0 | 0 | 0 |
| 0 | 0 | 0 |
| 0 | 0 | 0 |
| 0 | 0 | 0 |
| 0 | 0 | 0 |
| 0 | 0 | 0 |
| 1 | 1 | 1 |
| 0 | 0 | 0 |
| 0 | 1 | 0 |
| 0 | 1 | 0 |
| 0 | 1 | 0 |
| 1 | 1 | 1 |
| 0 | 1 | 0 |
| 0 | 0 | 0 |
| 0 | 0 | 0 |
| 0 | 0 | 0 |
| 0 | 0 | 0 |
| 0 | 0 | 0 |
| 0 | 0 | 0 |
| 0 | 0 | 0 |
| 0 | 0 | 0 |
| 0 | 0 | 0 |
| 1 | 1 | 1 |
| 0 | 0 | 0 |
| 0 | 0 | 0 |
| 0 | 0 | 0 |
| 0 | 0 | 0 |
| 0 | 1 | 0 |
| 0 | 0 | 0 |
| 0 | 0 | 0 |
| 1 | 1 | 1 |
| 0 | 1 | 0 |
| 1 | 1 | 1 |
| 0 | 1 | 0 |
| 0 | 1 | 0 |
| 0 | 0 | 0 |
| 0 | 0 | 0 |
| 0 | 0 | 0 |
| 0 | 0 | 0 |
| 0 | 1 | 0 |
| 0 | 0 | 0 |
| 1 | 1 | 1 |

| | | |
|---|---|---|
| 0 | 1 | 0 |
| 1 | 1 | 1 |
| 1 | 1 | 1 |
| 1 | 1 | 1 |
| 1 | 1 | 1 |
| 0 | 0 | 0 |
| 0 | 1 | 0 |
| 1 | 1 | 1 |
| 1 | 1 | 1 |
| 1 | 1 | 1 |
| 0 | 0 | 0 |
| 0 | 0 | 0 |
| 0 | 0 | 0 |
| 0 | 0 | 0 |
| 0 | 0 | 0 |
| 0 | 1 | 0 |
| 1 | 0 | 0 |
| 0 | 0 | 0 |
| 1 | 1 | 1 |
| 1 | 1 | 1 |
| 1 | 1 | 1 |
| 1 | 1 | 1 |
| 0 | 0 | 0 |
| 1 | 1 | 1 |
| 0 | 0 | 0 |
| 0 | 0 | 0 |
| 0 | 0 | 0 |
| 0 | 0 | 0 |
| 1 | 1 | 1 |
| 0 | 0 | 0 |
| 1 | 1 | 1 |
| 0 | 1 | 0 |
| 0 | 1 | 0 |
| 0 | 1 | 0 |
| 1 | 1 | 1 |
| 1 | 1 | 1 |
| 0 | 1 | 0 |
| 0 | 0 | 0 |
| 0 | 0 | 0 |
| 0 | 0 | 0 |
| 0 | 0 | 0 |
| 0 | 0 | 0 |
| 1 | 1 | 1 |
| 0 | 0 | 0 |
| 0 | 0 | 0 |
| 1 | 1 | 1 |
| 0 | 0 | 0 |
| 0 | 0 | 0 |
| 0 | 0 | 0 |
| 0 | 0 | 0 |
| 1 | 1 | 1 |
| 1 | 1 | 1 |
| 1 | 1 | 1 |
| 0 | 0 | 0 |

| | | |
|---|---|---|
| 0 | 0 | 0 |
| 1 | 1 | 1 |
| 0 | 0 | 0 |
| 0 | 1 | 0 |
| 1 | 0 | 0 |
| 1 | 0 | 0 |
| 1 | 0 | 0 |
| 1 | 1 | 1 |
| 0 | 0 | 0 |
| 1 | 1 | 1 |
| 0 | 0 | 0 |
| 0 | 0 | 0 |
| 0 | 0 | 0 |
| 0 | 0 | 0 |
| 1 | 1 | 1 |
| 1 | 1 | 1 |
| 0 | 0 | 0 |
| 0 | 0 | 0 |
| 0 | 1 | 0 |
| 1 | 1 | 1 |
| 0 | 0 | 0 |
| 1 | 0 | 0 |
| 1 | 0 | 0 |
| 1 | 1 | 1 |
| 0 | 0 | 0 |
| 1 | 0 | 0 |
| 0 | 0 | 0 |
| 0 | 0 | 0 |
| 0 | 0 | 0 |
| 1 | 1 | 1 |
| 0 | 0 | 0 |
| 0 | 0 | 0 |
| 0 | 0 | 0 |
| 1 | 1 | 1 |
| 0 | 0 | 0 |
| 1 | 1 | 1 |
| 0 | 0 | 0 |
| 0 | 0 | 0 |
| 0 | 0 | 0 |
| 0 | 0 | 0 |
| 0 | 0 | 0 |
| 1 | 1 | 1 |
| 0 | 0 | 0 |
| 0 | 0 | 0 |
| 0 | 0 | 0 |
| 0 | 0 | 0 |
| 0 | 0 | 0 |
| 0 | 0 | 0 |
| 0 | 0 | 0 |
| 0 | 0 | 0 |
| 0 | 0 | 0 |
| 0 | 0 | 0 |
| 1 | 1 | 1 |
| 0 | 0 | 0 |

| | | |
|---|---|---|
| 0 | 1 | 0 |
| 1 | 1 | 1 |
| 1 | 1 | 1 |
| 1 | 1 | 1 |
| 1 | 1 | 1 |
| 1 | 1 | 1 |
| 1 | 1 | 1 |
| 1 | 1 | 1 |
| 1 | 1 | 1 |
| 0 | 0 | 0 |
| 0 | 0 | 0 |
| 1 | 1 | 1 |
| 1 | 1 | 1 |
| 1 | 1 | 1 |
| 0 | 0 | 0 |
| 1 | 1 | 1 |
| 1 | 1 | 1 |
| 0 | 0 | 0 |
| 0 | 1 | 0 |
| 0 | 1 | 0 |
| 0 | 1 | 0 |
| 0 | 0 | 0 |
| 0 | 0 | 0 |
| 0 | 0 | 0 |
| 0 | 0 | 0 |
| 0 | 0 | 0 |
| 0 | 0 | 0 |
| 0 | 0 | 0 |
| 1 | 1 | 1 |
| 0 | 0 | 0 |
| 0 | 0 | 0 |
| 1 | 1 | 1 |
| 1 | 1 | 1 |
| 1 | 1 | 1 |
| 1 | 1 | 1 |
| 1 | 1 | 1 |
| 1 | 1 | 1 |
| 1 | 1 | 1 |
| 1 | 1 | 1 |
| 1 | 1 | 1 |
| 1 | 1 | 1 |
| 1 | 1 | 1 |
| 1 | 1 | 1 |
| 0 | 0 | 0 |
| 1 | 1 | 1 |
| 0 | 0 | 0 |
| 1 | 1 | 1 |
| 1 | 1 | 1 |
| 1 | 1 | 1 |
| 1 | 1 | 1 |
| 0 | 0 | 0 |
| 1 | 1 | 1 |
| 1 | 1 | 1 |
| 0 | 0 | 0 |

| | | |
|---|---|---|
| 0 | 0 | 0 |
| 0 | 0 | 0 |
| 0 | 0 | 0 |
| 0 | 0 | 0 |
| 1 | 1 | 1 |
| 0 | 1 | 0 |
| 1 | 1 | 1 |
| 0 | 0 | 0 |
| 0 | 0 | 0 |
| 0 | 0 | 0 |
| 0 | 0 | 0 |
| 1 | 1 | 1 |
| 1 | 1 | 1 |
| 1 | 1 | 1 |
| 1 | 1 | 1 |
| 0 | 0 | 0 |
| 0 | 0 | 0 |
| 0 | 1 | 0 |
| 0 | 1 | 0 |
| 1 | 1 | 1 |
| 1 | 1 | 1 |
| 0 | 0 | 0 |
| 1 | 1 | 1 |
| 0 | 1 | 0 |
| 0 | 0 | 0 |
| 0 | 0 | 0 |
| 0 | 0 | 0 |
| 0 | 0 | 0 |
| 1 | 1 | 1 |
| 1 | 1 | 1 |
| 1 | 1 | 1 |
| 0 | 0 | 0 |
| 1 | 1 | 1 |
| 1 | 1 | 1 |
| 1 | 1 | 1 |
| 1 | 1 | 1 |
| 1 | 1 | 1 |
| 1 | 1 | 1 |
| 1 | 1 | 1 |
| 1 | 1 | 1 |
| 1 | 1 | 1 |
| 1 | 1 | 1 |
| 0 | 0 | 0 |
| 0 | 1 | 0 |
| 0 | 1 | 0 |
| 1 | 1 | 1 |
| 0 | 1 | 0 |
| 1 | 1 | 1 |
| 1 | 1 | 1 |
| 0 | 0 | 0 |
| 1 | 1 | 1 |
| 1 | 1 | 1 |
| 1 | 1 | 1 |
| 1 | 1 | 1 |

| | | |
|---|---|---|
| 0 | 0 | 0 |
| 1 | 1 | 1 |
| 0 | 0 | 0 |
| 0 | 1 | 0 |
| 1 | 1 | 1 |
| 1 | 1 | 1 |
| 1 | 1 | 1 |
| 0 | 0 | 0 |
| 1 | 1 | 1 |
| 1 | 1 | 1 |
| 0 | 0 | 0 |
| 0 | 0 | 0 |
| 0 | 0 | 0 |
| 1 | 1 | 1 |
| 0 | 0 | 0 |
| 0 | 0 | 0 |
| 0 | 0 | 0 |
| 0 | 0 | 0 |
| 0 | 1 | 0 |
| 0 | 0 | 0 |
| 0 | 0 | 0 |
| 0 | 0 | 0 |
| 0 | 0 | 0 |
| 0 | 0 | 0 |
| 0 | 0 | 0 |
| 0 | 0 | 0 |
| 0 | 0 | 0 |
| 0 | 0 | 0 |
| 0 | 0 | 0 |
| 0 | 0 | 0 |
| 0 | 0 | 0 |
| 0 | 0 | 0 |
| 1 | 1 | 1 |
| 0 | 0 | 0 |
| 1 | 1 | 1 |
| 0 | 0 | 0 |
| 0 | 0 | 0 |
| 1 | 1 | 1 |
| 1 | 1 | 1 |
| 0 | 1 | 0 |
| 0 | 1 | 0 |
| 0 | 0 | 0 |
| 0 | 0 | 0 |
| 0 | 0 | 0 |
| 1 | 1 | 1 |
| 0 | 0 | 0 |
| 1 | 1 | 1 |
| 0 | 0 | 0 |
| 0 | 1 | 0 |
| 0 | 1 | 0 |
| 0 | 1 | 0 |
| 0 | 1 | 0 |
| 0 | 1 | 0 |
| 1 | 1 | 1 |

| | | |
|---|---|---|
| 1 | 1 | 1 |
| 1 | 1 | 1 |
| 0 | 1 | 0 |
| 1 | 1 | 1 |
| 1 | 1 | 1 |
| 0 | 0 | 0 |
| 0 | 0 | 0 |
| 1 | 1 | 1 |
| 0 | 0 | 0 |
| 0 | 0 | 0 |
| 1 | 1 | 1 |
| 0 | 1 | 0 |
| 1 | 1 | 1 |
| 1 | 1 | 1 |
| 1 | 1 | 1 |
| 0 | 0 | 0 |
| 1 | 1 | 1 |
| 1 | 1 | 1 |
| 0 | 0 | 0 |
| 0 | 0 | 0 |
| 0 | 0 | 0 |
| 1 | 1 | 1 |
| 1 | 1 | 1 |
| 0 | 0 | 0 |
| 1 | 1 | 1 |
| 0 | 0 | 0 |
| 0 | 0 | 0 |
| 1 | 1 | 1 |
| 1 | 1 | 1 |
| 1 | 1 | 1 |
| 0 | 0 | 0 |
| 1 | 0 | 0 |
| 0 | 0 | 0 |
| 0 | 0 | 0 |
| 0 | 0 | 0 |
| 0 | 0 | 0 |
| 0 | 0 | 0 |
| 1 | 1 | 1 |
| 0 | 0 | 0 |
| 0 | 1 | 0 |
| 0 | 0 | 0 |
| 0 | 0 | 0 |
| 0 | 0 | 0 |
| 0 | 0 | 0 |
| 1 | 1 | 1 |
| 0 | 1 | 0 |
| 0 | 1 | 0 |
| 0 | 0 | 0 |
| 0 | 0 | 0 |
| 0 | 0 | 0 |
| 1 | 1 | 1 |
| 0 | 1 | 0 |
| 1 | 1 | 1 |
| 1 | 1 | 1 |

| | | |
|---|---|---|
| 1 | 1 | 1 |
| 0 | 1 | 0 |
| 0 | 1 | 0 |
| 0 | 0 | 0 |
| 0 | 0 | 0 |
| 1 | 1 | 1 |
| 0 | 0 | 0 |
| 1 | 1 | 1 |
| 1 | 1 | 1 |
| 1 | 1 | 1 |
| 1 | 1 | 1 |
| 0 | 0 | 0 |
| 1 | 0 | 0 |
| 0 | 0 | 0 |
| 1 | 1 | 1 |
| 0 | 0 | 0 |
| 0 | 0 | 0 |
| 0 | 0 | 0 |
| 1 | 1 | 1 |
| 0 | 0 | 0 |
| 0 | 0 | 0 |
| 0 | 0 | 0 |
| 0 | 0 | 0 |
| 0 | 0 | 0 |
| 0 | 0 | 0 |
| 1 | 1 | 1 |
| 0 | 0 | 0 |
| 0 | 1 | 0 |
| 0 | 1 | 0 |
| 0 | 1 | 0 |
| 0 | 1 | 0 |
| 0 | 1 | 0 |
| 0 | 0 | 0 |
| 0 | 0 | 0 |
| 1 | 1 | 1 |
| 1 | 1 | 1 |
| 0 | 0 | 0 |
| 0 | 0 | 0 |
| 0 | 0 | 0 |
| 1 | 0 | 0 |
| 0 | 0 | 0 |
| 0 | 0 | 0 |
| 1 | 1 | 1 |
| 0 | 0 | 0 |
| 1 | 1 | 1 |
| 1 | 1 | 1 |
| 1 | 1 | 1 |
| 1 | 1 | 1 |
| 1 | 1 | 1 |
| 0 | 0 | 0 |
| 1 | 1 | 1 |
| 1 | 1 | 1 |
| 1 | 1 | 1 |
| 0 | 1 | 0 |

| | | |
|---|---|---|
| 1 | 1 | 1 |
| 1 | 1 | 1 |
| 1 | 1 | 1 |
| 0 | 0 | 0 |
| 1 | 1 | 1 |
| 1 | 1 | 1 |
| 0 | 1 | 0 |
| 0 | 0 | 0 |
| 1 | 1 | 1 |
| 1 | 1 | 1 |
| 1 | 1 | 1 |
| 1 | 1 | 1 |
| 1 | 1 | 1 |
| 0 | 1 | 0 |
| 0 | 1 | 0 |
| 0 | 1 | 0 |
| 0 | 0 | 0 |
| 0 | 0 | 0 |
| 0 | 1 | 0 |
| 1 | 1 | 1 |
| 0 | 1 | 0 |
| 1 | 1 | 1 |
| 0 | 1 | 0 |
| 1 | 1 | 1 |
| 1 | 1 | 1 |
| 1 | 1 | 1 |
| 1 | 1 | 1 |
| 0 | 0 | 0 |
| 0 | 0 | 0 |
| 1 | 1 | 1 |
| 1 | 1 | 1 |
| 0 | 0 | 0 |
| 0 | 1 | 0 |
| 0 | 0 | 0 |
| 0 | 0 | 0 |
| 0 | 0 | 0 |
| 0 | 0 | 0 |
| 0 | 0 | 0 |
| 1 | 1 | 1 |
| 1 | 1 | 1 |
| 1 | 1 | 1 |
| 0 | 1 | 0 |
| 1 | 1 | 1 |
| 1 | 1 | 1 |
| 0 | 0 | 0 |
| 0 | 0 | 0 |
| 0 | 0 | 0 |
| 1 | 1 | 1 |
| 1 | 1 | 1 |
| 1 | 1 | 1 |
| 1 | 1 | 1 |
| 0 | 0 | 0 |
| 0 | 0 | 0 |
| 0 | 0 | 0 |

| | | |
|---|---|---|
| 0 | 0 | 0 |
| 0 | 0 | 0 |
| 1 | 1 | 1 |
| 0 | 0 | 0 |
| 0 | 0 | 0 |
| 0 | 0 | 0 |
| 0 | 0 | 0 |
| 1 | 1 | 1 |
| 1 | 1 | 1 |
| 1 | 1 | 1 |
| 1 | 1 | 1 |
| 1 | 1 | 1 |
| 0 | 1 | 0 |
| 0 | 1 | 0 |
| 1 | 1 | 1 |
| 1 | 1 | 1 |
| 1 | 1 | 1 |
| 1 | 1 | 1 |
| 1 | 1 | 1 |
| 1 | 1 | 1 |
| 0 | 0 | 0 |
| 0 | 0 | 0 |
| 0 | 0 | 0 |
| 0 | 0 | 0 |
| 0 | 0 | 0 |
| 1 | 1 | 1 |
| 1 | 1 | 1 |
| 0 | 0 | 0 |
| 0 | 1 | 0 |
| 0 | 1 | 0 |
| 0 | 0 | 0 |
| 0 | 0 | 0 |
| 0 | 0 | 0 |
| 1 | 1 | 1 |
| 0 | 0 | 0 |
| 1 | 1 | 1 |
| 0 | 1 | 0 |
| 0 | 0 | 0 |
| 1 | 1 | 1 |
| 1 | 1 | 1 |
| 0 | 1 | 0 |
| 0 | 1 | 0 |
| 1 | 1 | 1 |
| 1 | 1 | 1 |
| 1 | 1 | 1 |
| 1 | 1 | 1 |
| 1 | 1 | 1 |
| 1 | 1 | 1 |
| 0 | 0 | 0 |
| 1 | 1 | 1 |
| 0 | 0 | 0 |
| 0 | 0 | 0 |
| 1 | 1 | 1 |
| 0 | 1 | 0 |

| | | |
|---|---|---|
| 0 | 0 | 0 |
| 1 | 1 | 1 |
| 1 | 1 | 1 |
| 1 | 1 | 1 |
| 1 | 1 | 1 |
| 0 | 0 | 0 |
| 0 | 0 | 0 |
| 1 | 1 | 1 |
| 0 | 1 | 0 |
| 1 | 1 | 1 |
| 1 | 1 | 1 |
| 0 | 0 | 0 |
| 0 | 1 | 0 |
| 1 | 1 | 1 |
| 0 | 1 | 0 |
| 1 | 1 | 1 |
| 0 | 0 | 0 |
| 0 | 0 | 0 |
| 1 | 1 | 1 |
| 1 | 1 | 1 |
| 1 | 1 | 1 |
| 1 | 1 | 1 |
| 1 | 1 | 1 |
| 1 | 1 | 1 |
| 0 | 0 | 0 |
| 1 | 1 | 1 |
| 1 | 1 | 1 |
| 1 | 1 | 1 |
| 1 | 1 | 1 |
| 1 | 1 | 1 |
| 0 | 1 | 0 |
| 0 | 0 | 0 |
| 0 | 0 | 0 |
| 0 | 1 | 0 |
| 1 | 1 | 1 |
| 0 | 0 | 0 |
| 0 | 0 | 0 |
| 0 | 0 | 0 |
| 0 | 0 | 0 |
| 0 | 0 | 0 |
| 0 | 0 | 0 |
| 0 | 0 | 0 |
| 0 | 0 | 0 |
| 0 | 0 | 0 |
| 0 | 0 | 0 |
| 0 | 0 | 0 |
| 0 | 0 | 0 |
| 0 | 0 | 0 |
| 1 | 0 | 0 |
| 1 | 1 | 1 |
| 1 | 0 | 0 |
| 1 | 1 | 1 |
| 0 | 0 | 0 |
| 1 | 1 | 1 |

| | | |
|---|---|---|
| 1 | 1 | 1 |
| 1 | 1 | 1 |
| 1 | 1 | 1 |
| 1 | 1 | 1 |
| 1 | 1 | 1 |
| 1 | 1 | 1 |
| 1 | 1 | 1 |
| 0 | 0 | 0 |
| 0 | 0 | 0 |
| 1 | 0 | 0 |
| 1 | 1 | 1 |
| 1 | 1 | 1 |
| 1 | 1 | 1 |
| 0 | 0 | 0 |
| 1 | 1 | 1 |
| 1 | 1 | 1 |
| 0 | 0 | 0 |
| 1 | 1 | 1 |
| 0 | 0 | 0 |
| 0 | 1 | 0 |
| 1 | 1 | 1 |
| 1 | 1 | 1 |
| 1 | 1 | 1 |
| 0 | 0 | 0 |
| 1 | 1 | 1 |
| 1 | 1 | 1 |
| 0 | 1 | 0 |
| 0 | 1 | 0 |
| 0 | 1 | 0 |
| 0 | 1 | 0 |
| 1 | 1 | 1 |
| 0 | 0 | 0 |
| 1 | 1 | 1 |
| 0 | 1 | 0 |
| 1 | 1 | 1 |
| 1 | 1 | 1 |
| 1 | 1 | 1 |
| 1 | 1 | 1 |
| 0 | 0 | 0 |
| 1 | 1 | 1 |
| 1 | 1 | 1 |
| 1 | 1 | 1 |
| 0 | 1 | 0 |
| 1 | 1 | 1 |
| 1 | 1 | 1 |
| 0 | 0 | 0 |
| 0 | 1 | 0 |
| 0 | 1 | 0 |
| 1 | 0 | 0 |
| 1 | 0 | 0 |
| 0 | 1 | 0 |
| 1 | 0 | 0 |
| 0 | 0 | 0 |
| 1 | 0 | 0 |

| | | |
|---|---|---|
| 1 | 0 | 0 |
| 1 | 0 | 0 |
| 1 | 0 | 0 |
| 1 | 0 | 0 |
| 0 | 0 | 0 |
| 0 | 0 | 0 |
| 0 | 0 | 0 |
| 0 | 0 | 0 |
| 1 | 0 | 0 |
| 1 | 0 | 0 |
| 1 | 0 | 0 |
| 1 | 0 | 0 |
| 1 | 0 | 0 |
| 0 | 0 | 0 |
| 1 | 0 | 0 |
| 0 | 0 | 0 |
| 1 | 0 | 0 |
| 0 | 0 | 0 |
| 1 | 0 | 0 |
| 0 | 0 | 0 |
| 0 | 0 | 0 |
| 1 | 0 | 0 |
| 1 | 0 | 0 |
| 0 | 0 | 0 |
| 0 | 0 | 0 |
| 1 | 0 | 0 |
| 0 | 0 | 0 |
| 0 | 0 | 0 |
| 0 | 0 | 0 |
| 1 | 0 | 0 |
| 1 | 0 | 0 |
| 0 | 0 | 0 |
| 1 | 0 | 0 |
| 1 | 0 | 0 |
| 1 | 0 | 0 |
| 1 | 0 | 0 |
| 1 | 0 | 0 |
| 1 | 0 | 0 |
| 1 | 0 | 0 |
| 1 | 0 | 0 |
| 1 | 0 | 0 |
| 1 | 0 | 0 |
| 1 | 0 | 0 |
| 0 | 0 | 0 |
| 1 | 1 | 1 |
| 1 | 1 | 1 |
| 0 | 0 | 0 |
| 0 | 0 | 0 |
| 0 | 1 | 0 |
| 0 | 0 | 0 |
| 0 | 0 | 0 |
| 0 | 0 | 0 |
| 1 | 1 | 1 |
| 1 | 1 | 1 |

| | | |
|---|---|---|
| 1 | 1 | 1 |
| 1 | 1 | 1 |
| 1 | 1 | 1 |
| 1 | 1 | 1 |
| 1 | 1 | 1 |
| 1 | 1 | 1 |
| 1 | 1 | 1 |
| 0 | 0 | 0 |
| 1 | 1 | 1 |
| 1 | 1 | 1 |
| 0 | 1 | 0 |
| 0 | 1 | 0 |
| 1 | 0 | 0 |
| 1 | 1 | 1 |
| 0 | 0 | 0 |
| 0 | 0 | 0 |
| 0 | 1 | 0 |
| 0 | 0 | 0 |
| 1 | 1 | 1 |
| 1 | 1 | 1 |
| 0 | 0 | 0 |
| 0 | 1 | 0 |
| 0 | 1 | 0 |
| 1 | 1 | 1 |
| 0 | 0 | 0 |
| 1 | 1 | 1 |
| 0 | 1 | 0 |
| 0 | 0 | 0 |
| 0 | 0 | 0 |
| 0 | 0 | 0 |
| 0 | 0 | 0 |
| 1 | 1 | 1 |
| 1 | 1 | 1 |
| 0 | 1 | 0 |
| 1 | 1 | 1 |
| 1 | 1 | 1 |
| 0 | 0 | 0 |
| 0 | 0 | 0 |
| 1 | 1 | 1 |
| 1 | 1 | 1 |
| 1 | 1 | 1 |
| 1 | 1 | 1 |
| 0 | 0 | 0 |
| 0 | 1 | 0 |
| 1 | 1 | 1 |
| 1 | 1 | 1 |
| 1 | 1 | 1 |
| 1 | 1 | 1 |
| 0 | 0 | 0 |
| 0 | 0 | 0 |
| 0 | 0 | 0 |
| 0 | 0 | 0 |
| 0 | 0 | 0 |
| 1 | 1 | 1 |

| | | |
|---|---|---|
| 0 | 0 | 0 |
| 0 | 0 | 0 |
| 0 | 1 | 0 |
| 0 | 0 | 0 |
| 1 | 1 | 1 |
| 0 | 1 | 0 |
| 0 | 1 | 0 |
| 1 | 1 | 1 |
| 1 | 1 | 1 |
| 0 | 0 | 0 |
| 1 | 1 | 1 |
| 0 | 1 | 0 |
| 1 | 1 | 1 |
| 0 | 1 | 0 |
| 0 | 1 | 0 |
| 0 | 1 | 0 |
| 0 | 1 | 0 |
| 1 | 1 | 1 |
| 1 | 1 | 1 |
| 0 | 1 | 0 |
| 0 | 1 | 0 |
| 0 | 1 | 0 |
| 0 | 1 | 0 |
| 0 | 1 | 0 |
| 0 | 1 | 0 |
| 0 | 1 | 0 |
| 0 | 1 | 0 |
| 0 | 1 | 0 |
| 0 | 1 | 0 |
| 0 | 1 | 0 |
| 1 | 1 | 1 |
| 1 | 1 | 1 |
| 0 | 0 | 0 |
| 1 | 1 | 1 |
| 1 | 1 | 1 |
| 1 | 1 | 1 |
| 1 | 1 | 1 |
| 0 | 1 | 0 |
| 1 | 1 | 1 |
| 1 | 1 | 1 |
| 1 | 1 | 1 |
| 1 | 1 | 1 |
| 1 | 1 | 1 |
| 0 | 1 | 0 |
| 0 | 1 | 0 |
| 1 | 1 | 1 |
| 1 | 1 | 1 |
| 0 | 0 | 0 |
| 0 | 0 | 0 |
| 0 | 0 | 0 |
| 0 | 0 | 0 |
| 0 | 0 | 0 |
| 1 | 1 | 1 |
| 0 | 0 | 0 |

| | | |
|---|---|---|
| 1 | 1 | 1 |
| 0 | 0 | 0 |
| 1 | 1 | 1 |
| 0 | 1 | 0 |
| 0 | 1 | 0 |
| 0 | 0 | 0 |
| 0 | 0 | 0 |
| 0 | 0 | 0 |
| 0 | 0 | 0 |
| 0 | 0 | 0 |
| 0 | 1 | 0 |
| 0 | 0 | 0 |
| 0 | 1 | 0 |
| 1 | 1 | 1 |
| 0 | 1 | 0 |
| 0 | 0 | 0 |
| 1 | 1 | 1 |
| 1 | 1 | 1 |
| 0 | 0 | 0 |
| 0 | 0 | 0 |
| 0 | 0 | 0 |
| 0 | 0 | 0 |
| 0 | 0 | 0 |
| 0 | 0 | 0 |
| 0 | 1 | 0 |
| 0 | 0 | 0 |
| 0 | 0 | 0 |
| 0 | 0 | 0 |
| 0 | 0 | 0 |
| 0 | 0 | 0 |
| 0 | 0 | 0 |
| 1 | 1 | 1 |
| 1 | 1 | 1 |
| 0 | 0 | 0 |
| 0 | 0 | 0 |
| 1 | 0 | 0 |
| 0 | 0 | 0 |
| 0 | 0 | 0 |
| 0 | 0 | 0 |
| 0 | 0 | 0 |
| 0 | 0 | 0 |
| 1 | 0 | 0 |
| 1 | 1 | 1 |
| 1 | 1 | 1 |
| 0 | 0 | 0 |
| 0 | 0 | 0 |
| 0 | 0 | 0 |
| 1 | 1 | 1 |
| 1 | 1 | 1 |
| 1 | 1 | 1 |
| 1 | 1 | 1 |
| 1 | 1 | 1 |
| 1 | 1 | 1 |
| 0 | 0 | 0 |

| | | |
|---|---|---|
| 1 | 1 | 1 |
| 1 | 1 | 1 |
| 1 | 1 | 1 |
| 1 | 1 | 1 |
| 0 | 0 | 0 |
| 0 | 0 | 0 |
| 1 | 0 | 0 |
| 1 | 0 | 0 |
| 0 | 1 | 0 |
| 1 | 1 | 1 |
| 1 | 1 | 1 |
| 0 | 0 | 0 |
| 1 | 1 | 1 |
| 0 | 0 | 0 |
| 0 | 0 | 0 |
| 0 | 0 | 0 |
| 0 | 0 | 0 |
| 0 | 0 | 0 |
| 0 | 0 | 0 |
| 0 | 0 | 0 |
| 0 | 0 | 0 |
| 0 | 0 | 0 |
| 0 | 0 | 0 |
| 0 | 1 | 0 |
| 0 | 1 | 0 |
| 0 | 1 | 0 |
| 1 | 1 | 1 |
| 1 | 1 | 1 |
| 1 | 1 | 1 |
| 1 | 1 | 1 |
| 1 | 1 | 1 |
| 0 | 0 | 0 |
| 1 | 1 | 1 |
| 0 | 0 | 0 |
| 0 | 0 | 0 |
| 0 | 0 | 0 |
| 0 | 0 | 0 |
| 1 | 1 | 1 |
| 1 | 1 | 1 |
| 0 | 1 | 0 |
| 1 | 1 | 1 |
| 1 | 1 | 1 |
| 1 | 1 | 1 |
| 0 | 1 | 0 |
| 1 | 1 | 1 |
| 0 | 0 | 0 |
| 1 | 1 | 1 |
| 1 | 1 | 1 |
| 1 | 1 | 1 |
| 0 | 0 | 0 |
| 0 | 0 | 0 |
| 0 | 0 | 0 |
| 0 | 1 | 0 |
| 0 | 1 | 0 |

| | | |
|---|---|---|
| 0 | 1 | 0 |
| 0 | 1 | 0 |
| 0 | 1 | 0 |
| 0 | 0 | 0 |
| 0 | 1 | 0 |
| 1 | 1 | 1 |
| 0 | 0 | 0 |
| 0 | 1 | 0 |
| 1 | 1 | 1 |
| 1 | 1 | 1 |
| 0 | 0 | 0 |
| 0 | 0 | 0 |
| 1 | 1 | 1 |
| 1 | 1 | 1 |
| 0 | 1 | 0 |
| 0 | 1 | 0 |
| 0 | 0 | 0 |
| 0 | 0 | 0 |
| 1 | 1 | 1 |
| 0 | 0 | 0 |
| 0 | 0 | 0 |
| 0 | 0 | 0 |
| 0 | 1 | 0 |
| 0 | 0 | 0 |
| 0 | 0 | 0 |
| 1 | 1 | 1 |
| 0 | 0 | 0 |
| 0 | 1 | 0 |
| 1 | 1 | 1 |
| 0 | 0 | 0 |
| 0 | 1 | 0 |
| 1 | 1 | 1 |
| 0 | 0 | 0 |
| 0 | 1 | 0 |
| 0 | 1 | 0 |
| 0 | 0 | 0 |
| 1 | 1 | 1 |
| 0 | 0 | 0 |
| 1 | 1 | 1 |
| 0 | 0 | 0 |
| 0 | 0 | 0 |
| 1 | 0 | 0 |
| 0 | 0 | 0 |
| 0 | 0 | 0 |
| 1 | 1 | 1 |
| 1 | 0 | 0 |
| 0 | 1 | 0 |
| 1 | 1 | 1 |
| 1 | 1 | 1 |
| 1 | 1 | 1 |
| 1 | 1 | 1 |
| 0 | 1 | 0 |
| 0 | 1 | 0 |
| 0 | 1 | 0 |

| | | |
|---|---|---|
| 0 | 1 | 0 |
| 0 | 1 | 0 |
| 0 | 1 | 0 |
| 0 | 0 | 0 |
| 1 | 0 | 0 |
| 0 | 0 | 0 |
| 1 | 1 | 1 |
| 1 | 0 | 0 |
| 0 | 0 | 0 |
| 0 | 0 | 0 |
| 1 | 1 | 1 |
| 0 | 0 | 0 |
| 0 | 0 | 0 |
| 0 | 0 | 0 |
| 1 | 0 | 0 |
| 0 | 0 | 0 |
| 0 | 0 | 0 |
| 0 | 1 | 0 |
| 0 | 0 | 0 |
| 0 | 0 | 0 |
| 0 | 0 | 0 |
| 0 | 0 | 0 |
| 1 | 1 | 1 |
| 0 | 0 | 0 |
| 0 | 0 | 0 |
| 0 | 0 | 0 |
| 0 | 0 | 0 |
| 0 | 0 | 0 |
| 0 | 0 | 0 |
| 0 | 0 | 0 |
| 0 | 0 | 0 |
| 0 | 0 | 0 |
| 0 | 0 | 0 |
| 0 | 1 | 0 |
| 0 | 1 | 0 |
| 0 | 0 | 0 |
| 1 | 1 | 1 |
| 0 | 0 | 0 |
| 0 | 0 | 0 |
| 0 | 0 | 0 |
| 0 | 0 | 0 |
| 1 | 1 | 1 |
| 0 | 0 | 0 |
| 0 | 0 | 0 |
| 1 | 1 | 1 |
| 0 | 1 | 0 |
| 1 | 1 | 1 |
| 1 | 1 | 1 |
| 1 | 1 | 1 |
| 0 | 1 | 0 |
| 1 | 1 | 1 |
| 1 | 1 | 1 |
| 1 | 1 | 1 |
| 1 | 1 | 1 |

| | | |
|---|---|---|
| 1 | 1 | 1 |
| 0 | 0 | 0 |
| 1 | 1 | 1 |
| 1 | 1 | 1 |
| 1 | 1 | 1 |
| 1 | 1 | 1 |
| 1 | 1 | 1 |
| 1 | 1 | 1 |
| 1 | 1 | 1 |
| 0 | 1 | 0 |
| 1 | 1 | 1 |
| 0 | 0 | 0 |
| 1 | 0 | 0 |
| 0 | 1 | 0 |
| 0 | 0 | 0 |
| 0 | 1 | 0 |
| 0 | 1 | 0 |
| 1 | 1 | 1 |
| 1 | 1 | 1 |
| 1 | 1 | 1 |
| 0 | 0 | 0 |
| 0 | 1 | 0 |
| 0 | 0 | 0 |
| 0 | 0 | 0 |
| 1 | 1 | 1 |
| 0 | 0 | 0 |
| 1 | 1 | 1 |
| 1 | 1 | 1 |
| 1 | 1 | 1 |
| 1 | 1 | 1 |
| 1 | 1 | 1 |
| 0 | 0 | 0 |
| 1 | 1 | 1 |
| 0 | 0 | 0 |
| 1 | 1 | 1 |
| 0 | 1 | 0 |
| 0 | 1 | 0 |
| 0 | 1 | 0 |
| 1 | 1 | 1 |
| 0 | 0 | 0 |
| 1 | 1 | 1 |
| 1 | 1 | 1 |
| 0 | 1 | 0 |
| 0 | 0 | 0 |
| 1 | 1 | 1 |
| 1 | 1 | 1 |
| 1 | 1 | 1 |
| 0 | 0 | 0 |
| 0 | 0 | 0 |
| 1 | 1 | 1 |
| 1 | 1 | 1 |
| 0 | 0 | 0 |
| 0 | 0 | 0 |
| 0 | 0 | 0 |

| | | |
|---|---|---|
| 0 | 0 | 0 |
| 1 | 1 | 1 |
| 0 | 0 | 0 |
| 0 | 0 | 0 |
| 0 | 0 | 0 |
| 1 | 1 | 1 |
| 1 | 1 | 1 |
| 0 | 1 | 0 |
| 1 | 1 | 1 |
| 0 | 0 | 0 |
| 1 | 1 | 1 |
| 0 | 1 | 0 |
| 1 | 1 | 1 |
| 1 | 1 | 1 |
| 0 | 0 | 0 |
| 0 | 1 | 0 |
| 1 | 1 | 1 |
| 0 | 0 | 0 |
| 1 | 1 | 1 |
| 0 | 0 | 0 |
| 0 | 0 | 0 |
| 1 | 1 | 1 |
| 0 | 0 | 0 |
| 1 | 1 | 1 |
| 0 | 0 | 0 |
| 0 | 0 | 0 |
| 1 | 1 | 1 |
| 0 | 1 | 0 |
| 0 | 0 | 0 |
| 1 | 1 | 1 |
| 1 | 1 | 1 |
| 1 | 1 | 1 |
| 1 | 1 | 1 |
| 0 | 1 | 0 |
| 0 | 0 | 0 |
| 0 | 0 | 0 |
| 1 | 1 | 1 |
| 0 | 0 | 0 |
| 0 | 0 | 0 |
| 0 | 0 | 0 |
| 0 | 0 | 0 |
| 0 | 0 | 0 |
| 0 | 0 | 0 |
| 0 | 1 | 0 |
| 0 | 0 | 0 |
| 1 | 1 | 1 |
| 0 | 1 | 0 |
| 0 | 0 | 0 |
| 0 | 0 | 0 |
| 0 | 0 | 0 |
| 0 | 0 | 0 |
| 0 | 0 | 0 |
| 1 | 1 | 1 |
| 1 | 1 | 1 |

| | | |
|---|---|---|
| 1 | 1 | 1 |
| 0 | 0 | 0 |
| 0 | 0 | 0 |
| 0 | 0 | 0 |
| 0 | 0 | 0 |
| 0 | 0 | 0 |
| 0 | 0 | 0 |
| 1 | 0 | 0 |
| 1 | 0 | 0 |
| 1 | 1 | 1 |
| 0 | 0 | 0 |
| 0 | 0 | 0 |
| 1 | 1 | 1 |
| 0 | 0 | 0 |
| 1 | 1 | 1 |
| 0 | 0 | 0 |
| 1 | 1 | 1 |
| 1 | 1 | 1 |
| 0 | 1 | 0 |
| 0 | 1 | 0 |
| 0 | 1 | 0 |
| 1 | 1 | 1 |
| 0 | 1 | 0 |
| 0 | 0 | 0 |
| 0 | 0 | 0 |
| 1 | 1 | 1 |
| 1 | 1 | 1 |
| 1 | 1 | 1 |
| 1 | 1 | 1 |
| 0 | 0 | 0 |
| 0 | 0 | 0 |
| 0 | 0 | 0 |
| 0 | 0 | 0 |
| 1 | 1 | 1 |
| 0 | 0 | 0 |
| 0 | 0 | 0 |
| 0 | 0 | 0 |
| 0 | 0 | 0 |
| 0 | 0 | 0 |
| 1 | 1 | 1 |
| 1 | 1 | 1 |
| 0 | 0 | 0 |
| 1 | 1 | 1 |
| 1 | 1 | 1 |
| 0 | 0 | 0 |
| 1 | 0 | 0 |
| 1 | 1 | 1 |
| 0 | 1 | 0 |
| 1 | 0 | 0 |
| 0 | 1 | 0 |
| 0 | 0 | 0 |
| 1 | 1 | 1 |
| 0 | 0 | 0 |
| 0 | 0 | 0 |

| | | |
|---|---|---|
| 0 | 1 | 0 |
| 1 | 1 | 1 |
| 0 | 0 | 0 |
| 0 | 0 | 0 |
| 1 | 1 | 1 |
| 1 | 1 | 1 |
| 1 | 1 | 1 |
| 1 | 1 | 1 |
| 1 | 1 | 1 |
| 1 | 1 | 1 |
| 1 | 1 | 1 |
| 1 | 1 | 1 |
| 0 | 0 | 0 |
| 1 | 1 | 1 |
| 1 | 1 | 1 |
| 1 | 1 | 1 |
| 1 | 1 | 1 |
| 1 | 1 | 1 |
| 1 | 1 | 1 |
| 1 | 1 | 1 |
| 1 | 1 | 1 |
| 1 | 1 | 1 |
| 1 | 1 | 1 |
| 1 | 1 | 1 |
| 1 | 1 | 1 |
| 1 | 1 | 1 |
| 0 | 1 | 0 |
| 1 | 1 | 1 |
| 1 | 1 | 1 |
| 1 | 1 | 1 |
| 1 | 1 | 1 |
| 0 | 0 | 0 |
| 1 | 1 | 1 |
| 0 | 1 | 0 |
| 1 | 1 | 1 |
| 1 | 1 | 1 |
| 0 | 1 | 0 |
| 1 | 1 | 1 |
| 1 | 1 | 1 |
| 1 | 1 | 1 |
| 0 | 0 | 0 |
| 0 | 0 | 0 |
| 1 | 1 | 1 |
| 1 | 1 | 1 |
| 1 | 1 | 1 |
| 1 | 0 | 0 |
| 1 | 0 | 0 |
| 1 | 1 | 1 |
| 0 | 0 | 0 |
| 0 | 0 | 0 |
| 1 | 1 | 1 |
| 0 | 1 | 0 |
| 0 | 0 | 0 |
| 1 | 1 | 1 |

| | | |
|---|---|---|
| 0 | 0 | 0 |
| 0 | 0 | 0 |
| 0 | 0 | 0 |
| 1 | 1 | 1 |
| 1 | 1 | 1 |
| 1 | 1 | 1 |
| 1 | 1 | 1 |
| 0 | 0 | 0 |
| 0 | 0 | 0 |
| 0 | 0 | 0 |
| 0 | 0 | 0 |
| 1 | 1 | 1 |
| 0 | 0 | 0 |
| 1 | 1 | 1 |
| 1 | 1 | 1 |
| 1 | 1 | 1 |
| 1 | 1 | 1 |
| 0 | 0 | 0 |
| 1 | 1 | 1 |
| 1 | 1 | 1 |
| 1 | 1 | 1 |
| 1 | 1 | 1 |
| 0 | 1 | 0 |
| 1 | 1 | 1 |
| 1 | 1 | 1 |
| 0 | 1 | 0 |
| 1 | 1 | 1 |
| 1 | 1 | 1 |
| 1 | 1 | 1 |
| 1 | 1 | 1 |
| 0 | 0 | 0 |
| 0 | 0 | 0 |
| 1 | 1 | 1 |
| 1 | 1 | 1 |
| 1 | 1 | 1 |
| 1 | 1 | 1 |
| 1 | 1 | 1 |
| 1 | 1 | 1 |
| 1 | 1 | 1 |
| 1 | 1 | 1 |
| 1 | 1 | 1 |
| 0 | 0 | 0 |
| 1 | 1 | 1 |
| 1 | 1 | 1 |
| 0 | 0 | 0 |
| 0 | 0 | 0 |
| 1 | 1 | 1 |
| 1 | 1 | 1 |
| 0 | 0 | 0 |
| 0 | 0 | 0 |
| 0 | 0 | 0 |
| 0 | 0 | 0 |
| 0 | 0 | 0 |
| 0 | 0 | 0 |

| | | |
|---|---|---|
| 1 | 1 | 1 |
| 1 | 1 | 1 |
| 1 | 1 | 1 |
| 1 | 1 | 1 |
| 1 | 1 | 1 |
| 0 | 0 | 0 |
| 0 | 0 | 0 |
| 0 | 0 | 0 |
| 0 | 0 | 0 |
| 0 | 0 | 0 |
| 0 | 1 | 0 |
| 0 | 0 | 0 |
| 0 | 0 | 0 |
| 0 | 0 | 0 |
| 0 | 1 | 0 |
| 0 | 0 | 0 |
| 0 | 1 | 0 |
| 0 | 1 | 0 |
| 0 | 0 | 0 |
| 0 | 0 | 0 |
| 0 | 0 | 0 |
| 0 | 0 | 0 |
| 1 | 0 | 0 |
| 0 | 0 | 0 |
| 0 | 0 | 0 |
| 0 | 0 | 0 |
| 0 | 0 | 0 |
| 1 | 1 | 1 |
| 1 | 1 | 1 |
| 1 | 1 | 1 |
| 1 | 1 | 1 |
| 1 | 1 | 1 |
| 0 | 0 | 0 |
| 0 | 1 | 0 |
| 0 | 0 | 0 |
| 0 | 0 | 0 |
| 0 | 0 | 0 |
| 0 | 0 | 0 |
| 0 | 0 | 0 |
| 0 | 0 | 0 |
| 0 | 0 | 0 |
| 0 | 0 | 0 |
| 0 | 0 | 0 |
| 1 | 1 | 1 |
| 1 | 1 | 1 |
| 1 | 1 | 1 |
| 1 | 1 | 1 |
| 0 | 0 | 0 |
| 0 | 0 | 0 |
| 0 | 0 | 0 |
| 1 | 1 | 1 |
| 1 | 1 | 1 |
| 0 | 0 | 0 |
| 1 | 1 | 1 |

| | | |
|---|---|---|
| 1 | 1 | 1 |
| 0 | 0 | 0 |
| 0 | 1 | 0 |
| 0 | 1 | 0 |
| 1 | 1 | 1 |
| 1 | 1 | 1 |
| 0 | 1 | 0 |
| 1 | 1 | 1 |
| 0 | 1 | 0 |
| 1 | 1 | 1 |
| 1 | 1 | 1 |
| 1 | 1 | 1 |
| 0 | 0 | 0 |
| 0 | 0 | 0 |
| 0 | 0 | 0 |
| 0 | 0 | 0 |
| 0 | 0 | 0 |
| 0 | 0 | 0 |
| 0 | 0 | 0 |
| 0 | 1 | 0 |
| 0 | 0 | 0 |
| 0 | 0 | 0 |
| 0 | 0 | 0 |
| 0 | 0 | 0 |
| 0 | 0 | 0 |
| 1 | 1 | 1 |
| 1 | 1 | 1 |
| 0 | 0 | 0 |
| 0 | 1 | 0 |
| 0 | 0 | 0 |
| 0 | 0 | 0 |
| 1 | 1 | 1 |
| 0 | 0 | 0 |
| 1 | 1 | 1 |
| 1 | 1 | 1 |
| 1 | 1 | 1 |
| 1 | 0 | 0 |
| 1 | 1 | 1 |
| 1 | 1 | 1 |
| 1 | 1 | 1 |
| 0 | 0 | 0 |
| 1 | 1 | 1 |
| 1 | 1 | 1 |
| 0 | 0 | 0 |
| 0 | 0 | 0 |
| 1 | 1 | 1 |
| 0 | 0 | 0 |
| 0 | 0 | 0 |
| 1 | 1 | 1 |
| 1 | 1 | 1 |
| 0 | 0 | 0 |
| 1 | 1 | 1 |
| 1 | 1 | 1 |
| 1 | 1 | 1 |

| | | |
|---|---|---|
| 1 | 1 | 1 |
| 1 | 1 | 1 |
| 1 | 1 | 1 |
| 1 | 1 | 1 |
| 0 | 0 | 0 |
| 0 | 0 | 0 |
| 1 | 1 | 1 |
| 1 | 1 | 1 |
| 1 | 1 | 1 |
| 0 | 1 | 0 |
| 1 | 1 | 1 |
| 0 | 0 | 0 |
| 0 | 0 | 0 |
| 1 | 1 | 1 |
| 1 | 1 | 1 |
| 0 | 0 | 0 |
| 0 | 1 | 0 |
| 1 | 1 | 1 |
| 0 | 0 | 0 |
| 1 | 1 | 1 |
| 1 | 1 | 1 |
| 1 | 1 | 1 |
| 1 | 1 | 1 |
| 0 | 0 | 0 |
| 1 | 1 | 1 |
| 0 | 0 | 0 |
| 1 | 1 | 1 |
| 1 | 1 | 1 |
| 0 | 0 | 0 |
| 0 | 0 | 0 |
| 0 | 0 | 0 |
| 0 | 0 | 0 |
| 0 | 0 | 0 |
| 0 | 0 | 0 |
| 1 | 1 | 1 |
| 1 | 1 | 1 |
| 1 | 1 | 1 |
| 0 | 0 | 0 |
| 1 | 1 | 1 |
| 1 | 1 | 1 |
| 0 | 0 | 0 |
| 0 | 0 | 0 |
| 0 | 0 | 0 |
| 1 | 1 | 1 |
| 0 | 0 | 0 |
| 0 | 1 | 0 |
| 0 | 1 | 0 |
| 0 | 0 | 0 |
| 1 | 1 | 1 |
| 1 | 1 | 1 |
| 1 | 1 | 1 |
| 0 | 0 | 0 |
| 1 | 1 | 1 |
| 0 | 0 | 0 |

| | | |
|---|---|---|
| 1 | 1 | 1 |
| 0 | 0 | 0 |
| 0 | 0 | 0 |
| 0 | 1 | 0 |
| 1 | 1 | 1 |
| 1 | 1 | 1 |
| 0 | 0 | 0 |
| 1 | 1 | 1 |
| 0 | 1 | 0 |
| 0 | 1 | 0 |
| 1 | 1 | 1 |
| 1 | 1 | 1 |
| 1 | 1 | 1 |
| 1 | 1 | 1 |
| 1 | 1 | 1 |
| 1 | 1 | 1 |
| 1 | 1 | 1 |
| 1 | 1 | 1 |
| 1 | 1 | 1 |
| 1 | 1 | 1 |
| 1 | 1 | 1 |
| 0 | 0 | 0 |
| 0 | 1 | 0 |
| 0 | 1 | 0 |
| 1 | 1 | 1 |
| 1 | 1 | 1 |
| 0 | 0 | 0 |
| 1 | 1 | 1 |
| 0 | 0 | 0 |
| 0 | 1 | 0 |
| 1 | 1 | 1 |
| 0 | 0 | 0 |
| 0 | 0 | 0 |
| 1 | 0 | 0 |
| 1 | 1 | 1 |
| 1 | 1 | 1 |
| 1 | 1 | 1 |
| 1 | 1 | 1 |
| 0 | 0 | 0 |
| 0 | 0 | 0 |
| 0 | 0 | 0 |
| 1 | 1 | 1 |
| 1 | 1 | 1 |
| 0 | 1 | 0 |
| 0 | 1 | 0 |
| 0 | 0 | 0 |
| 1 | 1 | 1 |
| 0 | 0 | 0 |
| 1 | 1 | 1 |
| 0 | 0 | 0 |
| 0 | 0 | 0 |
| 0 | 0 | 0 |
| 1 | 1 | 1 |
| 1 | 1 | 1 |

| | | |
|---|---|---|
| 0 | 0 | 0 |
| 0 | 0 | 0 |
| 1 | 1 | 1 |
| 1 | 1 | 1 |
| 1 | 1 | 1 |
| 1 | 1 | 1 |
| 1 | 1 | 1 |
| 1 | 1 | 1 |
| 1 | 1 | 1 |
| 1 | 1 | 1 |
| 1 | 1 | 1 |
| 1 | 1 | 1 |
| 1 | 1 | 1 |
| 1 | 1 | 1 |
| 1 | 1 | 1 |
| 0 | 0 | 0 |
| 0 | 0 | 0 |
| 0 | 0 | 0 |
| 1 | 1 | 1 |
| 1 | 1 | 1 |
| 0 | 1 | 0 |
| 0 | 0 | 0 |
| 0 | 0 | 0 |
| 0 | 0 | 0 |
| 0 | 0 | 0 |
| 1 | 1 | 1 |
| 1 | 1 | 1 |
| 1 | 1 | 1 |
| 1 | 1 | 1 |
| 1 | 1 | 1 |
| 0 | 0 | 0 |
| 0 | 1 | 0 |
| 0 | 0 | 0 |
| 1 | 1 | 1 |
| 1 | 0 | 0 |
| 1 | 0 | 0 |
| 0 | 0 | 0 |
| 0 | 0 | 0 |
| 0 | 1 | 0 |
| 1 | 1 | 1 |
| 1 | 0 | 0 |
| 0 | 0 | 0 |
| 1 | 1 | 1 |
| 0 | 1 | 0 |
| 0 | 0 | 0 |
| 1 | 0 | 0 |
| 0 | 0 | 0 |
| 0 | 0 | 0 |
| 1 | 1 | 1 |
| 0 | 1 | 0 |
| 0 | 0 | 0 |
| 0 | 0 | 0 |
| 1 | 1 | 1 |
| 0 | 0 | 0 |

| | | |
|---|---|---|
| 0 | 1 | 0 |
| 0 | 0 | 0 |
| 1 | 0 | 0 |
| 0 | 0 | 0 |
| 1 | 1 | 1 |
| 1 | 1 | 1 |
| 1 | 1 | 1 |
| 0 | 1 | 0 |
| 0 | 0 | 0 |
| 1 | 1 | 1 |
| 0 | 1 | 0 |
| 0 | 1 | 0 |
| 0 | 0 | 0 |
| 0 | 0 | 0 |
| 0 | 0 | 0 |
| 1 | 1 | 1 |
| 0 | 0 | 0 |
| 0 | 1 | 0 |
| 0 | 0 | 0 |
| 0 | 0 | 0 |
| 0 | 0 | 0 |
| 0 | 0 | 0 |
| 0 | 0 | 0 |
| 0 | 0 | 0 |
| 0 | 1 | 0 |
| 0 | 1 | 0 |
| 0 | 0 | 0 |
| 0 | 0 | 0 |
| 0 | 0 | 0 |
| 0 | 0 | 0 |
| 1 | 1 | 1 |
| 1 | 1 | 1 |
| 1 | 1 | 1 |
| 1 | 1 | 1 |
| 1 | 1 | 1 |
| 0 | 0 | 0 |
| 0 | 0 | 0 |
| 0 | 1 | 0 |
| 1 | 1 | 1 |
| 1 | 1 | 1 |
| 0 | 1 | 0 |
| 1 | 1 | 1 |
| 1 | 1 | 1 |
| 0 | 0 | 0 |
| 0 | 0 | 0 |
| 0 | 0 | 0 |
| 1 | 1 | 1 |
| 0 | 0 | 0 |
| 0 | 0 | 0 |
| 0 | 0 | 0 |
| 1 | 1 | 1 |
| 1 | 1 | 1 |
| 0 | 1 | 0 |
| 0 | 1 | 0 |

| | | |
|---|---|---|
| 0 | 1 | 0 |
| 1 | 1 | 1 |
| 1 | 0 | 0 |
| 1 | 1 | 1 |
| 0 | 0 | 0 |
| 1 | 0 | 0 |
| 0 | 0 | 0 |
| 0 | 0 | 0 |
| 0 | 0 | 0 |
| 0 | 0 | 0 |
| 0 | 0 | 0 |
| 0 | 0 | 0 |
| 0 | 1 | 0 |
| 0 | 0 | 0 |
| 0 | 0 | 0 |
| 1 | 1 | 1 |
| 1 | 1 | 1 |
| 1 | 1 | 1 |
| 0 | 0 | 0 |
| 0 | 0 | 0 |
| 0 | 0 | 0 |
| 0 | 0 | 0 |
| 0 | 1 | 0 |
| 0 | 1 | 0 |
| 1 | 1 | 1 |
| 1 | 1 | 1 |
| 1 | 1 | 1 |
| 0 | 0 | 0 |
| 0 | 0 | 0 |
| 0 | 0 | 0 |
| 0 | 0 | 0 |
| 0 | 0 | 0 |
| 0 | 0 | 0 |
| 0 | 1 | 0 |
| 1 | 1 | 1 |
| 0 | 0 | 0 |
| 1 | 1 | 1 |
| 1 | 1 | 1 |
| 1 | 1 | 1 |
| 1 | 1 | 1 |
| 0 | 0 | 0 |
| 1 | 1 | 1 |
| 0 | 0 | 0 |
| 0 | 1 | 0 |
| 1 | 1 | 1 |
| 1 | 1 | 1 |
| 1 | 1 | 1 |
| 0 | 0 | 0 |
| 1 | 1 | 1 |
| 1 | 1 | 1 |
| 1 | 1 | 1 |
| 1 | 1 | 1 |
| 1 | 1 | 1 |
| 0 | 1 | 0 |

| | | |
|---|---|---|
| 0 | 1 | 0 |
| 1 | 1 | 1 |
| 1 | 1 | 1 |
| 0 | 1 | 0 |
| 1 | 1 | 1 |
| 0 | 1 | 0 |
| 0 | 1 | 0 |
| 0 | 1 | 0 |
| 0 | 1 | 0 |
| 0 | 1 | 0 |
| 0 | 1 | 0 |
| 0 | 0 | 0 |
| 0 | 1 | 0 |
| 0 | 1 | 0 |
| 0 | 1 | 0 |
| 0 | 1 | 0 |
| 0 | 1 | 0 |
| 0 | 0 | 0 |
| 0 | 0 | 0 |
| 0 | 0 | 0 |
| 0 | 0 | 0 |
| 0 | 1 | 0 |
| 0 | 0 | 0 |
| 1 | 1 | 1 |
| 1 | 1 | 1 |
| 1 | 1 | 1 |
| 1 | 1 | 1 |
| 0 | 1 | 0 |
| 0 | 0 | 0 |
| 0 | 0 | 0 |
| 1 | 1 | 1 |
| 1 | 1 | 1 |
| 0 | 0 | 0 |
| 1 | 1 | 1 |
| 0 | 0 | 0 |
| 0 | 0 | 0 |
| 0 | 0 | 0 |
| 1 | 1 | 1 |
| 0 | 0 | 0 |
| 1 | 1 | 1 |
| 0 | 0 | 0 |
| 1 | 1 | 1 |
| 0 | 0 | 0 |
| 1 | 0 | 0 |
| 1 | 0 | 0 |
| 1 | 0 | 0 |
| 1 | 0 | 0 |
| 0 | 1 | 0 |
| 0 | 1 | 0 |
| 0 | 0 | 0 |
| 1 | 0 | 0 |
| 0 | 0 | 0 |
| 1 | 0 | 0 |
| 1 | 1 | 1 |

| | | |
|---|---|---|
| 0 | 0 | 0 |
| 0 | 0 | 0 |
| 0 | 0 | 0 |
| 0 | 0 | 0 |
| 0 | 0 | 0 |
| 1 | 1 | 1 |
| 0 | 0 | 0 |
| 0 | 0 | 0 |
| 1 | 1 | 1 |
| 0 | 0 | 0 |
| 0 | 0 | 0 |
| 0 | 0 | 0 |
| 0 | 1 | 0 |
| 0 | 1 | 0 |
| 0 | 1 | 0 |
| 0 | 1 | 0 |
| 0 | 1 | 0 |
| 0 | 1 | 0 |
| 0 | 1 | 0 |
| 0 | 1 | 0 |
| 0 | 1 | 0 |
| 0 | 1 | 0 |
| 0 | 1 | 0 |
| 0 | 1 | 0 |
| 0 | 1 | 0 |
| 0 | 1 | 0 |
| 0 | 1 | 0 |
| 0 | 1 | 0 |
| 0 | 1 | 0 |
| 0 | 1 | 0 |
| 0 | 1 | 0 |
| 0 | 1 | 0 |
| 0 | 1 | 0 |
| 0 | 1 | 0 |
| 0 | 1 | 0 |
| 0 | 1 | 0 |
| 0 | 1 | 0 |
| 0 | 1 | 0 |
| 0 | 1 | 0 |
| 0 | 1 | 0 |
| 0 | 1 | 0 |
| 0 | 1 | 0 |
| 0 | 1 | 0 |
| 0 | 1 | 0 |
| 0 | 0 | 0 |
| 1 | 1 | 1 |
| 1 | 1 | 1 |
| 1 | 1 | 1 |
| 1 | 1 | 1 |
| 0 | 0 | 0 |
| 1 | 1 | 1 |
| 1 | 1 | 1 |
| 0 | 1 | 0 |

| | | |
|---|---|---|
| 1 | 1 | 1 |
| 1 | 1 | 1 |
| 1 | 1 | 1 |
| 1 | 1 | 1 |
| 0 | 1 | 0 |
| 0 | 1 | 0 |
| 0 | 1 | 0 |
| 0 | 0 | 0 |
| 0 | 0 | 0 |
| 1 | 1 | 1 |
| 0 | 1 | 0 |
| 1 | 1 | 1 |
| 1 | 1 | 1 |
| 1 | 1 | 1 |
| 0 | 0 | 0 |
| 0 | 0 | 0 |
| 0 | 0 | 0 |
| 1 | 1 | 1 |
| 1 | 0 | 0 |
| 1 | 1 | 1 |
| 0 | 0 | 0 |
| 0 | 0 | 0 |
| 0 | 0 | 0 |
| 0 | 0 | 0 |
| 0 | 0 | 0 |
| 0 | 0 | 0 |
| 0 | 1 | 0 |
| 1 | 1 | 1 |
| 0 | 0 | 0 |
| 1 | 1 | 1 |
| 1 | 1 | 1 |
| 1 | 1 | 1 |
| 0 | 0 | 0 |
| 1 | 1 | 1 |
| 1 | 1 | 1 |
| 1 | 1 | 1 |
| 0 | 0 | 0 |
| 0 | 0 | 0 |
| 0 | 0 | 0 |
| 1 | 1 | 1 |
| 1 | 1 | 1 |
| 1 | 1 | 1 |
| 1 | 1 | 1 |
| 1 | 1 | 1 |
| 0 | 1 | 0 |
| 0 | 0 | 0 |
| 1 | 1 | 1 |
| 0 | 1 | 0 |
| 1 | 1 | 1 |
| 0 | 1 | 0 |
| 1 | 1 | 1 |
| 1 | 1 | 1 |
| 0 | 1 | 0 |
| 1 | 1 | 1 |

| | | |
|---|---|---|
| 1 | 1 | 1 |
| 0 | 1 | 0 |
| 1 | 1 | 1 |
| 0 | 0 | 0 |
| 1 | 1 | 1 |
| 1 | 1 | 1 |
| 1 | 1 | 1 |
| 0 | 0 | 0 |
| 0 | 0 | 0 |
| 1 | 1 | 1 |
| 1 | 1 | 1 |
| 1 | 1 | 1 |
| 0 | 0 | 0 |
| 0 | 1 | 0 |
| 0 | 1 | 0 |
| 0 | 1 | 0 |
| 0 | 0 | 0 |
| 1 | 1 | 1 |
| 1 | 1 | 1 |
| 1 | 1 | 1 |
| 1 | 1 | 1 |
| 1 | 1 | 1 |
| 1 | 1 | 1 |
| 1 | 1 | 1 |
| 0 | 1 | 0 |
| 1 | 1 | 1 |
| 0 | 1 | 0 |
| 0 | 0 | 0 |
| 0 | 0 | 0 |
| 0 | 0 | 0 |
| 0 | 0 | 0 |
| 0 | 0 | 0 |
| 0 | 0 | 0 |
| 0 | 1 | 0 |
| 0 | 0 | 0 |
| 0 | 0 | 0 |
| 0 | 0 | 0 |
| 1 | 1 | 1 |
| 1 | 1 | 1 |
| 1 | 1 | 1 |
| 1 | 1 | 1 |
| 0 | 1 | 0 |
| 0 | 0 | 0 |
| 0 | 0 | 0 |
| 1 | 1 | 1 |
| 1 | 1 | 1 |
| 0 | 0 | 0 |
| 1 | 1 | 1 |
| 1 | 1 | 1 |
| 1 | 1 | 1 |
| 1 | 1 | 1 |
| 1 | 1 | 1 |
| 0 | 0 | 0 |
| 0 | 0 | 0 |

| | | |
|---|---|---|
| 0 | 0 | 0 |
| 0 | 0 | 0 |
| 0 | 0 | 0 |
| 0 | 0 | 0 |
| 0 | 0 | 0 |
| 0 | 0 | 0 |
| 0 | 0 | 0 |
| 0 | 0 | 0 |
| 1 | 1 | 1 |
| 0 | 0 | 0 |
| 1 | 1 | 1 |
| 1 | 1 | 1 |
| 1 | 1 | 1 |
| 1 | 1 | 1 |
| 0 | 0 | 0 |
| 1 | 1 | 1 |
| 1 | 1 | 1 |
| 1 | 1 | 1 |
| 1 | 1 | 1 |
| 0 | 0 | 0 |
| 0 | 0 | 0 |
| 1 | 1 | 1 |
| 0 | 0 | 0 |
| 0 | 0 | 0 |
| 0 | 0 | 0 |
| 0 | 0 | 0 |
| 0 | 0 | 0 |
| 0 | 1 | 0 |
| 0 | 1 | 0 |
| 0 | 1 | 0 |
| 0 | 0 | 0 |
| 1 | 1 | 1 |
| 1 | 1 | 1 |
| 1 | 1 | 1 |
| 1 | 1 | 1 |
| 1 | 1 | 1 |
| 1 | 1 | 1 |
| 1 | 1 | 1 |
| 1 | 1 | 1 |
| 1 | 1 | 1 |
| 0 | 1 | 0 |
| 0 | 0 | 0 |
| 1 | 1 | 1 |
| 1 | 1 | 1 |
| 1 | 1 | 1 |
| 0 | 1 | 0 |
| 1 | 1 | 1 |
| 1 | 1 | 1 |
| 0 | 0 | 0 |
| 0 | 1 | 0 |
| 1 | 1 | 1 |
| 1 | 1 | 1 |
| 0 | 0 | 0 |
| 0 | 0 | 0 |

| | | |
|---|---|---|
| 1 | 1 | 1 |
| 1 | 1 | 1 |
| 0 | 1 | 0 |
| 1 | 0 | 0 |
| 0 | 0 | 0 |
| 0 | 0 | 0 |
| 0 | 1 | 0 |
| 0 | 0 | 0 |
| 1 | 1 | 1 |
| 1 | 1 | 1 |
| 0 | 0 | 0 |
| 1 | 0 | 0 |
| 0 | 0 | 0 |
| 0 | 0 | 0 |
| 0 | 0 | 0 |
| 0 | 0 | 0 |
| 1 | 1 | 1 |
| 1 | 1 | 1 |
| 0 | 0 | 0 |
| 0 | 0 | 0 |
| 0 | 0 | 0 |
| 0 | 0 | 0 |
| 0 | 0 | 0 |
| 1 | 1 | 1 |
| 1 | 1 | 1 |
| 0 | 0 | 0 |
| 1 | 1 | 1 |
| 0 | 0 | 0 |
| 0 | 0 | 0 |
| 0 | 0 | 0 |
| 0 | 1 | 0 |
| 0 | 0 | 0 |
| 0 | 0 | 0 |
| 1 | 1 | 1 |
| 1 | 1 | 1 |
| 0 | 0 | 0 |
| 0 | 0 | 0 |
| 1 | 0 | 0 |
| 0 | 0 | 0 |
| 0 | 0 | 0 |
| 0 | 0 | 0 |
| 0 | 0 | 0 |
| 0 | 1 | 0 |
| 1 | 1 | 1 |
| 0 | 0 | 0 |
| 0 | 1 | 0 |
| 0 | 0 | 0 |
| 1 | 1 | 1 |
| 1 | 1 | 1 |
| 1 | 1 | 1 |
| 1 | 1 | 1 |
| 1 | 1 | 1 |
| 0 | 1 | 0 |
| 1 | 1 | 1 |

| | | |
|---|---|---|
| 1 | 1 | 1 |
| 1 | 1 | 1 |
| 1 | 1 | 1 |
| 0 | 1 | 0 |
| 1 | 1 | 1 |
| 1 | 1 | 1 |
| 0 | 1 | 0 |
| 0 | 0 | 0 |
| 1 | 1 | 1 |
| 0 | 0 | 0 |
| 0 | 0 | 0 |
| 0 | 0 | 0 |
| 0 | 1 | 0 |
| 1 | 1 | 1 |
| 0 | 0 | 0 |
| 0 | 1 | 0 |
| 0 | 1 | 0 |
| 0 | 1 | 0 |
| 0 | 1 | 0 |
| 0 | 1 | 0 |
| 1 | 1 | 1 |
| 1 | 1 | 1 |
| 1 | 1 | 1 |
| 0 | 1 | 0 |
| 1 | 1 | 1 |
| 1 | 1 | 1 |
| 1 | 1 | 1 |
| 1 | 0 | 0 |
| 1 | 1 | 1 |
| 0 | 0 | 0 |
| 0 | 0 | 0 |
| 1 | 1 | 1 |
| 1 | 1 | 1 |
| 0 | 1 | 0 |
| 0 | 1 | 0 |
| 0 | 0 | 0 |
| 1 | 1 | 1 |
| 1 | 1 | 1 |
| 1 | 1 | 1 |
| 1 | 1 | 1 |
| 1 | 1 | 1 |
| 0 | 0 | 0 |
| 0 | 0 | 0 |
| 0 | 0 | 0 |
| 1 | 1 | 1 |
| 1 | 1 | 1 |
| 1 | 1 | 1 |
| 1 | 1 | 1 |
| 1 | 1 | 1 |
| 1 | 1 | 1 |
| 1 | 1 | 1 |
| 1 | 1 | 1 |
| 1 | 1 | 1 |
| 0 | 0 | 0 |

| | | |
|---|---|---|
| 1 | 1 | 1 |
| 1 | 1 | 1 |
| 1 | 1 | 1 |
| 1 | 1 | 1 |
| 1 | 1 | 1 |
| 1 | 1 | 1 |
| 1 | 1 | 1 |
| 0 | 1 | 0 |
| 0 | 0 | 0 |
| 0 | 0 | 0 |
| 1 | 1 | 1 |
| 0 | 1 | 0 |
| 0 | 1 | 0 |
| 0 | 0 | 0 |
| 1 | 1 | 1 |
| 1 | 1 | 1 |
| 1 | 1 | 1 |
| 0 | 0 | 0 |
| 0 | 0 | 0 |
| 0 | 0 | 0 |
| 0 | 0 | 0 |
| 0 | 1 | 0 |
| 1 | 1 | 1 |
| 1 | 1 | 1 |
| 1 | 1 | 1 |
| 0 | 0 | 0 |
| 1 | 1 | 1 |
| 0 | 1 | 0 |
| 1 | 1 | 1 |
| 1 | 1 | 1 |
| 1 | 1 | 1 |
| 1 | 1 | 1 |
| 1 | 1 | 1 |
| 1 | 1 | 1 |
| 1 | 1 | 1 |
| 1 | 1 | 1 |
| 1 | 1 | 1 |
| 1 | 1 | 1 |
| 1 | 1 | 1 |
| 0 | 0 | 0 |
| 1 | 1 | 1 |
| 1 | 1 | 1 |
| 0 | 0 | 0 |
| 1 | 1 | 1 |
| 1 | 1 | 1 |
| 1 | 1 | 1 |
| 0 | 0 | 0 |
| 0 | 0 | 0 |
| 1 | 1 | 1 |
| 1 | 1 | 1 |
| 0 | 0 | 0 |
| 1 | 1 | 1 |
| 0 | 1 | 0 |
| 1 | 1 | 1 |

| | | |
|---|---|---|
| 1 | 1 | 1 |
| 1 | 1 | 1 |
| 1 | 1 | 1 |
| 1 | 1 | 1 |
| 1 | 1 | 1 |
| 1 | 1 | 1 |
| 1 | 1 | 1 |
| 0 | 1 | 0 |
| 1 | 1 | 1 |
| 1 | 1 | 1 |
| 1 | 1 | 1 |
| 1 | 1 | 1 |
| 1 | 1 | 1 |
| 0 | 0 | 0 |
| 0 | 0 | 0 |
| 0 | 0 | 0 |
| 0 | 0 | 0 |
| 0 | 1 | 0 |
| 1 | 1 | 1 |
| 0 | 0 | 0 |
| 1 | 1 | 1 |
| 1 | 0 | 0 |
| 1 | 1 | 1 |
| 1 | 1 | 1 |
| 1 | 1 | 1 |
| 0 | 0 | 0 |
| 0 | 1 | 0 |
| 1 | 1 | 1 |
| 1 | 1 | 1 |
| 0 | 0 | 0 |
| 0 | 0 | 0 |
| 1 | 1 | 1 |
| 1 | 1 | 1 |
| 1 | 1 | 1 |
| 1 | 1 | 1 |
| 1 | 1 | 1 |
| 1 | 1 | 1 |
| 1 | 1 | 1 |
| 0 | 0 | 0 |
| 1 | 1 | 1 |
| 1 | 1 | 1 |
| 0 | 0 | 0 |
| 0 | 0 | 0 |
| 0 | 0 | 0 |
| 0 | 0 | 0 |
| 1 | 1 | 1 |
| 0 | 0 | 0 |
| 1 | 1 | 1 |
| 1 | 1 | 1 |
| 1 | 1 | 1 |
| 0 | 1 | 0 |
| 1 | 0 | 0 |
| 1 | 0 | 0 |
| 1 | 0 | 0 |

| | | |
|---|---|---|
| 1 | 1 | 1 |
| 1 | 1 | 1 |
| 0 | 0 | 0 |
| 1 | 1 | 1 |
| 1 | 1 | 1 |
| 1 | 1 | 1 |
| 1 | 1 | 1 |
| 0 | 1 | 0 |
| 1 | 1 | 1 |
| 1 | 1 | 1 |
| 1 | 1 | 1 |
| 0 | 1 | 0 |
| 0 | 1 | 0 |
| 0 | 1 | 0 |
| 0 | 1 | 0 |
| 0 | 1 | 0 |
| 0 | 1 | 0 |
| 0 | 1 | 0 |
| 0 | 1 | 0 |
| 0 | 1 | 0 |
| 0 | 1 | 0 |
| 0 | 1 | 0 |
| 0 | 1 | 0 |
| 0 | 1 | 0 |
| 0 | 1 | 0 |
| 0 | 1 | 0 |
| 0 | 1 | 0 |
| 0 | 1 | 0 |
| 1 | 1 | 1 |
| 1 | 1 | 1 |
| 0 | 0 | 0 |
| 1 | 1 | 1 |
| 0 | 0 | 0 |
| 0 | 0 | 0 |
| 0 | 0 | 0 |
| 0 | 0 | 0 |
| 1 | 1 | 1 |
| 0 | 0 | 0 |
| 1 | 1 | 1 |
| 1 | 1 | 1 |
| 0 | 0 | 0 |
| 0 | 0 | 0 |
| 0 | 0 | 0 |
| 1 | 1 | 1 |
| 0 | 0 | 0 |
| 0 | 0 | 0 |
| 1 | 1 | 1 |
| 1 | 1 | 1 |
| 1 | 1 | 1 |
| 1 | 1 | 1 |
| 0 | 0 | 0 |
| 1 | 1 | 1 |
| 1 | 1 | 1 |
| 1 | 1 | 1 |

| | | |
|---|---|---|
| 1 | 1 | 1 |
| 1 | 1 | 1 |
| 1 | 1 | 1 |
| 1 | 1 | 1 |
| 1 | 1 | 1 |
| 0 | 0 | 0 |
| 1 | 1 | 1 |
| 0 | 0 | 0 |
| 0 | 0 | 0 |
| 0 | 0 | 0 |
| 1 | 1 | 1 |
| 0 | 0 | 0 |
| 1 | 1 | 1 |
| 1 | 1 | 1 |
| 1 | 1 | 1 |
| 1 | 0 | 0 |
| 1 | 1 | 1 |
| 0 | 0 | 0 |
| 1 | 1 | 1 |
| 0 | 0 | 0 |
| 1 | 1 | 1 |
| 0 | 0 | 0 |
| 0 | 0 | 0 |
| 0 | 0 | 0 |
| 1 | 1 | 1 |
| 0 | 1 | 0 |
| 1 | 1 | 1 |
| 1 | 1 | 1 |
| 0 | 0 | 0 |
| 1 | 1 | 1 |
| 1 | 1 | 1 |
| 1 | 1 | 1 |
| 1 | 1 | 1 |
| 1 | 1 | 1 |
| 0 | 0 | 0 |
| 0 | 0 | 0 |
| 1 | 1 | 1 |
| 0 | 0 | 0 |
| 0 | 0 | 0 |
| 0 | 1 | 0 |
| 0 | 0 | 0 |
| 0 | 0 | 0 |
| 0 | 1 | 0 |
| 0 | 1 | 0 |
| 1 | 1 | 1 |
| 0 | 1 | 0 |
| 1 | 1 | 1 |
| 1 | 1 | 1 |
| 1 | 1 | 1 |
| 1 | 1 | 1 |
| 1 | 1 | 1 |
| 1 | 1 | 1 |
| 1 | 1 | 1 |
| 1 | 1 | 1 |

| | | |
|---|---|---|
| 1 | 1 | 1 |
| 0 | 0 | 0 |
| 0 | 1 | 0 |
| 1 | 1 | 1 |
| 0 | 0 | 0 |
| 1 | 1 | 1 |
| 1 | 1 | 1 |
| 1 | 1 | 1 |
| 0 | 0 | 0 |
| 0 | 0 | 0 |
| 1 | 1 | 1 |
| 0 | 0 | 0 |
| 0 | 0 | 0 |
| 1 | 1 | 1 |
| 1 | 1 | 1 |
| 0 | 0 | 0 |
| 0 | 1 | 0 |
| 0 | 0 | 0 |
| 0 | 0 | 0 |
| 1 | 1 | 1 |
| 0 | 0 | 0 |
| 0 | 0 | 0 |
| 0 | 0 | 0 |
| 1 | 1 | 1 |
| 0 | 1 | 0 |
| 0 | 0 | 0 |
| 1 | 1 | 1 |
| 1 | 1 | 1 |
| 1 | 1 | 1 |
| 1 | 1 | 1 |
| 0 | 0 | 0 |
| 1 | 1 | 1 |
| 0 | 0 | 0 |
| 1 | 1 | 1 |
| 0 | 0 | 0 |
| 0 | 0 | 0 |
| 1 | 1 | 1 |
| 1 | 1 | 1 |
| 1 | 1 | 1 |
| 1 | 1 | 1 |
| 1 | 1 | 1 |
| 1 | 1 | 1 |
| 1 | 1 | 1 |
| 1 | 1 | 1 |
| 1 | 1 | 1 |
| 1 | 1 | 1 |
| 1 | 0 | 0 |
| 1 | 1 | 1 |
| 1 | 1 | 1 |
| 1 | 1 | 1 |
| 1 | 1 | 1 |
| 1 | 1 | 1 |
| 1 | 1 | 1 |
| 0 | 0 | 0 |

| | | |
|---|---|---|
| 1 | 1 | 1 |
| 1 | 1 | 1 |
| 1 | 1 | 1 |
| 1 | 1 | 1 |
| 0 | 0 | 0 |
| 1 | 1 | 1 |
| 0 | 1 | 0 |
| 0 | 1 | 0 |
| 0 | 1 | 0 |
| 1 | 1 | 1 |
| 0 | 0 | 0 |
| 1 | 1 | 1 |
| 1 | 1 | 1 |
| 1 | 1 | 1 |
| 0 | 0 | 0 |
| 0 | 0 | 0 |
| 0 | 0 | 0 |
| 0 | 0 | 0 |
| 0 | 1 | 0 |
| 0 | 0 | 0 |
| 0 | 0 | 0 |
| 0 | 0 | 0 |
| 1 | 1 | 1 |
| 0 | 0 | 0 |
| 0 | 0 | 0 |
| 1 | 1 | 1 |
| 0 | 0 | 0 |
| 0 | 0 | 0 |
| 0 | 0 | 0 |
| 0 | 0 | 0 |
| 0 | 0 | 0 |
| 1 | 1 | 1 |
| 1 | 1 | 1 |
| 0 | 1 | 0 |
| 1 | 1 | 1 |
| 1 | 1 | 1 |
| 1 | 1 | 1 |
| 0 | 0 | 0 |
| 0 | 0 | 0 |
| 0 | 1 | 0 |
| 1 | 1 | 1 |
| 0 | 0 | 0 |
| 0 | 0 | 0 |
| 0 | 1 | 0 |
| 0 | 0 | 0 |
| 1 | 1 | 1 |
| 1 | 1 | 1 |
| 1 | 1 | 1 |
| 0 | 0 | 0 |
| 0 | 0 | 0 |
| 0 | 0 | 0 |
| 1 | 1 | 1 |
| 0 | 1 | 0 |
| 0 | 0 | 0 |

| | | |
|---|---|---|
| 1 | 1 | 1 |
| 0 | 0 | 0 |
| 1 | 1 | 1 |
| 0 | 0 | 0 |
| 0 | 0 | 0 |
| 0 | 1 | 0 |
| 0 | 0 | 0 |
| 0 | 0 | 0 |
| 1 | 1 | 1 |
| 1 | 1 | 1 |
| 1 | 1 | 1 |
| 1 | 1 | 1 |
| 1 | 1 | 1 |
| 1 | 1 | 1 |
| 1 | 1 | 1 |
| 1 | 1 | 1 |
| 1 | 1 | 1 |
| 0 | 1 | 0 |
| 1 | 1 | 1 |
| 1 | 1 | 1 |
| 0 | 0 | 0 |
| 0 | 1 | 0 |
| 0 | 1 | 0 |
| 0 | 1 | 0 |
| 0 | 1 | 0 |
| 0 | 0 | 0 |
| 0 | 0 | 0 |
| 1 | 1 | 1 |
| 1 | 1 | 1 |
| 1 | 1 | 1 |
| 1 | 1 | 1 |
| 1 | 1 | 1 |
| 1 | 1 | 1 |
| 0 | 1 | 0 |
| 1 | 1 | 1 |
| 1 | 1 | 1 |
| 1 | 1 | 1 |
| 1 | 1 | 1 |
| 0 | 1 | 0 |
| 1 | 1 | 1 |
| 1 | 1 | 1 |
| 0 | 1 | 0 |
| 1 | 1 | 1 |
| 0 | 0 | 0 |
| 1 | 1 | 1 |
| 0 | 0 | 0 |
| 0 | 0 | 0 |
| 0 | 0 | 0 |
| 0 | 0 | 0 |
| 0 | 0 | 0 |
| 0 | 0 | 0 |
| 0 | 1 | 0 |
| 1 | 1 | 1 |
| 1 | 1 | 1 |

| | | |
|---|---|---|
| 1 | 1 | 1 |
| 1 | 1 | 1 |
| 1 | 1 | 1 |
| 0 | 0 | 0 |
| 1 | 1 | 1 |
| 0 | 0 | 0 |
| 1 | 1 | 1 |
| 1 | 1 | 1 |
| 0 | 0 | 0 |
| 1 | 1 | 1 |
| 1 | 1 | 1 |
| 0 | 1 | 0 |
| 1 | 1 | 1 |
| 1 | 1 | 1 |
| 1 | 1 | 1 |
| 0 | 0 | 0 |
| 0 | 0 | 0 |
| 0 | 0 | 0 |
| 0 | 0 | 0 |
| 0 | 0 | 0 |
| 1 | 1 | 1 |
| 1 | 1 | 1 |
| 1 | 1 | 1 |
| 1 | 1 | 1 |
| 0 | 0 | 0 |
| 1 | 1 | 1 |
| 1 | 1 | 1 |
| 1 | 1 | 1 |
| 0 | 0 | 0 |
| 1 | 1 | 1 |
| 0 | 0 | 0 |
| 0 | 0 | 0 |
| 0 | 0 | 0 |
| 0 | 0 | 0 |
| 0 | 0 | 0 |
| 1 | 1 | 1 |
| 0 | 0 | 0 |
| 0 | 0 | 0 |
| 0 | 0 | 0 |
| 0 | 0 | 0 |
| 0 | 0 | 0 |
| 0 | 0 | 0 |
| 0 | 0 | 0 |
| 0 | 0 | 0 |
| 0 | 0 | 0 |
| 0 | 0 | 0 |
| 0 | 0 | 0 |
| 0 | 0 | 0 |
| 1 | 1 | 1 |
| 0 | 0 | 0 |
| 1 | 1 | 1 |
| 1 | 1 | 1 |
| 0 | 1 | 0 |
| 0 | 0 | 0 |

| | | |
|---|---|---|
| 0 | 0 | 0 |
| 0 | 1 | 0 |
| 1 | 1 | 1 |
| 1 | 1 | 1 |
| 1 | 1 | 1 |
| 0 | 0 | 0 |
| 1 | 1 | 1 |
| 0 | 1 | 0 |
| 1 | 1 | 1 |
| 0 | 1 | 0 |
| 0 | 1 | 0 |
| 0 | 1 | 0 |
| 1 | 1 | 1 |
| 0 | 0 | 0 |
| 0 | 1 | 0 |
| 1 | 1 | 1 |
| 1 | 1 | 1 |
| 0 | 0 | 0 |
| 1 | 1 | 1 |
| 1 | 1 | 1 |
| 1 | 1 | 1 |
| 1 | 1 | 1 |
| 0 | 0 | 0 |
| 1 | 1 | 1 |
| 1 | 1 | 1 |
| 0 | 0 | 0 |
| 0 | 0 | 0 |
| 0 | 0 | 0 |
| 1 | 1 | 1 |
| 0 | 1 | 0 |
| 1 | 1 | 1 |
| 1 | 1 | 1 |
| 1 | 1 | 1 |
| 1 | 1 | 1 |
| 1 | 1 | 1 |
| 1 | 1 | 1 |
| 1 | 1 | 1 |
| 1 | 1 | 1 |
| 1 | 1 | 1 |
| 1 | 1 | 1 |
| 1 | 1 | 1 |
| 1 | 1 | 1 |
| 1 | 1 | 1 |
| 1 | 1 | 1 |
| 1 | 1 | 1 |
| 1 | 1 | 1 |
| 1 | 1 | 1 |
| 1 | 1 | 1 |
| 1 | 0 | 0 |
| 1 | 1 | 1 |
| 1 | 1 | 1 |
| 1 | 1 | 1 |
| 1 | 1 | 1 |
| 1 | 1 | 1 |

| | | |
|---|---|---|
| 1 | 1 | 1 |
| 0 | 0 | 0 |
| 1 | 1 | 1 |
| 1 | 1 | 1 |
| 1 | 1 | 1 |
| 0 | 0 | 0 |
| 0 | 0 | 0 |
| 0 | 0 | 0 |
| 1 | 1 | 1 |
| 1 | 1 | 1 |
| 0 | 1 | 0 |
| 1 | 1 | 1 |
| 1 | 1 | 1 |
| 1 | 1 | 1 |
| 0 | 0 | 0 |
| 0 | 0 | 0 |
| 0 | 0 | 0 |
| 1 | 1 | 1 |
| 0 | 0 | 0 |
| 0 | 0 | 0 |
| 0 | 0 | 0 |
| 0 | 0 | 0 |
| 0 | 0 | 0 |
| 0 | 0 | 0 |
| 0 | 1 | 0 |
| 1 | 1 | 1 |
| 1 | 1 | 1 |
| 0 | 1 | 0 |
| 0 | 1 | 0 |
| 1 | 1 | 1 |
| 1 | 1 | 1 |
| 1 | 1 | 1 |
| 0 | 0 | 0 |
| 1 | 1 | 1 |
| 0 | 1 | 0 |
| 0 | 0 | 0 |
| 0 | 0 | 0 |
| 0 | 0 | 0 |
| 0 | 0 | 0 |
| 1 | 1 | 1 |
| 0 | 1 | 0 |
| 1 | 1 | 1 |
| 1 | 1 | 1 |
| 0 | 0 | 0 |
| 0 | 0 | 0 |
| 0 | 0 | 0 |
| 0 | 1 | 0 |
| 1 | 1 | 1 |
| 0 | 0 | 0 |
| 1 | 1 | 1 |
| 1 | 1 | 1 |
| 0 | 0 | 0 |
| 1 | 1 | 1 |
| 0 | 1 | 0 |

| | | |
|---|---|---|
| 1 | 1 | 1 |
| 0 | 1 | 0 |
| 1 | 1 | 1 |
| 1 | 1 | 1 |
| 1 | 1 | 1 |
| 1 | 1 | 1 |
| 0 | 1 | 0 |
| 1 | 1 | 1 |
| 0 | 0 | 0 |
| 0 | 0 | 0 |
| 1 | 1 | 1 |
| 0 | 0 | 0 |
| 0 | 0 | 0 |
| 0 | 0 | 0 |
| 1 | 1 | 1 |
| 1 | 1 | 1 |
| 0 | 0 | 0 |
| 0 | 0 | 0 |
| 0 | 0 | 0 |
| 1 | 1 | 1 |
| 1 | 1 | 1 |
| 1 | 1 | 1 |
| 0 | 0 | 0 |
| 1 | 1 | 1 |
| 0 | 0 | 0 |
| 1 | 1 | 1 |
| 1 | 1 | 1 |
| 1 | 1 | 1 |
| 1 | 1 | 1 |
| 0 | 0 | 0 |
| 0 | 0 | 0 |
| 0 | 0 | 0 |
| 1 | 1 | 1 |
| 0 | 1 | 0 |
| 1 | 1 | 1 |
| 1 | 1 | 1 |
| 1 | 1 | 1 |
| 0 | 1 | 0 |
| 0 | 1 | 0 |
| 0 | 1 | 0 |
| 0 | 0 | 0 |
| 1 | 1 | 1 |
| 1 | 1 | 1 |
| 1 | 1 | 1 |
| 1 | 1 | 1 |
| 0 | 0 | 0 |
| 1 | 0 | 0 |
| 0 | 0 | 0 |
| 1 | 1 | 1 |
| 0 | 0 | 0 |
| 0 | 0 | 0 |
| 0 | 0 | 0 |
| 0 | 1 | 0 |
| 1 | 0 | 0 |

| | | |
|---|---|---|
| 1 | 0 | 0 |
| 0 | 0 | 0 |
| 0 | 1 | 0 |
| 1 | 1 | 1 |
| 0 | 1 | 0 |
| 0 | 1 | 0 |
| 0 | 1 | 0 |
| 0 | 0 | 0 |
| 0 | 0 | 0 |
| 0 | 1 | 0 |
| 0 | 0 | 0 |
| 1 | 1 | 1 |
| 1 | 1 | 1 |
| 1 | 1 | 1 |
| 0 | 1 | 0 |
| 1 | 1 | 1 |
| 1 | 1 | 1 |
| 1 | 1 | 1 |
| 1 | 1 | 1 |
| 0 | 1 | 0 |
| 1 | 1 | 1 |
| 1 | 1 | 1 |
| 1 | 1 | 1 |
| 1 | 1 | 1 |
| 0 | 1 | 0 |
| 0 | 1 | 0 |
| 0 | 1 | 0 |
| 0 | 1 | 0 |
| 0 | 0 | 0 |
| 0 | 1 | 0 |
| 0 | 0 | 0 |
| 1 | 1 | 1 |
| 0 | 0 | 0 |
| 0 | 0 | 0 |
| 0 | 1 | 0 |
| 0 | 0 | 0 |
| 0 | 0 | 0 |
| 1 | 1 | 1 |
| 0 | 1 | 0 |
| 0 | 1 | 0 |
| 1 | 1 | 1 |
| 0 | 0 | 0 |
| 0 | 0 | 0 |
| 1 | 1 | 1 |
| 1 | 1 | 1 |
| 0 | 0 | 0 |
| 0 | 0 | 0 |
| 0 | 0 | 0 |
| 0 | 0 | 0 |
| 0 | 1 | 0 |
| 0 | 0 | 0 |
| 0 | 0 | 0 |
| 1 | 1 | 1 |
| 1 | 1 | 1 |

| | | |
|---|---|---|
| 1 | 1 | 1 |
| 1 | 1 | 1 |
| 1 | 1 | 1 |
| 1 | 1 | 1 |
| 0 | 0 | 0 |
| 0 | 0 | 0 |
| 0 | 0 | 0 |
| 0 | 0 | 0 |
| 1 | 1 | 1 |
| 0 | 0 | 0 |
| 0 | 0 | 0 |
| 0 | 0 | 0 |
| 0 | 0 | 0 |
| 1 | 1 | 1 |
| 1 | 0 | 0 |
| 0 | 1 | 0 |
| 1 | 1 | 1 |
| 1 | 1 | 1 |
| 1 | 1 | 1 |
| 0 | 1 | 0 |
| 1 | 1 | 1 |
| 1 | 1 | 1 |
| 1 | 1 | 1 |
| 1 | 1 | 1 |
| 1 | 1 | 1 |
| 0 | 0 | 0 |
| 0 | 0 | 0 |
| 0 | 0 | 0 |
| 0 | 0 | 0 |
| 0 | 1 | 0 |
| 0 | 1 | 0 |
| 0 | 1 | 0 |
| 1 | 1 | 1 |
| 0 | 1 | 0 |
| 0 | 1 | 0 |
| 1 | 1 | 1 |
| 1 | 1 | 1 |
| 1 | 1 | 1 |
| 1 | 1 | 1 |
| 1 | 1 | 1 |
| 1 | 1 | 1 |
| 1 | 1 | 1 |
| 1 | 1 | 1 |
| 1 | 1 | 1 |
| 1 | 1 | 1 |
| 1 | 1 | 1 |
| 1 | 1 | 1 |
| 1 | 1 | 1 |
| 1 | 1 | 1 |
| 1 | 1 | 1 |
| 1 | 1 | 1 |
| 1 | 1 | 1 |
| 1 | 1 | 1 |
| 1 | 1 | 1 |
| 0 | 0 | 0 |

| | | |
|---|---|---|
| 1 | 1 | 1 |
| 1 | 1 | 1 |
| 1 | 1 | 1 |
| 0 | 1 | 0 |
| 0 | 1 | 0 |
| 0 | 1 | 0 |
| 0 | 1 | 0 |
| 0 | 1 | 0 |
| 1 | 1 | 1 |
| 1 | 1 | 1 |
| 1 | 1 | 1 |
| 1 | 1 | 1 |
| 1 | 1 | 1 |
| 1 | 1 | 1 |
| 1 | 1 | 1 |
| 1 | 1 | 1 |
| 1 | 1 | 1 |
| 1 | 1 | 1 |
| 1 | 1 | 1 |
| 1 | 1 | 1 |
| 1 | 1 | 1 |
| 1 | 1 | 1 |
| 1 | 1 | 1 |
| 1 | 1 | 1 |
| 1 | 1 | 1 |
| 1 | 1 | 1 |
| 1 | 1 | 1 |
| 1 | 1 | 1 |
| 1 | 1 | 1 |
| 1 | 1 | 1 |
| 1 | 1 | 1 |
| 1 | 1 | 1 |
| 1 | 1 | 1 |
| 0 | 1 | 0 |
| 1 | 1 | 1 |
| 1 | 1 | 1 |
| 1 | 1 | 1 |
| 1 | 1 | 1 |
| 1 | 1 | 1 |
| 1 | 1 | 1 |
| 0 | 1 | 0 |
| 1 | 1 | 1 |
| 1 | 1 | 1 |
| 1 | 1 | 1 |
| 0 | 1 | 0 |
| 1 | 0 | 0 |
| 0 | 0 | 0 |
| 1 | 1 | 1 |
| 0 | 1 | 0 |
| 1 | 1 | 1 |
| 1 | 1 | 1 |
| 0 | 1 | 0 |
| 0 | 1 | 0 |
| 0 | 1 | 0 |

| | | |
|---|---|---|
| 0 | 1 | 0 |
| 1 | 1 | 1 |
| 0 | 0 | 0 |
| 1 | 1 | 1 |
| 1 | 1 | 1 |
| 1 | 1 | 1 |
| 0 | 0 | 0 |
| 1 | 1 | 1 |
| 1 | 1 | 1 |
| 0 | 1 | 0 |
| 0 | 1 | 0 |
| 0 | 0 | 0 |
| 1 | 1 | 1 |
| 0 | 0 | 0 |
| 0 | 0 | 0 |
| 0 | 0 | 0 |
| 0 | 0 | 0 |
| 0 | 0 | 0 |
| 0 | 0 | 0 |
| 0 | 0 | 0 |
| 0 | 0 | 0 |

| organism_scientific | evo_cons | cluster_id | pdb_id | chain_id | resi | resn | change | uniprot_id |
|---|---|---|---|---|---|---|---|---|
| HOMO SAPIENS | 5 | 87 | 3uzw | A | 106 | LEU | p.Leu106Phe | P51857 |
| HOMO SAPIENS | 3 | 87 | 3uzw | A | 133 | PRO | p.Pro133Arg | P51857 |
| HOMO SAPIENS | 0 | 87 | 3uzw | A | 198 | PRO | p.Pro198Leu | P51857 |
| HOMO SAPIENS | 7 | 87 | 3uzw | A | 261 | ARG | p.Arg261Cys | P51857 |
| HOMO SAPIENS | 0 | 4693 | 3mk1 | A | 3 | PRO | p.Pro25Leu | P05187 |
| HOMO SAPIENS | 8 | 4830 | 2h64 | B | 101 | CYS | p.Cys124Arg | P36894 |
| HOMO SAPIENS | 2 | 4830 | 2h64 | B | 116 | PRO | p.Pro139Ser | P36894 |
| HOMO SAPIENS | 0 | 6937 | 3dar | A | 186 | MET | p.Met186Thr | P21802 |
| HOMO SAPIENS | 3 | 5 | 1aut | C | 30 | GLN | p.Gln226His | P04070 |
| HOMO SAPIENS | 1 | 5 | 1aut | C | 68 | LEU | p.Leu265Phe | P04070 |
| HOMO SAPIENS | 0 | 5 | 1aut | C | 75 | ARG | p.Arg272Cys | P04070 |
| HOMO SAPIENS | 6 | 5 | 1aut | C | 92 | PRO | p.Pro289Leu | P04070 |
| HOMO SAPIENS | 0 | 5 | 1aut | C | 104 | ALA | p.Ala301Val | P04070 |
| HOMO SAPIENS | 1 | 5 | 1aut | C | 112 | ALA | p.Ala309Thr | P04070 |
| HOMO SAPIENS | 2 | 5 | 1aut | C | 115 | SER | p.Ser312Leu | P04070 |
| HOMO SAPIENS | 1 | 5 | 1aut | C | 133 | GLY | p.Gly334Ser | P04070 |
| HOMO SAPIENS | 0 | 5 | 1aut | C | 147 | ARG | p.Arg348Ter | P04070 |
| HOMO SAPIENS | 8 | 5 | 1aut | C | 194 | ASP | p.Asp401Asn | P04070 |
| HOMO SAPIENS | 7 | 5 | 1aut | C | 237 | TRP | p.Trp444Cys | P04070 |
| HOMO SAPIENS | 0 | 5 | 1aut | C | 238 | ILE | p.Ile445Met | P04070 |
| HOMO SAPIENS | 8 | 43 | 1aiy | F | 10 | HIS | p.His34Asp | P01308 |
| HOMO SAPIENS | 2 | 43 | 1aiy | F | 22 | ARG | p.Arg46Gln | P01308 |
| HOMO SAPIENS | 4 | 43 | 1aiy | F | 23 | GLY | p.Gly47Val | P01308 |
| HOMO SAPIENS | 7 | 43 | 1aiy | F | 24 | PHE | p.Phe48Ser | P01308 |
| HOMO SAPIENS | 5 | 24 | 2fg8 | A | 33 | THR | p.Thr30Ile | P02792 |
| HOMO SAPIENS | 4 | 24 | 2fg8 | A | 99 | ALA | p.Ala96Thr | P02792 |
| HOMO SAPIENS | 5 | 24 | 2fg8 | A | 107 | GLU | p.Glu104Ter | P02792 |
| HOMO SAPIENS | 3 | 5409 | 5bnq | A | 199 | MET | p.Met199Lys | O14788 |
| HOMO SAPIENS | -1 | 839 | 3swc | A | 959 | GLN | p.Gln990His | Q9H9B1 |
| HOMO SAPIENS | -1 | 839 | 3swc | A | 968 | ALA | p.Ala999Val | Q9H9B1 |
| HOMO SAPIENS | -1 | 839 | 3swc | A | 973 | SER | p.Ser1004Ile | Q9H9B1 |
| HOMO SAPIENS | 8 | 17 | 1eou | A | 18 | LYS | p.Lys18Glu | P00918 |
| HOMO SAPIENS | 2 | 17 | 1eou | A | 107 | HIS | p.His107Tyr | P00918 |
| HOMO SAPIENS | 1 | 17 | 1eou | A | 236 | PRO | p.Pro236His | P00918 |
| HOMO SAPIENS | 6 | 17 | 1eou | A | 252 | ASN | p.Asn252Asp | P00918 |
| HOMO SAPIENS | 2 | 9140 | 5c4v | A | 445 | ARG | p.Arg445Ter | Q13485 |
| HOMO SAPIENS | 7 | 9140 | 5c4v | A | 496 | ARG | p.Arg496Cys | Q13485 |
| HOMO SAPIENS | -1 | 42 | 4pwi | A | 6 | GLY | p.Gly26Ser | P02766 |
| HOMO SAPIENS | 6 | 42 | 4pwi | A | 10 | CYS | p.Cys30Arg | P02766 |
| HOMO SAPIENS | 8 | 42 | 4pwi | A | 12 | LEU | p.Leu32Pro | P02766 |
| HOMO SAPIENS | 9 | 42 | 4pwi | A | 18 | ASP | p.Asp38Gly | P02766 |
| HOMO SAPIENS | 8 | 42 | 4pwi | A | 20 | VAL | p.Val40Ile | P02766 |
| HOMO SAPIENS | 4 | 42 | 4pwi | A | 30 | MET | p.Val50Ala | P02766 |
| HOMO SAPIENS | 4 | 42 | 4pwi | A | 30 | MET | p.Val50Leu | P02766 |
| HOMO SAPIENS | 6 | 42 | 4pwi | A | 33 | PHE | p.Phe53Ile | P02766 |
| HOMO SAPIENS | 7 | 42 | 4pwi | A | 36 | ALA | p.Ala56Pro | P02766 |
| HOMO SAPIENS | 7 | 42 | 4pwi | A | 42 | GLU | p.Glu62Gly | P02766 |
| HOMO SAPIENS | 8 | 42 | 4pwi | A | 44 | PHE | p.Phe64Ser | P02766 |
| HOMO SAPIENS | 4 | 42 | 4pwi | A | 47 | GLY | p.Gly67Ala | P02766 |
| HOMO SAPIENS | 6 | 42 | 4pwi | A | 49 | THR | p.Thr69Ala | P02766 |
| HOMO SAPIENS | 6 | 42 | 4pwi | A | 50 | SER | p.Ser70Ile | P02766 |
| HOMO SAPIENS | 6 | 42 | 4pwi | A | 50 | SER | p.Ser70Arg | P02766 |
| HOMO SAPIENS | 0 | 42 | 4pwi | A | 53 | GLY | p.Gly73Glu | P02766 |

| | | | | | | | | |
|---|---|---|---|---|---|---|---|---|
| HOMO SAPIENS | 7 | 42 | 4pwi | A | 55 | LEU | p.Leu75Pro | P02766 |
| HOMO SAPIENS | 6 | 42 | 4pwi | A | 58 | LEU | p.Leu78His | P02766 |
| HOMO SAPIENS | 7 | 42 | 4pwi | A | 61 | GLU | p.Glu81Lys | P02766 |
| HOMO SAPIENS | 8 | 42 | 4pwi | A | 64 | PHE | p.Phe84Leu | P02766 |
| HOMO SAPIENS | 8 | 42 | 4pwi | A | 68 | ILE | p.Ile88Leu | P02766 |
| HOMO SAPIENS | 8 | 42 | 4pwi | A | 69 | TYR | p.Tyr89His | P02766 |
| HOMO SAPIENS | 4 | 42 | 4pwi | A | 71 | VAL | p.Val91Ala | P02766 |
| HOMO SAPIENS | 6 | 42 | 4pwi | A | 77 | SER | p.Ser97Tyr | P02766 |
| HOMO SAPIENS | 4 | 42 | 4pwi | A | 84 | ILE | p.Ile104Ser | P02766 |
| HOMO SAPIENS | 8 | 42 | 4pwi | A | 89 | GLU | p.Glu109Gln | P02766 |
| HOMO SAPIENS | 8 | 42 | 4pwi | A | 97 | ALA | p.Ala117Gly | P02766 |
| HOMO SAPIENS | 7 | 42 | 4pwi | A | 99 | ASP | p.Asp119Asn | P02766 |
| HOMO SAPIENS | 9 | 42 | 4pwi | A | 107 | ILE | p.Ile127Val | P02766 |
| HOMO SAPIENS | 9 | 42 | 4pwi | A | 111 | LEU | p.Leu131Met | P02766 |
| HOMO SAPIENS | 7 | 42 | 4pwi | A | 114 | TYR | p.Tyr134Cys | P02766 |
| HOMO SAPIENS | 7 | 42 | 4pwi | A | 114 | TYR | p.Tyr134His | P02766 |
| HOMO SAPIENS | 7 | 42 | 4pwi | A | 122 | VAL | p.Val142Ile | P02766 |
| HOMO SAPIENS | -1 | 42 | 4pwi | A | 126 | LYS | p.Lys146Arg | P02766 |
| HOMO SAPIENS | -1 | 42 | 4pwi | A | 127 | GLU | p.Glu147Ter | P02766 |
| HOMO SAPIENS | -1 | 1312 | 4dce | A | 1275 | ARG | p.Arg1275Gln | Q9UM73 |
| HOMO SAPIENS | -1 | 1312 | 4dce | A | 1280 | ALA | p.Ala1280Val | Q9UM73 |
| HOMO SAPIENS | 1 | 20 | 1y0x | X | 234 | ALA | p.Ala234Thr | P10828 |
| HOMO SAPIENS | 0 | 20 | 1y0x | X | 243 | ARG | p.Arg243Trp | P10828 |
| HOMO SAPIENS | 3 | 20 | 1y0x | X | 316 | ARG | p.Arg316His | P10828 |
| HOMO SAPIENS | 1 | 20 | 1y0x | X | 317 | ALA | p.Ala317Thr | P10828 |
| HOMO SAPIENS | 2 | 20 | 1y0x | X | 320 | ARG | p.Arg320Cys | P10828 |
| HOMO SAPIENS | 2 | 20 | 1y0x | X | 320 | ARG | p.Arg320His | P10828 |
| HOMO SAPIENS | 0 | 20 | 1y0x | X | 332 | GLY | p.Gly332Arg | P10828 |
| HOMO SAPIENS | 1 | 20 | 1y0x | X | 338 | ARG | p.Arg338Trp | P10828 |
| HOMO SAPIENS | 0 | 20 | 1y0x | X | 340 | GLN | p.Gln340His | P10828 |
| HOMO SAPIENS | 0 | 20 | 1y0x | X | 345 | GLY | p.Gly345Val | P10828 |
| HOMO SAPIENS | 0 | 20 | 1y0x | X | 345 | GLY | p.Gly345Ser | P10828 |
| HOMO SAPIENS | 0 | 20 | 1y0x | X | 347 | GLY | p.Gly347Glu | P10828 |
| HOMO SAPIENS | 1 | 20 | 1y0x | X | 438 | ARG | p.Arg438His | P10828 |
| HOMO SAPIENS | 0 | 20 | 1y0x | X | 442 | MET | p.Met442Val | P10828 |
| HOMO SAPIENS | 0 | 20 | 1y0x | X | 443 | LYS | p.Lys443Glu | P10828 |
| HOMO SAPIENS | 0 | 20 | 1y0x | X | 446 | CYS | p.Cys446Arg | P10828 |
| HOMO SAPIENS | 2 | 20 | 1y0x | X | 453 | PRO | p.Pro453His | P10828 |
| HOMO SAPIENS | 2 | 20 | 1y0x | X | 453 | PRO | p.Pro453Thr | P10828 |
| HOMO SAPIENS | 3 | 20 | 1y0x | X | 459 | PHE | p.Phe459Cys | P10828 |
| HOMO SAPIENS | 0 | 4409 | 3qb7 | D | 87 | GLU | p.Glu109Lys | P31785 |
| HOMO SAPIENS | 1 | 4409 | 3qb7 | D | 92 | GLY | p.Gly114Asp | P31785 |
| HOMO SAPIENS | 1 | 4409 | 3qb7 | D | 93 | CYS | p.Cys115Arg | P31785 |
| HOMO SAPIENS | 0 | 4409 | 3qb7 | D | 131 | ILE | p.Ile153Asn | P31785 |
| HOMO SAPIENS | 9 | 4409 | 3qb7 | D | 200 | ARG | p.Arg222Cys | P31785 |
| HOMO SAPIENS | 0 | 3891 | 3ije | B | 33 | LEU | p.Leu59Pro | P05106 |
| HOMO SAPIENS | 0 | 3891 | 3ije | B | 117 | LEU | p.Leu143Trp | P05106 |
| HOMO SAPIENS | 3 | 3891 | 3ije | B | 119 | ASP | p.Asp145Tyr | P05106 |
| HOMO SAPIENS | 2 | 3891 | 3ije | B | 143 | ARG | p.Arg169Gln | P05106 |
| HOMO SAPIENS | 2 | 3891 | 3ije | B | 214 | ARG | p.Arg240Gln | P05106 |
| HOMO SAPIENS | 2 | 3891 | 3ije | B | 214 | ARG | p.Arg240Trp | P05106 |
| HOMO SAPIENS | 2 | 3891 | 3ije | B | 374 | CYS | p.Cys400Tyr | P05106 |
| HOMO SAPIENS | 1 | 3891 | 3ije | B | 407 | PRO | p.Pro433Ala | P05106 |
| HOMO SAPIENS | 0 | 3891 | 3ije | B | 489 | ARG | p.Arg515Gln | P05106 |

| | | | | | | | | |
|---|---|---|---|---|---|---|---|---|
| HOMO SAPIENS | -1 | 9613 | 5mke | B | 28 | ARG | p.Arg28Pro | Q13563 |
| HOMO SAPIENS | -1 | 9613 | 5mke | B | 190 | ALA | p.Ala190Thr | Q13563 |
| HOMO SAPIENS | 4 | 9613 | 5mke | B | 262 | VAL | p.Val262Met | Q13563 |
| HOMO SAPIENS | 0 | 9613 | 5mke | B | 482 | PHE | p.Phe482Cys | Q13563 |
| HOMO SAPIENS | 4 | 9613 | 5mke | B | 511 | ASP | p.Asp511Val | Q13563 |
| HOMO SAPIENS | 5 | 9613 | 5mke | B | 516 | VAL | p.Val516Leu | Q13563 |
| HOMO SAPIENS | -1 | 9613 | 5mke | B | 807 | ARG | p.Arg807Gln | Q13563 |
| HOMO SAPIENS | 4 | 2428 | 1olh | C | 337 | ARG | p.Arg337Cys | P04637 |
| HOMO SAPIENS | 4 | 2428 | 1olh | C | 337 | ARG | p.Arg337His | P04637 |
| HOMO SAPIENS | 8 | 2428 | 1olh | C | 338 | PHE | p.Phe338Leu | P04637 |
| HOMO SAPIENS | 8 | 2428 | 1olh | C | 339 | GLU | p.Glu339Lys | P04637 |
| HOMO SAPIENS | 7 | 2428 | 1olh | C | 344 | LEU | p.Leu344Pro | P04637 |
| HOMO SAPIENS | 0 | 2428 | 1olh | C | 360 | GLY | p.Gly360Ala | P04637 |
| HOMO SAPIENS | 7 | 4867 | 1mox | B | 404 | GLY | p.Gly428Asp | P00533 |
| HOMO SAPIENS | 5 | 4867 | 1mox | B | 497 | ARG | p.Arg521Lys | P00533 |
| HOMO SAPIENS | 1 | 4630 | 1fu1 | A | 43 | TRP | p.Trp43Arg | Q13426 |
| HOMO SAPIENS | 2 | 4630 | 1fu1 | A | 161 | ARG | p.Arg161Ter | Q13426 |
| HOMO SAPIENS | 2 | 15 | 1lfy | B | 2 | HIS | p.His3Gln | P68871 |
| HOMO SAPIENS | 2 | 15 | 1lfy | B | 6 | GLU | p.Glu7Ala | P68871 |
| HOMO SAPIENS | 2 | 15 | 1lfy | B | 6 | GLU | p.Glu7Lys | P68871 |
| HOMO SAPIENS | 4 | 15 | 1lfy | B | 9 | SER | p.Ser10Cys | P68871 |
| HOMO SAPIENS | 6 | 15 | 1lfy | B | 15 | TRP | p.Trp16Ter | P68871 |
| HOMO SAPIENS | 3 | 15 | 1lfy | B | 17 | LYS | p.Lys18Glu | P68871 |
| HOMO SAPIENS | 6 | 15 | 1lfy | B | 19 | ASN | p.Asn20Ser | P68871 |
| HOMO SAPIENS | 6 | 15 | 1lfy | B | 22 | GLU | p.Glu23Ala | P68871 |
| HOMO SAPIENS | 6 | 15 | 1lfy | B | 22 | GLU | p.Glu23Lys | P68871 |
| HOMO SAPIENS | 8 | 15 | 1lfy | B | 26 | GLU | p.Glu27Lys | P68871 |
| HOMO SAPIENS | 8 | 15 | 1lfy | B | 28 | LEU | p.Leu29Pro | P68871 |
| HOMO SAPIENS | 8 | 15 | 1lfy | B | 30 | ARG | p.Arg31Ser | P68871 |
| HOMO SAPIENS | 8 | 15 | 1lfy | B | 30 | ARG | p.Arg31Lys | P68871 |
| HOMO SAPIENS | 6 | 15 | 1lfy | B | 39 | GLN | p.Gln40Ter | P68871 |
| HOMO SAPIENS | 2 | 15 | 1lfy | B | 43 | GLU | p.Glu44Gln | P68871 |
| HOMO SAPIENS | 6 | 15 | 1lfy | B | 57 | ASN | p.Asn58Lys | P68871 |
| HOMO SAPIENS | 6 | 15 | 1lfy | B | 59 | LYS | p.Lys60Asn | P68871 |
| HOMO SAPIENS | 8 | 15 | 1lfy | B | 61 | LYS | p.Lys62Glu | P68871 |
| HOMO SAPIENS | 4 | 15 | 1lfy | B | 73 | ASP | p.Asp74Tyr | P68871 |
| HOMO SAPIENS | 4 | 15 | 1lfy | B | 87 | THR | p.Thr88Ile | P68871 |
| HOMO SAPIENS | 4 | 15 | 1lfy | B | 87 | THR | p.Thr88Pro | P68871 |
| HOMO SAPIENS | 5 | 15 | 1lfy | B | 90 | GLU | p.Glu91Lys | P68871 |
| HOMO SAPIENS | 6 | 15 | 1lfy | B | 91 | LEU | p.Leu92Pro | P68871 |
| HOMO SAPIENS | 5 | 15 | 1lfy | B | 97 | HIS | p.His98Gln | P68871 |
| HOMO SAPIENS | 6 | 15 | 1lfy | B | 110 | LEU | p.Leu111Pro | P68871 |
| HOMO SAPIENS | 6 | 15 | 1lfy | B | 113 | VAL | p.Val114Glu | P68871 |
| HOMO SAPIENS | 2 | 15 | 1lfy | B | 119 | GLY | p.Gly120Ala | P68871 |
| HOMO SAPIENS | 4 | 15 | 1lfy | B | 121 | GLU | p.Glu122Gln | P68871 |
| HOMO SAPIENS | 5 | 15 | 1lfy | B | 134 | VAL | p.Val135Glu | P68871 |
| HOMO SAPIENS | 3 | 15 | 1lfy | B | 143 | HIS | p.His144Gln | P68871 |
| HOMO SAPIENS | 4 | 91 | 5d9k | A | 75 | GLY | p.Gly75Val | P51812 |
| HOMO SAPIENS | 5 | 91 | 5d9k | A | 82 | VAL | p.Val82Phe | P51812 |
| HOMO SAPIENS | 0 | 91 | 5d9k | A | 114 | ARG | p.Arg114Trp | P51812 |
| HOMO SAPIENS | 4 | 91 | 5d9k | A | 189 | ILE | p.Ile189Lys | P51812 |
| HOMO SAPIENS | -1 | 91 | 5d9k | A | 227 | SER | p.Ser227Ala | P51812 |
| HOMO SAPIENS | 1 | 91 | 5d9k | A | 268 | PHE | p.Phe268Ser | P51812 |
| HOMO SAPIENS | 0 | 954 | 2vt0 | A | 15 | VAL | p.Val54Leu | P04062 |

| | | | | | | | | |
|---|---|---|---|---|---|---|---|---|
| HOMO SAPIENS | 5 | 954 | 2vt0 | A | 46 | GLY | p.Gly85Glu | P04062 |
| HOMO SAPIENS | 5 | 954 | 2vt0 | A | 48 | ARG | p.Arg87Trp | P04062 |
| HOMO SAPIENS | 6 | 954 | 2vt0 | A | 79 | LYS | p.Lys118Asn | P04062 |
| HOMO SAPIENS | 0 | 954 | 2vt0 | A | 120 | ARG | p.Arg159Gln | P04062 |
| HOMO SAPIENS | 0 | 954 | 2vt0 | A | 120 | ARG | p.Arg159Trp | P04062 |
| HOMO SAPIENS | 0 | 954 | 2vt0 | A | 122 | PRO | p.Pro161Ser | P04062 |
| HOMO SAPIENS | 5 | 954 | 2vt0 | A | 131 | ARG | p.Arg170Leu | P04062 |
| HOMO SAPIENS | 5 | 954 | 2vt0 | A | 135 | TYR | p.Tyr174Cys | P04062 |
| HOMO SAPIENS | 0 | 954 | 2vt0 | A | 157 | LYS | p.Lys196Gln | P04062 |
| HOMO SAPIENS | 5 | 954 | 2vt0 | A | 163 | ARG | p.Arg202Gln | P04062 |
| HOMO SAPIENS | 5 | 954 | 2vt0 | A | 170 | ARG | p.Arg209Cys | P04062 |
| HOMO SAPIENS | 5 | 954 | 2vt0 | A | 188 | ASN | p.Asn227Ser | P04062 |
| HOMO SAPIENS | 3 | 954 | 2vt0 | A | 196 | SER | p.Ser235Pro | P04062 |
| HOMO SAPIENS | 5 | 954 | 2vt0 | A | 202 | GLY | p.Gly241Arg | P04062 |
| HOMO SAPIENS | 0 | 954 | 2vt0 | A | 212 | TYR | p.Tyr251His | P04062 |
| HOMO SAPIENS | 0 | 954 | 2vt0 | A | 213 | PHE | p.Phe252Ile | P04062 |
| HOMO SAPIENS | 5 | 954 | 2vt0 | A | 216 | PHE | p.Phe255Tyr | P04062 |
| HOMO SAPIENS | 6 | 954 | 2vt0 | A | 251 | PHE | p.Phe290Leu | P04062 |
| HOMO SAPIENS | 5 | 954 | 2vt0 | A | 255 | HIS | p.His294Gln | P04062 |
| HOMO SAPIENS | 6 | 954 | 2vt0 | A | 257 | ARG | p.Arg296Gln | P04062 |
| HOMO SAPIENS | 5 | 954 | 2vt0 | A | 289 | PRO | p.Pro328Leu | P04062 |
| HOMO SAPIENS | 0 | 954 | 2vt0 | A | 309 | ALA | p.Ala348Val | P04062 |
| HOMO SAPIENS | 0 | 954 | 2vt0 | A | 311 | HIS | p.His350Arg | P04062 |
| HOMO SAPIENS | 6 | 954 | 2vt0 | A | 312 | TRP | p.Trp351Cys | P04062 |
| HOMO SAPIENS | 5 | 954 | 2vt0 | A | 323 | THR | p.Thr362Ile | P04062 |
| HOMO SAPIENS | 5 | 954 | 2vt0 | A | 325 | GLY | p.Gly364Arg | P04062 |
| HOMO SAPIENS | 6 | 954 | 2vt0 | A | 342 | CYS | p.Cys381Gly | P04062 |
| HOMO SAPIENS | 5 | 954 | 2vt0 | A | 353 | ARG | p.Arg392Gly | P04062 |
| HOMO SAPIENS | 3 | 954 | 2vt0 | A | 364 | SER | p.Ser403Thr | P04062 |
| HOMO SAPIENS | 5 | 954 | 2vt0 | A | 369 | THR | p.Thr408Met | P04062 |
| HOMO SAPIENS | 3 | 954 | 2vt0 | A | 370 | ASN | p.Asn409Ser | P04062 |
| HOMO SAPIENS | 0 | 954 | 2vt0 | A | 371 | LEU | p.Leu410Val | P04062 |
| HOMO SAPIENS | 0 | 954 | 2vt0 | A | 375 | VAL | p.Val414Leu | P04062 |
| HOMO SAPIENS | 0 | 954 | 2vt0 | A | 377 | GLY | p.Gly416Ser | P04062 |
| HOMO SAPIENS | 8 | 954 | 2vt0 | A | 398 | VAL | p.Val437Phe | P04062 |
| HOMO SAPIENS | 0 | 954 | 2vt0 | A | 401 | PRO | p.Pro440Leu | P04062 |
| HOMO SAPIENS | 3 | 954 | 2vt0 | A | 409 | ASP | p.Asp448His | P04062 |
| HOMO SAPIENS | 3 | 954 | 2vt0 | A | 409 | ASP | p.Asp448Val | P04062 |
| HOMO SAPIENS | 0 | 954 | 2vt0 | A | 415 | PRO | p.Pro454Arg | P04062 |
| HOMO SAPIENS | 5 | 954 | 2vt0 | A | 444 | LEU | p.Leu483Arg | P04062 |
| HOMO SAPIENS | 5 | 954 | 2vt0 | A | 456 | ALA | p.Ala495Pro | P04062 |
| HOMO SAPIENS | 5 | 954 | 2vt0 | A | 463 | ARG | p.Arg502Cys | P04062 |
| HOMO SAPIENS | 5 | 954 | 2vt0 | A | 463 | ARG | p.Arg502His | P04062 |
| HOMO SAPIENS | 5 | 954 | 2vt0 | A | 478 | GLY | p.Gly517Ser | P04062 |
| HOMO SAPIENS | 5 | 954 | 2vt0 | A | 496 | ARG | p.Arg535Cys | P04062 |
| HOMO SAPIENS | 5 | 954 | 2vt0 | A | 496 | ARG | p.Arg535His | P04062 |
| HOMO SAPIENS | 1 | 9037 | 2rqb | A | 946 | ALA | p.Ala946Thr | Q9BYX4 |
| HOMO SAPIENS | 0 | 253 | 5E+092 | A | 308 | LEU | p.Leu308Pro | P37173 |
| HOMO SAPIENS | 0 | 253 | 5E+092 | A | 315 | THR | p.Thr315Met | P37173 |
| HOMO SAPIENS | 0 | 253 | 5E+092 | A | 328 | HIS | p.His328Gln | P37173 |
| HOMO SAPIENS | 0 | 253 | 5E+092 | A | 336 | TYR | p.Tyr336Asn | P37173 |
| HOMO SAPIENS | 0 | 253 | 5E+092 | A | 348 | ARG | p.Arg348Cys | P37173 |
| HOMO SAPIENS | 0 | 253 | 5E+092 | A | 355 | ALA | p.Ala355Pro | P37173 |
| HOMO SAPIENS | 1 | 253 | 5E+092 | A | 356 | ARG | p.Arg356Gly | P37173 |

| | | | | | | | | |
|---|---|---|---|---|---|---|---|---|
| HOMO SAPIENS | 0 | 253 | 5E+092 | A | 357 | GLY | p.Gly357Trp | P37173 |
| HOMO SAPIENS | 1 | 253 | 5E+092 | A | 403 | ARG | p.Arg403His | P37173 |
| HOMO SAPIENS | 5 | 253 | 5E+092 | A | 449 | SER | p.Ser449Phe | P37173 |
| HOMO SAPIENS | 0 | 253 | 5E+092 | A | 460 | ARG | p.Arg460Cys | P37173 |
| HOMO SAPIENS | 0 | 253 | 5E+092 | A | 460 | ARG | p.Arg460His | P37173 |
| HOMO SAPIENS | 2 | 253 | 5E+092 | A | 526 | GLU | p.Glu526Gln | P37173 |
| HOMO SAPIENS | 5 | 253 | 5E+092 | A | 528 | ARG | p.Arg528Cys | P37173 |
| HOMO SAPIENS | 5 | 253 | 5E+092 | A | 528 | ARG | p.Arg528His | P37173 |
| HOMO SAPIENS | 6 | 158 | 5amc | A | 231 | ARG | p.Arg260Leu | P12821 |
| HOMO SAPIENS | 7 | 158 | 5amc | A | 237 | TYR | p.Tyr266Ter | P12821 |
| HOMO SAPIENS | 6 | 158 | 5amc | A | 325 | GLY | p.Gly354Arg | P12821 |
| HOMO SAPIENS | 0 | 158 | 5amc | A | 467 | ARG | p.Arg496Ter | P12821 |
| HOMO SAPIENS | 4 | 158 | 5amc | A | 479 | ARG | p.Arg508Ter | P12821 |
| HOMO SAPIENS | 0 | 158 | 5amc | A | 563 | ASP | p.Asp592Gly | P12821 |
| HOMO SAPIENS | -1 | 33062 | 1gr3 | A | 545 | GLY | p.Gly545Arg | Q03692 |
| HOMO SAPIENS | 5 | 33062 | 1gr3 | A | 591 | CYS | p.Cys591Arg | Q03692 |
| HOMO SAPIENS | 2 | 33062 | 1gr3 | A | 595 | GLY | p.Gly595Glu | Q03692 |
| HOMO SAPIENS | 0 | 33062 | 1gr3 | A | 597 | TYR | p.Tyr597Cys | Q03692 |
| HOMO SAPIENS | 5 | 33062 | 1gr3 | A | 598 | TYR | p.Tyr598Asp | Q03692 |
| HOMO SAPIENS | 7 | 33062 | 1gr3 | A | 600 | SER | p.Ser600Pro | Q03692 |
| HOMO SAPIENS | 1 | 33062 | 1gr3 | A | 614 | LEU | p.Leu614Pro | Q03692 |
| HOMO SAPIENS | 6 | 33062 | 1gr3 | A | 651 | TRP | p.Trp651Arg | Q03692 |
| HOMO SAPIENS | 7 | 33062 | 1gr3 | A | 671 | SER | p.Ser671Pro | Q03692 |
| HOMO SAPIENS | -1 | 38257 | 2l2i | B | 87 | PRO | p.Pro87Ala | Q13351 |
| HOMO SAPIENS | 2 | 2036 | 1ghq | A | 231 | GLY | p.Gly1224Asp | P01024 |
| HOMO SAPIENS | 5 | 3594 | 4cde | F | 388 | TYR | p.Tyr412Cys | P53634 |
| HOMO SAPIENS | 9 | 3594 | 4cde | F | 405 | TRP | p.Trp429Cys | P53634 |
| HOMO SAPIENS | 0 | 3594 | 4cde | F | 429 | ILE | p.Ile453Val | P53634 |
| HOMO SAPIENS | 7 | 18 | 1jkb | A | 34 | TRP | p.Trp52Ter | P61626 |
| HOMO SAPIENS | 9 | 18 | 1jkb | A | 70 | THR | p.Thr88Asn | P61626 |
| HOMO SAPIENS | 8 | 18 | 1jkb | A | 121 | VAL | p.Val139Ile | P61626 |
| HOMO SAPIENS | 2 | 414 | 3fj6 | A | 136 | ARG | p.Arg135Cys | Q02127 |
| HOMO SAPIENS | 1 | 414 | 3fj6 | A | 153 | GLY | p.Gly152Arg | Q02127 |
| HOMO SAPIENS | 0 | 414 | 3fj6 | A | 200 | ARG | p.Arg199Cys | Q02127 |
| HOMO SAPIENS | 0 | 414 | 3fj6 | A | 203 | GLY | p.Gly202Ala | Q02127 |
| HOMO SAPIENS | 0 | 414 | 3fj6 | A | 245 | ARG | p.Arg244Trp | Q02127 |
| HOMO SAPIENS | 0 | 414 | 3fj6 | A | 342 | ALA | p.Ala341Val | Q02127 |
| HOMO SAPIENS | 1 | 414 | 3fj6 | A | 347 | ARG | p.Arg346Trp | Q02127 |
| HOMO SAPIENS | 6 | 103 | 1u3w | A | 77 | GLY | p.Gly78Ter | P00326 |
| HOMO SAPIENS | -1 | 8369 | 4ux8 | F | 16 | ARG | p.Arg93Trp | P39905 |
| HOMO SAPIENS | 0 | 366 | 2zjc | A | 32 | GLY | p.Arg108Trp | P01375 |
| HOMO SAPIENS | 0 | 1457 | 3v9i | A | 352 | LEU | p.Ser352Leu | P30038 |
| HOMO SAPIENS | 0 | 1457 | 3v9i | A | 470 | VAL | p.Val470Ile | P30038 |
| HOMO SAPIENS | 0 | 1457 | 3v9i | A | 473 | THR | p.Thr473Ala | P30038 |
| HOMO SAPIENS | 1 | 2 | 4wo5 | A | 462 | ARG | p.Arg462Ile | P15056 |
| HOMO SAPIENS | 3 | 2 | 4wo5 | A | 463 | ILE | p.Ile463Ser | P15056 |
| HOMO SAPIENS | 1 | 2 | 4wo5 | A | 466 | GLY | p.Gly466Val | P15056 |
| HOMO SAPIENS | 1 | 2 | 4wo5 | A | 468 | PHE | p.Phe468Ser | P15056 |
| HOMO SAPIENS | 2 | 2 | 4wo5 | A | 469 | GLY | p.Gly469Ala | P15056 |
| HOMO SAPIENS | 2 | 2 | 4wo5 | A | 469 | GLY | p.Gly469Arg | P15056 |
| HOMO SAPIENS | 2 | 2 | 4wo5 | A | 485 | LEU | p.Leu485Phe | P15056 |
| HOMO SAPIENS | 0 | 2 | 4wo5 | A | 501 | GLU | p.Glu501Gly | P15056 |
| HOMO SAPIENS | 0 | 2 | 4wo5 | A | 501 | GLU | p.Glu501Lys | P15056 |
| HOMO SAPIENS | 7 | 2 | 4wo5 | A | 581 | ASN | p.Asn581Asp | P15056 |

| | | | | | | | | |
|---|---|---|---|---|---|---|---|---|
| HOMO SAPIENS | 8 | 2 | 4wo5 | A | 594 | ASP | p.Asp594Gly | P15056 |
| HOMO SAPIENS | 0 | 2 | 4wo5 | A | 597 | LEU | p.Leu597Arg | P15056 |
| HOMO SAPIENS | 0 | 2 | 4wo5 | A | 597 | LEU | p.Leu597Val | P15056 |
| HOMO SAPIENS | 0 | 2 | 4wo5 | A | 600 | VAL | p.Val600Glu | P15056 |
| HOMO SAPIENS | 2 | 2 | 4wo5 | A | 601 | LYS | p.Lys601Glu | P15056 |
| HOMO SAPIENS | 8 | 2 | 4wo5 | A | 638 | ASP | p.Asp638Glu | P15056 |
| HOMO SAPIENS | -1 | 30366 | 5bqd | B | 19 | ALA | p.Ala19Ser | Q99593 |
| HOMO SAPIENS | -1 | 30366 | 5bqd | B | 23 | PRO | p.Pro23His | Q99593 |
| HOMO SAPIENS | 1 | 30366 | 5bqd | B | 49 | GLN | p.Gln49Lys | Q99593 |
| HOMO SAPIENS | 2 | 30366 | 5bqd | B | 54 | ILE | p.Ile54Thr | Q99593 |
| HOMO SAPIENS | 8 | 30366 | 5bqd | B | 80 | GLY | p.Gly80Arg | Q99593 |
| HOMO SAPIENS | 6 | 30366 | 5bqd | B | 106 | ILE | p.Ile106Val | Q99593 |
| HOMO SAPIENS | 7 | 30366 | 5bqd | B | 111 | ASP | p.Asp111Tyr | Q99593 |
| HOMO SAPIENS | -1 | 30366 | 5bqd | B | 237 | ARG | p.Arg237Gln | Q99593 |
| HOMO SAPIENS | -1 | 30366 | 5bqd | B | 237 | ARG | p.Arg237Trp | Q99593 |
| HOMO SAPIENS | 5 | 4789 | 1n4m | A | 567 | SER | p.Ser567Leu | P06400 |
| HOMO SAPIENS | 0 | 4789 | 1n4m | A | 658 | ALA | p.Ala658Asp | P06400 |
| HOMO SAPIENS | 7 | 4789 | 1n4m | A | 661 | ARG | p.Arg661Trp | P06400 |
| HOMO SAPIENS | 6 | 4789 | 1n4m | A | 668 | ARG | p.Arg668His | P06400 |
| HOMO SAPIENS | 8 | 4789 | 1n4m | A | 697 | ASP | p.Asp697Glu | P06400 |
| HOMO SAPIENS | 6 | 4789 | 1n4m | A | 712 | CYS | p.Cys712Arg | P06400 |
| HOMO SAPIENS | 1 | 1798 | 2r5g | A | 101 | HIS | p.His101Gln | O15247 |
| HOMO SAPIENS | 0 | 1798 | 2r5g | A | 160 | PRO | p.Pro160Ala | O15247 |
| HOMO SAPIENS | 2 | 22 | 4x0t | A | 82 | ARG | p.Arg110Ter | P49419 |
| HOMO SAPIENS | 0 | 22 | 4x0t | A | 149 | ALA | p.Ala177Pro | P49419 |
| HOMO SAPIENS | 4 | 22 | 4x0t | A | 164 | THR | p.Thr192Met | P49419 |
| HOMO SAPIENS | 9 | 22 | 4x0t | A | 167 | ASN | p.Asn195Ser | P49419 |
| HOMO SAPIENS | 0 | 22 | 4x0t | A | 175 | TRP | p.Trp203Gly | P49419 |
| HOMO SAPIENS | 0 | 22 | 4x0t | A | 194 | THR | p.Thr222Ala | P49419 |
| HOMO SAPIENS | 0 | 22 | 4x0t | A | 273 | ASN | p.Asn301Ile | P49419 |
| HOMO SAPIENS | 0 | 22 | 4x0t | A | 307 | ARG | p.Arg335Gln | P49419 |
| HOMO SAPIENS | 3 | 22 | 4x0t | A | 313 | SER | p.Ser341Asn | P49419 |
| HOMO SAPIENS | 0 | 22 | 4x0t | A | 329 | ILE | p.Ile357Val | P49419 |
| HOMO SAPIENS | 1 | 22 | 4x0t | A | 380 | TYR | p.Tyr408Ter | P49419 |
| HOMO SAPIENS | 9 | 22 | 4x0t | A | 399 | GLU | p.Glu427Gln | P49419 |
| HOMO SAPIENS | 0 | 22 | 4x0t | A | 406 | TYR | p.Tyr434Cys | P49419 |
| HOMO SAPIENS | 0 | 22 | 4x0t | A | 431 | ILE | p.Ile459Phe | P49419 |
| HOMO SAPIENS | 4 | 22 | 4x0t | A | 477 | GLY | p.Gly505Arg | P49419 |
| HOMO SAPIENS | 0 | 22 | 4x0t | A | 495 | THR | p.Thr523Ala | P49419 |
| HOMO SAPIENS | 8 | 11042 | 1y6b | A | 1145 | PRO | p.Pro1147Ser | P35968 |
| HOMO SAPIENS | -1 | 3867 | 2xij | A | 18 | GLN | p.Gln18Ter | P22033 |
| HOMO SAPIENS | -1 | 3867 | 2xij | A | 31 | ARG | p.Arg31Ter | P22033 |
| HOMO SAPIENS | 2 | 3867 | 2xij | A | 93 | ARG | p.Arg93His | P22033 |
| HOMO SAPIENS | 0 | 3867 | 2xij | A | 94 | GLY | p.Gly94Arg | P22033 |
| HOMO SAPIENS | 2 | 3867 | 2xij | A | 95 | PRO | p.Pro95Leu | P22033 |
| HOMO SAPIENS | 2 | 3867 | 2xij | A | 105 | TRP | p.Trp105Arg | P22033 |
| HOMO SAPIENS | 2 | 3867 | 2xij | A | 108 | ARG | p.Arg108Cys | P22033 |
| HOMO SAPIENS | 3 | 3867 | 2xij | A | 154 | ARG | p.Arg154His | P22033 |
| HOMO SAPIENS | 3 | 3867 | 2xij | A | 156 | ASP | p.Asp156Val | P22033 |
| HOMO SAPIENS | 0 | 3867 | 2xij | A | 189 | ASN | p.Asn189Ile | P22033 |
| HOMO SAPIENS | 0 | 3867 | 2xij | A | 191 | ALA | p.Ala191Glu | P22033 |
| HOMO SAPIENS | 0 | 3867 | 2xij | A | 203 | GLY | p.Gly203Arg | P22033 |
| HOMO SAPIENS | 2 | 3867 | 2xij | A | 215 | GLY | p.Gly215Ser | P22033 |
| HOMO SAPIENS | 0 | 3867 | 2xij | A | 219 | ASN | p.Asn219Tyr | P22033 |

| | | | | | | | | |
|---|---|---|---|---|---|---|---|---|
| HOMO SAPIENS | 3 | 3867 | 2xij | A | 228 | ARG | p.Arg228Ter | P22033 |
| HOMO SAPIENS | 1 | 3867 | 2xij | A | 297 | THR | p.Thr297Ile | P22033 |
| HOMO SAPIENS | 0 | 3867 | 2xij | A | 326 | ARG | p.Arg326Lys | P22033 |
| HOMO SAPIENS | 3 | 3867 | 2xij | A | 369 | ARG | p.Arg369His | P22033 |
| HOMO SAPIENS | 3 | 3867 | 2xij | A | 369 | ARG | p.Arg369Cys | P22033 |
| HOMO SAPIENS | 0 | 3867 | 2xij | A | 370 | THR | p.Thr370Pro | P22033 |
| HOMO SAPIENS | 0 | 3867 | 2xij | A | 377 | ALA | p.Ala377Glu | P22033 |
| HOMO SAPIENS | 2 | 3867 | 2xij | A | 426 | GLY | p.Gly426Ala | P22033 |
| HOMO SAPIENS | 2 | 3867 | 2xij | A | 427 | GLY | p.Gly427Asp | P22033 |
| HOMO SAPIENS | 1 | 3867 | 2xij | A | 499 | THR | p.Ala499Thr | P22033 |
| HOMO SAPIENS | 2 | 3867 | 2xij | A | 532 | ARG | p.Arg532His | P22033 |
| HOMO SAPIENS | 3 | 3867 | 2xij | A | 555 | ALA | p.Ala555Thr | P22033 |
| HOMO SAPIENS | 0 | 3867 | 2xij | A | 573 | PHE | p.Phe573Ser | P22033 |
| HOMO SAPIENS | 3 | 3867 | 2xij | A | 623 | GLY | p.Gly623Arg | P22033 |
| HOMO SAPIENS | 5 | 3867 | 2xij | A | 671 | VAL | p.Ile671Val | P22033 |
| HOMO SAPIENS | 4 | 3867 | 2xij | A | 694 | ARG | p.Arg694Trp | P22033 |
| HOMO SAPIENS | 1 | 3867 | 2xij | A | 700 | MET | p.Met700Lys | P22033 |
| HOMO SAPIENS | 8 | 3867 | 2xij | A | 703 | GLY | p.Gly703Arg | P22033 |
| HOMO SAPIENS | 0 | 3867 | 2xij | A | 717 | GLY | p.Gly717Val | P22033 |
| HOMO SAPIENS | 1 | 3867 | 2xij | A | 727 | ARG | p.Arg727Ter | P22033 |
| HOMO SAPIENS | -1 | 52 | 3k91 | B | 72 | GLY | p.Gly73Ser | P00441 |
| HOMO SAPIENS | 6 | 52 | 3k91 | B | 96 | ASP | p.Asp97Asn | P00441 |
| HOMO SAPIENS | 5 | 52 | 3k91 | B | 112 | ILE | p.Ile113Thr | P00441 |
| HOMO SAPIENS | -1 | 52 | 3k91 | B | 134 | SER | p.Ser135Asn | P00441 |
| HOMO SAPIENS | 0 | 3950 | 4fom | A | 296 | ASN | p.Asn296His | Q9NQS3 |
| HOMO SAPIENS | 1 | 10 | 1de4 | A | 31 | VAL | p.Val53Met | Q30201 |
| HOMO SAPIENS | 0 | 10 | 1de4 | A | 37 | VAL | p.Val59Met | Q30201 |
| HOMO SAPIENS | 0 | 10 | 1de4 | A | 71 | GLY | p.Gly93Arg | Q30201 |
| HOMO SAPIENS | 0 | 10 | 1de4 | A | 83 | ILE | p.Ile105Thr | Q30201 |
| HOMO SAPIENS | 0 | 10 | 1de4 | A | 105 | GLN | p.Gln127His | Q30201 |
| HOMO SAPIENS | 1 | 10 | 1de4 | A | 146 | GLU | p.Glu168Ter | Q30201 |
| HOMO SAPIENS | 2 | 10 | 1de4 | A | 261 | GLN | p.Gln283Pro | Q30201 |
| HOMO SAPIENS | 0 | 8038 | 4x7n | A | 586 | ILE | p.Ile586Leu | Q9NZJ5 |
| HOMO SAPIENS | 0 | 8038 | 4x7n | A | 588 | ARG | p.Arg588Gln | Q9NZJ5 |
| HOMO SAPIENS | 0 | 35175 | 5ugw | A | 1062 | ALA | p.Ala1062Thr | O14746 |
| HOMO SAPIENS | 2 | 35175 | 5ugw | A | 1090 | VAL | p.Val1090Met | O14746 |
| HOMO SAPIENS | 0 | 1722 | 4z3k | C | 147 | ARG | p.Arg150Gly | P35270 |
| HOMO SAPIENS | 0 | 1722 | 4z3k | C | 160 | PRO | p.Pro163Leu | P35270 |
| HOMO SAPIENS | 0 | 1722 | 4z3k | C | 207 | ARG | p.Arg210Trp | P35270 |
| HOMO SAPIENS | 0 | 1722 | 4z3k | C | 216 | ARG | p.Arg219Ter | P35270 |
| HOMO SAPIENS | 0 | 1722 | 4z3k | C | 248 | LYS | p.Lys251Ter | P35270 |
| HOMO SAPIENS | 1 | 2832 | 4x8v | L | 115 | HIS | p.His175Gln | P08709 |
| HOMO SAPIENS | 0 | 825 | 1emh | A | 242 | PHE | p.Phe251Ser | P13051 |
| HOMO SAPIENS | 0 | 122 | 5uwk | A | 109 | ARG | p.Arg109Ter | P45452 |
| HOMO SAPIENS | 1 | 122 | 5uwk | A | 170 | LYS | p.Lys170Thr | P45452 |
| HOMO SAPIENS | 9 | 122 | 5uwk | A | 232 | HIS | p.His232Asn | P45452 |
| HOMO SAPIENS | 5 | 122 | 5uwk | A | 257 | ASP | p.Asp257Val | P45452 |
| HOMO SAPIENS | 9 | 1074 | 3p5i | A | 264 | TRP | p.Trp264Arg | Q9UJ71 |
| HOMO SAPIENS | 0 | 146 | 3r3g | A | 14C | GLU | p.Glu352Lys | P00734 |
| HOMO SAPIENS | 3 | 9522 | 1wyq | A | 83 | LEU | p.Leu253Pro | O15020 |
| HOMO SAPIENS | 8 | 67 | 2vgb | A | 130 | SER | p.Ser130Tyr | P30613 |
| HOMO SAPIENS | 8 | 67 | 2vgb | A | 163 | ARG | p.Arg163Cys | P30613 |
| HOMO SAPIENS | 4 | 67 | 2vgb | A | 241 | GLU | p.Glu241Ter | P30613 |
| HOMO SAPIENS | 9 | 67 | 2vgb | A | 384 | THR | p.Thr384Met | P30613 |

| | | | | | | | | |
|---|---|---|---|---|---|---|---|---|
| HOMO SAPIENS | 0 | 67 | 2vgb | A | 421 | GLN | p.Gln421Lys | P30613 |
| HOMO SAPIENS | 6 | 67 | 2vgb | A | 479 | ARG | p.Arg479His | P30613 |
| HOMO SAPIENS | 8 | 67 | 2vgb | A | 510 | ARG | p.Arg510Gln | P30613 |
| HOMO SAPIENS | 4 | 67 | 2vgb | A | 569 | ARG | p.Arg569Trp | P30613 |
| HOMO SAPIENS | 0 | 9088 | 5v44 | A | 232 | ASN | p.Asn232Lys | Q9NZJ4 |
| HOMO SAPIENS | 4 | 16 | 3lbn | A | 12 | GLY | p.Gly12Val | P01112 |
| HOMO SAPIENS | 4 | 16 | 3lbn | A | 12 | GLY | p.Gly12Ser | P01112 |
| HOMO SAPIENS | 4 | 16 | 3lbn | A | 13 | GLY | p.Gly13Cys | P01112 |
| HOMO SAPIENS | 4 | 16 | 3lbn | A | 13 | GLY | p.Gly13Asp | P01112 |
| HOMO SAPIENS | 5 | 16 | 3lbn | A | 22 | GLN | p.Gln22Lys | P01112 |
| HOMO SAPIENS | 5 | 16 | 3lbn | A | 58 | THR | p.Thr58Ile | P01112 |
| HOMO SAPIENS | 3 | 16 | 3lbn | A | 61 | GLN | p.Gln61Lys | P01112 |
| HOMO SAPIENS | 1 | 16 | 3lbn | A | 63 | GLU | p.Glu63Lys | P01112 |
| HOMO SAPIENS | 9 | 16 | 3lbn | A | 117 | LYS | p.Lys117Arg | P01112 |
| HOMO SAPIENS | 6 | 16 | 3lbn | A | 123 | ARG | p.Arg123Cys | P01112 |
| HOMO SAPIENS | 1 | 16 | 3lbn | A | 138 | GLY | p.Gly138Ser | P01112 |
| HOMO SAPIENS | 9 | 16 | 3lbn | A | 146 | ALA | p.Ala146Thr | P01112 |
| HOMO SAPIENS | 9 | 16 | 3lbn | A | 146 | ALA | p.Ala146Val | P01112 |
| HOMO SAPIENS | 1 | 359 | 5nvy | C | 63 | LEU | p.Leu63Pro | P40337 |
| HOMO SAPIENS | 1 | 359 | 5nvy | C | 64 | ARG | p.Arg64Pro | P40337 |
| HOMO SAPIENS | 1 | 359 | 5nvy | C | 70 | GLU | p.Glu70Lys | P40337 |
| HOMO SAPIENS | 0 | 359 | 5nvy | C | 84 | VAL | p.Val84Leu | P40337 |
| HOMO SAPIENS | 1 | 359 | 5nvy | C | 88 | TRP | p.Trp88Ser | P40337 |
| HOMO SAPIENS | 1 | 359 | 5nvy | C | 93 | GLY | p.Gly93Cys | P40337 |
| HOMO SAPIENS | 1 | 359 | 5nvy | C | 98 | TYR | p.Tyr98His | P40337 |
| HOMO SAPIENS | 1 | 359 | 5nvy | C | 111 | SER | p.Ser111Arg | P40337 |
| HOMO SAPIENS | 1 | 359 | 5nvy | C | 112 | TYR | p.Tyr112His | P40337 |
| HOMO SAPIENS | 1 | 359 | 5nvy | C | 126 | ASP | p.Asp126Tyr | P40337 |
| HOMO SAPIENS | 1 | 359 | 5nvy | C | 130 | VAL | p.Val130Leu | P40337 |
| HOMO SAPIENS | 1 | 359 | 5nvy | C | 135 | LEU | p.Leu135Phe | P40337 |
| HOMO SAPIENS | 9 | 359 | 5nvy | C | 161 | ARG | p.Arg161Gly | P40337 |
| HOMO SAPIENS | 7 | 359 | 5nvy | C | 162 | CYS | p.Cys162Phe | P40337 |
| HOMO SAPIENS | 4 | 359 | 5nvy | C | 163 | LEU | p.Leu163Pro | P40337 |
| HOMO SAPIENS | 3 | 359 | 5nvy | C | 164 | GLN | p.Gln164Arg | P40337 |
| HOMO SAPIENS | 9 | 359 | 5nvy | C | 166 | VAL | p.Val166Phe | P40337 |
| HOMO SAPIENS | 7 | 359 | 5nvy | C | 167 | ARG | p.Arg167Gly | P40337 |
| HOMO SAPIENS | 7 | 359 | 5nvy | C | 167 | ARG | p.Arg167Gln | P40337 |
| HOMO SAPIENS | 1 | 359 | 5nvy | C | 183 | SER | p.Ser183Ter | P40337 |
| HOMO SAPIENS | 2 | 359 | 5nvy | C | 188 | LEU | p.Leu188Val | P40337 |
| HOMO SAPIENS | 1 | 359 | 5nvy | C | 191 | HIS | p.His191Asp | P40337 |
| HOMO SAPIENS | 1 | 359 | 5nvy | C | 192 | PRO | p.Pro192Ser | P40337 |
| HOMO SAPIENS | 1 | 359 | 5nvy | C | 200 | ARG | p.Arg200Trp | P40337 |
| HOMO SAPIENS | -1 | 359 | 5nvy | C | 211 | MET | p.Met211Leu | P40337 |
| HOMO SAPIENS | 0 | 14099 | 4btm | A | 282 | ARG | p.Arg258Gln | Q5TCY1 |
| HOMO SAPIENS | 0 | 696 | 5h1b | A | 150 | ARG | p.Arg150Gln | Q06609 |
| HOMO SAPIENS | -1 | 208 | 2bxl | A | 1 | ASP | p.Asp25Val | P02768 |
| HOMO SAPIENS | 0 | 208 | 2bxl | A | 269 | ASP | p.Asp293Gly | P02768 |
| HOMO SAPIENS | 0 | 208 | 2bxl | A | 570 | GLU | p.Glu594Lys | P02768 |
| HOMO SAPIENS | 0 | 208 | 2bxl | A | 573 | LYS | p.Lys597Glu | P02768 |
| HOMO SAPIENS | -1 | 12813 | 2uzx | B | 35 | MET | p.Met35Leu | P08581 |
| HOMO SAPIENS | 1 | 12813 | 2uzx | B | 156 | SER | p.Ser156Leu | P08581 |
| HOMO SAPIENS | 0 | 12813 | 2uzx | B | 168 | GLU | p.Glu168Asp | P08581 |
| HOMO SAPIENS | 2 | 12813 | 2uzx | B | 211 | LEU | p.Leu211Trp | P08581 |
| HOMO SAPIENS | 2 | 12813 | 2uzx | B | 218 | ARG | p.Arg218Ser | P08581 |

| | | | | | | | | |
|---|---|---|---|---|---|---|---|---|
| HOMO SAPIENS | 1 | 12813 | 2uzx | B | 238 | LEU | p.Leu238Ser | P08581 |
| HOMO SAPIENS | 4 | 12813 | 2uzx | B | 320 | ALA | p.Ala320Val | P08581 |
| HOMO SAPIENS | 1 | 12813 | 2uzx | B | 375 | ASN | p.Asn375Ser | P08581 |
| HOMO SAPIENS | 0 | 12813 | 2uzx | B | 575 | LEU | p.Leu575Ile | P08581 |
| HOMO SAPIENS | 1 | 10780 | 4ejx | A | 348 | ARG | p.Arg367Cys | P00450 |
| HOMO SAPIENS | 2 | 10780 | 4ejx | A | 525 | ASP | p.Asp544Glu | P00450 |
| HOMO SAPIENS | 1 | 1407 | 5jn5 | A | 88 | ILE | p.Ile88Val | P36871 |
| HOMO SAPIENS | 7 | 1407 | 5jn5 | A | 115 | THR | p.Thr115Ala | P36871 |
| HOMO SAPIENS | 3 | 1407 | 5jn5 | A | 121 | GLY | p.Gly121Arg | P36871 |
| HOMO SAPIENS | 2 | 1407 | 5jn5 | A | 221 | ARG | p.Arg221Cys | P36871 |
| HOMO SAPIENS | 6 | 1407 | 5jn5 | A | 263 | TYR | p.Asp263Tyr | P36871 |
| HOMO SAPIENS | 3 | 1407 | 5jn5 | A | 420 | TYR | p.Tyr420His | P36871 |
| HOMO SAPIENS | 0 | 1407 | 5jn5 | A | 501 | VAL | p.Val501Ile | P36871 |
| HOMO SAPIENS | -1 | 1878 | 1yow | A | 255 | ARG | p.Arg255Leu | Q13285 |
| HOMO SAPIENS | 6 | 1878 | 1yow | A | 293 | ASP | p.Asp293Asn | Q13285 |
| HOMO SAPIENS | 1 | 1878 | 1yow | A | 437 | LEU | p.Leu437Gln | Q13285 |
| HOMO SAPIENS | 0 | 3071 | 2wpm | E | 145 | ARG | p.Arg191His | P00740 |
| HOMO SAPIENS | 0 | 3071 | 2wpm | E | 145 | ARG | p.Arg191Cys | P00740 |
| HOMO SAPIENS | 3 | 589 | 3ggc | B | 15 | GLY | p.Gly16Ser | P00492 |
| HOMO SAPIENS | 4 | 589 | 3ggc | B | 40 | LEU | p.Leu41Pro | P00492 |
| HOMO SAPIENS | 2 | 589 | 3ggc | B | 50 | ARG | p.Arg51Gly | P00492 |
| HOMO SAPIENS | 5 | 589 | 3ggc | B | 51 | ASP | p.Asp52Gly | P00492 |
| HOMO SAPIENS | 0 | 589 | 3ggc | B | 57 | GLY | p.Gly58Arg | P00492 |
| HOMO SAPIENS | 6 | 589 | 3ggc | B | 69 | GLY | p.Gly70Glu | P00492 |
| HOMO SAPIENS | 1 | 589 | 3ggc | B | 73 | PHE | p.Phe74Leu | P00492 |
| HOMO SAPIENS | 0 | 589 | 3ggc | B | 77 | LEU | p.Leu78Val | P00492 |
| HOMO SAPIENS | 3 | 589 | 3ggc | B | 79 | ASP | p.Asp80Val | P00492 |
| HOMO SAPIENS | 0 | 589 | 3ggc | B | 103 | SER | p.Ser104Arg | P00492 |
| HOMO SAPIENS | 0 | 589 | 3ggc | B | 109 | SER | p.Ser110Leu | P00492 |
| HOMO SAPIENS | 1 | 589 | 3ggc | B | 129 | VAL | p.Val130Asp | P00492 |
| HOMO SAPIENS | 4 | 589 | 3ggc | B | 131 | ILE | p.Ile132Met | P00492 |
| HOMO SAPIENS | 7 | 589 | 3ggc | B | 142 | MET | p.Met143Lys | P00492 |
| HOMO SAPIENS | 0 | 589 | 3ggc | B | 160 | ALA | p.Ala161Ser | P00492 |
| HOMO SAPIENS | 6 | 589 | 3ggc | B | 167 | THR | p.Thr168Ile | P00492 |
| HOMO SAPIENS | 0 | 589 | 3ggc | B | 169 | ARG | p.Arg170Ter | P00492 |
| HOMO SAPIENS | 5 | 589 | 3ggc | B | 175 | PRO | p.Pro176Leu | P00492 |
| HOMO SAPIENS | 8 | 589 | 3ggc | B | 193 | ASP | p.Asp194Glu | P00492 |
| HOMO SAPIENS | 8 | 589 | 3ggc | B | 193 | ASP | p.Asp194Asn | P00492 |
| HOMO SAPIENS | 8 | 589 | 3ggc | B | 198 | PHE | p.Phe199Val | P00492 |
| HOMO SAPIENS | 8 | 589 | 3ggc | B | 200 | ASP | p.Asp201Gly | P00492 |
| HOMO SAPIENS | 0 | 4269 | 2pla | B | 174 | GLU | p.Glu174Lys | Q8N335 |
| HOMO SAPIENS | 1 | 4269 | 2pla | B | 220 | ARG | p.Arg220His | Q8N335 |
| HOMO SAPIENS | 4 | 4269 | 2pla | B | 280 | ALA | p.Ala280Val | Q8N335 |
| HOMO SAPIENS | 0 | 2712 | 1ii4 | E | 252 | TRP | p.Ser252Leu | P21802 |
| HOMO SAPIENS | 0 | 2712 | 1ii4 | E | 253 | PRO | p.Pro253Arg | P21802 |
| HOMO SAPIENS | 5 | 2712 | 1ii4 | E | 267 | SER | p.Ser267Pro | P21802 |
| HOMO SAPIENS | 0 | 2712 | 1ii4 | E | 278 | CYS | p.Cys278Phe | P21802 |
| HOMO SAPIENS | 0 | 2712 | 1ii4 | E | 289 | GLN | p.Gln289Pro | P21802 |
| HOMO SAPIENS | 0 | 2712 | 1ii4 | E | 290 | TRP | p.Trp290Cys | P21802 |
| HOMO SAPIENS | 0 | 2712 | 1ii4 | E | 290 | TRP | p.Trp290Arg | P21802 |
| HOMO SAPIENS | 0 | 2712 | 1ii4 | E | 292 | LYS | p.Lys292Glu | P21802 |
| HOMO SAPIENS | 0 | 2712 | 1ii4 | E | 315 | ALA | p.Ala315Ser | P21802 |
| HOMO SAPIENS | 0 | 2712 | 1ii4 | E | 321 | ASP | p.Asp321Ala | P21802 |
| HOMO SAPIENS | 1 | 2712 | 1ii4 | E | 328 | TYR | p.Tyr328Cys | P21802 |

| | | | | | | | | |
|---|---|---|---|---|---|---|---|---|
| HOMO SAPIENS | 9 | 2712 | 1ii4 | E | 340 | TYR | p.Tyr340His | P21802 |
| HOMO SAPIENS | 1 | 2712 | 1ii4 | E | 341 | THR | p.Thr341Pro | P21802 |
| HOMO SAPIENS | 0 | 2712 | 1ii4 | E | 342 | CYS | p.Cys342Arg | P21802 |
| HOMO SAPIENS | 0 | 2712 | 1ii4 | E | 342 | CYS | p.Cys342Trp | P21802 |
| HOMO SAPIENS | 0 | 2712 | 1ii4 | E | 344 | ALA | p.Ala344Gly | P21802 |
| HOMO SAPIENS | 0 | 2712 | 1ii4 | E | 347 | SER | p.Ser347Cys | P21802 |
| HOMO SAPIENS | 0 | 2712 | 1ii4 | E | 351 | SER | p.Ser351Cys | P21802 |
| HOMO SAPIENS | 1 | 2712 | 1ii4 | E | 354 | SER | p.Ser354Cys | P21802 |
| HOMO SAPIENS | 2 | 53 | 4pj1 | H | 5 | ASP | p.Asp29Gly | P10809 |
| HOMO SAPIENS | 0 | 53 | 4pj1 | H | 74 | VAL | p.Val98Ile | P10809 |
| HOMO SAPIENS | 0 | 29968 | 2fdb | M | 71 | LYS | p.Lys89Glu | P55075 |
| HOMO SAPIENS | 2 | 29968 | 2fdb | M | 98 | ARG | p.Arg116Gly | P55075 |
| HOMO SAPIENS | 2 | 29968 | 2fdb | M | 122 | GLY | p.Gly140Ser | P55075 |
| HOMO SAPIENS | 8 | 5061 | 5f85 | B | 60 | GLY | p.Gly106Ser | P00740 |
| HOMO SAPIENS | 0 | 35944 | 1h6h | A | 85 | THR | p.Thr85Asn | Q15080 |
| HOMO SAPIENS | 8 | 35944 | 1h6h | A | 105 | ARG | p.Arg105Gln | Q15080 |
| HOMO SAPIENS | 7 | 12016 | 1hj7 | A | 295 | ASN | p.Asn316Ser | P01130 |
| HOMO SAPIENS | 5 | 12016 | 1hj7 | A | 302 | GLY | p.Gly323Ser | P01130 |
| HOMO SAPIENS | 6 | 12016 | 1hj7 | A | 319 | CYS | p.Cys340Trp | P01130 |
| HOMO SAPIENS | 7 | 12016 | 1hj7 | A | 329 | ARG | p.Arg350Ter | P01130 |
| HOMO SAPIENS | 5 | 12016 | 1hj7 | A | 331 | CYS | p.Cys352Ter | P01130 |
| HOMO SAPIENS | 2 | 12016 | 1hj7 | A | 345 | GLN | p.Gln366Arg | P01130 |
| HOMO SAPIENS | 2 | 1481 | 2i07 | A | 239 | TYR | p.Tyr261Ter | P01024 |
| HOMO SAPIENS | -1 | 1481 | 2i07 | A | 292 | LEU | p.Pro314Leu | P01024 |
| HOMO SAPIENS | 2 | 1481 | 2i07 | A | 326 | ILE | p.Ile348Val | P01024 |
| HOMO SAPIENS | 2 | 1481 | 2i07 | A | 413 | GLU | p.Glu435Lys | P01024 |
| HOMO SAPIENS | 4 | 1481 | 2i07 | A | 447 | GLU | p.Glu469Asp | P01024 |
| HOMO SAPIENS | 2 | 1481 | 2i07 | A | 597 | VAL | p.Val619Met | P01024 |
| HOMO SAPIENS | 5 | 1481 | 2i07 | A | 603 | ILE | p.Ile625Phe | P01024 |
| HOMO SAPIENS | 0 | 1481 | 2i07 | A | 611 | LYS | p.Lys633Arg | P01024 |
| HOMO SAPIENS | 0 | 11123 | 3c5n | A | 315 | GLY | p.Gly368Trp | O00294 |
| HOMO SAPIENS | 8 | 11123 | 3c5n | A | 329 | PHE | p.Phe382Ser | O00294 |
| HOMO SAPIENS | 5 | 11123 | 3c5n | A | 429 | ARG | p.Arg482Trp | O00294 |
| HOMO SAPIENS | 8 | 11123 | 3c5n | A | 438 | PHE | p.Phe491Leu | O00294 |
| HOMO SAPIENS | 3 | 4398 | 3fx0 | B | 1288 | ALA | p.Ala288Gly | Q9Y6K9 |
| HOMO SAPIENS | 7 | 4398 | 3fx0 | B | 1315 | GLU | p.Glu315Ala | Q9Y6K9 |
| HOMO SAPIENS | 8 | 4398 | 3fx0 | B | 1319 | ARG | p.Arg319Gln | Q9Y6K9 |
| HOMO SAPIENS | 6 | 3989 | 3qd6 | S | 83 | CYS | p.Cys83Arg | P25942 |
| HOMO SAPIENS | 1 | 3989 | 3qd6 | S | 124 | SER | p.Ser124Leu | P25942 |
| HOMO SAPIENS | 2 | 24003 | 2m0v | A | 225 | GLU | p.Glu225Lys | O14745 |
| HOMO SAPIENS | 2 | 46 | 4aqd | A | 70 | ASP | p.Asp98Gly | P06276 |
| HOMO SAPIENS | 4 | 46 | 4aqd | A | 115 | GLY | p.Gly143Asp | P06276 |
| HOMO SAPIENS | 3 | 46 | 4aqd | A | 184 | ALA | p.Ala212Val | P06276 |
| HOMO SAPIENS | 0 | 46 | 4aqd | A | 225 | GLY | p.Gly253Ter | P06276 |
| HOMO SAPIENS | 2 | 46 | 4aqd | A | 243 | THR | p.Thr271Met | P06276 |
| HOMO SAPIENS | 2 | 46 | 4aqd | A | 255 | GLU | p.Glu283Asp | P06276 |
| HOMO SAPIENS | 5 | 46 | 4aqd | A | 267 | LYS | p.Lys295Arg | P06276 |
| HOMO SAPIENS | 3 | 46 | 4aqd | A | 271 | GLU | p.Glu299Ter | P06276 |
| HOMO SAPIENS | 2 | 46 | 4aqd | A | 307 | LEU | p.Leu335Pro | P06276 |
| HOMO SAPIENS | 1 | 46 | 4aqd | A | 330 | LEU | p.Leu358Ile | P06276 |
| HOMO SAPIENS | 0 | 46 | 4aqd | A | 497 | GLU | p.Glu525Val | P06276 |
| HOMO SAPIENS | 3 | 786 | 1ols | B | 51 | GLY | p.Gly101Asp | P21953 |
| HOMO SAPIENS | 2 | 786 | 1ols | B | 87 | ALA | p.Ala137Val | P21953 |
| HOMO SAPIENS | 0 | 786 | 1ols | B | 110 | ILE | p.Ile160Ser | P21953 |

| | | | | | | | | |
|---|---|---|---|---|---|---|---|---|
| HOMO SAPIENS | 4 | 786 | 1ols | B | 120 | ARG | p.Arg170Cys | P21953 |
| HOMO SAPIENS | 2 | 786 | 1ols | B | 126 | ASN | p.Asn176Tyr | P21953 |
| HOMO SAPIENS | 0 | 786 | 1ols | B | 133 | ARG | p.Arg183Gly | P21953 |
| HOMO SAPIENS | 0 | 786 | 1ols | B | 133 | ARG | p.Arg183Pro | P21953 |
| HOMO SAPIENS | 0 | 786 | 1ols | B | 135 | PRO | p.Pro185Leu | P21953 |
| HOMO SAPIENS | 3 | 786 | 1ols | B | 156 | HIS | p.His206Tyr | P21953 |
| HOMO SAPIENS | 0 | 786 | 1ols | B | 228 | GLY | p.Gly278Ser | P21953 |
| HOMO SAPIENS | 1 | 786 | 1ols | B | 235 | ARG | p.Arg285Ter | P21953 |
| HOMO SAPIENS | 6 | 786 | 1ols | B | 274 | ARG | p.Arg324Ter | P21953 |
| HOMO SAPIENS | 1 | 831 | 4zt1 | B | 48 | VAL | p.Val202Ile | P12830 |
| HOMO SAPIENS | 6 | 831 | 4zt1 | B | 70 | ARG | p.Arg224Cys | P12830 |
| HOMO SAPIENS | 5 | 831 | 4zt1 | B | 79 | HIS | p.His233Gln | P12830 |
| HOMO SAPIENS | 4 | 831 | 4zt1 | B | 120 | GLY | p.Gly274Ser | P12830 |
| HOMO SAPIENS | 7 | 831 | 4zt1 | B | 187 | TYR | p.Tyr341Ter | P12830 |
| HOMO SAPIENS | 0 | 8497 | 3sla | A | 287 | ASN | p.Asn287Ser | P35222 |
| HOMO SAPIENS | 7 | 23986 | 2ke1 | A | 301 | VAL | p.Val301Met | O43918 |
| HOMO SAPIENS | 0 | 4967 | 1rj7 | B | 334 | ARG | p.Arg334His | Q92838 |
| HOMO SAPIENS | 0 | 4967 | 1rj7 | B | 338 | THR | p.Thr338Met | Q92838 |
| HOMO SAPIENS | 3 | 4967 | 1rj7 | B | 349 | ALA | p.Ala349Thr | Q92838 |
| HOMO SAPIENS | 2 | 4967 | 1rj7 | B | 358 | GLN | p.Gln358Glu | Q92838 |
| HOMO SAPIENS | 1 | 4967 | 1rj7 | B | 375 | LYS | p.Lys375Glu | Q92838 |
| HOMO SAPIENS | 9 | 7623 | 1d4t | A | 32 | ARG | p.Arg32Thr | O60880 |
| HOMO SAPIENS | 6 | 7623 | 1d4t | A | 55 | ARG | p.Arg55Leu | O60880 |
| HOMO SAPIENS | 0 | 7623 | 1d4t | A | 68 | THR | p.Thr68Ile | O60880 |
| HOMO SAPIENS | 6 | 705 | 3idh | A | 59 | PRO | p.Pro59Ser | P35557 |
| HOMO SAPIENS | 5 | 705 | 3idh | A | 124 | ASP | p.Asp124Asn | P35557 |
| HOMO SAPIENS | 4 | 705 | 3idh | A | 131 | SER | p.Ser131Pro | P35557 |
| HOMO SAPIENS | 0 | 705 | 3idh | A | 147 | GLY | p.Gly147Asp | P35557 |
| HOMO SAPIENS | 5 | 705 | 3idh | A | 186 | ARG | p.Arg186Ter | P35557 |
| HOMO SAPIENS | 0 | 705 | 3idh | A | 202 | MET | p.Met202Thr | P35557 |
| HOMO SAPIENS | 2 | 705 | 3idh | A | 210 | MET | p.Met210Lys | P35557 |
| HOMO SAPIENS | 1 | 705 | 3idh | A | 215 | TYR | p.Tyr215Ter | P35557 |
| HOMO SAPIENS | 0 | 705 | 3idh | A | 226 | VAL | p.Val226Met | P35557 |
| HOMO SAPIENS | 9 | 705 | 3idh | A | 228 | THR | p.Thr228Met | P35557 |
| HOMO SAPIENS | 9 | 705 | 3idh | A | 256 | GLU | p.Glu256Lys | P35557 |
| HOMO SAPIENS | 2 | 705 | 3idh | A | 261 | GLY | p.Gly261Arg | P35557 |
| HOMO SAPIENS | 6 | 705 | 3idh | A | 279 | GLU | p.Glu279Gln | P35557 |
| HOMO SAPIENS | 7 | 705 | 3idh | A | 299 | GLY | p.Gly299Arg | P35557 |
| HOMO SAPIENS | 2 | 705 | 3idh | A | 318 | GLY | p.Gly318Trp | P35557 |
| HOMO SAPIENS | 4 | 705 | 3idh | A | 378 | ALA | p.Ala378Thr | P35557 |
| HOMO SAPIENS | 0 | 705 | 3idh | A | 387 | ALA | p.Ala387Glu | P35557 |
| HOMO SAPIENS | 5 | 705 | 3idh | A | 414 | LYS | p.Lys414Glu | P35557 |
| HOMO SAPIENS | 0 | 705 | 3idh | A | 455 | VAL | p.Val455Met | P35557 |
| HOMO SAPIENS | -1 | 705 | 3idh | A | 462 | MET | p.Met462Ile | P35557 |
| HOMO SAPIENS | 1 | 509 | 4nef | A | 22 | LEU | p.Leu22Val | P41181 |
| HOMO SAPIENS | 4 | 509 | 4nef | A | 57 | GLN | p.Gln57Pro | P41181 |
| HOMO SAPIENS | 8 | 509 | 4nef | A | 64 | GLY | p.Gly64Arg | P41181 |
| HOMO SAPIENS | 7 | 509 | 4nef | A | 68 | ASN | p.Asn68Ser | P41181 |
| HOMO SAPIENS | 1 | 509 | 4nef | A | 100 | GLY | p.Gly100Val | P41181 |
| HOMO SAPIENS | 0 | 509 | 4nef | A | 125 | THR | p.Thr125Met | P41181 |
| HOMO SAPIENS | 0 | 509 | 4nef | A | 126 | THR | p.Thr126Met | P41181 |
| HOMO SAPIENS | 4 | 509 | 4nef | A | 146 | PHE | p.Phe146Leu | P41181 |
| HOMO SAPIENS | 0 | 509 | 4nef | A | 175 | GLY | p.Gly175Arg | P41181 |
| HOMO SAPIENS | 5 | 509 | 4nef | A | 181 | CYS | p.Cys181Trp | P41181 |

| | | | | | | | | | |
|---|---|---|---|---|---|---|---|---|---|
| HOMO SAPIENS | 9 | 509 | 4nef | A | | 187 | ARG | p.Arg187Cys | P41181 |
| HOMO SAPIENS | 0 | 509 | 4nef | A | | 190 | ALA | p.Ala190Thr | P41181 |
| HOMO SAPIENS | 0 | 509 | 4nef | A | | 216 | SER | p.Ser216Pro | P41181 |
| HOMO SAPIENS | 4 | 2242 | 1z70 | X | | 1155 | SER | p.Ser155Pro | Q8NBK3 |
| HOMO SAPIENS | 3 | 2242 | 1z70 | X | | 1218 | CYS | p.Cys218Tyr | Q8NBK3 |
| HOMO SAPIENS | 0 | 2242 | 1z70 | X | | 1279 | ALA | p.Ala279Val | Q8NBK3 |
| HOMO SAPIENS | -1 | 2242 | 1z70 | X | | 1336 | OCS | p.Cys336Arg | Q8NBK3 |
| HOMO SAPIENS | 4 | 2242 | 1z70 | X | | 1345 | ARG | p.Arg345Cys | Q8NBK3 |
| HOMO SAPIENS | 2 | 2242 | 1z70 | X | | 1348 | ALA | p.Ala348Pro | Q8NBK3 |
| HOMO SAPIENS | 5 | 2242 | 1z70 | X | | 1349 | ARG | p.Arg349Gln | Q8NBK3 |
| HOMO SAPIENS | 5 | 2242 | 1z70 | X | | 1349 | ARG | p.Arg349Trp | Q8NBK3 |
| HOMO SAPIENS | 3 | 2242 | 1z70 | X | | 1359 | SER | p.Ser359Ter | Q8NBK3 |
| HOMO SAPIENS | 5 | 8269 | 5wrr | B | | 136 | ARG | p.Arg136Ter | Q96MK3 |
| HOMO SAPIENS | 1 | 8269 | 5wrr | B | | 332 | LYS | p.Asn332Lys | Q96MK3 |
| HOMO SAPIENS | 3 | 8269 | 5wrr | B | | 478 | ARG | p.Arg478Ter | Q96MK3 |
| HOMO SAPIENS | 5 | 8269 | 5wrr | B | | 504 | THR | p.Thr504Ile | Q96MK3 |
| HOMO SAPIENS | 5 | 1411 | 4h1s | B | | 221 | SER | p.Ser221Ter | P21589 |
| HOMO SAPIENS | 1 | 1411 | 4h1s | B | | 358 | CYS | p.Cys358Tyr | P21589 |
| HOMO SAPIENS | 2 | 23 | 5vvd | B | | 298 | GLU | p.Glu298Asp | P29474 |
| HOMO SAPIENS | 0 | 19 | 5jlh | E | | 88 | THR | p.Thr89Ile | P63261 |
| HOMO SAPIENS | 6 | 19 | 5jlh | E | | 117 | LYS | p.Lys118Met | P63261 |
| HOMO SAPIENS | 5 | 19 | 5jlh | E | | 263 | PRO | p.Pro264Leu | P63261 |
| HOMO SAPIENS | 1 | 19 | 5jlh | E | | 277 | THR | p.Thr278Ile | P63261 |
| HOMO SAPIENS | 6 | 19 | 5jlh | E | | 331 | PRO | p.Pro332Ala | P63261 |
| HOMO SAPIENS | 5 | 19 | 5jlh | E | | 369 | VAL | p.Val370Ala | P63261 |
| HOMO SAPIENS | 0 | 14306 | 4w4o | C | | 92 | ARG | p.Arg92Ter | P12314 |
| HOMO SAPIENS | 3 | 603 | 1nuh | A | | 4 | THR | p.Thr5Ile | P06744 |
| HOMO SAPIENS | 2 | 603 | 1nuh | A | | 19 | HIS | p.His20Pro | P06744 |
| HOMO SAPIENS | 8 | 603 | 1nuh | A | | 158 | GLY | p.Gly159Ser | P06744 |
| HOMO SAPIENS | 6 | 603 | 1nuh | A | | 223 | THR | p.Thr224Met | P06744 |
| HOMO SAPIENS | 0 | 603 | 1nuh | A | | 338 | LEU | p.Leu339Pro | P06744 |
| HOMO SAPIENS | 6 | 603 | 1nuh | A | | 342 | GLN | p.Gln343Arg | P06744 |
| HOMO SAPIENS | 6 | 603 | 1nuh | A | | 374 | THR | p.Thr375Arg | P06744 |
| HOMO SAPIENS | 4 | 603 | 1nuh | A | | 524 | ILE | p.Ile525Thr | P06744 |
| HOMO SAPIENS | 4 | 603 | 1nuh | A | | 538 | ASP | p.Asp539Asn | P06744 |
| HOMO SAPIENS | 1 | 3107 | 3kzj | A | | 1197 | VAL | p.Val1197Ala | P08603 |
| HOMO SAPIENS | 9 | 1748 | 2noh | A | | 46 | ARG | p.Arg46Gln | O15527 |
| HOMO SAPIENS | 2 | 11750 | 3s4y | A | | 33 | ARG | p.Arg33His | Q9H3S4 |
| HOMO SAPIENS | 2 | 11750 | 3s4y | A | | 40 | LEU | p.Leu40Pro | Q9H3S4 |
| HOMO SAPIENS | 4 | 11750 | 3s4y | A | | 142 | LEU | p.Leu142Phe | Q9H3S4 |
| HOMO SAPIENS | 5 | 11750 | 3s4y | A | | 219 | ASN | p.Asn219Ser | Q9H3S4 |
| HOMO SAPIENS | 1 | 2591 | 5vnj | A | | 382 | PHE | p.Phe382Leu | Q15436 |
| HOMO SAPIENS | 2 | 2591 | 5vnj | A | | 702 | MET | p.Met702Val | Q15436 |
| HOMO SAPIENS | 1 | 2426 | 5m8t | A | | 87 | ARG | p.Arg87Trp | P17643 |
| HOMO SAPIENS | 1 | 2426 | 5m8t | A | | 166 | SER | p.Ser166Ter | P17643 |
| HOMO SAPIENS | 0 | 2426 | 5m8t | A | | 356 | ARG | p.Arg356Gln | P17643 |
| HOMO SAPIENS | 2 | 2426 | 5m8t | A | | 374 | SER | p.Arg374Ter | P17643 |
| HOMO SAPIENS | 6 | 2426 | 5m8t | A | | 378 | ASN | p.Asn378Ser | P17643 |
| HOMO SAPIENS | 6 | 2426 | 5m8t | A | | 382 | LEU | p.Leu382Pro | P17643 |
| HOMO SAPIENS | 7 | 10097 | 4n6v | | 1 | 46 | GLN | p.Gln46Ter | P04080 |
| HOMO SAPIENS | 3 | 10097 | 4n6v | | 1 | 65 | VAL | p.Val65Ile | P04080 |
| HOMO SAPIENS | 8 | 10097 | 4n6v | | 1 | 68 | ARG | p.Arg68Ter | P04080 |
| HOMO SAPIENS | 2 | 859 | 4z9a | A | | 105 | GLN | p.Gln106Arg | P24666 |
| HOMO SAPIENS | 8 | 1420 | 5nan | F | | 95 | SER | p.Ser95Leu | Q96PD4 |

| | | | | | | | | |
|---|---|---|---|---|---|---|---|---|
| HOMO SAPIENS | 6 | 1420 | 5nan | F | 126 | GLU | p.Glu126Gly | Q96PD4 |
| HOMO SAPIENS | 8 | 1420 | 5nan | F | 155 | VAL | p.Val155Ile | Q96PD4 |
| HOMO SAPIENS | 6 | 1420 | 5nan | F | 157 | PRO | p.Pro157Ser | Q96PD4 |
| HOMO SAPIENS | -1 | 1420 | 5nan | F | 161 | HIS | p.His161Arg | Q96PD4 |
| HOMO SAPIENS | 3 | 288 | 5i96 | B | 140 | GLN | p.Arg140Gly | P48735 |
| HOMO SAPIENS | 3 | 288 | 5i96 | B | 140 | GLN | p.Arg140Gln | P48735 |
| HOMO SAPIENS | 2 | 288 | 5i96 | B | 261 | ARG | p.Arg261His | P48735 |
| HOMO SAPIENS | 2 | 288 | 5i96 | B | 435 | THR | p.Thr435Met | P48735 |
| HOMO SAPIENS | 9 | 120 | 1tup | B | 107 | TYR | p.Tyr107His | P04637 |
| HOMO SAPIENS | 8 | 120 | 1tup | B | 110 | ARG | p.Arg110Leu | P04637 |
| HOMO SAPIENS | 5 | 120 | 1tup | B | 120 | LYS | p.Lys120Glu | P04637 |
| HOMO SAPIENS | 4 | 120 | 1tup | B | 133 | MET | p.Met133Thr | P04637 |
| HOMO SAPIENS | 3 | 120 | 1tup | B | 134 | PHE | p.Phe134Cys | P04637 |
| HOMO SAPIENS | 8 | 120 | 1tup | B | 136 | GLN | p.Gln136His | P04637 |
| HOMO SAPIENS | 9 | 120 | 1tup | B | 138 | ALA | p.Ala138Pro | P04637 |
| HOMO SAPIENS | 9 | 120 | 1tup | B | 151 | PRO | p.Pro151Ser | P04637 |
| HOMO SAPIENS | 9 | 120 | 1tup | B | 152 | PRO | p.Pro152Leu | P04637 |
| HOMO SAPIENS | 9 | 120 | 1tup | B | 158 | ARG | p.Arg158Cys | P04637 |
| HOMO SAPIENS | 9 | 120 | 1tup | B | 158 | ARG | p.Arg158His | P04637 |
| HOMO SAPIENS | 9 | 120 | 1tup | B | 175 | ARG | p.Arg175Cys | P04637 |
| HOMO SAPIENS | 9 | 120 | 1tup | B | 175 | ARG | p.Arg175Leu | P04637 |
| HOMO SAPIENS | 8 | 120 | 1tup | B | 181 | ARG | p.Arg181Leu | P04637 |
| HOMO SAPIENS | 9 | 120 | 1tup | B | 189 | ALA | p.Ala189Val | P04637 |
| HOMO SAPIENS | 9 | 120 | 1tup | B | 196 | ARG | p.Arg196Gly | P04637 |
| HOMO SAPIENS | 9 | 120 | 1tup | B | 213 | ARG | p.Arg213Gln | P04637 |
| HOMO SAPIENS | 9 | 120 | 1tup | B | 213 | ARG | p.Arg213Ter | P04637 |
| HOMO SAPIENS | 9 | 120 | 1tup | B | 214 | HIS | p.His214Gln | P04637 |
| HOMO SAPIENS | 9 | 120 | 1tup | B | 220 | TYR | p.Tyr220Cys | P04637 |
| HOMO SAPIENS | 0 | 120 | 1tup | B | 234 | TYR | p.Tyr234Cys | P04637 |
| HOMO SAPIENS | 9 | 120 | 1tup | B | 237 | MET | p.Met237Ile | P04637 |
| HOMO SAPIENS | 0 | 120 | 1tup | B | 238 | CYS | p.Cys238Tyr | P04637 |
| HOMO SAPIENS | 9 | 120 | 1tup | B | 241 | SER | p.Ser241Phe | P04637 |
| HOMO SAPIENS | 9 | 120 | 1tup | B | 242 | CYS | p.Cys242Ser | P04637 |
| HOMO SAPIENS | 9 | 120 | 1tup | B | 245 | GLY | p.Gly245Cys | P04637 |
| HOMO SAPIENS | 9 | 120 | 1tup | B | 245 | GLY | p.Gly245Asp | P04637 |
| HOMO SAPIENS | 9 | 120 | 1tup | B | 248 | ARG | p.Arg248Leu | P04637 |
| HOMO SAPIENS | 9 | 120 | 1tup | B | 248 | ARG | p.Arg248Trp | P04637 |
| HOMO SAPIENS | 9 | 120 | 1tup | B | 251 | ILE | p.Ile251Ser | P04637 |
| HOMO SAPIENS | 9 | 120 | 1tup | B | 252 | LEU | p.Leu252Pro | P04637 |
| HOMO SAPIENS | 9 | 120 | 1tup | B | 258 | GLU | p.Glu258Lys | P04637 |
| HOMO SAPIENS | 8 | 120 | 1tup | B | 272 | VAL | p.Val272Leu | P04637 |
| HOMO SAPIENS | 9 | 120 | 1tup | B | 273 | ARG | p.Arg273Cys | P04637 |
| HOMO SAPIENS | 9 | 120 | 1tup | B | 273 | ARG | p.Arg273His | P04637 |
| HOMO SAPIENS | 9 | 120 | 1tup | B | 277 | CYS | p.Cys277Phe | P04637 |
| HOMO SAPIENS | 9 | 120 | 1tup | B | 280 | ARG | p.Arg280Ile | P04637 |
| HOMO SAPIENS | 9 | 120 | 1tup | B | 282 | ARG | p.Arg282Trp | P04637 |
| HOMO SAPIENS | 9 | 120 | 1tup | B | 283 | ARG | p.Arg283His | P04637 |
| HOMO SAPIENS | 9 | 120 | 1tup | B | 285 | GLU | p.Glu285Val | P04637 |
| HOMO SAPIENS | -1 | 120 | 1tup | B | 292 | LYS | p.Lys292Ile | P04637 |
| HOMO SAPIENS | -1 | 120 | 1tup | B | 298 | GLU | p.Glu298Lys | P04637 |
| HOMO SAPIENS | -1 | 2791 | 4ygb | A | 39 | VAL | p.Val39Ala | P49257 |
| HOMO SAPIENS | 2 | 106 | 4x6j | A | 32 | GLY | p.Gly146Arg | P43235 |
| HOMO SAPIENS | 0 | 106 | 4x6j | A | 76 | GLN | p.Gln190Ter | P43235 |
| HOMO SAPIENS | 7 | 106 | 4x6j | A | 198 | ARG | p.Arg312Ter | P43235 |

| | | | | | | | | |
|---|---|---|---|---|---|---|---|---|
| HOMO SAPIENS | 5 | 490 | 2hlo | B | 172 | LEU | p.Leu202Gln | P02675 |
| HOMO SAPIENS | 0 | 490 | 2hlo | B | 353 | LEU | p.Leu383Arg | P02675 |
| HOMO SAPIENS | 4 | 490 | 2hlo | B | 400 | GLY | p.Gly430Asp | P02675 |
| HOMO SAPIENS | 7 | 490 | 2hlo | B | 448 | ARG | p.Arg478Lys | P02675 |
| HOMO SAPIENS | 1 | 289 | 4p13 | B | 18 | GLU | p.Glu43Lys | P11310 |
| HOMO SAPIENS | 2 | 289 | 4p13 | B | 20 | GLN | p.Gln45Arg | P11310 |
| HOMO SAPIENS | 5 | 289 | 4p13 | B | 28 | ARG | p.Arg53Cys | P11310 |
| HOMO SAPIENS | 1 | 289 | 4p13 | B | 42 | TYR | p.Tyr67His | P11310 |
| HOMO SAPIENS | 1 | 289 | 4p13 | B | 53 | ILE | p.Ile78Thr | P11310 |
| HOMO SAPIENS | 1 | 289 | 4p13 | B | 59 | LEU | p.Leu84Phe | P11310 |
| HOMO SAPIENS | 2 | 289 | 4p13 | B | 96 | THR | p.Thr121Ile | P11310 |
| HOMO SAPIENS | 0 | 289 | 4p13 | B | 124 | MET | p.Met149Ile | P11310 |
| HOMO SAPIENS | 5 | 289 | 4p13 | B | 143 | ASP | p.Asp168Ala | P11310 |
| HOMO SAPIENS | 7 | 289 | 4p13 | B | 168 | THR | p.Thr193Ala | P11310 |
| HOMO SAPIENS | 1 | 289 | 4p13 | B | 170 | GLY | p.Gly195Arg | P11310 |
| HOMO SAPIENS | 2 | 289 | 4p13 | B | 181 | ARG | p.Arg206Cys | P11310 |
| HOMO SAPIENS | 2 | 289 | 4p13 | B | 181 | ARG | p.Arg206His | P11310 |
| HOMO SAPIENS | 8 | 289 | 4p13 | B | 203 | THR | p.Thr228Ile | P11310 |
| HOMO SAPIENS | 2 | 289 | 4p13 | B | 208 | ILE | p.Ile233Asn | P11310 |
| HOMO SAPIENS | 1 | 289 | 4p13 | B | 219 | CYS | p.Cys244Arg | P11310 |
| HOMO SAPIENS | 6 | 289 | 4p13 | B | 220 | SER | p.Ser245Leu | P11310 |
| HOMO SAPIENS | 5 | 289 | 4p13 | B | 228 | GLU | p.Glu253Lys | P11310 |
| HOMO SAPIENS | 2 | 289 | 4p13 | B | 241 | ASP | p.Asp266Gly | P11310 |
| HOMO SAPIENS | 1 | 289 | 4p13 | B | 242 | GLY | p.Gly267Arg | P11310 |
| HOMO SAPIENS | 7 | 289 | 4p13 | B | 256 | ARG | p.Arg281Thr | P11310 |
| HOMO SAPIENS | 4 | 289 | 4p13 | B | 269 | ARG | p.Arg294Thr | P11310 |
| HOMO SAPIENS | 4 | 289 | 4p13 | B | 304 | GLU | p.Lys329Glu | P11310 |
| HOMO SAPIENS | 1 | 289 | 4p13 | B | 324 | ARG | p.Arg349Ter | P11310 |
| HOMO SAPIENS | 3 | 289 | 4p13 | B | 326 | THR | p.Thr351Ile | P11310 |
| HOMO SAPIENS | 7 | 289 | 4p13 | B | 350 | ILE | p.Ile375Thr | P11310 |
| HOMO SAPIENS | 3 | 289 | 4p13 | B | 359 | GLU | p.Glu384Ter | P11310 |
| HOMO SAPIENS | 1 | 4563 | 3i2b | A | 16 | ARG | p.Arg16Cys | Q03393 |
| HOMO SAPIENS | 5 | 4563 | 3i2b | A | 25 | ARG | p.Arg25Gln | Q03393 |
| HOMO SAPIENS | 1 | 4563 | 3i2b | A | 47 | ASN | p.Asn47Asp | Q03393 |
| HOMO SAPIENS | 8 | 4563 | 3i2b | A | 52 | ASN | p.Asn52Ser | Q03393 |
| HOMO SAPIENS | 1 | 4563 | 3i2b | A | 56 | VAL | p.Val56Met | Q03393 |
| HOMO SAPIENS | 1 | 4563 | 3i2b | A | 87 | PRO | p.Pro87Ser | Q03393 |
| HOMO SAPIENS | 1 | 4563 | 3i2b | A | 96 | ASP | p.Asp96Asn | Q03393 |
| HOMO SAPIENS | 0 | 4563 | 3i2b | A | 99 | TYR | p.Tyr99Ter | Q03393 |
| HOMO SAPIENS | 2 | 4563 | 3i2b | A | 116 | ASP | p.Asp116Gly | Q03393 |
| HOMO SAPIENS | 2 | 41 | 3exx | A | 1 | GLY | p.Gly90Cys | P01308 |
| HOMO SAPIENS | 5 | 41 | 3exx | A | 3 | VAL | p.Val92Leu | P01308 |
| HOMO SAPIENS | 4 | 41 | 3exx | A | 7 | CYS | p.Cys96Ser | P01308 |
| HOMO SAPIENS | 9 | 41 | 3exx | A | 19 | TYR | p.Tyr108Cys | P01308 |
| HOMO SAPIENS | 9 | 131 | 4mjo | D | 118 | ASP | p.Asp119Asn | P09467 |
| HOMO SAPIENS | 0 | 131 | 4mjo | D | 163 | GLY | p.Gly164Ser | P09467 |
| HOMO SAPIENS | 0 | 131 | 4mjo | D | 176 | ALA | p.Ala177Asp | P09467 |
| HOMO SAPIENS | 5 | 131 | 4mjo | D | 193 | PHE | p.Phe194Ser | P09467 |
| HOMO SAPIENS | 4 | 131 | 4mjo | D | 217 | ARG | p.Arg218Lys | P09467 |
| HOMO SAPIENS | 4 | 131 | 4mjo | D | 223 | VAL | p.Val224Ile | P09467 |
| HOMO SAPIENS | 4 | 131 | 4mjo | D | 232 | PHE | p.Phe233Ile | P09467 |
| HOMO SAPIENS | 5 | 131 | 4mjo | D | 243 | ARG | p.Arg244Trp | P09467 |
| HOMO SAPIENS | 0 | 131 | 4mjo | D | 283 | PRO | p.Pro284Arg | P09467 |
| HOMO SAPIENS | 0 | 131 | 4mjo | D | 288 | MET | p.Met289Leu | P09467 |

| | | | | | | | | |
|---|---|---|---|---|---|---|---|---|
| HOMO SAPIENS | 0 | 131 | 4mjo | D | 323 | ASP | p.Asp324Asn | P09467 |
| HOMO SAPIENS | -1 | 13122 | 3hfc | A | 583 | ARG | p.Arg583Cys | P51787 |
| HOMO SAPIENS | 1 | 13122 | 3hfc | A | 587 | THR | p.Thr587Met | P51787 |
| HOMO SAPIENS | 0 | 13122 | 3hfc | A | 589 | GLY | p.Gly589Asp | P51787 |
| HOMO SAPIENS | 4 | 3002 | 3nbq | B | 94 | ARG | p.Arg94Cys | Q16831 |
| HOMO SAPIENS | 1 | 3616 | 3dbp | A | 97 | ILE | p.Ile286Asn | P11172 |
| HOMO SAPIENS | 0 | 3616 | 3dbp | A | 240 | GLY | p.Gly429Arg | P11172 |
| HOMO SAPIENS | 0 | 3616 | 3dbp | A | 257 | ILE | p.Ile446Val | P11172 |
| HOMO SAPIENS | 1 | 584 | 1drf | A | 79 | LEU | p.Leu80Phe | P00374 |
| HOMO SAPIENS | 3 | 1609 | 3qa2 | B | 49 | VAL | p.Val49Ile | P50053 |
| HOMO SAPIENS | 0 | 20560 | 5mug | A | 101 | HIS | p.His101Gln | P49638 |
| HOMO SAPIENS | 2 | 20560 | 5mug | A | 120 | ALA | p.Ala120Thr | P49638 |
| HOMO SAPIENS | 3 | 20560 | 5mug | A | 192 | ARG | p.Arg192His | P49638 |
| HOMO SAPIENS | 3 | 20560 | 5mug | A | 221 | ARG | p.Arg221Trp | P49638 |
| HOMO SAPIENS | 4 | 256 | 3vtv | A | 65 | THR | p.Glu50Lys | Q9GZQ8 |
| HOMO SAPIENS | 4 | 256 | 3vtv | A | 113 | VAL | p.Met98Lys | Q9GZQ8 |
| HOMO SAPIENS | 4 | 5397 | 1m61 | A | 192 | ARG | p.Arg192Trp | P43403 |
| HOMO SAPIENS | 5 | 39881 | 1aut | L | 107 | HIS | p.His149Pro | P04070 |
| HOMO SAPIENS | 5 | 4010 | 4uf8 | I | 33 | ALA | p.Ala33Val | Q13501 |
| HOMO SAPIENS | 3 | 313 | 5w49 | A | 98 | PRO | p.Pro98Leu | P23526 |
| HOMO SAPIENS | 2 | 313 | 5w49 | A | 143 | TYR | p.Tyr143Cys | P23526 |
| HOMO SAPIENS | -1 | 1643 | 2m8t | A | 102 | PRO | p.Pro102Leu | P04156 |
| HOMO SAPIENS | -1 | 1643 | 2m8t | A | 105 | PRO | p.Pro105Leu | P04156 |
| HOMO SAPIENS | -1 | 1643 | 2m8t | A | 117 | ALA | p.Ala117Val | P04156 |
| HOMO SAPIENS | 5 | 1643 | 2m8t | A | 129 | MET | p.Met129Val | P04156 |
| HOMO SAPIENS | 5 | 1643 | 2m8t | A | 131 | GLY | p.Gly131Val | P04156 |
| HOMO SAPIENS | 2 | 1643 | 2m8t | A | 142 | GLY | p.Gly142Ser | P04156 |
| HOMO SAPIENS | 3 | 1643 | 2m8t | A | 171 | ASN | p.Asn171Ser | P04156 |
| HOMO SAPIENS | 9 | 1643 | 2m8t | A | 178 | ASP | p.Asp178Asn | P04156 |
| HOMO SAPIENS | 9 | 1643 | 2m8t | A | 180 | VAL | p.Val180Ile | P04156 |
| HOMO SAPIENS | 9 | 1643 | 2m8t | A | 183 | THR | p.Thr183Ala | P04156 |
| HOMO SAPIENS | 9 | 1643 | 2m8t | A | 187 | HIS | p.His187Arg | P04156 |
| HOMO SAPIENS | 8 | 1643 | 2m8t | A | 198 | PHE | p.Phe198Ser | P04156 |
| HOMO SAPIENS | 9 | 1643 | 2m8t | A | 200 | GLU | p.Glu200Lys | P04156 |
| HOMO SAPIENS | 9 | 1643 | 2m8t | A | 208 | ARG | p.Arg208His | P04156 |
| HOMO SAPIENS | 3 | 1643 | 2m8t | A | 210 | VAL | p.Val210Ile | P04156 |
| HOMO SAPIENS | 6 | 1643 | 2m8t | A | 217 | GLN | p.Gln217Arg | P04156 |
| HOMO SAPIENS | 1 | 29 | 5kos | A | 151 | GLY | p.Gly217Arg | P00797 |
| HOMO SAPIENS | 1 | 29 | 5kos | A | 164 | ARG | p.Arg230Lys | P00797 |
| HOMO SAPIENS | 3 | 504 | 4zkv | B | 37 | ARG | p.Arg37Pro | P49773 |
| HOMO SAPIENS | 2 | 504 | 4zkv | B | 62 | GLN | p.Gln62Ter | P49773 |
| HOMO SAPIENS | 2 | 504 | 4zkv | B | 84 | CYS | p.Cys84Arg | P49773 |
| HOMO SAPIENS | 4 | 504 | 4zkv | B | 93 | GLY | p.Gly93Asp | P49773 |
| HOMO SAPIENS | 9 | 504 | 4zkv | B | 112 | HIS | p.His112Asn | P49773 |
| HOMO SAPIENS | 0 | 74 | 5vwo | C | 54 | ASN | p.Asn54Lys | P04181 |
| HOMO SAPIENS | 0 | 74 | 5vwo | C | 55 | TYR | p.Tyr55His | P04181 |
| HOMO SAPIENS | 2 | 74 | 5vwo | C | 90 | GLN | p.Gln90Glu | P04181 |
| HOMO SAPIENS | 6 | 74 | 5vwo | C | 91 | GLY | p.Gly91Glu | P04181 |
| HOMO SAPIENS | 2 | 74 | 5vwo | C | 93 | CYS | p.Cys93Phe | P04181 |
| HOMO SAPIENS | 1 | 74 | 5vwo | C | 154 | ARG | p.Arg154Leu | P04181 |
| HOMO SAPIENS | 0 | 74 | 5vwo | C | 180 | ARG | p.Arg180Thr | P04181 |
| HOMO SAPIENS | 0 | 74 | 5vwo | C | 199 | PRO | p.Pro199Gln | P04181 |
| HOMO SAPIENS | 4 | 74 | 5vwo | C | 209 | TYR | p.Tyr209Ter | P04181 |
| HOMO SAPIENS | 0 | 74 | 5vwo | C | 226 | ALA | p.Ala226Val | P04181 |

| | | | | | | | | |
|---|---|---|---|---|---|---|---|---|
| HOMO SAPIENS | 5 | 74 | 5vwo | C | 233 | GLN | p.Gln233Arg | P04181 |
| HOMO SAPIENS | 5 | 74 | 5vwo | C | 241 | PRO | p.Pro241Leu | P04181 |
| HOMO SAPIENS | 2 | 74 | 5vwo | C | 245 | TYR | p.Tyr245Cys | P04181 |
| HOMO SAPIENS | 6 | 74 | 5vwo | C | 250 | ARG | p.Arg250Pro | P04181 |
| HOMO SAPIENS | 6 | 74 | 5vwo | C | 250 | ARG | p.Arg250Ter | P04181 |
| HOMO SAPIENS | 8 | 74 | 5vwo | C | 267 | THR | p.Thr267Ile | P04181 |
| HOMO SAPIENS | 0 | 74 | 5vwo | C | 270 | ALA | p.Ala270Pro | P04181 |
| HOMO SAPIENS | 0 | 74 | 5vwo | C | 271 | ARG | p.Arg271Lys | P04181 |
| HOMO SAPIENS | 1 | 74 | 5vwo | C | 299 | TYR | p.Tyr299Ter | P04181 |
| HOMO SAPIENS | 0 | 74 | 5vwo | C | 318 | GLU | p.Glu318Lys | P04181 |
| HOMO SAPIENS | 1 | 74 | 5vwo | C | 319 | HIS | p.His319Tyr | P04181 |
| HOMO SAPIENS | 1 | 74 | 5vwo | C | 331 | ARG | p.Arg331Ter | P04181 |
| HOMO SAPIENS | 0 | 74 | 5vwo | C | 332 | VAL | p.Val332Met | P04181 |
| HOMO SAPIENS | 1 | 74 | 5vwo | C | 353 | GLY | p.Gly353Asp | P04181 |
| HOMO SAPIENS | 2 | 74 | 5vwo | C | 375 | GLY | p.Gly375Ala | P04181 |
| HOMO SAPIENS | 0 | 74 | 5vwo | C | 394 | CYS | p.Cys394Arg | P04181 |
| HOMO SAPIENS | 0 | 74 | 5vwo | C | 402 | LEU | p.Leu402Pro | P04181 |
| HOMO SAPIENS | 0 | 74 | 5vwo | C | 436 | ILE | p.Ile436Asn | P04181 |
| HOMO SAPIENS | 7 | 24356 | 4az0 | A | 104 | VAL | p.Val132Met | P10619 |
| HOMO SAPIENS | 4 | 24356 | 4az0 | A | 221 | TYR | p.Tyr249Asn | P10619 |
| HOMO SAPIENS | 3 | 245 | 1rsz | A | 24 | ARG | p.Arg24Ter | P00491 |
| HOMO SAPIENS | 3 | 245 | 1rsz | A | 51 | SER | p.Gly51Ser | P00491 |
| HOMO SAPIENS | 1 | 245 | 1rsz | A | 58 | ARG | p.Arg58Ter | P00491 |
| HOMO SAPIENS | 4 | 245 | 1rsz | A | 89 | GLU | p.Glu89Lys | P00491 |
| HOMO SAPIENS | 6 | 245 | 1rsz | A | 128 | ASP | p.Asp128Gly | P00491 |
| HOMO SAPIENS | 0 | 245 | 1rsz | A | 174 | ALA | p.Ala174Pro | P00491 |
| HOMO SAPIENS | 8 | 245 | 1rsz | A | 192 | TYR | p.Tyr192Cys | P00491 |
| HOMO SAPIENS | 8 | 245 | 1rsz | A | 234 | ARG | p.Arg234Pro | P00491 |
| HOMO SAPIENS | 0 | 37 | 5tby | E | 22 | GLU | p.Glu22Lys | P10916 |
| HOMO SAPIENS | 1 | 37 | 5tby | E | 86 | PHE | p.Phe86Ser | P10916 |
| HOMO SAPIENS | 9 | 71 | 5m9j | A | 12 | GLN | p.Gln36Leu | P03950 |
| HOMO SAPIENS | 3 | 71 | 5m9j | A | 17 | LYS | p.Lys41Glu | P03950 |
| HOMO SAPIENS | 3 | 71 | 5m9j | A | 17 | LYS | p.Lys41Ile | P03950 |
| HOMO SAPIENS | 4 | 71 | 5m9j | A | 28 | SER | p.Ser52Asn | P03950 |
| HOMO SAPIENS | 7 | 71 | 5m9j | A | 31 | ARG | p.Arg55Lys | P03950 |
| HOMO SAPIENS | 9 | 71 | 5m9j | A | 39 | CYS | p.Cys63Trp | P03950 |
| HOMO SAPIENS | 9 | 71 | 5m9j | A | 40 | LYS | p.Lys64Ile | P03950 |
| HOMO SAPIENS | 2 | 71 | 5m9j | A | 60 | GLU | p.Lys84Glu | P03950 |
| HOMO SAPIENS | 0 | 71 | 5m9j | A | 112 | PRO | p.Pro136Leu | P03950 |
| HOMO SAPIENS | 9 | 71 | 5m9j | A | 113 | VAL | p.Val137Ile | P03950 |
| HOMO SAPIENS | 3 | 326 | 5dqi | A | 209 | GLY | p.Gly209Val | Q9Y253 |
| HOMO SAPIENS | 1 | 326 | 5dqi | A | 242 | SER | p.Ser242Ter | Q9Y253 |
| HOMO SAPIENS | 1 | 326 | 5dqi | A | 303 | ARG | p.Arg303Ter | Q9Y253 |
| HOMO SAPIENS | 1 | 8203 | 3iu3 | K | 132 | VAL | p.Val153Ile | P01589 |
| HOMO SAPIENS | 0 | 4995 | 5ffo | H | 234 | THR | p.Thr263Ile | P01137 |
| HOMO SAPIENS | 5 | 38609 | 2ruk | A | 49 | ASP | p.Asp49His | P04637 |
| HOMO SAPIENS | 5 | 38609 | 2ruk | A | 58 | PRO | p.Pro58Arg | P04637 |
| HOMO SAPIENS | 0 | 1607 | 6pah | A | 117 | THR | p.Thr117Ile | P00439 |
| HOMO SAPIENS | 3 | 1607 | 6pah | A | 145 | ASP | p.Asp145Val | P00439 |
| HOMO SAPIENS | 5 | 1607 | 6pah | A | 158 | ARG | p.Arg158Gln | P00439 |
| HOMO SAPIENS | 5 | 1607 | 6pah | A | 158 | ARG | p.Arg158Trp | P00439 |
| HOMO SAPIENS | 5 | 1607 | 6pah | A | 169 | ARG | p.Arg169His | P00439 |
| HOMO SAPIENS | 5 | 1607 | 6pah | A | 169 | ARG | p.Arg169Cys | P00439 |
| HOMO SAPIENS | 3 | 1607 | 6pah | A | 176 | ARG | p.Arg176Leu | P00439 |

| | | | | | | | | |
|---|---|---|---|---|---|---|---|---|
| HOMO SAPIENS | 3 | 1607 | 6pah | A | 177 | VAL | p.Val177Leu | P00439 |
| HOMO SAPIENS | 1 | 1607 | 6pah | A | 194 | LEU | p.Leu194Pro | P00439 |
| HOMO SAPIENS | 1 | 1607 | 6pah | A | 204 | TYR | p.Tyr204Cys | P00439 |
| HOMO SAPIENS | 1 | 1607 | 6pah | A | 204 | TYR | p.Tyr204Ter | P00439 |
| HOMO SAPIENS | 2 | 1607 | 6pah | A | 230 | VAL | p.Val230Ile | P00439 |
| HOMO SAPIENS | 4 | 1607 | 6pah | A | 232 | GLN | p.Gln232Ter | P00439 |
| HOMO SAPIENS | 4 | 1607 | 6pah | A | 241 | ARG | p.Arg241Cys | P00439 |
| HOMO SAPIENS | 4 | 1607 | 6pah | A | 241 | ARG | p.Arg241His | P00439 |
| HOMO SAPIENS | 5 | 1607 | 6pah | A | 243 | ARG | p.Arg243Gln | P00439 |
| HOMO SAPIENS | 5 | 1607 | 6pah | A | 243 | ARG | p.Arg243Ter | P00439 |
| HOMO SAPIENS | 7 | 1607 | 6pah | A | 249 | LEU | p.Leu249Phe | P00439 |
| HOMO SAPIENS | 5 | 1607 | 6pah | A | 252 | ARG | p.Arg252Gly | P00439 |
| HOMO SAPIENS | 5 | 1607 | 6pah | A | 261 | ARG | p.Arg261Ter | P00439 |
| HOMO SAPIENS | 4 | 1607 | 6pah | A | 267 | GLN | p.Gln267Leu | P00439 |
| HOMO SAPIENS | 1 | 1607 | 6pah | A | 272 | GLY | p.Gly272Ter | P00439 |
| HOMO SAPIENS | 2 | 1607 | 6pah | A | 277 | TYR | p.Tyr277Asp | P00439 |
| HOMO SAPIENS | 7 | 1607 | 6pah | A | 280 | GLU | p.Glu280Lys | P00439 |
| HOMO SAPIENS | 8 | 1607 | 6pah | A | 281 | PRO | p.Pro281Leu | P00439 |
| HOMO SAPIENS | 0 | 1607 | 6pah | A | 299 | PHE | p.Phe299Cys | P00439 |
| HOMO SAPIENS | 5 | 1607 | 6pah | A | 300 | ALA | p.Ala300Ser | P00439 |
| HOMO SAPIENS | 5 | 1607 | 6pah | A | 309 | ALA | p.Ala309Asp | P00439 |
| HOMO SAPIENS | 4 | 1607 | 6pah | A | 320 | LYS | p.Lys320Asn | P00439 |
| HOMO SAPIENS | 0 | 1607 | 6pah | A | 345 | ALA | p.Ala345Ser | P00439 |
| HOMO SAPIENS | 0 | 1607 | 6pah | A | 348 | LEU | p.Leu348Val | P00439 |
| HOMO SAPIENS | 6 | 1607 | 6pah | A | 356 | TYR | p.Tyr356Ter | P00439 |
| HOMO SAPIENS | 2 | 1607 | 6pah | A | 380 | THR | p.Thr380Met | P00439 |
| HOMO SAPIENS | 0 | 1607 | 6pah | A | 386 | TYR | p.Tyr386Cys | P00439 |
| HOMO SAPIENS | 5 | 1607 | 6pah | A | 390 | GLU | p.Glu390Gly | P00439 |
| HOMO SAPIENS | 5 | 1607 | 6pah | A | 403 | ALA | p.Ala403Val | P00439 |
| HOMO SAPIENS | 4 | 1607 | 6pah | A | 408 | ARG | p.Arg408Trp | P00439 |
| HOMO SAPIENS | 3 | 1607 | 6pah | A | 413 | ARG | p.Arg413Pro | P00439 |
| HOMO SAPIENS | 4 | 1607 | 6pah | A | 414 | TYR | p.Tyr414Cys | P00439 |
| HOMO SAPIENS | 4 | 1607 | 6pah | A | 414 | TYR | p.Tyr414His | P00439 |
| HOMO SAPIENS | 4 | 1607 | 6pah | A | 415 | ASP | p.Asp415Asn | P00439 |
| HOMO SAPIENS | -1 | 676 | 5dc5 | A | 7 | PRO | p.Pro7Leu | Q9BY41 |
| HOMO SAPIENS | 0 | 676 | 5dc5 | A | 311 | THR | p.Thr311Met | Q9BY41 |
| HOMO SAPIENS | 1 | 124 | 5ftk | E | 95 | ARG | p.Arg95Gly | P55072 |
| HOMO SAPIENS | 1 | 124 | 5ftk | E | 155 | ARG | p.Arg155Cys | P55072 |
| HOMO SAPIENS | 1 | 124 | 5ftk | E | 155 | ARG | p.Arg155His | P55072 |
| HOMO SAPIENS | 1 | 124 | 5ftk | E | 159 | ARG | p.Arg159His | P55072 |
| HOMO SAPIENS | 1 | 124 | 5ftk | E | 159 | ARG | p.Arg159Gly | P55072 |
| HOMO SAPIENS | 0 | 124 | 5ftk | E | 191 | ARG | p.Arg191Gln | P55072 |
| HOMO SAPIENS | 0 | 124 | 5ftk | E | 232 | ALA | p.Ala232Glu | P55072 |
| HOMO SAPIENS | 5 | 5442 | 1fgg | B | 223 | GLY | p.Gly223Ser | O94766 |
| HOMO SAPIENS | 0 | 5442 | 1fgg | B | 277 | ARG | p.Arg277Gln | O94766 |
| HOMO SAPIENS | 0 | 9072 | 2gys | B | 225 | GLU | p.Glu249Gln | P32927 |
| HOMO SAPIENS | 0 | 9072 | 2gys | B | 247 | SER | p.Ser271Ter | P32927 |
| HOMO SAPIENS | 0 | 9072 | 2gys | B | 294 | ASP | p.Asp312Asn | P32927 |
| HOMO SAPIENS | 0 | 1484 | 4zrj | A | 62 | PHE | p.Phe62Ser | P35240 |
| HOMO SAPIENS | 3 | 1484 | 4zrj | A | 205 | MET | p.Met205Val | P35240 |
| HOMO SAPIENS | 3 | 1484 | 4zrj | A | 301 | ILE | p.Ile301Met | P35240 |
| HOMO SAPIENS | 4 | 40923 | 2enq | A | 68 | ILE | p.Ile391Met | P42336 |
| HOMO SAPIENS | 1 | 6179 | 5ofv | A | 125 | THR | p.Thr125Met | O43175 |
| HOMO SAPIENS | 0 | 6179 | 5ofv | A | 135 | ARG | p.Arg135Trp | O43175 |

| | | | | | | | | |
|---|---|---|---|---|---|---|---|---|
| HOMO SAPIENS | 1 | 6179 | 5ofv | A | 163 | ARG | p.Arg163Gln | O43175 |
| HOMO SAPIENS | 9 | 6179 | 5ofv | A | 261 | VAL | p.Val261Met | O43175 |
| HOMO SAPIENS | 7 | 34118 | 2i5o | A | 24 | MET | p.Met647Leu | Q9Y253 |
| HOMO SAPIENS | 8 | 3234 | 5c84 | A | 311 | LYS | p.Lys311Asn | P98170 |
| HOMO SAPIENS | 3 | 3234 | 5c84 | A | 321 | ALA | p.Ala321Gly | P98170 |
| HOMO SAPIENS | -1 | 15129 | 1zy8 | K | 1 | GLY | p.Gly54Ser | O00330 |
| HOMO SAPIENS | -1 | 15129 | 1zy8 | K | 48 | THR | p.Thr101Ala | O00330 |
| HOMO SAPIENS | -1 | 15129 | 1zy8 | K | 105 | GLU | p.Glu158Asp | O00330 |
| HOMO SAPIENS | 8 | 15129 | 1zy8 | K | 136 | ARG | p.Arg189His | O00330 |
| HOMO SAPIENS | -1 | 15129 | 1zy8 | K | 195 | GLN | p.Gln248Ter | O00330 |
| HOMO SAPIENS | 8 | 202 | 4bzb | A | 123 | HIS | p.His123Pro | Q9Y3Z3 |
| HOMO SAPIENS | 4 | 202 | 4bzb | A | 145 | ARG | p.Arg145Ter | Q9Y3Z3 |
| HOMO SAPIENS | 6 | 202 | 4bzb | A | 164 | ARG | p.Arg164Ter | Q9Y3Z3 |
| HOMO SAPIENS | 3 | 202 | 4bzb | A | 201 | ILE | p.Ile201Asn | Q9Y3Z3 |
| HOMO SAPIENS | 0 | 202 | 4bzb | A | 209 | GLY | p.Gly209Ser | Q9Y3Z3 |
| HOMO SAPIENS | 0 | 202 | 4bzb | A | 254 | MET | p.Met254Val | Q9Y3Z3 |
| HOMO SAPIENS | 5 | 202 | 4bzb | A | 531 | ARG | p.Arg531Ser | Q9Y3Z3 |
| HOMO SAPIENS | 6 | 202 | 4bzb | A | 548 | GLN | p.Gln548Ter | Q9Y3Z3 |
| HOMO SAPIENS | 3 | 30947 | 5lgd | A | 90 | PRO | p.Pro90Ser | P16671 |
| HOMO SAPIENS | 6 | 30947 | 5lgd | A | 254 | PHE | p.Phe254Leu | P16671 |
| HOMO SAPIENS | 0 | 30947 | 5lgd | A | 413 | ILE | p.Ile413Leu | P16671 |
| HOMO SAPIENS | 2 | 3906 | 5jtc | A | 735 | ARG | p.Arg735Trp | Q12797 |
| HOMO SAPIENS | 1 | 45 | 5uyx | C | 146 | GLU | p.Glu146Val | P48643 |
| HOMO SAPIENS | 1 | 45 | 5uyx | C | 147 | HIS | p.His147Arg | P48643 |
| HOMO SAPIENS | 0 | 45 | 5uyx | C | 362 | ILE | p.Ile362Met | P48643 |
| HOMO SAPIENS | 8 | 39596 | 1mhn | A | 95 | GLY | p.Gly95Arg | Q16637 |
| HOMO SAPIENS | 4 | 39596 | 1mhn | A | 111 | ALA | p.Ala111Gly | Q16637 |
| HOMO SAPIENS | 9 | 39596 | 1mhn | A | 116 | ILE | p.Ile116Phe | Q16637 |
| HOMO SAPIENS | 1 | 39596 | 1mhn | A | 136 | GLN | p.Gln136Glu | Q16637 |
| HOMO SAPIENS | 1 | 6408 | 5xjy | A | 85 | PRO | p.Pro85Leu | O95477 |
| HOMO SAPIENS | 0 | 6408 | 5xjy | A | 219 | ARG | p.Arg219Lys | O95477 |
| HOMO SAPIENS | 0 | 6408 | 5xjy | A | 230 | ARG | p.Arg230Cys | O95477 |
| HOMO SAPIENS | 1 | 6408 | 5xjy | A | 281 | MET | p.Met281Thr | O95477 |
| HOMO SAPIENS | -1 | 6408 | 5xjy | A | 343 | ALA | p.Ala343Gly | O95477 |
| HOMO SAPIENS | 1 | 6408 | 5xjy | A | 399 | VAL | p.Val399Ala | O95477 |
| HOMO SAPIENS | 1 | 6408 | 5xjy | A | 496 | ARG | p.Arg496Trp | O95477 |
| HOMO SAPIENS | 1 | 6408 | 5xjy | A | 590 | TRP | p.Trp590Ser | O95477 |
| HOMO SAPIENS | 1 | 6408 | 5xjy | A | 638 | ARG | p.Arg638Gln | O95477 |
| HOMO SAPIENS | 4 | 6408 | 5xjy | A | 697 | ALA | p.Ala697Thr | O95477 |
| HOMO SAPIENS | 1 | 6408 | 5xjy | A | 771 | VAL | p.Val771Met | O95477 |
| HOMO SAPIENS | 1 | 6408 | 5xjy | A | 774 | THR | p.Thr774Pro | O95477 |
| HOMO SAPIENS | 1 | 6408 | 5xjy | A | 776 | LYS | p.Lys776Asn | O95477 |
| HOMO SAPIENS | 0 | 6408 | 5xjy | A | 825 | VAL | p.Val825Ile | O95477 |
| HOMO SAPIENS | -1 | 6408 | 5xjy | A | 868 | GLU | p.Glu868Lys | O95477 |
| HOMO SAPIENS | -1 | 6408 | 5xjy | A | 883 | ILE | p.Ile883Met | O95477 |
| HOMO SAPIENS | 0 | 6408 | 5xjy | A | 935 | ASN | p.Asn935Ser | O95477 |
| HOMO SAPIENS | 0 | 6408 | 5xjy | A | 935 | ASN | p.Asn935His | O95477 |
| HOMO SAPIENS | 0 | 6408 | 5xjy | A | 937 | ALA | p.Ala937Val | O95477 |
| HOMO SAPIENS | 0 | 6408 | 5xjy | A | 1018 | ASP | p.Asp1018Gly | O95477 |
| HOMO SAPIENS | 0 | 6408 | 5xjy | A | 1096 | VAL | p.Val1096Ile | O95477 |
| HOMO SAPIENS | 0 | 6408 | 5xjy | A | 1099 | ASP | p.Asp1099Tyr | O95477 |
| HOMO SAPIENS | 0 | 6408 | 5xjy | A | 1172 | GLU | p.Glu1172Asp | O95477 |
| HOMO SAPIENS | 0 | 6408 | 5xjy | A | 1181 | SER | p.Ser1181Phe | O95477 |
| HOMO SAPIENS | 0 | 6408 | 5xjy | A | 1182 | ALA | p.Ala1182Thr | O95477 |

| | | | | | | | | |
|---|---|---|---|---|---|---|---|---|
| HOMO SAPIENS | -1 | 6408 | 5xjy | A | 1289 | ASP | p.Asp1289Asn | O95477 |
| HOMO SAPIENS | 0 | 6408 | 5xjy | A | 1399 | THR | p.Thr1399Ser | O95477 |
| HOMO SAPIENS | 1 | 6408 | 5xjy | A | 1408 | LEU | p.Leu1408Phe | O95477 |
| HOMO SAPIENS | 0 | 6408 | 5xjy | A | 1587 | LYS | p.Lys1587Arg | O95477 |
| HOMO SAPIENS | 1 | 6408 | 5xjy | A | 1674 | VAL | p.Val1674Ile | O95477 |
| HOMO SAPIENS | 1 | 6408 | 5xjy | A | 1680 | ARG | p.Arg1680Trp | O95477 |
| HOMO SAPIENS | 1 | 6408 | 5xjy | A | 1680 | ARG | p.Arg1680Gln | O95477 |
| HOMO SAPIENS | 0 | 6408 | 5xjy | A | 1800 | ASN | p.Asn1800His | O95477 |
| HOMO SAPIENS | -1 | 6408 | 5xjy | A | 1925 | ARG | p.Arg1925Gln | O95477 |
| HOMO SAPIENS | 0 | 6408 | 5xjy | A | 2009 | PHE | p.Phe2009Ser | O95477 |
| HOMO SAPIENS | -1 | 6408 | 5xjy | A | 2243 | ASP | p.Asp2243Glu | O95477 |
| HOMO SAPIENS | 4 | 23760 | 2mut | A | 231 | LEU | p.Phe231Leu | P07992 |
| HOMO SAPIENS | 6 | 1460 | 2hre | B | 96 | ARG | p.Arg96Gln | P22830 |
| HOMO SAPIENS | 8 | 1460 | 2hre | B | 129 | GLY | p.Gly129Arg | P22830 |
| HOMO SAPIENS | 9 | 1460 | 2hre | B | 172 | GLU | p.Glu172Lys | P22830 |
| HOMO SAPIENS | 4 | 1460 | 2hre | B | 285 | GLN | p.Gln285Arg | P22830 |
| HOMO SAPIENS | 5 | 1460 | 2hre | B | 362 | VAL | p.Val362Gly | P22830 |
| HOMO SAPIENS | 6 | 1460 | 2hre | B | 411 | CYS | p.Cys411Gly | P22830 |
| HOMO SAPIENS | 6 | 1460 | 2hre | B | 417 | PHE | p.Phe417Ser | P22830 |
| HOMO SAPIENS | 0 | 2210 | 5bug | A | 19 | ASP | p.Asp19Asn | P60484 |
| HOMO SAPIENS | 0 | 2210 | 5bug | A | 35 | MET | p.Met35Arg | P60484 |
| HOMO SAPIENS | 1 | 2210 | 5bug | A | 61 | HIS | p.His61Asp | P60484 |
| HOMO SAPIENS | 0 | 2210 | 5bug | A | 68 | TYR | p.Tyr68His | P60484 |
| HOMO SAPIENS | 8 | 2210 | 5bug | A | 70 | LEU | p.Leu70Pro | P60484 |
| HOMO SAPIENS | 6 | 2210 | 5bug | A | 93 | HIS | p.His93Arg | P60484 |
| HOMO SAPIENS | 1 | 2210 | 5bug | A | 112 | LEU | p.Leu112Pro | P60484 |
| HOMO SAPIENS | 0 | 2210 | 5bug | A | 121 | ALA | p.Ala121Gly | P60484 |
| HOMO SAPIENS | 3 | 2210 | 5bug | A | 123 | HIS | p.His123Arg | P60484 |
| HOMO SAPIENS | 7 | 2210 | 5bug | A | 124 | CYS | p.Cys124Arg | P60484 |
| HOMO SAPIENS | 9 | 2210 | 5bug | A | 129 | GLY | p.Gly129Glu | P60484 |
| HOMO SAPIENS | 9 | 2210 | 5bug | A | 130 | ARG | p.Arg130Gln | P60484 |
| HOMO SAPIENS | 9 | 2210 | 5bug | A | 130 | ARG | p.Arg130Gly | P60484 |
| HOMO SAPIENS | 0 | 2210 | 5bug | A | 132 | GLY | p.Gly132Val | P60484 |
| HOMO SAPIENS | 0 | 2210 | 5bug | A | 170 | SER | p.Ser170Arg | P60484 |
| HOMO SAPIENS | 1 | 2210 | 5bug | A | 173 | ARG | p.Arg173His | P60484 |
| HOMO SAPIENS | 0 | 2210 | 5bug | A | 217 | VAL | p.Val217Ile | P60484 |
| HOMO SAPIENS | 0 | 2210 | 5bug | A | 241 | PHE | p.Phe241Ser | P60484 |
| HOMO SAPIENS | 0 | 2210 | 5bug | A | 250 | CYS | p.Cys250Ter | P60484 |
| HOMO SAPIENS | 1 | 2210 | 5bug | A | 252 | ASP | p.Asp252Gly | P60484 |
| HOMO SAPIENS | 3 | 123 | 2ibu | A | 93 | ASN | p.Asn93Ser | P24752 |
| HOMO SAPIENS | 0 | 123 | 2ibu | A | 145 | GLN | p.Gln145Glu | P24752 |
| HOMO SAPIENS | 0 | 123 | 2ibu | A | 152 | GLY | p.Gly152Ala | P24752 |
| HOMO SAPIENS | 2 | 123 | 2ibu | A | 158 | ASN | p.Asn158Asp | P24752 |
| HOMO SAPIENS | 2 | 123 | 2ibu | A | 158 | ASN | p.Asn158Ser | P24752 |
| HOMO SAPIENS | 5 | 123 | 2ibu | A | 183 | GLY | p.Gly183Arg | P24752 |
| HOMO SAPIENS | 3 | 123 | 2ibu | A | 272 | GLN | p.Gln272Ter | P24752 |
| HOMO SAPIENS | 3 | 123 | 2ibu | A | 312 | ILE | p.Ile312Thr | P24752 |
| HOMO SAPIENS | 1 | 123 | 2ibu | A | 333 | ALA | p.Ala333Pro | P24752 |
| HOMO SAPIENS | 5 | 123 | 2ibu | A | 379 | GLY | p.Gly379Val | P24752 |
| HOMO SAPIENS | 0 | 123 | 2ibu | A | 380 | ALA | p.Ala380Thr | P24752 |
| HOMO SAPIENS | 2 | 369 | 5nrw | A | 1175 | TYR | p.Tyr1175Cys | Q92793 |
| HOMO SAPIENS | 2 | 830 | 2vc2 | A | 116 | LEU | p.Leu147Val | P08514 |
| HOMO SAPIENS | 0 | 830 | 2vc2 | A | 183 | LEU | p.Leu214Pro | P08514 |
| HOMO SAPIENS | 4 | 830 | 2vc2 | A | 242 | GLY | p.Gly273Asp | P08514 |

| | | | | | | | | |
|---|---|---|---|---|---|---|---|---|
| HOMO SAPIENS | 6 | 830 | 2vc2 | A | 324 | GLU | p.Glu355Lys | P08514 |
| HOMO SAPIENS | -1 | 10353 | 5ejc | E | 989 | ALA | p.Ala989Ser | Q92574 |
| HOMO SAPIENS | 0 | 15448 | 3lgg | A | 21 | GLY | p.Gly47Arg | Q9NZK5 |
| HOMO SAPIENS | 0 | 15448 | 3lgg | A | 21 | GLY | p.Gly47Val | Q9NZK5 |
| HOMO SAPIENS | 9 | 15448 | 3lgg | A | 86 | HIS | p.His112Gln | Q9NZK5 |
| HOMO SAPIENS | 0 | 15448 | 3lgg | A | 143 | ARG | p.Arg169Gln | Q9NZK5 |
| HOMO SAPIENS | 3 | 15448 | 3lgg | A | 221 | ALA | p.Ala247Val | Q9NZK5 |
| HOMO SAPIENS | 0 | 15448 | 3lgg | A | 225 | PRO | p.Pro251Leu | Q9NZK5 |
| HOMO SAPIENS | 0 | 15448 | 3lgg | A | 283 | MET | p.Met309Ile | Q9NZK5 |
| HOMO SAPIENS | 2 | 15448 | 3lgg | A | 334 | THR | p.Thr360Ala | Q9NZK5 |
| HOMO SAPIENS | 7 | 15448 | 3lgg | A | 357 | GLY | p.Gly383Ser | Q9NZK5 |
| HOMO SAPIENS | 0 | 15448 | 3lgg | A | 427 | TYR | p.Tyr453Cys | Q9NZK5 |
| HOMO SAPIENS | 0 | 872 | 2y3i | A | 87 | LEU | p.Leu88Pro | P00558 |
| HOMO SAPIENS | 1 | 872 | 2y3i | A | 163 | ASP | p.Asp164Val | P00558 |
| HOMO SAPIENS | 4 | 872 | 2y3i | A | 205 | ARG | p.Arg206Pro | P00558 |
| HOMO SAPIENS | 4 | 872 | 2y3i | A | 267 | ASP | p.Asp268Asn | P00558 |
| HOMO SAPIENS | 8 | 872 | 2y3i | A | 284 | ASP | p.Asp285Val | P00558 |
| HOMO SAPIENS | 4 | 8 | 5jco | B | 188 | ILE | p.Ile188Leu | Q71U36 |
| HOMO SAPIENS | 7 | 8 | 5jco | B | 263 | PRO | p.Pro263Thr | Q71U36 |
| HOMO SAPIENS | 6 | 8 | 5jco | B | 264 | ARG | p.Arg264Cys | Q71U36 |
| HOMO SAPIENS | 7 | 8 | 5jco | B | 402 | ARG | p.Arg402His | Q71U36 |
| HOMO SAPIENS | 7 | 8 | 5jco | B | 419 | SER | p.Ser419Leu | Q71U36 |
| HOMO SAPIENS | -1 | 880 | 4nwf | A | 2 | THR | p.Thr2Ile | Q06124 |
| HOMO SAPIENS | 2 | 880 | 4nwf | A | 58 | ASN | p.Asn58Ser | Q06124 |
| HOMO SAPIENS | 4 | 880 | 4nwf | A | 61 | ASP | p.Asp61Gly | Q06124 |
| HOMO SAPIENS | 2 | 880 | 4nwf | A | 62 | TYR | p.Tyr62Asp | Q06124 |
| HOMO SAPIENS | 3 | 880 | 4nwf | A | 63 | TYR | p.Tyr63Cys | Q06124 |
| HOMO SAPIENS | 2 | 880 | 4nwf | A | 69 | GLU | p.Glu69Gln | Q06124 |
| HOMO SAPIENS | 2 | 880 | 4nwf | A | 72 | ALA | p.Ala72Gly | Q06124 |
| HOMO SAPIENS | 2 | 880 | 4nwf | A | 72 | ALA | p.Ala72Ser | Q06124 |
| HOMO SAPIENS | 3 | 880 | 4nwf | A | 73 | THR | p.Thr73Ile | Q06124 |
| HOMO SAPIENS | 5 | 880 | 4nwf | A | 76 | GLU | p.Glu76Ala | Q06124 |
| HOMO SAPIENS | 5 | 880 | 4nwf | A | 76 | GLU | p.Glu76Lys | Q06124 |
| HOMO SAPIENS | 3 | 880 | 4nwf | A | 79 | GLN | p.Gln79Arg | Q06124 |
| HOMO SAPIENS | 5 | 880 | 4nwf | A | 152 | ARG | p.Arg152His | Q06124 |
| HOMO SAPIENS | 2 | 880 | 4nwf | A | 268 | GLY | p.Gly268Cys | Q06124 |
| HOMO SAPIENS | 9 | 880 | 4nwf | A | 276 | LYS | p.Lys276Arg | Q06124 |
| HOMO SAPIENS | 9 | 880 | 4nwf | A | 279 | TYR | p.Tyr279Cys | Q06124 |
| HOMO SAPIENS | 5 | 880 | 4nwf | A | 285 | PHE | p.Phe285Ser | Q06124 |
| HOMO SAPIENS | 0 | 880 | 4nwf | A | 308 | ASP | p.Asn308Ser | Q06124 |
| HOMO SAPIENS | 0 | 880 | 4nwf | A | 308 | ASP | p.Asn308Asp | Q06124 |
| HOMO SAPIENS | 5 | 880 | 4nwf | A | 392 | ALA | p.Ala392Thr | Q06124 |
| HOMO SAPIENS | 1 | 880 | 4nwf | A | 411 | THR | p.Thr415Met | Q06124 |
| HOMO SAPIENS | 0 | 880 | 4nwf | A | 461 | ALA | p.Ala465Thr | Q06124 |
| HOMO SAPIENS | 0 | 880 | 4nwf | A | 464 | GLY | p.Gly468Ala | Q06124 |
| HOMO SAPIENS | 1 | 880 | 4nwf | A | 491 | PRO | p.Pro495Ser | Q06124 |
| HOMO SAPIENS | 1 | 880 | 4nwf | A | 502 | SER | p.Ser506Thr | Q06124 |
| HOMO SAPIENS | 8 | 880 | 4nwf | A | 510 | GLN | p.Gln514Pro | Q06124 |
| HOMO SAPIENS | 4 | 1872 | 3wf1 | D | 109 | ARG | p.Arg109Trp | P16278 |
| HOMO SAPIENS | 0 | 1872 | 3wf1 | D | 521 | ARG | p.Arg521Cys | P16278 |
| HOMO SAPIENS | 5 | 910 | 5xte | A | 45 | SER | p.Ser45Phe | O14949 |
| HOMO SAPIENS | 2 | 9017 | 5f0k | A | 382 | LYS | p.Lys382Arg | Q96QK1 |
| HOMO SAPIENS | 1 | 710 | 4lsj | A | 559 | ILE | p.Ile559Asn | P04150 |
| HOMO SAPIENS | 5 | 710 | 4lsj | A | 571 | VAL | p.Val571Ala | P04150 |

| | | | | | | | | |
|---|---|---|---|---|---|---|---|---|
| HOMO SAPIENS | 0 | 710 | 4lsj | A | 641 | ASP | p.Asp641Val | P04150 |
| HOMO SAPIENS | 1 | 710 | 4lsj | A | 678 | ASP | p.Asp678Glu | P04150 |
| HOMO SAPIENS | 2 | 710 | 4lsj | A | 679 | GLY | p.Gly679Ser | P04150 |
| HOMO SAPIENS | 1 | 710 | 4lsj | A | 737 | PHE | p.Phe737Leu | P04150 |
| HOMO SAPIENS | 1 | 710 | 4lsj | A | 747 | ILE | p.Ile747Met | P04150 |
| HOMO SAPIENS | 1 | 710 | 4lsj | A | 753 | LEU | p.Leu753Phe | P04150 |
| HOMO SAPIENS | 1 | 710 | 4lsj | A | 773 | LEU | p.Leu773Pro | P04150 |
| HOMO SAPIENS | 1 | 2145 | 1av1 | B | 71 | PHE | p.Phe95Tyr | P02647 |
| HOMO SAPIENS | 1 | 2145 | 1av1 | B | 128 | GLU | p.Glu152Lys | P02647 |
| HOMO SAPIENS | 1 | 2145 | 1av1 | B | 136 | GLU | p.Glu160Lys | P02647 |
| HOMO SAPIENS | 2 | 2145 | 1av1 | B | 143 | PRO | p.Pro167Arg | P02647 |
| HOMO SAPIENS | 6 | 2145 | 1av1 | B | 164 | ALA | p.Ala188Ser | P02647 |
| HOMO SAPIENS | 7 | 2145 | 1av1 | B | 173 | ARG | p.Arg197Cys | P02647 |
| HOMO SAPIENS | 2 | 2145 | 1av1 | B | 198 | GLU | p.Glu222Lys | P02647 |
| HOMO SAPIENS | -1 | 3432 | 2beu | A | 1 | SER | p.Ser46Ter | P12694 |
| HOMO SAPIENS | 5 | 3432 | 2beu | A | 79 | ARG | p.Arg124Trp | P12694 |
| HOMO SAPIENS | 7 | 3432 | 2beu | A | 114 | ARG | p.Arg159Gln | P12694 |
| HOMO SAPIENS | 6 | 3432 | 2beu | A | 166 | THR | p.Thr211Met | P12694 |
| HOMO SAPIENS | 4 | 3432 | 2beu | A | 175 | ALA | p.Ala220Val | P12694 |
| HOMO SAPIENS | 5 | 3432 | 2beu | A | 204 | GLY | p.Gly249Ser | P12694 |
| HOMO SAPIENS | 4 | 3432 | 2beu | A | 209 | ALA | p.Ala254Asp | P12694 |
| HOMO SAPIENS | 6 | 3432 | 2beu | A | 220 | ARG | p.Arg265Trp | P12694 |
| HOMO SAPIENS | 6 | 3432 | 2beu | A | 234 | TYR | p.Tyr279Cys | P12694 |
| HOMO SAPIENS | 5 | 3432 | 2beu | A | 240 | ALA | p.Ala285Pro | P12694 |
| HOMO SAPIENS | 4 | 3432 | 2beu | A | 245 | GLY | p.Gly290Arg | P12694 |
| HOMO SAPIENS | 5 | 3432 | 2beu | A | 269 | ARG | p.Arg314Ter | P12694 |
| HOMO SAPIENS | 3 | 3432 | 2beu | A | 301 | ARG | p.Arg346Cys | P12694 |
| HOMO SAPIENS | 5 | 3432 | 2beu | A | 359 | ASN | p.Asn404Ser | P12694 |
| HOMO SAPIENS | 4 | 3432 | 2beu | A | 393 | TYR | p.Tyr438Asn | P12694 |
| HOMO SAPIENS | 8 | 4541 | 1rr8 | C | 533 | ASP | p.Asp533Gly | P11387 |
| HOMO SAPIENS | 2 | 3247 | 1tzn | d | 1105 | GLY | p.Gly105Asp | P58335 |
| HOMO SAPIENS | 3 | 3247 | 1tzn | d | 1111 | ARG | p.Arg111Cys | P58335 |
| HOMO SAPIENS | 2 | 3247 | 1tzn | d | 1189 | ILE | p.Ile189Thr | P58335 |
| HOMO SAPIENS | 9 | 467 | 1suv | C | 58 | ASP | p.Asp77Asn | P02787 |
| HOMO SAPIENS | 2 | 467 | 1suv | C | 277 | ASP | p.Asp296Gly | P02787 |
| HOMO SAPIENS | 9 | 6485 | 1xwd | B | 51 | CYS | p.Cys69Gly | P01225 |
| HOMO SAPIENS | 1 | 694 | 1wuu | A | 28 | PRO | p.Pro28Thr | P51570 |
| HOMO SAPIENS | 0 | 694 | 1wuu | A | 32 | VAL | p.Val32Met | P51570 |
| HOMO SAPIENS | 0 | 694 | 1wuu | A | 198 | ALA | p.Ala198Val | P51570 |
| HOMO SAPIENS | 0 | 143 | 4nkx | A | 93 | PHE | p.Phe93Cys | P05093 |
| HOMO SAPIENS | 3 | 143 | 4nkx | A | 96 | ARG | p.Arg96Trp | P05093 |
| HOMO SAPIENS | 3 | 143 | 4nkx | A | 96 | ARG | p.Arg96Gln | P05093 |
| HOMO SAPIENS | 0 | 143 | 4nkx | A | 106 | SER | p.Ser106Pro | P05093 |
| HOMO SAPIENS | 1 | 143 | 4nkx | A | 114 | PHE | p.Phe114Val | P05093 |
| HOMO SAPIENS | 1 | 143 | 4nkx | A | 116 | ASP | p.Asp116Val | P05093 |
| HOMO SAPIENS | 3 | 143 | 4nkx | A | 125 | ARG | p.Arg125Gln | P05093 |
| HOMO SAPIENS | 0 | 143 | 4nkx | A | 201 | TYR | p.Tyr201Asp | P05093 |
| HOMO SAPIENS | 0 | 143 | 4nkx | A | 239 | ARG | p.Arg239Ter | P05093 |
| HOMO SAPIENS | 0 | 143 | 4nkx | A | 329 | TYR | p.Tyr329Asp | P05093 |
| HOMO SAPIENS | 1 | 143 | 4nkx | A | 342 | PRO | p.Pro342Thr | P05093 |
| HOMO SAPIENS | 2 | 143 | 4nkx | A | 347 | ARG | p.Arg347His | P05093 |
| HOMO SAPIENS | 2 | 143 | 4nkx | A | 347 | ARG | p.Arg347Cys | P05093 |
| HOMO SAPIENS | 1 | 143 | 4nkx | A | 358 | ARG | p.Arg358Gln | P05093 |
| HOMO SAPIENS | 0 | 143 | 4nkx | A | 362 | ARG | p.Arg362Cys | P05093 |

| | | | | | | | | |
|---|---|---|---|---|---|---|---|---|
| HOMO SAPIENS | 1 | 143 | 4nkx | A | 406 | TRP | p.Trp406Arg | P05093 |
| HOMO SAPIENS | 0 | 143 | 4nkx | A | 428 | PRO | p.Pro428Leu | P05093 |
| HOMO SAPIENS | 6 | 1441 | 1uuh | A | 46 | ARG | p.Arg46Gly | P16070 |
| HOMO SAPIENS | 3 | 82 | 4wlv | D | 37 | GLY | p.Gly37Arg | P40926 |
| HOMO SAPIENS | 0 | 82 | 4wlv | D | 133 | PRO | p.Pro133Leu | P40926 |
| HOMO SAPIENS | 2 | 392 | 5xtd | A | 56 | SER | p.Ser56Pro | P49821 |
| HOMO SAPIENS | 2 | 392 | 5xtd | A | 59 | ARG | p.Arg59Ter | P49821 |
| HOMO SAPIENS | 2 | 392 | 5xtd | A | 74 | ASP | p.Asp74Asn | P49821 |
| HOMO SAPIENS | 4 | 392 | 5xtd | A | 116 | ASN | p.Asn116Lys | P49821 |
| HOMO SAPIENS | 4 | 392 | 5xtd | A | 117 | ALA | p.Ala117Thr | P49821 |
| HOMO SAPIENS | 5 | 392 | 5xtd | A | 122 | PRO | p.Pro122Leu | P49821 |
| HOMO SAPIENS | 3 | 392 | 5xtd | A | 160 | GLY | p.Gly160Glu | P49821 |
| HOMO SAPIENS | 4 | 392 | 5xtd | A | 214 | GLU | p.Glu214Lys | P49821 |
| HOMO SAPIENS | 2 | 392 | 5xtd | A | 357 | MET | p.Met357Thr | P49821 |
| HOMO SAPIENS | 4 | 392 | 5xtd | A | 423 | THR | p.Thr423Met | P49821 |
| HOMO SAPIENS | 0 | 22593 | 3pdf | A | 8 | TYR | p.Tyr32Ter | P53634 |
| HOMO SAPIENS | 0 | 22593 | 3pdf | A | 15 | TRP | p.Trp39Ser | P53634 |
| HOMO SAPIENS | 0 | 22593 | 3pdf | A | 103 | HIS | p.His127Pro | P53634 |
| HOMO SAPIENS | 9 | 22593 | 3pdf | A | 228 | GLN | p.Gln252Leu | P53634 |
| HOMO SAPIENS | 3 | 22593 | 3pdf | A | 262 | GLN | p.Gln286Arg | P53634 |
| HOMO SAPIENS | 4 | 22593 | 3pdf | A | 277 | GLY | p.Gly301Ser | P53634 |
| HOMO SAPIENS | 0 | 22593 | 3pdf | A | 323 | TYR | p.Tyr347Cys | P53634 |
| HOMO SAPIENS | 5 | 3461 | 4c2a | A | 1306 | GLN | p.Arg1306Trp | P04275 |
| HOMO SAPIENS | 5 | 3461 | 4c2a | A | 1374 | ARG | p.Arg1374Cys | P04275 |
| HOMO SAPIENS | 5 | 3461 | 4c2a | A | 1379 | ARG | p.Arg1379Cys | P04275 |
| HOMO SAPIENS | 4 | 3461 | 4c2a | A | 1380 | ILE | p.Ile1380Val | P04275 |
| HOMO SAPIENS | 0 | 3461 | 4c2a | A | 1381 | ALA | p.Thr1381Ala | P04275 |
| HOMO SAPIENS | 5 | 3461 | 4c2a | A | 1399 | ARG | p.Arg1399His | P04275 |
| HOMO SAPIENS | 5 | 3461 | 4c2a | A | 1435 | ASN | p.Asn1435Ser | P04275 |
| HOMO SAPIENS | 0 | 4533 | 5uj7 | C | 174 | TYR | p.Tyr174Cys | O43929 |
| HOMO SAPIENS | 0 | 4533 | 5uj7 | C | 202 | LEU | p.Leu202Val | O43929 |
| HOMO SAPIENS | 0 | 1374 | 3j07 | O | 39 | PRO | p.Pro39Leu | P02511 |
| HOMO SAPIENS | 0 | 1374 | 3j07 | O | 107 | ARG | p.Arg107His | P02511 |
| HOMO SAPIENS | 0 | 1374 | 3j07 | O | 157 | ARG | p.Arg157His | P02511 |
| HOMO SAPIENS | -1 | 942 | 1s9i | B | 57 | PHE | p.Phe57Cys | P36507 |
| HOMO SAPIENS | -1 | 942 | 1s9i | B | 57 | PHE | p.Phe57Val | P36507 |
| HOMO SAPIENS | 2 | 942 | 1s9i | B | 128 | PRO | p.Pro128Gln | P36507 |
| HOMO SAPIENS | 1 | 942 | 1s9i | B | 134 | TYR | p.Tyr134His | P36507 |
| HOMO SAPIENS | 1 | 942 | 1s9i | B | 188 | HIS | p.His188Gln | P36507 |
| HOMO SAPIENS | 0 | 942 | 1s9i | B | 262 | VAL | p.Val262Phe | P36507 |
| HOMO SAPIENS | 0 | 942 | 1s9i | B | 263 | GLY | p.Gly263Arg | P36507 |
| HOMO SAPIENS | -1 | 942 | 1s9i | B | 298 | PRO | p.Pro298Leu | P36507 |
| HOMO SAPIENS | -1 | 942 | 1s9i | B | 305 | VAL | p.Val305Ile | P36507 |
| HOMO SAPIENS | 0 | 3245 | 1fbv | A | 367 | GLN | p.Gln367Pro | P22681 |
| HOMO SAPIENS | 0 | 3245 | 1fbv | A | 371 | TYR | p.Tyr371His | P22681 |
| HOMO SAPIENS | 1 | 3245 | 1fbv | A | 382 | LYS | p.Lys382Glu | P22681 |
| HOMO SAPIENS | 2 | 3245 | 1fbv | A | 390 | ASP | p.Asp390Tyr | P22681 |
| HOMO SAPIENS | 9 | 3245 | 1fbv | A | 396 | CYS | p.Cys396Arg | P22681 |
| HOMO SAPIENS | 7 | 3245 | 1fbv | A | 420 | ARG | p.Arg420Gln | P22681 |
| HOMO SAPIENS | 1 | 3245 | 1fbv | A | 429 | ILE | p.Ile429Met | P22681 |
| HOMO SAPIENS | 0 | 2604 | 2pbd | P | 111 | LEU | p.Leu112Met | P07737 |
| HOMO SAPIENS | 3 | 2604 | 2pbd | P | 116 | GLU | p.Glu117Gly | P07737 |
| HOMO SAPIENS | 4 | 2013 | 3b7w | A | 472 | ARG | p.Arg472Gly | Q08AH3 |
| HOMO SAPIENS | 0 | 24346 | 3sh5 | A | 85 | ARG | p.Arg4281His | P98160 |

| | | | | | | | | |
|---|---|---|---|---|---|---|---|---|
| HOMO SAPIENS | 0 | 24346 | 3sh5 | A | 122 | ARG | p.Arg4318Trp | P98160 |
| HOMO SAPIENS | 0 | 24346 | 3sh5 | A | 135 | SER | p.Ser4331Asn | P98160 |
| HOMO SAPIENS | 0 | 24346 | 3sh5 | A | 136 | VAL | p.Val4332Ile | P98160 |
| HOMO SAPIENS | 0 | 707 | 4p2g | A | 26 | MET | p.Met26Ile | Q99497 |
| HOMO SAPIENS | 5 | 707 | 4p2g | A | 39 | ALA | p.Ala39Ser | Q99497 |
| HOMO SAPIENS | 5 | 707 | 4p2g | A | 64 | GLU | p.Glu64Asp | Q99497 |
| HOMO SAPIENS | 4 | 707 | 4p2g | A | 149 | ASP | p.Asp149Ala | Q99497 |
| HOMO SAPIENS | 6 | 707 | 4p2g | A | 163 | GLU | p.Glu163Lys | Q99497 |
| HOMO SAPIENS | 6 | 707 | 4p2g | A | 166 | LEU | p.Leu166Pro | Q99497 |
| HOMO SAPIENS | 3 | 5121 | 4xss | E | 31 | GLY | p.Gly58Arg | P06213 |
| HOMO SAPIENS | 2 | 5121 | 4xss | E | 119 | ILE | p.Ile146Met | P06213 |
| HOMO SAPIENS | 8 | 5121 | 4xss | E | 209 | HIS | p.His236Arg | P06213 |
| HOMO SAPIENS | 2 | 5121 | 4xss | E | 234 | ASP | p.Asp261Glu | P06213 |
| HOMO SAPIENS | 1 | 5121 | 4xss | E | 276 | GLN | p.Gln303His | P06213 |
| HOMO SAPIENS | 1 | 27790 | 5erp | B | 140 | THR | p.Thr275Met | Q02487 |
| HOMO SAPIENS | 0 | 27790 | 5erp | B | 150 | ILE | p.Ile285Thr | Q02487 |
| HOMO SAPIENS | 1 | 27790 | 5erp | B | 205 | THR | p.Thr340Ala | Q02487 |
| HOMO SAPIENS | 1 | 27790 | 5erp | B | 223 | THR | p.Thr358Ile | Q02487 |
| HOMO SAPIENS | 0 | 27790 | 5erp | B | 301 | GLY | p.Gly436Ala | Q02487 |
| HOMO SAPIENS | -1 | 27790 | 5erp | B | 315 | ARG | p.Arg450Ser | Q02487 |
| HOMO SAPIENS | 0 | 27790 | 5erp | B | 369 | SER | p.Ser504Gly | Q02487 |
| HOMO SAPIENS | 0 | 27790 | 5erp | B | 385 | ILE | p.Ile520Thr | Q02487 |
| HOMO SAPIENS | 0 | 27790 | 5erp | B | 461 | ALA | p.Ala596Val | Q02487 |
| HOMO SAPIENS | 0 | 27790 | 5erp | B | 503 | GLN | p.Gln638His | Q02487 |
| HOMO SAPIENS | 5 | 6599 | 4nsc | A | 129 | ARG | p.Arg129Leu | Q9BPX6 |
| HOMO SAPIENS | -1 | 6599 | 4nsc | A | 183 | GLN | p.Gln183Ter | Q9BPX6 |
| HOMO SAPIENS | 0 | 20420 | 3ghm | A | 398 | ARG | p.Arg398His | Q76LX8 |
| HOMO SAPIENS | 0 | 20420 | 3ghm | A | 448 | GLN | p.Gln448Glu | Q76LX8 |
| HOMO SAPIENS | 1 | 20420 | 3ghm | A | 449 | GLN | p.Gln449Ter | Q76LX8 |
| HOMO SAPIENS | 0 | 20420 | 3ghm | A | 456 | GLN | p.Gln456His | Q76LX8 |
| HOMO SAPIENS | 1 | 20420 | 3ghm | A | 528 | ARG | p.Arg528Gly | Q76LX8 |
| HOMO SAPIENS | 0 | 20420 | 3ghm | A | 618 | PRO | p.Pro618Ala | Q76LX8 |
| HOMO SAPIENS | 0 | 20420 | 3ghm | A | 625 | ARG | p.Arg625His | Q76LX8 |
| HOMO SAPIENS | 0 | 275 | 4rmz | A | 293 | GLN | p.Gln293Ter | Q9NWZ3 |
| HOMO SAPIENS | 0 | 275 | 4rmz | A | 428 | ALA | p.Ala428Thr | Q9NWZ3 |
| HOMO SAPIENS | 2 | 10458 | 3puf | B | 138 | LEU | p.Leu138Phe | Q5TBB1 |
| HOMO SAPIENS | 6 | 10458 | 3puf | B | 177 | ALA | p.Ala177Thr | Q5TBB1 |
| HOMO SAPIENS | -1 | 10458 | 3puf | B | 185 | VAL | p.Val185Gly | Q5TBB1 |
| HOMO SAPIENS | 0 | 26648 | 2n90 | A | 36 | ARG | p.Arg444Gln | P04629 |
| HOMO SAPIENS | 2 | 977 | 1dnw | D | 166 | ALA | p.Ala332Val | P05164 |
| HOMO SAPIENS | 4 | 977 | 1dnw | D | 333 | ARG | p.Arg499Cys | P05164 |
| HOMO SAPIENS | 1 | 977 | 1dnw | D | 403 | ARG | p.Arg569Trp | P05164 |
| HOMO SAPIENS | 0 | 25704 | 3ajb | A | 82 | GLN | p.Gln82Arg | P56589 |
| HOMO SAPIENS | 3 | 255 | 3t1p | A | 38 | TYR | p.Tyr62Ter | P01009 |
| HOMO SAPIENS | 3 | 255 | 3t1p | A | 39 | ARG | p.Arg63Cys | P01009 |
| HOMO SAPIENS | 0 | 255 | 3t1p | A | 101 | ARG | p.Arg125His | P01009 |
| HOMO SAPIENS | 0 | 255 | 3t1p | A | 212 | GLN | p.Gln236Arg | P01009 |
| HOMO SAPIENS | 0 | 255 | 3t1p | A | 213 | VAL | p.Val237Ala | P01009 |
| HOMO SAPIENS | 1 | 255 | 3t1p | A | 269 | HIS | p.His293Gln | P01009 |
| HOMO SAPIENS | 8 | 255 | 3t1p | A | 341 | ASP | p.Asp365Asn | P01009 |
| HOMO SAPIENS | 9 | 255 | 3t1p | A | 342 | GLU | p.Glu366Lys | P01009 |
| HOMO SAPIENS | 0 | 255 | 3t1p | A | 369 | PRO | p.Pro393Ser | P01009 |
| HOMO SAPIENS | 0 | 255 | 3t1p | A | 376 | GLU | p.Glu400Asp | P01009 |
| HOMO SAPIENS | -1 | 5175 | 3zxi | A | 35 | ALA | p.Ala35Asp | Q9Y2Z4 |

| | | | | | | | | | |
|---|---|---|---|---|---|---|---|---|---|
| HOMO SAPIENS | 0 | 5175 | 3zxi | A | | 52 | PHE | p.Phe52Leu | Q9Y2Z4 |
| HOMO SAPIENS | -1 | 5175 | 3zxi | A | | 68 | GLY | p.Gly68Ser | Q9Y2Z4 |
| HOMO SAPIENS | 0 | 5175 | 3zxi | A | | 179 | LYS | p.Lys179Gln | Q9Y2Z4 |
| HOMO SAPIENS | 0 | 5175 | 3zxi | A | | 191 | GLY | p.Gly191Val | Q9Y2Z4 |
| HOMO SAPIENS | 0 | 5175 | 3zxi | A | | 251 | ILE | p.Ile251Val | Q9Y2Z4 |
| HOMO SAPIENS | 2 | 5175 | 3zxi | A | | 295 | ASP | p.Asp295Asn | Q9Y2Z4 |
| HOMO SAPIENS | 0 | 5175 | 3zxi | A | | 311 | ASP | p.Asp311Glu | Q9Y2Z4 |
| HOMO SAPIENS | 0 | 5175 | 3zxi | A | | 360 | ARG | p.Arg360Ter | Q9Y2Z4 |
| HOMO SAPIENS | 1 | 3016 | 3exg | | 3 | 4 | ASP | p.Asp33His | P08559 |
| HOMO SAPIENS | 1 | 3016 | 3exg | | 3 | 176 | PHE | p.Phe205Leu | P08559 |
| HOMO SAPIENS | 1 | 3016 | 3exg | | 3 | 214 | TYR | p.Tyr243Asn | P08559 |
| HOMO SAPIENS | 1 | 3016 | 3exg | | 3 | 229 | ASP | p.Asp258Ala | P08559 |
| HOMO SAPIENS | 0 | 3016 | 3exg | | 3 | 233 | VAL | p.Val262Ile | P08559 |
| HOMO SAPIENS | 1 | 3016 | 3exg | | 3 | 234 | ARG | p.Arg263Gly | P08559 |
| HOMO SAPIENS | 0 | 3016 | 3exg | | 3 | 253 | MET | p.Met282Leu | P08559 |
| HOMO SAPIENS | 7 | 3016 | 3exg | | 3 | 259 | ARG | p.Arg288His | P08559 |
| HOMO SAPIENS | -1 | 3016 | 3exg | | 3 | 273 | ARG | p.Arg302Cys | P08559 |
| HOMO SAPIENS | 2 | 3016 | 3exg | | 3 | 286 | ASP | p.Asp315Asn | P08559 |
| HOMO SAPIENS | 1 | 3016 | 3exg | | 3 | 349 | ARG | p.Arg378His | P08559 |
| HOMO SAPIENS | 5 | 6729 | 4a0l | E | | 572 | ARG | p.Arg572Cys | Q13620 |
| HOMO SAPIENS | 0 | 1067 | 4kyo | A | | 9 | THR | p.Thr9Asn | P21549 |
| HOMO SAPIENS | 0 | 1067 | 4kyo | A | | 11 | PRO | p.Pro11Leu | P21549 |
| HOMO SAPIENS | 0 | 1067 | 4kyo | A | | 22 | ASN | p.Asn22Ser | P21549 |
| HOMO SAPIENS | 4 | 1067 | 4kyo | A | | 25 | LEU | p.Leu25Arg | P21549 |
| HOMO SAPIENS | 7 | 1067 | 4kyo | A | | 36 | ARG | p.Arg36His | P21549 |
| HOMO SAPIENS | 3 | 1067 | 4kyo | A | | 41 | GLY | p.Gly41Arg | P21549 |
| HOMO SAPIENS | 0 | 1067 | 4kyo | A | | 42 | GLY | p.Gly42Glu | P21549 |
| HOMO SAPIENS | 3 | 1067 | 4kyo | A | | 56 | ILE | p.Ile56Asn | P21549 |
| HOMO SAPIENS | 3 | 1067 | 4kyo | A | | 59 | GLU | p.Glu59Lys | P21549 |
| HOMO SAPIENS | 3 | 1067 | 4kyo | A | | 66 | TYR | p.Tyr66Ter | P21549 |
| HOMO SAPIENS | 3 | 1067 | 4kyo | A | | 111 | ARG | p.Arg111Leu | P21549 |
| HOMO SAPIENS | 7 | 1067 | 4kyo | A | | 118 | ARG | p.Arg118His | P21549 |
| HOMO SAPIENS | 3 | 1067 | 4kyo | A | | 122 | ARG | p.Arg122Ter | P21549 |
| HOMO SAPIENS | 3 | 1067 | 4kyo | A | | 124 | HIS | p.His124Pro | P21549 |
| HOMO SAPIENS | 0 | 1067 | 4kyo | A | | 152 | PHE | p.Phe152Ile | P21549 |
| HOMO SAPIENS | 1 | 1067 | 4kyo | A | | 156 | GLY | p.Gly156Arg | P21549 |
| HOMO SAPIENS | 0 | 1067 | 4kyo | A | | 161 | GLY | p.Gly161Arg | P21549 |
| HOMO SAPIENS | 3 | 1067 | 4kyo | A | | 166 | LEU | p.Leu166Pro | P21549 |
| HOMO SAPIENS | 5 | 1067 | 4kyo | A | | 170 | GLY | p.Gly170Arg | P21549 |
| HOMO SAPIENS | 5 | 1067 | 4kyo | A | | 186 | ALA | p.Ala186Val | P21549 |
| HOMO SAPIENS | 3 | 1067 | 4kyo | A | | 187 | SER | p.Ser187Phe | P21549 |
| HOMO SAPIENS | 3 | 1067 | 4kyo | A | | 197 | ARG | p.Arg197Pro | P21549 |
| HOMO SAPIENS | 5 | 1067 | 4kyo | A | | 201 | ASP | p.Asp201Glu | P21549 |
| HOMO SAPIENS | 0 | 1067 | 4kyo | A | | 202 | ILE | p.Ile202Asn | P21549 |
| HOMO SAPIENS | 1 | 1067 | 4kyo | A | | 205 | SER | p.Ser205Pro | P21549 |
| HOMO SAPIENS | 1 | 1067 | 4kyo | A | | 205 | SER | p.Ser205Ter | P21549 |
| HOMO SAPIENS | 2 | 1067 | 4kyo | A | | 216 | GLY | p.Gly216Arg | P21549 |
| HOMO SAPIENS | 5 | 1067 | 4kyo | A | | 233 | ARG | p.Arg233Cys | P21549 |
| HOMO SAPIENS | 5 | 1067 | 4kyo | A | | 244 | ILE | p.Ile244Thr | P21549 |
| HOMO SAPIENS | 5 | 1067 | 4kyo | A | | 244 | ILE | p.Ile244Met | P21549 |
| HOMO SAPIENS | 4 | 1067 | 4kyo | A | | 253 | CYS | p.Cys253Arg | P21549 |
| HOMO SAPIENS | 7 | 1067 | 4kyo | A | | 274 | GLU | p.Glu274Asp | P21549 |
| HOMO SAPIENS | 7 | 1067 | 4kyo | A | | 279 | ILE | p.Ile279Thr | P21549 |
| HOMO SAPIENS | 6 | 1067 | 4kyo | A | | 295 | ALA | p.Ala295Thr | P21549 |

| | | | | | | | | |
|---|---|---|---|---|---|---|---|---|
| HOMO SAPIENS | 5 | 1067 | 4kyo | A | 340 | ILE | p.Ile340Met | P21549 |
| HOMO SAPIENS | 1 | 790 | 2ric | C | 270 | SER | p.Ser290Thr | P35247 |
| HOMO SAPIENS | 0 | 790 | 2ric | C | 289 | GLU | p.Glu309Lys | P35247 |
| HOMO SAPIENS | 1 | 212 | 5sxe | A | 545 | GLU | p.Glu545Lys | P42336 |
| HOMO SAPIENS | 3 | 212 | 5sxe | A | 546 | GLN | p.Gln546Glu | P42336 |
| HOMO SAPIENS | 2 | 212 | 5sxe | A | 1047 | HIS | p.His1047Leu | P42336 |
| HOMO SAPIENS | 7 | 152 | 2zqq | F | 197 | ARG | p.Arg197Ter | Q13825 |
| HOMO SAPIENS | 0 | 152 | 2zqq | F | 248 | ALA | p.Ala248Thr | Q13825 |
| HOMO SAPIENS | 0 | 152 | 2zqq | F | 275 | ALA | p.Ala275Val | Q13825 |
| HOMO SAPIENS | 5 | 152 | 2zqq | F | 331 | LYS | p.Lys331Ter | Q13825 |
| HOMO SAPIENS | 6 | 17801 | 2j8l | A | 283 | PHE | p.Phe301Leu | P03951 |
| HOMO SAPIENS | 3 | 17801 | 2j8l | A | 308 | ARG | p.Arg326Cys | P03951 |
| HOMO SAPIENS | 0 | 17801 | 2j8l | A | 321 | CYS | p.Cys339Phe | P03951 |
| HOMO SAPIENS | 3 | 17801 | 2j8l | A | 351 | TYR | p.Tyr369Ter | P03951 |
| HOMO SAPIENS | 7 | 1802 | 5db0 | A | 22 | LEU | p.Leu22Arg | O00255 |
| HOMO SAPIENS | 7 | 1802 | 5db0 | A | 26 | GLU | p.Glu26Lys | O00255 |
| HOMO SAPIENS | 7 | 1802 | 5db0 | A | 139 | HIS | p.His139Asp | O00255 |
| HOMO SAPIENS | 0 | 1802 | 5db0 | A | 171 | ARG | p.Arg176Gln | O00255 |
| HOMO SAPIENS | 0 | 1802 | 5db0 | A | 184 | VAL | p.Val189Glu | O00255 |
| HOMO SAPIENS | 7 | 1802 | 5db0 | A | 255 | GLU | p.Glu260Lys | O00255 |
| HOMO SAPIENS | 4 | 1802 | 5db0 | A | 418 | ASP | p.Asp423Asn | O00255 |
| HOMO SAPIENS | 0 | 1802 | 5db0 | A | 436 | TRP | p.Trp441Arg | O00255 |
| HOMO SAPIENS | -1 | 1802 | 5db0 | A | 541 | ALA | p.Thr546Ala | O00255 |
| HOMO SAPIENS | 4 | 1802 | 5db0 | A | 552 | THR | p.Thr557Ser | O00255 |
| HOMO SAPIENS | 0 | 3450 | 2yuz | A | 36 | PRO | p.Pro461Gln | Q00872 |
| HOMO SAPIENS | 1 | 3450 | 2yuz | A | 56 | GLN | p.His481Gln | Q00872 |
| HOMO SAPIENS | 2 | 3450 | 2yuz | A | 68 | ALA | p.Ala493Val | Q00872 |
| HOMO SAPIENS | 0 | 3211 | 3rqg | A | 72 | VAL | p.Val72Leu | Q9BUL8 |
| HOMO SAPIENS | 2 | 4168 | 4q4f | A | 121 | ARG | p.Arg121Ter | Q14108 |
| HOMO SAPIENS | 2 | 4168 | 4q4f | A | 121 | ARG | p.Arg121Gln | Q14108 |
| HOMO SAPIENS | 2 | 4168 | 4q4f | A | 159 | MET | p.Met159Val | Q14108 |
| HOMO SAPIENS | 4 | 4168 | 4q4f | A | 363 | HIS | p.His363Asn | Q14108 |
| HOMO SAPIENS | 2 | 4168 | 4q4f | A | 396 | VAL | p.Val396Ile | Q14108 |
| HOMO SAPIENS | 0 | 1448 | 1y97 | A | 87 | ARG | p.Arg130Ter | Q9BQ50 |
| HOMO SAPIENS | 0 | 3444 | 4iwp | A | 151 | SER | p.Ser151Phe | Q9UHD2 |
| HOMO SAPIENS | 0 | 3444 | 4iwp | A | 401 | LYS | p.Lys401Glu | Q9UHD2 |
| HOMO SAPIENS | 0 | 116 | 1zy8 | B | 194 | GLY | p.Gly229Cys | P09622 |
| HOMO SAPIENS | 0 | 116 | 1zy8 | B | 255 | ASP | p.Asp290Gly | P09622 |
| HOMO SAPIENS | 2 | 116 | 1zy8 | B | 326 | MET | p.Met361Val | P09622 |
| HOMO SAPIENS | 0 | 116 | 1zy8 | B | 340 | GLU | p.Glu375Lys | P09622 |
| HOMO SAPIENS | 2 | 116 | 1zy8 | B | 358 | ILE | p.Ile393Thr | P09622 |
| HOMO SAPIENS | 1 | 116 | 1zy8 | B | 453 | PRO | p.Pro488Leu | P09622 |
| HOMO SAPIENS | 7 | 127 | 4qo7 | B | 160 | SER | p.Ser161Arg | P00338 |
| HOMO SAPIENS | 0 | 2025 | 1wrl | A | 8 | ALA | p.Ala8Val | P63316 |
| HOMO SAPIENS | 8 | 2025 | 1wrl | A | 24 | PHE | p.Phe24Leu | P63316 |
| HOMO SAPIENS | 0 | 2025 | 1wrl | A | 84 | SER | p.Cys84Tyr | P63316 |
| HOMO SAPIENS | 8 | 3791 | 1mhd | B | 93 | ARG | p.Arg93Ter | P84022 |
| HOMO SAPIENS | 0 | 1575 | 4phc | A | 205 | GLY | p.Gly205Asp | P12081 |
| HOMO SAPIENS | 0 | 1575 | 4phc | A | 454 | TYR | p.Tyr454Ser | P12081 |
| HOMO SAPIENS | 0 | 39620 | 1ozs | B | 135 | ARG | p.Arg136Gln | P19429 |
| HOMO SAPIENS | 5 | 39620 | 1ozs | B | 140 | ARG | p.Arg141Trp | P19429 |
| HOMO SAPIENS | 5 | 39620 | 1ozs | B | 142 | THR | p.Thr143Asn | P19429 |
| HOMO SAPIENS | 5 | 39620 | 1ozs | B | 143 | LEU | p.Leu144Gln | P19429 |
| HOMO SAPIENS | 5 | 39620 | 1ozs | B | 144 | ARG | p.Arg145Gly | P19429 |

| | | | | | | | | |
|---|---|---|---|---|---|---|---|---|
| HOMO SAPIENS | 5 | 39620 | 1ozs | B | 144 | ARG | p.Arg145Gln | P19429 |
| HOMO SAPIENS | -1 | 20934 | 4nuv | D | 42 | GLY | p.Gly42Asp | Q16570 |
| HOMO SAPIENS | -1 | 40565 | 5lcw | S | 4 | VAL | p.Val4Met | O60566 |
| HOMO SAPIENS | 0 | 40565 | 5lcw | S | 40 | THR | p.Thr40Met | O60566 |
| HOMO SAPIENS | 4 | 40565 | 5lcw | S | 198 | ALA | p.Glu390Asp | O60566 |
| HOMO SAPIENS | 4 | 758 | 4oiu | A | 726 | ARG | p.Arg727Leu | P10275 |
| HOMO SAPIENS | 4 | 758 | 4oiu | A | 730 | VAL | p.Val731Met | P10275 |
| HOMO SAPIENS | 2 | 758 | 4oiu | A | 798 | GLN | p.Gln799Glu | P10275 |
| HOMO SAPIENS | 3 | 758 | 4oiu | A | 855 | ARG | p.Arg856His | P10275 |
| HOMO SAPIENS | 1 | 758 | 4oiu | A | 886 | MET | p.Met887Val | P10275 |
| HOMO SAPIENS | 0 | 3303 | 4ie0 | A | 134 | ALA | p.Ala134Thr | Q9C0B1 |
| HOMO SAPIENS | 9 | 3303 | 4ie0 | A | 316 | ARG | p.Arg316Gln | Q9C0B1 |
| HOMO SAPIENS | 1 | 41502 | 3fgh | A | 136 | PRO | p.Pro178Leu | Q00059 |
| HOMO SAPIENS | 5 | 175 | 3ceh | B | 58 | THR | p.Thr59Lys | P06737 |
| HOMO SAPIENS | 8 | 175 | 3ceh | B | 93 | ARG | p.Arg94Ter | P06737 |
| HOMO SAPIENS | 5 | 175 | 3ceh | B | 221 | VAL | p.Val222Ile | P06737 |
| HOMO SAPIENS | 0 | 175 | 3ceh | B | 338 | ASN | p.Asn339Ser | P06737 |
| HOMO SAPIENS | 1 | 175 | 3ceh | B | 372 | PHE | p.Phe373Leu | P06737 |
| HOMO SAPIENS | 9 | 175 | 3ceh | B | 376 | ASN | p.Asn377Lys | P06737 |
| HOMO SAPIENS | 8 | 175 | 3ceh | B | 398 | ARG | p.Arg399Ter | P06737 |
| HOMO SAPIENS | 5 | 175 | 3ceh | B | 424 | ARG | p.Arg425Pro | P06737 |
| HOMO SAPIENS | 9 | 175 | 3ceh | B | 455 | VAL | p.Val456Met | P06737 |
| HOMO SAPIENS | 8 | 175 | 3ceh | B | 490 | ARG | p.Arg491Cys | P06737 |
| HOMO SAPIENS | 8 | 175 | 3ceh | B | 631 | ASN | p.Asn632Ile | P06737 |
| HOMO SAPIENS | 9 | 175 | 3ceh | B | 648 | TYR | p.Tyr649Ter | P06737 |
| HOMO SAPIENS | 9 | 175 | 3ceh | B | 672 | GLU | p.Glu673Lys | P06737 |
| HOMO SAPIENS | 9 | 175 | 3ceh | B | 674 | SER | p.Ser675Leu | P06737 |
| HOMO SAPIENS | 0 | 175 | 3ceh | B | 805 | ILE | p.Ile806Leu | P06737 |
| HOMO SAPIENS | 0 | 175 | 3ceh | B | 820 | TYR | p.Tyr821His | P06737 |
| HOMO SAPIENS | 1 | 6750 | 4fdj | A | 47 | GLY | p.Gly47Arg | P34059 |
| HOMO SAPIENS | 9 | 6750 | 4fdj | A | 60 | ASP | p.Asp60Asn | P34059 |
| HOMO SAPIENS | 5 | 6750 | 4fdj | A | 67 | LEU | p.Leu67Met | P34059 |
| HOMO SAPIENS | 0 | 6750 | 4fdj | A | 69 | PHE | p.Phe69Val | P34059 |
| HOMO SAPIENS | 3 | 6750 | 4fdj | A | 94 | ARG | p.Arg94Gly | P34059 |
| HOMO SAPIENS | 0 | 6750 | 4fdj | A | 113 | ILE | p.Ile113Phe | P34059 |
| HOMO SAPIENS | 0 | 6750 | 4fdj | A | 138 | VAL | p.Val138Ala | P34059 |
| HOMO SAPIENS | 0 | 6750 | 4fdj | A | 139 | GLY | p.Gly139Ser | P34059 |
| HOMO SAPIENS | 0 | 6750 | 4fdj | A | 151 | PRO | p.Pro151Thr | P34059 |
| HOMO SAPIENS | 0 | 6750 | 4fdj | A | 155 | GLY | p.Gly155Arg | P34059 |
| HOMO SAPIENS | 0 | 6750 | 4fdj | A | 159 | TRP | p.Trp159Ter | P34059 |
| HOMO SAPIENS | 0 | 6750 | 4fdj | A | 162 | SER | p.Ser162Phe | P34059 |
| HOMO SAPIENS | 0 | 6750 | 4fdj | A | 200 | THR | p.Thr200Met | P34059 |
| HOMO SAPIENS | 0 | 6750 | 4fdj | A | 204 | ASN | p.Asn204Lys | P34059 |
| HOMO SAPIENS | 0 | 6750 | 4fdj | A | 231 | ALA | p.Ala231Gly | P34059 |
| HOMO SAPIENS | 2 | 6750 | 4fdj | A | 235 | THR | p.Thr235Lys | P34059 |
| HOMO SAPIENS | 1 | 6750 | 4fdj | A | 259 | ARG | p.Arg259Gln | P34059 |
| HOMO SAPIENS | 0 | 6750 | 4fdj | A | 287 | SER | p.Ser287Leu | P34059 |
| HOMO SAPIENS | 0 | 6750 | 4fdj | A | 291 | ALA | p.Ala291Thr | P34059 |
| HOMO SAPIENS | 3 | 6750 | 4fdj | A | 301 | GLY | p.Gly301Cys | P34059 |
| HOMO SAPIENS | 0 | 6750 | 4fdj | A | 312 | THR | p.Thr312Ser | P34059 |
| HOMO SAPIENS | 0 | 6750 | 4fdj | A | 318 | MET | p.Met318Arg | P34059 |
| HOMO SAPIENS | 0 | 6750 | 4fdj | A | 340 | GLY | p.Gly340Asp | P34059 |
| HOMO SAPIENS | 0 | 6750 | 4fdj | A | 386 | ARG | p.Arg386Cys | P34059 |
| HOMO SAPIENS | 0 | 6750 | 4fdj | A | 391 | MET | p.Met391Val | P34059 |

| | | | | | | | | |
|---|---|---|---|---|---|---|---|---|
| HOMO SAPIENS | 0 | 6750 | 4fdj | A | 392 | ALA | p.Ala392Val | P34059 |
| HOMO SAPIENS | 0 | 6750 | 4fdj | A | 393 | ALA | p.Ala393Ser | P34059 |
| HOMO SAPIENS | 0 | 6750 | 4fdj | A | 459 | ALA | p.Ala459Val | P34059 |
| HOMO SAPIENS | 0 | 6750 | 4fdj | A | 473 | GLN | p.Gln473Ter | P34059 |
| HOMO SAPIENS | 2 | 6750 | 4fdj | A | 480 | VAL | p.Val480Phe | P34059 |
| HOMO SAPIENS | 0 | 6750 | 4fdj | A | 487 | ASN | p.Asn487Ser | P34059 |
| HOMO SAPIENS | 0 | 6750 | 4fdj | A | 488 | VAL | p.Val488Met | P34059 |
| HOMO SAPIENS | 0 | 6750 | 4fdj | A | 520 | TRP | p.Trp520Ter | P34059 |
| HOMO SAPIENS | 4 | 1088 | 2p3u | A | 102 | GLU | p.Glu142Lys | P00742 |
| HOMO SAPIENS | 6 | 370 | 1nr1 | D | 73 | ASP | p.Asp126His | P00367 |
| HOMO SAPIENS | 3 | 370 | 1nr1 | D | 221 | ARG | p.Arg274Cys | P00367 |
| HOMO SAPIENS | 2 | 370 | 1nr1 | D | 265 | ARG | p.Arg318Lys | P00367 |
| HOMO SAPIENS | 1 | 370 | 1nr1 | D | 269 | ARG | p.Arg322His | P00367 |
| HOMO SAPIENS | 1 | 370 | 1nr1 | D | 296 | GLU | p.Glu349Ala | P00367 |
| HOMO SAPIENS | 1 | 370 | 1nr1 | D | 445 | SER | p.Ser498Leu | P00367 |
| HOMO SAPIENS | 1 | 370 | 1nr1 | D | 446 | GLY | p.Gly499Asp | P00367 |
| HOMO SAPIENS | 1 | 370 | 1nr1 | D | 446 | GLY | p.Gly499Ser | P00367 |
| HOMO SAPIENS | 1 | 370 | 1nr1 | D | 448 | SER | p.Ser501Pro | P00367 |
| HOMO SAPIENS | 2 | 370 | 1nr1 | D | 454 | HIS | p.His507Tyr | P00367 |
| HOMO SAPIENS | 1 | 7720 | 2he0 | A | 147 | HIS | p.His2018Gln | P46531 |
| HOMO SAPIENS | 2 | 11844 | 5l0v | A | 218 | ARG | p.Arg218Ter | Q8NBL1 |
| HOMO SAPIENS | 2 | 11844 | 5l0v | A | 279 | ARG | p.Arg279Trp | Q8NBL1 |
| HOMO SAPIENS | 9 | 1485 | 5va0 | A | 477 | ARG | p.Arg477His | P04150 |
| HOMO SAPIENS | 0 | 235 | 2fun | B | 2248 | ARG | p.Arg248Trp | Q14790 |
| HOMO SAPIENS | 4 | 235 | 2fun | B | 2285 | HIS | p.Asp285His | Q14790 |
| HOMO SAPIENS | 0 | 4822 | 5y3n | A | 307 | ARG | p.Arg307Gly | Q12931 |
| HOMO SAPIENS | 8 | 32928 | 4m6w | A | 1886 | HIS | p.His1886Tyr | Q8IYD8 |
| HOMO SAPIENS | 0 | 2711 | 4y5v | F | 22 | GLY | p.Gly46Glu | P19235 |
| HOMO SAPIENS | -1 | 2711 | 4y5v | F | 166 | ALA | p.Ala190Ser | P19235 |
| HOMO SAPIENS | 4 | 5161 | 5bpu | A | 42 | HIS | p.His42Arg | Q00604 |
| HOMO SAPIENS | 0 | 5161 | 5bpu | A | 44 | TYR | p.Tyr44Cys | Q00604 |
| HOMO SAPIENS | 0 | 5161 | 5bpu | A | 45 | VAL | p.Val45Glu | Q00604 |
| HOMO SAPIENS | 0 | 5161 | 5bpu | A | 60 | VAL | p.Val60Glu | Q00604 |
| HOMO SAPIENS | 0 | 5161 | 5bpu | A | 61 | LEU | p.Leu61Phe | Q00604 |
| HOMO SAPIENS | 7 | 5161 | 5bpu | A | 69 | CYS | p.Cys69Ser | Q00604 |
| HOMO SAPIENS | 0 | 5161 | 5bpu | A | 75 | SER | p.Ser75Cys | Q00604 |
| HOMO SAPIENS | 0 | 5161 | 5bpu | A | 90 | ARG | p.Arg90Pro | Q00604 |
| HOMO SAPIENS | 9 | 5161 | 5bpu | A | 96 | CYS | p.Cys96Tyr | Q00604 |
| HOMO SAPIENS | 9 | 5161 | 5bpu | A | 96 | CYS | p.Cys96Trp | Q00604 |
| HOMO SAPIENS | 0 | 5161 | 5bpu | A | 101 | SER | p.Ser101Phe | Q00604 |
| HOMO SAPIENS | 0 | 5161 | 5bpu | A | 105 | ALA | p.Ala105Thr | Q00604 |
| HOMO SAPIENS | 0 | 5161 | 5bpu | A | 110 | CYS | p.Cys110Gly | Q00604 |
| HOMO SAPIENS | 0 | 5161 | 5bpu | A | 121 | ARG | p.Arg121Trp | Q00604 |
| HOMO SAPIENS | 0 | 5161 | 5bpu | A | 121 | ARG | p.Arg121Leu | Q00604 |
| HOMO SAPIENS | 0 | 5161 | 5bpu | A | 124 | LEU | p.Leu124Phe | Q00604 |
| HOMO SAPIENS | 8 | 5161 | 5bpu | A | 128 | CYS | p.Cys128Ter | Q00604 |
| HOMO SAPIENS | 1 | 13944 | 5xqh | A | 238 | ARG | p.Arg238His | Q9GZN7 |
| HOMO SAPIENS | -1 | 7508 | 5jl6 | A | 39 | TRP | p.Trp39Arg | P11511 |
| HOMO SAPIENS | 0 | 7508 | 5jl6 | A | 201 | THR | p.Thr201Met | P11511 |
| HOMO SAPIENS | 0 | 7508 | 5jl6 | A | 210 | GLU | p.Glu210Lys | P11511 |
| HOMO SAPIENS | 1 | 7508 | 5jl6 | A | 264 | ARG | p.Arg264Cys | P11511 |
| HOMO SAPIENS | 2 | 7508 | 5jl6 | A | 365 | ARG | p.Arg365Gln | P11511 |
| HOMO SAPIENS | 7 | 7508 | 5jl6 | A | 375 | ARG | p.Arg375Cys | P11511 |
| HOMO SAPIENS | 8 | 7508 | 5jl6 | A | 435 | ARG | p.Arg435Cys | P11511 |

| | | | | | | | | |
|---|---|---|---|---|---|---|---|---|
| HOMO SAPIENS | 9 | 7508 | 5jl6 | A | 437 | CYS | p.Cys437Tyr | P11511 |
| HOMO SAPIENS | -1 | 30206 | 5l95 | B | 246 | ARG | p.Arg246Ter | Q9GZZ9 |
| HOMO SAPIENS | 4 | 875 | 3efh | B | 43 | GLU | p.Glu43Asp | P60891 |
| HOMO SAPIENS | 5 | 875 | 3efh | B | 65 | ASP | p.Asp65Asn | P60891 |
| HOMO SAPIENS | 0 | 875 | 3efh | B | 87 | ALA | p.Ala87Thr | P60891 |
| HOMO SAPIENS | 2 | 875 | 3efh | B | 115 | MET | p.Met115Thr | P60891 |
| HOMO SAPIENS | 8 | 875 | 3efh | B | 133 | GLN | p.Gln133Pro | P60891 |
| HOMO SAPIENS | 8 | 875 | 3efh | B | 152 | LEU | p.Leu152Pro | P60891 |
| HOMO SAPIENS | 7 | 875 | 3efh | B | 290 | ILE | p.Ile290Thr | P60891 |
| HOMO SAPIENS | 6 | 875 | 3efh | B | 306 | GLY | p.Gly306Arg | P60891 |
| HOMO SAPIENS | 0 | 7978 | 5edm | A | 122 | MET | p.Thr165Met | P00734 |
| HOMO SAPIENS | -1 | 7978 | 5edm | A | 257 | ARG | p.Arg314Cys | P00734 |
| HOMO SAPIENS | 0 | 1716 | 1b98 | M | 126 | ARG | p.Arg206Gln | P34130 |
| HOMO SAPIENS | 0 | 1716 | 1b98 | M | 126 | ARG | p.Arg206Trp | P34130 |
| HOMO SAPIENS | 9 | 3960 | 2qyp | B | 28 | ILE | p.Ile338Val | P07602 |
| HOMO SAPIENS | 1 | 3960 | 2qyp | B | 39 | LEU | p.Leu349Pro | P07602 |
| HOMO SAPIENS | 0 | 31414 | 5u1d | A | 333 | ILE | p.Ile393Val | Q03518 |
| HOMO SAPIENS | 0 | 31414 | 5u1d | A | 637 | ASP | p.Asp697Gly | Q03518 |
| HOMO SAPIENS | 0 | 31414 | 5u1d | A | 659 | ARG | p.Arg719Gln | Q03518 |
| HOMO SAPIENS | 3 | 119 | 2pho | A | 11 | ILE | p.Ile11Thr | P05089 |
| HOMO SAPIENS | 8 | 119 | 2pho | A | 21 | ARG | p.Arg21Ter | P05089 |
| HOMO SAPIENS | 0 | 119 | 2pho | A | 122 | TRP | p.Trp122Ter | P05089 |
| HOMO SAPIENS | 6 | 119 | 2pho | A | 138 | GLY | p.Gly138Val | P05089 |
| HOMO SAPIENS | 5 | 119 | 2pho | A | 290 | THR | p.Thr290Ser | P05089 |
| HOMO SAPIENS | 5 | 119 | 2pho | A | 291 | ARG | p.Arg291Ter | P05089 |
| HOMO SAPIENS | 0 | 3732 | 2i99 | A | 247 | TYR | p.Tyr247Ter | Q14894 |
| HOMO SAPIENS | -1 | 1382 | 4ffx | A | 2 | ALA | p.Ala2Val | P30566 |
| HOMO SAPIENS | 1 | 1382 | 4ffx | A | 72 | ILE | p.Ile72Met | P30566 |
| HOMO SAPIENS | 1 | 1382 | 4ffx | A | 100 | PRO | p.Pro100Ala | P30566 |
| HOMO SAPIENS | 0 | 1382 | 4ffx | A | 114 | TYR | p.Tyr114His | P30566 |
| HOMO SAPIENS | 1 | 1382 | 4ffx | A | 141 | ARG | p.Arg141Trp | P30566 |
| HOMO SAPIENS | 0 | 1382 | 4ffx | A | 147 | LYS | p.Lys147Met | P30566 |
| HOMO SAPIENS | 3 | 1382 | 4ffx | A | 190 | ARG | p.Arg190Gln | P30566 |
| HOMO SAPIENS | 3 | 1382 | 4ffx | A | 190 | ARG | p.Arg190Ter | P30566 |
| HOMO SAPIENS | 0 | 1382 | 4ffx | A | 217 | HIS | p.His217Asp | P30566 |
| HOMO SAPIENS | 2 | 1382 | 4ffx | A | 246 | LYS | p.Lys246Glu | P30566 |
| HOMO SAPIENS | 5 | 1382 | 4ffx | A | 303 | ARG | p.Arg303Cys | P30566 |
| HOMO SAPIENS | 1 | 1382 | 4ffx | A | 318 | PRO | p.Pro318Leu | P30566 |
| HOMO SAPIENS | 6 | 1382 | 4ffx | A | 332 | ASP | p.Asp332His | P30566 |
| HOMO SAPIENS | 6 | 1382 | 4ffx | A | 337 | ARG | p.Arg337Ter | P30566 |
| HOMO SAPIENS | 1 | 1382 | 4ffx | A | 396 | ARG | p.Arg396His | P30566 |
| HOMO SAPIENS | 1 | 1382 | 4ffx | A | 426 | ARG | p.Arg426His | P30566 |
| HOMO SAPIENS | 0 | 1382 | 4ffx | A | 438 | SER | p.Ser438Pro | P30566 |
| HOMO SAPIENS | 1 | 1382 | 4ffx | A | 447 | SER | p.Ser447Pro | P30566 |
| HOMO SAPIENS | 2 | 1382 | 4ffx | A | 448 | SER | p.Ser448Pro | P30566 |
| HOMO SAPIENS | 2 | 1382 | 4ffx | A | 450 | THR | p.Thr450Ser | P30566 |
| HOMO SAPIENS | 0 | 6212 | 1ao3 | A | 46 | SER | p.Ser1731Thr | P04275 |
| HOMO SAPIENS | 0 | 1711 | 3f9c | A | 279 | VAL | p.Val279Phe | Q13093 |
| HOMO SAPIENS | 0 | 6112 | 4fvl | A | 312 | HIS | p.His312Arg | P45452 |
| HOMO SAPIENS | 0 | 6112 | 4fvl | A | 323 | THR | p.Thr323Met | P45452 |
| HOMO SAPIENS | 0 | 6112 | 4fvl | A | 390 | ASP | p.Asp390Gly | P45452 |
| HOMO SAPIENS | 0 | 6112 | 4fvl | A | 458 | ARG | p.Arg458Cys | P45452 |
| HOMO SAPIENS | 2 | 6180 | 5ggn | A | 97 | ARG | p.Arg97Trp | Q8WZA1 |
| HOMO SAPIENS | 2 | 6180 | 5ggn | A | 215 | ARG | p.Arg215Ter | Q8WZA1 |

| | | | | | | | | |
|---|---|---|---|---|---|---|---|---|
| HOMO SAPIENS | -1 | 5526 | 4ak3 | A | 11 | ASP | p.Asp1232Asn | P02461 |
| HOMO SAPIENS | 6 | 5526 | 4ak3 | A | 38 | ALA | p.Ala1259Thr | P02461 |
| HOMO SAPIENS | 0 | 5526 | 4ak3 | A | 92 | LYS | p.Lys1313Arg | P02461 |
| HOMO SAPIENS | 0 | 5526 | 4ak3 | A | 132 | GLN | p.His1353Gln | P02461 |
| HOMO SAPIENS | 4 | 16616 | 1hci | B | 433 | SER | p.Ser433Leu | P35609 |
| HOMO SAPIENS | 1 | 16616 | 1hci | B | 475 | ASP | p.Asp475Asn | P35609 |
| HOMO SAPIENS | 1 | 16616 | 1hci | B | 476 | ALA | p.Ala476Ser | P35609 |
| HOMO SAPIENS | 2 | 16616 | 1hci | B | 507 | MET | p.Met507Val | P35609 |
| HOMO SAPIENS | 1 | 16616 | 1hci | B | 604 | MET | p.Met604Val | P35609 |
| HOMO SAPIENS | 2 | 16616 | 1hci | B | 640 | ARG | p.Arg640His | P35609 |
| HOMO SAPIENS | 1 | 16616 | 1hci | B | 721 | ARG | p.Arg721Cys | P35609 |
| HOMO SAPIENS | -1 | 3402 | 5iaa | B | 57 | MET | p.Met57Val | Q9GZZ9 |
| HOMO SAPIENS | 7 | 3250 | 3irq | B | 193 | PRO | p.Pro193Ala | P55265 |
| HOMO SAPIENS | 2 | 15200 | 2wns | A | 30 | SER | p.Ser30Gly | P11172 |
| HOMO SAPIENS | 0 | 15200 | 2wns | A | 96 | ARG | p.Arg96Gly | P11172 |
| HOMO SAPIENS | 0 | 15200 | 2wns | A | 109 | VAL | p.Val109Gly | P11172 |
| HOMO SAPIENS | 2 | 15759 | 4x4w | B | 140 | LEU | p.Leu166Ser | Q96Q11 |
| HOMO SAPIENS | 0 | 15759 | 4x4w | B | 197 | ILE | p.Ile223Thr | Q96Q11 |
| HOMO SAPIENS | 0 | 3156 | 3w03 | B | 14 | ALA | p.Ala14Thr | Q9H9Q4 |
| HOMO SAPIENS | 0 | 3156 | 3w03 | B | 57 | ARG | p.Arg57Gly | Q9H9Q4 |
| HOMO SAPIENS | 0 | 3156 | 3w03 | B | 123 | CYS | p.Cys123Arg | Q9H9Q4 |
| HOMO SAPIENS | 0 | 11693 | 3ojy | A | 63 | GLN | p.Gln93Lys | P07357 |
| HOMO SAPIENS | 0 | 1421 | 2qc8 | E | 275 | GLU | p.Glu275Asp | P15104 |
| HOMO SAPIENS | 9 | 1421 | 2qc8 | E | 324 | ARG | p.Arg324Cys | P15104 |
| HOMO SAPIENS | 6 | 1421 | 2qc8 | E | 341 | ARG | p.Arg341Cys | P15104 |
| HOMO SAPIENS | 6 | 2850 | 3qvb | A | 83 | THR | p.Thr83Met | P46976 |
| HOMO SAPIENS | 8 | 2850 | 3qvb | A | 102 | ASP | p.Asp102His | P46976 |
| HOMO SAPIENS | 1 | 24314 | 4m9p | A | 480 | PRO | p.Pro480Leu | P21333 |
| HOMO SAPIENS | 0 | 24314 | 4m9p | A | 528 | VAL | p.Val528Met | P21333 |
| HOMO SAPIENS | 0 | 24314 | 4m9p | A | 590 | VAL | p.Val590Ile | P21333 |
| HOMO SAPIENS | 0 | 24314 | 4m9p | A | 605 | ASP | p.Asp605Asn | P21333 |
| HOMO SAPIENS | 1 | 24314 | 4m9p | A | 637 | PRO | p.Pro637Gln | P21333 |
| HOMO SAPIENS | 0 | 24314 | 4m9p | A | 656 | LEU | p.Leu656Phe | P21333 |
| HOMO SAPIENS | 2 | 24314 | 4m9p | A | 711 | VAL | p.Val711Asp | P21333 |
| HOMO SAPIENS | 0 | 4951 | 5joe | A | 40 | THR | p.Thr9621Ser | Q8WZ42 |
| HOMO SAPIENS | 1 | 4951 | 5joe | A | 41 | GLU | p.Glu9622Gln | Q8WZ42 |
| HOMO SAPIENS | 0 | 4951 | 5joe | A | 82 | ALA | p.Ala9663Ser | Q8WZ42 |
| HOMO SAPIENS | 4 | 4951 | 5joe | A | 89 | ILE | p.Ile9670Asn | Q8WZ42 |
| HOMO SAPIENS | 0 | 9664 | 3c6m | A | 56 | GLY | p.Gly56Ser | P52788 |
| HOMO SAPIENS | 0 | 9664 | 3c6m | A | 132 | VAL | p.Val132Gly | P52788 |
| HOMO SAPIENS | 1 | 9664 | 3c6m | A | 190 | THR | p.Thr190Ile | P52788 |
| HOMO SAPIENS | -1 | 12118 | 5hcd | B | 19 | GLN | p.Gln19Ter | P01031 |
| HOMO SAPIENS | 0 | 1583 | 2b4x | L | 7 | ILE | p.Ile39Asn | P01008 |
| HOMO SAPIENS | 0 | 1583 | 2b4x | L | 24 | ARG | p.Arg56Cys | P01008 |
| HOMO SAPIENS | 0 | 1583 | 2b4x | L | 47 | ARG | p.Arg79Cys | P01008 |
| HOMO SAPIENS | 0 | 1583 | 2b4x | L | 47 | ARG | p.Arg79His | P01008 |
| HOMO SAPIENS | 1 | 1583 | 2b4x | L | 95 | CYS | p.Cys127Arg | P01008 |
| HOMO SAPIENS | 1 | 1583 | 2b4x | L | 99 | LEU | p.Leu131Phe | P01008 |
| HOMO SAPIENS | 0 | 1583 | 2b4x | L | 116 | SER | p.Ser148Pro | P01008 |
| HOMO SAPIENS | 0 | 1583 | 2b4x | L | 129 | ARG | p.Arg161Gln | P01008 |
| HOMO SAPIENS | 0 | 1583 | 2b4x | L | 135 | ASN | p.Asn167Thr | P01008 |
| HOMO SAPIENS | 2 | 1583 | 2b4x | L | 187 | ASN | p.Asn219Asp | P01008 |
| HOMO SAPIENS | 0 | 1583 | 2b4x | L | 191 | SER | p.Ser223Pro | P01008 |
| HOMO SAPIENS | 2 | 1583 | 2b4x | L | 349 | SER | p.Ser381Pro | P01008 |

| | | | | | | | | |
|---|---|---|---|---|---|---|---|---|
| HOMO SAPIENS | 0 | 1583 | 2b4x | L | 382 | ALA | p.Ala414Thr | P01008 |
| HOMO SAPIENS | 0 | 1583 | 2b4x | L | 387 | ALA | p.Ala419Val | P01008 |
| HOMO SAPIENS | 0 | 1583 | 2b4x | L | 392 | GLY | p.Gly424Asp | P01008 |
| HOMO SAPIENS | 0 | 1583 | 2b4x | L | 393 | ARG | p.Arg425His | P01008 |
| HOMO SAPIENS | 0 | 1583 | 2b4x | L | 393 | ARG | p.Arg425Cys | P01008 |
| HOMO SAPIENS | 0 | 1583 | 2b4x | L | 394 | SER | p.Ser426Leu | P01008 |
| HOMO SAPIENS | 0 | 1583 | 2b4x | L | 404 | ALA | p.Ala436Thr | P01008 |
| HOMO SAPIENS | 2 | 1583 | 2b4x | L | 407 | PRO | p.Pro439Leu | P01008 |
| HOMO SAPIENS | 2 | 1583 | 2b4x | L | 429 | PRO | p.Pro461Leu | P01008 |
| HOMO SAPIENS | 0 | 11647 | 2a1t | S | 78 | ALA | p.Ala78Ser | P38117 |
| HOMO SAPIENS | 0 | 11647 | 2a1t | S | 98 | ARG | p.Arg98Cys | P38117 |
| HOMO SAPIENS | 8 | 11647 | 2a1t | S | 128 | ASP | p.Asp128Asn | P38117 |
| HOMO SAPIENS | 5 | 11647 | 2a1t | S | 154 | THR | p.Thr154Met | P38117 |
| HOMO SAPIENS | 6 | 11647 | 2a1t | S | 164 | ARG | p.Arg164Gln | P38117 |
| HOMO SAPIENS | 5 | 11647 | 2a1t | S | 174 | ARG | p.Arg174His | P38117 |
| HOMO SAPIENS | -1 | 11200 | 4tx5 | A | 4 | ILE | p.Ile59Val | Q9NR28 |
| HOMO SAPIENS | 0 | 11200 | 4tx5 | A | 169 | GLY | p.Gly224Arg | Q9NR28 |
| HOMO SAPIENS | 3 | 417 | 2vr2 | A | 6 | ARG | p.Arg6Gln | Q14117 |
| HOMO SAPIENS | 1 | 417 | 2vr2 | A | 334 | GLN | p.Gln334Arg | Q14117 |
| HOMO SAPIENS | 2 | 417 | 2vr2 | A | 360 | TRP | p.Trp360Arg | Q14117 |
| HOMO SAPIENS | 0 | 417 | 2vr2 | A | 379 | SER | p.Ser379Arg | Q14117 |
| HOMO SAPIENS | 0 | 417 | 2vr2 | A | 435 | GLY | p.Gly435Arg | Q14117 |
| HOMO SAPIENS | 0 | 12498 | 3eo3 | B | 423 | ILE | p.Ile423Val | Q9Y223 |
| HOMO SAPIENS | 5 | 12498 | 3eo3 | B | 572 | VAL | p.Val572Leu | Q9Y223 |
| HOMO SAPIENS | 0 | 12498 | 3eo3 | B | 576 | GLY | p.Gly576Glu | Q9Y223 |
| HOMO SAPIENS | 0 | 12498 | 3eo3 | B | 587 | ILE | p.Ile587Thr | Q9Y223 |
| HOMO SAPIENS | 0 | 12498 | 3eo3 | B | 615 | SER | p.Ser615Ter | Q9Y223 |
| HOMO SAPIENS | 1 | 12498 | 3eo3 | B | 631 | ALA | p.Ala631Val | Q9Y223 |
| HOMO SAPIENS | 1 | 12498 | 3eo3 | B | 669 | GLY | p.Gly669Arg | Q9Y223 |
| HOMO SAPIENS | 1 | 12498 | 3eo3 | B | 696 | VAL | p.Val696Met | Q9Y223 |
| HOMO SAPIENS | 2 | 12498 | 3eo3 | B | 712 | MET | p.Met712Thr | Q9Y223 |
| HOMO SAPIENS | 0 | 1568 | 1r74 | A | 49 | LEU | p.Leu50Pro | Q14749 |
| HOMO SAPIENS | 0 | 1568 | 1r74 | A | 176 | HIS | p.His177Asn | Q14749 |
| HOMO SAPIENS | 3 | 147 | 5eig | A | 67 | THR | p.Thr67Ile | P32929 |
| HOMO SAPIENS | 3 | 147 | 5eig | A | 240 | GLN | p.Gln240Glu | P32929 |
| HOMO SAPIENS | -1 | 147 | 5eig | A | 403 | SER | p.Ser403Ile | P32929 |
| HOMO SAPIENS | 8 | 20036 | 2lsr | A | 31 | ARG | p.Arg31Ter | Q9Y6N9 |
| HOMO SAPIENS | -1 | 41382 | 4uis | B | 431 | UNK | p.Ala431Glu | P49768 |
| HOMO SAPIENS | -1 | 8536 | 3i6w | B | 85 | PRO | p.Pro85Leu | O96017 |
| HOMO SAPIENS | 0 | 8536 | 3i6w | B | 93 | TRP | p.Trp93Ter | O96017 |
| HOMO SAPIENS | 0 | 8536 | 3i6w | B | 95 | ARG | p.Arg95Ter | O96017 |
| HOMO SAPIENS | 0 | 8536 | 3i6w | B | 117 | ARG | p.Arg117Gly | O96017 |
| HOMO SAPIENS | 0 | 8536 | 3i6w | B | 119 | LYS | p.Lys119Arg | O96017 |
| HOMO SAPIENS | 0 | 8536 | 3i6w | B | 137 | ARG | p.Arg137Gln | O96017 |
| HOMO SAPIENS | 0 | 8536 | 3i6w | B | 137 | ARG | p.Arg137Ter | O96017 |
| HOMO SAPIENS | 0 | 8536 | 3i6w | B | 145 | ARG | p.Arg145Trp | O96017 |
| HOMO SAPIENS | 0 | 8536 | 3i6w | B | 160 | ILE | p.Ile160Met | O96017 |
| HOMO SAPIENS | 0 | 41322 | 4bxl | C | 46 | GLU | p.Glu46Lys | P37840 |
| HOMO SAPIENS | 5 | 41322 | 4bxl | C | 53 | ALA | p.Ala53Thr | P37840 |
| HOMO SAPIENS | 2 | 24966 | 3hve | A | 15 | ARG | p.Arg15Ser | Q9H2C0 |
| HOMO SAPIENS | 1 | 24966 | 3hve | A | 138 | ARG | p.Arg138His | Q9H2C0 |
| HOMO SAPIENS | 0 | 24966 | 3hve | A | 201 | ARG | p.Arg201Ter | Q9H2C0 |
| HOMO SAPIENS | 0 | 11680 | 5an9 | J | 8 | ASN | p.Asn8Lys | Q9Y3A5 |
| HOMO SAPIENS | 1 | 11680 | 5an9 | J | 33 | LYS | p.Lys33Arg | Q9Y3A5 |

| | | | | | | | | |
|---|---|---|---|---|---|---|---|---|
| HOMO SAPIENS | 1 | 11680 | 5an9 | J | 126 | ARG | p.Arg126Thr | Q9Y3A5 |
| HOMO SAPIENS | 1 | 11680 | 5an9 | J | 130 | VAL | p.Val130Met | Q9Y3A5 |
| HOMO SAPIENS | 1 | 11680 | 5an9 | J | 169 | ARG | p.Arg169Cys | Q9Y3A5 |
| HOMO SAPIENS | 1 | 11680 | 5an9 | J | 218 | ARG | p.Arg218Ter | Q9Y3A5 |
| HOMO SAPIENS | 4 | 978 | 3zs0 | B | 7 | TYR | p.Tyr173Cys | P05164 |
| HOMO SAPIENS | 1 | 978 | 3zs0 | B | 85 | MET | p.Met251Thr | P05164 |
| HOMO SAPIENS | 4 | 5140 | 2l4r | A | 52 | CYS | p.Cys52Ter | Q12809 |
| HOMO SAPIENS | 4 | 5140 | 2l4r | A | 63 | PRO | p.Pro63His | Q12809 |
| HOMO SAPIENS | 0 | 5140 | 2l4r | A | 100 | ARG | p.Arg100Gly | Q12809 |
| HOMO SAPIENS | 4 | 5140 | 2l4r | A | 129 | PHE | p.Phe129Leu | Q12809 |
| HOMO SAPIENS | 0 | 25836 | 2gy7 | B | 103 | GLU | p.Glu103Asp | Q02763 |
| HOMO SAPIENS | 0 | 25836 | 2gy7 | B | 148 | ILE | p.Ile148Thr | Q02763 |
| HOMO SAPIENS | 0 | 25836 | 2gy7 | B | 226 | ALA | p.Ala226Val | Q02763 |
| HOMO SAPIENS | 0 | 25836 | 2gy7 | B | 391 | THR | p.Thr391Ile | Q02763 |
| HOMO SAPIENS | 4 | 12563 | 5kwy | D | 20 | GLU | p.Glu20Ter | P61916 |
| HOMO SAPIENS | 2 | 12563 | 5kwy | D | 30 | VAL | p.Val30Met | P61916 |
| HOMO SAPIENS | 4 | 12563 | 5kwy | D | 39 | VAL | p.Val39Met | P61916 |
| HOMO SAPIENS | 8 | 12563 | 5kwy | D | 67 | SER | p.Ser67Pro | P61916 |
| HOMO SAPIENS | 8 | 12563 | 5kwy | D | 99 | CYS | p.Cys99Arg | P61916 |
| HOMO SAPIENS | 2 | 12563 | 5kwy | D | 118 | GLU | p.Glu118Ter | P61916 |
| HOMO SAPIENS | 2 | 12563 | 5kwy | D | 120 | PRO | p.Pro120Ser | P61916 |
| HOMO SAPIENS | -1 | 1679 | 1pmk | A | -2 | VAL | p.Val374Phe | P00747 |
| HOMO SAPIENS | -1 | 5315 | 2w2n | P | 53 | ALA | p.Ala53Val | Q8NBP7 |
| HOMO SAPIENS | 8 | 13496 | 2yad | A | 138 | THR | p.Asn138Thr | P11686 |
| HOMO SAPIENS | -1 | 13496 | 2yad | A | 173 | ARG | p.Arg167Gln | P11686 |
| HOMO SAPIENS | 2 | 13496 | 2yad | A | 186 | SER | p.Asn186Ser | P11686 |
| HOMO SAPIENS | 0 | 4426 | 3qd6 | C | 123 | ALA | p.Ala123Glu | P29965 |
| HOMO SAPIENS | 1 | 4426 | 3qd6 | C | 140 | TRP | p.Trp140Gly | P29965 |
| HOMO SAPIENS | 0 | 4426 | 3qd6 | C | 155 | LEU | p.Leu155Pro | P29965 |
| HOMO SAPIENS | 6 | 4426 | 3qd6 | C | 227 | GLY | p.Gly227Val | P29965 |
| HOMO SAPIENS | 0 | 4426 | 3qd6 | C | 235 | ALA | p.Ala235Pro | P29965 |
| HOMO SAPIENS | 0 | 10734 | 3dyd | B | 45 | PRO | p.Pro45Ser | P17735 |
| HOMO SAPIENS | -1 | 10734 | 3dyd | B | 57 | ARG | p.Arg57Ter | P17735 |
| HOMO SAPIENS | 8 | 10734 | 3dyd | B | 417 | ARG | p.Arg417Ter | P17735 |
| HOMO SAPIENS | 0 | 6956 | 4pzs | A | 1600 | LEU | p.Leu1600Phe | Q09472 |
| HOMO SAPIENS | 0 | 9370 | 3hfv | A | 21 | SER | p.Ser57Cys | O95363 |
| HOMO SAPIENS | 1 | 9370 | 3hfv | A | 25 | ASP | p.Asp61Glu | O95363 |
| HOMO SAPIENS | 1 | 9370 | 3hfv | A | 130 | ALA | p.Ala166Thr | O95363 |
| HOMO SAPIENS | 5 | 9370 | 3hfv | A | 148 | ASP | p.Asp184Asn | O95363 |
| HOMO SAPIENS | 0 | 9370 | 3hfv | A | 187 | ARG | p.Arg223Cys | O95363 |
| HOMO SAPIENS | 0 | 9370 | 3hfv | A | 210 | THR | p.Thr246Met | O95363 |
| HOMO SAPIENS | 1 | 9370 | 3hfv | A | 244 | ASN | p.Asn280Ser | O95363 |
| HOMO SAPIENS | 2 | 9370 | 3hfv | A | 288 | TYR | p.Tyr324Cys | O95363 |
| HOMO SAPIENS | 5 | 9370 | 3hfv | A | 293 | ILE | p.Ile329Thr | O95363 |
| HOMO SAPIENS | 1 | 9370 | 3hfv | A | 355 | ASP | p.Asp391Val | O95363 |
| HOMO SAPIENS | 1 | 9370 | 3hfv | A | 371 | THR | p.Thr407Met | O95363 |
| HOMO SAPIENS | 1 | 9370 | 3hfv | A | 383 | ARG | p.Arg419Cys | O95363 |
| HOMO SAPIENS | 1 | 9370 | 3hfv | A | 387 | ARG | p.Arg423Gln | O95363 |
| HOMO SAPIENS | 0 | 14401 | 1q3x | A | 371 | ASP | p.Asp371Tyr | O00187 |
| HOMO SAPIENS | 0 | 14401 | 1q3x | A | 377 | VAL | p.Val377Ala | O00187 |
| HOMO SAPIENS | 0 | 14401 | 1q3x | A | 439 | ARG | p.Arg439His | O00187 |
| HOMO SAPIENS | 0 | 14401 | 1q3x | A | 577 | GLN | p.Gln577His | O00187 |
| HOMO SAPIENS | 2 | 18270 | 3teq | A | 429 | ARG | p.Arg429Cys | Q13586 |
| HOMO SAPIENS | -1 | 16081 | 4mqt | A | 231 | VAL | p.Val231Ile | P08172 |

| | | | | | | | | |
|---|---|---|---|---|---|---|---|---|
| HOMO SAPIENS | 9 | 6607 | 3whu | B | 81 | ASP | p.Asp81His | Q8NI22 |
| HOMO SAPIENS | 9 | 6607 | 3whu | B | 129 | ASP | p.Asp129Glu | Q8NI22 |
| HOMO SAPIENS | 9 | 6607 | 3whu | B | 136 | ILE | p.Ile136Thr | Q8NI22 |
| HOMO SAPIENS | 0 | 7208 | 2e9w | D | 11 | ASN | p.Asn36Ser | P21583 |
| HOMO SAPIENS | 0 | 24566 | 4bdx | A | 10 | LEU | p.Leu140Val | P00748 |
| HOMO SAPIENS | 5 | 24566 | 4bdx | A | 77 | PRO | p.Ala207Pro | P00748 |
| HOMO SAPIENS | 7 | 606 | 3qnf | A | 319 | MET | p.Met319Val | Q9NZ08 |
| HOMO SAPIENS | 8 | 4097 | 1ydl | A | 21 | LEU | p.Leu21Pro | Q6ZYL4 |
| HOMO SAPIENS | 6 | 4097 | 1ydl | A | 56 | ARG | p.Arg56Ter | Q6ZYL4 |
| HOMO SAPIENS | 8 | 36691 | 1dqb | A | 42 | ARG | p.Arg403Lys | P07204 |
| HOMO SAPIENS | 0 | 5642 | 3j7y | X | 59 | ARG | p.Arg59Pro | Q13084 |
| HOMO SAPIENS | 1 | 5642 | 3j7y | X | 204 | VAL | p.Val204Met | Q13084 |
| HOMO SAPIENS | 1 | 1119 | 3nwu | B | 252 | ALA | p.Ala252Thr | Q92743 |
| HOMO SAPIENS | 3 | 1119 | 3nwu | B | 274 | ARG | p.Arg274Gln | Q92743 |
| HOMO SAPIENS | 9 | 1119 | 3nwu | B | 297 | VAL | p.Val297Met | Q92743 |
| HOMO SAPIENS | -1 | 1119 | 3nwu | B | 302 | ARG | p.Arg302Ter | Q92743 |
| HOMO SAPIENS | 9 | 1119 | 3nwu | B | 321 | ALA | p.Ala321Thr | Q92743 |
| HOMO SAPIENS | -1 | 1119 | 3nwu | B | 370 | ARG | p.Arg370Ter | Q92743 |
| HOMO SAPIENS | 0 | 416 | 4ycu | B | 105 | LEU | p.Leu105His | Q15046 |
| HOMO SAPIENS | 0 | 416 | 4ycu | B | 145 | TYR | p.Tyr145His | Q15046 |
| HOMO SAPIENS | 0 | 416 | 4ycu | B | 200 | PRO | p.Pro200Leu | Q15046 |
| HOMO SAPIENS | 0 | 416 | 4ycu | B | 201 | TYR | p.Tyr201His | Q15046 |
| HOMO SAPIENS | 0 | 416 | 4ycu | B | 420 | ARG | p.Arg420Cys | Q15046 |
| HOMO SAPIENS | 9 | 3297 | 4fpb | A | 680 | THR | p.Thr680Ser | P19367 |
| HOMO SAPIENS | 0 | 6771 | 2zt6 | A | 201 | ALA | p.Ala255Val | P41250 |
| HOMO SAPIENS | 0 | 6771 | 2zt6 | A | 202 | ASP | p.Asp256His | P41250 |
| HOMO SAPIENS | 0 | 6771 | 2zt6 | A | 334 | ARG | p.Arg388Gln | P41250 |
| HOMO SAPIENS | -1 | 6771 | 2zt6 | A | 439 | ASN | p.Asn493Ser | P41250 |
| HOMO SAPIENS | -1 | 6771 | 2zt6 | A | 500 | ASP | p.Asp554Asn | P41250 |
| HOMO SAPIENS | 1 | 6771 | 2zt6 | A | 581 | SER | p.Ser635Leu | P41250 |
| HOMO SAPIENS | -1 | 6771 | 2zt6 | A | 684 | GLU | p.Glu738Lys | P41250 |
| HOMO SAPIENS | 0 | 2687 | 4grw | B | 276 | VAL | p.Val298Phe | P29460 |
| HOMO SAPIENS | 6 | 913 | 1m75 | B | 45 | ASP | p.Asp57Glu | Q16836 |
| HOMO SAPIENS | 2 | 913 | 1m75 | B | 74 | PRO | p.Leu86Pro | Q16836 |
| HOMO SAPIENS | 3 | 913 | 1m75 | B | 214 | TYR | p.Tyr226His | Q16836 |
| HOMO SAPIENS | 3 | 913 | 1m75 | B | 224 | ARG | p.Arg236Ter | Q16836 |
| HOMO SAPIENS | 4 | 913 | 1m75 | B | 246 | PRO | p.Pro258Leu | Q16836 |
| HOMO SAPIENS | 2 | 4713 | 4djn | B | 40 | ARG | p.Arg40Cys | Q7LG56 |
| HOMO SAPIENS | 1 | 4713 | 4djn | B | 41 | ARG | p.Arg41Pro | Q7LG56 |
| HOMO SAPIENS | 2 | 4713 | 4djn | B | 110 | ARG | p.Arg110His | Q7LG56 |
| HOMO SAPIENS | 2 | 4713 | 4djn | B | 121 | ARG | p.Arg121His | Q7LG56 |
| HOMO SAPIENS | 9 | 4713 | 4djn | B | 131 | GLU | p.Glu131Lys | Q7LG56 |
| HOMO SAPIENS | 2 | 4713 | 4djn | B | 182 | THR | p.Thr182Ala | Q7LG56 |
| HOMO SAPIENS | 6 | 4713 | 4djn | B | 194 | GLU | p.Glu194Lys | Q7LG56 |
| HOMO SAPIENS | 2 | 4713 | 4djn | B | 211 | ARG | p.Arg211Lys | Q7LG56 |
| HOMO SAPIENS | 0 | 4713 | 4djn | B | 224 | ILE | p.Ile224Ser | Q7LG56 |
| HOMO SAPIENS | 3 | 4713 | 4djn | B | 229 | GLY | p.Gly229Val | Q7LG56 |
| HOMO SAPIENS | 2 | 4713 | 4djn | B | 236 | CYS | p.Cys236Phe | Q7LG56 |
| HOMO SAPIENS | 0 | 4713 | 4djn | B | 273 | GLY | p.Gly273Ser | Q7LG56 |
| HOMO SAPIENS | -1 | 4713 | 4djn | B | 282 | MSE | p.Met282Ile | Q7LG56 |
| HOMO SAPIENS | 3 | 4713 | 4djn | B | 284 | GLN | p.Gln284Ter | Q7LG56 |
| HOMO SAPIENS | 3 | 4713 | 4djn | B | 305 | ALA | p.Ala305Ser | Q7LG56 |
| HOMO SAPIENS | -1 | 8974 | 5n7i | A | 500 | ARG | p.Arg500Ter | Q8N4S9 |
| HOMO SAPIENS | 0 | 17072 | 3fxi | B | 299 | ASP | p.Asp299Gly | O00206 |

| | | | | | | | | |
|---|---|---|---|---|---|---|---|---|
| HOMO SAPIENS | 1 | 14355 | 2wwk | T | 55 | ILE | p.Ile34306Asn | Q8WZ42 |
| HOMO SAPIENS | 0 | 14355 | 2wwk | T | 64 | LEU | p.Leu34315Pro | Q8WZ42 |
| HOMO SAPIENS | 2 | 17797 | 1xu1 | S | 72 | ARG | p.Arg72His | O14836 |
| HOMO SAPIENS | -1 | 10836 | 4ztu | A | 39 | GLN | p.Gln39Arg | P54098 |
| HOMO SAPIENS | -1 | 10836 | 4ztu | A | 42 | ARG | p.Arg42Gln | P54098 |
| HOMO SAPIENS | -1 | 10836 | 4ztu | A | 45 | GLN | p.Gln45Arg | P54098 |
| HOMO SAPIENS | -1 | 10836 | 4ztu | A | 52 | GLN | p.Gln52Glu | P54098 |
| HOMO SAPIENS | 0 | 10836 | 4ztu | A | 88 | PHE | p.Phe88Leu | P54098 |
| HOMO SAPIENS | 1 | 10836 | 4ztu | A | 194 | ALA | p.Ala194Val | P54098 |
| HOMO SAPIENS | 1 | 10836 | 4ztu | A | 227 | ARG | p.Arg227Trp | P54098 |
| HOMO SAPIENS | 0 | 10836 | 4ztu | A | 232 | ARG | p.Arg232His | P54098 |
| HOMO SAPIENS | 1 | 10836 | 4ztu | A | 285 | GLN | p.Gln285His | P54098 |
| HOMO SAPIENS | 1 | 10836 | 4ztu | A | 434 | TYR | p.Tyr434His | P54098 |
| HOMO SAPIENS | -1 | 10836 | 4ztu | A | 511 | SER | p.Ser511Asn | P54098 |
| HOMO SAPIENS | 0 | 10836 | 4ztu | A | 533 | CYS | p.Cys533Trp | P54098 |
| HOMO SAPIENS | 1 | 10836 | 4ztu | A | 546 | ARG | p.Arg546Cys | P54098 |
| HOMO SAPIENS | 1 | 10836 | 4ztu | A | 587 | PRO | p.Pro587Leu | P54098 |
| HOMO SAPIENS | 0 | 10836 | 4ztu | A | 603 | MET | p.Met603Thr | P54098 |
| HOMO SAPIENS | 0 | 10836 | 4ztu | A | 623 | LEU | p.Leu623Trp | P54098 |
| HOMO SAPIENS | -1 | 10836 | 4ztu | A | 627 | ARG | p.Arg627Trp | P54098 |
| HOMO SAPIENS | -1 | 10836 | 4ztu | A | 628 | ARG | p.Arg628Pro | P54098 |
| HOMO SAPIENS | 0 | 10836 | 4ztu | A | 635 | PRO | p.Pro635Leu | P54098 |
| HOMO SAPIENS | 0 | 10836 | 4ztu | A | 662 | GLU | p.Glu662Lys | P54098 |
| HOMO SAPIENS | -1 | 10836 | 4ztu | A | 674 | GLY | p.Gly674Asp | P54098 |
| HOMO SAPIENS | -1 | 10836 | 4ztu | A | 722 | ARG | p.Arg722His | P54098 |
| HOMO SAPIENS | -1 | 10836 | 4ztu | A | 737 | GLY | p.Gly737Arg | P54098 |
| HOMO SAPIENS | 0 | 10836 | 4ztu | A | 748 | TRP | p.Trp748Ser | P54098 |
| HOMO SAPIENS | 0 | 10836 | 4ztu | A | 749 | PHE | p.Phe749Ser | P54098 |
| HOMO SAPIENS | 1 | 10836 | 4ztu | A | 807 | ARG | p.Arg807Cys | P54098 |
| HOMO SAPIENS | 1 | 10836 | 4ztu | A | 831 | TYR | p.Tyr831Cys | P54098 |
| HOMO SAPIENS | 8 | 10836 | 4ztu | A | 848 | GLY | p.Gly848Ser | P54098 |
| HOMO SAPIENS | 1 | 10836 | 4ztu | A | 852 | ARG | p.Arg852Cys | P54098 |
| HOMO SAPIENS | 9 | 10836 | 4ztu | A | 853 | ARG | p.Arg853Trp | P54098 |
| HOMO SAPIENS | 1 | 10836 | 4ztu | A | 864 | ASN | p.Asn864Ser | P54098 |
| HOMO SAPIENS | 1 | 10836 | 4ztu | A | 879 | GLN | p.Gln879Arg | P54098 |
| HOMO SAPIENS | 1 | 10836 | 4ztu | A | 914 | THR | p.Thr914Pro | P54098 |
| HOMO SAPIENS | 6 | 10836 | 4ztu | A | 932 | HIS | p.His932Tyr | P54098 |
| HOMO SAPIENS | 2 | 10836 | 4ztu | A | 944 | GLU | p.Glu944Lys | P54098 |
| HOMO SAPIENS | 6 | 10836 | 4ztu | A | 955 | TYR | p.Tyr955Cys | P54098 |
| HOMO SAPIENS | 1 | 10836 | 4ztu | A | 957 | ALA | p.Ala957Ser | P54098 |
| HOMO SAPIENS | 1 | 10836 | 4ztu | A | 957 | ALA | p.Ala957Val | P54098 |
| HOMO SAPIENS | 1 | 10836 | 4ztu | A | 964 | ARG | p.Arg964Cys | P54098 |
| HOMO SAPIENS | 1 | 10836 | 4ztu | A | 966 | LEU | p.Leu966Arg | P54098 |
| HOMO SAPIENS | 0 | 10836 | 4ztu | A | 1051 | GLY | p.Gly1051Arg | P54098 |
| HOMO SAPIENS | 1 | 10836 | 4ztu | A | 1073 | PRO | p.Pro1073Leu | P54098 |
| HOMO SAPIENS | 1 | 10836 | 4ztu | A | 1096 | ARG | p.Arg1096Cys | P54098 |
| HOMO SAPIENS | 1 | 10836 | 4ztu | A | 1096 | ARG | p.Arg1096His | P54098 |
| HOMO SAPIENS | 0 | 10836 | 4ztu | A | 1116 | MET | p.Met1116Thr | P54098 |
| HOMO SAPIENS | 1 | 10836 | 4ztu | A | 1142 | ARG | p.Arg1142Trp | P54098 |
| HOMO SAPIENS | 1 | 10836 | 4ztu | A | 1146 | ARG | p.Arg1146Cys | P54098 |
| HOMO SAPIENS | 1 | 10836 | 4ztu | A | 1157 | ASN | p.Asn1157Ser | P54098 |
| HOMO SAPIENS | -1 | 10836 | 4ztu | A | 1234 | ARG | p.Arg1234Gln | P54098 |
| HOMO SAPIENS | -1 | 10836 | 4ztu | A | 1236 | GLN | p.Gln1236His | P54098 |
| HOMO SAPIENS | 5 | 2666 | 1jph | A | 62 | PRO | p.Pro62Leu | P06132 |

| | | | | | | | | |
|---|---|---|---|---|---|---|---|---|
| HOMO SAPIENS | 5 | 2666 | 1jph | A | 116 | GLN | p.Gln116Ter | P06132 |
| HOMO SAPIENS | 0 | 2666 | 1jph | A | 165 | MET | p.Met165Arg | P06132 |
| HOMO SAPIENS | 6 | 2666 | 1jph | A | 167 | GLU | p.Glu167Lys | P06132 |
| HOMO SAPIENS | 0 | 2666 | 1jph | A | 195 | LEU | p.Leu195Phe | P06132 |
| HOMO SAPIENS | 7 | 2666 | 1jph | A | 253 | LEU | p.Leu253Gln | P06132 |
| HOMO SAPIENS | 0 | 2666 | 1jph | A | 281 | GLY | p.Gly281Glu | P06132 |
| HOMO SAPIENS | 9 | 2666 | 1jph | A | 304 | ASN | p.Asn304Lys | P06132 |
| HOMO SAPIENS | 8 | 2666 | 1jph | A | 311 | TYR | p.Tyr311Cys | P06132 |
| HOMO SAPIENS | 5 | 2666 | 1jph | A | 332 | ARG | p.Arg332His | P06132 |
| HOMO SAPIENS | 1 | 889 | 1m51 | A | 184 | LEU | p.Val184Leu | P35558 |
| HOMO SAPIENS | 0 | 889 | 1m51 | A | 267 | VAL | p.Ile267Val | P35558 |
| HOMO SAPIENS | 2 | 889 | 1m51 | A | 276 | GLU | p.Glu276Lys | P35558 |
| HOMO SAPIENS | 6 | 889 | 1m51 | A | 368 | VAL | p.Val368Ile | P35558 |
| HOMO SAPIENS | 0 | 889 | 1m51 | A | 579 | SER | p.Ser579Arg | P35558 |
| HOMO SAPIENS | 1 | 4084 | 2gjx | A | 28 | GLN | p.Gln28Ter | P06865 |
| HOMO SAPIENS | 0 | 4084 | 2gjx | A | 39 | LEU | p.Leu39Arg | P06865 |
| HOMO SAPIENS | 0 | 4084 | 2gjx | A | 127 | LEU | p.Leu127Arg | P06865 |
| HOMO SAPIENS | 6 | 4084 | 2gjx | A | 137 | ARG | p.Arg137Ter | P06865 |
| HOMO SAPIENS | 6 | 4084 | 2gjx | A | 170 | ARG | p.Arg170Gln | P06865 |
| HOMO SAPIENS | 9 | 4084 | 2gjx | A | 178 | ARG | p.Arg178Cys | P06865 |
| HOMO SAPIENS | 9 | 4084 | 2gjx | A | 178 | ARG | p.Arg178His | P06865 |
| HOMO SAPIENS | 5 | 4084 | 2gjx | A | 180 | TYR | p.Tyr180His | P06865 |
| HOMO SAPIENS | 5 | 4084 | 2gjx | A | 180 | TYR | p.Tyr180Ter | P06865 |
| HOMO SAPIENS | 1 | 4084 | 2gjx | A | 197 | LYS | p.Lys197Thr | P06865 |
| HOMO SAPIENS | 0 | 4084 | 2gjx | A | 200 | VAL | p.Val200Met | P06865 |
| HOMO SAPIENS | 9 | 4084 | 2gjx | A | 204 | HIS | p.His204Arg | P06865 |
| HOMO SAPIENS | 5 | 4084 | 2gjx | A | 210 | SER | p.Ser210Phe | P06865 |
| HOMO SAPIENS | 0 | 4084 | 2gjx | A | 211 | PHE | p.Phe211Ser | P06865 |
| HOMO SAPIENS | 2 | 4084 | 2gjx | A | 250 | GLY | p.Gly250Asp | P06865 |
| HOMO SAPIENS | 9 | 4084 | 2gjx | A | 258 | ASP | p.Asp258His | P06865 |
| HOMO SAPIENS | 0 | 4084 | 2gjx | A | 269 | GLY | p.Gly269Ser | P06865 |
| HOMO SAPIENS | 0 | 4084 | 2gjx | A | 301 | MET | p.Met301Arg | P06865 |
| HOMO SAPIENS | 3 | 4084 | 2gjx | A | 393 | ARG | p.Arg393Ter | P06865 |
| HOMO SAPIENS | 3 | 4084 | 2gjx | A | 393 | ARG | p.Arg393Pro | P06865 |
| HOMO SAPIENS | 0 | 4084 | 2gjx | A | 399 | ASN | p.Asn399Asp | P06865 |
| HOMO SAPIENS | 2 | 4084 | 2gjx | A | 420 | TRP | p.Trp420Cys | P06865 |
| HOMO SAPIENS | 1 | 4084 | 2gjx | A | 434 | PHE | p.Phe434Leu | P06865 |
| HOMO SAPIENS | 0 | 4084 | 2gjx | A | 454 | GLY | p.Gly454Ser | P06865 |
| HOMO SAPIENS | 1 | 4084 | 2gjx | A | 474 | TRP | p.Trp474Cys | P06865 |
| HOMO SAPIENS | 0 | 4084 | 2gjx | A | 479 | ALA | p.Ala479Thr | P06865 |
| HOMO SAPIENS | 5 | 4084 | 2gjx | A | 485 | TRP | p.Trp485Arg | P06865 |
| HOMO SAPIENS | 4 | 4084 | 2gjx | A | 499 | ARG | p.Arg499Cys | P06865 |
| HOMO SAPIENS | 4 | 4084 | 2gjx | A | 499 | ARG | p.Arg499His | P06865 |
| HOMO SAPIENS | 2 | 4084 | 2gjx | A | 504 | ARG | p.Arg504Cys | P06865 |
| HOMO SAPIENS | 2 | 4084 | 2gjx | A | 504 | ARG | p.Arg504His | P06865 |
| HOMO SAPIENS | -1 | 1943 | 1xmj | A | 417 | ASN | p.Asn417Lys | P13569 |
| HOMO SAPIENS | -1 | 1943 | 1xmj | A | 422 | SER | p.Ser422Phe | P13569 |
| HOMO SAPIENS | 1 | 1943 | 1xmj | A | 438 | THR | p.Thr438Ala | P13569 |
| HOMO SAPIENS | 0 | 1943 | 1xmj | A | 455 | ALA | p.Ala455Glu | P13569 |
| HOMO SAPIENS | 4 | 1943 | 1xmj | A | 456 | VAL | p.Val456Ala | P13569 |
| HOMO SAPIENS | 9 | 1943 | 1xmj | A | 458 | GLY | p.Gly458Val | P13569 |
| HOMO SAPIENS | 8 | 1943 | 1xmj | A | 466 | SER | p.Ser466Ter | P13569 |
| HOMO SAPIENS | 0 | 1943 | 1xmj | A | 467 | LEU | p.Leu467Pro | P13569 |
| HOMO SAPIENS | 1 | 1943 | 1xmj | A | 470 | MET | p.Val470Met | P13569 |

| | | | | | | | | |
|---|---|---|---|---|---|---|---|---|
| HOMO SAPIENS | 5 | 1943 | 1xmj | A | 480 | GLY | p.Gly480Cys | P13569 |
| HOMO SAPIENS | 3 | 1943 | 1xmj | A | 492 | SER | p.Ser492Phe | P13569 |
| HOMO SAPIENS | 8 | 1943 | 1xmj | A | 493 | GLN | p.Gln493Ter | P13569 |
| HOMO SAPIENS | 5 | 1943 | 1xmj | A | 506 | ILE | p.Ile506Met | P13569 |
| HOMO SAPIENS | 3 | 1943 | 1xmj | A | 520 | VAL | p.Val520Phe | P13569 |
| HOMO SAPIENS | 1 | 1943 | 1xmj | A | 528 | GLU | p.Glu528Asp | P13569 |
| HOMO SAPIENS | 1 | 1943 | 1xmj | A | 534 | ALA | p.Ala534Glu | P13569 |
| HOMO SAPIENS | 4 | 1943 | 1xmj | A | 542 | GLY | p.Gly542Ter | P13569 |
| HOMO SAPIENS | 6 | 1943 | 1xmj | A | 549 | SER | p.Ser549Asn | P13569 |
| HOMO SAPIENS | 6 | 1943 | 1xmj | A | 549 | SER | p.Ser549Arg | P13569 |
| HOMO SAPIENS | 6 | 1943 | 1xmj | A | 549 | SER | p.Ser549Arg | P13569 |
| HOMO SAPIENS | 6 | 1943 | 1xmj | A | 551 | GLY | p.Gly551Asp | P13569 |
| HOMO SAPIENS | 6 | 1943 | 1xmj | A | 551 | GLY | p.Gly551Ser | P13569 |
| HOMO SAPIENS | 6 | 1943 | 1xmj | A | 553 | GLN | p.Arg553Gln | P13569 |
| HOMO SAPIENS | 6 | 1943 | 1xmj | A | 553 | GLN | p.Arg553Ter | P13569 |
| HOMO SAPIENS | 0 | 1943 | 1xmj | A | 559 | ALA | p.Ala559Thr | P13569 |
| HOMO SAPIENS | 7 | 1943 | 1xmj | A | 560 | ARG | p.Arg560Lys | P13569 |
| HOMO SAPIENS | 4 | 1943 | 1xmj | A | 562 | VAL | p.Val562Ile | P13569 |
| HOMO SAPIENS | 3 | 1943 | 1xmj | A | 563 | TYR | p.Tyr563Asn | P13569 |
| HOMO SAPIENS | 3 | 1943 | 1xmj | A | 569 | TYR | p.Tyr569Asp | P13569 |
| HOMO SAPIENS | 3 | 1943 | 1xmj | A | 585 | GLU | p.Glu585Ter | P13569 |
| HOMO SAPIENS | 1 | 1943 | 1xmj | A | 618 | ILE | p.Ile618Thr | P13569 |
| HOMO SAPIENS | 9 | 1943 | 1xmj | A | 622 | GLY | p.Gly622Asp | P13569 |
| HOMO SAPIENS | -1 | 1943 | 1xmj | A | 648 | ASP | p.Asp648Val | P13569 |
| HOMO SAPIENS | 0 | 1943 | 1xmj | A | 673 | GLY | p.Gly673Ter | P13569 |
| HOMO SAPIENS | 5 | 5111 | 4rec | A | 749 | ARG | p.Arg749Ter | Q9Y2M0 |
| HOMO SAPIENS | -1 | 4580 | 4l0d | B | 2 | GLY | p.Pro2Leu | P35520 |
| HOMO SAPIENS | -1 | 4580 | 4l0d | B | 12 | PRO | p.Pro12Ser | P35520 |
| HOMO SAPIENS | -1 | 4580 | 4l0d | B | 18 | ARG | p.Arg18Cys | P35520 |
| HOMO SAPIENS | 1 | 4580 | 4l0d | B | 49 | PRO | p.Pro49Leu | P35520 |
| HOMO SAPIENS | 2 | 4580 | 4l0d | B | 70 | PRO | p.Pro70Leu | P35520 |
| HOMO SAPIENS | 1 | 4580 | 4l0d | B | 102 | LYS | p.Lys102Gln | P35520 |
| HOMO SAPIENS | 7 | 4580 | 4l0d | B | 114 | ALA | p.Ala114Val | P35520 |
| HOMO SAPIENS | 5 | 4580 | 4l0d | B | 116 | GLY | p.Gly116Arg | P35520 |
| HOMO SAPIENS | 8 | 4580 | 4l0d | B | 121 | ARG | p.Arg121Cys | P35520 |
| HOMO SAPIENS | 8 | 4580 | 4l0d | B | 121 | ARG | p.Arg121His | P35520 |
| HOMO SAPIENS | 5 | 4580 | 4l0d | B | 125 | ARG | p.Arg125Gln | P35520 |
| HOMO SAPIENS | 4 | 4580 | 4l0d | B | 134 | GLY | p.Gly134Arg | P35520 |
| HOMO SAPIENS | 4 | 4580 | 4l0d | B | 139 | GLY | p.Gly139Arg | P35520 |
| HOMO SAPIENS | 0 | 4580 | 4l0d | B | 145 | PRO | p.Pro145Leu | P35520 |
| HOMO SAPIENS | 0 | 4580 | 4l0d | B | 148 | GLY | p.Gly148Arg | P35520 |
| HOMO SAPIENS | 0 | 4580 | 4l0d | B | 168 | VAL | p.Val168Met | P35520 |
| HOMO SAPIENS | 2 | 4580 | 4l0d | B | 191 | THR | p.Thr191Met | P35520 |
| HOMO SAPIENS | 5 | 4580 | 4l0d | B | 234 | ASP | p.Asp234Asn | P35520 |
| HOMO SAPIENS | 0 | 4580 | 4l0d | B | 257 | THR | p.Thr257Met | P35520 |
| HOMO SAPIENS | 4 | 4580 | 4l0d | B | 262 | THR | p.Thr262Met | P35520 |
| HOMO SAPIENS | 0 | 4580 | 4l0d | B | 278 | ILE | p.Ile278Thr | P35520 |
| HOMO SAPIENS | 2 | 4580 | 4l0d | B | 307 | GLY | p.Gly307Ser | P35520 |
| HOMO SAPIENS | 4 | 4580 | 4l0d | B | 320 | VAL | p.Val320Ala | P35520 |
| HOMO SAPIENS | 2 | 4580 | 4l0d | B | 336 | ARG | p.Arg336Cys | P35520 |
| HOMO SAPIENS | 2 | 4580 | 4l0d | B | 336 | ARG | p.Arg336His | P35520 |
| HOMO SAPIENS | 0 | 4580 | 4l0d | B | 351 | GLY | p.Gly351Arg | P35520 |
| HOMO SAPIENS | 0 | 4580 | 4l0d | B | 353 | THR | p.Thr353Met | P35520 |
| HOMO SAPIENS | 0 | 4580 | 4l0d | B | 371 | VAL | p.Val371Met | P35520 |

| | | | | | | | | |
|---|---|---|---|---|---|---|---|---|
| HOMO SAPIENS | 2 | 4580 | 4l0d | B | 379 | ARG | p.Arg379Gln | P35520 |
| HOMO SAPIENS | 2 | 4580 | 4l0d | B | 379 | ARG | p.Arg379Trp | P35520 |
| HOMO SAPIENS | 2 | 4580 | 4l0d | B | 384 | LYS | p.Lys384Glu | P35520 |
| HOMO SAPIENS | 0 | 4580 | 4l0d | B | 422 | PRO | p.Pro422Leu | P35520 |
| HOMO SAPIENS | 0 | 4580 | 4l0d | B | 444 | ASP | p.Asp444Asn | P35520 |
| HOMO SAPIENS | 0 | 4580 | 4l0d | B | 466 | SER | p.Ser466Leu | P35520 |
| HOMO SAPIENS | 0 | 4580 | 4l0d | B | 539 | LEU | p.Leu539Ser | P35520 |
| HOMO SAPIENS | 0 | 4580 | 4l0d | B | 548 | ARG | p.Arg548Gln | P35520 |
| HOMO SAPIENS | 1 | 6536 | 3ulu | A | 412 | LEU | p.Leu412Phe | O15455 |
| HOMO SAPIENS | 1 | 28493 | 3ii6 | X | 814 | ARG | p.Arg814Ter | P49917 |
| HOMO SAPIENS | 1 | 28493 | 3ii6 | X | 857 | ALA | p.Ala857Thr | P49917 |
| HOMO SAPIENS | 9 | 3543 | 2q51 | A | 24 | GLU | p.Glu24Gly | P45381 |
| HOMO SAPIENS | 1 | 3543 | 2q51 | A | 27 | GLY | p.Gly27Arg | P45381 |
| HOMO SAPIENS | 9 | 3543 | 2q51 | A | 71 | ARG | p.Arg71His | P45381 |
| HOMO SAPIENS | -1 | 3543 | 2q51 | A | 82 | MSE | p.Met82Thr | P45381 |
| HOMO SAPIENS | 0 | 3543 | 2q51 | A | 143 | ILE | p.Ile143Phe | P45381 |
| HOMO SAPIENS | 1 | 3543 | 2q51 | A | 152 | CYS | p.Cys152Arg | P45381 |
| HOMO SAPIENS | 2 | 3543 | 2q51 | A | 218 | CYS | p.Cys218Ter | P45381 |
| HOMO SAPIENS | 1 | 3543 | 2q51 | A | 231 | TYR | p.Tyr231Cys | P45381 |
| HOMO SAPIENS | 1 | 3543 | 2q51 | A | 231 | TYR | p.Tyr231Ter | P45381 |
| HOMO SAPIENS | 2 | 3543 | 2q51 | A | 274 | GLY | p.Gly274Arg | P45381 |
| HOMO SAPIENS | 6 | 3543 | 2q51 | A | 285 | GLU | p.Glu285Ala | P45381 |
| HOMO SAPIENS | 5 | 3543 | 2q51 | A | 287 | ALA | p.Ala287Thr | P45381 |
| HOMO SAPIENS | 2 | 3543 | 2q51 | A | 305 | ALA | p.Ala305Glu | P45381 |
| HOMO SAPIENS | 2 | 24687 | 1hfi | A | 25 | SER | p.Ser890Ile | P08603 |
| HOMO SAPIENS | 0 | 1508 | 3wn5 | C | 45 | LEU | p.Leu66Arg | P08637 |
| HOMO SAPIENS | 0 | 1508 | 3wn5 | C | 155 | VAL | p.Phe176Val | P08637 |
| HOMO SAPIENS | 0 | 750 | 5tpa | A | 54 | VAL | p.Val452Met | Q12879 |
| HOMO SAPIENS | 9 | 750 | 5tpa | A | 120 | ARG | p.Arg518Cys | Q12879 |
| HOMO SAPIENS | 0 | 750 | 5tpa | A | 199 | ALA | p.Ala716Thr | Q12879 |
| HOMO SAPIENS | 4 | 1363 | 3hg2 | B | 40 | PRO | p.Pro40Ser | P06280 |
| HOMO SAPIENS | 3 | 1363 | 3hg2 | B | 65 | SER | p.Ser65Thr | P06280 |
| HOMO SAPIENS | 4 | 1363 | 3hg2 | B | 66 | GLU | p.Glu66Gln | P06280 |
| HOMO SAPIENS | 2 | 1363 | 3hg2 | B | 80 | GLY | p.Gly80Asp | P06280 |
| HOMO SAPIENS | 3 | 1363 | 3hg2 | B | 112 | ARG | p.Arg112Cys | P06280 |
| HOMO SAPIENS | 3 | 1363 | 3hg2 | B | 112 | ARG | p.Arg112His | P06280 |
| HOMO SAPIENS | 0 | 1363 | 3hg2 | B | 139 | ASN | p.Asn139Ser | P06280 |
| HOMO SAPIENS | 7 | 1363 | 3hg2 | B | 156 | ALA | p.Ala156Thr | P06280 |
| HOMO SAPIENS | 5 | 1363 | 3hg2 | B | 162 | TRP | p.Trp162Arg | P06280 |
| HOMO SAPIENS | 5 | 1363 | 3hg2 | B | 207 | TYR | p.Tyr207His | P06280 |
| HOMO SAPIENS | 0 | 1363 | 3hg2 | B | 215 | ASN | p.Asn215Ser | P06280 |
| HOMO SAPIENS | 0 | 1363 | 3hg2 | B | 264 | ASP | p.Asp264Val | P06280 |
| HOMO SAPIENS | 5 | 1363 | 3hg2 | B | 266 | ASP | p.Asp266Val | P06280 |
| HOMO SAPIENS | 2 | 1363 | 3hg2 | B | 269 | VAL | p.Val269Ala | P06280 |
| HOMO SAPIENS | 3 | 1363 | 3hg2 | B | 272 | ASN | p.Asn272Ser | P06280 |
| HOMO SAPIENS | 3 | 1363 | 3hg2 | B | 279 | GLN | p.Gln279Glu | P06280 |
| HOMO SAPIENS | 0 | 1363 | 3hg2 | B | 289 | ILE | p.Ile289Val | P06280 |
| HOMO SAPIENS | 0 | 1363 | 3hg2 | B | 296 | MET | p.Met296Val | P06280 |
| HOMO SAPIENS | 0 | 1363 | 3hg2 | B | 296 | MET | p.Met296Ile | P06280 |
| HOMO SAPIENS | 0 | 1363 | 3hg2 | B | 297 | SER | p.Ser297Phe | P06280 |
| HOMO SAPIENS | 2 | 1363 | 3hg2 | B | 301 | ARG | p.Arg301Gln | P06280 |
| HOMO SAPIENS | 3 | 1363 | 3hg2 | B | 327 | GLN | p.Gln327Lys | P06280 |
| HOMO SAPIENS | 0 | 1363 | 3hg2 | B | 328 | GLY | p.Gly328Arg | P06280 |
| HOMO SAPIENS | 0 | 1363 | 3hg2 | B | 328 | GLY | p.Gly328Ala | P06280 |

| | | | | | | | | |
|---|---|---|---|---|---|---|---|---|
| HOMO SAPIENS | 2 | 1363 | 3hg2 | B | 331 | LEU | p.Leu331Phe | P06280 |
| HOMO SAPIENS | 0 | 1363 | 3hg2 | B | 342 | ARG | p.Arg342Gln | P06280 |
| HOMO SAPIENS | 1 | 1363 | 3hg2 | B | 361 | GLY | p.Gly361Arg | P06280 |
| HOMO SAPIENS | 1 | 1363 | 3hg2 | B | 363 | ARG | p.Arg363His | P06280 |
| HOMO SAPIENS | 1 | 1363 | 3hg2 | B | 385 | THR | p.Thr385Ala | P06280 |
| HOMO SAPIENS | 1 | 1363 | 3hg2 | B | 399 | TRP | p.Trp399Ser | P06280 |
| HOMO SAPIENS | 1 | 1363 | 3hg2 | B | 410 | THR | p.Thr410Ala | P06280 |
| HOMO SAPIENS | 2 | 3904 | 3caa | B | 385 | MET | p.Met414Val | P01011 |
| HOMO SAPIENS | 0 | 558 | 5eub | A | 56 | VAL | p.Val56Ile | Q13126 |
| HOMO SAPIENS | 0 | 558 | 5eub | A | 189 | TRP | p.Trp189Leu | Q13126 |
| HOMO SAPIENS | 0 | 558 | 5eub | A | 218 | ALA | p.Ala218Gly | Q13126 |
| HOMO SAPIENS | 1 | 3865 | 1cb5 | B | 443 | ILE | p.Ile443Val | Q13867 |
| HOMO SAPIENS | 2 | 14329 | 4m92 | A | 24 | ILE | p.Ile65Val | Q13454 |
| HOMO SAPIENS | 1 | 7381 | 5kz5 | k | 50 | GLY | p.Gly50Glu | Q9H1K1 |
| HOMO SAPIENS | 0 | 1102 | 4fdh | E | 86 | PRO | p.Pro86Ala | P19099 |
| HOMO SAPIENS | 0 | 1102 | 4fdh | E | 118 | ALA | p.Ala118Thr | P19099 |
| HOMO SAPIENS | 0 | 1102 | 4fdh | E | 159 | PRO | p.Pro159Leu | P19099 |
| HOMO SAPIENS | 0 | 1102 | 4fdh | E | 173 | LYS | p.Lys173Arg | P19099 |
| HOMO SAPIENS | 0 | 1102 | 4fdh | E | 181 | ARG | p.Arg181Trp | P19099 |
| HOMO SAPIENS | 0 | 1102 | 4fdh | E | 185 | THR | p.Thr185Ile | P19099 |
| HOMO SAPIENS | 0 | 1102 | 4fdh | E | 198 | GLU | p.Glu198Asp | P19099 |
| HOMO SAPIENS | 0 | 1102 | 4fdh | E | 214 | HIS | p.His214Asp | P19099 |
| HOMO SAPIENS | 0 | 1102 | 4fdh | E | 251 | LYS | p.Lys251Arg | P19099 |
| HOMO SAPIENS | 0 | 1102 | 4fdh | E | 255 | GLU | p.Glu255Ter | P19099 |
| HOMO SAPIENS | 0 | 1102 | 4fdh | E | 281 | ASN | p.Asn281Ser | P19099 |
| HOMO SAPIENS | 0 | 1102 | 4fdh | E | 320 | ALA | p.Ala320Val | P19099 |
| HOMO SAPIENS | 0 | 1102 | 4fdh | E | 339 | ILE | p.Ile339Thr | P19099 |
| HOMO SAPIENS | 0 | 1102 | 4fdh | E | 341 | ARG | p.Arg341Cys | P19099 |
| HOMO SAPIENS | 2 | 1102 | 4fdh | E | 374 | ARG | p.Arg374Trp | P19099 |
| HOMO SAPIENS | 0 | 1102 | 4fdh | E | 386 | VAL | p.Val386Ala | P19099 |
| HOMO SAPIENS | 0 | 1102 | 4fdh | E | 390 | LEU | p.Leu390Phe | P19099 |
| HOMO SAPIENS | -1 | 1102 | 4fdh | E | 435 | GLY | p.Gly435Ser | P19099 |
| HOMO SAPIENS | 0 | 1102 | 4fdh | E | 461 | LEU | p.Leu461Pro | P19099 |
| HOMO SAPIENS | 0 | 1102 | 4fdh | E | 498 | THR | p.Thr498Ala | P19099 |
| HOMO SAPIENS | 6 | 10292 | 2of5 | F | 128 | GLY | p.Gly128Arg | P78560 |
| HOMO SAPIENS | 6 | 10292 | 2of5 | F | 170 | ARG | p.Arg170Cys | P78560 |
| HOMO SAPIENS | -1 | 18965 | 3ghg | A | 16 | ARG | p.Arg35Cys | P02671 |
| HOMO SAPIENS | -1 | 18965 | 3ghg | A | 16 | ARG | p.Arg35His | P02671 |
| HOMO SAPIENS | -1 | 18965 | 3ghg | A | 283 | PRO | p.Pro302Ala | P02671 |
| HOMO SAPIENS | -1 | 18965 | 3ghg | A | 312 | THR | p.Thr331Ala | P02671 |
| HOMO SAPIENS | -1 | 18965 | 3ghg | A | 381 | SER | p.Ser400Phe | P02671 |
| HOMO SAPIENS | -1 | 18965 | 3ghg | A | 463 | VAL | p.Val482Met | P02671 |
| HOMO SAPIENS | -1 | 18965 | 3ghg | A | 526 | GLU | p.Glu545Val | P02671 |
| HOMO SAPIENS | -1 | 18965 | 3ghg | A | 554 | ARG | p.Arg573Leu | P02671 |
| HOMO SAPIENS | 8 | 3023 | 4ttb | A | 101 | ARG | p.Arg101Trp | Q6PHW0 |
| HOMO SAPIENS | 2 | 3023 | 4ttb | A | 116 | ILE | p.Ile116Thr | Q6PHW0 |
| HOMO SAPIENS | 0 | 3023 | 4ttb | A | 220 | ALA | p.Ala220Thr | Q6PHW0 |
| HOMO SAPIENS | 2 | 3023 | 4ttb | A | 271 | GLU | p.Glu271Lys | Q6PHW0 |
| HOMO SAPIENS | 3 | 33226 | 1ya5 | T | 11 | SER | p.Ser11Ter | O15273 |
| HOMO SAPIENS | 6 | 33226 | 1ya5 | T | 64 | SER | p.Ser64Leu | O15273 |
| HOMO SAPIENS | 6 | 33226 | 1ya5 | T | 70 | ARG | p.Arg70Gln | O15273 |
| HOMO SAPIENS | 3 | 33226 | 1ya5 | T | 75 | GLY | p.Gly75Ser | O15273 |
| HOMO SAPIENS | 0 | 33226 | 1ya5 | T | 87 | ARG | p.Arg87Trp | O15273 |
| HOMO SAPIENS | 2 | 5667 | 3dax | A | 398 | PRO | p.Pro398Ala | P22680 |

| | | | | | | | | |
|---|---|---|---|---|---|---|---|---|
| HOMO SAPIENS | 0 | 12861 | 5mya | A | 464 | ASN | p.Asn464His | Q02763 |
| HOMO SAPIENS | 2 | 12861 | 5mya | A | 486 | VAL | p.Val486Ile | Q02763 |
| HOMO SAPIENS | 0 | 12861 | 5mya | A | 522 | ARG | p.Arg522His | Q02763 |
| HOMO SAPIENS | 2 | 12861 | 5mya | A | 600 | VAL | p.Val600Leu | Q02763 |
| HOMO SAPIENS | 0 | 12861 | 5mya | A | 634 | LEU | p.Leu634Phe | Q02763 |
| HOMO SAPIENS | 0 | 12861 | 5mya | A | 684 | HIS | p.His684Gln | Q02763 |
| HOMO SAPIENS | 0 | 12861 | 5mya | A | 724 | ALA | p.Ala724Thr | Q02763 |
| HOMO SAPIENS | 0 | 7743 | 3qzw | G | 90 | GLY | p.Gly111Ser | P01732 |
| HOMO SAPIENS | 2 | 9498 | 3kci | A | 4267 | ASP | p.Asp4267Glu | O95714 |
| HOMO SAPIENS | 5 | 201 | 3ips | B | 415 | ARG | p.Arg415Gln | Q13133 |
| HOMO SAPIENS | 3 | 229 | 2q50 | A | 68 | LEU | p.Leu68Pro | Q9UBQ7 |
| HOMO SAPIENS | 2 | 229 | 2q50 | A | 99 | ARG | p.Arg99Ter | Q9UBQ7 |
| HOMO SAPIENS | 3 | 229 | 2q50 | A | 160 | GLY | p.Gly160Arg | Q9UBQ7 |
| HOMO SAPIENS | 5 | 229 | 2q50 | A | 165 | GLY | p.Gly165Asp | Q9UBQ7 |
| HOMO SAPIENS | 4 | 229 | 2q50 | A | 302 | ARG | p.Arg302Cys | Q9UBQ7 |
| HOMO SAPIENS | 4 | 229 | 2q50 | A | 302 | ARG | p.Arg302His | Q9UBQ7 |
| HOMO SAPIENS | -1 | 229 | 2q50 | A | 322 | MSE | p.Met322Arg | Q9UBQ7 |
| HOMO SAPIENS | -1 | 4519 | 5clw | A | 40 | PRO | p.Pro40Thr | Q04446 |
| HOMO SAPIENS | 1 | 4519 | 5clw | A | 59 | ILE | p.Ile59Thr | Q04446 |
| HOMO SAPIENS | 0 | 4519 | 5clw | A | 190 | ARG | p.Arg190Gly | Q04446 |
| HOMO SAPIENS | 1 | 4519 | 5clw | A | 224 | LEU | p.Leu224Pro | Q04446 |
| HOMO SAPIENS | 0 | 4519 | 5clw | A | 236 | GLN | p.Gln236His | Q04446 |
| HOMO SAPIENS | 1 | 4519 | 5clw | A | 254 | THR | p.Thr254Ala | Q04446 |
| HOMO SAPIENS | 0 | 4519 | 5clw | A | 257 | PHE | p.Phe257Leu | Q04446 |
| HOMO SAPIENS | 1 | 4519 | 5clw | A | 262 | ARG | p.Arg262Cys | Q04446 |
| HOMO SAPIENS | 3 | 4519 | 5clw | A | 329 | TYR | p.Tyr329Ser | Q04446 |
| HOMO SAPIENS | 0 | 4519 | 5clw | A | 507 | THR | p.Thr507Ala | Q04446 |
| HOMO SAPIENS | 0 | 4519 | 5clw | A | 515 | ARG | p.Arg515Cys | Q04446 |
| HOMO SAPIENS | 0 | 4519 | 5clw | A | 515 | ARG | p.Arg515His | Q04446 |
| HOMO SAPIENS | 0 | 4519 | 5clw | A | 524 | ARG | p.Arg524Gln | Q04446 |
| HOMO SAPIENS | 0 | 4519 | 5clw | A | 524 | ARG | p.Arg524Ter | Q04446 |
| HOMO SAPIENS | 0 | 4519 | 5clw | A | 545 | HIS | p.His545Arg | Q04446 |
| HOMO SAPIENS | 2 | 4519 | 5clw | A | 548 | TRP | p.Trp548Ter | Q04446 |
| HOMO SAPIENS | 0 | 4519 | 5clw | A | 592 | GLU | p.Glu592Ter | Q04446 |
| HOMO SAPIENS | 0 | 4519 | 5clw | A | 628 | HIS | p.His628Arg | Q04446 |
| HOMO SAPIENS | 0 | 4519 | 5clw | A | 637 | ARG | p.Arg637Ter | Q04446 |
| HOMO SAPIENS | 0 | 4519 | 5clw | A | 673 | ALA | p.Ala673Thr | Q04446 |
| HOMO SAPIENS | 1 | 10405 | 3bg3 | D | 507 | VAL | p.Val507Ile | P11498 |
| HOMO SAPIENS | 0 | 10405 | 3bg3 | D | 583 | ARG | p.Arg583Leu | P11498 |
| HOMO SAPIENS | 1 | 10405 | 3bg3 | D | 589 | LYS | p.Lys589Asn | P11498 |
| HOMO SAPIENS | 6 | 10405 | 3bg3 | D | 610 | ALA | p.Ala610Thr | P11498 |
| HOMO SAPIENS | 2 | 10405 | 3bg3 | D | 631 | ARG | p.Arg631Gln | P11498 |
| HOMO SAPIENS | 1 | 10405 | 3bg3 | D | 732 | ARG | p.Arg732Gln | P11498 |
| HOMO SAPIENS | 6 | 10405 | 3bg3 | D | 743 | MET | p.Met743Ile | P11498 |
| HOMO SAPIENS | 0 | 10405 | 3bg3 | D | 750 | THR | p.Thr750Met | P11498 |
| HOMO SAPIENS | -1 | 13751 | 2o8d | B | 350 | ALA | p.Ala350Val | P52701 |
| HOMO SAPIENS | -1 | 13751 | 2o8d | B | 352 | VAL | p.Val352Ile | P52701 |
| HOMO SAPIENS | -1 | 13751 | 2o8d | B | 355 | GLY | p.Gly355Ser | P52701 |
| HOMO SAPIENS | 0 | 13751 | 2o8d | B | 385 | ARG | p.Arg385Trp | P52701 |
| HOMO SAPIENS | 0 | 13751 | 2o8d | B | 390 | ASP | p.Asp390Glu | P52701 |
| HOMO SAPIENS | 0 | 13751 | 2o8d | B | 396 | LEU | p.Leu396Val | P52701 |
| HOMO SAPIENS | 0 | 13751 | 2o8d | B | 449 | LEU | p.Leu449Pro | P52701 |
| HOMO SAPIENS | 0 | 13751 | 2o8d | B | 468 | ARG | p.Arg468His | P52701 |
| HOMO SAPIENS | 0 | 13751 | 2o8d | B | 503 | SER | p.Ser503Cys | P52701 |

| | | | | | | | | |
|---|---|---|---|---|---|---|---|---|
| HOMO SAPIENS | 1 | 13751 | 2o8d | B | 509 | VAL | p.Val509Ala | P52701 |
| HOMO SAPIENS | 0 | 13751 | 2o8d | B | 675 | GLU | p.Glu675Asp | P52701 |
| HOMO SAPIENS | 0 | 13751 | 2o8d | B | 795 | ILE | p.Ile795Thr | P52701 |
| HOMO SAPIENS | 0 | 13751 | 2o8d | B | 850 | TYR | p.Tyr850Ter | P52701 |
| HOMO SAPIENS | 0 | 13751 | 2o8d | B | 878 | VAL | p.Val878Ala | P52701 |
| HOMO SAPIENS | 0 | 13751 | 2o8d | B | 911 | ARG | p.Arg911Ter | P52701 |
| HOMO SAPIENS | 0 | 13751 | 2o8d | B | 922 | ARG | p.Arg922Ter | P52701 |
| HOMO SAPIENS | 0 | 13751 | 2o8d | B | 969 | TYR | p.Tyr969Cys | P52701 |
| HOMO SAPIENS | 0 | 13751 | 2o8d | B | 1005 | ARG | p.Arg1005Ter | P52701 |
| HOMO SAPIENS | 0 | 13751 | 2o8d | B | 1035 | ARG | p.Arg1035Ter | P52701 |
| HOMO SAPIENS | 1 | 13751 | 2o8d | B | 1048 | GLN | p.Gln1048Ter | P52701 |
| HOMO SAPIENS | 1 | 13751 | 2o8d | B | 1051 | VAL | p.Val1051Ile | P52701 |
| HOMO SAPIENS | 0 | 13751 | 2o8d | B | 1054 | ILE | p.Ile1054Phe | P52701 |
| HOMO SAPIENS | 0 | 13751 | 2o8d | B | 1054 | ILE | p.Ile1054Met | P52701 |
| HOMO SAPIENS | 0 | 13751 | 2o8d | B | 1163 | GLU | p.Glu1163Val | P52701 |
| HOMO SAPIENS | 0 | 13751 | 2o8d | B | 1237 | GLU | p.Glu1237Asp | P52701 |
| HOMO SAPIENS | 0 | 13751 | 2o8d | B | 1242 | ARG | p.Arg1242His | P52701 |
| HOMO SAPIENS | -1 | 13751 | 2o8d | B | 1278 | PRO | p.Pro1278Thr | P52701 |
| HOMO SAPIENS | 0 | 13751 | 2o8d | B | 1304 | ARG | p.Arg1304Lys | P52701 |
| HOMO SAPIENS | 0 | 13751 | 2o8d | B | 1317 | HIS | p.His1317Gln | P52701 |
| HOMO SAPIENS | 0 | 13751 | 2o8d | B | 1329 | SER | p.Ser1329Leu | P52701 |
| HOMO SAPIENS | 3 | 13273 | 4g8k | A | 265 | GLU | p.Glu265Ter | Q05823 |
| HOMO SAPIENS | -1 | 11088 | 5uj7 | A | 666 | ARG | p.Arg666Trp | Q13415 |
| HOMO SAPIENS | 4 | 11088 | 5uj7 | A | 720 | ARG | p.Arg720Gln | Q13415 |
| HOMO SAPIENS | 0 | 11088 | 5uj7 | A | 816 | MET | p.Met816Thr | Q13415 |
| HOMO SAPIENS | 1 | 4352 | 5va1 | A | 421 | THR | p.Thr421Met | Q12809 |
| HOMO SAPIENS | 0 | 4352 | 5va1 | A | 451 | PRO | p.Pro451Leu | Q12809 |
| HOMO SAPIENS | 0 | 4352 | 5va1 | A | 470 | ASN | p.Asn470Asp | Q12809 |
| HOMO SAPIENS | 0 | 4352 | 5va1 | A | 490 | ALA | p.Ala490Thr | Q12809 |
| HOMO SAPIENS | 0 | 4352 | 5va1 | A | 491 | VAL | p.Val491Ile | Q12809 |
| HOMO SAPIENS | 1 | 4352 | 5va1 | A | 533 | VAL | p.Val533Gly | Q12809 |
| HOMO SAPIENS | 0 | 4352 | 5va1 | A | 552 | LEU | p.Leu552Ser | Q12809 |
| HOMO SAPIENS | 0 | 4352 | 5va1 | A | 558 | ALA | p.Ala558Pro | Q12809 |
| HOMO SAPIENS | 1 | 4352 | 5va1 | A | 561 | ALA | p.Ala561Val | Q12809 |
| HOMO SAPIENS | 0 | 4352 | 5va1 | A | 572 | GLY | p.Gly572Arg | Q12809 |
| HOMO SAPIENS | -1 | 4352 | 5va1 | A | 582 | ARG | p.Arg582Cys | Q12809 |
| HOMO SAPIENS | 0 | 4352 | 5va1 | A | 584 | GLY | p.Gly584Ser | Q12809 |
| HOMO SAPIENS | 0 | 4352 | 5va1 | A | 588 | ASN | p.Asn588Lys | Q12809 |
| HOMO SAPIENS | 0 | 4352 | 5va1 | A | 593 | ILE | p.Ile593Arg | Q12809 |
| HOMO SAPIENS | 1 | 4352 | 5va1 | A | 627 | PHE | p.Phe627Leu | Q12809 |
| HOMO SAPIENS | 0 | 4352 | 5va1 | A | 630 | VAL | p.Val630Leu | Q12809 |
| HOMO SAPIENS | 0 | 4352 | 5va1 | A | 652 | TYR | p.Tyr652Ter | Q12809 |
| HOMO SAPIENS | 0 | 4352 | 5va1 | A | 711 | ILE | p.Ile711Val | Q12809 |
| HOMO SAPIENS | 0 | 4352 | 5va1 | A | 735 | SER | p.Ser735Leu | Q12809 |
| HOMO SAPIENS | 0 | 4352 | 5va1 | A | 744 | ARG | p.Arg744Ter | Q12809 |
| HOMO SAPIENS | 0 | 4352 | 5va1 | A | 784 | ARG | p.Arg784Trp | Q12809 |
| HOMO SAPIENS | 1 | 4352 | 5va1 | A | 820 | GLY | p.Gly820Arg | Q12809 |
| HOMO SAPIENS | 5 | 4352 | 5va1 | A | 822 | VAL | p.Val822Met | Q12809 |
| HOMO SAPIENS | 1 | 4352 | 5va1 | A | 863 | ARG | p.Arg863Ter | Q12809 |
| HOMO SAPIENS | 5 | 303 | 2obv | A | 55 | ALA | p.Ala55Asp | Q00266 |
| HOMO SAPIENS | 7 | 303 | 2obv | A | 259 | ALA | p.Ala259Gly | Q00266 |
| HOMO SAPIENS | 9 | 303 | 2obv | A | 264 | ARG | p.Arg264His | Q00266 |
| HOMO SAPIENS | 9 | 303 | 2obv | A | 264 | ARG | p.Arg264Cys | Q00266 |
| HOMO SAPIENS | 0 | 303 | 2obv | A | 305 | LEU | p.Leu305Pro | Q00266 |

| | | | | | | | | |
|---|---|---|---|---|---|---|---|---|
| HOMO SAPIENS | 7 | 303 | 2obv | A | 322 | ILE | p.Ile322Met | Q00266 |
| HOMO SAPIENS | 0 | 303 | 2obv | A | 336 | GLY | p.Gly336Arg | Q00266 |
| HOMO SAPIENS | 0 | 303 | 2obv | A | 357 | PRO | p.Pro357Leu | Q00266 |
| HOMO SAPIENS | 0 | 2077 | 4zxb | E | 462 | ASN | p.Asn489Ser | P06213 |
| HOMO SAPIENS | -1 | 2077 | 4zxb | E | 519 | ASP | p.Asp546Glu | P06213 |
| HOMO SAPIENS | 2 | 2077 | 4zxb | E | 819 | THR | p.Thr858Ala | P06213 |
| HOMO SAPIENS | 0 | 2077 | 4zxb | E | 907 | ASP | p.Asp946Glu | P06213 |
| HOMO SAPIENS | 7 | 3274 | 1wsv | B | 14 | HIS | p.His42Arg | P48728 |
| HOMO SAPIENS | 0 | 3274 | 1wsv | B | 19 | GLY | p.Gly47Arg | P48728 |
| HOMO SAPIENS | 4 | 3274 | 1wsv | B | 45 | ARG | p.Arg73Cys | P48728 |
| HOMO SAPIENS | 1 | 3274 | 1wsv | B | 49 | SER | p.Ser77Leu | P48728 |
| HOMO SAPIENS | 3 | 3274 | 1wsv | B | 241 | GLY | p.Gly269Asp | P48728 |
| HOMO SAPIENS | 7 | 3274 | 1wsv | B | 248 | ASP | p.Asp276His | P48728 |
| HOMO SAPIENS | 0 | 3274 | 1wsv | B | 272 | MET | p.Met300Val | P48728 |
| HOMO SAPIENS | 4 | 3274 | 1wsv | B | 292 | ARG | p.Arg320His | P48728 |
| HOMO SAPIENS | 0 | 3274 | 1wsv | B | 354 | ARG | p.Arg382Gln | P48728 |
| HOMO SAPIENS | -1 | 435 | 3uyt | A | 44 | THR | p.Thr44Ala | P48730 |
| HOMO SAPIENS | 1 | 4698 | 5ife | C | 666 | LYS | p.Lys666Asn | O75533 |
| HOMO SAPIENS | 1 | 16917 | 1apy | C | 37 | GLY | p.Gly60Asp | P20933 |
| HOMO SAPIENS | 0 | 16917 | 1apy | C | 49 | SER | p.Ser72Pro | P20933 |
| HOMO SAPIENS | 0 | 16917 | 1apy | C | 84 | ARG | p.Arg107Ter | P20933 |
| HOMO SAPIENS | 0 | 16917 | 1apy | C | 126 | SER | p.Thr149Ser | P20933 |
| HOMO SAPIENS | 0 | 16917 | 1apy | C | 135 | TRP | p.Trp158Ter | P20933 |
| HOMO SAPIENS | 0 | 16917 | 1apy | C | 140 | CYS | p.Cys163Ser | P20933 |
| HOMO SAPIENS | 0 | 16917 | 1apy | C | 145 | TRP | p.Trp168Ter | P20933 |
| HOMO SAPIENS | 0 | 10413 | 4frw | A | 185 | THR | p.Thr185Met | Q96NY8 |
| HOMO SAPIENS | 0 | 5217 | 5n6w | G | 96 | TRP | p.Trp96Arg | O15537 |
| HOMO SAPIENS | 1 | 5217 | 5n6w | G | 102 | ARG | p.Arg102Trp | O15537 |
| HOMO SAPIENS | 0 | 5217 | 5n6w | G | 109 | GLY | p.Gly109Arg | O15537 |
| HOMO SAPIENS | -1 | 5217 | 5n6w | G | 223 | CYS | p.Cys223Arg | O15537 |
| HOMO SAPIENS | 0 | 9485 | 4z6y | E | 138 | ALA | p.Ala138Asp | Q9P0N9 |
| HOMO SAPIENS | 3 | 4504 | 4mwt | B | 411 | GLY | p.Gly439Ser | P10619 |
| HOMO SAPIENS | 3 | 4504 | 4mwt | B | 412 | PHE | p.Phe440Val | P10619 |
| HOMO SAPIENS | 6 | 6391 | 4kxy | A | 318 | ARG | p.Arg318Cys | P29401 |
| HOMO SAPIENS | 6 | 19546 | 2bx6 | A | 87 | THR | p.Thr87Ile | O75695 |
| HOMO SAPIENS | 1 | 19546 | 2bx6 | A | 118 | ARG | p.Arg118His | O75695 |
| HOMO SAPIENS | 1 | 19546 | 2bx6 | A | 201 | ILE | p.Ile201Val | O75695 |
| HOMO SAPIENS | 1 | 19546 | 2bx6 | A | 282 | ARG | p.Arg282Trp | O75695 |
| HOMO SAPIENS | 1 | 5265 | 5usr | H | 68 | ARG | p.Arg68Leu | Q9HD34 |
| HOMO SAPIENS | 2 | 1012 | 4nyi | S | 569 | LEU | p.Leu569Val | Q07889 |
| HOMO SAPIENS | 2 | 1012 | 4nyi | S | 655 | PRO | p.Pro655Leu | Q07889 |
| HOMO SAPIENS | 4 | 1012 | 4nyi | S | 663 | ILE | p.Ile663Met | Q07889 |
| HOMO SAPIENS | 2 | 1012 | 4nyi | S | 702 | TYR | p.Tyr702Cys | Q07889 |
| HOMO SAPIENS | 2 | 1012 | 4nyi | S | 708 | ALA | p.Ala708Thr | Q07889 |
| HOMO SAPIENS | 4 | 1012 | 4nyi | S | 733 | ILE | p.Ile733Phe | Q07889 |
| HOMO SAPIENS | 1 | 1848 | 3g1n | A | 4013 | ARG | p.Arg4013Trp | Q7Z6Z7 |
| HOMO SAPIENS | 1 | 1848 | 3g1n | A | 4187 | ARG | p.Arg4187Cys | Q7Z6Z7 |
| HOMO SAPIENS | 1 | 77 | 4pod | A | 41 | CYS | p.Cys79Tyr | P60174 |
| HOMO SAPIENS | 9 | 77 | 4pod | A | 104 | GLU | p.Glu142Asp | P60174 |
| HOMO SAPIENS | 1 | 77 | 4pod | A | 122 | GLY | p.Gly160Arg | P60174 |
| HOMO SAPIENS | -1 | 8672 | 3thz | A | 8 | THR | p.Thr8Met | P43246 |
| HOMO SAPIENS | 0 | 8672 | 3thz | A | 19 | PHE | p.Phe19Leu | P43246 |
| HOMO SAPIENS | 0 | 8672 | 3thz | A | 23 | PHE | p.Phe23Leu | P43246 |
| HOMO SAPIENS | 0 | 8672 | 3thz | A | 96 | ARG | p.Arg96His | P43246 |

| | | | | | | | | |
|---|---|---|---|---|---|---|---|---|
| HOMO SAPIENS | 0 | 8672 | 3thz | A | 102 | VAL | p.Val102Ile | P43246 |
| HOMO SAPIENS | 0 | 8672 | 3thz | A | 106 | ARG | p.Arg106Lys | P43246 |
| HOMO SAPIENS | 0 | 8672 | 3thz | A | 127 | ASN | p.Asn127Ser | P43246 |
| HOMO SAPIENS | 0 | 8672 | 3thz | A | 128 | LEU | p.Leu128Val | P43246 |
| HOMO SAPIENS | 0 | 8672 | 3thz | A | 163 | VAL | p.Val163Gly | P43246 |
| HOMO SAPIENS | 0 | 8672 | 3thz | A | 169 | ILE | p.Ile169Val | P43246 |
| HOMO SAPIENS | 0 | 8672 | 3thz | A | 315 | GLY | p.Gly315Asp | P43246 |
| HOMO SAPIENS | -1 | 8672 | 3thz | A | 322 | GLY | p.Gly322Asp | P43246 |
| HOMO SAPIENS | 0 | 8672 | 3thz | A | 331 | ASN | p.Asn331Asp | P43246 |
| HOMO SAPIENS | 1 | 8672 | 3thz | A | 390 | LEU | p.Leu390Phe | P43246 |
| HOMO SAPIENS | 0 | 8672 | 3thz | A | 419 | GLN | p.Gln419Lys | P43246 |
| HOMO SAPIENS | 0 | 8672 | 3thz | A | 447 | PHE | p.Phe447Leu | P43246 |
| HOMO SAPIENS | 0 | 8672 | 3thz | A | 524 | ARG | p.Arg524Pro | P43246 |
| HOMO SAPIENS | 0 | 8672 | 3thz | A | 539 | PHE | p.Phe539Leu | P43246 |
| HOMO SAPIENS | -1 | 8672 | 3thz | A | 546 | LYS | p.Lys546Asn | P43246 |
| HOMO SAPIENS | 0 | 8672 | 3thz | A | 564 | THR | p.Thr564Ala | P43246 |
| HOMO SAPIENS | 0 | 8672 | 3thz | A | 577 | ILE | p.Ile577Thr | P43246 |
| HOMO SAPIENS | 0 | 8672 | 3thz | A | 588 | TYR | p.Tyr588Ter | P43246 |
| HOMO SAPIENS | 0 | 8672 | 3thz | A | 622 | PRO | p.Pro622Leu | P43246 |
| HOMO SAPIENS | 0 | 8672 | 3thz | A | 629 | GLN | p.Gln629Arg | P43246 |
| HOMO SAPIENS | 0 | 8672 | 3thz | A | 636 | ALA | p.Ala636Pro | P43246 |
| HOMO SAPIENS | 2 | 8672 | 3thz | A | 642 | VAL | p.Val642Ile | P43246 |
| HOMO SAPIENS | 1 | 8672 | 3thz | A | 654 | ASP | p.Asp654Glu | P43246 |
| HOMO SAPIENS | 0 | 8672 | 3thz | A | 662 | GLN | p.Gln662His | P43246 |
| HOMO SAPIENS | 2 | 8672 | 3thz | A | 681 | GLN | p.Gln681His | P43246 |
| HOMO SAPIENS | 0 | 8672 | 3thz | A | 707 | CYS | p.Cys707Ser | P43246 |
| HOMO SAPIENS | 0 | 8672 | 3thz | A | 809 | GLU | p.Glu809Lys | P43246 |
| HOMO SAPIENS | 0 | 8672 | 3thz | A | 813 | MET | p.Met813Val | P43246 |
| HOMO SAPIENS | 1 | 8672 | 3thz | A | 834 | ALA | p.Ala834Thr | P43246 |
| HOMO SAPIENS | 0 | 8672 | 3thz | A | 849 | LEU | p.Leu849Val | P43246 |
| HOMO SAPIENS | -1 | 8672 | 3thz | A | 860 | SER | p.Ser860Leu | P43246 |
| HOMO SAPIENS | -1 | 8672 | 3thz | A | 934 | THR | p.Thr934Lys | P43246 |
| HOMO SAPIENS | 2 | 292 | 1i3k | A | 34 | ASN | p.Asn34Ser | Q14376 |
| HOMO SAPIENS | 7 | 292 | 1i3k | A | 90 | GLY | p.Gly90Glu | Q14376 |
| HOMO SAPIENS | 2 | 292 | 1i3k | A | 103 | ASP | p.Asp103Gly | Q14376 |
| HOMO SAPIENS | 2 | 292 | 1i3k | A | 136 | TYR | p.Tyr136Ter | Q14376 |
| HOMO SAPIENS | 0 | 292 | 1i3k | A | 169 | ARG | p.Arg169Trp | Q14376 |
| HOMO SAPIENS | 0 | 292 | 1i3k | A | 183 | LEU | p.Leu183Pro | Q14376 |
| HOMO SAPIENS | 0 | 292 | 1i3k | A | 302 | GLY | p.Gly302Asp | Q14376 |
| HOMO SAPIENS | 0 | 292 | 1i3k | A | 313 | LEU | p.Leu313Met | Q14376 |
| HOMO SAPIENS | 1 | 2084 | 5vc8 | A | 295 | GLN | p.Gln295Leu | O96028 |
| HOMO SAPIENS | -1 | 10157 | 5wrw | A | 7 | GLY | p.Gly7Ala | O76094 |
| HOMO SAPIENS | -1 | 10157 | 5wrw | A | 7 | GLY | p.Gly7Arg | O76094 |
| HOMO SAPIENS | -1 | 10157 | 5wrw | A | 8 | GLY | p.Gly8Ala | O76094 |
| HOMO SAPIENS | 0 | 3113 | 3in5 | A | 29 | GLU | p.Glu29Lys | Q9UBT6 |
| HOMO SAPIENS | 5 | 4546 | 5in3 | B | 44 | VAL | p.Val44Leu | P07902 |
| HOMO SAPIENS | -1 | 4546 | 5in3 | B | 51 | ARG | p.Arg51Leu | P07902 |
| HOMO SAPIENS | -1 | 4546 | 5in3 | B | 62 | LEU | p.Leu62Met | P07902 |
| HOMO SAPIENS | 4 | 4546 | 5in3 | B | 66 | PRO | p.Pro66His | P07902 |
| HOMO SAPIENS | 2 | 4546 | 5in3 | B | 74 | LEU | p.Leu74Pro | P07902 |
| HOMO SAPIENS | 5 | 4546 | 5in3 | B | 97 | ASN | p.Asn97Ser | P07902 |
| HOMO SAPIENS | 1 | 4546 | 5in3 | B | 135 | SER | p.Ser135Leu | P07902 |
| HOMO SAPIENS | 5 | 4546 | 5in3 | B | 138 | THR | p.Thr138Met | P07902 |
| HOMO SAPIENS | 7 | 4546 | 5in3 | B | 139 | LEU | p.Leu139Pro | P07902 |

| | | | | | | | | |
|---|---|---|---|---|---|---|---|---|
| HOMO SAPIENS | 2 | 4546 | 5in3 | B | 142 | MET | p.Met142Lys | P07902 |
| HOMO SAPIENS | 1 | 4546 | 5in3 | B | 143 | SER | p.Ser143Leu | P07902 |
| HOMO SAPIENS | 2 | 4546 | 5in3 | B | 148 | ARG | p.Arg148Gly | P07902 |
| HOMO SAPIENS | 2 | 4546 | 5in3 | B | 148 | ARG | p.Arg148Gln | P07902 |
| HOMO SAPIENS | 5 | 4546 | 5in3 | B | 183 | PRO | p.Pro183Thr | P07902 |
| HOMO SAPIENS | 5 | 4546 | 5in3 | B | 185 | PRO | p.Pro185His | P07902 |
| HOMO SAPIENS | 5 | 4546 | 5in3 | B | 186 | HIS | p.His186Tyr | P07902 |
| HOMO SAPIENS | 7 | 4546 | 5in3 | B | 188 | GLN | p.Gln188Arg | P07902 |
| HOMO SAPIENS | 5 | 4546 | 5in3 | B | 195 | LEU | p.Leu195Pro | P07902 |
| HOMO SAPIENS | 4 | 4546 | 5in3 | B | 201 | ARG | p.Arg201His | P07902 |
| HOMO SAPIENS | 4 | 4546 | 5in3 | B | 201 | ARG | p.Arg201Cys | P07902 |
| HOMO SAPIENS | 4 | 4546 | 5in3 | B | 203 | GLU | p.Glu203Lys | P07902 |
| HOMO SAPIENS | 2 | 4546 | 5in3 | B | 209 | TYR | p.Tyr209Cys | P07902 |
| HOMO SAPIENS | 5 | 4546 | 5in3 | B | 223 | ARG | p.Arg223Ser | P07902 |
| HOMO SAPIENS | 5 | 4546 | 5in3 | B | 226 | LEU | p.Leu226Pro | P07902 |
| HOMO SAPIENS | 5 | 4546 | 5in3 | B | 226 | LEU | p.Leu226Val | P07902 |
| HOMO SAPIENS | 2 | 4546 | 5in3 | B | 227 | LEU | p.Leu227Pro | P07902 |
| HOMO SAPIENS | 8 | 4546 | 5in3 | B | 258 | ARG | p.Arg258Cys | P07902 |
| HOMO SAPIENS | 1 | 4546 | 5in3 | B | 265 | PRO | p.Pro265Ala | P07902 |
| HOMO SAPIENS | 8 | 4546 | 5in3 | B | 268 | THR | p.Thr268Asn | P07902 |
| HOMO SAPIENS | 7 | 4546 | 5in3 | B | 278 | ILE | p.Ile278Asn | P07902 |
| HOMO SAPIENS | 5 | 4546 | 5in3 | B | 285 | LYS | p.Lys285Asn | P07902 |
| HOMO SAPIENS | 1 | 4546 | 5in3 | B | 319 | HIS | p.His319Gln | P07902 |
| HOMO SAPIENS | 4 | 4546 | 5in3 | B | 321 | HIS | p.His321Tyr | P07902 |
| HOMO SAPIENS | 4 | 4546 | 5in3 | B | 325 | PRO | p.Pro325Leu | P07902 |
| HOMO SAPIENS | 2 | 4546 | 5in3 | B | 328 | ARG | p.Arg328His | P07902 |
| HOMO SAPIENS | 2 | 4546 | 5in3 | B | 329 | SER | p.Ser329Phe | P07902 |
| HOMO SAPIENS | 5 | 4546 | 5in3 | B | 334 | LYS | p.Lys334Arg | P07902 |
| HOMO SAPIENS | 6 | 1290 | 5uav | C | 119 | ARG | p.Arg119Gly | P32322 |
| HOMO SAPIENS | 6 | 1290 | 5uav | C | 119 | ARG | p.Arg119His | P32322 |
| HOMO SAPIENS | 3 | 1290 | 5uav | C | 251 | ARG | p.Arg251His | P32322 |
| HOMO SAPIENS | 7 | 1290 | 5uav | C | 266 | ARG | p.Arg266Gln | P32322 |
| HOMO SAPIENS | 0 | 9618 | 2r7e | A | 22 | GLY | p.Gly41Cys | P00451 |
| HOMO SAPIENS | 2 | 9618 | 2r7e | A | 113 | GLU | p.Glu132Asp | P00451 |
| HOMO SAPIENS | -1 | 9618 | 2r7e | A | 336 | ARG | p.Arg355Ter | P00451 |
| HOMO SAPIENS | 0 | 9618 | 2r7e | A | 372 | ARG | p.Arg391His | P00451 |
| HOMO SAPIENS | 0 | 9618 | 2r7e | A | 373 | SER | p.Ser392Leu | P00451 |
| HOMO SAPIENS | 0 | 9618 | 2r7e | A | 373 | SER | p.Ser392Pro | P00451 |
| HOMO SAPIENS | 3 | 9618 | 2r7e | A | 386 | ILE | p.Ile405Ser | P00451 |
| HOMO SAPIENS | 6 | 9618 | 2r7e | A | 390 | GLU | p.Glu409Gly | P00451 |
| HOMO SAPIENS | 2 | 9618 | 2r7e | A | 412 | LEU | p.Leu431Phe | P00451 |
| HOMO SAPIENS | 5 | 9618 | 2r7e | A | 425 | LYS | p.Lys444Arg | P00451 |
| HOMO SAPIENS | 0 | 9618 | 2r7e | A | 484 | ARG | p.Arg503His | P00451 |
| HOMO SAPIENS | 2 | 9618 | 2r7e | A | 531 | ARG | p.Arg550His | P00451 |
| HOMO SAPIENS | 3 | 9618 | 2r7e | A | 535 | SER | p.Ser554Gly | P00451 |
| HOMO SAPIENS | 2 | 9618 | 2r7e | A | 593 | ARG | p.Arg612Cys | P00451 |
| HOMO SAPIENS | 0 | 9618 | 2r7e | A | 704 | ALA | p.Ala723Thr | P00451 |
| HOMO SAPIENS | 0 | 9618 | 2r7e | A | 720 | GLU | p.Glu739Lys | P00451 |
| HOMO SAPIENS | 1 | 13085 | 3ud1 | A | 1075 | ILE | p.Thr1075Ile | P16157 |
| HOMO SAPIENS | -1 | 3289 | 2fy4 | A | 2 | ALA | p.Ala120Thr | P28329 |
| HOMO SAPIENS | 7 | 3289 | 2fy4 | A | 18 | VAL | p.Val136Met | P28329 |
| HOMO SAPIENS | 5 | 3289 | 2fy4 | A | 92 | LEU | p.Leu210Pro | P28329 |
| HOMO SAPIENS | 2 | 3289 | 2fy4 | A | 93 | PRO | p.Pro211Ala | P28329 |
| HOMO SAPIENS | 4 | 3289 | 2fy4 | A | 131 | LEU | p.Leu249Val | P28329 |

| | | | | | | | | |
|---|---|---|---|---|---|---|---|---|
| HOMO SAPIENS | 2 | 3289 | 2fy4 | A | 181 | PRO | p.Pro299Leu | P28329 |
| HOMO SAPIENS | 0 | 3289 | 2fy4 | A | 218 | ILE | p.Ile336Thr | P28329 |
| HOMO SAPIENS | 2 | 3289 | 2fy4 | A | 261 | ASP | p.Asp379His | P28329 |
| HOMO SAPIENS | 0 | 3289 | 2fy4 | A | 302 | ARG | p.Arg420Cys | P28329 |
| HOMO SAPIENS | 5 | 3289 | 2fy4 | A | 323 | GLU | p.Glu441Lys | P28329 |
| HOMO SAPIENS | 0 | 3289 | 2fy4 | A | 343 | MET | p.Val461Met | P28329 |
| HOMO SAPIENS | 2 | 3289 | 2fy4 | A | 364 | ARG | p.Arg482Gly | P28329 |
| HOMO SAPIENS | 3 | 3289 | 2fy4 | A | 380 | SER | p.Ser498Leu | P28329 |
| HOMO SAPIENS | 6 | 3289 | 2fy4 | A | 388 | VAL | p.Val506Leu | P28329 |
| HOMO SAPIENS | 7 | 3289 | 2fy4 | A | 429 | HIS | p.His547Gln | P28329 |
| HOMO SAPIENS | 8 | 3289 | 2fy4 | A | 442 | ARG | p.Arg560His | P28329 |
| HOMO SAPIENS | 3 | 3289 | 2fy4 | A | 443 | ARG | p.Arg561Gln | P28329 |
| HOMO SAPIENS | 0 | 4137 | 3hhr | B | 38 | CYS | p.Cys56Ter | P10912 |
| HOMO SAPIENS | 0 | 4137 | 3hhr | B | 43 | ARG | p.Arg61Ter | P10912 |
| HOMO SAPIENS | 0 | 4137 | 3hhr | B | 44 | GLU | p.Glu62Lys | P10912 |
| HOMO SAPIENS | 0 | 4137 | 3hhr | B | 96 | PHE | p.Phe114Ser | P10912 |
| HOMO SAPIENS | 5 | 4137 | 3hhr | B | 131 | PRO | p.Pro149Gln | P10912 |
| HOMO SAPIENS | 0 | 4137 | 3hhr | B | 144 | VAL | p.Val162Ile | P10912 |
| HOMO SAPIENS | 0 | 4137 | 3hhr | B | 150 | HIS | p.His168Gln | P10912 |
| HOMO SAPIENS | 0 | 4137 | 3hhr | B | 152 | ASP | p.Asp170His | P10912 |
| HOMO SAPIENS | 0 | 4137 | 3hhr | B | 153 | ILE | p.Ile171Thr | P10912 |
| HOMO SAPIENS | 0 | 4137 | 3hhr | B | 154 | GLN | p.Gln172Pro | P10912 |
| HOMO SAPIENS | 0 | 4137 | 3hhr | B | 155 | VAL | p.Val173Gly | P10912 |
| HOMO SAPIENS | 0 | 4137 | 3hhr | B | 217 | ARG | p.Arg235Ter | P10912 |
| HOMO SAPIENS | 5 | 4137 | 3hhr | B | 224 | GLU | p.Glu242Asp | P10912 |
| HOMO SAPIENS | 5 | 4137 | 3hhr | B | 224 | GLU | p.Glu242Ter | P10912 |
| HOMO SAPIENS | 6 | 2493 | 5bpb | A | 69 | HIS | p.His69Tyr | Q9ULV1 |
| HOMO SAPIENS | 6 | 2493 | 5bpb | A | 105 | MET | p.Met105Val | Q9ULV1 |
| HOMO SAPIENS | -1 | 2493 | 5bpb | A | 168 | PRO | p.Pro168Ser | Q9ULV1 |
| HOMO SAPIENS | 0 | 32123 | 4ccz | A | 314 | ASP | p.Asp314Asn | Q99707 |
| HOMO SAPIENS | 0 | 32123 | 4ccz | A | 345 | VAL | p.Val345Phe | Q99707 |
| HOMO SAPIENS | 1 | 32123 | 4ccz | A | 585 | ARG | p.Arg585Ter | Q99707 |
| HOMO SAPIENS | 4 | 4618 | 2ag4 | A | 26 | PRO | p.Pro55Leu | P17900 |
| HOMO SAPIENS | 4 | 4618 | 2ag4 | A | 30 | VAL | p.Ile59Val | P17900 |
| HOMO SAPIENS | 2 | 4618 | 2ag4 | A | 40 | VAL | p.Met69Val | P17900 |
| HOMO SAPIENS | 2 | 4618 | 2ag4 | A | 124 | VAL | p.Val153Ala | P17900 |
| HOMO SAPIENS | 8 | 4618 | 2ag4 | A | 140 | ARG | p.Arg169Pro | P17900 |
| HOMO SAPIENS | 5 | 1082 | 1hk0 | X | 14 | ARG | p.Arg15Cys | P07320 |
| HOMO SAPIENS | 7 | 1082 | 1hk0 | X | 23 | PRO | p.Pro24Thr | P07320 |
| HOMO SAPIENS | 6 | 1082 | 1hk0 | X | 36 | ARG | p.Arg37Ser | P07320 |
| HOMO SAPIENS | 5 | 1082 | 1hk0 | X | 55 | TYR | p.Tyr56Ter | P07320 |
| HOMO SAPIENS | 3 | 1082 | 1hk0 | X | 58 | ARG | p.Arg59His | P07320 |
| HOMO SAPIENS | 7 | 1082 | 1hk0 | X | 88 | HIS | p.His88Gln | P07320 |
| HOMO SAPIENS | 6 | 1082 | 1hk0 | X | 95 | ARG | p.Arg95Ser | P07320 |
| HOMO SAPIENS | 5 | 1082 | 1hk0 | X | 126 | VAL | p.Val126Met | P07320 |
| HOMO SAPIENS | 8 | 1082 | 1hk0 | X | 157 | TRP | p.Trp157Ter | P07320 |
| HOMO SAPIENS | 0 | 22879 | 3j0a | B | 644 | ILE | p.Ile644Phe | O60602 |
| HOMO SAPIENS | 8 | 6 | 2o23 | B | 122 | LEU | p.Leu122Val | Q99714 |
| HOMO SAPIENS | 1 | 6 | 2o23 | B | 130 | ARG | p.Arg130Cys | Q99714 |
| HOMO SAPIENS | 1 | 6 | 2o23 | B | 247 | ASN | p.Asn247Ser | Q99714 |
| HOMO SAPIENS | 0 | 6 | 2o23 | B | 249 | GLU | p.Glu249Gln | Q99714 |
| HOMO SAPIENS | 1 | 12910 | 4hl4 | A | 67 | ARG | p.Arg67Ter | Q96BZ9 |
| HOMO SAPIENS | 8 | 16105 | 1lmj | A | 7 | GLU | p.Glu1073Lys | P35555 |
| HOMO SAPIENS | 8 | 16105 | 1lmj | A | 8 | CYS | p.Cys1074Arg | P35555 |

| | | | | | | | | |
|---|---|---|---|---|---|---|---|---|
| HOMO SAPIENS | 9 | 16105 | 1lmj | A | 51 | CYS | p.Cys1117Tyr | P35555 |
| HOMO SAPIENS | 9 | 16105 | 1lmj | A | 63 | CYS | p.Cys1129Tyr | P35555 |
| HOMO SAPIENS | 9 | 16105 | 1lmj | A | 65 | ASN | p.Asn1131Tyr | P35555 |
| HOMO SAPIENS | 4 | 16105 | 1lmj | A | 71 | ARG | p.Arg1137Pro | P35555 |
| HOMO SAPIENS | 1 | 16105 | 1lmj | A | 82 | PRO | p.Pro1148Ala | P35555 |
| HOMO SAPIENS | 0 | 4795 | 3s5f | A | 106 | LEU | p.Leu106Ser | Q16595 |
| HOMO SAPIENS | 0 | 4795 | 3s5f | A | 130 | GLY | p.Gly130Val | Q16595 |
| HOMO SAPIENS | 0 | 4795 | 3s5f | A | 154 | ILE | p.Ile154Phe | Q16595 |
| HOMO SAPIENS | -1 | 4795 | 3s5f | A | 209 | ASP | p.Asp209Gly | Q16595 |
| HOMO SAPIENS | -1 | 2884 | 4lli | A | 1655 | ASP | p.Asp1655Glu | Q9Y4I1 |
| HOMO SAPIENS | 0 | 1786 | 7pck | C | 56P | TYR | p.Tyr71Ter | P43235 |
| HOMO SAPIENS | 0 | 1786 | 7pck | C | 64P | GLY | p.Gly79Glu | P43235 |
| HOMO SAPIENS | 7 | 38286 | 2l53 | B | 1914 | ARG | p.Arg1914Gly | Q14524 |
| HOMO SAPIENS | 1 | 38286 | 2l53 | B | 1924 | ALA | p.Ala1924Thr | Q14524 |
| HOMO SAPIENS | 0 | 31747 | 3ksy | A | 25 | PRO | p.Pro25Ser | Q07889 |
| HOMO SAPIENS | 3 | 31747 | 3ksy | A | 37 | THR | p.Thr37Ala | Q07889 |
| HOMO SAPIENS | 0 | 31747 | 3ksy | A | 54 | GLN | p.Gln54His | Q07889 |
| HOMO SAPIENS | -1 | 31747 | 3ksy | A | 190 | ASP | p.Asp190Glu | Q07889 |
| HOMO SAPIENS | 2 | 31747 | 3ksy | A | 266 | THR | p.Thr266Lys | Q07889 |
| HOMO SAPIENS | 1 | 31747 | 3ksy | A | 269 | MET | p.Met269Arg | Q07889 |
| HOMO SAPIENS | 1 | 31747 | 3ksy | A | 340 | PRO | p.Pro340Ser | Q07889 |
| HOMO SAPIENS | 1 | 31747 | 3ksy | A | 423 | ASN | p.Asn423Lys | Q07889 |
| HOMO SAPIENS | 1 | 31747 | 3ksy | A | 424 | GLU | p.Glu424Lys | Q07889 |
| HOMO SAPIENS | 1 | 31747 | 3ksy | A | 432 | TRP | p.Trp432Arg | Q07889 |
| HOMO SAPIENS | 1 | 31747 | 3ksy | A | 552 | ARG | p.Arg552Gly | Q07889 |
| HOMO SAPIENS | 1 | 31747 | 3ksy | A | 552 | ARG | p.Arg552Ser | Q07889 |
| HOMO SAPIENS | 3 | 7529 | 6jdw | A | 93 | ILE | p.Ile93Met | P50440 |
| HOMO SAPIENS | 2 | 7529 | 6jdw | A | 110 | GLN | p.Gln110His | P50440 |
| HOMO SAPIENS | 5 | 7529 | 6jdw | A | 149 | TRP | p.Trp149Ter | P50440 |
| HOMO SAPIENS | 0 | 7529 | 6jdw | A | 169 | ARG | p.Arg169Ter | P50440 |
| HOMO SAPIENS | 0 | 7529 | 6jdw | A | 260 | ARG | p.Arg260Ter | P50440 |
| HOMO SAPIENS | 3 | 7529 | 6jdw | A | 344 | ILE | p.Ile344Val | P50440 |
| HOMO SAPIENS | 5 | 7529 | 6jdw | A | 418 | LEU | p.Leu418Ile | P50440 |
| HOMO SAPIENS | 1 | 14788 | 1eak | A | 101 | ARG | p.Arg101His | P08253 |
| HOMO SAPIENS | 0 | 14788 | 1eak | A | 383 | ASP | p.Asp383Glu | P08253 |
| HOMO SAPIENS | 7 | 14788 | 1eak | A | 404 | GLN | p.Glu404Lys | P08253 |
| HOMO SAPIENS | 2 | 14788 | 1eak | A | 447 | ALA | p.Ala447Val | P08253 |
| HOMO SAPIENS | 0 | 1255 | 5xtc | p | 8 | PRO | p.Pro8His | Q9Y6M9 |
| HOMO SAPIENS | 0 | 1255 | 5xtc | p | 146 | PRO | p.Pro146Ser | Q9Y6M9 |
| HOMO SAPIENS | 3 | 6721 | 1a22 | A | 25 | PHE | p.Phe51Tyr | P01241 |
| HOMO SAPIENS | 2 | 6721 | 1a22 | A | 95 | SER | p.Ser121Arg | P01241 |
| HOMO SAPIENS | 2 | 6721 | 1a22 | A | 110 | VAL | p.Val136Ile | P01241 |
| HOMO SAPIENS | 0 | 1380 | 5i6z | A | 605 | LYS | p.Lys605Asn | P31645 |
| HOMO SAPIENS | 0 | 941 | 1qki | D | 48 | ILE | p.Ile48Thr | P11413 |
| HOMO SAPIENS | 3 | 941 | 1qki | D | 68 | VAL | p.Val68Met | P11413 |
| HOMO SAPIENS | 2 | 941 | 1qki | D | 126 | ASN | p.Asn126Asp | P11413 |
| HOMO SAPIENS | 8 | 941 | 1qki | D | 163 | GLY | p.Gly163Ser | P11413 |
| HOMO SAPIENS | 6 | 941 | 1qki | D | 165 | ASN | p.Asn165Asp | P11413 |
| HOMO SAPIENS | 4 | 941 | 1qki | D | 181 | ASP | p.Asp181Val | P11413 |
| HOMO SAPIENS | 4 | 941 | 1qki | D | 182 | ARG | p.Arg182Trp | P11413 |
| HOMO SAPIENS | 5 | 941 | 1qki | D | 188 | SER | p.Ser188Phe | P11413 |
| HOMO SAPIENS | 9 | 941 | 1qki | D | 198 | ARG | p.Arg198Cys | P11413 |
| HOMO SAPIENS | 8 | 941 | 1qki | D | 213 | VAL | p.Val213Leu | P11413 |
| HOMO SAPIENS | 5 | 941 | 1qki | D | 282 | ASP | p.Asp282His | P11413 |

| | | | | | | | | |
|---|---|---|---|---|---|---|---|---|
| HOMO SAPIENS | 7 | 941 | 1qki | D | 285 | ARG | p.Arg285His | P11413 |
| HOMO SAPIENS | 5 | 941 | 1qki | D | 291 | VAL | p.Val291Met | P11413 |
| HOMO SAPIENS | 8 | 941 | 1qki | D | 323 | LEU | p.Leu323Pro | P11413 |
| HOMO SAPIENS | 3 | 941 | 1qki | D | 335 | ALA | p.Ala335Thr | P11413 |
| HOMO SAPIENS | 6 | 941 | 1qki | D | 350 | ASP | p.Asp350His | P11413 |
| HOMO SAPIENS | 5 | 941 | 1qki | D | 393 | ARG | p.Arg393His | P11413 |
| HOMO SAPIENS | 5 | 941 | 1qki | D | 454 | ARG | p.Arg454Cys | P11413 |
| HOMO SAPIENS | 2 | 941 | 1qki | D | 459 | LEU | p.Arg459Leu | P11413 |
| HOMO SAPIENS | 4 | 941 | 1qki | D | 463 | ARG | p.Arg463His | P11413 |
| HOMO SAPIENS | 4 | 2483 | 1tij | A | 68 | LEU | p.Leu94Gln | P01034 |
| HOMO SAPIENS | 2 | 20632 | 3v5b | A | 331 | ARG | p.Arg331Pro | P02545 |
| HOMO SAPIENS | 2 | 20632 | 3v5b | A | 374 | HIS | p.His374Gln | P02545 |
| HOMO SAPIENS | 4 | 13929 | 5xtd | J | 75 | ARG | p.Arg75His | Q16795 |
| HOMO SAPIENS | 3 | 13929 | 5xtd | J | 243 | VAL | p.Val243Ala | Q16795 |
| HOMO SAPIENS | 1 | 13929 | 5xtd | J | 294 | PRO | p.Pro294Leu | Q16795 |
| HOMO SAPIENS | 2 | 13929 | 5xtd | J | 321 | ARG | p.Arg321Pro | Q16795 |
| HOMO SAPIENS | 0 | 9411 | 5thl | A | 41 | ARG | p.Gly41Arg | P54577 |
| HOMO SAPIENS | 0 | 9411 | 5thl | A | 196 | GLU | p.Glu196Lys | P54577 |
| HOMO SAPIENS | 0 | 9411 | 5thl | A | 316 | ALA | p.Ala316Ser | P54577 |
| HOMO SAPIENS | 0 | 9411 | 5thl | A | 339 | ALA | p.Ala339Thr | P54577 |
| HOMO SAPIENS | 4 | 16273 | 2e8n | A | 46 | THR | p.Thr940Ile | P29317 |
| HOMO SAPIENS | 7 | 16273 | 2e8n | A | 54 | GLY | p.Gly948Trp | P29317 |
| HOMO SAPIENS | 6 | 16273 | 2e8n | A | 65 | ALA | p.Ala959Thr | P29317 |
| HOMO SAPIENS | 7 | 30387 | 2k9y | A | 541 | VAL | p.Val541Met | P29317 |
| HOMO SAPIENS | 0 | 6961 | 4nha | A | 843 | ILE | p.Ile853Val | Q9UPY3 |
| HOMO SAPIENS | -1 | 6961 | 4nha | A | 862 | ALA | p.Ala872Thr | Q9UPY3 |
| HOMO SAPIENS | 3 | 15708 | 4pzg | A | 16 | ARG | p.Arg16Ter | Q9Y5X0 |
| HOMO SAPIENS | 4 | 15708 | 4pzg | A | 51 | ARG | p.Arg51Gln | Q9Y5X0 |
| HOMO SAPIENS | 3 | 30166 | 4o00 | A | 44 | ARG | p.Arg14257Ter | Q8WZ42 |
| HOMO SAPIENS | 2 | 30166 | 4o00 | A | 54 | ARG | p.Arg14267His | Q8WZ42 |
| HOMO SAPIENS | 8 | 11158 | 4hca | A | 275 | TRP | p.Trp274Arg | P23771 |
| HOMO SAPIENS | 2 | 4328 | 4e0s | A | 1476 | ARG | p.Arg1476Ter | P01031 |
| HOMO SAPIENS | 0 | 11462 | 1dx5 | I | 455 | ALA | p.Ala473Val | P07204 |
| HOMO SAPIENS | -1 | 11150 | 2fic | A | 45 | LYS | p.Lys35Asn | O00499 |
| HOMO SAPIENS | 6 | 11150 | 2fic | A | 161 | ASP | p.Asp151Asn | O00499 |
| HOMO SAPIENS | 8 | 27382 | 4mqv | B | 17 | ARG | p.Arg17Ter | Q9NZC9 |
| HOMO SAPIENS | -1 | 37991 | 1q0p | A | 231 | GLN | p.Gln256Ter | P00751 |
| HOMO SAPIENS | 1 | 443 | 5xte | U | 215 | LEU | p.Leu215Phe | P08574 |
| HOMO SAPIENS | 8 | 21471 | 5y4e | A | 1461 | ASN | p.Thr692Ala | Q01484 |
| HOMO SAPIENS | -1 | 35443 | 3um3 | B | 129 | ASP | p.Asp129Val | Q9H444 |
| HOMO SAPIENS | -1 | 35443 | 3um3 | B | 161 | GLU | p.Glu161Lys | Q9H444 |
| HOMO SAPIENS | 6 | 28781 | 2n2a | B | 111 | ILE | p.Ile654Val | P04626 |
| HOMO SAPIENS | 7 | 28781 | 2n2a | B | 112 | ILE | p.Ile655Val | P04626 |
| HOMO SAPIENS | -1 | 3346 | 4ztz | C | 61 | GLY | p.Gly61Arg | Q9UHN1 |
| HOMO SAPIENS | 4 | 3346 | 4ztz | C | 103 | GLY | p.Gly103Ser | Q9UHN1 |
| HOMO SAPIENS | 4 | 3346 | 4ztz | C | 133 | HIS | p.His133Tyr | Q9UHN1 |
| HOMO SAPIENS | -1 | 3346 | 4ztz | C | 153 | LEU | p.Leu153Val | Q9UHN1 |
| HOMO SAPIENS | -1 | 3346 | 4ztz | C | 169 | ALA | p.Ala169Thr | Q9UHN1 |
| HOMO SAPIENS | 7 | 3346 | 4ztz | C | 369 | ARG | p.Arg369Gly | Q9UHN1 |
| HOMO SAPIENS | 4 | 3346 | 4ztz | C | 386 | ASP | p.Asp386Glu | Q9UHN1 |
| HOMO SAPIENS | 4 | 3346 | 4ztz | C | 451 | GLY | p.Gly451Glu | Q9UHN1 |
| HOMO SAPIENS | 4 | 2131 | 5fo9 | E | 1549 | LEU | p.Leu1549Met | P01024 |
| HOMO SAPIENS | 2 | 2131 | 5fo9 | E | 1619 | SER | p.Ser1619Arg | P01024 |
| HOMO SAPIENS | 3 | 12662 | 4jhp | C | 75 | ILE | p.Ile75Val | Q92834 |

| | | | | | | | | |
|---|---|---|---|---|---|---|---|---|
| HOMO SAPIENS | 1 | 12662 | 4jhp | C | 173 | GLY | p.Gly173Arg | Q92834 |
| HOMO SAPIENS | 0 | 12662 | 4jhp | C | 184 | GLN | p.Gln184His | Q92834 |
| HOMO SAPIENS | 3 | 12662 | 4jhp | C | 262 | ALA | p.Ala262Gly | Q92834 |
| HOMO SAPIENS | 3 | 12662 | 4jhp | C | 302 | CYS | p.Cys302Tyr | Q92834 |
| HOMO SAPIENS | 5 | 1093 | 2jg9 | A | 186 | GLN | p.Gln208Ter | P02745 |
| HOMO SAPIENS | 0 | 13181 | 4r4l | A | 12 | MET | p.Met13Ile | Q13976 |
| HOMO SAPIENS | 8 | 25939 | 4iyp | A | 106 | ARG | p.Arg106Gln | P78318 |
| HOMO SAPIENS | 1 | 217 | 2xsx | A | 71 | SER | p.Asn71Ser | P13929 |
| HOMO SAPIENS | 0 | 217 | 2xsx | A | 76 | ALA | p.Ala76Pro | P13929 |
| HOMO SAPIENS | 1 | 217 | 2xsx | A | 85 | ALA | p.Val85Ala | P13929 |
| HOMO SAPIENS | 8 | 217 | 2xsx | A | 156 | GLY | p.Gly156Asp | P13929 |
| HOMO SAPIENS | 5 | 217 | 2xsx | A | 332 | VAL | p.Val332Ile | P13929 |
| HOMO SAPIENS | 7 | 217 | 2xsx | A | 374 | GLY | p.Gly374Glu | P13929 |
| HOMO SAPIENS | 1 | 5508 | 3szm | A | 682 | THR | p.Thr682Asn | Q8NEM0 |
| HOMO SAPIENS | 0 | 5508 | 3szm | A | 715 | TRP | p.Trp715Ter | Q8NEM0 |
| HOMO SAPIENS | 1 | 5508 | 3szm | A | 761 | VAL | p.Ala761Val | Q8NEM0 |
| HOMO SAPIENS | 1 | 5508 | 3szm | A | 828 | PRO | p.Pro828Ser | Q8NEM0 |
| HOMO SAPIENS | 4 | 5508 | 3szm | A | 829 | GLU | p.Glu829Asp | Q8NEM0 |
| HOMO SAPIENS | 2 | 16353 | 2q0z | X | 171 | GLY | p.Gly1978Ala | O75643 |
| HOMO SAPIENS | 2 | 16353 | 2q0z | X | 188 | ALA | p.Ala1995Thr | O75643 |
| HOMO SAPIENS | 0 | 4127 | 3swr | A | 860 | GLY | p.Gly860Arg | P26358 |
| HOMO SAPIENS | -1 | 4127 | 3swr | A | 1122 | SER | p.Ser1122Cys | P26358 |
| HOMO SAPIENS | 0 | 4127 | 3swr | A | 1216 | ASP | p.Asp1216Glu | P26358 |
| HOMO SAPIENS | 0 | 6382 | 3o96 | A | 17 | GLU | p.Glu17Lys | P31749 |
| HOMO SAPIENS | 6 | 14314 | 5xtc | Q | 71 | PRO | p.Pro71Leu | O75306 |
| HOMO SAPIENS | 2 | 5886 | 4fao | E | 22 | ARG | p.Arg40His | Q13705 |
| HOMO SAPIENS | 1 | 2534 | 3mp4 | A | 37 | GLU | p.Glu37Ter | P35914 |
| HOMO SAPIENS | 3 | 2534 | 3mp4 | A | 41 | MET | p.Arg41Gln | P35914 |
| HOMO SAPIENS | 1 | 2534 | 3mp4 | A | 70 | VAL | p.Val70Leu | P35914 |
| HOMO SAPIENS | 2 | 2196 | 3ifq | A | 142 | ARG | p.Arg142His | P14923 |
| HOMO SAPIENS | 2 | 2196 | 3ifq | A | 153 | ASP | p.Asp153Glu | P14923 |
| HOMO SAPIENS | 3 | 2196 | 3ifq | A | 176 | ARG | p.Arg176Gln | P14923 |
| HOMO SAPIENS | 3 | 2196 | 3ifq | A | 265 | ARG | p.Arg265His | P14923 |
| HOMO SAPIENS | 2 | 2196 | 3ifq | A | 301 | GLU | p.Glu301Gly | P14923 |
| HOMO SAPIENS | 0 | 2196 | 3ifq | A | 456 | VAL | p.Val456Ile | P14923 |
| HOMO SAPIENS | 3 | 2196 | 3ifq | A | 603 | VAL | p.Val603Leu | P14923 |
| HOMO SAPIENS | 0 | 2196 | 3ifq | A | 648 | VAL | p.Val648Ile | P14923 |
| HOMO SAPIENS | 1 | 8746 | 1cgj | I | 32 | PRO | p.Pro55Ser | P00995 |
| HOMO SAPIENS | 1 | 8746 | 1cgj | I | 44 | ARG | p.Arg67His | P00995 |
| HOMO SAPIENS | 1 | 8746 | 1cgj | I | 44 | ARG | p.Arg67Cys | P00995 |
| HOMO SAPIENS | 3 | 3212 | 1syq | A | 105 | ARG | p.Arg105Ter | P18206 |
| HOMO SAPIENS | 3 | 3212 | 1syq | A | 126 | PHE | p.Phe126Leu | P18206 |
| HOMO SAPIENS | 4 | 10028 | 3eto | A | 1609 | ARG | p.Arg1608His | P46531 |
| HOMO SAPIENS | 1 | 10028 | 3eto | A | 1672 | VAL | p.Val1671Ile | P46531 |
| HOMO SAPIENS | 5 | 10028 | 3eto | A | 1692 | GLN | p.Gln1691His | P46531 |
| HOMO SAPIENS | 3 | 3749 | 5xtc | Z | 70 | GLY | p.Gly70Ter | O43676 |
| HOMO SAPIENS | 2 | 17039 | 3brv | B | 57 | GLU | p.Glu57Lys | Q9Y6K9 |
| HOMO SAPIENS | 8 | 3589 | 2fh3 | A | 433 | VAL | p.Val460Leu | P06396 |
| HOMO SAPIENS | 1 | 3589 | 2fh3 | A | 454 | ARG | p.Arg481Cys | P06396 |
| HOMO SAPIENS | 4 | 3589 | 2fh3 | A | 498 | ARG | p.Arg525His | P06396 |
| HOMO SAPIENS | 2 | 3589 | 2fh3 | A | 520 | TYR | p.Tyr547Cys | P06396 |
| HOMO SAPIENS | 1 | 3589 | 2fh3 | A | 529 | GLY | p.Gly556Arg | P06396 |
| HOMO SAPIENS | 2 | 3589 | 2fh3 | A | 536 | THR | p.Thr563Ser | P06396 |
| HOMO SAPIENS | 2 | 3589 | 2fh3 | A | 579 | VAL | p.Val606Met | P06396 |

| | | | | | | | | |
|---|---|---|---|---|---|---|---|---|
| HOMO SAPIENS | 1 | 3589 | 2fh3 | A | 589 | THR | p.Thr616Met | P06396 |
| HOMO SAPIENS | 5 | 3589 | 2fh3 | A | 668 | THR | p.Thr695Met | P06396 |
| HOMO SAPIENS | 0 | 148 | 1lcl | A | 130 | ILE | p.Ile130Ser | Q05315 |
| HOMO SAPIENS | -1 | 24727 | 4fj3 | P | 237 | ALA | p.Ala237Thr | P04049 |
| HOMO SAPIENS | 0 | 24727 | 4fj3 | P | 257 | SER | p.Ser257Leu | P04049 |
| HOMO SAPIENS | 0 | 24727 | 4fj3 | P | 261 | PRO | p.Pro261Ala | P04049 |
| HOMO SAPIENS | 0 | 2949 | 5xtb | B | 77 | ARG | p.Arg77Gly | O00217 |
| HOMO SAPIENS | 0 | 2949 | 5xtb | B | 79 | PRO | p.Pro79Leu | O00217 |
| HOMO SAPIENS | 0 | 2949 | 5xtb | B | 100 | ALA | p.Ala100Val | O00217 |
| HOMO SAPIENS | 1 | 2949 | 5xtb | B | 115 | LYS | p.Lys115Glu | O00217 |
| HOMO SAPIENS | 0 | 2949 | 5xtb | B | 138 | ARG | p.Arg138His | O00217 |
| HOMO SAPIENS | 1 | 3959 | 5k5s | B | 25 | ARG | p.Arg25Ter | P41180 |
| HOMO SAPIENS | 4 | 3959 | 5k5s | B | 39 | PRO | p.Pro39Ala | P41180 |
| HOMO SAPIENS | 0 | 3959 | 5k5s | B | 47 | LYS | p.Lys47Asn | P41180 |
| HOMO SAPIENS | 2 | 3959 | 5k5s | B | 62 | ARG | p.Arg62Met | P41180 |
| HOMO SAPIENS | 2 | 3959 | 5k5s | B | 66 | ARG | p.Arg66Cys | P41180 |
| HOMO SAPIENS | 4 | 3959 | 5k5s | B | 90 | ASN | p.Asn90Thr | P41180 |
| HOMO SAPIENS | 1 | 3959 | 5k5s | B | 116 | ALA | p.Ala116Thr | P41180 |
| HOMO SAPIENS | 1 | 3959 | 5k5s | B | 118 | ASN | p.Asn118Lys | P41180 |
| HOMO SAPIENS | -1 | 3959 | 5k5s | B | 125 | LEU | p.Leu125Pro | P41180 |
| HOMO SAPIENS | -1 | 3959 | 5k5s | B | 127 | GLU | p.Glu127Ala | P41180 |
| HOMO SAPIENS | -1 | 3959 | 5k5s | B | 128 | PHE | p.Phe128Leu | P41180 |
| HOMO SAPIENS | -1 | 3959 | 5k5s | B | 131 | CYS | p.Cys131Trp | P41180 |
| HOMO SAPIENS | 1 | 3959 | 5k5s | B | 138 | THR | p.Thr138Met | P41180 |
| HOMO SAPIENS | 0 | 3959 | 5k5s | B | 143 | GLY | p.Gly143Glu | P41180 |
| HOMO SAPIENS | 5 | 3959 | 5k5s | B | 151 | THR | p.Thr151Met | P41180 |
| HOMO SAPIENS | 4 | 3959 | 5k5s | B | 180 | PHE | p.Phe180Cys | P41180 |
| HOMO SAPIENS | 0 | 3959 | 5k5s | B | 185 | ARG | p.Arg185Gln | P41180 |
| HOMO SAPIENS | 0 | 3959 | 5k5s | B | 185 | ARG | p.Arg185Ter | P41180 |
| HOMO SAPIENS | 2 | 3959 | 5k5s | B | 191 | GLU | p.Glu191Lys | P41180 |
| HOMO SAPIENS | 2 | 3959 | 5k5s | B | 227 | ARG | p.Arg227Leu | P41180 |
| HOMO SAPIENS | 2 | 3959 | 5k5s | B | 250 | GLU | p.Glu250Lys | P41180 |
| HOMO SAPIENS | 6 | 3959 | 5k5s | B | 297 | GLU | p.Glu297Lys | P41180 |
| HOMO SAPIENS | 1 | 3959 | 5k5s | B | 336 | LYS | p.Lys336Asn | P41180 |
| HOMO SAPIENS | 6 | 3959 | 5k5s | B | 429 | HIS | p.His429Tyr | P41180 |
| HOMO SAPIENS | 0 | 3959 | 5k5s | B | 445 | THR | p.Thr445Ala | P41180 |
| HOMO SAPIENS | 3 | 3959 | 5k5s | B | 509 | GLY | p.Gly509Arg | P41180 |
| HOMO SAPIENS | 1 | 3959 | 5k5s | B | 553 | GLY | p.Gly553Arg | P41180 |
| HOMO SAPIENS | 3 | 3959 | 5k5s | B | 592 | ASN | p.Asn592Ser | P41180 |
| HOMO SAPIENS | -1 | 3959 | 5k5s | B | 604 | GLU | p.Glu604Lys | P41180 |
| HOMO SAPIENS | 4 | 2564 | 5l0c | C | 975 | ARG | p.Arg975Trp | P18206 |
| HOMO SAPIENS | 4 | 2564 | 5l0c | C | 975 | ARG | p.Arg975Leu | P18206 |
| HOMO SAPIENS | 1 | 5695 | 5i8r | A | 294 | ARG | p.Arg294Gln | P17405 |
| HOMO SAPIENS | 0 | 5695 | 5i8r | A | 485 | ALA | p.Ala485Val | P17405 |
| HOMO SAPIENS | 1 | 5695 | 5i8r | A | 506 | GLY | p.Gly506Arg | P17405 |
| HOMO SAPIENS | -1 | 1978 | 3avs | A | 1060 | HIS | p.His1060Leu | O15550 |
| HOMO SAPIENS | 0 | 11173 | 4v0c | B | 362 | LYS | p.Lys362Arg | P51787 |
| HOMO SAPIENS | 3 | 11173 | 4v0c | B | 381 | CYS | p.Cys381Tyr | P51787 |
| HOMO SAPIENS | 7 | 11173 | 4v0c | B | 518 | ARG | p.Arg518Gly | P51787 |
| HOMO SAPIENS | 2 | 11173 | 4v0c | B | 525 | ALA | p.Ala525Thr | P51787 |
| HOMO SAPIENS | 0 | 11173 | 4v0c | B | 530 | GLN | p.Gln530Ter | P51787 |
| HOMO SAPIENS | -1 | 11173 | 4v0c | B | 536 | TYR | p.Tyr536Ter | P51787 |
| HOMO SAPIENS | 2 | 1113 | 5xtd | N | 60 | ARG | p.Arg60Ter | Q9UI09 |
| HOMO SAPIENS | 3 | 6061 | 2mmz | A | 148 | LEU | p.Leu148Ser | Q86SX6 |

| | | | | | | | | |
|---|---|---|---|---|---|---|---|---|
| HOMO SAPIENS | -1 | 26081 | 3eu7 | A | 837 | GLU | p.Glu837Ter | Q86YC2 |
| HOMO SAPIENS | 0 | 26081 | 3eu7 | A | 864 | PRO | p.Pro864Ser | Q86YC2 |
| HOMO SAPIENS | 5 | 26081 | 3eu7 | A | 891 | CYS | p.Cys891Trp | Q86YC2 |
| HOMO SAPIENS | 0 | 26081 | 3eu7 | A | 932 | VAL | p.Val932Met | Q86YC2 |
| HOMO SAPIENS | 5 | 26081 | 3eu7 | A | 961 | LEU | p.Leu961Met | Q86YC2 |
| HOMO SAPIENS | 5 | 26081 | 3eu7 | A | 1038 | TRP | p.Trp1038Ter | Q86YC2 |
| HOMO SAPIENS | -1 | 26081 | 3eu7 | A | 1086 | ARG | p.Arg1086Ter | Q86YC2 |
| HOMO SAPIENS | 0 | 26081 | 3eu7 | A | 1122 | ASP | p.Asp1122Glu | Q86YC2 |
| HOMO SAPIENS | 5 | 26081 | 3eu7 | A | 1158 | HIS | p.His1158Gln | Q86YC2 |
| HOMO SAPIENS | 5 | 26081 | 3eu7 | A | 1183 | TYR | p.Tyr1183Ter | Q86YC2 |
| HOMO SAPIENS | 4 | 199 | 1qo5 | C | 45 | ARG | p.Arg46Trp | P05062 |
| HOMO SAPIENS | 5 | 199 | 1qo5 | C | 147 | TRP | p.Trp148Arg | P05062 |
| HOMO SAPIENS | 0 | 199 | 1qo5 | C | 149 | ALA | p.Ala150Pro | P05062 |
| HOMO SAPIENS | 0 | 199 | 1qo5 | C | 174 | ALA | p.Ala175Asp | P05062 |
| HOMO SAPIENS | 0 | 199 | 1qo5 | C | 334 | ASN | p.Asn335Lys | P05062 |
| HOMO SAPIENS | 0 | 199 | 1qo5 | C | 337 | ALA | p.Ala338Val | P05062 |
| HOMO SAPIENS | 5 | 199 | 1qo5 | C | 342 | TYR | p.Tyr343His | P05062 |
| HOMO SAPIENS | 0 | 6910 | 4zht | B | 59 | ARG | p.Arg59Ter | Q9Y223 |
| HOMO SAPIENS | 4 | 6910 | 4zht | B | 129 | ARG | p.Arg129Ter | Q9Y223 |
| HOMO SAPIENS | 4 | 6910 | 4zht | B | 129 | ARG | p.Arg129Gln | Q9Y223 |
| HOMO SAPIENS | 1 | 6910 | 4zht | B | 162 | ARG | p.Arg162Cys | Q9Y223 |
| HOMO SAPIENS | 0 | 6910 | 4zht | B | 176 | ASP | p.Asp176Val | Q9Y223 |
| HOMO SAPIENS | 0 | 6910 | 4zht | B | 216 | VAL | p.Val216Ala | Q9Y223 |
| HOMO SAPIENS | 0 | 6910 | 4zht | B | 246 | ARG | p.Arg246Trp | Q9Y223 |
| HOMO SAPIENS | 0 | 6910 | 4zht | B | 263 | ARG | p.Arg263Leu | Q9Y223 |
| HOMO SAPIENS | 0 | 6910 | 4zht | B | 266 | ARG | p.Arg266Gln | Q9Y223 |
| HOMO SAPIENS | 0 | 6910 | 4zht | B | 266 | ARG | p.Arg266Trp | Q9Y223 |
| HOMO SAPIENS | 0 | 6910 | 4zht | B | 277 | ARG | p.Arg277Cys | Q9Y223 |
| HOMO SAPIENS | 0 | 6910 | 4zht | B | 378 | ASP | p.Asp378Tyr | Q9Y223 |
| HOMO SAPIENS | 3 | 211 | 5kxi | C | 262 | VAL | p.Val287Leu | P17787 |
| HOMO SAPIENS | 0 | 211 | 5kxi | C | 369 | PHE | p.Phe478Leu | P17787 |
| HOMO SAPIENS | 0 | 2454 | 5iy9 | V | null | PHE | p.Phe99Ser | P19447 |
| HOMO SAPIENS | 0 | 2454 | 5iy9 | V | null | ARG | p.Arg109Ter | P19447 |
| HOMO SAPIENS | 0 | 2454 | 5iy9 | V | null | THR | p.Thr119Pro | P19447 |
| HOMO SAPIENS | 0 | 2454 | 5iy9 | V | 425 | ARG | p.Arg425Ter | P19447 |
| HOMO SAPIENS | 1 | 2454 | 5iy9 | V | 545 | GLN | p.Gln545Ter | P19447 |
| HOMO SAPIENS | -1 | 3805 | 3rbq | A | 57 | LYS | p.Lys57Ter | Q13432 |
| HOMO SAPIENS | 2 | 2649 | 4ido | A | 43 | ASP | p.Asp43Glu | Q8WXF7 |
| HOMO SAPIENS | 2 | 2649 | 4ido | A | 157 | LEU | p.Leu157Trp | Q8WXF7 |
| HOMO SAPIENS | 2 | 2649 | 4ido | A | 193 | PHE | p.Phe193Cys | Q8WXF7 |
| HOMO SAPIENS | 7 | 2649 | 4ido | A | 217 | ARG | p.Arg217Gln | Q8WXF7 |
| HOMO SAPIENS | 4 | 2649 | 4ido | A | 239 | ARG | p.Arg239Cys | Q8WXF7 |
| HOMO SAPIENS | 3 | 2649 | 4ido | A | 258 | HIS | p.His258Arg | Q8WXF7 |
| HOMO SAPIENS | 4 | 2649 | 4ido | A | 259 | SER | p.Ser259Tyr | Q8WXF7 |
| HOMO SAPIENS | 4 | 2649 | 4ido | A | 408 | MET | p.Met408Val | Q8WXF7 |
| HOMO SAPIENS | 2 | 2649 | 4ido | A | 415 | ARG | p.Arg415Trp | Q8WXF7 |
| HOMO SAPIENS | 3 | 2649 | 4ido | A | 416 | ARG | p.Arg416Cys | Q8WXF7 |
| HOMO SAPIENS | 5 | 2203 | 3wff | A | 810 | LEU | p.Ser810Leu | P08235 |
| HOMO SAPIENS | 4 | 12648 | 5ep6 | B | 696 | GLU | p.Glu696Lys | Q9UHD2 |
| HOMO SAPIENS | 0 | 30622 | 2pe4 | A | 90 | GLU | p.Glu90Asp | Q12794 |
| HOMO SAPIENS | 0 | 30622 | 2pe4 | A | 256 | GLY | p.Gly256Arg | Q12794 |
| HOMO SAPIENS | 1 | 30622 | 2pe4 | A | 268 | GLU | p.Glu268Lys | Q12794 |
| HOMO SAPIENS | 0 | 30622 | 2pe4 | A | 392 | THR | p.Thr392Met | Q12794 |
| HOMO SAPIENS | 5 | 5411 | 1lya | A | 93 | SER | p.Ser157Leu | P07339 |

| | | | | | | | | |
|---|---|---|---|---|---|---|---|---|
| HOMO SAPIENS | 3 | 5743 | 2uzp | A | 189 | ASN | p.Asn189Lys | P05129 |
| HOMO SAPIENS | 1 | 12529 | 3kfd | J | 21 | ASN | p.Asn45Ser | P36897 |
| HOMO SAPIENS | 1 | 1063 | 2e1q | B | 149 | ARG | p.Arg149Cys | P47989 |
| HOMO SAPIENS | -1 | 1063 | 2e1q | B | 172 | GLY | p.Gly172Arg | P47989 |
| HOMO SAPIENS | 4 | 1063 | 2e1q | B | 228 | ARG | p.Arg228Ter | P47989 |
| HOMO SAPIENS | 5 | 1063 | 2e1q | B | 395 | LYS | p.Lys395Met | P47989 |
| HOMO SAPIENS | 2 | 1063 | 2e1q | B | 646 | ILE | p.Ile646Val | P47989 |
| HOMO SAPIENS | 4 | 1063 | 2e1q | B | 703 | ILE | p.Ile703Val | P47989 |
| HOMO SAPIENS | 8 | 2525 | 1kqn | F | 18 | ASN | p.Asn18Ser | Q9HAN9 |
| HOMO SAPIENS | 0 | 2525 | 1kqn | F | 151 | VAL | p.Val151Phe | Q9HAN9 |
| HOMO SAPIENS | 0 | 2525 | 1kqn | F | 153 | LEU | p.Leu153Val | Q9HAN9 |
| HOMO SAPIENS | 0 | 2525 | 1kqn | F | 169 | TRP | p.Trp169Ter | Q9HAN9 |
| HOMO SAPIENS | 0 | 2525 | 1kqn | F | 207 | ARG | p.Arg207Trp | Q9HAN9 |
| HOMO SAPIENS | 0 | 2525 | 1kqn | F | 273 | ASN | p.Asn273Asp | Q9HAN9 |
| HOMO SAPIENS | 0 | 12299 | 5duq | A | 432 | VAL | p.Val454Glu | P05155 |
| HOMO SAPIENS | 0 | 12299 | 5duq | A | 436 | ALA | p.Ala458Thr | P05155 |
| HOMO SAPIENS | 0 | 12299 | 5duq | A | 443 | ALA | p.Ala465Val | P05155 |
| HOMO SAPIENS | 0 | 12299 | 5duq | A | 444 | ARG | p.Arg466His | P05155 |
| HOMO SAPIENS | 0 | 12299 | 5duq | A | 458 | MET | p.Val480Met | P05155 |
| HOMO SAPIENS | 0 | 327 | 1uhp | A | 36 | ASP | p.Asp274Asn | Q9UPQ7 |
| HOMO SAPIENS | 0 | 64 | 2mo5 | A | 54 | ALA | p.Ala55Thr | P12104 |
| HOMO SAPIENS | 2 | 35969 | 1jrh | I | 60 | CYS | p.Cys77Tyr | P15260 |
| HOMO SAPIENS | 4 | 35969 | 1jrh | I | 70 | ILE | p.Ile87Thr | P15260 |
| HOMO SAPIENS | 2 | 1068 | 1mu7 | A | 304 | ARG | p.Arg304Gln | Q9NUW8 |
| HOMO SAPIENS | 8 | 1068 | 1mu7 | A | 493 | HIS | p.His493Arg | Q9NUW8 |
| HOMO SAPIENS | 0 | 32360 | 2kqp | A | 31 | ARG | p.Arg55Cys | P01308 |
| HOMO SAPIENS | 0 | 32360 | 2kqp | A | 65 | ARG | p.Arg89His | P01308 |
| HOMO SAPIENS | 0 | 32360 | 2kqp | A | 65 | ARG | p.Arg89Cys | P01308 |
| HOMO SAPIENS | 6 | 10805 | 3nks | A | 12 | ILE | p.Ile12Thr | P50336 |
| HOMO SAPIENS | 0 | 10805 | 3nks | A | 20 | HIS | p.His20Pro | P50336 |
| HOMO SAPIENS | 0 | 10805 | 3nks | A | 59 | ARG | p.Arg59Trp | P50336 |
| HOMO SAPIENS | 0 | 10805 | 3nks | A | 168 | ARG | p.Arg168Cys | P50336 |
| HOMO SAPIENS | 0 | 10805 | 3nks | A | 168 | ARG | p.Arg168His | P50336 |
| HOMO SAPIENS | 2 | 10805 | 3nks | A | 232 | GLY | p.Gly232Arg | P50336 |
| HOMO SAPIENS | 3 | 10805 | 3nks | A | 256 | PRO | p.Pro256Arg | P50336 |
| HOMO SAPIENS | 0 | 10805 | 3nks | A | 304 | ARG | p.Arg304His | P50336 |
| HOMO SAPIENS | 0 | 10805 | 3nks | A | 349 | ASP | p.Asp349Ala | P50336 |
| HOMO SAPIENS | 9 | 7008 | 3u2l | A | 125 | GLN | p.Gln125Ter | P55789 |
| HOMO SAPIENS | 4 | 7008 | 3u2l | A | 172 | HIS | p.His172Asn | P55789 |
| HOMO SAPIENS | 4 | 7008 | 3u2l | A | 194 | ARG | p.Arg194His | P55789 |
| HOMO SAPIENS | 3 | 7008 | 3u2l | A | 198 | GLY | p.Gly198Ser | P55789 |
| HOMO SAPIENS | 0 | 20821 | 3e0g | A | 65 | HIS | p.His116Tyr | P42702 |
| HOMO SAPIENS | 3 | 1151 | 5l9v | A | 278 | ASP | p.Asp278Asn | Q9GZT9 |
| HOMO SAPIENS | 4 | 1151 | 5l9v | A | 317 | CYS | p.Pro317Arg | Q9GZT9 |
| HOMO SAPIENS | 5 | 1151 | 5l9v | A | 371 | ARG | p.Arg371His | Q9GZT9 |
| HOMO SAPIENS | 2 | 3935 | 1gwb | A | 36 | ARG | p.Arg36Gly | P07359 |
| HOMO SAPIENS | 1 | 3935 | 1gwb | A | 73 | LEU | p.Leu73Phe | P07359 |
| HOMO SAPIENS | 1 | 3935 | 1gwb | A | 172 | ALA | p.Ala172Val | P07359 |
| HOMO SAPIENS | 0 | 3935 | 1gwb | A | 249 | GLY | p.Gly249Val | P07359 |
| HOMO SAPIENS | 0 | 3935 | 1gwb | A | 255 | MET | p.Met255Val | P07359 |
| HOMO SAPIENS | 2 | 14541 | 5cuz | A | 138 | GLU | p.Glu138Lys | Q9H3L0 |
| HOMO SAPIENS | 2 | 14541 | 5cuz | A | 143 | SER | p.Ser143Ile | Q9H3L0 |
| HOMO SAPIENS | 4 | 14541 | 5cuz | A | 182 | THR | p.Thr182Asn | Q9H3L0 |
| HOMO SAPIENS | 6 | 14541 | 5cuz | A | 206 | ASN | p.Asn206Ile | Q9H3L0 |

| | | | | | | | | |
|---|---|---|---|---|---|---|---|---|
| HOMO SAPIENS | 0 | 14541 | 5cuz | A | 249 | TYR | p.Tyr249Cys | Q9H3L0 |
| HOMO SAPIENS | 4 | 14541 | 5cuz | A | 250 | ARG | p.Arg250Ter | Q9H3L0 |
| HOMO SAPIENS | 2 | 14541 | 5cuz | A | 259 | LEU | p.Leu259Pro | Q9H3L0 |
| HOMO SAPIENS | 0 | 4206 | 4fao | D | 30 | PRO | p.Pro30Ser | P37023 |
| HOMO SAPIENS | 1 | 4206 | 4fao | D | 50 | TRP | p.Trp50Cys | P37023 |
| HOMO SAPIENS | 0 | 16553 | 3v30 | A | 251 | GLN | p.Gln251Glu | O14593 |
| HOMO SAPIENS | 0 | 5012 | 5hbf | A | 442 | ALA | p.Ala442Gly | Q13231 |
| HOMO SAPIENS | 5 | 4844 | 5tdc | A | 136 | HIS | p.His136Arg | Q8IWV7 |
| HOMO SAPIENS | 0 | 15435 | 4cgz | A | 643 | ARG | p.Arg643His | P54132 |
| HOMO SAPIENS | 1 | 15435 | 4cgz | A | 672 | GLN | p.Gln672Arg | P54132 |
| HOMO SAPIENS | 1 | 15435 | 4cgz | A | 700 | GLN | p.Gln700Ter | P54132 |
| HOMO SAPIENS | 0 | 15435 | 4cgz | A | 707 | PRO | p.Pro707Ser | P54132 |
| HOMO SAPIENS | 0 | 15435 | 4cgz | A | 755 | LYS | p.Lys755Glu | P54132 |
| HOMO SAPIENS | 0 | 15435 | 4cgz | A | 881 | TRP | p.Trp881Ter | P54132 |
| HOMO SAPIENS | 2 | 15435 | 4cgz | A | 899 | ARG | p.Arg899Ter | P54132 |
| HOMO SAPIENS | 0 | 15435 | 4cgz | A | 1036 | CYS | p.Cys1036Phe | P54132 |
| HOMO SAPIENS | 0 | 15435 | 4cgz | A | 1055 | CYS | p.Cys1055Ser | P54132 |
| HOMO SAPIENS | 0 | 15435 | 4cgz | A | 1139 | ARG | p.Arg1139Ter | P54132 |
| HOMO SAPIENS | -1 | 15435 | 4cgz | A | 1198 | VAL | p.Val1198Met | P54132 |
| HOMO SAPIENS | -1 | 15435 | 4cgz | A | 1205 | VAL | p.Val1205Ile | P54132 |
| HOMO SAPIENS | 0 | 15435 | 4cgz | A | 1251 | GLU | p.Glu1251Gln | P54132 |
| HOMO SAPIENS | 0 | 11185 | 1kb4 | A | 30 | ARG | p.Arg30Ter | P11473 |
| HOMO SAPIENS | 5 | 11185 | 1kb4 | A | 33 | GLY | p.Gly33Asp | P11473 |
| HOMO SAPIENS | 5 | 11185 | 1kb4 | A | 46 | GLY | p.Gly46Asp | P11473 |
| HOMO SAPIENS | 6 | 11185 | 1kb4 | A | 50 | ARG | p.Arg50Gln | P11473 |
| HOMO SAPIENS | 9 | 11185 | 1kb4 | A | 73 | ARG | p.Arg73Gln | P11473 |
| HOMO SAPIENS | 9 | 11185 | 1kb4 | A | 80 | ARG | p.Arg80Gln | P11473 |
| HOMO SAPIENS | 0 | 5382 | 3fl7 | A | 41 | LEU | p.Leu41Val | P29317 |
| HOMO SAPIENS | 2 | 5382 | 3fl7 | A | 175 | ARG | p.Arg175Cys | P29317 |
| HOMO SAPIENS | 1 | 5382 | 3fl7 | A | 195 | ARG | p.Arg195Cys | P29317 |
| HOMO SAPIENS | 0 | 5382 | 3fl7 | A | 232 | ASP | p.Asp232Gly | P29317 |
| HOMO SAPIENS | 0 | 5382 | 3fl7 | A | 277 | SER | p.Ser277Leu | P29317 |
| HOMO SAPIENS | 0 | 5382 | 3fl7 | A | 315 | ARG | p.Arg315Gln | P29317 |
| HOMO SAPIENS | 7 | 5382 | 3fl7 | A | 350 | PRO | p.Pro350Thr | P29317 |
| HOMO SAPIENS | 0 | 5382 | 3fl7 | A | 391 | GLY | p.Gly391Arg | P29317 |
| HOMO SAPIENS | 0 | 5382 | 3fl7 | A | 418 | VAL | p.Val418Ile | P29317 |
| HOMO SAPIENS | 0 | 5382 | 3fl7 | A | 511 | THR | p.Thr511Met | P29317 |
| HOMO SAPIENS | 2 | 29168 | 4p22 | B | 144 | VAL | p.Val144Ile | P22314 |
| HOMO SAPIENS | 5 | 4348 | 1bf5 | A | 241 | ARG | p.Arg241Gln | P42224 |
| HOMO SAPIENS | 2 | 4348 | 1bf5 | A | 266 | VAL | p.Val266Ile | P42224 |
| HOMO SAPIENS | 4 | 4348 | 1bf5 | A | 320 | GLU | p.Glu320Gln | P42224 |
| HOMO SAPIENS | 1 | 4348 | 1bf5 | A | 372 | ASN | p.Asn372Lys | P42224 |
| HOMO SAPIENS | 1 | 4348 | 1bf5 | A | 447 | ASP | p.Asp447Glu | P42224 |
| HOMO SAPIENS | 2 | 4348 | 1bf5 | A | 463 | GLN | p.Gln463His | P42224 |
| HOMO SAPIENS | 4 | 4348 | 1bf5 | A | 600 | LEU | p.Leu600Pro | P42224 |
| HOMO SAPIENS | 0 | 4348 | 1bf5 | A | 706 | LEU | p.Leu706Ser | P42224 |
| HOMO SAPIENS | 4 | 4991 | 4bej | B | 141 | ASN | p.Asn141Ser | O00429 |
| HOMO SAPIENS | 0 | 4991 | 4bej | B | 395 | ALA | p.Ala395Asp | O00429 |
| HOMO SAPIENS | 2 | 11792 | 1fvo | A | 40 | ARG | p.Arg40Cys | P00480 |
| HOMO SAPIENS | 2 | 11792 | 1fvo | A | 40 | ARG | p.Arg40His | P00480 |
| HOMO SAPIENS | 0 | 11792 | 1fvo | A | 45 | LEU | p.Leu45Pro | P00480 |
| HOMO SAPIENS | 1 | 11792 | 1fvo | A | 47 | ASN | p.Asn47Ile | P00480 |
| HOMO SAPIENS | 0 | 11792 | 1fvo | A | 50 | GLY | p.Gly50Arg | P00480 |
| HOMO SAPIENS | 5 | 11792 | 1fvo | A | 87 | GLU | p.Glu87Lys | P00480 |

| | | | | | | | | |
|---|---|---|---|---|---|---|---|---|
| HOMO SAPIENS | 3 | 11792 | 1fvo | A | 111 | LEU | p.Leu111Pro | P00480 |
| HOMO SAPIENS | 1 | 11792 | 1fvo | A | 125 | THR | p.Thr125Met | P00480 |
| HOMO SAPIENS | 3 | 11792 | 1fvo | A | 129 | ARG | p.Arg129His | P00480 |
| HOMO SAPIENS | 9 | 11792 | 1fvo | A | 141 | ARG | p.Arg141Pro | P00480 |
| HOMO SAPIENS | 0 | 11792 | 1fvo | A | 206 | MET | p.Met206Arg | P00480 |
| HOMO SAPIENS | 0 | 11792 | 1fvo | A | 207 | SER | p.Ser207Arg | P00480 |
| HOMO SAPIENS | 0 | 11792 | 1fvo | A | 216 | GLN | p.Gln216Glu | P00480 |
| HOMO SAPIENS | 1 | 11792 | 1fvo | A | 225 | PRO | p.Pro225Leu | P00480 |
| HOMO SAPIENS | 0 | 11792 | 1fvo | A | 270 | ARG | p.Gln270Arg | P00480 |
| HOMO SAPIENS | 0 | 11792 | 1fvo | A | 270 | ARG | p.Gln270Ter | P00480 |
| HOMO SAPIENS | 2 | 11792 | 1fvo | A | 277 | ARG | p.Arg277Trp | P00480 |
| HOMO SAPIENS | 0 | 3721 | 4r3z | B | 2 | ASP | p.Asp2Gly | P54136 |
| HOMO SAPIENS | 0 | 3721 | 4r3z | B | 456 | SER | p.Ser456Leu | P54136 |
| HOMO SAPIENS | 5 | 3721 | 4r3z | B | 512 | ARG | p.Arg512Gln | P54136 |
| HOMO SAPIENS | 3 | 5749 | 3hl2 | A | 239 | ALA | p.Ala239Thr | Q9HD40 |
| HOMO SAPIENS | 1 | 5749 | 3hl2 | A | 334 | TYR | p.Tyr334Cys | Q9HD40 |
| HOMO SAPIENS | 1 | 5749 | 3hl2 | A | 452 | LYS | p.Lys452Asn | Q9HD40 |
| HOMO SAPIENS | -1 | 5749 | 3hl2 | A | 465 | ARG | p.Arg465Leu | Q9HD40 |
| HOMO SAPIENS | -1 | 5749 | 3hl2 | A | 489 | ASP | p.Asp489Gly | Q9HD40 |
| HOMO SAPIENS | 5 | 1721 | 3k15 | A | 1652 | MET | p.Met1652Ile | P38398 |
| HOMO SAPIENS | 5 | 1721 | 3k15 | A | 1652 | MET | p.Met1652Lys | P38398 |
| HOMO SAPIENS | 5 | 1721 | 3k15 | A | 1662 | PHE | p.Phe1662Cys | P38398 |
| HOMO SAPIENS | 5 | 1721 | 3k15 | A | 1664 | LEU | p.Leu1664Pro | P38398 |
| HOMO SAPIENS | 0 | 1721 | 3k15 | A | 1665 | VAL | p.Val1665Met | P38398 |
| HOMO SAPIENS | 5 | 1721 | 3k15 | A | 1690 | LYS | p.Lys1690Gln | P38398 |
| HOMO SAPIENS | 5 | 1721 | 3k15 | A | 1692 | ASP | p.Asp1692Asn | P38398 |
| HOMO SAPIENS | 5 | 1721 | 3k15 | A | 1699 | ARG | p.Arg1699Gln | P38398 |
| HOMO SAPIENS | 5 | 1721 | 3k15 | A | 1699 | ARG | p.Arg1699Trp | P38398 |
| HOMO SAPIENS | 3 | 1721 | 3k15 | A | 1706 | GLY | p.Gly1706Glu | P38398 |
| HOMO SAPIENS | 5 | 1721 | 3k15 | A | 1720 | THR | p.Thr1720Ala | P38398 |
| HOMO SAPIENS | 5 | 1721 | 3k15 | A | 1751 | ARG | p.Arg1751Pro | P38398 |
| HOMO SAPIENS | 4 | 1721 | 3k15 | A | 1775 | MET | p.Met1775Arg | P38398 |
| HOMO SAPIENS | 2 | 1721 | 3k15 | A | 1788 | GLY | p.Gly1788Val | P38398 |
| HOMO SAPIENS | 5 | 1721 | 3k15 | A | 1794 | GLU | p.Glu1794Asp | P38398 |
| HOMO SAPIENS | 5 | 1721 | 3k15 | A | 1804 | VAL | p.Val1804Asp | P38398 |
| HOMO SAPIENS | 3 | 1721 | 3k15 | A | 1815 | TRP | p.Trp1815Ter | P38398 |
| HOMO SAPIENS | 1 | 1721 | 3k15 | A | 1819 | ASN | p.Asn1819Ser | P38398 |
| HOMO SAPIENS | 5 | 1721 | 3k15 | A | 1856 | PRO | p.Pro1856Ser | P38398 |
| HOMO SAPIENS | 5 | 1721 | 3k15 | A | 1858 | ILE | p.Ile1858Leu | P38398 |
| HOMO SAPIENS | 5 | 1721 | 3k15 | A | 1858 | ILE | p.Ile1858Thr | P38398 |
| HOMO SAPIENS | 5 | 1721 | 3k15 | A | 1859 | PRO | p.Pro1859Arg | P38398 |
| HOMO SAPIENS | 2 | 22086 | 3kq4 | B | 1297 | PRO | p.Pro1297Leu | O60494 |
| HOMO SAPIENS | 1 | 12135 | 2k9j | B | 724 | ARG | p.Arg750Ter | P05106 |
| HOMO SAPIENS | -1 | 24027 | 2qtl | A | 175 | LEU | p.Ser202Leu | Q9UBK8 |
| HOMO SAPIENS | 0 | 24027 | 2qtl | A | 257 | SER | p.Ser284Thr | Q9UBK8 |
| HOMO SAPIENS | 0 | 24027 | 2qtl | A | 333 | LEU | p.Leu360Val | Q9UBK8 |
| HOMO SAPIENS | 0 | 24027 | 2qtl | A | 350 | LYS | p.Lys377Arg | Q9UBK8 |
| HOMO SAPIENS | 0 | 24027 | 2qtl | A | 415 | ARG | p.Arg442Cys | Q9UBK8 |
| HOMO SAPIENS | 2 | 24027 | 2qtl | A | 450 | PRO | p.Pro477Arg | Q9UBK8 |
| HOMO SAPIENS | 4 | 24027 | 2qtl | A | 487 | GLY | p.Gly514Arg | Q9UBK8 |
| HOMO SAPIENS | 1 | 24027 | 2qtl | A | 595 | HIS | p.His622Tyr | Q9UBK8 |
| HOMO SAPIENS | 0 | 5758 | 3di2 | B | 46 | THR | p.Thr66Ile | P16871 |
| HOMO SAPIENS | 1 | 5758 | 3di2 | B | 118 | VAL | p.Ile138Val | P16871 |
| HOMO SAPIENS | 4 | 848 | 5oa3 | Y | 79 | LEU | p.Leu79Val | P62847 |

| | | | | | | | | |
|---|---|---|---|---|---|---|---|---|
| HOMO SAPIENS | 2 | 848 | 5oa3 | Y | 124 | ASN | p.Asn124Ser | P62847 |
| HOMO SAPIENS | 0 | 81 | 3s5o | A | 36 | ALA | p.Ala36Glu | Q86XE5 |
| HOMO SAPIENS | 0 | 81 | 3s5o | A | 39 | TYR | p.Tyr39Ter | Q86XE5 |
| HOMO SAPIENS | 0 | 81 | 3s5o | A | 45 | PRO | p.Pro45Leu | Q86XE5 |
| HOMO SAPIENS | 0 | 81 | 3s5o | A | 70 | ARG | p.Arg70Ter | Q86XE5 |
| HOMO SAPIENS | 0 | 81 | 3s5o | A | 70 | ARG | p.Arg70Pro | Q86XE5 |
| HOMO SAPIENS | 1 | 81 | 3s5o | A | 97 | ARG | p.Arg97Cys | Q86XE5 |
| HOMO SAPIENS | 4 | 81 | 3s5o | A | 113 | GLU | p.Glu113Lys | Q86XE5 |
| HOMO SAPIENS | 1 | 81 | 3s5o | A | 190 | PRO | p.Pro190Leu | Q86XE5 |
| HOMO SAPIENS | 0 | 81 | 3s5o | A | 245 | VAL | p.Val245Phe | Q86XE5 |
| HOMO SAPIENS | 0 | 81 | 3s5o | A | 257 | CYS | p.Cys257Gly | Q86XE5 |
| HOMO SAPIENS | 0 | 81 | 3s5o | A | 287 | GLY | p.Gly287Val | Q86XE5 |
| HOMO SAPIENS | 7 | 81 | 3s5o | A | 303 | ARG | p.Arg303Cys | Q86XE5 |
| HOMO SAPIENS | 0 | 81 | 3s5o | A | 325 | GLY | p.Gly325Ser | Q86XE5 |
| HOMO SAPIENS | -1 | 622 | 5era | A | 12 | GLY | p.Gly12Arg | P29033 |
| HOMO SAPIENS | -1 | 622 | 5era | A | 12 | GLY | p.Gly12Asp | P29033 |
| HOMO SAPIENS | -1 | 622 | 5era | A | 17 | SER | p.Ser17Phe | P29033 |
| HOMO SAPIENS | 5 | 622 | 5era | A | 24 | TRP | p.Trp24Ter | P29033 |
| HOMO SAPIENS | 0 | 622 | 5era | A | 32 | ARG | p.Arg32Cys | P29033 |
| HOMO SAPIENS | 5 | 622 | 5era | A | 34 | MET | p.Met34Thr | P29033 |
| HOMO SAPIENS | 5 | 622 | 5era | A | 36 | LEU | p.Leu36Pro | P29033 |
| HOMO SAPIENS | 5 | 622 | 5era | A | 37 | VAL | p.Val37Ile | P29033 |
| HOMO SAPIENS | 5 | 622 | 5era | A | 40 | ALA | p.Ala40Glu | P29033 |
| HOMO SAPIENS | 5 | 622 | 5era | A | 44 | TRP | p.Trp44Cys | P29033 |
| HOMO SAPIENS | 5 | 622 | 5era | A | 45 | GLY | p.Gly45Glu | P29033 |
| HOMO SAPIENS | 5 | 622 | 5era | A | 47 | GLU | p.Glu47Ter | P29033 |
| HOMO SAPIENS | 5 | 622 | 5era | A | 50 | ASP | p.Asp50Asn | P29033 |
| HOMO SAPIENS | 5 | 622 | 5era | A | 54 | ASN | p.Asn54Lys | P29033 |
| HOMO SAPIENS | 5 | 622 | 5era | A | 57 | GLN | p.Gln57Ter | P29033 |
| HOMO SAPIENS | 5 | 622 | 5era | A | 59 | GLY | p.Gly59Ala | P29033 |
| HOMO SAPIENS | 5 | 622 | 5era | A | 59 | GLY | p.Gly59Ser | P29033 |
| HOMO SAPIENS | 5 | 622 | 5era | A | 66 | ASP | p.Asp66His | P29033 |
| HOMO SAPIENS | 5 | 622 | 5era | A | 68 | TYR | p.Tyr68Cys | P29033 |
| HOMO SAPIENS | 5 | 622 | 5era | A | 70 | PRO | p.Pro70Ala | P29033 |
| HOMO SAPIENS | 5 | 622 | 5era | A | 73 | HIS | p.His73Arg | P29033 |
| HOMO SAPIENS | 5 | 622 | 5era | A | 75 | ARG | p.Arg75Trp | P29033 |
| HOMO SAPIENS | 5 | 622 | 5era | A | 75 | ARG | p.Arg75Gln | P29033 |
| HOMO SAPIENS | 2 | 622 | 5era | A | 77 | TRP | p.Trp77Arg | P29033 |
| HOMO SAPIENS | 2 | 622 | 5era | A | 77 | TRP | p.Trp77Ter | P29033 |
| HOMO SAPIENS | 0 | 622 | 5era | A | 80 | GLN | p.Gln80Ter | P29033 |
| HOMO SAPIENS | 0 | 622 | 5era | A | 80 | GLN | p.Gln80Pro | P29033 |
| HOMO SAPIENS | 5 | 622 | 5era | A | 81 | LEU | p.Leu81Val | P29033 |
| HOMO SAPIENS | 5 | 622 | 5era | A | 82 | ILE | p.Ile82Met | P29033 |
| HOMO SAPIENS | 5 | 622 | 5era | A | 83 | PHE | p.Phe83Leu | P29033 |
| HOMO SAPIENS | 5 | 622 | 5era | A | 84 | VAL | p.Val84Leu | P29033 |
| HOMO SAPIENS | 5 | 622 | 5era | A | 90 | LEU | p.Leu90Pro | P29033 |
| HOMO SAPIENS | 5 | 622 | 5era | A | 95 | VAL | p.Val95Met | P29033 |
| HOMO SAPIENS | -1 | 622 | 5era | A | 100 | HIS | p.His100Tyr | P29033 |
| HOMO SAPIENS | -1 | 622 | 5era | A | 113 | SER | p.Ser113Arg | P29033 |
| HOMO SAPIENS | -1 | 622 | 5era | A | 119 | GLU | p.Glu119Lys | P29033 |
| HOMO SAPIENS | -1 | 622 | 5era | A | 122 | LYS | p.Lys122Ile | P29033 |
| HOMO SAPIENS | -1 | 622 | 5era | A | 123 | THR | p.Thr123Asn | P29033 |
| HOMO SAPIENS | -1 | 622 | 5era | A | 124 | GLN | p.Gln124Ter | P29033 |
| HOMO SAPIENS | 5 | 622 | 5era | A | 139 | SER | p.Ser139Asn | P29033 |

| | | | | | | | | |
|---|---|---|---|---|---|---|---|---|
| HOMO SAPIENS | 5 | 622 | 5era | A | 143 | ARG | p.Arg143Trp | P29033 |
| HOMO SAPIENS | 5 | 622 | 5era | A | 143 | ARG | p.Arg143Gln | P29033 |
| HOMO SAPIENS | 0 | 622 | 5era | A | 147 | GLU | p.Glu147Lys | P29033 |
| HOMO SAPIENS | 7 | 622 | 5era | A | 155 | TYR | p.Tyr155Ter | P29033 |
| HOMO SAPIENS | 2 | 622 | 5era | A | 159 | ASP | p.Asp159Val | P29033 |
| HOMO SAPIENS | 5 | 622 | 5era | A | 163 | MET | p.Met163Val | P29033 |
| HOMO SAPIENS | 5 | 622 | 5era | A | 165 | ARG | p.Arg165Trp | P29033 |
| HOMO SAPIENS | 5 | 622 | 5era | A | 167 | VAL | p.Val167Met | P29033 |
| HOMO SAPIENS | 5 | 622 | 5era | A | 169 | CYS | p.Cys169Tyr | P29033 |
| HOMO SAPIENS | 5 | 622 | 5era | A | 179 | ASP | p.Asp179Asn | P29033 |
| HOMO SAPIENS | 0 | 622 | 5era | A | 199 | SER | p.Ser199Phe | P29033 |
| HOMO SAPIENS | 0 | 622 | 5era | A | 202 | CYS | p.Cys202Phe | P29033 |
| HOMO SAPIENS | 5 | 622 | 5era | A | 203 | ILE | p.Ile203Thr | P29033 |
| HOMO SAPIENS | 5 | 622 | 5era | A | 206 | ASN | p.Asn206Ser | P29033 |
| HOMO SAPIENS | 0 | 8103 | 5dot | A | 163 | GLY | p.Gly163Ter | P31327 |
| HOMO SAPIENS | 0 | 8103 | 5dot | A | 337 | HIS | p.His337Arg | P31327 |
| HOMO SAPIENS | 3 | 8103 | 5dot | A | 344 | THR | p.Thr344Ala | P31327 |
| HOMO SAPIENS | 1 | 8103 | 5dot | A | 405 | THR | p.Thr405Asn | P31327 |
| HOMO SAPIENS | 5 | 8103 | 5dot | A | 544 | THR | p.Thr544Met | P31327 |
| HOMO SAPIENS | 5 | 8103 | 5dot | A | 716 | ASN | p.Asn716Lys | P31327 |
| HOMO SAPIENS | -1 | 8103 | 5dot | A | 780 | ARG | p.Arg780His | P31327 |
| HOMO SAPIENS | 0 | 8103 | 5dot | A | 982 | GLY | p.Gly982Asp | P31327 |
| HOMO SAPIENS | 3 | 8103 | 5dot | A | 1119 | ALA | p.Ala1119Ser | P31327 |
| HOMO SAPIENS | 1 | 8103 | 5dot | A | 1376 | GLY | p.Gly1376Ser | P31327 |
| HOMO SAPIENS | 4 | 6833 | 1l8l | B | 27 | ARG | p.Arg27Ser | P78330 |
| HOMO SAPIENS | 3 | 6833 | 1l8l | B | 32 | ASP | p.Asp32Gly | P78330 |
| HOMO SAPIENS | 3 | 6833 | 1l8l | B | 32 | ASP | p.Asp32Asn | P78330 |
| HOMO SAPIENS | 1 | 6833 | 1l8l | B | 49 | ARG | p.Arg49Trp | P78330 |
| HOMO SAPIENS | 0 | 6833 | 1l8l | B | 52 | MET | p.Met52Thr | P78330 |
| HOMO SAPIENS | 0 | 6833 | 1l8l | B | 83 | GLN | p.Gln83His | P78330 |
| HOMO SAPIENS | 6 | 810 | 2iw2 | A | 185 | ARG | p.Arg184Gln | P12955 |
| HOMO SAPIENS | 0 | 810 | 2iw2 | A | 213 | ALA | p.Ala212Pro | P12955 |
| HOMO SAPIENS | 1 | 810 | 2iw2 | A | 266 | ARG | p.Arg265Ter | P12955 |
| HOMO SAPIENS | 9 | 810 | 2iw2 | A | 277 | ASP | p.Asp276Asn | P12955 |
| HOMO SAPIENS | 0 | 810 | 2iw2 | A | 279 | GLY | p.Gly278Asp | P12955 |
| HOMO SAPIENS | 9 | 810 | 2iw2 | A | 378 | HIS | p.His377Gln | P12955 |
| HOMO SAPIENS | 0 | 810 | 2iw2 | A | 389 | ARG | p.Arg388His | P12955 |
| HOMO SAPIENS | 0 | 810 | 2iw2 | A | 436 | PHE | p.Leu435Phe | P12955 |
| HOMO SAPIENS | 0 | 810 | 2iw2 | A | 449 | GLY | p.Gly448Arg | P12955 |
| HOMO SAPIENS | -1 | 1530 | 5fwj | A | 388 | ALA | p.Ala388Pro | P41229 |
| HOMO SAPIENS | 4 | 1530 | 5fwj | A | 451 | SER | p.Ser451Arg | P41229 |
| HOMO SAPIENS | 3 | 1530 | 5fwj | A | 731 | LEU | p.Leu731Phe | P41229 |
| HOMO SAPIENS | 3 | 578 | 5net | B | 123 | ALA | p.Ala143Thr | P18564 |
| HOMO SAPIENS | 6 | 578 | 5net | B | 176 | PRO | p.Pro196Thr | P18564 |
| HOMO SAPIENS | 3 | 578 | 5net | B | 280 | GLU | p.Glu300Lys | P18564 |
| HOMO SAPIENS | -1 | 5247 | 4j15 | A | 274 | ALA | p.Ala274Val | P14868 |
| HOMO SAPIENS | -1 | 5247 | 4j15 | A | 280 | HIS | p.His280Leu | P14868 |
| HOMO SAPIENS | 2 | 5247 | 4j15 | A | 460 | ARG | p.Arg460His | P14868 |
| HOMO SAPIENS | 5 | 5247 | 4j15 | A | 494 | ARG | p.Arg494Cys | P14868 |
| HOMO SAPIENS | 7 | 2566 | 1b55 | A | 28 | ARG | p.Arg28His | Q06187 |
| HOMO SAPIENS | 5 | 2566 | 1b55 | A | 33 | THR | p.Thr33Pro | Q06187 |
| HOMO SAPIENS | 0 | 2566 | 1b55 | A | 113 | VAL | p.Val113Asp | Q06187 |
| HOMO SAPIENS | 0 | 356 | 5a2s | A | 786 | ALA | p.Ala786Thr | P56524 |
| HOMO SAPIENS | 5 | 2960 | 4cak | A | 553 | ARG | p.Arg584Ter | P08514 |

| | | | | | | | | |
|---|---|---|---|---|---|---|---|---|
| HOMO SAPIENS | 5 | 2960 | 4cak | A | 565 | ILE | p.Ile596Thr | P08514 |
| HOMO SAPIENS | -1 | 2960 | 4cak | A | 841 | LEU | p.Leu872Met | P08514 |
| HOMO SAPIENS | -1 | 2960 | 4cak | A | 843 | ILE | p.Ile874Ser | P08514 |
| HOMO SAPIENS | 0 | 5054 | 4k8r | B | 216 | PHE | p.Phe216Leu | Q8NBP7 |
| HOMO SAPIENS | 1 | 5054 | 4k8r | B | 417 | HIS | p.His417Gln | Q8NBP7 |
| HOMO SAPIENS | 1 | 5054 | 4k8r | B | 425 | ASN | p.Asn425Ser | Q8NBP7 |
| HOMO SAPIENS | 1 | 5054 | 4k8r | B | 443 | ALA | p.Ala443Thr | Q8NBP7 |
| HOMO SAPIENS | 0 | 27283 | 3n5n | X | 71 | HIS | p.His82Arg | Q9UIF7 |
| HOMO SAPIENS | 1 | 27283 | 3n5n | X | 83 | ARG | p.Arg94Ter | Q9UIF7 |
| HOMO SAPIENS | 8 | 27283 | 3n5n | X | 89 | TRP | p.Trp100Ter | Q9UIF7 |
| HOMO SAPIENS | 8 | 27283 | 3n5n | X | 90 | TYR | p.Tyr101Ter | Q9UIF7 |
| HOMO SAPIENS | 7 | 27283 | 3n5n | X | 95 | ARG | p.Arg106Trp | Q9UIF7 |
| HOMO SAPIENS | 7 | 27283 | 3n5n | X | 117 | TRP | p.Trp128Arg | Q9UIF7 |
| HOMO SAPIENS | 0 | 27283 | 3n5n | X | 137 | GLY | p.Gly148Arg | Q9UIF7 |
| HOMO SAPIENS | 5 | 27283 | 3n5n | X | 142 | TRP | p.Trp153Ter | Q9UIF7 |
| HOMO SAPIENS | 0 | 27283 | 3n5n | X | 165 | TYR | p.Tyr176Cys | Q9UIF7 |
| HOMO SAPIENS | 0 | 27283 | 3n5n | X | 168 | ARG | p.Arg179His | Q9UIF7 |
| HOMO SAPIENS | 0 | 27283 | 3n5n | X | 168 | ARG | p.Arg179Cys | Q9UIF7 |
| HOMO SAPIENS | 0 | 27283 | 3n5n | X | 196 | GLN | p.Gln207Ter | Q9UIF7 |
| HOMO SAPIENS | 8 | 27283 | 3n5n | X | 201 | VAL | p.Val212Met | Q9UIF7 |
| HOMO SAPIENS | 9 | 27283 | 3n5n | X | 227 | ARG | p.Arg238Trp | Q9UIF7 |
| HOMO SAPIENS | 9 | 27283 | 3n5n | X | 231 | ARG | p.Arg242Cys | Q9UIF7 |
| HOMO SAPIENS | 9 | 27283 | 3n5n | X | 231 | ARG | p.Arg242His | Q9UIF7 |
| HOMO SAPIENS | 3 | 27283 | 3n5n | X | 232 | VAL | p.Val243Phe | Q9UIF7 |
| HOMO SAPIENS | 3 | 27283 | 3n5n | X | 246 | GLN | p.Gln257Ter | Q9UIF7 |
| HOMO SAPIENS | 9 | 27283 | 3n5n | X | 272 | GLY | p.Gly283Glu | Q9UIF7 |
| HOMO SAPIENS | 9 | 27283 | 3n5n | X | 281 | PRO | p.Pro292Leu | Q9UIF7 |
| HOMO SAPIENS | -1 | 27283 | 3n5n | X | 314 | ASP | p.Asp325Glu | Q9UIF7 |
| HOMO SAPIENS | 0 | 27283 | 3n5n | X | 324 | GLN | p.Gln335His | Q9UIF7 |
| HOMO SAPIENS | 0 | 27283 | 3n5n | X | 324 | GLN | p.Gln335Arg | Q9UIF7 |
| HOMO SAPIENS | 0 | 27283 | 3n5n | X | 324 | GLN | p.Gln335Ter | Q9UIF7 |
| HOMO SAPIENS | -1 | 20852 | 4p60 | B | 243 | ARG | p.Arg243Lys | O75746 |
| HOMO SAPIENS | -1 | 1875 | 5dd2 | G | 152 | VAL | p.Val179Leu | P06396 |
| HOMO SAPIENS | 9 | 2126 | 3n1p | B | 46 | PRO | p.Pro46Leu | Q14623 |
| HOMO SAPIENS | 8 | 2126 | 3n1p | B | 76 | GLU | p.Glu76Ala | Q14623 |
| HOMO SAPIENS | 9 | 2126 | 3n1p | B | 77 | ARG | p.Arg77Ser | Q14623 |
| HOMO SAPIENS | 6 | 2126 | 3n1p | B | 95 | GLU | p.Glu95Lys | Q14623 |
| HOMO SAPIENS | 9 | 2126 | 3n1p | B | 100 | ASP | p.Asp100Glu | Q14623 |
| HOMO SAPIENS | 9 | 2126 | 3n1p | B | 131 | GLU | p.Glu131Lys | Q14623 |
| HOMO SAPIENS | 0 | 2126 | 3n1p | B | 190 | VAL | p.Val190Ala | Q14623 |
| HOMO SAPIENS | 5 | 40410 | 3ioz | B | 70 | GLN | p.Gln70Leu | P20963 |
| HOMO SAPIENS | 1 | 5754 | 5lkz | A | 1108 | THR | p.Tyr1191Ter | Q09472 |
| HOMO SAPIENS | 3 | 8159 | 1tiu | A | 3 | GLU | p.Lys12679Glu | Q8WZ42 |
| HOMO SAPIENS | 4 | 3275 | 4ldq | A | 210 | ARG | p.Arg210Gln | Q9BV20 |
| HOMO SAPIENS | 0 | 8143 | 4miv | B | 66 | SER | p.Ser66Trp | P51688 |
| HOMO SAPIENS | 1 | 8143 | 4miv | B | 74 | ARG | p.Arg74Cys | P51688 |
| HOMO SAPIENS | 0 | 8143 | 4miv | B | 128 | PRO | p.Pro128Leu | P51688 |
| HOMO SAPIENS | 1 | 8143 | 4miv | B | 150 | ARG | p.Arg150Gln | P51688 |
| HOMO SAPIENS | 0 | 8143 | 4miv | B | 191 | GLY | p.Gly191Arg | P51688 |
| HOMO SAPIENS | 0 | 8143 | 4miv | B | 206 | ARG | p.Arg206Pro | P51688 |
| HOMO SAPIENS | 0 | 8143 | 4miv | B | 225 | PHE | p.Phe225Leu | P51688 |
| HOMO SAPIENS | 0 | 8143 | 4miv | B | 233 | ARG | p.Arg233Ter | P51688 |
| HOMO SAPIENS | 0 | 8143 | 4miv | B | 234 | ALA | p.Ala234Gly | P51688 |
| HOMO SAPIENS | 0 | 8143 | 4miv | B | 245 | ARG | p.Arg245His | P51688 |

| | | | | | | | | |
|---|---|---|---|---|---|---|---|---|
| HOMO SAPIENS | 0 | 8143 | 4miv | B | 293 | PRO | p.Pro293Ser | P51688 |
| HOMO SAPIENS | 1 | 8143 | 4miv | B | 298 | SER | p.Ser298Pro | P51688 |
| HOMO SAPIENS | 0 | 8143 | 4miv | B | 361 | VAL | p.Val361Ile | P51688 |
| HOMO SAPIENS | 0 | 8143 | 4miv | B | 369 | GLU | p.Glu369Lys | P51688 |
| HOMO SAPIENS | 0 | 8143 | 4miv | B | 372 | MET | p.Met372Ile | P51688 |
| HOMO SAPIENS | 1 | 8143 | 4miv | B | 380 | GLN | p.Gln380Arg | P51688 |
| HOMO SAPIENS | 0 | 8143 | 4miv | B | 387 | VAL | p.Val387Met | P51688 |
| HOMO SAPIENS | 0 | 8143 | 4miv | B | 394 | MET | p.Met394Ile | P51688 |
| HOMO SAPIENS | 0 | 8143 | 4miv | B | 433 | ARG | p.Arg433Gln | P51688 |
| HOMO SAPIENS | 1 | 8143 | 4miv | B | 446 | HIS | p.His446Arg | P51688 |
| HOMO SAPIENS | 5 | 8143 | 4miv | B | 447 | GLU | p.Glu447Lys | P51688 |
| HOMO SAPIENS | 2 | 8143 | 4miv | B | 456 | ARG | p.Arg456His | P51688 |
| HOMO SAPIENS | 1 | 21880 | 2oo9 | B | 857 | LEU | p.Leu857Phe | P22681 |
| HOMO SAPIENS | 0 | 9467 | 1s1d | A | 154 | LEU | p.Leu224Pro | Q8WVQ1 |
| HOMO SAPIENS | 0 | 9467 | 1s1d | A | 156 | VAL | p.Val226Met | Q8WVQ1 |
| HOMO SAPIENS | 0 | 9467 | 1s1d | A | 229 | PRO | p.Pro299Leu | Q8WVQ1 |
| HOMO SAPIENS | 4 | 9467 | 1s1d | A | 230 | ARG | p.Arg300Cys | Q8WVQ1 |
| HOMO SAPIENS | 4 | 9467 | 1s1d | A | 230 | ARG | p.Arg300His | Q8WVQ1 |
| HOMO SAPIENS | 0 | 9467 | 1s1d | A | 253 | ALA | p.Ala323Thr | Q8WVQ1 |
| HOMO SAPIENS | 2 | 9467 | 1s1d | A | 305 | MET | p.Met375Leu | Q8WVQ1 |
| HOMO SAPIENS | -1 | 2361 | 5llm | A | 5 | ASN | p.Asn5Lys | P43003 |
| HOMO SAPIENS | 0 | 2361 | 5llm | A | 71 | TYR | p.Tyr71Cys | P43003 |
| HOMO SAPIENS | -1 | 2361 | 5llm | A | 211 | LYS | p.Glu219Asp | Q15758 |
| HOMO SAPIENS | 8 | 2361 | 5llm | A | 270 | PRO | p.Pro290Arg | P43003 |
| HOMO SAPIENS | 1 | 2361 | 5llm | A | 321 | THR | p.Ala329Thr | Q15758 |
| HOMO SAPIENS | 1 | 2361 | 5llm | A | 365 | ARG | p.Arg385His | P43003 |
| HOMO SAPIENS | 1 | 2361 | 5llm | A | 479 | ARG | p.Arg499Gln | P43003 |
| HOMO SAPIENS | 3 | 5412 | 4obz | D | 165 | PHE | p.Phe229Ile | P07339 |
| HOMO SAPIENS | 2 | 5412 | 4obz | D | 187 | ASP | p.Asp251Asn | P07339 |
| HOMO SAPIENS | 0 | 5412 | 4obz | D | 218 | GLY | p.Gly282Arg | P07339 |
| HOMO SAPIENS | 0 | 5412 | 4obz | D | 245 | ARG | p.Arg309His | P07339 |
| HOMO SAPIENS | 2 | 5412 | 4obz | D | 319 | TRP | p.Trp383Cys | P07339 |
| HOMO SAPIENS | 0 | 3544 | 5h64 | A | 277 | MET | p.Met277Ile | P42345 |
| HOMO SAPIENS | 0 | 14230 | 2kbx | A | 145 | ARG | p.Arg145Ser | Q13418 |
| HOMO SAPIENS | 2 | 34970 | 4xhz | A | 834 | ARG | p.Arg855Trp | Q96QU1 |
| HOMO SAPIENS | 4 | 34970 | 4xhz | A | 854 | SER | p.Ser875Ter | Q96QU1 |
| HOMO SAPIENS | 1 | 34970 | 4xhz | A | 908 | ARG | p.Arg929Gln | Q96QU1 |
| HOMO SAPIENS | 1 | 34970 | 4xhz | A | 941 | ARG | p.Arg962Cys | Q96QU1 |
| HOMO SAPIENS | 0 | 34970 | 4xhz | A | 970 | ARG | p.Arg991Ter | Q96QU1 |
| HOMO SAPIENS | 0 | 34970 | 4xhz | A | 976 | GLU | p.Glu997Gly | Q96QU1 |
| HOMO SAPIENS | 0 | 25734 | 4asc | A | 345 | SER | p.Asn345Ser | Q2TBA0 |
| HOMO SAPIENS | 0 | 25734 | 4asc | A | 514 | VAL | p.Val514Met | Q2TBA0 |
| HOMO SAPIENS | 6 | 25734 | 4asc | A | 528 | GLU | p.Glu528Lys | Q2TBA0 |
| HOMO SAPIENS | 0 | 25734 | 4asc | A | 580 | LEU | p.Leu580Ile | Q2TBA0 |
| HOMO SAPIENS | 8 | 2356 | 3kse | D | 22 | LYS | p.Lys22Ter | P01040 |
| HOMO SAPIENS | 1 | 2356 | 3kse | D | 58 | ARG | p.Arg58Ter | P01040 |
| HOMO SAPIENS | 2 | 6000 | 1jqd | A | 60 | GLY | p.Gly60Asp | P50135 |
| HOMO SAPIENS | 9 | 6000 | 1jqd | A | 208 | LEU | p.Leu208Pro | P50135 |
| HOMO SAPIENS | 0 | 14741 | 5eh1 | A | 58 | THR | p.Thr58Arg | P38484 |
| HOMO SAPIENS | 0 | 14741 | 5eh1 | A | 168 | THR | p.Thr168Asn | P38484 |
| HOMO SAPIENS | 1 | 4814 | 4kxk | D | 427 | ARG | p.Arg427Ter | P50542 |
| HOMO SAPIENS | 0 | 188 | 2xt3 | A | 49 | ASP | p.Asp52Asn | Q2M1P5 |
| HOMO SAPIENS | 3 | 188 | 2xt3 | A | 142 | TYR | p.Tyr145Ser | Q2M1P5 |
| HOMO SAPIENS | 9 | 8681 | 5tby | D | 57 | ALA | p.Ala57Asp | P08590 |

| | | | | | | | | |
|---|---|---|---|---|---|---|---|---|
| HOMO SAPIENS | 3 | 8681 | 5tby | D | 62 | ASP | p.Asp62Asn | P08590 |
| HOMO SAPIENS | 0 | 8681 | 5tby | D | 65 | PRO | p.Pro65Arg | P08590 |
| HOMO SAPIENS | 7 | 8681 | 5tby | D | 79 | VAL | p.Val79Ile | P08590 |
| HOMO SAPIENS | 0 | 8681 | 5tby | D | 143 | GLU | p.Glu143Lys | P08590 |
| HOMO SAPIENS | 0 | 8681 | 5tby | D | 149 | MET | p.Met149Val | P08590 |
| HOMO SAPIENS | 3 | 8681 | 5tby | D | 154 | ARG | p.Arg154His | P08590 |
| HOMO SAPIENS | 6 | 30984 | 4xwh | A | 140 | TYR | p.Tyr140Cys | P54802 |
| HOMO SAPIENS | 4 | 30984 | 4xwh | A | 141 | SER | p.Ser141Thr | P54802 |
| HOMO SAPIENS | 2 | 30984 | 4xwh | A | 234 | ARG | p.Arg234Cys | P54802 |
| HOMO SAPIENS | 2 | 30984 | 4xwh | A | 336 | GLU | p.Glu336Ter | P54802 |
| HOMO SAPIENS | 6 | 30984 | 4xwh | A | 482 | ARG | p.Arg482Trp | P54802 |
| HOMO SAPIENS | 2 | 30984 | 4xwh | A | 521 | PRO | p.Pro521Leu | P54802 |
| HOMO SAPIENS | 0 | 30984 | 4xwh | A | 565 | ARG | p.Arg565Gln | P54802 |
| HOMO SAPIENS | 0 | 30984 | 4xwh | A | 565 | ARG | p.Arg565Trp | P54802 |
| HOMO SAPIENS | 6 | 30984 | 4xwh | A | 626 | ARG | p.Arg626Ter | P54802 |
| HOMO SAPIENS | 6 | 30984 | 4xwh | A | 643 | ARG | p.Arg643His | P54802 |
| HOMO SAPIENS | 6 | 30984 | 4xwh | A | 643 | ARG | p.Arg643Cys | P54802 |
| HOMO SAPIENS | 2 | 30984 | 4xwh | A | 674 | ARG | p.Arg674His | P54802 |
| HOMO SAPIENS | 2 | 30984 | 4xwh | A | 737 | ARG | p.Arg737Gly | P54802 |
| HOMO SAPIENS | 5 | 10545 | 1oqe | N | 6 | PRO | p.Pro21Arg | Q96RJ3 |
| HOMO SAPIENS | 1 | 16995 | 5d3a | B | 12 | MET | p.Met947Arg | Q7Z4S6 |
| HOMO SAPIENS | 1 | 16995 | 5d3a | B | 12 | MET | p.Met947Val | Q7Z4S6 |
| HOMO SAPIENS | 2 | 16995 | 5d3a | B | 19 | ARG | p.Arg954Gln | Q7Z4S6 |
| HOMO SAPIENS | 2 | 16995 | 5d3a | B | 19 | ARG | p.Arg954Trp | Q7Z4S6 |
| HOMO SAPIENS | 0 | 16995 | 5d3a | B | 75 | ILE | p.Ile1010Thr | Q7Z4S6 |
| HOMO SAPIENS | 0 | 17091 | 2yin | B | 1324 | GLN | p.Gln1324Ter | Q92608 |
| HOMO SAPIENS | 5 | 27606 | 1lm7 | B | 2366 | ARG | p.Arg2366Cys | P15924 |
| HOMO SAPIENS | 2 | 15731 | 1bi7 | B | 24 | ARG | p.Arg24Pro | P42771 |
| HOMO SAPIENS | 0 | 15731 | 1bi7 | B | 28 | VAL | p.Val28Gly | P42771 |
| HOMO SAPIENS | 0 | 15731 | 1bi7 | B | 49 | ILE | p.Ile49Thr | P42771 |
| HOMO SAPIENS | 0 | 15731 | 1bi7 | B | 53 | MET | p.Met53Ile | P42771 |
| HOMO SAPIENS | 2 | 15731 | 1bi7 | B | 56 | SER | p.Ser56Ile | P42771 |
| HOMO SAPIENS | 1 | 15731 | 1bi7 | B | 57 | ALA | p.Ala57Val | P42771 |
| HOMO SAPIENS | 4 | 15731 | 1bi7 | B | 59 | VAL | p.Val59Gly | P42771 |
| HOMO SAPIENS | 9 | 15731 | 1bi7 | B | 89 | GLY | p.Gly89Asp | P42771 |
| HOMO SAPIENS | 0 | 15731 | 1bi7 | B | 101 | GLY | p.Gly101Trp | P42771 |
| HOMO SAPIENS | 0 | 15731 | 1bi7 | B | 114 | PRO | p.Pro114Ser | P42771 |
| HOMO SAPIENS | 9 | 15731 | 1bi7 | B | 123 | HIS | p.His123Gln | P42771 |
| HOMO SAPIENS | 8 | 15731 | 1bi7 | B | 126 | VAL | p.Val126Asp | P42771 |
| HOMO SAPIENS | 7 | 15731 | 1bi7 | B | 127 | ALA | p.Ala127Ser | P42771 |
| HOMO SAPIENS | -1 | 15731 | 1bi7 | B | 148 | ALA | p.Ala148Thr | P42771 |
| HOMO SAPIENS | -1 | 19858 | 3jbk | M | 866 | ALA | p.Ala934Val | P18206 |
| HOMO SAPIENS | 0 | 19484 | 3ee6 | B | 62 | SER | p.Ser62Leu | O14773 |
| HOMO SAPIENS | 1 | 19484 | 3ee6 | B | 66 | GLN | p.Gln66Ter | O14773 |
| HOMO SAPIENS | 0 | 19484 | 3ee6 | B | 100 | GLN | p.Gln100Arg | O14773 |
| HOMO SAPIENS | 0 | 19484 | 3ee6 | B | 104 | LEU | p.Leu104Ter | O14773 |
| HOMO SAPIENS | 0 | 19484 | 3ee6 | B | 127 | ARG | p.Arg127Gln | O14773 |
| HOMO SAPIENS | 0 | 19484 | 3ee6 | B | 127 | ARG | p.Arg127Ter | O14773 |
| HOMO SAPIENS | -1 | 19484 | 3ee6 | B | 185 | ARG | p.Arg185Cys | O14773 |
| HOMO SAPIENS | 0 | 19484 | 3ee6 | B | 206 | ARG | p.Arg206Cys | O14773 |
| HOMO SAPIENS | 0 | 19484 | 3ee6 | B | 208 | ARG | p.Arg208Ter | O14773 |
| HOMO SAPIENS | 0 | 19484 | 3ee6 | B | 214 | GLN | p.Gln214Ter | O14773 |
| HOMO SAPIENS | 1 | 19484 | 3ee6 | B | 276 | ASP | p.Asp276Val | O14773 |
| HOMO SAPIENS | 0 | 19484 | 3ee6 | B | 284 | GLY | p.Gly284Val | O14773 |

| | | | | | | | | |
|---|---|---|---|---|---|---|---|---|
| HOMO SAPIENS | 0 | 19484 | 3ee6 | B | 286 | ASN | p.Asn286Ser | O14773 |
| HOMO SAPIENS | 0 | 19484 | 3ee6 | B | 339 | ARG | p.Arg339Gly | O14773 |
| HOMO SAPIENS | 2 | 19484 | 3ee6 | B | 345 | MET | p.Met345Leu | O14773 |
| HOMO SAPIENS | 0 | 19484 | 3ee6 | B | 365 | CYS | p.Cys365Arg | O14773 |
| HOMO SAPIENS | 0 | 19484 | 3ee6 | B | 365 | CYS | p.Cys365Tyr | O14773 |
| HOMO SAPIENS | 0 | 19484 | 3ee6 | B | 373 | GLN | p.Gln373Glu | O14773 |
| HOMO SAPIENS | 6 | 19484 | 3ee6 | B | 418 | ARG | p.Arg418Gln | O14773 |
| HOMO SAPIENS | 7 | 19484 | 3ee6 | B | 422 | GLN | p.Gln422His | O14773 |
| HOMO SAPIENS | 0 | 19484 | 3ee6 | B | 447 | ARG | p.Arg447His | O14773 |
| HOMO SAPIENS | 0 | 19484 | 3ee6 | B | 466 | VAL | p.Val466Met | O14773 |
| HOMO SAPIENS | 0 | 4654 | 5t5w | A | 47 | LYS | p.Lys47Glu | Q08334 |
| HOMO SAPIENS | 0 | 4654 | 5t5w | A | 141 | GLU | p.Glu141Ter | Q08334 |
| HOMO SAPIENS | -1 | 18815 | 4wj7 | A | 53 | VAL | p.Val53Ile | Q9BSQ5 |
| HOMO SAPIENS | 0 | 18815 | 4wj7 | A | 120 | VAL | p.Val120Ile | Q9BSQ5 |
| HOMO SAPIENS | 0 | 18815 | 4wj7 | A | 198 | LEU | p.Leu198Arg | Q9BSQ5 |
| HOMO SAPIENS | 0 | 4602 | 1mjd | A | 47 | SER | p.Ser47Arg | O43602 |
| HOMO SAPIENS | 6 | 4602 | 1mjd | A | 59 | ARG | p.Arg59Leu | O43602 |
| HOMO SAPIENS | 7 | 4602 | 1mjd | A | 62 | ASP | p.Asp62Asn | O43602 |
| HOMO SAPIENS | 7 | 4602 | 1mjd | A | 71 | ALA | p.Ala71Ser | O43602 |
| HOMO SAPIENS | 7 | 4602 | 1mjd | A | 78 | ARG | p.Arg78Leu | O43602 |
| HOMO SAPIENS | 7 | 4602 | 1mjd | A | 89 | ARG | p.Arg89Gly | O43602 |
| HOMO SAPIENS | 8 | 4602 | 1mjd | A | 104 | ILE | p.Ile104Phe | O43602 |
| HOMO SAPIENS | 7 | 4602 | 1mjd | A | 125 | TYR | p.Tyr125His | O43602 |
| HOMO SAPIENS | 5 | 34467 | 1n0w | B | 1529 | GLY | p.Gly1529Arg | P51587 |
| HOMO SAPIENS | 8 | 5738 | 1u31 | A | 130 | ASN | p.Asn1009Lys | Q13423 |
| HOMO SAPIENS | 0 | 5760 | 2i6s | A | 298 | GLU | p.Glu318Asp | P06681 |
| HOMO SAPIENS | 0 | 5760 | 2i6s | A | 573 | ARG | p.Arg593Gln | P06681 |
| HOMO SAPIENS | 0 | 5760 | 2i6s | A | 592 | SER | p.Ser612Asn | P06681 |
| HOMO SAPIENS | 0 | 5760 | 2i6s | A | 621 | VAL | p.Val641Ala | P06681 |
| HOMO SAPIENS | 1 | 10977 | 2fby | A | 114 | VAL | p.Val114Ile | Q14191 |
| HOMO SAPIENS | 0 | 10977 | 2fby | A | 135 | LYS | p.Lys135Glu | Q14191 |
| HOMO SAPIENS | -1 | 1175 | 1qrk | A | 34 | VAL | p.Val35Leu | P00488 |
| HOMO SAPIENS | 4 | 1175 | 1qrk | A | 564 | PRO | p.Pro565Leu | P00488 |
| HOMO SAPIENS | 0 | 1175 | 1qrk | A | 650 | VAL | p.Val651Ile | P00488 |
| HOMO SAPIENS | 0 | 1175 | 1qrk | A | 651 | GLU | p.Gln652Glu | P00488 |
| HOMO SAPIENS | 3 | 909 | 4h0w | A | 375 | GLU | p.Glu394Lys | P02787 |
| HOMO SAPIENS | 0 | 909 | 4h0w | A | 458 | ALA | p.Ala477Pro | P02787 |
| HOMO SAPIENS | 6 | 909 | 4h0w | A | 515 | TYR | p.Tyr534Ter | P02787 |
| HOMO SAPIENS | 7 | 909 | 4h0w | A | 570 | PRO | p.Pro589Ser | P02787 |
| HOMO SAPIENS | 3 | 909 | 4h0w | A | 652 | GLY | p.Gly671Glu | P02787 |
| HOMO SAPIENS | 5 | 1964 | 2car | A | 151 | TRP | p.Trp151Ter | Q9BY32 |
| HOMO SAPIENS | 7 | 1964 | 2car | A | 178 | ARG | p.Arg178Cys | Q9BY32 |
| HOMO SAPIENS | 0 | 144 | 2n0a | J | 30 | ALA | p.Ala30Pro | P37840 |
| HOMO SAPIENS | 1 | 12288 | 4q58 | D | 1225 | ARG | p.Arg1225His | P16144 |
| HOMO SAPIENS | 2 | 12288 | 4q58 | D | 1281 | ARG | p.Arg1281Trp | P16144 |
| HOMO SAPIENS | 2 | 12288 | 4q58 | D | 1287 | ASN | p.Asn1287Lys | P16144 |
| HOMO SAPIENS | 0 | 5085 | 3s94 | A | 360 | ARG | p.Arg360His | O75581 |
| HOMO SAPIENS | 0 | 5085 | 3s94 | A | 433 | ASN | p.Asn433Ser | O75581 |
| HOMO SAPIENS | 2 | 5085 | 3s94 | A | 611 | ARG | p.Arg611Cys | O75581 |
| HOMO SAPIENS | 0 | 15452 | 4dur | A | 19 | LYS | p.Lys38Glu | P00747 |
| HOMO SAPIENS | 2 | 15452 | 4dur | A | 216 | ARG | p.Arg235His | P00747 |
| HOMO SAPIENS | 7 | 8406 | 4wo1 | D | 28 | VAL | p.Val55Leu | O43914 |
| HOMO SAPIENS | 2 | 895 | 5xte | K | 183 | ARG | p.Arg183Trp | P22695 |
| HOMO SAPIENS | 2 | 4191 | 4wnd | A | 120 | ARG | p.Arg127Ter | P81274 |

| | | | | | | | | |
|---|---|---|---|---|---|---|---|---|
| HOMO SAPIENS | 2 | 4191 | 4wnd | A | 120 | ARG | p.Arg127Gln | P81274 |
| HOMO SAPIENS | 1 | 4191 | 4wnd | A | 349 | GLY | p.Gly356Arg | P81274 |
| HOMO SAPIENS | 0 | 41734 | 2kbi | A | 1777 | VAL | p.Val1777Met | Q14524 |
| HOMO SAPIENS | 1 | 41734 | 2kbi | A | 1784 | GLU | p.Glu1784Lys | Q14524 |
| HOMO SAPIENS | 5 | 41734 | 2kbi | A | 1795 | TYR | p.Tyr1795Cys | Q14524 |
| HOMO SAPIENS | 5 | 41734 | 2kbi | A | 1795 | TYR | p.Tyr1795His | Q14524 |
| HOMO SAPIENS | 7 | 41734 | 2kbi | A | 1826 | ARG | p.Arg1826Cys | Q14524 |
| HOMO SAPIENS | 3 | 2119 | 1pl0 | A | 426 | LYS | p.Lys426Arg | P31939 |
| HOMO SAPIENS | 5 | 3475 | 5u62 | C | 50 | LEU | p.Leu50Ser | Q15910 |
| HOMO SAPIENS | 3 | 379 | 4uez | A | 200 | TYR | p.Tyr200Ter | P15085 |
| HOMO SAPIENS | 0 | 8195 | 5xaw | B | 67 | ARG | p.Arg67Trp | Q8IWT1 |
| HOMO SAPIENS | 0 | 8195 | 5xaw | B | 94 | THR | p.Thr94Met | Q8IWT1 |
| HOMO SAPIENS | 1 | 5711 | 1fo3 | A | 265 | ARG | p.Arg265His | Q9UKM7 |
| HOMO SAPIENS | 4 | 5711 | 1fo3 | A | 334 | ARG | p.Arg334Cys | Q9UKM7 |
| HOMO SAPIENS | 1 | 5711 | 1fo3 | A | 478 | LYS | p.Lys478Arg | Q9UKM7 |
| HOMO SAPIENS | 1 | 5711 | 1fo3 | A | 682 | ASP | p.Asp682Glu | Q9UKM7 |
| HOMO SAPIENS | 0 | 170 | 2n3y | A | 41 | GLY | p.Gly42Ser | P99999 |
| HOMO SAPIENS | 0 | 4122 | 3w81 | A | 33 | HIS | p.His33Gln | P35475 |
| HOMO SAPIENS | 0 | 4122 | 3w81 | A | 64 | TYR | p.Tyr64Ter | P35475 |
| HOMO SAPIENS | 0 | 4122 | 3w81 | A | 70 | GLN | p.Gln70Ter | P35475 |
| HOMO SAPIENS | 0 | 4122 | 3w81 | A | 75 | ALA | p.Ala75Thr | P35475 |
| HOMO SAPIENS | 0 | 4122 | 3w81 | A | 79 | ALA | p.Ala79Val | P35475 |
| HOMO SAPIENS | 0 | 4122 | 3w81 | A | 79 | ALA | p.Ala79Thr | P35475 |
| HOMO SAPIENS | 0 | 4122 | 3w81 | A | 89 | ARG | p.Arg89Gln | P35475 |
| HOMO SAPIENS | 0 | 4122 | 3w81 | A | 99 | THR | p.Thr99Ile | P35475 |
| HOMO SAPIENS | 0 | 4122 | 3w81 | A | 105 | ARG | p.Arg105Gln | P35475 |
| HOMO SAPIENS | 0 | 4122 | 3w81 | A | 238 | LEU | p.Leu238Gln | P35475 |
| HOMO SAPIENS | 0 | 4122 | 3w81 | A | 300 | ALA | p.Ala300Thr | P35475 |
| HOMO SAPIENS | 0 | 4122 | 3w81 | A | 322 | VAL | p.Val322Glu | P35475 |
| HOMO SAPIENS | 0 | 4122 | 3w81 | A | 327 | ALA | p.Ala327Pro | P35475 |
| HOMO SAPIENS | 0 | 4122 | 3w81 | A | 346 | LEU | p.Leu346Arg | P35475 |
| HOMO SAPIENS | 0 | 4122 | 3w81 | A | 349 | ASP | p.Asp349Asn | P35475 |
| HOMO SAPIENS | 0 | 4122 | 3w81 | A | 361 | ALA | p.Ala361Thr | P35475 |
| HOMO SAPIENS | 0 | 4122 | 3w81 | A | 366 | THR | p.Thr366Pro | P35475 |
| HOMO SAPIENS | 0 | 4122 | 3w81 | A | 402 | TRP | p.Trp402Ter | P35475 |
| HOMO SAPIENS | 0 | 4122 | 3w81 | A | 409 | GLY | p.Gly409Arg | P35475 |
| HOMO SAPIENS | 0 | 4122 | 3w81 | A | 454 | VAL | p.Val454Ile | P35475 |
| HOMO SAPIENS | 1 | 4122 | 3w81 | A | 490 | LEU | p.Leu490Pro | P35475 |
| HOMO SAPIENS | 1 | 4122 | 3w81 | A | 492 | ARG | p.Arg492Pro | P35475 |
| HOMO SAPIENS | 0 | 4122 | 3w81 | A | 533 | PRO | p.Pro533Arg | P35475 |
| HOMO SAPIENS | 0 | 4122 | 3w81 | A | 591 | ALA | p.Ala591Thr | P35475 |
| HOMO SAPIENS | 7 | 4122 | 3w81 | A | 619 | ARG | p.Arg619Gly | P35475 |
| HOMO SAPIENS | 0 | 15325 | 3wvz | B | 54 | VAL | p.Val54Leu | Q53FT3 |
| HOMO SAPIENS | 3 | 3228 | 5iz8 | A | 421 | TRP | p.Trp421Ter | P25054 |
| HOMO SAPIENS | 2 | 3228 | 5iz8 | A | 437 | PRO | p.Pro437Arg | P25054 |
| HOMO SAPIENS | 3 | 3228 | 5iz8 | A | 499 | ARG | p.Arg499Ter | P25054 |
| HOMO SAPIENS | -1 | 29229 | 5buo | A | 539 | SER | p.Ser614Gly | P05067 |
| HOMO SAPIENS | -1 | 34931 | 4n7v | C | 54 | SER | p.Ser54Leu | O94986 |
| HOMO SAPIENS | 0 | 2450 | 5kzx | A | 123 | GLY | p.Gly123Glu | P10253 |
| HOMO SAPIENS | 1 | 2450 | 5kzx | A | 190 | ARG | p.Arg190His | P10253 |
| HOMO SAPIENS | -1 | 2450 | 5kzx | A | 199 | HIS | p.His199Arg | P10253 |
| HOMO SAPIENS | 0 | 2450 | 5kzx | A | 219 | GLY | p.Gly219Arg | P10253 |
| HOMO SAPIENS | 0 | 2450 | 5kzx | A | 220 | VAL | p.Val220Leu | P10253 |
| HOMO SAPIENS | 0 | 2450 | 5kzx | A | 222 | VAL | p.Val222Met | P10253 |

| | | | | | | | | |
|---|---|---|---|---|---|---|---|---|
| HOMO SAPIENS | 0 | 2450 | 5kzx | A | 237 | ALA | p.Ala237Val | P10253 |
| HOMO SAPIENS | 0 | 2450 | 5kzx | A | 251 | SER | p.Ser251Leu | P10253 |
| HOMO SAPIENS | 0 | 2450 | 5kzx | A | 254 | SER | p.Ser254Leu | P10253 |
| HOMO SAPIENS | 0 | 2450 | 5kzx | A | 262 | GLU | p.Glu262Lys | P10253 |
| HOMO SAPIENS | 0 | 2450 | 5kzx | A | 293 | GLY | p.Gly293Arg | P10253 |
| HOMO SAPIENS | 0 | 2450 | 5kzx | A | 299 | LEU | p.Leu299Arg | P10253 |
| HOMO SAPIENS | 0 | 2450 | 5kzx | A | 309 | GLY | p.Gly309Arg | P10253 |
| HOMO SAPIENS | 0 | 2450 | 5kzx | A | 318 | MET | p.Met318Thr | P10253 |
| HOMO SAPIENS | 0 | 2450 | 5kzx | A | 355 | LEU | p.Leu355Pro | P10253 |
| HOMO SAPIENS | 1 | 2450 | 5kzx | A | 361 | PRO | p.Pro361Leu | P10253 |
| HOMO SAPIENS | 4 | 2450 | 5kzx | A | 408 | MET | p.Met408Val | P10253 |
| HOMO SAPIENS | 1 | 2450 | 5kzx | A | 429 | GLN | p.Gln429Glu | P10253 |
| HOMO SAPIENS | 4 | 2450 | 5kzx | A | 437 | ARG | p.Arg437Cys | P10253 |
| HOMO SAPIENS | 0 | 2450 | 5kzx | A | 439 | MET | p.Met439Lys | P10253 |
| HOMO SAPIENS | 1 | 2450 | 5kzx | A | 448 | SER | p.Ser448Thr | P10253 |
| HOMO SAPIENS | 0 | 2450 | 5kzx | A | 451 | PRO | p.Pro451Arg | P10253 |
| HOMO SAPIENS | 1 | 2450 | 5kzx | A | 459 | ASP | p.Asp459His | P10253 |
| HOMO SAPIENS | 8 | 2450 | 5kzx | A | 481 | TRP | p.Trp481Arg | P10253 |
| HOMO SAPIENS | 1 | 2450 | 5kzx | A | 489 | ASP | p.Asp489Asn | P10253 |
| HOMO SAPIENS | 0 | 2450 | 5kzx | A | 521 | GLU | p.Glu521Lys | P10253 |
| HOMO SAPIENS | 0 | 2450 | 5kzx | A | 529 | SER | p.Ser529Val | P10253 |
| HOMO SAPIENS | 0 | 2450 | 5kzx | A | 545 | PRO | p.Pro545Leu | P10253 |
| HOMO SAPIENS | 0 | 2450 | 5kzx | A | 552 | LEU | p.Leu552Pro | P10253 |
| HOMO SAPIENS | 0 | 2450 | 5kzx | A | 576 | GLY | p.Gly576Ser | P10253 |
| HOMO SAPIENS | 9 | 2450 | 5kzx | A | 600 | ARG | p.Arg600His | P10253 |
| HOMO SAPIENS | 8 | 2450 | 5kzx | A | 615 | GLY | p.Gly615Arg | P10253 |
| HOMO SAPIENS | 3 | 2450 | 5kzx | A | 619 | SER | p.Ser619Asn | P10253 |
| HOMO SAPIENS | 0 | 2450 | 5kzx | A | 638 | GLY | p.Gly638Trp | P10253 |
| HOMO SAPIENS | 0 | 2450 | 5kzx | A | 643 | GLY | p.Gly643Arg | P10253 |
| HOMO SAPIENS | 9 | 2450 | 5kzx | A | 645 | ASP | p.Asp645Glu | P10253 |
| HOMO SAPIENS | 9 | 2450 | 5kzx | A | 645 | ASP | p.Asp645His | P10253 |
| HOMO SAPIENS | 0 | 2450 | 5kzx | A | 648 | GLY | p.Gly648Ser | P10253 |
| HOMO SAPIENS | 5 | 2450 | 5kzx | A | 660 | ARG | p.Arg660His | P10253 |
| HOMO SAPIENS | 8 | 2450 | 5kzx | A | 672 | ARG | p.Arg672Gln | P10253 |
| HOMO SAPIENS | 0 | 2450 | 5kzx | A | 702 | ARG | p.Arg702Leu | P10253 |
| HOMO SAPIENS | 3 | 2450 | 5kzx | A | 725 | ARG | p.Arg725Trp | P10253 |
| HOMO SAPIENS | 0 | 2450 | 5kzx | A | 746 | TRP | p.Trp746Cys | P10253 |
| HOMO SAPIENS | 0 | 2450 | 5kzx | A | 780 | ILE | p.Val780Ile | P10253 |
| HOMO SAPIENS | 1 | 2450 | 5kzx | A | 816 | VAL | p.Val816Ile | P10253 |
| HOMO SAPIENS | 0 | 2450 | 5kzx | A | 838 | GLN | p.Gln838Ter | P10253 |
| HOMO SAPIENS | 2 | 2450 | 5kzx | A | 854 | ARG | p.Arg854Ter | P10253 |
| HOMO SAPIENS | 3 | 2450 | 5kzx | A | 870 | ARG | p.Arg870Ter | P10253 |
| HOMO SAPIENS | 0 | 2450 | 5kzx | A | 890 | VAL | p.Val890Leu | P10253 |
| HOMO SAPIENS | 6 | 1882 | 4km6 | A | 175 | CYS | p.Cys175Ter | P15328 |
| HOMO SAPIENS | 3 | 14767 | 1lm5 | A | 2628 | LEU | p.Leu2628Pro | P15924 |
| HOMO SAPIENS | 5 | 14767 | 1lm5 | A | 2655 | ALA | p.Ala2655Gly | P15924 |
| HOMO SAPIENS | 3 | 14767 | 1lm5 | A | 2731 | TYR | p.Tyr2731His | P15924 |
| HOMO SAPIENS | 1 | 14767 | 1lm5 | A | 2767 | THR | p.Thr2767Asn | P15924 |
| HOMO SAPIENS | -1 | 14767 | 1lm5 | A | 2819 | MET | p.Met2819Leu | P15924 |
| HOMO SAPIENS | -1 | 11246 | 4y5o | A | 289 | SER | p.Ser289Asn | Q9BSQ5 |
| HOMO SAPIENS | 0 | 7380 | 5kz5 | I | 60 | ARG | p.Arg60His | Q16595 |
| HOMO SAPIENS | 0 | 7380 | 5kz5 | I | 76 | MET | p.Met76Val | Q16595 |
| HOMO SAPIENS | 2 | 22187 | 3fby | A | 328 | CYS | p.Cys328Arg | P49747 |
| HOMO SAPIENS | 2 | 22187 | 3fby | A | 342 | ASP | p.Asp342Tyr | P49747 |

| | | | | | | | | |
|---|---|---|---|---|---|---|---|---|
| HOMO SAPIENS | 2 | 22187 | 3fby | A | 348 | CYS | p.Cys348Arg | P49747 |
| HOMO SAPIENS | 2 | 22187 | 3fby | A | 386 | ASN | p.Asn386Asp | P49747 |
| HOMO SAPIENS | 2 | 22187 | 3fby | A | 453 | ASN | p.Asn453Ser | P49747 |
| HOMO SAPIENS | 2 | 22187 | 3fby | A | 468 | CYS | p.Cys468Tyr | P49747 |
| HOMO SAPIENS | 3 | 22187 | 3fby | A | 472 | ASP | p.Asp472Tyr | P49747 |
| HOMO SAPIENS | 3 | 22187 | 3fby | A | 473 | ASP | p.Asp473Gly | P49747 |
| HOMO SAPIENS | 2 | 22187 | 3fby | A | 523 | ASN | p.Asn523Lys | P49747 |
| HOMO SAPIENS | 1 | 22187 | 3fby | A | 718 | ARG | p.Arg718Trp | P49747 |
| HOMO SAPIENS | 1 | 22187 | 3fby | A | 719 | GLY | p.Gly719Asp | P49747 |
| HOMO SAPIENS | 0 | 22187 | 3fby | A | 756 | GLN | p.Gln756Arg | P49747 |
| HOMO SAPIENS | -1 | 2784 | 5kvh | A | 88 | MET | p.Met88Val | O95831 |
| HOMO SAPIENS | -1 | 2784 | 5kvh | A | 96 | ASN | p.Asn96Ser | O95831 |
| HOMO SAPIENS | 0 | 2784 | 5kvh | A | 282 | THR | p.Thr282Met | O95831 |
| HOMO SAPIENS | 1 | 2784 | 5kvh | A | 366 | ASN | p.Asn366Ser | O95831 |
| HOMO SAPIENS | 3 | 2784 | 5kvh | A | 463 | ARG | p.Arg463Ile | O95831 |
| HOMO SAPIENS | 0 | 2784 | 5kvh | A | 498 | VAL | p.Val498Met | O95831 |
| HOMO SAPIENS | -1 | 2784 | 5kvh | A | 549 | ALA | p.Ala549Val | O95831 |
| HOMO SAPIENS | 0 | 38316 | 2lnk | C | 1933 | ARG | p.Arg1933Ter | P35579 |
| HOMO SAPIENS | 0 | 33808 | 2miu | A | 29 | VAL | p.Val29Met | Q14192 |
| HOMO SAPIENS | 1 | 33808 | 2miu | A | 37 | ALA | p.Ala37Ser | Q14192 |
| HOMO SAPIENS | 2 | 20994 | 2hey | R | 65 | ARG | p.Arg65Cys | P43489 |
| HOMO SAPIENS | -1 | 5687 | 2xwj | I | 7 | ARG | p.Arg32Gln | P00751 |
| HOMO SAPIENS | -1 | 5687 | 2xwj | I | 7 | ARG | p.Arg32Trp | P00751 |
| HOMO SAPIENS | 0 | 5687 | 2xwj | I | 49 | ARG | p.Arg74His | P00751 |
| HOMO SAPIENS | 0 | 5687 | 2xwj | I | 177 | ARG | p.Arg202Gly | P00751 |
| HOMO SAPIENS | 0 | 5687 | 2xwj | I | 217 | ILE | p.Ile242Leu | P00751 |
| HOMO SAPIENS | -1 | 5687 | 2xwj | I | 227 | GLY | p.Gly252Ser | P00751 |
| HOMO SAPIENS | 0 | 5687 | 2xwj | I | 483 | HIS | p.His508Gln | P00751 |
| HOMO SAPIENS | 0 | 5687 | 2xwj | I | 508 | LYS | p.Lys533Arg | P00751 |
| HOMO SAPIENS | 0 | 5687 | 2xwj | I | 540 | LYS | p.Lys565Glu | P00751 |
| HOMO SAPIENS | 0 | 5687 | 2xwj | I | 541 | GLU | p.Glu566Ala | P00751 |
| HOMO SAPIENS | 2 | 953 | 3d4j | A | 249 | PHE | p.Phe249Ser | P53602 |
| HOMO SAPIENS | 1 | 953 | 3d4j | A | 292 | ASN | p.Asn292Ser | P53602 |
| HOMO SAPIENS | 0 | 3300 | 1s9c | A | 140 | ASN | p.Asn457Tyr | P51659 |
| HOMO SAPIENS | 2 | 3300 | 1s9c | A | 167 | ILE | p.Ile484Val | P51659 |
| HOMO SAPIENS | 0 | 3300 | 1s9c | A | 174 | ALA | p.Ala491Thr | P51659 |
| HOMO SAPIENS | 8 | 3300 | 1s9c | A | 193 | ASP | p.Asp510Asn | P51659 |
| HOMO SAPIENS | 3 | 3300 | 1s9c | A | 194 | TRP | p.Trp511Arg | P51659 |
| HOMO SAPIENS | 0 | 3300 | 1s9c | A | 242 | VAL | p.Ile559Val | P51659 |
| HOMO SAPIENS | 0 | 10810 | 5i25 | A | 5 | GLN | p.Gln23Ter | P03951 |
| HOMO SAPIENS | 0 | 10810 | 5i25 | A | 55 | TRP | p.Trp73Ter | P03951 |
| HOMO SAPIENS | 0 | 10810 | 5i25 | A | 91 | ALA | p.Ala109Thr | P03951 |
| HOMO SAPIENS | 0 | 10810 | 5i25 | A | 117 | GLU | p.Glu135Ter | P03951 |
| HOMO SAPIENS | 0 | 10810 | 5i25 | A | 118 | CYS | p.Cys136Ter | P03951 |
| HOMO SAPIENS | 0 | 10810 | 5i25 | A | 128 | CYS | p.Cys146Ter | P03951 |
| HOMO SAPIENS | 0 | 10810 | 5i25 | A | 210 | ARG | p.Arg228Ter | P03951 |
| HOMO SAPIENS | 0 | 10810 | 5i25 | A | 226 | GLN | p.Gln244Ter | P03951 |
| HOMO SAPIENS | 0 | 10810 | 5i25 | A | 233 | GLN | p.Gln251Ter | P03951 |
| HOMO SAPIENS | 0 | 16668 | 3lbx | A | 28 | ARG | p.Arg28Leu | P02549 |
| HOMO SAPIENS | 0 | 16668 | 3lbx | A | 28 | ARG | p.Arg28Cys | P02549 |
| HOMO SAPIENS | 1 | 16668 | 3lbx | A | 41 | ARG | p.Arg41Trp | P02549 |
| HOMO SAPIENS | 1 | 16668 | 3lbx | A | 45 | ARG | p.Arg45Ser | P02549 |
| HOMO SAPIENS | 2 | 16668 | 3lbx | A | 48 | LYS | p.Lys48Arg | P02549 |
| HOMO SAPIENS | 1 | 16668 | 3lbx | A | 152 | ASP | p.Asp152Asn | P02549 |

| | | | | | | | | |
|---|---|---|---|---|---|---|---|---|
| HOMO SAPIENS | 0 | 226 | 2ble | A | 256 | ILE | p.Phe256Ile | P36959 |
| HOMO SAPIENS | 2 | 36808 | 4c7n | A | 3 | SER | p.Ser357Pro | O75030 |
| HOMO SAPIENS | 0 | 4092 | 5jnx | A | 113 | CYS | p.Cys113Arg | O15118 |
| HOMO SAPIENS | 4 | 4092 | 5jnx | A | 215 | HIS | p.His215Arg | O15118 |
| HOMO SAPIENS | 6 | 4092 | 5jnx | A | 237 | PRO | p.Pro237Ser | O15118 |
| HOMO SAPIENS | 0 | 4092 | 5jnx | A | 363 | ALA | p.Ala363Val | O15118 |
| HOMO SAPIENS | 0 | 4092 | 5jnx | A | 378 | VAL | p.Val378Ala | O15118 |
| HOMO SAPIENS | 8 | 4092 | 5jnx | A | 404 | ARG | p.Arg404Gln | O15118 |
| HOMO SAPIENS | 8 | 4092 | 5jnx | A | 411 | ARG | p.Arg411Gln | O15118 |
| HOMO SAPIENS | 2 | 4092 | 5jnx | A | 474 | PRO | p.Pro474Leu | O15118 |
| HOMO SAPIENS | 8 | 4092 | 5jnx | A | 511 | THR | p.Thr511Met | O15118 |
| HOMO SAPIENS | 0 | 4092 | 5jnx | A | 642 | MET | p.Met642Ile | O15118 |
| HOMO SAPIENS | 2 | 4092 | 5jnx | A | 664 | VAL | p.Val664Met | O15118 |
| HOMO SAPIENS | 0 | 4092 | 5jnx | A | 734 | SER | p.Ser734Ile | O15118 |
| HOMO SAPIENS | -1 | 4092 | 5jnx | A | 775 | GLN | p.Gln775Pro | O15118 |
| HOMO SAPIENS | 0 | 4092 | 5jnx | A | 858 | ILE | p.Ile858Val | O15118 |
| HOMO SAPIENS | 0 | 4092 | 5jnx | A | 889 | VAL | p.Val889Met | O15118 |
| HOMO SAPIENS | 0 | 4092 | 5jnx | A | 910 | GLY | p.Gly910Ser | O15118 |
| HOMO SAPIENS | 0 | 4092 | 5jnx | A | 911 | GLY | p.Gly911Ser | O15118 |
| HOMO SAPIENS | 0 | 4092 | 5jnx | A | 928 | GLN | p.Gln928Pro | O15118 |
| HOMO SAPIENS | 0 | 4092 | 5jnx | A | 931 | ASN | p.Asn931Lys | O15118 |
| HOMO SAPIENS | 0 | 4092 | 5jnx | A | 940 | SER | p.Ser940Leu | O15118 |
| HOMO SAPIENS | 0 | 4092 | 5jnx | A | 950 | VAL | p.Val950Met | O15118 |
| HOMO SAPIENS | 0 | 4092 | 5jnx | A | 954 | SER | p.Ser954Leu | O15118 |
| HOMO SAPIENS | 0 | 4092 | 5jnx | A | 958 | ARG | p.Arg958Gln | O15118 |
| HOMO SAPIENS | 0 | 4092 | 5jnx | A | 958 | ARG | p.Arg958Ter | O15118 |
| HOMO SAPIENS | -1 | 4092 | 5jnx | A | 978 | ARG | p.Arg978Cys | O15118 |
| HOMO SAPIENS | 0 | 4092 | 5jnx | A | 992 | GLY | p.Gly992Trp | O15118 |
| HOMO SAPIENS | 0 | 4092 | 5jnx | A | 1007 | PRO | p.Pro1007Ala | O15118 |
| HOMO SAPIENS | 0 | 4092 | 5jnx | A | 1019 | TYR | p.Tyr1019Cys | O15118 |
| HOMO SAPIENS | 2 | 4092 | 5jnx | A | 1035 | ALA | p.Ala1035Val | O15118 |
| HOMO SAPIENS | 2 | 4092 | 5jnx | A | 1054 | ALA | p.Ala1054Thr | O15118 |
| HOMO SAPIENS | 2 | 4092 | 5jnx | A | 1061 | ILE | p.Ile1061Thr | O15118 |
| HOMO SAPIENS | 0 | 4092 | 5jnx | A | 1073 | GLY | p.Gly1073Ser | O15118 |
| HOMO SAPIENS | 2 | 4092 | 5jnx | A | 1088 | TYR | p.Tyr1088Cys | O15118 |
| HOMO SAPIENS | 2 | 4092 | 5jnx | A | 1142 | MET | p.Met1142Thr | O15118 |
| HOMO SAPIENS | 2 | 4092 | 5jnx | A | 1156 | ASN | p.Asn1156Ser | O15118 |
| HOMO SAPIENS | 0 | 4092 | 5jnx | A | 1179 | MET | p.Met1179Val | O15118 |
| HOMO SAPIENS | 0 | 4092 | 5jnx | A | 1186 | ARG | p.Arg1186His | O15118 |
| HOMO SAPIENS | 0 | 4092 | 5jnx | A | 1187 | ALA | p.Ala1187Gly | O15118 |
| HOMO SAPIENS | 2 | 4092 | 5jnx | A | 1189 | GLU | p.Glu1189Gly | O15118 |
| HOMO SAPIENS | 2 | 4092 | 5jnx | A | 1200 | SER | p.Ser1200Gly | O15118 |
| HOMO SAPIENS | 2 | 4092 | 5jnx | A | 1205 | THR | p.Thr1205Lys | O15118 |
| HOMO SAPIENS | 0 | 4092 | 5jnx | A | 1213 | LEU | p.Leu1213Phe | O15118 |
| HOMO SAPIENS | -1 | 4092 | 5jnx | A | 1266 | ARG | p.Arg1266Gln | O15118 |
| HOMO SAPIENS | 9 | 16498 | 4i6o | A | 64 | ARG | p.Arg735Trp | P01024 |
| HOMO SAPIENS | 1 | 21902 | 3un9 | C | 751 | ARG | p.Arg751His | Q86UT6 |
| HOMO SAPIENS | 7 | 20391 | 2h32 | B | 105 | PRO | p.Pro142Leu | P15814 |
| HOMO SAPIENS | 6 | 1635 | 4x45 | A | 65 | ASP | p.Asp65Val | P07741 |
| HOMO SAPIENS | 1 | 1635 | 4x45 | A | 98 | TRP | p.Trp98Ter | P07741 |
| HOMO SAPIENS | 0 | 1635 | 4x45 | A | 110 | LEU | p.Leu110Pro | P07741 |
| HOMO SAPIENS | 7 | 1635 | 4x45 | A | 136 | MET | p.Met136Thr | P07741 |
| HOMO SAPIENS | 1 | 1635 | 4x45 | A | 150 | VAL | p.Val150Phe | P07741 |
| HOMO SAPIENS | 4 | 34810 | 2rpc | A | 16 | CYS | p.Cys253Ser | O60481 |

| | | | | | | | | |
|---|---|---|---|---|---|---|---|---|
| HOMO SAPIENS | 1 | 34810 | 2rpc | A | 18 | TRP | p.Trp255Gly | O60481 |
| HOMO SAPIENS | 1 | 14359 | 4hxi | A | 78 | MET | p.Met78Val | Q9UH77 |
| HOMO SAPIENS | 0 | 5342 | 4d9t | A | 628 | GLU | p.Glu628Asp | P51812 |
| HOMO SAPIENS | -1 | 5342 | 4d9t | A | 723 | ARG | p.Arg723His | P51812 |
| HOMO SAPIENS | -1 | 5342 | 4d9t | A | 729 | ARG | p.Arg729Gln | P51812 |
| HOMO SAPIENS | 1 | 9619 | 3j2s | B | 1781 | ARG | p.Arg1800Cys | P00451 |
| HOMO SAPIENS | 0 | 9619 | 3j2s | B | 1823 | MET | p.Met1842Ile | P00451 |
| HOMO SAPIENS | 1 | 9619 | 3j2s | B | 1825 | PRO | p.Pro1844Ser | P00451 |
| HOMO SAPIENS | 0 | 9619 | 3j2s | B | 1826 | THR | p.Thr1845Pro | P00451 |
| HOMO SAPIENS | 0 | 9619 | 3j2s | B | 1834 | ALA | p.Ala1853Val | P00451 |
| HOMO SAPIENS | 0 | 9619 | 3j2s | B | 1846 | ASP | p.Asp1865Asn | P00451 |
| HOMO SAPIENS | 1 | 9619 | 3j2s | B | 1848 | HIS | p.His1867Arg | P00451 |
| HOMO SAPIENS | 0 | 9619 | 3j2s | B | 1854 | PRO | p.Pro1873Arg | P00451 |
| HOMO SAPIENS | 1 | 9619 | 3j2s | B | 1885 | GLU | p.Glu1904Lys | P00451 |
| HOMO SAPIENS | 0 | 9619 | 3j2s | B | 1922 | ASN | p.Asn1941Ser | P00451 |
| HOMO SAPIENS | 1 | 9619 | 3j2s | B | 1941 | ARG | p.Arg1960Gln | P00451 |
| HOMO SAPIENS | 4 | 9619 | 3j2s | B | 2105 | TYR | p.Tyr2124Cys | P00451 |
| HOMO SAPIENS | 6 | 9619 | 3j2s | B | 2150 | ARG | p.Arg2169His | P00451 |
| HOMO SAPIENS | 0 | 9619 | 3j2s | B | 2159 | ARG | p.Arg2178Cys | P00451 |
| HOMO SAPIENS | 0 | 9619 | 3j2s | B | 2159 | ARG | p.Arg2178His | P00451 |
| HOMO SAPIENS | 2 | 9619 | 3j2s | B | 2238 | MET | p.Met2257Val | P00451 |
| HOMO SAPIENS | 2 | 9619 | 3j2s | B | 2300 | PRO | p.Pro2319Ser | P00451 |
| HOMO SAPIENS | 1 | 667 | 1z2m | A | 127 | GLU | p.Glu127Ter | P05161 |
| HOMO SAPIENS | 0 | 15717 | 2nn6 | G | 30 | GLY | p.Gly31Ala | Q9NQT5 |
| HOMO SAPIENS | -1 | 15717 | 2nn6 | G | 55 | ASN | p.Asn56His | Q9NQT5 |
| HOMO SAPIENS | 7 | 15717 | 2nn6 | G | 131 | ASP | p.Asp132Ala | Q9NQT5 |
| HOMO SAPIENS | 7 | 784 | 3rkq | A | 161 | ARG | p.Arg161Pro | P52952 |
| HOMO SAPIENS | 6 | 784 | 3rkq | A | 178 | THR | p.Thr178Met | P52952 |
| HOMO SAPIENS | 7 | 784 | 3rkq | A | 183 | LYS | p.Lys183Glu | P52952 |
| HOMO SAPIENS | 8 | 784 | 3rkq | A | 190 | ARG | p.Arg190Cys | P52952 |
| HOMO SAPIENS | 6 | 385 | 30000 | C | 70 | ALA | p.Ala70Thr | P07954 |
| HOMO SAPIENS | 2 | 385 | 30000 | C | 101 | ARG | p.Arg101Pro | P07954 |
| HOMO SAPIENS | 2 | 385 | 30000 | C | 174 | PRO | p.Pro174Arg | P07954 |
| HOMO SAPIENS | 6 | 385 | 30000 | C | 187 | SER | p.Ser187Ter | P07954 |
| HOMO SAPIENS | 9 | 385 | 30000 | C | 233 | ARG | p.Arg233His | P07954 |
| HOMO SAPIENS | 3 | 385 | 30000 | C | 273 | ALA | p.Ala273Thr | P07954 |
| HOMO SAPIENS | 4 | 385 | 30000 | C | 334 | SER | p.Ser334Arg | P07954 |
| HOMO SAPIENS | 6 | 385 | 30000 | C | 340 | ASN | p.Asn340Lys | P07954 |
| HOMO SAPIENS | 8 | 385 | 30000 | C | 343 | ARG | p.Arg343Ter | P07954 |
| HOMO SAPIENS | 3 | 385 | 30000 | C | 350 | ARG | p.Arg350Trp | P07954 |
| HOMO SAPIENS | 4 | 385 | 30000 | C | 376 | GLN | p.Gln376Pro | P07954 |
| HOMO SAPIENS | 6 | 385 | 30000 | C | 434 | CYS | p.Cys434Ter | P07954 |
| HOMO SAPIENS | 0 | 3316 | 4f91 | B | 934 | THR | p.Thr934Ala | O75643 |
| HOMO SAPIENS | 0 | 3316 | 4f91 | B | 1002 | ASN | p.Asn1002Ser | O75643 |
| HOMO SAPIENS | 0 | 3316 | 4f91 | B | 1087 | SER | p.Ser1087Leu | O75643 |
| HOMO SAPIENS | 4 | 3316 | 4f91 | B | 1389 | VAL | p.Val1389Ile | O75643 |
| HOMO SAPIENS | 0 | 619 | 5hbt | B | 195 | ASP | p.Asp240His | P02708 |
| HOMO SAPIENS | 1 | 619 | 5hbt | B | 209 | ARG | p.Arg254Leu | P02708 |
| HOMO SAPIENS | 0 | 18171 | 2y25 | B | 1408 | ASP | p.Asp1408Asn | P52179 |
| HOMO SAPIENS | 0 | 18171 | 2y25 | B | 1453 | MET | p.Met1453Thr | P52179 |
| HOMO SAPIENS | 0 | 18171 | 2y25 | B | 1459 | ILE | p.Ile1459Lys | P52179 |
| HOMO SAPIENS | 0 | 18171 | 2y25 | B | 1558 | PHE | p.Phe1558Leu | P52179 |
| HOMO SAPIENS | 0 | 18171 | 2y25 | B | 1573 | ARG | p.Arg1573Gln | P52179 |
| HOMO SAPIENS | 7 | 18171 | 2y25 | B | 1663 | VAL | p.Val1663Met | P52179 |

| | | | | | | | | |
|---|---|---|---|---|---|---|---|---|
| HOMO SAPIENS | 8 | 14262 | 2fcw | B | 117 | SER | p.Ser138Ter | P01130 |
| HOMO SAPIENS | 8 | 14262 | 2fcw | B | 118 | ASP | p.Asp139Glu | P01130 |
| HOMO SAPIENS | 9 | 14262 | 2fcw | B | 119 | GLU | p.Glu140Lys | P01130 |
| HOMO SAPIENS | 7 | 14262 | 2fcw | B | 122 | CYS | p.Cys143Ter | P01130 |
| HOMO SAPIENS | 6 | 14262 | 2fcw | B | 134 | CYS | p.Cys155Ter | P01130 |
| HOMO SAPIENS | 7 | 14262 | 2fcw | B | 147 | ASP | p.Asp168Glu | P01130 |
| HOMO SAPIENS | 4 | 2624 | 3b7b | A | 855 | VAL | p.Val886Met | Q9H9B1 |
| HOMO SAPIENS | 7 | 2624 | 3b7b | A | 868 | ILE | p.Ile899Val | Q9H9B1 |
| HOMO SAPIENS | 0 | 2624 | 3b7b | A | 892 | GLU | p.Glu923Lys | Q9H9B1 |
| HOMO SAPIENS | 0 | 4791 | 5ctd | B | 68 | LEU | p.Pro549Ala | Q14055 |
| HOMO SAPIENS | 5 | 460 | 4ui1 | D | 80 | CYS | p.Cys80Arg | Q6ZVN8 |
| HOMO SAPIENS | 3 | 460 | 4ui1 | D | 101 | LEU | p.Leu101Pro | Q6ZVN8 |
| HOMO SAPIENS | 0 | 40782 | 1sy6 | A | 47 | LYS | p.Lys69Ter | P09693 |
| HOMO SAPIENS | -1 | 1092 | 2idx | A | 62 | THR | p.Thr62Met | Q96EY8 |
| HOMO SAPIENS | 6 | 1092 | 2idx | A | 183 | HIS | p.His183Leu | Q96EY8 |
| HOMO SAPIENS | 7 | 1092 | 2idx | A | 186 | ARG | p.Arg186Trp | Q96EY8 |
| HOMO SAPIENS | 6 | 1092 | 2idx | A | 190 | ARG | p.Arg190Cys | Q96EY8 |
| HOMO SAPIENS | 6 | 1092 | 2idx | A | 190 | ARG | p.Arg190His | Q96EY8 |
| HOMO SAPIENS | 7 | 1092 | 2idx | A | 191 | ARG | p.Arg191Trp | Q96EY8 |
| HOMO SAPIENS | 7 | 1092 | 2idx | A | 191 | ARG | p.Arg191Gln | Q96EY8 |
| HOMO SAPIENS | 6 | 1092 | 2idx | A | 195 | ARG | p.Arg195His | Q96EY8 |
| HOMO SAPIENS | 1 | 1092 | 2idx | A | 234 | GLN | p.Gln234Ter | Q96EY8 |
| HOMO SAPIENS | 0 | 1092 | 2idx | A | 239 | MET | p.Met239Lys | Q96EY8 |
| HOMO SAPIENS | -1 | 6418 | 5fur | G | 11 | THR | p.Thr11Ile | P21675 |
| HOMO SAPIENS | -1 | 6418 | 5fur | G | 290 | LEU | p.Leu269Val | P21675 |
| HOMO SAPIENS | 0 | 6100 | 1wiq | A | 240 | ARG | p.Arg265Trp | P01730 |
| HOMO SAPIENS | 0 | 13757 | 4ye6 | A | 45 | GLY | p.Gly45Val | P47897 |
| HOMO SAPIENS | 1 | 13757 | 4ye6 | A | 57 | TYR | p.Tyr57His | P47897 |
| HOMO SAPIENS | 8 | 13757 | 4ye6 | A | 403 | ARG | p.Arg403Trp | P47897 |
| HOMO SAPIENS | 7 | 13757 | 4ye6 | A | 524 | ARG | p.Arg524Trp | P47897 |
| HOMO SAPIENS | 6 | 13757 | 4ye6 | A | 690 | ARG | p.Arg690Cys | P47897 |
| HOMO SAPIENS | 0 | 26958 | 2lzl | A | 370 | GLY | p.Gly370Cys | P22607 |
| HOMO SAPIENS | 0 | 26958 | 2lzl | A | 371 | SER | p.Ser371Cys | P22607 |
| HOMO SAPIENS | 0 | 26958 | 2lzl | A | 373 | TYR | p.Tyr373Cys | P22607 |
| HOMO SAPIENS | 0 | 26958 | 2lzl | A | 380 | GLY | p.Gly380Arg | P22607 |
| HOMO SAPIENS | 0 | 26958 | 2lzl | A | 384 | PHE | p.Phe384Leu | P22607 |
| HOMO SAPIENS | 5 | 26958 | 2lzl | A | 391 | ALA | p.Ala391Glu | P22607 |
| HOMO SAPIENS | 6 | 11325 | 2wa6 | A | 121 | TYR | p.Tyr121Phe | O75369 |
| HOMO SAPIENS | 2 | 11325 | 2wa6 | A | 161 | PHE | p.Phe161Cys | O75369 |
| HOMO SAPIENS | 0 | 11325 | 2wa6 | A | 171 | LEU | p.Leu171Arg | O75369 |
| HOMO SAPIENS | 2 | 11325 | 2wa6 | A | 173 | ALA | p.Ala173Val | O75369 |
| HOMO SAPIENS | 0 | 11325 | 2wa6 | A | 202 | MET | p.Met202Val | O75369 |
| HOMO SAPIENS | 3 | 11325 | 2wa6 | A | 227 | GLU | p.Glu227Lys | O75369 |
| HOMO SAPIENS | 0 | 11325 | 2wa6 | A | 234 | LEU | p.Leu234Val | O75369 |
| HOMO SAPIENS | 2 | 11325 | 2wa6 | A | 235 | SER | p.Ser235Pro | O75369 |
| HOMO SAPIENS | 9 | 7240 | 3tgx | A | 137 | TYR | p.Tyr156Cys | Q9HBE5 |
| HOMO SAPIENS | 0 | 35479 | 4elj | A | 307 | THR | p.Thr307Ile | P06400 |
| HOMO SAPIENS | 0 | 35479 | 4elj | A | 310 | GLY | p.Gly310Glu | P06400 |
| HOMO SAPIENS | 0 | 35479 | 4elj | A | 377 | THR | p.Thr377Ser | P06400 |
| HOMO SAPIENS | 9 | 36706 | 1he7 | A | 359 | TYR | p.Tyr359Cys | P04629 |
| HOMO SAPIENS | 4 | 6538 | 4d0e | A | 504 | ARG | p.Arg504His | P46531 |
| HOMO SAPIENS | 1 | 26518 | 5fm5 | M | 600 | GLU | p.Glu600Val | P52179 |
| HOMO SAPIENS | -1 | 26518 | 5fm5 | M | 651 | ARG | p.Arg651Gln | P52179 |
| HOMO SAPIENS | -1 | 26518 | 5fm5 | M | 660 | PRO | p.Pro660Ala | P52179 |

| | | | | | | | | |
|---|---|---|---|---|---|---|---|---|
| HOMO SAPIENS | -1 | 26518 | 5fm5 | M | 688 | THR | p.Thr688Ser | P52179 |
| HOMO SAPIENS | -1 | 26518 | 5fm5 | M | 704 | GLU | p.Glu704Lys | P52179 |
| HOMO SAPIENS | -1 | 26518 | 5fm5 | M | 727 | THR | p.Thr727Ala | P52179 |
| HOMO SAPIENS | 2 | 27054 | 4rhp | B | 209 | LEU | p.Leu209Val | O75208 |
| HOMO SAPIENS | -1 | 27054 | 4rhp | B | 288 | LYS | p.Lys288Asn | O75208 |
| HOMO SAPIENS | 5 | 2384 | 5fqd | B | 419 | ARG | p.Arg419Ter | Q96SW2 |
| HOMO SAPIENS | 4 | 5154 | 2hl3 | A | 56 | LYS | p.Lys56Arg | Q14203 |
| HOMO SAPIENS | 0 | 5154 | 2hl3 | A | 59 | GLY | p.Gly59Ser | Q14203 |
| HOMO SAPIENS | 0 | 5154 | 2hl3 | A | 71 | GLY | p.Gly71Ala | Q14203 |
| HOMO SAPIENS | 0 | 5154 | 2hl3 | A | 71 | GLY | p.Gly71Arg | Q14203 |
| HOMO SAPIENS | 7 | 5154 | 2hl3 | A | 72 | THR | p.Thr72Pro | Q14203 |
| HOMO SAPIENS | 2 | 5154 | 2hl3 | A | 74 | GLN | p.Gln74Pro | Q14203 |
| HOMO SAPIENS | 2 | 20039 | 2l1g | A | 9 | GLY | p.Gly9Cys | Q9NVV9 |
| HOMO SAPIENS | 5 | 20039 | 2l1g | A | 81 | PHE | p.Phe81Leu | Q9NVV9 |
| HOMO SAPIENS | -1 | 97 | 5uz7 | A | 201 | ARG | p.Arg201His | P63092 |
| HOMO SAPIENS | -1 | 97 | 5uz7 | A | 201 | ARG | p.Arg201Cys | P63092 |
| HOMO SAPIENS | 0 | 3257 | 4b7w | B | 278 | GLU | p.Glu278Asp | P41235 |
| HOMO SAPIENS | 1 | 3257 | 4b7w | B | 331 | ARG | p.Arg331Cys | P41235 |
| HOMO SAPIENS | 0 | 2814 | 4qgk | B | 10 | GLN | p.Gln10Ter | P51648 |
| HOMO SAPIENS | 0 | 2814 | 4qgk | B | 177 | ARG | p.Arg177Ter | P51648 |
| HOMO SAPIENS | 9 | 2814 | 4qgk | B | 184 | THR | p.Thr184Met | P51648 |
| HOMO SAPIENS | 1 | 2814 | 4qgk | B | 188 | ALA | p.Ala188Val | P51648 |
| HOMO SAPIENS | 0 | 2814 | 4qgk | B | 214 | CYS | p.Cys214Tyr | P51648 |
| HOMO SAPIENS | 0 | 2814 | 4qgk | B | 245 | ASP | p.Asp245Asn | P51648 |
| HOMO SAPIENS | 0 | 2814 | 4qgk | B | 266 | LYS | p.Lys266Asn | P51648 |
| HOMO SAPIENS | 0 | 2814 | 4qgk | B | 315 | PRO | p.Pro315Ser | P51648 |
| HOMO SAPIENS | 0 | 2814 | 4qgk | B | 386 | ASN | p.Asn386Ser | P51648 |
| HOMO SAPIENS | -1 | 5486 | 5u6o | B | 47 | GLY | p.Gly47Val | O60741 |
| HOMO SAPIENS | 1 | 21475 | 3pdy | A | 679 | THR | p.Thr789Ile | Q15149 |
| HOMO SAPIENS | 0 | 32890 | 4djc | B | 1501 | LEU | p.Leu1501Val | Q14524 |
| HOMO SAPIENS | -1 | 32890 | 4djc | B | 1512 | ARG | p.Arg1512Trp | Q14524 |
| HOMO SAPIENS | 0 | 26683 | 2xze | B | 38 | ARG | p.Arg38Cys | O95630 |
| HOMO SAPIENS | 0 | 26683 | 2xze | B | 100 | PHE | p.Phe100Tyr | O95630 |
| HOMO SAPIENS | -1 | 23175 | 5dka | A | 163 | SER | p.Ser163Leu | Q96EQ8 |
| HOMO SAPIENS | -1 | 23175 | 5dka | A | 174 | ARG | p.Arg174Cys | Q96EQ8 |
| HOMO SAPIENS | 6 | 21177 | 3k1r | A | 155 | ARG | p.Arg155Ter | Q9Y6N9 |
| HOMO SAPIENS | 0 | 1237 | 2wwm | O | 40 | PRO | p.Pro40Arg | O75147 |
| HOMO SAPIENS | 1 | 2543 | 5ewl | B | 64 | VAL | p.Val64Leu | Q13224 |
| HOMO SAPIENS | 2 | 2543 | 5ewl | B | 271 | ALA | p.Ala271Glu | Q13224 |
| HOMO SAPIENS | 3 | 34262 | 2mng | A | 613 | PHE | p.Phe613Leu | Q9Y3Q4 |
| HOMO SAPIENS | 7 | 34262 | 2mng | A | 672 | SER | p.Ser672Arg | Q9Y3Q4 |
| HOMO SAPIENS | 4 | 1667 | 1f05 | A | 321 | LYS | p.Lys321Met | P37837 |
| HOMO SAPIENS | 0 | 25179 | 2knc | B | 752 | SER | p.Ser778Pro | P05106 |
| HOMO SAPIENS | 0 | 2185 | 2ig7 | A | 50 | TYR | p.Tyr50Cys | Q9Y259 |
| HOMO SAPIENS | 0 | 2185 | 2ig7 | A | 224 | ASN | p.Asn224His | Q9Y259 |
| HOMO SAPIENS | 0 | 2185 | 2ig7 | A | 270 | TYR | p.Tyr270Ter | Q9Y259 |
| HOMO SAPIENS | 0 | 2185 | 2ig7 | A | 328 | GLN | p.Gln328Arg | Q9Y259 |
| HOMO SAPIENS | 3 | 2201 | 2r0o | A | 179 | PRO | p.Pro179Leu | O43707 |
| HOMO SAPIENS | 3 | 2201 | 2r0o | A | 255 | GLU | p.Lys255Glu | O43707 |
| HOMO SAPIENS | 3 | 2201 | 2r0o | A | 259 | THR | p.Thr259Ile | O43707 |
| HOMO SAPIENS | 4 | 2201 | 2r0o | A | 262 | SER | p.Ser262Pro | O43707 |
| HOMO SAPIENS | -1 | 31221 | 2wvr | C | 44 | THR | p.Thr44Pro | Q9H211 |
| HOMO SAPIENS | -1 | 31221 | 2wvr | C | 135 | ALA | p.Ala135Val | Q9H211 |
| HOMO SAPIENS | -1 | 31221 | 2wvr | C | 456 | GLU | p.Glu456Ala | Q9H211 |

| | | | | | | | | |
|---|---|---|---|---|---|---|---|---|
| HOMO SAPIENS | -1 | 31221 | 2wvr | C | 462 | ARG | p.Arg462Gln | Q9H211 |
| HOMO SAPIENS | -1 | 31221 | 2wvr | C | 468 | GLU | p.Glu468Lys | Q9H211 |
| HOMO SAPIENS | -1 | 31221 | 2wvr | C | 537 | ALA | p.Ala537Val | Q9H211 |
| HOMO SAPIENS | 5 | 2576 | 5o3l | C | 363 | VAL | p.Val680Ile | P10636 |
| HOMO SAPIENS | 5 | 1345 | 1czt | A | 37 | ARG | p.Arg2102Cys | P12259 |
| HOMO SAPIENS | 1 | 1345 | 1czt | A | 83 | MET | p.Met2148Thr | P12259 |
| HOMO SAPIENS | 1 | 1345 | 1czt | A | 157 | ASP | p.Asp2222Gly | P12259 |
| HOMO SAPIENS | 8 | 35985 | 1lo1 | A | 110 | ALA | p.Ala110Val | O95718 |
| HOMO SAPIENS | 1 | 617 | 2ocp | B | 44 | GLU | p.Glu44Lys | Q16854 |
| HOMO SAPIENS | 5 | 617 | 2ocp | B | 46 | ASN | p.Asn46Ser | Q16854 |
| HOMO SAPIENS | 0 | 617 | 2ocp | B | 59 | THR | p.Thr59Met | Q16854 |
| HOMO SAPIENS | 7 | 617 | 2ocp | B | 142 | ARG | p.Arg142Lys | Q16854 |
| HOMO SAPIENS | 3 | 617 | 2ocp | B | 154 | ASN | p.Asn154Lys | Q16854 |
| HOMO SAPIENS | 3 | 617 | 2ocp | B | 170 | GLN | p.Gln170Arg | Q16854 |
| HOMO SAPIENS | 3 | 617 | 2ocp | B | 227 | GLU | p.Glu227Lys | Q16854 |
| HOMO SAPIENS | 3 | 617 | 2ocp | B | 233 | LYS | p.Lys233Glu | Q16854 |
| HOMO SAPIENS | 3 | 617 | 2ocp | B | 255 | ASP | p.Asp255Tyr | Q16854 |
| HOMO SAPIENS | -1 | 38975 | 5itz | D | 421 | GLN | p.Gln421His | Q9HC77 |
| HOMO SAPIENS | 2 | 6235 | 2f8v | A | 15 | VAL | p.Val15Ile | Q8WZ42 |
| HOMO SAPIENS | 0 | 6235 | 2f8v | A | 54 | VAL | p.Val54Met | Q8WZ42 |
| HOMO SAPIENS | 0 | 6235 | 2f8v | A | 59 | SER | p.Ser59Arg | Q8WZ42 |
| HOMO SAPIENS | 0 | 6235 | 2f8v | A | 60 | ASP | p.Asp60Tyr | Q8WZ42 |
| HOMO SAPIENS | 0 | 12630 | 1ext | B | 12 | ASP | p.Asp41Glu | P19438 |
| HOMO SAPIENS | 3 | 12630 | 1ext | B | 30 | CYS | p.Cys59Arg | P19438 |
| HOMO SAPIENS | 3 | 12630 | 1ext | B | 30 | CYS | p.Cys59Ser | P19438 |
| HOMO SAPIENS | 5 | 12630 | 1ext | B | 33 | CYS | p.Cys62Tyr | P19438 |
| HOMO SAPIENS | 5 | 12630 | 1ext | B | 33 | CYS | p.Cys62Gly | P19438 |
| HOMO SAPIENS | 1 | 12630 | 1ext | B | 46 | PRO | p.Pro75Leu | P19438 |
| HOMO SAPIENS | 3 | 12630 | 1ext | B | 50 | THR | p.Thr79Met | P19438 |
| HOMO SAPIENS | 6 | 12630 | 1ext | B | 52 | CYS | p.Cys81Phe | P19438 |
| HOMO SAPIENS | 6 | 12630 | 1ext | B | 70 | CYS | p.Cys99Ser | P19438 |
| HOMO SAPIENS | 5 | 12630 | 1ext | B | 88 | CYS | p.Cys117Arg | P19438 |
| HOMO SAPIENS | 5 | 12630 | 1ext | B | 88 | CYS | p.Cys117Tyr | P19438 |
| HOMO SAPIENS | -1 | 12630 | 1ext | B | 170 | ILE | p.Ile199Asn | P19438 |
| HOMO SAPIENS | 0 | 994 | 1aos | B | 95 | ARG | p.Arg95Cys | P04424 |
| HOMO SAPIENS | 0 | 994 | 1aos | B | 111 | ARG | p.Arg111Gln | P04424 |
| HOMO SAPIENS | 0 | 994 | 1aos | B | 131 | THR | p.Thr131Met | P04424 |
| HOMO SAPIENS | 6 | 994 | 1aos | B | 178 | VAL | p.Val178Met | P04424 |
| HOMO SAPIENS | 9 | 994 | 1aos | B | 182 | ARG | p.Arg182Gln | P04424 |
| HOMO SAPIENS | 9 | 994 | 1aos | B | 186 | ARG | p.Arg186Trp | P04424 |
| HOMO SAPIENS | 8 | 994 | 1aos | B | 193 | ARG | p.Arg193Gln | P04424 |
| HOMO SAPIENS | 5 | 994 | 1aos | B | 217 | ARG | p.Arg217Ter | P04424 |
| HOMO SAPIENS | 0 | 994 | 1aos | B | 286 | GLN | p.Gln286Arg | P04424 |
| HOMO SAPIENS | 5 | 994 | 1aos | B | 379 | ARG | p.Arg379Cys | P04424 |
| HOMO SAPIENS | 4 | 16066 | 5mv9 | A | 1719 | TYR | p.Tyr1719Cys | Q13402 |
| HOMO SAPIENS | -1 | 16066 | 5mv9 | A | 1739 | ARG | p.Arg1739Ter | Q13402 |
| HOMO SAPIENS | 4 | 16066 | 5mv9 | A | 1775 | ILE | p.Ile1775Thr | Q13402 |
| HOMO SAPIENS | 2 | 16066 | 5mv9 | A | 1786 | SER | p.Ser1786Thr | Q13402 |
| HOMO SAPIENS | 4 | 16066 | 5mv9 | A | 1798 | GLN | p.Gln1798Ter | Q13402 |
| HOMO SAPIENS | 0 | 16066 | 5mv9 | A | 1858 | LEU | p.Leu1858Pro | Q13402 |
| HOMO SAPIENS | 6 | 16066 | 5mv9 | A | 1873 | ARG | p.Arg1873Trp | Q13402 |
| HOMO SAPIENS | 6 | 16066 | 5mv9 | A | 1873 | ARG | p.Arg1873Leu | Q13402 |
| HOMO SAPIENS | 6 | 16066 | 5mv9 | A | 1883 | ARG | p.Arg1883Gln | Q13402 |
| HOMO SAPIENS | 4 | 16066 | 5mv9 | A | 1887 | PRO | p.Pro1887Leu | Q13402 |

| | | | | | | | | |
|---|---|---|---|---|---|---|---|---|
| HOMO SAPIENS | 5 | 16066 | 5mv9 | A | 1942 | GLY | p.Gly1942Ter | Q13402 |
| HOMO SAPIENS | 5 | 16066 | 5mv9 | A | 1954 | LEU | p.Leu1954Ile | Q13402 |
| HOMO SAPIENS | 5 | 16066 | 5mv9 | A | 1956 | VAL | p.Val1956Ile | Q13402 |
| HOMO SAPIENS | 5 | 16066 | 5mv9 | A | 1967 | ARG | p.Arg1967Ter | Q13402 |
| HOMO SAPIENS | 5 | 16066 | 5mv9 | A | 1990 | GLN | p.Gln1990Ter | Q13402 |
| HOMO SAPIENS | 2 | 16066 | 5mv9 | A | 2005 | LYS | p.Lys2005Glu | Q13402 |
| HOMO SAPIENS | 1 | 16066 | 5mv9 | A | 2072 | VAL | p.Val2072Ile | Q13402 |
| HOMO SAPIENS | 4 | 16066 | 5mv9 | A | 2109 | THR | p.Thr2109Ile | Q13402 |
| HOMO SAPIENS | 4 | 16066 | 5mv9 | A | 2142 | ASP | p.Asp2142Asn | Q13402 |
| HOMO SAPIENS | -1 | 16066 | 5mv9 | A | 2214 | GLY | p.Gly2214Ser | Q13402 |
| HOMO SAPIENS | 7 | 27942 | 1deb | A | 24 | ARG | p.Arg24Ter | P25054 |
| HOMO SAPIENS | 0 | 876 | 5hnr | B | 12 | PHE | p.Phe12Leu | P13716 |
| HOMO SAPIENS | 0 | 876 | 5hnr | B | 59 | LYS | p.Lys59Asn | P13716 |
| HOMO SAPIENS | -1 | 876 | 5hnr | B | 133 | GLY | p.Gly133Arg | P13716 |
| HOMO SAPIENS | 2 | 876 | 5hnr | B | 240 | ARG | p.Arg240Trp | P13716 |
| HOMO SAPIENS | 0 | 876 | 5hnr | B | 274 | ALA | p.Ala274Thr | P13716 |
| HOMO SAPIENS | 0 | 876 | 5hnr | B | 275 | VAL | p.Val275Met | P13716 |
| HOMO SAPIENS | 0 | 1142 | 3qe2 | A | 503 | VAL | p.Ala500Val | P16435 |
| HOMO SAPIENS | 8 | 14358 | 4dkd | C | 32 | VAL | p.Val32Gly | P07333 |
| HOMO SAPIENS | 0 | 14358 | 4dkd | C | 60 | LEU | p.Leu60Val | P07333 |
| HOMO SAPIENS | 7 | 14358 | 4dkd | C | 75 | THR | p.Thr75Ile | P07333 |
| HOMO SAPIENS | -1 | 14358 | 4dkd | C | 90 | PRO | p.Pro90Thr | P07333 |
| HOMO SAPIENS | 0 | 14358 | 4dkd | C | 106 | ARG | p.Arg106Trp | P07333 |
| HOMO SAPIENS | 0 | 14358 | 4dkd | C | 241 | ASN | p.Asn241Asp | P07333 |
| HOMO SAPIENS | 0 | 14358 | 4dkd | C | 245 | ALA | p.Ala245Ser | P07333 |
| HOMO SAPIENS | 0 | 14358 | 4dkd | C | 255 | ASN | p.Asn255Ile | P07333 |
| HOMO SAPIENS | 0 | 14358 | 4dkd | C | 268 | VAL | p.Val268Ile | P07333 |
| HOMO SAPIENS | 0 | 14358 | 4dkd | C | 279 | VAL | p.Val279Met | P07333 |
| HOMO SAPIENS | 0 | 14358 | 4dkd | C | 294 | ARG | p.Arg294Gln | P07333 |
| HOMO SAPIENS | 5 | 8968 | 3ppy | A | 1565 | VAL | p.Val1565Leu | P04275 |
| HOMO SAPIENS | 4 | 8968 | 3ppy | A | 1584 | TYR | p.Tyr1584Cys | P04275 |
| HOMO SAPIENS | 5 | 5341 | 3k3n | A | 279 | PHE | p.Phe315Ser | Q9UPP1 |
| HOMO SAPIENS | 0 | 29463 | 1q7l | A | 197 | ARG | p.Arg197Trp | Q03154 |
| HOMO SAPIENS | 0 | 33761 | 2leh | B | 30 | ASP | p.Asp30Asn | Q16637 |
| HOMO SAPIENS | 6 | 33761 | 2leh | B | 44 | ASP | p.Asp44Val | Q16637 |
| HOMO SAPIENS | -1 | 12249 | 2pnr | C | 224 | GLU | p.Glu310Lys | P10515 |
| HOMO SAPIENS | -1 | 12249 | 2pnr | C | 232 | VAL | p.Val318Ala | P10515 |
| HOMO SAPIENS | -1 | 7 | 5whk | B | -9 | ALA | p.Ala11Pro | P61769 |
| HOMO SAPIENS | 8 | 20298 | 4euw | A | 154 | PHE | p.Phe154Leu | P48436 |
| HOMO SAPIENS | 9 | 20298 | 4euw | A | 158 | ALA | p.Ala158Thr | P48436 |
| HOMO SAPIENS | 5 | 20298 | 4euw | A | 165 | HIS | p.His165Tyr | P48436 |
| HOMO SAPIENS | 3 | 20298 | 4euw | A | 169 | HIS | p.His169Gln | P48436 |
| HOMO SAPIENS | 0 | 20298 | 4euw | A | 173 | LYS | p.Lys173Glu | P48436 |
| HOMO SAPIENS | 1 | 16465 | 2jgx | A | 402 | TYR | p.Tyr402His | P08603 |
| HOMO SAPIENS | 0 | 8993 | 3v8o | A | 659 | LEU | p.Leu659Arg | Q14315 |
| HOMO SAPIENS | 6 | 8993 | 3v8o | A | 693 | ASP | p.Asp693Ala | Q14315 |
| HOMO SAPIENS | 0 | 8993 | 3v8o | A | 709 | ALA | p.Ala709Thr | Q14315 |
| HOMO SAPIENS | 2 | 3852 | 5es4 | B | 106 | ASP | p.Asp128Asn | P05107 |
| HOMO SAPIENS | 2 | 3852 | 5es4 | B | 116 | SER | p.Ser138Pro | P05107 |
| HOMO SAPIENS | 1 | 3852 | 5es4 | B | 127 | LEU | p.Leu149Pro | P05107 |
| HOMO SAPIENS | 0 | 3852 | 5es4 | B | 147 | GLY | p.Gly169Arg | P05107 |
| HOMO SAPIENS | 3 | 3852 | 5es4 | B | 156 | PRO | p.Pro178Leu | P05107 |
| HOMO SAPIENS | 0 | 3852 | 5es4 | B | 174 | LYS | p.Lys196Thr | P05107 |
| HOMO SAPIENS | 2 | 3852 | 5es4 | B | 251 | GLY | p.Gly273Arg | P05107 |

| | | | | | | | | |
|---|---|---|---|---|---|---|---|---|
| HOMO SAPIENS | 2 | 3852 | 5es4 | B | 329 | ASN | p.Asn351Ser | P05107 |
| HOMO SAPIENS | 2 | 3852 | 5es4 | B | 571 | ARG | p.Arg593Cys | P05107 |
| HOMO SAPIENS | 7 | 5962 | 5fn2 | B | 146 | MET | p.Met146Leu | P49768 |
| HOMO SAPIENS | 3 | 5962 | 5fn2 | B | 166 | LEU | p.Leu166Pro | P49768 |
| HOMO SAPIENS | 1 | 5962 | 5fn2 | B | 233 | MET | p.Met233Leu | P49768 |
| HOMO SAPIENS | 1 | 5962 | 5fn2 | B | 261 | VAL | p.Val261Gly | P49768 |
| HOMO SAPIENS | 1 | 5962 | 5fn2 | B | 266 | GLY | p.Gly266Ser | P49768 |
| HOMO SAPIENS | -1 | 5962 | 5fn2 | B | 333 | ASP | p.Asp333Gly | P49768 |
| HOMO SAPIENS | 6 | 1918 | 3dlx | A | 58 | THR | p.Thr58Met | P55809 |
| HOMO SAPIENS | 5 | 1918 | 3dlx | A | 133 | VAL | p.Val133Glu | P55809 |
| HOMO SAPIENS | 5 | 1918 | 3dlx | A | 219 | GLY | p.Gly219Glu | P55809 |
| HOMO SAPIENS | 0 | 1918 | 3dlx | A | 221 | VAL | p.Val221Met | P55809 |
| HOMO SAPIENS | 5 | 1918 | 3dlx | A | 283 | SER | p.Ser283Ter | P55809 |
| HOMO SAPIENS | 4 | 1918 | 3dlx | A | 324 | GLY | p.Gly324Glu | P55809 |
| HOMO SAPIENS | 0 | 1918 | 3dlx | A | 456 | CYS | p.Cys456Phe | P55809 |
| HOMO SAPIENS | 0 | 9010 | 2wfr | A | 162 | LEU | p.Leu162Pro | O43323 |
| HOMO SAPIENS | 5 | 3770 | 5n38 | A | 56 | VAL | p.Val56Glu | O60260 |
| HOMO SAPIENS | 1 | 3770 | 5n38 | A | 161 | LYS | p.Lys161Asn | O60260 |
| HOMO SAPIENS | 1 | 3770 | 5n38 | A | 167 | SER | p.Ser167Asn | O60260 |
| HOMO SAPIENS | 1 | 3770 | 5n38 | A | 211 | LYS | p.Lys211Asn | O60260 |
| HOMO SAPIENS | 1 | 3770 | 5n38 | A | 212 | CYS | p.Cys212Tyr | O60260 |
| HOMO SAPIENS | 1 | 3770 | 5n38 | A | 275 | ARG | p.Arg275Trp | O60260 |
| HOMO SAPIENS | 2 | 3770 | 5n38 | A | 334 | ARG | p.Arg334Cys | O60260 |
| HOMO SAPIENS | 1 | 3770 | 5n38 | A | 366 | ARG | p.Arg366Trp | O60260 |
| HOMO SAPIENS | -1 | 3770 | 5n38 | A | 380 | VAL | p.Val380Leu | O60260 |
| HOMO SAPIENS | 1 | 3770 | 5n38 | A | 394 | ASP | p.Asp394Asn | O60260 |
| HOMO SAPIENS | 1 | 3770 | 5n38 | A | 430 | GLY | p.Gly430Asp | O60260 |
| HOMO SAPIENS | -1 | 2467 | 4xx1 | A | 10 | PRO | p.Pro34Leu | P04180 |
| HOMO SAPIENS | 4 | 2467 | 4xx1 | A | 83 | TYR | p.Tyr107Ter | P04180 |
| HOMO SAPIENS | 4 | 2467 | 4xx1 | A | 93 | ALA | p.Ala117Thr | P04180 |
| HOMO SAPIENS | 4 | 2467 | 4xx1 | A | 123 | THR | p.Thr147Ile | P04180 |
| HOMO SAPIENS | 4 | 2467 | 4xx1 | A | 135 | ARG | p.Arg159Trp | P04180 |
| HOMO SAPIENS | 3 | 2467 | 4xx1 | A | 158 | ARG | p.Arg182Cys | P04180 |
| HOMO SAPIENS | 0 | 2467 | 4xx1 | A | 209 | LEU | p.Leu233Pro | P04180 |
| HOMO SAPIENS | 1 | 2467 | 4xx1 | A | 228 | ASN | p.Asn252Lys | P04180 |
| HOMO SAPIENS | 4 | 2467 | 4xx1 | A | 252 | MET | p.Met276Lys | P04180 |
| HOMO SAPIENS | 3 | 2467 | 4xx1 | A | 293 | MET | p.Met317Ile | P04180 |
| HOMO SAPIENS | 5 | 2467 | 4xx1 | A | 321 | THR | p.Thr345Met | P04180 |
| HOMO SAPIENS | 9 | 2467 | 4xx1 | A | 347 | THR | p.Thr371Met | P04180 |
| HOMO SAPIENS | 1 | 5834 | 3qrf | G | 371 | PHE | p.Phe371Cys | Q9BZS1 |
| HOMO SAPIENS | 1 | 5834 | 3qrf | G | 373 | PHE | p.Phe373Ala | Q9BZS1 |
| HOMO SAPIENS | 9 | 5834 | 3qrf | G | 384 | ALA | p.Ala384Thr | Q9BZS1 |
| HOMO SAPIENS | 6 | 5834 | 3qrf | G | 397 | ARG | p.Arg397Trp | Q9BZS1 |
| HOMO SAPIENS | -1 | 11869 | 4km8 | A | 15 | PRO | p.Pro15Leu | Q9UMX1 |
| HOMO SAPIENS | 5 | 262 | 4kbo | A | 126 | ARG | p.Arg126Trp | P38646 |
| HOMO SAPIENS | 3 | 262 | 4kbo | A | 128 | TYR | p.Tyr128Cys | P38646 |
| HOMO SAPIENS | -1 | 11347 | 4uf9 | D | 117 | ALA | p.Ala117Val | Q13501 |
| HOMO SAPIENS | 0 | 2173 | 4ah6 | A | 45 | SER | p.Ser45Gly | Q6PI48 |
| HOMO SAPIENS | 2 | 2173 | 4ah6 | A | 152 | CYS | p.Cys152Phe | Q6PI48 |
| HOMO SAPIENS | 4 | 2173 | 4ah6 | A | 179 | ARG | p.Arg179His | Q6PI48 |
| HOMO SAPIENS | 0 | 2173 | 4ah6 | A | 196 | LYS | p.Lys196Arg | Q6PI48 |
| HOMO SAPIENS | 8 | 2173 | 4ah6 | A | 263 | ARG | p.Arg263Gln | Q6PI48 |
| HOMO SAPIENS | 1 | 2173 | 4ah6 | A | 338 | GLY | p.Gly338Glu | Q6PI48 |
| HOMO SAPIENS | 0 | 2173 | 4ah6 | A | 425 | GLU | p.Glu425Ter | Q6PI48 |

| | | | | | | | | |
|---|---|---|---|---|---|---|---|---|
| HOMO SAPIENS | 1 | 2173 | 4ah6 | A | 548 | LEU | p.Leu548Met | Q6PI48 |
| HOMO SAPIENS | 0 | 2173 | 4ah6 | A | 609 | ARG | p.Arg609Trp | Q6PI48 |
| HOMO SAPIENS | 2 | 2173 | 4ah6 | A | 613 | LEU | p.Leu613Phe | Q6PI48 |
| HOMO SAPIENS | 0 | 2173 | 4ah6 | A | 626 | LEU | p.Leu626Gln | Q6PI48 |
| HOMO SAPIENS | 0 | 2173 | 4ah6 | A | 626 | LEU | p.Leu626Val | Q6PI48 |
| HOMO SAPIENS | 7 | 3203 | 2cn1 | A | 87 | ASP | p.Asp137Val | Q9H0P0 |
| HOMO SAPIENS | 8 | 3203 | 2cn1 | A | 166 | GLN | p.Gln216Ter | Q9H0P0 |
| HOMO SAPIENS | 8 | 3203 | 2cn1 | A | 179 | ASN | p.Asn229Ser | Q9H0P0 |
| HOMO SAPIENS | 7 | 3203 | 2cn1 | A | 230 | GLY | p.Gly280Arg | Q9H0P0 |
| HOMO SAPIENS | 2 | 3424 | 2a1t | R | 116 | GLY | p.Gly116Arg | P13804 |
| HOMO SAPIENS | 2 | 3424 | 2a1t | R | 157 | VAL | p.Val157Gly | P13804 |
| HOMO SAPIENS | 2 | 3424 | 2a1t | R | 171 | THR | p.Thr171Ile | P13804 |
| HOMO SAPIENS | 0 | 3424 | 2a1t | R | 178 | THR | p.Thr178Arg | P13804 |
| HOMO SAPIENS | 0 | 3424 | 2a1t | R | 194 | VAL | p.Val194Met | P13804 |
| HOMO SAPIENS | 1 | 3424 | 2a1t | R | 209 | ARG | p.Arg209Ter | P13804 |
| HOMO SAPIENS | 6 | 3424 | 2a1t | R | 266 | THR | p.Thr266Met | P13804 |
| HOMO SAPIENS | 0 | 27598 | 1g1c | A | 26 | ARG | p.Arg2098Gly | Q8WZ42 |
| HOMO SAPIENS | 1 | 27598 | 1g1c | A | 46 | ILE | p.Ile2118Thr | Q8WZ42 |
| HOMO SAPIENS | 0 | 32885 | 4bsk | A | 149 | ASN | p.Asn149Asp | P35916 |
| HOMO SAPIENS | -1 | 8074 | 3ghg | B | 17 | ARG | p.Arg47Ter | P02675 |
| HOMO SAPIENS | -1 | 447 | 5g4p | A | 190 | ALA | p.Ala190Thr | P84074 |
| HOMO SAPIENS | 0 | 10761 | 5u1d | B | 379 | VAL | p.Val379Ile | Q03519 |
| HOMO SAPIENS | 0 | 10761 | 5u1d | B | 665 | THR | p.Thr665Ala | Q03519 |
| HOMO SAPIENS | 5 | 29251 | 5fn7 | A | 79 | ILE | p.Ile294Leu | P08575 |
| HOMO SAPIENS | -1 | 27727 | 4ky9 | A | 56 | LYS | p.Lys56Glu | P02730 |
| HOMO SAPIENS | 2 | 27727 | 4ky9 | A | 72 | GLU | p.Glu72Asp | P02730 |
| HOMO SAPIENS | 2 | 27727 | 4ky9 | A | 90 | GLU | p.Glu90Lys | P02730 |
| HOMO SAPIENS | 1 | 27727 | 4ky9 | A | 130 | GLY | p.Gly130Arg | P02730 |
| HOMO SAPIENS | 2 | 27727 | 4ky9 | A | 150 | ARG | p.Arg150Ter | P02730 |
| HOMO SAPIENS | 0 | 27727 | 4ky9 | A | 327 | PRO | p.Pro327Arg | P02730 |
| HOMO SAPIENS | 2 | 13202 | 2fxo | B | 870 | ARG | p.Arg870His | P12883 |
| HOMO SAPIENS | 2 | 13202 | 2fxo | B | 870 | ARG | p.Arg870Cys | P12883 |
| HOMO SAPIENS | 4 | 13202 | 2fxo | B | 904 | ARG | p.Arg904Cys | P12883 |
| HOMO SAPIENS | 4 | 13202 | 2fxo | B | 906 | ASP | p.Asp906Gly | P12883 |
| HOMO SAPIENS | 2 | 13202 | 2fxo | B | 908 | LEU | p.Leu908Val | P12883 |
| HOMO SAPIENS | 2 | 13202 | 2fxo | B | 923 | ASN | p.Asn923Lys | P12883 |
| HOMO SAPIENS | 7 | 13202 | 2fxo | B | 935 | GLU | p.Glu935Lys | P12883 |
| HOMO SAPIENS | 7 | 13202 | 2fxo | B | 935 | GLU | p.Glu935Val | P12883 |
| HOMO SAPIENS | 8 | 13202 | 2fxo | B | 949 | GLU | p.Glu949Lys | P12883 |
| HOMO SAPIENS | 0 | 13200 | 2a1s | A | 349 | ARG | p.Arg349Trp | O95453 |
| HOMO SAPIENS | 0 | 13200 | 2a1s | A | 421 | LYS | p.Lys421Arg | O95453 |
| HOMO SAPIENS | 6 | 3196 | 5xtb | T | 115 | CYS | p.Cys115Tyr | O75380 |
| HOMO SAPIENS | 0 | 1108 | 5ukf | A | 89 | ARG | p.Arg89Gln | Q99986 |
| HOMO SAPIENS | 0 | 1108 | 5ukf | A | 119 | HIS | p.His119Arg | Q99986 |
| HOMO SAPIENS | 0 | 1108 | 5ukf | A | 133 | ARG | p.Arg133Cys | Q99986 |
| HOMO SAPIENS | 0 | 1108 | 5ukf | A | 236 | VAL | p.Val236Leu | Q99986 |
| HOMO SAPIENS | 0 | 1108 | 5ukf | A | 321 | ARG | p.Arg321Cys | Q99986 |
| HOMO SAPIENS | -1 | 1108 | 5ukf | A | 358 | ARG | p.Arg358Ter | Q99986 |
| HOMO SAPIENS | 0 | 5842 | 4yli | C | 204 | GLY | p.Gly204Ser | Q9BWP8 |
| HOMO SAPIENS | -1 | 31362 | 4wqo | A | 25 | PRO | p.Pro25Leu | P40337 |
| HOMO SAPIENS | 2 | 10107 | 2j8h | A | 34 | VAL | p.Val31886Ile | Q8WZ42 |
| HOMO SAPIENS | 8 | 20190 | 1wm5 | A | 77 | ARG | p.Arg77Gln | P19878 |
| HOMO SAPIENS | 1 | 20190 | 1wm5 | A | 100 | GLN | p.Gln100Ter | P19878 |
| HOMO SAPIENS | 0 | 20190 | 1wm5 | A | 102 | ARG | p.Arg102Ter | P19878 |

| | | | | | | | | | |
|---|---|---|---|---|---|---|---|---|---|
| HOMO SAPIENS | 0 | 20190 | 1wm5 | A | | 128 | ALA | p.Ala128Val | P19878 |
| HOMO SAPIENS | 0 | 20190 | 1wm5 | A | | 181 | LYS | p.Lys181Arg | P19878 |
| HOMO SAPIENS | 7 | 1278 | 5a2q | a | | 33 | ASP | p.Asp33Asn | P62854 |
| HOMO SAPIENS | 3 | 19515 | 2obd | A | | 11 | ILE | p.Ile28Met | P11597 |
| HOMO SAPIENS | 3 | 19515 | 2obd | A | | 89 | VAL | p.Val106Met | P11597 |
| HOMO SAPIENS | 3 | 19515 | 2obd | A | | 137 | ARG | p.Arg154Trp | P11597 |
| HOMO SAPIENS | 3 | 19515 | 2obd | A | | 139 | ASP | p.Asp156Asn | P11597 |
| HOMO SAPIENS | 3 | 19515 | 2obd | A | | 204 | SER | p.Ser221Arg | P11597 |
| HOMO SAPIENS | 0 | 19515 | 2obd | A | | 297 | GLU | p.Glu314Lys | P11597 |
| HOMO SAPIENS | 0 | 19515 | 2obd | A | | 314 | GLY | p.Gly331Ser | P11597 |
| HOMO SAPIENS | 0 | 19515 | 2obd | A | | 332 | SER | p.Ser349Tyr | P11597 |
| HOMO SAPIENS | 0 | 19515 | 2obd | A | | 368 | VAL | p.Val385Met | P11597 |
| HOMO SAPIENS | 0 | 19515 | 2obd | A | | 373 | ALA | p.Ala390Pro | P11597 |
| HOMO SAPIENS | 3 | 19515 | 2obd | A | | 405 | ILE | p.Val422Ile | P11597 |
| HOMO SAPIENS | 0 | 19515 | 2obd | A | | 451 | ARG | p.Arg468Gln | P11597 |
| HOMO SAPIENS | 0 | 19515 | 2obd | A | | 451 | ARG | p.Arg468Ter | P11597 |
| HOMO SAPIENS | 0 | 6626 | 3hop | A | | 39 | ALA | p.Ala39Gly | P21333 |
| HOMO SAPIENS | 1 | 6626 | 3hop | A | | 82 | GLU | p.Glu82Val | P21333 |
| HOMO SAPIENS | 1 | 6626 | 3hop | A | | 128 | ALA | p.Ala128Val | P21333 |
| HOMO SAPIENS | 2 | 6626 | 3hop | A | | 203 | ASP | p.Asp203Tyr | P21333 |
| HOMO SAPIENS | 1 | 6626 | 3hop | A | | 207 | PRO | p.Pro207Leu | P21333 |
| HOMO SAPIENS | 2 | 6626 | 3hop | A | | 210 | CYS | p.Cys210Phe | P21333 |
| HOMO SAPIENS | 7 | 6626 | 3hop | A | | 254 | GLU | p.Glu254Lys | P21333 |
| HOMO SAPIENS | 4 | 4210 | 2avg | A | | 8 | VAL | p.Val158Met | Q14896 |
| HOMO SAPIENS | 6 | 4210 | 2avg | A | | 15 | GLU | p.Glu165Asp | Q14896 |
| HOMO SAPIENS | 4 | 4210 | 2avg | A | | 27 | ARG | p.Arg177His | Q14896 |
| HOMO SAPIENS | 0 | 4210 | 2avg | A | | 28 | VAL | p.Val178Met | Q14896 |
| HOMO SAPIENS | 6 | 4210 | 2avg | A | | 39 | VAL | p.Val189Ile | Q14896 |
| HOMO SAPIENS | 2 | 4210 | 2avg | A | | 52 | LYS | p.Lys202Gln | Q14896 |
| HOMO SAPIENS | 4 | 4210 | 2avg | A | | 67 | SER | p.Ser217Gly | Q14896 |
| HOMO SAPIENS | 6 | 4210 | 2avg | A | | 69 | VAL | p.Val219Leu | Q14896 |
| HOMO SAPIENS | 2 | 4210 | 2avg | A | | 86 | GLY | p.Ser236Gly | Q14896 |
| HOMO SAPIENS | 4 | 4210 | 2avg | A | | 108 | GLU | p.Glu258Lys | Q14896 |
| HOMO SAPIENS | 0 | 22237 | 1w6k | A | | 588 | GLY | p.Gly588Ser | P48449 |
| HOMO SAPIENS | 0 | 10765 | 4a5w | B | | 98 | GLU | p.Glu98Ala | P13671 |
| HOMO SAPIENS | 0 | 10765 | 4a5w | B | | 596 | ARG | p.Arg596Ter | P13671 |
| HOMO SAPIENS | -1 | 15515 | 3hx3 | A | | 11 | MSE | p.Met10Lys | P12271 |
| HOMO SAPIENS | 8 | 15515 | 3hx3 | A | | 64 | ARG | p.Arg64Gln | P12271 |
| HOMO SAPIENS | 0 | 15515 | 3hx3 | A | | 182 | PHE | p.Phe182Cys | P12271 |
| HOMO SAPIENS | 0 | 15515 | 3hx3 | A | | 216 | ARG | p.Arg216Gln | P12271 |
| HOMO SAPIENS | -1 | 15515 | 3hx3 | A | | 226 | MSE | p.Met226Lys | P12271 |
| HOMO SAPIENS | 0 | 15515 | 3hx3 | A | | 234 | TRP | p.Arg234Trp | P12271 |
| HOMO SAPIENS | 0 | 15515 | 3hx3 | A | | 273 | LEU | p.Leu273Phe | P12271 |
| HOMO SAPIENS | 0 | 15515 | 3hx3 | A | | 292 | THR | p.Thr292Met | P12271 |
| HOMO SAPIENS | 0 | 5152 | 2www | D | | 95 | GLN | p.Gln95Ter | Q8IVH4 |
| HOMO SAPIENS | 0 | 5152 | 2www | D | | 133 | GLN | p.Gln133Ter | Q8IVH4 |
| HOMO SAPIENS | 0 | 5152 | 2www | D | | 145 | ARG | p.Arg145Ter | Q8IVH4 |
| HOMO SAPIENS | 0 | 5152 | 2www | D | | 207 | TYR | p.Tyr207Cys | Q8IVH4 |
| HOMO SAPIENS | 8 | 5152 | 2www | D | | 330 | ARG | p.Arg330Ter | Q8IVH4 |
| HOMO SAPIENS | 0 | 5152 | 2www | D | | 363 | GLN | p.Gln363His | Q8IVH4 |
| HOMO SAPIENS | 1 | 1626 | 1ilr | | 1 | 105 | SER | p.Ser130Arg | P18510 |
| HOMO SAPIENS | -1 | 929 | 5loh | A | | 12 | GLY | p.Gly12Arg | Q96GX5 |
| HOMO SAPIENS | 9 | 3206 | 1x63 | A | | 21 | CYS | p.Cys104Arg | Q13642 |
| HOMO SAPIENS | 8 | 3206 | 1x63 | A | | 39 | TRP | p.Trp122Ser | Q13642 |

| | | | | | | | | |
|---|---|---|---|---|---|---|---|---|
| HOMO SAPIENS | 9 | 3206 | 1x63 | A | 40 | HIS | p.His123Tyr | Q13642 |
| HOMO SAPIENS | 9 | 3206 | 1x63 | A | 40 | HIS | p.His123Leu | Q13642 |
| HOMO SAPIENS | 9 | 3206 | 1x63 | A | 40 | HIS | p.His123Gln | Q13642 |
| HOMO SAPIENS | 3 | 3206 | 1x63 | A | 49 | CYS | p.Cys132Phe | Q13642 |
| HOMO SAPIENS | 3 | 3206 | 1x63 | A | 67 | CYS | p.Cys150Tyr | Q13642 |
| HOMO SAPIENS | 3 | 3206 | 1x63 | A | 70 | CYS | p.Cys153Arg | Q13642 |
| HOMO SAPIENS | 3 | 3206 | 1x63 | A | 70 | CYS | p.Cys153Tyr | Q13642 |
| HOMO SAPIENS | 5 | 1052 | 3ecs | E | 208 | ASN | p.Asn208Tyr | Q14232 |
| HOMO SAPIENS | 2 | 1052 | 3ecs | E | 275 | TYR | p.Tyr275Cys | Q14232 |
| HOMO SAPIENS | 0 | 24253 | 5i35 | A | 271 | ARG | p.Arg271Cys | Q8NI60 |
| HOMO SAPIENS | 0 | 24253 | 5i35 | A | 272 | GLY | p.Gly272Asp | Q8NI60 |
| HOMO SAPIENS | 0 | 24253 | 5i35 | A | 299 | ARG | p.Arg299Trp | Q8NI60 |
| HOMO SAPIENS | 1 | 24253 | 5i35 | A | 339 | ALA | p.Ala339Thr | Q8NI60 |
| HOMO SAPIENS | 0 | 24253 | 5i35 | A | 348 | ARG | p.Arg348Ter | Q8NI60 |
| HOMO SAPIENS | 0 | 24253 | 5i35 | A | 395 | ILE | p.Ile395Met | Q8NI60 |
| HOMO SAPIENS | 0 | 24253 | 5i35 | A | 396 | ASP | p.Asp396Glu | Q8NI60 |
| HOMO SAPIENS | 0 | 24253 | 5i35 | A | 480 | PHE | p.Phe480Leu | Q8NI60 |
| HOMO SAPIENS | 2 | 24253 | 5i35 | A | 511 | THR | p.Thr511Met | Q8NI60 |
| HOMO SAPIENS | 0 | 24253 | 5i35 | A | 514 | TYR | p.Tyr514Cys | Q8NI60 |
| HOMO SAPIENS | 0 | 24253 | 5i35 | A | 538 | ARG | p.Arg538Gln | Q8NI60 |
| HOMO SAPIENS | 0 | 24253 | 5i35 | A | 549 | GLY | p.Gly549Ser | Q8NI60 |
| HOMO SAPIENS | 0 | 24253 | 5i35 | A | 551 | GLU | p.Glu551Lys | Q8NI60 |
| HOMO SAPIENS | -1 | 24253 | 5i35 | A | 555 | MSE | p.Met555Ile | Q8NI60 |
| HOMO SAPIENS | 0 | 24253 | 5i35 | A | 568 | GLU | p.Glu568Ter | Q8NI60 |
| HOMO SAPIENS | 0 | 24253 | 5i35 | A | 602 | PRO | p.Pro602Arg | Q8NI60 |
| HOMO SAPIENS | -1 | 15505 | 4goj | C | 22 | GLY | p.Gly22Val | Q13432 |
| HOMO SAPIENS | 5 | 23902 | 2lrr | A | 765 | HIS | p.His765Gln | P38935 |
| HOMO SAPIENS | 1 | 29650 | 2vj2 | A | 234 | CYS | p.Cys234Tyr | P78504 |
| HOMO SAPIENS | 0 | 29650 | 2vj2 | A | 255 | TYR | p.Tyr255Ter | P78504 |
| HOMO SAPIENS | 3 | 29650 | 2vj2 | A | 274 | GLY | p.Gly274Asp | P78504 |
| HOMO SAPIENS | 1 | 105 | 5he0 | B | 77 | GLY | p.Gly77Ser | P62873 |
| HOMO SAPIENS | 2 | 105 | 5he0 | B | 80 | ILE | p.Ile80Thr | P62873 |
| HOMO SAPIENS | 3 | 4587 | 3rgh | A | 159 | ASP | p.Asp1159Ala | P21333 |
| HOMO SAPIENS | 3 | 4587 | 3rgh | A | 184 | ASP | p.Asp1184Glu | P21333 |
| HOMO SAPIENS | 2 | 4587 | 3rgh | A | 186 | SER | p.Ser1186Leu | P21333 |
| HOMO SAPIENS | 4 | 4587 | 3rgh | A | 188 | ALA | p.Ala1188Thr | P21333 |
| HOMO SAPIENS | 2 | 4587 | 3rgh | A | 199 | SER | p.Ser1199Leu | P21333 |
| HOMO SAPIENS | 0 | 20283 | 4n3w | A | 1278 | GLU | p.Glu1278Lys | Q92793 |
| HOMO SAPIENS | 1 | 14294 | 2d9q | B | 208 | MET | p.Met231Thr | Q99062 |
| HOMO SAPIENS | 6 | 14294 | 2d9q | B | 297 | ASP | p.Asp320Asn | Q99062 |
| HOMO SAPIENS | 0 | 11642 | 3bpt | A | 66 | ARG | p.Arg66Trp | Q6NVY1 |
| HOMO SAPIENS | 0 | 11642 | 3bpt | A | 122 | TYR | p.Tyr122Cys | Q6NVY1 |
| HOMO SAPIENS | 0 | 11642 | 3bpt | A | 345 | GLY | p.Gly345Ser | Q6NVY1 |
| HOMO SAPIENS | 1 | 3942 | 3sby | A | 60 | ASP | p.Asp60His | Q9Y4U1 |
| HOMO SAPIENS | 0 | 3942 | 3sby | A | 111 | ARG | p.Arg111Ter | Q9Y4U1 |
| HOMO SAPIENS | 1 | 3942 | 3sby | A | 116 | LEU | p.Leu116Pro | Q9Y4U1 |
| HOMO SAPIENS | 1 | 3942 | 3sby | A | 130 | TYR | p.Tyr130Cys | Q9Y4U1 |
| HOMO SAPIENS | 1 | 3942 | 3sby | A | 132 | ARG | p.Arg132Ter | Q9Y4U1 |
| HOMO SAPIENS | 1 | 3942 | 3sby | A | 147 | GLY | p.Gly147Ala | Q9Y4U1 |
| HOMO SAPIENS | 1 | 3942 | 3sby | A | 161 | ARG | p.Arg161Gln | Q9Y4U1 |
| HOMO SAPIENS | 1 | 3942 | 3sby | A | 161 | ARG | p.Arg161Ter | Q9Y4U1 |
| HOMO SAPIENS | 1 | 3942 | 3sby | A | 203 | TRP | p.Trp203Ter | Q9Y4U1 |
| HOMO SAPIENS | 1 | 3942 | 3sby | A | 203 | TRP | p.Trp203Ter | Q9Y4U1 |
| HOMO SAPIENS | 1 | 3942 | 3sby | A | 205 | TYR | p.Tyr205Ter | Q9Y4U1 |

| | | | | | | | | | |
|---|---|---|---|---|---|---|---|---|---|
| HOMO SAPIENS | 0 | 38340 | 2mbh | B | | 39 | MET | p.Met39Leu | Q13351 |
| HOMO SAPIENS | 2 | 7360 | 4k30 | A | | 430 | LEU | p.Leu430Pro | Q8N159 |
| HOMO SAPIENS | 2 | 7360 | 4k30 | A | | 433 | GLU | p.Glu433Asp | Q8N159 |
| HOMO SAPIENS | 7 | 7360 | 4k30 | A | | 484 | TRP | p.Trp484Arg | Q8N159 |
| HOMO SAPIENS | 1 | 41903 | 2xs6 | A | | 151 | PRO | p.Pro151Ser | O00459 |
| HOMO SAPIENS | 0 | 172 | 5glh | A | | 183 | ALA | p.Ala183Gly | P24530 |
| HOMO SAPIENS | 0 | 172 | 5glh | A | | 201 | ARG | p.Arg201Ter | P24530 |
| HOMO SAPIENS | 0 | 172 | 5glh | A | | 244 | THR | p.Thr244Met | P24530 |
| HOMO SAPIENS | 0 | 172 | 5glh | A | | 253 | ARG | p.Arg253Ter | P24530 |
| HOMO SAPIENS | 9 | 172 | 5glh | A | | 1017 | ILE | p.Leu17Phe | P00720 |
| HOMO SAPIENS | -1 | 3342 | 1ljm | A | | 1058 | HIS | p.His58Asn | Q01196 |
| HOMO SAPIENS | 4 | 3342 | 1ljm | A | | 1174 | ARG | p.Arg174Gln | Q01196 |
| HOMO SAPIENS | 5 | 25607 | 1p22 | C | | 32 | ASP | p.Asp32Ala | P35222 |
| HOMO SAPIENS | 5 | 25607 | 1p22 | C | | 32 | ASP | p.Asp32Tyr | P35222 |
| HOMO SAPIENS | -1 | 25607 | 1p22 | C | | 33 | SEP | p.Ser33Phe | P35222 |
| HOMO SAPIENS | 5 | 25607 | 1p22 | C | | 34 | GLY | p.Gly34Glu | P35222 |
| HOMO SAPIENS | -1 | 25607 | 1p22 | C | | 37 | SEP | p.Ser37Cys | P35222 |
| HOMO SAPIENS | -1 | 25607 | 1p22 | C | | 41 | THR | p.Thr41Ile | P35222 |
| HOMO SAPIENS | 1 | 48 | 4uow | | 9 | 60 | GLN | p.Arg34111His | Q8WZ42 |
| HOMO SAPIENS | 0 | 48 | 4uow | | 9 | 64 | LEU | p.Val34115Ala | Q8WZ42 |
| HOMO SAPIENS | 0 | 1127 | 5c01 | A | | 638 | ARG | p.Arg638Ter | P29597 |
| HOMO SAPIENS | 0 | 1127 | 5c01 | A | | 684 | ILE | p.Ile684Ser | P29597 |
| HOMO SAPIENS | 0 | 1127 | 5c01 | A | | 703 | ARG | p.Arg703Trp | P29597 |
| HOMO SAPIENS | 5 | 16389 | 2aq0 | B | | 128 | ALA | p.Ala860Asp | Q92889 |
| HOMO SAPIENS | 2 | 16389 | 2aq0 | B | | 141 | ILE | p.Ile873Val | Q92889 |
| HOMO SAPIENS | 5 | 16389 | 2aq0 | B | | 143 | GLU | p.Glu875Gly | Q92889 |
| HOMO SAPIENS | 0 | 16389 | 2aq0 | B | | 180 | GLY | p.Gly912Arg | Q92889 |
| HOMO SAPIENS | 0 | 15816 | 2xrc | A | | 89 | THR | p.Thr107Ala | P05156 |
| HOMO SAPIENS | 0 | 15816 | 2xrc | A | | 133 | ASN | p.Asn151Ser | P05156 |
| HOMO SAPIENS | 0 | 15816 | 2xrc | A | | 185 | THR | p.Thr203Ile | P05156 |
| HOMO SAPIENS | 0 | 15816 | 2xrc | A | | 225 | GLY | p.Gly243Asp | P05156 |
| HOMO SAPIENS | 0 | 15816 | 2xrc | A | | 243 | GLY | p.Gly261Ala | P05156 |
| HOMO SAPIENS | 0 | 15816 | 2xrc | A | | 269 | GLY | p.Gly287Arg | P05156 |
| HOMO SAPIENS | -1 | 15816 | 2xrc | A | | 288 | ILE | p.Ile306Val | P05156 |
| HOMO SAPIENS | -1 | 15816 | 2xrc | A | | 388 | ARG | p.Arg406His | P05156 |
| HOMO SAPIENS | 7 | 15816 | 2xrc | A | | 398 | ILE | p.Ile416Leu | P05156 |
| HOMO SAPIENS | 9 | 15816 | 2xrc | A | | 400 | HIS | p.His418Leu | P05156 |
| HOMO SAPIENS | 0 | 15816 | 2xrc | A | | 423 | LYS | p.Lys441Arg | P05156 |
| HOMO SAPIENS | -1 | 15816 | 2xrc | A | | 530 | GLU | p.Glu548Gln | P05156 |
| HOMO SAPIENS | 0 | 15816 | 2xrc | A | | 535 | PRO | p.Pro553Ala | P05156 |
| HOMO SAPIENS | 0 | 15816 | 2xrc | A | | 536 | GLU | p.Glu554Gly | P05156 |
| HOMO SAPIENS | 1 | 12385 | 4zlp | B | | 1450 | ALA | p.Ala1450Pro | Q9UM47 |
| HOMO SAPIENS | 7 | 12385 | 4zlp | B | | 1482 | GLY | p.Gly1482Arg | Q9UM47 |
| HOMO SAPIENS | 2 | 12385 | 4zlp | B | | 1518 | LEU | p.Leu1518Met | Q9UM47 |
| HOMO SAPIENS | 0 | 12385 | 4zlp | B | | 1560 | ARG | p.Arg1560Pro | Q9UM47 |
| HOMO SAPIENS | 0 | 1227 | 4a04 | A | | 148 | PHE | p.Phe148Tyr | O43435 |
| HOMO SAPIENS | 4 | 1227 | 4a04 | A | | 194 | HIS | p.His194Gln | O43435 |
| HOMO SAPIENS | 2 | 8017 | 3ei4 | B | | 244 | LYS | p.Lys244Glu | Q92466 |
| HOMO SAPIENS | 1 | 8017 | 3ei4 | B | | 273 | ARG | p.Arg273His | Q92466 |
| HOMO SAPIENS | 3 | 36112 | 2k8f | B | | 9 | SER | p.Ser9Arg | P04637 |
| HOMO SAPIENS | 2 | 12880 | 5ggf | A | | 265 | ARG | p.Arg265His | Q8WZA1 |
| HOMO SAPIENS | 2 | 12880 | 5ggf | A | | 287 | ILE | p.Ile287Ser | Q8WZA1 |
| HOMO SAPIENS | 2 | 12880 | 5ggf | A | | 311 | ARG | p.Arg311Gln | Q8WZA1 |
| HOMO SAPIENS | 2 | 12880 | 5ggf | A | | 425 | TRP | p.Trp425Ser | Q8WZA1 |

| | | | | | | | | |
|---|---|---|---|---|---|---|---|---|
| HOMO SAPIENS | 2 | 12880 | 5ggf | A | 442 | ARG | p.Arg442Cys | Q8WZA1 |
| HOMO SAPIENS | 2 | 12880 | 5ggf | A | 448 | GLY | p.Gly448Arg | Q8WZA1 |
| HOMO SAPIENS | 0 | 12880 | 5ggf | A | 490 | CYS | p.Cys490Tyr | Q8WZA1 |
| HOMO SAPIENS | 0 | 12880 | 5ggf | A | 493 | PRO | p.Pro493Arg | Q8WZA1 |
| HOMO SAPIENS | 3 | 12880 | 5ggf | A | 550 | SER | p.Ser550Asn | Q8WZA1 |
| HOMO SAPIENS | 1 | 12880 | 5ggf | A | 556 | ASP | p.Asp556Asn | Q8WZA1 |
| HOMO SAPIENS | 0 | 12880 | 5ggf | A | 580 | ARG | p.Arg580Ter | Q8WZA1 |
| HOMO SAPIENS | 2 | 12880 | 5ggf | A | 605 | ARG | p.Arg605Pro | Q8WZA1 |
| HOMO SAPIENS | 0 | 12880 | 5ggf | A | 623 | VAL | p.Met623Val | Q8WZA1 |
| HOMO SAPIENS | 1 | 10187 | 4qw2 | A | 145 | ALA | p.Ala145Ser | Q06787 |
| HOMO SAPIENS | 1 | 10187 | 4qw2 | A | 174 | ASN | p.Asn174Lys | Q06787 |
| HOMO SAPIENS | 0 | 24002 | 2m0t | A | 110 | LEU | p.Leu110Val | O14745 |
| HOMO SAPIENS | 9 | 3126 | 4mjh | A | 99 | LEU | p.Leu99Met | P04792 |
| HOMO SAPIENS | 1 | 3126 | 4mjh | A | 127 | ARG | p.Arg127Trp | P04792 |
| HOMO SAPIENS | 1 | 3126 | 4mjh | A | 127 | ARG | p.Arg127Leu | P04792 |
| HOMO SAPIENS | 1 | 3126 | 4mjh | A | 135 | SER | p.Ser135Phe | P04792 |
| HOMO SAPIENS | 2 | 3126 | 4mjh | A | 136 | ARG | p.Arg136Trp | P04792 |
| HOMO SAPIENS | 1 | 3126 | 4mjh | A | 140 | ARG | p.Arg140Gly | P04792 |
| HOMO SAPIENS | 0 | 3126 | 4mjh | A | 151 | THR | p.Thr151Ile | P04792 |
| HOMO SAPIENS | -1 | 15780 | 2jlp | A | 213 | ARG | p.Arg231Gly | P08294 |
| HOMO SAPIENS | 8 | 3578 | 1apz | D | 203 | GLY | p.Gly226Asp | P20933 |
| HOMO SAPIENS | 1 | 3578 | 1apz | D | 279 | GLY | p.Gly302Arg | P20933 |
| HOMO SAPIENS | 0 | 3578 | 1apz | D | 283 | CYS | p.Cys306Arg | P20933 |
| HOMO SAPIENS | 5 | 1335 | 5xtd | C | 115 | ARG | p.Arg115Cys | O75251 |
| HOMO SAPIENS | 0 | 1335 | 5xtd | C | 122 | VAL | p.Val122Met | O75251 |
| HOMO SAPIENS | 2 | 5063 | 2le4 | A | 37 | ARG | p.Arg74Pro | P48431 |
| HOMO SAPIENS | 1 | 25870 | 2o10 | A | 44 | LEU | p.Leu44Pro | P50461 |
| HOMO SAPIENS | 4 | 25870 | 2o10 | A | 58 | CYS | p.Cys58Gly | P50461 |
| HOMO SAPIENS | 0 | 18748 | 5fii | C | 39 | PHE | p.Phe39Leu | P00439 |
| HOMO SAPIENS | 6 | 18748 | 5fii | C | 48 | LEU | p.Leu48Ser | P00439 |
| HOMO SAPIENS | 2 | 18748 | 5fii | C | 65 | ILE | p.Ile65Asn | P00439 |
| HOMO SAPIENS | 3 | 18748 | 5fii | C | 95 | ILE | p.Ile95Phe | P00439 |
| HOMO SAPIENS | 1 | 14261 | 2vay | B | 1539 | ARG | p.Arg1539Cys | Q13698 |
| HOMO SAPIENS | 0 | 2494 | 4dx1 | A | 183 | GLU | p.Glu183Asp | Q9HBA0 |
| HOMO SAPIENS | 0 | 2494 | 4dx1 | A | 217 | ALA | p.Ala217Ser | Q9HBA0 |
| HOMO SAPIENS | 0 | 2494 | 4dx1 | A | 217 | ALA | p.Ala217Val | Q9HBA0 |
| HOMO SAPIENS | 4 | 2494 | 4dx1 | A | 224 | ARG | p.Arg224Trp | Q9HBA0 |
| HOMO SAPIENS | 4 | 2494 | 4dx1 | A | 254 | VAL | p.Val254Leu | Q9HBA0 |
| HOMO SAPIENS | 0 | 2494 | 4dx1 | A | 269 | ARG | p.Arg269Cys | Q9HBA0 |
| HOMO SAPIENS | 0 | 2494 | 4dx1 | A | 269 | ARG | p.Arg269His | Q9HBA0 |
| HOMO SAPIENS | 0 | 2494 | 4dx1 | A | 278 | GLU | p.Glu278Lys | Q9HBA0 |
| HOMO SAPIENS | 0 | 2494 | 4dx1 | A | 315 | ARG | p.Arg315Trp | Q9HBA0 |
| HOMO SAPIENS | 0 | 2494 | 4dx1 | A | 316 | ARG | p.Arg316Cys | Q9HBA0 |
| HOMO SAPIENS | 0 | 2494 | 4dx1 | A | 331 | ILE | p.Ile331Phe | Q9HBA0 |
| HOMO SAPIENS | 0 | 2494 | 4dx1 | A | 333 | ASP | p.Asp333Gly | Q9HBA0 |
| HOMO SAPIENS | 1 | 2494 | 4dx1 | A | 365 | VAL | p.Val365Met | Q9HBA0 |
| HOMO SAPIENS | 0 | 23998 | 2l1r | B | 161 | ARG | p.Arg162Trp | P19429 |
| HOMO SAPIENS | 0 | 8354 | 4f0x | D | 187 | SER | p.Ser187Ter | O95822 |
| HOMO SAPIENS | 2 | 8354 | 4f0x | D | 259 | GLY | p.Gly259Ala | O95822 |
| HOMO SAPIENS | 2 | 8354 | 4f0x | D | 284 | THR | p.Thr284Ala | O95822 |
| HOMO SAPIENS | 3 | 8354 | 4f0x | D | 309 | LYS | p.Lys309Gln | O95822 |
| HOMO SAPIENS | 0 | 21182 | 4apo | A | 228 | GLN | p.Gln228Lys | O00170 |
| HOMO SAPIENS | 1 | 21182 | 4apo | A | 307 | GLN | p.Gln307Arg | O00170 |
| HOMO SAPIENS | 0 | 2287 | 5of4 | B | 112 | ARG | p.Arg112His | P18074 |

| | | | | | | | | |
|---|---|---|---|---|---|---|---|---|
| HOMO SAPIENS | 0 | 2287 | 5of4 | B | 312 | ASP | p.Asp312Asn | P18074 |
| HOMO SAPIENS | 3 | 2287 | 5of4 | B | 461 | LEU | p.Leu461Val | P18074 |
| HOMO SAPIENS | 0 | 2287 | 5of4 | B | 485 | LEU | p.Leu485Pro | P18074 |
| HOMO SAPIENS | 1 | 2287 | 5of4 | B | 541 | SER | p.Ser541Arg | P18074 |
| HOMO SAPIENS | 0 | 2287 | 5of4 | B | 616 | ARG | p.Arg616Pro | P18074 |
| HOMO SAPIENS | 0 | 2287 | 5of4 | B | 616 | ARG | p.Arg616Trp | P18074 |
| HOMO SAPIENS | 1 | 2287 | 5of4 | B | 658 | ARG | p.Arg658Cys | P18074 |
| HOMO SAPIENS | 1 | 2287 | 5of4 | B | 681 | ASP | p.Asp681Asn | P18074 |
| HOMO SAPIENS | 1 | 2287 | 5of4 | B | 683 | ARG | p.Arg683Trp | P18074 |
| HOMO SAPIENS | 0 | 2287 | 5of4 | B | 713 | GLY | p.Gly713Arg | P18074 |
| HOMO SAPIENS | 0 | 2287 | 5of4 | B | 717 | ALA | p.Ala717Gly | P18074 |
| HOMO SAPIENS | 0 | 2287 | 5of4 | B | 722 | ARG | p.Arg722Trp | P18074 |
| HOMO SAPIENS | 0 | 2287 | 5of4 | B | 725 | ALA | p.Ala725Pro | P18074 |
| HOMO SAPIENS | 4 | 12188 | 4its | A | 122 | ARG | p.Arg122Gly | Q9Y606 |
| HOMO SAPIENS | 4 | 12188 | 4its | A | 133 | ASP | p.Asp133Asn | Q9Y606 |
| HOMO SAPIENS | 4 | 12188 | 4its | A | 144 | ARG | p.Arg144Trp | Q9Y606 |
| HOMO SAPIENS | 0 | 31529 | 1kgd | A | 761 | ARG | p.Arg766Gln | O14936 |
| HOMO SAPIENS | 1 | 31529 | 1kgd | A | 824 | GLN | p.Gln829Arg | O14936 |
| HOMO SAPIENS | 0 | 10906 | 2h63 | C | 18 | ARG | p.Arg18Ter | P53004 |
| HOMO SAPIENS | 9 | 27767 | 4wyq | A | 280 | LEU | p.Leu278Phe | Q9UPY3 |
| HOMO SAPIENS | 0 | 27767 | 4wyq | A | 297 | SER | p.Ser295Cys | Q9UPY3 |
| HOMO SAPIENS | 1 | 27767 | 4wyq | A | 334 | TYR | p.Tyr332Ter | Q9UPY3 |
| HOMO SAPIENS | 7 | 7775 | 5sv7 | C | 136 | SER | p.Ser136Cys | Q9NZJ5 |
| HOMO SAPIENS | 2 | 7775 | 5sv7 | C | 166 | GLN | p.Gln166Arg | Q9NZJ5 |
| HOMO SAPIENS | 2 | 7775 | 5sv7 | C | 332 | GLU | p.Glu332Ter | Q9NZJ5 |
| HOMO SAPIENS | 0 | 21449 | 2e9w | B | 84 | THR | p.Thr84Met | P10721 |
| HOMO SAPIENS | 0 | 21449 | 2e9w | B | 168 | ALA | p.Ala168Glu | P10721 |
| HOMO SAPIENS | 0 | 21449 | 2e9w | B | 178 | ALA | p.Ala178Thr | P10721 |
| HOMO SAPIENS | 0 | 21449 | 2e9w | B | 293 | ASN | p.Asn293Ser | P10721 |
| HOMO SAPIENS | -1 | 21449 | 2e9w | B | 318 | MET | p.Met318Leu | P10721 |
| HOMO SAPIENS | 0 | 21449 | 2e9w | B | 374 | VAL | p.Val374Ile | P10721 |
| HOMO SAPIENS | 6 | 3010 | 3flg | A | 66 | SER | p.Ser270Pro | Q9UBC3 |
| HOMO SAPIENS | 1 | 33918 | 2xzz | A | 760 | ARG | p.Arg760Ter | P22735 |
| HOMO SAPIENS | 2 | 1254 | 5xtc | w | 65 | ASN | p.Asn65Ser | O95299 |
| HOMO SAPIENS | 1 | 1254 | 5xtc | w | 238 | GLU | p.Glu238Lys | O95299 |
| HOMO SAPIENS | 3 | 10245 | 3siv | B | 194 | ALA | p.Ala194Glu | Q8WWY3 |
| HOMO SAPIENS | 4 | 10245 | 3siv | B | 216 | ALA | p.Ala216Pro | Q8WWY3 |
| HOMO SAPIENS | 0 | 2670 | 3dwb | A | 341 | THR | p.Thr341Ile | P42892 |
| HOMO SAPIENS | 1 | 2670 | 3dwb | A | 462 | SER | p.Ser462Asn | P42892 |
| HOMO SAPIENS | 1 | 2670 | 3dwb | A | 754 | ARG | p.Arg754Cys | P42892 |
| HOMO SAPIENS | 1 | 2961 | 5xtb | O | 40 | VAL | p.Val40Met | P19404 |
| HOMO SAPIENS | 5 | 2961 | 5xtb | O | 172 | ILE | p.Ile172Thr | P19404 |
| HOMO SAPIENS | 5 | 2961 | 5xtb | O | 190 | ASP | p.Asp190Gly | P19404 |
| HOMO SAPIENS | 5 | 29655 | 2xdg | A | 57 | ALA | p.Ala57Thr | Q02643 |
| HOMO SAPIENS | 4 | 29655 | 2xdg | A | 72 | GLU | p.Glu72Ter | Q02643 |
| HOMO SAPIENS | 0 | 29655 | 2xdg | A | 121 | GLU | p.Glu121Asp | Q02643 |
| HOMO SAPIENS | -1 | 17935 | 1hyn | P | 38 | ASP | p.Asp38Ala | P02730 |
| HOMO SAPIENS | 7 | 12825 | 1g82 | D | 99 | SER | p.Ser99Asn | P31371 |
| HOMO SAPIENS | 6 | 24927 | 1jm7 | B | 85 | VAL | p.Val85Leu | Q99728 |
| HOMO SAPIENS | 1 | 24927 | 1jm7 | B | 112 | ARG | p.Arg112Ter | Q99728 |
| HOMO SAPIENS | 0 | 24927 | 1jm7 | B | 116 | HIS | p.His116Tyr | Q99728 |
| HOMO SAPIENS | 7 | 92 | 5hde | A | 61 | LYS | p.Lys61Arg | Q05209 |
| HOMO SAPIENS | 0 | 17246 | 1szb | A | 84 | ARG | p.Arg99Gln | O00187 |
| HOMO SAPIENS | 1 | 17246 | 1szb | A | 105 | ASP | p.Asp120Gly | O00187 |

| | | | | | | | | |
|---|---|---|---|---|---|---|---|---|
| HOMO SAPIENS | 0 | 17246 | 1szb | A | 111 | PRO | p.Pro126Leu | O00187 |
| HOMO SAPIENS | 9 | 17246 | 1szb | A | 141 | CYS | p.Cys156Tyr | O00187 |
| HOMO SAPIENS | 0 | 32473 | 1b72 | A | 207 | ARG | p.Arg207Cys | P14653 |
| HOMO SAPIENS | 0 | 1920 | 4yzf | B | 488 | VAL | p.Val488Met | P02730 |
| HOMO SAPIENS | 0 | 1920 | 4yzf | B | 589 | ARG | p.Arg589Cys | P02730 |
| HOMO SAPIENS | 0 | 1920 | 4yzf | B | 589 | ARG | p.Arg589His | P02730 |
| HOMO SAPIENS | 0 | 1920 | 4yzf | B | 602 | ARG | p.Arg602Pro | P02730 |
| HOMO SAPIENS | 0 | 1920 | 4yzf | B | 613 | SER | p.Ser613Phe | P02730 |
| HOMO SAPIENS | -1 | 1920 | 4yzf | B | 646 | ARG | p.Arg646Gln | P02730 |
| HOMO SAPIENS | -1 | 1920 | 4yzf | B | 646 | ARG | p.Arg646Trp | P02730 |
| HOMO SAPIENS | 0 | 1920 | 4yzf | B | 701 | GLY | p.Gly701Asp | P02730 |
| HOMO SAPIENS | 0 | 1920 | 4yzf | B | 760 | ARG | p.Arg760Gln | P02730 |
| HOMO SAPIENS | 0 | 1920 | 4yzf | B | 771 | GLY | p.Gly771Asp | P02730 |
| HOMO SAPIENS | 0 | 1920 | 4yzf | B | 773 | SER | p.Ser773Pro | P02730 |
| HOMO SAPIENS | 0 | 1920 | 4yzf | B | 837 | THR | p.Thr837Ala | P02730 |
| HOMO SAPIENS | 0 | 1920 | 4yzf | B | 854 | PRO | p.Pro854Leu | P02730 |
| HOMO SAPIENS | 0 | 1920 | 4yzf | B | 858 | ALA | p.Ala858Asp | P02730 |
| HOMO SAPIENS | 0 | 1920 | 4yzf | B | 862 | VAL | p.Val862Ile | P02730 |
| HOMO SAPIENS | 0 | 1920 | 4yzf | B | 868 | PRO | p.Pro868Leu | P02730 |
| HOMO SAPIENS | 0 | 1920 | 4yzf | B | 870 | ARG | p.Arg870Trp | P02730 |
| HOMO SAPIENS | 0 | 9863 | 2juj | A | 54 | VAL | p.Val904Ile | P22681 |
| HOMO SAPIENS | 0 | 21934 | 2na7 | A | 184 | ILE | p.Ile184Val | P25445 |
| HOMO SAPIENS | -1 | 6746 | 1lw3 | A | 3 | THR | p.Lys3Thr | Q13614 |
| HOMO SAPIENS | -1 | 6746 | 1lw3 | A | 44 | VAL | p.Val44Ile | Q13614 |
| HOMO SAPIENS | 3 | 6746 | 1lw3 | A | 183 | ALA | p.Ala183Ser | Q13614 |
| HOMO SAPIENS | 4 | 6746 | 1lw3 | A | 482 | GLN | p.Gln482Ter | Q13614 |
| HOMO SAPIENS | 4 | 6746 | 1lw3 | A | 502 | GLU | p.Glu502Gln | Q13614 |
| HOMO SAPIENS | 3 | 6746 | 1lw3 | A | 545 | ASN | p.Asn545Ser | Q13614 |
| HOMO SAPIENS | 0 | 6746 | 1lw3 | A | 586 | ARG | p.Arg586Trp | Q13614 |
| HOMO SAPIENS | -1 | 6746 | 1lw3 | A | 602 | ALA | p.Ala602Gly | Q13614 |
| HOMO SAPIENS | 1 | 319 | 4ch9 | B | 336 | ARG | p.Arg336Ile | Q9UH77 |
| HOMO SAPIENS | 3 | 319 | 4ch9 | B | 340 | ALA | p.Ala340Val | Q9UH77 |
| HOMO SAPIENS | 0 | 319 | 4ch9 | B | 384 | ARG | p.Arg384Gln | Q9UH77 |
| HOMO SAPIENS | 5 | 319 | 4ch9 | B | 426 | PRO | p.Pro426Leu | Q9UH77 |
| HOMO SAPIENS | 3 | 319 | 4ch9 | B | 431 | ARG | p.Arg431Gln | Q9UH77 |
| HOMO SAPIENS | 2 | 319 | 4ch9 | B | 575 | ARG | p.Arg575Trp | Q9UH77 |
| HOMO SAPIENS | 0 | 574 | 2qkk | A | 142 | VAL | p.Val142Ile | O60930 |
| HOMO SAPIENS | 0 | 24021 | 2n8y | A | 752 | ARG | p.Arg752Gln | P12814 |
| HOMO SAPIENS | 0 | 18693 | 2y7j | B | 44 | ARG | p.Arg44Ter | P15735 |
| HOMO SAPIENS | 0 | 18693 | 2y7j | B | 106 | VAL | p.Val106Glu | P15735 |
| HOMO SAPIENS | 3 | 18693 | 2y7j | B | 189 | GLY | p.Gly189Glu | P15735 |
| HOMO SAPIENS | 0 | 17087 | 2ifg | A | 237 | THR | p.Thr237Met | P04629 |
| HOMO SAPIENS | 0 | 7980 | 1qub | A | 247 | VAL | p.Val266Leu | P02749 |
| HOMO SAPIENS | -1 | 2205 | 1h7u | A | 18 | ILE | p.Ile18Val | P54278 |
| HOMO SAPIENS | -1 | 2205 | 1h7u | A | 20 | ARG | p.Arg20Gln | P54278 |
| HOMO SAPIENS | 9 | 2205 | 1h7u | A | 46 | SER | p.Ser46Ile | P54278 |
| HOMO SAPIENS | 0 | 2205 | 1h7u | A | 60 | ASP | p.Asp60Glu | P54278 |
| HOMO SAPIENS | 0 | 2205 | 1h7u | A | 80 | PHE | p.Phe80Leu | P54278 |
| HOMO SAPIENS | 0 | 2205 | 1h7u | A | 118 | SER | p.Ser118Arg | P54278 |
| HOMO SAPIENS | 0 | 2205 | 1h7u | A | 126 | HIS | p.His126Gln | P54278 |
| HOMO SAPIENS | 0 | 2205 | 1h7u | A | 128 | SER | p.Ser128Leu | P54278 |
| HOMO SAPIENS | 1 | 2205 | 1h7u | A | 134 | ARG | p.Arg134Ter | P54278 |
| HOMO SAPIENS | 0 | 2205 | 1h7u | A | 166 | LEU | p.Leu166Pro | P54278 |
| HOMO SAPIENS | 1 | 2205 | 1h7u | A | 191 | TYR | p.Tyr191Cys | P54278 |

| | | | | | | | | |
|---|---|---|---|---|---|---|---|---|
| HOMO SAPIENS | 0 | 2205 | 1h7u | A | 211 | ARG | p.Arg211Ter | P54278 |
| HOMO SAPIENS | 0 | 2205 | 1h7u | A | 233 | GLN | p.Gln233Ter | P54278 |
| HOMO SAPIENS | 0 | 2205 | 1h7u | A | 255 | TYR | p.Tyr255Ter | P54278 |
| HOMO SAPIENS | 0 | 2205 | 1h7u | A | 277 | THR | p.Thr277Lys | P54278 |
| HOMO SAPIENS | 0 | 2205 | 1h7u | A | 301 | LYS | p.Lys301Asn | P54278 |
| HOMO SAPIENS | 0 | 2205 | 1h7u | A | 312 | MET | p.Met312Ile | P54278 |
| HOMO SAPIENS | 0 | 2205 | 1h7u | A | 312 | MET | p.Met312Thr | P54278 |
| HOMO SAPIENS | 0 | 2205 | 1h7u | A | 315 | ARG | p.Arg315Ter | P54278 |
| HOMO SAPIENS | 0 | 2205 | 1h7u | A | 318 | TYR | p.Tyr318Cys | P54278 |
| HOMO SAPIENS | 0 | 2205 | 1h7u | A | 335 | ASN | p.Asn335Ile | P54278 |
| HOMO SAPIENS | 0 | 2205 | 1h7u | A | 360 | ILE | p.Ile360Met | P54278 |
| HOMO SAPIENS | 6 | 20754 | 5mpo | B | 7 | VAL | p.Val7Phe | O96033 |
| HOMO SAPIENS | 4 | 20754 | 5mpo | B | 64 | VAL | p.Val64Ile | O96033 |
| HOMO SAPIENS | 5 | 34944 | 4qam | B | 1114 | ASP | p.Asp1114Gly | Q96KN7 |
| HOMO SAPIENS | 5 | 34944 | 4qam | B | 1240 | GLY | p.Gly1240Glu | Q96KN7 |
| HOMO SAPIENS | 9 | 40496 | 3vnn | A | 278 | ARG | p.Arg278His | P49917 |
| HOMO SAPIENS | 6 | 30131 | 4dt7 | E | 40 | PRO | p.Pro210Leu | P04070 |
| HOMO SAPIENS | -1 | 30131 | 4dt7 | E | 50 | ARG | p.Arg220Trp | P04070 |
| HOMO SAPIENS | 7 | 15311 | 2pqt | A | 64 | ARG | p.Arg64Trp | P18440 |
| HOMO SAPIENS | 2 | 15311 | 2pqt | A | 149 | VAL | p.Val149Ile | P18440 |
| HOMO SAPIENS | -1 | 19581 | 3hy8 | A | 33 | ASP | p.Asp33Val | Q9NVS9 |
| HOMO SAPIENS | 5 | 19581 | 3hy8 | A | 116 | ARG | p.Arg116Gln | Q9NVS9 |
| HOMO SAPIENS | 4 | 19581 | 3hy8 | A | 225 | ARG | p.Arg225His | Q9NVS9 |
| HOMO SAPIENS | 2 | 19581 | 3hy8 | A | 229 | TRP | p.Arg229Trp | Q9NVS9 |
| HOMO SAPIENS | 2 | 19581 | 3hy8 | A | 229 | TRP | p.Arg229Gln | Q9NVS9 |
| HOMO SAPIENS | 0 | 10266 | 3t1i | A | 41 | ASP | p.Asp41His | P49959 |
| HOMO SAPIENS | 1 | 10266 | 3t1i | A | 305 | ARG | p.Arg305Trp | P49959 |
| HOMO SAPIENS | 0 | 10266 | 3t1i | A | 308 | PHE | p.Phe308Leu | P49959 |
| HOMO SAPIENS | 0 | 10266 | 3t1i | A | 334 | SER | p.Ser334Arg | P49959 |
| HOMO SAPIENS | 1 | 1600 | 3rbf | A | 61 | GLU | p.Glu61Asp | P20711 |
| HOMO SAPIENS | 0 | 1600 | 3rbf | A | 91 | ALA | p.Ala91Val | P20711 |
| HOMO SAPIENS | -1 | 1600 | 3rbf | A | 102 | GLY | p.Gly102Ser | P20711 |
| HOMO SAPIENS | 3 | 1600 | 3rbf | A | 147 | SER | p.Ser147Arg | P20711 |
| HOMO SAPIENS | 1 | 1600 | 3rbf | A | 210 | PRO | p.Pro210Leu | P20711 |
| HOMO SAPIENS | 0 | 1600 | 3rbf | A | 217 | MET | p.Met217Val | P20711 |
| HOMO SAPIENS | 4 | 1600 | 3rbf | A | 250 | SER | p.Ser250Phe | P20711 |
| HOMO SAPIENS | 2 | 1600 | 3rbf | A | 275 | ALA | p.Ala275Thr | P20711 |
| HOMO SAPIENS | 1 | 1600 | 3rbf | A | 309 | PHE | p.Phe309Leu | P20711 |
| HOMO SAPIENS | -1 | 1600 | 3rbf | A | 347 | ARG | p.Arg347Gln | P20711 |
| HOMO SAPIENS | 0 | 1600 | 3rbf | A | 462 | ARG | p.Arg462Gln | P20711 |
| HOMO SAPIENS | 6 | 20830 | 5y4f | A | 714 | THR | p.Thr715Ile | Q01484 |
| HOMO SAPIENS | 7 | 20830 | 5y4f | A | 719 | ASP | p.Asp720Glu | Q01484 |
| HOMO SAPIENS | 0 | 20830 | 5y4f | A | 814 | VAL | p.Val815Ile | Q01484 |
| HOMO SAPIENS | -1 | 302 | 2c9y | A | 9 | GLU | p.Glu9Ter | P54819 |
| HOMO SAPIENS | 0 | 302 | 2c9y | A | 103 | ARG | p.Arg103Trp | P54819 |
| HOMO SAPIENS | 3 | 302 | 2c9y | A | 165 | ASP | p.Asp165Gly | P54819 |
| HOMO SAPIENS | 0 | 302 | 2c9y | A | 182 | ALA | p.Ala182Asp | P54819 |
| HOMO SAPIENS | 2 | 302 | 2c9y | A | 186 | ARG | p.Arg186Cys | P54819 |
| HOMO SAPIENS | 0 | 302 | 2c9y | A | 209 | ALA | p.Ala209Thr | P54819 |
| HOMO SAPIENS | -1 | 16920 | 1jlj | B | 10 | ASN | p.Asn10Tyr | Q9NQX3 |
| HOMO SAPIENS | 9 | 8462 | 1zbq | B | 16 | GLY | p.Gly16Ser | P51659 |
| HOMO SAPIENS | 5 | 8462 | 1zbq | B | 19 | ALA | p.Ala19Gly | P51659 |
| HOMO SAPIENS | 1 | 8462 | 1zbq | B | 34 | ALA | p.Ala34Val | P51659 |
| HOMO SAPIENS | 1 | 8462 | 1zbq | B | 106 | ARG | p.Arg106His | P51659 |

| | | | | | | | | |
|---|---|---|---|---|---|---|---|---|
| HOMO SAPIENS | 0 | 8462 | 1zbq | B | 140 | LYS | p.Lys140Asn | P51659 |
| HOMO SAPIENS | 0 | 8462 | 1zbq | B | 292 | THR | p.Thr292Ser | P51659 |
| HOMO SAPIENS | 0 | 4375 | 4ndi | A | 199 | ARG | p.Arg213His | Q7Z2E3 |
| HOMO SAPIENS | 3 | 4375 | 4ndi | A | 201 | HIS | p.His215Arg | Q7Z2E3 |
| HOMO SAPIENS | 0 | 4375 | 4ndi | A | 206 | PRO | p.Pro220Leu | Q7Z2E3 |
| HOMO SAPIENS | 2 | 4375 | 4ndi | A | 223 | LEU | p.Leu237Phe | Q7Z2E3 |
| HOMO SAPIENS | 0 | 4375 | 4ndi | A | 245 | ARG | p.Arg259His | Q7Z2E3 |
| HOMO SAPIENS | 0 | 4375 | 4ndi | A | 263 | VAL | p.Val277Gly | Q7Z2E3 |
| HOMO SAPIENS | 0 | 4375 | 4ndi | A | 279 | TRP | p.Trp293Ter | Q7Z2E3 |
| HOMO SAPIENS | 0 | 27004 | 4ccg | Y | 321 | ASP | p.Asp321Glu | Q9NW38 |
| HOMO SAPIENS | -1 | 27004 | 4ccg | Y | 372 | GLY | p.Gly372Ala | Q9NW38 |
| HOMO SAPIENS | -1 | 6383 | 5fai | A | 34 | ALA | p.Ala34Gly | Q92979 |
| HOMO SAPIENS | 8 | 6383 | 5fai | A | 86 | ASP | p.Asp86Gly | Q92979 |
| HOMO SAPIENS | 2 | 24585 | 4mi5 | A | 684 | ARG | p.Arg679Cys | Q15910 |
| HOMO SAPIENS | 0 | 15967 | 3nhb | A | 723 | ARG | p.Arg723Gln | Q9NP58 |
| HOMO SAPIENS | 5 | 18152 | 1oey | A | 389 | HIS | p.His389Gln | P19878 |
| HOMO SAPIENS | 5 | 18152 | 1oey | A | 395 | ARG | p.Arg395Trp | P19878 |
| HOMO SAPIENS | -1 | 1044 | 1li1 | F | 2 | VAL | p.Val1486Ile | P08572 |
| HOMO SAPIENS | 6 | 1044 | 1li1 | F | 37 | GLU | p.Glu1521Lys | P08572 |
| HOMO SAPIENS | 5 | 1044 | 1li1 | F | 157 | SER | p.Ser1639Leu | P08572 |
| HOMO SAPIENS | 5 | 1044 | 1li1 | F | 179 | ARG | p.Arg1662His | P08572 |
| HOMO SAPIENS | 5 | 1044 | 1li1 | F | 180 | GLY | p.Gly1663Arg | P08572 |
| HOMO SAPIENS | 6 | 20076 | 4uyi | A | 671 | LEU | p.Leu671Ser | Q8IY92 |
| HOMO SAPIENS | 0 | 19588 | 1jmj | A | 189 | ARG | p.Arg208His | P05546 |
| HOMO SAPIENS | -1 | 38023 | 1ry7 | B | 65 | GLY | p.Gly65Arg | P22607 |
| HOMO SAPIENS | -1 | 38023 | 1ry7 | B | 139 | ASP | p.Asp139Glu | P22607 |
| HOMO SAPIENS | 0 | 38023 | 1ry7 | B | 248 | ARG | p.Arg248Cys | P22607 |
| HOMO SAPIENS | 0 | 38023 | 1ry7 | B | 249 | SER | p.Ser249Cys | P22607 |
| HOMO SAPIENS | 0 | 38023 | 1ry7 | B | 250 | PRO | p.Pro250Arg | P22607 |
| HOMO SAPIENS | 0 | 38023 | 1ry7 | B | 322 | GLU | p.Glu322Lys | P22607 |
| HOMO SAPIENS | 5 | 16884 | 5eli | A | 33 | GLN | p.Gln33Ter | Q9NZC2 |
| HOMO SAPIENS | 0 | 16884 | 5eli | A | 44 | TRP | p.Trp44Ter | Q9NZC2 |
| HOMO SAPIENS | 0 | 16884 | 5eli | A | 47 | ARG | p.Arg47His | Q9NZC2 |
| HOMO SAPIENS | 0 | 16884 | 5eli | A | 66 | THR | p.Thr66Met | Q9NZC2 |
| HOMO SAPIENS | 0 | 16884 | 5eli | A | 78 | TRP | p.Trp78Ter | Q9NZC2 |
| HOMO SAPIENS | 1 | 16884 | 5eli | A | 96 | THR | p.Thr96Lys | Q9NZC2 |
| HOMO SAPIENS | 9 | 16884 | 5eli | A | 126 | VAL | p.Val126Gly | Q9NZC2 |
| HOMO SAPIENS | -1 | 12267 | 3w5o | A | 3 | ALA | p.Ala3Val | P49917 |
| HOMO SAPIENS | 1 | 12267 | 3w5o | A | 9 | THR | p.Thr9Ile | P49917 |
| HOMO SAPIENS | 2 | 12267 | 3w5o | A | 188 | ARG | p.Arg188Gln | P49917 |
| HOMO SAPIENS | 0 | 12267 | 3w5o | A | 469 | GLY | p.Gly469Glu | P49917 |
| HOMO SAPIENS | 1 | 12267 | 3w5o | A | 580 | ARG | p.Arg580Ter | P49917 |
| HOMO SAPIENS | 0 | 12267 | 3w5o | A | 600 | GLU | p.Glu600Lys | P49917 |
| HOMO SAPIENS | 9 | 2839 | 2hw5 | B | 54 | ARG | p.Arg54Cys | P30084 |
| HOMO SAPIENS | 9 | 2839 | 2hw5 | B | 59 | ASN | p.Asn59Ser | P30084 |
| HOMO SAPIENS | 0 | 2839 | 2hw5 | B | 273 | LYS | p.Lys273Glu | P30084 |
| HOMO SAPIENS | 1 | 12266 | 2o2k | B | 939 | GLY | p.Gly939Cys | Q99707 |
| HOMO SAPIENS | 5 | 12266 | 2o2k | B | 1164 | ARG | p.Arg1164His | Q99707 |
| HOMO SAPIENS | 1 | 12266 | 2o2k | B | 1166 | LEU | p.Leu1166Met | Q99707 |
| HOMO SAPIENS | 5 | 12266 | 2o2k | B | 1173 | PRO | p.Pro1173Leu | Q99707 |
| HOMO SAPIENS | 5 | 12266 | 2o2k | B | 1222 | ASN | p.Asn1222Ser | Q99707 |
| HOMO SAPIENS | 0 | 30891 | 3ch4 | B | 138 | ARG | p.Arg138Ter | Q15126 |
| HOMO SAPIENS | -1 | 4433 | 1qlz | A | 54 | GLY | p.Gly54Ser | P04156 |
| HOMO SAPIENS | 0 | 22568 | 3cxl | A | 20 | LEU | p.Leu20Phe | P15882 |

| | | | | | | | | |
|---|---|---|---|---|---|---|---|---|
| HOMO SAPIENS | 0 | 22568 | 3cxl | A | 126 | ILE | p.Ile126Met | P15882 |
| HOMO SAPIENS | 0 | 22568 | 3cxl | A | 141 | PRO | p.Pro141Leu | P15882 |
| HOMO SAPIENS | 0 | 22568 | 3cxl | A | 143 | TYR | p.Tyr143His | P15882 |
| HOMO SAPIENS | 0 | 22568 | 3cxl | A | 223 | ALA | p.Ala223Val | P15882 |
| HOMO SAPIENS | 0 | 22568 | 3cxl | A | 228 | GLY | p.Gly228Ser | P15882 |
| HOMO SAPIENS | 0 | 22568 | 3cxl | A | 252 | PRO | p.Pro252Gln | P15882 |
| HOMO SAPIENS | 0 | 22568 | 3cxl | A | 313 | GLU | p.Glu313Lys | P15882 |
| HOMO SAPIENS | 3 | 2668 | 5eqg | A | 34 | ASN | p.Asn34Ile | P11166 |
| HOMO SAPIENS | 3 | 2668 | 5eqg | A | 77 | MET | p.Met77Val | P11166 |
| HOMO SAPIENS | 6 | 2668 | 5eqg | A | 91 | GLY | p.Gly91Asp | P11166 |
| HOMO SAPIENS | 6 | 2668 | 5eqg | A | 92 | ARG | p.Arg92Trp | P11166 |
| HOMO SAPIENS | 6 | 2668 | 5eqg | A | 93 | ARG | p.Arg93Trp | P11166 |
| HOMO SAPIENS | 0 | 2668 | 5eqg | A | 95 | SER | p.Ser95Ile | P11166 |
| HOMO SAPIENS | 3 | 2668 | 5eqg | A | 126 | ARG | p.Arg126His | P11166 |
| HOMO SAPIENS | 3 | 2668 | 5eqg | A | 126 | ARG | p.Arg126Cys | P11166 |
| HOMO SAPIENS | 4 | 2668 | 5eqg | A | 232 | ARG | p.Arg232Cys | P11166 |
| HOMO SAPIENS | 3 | 2668 | 5eqg | A | 249 | ARG | p.Arg249Gln | P11166 |
| HOMO SAPIENS | 3 | 2668 | 5eqg | A | 255 | LYS | p.Lys255Arg | P11166 |
| HOMO SAPIENS | 0 | 2668 | 5eqg | A | 275 | ALA | p.Ala275Thr | P11166 |
| HOMO SAPIENS | 4 | 2668 | 5eqg | A | 299 | GLU | p.Glu299Lys | P11166 |
| HOMO SAPIENS | 6 | 2668 | 5eqg | A | 314 | GLY | p.Gly314Ser | P11166 |
| HOMO SAPIENS | 7 | 2668 | 5eqg | A | 315 | ILE | p.Ile315Met | P11166 |
| HOMO SAPIENS | -1 | 2668 | 5eqg | A | 458 | ARG | p.Arg458Trp | P11166 |
| HOMO SAPIENS | -1 | 2668 | 5eqg | A | 468 | ARG | p.Arg468Trp | P11166 |
| HOMO SAPIENS | 3 | 2291 | 2amy | A | 31 | PHE | p.Phe31Leu | O15305 |
| HOMO SAPIENS | 7 | 2291 | 2amy | A | 32 | LEU | p.Leu32Arg | O15305 |
| HOMO SAPIENS | 7 | 2291 | 2amy | A | 65 | ASP | p.Asp65Tyr | O15305 |
| HOMO SAPIENS | 1 | 2291 | 2amy | A | 108 | ALA | p.Ala108Val | O15305 |
| HOMO SAPIENS | 3 | 2291 | 2amy | A | 113 | PRO | p.Pro113Leu | O15305 |
| HOMO SAPIENS | 3 | 2291 | 2amy | A | 113 | PRO | p.Pro113Ser | O15305 |
| HOMO SAPIENS | 6 | 2291 | 2amy | A | 117 | GLY | p.Gly117Arg | O15305 |
| HOMO SAPIENS | 6 | 2291 | 2amy | A | 119 | PHE | p.Phe119Leu | O15305 |
| HOMO SAPIENS | 6 | 2291 | 2amy | A | 123 | ARG | p.Arg123Gln | O15305 |
| HOMO SAPIENS | 0 | 2291 | 2amy | A | 129 | VAL | p.Val129Met | O15305 |
| HOMO SAPIENS | 7 | 2291 | 2amy | A | 139 | GLU | p.Glu139Lys | O15305 |
| HOMO SAPIENS | 5 | 2291 | 2amy | A | 141 | ARG | p.Arg141His | O15305 |
| HOMO SAPIENS | 7 | 2291 | 2amy | A | 144 | PHE | p.Phe144Leu | O15305 |
| HOMO SAPIENS | 6 | 2291 | 2amy | A | 148 | ASP | p.Asp148Asn | O15305 |
| HOMO SAPIENS | 6 | 2291 | 2amy | A | 153 | ILE | p.Ile153Thr | O15305 |
| HOMO SAPIENS | 2 | 2291 | 2amy | A | 157 | PHE | p.Phe157Ser | O15305 |
| HOMO SAPIENS | 1 | 2291 | 2amy | A | 162 | ARG | p.Arg162Trp | O15305 |
| HOMO SAPIENS | 6 | 2291 | 2amy | A | 188 | ASP | p.Asp188Gly | O15305 |
| HOMO SAPIENS | 5 | 2291 | 2amy | A | 194 | ARG | p.Arg194Ter | O15305 |
| HOMO SAPIENS | 0 | 2291 | 2amy | A | 207 | PHE | p.Phe207Ser | O15305 |
| HOMO SAPIENS | 5 | 2291 | 2amy | A | 208 | GLY | p.Gly208Ala | O15305 |
| HOMO SAPIENS | 1 | 2291 | 2amy | A | 216 | ASN | p.Asn216Ile | O15305 |
| HOMO SAPIENS | 5 | 2291 | 2amy | A | 218 | HIS | p.His218Leu | O15305 |
| HOMO SAPIENS | 6 | 2291 | 2amy | A | 223 | ASP | p.Asp223Glu | O15305 |
| HOMO SAPIENS | 6 | 2291 | 2amy | A | 226 | THR | p.Thr226Ser | O15305 |
| HOMO SAPIENS | 5 | 2291 | 2amy | A | 231 | VAL | p.Val231Met | O15305 |
| HOMO SAPIENS | 1 | 2291 | 2amy | A | 237 | THR | p.Thr237Met | O15305 |
| HOMO SAPIENS | 1 | 2291 | 2amy | A | 241 | CYS | p.Cys241Ser | O15305 |
| HOMO SAPIENS | 5 | 35907 | 1d6g | A | 21 | GLY | p.Gly21Arg | P32238 |
| HOMO SAPIENS | -1 | 33209 | 1x9d | A | 197 | PRO | p.Pro197Leu | Q9UKM7 |

| | | | | | | | | |
|---|---|---|---|---|---|---|---|---|
| HOMO SAPIENS | -1 | 33209 | 1x9d | A | 201 | PRO | p.Pro201Leu | Q9UKM7 |
| HOMO SAPIENS | 6 | 21305 | 4yep | B | 1326 | ARG | p.Arg1326Ter | P24043 |
| HOMO SAPIENS | 6 | 21305 | 4yep | B | 1350 | ARG | p.Arg1350Ter | P24043 |
| HOMO SAPIENS | 0 | 21553 | 3qyb | A | 1120 | VAL | p.Val1119Ala | O60343 |
| HOMO SAPIENS | 2 | 21553 | 3qyb | A | 1148 | THR | p.Thr1147Met | O60343 |
| HOMO SAPIENS | -1 | 30645 | 3m0c | C | -17 | TRP | p.Trp4Ter | P01130 |
| HOMO SAPIENS | -1 | 30645 | 3m0c | C | -17 | TRP | p.Trp4Ter | P01130 |
| HOMO SAPIENS | -1 | 30645 | 3m0c | C | 12 | GLN | p.Gln33Ter | P01130 |
| HOMO SAPIENS | -1 | 30645 | 3m0c | C | 15 | ASP | p.Asp36Glu | P01130 |
| HOMO SAPIENS | -1 | 30645 | 3m0c | C | 28 | SER | p.Ser49Arg | P01130 |
| HOMO SAPIENS | -1 | 30645 | 3m0c | C | 41 | THR | p.Thr62Met | P01130 |
| HOMO SAPIENS | -1 | 30645 | 3m0c | C | 69 | ASP | p.Asp90Asn | P01130 |
| HOMO SAPIENS | -1 | 30645 | 3m0c | C | 76 | ASN | p.Asn97Lys | P01130 |
| HOMO SAPIENS | -1 | 30645 | 3m0c | C | 78 | SER | p.Ser99Ter | P01130 |
| HOMO SAPIENS | -1 | 30645 | 3m0c | C | 81 | GLN | p.Gln102Ter | P01130 |
| HOMO SAPIENS | -1 | 30645 | 3m0c | C | 84 | PRO | p.Pro105Ser | P01130 |
| HOMO SAPIENS | -1 | 30645 | 3m0c | C | 194 | ARG | p.Arg215Cys | P01130 |
| HOMO SAPIENS | -1 | 30645 | 3m0c | C | 198 | GLY | p.Gly219Asp | P01130 |
| HOMO SAPIENS | -1 | 30645 | 3m0c | C | 201 | CYS | p.Cys222Tyr | P01130 |
| HOMO SAPIENS | -1 | 30645 | 3m0c | C | 201 | CYS | p.Cys222Ter | P01130 |
| HOMO SAPIENS | -1 | 30645 | 3m0c | C | 206 | ASP | p.Asp227Glu | P01130 |
| HOMO SAPIENS | -1 | 30645 | 3m0c | C | 249 | CYS | p.Cys270Ter | P01130 |
| HOMO SAPIENS | -1 | 30645 | 3m0c | C | 250 | VAL | p.Val271Ile | P01130 |
| HOMO SAPIENS | 0 | 30645 | 3m0c | C | 255 | CYS | p.Cys276Tyr | P01130 |
| HOMO SAPIENS | 0 | 30645 | 3m0c | C | 255 | CYS | p.Cys276Ter | P01130 |
| HOMO SAPIENS | 0 | 30645 | 3m0c | C | 256 | GLU | p.Glu277Lys | P01130 |
| HOMO SAPIENS | 0 | 30645 | 3m0c | C | 265 | SER | p.Ser286Arg | P01130 |
| HOMO SAPIENS | 0 | 30645 | 3m0c | C | 284 | TRP | p.Trp305Ter | P01130 |
| HOMO SAPIENS | 0 | 30645 | 3m0c | C | 292 | CYS | p.Cys313Ter | P01130 |
| HOMO SAPIENS | 1 | 30645 | 3m0c | C | 385 | ARG | p.Arg406Trp | P01130 |
| HOMO SAPIENS | 0 | 30645 | 3m0c | C | 395 | ARG | p.Arg416Gln | P01130 |
| HOMO SAPIENS | 0 | 30645 | 3m0c | C | 398 | TYR | p.Tyr419Ter | P01130 |
| HOMO SAPIENS | 1 | 30645 | 3m0c | C | 415 | VAL | p.Val436Ala | P01130 |
| HOMO SAPIENS | 1 | 30645 | 3m0c | C | 424 | ASP | p.Asp445Glu | P01130 |
| HOMO SAPIENS | 1 | 30645 | 3m0c | C | 425 | LEU | p.Leu446Val | P01130 |
| HOMO SAPIENS | -1 | 30645 | 3m0c | C | 440 | GLY | p.Gly461Cys | P01130 |
| HOMO SAPIENS | -1 | 30645 | 3m0c | C | 441 | VAL | p.Val462Ile | P01130 |
| HOMO SAPIENS | 1 | 30645 | 3m0c | C | 447 | VAL | p.Val468Ile | P01130 |
| HOMO SAPIENS | 1 | 30645 | 3m0c | C | 489 | THR | p.Thr510Met | P01130 |
| HOMO SAPIENS | 0 | 30645 | 3m0c | C | 524 | GLY | p.Gly545Glu | P01130 |
| HOMO SAPIENS | 1 | 30645 | 3m0c | C | 615 | GLY | p.Gly636Ser | P01130 |
| HOMO SAPIENS | 1 | 30645 | 3m0c | C | 625 | LEU | p.Leu646Ile | P01130 |
| HOMO SAPIENS | 1 | 30645 | 3m0c | C | 630 | ASP | p.Asp651Gly | P01130 |
| HOMO SAPIENS | 3 | 30645 | 3m0c | C | 646 | CYS | p.Cys667Tyr | P01130 |
| HOMO SAPIENS | 4 | 30645 | 3m0c | C | 654 | GLY | p.Gly675Ser | P01130 |
| HOMO SAPIENS | 6 | 30645 | 3m0c | C | 660 | CYS | p.Cys681Ter | P01130 |
| HOMO SAPIENS | 5 | 30645 | 3m0c | C | 676 | ALA | p.Ala697Pro | P01130 |
| HOMO SAPIENS | -1 | 30645 | 3m0c | C | 749 | GLN | p.Gln770Ter | P01130 |
| HOMO SAPIENS | -1 | 41323 | 4bxo | A | 1812 | PRO | p.Pro1812Ala | Q8IYD8 |
| HOMO SAPIENS | 5 | 8335 | 3h0t | C | 12 | GLY | p.Gly71Asp | P81172 |
| HOMO SAPIENS | 6 | 609 | 1bhg | B | 148 | PRO | p.Pro148Ser | P08236 |
| HOMO SAPIENS | 1 | 609 | 1bhg | B | 176 | LEU | p.Leu176Phe | P08236 |
| HOMO SAPIENS | 6 | 609 | 1bhg | B | 216 | ARG | p.Arg216Trp | P08236 |
| HOMO SAPIENS | 0 | 609 | 1bhg | B | 350 | LYS | p.Lys350Asn | P08236 |

| | | | | | | | | |
|---|---|---|---|---|---|---|---|---|
| HOMO SAPIENS | 1 | 609 | 1bhg | B | 354 | ALA | p.Ala354Val | P08236 |
| HOMO SAPIENS | 1 | 609 | 1bhg | B | 357 | ARG | p.Arg357Ter | P08236 |
| HOMO SAPIENS | 0 | 609 | 1bhg | B | 382 | ARG | p.Arg382Cys | P08236 |
| HOMO SAPIENS | 0 | 609 | 1bhg | B | 495 | TYR | p.Tyr495Cys | P08236 |
| HOMO SAPIENS | 2 | 609 | 1bhg | B | 507 | TRP | p.Trp507Ter | P08236 |
| HOMO SAPIENS | 0 | 609 | 1bhg | B | 577 | ARG | p.Arg577Leu | P08236 |
| HOMO SAPIENS | 1 | 609 | 1bhg | B | 611 | ARG | p.Arg611Trp | P08236 |
| HOMO SAPIENS | 0 | 609 | 1bhg | B | 619 | ALA | p.Ala619Val | P08236 |
| HOMO SAPIENS | 0 | 609 | 1bhg | B | 627 | TRP | p.Trp627Cys | P08236 |
| HOMO SAPIENS | 5 | 3422 | 1x9n | A | 566 | GLU | p.Glu566Lys | P18858 |
| HOMO SAPIENS | 2 | 3422 | 1x9n | A | 771 | ARG | p.Arg771Trp | P18858 |
| HOMO SAPIENS | 0 | 3279 | 5tby | B | 797 | ALA | p.Ala797Thr | P12883 |
| HOMO SAPIENS | -1 | 3279 | 5tby | B | 982 | MET | p.Met982Thr | P12883 |
| HOMO SAPIENS | -1 | 3279 | 5tby | B | 992 | LEU | p.Leu992Met | P12883 |
| HOMO SAPIENS | -1 | 3279 | 5tby | B | 1004 | GLN | p.Gln1004Glu | P12883 |
| HOMO SAPIENS | -1 | 3279 | 5tby | B | 1013 | GLU | p.Glu1013Lys | P12883 |
| HOMO SAPIENS | -1 | 3279 | 5tby | B | 1032 | ASP | p.Asp1032Asn | P12883 |
| HOMO SAPIENS | -1 | 3279 | 5tby | B | 1053 | ARG | p.Arg1053Gln | P12883 |
| HOMO SAPIENS | -1 | 3279 | 5tby | B | 1117 | GLU | p.Glu1117Ter | P12883 |
| HOMO SAPIENS | -1 | 3279 | 5tby | B | 1193 | ARG | p.Arg1193His | P12883 |
| HOMO SAPIENS | -1 | 3279 | 5tby | B | 1205 | GLU | p.Glu1205Lys | P12883 |
| HOMO SAPIENS | -1 | 3279 | 5tby | B | 1207 | ILE | p.Ile1207Met | P12883 |
| HOMO SAPIENS | -1 | 3279 | 5tby | B | 1250 | ARG | p.Arg1250Pro | P12883 |
| HOMO SAPIENS | -1 | 3279 | 5tby | B | 1259 | HIS | p.His1259Gln | P12883 |
| HOMO SAPIENS | -1 | 3279 | 5tby | B | 1272 | ASP | p.Asp1272Asn | P12883 |
| HOMO SAPIENS | -1 | 3279 | 5tby | B | 1295 | GLU | p.Glu1295Lys | P12883 |
| HOMO SAPIENS | -1 | 3279 | 5tby | B | 1335 | SER | p.Ser1335Leu | P12883 |
| HOMO SAPIENS | -1 | 3279 | 5tby | B | 1399 | GLU | p.Glu1399Asp | P12883 |
| HOMO SAPIENS | -1 | 3279 | 5tby | B | 1413 | SER | p.Ser1413Ter | P12883 |
| HOMO SAPIENS | -1 | 3279 | 5tby | B | 1459 | LYS | p.Lys1459Asn | P12883 |
| HOMO SAPIENS | -1 | 3279 | 5tby | B | 1467 | LEU | p.Leu1467Val | P12883 |
| HOMO SAPIENS | -1 | 3279 | 5tby | B | 1491 | SER | p.Ser1491Cys | P12883 |
| HOMO SAPIENS | -1 | 3279 | 5tby | B | 1496 | GLU | p.Glu1496Ala | P12883 |
| HOMO SAPIENS | -1 | 3279 | 5tby | B | 1500 | ARG | p.Arg1500Pro | P12883 |
| HOMO SAPIENS | -1 | 3279 | 5tby | B | 1606 | ARG | p.Arg1606His | P12883 |
| HOMO SAPIENS | -1 | 3279 | 5tby | B | 1664 | ASN | p.Asn1664Lys | P12883 |
| HOMO SAPIENS | -1 | 3279 | 5tby | B | 1712 | ARG | p.Arg1712Trp | P12883 |
| HOMO SAPIENS | -1 | 3279 | 5tby | B | 1769 | LEU | p.Leu1769Met | P12883 |
| HOMO SAPIENS | -1 | 3279 | 5tby | B | 1808 | GLY | p.Gly1808Ser | P12883 |
| HOMO SAPIENS | -1 | 3279 | 5tby | B | 1818 | ARG | p.Arg1818Trp | P12883 |
| HOMO SAPIENS | -1 | 3279 | 5tby | B | 1820 | ARG | p.Arg1820Trp | P12883 |
| HOMO SAPIENS | -1 | 3279 | 5tby | B | 1844 | GLU | p.Glu1844Lys | P12883 |
| HOMO SAPIENS | -1 | 3279 | 5tby | B | 1845 | ARG | p.Arg1845Trp | P12883 |
| HOMO SAPIENS | -1 | 3279 | 5tby | B | 1854 | THR | p.Thr1854Met | P12883 |
| HOMO SAPIENS | -1 | 3279 | 5tby | B | 1909 | ARG | p.Arg1909Trp | P12883 |
| HOMO SAPIENS | 2 | 2260 | 4kl1 | A | 553 | ASP | p.Asp553Asn | Q9Y3Q4 |
| HOMO SAPIENS | 0 | 40570 | 5lst | A | 522 | ARG | p.Arg522Cys | O94761 |
| HOMO SAPIENS | 0 | 40570 | 5lst | A | 522 | ARG | p.Arg522His | O94761 |
| HOMO SAPIENS | 0 | 40570 | 5lst | A | 591 | PRO | p.Pro591Leu | O94761 |
| HOMO SAPIENS | 0 | 40570 | 5lst | A | 793 | PRO | p.Pro793Ser | O94761 |
| HOMO SAPIENS | 4 | 40570 | 5lst | A | 799 | VAL | p.Val799Met | O94761 |
| HOMO SAPIENS | 0 | 40570 | 5lst | A | 1004 | ARG | p.Arg1004Trp | O94761 |
| HOMO SAPIENS | 0 | 40570 | 5lst | A | 1005 | ARG | p.Arg1005Gln | O94761 |
| HOMO SAPIENS | 0 | 40570 | 5lst | A | 1021 | ARG | p.Arg1021Gln | O94761 |

| | | | | | | | | |
|---|---|---|---|---|---|---|---|---|
| HOMO SAPIENS | 0 | 40570 | 5lst | A | 1043 | LEU | p.Leu1043Pro | O94761 |
| HOMO SAPIENS | 0 | 40570 | 5lst | A | 1045 | ALA | p.Ala1045Thr | O94761 |
| HOMO SAPIENS | 0 | 40570 | 5lst | A | 1105 | GLY | p.Gly1105Asp | O94761 |
| HOMO SAPIENS | 0 | 40570 | 5lst | A | 1105 | GLY | p.Gly1105Ser | O94761 |
| HOMO SAPIENS | -1 | 40570 | 5lst | A | 1113 | GLY | p.Gly1113Arg | O94761 |
| HOMO SAPIENS | -1 | 18865 | 1gxc | A | 64 | GLU | p.Glu64Lys | O96017 |
| HOMO SAPIENS | -1 | 11673 | 3dzu | D | 85 | PRO | p.Pro113Gln | P37231 |
| HOMO SAPIENS | 0 | 19383 | 3b96 | A | 63 | LYS | p.Lys103Arg | P49748 |
| HOMO SAPIENS | 4 | 19383 | 3b96 | A | 140 | ALA | p.Ala180Thr | P49748 |
| HOMO SAPIENS | 1 | 19383 | 3b96 | A | 145 | GLY | p.Gly185Ser | P49748 |
| HOMO SAPIENS | 0 | 19383 | 3b96 | A | 220 | THR | p.Thr260Met | P49748 |
| HOMO SAPIENS | 5 | 19383 | 3b96 | A | 243 | VAL | p.Val283Ala | P49748 |
| HOMO SAPIENS | 0 | 19383 | 3b96 | A | 249 | GLY | p.Gly289Arg | P49748 |
| HOMO SAPIENS | 0 | 19383 | 3b96 | A | 254 | GLY | p.Gly294Glu | P49748 |
| HOMO SAPIENS | 7 | 19383 | 3b96 | A | 326 | ARG | p.Arg366Cys | P49748 |
| HOMO SAPIENS | 7 | 19383 | 3b96 | A | 326 | ARG | p.Arg366His | P49748 |
| HOMO SAPIENS | 5 | 19383 | 3b96 | A | 342 | LYS | p.Lys382Gln | P49748 |
| HOMO SAPIENS | 0 | 19383 | 3b96 | A | 369 | THR | p.Thr409Met | P49748 |
| HOMO SAPIENS | 5 | 19383 | 3b96 | A | 376 | ALA | p.Ala416Ser | P49748 |
| HOMO SAPIENS | 0 | 19383 | 3b96 | A | 385 | ALA | p.Ala425Thr | P49748 |
| HOMO SAPIENS | 5 | 19383 | 3b96 | A | 399 | GLY | p.Gly439Asp | P49748 |
| HOMO SAPIENS | 7 | 19383 | 3b96 | A | 401 | GLY | p.Gly441Asp | P49748 |
| HOMO SAPIENS | 9 | 19383 | 3b96 | A | 413 | ARG | p.Arg453Gln | P49748 |
| HOMO SAPIENS | 2 | 19383 | 3b96 | A | 418 | PHE | p.Phe458Leu | P49748 |
| HOMO SAPIENS | 6 | 19383 | 3b96 | A | 419 | ARG | p.Arg459Trp | P49748 |
| HOMO SAPIENS | 6 | 19383 | 3b96 | A | 419 | ARG | p.Arg459Gln | P49748 |
| HOMO SAPIENS | -1 | 19383 | 3b96 | A | 471 | ARG | p.Arg511Gln | P49748 |
| HOMO SAPIENS | 0 | 19383 | 3b96 | A | 573 | ARG | p.Arg613Trp | P49748 |
| HOMO SAPIENS | 0 | 19383 | 3b96 | A | 575 | ARG | p.Arg615Gln | P49748 |
| HOMO SAPIENS | 0 | 22286 | 3k2s | A | 3 | PRO | p.Pro27His | P02647 |
| HOMO SAPIENS | 0 | 22286 | 3k2s | A | 4 | PRO | p.Pro28Arg | P02647 |
| HOMO SAPIENS | 0 | 22286 | 3k2s | A | 10 | ARG | p.Arg34Leu | P02647 |
| HOMO SAPIENS | 0 | 22286 | 3k2s | A | 26 | GLY | p.Gly50Arg | P02647 |
| HOMO SAPIENS | 0 | 22286 | 3k2s | A | 37 | ALA | p.Ala61Thr | P02647 |
| HOMO SAPIENS | 1 | 1622 | 2qrv | D | 631 | ARG | p.Arg635Trp | Q9Y6K1 |
| HOMO SAPIENS | 1 | 1622 | 2qrv | D | 900 | PRO | p.Pro904Leu | Q9Y6K1 |
| HOMO SAPIENS | 6 | 40796 | 1v18 | B | 1509 | LEU | p.Leu1509Val | P25054 |
| HOMO SAPIENS | 1 | 22375 | 4pki | G | 1214 | LYS | p.Asp214Asn | P28289 |
| HOMO SAPIENS | 1 | 16307 | 5mv8 | B | 483 | ALA | p.Ala483Ser | Q9Y6N9 |
| HOMO SAPIENS | 6 | 16307 | 5mv8 | B | 506 | ASN | p.Asn506Lys | Q9Y6N9 |
| HOMO SAPIENS | 4 | 16307 | 5mv8 | B | 519 | ASP | p.Glu519Asp | Q9Y6N9 |
| HOMO SAPIENS | 0 | 16307 | 5mv8 | B | 530 | GLY | p.Gly530Arg | Q9Y6N9 |
| HOMO SAPIENS | 4 | 36221 | 4cvo | A | 127 | VAL | p.Val127Ile | Q03468 |
| HOMO SAPIENS | 7 | 36221 | 4cvo | A | 134 | ARG | p.Arg134Trp | Q03468 |
| HOMO SAPIENS | 0 | 36221 | 4cvo | A | 156 | GLN | p.Gln156Ter | Q03468 |
| HOMO SAPIENS | 0 | 10829 | 4a11 | B | 13 | GLU | p.Glu13Ter | Q13216 |
| HOMO SAPIENS | 1 | 10829 | 4a11 | B | 100 | TYR | p.Tyr100Ter | Q13216 |
| HOMO SAPIENS | 0 | 10829 | 4a11 | B | 160 | ALA | p.Ala160Val | Q13216 |
| HOMO SAPIENS | 1 | 6759 | 5lev | A | 29 | ILE | p.Ile29Phe | Q9H3H5 |
| HOMO SAPIENS | 0 | 6759 | 5lev | A | 108 | MET | p.Met108Ile | Q9H3H5 |
| HOMO SAPIENS | 2 | 6759 | 5lev | A | 117 | VAL | p.Val117Ile | Q9H3H5 |
| HOMO SAPIENS | 1 | 6759 | 5lev | A | 170 | TYR | p.Tyr170Cys | Q9H3H5 |
| HOMO SAPIENS | 0 | 6759 | 5lev | A | 264 | GLY | p.Val264Gly | Q9H3H5 |
| HOMO SAPIENS | 0 | 6759 | 5lev | A | 332 | PHE | p.Phe332Val | Q9H3H5 |

| | | | | | | | | |
|---|---|---|---|---|---|---|---|---|
| HOMO SAPIENS | 7 | 6759 | 5lev | A | 393 | ILE | p.Ile393Val | Q9H3H5 |
| HOMO SAPIENS | -1 | 33473 | 5m5e | C | 9 | ASN | p.Asn31Ser | P31785 |
| HOMO SAPIENS | 0 | 26155 | 1ic8 | B | 98 | ALA | p.Ala98Val | P20823 |
| HOMO SAPIENS | 0 | 15329 | 4iqr | B | 127 | ARG | p.Arg136Trp | P41235 |
| HOMO SAPIENS | 0 | 20725 | 4bsj | A | 350 | GLU | p.Glu350Val | P35916 |
| HOMO SAPIENS | 0 | 20725 | 4bsj | A | 494 | THR | p.Thr494Ala | P35916 |
| HOMO SAPIENS | 0 | 20725 | 4bsj | A | 527 | ASN | p.Asn527Ser | P35916 |
| HOMO SAPIENS | 6 | 29959 | 1xiw | B | 47 | ARG | p.Arg68Ter | P04234 |
| HOMO SAPIENS | 2 | 29959 | 1xiw | B | 72 | CYS | p.Cys93Ter | P04234 |
| HOMO SAPIENS | 3 | 20242 | 5tjc | A | 102 | ARG | p.Arg102Ter | Q9GZU1 |
| HOMO SAPIENS | 3 | 20242 | 5tjc | A | 232 | THR | p.Thr232Pro | Q9GZU1 |
| HOMO SAPIENS | 7 | 9590 | 4ap8 | A | 50 | THR | p.Thr50Ala | O96007 |
| HOMO SAPIENS | 4 | 9590 | 4ap8 | A | 168 | GLU | p.Glu168Lys | O96007 |
| HOMO SAPIENS | -1 | 4297 | 5do7 | B | 19 | ASP | p.Asp19His | Q9H221 |
| HOMO SAPIENS | -1 | 4297 | 5do7 | B | 54 | TYR | p.Tyr54Cys | Q9H221 |
| HOMO SAPIENS | -1 | 4297 | 5do7 | B | 80 | CYS | p.Cys80Tyr | Q9H221 |
| HOMO SAPIENS | 5 | 4297 | 5do7 | B | 107 | SER | p.Ser107Ter | Q9H221 |
| HOMO SAPIENS | 1 | 4297 | 5do7 | B | 164 | ARG | p.Arg164Pro | Q9H221 |
| HOMO SAPIENS | 0 | 4297 | 5do7 | B | 210 | VAL | p.Val210Met | Q9H221 |
| HOMO SAPIENS | 0 | 4297 | 5do7 | B | 231 | PRO | p.Pro231Thr | Q9H221 |
| HOMO SAPIENS | 0 | 4297 | 5do7 | B | 263 | ARG | p.Arg263Gln | Q9H221 |
| HOMO SAPIENS | 0 | 4297 | 5do7 | B | 361 | TRP | p.Trp361Ter | Q9H221 |
| HOMO SAPIENS | 0 | 4297 | 5do7 | B | 400 | THR | p.Thr400Lys | Q9H221 |
| HOMO SAPIENS | 0 | 4297 | 5do7 | B | 549 | ALA | p.Ala549Pro | Q9H221 |
| HOMO SAPIENS | 0 | 4297 | 5do7 | B | 596 | LEU | p.Leu596Arg | Q9H221 |
| HOMO SAPIENS | 0 | 4297 | 5do7 | B | 632 | VAL | p.Val632Ala | Q9H221 |
| HOMO SAPIENS | 0 | 4297 | 5do7 | B | 658 | TYR | p.Tyr658Ter | Q9H221 |
| HOMO SAPIENS | 2 | 9356 | 4igg | B | 206 | GLN | p.Gln206His | P35221 |
| HOMO SAPIENS | 6 | 2179 | 2wk6 | A | 44 | ARG | p.Arg44Gln | P19971 |
| HOMO SAPIENS | 6 | 2179 | 2wk6 | A | 81 | ARG | p.Arg81Gln | P19971 |
| HOMO SAPIENS | 6 | 2179 | 2wk6 | A | 145 | GLY | p.Gly145Arg | P19971 |
| HOMO SAPIENS | 7 | 2179 | 2wk6 | A | 146 | ARG | p.Arg146His | P19971 |
| HOMO SAPIENS | 7 | 2179 | 2wk6 | A | 153 | GLY | p.Gly153Ser | P19971 |
| HOMO SAPIENS | 0 | 2179 | 2wk6 | A | 173 | MET | p.Met173Ile | P19971 |
| HOMO SAPIENS | 6 | 2179 | 2wk6 | A | 202 | ARG | p.Arg202Lys | P19971 |
| HOMO SAPIENS | 6 | 2179 | 2wk6 | A | 208 | VAL | p.Val208Met | P19971 |
| HOMO SAPIENS | 5 | 2179 | 2wk6 | A | 222 | LYS | p.Lys222Arg | P19971 |
| HOMO SAPIENS | 5 | 2179 | 2wk6 | A | 289 | GLU | p.Glu289Ala | P19971 |
| HOMO SAPIENS | 2 | 2179 | 2wk6 | A | 311 | GLY | p.Gly311Arg | P19971 |
| HOMO SAPIENS | 2 | 2179 | 2wk6 | A | 376 | ARG | p.Arg376Pro | P19971 |
| HOMO SAPIENS | 2 | 2179 | 2wk6 | A | 465 | ALA | p.Ala465Thr | P19971 |
| HOMO SAPIENS | -1 | 11856 | 3orh | A | 3 | ALA | p.Ala3Ser | Q14353 |
| HOMO SAPIENS | 1 | 11856 | 3orh | A | 20 | TRP | p.Trp20Ser | Q14353 |
| HOMO SAPIENS | 1 | 11856 | 3orh | A | 50 | MET | p.Met50Leu | Q14353 |
| HOMO SAPIENS | 1 | 11856 | 3orh | A | 110 | VAL | p.Val110Phe | Q14353 |
| HOMO SAPIENS | 8 | 11856 | 3orh | A | 164 | GLY | p.Gly164Asp | Q14353 |
| HOMO SAPIENS | 1 | 11856 | 3orh | A | 169 | CYS | p.Cys169Tyr | Q14353 |
| HOMO SAPIENS | 1 | 11856 | 3orh | A | 174 | TRP | p.Trp174Ter | Q14353 |
| HOMO SAPIENS | 2 | 11856 | 3orh | A | 196 | ALA | p.Ala196Val | Q14353 |
| HOMO SAPIENS | 1 | 11856 | 3orh | A | 209 | THR | p.Thr209Met | Q14353 |
| HOMO SAPIENS | 1 | 11856 | 3orh | A | 219 | ASP | p.Asp219Asn | Q14353 |
| HOMO SAPIENS | 1 | 11856 | 3orh | A | 224 | ALA | p.Ala224Thr | Q14353 |
| HOMO SAPIENS | 1 | 13244 | 2m6i | D | 218 | ARG | p.Arg246Leu | P23415 |
| HOMO SAPIENS | 1 | 13244 | 2m6i | D | 244 | ILE | p.Ile272Asn | P23415 |

| | | | | | | | | |
|---|---|---|---|---|---|---|---|---|
| HOMO SAPIENS | 5 | 13244 | 2m6i | D | 250 | PRO | p.Pro278Thr | P23415 |
| HOMO SAPIENS | 6 | 13244 | 2m6i | D | 266 | GLN | p.Gln294His | P23415 |
| HOMO SAPIENS | 1 | 13244 | 2m6i | D | 271 | ARG | p.Arg299Leu | P23415 |
| HOMO SAPIENS | 1 | 13244 | 2m6i | D | 276 | LYS | p.Lys304Glu | P23415 |
| HOMO SAPIENS | 1 | 13244 | 2m6i | D | 279 | TYR | p.Tyr307Cys | P23415 |
| HOMO SAPIENS | 0 | 13244 | 2m6i | D | 316 | GLY | p.Gly378Ser | P23415 |
| HOMO SAPIENS | 1 | 13244 | 2m6i | D | 400 | ARG | p.Arg428His | P23415 |
| HOMO SAPIENS | 3 | 1967 | 2i6b | A | 301 | ALA | p.Ala318Glu | P55263 |
| HOMO SAPIENS | 4 | 11096 | 1h8b | A | 61 | ALA | p.Ala882Thr | P35609 |
| HOMO SAPIENS | 0 | 15423 | 5ngz | A | 2 | GLN | p.Gln2Glu | Q9NPD8 |
| HOMO SAPIENS | 3 | 30475 | 3lpw | A | 21 | SER | p.Ser22894Tyr | Q8WZ42 |
| HOMO SAPIENS | 6 | 32687 | 2arf | A | 1041 | ARG | p.Arg1041Trp | P35670 |
| HOMO SAPIENS | 8 | 32687 | 2arf | A | 1061 | GLY | p.Gly1061Glu | P35670 |
| HOMO SAPIENS | 6 | 32687 | 2arf | A | 1064 | GLU | p.Glu1064Ala | P35670 |
| HOMO SAPIENS | 2 | 32687 | 2arf | A | 1069 | HIS | p.His1069Gln | P35670 |
| HOMO SAPIENS | 2 | 32687 | 2arf | A | 1088 | LEU | p.Leu1088Ter | P35670 |
| HOMO SAPIENS | 0 | 32687 | 2arf | A | 1099 | GLY | p.Gly1099Ser | P35670 |
| HOMO SAPIENS | 4 | 32687 | 2arf | A | 1106 | VAL | p.Val1106Asp | P35670 |
| HOMO SAPIENS | 2 | 32687 | 2arf | A | 1142 | GLN | p.Gln1142His | P35670 |
| HOMO SAPIENS | 8 | 32687 | 2arf | A | 1148 | ILE | p.Ile1148Thr | P35670 |
| HOMO SAPIENS | 6 | 32687 | 2arf | A | 1151 | ARG | p.Arg1151Cys | P35670 |
| HOMO SAPIENS | 4 | 32687 | 2arf | A | 1169 | MET | p.Met1169Val | P35670 |
| HOMO SAPIENS | 4 | 32687 | 2arf | A | 1173 | GLU | p.Glu1173Lys | P35670 |
| HOMO SAPIENS | 2 | 32687 | 2arf | A | 1176 | GLY | p.Gly1176Arg | P35670 |
| HOMO SAPIENS | 3 | 15544 | 5uq3 | C | 575 | PRO | p.Pro69Leu | P46527 |
| HOMO SAPIENS | 6 | 15544 | 5uq3 | C | 592 | GLU | p.Glu86Asp | P46527 |
| HOMO SAPIENS | 6 | 15544 | 5uq3 | C | 599 | ARG | p.Arg93Trp | P46527 |
| HOMO SAPIENS | -1 | 15544 | 5uq3 | C | 615 | VAL | p.Val109Gly | P46527 |
| HOMO SAPIENS | -1 | 15544 | 5uq3 | C | 631 | SER | p.Ser125Pro | P46527 |
| HOMO SAPIENS | -1 | 15544 | 5uq3 | C | 639 | PRO | p.Pro133Ser | P46527 |
| HOMO SAPIENS | 2 | 2703 | 4wxq | A | 245 | CYS | p.Cys245Tyr | Q99972 |
| HOMO SAPIENS | 0 | 2703 | 4wxq | A | 252 | GLY | p.Gly252Arg | Q99972 |
| HOMO SAPIENS | 1 | 2703 | 4wxq | A | 337 | GLN | p.Gln337Arg | Q99972 |
| HOMO SAPIENS | 5 | 2703 | 4wxq | A | 347 | TYR | p.Tyr347Ter | Q99972 |
| HOMO SAPIENS | 3 | 2703 | 4wxq | A | 364 | GLY | p.Gly364Val | Q99972 |
| HOMO SAPIENS | 2 | 2703 | 4wxq | A | 367 | GLY | p.Gly367Arg | Q99972 |
| HOMO SAPIENS | 2 | 2703 | 4wxq | A | 368 | GLN | p.Gln368Ter | Q99972 |
| HOMO SAPIENS | 6 | 2703 | 4wxq | A | 370 | PRO | p.Pro370Leu | Q99972 |
| HOMO SAPIENS | 1 | 2703 | 4wxq | A | 380 | ASP | p.Asp380His | Q99972 |
| HOMO SAPIENS | 8 | 2703 | 4wxq | A | 399 | GLY | p.Gly399Val | Q99972 |
| HOMO SAPIENS | 0 | 2703 | 4wxq | A | 423 | LYS | p.Lys423Glu | Q99972 |
| HOMO SAPIENS | 3 | 2703 | 4wxq | A | 433 | CYS | p.Cys433Arg | Q99972 |
| HOMO SAPIENS | 3 | 2703 | 4wxq | A | 437 | TYR | p.Tyr437His | Q99972 |
| HOMO SAPIENS | 2 | 2703 | 4wxq | A | 477 | ILE | p.Ile477Ser | Q99972 |
| HOMO SAPIENS | 3 | 2703 | 4wxq | A | 480 | ASN | p.Asn480Lys | Q99972 |
| HOMO SAPIENS | -1 | 34100 | 2fot | C | 1171 | GLN | p.Gln1173Pro | Q13813 |
| HOMO SAPIENS | 0 | 8773 | 1kw2 | A | 432 | ASP | p.Asp432Glu | P02774 |
| HOMO SAPIENS | 0 | 8773 | 1kw2 | A | 436 | LYS | p.Lys436Thr | P02774 |
| HOMO SAPIENS | 0 | 8402 | 4dzo | B | 649 | TYR | p.Tyr649Ter | Q9Y6D9 |
| HOMO SAPIENS | 0 | 39962 | 1nst | A | 611 | GLY | p.Gly611Ser | P52848 |
| HOMO SAPIENS | 2 | 22578 | 3dof | B | 45 | MET | p.Met45Arg | Q9Y2Y0 |
| HOMO SAPIENS | 1 | 23290 | 5j1s | A | 292 | ILE | p.Ile292Thr | O14656 |
| HOMO SAPIENS | 4 | 9505 | 4j1y | A | 534 | ARG | p.Arg534Trp | P09871 |
| HOMO SAPIENS | 9 | 260 | 2h57 | A | 31 | THR | p.Thr31Met | Q9H0F7 |

| | | | | | | | | |
|---|---|---|---|---|---|---|---|---|
| HOMO SAPIENS | 0 | 260 | 2h57 | A | 91 | ILE | p.Ile91Thr | Q9H0F7 |
| HOMO SAPIENS | 0 | 260 | 2h57 | A | 121 | ARG | p.Arg121His | Q9H0F7 |
| HOMO SAPIENS | 0 | 260 | 2h57 | A | 122 | ARG | p.Arg122Ter | Q9H0F7 |
| HOMO SAPIENS | 2 | 260 | 2h57 | A | 169 | GLY | p.Gly169Ala | Q9H0F7 |
| HOMO SAPIENS | 4 | 260 | 2h57 | A | 170 | LEU | p.Leu170Trp | Q9H0F7 |
| HOMO SAPIENS | 4 | 34023 | 2cr3 | A | 18 | GLY | p.Gly48Ser | P11362 |
| HOMO SAPIENS | 0 | 37667 | 3ii7 | A | 420 | ARG | p.Arg420Cys | Q8IXQ5 |
| HOMO SAPIENS | 0 | 41720 | 2k9u | A | 68 | ARG | p.Arg2364His | Q14315 |
| HOMO SAPIENS | 1 | 29250 | 5fmv | A | 568 | HIS | p.His568Gln | P08575 |
| HOMO SAPIENS | 1 | 1280 | 1fb1 | A | 69 | PRO | p.Pro69Leu | P30793 |
| HOMO SAPIENS | 3 | 1280 | 1fb1 | A | 88 | ARG | p.Arg88Trp | P30793 |
| HOMO SAPIENS | 2 | 1280 | 1fb1 | A | 108 | GLY | p.Gly108Asp | P30793 |
| HOMO SAPIENS | 4 | 1280 | 1fb1 | A | 134 | ASP | p.Asp134Val | P30793 |
| HOMO SAPIENS | 6 | 1280 | 1fb1 | A | 135 | ILE | p.Ile135Lys | P30793 |
| HOMO SAPIENS | 9 | 1280 | 1fb1 | A | 144 | HIS | p.His144Pro | P30793 |
| HOMO SAPIENS | 9 | 1280 | 1fb1 | A | 184 | ARG | p.Arg184His | P30793 |
| HOMO SAPIENS | 1 | 1280 | 1fb1 | A | 196 | ALA | p.Ala196Ser | P30793 |
| HOMO SAPIENS | 3 | 1280 | 1fb1 | A | 201 | GLY | p.Gly201Glu | P30793 |
| HOMO SAPIENS | 2 | 1280 | 1fb1 | A | 204 | VAL | p.Val204Ile | P30793 |
| HOMO SAPIENS | 2 | 1280 | 1fb1 | A | 211 | MET | p.Met211Ile | P30793 |
| HOMO SAPIENS | 0 | 1280 | 1fb1 | A | 221 | MET | p.Met221Thr | P30793 |
| HOMO SAPIENS | 3 | 1280 | 1fb1 | A | 224 | LYS | p.Lys224Arg | P30793 |
| HOMO SAPIENS | 2 | 1280 | 1fb1 | A | 249 | ARG | p.Arg249Ser | P30793 |
| HOMO SAPIENS | 8 | 41352 | 4k12 | A | 47 | ILE | p.Ile551Thr | P08603 |
| HOMO SAPIENS | 9 | 20703 | 5nir | A | 57 | CYS | p.Cys57Tyr | P02458 |
| HOMO SAPIENS | 9 | 20703 | 5nir | A | 64 | CYS | p.Cys64Ter | P02458 |
| HOMO SAPIENS | 0 | 20087 | 4cmn | A | 299 | MET | p.Met299Ile | Q01968 |
| HOMO SAPIENS | 1 | 20819 | 3aaf | A | 979 | ILE | p.Ile979Met | Q14191 |
| HOMO SAPIENS | 1 | 20819 | 3aaf | A | 984 | LEU | p.Leu984Ile | Q14191 |
| HOMO SAPIENS | 3 | 20819 | 3aaf | A | 987 | ARG | p.Arg987Ter | Q14191 |
| HOMO SAPIENS | -1 | 20819 | 3aaf | A | 1074 | PHE | p.Leu1074Phe | Q14191 |
| HOMO SAPIENS | 0 | 3619 | 3dem | A | 206 | ASN | p.Asn225Ser | P48740 |
| HOMO SAPIENS | 1 | 3619 | 3dem | A | 269 | ARG | p.Arg288Leu | P48740 |
| HOMO SAPIENS | 0 | 3105 | 3ooi | A | 164 | TYR | p.Tyr2015His | Q96L73 |
| HOMO SAPIENS | 6 | 15828 | 2lkx | A | 16 | LEU | p.Leu100Gln | Q99697 |
| HOMO SAPIENS | 8 | 15828 | 2lkx | A | 24 | ARG | p.Arg108His | Q99697 |
| HOMO SAPIENS | 2 | 15828 | 2lkx | A | 30 | THR | p.Thr114Pro | Q99697 |
| HOMO SAPIENS | 8 | 15828 | 2lkx | A | 45 | VAL | p.Val129Leu | Q99697 |
| HOMO SAPIENS | 8 | 15828 | 2lkx | A | 46 | ARG | p.Arg130Trp | Q99697 |
| HOMO SAPIENS | 9 | 15828 | 2lkx | A | 53 | ARG | p.Arg137Pro | Q99697 |
| HOMO SAPIENS | 9 | 13017 | 4irq | A | 141 | ARG | p.Arg141Trp | Q9UBV7 |
| HOMO SAPIENS | 0 | 13017 | 4irq | A | 186 | ALA | p.Ala186Asp | Q9UBV7 |
| HOMO SAPIENS | 0 | 13017 | 4irq | A | 206 | LEU | p.Leu206Pro | Q9UBV7 |
| HOMO SAPIENS | 1 | 13017 | 4irq | A | 214 | CYS | p.Cys214Tyr | Q9UBV7 |
| HOMO SAPIENS | 0 | 13017 | 4irq | A | 270 | ARG | p.Arg270Cys | Q9UBV7 |
| HOMO SAPIENS | -1 | 13053 | 4mqw | X | 307 | ALA | p.Ala307Thr | P23945 |
| HOMO SAPIENS | 0 | 19406 | 4b3g | A | 43 | ARG | p.Arg43Ter | P38935 |
| HOMO SAPIENS | 0 | 19406 | 4b3g | A | 46 | CYS | p.Cys46Ter | P38935 |
| HOMO SAPIENS | 0 | 19406 | 4b3g | A | 75 | ALA | p.Ala75Thr | P38935 |
| HOMO SAPIENS | 0 | 19406 | 4b3g | A | 201 | LEU | p.Leu201Ser | P38935 |
| HOMO SAPIENS | 3 | 19406 | 4b3g | A | 213 | HIS | p.His213Arg | P38935 |
| HOMO SAPIENS | 0 | 19406 | 4b3g | A | 275 | ILE | p.Ile275Val | P38935 |
| HOMO SAPIENS | -1 | 19406 | 4b3g | A | 320 | ARG | p.Arg320Ter | P38935 |
| HOMO SAPIENS | 0 | 19406 | 4b3g | A | 361 | LEU | p.Leu361Pro | P38935 |

| | | | | | | | | |
|---|---|---|---|---|---|---|---|---|
| HOMO SAPIENS | 0 | 19406 | 4b3g | A | 369 | PHE | p.Phe369Leu | P38935 |
| HOMO SAPIENS | 0 | 19406 | 4b3g | A | 493 | THR | p.Thr493Ile | P38935 |
| HOMO SAPIENS | 0 | 19406 | 4b3g | A | 496 | CYS | p.Cys496Ter | P38935 |
| HOMO SAPIENS | 0 | 19406 | 4b3g | A | 514 | GLU | p.Glu514Lys | P38935 |
| HOMO SAPIENS | 0 | 19406 | 4b3g | A | 528 | ALA | p.Ala528Thr | P38935 |
| HOMO SAPIENS | 0 | 19406 | 4b3g | A | 531 | PRO | p.Pro531Thr | P38935 |
| HOMO SAPIENS | 0 | 19406 | 4b3g | A | 580 | VAL | p.Val580Ile | P38935 |
| HOMO SAPIENS | 6 | 19406 | 4b3g | A | 603 | ARG | p.Arg603His | P38935 |
| HOMO SAPIENS | 0 | 21501 | 5jde | A | 40 | SER | p.Ser2831Asn | Q8WZ42 |
| HOMO SAPIENS | 5 | 21501 | 5jde | A | 189 | ALA | p.Ala2980Thr | Q8WZ42 |
| HOMO SAPIENS | 6 | 15008 | 3mop | H | 12 | ARG | p.Arg12Cys | Q9NWZ3 |
| HOMO SAPIENS | 8 | 17455 | 3jtc | C | 15 | ARG | p.Arg57Trp | P04070 |
| HOMO SAPIENS | -1 | 17455 | 3jtc | C | 20 | CGU | p.Glu62Ala | P04070 |
| HOMO SAPIENS | 0 | 27795 | 5gox | B | 625 | TYR | p.Tyr625Ter | Q92878 |
| HOMO SAPIENS | 0 | 27795 | 5gox | B | 653 | SER | p.Ser653Ter | Q92878 |
| HOMO SAPIENS | -1 | 6558 | 5nug | A | 137 | VAL | p.Val137Ile | Q14204 |
| HOMO SAPIENS | -1 | 6558 | 5nug | A | 567 | ARG | p.Arg567His | Q14204 |
| HOMO SAPIENS | -1 | 6558 | 5nug | A | 1140 | THR | p.Thr1140Met | Q14204 |
| HOMO SAPIENS | -1 | 6558 | 5nug | A | 1991 | ASP | p.Asp1991Asn | Q14204 |
| HOMO SAPIENS | -1 | 6558 | 5nug | A | 2401 | LYS | p.Lys2401Asn | Q14204 |
| HOMO SAPIENS | 0 | 6558 | 5nug | A | 2488 | ARG | p.Arg2488His | Q14204 |
| HOMO SAPIENS | 0 | 6558 | 5nug | A | 3907 | HIS | p.His3907Gln | Q14204 |
| HOMO SAPIENS | 0 | 6558 | 5nug | A | 3981 | THR | p.Thr3981Arg | Q14204 |
| HOMO SAPIENS | 0 | 6558 | 5nug | A | 4394 | THR | p.Thr4394Met | Q14204 |
| HOMO SAPIENS | 0 | 6558 | 5nug | A | 4573 | ASN | p.Asn4573Ser | Q14204 |
| HOMO SAPIENS | 0 | 6558 | 5nug | A | 4603 | SER | p.Ser4603Ile | Q14204 |
| HOMO SAPIENS | 1 | 4244 | 1umk | A | 57 | ARG | p.Arg58Gln | P00387 |
| HOMO SAPIENS | 1 | 4244 | 1umk | A | 72 | LEU | p.Leu73Pro | P00387 |
| HOMO SAPIENS | 1 | 4244 | 1umk | A | 105 | VAL | p.Val106Met | P00387 |
| HOMO SAPIENS | 0 | 4244 | 1umk | A | 116 | THR | p.Thr117Ser | P00387 |
| HOMO SAPIENS | 8 | 4244 | 1umk | A | 127 | SER | p.Ser128Pro | P00387 |
| HOMO SAPIENS | 0 | 4244 | 1umk | A | 148 | LEU | p.Leu149Pro | P00387 |
| HOMO SAPIENS | 0 | 4244 | 1umk | A | 159 | ARG | p.Arg160Ter | P00387 |
| HOMO SAPIENS | 0 | 4244 | 1umk | A | 203 | CYS | p.Cys204Arg | P00387 |
| HOMO SAPIENS | 0 | 4244 | 1umk | A | 203 | CYS | p.Cys204Tyr | P00387 |
| HOMO SAPIENS | 2 | 4244 | 1umk | A | 239 | ASP | p.Asp240Gly | P00387 |
| HOMO SAPIENS | 0 | 4244 | 1umk | A | 291 | GLY | p.Gly292Asp | P00387 |
| HOMO SAPIENS | 4 | 34495 | 1s35 | A | 1151 | ASP | p.Asn1151Asp | P11277 |
| HOMO SAPIENS | 3 | 30110 | 3zqs | A | 224 | THR | p.Thr224Ala | Q9NW38 |
| HOMO SAPIENS | 3 | 17588 | 5dse | B | 53 | LEU | p.Leu53Pro | Q9BYI3 |
| HOMO SAPIENS | 0 | 1564 | 4yqf | B | 72 | HIS | p.His347Gln | Q9UHD8 |
| HOMO SAPIENS | 0 | 17673 | 4jnw | A | 32 | ARG | p.Arg32186His | Q8WZ42 |
| HOMO SAPIENS | 3 | 17673 | 4jnw | A | 140 | ILE | p.Ile32294Val | Q8WZ42 |
| HOMO SAPIENS | 5 | 17673 | 4jnw | A | 269 | ALA | p.Ala32423Thr | Q8WZ42 |
| HOMO SAPIENS | 3 | 12858 | 3s97 | C | 302 | LEU | p.Leu302Pro | Q12860 |
| HOMO SAPIENS | 0 | 6147 | 4nn0 | A | 114 | ARG | p.Arg114Gln | Q9BXJ0 |
| HOMO SAPIENS | 0 | 7549 | 1tr2 | A | 277 | LEU | p.Leu277Met | P18206 |
| HOMO SAPIENS | 1 | 7549 | 1tr2 | A | 413 | ALA | p.Ala413Thr | P18206 |
| HOMO SAPIENS | 1 | 7549 | 1tr2 | A | 519 | ILE | p.Ile519Leu | P18206 |
| HOMO SAPIENS | 5 | 7549 | 1tr2 | A | 547 | ARG | p.Arg547Ter | P18206 |
| HOMO SAPIENS | 2 | 7549 | 1tr2 | A | 636 | HIS | p.His636Arg | P18206 |
| HOMO SAPIENS | 2 | 7549 | 1tr2 | A | 824 | ILE | p.Ile824Met | P18206 |
| HOMO SAPIENS | 2 | 13900 | 3gro | A | 45 | CYS | p.Cys45Tyr | P50897 |
| HOMO SAPIENS | 2 | 13900 | 3gro | A | 75 | THR | p.Thr75Pro | P50897 |

| | | | | | | | | |
|---|---|---|---|---|---|---|---|---|
| HOMO SAPIENS | 2 | 13900 | 3gro | A | 79 | ASP | p.Asp79Gly | P50897 |
| HOMO SAPIENS | 5 | 13900 | 3gro | A | 91 | GLN | p.Gln91Pro | P50897 |
| HOMO SAPIENS | 5 | 13900 | 3gro | A | 96 | CYS | p.Cys96Tyr | P50897 |
| HOMO SAPIENS | 0 | 13900 | 3gro | A | 104 | LYS | p.Lys104Ter | P50897 |
| HOMO SAPIENS | 2 | 13900 | 3gro | A | 108 | GLY | p.Gly108Arg | P50897 |
| HOMO SAPIENS | 2 | 13900 | 3gro | A | 109 | TYR | p.Tyr109Asp | P50897 |
| HOMO SAPIENS | 0 | 13900 | 3gro | A | 122 | ARG | p.Arg122Trp | P50897 |
| HOMO SAPIENS | 0 | 13900 | 3gro | A | 138 | SER | p.Ser138Leu | P50897 |
| HOMO SAPIENS | 5 | 13900 | 3gro | A | 151 | ARG | p.Arg151Ter | P50897 |
| HOMO SAPIENS | 7 | 13900 | 3gro | A | 164 | ARG | p.Arg164Ter | P50897 |
| HOMO SAPIENS | 0 | 13900 | 3gro | A | 177 | GLN | p.Gln177Glu | P50897 |
| HOMO SAPIENS | 0 | 13900 | 3gro | A | 181 | VAL | p.Val181Leu | P50897 |
| HOMO SAPIENS | 0 | 13900 | 3gro | A | 184 | GLU | p.Glu184Lys | P50897 |
| HOMO SAPIENS | 5 | 13900 | 3gro | A | 186 | TRP | p.Trp186Ter | P50897 |
| HOMO SAPIENS | 0 | 13900 | 3gro | A | 219 | LEU | p.Leu219Gln | P50897 |
| HOMO SAPIENS | 0 | 13900 | 3gro | A | 222 | LEU | p.Leu222Pro | P50897 |
| HOMO SAPIENS | 5 | 13900 | 3gro | A | 296 | TRP | p.Trp296Ter | P50897 |
| HOMO SAPIENS | 6 | 35028 | 5mwb | A | 444 | CYS | p.Cys444Tyr | Q04721 |
| HOMO SAPIENS | 0 | 35028 | 5mwb | A | 514 | ASN | p.Asn514Ser | Q04721 |
| HOMO SAPIENS | 5 | 1483 | 4gl2 | A | 822 | ARG | p.Arg822Gln | Q9BYX4 |
| HOMO SAPIENS | 0 | 1483 | 4gl2 | A | 843 | ARG | p.His843Arg | Q9BYX4 |
| HOMO SAPIENS | -1 | 28838 | 3cl3 | D | 153 | LEU | p.Leu153Arg | Q9Y6K9 |
| HOMO SAPIENS | 2 | 22563 | 4zel | B | 79 | ARG | p.Arg79Trp | P09172 |
| HOMO SAPIENS | 2 | 22563 | 4zel | B | 101 | VAL | p.Val101Met | P09172 |
| HOMO SAPIENS | 2 | 22563 | 4zel | B | 114 | ASP | p.Asp114Glu | P09172 |
| HOMO SAPIENS | 0 | 22563 | 4zel | B | 201 | ASN | p.Asn201Ser | P09172 |
| HOMO SAPIENS | 2 | 22563 | 4zel | B | 211 | ALA | p.Ala211Thr | P09172 |
| HOMO SAPIENS | -1 | 22563 | 4zel | B | 290 | ASP | p.Asp290Asn | P09172 |
| HOMO SAPIENS | 2 | 22563 | 4zel | B | 318 | ALA | p.Ala318Ser | P09172 |
| HOMO SAPIENS | 2 | 22563 | 4zel | B | 345 | ASP | p.Asp345Asn | P09172 |
| HOMO SAPIENS | 2 | 22563 | 4zel | B | 362 | ALA | p.Ala362Glu | P09172 |
| HOMO SAPIENS | 2 | 22563 | 4zel | B | 549 | ARG | p.Arg549Cys | P09172 |
| HOMO SAPIENS | 9 | 9780 | 1y6k | R | 36 | TYR | p.Tyr57Cys | Q13651 |
| HOMO SAPIENS | 1 | 9780 | 1y6k | R | 40 | LEU | p.Leu61Val | Q13651 |
| HOMO SAPIENS | 1 | 9780 | 1y6k | R | 63 | THR | p.Thr84Ile | Q13651 |
| HOMO SAPIENS | 9 | 9780 | 1y6k | R | 80 | ARG | p.Arg101Trp | Q13651 |
| HOMO SAPIENS | 0 | 9780 | 1y6k | R | 120 | GLY | p.Gly141Arg | Q13651 |
| HOMO SAPIENS | 0 | 9780 | 1y6k | R | 138 | SER | p.Ser159Gly | Q13651 |
| HOMO SAPIENS | 0 | 9780 | 1y6k | R | 203 | ILE | p.Ile224Val | Q13651 |
| HOMO SAPIENS | 0 | 16019 | 3rep | A | 211 | ARG | p.Arg211Cys | Q13418 |
| HOMO SAPIENS | 3 | 7879 | 5ec3 | A | 33 | ALA | p.Ala33Thr | P32754 |
| HOMO SAPIENS | 4 | 7879 | 5ec3 | A | 113 | ARG | p.Arg113Gln | P32754 |
| HOMO SAPIENS | 3 | 7879 | 5ec3 | A | 340 | VAL | p.Val340Leu | P32754 |
| HOMO SAPIENS | 0 | 8904 | 1qk9 | A | 6 | LYS | p.Lys82Arg | P51608 |
| HOMO SAPIENS | 4 | 8904 | 1qk9 | A | 24 | LEU | p.Leu100Val | P51608 |
| HOMO SAPIENS | 7 | 8904 | 1qk9 | A | 30 | ARG | p.Arg106Trp | P51608 |
| HOMO SAPIENS | 6 | 8904 | 1qk9 | A | 57 | ARG | p.Arg133Cys | P51608 |
| HOMO SAPIENS | 3 | 8904 | 1qk9 | A | 61 | GLU | p.Glu137Gly | P51608 |
| HOMO SAPIENS | 2 | 8904 | 1qk9 | A | 64 | ALA | p.Ala140Val | P51608 |
| HOMO SAPIENS | 5 | 8904 | 1qk9 | A | 79 | PHE | p.Phe155Ser | P51608 |
| HOMO SAPIENS | 3 | 8904 | 1qk9 | A | 82 | THR | p.Thr158Met | P51608 |
| HOMO SAPIENS | 1 | 23121 | 3e9k | A | 198 | THR | p.Thr198Ala | Q16719 |
| HOMO SAPIENS | -1 | 26090 | 3tje | F | 19 | LEU | p.Leu35Met | P25445 |
| HOMO SAPIENS | 8 | 26090 | 3tje | F | 106 | THR | p.Thr122Ile | P25445 |

| | | | | | | | | |
|---|---|---|---|---|---|---|---|---|
| HOMO SAPIENS | 2 | 36404 | 5cqr | A | 765 | GLY | p.Gly765Glu | O95163 |
| HOMO SAPIENS | 2 | 36404 | 5cqr | A | 816 | ILE | p.Ile816Leu | O95163 |
| HOMO SAPIENS | 2 | 36404 | 5cqr | A | 830 | ILE | p.Ile830Met | O95163 |
| HOMO SAPIENS | 5 | 36404 | 5cqr | A | 939 | TYR | p.Tyr939Ter | O95163 |
| HOMO SAPIENS | 2 | 36404 | 5cqr | A | 952 | LYS | p.Lys952Ile | O95163 |
| HOMO SAPIENS | 5 | 36404 | 5cqr | A | 1013 | GLY | p.Gly1013Ser | O95163 |
| HOMO SAPIENS | 5 | 36404 | 5cqr | A | 1072 | CYS | p.Cys1072Ser | O95163 |
| HOMO SAPIENS | -1 | 36404 | 5cqr | A | 1158 | PRO | p.Pro1158Leu | O95163 |
| HOMO SAPIENS | 1 | 34149 | 2k40 | A | 18 | ASN | p.Asn125Ser | Q9UBX0 |
| HOMO SAPIENS | 0 | 34149 | 2k40 | A | 42 | LEU | p.Glu149Lys | Q9UBX0 |
| HOMO SAPIENS | 9 | 34149 | 2k40 | A | 53 | ARG | p.Arg160Cys | Q9UBX0 |
| HOMO SAPIENS | 0 | 34149 | 2k40 | A | 63 | SER | p.Ser170Leu | Q9UBX0 |
| HOMO SAPIENS | 2 | 2468 | 3vp6 | A | 484 | LEU | p.Leu484Met | Q99259 |
| HOMO SAPIENS | 0 | 2468 | 3vp6 | A | 532 | ARG | p.Arg532Gln | Q99259 |
| HOMO SAPIENS | 5 | 1253 | 5xtd | S | 8 | GLY | p.Gly8Arg | O15239 |
| HOMO SAPIENS | 7 | 1253 | 5xtd | S | 32 | GLY | p.Gly32Arg | O15239 |
| HOMO SAPIENS | 7 | 1253 | 5xtd | S | 37 | ARG | p.Arg37Ser | O15239 |
| HOMO SAPIENS | -1 | 13292 | 3u1k | A | 121 | VAL | p.Ile121Val | Q8TCS8 |
| HOMO SAPIENS | 2 | 13292 | 3u1k | A | 165 | PRO | p.Pro165Ser | Q8TCS8 |
| HOMO SAPIENS | 2 | 13292 | 3u1k | A | 192 | ARG | p.Arg192Gln | Q8TCS8 |
| HOMO SAPIENS | 1 | 13292 | 3u1k | A | 315 | LYS | p.Lys315Arg | Q8TCS8 |
| HOMO SAPIENS | 4 | 13292 | 3u1k | A | 509 | VAL | p.Val509Ile | Q8TCS8 |
| HOMO SAPIENS | 7 | 13292 | 3u1k | A | 531 | THR | p.Thr531Arg | Q8TCS8 |
| HOMO SAPIENS | 0 | 13292 | 3u1k | A | 590 | ASN | p.Asn590Asp | Q8TCS8 |
| HOMO SAPIENS | 0 | 27048 | 4p5w | B | 184 | ARG | p.Arg184Ter | Q9UJS0 |
| HOMO SAPIENS | 0 | 27048 | 4p5w | B | 225 | SER | p.Ser225Ter | Q9UJS0 |
| HOMO SAPIENS | -1 | 27048 | 4p5w | B | 631 | ALA | p.Arg360Ter | Q9UJS0 |
| HOMO SAPIENS | 0 | 24240 | 4mdr | A | 318 | ARG | p.Arg318Ter | O00189 |
| HOMO SAPIENS | 0 | 24240 | 4mdr | A | 338 | ARG | p.Arg338Ter | O00189 |
| HOMO SAPIENS | 0 | 13429 | 4cg4 | A | 479 | PHE | p.Phe479Leu | O15553 |
| HOMO SAPIENS | 0 | 13429 | 4cg4 | A | 487 | VAL | p.Val487Leu | O15553 |
| HOMO SAPIENS | 3 | 13429 | 4cg4 | A | 653 | ARG | p.Arg653His | O15553 |
| HOMO SAPIENS | 1 | 13429 | 4cg4 | A | 680 | MET | p.Met680Ile | O15553 |
| HOMO SAPIENS | 1 | 13429 | 4cg4 | A | 688 | TYR | p.Tyr688Ter | O15553 |
| HOMO SAPIENS | 0 | 13429 | 4cg4 | A | 694 | MET | p.Met694Ile | O15553 |
| HOMO SAPIENS | 0 | 13429 | 4cg4 | A | 694 | MET | p.Met694Val | O15553 |
| HOMO SAPIENS | 1 | 13429 | 4cg4 | A | 726 | VAL | p.Val726Ala | O15553 |
| HOMO SAPIENS | 0 | 13429 | 4cg4 | A | 761 | ARG | p.Arg761His | O15553 |
| HOMO SAPIENS | 0 | 2079 | 4okh | A | 691 | GLY | p.Gly691Arg | P20807 |
| HOMO SAPIENS | 2 | 2079 | 4okh | A | 748 | ARG | p.Arg748Ter | P20807 |
| HOMO SAPIENS | 0 | 2079 | 4okh | A | 763 | TYR | p.Tyr763Cys | P20807 |
| HOMO SAPIENS | 1 | 2079 | 4okh | A | 769 | ARG | p.Arg769Gln | P20807 |
| HOMO SAPIENS | 9 | 2079 | 4okh | A | 777 | ILE | p.Ile777Val | P20807 |
| HOMO SAPIENS | 8 | 2079 | 4okh | A | 778 | ASP | p.Asp778His | P20807 |
| HOMO SAPIENS | 5 | 2079 | 4okh | A | 780 | ASP | p.Asp780His | P20807 |
| HOMO SAPIENS | 1 | 1200 | 2gi7 | A | 142 | ARG | p.Arg162Trp | Q9HCN6 |
| HOMO SAPIENS | 4 | 9199 | 3ezq | C | 260 | ASP | p.Asp260Val | P25445 |
| HOMO SAPIENS | 0 | 36522 | 5ms9 | A | 122 | ARG | p.Arg122Cys | P35555 |
| HOMO SAPIENS | 0 | 36522 | 5ms9 | A | 240 | ARG | p.Arg240Cys | P35555 |
| HOMO SAPIENS | 0 | 17448 | 3g2f | A | 225 | ARG | p.Arg225His | Q13873 |
| HOMO SAPIENS | 0 | 17448 | 3g2f | A | 347 | CYS | p.Cys347Tyr | Q13873 |
| HOMO SAPIENS | 3 | 17448 | 3g2f | A | 485 | ASP | p.Asp485Gly | Q13873 |
| HOMO SAPIENS | 7 | 17448 | 3g2f | A | 491 | ARG | p.Arg491Gln | Q13873 |
| HOMO SAPIENS | 7 | 17448 | 3g2f | A | 491 | ARG | p.Arg491Trp | Q13873 |

| | | | | | | | | |
|---|---|---|---|---|---|---|---|---|
| HOMO SAPIENS | 0 | 4861 | 4ux8 | A | 56 | LEU | p.Leu56Met | P07949 |
| HOMO SAPIENS | 1 | 4861 | 4ux8 | A | 88 | ILE | p.Ile88Leu | P07949 |
| HOMO SAPIENS | 0 | 4861 | 4ux8 | A | 114 | ARG | p.Arg114His | P07949 |
| HOMO SAPIENS | 5 | 4861 | 4ux8 | A | 180 | ARG | p.Arg180Gln | P07949 |
| HOMO SAPIENS | 0 | 4861 | 4ux8 | A | 208 | GLU | p.Glu208Asp | P07949 |
| HOMO SAPIENS | 0 | 4861 | 4ux8 | A | 373 | ALA | p.Ala373Val | P07949 |
| HOMO SAPIENS | 0 | 4861 | 4ux8 | A | 375 | LEU | p.Leu375Gln | P07949 |
| HOMO SAPIENS | -1 | 4861 | 4ux8 | A | 386 | ALA | p.Ala386Val | P07949 |
| HOMO SAPIENS | 0 | 4861 | 4ux8 | A | 446 | GLY | p.Gly446Arg | P07949 |
| HOMO SAPIENS | 0 | 4861 | 4ux8 | A | 480 | GLU | p.Glu480Lys | P07949 |
| HOMO SAPIENS | 0 | 4861 | 4ux8 | A | 489 | ASP | p.Asp489Asn | P07949 |
| HOMO SAPIENS | -1 | 4861 | 4ux8 | A | 510 | ALA | p.Ala510Val | P07949 |
| HOMO SAPIENS | -1 | 4861 | 4ux8 | A | 533 | GLY | p.Gly533Cys | P07949 |
| HOMO SAPIENS | -1 | 4861 | 4ux8 | A | 567 | ASP | p.Asp567Asn | P07949 |
| HOMO SAPIENS | -1 | 4861 | 4ux8 | A | 609 | CYS | p.Cys609Tyr | P07949 |
| HOMO SAPIENS | -1 | 4861 | 4ux8 | A | 609 | CYS | p.Cys609Gly | P07949 |
| HOMO SAPIENS | -1 | 4861 | 4ux8 | A | 611 | CYS | p.Cys611Trp | P07949 |
| HOMO SAPIENS | -1 | 4861 | 4ux8 | A | 618 | CYS | p.Cys618Gly | P07949 |
| HOMO SAPIENS | -1 | 4861 | 4ux8 | A | 618 | CYS | p.Cys618Phe | P07949 |
| HOMO SAPIENS | -1 | 4861 | 4ux8 | A | 620 | CYS | p.Cys620Gly | P07949 |
| HOMO SAPIENS | -1 | 4861 | 4ux8 | A | 620 | CYS | p.Cys620Ser | P07949 |
| HOMO SAPIENS | -1 | 4861 | 4ux8 | A | 620 | CYS | p.Cys620Trp | P07949 |
| HOMO SAPIENS | -1 | 4861 | 4ux8 | A | 623 | GLU | p.Glu623Lys | P07949 |
| HOMO SAPIENS | -1 | 4861 | 4ux8 | A | 631 | ASP | p.Asp631Gly | P07949 |
| HOMO SAPIENS | -1 | 4861 | 4ux8 | A | 631 | ASP | p.Asp631Glu | P07949 |
| HOMO SAPIENS | -1 | 4861 | 4ux8 | A | 634 | CYS | p.Cys634Gly | P07949 |
| HOMO SAPIENS | -1 | 4861 | 4ux8 | A | 634 | CYS | p.Cys634Phe | P07949 |
| HOMO SAPIENS | -1 | 4861 | 4ux8 | A | 634 | CYS | p.Cys634Trp | P07949 |
| HOMO SAPIENS | 0 | 36308 | 4nkq | B | 123 | PRO | p.Pro142Leu | P15509 |
| HOMO SAPIENS | 0 | 36308 | 4nkq | B | 145 | ARG | p.Arg164Gln | P15509 |
| HOMO SAPIENS | -1 | 36308 | 4nkq | B | 177 | GLY | p.Gly196Arg | P15509 |
| HOMO SAPIENS | 0 | 36006 | 1oqa | A | 107 | HIS | p.His1860Pro | P38398 |
| HOMO SAPIENS | 7 | 15394 | 2ozo | A | 465 | ARG | p.Arg465His | P43403 |
| HOMO SAPIENS | 7 | 15394 | 2ozo | A | 465 | ARG | p.Arg465Cys | P43403 |
| HOMO SAPIENS | 4 | 15394 | 2ozo | A | 518 | SER | p.Ser518Arg | P43403 |
| HOMO SAPIENS | 1 | 20813 | 3o5z | A | 19 | ARG | p.Arg19Trp | O00459 |
| HOMO SAPIENS | 2 | 4329 | 2bb5 | A | 5 | ILE | p.Ile23Val | P20062 |
| HOMO SAPIENS | 4 | 4329 | 2bb5 | A | 59 | LYS | p.Lys77Met | P20062 |
| HOMO SAPIENS | 4 | 4329 | 2bb5 | A | 197 | ARG | p.Arg215Trp | P20062 |
| HOMO SAPIENS | 1 | 4329 | 2bb5 | A | 241 | PRO | p.Arg259Pro | P20062 |
| HOMO SAPIENS | 2 | 4329 | 2bb5 | A | 315 | THR | p.Thr333Lys | P20062 |
| HOMO SAPIENS | 5 | 4329 | 2bb5 | A | 330 | SER | p.Ser348Phe | P20062 |
| HOMO SAPIENS | 4 | 4329 | 2bb5 | A | 358 | LEU | p.Leu376Ser | P20062 |
| HOMO SAPIENS | 5 | 4329 | 2bb5 | A | 381 | ARG | p.Arg399Gln | P20062 |
| HOMO SAPIENS | 4 | 18294 | 4r30 | A | 163 | ASN | p.Asn163Asp | O95278 |
| HOMO SAPIENS | 6 | 18294 | 4r30 | A | 227 | ALA | p.Ala227Val | O95278 |
| HOMO SAPIENS | 4 | 18294 | 4r30 | A | 241 | ARG | p.Arg241Ter | O95278 |
| HOMO SAPIENS | 4 | 18294 | 4r30 | A | 325 | ARG | p.Arg325His | O95278 |
| HOMO SAPIENS | 0 | 2632 | 4xem | A | 22 | GLU | p.Glu22Lys | P49588 |
| HOMO SAPIENS | 0 | 2632 | 4xem | A | 275 | GLY | p.Gly275Asp | P49588 |
| HOMO SAPIENS | 1 | 2632 | 4xem | A | 329 | ARG | p.Arg329His | P49588 |
| HOMO SAPIENS | 0 | 11431 | 5t4n | A | 284 | GLN | p.Gln305Glu | Q96QU1 |
| HOMO SAPIENS | 0 | 11431 | 5t4n | A | 315 | ARG | p.Arg336Ter | Q96QU1 |
| HOMO SAPIENS | 0 | 11431 | 5t4n | A | 322 | ARG | p.Arg343Lys | Q96QU1 |

| | | | | | | | | |
|---|---|---|---|---|---|---|---|---|
| HOMO SAPIENS | 1 | 11431 | 5t4n | A | 326 | LEU | p.Leu347Phe | Q96QU1 |
| HOMO SAPIENS | 0 | 11431 | 5t4n | A | 373 | TYR | p.Tyr394Cys | Q96QU1 |
| HOMO SAPIENS | 0 | 11431 | 5t4n | A | 398 | ASN | p.Asn419Ser | Q96QU1 |
| HOMO SAPIENS | 0 | 11431 | 5t4n | A | 414 | ALA | p.Asp435Ala | Q96QU1 |
| HOMO SAPIENS | 0 | 11431 | 5t4n | A | 433 | VAL | p.Val454Phe | Q96QU1 |
| HOMO SAPIENS | 1 | 11431 | 5t4n | A | 507 | VAL | p.Val528Asp | Q96QU1 |
| HOMO SAPIENS | 0 | 11431 | 5t4n | A | 547 | ALA | p.Ala568Ser | Q96QU1 |
| HOMO SAPIENS | 3 | 3194 | 5xtd | L | 84 | ARG | p.Arg84His | O43181 |
| HOMO SAPIENS | 1 | 3194 | 5xtd | L | 130 | THR | p.Thr130Ser | O43181 |
| HOMO SAPIENS | 8 | 3194 | 5xtd | L | 136 | SER | p.Ser136Ala | O43181 |
| HOMO SAPIENS | 0 | 40749 | 1nun | A | 106 | CYS | p.Cys106Phe | O15520 |
| HOMO SAPIENS | 9 | 40749 | 1nun | A | 156 | ILE | p.Ile156Arg | O15520 |
| HOMO SAPIENS | 1 | 40749 | 1nun | A | 193 | ARG | p.Arg193Ter | O15520 |
| HOMO SAPIENS | 0 | 40749 | 1nun | A | 207 | HIS | p.His207Pro | O15520 |
| HOMO SAPIENS | 0 | 7026 | 3n1f | D | 875 | SER | p.Ser875Gly | Q4KMG0 |
| HOMO SAPIENS | 6 | 23802 | 3rzv | A | 424 | ARG | p.Arg424Ter | O95630 |
| HOMO SAPIENS | 0 | 36720 | 1jm7 | A | 12 | GLN | p.Gln12Ter | P38398 |
| HOMO SAPIENS | 3 | 36720 | 1jm7 | A | 23 | GLU | p.Glu23Asp | P38398 |
| HOMO SAPIENS | 1 | 36720 | 1jm7 | A | 45 | LYS | p.Lys45Gln | P38398 |
| HOMO SAPIENS | 8 | 36720 | 1jm7 | A | 47 | CYS | p.Cys47Ter | P38398 |
| HOMO SAPIENS | 0 | 36720 | 1jm7 | A | 59 | SER | p.Ser59Ter | P38398 |
| HOMO SAPIENS | 8 | 36720 | 1jm7 | A | 61 | CYS | p.Cys61Gly | P38398 |
| HOMO SAPIENS | 8 | 36720 | 1jm7 | A | 63 | LEU | p.Leu63Ter | P38398 |
| HOMO SAPIENS | 7 | 36720 | 1jm7 | A | 64 | CYS | p.Cys64Gly | P38398 |
| HOMO SAPIENS | 0 | 36720 | 1jm7 | A | 101 | TYR | p.Tyr101Ter | P38398 |
| HOMO SAPIENS | 0 | 36720 | 1jm7 | A | null | TYR | p.Tyr105Cys | P38398 |
| HOMO SAPIENS | 0 | 28500 | 3kuz | A | 1217 | CYS | p.Cys1217Arg | O60486 |
| HOMO SAPIENS | 5 | 8097 | 3p56 | C | 69 | ARG | p.Arg69Trp | Q8TDP1 |
| HOMO SAPIENS | 5 | 8097 | 3p56 | C | 143 | LYS | p.Lys143Ile | Q8TDP1 |
| HOMO SAPIENS | 2 | 30073 | 3nmz | A | 380 | ARG | p.Arg380Gln | P25054 |
| HOMO SAPIENS | 1 | 22573 | 1cjy | A | 111 | SER | p.Ser111Pro | P47712 |
| HOMO SAPIENS | 0 | 22573 | 1cjy | A | 485 | ARG | p.Arg485His | P47712 |
| HOMO SAPIENS | 2 | 965 | 3E+077 | A | 99 | ALA | p.Ala99Val | Q9Y617 |
| HOMO SAPIENS | 2 | 965 | 3E+077 | A | 100 | ASP | p.Asp100Ala | Q9Y617 |
| HOMO SAPIENS | 4 | 1503 | 3kxc | A | 62 | ARG | p.Arg62Trp | O43617 |
| HOMO SAPIENS | 6 | 9418 | 1hdr | A | 22 | GLY | p.Gly23Asp | P09417 |
| HOMO SAPIENS | 2 | 9418 | 1hdr | A | 35 | TRP | p.Trp36Arg | P09417 |
| HOMO SAPIENS | 2 | 9418 | 1hdr | A | 107 | TRP | p.Trp108Gly | P09417 |
| HOMO SAPIENS | 6 | 9418 | 1hdr | A | 149 | TYR | p.Tyr150Cys | P09417 |
| HOMO SAPIENS | -1 | 12079 | 3tnu | B | 373 | GLY | p.Gly373Ala | P13647 |
| HOMO SAPIENS | 1 | 12079 | 3tnu | B | 463 | LEU | p.Leu463Pro | P13647 |
| HOMO SAPIENS | 1 | 12079 | 3tnu | B | 471 | ARG | p.Arg471Cys | P13647 |
| HOMO SAPIENS | 0 | 8135 | 3rbn | B | 89 | SER | p.Ser556Ile | P40692 |
| HOMO SAPIENS | 0 | 8135 | 3rbn | B | 111 | GLU | p.Glu578Gly | P40692 |
| HOMO SAPIENS | 2 | 8135 | 3rbn | B | 114 | PRO | p.Pro581Leu | P40692 |
| HOMO SAPIENS | 0 | 8135 | 3rbn | B | 133 | GLU | p.Glu600Gly | P40692 |
| HOMO SAPIENS | 2 | 8135 | 3rbn | B | 136 | PRO | p.Pro603Arg | P40692 |
| HOMO SAPIENS | 2 | 8135 | 3rbn | B | 136 | PRO | p.Pro603Leu | P40692 |
| HOMO SAPIENS | 7 | 8135 | 3rbn | B | 140 | LEU | p.Leu607His | P40692 |
| HOMO SAPIENS | 2 | 8135 | 3rbn | B | 151 | LYS | p.Lys618Ala | P40692 |
| HOMO SAPIENS | 2 | 8135 | 3rbn | B | 151 | LYS | p.Lys618Thr | P40692 |
| HOMO SAPIENS | 5 | 8135 | 3rbn | B | 181 | PRO | p.Pro648Ser | P40692 |
| HOMO SAPIENS | 2 | 8135 | 3rbn | B | 188 | ILE | p.Ile655Val | P40692 |
| HOMO SAPIENS | 2 | 8135 | 3rbn | B | 214 | ALA | p.Ala681Thr | P40692 |

| | | | | | | | | |
|---|---|---|---|---|---|---|---|---|
| HOMO SAPIENS | 2 | 8135 | 3rbn | B | 220 | ARG | p.Arg687Trp | P40692 |
| HOMO SAPIENS | 2 | 8135 | 3rbn | B | 222 | GLN | p.Gln689Arg | P40692 |
| HOMO SAPIENS | -1 | 8135 | 3rbn | B | 236 | GLN | p.Gln701Ter | P40692 |
| HOMO SAPIENS | 7 | 8135 | 3rbn | B | 251 | HIS | p.His718Tyr | P40692 |
| HOMO SAPIENS | 0 | 8135 | 3rbn | B | 284 | LYS | p.Lys751Arg | P40692 |
| HOMO SAPIENS | 1 | 21998 | 5d68 | A | 355 | LEU | p.Leu355Phe | O00522 |
| HOMO SAPIENS | 3 | 21998 | 5d68 | A | 380 | THR | p.Thr380Met | O00522 |
| HOMO SAPIENS | 0 | 21998 | 5d68 | A | 415 | ILE | p.Ile415Met | O00522 |
| HOMO SAPIENS | 0 | 33789 | 2m0c | A | 13 | ARG | p.Arg218Gln | Q9H161 |
| HOMO SAPIENS | 7 | 33789 | 2m0c | A | 52 | ARG | p.Arg257Thr | Q9H161 |
| HOMO SAPIENS | 0 | 33789 | 2m0c | A | 67 | ARG | p.Arg272Pro | Q9H161 |
| HOMO SAPIENS | 6 | 8263 | 2osg | A | 51 | GLY | p.Gly355Arg | Q9UDY2 |
| HOMO SAPIENS | 5 | 8263 | 2osg | A | 60 | THR | p.Thr364Met | Q9UDY2 |
| HOMO SAPIENS | 1 | 2475 | 5E+026 | B | 234 | THR | p.Thr234Ala | Q9BZ23 |
| HOMO SAPIENS | 2 | 2475 | 5E+026 | B | 264 | ARG | p.Arg264Trp | Q9BZ23 |
| HOMO SAPIENS | 4 | 2475 | 5E+026 | B | 286 | ARG | p.Arg286Cys | Q9BZ23 |
| HOMO SAPIENS | 0 | 2475 | 5E+026 | B | 357 | ARG | p.Arg357Trp | Q9BZ23 |
| HOMO SAPIENS | 2 | 2475 | 5E+026 | B | 481 | ARG | p.Arg481Ter | Q9BZ23 |
| HOMO SAPIENS | 8 | 2475 | 5E+026 | B | 521 | GLY | p.Gly521Arg | Q9BZ23 |
| HOMO SAPIENS | 2 | 2475 | 5E+026 | B | 528 | THR | p.Thr528Met | Q9BZ23 |
| HOMO SAPIENS | 1 | 3742 | 5xtd | E | 108 | HIS | p.His134Tyr | P56556 |
| HOMO SAPIENS | 1 | 16725 | 3g04 | C | 36 | ASP | p.Asp36His | P16473 |
| HOMO SAPIENS | 3 | 16725 | 3g04 | C | 52 | PRO | p.Pro52Thr | P16473 |
| HOMO SAPIENS | 1 | 16725 | 3g04 | C | 162 | PRO | p.Pro162Ala | P16473 |
| HOMO SAPIENS | -1 | 12366 | 5d2d | C | 750 | PRO | p.Pro750Leu | P13569 |
| HOMO SAPIENS | 0 | 12366 | 5d2d | C | 764 | ARG | p.Arg764Ter | P13569 |
| HOMO SAPIENS | 0 | 34845 | 2y23 | A | 1151 | ASP | p.Asp1151Glu | P52179 |
| HOMO SAPIENS | 0 | 34845 | 2y23 | A | 1180 | ASP | p.Asp1180Ala | P52179 |
| HOMO SAPIENS | 6 | 36740 | 3lbx | B | 2053 | ALA | p.Ala2053Pro | P11277 |
| HOMO SAPIENS | 0 | 19289 | 3ihj | A | 404 | ARG | p.Arg404Ter | Q8TD30 |
| HOMO SAPIENS | 5 | 1034 | 5xtd | Q | 228 | ARG | p.Arg228Gln | O75306 |
| HOMO SAPIENS | 5 | 1034 | 5xtd | Q | 229 | PRO | p.Pro229Gln | O75306 |
| HOMO SAPIENS | 4 | 1034 | 5xtd | Q | 323 | ARG | p.Arg323Gln | O75306 |
| HOMO SAPIENS | 4 | 1034 | 5xtd | Q | 352 | PRO | p.Pro352Ala | O75306 |
| HOMO SAPIENS | 5 | 1034 | 5xtd | Q | 413 | SER | p.Ser413Pro | O75306 |
| HOMO SAPIENS | 1 | 2772 | 2vaf | A | 1047 | THR | p.Thr66Ala | O14958 |
| HOMO SAPIENS | 5 | 2772 | 2vaf | A | 1057 | VAL | p.Val76Met | O14958 |
| HOMO SAPIENS | 0 | 2772 | 2vaf | A | 1148 | LEU | p.Leu167His | O14958 |
| HOMO SAPIENS | 1 | 2772 | 2vaf | A | 1149 | ILE | p.Ile168Thr | O14958 |
| HOMO SAPIENS | 8 | 2772 | 2vaf | A | 1225 | HIS | p.His244Arg | O14958 |
| HOMO SAPIENS | 8 | 2772 | 2vaf | A | 1225 | HIS | p.His244Tyr | O14958 |
| HOMO SAPIENS | 7 | 2772 | 2vaf | A | 1251 | ILE | p.Ile270Thr | O14958 |
| HOMO SAPIENS | 3 | 2772 | 2vaf | A | 1288 | ASP | p.Asp307His | O14958 |
| HOMO SAPIENS | 5 | 2772 | 2vaf | A | 1289 | PRO | p.Pro308Leu | O14958 |
| HOMO SAPIENS | 1 | 31885 | 2aex | A | 132 | ALA | p.Ala132Val | P36551 |
| HOMO SAPIENS | 1 | 31885 | 2aex | A | 272 | ASN | p.Asn272His | P36551 |
| HOMO SAPIENS | 8 | 31885 | 2aex | A | 279 | GLY | p.Gly279Arg | P36551 |
| HOMO SAPIENS | 5 | 31885 | 2aex | A | 294 | VAL | p.Val294Ile | P36551 |
| HOMO SAPIENS | 5 | 31885 | 2aex | A | 295 | HIS | p.His295Asp | P36551 |
| HOMO SAPIENS | 1 | 31885 | 2aex | A | 331 | ARG | p.Arg331Trp | P36551 |
| HOMO SAPIENS | 3 | 31885 | 2aex | A | 352 | ARG | p.Arg352Cys | P36551 |
| HOMO SAPIENS | 1 | 31885 | 2aex | A | 404 | LYS | p.Lys404Glu | P36551 |
| HOMO SAPIENS | 2 | 31885 | 2aex | A | 447 | ARG | p.Arg447Cys | P36551 |
| HOMO SAPIENS | 5 | 11721 | 2r3v | A | 20 | HIS | p.His20Pro | Q03426 |

| | | | | | | | | |
|---|---|---|---|---|---|---|---|---|
| HOMO SAPIENS | 1 | 11721 | 2r3v | A | 116 | TYR | p.Tyr116His | Q03426 |
| HOMO SAPIENS | 1 | 11721 | 2r3v | A | 148 | ALA | p.Ala148Thr | Q03426 |
| HOMO SAPIENS | 1 | 11721 | 2r3v | A | 167 | PRO | p.Pro167Leu | Q03426 |
| HOMO SAPIENS | 6 | 11721 | 2r3v | A | 203 | VAL | p.Val203Ala | Q03426 |
| HOMO SAPIENS | 2 | 11721 | 2r3v | A | 215 | ARG | p.Arg215Ter | Q03426 |
| HOMO SAPIENS | 2 | 11721 | 2r3v | A | 237 | THR | p.Thr237Ser | Q03426 |
| HOMO SAPIENS | 0 | 11721 | 2r3v | A | 268 | ILE | p.Ile268Thr | Q03426 |
| HOMO SAPIENS | 0 | 11721 | 2r3v | A | 268 | ILE | p.Ile268Val | Q03426 |
| HOMO SAPIENS | 0 | 11721 | 2r3v | A | 301 | ASN | p.Asn301Ser | Q03426 |
| HOMO SAPIENS | 6 | 11721 | 2r3v | A | 310 | VAL | p.Val310Met | Q03426 |
| HOMO SAPIENS | 5 | 11721 | 2r3v | A | 334 | ALA | p.Ala334Thr | Q03426 |
| HOMO SAPIENS | 2 | 11721 | 2r3v | A | 377 | VAL | p.Val377Ile | Q03426 |
| HOMO SAPIENS | 1 | 11721 | 2r3v | A | 386 | ASP | p.Asp386Asn | Q03426 |
| HOMO SAPIENS | 1 | 11721 | 2r3v | A | 388 | ARG | p.Arg388Ter | Q03426 |
| HOMO SAPIENS | 8 | 5010 | 2hgs | A | 125 | ARG | p.Arg125Cys | P48637 |
| HOMO SAPIENS | 1 | 5010 | 2hgs | A | 164 | ARG | p.Arg164Gln | P48637 |
| HOMO SAPIENS | 0 | 5010 | 2hgs | A | 219 | ASP | p.Asp219Gly | P48637 |
| HOMO SAPIENS | 7 | 5010 | 2hgs | A | 267 | ARG | p.Arg267Trp | P48637 |
| HOMO SAPIENS | 0 | 5010 | 2hgs | A | 283 | ARG | p.Arg283Cys | P48637 |
| HOMO SAPIENS | 0 | 6892 | 5erd | B | 7 | VAL | p.Val56Met | Q14126 |
| HOMO SAPIENS | 1 | 6892 | 5erd | B | 25 | HIS | p.His74Arg | Q14126 |
| HOMO SAPIENS | 0 | 6892 | 5erd | B | 40 | TYR | p.Tyr89Cys | Q14126 |
| HOMO SAPIENS | 0 | 6892 | 5erd | B | 97 | ARG | p.Arg146Leu | Q14126 |
| HOMO SAPIENS | 7 | 6892 | 5erd | B | 217 | ASN | p.Asn266Ser | Q14126 |
| HOMO SAPIENS | 0 | 6892 | 5erd | B | 243 | ARG | p.Arg292His | Q14126 |
| HOMO SAPIENS | 0 | 6892 | 5erd | B | 244 | ILE | p.Ile293Val | Q14126 |
| HOMO SAPIENS | 0 | 6892 | 5erd | B | 343 | VAL | p.Val392Ile | Q14126 |
| HOMO SAPIENS | 4 | 6892 | 5erd | B | 350 | GLU | p.Glu399Lys | Q14126 |
| HOMO SAPIENS | 0 | 6892 | 5erd | B | 410 | TYR | p.Tyr459Cys | Q14126 |
| HOMO SAPIENS | 0 | 6892 | 5erd | B | 466 | VAL | p.Val515Ile | Q14126 |
| HOMO SAPIENS | 0 | 6892 | 5erd | B | 482 | PHE | p.Phe531Cys | Q14126 |
| HOMO SAPIENS | 0 | 6892 | 5erd | B | 484 | VAL | p.Val533Ile | Q14126 |
| HOMO SAPIENS | 0 | 6892 | 5erd | B | 545 | LEU | p.Leu594Pro | Q14126 |
| HOMO SAPIENS | 0 | 1338 | 2ax4 | A | 58 | THR | p.Thr48Arg | O95340 |
| HOMO SAPIENS | 7 | 8675 | 2zkm | X | 385 | LYS | p.Lys385Arg | Q00722 |
| HOMO SAPIENS | 1 | 11769 | 2wya | B | 54 | VAL | p.Val54Met | P54868 |
| HOMO SAPIENS | 1 | 11769 | 2wya | B | 116 | ARG | p.Arg116Cys | P54868 |
| HOMO SAPIENS | 5 | 11769 | 2wya | B | 167 | TYR | p.Tyr167Cys | P54868 |
| HOMO SAPIENS | 1 | 11769 | 2wya | B | 174 | PHE | p.Phe174Leu | P54868 |
| HOMO SAPIENS | 0 | 11769 | 2wya | B | 212 | GLY | p.Gly212Arg | P54868 |
| HOMO SAPIENS | 1 | 11769 | 2wya | B | 288 | ARG | p.Arg288Ter | P54868 |
| HOMO SAPIENS | 0 | 11769 | 2wya | B | 375 | SER | p.Ser375Cys | P54868 |
| HOMO SAPIENS | 4 | 11769 | 2wya | B | 500 | ARG | p.Arg500His | P54868 |
| HOMO SAPIENS | 0 | 3102 | 2iwo | A | 1880 | ARG | p.Arg1908Lys | O75970 |
| HOMO SAPIENS | 6 | 14296 | 2m9w | A | 38 | GLY | p.Gly296Ser | P43694 |
| HOMO SAPIENS | 0 | 14296 | 2m9w | A | 58 | GLN | p.Gln316Glu | P43694 |
| HOMO SAPIENS | 2 | 704 | 5ekd | B | 239 | ARG | p.Arg239Ter | Q9UGM6 |
| HOMO SAPIENS | 2 | 704 | 5ekd | B | 264 | TYR | p.Tyr264Cys | Q9UGM6 |
| HOMO SAPIENS | 6 | 2228 | 1ey2 | A | 47 | SER | p.Ser47Ter | Q93099 |
| HOMO SAPIENS | 6 | 2228 | 1ey2 | A | 48 | ALA | p.Ala48Ser | Q93099 |
| HOMO SAPIENS | 6 | 2228 | 1ey2 | A | 53 | ARG | p.Arg53Gln | Q93099 |
| HOMO SAPIENS | 3 | 2228 | 1ey2 | A | 80 | HIS | p.Gln80His | Q93099 |
| HOMO SAPIENS | 0 | 2228 | 1ey2 | A | 120 | CYS | p.Cys120Trp | Q93099 |
| HOMO SAPIENS | 3 | 2228 | 1ey2 | A | 122 | ALA | p.Ala122Val | Q93099 |

| | | | | | | | | |
|---|---|---|---|---|---|---|---|---|
| HOMO SAPIENS | 3 | 2228 | 1ey2 | A | 161 | GLY | p.Gly161Arg | Q93099 |
| HOMO SAPIENS | 6 | 2228 | 1ey2 | A | 225 | ARG | p.Arg225His | Q93099 |
| HOMO SAPIENS | 0 | 2228 | 1ey2 | A | 230 | PRO | p.Pro230Ser | Q93099 |
| HOMO SAPIENS | 6 | 2228 | 1ey2 | A | 270 | GLY | p.Gly270Arg | Q93099 |
| HOMO SAPIENS | 6 | 2228 | 1ey2 | A | 330 | ARG | p.Arg330Ser | Q93099 |
| HOMO SAPIENS | -1 | 2228 | 1ey2 | A | 368 | MSE | p.Met368Val | Q93099 |
| HOMO SAPIENS | 6 | 2228 | 1ey2 | A | 371 | HIS | p.His371Arg | Q93099 |
| HOMO SAPIENS | 6 | 2228 | 1ey2 | A | 401 | GLU | p.Glu401Gln | Q93099 |
| HOMO SAPIENS | 5 | 34648 | 2k27 | A | 31 | ARG | p.Arg31His | Q06710 |
| HOMO SAPIENS | 7 | 34648 | 2k27 | A | 40 | GLN | p.Gln40Pro | Q06710 |
| HOMO SAPIENS | 5 | 34648 | 2k27 | A | 57 | CYS | p.Cys57Tyr | Q06710 |
| HOMO SAPIENS | 5 | 34648 | 2k27 | A | 62 | LEU | p.Leu62Arg | Q06710 |
| HOMO SAPIENS | 1 | 21399 | 3edu | A | 1819 | GLU | p.Glu1819Ter | P11277 |
| HOMO SAPIENS | 1 | 18153 | 1oey | J | 245 | TYR | p.Tyr245Ter | Q15080 |
| HOMO SAPIENS | 2 | 41567 | 3sw0 | X | 1050 | ASN | p.Asn1050Tyr | P08603 |
| HOMO SAPIENS | 5 | 41567 | 3sw0 | X | 1059 | ILE | p.Ile1059Thr | P08603 |
| HOMO SAPIENS | 0 | 36362 | 4wrm | A | 362 | HIS | p.His362Arg | P07333 |
| HOMO SAPIENS | -1 | 36362 | 4wrm | A | 413 | GLY | p.Gly413Ser | P07333 |
| HOMO SAPIENS | 0 | 31255 | 2w2j | A | 100 | SER | p.Ser100Pro | P35219 |
| HOMO SAPIENS | 0 | 31255 | 2w2j | A | 100 | SER | p.Ser100Leu | P35219 |
| HOMO SAPIENS | 5 | 31255 | 2w2j | A | 237 | ARG | p.Arg237Gln | P35219 |
| HOMO SAPIENS | -1 | 26450 | 1th1 | C | 1395 | SER | p.Ser1395Cys | P25054 |
| HOMO SAPIENS | -1 | 26450 | 1th1 | C | 1413 | MET | p.Met1413Val | P25054 |
| HOMO SAPIENS | -1 | 26450 | 1th1 | C | 1446 | ALA | p.Ala1446Thr | P25054 |
| HOMO SAPIENS | -1 | 26450 | 1th1 | C | 1454 | LYS | p.Lys1454Glu | P25054 |
| HOMO SAPIENS | 0 | 26450 | 1th1 | C | 1474 | ALA | p.Ala1474Thr | P25054 |
| HOMO SAPIENS | 1 | 981 | 5xtb | F | 14 | LEU | p.Leu14Val | O43678 |
| HOMO SAPIENS | 1 | 11069 | 3kq4 | A | 46 | SER | p.Ser46Leu | P27352 |
| HOMO SAPIENS | 1 | 11069 | 3kq4 | A | 97 | MET | p.Met97Thr | P27352 |
| HOMO SAPIENS | 1 | 11069 | 3kq4 | A | 255 | ASN | p.Asn255Ser | P27352 |
| HOMO SAPIENS | 1 | 11069 | 3kq4 | A | 330 | ASN | p.Asn330Lys | P27352 |
| HOMO SAPIENS | 0 | 41489 | 3erb | A | 53 | PRO | p.Pro73Leu | P06681 |
| HOMO SAPIENS | 6 | 8715 | 2bq8 | X | 23 | THR | p.Thr44Met | P13686 |
| HOMO SAPIENS | 0 | 8715 | 2bq8 | X | 88 | GLY | p.Gly109Arg | P13686 |
| HOMO SAPIENS | 6 | 8715 | 2bq8 | X | 127 | VAL | p.Val148Met | P13686 |
| HOMO SAPIENS | 2 | 8715 | 2bq8 | X | 179 | VAL | p.Val200Met | P13686 |
| HOMO SAPIENS | 6 | 8715 | 2bq8 | X | 251 | ARG | p.Arg272Cys | P13686 |
| HOMO SAPIENS | 7 | 8112 | 3p56 | A | 37 | GLY | p.Gly37Ser | O75792 |
| HOMO SAPIENS | -1 | 8112 | 3p56 | A | 202 | LEU | p.Leu202Ser | O75792 |
| HOMO SAPIENS | 1 | 8112 | 3p56 | A | 205 | ASP | p.Asp205Glu | O75792 |
| HOMO SAPIENS | 4 | 8112 | 3p56 | A | 212 | ASN | p.Asn212Ile | O75792 |
| HOMO SAPIENS | 0 | 8112 | 3p56 | A | 235 | ARG | p.Arg235Gln | O75792 |
| HOMO SAPIENS | 4 | 8112 | 3p56 | A | 240 | THR | p.Thr240Met | O75792 |
| HOMO SAPIENS | 0 | 8112 | 3p56 | A | 291 | ARG | p.Arg291His | O75792 |
| HOMO SAPIENS | 0 | 37603 | 2yqg | A | 10 | ARG | p.Arg49His | Q14126 |
| HOMO SAPIENS | 0 | 21178 | 3k1r | B | 458 | ASP | p.Asp458Val | Q495M9 |
| HOMO SAPIENS | -1 | 36828 | 4fq3 | B | 507 | GLY | p.Gly507Asp | P35637 |
| HOMO SAPIENS | 5 | 36828 | 4fq3 | B | 517 | HIS | p.His517Gln | P35637 |
| HOMO SAPIENS | 8 | 36828 | 4fq3 | B | 518 | ARG | p.Arg518Lys | P35637 |
| HOMO SAPIENS | 6 | 36828 | 4fq3 | B | 521 | ARG | p.Arg521Cys | P35637 |
| HOMO SAPIENS | 6 | 36828 | 4fq3 | B | 521 | ARG | p.Arg521His | P35637 |
| HOMO SAPIENS | 0 | 6363 | 3j7y | c | null | HIS | p.His13Arg | Q9H9J2 |
| HOMO SAPIENS | 2 | 6363 | 3j7y | c | 105 | ARG | p.Arg105Cys | Q9H9J2 |
| HOMO SAPIENS | 2 | 6363 | 3j7y | c | 138 | THR | p.Thr138Ile | Q9H9J2 |

| | | | | | | | | |
|---|---|---|---|---|---|---|---|---|
| HOMO SAPIENS | 1 | 6363 | 3j7y | c | 156 | LEU | p.Leu156Arg | Q9H9J2 |
| HOMO SAPIENS | 1 | 6782 | 5l46 | A | 109 | HIS | p.His109Arg | Q9UI17 |
| HOMO SAPIENS | 0 | 23951 | 5t76 | A | 175 | GLY | p.Gly931Ser | P49588 |
| HOMO SAPIENS | 8 | 12604 | 3p0l | A | 182 | ARG | p.Arg182Leu | P49675 |
| HOMO SAPIENS | 2 | 12604 | 3p0l | A | 187 | VAL | p.Val187Met | P49675 |
| HOMO SAPIENS | 2 | 12604 | 3p0l | A | 188 | ARG | p.Arg188Cys | P49675 |
| HOMO SAPIENS | 0 | 12604 | 3p0l | A | 193 | ARG | p.Arg193Ter | P49675 |
| HOMO SAPIENS | 9 | 12604 | 3p0l | A | 217 | ARG | p.Arg217Thr | P49675 |
| HOMO SAPIENS | 4 | 12604 | 3p0l | A | 218 | ALA | p.Ala218Val | P49675 |
| HOMO SAPIENS | 0 | 12604 | 3p0l | A | 250 | TRP | p.Trp250Ter | P49675 |
| HOMO SAPIENS | 1 | 12604 | 3p0l | A | 258 | GLN | p.Gln258Ter | P49675 |
| HOMO SAPIENS | -1 | 27765 | 4wkr | A | 4 | GLU | p.Glu4Lys | Q4G0J3 |
| HOMO SAPIENS | 2 | 3587 | 4kit | C | 2301 | PRO | p.Pro2301Thr | Q6P2Q9 |
| HOMO SAPIENS | 3 | 3587 | 4kit | C | 2304 | PHE | p.Phe2304Leu | Q6P2Q9 |
| HOMO SAPIENS | 3 | 3587 | 4kit | C | 2309 | HIS | p.His2309Pro | Q6P2Q9 |
| HOMO SAPIENS | 3 | 3587 | 4kit | C | 2310 | ARG | p.Arg2310Lys | Q6P2Q9 |
| HOMO SAPIENS | 0 | 9073 | 1n69 | B | 21 | ASN | p.Asn215His | P07602 |
| HOMO SAPIENS | 0 | 9073 | 1n69 | B | 23 | THR | p.Thr217Ile | P07602 |
| HOMO SAPIENS | 0 | 9073 | 1n69 | B | 47 | CYS | p.Cys241Ser | P07602 |
| HOMO SAPIENS | 0 | 475 | 2vd5 | A | 225 | THR | p.Thr225Met | Q09013 |
| HOMO SAPIENS | 1 | 9066 | 4mzv | A | 31 | ASN | p.Asn31Lys | P16422 |
| HOMO SAPIENS | 1 | 9066 | 4mzv | A | 47 | GLN | p.Gln47His | P16422 |
| HOMO SAPIENS | 3 | 9066 | 4mzv | A | 66 | CYS | p.Cys66Tyr | P16422 |
| HOMO SAPIENS | 1 | 9066 | 4mzv | A | 102 | SER | p.Ser102Gly | P16422 |
| HOMO SAPIENS | 5 | 9066 | 4mzv | A | 115 | THR | p.Met115Thr | P16422 |
| HOMO SAPIENS | 1 | 9066 | 4mzv | A | 138 | ARG | p.Arg138Ter | P16422 |
| HOMO SAPIENS | 1 | 9066 | 4mzv | A | 153 | ARG | p.Arg153Thr | P16422 |
| HOMO SAPIENS | 1 | 9066 | 4mzv | A | 165 | ALA | p.Ala165Thr | P16422 |
| HOMO SAPIENS | 8 | 1827 | 2nz2 | A | 14 | GLY | p.Gly14Ser | P00966 |
| HOMO SAPIENS | 5 | 1827 | 2nz2 | A | 18 | SER | p.Ser18Leu | P00966 |
| HOMO SAPIENS | 3 | 1827 | 2nz2 | A | 86 | ARG | p.Arg86Cys | P00966 |
| HOMO SAPIENS | 3 | 1827 | 2nz2 | A | 86 | ARG | p.Arg86His | P00966 |
| HOMO SAPIENS | 1 | 1827 | 2nz2 | A | 179 | TRP | p.Trp179Arg | P00966 |
| HOMO SAPIENS | 7 | 1827 | 2nz2 | A | 180 | SER | p.Ser180Asn | P00966 |
| HOMO SAPIENS | 7 | 1827 | 2nz2 | A | 191 | GLU | p.Glu191Lys | P00966 |
| HOMO SAPIENS | 4 | 1827 | 2nz2 | A | 256 | GLU | p.Glu256Lys | P00966 |
| HOMO SAPIENS | 4 | 1827 | 2nz2 | A | 263 | VAL | p.Val263Met | P00966 |
| HOMO SAPIENS | 5 | 1827 | 2nz2 | A | 265 | ARG | p.Arg265Cys | P00966 |
| HOMO SAPIENS | 5 | 1827 | 2nz2 | A | 268 | ILE | p.Ile268Met | P00966 |
| HOMO SAPIENS | 6 | 1827 | 2nz2 | A | 269 | VAL | p.Val269Met | P00966 |
| HOMO SAPIENS | 7 | 1827 | 2nz2 | A | 272 | ARG | p.Arg272Cys | P00966 |
| HOMO SAPIENS | 3 | 1827 | 2nz2 | A | 304 | ARG | p.Arg304Trp | P00966 |
| HOMO SAPIENS | 3 | 1827 | 2nz2 | A | 307 | ARG | p.Arg307Cys | P00966 |
| HOMO SAPIENS | 3 | 1827 | 2nz2 | A | 310 | LYS | p.Lys310Gln | P00966 |
| HOMO SAPIENS | 2 | 1827 | 2nz2 | A | 362 | GLY | p.Gly362Val | P00966 |
| HOMO SAPIENS | 2 | 1827 | 2nz2 | A | 363 | ARG | p.Arg363Trp | P00966 |
| HOMO SAPIENS | 2 | 1827 | 2nz2 | A | 363 | ARG | p.Arg363Gln | P00966 |
| HOMO SAPIENS | 1 | 1827 | 2nz2 | A | 390 | GLY | p.Gly390Arg | P00966 |
| HOMO SAPIENS | 0 | 29524 | 5vbn | B | 2165 | ARG | p.Arg2165His | Q07864 |
| HOMO SAPIENS | 2 | 29524 | 5vbn | B | 2255 | ILE | p.Ile2255Phe | Q07864 |
| HOMO SAPIENS | 2 | 29524 | 5vbn | B | 2274 | LEU | p.Leu2274Val | Q07864 |
| HOMO SAPIENS | 0 | 31897 | 3t0o | A | 184 | CYS | p.Cys184Arg | O00584 |
| HOMO SAPIENS | -1 | 31897 | 3t0o | A | 215 | GLU | p.Glu215Lys | O00584 |
| HOMO SAPIENS | 0 | 31897 | 3t0o | A | 236 | ARG | p.Arg236Trp | O00584 |

| | | | | | | | | |
|---|---|---|---|---|---|---|---|---|
| HOMO SAPIENS | 7 | 18490 | 4pcw | B | 31 | GLU | p.Glu32Ter | P20930 |
| HOMO SAPIENS | -1 | 39865 | 2h9e | L | 152 | GLY | p.Gly192Arg | P00742 |
| HOMO SAPIENS | 2 | 4309 | 2x0b | F | 174 | THR | p.Thr207Met | P01019 |
| HOMO SAPIENS | 3 | 4309 | 2x0b | F | 342 | ARG | p.Arg375Gln | P01019 |
| HOMO SAPIENS | 2 | 3094 | 1zll | B | 9 | ARG | p.Arg9Cys | P26678 |
| HOMO SAPIENS | 9 | 3094 | 1zll | B | 39 | LEU | p.Leu39Ter | P26678 |
| HOMO SAPIENS | 5 | 3094 | 1zll | B | 49 | VAL | p.Val49Met | P26678 |
| HOMO SAPIENS | -1 | 41498 | 3ffd | P | 8 | LEU | p.Leu44Pro | P12272 |
| HOMO SAPIENS | 1 | 20697 | 1igr | A | 407 | ARG | p.Arg437His | P08069 |
| HOMO SAPIENS | 0 | 20697 | 1igr | A | 416 | MET | p.Met446Val | P08069 |
| HOMO SAPIENS | -1 | 20697 | 1igr | A | 462 | VAL | p.Val492Ile | P08069 |
| HOMO SAPIENS | 0 | 3973 | 4cbz | A | 37 | LEU | p.Leu37Ser | P78504 |
| HOMO SAPIENS | 0 | 3973 | 4cbz | A | 45 | VAL | p.Val45Leu | P78504 |
| HOMO SAPIENS | 0 | 3973 | 4cbz | A | 134 | LEU | p.Leu134Met | P78504 |
| HOMO SAPIENS | -1 | 3973 | 4cbz | A | 146 | VAL | p.Val146Ile | P78504 |
| HOMO SAPIENS | 0 | 3973 | 4cbz | A | 176 | VAL | p.Val176Ile | P78504 |
| HOMO SAPIENS | 0 | 3973 | 4cbz | A | 184 | ARG | p.Arg184Cys | P78504 |
| HOMO SAPIENS | 6 | 1697 | 4hmy | S | 33 | ARG | p.Arg33Trp | Q96PC3 |
| HOMO SAPIENS | 2 | 23752 | 5j4o | A | 13 | ARG | p.Arg1611Ser | P02549 |
| HOMO SAPIENS | 0 | 23752 | 5j4o | A | 42 | ALA | p.Ala1640Gly | P02549 |
| HOMO SAPIENS | 0 | 23752 | 5j4o | A | 108 | HIS | p.His1706Gln | P02549 |
| HOMO SAPIENS | 6 | 23752 | 5j4o | A | 206 | LEU | p.Leu1804Phe | P02549 |
| HOMO SAPIENS | 8 | 531 | 2qz4 | A | 349 | GLY | p.Gly349Ser | Q9UQ90 |
| HOMO SAPIENS | 0 | 531 | 2qz4 | A | 486 | ARG | p.Arg486Gln | Q9UQ90 |
| HOMO SAPIENS | 1 | 531 | 2qz4 | A | 510 | ALA | p.Ala510Val | Q9UQ90 |
| HOMO SAPIENS | 0 | 531 | 2qz4 | A | 529 | ALA | p.Ala529Val | Q9UQ90 |
| HOMO SAPIENS | 0 | 531 | 2qz4 | A | 558 | LYS | p.Lys558Ter | Q9UQ90 |
| HOMO SAPIENS | 1 | 531 | 2qz4 | A | 559 | LYS | p.Lys559Ter | Q9UQ90 |
| HOMO SAPIENS | 2 | 7736 | 1fnh | A | 114 | TRP | p.Trp1834Arg | P02751 |
| HOMO SAPIENS | 0 | 7736 | 1fnh | A | 163 | LEU | p.Leu1883Arg | P02751 |
| HOMO SAPIENS | 3 | 20337 | 5wy5 | B | 264 | LEU | p.Leu264Phe | Q96MG7 |
| HOMO SAPIENS | 8 | 36096 | 2jku | A | 643 | GLY | p.Gly668Arg | P05165 |
| HOMO SAPIENS | 6 | 36096 | 2jku | A | 663 | CYS | p.Cys688Arg | P05165 |
| HOMO SAPIENS | 0 | 9850 | 3kbt | A | 1606 | TYR | p.Tyr1606Ter | P11277 |
| HOMO SAPIENS | 4 | 6876 | 4jkq | A | 217 | ILE | p.Ile217Val | Q92736 |
| HOMO SAPIENS | 5 | 6876 | 4jkq | A | 420 | ARG | p.Arg420Trp | Q92736 |
| HOMO SAPIENS | 2 | 27603 | 1itv | B | 62 | PRO | p.Arg574Pro | P14780 |
| HOMO SAPIENS | 4 | 27603 | 1itv | B | 156 | ARG | p.Arg668Gln | P14780 |
| HOMO SAPIENS | 2 | 22954 | 4f52 | E | 48 | ILE | p.Ile48Met | Q92990 |
| HOMO SAPIENS | 0 | 22954 | 4f52 | E | 91 | ASP | p.Asp91Asn | Q92990 |
| HOMO SAPIENS | 0 | 22954 | 4f52 | E | 146 | ALA | p.Ala146Thr | Q92990 |
| HOMO SAPIENS | 0 | 22954 | 4f52 | E | 181 | ILE | p.Ile181Lys | Q92990 |
| HOMO SAPIENS | 0 | 22954 | 4f52 | E | 194 | ASN | p.Asn194Ser | Q92990 |
| HOMO SAPIENS | 2 | 22954 | 4f52 | E | 254 | PRO | p.Pro254Arg | Q92990 |
| HOMO SAPIENS | 2 | 22954 | 4f52 | E | 300 | ILE | p.Ile300Thr | Q92990 |
| HOMO SAPIENS | 2 | 22954 | 4f52 | E | 336 | LEU | p.Leu336Ser | Q92990 |
| HOMO SAPIENS | 0 | 22954 | 4f52 | E | 353 | TYR | p.Tyr353His | Q92990 |
| HOMO SAPIENS | 0 | 22954 | 4f52 | E | 387 | MET | p.Met387Val | Q92990 |
| HOMO SAPIENS | 4 | 2946 | 5gl7 | A | 344 | TYR | p.Tyr344Cys | P56192 |
| HOMO SAPIENS | 0 | 2946 | 5gl7 | A | 370 | PHE | p.Phe370Leu | P56192 |
| HOMO SAPIENS | 0 | 2946 | 5gl7 | A | 393 | ALA | p.Ala393Thr | P56192 |
| HOMO SAPIENS | 0 | 2946 | 5gl7 | A | 523 | ILE | p.Ile523Thr | P56192 |
| HOMO SAPIENS | 0 | 2946 | 5gl7 | A | 567 | SER | p.Ser567Leu | P56192 |
| HOMO SAPIENS | 0 | 2946 | 5gl7 | A | 800 | PRO | p.Pro800Thr | P56192 |

| | | | | | | | | |
|---|---|---|---|---|---|---|---|---|
| HOMO SAPIENS | 3 | 34961 | 4wy4 | C | 44 | TYR | p.Tyr44His | O95721 |
| HOMO SAPIENS | 8 | 34961 | 4wy4 | C | 89 | GLU | p.Glu89Lys | O95721 |
| HOMO SAPIENS | 0 | 9427 | 4chl | B | 21 | ALA | p.Ala21Ser | O95571 |
| HOMO SAPIENS | 2 | 9427 | 4chl | B | 62 | ALA | p.Ala62Thr | O95571 |
| HOMO SAPIENS | 0 | 9427 | 4chl | B | 114 | ILE | p.Ile114Phe | O95571 |
| HOMO SAPIENS | 2 | 9427 | 4chl | B | 159 | ARG | p.Arg159His | O95571 |
| HOMO SAPIENS | 8 | 9427 | 4chl | B | 163 | ARG | p.Arg163Trp | O95571 |
| HOMO SAPIENS | 8 | 9427 | 4chl | B | 163 | ARG | p.Arg163Gln | O95571 |
| HOMO SAPIENS | 2 | 9427 | 4chl | B | 185 | LEU | p.Leu185Arg | O95571 |
| HOMO SAPIENS | 0 | 40917 | 2e8o | A | 11 | PRO | p.Pro26Leu | Q9Y3Z3 |
| HOMO SAPIENS | -1 | 39190 | 4gow | A | 556 | TYR | p.Tyr556Ter | P56696 |
| HOMO SAPIENS | 0 | 2399 | 2mkv | A | 8 | GLY | p.Gly10Ser | P54710 |
| HOMO SAPIENS | 1 | 2399 | 2mkv | A | 24 | VAL | p.Val26Ile | P54710 |
| HOMO SAPIENS | 4 | 2399 | 2mkv | A | 39 | GLY | p.Gly41Arg | P54710 |
| HOMO SAPIENS | 0 | 20801 | 2zej | B | 1441 | ARG | p.Arg1441Cys | Q5S007 |
| HOMO SAPIENS | 0 | 20801 | 2zej | B | 1441 | ARG | p.Arg1441His | Q5S007 |
| HOMO SAPIENS | 5 | 1019 | 5xtb | M | 62 | ARG | p.Arg62Gln | P28331 |
| HOMO SAPIENS | 1 | 1019 | 5xtb | M | 85 | ALA | p.Ala85Thr | P28331 |
| HOMO SAPIENS | 5 | 1019 | 5xtb | M | 231 | LEU | p.Leu231Val | P28331 |
| HOMO SAPIENS | 2 | 1019 | 5xtb | M | 241 | ARG | p.Arg241Trp | P28331 |
| HOMO SAPIENS | 1 | 1019 | 5xtb | M | 252 | ASP | p.Asp252Gly | P28331 |
| HOMO SAPIENS | 1 | 1019 | 5xtb | M | 408 | ARG | p.Arg408Cys | P28331 |
| HOMO SAPIENS | 2 | 1019 | 5xtb | M | 417 | ARG | p.Arg417Ser | P28331 |
| HOMO SAPIENS | 1 | 1019 | 5xtb | M | 557 | ARG | p.Arg557Ter | P28331 |
| HOMO SAPIENS | 0 | 1019 | 5xtb | M | 567 | ILE | p.Ile567Val | P28331 |
| HOMO SAPIENS | 1 | 1019 | 5xtb | M | 595 | THR | p.Thr595Ala | P28331 |
| HOMO SAPIENS | 1 | 1019 | 5xtb | M | 703 | ALA | p.Ala703Thr | P28331 |
| HOMO SAPIENS | -1 | 3437 | 5hyn | A | 185 | ASP | p.Asp185His | Q15910 |
| HOMO SAPIENS | -1 | 3437 | 5hyn | A | 482 | ALA | p.Ala482Thr | Q15910 |
| HOMO SAPIENS | 0 | 8201 | 2wfd | A | 373 | TYR | p.Tyr373Cys | Q9P2J5 |
| HOMO SAPIENS | 1 | 3590 | 3edi | A | 90 | VAL | p.Val238Ala | O43897 |
| HOMO SAPIENS | 6 | 4555 | 1ytq | A | 64 | ALA | p.Ala65Ser | P43320 |
| HOMO SAPIENS | 0 | 4555 | 1ytq | A | 194 | TRP | p.Trp195Gly | P43320 |
| HOMO SAPIENS | 0 | 1013 | 2nn6 | C | 272 | SER | p.Ser272Thr | Q96B26 |
| HOMO SAPIENS | -1 | 29026 | 5m73 | D | 530 | ILE | p.Ile530Asn | O76094 |
| HOMO SAPIENS | 8 | 2252 | 1d5v | A | 57 | SER | p.Ser125Leu | Q99958 |
| HOMO SAPIENS | 1 | 4337 | 3ecr | A | 29 | ARG | p.Arg26His | P08397 |
| HOMO SAPIENS | 2 | 4337 | 3ecr | A | 34 | ALA | p.Ala31Thr | P08397 |
| HOMO SAPIENS | 2 | 4337 | 3ecr | A | 37 | GLN | p.Gln34Lys | P08397 |
| HOMO SAPIENS | 1 | 4337 | 3ecr | A | 58 | ALA | p.Ala55Ser | P08397 |
| HOMO SAPIENS | 0 | 4337 | 3ecr | A | 84 | LEU | p.Leu81Pro | P08397 |
| HOMO SAPIENS | 0 | 4337 | 3ecr | A | 114 | GLY | p.Gly111Arg | P08397 |
| HOMO SAPIENS | 1 | 4337 | 3ecr | A | 119 | ARG | p.Arg116Trp | P08397 |
| HOMO SAPIENS | 1 | 4337 | 3ecr | A | 152 | ARG | p.Arg149Gln | P08397 |
| HOMO SAPIENS | 1 | 4337 | 3ecr | A | 170 | ARG | p.Arg167Trp | P08397 |
| HOMO SAPIENS | 1 | 4337 | 3ecr | A | 176 | ARG | p.Arg173Gln | P08397 |
| HOMO SAPIENS | 0 | 4337 | 3ecr | A | 180 | LEU | p.Leu177Arg | P08397 |
| HOMO SAPIENS | 0 | 4337 | 3ecr | A | 181 | ASP | p.Asp178Asn | P08397 |
| HOMO SAPIENS | 1 | 4337 | 3ecr | A | 198 | ARG | p.Arg195Cys | P08397 |
| HOMO SAPIENS | 0 | 4337 | 3ecr | A | 201 | TRP | p.Trp198Ter | P08397 |
| HOMO SAPIENS | 1 | 4337 | 3ecr | A | 204 | ARG | p.Arg201Trp | P08397 |
| HOMO SAPIENS | 1 | 4337 | 3ecr | A | 219 | GLY | p.Gly216Asp | P08397 |
| HOMO SAPIENS | 1 | 4337 | 3ecr | A | 226 | GLU | p.Glu223Lys | P08397 |
| HOMO SAPIENS | 0 | 4337 | 3ecr | A | 248 | LEU | p.Leu245Arg | P08397 |

| | | | | | | | | |
|---|---|---|---|---|---|---|---|---|
| HOMO SAPIENS | 1 | 4337 | 3ecr | A | 250 | CYS | p.Cys247Arg | P08397 |
| HOMO SAPIENS | 0 | 4337 | 3ecr | A | 253 | GLU | p.Glu250Lys | P08397 |
| HOMO SAPIENS | 0 | 4337 | 3ecr | A | 255 | ALA | p.Ala252Thr | P08397 |
| HOMO SAPIENS | 0 | 4337 | 3ecr | A | 255 | ALA | p.Ala252Val | P08397 |
| HOMO SAPIENS | 0 | 4337 | 3ecr | A | 259 | HIS | p.His256Asn | P08397 |
| HOMO SAPIENS | 0 | 4337 | 3ecr | A | 324 | ARG | p.Arg321His | P08397 |
| HOMO SAPIENS | -1 | 4337 | 3ecr | A | 362 | ASP | p.Asp359Asn | P08397 |
| HOMO SAPIENS | 1 | 1204 | 5lwv | A | 854 | ILE | p.Ala864Thr | O15294 |
| HOMO SAPIENS | 1 | 25016 | 1m4u | A | 35 | PRO | p.Pro35Arg | Q13253 |
| HOMO SAPIENS | 1 | 25016 | 1m4u | A | 35 | PRO | p.Pro35Ser | Q13253 |
| HOMO SAPIENS | 1 | 25016 | 1m4u | A | 167 | ARG | p.Arg167Gly | Q13253 |
| HOMO SAPIENS | 1 | 25016 | 1m4u | A | 184 | CYS | p.Cys184Tyr | Q13253 |
| HOMO SAPIENS | 0 | 25016 | 1m4u | A | 189 | GLY | p.Gly189Cys | Q13253 |
| HOMO SAPIENS | 1 | 25016 | 1m4u | A | 204 | ARG | p.Arg204Leu | Q13253 |
| HOMO SAPIENS | 1 | 25016 | 1m4u | A | 205 | TRP | p.Trp205Cys | Q13253 |
| HOMO SAPIENS | 1 | 25016 | 1m4u | A | 217 | TRP | p.Trp217Gly | Q13253 |
| HOMO SAPIENS | 0 | 25016 | 1m4u | A | 222 | TYR | p.Tyr222Cys | Q13253 |
| HOMO SAPIENS | 0 | 25016 | 1m4u | A | 222 | TYR | p.Tyr222Asp | Q13253 |
| HOMO SAPIENS | 1 | 25016 | 1m4u | A | 223 | PRO | p.Pro223Leu | Q13253 |
| HOMO SAPIENS | 0 | 1394 | 1um7 | A | 95 | ALA | p.Ala463Val | Q92796 |
| HOMO SAPIENS | 0 | 21294 | 4jpz | B | 1788 | GLU | p.Glu1788Asp | Q99250 |
| HOMO SAPIENS | 9 | 24068 | 2db6 | A | 47 | ARG | p.Arg119Cys | Q96MF2 |
| HOMO SAPIENS | 5 | 1869 | 4omt | A | 39 | ARG | p.Arg39Leu | P08237 |
| HOMO SAPIENS | 2 | 1869 | 4omt | A | 100 | ARG | p.Arg100Gln | P08237 |
| HOMO SAPIENS | 7 | 1869 | 4omt | A | 543 | ASP | p.Asp543Ala | P08237 |
| HOMO SAPIENS | 1 | 1869 | 4omt | A | 686 | TRP | p.Trp686Cys | P08237 |
| HOMO SAPIENS | 1 | 1869 | 4omt | A | 696 | ARG | p.Arg696His | P08237 |
| HOMO SAPIENS | 0 | 12163 | 4wxx | A | 495 | TYR | p.Tyr495Cys | P26358 |
| HOMO SAPIENS | 0 | 12163 | 4wxx | A | 528 | ILE | p.Ile528Met | P26358 |
| HOMO SAPIENS | 0 | 12391 | 1j1d | C | 36 | LYS | p.Lys36Gln | P19429 |
| HOMO SAPIENS | 0 | 12391 | 1j1d | C | 82 | PRO | p.Pro82Ser | P19429 |
| HOMO SAPIENS | 7 | 12391 | 1j1d | C | 102 | ALA | p.Ala102Thr | P19429 |
| HOMO SAPIENS | 1 | 5000 | 4cca | A | 17 | GLY | p.Gly17Arg | Q15833 |
| HOMO SAPIENS | 0 | 5000 | 4cca | A | 205 | VAL | p.Val205Ile | Q15833 |
| HOMO SAPIENS | 0 | 5000 | 4cca | A | 209 | LEU | p.Leu209Pro | Q15833 |
| HOMO SAPIENS | 1 | 5000 | 4cca | A | 459 | ARG | p.Arg459Trp | Q15833 |
| HOMO SAPIENS | 7 | 5000 | 4cca | A | 477 | PRO | p.Pro477Leu | Q15833 |
| HOMO SAPIENS | -1 | 5000 | 4cca | A | 526 | ILE | p.Ile526Val | Q15833 |
| HOMO SAPIENS | 0 | 5000 | 4cca | A | 541 | GLY | p.Gly541Ser | Q15833 |
| HOMO SAPIENS | 0 | 5000 | 4cca | A | 555 | ARG | p.Arg555Gly | Q15833 |
| HOMO SAPIENS | 5 | 28551 | 3vcm | P | 26 | ARG | p.Arg49Ter | P00797 |
| HOMO SAPIENS | 6 | 36453 | 5dzz | A | 2070 | ASP | p.Asp2070His | P15924 |
| HOMO SAPIENS | 0 | 36453 | 5dzz | A | 2166 | ARG | p.Arg2166Ter | P15924 |
| HOMO SAPIENS | 0 | 20268 | 2da6 | A | 96 | ASN | p.Asn321Ser | P35680 |
| HOMO SAPIENS | 1 | 1405 | 3iar | A | 8 | ASP | p.Asp8Asn | P00813 |
| HOMO SAPIENS | 8 | 1405 | 3iar | A | 15 | HIS | p.His15Asp | P00813 |
| HOMO SAPIENS | 6 | 1405 | 3iar | A | 74 | GLY | p.Gly74Val | P00813 |
| HOMO SAPIENS | 6 | 1405 | 3iar | A | 80 | LYS | p.Lys80Arg | P00813 |
| HOMO SAPIENS | 5 | 1405 | 3iar | A | 97 | TYR | p.Tyr97Cys | P00813 |
| HOMO SAPIENS | 5 | 1405 | 3iar | A | 101 | ARG | p.Arg101Leu | P00813 |
| HOMO SAPIENS | 5 | 1405 | 3iar | A | 101 | ARG | p.Arg101Trp | P00813 |
| HOMO SAPIENS | 5 | 1405 | 3iar | A | 106 | LEU | p.Leu106Val | P00813 |
| HOMO SAPIENS | 0 | 1405 | 3iar | A | 107 | LEU | p.Leu107Pro | P00813 |
| HOMO SAPIENS | 9 | 1405 | 3iar | A | 156 | ARG | p.Arg156Cys | P00813 |

| | | | | | | | | |
|---|---|---|---|---|---|---|---|---|
| HOMO SAPIENS | 6 | 1405 | 3iar | A | 211 | ARG | p.Arg211His | P00813 |
| HOMO SAPIENS | 0 | 1405 | 3iar | A | 216 | GLY | p.Gly216Arg | P00813 |
| HOMO SAPIENS | 0 | 1405 | 3iar | A | 291 | SER | p.Ser291Leu | P00813 |
| HOMO SAPIENS | 6 | 1405 | 3iar | A | 304 | LEU | p.Leu304Arg | P00813 |
| HOMO SAPIENS | 0 | 1405 | 3iar | A | 329 | ALA | p.Ala329Val | P00813 |
| HOMO SAPIENS | -1 | 31340 | 4atu | I | 186 | ARG | p.Arg186His | O43602 |
| HOMO SAPIENS | -1 | 31340 | 4atu | I | 192 | ARG | p.Arg192Trp | O43602 |
| HOMO SAPIENS | -1 | 31340 | 4atu | I | 196 | ARG | p.Arg196His | O43602 |
| HOMO SAPIENS | -1 | 31340 | 4atu | I | 203 | THR | p.Thr203Arg | O43602 |
| HOMO SAPIENS | -1 | 31340 | 4atu | I | 286 | ALA | p.Ala286Ser | O43602 |
| HOMO SAPIENS | 4 | 30734 | 1fsu | A | 95 | ARG | p.Arg95Gln | P15848 |
| HOMO SAPIENS | 0 | 30734 | 1fsu | A | 117 | CYS | p.Cys117Arg | P15848 |
| HOMO SAPIENS | 0 | 30734 | 1fsu | A | 137 | GLY | p.Gly137Val | P15848 |
| HOMO SAPIENS | 0 | 30734 | 1fsu | A | 210 | TYR | p.Tyr210Cys | P15848 |
| HOMO SAPIENS | 0 | 30734 | 1fsu | A | 236 | LEU | p.Leu236Pro | P15848 |
| HOMO SAPIENS | 1 | 30734 | 1fsu | A | 315 | ARG | p.Arg315Gln | P15848 |
| HOMO SAPIENS | 0 | 30734 | 1fsu | A | 358 | VAL | p.Val358Met | P15848 |
| HOMO SAPIENS | 2 | 30734 | 1fsu | A | 376 | VAL | p.Val376Met | P15848 |
| HOMO SAPIENS | 0 | 30734 | 1fsu | A | 384 | SER | p.Ser384Asn | P15848 |
| HOMO SAPIENS | 0 | 30734 | 1fsu | A | 393 | HIS | p.His393Pro | P15848 |
| HOMO SAPIENS | 0 | 30734 | 1fsu | A | 405 | CYS | p.Cys405Tyr | P15848 |
| HOMO SAPIENS | 0 | 30734 | 1fsu | A | 458 | ASN | p.Asn458Ser | P15848 |
| HOMO SAPIENS | 1 | 30734 | 1fsu | A | 484 | ARG | p.Arg484Gly | P15848 |
| HOMO SAPIENS | 0 | 38072 | 1wmv | A | 51 | PRO | p.Pro98Leu | Q9NZC7 |
| HOMO SAPIENS | -1 | 5674 | 1j1e | C | 185 | ASN | p.Asn185Lys | P19429 |
| HOMO SAPIENS | -1 | 5674 | 1j1e | C | 192 | ARG | p.Arg192His | P19429 |
| HOMO SAPIENS | -1 | 5674 | 1j1e | C | 196 | ASP | p.Asp196Asn | P19429 |
| HOMO SAPIENS | -1 | 5674 | 1j1e | C | 206 | LYS | p.Lys206Gln | P19429 |
| HOMO SAPIENS | 0 | 38067 | 1wlp | A | 13 | PRO | p.Pro156Gln | P13498 |
| HOMO SAPIENS | -1 | 20441 | 3vfd | A | 289 | HIS | p.His289Tyr | Q9UBP0 |
| HOMO SAPIENS | 2 | 20441 | 3vfd | A | 344 | ILE | p.Ile344Lys | Q9UBP0 |
| HOMO SAPIENS | 0 | 20441 | 3vfd | A | 362 | SER | p.Ser362Cys | Q9UBP0 |
| HOMO SAPIENS | 1 | 20441 | 3vfd | A | 386 | ASN | p.Asn386Ser | Q9UBP0 |
| HOMO SAPIENS | 9 | 20441 | 3vfd | A | 441 | ASP | p.Asp441Gly | Q9UBP0 |
| HOMO SAPIENS | -1 | 20441 | 3vfd | A | 448 | CYS | p.Cys448Tyr | Q9UBP0 |
| HOMO SAPIENS | 2 | 20441 | 3vfd | A | 500 | ARG | p.Arg499Cys | Q9UBP0 |
| HOMO SAPIENS | 1 | 20441 | 3vfd | A | 563 | ARG | p.Arg562Gly | Q9UBP0 |
| HOMO SAPIENS | 3 | 18471 | 4dx8 | J | 26 | ARG | p.Arg26Gln | O00522 |
| HOMO SAPIENS | -1 | 18471 | 4dx8 | J | 101 | GLY | p.Gly101Val | O00522 |
| HOMO SAPIENS | -1 | 34735 | 2ly4 | B | 83 | ALA | p.Ala83Glu | P04637 |
| HOMO SAPIENS | 0 | 31128 | 4cvh | A | 116 | ARG | p.Arg116Cys | A4D126 |
| HOMO SAPIENS | 8 | 31128 | 4cvh | A | 215 | GLN | p.Gln215Ter | A4D126 |
| HOMO SAPIENS | 0 | 31128 | 4cvh | A | 268 | ARG | p.Arg268Ter | A4D126 |
| HOMO SAPIENS | 0 | 31128 | 4cvh | A | 316 | THR | p.Thr316Lys | A4D126 |
| HOMO SAPIENS | 1 | 7164 | 3mt5 | A | 369 | ASP | p.Asp434Gly | Q12791 |
| HOMO SAPIENS | 2 | 26812 | 2wtk | F | 49 | TYR | p.Tyr49Asp | Q15831 |
| HOMO SAPIENS | 0 | 26812 | 2wtk | F | 67 | LEU | p.Leu67Pro | Q15831 |
| HOMO SAPIENS | 4 | 26812 | 2wtk | F | 104 | ARG | p.Arg104Ser | Q15831 |
| HOMO SAPIENS | 0 | 26812 | 2wtk | F | 135 | GLY | p.Gly135Arg | Q15831 |
| HOMO SAPIENS | 0 | 26812 | 2wtk | F | 163 | GLY | p.Gly163Asp | Q15831 |
| HOMO SAPIENS | 2 | 26812 | 2wtk | F | 194 | ALA | p.Asp194Asn | Q15831 |
| HOMO SAPIENS | 2 | 1357 | 5xtd | W | 57 | ARG | p.Arg57His | Q9P0J0 |
| HOMO SAPIENS | 0 | 28086 | 4nn2 | A | 229 | HIS | p.His229Arg | Q8IWS0 |
| HOMO SAPIENS | 0 | 28086 | 4nn2 | A | 234 | LYS | p.Lys234Glu | Q8IWS0 |

| | | | | | | | | |
|---|---|---|---|---|---|---|---|---|
| HOMO SAPIENS | 0 | 28086 | 4nn2 | A | 257 | ARG | p.Arg257Gly | Q8IWS0 |
| HOMO SAPIENS | 5 | 31182 | 2a1x | A | 176 | GLN | p.Gln176Lys | O14832 |
| HOMO SAPIENS | 1 | 31182 | 2a1x | A | 204 | GLY | p.Gly204Ser | O14832 |
| HOMO SAPIENS | 0 | 31182 | 2a1x | A | 269 | ASN | p.Asn269His | O14832 |
| HOMO SAPIENS | 3 | 31182 | 2a1x | A | 275 | ARG | p.Arg275Gln | O14832 |
| HOMO SAPIENS | 3 | 31182 | 2a1x | A | 275 | ARG | p.Arg275Trp | O14832 |
| HOMO SAPIENS | 4 | 2823 | 5xtb | P | 145 | THR | p.Thr145Ile | O75489 |
| HOMO SAPIENS | 5 | 2823 | 5xtb | P | 199 | ARG | p.Arg199Trp | O75489 |
| HOMO SAPIENS | 0 | 8645 | 2h5g | A | 372 | SER | p.Ser372Tyr | P54886 |
| HOMO SAPIENS | 3 | 8645 | 2h5g | A | 715 | ASP | p.Asp715His | P54886 |
| HOMO SAPIENS | 0 | 8645 | 2h5g | A | 765 | ARG | p.Arg765Gln | P54886 |
| HOMO SAPIENS | 0 | 8645 | 2h5g | A | 782 | TYR | p.Tyr782Cys | P54886 |
| HOMO SAPIENS | 0 | 8645 | 2h5g | A | 784 | HIS | p.His784Tyr | P54886 |
| HOMO SAPIENS | 1 | 13992 | 2cue | A | 39 | ARG | p.Arg242Thr | P26367 |
| HOMO SAPIENS | 9 | 13992 | 2cue | A | 55 | PHE | p.Phe258Ser | P26367 |
| HOMO SAPIENS | 0 | 30418 | 2xsw | A | 303 | VAL | p.Val303Met | Q9NRR6 |
| HOMO SAPIENS | 0 | 30418 | 2xsw | A | 315 | PRO | p.Pro315Leu | Q9NRR6 |
| HOMO SAPIENS | 0 | 30418 | 2xsw | A | 341 | GLY | p.Gly341Ser | Q9NRR6 |
| HOMO SAPIENS | 0 | 30418 | 2xsw | A | 378 | ARG | p.Arg378Cys | Q9NRR6 |
| HOMO SAPIENS | 0 | 30418 | 2xsw | A | 435 | ARG | p.Arg435Gln | Q9NRR6 |
| HOMO SAPIENS | 0 | 30418 | 2xsw | A | 515 | ARG | p.Arg515Trp | Q9NRR6 |
| HOMO SAPIENS | 1 | 30418 | 2xsw | A | 526 | PRO | p.Pro526Leu | Q9NRR6 |
| HOMO SAPIENS | 0 | 30418 | 2xsw | A | 563 | ARG | p.Arg563His | Q9NRR6 |
| HOMO SAPIENS | 2 | 30418 | 2xsw | A | 585 | ARG | p.Arg585His | Q9NRR6 |
| HOMO SAPIENS | 2 | 11167 | 4y99 | B | 205 | ARG | p.Arg215Leu | P45379 |
| HOMO SAPIENS | 5 | 11167 | 4y99 | B | 244 | GLU | p.Glu254Asp | P45379 |
| HOMO SAPIENS | 1 | 11167 | 4y99 | B | 253 | LYS | p.Lys263Arg | P45379 |
| HOMO SAPIENS | 8 | 11167 | 4y99 | B | 265 | ARG | p.Arg275Gln | P45379 |
| HOMO SAPIENS | 3 | 11167 | 4y99 | B | 270 | ASP | p.Asp280Asn | P45379 |
| HOMO SAPIENS | -1 | 11167 | 4y99 | B | 278 | ARG | p.Arg288Pro | P45379 |
| HOMO SAPIENS | -1 | 11167 | 4y99 | B | 286 | ARG | p.Arg296Cys | P45379 |
| HOMO SAPIENS | -1 | 11167 | 4y99 | B | 286 | ARG | p.Arg296His | P45379 |
| HOMO SAPIENS | 0 | 15993 | 4po6 | A | 197 | ARG | p.Arg197His | P29597 |
| HOMO SAPIENS | -1 | 15993 | 4po6 | A | 362 | VAL | p.Val362Phe | P29597 |
| HOMO SAPIENS | -1 | 15993 | 4po6 | A | 363 | GLY | p.Gly363Ser | P29597 |
| HOMO SAPIENS | 0 | 5646 | 5uak | A | 7 | GLU | p.Glu7Ter | P13569 |
| HOMO SAPIENS | 0 | 5646 | 5uak | A | 27 | GLY | p.Gly27Ter | P13569 |
| HOMO SAPIENS | 0 | 5646 | 5uak | A | 46 | ALA | p.Ala46Asp | P13569 |
| HOMO SAPIENS | 0 | 5646 | 5uak | A | 60 | GLU | p.Glu60Ter | P13569 |
| HOMO SAPIENS | 0 | 5646 | 5uak | A | 67 | PRO | p.Pro67Leu | P13569 |
| HOMO SAPIENS | 0 | 5646 | 5uak | A | 74 | ARG | p.Arg74Trp | P13569 |
| HOMO SAPIENS | 0 | 5646 | 5uak | A | 75 | ARG | p.Arg75Ter | P13569 |
| HOMO SAPIENS | 0 | 5646 | 5uak | A | 85 | GLY | p.Gly85Glu | P13569 |
| HOMO SAPIENS | 0 | 5646 | 5uak | A | 91 | GLY | p.Gly91Arg | P13569 |
| HOMO SAPIENS | 0 | 5646 | 5uak | A | 92 | GLU | p.Glu92Lys | P13569 |
| HOMO SAPIENS | 0 | 5646 | 5uak | A | 98 | GLN | p.Gln98Arg | P13569 |
| HOMO SAPIENS | 0 | 5646 | 5uak | A | 98 | GLN | p.Gln98Ter | P13569 |
| HOMO SAPIENS | 0 | 5646 | 5uak | A | 109 | TYR | p.Tyr109Cys | P13569 |
| HOMO SAPIENS | 0 | 5646 | 5uak | A | 110 | ASP | p.Asp110His | P13569 |
| HOMO SAPIENS | 0 | 5646 | 5uak | A | 117 | ARG | p.Arg117Cys | P13569 |
| HOMO SAPIENS | 0 | 5646 | 5uak | A | 117 | ARG | p.Arg117His | P13569 |
| HOMO SAPIENS | 0 | 5646 | 5uak | A | 125 | ILE | p.Ile125Thr | P13569 |
| HOMO SAPIENS | 0 | 5646 | 5uak | A | 148 | ILE | p.Ile148Thr | P13569 |
| HOMO SAPIENS | 0 | 5646 | 5uak | A | 170 | ARG | p.Arg170His | P13569 |

| | | | | | | | | |
|---|---|---|---|---|---|---|---|---|
| HOMO SAPIENS | 0 | 5646 | 5uak | A | 178 | GLY | p.Gly178Arg | P13569 |
| HOMO SAPIENS | 0 | 5646 | 5uak | A | 206 | LEU | p.Leu206Trp | P13569 |
| HOMO SAPIENS | 0 | 5646 | 5uak | A | 373 | ASP | p.Asp373Asn | P13569 |
| HOMO SAPIENS | -1 | 5646 | 5uak | A | 709 | ARG | p.Arg709Ter | P13569 |
| HOMO SAPIENS | -1 | 5646 | 5uak | A | 732 | LEU | p.Leu732Ter | P13569 |
| HOMO SAPIENS | -1 | 5646 | 5uak | A | 785 | ARG | p.Arg785Ter | P13569 |
| HOMO SAPIENS | -1 | 5646 | 5uak | A | 831 | GLU | p.Glu831Ter | P13569 |
| HOMO SAPIENS | -1 | 5646 | 5uak | A | 836 | ASP | p.Asp836Tyr | P13569 |
| HOMO SAPIENS | 0 | 5646 | 5uak | A | 851 | ARG | p.Arg851Ter | P13569 |
| HOMO SAPIENS | 0 | 5646 | 5uak | A | 913 | TYR | p.Tyr913Cys | P13569 |
| HOMO SAPIENS | 0 | 5646 | 5uak | A | 913 | TYR | p.Tyr913Ter | P13569 |
| HOMO SAPIENS | 0 | 5646 | 5uak | A | 945 | SER | p.Ser945Leu | P13569 |
| HOMO SAPIENS | 0 | 5646 | 5uak | A | 949 | HIS | p.His949Tyr | P13569 |
| HOMO SAPIENS | 0 | 5646 | 5uak | A | 970 | GLY | p.Gly970Asp | P13569 |
| HOMO SAPIENS | 0 | 5646 | 5uak | A | 1027 | ILE | p.Ile1027Thr | P13569 |
| HOMO SAPIENS | 0 | 5646 | 5uak | A | 1032 | TYR | p.Tyr1032Cys | P13569 |
| HOMO SAPIENS | 0 | 5646 | 5uak | A | 1061 | GLY | p.Gly1061Arg | P13569 |
| HOMO SAPIENS | 1 | 5646 | 5uak | A | 1065 | LEU | p.Leu1065Pro | P13569 |
| HOMO SAPIENS | 1 | 5646 | 5uak | A | 1066 | ARG | p.Arg1066Cys | P13569 |
| HOMO SAPIENS | 1 | 5646 | 5uak | A | 1066 | ARG | p.Arg1066His | P13569 |
| HOMO SAPIENS | 1 | 5646 | 5uak | A | 1067 | ALA | p.Ala1067Thr | P13569 |
| HOMO SAPIENS | 0 | 5646 | 5uak | A | 1069 | GLY | p.Gly1069Arg | P13569 |
| HOMO SAPIENS | 0 | 5646 | 5uak | A | 1070 | ARG | p.Arg1070Trp | P13569 |
| HOMO SAPIENS | 1 | 5646 | 5uak | A | 1071 | GLN | p.Gln1071Pro | P13569 |
| HOMO SAPIENS | 0 | 5646 | 5uak | A | 1074 | PHE | p.Phe1074Leu | P13569 |
| HOMO SAPIENS | 0 | 5646 | 5uak | A | 1077 | LEU | p.Leu1077Pro | P13569 |
| HOMO SAPIENS | 0 | 5646 | 5uak | A | 1089 | TRP | p.Trp1089Ter | P13569 |
| HOMO SAPIENS | 0 | 5646 | 5uak | A | 1092 | TYR | p.Tyr1092Ter | P13569 |
| HOMO SAPIENS | 0 | 5646 | 5uak | A | 1101 | MET | p.Met1101Lys | P13569 |
| HOMO SAPIENS | 0 | 5646 | 5uak | A | 1156 | LEU | p.Leu1156Phe | P13569 |
| HOMO SAPIENS | 0 | 5646 | 5uak | A | 1158 | ARG | p.Arg1158Ter | P13569 |
| HOMO SAPIENS | 0 | 5646 | 5uak | A | 1162 | ARG | p.Arg1162Leu | P13569 |
| HOMO SAPIENS | 0 | 5646 | 5uak | A | 1162 | ARG | p.Arg1162Ter | P13569 |
| HOMO SAPIENS | -1 | 5646 | 5uak | A | 1476 | GLN | p.Gln1476Ter | P13569 |
| HOMO SAPIENS | 0 | 22763 | 1imv | A | 52 | MET | p.Thr72Met | P36955 |
| HOMO SAPIENS | 0 | 22763 | 1imv | A | 301 | TYR | p.Tyr321Ter | P36955 |
| HOMO SAPIENS | 0 | 23179 | 5fql | A | 160 | PRO | p.Pro160Arg | P22304 |
| HOMO SAPIENS | 0 | 23179 | 5fql | A | 172 | ARG | p.Arg172Ter | P22304 |
| HOMO SAPIENS | 0 | 23179 | 5fql | A | 214 | THR | p.Thr214Met | P22304 |
| HOMO SAPIENS | 0 | 23179 | 5fql | A | 309 | THR | p.Thr309Ala | P22304 |
| HOMO SAPIENS | 0 | 23179 | 5fql | A | 333 | SER | p.Ser333Leu | P22304 |
| HOMO SAPIENS | 0 | 23179 | 5fql | A | 443 | ARG | p.Arg443Ter | P22304 |
| HOMO SAPIENS | 0 | 23179 | 5fql | A | 468 | ARG | p.Arg468Leu | P22304 |
| HOMO SAPIENS | 0 | 23179 | 5fql | A | 468 | ARG | p.Arg468Trp | P22304 |
| HOMO SAPIENS | 0 | 23179 | 5fql | A | 502 | TRP | p.Trp502Ser | P22304 |
| HOMO SAPIENS | 9 | 20989 | 1nfi | F | 185 | THR | p.Thr185Met | P25963 |
| HOMO SAPIENS | 4 | 20989 | 1nfi | F | 194 | GLY | p.Gly194Ala | P25963 |
| HOMO SAPIENS | 0 | 11538 | 3eab | A | 162 | VAL | p.Val162Ile | Q9UBP0 |
| HOMO SAPIENS | 0 | 33399 | 2p39 | A | 71 | SER | p.Ser71Gly | Q9GZV9 |
| HOMO SAPIENS | 3 | 33399 | 2p39 | A | 96 | MET | p.Met96Thr | Q9GZV9 |
| HOMO SAPIENS | 0 | 33399 | 2p39 | A | 129 | SER | p.Ser129Phe | Q9GZV9 |
| HOMO SAPIENS | -1 | 33399 | 2p39 | A | 172 | PRO | p.Pro172Leu | Q9GZV9 |
| HOMO SAPIENS | -1 | 33399 | 2p39 | A | 176 | ARG | p.Arg176Gln | Q9GZV9 |
| HOMO SAPIENS | -1 | 33399 | 2p39 | A | 179 | ARG | p.Arg179Trp | Q9GZV9 |

| | | | | | | | | |
|---|---|---|---|---|---|---|---|---|
| HOMO SAPIENS | -1 | 33399 | 2p39 | A | 179 | ARG | p.Arg179Gln | Q9GZV9 |
| HOMO SAPIENS | 0 | 16217 | 2dne | A | 79 | ILE | p.Ile163Leu | P10515 |
| HOMO SAPIENS | 8 | 7666 | 4kbq | A | 65 | ASN | p.Asn65Ser | Q9UNE7 |
| HOMO SAPIENS | 1 | 6407 | 1ikt | A | 71 | THR | p.Thr687Ile | P51659 |
| HOMO SAPIENS | 0 | 6407 | 1ikt | A | 112 | MET | p.Met728Val | P51659 |
| HOMO SAPIENS | 0 | 5849 | 4ihb | A | 67 | VAL | p.Val67Asp | O75923 |
| HOMO SAPIENS | 6 | 17703 | 5fcn | A | 20 | HIS | p.His99Gln | Q66GS9 |
| HOMO SAPIENS | 8 | 17703 | 5fcn | A | 33 | ARG | p.Arg112His | Q66GS9 |
| HOMO SAPIENS | 0 | 34095 | 2E+073 | A | 101 | HIS | p.His101Tyr | P05129 |
| HOMO SAPIENS | -1 | 41358 | 4liq | E | 507 | THR | p.Thr507Met | P07333 |
| HOMO SAPIENS | -1 | 21290 | 1fch | A | 235 | MET | p.Met272Arg | P50542 |
| HOMO SAPIENS | 5 | 23431 | 6pax | A | 23 | ARG | p.Arg26Gly | P26367 |
| HOMO SAPIENS | 1 | 23431 | 6pax | A | 61 | GLY | p.Gly64Val | P26367 |
| HOMO SAPIENS | 1 | 23431 | 6pax | A | 65 | PRO | p.Pro68Ser | P26367 |
| HOMO SAPIENS | 1 | 23431 | 6pax | A | 116 | SER | p.Ser119Arg | P26367 |
| HOMO SAPIENS | 9 | 23431 | 6pax | A | 123 | VAL | p.Val126Asp | P26367 |
| HOMO SAPIENS | 9 | 23431 | 6pax | A | 125 | ARG | p.Arg128Cys | P26367 |
| HOMO SAPIENS | 9 | 33003 | 4zrj | B | 535 | LEU | p.Leu535Pro | P35240 |
| HOMO SAPIENS | 7 | 33003 | 4zrj | B | 538 | GLN | p.Gln538Pro | P35240 |
| HOMO SAPIENS | 0 | 33003 | 4zrj | B | 585 | TRP | p.Ala585Thr | P35240 |
| HOMO SAPIENS | 0 | 3800 | 2nzj | A | 193 | VAL | p.Val193Met | O75628 |

| gene_name | max_cons | max_bs_id | max_pdb_id | max_chain_id |
|---|---|---|---|---|
| AKR1D1 | None | None | None | None |
| AKR1D1 | None | None | None | None |
| AKR1D1 | None | None | None | None |
| AKR1D1 | None | None | None | None |
| ALPP | None | None | None | None |
| BMPR1A | None | None | None | None |
| BMPR1A | None | None | None | None |
| FGFR2 | None | None | None | None |
| PROC | None | None | None | None |
| PROC | None | None | None | None |
| PROC | None | None | None | None |
| PROC | None | None | None | None |
| PROC | None | None | None | None |
| PROC | None | None | None | None |
| PROC | None | None | None | None |
| PROC | None | None | None | None |
| PROC | None | None | None | None |
| PROC | None | None | None | None |
| PROC | None | None | None | None |
| PROC | None | None | None | None |
| INS | 1 | 2 | 3zs2 | J |
| INS | 0.85 | 1 | 4ajx | D |
| INS | 1 | 1 | 3zs2 | J |
| INS | 1 | 1 | 3zs2 | J |
| FTL | 0.627 | 11 | 2fg8 | B |
| FTL | None | None | None | None |
| FTL | None | None | None | None |
| TNFSF11 | None | None | None | None |
| EHMT1 | None | None | None | None |
| EHMT1 | None | None | None | None |
| EHMT1 | None | None | None | None |
| CA2 | 0.939 | 1 | 4idr | X |
| CA2 | 0.939 | 1 | 4idr | X |
| CA2 | 0.902 | 5 | 4idr | X |
| CA2 | 0.902 | 5 | 4idr | X |
| SMAD4 | None | None | None | None |
| SMAD4 | None | None | None | None |
| TTR | 0.691 | 12 | 1ttb | B |
| TTR | 0.961 | 7 | 3grb | A |
| TTR | 0.891 | 4 | 3dk2 | B |
| TTR | 0.947 | 3 | 1tta | A |
| TTR | 0.923 | 1 | 3grb | B |
| TTR | 0.891 | 1 | 4y9b | B |
| TTR | 0.891 | 1 | 4y9b | B |
| TTR | 0.882 | 15 | 3grb | A |
| TTR | 0.909 | 5 | 4tl4 | A |
| TTR | 0.836 | 7 | 1z7j | B |
| TTR | 0.882 | 15 | 3grb | A |
| TTR | 0.891 | 5 | 3i9i | B |
| TTR | 0.902 | 9 | 3m1o | B |
| TTR | 0.902 | 9 | 3m1o | B |
| TTR | 0.902 | 9 | 3m1o | B |
| TTR | 0.636 | 16 | 1thc | A |

| | | | | |
|---|---|---|---|---|
| TTR | 0.902 | 9 | 3m1o | B |
| TTR | 0.836 | 7 | 1bze | A |
| TTR | 0.846 | 16 | 3gps | A |
| TTR | 0.846 | 16 | 3gps | A |
| TTR | 0.909 | 5 | 4tl4 | A |
| TTR | 0.909 | 5 | 4tl4 | A |
| TTR | 0.673 | 15 | 3dk2 | B |
| TTR | 0.945 | 4 | 4deu | A |
| TTR | 0.941 | 3 | 3did | B |
| TTR | 0.941 | 3 | 3did | B |
| TTR | 0.868 | 6 | 4l1t | A |
| TTR | 0.882 | 19 | 3gps | B |
| TTR | 0.891 | 4 | 3dk2 | B |
| TTR | 0.941 | 3 | 3did | B |
| TTR | 0.923 | 1 | 4hiq | A |
| TTR | 0.923 | 1 | 4hiq | A |
| TTR | 0.961 | 7 | 3grb | A |
| TTR | 0.839 | 7 | 1tsh | B |
| TTR | 0.891 | 6 | 1ttb | B |
| ALK | 0.682 | 1 | 4cd0 | A |
| ALK | None | None | None | None |
| THRB | None | None | None | None |
| THRB | None | None | None | None |
| THRB | None | None | None | None |
| THRB | None | None | None | None |
| THRB | None | None | None | None |
| THRB | None | None | None | None |
| THRB | None | None | None | None |
| THRB | None | None | None | None |
| THRB | None | None | None | None |
| THRB | None | None | None | None |
| THRB | None | None | None | None |
| THRB | None | None | None | None |
| THRB | None | None | None | None |
| THRB | None | None | None | None |
| THRB | None | None | None | None |
| THRB | None | None | None | None |
| THRB | None | None | None | None |
| THRB | None | None | None | None |
| THRB | None | None | None | None |
| IL2RG | None | None | None | None |
| IL2RG | None | None | None | None |
| IL2RG | None | None | None | None |
| IL2RG | None | None | None | None |
| IL2RG | None | None | None | None |
| ITGB3 | None | None | None | None |
| ITGB3 | None | None | None | None |
| ITGB3 | None | None | None | None |
| ITGB3 | None | None | None | None |
| ITGB3 | None | None | None | None |
| ITGB3 | None | None | None | None |
| ITGB3 | None | None | None | None |
| ITGB3 | None | None | None | None |
| ITGB3 | None | None | None | None |

| | | | | |
|---|---|---|---|---|
| PKD2 | None | None | None | None |
| PKD2 | None | None | None | None |
| PKD2 | None | None | None | None |
| PKD2 | None | None | None | None |
| PKD2 | None | None | None | None |
| PKD2 | None | None | None | None |
| PKD2 | None | None | None | None |
| TP53 | None | None | None | None |
| TP53 | None | None | None | None |
| TP53 | None | None | None | None |
| TP53 | None | None | None | None |
| TP53 | None | None | None | None |
| TP53 | None | None | None | None |
| EGFR | None | None | None | None |
| EGFR | None | None | None | None |
| XRCC4 | None | None | None | None |
| XRCC4 | None | None | None | None |
| HBB | None | None | None | None |
| HBB | None | None | None | None |
| HBB | None | None | None | None |
| HBB | None | None | None | None |
| HBB | None | None | None | None |
| HBB | None | None | None | None |
| HBB | None | None | None | None |
| HBB | None | None | None | None |
| HBB | None | None | None | None |
| HBB | None | None | None | None |
| HBB | None | None | None | None |
| HBB | None | None | None | None |
| HBB | None | None | None | None |
| HBB | None | None | None | None |
| HBB | None | None | None | None |
| HBB | None | None | None | None |
| HBB | None | None | None | None |
| HBB | None | None | None | None |
| HBB | None | None | None | None |
| HBB | None | None | None | None |
| HBB | None | None | None | None |
| HBB | None | None | None | None |
| HBB | None | None | None | None |
| HBB | None | None | None | None |
| HBB | None | None | None | None |
| HBB | 0.6 | 1 | 2w72 | D |
| HBB | None | None | None | None |
| HBB | None | None | None | None |
| HBB | None | None | None | None |
| HBB | 0.733 | 1 | 1yff | D |
| HBB | 0.692 | 2 | 1buw | B |
| RPS6KA3 | None | None | None | None |
| RPS6KA3 | None | None | None | None |
| RPS6KA3 | None | None | None | None |
| RPS6KA3 | None | None | None | None |
| RPS6KA3 | None | None | None | None |
| RPS6KA3 | None | None | None | None |
| GBA | 1 | 15 | 3gxd | B |

| | | | | | |
|---|---|---|---|---|---|
| GBA | | 1 | 55 | 3gxd | B |
| GBA | | 1 | 15 | 3gxd | B |
| GBA | | 0.75 | 45 | 3ril | A |
| GBA | None | None | | None | None |
| GBA | None | None | | None | None |
| GBA | | 1 | 12 | 1ogs | A |
| GBA | | 0.75 | 122 | 3gxi | A |
| GBA | | 1 | 2 | 1ogs | A |
| GBA | | 1 | 11 | 1ogs | A |
| GBA | None | None | | None | None |
| GBA | | 0.75 | 45 | 3ril | A |
| GBA | | 1 | 18 | 1ogs | A |
| GBA | | 1 | 3 | 1ogs | A |
| GBA | None | None | | None | None |
| GBA | | 1 | 2 | 1ogs | A |
| GBA | None | None | | None | None |
| GBA | | 0.75 | 76 | 3gxm | C |
| GBA | | 1 | 3 | 1ogs | A |
| GBA | | 1 | 3 | 1ogs | A |
| GBA | | 0.75 | 62 | 3ril | A |
| GBA | | 1 | 28 | 1ogs | A |
| GBA | None | None | | None | None |
| GBA | None | None | | None | None |
| GBA | None | None | | None | None |
| GBA | None | None | | None | None |
| GBA | None | None | | None | None |
| GBA | None | None | | None | None |
| GBA | | 0.75 | 49 | 3ril | A |
| GBA | None | None | | None | None |
| GBA | | 0.667 | 60 | 1ogs | A |
| GBA | None | None | | None | None |
| GBA | None | None | | None | None |
| GBA | | 0.75 | 21 | 3gxi | A |
| GBA | None | None | | None | None |
| GBA | None | None | | None | None |
| GBA | | 0.75 | 25 | 3keh | B |
| GBA | | 0.75 | 70 | 3ril | A |
| GBA | | 0.75 | 70 | 3ril | A |
| GBA | | 1 | 15 | 3gxd | B |
| GBA | | 1 | 44 | 3gxd | B |
| GBA | None | None | | None | None |
| GBA | | 1 | 44 | 3gxd | B |
| GBA | | 1 | 44 | 3gxd | B |
| GBA | None | None | | None | None |
| GBA | | 0.75 | 94 | 3gxi | A |
| GBA | | 0.75 | 94 | 3gxi | A |
| IFIH1 | None | None | | None | None |
| TGFBR2 | None | None | | None | None |
| TGFBR2 | None | None | | None | None |
| TGFBR2 | None | None | | None | None |
| TGFBR2 | None | None | | None | None |
| TGFBR2 | None | None | | None | None |
| TGFBR2 | None | None | | None | None |
| TGFBR2 | None | None | | None | None |

| | | | | | |
|---|---|---|---|---|---|
| TGFBR2 | None | None | None | None | |
| TGFBR2 | None | None | None | None | |
| TGFBR2 | 0.742 | | 1 | 5E+091 | A |
| TGFBR2 | None | None | None | None | |
| TGFBR2 | None | None | None | None | |
| TGFBR2 | None | None | None | None | |
| TGFBR2 | 0.676 | | 1 | 5e8v | A |
| TGFBR2 | 0.676 | | 1 | 5e8v | A |
| ACE | 0.684 | | 17 | 3nxq | A |
| ACE | None | None | None | None | |
| ACE | None | None | None | None | |
| ACE | None | None | None | None | |
| ACE | None | None | None | None | |
| ACE | None | None | None | None | |
| COL10A1 | None | None | None | None | |
| COL10A1 | None | None | None | None | |
| COL10A1 | None | None | None | None | |
| COL10A1 | None | None | None | None | |
| COL10A1 | None | None | None | None | |
| COL10A1 | None | None | None | None | |
| COL10A1 | None | None | None | None | |
| COL10A1 | None | None | None | None | |
| COL10A1 | None | None | None | None | |
| KLF1 | None | None | None | None | |
| C3 | None | None | None | None | |
| CTSC | None | None | None | None | |
| CTSC | 0.934 | | 1 | 2djg | C |
| CTSC | None | None | None | None | |
| LYZ | 0.877 | | 3 | 1bb5 | A |
| LYZ | 0.923 | | 1 | 1lz4 | A |
| LYZ | 0.852 | | 2 | 1ubz | A |
| DHODH | None | None | None | None | |
| DHODH | None | None | None | None | |
| DHODH | None | None | None | None | |
| DHODH | None | None | None | None | |
| DHODH | None | None | None | None | |
| DHODH | None | None | None | None | |
| DHODH | None | None | None | None | |
| ADH1C | None | None | None | None | |
| GDNF | None | None | None | None | |
| TNF | None | None | None | None | |
| ALDH4A1 | 0.636 | | 3 | 3v9h | A |
| ALDH4A1 | None | None | None | None | |
| ALDH4A1 | None | None | None | None | |
| BRAF | None | None | None | None | |
| BRAF | None | None | None | None | |
| BRAF | 0.752 | | 2 | 3psb | A |
| BRAF | 0.752 | | 2 | 3psb | A |
| BRAF | 0.737 | | 1 | 3pri | A |
| BRAF | 0.737 | | 1 | 3pri | A |
| BRAF | 0.737 | | 1 | 3pri | A |
| BRAF | 0.737 | | 1 | 3prf | A |
| BRAF | 0.737 | | 1 | 3prf | A |
| BRAF | 0.752 | | 2 | 3psb | A |

| | | | | |
|---|---|---|---|---|
| BRAF | 0.752 | 2 | 3psb | A |
| BRAF | 0.752 | 2 | 3psb | A |
| BRAF | 0.752 | 2 | 3psb | A |
| BRAF | None | None | None | None |
| BRAF | 0.663 | 1 | 4ehe | B |
| BRAF | 0.734 | 1 | 5csx | A |
| TBX5 | None | None | None | None |
| TBX5 | None | None | None | None |
| TBX5 | None | None | None | None |
| TBX5 | None | None | None | None |
| TBX5 | None | None | None | None |
| TBX5 | None | None | None | None |
| TBX5 | None | None | None | None |
| TBX5 | None | None | None | None |
| TBX5 | None | None | None | None |
| RB1 | None | None | None | None |
| RB1 | None | None | None | None |
| RB1 | None | None | None | None |
| RB1 | None | None | None | None |
| RB1 | None | None | None | None |
| RB1 | None | None | None | None |
| CLIC2 | None | None | None | None |
| CLIC2 | None | None | None | None |
| ALDH7A1 | None | None | None | None |
| ALDH7A1 | None | None | None | None |
| ALDH7A1 | None | None | None | None |
| ALDH7A1 | None | None | None | None |
| ALDH7A1 | None | None | None | None |
| ALDH7A1 | None | None | None | None |
| ALDH7A1 | None | None | None | None |
| ALDH7A1 | None | None | None | None |
| ALDH7A1 | None | None | None | None |
| ALDH7A1 | None | None | None | None |
| ALDH7A1 | None | None | None | None |
| ALDH7A1 | None | None | None | None |
| ALDH7A1 | None | None | None | None |
| ALDH7A1 | None | None | None | None |
| ALDH7A1 | 0.632 | 1 | 4zuk | A |
| ALDH7A1 | None | None | None | None |
| KDR | None | None | None | None |
| MUT | None | None | None | None |
| MUT | None | None | None | None |
| MUT | None | None | None | None |
| MUT | None | None | None | None |
| MUT | None | None | None | None |
| MUT | None | None | None | None |
| MUT | None | None | None | None |
| MUT | None | None | None | None |
| MUT | None | None | None | None |
| MUT | None | None | None | None |
| MUT | None | None | None | None |
| MUT | None | None | None | None |
| MUT | None | None | None | None |
| MUT | None | None | None | None |

| | | | | |
|---|---|---|---|---|
| MUT | None | None | None | None |
| MUT | None | None | None | None |
| MUT | 0.6 | 1 | 3bic | A |
| MUT | 0.6 | 2 | 2xiq | A |
| MUT | 0.6 | 2 | 2xiq | A |
| MUT | None | None | None | None |
| MUT | None | None | None | None |
| MUT | None | None | None | None |
| MUT | None | None | None | None |
| MUT | None | None | None | None |
| MUT | None | None | None | None |
| MUT | None | None | None | None |
| MUT | None | None | None | None |
| MUT | None | None | None | None |
| MUT | None | None | None | None |
| MUT | None | None | None | None |
| MUT | None | None | None | None |
| MUT | None | None | None | None |
| MUT | None | None | None | None |
| MUT | None | None | None | None |
| SOD1 | 0.778 | 1 | 2gbu | A |
| SOD1 | 0.727 | 4 | 4xcr | A |
| SOD1 | None | None | None | None |
| SOD1 | 0.621 | 12 | 1hl5 | H |
| NECTIN3 | None | None | None | None |
| HFE | None | None | None | None |
| HFE | None | None | None | None |
| HFE | None | None | None | None |
| HFE | None | None | None | None |
| HFE | None | None | None | None |
| HFE | None | None | None | None |
| HFE | None | None | None | None |
| EIF2AK3 | None | None | None | None |
| EIF2AK3 | None | None | None | None |
| TERT | None | None | None | None |
| TERT | None | None | None | None |
| SPR | None | None | None | None |
| SPR | None | None | None | None |
| SPR | None | None | None | None |
| SPR | None | None | None | None |
| SPR | None | None | None | None |
| F7 | 0.625 | 2 | 1klj | L |
| UNG | None | None | None | None |
| MMP13 | None | None | None | None |
| MMP13 | 0.786 | 1 | 3tvc | A |
| MMP13 | 0.766 | 1 | 3zxh | B |
| MMP13 | 0.766 | 1 | 3zxh | B |
| CD207 | 0.621 | 1 | 3kqg | B |
| F2 | 1 | 2 | 1nrq | L |
| SPTBN2 | None | None | None | None |
| PKLR | None | None | None | None |
| PKLR | None | None | None | None |
| PKLR | None | None | None | None |
| PKLR | None | None | None | None |

| | | | | |
|---|---|---|---|---|
| PKLR | None | None | None | None |
| PKLR | None | None | None | None |
| PKLR | None | None | None | None |
| PKLR | None | None | None | None |
| SACS | None | None | None | None |
| HRAS | None | None | None | None |
| HRAS | None | None | None | None |
| HRAS | None | None | None | None |
| HRAS | None | None | None | None |
| HRAS | 0.664 | 1 | 2lwi | A |
| HRAS | 0.742 | 1 | 1ctq | A |
| HRAS | 0.69 | 1 | 1xd2 | A |
| HRAS | None | None | None | None |
| HRAS | None | None | None | None |
| HRAS | None | None | None | None |
| HRAS | None | None | None | None |
| HRAS | 0.664 | 1 | 2lwi | A |
| HRAS | 0.664 | 1 | 2lwi | A |
| VHL | None | None | None | None |
| VHL | None | None | None | None |
| VHL | None | None | None | None |
| VHL | None | None | None | None |
| VHL | None | None | None | None |
| VHL | None | None | None | None |
| VHL | None | None | None | None |
| VHL | None | None | None | None |
| VHL | None | None | None | None |
| VHL | None | None | None | None |
| VHL | None | None | None | None |
| VHL | None | None | None | None |
| VHL | None | None | None | None |
| VHL | None | None | None | None |
| VHL | None | None | None | None |
| VHL | None | None | None | None |
| VHL | None | None | None | None |
| VHL | None | None | None | None |
| VHL | None | None | None | None |
| VHL | None | None | None | None |
| VHL | None | None | None | None |
| VHL | None | None | None | None |
| VHL | None | None | None | None |
| VHL | None | None | None | None |
| VHL | None | None | None | None |
| TTBK1 | None | None | None | None |
| RAD51 | None | None | None | None |
| ALB | None | None | None | None |
| ALB | None | None | None | None |
| ALB | None | None | None | None |
| ALB | None | None | None | None |
| MET | None | None | None | None |
| MET | None | None | None | None |
| MET | None | None | None | None |
| MET | None | None | None | None |
| MET | None | None | None | None |

| | | | | |
|---|---|---|---|---|
| MET | None | None | None | None |
| MET | None | None | None | None |
| MET | None | None | None | None |
| MET | None | None | None | None |
| CP | None | None | None | None |
| CP | None | None | None | None |
| PGM1 | None | None | None | None |
| PGM1 | None | None | None | None |
| PGM1 | None | None | None | None |
| PGM1 | None | None | None | None |
| PGM1 | None | None | None | None |
| PGM1 | None | None | None | None |
| PGM1 | None | None | None | None |
| NR5A1 | None | None | None | None |
| NR5A1 | None | None | None | None |
| NR5A1 | None | None | None | None |
| F9 | None | None | None | None |
| F9 | None | None | None | None |
| HPRT1 | None | None | None | None |
| HPRT1 | None | None | None | None |
| HPRT1 | None | None | None | None |
| HPRT1 | None | None | None | None |
| HPRT1 | None | None | None | None |
| HPRT1 | 0.871 | 3 | 4rac | B |
| HPRT1 | 0.806 | 2 | 4ran | D |
| HPRT1 | None | None | None | None |
| HPRT1 | None | None | None | None |
| HPRT1 | None | None | None | None |
| HPRT1 | None | None | None | None |
| HPRT1 | None | None | None | None |
| HPRT1 | None | None | None | None |
| HPRT1 | 0.703 | 4 | 5w8v | A |
| HPRT1 | None | None | None | None |
| HPRT1 | None | None | None | None |
| HPRT1 | None | None | None | None |
| HPRT1 | None | None | None | None |
| HPRT1 | 0.871 | 3 | 4rac | B |
| HPRT1 | 0.871 | 3 | 4rac | B |
| HPRT1 | 0.765 | 2 | 4rao | D |
| HPRT1 | 0.774 | 2 | 4rac | B |
| GPD1L | None | None | None | None |
| GPD1L | None | None | None | None |
| GPD1L | None | None | None | None |
| FGFR2 | None | None | None | None |
| FGFR2 | None | None | None | None |
| FGFR2 | None | None | None | None |
| FGFR2 | None | None | None | None |
| FGFR2 | None | None | None | None |
| FGFR2 | None | None | None | None |
| FGFR2 | None | None | None | None |
| FGFR2 | None | None | None | None |
| FGFR2 | None | None | None | None |
| FGFR2 | None | None | None | None |
| FGFR2 | None | None | None | None |

| | | | | |
|---|---|---|---|---|
| FGFR2 | None | None | None | None |
| FGFR2 | None | None | None | None |
| FGFR2 | None | None | None | None |
| FGFR2 | None | None | None | None |
| FGFR2 | None | None | None | None |
| FGFR2 | None | None | None | None |
| FGFR2 | None | None | None | None |
| FGFR2 | None | None | None | None |
| HSPD1 | None | None | None | None |
| HSPD1 | None | None | None | None |
| FGF8 | None | None | None | None |
| FGF8 | None | None | None | None |
| FGF8 | None | None | None | None |
| F9 | None | None | None | None |
| NCF4 | None | None | None | None |
| NCF4 | None | None | None | None |
| LDLR | None | None | None | None |
| LDLR | None | None | None | None |
| LDLR | None | None | None | None |
| LDLR | None | None | None | None |
| LDLR | None | None | None | None |
| LDLR | None | None | None | None |
| C3 | None | None | None | None |
| C3 | None | None | None | None |
| C3 | None | None | None | None |
| C3 | None | None | None | None |
| C3 | None | None | None | None |
| C3 | None | None | None | None |
| C3 | None | None | None | None |
| C3 | None | None | None | None |
| TULP1 | None | None | None | None |
| TULP1 | None | None | None | None |
| TULP1 | None | None | None | None |
| TULP1 | None | None | None | None |
| IKBKG | None | None | None | None |
| IKBKG | None | None | None | None |
| IKBKG | None | None | None | None |
| CD40 | None | None | None | None |
| CD40 | None | None | None | None |
| SLC9A3R1 | None | None | None | None |
| BCHE | None | None | None | None |
| BCHE | None | None | None | None |
| BCHE | None | None | None | None |
| BCHE | None | None | None | None |
| BCHE | None | None | None | None |
| BCHE | None | None | None | None |
| BCHE | 0.63 | 4 | 2xqg | A |
| BCHE | None | None | None | None |
| BCHE | None | None | None | None |
| BCHE | None | None | None | None |
| BCHE | None | None | None | None |
| BCKDHB | 0.833 | 1 | 1v11 | B |
| BCKDHB | None | None | None | None |
| BCKDHB | None | None | None | None |

| | | | | |
|---|---|---|---|---|
| BCKDHB | 0.667 | 11 | 2bfd | B |
| BCKDHB | 0.833 | 10 | 1olx | B |
| BCKDHB | 0.833 | 1 | 1v11 | B |
| BCKDHB | 0.833 | 1 | 1v11 | B |
| BCKDHB | 0.625 | 2 | 1x7z | B |
| BCKDHB | None | None | None | None |
| BCKDHB | None | None | None | None |
| BCKDHB | None | None | None | None |
| BCKDHB | 0.857 | 1 | 2bff | B |
| CDH1 | None | None | None | None |
| CDH1 | None | None | None | None |
| CDH1 | None | None | None | None |
| CDH1 | None | None | None | None |
| CDH1 | None | None | None | None |
| CTNNB1 | None | None | None | None |
| AIRE | None | None | None | None |
| EDA | 1 | 1 | 1rj8 | A |
| EDA | None | None | None | None |
| EDA | None | None | None | None |
| EDA | None | None | None | None |
| EDA | None | None | None | None |
| SH2D1A | None | None | None | None |
| SH2D1A | None | None | None | None |
| SH2D1A | None | None | None | None |
| GCK | None | None | None | None |
| GCK | None | None | None | None |
| GCK | None | None | None | None |
| GCK | None | None | None | None |
| GCK | None | None | None | None |
| GCK | 0.611 | 13 | 3h1v | X |
| GCK | None | None | None | None |
| GCK | None | None | None | None |
| GCK | None | None | None | None |
| GCK | 0.778 | 5 | 3id8 | A |
| GCK | 0.833 | 1 | 4dhy | A |
| GCK | None | None | None | None |
| GCK | None | None | None | None |
| GCK | 0.611 | 6 | 3id8 | A |
| GCK | None | None | None | None |
| GCK | None | None | None | None |
| GCK | None | None | None | None |
| GCK | None | None | None | None |
| GCK | None | None | None | None |
| GCK | None | None | None | None |
| AQP2 | None | None | None | None |
| AQP2 | None | None | None | None |
| AQP2 | None | None | None | None |
| AQP2 | None | None | None | None |
| AQP2 | None | None | None | None |
| AQP2 | None | None | None | None |
| AQP2 | None | None | None | None |
| AQP2 | None | None | None | None |
| AQP2 | None | None | None | None |
| AQP2 | None | None | None | None |

| | | | | |
|---|---|---|---|---|
| AQP2 | None | None | None | None |
| AQP2 | None | None | None | None |
| AQP2 | None | None | None | None |
| SUMF1 | 0.75 | 6 | 1z70 | X |
| SUMF1 | None | None | None | None |
| SUMF1 | None | None | None | None |
| SUMF1 | 0.625 | 18 | 2aij | X |
| SUMF1 | 0.875 | 35 | 2afy | X |
| SUMF1 | 0.667 | 19 | 1y1i | X |
| SUMF1 | 0.75 | 17 | 1y1j | X |
| SUMF1 | 0.75 | 17 | 1y1j | X |
| SUMF1 | 0.625 | 27 | 1y1g | X |
| FAM20A | None | None | None | None |
| FAM20A | None | None | None | None |
| FAM20A | None | None | None | None |
| FAM20A | None | None | None | None |
| NT5E | None | None | None | None |
| NT5E | None | None | None | None |
| NOS3 | None | None | None | None |
| ACTG1 | None | None | None | None |
| ACTG1 | None | None | None | None |
| ACTG1 | None | None | None | None |
| ACTG1 | None | None | None | None |
| ACTG1 | None | None | None | None |
| ACTG1 | None | None | None | None |
| FCGR1A | None | None | None | None |
| GPI | None | None | None | None |
| GPI | None | None | None | None |
| GPI | 1 | 3 | 1jiq | A |
| GPI | None | None | None | None |
| GPI | 0.696 | 22 | 1iat | A |
| GPI | None | None | None | None |
| GPI | 0.696 | 27 | 1iat | A |
| GPI | None | None | None | None |
| GPI | None | None | None | None |
| CFH | None | None | None | None |
| OGG1 | 0.895 | 13 | 1m3h | A |
| TPK1 | None | None | None | None |
| TPK1 | None | None | None | None |
| TPK1 | None | None | None | None |
| TPK1 | None | None | None | None |
| SEC23A | None | None | None | None |
| SEC23A | None | None | None | None |
| TYRP1 | None | None | None | None |
| TYRP1 | None | None | None | None |
| TYRP1 | None | None | None | None |
| TYRP1 | None | None | None | None |
| TYRP1 | None | None | None | None |
| TYRP1 | None | None | None | None |
| CSTB | None | None | None | None |
| CSTB | None | None | None | None |
| CSTB | None | None | None | None |
| ACP1 | None | None | None | None |
| IL17F | None | None | None | None |

| | | | | |
|---|---|---|---|---|
| IL17F | None | None | None | None |
| IL17F | None | None | None | None |
| IL17F | None | None | None | None |
| IL17F | None | None | None | None |
| IDH2 | None | None | None | None |
| IDH2 | None | None | None | None |
| IDH2 | None | None | None | None |
| IDH2 | None | None | None | None |
| TP53 | 0.925 | 11 | 2ocj | A |
| TP53 | 0.8 | 16 | 2ahi | D |
| TP53 | 0.65 | 49 | 4ibq | A |
| TP53 | 0.725 | 27 | 5g4o | A |
| TP53 | None | None | None | None |
| TP53 | 0.9 | 1 | 4agp | A |
| TP53 | 0.9 | 1 | 4agp | A |
| TP53 | 0.925 | 10 | 4hje | A |
| TP53 | 0.7 | 29 | 3d05 | A |
| TP53 | 0.875 | 7 | 3kmd | B |
| TP53 | 0.875 | 7 | 3kmd | B |
| TP53 | 0.925 | 6 | 3kz8 | B |
| TP53 | 0.925 | 6 | 3kz8 | B |
| TP53 | 0.609 | 22 | 3q01 | B |
| TP53 | 0.925 | 5 | 3kmd | B |
| TP53 | 0.925 | 5 | 3kmd | B |
| TP53 | 0.925 | 8 | 3kz8 | B |
| TP53 | 0.925 | 8 | 3kz8 | B |
| TP53 | 0.925 | 1 | 3kmd | B |
| TP53 | 0.9 | 11 | 2pcx | A |
| TP53 | 0.9 | 1 | 5aol | A |
| TP53 | 0.85 | 5 | 5lgy | A |
| TP53 | 0.9 | 1 | 3q06 | B |
| TP53 | 0.725 | 19 | 5lgy | A |
| TP53 | 0.775 | 14 | 4mzi | A |
| TP53 | 0.825 | 8 | 1kzy | A |
| TP53 | 0.825 | 8 | 1kzy | A |
| TP53 | 0.8 | 11 | 4ibt | A |
| TP53 | 0.8 | 11 | 4ibt | A |
| TP53 | 0.725 | 15 | 5lgy | A |
| TP53 | 0.725 | 29 | 2h1l | O |
| TP53 | 0.875 | 7 | 3kmd | B |
| TP53 | 0.725 | 15 | 5lgy | A |
| TP53 | 0.8 | 10 | 4agp | A |
| TP53 | 0.8 | 10 | 4agp | A |
| TP53 | 0.8 | 10 | 4agp | A |
| TP53 | 0.8 | 10 | 4agp | A |
| TP53 | 0.675 | 35 | 4ibq | A |
| TP53 | None | None | None | None |
| TP53 | 0.675 | 17 | 3kz8 | B |
| TP53 | None | None | None | None |
| TP53 | None | None | None | None |
| LMAN1 | None | None | None | None |
| CTSK | 0.75 | 2 | 4n79 | A |
| CTSK | None | None | None | None |
| CTSK | None | None | None | None |

| | | | | |
|---|---|---|---|---|
| FGB | None | None | None | None |
| FGB | None | None | None | None |
| FGB | 0.688 | 1 | 1fzf | E |
| FGB | None | None | None | None |
| ACADM | None | None | None | None |
| ACADM | None | None | None | None |
| ACADM | None | None | None | None |
| ACADM | None | None | None | None |
| ACADM | None | None | None | None |
| ACADM | None | None | None | None |
| ACADM | None | None | None | None |
| ACADM | None | None | None | None |
| ACADM | None | None | None | None |
| ACADM | None | None | None | None |
| ACADM | None | None | None | None |
| ACADM | None | None | None | None |
| ACADM | None | None | None | None |
| ACADM | None | None | None | None |
| ACADM | None | None | None | None |
| ACADM | None | None | None | None |
| ACADM | None | None | None | None |
| ACADM | None | None | None | None |
| ACADM | None | None | None | None |
| ACADM | None | None | None | None |
| ACADM | None | None | None | None |
| ACADM | None | None | None | None |
| ACADM | None | None | None | None |
| ACADM | None | None | None | None |
| ACADM | None | None | None | None |
| ACADM | None | None | None | None |
| ACADM | None | None | None | None |
| PTS | None | None | None | None |
| PTS | None | None | None | None |
| PTS | None | None | None | None |
| PTS | 0.778 | 1 | 3i2b | D |
| PTS | None | None | None | None |
| PTS | None | None | None | None |
| PTS | None | None | None | None |
| PTS | None | None | None | None |
| PTS | None | None | None | None |
| INS | 0.75 | 2 | 3zqr | E |
| INS | 1 | 5 | 2l1z | A |
| INS | 1 | 5 | 2l1z | A |
| INS | 1 | 1 | 1his | A |
| FBP1 | 0.71 | 26 | 2y5l | C |
| FBP1 | None | None | None | None |
| FBP1 | None | None | None | None |
| FBP1 | 0.7 | 10 | 2y5k | A |
| FBP1 | None | None | None | None |
| FBP1 | 0.733 | 10 | 3kc1 | A |
| FBP1 | 0.633 | 43 | 2jjk | A |
| FBP1 | 0.8 | 11 | 2y5k | A |
| FBP1 | None | None | None | None |
| FBP1 | None | None | None | None |

| | | | | |
|---|---|---|---|---|
| FBP1 | 0.643 | 7 | 4mjo | B |
| KCNQ1 | None | None | None | None |
| KCNQ1 | None | None | None | None |
| KCNQ1 | None | None | None | None |
| UPP1 | 0.706 | 4 | 3euf | A |
| UMPS | 1 | 3 | 2qcl | A |
| UMPS | None | None | None | None |
| UMPS | None | None | None | None |
| DHFR | None | None | None | None |
| KHK | 0.667 | 2 | 2hw1 | A |
| TTPA | None | None | None | None |
| TTPA | None | None | None | None |
| TTPA | None | None | None | None |
| TTPA | None | None | None | None |
| MAP1LC3B | None | None | None | None |
| MAP1LC3B | None | None | None | None |
| ZAP70 | None | None | None | None |
| PROC | None | None | None | None |
| SQSTM1 | None | None | None | None |
| AHCY | None | None | None | None |
| AHCY | None | None | None | None |
| PRNP | None | None | None | None |
| PRNP | None | None | None | None |
| PRNP | None | None | None | None |
| PRNP | None | None | None | None |
| PRNP | None | None | None | None |
| PRNP | None | None | None | None |
| PRNP | None | None | None | None |
| PRNP | None | None | None | None |
| PRNP | None | None | None | None |
| PRNP | None | None | None | None |
| PRNP | None | None | None | None |
| PRNP | None | None | None | None |
| PRNP | None | None | None | None |
| PRNP | None | None | None | None |
| PRNP | None | None | None | None |
| PRNP | None | None | None | None |
| REN | None | None | None | None |
| REN | None | None | None | None |
| HINT1 | None | None | None | None |
| HINT1 | 0.897 | 1 | 5km4 | A |
| HINT1 | None | None | None | None |
| HINT1 | 0.667 | 11 | 5ed6 | B |
| HINT1 | 0.923 | 1 | 5ipc | A |
| OAT | None | None | None | None |
| OAT | None | None | None | None |
| OAT | None | None | None | None |
| OAT | None | None | None | None |
| OAT | None | None | None | None |
| OAT | None | None | None | None |
| OAT | None | None | None | None |
| OAT | None | None | None | None |
| OAT | None | None | None | None |
| OAT | None | None | None | None |

| | | | | |
|---|---|---|---|---|
| OAT | 0.675 | 2 | 2byl | B |
| OAT | None | None | None | None |
| OAT | None | None | None | None |
| OAT | None | None | None | None |
| OAT | None | None | None | None |
| OAT | 0.758 | 1 | 2byl | A |
| OAT | None | None | None | None |
| OAT | None | None | None | None |
| OAT | None | None | None | None |
| OAT | None | None | None | None |
| OAT | None | None | None | None |
| OAT | None | None | None | None |
| OAT | None | None | None | None |
| OAT | None | None | None | None |
| OAT | None | None | None | None |
| OAT | None | None | None | None |
| OAT | None | None | None | None |
| OAT | None | None | None | None |
| CTSA | None | None | None | None |
| CTSA | None | None | None | None |
| PNP | None | None | None | None |
| PNP | None | None | None | None |
| PNP | None | None | None | None |
| PNP | 0.706 | 5 | 4ear | C |
| PNP | None | None | None | None |
| PNP | None | None | None | None |
| PNP | 0.933 | 1 | 3ggs | C |
| PNP | 0.833 | 1 | 3d1v | A |
| MYL2 | None | None | None | None |
| MYL2 | None | None | None | None |
| ANG | None | None | None | None |
| ANG | None | None | None | None |
| ANG | None | None | None | None |
| ANG | None | None | None | None |
| ANG | 0.761 | 1 | 5eop | A |
| ANG | None | None | None | None |
| ANG | None | None | None | None |
| ANG | None | None | None | None |
| ANG | None | None | None | None |
| ANG | None | None | None | None |
| POLH | None | None | None | None |
| POLH | None | None | None | None |
| POLH | None | None | None | None |
| IL2RA | None | None | None | None |
| TGFB1 | None | None | None | None |
| TP53 | None | None | None | None |
| TP53 | None | None | None | None |
| PAH | None | None | None | None |
| PAH | None | None | None | None |
| PAH | None | None | None | None |
| PAH | None | None | None | None |
| PAH | None | None | None | None |
| PAH | None | None | None | None |
| PAH | None | None | None | None |

| | | | | |
|---|---|---|---|---|
| PAH | None | None | None | None |
| PAH | None | None | None | None |
| PAH | None | None | None | None |
| PAH | None | None | None | None |
| PAH | None | None | None | None |
| PAH | None | None | None | None |
| PAH | None | None | None | None |
| PAH | None | None | None | None |
| PAH | None | None | None | None |
| PAH | None | None | None | None |
| PAH | None | None | None | None |
| PAH | None | None | None | None |
| PAH | None | None | None | None |
| PAH | None | None | None | None |
| PAH | None | None | None | None |
| PAH | None | None | None | None |
| PAH | 0.68 | 6 | 2pah | A |
| PAH | 0.76 | 1 | 1tdw | A |
| PAH | None | None | None | None |
| PAH | None | None | None | None |
| PAH | None | None | None | None |
| PAH | None | None | None | None |
| PAH | None | None | None | None |
| PAH | None | None | None | None |
| PAH | None | None | None | None |
| PAH | 0.64 | 2 | 1lrm | A |
| PAH | None | None | None | None |
| PAH | None | None | None | None |
| PAH | None | None | None | None |
| PAH | None | None | None | None |
| PAH | None | None | None | None |
| PAH | None | None | None | None |
| PAH | None | None | None | None |
| PAH | None | None | None | None |
| HDAC8 | None | None | None | None |
| HDAC8 | 0.732 | 3 | 3rqd | A |
| VCP | None | None | None | None |
| VCP | None | None | None | None |
| VCP | None | None | None | None |
| VCP | None | None | None | None |
| VCP | None | None | None | None |
| VCP | None | None | None | None |
| VCP | None | None | None | None |
| B3GAT3 | None | None | None | None |
| B3GAT3 | None | None | None | None |
| CSF2RB | None | None | None | None |
| CSF2RB | None | None | None | None |
| CSF2RB | None | None | None | None |
| NF2 | None | None | None | None |
| NF2 | None | None | None | None |
| NF2 | None | None | None | None |
| PIK3CA | None | None | None | None |
| PHGDH | None | None | None | None |
| PHGDH | None | None | None | None |

| | | | | |
|---|---|---|---|---|
| PHGDH | None | None | None | None |
| PHGDH | 0.66 | | 1 5n53 | A |
| POLH | None | None | None | None |
| XIAP | None | None | None | None |
| XIAP | None | None | None | None |
| PDHX | None | None | None | None |
| PDHX | None | None | None | None |
| PDHX | None | None | None | None |
| PDHX | None | None | None | None |
| PDHX | None | None | None | None |
| SAMHD1 | None | None | None | None |
| SAMHD1 | None | None | None | None |
| SAMHD1 | None | None | None | None |
| SAMHD1 | None | None | None | None |
| SAMHD1 | None | None | None | None |
| SAMHD1 | None | None | None | None |
| SAMHD1 | None | None | None | None |
| SAMHD1 | None | None | None | None |
| CD36 | None | None | None | None |
| CD36 | None | None | None | None |
| CD36 | None | None | None | None |
| ASPH | None | None | None | None |
| CCT5 | None | None | None | None |
| CCT5 | None | None | None | None |
| CCT5 | None | None | None | None |
| SMN1 | None | None | None | None |
| SMN1 | None | None | None | None |
| SMN1 | None | None | None | None |
| SMN1 | None | None | None | None |
| ABCA1 | None | None | None | None |
| ABCA1 | None | None | None | None |
| ABCA1 | None | None | None | None |
| ABCA1 | None | None | None | None |
| ABCA1 | None | None | None | None |
| ABCA1 | None | None | None | None |
| ABCA1 | None | None | None | None |
| ABCA1 | None | None | None | None |
| ABCA1 | None | None | None | None |
| ABCA1 | None | None | None | None |
| ABCA1 | None | None | None | None |
| ABCA1 | None | None | None | None |
| ABCA1 | None | None | None | None |
| ABCA1 | None | None | None | None |
| ABCA1 | None | None | None | None |
| ABCA1 | None | None | None | None |
| ABCA1 | None | None | None | None |
| ABCA1 | None | None | None | None |
| ABCA1 | None | None | None | None |
| ABCA1 | None | None | None | None |
| ABCA1 | None | None | None | None |
| ABCA1 | None | None | None | None |
| ABCA1 | None | None | None | None |
| ABCA1 | None | None | None | None |
| ABCA1 | None | None | None | None |

| | | | | |
|---|---|---|---|---|
| ABCA1 | None | None | None | None |
| ABCA1 | None | None | None | None |
| ABCA1 | None | None | None | None |
| ABCA1 | None | None | None | None |
| ABCA1 | None | None | None | None |
| ABCA1 | None | None | None | None |
| ABCA1 | None | None | None | None |
| ABCA1 | None | None | None | None |
| ABCA1 | None | None | None | None |
| ABCA1 | None | None | None | None |
| ABCA1 | None | None | None | None |
| ERCC1 | None | None | None | None |
| FECH | None | None | None | None |
| FECH | 0.6 | 60 | 2hre | A |
| FECH | 0.636 | 20 | 2po7 | A |
| FECH | None | None | None | None |
| FECH | None | None | None | None |
| FECH | 0.667 | 10 | 2po5 | A |
| FECH | None | None | None | None |
| PTEN | None | None | None | None |
| PTEN | None | None | None | None |
| PTEN | None | None | None | None |
| PTEN | None | None | None | None |
| PTEN | None | None | None | None |
| PTEN | None | None | None | None |
| PTEN | None | None | None | None |
| PTEN | None | None | None | None |
| PTEN | None | None | None | None |
| PTEN | None | None | None | None |
| PTEN | None | None | None | None |
| PTEN | None | None | None | None |
| PTEN | None | None | None | None |
| PTEN | None | None | None | None |
| PTEN | None | None | None | None |
| PTEN | None | None | None | None |
| PTEN | None | None | None | None |
| PTEN | None | None | None | None |
| PTEN | None | None | None | None |
| PTEN | None | None | None | None |
| ACAT1 | 0.667 | 1 | 2ib9 | C |
| ACAT1 | None | None | None | None |
| ACAT1 | None | None | None | None |
| ACAT1 | 0.667 | 1 | 2ib9 | C |
| ACAT1 | 0.667 | 1 | 2ib9 | C |
| ACAT1 | None | None | None | None |
| ACAT1 | 0.833 | 6 | 2ib7 | C |
| ACAT1 | None | None | None | None |
| ACAT1 | None | None | None | None |
| ACAT1 | None | None | None | None |
| ACAT1 | 0.703 | 16 | 2ibu | A |
| CREBBP | None | None | None | None |
| ITGA2B | None | None | None | None |
| ITGA2B | None | None | None | None |
| ITGA2B | None | None | None | None |

| | | | | |
|---|---|---|---|---|
| ITGA2B | None | None | None | None |
| TSC1 | None | None | None | None |
| ADA2 | None | None | None | None |
| ADA2 | None | None | None | None |
| ADA2 | 0.75 | 1 | 3lgd | A |
| ADA2 | None | None | None | None |
| ADA2 | None | None | None | None |
| ADA2 | None | None | None | None |
| ADA2 | None | None | None | None |
| ADA2 | None | None | None | None |
| ADA2 | 1 | 1 | 3lgd | B |
| ADA2 | None | None | None | None |
| PGK1 | None | None | None | None |
| PGK1 | None | None | None | None |
| PGK1 | None | None | None | None |
| PGK1 | None | None | None | None |
| PGK1 | None | None | None | None |
| TUBA1A | None | None | None | None |
| TUBA1A | None | None | None | None |
| TUBA1A | None | None | None | None |
| TUBA1A | None | None | None | None |
| TUBA1A | None | None | None | None |
| PTPN11 | None | None | None | None |
| PTPN11 | None | None | None | None |
| PTPN11 | None | None | None | None |
| PTPN11 | None | None | None | None |
| PTPN11 | None | None | None | None |
| PTPN11 | None | None | None | None |
| PTPN11 | None | None | None | None |
| PTPN11 | None | None | None | None |
| PTPN11 | None | None | None | None |
| PTPN11 | None | None | None | None |
| PTPN11 | None | None | None | None |
| PTPN11 | None | None | None | None |
| PTPN11 | None | None | None | None |
| PTPN11 | None | None | None | None |
| PTPN11 | 0.621 | 2 | 3b7o | A |
| PTPN11 | 0.776 | 1 | 3zm1 | A |
| PTPN11 | None | None | None | None |
| PTPN11 | None | None | None | None |
| PTPN11 | None | None | None | None |
| PTPN11 | None | None | None | None |
| PTPN11 | None | None | None | None |
| PTPN11 | 0.776 | 1 | 3zm1 | A |
| PTPN11 | 0.621 | 6 | 3zm1 | A |
| PTPN11 | None | None | None | None |
| PTPN11 | None | None | None | None |
| PTPN11 | 0.621 | 6 | 3zm1 | A |
| GLB1 | None | None | None | None |
| GLB1 | None | None | None | None |
| UQCRQ | None | None | None | None |
| VPS35 | None | None | None | None |
| NR3C1 | None | None | None | None |
| NR3C1 | None | None | None | None |

| | | | | |
|---|---|---|---|---|
| NR3C1 | None | None | None | None |
| NR3C1 | None | None | None | None |
| NR3C1 | None | None | None | None |
| NR3C1 | None | None | None | None |
| NR3C1 | None | None | None | None |
| NR3C1 | None | None | None | None |
| NR3C1 | None | None | None | None |
| APOA1 | None | None | None | None |
| APOA1 | None | None | None | None |
| APOA1 | None | None | None | None |
| APOA1 | None | None | None | None |
| APOA1 | None | None | None | None |
| APOA1 | None | None | None | None |
| APOA1 | None | None | None | None |
| BCKDHA | None | None | None | None |
| BCKDHA | None | None | None | None |
| BCKDHA | 0.963 | 1 | 1ols | A |
| BCKDHA | 0.696 | 3 | 1v1r | A |
| BCKDHA | 0.625 | 16 | 2beu | A |
| BCKDHA | None | None | None | None |
| BCKDHA | 0.652 | 11 | 1v1r | A |
| BCKDHA | 1 | 1 | 2beu | A |
| BCKDHA | 0.84 | 2 | 2j9f | A |
| BCKDHA | None | None | None | None |
| BCKDHA | 0.609 | 90 | 1v1r | A |
| BCKDHA | 0.652 | 25 | 1v1r | A |
| BCKDHA | None | None | None | None |
| BCKDHA | 0.625 | 60 | 2beu | A |
| BCKDHA | 0.625 | 28 | 2beu | A |
| TOP1 | None | None | None | None |
| ANTXR2 | None | None | None | None |
| ANTXR2 | None | None | None | None |
| ANTXR2 | None | None | None | None |
| TF | 0.625 | 35 | 5dyh | A |
| TF | None | None | None | None |
| FSHB | None | None | None | None |
| GALK1 | None | None | None | None |
| GALK1 | None | None | None | None |
| GALK1 | None | None | None | None |
| CYP17A1 | None | None | None | None |
| CYP17A1 | None | None | None | None |
| CYP17A1 | None | None | None | None |
| CYP17A1 | None | None | None | None |
| CYP17A1 | None | None | None | None |
| CYP17A1 | None | None | None | None |
| CYP17A1 | None | None | None | None |
| CYP17A1 | None | None | None | None |
| CYP17A1 | None | None | None | None |
| CYP17A1 | None | None | None | None |
| CYP17A1 | None | None | None | None |
| CYP17A1 | None | None | None | None |
| CYP17A1 | None | None | None | None |
| CYP17A1 | None | None | None | None |
| CYP17A1 | None | None | None | None |

| | | | | |
|---|---|---|---|---|
| CYP17A1 | None | None | None | None |
| CYP17A1 | None | None | None | None |
| CD44 | 0.667 | 24 | 1uuh | A |
| MDH2 | None | None | None | None |
| MDH2 | None | None | None | None |
| NDUFV1 | None | None | None | None |
| NDUFV1 | None | None | None | None |
| NDUFV1 | None | None | None | None |
| NDUFV1 | None | None | None | None |
| NDUFV1 | None | None | None | None |
| NDUFV1 | None | None | None | None |
| NDUFV1 | None | None | None | None |
| NDUFV1 | None | None | None | None |
| NDUFV1 | None | None | None | None |
| NDUFV1 | None | None | None | None |
| CTSC | None | None | None | None |
| CTSC | None | None | None | None |
| CTSC | None | None | None | None |
| CTSC | None | None | None | None |
| CTSC | 0.643 | 1 | 4oel | A |
| CTSC | None | None | None | None |
| CTSC | None | None | None | None |
| VWF | None | None | None | None |
| VWF | None | None | None | None |
| VWF | None | None | None | None |
| VWF | None | None | None | None |
| VWF | None | None | None | None |
| VWF | None | None | None | None |
| VWF | None | None | None | None |
| ORC4 | None | None | None | None |
| ORC4 | None | None | None | None |
| CRYAB | None | None | None | None |
| CRYAB | None | None | None | None |
| CRYAB | None | None | None | None |
| MAP2K2 | None | None | None | None |
| MAP2K2 | None | None | None | None |
| MAP2K2 | None | None | None | None |
| MAP2K2 | None | None | None | None |
| MAP2K2 | None | None | None | None |
| MAP2K2 | None | None | None | None |
| MAP2K2 | None | None | None | None |
| MAP2K2 | None | None | None | None |
| MAP2K2 | None | None | None | None |
| CBL | None | None | None | None |
| CBL | None | None | None | None |
| CBL | None | None | None | None |
| CBL | None | None | None | None |
| CBL | None | None | None | None |
| CBL | None | None | None | None |
| CBL | None | None | None | None |
| PFN1 | None | None | None | None |
| PFN1 | 0.75 | 2 | 1fik | A |
| ACSM2A | None | None | None | None |
| HSPG2 | None | None | None | None |

| | | | | |
|---|---|---|---|---|
| HSPG2 | None | None | None | None |
| HSPG2 | None | None | None | None |
| HSPG2 | None | None | None | None |
| PARK7 | None | None | None | None |
| PARK7 | 0.769 | 4 | 1pe0 | A |
| PARK7 | 0.808 | 2 | 1pe0 | A |
| PARK7 | 0.667 | 8 | 4rky | A |
| PARK7 | 0.889 | 1 | 3cz9 | A |
| PARK7 | 0.75 | 2 | 1pdw | B |
| INSR | None | None | None | None |
| INSR | None | None | None | None |
| INSR | None | None | None | None |
| INSR | None | None | None | None |
| INSR | None | None | None | None |
| DSC2 | None | None | None | None |
| DSC2 | None | None | None | None |
| DSC2 | None | None | None | None |
| DSC2 | None | None | None | None |
| DSC2 | None | None | None | None |
| DSC2 | None | None | None | None |
| DSC2 | None | None | None | None |
| DSC2 | None | None | None | None |
| DSC2 | None | None | None | None |
| DSC2 | None | None | None | None |
| MICU1 | None | None | None | None |
| MICU1 | None | None | None | None |
| ADAMTS13 | None | None | None | None |
| ADAMTS13 | None | None | None | None |
| ADAMTS13 | None | None | None | None |
| ADAMTS13 | None | None | None | None |
| ADAMTS13 | None | None | None | None |
| ADAMTS13 | None | None | None | None |
| ADAMTS13 | None | None | None | None |
| IRAK4 | None | None | None | None |
| IRAK4 | None | None | None | None |
| RNASEH2B | None | None | None | None |
| RNASEH2B | None | None | None | None |
| RNASEH2B | None | None | None | None |
| NTRK1 | None | None | None | None |
| MPO | 0.75 | 3 | 1d5l | C |
| MPO | 0.6 | 25 | 1myp | D |
| MPO | None | None | None | None |
| PEX3 | None | None | None | None |
| SERPINA1 | None | None | None | None |
| SERPINA1 | None | None | None | None |
| SERPINA1 | None | None | None | None |
| SERPINA1 | None | None | None | None |
| SERPINA1 | None | None | None | None |
| SERPINA1 | None | None | None | None |
| SERPINA1 | None | None | None | None |
| SERPINA1 | 0.6 | 1 | 1ezx | A |
| SERPINA1 | None | None | None | None |
| SERPINA1 | None | None | None | None |
| YARS2 | None | None | None | None |

| | | | | |
|---|---|---|---|---|
| YARS2 | None | None | None | None |
| YARS2 | None | None | None | None |
| YARS2 | None | None | None | None |
| YARS2 | None | None | None | None |
| YARS2 | 0.781 | 1 | 2pid | B |
| YARS2 | None | None | None | None |
| YARS2 | None | None | None | None |
| YARS2 | None | None | None | None |
| PDHA1 | None | None | None | None |
| PDHA1 | None | None | None | None |
| PDHA1 | None | None | None | None |
| PDHA1 | None | None | None | None |
| PDHA1 | None | None | None | None |
| PDHA1 | None | None | None | None |
| PDHA1 | None | None | None | None |
| PDHA1 | 0.96 | 1 | 3exh | C |
| PDHA1 | None | None | None | None |
| PDHA1 | None | None | None | None |
| PDHA1 | None | None | None | None |
| CUL4B | None | None | None | None |
| AGXT | None | None | None | None |
| AGXT | None | None | None | None |
| AGXT | None | None | None | None |
| AGXT | 0.692 | 3 | 2yob | A |
| AGXT | 0.786 | 7 | 4kxk | C |
| AGXT | None | None | None | None |
| AGXT | None | None | None | None |
| AGXT | None | None | None | None |
| AGXT | None | None | None | None |
| AGXT | 0.611 | 4 | 5hhy | B |
| AGXT | None | None | None | None |
| AGXT | None | None | None | None |
| AGXT | None | None | None | None |
| AGXT | None | None | None | None |
| AGXT | None | None | None | None |
| AGXT | 0.684 | 1 | 1h0c | A |
| AGXT | None | None | None | None |
| AGXT | None | None | None | None |
| AGXT | None | None | None | None |
| AGXT | 0.75 | 2 | 5luc | B |
| AGXT | 0.722 | 3 | 5hhy | B |
| AGXT | None | None | None | None |
| AGXT | None | None | None | None |
| AGXT | None | None | None | None |
| AGXT | 0.778 | 1 | 5hhy | B |
| AGXT | 0.778 | 1 | 5hhy | B |
| AGXT | None | None | None | None |
| AGXT | 0.692 | 7 | 4kyo | A |
| AGXT | None | None | None | None |
| AGXT | None | None | None | None |
| AGXT | 0.615 | 13 | 3r9a | A |
| AGXT | 0.786 | 7 | 4kxk | C |
| AGXT | 0.846 | 1 | 4kxk | A |
| AGXT | 0.615 | 36 | 3r9a | A |

| | | | | |
|---|---|---|---|---|
| AGXT | None | None | None | None |
| SFTPD | None | None | None | None |
| SFTPD | None | None | None | None |
| PIK3CA | None | None | None | None |
| PIK3CA | None | None | None | None |
| PIK3CA | None | None | None | None |
| AUH | None | None | None | None |
| AUH | None | None | None | None |
| AUH | None | None | None | None |
| AUH | None | None | None | None |
| F11 | None | None | None | None |
| F11 | None | None | None | None |
| F11 | None | None | None | None |
| F11 | None | None | None | None |
| MEN1 | 0.6 | 81 | 5db0 | A |
| MEN1 | 0.8 | 54 | 4i80 | A |
| MEN1 | 0.8 | 84 | 5db0 | A |
| MEN1 | 0.8 | 30 | 5ddb | A |
| MEN1 | 1 | 2 | 4i80 | A |
| MEN1 | 0.8 | 17 | 5db0 | A |
| MEN1 | 0.6 | 17 | 5ddb | A |
| MEN1 | 0.6 | 31 | 3u88 | A |
| MEN1 | None | None | None | None |
| MEN1 | 0.6 | 56 | 5ddb | A |
| MYBPC1 | None | None | None | None |
| MYBPC1 | None | None | None | None |
| MYBPC1 | None | None | None | None |
| PDCD10 | None | None | None | None |
| SCARB2 | None | None | None | None |
| SCARB2 | None | None | None | None |
| SCARB2 | None | None | None | None |
| SCARB2 | None | None | None | None |
| SCARB2 | None | None | None | None |
| TREX2 | None | None | None | None |
| TBK1 | None | None | None | None |
| TBK1 | None | None | None | None |
| DLD | None | None | None | None |
| DLD | None | None | None | None |
| DLD | None | None | None | None |
| DLD | None | None | None | None |
| DLD | None | None | None | None |
| DLD | None | None | None | None |
| LDHA | 0.802 | 1 | 4qt0 | A |
| TNNC1 | None | None | None | None |
| TNNC1 | None | None | None | None |
| TNNC1 | None | None | None | None |
| SMAD3 | None | None | None | None |
| HARS | None | None | None | None |
| HARS | None | None | None | None |
| TNNI3 | None | None | None | None |
| TNNI3 | None | None | None | None |
| TNNI3 | None | None | None | None |
| TNNI3 | None | None | None | None |
| TNNI3 | None | None | None | None |

| | | | | |
|---|---|---|---|---|
| TNNI3 | None | None | None | None |
| ACKR1 | None | None | None | None |
| BUB1B | None | None | None | None |
| BUB1B | None | None | None | None |
| BUB1B | None | None | None | None |
| AR | None | None | None | None |
| AR | None | None | None | None |
| AR | None | None | None | None |
| AR | None | None | None | None |
| AR | None | None | None | None |
| FTO | None | None | None | None |
| FTO | None | None | None | None |
| TFAM | None | None | None | None |
| PYGL | None | None | None | None |
| PYGL | 0.778 | 3 | 1fa9 | A |
| PYGL | None | None | None | None |
| PYGL | 0.722 | 2 | 1fa9 | A |
| PYGL | None | None | None | None |
| PYGL | 0.611 | 25 | 3ceh | B |
| PYGL | 0.611 | 32 | 1l7x | B |
| PYGL | None | None | None | None |
| PYGL | 0.722 | 2 | 1fc0 | B |
| PYGL | 0.778 | 3 | 1fa9 | A |
| PYGL | None | None | None | None |
| PYGL | 0.722 | 12 | 2ati | A |
| PYGL | 0.722 | 1 | 3ceh | B |
| PYGL | 0.611 | 11 | 3ceh | B |
| PYGL | 0.611 | 18 | 1fc0 | B |
| PYGL | 0.722 | 2 | 1fc0 | B |
| GALNS | None | None | None | None |
| GALNS | None | None | None | None |
| GALNS | None | None | None | None |
| GALNS | None | None | None | None |
| GALNS | None | None | None | None |
| GALNS | None | None | None | None |
| GALNS | None | None | None | None |
| GALNS | None | None | None | None |
| GALNS | None | None | None | None |
| GALNS | None | None | None | None |
| GALNS | None | None | None | None |
| GALNS | None | None | None | None |
| GALNS | None | None | None | None |
| GALNS | None | None | None | None |
| GALNS | None | None | None | None |
| GALNS | None | None | None | None |
| GALNS | None | None | None | None |
| GALNS | None | None | None | None |
| GALNS | None | None | None | None |
| GALNS | None | None | None | None |
| GALNS | None | None | None | None |
| GALNS | None | None | None | None |
| GALNS | None | None | None | None |
| GALNS | None | None | None | None |
| GALNS | None | None | None | None |
| GALNS | None | None | None | None |

| | | | | |
|---|---|---|---|---|
| GALNS | None | None | None | None |
| GALNS | None | None | None | None |
| GALNS | None | None | None | None |
| GALNS | None | None | None | None |
| GALNS | None | None | None | None |
| GALNS | None | None | None | None |
| GALNS | None | None | None | None |
| GALNS | None | None | None | None |
| F10 | None | None | None | None |
| GLUD1 | None | None | None | None |
| GLUD1 | None | None | None | None |
| GLUD1 | None | None | None | None |
| GLUD1 | None | None | None | None |
| GLUD1 | None | None | None | None |
| GLUD1 | None | None | None | None |
| GLUD1 | None | None | None | None |
| GLUD1 | None | None | None | None |
| GLUD1 | None | None | None | None |
| GLUD1 | None | None | None | None |
| NOTCH1 | None | None | None | None |
| POGLUT1 | None | None | None | None |
| POGLUT1 | None | None | None | None |
| NR3C1 | None | None | None | None |
| CASP8 | None | None | None | None |
| CASP8 | None | None | None | None |
| TRAP1 | None | None | None | None |
| FANCM | None | None | None | None |
| EPOR | None | None | None | None |
| EPOR | None | None | None | None |
| NDP | None | None | None | None |
| NDP | None | None | None | None |
| NDP | None | None | None | None |
| NDP | None | None | None | None |
| NDP | None | None | None | None |
| NDP | None | None | None | None |
| NDP | None | None | None | None |
| NDP | None | None | None | None |
| NDP | None | None | None | None |
| NDP | None | None | None | None |
| NDP | None | None | None | None |
| NDP | None | None | None | None |
| NDP | None | None | None | None |
| NDP | None | None | None | None |
| NDP | None | None | None | None |
| NDP | None | None | None | None |
| NDP | None | None | None | None |
| ROGDI | None | None | None | None |
| CYP19A1 | None | None | None | None |
| CYP19A1 | None | None | None | None |
| CYP19A1 | None | None | None | None |
| CYP19A1 | None | None | None | None |
| CYP19A1 | None | None | None | None |
| CYP19A1 | None | None | None | None |
| CYP19A1 | None | None | None | None |

| | | | | |
|---|---|---|---|---|
| CYP19A1 | None | None | None | None |
| UBA5 | None | None | None | None |
| PRPS1 | None | None | None | None |
| PRPS1 | None | None | None | None |
| PRPS1 | None | None | None | None |
| PRPS1 | None | None | None | None |
| PRPS1 | None | None | None | None |
| PRPS1 | None | None | None | None |
| PRPS1 | None | None | None | None |
| PRPS1 | None | None | None | None |
| F2 | None | None | None | None |
| F2 | None | None | None | None |
| NTF4 | None | None | None | None |
| NTF4 | None | None | None | None |
| PSAP | None | None | None | None |
| PSAP | None | None | None | None |
| TAP1 | None | None | None | None |
| TAP1 | None | None | None | None |
| TAP1 | None | None | None | None |
| ARG1 | None | None | None | None |
| ARG1 | 0.714 | 1 | 2pha | A |
| ARG1 | None | None | None | None |
| ARG1 | None | None | None | None |
| ARG1 | None | None | None | None |
| ARG1 | None | None | None | None |
| CRYM | None | None | None | None |
| ADSL | None | None | None | None |
| ADSL | None | None | None | None |
| ADSL | None | None | None | None |
| ADSL | None | None | None | None |
| ADSL | None | None | None | None |
| ADSL | None | None | None | None |
| ADSL | None | None | None | None |
| ADSL | None | None | None | None |
| ADSL | None | None | None | None |
| ADSL | None | None | None | None |
| ADSL | None | None | None | None |
| ADSL | None | None | None | None |
| ADSL | 0.647 | 1 | 2vd6 | D |
| ADSL | None | None | None | None |
| ADSL | None | None | None | None |
| ADSL | None | None | None | None |
| ADSL | None | None | None | None |
| ADSL | None | None | None | None |
| ADSL | None | None | None | None |
| ADSL | None | None | None | None |
| VWF | None | None | None | None |
| PLA2G7 | None | None | None | None |
| MMP13 | None | None | None | None |
| MMP13 | None | None | None | None |
| MMP13 | None | None | None | None |
| MMP13 | None | None | None | None |
| POMGNT1 | None | None | None | None |
| POMGNT1 | None | None | None | None |

| | | | | |
|---|---|---|---|---|
| COL3A1 | None | None | None | None |
| COL3A1 | None | None | None | None |
| COL3A1 | None | None | None | None |
| COL3A1 | None | None | None | None |
| ACTN2 | None | None | None | None |
| ACTN2 | None | None | None | None |
| ACTN2 | None | None | None | None |
| ACTN2 | None | None | None | None |
| ACTN2 | None | None | None | None |
| ACTN2 | None | None | None | None |
| ACTN2 | None | None | None | None |
| UBA5 | None | None | None | None |
| ADAR | 0.75 | 1 | 1qbj | B |
| UMPS | None | None | None | None |
| UMPS | None | None | None | None |
| UMPS | None | None | None | None |
| TRNT1 | None | None | None | None |
| TRNT1 | None | None | None | None |
| NHEJ1 | None | None | None | None |
| NHEJ1 | None | None | None | None |
| NHEJ1 | None | None | None | None |
| C8A | None | None | None | None |
| GLUL | None | None | None | None |
| GLUL | None | None | None | None |
| GLUL | None | None | None | None |
| GYG1 | 0.615 | 45 | 3rmv | A |
| GYG1 | 0.923 | 1 | 3u2u | B |
| FLNA | None | None | None | None |
| FLNA | None | None | None | None |
| FLNA | None | None | None | None |
| FLNA | None | None | None | None |
| FLNA | None | None | None | None |
| FLNA | None | None | None | None |
| FLNA | None | None | None | None |
| TTN | None | None | None | None |
| TTN | None | None | None | None |
| TTN | None | None | None | None |
| TTN | None | None | None | None |
| SMS | None | None | None | None |
| SMS | None | None | None | None |
| SMS | None | None | None | None |
| C5 | None | None | None | None |
| SERPINC1 | None | None | None | None |
| SERPINC1 | None | None | None | None |
| SERPINC1 | None | None | None | None |
| SERPINC1 | None | None | None | None |
| SERPINC1 | None | None | None | None |
| SERPINC1 | None | None | None | None |
| SERPINC1 | None | None | None | None |
| SERPINC1 | None | None | None | None |
| SERPINC1 | None | None | None | None |
| SERPINC1 | None | None | None | None |
| SERPINC1 | None | None | None | None |
| SERPINC1 | None | None | None | None |

| | | | | |
|---|---|---|---|---|
| SERPINC1 | None | None | None | None |
| SERPINC1 | None | None | None | None |
| SERPINC1 | None | None | None | None |
| SERPINC1 | None | None | None | None |
| SERPINC1 | None | None | None | None |
| SERPINC1 | None | None | None | None |
| SERPINC1 | None | None | None | None |
| SERPINC1 | None | None | None | None |
| SERPINC1 | None | None | None | None |
| ETFB | None | None | None | None |
| ETFB | None | None | None | None |
| ETFB | None | None | None | None |
| ETFB | None | None | None | None |
| ETFB | None | None | None | None |
| ETFB | None | None | None | None |
| DIABLO | None | None | None | None |
| DIABLO | None | None | None | None |
| DPYS | None | None | None | None |
| DPYS | None | None | None | None |
| DPYS | None | None | None | None |
| DPYS | None | None | None | None |
| DPYS | None | None | None | None |
| GNE | None | None | None | None |
| GNE | None | None | None | None |
| GNE | None | None | None | None |
| GNE | None | None | None | None |
| GNE | None | None | None | None |
| GNE | None | None | None | None |
| GNE | None | None | None | None |
| GNE | None | None | None | None |
| GNE | None | None | None | None |
| GNMT | None | None | None | None |
| GNMT | None | None | None | None |
| CTH | None | None | None | None |
| CTH | None | None | None | None |
| CTH | None | None | None | None |
| USH1C | None | None | None | None |
| PSEN1 | None | None | None | None |
| CHEK2 | None | None | None | None |
| CHEK2 | None | None | None | None |
| CHEK2 | None | None | None | None |
| CHEK2 | None | None | None | None |
| CHEK2 | None | None | None | None |
| CHEK2 | None | None | None | None |
| CHEK2 | None | None | None | None |
| CHEK2 | None | None | None | None |
| CHEK2 | None | None | None | None |
| SNCA | None | None | None | None |
| SNCA | None | None | None | None |
| GAN | None | None | None | None |
| GAN | None | None | None | None |
| GAN | None | None | None | None |
| SBDS | None | None | None | None |
| SBDS | None | None | None | None |

| | | | | |
|---|---|---|---|---|
| SBDS | None | None | None | None |
| SBDS | None | None | None | None |
| SBDS | None | None | None | None |
| SBDS | None | None | None | None |
| MPO | | 1 | 3 | 3f9p | A |
| MPO | | 0.6 | 12 | 5mfa | A |
| KCNH2 | None | None | None | None |
| KCNH2 | None | None | None | None |
| KCNH2 | | 0.75 | 1 | 4hp9 | A |
| KCNH2 | None | None | None | None |
| TEK | None | None | None | None |
| TEK | None | None | None | None |
| TEK | None | None | None | None |
| TEK | None | None | None | None |
| NPC2 | None | None | None | None |
| NPC2 | None | None | None | None |
| NPC2 | None | None | None | None |
| NPC2 | None | None | None | None |
| NPC2 | None | None | None | None |
| NPC2 | None | None | None | None |
| NPC2 | None | None | None | None |
| PLG | None | None | None | None |
| PCSK9 | None | None | None | None |
| SFTPC | None | None | None | None |
| SFTPC | None | None | None | None |
| SFTPC | None | None | None | None |
| CD40LG | None | None | None | None |
| CD40LG | None | None | None | None |
| CD40LG | None | None | None | None |
| CD40LG | None | None | None | None |
| CD40LG | None | None | None | None |
| TAT | None | None | None | None |
| TAT | None | None | None | None |
| TAT | None | None | None | None |
| EP300 | None | None | None | None |
| FARS2 | None | None | None | None |
| FARS2 | None | None | None | None |
| FARS2 | None | None | None | None |
| FARS2 | None | None | None | None |
| FARS2 | None | None | None | None |
| FARS2 | None | None | None | None |
| FARS2 | None | None | None | None |
| FARS2 | None | None | None | None |
| FARS2 | None | None | None | None |
| FARS2 | None | None | None | None |
| FARS2 | None | None | None | None |
| FARS2 | None | None | None | None |
| FARS2 | None | None | None | None |
| MASP2 | None | None | None | None |
| MASP2 | None | None | None | None |
| MASP2 | None | None | None | None |
| MASP2 | None | None | None | None |
| STIM1 | None | None | None | None |
| CHRM2 | None | None | None | None |

| | | | | |
|---|---|---|---|---|
| MCFD2 | None | None | None | None |
| MCFD2 | None | None | None | None |
| MCFD2 | 0.623 | 1 | 3wnx | B |
| KITLG | None | None | None | None |
| F12 | None | None | None | None |
| F12 | None | None | None | None |
| ERAP1 | None | None | None | None |
| GTF2H5 | None | None | None | None |
| GTF2H5 | None | None | None | None |
| THBD | None | None | None | None |
| MRPL28 | None | None | None | None |
| MRPL28 | None | None | None | None |
| HTRA1 | None | None | None | None |
| HTRA1 | None | None | None | None |
| HTRA1 | None | None | None | None |
| HTRA1 | None | None | None | None |
| HTRA1 | None | None | None | None |
| HTRA1 | None | None | None | None |
| KARS | None | None | None | None |
| KARS | None | None | None | None |
| KARS | None | None | None | None |
| KARS | None | None | None | None |
| KARS | None | None | None | None |
| HK1 | 0.722 | 11 | 4fpb | B |
| GARS | None | None | None | None |
| GARS | None | None | None | None |
| GARS | None | None | None | None |
| GARS | None | None | None | None |
| GARS | None | None | None | None |
| GARS | None | None | None | None |
| GARS | None | None | None | None |
| IL12B | None | None | None | None |
| HADH | None | None | None | None |
| HADH | None | None | None | None |
| HADH | None | None | None | None |
| HADH | None | None | None | None |
| HADH | None | None | None | None |
| RRM2B | None | None | None | None |
| RRM2B | None | None | None | None |
| RRM2B | None | None | None | None |
| RRM2B | None | None | None | None |
| RRM2B | None | None | None | None |
| RRM2B | None | None | None | None |
| RRM2B | None | None | None | None |
| RRM2B | None | None | None | None |
| RRM2B | None | None | None | None |
| RRM2B | None | None | None | None |
| RRM2B | None | None | None | None |
| RRM2B | None | None | None | None |
| RRM2B | None | None | None | None |
| RRM2B | None | None | None | None |
| RRM2B | None | None | None | None |
| MARVELD2 | None | None | None | None |
| TLR4 | None | None | None | None |

| | | | | |
|---|---|---|---|---|
| TTN | None | None | None | None |
| TTN | None | None | None | None |
| TNFRSF13B | None | None | None | None |
| POLG | None | None | None | None |
| POLG | None | None | None | None |
| POLG | None | None | None | None |
| POLG | None | None | None | None |
| POLG | None | None | None | None |
| POLG | None | None | None | None |
| POLG | None | None | None | None |
| POLG | None | None | None | None |
| POLG | None | None | None | None |
| POLG | None | None | None | None |
| POLG | None | None | None | None |
| POLG | None | None | None | None |
| POLG | None | None | None | None |
| POLG | None | None | None | None |
| POLG | None | None | None | None |
| POLG | None | None | None | None |
| POLG | None | None | None | None |
| POLG | None | None | None | None |
| POLG | None | None | None | None |
| POLG | None | None | None | None |
| POLG | None | None | None | None |
| POLG | None | None | None | None |
| POLG | None | None | None | None |
| POLG | None | None | None | None |
| POLG | None | None | None | None |
| POLG | None | None | None | None |
| POLG | None | None | None | None |
| POLG | None | None | None | None |
| POLG | None | None | None | None |
| POLG | None | None | None | None |
| POLG | None | None | None | None |
| POLG | None | None | None | None |
| POLG | None | None | None | None |
| POLG | None | None | None | None |
| POLG | None | None | None | None |
| POLG | None | None | None | None |
| POLG | None | None | None | None |
| POLG | None | None | None | None |
| POLG | None | None | None | None |
| POLG | None | None | None | None |
| POLG | None | None | None | None |
| POLG | None | None | None | None |
| POLG | None | None | None | None |
| POLG | None | None | None | None |
| POLG | None | None | None | None |
| POLG | None | None | None | None |
| POLG | None | None | None | None |
| POLG | None | None | None | None |
| POLG | None | None | None | None |
| POLG | None | None | None | None |
| POLG | None | None | None | None |
| POLG | None | None | None | None |
| UROD | 0.632 | 24 | 1jpk | A |

| | | | | |
|---|---|---|---|---|
| UROD | None | None | None | None |
| UROD | None | None | None | None |
| UROD | None | None | None | None |
| UROD | None | None | None | None |
| UROD | 0.632 | 13 | 3gvw | A |
| UROD | 0.789 | 6 | 3gvw | A |
| UROD | 0.842 | 2 | 3gvv | A |
| UROD | 0.737 | 31 | 1r3q | A |
| UROD | 0.667 | 14 | 2q71 | A |
| PCK1 | None | None | None | None |
| PCK1 | None | None | None | None |
| PCK1 | None | None | None | None |
| PCK1 | None | None | None | None |
| PCK1 | None | None | None | None |
| HEXA | None | None | None | None |
| HEXA | None | None | None | None |
| HEXA | None | None | None | None |
| HEXA | None | None | None | None |
| HEXA | None | None | None | None |
| HEXA | None | None | None | None |
| HEXA | None | None | None | None |
| HEXA | None | None | None | None |
| HEXA | None | None | None | None |
| HEXA | None | None | None | None |
| HEXA | None | None | None | None |
| HEXA | None | None | None | None |
| HEXA | None | None | None | None |
| HEXA | None | None | None | None |
| HEXA | None | None | None | None |
| HEXA | None | None | None | None |
| HEXA | None | None | None | None |
| HEXA | None | None | None | None |
| HEXA | None | None | None | None |
| HEXA | None | None | None | None |
| HEXA | None | None | None | None |
| HEXA | None | None | None | None |
| HEXA | None | None | None | None |
| HEXA | None | None | None | None |
| HEXA | None | None | None | None |
| HEXA | None | None | None | None |
| HEXA | None | None | None | None |
| HEXA | None | None | None | None |
| HEXA | None | None | None | None |
| HEXA | None | None | None | None |
| HEXA | None | None | None | None |
| CFTR | None | None | None | None |
| CFTR | None | None | None | None |
| CFTR | None | None | None | None |
| CFTR | None | None | None | None |
| CFTR | None | None | None | None |
| CFTR | None | None | None | None |
| CFTR | None | None | None | None |
| CFTR | None | None | None | None |
| CFTR | None | None | None | None |

| | | | | |
|---|---|---|---|---|
| CFTR | None | None | None | None |
| CFTR | None | None | None | None |
| CFTR | None | None | None | None |
| CFTR | None | None | None | None |
| CFTR | None | None | None | None |
| CFTR | None | None | None | None |
| CFTR | None | None | None | None |
| CFTR | None | None | None | None |
| CFTR | None | None | None | None |
| CFTR | None | None | None | None |
| CFTR | None | None | None | None |
| CFTR | None | None | None | None |
| CFTR | None | None | None | None |
| CFTR | None | None | None | None |
| CFTR | None | None | None | None |
| CFTR | None | None | None | None |
| CFTR | None | None | None | None |
| CFTR | None | None | None | None |
| CFTR | None | None | None | None |
| CFTR | None | None | None | None |
| CFTR | None | None | None | None |
| CFTR | None | None | None | None |
| CFTR | None | None | None | None |
| CFTR | None | None | None | None |
| CFTR | None | None | None | None |
| FAN1 | None | None | None | None |
| CBS | None | None | None | None |
| CBS | None | None | None | None |
| CBS | None | None | None | None |
| CBS | None | None | None | None |
| CBS | None | None | None | None |
| CBS | None | None | None | None |
| CBS | None | None | None | None |
| CBS | None | None | None | None |
| CBS | None | None | None | None |
| CBS | None | None | None | None |
| CBS | None | None | None | None |
| CBS | None | None | None | None |
| CBS | None | None | None | None |
| CBS | None | None | None | None |
| CBS | None | None | None | None |
| CBS | None | None | None | None |
| CBS | None | None | None | None |
| CBS | None | None | None | None |
| CBS | 0.698 | 2 | 1m54 | B |
| CBS | None | None | None | None |
| CBS | None | None | None | None |
| CBS | None | None | None | None |
| CBS | None | None | None | None |
| CBS | None | None | None | None |
| CBS | None | None | None | None |
| CBS | None | None | None | None |
| CBS | None | None | None | None |
| CBS | None | None | None | None |

| | | | | |
|---|---|---|---|---|
| CBS | 0.603 | 3 | 4coo | A |
| CBS | 0.603 | 3 | 4coo | A |
| CBS | None | None | None | None |
| CBS | None | None | None | None |
| CBS | None | None | None | None |
| CBS | None | None | None | None |
| CBS | None | None | None | None |
| CBS | None | None | None | None |
| TLR3 | None | None | None | None |
| LIG4 | None | None | None | None |
| LIG4 | None | None | None | None |
| ASPA | 0.769 | 1 | 2i3c | B |
| ASPA | 0.769 | 1 | 2i3c | B |
| ASPA | None | None | None | None |
| ASPA | None | None | None | None |
| ASPA | None | None | None | None |
| ASPA | None | None | None | None |
| ASPA | None | None | None | None |
| ASPA | None | None | None | None |
| ASPA | None | None | None | None |
| ASPA | None | None | None | None |
| ASPA | None | None | None | None |
| ASPA | None | None | None | None |
| ASPA | None | None | None | None |
| CFH | None | None | None | None |
| FCGR3A | None | None | None | None |
| FCGR3A | None | None | None | None |
| GRIN2A | None | None | None | None |
| GRIN2A | 0.658 | 1 | 5i2n | A |
| GRIN2A | None | None | None | None |
| GLA | None | None | None | None |
| GLA | None | None | None | None |
| GLA | None | None | None | None |
| GLA | None | None | None | None |
| GLA | None | None | None | None |
| GLA | None | None | None | None |
| GLA | None | None | None | None |
| GLA | 0.647 | 2 | 3s5y | A |
| GLA | None | None | None | None |
| GLA | None | None | None | None |
| GLA | None | None | None | None |
| GLA | 0.75 | 3 | 3gxn | B |
| GLA | 0.75 | 3 | 3gxn | B |
| GLA | None | None | None | None |
| GLA | None | None | None | None |
| GLA | None | None | None | None |
| GLA | None | None | None | None |
| GLA | None | None | None | None |
| GLA | None | None | None | None |
| GLA | None | None | None | None |
| GLA | None | None | None | None |
| GLA | None | None | None | None |
| GLA | None | None | None | None |
| GLA | None | None | None | None |

| | | | | |
|---|---|---|---|---|
| GLA | None | None | None | None |
| GLA | None | None | None | None |
| GLA | None | None | None | None |
| GLA | None | None | None | None |
| GLA | None | None | None | None |
| GLA | None | None | None | None |
| GLA | None | None | None | None |
| SERPINA3 | None | None | None | None |
| MTAP | None | None | None | None |
| MTAP | None | None | None | None |
| MTAP | None | None | None | None |
| BLMH | None | None | None | None |
| TUSC3 | None | None | None | None |
| ISCU | None | None | None | None |
| CYP11B2 | None | None | None | None |
| CYP11B2 | None | None | None | None |
| CYP11B2 | None | None | None | None |
| CYP11B2 | None | None | None | None |
| CYP11B2 | None | None | None | None |
| CYP11B2 | None | None | None | None |
| CYP11B2 | None | None | None | None |
| CYP11B2 | None | None | None | None |
| CYP11B2 | None | None | None | None |
| CYP11B2 | None | None | None | None |
| CYP11B2 | None | None | None | None |
| CYP11B2 | None | None | None | None |
| CYP11B2 | None | None | None | None |
| CYP11B2 | None | None | None | None |
| CYP11B2 | None | None | None | None |
| CYP11B2 | None | None | None | None |
| CYP11B2 | None | None | None | None |
| CYP11B2 | None | None | None | None |
| CYP11B2 | None | None | None | None |
| CRADD | None | None | None | None |
| CRADD | None | None | None | None |
| FGA | None | None | None | None |
| FGA | None | None | None | None |
| FGA | None | None | None | None |
| FGA | None | None | None | None |
| FGA | None | None | None | None |
| FGA | None | None | None | None |
| FGA | None | None | None | None |
| FGA | None | None | None | None |
| IYD | 1 | 1 | 4ttb | A |
| IYD | None | None | None | None |
| IYD | None | None | None | None |
| IYD | None | None | None | None |
| TCAP | None | None | None | None |
| TCAP | None | None | None | None |
| TCAP | None | None | None | None |
| TCAP | None | None | None | None |
| TCAP | None | None | None | None |
| CYP7A1 | None | None | None | None |

| | | | | |
|---|---|---|---|---|
| TEK | None | None | None | None |
| TEK | None | None | None | None |
| TEK | None | None | None | None |
| TEK | None | None | None | None |
| TEK | None | None | None | None |
| TEK | None | None | None | None |
| TEK | None | None | None | None |
| CD8A | None | None | None | None |
| HERC2 | None | None | None | None |
| NR1H3 | None | None | None | None |
| GRHPR | None | None | None | None |
| GRHPR | None | None | None | None |
| GRHPR | None | None | None | None |
| GRHPR | None | None | None | None |
| GRHPR | 0.697 | 1 | 2wwr | D |
| GRHPR | 0.697 | 1 | 2wwr | D |
| GRHPR | None | None | None | None |
| GBE1 | None | None | None | None |
| GBE1 | None | None | None | None |
| GBE1 | None | None | None | None |
| GBE1 | None | None | None | None |
| GBE1 | None | None | None | None |
| GBE1 | None | None | None | None |
| GBE1 | None | None | None | None |
| GBE1 | None | None | None | None |
| GBE1 | None | None | None | None |
| GBE1 | None | None | None | None |
| GBE1 | None | None | None | None |
| GBE1 | None | None | None | None |
| GBE1 | None | None | None | None |
| GBE1 | None | None | None | None |
| GBE1 | None | None | None | None |
| GBE1 | None | None | None | None |
| GBE1 | None | None | None | None |
| GBE1 | None | None | None | None |
| GBE1 | None | None | None | None |
| PC | None | None | None | None |
| PC | None | None | None | None |
| PC | None | None | None | None |
| PC | None | None | None | None |
| PC | None | None | None | None |
| PC | None | None | None | None |
| PC | None | None | None | None |
| PC | None | None | None | None |
| MSH6 | None | None | None | None |
| MSH6 | None | None | None | None |
| MSH6 | None | None | None | None |
| MSH6 | None | None | None | None |
| MSH6 | None | None | None | None |
| MSH6 | None | None | None | None |
| MSH6 | None | None | None | None |
| MSH6 | None | None | None | None |
| MSH6 | None | None | None | None |

| | | | | |
|---|---|---|---|---|
| MSH6 | None | None | None | None |
| MSH6 | None | None | None | None |
| MSH6 | None | None | None | None |
| MSH6 | None | None | None | None |
| MSH6 | None | None | None | None |
| MSH6 | None | None | None | None |
| MSH6 | None | None | None | None |
| MSH6 | None | None | None | None |
| MSH6 | None | None | None | None |
| MSH6 | None | None | None | None |
| MSH6 | None | None | None | None |
| MSH6 | None | None | None | None |
| MSH6 | None | None | None | None |
| MSH6 | None | None | None | None |
| MSH6 | None | None | None | None |
| MSH6 | None | None | None | None |
| MSH6 | None | None | None | None |
| MSH6 | None | None | None | None |
| MSH6 | None | None | None | None |
| MSH6 | None | None | None | None |
| MSH6 | None | None | None | None |
| RNASEL | None | None | None | None |
| ORC1 | None | None | None | None |
| ORC1 | None | None | None | None |
| ORC1 | None | None | None | None |
| KCNH2 | None | None | None | None |
| KCNH2 | None | None | None | None |
| KCNH2 | None | None | None | None |
| KCNH2 | None | None | None | None |
| KCNH2 | None | None | None | None |
| KCNH2 | None | None | None | None |
| KCNH2 | None | None | None | None |
| KCNH2 | None | None | None | None |
| KCNH2 | None | None | None | None |
| KCNH2 | None | None | None | None |
| KCNH2 | None | None | None | None |
| KCNH2 | None | None | None | None |
| KCNH2 | None | None | None | None |
| KCNH2 | None | None | None | None |
| KCNH2 | None | None | None | None |
| KCNH2 | None | None | None | None |
| KCNH2 | None | None | None | None |
| KCNH2 | None | None | None | None |
| KCNH2 | None | None | None | None |
| KCNH2 | None | None | None | None |
| KCNH2 | None | None | None | None |
| KCNH2 | None | None | None | None |
| KCNH2 | None | None | None | None |
| KCNH2 | None | None | None | None |
| MAT1A | None | None | None | None |
| MAT1A | None | None | None | None |
| MAT1A | None | None | None | None |
| MAT1A | None | None | None | None |
| MAT1A | None | None | None | None |

| | | | | |
|---|---|---|---|---|
| MAT1A | None | None | None | None |
| MAT1A | None | None | None | None |
| MAT1A | None | None | None | None |
| INSR | None | None | None | None |
| INSR | None | None | None | None |
| INSR | None | None | None | None |
| INSR | None | None | None | None |
| AMT | None | None | None | None |
| AMT | None | None | None | None |
| AMT | None | None | None | None |
| AMT | None | None | None | None |
| AMT | None | None | None | None |
| AMT | None | None | None | None |
| AMT | None | None | None | None |
| AMT | None | None | None | None |
| AMT | None | None | None | None |
| CSNK1D | None | None | None | None |
| SF3B1 | None | None | None | None |
| AGA | None | None | None | None |
| AGA | None | None | None | None |
| AGA | None | None | None | None |
| AGA | None | None | None | None |
| AGA | None | None | None | None |
| AGA | None | None | None | None |
| AGA | None | None | None | None |
| NECTIN4 | None | None | None | None |
| RS1 | None | None | None | None |
| RS1 | None | None | None | None |
| RS1 | None | None | None | None |
| RS1 | None | None | None | None |
| TBC1D7 | None | None | None | None |
| CTSA | None | None | None | None |
| CTSA | None | None | None | None |
| TKT | None | None | None | None |
| RP2 | None | None | None | None |
| RP2 | None | None | None | None |
| RP2 | None | None | None | None |
| RP2 | None | None | None | None |
| LYRM4 | None | None | None | None |
| SOS1 | None | None | None | None |
| SOS1 | None | None | None | None |
| SOS1 | None | None | None | None |
| SOS1 | None | None | None | None |
| SOS1 | None | None | None | None |
| SOS1 | None | None | None | None |
| HUWE1 | None | None | None | None |
| HUWE1 | None | None | None | None |
| TPI1 | None | None | None | None |
| TPI1 | 0.886 | 1 | 4unk | B |
| TPI1 | None | None | None | None |
| MSH2 | None | None | None | None |
| MSH2 | None | None | None | None |
| MSH2 | None | None | None | None |
| MSH2 | None | None | None | None |

| | | | | |
|---|---|---|---|---|
| MSH2 | None | None | None | None |
| MSH2 | None | None | None | None |
| MSH2 | None | None | None | None |
| MSH2 | None | None | None | None |
| MSH2 | None | None | None | None |
| MSH2 | None | None | None | None |
| MSH2 | None | None | None | None |
| MSH2 | None | None | None | None |
| MSH2 | None | None | None | None |
| MSH2 | None | None | None | None |
| MSH2 | None | None | None | None |
| MSH2 | None | None | None | None |
| MSH2 | None | None | None | None |
| MSH2 | None | None | None | None |
| MSH2 | None | None | None | None |
| MSH2 | None | None | None | None |
| MSH2 | None | None | None | None |
| MSH2 | None | None | None | None |
| MSH2 | None | None | None | None |
| MSH2 | None | None | None | None |
| MSH2 | None | None | None | None |
| MSH2 | None | None | None | None |
| MSH2 | None | None | None | None |
| MSH2 | None | None | None | None |
| MSH2 | None | None | None | None |
| MSH2 | None | None | None | None |
| MSH2 | None | None | None | None |
| MSH2 | None | None | None | None |
| MSH2 | None | None | None | None |
| MSH2 | None | None | None | None |
| MSH2 | None | None | None | None |
| MSH2 | None | None | None | None |
| GALE | None | None | None | None |
| GALE | 0.78 | 1 | 1i3n | A |
| GALE | None | None | None | None |
| GALE | None | None | None | None |
| GALE | None | None | None | None |
| GALE | None | None | None | None |
| GALE | None | None | None | None |
| GALE | None | None | None | None |
| NSD2 | None | None | None | None |
| SRP72 | None | None | None | None |
| SRP72 | None | None | None | None |
| SRP72 | None | None | None | None |
| POLK | None | None | None | None |
| GALT | None | None | None | None |
| GALT | None | None | None | None |
| GALT | None | None | None | None |
| GALT | None | None | None | None |
| GALT | None | None | None | None |
| GALT | None | None | None | None |
| GALT | None | None | None | None |
| GALT | None | None | None | None |
| GALT | None | None | None | None |

| | | | | |
|---|---|---|---|---|
| GALT | None | None | None | None |
| GALT | None | None | None | None |
| GALT | None | None | None | None |
| GALT | None | None | None | None |
| GALT | None | None | None | None |
| GALT | None | None | None | None |
| GALT | None | None | None | None |
| GALT | None | None | None | None |
| GALT | None | None | None | None |
| GALT | None | None | None | None |
| GALT | None | None | None | None |
| GALT | None | None | None | None |
| GALT | None | None | None | None |
| GALT | None | None | None | None |
| GALT | None | None | None | None |
| GALT | None | None | None | None |
| GALT | None | None | None | None |
| GALT | None | None | None | None |
| GALT | None | None | None | None |
| GALT | None | None | None | None |
| GALT | None | None | None | None |
| GALT | None | None | None | None |
| GALT | None | None | None | None |
| GALT | None | None | None | None |
| GALT | None | None | None | None |
| GALT | None | None | None | None |
| GALT | None | None | None | None |
| GALT | None | None | None | None |
| PYCR1 | 0.75 | 11 | 5uav | A |
| PYCR1 | 0.75 | 11 | 5uav | A |
| PYCR1 | None | None | None | None |
| PYCR1 | None | None | None | None |
| F8 | None | None | None | None |
| F8 | None | None | None | None |
| F8 | None | None | None | None |
| F8 | None | None | None | None |
| F8 | None | None | None | None |
| F8 | None | None | None | None |
| F8 | None | None | None | None |
| F8 | None | None | None | None |
| F8 | None | None | None | None |
| F8 | None | None | None | None |
| F8 | None | None | None | None |
| F8 | None | None | None | None |
| F8 | None | None | None | None |
| F8 | None | None | None | None |
| F8 | None | None | None | None |
| F8 | None | None | None | None |
| ANK1 | None | None | None | None |
| CHAT | None | None | None | None |
| CHAT | None | None | None | None |
| CHAT | 0.615 | 8 | 2fy5 | A |
| CHAT | None | None | None | None |
| CHAT | None | None | None | None |

| | | | | |
|---|---|---|---|---|
| CHAT | None | None | None | None |
| CHAT | None | None | None | None |
| CHAT | None | None | None | None |
| CHAT | None | None | None | None |
| CHAT | None | None | None | None |
| CHAT | None | None | None | None |
| CHAT | None | None | None | None |
| CHAT | None | None | None | None |
| CHAT | None | None | None | None |
| CHAT | None | None | None | None |
| CHAT | None | None | None | None |
| CHAT | None | None | None | None |
| GHR | None | None | None | None |
| GHR | None | None | None | None |
| GHR | None | None | None | None |
| GHR | None | None | None | None |
| GHR | None | None | None | None |
| GHR | None | None | None | None |
| GHR | None | None | None | None |
| GHR | None | None | None | None |
| GHR | None | None | None | None |
| GHR | None | None | None | None |
| GHR | None | None | None | None |
| GHR | None | None | None | None |
| GHR | None | None | None | None |
| GHR | None | None | None | None |
| FZD4 | None | None | None | None |
| FZD4 | None | None | None | None |
| FZD4 | None | None | None | None |
| MTR | None | None | None | None |
| MTR | None | None | None | None |
| MTR | None | None | None | None |
| GM2A | None | None | None | None |
| GM2A | None | None | None | None |
| GM2A | None | None | None | None |
| GM2A | None | None | None | None |
| GM2A | None | None | None | None |
| CRYGD | None | None | None | None |
| CRYGD | None | None | None | None |
| CRYGD | None | None | None | None |
| CRYGD | None | None | None | None |
| CRYGD | None | None | None | None |
| CRYGD | None | None | None | None |
| CRYGD | None | None | None | None |
| CRYGD | None | None | None | None |
| CRYGD | None | None | None | None |
| TLR5 | None | None | None | None |
| HSD17B10 | 0.855 | 1 | 1u7t | B |
| HSD17B10 | None | None | None | None |
| HSD17B10 | None | None | None | None |
| HSD17B10 | None | None | None | None |
| TBC1D20 | None | None | None | None |
| FBN1 | None | None | None | None |
| FBN1 | None | None | None | None |

| | | | | |
|---|---|---|---|---|
| FBN1 | None | None | None | None |
| FBN1 | None | None | None | None |
| FBN1 | None | None | None | None |
| FBN1 | None | None | None | None |
| FBN1 | None | None | None | None |
| FXN | None | None | None | None |
| FXN | None | None | None | None |
| FXN | None | None | None | None |
| FXN | None | None | None | None |
| MYO5A | None | None | None | None |
| CTSK | None | None | None | None |
| CTSK | None | None | None | None |
| SCN5A | None | None | None | None |
| SCN5A | None | None | None | None |
| SOS1 | None | None | None | None |
| SOS1 | None | None | None | None |
| SOS1 | None | None | None | None |
| SOS1 | None | None | None | None |
| SOS1 | None | None | None | None |
| SOS1 | None | None | None | None |
| SOS1 | None | None | None | None |
| SOS1 | None | None | None | None |
| SOS1 | None | None | None | None |
| SOS1 | None | None | None | None |
| SOS1 | None | None | None | None |
| SOS1 | None | None | None | None |
| GATM | None | None | None | None |
| GATM | None | None | None | None |
| GATM | 0.75 | 4 | 4jdw | A |
| GATM | None | None | None | None |
| GATM | None | None | None | None |
| GATM | 0.75 | 6 | 6jdw | A |
| GATM | 0.75 | 11 | 6jdw | A |
| MMP2 | None | None | None | None |
| MMP2 | None | None | None | None |
| MMP2 | None | None | None | None |
| MMP2 | None | None | None | None |
| NDUFB9 | None | None | None | None |
| NDUFB9 | None | None | None | None |
| GH1 | None | None | None | None |
| GH1 | None | None | None | None |
| GH1 | None | None | None | None |
| SLC6A4 | None | None | None | None |
| G6PD | None | None | None | None |
| G6PD | None | None | None | None |
| G6PD | None | None | None | None |
| G6PD | None | None | None | None |
| G6PD | None | None | None | None |
| G6PD | None | None | None | None |
| G6PD | None | None | None | None |
| G6PD | None | None | None | None |
| G6PD | None | None | None | None |
| G6PD | None | None | None | None |
| G6PD | None | None | None | None |

| | | | | |
|---|---|---|---|---|
| G6PD | None | None | None | None |
| G6PD | None | None | None | None |
| G6PD | None | None | None | None |
| G6PD | None | None | None | None |
| G6PD | None | None | None | None |
| G6PD | None | None | None | None |
| G6PD | None | None | None | None |
| G6PD | None | None | None | None |
| G6PD | None | None | None | None |
| CST3 | None | None | None | None |
| LMNA | None | None | None | None |
| LMNA | None | None | None | None |
| NDUFA9 | None | None | None | None |
| NDUFA9 | None | None | None | None |
| NDUFA9 | None | None | None | None |
| NDUFA9 | None | None | None | None |
| YARS | None | None | None | None |
| YARS | None | None | None | None |
| YARS | None | None | None | None |
| YARS | None | None | None | None |
| EPHA2 | None | None | None | None |
| EPHA2 | None | None | None | None |
| EPHA2 | None | None | None | None |
| EPHA2 | None | None | None | None |
| DICER1 | None | None | None | None |
| DICER1 | None | None | None | None |
| SNX10 | None | None | None | None |
| SNX10 | None | None | None | None |
| TTN | None | None | None | None |
| TTN | None | None | None | None |
| GATA3 | None | None | None | None |
| C5 | None | None | None | None |
| THBD | None | None | None | None |
| BIN1 | None | None | None | None |
| BIN1 | None | None | None | None |
| SMARCAL1 | None | None | None | None |
| CFB | None | None | None | None |
| CYC1 | None | None | None | None |
| ANK2 | None | None | None | None |
| CHMP4B | None | None | None | None |
| CHMP4B | None | None | None | None |
| ERBB2 | None | None | None | None |
| ERBB2 | None | None | None | None |
| POLG2 | None | None | None | None |
| POLG2 | None | None | None | None |
| POLG2 | None | None | None | None |
| POLG2 | None | None | None | None |
| POLG2 | None | None | None | None |
| POLG2 | None | None | None | None |
| POLG2 | None | None | None | None |
| POLG2 | None | None | None | None |
| C3 | None | None | None | None |
| C3 | None | None | None | None |
| RPGR | None | None | None | None |

| | | | | |
|---|---|---|---|---|
| RPGR | None | None | None | None |
| RPGR | None | None | None | None |
| RPGR | None | None | None | None |
| RPGR | None | None | None | None |
| C1QA | None | None | None | None |
| PRKG1 | None | None | None | None |
| IGBP1 | None | None | None | None |
| ENO3 | None | None | None | None |
| ENO3 | None | None | None | None |
| ENO3 | None | None | None | None |
| ENO3 | None | None | None | None |
| ENO3 | None | None | None | None |
| ENO3 | 0.897 | 2 | 2xsx | B |
| MCPH1 | None | None | None | None |
| MCPH1 | None | None | None | None |
| MCPH1 | None | None | None | None |
| MCPH1 | None | None | None | None |
| MCPH1 | None | None | None | None |
| SNRNP200 | None | None | None | None |
| SNRNP200 | None | None | None | None |
| DNMT1 | None | None | None | None |
| DNMT1 | None | None | None | None |
| DNMT1 | None | None | None | None |
| AKT1 | None | None | None | None |
| NDUFS2 | None | None | None | None |
| ACVR2B | None | None | None | None |
| HMGCL | None | None | None | None |
| HMGCL | None | None | None | None |
| HMGCL | None | None | None | None |
| JUP | None | None | None | None |
| JUP | None | None | None | None |
| JUP | None | None | None | None |
| JUP | None | None | None | None |
| JUP | None | None | None | None |
| JUP | None | None | None | None |
| JUP | None | None | None | None |
| JUP | None | None | None | None |
| SPINK1 | None | None | None | None |
| SPINK1 | None | None | None | None |
| SPINK1 | None | None | None | None |
| VCL | None | None | None | None |
| VCL | None | None | None | None |
| NOTCH1 | None | None | None | None |
| NOTCH1 | None | None | None | None |
| NOTCH1 | None | None | None | None |
| NDUFB3 | None | None | None | None |
| IKBKG | None | None | None | None |
| GSN | None | None | None | None |
| GSN | None | None | None | None |
| GSN | None | None | None | None |
| GSN | None | None | None | None |
| GSN | None | None | None | None |
| GSN | None | None | None | None |
| GSN | None | None | None | None |

| | | | | |
|---|---|---|---|---|
| GSN | None | None | None | None |
| GSN | None | None | None | None |
| CLC | None | None | None | None |
| RAF1 | None | None | None | None |
| RAF1 | None | None | None | None |
| RAF1 | None | None | None | None |
| NDUFS8 | None | None | None | None |
| NDUFS8 | None | None | None | None |
| NDUFS8 | None | None | None | None |
| NDUFS8 | None | None | None | None |
| NDUFS8 | None | None | None | None |
| CASR | None | None | None | None |
| CASR | None | None | None | None |
| CASR | None | None | None | None |
| CASR | None | None | None | None |
| CASR | None | None | None | None |
| CASR | None | None | None | None |
| CASR | None | None | None | None |
| CASR | None | None | None | None |
| CASR | None | None | None | None |
| CASR | None | None | None | None |
| CASR | None | None | None | None |
| CASR | None | None | None | None |
| CASR | None | None | None | None |
| CASR | None | None | None | None |
| CASR | None | None | None | None |
| CASR | None | None | None | None |
| CASR | None | None | None | None |
| CASR | None | None | None | None |
| CASR | None | None | None | None |
| CASR | None | None | None | None |
| CASR | None | None | None | None |
| CASR | None | None | None | None |
| CASR | None | None | None | None |
| CASR | None | None | None | None |
| CASR | None | None | None | None |
| CASR | None | None | None | None |
| CASR | None | None | None | None |
| CASR | None | None | None | None |
| CASR | None | None | None | None |
| VCL | None | None | None | None |
| VCL | None | None | None | None |
| SMPD1 | None | None | None | None |
| SMPD1 | None | None | None | None |
| SMPD1 | None | None | None | None |
| KDM6A | None | None | None | None |
| KCNQ1 | None | None | None | None |
| KCNQ1 | None | None | None | None |
| KCNQ1 | None | None | None | None |
| KCNQ1 | None | None | None | None |
| KCNQ1 | None | None | None | None |
| KCNQ1 | None | None | None | None |
| NDUFA12 | None | None | None | None |
| GLRX5 | None | None | None | None |

| | | | | |
|---|---|---|---|---|
| PALB2 | None | None | None | None |
| PALB2 | None | None | None | None |
| PALB2 | None | None | None | None |
| PALB2 | None | None | None | None |
| PALB2 | None | None | None | None |
| PALB2 | None | None | None | None |
| PALB2 | None | None | None | None |
| PALB2 | None | None | None | None |
| PALB2 | None | None | None | None |
| PALB2 | None | None | None | None |
| ALDOB | None | None | None | None |
| ALDOB | 0.731 | 2 | 1xdm | A |
| ALDOB | 0.731 | 1 | 1qo5 | B |
| ALDOB | None | None | None | None |
| ALDOB | 0.692 | 7 | 1qo5 | C |
| ALDOB | None | None | None | None |
| ALDOB | None | None | None | None |
| GNE | None | None | None | None |
| GNE | None | None | None | None |
| GNE | None | None | None | None |
| GNE | None | None | None | None |
| GNE | None | None | None | None |
| GNE | None | None | None | None |
| GNE | None | None | None | None |
| GNE | None | None | None | None |
| GNE | None | None | None | None |
| GNE | None | None | None | None |
| GNE | None | None | None | None |
| GNE | None | None | None | None |
| CHRNB2 | None | None | None | None |
| CHRNB2 | None | None | None | None |
| ERCC3 | None | None | None | None |
| ERCC3 | None | None | None | None |
| ERCC3 | None | None | None | None |
| ERCC3 | None | None | None | None |
| ERCC3 | None | None | None | None |
| UNC119 | None | None | None | None |
| ATL1 | None | None | None | None |
| ATL1 | None | None | None | None |
| ATL1 | None | None | None | None |
| ATL1 | 0.8 | 1 | 3q5d | A |
| ATL1 | None | None | None | None |
| ATL1 | None | None | None | None |
| ATL1 | None | None | None | None |
| ATL1 | None | None | None | None |
| ATL1 | None | None | None | None |
| ATL1 | None | None | None | None |
| NR3C2 | 0.701 | 1 | 2oax | A |
| TBK1 | None | None | None | None |
| HYAL1 | None | None | None | None |
| HYAL1 | None | None | None | None |
| HYAL1 | None | None | None | None |
| HYAL1 | None | None | None | None |
| CTSD | 0.633 | 1 | 4obz | A |

| | | | | |
|---|---|---|---|---|
| PRKCG | None | None | None | None |
| TGFBR1 | None | None | None | None |
| XDH | None | None | None | None |
| XDH | None | None | None | None |
| XDH | None | None | None | None |
| XDH | None | None | None | None |
| XDH | None | None | None | None |
| XDH | None | None | None | None |
| NMNAT1 | 0.833 | 1 | 1kqn | E |
| NMNAT1 | None | None | None | None |
| NMNAT1 | None | None | None | None |
| NMNAT1 | None | None | None | None |
| NMNAT1 | None | None | None | None |
| NMNAT1 | None | None | None | None |
| SERPING1 | None | None | None | None |
| SERPING1 | None | None | None | None |
| SERPING1 | None | None | None | None |
| SERPING1 | None | None | None | None |
| SERPING1 | None | None | None | None |
| PDZRN3 | None | None | None | None |
| FABP2 | None | None | None | None |
| IFNGR1 | None | None | None | None |
| IFNGR1 | None | None | None | None |
| TDP1 | None | None | None | None |
| TDP1 | None | None | None | None |
| INS | None | None | None | None |
| INS | 0.769 | 1 | 2r34 | C |
| INS | 0.769 | 1 | 2r34 | C |
| PPOX | 0.857 | 1 | 4ivm | B |
| PPOX | None | None | None | None |
| PPOX | None | None | None | None |
| PPOX | None | None | None | None |
| PPOX | None | None | None | None |
| PPOX | None | None | None | None |
| PPOX | None | None | None | None |
| PPOX | None | None | None | None |
| PPOX | None | None | None | None |
| GFER | None | None | None | None |
| GFER | 0.769 | 4 | 3tk0 | A |
| GFER | None | None | None | None |
| GFER | None | None | None | None |
| LIFR | None | None | None | None |
| EGLN1 | None | None | None | None |
| EGLN1 | 0.889 | 6 | 5v18 | A |
| EGLN1 | 0.944 | 2 | 5a3u | C |
| GP1BA | None | None | None | None |
| GP1BA | None | None | None | None |
| GP1BA | None | None | None | None |
| GP1BA | None | None | None | None |
| GP1BA | None | None | None | None |
| MMADHC | None | None | None | None |
| MMADHC | None | None | None | None |
| MMADHC | None | None | None | None |
| MMADHC | None | None | None | None |

| | | | | |
|---|---|---|---|---|
| MMADHC | None | None | None | None |
| MMADHC | None | None | None | None |
| MMADHC | None | None | None | None |
| ACVRL1 | None | None | None | None |
| ACVRL1 | None | None | None | None |
| RFXANK | None | None | None | None |
| CHIT1 | None | None | None | None |
| UBR1 | None | None | None | None |
| BLM | None | None | None | None |
| BLM | None | None | None | None |
| BLM | None | None | None | None |
| BLM | None | None | None | None |
| BLM | None | None | None | None |
| BLM | None | None | None | None |
| BLM | None | None | None | None |
| BLM | None | None | None | None |
| BLM | None | None | None | None |
| BLM | None | None | None | None |
| BLM | None | None | None | None |
| BLM | None | None | None | None |
| BLM | None | None | None | None |
| VDR | None | None | None | None |
| VDR | None | None | None | None |
| VDR | None | None | None | None |
| VDR | None | None | None | None |
| VDR | None | None | None | None |
| VDR | None | None | None | None |
| EPHA2 | None | None | None | None |
| EPHA2 | None | None | None | None |
| EPHA2 | None | None | None | None |
| EPHA2 | None | None | None | None |
| EPHA2 | None | None | None | None |
| EPHA2 | None | None | None | None |
| EPHA2 | None | None | None | None |
| EPHA2 | None | None | None | None |
| EPHA2 | None | None | None | None |
| EPHA2 | None | None | None | None |
| UBA1 | None | None | None | None |
| STAT1 | None | None | None | None |
| STAT1 | None | None | None | None |
| STAT1 | None | None | None | None |
| STAT1 | None | None | None | None |
| STAT1 | None | None | None | None |
| STAT1 | None | None | None | None |
| STAT1 | None | None | None | None |
| STAT1 | None | None | None | None |
| DNM1L | None | None | None | None |
| DNM1L | None | None | None | None |
| OTC | None | None | None | None |
| OTC | None | None | None | None |
| OTC | None | None | None | None |
| OTC | None | None | None | None |
| OTC | None | None | None | None |
| OTC | None | None | None | None |

| | | | | |
|---|---|---|---|---|
| OTC | None | None | None | None |
| OTC | None | None | None | None |
| OTC | None | None | None | None |
| OTC | None | None | None | None |
| OTC | None | None | None | None |
| OTC | None | None | None | None |
| OTC | None | None | None | None |
| OTC | None | None | None | None |
| OTC | None | None | None | None |
| OTC | None | None | None | None |
| OTC | None | None | None | None |
| RARS | None | None | None | None |
| RARS | None | None | None | None |
| RARS | None | None | None | None |
| SEPSECS | None | None | None | None |
| SEPSECS | None | None | None | None |
| SEPSECS | None | None | None | None |
| SEPSECS | None | None | None | None |
| SEPSECS | None | None | None | None |
| BRCA1 | None | None | None | None |
| BRCA1 | None | None | None | None |
| BRCA1 | None | None | None | None |
| BRCA1 | None | None | None | None |
| BRCA1 | None | None | None | None |
| BRCA1 | None | None | None | None |
| BRCA1 | None | None | None | None |
| BRCA1 | None | None | None | None |
| BRCA1 | None | None | None | None |
| BRCA1 | None | None | None | None |
| BRCA1 | None | None | None | None |
| BRCA1 | None | None | None | None |
| BRCA1 | None | None | None | None |
| BRCA1 | None | None | None | None |
| BRCA1 | None | None | None | None |
| BRCA1 | None | None | None | None |
| BRCA1 | None | None | None | None |
| BRCA1 | None | None | None | None |
| BRCA1 | None | None | None | None |
| BRCA1 | None | None | None | None |
| BRCA1 | None | None | None | None |
| BRCA1 | None | None | None | None |
| CUBN | None | None | None | None |
| ITGB3 | None | None | None | None |
| MTRR | None | None | None | None |
| MTRR | None | None | None | None |
| MTRR | None | None | None | None |
| MTRR | None | None | None | None |
| MTRR | None | None | None | None |
| MTRR | None | None | None | None |
| MTRR | None | None | None | None |
| MTRR | None | None | None | None |
| IL7R | None | None | None | None |
| IL7R | None | None | None | None |
| RPS24 | None | None | None | None |

| | | | | |
|---|---|---|---|---|
| RPS24 | None | None | None | None |
| HOGA1 | None | None | None | None |
| HOGA1 | None | None | None | None |
| HOGA1 | None | None | None | None |
| HOGA1 | None | None | None | None |
| HOGA1 | None | None | None | None |
| HOGA1 | None | None | None | None |
| HOGA1 | 0.791 | 1 | 3s5n | A |
| HOGA1 | None | None | None | None |
| HOGA1 | None | None | None | None |
| HOGA1 | None | None | None | None |
| HOGA1 | None | None | None | None |
| HOGA1 | None | None | None | None |
| HOGA1 | None | None | None | None |
| GJB2 | None | None | None | None |
| GJB2 | None | None | None | None |
| GJB2 | None | None | None | None |
| GJB2 | None | None | None | None |
| GJB2 | None | None | None | None |
| GJB2 | None | None | None | None |
| GJB2 | None | None | None | None |
| GJB2 | None | None | None | None |
| GJB2 | None | None | None | None |
| GJB2 | None | None | None | None |
| GJB2 | None | None | None | None |
| GJB2 | None | None | None | None |
| GJB2 | None | None | None | None |
| GJB2 | None | None | None | None |
| GJB2 | None | None | None | None |
| GJB2 | None | None | None | None |
| GJB2 | None | None | None | None |
| GJB2 | None | None | None | None |
| GJB2 | None | None | None | None |
| GJB2 | None | None | None | None |
| GJB2 | None | None | None | None |
| GJB2 | None | None | None | None |
| GJB2 | None | None | None | None |
| GJB2 | None | None | None | None |
| GJB2 | None | None | None | None |
| GJB2 | None | None | None | None |
| GJB2 | None | None | None | None |
| GJB2 | None | None | None | None |
| GJB2 | None | None | None | None |
| GJB2 | None | None | None | None |
| GJB2 | None | None | None | None |
| GJB2 | None | None | None | None |
| GJB2 | None | None | None | None |
| GJB2 | None | None | None | None |
| GJB2 | None | None | None | None |
| GJB2 | None | None | None | None |
| GJB2 | None | None | None | None |
| GJB2 | None | None | None | None |
| GJB2 | None | None | None | None |
| GJB2 | None | None | None | None |

| | | | | |
|---|---|---|---|---|
| GJB2 | None | None | None | None |
| GJB2 | None | None | None | None |
| GJB2 | None | None | None | None |
| GJB2 | None | None | None | None |
| GJB2 | None | None | None | None |
| GJB2 | None | None | None | None |
| GJB2 | None | None | None | None |
| GJB2 | None | None | None | None |
| GJB2 | None | None | None | None |
| GJB2 | None | None | None | None |
| GJB2 | None | None | None | None |
| GJB2 | None | None | None | None |
| GJB2 | None | None | None | None |
| GJB2 | None | None | None | None |
| CPS1 | None | None | None | None |
| CPS1 | None | None | None | None |
| CPS1 | None | None | None | None |
| CPS1 | None | None | None | None |
| CPS1 | None | None | None | None |
| CPS1 | None | None | None | None |
| CPS1 | None | None | None | None |
| CPS1 | None | None | None | None |
| CPS1 | None | None | None | None |
| CPS1 | None | None | None | None |
| PSPH | None | None | None | None |
| PSPH | None | None | None | None |
| PSPH | None | None | None | None |
| PSPH | None | None | None | None |
| PSPH | None | None | None | None |
| PSPH | None | None | None | None |
| PEPD | None | None | None | None |
| PEPD | None | None | None | None |
| PEPD | None | None | None | None |
| PEPD | 0.643 | 2 | 5m4j | B |
| PEPD | 0.643 | 2 | 5m4j | B |
| PEPD | None | None | None | None |
| PEPD | None | None | None | None |
| PEPD | None | None | None | None |
| PEPD | None | None | None | None |
| KDM5C | None | None | None | None |
| KDM5C | None | None | None | None |
| KDM5C | None | None | None | None |
| ITGB6 | None | None | None | None |
| ITGB6 | None | None | None | None |
| ITGB6 | None | None | None | None |
| DARS | None | None | None | None |
| DARS | None | None | None | None |
| DARS | None | None | None | None |
| DARS | None | None | None | None |
| BTK | None | None | None | None |
| BTK | None | None | None | None |
| BTK | None | None | None | None |
| HDAC4 | None | None | None | None |
| ITGA2B | None | None | None | None |

| | | | | |
|---|---|---|---|---|
| ITGA2B | None | None | None | None |
| ITGA2B | None | None | None | None |
| ITGA2B | None | None | None | None |
| PCSK9 | None | None | None | None |
| PCSK9 | None | None | None | None |
| PCSK9 | None | None | None | None |
| PCSK9 | None | None | None | None |
| MUTYH | None | None | None | None |
| MUTYH | None | None | None | None |
| MUTYH | None | None | None | None |
| MUTYH | None | None | None | None |
| MUTYH | None | None | None | None |
| MUTYH | None | None | None | None |
| MUTYH | None | None | None | None |
| MUTYH | None | None | None | None |
| MUTYH | None | None | None | None |
| MUTYH | None | None | None | None |
| MUTYH | None | None | None | None |
| MUTYH | None | None | None | None |
| MUTYH | None | None | None | None |
| MUTYH | None | None | None | None |
| MUTYH | None | None | None | None |
| MUTYH | None | None | None | None |
| MUTYH | None | None | None | None |
| MUTYH | None | None | None | None |
| MUTYH | None | None | None | None |
| MUTYH | None | None | None | None |
| MUTYH | None | None | None | None |
| MUTYH | None | None | None | None |
| MUTYH | None | None | None | None |
| MUTYH | None | None | None | None |
| SLC25A12 | None | None | None | None |
| GSN | None | None | None | None |
| IHH | 0.909 | 1 | 3n1p | B |
| IHH | None | None | None | None |
| IHH | None | None | None | None |
| IHH | 0.818 | 2 | 3n1p | B |
| IHH | 0.727 | 12 | 3n1p | B |
| IHH | 0.636 | 15 | 3n1p | B |
| IHH | 0.818 | 7 | 3n1p | B |
| CD247 | None | None | None | None |
| EP300 | None | None | None | None |
| TTN | None | None | None | None |
| MRI1 | None | None | None | None |
| SGSH | None | None | None | None |
| SGSH | None | None | None | None |
| SGSH | None | None | None | None |
| SGSH | None | None | None | None |
| SGSH | None | None | None | None |
| SGSH | None | None | None | None |
| SGSH | None | None | None | None |
| SGSH | None | None | None | None |
| SGSH | None | None | None | None |
| SGSH | None | None | None | None |

| | | | | |
|---|---|---|---|---|
| SGSH | None | None | None | None |
| SGSH | None | None | None | None |
| SGSH | None | None | None | None |
| SGSH | None | None | None | None |
| SGSH | None | None | None | None |
| SGSH | None | None | None | None |
| SGSH | None | None | None | None |
| SGSH | None | None | None | None |
| SGSH | None | None | None | None |
| SGSH | None | None | None | None |
| SGSH | None | None | None | None |
| SGSH | None | None | None | None |
| CBL | None | None | None | None |
| CANT1 | None | None | None | None |
| CANT1 | None | None | None | None |
| CANT1 | None | None | None | None |
| CANT1 | None | None | None | None |
| CANT1 | None | None | None | None |
| CANT1 | None | None | None | None |
| CANT1 | None | None | None | None |
| SLC1A3 | None | None | None | None |
| SLC1A3 | None | None | None | None |
| SLC1A5 | None | None | None | None |
| SLC1A3 | None | None | None | None |
| SLC1A5 | None | None | None | None |
| SLC1A3 | None | None | None | None |
| SLC1A3 | None | None | None | None |
| CTSD | None | None | None | None |
| CTSD | None | None | None | None |
| CTSD | None | None | None | None |
| CTSD | None | None | None | None |
| CTSD | None | None | None | None |
| MTOR | None | None | None | None |
| ILK | None | None | None | None |
| PCDH15 | None | None | None | None |
| PCDH15 | None | None | None | None |
| PCDH15 | None | None | None | None |
| PCDH15 | None | None | None | None |
| PCDH15 | None | None | None | None |
| PCDH15 | None | None | None | None |
| KLHL40 | None | None | None | None |
| KLHL40 | None | None | None | None |
| KLHL40 | None | None | None | None |
| KLHL40 | None | None | None | None |
| CSTA | None | None | None | None |
| CSTA | None | None | None | None |
| HNMT | None | None | None | None |
| HNMT | None | None | None | None |
| IFNGR2 | None | None | None | None |
| IFNGR2 | None | None | None | None |
| PEX5 | None | None | None | None |
| KIF7 | None | None | None | None |
| KIF7 | None | None | None | None |
| MYL3 | None | None | None | None |

| | | | | |
|---|---|---|---|---|
| MYL3 | None | None | None | None |
| MYL3 | None | None | None | None |
| MYL3 | None | None | None | None |
| MYL3 | None | None | None | None |
| MYL3 | None | None | None | None |
| MYL3 | None | None | None | None |
| NAGLU | None | None | None | None |
| NAGLU | None | None | None | None |
| NAGLU | None | None | None | None |
| NAGLU | None | None | None | None |
| NAGLU | None | None | None | None |
| NAGLU | None | None | None | None |
| NAGLU | None | None | None | None |
| NAGLU | None | None | None | None |
| NAGLU | None | None | None | None |
| NAGLU | None | None | None | None |
| NAGLU | None | None | None | None |
| NAGLU | None | None | None | None |
| NAGLU | None | None | None | None |
| TNFRSF13C | None | None | None | None |
| KIF21A | None | None | None | None |
| KIF21A | None | None | None | None |
| KIF21A | None | None | None | None |
| KIF21A | None | None | None | None |
| KIF21A | None | None | None | None |
| DOCK2 | None | None | None | None |
| DSP | None | None | None | None |
| CDKN2A | None | None | None | None |
| CDKN2A | None | None | None | None |
| CDKN2A | None | None | None | None |
| CDKN2A | None | None | None | None |
| CDKN2A | None | None | None | None |
| CDKN2A | None | None | None | None |
| CDKN2A | None | None | None | None |
| CDKN2A | None | None | None | None |
| CDKN2A | None | None | None | None |
| CDKN2A | None | None | None | None |
| CDKN2A | None | None | None | None |
| CDKN2A | None | None | None | None |
| CDKN2A | None | None | None | None |
| CDKN2A | None | None | None | None |
| VCL | None | None | None | None |
| TPP1 | None | None | None | None |
| TPP1 | None | None | None | None |
| TPP1 | None | None | None | None |
| TPP1 | None | None | None | None |
| TPP1 | None | None | None | None |
| TPP1 | None | None | None | None |
| TPP1 | None | None | None | None |
| TPP1 | None | None | None | None |
| TPP1 | None | None | None | None |
| TPP1 | None | None | None | None |
| TPP1 | None | None | None | None |
| TPP1 | None | None | None | None |

| | | | | |
|---|---|---|---|---|
| TPP1 | None | None | None | None |
| TPP1 | None | None | None | None |
| TPP1 | None | None | None | None |
| TPP1 | None | None | None | None |
| TPP1 | None | None | None | None |
| TPP1 | None | None | None | None |
| TPP1 | None | None | None | None |
| TPP1 | None | None | None | None |
| TPP1 | None | None | None | None |
| TPP1 | None | None | None | None |
| IL10RB | None | None | None | None |
| IL10RB | None | None | None | None |
| CCM2 | None | None | None | None |
| CCM2 | None | None | None | None |
| CCM2 | None | None | None | None |
| DCX | None | None | None | None |
| DCX | None | None | None | None |
| DCX | None | None | None | None |
| DCX | None | None | None | None |
| DCX | None | None | None | None |
| DCX | None | None | None | None |
| DCX | None | None | None | None |
| DCX | None | None | None | None |
| BRCA2 | None | None | None | None |
| NNT | None | None | None | None |
| C2 | None | None | None | None |
| C2 | None | None | None | None |
| C2 | None | None | None | None |
| C2 | None | None | None | None |
| WRN | None | None | None | None |
| WRN | None | None | None | None |
| F13A1 | None | None | None | None |
| F13A1 | None | None | None | None |
| F13A1 | None | None | None | None |
| F13A1 | None | None | None | None |
| TF | None | None | None | None |
| TF | None | None | None | None |
| TF | None | None | None | None |
| TF | None | None | None | None |
| TF | None | None | None | None |
| ITPA | None | None | None | None |
| ITPA | None | None | None | None |
| SNCA | None | None | None | None |
| ITGB4 | None | None | None | None |
| ITGB4 | None | None | None | None |
| ITGB4 | None | None | None | None |
| LRP6 | None | None | None | None |
| LRP6 | None | None | None | None |
| LRP6 | None | None | None | None |
| PLG | None | None | None | None |
| PLG | 0.61 | 1 | 1i5k | A |
| TYROBP | None | None | None | None |
| UQCRC2 | None | None | None | None |
| GPSM2 | None | None | None | None |

| | | | | | |
|---|---|---|---|---|---|
| GPSM2 | None | None | | None | None |
| GPSM2 | None | None | | None | None |
| SCN5A | None | None | | None | None |
| SCN5A | None | None | | None | None |
| SCN5A | None | None | | None | None |
| SCN5A | None | None | | None | None |
| SCN5A | None | None | | None | None |
| ATIC | 0.6 | | 6 | 1p4r | A |
| EZH2 | None | None | | None | None |
| CPA1 | None | None | | None | None |
| SCN4B | None | None | | None | None |
| SCN4B | None | None | | None | None |
| MAN1B1 | None | None | | None | None |
| MAN1B1 | 0.6 | | 9 | 1fmi | A |
| MAN1B1 | None | None | | None | None |
| MAN1B1 | None | None | | None | None |
| CYCS | 0.74 | | 1 | 2n9i | A |
| IDUA | None | None | | None | None |
| IDUA | None | None | | None | None |
| IDUA | None | None | | None | None |
| IDUA | None | None | | None | None |
| IDUA | None | None | | None | None |
| IDUA | None | None | | None | None |
| IDUA | None | None | | None | None |
| IDUA | None | None | | None | None |
| IDUA | None | None | | None | None |
| IDUA | None | None | | None | None |
| IDUA | None | None | | None | None |
| IDUA | None | None | | None | None |
| IDUA | None | None | | None | None |
| IDUA | None | None | | None | None |
| IDUA | None | None | | None | None |
| IDUA | None | None | | None | None |
| IDUA | None | None | | None | None |
| IDUA | None | None | | None | None |
| IDUA | None | None | | None | None |
| IDUA | None | None | | None | None |
| IDUA | None | None | | None | None |
| IDUA | None | None | | None | None |
| IDUA | None | None | | None | None |
| IDUA | None | None | | None | None |
| IDUA | None | None | | None | None |
| HIKESHI | None | None | | None | None |
| APC | None | None | | None | None |
| APC | None | None | | None | None |
| APC | None | None | | None | None |
| APP | None | None | | None | None |
| CEP152 | None | None | | None | None |
| GAA | 0.609 | | 1 | 5kzx | A |
| GAA | 0.609 | | 1 | 5kzx | A |
| GAA | None | None | | None | None |
| GAA | None | None | | None | None |
| GAA | None | None | | None | None |
| GAA | 0.609 | | 1 | 5kzx | A |

| | | | | |
|---|---|---|---|---|
| GAA | 0.609 | 1 | 5kzx | A |
| GAA | 0.609 | 1 | 5kzx | A |
| GAA | 0.609 | 1 | 5kzx | A |
| GAA | 0.609 | 1 | 5kzx | A |
| GAA | 0.609 | 1 | 5kzx | A |
| GAA | 0.609 | 1 | 5kzx | A |
| GAA | 0.609 | 1 | 5kzx | A |
| GAA | 0.609 | 1 | 5kzx | A |
| GAA | 0.609 | 1 | 5kzx | A |
| GAA | 0.609 | 1 | 5kzx | A |
| GAA | 0.609 | 1 | 5kzx | A |
| GAA | 0.609 | 1 | 5kzx | A |
| GAA | 0.609 | 1 | 5kzx | A |
| GAA | 0.609 | 1 | 5kzx | A |
| GAA | 0.609 | 1 | 5kzx | A |
| GAA | 0.609 | 1 | 5kzx | A |
| GAA | 0.609 | 1 | 5kzx | A |
| GAA | 0.609 | 1 | 5kzx | A |
| GAA | 0.609 | 1 | 5kzx | A |
| GAA | 0.609 | 1 | 5kzx | A |
| GAA | 0.609 | 1 | 5kzx | A |
| GAA | 0.609 | 1 | 5kzx | A |
| GAA | 0.609 | 1 | 5kzx | A |
| GAA | 0.609 | 1 | 5kzx | A |
| GAA | 0.609 | 1 | 5kzx | A |
| GAA | 0.609 | 1 | 5kzx | A |
| GAA | 0.609 | 1 | 5kzx | A |
| GAA | 0.609 | 1 | 5kzx | A |
| GAA | 0.609 | 1 | 5kzx | A |
| GAA | 0.609 | 1 | 5kzx | A |
| GAA | 0.609 | 1 | 5kzx | A |
| GAA | 0.609 | 1 | 5kzx | A |
| GAA | 0.609 | 1 | 5kzx | A |
| GAA | 0.609 | 1 | 5kzx | A |
| GAA | 0.609 | 1 | 5kzx | A |
| GAA | 0.609 | 1 | 5kzx | A |
| GAA | 0.609 | 1 | 5kzx | A |
| GAA | 0.609 | 1 | 5kzx | A |
| GAA | 0.609 | 1 | 5kzx | A |
| GAA | 0.609 | 1 | 5kzx | A |
| GAA | 0.609 | 1 | 5kzx | A |
| GAA | 0.609 | 1 | 5kzx | A |
| GAA | None | None | None | None |
| FOLR1 | None | None | None | None |
| DSP | None | None | None | None |
| DSP | None | None | None | None |
| DSP | None | None | None | None |
| DSP | None | None | None | None |
| DSP | None | None | None | None |
| CCM2 | None | None | None | None |
| FXN | None | None | None | None |
| FXN | None | None | None | None |
| COMP | None | None | None | None |
| COMP | None | None | None | None |

| | | | | |
|---|---|---|---|---|
| COMP | None | None | None | None |
| COMP | None | None | None | None |
| COMP | None | None | None | None |
| COMP | None | None | None | None |
| COMP | None | None | None | None |
| COMP | None | None | None | None |
| COMP | None | None | None | None |
| COMP | None | None | None | None |
| COMP | None | None | None | None |
| COMP | None | None | None | None |
| AIFM1 | None | None | None | None |
| AIFM1 | None | None | None | None |
| AIFM1 | None | None | None | None |
| AIFM1 | None | None | None | None |
| AIFM1 | None | None | None | None |
| AIFM1 | None | None | None | None |
| AIFM1 | None | None | None | None |
| MYH9 | None | None | None | None |
| FHL2 | None | None | None | None |
| FHL2 | None | None | None | None |
| TNFRSF4 | None | None | None | None |
| CFB | None | None | None | None |
| CFB | None | None | None | None |
| CFB | None | None | None | None |
| CFB | None | None | None | None |
| CFB | None | None | None | None |
| CFB | None | None | None | None |
| CFB | None | None | None | None |
| CFB | None | None | None | None |
| CFB | None | None | None | None |
| CFB | None | None | None | None |
| MVD | None | None | None | None |
| MVD | None | None | None | None |
| HSD17B4 | None | None | None | None |
| HSD17B4 | None | None | None | None |
| HSD17B4 | None | None | None | None |
| HSD17B4 | None | None | None | None |
| HSD17B4 | None | None | None | None |
| HSD17B4 | None | None | None | None |
| F11 | None | None | None | None |
| F11 | None | None | None | None |
| F11 | None | None | None | None |
| F11 | None | None | None | None |
| F11 | None | None | None | None |
| F11 | None | None | None | None |
| F11 | None | None | None | None |
| F11 | None | None | None | None |
| F11 | None | None | None | None |
| SPTA1 | None | None | None | None |
| SPTA1 | None | None | None | None |
| SPTA1 | None | None | None | None |
| SPTA1 | None | None | None | None |
| SPTA1 | None | None | None | None |
| SPTA1 | None | None | None | None |

| | | | | |
|---|---|---|---|---|
| GMPR | None | None | None | None |
| MITF | None | None | None | None |
| NPC1 | None | None | None | None |
| NPC1 | None | None | None | None |
| NPC1 | None | None | None | None |
| NPC1 | None | None | None | None |
| NPC1 | None | None | None | None |
| NPC1 | None | None | None | None |
| NPC1 | None | None | None | None |
| NPC1 | None | None | None | None |
| NPC1 | None | None | None | None |
| NPC1 | None | None | None | None |
| NPC1 | None | None | None | None |
| NPC1 | None | None | None | None |
| NPC1 | None | None | None | None |
| NPC1 | None | None | None | None |
| NPC1 | None | None | None | None |
| NPC1 | None | None | None | None |
| NPC1 | None | None | None | None |
| NPC1 | None | None | None | None |
| NPC1 | None | None | None | None |
| NPC1 | None | None | None | None |
| NPC1 | None | None | None | None |
| NPC1 | None | None | None | None |
| NPC1 | None | None | None | None |
| NPC1 | None | None | None | None |
| NPC1 | None | None | None | None |
| NPC1 | None | None | None | None |
| NPC1 | None | None | None | None |
| NPC1 | None | None | None | None |
| NPC1 | None | None | None | None |
| NPC1 | None | None | None | None |
| NPC1 | None | None | None | None |
| NPC1 | None | None | None | None |
| NPC1 | None | None | None | None |
| NPC1 | None | None | None | None |
| NPC1 | None | None | None | None |
| NPC1 | None | None | None | None |
| NPC1 | None | None | None | None |
| NPC1 | None | None | None | None |
| NPC1 | None | None | None | None |
| NPC1 | None | None | None | None |
| NPC1 | None | None | None | None |
| NPC1 | None | None | None | None |
| NPC1 | None | None | None | None |
| NPC1 | None | None | None | None |
| C3 | None | None | None | None |
| NLRX1 | None | None | None | None |
| IGLL1 | None | None | None | None |
| APRT | 0.769 | 4 | 1zn9 | B |
| APRT | None | None | None | None |
| APRT | None | None | None | None |
| APRT | 0.923 | 1 | 1zn8 | A |
| APRT | None | None | None | None |
| ZIC3 | None | None | None | None |

| | | | | |
|---|---|---|---|---|
| ZIC3 | None | None | None | None |
| KLHL3 | None | None | None | None |
| RPS6KA3 | None | None | None | None |
| RPS6KA3 | None | None | None | None |
| RPS6KA3 | None | None | None | None |
| F8 | None | None | None | None |
| F8 | None | None | None | None |
| F8 | None | None | None | None |
| F8 | None | None | None | None |
| F8 | None | None | None | None |
| F8 | None | None | None | None |
| F8 | None | None | None | None |
| F8 | None | None | None | None |
| F8 | None | None | None | None |
| F8 | None | None | None | None |
| F8 | None | None | None | None |
| F8 | None | None | None | None |
| F8 | None | None | None | None |
| F8 | None | None | None | None |
| F8 | None | None | None | None |
| F8 | None | None | None | None |
| F8 | None | None | None | None |
| ISG15 | None | None | None | None |
| EXOSC3 | None | None | None | None |
| EXOSC3 | None | None | None | None |
| EXOSC3 | None | None | None | None |
| NKX2-5 | None | None | None | None |
| NKX2-5 | None | None | None | None |
| NKX2-5 | None | None | None | None |
| NKX2-5 | None | None | None | None |
| FH | None | None | None | None |
| FH | None | None | None | None |
| FH | None | None | None | None |
| FH | None | None | None | None |
| FH | 0.621 | 4 | 30000 | A |
| FH | None | None | None | None |
| FH | None | None | None | None |
| FH | None | None | None | None |
| FH | 0.64 | 6 | 5d6b | A |
| FH | None | None | None | None |
| FH | 0.64 | 1 | 5d6b | A |
| FH | 0.64 | 1 | 5d6b | A |
| SNRNP200 | None | None | None | None |
| SNRNP200 | None | None | None | None |
| SNRNP200 | None | None | None | None |
| SNRNP200 | None | None | None | None |
| CHRNA1 | None | None | None | None |
| CHRNA1 | None | None | None | None |
| MYOM1 | None | None | None | None |
| MYOM1 | None | None | None | None |
| MYOM1 | None | None | None | None |
| MYOM1 | None | None | None | None |
| MYOM1 | None | None | None | None |
| MYOM1 | None | None | None | None |

| | | | | |
|---|---|---|---|---|
| LDLR | None | None | None | None |
| LDLR | None | None | None | None |
| LDLR | None | None | None | None |
| LDLR | None | None | None | None |
| LDLR | None | None | None | None |
| LDLR | None | None | None | None |
| EHMT1 | None | None | None | None |
| EHMT1 | None | None | None | None |
| EHMT1 | None | None | None | None |
| COL9A2 | None | None | None | None |
| HFE2 | None | None | None | None |
| HFE2 | None | None | None | None |
| CD3G | None | None | None | None |
| MMAB | None | None | None | None |
| MMAB | None | None | None | None |
| MMAB | None | None | None | None |
| MMAB | 0.65 | 3 | 2idx | A |
| MMAB | 0.65 | 3 | 2idx | A |
| MMAB | 0.7 | 2 | 2idx | B |
| MMAB | 0.7 | 2 | 2idx | B |
| MMAB | None | None | None | None |
| MMAB | None | None | None | None |
| MMAB | None | None | None | None |
| TAF1 | None | None | None | None |
| TAF1 | None | None | None | None |
| CD4 | None | None | None | None |
| QARS | None | None | None | None |
| QARS | None | None | None | None |
| QARS | None | None | None | None |
| QARS | None | None | None | None |
| QARS | None | None | None | None |
| FGFR3 | None | None | None | None |
| FGFR3 | None | None | None | None |
| FGFR3 | None | None | None | None |
| FGFR3 | None | None | None | None |
| FGFR3 | None | None | None | None |
| FGFR3 | None | None | None | None |
| FLNB | None | None | None | None |
| FLNB | None | None | None | None |
| FLNB | None | None | None | None |
| FLNB | None | None | None | None |
| FLNB | None | None | None | None |
| FLNB | None | None | None | None |
| FLNB | None | None | None | None |
| FLNB | None | None | None | None |
| IL21R | None | None | None | None |
| RB1 | None | None | None | None |
| RB1 | None | None | None | None |
| RB1 | None | None | None | None |
| NTRK1 | None | None | None | None |
| NOTCH1 | None | None | None | None |
| MYOM1 | None | None | None | None |
| MYOM1 | None | None | None | None |
| MYOM1 | None | None | None | None |

| | | | | |
|---|---|---|---|---|
| MYOM1 | None | None | None | None |
| MYOM1 | None | None | None | None |
| MYOM1 | None | None | None | None |
| COQ9 | None | None | None | None |
| COQ9 | None | None | None | None |
| CRBN | None | None | None | None |
| DCTN1 | None | None | None | None |
| DCTN1 | None | None | None | None |
| DCTN1 | None | None | None | None |
| DCTN1 | None | None | None | None |
| DCTN1 | None | None | None | None |
| DCTN1 | None | None | None | None |
| THAP1 | None | None | None | None |
| THAP1 | None | None | None | None |
| GNAS | None | None | None | None |
| GNAS | None | None | None | None |
| HNF4A | None | None | None | None |
| HNF4A | None | None | None | None |
| ALDH3A2 | None | None | None | None |
| ALDH3A2 | None | None | None | None |
| ALDH3A2 | None | None | None | None |
| ALDH3A2 | None | None | None | None |
| ALDH3A2 | None | None | None | None |
| ALDH3A2 | None | None | None | None |
| ALDH3A2 | None | None | None | None |
| ALDH3A2 | None | None | None | None |
| ALDH3A2 | None | None | None | None |
| HCN1 | None | None | None | None |
| PLEC | None | None | None | None |
| SCN5A | None | None | None | None |
| SCN5A | None | None | None | None |
| STAMBP | None | None | None | None |
| STAMBP | None | None | None | None |
| RNF125 | None | None | None | None |
| RNF125 | None | None | None | None |
| USH1C | None | None | None | None |
| OBSL1 | None | None | None | None |
| GRIN2B | None | None | None | None |
| GRIN2B | None | None | None | None |
| HCN4 | None | None | None | None |
| HCN4 | None | None | None | None |
| TALDO1 | None | None | None | None |
| ITGB3 | None | None | None | None |
| CHKB | None | None | None | None |
| CHKB | None | None | None | None |
| CHKB | None | None | None | None |
| CHKB | None | None | None | None |
| ACTN4 | None | None | None | None |
| ACTN4 | None | None | None | None |
| ACTN4 | None | None | None | None |
| ACTN4 | None | None | None | None |
| CDT1 | None | None | None | None |
| CDT1 | None | None | None | None |
| CDT1 | None | None | None | None |

| | | | | |
|---|---|---|---|---|
| CDT1 | None | None | None | None |
| CDT1 | None | None | None | None |
| CDT1 | None | None | None | None |
| MAPT | None | None | None | None |
| F5 | None | None | None | None |
| F5 | None | None | None | None |
| F5 | None | None | None | None |
| ESRRB | None | None | None | None |
| DGUOK | None | None | None | None |
| DGUOK | None | None | None | None |
| DGUOK | None | None | None | None |
| DGUOK | None | None | None | None |
| DGUOK | 0.765 | 1 | 2ocp | A |
| DGUOK | None | None | None | None |
| DGUOK | None | None | None | None |
| DGUOK | None | None | None | None |
| DGUOK | None | None | None | None |
| CENPJ | None | None | None | None |
| TTN | None | None | None | None |
| TTN | None | None | None | None |
| TTN | None | None | None | None |
| TTN | None | None | None | None |
| TNFRSF1A | None | None | None | None |
| TNFRSF1A | None | None | None | None |
| TNFRSF1A | None | None | None | None |
| TNFRSF1A | None | None | None | None |
| TNFRSF1A | None | None | None | None |
| TNFRSF1A | None | None | None | None |
| TNFRSF1A | None | None | None | None |
| TNFRSF1A | None | None | None | None |
| TNFRSF1A | None | None | None | None |
| TNFRSF1A | None | None | None | None |
| TNFRSF1A | None | None | None | None |
| TNFRSF1A | None | None | None | None |
| ASL | None | None | None | None |
| ASL | 0.625 | 25 | 1aos | A |
| ASL | None | None | None | None |
| ASL | None | None | None | None |
| ASL | None | None | None | None |
| ASL | None | None | None | None |
| ASL | 0.625 | 1 | 1aos | A |
| ASL | None | None | None | None |
| ASL | None | None | None | None |
| ASL | None | None | None | None |
| MYO7A | None | None | None | None |
| MYO7A | None | None | None | None |
| MYO7A | None | None | None | None |
| MYO7A | None | None | None | None |
| MYO7A | None | None | None | None |
| MYO7A | None | None | None | None |
| MYO7A | None | None | None | None |
| MYO7A | None | None | None | None |
| MYO7A | None | None | None | None |
| MYO7A | None | None | None | None |

| | | | | |
|---|---|---|---|---|
| MYO7A | None | None | None | None |
| MYO7A | None | None | None | None |
| MYO7A | None | None | None | None |
| MYO7A | None | None | None | None |
| MYO7A | None | None | None | None |
| MYO7A | None | None | None | None |
| MYO7A | None | None | None | None |
| MYO7A | None | None | None | None |
| MYO7A | None | None | None | None |
| MYO7A | None | None | None | None |
| APC | None | None | None | None |
| ALAD | None | None | None | None |
| ALAD | None | None | None | None |
| ALAD | None | None | None | None |
| ALAD | None | None | None | None |
| ALAD | None | None | None | None |
| ALAD | None | None | None | None |
| POR | None | None | None | None |
| CSF1R | None | None | None | None |
| CSF1R | None | None | None | None |
| CSF1R | None | None | None | None |
| CSF1R | None | None | None | None |
| CSF1R | None | None | None | None |
| CSF1R | None | None | None | None |
| CSF1R | None | None | None | None |
| CSF1R | None | None | None | None |
| CSF1R | None | None | None | None |
| CSF1R | None | None | None | None |
| CSF1R | None | None | None | None |
| VWF | None | None | None | None |
| VWF | None | None | None | None |
| PHF8 | None | None | None | None |
| ACY1 | None | None | None | None |
| SMN1 | None | None | None | None |
| SMN1 | None | None | None | None |
| DLAT | None | None | None | None |
| DLAT | None | None | None | None |
| B2M | None | None | None | None |
| SOX9 | None | None | None | None |
| SOX9 | None | None | None | None |
| SOX9 | None | None | None | None |
| SOX9 | None | None | None | None |
| SOX9 | None | None | None | None |
| CFH | None | None | None | None |
| FLNC | None | None | None | None |
| FLNC | None | None | None | None |
| FLNC | None | None | None | None |
| ITGB2 | None | None | None | None |
| ITGB2 | None | None | None | None |
| ITGB2 | None | None | None | None |
| ITGB2 | None | None | None | None |
| ITGB2 | None | None | None | None |
| ITGB2 | None | None | None | None |
| ITGB2 | None | None | None | None |

| | | | | |
|---|---|---|---|---|
| ITGB2 | None | None | None | None |
| ITGB2 | None | None | None | None |
| PSEN1 | None | None | None | None |
| PSEN1 | None | None | None | None |
| PSEN1 | None | None | None | None |
| PSEN1 | None | None | None | None |
| PSEN1 | None | None | None | None |
| PSEN1 | None | None | None | None |
| OXCT1 | None | None | None | None |
| OXCT1 | None | None | None | None |
| OXCT1 | None | None | None | None |
| OXCT1 | None | None | None | None |
| OXCT1 | None | None | None | None |
| OXCT1 | None | None | None | None |
| OXCT1 | None | None | None | None |
| DHH | None | None | None | None |
| PRKN | None | None | None | None |
| PRKN | None | None | None | None |
| PRKN | None | None | None | None |
| PRKN | None | None | None | None |
| PRKN | None | None | None | None |
| PRKN | None | None | None | None |
| PRKN | None | None | None | None |
| PRKN | None | None | None | None |
| PRKN | None | None | None | None |
| PRKN | None | None | None | None |
| PRKN | None | None | None | None |
| LCAT | None | None | None | None |
| LCAT | None | None | None | None |
| LCAT | None | None | None | None |
| LCAT | None | None | None | None |
| LCAT | None | None | None | None |
| LCAT | None | None | None | None |
| LCAT | None | None | None | None |
| LCAT | None | None | None | None |
| LCAT | None | None | None | None |
| LCAT | None | None | None | None |
| LCAT | None | None | None | None |
| LCAT | None | None | None | None |
| FOXP3 | None | None | None | None |
| FOXP3 | None | None | None | None |
| FOXP3 | None | None | None | None |
| FOXP3 | None | None | None | None |
| SUFU | None | None | None | None |
| HSPA9 | None | None | None | None |
| HSPA9 | None | None | None | None |
| SQSTM1 | None | None | None | None |
| DARS2 | None | None | None | None |
| DARS2 | None | None | None | None |
| DARS2 | None | None | None | None |
| DARS2 | None | None | None | None |
| DARS2 | None | None | None | None |
| DARS2 | None | None | None | None |
| DARS2 | None | None | None | None |

| | | | | |
|---|---|---|---|---|
| DARS2 | None | None | None | None |
| DARS2 | None | None | None | None |
| DARS2 | None | None | None | None |
| DARS2 | None | None | None | None |
| DARS2 | None | None | None | None |
| NT5C3A | None | None | None | None |
| NT5C3A | None | None | None | None |
| NT5C3A | None | None | None | None |
| NT5C3A | 0.6 | 14 | 2vkq | A |
| ETFA | None | None | None | None |
| ETFA | None | None | None | None |
| ETFA | None | None | None | None |
| ETFA | None | None | None | None |
| ETFA | None | None | None | None |
| ETFA | None | None | None | None |
| ETFA | None | None | None | None |
| TTN | None | None | None | None |
| TTN | None | None | None | None |
| FLT4 | None | None | None | None |
| FGB | None | None | None | None |
| HPCA | None | None | None | None |
| TAP2 | None | None | None | None |
| TAP2 | None | None | None | None |
| PTPRC | None | None | None | None |
| SLC4A1 | None | None | None | None |
| SLC4A1 | None | None | None | None |
| SLC4A1 | None | None | None | None |
| SLC4A1 | None | None | None | None |
| SLC4A1 | None | None | None | None |
| SLC4A1 | None | None | None | None |
| MYH7 | None | None | None | None |
| MYH7 | None | None | None | None |
| MYH7 | None | None | None | None |
| MYH7 | None | None | None | None |
| MYH7 | None | None | None | None |
| MYH7 | None | None | None | None |
| MYH7 | None | None | None | None |
| MYH7 | None | None | None | None |
| MYH7 | None | None | None | None |
| PARN | None | None | None | None |
| PARN | None | None | None | None |
| NDUFS6 | None | None | None | None |
| VRK1 | None | None | None | None |
| VRK1 | None | None | None | None |
| VRK1 | None | None | None | None |
| VRK1 | None | None | None | None |
| VRK1 | None | None | None | None |
| VRK1 | None | None | None | None |
| COLEC11 | None | None | None | None |
| VHL | None | None | None | None |
| TTN | None | None | None | None |
| NCF2 | None | None | None | None |
| NCF2 | None | None | None | None |
| NCF2 | None | None | None | None |

| | | | | |
|---|---|---|---|---|
| NCF2 | None | None | None | None |
| NCF2 | None | None | None | None |
| RPS26 | None | None | None | None |
| CETP | None | None | None | None |
| CETP | None | None | None | None |
| CETP | None | None | None | None |
| CETP | None | None | None | None |
| CETP | None | None | None | None |
| CETP | None | None | None | None |
| CETP | None | None | None | None |
| CETP | None | None | None | None |
| CETP | None | None | None | None |
| CETP | None | None | None | None |
| CETP | None | None | None | None |
| CETP | None | None | None | None |
| CETP | None | None | None | None |
| FLNA | None | None | None | None |
| FLNA | None | None | None | None |
| FLNA | None | None | None | None |
| FLNA | None | None | None | None |
| FLNA | None | None | None | None |
| FLNA | None | None | None | None |
| FLNA | None | None | None | None |
| MYBPC3 | None | None | None | None |
| MYBPC3 | None | None | None | None |
| MYBPC3 | None | None | None | None |
| MYBPC3 | None | None | None | None |
| MYBPC3 | None | None | None | None |
| MYBPC3 | None | None | None | None |
| MYBPC3 | None | None | None | None |
| MYBPC3 | None | None | None | None |
| MYBPC3 | None | None | None | None |
| MYBPC3 | None | None | None | None |
| LSS | None | None | None | None |
| C6 | None | None | None | None |
| C6 | None | None | None | None |
| RLBP1 | None | None | None | None |
| RLBP1 | None | None | None | None |
| RLBP1 | None | None | None | None |
| RLBP1 | None | None | None | None |
| RLBP1 | None | None | None | None |
| RLBP1 | None | None | None | None |
| RLBP1 | None | None | None | None |
| RLBP1 | None | None | None | None |
| MMAA | None | None | None | None |
| MMAA | None | None | None | None |
| MMAA | None | None | None | None |
| MMAA | None | None | None | None |
| MMAA | None | None | None | None |
| MMAA | None | None | None | None |
| IL1RN | None | None | None | None |
| MASTL | None | None | None | None |
| FHL1 | None | None | None | None |
| FHL1 | None | None | None | None |

| | | | | |
|---|---|---|---|---|
| FHL1 | None | None | None | None |
| FHL1 | None | None | None | None |
| FHL1 | None | None | None | None |
| FHL1 | None | None | None | None |
| FHL1 | None | None | None | None |
| FHL1 | None | None | None | None |
| FHL1 | None | None | None | None |
| EIF2B1 | 0.636 | 1 | 3ecs | H |
| EIF2B1 | 0.636 | 1 | 3ecs | H |
| COQ8A | None | None | None | None |
| COQ8A | None | None | None | None |
| COQ8A | None | None | None | None |
| COQ8A | None | None | None | None |
| COQ8A | None | None | None | None |
| COQ8A | None | None | None | None |
| COQ8A | None | None | None | None |
| COQ8A | None | None | None | None |
| COQ8A | None | None | None | None |
| COQ8A | None | None | None | None |
| COQ8A | None | None | None | None |
| COQ8A | None | None | None | None |
| COQ8A | None | None | None | None |
| COQ8A | None | None | None | None |
| COQ8A | None | None | None | None |
| COQ8A | None | None | None | None |
| UNC119 | None | None | None | None |
| IGHMBP2 | None | None | None | None |
| JAG1 | None | None | None | None |
| JAG1 | None | None | None | None |
| JAG1 | None | None | None | None |
| GNB1 | None | None | None | None |
| GNB1 | None | None | None | None |
| FLNA | None | None | None | None |
| FLNA | None | None | None | None |
| FLNA | None | None | None | None |
| FLNA | None | None | None | None |
| FLNA | None | None | None | None |
| CREBBP | None | None | None | None |
| CSF3R | None | None | None | None |
| CSF3R | None | None | None | None |
| HIBCH | None | None | None | None |
| HIBCH | None | None | None | None |
| HIBCH | None | None | None | None |
| MMACHC | None | None | None | None |
| MMACHC | None | None | None | None |
| MMACHC | None | None | None | None |
| MMACHC | None | None | None | None |
| MMACHC | None | None | None | None |
| MMACHC | None | None | None | None |
| MMACHC | None | None | None | None |
| MMACHC | None | None | None | None |
| MMACHC | None | None | None | None |
| MMACHC | None | None | None | None |
| MMACHC | None | None | None | None |

| | | | | |
|---|---|---|---|---|
| KLF1 | None | None | None | None |
| NAGS | None | None | None | None |
| NAGS | None | None | None | None |
| NAGS | None | None | None | None |
| PIK3R2 | None | None | None | None |
| EDNRB | None | None | None | None |
| EDNRB | None | None | None | None |
| EDNRB | None | None | None | None |
| EDNRB | None | None | None | None |
| E | None | None | None | None |
| RUNX1 | None | None | None | None |
| RUNX1 | None | None | None | None |
| CTNNB1 | None | None | None | None |
| CTNNB1 | None | None | None | None |
| CTNNB1 | None | None | None | None |
| CTNNB1 | None | None | None | None |
| CTNNB1 | None | None | None | None |
| CTNNB1 | None | None | None | None |
| TTN | None | None | None | None |
| TTN | None | None | None | None |
| TYK2 | None | None | None | None |
| TYK2 | None | None | None | None |
| TYK2 | None | None | None | None |
| ERCC4 | None | None | None | None |
| ERCC4 | None | None | None | None |
| ERCC4 | None | None | None | None |
| ERCC4 | None | None | None | None |
| CFI | None | None | None | None |
| CFI | None | None | None | None |
| CFI | None | None | None | None |
| CFI | None | None | None | None |
| CFI | None | None | None | None |
| CFI | None | None | None | None |
| CFI | None | None | None | None |
| CFI | None | None | None | None |
| CFI | None | None | None | None |
| CFI | None | None | None | None |
| CFI | None | None | None | None |
| CFI | 0.691 | 2 | 5o32 | J |
| CFI | 0.691 | 2 | 5o32 | J |
| CFI | None | None | None | None |
| NOTCH3 | None | None | None | None |
| NOTCH3 | None | None | None | None |
| NOTCH3 | None | None | None | None |
| NOTCH3 | None | None | None | None |
| TBX1 | None | None | None | None |
| TBX1 | None | None | None | None |
| DDB2 | None | None | None | None |
| DDB2 | None | None | None | None |
| TP53 | None | None | None | None |
| POMGNT1 | None | None | None | None |
| POMGNT1 | None | None | None | None |
| POMGNT1 | None | None | None | None |
| POMGNT1 | None | None | None | None |

| | | | | |
|---|---|---|---|---|
| POMGNT1 | None | None | None | None |
| POMGNT1 | None | None | None | None |
| POMGNT1 | None | None | None | None |
| POMGNT1 | None | None | None | None |
| POMGNT1 | None | None | None | None |
| POMGNT1 | None | None | None | None |
| POMGNT1 | None | None | None | None |
| POMGNT1 | None | None | None | None |
| POMGNT1 | None | None | None | None |
| FMR1 | None | None | None | None |
| FMR1 | None | None | None | None |
| SLC9A3R1 | None | None | None | None |
| HSPB1 | None | None | None | None |
| HSPB1 | None | None | None | None |
| HSPB1 | None | None | None | None |
| HSPB1 | None | None | None | None |
| HSPB1 | None | None | None | None |
| HSPB1 | None | None | None | None |
| HSPB1 | None | None | None | None |
| SOD3 | None | None | None | None |
| AGA | 1 | 2 | 1apz | D |
| AGA | None | None | None | None |
| AGA | None | None | None | None |
| NDUFS7 | None | None | None | None |
| NDUFS7 | None | None | None | None |
| SOX2 | None | None | None | None |
| CSRP3 | None | None | None | None |
| CSRP3 | None | None | None | None |
| PAH | None | None | None | None |
| PAH | None | None | None | None |
| PAH | None | None | None | None |
| PAH | None | None | None | None |
| CACNA1S | None | None | None | None |
| TRPV4 | None | None | None | None |
| TRPV4 | None | None | None | None |
| TRPV4 | None | None | None | None |
| TRPV4 | None | None | None | None |
| TRPV4 | None | None | None | None |
| TRPV4 | None | None | None | None |
| TRPV4 | None | None | None | None |
| TRPV4 | None | None | None | None |
| TRPV4 | None | None | None | None |
| TRPV4 | None | None | None | None |
| TRPV4 | None | None | None | None |
| TRPV4 | None | None | None | None |
| TRPV4 | None | None | None | None |
| TNNI3 | None | None | None | None |
| MLYCD | None | None | None | None |
| MLYCD | None | None | None | None |
| MLYCD | None | None | None | None |
| MLYCD | None | None | None | None |
| AIP | None | None | None | None |
| AIP | None | None | None | None |
| ERCC2 | None | None | None | None |

| | | | | |
|---|---|---|---|---|
| ERCC2 | None | None | None | None |
| ERCC2 | None | None | None | None |
| ERCC2 | None | None | None | None |
| ERCC2 | None | None | None | None |
| ERCC2 | None | None | None | None |
| ERCC2 | None | None | None | None |
| ERCC2 | None | None | None | None |
| ERCC2 | None | None | None | None |
| ERCC2 | None | None | None | None |
| ERCC2 | None | None | None | None |
| ERCC2 | None | None | None | None |
| ERCC2 | None | None | None | None |
| ERCC2 | None | None | None | None |
| PUS1 | None | None | None | None |
| PUS1 | None | None | None | None |
| PUS1 | None | None | None | None |
| CASK | None | None | None | None |
| CASK | None | None | None | None |
| BLVRA | None | None | None | None |
| DICER1 | None | None | None | None |
| DICER1 | None | None | None | None |
| DICER1 | None | None | None | None |
| EIF2AK3 | None | None | None | None |
| EIF2AK3 | None | None | None | None |
| EIF2AK3 | None | None | None | None |
| KIT | None | None | None | None |
| KIT | None | None | None | None |
| KIT | None | None | None | None |
| KIT | None | None | None | None |
| KIT | None | None | None | None |
| KIT | None | None | None | None |
| DNMT3B | None | None | None | None |
| TGM1 | None | None | None | None |
| NDUFA10 | None | None | None | None |
| NDUFA10 | None | None | None | None |
| PRPF31 | None | None | None | None |
| PRPF31 | None | None | None | None |
| ECE1 | None | None | None | None |
| ECE1 | None | None | None | None |
| ECE1 | None | None | None | None |
| NDUFV2 | None | None | None | None |
| NDUFV2 | None | None | None | None |
| NDUFV2 | None | None | None | None |
| GHRHR | None | None | None | None |
| GHRHR | None | None | None | None |
| GHRHR | None | None | None | None |
| SLC4A1 | None | None | None | None |
| FGF9 | None | None | None | None |
| BARD1 | None | None | None | None |
| BARD1 | None | None | None | None |
| BARD1 | None | None | None | None |
| PTPN12 | 0.741 | 2 | 5j8r | D |
| MASP2 | None | None | None | None |
| MASP2 | None | None | None | None |

| | | | | |
|---|---|---|---|---|
| MASP2 | None | None | None | None |
| MASP2 | None | None | None | None |
| HOXB1 | None | None | None | None |
| SLC4A1 | None | None | None | None |
| SLC4A1 | None | None | None | None |
| SLC4A1 | None | None | None | None |
| SLC4A1 | None | None | None | None |
| SLC4A1 | None | None | None | None |
| SLC4A1 | None | None | None | None |
| SLC4A1 | None | None | None | None |
| SLC4A1 | None | None | None | None |
| SLC4A1 | None | None | None | None |
| SLC4A1 | None | None | None | None |
| SLC4A1 | None | None | None | None |
| SLC4A1 | None | None | None | None |
| SLC4A1 | None | None | None | None |
| SLC4A1 | None | None | None | None |
| SLC4A1 | None | None | None | None |
| SLC4A1 | None | None | None | None |
| SLC4A1 | None | None | None | None |
| SLC4A1 | None | None | None | None |
| CBL | None | None | None | None |
| FAS | None | None | None | None |
| MTMR2 | None | None | None | None |
| MTMR2 | None | None | None | None |
| MTMR2 | None | None | None | None |
| MTMR2 | None | None | None | None |
| MTMR2 | None | None | None | None |
| MTMR2 | None | None | None | None |
| MTMR2 | None | None | None | None |
| MTMR2 | None | None | None | None |
| KLHL3 | None | None | None | None |
| KLHL3 | None | None | None | None |
| KLHL3 | 0.667 | 1 | 4ch9 | B |
| KLHL3 | 0.733 | 1 | 5nkp | A |
| KLHL3 | None | None | None | None |
| KLHL3 | None | None | None | None |
| RNASEH1 | None | None | None | None |
| ACTN1 | None | None | None | None |
| PHKG2 | None | None | None | None |
| PHKG2 | None | None | None | None |
| PHKG2 | None | None | None | None |
| NTRK1 | None | None | None | None |
| APOH | None | None | None | None |
| PMS2 | None | None | None | None |
| PMS2 | None | None | None | None |
| PMS2 | 0.797 | 1 | 1ea6 | A |
| PMS2 | None | None | None | None |
| PMS2 | None | None | None | None |
| PMS2 | None | None | None | None |
| PMS2 | None | None | None | None |
| PMS2 | None | None | None | None |
| PMS2 | None | None | None | None |
| PMS2 | None | None | None | None |
| PMS2 | None | None | None | None |

| | | | | |
|---|---|---|---|---|
| PMS2 | None | None | None | None |
| PMS2 | None | None | None | None |
| PMS2 | None | None | None | None |
| PMS2 | None | None | None | None |
| PMS2 | None | None | None | None |
| PMS2 | None | None | None | None |
| PMS2 | None | None | None | None |
| PMS2 | None | None | None | None |
| PMS2 | None | None | None | None |
| PMS2 | None | None | None | None |
| PMS2 | None | None | None | None |
| MOCS2 | None | None | None | None |
| MOCS2 | None | None | None | None |
| RPGRIP1 | None | None | None | None |
| RPGRIP1 | None | None | None | None |
| LIG4 | None | None | None | None |
| PROC | None | None | None | None |
| PROC | None | None | None | None |
| NAT1 | None | None | None | None |
| NAT1 | None | None | None | None |
| PNPO | None | None | None | None |
| PNPO | None | None | None | None |
| PNPO | None | None | None | None |
| PNPO | None | None | None | None |
| PNPO | None | None | None | None |
| MRE11 | None | None | None | None |
| MRE11 | None | None | None | None |
| MRE11 | None | None | None | None |
| MRE11 | None | None | None | None |
| DDC | None | None | None | None |
| DDC | None | None | None | None |
| DDC | None | None | None | None |
| DDC | None | None | None | None |
| DDC | None | None | None | None |
| DDC | 0.909 | 7 | 3rch | B |
| DDC | 0.909 | 7 | 3rch | B |
| DDC | 0.643 | 4 | 3rbf | A |
| DDC | None | None | None | None |
| DDC | None | None | None | None |
| DDC | None | None | None | None |
| ANK2 | None | None | None | None |
| ANK2 | None | None | None | None |
| ANK2 | None | None | None | None |
| AK2 | None | None | None | None |
| AK2 | None | None | None | None |
| AK2 | None | None | None | None |
| AK2 | None | None | None | None |
| AK2 | None | None | None | None |
| AK2 | None | None | None | None |
| GPHN | None | None | None | None |
| HSD17B4 | 0.66 | 1 | 1zbq | A |
| HSD17B4 | 0.66 | 1 | 1zbq | A |
| HSD17B4 | None | None | None | None |
| HSD17B4 | None | None | None | None |

| | | | | |
|---|---|---|---|---|
| HSD17B4 | None | None | None | None |
| HSD17B4 | None | None | None | None |
| APTX | 0.6 | 5 | 4ndg | A |
| APTX | 0.6 | 5 | 4ndg | A |
| APTX | None | None | None | None |
| APTX | None | None | None | None |
| APTX | None | None | None | None |
| APTX | None | None | None | None |
| APTX | None | None | None | None |
| FANCL | None | None | None | None |
| FANCL | None | None | None | None |
| EMG1 | None | None | None | None |
| EMG1 | None | None | None | None |
| EZH2 | None | None | None | None |
| ABCB6 | None | None | None | None |
| NCF2 | None | None | None | None |
| NCF2 | None | None | None | None |
| COL4A2 | None | None | None | None |
| COL4A2 | 1 | 1 | 1li1 | F |
| COL4A2 | None | None | None | None |
| COL4A2 | None | None | None | None |
| COL4A2 | 0.75 | 16 | 1li1 | C |
| SLX4 | None | None | None | None |
| SERPIND1 | None | None | None | None |
| FGFR3 | None | None | None | None |
| FGFR3 | None | None | None | None |
| FGFR3 | None | None | None | None |
| FGFR3 | None | None | None | None |
| FGFR3 | None | None | None | None |
| FGFR3 | None | None | None | None |
| TREM2 | None | None | None | None |
| TREM2 | None | None | None | None |
| TREM2 | None | None | None | None |
| TREM2 | None | None | None | None |
| TREM2 | None | None | None | None |
| TREM2 | None | None | None | None |
| TREM2 | None | None | None | None |
| LIG4 | None | None | None | None |
| LIG4 | None | None | None | None |
| LIG4 | None | None | None | None |
| LIG4 | None | None | None | None |
| LIG4 | None | None | None | None |
| LIG4 | None | None | None | None |
| ECHS1 | None | None | None | None |
| ECHS1 | None | None | None | None |
| ECHS1 | None | None | None | None |
| MTR | None | None | None | None |
| MTR | None | None | None | None |
| MTR | None | None | None | None |
| MTR | None | None | None | None |
| MTR | None | None | None | None |
| PMVK | None | None | None | None |
| PRNP | None | None | None | None |
| CHN1 | None | None | None | None |

| | | | | |
|---|---|---|---|---|
| CHN1 | None | None | None | None |
| CHN1 | None | None | None | None |
| CHN1 | None | None | None | None |
| CHN1 | None | None | None | None |
| CHN1 | None | None | None | None |
| CHN1 | None | None | None | None |
| CHN1 | None | None | None | None |
| SLC2A1 | None | None | None | None |
| SLC2A1 | None | None | None | None |
| SLC2A1 | None | None | None | None |
| SLC2A1 | None | None | None | None |
| SLC2A1 | None | None | None | None |
| SLC2A1 | None | None | None | None |
| SLC2A1 | None | None | None | None |
| SLC2A1 | None | None | None | None |
| SLC2A1 | None | None | None | None |
| SLC2A1 | None | None | None | None |
| SLC2A1 | None | None | None | None |
| SLC2A1 | None | None | None | None |
| SLC2A1 | None | None | None | None |
| SLC2A1 | None | None | None | None |
| SLC2A1 | None | None | None | None |
| SLC2A1 | None | None | None | None |
| SLC2A1 | None | None | None | None |
| PMM2 | None | None | None | None |
| PMM2 | None | None | None | None |
| PMM2 | None | None | None | None |
| PMM2 | None | None | None | None |
| PMM2 | None | None | None | None |
| PMM2 | None | None | None | None |
| PMM2 | None | None | None | None |
| PMM2 | None | None | None | None |
| PMM2 | None | None | None | None |
| PMM2 | None | None | None | None |
| PMM2 | None | None | None | None |
| PMM2 | None | None | None | None |
| PMM2 | None | None | None | None |
| PMM2 | None | None | None | None |
| PMM2 | None | None | None | None |
| PMM2 | None | None | None | None |
| PMM2 | None | None | None | None |
| PMM2 | None | None | None | None |
| PMM2 | None | None | None | None |
| PMM2 | None | None | None | None |
| PMM2 | None | None | None | None |
| PMM2 | None | None | None | None |
| PMM2 | None | None | None | None |
| PMM2 | None | None | None | None |
| PMM2 | None | None | None | None |
| PMM2 | None | None | None | None |
| PMM2 | None | None | None | None |
| PMM2 | None | None | None | None |
| CCKAR | None | None | None | None |
| MAN1B1 | None | None | None | None |

| | | | | |
|---|---|---|---|---|
| MAN1B1 | None | None | None | None |
| LAMA2 | None | None | None | None |
| LAMA2 | None | None | None | None |
| TBC1D4 | None | None | None | None |
| TBC1D4 | None | None | None | None |
| LDLR | None | None | None | None |
| LDLR | None | None | None | None |
| LDLR | None | None | None | None |
| LDLR | None | None | None | None |
| LDLR | None | None | None | None |
| LDLR | None | None | None | None |
| LDLR | None | None | None | None |
| LDLR | None | None | None | None |
| LDLR | None | None | None | None |
| LDLR | None | None | None | None |
| LDLR | None | None | None | None |
| LDLR | None | None | None | None |
| LDLR | None | None | None | None |
| LDLR | None | None | None | None |
| LDLR | None | None | None | None |
| LDLR | None | None | None | None |
| LDLR | None | None | None | None |
| LDLR | None | None | None | None |
| LDLR | None | None | None | None |
| LDLR | None | None | None | None |
| LDLR | None | None | None | None |
| LDLR | None | None | None | None |
| LDLR | None | None | None | None |
| LDLR | None | None | None | None |
| LDLR | None | None | None | None |
| LDLR | None | None | None | None |
| LDLR | None | None | None | None |
| LDLR | None | None | None | None |
| LDLR | None | None | None | None |
| LDLR | None | None | None | None |
| LDLR | None | None | None | None |
| LDLR | None | None | None | None |
| LDLR | None | None | None | None |
| LDLR | None | None | None | None |
| LDLR | None | None | None | None |
| LDLR | None | None | None | None |
| LDLR | None | None | None | None |
| LDLR | None | None | None | None |
| LDLR | None | None | None | None |
| LDLR | None | None | None | None |
| LDLR | None | None | None | None |
| LDLR | None | None | None | None |
| LDLR | None | None | None | None |
| FANCM | None | None | None | None |
| HAMP | None | None | None | None |
| GUSB | None | None | None | None |
| GUSB | None | None | None | None |
| GUSB | None | None | None | None |
| GUSB | None | None | None | None |

| | | | | |
|---|---|---|---|---|
| GUSB | None | None | None | None |
| GUSB | None | None | None | None |
| GUSB | None | None | None | None |
| GUSB | None | None | None | None |
| GUSB | None | None | None | None |
| GUSB | None | None | None | None |
| GUSB | None | None | None | None |
| GUSB | None | None | None | None |
| GUSB | None | None | None | None |
| LIG1 | None | None | None | None |
| LIG1 | None | None | None | None |
| MYH7 | None | None | None | None |
| MYH7 | None | None | None | None |
| MYH7 | None | None | None | None |
| MYH7 | None | None | None | None |
| MYH7 | None | None | None | None |
| MYH7 | None | None | None | None |
| MYH7 | None | None | None | None |
| MYH7 | None | None | None | None |
| MYH7 | None | None | None | None |
| MYH7 | None | None | None | None |
| MYH7 | None | None | None | None |
| MYH7 | None | None | None | None |
| MYH7 | None | None | None | None |
| MYH7 | None | None | None | None |
| MYH7 | None | None | None | None |
| MYH7 | None | None | None | None |
| MYH7 | None | None | None | None |
| MYH7 | None | None | None | None |
| MYH7 | None | None | None | None |
| MYH7 | None | None | None | None |
| MYH7 | None | None | None | None |
| MYH7 | None | None | None | None |
| MYH7 | None | None | None | None |
| MYH7 | None | None | None | None |
| MYH7 | None | None | None | None |
| MYH7 | None | None | None | None |
| MYH7 | None | None | None | None |
| MYH7 | None | None | None | None |
| MYH7 | None | None | None | None |
| MYH7 | None | None | None | None |
| MYH7 | None | None | None | None |
| MYH7 | None | None | None | None |
| MYH7 | None | None | None | None |
| MYH7 | None | None | None | None |
| HCN4 | None | None | None | None |
| RECQL4 | None | None | None | None |
| RECQL4 | None | None | None | None |
| RECQL4 | None | None | None | None |
| RECQL4 | None | None | None | None |
| RECQL4 | None | None | None | None |
| RECQL4 | None | None | None | None |
| RECQL4 | None | None | None | None |
| RECQL4 | None | None | None | None |

| | | | | |
|---|---|---|---|---|
| RECQL4 | None | None | None | None |
| RECQL4 | None | None | None | None |
| RECQL4 | None | None | None | None |
| RECQL4 | None | None | None | None |
| RECQL4 | None | None | None | None |
| CHEK2 | None | None | None | None |
| PPARG | None | None | None | None |
| ACADVL | None | None | None | None |
| ACADVL | None | None | None | None |
| ACADVL | None | None | None | None |
| ACADVL | None | None | None | None |
| ACADVL | None | None | None | None |
| ACADVL | None | None | None | None |
| ACADVL | None | None | None | None |
| ACADVL | None | None | None | None |
| ACADVL | None | None | None | None |
| ACADVL | None | None | None | None |
| ACADVL | None | None | None | None |
| ACADVL | None | None | None | None |
| ACADVL | None | None | None | None |
| ACADVL | None | None | None | None |
| ACADVL | None | None | None | None |
| ACADVL | None | None | None | None |
| ACADVL | None | None | None | None |
| ACADVL | None | None | None | None |
| ACADVL | None | None | None | None |
| ACADVL | None | None | None | None |
| ACADVL | None | None | None | None |
| ACADVL | None | None | None | None |
| APOA1 | None | None | None | None |
| APOA1 | None | None | None | None |
| APOA1 | None | None | None | None |
| APOA1 | None | None | None | None |
| APOA1 | None | None | None | None |
| DNMT3A | None | None | None | None |
| DNMT3A | None | None | None | None |
| APC | None | None | None | None |
| TMOD1 | None | None | None | None |
| USH1C | None | None | None | None |
| USH1C | None | None | None | None |
| USH1C | None | None | None | None |
| USH1C | None | None | None | None |
| ERCC6 | None | None | None | None |
| ERCC6 | None | None | None | None |
| ERCC6 | None | None | None | None |
| ERCC8 | None | None | None | None |
| ERCC8 | None | None | None | None |
| ERCC8 | None | None | None | None |
| DPAGT1 | None | None | None | None |
| DPAGT1 | None | None | None | None |
| DPAGT1 | None | None | None | None |
| DPAGT1 | None | None | None | None |
| DPAGT1 | None | None | None | None |
| DPAGT1 | None | None | None | None |

| | | | | |
|---|---|---|---|---|
| DPAGT1 | None | None | None | None |
| IL2RG | None | None | None | None |
| HNF1A | None | None | None | None |
| HNF4A | None | None | None | None |
| FLT4 | None | None | None | None |
| FLT4 | None | None | None | None |
| FLT4 | None | None | None | None |
| CD3D | None | None | None | None |
| CD3D | None | None | None | None |
| MCOLN1 | None | None | None | None |
| MCOLN1 | None | None | None | None |
| MOCS2 | None | None | None | None |
| MOCS2 | None | None | None | None |
| ABCG8 | None | None | None | None |
| ABCG8 | None | None | None | None |
| ABCG8 | None | None | None | None |
| ABCG8 | None | None | None | None |
| ABCG8 | None | None | None | None |
| ABCG8 | None | None | None | None |
| ABCG8 | None | None | None | None |
| ABCG8 | None | None | None | None |
| ABCG8 | None | None | None | None |
| ABCG8 | None | None | None | None |
| ABCG8 | None | None | None | None |
| ABCG8 | None | None | None | None |
| ABCG8 | None | None | None | None |
| ABCG8 | None | None | None | None |
| CTNNA1 | None | None | None | None |
| TYMP | None | None | None | None |
| TYMP | None | None | None | None |
| TYMP | None | None | None | None |
| TYMP | None | None | None | None |
| TYMP | None | None | None | None |
| TYMP | None | None | None | None |
| TYMP | None | None | None | None |
| TYMP | None | None | None | None |
| TYMP | None | None | None | None |
| TYMP | None | None | None | None |
| TYMP | None | None | None | None |
| TYMP | None | None | None | None |
| TYMP | None | None | None | None |
| GAMT | None | None | None | None |
| GAMT | None | None | None | None |
| GAMT | None | None | None | None |
| GAMT | None | None | None | None |
| GAMT | None | None | None | None |
| GAMT | None | None | None | None |
| GAMT | None | None | None | None |
| GAMT | None | None | None | None |
| GAMT | None | None | None | None |
| GAMT | None | None | None | None |
| GAMT | None | None | None | None |
| GLRA1 | None | None | None | None |
| GLRA1 | None | None | None | None |

| | | | | |
|---|---|---|---|---|
| GLRA1 | None | None | None | None |
| GLRA1 | None | None | None | None |
| GLRA1 | None | None | None | None |
| GLRA1 | None | None | None | None |
| GLRA1 | None | None | None | None |
| GLRA1 | None | None | None | None |
| GLRA1 | None | None | None | None |
| ADK | None | None | None | None |
| ACTN2 | None | None | None | None |
| UBE2T | None | None | None | None |
| TTN | None | None | None | None |
| ATP7B | None | None | None | None |
| ATP7B | None | None | None | None |
| ATP7B | None | None | None | None |
| ATP7B | None | None | None | None |
| ATP7B | None | None | None | None |
| ATP7B | None | None | None | None |
| ATP7B | None | None | None | None |
| ATP7B | None | None | None | None |
| ATP7B | None | None | None | None |
| ATP7B | None | None | None | None |
| ATP7B | None | None | None | None |
| ATP7B | None | None | None | None |
| ATP7B | None | None | None | None |
| CDKN1B | None | None | None | None |
| CDKN1B | None | None | None | None |
| CDKN1B | None | None | None | None |
| CDKN1B | None | None | None | None |
| CDKN1B | None | None | None | None |
| CDKN1B | None | None | None | None |
| MYOC | None | None | None | None |
| MYOC | None | None | None | None |
| MYOC | None | None | None | None |
| MYOC | None | None | None | None |
| MYOC | None | None | None | None |
| MYOC | None | None | None | None |
| MYOC | None | None | None | None |
| MYOC | None | None | None | None |
| MYOC | None | None | None | None |
| MYOC | None | None | None | None |
| MYOC | None | None | None | None |
| MYOC | None | None | None | None |
| MYOC | None | None | None | None |
| MYOC | None | None | None | None |
| MYOC | None | None | None | None |
| SPTAN1 | None | None | None | None |
| GC | None | None | None | None |
| GC | None | None | None | None |
| MAD1L1 | None | None | None | None |
| NDST1 | None | None | None | None |
| ARL2BP | None | None | None | None |
| TOR1A | None | None | None | None |
| C1S | None | None | None | None |
| ARL6 | None | None | None | None |

| | | | | |
|---|---|---|---|---|
| ARL6 | None | None | None | None |
| ARL6 | None | None | None | None |
| ARL6 | None | None | None | None |
| ARL6 | None | None | None | None |
| ARL6 | None | None | None | None |
| FGFR1 | None | None | None | None |
| KLHL7 | None | None | None | None |
| FLNC | None | None | None | None |
| PTPRC | None | None | None | None |
| GCH1 | None | None | None | None |
| GCH1 | None | None | None | None |
| GCH1 | None | None | None | None |
| GCH1 | None | None | None | None |
| GCH1 | None | None | None | None |
| GCH1 | None | None | None | None |
| GCH1 | None | None | None | None |
| GCH1 | None | None | None | None |
| GCH1 | None | None | None | None |
| GCH1 | None | None | None | None |
| GCH1 | None | None | None | None |
| GCH1 | None | None | None | None |
| GCH1 | None | None | None | None |
| GCH1 | None | None | None | None |
| CFH | None | None | None | None |
| COL2A1 | None | None | None | None |
| COL2A1 | None | None | None | None |
| OCRL | None | None | None | None |
| WRN | None | None | None | None |
| WRN | None | None | None | None |
| WRN | None | None | None | None |
| WRN | None | None | None | None |
| MASP1 | None | None | None | None |
| MASP1 | None | None | None | None |
| NSD1 | None | None | None | None |
| PITX2 | None | None | None | None |
| PITX2 | None | None | None | None |
| PITX2 | None | None | None | None |
| PITX2 | None | None | None | None |
| PITX2 | None | None | None | None |
| PITX2 | None | None | None | None |
| B4GALT7 | 0.917 | 1 | 4irq | A |
| B4GALT7 | None | None | None | None |
| B4GALT7 | None | None | None | None |
| B4GALT7 | None | None | None | None |
| B4GALT7 | None | None | None | None |
| FSHR | None | None | None | None |
| IGHMBP2 | None | None | None | None |
| IGHMBP2 | None | None | None | None |
| IGHMBP2 | None | None | None | None |
| IGHMBP2 | None | None | None | None |
| IGHMBP2 | None | None | None | None |
| IGHMBP2 | None | None | None | None |
| IGHMBP2 | None | None | None | None |
| IGHMBP2 | None | None | None | None |

| | | | | |
|---|---|---|---|---|
| IGHMBP2 | None | None | None | None |
| IGHMBP2 | None | None | None | None |
| IGHMBP2 | None | None | None | None |
| IGHMBP2 | None | None | None | None |
| IGHMBP2 | None | None | None | None |
| IGHMBP2 | None | None | None | None |
| IGHMBP2 | None | None | None | None |
| IGHMBP2 | None | None | None | None |
| TTN | None | None | None | None |
| TTN | None | None | None | None |
| IRAK4 | None | None | None | None |
| PROC | None | None | None | None |
| PROC | None | None | None | None |
| RAD50 | None | None | None | None |
| RAD50 | None | None | None | None |
| DYNC1H1 | None | None | None | None |
| DYNC1H1 | None | None | None | None |
| DYNC1H1 | None | None | None | None |
| DYNC1H1 | None | None | None | None |
| DYNC1H1 | None | None | None | None |
| DYNC1H1 | None | None | None | None |
| DYNC1H1 | None | None | None | None |
| DYNC1H1 | None | None | None | None |
| DYNC1H1 | None | None | None | None |
| DYNC1H1 | None | None | None | None |
| DYNC1H1 | None | None | None | None |
| CYB5R3 | None | None | None | None |
| CYB5R3 | None | None | None | None |
| CYB5R3 | None | None | None | None |
| CYB5R3 | None | None | None | None |
| CYB5R3 | None | None | None | None |
| CYB5R3 | None | None | None | None |
| CYB5R3 | None | None | None | None |
| CYB5R3 | None | None | None | None |
| CYB5R3 | None | None | None | None |
| CYB5R3 | None | None | None | None |
| CYB5R3 | None | None | None | None |
| SPTB | None | None | None | None |
| FANCL | None | None | None | None |
| FAM126A | None | None | None | None |
| SEPT9 | None | None | None | None |
| TTN | None | None | None | None |
| TTN | None | None | None | None |
| TTN | None | None | None | None |
| CNTN1 | None | None | None | None |
| C1QTNF5 | None | None | None | None |
| VCL | None | None | None | None |
| VCL | None | None | None | None |
| VCL | None | None | None | None |
| VCL | None | None | None | None |
| VCL | None | None | None | None |
| VCL | None | None | None | None |
| PPT1 | None | None | None | None |
| PPT1 | None | None | None | None |

| | | | | |
|---|---|---|---|---|
| PPT1 | None | None | None | None |
| PPT1 | None | None | None | None |
| PPT1 | None | None | None | None |
| PPT1 | None | None | None | None |
| PPT1 | None | None | None | None |
| PPT1 | None | None | None | None |
| PPT1 | None | None | None | None |
| PPT1 | None | None | None | None |
| PPT1 | None | None | None | None |
| PPT1 | None | None | None | None |
| PPT1 | None | None | None | None |
| PPT1 | None | None | None | None |
| PPT1 | None | None | None | None |
| PPT1 | None | None | None | None |
| PPT1 | None | None | None | None |
| PPT1 | None | None | None | None |
| PPT1 | None | None | None | None |
| NOTCH2 | None | None | None | None |
| NOTCH2 | None | None | None | None |
| IFIH1 | None | None | None | None |
| IFIH1 | None | None | None | None |
| IKBKG | None | None | None | None |
| DBH | None | None | None | None |
| DBH | None | None | None | None |
| DBH | None | None | None | None |
| DBH | None | None | None | None |
| DBH | None | None | None | None |
| DBH | None | None | None | None |
| DBH | None | None | None | None |
| DBH | None | None | None | None |
| DBH | None | None | None | None |
| DBH | None | None | None | None |
| IL10RA | None | None | None | None |
| IL10RA | None | None | None | None |
| IL10RA | None | None | None | None |
| IL10RA | None | None | None | None |
| IL10RA | None | None | None | None |
| IL10RA | None | None | None | None |
| IL10RA | None | None | None | None |
| ILK | None | None | None | None |
| HPD | None | None | None | None |
| HPD | None | None | None | None |
| HPD | None | None | None | None |
| MECP2 | None | None | None | None |
| MECP2 | None | None | None | None |
| MECP2 | None | None | None | None |
| MECP2 | None | None | None | None |
| MECP2 | None | None | None | None |
| MECP2 | None | None | None | None |
| MECP2 | None | None | None | None |
| MECP2 | None | None | None | None |
| KYNU | None | None | None | None |
| FAS | None | None | None | None |
| FAS | None | None | None | None |

| | | | | |
|---|---|---|---|---|
| ELP1 | None | None | None | None |
| ELP1 | None | None | None | None |
| ELP1 | None | None | None | None |
| ELP1 | None | None | None | None |
| ELP1 | None | None | None | None |
| ELP1 | None | None | None | None |
| ELP1 | None | None | None | None |
| ELP1 | None | None | None | None |
| HESX1 | None | None | None | None |
| HESX1 | None | None | None | None |
| HESX1 | None | None | None | None |
| HESX1 | None | None | None | None |
| GAD1 | None | None | None | None |
| GAD1 | None | None | None | None |
| NDUFA1 | None | None | None | None |
| NDUFA1 | None | None | None | None |
| NDUFA1 | None | None | None | None |
| PNPT1 | None | None | None | None |
| PNPT1 | None | None | None | None |
| PNPT1 | None | None | None | None |
| PNPT1 | None | None | None | None |
| PNPT1 | None | None | None | None |
| PNPT1 | None | None | None | None |
| PNPT1 | None | None | None | None |
| SLC25A13 | None | None | None | None |
| SLC25A13 | None | None | None | None |
| SLC25A13 | None | None | None | None |
| AP4M1 | None | None | None | None |
| AP4M1 | None | None | None | None |
| MEFV | None | None | None | None |
| MEFV | None | None | None | None |
| MEFV | None | None | None | None |
| MEFV | None | None | None | None |
| MEFV | None | None | None | None |
| MEFV | None | None | None | None |
| MEFV | None | None | None | None |
| MEFV | None | None | None | None |
| MEFV | None | None | None | None |
| CAPN3 | None | None | None | None |
| CAPN3 | None | None | None | None |
| CAPN3 | None | None | None | None |
| CAPN3 | None | None | None | None |
| CAPN3 | None | None | None | None |
| CAPN3 | None | None | None | None |
| CAPN3 | None | None | None | None |
| GP6 | None | None | None | None |
| FAS | None | None | None | None |
| FBN1 | None | None | None | None |
| FBN1 | None | None | None | None |
| BMPR2 | None | None | None | None |
| BMPR2 | None | None | None | None |
| BMPR2 | None | None | None | None |
| BMPR2 | None | None | None | None |
| BMPR2 | None | None | None | None |

| | | | | |
|---|---|---|---|---|
| RET | None | None | None | None |
| RET | None | None | None | None |
| RET | None | None | None | None |
| RET | None | None | None | None |
| RET | None | None | None | None |
| RET | None | None | None | None |
| RET | None | None | None | None |
| RET | None | None | None | None |
| RET | None | None | None | None |
| RET | None | None | None | None |
| RET | None | None | None | None |
| RET | None | None | None | None |
| RET | None | None | None | None |
| RET | None | None | None | None |
| RET | None | None | None | None |
| RET | None | None | None | None |
| RET | None | None | None | None |
| RET | None | None | None | None |
| RET | None | None | None | None |
| RET | None | None | None | None |
| RET | None | None | None | None |
| RET | None | None | None | None |
| RET | None | None | None | None |
| RET | None | None | None | None |
| RET | None | None | None | None |
| RET | None | None | None | None |
| RET | None | None | None | None |
| RET | None | None | None | None |
| CSF2RA | None | None | None | None |
| CSF2RA | None | None | None | None |
| CSF2RA | None | None | None | None |
| BRCA1 | None | None | None | None |
| ZAP70 | 0.636 | 1 | 2ozo | A |
| ZAP70 | 0.636 | 1 | 2ozo | A |
| ZAP70 | None | None | None | None |
| PIK3R2 | None | None | None | None |
| TCN2 | None | None | None | None |
| TCN2 | None | None | None | None |
| TCN2 | None | None | None | None |
| TCN2 | None | None | None | None |
| TCN2 | None | None | None | None |
| TCN2 | None | None | None | None |
| TCN2 | None | None | None | None |
| TCN2 | None | None | None | None |
| EPM2A | None | None | None | None |
| EPM2A | None | None | None | None |
| EPM2A | None | None | None | None |
| EPM2A | None | None | None | None |
| AARS | None | None | None | None |
| AARS | None | None | None | None |
| AARS | None | None | None | None |
| PCDH15 | None | None | None | None |
| PCDH15 | None | None | None | None |
| PCDH15 | None | None | None | None |

| | | | | |
|---|---|---|---|---|
| PCDH15 | None | None | None | None |
| PCDH15 | None | None | None | None |
| PCDH15 | None | None | None | None |
| PCDH15 | None | None | None | None |
| PCDH15 | None | None | None | None |
| PCDH15 | None | None | None | None |
| PCDH15 | None | None | None | None |
| NDUFS4 | None | None | None | None |
| NDUFS4 | None | None | None | None |
| NDUFS4 | None | None | None | None |
| FGF10 | None | None | None | None |
| FGF10 | None | None | None | None |
| FGF10 | None | None | None | None |
| FGF10 | None | None | None | None |
| CDON | None | None | None | None |
| STAMBP | 0.909 | 2 | 3rzv | A |
| BRCA1 | None | None | None | None |
| BRCA1 | None | None | None | None |
| BRCA1 | None | None | None | None |
| BRCA1 | None | None | None | None |
| BRCA1 | None | None | None | None |
| BRCA1 | None | None | None | None |
| BRCA1 | None | None | None | None |
| BRCA1 | None | None | None | None |
| BRCA1 | None | None | None | None |
| BRCA1 | None | None | None | None |
| PLXNC1 | None | None | None | None |
| RNASEH2C | None | None | None | None |
| RNASEH2C | None | None | None | None |
| APC | None | None | None | None |
| PLA2G4A | None | None | None | None |
| PLA2G4A | None | None | None | None |
| PSAT1 | None | None | None | None |
| PSAT1 | None | None | None | None |
| TRAPPC3 | None | None | None | None |
| QDPR | None | None | None | None |
| QDPR | None | None | None | None |
| QDPR | None | None | None | None |
| QDPR | None | None | None | None |
| KRT5 | None | None | None | None |
| KRT5 | None | None | None | None |
| KRT5 | None | None | None | None |
| MLH1 | None | None | None | None |
| MLH1 | None | None | None | None |
| MLH1 | None | None | None | None |
| MLH1 | None | None | None | None |
| MLH1 | None | None | None | None |
| MLH1 | None | None | None | None |
| MLH1 | None | None | None | None |
| MLH1 | None | None | None | None |
| MLH1 | None | None | None | None |
| MLH1 | None | None | None | None |
| MLH1 | None | None | None | None |
| MLH1 | None | None | None | None |

| | | | | |
|---|---|---|---|---|
| MLH1 | None | None | None | None |
| MLH1 | None | None | None | None |
| MLH1 | None | None | None | None |
| MLH1 | None | None | None | None |
| MLH1 | None | None | None | None |
| KRIT1 | None | None | None | None |
| KRIT1 | None | None | None | None |
| KRIT1 | None | None | None | None |
| ALX4 | None | None | None | None |
| ALX4 | None | None | None | None |
| ALX4 | None | None | None | None |
| TJP2 | None | None | None | None |
| TJP2 | None | None | None | None |
| PANK2 | None | None | None | None |
| PANK2 | None | None | None | None |
| PANK2 | None | None | None | None |
| PANK2 | None | None | None | None |
| PANK2 | None | None | None | None |
| PANK2 | None | None | None | None |
| PANK2 | None | None | None | None |
| NDUFA6 | None | None | None | None |
| TSHR | None | None | None | None |
| TSHR | None | None | None | None |
| TSHR | None | None | None | None |
| CFTR | None | None | None | None |
| CFTR | None | None | None | None |
| MYOM1 | None | None | None | None |
| MYOM1 | None | None | None | None |
| SPTB | None | None | None | None |
| GPT2 | None | None | None | None |
| NDUFS2 | None | None | None | None |
| NDUFS2 | None | None | None | None |
| NDUFS2 | None | None | None | None |
| NDUFS2 | None | None | None | None |
| NDUFS2 | None | None | None | None |
| CASQ2 | None | None | None | None |
| CASQ2 | None | None | None | None |
| CASQ2 | None | None | None | None |
| CASQ2 | None | None | None | None |
| CASQ2 | None | None | None | None |
| CASQ2 | None | None | None | None |
| CASQ2 | None | None | None | None |
| CASQ2 | None | None | None | None |
| CASQ2 | None | None | None | None |
| CPOX | None | None | None | None |
| CPOX | None | None | None | None |
| CPOX | 0.857 | 2 | 2aex | A |
| CPOX | None | None | None | None |
| CPOX | None | None | None | None |
| CPOX | None | None | None | None |
| CPOX | None | None | None | None |
| CPOX | None | None | None | None |
| CPOX | None | None | None | None |
| MVK | None | None | None | None |

| | | | | |
|---|---|---|---|---|
| MVK | None | None | None | None |
| MVK | None | None | None | None |
| MVK | None | None | None | None |
| MVK | None | None | None | None |
| MVK | None | None | None | None |
| MVK | None | None | None | None |
| MVK | None | None | None | None |
| MVK | None | None | None | None |
| MVK | None | None | None | None |
| MVK | None | None | None | None |
| MVK | None | None | None | None |
| MVK | None | None | None | None |
| MVK | None | None | None | None |
| MVK | None | None | None | None |
| GSS | None | None | None | None |
| GSS | None | None | None | None |
| GSS | None | None | None | None |
| GSS | None | None | None | None |
| GSS | None | None | None | None |
| DSG2 | None | None | None | None |
| DSG2 | None | None | None | None |
| DSG2 | None | None | None | None |
| DSG2 | None | None | None | None |
| DSG2 | None | None | None | None |
| DSG2 | None | None | None | None |
| DSG2 | None | None | None | None |
| DSG2 | None | None | None | None |
| DSG2 | None | None | None | None |
| DSG2 | None | None | None | None |
| DSG2 | None | None | None | None |
| DSG2 | None | None | None | None |
| DSG2 | None | None | None | None |
| DSG2 | None | None | None | None |
| PAPSS2 | None | None | None | None |
| PLCB2 | None | None | None | None |
| HMGCS2 | None | None | None | None |
| HMGCS2 | None | None | None | None |
| HMGCS2 | 0.938 | 1 | 2wya | C |
| HMGCS2 | None | None | None | None |
| HMGCS2 | None | None | None | None |
| HMGCS2 | None | None | None | None |
| HMGCS2 | None | None | None | None |
| HMGCS2 | None | None | None | None |
| MPDZ | None | None | None | None |
| GATA4 | None | None | None | None |
| GATA4 | None | None | None | None |
| WARS2 | None | None | None | None |
| WARS2 | None | None | None | None |
| HGD | 1 | 47 | 1eyb | A |
| HGD | 1 | 47 | 1eyb | A |
| HGD | 1 | 70 | 1eyb | A |
| HGD | None | None | None | None |
| HGD | None | None | None | None |
| HGD | None | None | None | None |

| | | | | |
|---|---|---|---|---|
| HGD | None | None | None | None |
| HGD | None | None | None | None |
| HGD | None | None | None | None |
| HGD | 1 | 1 | 1ey2 | A |
| HGD | 1 | 9 | 1ey2 | A |
| HGD | None | None | None | None |
| HGD | 1 | 10 | 1eyb | A |
| HGD | 1 | 7 | 1ey2 | A |
| PAX8 | None | None | None | None |
| PAX8 | None | None | None | None |
| PAX8 | None | None | None | None |
| PAX8 | None | None | None | None |
| SPTB | None | None | None | None |
| NCF4 | None | None | None | None |
| CFH | None | None | None | None |
| CFH | None | None | None | None |
| CSF1R | None | None | None | None |
| CSF1R | None | None | None | None |
| CA8 | None | None | None | None |
| CA8 | None | None | None | None |
| CA8 | 0.748 | 7 | 2w2j | A |
| APC | None | None | None | None |
| APC | None | None | None | None |
| APC | None | None | None | None |
| APC | None | None | None | None |
| APC | None | None | None | None |
| NDUFA2 | None | None | None | None |
| GIF | None | None | None | None |
| GIF | None | None | None | None |
| GIF | None | None | None | None |
| GIF | None | None | None | None |
| C2 | None | None | None | None |
| ACP5 | None | None | None | None |
| ACP5 | None | None | None | None |
| ACP5 | None | None | None | None |
| ACP5 | None | None | None | None |
| ACP5 | None | None | None | None |
| RNASEH2A | None | None | None | None |
| RNASEH2A | None | None | None | None |
| RNASEH2A | None | None | None | None |
| RNASEH2A | None | None | None | None |
| RNASEH2A | None | None | None | None |
| RNASEH2A | None | None | None | None |
| RNASEH2A | None | None | None | None |
| DSG2 | None | None | None | None |
| USH1G | None | None | None | None |
| FUS | None | None | None | None |
| FUS | None | None | None | None |
| FUS | None | None | None | None |
| FUS | None | None | None | None |
| FUS | None | None | None | None |
| MRPL44 | None | None | None | None |
| MRPL44 | None | None | None | None |
| MRPL44 | None | None | None | None |

| | | | | |
|---|---|---|---|---|
| MRPL44 | None | None | None | None |
| DMGDH | None | None | None | None |
| AARS | None | None | None | None |
| STAR | None | None | None | None |
| STAR | None | None | None | None |
| STAR | None | None | None | None |
| STAR | None | None | None | None |
| STAR | None | None | None | None |
| STAR | None | None | None | None |
| STAR | None | None | None | None |
| STAR | None | None | None | None |
| LARP7 | None | None | None | None |
| PRPF8 | None | None | None | None |
| PRPF8 | None | None | None | None |
| PRPF8 | None | None | None | None |
| PRPF8 | None | None | None | None |
| PSAP | None | None | None | None |
| PSAP | None | None | None | None |
| PSAP | None | None | None | None |
| DMPK | None | None | None | None |
| EPCAM | None | None | None | None |
| EPCAM | None | None | None | None |
| EPCAM | None | None | None | None |
| EPCAM | None | None | None | None |
| EPCAM | None | None | None | None |
| EPCAM | None | None | None | None |
| EPCAM | None | None | None | None |
| EPCAM | None | None | None | None |
| ASS1 | None | None | None | None |
| ASS1 | None | None | None | None |
| ASS1 | None | None | None | None |
| ASS1 | None | None | None | None |
| ASS1 | None | None | None | None |
| ASS1 | 0.857 | 20 | 2nz2 | A |
| ASS1 | 0.857 | 9 | 2nz2 | A |
| ASS1 | None | None | None | None |
| ASS1 | None | None | None | None |
| ASS1 | None | None | None | None |
| ASS1 | 0.857 | 9 | 2nz2 | A |
| ASS1 | 0.857 | 9 | 2nz2 | A |
| ASS1 | 0.857 | 9 | 2nz2 | A |
| ASS1 | None | None | None | None |
| ASS1 | None | None | None | None |
| ASS1 | None | None | None | None |
| ASS1 | None | None | None | None |
| ASS1 | None | None | None | None |
| ASS1 | None | None | None | None |
| ASS1 | None | None | None | None |
| POLE | None | None | None | None |
| POLE | None | None | None | None |
| POLE | None | None | None | None |
| RNASET2 | None | None | None | None |
| RNASET2 | None | None | None | None |
| RNASET2 | None | None | None | None |

| | | | | |
|---|---|---|---|---|
| FLG | None | None | None | None |
| F10 | None | None | None | None |
| AGT | None | None | None | None |
| AGT | None | None | None | None |
| PLN | None | None | None | None |
| PLN | None | None | None | None |
| PLN | None | None | None | None |
| PTHLH | None | None | None | None |
| IGF1R | None | None | None | None |
| IGF1R | None | None | None | None |
| IGF1R | None | None | None | None |
| JAG1 | None | None | None | None |
| JAG1 | None | None | None | None |
| JAG1 | None | None | None | None |
| JAG1 | None | None | None | None |
| JAG1 | None | None | None | None |
| JAG1 | None | None | None | None |
| AP1S3 | None | None | None | None |
| SPTA1 | None | None | None | None |
| SPTA1 | None | None | None | None |
| SPTA1 | None | None | None | None |
| SPTA1 | None | None | None | None |
| SPG7 | None | None | None | None |
| SPG7 | None | None | None | None |
| SPG7 | None | None | None | None |
| SPG7 | None | None | None | None |
| SPG7 | None | None | None | None |
| SPG7 | None | None | None | None |
| FN1 | None | None | None | None |
| FN1 | None | None | None | None |
| NSMCE3 | None | None | None | None |
| PCCA | None | None | None | None |
| PCCA | None | None | None | None |
| SPTB | None | None | None | None |
| RYR2 | None | None | None | None |
| RYR2 | None | None | None | None |
| MMP9 | None | None | None | None |
| MMP9 | None | None | None | None |
| GLMN | None | None | None | None |
| GLMN | None | None | None | None |
| GLMN | None | None | None | None |
| GLMN | None | None | None | None |
| GLMN | None | None | None | None |
| GLMN | None | None | None | None |
| GLMN | None | None | None | None |
| GLMN | None | None | None | None |
| GLMN | None | None | None | None |
| GLMN | None | None | None | None |
| MARS | None | None | None | None |
| MARS | None | None | None | None |
| MARS | None | None | None | None |
| MARS | None | None | None | None |
| MARS | None | None | None | None |
| MARS | None | None | None | None |

| | | | | |
|---|---|---|---|---|
| SNAP29 | None | None | None | None |
| SNAP29 | None | None | None | None |
| ETHE1 | None | None | None | None |
| ETHE1 | None | None | None | None |
| ETHE1 | None | None | None | None |
| ETHE1 | None | None | None | None |
| ETHE1 | None | None | None | None |
| ETHE1 | None | None | None | None |
| ETHE1 | None | None | None | None |
| SAMHD1 | None | None | None | None |
| KCNQ4 | None | None | None | None |
| FXYD2 | None | None | None | None |
| FXYD2 | None | None | None | None |
| FXYD2 | None | None | None | None |
| LRRK2 | None | None | None | None |
| LRRK2 | None | None | None | None |
| NDUFS1 | None | None | None | None |
| NDUFS1 | None | None | None | None |
| NDUFS1 | None | None | None | None |
| NDUFS1 | None | None | None | None |
| NDUFS1 | None | None | None | None |
| NDUFS1 | None | None | None | None |
| NDUFS1 | None | None | None | None |
| NDUFS1 | None | None | None | None |
| NDUFS1 | None | None | None | None |
| NDUFS1 | None | None | None | None |
| NDUFS1 | None | None | None | None |
| EZH2 | None | None | None | None |
| EZH2 | None | None | None | None |
| LARS | None | None | None | None |
| TLL1 | None | None | None | None |
| CRYBB2 | None | None | None | None |
| CRYBB2 | None | None | None | None |
| EXOSC8 | None | None | None | None |
| SRP72 | None | None | None | None |
| FOXC2 | None | None | None | None |
| HMBS | None | None | None | None |
| HMBS | None | None | None | None |
| HMBS | None | None | None | None |
| HMBS | None | None | None | None |
| HMBS | None | None | None | None |
| HMBS | None | None | None | None |
| HMBS | None | None | None | None |
| HMBS | None | None | None | None |
| HMBS | None | None | None | None |
| HMBS | None | None | None | None |
| HMBS | None | None | None | None |
| HMBS | None | None | None | None |
| HMBS | None | None | None | None |
| HMBS | None | None | None | None |
| HMBS | None | None | None | None |
| HMBS | None | None | None | None |
| HMBS | None | None | None | None |
| HMBS | None | None | None | None |

| | | | | |
|---|---|---|---|---|
| HMBS | None | None | None | None |
| HMBS | None | None | None | None |
| HMBS | None | None | None | None |
| HMBS | None | None | None | None |
| HMBS | None | None | None | None |
| HMBS | None | None | None | None |
| HMBS | None | None | None | None |
| OGT | None | None | None | None |
| NOG | None | None | None | None |
| NOG | None | None | None | None |
| NOG | None | None | None | None |
| NOG | None | None | None | None |
| NOG | None | None | None | None |
| NOG | None | None | None | None |
| NOG | None | None | None | None |
| NOG | None | None | None | None |
| NOG | None | None | None | None |
| NOG | None | None | None | None |
| NOG | None | None | None | None |
| DLG3 | None | None | None | None |
| SCN2A | None | None | None | None |
| STAC3 | None | None | None | None |
| PFKM | None | None | None | None |
| PFKM | None | None | None | None |
| PFKM | None | None | None | None |
| PFKM | None | None | None | None |
| PFKM | None | None | None | None |
| DNMT1 | None | None | None | None |
| DNMT1 | None | None | None | None |
| TNNI3 | None | None | None | None |
| TNNI3 | None | None | None | None |
| TNNI3 | None | None | None | None |
| STXBP2 | None | None | None | None |
| STXBP2 | None | None | None | None |
| STXBP2 | None | None | None | None |
| STXBP2 | None | None | None | None |
| STXBP2 | None | None | None | None |
| STXBP2 | None | None | None | None |
| STXBP2 | None | None | None | None |
| STXBP2 | None | None | None | None |
| REN | None | None | None | None |
| DSP | None | None | None | None |
| DSP | None | None | None | None |
| HNF1B | None | None | None | None |
| ADA | None | None | None | None |
| ADA | None | None | None | None |
| ADA | 0.615 | 7 | 3iar | A |
| ADA | None | None | None | None |
| ADA | None | None | None | None |
| ADA | None | None | None | None |
| ADA | None | None | None | None |
| ADA | None | None | None | None |
| ADA | None | None | None | None |
| ADA | 0.615 | 1 | 3iar | A |

| | | | | |
|---|---|---|---|---|
| ADA | None | None | None | None |
| ADA | 0.615 | | 1 3iar | A |
| ADA | None | None | None | None |
| ADA | None | None | None | None |
| ADA | None | None | None | None |
| DCX | None | None | None | None |
| DCX | None | None | None | None |
| DCX | None | None | None | None |
| DCX | None | None | None | None |
| DCX | None | None | None | None |
| ARSB | None | None | None | None |
| ARSB | None | None | None | None |
| ARSB | None | None | None | None |
| ARSB | None | None | None | None |
| ARSB | None | None | None | None |
| ARSB | None | None | None | None |
| ARSB | None | None | None | None |
| ARSB | None | None | None | None |
| ARSB | None | None | None | None |
| ARSB | None | None | None | None |
| ARSB | None | None | None | None |
| ARSB | None | None | None | None |
| ARSB | None | None | None | None |
| WWOX | None | None | None | None |
| TNNI3 | None | None | None | None |
| TNNI3 | None | None | None | None |
| TNNI3 | None | None | None | None |
| TNNI3 | None | None | None | None |
| CYBA | None | None | None | None |
| SPAST | None | None | None | None |
| SPAST | None | None | None | None |
| SPAST | None | None | None | None |
| SPAST | None | None | None | None |
| SPAST | None | None | None | None |
| SPAST | None | None | None | None |
| SPAST | None | None | None | None |
| SPAST | None | None | None | None |
| KRIT1 | None | None | None | None |
| KRIT1 | None | None | None | None |
| TP53 | None | None | None | None |
| ISPD | None | None | None | None |
| ISPD | None | None | None | None |
| ISPD | None | None | None | None |
| ISPD | None | None | None | None |
| KCNMA1 | None | None | None | None |
| STK11 | None | None | None | None |
| STK11 | None | None | None | None |
| STK11 | None | None | None | None |
| STK11 | None | None | None | None |
| STK11 | None | None | None | None |
| STK11 | None | None | None | None |
| NDUFA13 | None | None | None | None |
| PHF6 | None | None | None | None |
| PHF6 | None | None | None | None |

| | | | | |
|---|---|---|---|---|
| PHF6 | None | None | None | None |
| PHYH | None | None | None | None |
| PHYH | None | None | None | None |
| PHYH | None | None | None | None |
| PHYH | None | None | None | None |
| PHYH | None | None | None | None |
| NDUFS3 | None | None | None | None |
| NDUFS3 | None | None | None | None |
| ALDH18A1 | None | None | None | None |
| ALDH18A1 | None | None | None | None |
| ALDH18A1 | None | None | None | None |
| ALDH18A1 | None | None | None | None |
| ALDH18A1 | None | None | None | None |
| PAX6 | None | None | None | None |
| PAX6 | None | None | None | None |
| INPP5E | None | None | None | None |
| INPP5E | None | None | None | None |
| INPP5E | None | None | None | None |
| INPP5E | None | None | None | None |
| INPP5E | None | None | None | None |
| INPP5E | None | None | None | None |
| INPP5E | None | None | None | None |
| INPP5E | None | None | None | None |
| INPP5E | None | None | None | None |
| TNNT2 | None | None | None | None |
| TNNT2 | None | None | None | None |
| TNNT2 | None | None | None | None |
| TNNT2 | None | None | None | None |
| TNNT2 | None | None | None | None |
| TNNT2 | None | None | None | None |
| TNNT2 | None | None | None | None |
| TNNT2 | None | None | None | None |
| TYK2 | None | None | None | None |
| TYK2 | None | None | None | None |
| TYK2 | None | None | None | None |
| CFTR | None | None | None | None |
| CFTR | None | None | None | None |
| CFTR | None | None | None | None |
| CFTR | None | None | None | None |
| CFTR | None | None | None | None |
| CFTR | None | None | None | None |
| CFTR | None | None | None | None |
| CFTR | None | None | None | None |
| CFTR | None | None | None | None |
| CFTR | None | None | None | None |
| CFTR | None | None | None | None |
| CFTR | None | None | None | None |
| CFTR | None | None | None | None |
| CFTR | None | None | None | None |
| CFTR | None | None | None | None |
| CFTR | None | None | None | None |
| CFTR | None | None | None | None |
| CFTR | None | None | None | None |
| CFTR | None | None | None | None |

| | | | | |
|---|---|---|---|---|
| CFTR | None | None | None | None |
| CFTR | None | None | None | None |
| CFTR | None | None | None | None |
| CFTR | None | None | None | None |
| CFTR | None | None | None | None |
| CFTR | None | None | None | None |
| CFTR | None | None | None | None |
| CFTR | None | None | None | None |
| CFTR | None | None | None | None |
| CFTR | None | None | None | None |
| CFTR | None | None | None | None |
| CFTR | None | None | None | None |
| CFTR | None | None | None | None |
| CFTR | None | None | None | None |
| CFTR | None | None | None | None |
| CFTR | None | None | None | None |
| CFTR | None | None | None | None |
| CFTR | None | None | None | None |
| CFTR | None | None | None | None |
| CFTR | None | None | None | None |
| CFTR | None | None | None | None |
| CFTR | None | None | None | None |
| CFTR | None | None | None | None |
| CFTR | None | None | None | None |
| CFTR | None | None | None | None |
| CFTR | None | None | None | None |
| CFTR | None | None | None | None |
| CFTR | None | None | None | None |
| CFTR | None | None | None | None |
| CFTR | None | None | None | None |
| CFTR | None | None | None | None |
| CFTR | None | None | None | None |
| CFTR | None | None | None | None |
| CFTR | None | None | None | None |
| SERPINF1 | None | None | None | None |
| SERPINF1 | None | None | None | None |
| IDS | None | None | None | None |
| IDS | None | None | None | None |
| IDS | None | None | None | None |
| IDS | None | None | None | None |
| IDS | None | None | None | None |
| IDS | None | None | None | None |
| IDS | None | None | None | None |
| IDS | None | None | None | None |
| IDS | None | None | None | None |
| NFKBIA | 0.7 | 1 | 1nfi | F |
| NFKBIA | None | None | None | None |
| SPAST | None | None | None | None |
| FGF23 | None | None | None | None |
| FGF23 | None | None | None | None |
| FGF23 | None | None | None | None |
| FGF23 | None | None | None | None |
| FGF23 | None | None | None | None |
| FGF23 | None | None | None | None |

| | | | | |
|---|---|---|---|---|
| FGF23 | None | None | None | None |
| DLAT | None | None | None | None |
| STUB1 | None | None | None | None |
| HSD17B4 | None | None | None | None |
| HSD17B4 | None | None | None | None |
| DYSF | None | None | None | None |
| CEP135 | None | None | None | None |
| CEP135 | None | None | None | None |
| PRKCG | None | None | None | None |
| CSF1R | None | None | None | None |
| PEX5 | None | None | None | None |
| PAX6 | None | None | None | None |
| PAX6 | None | None | None | None |
| PAX6 | None | None | None | None |
| PAX6 | None | None | None | None |
| PAX6 | None | None | None | None |
| PAX6 | None | None | None | None |
| NF2 | None | None | None | None |
| NF2 | None | None | None | None |
| NF2 | None | None | None | None |
| REM1 | None | None | None | None |

| source_id | classification | binding_site | binding_site_single_water | protein_binding_site |
|---|---|---|---|---|
| rs121918343 | pathogenic | 1 | 1 | 0 |
| rs267606649 | pathogenic | 1 | 1 | 1 |
| rs121918342 | pathogenic | 0 | 1 | 0 |
| rs267606650 | pathogenic | 1 | 1 | 1 |
| rs1130335 | benign | 1 | 1 | 1 |
| rs199476087 | pathogenic | 1 | 1 | 1 |
| rs772163112 | benign | 0 | 0 | 0 |
| rs755793 | benign | 0 | 0 | 0 |
| rs121918155 | pathogenic | 1 | 1 | 0 |
| rs121918156 | pathogenic | 1 | 1 | 0 |
| rs121918154 | pathogenic | 1 | 1 | 1 |
| rs121918151 | pathogenic | 1 | 1 | 1 |
| rs121918144 | pathogenic | 0 | 1 | 0 |
| rs121918146 | pathogenic | 1 | 1 | 1 |
| rs121918160 | pathogenic | 1 | 1 | 1 |
| rs121918150 | pathogenic | 1 | 1 | 1 |
| rs121918141 | pathogenic | 1 | 1 | 1 |
| rs142742242 | pathogenic | 1 | 1 | 1 |
| rs121918142 | pathogenic | 1 | 1 | 1 |
| rs121918157 | pathogenic | 1 | 1 | 1 |
| rs121918101 | pathogenic | 1 | 1 | 1 |
| rs121908260 | pathogenic | 1 | 1 | 1 |
| rs80356667 | pathogenic | 1 | 1 | 1 |
| rs80356668 | pathogenic | 1 | 1 | 1 |
| rs397514540 | pathogenic | 1 | 1 | 1 |
| rs104894685 | pathogenic | 0 | 1 | 0 |
| rs199869995 | pathogenic | 1 | 1 | 1 |
| rs121909072 | pathogenic | 0 | 0 | 0 |
| rs143669310 | benign | 1 | 1 | 1 |
| rs747347498 | benign | 0 | 0 | 0 |
| rs138283222 | benign | 0 | 0 | 0 |
| rs118203931 | pathogenic | 1 | 1 | 1 |
| rs118203933 | pathogenic | 1 | 1 | 0 |
| rs118203932 | pathogenic | 1 | 1 | 0 |
| rs2228063 | benign | 1 | 1 | 1 |
| rs377767360 | pathogenic | 0 | 0 | 0 |
| rs397518413 | pathogenic | 1 | 1 | 1 |
| rs1800458 | benign | 1 | 1 | 0 |
| rs121918083 | pathogenic | 1 | 1 | 1 |
| rs121918094 | pathogenic | 1 | 1 | 1 |
| rs121918098 | pathogenic | 1 | 1 | 1 |
| rs121918093 | pathogenic | 1 | 1 | 1 |
| rs79977247 | pathogenic | 1 | 1 | 1 |
| rs28933979 | pathogenic | 1 | 1 | 1 |
| rs121918068 | pathogenic | 1 | 1 | 1 |
| rs121918077 | pathogenic | 1 | 1 | 1 |
| rs11541796 | pathogenic | 1 | 1 | 1 |
| rs104894665 | pathogenic | 1 | 1 | 1 |
| rs121918090 | pathogenic | 1 | 1 | 1 |
| rs121918081 | pathogenic | 1 | 1 | 1 |
| rs121918080 | pathogenic | 1 | 1 | 1 |
| rs121918076 | pathogenic | 1 | 1 | 1 |
| rs121918097 | pathogenic | 1 | 1 | 1 |

| | | | | |
|---|---|---|---|---|
| rs121918079 | pathogenic | 1 | 1 | 1 |
| rs121918069 | pathogenic | 1 | 1 | 1 |
| rs121918086 | pathogenic | 1 | 1 | 1 |
| rs121918091 | pathogenic | 1 | 1 | 1 |
| rs121918085 | pathogenic | 1 | 1 | 1 |
| rs121918100 | pathogenic | 1 | 1 | 1 |
| rs121918084 | pathogenic | 1 | 1 | 1 |
| rs121918071 | pathogenic | 1 | 1 | 1 |
| rs121918072 | pathogenic | 1 | 1 | 1 |
| rs121918082 | pathogenic | 1 | 1 | 1 |
| rs121918087 | pathogenic | 1 | 1 | 1 |
| rs76410435 | benign | 1 | 1 | 1 |
| rs121918089 | pathogenic | 1 | 1 | 1 |
| rs121918073 | pathogenic | 1 | 1 | 1 |
| rs121918075 | pathogenic | 1 | 1 | 1 |
| rs121918088 | pathogenic | 1 | 1 | 1 |
| rs76992529 | pathogenic | 1 | 1 | 1 |
| rs536294863 | benign | 1 | 1 | 1 |
| rs730881162 | benign | 1 | 1 | 1 |
| rs113994087 | pathogenic | 1 | 1 | 1 |
| rs74716434 | benign | 1 | 1 | 1 |
| rs121918694 | pathogenic | 1 | 1 | 1 |
| rs121918707 | pathogenic | 1 | 1 | 1 |
| rs121918695 | pathogenic | 1 | 1 | 0 |
| rs121918690 | pathogenic | 1 | 1 | 0 |
| rs121918696 | pathogenic | 1 | 1 | 0 |
| rs121918693 | pathogenic | 1 | 1 | 0 |
| rs28999969 | pathogenic | 1 | 1 | 1 |
| rs121918697 | pathogenic | 1 | 1 | 1 |
| rs121918688 | pathogenic | 1 | 1 | 0 |
| rs28999970 | pathogenic | 1 | 1 | 0 |
| rs121918686 | pathogenic | 1 | 1 | 0 |
| rs28999971 | pathogenic | 1 | 1 | 1 |
| rs121918698 | pathogenic | 1 | 1 | 1 |
| rs121918691 | pathogenic | 1 | 1 | 0 |
| rs121918692 | pathogenic | 1 | 1 | 1 |
| rs121918703 | pathogenic | 1 | 1 | 1 |
| rs121918687 | pathogenic | 1 | 1 | 1 |
| rs28933408 | pathogenic | 1 | 1 | 1 |
| rs121918702 | pathogenic | 1 | 1 | 1 |
| rs17875899 | benign | 1 | 1 | 1 |
| rs111033620 | pathogenic | 1 | 1 | 1 |
| rs111033622 | pathogenic | 1 | 1 | 1 |
| rs111033621 | pathogenic | 1 | 1 | 1 |
| rs111033618 | pathogenic | 1 | 1 | 1 |
| rs5918 | benign | 0 | 0 | 0 |
| rs121918452 | pathogenic | 0 | 0 | 0 |
| rs121918445 | pathogenic | 1 | 1 | 1 |
| rs5917 | benign | 1 | 1 | 1 |
| rs121918444 | pathogenic | 1 | 1 | 1 |
| rs121918446 | pathogenic | 1 | 1 | 1 |
| rs121918449 | pathogenic | 0 | 0 | 0 |
| rs121918448 | benign | 0 | 0 | 0 |
| rs13306487 | benign | 1 | 1 | 1 |

| | | | | |
|---|---|---|---|---|
| rs1805044 | benign | 0 | 0 | 0 |
| rs117078377 | benign | 0 | 0 | 0 |
| rs138132026 | benign | 1 | 1 | 1 |
| rs75762896 | benign | 1 | 1 | 0 |
| rs121918043 | pathogenic | 1 | 1 | 0 |
| rs143581690 | benign | 1 | 1 | 0 |
| rs147654263 | benign | 0 | 0 | 0 |
| rs587782529 | pathogenic | 1 | 1 | 1 |
| rs121912664 | pathogenic | 1 | 1 | 1 |
| rs150293825 | benign | 1 | 1 | 1 |
| rs17882252 | benign | 1 | 1 | 1 |
| rs121912662 | pathogenic | 1 | 1 | 1 |
| rs35993958 | benign | 1 | 1 | 1 |
| rs606231253 | pathogenic | 1 | 1 | 1 |
| rs2227983 | benign | 1 | 1 | 1 |
| rs587779351 | pathogenic | 0 | 0 | 0 |
| rs779773463 | pathogenic | 1 | 1 | 1 |
| rs713040 | benign | 1 | 1 | 1 |
| rs334 | pathogenic | 1 | 1 | 1 |
| rs33930165 | pathogenic | 1 | 1 | 1 |
| rs33918131 | benign | 1 | 1 | 1 |
| rs63750783 | pathogenic | 1 | 1 | 1 |
| rs33986703 | pathogenic | 1 | 1 | 1 |
| rs33972047 | pathogenic | 1 | 1 | 1 |
| rs33936254 | pathogenic | 1 | 1 | 1 |
| rs33959855 | benign | 1 | 1 | 1 |
| rs33950507 | pathogenic | 1 | 1 | 1 |
| rs33916412 | pathogenic | 1 | 1 | 0 |
| rs1135071 | pathogenic | 1 | 1 | 1 |
| rs33960103 | pathogenic | 1 | 1 | 1 |
| rs11549407 | pathogenic | 1 | 1 | 1 |
| rs33922842 | pathogenic | 1 | 1 | 1 |
| rs35278874 | benign | 1 | 1 | 1 |
| rs34621955 | benign | 1 | 1 | 1 |
| rs33995148 | pathogenic | 1 | 1 | 1 |
| rs33945705 | benign | 1 | 1 | 1 |
| rs33993568 | benign | 1 | 1 | 1 |
| rs35553496 | benign | 1 | 1 | 1 |
| rs33913712 | pathogenic | 1 | 1 | 1 |
| rs33917785 | pathogenic | 1 | 1 | 1 |
| rs34515413 | benign | 1 | 1 | 1 |
| rs35256489 | pathogenic | 1 | 1 | 0 |
| rs34484056 | pathogenic | 1 | 1 | 1 |
| rs33947020 | benign | 1 | 1 | 1 |
| rs33946267 | pathogenic | 1 | 1 | 1 |
| rs33966761 | pathogenic | 1 | 1 | 1 |
| rs36020563 | benign | 1 | 1 | 1 |
| rs122454124 | pathogenic | 1 | 1 | 1 |
| rs122454126 | pathogenic | 1 | 1 | 0 |
| rs122454127 | pathogenic | 1 | 1 | 1 |
| rs122454130 | pathogenic | 1 | 1 | 1 |
| rs122454125 | pathogenic | 1 | 1 | 1 |
| rs122454131 | pathogenic | 1 | 1 | 0 |
| rs121908302 | pathogenic | 0 | 1 | 0 |

| | | | | |
|---|---|---|---|---|
| rs77829017 | pathogenic | 0 | 1 | 0 |
| rs1141814 | pathogenic | 0 | 1 | 0 |
| rs121908312 | pathogenic | 0 | 1 | 0 |
| rs79653797 | pathogenic | 1 | 1 | 0 |
| rs439898 | pathogenic | 1 | 1 | 0 |
| rs121908299 | pathogenic | 0 | 1 | 0 |
| rs80356763 | pathogenic | 1 | 1 | 1 |
| rs781152868 | pathogenic | 0 | 1 | 0 |
| rs121908297 | pathogenic | 0 | 1 | 0 |
| rs398123531 | benign | 0 | 1 | 0 |
| rs398123532 | pathogenic | 0 | 1 | 0 |
| rs364897 | pathogenic | 1 | 1 | 1 |
| rs1064644 | pathogenic | 0 | 1 | 0 |
| rs409652 | pathogenic | 1 | 1 | 1 |
| rs121908300 | pathogenic | 0 | 1 | 0 |
| rs381737 | pathogenic | 0 | 1 | 0 |
| rs74500255 | pathogenic | 0 | 1 | 0 |
| rs121908313 | pathogenic | 0 | 1 | 0 |
| rs367968666 | pathogenic | 0 | 1 | 0 |
| rs78973108 | pathogenic | 1 | 1 | 1 |
| rs121908298 | pathogenic | 1 | 1 | 1 |
| rs78396650 | pathogenic | 0 | 0 | 0 |
| rs78198234 | pathogenic | 1 | 1 | 0 |
| rs121908304 | pathogenic | 1 | 1 | 1 |
| rs76539814 | pathogenic | 1 | 1 | 1 |
| rs121908305 | pathogenic | 0 | 1 | 0 |
| rs121908306 | pathogenic | 1 | 1 | 0 |
| rs121908308 | pathogenic | 1 | 1 | 1 |
| rs121908307 | pathogenic | 0 | 1 | 0 |
| rs75548401 | benign | 0 | 1 | 0 |
| rs76763715 | pathogenic | 0 | 0 | 0 |
| rs121908314 | pathogenic | 0 | 1 | 0 |
| rs398123528 | pathogenic | 0 | 1 | 0 |
| rs121908311 | pathogenic | 0 | 0 | 0 |
| rs121908310 | pathogenic | 1 | 1 | 0 |
| rs74598136 | pathogenic | 0 | 1 | 0 |
| rs1064651 | pathogenic | 1 | 1 | 1 |
| rs77369218 | pathogenic | 1 | 1 | 1 |
| rs121908295 | pathogenic | 0 | 1 | 0 |
| rs421016 | pathogenic | 0 | 1 | 0 |
| rs368060 | pathogenic | 0 | 1 | 0 |
| rs80356771 | pathogenic | 1 | 1 | 1 |
| rs80356772 | pathogenic | 1 | 1 | 1 |
| rs121908301 | pathogenic | 0 | 0 | 0 |
| rs747506979 | pathogenic | 0 | 1 | 0 |
| rs75822236 | pathogenic | 0 | 1 | 0 |
| rs1990760 | benign | 1 | 1 | 0 |
| rs28934568 | pathogenic | 1 | 1 | 0 |
| rs34833812 | benign | 1 | 1 | 0 |
| rs193922666 | benign | 1 | 1 | 0 |
| rs104893812 | pathogenic | 1 | 1 | 0 |
| rs144701411 | benign | 0 | 0 | 0 |
| rs104893813 | pathogenic | 0 | 1 | 0 |
| rs199660234 | pathogenic | 1 | 1 | 0 |

| | | | | |
|---|---|---|---|---|
| rs104893814 | pathogenic | 1 | 1 | 0 |
| rs143095746 | benign | 1 | 1 | 1 |
| rs104893807 | pathogenic | 1 | 1 | 0 |
| rs104893811 | pathogenic | 0 | 0 | 0 |
| rs104893816 | pathogenic | 0 | 0 | 0 |
| rs121918714 | pathogenic | 0 | 1 | 0 |
| rs104893810 | pathogenic | 1 | 1 | 0 |
| rs104893815 | pathogenic | 1 | 1 | 0 |
| rs4303 | benign | 0 | 1 | 0 |
| rs121912704 | pathogenic | 0 | 1 | 0 |
| rs56394458 | benign | 1 | 1 | 0 |
| rs397514688 | pathogenic | 0 | 0 | 0 |
| rs367797185 | pathogenic | 1 | 1 | 0 |
| rs12709426 | benign | 0 | 1 | 0 |
| rs2228547 | benign | 0 | 0 | 0 |
| rs111033546 | pathogenic | 0 | 0 | 0 |
| rs111033553 | pathogenic | 0 | 0 | 0 |
| rs111033554 | pathogenic | 1 | 1 | 1 |
| rs111033544 | pathogenic | 1 | 1 | 1 |
| rs111033555 | pathogenic | 1 | 1 | 1 |
| rs111033545 | pathogenic | 1 | 1 | 1 |
| rs111033549 | pathogenic | 0 | 1 | 0 |
| rs111033552 | pathogenic | 1 | 1 | 1 |
| rs752204035 | benign | 0 | 0 | 0 |
| rs11569534 | benign | 0 | 0 | 0 |
| rs28937571 | pathogenic | 1 | 1 | 1 |
| rs104894215 | pathogenic | 1 | 1 | 1 |
| rs3888798 | benign | 1 | 1 | 1 |
| rs764263034 | benign | 1 | 1 | 1 |
| rs1800973 | benign | 1 | 1 | 1 |
| rs367623154 | benign | 1 | 1 | 1 |
| rs201230446 | pathogenic | 1 | 1 | 1 |
| rs267606766 | pathogenic | 0 | 1 | 0 |
| rs267606769 | pathogenic | 0 | 1 | 0 |
| rs267606767 | pathogenic | 1 | 1 | 0 |
| rs267606768 | pathogenic | 0 | 1 | 0 |
| rs61733129 | benign | 0 | 1 | 0 |
| rs201947120 | pathogenic | 1 | 1 | 1 |
| rs283413 | benign | 0 | 1 | 0 |
| rs36119840 | benign | 0 | 0 | 0 |
| rs281865419 | pathogenic | 1 | 1 | 1 |
| rs137852937 | pathogenic | 0 | 1 | 0 |
| rs2230709 | benign | 0 | 1 | 0 |
| rs6695033 | benign | 0 | 1 | 0 |
| rs180177032 | pathogenic | 1 | 1 | 1 |
| rs180177033 | pathogenic | 1 | 1 | 1 |
| rs121913351 | pathogenic | 1 | 1 | 1 |
| rs397507473 | pathogenic | 1 | 1 | 1 |
| rs121913355 | pathogenic | 1 | 1 | 1 |
| rs121913357 | pathogenic | 1 | 1 | 1 |
| rs180177036 | pathogenic | 1 | 1 | 1 |
| rs180177039 | pathogenic | 1 | 1 | 1 |
| rs180177038 | pathogenic | 1 | 1 | 1 |
| rs180177040 | pathogenic | 1 | 1 | 1 |

| | | | | |
|---|---|---|---|---|
| rs121913338 | pathogenic | 1 | 1 | 1 |
| rs121913366 | pathogenic | 1 | 1 | 1 |
| rs121913369 | pathogenic | 1 | 1 | 1 |
| rs113488022 | pathogenic | 1 | 1 | 1 |
| rs121913364 | pathogenic | 1 | 1 | 1 |
| rs180177042 | pathogenic | 1 | 1 | 1 |
| rs200461617 | benign | 0 | 0 | 0 |
| rs141609745 | benign | 0 | 0 | 0 |
| rs104894383 | pathogenic | 1 | 1 | 1 |
| rs104894384 | pathogenic | 1 | 1 | 1 |
| rs104894381 | pathogenic | 1 | 1 | 0 |
| rs147710408 | benign | 1 | 1 | 1 |
| rs77357563 | benign | 1 | 1 | 1 |
| rs104894378 | pathogenic | 0 | 0 | 0 |
| rs104894382 | pathogenic | 0 | 0 | 0 |
| rs137853292 | pathogenic | 1 | 1 | 1 |
| rs587778834 | pathogenic | 0 | 0 | 0 |
| rs137853294 | pathogenic | 1 | 1 | 1 |
| rs551747882 | benign | 0 | 0 | 0 |
| rs3092903 | benign | 1 | 1 | 1 |
| rs137853296 | pathogenic | 0 | 0 | 0 |
| rs398122917 | pathogenic | 0 | 1 | 0 |
| rs41304992 | benign | 0 | 0 | 0 |
| rs121912708 | pathogenic | 1 | 1 | 1 |
| rs140102105 | pathogenic | 1 | 1 | 1 |
| rs376917645 | pathogenic | 1 | 1 | 0 |
| rs372660425 | pathogenic | 1 | 1 | 0 |
| rs555896752 | pathogenic | 1 | 1 | 1 |
| rs777829351 | pathogenic | 1 | 1 | 0 |
| rs121912711 | pathogenic | 1 | 1 | 0 |
| rs754449549 | pathogenic | 0 | 1 | 0 |
| rs141701364 | benign | 0 | 1 | 0 |
| rs761011399 | benign | 1 | 1 | 1 |
| rs121912710 | benign | 1 | 1 | 0 |
| rs121912707 | pathogenic | 1 | 1 | 0 |
| rs747597620 | pathogenic | 0 | 1 | 0 |
| rs186558364 | pathogenic | 1 | 1 | 1 |
| rs556400964 | pathogenic | 1 | 1 | 1 |
| rs61757684 | benign | 1 | 1 | 1 |
| rs121917766 | pathogenic | 1 | 1 | 1 |
| rs121918248 | pathogenic | 0 | 0 | 0 |
| rs398123278 | pathogenic | 0 | 0 | 0 |
| rs121918251 | pathogenic | 1 | 1 | 1 |
| rs727504022 | pathogenic | 0 | 0 | 0 |
| rs190834116 | pathogenic | 1 | 1 | 1 |
| rs121918249 | pathogenic | 1 | 1 | 0 |
| rs121918257 | pathogenic | 1 | 1 | 0 |
| rs142068898 | benign | 0 | 0 | 0 |
| rs757000253 | pathogenic | 0 | 0 | 0 |
| rs200908035 | pathogenic | 1 | 1 | 0 |
| rs760782399 | pathogenic | 0 | 0 | 0 |
| rs778702777 | pathogenic | 0 | 0 | 0 |
| rs121918258 | pathogenic | 1 | 1 | 0 |
| rs121918256 | pathogenic | 0 | 0 | 0 |

| | | | | |
|---|---|---|---|---|
| rs200596762 | pathogenic | 1 | 1 | 0 |
| rs547709692 | pathogenic | 0 | 0 | 0 |
| rs758577372 | pathogenic | 0 | 1 | 0 |
| rs564069299 | pathogenic | 1 | 1 | 1 |
| rs772552898 | pathogenic | 1 | 1 | 1 |
| rs368790885 | pathogenic | 1 | 1 | 1 |
| rs121918250 | pathogenic | 0 | 0 | 0 |
| rs533755473 | pathogenic | 1 | 1 | 1 |
| rs753288303 | pathogenic | 1 | 1 | 1 |
| rs2229385 | benign | 0 | 0 | 0 |
| rs1141321 | benign | 0 | 0 | 0 |
| rs753564352 | pathogenic | 0 | 0 | 0 |
| rs775593146 | pathogenic | 0 | 0 | 0 |
| rs121918254 | pathogenic | 0 | 0 | 0 |
| rs8589 | benign | 1 | 1 | 0 |
| rs777758903 | pathogenic | 0 | 0 | 0 |
| rs140600746 | pathogenic | 1 | 1 | 0 |
| rs121918255 | pathogenic | 1 | 1 | 0 |
| rs121918252 | pathogenic | 0 | 0 | 0 |
| rs779990936 | pathogenic | 1 | 1 | 0 |
| rs121912455 | pathogenic | 1 | 1 | 1 |
| rs121912459 | pathogenic | 1 | 1 | 1 |
| rs74315452 | pathogenic | 1 | 1 | 1 |
| rs121912451 | pathogenic | 1 | 1 | 1 |
| rs79006549 | pathogenic | 0 | 0 | 0 |
| rs28934889 | benign | 0 | 1 | 0 |
| rs111033557 | benign | 1 | 1 | 1 |
| rs28934597 | pathogenic | 1 | 1 | 1 |
| rs28934596 | pathogenic | 1 | 1 | 1 |
| rs28934595 | pathogenic | 1 | 1 | 1 |
| rs146519482 | pathogenic | 1 | 1 | 1 |
| rs111033563 | pathogenic | 1 | 1 | 1 |
| rs75385605 | benign | 1 | 1 | 1 |
| rs121908569 | pathogenic | 1 | 1 | 1 |
| rs35719940 | benign | 0 | 0 | 0 |
| rs121918664 | pathogenic | 0 | 0 | 0 |
| rs104893665 | pathogenic | 1 | 1 | 1 |
| rs104893666 | pathogenic | 1 | 1 | 1 |
| rs200845457 | benign | 1 | 1 | 1 |
| rs779204655 | pathogenic | 1 | 1 | 1 |
| rs121917747 | pathogenic | 1 | 1 | 1 |
| rs6042 | benign | 1 | 1 | 1 |
| rs104894380 | pathogenic | 0 | 0 | 0 |
| rs369083541 | pathogenic | 1 | 1 | 1 |
| rs145243532 | benign | 1 | 1 | 1 |
| rs121909500 | pathogenic | 1 | 1 | 1 |
| rs140993310 | benign | 1 | 1 | 1 |
| rs200837270 | pathogenic | 1 | 1 | 1 |
| rs121918484 | pathogenic | 1 | 1 | 1 |
| rs121918306 | pathogenic | 0 | 0 | 0 |
| rs118204089 | pathogenic | 1 | 1 | 0 |
| rs118204083 | pathogenic | 1 | 1 | 0 |
| rs201953584 | pathogenic | 1 | 1 | 1 |
| rs74315362 | pathogenic | 1 | 1 | 1 |

| | | | | |
|---|---|---|---|---|
| rs118204084 | pathogenic | 1 | 1 | 0 |
| rs118204085 | pathogenic | 1 | 1 | 1 |
| rs113403872 | pathogenic | 1 | 1 | 0 |
| rs1052176 | benign | 1 | 1 | 1 |
| rs2031640 | benign | 0 | 0 | 0 |
| rs104894230 | pathogenic | 1 | 1 | 1 |
| rs104894229 | pathogenic | 1 | 1 | 1 |
| rs104894228 | pathogenic | 1 | 1 | 1 |
| rs104894226 | pathogenic | 1 | 1 | 1 |
| rs121917757 | pathogenic | 1 | 1 | 1 |
| rs121917758 | pathogenic | 1 | 1 | 1 |
| rs28933406 | pathogenic | 1 | 1 | 1 |
| rs121917756 | pathogenic | 1 | 1 | 1 |
| rs104894227 | pathogenic | 1 | 1 | 1 |
| rs369106578 | pathogenic | 1 | 1 | 1 |
| rs397517142 | benign | 1 | 1 | 1 |
| rs104894231 | pathogenic | 1 | 1 | 1 |
| rs121917759 | pathogenic | 1 | 1 | 1 |
| rs104893827 | pathogenic | 0 | 0 | 0 |
| rs104893826 | pathogenic | 1 | 1 | 1 |
| rs5030802 | pathogenic | 1 | 1 | 1 |
| rs5030827 | pathogenic | 1 | 1 | 0 |
| rs119103277 | pathogenic | 1 | 1 | 1 |
| rs5030808 | pathogenic | 0 | 1 | 0 |
| rs5030809 | pathogenic | 1 | 1 | 1 |
| rs765978945 | pathogenic | 1 | 1 | 1 |
| rs104893824 | pathogenic | 1 | 1 | 1 |
| rs104893831 | pathogenic | 0 | 1 | 0 |
| rs104893830 | pathogenic | 0 | 1 | 0 |
| rs119103278 | pathogenic | 0 | 1 | 0 |
| rs5030818 | pathogenic | 1 | 1 | 1 |
| rs397516444 | pathogenic | 1 | 1 | 1 |
| rs28940297 | pathogenic | 1 | 1 | 1 |
| rs267607170 | pathogenic | 1 | 1 | 1 |
| rs104893825 | pathogenic | 1 | 1 | 1 |
| rs5030820 | pathogenic | 0 | 1 | 0 |
| rs5030821 | pathogenic | 0 | 1 | 0 |
| rs5030823 | pathogenic | 1 | 1 | 1 |
| rs5030824 | pathogenic | 1 | 1 | 1 |
| rs28940301 | pathogenic | 0 | 1 | 0 |
| rs28940300 | pathogenic | 0 | 1 | 0 |
| rs28940298 | pathogenic | 0 | 1 | 0 |
| rs200019083 | benign | 0 | 0 | 0 |
| rs759748655 | pathogenic | 0 | 0 | 0 |
| rs121917739 | pathogenic | 1 | 1 | 1 |
| rs75353611 | pathogenic | 1 | 1 | 1 |
| rs79744198 | benign | 1 | 1 | 1 |
| rs79228041 | pathogenic | 1 | 1 | 1 |
| rs80106970 | benign | 1 | 1 | 1 |
| rs375353223 | benign | 0 | 0 | 0 |
| rs56311081 | benign | 0 | 0 | 0 |
| rs55985569 | benign | 1 | 1 | 1 |
| rs45483396 | benign | 0 | 0 | 0 |
| rs35284565 | benign | 1 | 1 | 1 |

| | | | | |
|---|---|---|---|---|
| rs34349517 | benign | 0 | 0 | 0 |
| rs35776110 | benign | 0 | 0 | 0 |
| rs33917957 | benign | 1 | 1 | 1 |
| rs587778445 | benign | 0 | 0 | 0 |
| rs34624984 | benign | 0 | 0 | 0 |
| rs701753 | benign | 0 | 0 | 0 |
| rs855314 | benign | 0 | 0 | 0 |
| rs121918371 | pathogenic | 1 | 1 | 0 |
| rs398122912 | pathogenic | 0 | 1 | 0 |
| rs1126728 | benign | 0 | 0 | 0 |
| rs587777404 | pathogenic | 1 | 1 | 0 |
| rs11208257 | benign | 0 | 0 | 0 |
| rs6676290 | benign | 0 | 0 | 0 |
| rs104894118 | pathogenic | 0 | 0 | 0 |
| rs121918655 | pathogenic | 1 | 1 | 1 |
| rs104894120 | pathogenic | 1 | 1 | 1 |
| rs137852238 | pathogenic | 0 | 0 | 0 |
| rs137852237 | pathogenic | 0 | 0 | 0 |
| rs137852499 | pathogenic | 1 | 1 | 1 |
| rs137852480 | pathogenic | 1 | 1 | 1 |
| rs137852494 | pathogenic | 1 | 1 | 1 |
| rs137852502 | pathogenic | 1 | 1 | 1 |
| rs137852500 | pathogenic | 1 | 1 | 1 |
| rs137852487 | pathogenic | 1 | 1 | 1 |
| rs137852481 | pathogenic | 1 | 1 | 1 |
| rs137852501 | pathogenic | 0 | 0 | 0 |
| rs137852478 | pathogenic | 1 | 1 | 1 |
| rs137852485 | pathogenic | 1 | 1 | 1 |
| rs137852482 | pathogenic | 1 | 1 | 0 |
| rs137852483 | pathogenic | 0 | 1 | 0 |
| rs137852477 | pathogenic | 1 | 1 | 0 |
| rs137852496 | pathogenic | 1 | 1 | 0 |
| rs137852484 | pathogenic | 0 | 1 | 0 |
| rs137852498 | pathogenic | 1 | 1 | 0 |
| rs137852497 | pathogenic | 1 | 1 | 0 |
| rs137852493 | pathogenic | 1 | 1 | 0 |
| rs137852504 | pathogenic | 1 | 1 | 1 |
| rs267606863 | pathogenic | 1 | 1 | 1 |
| rs137852486 | pathogenic | 1 | 1 | 1 |
| rs137852479 | pathogenic | 1 | 1 | 1 |
| rs112122950 | benign | 0 | 0 | 0 |
| rs72546648 | benign | 1 | 1 | 1 |
| rs72552291 | pathogenic | 0 | 0 | 0 |
| rs79184941 | pathogenic | 1 | 1 | 1 |
| rs77543610 | pathogenic | 1 | 1 | 1 |
| rs121918505 | pathogenic | 1 | 1 | 1 |
| rs776587763 | pathogenic | 1 | 1 | 1 |
| rs121918497 | pathogenic | 1 | 1 | 1 |
| rs121918499 | pathogenic | 1 | 1 | 1 |
| rs121918501 | pathogenic | 1 | 1 | 1 |
| rs121918500 | pathogenic | 0 | 0 | 0 |
| rs121918504 | pathogenic | 1 | 1 | 1 |
| rs121918510 | pathogenic | 1 | 1 | 1 |
| rs121918493 | pathogenic | 1 | 1 | 1 |

| | | | | |
|---|---|---|---|---|
| rs121918489 | pathogenic | 0 | 0 | 0 |
| rs121918495 | pathogenic | 1 | 1 | 1 |
| rs121918488 | pathogenic | 0 | 0 | 0 |
| rs121918496 | pathogenic | 0 | 0 | 0 |
| rs121918492 | pathogenic | 1 | 1 | 1 |
| rs121918494 | pathogenic | 1 | 1 | 1 |
| rs121918502 | pathogenic | 1 | 1 | 1 |
| rs121918490 | pathogenic | 1 | 1 | 1 |
| rs72466451 | pathogenic | 1 | 1 | 1 |
| rs66468541 | pathogenic | 0 | 0 | 0 |
| rs137852662 | pathogenic | 0 | 0 | 0 |
| rs137852663 | pathogenic | 0 | 0 | 0 |
| rs606231407 | pathogenic | 0 | 0 | 0 |
| rs137852233 | pathogenic | 1 | 1 | 1 |
| rs112306225 | benign | 0 | 0 | 0 |
| rs387906808 | pathogenic | 1 | 1 | 0 |
| rs730882094 | pathogenic | 1 | 1 | 1 |
| rs373869746 | pathogenic | 1 | 1 | 1 |
| rs777524402 | pathogenic | 1 | 1 | 1 |
| rs769737896 | pathogenic | 0 | 1 | 0 |
| rs13306515 | benign | 1 | 1 | 0 |
| rs746982741 | pathogenic | 0 | 0 | 0 |
| rs2230200 | benign | 1 | 1 | 1 |
| rs1047286 | benign | 0 | 0 | 0 |
| rs141737564 | benign | 0 | 0 | 0 |
| rs774826179 | benign | 0 | 0 | 0 |
| rs11569422 | benign | 1 | 1 | 1 |
| rs146613648 | benign | 0 | 0 | 0 |
| rs144432231 | benign | 0 | 0 | 0 |
| rs140655115 | benign | 0 | 0 | 0 |
| rs387906837 | pathogenic | 1 | 1 | 0 |
| rs121909076 | pathogenic | 1 | 1 | 0 |
| rs121909077 | pathogenic | 0 | 0 | 0 |
| rs121909074 | pathogenic | 0 | 0 | 0 |
| rs137853330 | pathogenic | 1 | 1 | 1 |
| rs137853331 | pathogenic | 1 | 1 | 1 |
| rs137853332 | pathogenic | 1 | 1 | 1 |
| rs28931586 | pathogenic | 1 | 1 | 0 |
| rs11569321 | benign | 1 | 1 | 1 |
| rs119486097 | benign | 0 | 0 | 0 |
| rs1799807 | pathogenic | 1 | 1 | 1 |
| rs201820739 | pathogenic | 1 | 1 | 0 |
| rs114706984 | pathogenic | 1 | 1 | 0 |
| rs140080572 | pathogenic | 1 | 1 | 0 |
| rs28933389 | pathogenic | 0 | 1 | 0 |
| rs16849700 | benign | 1 | 1 | 0 |
| rs115624085 | pathogenic | 1 | 1 | 1 |
| rs747196387 | pathogenic | 1 | 1 | 1 |
| rs104893684 | pathogenic | 1 | 1 | 0 |
| rs121918557 | pathogenic | 1 | 1 | 0 |
| rs121918556 | pathogenic | 1 | 1 | 1 |
| rs398124571 | pathogenic | 1 | 1 | 1 |
| rs776631396 | pathogenic | 0 | 1 | 0 |
| rs398124576 | pathogenic | 1 | 1 | 1 |

| | | | | |
|---|---|---|---|---|
| rs398124581 | pathogenic | 1 | 1 | 1 |
| rs398124582 | pathogenic | 1 | 1 | 1 |
| rs149766077 | pathogenic | 0 | 1 | 0 |
| rs79761867 | pathogenic | 0 | 1 | 0 |
| rs148905512 | pathogenic | 0 | 1 | 0 |
| rs121965004 | pathogenic | 1 | 1 | 1 |
| rs386834233 | pathogenic | 0 | 1 | 0 |
| rs398124598 | pathogenic | 0 | 1 | 0 |
| rs398124603 | pathogenic | 1 | 1 | 1 |
| rs546716073 | benign | 1 | 1 | 1 |
| rs200310662 | benign | 1 | 1 | 1 |
| rs115494727 | benign | 1 | 1 | 1 |
| rs781513008 | benign | 0 | 0 | 0 |
| rs587776398 | benign | 1 | 1 | 0 |
| rs35288908 | benign | 1 | 1 | 1 |
| rs150634562 | pathogenic | 1 | 1 | 0 |
| rs142948132 | pathogenic | 1 | 1 | 1 |
| rs132630321 | pathogenic | 1 | 1 | 1 |
| rs132630317 | pathogenic | 1 | 1 | 1 |
| rs132630320 | pathogenic | 0 | 0 | 0 |
| rs781394318 | pathogenic | 1 | 1 | 1 |
| rs111033624 | pathogenic | 1 | 1 | 1 |
| rs111033630 | pathogenic | 1 | 1 | 1 |
| rs111033627 | pathogenic | 1 | 1 | 1 |
| rs193922287 | pathogenic | 1 | 1 | 1 |
| rs759072800 | pathogenic | 0 | 1 | 0 |
| rs104894010 | pathogenic | 0 | 0 | 0 |
| rs193922296 | pathogenic | 0 | 0 | 0 |
| rs104894006 | pathogenic | 1 | 1 | 1 |
| rs193922311 | pathogenic | 1 | 1 | 0 |
| rs80356654 | pathogenic | 1 | 1 | 0 |
| rs144723656 | benign | 1 | 1 | 1 |
| rs148311934 | pathogenic | 1 | 1 | 0 |
| rs80356655 | pathogenic | 1 | 1 | 1 |
| rs769268803 | pathogenic | 1 | 1 | 0 |
| rs104894008 | pathogenic | 0 | 1 | 0 |
| rs104894005 | pathogenic | 1 | 1 | 1 |
| rs104894009 | pathogenic | 1 | 1 | 0 |
| rs193922340 | pathogenic | 0 | 0 | 0 |
| rs104894016 | pathogenic | 0 | 1 | 0 |
| rs193921338 | pathogenic | 0 | 0 | 0 |
| rs193922272 | pathogenic | 1 | 1 | 1 |
| rs104894012 | pathogenic | 1 | 1 | 0 |
| rs193922285 | pathogenic | 1 | 1 | 1 |
| rs104894336 | pathogenic | 1 | 1 | 1 |
| rs28931580 | pathogenic | 1 | 1 | 1 |
| rs104894326 | pathogenic | 1 | 1 | 1 |
| rs104894331 | pathogenic | 0 | 0 | 0 |
| rs104894338 | pathogenic | 0 | 0 | 0 |
| rs104894333 | pathogenic | 1 | 1 | 1 |
| rs104894330 | pathogenic | 1 | 1 | 1 |
| rs143886391 | benign | 1 | 1 | 1 |
| rs104894335 | pathogenic | 1 | 1 | 1 |
| rs104894337 | pathogenic | 0 | 0 | 0 |

| | | | | |
|---|---|---|---|---|
| rs104894328 | pathogenic | 0 | 0 | 0 |
| rs104894341 | pathogenic | 1 | 1 | 0 |
| rs104894329 | pathogenic | 0 | 0 | 0 |
| rs137852850 | pathogenic | 1 | 1 | 1 |
| rs137852854 | pathogenic | 0 | 1 | 0 |
| rs137852849 | pathogenic | 0 | 1 | 0 |
| rs137852848 | pathogenic | 1 | 1 | 1 |
| rs137852852 | pathogenic | 0 | 1 | 0 |
| rs137852853 | pathogenic | 0 | 1 | 0 |
| rs137852847 | pathogenic | 0 | 1 | 0 |
| rs137852846 | pathogenic | 0 | 1 | 0 |
| rs137852844 | pathogenic | 0 | 1 | 0 |
| rs144411158 | pathogenic | 0 | 0 | 0 |
| rs2302234 | benign | 0 | 0 | 0 |
| rs139620139 | pathogenic | 0 | 0 | 0 |
| rs146689929 | benign | 0 | 0 | 0 |
| rs373328681 | pathogenic | 1 | 1 | 0 |
| rs387906620 | pathogenic | 1 | 1 | 1 |
| rs1799983 | benign | 0 | 1 | 0 |
| rs28999111 | pathogenic | 1 | 1 | 1 |
| rs104894544 | pathogenic | 1 | 1 | 1 |
| rs104894546 | pathogenic | 1 | 1 | 1 |
| rs28999112 | pathogenic | 0 | 1 | 0 |
| rs104894545 | pathogenic | 1 | 1 | 1 |
| rs104894547 | pathogenic | 1 | 1 | 1 |
| rs74315310 | pathogenic | 0 | 0 | 0 |
| rs267606852 | pathogenic | 0 | 1 | 0 |
| rs137853586 | pathogenic | 0 | 1 | 0 |
| rs137853582 | pathogenic | 1 | 1 | 1 |
| rs61754634 | pathogenic | 1 | 1 | 1 |
| rs137853587 | pathogenic | 0 | 1 | 0 |
| rs267606851 | pathogenic | 1 | 1 | 1 |
| rs267606853 | pathogenic | 1 | 1 | 1 |
| rs137853584 | pathogenic | 1 | 1 | 1 |
| rs137853585 | pathogenic | 1 | 1 | 1 |
| rs460184 | pathogenic | 1 | 1 | 1 |
| rs104893751 | pathogenic | 0 | 1 | 0 |
| rs77358162 | benign | 0 | 0 | 0 |
| rs387906936 | pathogenic | 0 | 0 | 0 |
| rs769525399 | pathogenic | 0 | 0 | 0 |
| rs371271054 | pathogenic | 1 | 1 | 0 |
| rs118204000 | pathogenic | 0 | 0 | 0 |
| rs138568622 | pathogenic | 1 | 1 | 1 |
| rs34509359 | benign | 0 | 0 | 0 |
| rs104894130 | pathogenic | 1 | 1 | 0 |
| rs281865424 | pathogenic | 0 | 0 | 0 |
| rs121912778 | pathogenic | 1 | 1 | 0 |
| rs771055145 | pathogenic | 1 | 1 | 0 |
| rs776174514 | pathogenic | 1 | 1 | 0 |
| rs545986367 | pathogenic | 1 | 1 | 1 |
| rs570768038 | benign | 1 | 1 | 1 |
| rs74315442 | pathogenic | 1 | 1 | 1 |
| rs79716074 | benign | 1 | 1 | 1 |
| rs748486078 | pathogenic | 1 | 1 | 1 |

| | | | | |
|---|---|---|---|---|
| rs2397084 | benign | 1 | 1 | 1 |
| rs11465553 | benign | 1 | 1 | 1 |
| rs144576902 | benign | 1 | 1 | 1 |
| rs763780 | benign | 1 | 1 | 1 |
| rs267606870 | pathogenic | 1 | 1 | 0 |
| rs121913502 | pathogenic | 1 | 1 | 0 |
| rs118101777 | benign | 1 | 1 | 1 |
| rs118053940 | benign | 0 | 0 | 0 |
| rs368771578 | benign | 1 | 1 | 1 |
| rs11540654 | pathogenic | 1 | 1 | 1 |
| rs121912658 | pathogenic | 1 | 1 | 1 |
| rs28934873 | pathogenic | 0 | 1 | 0 |
| rs780442292 | pathogenic | 1 | 1 | 1 |
| rs758781593 | benign | 1 | 1 | 1 |
| rs28934875 | pathogenic | 1 | 1 | 1 |
| rs28934874 | pathogenic | 1 | 1 | 0 |
| rs587782705 | pathogenic | 1 | 1 | 1 |
| rs587780068 | pathogenic | 1 | 1 | 1 |
| rs587782144 | pathogenic | 1 | 1 | 1 |
| rs138729528 | pathogenic | 1 | 1 | 1 |
| rs28934578 | pathogenic | 1 | 1 | 1 |
| rs397514495 | pathogenic | 1 | 1 | 1 |
| rs121912665 | pathogenic | 1 | 1 | 1 |
| rs397516435 | pathogenic | 1 | 1 | 1 |
| rs587778720 | pathogenic | 1 | 1 | 1 |
| rs397516436 | pathogenic | 1 | 1 | 1 |
| rs587781386 | benign | 1 | 1 | 1 |
| rs121912666 | pathogenic | 1 | 1 | 0 |
| rs587780073 | pathogenic | 1 | 1 | 1 |
| rs587782664 | pathogenic | 1 | 1 | 1 |
| rs730882005 | pathogenic | 1 | 1 | 1 |
| rs28934573 | pathogenic | 1 | 1 | 1 |
| rs121912655 | pathogenic | 1 | 1 | 1 |
| rs28934575 | pathogenic | 1 | 1 | 1 |
| rs121912656 | pathogenic | 1 | 1 | 1 |
| rs11540652 | pathogenic | 1 | 1 | 1 |
| rs121912651 | pathogenic | 1 | 1 | 1 |
| rs730882027 | pathogenic | 1 | 1 | 1 |
| rs121912653 | pathogenic | 1 | 1 | 1 |
| rs121912652 | pathogenic | 1 | 1 | 1 |
| rs121912657 | pathogenic | 0 | 1 | 0 |
| rs121913343 | pathogenic | 1 | 1 | 1 |
| rs28934576 | pathogenic | 1 | 1 | 1 |
| rs763098116 | pathogenic | 1 | 1 | 1 |
| rs121912660 | pathogenic | 1 | 1 | 1 |
| rs28934574 | pathogenic | 1 | 1 | 1 |
| rs371409680 | pathogenic | 1 | 1 | 1 |
| rs121912667 | pathogenic | 1 | 1 | 1 |
| rs121912663 | pathogenic | 1 | 1 | 1 |
| rs201744589 | pathogenic | 0 | 0 | 0 |
| rs33926449 | benign | 0 | 0 | 0 |
| rs74315302 | pathogenic | 0 | 1 | 0 |
| rs202040269 | pathogenic | 1 | 1 | 1 |
| rs375958814 | pathogenic | 1 | 1 | 1 |

| | | | | |
|---|---|---|---|---|
| rs121909624 | pathogenic | 1 | 1 | 1 |
| rs121909621 | pathogenic | 1 | 1 | 1 |
| rs121909622 | pathogenic | 1 | 1 | 0 |
| rs4220 | benign | 1 | 1 | 1 |
| rs147559466 | pathogenic | 1 | 1 | 1 |
| rs757434857 | pathogenic | 1 | 1 | 1 |
| rs398123072 | pathogenic | 1 | 1 | 1 |
| rs121434280 | pathogenic | 0 | 1 | 0 |
| rs398123074 | pathogenic | 0 | 1 | 0 |
| rs762114560 | pathogenic | 1 | 1 | 1 |
| rs121434283 | pathogenic | 1 | 1 | 0 |
| rs121434277 | pathogenic | 0 | 1 | 0 |
| rs745844469 | pathogenic | 1 | 1 | 1 |
| rs121434279 | pathogenic | 1 | 1 | 0 |
| rs121434278 | pathogenic | 1 | 1 | 1 |
| rs373715782 | pathogenic | 1 | 1 | 1 |
| rs200724875 | pathogenic | 1 | 1 | 1 |
| rs149678400 | pathogenic | 0 | 1 | 0 |
| rs770273135 | pathogenic | 1 | 1 | 1 |
| rs121434276 | pathogenic | 0 | 1 | 0 |
| rs121434281 | pathogenic | 1 | 1 | 0 |
| rs768884003 | pathogenic | 1 | 1 | 1 |
| rs201375579 | pathogenic | 0 | 0 | 0 |
| rs121434274 | pathogenic | 1 | 1 | 0 |
| rs121434282 | pathogenic | 1 | 1 | 0 |
| rs779759347 | pathogenic | 0 | 1 | 0 |
| rs77931234 | pathogenic | 1 | 1 | 1 |
| rs148207467 | pathogenic | 1 | 1 | 1 |
| rs766140986 | pathogenic | 1 | 1 | 1 |
| rs121434275 | pathogenic | 1 | 1 | 1 |
| rs748110745 | pathogenic | 1 | 1 | 1 |
| rs104894274 | pathogenic | 1 | 1 | 1 |
| rs104894273 | pathogenic | 0 | 1 | 0 |
| rs104894278 | pathogenic | 1 | 1 | 1 |
| rs104894275 | pathogenic | 1 | 1 | 1 |
| rs104894277 | pathogenic | 1 | 1 | 1 |
| rs104894276 | pathogenic | 0 | 0 | 0 |
| rs104894280 | pathogenic | 1 | 1 | 1 |
| rs145882709 | pathogenic | 1 | 1 | 1 |
| rs104894279 | pathogenic | 1 | 1 | 1 |
| rs80356670 | pathogenic | 1 | 1 | 1 |
| rs121918102 | pathogenic | 1 | 1 | 1 |
| rs80356671 | pathogenic | 1 | 1 | 1 |
| rs80356672 | pathogenic | 1 | 1 | 1 |
| rs758609113 | pathogenic | 1 | 1 | 0 |
| rs121918188 | pathogenic | 0 | 1 | 0 |
| rs121918189 | pathogenic | 1 | 1 | 0 |
| rs121918191 | pathogenic | 1 | 1 | 1 |
| rs1769259 | benign | 1 | 1 | 1 |
| rs140222720 | benign | 1 | 1 | 1 |
| rs2297085 | benign | 1 | 1 | 1 |
| rs200679026 | pathogenic | 1 | 1 | 1 |
| rs121918192 | pathogenic | 0 | 1 | 0 |
| rs145771944 | benign | 0 | 1 | 0 |

| | | | | |
|---|---|---|---|---|
| rs201508880 | pathogenic | 0 | 1 | 0 |
| rs17221854 | pathogenic | 0 | 0 | 0 |
| rs120074189 | pathogenic | 1 | 1 | 1 |
| rs120074190 | pathogenic | 1 | 1 | 1 |
| rs758600283 | benign | 1 | 1 | 1 |
| rs200305064 | pathogenic | 1 | 1 | 1 |
| rs121917891 | pathogenic | 1 | 1 | 1 |
| rs3772809 | benign | 0 | 1 | 0 |
| rs387906619 | pathogenic | 1 | 1 | 0 |
| rs2304681 | benign | 0 | 1 | 0 |
| rs121917849 | pathogenic | 0 | 0 | 0 |
| rs143010236 | pathogenic | 0 | 0 | 0 |
| rs121917850 | pathogenic | 1 | 1 | 0 |
| rs35916840 | pathogenic | 1 | 1 | 0 |
| rs28939688 | pathogenic | 1 | 1 | 1 |
| rs11258194 | benign | 1 | 1 | 1 |
| rs199840952 | pathogenic | 1 | 1 | 1 |
| rs121918159 | pathogenic | 1 | 1 | 1 |
| rs200396166 | pathogenic | 0 | 0 | 0 |
| rs757966746 | pathogenic | 0 | 1 | 0 |
| rs121918608 | pathogenic | 0 | 1 | 0 |
| rs74315401 | pathogenic | 1 | 1 | 1 |
| rs11538758 | pathogenic | 0 | 0 | 0 |
| rs74315402 | pathogenic | 1 | 1 | 1 |
| rs1799990 | pathogenic | 1 | 1 | 1 |
| rs74315410 | pathogenic | 1 | 1 | 1 |
| rs150351644 | benign | 1 | 1 | 1 |
| rs16990018 | benign | 1 | 1 | 1 |
| rs74315403 | pathogenic | 1 | 1 | 1 |
| rs74315408 | pathogenic | 1 | 1 | 1 |
| rs74315411 | pathogenic | 1 | 1 | 1 |
| rs74315413 | pathogenic | 1 | 1 | 1 |
| rs74315405 | pathogenic | 1 | 1 | 1 |
| rs28933385 | pathogenic | 1 | 1 | 1 |
| rs74315412 | pathogenic | 1 | 1 | 1 |
| rs74315407 | pathogenic | 1 | 1 | 1 |
| rs74315406 | pathogenic | 1 | 1 | 1 |
| rs11571117 | benign | 1 | 1 | 1 |
| rs121917742 | pathogenic | 1 | 1 | 1 |
| rs149782619 | pathogenic | 1 | 1 | 1 |
| rs397514492 | pathogenic | 1 | 1 | 1 |
| rs397514489 | pathogenic | 0 | 1 | 0 |
| rs397514493 | pathogenic | 1 | 1 | 1 |
| rs373849532 | pathogenic | 1 | 1 | 1 |
| rs121965048 | pathogenic | 1 | 1 | 1 |
| rs121965037 | pathogenic | 1 | 1 | 1 |
| rs121965060 | pathogenic | 0 | 1 | 0 |
| rs386833603 | pathogenic | 1 | 1 | 1 |
| rs121965038 | pathogenic | 1 | 1 | 1 |
| rs121965039 | pathogenic | 1 | 1 | 1 |
| rs121965040 | pathogenic | 1 | 1 | 1 |
| rs267606925 | pathogenic | 1 | 1 | 1 |
| rs121965056 | pathogenic | 0 | 1 | 0 |
| rs121965059 | pathogenic | 0 | 0 | 0 |

| | | | | |
|---|---|---|---|---|
| rs386833615 | pathogenic | 1 | 1 | 0 |
| rs121965051 | pathogenic | 1 | 1 | 0 |
| rs121965046 | pathogenic | 0 | 1 | 0 |
| rs121965052 | pathogenic | 0 | 1 | 0 |
| rs386833617 | pathogenic | 0 | 1 | 0 |
| rs386833618 | pathogenic | 1 | 1 | 0 |
| rs121965041 | pathogenic | 0 | 1 | 0 |
| rs121965042 | pathogenic | 0 | 1 | 0 |
| rs121965057 | pathogenic | 1 | 1 | 1 |
| rs386833621 | pathogenic | 1 | 1 | 1 |
| rs121965049 | pathogenic | 1 | 1 | 1 |
| rs386833623 | pathogenic | 1 | 1 | 1 |
| rs121965047 | pathogenic | 1 | 1 | 1 |
| rs121965053 | pathogenic | 0 | 1 | 0 |
| rs121965045 | pathogenic | 0 | 1 | 0 |
| rs121965054 | pathogenic | 0 | 1 | 0 |
| rs121965043 | pathogenic | 0 | 1 | 0 |
| rs386833598 | pathogenic | 0 | 0 | 0 |
| rs137854545 | pathogenic | 1 | 1 | 1 |
| rs137854544 | pathogenic | 1 | 1 | 1 |
| rs104894455 | pathogenic | 0 | 1 | 0 |
| rs1049564 | benign | 0 | 1 | 0 |
| rs104894460 | pathogenic | 1 | 1 | 1 |
| rs104894453 | pathogenic | 1 | 1 | 1 |
| rs104894450 | pathogenic | 0 | 1 | 0 |
| rs104894454 | pathogenic | 0 | 1 | 0 |
| rs104894452 | pathogenic | 1 | 1 | 1 |
| rs104894451 | pathogenic | 0 | 1 | 0 |
| rs104894368 | pathogenic | 1 | 1 | 1 |
| rs730880950 | pathogenic | 1 | 1 | 1 |
| rs121909535 | pathogenic | 1 | 1 | 1 |
| rs121909537 | pathogenic | 1 | 1 | 1 |
| rs121909536 | pathogenic | 1 | 1 | 1 |
| rs121909542 | pathogenic | 1 | 1 | 1 |
| rs121909538 | pathogenic | 1 | 1 | 1 |
| rs121909539 | pathogenic | 1 | 1 | 1 |
| rs121909540 | pathogenic | 1 | 1 | 1 |
| rs17560 | benign | 1 | 1 | 1 |
| rs121909543 | pathogenic | 0 | 1 | 0 |
| rs121909544 | pathogenic | 1 | 1 | 1 |
| rs2307456 | benign | 0 | 1 | 0 |
| rs745778317 | pathogenic | 0 | 1 | 0 |
| rs759607901 | pathogenic | 0 | 1 | 0 |
| rs201188114 | benign | 0 | 0 | 0 |
| rs1800472 | benign | 0 | 0 | 0 |
| rs587780728 | benign | 1 | 1 | 1 |
| rs144386518 | benign | 1 | 1 | 1 |
| rs281865439 | pathogenic | 1 | 1 | 1 |
| rs140175796 | pathogenic | 1 | 1 | 1 |
| rs5030843 | pathogenic | 1 | 1 | 0 |
| rs75166491 | pathogenic | 1 | 1 | 0 |
| rs199475679 | pathogenic | 1 | 1 | 1 |
| rs281865440 | pathogenic | 1 | 1 | 1 |
| rs74486803 | pathogenic | 0 | 1 | 0 |

| | | | | |
|---|---|---|---|---|
| rs199475602 | pathogenic | 0 | 1 | 0 |
| rs5030844 | pathogenic | 1 | 1 | 0 |
| rs62514927 | pathogenic | 1 | 1 | 1 |
| rs62514928 | pathogenic | 1 | 1 | 1 |
| rs62516152 | pathogenic | 0 | 1 | 0 |
| rs62507348 | pathogenic | 1 | 1 | 1 |
| rs76687508 | pathogenic | 1 | 1 | 1 |
| rs62508730 | pathogenic | 1 | 1 | 1 |
| rs62508588 | pathogenic | 0 | 1 | 0 |
| rs5030846 | pathogenic | 0 | 1 | 0 |
| rs74503222 | pathogenic | 1 | 1 | 0 |
| rs5030847 | pathogenic | 1 | 1 | 0 |
| rs5030850 | pathogenic | 1 | 1 | 1 |
| rs778154939 | pathogenic | 0 | 1 | 0 |
| rs62514952 | pathogenic | 1 | 1 | 1 |
| rs78655458 | pathogenic | 1 | 1 | 1 |
| rs62508698 | pathogenic | 1 | 1 | 0 |
| rs5030851 | pathogenic | 1 | 1 | 0 |
| rs62642933 | pathogenic | 1 | 1 | 1 |
| rs5030853 | pathogenic | 1 | 1 | 1 |
| rs62642935 | pathogenic | 1 | 1 | 1 |
| rs199475615 | pathogenic | 0 | 0 | 0 |
| rs62516062 | pathogenic | 1 | 1 | 0 |
| rs62516092 | pathogenic | 0 | 0 | 0 |
| rs62516095 | pathogenic | 1 | 1 | 1 |
| rs62642937 | pathogenic | 1 | 1 | 1 |
| rs62516141 | pathogenic | 0 | 1 | 0 |
| rs5030856 | pathogenic | 1 | 1 | 1 |
| rs5030857 | pathogenic | 1 | 1 | 1 |
| rs5030858 | pathogenic | 1 | 1 | 1 |
| rs79931499 | pathogenic | 1 | 1 | 1 |
| rs5030860 | pathogenic | 1 | 1 | 1 |
| rs281865437 | pathogenic | 1 | 1 | 1 |
| rs62644499 | pathogenic | 1 | 1 | 1 |
| rs147689487 | benign | 0 | 0 | 0 |
| rs397515417 | pathogenic | 1 | 1 | 1 |
| rs121909332 | pathogenic | 1 | 1 | 1 |
| rs121909330 | pathogenic | 1 | 1 | 0 |
| rs121909329 | pathogenic | 1 | 1 | 0 |
| rs121909335 | pathogenic | 1 | 1 | 1 |
| rs387906789 | pathogenic | 1 | 1 | 1 |
| rs121909334 | pathogenic | 1 | 1 | 1 |
| rs121909331 | pathogenic | 1 | 1 | 1 |
| rs372487178 | pathogenic | 1 | 1 | 1 |
| rs387906937 | pathogenic | 1 | 1 | 0 |
| rs16845 | benign | 1 | 1 | 1 |
| rs672601313 | pathogenic | 1 | 1 | 1 |
| rs117805308 | benign | 0 | 0 | 0 |
| rs121434261 | pathogenic | 1 | 1 | 1 |
| rs141629512 | benign | 0 | 0 | 0 |
| rs199957176 | benign | 1 | 1 | 1 |
| rs2230461 | benign | 1 | 1 | 1 |
| rs764618040 | pathogenic | 1 | 1 | 1 |
| rs267606949 | pathogenic | 1 | 1 | 1 |

| | | | | |
|---|---|---|---|---|
| rs587777483 | pathogenic | 1 | 1 | 1 |
| rs267606947 | pathogenic | 1 | 1 | 1 |
| rs6941583 | benign | 0 | 0 | 0 |
| rs749597959 | benign | 1 | 1 | 1 |
| rs182340753 | benign | 1 | 1 | 1 |
| rs146678221 | benign | 0 | 0 | 0 |
| rs11539202 | benign | 0 | 0 | 0 |
| rs148723565 | benign | 0 | 0 | 0 |
| rs61752925 | benign | 1 | 1 | 1 |
| rs113309941 | pathogenic | 0 | 0 | 0 |
| rs121434520 | pathogenic | 1 | 1 | 1 |
| rs121434517 | pathogenic | 1 | 1 | 1 |
| rs267607027 | pathogenic | 1 | 1 | 1 |
| rs138603088 | pathogenic | 0 | 0 | 0 |
| rs121434516 | pathogenic | 1 | 1 | 0 |
| rs121434521 | pathogenic | 0 | 0 | 0 |
| rs145735112 | benign | 1 | 1 | 1 |
| rs121434519 | pathogenic | 1 | 1 | 1 |
| rs75326924 | pathogenic | 1 | 1 | 0 |
| rs142186404 | pathogenic | 0 | 0 | 0 |
| rs121918035 | pathogenic | 0 | 0 | 0 |
| rs374385878 | pathogenic | 1 | 1 | 0 |
| rs11557652 | benign | 0 | 0 | 0 |
| rs118203986 | pathogenic | 1 | 1 | 1 |
| rs141675330 | benign | 1 | 1 | 1 |
| rs104893927 | pathogenic | 0 | 0 | 0 |
| rs104893935 | pathogenic | 0 | 0 | 0 |
| rs104893933 | pathogenic | 1 | 1 | 0 |
| rs104893934 | pathogenic | 0 | 0 | 0 |
| rs145183203 | benign | 1 | 1 | 0 |
| rs2230806 | benign | 0 | 0 | 0 |
| rs9282541 | benign | 0 | 0 | 0 |
| rs559110055 | benign | 0 | 0 | 0 |
| rs200030513 | benign | 0 | 0 | 0 |
| rs9282543 | benign | 0 | 0 | 0 |
| rs147675550 | benign | 0 | 0 | 0 |
| rs137854496 | pathogenic | 0 | 0 | 0 |
| rs374190304 | benign | 0 | 0 | 0 |
| rs114620717 | benign | 0 | 0 | 0 |
| rs2066718 | benign | 0 | 0 | 0 |
| rs35819696 | benign | 0 | 0 | 0 |
| rs138880920 | benign | 0 | 0 | 0 |
| rs2066715 | benign | 0 | 0 | 0 |
| rs35207495 | benign | 0 | 0 | 0 |
| rs2066714 | benign | 0 | 0 | 0 |
| rs28937313 | pathogenic | 0 | 0 | 0 |
| rs28937314 | pathogenic | 0 | 0 | 0 |
| rs137854495 | pathogenic | 0 | 0 | 0 |
| rs140365800 | benign | 0 | 0 | 0 |
| rs13306073 | benign | 0 | 0 | 0 |
| rs28933692 | pathogenic | 0 | 0 | 0 |
| rs33918808 | benign | 0 | 0 | 0 |
| rs76881554 | benign | 0 | 0 | 0 |
| rs143180998 | benign | 0 | 0 | 0 |

| | | | | |
|---|---|---|---|---|
| rs137854500 | pathogenic | 0 | 0 | 0 |
| rs538614702 | benign | 0 | 0 | 0 |
| rs201879964 | benign | 0 | 0 | 0 |
| rs2230808 | benign | 0 | 0 | 0 |
| rs138422574 | benign | 0 | 0 | 0 |
| rs137854498 | pathogenic | 0 | 0 | 0 |
| rs150125857 | benign | 0 | 0 | 0 |
| rs146292819 | pathogenic | 0 | 0 | 0 |
| rs142688906 | benign | 0 | 0 | 0 |
| rs137854499 | pathogenic | 0 | 0 | 0 |
| rs34879708 | benign | 0 | 0 | 0 |
| rs121913028 | pathogenic | 1 | 1 | 1 |
| rs1041951 | benign | 1 | 1 | 0 |
| rs138714094 | benign | 1 | 1 | 0 |
| rs200650502 | benign | 0 | 1 | 0 |
| rs370708663 | pathogenic | 1 | 1 | 1 |
| rs118204040 | pathogenic | 0 | 1 | 0 |
| rs146899669 | pathogenic | 1 | 1 | 0 |
| rs118204039 | pathogenic | 0 | 1 | 0 |
| rs121909233 | pathogenic | 0 | 0 | 0 |
| rs121909225 | pathogenic | 1 | 1 | 0 |
| rs121909236 | pathogenic | 0 | 0 | 0 |
| rs398123317 | pathogenic | 0 | 0 | 0 |
| rs121909226 | pathogenic | 1 | 1 | 0 |
| rs121909238 | pathogenic | 1 | 1 | 0 |
| rs121909230 | pathogenic | 0 | 0 | 0 |
| rs121909237 | pathogenic | 0 | 0 | 0 |
| rs121909222 | pathogenic | 1 | 1 | 0 |
| rs121909223 | pathogenic | 1 | 1 | 0 |
| rs121909218 | pathogenic | 1 | 1 | 0 |
| rs121909229 | pathogenic | 1 | 1 | 0 |
| rs121909224 | pathogenic | 1 | 1 | 0 |
| rs121909241 | pathogenic | 0 | 0 | 0 |
| rs121909221 | pathogenic | 0 | 0 | 0 |
| rs121913294 | pathogenic | 0 | 0 | 0 |
| rs121909234 | pathogenic | 0 | 0 | 0 |
| rs121909240 | pathogenic | 0 | 0 | 0 |
| rs767623493 | benign | 0 | 0 | 0 |
| rs121909239 | pathogenic | 0 | 0 | 0 |
| rs120074145 | pathogenic | 1 | 1 | 1 |
| rs120074148 | pathogenic | 1 | 1 | 1 |
| rs762991875 | pathogenic | 0 | 0 | 0 |
| rs148639841 | pathogenic | 1 | 1 | 1 |
| rs199524907 | pathogenic | 1 | 1 | 1 |
| rs120074141 | pathogenic | 1 | 1 | 1 |
| rs120074144 | pathogenic | 0 | 1 | 0 |
| rs120074146 | pathogenic | 1 | 1 | 1 |
| rs120074147 | pathogenic | 0 | 1 | 0 |
| rs120074143 | pathogenic | 0 | 1 | 0 |
| rs120074140 | pathogenic | 1 | 1 | 0 |
| rs28937315 | pathogenic | 1 | 1 | 1 |
| rs76066357 | benign | 1 | 1 | 1 |
| rs137852911 | pathogenic | 0 | 1 | 0 |
| rs137852907 | pathogenic | 1 | 1 | 0 |

| | | | | |
|---|---|---|---|---|
| rs137852910 | pathogenic | 1 | 1 | 1 |
| rs537585211 | benign | 0 | 0 | 0 |
| rs202134424 | pathogenic | 0 | 0 | 0 |
| rs200930463 | pathogenic | 0 | 0 | 0 |
| rs587777241 | pathogenic | 1 | 1 | 0 |
| rs77563738 | pathogenic | 0 | 0 | 0 |
| rs750868279 | pathogenic | 1 | 1 | 0 |
| rs148936893 | pathogenic | 0 | 0 | 0 |
| rs146597836 | pathogenic | 0 | 0 | 0 |
| rs775440641 | pathogenic | 1 | 1 | 1 |
| rs770689762 | pathogenic | 0 | 1 | 0 |
| rs376785840 | pathogenic | 0 | 0 | 0 |
| rs137852531 | pathogenic | 0 | 0 | 0 |
| rs137852538 | pathogenic | 1 | 1 | 0 |
| rs137852529 | pathogenic | 0 | 1 | 0 |
| rs137852528 | pathogenic | 0 | 1 | 0 |
| rs137852535 | pathogenic | 1 | 1 | 0 |
| rs137853045 | pathogenic | 0 | 1 | 0 |
| rs137853046 | pathogenic | 1 | 1 | 1 |
| rs137853043 | pathogenic | 1 | 1 | 1 |
| rs137853044 | pathogenic | 1 | 1 | 1 |
| rs137853047 | pathogenic | 1 | 1 | 1 |
| rs267606990 | pathogenic | 1 | 1 | 1 |
| rs751437780 | pathogenic | 1 | 1 | 0 |
| rs121918461 | pathogenic | 1 | 1 | 1 |
| rs121918460 | pathogenic | 1 | 1 | 1 |
| rs121918459 | pathogenic | 1 | 1 | 0 |
| rs397507511 | pathogenic | 1 | 1 | 1 |
| rs121918454 | pathogenic | 1 | 1 | 1 |
| rs121918453 | pathogenic | 1 | 1 | 1 |
| rs121918462 | pathogenic | 1 | 1 | 0 |
| rs121918465 | pathogenic | 1 | 1 | 0 |
| rs121918464 | pathogenic | 1 | 1 | 0 |
| rs121918466 | pathogenic | 1 | 1 | 1 |
| rs397507521 | pathogenic | 1 | 1 | 1 |
| rs397507527 | pathogenic | 1 | 1 | 1 |
| rs777603059 | pathogenic | 0 | 1 | 0 |
| rs121918456 | pathogenic | 1 | 1 | 1 |
| rs121918463 | pathogenic | 1 | 1 | 1 |
| rs121918455 | pathogenic | 0 | 1 | 0 |
| rs28933386 | pathogenic | 0 | 1 | 0 |
| rs774356443 | benign | 1 | 1 | 1 |
| rs121918467 | pathogenic | 1 | 1 | 1 |
| rs121918468 | pathogenic | 1 | 1 | 1 |
| rs121918469 | pathogenic | 1 | 1 | 1 |
| rs397507539 | pathogenic | 1 | 1 | 0 |
| rs121918458 | pathogenic | 1 | 1 | 1 |
| rs121918470 | pathogenic | 1 | 1 | 1 |
| rs35289681 | benign | 0 | 1 | 0 |
| rs4302331 | benign | 0 | 1 | 0 |
| rs11544803 | pathogenic | 1 | 1 | 1 |
| rs116254156 | benign | 0 | 0 | 0 |
| rs104893909 | pathogenic | 1 | 1 | 1 |
| rs104893911 | pathogenic | 1 | 1 | 1 |

| | | | | |
|---|---|---|---|---|
| rs104893908 | pathogenic | 1 | 1 | 0 |
| rs258751 | benign | 1 | 1 | 1 |
| rs104893914 | pathogenic | 1 | 1 | 0 |
| rs121909727 | pathogenic | 1 | 1 | 1 |
| rs104893910 | pathogenic | 1 | 1 | 1 |
| rs121909726 | pathogenic | 1 | 1 | 1 |
| rs104893912 | pathogenic | 1 | 1 | 1 |
| rs138407155 | benign | 1 | 1 | 1 |
| rs574061789 | benign | 1 | 1 | 1 |
| rs121912718 | pathogenic | 1 | 1 | 1 |
| rs121912719 | pathogenic | 1 | 1 | 1 |
| rs140770089 | benign | 1 | 1 | 1 |
| rs28931573 | pathogenic | 1 | 1 | 1 |
| rs121912717 | pathogenic | 1 | 1 | 1 |
| rs376456598 | pathogenic | 0 | 0 | 0 |
| rs398123499 | pathogenic | 1 | 1 | 1 |
| rs773048903 | pathogenic | 1 | 1 | 0 |
| rs398123503 | pathogenic | 1 | 1 | 1 |
| rs375785084 | pathogenic | 0 | 1 | 0 |
| rs137852874 | pathogenic | 1 | 1 | 1 |
| rs373713279 | pathogenic | 1 | 1 | 1 |
| rs137852873 | pathogenic | 1 | 1 | 0 |
| rs367823977 | pathogenic | 1 | 1 | 1 |
| rs398123508 | pathogenic | 1 | 1 | 1 |
| rs137852871 | pathogenic | 1 | 1 | 1 |
| rs753698250 | pathogenic | 0 | 1 | 0 |
| rs182923857 | pathogenic | 1 | 1 | 1 |
| rs148090804 | benign | 1 | 1 | 1 |
| rs137852870 | pathogenic | 1 | 1 | 1 |
| rs267607131 | pathogenic | 1 | 1 | 0 |
| rs137852902 | pathogenic | 0 | 0 | 0 |
| rs61741646 | benign | 1 | 1 | 1 |
| rs137852905 | pathogenic | 0 | 0 | 0 |
| rs121918681 | pathogenic | 0 | 1 | 0 |
| rs8177238 | benign | 1 | 1 | 0 |
| rs5030776 | pathogenic | 1 | 1 | 1 |
| rs104894572 | pathogenic | 0 | 0 | 0 |
| rs104894576 | pathogenic | 0 | 0 | 0 |
| rs80084721 | pathogenic | 0 | 0 | 0 |
| rs104894146 | pathogenic | 0 | 1 | 0 |
| rs104894138 | pathogenic | 1 | 1 | 0 |
| rs104894153 | pathogenic | 1 | 1 | 0 |
| rs104894135 | pathogenic | 1 | 1 | 0 |
| rs104894147 | pathogenic | 1 | 1 | 0 |
| rs104894148 | pathogenic | 1 | 1 | 0 |
| rs104894154 | pathogenic | 1 | 1 | 0 |
| rs104894150 | pathogenic | 1 | 1 | 0 |
| rs104894136 | pathogenic | 1 | 1 | 0 |
| rs104894144 | pathogenic | 0 | 0 | 0 |
| rs104894137 | pathogenic | 0 | 0 | 0 |
| rs61754278 | pathogenic | 1 | 1 | 0 |
| rs104894149 | pathogenic | 1 | 1 | 0 |
| rs104894139 | pathogenic | 1 | 1 | 0 |
| rs104894142 | pathogenic | 0 | 0 | 0 |

| | | | | |
|---|---|---|---|---|
| rs104894143 | pathogenic | 0 | 0 | 0 |
| rs104894145 | pathogenic | 0 | 0 | 0 |
| rs121909545 | benign | 0 | 1 | 0 |
| rs782308462 | pathogenic | 1 | 1 | 1 |
| rs375002796 | pathogenic | 0 | 1 | 0 |
| rs201727685 | pathogenic | 0 | 0 | 0 |
| rs768050261 | pathogenic | 1 | 1 | 1 |
| rs780851340 | benign | 0 | 0 | 0 |
| rs139824241 | benign | 1 | 1 | 0 |
| rs757486575 | pathogenic | 1 | 1 | 0 |
| rs750831299 | pathogenic | 1 | 1 | 1 |
| rs767602961 | pathogenic | 1 | 1 | 1 |
| rs121913661 | pathogenic | 1 | 1 | 1 |
| rs777692271 | pathogenic | 1 | 1 | 1 |
| rs121913659 | pathogenic | 1 | 1 | 1 |
| rs587777533 | pathogenic | 1 | 1 | 1 |
| rs104894210 | pathogenic | 0 | 0 | 0 |
| rs104894216 | pathogenic | 1 | 1 | 1 |
| rs104894207 | pathogenic | 1 | 1 | 1 |
| rs104894208 | pathogenic | 0 | 1 | 0 |
| rs104894214 | pathogenic | 1 | 1 | 1 |
| rs104894211 | pathogenic | 1 | 1 | 1 |
| rs61749384 | pathogenic | 0 | 0 | 0 |
| rs61750071 | pathogenic | 1 | 1 | 1 |
| rs61750074 | pathogenic | 0 | 0 | 0 |
| rs11063988 | benign | 0 | 0 | 0 |
| rs216311 | benign | 0 | 0 | 0 |
| rs1800382 | benign | 1 | 1 | 1 |
| rs11063987 | benign | 1 | 1 | 1 |
| rs387906847 | pathogenic | 1 | 1 | 1 |
| rs73003466 | benign | 0 | 0 | 0 |
| rs149787233 | benign | 1 | 1 | 1 |
| rs144451841 | pathogenic | 1 | 1 | 1 |
| rs141638421 | pathogenic | 1 | 1 | 1 |
| rs121434497 | pathogenic | 0 | 0 | 0 |
| rs121434498 | pathogenic | 0 | 0 | 0 |
| rs267607230 | pathogenic | 1 | 1 | 0 |
| rs121434499 | pathogenic | 1 | 1 | 0 |
| rs546749336 | benign | 1 | 1 | 1 |
| rs138873805 | benign | 0 | 0 | 0 |
| rs730880522 | pathogenic | 0 | 0 | 0 |
| rs200371894 | benign | 0 | 0 | 0 |
| rs730880509 | benign | 0 | 0 | 0 |
| rs267606704 | pathogenic | 1 | 1 | 1 |
| rs267606706 | pathogenic | 0 | 0 | 0 |
| rs267606705 | pathogenic | 1 | 1 | 1 |
| rs267606707 | pathogenic | 0 | 0 | 0 |
| rs387906665 | pathogenic | 1 | 1 | 1 |
| rs267606708 | pathogenic | 1 | 1 | 1 |
| rs148368481 | benign | 0 | 0 | 0 |
| rs13204 | benign | 0 | 0 | 0 |
| rs140547520 | pathogenic | 1 | 1 | 1 |
| rs755234990 | benign | 1 | 1 | 0 |
| rs151184052 | benign | 0 | 0 | 0 |

| | | | | |
|---|---|---|---|---|
| rs3736358 | benign | 0 | 0 | 0 |
| rs3736360 | benign | 0 | 0 | 0 |
| rs1138469 | benign | 0 | 0 | 0 |
| rs74315351 | pathogenic | 0 | 1 | 0 |
| rs137853051 | pathogenic | 1 | 1 | 0 |
| rs74315353 | pathogenic | 0 | 1 | 0 |
| rs74315352 | pathogenic | 0 | 1 | 0 |
| rs74315354 | pathogenic | 1 | 1 | 1 |
| rs28938172 | pathogenic | 1 | 1 | 1 |
| rs52836744 | pathogenic | 0 | 0 | 0 |
| rs121913159 | pathogenic | 1 | 1 | 1 |
| rs121913145 | pathogenic | 1 | 1 | 1 |
| rs891087 | benign | 1 | 1 | 1 |
| rs9282757 | benign | 1 | 1 | 1 |
| rs397517404 | pathogenic | 1 | 1 | 1 |
| rs199918720 | benign | 1 | 1 | 0 |
| rs368299411 | benign | 1 | 1 | 0 |
| rs139399951 | benign | 1 | 1 | 0 |
| rs763981974 | benign | 1 | 1 | 0 |
| rs144242114 | benign | 0 | 0 | 0 |
| rs750965801 | benign | 0 | 0 | 0 |
| rs561310777 | benign | 0 | 0 | 0 |
| rs148185335 | benign | 0 | 0 | 0 |
| rs147742157 | benign | 0 | 0 | 0 |
| rs375664373 | pathogenic | 1 | 1 | 1 |
| rs777327250 | pathogenic | 0 | 0 | 0 |
| rs121908471 | pathogenic | 1 | 1 | 0 |
| rs2301612 | pathogenic | 1 | 1 | 0 |
| rs121908476 | pathogenic | 0 | 0 | 0 |
| rs36220239 | benign | 0 | 0 | 0 |
| rs121908473 | pathogenic | 0 | 0 | 0 |
| rs28647808 | benign | 0 | 0 | 0 |
| rs36090624 | benign | 0 | 0 | 0 |
| rs121908002 | pathogenic | 1 | 1 | 1 |
| rs4251545 | benign | 1 | 1 | 1 |
| rs78705382 | pathogenic | 0 | 0 | 0 |
| rs75184679 | pathogenic | 1 | 1 | 1 |
| rs74555752 | pathogenic | 0 | 0 | 0 |
| rs56320207 | benign | 1 | 1 | 1 |
| rs28730837 | pathogenic | 1 | 1 | 1 |
| rs119469014 | pathogenic | 1 | 1 | 1 |
| rs119468010 | pathogenic | 1 | 1 | 1 |
| rs35220041 | benign | 0 | 0 | 0 |
| rs762321137 | pathogenic | 1 | 1 | 1 |
| rs28931570 | pathogenic | 1 | 1 | 1 |
| rs709932 | benign | 1 | 1 | 1 |
| rs746531546 | benign | 1 | 1 | 1 |
| rs6647 | benign | 1 | 1 | 1 |
| rs141095970 | benign | 1 | 1 | 1 |
| rs143370956 | benign | 1 | 1 | 1 |
| rs28929474 | pathogenic | 1 | 1 | 1 |
| rs61761869 | pathogenic | 1 | 1 | 1 |
| rs1303 | benign | 1 | 1 | 1 |
| rs149447502 | benign | 0 | 0 | 0 |

| | | | | |
|---|---|---|---|---|
| rs267607180 | pathogenic | 1 | 1 | 1 |
| rs150304121 | benign | 0 | 0 | 0 |
| rs147630375 | benign | 1 | 1 | 1 |
| rs11539445 | benign | 1 | 1 | 1 |
| rs372098364 | pathogenic | 1 | 1 | 0 |
| rs760072286 | benign | 0 | 0 | 0 |
| rs745595833 | pathogenic | 1 | 1 | 1 |
| rs587777214 | pathogenic | 0 | 0 | 0 |
| rs150318528 | benign | 0 | 0 | 0 |
| rs137853254 | pathogenic | 1 | 1 | 1 |
| rs137853255 | pathogenic | 0 | 0 | 0 |
| rs137853253 | pathogenic | 0 | 0 | 0 |
| rs202166915 | benign | 0 | 0 | 0 |
| rs137853259 | pathogenic | 0 | 0 | 0 |
| rs2229137 | benign | 0 | 0 | 0 |
| rs137853258 | pathogenic | 1 | 1 | 0 |
| rs137853252 | pathogenic | 1 | 1 | 1 |
| rs137853256 | pathogenic | 0 | 0 | 0 |
| rs137853250 | pathogenic | 1 | 1 | 1 |
| rs121434615 | pathogenic | 0 | 0 | 0 |
| rs115014558 | benign | 1 | 1 | 1 |
| rs34116584 | benign | 1 | 1 | 1 |
| rs34885252 | benign | 1 | 1 | 1 |
| rs180177262 | pathogenic | 1 | 1 | 1 |
| rs180177162 | pathogenic | 1 | 1 | 1 |
| rs121908523 | pathogenic | 1 | 1 | 1 |
| rs180177170 | pathogenic | 1 | 1 | 1 |
| rs180177180 | pathogenic | 1 | 1 | 1 |
| rs767586362 | pathogenic | 1 | 1 | 1 |
| rs121908521 | pathogenic | 1 | 1 | 1 |
| rs180177203 | pathogenic | 1 | 1 | 1 |
| rs138025751 | pathogenic | 1 | 1 | 1 |
| rs180177210 | pathogenic | 0 | 1 | 0 |
| rs180177211 | pathogenic | 0 | 1 | 0 |
| rs121908524 | pathogenic | 1 | 1 | 0 |
| rs121908530 | pathogenic | 1 | 1 | 0 |
| rs180177227 | pathogenic | 0 | 1 | 0 |
| rs180177230 | pathogenic | 1 | 1 | 1 |
| rs121908529 | pathogenic | 0 | 1 | 0 |
| rs117195882 | benign | 1 | 1 | 0 |
| rs180177238 | pathogenic | 1 | 1 | 0 |
| rs34664134 | benign | 1 | 1 | 1 |
| rs180177246 | pathogenic | 0 | 1 | 0 |
| rs536352238 | pathogenic | 0 | 0 | 0 |
| rs121908520 | pathogenic | 1 | 1 | 0 |
| rs180177248 | pathogenic | 1 | 1 | 0 |
| rs180177252 | pathogenic | 1 | 1 | 1 |
| rs121908526 | pathogenic | 1 | 1 | 1 |
| rs121908525 | pathogenic | 1 | 1 | 1 |
| rs147106773 | benign | 1 | 1 | 1 |
| rs180177264 | pathogenic | 0 | 1 | 0 |
| rs146525143 | pathogenic | 1 | 1 | 1 |
| rs140992177 | benign | 1 | 1 | 1 |
| rs13408961 | benign | 0 | 1 | 0 |

| | | | | |
|---|---|---|---|---|
| rs4426527 | benign | 1 | 1 | 1 |
| rs3088308 | benign | 0 | 1 | 0 |
| rs4469829 | benign | 1 | 1 | 0 |
| rs104886003 | pathogenic | 1 | 1 | 1 |
| rs121913286 | pathogenic | 1 | 1 | 1 |
| rs121913279 | pathogenic | 1 | 1 | 1 |
| rs121434636 | pathogenic | 1 | 1 | 1 |
| rs755299132 | pathogenic | 1 | 1 | 1 |
| rs748318386 | pathogenic | 1 | 1 | 1 |
| rs387906757 | pathogenic | 1 | 1 | 1 |
| rs121965064 | pathogenic | 1 | 1 | 1 |
| rs28934608 | pathogenic | 0 | 0 | 0 |
| rs5967 | benign | 1 | 1 | 1 |
| rs773905328 | pathogenic | 0 | 0 | 0 |
| rs104894256 | pathogenic | 0 | 1 | 0 |
| rs28931612 | pathogenic | 0 | 1 | 0 |
| rs104894263 | pathogenic | 1 | 1 | 1 |
| rs607969 | benign | 1 | 1 | 0 |
| rs104894262 | pathogenic | 0 | 1 | 0 |
| rs104894268 | pathogenic | 1 | 1 | 1 |
| rs104894264 | pathogenic | 0 | 1 | 0 |
| rs104894259 | pathogenic | 1 | 1 | 0 |
| rs2959656 | benign | 0 | 0 | 0 |
| rs121913035 | pathogenic | 1 | 1 | 0 |
| rs139605474 | benign | 0 | 0 | 0 |
| rs3817552 | benign | 0 | 0 | 0 |
| rs150541115 | benign | 0 | 0 | 0 |
| rs562763010 | benign | 1 | 1 | 1 |
| rs200053119 | pathogenic | 0 | 0 | 0 |
| rs73826386 | benign | 0 | 0 | 0 |
| rs143655258 | benign | 0 | 0 | 0 |
| rs758857853 | pathogenic | 1 | 1 | 0 |
| rs2228380 | benign | 0 | 0 | 0 |
| rs141078733 | benign | 1 | 1 | 1 |
| rs55824172 | pathogenic | 1 | 1 | 1 |
| rs756751089 | pathogenic | 1 | 1 | 1 |
| rs121964990 | pathogenic | 0 | 0 | 0 |
| rs143555186 | benign | 0 | 0 | 0 |
| rs121964993 | pathogenic | 1 | 1 | 0 |
| rs121964992 | pathogenic | 1 | 1 | 1 |
| rs121964991 | pathogenic | 1 | 1 | 0 |
| rs121964988 | pathogenic | 1 | 1 | 1 |
| rs4687 | benign | 1 | 1 | 1 |
| rs267607125 | pathogenic | 1 | 1 | 1 |
| rs751682878 | benign | 1 | 1 | 1 |
| rs267607126 | pathogenic | 1 | 1 | 1 |
| rs768713596 | pathogenic | 0 | 0 | 0 |
| rs147288996 | benign | 0 | 0 | 0 |
| rs387906639 | pathogenic | 1 | 1 | 1 |
| rs730881069 | pathogenic | 1 | 1 | 1 |
| rs730881071 | pathogenic | 1 | 1 | 1 |
| rs397516348 | pathogenic | 0 | 0 | 0 |
| rs121917760 | pathogenic | 0 | 0 | 0 |
| rs104894724 | pathogenic | 1 | 1 | 1 |

| | | | | |
|---|---|---|---|---|
| rs397516349 | pathogenic | 1 | 1 | 1 |
| rs12075 | benign | 0 | 0 | 0 |
| rs147549987 | benign | 0 | 0 | 0 |
| rs56079734 | benign | 0 | 0 | 0 |
| rs1017842 | benign | 1 | 1 | 1 |
| rs137852593 | pathogenic | 1 | 1 | 1 |
| rs137852571 | pathogenic | 1 | 1 | 1 |
| rs137852591 | pathogenic | 1 | 1 | 1 |
| rs9332971 | pathogenic | 1 | 1 | 1 |
| rs755226547 | pathogenic | 1 | 1 | 1 |
| rs79206939 | benign | 0 | 0 | 0 |
| rs121918214 | pathogenic | 1 | 1 | 0 |
| rs757075712 | pathogenic | 1 | 1 | 0 |
| rs150483902 | benign | 1 | 1 | 1 |
| rs113993973 | pathogenic | 0 | 1 | 0 |
| rs946616 | benign | 0 | 0 | 0 |
| rs113993976 | pathogenic | 1 | 1 | 0 |
| rs151022264 | benign | 0 | 1 | 0 |
| rs113993977 | pathogenic | 1 | 1 | 0 |
| rs113993978 | pathogenic | 1 | 1 | 0 |
| rs2228499 | benign | 0 | 1 | 0 |
| rs113993979 | pathogenic | 1 | 1 | 0 |
| rs113993980 | pathogenic | 1 | 1 | 0 |
| rs113993983 | pathogenic | 0 | 1 | 0 |
| rs150547274 | pathogenic | 1 | 1 | 0 |
| rs113993984 | pathogenic | 1 | 1 | 0 |
| rs113993986 | pathogenic | 1 | 1 | 0 |
| rs34313873 | benign | 0 | 1 | 0 |
| rs113993988 | pathogenic | 1 | 1 | 0 |
| rs199638097 | pathogenic | 1 | 1 | 1 |
| rs118204447 | pathogenic | 1 | 1 | 1 |
| rs11862754 | benign | 0 | 0 | 0 |
| rs118204445 | pathogenic | 0 | 0 | 0 |
| rs118204441 | pathogenic | 0 | 0 | 0 |
| rs118204438 | pathogenic | 0 | 0 | 0 |
| rs118204436 | pathogenic | 0 | 0 | 0 |
| rs146093755 | pathogenic | 0 | 0 | 0 |
| rs781439830 | pathogenic | 0 | 0 | 0 |
| rs398123438 | pathogenic | 0 | 0 | 0 |
| rs398123439 | pathogenic | 0 | 0 | 0 |
| rs118204444 | pathogenic | 0 | 0 | 0 |
| rs7187889 | benign | 0 | 0 | 0 |
| rs118204435 | pathogenic | 1 | 1 | 0 |
| rs34745339 | benign | 0 | 0 | 0 |
| rs398123440 | pathogenic | 0 | 0 | 0 |
| rs118204442 | pathogenic | 0 | 0 | 0 |
| rs770053354 | pathogenic | 0 | 0 | 0 |
| rs118204448 | pathogenic | 0 | 0 | 0 |
| rs118204443 | pathogenic | 0 | 0 | 0 |
| rs118204446 | pathogenic | 0 | 0 | 0 |
| rs746756997 | pathogenic | 0 | 0 | 0 |
| rs267606838 | pathogenic | 0 | 0 | 0 |
| rs118204437 | pathogenic | 0 | 0 | 0 |
| rs398123429 | pathogenic | 0 | 0 | 0 |

| | | | | |
|---|---|---|---|---|
| rs398123430 | pathogenic | 0 | 0 | 0 |
| rs2303269 | benign | 0 | 0 | 0 |
| rs114703967 | benign | 0 | 0 | 0 |
| rs118204439 | pathogenic | 0 | 0 | 0 |
| rs151296605 | benign | 0 | 0 | 0 |
| rs118204440 | pathogenic | 0 | 0 | 0 |
| rs78127134 | benign | 0 | 0 | 0 |
| rs372893383 | pathogenic | 0 | 0 | 0 |
| rs61753266 | pathogenic | 1 | 1 | 1 |
| rs139579928 | benign | 1 | 1 | 1 |
| rs56275071 | pathogenic | 1 | 1 | 1 |
| rs121909736 | pathogenic | 1 | 1 | 0 |
| rs121909737 | pathogenic | 1 | 1 | 0 |
| rs121909735 | pathogenic | 1 | 1 | 0 |
| rs121909731 | pathogenic | 1 | 1 | 1 |
| rs121909734 | pathogenic | 1 | 1 | 1 |
| rs121909733 | pathogenic | 1 | 1 | 1 |
| rs121909732 | pathogenic | 1 | 1 | 1 |
| rs121909730 | pathogenic | 1 | 1 | 1 |
| rs202198360 | benign | 1 | 1 | 1 |
| rs587777294 | pathogenic | 1 | 1 | 0 |
| rs587777297 | pathogenic | 1 | 1 | 0 |
| rs104893913 | pathogenic | 1 | 1 | 0 |
| rs17860424 | pathogenic | 1 | 1 | 1 |
| rs1045485 | benign | 1 | 1 | 1 |
| rs13926 | benign | 0 | 0 | 0 |
| rs79343837 | benign | 0 | 0 | 0 |
| rs45516306 | benign | 1 | 1 | 1 |
| rs139849355 | benign | 1 | 1 | 1 |
| rs104894874 | pathogenic | 1 | 1 | 1 |
| rs104894870 | pathogenic | 1 | 1 | 1 |
| rs137852221 | pathogenic | 1 | 1 | 1 |
| rs104894869 | pathogenic | 1 | 1 | 1 |
| rs104894880 | pathogenic | 1 | 1 | 1 |
| rs104894872 | pathogenic | 1 | 1 | 1 |
| rs104894868 | pathogenic | 1 | 1 | 1 |
| rs104894867 | pathogenic | 1 | 1 | 1 |
| rs104894871 | pathogenic | 1 | 1 | 1 |
| rs104894877 | pathogenic | 1 | 1 | 1 |
| rs104894883 | pathogenic | 1 | 1 | 1 |
| rs104894875 | pathogenic | 1 | 1 | 1 |
| rs104894876 | pathogenic | 1 | 1 | 1 |
| rs104894878 | pathogenic | 1 | 1 | 1 |
| rs137852220 | pathogenic | 1 | 1 | 1 |
| rs28933684 | pathogenic | 1 | 1 | 1 |
| rs104894873 | pathogenic | 1 | 1 | 1 |
| rs138409264 | benign | 0 | 0 | 0 |
| rs2236722 | benign | 0 | 0 | 0 |
| rs28757184 | benign | 0 | 0 | 0 |
| rs121434538 | pathogenic | 0 | 0 | 0 |
| rs700519 | benign | 0 | 0 | 0 |
| rs80051519 | pathogenic | 0 | 0 | 0 |
| rs121434536 | pathogenic | 1 | 1 | 0 |
| rs121434534 | pathogenic | 1 | 1 | 0 |

| | | | | |
|---|---|---|---|---|
| rs78310315 | pathogenic | 1 | 1 | 0 |
| rs540839115 | pathogenic | 1 | 1 | 1 |
| rs80338731 | pathogenic | 1 | 1 | 1 |
| rs180177151 | pathogenic | 1 | 1 | 1 |
| rs180177152 | pathogenic | 0 | 0 | 0 |
| rs80338732 | pathogenic | 1 | 1 | 1 |
| rs80338675 | pathogenic | 1 | 1 | 1 |
| rs80338676 | pathogenic | 1 | 1 | 1 |
| rs180177153 | pathogenic | 0 | 0 | 0 |
| rs180177154 | pathogenic | 1 | 1 | 1 |
| rs5896 | benign | 1 | 1 | 1 |
| rs121918477 | pathogenic | 0 | 0 | 0 |
| rs121918428 | pathogenic | 1 | 1 | 1 |
| rs121918427 | pathogenic | 1 | 1 | 1 |
| rs544300820 | benign | 1 | 1 | 1 |
| rs121918110 | pathogenic | 1 | 1 | 1 |
| rs1057141 | benign | 0 | 0 | 0 |
| rs1135216 | benign | 1 | 1 | 1 |
| rs121917702 | pathogenic | 1 | 1 | 1 |
| rs28941474 | pathogenic | 0 | 0 | 0 |
| rs104893944 | pathogenic | 1 | 1 | 1 |
| rs104893947 | pathogenic | 1 | 1 | 0 |
| rs104893943 | pathogenic | 1 | 1 | 1 |
| rs104893942 | pathogenic | 1 | 1 | 1 |
| rs104893940 | pathogenic | 1 | 1 | 1 |
| rs34045013 | benign | 0 | 0 | 0 |
| rs143083947 | pathogenic | 0 | 0 | 0 |
| rs145786986 | benign | 0 | 0 | 0 |
| rs119450942 | pathogenic | 0 | 0 | 0 |
| rs374259530 | pathogenic | 0 | 0 | 0 |
| rs756210458 | pathogenic | 0 | 0 | 0 |
| rs11089991 | benign | 0 | 0 | 0 |
| rs28941471 | pathogenic | 0 | 1 | 0 |
| rs750614500 | pathogenic | 0 | 1 | 0 |
| rs199761158 | benign | 1 | 1 | 1 |
| rs119450944 | pathogenic | 1 | 1 | 1 |
| rs373458753 | pathogenic | 1 | 1 | 1 |
| rs202064195 | pathogenic | 1 | 1 | 1 |
| rs776496275 | pathogenic | 1 | 1 | 1 |
| rs761493155 | pathogenic | 1 | 1 | 1 |
| rs763542069 | pathogenic | 0 | 0 | 0 |
| rs119450941 | pathogenic | 0 | 0 | 0 |
| rs119450940 | pathogenic | 0 | 0 | 0 |
| rs777821034 | pathogenic | 1 | 1 | 1 |
| rs771121666 | pathogenic | 1 | 1 | 1 |
| rs372895468 | pathogenic | 1 | 1 | 1 |
| rs61750603 | pathogenic | 0 | 0 | 0 |
| rs76863441 | pathogenic | 0 | 0 | 0 |
| rs142601143 | benign | 1 | 1 | 1 |
| rs185832993 | benign | 1 | 1 | 0 |
| rs17860568 | benign | 0 | 0 | 0 |
| rs142064825 | benign | 0 | 0 | 0 |
| rs564076183 | benign | 0 | 0 | 0 |
| rs386834034 | pathogenic | 0 | 0 | 0 |

| | | | | |
|---|---|---|---|---|
| rs147663769 | benign | 0 | 0 | 0 |
| rs776478974 | benign | 0 | 0 | 0 |
| rs111840783 | benign | 0 | 0 | 0 |
| rs1516446 | benign | 1 | 1 | 1 |
| rs143749154 | benign | 1 | 1 | 1 |
| rs80257412 | benign | 0 | 0 | 0 |
| rs142943120 | benign | 0 | 0 | 0 |
| rs144553482 | benign | 0 | 0 | 0 |
| rs35997569 | benign | 0 | 0 | 0 |
| rs772909106 | pathogenic | 0 | 0 | 0 |
| rs149433837 | benign | 1 | 1 | 1 |
| rs532178791 | pathogenic | 0 | 0 | 0 |
| rs145588689 | pathogenic | 1 | 1 | 0 |
| rs17843776 | benign | 1 | 1 | 0 |
| rs121917890 | pathogenic | 1 | 1 | 1 |
| rs121917892 | pathogenic | 0 | 0 | 0 |
| rs606231289 | pathogenic | 0 | 0 | 0 |
| rs370011798 | pathogenic | 0 | 0 | 0 |
| rs34689457 | benign | 0 | 0 | 0 |
| rs118204451 | pathogenic | 0 | 0 | 0 |
| rs118204452 | pathogenic | 1 | 1 | 1 |
| rs652785 | benign | 0 | 0 | 0 |
| rs35378126 | benign | 0 | 0 | 0 |
| rs80358214 | pathogenic | 1 | 1 | 1 |
| rs80358215 | pathogenic | 0 | 1 | 0 |
| rs267606858 | pathogenic | 1 | 1 | 1 |
| rs143137713 | pathogenic | 1 | 1 | 1 |
| rs782168634 | benign | 0 | 0 | 0 |
| rs143873938 | benign | 0 | 0 | 0 |
| rs782776511 | benign | 0 | 0 | 0 |
| rs201093148 | benign | 0 | 0 | 0 |
| rs267606815 | pathogenic | 0 | 0 | 0 |
| rs137853311 | pathogenic | 0 | 0 | 0 |
| rs267606817 | pathogenic | 0 | 0 | 0 |
| rs72650006 | benign | 0 | 0 | 0 |
| rs727503638 | benign | 1 | 1 | 0 |
| rs189286381 | benign | 0 | 0 | 0 |
| rs376644553 | benign | 0 | 0 | 0 |
| rs121434610 | pathogenic | 1 | 1 | 1 |
| rs267607076 | pathogenic | 1 | 1 | 1 |
| rs201275730 | benign | 0 | 0 | 0 |
| rs121909587 | pathogenic | 0 | 0 | 0 |
| rs121909558 | pathogenic | 1 | 1 | 0 |
| rs28929469 | pathogenic | 1 | 1 | 0 |
| rs121909547 | pathogenic | 1 | 1 | 0 |
| rs121909552 | pathogenic | 1 | 1 | 0 |
| rs121909573 | pathogenic | 1 | 1 | 1 |
| rs121909567 | pathogenic | 1 | 1 | 0 |
| rs121909569 | pathogenic | 0 | 0 | 0 |
| rs121909563 | pathogenic | 1 | 1 | 0 |
| rs121909570 | pathogenic | 1 | 1 | 1 |
| rs121909571 | pathogenic | 1 | 1 | 0 |
| rs121909572 | pathogenic | 1 | 1 | 1 |
| rs121909565 | pathogenic | 1 | 1 | 1 |

| | | | | |
|---|---|---|---|---|
| rs121909557 | pathogenic | 1 | 1 | 1 |
| rs121909568 | pathogenic | 1 | 1 | 1 |
| rs121909566 | pathogenic | 1 | 1 | 1 |
| rs121909549 | pathogenic | 1 | 1 | 1 |
| rs121909554 | pathogenic | 1 | 1 | 1 |
| rs121909550 | pathogenic | 1 | 1 | 1 |
| rs121909546 | pathogenic | 1 | 1 | 0 |
| rs121909555 | pathogenic | 1 | 1 | 1 |
| rs121909564 | pathogenic | 1 | 1 | 1 |
| rs548046212 | pathogenic | 0 | 0 | 0 |
| rs147353781 | benign | 0 | 0 | 0 |
| rs104894678 | pathogenic | 1 | 1 | 1 |
| rs1130426 | benign | 0 | 0 | 0 |
| rs104894677 | pathogenic | 1 | 1 | 1 |
| rs141917423 | benign | 1 | 1 | 1 |
| rs574777883 | benign | 0 | 0 | 0 |
| rs150199226 | benign | 1 | 1 | 1 |
| rs199618701 | benign | 0 | 1 | 0 |
| rs121964923 | pathogenic | 0 | 0 | 0 |
| rs121964924 | pathogenic | 0 | 0 | 0 |
| rs201258823 | pathogenic | 0 | 1 | 0 |
| rs267606773 | pathogenic | 0 | 0 | 0 |
| rs35638832 | benign | 0 | 0 | 0 |
| rs121908632 | pathogenic | 1 | 1 | 1 |
| rs121908625 | pathogenic | 1 | 1 | 1 |
| rs748949603 | pathogenic | 0 | 0 | 0 |
| rs757523840 | pathogenic | 0 | 0 | 0 |
| rs62541771 | pathogenic | 0 | 0 | 0 |
| rs776384541 | pathogenic | 1 | 1 | 0 |
| rs121908627 | pathogenic | 1 | 1 | 0 |
| rs28937594 | pathogenic | 0 | 0 | 0 |
| rs121907888 | pathogenic | 0 | 0 | 0 |
| rs121907889 | pathogenic | 0 | 1 | 0 |
| rs28941785 | pathogenic | 1 | 1 | 1 |
| rs28941786 | pathogenic | 1 | 1 | 1 |
| rs1021737 | benign | 0 | 0 | 0 |
| rs121908370 | pathogenic | 1 | 1 | 1 |
| rs63750083 | pathogenic | 0 | 0 | 0 |
| rs17883862 | benign | 0 | 0 | 0 |
| rs587782070 | pathogenic | 1 | 1 | 1 |
| rs587781269 | pathogenic | 1 | 1 | 1 |
| rs28909982 | pathogenic | 1 | 1 | 1 |
| rs373648967 | benign | 1 | 1 | 1 |
| rs368570187 | benign | 1 | 1 | 1 |
| rs730881701 | pathogenic | 1 | 1 | 1 |
| rs137853007 | pathogenic | 0 | 0 | 0 |
| rs575910805 | benign | 0 | 0 | 0 |
| rs104893875 | pathogenic | 1 | 1 | 1 |
| rs104893877 | pathogenic | 1 | 1 | 1 |
| rs119485093 | pathogenic | 1 | 1 | 1 |
| rs119485092 | pathogenic | 0 | 0 | 0 |
| rs119485090 | pathogenic | 0 | 0 | 0 |
| rs28942099 | pathogenic | 1 | 1 | 1 |
| rs373730800 | pathogenic | 1 | 1 | 1 |

| | | | | |
|---|---|---|---|---|
| rs113993995 | pathogenic | 0 | 0 | 0 |
| rs201070132 | pathogenic | 0 | 0 | 0 |
| rs113993996 | pathogenic | 0 | 0 | 0 |
| rs113993998 | pathogenic | 0 | 0 | 0 |
| rs78950939 | pathogenic | 1 | 1 | 1 |
| rs56378716 | pathogenic | 1 | 1 | 1 |
| rs754921704 | benign | 0 | 1 | 0 |
| rs766379103 | pathogenic | 1 | 1 | 0 |
| rs121912515 | pathogenic | 0 | 1 | 0 |
| rs764831888 | benign | 1 | 1 | 0 |
| rs572527340 | benign | 0 | 0 | 0 |
| rs35969327 | benign | 0 | 0 | 0 |
| rs35814893 | benign | 0 | 0 | 0 |
| rs34032300 | benign | 0 | 0 | 0 |
| rs80358260 | pathogenic | 0 | 0 | 0 |
| rs151220873 | benign | 0 | 0 | 0 |
| rs80358261 | pathogenic | 1 | 1 | 0 |
| rs11694 | pathogenic | 0 | 0 | 0 |
| rs80358264 | pathogenic | 0 | 0 | 0 |
| rs80358266 | pathogenic | 1 | 1 | 0 |
| rs104894458 | pathogenic | 1 | 1 | 1 |
| rs121918028 | pathogenic | 0 | 0 | 0 |
| rs11583680 | pathogenic | 0 | 0 | 0 |
| rs4715 | benign | 1 | 1 | 1 |
| rs34957318 | benign | 0 | 0 | 0 |
| rs1124 | benign | 1 | 1 | 1 |
| rs104894778 | pathogenic | 1 | 1 | 1 |
| rs104894777 | pathogenic | 1 | 1 | 1 |
| rs104894769 | pathogenic | 1 | 1 | 1 |
| rs104894768 | pathogenic | 1 | 1 | 1 |
| rs104894771 | pathogenic | 1 | 1 | 1 |
| rs575372249 | benign | 1 | 1 | 1 |
| rs118203914 | pathogenic | 0 | 0 | 0 |
| rs118203916 | pathogenic | 1 | 1 | 0 |
| rs140154690 | benign | 0 | 0 | 0 |
| rs34382405 | benign | 0 | 0 | 0 |
| rs73718082 | benign | 0 | 0 | 0 |
| rs764932724 | benign | 0 | 0 | 0 |
| rs554931092 | pathogenic | 1 | 1 | 0 |
| rs202060864 | pathogenic | 0 | 0 | 0 |
| rs116567033 | benign | 0 | 0 | 0 |
| rs11243011 | benign | 0 | 0 | 0 |
| rs142073519 | pathogenic | 0 | 0 | 0 |
| rs397514611 | pathogenic | 1 | 1 | 0 |
| rs397514612 | pathogenic | 0 | 0 | 0 |
| rs372054960 | pathogenic | 0 | 0 | 0 |
| rs775690041 | pathogenic | 0 | 0 | 0 |
| rs148921184 | pathogenic | 0 | 0 | 0 |
| rs12711521 | benign | 0 | 0 | 0 |
| rs2273346 | benign | 0 | 0 | 0 |
| rs12085877 | benign | 1 | 1 | 1 |
| rs144471433 | benign | 1 | 1 | 1 |
| rs397514671 | pathogenic | 1 | 1 | 1 |
| rs76394680 | benign | 0 | 0 | 0 |

| | | | | |
|---|---|---|---|---|
| rs78289603 | pathogenic | 1 | 1 | 1 |
| rs137852913 | pathogenic | 1 | 1 | 1 |
| rs137852914 | pathogenic | 1 | 1 | 1 |
| rs121918653 | pathogenic | 1 | 1 | 1 |
| rs35515200 | benign | 1 | 1 | 0 |
| rs17876030 | benign | 1 | 1 | 1 |
| rs556623443 | benign | 1 | 1 | 1 |
| rs121434365 | pathogenic | 1 | 1 | 1 |
| rs121434364 | pathogenic | 0 | 0 | 0 |
| rs41400249 | benign | 0 | 0 | 0 |
| rs149440376 | benign | 1 | 1 | 1 |
| rs181590179 | benign | 0 | 0 | 0 |
| rs113993968 | pathogenic | 0 | 0 | 0 |
| rs587776445 | pathogenic | 1 | 1 | 1 |
| rs113993969 | pathogenic | 1 | 1 | 1 |
| rs113993970 | pathogenic | 1 | 1 | 1 |
| rs587776449 | pathogenic | 1 | 1 | 1 |
| rs113993971 | pathogenic | 1 | 1 | 1 |
| rs267607194 | pathogenic | 1 | 1 | 1 |
| rs397514745 | pathogenic | 0 | 0 | 0 |
| rs201650281 | pathogenic | 0 | 0 | 0 |
| rs150529876 | benign | 1 | 1 | 1 |
| rs77573084 | benign | 0 | 0 | 0 |
| rs398122379 | pathogenic | 1 | 1 | 0 |
| rs765478968 | benign | 0 | 0 | 0 |
| rs201399681 | benign | 0 | 0 | 0 |
| rs17159287 | benign | 0 | 0 | 0 |
| rs538571144 | benign | 0 | 0 | 0 |
| rs137852647 | pathogenic | 0 | 0 | 0 |
| rs201358272 | pathogenic | 1 | 1 | 1 |
| rs181251337 | benign | 0 | 0 | 0 |
| rs3213119 | benign | 0 | 0 | 0 |
| rs137853102 | pathogenic | 1 | 1 | 0 |
| rs4956145 | benign | 0 | 0 | 0 |
| rs146036912 | pathogenic | 1 | 1 | 1 |
| rs375717077 | pathogenic | 1 | 1 | 1 |
| rs137853103 | pathogenic | 1 | 1 | 1 |
| rs776184830 | pathogenic | 1 | 1 | 1 |
| rs200273673 | pathogenic | 1 | 1 | 1 |
| rs267607025 | pathogenic | 1 | 1 | 1 |
| rs267607024 | pathogenic | 1 | 1 | 1 |
| rs515726188 | pathogenic | 1 | 1 | 1 |
| rs147315735 | benign | 0 | 0 | 0 |
| rs121918308 | pathogenic | 1 | 1 | 0 |
| rs515726195 | pathogenic | 0 | 0 | 0 |
| rs515726196 | pathogenic | 1 | 1 | 0 |
| rs121918311 | pathogenic | 0 | 1 | 0 |
| rs121918309 | pathogenic | 0 | 0 | 0 |
| rs387906891 | pathogenic | 0 | 0 | 0 |
| rs182614164 | pathogenic | 1 | 1 | 0 |
| rs121918307 | pathogenic | 0 | 0 | 0 |
| rs201440849 | benign | 0 | 0 | 0 |
| rs118203957 | pathogenic | 0 | 0 | 0 |
| rs4986790 | benign | 0 | 0 | 0 |

| | | | | |
|---|---|---|---|---|
| rs281864928 | pathogenic | 1 | 1 | 0 |
| rs267607156 | pathogenic | 0 | 0 | 0 |
| rs55916807 | benign | 0 | 0 | 0 |
| rs749750052 | benign | 0 | 0 | 0 |
| rs74382477 | benign | 0 | 0 | 0 |
| rs201016638 | benign | 0 | 0 | 0 |
| rs376683989 | benign | 0 | 0 | 0 |
| rs144439703 | pathogenic | 0 | 0 | 0 |
| rs779122022 | benign | 0 | 0 | 0 |
| rs121918056 | pathogenic | 0 | 0 | 0 |
| rs113994093 | pathogenic | 1 | 1 | 1 |
| rs141367015 | pathogenic | 0 | 0 | 0 |
| rs775538075 | pathogenic | 0 | 0 | 0 |
| rs121918055 | pathogenic | 0 | 0 | 0 |
| rs754246294 | benign | 1 | 1 | 1 |
| rs2307447 | benign | 1 | 1 | 1 |
| rs113994096 | pathogenic | 1 | 1 | 1 |
| rs367610201 | pathogenic | 1 | 1 | 1 |
| rs758438414 | pathogenic | 1 | 1 | 0 |
| rs121918046 | pathogenic | 0 | 0 | 0 |
| rs201871736 | benign | 0 | 0 | 0 |
| rs773994204 | benign | 0 | 0 | 0 |
| rs2307450 | benign | 1 | 1 | 1 |
| rs200257554 | benign | 0 | 0 | 0 |
| rs185645212 | benign | 0 | 0 | 0 |
| rs121918054 | pathogenic | 0 | 0 | 0 |
| rs113994097 | pathogenic | 0 | 0 | 0 |
| rs202037973 | pathogenic | 0 | 0 | 0 |
| rs769827124 | pathogenic | 1 | 1 | 0 |
| rs41549716 | benign | 1 | 1 | 1 |
| rs113994098 | pathogenic | 1 | 1 | 0 |
| rs144500145 | pathogenic | 1 | 1 | 0 |
| rs121918053 | pathogenic | 1 | 1 | 0 |
| rs121918050 | pathogenic | 1 | 1 | 0 |
| rs368587966 | pathogenic | 0 | 0 | 0 |
| rs139590686 | pathogenic | 0 | 0 | 0 |
| rs121918048 | pathogenic | 1 | 1 | 0 |
| rs768653086 | pathogenic | 1 | 1 | 0 |
| rs113994099 | pathogenic | 1 | 1 | 0 |
| rs121918051 | pathogenic | 1 | 1 | 0 |
| rs753160398 | pathogenic | 1 | 1 | 0 |
| rs201477273 | pathogenic | 1 | 1 | 0 |
| rs142347031 | pathogenic | 0 | 0 | 0 |
| rs121918049 | pathogenic | 0 | 0 | 0 |
| rs267606959 | pathogenic | 1 | 1 | 0 |
| rs201732356 | pathogenic | 0 | 0 | 0 |
| rs368435864 | pathogenic | 0 | 0 | 0 |
| rs764036283 | pathogenic | 0 | 0 | 0 |
| rs2307442 | benign | 0 | 0 | 0 |
| rs2307440 | benign | 0 | 0 | 0 |
| rs548076633 | pathogenic | 0 | 1 | 0 |
| rs750792237 | benign | 0 | 0 | 0 |
| rs3087374 | benign | 0 | 0 | 0 |
| rs121918060 | pathogenic | 0 | 1 | 0 |

| | | | | |
|---|---|---|---|---|
| rs397514765 | pathogenic | 1 | 1 | 1 |
| rs121918063 | pathogenic | 0 | 0 | 0 |
| rs121918058 | pathogenic | 1 | 1 | 1 |
| rs121918064 | pathogenic | 0 | 0 | 0 |
| rs36033115 | benign | 0 | 1 | 0 |
| rs121918057 | pathogenic | 0 | 1 | 0 |
| rs121918065 | pathogenic | 1 | 1 | 1 |
| rs121918061 | pathogenic | 1 | 1 | 1 |
| rs121918066 | pathogenic | 0 | 1 | 0 |
| rs707555 | benign | 0 | 1 | 0 |
| rs8192708 | benign | 0 | 1 | 0 |
| rs11552145 | benign | 0 | 0 | 0 |
| rs1804160 | benign | 0 | 1 | 0 |
| rs34103538 | benign | 0 | 1 | 0 |
| rs751393950 | pathogenic | 1 | 1 | 1 |
| rs121907979 | pathogenic | 1 | 1 | 1 |
| rs121907975 | pathogenic | 1 | 1 | 1 |
| rs121907962 | pathogenic | 1 | 1 | 1 |
| rs121907957 | pathogenic | 0 | 0 | 0 |
| rs121907953 | pathogenic | 1 | 1 | 1 |
| rs28941770 | pathogenic | 1 | 1 | 1 |
| rs28941771 | pathogenic | 1 | 1 | 1 |
| rs121907969 | pathogenic | 1 | 1 | 1 |
| rs121907973 | pathogenic | 1 | 1 | 1 |
| rs1800429 | pathogenic | 0 | 0 | 0 |
| rs121907976 | pathogenic | 1 | 1 | 0 |
| rs121907961 | pathogenic | 0 | 0 | 0 |
| rs121907974 | pathogenic | 0 | 0 | 0 |
| rs121907959 | pathogenic | 0 | 0 | 0 |
| rs121907971 | pathogenic | 1 | 1 | 0 |
| rs121907954 | pathogenic | 0 | 0 | 0 |
| rs121907977 | pathogenic | 0 | 0 | 0 |
| rs121907963 | pathogenic | 1 | 1 | 0 |
| rs370266293 | pathogenic | 1 | 1 | 0 |
| rs1800430 | benign | 0 | 0 | 0 |
| rs121907958 | pathogenic | 0 | 0 | 0 |
| rs202173526 | pathogenic | 0 | 0 | 0 |
| rs121907978 | pathogenic | 0 | 0 | 0 |
| rs121907981 | pathogenic | 0 | 1 | 0 |
| rs145012038 | benign | 1 | 1 | 1 |
| rs121907968 | pathogenic | 1 | 1 | 1 |
| rs121907966 | pathogenic | 1 | 1 | 1 |
| rs121907956 | pathogenic | 1 | 1 | 1 |
| rs28942071 | pathogenic | 1 | 1 | 1 |
| rs121907955 | pathogenic | 1 | 1 | 1 |
| rs4727853 | benign | 0 | 0 | 0 |
| rs201880593 | benign | 0 | 0 | 0 |
| rs201434579 | benign | 1 | 1 | 1 |
| rs74551128 | pathogenic | 1 | 1 | 0 |
| rs193922500 | pathogenic | 0 | 0 | 0 |
| rs121909009 | pathogenic | 1 | 1 | 0 |
| rs121908805 | pathogenic | 1 | 1 | 0 |
| rs139573311 | pathogenic | 0 | 0 | 0 |
| rs213950 | benign | 0 | 0 | 0 |

| | | | | |
|---|---|---|---|---|
| rs79282516 | pathogenic | 0 | 0 | 0 |
| rs121909017 | pathogenic | 1 | 1 | 1 |
| rs77101217 | pathogenic | 1 | 1 | 1 |
| rs1800092 | pathogenic | 0 | 1 | 0 |
| rs77646904 | pathogenic | 0 | 0 | 0 |
| rs1800095 | benign | 1 | 1 | 1 |
| rs387906368 | pathogenic | 1 | 1 | 1 |
| rs113993959 | pathogenic | 1 | 1 | 1 |
| rs121908755 | pathogenic | 1 | 1 | 1 |
| rs121909005 | pathogenic | 1 | 1 | 1 |
| rs121908757 | pathogenic | 1 | 1 | 1 |
| rs75527207 | pathogenic | 1 | 1 | 0 |
| rs121909013 | pathogenic | 1 | 1 | 0 |
| rs121909044 | pathogenic | 1 | 1 | 0 |
| rs74597325 | pathogenic | 1 | 1 | 0 |
| rs75549581 | pathogenic | 0 | 0 | 0 |
| rs80055610 | pathogenic | 1 | 1 | 1 |
| rs1800097 | pathogenic | 0 | 0 | 0 |
| rs121909006 | pathogenic | 0 | 1 | 0 |
| rs397508276 | pathogenic | 0 | 0 | 0 |
| rs397508296 | pathogenic | 1 | 1 | 1 |
| rs139468767 | pathogenic | 0 | 0 | 0 |
| rs121908759 | pathogenic | 1 | 1 | 0 |
| rs121909033 | pathogenic | 1 | 1 | 0 |
| rs397508331 | pathogenic | 0 | 0 | 0 |
| rs387907279 | pathogenic | 1 | 1 | 1 |
| rs546530618 | benign | 0 | 0 | 0 |
| rs558259739 | benign | 0 | 0 | 0 |
| rs201827340 | benign | 0 | 0 | 0 |
| rs148865119 | pathogenic | 1 | 1 | 0 |
| rs2229413 | benign | 1 | 1 | 1 |
| rs34040148 | benign | 1 | 1 | 1 |
| rs121964964 | pathogenic | 1 | 1 | 1 |
| rs760214620 | pathogenic | 1 | 1 | 1 |
| rs775992753 | pathogenic | 1 | 1 | 1 |
| rs770095972 | pathogenic | 1 | 1 | 1 |
| rs781444670 | pathogenic | 1 | 1 | 1 |
| rs147474549 | pathogenic | 0 | 0 | 0 |
| rs121964965 | pathogenic | 0 | 1 | 0 |
| rs121964963 | pathogenic | 1 | 1 | 1 |
| rs755952006 | pathogenic | 1 | 1 | 1 |
| rs121964970 | pathogenic | 1 | 1 | 1 |
| rs121964973 | pathogenic | 1 | 1 | 1 |
| rs773734233 | pathogenic | 1 | 1 | 1 |
| rs758236584 | pathogenic | 1 | 1 | 1 |
| rs149119723 | pathogenic | 1 | 1 | 0 |
| rs5742905 | pathogenic | 0 | 0 | 0 |
| rs121964962 | pathogenic | 1 | 1 | 1 |
| rs781567152 | pathogenic | 1 | 1 | 0 |
| rs398123151 | pathogenic | 1 | 1 | 1 |
| rs760417941 | pathogenic | 1 | 1 | 1 |
| rs774926464 | pathogenic | 0 | 0 | 0 |
| rs121964972 | pathogenic | 1 | 1 | 0 |
| rs372010465 | pathogenic | 1 | 1 | 1 |

| | | | | |
|---|---|---|---|---|
| rs763036586 | pathogenic | 1 | 1 | 1 |
| rs769080151 | pathogenic | 1 | 1 | 1 |
| rs121964967 | pathogenic | 1 | 1 | 1 |
| rs28934892 | pathogenic | 1 | 1 | 0 |
| rs28934891 | pathogenic | 1 | 1 | 1 |
| rs121964971 | pathogenic | 1 | 1 | 1 |
| rs121964968 | pathogenic | 1 | 1 | 1 |
| rs150828989 | benign | 1 | 1 | 1 |
| rs3775291 | benign | 1 | 1 | 0 |
| rs104894419 | pathogenic | 1 | 1 | 1 |
| rs2232642 | benign | 0 | 0 | 0 |
| rs104894551 | pathogenic | 1 | 1 | 0 |
| rs766328537 | pathogenic | 0 | 1 | 0 |
| rs104894553 | pathogenic | 1 | 1 | 0 |
| rs80099330 | benign | 0 | 0 | 0 |
| rs199565861 | pathogenic | 0 | 0 | 0 |
| rs104894548 | pathogenic | 0 | 0 | 0 |
| rs104894549 | pathogenic | 0 | 0 | 0 |
| rs104894550 | pathogenic | 1 | 1 | 1 |
| rs12948217 | pathogenic | 1 | 1 | 1 |
| rs761064915 | pathogenic | 0 | 0 | 0 |
| rs28940279 | pathogenic | 1 | 1 | 1 |
| rs774323189 | pathogenic | 1 | 1 | 1 |
| rs28940574 | pathogenic | 0 | 0 | 0 |
| rs515299 | benign | 1 | 1 | 1 |
| rs10127939 | benign | 1 | 1 | 1 |
| rs396991 | benign | 1 | 1 | 1 |
| rs145956175 | benign | 0 | 1 | 0 |
| rs747838255 | pathogenic | 1 | 1 | 1 |
| rs762659685 | pathogenic | 1 | 1 | 0 |
| rs104894831 | pathogenic | 0 | 1 | 0 |
| rs104894848 | pathogenic | 1 | 1 | 0 |
| rs104894833 | pathogenic | 0 | 1 | 0 |
| rs781838005 | pathogenic | 0 | 1 | 0 |
| rs104894834 | pathogenic | 1 | 1 | 0 |
| rs372966991 | pathogenic | 1 | 1 | 0 |
| rs138886989 | benign | 1 | 1 | 1 |
| rs28935195 | pathogenic | 0 | 1 | 0 |
| rs28935196 | pathogenic | 0 | 1 | 0 |
| rs372416832 | pathogenic | 1 | 1 | 1 |
| rs28935197 | pathogenic | 1 | 1 | 0 |
| rs28935486 | pathogenic | 1 | 1 | 0 |
| rs28935487 | pathogenic | 1 | 1 | 0 |
| rs28935488 | pathogenic | 1 | 1 | 1 |
| rs28935495 | pathogenic | 1 | 1 | 1 |
| rs28935485 | pathogenic | 1 | 1 | 1 |
| rs140329381 | pathogenic | 0 | 1 | 0 |
| rs104894830 | pathogenic | 0 | 1 | 0 |
| rs104894846 | pathogenic | 0 | 1 | 0 |
| rs28935489 | pathogenic | 0 | 1 | 0 |
| rs104894828 | pathogenic | 1 | 1 | 0 |
| rs28935491 | pathogenic | 1 | 1 | 0 |
| rs104894832 | pathogenic | 1 | 1 | 0 |
| rs28935492 | pathogenic | 1 | 1 | 0 |

| | | | | |
|---|---|---|---|---|
| rs730880437 | benign | 1 | 1 | 0 |
| rs28935493 | pathogenic | 0 | 1 | 0 |
| rs28935494 | pathogenic | 1 | 1 | 1 |
| rs111422676 | pathogenic | 1 | 1 | 1 |
| rs397515869 | benign | 1 | 1 | 0 |
| rs782449839 | pathogenic | 0 | 1 | 0 |
| rs104894852 | pathogenic | 1 | 1 | 1 |
| rs116929575 | pathogenic | 1 | 1 | 1 |
| rs7023954 | benign | 0 | 0 | 0 |
| rs184520335 | benign | 1 | 1 | 1 |
| rs143080527 | benign | 0 | 0 | 0 |
| rs1050565 | benign | 0 | 0 | 0 |
| rs11545035 | benign | 0 | 0 | 0 |
| rs267607190 | pathogenic | 1 | 1 | 1 |
| rs200283987 | benign | 0 | 0 | 0 |
| rs372556807 | benign | 1 | 1 | 0 |
| rs563073392 | benign | 0 | 0 | 0 |
| rs4539 | benign | 0 | 0 | 0 |
| rs28931609 | pathogenic | 1 | 1 | 1 |
| rs121912978 | pathogenic | 1 | 1 | 1 |
| rs104894072 | pathogenic | 1 | 1 | 1 |
| rs551933154 | benign | 1 | 1 | 1 |
| rs752962897 | benign | 1 | 1 | 0 |
| rs121912977 | pathogenic | 1 | 1 | 0 |
| rs4537 | benign | 1 | 1 | 1 |
| rs201830462 | benign | 0 | 0 | 0 |
| rs4544 | benign | 0 | 0 | 0 |
| rs562670189 | benign | 0 | 0 | 0 |
| rs4538 | benign | 0 | 1 | 0 |
| rs61757294 | pathogenic | 0 | 0 | 0 |
| rs5313 | benign | 0 | 0 | 0 |
| rs4545 | benign | 0 | 0 | 0 |
| rs72554627 | pathogenic | 1 | 1 | 1 |
| rs72554626 | pathogenic | 1 | 1 | 1 |
| rs387906861 | pathogenic | 1 | 1 | 1 |
| rs749655461 | pathogenic | 1 | 1 | 1 |
| rs121909606 | pathogenic | 1 | 1 | 1 |
| rs121909607 | pathogenic | 1 | 1 | 1 |
| rs200203992 | benign | 0 | 0 | 0 |
| rs6050 | benign | 0 | 0 | 0 |
| rs184635235 | benign | 0 | 0 | 0 |
| rs139146037 | benign | 0 | 0 | 0 |
| rs121909612 | pathogenic | 0 | 0 | 0 |
| rs78506343 | pathogenic | 0 | 0 | 0 |
| rs121918138 | pathogenic | 1 | 1 | 1 |
| rs121918139 | pathogenic | 1 | 1 | 1 |
| rs121918140 | pathogenic | 0 | 0 | 0 |
| rs36063028 | benign | 1 | 1 | 1 |
| rs45495192 | benign | 1 | 1 | 1 |
| rs45458802 | benign | 1 | 1 | 1 |
| rs552865793 | pathogenic | 1 | 1 | 1 |
| rs753744791 | pathogenic | 1 | 1 | 1 |
| rs777518512 | benign | 1 | 1 | 1 |
| rs142708991 | benign | 1 | 1 | 0 |

| | | | | |
|---|---|---|---|---|
| rs746069957 | benign | 0 | 0 | 0 |
| rs1334811 | benign | 0 | 0 | 0 |
| rs138894008 | benign | 1 | 1 | 0 |
| rs35030851 | benign | 1 | 1 | 0 |
| rs35378598 | benign | 0 | 0 | 0 |
| rs35818764 | benign | 1 | 1 | 1 |
| rs4631561 | benign | 1 | 1 | 1 |
| rs121918660 | pathogenic | 0 | 0 | 0 |
| rs373175587 | benign | 0 | 0 | 0 |
| rs61731956 | pathogenic | 1 | 1 | 1 |
| rs180177305 | pathogenic | 0 | 0 | 0 |
| rs119490108 | pathogenic | 0 | 0 | 0 |
| rs180177312 | pathogenic | 1 | 1 | 0 |
| rs180177314 | pathogenic | 1 | 1 | 0 |
| rs180177322 | pathogenic | 1 | 1 | 1 |
| rs180177323 | pathogenic | 1 | 1 | 1 |
| rs180177325 | pathogenic | 0 | 0 | 0 |
| rs35196441 | benign | 0 | 0 | 0 |
| rs28763904 | benign | 0 | 0 | 0 |
| rs2229519 | benign | 0 | 0 | 0 |
| rs137852886 | pathogenic | 0 | 0 | 0 |
| rs137852892 | pathogenic | 0 | 0 | 0 |
| rs770427750 | pathogenic | 0 | 0 | 0 |
| rs137852887 | pathogenic | 0 | 0 | 0 |
| rs137852893 | pathogenic | 0 | 0 | 0 |
| rs80338671 | pathogenic | 1 | 1 | 0 |
| rs2228389 | benign | 0 | 0 | 0 |
| rs80338672 | pathogenic | 0 | 0 | 0 |
| rs201958741 | pathogenic | 0 | 0 | 0 |
| rs80338673 | pathogenic | 0 | 0 | 0 |
| rs137852888 | pathogenic | 0 | 0 | 0 |
| rs137852889 | pathogenic | 0 | 0 | 0 |
| rs137852894 | pathogenic | 0 | 0 | 0 |
| rs137852890 | pathogenic | 0 | 0 | 0 |
| rs137852891 | pathogenic | 0 | 0 | 0 |
| rs766935302 | pathogenic | 0 | 0 | 0 |
| rs193074572 | benign | 0 | 0 | 0 |
| rs555210692 | benign | 1 | 1 | 0 |
| rs119103242 | pathogenic | 0 | 1 | 0 |
| rs143725101 | benign | 0 | 1 | 0 |
| rs28940589 | pathogenic | 1 | 1 | 1 |
| rs113994145 | pathogenic | 0 | 0 | 0 |
| rs773872994 | benign | 0 | 0 | 0 |
| rs28940590 | pathogenic | 1 | 1 | 0 |
| rs145400751 | benign | 1 | 1 | 1 |
| rs587782331 | benign | 0 | 0 | 0 |
| rs730881787 | benign | 0 | 0 | 0 |
| rs587778531 | benign | 0 | 0 | 0 |
| rs781652274 | benign | 0 | 0 | 0 |
| rs55882234 | benign | 0 | 0 | 0 |
| rs2020908 | benign | 0 | 0 | 0 |
| rs63750741 | pathogenic | 1 | 1 | 1 |
| rs41295268 | benign | 1 | 1 | 1 |
| rs63750897 | benign | 1 | 1 | 0 |

| | | | | |
|---|---|---|---|---|
| rs63751005 | benign | 1 | 1 | 0 |
| rs587779223 | benign | 0 | 0 | 0 |
| rs202127474 | benign | 0 | 0 | 0 |
| rs374230313 | benign | 0 | 0 | 0 |
| rs2020912 | benign | 0 | 0 | 0 |
| rs63751017 | pathogenic | 0 | 0 | 0 |
| rs587779246 | pathogenic | 0 | 0 | 0 |
| rs63749919 | pathogenic | 0 | 0 | 0 |
| rs63750563 | pathogenic | 1 | 1 | 1 |
| rs63749999 | pathogenic | 0 | 0 | 0 |
| rs200492211 | pathogenic | 0 | 0 | 0 |
| rs576269342 | benign | 0 | 0 | 0 |
| rs267608075 | benign | 0 | 0 | 0 |
| rs149605979 | benign | 0 | 0 | 0 |
| rs63750252 | benign | 0 | 0 | 0 |
| rs754289472 | benign | 0 | 0 | 0 |
| rs63750119 | pathogenic | 0 | 0 | 0 |
| rs587782109 | benign | 0 | 0 | 0 |
| rs34625968 | benign | 1 | 1 | 1 |
| rs764786814 | benign | 1 | 1 | 1 |
| rs199594809 | benign | 1 | 1 | 1 |
| rs74315364 | pathogenic | 1 | 1 | 1 |
| rs201253919 | pathogenic | 0 | 0 | 0 |
| rs387906828 | pathogenic | 1 | 1 | 1 |
| rs34521609 | benign | 1 | 1 | 1 |
| rs199472894 | pathogenic | 0 | 0 | 0 |
| rs199472902 | pathogenic | 0 | 0 | 0 |
| rs121912505 | pathogenic | 0 | 0 | 0 |
| rs28928905 | pathogenic | 0 | 0 | 0 |
| rs374376640 | benign | 0 | 0 | 0 |
| rs202102276 | benign | 0 | 0 | 0 |
| rs199472918 | pathogenic | 0 | 0 | 0 |
| rs121912516 | pathogenic | 1 | 1 | 1 |
| rs121912504 | pathogenic | 0 | 0 | 0 |
| rs9333649 | pathogenic | 0 | 0 | 0 |
| rs121912508 | pathogenic | 0 | 0 | 0 |
| rs199473428 | pathogenic | 1 | 1 | 1 |
| rs104894021 | pathogenic | 0 | 0 | 0 |
| rs28928904 | pathogenic | 1 | 1 | 1 |
| rs199473039 | benign | 1 | 1 | 1 |
| rs199472958 | pathogenic | 1 | 1 | 1 |
| rs1137617 | pathogenic | 1 | 1 | 1 |
| rs199473532 | pathogenic | 1 | 1 | 1 |
| rs199472988 | pathogenic | 0 | 0 | 0 |
| rs189014161 | pathogenic | 0 | 0 | 0 |
| rs12720441 | pathogenic | 0 | 0 | 0 |
| rs199473001 | pathogenic | 1 | 1 | 0 |
| rs121912506 | pathogenic | 1 | 1 | 0 |
| rs773724817 | pathogenic | 1 | 1 | 1 |
| rs118204002 | pathogenic | 1 | 1 | 1 |
| rs138556525 | pathogenic | 1 | 1 | 1 |
| rs72558181 | pathogenic | 1 | 1 | 1 |
| rs118204005 | pathogenic | 1 | 1 | 1 |
| rs118204004 | pathogenic | 0 | 0 | 0 |

| | | | | |
|---|---|---|---|---|
| rs118204001 | pathogenic | 1 | 1 | 1 |
| rs118204006 | pathogenic | 1 | 1 | 1 |
| rs118204003 | pathogenic | 0 | 0 | 0 |
| rs121913147 | pathogenic | 1 | 1 | 0 |
| rs2229429 | benign | 1 | 1 | 0 |
| rs182552223 | benign | 0 | 0 | 0 |
| rs146588336 | benign | 0 | 0 | 0 |
| rs121964983 | pathogenic | 0 | 1 | 0 |
| rs121964982 | pathogenic | 0 | 0 | 0 |
| rs386833679 | pathogenic | 0 | 0 | 0 |
| rs386833680 | pathogenic | 0 | 0 | 0 |
| rs121964981 | pathogenic | 1 | 1 | 0 |
| rs121964984 | pathogenic | 0 | 0 | 0 |
| rs144971200 | benign | 0 | 0 | 0 |
| rs121964985 | pathogenic | 1 | 1 | 0 |
| rs141246107 | benign | 0 | 0 | 0 |
| rs104894561 | pathogenic | 1 | 1 | 0 |
| rs377023736 | pathogenic | 0 | 0 | 0 |
| rs121964907 | pathogenic | 1 | 1 | 1 |
| rs121964909 | pathogenic | 1 | 1 | 1 |
| rs765070743 | pathogenic | 0 | 0 | 0 |
| rs2228119 | benign | 0 | 0 | 0 |
| rs745976989 | pathogenic | 1 | 1 | 1 |
| rs121964904 | pathogenic | 0 | 0 | 0 |
| rs386833430 | pathogenic | 1 | 1 | 1 |
| rs267606992 | pathogenic | 0 | 0 | 0 |
| rs61752063 | pathogenic | 0 | 0 | 0 |
| rs61752067 | pathogenic | 1 | 1 | 1 |
| rs104894934 | pathogenic | 0 | 0 | 0 |
| rs104894929 | pathogenic | 1 | 1 | 1 |
| rs80189640 | benign | 0 | 0 | 0 |
| rs137854547 | pathogenic | 1 | 1 | 1 |
| rs137854540 | pathogenic | 1 | 1 | 1 |
| rs782092363 | pathogenic | 1 | 1 | 1 |
| rs147783915 | benign | 0 | 0 | 0 |
| rs28933687 | pathogenic | 1 | 1 | 1 |
| rs149503319 | benign | 0 | 0 | 0 |
| rs1805147 | benign | 0 | 0 | 0 |
| rs587777218 | pathogenic | 1 | 1 | 1 |
| rs200786705 | benign | 0 | 0 | 0 |
| rs56219475 | benign | 0 | 0 | 0 |
| rs587781172 | benign | 0 | 0 | 0 |
| rs757094189 | pathogenic | 0 | 0 | 0 |
| rs140811086 | benign | 1 | 1 | 1 |
| rs574088829 | pathogenic | 1 | 1 | 1 |
| rs121918525 | pathogenic | 0 | 0 | 0 |
| rs121918527 | pathogenic | 0 | 0 | 0 |
| rs121964848 | pathogenic | 0 | 1 | 0 |
| rs121964845 | pathogenic | 1 | 1 | 1 |
| rs121964846 | pathogenic | 0 | 1 | 0 |
| rs17217716 | benign | 1 | 1 | 1 |
| rs141711342 | benign | 0 | 0 | 0 |
| rs372619120 | benign | 0 | 0 | 0 |
| rs63750002 | benign | 1 | 1 | 1 |

| | | | | |
|---|---|---|---|---|
| rs193922373 | benign | 0 | 0 | 0 |
| rs41295286 | benign | 0 | 0 | 0 |
| rs17217772 | benign | 0 | 0 | 0 |
| rs145649774 | benign | 0 | 0 | 0 |
| rs63750214 | pathogenic | 0 | 0 | 0 |
| rs63750716 | benign | 0 | 0 | 0 |
| rs202026056 | benign | 0 | 0 | 0 |
| rs4987188 | benign | 1 | 1 | 0 |
| rs267607938 | benign | 0 | 0 | 0 |
| rs17224367 | benign | 0 | 0 | 0 |
| rs63750006 | benign | 0 | 0 | 0 |
| rs587781373 | benign | 0 | 0 | 0 |
| rs63751207 | pathogenic | 1 | 1 | 0 |
| rs730881759 | benign | 0 | 0 | 0 |
| rs372350768 | benign | 1 | 1 | 1 |
| rs55778204 | benign | 1 | 1 | 0 |
| rs63749910 | benign | 0 | 0 | 0 |
| rs63750844 | benign | 0 | 0 | 0 |
| rs28929483 | pathogenic | 0 | 0 | 0 |
| rs61756468 | benign | 0 | 0 | 0 |
| rs63750875 | pathogenic | 1 | 1 | 0 |
| rs776528054 | benign | 1 | 1 | 0 |
| rs751939698 | benign | 0 | 0 | 0 |
| rs587780685 | benign | 0 | 0 | 0 |
| rs730881763 | benign | 1 | 1 | 0 |
| rs373226409 | benign | 0 | 0 | 0 |
| rs202145681 | benign | 1 | 1 | 1 |
| rs63749841 | benign | 1 | 1 | 1 |
| rs63750757 | benign | 1 | 1 | 1 |
| rs587778527 | benign | 1 | 1 | 1 |
| rs63750849 | benign | 0 | 0 | 0 |
| rs587779969 | benign | 0 | 0 | 0 |
| rs121908046 | pathogenic | 1 | 1 | 1 |
| rs28940882 | pathogenic | 1 | 1 | 0 |
| rs28940883 | pathogenic | 1 | 1 | 1 |
| rs727503943 | pathogenic | 1 | 1 | 1 |
| rs137853859 | pathogenic | 1 | 1 | 1 |
| rs121908045 | pathogenic | 1 | 1 | 0 |
| rs137853861 | pathogenic | 1 | 1 | 0 |
| rs3180383 | pathogenic | 0 | 1 | 0 |
| rs144431814 | benign | 0 | 0 | 0 |
| rs139502866 | benign | 0 | 0 | 0 |
| rs17524437 | benign | 0 | 0 | 0 |
| rs760960779 | benign | 0 | 0 | 0 |
| rs148960463 | pathogenic | 0 | 0 | 0 |
| rs111033647 | pathogenic | 1 | 1 | 1 |
| rs111033648 | pathogenic | 0 | 0 | 0 |
| rs1800461 | benign | 0 | 0 | 0 |
| rs111033656 | pathogenic | 0 | 0 | 0 |
| rs111033663 | pathogenic | 1 | 1 | 0 |
| rs111033669 | pathogenic | 1 | 1 | 1 |
| rs111033690 | pathogenic | 0 | 0 | 0 |
| rs111033686 | pathogenic | 0 | 0 | 0 |
| rs111033687 | pathogenic | 0 | 0 | 0 |

| | | | | |
|---|---|---|---|---|
| rs111033695 | pathogenic | 0 | 0 | 0 |
| rs111033697 | pathogenic | 0 | 0 | 0 |
| rs111033693 | pathogenic | 0 | 0 | 0 |
| rs111033694 | pathogenic | 0 | 0 | 0 |
| rs111033721 | pathogenic | 1 | 1 | 1 |
| rs111033722 | pathogenic | 0 | 0 | 0 |
| rs111033725 | pathogenic | 1 | 1 | 0 |
| rs75391579 | pathogenic | 1 | 1 | 1 |
| rs111033728 | pathogenic | 1 | 1 | 1 |
| rs111033735 | pathogenic | 1 | 1 | 1 |
| rs111033739 | pathogenic | 1 | 1 | 1 |
| rs111033736 | pathogenic | 0 | 0 | 0 |
| rs111033744 | pathogenic | 1 | 1 | 1 |
| rs111033750 | pathogenic | 1 | 1 | 1 |
| rs111033752 | pathogenic | 0 | 0 | 0 |
| rs111033751 | pathogenic | 0 | 0 | 0 |
| rs111033846 | pathogenic | 0 | 0 | 0 |
| rs368166217 | pathogenic | 0 | 0 | 0 |
| rs111033764 | pathogenic | 0 | 0 | 0 |
| rs111033847 | benign | 0 | 0 | 0 |
| rs111033778 | pathogenic | 0 | 0 | 0 |
| rs111033773 | pathogenic | 0 | 0 | 0 |
| rs111033792 | pathogenic | 0 | 0 | 0 |
| rs367543266 | pathogenic | 1 | 1 | 1 |
| rs111033794 | pathogenic | 0 | 0 | 0 |
| rs111033802 | pathogenic | 0 | 0 | 0 |
| rs111033803 | pathogenic | 0 | 0 | 0 |
| rs111033809 | pathogenic | 1 | 1 | 0 |
| rs121918376 | pathogenic | 0 | 1 | 0 |
| rs121918377 | pathogenic | 0 | 1 | 0 |
| rs121918378 | pathogenic | 1 | 1 | 1 |
| rs121918374 | pathogenic | 1 | 1 | 1 |
| rs137852379 | pathogenic | 0 | 0 | 0 |
| rs137852388 | pathogenic | 1 | 1 | 1 |
| rs137852368 | pathogenic | 0 | 0 | 0 |
| rs28935499 | pathogenic | 0 | 0 | 0 |
| rs28933668 | pathogenic | 0 | 0 | 0 |
| rs28933669 | pathogenic | 0 | 0 | 0 |
| rs28933670 | pathogenic | 0 | 0 | 0 |
| rs28933671 | pathogenic | 0 | 0 | 0 |
| rs28933672 | pathogenic | 0 | 0 | 0 |
| rs28937272 | pathogenic | 0 | 0 | 0 |
| rs35383156 | benign | 0 | 0 | 0 |
| rs137852418 | pathogenic | 0 | 0 | 0 |
| rs137852419 | pathogenic | 0 | 0 | 0 |
| rs137852428 | pathogenic | 0 | 0 | 0 |
| rs137852436 | pathogenic | 0 | 0 | 0 |
| rs28937285 | pathogenic | 0 | 0 | 0 |
| rs35213384 | benign | 1 | 1 | 1 |
| rs3810950 | benign | 0 | 0 | 0 |
| rs201479289 | pathogenic | 0 | 0 | 0 |
| rs121912820 | pathogenic | 0 | 1 | 0 |
| rs121912815 | pathogenic | 0 | 1 | 0 |
| rs115510708 | benign | 0 | 0 | 0 |

| | | | | |
|---|---|---|---|---|
| rs868749 | benign | 0 | 0 | 0 |
| rs121912823 | pathogenic | 0 | 0 | 0 |
| rs115212829 | benign | 0 | 0 | 0 |
| rs121912822 | pathogenic | 0 | 0 | 0 |
| rs121912816 | pathogenic | 0 | 1 | 0 |
| rs4838544 | benign | 0 | 0 | 0 |
| rs121912818 | pathogenic | 0 | 0 | 0 |
| rs121912821 | pathogenic | 0 | 0 | 0 |
| rs121912817 | pathogenic | 0 | 0 | 0 |
| rs8178992 | benign | 0 | 0 | 0 |
| rs121912819 | pathogenic | 1 | 1 | 0 |
| rs80097077 | benign | 1 | 1 | 0 |
| rs121909359 | pathogenic | 1 | 1 | 0 |
| rs121909358 | pathogenic | 1 | 1 | 1 |
| rs121909361 | pathogenic | 1 | 1 | 1 |
| rs121909357 | pathogenic | 0 | 1 | 0 |
| rs121909365 | pathogenic | 1 | 1 | 1 |
| rs6413484 | pathogenic | 1 | 1 | 1 |
| rs121909373 | pathogenic | 1 | 1 | 1 |
| rs121909366 | pathogenic | 1 | 1 | 1 |
| rs121909367 | pathogenic | 1 | 1 | 1 |
| rs121909368 | pathogenic | 1 | 1 | 1 |
| rs121909369 | pathogenic | 0 | 0 | 0 |
| rs121909363 | pathogenic | 1 | 1 | 1 |
| rs45588036 | pathogenic | 1 | 1 | 1 |
| rs121909364 | pathogenic | 1 | 1 | 1 |
| rs80358282 | benign | 0 | 0 | 0 |
| rs80358284 | pathogenic | 1 | 1 | 1 |
| rs61735303 | benign | 0 | 0 | 0 |
| rs2229274 | benign | 0 | 0 | 0 |
| rs145006491 | benign | 0 | 0 | 0 |
| rs121913580 | pathogenic | 1 | 1 | 0 |
| rs730882196 | pathogenic | 0 | 0 | 0 |
| rs153477 | benign | 0 | 0 | 0 |
| rs153478 | benign | 1 | 1 | 0 |
| rs61740602 | benign | 1 | 1 | 0 |
| rs104893892 | pathogenic | 1 | 1 | 0 |
| rs121909595 | pathogenic | 1 | 1 | 1 |
| rs28931605 | pathogenic | 1 | 1 | 1 |
| rs121909597 | pathogenic | 0 | 1 | 0 |
| rs202233735 | pathogenic | 0 | 1 | 0 |
| rs121909596 | pathogenic | 1 | 1 | 1 |
| rs75156162 | benign | 0 | 1 | 0 |
| rs2305430 | benign | 1 | 1 | 1 |
| rs150318966 | benign | 0 | 0 | 0 |
| rs121909598 | pathogenic | 0 | 1 | 0 |
| rs5744175 | benign | 0 | 0 | 0 |
| rs28935476 | pathogenic | 1 | 1 | 1 |
| rs28935475 | pathogenic | 1 | 1 | 1 |
| rs122461163 | pathogenic | 1 | 1 | 1 |
| rs62626305 | pathogenic | 1 | 1 | 1 |
| rs587777157 | pathogenic | 0 | 0 | 0 |
| rs137854478 | pathogenic | 1 | 1 | 0 |
| rs137854465 | pathogenic | 0 | 0 | 0 |

| | | | | |
|---|---|---|---|---|
| rs137854470 | pathogenic | 0 | 0 | 0 |
| rs137854482 | pathogenic | 0 | 0 | 0 |
| rs137854473 | pathogenic | 0 | 0 | 0 |
| rs137854456 | pathogenic | 0 | 0 | 0 |
| rs140598 | benign | 0 | 0 | 0 |
| rs104894105 | pathogenic | 1 | 1 | 1 |
| rs104894107 | pathogenic | 1 | 1 | 1 |
| rs104894106 | pathogenic | 1 | 1 | 1 |
| rs74621026 | benign | 1 | 1 | 1 |
| rs2290332 | benign | 0 | 0 | 0 |
| rs780202604 | pathogenic | 1 | 1 | 1 |
| rs74315305 | pathogenic | 1 | 1 | 0 |
| rs766751878 | pathogenic | 1 | 1 | 1 |
| rs137854603 | pathogenic | 1 | 1 | 1 |
| rs139592595 | benign | 1 | 1 | 0 |
| rs150565592 | benign | 1 | 1 | 1 |
| rs763520126 | benign | 0 | 0 | 0 |
| rs55980502 | benign | 0 | 0 | 0 |
| rs137852812 | pathogenic | 0 | 0 | 0 |
| rs137852813 | pathogenic | 0 | 0 | 0 |
| rs190222208 | benign | 0 | 0 | 0 |
| rs138459502 | pathogenic | 0 | 0 | 0 |
| rs730881041 | pathogenic | 0 | 0 | 0 |
| rs267607080 | pathogenic | 0 | 0 | 0 |
| rs137852814 | pathogenic | 0 | 0 | 0 |
| rs267607079 | pathogenic | 0 | 0 | 0 |
| rs192378417 | benign | 0 | 0 | 0 |
| rs1288775 | benign | 0 | 0 | 0 |
| rs80338737 | pathogenic | 1 | 1 | 1 |
| rs397514708 | pathogenic | 0 | 0 | 0 |
| rs775933965 | pathogenic | 0 | 0 | 0 |
| rs747608698 | benign | 0 | 1 | 0 |
| rs1145086 | benign | 0 | 1 | 0 |
| rs121912953 | pathogenic | 1 | 1 | 0 |
| rs243849 | benign | 0 | 0 | 0 |
| rs121912955 | pathogenic | 1 | 1 | 1 |
| rs17859943 | benign | 0 | 1 | 0 |
| rs755557988 | benign | 1 | 1 | 1 |
| rs10195 | benign | 0 | 0 | 0 |
| rs61735357 | benign | 1 | 1 | 1 |
| rs137882374 | benign | 0 | 0 | 0 |
| rs5388 | benign | 0 | 0 | 0 |
| rs6352 | benign | 0 | 0 | 0 |
| rs76645461 | pathogenic | 1 | 1 | 0 |
| rs1050828 | pathogenic | 0 | 0 | 0 |
| rs1050829 | pathogenic | 0 | 0 | 0 |
| rs137852314 | pathogenic | 0 | 0 | 0 |
| rs137852331 | pathogenic | 0 | 0 | 0 |
| rs5030872 | pathogenic | 0 | 0 | 0 |
| rs267606836 | pathogenic | 0 | 0 | 0 |
| rs5030868 | pathogenic | 0 | 0 | 0 |
| rs137852330 | pathogenic | 0 | 1 | 0 |
| rs137852326 | pathogenic | 1 | 1 | 1 |
| rs137852318 | pathogenic | 1 | 1 | 1 |

| | | | | |
|---|---|---|---|---|
| rs74575103 | benign | 0 | 0 | 0 |
| rs137852327 | pathogenic | 1 | 1 | 1 |
| rs76723693 | pathogenic | 0 | 0 | 0 |
| rs5030869 | pathogenic | 0 | 0 | 0 |
| rs34193178 | pathogenic | 1 | 1 | 1 |
| rs137852316 | pathogenic | 1 | 1 | 0 |
| rs398123546 | pathogenic | 1 | 1 | 1 |
| rs72554665 | pathogenic | 0 | 0 | 0 |
| rs72554664 | pathogenic | 0 | 0 | 0 |
| rs28939068 | pathogenic | 1 | 1 | 1 |
| rs59301204 | pathogenic | 1 | 1 | 1 |
| rs143715750 | benign | 1 | 1 | 1 |
| rs35263902 | pathogenic | 1 | 1 | 1 |
| rs202214518 | benign | 1 | 1 | 1 |
| rs34076756 | benign | 0 | 0 | 0 |
| rs199592341 | pathogenic | 1 | 1 | 0 |
| rs121908833 | pathogenic | 1 | 1 | 1 |
| rs121908834 | pathogenic | 1 | 1 | 0 |
| rs138454151 | benign | 0 | 0 | 0 |
| rs141323599 | benign | 0 | 0 | 0 |
| rs137853200 | pathogenic | 0 | 0 | 0 |
| rs137853199 | pathogenic | 0 | 0 | 0 |
| rs139787163 | benign | 0 | 0 | 0 |
| rs61731097 | benign | 1 | 1 | 1 |
| rs61729795 | benign | 0 | 0 | 0 |
| rs149242330 | benign | 0 | 0 | 0 |
| rs587777490 | pathogenic | 1 | 1 | 1 |
| rs398123011 | pathogenic | 1 | 1 | 0 |
| rs775186117 | pathogenic | 0 | 0 | 0 |
| rs72677237 | benign | 0 | 0 | 0 |
| rs104894163 | pathogenic | 1 | 1 | 1 |
| rs121909588 | pathogenic | 0 | 0 | 0 |
| rs1042579 | benign | 0 | 0 | 0 |
| rs121909273 | pathogenic | 0 | 0 | 0 |
| rs121909274 | pathogenic | 0 | 0 | 0 |
| rs119473034 | pathogenic | 1 | 1 | 1 |
| rs398123065 | pathogenic | 0 | 0 | 0 |
| rs587777042 | pathogenic | 1 | 1 | 0 |
| rs761413864 | pathogenic | 1 | 1 | 1 |
| rs118203965 | pathogenic | 0 | 0 | 0 |
| rs118203966 | pathogenic | 0 | 0 | 0 |
| rs1801201 | benign | 1 | 1 | 0 |
| rs1136201 | benign | 1 | 1 | 1 |
| rs144148008 | benign | 0 | 0 | 0 |
| rs200118378 | benign | 1 | 1 | 1 |
| rs112984118 | benign | 1 | 1 | 1 |
| rs149446102 | benign | 1 | 1 | 1 |
| rs1427463 | benign | 0 | 0 | 0 |
| rs201936720 | benign | 1 | 1 | 1 |
| rs61751983 | benign | 1 | 1 | 1 |
| rs104894632 | pathogenic | 1 | 1 | 1 |
| rs149202905 | benign | 0 | 0 | 0 |
| rs2230210 | benign | 1 | 1 | 1 |
| rs111631988 | benign | 0 | 0 | 0 |

| | | | | |
|---|---|---|---|---|
| rs137852550 | pathogenic | 0 | 0 | 0 |
| rs5963403 | benign | 0 | 0 | 0 |
| rs138018739 | benign | 0 | 0 | 0 |
| rs62640590 | benign | 0 | 0 | 0 |
| rs121909581 | pathogenic | 1 | 1 | 1 |
| rs202017913 | benign | 1 | 1 | 1 |
| rs143163513 | benign | 0 | 0 | 0 |
| rs238238 | benign | 0 | 1 | 0 |
| rs143945974 | benign | 0 | 0 | 0 |
| rs238239 | benign | 1 | 1 | 1 |
| rs121918403 | pathogenic | 0 | 1 | 0 |
| rs61735456 | benign | 0 | 1 | 0 |
| rs121918404 | pathogenic | 1 | 1 | 0 |
| rs12674488 | benign | 0 | 0 | 0 |
| rs201599657 | pathogenic | 0 | 0 | 0 |
| rs1057090 | benign | 0 | 0 | 0 |
| rs1057091 | benign | 0 | 0 | 0 |
| rs587783740 | benign | 0 | 0 | 0 |
| rs75956769 | benign | 0 | 0 | 0 |
| rs201691299 | benign | 0 | 0 | 0 |
| rs62621087 | benign | 0 | 0 | 0 |
| rs201308454 | benign | 0 | 0 | 0 |
| rs145657360 | benign | 0 | 0 | 0 |
| rs121434592 | pathogenic | 1 | 1 | 0 |
| rs375650413 | benign | 0 | 0 | 0 |
| rs121434437 | pathogenic | 1 | 1 | 0 |
| rs763494292 | pathogenic | 0 | 0 | 0 |
| rs121964997 | pathogenic | 1 | 1 | 0 |
| rs121964996 | pathogenic | 0 | 0 | 0 |
| rs41283425 | benign | 1 | 1 | 1 |
| rs376289528 | benign | 1 | 1 | 1 |
| rs144171604 | benign | 1 | 1 | 1 |
| rs782440692 | pathogenic | 1 | 1 | 1 |
| rs782058451 | pathogenic | 1 | 1 | 1 |
| rs78437817 | benign | 0 | 0 | 0 |
| rs200327969 | benign | 1 | 1 | 1 |
| rs143043662 | benign | 0 | 0 | 0 |
| rs111966833 | benign | 1 | 1 | 1 |
| rs35523678 | benign | 0 | 0 | 0 |
| rs515726208 | pathogenic | 0 | 0 | 0 |
| rs397517239 | pathogenic | 1 | 1 | 1 |
| rs148966602 | benign | 1 | 1 | 1 |
| rs76371972 | benign | 1 | 1 | 1 |
| rs2229968 | benign | 1 | 1 | 1 |
| rs61751538 | benign | 1 | 1 | 1 |
| rs200800978 | pathogenic | 1 | 1 | 1 |
| rs148695964 | benign | 1 | 1 | 1 |
| rs140042418 | benign | 1 | 1 | 1 |
| rs116185403 | benign | 0 | 0 | 0 |
| rs142828669 | benign | 1 | 1 | 1 |
| rs139832048 | benign | 0 | 1 | 0 |
| rs58750568 | benign | 0 | 1 | 0 |
| rs77681311 | benign | 0 | 0 | 0 |
| rs151208452 | benign | 0 | 0 | 0 |

| | | | | |
|---|---|---|---|---|
| rs76463933 | benign | 0 | 0 | 0 |
| rs144434647 | benign | 0 | 0 | 0 |
| rs117361485 | benign | 0 | 0 | 0 |
| rs587777588 | pathogenic | 0 | 0 | 0 |
| rs80338796 | pathogenic | 0 | 0 | 0 |
| rs121434594 | pathogenic | 0 | 0 | 0 |
| rs146766138 | pathogenic | 1 | 1 | 1 |
| rs28939679 | pathogenic | 1 | 1 | 1 |
| rs748754134 | pathogenic | 1 | 1 | 0 |
| rs764276946 | pathogenic | 1 | 1 | 1 |
| rs111033588 | pathogenic | 1 | 1 | 1 |
| rs201633414 | pathogenic | 1 | 1 | 1 |
| rs121909262 | pathogenic | 1 | 1 | 1 |
| rs104893702 | pathogenic | 1 | 1 | 1 |
| rs121909265 | pathogenic | 1 | 1 | 1 |
| rs121909266 | pathogenic | 1 | 1 | 1 |
| rs193922439 | pathogenic | 1 | 1 | 0 |
| rs104893691 | pathogenic | 1 | 1 | 1 |
| rs104893695 | pathogenic | 0 | 0 | 0 |
| rs104893708 | pathogenic | 1 | 1 | 1 |
| rs121909260 | pathogenic | 0 | 0 | 0 |
| rs104893696 | pathogenic | 0 | 0 | 0 |
| rs121909267 | pathogenic | 0 | 0 | 0 |
| rs121909263 | pathogenic | 0 | 0 | 0 |
| rs121909264 | pathogenic | 0 | 0 | 0 |
| rs104893694 | pathogenic | 1 | 1 | 1 |
| rs121909268 | pathogenic | 1 | 1 | 1 |
| rs104893689 | pathogenic | 1 | 1 | 0 |
| rs104893707 | pathogenic | 1 | 1 | 0 |
| rs104893697 | pathogenic | 1 | 1 | 1 |
| rs28936684 | pathogenic | 1 | 1 | 1 |
| rs62269092 | benign | 1 | 1 | 0 |
| rs121909259 | pathogenic | 1 | 1 | 0 |
| rs548403340 | benign | 1 | 1 | 0 |
| rs142818334 | benign | 0 | 0 | 0 |
| rs12493789 | benign | 1 | 1 | 1 |
| rs193922423 | pathogenic | 1 | 1 | 0 |
| rs104893719 | pathogenic | 1 | 1 | 1 |
| rs117375173 | benign | 1 | 1 | 0 |
| rs104893712 | pathogenic | 0 | 0 | 0 |
| rs121917776 | pathogenic | 1 | 1 | 1 |
| rs767325003 | benign | 1 | 1 | 1 |
| rs35824453 | benign | 0 | 0 | 0 |
| rs141641266 | benign | 0 | 0 | 0 |
| rs1050239 | benign | 0 | 0 | 0 |
| rs141303384 | benign | 0 | 0 | 0 |
| rs12720458 | pathogenic | 0 | 0 | 0 |
| rs368507376 | pathogenic | 1 | 1 | 1 |
| rs17215500 | pathogenic | 1 | 1 | 1 |
| rs120074188 | pathogenic | 1 | 1 | 1 |
| rs397508097 | pathogenic | 1 | 1 | 1 |
| rs138551008 | pathogenic | 0 | 0 | 0 |
| rs387907139 | pathogenic | 1 | 1 | 1 |
| rs765487627 | pathogenic | 0 | 0 | 0 |

| | | | | |
|---|---|---|---|---|
| rs587778587 | benign | 0 | 0 | 0 |
| rs45568339 | benign | 0 | 0 | 0 |
| rs78179744 | benign | 0 | 0 | 0 |
| rs45624036 | benign | 0 | 0 | 0 |
| rs61755166 | benign | 0 | 0 | 0 |
| rs180177132 | pathogenic | 1 | 1 | 1 |
| rs587776527 | pathogenic | 0 | 0 | 0 |
| rs373783514 | benign | 0 | 0 | 0 |
| rs730881896 | benign | 0 | 0 | 0 |
| rs118203998 | pathogenic | 0 | 0 | 0 |
| rs41281039 | pathogenic | 1 | 1 | 1 |
| rs118204430 | pathogenic | 1 | 1 | 1 |
| rs1800546 | pathogenic | 1 | 1 | 0 |
| rs76917243 | pathogenic | 0 | 1 | 0 |
| rs78340951 | pathogenic | 0 | 1 | 0 |
| rs77718928 | pathogenic | 0 | 0 | 0 |
| rs369586696 | pathogenic | 0 | 1 | 0 |
| rs745517517 | pathogenic | 1 | 1 | 0 |
| rs372872777 | pathogenic | 0 | 0 | 0 |
| rs748704459 | pathogenic | 0 | 0 | 0 |
| rs769215411 | pathogenic | 0 | 0 | 0 |
| rs139425890 | pathogenic | 0 | 0 | 0 |
| rs779694939 | pathogenic | 1 | 1 | 0 |
| rs773729410 | pathogenic | 0 | 0 | 0 |
| rs121908623 | pathogenic | 1 | 1 | 1 |
| rs121908622 | pathogenic | 1 | 1 | 1 |
| rs121908621 | pathogenic | 1 | 1 | 1 |
| rs762106720 | pathogenic | 1 | 1 | 0 |
| rs199877522 | pathogenic | 0 | 0 | 0 |
| rs74315291 | pathogenic | 1 | 1 | 1 |
| rs79137415 | benign | 0 | 0 | 0 |
| rs121913045 | pathogenic | 0 | 0 | 0 |
| rs34295337 | pathogenic | 0 | 0 | 0 |
| rs121913046 | pathogenic | 0 | 0 | 0 |
| rs121913047 | pathogenic | 0 | 0 | 0 |
| rs121913048 | pathogenic | 0 | 0 | 0 |
| rs267607166 | pathogenic | 0 | 0 | 0 |
| rs17850684 | benign | 1 | 1 | 1 |
| rs119476051 | pathogenic | 1 | 1 | 1 |
| rs17850683 | benign | 1 | 1 | 1 |
| rs119476049 | pathogenic | 1 | 1 | 1 |
| rs119476046 | pathogenic | 1 | 1 | 1 |
| rs119476048 | pathogenic | 0 | 0 | 0 |
| rs119476047 | pathogenic | 1 | 1 | 1 |
| rs28939094 | pathogenic | 1 | 1 | 1 |
| rs119476050 | pathogenic | 0 | 0 | 0 |
| rs387906941 | pathogenic | 1 | 1 | 1 |
| rs41511344 | pathogenic | 1 | 1 | 0 |
| rs748112833 | pathogenic | 1 | 1 | 1 |
| rs74342080 | benign | 1 | 1 | 0 |
| rs116482870 | benign | 0 | 0 | 0 |
| rs104893743 | pathogenic | 0 | 0 | 0 |
| rs117179004 | benign | 0 | 0 | 0 |
| rs587779409 | pathogenic | 1 | 1 | 1 |

| | | | | |
|---|---|---|---|---|
| rs3745406 | benign | 1 | 1 | 0 |
| rs387906696 | pathogenic | 0 | 0 | 0 |
| rs72549369 | pathogenic | 1 | 1 | 0 |
| rs45523133 | benign | 0 | 0 | 0 |
| rs119460972 | pathogenic | 0 | 1 | 0 |
| rs34929837 | benign | 0 | 1 | 0 |
| rs17323225 | benign | 0 | 1 | 0 |
| rs17011368 | benign | 0 | 0 | 0 |
| rs748902766 | pathogenic | 1 | 1 | 1 |
| rs387907292 | pathogenic | 0 | 0 | 0 |
| rs387907293 | pathogenic | 1 | 1 | 0 |
| rs371526758 | pathogenic | 1 | 1 | 1 |
| rs142968179 | pathogenic | 1 | 1 | 1 |
| rs387907291 | pathogenic | 1 | 1 | 0 |
| rs121907949 | pathogenic | 0 | 0 | 0 |
| rs121907947 | pathogenic | 0 | 0 | 0 |
| rs121907950 | pathogenic | 1 | 1 | 1 |
| rs121907948 | pathogenic | 1 | 1 | 1 |
| rs4926 | benign | 0 | 0 | 0 |
| rs748809996 | pathogenic | 0 | 0 | 0 |
| rs1799883 | benign | 1 | 1 | 0 |
| rs104893974 | pathogenic | 0 | 1 | 0 |
| rs104893973 | pathogenic | 0 | 1 | 0 |
| rs34452707 | benign | 0 | 0 | 0 |
| rs119467003 | pathogenic | 1 | 1 | 1 |
| rs121908261 | pathogenic | 0 | 1 | 0 |
| rs28933985 | pathogenic | 1 | 1 | 1 |
| rs80356669 | pathogenic | 1 | 1 | 1 |
| rs28936677 | pathogenic | 1 | 1 | 0 |
| rs121918326 | pathogenic | 0 | 0 | 0 |
| rs121918324 | pathogenic | 1 | 1 | 0 |
| rs121918325 | pathogenic | 1 | 1 | 0 |
| rs41270025 | pathogenic | 1 | 1 | 0 |
| rs121918323 | pathogenic | 0 | 0 | 0 |
| rs12735723 | pathogenic | 1 | 1 | 0 |
| rs36013429 | benign | 0 | 0 | 0 |
| rs28936676 | pathogenic | 1 | 1 | 0 |
| rs771809901 | pathogenic | 1 | 1 | 1 |
| rs758350837 | benign | 1 | 1 | 0 |
| rs121908192 | pathogenic | 1 | 1 | 1 |
| rs147568500 | pathogenic | 1 | 1 | 1 |
| rs3729734 | benign | 0 | 0 | 0 |
| rs371536347 | benign | 1 | 1 | 1 |
| rs80358193 | pathogenic | 1 | 1 | 1 |
| rs119476044 | pathogenic | 1 | 1 | 1 |
| rs111292798 | benign | 1 | 1 | 1 |
| rs121908063 | pathogenic | 1 | 1 | 0 |
| rs121908065 | pathogenic | 1 | 1 | 1 |
| rs121908062 | pathogenic | 1 | 1 | 1 |
| rs121908064 | pathogenic | 1 | 1 | 1 |
| rs61746421 | benign | 1 | 1 | 1 |
| rs34886916 | benign | 1 | 1 | 1 |
| rs118204045 | pathogenic | 0 | 0 | 0 |
| rs138607412 | benign | 1 | 1 | 1 |

| | | | | |
|---|---|---|---|---|
| rs118204046 | pathogenic | 0 | 0 | 0 |
| rs118204048 | pathogenic | 0 | 0 | 0 |
| rs118204044 | pathogenic | 0 | 0 | 0 |
| rs149664056 | benign | 0 | 0 | 0 |
| rs121909285 | pathogenic | 0 | 0 | 0 |
| rs1802498 | benign | 0 | 0 | 0 |
| rs1065761 | benign | 0 | 0 | 0 |
| rs119477054 | pathogenic | 0 | 0 | 0 |
| rs12720097 | benign | 0 | 0 | 0 |
| rs747281324 | pathogenic | 1 | 1 | 0 |
| rs367543028 | pathogenic | 1 | 1 | 0 |
| rs146077918 | benign | 0 | 0 | 0 |
| rs142551229 | benign | 0 | 0 | 0 |
| rs367543039 | pathogenic | 0 | 0 | 0 |
| rs587779884 | pathogenic | 1 | 1 | 0 |
| rs137853153 | pathogenic | 0 | 0 | 0 |
| rs367543029 | pathogenic | 0 | 0 | 0 |
| rs587783037 | pathogenic | 1 | 1 | 0 |
| rs142928725 | benign | 0 | 0 | 0 |
| rs28385141 | benign | 0 | 0 | 0 |
| rs587779888 | pathogenic | 0 | 0 | 0 |
| rs121909801 | pathogenic | 1 | 1 | 1 |
| rs121909790 | pathogenic | 1 | 1 | 0 |
| rs121909797 | pathogenic | 1 | 1 | 0 |
| rs121909794 | pathogenic | 1 | 1 | 1 |
| rs121909791 | pathogenic | 1 | 1 | 1 |
| rs121909793 | pathogenic | 1 | 1 | 1 |
| rs147977279 | benign | 1 | 1 | 1 |
| rs147352564 | benign | 0 | 0 | 0 |
| rs368758045 | benign | 1 | 1 | 1 |
| rs114498261 | benign | 1 | 1 | 0 |
| rs35484156 | benign | 0 | 0 | 0 |
| rs139176878 | benign | 0 | 0 | 0 |
| rs11543934 | benign | 0 | 0 | 0 |
| rs34192549 | benign | 0 | 0 | 0 |
| rs568736295 | benign | 0 | 0 | 0 |
| rs55747232 | benign | 0 | 0 | 0 |
| rs781887272 | benign | 0 | 0 | 0 |
| rs146273341 | benign | 0 | 0 | 0 |
| rs41473544 | benign | 0 | 0 | 0 |
| rs137852680 | pathogenic | 0 | 0 | 0 |
| rs55891000 | benign | 0 | 0 | 0 |
| rs140351189 | benign | 0 | 0 | 0 |
| rs137852679 | pathogenic | 1 | 1 | 0 |
| rs137852678 | pathogenic | 0 | 0 | 0 |
| rs137852677 | pathogenic | 1 | 1 | 1 |
| rs138620818 | benign | 0 | 0 | 0 |
| rs121908531 | pathogenic | 0 | 0 | 0 |
| rs72554307 | pathogenic | 0 | 0 | 0 |
| rs72554308 | pathogenic | 0 | 0 | 0 |
| rs72554312 | pathogenic | 1 | 1 | 0 |
| rs67939655 | pathogenic | 0 | 0 | 0 |
| rs67486158 | pathogenic | 0 | 0 | 0 |
| rs72554338 | pathogenic | 1 | 1 | 1 |

| | | | | |
|---|---|---|---|---|
| rs1800324 | pathogenic | 1 | 1 | 1 |
| rs72554356 | pathogenic | 1 | 1 | 1 |
| rs66656800 | pathogenic | 1 | 1 | 1 |
| rs68026851 | pathogenic | 1 | 1 | 1 |
| rs72558412 | pathogenic | 0 | 0 | 0 |
| rs72558415 | pathogenic | 0 | 0 | 0 |
| rs72558423 | pathogenic | 0 | 0 | 0 |
| rs67120076 | pathogenic | 1 | 1 | 1 |
| rs1800328 | pathogenic | 1 | 1 | 1 |
| rs72558451 | pathogenic | 1 | 1 | 1 |
| rs72558454 | pathogenic | 1 | 1 | 1 |
| rs672601372 | pathogenic | 0 | 0 | 0 |
| rs139644798 | pathogenic | 0 | 0 | 0 |
| rs369398935 | pathogenic | 0 | 0 | 0 |
| rs267607035 | pathogenic | 0 | 0 | 0 |
| rs267607036 | pathogenic | 1 | 1 | 1 |
| rs2302566 | benign | 1 | 1 | 0 |
| rs115959591 | benign | 0 | 0 | 0 |
| rs773876739 | pathogenic | 0 | 0 | 0 |
| rs1799967 | benign | 0 | 0 | 0 |
| rs80356968 | benign | 0 | 0 | 0 |
| rs28897695 | benign | 1 | 1 | 1 |
| rs80357314 | benign | 0 | 0 | 0 |
| rs80357169 | benign | 0 | 0 | 0 |
| rs397507239 | benign | 1 | 1 | 1 |
| rs80187739 | pathogenic | 1 | 1 | 1 |
| rs41293459 | pathogenic | 1 | 1 | 1 |
| rs55770810 | pathogenic | 1 | 1 | 1 |
| rs80356860 | pathogenic | 0 | 0 | 0 |
| rs56195342 | benign | 0 | 0 | 0 |
| rs80357442 | benign | 0 | 0 | 0 |
| rs41293463 | pathogenic | 1 | 1 | 1 |
| rs80357069 | pathogenic | 0 | 0 | 0 |
| rs397509275 | benign | 0 | 0 | 0 |
| rs80356920 | benign | 0 | 0 | 0 |
| rs397509284 | pathogenic | 0 | 0 | 0 |
| rs80357286 | benign | 0 | 0 | 0 |
| rs80357274 | benign | 0 | 0 | 0 |
| rs765656957 | benign | 0 | 0 | 0 |
| rs755427809 | benign | 0 | 0 | 0 |
| rs80357322 | benign | 0 | 0 | 0 |
| rs121434430 | pathogenic | 0 | 0 | 0 |
| rs121918450 | pathogenic | 1 | 1 | 1 |
| rs1532268 | benign | 0 | 0 | 0 |
| rs2303080 | benign | 0 | 0 | 0 |
| rs10064631 | benign | 0 | 0 | 0 |
| rs162036 | benign | 0 | 0 | 0 |
| rs2287780 | benign | 0 | 0 | 0 |
| rs16879334 | benign | 0 | 0 | 0 |
| rs137853061 | pathogenic | 1 | 1 | 0 |
| rs10380 | benign | 0 | 0 | 0 |
| rs1494558 | benign | 0 | 0 | 0 |
| rs1494555 | benign | 0 | 0 | 0 |
| rs192994060 | benign | 0 | 0 | 0 |

| | | | | |
|---|---|---|---|---|
| rs147128920 | benign | 1 | 1 | 0 |
| rs201803986 | pathogenic | 0 | 0 | 0 |
| rs746419489 | pathogenic | 0 | 1 | 0 |
| rs764396564 | pathogenic | 0 | 0 | 0 |
| rs758304537 | pathogenic | 0 | 0 | 0 |
| rs267606763 | pathogenic | 0 | 0 | 0 |
| rs267606762 | pathogenic | 0 | 0 | 0 |
| rs150702945 | pathogenic | 1 | 1 | 1 |
| rs202047589 | pathogenic | 0 | 0 | 0 |
| rs755562733 | pathogenic | 0 | 0 | 0 |
| rs267606764 | pathogenic | 1 | 1 | 1 |
| rs138207257 | pathogenic | 1 | 1 | 0 |
| rs149150736 | pathogenic | 1 | 1 | 1 |
| rs777046879 | pathogenic | 0 | 0 | 0 |
| rs104894408 | pathogenic | 1 | 1 | 1 |
| rs1801002 | pathogenic | 1 | 1 | 1 |
| rs28929485 | pathogenic | 1 | 1 | 1 |
| rs104894396 | pathogenic | 1 | 1 | 1 |
| rs371024165 | pathogenic | 0 | 0 | 0 |
| rs35887622 | pathogenic | 1 | 1 | 1 |
| rs587783644 | pathogenic | 1 | 1 | 1 |
| rs72474224 | pathogenic | 1 | 1 | 1 |
| rs111033296 | pathogenic | 1 | 1 | 1 |
| rs104894407 | pathogenic | 1 | 1 | 1 |
| rs72561723 | pathogenic | 1 | 1 | 1 |
| rs104894398 | pathogenic | 1 | 1 | 1 |
| rs28931594 | pathogenic | 1 | 1 | 1 |
| rs104894412 | pathogenic | 1 | 1 | 1 |
| rs111033297 | pathogenic | 1 | 1 | 1 |
| rs104894404 | pathogenic | 1 | 1 | 1 |
| rs104894410 | pathogenic | 1 | 1 | 1 |
| rs104894403 | pathogenic | 1 | 1 | 1 |
| rs397516870 | benign | 0 | 0 | 0 |
| rs200023879 | pathogenic | 1 | 1 | 1 |
| rs121912968 | pathogenic | 0 | 0 | 0 |
| rs104894402 | pathogenic | 1 | 1 | 1 |
| rs28931593 | pathogenic | 1 | 1 | 1 |
| rs104894397 | pathogenic | 0 | 0 | 0 |
| rs80338944 | pathogenic | 0 | 0 | 0 |
| rs199883710 | pathogenic | 0 | 0 | 0 |
| rs727504302 | pathogenic | 0 | 0 | 0 |
| rs145216882 | benign | 1 | 1 | 1 |
| rs781534323 | pathogenic | 1 | 1 | 1 |
| rs111033218 | benign | 1 | 1 | 1 |
| rs104894409 | pathogenic | 0 | 0 | 0 |
| rs80338945 | pathogenic | 1 | 1 | 1 |
| rs111033299 | pathogenic | 0 | 0 | 0 |
| rs143343083 | pathogenic | 1 | 1 | 1 |
| rs80338946 | pathogenic | 0 | 0 | 0 |
| rs150529554 | pathogenic | 0 | 0 | 0 |
| rs111033295 | pathogenic | 0 | 0 | 0 |
| rs111033188 | benign | 0 | 0 | 0 |
| rs397516874 | pathogenic | 0 | 0 | 0 |
| rs76434661 | pathogenic | 0 | 0 | 0 |

| | | | | |
|---|---|---|---|---|
| rs80338948 | pathogenic | 0 | 0 | 0 |
| rs104894401 | pathogenic | 0 | 0 | 0 |
| rs767178508 | pathogenic | 0 | 0 | 0 |
| rs772264564 | pathogenic | 0 | 0 | 0 |
| rs28931592 | pathogenic | 0 | 0 | 0 |
| rs80338949 | pathogenic | 1 | 1 | 1 |
| rs376898963 | benign | 1 | 1 | 1 |
| rs111033360 | benign | 0 | 0 | 0 |
| rs774518779 | pathogenic | 1 | 1 | 1 |
| rs28931595 | pathogenic | 1 | 1 | 1 |
| rs771748289 | pathogenic | 0 | 0 | 0 |
| rs104894406 | pathogenic | 0 | 0 | 0 |
| rs76838169 | benign | 0 | 0 | 0 |
| rs111033294 | pathogenic | 0 | 0 | 0 |
| rs200214298 | pathogenic | 0 | 0 | 0 |
| rs28940283 | pathogenic | 0 | 1 | 0 |
| rs1047883 | benign | 0 | 0 | 0 |
| rs138254425 | benign | 0 | 0 | 0 |
| rs121912592 | pathogenic | 1 | 1 | 0 |
| rs369061090 | pathogenic | 1 | 1 | 0 |
| rs758724746 | pathogenic | 0 | 1 | 0 |
| rs121912595 | pathogenic | 0 | 0 | 0 |
| rs76340296 | benign | 0 | 0 | 0 |
| rs140578009 | benign | 1 | 1 | 1 |
| rs74445297 | benign | 0 | 0 | 0 |
| rs78599516 | benign | 1 | 1 | 0 |
| rs104894035 | pathogenic | 1 | 1 | 0 |
| rs79451216 | benign | 0 | 0 | 0 |
| rs104894036 | pathogenic | 0 | 0 | 0 |
| rs73343757 | benign | 0 | 0 | 0 |
| rs121917722 | pathogenic | 1 | 1 | 1 |
| rs747700126 | pathogenic | 0 | 1 | 0 |
| rs121917725 | pathogenic | 1 | 1 | 1 |
| rs121917721 | pathogenic | 1 | 1 | 1 |
| rs121917723 | pathogenic | 1 | 1 | 0 |
| rs17569 | benign | 1 | 1 | 1 |
| rs2230062 | benign | 1 | 1 | 1 |
| rs17570 | benign | 0 | 0 | 0 |
| rs121917724 | pathogenic | 0 | 1 | 0 |
| rs199422235 | pathogenic | 0 | 0 | 0 |
| rs199422237 | pathogenic | 1 | 1 | 1 |
| rs199422234 | pathogenic | 1 | 1 | 0 |
| rs140015315 | pathogenic | 1 | 1 | 1 |
| rs730880298 | pathogenic | 1 | 1 | 1 |
| rs779692470 | pathogenic | 0 | 0 | 0 |
| rs369152939 | pathogenic | 0 | 0 | 0 |
| rs527236040 | pathogenic | 0 | 0 | 0 |
| rs587776985 | pathogenic | 1 | 1 | 1 |
| rs147077598 | pathogenic | 1 | 1 | 1 |
| rs128620185 | pathogenic | 1 | 1 | 1 |
| rs128620189 | pathogenic | 0 | 0 | 0 |
| rs128621190 | pathogenic | 0 | 0 | 0 |
| rs61754648 | benign | 0 | 0 | 0 |
| rs137852906 | pathogenic | 0 | 0 | 0 |

| | | | | |
|---|---|---|---|---|
| rs76811038 | pathogenic | 0 | 0 | 0 |
| rs149468422 | benign | 0 | 0 | 0 |
| rs5911 | benign | 0 | 0 | 0 |
| rs28942112 | pathogenic | 1 | 1 | 1 |
| rs143275858 | pathogenic | 1 | 1 | 0 |
| rs28362261 | benign | 0 | 0 | 0 |
| rs28362263 | benign | 1 | 1 | 1 |
| rs558707786 | pathogenic | 0 | 0 | 0 |
| rs138775799 | pathogenic | 0 | 0 | 0 |
| rs748170941 | pathogenic | 0 | 0 | 0 |
| rs121908380 | pathogenic | 0 | 0 | 0 |
| rs765123255 | pathogenic | 1 | 1 | 1 |
| rs730881832 | pathogenic | 0 | 1 | 0 |
| rs587782165 | benign | 1 | 1 | 1 |
| rs762307622 | pathogenic | 0 | 0 | 0 |
| rs34612342 | pathogenic | 1 | 1 | 0 |
| rs143353451 | pathogenic | 0 | 1 | 0 |
| rs747993448 | pathogenic | 0 | 1 | 0 |
| rs376561094 | pathogenic | 0 | 0 | 0 |
| rs776487884 | pathogenic | 1 | 1 | 0 |
| rs34126013 | pathogenic | 1 | 1 | 0 |
| rs200495564 | pathogenic | 1 | 1 | 0 |
| rs140342925 | pathogenic | 1 | 1 | 0 |
| rs587780749 | pathogenic | 1 | 1 | 0 |
| rs773087549 | pathogenic | 1 | 1 | 0 |
| rs730881833 | pathogenic | 0 | 0 | 0 |
| rs374950566 | pathogenic | 1 | 1 | 0 |
| rs587780752 | benign | 0 | 0 | 0 |
| rs3219489 | benign | 0 | 0 | 0 |
| rs199742231 | benign | 0 | 0 | 0 |
| rs587780082 | pathogenic | 0 | 0 | 0 |
| rs35881803 | benign | 0 | 0 | 0 |
| rs41305623 | benign | 1 | 1 | 1 |
| rs121917856 | pathogenic | 1 | 1 | 1 |
| rs576284122 | benign | 0 | 0 | 0 |
| rs142036701 | benign | 0 | 0 | 0 |
| rs121917852 | pathogenic | 1 | 1 | 1 |
| rs121917854 | pathogenic | 1 | 1 | 1 |
| rs121917853 | pathogenic | 1 | 1 | 1 |
| rs121917857 | pathogenic | 0 | 1 | 0 |
| rs193922740 | pathogenic | 1 | 1 | 1 |
| rs565779970 | pathogenic | 1 | 1 | 1 |
| rs6723526 | benign | 0 | 0 | 0 |
| rs141094096 | pathogenic | 1 | 1 | 1 |
| rs104894637 | pathogenic | 0 | 0 | 0 |
| rs104894636 | pathogenic | 0 | 0 | 0 |
| rs104894642 | pathogenic | 0 | 0 | 0 |
| rs104894638 | pathogenic | 0 | 0 | 0 |
| rs753666460 | pathogenic | 0 | 0 | 0 |
| rs104894643 | pathogenic | 1 | 1 | 0 |
| rs34520362 | pathogenic | 0 | 0 | 0 |
| rs374621913 | pathogenic | 0 | 0 | 0 |
| rs113641837 | benign | 0 | 0 | 0 |
| rs104894635 | pathogenic | 0 | 0 | 0 |

| | | | | |
|---|---|---|---|---|
| rs143947056 | pathogenic | 0 | 0 | 0 |
| rs138504221 | pathogenic | 0 | 0 | 0 |
| rs9894254 | benign | 0 | 0 | 0 |
| rs104894640 | pathogenic | 1 | 1 | 1 |
| rs58786455 | benign | 0 | 0 | 0 |
| rs144143780 | pathogenic | 0 | 0 | 0 |
| rs62620232 | benign | 0 | 0 | 0 |
| rs34297805 | benign | 1 | 1 | 1 |
| rs104894641 | pathogenic | 0 | 0 | 0 |
| rs148876719 | benign | 0 | 0 | 0 |
| rs104894639 | pathogenic | 0 | 0 | 0 |
| rs7503034 | benign | 0 | 0 | 0 |
| rs201631570 | benign | 0 | 0 | 0 |
| rs150181226 | pathogenic | 0 | 0 | 0 |
| rs377546036 | pathogenic | 0 | 0 | 0 |
| rs267606700 | pathogenic | 0 | 0 | 0 |
| rs267606701 | pathogenic | 1 | 1 | 0 |
| rs267606699 | pathogenic | 1 | 1 | 0 |
| rs9903215 | benign | 0 | 0 | 0 |
| rs139486406 | benign | 0 | 0 | 0 |
| rs143791614 | benign | 0 | 0 | 0 |
| rs573483474 | benign | 0 | 0 | 0 |
| rs2032892 | benign | 0 | 0 | 0 |
| rs137852619 | pathogenic | 0 | 0 | 0 |
| rs200243548 | benign | 0 | 0 | 0 |
| rs115702388 | benign | 0 | 0 | 0 |
| rs138085358 | pathogenic | 0 | 0 | 0 |
| rs121912789 | pathogenic | 1 | 1 | 1 |
| rs763407972 | pathogenic | 1 | 1 | 1 |
| rs147278302 | benign | 0 | 0 | 0 |
| rs374540411 | benign | 0 | 0 | 0 |
| rs121912790 | pathogenic | 1 | 1 | 0 |
| rs752049917 | benign | 0 | 0 | 0 |
| rs139640491 | benign | 1 | 1 | 1 |
| rs138010738 | benign | 1 | 1 | 0 |
| rs201328768 | pathogenic | 1 | 1 | 0 |
| rs2135720 | benign | 1 | 1 | 0 |
| rs201816080 | benign | 1 | 1 | 0 |
| rs754391973 | pathogenic | 0 | 0 | 0 |
| rs139087859 | benign | 0 | 0 | 0 |
| rs6805421 | benign | 0 | 0 | 0 |
| rs146266900 | benign | 0 | 0 | 0 |
| rs397509419 | pathogenic | 0 | 0 | 0 |
| rs149980411 | benign | 0 | 0 | 0 |
| rs747711488 | pathogenic | 1 | 1 | 1 |
| rs149474339 | pathogenic | 1 | 1 | 1 |
| rs758252808 | pathogenic | 1 | 1 | 0 |
| rs745756308 | pathogenic | 0 | 0 | 0 |
| rs4986958 | benign | 0 | 0 | 0 |
| rs74315444 | pathogenic | 0 | 0 | 0 |
| rs61752137 | pathogenic | 0 | 0 | 0 |
| rs8179065 | benign | 1 | 1 | 1 |
| rs758361736 | pathogenic | 1 | 1 | 0 |
| rs139794067 | pathogenic | 1 | 1 | 1 |

| | | | | |
|---|---|---|---|---|
| rs730880954 | pathogenic | 1 | 1 | 1 |
| rs730880955 | pathogenic | 1 | 1 | 1 |
| rs150634297 | pathogenic | 0 | 0 | 0 |
| rs104893750 | pathogenic | 1 | 1 | 1 |
| rs104893748 | pathogenic | 1 | 1 | 1 |
| rs104893749 | pathogenic | 1 | 1 | 1 |
| rs753520553 | pathogenic | 1 | 1 | 0 |
| rs61737294 | benign | 0 | 0 | 0 |
| rs104894601 | pathogenic | 0 | 0 | 0 |
| rs376090795 | pathogenic | 0 | 0 | 0 |
| rs104894596 | pathogenic | 0 | 0 | 0 |
| rs104894595 | pathogenic | 0 | 0 | 0 |
| rs104894598 | pathogenic | 0 | 0 | 0 |
| rs104894597 | pathogenic | 0 | 0 | 0 |
| rs104894591 | pathogenic | 0 | 0 | 0 |
| rs104894593 | pathogenic | 0 | 0 | 0 |
| rs104894594 | pathogenic | 0 | 0 | 0 |
| rs104894590 | pathogenic | 0 | 0 | 0 |
| rs86312 | benign | 0 | 0 | 0 |
| rs77874543 | benign | 0 | 0 | 0 |
| rs121912590 | pathogenic | 1 | 1 | 1 |
| rs121912589 | pathogenic | 1 | 1 | 1 |
| rs121912586 | pathogenic | 1 | 1 | 1 |
| rs121912585 | pathogenic | 1 | 1 | 1 |
| rs121912587 | pathogenic | 1 | 1 | 1 |
| rs762909359 | pathogenic | 0 | 0 | 0 |
| rs28931610 | pathogenic | 0 | 0 | 0 |
| rs104894097 | pathogenic | 1 | 1 | 1 |
| rs775176191 | benign | 0 | 0 | 0 |
| rs199907548 | pathogenic | 0 | 0 | 0 |
| rs104894095 | pathogenic | 1 | 1 | 1 |
| rs104894109 | pathogenic | 1 | 1 | 1 |
| rs372266620 | benign | 1 | 1 | 1 |
| rs104894099 | pathogenic | 0 | 0 | 0 |
| rs137854599 | pathogenic | 1 | 1 | 1 |
| rs104894094 | pathogenic | 0 | 0 | 0 |
| rs104894104 | pathogenic | 0 | 0 | 0 |
| rs6413463 | benign | 1 | 1 | 1 |
| rs104894098 | pathogenic | 0 | 0 | 0 |
| rs6413464 | benign | 0 | 0 | 0 |
| rs3731249 | benign | 0 | 0 | 0 |
| rs16931179 | benign | 0 | 0 | 0 |
| rs2734715 | benign | 0 | 0 | 0 |
| rs759080581 | pathogenic | 0 | 0 | 0 |
| rs1800746 | benign | 0 | 0 | 0 |
| rs202189057 | pathogenic | 0 | 0 | 0 |
| rs121908204 | pathogenic | 0 | 0 | 0 |
| rs756564767 | pathogenic | 0 | 0 | 0 |
| rs34758634 | benign | 1 | 1 | 1 |
| rs28940573 | pathogenic | 0 | 0 | 0 |
| rs119455955 | pathogenic | 1 | 1 | 0 |
| rs752164603 | pathogenic | 1 | 1 | 0 |
| rs763162812 | pathogenic | 0 | 0 | 0 |
| rs119455957 | pathogenic | 0 | 0 | 0 |

| | | | | |
|---|---|---|---|---|
| rs119455958 | pathogenic | 1 | 1 | 0 |
| rs750428882 | pathogenic | 0 | 0 | 0 |
| rs141482368 | benign | 0 | 0 | 0 |
| rs119455953 | pathogenic | 0 | 0 | 0 |
| rs119455954 | pathogenic | 0 | 0 | 0 |
| rs112812685 | benign | 1 | 1 | 0 |
| rs138744051 | benign | 1 | 1 | 0 |
| rs121908200 | pathogenic | 0 | 0 | 0 |
| rs119455956 | pathogenic | 0 | 0 | 0 |
| rs78970155 | benign | 0 | 0 | 0 |
| rs2834167 | benign | 0 | 0 | 0 |
| rs387907326 | pathogenic | 1 | 1 | 1 |
| rs2107732 | benign | 0 | 0 | 0 |
| rs11552377 | benign | 0 | 0 | 0 |
| rs137852843 | pathogenic | 0 | 0 | 0 |
| rs104894783 | pathogenic | 0 | 0 | 0 |
| rs122457137 | pathogenic | 1 | 1 | 1 |
| rs104894779 | pathogenic | 1 | 1 | 1 |
| rs104894786 | pathogenic | 0 | 0 | 0 |
| rs104894784 | pathogenic | 0 | 0 | 0 |
| rs104894785 | pathogenic | 0 | 0 | 0 |
| rs201870761 | pathogenic | 0 | 0 | 0 |
| rs104894781 | pathogenic | 0 | 0 | 0 |
| rs28897728 | benign | 0 | 0 | 0 |
| rs370273690 | pathogenic | 1 | 1 | 0 |
| rs9332739 | benign | 0 | 0 | 0 |
| rs150878060 | benign | 0 | 0 | 0 |
| rs573509224 | benign | 0 | 0 | 0 |
| rs36221133 | benign | 0 | 0 | 0 |
| rs2230009 | benign | 0 | 0 | 0 |
| rs267607008 | pathogenic | 0 | 0 | 0 |
| rs5985 | benign | 1 | 1 | 1 |
| rs5982 | benign | 1 | 1 | 1 |
| rs5987 | benign | 0 | 0 | 0 |
| rs5988 | benign | 0 | 0 | 0 |
| rs121918680 | pathogenic | 1 | 1 | 0 |
| rs121918679 | pathogenic | 1 | 1 | 0 |
| rs7845 | benign | 1 | 1 | 0 |
| rs1049296 | benign | 1 | 1 | 1 |
| rs121918677 | pathogenic | 1 | 1 | 0 |
| rs200086262 | pathogenic | 1 | 1 | 0 |
| rs746930990 | pathogenic | 1 | 1 | 0 |
| rs104893878 | pathogenic | 1 | 1 | 1 |
| rs121912468 | pathogenic | 1 | 1 | 1 |
| rs121912467 | pathogenic | 1 | 1 | 1 |
| rs145351926 | benign | 1 | 1 | 1 |
| rs141212743 | pathogenic | 1 | 1 | 1 |
| rs397515473 | pathogenic | 1 | 1 | 0 |
| rs121918313 | pathogenic | 0 | 1 | 0 |
| rs73015965 | pathogenic | 0 | 0 | 0 |
| rs121918030 | pathogenic | 0 | 1 | 0 |
| rs77782321 | benign | 0 | 0 | 0 |
| rs374661051 | pathogenic | 1 | 1 | 1 |
| rs267606854 | pathogenic | 0 | 0 | 0 |

| | | | | |
|---|---|---|---|---|
| rs41279678 | benign | 0 | 0 | 0 |
| rs61754640 | benign | 0 | 0 | 0 |
| rs199473314 | pathogenic | 0 | 0 | 0 |
| rs137854601 | pathogenic | 0 | 0 | 0 |
| rs137854614 | pathogenic | 0 | 0 | 0 |
| rs137854615 | pathogenic | 0 | 0 | 0 |
| rs199473635 | pathogenic | 0 | 0 | 0 |
| rs121434478 | pathogenic | 1 | 1 | 1 |
| rs775407864 | pathogenic | 1 | 1 | 1 |
| rs12706927 | benign | 0 | 1 | 0 |
| rs775614261 | benign | 0 | 0 | 0 |
| rs768926261 | benign | 0 | 0 | 0 |
| rs114057640 | benign | 0 | 0 | 0 |
| rs387906886 | pathogenic | 1 | 1 | 0 |
| rs138658585 | benign | 0 | 0 | 0 |
| rs4880091 | benign | 0 | 0 | 0 |
| rs121918552 | pathogenic | 1 | 1 | 1 |
| rs10794537 | benign | 0 | 0 | 0 |
| rs121965022 | pathogenic | 1 | 1 | 0 |
| rs121965020 | pathogenic | 0 | 0 | 0 |
| rs758452450 | pathogenic | 0 | 0 | 0 |
| rs747981483 | benign | 0 | 0 | 0 |
| rs58037052 | benign | 0 | 0 | 0 |
| rs121965029 | pathogenic | 1 | 1 | 0 |
| rs147490060 | benign | 0 | 0 | 0 |
| rs3755955 | benign | 0 | 0 | 0 |
| rs148789453 | pathogenic | 0 | 0 | 0 |
| rs121965030 | pathogenic | 0 | 0 | 0 |
| rs76722191 | benign | 0 | 0 | 0 |
| rs199801029 | pathogenic | 0 | 0 | 0 |
| rs121965033 | pathogenic | 0 | 0 | 0 |
| rs368454909 | pathogenic | 1 | 1 | 0 |
| rs6831280 | benign | 0 | 0 | 0 |
| rs121965024 | pathogenic | 0 | 0 | 0 |
| rs121965019 | pathogenic | 0 | 0 | 0 |
| rs11934801 | benign | 0 | 0 | 0 |
| rs73066479 | benign | 0 | 0 | 0 |
| rs121965027 | pathogenic | 0 | 0 | 0 |
| rs121965026 | pathogenic | 1 | 1 | 0 |
| rs121965021 | pathogenic | 0 | 0 | 0 |
| rs398123257 | benign | 0 | 0 | 0 |
| rs121965031 | pathogenic | 0 | 0 | 0 |
| rs202003795 | pathogenic | 0 | 0 | 0 |
| rs559510809 | pathogenic | 0 | 0 | 0 |
| rs762936223 | benign | 0 | 0 | 0 |
| rs137854580 | pathogenic | 1 | 1 | 1 |
| rs112263157 | benign | 0 | 0 | 0 |
| rs2289181 | benign | 0 | 0 | 0 |
| rs138034915 | benign | 0 | 1 | 0 |
| rs528367092 | pathogenic | 0 | 1 | 0 |
| rs1042393 | benign | 0 | 0 | 0 |
| rs370950728 | pathogenic | 0 | 1 | 0 |
| rs530478036 | benign | 0 | 1 | 0 |
| rs374569672 | benign | 0 | 1 | 0 |

| | | | | |
|---|---|---|---|---|
| rs121907944 | pathogenic | 0 | 1 | 0 |
| rs200856561 | pathogenic | 0 | 1 | 0 |
| rs577915581 | pathogenic | 0 | 1 | 0 |
| rs201896815 | pathogenic | 0 | 1 | 0 |
| rs121907945 | pathogenic | 0 | 1 | 0 |
| rs121907940 | pathogenic | 0 | 1 | 0 |
| rs543300039 | pathogenic | 0 | 1 | 0 |
| rs121907936 | pathogenic | 0 | 1 | 0 |
| rs766074609 | pathogenic | 0 | 1 | 0 |
| rs755253527 | pathogenic | 0 | 1 | 0 |
| rs560575383 | pathogenic | 0 | 1 | 0 |
| rs528369909 | benign | 0 | 1 | 0 |
| rs770610356 | pathogenic | 0 | 1 | 0 |
| rs747610090 | pathogenic | 0 | 1 | 0 |
| rs145712232 | benign | 0 | 1 | 0 |
| rs7215458 | benign | 1 | 1 | 0 |
| rs535644999 | pathogenic | 0 | 1 | 0 |
| rs772883420 | pathogenic | 1 | 1 | 0 |
| rs398123169 | pathogenic | 0 | 1 | 0 |
| rs121907937 | pathogenic | 0 | 1 | 0 |
| rs121907941 | pathogenic | 1 | 1 | 0 |
| rs121907942 | pathogenic | 0 | 1 | 0 |
| rs779556619 | pathogenic | 0 | 1 | 0 |
| rs1800307 | benign | 0 | 1 | 0 |
| rs377544304 | pathogenic | 1 | 1 | 0 |
| rs549029029 | pathogenic | 0 | 1 | 0 |
| rs753269119 | pathogenic | 1 | 1 | 1 |
| rs757617999 | pathogenic | 0 | 1 | 0 |
| rs28937909 | pathogenic | 0 | 1 | 0 |
| rs28940868 | pathogenic | 0 | 1 | 0 |
| rs368438393 | pathogenic | 0 | 1 | 0 |
| rs536906561 | pathogenic | 0 | 1 | 0 |
| rs374143224 | pathogenic | 0 | 1 | 0 |
| rs778418246 | pathogenic | 0 | 1 | 0 |
| rs398123172 | pathogenic | 0 | 1 | 0 |
| rs121907938 | pathogenic | 0 | 1 | 0 |
| rs1800312 | pathogenic | 0 | 1 | 0 |
| rs1126690 | benign | 0 | 1 | 0 |
| rs1800314 | benign | 0 | 1 | 0 |
| rs369532274 | pathogenic | 0 | 1 | 0 |
| rs121907943 | pathogenic | 0 | 1 | 0 |
| rs780321415 | pathogenic | 0 | 1 | 0 |
| rs377286472 | benign | 0 | 1 | 0 |
| rs121918406 | pathogenic | 1 | 1 | 1 |
| rs147484870 | benign | 0 | 0 | 0 |
| rs193922671 | pathogenic | 0 | 0 | 0 |
| rs201397978 | benign | 1 | 1 | 1 |
| rs34884895 | benign | 1 | 1 | 1 |
| rs138329459 | benign | 0 | 0 | 0 |
| rs2289366 | benign | 1 | 1 | 1 |
| rs141858334 | benign | 1 | 1 | 1 |
| rs59907886 | benign | 1 | 1 | 1 |
| rs137852653 | pathogenic | 0 | 0 | 0 |
| rs137852652 | pathogenic | 0 | 0 | 0 |

| | | | | |
|---|---|---|---|---|
| rs137852656 | pathogenic | 0 | 0 | 0 |
| rs61739916 | benign | 0 | 0 | 0 |
| rs28936668 | pathogenic | 0 | 0 | 0 |
| rs137852651 | pathogenic | 0 | 0 | 0 |
| rs137852650 | pathogenic | 0 | 0 | 0 |
| rs28936669 | pathogenic | 0 | 0 | 0 |
| rs137852654 | pathogenic | 0 | 0 | 0 |
| rs28936368 | pathogenic | 1 | 1 | 0 |
| rs137852655 | pathogenic | 0 | 0 | 0 |
| rs61752496 | benign | 0 | 0 | 0 |
| rs750098055 | benign | 0 | 0 | 0 |
| rs764149793 | benign | 0 | 0 | 0 |
| rs724160017 | pathogenic | 0 | 1 | 0 |
| rs724160019 | pathogenic | 0 | 0 | 0 |
| rs202219398 | pathogenic | 1 | 1 | 0 |
| rs724160023 | pathogenic | 1 | 1 | 0 |
| rs761953453 | pathogenic | 0 | 0 | 0 |
| rs80338835 | pathogenic | 1 | 1 | 1 |
| rs56284940 | benign | 0 | 0 | 0 |
| rs74575749 | benign | 0 | 0 | 0 |
| rs587777075 | pathogenic | 1 | 1 | 1 |
| rs641153 | benign | 0 | 0 | 0 |
| rs12614 | benign | 0 | 0 | 0 |
| rs117314762 | benign | 0 | 0 | 0 |
| rs537478097 | benign | 1 | 1 | 1 |
| rs144812066 | benign | 0 | 0 | 0 |
| rs4151651 | benign | 0 | 0 | 0 |
| rs138207668 | benign | 1 | 1 | 1 |
| rs149101394 | benign | 1 | 1 | 1 |
| rs4151659 | benign | 1 | 1 | 1 |
| rs45484591 | benign | 1 | 1 | 1 |
| rs761991070 | pathogenic | 0 | 0 | 0 |
| rs755948940 | pathogenic | 0 | 0 | 0 |
| rs137853097 | pathogenic | 0 | 0 | 0 |
| rs775297352 | benign | 0 | 0 | 0 |
| rs28943591 | benign | 0 | 0 | 0 |
| rs191468413 | pathogenic | 1 | 1 | 0 |
| rs11539471 | benign | 1 | 1 | 1 |
| rs11205 | benign | 0 | 0 | 0 |
| rs768409400 | pathogenic | 0 | 0 | 0 |
| rs762013077 | pathogenic | 0 | 0 | 0 |
| rs768474112 | pathogenic | 0 | 0 | 0 |
| rs121965063 | pathogenic | 1 | 1 | 1 |
| rs143648758 | pathogenic | 0 | 0 | 0 |
| rs121965066 | pathogenic | 0 | 0 | 0 |
| rs752907087 | pathogenic | 0 | 0 | 0 |
| rs747702749 | pathogenic | 0 | 0 | 0 |
| rs754087775 | pathogenic | 0 | 0 | 0 |
| rs121918641 | pathogenic | 1 | 1 | 1 |
| rs121918642 | pathogenic | 1 | 1 | 1 |
| rs121918640 | pathogenic | 1 | 1 | 1 |
| rs121918637 | pathogenic | 1 | 1 | 1 |
| rs121918644 | pathogenic | 1 | 1 | 1 |
| rs16840544 | benign | 0 | 0 | 0 |

| | | | | |
|---|---|---|---|---|
| rs1042391 | benign | 1 | 1 | 1 |
| rs104893744 | pathogenic | 1 | 1 | 1 |
| rs120074136 | pathogenic | 0 | 0 | 0 |
| rs1805081 | benign | 1 | 1 | 0 |
| rs80358251 | benign | 0 | 0 | 0 |
| rs774588125 | benign | 0 | 0 | 0 |
| rs120074134 | pathogenic | 0 | 0 | 0 |
| rs139751448 | pathogenic | 0 | 0 | 0 |
| rs77080672 | benign | 0 | 0 | 0 |
| rs372445155 | pathogenic | 1 | 1 | 0 |
| rs13381670 | benign | 0 | 0 | 0 |
| rs1788799 | benign | 0 | 0 | 0 |
| rs376213990 | pathogenic | 0 | 0 | 0 |
| rs757475924 | pathogenic | 0 | 0 | 0 |
| rs80358253 | pathogenic | 0 | 0 | 0 |
| rs1805082 | benign | 0 | 0 | 0 |
| rs120074130 | pathogenic | 0 | 0 | 0 |
| rs768999208 | pathogenic | 0 | 0 | 0 |
| rs34302553 | benign | 0 | 0 | 0 |
| rs28940897 | pathogenic | 0 | 0 | 0 |
| rs1140458 | benign | 0 | 0 | 0 |
| rs143124972 | pathogenic | 0 | 0 | 0 |
| rs120074135 | pathogenic | 0 | 0 | 0 |
| rs543206298 | pathogenic | 0 | 0 | 0 |
| rs120074132 | pathogenic | 0 | 0 | 0 |
| rs759826138 | pathogenic | 0 | 0 | 0 |
| rs28942108 | pathogenic | 0 | 0 | 0 |
| rs80358254 | pathogenic | 0 | 0 | 0 |
| rs80358257 | pathogenic | 0 | 0 | 0 |
| rs781261962 | pathogenic | 0 | 0 | 0 |
| rs28942107 | pathogenic | 0 | 0 | 0 |
| rs80358258 | pathogenic | 0 | 0 | 0 |
| rs80358259 | pathogenic | 1 | 1 | 0 |
| rs141440861 | benign | 0 | 0 | 0 |
| rs28942106 | pathogenic | 0 | 0 | 0 |
| rs778878523 | pathogenic | 0 | 0 | 0 |
| rs28942105 | pathogenic | 0 | 0 | 0 |
| rs61731969 | benign | 0 | 0 | 0 |
| rs200444084 | pathogenic | 0 | 0 | 0 |
| rs113371321 | pathogenic | 0 | 0 | 0 |
| rs369098773 | pathogenic | 0 | 0 | 0 |
| rs35248744 | benign | 0 | 0 | 0 |
| rs758902805 | pathogenic | 0 | 0 | 0 |
| rs120074131 | pathogenic | 0 | 0 | 0 |
| rs1805084 | benign | 0 | 0 | 0 |
| rs117793540 | benign | 0 | 0 | 0 |
| rs530058063 | benign | 0 | 0 | 0 |
| rs1064422 | pathogenic | 1 | 1 | 1 |
| rs104894506 | pathogenic | 1 | 1 | 1 |
| rs104894507 | pathogenic | 1 | 1 | 1 |
| rs104894508 | pathogenic | 1 | 1 | 0 |
| rs28999113 | pathogenic | 1 | 1 | 0 |
| rs281860266 | pathogenic | 0 | 0 | 0 |
| rs104894961 | pathogenic | 1 | 1 | 0 |

| | | | | |
|---|---|---|---|---|
| rs122463168 | pathogenic | 0 | 0 | 0 |
| rs199469624 | pathogenic | 1 | 1 | 1 |
| rs150107747 | benign | 0 | 0 | 0 |
| rs144984628 | pathogenic | 0 | 0 | 0 |
| rs28935171 | pathogenic | 0 | 0 | 0 |
| rs137852443 | pathogenic | 1 | 1 | 0 |
| rs28933674 | pathogenic | 1 | 1 | 1 |
| rs28933675 | pathogenic | 1 | 1 | 1 |
| rs28933676 | pathogenic | 1 | 1 | 1 |
| rs28933677 | pathogenic | 1 | 1 | 1 |
| rs28933678 | pathogenic | 0 | 0 | 0 |
| rs28933679 | pathogenic | 1 | 1 | 1 |
| rs28933680 | pathogenic | 0 | 0 | 0 |
| rs28933681 | pathogenic | 0 | 0 | 0 |
| rs28933682 | pathogenic | 0 | 0 | 0 |
| rs28937294 | pathogenic | 1 | 1 | 0 |
| rs137852459 | pathogenic | 0 | 0 | 0 |
| rs137852461 | pathogenic | 0 | 0 | 0 |
| rs137852464 | pathogenic | 1 | 1 | 1 |
| rs137852465 | pathogenic | 1 | 1 | 1 |
| rs1800297 | benign | 1 | 1 | 0 |
| rs137852374 | pathogenic | 1 | 1 | 1 |
| rs672601312 | pathogenic | 1 | 1 | 1 |
| rs387907196 | pathogenic | 1 | 1 | 1 |
| rs148348866 | benign | 0 | 0 | 0 |
| rs141138948 | pathogenic | 1 | 1 | 1 |
| rs137852685 | pathogenic | 1 | 1 | 0 |
| rs104893900 | pathogenic | 1 | 1 | 0 |
| rs137852686 | pathogenic | 1 | 1 | 1 |
| rs104893906 | pathogenic | 1 | 1 | 0 |
| rs587782207 | pathogenic | 1 | 1 | 1 |
| rs75086406 | pathogenic | 0 | 0 | 0 |
| rs199822819 | pathogenic | 0 | 0 | 0 |
| rs398123166 | pathogenic | 1 | 1 | 1 |
| rs121913123 | pathogenic | 1 | 1 | 1 |
| rs772190176 | pathogenic | 0 | 1 | 0 |
| rs587782216 | pathogenic | 1 | 1 | 1 |
| rs398123159 | pathogenic | 1 | 1 | 1 |
| rs121913122 | pathogenic | 1 | 1 | 1 |
| rs755436052 | pathogenic | 1 | 1 | 1 |
| rs200796606 | pathogenic | 1 | 1 | 1 |
| rs2070080 | pathogenic | 0 | 1 | 0 |
| rs149616320 | benign | 0 | 0 | 0 |
| rs143529458 | benign | 0 | 0 | 0 |
| rs267607077 | pathogenic | 1 | 1 | 1 |
| rs143898031 | benign | 1 | 1 | 0 |
| rs148304857 | benign | 1 | 1 | 1 |
| rs137852809 | pathogenic | 0 | 0 | 0 |
| rs3765623 | benign | 0 | 0 | 0 |
| rs16944397 | benign | 0 | 0 | 0 |
| rs730880167 | benign | 0 | 0 | 0 |
| rs187108957 | benign | 0 | 0 | 0 |
| rs117342470 | benign | 0 | 0 | 0 |
| rs751200138 | pathogenic | 1 | 1 | 1 |

| | | | | |
|---|---|---|---|---|
| rs755799528 | pathogenic | 0 | 0 | 0 |
| rs537484504 | benign | 0 | 0 | 0 |
| rs748944640 | pathogenic | 0 | 0 | 0 |
| rs199774121 | pathogenic | 0 | 0 | 0 |
| rs766094434 | pathogenic | 1 | 1 | 0 |
| rs777321035 | pathogenic | 1 | 1 | 1 |
| rs146269774 | benign | 0 | 1 | 0 |
| rs144085805 | benign | 0 | 1 | 0 |
| rs367643028 | benign | 0 | 0 | 0 |
| rs42524 | benign | 0 | 0 | 0 |
| rs28940586 | pathogenic | 0 | 0 | 0 |
| rs74315327 | pathogenic | 1 | 1 | 1 |
| rs199676861 | pathogenic | 0 | 0 | 0 |
| rs115802744 | benign | 1 | 1 | 0 |
| rs752866643 | pathogenic | 1 | 1 | 1 |
| rs28941784 | pathogenic | 1 | 1 | 1 |
| rs398124434 | pathogenic | 1 | 1 | 1 |
| rs756414548 | pathogenic | 1 | 1 | 1 |
| rs376128990 | pathogenic | 1 | 1 | 1 |
| rs746219370 | pathogenic | 1 | 1 | 1 |
| rs756195708 | pathogenic | 1 | 1 | 1 |
| rs369296618 | pathogenic | 1 | 1 | 1 |
| rs9593 | benign | 1 | 1 | 1 |
| rs139510844 | benign | 0 | 0 | 0 |
| rs28382158 | benign | 0 | 0 | 0 |
| rs28919570 | pathogenic | 0 | 0 | 0 |
| rs587777331 | pathogenic | 0 | 0 | 0 |
| rs587777333 | pathogenic | 0 | 0 | 0 |
| rs587777332 | pathogenic | 0 | 0 | 0 |
| rs774980346 | pathogenic | 0 | 0 | 0 |
| rs370934093 | pathogenic | 0 | 0 | 0 |
| rs121913479 | pathogenic | 1 | 1 | 1 |
| rs121913484 | pathogenic | 0 | 0 | 0 |
| rs121913485 | pathogenic | 1 | 1 | 1 |
| rs28931614 | pathogenic | 1 | 1 | 1 |
| rs17881656 | benign | 1 | 1 | 1 |
| rs28931615 | pathogenic | 1 | 1 | 1 |
| rs147846832 | benign | 0 | 0 | 0 |
| rs80356506 | pathogenic | 0 | 0 | 0 |
| rs80356494 | pathogenic | 0 | 0 | 0 |
| rs121908894 | pathogenic | 0 | 0 | 0 |
| rs121908895 | pathogenic | 0 | 0 | 0 |
| rs80356508 | pathogenic | 0 | 0 | 0 |
| rs80356507 | pathogenic | 0 | 0 | 0 |
| rs121908896 | pathogenic | 0 | 0 | 0 |
| rs745372589 | pathogenic | 1 | 1 | 0 |
| rs183898408 | benign | 0 | 0 | 0 |
| rs200844292 | benign | 0 | 0 | 0 |
| rs146897002 | benign | 0 | 0 | 0 |
| rs121964869 | pathogenic | 1 | 1 | 1 |
| rs201768800 | benign | 1 | 1 | 1 |
| rs9807556 | benign | 1 | 1 | 1 |
| rs184721031 | benign | 0 | 0 | 0 |
| rs201104206 | benign | 0 | 0 | 0 |

| | | | | |
|---|---|---|---|---|
| rs188677538 | benign | 0 | 0 | 0 |
| rs149528866 | benign | 0 | 0 | 0 |
| rs115382168 | benign | 0 | 0 | 0 |
| rs78846023 | benign | 1 | 1 | 1 |
| rs61730662 | benign | 0 | 0 | 0 |
| rs121918368 | pathogenic | 0 | 0 | 0 |
| rs566433112 | pathogenic | 1 | 1 | 1 |
| rs121909342 | pathogenic | 0 | 0 | 0 |
| rs67586389 | pathogenic | 0 | 0 | 0 |
| rs72466485 | pathogenic | 0 | 0 | 0 |
| rs72466486 | pathogenic | 1 | 1 | 1 |
| rs72466487 | pathogenic | 1 | 1 | 1 |
| rs267607112 | pathogenic | 0 | 0 | 0 |
| rs118204013 | pathogenic | 0 | 0 | 0 |
| rs121913495 | pathogenic | 0 | 0 | 0 |
| rs11554273 | pathogenic | 0 | 0 | 0 |
| rs193922477 | pathogenic | 1 | 1 | 1 |
| rs193922479 | pathogenic | 1 | 1 | 1 |
| rs72547554 | pathogenic | 0 | 0 | 0 |
| rs72547561 | pathogenic | 1 | 1 | 1 |
| rs72547562 | pathogenic | 1 | 1 | 0 |
| rs115977487 | benign | 1 | 1 | 0 |
| rs72547564 | pathogenic | 0 | 0 | 0 |
| rs72547568 | pathogenic | 1 | 1 | 0 |
| rs72547569 | pathogenic | 1 | 1 | 1 |
| rs72547571 | pathogenic | 0 | 0 | 0 |
| rs72547575 | pathogenic | 1 | 1 | 1 |
| rs544994462 | benign | 0 | 0 | 0 |
| rs190470017 | benign | 0 | 0 | 0 |
| rs199473266 | pathogenic | 1 | 1 | 1 |
| rs137854602 | pathogenic | 0 | 0 | 0 |
| rs143739249 | pathogenic | 0 | 0 | 0 |
| rs397514697 | pathogenic | 1 | 1 | 1 |
| rs373764886 | pathogenic | 0 | 0 | 0 |
| rs370242930 | pathogenic | 0 | 0 | 0 |
| rs377145777 | pathogenic | 1 | 1 | 1 |
| rs191256186 | benign | 0 | 0 | 0 |
| rs150070901 | benign | 1 | 1 | 1 |
| rs138098032 | benign | 0 | 0 | 0 |
| rs117731813 | benign | 1 | 1 | 1 |
| rs104894488 | pathogenic | 1 | 1 | 1 |
| rs11302 | benign | 0 | 1 | 0 |
| rs121918447 | pathogenic | 1 | 1 | 1 |
| rs138205828 | benign | 1 | 1 | 0 |
| rs149858290 | benign | 0 | 0 | 0 |
| rs750764003 | pathogenic | 0 | 1 | 0 |
| rs141381896 | benign | 0 | 0 | 0 |
| rs149027682 | benign | 0 | 0 | 0 |
| rs121908415 | pathogenic | 0 | 0 | 0 |
| rs121908416 | pathogenic | 0 | 0 | 0 |
| rs121908417 | pathogenic | 0 | 0 | 0 |
| rs587780306 | benign | 0 | 0 | 0 |
| rs3218725 | benign | 0 | 0 | 0 |
| rs3218729 | benign | 0 | 0 | 0 |

| | | | | |
|---|---|---|---|---|
| rs387906917 | pathogenic | 0 | 0 | 0 |
| rs200652608 | pathogenic | 0 | 0 | 0 |
| rs3218721 | benign | 0 | 0 | 0 |
| rs63750869 | pathogenic | 1 | 1 | 1 |
| rs118203910 | pathogenic | 0 | 1 | 0 |
| rs9332701 | benign | 1 | 1 | 0 |
| rs6027 | benign | 0 | 0 | 0 |
| rs121909110 | pathogenic | 0 | 1 | 0 |
| rs762550967 | pathogenic | 1 | 1 | 0 |
| rs763615602 | pathogenic | 1 | 1 | 0 |
| rs144907403 | benign | 0 | 0 | 0 |
| rs104893631 | pathogenic | 1 | 1 | 0 |
| rs144181978 | pathogenic | 1 | 1 | 0 |
| rs74874677 | benign | 1 | 1 | 1 |
| rs104893632 | pathogenic | 0 | 1 | 0 |
| rs371342464 | benign | 1 | 1 | 1 |
| rs104893633 | pathogenic | 1 | 1 | 0 |
| rs201088712 | pathogenic | 0 | 0 | 0 |
| rs201857541 | benign | 1 | 1 | 1 |
| rs139517732 | pathogenic | 1 | 1 | 0 |
| rs191057824 | benign | 1 | 1 | 0 |
| rs35683768 | benign | 1 | 1 | 0 |
| rs104895271 | pathogenic | 0 | 0 | 0 |
| rs104895217 | pathogenic | 1 | 1 | 1 |
| rs104895223 | pathogenic | 1 | 1 | 1 |
| rs104895218 | pathogenic | 1 | 1 | 1 |
| rs104895225 | pathogenic | 1 | 1 | 1 |
| rs4149637 | benign | 1 | 1 | 1 |
| rs104895219 | pathogenic | 1 | 1 | 1 |
| rs104895220 | pathogenic | 1 | 1 | 1 |
| rs104895228 | pathogenic | 1 | 1 | 1 |
| rs104895221 | pathogenic | 1 | 1 | 1 |
| rs104895222 | pathogenic | 1 | 1 | 1 |
| rs104895247 | pathogenic | 0 | 0 | 0 |
| rs28940585 | pathogenic | 0 | 0 | 0 |
| rs561367199 | pathogenic | 1 | 1 | 1 |
| rs143793815 | benign | 0 | 0 | 0 |
| rs28941473 | pathogenic | 1 | 1 | 1 |
| rs751590073 | pathogenic | 1 | 1 | 1 |
| rs111407265 | benign | 1 | 1 | 1 |
| rs373697663 | pathogenic | 1 | 1 | 1 |
| rs369879957 | pathogenic | 0 | 0 | 0 |
| rs28941472 | pathogenic | 1 | 1 | 1 |
| rs28940287 | pathogenic | 1 | 1 | 1 |
| rs77625410 | benign | 0 | 0 | 0 |
| rs111033477 | benign | 0 | 0 | 0 |
| rs115123584 | benign | 0 | 0 | 0 |
| rs374576916 | benign | 0 | 0 | 0 |
| rs397516317 | pathogenic | 0 | 0 | 0 |
| rs368657015 | pathogenic | 0 | 0 | 0 |
| rs397516321 | pathogenic | 0 | 0 | 0 |
| rs397516322 | pathogenic | 0 | 0 | 0 |
| rs111033215 | pathogenic | 0 | 0 | 0 |
| rs199606180 | pathogenic | 0 | 0 | 0 |

| | | | | |
|---|---|---|---|---|
| rs111033192 | pathogenic | 0 | 0 | 0 |
| rs948962 | benign | 1 | 1 | 1 |
| rs142293185 | benign | 0 | 0 | 0 |
| rs376764423 | pathogenic | 0 | 0 | 0 |
| rs773844428 | pathogenic | 0 | 0 | 0 |
| rs186644871 | benign | 0 | 0 | 0 |
| rs200313391 | benign | 0 | 0 | 0 |
| rs377670513 | pathogenic | 0 | 0 | 0 |
| rs1132036 | benign | 0 | 0 | 0 |
| rs111033231 | benign | 0 | 0 | 0 |
| rs145945630 | pathogenic | 0 | 0 | 0 |
| rs121912984 | pathogenic | 1 | 1 | 1 |
| rs1800435 | benign | 0 | 1 | 0 |
| rs121912980 | pathogenic | 0 | 0 | 0 |
| rs121912982 | pathogenic | 1 | 1 | 1 |
| rs121912983 | pathogenic | 0 | 0 | 0 |
| rs121912981 | pathogenic | 0 | 0 | 0 |
| rs1057868 | benign | 0 | 0 | 0 |
| rs56048668 | benign | 0 | 0 | 0 |
| rs55865465 | benign | 0 | 0 | 0 |
| rs748096324 | benign | 1 | 1 | 0 |
| rs150475750 | benign | 0 | 0 | 0 |
| rs143025739 | benign | 1 | 1 | 0 |
| rs79702016 | benign | 1 | 1 | 0 |
| rs41338945 | benign | 0 | 0 | 0 |
| rs146406037 | benign | 1 | 1 | 1 |
| rs143274491 | benign | 0 | 0 | 0 |
| rs3829986 | benign | 0 | 0 | 0 |
| rs149168939 | benign | 0 | 0 | 0 |
| rs1800385 | benign | 0 | 0 | 0 |
| rs1800386 | pathogenic | 0 | 0 | 0 |
| rs121918524 | pathogenic | 1 | 1 | 1 |
| rs121912700 | pathogenic | 0 | 0 | 0 |
| rs104893930 | pathogenic | 1 | 1 | 1 |
| rs104893931 | pathogenic | 0 | 0 | 0 |
| rs116125936 | benign | 0 | 0 | 0 |
| rs627441 | benign | 0 | 0 | 0 |
| rs104894481 | pathogenic | 0 | 0 | 0 |
| rs137853129 | pathogenic | 0 | 0 | 0 |
| rs137853130 | pathogenic | 1 | 1 | 0 |
| rs28940282 | pathogenic | 0 | 0 | 0 |
| rs2229989 | benign | 0 | 0 | 0 |
| rs104894647 | pathogenic | 0 | 0 | 0 |
| rs1061170 | benign | 1 | 1 | 1 |
| rs576402053 | benign | 0 | 0 | 0 |
| rs34972246 | benign | 0 | 0 | 0 |
| rs192725607 | benign | 0 | 0 | 0 |
| rs137852615 | pathogenic | 1 | 1 | 0 |
| rs137852617 | pathogenic | 1 | 1 | 1 |
| rs137852611 | pathogenic | 0 | 0 | 0 |
| rs137852612 | pathogenic | 0 | 0 | 0 |
| rs137852614 | pathogenic | 1 | 1 | 1 |
| rs137852610 | pathogenic | 1 | 1 | 1 |
| rs137852618 | pathogenic | 1 | 1 | 1 |

| | | | | |
|---|---|---|---|---|
| rs137852613 | pathogenic | 1 | 1 | 0 |
| rs137852609 | pathogenic | 0 | 0 | 0 |
| rs63750306 | pathogenic | 1 | 1 | 1 |
| rs63750265 | pathogenic | 1 | 1 | 1 |
| rs63751287 | pathogenic | 1 | 1 | 1 |
| rs199723282 | benign | 0 | 0 | 0 |
| rs121917807 | pathogenic | 0 | 0 | 0 |
| rs121917809 | pathogenic | 0 | 0 | 0 |
| rs75134564 | pathogenic | 0 | 0 | 0 |
| rs267606930 | pathogenic | 1 | 1 | 0 |
| rs121909302 | pathogenic | 0 | 0 | 0 |
| rs121909303 | pathogenic | 0 | 0 | 0 |
| rs121909299 | pathogenic | 0 | 0 | 0 |
| rs121909301 | pathogenic | 0 | 0 | 0 |
| rs121909300 | pathogenic | 1 | 1 | 0 |
| rs111033589 | pathogenic | 0 | 0 | 0 |
| rs137853059 | pathogenic | 0 | 0 | 0 |
| rs137853057 | pathogenic | 1 | 1 | 1 |
| rs1801474 | benign | 0 | 0 | 0 |
| rs137853060 | pathogenic | 1 | 1 | 1 |
| rs137853058 | pathogenic | 1 | 1 | 1 |
| rs34424986 | pathogenic | 0 | 0 | 0 |
| rs199657839 | benign | 1 | 1 | 1 |
| rs56092260 | benign | 0 | 0 | 0 |
| rs1801582 | benign | 1 | 1 | 1 |
| rs1801334 | benign | 1 | 1 | 1 |
| rs191486604 | pathogenic | 0 | 0 | 0 |
| rs121908051 | pathogenic | 0 | 0 | 0 |
| rs121908055 | pathogenic | 1 | 1 | 0 |
| rs28940886 | pathogenic | 0 | 0 | 0 |
| rs121908050 | pathogenic | 0 | 0 | 0 |
| rs28940887 | pathogenic | 0 | 0 | 0 |
| rs387906300 | pathogenic | 0 | 0 | 0 |
| rs28942087 | pathogenic | 0 | 0 | 0 |
| rs121908049 | pathogenic | 1 | 1 | 0 |
| rs121908054 | pathogenic | 1 | 1 | 0 |
| rs121908048 | pathogenic | 0 | 0 | 0 |
| rs28940888 | pathogenic | 1 | 1 | 1 |
| rs121908053 | pathogenic | 1 | 1 | 0 |
| rs122467169 | pathogenic | 1 | 1 | 1 |
| rs122467172 | pathogenic | 1 | 1 | 1 |
| rs122467170 | pathogenic | 1 | 1 | 1 |
| rs28935477 | pathogenic | 1 | 1 | 0 |
| rs28942088 | pathogenic | 0 | 0 | 0 |
| rs751478142 | pathogenic | 1 | 1 | 0 |
| rs765368797 | pathogenic | 0 | 0 | 0 |
| rs147810437 | benign | 0 | 0 | 0 |
| rs121918209 | pathogenic | 1 | 1 | 1 |
| rs121918208 | pathogenic | 1 | 1 | 1 |
| rs121918210 | pathogenic | 1 | 1 | 1 |
| rs35515638 | benign | 1 | 1 | 1 |
| rs121918207 | pathogenic | 1 | 1 | 1 |
| rs141298312 | benign | 0 | 0 | 0 |
| rs121918211 | pathogenic | 0 | 0 | 0 |

| | | | | |
|---|---|---|---|---|
| rs149660059 | benign | 0 | 0 | 0 |
| rs200670286 | pathogenic | 1 | 1 | 1 |
| rs121918212 | pathogenic | 1 | 1 | 1 |
| rs121918213 | pathogenic | 1 | 1 | 1 |
| rs121918205 | pathogenic | 1 | 1 | 1 |
| rs104894025 | pathogenic | 1 | 1 | 1 |
| rs104894026 | pathogenic | 1 | 1 | 1 |
| rs104894028 | pathogenic | 0 | 1 | 0 |
| rs104894029 | pathogenic | 0 | 1 | 0 |
| rs119458971 | pathogenic | 0 | 0 | 0 |
| rs119458969 | pathogenic | 1 | 1 | 1 |
| rs1801591 | benign | 1 | 1 | 1 |
| rs140169311 | benign | 0 | 0 | 0 |
| rs753363278 | benign | 1 | 1 | 1 |
| rs199763682 | pathogenic | 0 | 0 | 0 |
| rs119458970 | pathogenic | 1 | 1 | 1 |
| rs763722868 | pathogenic | 0 | 0 | 0 |
| rs56404770 | benign | 0 | 0 | 0 |
| rs34221241 | benign | 0 | 0 | 0 |
| rs121909625 | pathogenic | 0 | 0 | 0 |
| rs550921485 | pathogenic | 0 | 0 | 0 |
| rs1800454 | benign | 0 | 0 | 0 |
| rs241447 | benign | 0 | 0 | 0 |
| rs2230606 | benign | 1 | 1 | 0 |
| rs5036 | pathogenic | 1 | 1 | 1 |
| rs13306788 | benign | 0 | 0 | 0 |
| rs28929480 | pathogenic | 0 | 0 | 0 |
| rs121912749 | pathogenic | 0 | 0 | 0 |
| rs56361140 | pathogenic | 0 | 0 | 0 |
| rs28931583 | pathogenic | 1 | 1 | 1 |
| rs36211715 | pathogenic | 1 | 1 | 1 |
| rs138049878 | pathogenic | 1 | 1 | 1 |
| rs727503253 | pathogenic | 1 | 1 | 1 |
| rs267606908 | pathogenic | 0 | 0 | 0 |
| rs121913631 | pathogenic | 1 | 1 | 1 |
| rs36211716 | benign | 1 | 1 | 1 |
| rs121913639 | pathogenic | 1 | 1 | 1 |
| rs730880761 | pathogenic | 1 | 1 | 1 |
| rs121913629 | pathogenic | 1 | 1 | 1 |
| rs754368658 | pathogenic | 0 | 0 | 0 |
| rs777090017 | pathogenic | 0 | 0 | 0 |
| rs267606913 | pathogenic | 1 | 1 | 1 |
| rs773138218 | pathogenic | 0 | 0 | 0 |
| rs371295780 | pathogenic | 0 | 0 | 0 |
| rs387906830 | pathogenic | 1 | 1 | 0 |
| rs771364038 | pathogenic | 0 | 1 | 0 |
| rs772731615 | pathogenic | 0 | 0 | 0 |
| rs137853063 | pathogenic | 0 | 0 | 0 |
| rs387907076 | pathogenic | 0 | 1 | 0 |
| rs35460768 | benign | 0 | 0 | 0 |
| rs2278196 | benign | 1 | 1 | 1 |
| rs119103275 | pathogenic | 0 | 0 | 0 |
| rs119103276 | pathogenic | 1 | 1 | 1 |
| rs374402066 | pathogenic | 1 | 1 | 1 |

| | | | | |
|---|---|---|---|---|
| rs119103274 | pathogenic | 0 | 0 | 0 |
| rs2274064 | benign | 0 | 0 | 0 |
| rs267607023 | pathogenic | 1 | 1 | 0 |
| rs142117489 | benign | 1 | 1 | 0 |
| rs371258270 | benign | 0 | 0 | 0 |
| rs34716057 | benign | 0 | 0 | 0 |
| rs201778606 | benign | 0 | 0 | 0 |
| rs201438792 | benign | 0 | 0 | 0 |
| rs140547417 | benign | 0 | 0 | 0 |
| rs5881 | benign | 0 | 0 | 0 |
| rs752298084 | benign | 0 | 0 | 0 |
| rs34855278 | benign | 0 | 0 | 0 |
| rs5880 | benign | 1 | 1 | 0 |
| rs5882 | benign | 0 | 0 | 0 |
| rs1800777 | benign | 0 | 0 | 0 |
| rs577301331 | benign | 0 | 0 | 0 |
| rs137853313 | pathogenic | 0 | 0 | 0 |
| rs28935169 | pathogenic | 0 | 0 | 0 |
| rs137853315 | pathogenic | 0 | 0 | 0 |
| rs137853314 | pathogenic | 0 | 0 | 0 |
| rs28935469 | pathogenic | 0 | 0 | 0 |
| rs137853318 | pathogenic | 0 | 0 | 0 |
| rs28935470 | pathogenic | 0 | 0 | 0 |
| rs3729986 | benign | 0 | 0 | 0 |
| rs730880619 | pathogenic | 0 | 0 | 0 |
| rs201012766 | benign | 0 | 0 | 0 |
| rs727503218 | pathogenic | 0 | 0 | 0 |
| rs11570052 | benign | 0 | 0 | 0 |
| rs730880623 | pathogenic | 0 | 0 | 0 |
| rs138753870 | benign | 0 | 0 | 0 |
| rs397516068 | pathogenic | 0 | 0 | 0 |
| rs3729989 | benign | 1 | 1 | 1 |
| rs397516074 | pathogenic | 0 | 0 | 0 |
| rs561449819 | pathogenic | 0 | 0 | 0 |
| rs121917779 | benign | 0 | 0 | 0 |
| rs142881576 | pathogenic | 0 | 0 | 0 |
| rs77384282 | benign | 0 | 0 | 0 |
| rs201865787 | benign | 0 | 0 | 0 |
| rs142244640 | benign | 0 | 0 | 0 |
| rs200488706 | benign | 0 | 0 | 0 |
| rs137853291 | pathogenic | 1 | 1 | 0 |
| rs28933990 | pathogenic | 1 | 1 | 0 |
| rs140570189 | benign | 0 | 0 | 0 |
| rs201588178 | benign | 0 | 0 | 0 |
| rs104893846 | pathogenic | 0 | 0 | 0 |
| rs754545360 | pathogenic | 0 | 1 | 0 |
| rs104893851 | pathogenic | 0 | 1 | 0 |
| rs104893849 | pathogenic | 0 | 0 | 0 |
| rs571038432 | pathogenic | 1 | 1 | 0 |
| rs2270655 | benign | 1 | 1 | 1 |
| rs315952 | benign | 0 | 0 | 0 |
| rs151176548 | benign | 0 | 0 | 0 |
| rs122459147 | pathogenic | 1 | 1 | 0 |
| rs122458140 | pathogenic | 0 | 0 | 0 |

| | | | | |
|---|---|---|---|---|
| rs122458142 | pathogenic | 1 | 1 | 0 |
| rs267606812 | pathogenic | 1 | 1 | 0 |
| rs267606813 | pathogenic | 1 | 1 | 0 |
| rs122458143 | pathogenic | 0 | 0 | 0 |
| rs122459146 | pathogenic | 0 | 0 | 0 |
| rs122458144 | pathogenic | 0 | 0 | 0 |
| rs122458145 | pathogenic | 0 | 0 | 0 |
| rs113994007 | pathogenic | 1 | 1 | 0 |
| rs758746181 | pathogenic | 1 | 1 | 1 |
| rs145034527 | pathogenic | 1 | 1 | 0 |
| rs119468006 | pathogenic | 1 | 1 | 0 |
| rs201908721 | pathogenic | 0 | 0 | 0 |
| rs755933881 | pathogenic | 1 | 1 | 0 |
| rs771578775 | pathogenic | 0 | 0 | 0 |
| rs17849927 | benign | 0 | 0 | 0 |
| rs139133094 | benign | 0 | 0 | 0 |
| rs12593 | benign | 0 | 0 | 0 |
| rs578189699 | pathogenic | 1 | 1 | 0 |
| rs119468008 | pathogenic | 0 | 0 | 0 |
| rs768524626 | benign | 0 | 0 | 0 |
| rs119468009 | pathogenic | 0 | 0 | 0 |
| rs119468004 | pathogenic | 0 | 0 | 0 |
| rs199874519 | pathogenic | 0 | 0 | 0 |
| rs376478331 | pathogenic | 0 | 0 | 0 |
| rs61995958 | pathogenic | 0 | 0 | 0 |
| rs199714731 | benign | 0 | 0 | 0 |
| rs149185954 | benign | 0 | 0 | 0 |
| rs121918353 | pathogenic | 0 | 0 | 0 |
| rs1131695 | benign | 0 | 0 | 0 |
| rs28939668 | pathogenic | 0 | 0 | 0 |
| rs758432471 | pathogenic | 1 | 1 | 1 |
| rs752746786 | pathogenic | 1 | 1 | 1 |
| rs28935471 | pathogenic | 0 | 0 | 0 |
| rs80338837 | pathogenic | 0 | 0 | 0 |
| rs137853312 | pathogenic | 0 | 0 | 0 |
| rs28935472 | pathogenic | 0 | 0 | 0 |
| rs28935473 | pathogenic | 0 | 0 | 0 |
| rs267606752 | pathogenic | 0 | 0 | 0 |
| rs3917973 | benign | 0 | 0 | 0 |
| rs3918018 | benign | 0 | 0 | 0 |
| rs757976755 | pathogenic | 0 | 0 | 0 |
| rs121918329 | pathogenic | 1 | 1 | 0 |
| rs770114459 | pathogenic | 0 | 0 | 0 |
| rs6662272 | benign | 0 | 0 | 0 |
| rs121918242 | pathogenic | 1 | 1 | 0 |
| rs121918240 | pathogenic | 1 | 1 | 0 |
| rs200094982 | pathogenic | 0 | 0 | 0 |
| rs121918241 | pathogenic | 0 | 0 | 0 |
| rs140522266 | pathogenic | 1 | 1 | 0 |
| rs121918243 | pathogenic | 1 | 1 | 0 |
| rs370596113 | pathogenic | 1 | 1 | 0 |
| rs398124295 | pathogenic | 1 | 1 | 0 |
| rs587776889 | pathogenic | 1 | 1 | 0 |
| rs747527726 | pathogenic | 1 | 1 | 0 |

| | | | | |
|---|---|---|---|---|
| rs112631212 | benign | 0 | 0 | 0 |
| rs104894605 | pathogenic | 0 | 0 | 0 |
| rs104894607 | pathogenic | 0 | 0 | 0 |
| rs104894606 | pathogenic | 0 | 0 | 0 |
| rs143873873 | benign | 0 | 0 | 0 |
| rs104894388 | pathogenic | 1 | 1 | 0 |
| rs104894391 | pathogenic | 1 | 1 | 0 |
| rs5350 | benign | 1 | 1 | 1 |
| rs104894390 | pathogenic | 1 | 1 | 1 |
| rs5346 | benign | 1 | 1 | 1 |
| rs121912500 | pathogenic | 0 | 0 | 0 |
| rs74315450 | pathogenic | 1 | 1 | 0 |
| rs121913396 | pathogenic | 1 | 1 | 1 |
| rs28931588 | pathogenic | 1 | 1 | 1 |
| rs121913400 | pathogenic | 0 | 0 | 0 |
| rs28931589 | pathogenic | 1 | 1 | 1 |
| rs121913403 | pathogenic | 0 | 0 | 0 |
| rs121913413 | pathogenic | 0 | 0 | 0 |
| rs760107623 | pathogenic | 1 | 1 | 0 |
| rs16866378 | benign | 1 | 1 | 0 |
| rs201025290 | pathogenic | 1 | 1 | 0 |
| rs12720356 | benign | 1 | 1 | 0 |
| rs55882956 | benign | 0 | 1 | 0 |
| rs4986933 | benign | 1 | 1 | 0 |
| rs2020957 | benign | 1 | 1 | 1 |
| rs1800124 | benign | 1 | 1 | 1 |
| rs150077735 | benign | 1 | 1 | 1 |
| rs201419000 | benign | 0 | 0 | 0 |
| rs772044176 | pathogenic | 0 | 0 | 0 |
| rs138346388 | benign | 0 | 0 | 0 |
| rs121964916 | pathogenic | 0 | 0 | 0 |
| rs112534524 | pathogenic | 0 | 0 | 0 |
| rs182078921 | benign | 1 | 1 | 1 |
| rs113273712 | benign | 0 | 0 | 0 |
| rs74817407 | benign | 1 | 1 | 1 |
| rs61733901 | benign | 1 | 1 | 1 |
| rs121964912 | pathogenic | 1 | 1 | 1 |
| rs41278047 | benign | 1 | 1 | 1 |
| rs7437875 | benign | 1 | 1 | 1 |
| rs113460688 | benign | 1 | 1 | 1 |
| rs754572081 | benign | 1 | 1 | 1 |
| rs201082692 | benign | 1 | 1 | 1 |
| rs202085803 | benign | 1 | 1 | 1 |
| rs141320511 | benign | 0 | 0 | 0 |
| rs78501403 | benign | 0 | 0 | 0 |
| rs28939675 | pathogenic | 0 | 0 | 0 |
| rs74315522 | pathogenic | 0 | 0 | 0 |
| rs121434639 | pathogenic | 1 | 1 | 0 |
| rs121434640 | pathogenic | 1 | 1 | 1 |
| rs757282628 | benign | 1 | 1 | 1 |
| rs386834010 | pathogenic | 0 | 0 | 0 |
| rs200863680 | pathogenic | 0 | 0 | 0 |
| rs193919336 | pathogenic | 1 | 1 | 0 |
| rs386834011 | pathogenic | 0 | 0 | 0 |

| | | | | |
|---|---|---|---|---|
| rs28940869 | pathogenic | 0 | 0 | 0 |
| rs386834014 | pathogenic | 1 | 1 | 0 |
| rs267606960 | pathogenic | 0 | 0 | 0 |
| rs28942068 | pathogenic | 0 | 0 | 0 |
| rs193919335 | pathogenic | 0 | 0 | 0 |
| rs74374973 | pathogenic | 0 | 0 | 0 |
| rs386834018 | pathogenic | 0 | 0 | 0 |
| rs267606962 | pathogenic | 0 | 0 | 0 |
| rs6659553 | benign | 0 | 0 | 0 |
| rs29281 | benign | 0 | 0 | 0 |
| rs782265090 | benign | 0 | 0 | 0 |
| rs35910969 | pathogenic | 0 | 0 | 0 |
| rs121909113 | pathogenic | 1 | 1 | 1 |
| rs29001571 | pathogenic | 1 | 1 | 1 |
| rs587781250 | pathogenic | 1 | 1 | 1 |
| rs28939680 | pathogenic | 1 | 1 | 1 |
| rs28939681 | pathogenic | 1 | 1 | 1 |
| rs121909112 | pathogenic | 1 | 1 | 1 |
| rs28937568 | pathogenic | 1 | 1 | 1 |
| rs1799895 | pathogenic | 0 | 0 | 0 |
| rs386833431 | pathogenic | 1 | 1 | 1 |
| rs121964905 | pathogenic | 1 | 1 | 1 |
| rs121964906 | pathogenic | 1 | 1 | 1 |
| rs11551664 | pathogenic | 1 | 1 | 1 |
| rs104894705 | pathogenic | 1 | 1 | 1 |
| rs104893805 | pathogenic | 0 | 0 | 0 |
| rs104894205 | pathogenic | 0 | 0 | 0 |
| rs104894204 | pathogenic | 1 | 1 | 0 |
| rs62642926 | pathogenic | 0 | 0 | 0 |
| rs5030841 | pathogenic | 1 | 1 | 1 |
| rs75193786 | pathogenic | 1 | 1 | 1 |
| rs62508682 | pathogenic | 0 | 0 | 0 |
| rs3850625 | benign | 1 | 1 | 1 |
| rs141908793 | benign | 0 | 0 | 0 |
| rs187864727 | benign | 0 | 0 | 0 |
| rs548909101 | benign | 0 | 0 | 0 |
| rs3825394 | benign | 0 | 1 | 0 |
| rs143548402 | benign | 0 | 0 | 0 |
| rs267607146 | pathogenic | 0 | 0 | 0 |
| rs267607144 | pathogenic | 0 | 0 | 0 |
| rs267607148 | pathogenic | 1 | 1 | 1 |
| rs267607143 | pathogenic | 0 | 0 | 0 |
| rs267607145 | pathogenic | 0 | 0 | 0 |
| rs121912636 | pathogenic | 1 | 1 | 1 |
| rs121912634 | pathogenic | 1 | 1 | 1 |
| rs570675468 | benign | 0 | 0 | 0 |
| rs368861241 | pathogenic | 1 | 1 | 1 |
| rs104894528 | pathogenic | 1 | 1 | 1 |
| rs77493891 | benign | 0 | 0 | 0 |
| rs62640903 | benign | 0 | 0 | 0 |
| rs149926091 | benign | 1 | 1 | 1 |
| rs641081 | benign | 0 | 0 | 0 |
| rs4930199 | benign | 0 | 0 | 0 |
| rs121913020 | pathogenic | 0 | 0 | 0 |

| | | | | |
|---|---|---|---|---|
| rs1799793 | benign | 0 | 0 | 0 |
| rs121913016 | pathogenic | 0 | 0 | 0 |
| rs121913025 | pathogenic | 0 | 0 | 0 |
| rs121913019 | pathogenic | 0 | 0 | 0 |
| rs376556895 | pathogenic | 1 | 1 | 1 |
| rs121913024 | pathogenic | 1 | 1 | 1 |
| rs121913021 | pathogenic | 0 | 0 | 0 |
| rs121913023 | pathogenic | 0 | 0 | 0 |
| rs41556519 | pathogenic | 0 | 0 | 0 |
| rs121913022 | pathogenic | 0 | 0 | 0 |
| rs144564120 | pathogenic | 0 | 0 | 0 |
| rs121913026 | pathogenic | 1 | 1 | 1 |
| rs121913018 | pathogenic | 1 | 1 | 1 |
| rs142954643 | benign | 0 | 0 | 0 |
| rs76655496 | benign | 0 | 0 | 0 |
| rs104894371 | pathogenic | 1 | 1 | 0 |
| rs137964936 | benign | 0 | 0 | 0 |
| rs759736132 | pathogenic | 0 | 0 | 0 |
| rs387906595 | pathogenic | 1 | 1 | 0 |
| rs768248216 | pathogenic | 0 | 0 | 0 |
| rs548231008 | benign | 0 | 0 | 0 |
| rs757094384 | benign | 0 | 0 | 0 |
| rs867529 | benign | 0 | 0 | 0 |
| rs13045 | benign | 1 | 1 | 1 |
| rs121908570 | pathogenic | 0 | 0 | 0 |
| rs201872586 | benign | 1 | 1 | 0 |
| rs149092990 | benign | 0 | 0 | 0 |
| rs115585711 | benign | 0 | 0 | 0 |
| rs137909416 | benign | 1 | 1 | 0 |
| rs143388949 | benign | 1 | 1 | 1 |
| rs73137716 | benign | 1 | 1 | 1 |
| rs121908947 | pathogenic | 1 | 1 | 1 |
| rs398122904 | pathogenic | 1 | 1 | 1 |
| rs35715497 | benign | 1 | 1 | 0 |
| rs35462421 | benign | 0 | 0 | 0 |
| rs119475043 | pathogenic | 0 | 0 | 0 |
| rs119475042 | pathogenic | 0 | 0 | 0 |
| rs1076669 | benign | 1 | 1 | 0 |
| rs141146885 | benign | 0 | 0 | 0 |
| rs3026906 | pathogenic | 0 | 0 | 0 |
| rs75362221 | benign | 1 | 1 | 1 |
| rs138472116 | benign | 1 | 1 | 1 |
| rs371040282 | pathogenic | 1 | 1 | 1 |
| rs4988496 | benign | 0 | 0 | 0 |
| rs121918117 | pathogenic | 1 | 1 | 1 |
| rs4988498 | benign | 0 | 0 | 0 |
| rs5035 | benign | 0 | 0 | 0 |
| rs121918322 | pathogenic | 1 | 1 | 1 |
| rs370359540 | benign | 0 | 0 | 0 |
| rs758972589 | pathogenic | 0 | 0 | 0 |
| rs144856889 | benign | 0 | 0 | 0 |
| rs121434623 | pathogenic | 1 | 1 | 1 |
| rs61735600 | benign | 0 | 0 | 0 |
| rs72550870 | benign | 1 | 1 | 1 |

| | | | | |
|---|---|---|---|---|
| rs56392418 | benign | 0 | 0 | 0 |
| rs41307788 | benign | 1 | 1 | 1 |
| rs387907239 | pathogenic | 1 | 1 | 0 |
| rs28931584 | pathogenic | 0 | 0 | 0 |
| rs121912745 | pathogenic | 0 | 0 | 0 |
| rs121912744 | pathogenic | 0 | 0 | 0 |
| rs121912754 | pathogenic | 1 | 1 | 1 |
| rs121912746 | pathogenic | 0 | 0 | 0 |
| rs121912757 | pathogenic | 0 | 0 | 0 |
| rs121912758 | pathogenic | 0 | 0 | 0 |
| rs121912748 | pathogenic | 0 | 0 | 0 |
| rs121912755 | pathogenic | 0 | 0 | 0 |
| rs121912741 | pathogenic | 0 | 0 | 0 |
| rs121912753 | pathogenic | 0 | 0 | 0 |
| rs121912750 | pathogenic | 0 | 0 | 0 |
| rs2285644 | pathogenic | 0 | 0 | 0 |
| rs121912751 | pathogenic | 0 | 0 | 0 |
| rs5026 | benign | 0 | 0 | 0 |
| rs121912759 | pathogenic | 0 | 0 | 0 |
| rs28931585 | pathogenic | 0 | 0 | 0 |
| rs17122769 | benign | 0 | 0 | 0 |
| rs28362322 | benign | 1 | 1 | 1 |
| rs3824874 | benign | 0 | 0 | 0 |
| rs146004831 | benign | 0 | 0 | 0 |
| rs142155860 | benign | 0 | 0 | 0 |
| rs121434404 | pathogenic | 0 | 0 | 0 |
| rs61735578 | benign | 1 | 1 | 1 |
| rs558018 | benign | 0 | 0 | 0 |
| rs61735577 | benign | 1 | 1 | 1 |
| rs76784113 | benign | 0 | 0 | 0 |
| rs199469640 | pathogenic | 0 | 0 | 0 |
| rs199469628 | pathogenic | 1 | 1 | 0 |
| rs199469629 | pathogenic | 0 | 1 | 0 |
| rs387907156 | pathogenic | 0 | 1 | 0 |
| rs199469643 | pathogenic | 0 | 0 | 0 |
| rs199469646 | pathogenic | 0 | 0 | 0 |
| rs766294940 | pathogenic | 0 | 0 | 0 |
| rs387907347 | pathogenic | 0 | 0 | 0 |
| rs137853590 | pathogenic | 1 | 1 | 1 |
| rs137853589 | pathogenic | 1 | 1 | 1 |
| rs137853588 | pathogenic | 1 | 1 | 1 |
| rs55909005 | benign | 0 | 0 | 0 |
| rs4581 | benign | 0 | 0 | 0 |
| rs63750123 | benign | 0 | 0 | 0 |
| rs10254120 | benign | 0 | 0 | 0 |
| rs121434629 | pathogenic | 1 | 1 | 0 |
| rs200313585 | benign | 0 | 0 | 0 |
| rs143162541 | benign | 1 | 1 | 0 |
| rs760615315 | benign | 0 | 0 | 0 |
| rs768488890 | benign | 1 | 1 | 1 |
| rs116373169 | benign | 1 | 1 | 0 |
| rs63750871 | pathogenic | 1 | 1 | 0 |
| rs116349687 | benign | 0 | 0 | 0 |
| rs375289386 | benign | 0 | 0 | 0 |

| | | | | |
|---|---|---|---|---|
| rs760228510 | pathogenic | 1 | 1 | 1 |
| rs587779343 | pathogenic | 1 | 1 | 1 |
| rs573125799 | benign | 0 | 0 | 0 |
| rs1805322 | benign | 0 | 0 | 0 |
| rs267608153 | pathogenic | 0 | 0 | 0 |
| rs139194813 | benign | 0 | 0 | 0 |
| rs530021751 | benign | 0 | 0 | 0 |
| rs200640585 | pathogenic | 0 | 0 | 0 |
| rs139438201 | benign | 0 | 0 | 0 |
| rs200513014 | benign | 0 | 0 | 0 |
| rs567102013 | benign | 0 | 0 | 0 |
| rs121908608 | pathogenic | 0 | 0 | 0 |
| rs121908606 | pathogenic | 0 | 0 | 0 |
| rs17103671 | benign | 0 | 0 | 0 |
| rs34725281 | benign | 0 | 0 | 0 |
| rs104894421 | pathogenic | 1 | 1 | 0 |
| rs121918145 | pathogenic | 1 | 1 | 1 |
| rs121918152 | pathogenic | 1 | 1 | 1 |
| rs56379106 | benign | 0 | 0 | 0 |
| rs4987076 | benign | 0 | 0 | 0 |
| rs370243877 | pathogenic | 0 | 0 | 0 |
| rs17679445 | benign | 1 | 1 | 1 |
| rs550423482 | pathogenic | 1 | 1 | 1 |
| rs104894629 | pathogenic | 1 | 1 | 1 |
| rs773450573 | pathogenic | 1 | 1 | 1 |
| rs116679717 | benign | 0 | 0 | 0 |
| rs372000848 | pathogenic | 0 | 0 | 0 |
| rs138595973 | benign | 0 | 0 | 0 |
| rs115244417 | benign | 0 | 0 | 0 |
| rs11575292 | benign | 0 | 0 | 0 |
| rs137853211 | pathogenic | 1 | 1 | 1 |
| rs137853207 | pathogenic | 0 | 0 | 0 |
| rs137853210 | pathogenic | 1 | 1 | 1 |
| rs6262 | benign | 0 | 0 | 0 |
| rs6263 | benign | 0 | 1 | 0 |
| rs137853208 | pathogenic | 0 | 1 | 0 |
| rs137853212 | pathogenic | 1 | 1 | 0 |
| rs137853209 | pathogenic | 1 | 1 | 1 |
| rs201951824 | pathogenic | 0 | 0 | 0 |
| rs11575542 | benign | 0 | 0 | 0 |
| rs199681788 | benign | 0 | 1 | 0 |
| rs748835902 | benign | 1 | 1 | 1 |
| rs753719949 | benign | 0 | 0 | 0 |
| rs267606647 | pathogenic | 0 | 0 | 0 |
| rs267606648 | pathogenic | 1 | 1 | 1 |
| rs267606643 | pathogenic | 0 | 1 | 0 |
| rs559947967 | pathogenic | 1 | 1 | 0 |
| rs267606645 | pathogenic | 1 | 1 | 1 |
| rs12116440 | benign | 0 | 0 | 0 |
| rs121908539 | pathogenic | 0 | 0 | 0 |
| rs137853096 | pathogenic | 1 | 1 | 0 |
| rs148363262 | benign | 1 | 1 | 0 |
| rs587777442 | pathogenic | 0 | 0 | 0 |
| rs25640 | benign | 1 | 1 | 1 |

| | | | | |
|---|---|---|---|---|
| rs28943589 | benign | 0 | 0 | 0 |
| rs1143650 | benign | 1 | 1 | 1 |
| rs150886026 | benign | 0 | 1 | 0 |
| rs121908133 | pathogenic | 1 | 1 | 0 |
| rs121908131 | pathogenic | 0 | 0 | 0 |
| rs754477154 | pathogenic | 0 | 0 | 0 |
| rs747334007 | benign | 0 | 0 | 0 |
| rs121908132 | pathogenic | 0 | 0 | 0 |
| rs104894103 | pathogenic | 1 | 1 | 0 |
| rs140088149 | benign | 0 | 0 | 0 |
| rs149803148 | benign | 0 | 0 | 0 |
| rs11064480 | benign | 0 | 0 | 0 |
| rs74435397 | pathogenic | 1 | 1 | 1 |
| rs587783626 | pathogenic | 1 | 1 | 1 |
| rs148211042 | pathogenic | 1 | 1 | 1 |
| rs17849502 | benign | 0 | 0 | 0 |
| rs13306575 | pathogenic | 1 | 1 | 1 |
| rs201426733 | benign | 0 | 0 | 0 |
| rs182278784 | benign | 1 | 1 | 1 |
| rs77786415 | benign | 1 | 1 | 1 |
| rs200192119 | benign | 1 | 1 | 1 |
| rs12877501 | benign | 1 | 1 | 1 |
| rs77985244 | benign | 1 | 1 | 1 |
| rs5907 | pathogenic | 1 | 1 | 0 |
| rs2305178 | benign | 0 | 0 | 0 |
| rs3135867 | benign | 0 | 0 | 0 |
| rs121913482 | pathogenic | 1 | 1 | 1 |
| rs121913483 | pathogenic | 1 | 1 | 1 |
| rs4647924 | pathogenic | 1 | 1 | 1 |
| rs121913111 | pathogenic | 1 | 1 | 1 |
| rs104894002 | pathogenic | 0 | 0 | 0 |
| rs104894001 | pathogenic | 0 | 0 | 0 |
| rs75932628 | benign | 0 | 0 | 0 |
| rs201258663 | pathogenic | 0 | 0 | 0 |
| rs104893998 | pathogenic | 0 | 0 | 0 |
| rs2234253 | benign | 1 | 1 | 0 |
| rs121908402 | pathogenic | 1 | 1 | 0 |
| rs1805389 | benign | 0 | 0 | 0 |
| rs1805388 | benign | 0 | 0 | 0 |
| rs748385144 | pathogenic | 0 | 0 | 0 |
| rs104894420 | pathogenic | 0 | 0 | 0 |
| rs104894418 | pathogenic | 0 | 0 | 0 |
| rs61731910 | benign | 0 | 0 | 0 |
| rs758723288 | pathogenic | 1 | 1 | 0 |
| rs201865375 | pathogenic | 1 | 1 | 0 |
| rs565090080 | pathogenic | 1 | 1 | 1 |
| rs113042166 | benign | 0 | 0 | 0 |
| rs61736326 | benign | 1 | 1 | 1 |
| rs12030699 | benign | 1 | 1 | 1 |
| rs121913578 | pathogenic | 1 | 1 | 1 |
| rs61739582 | benign | 0 | 0 | 0 |
| rs745983207 | pathogenic | 0 | 0 | 0 |
| rs763524380 | benign | 0 | 0 | 0 |
| rs121912792 | pathogenic | 0 | 0 | 0 |

| | | | | |
|---|---|---|---|---|
| rs121912793 | pathogenic | 0 | 0 | 0 |
| rs387906599 | pathogenic | 0 | 0 | 0 |
| rs121912794 | pathogenic | 0 | 1 | 0 |
| rs121912795 | pathogenic | 0 | 0 | 0 |
| rs121912796 | pathogenic | 1 | 1 | 0 |
| rs121912797 | pathogenic | 0 | 1 | 0 |
| rs121912798 | pathogenic | 1 | 1 | 1 |
| rs80359812 | pathogenic | 1 | 1 | 0 |
| rs776583130 | benign | 1 | 1 | 0 |
| rs80359814 | pathogenic | 1 | 1 | 0 |
| rs202060209 | pathogenic | 1 | 1 | 0 |
| rs267607061 | pathogenic | 1 | 1 | 0 |
| rs267607060 | pathogenic | 0 | 0 | 0 |
| rs80359816 | pathogenic | 1 | 1 | 0 |
| rs80359818 | pathogenic | 1 | 1 | 0 |
| rs387907313 | pathogenic | 0 | 0 | 0 |
| rs587784395 | benign | 0 | 0 | 0 |
| rs5811 | benign | 0 | 0 | 0 |
| rs121909740 | pathogenic | 1 | 1 | 0 |
| rs148518827 | benign | 1 | 1 | 0 |
| rs121909739 | pathogenic | 1 | 1 | 0 |
| rs780528770 | benign | 1 | 1 | 0 |
| rs13306758 | pathogenic | 0 | 0 | 0 |
| rs267607059 | pathogenic | 0 | 0 | 0 |
| rs61730638 | benign | 0 | 0 | 0 |
| rs104894533 | pathogenic | 0 | 0 | 0 |
| rs104894527 | pathogenic | 0 | 0 | 0 |
| rs200503569 | pathogenic | 1 | 1 | 1 |
| rs80338700 | pathogenic | 1 | 1 | 1 |
| rs765433263 | pathogenic | 1 | 1 | 1 |
| rs104894530 | pathogenic | 1 | 1 | 1 |
| rs80338701 | pathogenic | 1 | 1 | 1 |
| rs141498002 | pathogenic | 1 | 1 | 1 |
| rs104894525 | pathogenic | 0 | 0 | 0 |
| rs80338703 | pathogenic | 0 | 0 | 0 |
| rs28936415 | pathogenic | 1 | 1 | 0 |
| rs150719105 | pathogenic | 0 | 0 | 0 |
| rs148032587 | pathogenic | 0 | 0 | 0 |
| rs150577656 | pathogenic | 0 | 0 | 0 |
| rs190521996 | pathogenic | 0 | 0 | 0 |
| rs104894526 | pathogenic | 1 | 1 | 1 |
| rs80338704 | pathogenic | 0 | 0 | 0 |
| rs199562225 | pathogenic | 1 | 1 | 1 |
| rs532870929 | pathogenic | 0 | 0 | 0 |
| rs398123309 | pathogenic | 0 | 0 | 0 |
| rs78290141 | pathogenic | 0 | 0 | 0 |
| rs80338705 | pathogenic | 0 | 0 | 0 |
| rs104894531 | pathogenic | 1 | 1 | 1 |
| rs80338706 | pathogenic | 0 | 0 | 0 |
| rs80338707 | pathogenic | 0 | 0 | 0 |
| rs80338708 | pathogenic | 0 | 0 | 0 |
| rs80338709 | pathogenic | 0 | 0 | 0 |
| rs104893833 | benign | 0 | 0 | 0 |
| rs61744585 | benign | 0 | 0 | 0 |

| | | | | |
|---|---|---|---|---|
| rs73569515 | benign | 0 | 0 | 0 |
| rs398123373 | pathogenic | 0 | 0 | 0 |
| rs756854513 | pathogenic | 0 | 0 | 0 |
| rs58232698 | benign | 0 | 0 | 0 |
| rs9600455 | benign | 0 | 0 | 0 |
| rs201016593 | pathogenic | 0 | 0 | 0 |
| rs756039188 | pathogenic | 0 | 0 | 0 |
| rs121908024 | pathogenic | 0 | 0 | 0 |
| rs373144619 | benign | 0 | 0 | 0 |
| rs138078086 | benign | 0 | 0 | 0 |
| rs376207800 | pathogenic | 0 | 0 | 0 |
| rs749038326 | pathogenic | 1 | 1 | 0 |
| rs372845091 | benign | 0 | 0 | 0 |
| rs377271627 | pathogenic | 0 | 0 | 0 |
| rs563390335 | pathogenic | 0 | 0 | 0 |
| rs13306510 | benign | 0 | 0 | 0 |
| rs764042910 | pathogenic | 0 | 0 | 0 |
| rs201384282 | benign | 1 | 1 | 1 |
| rs730882086 | pathogenic | 1 | 1 | 1 |
| rs756613387 | pathogenic | 1 | 1 | 1 |
| rs121908028 | benign | 1 | 1 | 0 |
| rs773328511 | benign | 0 | 0 | 0 |
| rs749220643 | pathogenic | 1 | 1 | 0 |
| rs730882089 | pathogenic | 0 | 0 | 0 |
| rs146651743 | benign | 0 | 0 | 0 |
| rs148698650 | pathogenic | 0 | 0 | 0 |
| rs140241383 | benign | 0 | 0 | 0 |
| rs757252110 | pathogenic | 0 | 0 | 0 |
| rs13306512 | pathogenic | 1 | 1 | 1 |
| rs121908043 | pathogenic | 1 | 1 | 1 |
| rs773658037 | pathogenic | 0 | 0 | 0 |
| rs774439908 | pathogenic | 0 | 0 | 0 |
| rs779732323 | pathogenic | 1 | 1 | 0 |
| rs749780672 | pathogenic | 0 | 0 | 0 |
| rs375651668 | benign | 1 | 1 | 1 |
| rs193922568 | pathogenic | 1 | 1 | 0 |
| rs750363970 | benign | 1 | 1 | 0 |
| rs5932 | benign | 0 | 0 | 0 |
| rs755154048 | pathogenic | 1 | 1 | 0 |
| rs759876319 | pathogenic | 0 | 0 | 0 |
| rs764550980 | pathogenic | 1 | 1 | 0 |
| rs779940524 | pathogenic | 0 | 0 | 0 |
| rs563382937 | pathogenic | 1 | 1 | 1 |
| rs28942083 | pathogenic | 0 | 1 | 0 |
| rs770744861 | pathogenic | 0 | 1 | 0 |
| rs121908031 | pathogenic | 0 | 1 | 0 |
| rs776217028 | pathogenic | 0 | 1 | 0 |
| rs763625913 | pathogenic | 0 | 0 | 0 |
| rs3736772 | benign | 0 | 0 | 0 |
| rs104894696 | benign | 1 | 1 | 1 |
| rs121918177 | pathogenic | 0 | 1 | 0 |
| rs121918181 | pathogenic | 1 | 1 | 0 |
| rs121918174 | pathogenic | 0 | 1 | 0 |
| rs121918182 | pathogenic | 0 | 0 | 0 |

| | | | | |
|---|---|---|---|---|
| rs121918175 | pathogenic | 1 | 1 | 1 |
| rs121918185 | pathogenic | 1 | 1 | 1 |
| rs121918173 | pathogenic | 1 | 1 | 0 |
| rs121918178 | pathogenic | 1 | 1 | 1 |
| rs121918179 | pathogenic | 1 | 1 | 1 |
| rs121918183 | pathogenic | 0 | 0 | 0 |
| rs121918176 | pathogenic | 1 | 1 | 1 |
| rs121918172 | pathogenic | 0 | 1 | 0 |
| rs121918184 | pathogenic | 0 | 0 | 0 |
| rs121434560 | pathogenic | 1 | 1 | 0 |
| rs121434561 | pathogenic | 1 | 1 | 0 |
| rs3218716 | pathogenic | 1 | 1 | 1 |
| rs145532615 | benign | 0 | 0 | 0 |
| rs149840927 | pathogenic | 0 | 0 | 0 |
| rs730880762 | pathogenic | 0 | 0 | 0 |
| rs730880764 | pathogenic | 0 | 0 | 0 |
| rs730880765 | pathogenic | 0 | 0 | 0 |
| rs587782962 | pathogenic | 0 | 0 | 0 |
| rs141735183 | pathogenic | 0 | 0 | 0 |
| rs397516187 | pathogenic | 0 | 0 | 0 |
| rs727505026 | pathogenic | 0 | 0 | 0 |
| rs529700838 | pathogenic | 0 | 0 | 0 |
| rs540263945 | pathogenic | 0 | 0 | 0 |
| rs149103761 | benign | 0 | 0 | 0 |
| rs730880906 | pathogenic | 0 | 0 | 0 |
| rs730880785 | pathogenic | 0 | 0 | 0 |
| rs397516199 | pathogenic | 0 | 0 | 0 |
| rs730880725 | benign | 0 | 0 | 0 |
| rs730880796 | pathogenic | 0 | 0 | 0 |
| rs201307101 | benign | 0 | 0 | 0 |
| rs397516214 | pathogenic | 0 | 0 | 0 |
| rs3729823 | benign | 0 | 0 | 0 |
| rs397516218 | pathogenic | 0 | 0 | 0 |
| rs121913647 | pathogenic | 0 | 0 | 0 |
| rs373514686 | pathogenic | 0 | 0 | 0 |
| rs763538103 | pathogenic | 0 | 0 | 0 |
| rs121913650 | pathogenic | 0 | 0 | 0 |
| rs139222507 | pathogenic | 0 | 0 | 0 |
| rs369940645 | pathogenic | 0 | 0 | 0 |
| rs763073072 | pathogenic | 0 | 0 | 0 |
| rs145734640 | pathogenic | 0 | 0 | 0 |
| rs730880821 | pathogenic | 0 | 0 | 0 |
| rs28933098 | pathogenic | 0 | 0 | 0 |
| rs372381770 | pathogenic | 0 | 0 | 0 |
| rs146796870 | benign | 0 | 0 | 0 |
| rs104894485 | pathogenic | 1 | 1 | 1 |
| rs35407712 | benign | 0 | 0 | 0 |
| rs35842750 | benign | 0 | 0 | 0 |
| rs2721191 | benign | 0 | 0 | 0 |
| rs35098923 | benign | 0 | 0 | 0 |
| rs34293591 | benign | 1 | 1 | 0 |
| rs36023964 | benign | 0 | 0 | 0 |
| rs4251691 | benign | 0 | 0 | 0 |
| rs34666647 | benign | 0 | 0 | 0 |

| | | | | |
|---|---|---|---|---|
| rs4925828 | benign | 0 | 0 | 0 |
| rs35348691 | benign | 0 | 0 | 0 |
| rs36078464 | benign | 0 | 0 | 0 |
| rs34915097 | benign | 0 | 0 | 0 |
| rs35101495 | benign | 0 | 0 | 0 |
| rs141568342 | pathogenic | 0 | 0 | 0 |
| rs1800571 | pathogenic | 0 | 0 | 0 |
| rs140566084 | benign | 1 | 1 | 1 |
| rs727503791 | pathogenic | 1 | 1 | 0 |
| rs545215807 | pathogenic | 0 | 0 | 0 |
| rs113994168 | pathogenic | 0 | 0 | 0 |
| rs113994167 | pathogenic | 0 | 0 | 0 |
| rs200788251 | pathogenic | 0 | 0 | 0 |
| rs200573371 | pathogenic | 0 | 0 | 0 |
| rs771874163 | pathogenic | 1 | 1 | 1 |
| rs112406105 | pathogenic | 1 | 1 | 1 |
| rs118204015 | pathogenic | 1 | 1 | 1 |
| rs113994169 | pathogenic | 0 | 0 | 0 |
| rs118204018 | pathogenic | 0 | 1 | 0 |
| rs138834083 | pathogenic | 0 | 0 | 0 |
| rs533055438 | pathogenic | 1 | 1 | 1 |
| rs2309689 | pathogenic | 1 | 1 | 1 |
| rs138058572 | pathogenic | 1 | 1 | 1 |
| rs118204017 | pathogenic | 1 | 1 | 0 |
| rs766742117 | pathogenic | 1 | 1 | 1 |
| rs751995154 | pathogenic | 1 | 1 | 1 |
| rs200771970 | pathogenic | 0 | 0 | 0 |
| rs118204014 | pathogenic | 0 | 0 | 0 |
| rs148584617 | pathogenic | 1 | 1 | 1 |
| rs121912720 | pathogenic | 0 | 0 | 0 |
| rs121912721 | pathogenic | 0 | 0 | 0 |
| rs28929476 | pathogenic | 0 | 0 | 0 |
| rs28931574 | pathogenic | 1 | 1 | 1 |
| rs12718465 | benign | 0 | 0 | 0 |
| rs144689354 | pathogenic | 0 | 0 | 0 |
| rs149095705 | pathogenic | 0 | 0 | 0 |
| rs572839648 | benign | 1 | 1 | 1 |
| rs121909715 | pathogenic | 1 | 1 | 0 |
| rs34077456 | benign | 1 | 1 | 1 |
| rs397517876 | benign | 0 | 0 | 0 |
| rs1064074 | benign | 0 | 0 | 0 |
| rs142751309 | benign | 0 | 0 | 0 |
| rs116275562 | benign | 0 | 0 | 0 |
| rs148095899 | benign | 0 | 0 | 0 |
| rs751838040 | pathogenic | 0 | 0 | 0 |
| rs121434324 | pathogenic | 1 | 1 | 1 |
| rs143367518 | pathogenic | 0 | 0 | 0 |
| rs121434325 | pathogenic | 0 | 0 | 0 |
| rs397515328 | pathogenic | 0 | 0 | 0 |
| rs376039938 | pathogenic | 0 | 0 | 0 |
| rs387907243 | pathogenic | 0 | 0 | 0 |
| rs28934876 | pathogenic | 0 | 0 | 0 |
| rs387907245 | pathogenic | 0 | 0 | 0 |
| rs138544311 | benign | 0 | 0 | 0 |

| | | | | |
|---|---|---|---|---|
| rs643788 | benign | 0 | 0 | 0 |
| rs749977756 | benign | 0 | 0 | 0 |
| rs1800574 | benign | 0 | 0 | 0 |
| rs137853336 | pathogenic | 1 | 1 | 1 |
| rs111916034 | benign | 0 | 0 | 0 |
| rs307826 | benign | 0 | 0 | 0 |
| rs35874891 | benign | 0 | 0 | 0 |
| rs111033580 | pathogenic | 1 | 1 | 1 |
| rs111033581 | pathogenic | 1 | 1 | 1 |
| rs121908373 | pathogenic | 0 | 0 | 0 |
| rs767122713 | pathogenic | 0 | 0 | 0 |
| rs2233213 | benign | 0 | 0 | 0 |
| rs121908605 | pathogenic | 1 | 1 | 0 |
| rs11887534 | benign | 0 | 0 | 0 |
| rs4148211 | benign | 0 | 0 | 0 |
| rs80025980 | benign | 0 | 0 | 0 |
| rs137854891 | pathogenic | 1 | 1 | 0 |
| rs74358901 | benign | 0 | 0 | 0 |
| rs9282574 | benign | 0 | 0 | 0 |
| rs137852993 | pathogenic | 0 | 0 | 0 |
| rs137852990 | pathogenic | 0 | 0 | 0 |
| rs137852987 | pathogenic | 0 | 0 | 0 |
| rs4148217 | benign | 0 | 0 | 0 |
| rs140690030 | benign | 0 | 0 | 0 |
| rs137852992 | pathogenic | 0 | 0 | 0 |
| rs6544718 | benign | 0 | 0 | 0 |
| rs137852989 | pathogenic | 0 | 0 | 0 |
| rs150893072 | benign | 1 | 1 | 1 |
| rs28931613 | pathogenic | 1 | 1 | 1 |
| rs143789597 | benign | 1 | 1 | 1 |
| rs121913037 | pathogenic | 1 | 1 | 0 |
| rs188802138 | pathogenic | 1 | 1 | 0 |
| rs121913038 | pathogenic | 1 | 1 | 0 |
| rs146835206 | benign | 0 | 0 | 0 |
| rs121913041 | pathogenic | 1 | 1 | 1 |
| rs121913039 | pathogenic | 1 | 1 | 1 |
| rs149977726 | pathogenic | 1 | 1 | 0 |
| rs121913036 | pathogenic | 0 | 0 | 0 |
| rs121913040 | pathogenic | 0 | 0 | 0 |
| rs765007158 | benign | 0 | 0 | 0 |
| rs112723255 | benign | 0 | 0 | 0 |
| rs574164748 | benign | 0 | 0 | 0 |
| rs80338734 | pathogenic | 1 | 1 | 0 |
| rs104894694 | pathogenic | 1 | 1 | 0 |
| rs753198836 | pathogenic | 0 | 0 | 0 |
| rs760101382 | pathogenic | 0 | 0 | 0 |
| rs121909272 | pathogenic | 1 | 1 | 0 |
| rs370421531 | pathogenic | 0 | 0 | 0 |
| rs565109128 | benign | 0 | 0 | 0 |
| rs17851582 | benign | 0 | 0 | 0 |
| rs753228876 | benign | 0 | 0 | 0 |
| rs141471799 | benign | 0 | 0 | 0 |
| rs281864916 | pathogenic | 1 | 1 | 0 |
| rs121918409 | pathogenic | 1 | 1 | 1 |

| | | | | |
|---|---|---|---|---|
| rs121918413 | pathogenic | 1 | 1 | 1 |
| rs121918411 | pathogenic | 1 | 1 | 1 |
| rs121918408 | pathogenic | 1 | 1 | 1 |
| rs121918412 | pathogenic | 1 | 1 | 1 |
| rs121918410 | pathogenic | 1 | 1 | 1 |
| rs116474260 | benign | 1 | 1 | 1 |
| rs281864919 | pathogenic | 1 | 1 | 1 |
| rs397514452 | pathogenic | 1 | 1 | 0 |
| rs747843638 | benign | 0 | 0 | 0 |
| rs774357609 | pathogenic | 1 | 1 | 1 |
| rs201804005 | benign | 0 | 0 | 0 |
| rs746485916 | pathogenic | 0 | 0 | 0 |
| rs764131178 | pathogenic | 0 | 0 | 0 |
| rs374094065 | pathogenic | 0 | 0 | 0 |
| rs76151636 | pathogenic | 1 | 1 | 0 |
| rs753250853 | pathogenic | 0 | 0 | 0 |
| rs761632029 | pathogenic | 0 | 0 | 0 |
| rs775541743 | pathogenic | 0 | 0 | 0 |
| rs778749563 | pathogenic | 0 | 0 | 0 |
| rs60431989 | pathogenic | 1 | 1 | 0 |
| rs755554442 | pathogenic | 1 | 1 | 0 |
| rs749085322 | pathogenic | 0 | 0 | 0 |
| rs756029120 | pathogenic | 1 | 1 | 0 |
| rs137853279 | pathogenic | 1 | 1 | 0 |
| rs777354267 | pathogenic | 1 | 1 | 1 |
| rs142940486 | benign | 1 | 1 | 1 |
| rs140167393 | benign | 1 | 1 | 1 |
| rs2066827 | benign | 0 | 0 | 0 |
| rs765681672 | benign | 0 | 0 | 0 |
| rs137985549 | benign | 0 | 0 | 0 |
| rs74315340 | pathogenic | 1 | 1 | 1 |
| rs74315341 | pathogenic | 0 | 0 | 0 |
| rs74315335 | pathogenic | 0 | 0 | 0 |
| rs61730974 | benign | 0 | 0 | 0 |
| rs121909193 | pathogenic | 0 | 0 | 0 |
| rs74315334 | pathogenic | 0 | 0 | 0 |
| rs74315329 | pathogenic | 0 | 0 | 0 |
| rs74315330 | pathogenic | 0 | 0 | 0 |
| rs121909194 | pathogenic | 0 | 0 | 0 |
| rs28936694 | pathogenic | 0 | 0 | 0 |
| rs74315336 | pathogenic | 0 | 0 | 0 |
| rs74315338 | pathogenic | 1 | 1 | 1 |
| rs74315328 | pathogenic | 0 | 0 | 0 |
| rs74315331 | pathogenic | 0 | 0 | 0 |
| rs74315332 | pathogenic | 0 | 0 | 0 |
| rs141696651 | benign | 0 | 0 | 0 |
| rs7041 | benign | 0 | 0 | 0 |
| rs4588 | benign | 0 | 0 | 0 |
| rs121908981 | pathogenic | 1 | 1 | 1 |
| rs606231459 | pathogenic | 0 | 0 | 0 |
| rs398123053 | pathogenic | 0 | 0 | 0 |
| rs774259802 | benign | 0 | 0 | 0 |
| rs121909582 | pathogenic | 1 | 1 | 1 |
| rs104893680 | pathogenic | 1 | 1 | 1 |

| | | | | |
|---|---|---|---|---|
| rs137854907 | pathogenic | 0 | 0 | 0 |
| rs765715798 | pathogenic | 0 | 0 | 0 |
| rs104893678 | pathogenic | 0 | 0 | 0 |
| rs104893679 | pathogenic | 0 | 1 | 0 |
| rs104893681 | pathogenic | 1 | 1 | 0 |
| rs121909640 | pathogenic | 1 | 1 | 0 |
| rs780705654 | pathogenic | 0 | 0 | 0 |
| rs201672146 | benign | 1 | 1 | 0 |
| rs12136658 | benign | 0 | 0 | 0 |
| rs56127440 | benign | 0 | 0 | 0 |
| rs104894433 | pathogenic | 1 | 1 | 1 |
| rs104894435 | pathogenic | 0 | 1 | 0 |
| rs104894437 | pathogenic | 1 | 1 | 1 |
| rs104894441 | pathogenic | 1 | 1 | 1 |
| rs104894440 | pathogenic | 1 | 1 | 1 |
| rs104894445 | pathogenic | 1 | 1 | 1 |
| rs104894436 | pathogenic | 0 | 0 | 0 |
| rs104894438 | pathogenic | 0 | 0 | 0 |
| rs200891969 | pathogenic | 1 | 1 | 0 |
| rs104894443 | pathogenic | 1 | 1 | 1 |
| rs104894434 | pathogenic | 1 | 1 | 1 |
| rs41298442 | pathogenic | 1 | 1 | 1 |
| rs104894442 | pathogenic | 1 | 1 | 1 |
| rs35453854 | benign | 0 | 0 | 0 |
| rs121912898 | pathogenic | 0 | 0 | 0 |
| rs121912897 | pathogenic | 1 | 1 | 0 |
| rs138260625 | benign | 0 | 0 | 0 |
| rs147802438 | benign | 1 | 1 | 0 |
| rs146443310 | benign | 1 | 1 | 0 |
| rs747319628 | pathogenic | 1 | 1 | 0 |
| rs1801195 | benign | 0 | 0 | 0 |
| rs145045341 | benign | 1 | 1 | 0 |
| rs116001173 | benign | 1 | 1 | 0 |
| rs587784175 | pathogenic | 0 | 0 | 0 |
| rs104893857 | pathogenic | 0 | 0 | 0 |
| rs104893862 | pathogenic | 1 | 1 | 0 |
| rs104893858 | pathogenic | 1 | 1 | 1 |
| rs121909249 | pathogenic | 0 | 1 | 0 |
| rs121909248 | pathogenic | 1 | 1 | 1 |
| rs104893859 | pathogenic | 1 | 1 | 0 |
| rs187063864 | pathogenic | 1 | 1 | 0 |
| rs121917817 | pathogenic | 1 | 1 | 0 |
| rs121917818 | pathogenic | 0 | 0 | 0 |
| rs753594601 | pathogenic | 0 | 0 | 0 |
| rs28937869 | pathogenic | 1 | 1 | 0 |
| rs6165 | benign | 0 | 0 | 0 |
| rs200089714 | pathogenic | 1 | 1 | 0 |
| rs372000714 | pathogenic | 0 | 0 | 0 |
| rs2228206 | benign | 0 | 0 | 0 |
| rs560096 | benign | 0 | 0 | 0 |
| rs137852666 | pathogenic | 0 | 0 | 0 |
| rs10896380 | benign | 0 | 0 | 0 |
| rs773690764 | pathogenic | 0 | 0 | 0 |
| rs201060167 | pathogenic | 0 | 0 | 0 |

| | | | | |
|---|---|---|---|---|
| rs137852670 | pathogenic | 0 | 0 | 0 |
| rs780594709 | pathogenic | 0 | 0 | 0 |
| rs145226920 | pathogenic | 0 | 0 | 0 |
| rs137852665 | pathogenic | 0 | 0 | 0 |
| rs724159960 | pathogenic | 0 | 0 | 0 |
| rs756985703 | pathogenic | 0 | 0 | 0 |
| rs137852667 | pathogenic | 0 | 0 | 0 |
| rs151079750 | pathogenic | 1 | 1 | 0 |
| rs2306636 | benign | 0 | 0 | 0 |
| rs72647885 | benign | 1 | 1 | 0 |
| rs377584435 | pathogenic | 1 | 1 | 1 |
| rs757583846 | pathogenic | 1 | 1 | 0 |
| rs121918148 | pathogenic | 0 | 0 | 0 |
| rs149201802 | benign | 0 | 0 | 0 |
| rs587781904 | pathogenic | 1 | 1 | 1 |
| rs143421325 | benign | 0 | 0 | 0 |
| rs587781253 | pathogenic | 0 | 0 | 0 |
| rs587780331 | benign | 0 | 0 | 0 |
| rs151001016 | benign | 0 | 0 | 0 |
| rs150888094 | benign | 0 | 0 | 0 |
| rs149581331 | benign | 0 | 0 | 0 |
| rs774840535 | benign | 0 | 0 | 0 |
| rs138428684 | benign | 0 | 0 | 0 |
| rs149300055 | benign | 0 | 0 | 0 |
| rs571005578 | benign | 0 | 0 | 0 |
| rs200731839 | benign | 0 | 0 | 0 |
| rs121965007 | pathogenic | 0 | 0 | 0 |
| rs121965013 | pathogenic | 0 | 1 | 0 |
| rs121965009 | pathogenic | 0 | 0 | 0 |
| rs1800457 | benign | 1 | 1 | 0 |
| rs121965006 | pathogenic | 1 | 1 | 0 |
| rs121965008 | pathogenic | 0 | 0 | 0 |
| rs61732609 | pathogenic | 0 | 0 | 0 |
| rs121965011 | pathogenic | 0 | 0 | 0 |
| rs121965015 | pathogenic | 0 | 0 | 0 |
| rs121965018 | pathogenic | 1 | 1 | 0 |
| rs121965016 | pathogenic | 0 | 0 | 0 |
| rs77806 | benign | 0 | 0 | 0 |
| rs149731356 | benign | 0 | 0 | 0 |
| rs72549407 | pathogenic | 0 | 0 | 0 |
| rs200363929 | benign | 0 | 0 | 0 |
| rs376403708 | benign | 1 | 1 | 0 |
| rs56376197 | benign | 1 | 1 | 1 |
| rs200237973 | benign | 1 | 1 | 1 |
| rs548330075 | benign | 0 | 0 | 0 |
| rs148291350 | benign | 0 | 0 | 0 |
| rs71579353 | pathogenic | 0 | 0 | 0 |
| rs146278697 | benign | 0 | 0 | 0 |
| rs141033098 | benign | 0 | 0 | 0 |
| rs397517234 | pathogenic | 0 | 0 | 0 |
| rs71579374 | benign | 0 | 0 | 0 |
| rs140381835 | benign | 0 | 0 | 0 |
| rs137852702 | pathogenic | 0 | 0 | 0 |
| rs137852696 | pathogenic | 1 | 1 | 1 |

| | | | | |
|---|---|---|---|---|
| rs137852697 | pathogenic | 0 | 0 | 0 |
| rs386833639 | pathogenic | 0 | 0 | 0 |
| rs386833640 | pathogenic | 0 | 0 | 0 |
| rs386833641 | benign | 0 | 0 | 0 |
| rs137852701 | pathogenic | 0 | 0 | 0 |
| rs386833642 | pathogenic | 0 | 0 | 0 |
| rs137852695 | pathogenic | 0 | 0 | 0 |
| rs386833646 | pathogenic | 0 | 0 | 0 |
| rs137852700 | pathogenic | 1 | 1 | 0 |
| rs386833649 | pathogenic | 1 | 1 | 0 |
| rs386833650 | pathogenic | 0 | 0 | 0 |
| rs148412181 | pathogenic | 0 | 0 | 0 |
| rs386833655 | pathogenic | 0 | 0 | 0 |
| rs386833656 | pathogenic | 1 | 1 | 0 |
| rs137852698 | pathogenic | 0 | 0 | 0 |
| rs386833661 | pathogenic | 0 | 0 | 0 |
| rs386833670 | pathogenic | 0 | 0 | 0 |
| rs111033632 | pathogenic | 1 | 1 | 1 |
| rs34308573 | benign | 1 | 1 | 1 |
| rs376048533 | pathogenic | 1 | 1 | 0 |
| rs3747517 | benign | 1 | 1 | 0 |
| rs137853328 | pathogenic | 0 | 0 | 0 |
| rs77273740 | benign | 0 | 0 | 0 |
| rs267606760 | pathogenic | 0 | 0 | 0 |
| rs77576840 | pathogenic | 0 | 0 | 0 |
| rs45465204 | benign | 1 | 1 | 0 |
| rs5320 | benign | 0 | 0 | 0 |
| rs5324 | benign | 0 | 0 | 0 |
| rs4531 | benign | 0 | 0 | 0 |
| rs267606761 | pathogenic | 0 | 0 | 0 |
| rs75215331 | pathogenic | 0 | 0 | 0 |
| rs6271 | benign | 0 | 0 | 0 |
| rs201643277 | pathogenic | 0 | 0 | 0 |
| rs4252250 | benign | 0 | 0 | 0 |
| rs137853580 | pathogenic | 0 | 0 | 0 |
| rs368287711 | pathogenic | 0 | 0 | 0 |
| rs137853579 | pathogenic | 0 | 0 | 0 |
| rs3135932 | benign | 0 | 0 | 0 |
| rs2228055 | benign | 0 | 0 | 0 |
| rs140322345 | benign | 1 | 1 | 1 |
| rs1154510 | pathogenic | 0 | 0 | 0 |
| rs11833399 | benign | 0 | 0 | 0 |
| rs36023382 | benign | 1 | 1 | 1 |
| rs61754444 | benign | 0 | 0 | 0 |
| rs28935168 | pathogenic | 0 | 0 | 0 |
| rs28934907 | pathogenic | 0 | 0 | 0 |
| rs28934904 | pathogenic | 1 | 1 | 0 |
| rs61748392 | pathogenic | 1 | 1 | 0 |
| rs28934908 | pathogenic | 0 | 0 | 0 |
| rs28934905 | pathogenic | 0 | 0 | 0 |
| rs28934906 | pathogenic | 1 | 1 | 0 |
| rs606231307 | pathogenic | 0 | 0 | 0 |
| rs9333296 | benign | 0 | 0 | 0 |
| rs3218614 | benign | 0 | 0 | 0 |

| | | | | |
|---|---|---|---|---|
| rs2230792 | benign | 0 | 0 | 0 |
| rs2230793 | benign | 0 | 0 | 0 |
| rs2230794 | benign | 0 | 0 | 0 |
| rs749052963 | pathogenic | 0 | 0 | 0 |
| rs2230798 | benign | 0 | 0 | 0 |
| rs2230795 | benign | 0 | 0 | 0 |
| rs3204145 | benign | 0 | 0 | 0 |
| rs1538660 | benign | 0 | 0 | 0 |
| rs9878928 | benign | 0 | 0 | 0 |
| rs104893742 | pathogenic | 0 | 0 | 0 |
| rs28936702 | pathogenic | 0 | 0 | 0 |
| rs28936703 | pathogenic | 0 | 0 | 0 |
| rs769392 | benign | 0 | 0 | 0 |
| rs769402 | benign | 1 | 1 | 1 |
| rs104894884 | pathogenic | 1 | 1 | 1 |
| rs1801316 | benign | 1 | 1 | 1 |
| rs104894885 | pathogenic | 1 | 1 | 1 |
| rs782572 | benign | 0 | 0 | 0 |
| rs151166046 | pathogenic | 0 | 0 | 0 |
| rs772153760 | pathogenic | 1 | 1 | 0 |
| rs35405862 | benign | 0 | 0 | 0 |
| rs146571352 | pathogenic | 0 | 1 | 0 |
| rs374698153 | pathogenic | 0 | 0 | 0 |
| rs7594497 | benign | 0 | 0 | 0 |
| rs80338716 | pathogenic | 0 | 0 | 0 |
| rs80338719 | pathogenic | 1 | 1 | 1 |
| rs80338721 | pathogenic | 0 | 0 | 0 |
| rs730882249 | pathogenic | 0 | 0 | 0 |
| rs146262009 | pathogenic | 0 | 0 | 0 |
| rs104895083 | pathogenic | 1 | 1 | 1 |
| rs104895100 | benign | 1 | 1 | 1 |
| rs104895085 | pathogenic | 0 | 0 | 0 |
| rs28940580 | pathogenic | 0 | 0 | 0 |
| rs104895098 | pathogenic | 0 | 0 | 0 |
| rs28940578 | pathogenic | 0 | 0 | 0 |
| rs61752717 | pathogenic | 0 | 0 | 0 |
| rs28940579 | pathogenic | 0 | 0 | 0 |
| rs104895097 | pathogenic | 0 | 0 | 0 |
| rs140425651 | benign | 1 | 1 | 0 |
| rs768090444 | pathogenic | 0 | 0 | 0 |
| rs764459544 | pathogenic | 0 | 0 | 0 |
| rs80338802 | pathogenic | 1 | 1 | 1 |
| rs149969786 | pathogenic | 1 | 1 | 0 |
| rs115311625 | pathogenic | 1 | 1 | 1 |
| rs778768583 | pathogenic | 1 | 1 | 1 |
| rs892090 | benign | 0 | 0 | 0 |
| rs28929498 | pathogenic | 1 | 1 | 1 |
| rs137854467 | pathogenic | 0 | 0 | 0 |
| rs137854480 | pathogenic | 0 | 0 | 0 |
| rs148682262 | benign | 0 | 0 | 0 |
| rs137852744 | pathogenic | 0 | 0 | 0 |
| rs137852745 | pathogenic | 0 | 1 | 0 |
| rs137852749 | pathogenic | 0 | 1 | 0 |
| rs137852746 | pathogenic | 0 | 1 | 0 |

| | | | | |
|---|---|---|---|---|
| rs145633958 | benign | 0 | 0 | 0 |
| rs141679950 | benign | 0 | 0 | 0 |
| rs76397662 | benign | 1 | 1 | 1 |
| rs370736139 | benign | 0 | 0 | 0 |
| rs781750106 | benign | 0 | 0 | 0 |
| rs546866208 | benign | 0 | 0 | 0 |
| rs142338976 | benign | 0 | 0 | 0 |
| rs115272158 | benign | 0 | 0 | 0 |
| rs115423919 | benign | 0 | 0 | 0 |
| rs537874538 | benign | 0 | 0 | 0 |
| rs9282834 | benign | 0 | 0 | 0 |
| rs201745826 | benign | 0 | 0 | 0 |
| rs75873440 | pathogenic | 0 | 0 | 0 |
| rs147219360 | benign | 0 | 0 | 0 |
| rs77939446 | pathogenic | 0 | 0 | 0 |
| rs77558292 | pathogenic | 0 | 0 | 0 |
| rs80069458 | pathogenic | 0 | 0 | 0 |
| rs76262710 | pathogenic | 0 | 0 | 0 |
| rs79781594 | pathogenic | 0 | 0 | 0 |
| rs77316810 | pathogenic | 0 | 0 | 0 |
| rs77503355 | pathogenic | 0 | 0 | 0 |
| rs79890926 | pathogenic | 0 | 0 | 0 |
| rs377767402 | benign | 0 | 0 | 0 |
| rs121913308 | pathogenic | 0 | 0 | 0 |
| rs55846256 | benign | 0 | 0 | 0 |
| rs75076352 | pathogenic | 0 | 0 | 0 |
| rs75996173 | pathogenic | 0 | 0 | 0 |
| rs77709286 | pathogenic | 0 | 0 | 0 |
| rs151058706 | benign | 0 | 0 | 0 |
| rs138964358 | benign | 0 | 0 | 0 |
| rs137852353 | pathogenic | 1 | 1 | 0 |
| rs201196020 | benign | 0 | 0 | 0 |
| rs137853201 | pathogenic | 1 | 1 | 1 |
| rs113994174 | pathogenic | 1 | 1 | 1 |
| rs104893674 | pathogenic | 1 | 1 | 1 |
| rs779519249 | benign | 1 | 1 | 1 |
| rs9606756 | benign | 0 | 0 | 0 |
| rs75680863 | benign | 1 | 1 | 1 |
| rs35838082 | benign | 0 | 0 | 0 |
| rs1801198 | benign | 0 | 0 | 0 |
| rs117458738 | benign | 1 | 1 | 0 |
| rs9621049 | benign | 1 | 1 | 0 |
| rs1131603 | benign | 1 | 1 | 0 |
| rs4820889 | benign | 0 | 0 | 0 |
| rs777767978 | pathogenic | 1 | 1 | 0 |
| rs147399860 | benign | 1 | 1 | 1 |
| rs104893950 | pathogenic | 0 | 0 | 0 |
| rs777308612 | benign | 0 | 0 | 0 |
| rs151091410 | benign | 0 | 0 | 0 |
| rs11537667 | benign | 0 | 0 | 0 |
| rs267606621 | pathogenic | 1 | 1 | 0 |
| rs143058902 | benign | 0 | 0 | 0 |
| rs370261904 | pathogenic | 0 | 0 | 0 |
| rs141169746 | benign | 0 | 0 | 0 |

| | | | | |
|---|---|---|---|---|
| rs111033436 | benign | 0 | 0 | 0 |
| rs148533341 | benign | 0 | 0 | 0 |
| rs143827620 | benign | 0 | 0 | 0 |
| rs4935502 | benign | 0 | 0 | 0 |
| rs61735473 | benign | 0 | 0 | 0 |
| rs267606932 | pathogenic | 1 | 1 | 0 |
| rs61730754 | benign | 0 | 0 | 0 |
| rs374491359 | pathogenic | 1 | 1 | 1 |
| rs761800077 | benign | 1 | 1 | 1 |
| rs760391381 | benign | 1 | 1 | 1 |
| rs104893885 | pathogenic | 0 | 0 | 0 |
| rs104893886 | pathogenic | 1 | 1 | 1 |
| rs104893884 | pathogenic | 0 | 0 | 0 |
| rs147715509 | benign | 0 | 0 | 0 |
| rs115533243 | benign | 0 | 0 | 0 |
| rs397509390 | pathogenic | 0 | 1 | 0 |
| rs80357134 | pathogenic | 0 | 0 | 0 |
| rs766004110 | benign | 1 | 1 | 1 |
| rs769650474 | benign | 0 | 0 | 0 |
| rs398122635 | benign | 1 | 1 | 0 |
| rs199522616 | pathogenic | 0 | 0 | 0 |
| rs28897672 | pathogenic | 1 | 1 | 0 |
| rs80357086 | pathogenic | 1 | 1 | 1 |
| rs80357064 | pathogenic | 1 | 1 | 0 |
| rs80356936 | benign | 0 | 0 | 0 |
| rs28897673 | benign | 0 | 0 | 0 |
| rs146747324 | benign | 0 | 0 | 0 |
| rs78635798 | pathogenic | 0 | 0 | 0 |
| rs75146158 | pathogenic | 1 | 1 | 1 |
| rs587782886 | benign | 0 | 0 | 0 |
| rs121434634 | pathogenic | 0 | 0 | 0 |
| rs121434635 | pathogenic | 0 | 0 | 0 |
| rs587777778 | pathogenic | 0 | 0 | 0 |
| rs118203967 | pathogenic | 0 | 0 | 0 |
| rs751375244 | pathogenic | 1 | 1 | 1 |
| rs104893863 | pathogenic | 1 | 1 | 0 |
| rs104893865 | pathogenic | 0 | 0 | 0 |
| rs104893864 | pathogenic | 1 | 1 | 1 |
| rs104893866 | pathogenic | 1 | 1 | 0 |
| rs561738497 | benign | 0 | 0 | 0 |
| rs57599352 | pathogenic | 1 | 1 | 1 |
| rs267607448 | pathogenic | 1 | 1 | 1 |
| rs63751596 | pathogenic | 0 | 0 | 0 |
| rs63751612 | benign | 0 | 0 | 0 |
| rs63751684 | benign | 0 | 0 | 0 |
| rs267607861 | benign | 0 | 0 | 0 |
| rs35831931 | benign | 0 | 0 | 0 |
| rs63750876 | benign | 0 | 0 | 0 |
| rs41295284 | benign | 0 | 0 | 0 |
| rs35502531 | benign | 0 | 0 | 0 |
| rs63750449 | benign | 0 | 0 | 0 |
| rs63750899 | pathogenic | 0 | 0 | 0 |
| rs55907433 | benign | 0 | 0 | 0 |
| rs63750217 | pathogenic | 0 | 0 | 0 |

| | | | | |
|---|---|---|---|---|
| rs63751275 | pathogenic | 0 | 0 | 0 |
| rs63750702 | benign | 1 | 1 | 1 |
| rs63750114 | benign | 0 | 0 | 0 |
| rs2020873 | benign | 0 | 0 | 0 |
| rs140195825 | benign | 1 | 1 | 1 |
| rs182763805 | benign | 0 | 0 | 0 |
| rs533696377 | benign | 0 | 0 | 0 |
| rs41278788 | benign | 0 | 0 | 0 |
| rs104894193 | pathogenic | 1 | 1 | 0 |
| rs3824915 | benign | 1 | 1 | 1 |
| rs104894196 | pathogenic | 0 | 0 | 0 |
| rs78681604 | benign | 1 | 1 | 1 |
| rs77321498 | benign | 1 | 1 | 1 |
| rs137852965 | pathogenic | 0 | 0 | 0 |
| rs137852961 | pathogenic | 0 | 0 | 0 |
| rs137852962 | pathogenic | 1 | 1 | 0 |
| rs753376100 | pathogenic | 0 | 0 | 0 |
| rs137852968 | pathogenic | 1 | 1 | 1 |
| rs137852959 | pathogenic | 1 | 1 | 0 |
| rs137852967 | pathogenic | 0 | 0 | 0 |
| rs113437301 | benign | 1 | 1 | 1 |
| rs61747482 | benign | 1 | 1 | 1 |
| rs2234919 | benign | 1 | 1 | 0 |
| rs121908863 | pathogenic | 0 | 0 | 0 |
| rs140455771 | pathogenic | 0 | 0 | 0 |
| rs121908810 | pathogenic | 1 | 1 | 1 |
| rs143879853 | benign | 0 | 0 | 0 |
| rs188319622 | benign | 0 | 0 | 0 |
| rs121918645 | pathogenic | 0 | 0 | 0 |
| rs115352435 | pathogenic | 0 | 0 | 0 |
| rs121434427 | pathogenic | 1 | 1 | 1 |
| rs121434428 | pathogenic | 0 | 0 | 0 |
| rs35086265 | benign | 1 | 1 | 1 |
| rs11576415 | benign | 0 | 0 | 0 |
| rs121434429 | pathogenic | 1 | 1 | 1 |
| rs4074536 | benign | 1 | 1 | 1 |
| rs10801999 | benign | 0 | 0 | 0 |
| rs121434550 | pathogenic | 0 | 0 | 0 |
| rs148057999 | pathogenic | 0 | 0 | 0 |
| rs28730716 | benign | 0 | 0 | 0 |
| rs142036299 | benign | 0 | 0 | 0 |
| rs762381137 | pathogenic | 0 | 0 | 0 |
| rs121434549 | pathogenic | 0 | 0 | 0 |
| rs139228801 | pathogenic | 0 | 0 | 0 |
| rs147219463 | benign | 0 | 0 | 0 |
| rs1131857 | benign | 0 | 0 | 0 |
| rs121917874 | pathogenic | 0 | 1 | 0 |
| rs2228056 | benign | 0 | 0 | 0 |
| rs121917870 | pathogenic | 0 | 0 | 0 |
| rs121917866 | pathogenic | 0 | 0 | 0 |
| rs11921054 | benign | 0 | 0 | 0 |
| rs121917868 | pathogenic | 0 | 0 | 0 |
| rs28931603 | pathogenic | 0 | 0 | 0 |
| rs104895295 | pathogenic | 1 | 1 | 0 |

| | | | | |
|---|---|---|---|---|
| rs104895382 | pathogenic | 0 | 0 | 0 |
| rs104895298 | pathogenic | 0 | 0 | 0 |
| rs104895300 | pathogenic | 0 | 0 | 0 |
| rs104895332 | pathogenic | 0 | 0 | 0 |
| rs758026399 | pathogenic | 1 | 1 | 1 |
| rs104895366 | pathogenic | 0 | 0 | 0 |
| rs104895304 | pathogenic | 0 | 0 | 0 |
| rs759997079 | pathogenic | 0 | 0 | 0 |
| rs121917789 | pathogenic | 0 | 0 | 0 |
| rs104895319 | pathogenic | 0 | 0 | 0 |
| rs104895317 | pathogenic | 1 | 1 | 0 |
| rs28934897 | pathogenic | 0 | 0 | 0 |
| rs104895380 | benign | 0 | 0 | 0 |
| rs104895360 | pathogenic | 0 | 0 | 0 |
| rs28936396 | pathogenic | 1 | 1 | 0 |
| rs121909307 | pathogenic | 1 | 1 | 1 |
| rs28938472 | pathogenic | 1 | 1 | 0 |
| rs121909308 | pathogenic | 1 | 1 | 0 |
| rs121909309 | pathogenic | 0 | 0 | 0 |
| rs121913013 | benign | 1 | 1 | 1 |
| rs201855245 | benign | 1 | 1 | 1 |
| rs2230232 | benign | 1 | 1 | 0 |
| rs113451409 | benign | 0 | 0 | 0 |
| rs121913011 | pathogenic | 1 | 1 | 0 |
| rs185821167 | pathogenic | 1 | 1 | 0 |
| rs2230234 | benign | 0 | 0 | 0 |
| rs193922639 | benign | 1 | 1 | 0 |
| rs774863785 | pathogenic | 1 | 1 | 0 |
| rs576404380 | pathogenic | 1 | 1 | 0 |
| rs2230235 | benign | 1 | 1 | 0 |
| rs200484060 | pathogenic | 1 | 1 | 0 |
| rs199761749 | benign | 0 | 0 | 0 |
| rs199681901 | benign | 0 | 0 | 0 |
| rs121908951 | pathogenic | 0 | 0 | 0 |
| rs769251460 | benign | 0 | 0 | 0 |
| rs28937320 | pathogenic | 0 | 0 | 0 |
| rs200607527 | pathogenic | 0 | 0 | 0 |
| rs137852640 | pathogenic | 1 | 1 | 1 |
| rs137852636 | pathogenic | 1 | 1 | 1 |
| rs137852638 | pathogenic | 0 | 0 | 0 |
| rs142637231 | pathogenic | 0 | 0 | 0 |
| rs151328418 | benign | 0 | 1 | 0 |
| rs137852639 | pathogenic | 1 | 1 | 1 |
| rs34605667 | benign | 0 | 0 | 0 |
| rs104894073 | pathogenic | 1 | 1 | 0 |
| rs56298569 | pathogenic | 0 | 0 | 0 |
| rs757600616 | pathogenic | 0 | 0 | 0 |
| rs139194636 | pathogenic | 0 | 0 | 0 |
| rs369517993 | pathogenic | 1 | 1 | 1 |
| rs138846036 | benign | 1 | 1 | 1 |
| rs200808744 | pathogenic | 1 | 1 | 1 |
| rs2255543 | benign | 0 | 0 | 0 |
| rs149165166 | pathogenic | 0 | 0 | 0 |
| rs544956641 | pathogenic | 1 | 1 | 1 |

| | | | | |
|---|---|---|---|---|
| rs28941783 | pathogenic | 0 | 0 | 0 |
| rs562853291 | pathogenic | 1 | 1 | 1 |
| rs28942100 | pathogenic | 0 | 0 | 0 |
| rs120074174 | pathogenic | 1 | 1 | 1 |
| rs120074171 | pathogenic | 1 | 1 | 1 |
| rs120074173 | pathogenic | 0 | 0 | 0 |
| rs120074172 | pathogenic | 1 | 1 | 1 |
| rs767159114 | pathogenic | 1 | 1 | 1 |
| rs104893657 | pathogenic | 0 | 0 | 0 |
| rs104893656 | pathogenic | 0 | 0 | 0 |
| rs104893659 | pathogenic | 0 | 0 | 0 |
| rs104893658 | pathogenic | 0 | 0 | 0 |
| rs200386310 | pathogenic | 0 | 0 | 0 |
| rs2072712 | benign | 0 | 0 | 0 |
| rs35274867 | benign | 0 | 0 | 0 |
| rs35343172 | benign | 0 | 0 | 0 |
| rs10079250 | benign | 0 | 0 | 0 |
| rs34951517 | benign | 0 | 0 | 0 |
| rs267606695 | pathogenic | 0 | 0 | 0 |
| rs149391728 | benign | 0 | 0 | 0 |
| rs387906598 | pathogenic | 1 | 1 | 0 |
| rs137854578 | pathogenic | 0 | 0 | 0 |
| rs141519952 | benign | 0 | 0 | 0 |
| rs146572883 | benign | 0 | 0 | 0 |
| rs111866410 | benign | 0 | 0 | 0 |
| rs139387758 | benign | 1 | 1 | 1 |
| rs35672074 | benign | 1 | 1 | 1 |
| rs121434322 | pathogenic | 0 | 0 | 0 |
| rs150884181 | benign | 0 | 0 | 0 |
| rs35867471 | benign | 0 | 0 | 0 |
| rs2867802 | benign | 0 | 0 | 0 |
| rs137902889 | benign | 0 | 0 | 0 |
| rs369804864 | pathogenic | 0 | 0 | 0 |
| rs781050795 | pathogenic | 0 | 0 | 0 |
| rs2305799 | benign | 0 | 0 | 0 |
| rs2229531 | benign | 0 | 0 | 0 |
| rs147025508 | benign | 0 | 0 | 0 |
| rs76857106 | pathogenic | 0 | 0 | 0 |
| rs7247284 | benign | 0 | 0 | 0 |
| rs62619782 | benign | 0 | 0 | 0 |
| rs377244188 | pathogenic | 0 | 0 | 0 |
| rs75718910 | pathogenic | 0 | 0 | 0 |
| rs79843600 | pathogenic | 0 | 0 | 0 |
| rs75037667 | pathogenic | 1 | 1 | 1 |
| rs121913006 | pathogenic | 0 | 0 | 0 |
| rs397517925 | pathogenic | 1 | 1 | 1 |
| rs267606831 | pathogenic | 1 | 1 | 1 |
| rs121909667 | pathogenic | 1 | 1 | 1 |
| rs121909669 | pathogenic | 1 | 1 | 1 |
| rs121909668 | pathogenic | 1 | 1 | 1 |
| rs121909671 | pathogenic | 1 | 1 | 1 |
| rs534596036 | benign | 0 | 0 | 0 |
| rs144037629 | pathogenic | 0 | 0 | 0 |
| rs11546406 | benign | 0 | 0 | 0 |

| | | | | |
|---|---|---|---|---|
| rs143697995 | pathogenic | 0 | 0 | 0 |
| rs121908331 | pathogenic | 0 | 0 | 0 |
| rs149377346 | benign | 0 | 0 | 0 |
| rs104894086 | pathogenic | 1 | 1 | 0 |
| rs104894089 | pathogenic | 0 | 0 | 0 |
| rs104894090 | pathogenic | 1 | 1 | 0 |
| rs387907235 | pathogenic | 0 | 0 | 0 |
| rs137852689 | pathogenic | 0 | 0 | 0 |
| rs137852690 | pathogenic | 1 | 1 | 0 |
| rs104894087 | pathogenic | 0 | 0 | 0 |
| rs104894085 | pathogenic | 0 | 0 | 0 |
| rs79383654 | benign | 0 | 0 | 0 |
| rs121434239 | pathogenic | 0 | 0 | 0 |
| rs121434240 | pathogenic | 0 | 0 | 0 |
| rs121434236 | pathogenic | 0 | 0 | 0 |
| rs121434238 | pathogenic | 1 | 1 | 1 |
| rs121918107 | pathogenic | 1 | 1 | 1 |
| rs121918103 | pathogenic | 1 | 1 | 1 |
| rs121918104 | pathogenic | 1 | 1 | 1 |
| rs61747614 | benign | 1 | 1 | 1 |
| rs555329870 | benign | 0 | 0 | 0 |
| rs186094931 | benign | 0 | 0 | 0 |
| rs267606785 | pathogenic | 0 | 0 | 0 |
| rs34474955 | benign | 0 | 0 | 0 |
| rs1126497 | benign | 1 | 1 | 1 |
| rs397514661 | pathogenic | 0 | 0 | 0 |
| rs189732445 | benign | 0 | 0 | 0 |
| rs146685071 | benign | 0 | 0 | 0 |
| rs121908636 | pathogenic | 1 | 1 | 0 |
| rs121908643 | pathogenic | 0 | 0 | 0 |
| rs121908644 | pathogenic | 1 | 1 | 1 |
| rs575001023 | pathogenic | 1 | 1 | 1 |
| rs121908646 | pathogenic | 1 | 1 | 0 |
| rs121908638 | pathogenic | 1 | 1 | 1 |
| rs777828000 | pathogenic | 1 | 1 | 1 |
| rs74923032 | benign | 0 | 0 | 0 |
| rs192838388 | pathogenic | 1 | 1 | 1 |
| rs148918985 | pathogenic | 1 | 1 | 1 |
| rs773540764 | benign | 1 | 1 | 1 |
| rs370595480 | pathogenic | 1 | 1 | 1 |
| rs762387914 | pathogenic | 1 | 1 | 1 |
| rs121908642 | pathogenic | 1 | 1 | 1 |
| rs183276875 | pathogenic | 1 | 1 | 1 |
| rs121908648 | pathogenic | 1 | 1 | 1 |
| rs121908647 | pathogenic | 1 | 1 | 1 |
| rs121908640 | pathogenic | 1 | 1 | 1 |
| rs771937610 | pathogenic | 1 | 1 | 1 |
| rs121908641 | pathogenic | 1 | 1 | 1 |
| rs5745068 | benign | 0 | 0 | 0 |
| rs73155056 | benign | 1 | 1 | 1 |
| rs148788180 | benign | 0 | 0 | 0 |
| rs121918137 | pathogenic | 0 | 0 | 0 |
| rs41269593 | benign | 0 | 0 | 0 |
| rs11159 | benign | 1 | 1 | 0 |

| | | | | |
|---|---|---|---|---|
| rs114733570 | pathogenic | 1 | 1 | 0 |
| rs3211783 | benign | 0 | 0 | 0 |
| rs4762 | benign | 0 | 0 | 0 |
| rs74315283 | pathogenic | 0 | 0 | 0 |
| rs111033559 | pathogenic | 0 | 0 | 0 |
| rs111033560 | pathogenic | 1 | 1 | 1 |
| rs749962743 | benign | 0 | 0 | 0 |
| rs267606986 | pathogenic | 0 | 0 | 0 |
| rs34516635 | benign | 0 | 0 | 0 |
| rs117440569 | benign | 0 | 0 | 0 |
| rs61740877 | benign | 0 | 0 | 0 |
| rs121918352 | pathogenic | 0 | 0 | 0 |
| rs183974372 | benign | 0 | 0 | 0 |
| rs140511204 | benign | 1 | 1 | 0 |
| rs6040067 | benign | 0 | 0 | 0 |
| rs199674138 | benign | 0 | 0 | 0 |
| rs121918350 | pathogenic | 0 | 0 | 0 |
| rs138292988 | benign | 1 | 1 | 1 |
| rs145288947 | benign | 0 | 0 | 0 |
| rs369714529 | benign | 0 | 0 | 0 |
| rs200945419 | benign | 0 | 0 | 0 |
| rs116959874 | benign | 0 | 0 | 0 |
| rs141659620 | pathogenic | 1 | 1 | 0 |
| rs111475461 | benign | 0 | 0 | 0 |
| rs61755320 | pathogenic | 0 | 0 | 0 |
| rs748600162 | pathogenic | 1 | 1 | 1 |
| rs369227537 | pathogenic | 1 | 1 | 1 |
| rs372981030 | pathogenic | 1 | 1 | 1 |
| rs137854486 | pathogenic | 0 | 0 | 0 |
| rs137854487 | pathogenic | 0 | 0 | 0 |
| rs199905054 | pathogenic | 0 | 0 | 0 |
| rs771438170 | pathogenic | 0 | 0 | 0 |
| rs774949844 | pathogenic | 0 | 0 | 0 |
| rs4899145 | benign | 0 | 0 | 0 |
| rs200642525 | benign | 0 | 0 | 0 |
| rs190140598 | pathogenic | 0 | 0 | 0 |
| rs2250889 | benign | 0 | 0 | 0 |
| rs17577 | benign | 0 | 0 | 0 |
| rs142032681 | benign | 0 | 0 | 0 |
| rs144577963 | benign | 0 | 0 | 0 |
| rs61754623 | benign | 0 | 0 | 0 |
| rs139873200 | benign | 0 | 0 | 0 |
| rs532530906 | benign | 0 | 0 | 0 |
| rs145762716 | benign | 0 | 0 | 0 |
| rs34062020 | benign | 0 | 0 | 0 |
| rs35258161 | benign | 0 | 0 | 0 |
| rs149792649 | benign | 0 | 0 | 0 |
| rs78101774 | benign | 0 | 0 | 0 |
| rs766466297 | pathogenic | 0 | 1 | 0 |
| rs140467171 | pathogenic | 0 | 0 | 0 |
| rs141340466 | pathogenic | 0 | 0 | 0 |
| rs201555303 | pathogenic | 0 | 0 | 0 |
| rs143592405 | pathogenic | 0 | 0 | 0 |
| rs781249411 | pathogenic | 0 | 0 | 0 |

| | | | | |
|---|---|---|---|---|
| rs116644127 | benign | 0 | 0 | 0 |
| rs151146863 | benign | 1 | 1 | 1 |
| rs116440799 | benign | 0 | 0 | 0 |
| rs138958351 | benign | 0 | 0 | 0 |
| rs143319354 | benign | 0 | 0 | 0 |
| rs768669208 | pathogenic | 1 | 1 | 1 |
| rs28940289 | pathogenic | 1 | 1 | 0 |
| rs745656120 | pathogenic | 1 | 1 | 0 |
| rs387906987 | pathogenic | 0 | 0 | 0 |
| rs147240777 | benign | 0 | 0 | 0 |
| rs140945833 | benign | 0 | 0 | 0 |
| rs146614981 | benign | 0 | 0 | 0 |
| rs149878562 | benign | 0 | 0 | 0 |
| rs28938168 | pathogenic | 0 | 0 | 0 |
| rs33939927 | pathogenic | 1 | 1 | 1 |
| rs34995376 | pathogenic | 1 | 1 | 1 |
| rs201034481 | pathogenic | 1 | 1 | 1 |
| rs774069983 | benign | 0 | 0 | 0 |
| rs199422226 | pathogenic | 1 | 1 | 0 |
| rs199422225 | pathogenic | 1 | 1 | 1 |
| rs199422224 | pathogenic | 1 | 1 | 0 |
| rs149271416 | pathogenic | 1 | 1 | 0 |
| rs1801318 | benign | 1 | 1 | 1 |
| rs372691318 | pathogenic | 1 | 1 | 1 |
| rs147685849 | benign | 0 | 0 | 0 |
| rs387907199 | pathogenic | 0 | 0 | 0 |
| rs767122069 | pathogenic | 0 | 0 | 0 |
| rs2302427 | benign | 0 | 0 | 0 |
| rs201135441 | benign | 0 | 0 | 0 |
| rs201861847 | pathogenic | 0 | 0 | 0 |
| rs137852952 | pathogenic | 0 | 0 | 0 |
| rs16986560 | benign | 0 | 0 | 0 |
| rs147344332 | pathogenic | 1 | 1 | 1 |
| rs36027220 | pathogenic | 0 | 0 | 0 |
| rs192401229 | benign | 0 | 0 | 0 |
| rs121909106 | pathogenic | 1 | 1 | 0 |
| rs118204103 | pathogenic | 1 | 1 | 0 |
| rs118204104 | pathogenic | 1 | 1 | 0 |
| rs118204105 | pathogenic | 1 | 1 | 0 |
| rs118204106 | pathogenic | 0 | 0 | 0 |
| rs118204119 | pathogenic | 0 | 0 | 0 |
| rs118204107 | pathogenic | 0 | 0 | 0 |
| rs118204094 | pathogenic | 0 | 0 | 0 |
| rs118204098 | pathogenic | 1 | 1 | 0 |
| rs118204101 | pathogenic | 1 | 1 | 0 |
| rs118204096 | pathogenic | 1 | 1 | 0 |
| rs118204108 | pathogenic | 0 | 0 | 0 |
| rs536814318 | pathogenic | 0 | 0 | 0 |
| rs34413634 | pathogenic | 1 | 1 | 0 |
| rs118204100 | pathogenic | 0 | 0 | 0 |
| rs118204109 | pathogenic | 0 | 0 | 0 |
| rs118204116 | pathogenic | 1 | 1 | 0 |
| rs118204110 | pathogenic | 0 | 0 | 0 |
| rs118204099 | pathogenic | 0 | 0 | 0 |

| | | | | |
|---|---|---|---|---|
| rs118204111 | pathogenic | 0 | 0 | 0 |
| rs118204112 | pathogenic | 0 | 0 | 0 |
| rs118204113 | pathogenic | 0 | 0 | 0 |
| rs118204114 | pathogenic | 0 | 0 | 0 |
| rs118204115 | pathogenic | 0 | 0 | 0 |
| rs150428209 | benign | 0 | 0 | 0 |
| rs144949995 | benign | 0 | 0 | 0 |
| rs190023981 | benign | 0 | 0 | 0 |
| rs104894611 | pathogenic | 1 | 1 | 1 |
| rs28937580 | pathogenic | 1 | 1 | 1 |
| rs121908949 | pathogenic | 0 | 0 | 0 |
| rs104894612 | pathogenic | 1 | 1 | 1 |
| rs104894609 | pathogenic | 1 | 1 | 1 |
| rs104894610 | pathogenic | 1 | 1 | 1 |
| rs104894615 | pathogenic | 0 | 0 | 0 |
| rs104894603 | pathogenic | 0 | 0 | 0 |
| rs104894602 | pathogenic | 1 | 1 | 1 |
| rs121908948 | pathogenic | 1 | 1 | 1 |
| rs104894608 | pathogenic | 1 | 1 | 1 |
| rs371545673 | benign | 0 | 0 | 0 |
| rs199925238 | benign | 0 | 0 | 0 |
| rs146313451 | benign | 0 | 0 | 0 |
| rs121918193 | pathogenic | 1 | 1 | 0 |
| rs2228500 | benign | 0 | 0 | 0 |
| rs121918194 | pathogenic | 1 | 1 | 0 |
| rs121918196 | pathogenic | 0 | 0 | 0 |
| rs41291971 | benign | 0 | 0 | 0 |
| rs199473690 | pathogenic | 0 | 0 | 0 |
| rs2228613 | benign | 0 | 0 | 0 |
| rs267607130 | pathogenic | 1 | 1 | 1 |
| rs77615401 | benign | 1 | 1 | 1 |
| rs374618872 | benign | 1 | 1 | 1 |
| rs146165014 | benign | 0 | 0 | 0 |
| rs144586070 | benign | 0 | 0 | 0 |
| rs121918541 | pathogenic | 0 | 0 | 0 |
| rs142105943 | benign | 0 | 0 | 0 |
| rs121918540 | pathogenic | 0 | 0 | 0 |
| rs6791 | benign | 0 | 0 | 0 |
| rs61736587 | pathogenic | 0 | 0 | 0 |
| rs61736586 | benign | 0 | 0 | 0 |
| rs121917741 | pathogenic | 1 | 1 | 1 |
| rs41302885 | benign | 0 | 0 | 0 |
| rs141026028 | benign | 0 | 0 | 0 |
| rs193922493 | pathogenic | 0 | 0 | 0 |
| rs73598374 | benign | 0 | 0 | 0 |
| rs121908725 | pathogenic | 1 | 1 | 0 |
| rs199422328 | pathogenic | 0 | 1 | 0 |
| rs11555566 | benign | 1 | 1 | 1 |
| rs267606634 | pathogenic | 0 | 0 | 0 |
| rs121908714 | pathogenic | 1 | 1 | 0 |
| rs121908717 | pathogenic | 1 | 1 | 0 |
| rs267606635 | pathogenic | 1 | 1 | 0 |
| rs121908739 | pathogenic | 0 | 0 | 0 |
| rs121908735 | pathogenic | 0 | 1 | 0 |

| | | | | |
|---|---|---|---|---|
| rs121908716 | pathogenic | 0 | 1 | 0 |
| rs121908723 | pathogenic | 0 | 1 | 0 |
| rs121908721 | pathogenic | 0 | 0 | 0 |
| rs199422327 | pathogenic | 0 | 0 | 0 |
| rs121908715 | pathogenic | 0 | 0 | 0 |
| rs587783563 | pathogenic | 0 | 0 | 0 |
| rs104894780 | pathogenic | 0 | 0 | 0 |
| rs56030372 | pathogenic | 1 | 1 | 1 |
| rs104894782 | pathogenic | 1 | 1 | 1 |
| rs149495971 | benign | 0 | 0 | 0 |
| rs118203942 | pathogenic | 1 | 1 | 0 |
| rs118203939 | pathogenic | 0 | 0 | 0 |
| rs118203938 | pathogenic | 0 | 0 | 0 |
| rs118203943 | pathogenic | 0 | 0 | 0 |
| rs118203940 | pathogenic | 0 | 0 | 0 |
| rs727503809 | pathogenic | 0 | 0 | 0 |
| rs1065757 | benign | 0 | 0 | 0 |
| rs1071598 | benign | 0 | 0 | 0 |
| rs25414 | benign | 0 | 0 | 0 |
| rs118203944 | pathogenic | 0 | 0 | 0 |
| rs118203941 | pathogenic | 0 | 0 | 0 |
| rs144222167 | benign | 0 | 0 | 0 |
| rs201101343 | pathogenic | 0 | 0 | 0 |
| rs144601717 | benign | 0 | 0 | 0 |
| rs267607129 | pathogenic | 0 | 0 | 0 |
| rs104894729 | pathogenic | 0 | 0 | 0 |
| rs104894727 | pathogenic | 0 | 0 | 0 |
| rs104894725 | pathogenic | 0 | 0 | 0 |
| rs104894515 | pathogenic | 1 | 1 | 1 |
| rs74688377 | benign | 0 | 0 | 0 |
| rs121908513 | pathogenic | 1 | 1 | 0 |
| rs121908509 | pathogenic | 0 | 0 | 0 |
| rs121908514 | pathogenic | 1 | 1 | 0 |
| rs121908512 | pathogenic | 1 | 1 | 0 |
| rs121908510 | pathogenic | 0 | 0 | 0 |
| rs121908511 | pathogenic | 1 | 1 | 1 |
| rs121908518 | pathogenic | 1 | 1 | 1 |
| rs34358665 | benign | 0 | 0 | 0 |
| rs147568834 | benign | 0 | 0 | 0 |
| rs201717599 | benign | 0 | 0 | 0 |
| rs61744487 | benign | 0 | 0 | 0 |
| rs370627877 | pathogenic | 1 | 1 | 1 |
| rs368593151 | pathogenic | 1 | 1 | 1 |
| rs114363936 | benign | 0 | 0 | 0 |
| rs137853333 | pathogenic | 0 | 0 | 0 |
| rs137853080 | pathogenic | 1 | 1 | 1 |
| rs137853077 | pathogenic | 1 | 1 | 1 |
| rs780749732 | benign | 1 | 1 | 1 |
| rs137853081 | pathogenic | 1 | 1 | 0 |
| rs137853078 | pathogenic | 0 | 0 | 0 |
| rs121913315 | pathogenic | 1 | 1 | 1 |
| rs752513525 | pathogenic | 1 | 1 | 1 |
| rs104894918 | pathogenic | 0 | 0 | 0 |
| rs104894917 | pathogenic | 0 | 0 | 0 |

| | | | | |
|---|---|---|---|---|
| rs104894919 | pathogenic | 0 | 0 | 0 |
| rs28939672 | pathogenic | 0 | 0 | 0 |
| rs104894173 | pathogenic | 0 | 0 | 0 |
| rs104894179 | pathogenic | 0 | 0 | 0 |
| rs104894174 | pathogenic | 1 | 1 | 0 |
| rs104894178 | pathogenic | 1 | 1 | 0 |
| rs28939714 | pathogenic | 1 | 1 | 1 |
| rs104894270 | pathogenic | 1 | 1 | 1 |
| rs3765571 | benign | 0 | 0 | 0 |
| rs752669339 | pathogenic | 1 | 1 | 1 |
| rs537043237 | pathogenic | 1 | 1 | 1 |
| rs774047299 | pathogenic | 1 | 1 | 1 |
| rs121434583 | pathogenic | 1 | 1 | 1 |
| rs121907927 | pathogenic | 0 | 0 | 0 |
| rs121907925 | pathogenic | 0 | 1 | 0 |
| rs746212325 | pathogenic | 0 | 0 | 0 |
| rs754637179 | pathogenic | 0 | 0 | 0 |
| rs780882740 | pathogenic | 0 | 0 | 0 |
| rs121918130 | pathogenic | 0 | 0 | 0 |
| rs121918129 | pathogenic | 0 | 0 | 0 |
| rs13297509 | pathogenic | 0 | 0 | 0 |
| rs746867724 | pathogenic | 0 | 0 | 0 |
| rs121918128 | pathogenic | 0 | 0 | 0 |
| rs752300607 | pathogenic | 0 | 0 | 0 |
| rs121964860 | pathogenic | 0 | 0 | 0 |
| rs45466197 | pathogenic | 1 | 1 | 1 |
| rs3730238 | benign | 1 | 1 | 1 |
| rs730881125 | pathogenic | 1 | 1 | 1 |
| rs121964861 | pathogenic | 1 | 1 | 1 |
| rs397516484 | pathogenic | 0 | 0 | 0 |
| rs367785431 | pathogenic | 0 | 0 | 0 |
| rs141121678 | pathogenic | 0 | 0 | 0 |
| rs12720263 | benign | 0 | 0 | 0 |
| rs2304256 | benign | 0 | 0 | 0 |
| rs2304255 | benign | 0 | 0 | 0 |
| rs121909045 | pathogenic | 0 | 0 | 0 |
| rs397508796 | pathogenic | 0 | 0 | 0 |
| rs151020603 | pathogenic | 0 | 0 | 0 |
| rs77284892 | pathogenic | 0 | 0 | 0 |
| rs368505753 | pathogenic | 0 | 0 | 0 |
| rs115545701 | pathogenic | 0 | 0 | 0 |
| rs121908749 | pathogenic | 0 | 0 | 0 |
| rs75961395 | pathogenic | 0 | 0 | 0 |
| rs121908750 | pathogenic | 0 | 0 | 0 |
| rs121908751 | pathogenic | 0 | 0 | 0 |
| rs397508464 | pathogenic | 0 | 0 | 0 |
| rs397508461 | pathogenic | 0 | 0 | 0 |
| rs121909031 | pathogenic | 0 | 0 | 0 |
| rs113993958 | pathogenic | 0 | 0 | 0 |
| rs77834169 | pathogenic | 0 | 0 | 0 |
| rs78655421 | pathogenic | 0 | 0 | 0 |
| rs141723617 | benign | 0 | 0 | 0 |
| rs35516286 | benign | 0 | 0 | 0 |
| rs1800079 | pathogenic | 1 | 1 | 0 |

| | | | | |
|---|---|---|---|---|
| rs80282562 | pathogenic | 0 | 0 | 0 |
| rs121908752 | pathogenic | 0 | 0 | 0 |
| rs556880586 | pathogenic | 0 | 0 | 0 |
| rs121908760 | pathogenic | 0 | 0 | 0 |
| rs397508350 | pathogenic | 0 | 0 | 0 |
| rs374946172 | pathogenic | 0 | 0 | 0 |
| rs397508387 | pathogenic | 0 | 0 | 0 |
| rs201386642 | pathogenic | 0 | 0 | 0 |
| rs121909012 | pathogenic | 0 | 0 | 0 |
| rs121909008 | pathogenic | 0 | 0 | 0 |
| rs149790377 | pathogenic | 0 | 0 | 0 |
| rs397508442 | pathogenic | 0 | 0 | 0 |
| rs121909035 | pathogenic | 0 | 0 | 0 |
| rs386134230 | pathogenic | 0 | 0 | 0 |
| rs1800112 | benign | 0 | 0 | 0 |
| rs144055758 | pathogenic | 0 | 0 | 0 |
| rs142394380 | pathogenic | 0 | 0 | 0 |
| rs121909036 | pathogenic | 0 | 0 | 0 |
| rs78194216 | pathogenic | 0 | 0 | 0 |
| rs121909019 | pathogenic | 0 | 0 | 0 |
| rs121909020 | pathogenic | 0 | 0 | 0 |
| rs200321110 | pathogenic | 0 | 0 | 0 |
| rs202179988 | pathogenic | 0 | 0 | 0 |
| rs121909037 | pathogenic | 0 | 0 | 0 |
| rs186045772 | pathogenic | 0 | 0 | 0 |
| rs139304906 | pathogenic | 0 | 0 | 0 |
| rs78802634 | pathogenic | 0 | 0 | 0 |
| rs121908761 | pathogenic | 0 | 0 | 0 |
| rs36210737 | pathogenic | 0 | 0 | 0 |
| rs139729994 | pathogenic | 0 | 0 | 0 |
| rs79850223 | pathogenic | 0 | 0 | 0 |
| rs1800120 | benign | 0 | 0 | 0 |
| rs74767530 | pathogenic | 0 | 0 | 0 |
| rs374705585 | pathogenic | 0 | 0 | 0 |
| rs1136287 | benign | 0 | 0 | 0 |
| rs6828 | benign | 0 | 0 | 0 |
| rs104894856 | pathogenic | 0 | 0 | 0 |
| rs104894860 | pathogenic | 0 | 0 | 0 |
| rs61736892 | pathogenic | 0 | 0 | 0 |
| rs145807417 | benign | 0 | 0 | 0 |
| rs104894853 | pathogenic | 0 | 0 | 0 |
| rs199422227 | pathogenic | 0 | 0 | 0 |
| rs113993946 | pathogenic | 0 | 0 | 0 |
| rs199422231 | pathogenic | 0 | 0 | 0 |
| rs199422228 | pathogenic | 0 | 0 | 0 |
| rs142195196 | benign | 0 | 1 | 0 |
| rs148656104 | benign | 1 | 1 | 1 |
| rs141944844 | benign | 0 | 0 | 0 |
| rs104894342 | pathogenic | 0 | 0 | 0 |
| rs104894343 | pathogenic | 0 | 0 | 0 |
| rs104894344 | pathogenic | 0 | 0 | 0 |
| rs573322878 | benign | 0 | 0 | 0 |
| rs104894347 | pathogenic | 0 | 0 | 0 |
| rs28937882 | pathogenic | 0 | 0 | 0 |

| | | | | |
|---|---|---|---|---|
| rs193922702 | pathogenic | 0 | 0 | 0 |
| rs200786104 | benign | 0 | 0 | 0 |
| rs690016544 | pathogenic | 1 | 1 | 1 |
| rs28943592 | benign | 0 | 0 | 0 |
| rs28943594 | benign | 0 | 0 | 0 |
| rs121908957 | pathogenic | 0 | 0 | 0 |
| rs115408010 | benign | 1 | 1 | 1 |
| rs77591659 | benign | 0 | 0 | 0 |
| rs121918511 | pathogenic | 0 | 0 | 0 |
| rs538359948 | benign | 0 | 0 | 0 |
| rs76708142 | benign | 0 | 0 | 0 |
| rs121907913 | pathogenic | 1 | 1 | 0 |
| rs121907920 | pathogenic | 0 | 0 | 0 |
| rs121907923 | pathogenic | 1 | 1 | 0 |
| rs121907928 | pathogenic | 1 | 1 | 0 |
| rs121907919 | pathogenic | 1 | 1 | 0 |
| rs121907918 | pathogenic | 1 | 1 | 0 |
| rs74315493 | pathogenic | 1 | 1 | 1 |
| rs74315494 | pathogenic | 1 | 1 | 1 |
| rs145446060 | benign | 1 | 1 | 1 |
| rs550409143 | benign | 1 | 1 | 0 |

| compound_binding_site | cofactor_binding_site | nucleic_binding_site | ion_binding_site |
|---|---|---|---|
| 1 | 0 | 0 | 0 |
| 1 | 0 | 0 | 0 |
| 0 | 0 | 0 | 0 |
| 1 | 0 | 0 | 0 |
| 0 | 0 | 0 | 0 |
| 0 | 0 | 0 | 0 |
| 0 | 0 | 0 | 0 |
| 0 | 0 | 0 | 0 |
| 1 | 0 | 0 | 1 |
| 1 | 0 | 0 | 0 |
| 0 | 0 | 0 | 1 |
| 1 | 0 | 1 | 0 |
| 0 | 0 | 0 | 0 |
| 0 | 0 | 0 | 0 |
| 0 | 0 | 0 | 0 |
| 0 | 0 | 0 | 0 |
| 0 | 0 | 0 | 0 |
| 1 | 0 | 0 | 1 |
| 1 | 0 | 1 | 0 |
| 0 | 0 | 0 | 0 |
| 1 | 0 | 0 | 1 |
| 1 | 0 | 0 | 0 |
| 1 | 0 | 0 | 0 |
| 1 | 0 | 0 | 0 |
| 0 | 0 | 0 | 0 |
| 0 | 0 | 0 | 0 |
| 1 | 0 | 0 | 1 |
| 0 | 0 | 0 | 0 |
| 0 | 0 | 0 | 0 |
| 0 | 0 | 0 | 0 |
| 0 | 0 | 0 | 0 |
| 1 | 0 | 0 | 0 |
| 1 | 0 | 0 | 1 |
| 1 | 0 | 0 | 0 |
| 0 | 0 | 0 | 0 |
| 0 | 0 | 0 | 0 |
| 0 | 0 | 0 | 0 |
| 1 | 0 | 0 | 1 |
| 1 | 0 | 0 | 1 |
| 1 | 0 | 0 | 1 |
| 1 | 0 | 0 | 0 |
| 1 | 0 | 0 | 0 |
| 1 | 0 | 0 | 0 |
| 1 | 0 | 0 | 0 |
| 1 | 0 | 0 | 0 |
| 1 | 0 | 0 | 0 |
| 1 | 0 | 0 | 0 |
| 1 | 0 | 0 | 0 |
| 1 | 0 | 0 | 0 |
| 1 | 0 | 0 | 0 |
| 1 | 0 | 0 | 0 |
| 1 | 0 | 0 | 0 |
| 1 | 0 | 0 | 0 |

| | | | |
|---|---|---|---|
| 1 | 0 | 0 | 1 |
| 1 | 0 | 0 | 0 |
| 1 | 0 | 0 | 0 |
| 1 | 0 | 0 | 0 |
| 0 | 0 | 0 | 0 |
| 1 | 0 | 0 | 0 |
| 0 | 0 | 0 | 0 |
| 1 | 0 | 0 | 0 |
| 1 | 0 | 0 | 0 |
| 1 | 0 | 0 | 1 |
| 1 | 0 | 0 | 0 |
| 1 | 0 | 0 | 0 |
| 1 | 0 | 0 | 0 |
| 1 | 0 | 0 | 0 |
| 1 | 0 | 0 | 0 |
| 1 | 0 | 0 | 0 |
| 1 | 0 | 0 | 0 |
| 1 | 0 | 0 | 1 |
| 1 | 0 | 0 | 1 |
| 1 | 1 | 0 | 1 |
| 1 | 0 | 0 | 0 |
| 1 | 0 | 0 | 0 |
| 1 | 0 | 0 | 0 |
| 1 | 0 | 0 | 0 |
| 1 | 1 | 0 | 0 |
| 1 | 0 | 0 | 0 |
| 1 | 0 | 0 | 0 |
| 1 | 0 | 0 | 0 |
| 1 | 0 | 0 | 0 |
| 1 | 0 | 0 | 0 |
| 1 | 1 | 0 | 0 |
| 1 | 1 | 0 | 0 |
| 1 | 1 | 0 | 0 |
| 1 | 1 | 0 | 0 |
| 1 | 1 | 0 | 0 |
| 1 | 0 | 0 | 0 |
| 1 | 1 | 0 | 0 |
| 1 | 0 | 0 | 0 |
| 1 | 0 | 0 | 0 |
| 1 | 1 | 0 | 0 |
| 0 | 0 | 0 | 0 |
| 0 | 0 | 0 | 0 |
| 0 | 0 | 0 | 0 |
| 0 | 0 | 0 | 0 |
| 0 | 0 | 0 | 0 |
| 0 | 0 | 0 | 0 |
| 0 | 0 | 0 | 0 |
| 1 | 0 | 0 | 1 |
| 0 | 0 | 0 | 0 |
| 1 | 0 | 0 | 0 |
| 1 | 0 | 0 | 0 |
| 0 | 0 | 0 | 0 |
| 0 | 0 | 0 | 0 |
| 0 | 0 | 0 | 0 |

| | | | |
|---|---|---|---|
| 0 | 0 | 0 | 0 |
| 0 | 0 | 0 | 0 |
| 0 | 0 | 0 | 0 |
| 1 | 0 | 0 | 0 |
| 1 | 0 | 0 | 0 |
| 1 | 0 | 0 | 0 |
| 0 | 0 | 0 | 0 |
| 0 | 0 | 1 | 0 |
| 0 | 0 | 1 | 0 |
| 0 | 0 | 1 | 0 |
| 0 | 0 | 1 | 0 |
| 0 | 0 | 1 | 0 |
| 0 | 0 | 0 | 0 |
| 0 | 0 | 0 | 0 |
| 1 | 0 | 0 | 0 |
| 0 | 0 | 0 | 0 |
| 0 | 0 | 0 | 0 |
| 1 | 0 | 0 | 0 |
| 1 | 0 | 0 | 0 |
| 1 | 0 | 0 | 0 |
| 1 | 0 | 0 | 0 |
| 1 | 1 | 0 | 0 |
| 1 | 0 | 0 | 0 |
| 1 | 0 | 0 | 0 |
| 0 | 0 | 0 | 0 |
| 0 | 0 | 0 | 0 |
| 1 | 0 | 0 | 0 |
| 1 | 1 | 0 | 0 |
| 1 | 0 | 0 | 0 |
| 1 | 0 | 0 | 0 |
| 1 | 1 | 0 | 0 |
| 1 | 1 | 0 | 0 |
| 1 | 1 | 0 | 0 |
| 1 | 1 | 0 | 0 |
| 1 | 1 | 0 | 0 |
| 1 | 1 | 0 | 0 |
| 1 | 1 | 0 | 0 |
| 1 | 1 | 0 | 0 |
| 1 | 1 | 0 | 0 |
| 1 | 1 | 0 | 0 |
| 1 | 1 | 0 | 0 |
| 1 | 1 | 0 | 0 |
| 1 | 0 | 0 | 0 |
| 1 | 0 | 0 | 0 |
| 1 | 0 | 0 | 0 |
| 1 | 1 | 0 | 0 |
| 1 | 0 | 0 | 0 |
| 1 | 1 | 1 | 0 |
| 1 | 1 | 1 | 0 |
| 1 | 1 | 0 | 0 |
| 1 | 1 | 0 | 0 |
| 1 | 0 | 0 | 0 |
| 1 | 0 | 0 | 0 |
| 0 | 0 | 0 | 0 |

| | | | |
|---|---|---|---|
| 0 | 0 | 0 | 0 |
| 0 | 0 | 0 | 0 |
| 0 | 0 | 0 | 0 |
| 1 | 0 | 0 | 0 |
| 1 | 0 | 0 | 0 |
| 0 | 0 | 0 | 0 |
| 0 | 0 | 0 | 0 |
| 0 | 0 | 0 | 0 |
| 0 | 0 | 0 | 0 |
| 0 | 0 | 0 | 0 |
| 0 | 0 | 0 | 0 |
| 0 | 0 | 0 | 0 |
| 0 | 0 | 0 | 0 |
| 0 | 0 | 0 | 0 |
| 0 | 0 | 0 | 0 |
| 0 | 0 | 0 | 0 |
| 0 | 0 | 0 | 0 |
| 0 | 0 | 0 | 0 |
| 0 | 0 | 0 | 0 |
| 0 | 0 | 0 | 0 |
| 0 | 0 | 0 | 0 |
| 0 | 0 | 0 | 0 |
| 1 | 0 | 0 | 0 |
| 1 | 0 | 0 | 0 |
| 0 | 0 | 0 | 0 |
| 0 | 0 | 0 | 0 |
| 1 | 0 | 0 | 0 |
| 0 | 0 | 0 | 0 |
| 0 | 0 | 0 | 0 |
| 0 | 0 | 0 | 0 |
| 0 | 0 | 0 | 0 |
| 0 | 0 | 0 | 0 |
| 0 | 0 | 0 | 0 |
| 0 | 0 | 0 | 0 |
| 1 | 0 | 0 | 0 |
| 0 | 0 | 0 | 0 |
| 0 | 0 | 0 | 0 |
| 0 | 0 | 0 | 0 |
| 0 | 0 | 0 | 0 |
| 0 | 0 | 0 | 0 |
| 0 | 0 | 0 | 0 |
| 0 | 0 | 0 | 0 |
| 0 | 0 | 0 | 0 |
| 0 | 0 | 0 | 0 |
| 0 | 0 | 0 | 0 |
| 0 | 0 | 0 | 0 |
| 0 | 0 | 1 | 0 |
| 1 | 0 | 0 | 0 |
| 1 | 0 | 0 | 0 |
| 1 | 1 | 0 | 0 |
| 1 | 0 | 0 | 0 |
| 0 | 0 | 0 | 0 |
| 0 | 0 | 0 | 0 |
| 1 | 0 | 0 | 0 |

| | | | |
|---|---|---|---|
| 1 | 0 | 0 | 0 |
| 1 | 0 | 0 | 0 |
| 1 | 0 | 0 | 0 |
| 0 | 0 | 0 | 0 |
| 0 | 0 | 0 | 0 |
| 0 | 0 | 0 | 0 |
| 1 | 0 | 0 | 0 |
| 1 | 0 | 0 | 0 |
| 0 | 0 | 0 | 0 |
| 0 | 0 | 0 | 0 |
| 1 | 0 | 0 | 0 |
| 0 | 0 | 0 | 0 |
| 1 | 0 | 0 | 0 |
| 0 | 0 | 0 | 0 |
| 0 | 0 | 0 | 0 |
| 0 | 0 | 0 | 0 |
| 0 | 0 | 0 | 0 |
| 0 | 0 | 0 | 0 |
| 0 | 0 | 0 | 0 |
| 0 | 0 | 0 | 0 |
| 0 | 0 | 0 | 0 |
| 0 | 0 | 0 | 0 |
| 0 | 0 | 0 | 0 |
| 0 | 0 | 0 | 0 |
| 0 | 0 | 0 | 0 |
| 0 | 0 | 0 | 0 |
| 1 | 0 | 0 | 0 |
| 1 | 0 | 0 | 0 |
| 1 | 0 | 0 | 0 |
| 1 | 0 | 0 | 0 |
| 1 | 0 | 0 | 0 |
| 1 | 0 | 0 | 0 |
| 0 | 0 | 0 | 0 |
| 0 | 0 | 0 | 0 |
| 1 | 0 | 0 | 0 |
| 0 | 0 | 0 | 0 |
| 0 | 0 | 0 | 0 |
| 0 | 0 | 0 | 0 |
| 0 | 0 | 0 | 0 |
| 0 | 0 | 0 | 0 |
| 0 | 0 | 0 | 0 |
| 0 | 0 | 0 | 0 |
| 0 | 0 | 0 | 0 |
| 0 | 0 | 0 | 0 |
| 1 | 1 | 0 | 0 |
| 1 | 1 | 0 | 0 |
| 1 | 1 | 0 | 1 |
| 1 | 1 | 0 | 1 |
| 1 | 1 | 0 | 1 |
| 1 | 1 | 0 | 1 |
| 1 | 1 | 0 | 0 |
| 1 | 1 | 0 | 1 |
| 1 | 1 | 0 | 1 |
| 1 | 1 | 0 | 1 |

| | | | |
|---|---|---|---|
| 1 | 1 | 0 | 1 |
| 1 | 1 | 0 | 1 |
| 1 | 1 | 0 | 1 |
| 1 | 1 | 0 | 1 |
| 1 | 1 | 0 | 1 |
| 1 | 0 | 0 | 0 |
| 0 | 0 | 0 | 0 |
| 0 | 0 | 0 | 0 |
| 0 | 0 | 0 | 0 |
| 1 | 0 | 0 | 0 |
| 0 | 0 | 1 | 0 |
| 0 | 0 | 0 | 0 |
| 0 | 0 | 0 | 0 |
| 0 | 0 | 0 | 0 |
| 0 | 0 | 0 | 0 |
| 0 | 0 | 0 | 0 |
| 0 | 0 | 0 | 0 |
| 0 | 0 | 0 | 0 |
| 0 | 0 | 0 | 0 |
| 0 | 0 | 0 | 0 |
| 0 | 0 | 0 | 0 |
| 0 | 0 | 0 | 0 |
| 0 | 0 | 0 | 0 |
| 0 | 0 | 0 | 0 |
| 0 | 0 | 0 | 0 |
| 1 | 1 | 0 | 0 |
| 1 | 1 | 0 | 0 |
| 1 | 1 | 0 | 0 |
| 0 | 1 | 0 | 0 |
| 1 | 0 | 0 | 0 |
| 0 | 0 | 0 | 0 |
| 0 | 0 | 0 | 0 |
| 0 | 0 | 0 | 0 |
| 1 | 0 | 0 | 0 |
| 1 | 1 | 0 | 0 |
| 0 | 0 | 0 | 0 |
| 0 | 0 | 0 | 0 |
| 0 | 0 | 0 | 0 |
| 0 | 0 | 0 | 0 |
| 1 | 0 | 0 | 0 |
| 0 | 0 | 0 | 0 |
| 0 | 0 | 0 | 0 |
| 0 | 0 | 0 | 0 |
| 0 | 0 | 0 | 0 |
| 0 | 0 | 0 | 0 |
| 0 | 1 | 0 | 0 |
| 0 | 1 | 0 | 0 |
| 0 | 0 | 0 | 0 |
| 0 | 0 | 0 | 0 |
| 0 | 1 | 0 | 0 |
| 0 | 0 | 0 | 0 |
| 0 | 0 | 0 | 0 |
| 0 | 1 | 0 | 0 |
| 0 | 0 | 0 | 0 |

| | | | |
|---|---|---|---|
| 1 | 1 | 0 | 0 |
| 0 | 0 | 0 | 0 |
| 0 | 0 | 0 | 0 |
| 0 | 0 | 0 | 0 |
| 0 | 0 | 0 | 0 |
| 0 | 0 | 0 | 0 |
| 0 | 0 | 0 | 0 |
| 0 | 0 | 0 | 0 |
| 0 | 0 | 0 | 0 |
| 0 | 0 | 0 | 0 |
| 0 | 0 | 0 | 0 |
| 0 | 0 | 0 | 0 |
| 0 | 0 | 0 | 0 |
| 0 | 0 | 0 | 0 |
| 0 | 1 | 0 | 0 |
| 0 | 0 | 0 | 0 |
| 0 | 1 | 0 | 0 |
| 0 | 1 | 0 | 0 |
| 0 | 0 | 0 | 0 |
| 0 | 1 | 0 | 0 |
| 1 | 0 | 0 | 1 |
| 1 | 1 | 0 | 0 |
| 1 | 0 | 0 | 0 |
| 1 | 0 | 0 | 1 |
| 0 | 0 | 0 | 0 |
| 0 | 0 | 0 | 0 |
| 0 | 0 | 0 | 0 |
| 0 | 0 | 0 | 0 |
| 1 | 0 | 0 | 0 |
| 0 | 0 | 0 | 0 |
| 1 | 0 | 0 | 0 |
| 0 | 0 | 0 | 0 |
| 0 | 0 | 0 | 0 |
| 0 | 0 | 0 | 0 |
| 0 | 0 | 0 | 0 |
| 0 | 0 | 0 | 0 |
| 0 | 0 | 0 | 0 |
| 1 | 1 | 0 | 0 |
| 1 | 1 | 0 | 0 |
| 1 | 1 | 0 | 0 |
| 0 | 0 | 0 | 0 |
| 0 | 0 | 0 | 0 |
| 0 | 0 | 0 | 0 |
| 1 | 0 | 0 | 0 |
| 1 | 0 | 0 | 1 |
| 1 | 0 | 0 | 1 |
| 1 | 0 | 0 | 0 |
| 1 | 0 | 0 | 1 |
| 1 | 0 | 0 | 0 |
| 0 | 0 | 0 | 0 |
| 1 | 1 | 0 | 0 |
| 1 | 1 | 0 | 0 |
| 0 | 0 | 0 | 0 |
| 0 | 0 | 0 | 0 |

| | | | |
|---|---|---|---|
| 1 | 0 | 0 | 0 |
| 1 | 0 | 0 | 0 |
| 1 | 0 | 0 | 0 |
| 0 | 0 | 0 | 0 |
| 0 | 0 | 0 | 0 |
| 1 | 1 | 0 | 1 |
| 1 | 1 | 0 | 1 |
| 1 | 1 | 0 | 1 |
| 1 | 1 | 0 | 1 |
| 1 | 1 | 0 | 1 |
| 1 | 1 | 0 | 1 |
| 1 | 1 | 0 | 1 |
| 1 | 1 | 0 | 0 |
| 1 | 1 | 0 | 0 |
| 1 | 1 | 1 | 1 |
| 0 | 0 | 0 | 0 |
| 1 | 1 | 0 | 0 |
| 1 | 1 | 0 | 0 |
| 0 | 0 | 0 | 0 |
| 0 | 0 | 0 | 0 |
| 0 | 0 | 0 | 0 |
| 1 | 0 | 0 | 0 |
| 1 | 0 | 0 | 0 |
| 0 | 0 | 0 | 0 |
| 1 | 0 | 0 | 0 |
| 1 | 0 | 0 | 0 |
| 1 | 0 | 0 | 0 |
| 0 | 0 | 0 | 0 |
| 0 | 0 | 0 | 0 |
| 0 | 0 | 0 | 0 |
| 0 | 0 | 0 | 0 |
| 0 | 0 | 0 | 0 |
| 0 | 0 | 0 | 0 |
| 0 | 0 | 0 | 0 |
| 0 | 0 | 0 | 0 |
| 0 | 0 | 0 | 0 |
| 0 | 0 | 0 | 0 |
| 0 | 0 | 0 | 0 |
| 0 | 0 | 0 | 0 |
| 0 | 0 | 0 | 0 |
| 0 | 0 | 0 | 0 |
| 0 | 0 | 0 | 0 |
| 0 | 0 | 0 | 0 |
| 0 | 0 | 0 | 0 |
| 0 | 0 | 0 | 0 |
| 1 | 0 | 0 | 0 |
| 1 | 0 | 0 | 0 |
| 1 | 0 | 0 | 0 |
| 1 | 0 | 0 | 0 |
| 0 | 0 | 0 | 0 |
| 0 | 0 | 0 | 0 |
| 0 | 0 | 0 | 0 |
| 0 | 0 | 0 | 0 |
| 0 | 0 | 0 | 0 |

| | | | |
|---|---|---|---|
| 0 | 0 | 0 | 0 |
| 0 | 0 | 0 | 0 |
| 0 | 0 | 0 | 0 |
| 0 | 0 | 0 | 0 |
| 0 | 0 | 0 | 0 |
| 0 | 0 | 0 | 0 |
| 0 | 0 | 0 | 0 |
| 1 | 0 | 0 | 1 |
| 0 | 0 | 0 | 0 |
| 0 | 0 | 0 | 0 |
| 1 | 0 | 0 | 0 |
| 0 | 0 | 0 | 0 |
| 0 | 0 | 0 | 0 |
| 0 | 0 | 0 | 0 |
| 0 | 0 | 0 | 0 |
| 1 | 0 | 0 | 0 |
| 0 | 0 | 0 | 0 |
| 0 | 0 | 0 | 0 |
| 0 | 0 | 0 | 0 |
| 0 | 0 | 0 | 0 |
| 0 | 0 | 0 | 0 |
| 0 | 0 | 0 | 0 |
| 0 | 0 | 0 | 0 |
| 1 | 1 | 0 | 1 |
| 0 | 0 | 0 | 0 |
| 0 | 0 | 0 | 0 |
| 0 | 0 | 0 | 0 |
| 1 | 1 | 0 | 1 |
| 1 | 1 | 0 | 0 |
| 0 | 0 | 0 | 0 |
| 1 | 0 | 0 | 0 |
| 1 | 1 | 0 | 0 |
| 0 | 0 | 0 | 0 |
| 1 | 0 | 0 | 0 |
| 1 | 1 | 0 | 0 |
| 1 | 0 | 0 | 0 |
| 1 | 1 | 0 | 1 |
| 1 | 1 | 0 | 1 |
| 1 | 0 | 0 | 0 |
| 0 | 0 | 0 | 0 |
| 0 | 0 | 0 | 0 |
| 0 | 0 | 0 | 0 |
| 0 | 0 | 0 | 0 |
| 0 | 0 | 0 | 0 |
| 0 | 0 | 0 | 0 |
| 0 | 0 | 0 | 0 |
| 0 | 0 | 0 | 0 |
| 0 | 0 | 0 | 0 |
| 0 | 0 | 0 | 0 |
| 0 | 0 | 0 | 0 |
| 0 | 0 | 0 | 0 |
| 0 | 0 | 0 | 0 |
| 0 | 0 | 0 | 0 |
| 0 | 0 | 0 | 0 |

| | | | |
|---|---|---|---|
| 0 | 0 | 0 | 0 |
| 0 | 0 | 0 | 0 |
| 0 | 0 | 0 | 0 |
| 0 | 0 | 0 | 0 |
| 0 | 0 | 0 | 0 |
| 0 | 0 | 0 | 0 |
| 0 | 0 | 0 | 0 |
| 0 | 0 | 0 | 0 |
| 0 | 0 | 0 | 0 |
| 0 | 0 | 0 | 0 |
| 0 | 0 | 0 | 0 |
| 0 | 0 | 0 | 0 |
| 0 | 0 | 0 | 0 |
| 1 | 1 | 0 | 0 |
| 0 | 0 | 0 | 0 |
| 1 | 0 | 0 | 0 |
| 0 | 0 | 0 | 1 |
| 0 | 0 | 0 | 0 |
| 0 | 0 | 0 | 0 |
| 0 | 0 | 0 | 0 |
| 0 | 0 | 0 | 0 |
| 0 | 0 | 0 | 0 |
| 0 | 0 | 0 | 0 |
| 0 | 0 | 0 | 0 |
| 0 | 0 | 0 | 0 |
| 0 | 0 | 0 | 0 |
| 0 | 0 | 0 | 0 |
| 0 | 0 | 0 | 0 |
| 0 | 0 | 0 | 0 |
| 0 | 0 | 0 | 0 |
| 1 | 0 | 0 | 0 |
| 1 | 0 | 0 | 0 |
| 0 | 0 | 0 | 0 |
| 0 | 0 | 0 | 0 |
| 0 | 0 | 0 | 0 |
| 0 | 0 | 0 | 0 |
| 0 | 0 | 0 | 0 |
| 0 | 0 | 0 | 0 |
| 0 | 0 | 0 | 0 |
| 0 | 0 | 0 | 0 |
| 1 | 0 | 0 | 0 |
| 1 | 0 | 0 | 0 |
| 1 | 0 | 0 | 0 |
| 1 | 0 | 0 | 0 |
| 0 | 0 | 0 | 0 |
| 1 | 0 | 0 | 0 |
| 0 | 0 | 0 | 0 |
| 0 | 0 | 0 | 0 |
| 1 | 0 | 0 | 0 |
| 1 | 0 | 0 | 0 |
| 0 | 0 | 0 | 0 |
| 0 | 0 | 0 | 0 |
| 0 | 0 | 0 | 0 |
| 0 | 0 | 0 | 0 |

| | | | |
|---|---|---|---|
| 0 | 0 | 0 | 0 |
| 0 | 0 | 0 | 0 |
| 0 | 0 | 0 | 0 |
| 0 | 0 | 0 | 0 |
| 0 | 0 | 0 | 0 |
| 0 | 0 | 0 | 0 |
| 0 | 0 | 0 | 0 |
| 0 | 0 | 0 | 0 |
| 0 | 0 | 0 | 0 |
| 0 | 0 | 0 | 0 |
| 0 | 0 | 0 | 0 |
| 0 | 0 | 0 | 0 |
| 0 | 0 | 0 | 0 |
| 0 | 0 | 0 | 0 |
| 0 | 0 | 0 | 0 |
| 1 | 0 | 0 | 1 |
| 0 | 0 | 0 | 0 |
| 0 | 0 | 0 | 0 |
| 0 | 0 | 0 | 0 |
| 0 | 0 | 0 | 0 |
| 0 | 0 | 0 | 0 |
| 1 | 1 | 0 | 0 |
| 1 | 1 | 0 | 0 |
| 1 | 0 | 0 | 0 |
| 1 | 0 | 0 | 0 |
| 0 | 0 | 0 | 0 |
| 0 | 0 | 0 | 0 |
| 0 | 0 | 0 | 0 |
| 0 | 0 | 0 | 0 |
| 1 | 0 | 0 | 0 |
| 1 | 0 | 0 | 0 |
| 1 | 0 | 0 | 0 |
| 1 | 1 | 0 | 0 |
| 1 | 1 | 0 | 0 |
| 1 | 0 | 0 | 0 |
| 0 | 0 | 0 | 0 |
| 0 | 0 | 0 | 0 |
| 0 | 1 | 0 | 0 |
| 0 | 0 | 0 | 0 |
| 0 | 0 | 0 | 0 |
| 0 | 0 | 0 | 0 |
| 1 | 1 | 0 | 0 |
| 1 | 0 | 0 | 0 |
| 1 | 0 | 0 | 0 |
| 0 | 0 | 0 | 0 |
| 0 | 0 | 0 | 0 |
| 0 | 0 | 0 | 0 |
| 0 | 0 | 0 | 0 |
| 0 | 0 | 0 | 0 |
| 0 | 0 | 0 | 0 |
| 0 | 0 | 0 | 0 |
| 0 | 0 | 0 | 0 |
| 0 | 0 | 0 | 0 |
| 0 | 0 | 0 | 0 |

| | | | |
|---|---|---|---|
| 0 | 0 | 0 | 0 |
| 1 | 0 | 0 | 0 |
| 0 | 0 | 0 | 0 |
| 0 | 0 | 0 | 0 |
| 0 | 0 | 0 | 0 |
| 0 | 0 | 0 | 0 |
| 1 | 0 | 0 | 0 |
| 0 | 0 | 0 | 0 |
| 0 | 0 | 0 | 0 |
| 0 | 0 | 0 | 0 |
| 0 | 0 | 0 | 0 |
| 0 | 0 | 0 | 0 |
| 0 | 0 | 0 | 0 |
| 0 | 0 | 0 | 0 |
| 0 | 0 | 0 | 0 |
| 0 | 0 | 0 | 0 |
| 0 | 1 | 0 | 0 |
| 0 | 0 | 0 | 0 |
| 0 | 0 | 0 | 0 |
| 0 | 0 | 0 | 0 |
| 1 | 0 | 0 | 0 |
| 0 | 0 | 0 | 0 |
| 0 | 0 | 0 | 0 |
| 1 | 1 | 0 | 0 |
| 0 | 0 | 0 | 0 |
| 0 | 0 | 0 | 0 |
| 0 | 0 | 0 | 0 |
| 0 | 0 | 0 | 0 |
| 1 | 0 | 0 | 0 |
| 0 | 0 | 0 | 0 |
| 0 | 0 | 0 | 0 |
| 0 | 0 | 0 | 0 |
| 0 | 0 | 0 | 0 |
| 0 | 0 | 0 | 0 |
| 0 | 0 | 0 | 0 |
| 1 | 0 | 0 | 0 |
| 0 | 0 | 0 | 0 |
| 0 | 0 | 0 | 0 |
| 0 | 0 | 0 | 0 |
| 0 | 0 | 0 | 0 |
| 1 | 1 | 0 | 0 |
| 0 | 0 | 0 | 0 |
| 0 | 0 | 0 | 0 |
| 0 | 0 | 0 | 0 |
| 0 | 0 | 0 | 0 |
| 0 | 0 | 0 | 0 |
| 1 | 0 | 0 | 0 |
| 1 | 0 | 0 | 1 |
| 1 | 0 | 0 | 0 |
| 0 | 0 | 0 | 0 |
| 0 | 0 | 0 | 0 |
| 0 | 0 | 0 | 0 |
| 1 | 0 | 0 | 0 |
| 1 | 0 | 0 | 0 |

| | | | |
|---|---|---|---|
| 1 | 0 | 0 | 0 |
| 0 | 0 | 0 | 0 |
| 0 | 0 | 0 | 0 |
| 0 | 0 | 0 | 0 |
| 1 | 1 | 0 | 0 |
| 1 | 1 | 0 | 0 |
| 0 | 0 | 0 | 0 |
| 0 | 0 | 0 | 0 |
| 1 | 0 | 0 | 0 |
| 1 | 0 | 0 | 0 |
| 1 | 0 | 1 | 0 |
| 0 | 0 | 0 | 0 |
| 0 | 0 | 1 | 0 |
| 0 | 0 | 1 | 0 |
| 0 | 0 | 1 | 0 |
| 1 | 0 | 0 | 0 |
| 1 | 0 | 0 | 0 |
| 0 | 0 | 0 | 0 |
| 0 | 0 | 0 | 0 |
| 1 | 0 | 0 | 1 |
| 1 | 0 | 0 | 1 |
| 1 | 0 | 0 | 0 |
| 0 | 0 | 0 | 0 |
| 0 | 0 | 0 | 0 |
| 0 | 0 | 0 | 0 |
| 0 | 0 | 0 | 0 |
| 0 | 0 | 0 | 0 |
| 1 | 0 | 0 | 0 |
| 0 | 0 | 0 | 0 |
| 0 | 0 | 0 | 1 |
| 0 | 0 | 0 | 1 |
| 0 | 0 | 1 | 1 |
| 0 | 0 | 1 | 1 |
| 0 | 0 | 1 | 1 |
| 0 | 0 | 1 | 1 |
| 0 | 0 | 1 | 0 |
| 0 | 0 | 1 | 0 |
| 0 | 0 | 0 | 0 |
| 0 | 0 | 0 | 0 |
| 0 | 0 | 0 | 0 |
| 0 | 0 | 0 | 0 |
| 0 | 0 | 1 | 0 |
| 0 | 0 | 1 | 0 |
| 0 | 0 | 1 | 0 |
| 0 | 0 | 1 | 0 |
| 0 | 0 | 1 | 0 |
| 0 | 0 | 1 | 0 |
| 0 | 0 | 1 | 0 |
| 0 | 0 | 1 | 0 |
| 0 | 0 | 0 | 0 |
| 0 | 0 | 0 | 0 |
| 0 | 0 | 0 | 0 |
| 0 | 0 | 0 | 0 |
| 1 | 0 | 0 | 0 |

| | | | |
|---|---|---|---|
| 0 | 0 | 0 | 0 |
| 0 | 0 | 0 | 0 |
| 0 | 0 | 0 | 0 |
| 1 | 0 | 0 | 0 |
| 0 | 0 | 0 | 0 |
| 0 | 0 | 0 | 0 |
| 0 | 0 | 0 | 0 |
| 0 | 0 | 0 | 0 |
| 0 | 0 | 0 | 0 |
| 0 | 0 | 0 | 0 |
| 1 | 1 | 0 | 0 |
| 0 | 0 | 0 | 0 |
| 1 | 1 | 0 | 0 |
| 1 | 1 | 0 | 0 |
| 0 | 1 | 0 | 0 |
| 0 | 0 | 0 | 0 |
| 0 | 0 | 0 | 0 |
| 0 | 0 | 0 | 0 |
| 0 | 0 | 0 | 0 |
| 0 | 0 | 0 | 0 |
| 1 | 1 | 0 | 0 |
| 0 | 1 | 0 | 0 |
| 0 | 0 | 0 | 0 |
| 0 | 1 | 0 | 0 |
| 1 | 1 | 0 | 0 |
| 0 | 0 | 0 | 0 |
| 0 | 0 | 0 | 0 |
| 1 | 1 | 0 | 0 |
| 1 | 1 | 0 | 0 |
| 0 | 1 | 0 | 0 |
| 0 | 0 | 0 | 0 |
| 0 | 0 | 0 | 0 |
| 0 | 0 | 0 | 0 |
| 0 | 0 | 0 | 0 |
| 0 | 0 | 0 | 0 |
| 0 | 0 | 0 | 0 |
| 0 | 0 | 0 | 0 |
| 0 | 0 | 0 | 0 |
| 0 | 0 | 0 | 0 |
| 0 | 0 | 0 | 0 |
| 1 | 0 | 0 | 0 |
| 1 | 0 | 0 | 0 |
| 1 | 0 | 0 | 0 |
| 1 | 0 | 0 | 0 |
| 1 | 1 | 0 | 1 |
| 0 | 0 | 0 | 0 |
| 1 | 0 | 0 | 0 |
| 0 | 0 | 0 | 0 |
| 1 | 0 | 0 | 0 |
| 0 | 0 | 0 | 0 |
| 1 | 0 | 0 | 0 |
| 1 | 0 | 0 | 0 |
| 0 | 0 | 0 | 0 |
| 0 | 0 | 0 | 0 |

| | | | |
|---|---|---|---|
| 0 | 0 | 0 | 0 |
| 0 | 0 | 0 | 0 |
| 0 | 0 | 0 | 0 |
| 0 | 0 | 0 | 0 |
| 1 | 0 | 0 | 0 |
| 1 | 0 | 0 | 0 |
| 1 | 0 | 0 | 0 |
| 0 | 0 | 0 | 0 |
| 1 | 1 | 0 | 0 |
| 0 | 0 | 0 | 0 |
| 0 | 0 | 0 | 0 |
| 0 | 0 | 0 | 0 |
| 1 | 1 | 0 | 0 |
| 1 | 0 | 0 | 0 |
| 0 | 0 | 0 | 0 |
| 0 | 0 | 0 | 0 |
| 1 | 1 | 0 | 0 |
| 0 | 0 | 0 | 0 |
| 0 | 0 | 0 | 0 |
| 0 | 0 | 0 | 0 |
| 0 | 0 | 0 | 0 |
| 0 | 0 | 0 | 0 |
| 0 | 0 | 0 | 0 |
| 0 | 0 | 0 | 0 |
| 1 | 0 | 0 | 0 |
| 0 | 0 | 0 | 0 |
| 1 | 0 | 0 | 0 |
| 0 | 0 | 0 | 0 |
| 0 | 0 | 0 | 0 |
| 0 | 0 | 0 | 0 |
| 1 | 0 | 0 | 0 |
| 1 | 0 | 0 | 0 |
| 0 | 0 | 0 | 0 |
| 0 | 0 | 0 | 0 |
| 1 | 0 | 0 | 0 |
| 0 | 0 | 0 | 0 |
| 0 | 0 | 0 | 0 |
| 1 | 0 | 0 | 0 |
| 0 | 0 | 0 | 0 |
| 1 | 0 | 0 | 0 |
| 1 | 0 | 0 | 0 |
| 0 | 0 | 0 | 0 |
| 0 | 0 | 0 | 0 |
| 1 | 1 | 0 | 0 |
| 1 | 0 | 0 | 0 |
| 1 | 1 | 0 | 0 |
| 0 | 0 | 0 | 0 |
| 0 | 0 | 0 | 0 |
| 0 | 0 | 0 | 0 |
| 0 | 0 | 0 | 0 |
| 1 | 1 | 0 | 0 |
| 0 | 0 | 0 | 0 |
| 0 | 0 | 0 | 0 |
| 0 | 0 | 0 | 0 |

| | | | |
|---|---|---|---|
| 0 | 1 | 0 | 0 |
| 1 | 0 | 0 | 0 |
| 0 | 0 | 0 | 0 |
| 0 | 0 | 0 | 0 |
| 0 | 0 | 0 | 0 |
| 1 | 1 | 0 | 0 |
| 0 | 0 | 0 | 0 |
| 0 | 0 | 0 | 0 |
| 0 | 0 | 0 | 0 |
| 1 | 0 | 0 | 0 |
| 1 | 0 | 0 | 0 |
| 0 | 0 | 0 | 0 |
| 0 | 0 | 0 | 0 |
| 0 | 0 | 0 | 0 |
| 0 | 0 | 0 | 0 |
| 0 | 0 | 0 | 0 |
| 0 | 0 | 0 | 0 |
| 0 | 0 | 0 | 0 |
| 0 | 0 | 0 | 0 |
| 0 | 0 | 0 | 0 |
| 0 | 0 | 0 | 0 |
| 0 | 0 | 0 | 0 |
| 0 | 0 | 0 | 0 |
| 1 | 1 | 0 | 0 |
| 0 | 0 | 0 | 0 |
| 0 | 0 | 0 | 0 |
| 1 | 1 | 1 | 0 |
| 0 | 0 | 0 | 0 |
| 0 | 0 | 0 | 0 |
| 0 | 0 | 0 | 0 |
| 1 | 1 | 1 | 0 |
| 1 | 0 | 0 | 0 |
| 1 | 0 | 0 | 0 |
| 1 | 0 | 0 | 0 |
| 1 | 0 | 0 | 0 |
| 1 | 1 | 1 | 0 |
| 1 | 1 | 1 | 0 |
| 0 | 0 | 0 | 0 |
| 0 | 0 | 0 | 0 |
| 1 | 1 | 1 | 0 |
| 0 | 0 | 0 | 0 |
| 0 | 0 | 0 | 0 |
| 0 | 0 | 0 | 0 |
| 0 | 0 | 0 | 0 |
| 0 | 0 | 0 | 0 |
| 0 | 0 | 0 | 0 |
| 0 | 0 | 0 | 0 |
| 0 | 0 | 0 | 0 |
| 1 | 0 | 0 | 0 |
| 1 | 0 | 0 | 0 |
| 1 | 0 | 0 | 0 |
| 0 | 0 | 0 | 0 |
| 0 | 0 | 0 | 0 |
| 0 | 0 | 0 | 0 |

| | | | |
|---|---|---|---|
| 0 | 0 | 0 | 0 |
| 1 | 0 | 0 | 0 |
| 0 | 0 | 0 | 0 |
| 0 | 0 | 0 | 0 |
| 0 | 0 | 0 | 0 |
| 0 | 0 | 0 | 0 |
| 0 | 0 | 0 | 0 |
| 0 | 0 | 0 | 0 |
| 0 | 0 | 0 | 0 |
| 0 | 0 | 0 | 0 |
| 1 | 1 | 0 | 0 |
| 1 | 1 | 0 | 0 |
| 0 | 0 | 0 | 0 |
| 0 | 0 | 0 | 0 |
| 0 | 0 | 0 | 0 |
| 1 | 0 | 0 | 0 |
| 1 | 0 | 0 | 0 |
| 1 | 1 | 0 | 1 |
| 0 | 0 | 0 | 0 |
| 0 | 0 | 0 | 0 |
| 0 | 0 | 0 | 0 |
| 0 | 0 | 0 | 0 |
| 1 | 0 | 0 | 1 |
| 0 | 0 | 0 | 0 |
| 0 | 0 | 0 | 0 |
| 1 | 0 | 0 | 0 |
| 0 | 0 | 0 | 0 |
| 0 | 0 | 0 | 0 |
| 0 | 0 | 0 | 0 |
| 0 | 0 | 0 | 0 |
| 0 | 0 | 0 | 0 |
| 0 | 0 | 0 | 0 |
| 0 | 0 | 0 | 0 |
| 0 | 0 | 0 | 0 |
| 0 | 0 | 0 | 0 |
| 1 | 0 | 0 | 0 |
| 1 | 0 | 0 | 0 |
| 1 | 1 | 0 | 0 |
| 1 | 1 | 0 | 0 |
| 0 | 0 | 0 | 0 |
| 0 | 0 | 0 | 0 |
| 1 | 0 | 0 | 0 |
| 0 | 0 | 0 | 0 |
| 1 | 0 | 0 | 0 |
| 1 | 1 | 0 | 0 |
| 0 | 0 | 0 | 0 |
| 0 | 0 | 0 | 0 |
| 0 | 0 | 0 | 0 |
| 0 | 0 | 0 | 0 |
| 0 | 0 | 0 | 0 |
| 0 | 0 | 0 | 0 |
| 1 | 0 | 0 | 0 |
| 0 | 0 | 0 | 0 |
| 0 | 0 | 0 | 0 |

| | | | |
|---|---|---|---|
| 0 | 0 | 0 | 0 |
| 1 | 1 | 0 | 0 |
| 0 | 0 | 0 | 0 |
| 1 | 0 | 0 | 0 |
| 1 | 0 | 0 | 0 |
| 0 | 0 | 0 | 0 |
| 0 | 0 | 0 | 0 |
| 0 | 0 | 0 | 0 |
| 0 | 0 | 0 | 0 |
| 0 | 0 | 0 | 0 |
| 0 | 0 | 0 | 0 |
| 1 | 1 | 0 | 0 |
| 1 | 1 | 0 | 1 |
| 0 | 0 | 0 | 0 |
| 0 | 1 | 0 | 0 |
| 0 | 0 | 0 | 0 |
| 0 | 0 | 0 | 0 |
| 0 | 0 | 0 | 0 |
| 0 | 0 | 0 | 0 |
| 0 | 0 | 0 | 0 |
| 0 | 0 | 0 | 0 |
| 1 | 0 | 0 | 0 |
| 0 | 0 | 0 | 0 |
| 0 | 0 | 0 | 0 |
| 0 | 0 | 0 | 0 |
| 0 | 0 | 0 | 0 |
| 0 | 0 | 0 | 0 |
| 1 | 0 | 0 | 0 |
| 0 | 0 | 0 | 0 |
| 0 | 0 | 0 | 0 |
| 0 | 0 | 0 | 0 |
| 0 | 0 | 0 | 0 |
| 0 | 0 | 0 | 0 |
| 0 | 0 | 0 | 0 |
| 0 | 0 | 0 | 0 |
| 0 | 0 | 0 | 0 |
| 0 | 0 | 0 | 0 |
| 0 | 0 | 0 | 0 |
| 0 | 0 | 0 | 0 |
| 0 | 0 | 0 | 0 |
| 0 | 0 | 0 | 0 |
| 0 | 0 | 0 | 0 |
| 0 | 0 | 0 | 0 |
| 0 | 0 | 0 | 0 |
| 0 | 0 | 0 | 0 |
| 0 | 0 | 0 | 0 |
| 0 | 0 | 0 | 0 |
| 0 | 0 | 0 | 0 |
| 0 | 0 | 0 | 0 |
| 0 | 0 | 0 | 0 |
| 0 | 0 | 0 | 0 |
| 0 | 0 | 0 | 0 |
| 0 | 0 | 0 | 0 |
| 0 | 0 | 0 | 0 |

| | | | |
|---|---|---|---|
| 0 | 0 | 0 | 0 |
| 0 | 0 | 0 | 0 |
| 0 | 0 | 0 | 0 |
| 0 | 0 | 0 | 0 |
| 0 | 0 | 0 | 0 |
| 0 | 0 | 0 | 0 |
| 0 | 0 | 0 | 0 |
| 0 | 0 | 0 | 0 |
| 0 | 0 | 0 | 0 |
| 0 | 0 | 0 | 0 |
| 0 | 0 | 0 | 0 |
| 0 | 0 | 0 | 0 |
| 1 | 0 | 0 | 0 |
| 1 | 1 | 0 | 0 |
| 0 | 0 | 0 | 0 |
| 1 | 0 | 0 | 0 |
| 0 | 0 | 0 | 0 |
| 1 | 0 | 0 | 0 |
| 0 | 0 | 0 | 0 |
| 0 | 0 | 0 | 0 |
| 1 | 0 | 0 | 0 |
| 0 | 0 | 0 | 0 |
| 0 | 0 | 0 | 0 |
| 1 | 0 | 0 | 0 |
| 1 | 0 | 0 | 0 |
| 0 | 0 | 0 | 0 |
| 0 | 0 | 0 | 0 |
| 1 | 0 | 0 | 0 |
| 1 | 0 | 0 | 0 |
| 1 | 0 | 0 | 0 |
| 1 | 0 | 0 | 0 |
| 1 | 0 | 0 | 0 |
| 0 | 0 | 0 | 0 |
| 0 | 0 | 0 | 0 |
| 0 | 0 | 0 | 0 |
| 0 | 0 | 0 | 0 |
| 0 | 0 | 0 | 0 |
| 0 | 0 | 0 | 0 |
| 0 | 0 | 0 | 0 |
| 0 | 0 | 0 | 0 |
| 0 | 0 | 0 | 0 |
| 0 | 0 | 0 | 0 |
| 0 | 0 | 0 | 0 |
| 0 | 0 | 0 | 0 |
| 1 | 1 | 0 | 0 |
| 0 | 0 | 0 | 0 |
| 0 | 0 | 0 | 0 |
| 0 | 0 | 0 | 0 |
| 0 | 0 | 0 | 0 |
| 1 | 1 | 0 | 0 |
| 1 | 0 | 0 | 0 |
| 0 | 0 | 0 | 0 |
| 0 | 0 | 0 | 0 |
| 0 | 0 | 0 | 1 |

| | | | |
|---|---|---|---|
| 0 | 0 | 0 | 0 |
| 0 | 0 | 0 | 0 |
| 0 | 0 | 0 | 0 |
| 0 | 0 | 0 | 0 |
| 1 | 0 | 0 | 1 |
| 0 | 0 | 0 | 0 |
| 1 | 0 | 0 | 0 |
| 0 | 0 | 0 | 0 |
| 0 | 0 | 0 | 0 |
| 0 | 0 | 0 | 0 |
| 0 | 0 | 0 | 0 |
| 0 | 0 | 0 | 0 |
| 0 | 0 | 0 | 0 |
| 1 | 0 | 0 | 0 |
| 0 | 0 | 0 | 0 |
| 0 | 0 | 0 | 0 |
| 1 | 1 | 0 | 0 |
| 0 | 0 | 0 | 0 |
| 1 | 0 | 0 | 0 |
| 1 | 0 | 0 | 0 |
| 0 | 0 | 0 | 0 |
| 0 | 0 | 0 | 0 |
| 0 | 0 | 0 | 0 |
| 1 | 0 | 0 | 0 |
| 1 | 0 | 0 | 0 |
| 1 | 0 | 0 | 0 |
| 1 | 0 | 0 | 0 |
| 1 | 0 | 0 | 0 |
| 1 | 0 | 0 | 0 |
| 1 | 0 | 0 | 0 |
| 1 | 0 | 0 | 0 |
| 1 | 0 | 0 | 0 |
| 1 | 0 | 0 | 0 |
| 1 | 0 | 0 | 0 |
| 0 | 0 | 0 | 0 |
| 0 | 0 | 0 | 0 |
| 0 | 0 | 0 | 0 |
| 1 | 0 | 0 | 0 |
| 1 | 0 | 0 | 0 |
| 0 | 0 | 0 | 0 |
| 0 | 0 | 0 | 0 |
| 1 | 0 | 0 | 0 |
| 0 | 0 | 0 | 0 |
| 1 | 0 | 0 | 0 |
| 1 | 0 | 0 | 0 |
| 1 | 0 | 0 | 0 |
| 1 | 0 | 0 | 0 |
| 1 | 0 | 0 | 0 |
| 0 | 0 | 0 | 0 |
| 0 | 0 | 0 | 0 |
| 1 | 0 | 0 | 0 |
| 0 | 0 | 0 | 0 |
| 1 | 0 | 0 | 0 |
| 1 | 0 | 0 | 0 |

| | | | |
|---|---|---|---|
| 1 | 0 | 0 | 0 |
| 1 | 0 | 0 | 0 |
| 1 | 0 | 0 | 0 |
| 1 | 0 | 0 | 0 |
| 1 | 0 | 0 | 0 |
| 1 | 0 | 0 | 0 |
| 0 | 0 | 0 | 0 |
| 1 | 0 | 0 | 0 |
| 1 | 0 | 0 | 0 |
| 1 | 0 | 0 | 0 |
| 1 | 0 | 0 | 0 |
| 1 | 0 | 0 | 0 |
| 1 | 0 | 0 | 0 |
| 1 | 0 | 0 | 0 |
| 0 | 0 | 0 | 0 |
| 0 | 0 | 0 | 0 |
| 1 | 1 | 0 | 0 |
| 1 | 1 | 0 | 0 |
| 0 | 0 | 0 | 0 |
| 0 | 0 | 0 | 0 |
| 0 | 0 | 0 | 0 |
| 1 | 1 | 0 | 1 |
| 0 | 0 | 0 | 0 |
| 0 | 0 | 0 | 0 |
| 0 | 0 | 0 | 0 |
| 0 | 0 | 0 | 0 |
| 0 | 0 | 0 | 0 |
| 0 | 0 | 0 | 0 |
| 0 | 0 | 0 | 0 |
| 1 | 0 | 1 | 0 |
| 0 | 0 | 0 | 0 |
| 0 | 0 | 0 | 0 |
| 0 | 0 | 0 | 0 |
| 0 | 0 | 0 | 0 |
| 0 | 0 | 0 | 0 |
| 0 | 0 | 0 | 0 |
| 0 | 0 | 0 | 0 |
| 0 | 0 | 0 | 0 |
| 0 | 0 | 0 | 0 |
| 0 | 0 | 0 | 0 |
| 1 | 1 | 0 | 0 |
| 1 | 1 | 0 | 0 |
| 1 | 0 | 0 | 0 |
| 1 | 1 | 0 | 0 |
| 1 | 0 | 0 | 0 |
| 1 | 1 | 0 | 0 |
| 1 | 0 | 0 | 0 |
| 1 | 0 | 0 | 0 |
| 0 | 0 | 0 | 0 |
| 0 | 0 | 0 | 0 |
| 1 | 0 | 0 | 0 |
| 1 | 0 | 0 | 0 |
| 1 | 0 | 0 | 0 |
| 0 | 0 | 0 | 0 |

| | | | |
|---|---|---|---|
| 0 | 0 | 0 | 0 |
| 0 | 0 | 0 | 0 |
| 0 | 0 | 0 | 0 |
| 1 | 1 | 1 | 0 |
| 0 | 0 | 0 | 0 |
| 0 | 0 | 0 | 0 |
| 0 | 0 | 0 | 0 |
| 0 | 0 | 0 | 0 |
| 0 | 1 | 0 | 0 |
| 0 | 1 | 0 | 0 |
| 0 | 1 | 0 | 0 |
| 0 | 0 | 0 | 0 |
| 0 | 0 | 0 | 0 |
| 0 | 0 | 0 | 0 |
| 1 | 0 | 0 | 0 |
| 1 | 0 | 0 | 0 |
| 0 | 0 | 0 | 0 |
| 0 | 0 | 0 | 0 |
| 1 | 0 | 0 | 0 |
| 0 | 0 | 0 | 0 |
| 1 | 0 | 0 | 0 |
| 1 | 0 | 0 | 0 |
| 0 | 0 | 0 | 0 |
| 0 | 0 | 0 | 0 |
| 0 | 0 | 0 | 0 |
| 0 | 0 | 0 | 0 |
| 0 | 0 | 0 | 0 |
| 0 | 0 | 1 | 0 |
| 0 | 0 | 0 | 0 |
| 0 | 1 | 0 | 0 |
| 0 | 0 | 0 | 0 |
| 0 | 0 | 0 | 0 |
| 0 | 0 | 0 | 0 |
| 0 | 0 | 0 | 0 |
| 0 | 0 | 0 | 0 |
| 0 | 0 | 0 | 0 |
| 1 | 0 | 0 | 0 |
| 1 | 0 | 0 | 0 |
| 1 | 0 | 0 | 0 |
| 0 | 0 | 0 | 0 |
| 0 | 0 | 0 | 0 |
| 0 | 0 | 0 | 0 |
| 0 | 0 | 0 | 0 |
| 0 | 0 | 0 | 0 |
| 0 | 0 | 0 | 0 |
| 0 | 0 | 0 | 0 |
| 0 | 0 | 0 | 0 |
| 0 | 0 | 0 | 1 |
| 0 | 0 | 0 | 0 |
| 0 | 0 | 0 | 0 |
| 0 | 0 | 0 | 0 |
| 0 | 0 | 0 | 0 |
| 1 | 1 | 0 | 0 |
| 0 | 0 | 0 | 0 |

| | | | |
|---|---|---|---|
| 0 | 0 | 0 | 0 |
| 0 | 0 | 0 | 0 |
| 0 | 0 | 0 | 0 |
| 0 | 0 | 0 | 0 |
| 1 | 0 | 0 | 0 |
| 0 | 0 | 0 | 0 |
| 0 | 0 | 0 | 0 |
| 0 | 0 | 0 | 0 |
| 0 | 0 | 0 | 0 |
| 0 | 0 | 0 | 0 |
| 0 | 0 | 0 | 0 |
| 0 | 0 | 0 | 0 |
| 0 | 0 | 0 | 0 |
| 0 | 0 | 0 | 0 |
| 0 | 0 | 0 | 0 |
| 0 | 0 | 0 | 0 |
| 0 | 0 | 0 | 0 |
| 0 | 0 | 0 | 0 |
| 0 | 0 | 0 | 0 |
| 0 | 0 | 0 | 0 |
| 0 | 0 | 0 | 0 |
| 0 | 0 | 0 | 0 |
| 0 | 0 | 0 | 0 |
| 0 | 0 | 0 | 0 |
| 0 | 0 | 0 | 0 |
| 0 | 0 | 0 | 0 |
| 0 | 0 | 0 | 0 |
| 0 | 0 | 0 | 0 |
| 0 | 0 | 0 | 0 |
| 0 | 0 | 0 | 0 |
| 0 | 0 | 0 | 0 |
| 0 | 0 | 0 | 0 |
| 0 | 0 | 0 | 0 |
| 0 | 0 | 0 | 0 |
| 0 | 0 | 0 | 0 |
| 0 | 0 | 0 | 0 |
| 0 | 0 | 0 | 0 |
| 0 | 0 | 0 | 0 |
| 0 | 0 | 0 | 0 |
| 0 | 0 | 0 | 0 |
| 0 | 0 | 0 | 0 |
| 1 | 1 | 0 | 0 |
| 0 | 0 | 0 | 0 |
| 0 | 0 | 0 | 0 |
| 0 | 0 | 0 | 0 |
| 0 | 0 | 0 | 0 |
| 1 | 0 | 0 | 0 |
| 1 | 0 | 0 | 0 |
| 0 | 0 | 0 | 0 |
| 1 | 0 | 0 | 0 |
| 0 | 0 | 0 | 0 |
| 0 | 0 | 0 | 0 |
| 1 | 0 | 0 | 0 |
| 1 | 0 | 0 | 0 |
| 0 | 0 | 0 | 0 |

| | | | |
|---|---|---|---|
| 0 | 0 | 0 | 0 |
| 0 | 0 | 0 | 0 |
| 0 | 0 | 0 | 0 |
| 0 | 0 | 0 | 0 |
| 1 | 0 | 0 | 0 |
| 0 | 0 | 0 | 0 |
| 0 | 0 | 0 | 0 |
| 0 | 0 | 0 | 0 |
| 0 | 0 | 0 | 0 |
| 0 | 0 | 0 | 0 |
| 0 | 0 | 0 | 0 |
| 0 | 0 | 0 | 0 |
| 0 | 0 | 0 | 0 |
| 0 | 0 | 0 | 0 |
| 0 | 0 | 0 | 0 |
| 1 | 1 | 0 | 1 |
| 0 | 0 | 0 | 0 |
| 0 | 0 | 0 | 0 |
| 0 | 0 | 0 | 0 |
| 0 | 0 | 0 | 0 |
| 0 | 0 | 0 | 0 |
| 0 | 0 | 0 | 0 |
| 0 | 0 | 0 | 0 |
| 0 | 0 | 0 | 0 |
| 0 | 0 | 0 | 0 |
| 0 | 0 | 0 | 0 |
| 0 | 0 | 0 | 0 |
| 1 | 0 | 0 | 0 |
| 1 | 0 | 0 | 0 |
| 0 | 0 | 0 | 0 |
| 1 | 1 | 0 | 0 |
| 0 | 0 | 0 | 0 |
| 0 | 0 | 0 | 0 |
| 0 | 0 | 0 | 0 |
| 1 | 1 | 0 | 0 |
| 1 | 1 | 0 | 0 |
| 0 | 0 | 0 | 0 |
| 1 | 0 | 0 | 0 |
| 0 | 0 | 0 | 0 |
| 1 | 1 | 0 | 0 |
| 0 | 1 | 0 | 0 |
| 0 | 0 | 0 | 0 |
| 0 | 0 | 0 | 0 |
| 0 | 0 | 0 | 0 |
| 1 | 1 | 0 | 0 |
| 1 | 1 | 0 | 0 |
| 1 | 1 | 0 | 0 |
| 0 | 0 | 0 | 0 |
| 0 | 0 | 0 | 0 |
| 0 | 0 | 0 | 0 |
| 0 | 0 | 0 | 0 |
| 0 | 0 | 0 | 0 |
| 0 | 0 | 0 | 0 |
| 0 | 0 | 0 | 0 |

| | | | |
|---|---|---|---|
| 0 | 0 | 0 | 0 |
| 0 | 0 | 0 | 0 |
| 1 | 0 | 0 | 0 |
| 1 | 0 | 0 | 0 |
| 0 | 0 | 0 | 0 |
| 1 | 0 | 0 | 0 |
| 0 | 0 | 0 | 0 |
| 0 | 0 | 0 | 0 |
| 0 | 0 | 0 | 0 |
| 0 | 1 | 0 | 0 |
| 0 | 0 | 0 | 0 |
| 0 | 0 | 0 | 0 |
| 0 | 0 | 0 | 0 |
| 0 | 0 | 0 | 0 |
| 0 | 0 | 0 | 0 |
| 0 | 0 | 0 | 0 |
| 0 | 0 | 0 | 0 |
| 1 | 0 | 0 | 0 |
| 0 | 0 | 0 | 0 |
| 1 | 0 | 0 | 0 |
| 0 | 0 | 0 | 0 |
| 1 | 0 | 0 | 0 |
| 0 | 0 | 0 | 0 |
| 1 | 0 | 0 | 0 |
| 0 | 0 | 0 | 0 |
| 0 | 0 | 0 | 0 |
| 0 | 0 | 0 | 0 |
| 0 | 0 | 0 | 0 |
| 0 | 0 | 0 | 0 |
| 0 | 0 | 0 | 0 |
| 0 | 0 | 0 | 0 |
| 1 | 0 | 0 | 0 |
| 0 | 0 | 0 | 0 |
| 0 | 0 | 0 | 0 |
| 0 | 0 | 0 | 0 |
| 0 | 0 | 0 | 0 |
| 0 | 0 | 0 | 0 |
| 0 | 0 | 0 | 0 |
| 1 | 1 | 0 | 0 |
| 1 | 1 | 0 | 0 |
| 1 | 1 | 0 | 0 |
| 1 | 1 | 0 | 0 |
| 1 | 1 | 0 | 0 |
| 1 | 0 | 0 | 0 |
| 1 | 0 | 0 | 0 |
| 1 | 0 | 0 | 0 |
| 0 | 0 | 0 | 0 |
| 0 | 0 | 0 | 0 |
| 0 | 0 | 0 | 0 |
| 0 | 0 | 0 | 0 |
| 0 | 0 | 0 | 0 |
| 0 | 0 | 0 | 0 |
| 0 | 0 | 0 | 0 |
| 0 | 0 | 0 | 0 |

| | | | |
|---|---|---|---|
| 0 | 0 | 0 | 0 |
| 0 | 0 | 0 | 0 |
| 0 | 0 | 0 | 0 |
| 0 | 0 | 0 | 0 |
| 0 | 0 | 0 | 0 |
| 1 | 0 | 0 | 0 |
| 1 | 0 | 0 | 0 |
| 0 | 0 | 0 | 0 |
| 0 | 0 | 0 | 0 |
| 0 | 0 | 0 | 0 |
| 0 | 0 | 0 | 0 |
| 1 | 0 | 0 | 0 |
| 0 | 0 | 1 | 0 |
| 0 | 0 | 0 | 0 |
| 0 | 0 | 0 | 0 |
| 0 | 0 | 0 | 0 |
| 1 | 0 | 0 | 0 |
| 0 | 0 | 0 | 0 |
| 1 | 0 | 0 | 0 |
| 1 | 0 | 0 | 0 |
| 0 | 0 | 0 | 0 |
| 1 | 0 | 0 | 0 |
| 1 | 1 | 0 | 0 |
| 0 | 0 | 0 | 0 |
| 1 | 1 | 0 | 0 |
| 1 | 1 | 0 | 0 |
| 1 | 1 | 0 | 0 |
| 0 | 0 | 0 | 0 |
| 1 | 0 | 0 | 0 |
| 0 | 0 | 0 | 0 |
| 0 | 0 | 0 | 0 |
| 0 | 0 | 0 | 0 |
| 0 | 0 | 0 | 0 |
| 0 | 0 | 0 | 0 |
| 0 | 0 | 0 | 0 |
| 0 | 0 | 0 | 0 |
| 0 | 0 | 0 | 0 |
| 0 | 0 | 0 | 0 |
| 0 | 0 | 0 | 0 |
| 0 | 0 | 0 | 0 |
| 0 | 0 | 0 | 0 |
| 0 | 0 | 0 | 0 |
| 0 | 0 | 0 | 0 |
| 0 | 0 | 0 | 0 |
| 0 | 0 | 0 | 0 |
| 0 | 0 | 0 | 0 |
| 0 | 0 | 0 | 0 |
| 0 | 0 | 0 | 0 |
| 0 | 0 | 0 | 0 |
| 0 | 0 | 0 | 0 |
| 0 | 0 | 0 | 0 |
| 0 | 0 | 0 | 0 |
| 0 | 0 | 0 | 0 |
| 0 | 0 | 0 | 0 |

| | | | |
|---|---|---|---|
| 0 | 0 | 0 | 0 |
| 0 | 0 | 0 | 0 |
| 0 | 0 | 0 | 0 |
| 0 | 0 | 0 | 0 |
| 0 | 0 | 0 | 0 |
| 0 | 0 | 0 | 0 |
| 0 | 0 | 0 | 0 |
| 0 | 0 | 0 | 0 |
| 0 | 0 | 0 | 0 |
| 0 | 0 | 0 | 0 |
| 0 | 1 | 0 | 0 |
| 0 | 1 | 0 | 0 |
| 0 | 1 | 0 | 0 |
| 0 | 1 | 0 | 0 |
| 0 | 0 | 0 | 0 |
| 0 | 1 | 0 | 0 |
| 0 | 1 | 0 | 0 |
| 1 | 1 | 0 | 0 |
| 0 | 1 | 0 | 0 |
| 1 | 0 | 0 | 0 |
| 1 | 1 | 0 | 0 |
| 1 | 1 | 0 | 0 |
| 0 | 0 | 1 | 1 |
| 0 | 0 | 0 | 0 |
| 1 | 0 | 0 | 0 |
| 0 | 0 | 0 | 0 |
| 0 | 0 | 0 | 0 |
| 0 | 0 | 0 | 0 |
| 0 | 0 | 0 | 0 |
| 0 | 0 | 0 | 0 |
| 0 | 0 | 0 | 0 |
| 0 | 0 | 0 | 0 |
| 0 | 0 | 0 | 0 |
| 0 | 0 | 0 | 0 |
| 0 | 0 | 0 | 0 |
| 0 | 0 | 0 | 0 |
| 0 | 0 | 0 | 0 |
| 0 | 0 | 0 | 0 |
| 0 | 0 | 0 | 0 |
| 0 | 0 | 0 | 0 |
| 1 | 0 | 0 | 0 |
| 0 | 0 | 0 | 0 |
| 1 | 0 | 0 | 0 |
| 1 | 0 | 0 | 0 |
| 0 | 0 | 0 | 0 |
| 1 | 0 | 0 | 0 |
| 0 | 0 | 0 | 0 |
| 0 | 0 | 0 | 0 |
| 0 | 0 | 0 | 0 |
| 0 | 0 | 0 | 0 |
| 0 | 0 | 0 | 0 |
| 0 | 0 | 0 | 0 |
| 1 | 1 | 0 | 0 |
| 1 | 1 | 0 | 0 |

| | | | |
|---|---|---|---|
| 1 | 1 | 0 | 0 |
| 0 | 0 | 0 | 0 |
| 0 | 0 | 0 | 0 |
| 0 | 0 | 0 | 0 |
| 0 | 0 | 0 | 0 |
| 0 | 0 | 0 | 0 |
| 0 | 1 | 0 | 0 |
| 0 | 0 | 0 | 0 |
| 0 | 0 | 0 | 0 |
| 0 | 1 | 0 | 0 |
| 1 | 0 | 0 | 0 |
| 0 | 0 | 0 | 0 |
| 0 | 0 | 1 | 0 |
| 0 | 0 | 1 | 0 |
| 0 | 0 | 0 | 0 |
| 0 | 0 | 0 | 0 |
| 0 | 0 | 0 | 0 |
| 0 | 1 | 0 | 0 |
| 0 | 0 | 0 | 0 |
| 0 | 0 | 0 | 0 |
| 1 | 0 | 0 | 0 |
| 0 | 0 | 0 | 1 |
| 0 | 0 | 0 | 0 |
| 0 | 0 | 0 | 0 |
| 0 | 0 | 0 | 0 |
| 0 | 0 | 0 | 0 |
| 0 | 0 | 0 | 0 |
| 0 | 0 | 0 | 0 |
| 0 | 0 | 0 | 0 |
| 0 | 0 | 0 | 0 |
| 0 | 0 | 0 | 0 |
| 0 | 0 | 0 | 0 |
| 0 | 0 | 0 | 0 |
| 0 | 0 | 0 | 0 |
| 0 | 0 | 0 | 0 |
| 0 | 0 | 0 | 0 |
| 1 | 0 | 0 | 0 |
| 0 | 0 | 0 | 0 |
| 1 | 0 | 0 | 0 |
| 0 | 0 | 0 | 0 |
| 0 | 0 | 0 | 0 |
| 0 | 0 | 0 | 0 |
| 0 | 0 | 0 | 0 |
| 0 | 0 | 0 | 0 |
| 0 | 0 | 0 | 0 |
| 0 | 0 | 0 | 0 |
| 0 | 0 | 0 | 0 |
| 0 | 0 | 0 | 0 |
| 0 | 0 | 0 | 0 |
| 1 | 0 | 0 | 0 |
| 0 | 0 | 0 | 0 |
| 0 | 0 | 0 | 0 |
| 0 | 0 | 0 | 0 |
| 0 | 0 | 0 | 0 |

| | | | |
|---|---|---|---|
| 0 | 0 | 0 | 0 |
| 0 | 0 | 0 | 0 |
| 0 | 0 | 0 | 0 |
| 0 | 0 | 0 | 0 |
| 0 | 0 | 0 | 0 |
| 0 | 0 | 0 | 0 |
| 0 | 0 | 0 | 0 |
| 0 | 0 | 0 | 0 |
| 0 | 0 | 0 | 0 |
| 0 | 0 | 0 | 0 |
| 0 | 0 | 0 | 0 |
| 0 | 0 | 0 | 0 |
| 0 | 0 | 1 | 0 |
| 1 | 0 | 0 | 0 |
| 1 | 0 | 0 | 0 |
| 0 | 0 | 0 | 0 |
| 0 | 0 | 0 | 0 |
| 0 | 0 | 0 | 0 |
| 0 | 0 | 0 | 0 |
| 0 | 0 | 0 | 0 |
| 0 | 0 | 0 | 0 |
| 0 | 0 | 0 | 0 |
| 0 | 0 | 0 | 0 |
| 1 | 1 | 0 | 1 |
| 0 | 0 | 0 | 0 |
| 1 | 1 | 0 | 0 |
| 1 | 1 | 0 | 1 |
| 0 | 0 | 0 | 0 |
| 0 | 0 | 0 | 0 |
| 0 | 0 | 0 | 0 |
| 0 | 0 | 0 | 0 |
| 0 | 0 | 0 | 0 |
| 0 | 0 | 0 | 0 |
| 0 | 0 | 0 | 0 |
| 0 | 0 | 0 | 0 |
| 1 | 0 | 0 | 0 |
| 0 | 0 | 0 | 0 |
| 0 | 0 | 0 | 0 |
| 0 | 0 | 0 | 0 |
| 0 | 0 | 0 | 0 |
| 0 | 0 | 0 | 0 |
| 0 | 0 | 0 | 0 |
| 1 | 0 | 0 | 0 |
| 1 | 0 | 0 | 0 |
| 1 | 0 | 0 | 0 |
| 1 | 0 | 0 | 0 |
| 1 | 0 | 0 | 0 |
| 0 | 0 | 0 | 0 |
| 0 | 0 | 0 | 0 |
| 1 | 0 | 0 | 0 |
| 1 | 0 | 0 | 0 |
| 0 | 0 | 0 | 0 |
| 1 | 0 | 0 | 0 |
| 1 | 0 | 0 | 0 |

| | | | |
|---|---|---|---|
| 0 | 0 | 0 | 0 |
| 1 | 0 | 0 | 0 |
| 0 | 0 | 0 | 0 |
| 0 | 0 | 0 | 0 |
| 0 | 0 | 0 | 0 |
| 0 | 0 | 0 | 0 |
| 1 | 0 | 0 | 0 |
| 0 | 0 | 0 | 0 |
| 0 | 0 | 0 | 0 |
| 0 | 0 | 0 | 0 |
| 0 | 0 | 0 | 0 |
| 1 | 1 | 0 | 0 |
| 0 | 0 | 0 | 0 |
| 0 | 0 | 0 | 0 |
| 0 | 0 | 0 | 0 |
| 0 | 0 | 0 | 0 |
| 0 | 0 | 0 | 0 |
| 0 | 0 | 0 | 0 |
| 0 | 0 | 0 | 0 |
| 0 | 0 | 0 | 0 |
| 0 | 0 | 0 | 0 |
| 0 | 0 | 0 | 0 |
| 0 | 0 | 0 | 0 |
| 0 | 0 | 0 | 0 |
| 0 | 0 | 0 | 0 |
| 0 | 0 | 0 | 0 |
| 0 | 0 | 0 | 0 |
| 0 | 0 | 0 | 0 |
| 1 | 1 | 0 | 0 |
| 1 | 0 | 0 | 0 |
| 0 | 0 | 0 | 0 |
| 0 | 0 | 0 | 0 |
| 0 | 0 | 0 | 0 |
| 0 | 0 | 0 | 0 |
| 0 | 0 | 0 | 0 |
| 0 | 0 | 0 | 0 |
| 0 | 0 | 0 | 0 |
| 0 | 0 | 0 | 0 |
| 0 | 0 | 0 | 0 |
| 0 | 0 | 0 | 0 |
| 0 | 0 | 0 | 0 |
| 0 | 0 | 0 | 0 |
| 0 | 0 | 0 | 0 |
| 0 | 0 | 0 | 0 |
| 0 | 0 | 0 | 0 |
| 0 | 0 | 0 | 0 |
| 0 | 0 | 0 | 0 |
| 0 | 0 | 0 | 0 |
| 0 | 0 | 0 | 0 |
| 0 | 0 | 0 | 0 |
| 0 | 0 | 0 | 0 |
| 0 | 0 | 0 | 0 |
| 0 | 0 | 1 | 0 |
| 0 | 0 | 0 | 0 |

| | | | |
|---|---|---|---|
| 0 | 0 | 0 | 0 |
| 0 | 0 | 0 | 0 |
| 0 | 0 | 0 | 0 |
| 0 | 0 | 0 | 0 |
| 1 | 0 | 0 | 0 |
| 0 | 0 | 0 | 0 |
| 0 | 0 | 0 | 0 |
| 1 | 1 | 0 | 0 |
| 0 | 0 | 0 | 0 |
| 1 | 1 | 0 | 0 |
| 0 | 0 | 0 | 0 |
| 0 | 0 | 0 | 0 |
| 0 | 0 | 0 | 0 |
| 0 | 0 | 0 | 0 |
| 0 | 0 | 0 | 0 |
| 0 | 0 | 0 | 0 |
| 1 | 0 | 0 | 0 |
| 0 | 0 | 0 | 0 |
| 0 | 0 | 0 | 0 |
| 1 | 0 | 0 | 0 |
| 1 | 0 | 0 | 0 |
| 0 | 0 | 0 | 0 |
| 0 | 0 | 0 | 0 |
| 0 | 0 | 0 | 0 |
| 0 | 0 | 0 | 0 |
| 0 | 0 | 0 | 0 |
| 1 | 0 | 0 | 0 |
| 0 | 0 | 0 | 0 |
| 0 | 0 | 0 | 0 |
| 1 | 0 | 0 | 0 |
| 0 | 0 | 0 | 0 |
| 0 | 0 | 0 | 0 |
| 0 | 0 | 0 | 0 |
| 1 | 1 | 0 | 0 |
| 0 | 0 | 0 | 0 |
| 0 | 0 | 0 | 0 |
| 0 | 0 | 0 | 0 |
| 0 | 0 | 0 | 0 |
| 0 | 1 | 0 | 0 |
| 0 | 0 | 0 | 0 |
| 0 | 0 | 0 | 0 |
| 0 | 0 | 0 | 0 |
| 0 | 0 | 0 | 0 |
| 1 | 1 | 0 | 0 |
| 0 | 0 | 0 | 0 |
| 0 | 0 | 0 | 0 |
| 0 | 0 | 0 | 0 |
| 0 | 0 | 0 | 0 |
| 0 | 0 | 0 | 0 |
| 0 | 0 | 0 | 0 |
| 0 | 0 | 0 | 0 |
| 1 | 0 | 0 | 0 |
| 0 | 0 | 0 | 0 |
| 0 | 0 | 0 | 0 |

| | | | |
|---|---|---|---|
| 1 | 0 | 0 | 1 |
| 1 | 0 | 0 | 1 |
| 1 | 0 | 0 | 1 |
| 0 | 0 | 0 | 0 |
| 0 | 0 | 0 | 0 |
| 0 | 0 | 0 | 0 |
| 1 | 0 | 0 | 0 |
| 0 | 0 | 0 | 0 |
| 0 | 0 | 0 | 0 |
| 0 | 0 | 0 | 0 |
| 0 | 0 | 1 | 0 |
| 0 | 0 | 0 | 0 |
| 0 | 0 | 0 | 0 |
| 0 | 0 | 0 | 0 |
| 0 | 0 | 0 | 0 |
| 0 | 0 | 0 | 0 |
| 1 | 0 | 0 | 0 |
| 0 | 0 | 0 | 0 |
| 0 | 0 | 0 | 0 |
| 0 | 0 | 0 | 0 |
| 0 | 0 | 0 | 0 |
| 0 | 0 | 0 | 0 |
| 0 | 0 | 0 | 0 |
| 1 | 1 | 0 | 0 |
| 0 | 0 | 0 | 0 |
| 0 | 0 | 0 | 0 |
| 0 | 0 | 0 | 0 |
| 0 | 0 | 0 | 0 |
| 0 | 0 | 0 | 0 |
| 0 | 0 | 0 | 0 |
| 0 | 0 | 0 | 0 |
| 0 | 0 | 0 | 0 |
| 0 | 1 | 0 | 0 |
| 0 | 0 | 0 | 0 |
| 1 | 1 | 0 | 0 |
| 0 | 0 | 0 | 0 |
| 0 | 0 | 0 | 0 |
| 0 | 0 | 0 | 0 |
| 0 | 0 | 0 | 0 |
| 0 | 0 | 0 | 0 |
| 0 | 0 | 0 | 0 |
| 1 | 0 | 0 | 1 |
| 0 | 0 | 0 | 0 |
| 1 | 0 | 0 | 1 |
| 0 | 0 | 0 | 0 |
| 1 | 0 | 0 | 1 |
| 0 | 0 | 0 | 0 |
| 0 | 0 | 0 | 0 |
| 0 | 0 | 0 | 0 |
| 1 | 0 | 0 | 0 |
| 0 | 0 | 0 | 0 |
| 0 | 0 | 0 | 0 |
| 0 | 0 | 0 | 0 |
| 0 | 0 | 0 | 0 |

| | | | |
|---|---|---|---|
| 0 | 0 | 0 | 0 |
| 0 | 0 | 0 | 0 |
| 0 | 0 | 0 | 0 |
| 0 | 0 | 0 | 0 |
| 0 | 0 | 0 | 0 |
| 0 | 0 | 0 | 0 |
| 0 | 0 | 0 | 0 |
| 0 | 0 | 0 | 0 |
| 0 | 0 | 0 | 0 |
| 0 | 0 | 0 | 0 |
| 0 | 0 | 0 | 0 |
| 0 | 0 | 0 | 0 |
| 0 | 0 | 0 | 0 |
| 0 | 0 | 0 | 0 |
| 0 | 0 | 0 | 0 |
| 0 | 0 | 0 | 0 |
| 0 | 0 | 0 | 0 |
| 0 | 0 | 0 | 0 |
| 0 | 0 | 1 | 0 |
| 0 | 0 | 0 | 0 |
| 0 | 0 | 0 | 0 |
| 0 | 0 | 0 | 0 |
| 0 | 0 | 0 | 0 |
| 0 | 0 | 0 | 0 |
| 0 | 0 | 0 | 0 |
| 0 | 0 | 0 | 0 |
| 0 | 0 | 0 | 0 |
| 0 | 0 | 0 | 0 |
| 0 | 0 | 1 | 0 |
| 0 | 0 | 0 | 0 |
| 0 | 0 | 1 | 0 |
| 1 | 0 | 0 | 0 |
| 1 | 1 | 1 | 0 |
| 1 | 1 | 1 | 0 |
| 0 | 0 | 0 | 0 |
| 0 | 0 | 0 | 0 |
| 0 | 1 | 0 | 0 |
| 1 | 1 | 1 | 0 |
| 1 | 1 | 1 | 0 |
| 0 | 0 | 1 | 0 |
| 0 | 0 | 1 | 0 |
| 1 | 0 | 0 | 0 |
| 0 | 0 | 0 | 0 |
| 0 | 0 | 0 | 0 |
| 1 | 1 | 0 | 0 |
| 0 | 0 | 0 | 0 |
| 0 | 0 | 0 | 0 |
| 0 | 0 | 0 | 0 |
| 0 | 0 | 0 | 0 |
| 0 | 0 | 0 | 0 |
| 0 | 0 | 0 | 0 |
| 0 | 0 | 0 | 0 |
| 0 | 0 | 0 | 0 |
| 0 | 0 | 0 | 0 |

| | | | |
|---|---|---|---|
| 0 | 0 | 0 | 0 |
| 0 | 0 | 0 | 0 |
| 0 | 0 | 0 | 0 |
| 0 | 0 | 0 | 0 |
| 0 | 0 | 0 | 0 |
| 0 | 0 | 0 | 0 |
| 0 | 0 | 0 | 0 |
| 0 | 0 | 0 | 0 |
| 0 | 0 | 0 | 0 |
| 0 | 0 | 0 | 0 |
| 0 | 0 | 0 | 0 |
| 0 | 0 | 0 | 0 |
| 0 | 0 | 0 | 0 |
| 0 | 0 | 0 | 0 |
| 0 | 0 | 0 | 0 |
| 0 | 0 | 0 | 0 |
| 0 | 0 | 0 | 0 |
| 0 | 0 | 0 | 0 |
| 0 | 0 | 0 | 0 |
| 1 | 0 | 0 | 0 |
| 1 | 0 | 0 | 0 |
| 0 | 0 | 0 | 0 |
| 0 | 0 | 0 | 0 |
| 0 | 0 | 0 | 0 |
| 0 | 0 | 0 | 0 |
| 1 | 0 | 0 | 0 |
| 0 | 0 | 0 | 0 |
| 0 | 0 | 0 | 0 |
| 0 | 0 | 0 | 0 |
| 1 | 0 | 0 | 0 |
| 0 | 0 | 0 | 0 |
| 0 | 0 | 0 | 0 |
| 1 | 0 | 0 | 0 |
| 1 | 0 | 0 | 0 |
| 0 | 0 | 0 | 0 |
| 0 | 0 | 0 | 0 |
| 0 | 0 | 0 | 0 |
| 0 | 0 | 0 | 0 |
| 0 | 0 | 0 | 0 |
| 0 | 0 | 0 | 0 |
| 0 | 0 | 0 | 0 |
| 0 | 0 | 0 | 0 |
| 0 | 0 | 0 | 0 |
| 0 | 0 | 0 | 0 |
| 0 | 0 | 0 | 0 |
| 0 | 0 | 0 | 0 |
| 0 | 0 | 0 | 0 |
| 0 | 1 | 0 | 0 |
| 0 | 1 | 0 | 0 |
| 0 | 0 | 0 | 0 |
| 0 | 1 | 0 | 0 |
| 0 | 1 | 0 | 0 |
| 0 | 0 | 0 | 0 |
| 0 | 0 | 0 | 0 |

| | | | |
|---|---|---|---|
| 0 | 0 | 0 | 0 |
| 0 | 0 | 0 | 0 |
| 1 | 1 | 0 | 1 |
| 0 | 0 | 0 | 0 |
| 0 | 0 | 0 | 0 |
| 0 | 0 | 0 | 0 |
| 0 | 0 | 0 | 0 |
| 0 | 1 | 0 | 0 |
| 1 | 1 | 0 | 0 |
| 1 | 1 | 0 | 0 |
| 1 | 1 | 0 | 0 |
| 1 | 1 | 0 | 0 |
| 1 | 1 | 0 | 0 |
| 0 | 1 | 0 | 0 |
| 0 | 1 | 0 | 0 |
| 0 | 0 | 0 | 0 |
| 0 | 0 | 0 | 0 |
| 0 | 0 | 0 | 0 |
| 0 | 0 | 0 | 0 |
| 0 | 0 | 0 | 0 |
| 0 | 0 | 0 | 0 |
| 0 | 0 | 0 | 0 |
| 0 | 1 | 0 | 0 |
| 0 | 1 | 0 | 0 |
| 0 | 0 | 0 | 0 |
| 0 | 0 | 1 | 0 |
| 0 | 0 | 0 | 0 |
| 0 | 0 | 0 | 0 |
| 0 | 0 | 0 | 0 |
| 0 | 1 | 0 | 0 |
| 0 | 0 | 0 | 0 |
| 0 | 0 | 0 | 0 |
| 0 | 0 | 0 | 0 |
| 0 | 0 | 0 | 0 |
| 0 | 0 | 0 | 0 |
| 0 | 0 | 0 | 0 |
| 0 | 0 | 0 | 0 |
| 0 | 0 | 0 | 0 |
| 0 | 0 | 0 | 0 |
| 1 | 0 | 0 | 0 |
| 1 | 1 | 0 | 0 |
| 1 | 0 | 0 | 0 |
| 0 | 0 | 0 | 0 |
| 0 | 1 | 0 | 0 |
| 1 | 1 | 0 | 0 |
| 0 | 1 | 0 | 0 |
| 0 | 0 | 0 | 0 |
| 1 | 0 | 0 | 0 |
| 0 | 1 | 0 | 0 |
| 0 | 0 | 0 | 0 |
| 0 | 0 | 0 | 0 |
| 0 | 0 | 0 | 0 |
| 1 | 1 | 0 | 0 |
| 0 | 0 | 0 | 0 |

| | | | |
|---|---|---|---|
| 0 | 0 | 0 | 0 |
| 0 | 0 | 0 | 0 |
| 1 | 0 | 0 | 0 |
| 1 | 1 | 0 | 0 |
| 1 | 1 | 0 | 0 |
| 0 | 0 | 0 | 0 |
| 0 | 0 | 0 | 0 |
| 0 | 0 | 0 | 0 |
| 1 | 0 | 0 | 0 |
| 0 | 0 | 0 | 0 |
| 0 | 0 | 0 | 0 |
| 1 | 0 | 0 | 1 |
| 0 | 0 | 0 | 0 |
| 1 | 0 | 0 | 0 |
| 0 | 0 | 0 | 0 |
| 0 | 0 | 0 | 0 |
| 0 | 0 | 0 | 0 |
| 0 | 0 | 0 | 0 |
| 0 | 0 | 0 | 0 |
| 0 | 0 | 0 | 0 |
| 0 | 0 | 0 | 0 |
| 1 | 0 | 0 | 1 |
| 1 | 0 | 0 | 0 |
| 0 | 0 | 0 | 0 |
| 0 | 0 | 0 | 0 |
| 0 | 0 | 0 | 0 |
| 0 | 0 | 0 | 0 |
| 0 | 0 | 0 | 0 |
| 1 | 0 | 0 | 0 |
| 1 | 0 | 0 | 0 |
| 0 | 0 | 0 | 0 |
| 0 | 0 | 0 | 0 |
| 0 | 0 | 0 | 0 |
| 0 | 0 | 0 | 0 |
| 0 | 0 | 0 | 0 |
| 0 | 0 | 0 | 0 |
| 1 | 0 | 0 | 0 |
| 0 | 0 | 0 | 0 |
| 0 | 0 | 0 | 0 |
| 1 | 0 | 0 | 0 |
| 1 | 0 | 0 | 0 |
| 1 | 0 | 0 | 0 |
| 1 | 0 | 0 | 0 |
| 0 | 0 | 0 | 0 |
| 0 | 0 | 0 | 0 |
| 1 | 0 | 0 | 0 |
| 0 | 0 | 0 | 0 |
| 0 | 0 | 0 | 0 |
| 0 | 0 | 0 | 0 |
| 0 | 0 | 0 | 0 |
| 0 | 0 | 0 | 0 |
| 1 | 0 | 0 | 0 |
| 1 | 0 | 0 | 0 |
| 1 | 0 | 0 | 0 |

| | | | |
|---|---|---|---|
| 1 | 0 | 0 | 0 |
| 0 | 0 | 0 | 0 |
| 0 | 0 | 0 | 0 |
| 0 | 0 | 0 | 0 |
| 1 | 0 | 0 | 0 |
| 0 | 0 | 0 | 0 |
| 1 | 0 | 0 | 0 |
| 0 | 0 | 0 | 0 |
| 0 | 0 | 0 | 0 |
| 0 | 0 | 0 | 0 |
| 0 | 0 | 0 | 0 |
| 0 | 0 | 0 | 0 |
| 0 | 0 | 0 | 0 |
| 0 | 0 | 0 | 0 |
| 0 | 0 | 0 | 0 |
| 1 | 0 | 0 | 0 |
| 0 | 0 | 0 | 0 |
| 0 | 0 | 0 | 0 |
| 0 | 0 | 0 | 0 |
| 0 | 0 | 0 | 0 |
| 0 | 0 | 0 | 0 |
| 0 | 0 | 0 | 0 |
| 1 | 0 | 0 | 0 |
| 1 | 0 | 0 | 0 |
| 0 | 0 | 0 | 0 |
| 0 | 0 | 0 | 0 |
| 0 | 0 | 0 | 0 |
| 0 | 0 | 0 | 0 |
| 0 | 0 | 0 | 0 |
| 0 | 0 | 0 | 0 |
| 0 | 0 | 0 | 0 |
| 0 | 0 | 0 | 0 |
| 1 | 0 | 0 | 0 |
| 0 | 0 | 0 | 0 |
| 0 | 0 | 0 | 0 |
| 0 | 0 | 0 | 0 |
| 0 | 0 | 0 | 0 |
| 0 | 0 | 0 | 0 |
| 0 | 0 | 0 | 0 |
| 0 | 0 | 0 | 0 |
| 0 | 0 | 0 | 0 |
| 0 | 0 | 0 | 0 |
| 0 | 0 | 0 | 0 |
| 0 | 0 | 0 | 0 |
| 0 | 1 | 0 | 0 |
| 0 | 0 | 0 | 0 |
| 0 | 0 | 0 | 0 |
| 0 | 0 | 0 | 0 |
| 0 | 0 | 0 | 0 |
| 0 | 0 | 0 | 0 |
| 0 | 0 | 0 | 0 |
| 0 | 0 | 0 | 0 |
| 0 | 0 | 0 | 0 |
| 1 | 0 | 0 | 0 |

| | | | |
|---|---|---|---|
| 0 | 0 | 0 | 0 |
| 0 | 0 | 0 | 0 |
| 0 | 0 | 0 | 0 |
| 0 | 0 | 0 | 0 |
| 0 | 0 | 0 | 0 |
| 0 | 0 | 0 | 0 |
| 0 | 0 | 0 | 0 |
| 0 | 0 | 0 | 0 |
| 0 | 0 | 0 | 0 |
| 1 | 0 | 0 | 0 |
| 0 | 0 | 0 | 0 |
| 0 | 0 | 0 | 0 |
| 1 | 1 | 0 | 0 |
| 1 | 1 | 0 | 0 |
| 1 | 1 | 0 | 0 |
| 1 | 1 | 0 | 0 |
| 0 | 0 | 0 | 0 |
| 0 | 0 | 0 | 0 |
| 0 | 0 | 0 | 0 |
| 0 | 0 | 0 | 0 |
| 0 | 0 | 0 | 0 |
| 0 | 0 | 0 | 0 |
| 0 | 0 | 0 | 0 |
| 0 | 0 | 0 | 0 |
| 0 | 0 | 0 | 0 |
| 0 | 0 | 0 | 1 |
| 0 | 0 | 0 | 0 |
| 0 | 0 | 0 | 0 |
| 0 | 0 | 0 | 0 |
| 0 | 0 | 0 | 0 |
| 0 | 0 | 0 | 0 |
| 0 | 0 | 0 | 0 |
| 0 | 0 | 0 | 0 |
| 0 | 0 | 0 | 0 |
| 0 | 0 | 0 | 0 |
| 0 | 0 | 0 | 0 |
| 0 | 0 | 0 | 0 |
| 1 | 1 | 0 | 0 |
| 0 | 0 | 0 | 0 |
| 0 | 0 | 0 | 0 |
| 1 | 1 | 0 | 0 |
| 0 | 0 | 0 | 0 |
| 0 | 0 | 0 | 0 |
| 1 | 0 | 0 | 1 |
| 0 | 0 | 0 | 0 |
| 0 | 0 | 0 | 0 |
| 0 | 0 | 0 | 0 |
| 0 | 0 | 0 | 0 |
| 0 | 0 | 0 | 0 |
| 0 | 0 | 0 | 0 |
| 0 | 0 | 0 | 0 |
| 0 | 0 | 1 | 0 |
| 0 | 0 | 1 | 0 |
| 0 | 0 | 1 | 0 |

| | | | |
|---|---|---|---|
| 0 | 0 | 1 | 0 |
| 0 | 0 | 0 | 0 |
| 0 | 0 | 0 | 0 |
| 0 | 0 | 0 | 0 |
| 0 | 0 | 0 | 0 |
| 0 | 0 | 0 | 0 |
| 0 | 0 | 0 | 0 |
| 0 | 0 | 0 | 0 |
| 0 | 0 | 0 | 0 |
| 0 | 0 | 0 | 0 |
| 0 | 0 | 0 | 0 |
| 0 | 0 | 0 | 0 |
| 0 | 0 | 0 | 0 |
| 0 | 0 | 0 | 0 |
| 0 | 0 | 0 | 0 |
| 0 | 0 | 0 | 0 |
| 0 | 0 | 0 | 0 |
| 0 | 0 | 0 | 0 |
| 0 | 0 | 0 | 0 |
| 0 | 0 | 0 | 0 |
| 0 | 0 | 0 | 0 |
| 0 | 0 | 0 | 0 |
| 0 | 0 | 0 | 0 |
| 0 | 1 | 0 | 1 |
| 0 | 0 | 0 | 0 |
| 0 | 0 | 0 | 0 |
| 0 | 0 | 0 | 0 |
| 0 | 0 | 0 | 0 |
| 0 | 0 | 0 | 0 |
| 0 | 0 | 0 | 0 |
| 0 | 0 | 0 | 0 |
| 0 | 0 | 0 | 0 |
| 0 | 0 | 0 | 0 |
| 0 | 0 | 0 | 0 |
| 0 | 0 | 0 | 0 |
| 0 | 0 | 0 | 0 |
| 0 | 0 | 0 | 0 |
| 0 | 0 | 0 | 0 |
| 0 | 0 | 0 | 0 |
| 0 | 0 | 0 | 0 |
| 0 | 0 | 0 | 0 |
| 0 | 0 | 0 | 0 |
| 0 | 0 | 0 | 0 |
| 0 | 0 | 0 | 0 |
| 0 | 0 | 0 | 0 |
| 0 | 0 | 0 | 0 |
| 1 | 0 | 0 | 0 |
| 1 | 0 | 0 | 0 |
| 0 | 0 | 0 | 0 |
| 0 | 1 | 0 | 0 |
| 1 | 1 | 0 | 1 |
| 1 | 1 | 0 | 1 |
| 1 | 1 | 0 | 1 |
| 0 | 0 | 0 | 0 |

| | | | |
|---|---|---|---|
| 1 | 1 | 0 | 0 |
| 0 | 0 | 0 | 0 |
| 0 | 0 | 0 | 0 |
| 0 | 0 | 0 | 0 |
| 0 | 0 | 0 | 0 |
| 0 | 0 | 0 | 0 |
| 0 | 0 | 0 | 0 |
| 0 | 0 | 0 | 0 |
| 0 | 0 | 0 | 0 |
| 0 | 0 | 0 | 0 |
| 0 | 0 | 0 | 0 |
| 0 | 1 | 0 | 0 |
| 0 | 0 | 0 | 0 |
| 0 | 0 | 0 | 0 |
| 0 | 1 | 0 | 0 |
| 0 | 0 | 0 | 0 |
| 1 | 1 | 0 | 0 |
| 0 | 0 | 0 | 0 |
| 0 | 0 | 0 | 0 |
| 1 | 0 | 0 | 0 |
| 0 | 0 | 0 | 0 |
| 0 | 0 | 0 | 0 |
| 0 | 0 | 0 | 0 |
| 0 | 0 | 0 | 0 |
| 0 | 0 | 0 | 0 |
| 0 | 0 | 0 | 0 |
| 0 | 0 | 0 | 0 |
| 0 | 0 | 0 | 0 |
| 0 | 0 | 0 | 0 |
| 0 | 0 | 0 | 0 |
| 0 | 0 | 0 | 0 |
| 0 | 0 | 0 | 0 |
| 0 | 0 | 0 | 0 |
| 1 | 0 | 0 | 0 |
| 0 | 0 | 0 | 0 |
| 0 | 1 | 0 | 0 |
| 0 | 0 | 0 | 0 |
| 0 | 0 | 0 | 0 |
| 0 | 0 | 0 | 0 |
| 0 | 0 | 0 | 0 |
| 0 | 0 | 0 | 0 |
| 0 | 0 | 0 | 0 |
| 0 | 0 | 0 | 0 |
| 0 | 0 | 0 | 0 |
| 0 | 0 | 0 | 0 |
| 0 | 0 | 0 | 0 |
| 0 | 0 | 0 | 0 |
| 0 | 0 | 0 | 0 |
| 1 | 1 | 0 | 0 |
| 0 | 0 | 0 | 0 |
| 0 | 0 | 0 | 0 |
| 0 | 0 | 0 | 0 |
| 0 | 0 | 0 | 0 |
| 0 | 0 | 0 | 0 |

| | | | |
|---|---|---|---|
| 0 | 0 | 0 | 0 |
| 0 | 0 | 0 | 0 |
| 0 | 0 | 0 | 0 |
| 0 | 0 | 0 | 0 |
| 0 | 0 | 0 | 0 |
| 0 | 0 | 0 | 0 |
| 0 | 0 | 0 | 0 |
| 0 | 1 | 0 | 0 |
| 0 | 0 | 0 | 0 |
| 0 | 0 | 0 | 0 |
| 0 | 0 | 0 | 0 |
| 0 | 0 | 0 | 0 |
| 0 | 0 | 1 | 0 |
| 0 | 0 | 0 | 0 |
| 0 | 0 | 1 | 0 |
| 0 | 0 | 1 | 0 |
| 0 | 0 | 0 | 0 |
| 0 | 0 | 0 | 0 |
| 0 | 0 | 0 | 0 |
| 0 | 0 | 0 | 0 |
| 0 | 1 | 0 | 0 |
| 0 | 1 | 0 | 0 |
| 0 | 0 | 0 | 0 |
| 0 | 0 | 0 | 0 |
| 0 | 1 | 0 | 0 |
| 0 | 0 | 0 | 0 |
| 0 | 0 | 0 | 0 |
| 0 | 1 | 0 | 0 |
| 0 | 0 | 0 | 0 |
| 0 | 0 | 0 | 0 |
| 0 | 0 | 0 | 0 |
| 0 | 0 | 0 | 0 |
| 1 | 1 | 0 | 0 |
| 1 | 1 | 0 | 0 |
| 1 | 1 | 0 | 0 |
| 0 | 1 | 0 | 0 |
| 1 | 0 | 0 | 0 |
| 0 | 1 | 0 | 0 |
| 1 | 1 | 0 | 0 |
| 0 | 0 | 0 | 0 |
| 0 | 0 | 0 | 0 |
| 0 | 0 | 0 | 0 |
| 0 | 0 | 0 | 0 |
| 0 | 0 | 0 | 0 |
| 0 | 0 | 0 | 0 |
| 0 | 0 | 0 | 0 |
| 0 | 0 | 0 | 0 |
| 0 | 0 | 0 | 0 |
| 0 | 0 | 0 | 0 |
| 1 | 1 | 0 | 0 |
| 1 | 1 | 0 | 0 |
| 0 | 0 | 0 | 0 |
| 0 | 0 | 0 | 0 |
| 0 | 0 | 0 | 0 |

| | | | |
|---|---|---|---|
| 0 | 0 | 0 | 0 |
| 0 | 0 | 0 | 0 |
| 0 | 0 | 0 | 0 |
| 0 | 0 | 0 | 0 |
| 0 | 0 | 0 | 0 |
| 0 | 0 | 0 | 0 |
| 1 | 1 | 0 | 0 |
| 1 | 1 | 0 | 0 |
| 0 | 0 | 0 | 0 |
| 0 | 0 | 0 | 0 |
| 0 | 0 | 0 | 0 |
| 0 | 0 | 0 | 0 |
| 0 | 0 | 0 | 0 |
| 0 | 0 | 0 | 0 |
| 0 | 0 | 0 | 0 |
| 0 | 0 | 0 | 0 |
| 0 | 0 | 0 | 0 |
| 0 | 0 | 0 | 0 |
| 0 | 0 | 0 | 0 |
| 0 | 0 | 0 | 0 |
| 0 | 0 | 0 | 0 |
| 0 | 0 | 0 | 0 |
| 0 | 0 | 0 | 0 |
| 0 | 0 | 0 | 0 |
| 0 | 0 | 0 | 0 |
| 0 | 0 | 0 | 0 |
| 0 | 0 | 0 | 0 |
| 1 | 1 | 0 | 0 |
| 0 | 0 | 0 | 0 |
| 0 | 0 | 0 | 0 |
| 0 | 0 | 0 | 0 |
| 0 | 0 | 0 | 0 |
| 0 | 0 | 0 | 0 |
| 0 | 0 | 0 | 0 |
| 0 | 0 | 0 | 0 |
| 0 | 0 | 0 | 0 |
| 0 | 0 | 0 | 0 |
| 0 | 0 | 0 | 0 |
| 0 | 0 | 0 | 0 |
| 0 | 0 | 0 | 0 |
| 0 | 0 | 0 | 0 |
| 0 | 0 | 0 | 0 |
| 0 | 0 | 0 | 0 |
| 0 | 0 | 0 | 0 |
| 0 | 0 | 0 | 0 |
| 0 | 0 | 0 | 0 |
| 0 | 0 | 0 | 0 |
| 0 | 0 | 0 | 0 |
| 0 | 0 | 0 | 0 |
| 0 | 0 | 0 | 0 |
| 0 | 0 | 0 | 0 |
| 0 | 0 | 0 | 0 |
| 0 | 0 | 0 | 0 |
| 0 | 0 | 0 | 0 |

| | | | |
|---|---|---|---|
| 0 | 0 | 0 | 0 |
| 0 | 0 | 0 | 0 |
| 0 | 0 | 0 | 0 |
| 0 | 0 | 0 | 0 |
| 0 | 0 | 0 | 0 |
| 0 | 0 | 0 | 0 |
| 0 | 0 | 0 | 0 |
| 0 | 0 | 0 | 0 |
| 0 | 0 | 0 | 0 |
| 0 | 0 | 0 | 0 |
| 1 | 0 | 0 | 0 |
| 1 | 1 | 0 | 0 |
| 1 | 0 | 0 | 0 |
| 0 | 0 | 0 | 0 |
| 0 | 0 | 0 | 0 |
| 0 | 0 | 0 | 0 |
| 0 | 0 | 0 | 0 |
| 0 | 0 | 0 | 0 |
| 0 | 0 | 0 | 0 |
| 0 | 0 | 0 | 0 |
| 0 | 0 | 0 | 0 |
| 0 | 0 | 0 | 0 |
| 0 | 0 | 0 | 0 |
| 0 | 0 | 0 | 0 |
| 0 | 0 | 0 | 0 |
| 0 | 0 | 0 | 0 |
| 0 | 0 | 0 | 0 |
| 0 | 0 | 0 | 0 |
| 0 | 0 | 0 | 0 |
| 0 | 0 | 0 | 0 |
| 0 | 0 | 0 | 0 |
| 0 | 1 | 0 | 0 |
| 0 | 0 | 0 | 0 |
| 0 | 0 | 0 | 0 |
| 1 | 0 | 0 | 0 |
| 1 | 0 | 0 | 0 |
| 1 | 0 | 0 | 0 |
| 0 | 0 | 0 | 0 |
| 0 | 0 | 0 | 0 |
| 0 | 0 | 0 | 0 |
| 0 | 0 | 0 | 0 |
| 0 | 0 | 0 | 0 |
| 0 | 0 | 0 | 0 |
| 0 | 0 | 0 | 0 |
| 0 | 0 | 0 | 0 |
| 0 | 0 | 0 | 0 |
| 0 | 0 | 0 | 0 |
| 0 | 0 | 0 | 0 |
| 0 | 0 | 0 | 0 |
| 0 | 0 | 0 | 0 |
| 0 | 0 | 0 | 0 |
| 0 | 0 | 0 | 0 |
| 0 | 0 | 0 | 1 |
| 0 | 0 | 0 | 0 |

| | | | |
|---|---|---|---|
| 0 | 0 | 0 | 0 |
| 0 | 0 | 0 | 0 |
| 0 | 0 | 0 | 0 |
| 0 | 0 | 0 | 0 |
| 0 | 0 | 0 | 0 |
| 0 | 0 | 0 | 0 |
| 0 | 0 | 0 | 0 |
| 0 | 0 | 0 | 0 |
| 0 | 0 | 0 | 0 |
| 0 | 0 | 0 | 0 |
| 0 | 0 | 0 | 0 |
| 1 | 0 | 0 | 0 |
| 0 | 0 | 0 | 0 |
| 0 | 0 | 0 | 0 |
| 0 | 0 | 1 | 0 |
| 0 | 0 | 0 | 0 |
| 0 | 0 | 0 | 0 |
| 0 | 0 | 0 | 0 |
| 0 | 0 | 0 | 0 |
| 0 | 0 | 0 | 0 |
| 0 | 0 | 0 | 0 |
| 0 | 0 | 0 | 0 |
| 0 | 0 | 0 | 0 |
| 0 | 0 | 0 | 0 |
| 0 | 0 | 0 | 0 |
| 0 | 0 | 0 | 0 |
| 0 | 0 | 0 | 0 |
| 0 | 0 | 0 | 0 |
| 0 | 0 | 0 | 0 |
| 0 | 0 | 0 | 0 |
| 0 | 0 | 0 | 0 |
| 0 | 0 | 0 | 0 |
| 0 | 0 | 0 | 0 |
| 1 | 0 | 0 | 0 |
| 0 | 0 | 0 | 0 |
| 1 | 0 | 0 | 1 |
| 0 | 0 | 0 | 0 |
| 0 | 0 | 0 | 0 |
| 0 | 0 | 0 | 0 |
| 0 | 0 | 0 | 0 |
| 0 | 0 | 0 | 0 |
| 0 | 0 | 0 | 0 |
| 0 | 0 | 0 | 0 |
| 0 | 1 | 0 | 0 |
| 0 | 0 | 0 | 0 |
| 0 | 0 | 0 | 0 |
| 0 | 0 | 0 | 0 |
| 0 | 0 | 0 | 0 |
| 0 | 0 | 0 | 0 |
| 0 | 0 | 0 | 0 |
| 0 | 0 | 0 | 0 |
| 0 | 0 | 0 | 0 |
| 0 | 0 | 0 | 0 |
| 0 | 0 | 0 | 0 |

| | | | |
|---|---|---|---|
| 0 | 0 | 0 | 0 |
| 0 | 0 | 0 | 0 |
| 0 | 0 | 0 | 0 |
| 0 | 0 | 0 | 0 |
| 0 | 0 | 0 | 0 |
| 0 | 1 | 0 | 0 |
| 0 | 0 | 0 | 0 |
| 0 | 0 | 0 | 0 |
| 0 | 0 | 0 | 0 |
| 0 | 0 | 0 | 0 |
| 0 | 0 | 0 | 0 |
| 0 | 0 | 0 | 0 |
| 0 | 0 | 0 | 0 |
| 0 | 0 | 0 | 0 |
| 0 | 0 | 0 | 0 |
| 0 | 1 | 0 | 0 |
| 1 | 1 | 0 | 0 |
| 0 | 0 | 1 | 0 |
| 0 | 0 | 0 | 0 |
| 0 | 0 | 0 | 0 |
| 0 | 0 | 0 | 0 |
| 0 | 0 | 0 | 0 |
| 0 | 0 | 0 | 0 |
| 0 | 0 | 0 | 0 |
| 0 | 0 | 0 | 0 |
| 0 | 0 | 0 | 0 |
| 0 | 0 | 0 | 0 |
| 1 | 0 | 0 | 0 |
| 0 | 0 | 0 | 0 |
| 0 | 0 | 0 | 0 |
| 0 | 0 | 1 | 0 |
| 0 | 0 | 0 | 0 |
| 0 | 0 | 0 | 0 |
| 0 | 0 | 0 | 0 |
| 0 | 0 | 0 | 0 |
| 0 | 0 | 0 | 0 |
| 0 | 0 | 0 | 0 |
| 1 | 1 | 0 | 0 |
| 0 | 0 | 0 | 0 |
| 0 | 0 | 0 | 0 |
| 0 | 0 | 0 | 0 |
| 1 | 0 | 0 | 0 |
| 1 | 0 | 0 | 0 |
| 0 | 0 | 0 | 0 |
| 0 | 0 | 0 | 0 |
| 0 | 0 | 0 | 0 |
| 0 | 0 | 0 | 0 |
| 0 | 0 | 0 | 0 |
| 0 | 0 | 0 | 0 |
| 0 | 0 | 0 | 0 |
| 0 | 0 | 0 | 0 |
| 0 | 0 | 0 | 0 |
| 0 | 0 | 0 | 0 |
| 0 | 0 | 0 | 0 |

| | | | |
|---|---|---|---|
| 0 | 0 | 0 | 0 |
| 0 | 0 | 0 | 0 |
| 0 | 0 | 0 | 0 |
| 0 | 0 | 0 | 0 |
| 0 | 0 | 0 | 0 |
| 0 | 0 | 0 | 0 |
| 0 | 0 | 0 | 0 |
| 0 | 0 | 0 | 0 |
| 0 | 0 | 0 | 0 |
| 0 | 0 | 0 | 0 |
| 0 | 0 | 0 | 0 |
| 0 | 0 | 0 | 0 |
| 1 | 0 | 0 | 0 |
| 0 | 0 | 0 | 0 |
| 0 | 0 | 0 | 0 |
| 0 | 0 | 0 | 0 |
| 0 | 0 | 0 | 0 |
| 0 | 0 | 0 | 0 |
| 0 | 0 | 0 | 0 |
| 0 | 0 | 0 | 0 |
| 0 | 0 | 0 | 0 |
| 0 | 0 | 0 | 0 |
| 0 | 0 | 0 | 0 |
| 1 | 0 | 0 | 0 |
| 0 | 0 | 0 | 0 |
| 0 | 0 | 0 | 0 |
| 0 | 0 | 0 | 0 |
| 1 | 1 | 0 | 1 |
| 0 | 0 | 0 | 0 |
| 0 | 0 | 0 | 0 |
| 0 | 0 | 0 | 0 |
| 0 | 0 | 0 | 0 |
| 0 | 0 | 0 | 0 |
| 0 | 0 | 0 | 0 |
| 0 | 0 | 0 | 0 |
| 0 | 0 | 0 | 0 |
| 0 | 0 | 0 | 0 |
| 0 | 0 | 0 | 0 |
| 0 | 0 | 0 | 0 |
| 0 | 0 | 0 | 0 |
| 0 | 0 | 0 | 0 |
| 0 | 0 | 0 | 0 |
| 0 | 0 | 0 | 0 |
| 0 | 0 | 0 | 0 |
| 0 | 0 | 0 | 0 |
| 0 | 0 | 0 | 0 |
| 0 | 0 | 0 | 0 |
| 0 | 0 | 0 | 0 |
| 0 | 0 | 0 | 0 |
| 0 | 0 | 0 | 0 |
| 0 | 0 | 0 | 0 |
| 0 | 0 | 0 | 0 |
| 0 | 0 | 0 | 0 |
| 0 | 0 | 0 | 0 |

| | | | |
|---|---|---|---|
| 0 | 0 | 0 | 0 |
| 0 | 0 | 0 | 0 |
| 0 | 0 | 0 | 0 |
| 0 | 0 | 0 | 0 |
| 0 | 0 | 0 | 0 |
| 0 | 0 | 0 | 0 |
| 0 | 0 | 0 | 0 |
| 0 | 0 | 0 | 0 |
| 1 | 0 | 0 | 0 |
| 1 | 0 | 0 | 0 |
| 1 | 0 | 0 | 0 |
| 0 | 0 | 0 | 0 |
| 1 | 0 | 0 | 0 |
| 0 | 0 | 0 | 0 |
| 0 | 0 | 0 | 0 |
| 1 | 0 | 0 | 0 |
| 0 | 0 | 0 | 0 |
| 0 | 0 | 0 | 0 |
| 0 | 0 | 0 | 0 |
| 0 | 0 | 0 | 0 |
| 0 | 0 | 0 | 0 |
| 0 | 0 | 0 | 0 |
| 0 | 0 | 0 | 0 |
| 0 | 0 | 0 | 0 |
| 0 | 0 | 0 | 0 |
| 0 | 0 | 0 | 0 |
| 0 | 0 | 0 | 0 |
| 1 | 0 | 0 | 0 |
| 1 | 0 | 0 | 0 |
| 0 | 0 | 0 | 0 |
| 0 | 0 | 0 | 0 |
| 0 | 0 | 0 | 0 |
| 1 | 0 | 0 | 0 |
| 0 | 0 | 0 | 0 |
| 0 | 0 | 0 | 0 |
| 0 | 0 | 0 | 0 |
| 0 | 0 | 0 | 0 |
| 0 | 0 | 0 | 0 |
| 0 | 0 | 0 | 0 |
| 0 | 0 | 0 | 0 |
| 1 | 0 | 0 | 0 |
| 1 | 0 | 0 | 0 |
| 0 | 0 | 0 | 0 |
| 0 | 0 | 0 | 0 |
| 0 | 0 | 0 | 0 |
| 0 | 0 | 0 | 0 |
| 0 | 0 | 0 | 0 |
| 0 | 0 | 0 | 0 |
| 0 | 0 | 0 | 0 |
| 0 | 0 | 0 | 0 |
| 0 | 0 | 0 | 0 |
| 0 | 0 | 0 | 0 |
| 0 | 0 | 0 | 0 |
| 0 | 0 | 0 | 0 |

| | | | |
|---|---|---|---|
| 0 | 0 | 0 | 0 |
| 0 | 0 | 0 | 0 |
| 0 | 0 | 0 | 0 |
| 0 | 0 | 0 | 0 |
| 0 | 0 | 0 | 0 |
| 0 | 0 | 0 | 0 |
| 0 | 0 | 0 | 0 |
| 0 | 0 | 0 | 0 |
| 0 | 0 | 0 | 0 |
| 0 | 0 | 0 | 0 |
| 1 | 0 | 0 | 0 |
| 0 | 0 | 0 | 0 |
| 1 | 0 | 0 | 0 |
| 0 | 0 | 0 | 0 |
| 0 | 0 | 0 | 0 |
| 0 | 0 | 0 | 0 |
| 0 | 0 | 0 | 0 |
| 0 | 1 | 0 | 0 |
| 0 | 0 | 0 | 0 |
| 0 | 0 | 0 | 0 |
| 0 | 0 | 0 | 0 |
| 0 | 0 | 0 | 0 |
| 1 | 0 | 0 | 0 |
| 0 | 0 | 0 | 0 |
| 0 | 0 | 0 | 0 |
| 1 | 0 | 0 | 0 |
| 1 | 0 | 0 | 0 |
| 1 | 0 | 0 | 0 |
| 0 | 0 | 0 | 0 |
| 0 | 0 | 0 | 0 |
| 0 | 0 | 0 | 0 |
| 0 | 0 | 0 | 0 |
| 0 | 0 | 0 | 0 |
| 0 | 0 | 0 | 0 |
| 0 | 0 | 0 | 0 |
| 0 | 0 | 0 | 0 |
| 0 | 0 | 0 | 0 |
| 0 | 0 | 0 | 0 |
| 0 | 0 | 0 | 0 |
| 0 | 0 | 0 | 0 |
| 1 | 1 | 0 | 0 |
| 0 | 0 | 0 | 0 |
| 0 | 0 | 0 | 0 |
| 0 | 0 | 0 | 0 |
| 0 | 0 | 0 | 0 |
| 0 | 0 | 0 | 0 |
| 0 | 0 | 0 | 0 |
| 1 | 0 | 0 | 0 |
| 0 | 0 | 0 | 0 |
| 0 | 0 | 0 | 0 |
| 0 | 0 | 0 | 0 |
| 0 | 0 | 0 | 0 |
| 0 | 0 | 0 | 0 |
| 0 | 0 | 0 | 0 |

| | | | |
|---|---|---|---|
| 1 | 0 | 0 | 0 |
| 0 | 0 | 0 | 0 |
| 1 | 0 | 0 | 0 |
| 0 | 0 | 0 | 0 |
| 0 | 0 | 0 | 0 |
| 0 | 0 | 0 | 0 |
| 0 | 0 | 0 | 0 |
| 0 | 0 | 0 | 0 |
| 1 | 1 | 0 | 0 |
| 0 | 0 | 0 | 0 |
| 1 | 1 | 0 | 0 |
| 1 | 1 | 0 | 0 |
| 0 | 0 | 0 | 0 |
| 1 | 1 | 0 | 0 |
| 0 | 0 | 0 | 0 |
| 0 | 0 | 0 | 0 |
| 0 | 0 | 0 | 0 |
| 0 | 0 | 0 | 0 |
| 0 | 0 | 0 | 0 |
| 0 | 0 | 0 | 0 |
| 1 | 0 | 0 | 0 |
| 0 | 0 | 0 | 0 |
| 0 | 0 | 0 | 0 |
| 0 | 0 | 0 | 0 |
| 1 | 0 | 0 | 0 |
| 0 | 0 | 0 | 0 |
| 1 | 0 | 0 | 0 |
| 1 | 0 | 0 | 0 |
| 1 | 1 | 0 | 0 |
| 0 | 0 | 0 | 0 |
| 1 | 1 | 0 | 0 |
| 1 | 1 | 0 | 0 |
| 1 | 1 | 0 | 0 |
| 0 | 0 | 0 | 0 |
| 1 | 1 | 0 | 0 |
| 0 | 0 | 0 | 0 |
| 1 | 0 | 0 | 0 |
| 0 | 0 | 0 | 0 |
| 1 | 1 | 0 | 0 |
| 1 | 1 | 0 | 0 |
| 1 | 0 | 0 | 0 |
| 0 | 0 | 0 | 0 |
| 0 | 0 | 0 | 0 |
| 1 | 0 | 0 | 0 |
| 0 | 0 | 0 | 0 |
| 0 | 0 | 0 | 0 |
| 1 | 0 | 0 | 0 |
| 0 | 0 | 0 | 0 |
| 1 | 0 | 0 | 0 |
| 0 | 0 | 0 | 0 |
| 0 | 0 | 0 | 0 |
| 0 | 0 | 0 | 0 |
| 0 | 0 | 0 | 0 |
| 0 | 0 | 0 | 0 |

| | | | |
|---|---|---|---|
| 0 | 0 | 0 | 0 |
| 0 | 0 | 0 | 0 |
| 0 | 0 | 0 | 0 |
| 0 | 0 | 0 | 0 |
| 0 | 0 | 0 | 0 |
| 0 | 0 | 0 | 0 |
| 0 | 0 | 0 | 0 |
| 0 | 0 | 0 | 0 |
| 0 | 0 | 0 | 0 |
| 1 | 1 | 0 | 0 |
| 1 | 1 | 0 | 0 |
| 0 | 0 | 0 | 0 |
| 0 | 0 | 0 | 0 |
| 0 | 0 | 0 | 0 |
| 1 | 0 | 1 | 0 |
| 0 | 0 | 0 | 0 |
| 0 | 0 | 0 | 0 |
| 1 | 0 | 1 | 0 |
| 0 | 0 | 0 | 0 |
| 0 | 0 | 0 | 0 |
| 0 | 0 | 0 | 0 |
| 0 | 0 | 1 | 0 |
| 1 | 0 | 1 | 0 |
| 0 | 0 | 1 | 0 |
| 0 | 0 | 1 | 0 |
| 0 | 0 | 1 | 1 |
| 0 | 0 | 1 | 1 |
| 0 | 0 | 0 | 0 |
| 0 | 0 | 0 | 0 |
| 0 | 0 | 0 | 0 |
| 0 | 0 | 0 | 0 |
| 0 | 0 | 0 | 0 |
| 0 | 0 | 0 | 0 |
| 0 | 0 | 0 | 0 |
| 0 | 0 | 0 | 0 |
| 0 | 0 | 0 | 0 |
| 0 | 0 | 0 | 0 |
| 0 | 0 | 0 | 0 |
| 0 | 0 | 0 | 0 |
| 0 | 0 | 0 | 0 |
| 0 | 0 | 0 | 0 |
| 0 | 0 | 0 | 0 |
| 0 | 0 | 0 | 0 |
| 0 | 0 | 1 | 0 |
| 0 | 0 | 0 | 0 |
| 0 | 0 | 0 | 0 |
| 0 | 0 | 0 | 0 |
| 0 | 0 | 0 | 0 |
| 0 | 0 | 0 | 0 |
| 0 | 0 | 0 | 0 |
| 1 | 0 | 0 | 0 |
| 0 | 0 | 0 | 0 |
| 0 | 0 | 0 | 0 |
| 0 | 0 | 0 | 0 |

| | | | |
|---|---|---|---|
| 0 | 0 | 0 | 0 |
| 0 | 0 | 0 | 0 |
| 0 | 0 | 0 | 0 |
| 1 | 0 | 0 | 0 |
| 0 | 0 | 0 | 0 |
| 0 | 0 | 0 | 0 |
| 0 | 0 | 0 | 0 |
| 0 | 0 | 0 | 0 |
| 1 | 0 | 0 | 0 |
| 1 | 0 | 0 | 0 |
| 0 | 0 | 0 | 0 |
| 0 | 0 | 0 | 0 |
| 0 | 0 | 0 | 0 |
| 0 | 0 | 0 | 0 |
| 0 | 0 | 0 | 0 |
| 0 | 0 | 0 | 0 |
| 0 | 0 | 1 | 0 |
| 0 | 0 | 0 | 0 |
| 0 | 0 | 0 | 0 |
| 0 | 0 | 0 | 0 |
| 0 | 0 | 0 | 0 |
| 0 | 0 | 0 | 0 |
| 0 | 0 | 0 | 0 |
| 0 | 0 | 0 | 0 |
| 0 | 0 | 0 | 0 |
| 0 | 0 | 0 | 0 |
| 0 | 0 | 0 | 0 |
| 0 | 0 | 0 | 0 |
| 0 | 0 | 0 | 0 |
| 0 | 0 | 0 | 0 |
| 0 | 0 | 0 | 0 |
| 0 | 0 | 0 | 0 |
| 0 | 0 | 0 | 0 |
| 0 | 0 | 0 | 0 |
| 0 | 0 | 0 | 0 |
| 0 | 0 | 0 | 0 |
| 0 | 0 | 0 | 0 |
| 0 | 0 | 0 | 0 |
| 0 | 0 | 0 | 0 |
| 0 | 0 | 0 | 0 |
| 0 | 0 | 0 | 0 |
| 0 | 0 | 0 | 0 |
| 0 | 0 | 0 | 0 |
| 0 | 0 | 0 | 0 |
| 0 | 0 | 0 | 0 |
| 0 | 0 | 0 | 0 |
| 0 | 0 | 0 | 0 |
| 0 | 0 | 0 | 0 |
| 0 | 0 | 0 | 0 |
| 0 | 1 | 0 | 0 |
| 0 | 0 | 0 | 0 |
| 0 | 0 | 0 | 0 |
| 0 | 0 | 0 | 0 |
| 0 | 0 | 0 | 0 |

| | | | |
|---|---|---|---|
| 0 | 0 | 1 | 0 |
| 0 | 0 | 0 | 0 |
| 0 | 0 | 0 | 0 |
| 0 | 0 | 0 | 0 |
| 0 | 0 | 0 | 0 |
| 0 | 0 | 0 | 0 |
| 0 | 0 | 0 | 0 |
| 1 | 0 | 0 | 0 |
| 0 | 0 | 0 | 0 |
| 0 | 0 | 0 | 0 |
| 0 | 0 | 0 | 0 |
| 1 | 0 | 0 | 0 |
| 0 | 0 | 0 | 0 |
| 0 | 0 | 0 | 0 |
| 0 | 0 | 0 | 0 |
| 0 | 0 | 0 | 0 |
| 0 | 0 | 0 | 0 |
| 0 | 0 | 0 | 0 |
| 0 | 0 | 0 | 0 |
| 0 | 0 | 0 | 0 |
| 0 | 0 | 0 | 0 |
| 0 | 0 | 0 | 0 |
| 0 | 0 | 0 | 0 |
| 0 | 0 | 0 | 0 |
| 0 | 0 | 0 | 0 |
| 0 | 0 | 0 | 0 |
| 0 | 0 | 0 | 0 |
| 0 | 0 | 0 | 0 |
| 0 | 0 | 0 | 0 |
| 0 | 0 | 0 | 0 |
| 0 | 0 | 0 | 0 |
| 0 | 0 | 0 | 0 |
| 0 | 0 | 0 | 0 |
| 0 | 0 | 0 | 0 |
| 0 | 0 | 0 | 0 |
| 0 | 0 | 0 | 0 |
| 0 | 0 | 0 | 0 |
| 0 | 0 | 0 | 0 |
| 0 | 0 | 0 | 0 |
| 0 | 0 | 0 | 0 |
| 0 | 0 | 0 | 0 |
| 0 | 0 | 0 | 0 |
| 0 | 0 | 0 | 0 |
| 0 | 0 | 0 | 0 |
| 0 | 0 | 0 | 0 |
| 0 | 0 | 0 | 0 |
| 0 | 0 | 0 | 0 |
| 0 | 0 | 0 | 0 |
| 0 | 0 | 0 | 0 |
| 0 | 0 | 0 | 0 |
| 0 | 0 | 0 | 0 |
| 0 | 0 | 0 | 0 |
| 0 | 0 | 0 | 0 |
| 0 | 0 | 0 | 0 |

| | | | |
|---|---|---|---|
| 0 | 0 | 0 | 0 |
| 0 | 0 | 0 | 0 |
| 0 | 0 | 0 | 0 |
| 0 | 0 | 0 | 0 |
| 0 | 0 | 0 | 0 |
| 0 | 0 | 0 | 0 |
| 0 | 0 | 0 | 0 |
| 0 | 0 | 0 | 0 |
| 0 | 0 | 0 | 0 |
| 0 | 0 | 0 | 0 |
| 0 | 0 | 0 | 0 |
| 0 | 0 | 0 | 0 |
| 0 | 0 | 0 | 0 |
| 0 | 0 | 0 | 0 |
| 0 | 0 | 0 | 0 |
| 0 | 0 | 0 | 0 |
| 0 | 0 | 0 | 0 |
| 0 | 0 | 0 | 0 |
| 0 | 0 | 0 | 1 |
| 0 | 1 | 0 | 1 |
| 0 | 0 | 0 | 0 |
| 0 | 0 | 0 | 0 |
| 0 | 0 | 0 | 0 |
| 0 | 0 | 0 | 0 |
| 0 | 0 | 0 | 0 |
| 1 | 0 | 0 | 0 |
| 1 | 0 | 0 | 0 |
| 0 | 0 | 0 | 0 |
| 0 | 0 | 0 | 0 |
| 0 | 0 | 0 | 0 |
| 0 | 0 | 0 | 0 |
| 0 | 0 | 0 | 0 |
| 0 | 0 | 0 | 0 |
| 1 | 0 | 0 | 1 |
| 1 | 0 | 0 | 0 |
| 1 | 0 | 0 | 1 |
| 1 | 0 | 0 | 0 |
| 0 | 0 | 0 | 0 |
| 0 | 0 | 0 | 0 |
| 0 | 0 | 0 | 0 |
| 0 | 0 | 0 | 0 |
| 1 | 0 | 0 | 0 |
| 1 | 0 | 0 | 1 |
| 1 | 0 | 0 | 0 |
| 0 | 0 | 0 | 0 |
| 0 | 0 | 0 | 0 |
| 0 | 0 | 0 | 0 |
| 0 | 0 | 0 | 0 |
| 0 | 0 | 0 | 0 |
| 1 | 0 | 0 | 0 |
| 0 | 0 | 0 | 0 |
| 0 | 0 | 0 | 0 |
| 0 | 0 | 0 | 0 |
| 0 | 0 | 0 | 0 |

| | | | |
|---|---|---|---|
| 0 | 0 | 0 | 0 |
| 0 | 0 | 0 | 0 |
| 0 | 0 | 0 | 0 |
| 1 | 0 | 0 | 0 |
| 1 | 0 | 0 | 0 |
| 0 | 0 | 0 | 0 |
| 1 | 0 | 0 | 0 |
| 0 | 0 | 0 | 0 |
| 0 | 0 | 0 | 0 |
| 0 | 0 | 0 | 0 |
| 0 | 0 | 0 | 0 |
| 1 | 0 | 1 | 0 |
| 0 | 0 | 0 | 0 |
| 0 | 0 | 0 | 0 |
| 0 | 0 | 0 | 0 |
| 0 | 0 | 1 | 0 |
| 0 | 0 | 0 | 0 |
| 0 | 0 | 0 | 0 |
| 0 | 0 | 0 | 0 |
| 0 | 0 | 1 | 0 |
| 1 | 0 | 1 | 0 |
| 1 | 0 | 0 | 0 |
| 1 | 0 | 0 | 0 |
| 1 | 0 | 0 | 0 |
| 0 | 0 | 1 | 0 |
| 0 | 0 | 0 | 0 |
| 1 | 0 | 0 | 0 |
| 0 | 0 | 0 | 0 |
| 0 | 0 | 0 | 0 |
| 0 | 0 | 0 | 0 |
| 0 | 0 | 0 | 0 |
| 0 | 0 | 0 | 0 |
| 0 | 0 | 0 | 0 |
| 0 | 0 | 0 | 0 |
| 0 | 0 | 0 | 0 |
| 0 | 0 | 0 | 0 |
| 0 | 0 | 0 | 1 |
| 0 | 0 | 0 | 1 |
| 0 | 0 | 0 | 1 |
| 0 | 0 | 0 | 0 |
| 0 | 0 | 0 | 0 |
| 0 | 0 | 0 | 0 |
| 0 | 0 | 0 | 0 |
| 0 | 0 | 0 | 0 |
| 0 | 0 | 0 | 0 |
| 0 | 0 | 0 | 0 |
| 0 | 0 | 0 | 0 |
| 0 | 0 | 0 | 0 |
| 0 | 0 | 0 | 0 |
| 0 | 0 | 0 | 0 |
| 0 | 0 | 0 | 0 |
| 0 | 0 | 0 | 0 |
| 0 | 0 | 0 | 0 |
| 0 | 0 | 0 | 0 |

| | | | |
|---|---|---|---|
| 0 | 0 | 0 | 0 |
| 0 | 0 | 0 | 0 |
| 0 | 0 | 0 | 0 |
| 0 | 0 | 0 | 0 |
| 0 | 0 | 0 | 0 |
| 0 | 0 | 0 | 0 |
| 0 | 0 | 0 | 0 |
| 0 | 0 | 0 | 0 |
| 0 | 0 | 0 | 0 |
| 0 | 0 | 0 | 0 |
| 0 | 0 | 0 | 0 |
| 0 | 0 | 0 | 0 |
| 0 | 0 | 0 | 0 |
| 0 | 0 | 0 | 0 |
| 0 | 0 | 0 | 0 |
| 0 | 0 | 0 | 0 |
| 0 | 1 | 0 | 0 |
| 0 | 1 | 0 | 0 |
| 0 | 0 | 0 | 0 |
| 0 | 0 | 0 | 0 |
| 0 | 0 | 0 | 0 |
| 0 | 0 | 0 | 0 |
| 0 | 0 | 0 | 0 |
| 0 | 0 | 0 | 0 |
| 0 | 0 | 0 | 0 |
| 0 | 0 | 0 | 0 |
| 0 | 0 | 0 | 0 |
| 0 | 0 | 0 | 0 |
| 0 | 0 | 0 | 0 |
| 0 | 0 | 0 | 0 |
| 0 | 0 | 0 | 0 |
| 1 | 0 | 0 | 0 |
| 0 | 0 | 0 | 0 |
| 0 | 0 | 0 | 0 |
| 0 | 0 | 0 | 0 |
| 0 | 0 | 0 | 0 |
| 0 | 0 | 0 | 0 |
| 0 | 0 | 0 | 0 |
| 0 | 0 | 0 | 0 |
| 0 | 0 | 0 | 0 |
| 0 | 0 | 0 | 0 |
| 0 | 0 | 0 | 0 |
| 0 | 0 | 0 | 0 |
| 0 | 0 | 0 | 0 |
| 0 | 0 | 0 | 0 |
| 0 | 0 | 0 | 0 |
| 1 | 1 | 0 | 0 |
| 0 | 0 | 0 | 0 |
| 0 | 0 | 0 | 0 |
| 0 | 0 | 0 | 0 |
| 0 | 0 | 0 | 0 |
| 0 | 0 | 0 | 0 |
| 1 | 0 | 0 | 0 |
| 0 | 0 | 0 | 0 |

| | | | |
|---|---|---|---|
| 0 | 0 | 0 | 0 |
| 0 | 0 | 0 | 0 |
| 0 | 0 | 0 | 0 |
| 0 | 0 | 0 | 0 |
| 0 | 0 | 0 | 0 |
| 0 | 0 | 0 | 0 |
| 1 | 0 | 0 | 0 |
| 0 | 0 | 0 | 0 |
| 0 | 0 | 0 | 0 |
| 0 | 0 | 0 | 0 |
| 0 | 0 | 0 | 0 |
| 0 | 0 | 0 | 0 |
| 0 | 0 | 0 | 0 |
| 0 | 0 | 0 | 0 |
| 0 | 0 | 0 | 0 |
| 0 | 0 | 0 | 0 |
| 0 | 0 | 0 | 0 |
| 0 | 0 | 0 | 0 |
| 0 | 0 | 0 | 0 |
| 0 | 0 | 0 | 0 |
| 0 | 0 | 0 | 0 |
| 0 | 0 | 0 | 0 |
| 0 | 0 | 0 | 0 |
| 0 | 0 | 0 | 0 |
| 0 | 0 | 0 | 0 |
| 0 | 0 | 0 | 0 |
| 0 | 0 | 0 | 0 |
| 0 | 0 | 0 | 0 |
| 0 | 0 | 0 | 0 |
| 0 | 0 | 0 | 0 |
| 0 | 0 | 0 | 0 |
| 0 | 0 | 0 | 0 |
| 0 | 0 | 0 | 0 |
| 0 | 0 | 0 | 0 |
| 0 | 0 | 0 | 0 |
| 0 | 0 | 0 | 0 |
| 0 | 0 | 0 | 0 |
| 0 | 0 | 0 | 0 |
| 0 | 0 | 0 | 0 |
| 0 | 0 | 0 | 0 |
| 0 | 0 | 0 | 0 |
| 0 | 0 | 0 | 0 |
| 0 | 0 | 0 | 0 |
| 0 | 0 | 0 | 0 |
| 0 | 0 | 0 | 0 |
| 0 | 0 | 0 | 0 |
| 0 | 0 | 0 | 0 |
| 0 | 0 | 0 | 0 |
| 0 | 0 | 0 | 0 |
| 0 | 0 | 0 | 0 |
| 0 | 0 | 0 | 0 |
| 0 | 0 | 0 | 0 |
| 0 | 0 | 0 | 0 |
| 0 | 0 | 0 | 0 |

| | | | |
|---|---|---|---|
| 0 | 0 | 0 | 0 |
| 0 | 0 | 0 | 0 |
| 0 | 0 | 0 | 0 |
| 0 | 0 | 0 | 0 |
| 0 | 0 | 0 | 0 |
| 0 | 0 | 0 | 0 |
| 0 | 0 | 0 | 0 |
| 0 | 0 | 0 | 0 |
| 0 | 0 | 0 | 0 |
| 0 | 0 | 0 | 0 |
| 0 | 0 | 0 | 0 |
| 0 | 0 | 0 | 0 |
| 0 | 0 | 0 | 0 |
| 0 | 0 | 0 | 0 |
| 0 | 0 | 0 | 0 |
| 0 | 0 | 0 | 0 |
| 0 | 0 | 0 | 0 |
| 0 | 0 | 0 | 0 |
| 0 | 0 | 0 | 0 |
| 0 | 0 | 0 | 0 |
| 0 | 0 | 0 | 0 |
| 0 | 0 | 0 | 0 |
| 0 | 0 | 0 | 0 |
| 0 | 0 | 0 | 0 |
| 1 | 1 | 0 | 0 |
| 0 | 0 | 0 | 0 |
| 0 | 0 | 0 | 0 |
| 0 | 0 | 0 | 0 |
| 0 | 0 | 0 | 0 |
| 0 | 0 | 0 | 0 |
| 0 | 0 | 0 | 0 |
| 0 | 0 | 0 | 0 |
| 0 | 0 | 0 | 0 |
| 0 | 0 | 0 | 0 |
| 0 | 0 | 0 | 0 |
| 1 | 0 | 0 | 0 |
| 1 | 0 | 0 | 1 |
| 1 | 0 | 0 | 0 |
| 0 | 0 | 0 | 0 |
| 1 | 0 | 0 | 0 |
| 1 | 1 | 0 | 0 |
| 1 | 1 | 0 | 0 |
| 0 | 0 | 0 | 0 |
| 0 | 0 | 0 | 0 |
| 0 | 0 | 0 | 0 |
| 0 | 0 | 0 | 0 |
| 0 | 0 | 0 | 0 |
| 0 | 0 | 0 | 0 |
| 0 | 0 | 0 | 0 |
| 0 | 0 | 0 | 0 |
| 0 | 0 | 0 | 0 |
| 0 | 0 | 0 | 0 |
| 0 | 0 | 0 | 0 |
| 0 | 0 | 0 | 0 |

| | | | |
|---|---|---|---|
| 0 | 0 | 0 | 0 |
| 0 | 0 | 0 | 0 |
| 0 | 0 | 0 | 0 |
| 0 | 0 | 0 | 0 |
| 0 | 0 | 0 | 0 |
| 0 | 0 | 0 | 0 |
| 0 | 0 | 0 | 0 |
| 0 | 0 | 0 | 0 |
| 0 | 0 | 0 | 0 |
| 0 | 0 | 0 | 0 |
| 0 | 0 | 0 | 0 |
| 0 | 0 | 0 | 0 |
| 0 | 0 | 0 | 0 |
| 1 | 0 | 0 | 0 |
| 0 | 0 | 0 | 0 |
| 0 | 0 | 0 | 0 |
| 1 | 1 | 0 | 0 |
| 0 | 0 | 0 | 0 |
| 0 | 0 | 0 | 0 |
| 0 | 0 | 0 | 0 |
| 0 | 0 | 0 | 0 |
| 0 | 0 | 0 | 0 |
| 0 | 0 | 0 | 0 |
| 1 | 0 | 0 | 0 |
| 0 | 0 | 0 | 0 |
| 0 | 0 | 0 | 0 |
| 0 | 0 | 0 | 0 |
| 0 | 0 | 0 | 0 |
| 0 | 0 | 0 | 0 |
| 0 | 0 | 0 | 0 |
| 0 | 0 | 0 | 0 |
| 1 | 0 | 0 | 0 |
| 0 | 0 | 0 | 0 |
| 0 | 0 | 0 | 0 |
| 0 | 0 | 0 | 0 |
| 0 | 0 | 0 | 0 |
| 0 | 0 | 0 | 0 |
| 0 | 0 | 0 | 0 |
| 0 | 0 | 0 | 0 |
| 0 | 0 | 0 | 0 |
| 0 | 0 | 0 | 0 |
| 0 | 0 | 0 | 0 |
| 0 | 0 | 0 | 0 |
| 0 | 0 | 0 | 0 |
| 0 | 0 | 0 | 0 |
| 0 | 0 | 0 | 0 |
| 0 | 0 | 0 | 0 |
| 0 | 0 | 0 | 0 |
| 0 | 0 | 0 | 0 |
| 0 | 0 | 0 | 0 |
| 0 | 0 | 0 | 0 |
| 0 | 0 | 0 | 0 |
| 0 | 0 | 0 | 0 |
| 0 | 0 | 0 | 0 |

| | | | |
|---|---|---|---|
| 0 | 0 | 0 | 0 |
| 0 | 0 | 0 | 0 |
| 0 | 0 | 0 | 0 |
| 0 | 0 | 0 | 0 |
| 0 | 0 | 0 | 0 |
| 0 | 0 | 0 | 0 |
| 0 | 0 | 0 | 0 |
| 0 | 0 | 0 | 0 |
| 0 | 0 | 0 | 0 |
| 0 | 0 | 0 | 0 |
| 0 | 0 | 0 | 0 |
| 0 | 0 | 0 | 0 |
| 0 | 0 | 0 | 0 |
| 0 | 0 | 0 | 0 |
| 0 | 0 | 0 | 0 |
| 0 | 0 | 0 | 0 |
| 0 | 0 | 0 | 0 |
| 1 | 0 | 0 | 0 |
| 0 | 0 | 0 | 0 |
| 0 | 0 | 0 | 0 |
| 0 | 0 | 0 | 0 |
| 0 | 0 | 0 | 0 |
| 0 | 0 | 0 | 0 |
| 0 | 0 | 0 | 0 |
| 1 | 0 | 0 | 0 |
| 0 | 0 | 0 | 0 |
| 0 | 0 | 0 | 0 |
| 0 | 0 | 0 | 0 |
| 0 | 0 | 0 | 0 |
| 0 | 0 | 0 | 0 |
| 0 | 0 | 0 | 0 |
| 0 | 0 | 0 | 0 |
| 0 | 0 | 0 | 0 |
| 0 | 0 | 0 | 0 |
| 0 | 0 | 0 | 0 |
| 0 | 0 | 0 | 0 |
| 0 | 0 | 0 | 0 |
| 0 | 0 | 0 | 0 |
| 0 | 0 | 0 | 0 |
| 0 | 0 | 0 | 0 |
| 0 | 0 | 0 | 0 |
| 0 | 0 | 0 | 0 |
| 0 | 0 | 0 | 0 |
| 0 | 0 | 0 | 0 |
| 0 | 0 | 0 | 0 |
| 0 | 0 | 0 | 0 |
| 0 | 0 | 0 | 0 |
| 0 | 0 | 0 | 0 |
| 0 | 0 | 0 | 0 |
| 0 | 0 | 0 | 0 |
| 0 | 0 | 0 | 0 |
| 0 | 0 | 0 | 0 |
| 0 | 0 | 0 | 0 |
| 0 | 0 | 0 | 0 |

| | | | |
|---|---|---|---|
| 0 | 0 | 0 | 0 |
| 0 | 0 | 0 | 0 |
| 0 | 0 | 0 | 0 |
| 0 | 0 | 0 | 0 |
| 0 | 0 | 0 | 0 |
| 0 | 0 | 0 | 0 |
| 0 | 0 | 0 | 0 |
| 0 | 0 | 0 | 0 |
| 0 | 0 | 0 | 0 |
| 0 | 0 | 0 | 0 |
| 0 | 0 | 0 | 0 |
| 0 | 0 | 0 | 0 |
| 0 | 0 | 0 | 0 |
| 0 | 0 | 0 | 0 |
| 1 | 1 | 0 | 0 |
| 1 | 1 | 0 | 0 |
| 0 | 0 | 0 | 0 |
| 0 | 0 | 0 | 0 |
| 0 | 0 | 0 | 0 |
| 0 | 0 | 0 | 0 |
| 0 | 0 | 0 | 0 |
| 0 | 0 | 0 | 0 |
| 0 | 0 | 0 | 0 |
| 0 | 0 | 0 | 0 |
| 0 | 0 | 0 | 0 |
| 0 | 0 | 0 | 0 |
| 0 | 0 | 0 | 0 |
| 1 | 0 | 0 | 0 |
| 0 | 0 | 0 | 0 |
| 1 | 0 | 0 | 0 |
| 0 | 0 | 0 | 0 |
| 0 | 0 | 0 | 0 |
| 0 | 0 | 0 | 0 |
| 0 | 0 | 0 | 0 |
| 0 | 0 | 0 | 0 |
| 0 | 0 | 0 | 0 |
| 1 | 0 | 0 | 0 |
| 0 | 0 | 0 | 0 |
| 0 | 0 | 0 | 0 |
| 0 | 0 | 0 | 0 |
| 0 | 0 | 0 | 0 |
| 0 | 0 | 0 | 0 |
| 0 | 0 | 0 | 0 |
| 0 | 0 | 0 | 0 |
| 0 | 0 | 0 | 0 |
| 0 | 0 | 0 | 0 |
| 0 | 0 | 0 | 0 |
| 0 | 0 | 0 | 0 |
| 0 | 0 | 0 | 0 |
| 0 | 0 | 0 | 0 |
| 0 | 0 | 0 | 0 |
| 0 | 0 | 0 | 0 |
| 0 | 0 | 0 | 0 |
| 0 | 0 | 0 | 0 |

| | | | |
|---|---|---|---|
| 0 | 0 | 0 | 0 |
| 0 | 0 | 0 | 0 |
| 0 | 0 | 0 | 0 |
| 0 | 0 | 0 | 0 |
| 0 | 0 | 0 | 0 |
| 0 | 0 | 0 | 0 |
| 0 | 0 | 0 | 0 |
| 0 | 0 | 0 | 0 |
| 0 | 0 | 0 | 0 |
| 0 | 0 | 0 | 0 |
| 0 | 0 | 0 | 0 |
| 0 | 0 | 0 | 0 |
| 0 | 0 | 0 | 0 |
| 0 | 0 | 0 | 0 |
| 0 | 0 | 0 | 0 |
| 0 | 0 | 0 | 0 |
| 0 | 0 | 0 | 0 |
| 0 | 0 | 0 | 0 |
| 0 | 0 | 0 | 0 |
| 0 | 0 | 0 | 0 |
| 0 | 0 | 0 | 0 |
| 0 | 0 | 0 | 0 |
| 0 | 0 | 0 | 0 |
| 0 | 0 | 0 | 0 |
| 0 | 0 | 0 | 0 |
| 0 | 0 | 0 | 0 |
| 0 | 0 | 0 | 0 |
| 0 | 0 | 0 | 0 |
| 0 | 0 | 0 | 0 |
| 0 | 0 | 0 | 0 |
| 0 | 0 | 0 | 0 |
| 0 | 0 | 0 | 0 |
| 0 | 0 | 0 | 0 |
| 0 | 0 | 0 | 0 |
| 0 | 0 | 0 | 0 |
| 0 | 0 | 0 | 0 |
| 0 | 0 | 0 | 0 |
| 0 | 0 | 0 | 0 |
| 0 | 0 | 0 | 0 |
| 0 | 0 | 0 | 0 |
| 0 | 0 | 0 | 0 |
| 0 | 0 | 0 | 0 |
| 0 | 0 | 0 | 0 |
| 0 | 0 | 0 | 0 |
| 0 | 0 | 0 | 0 |
| 0 | 0 | 0 | 0 |
| 0 | 0 | 0 | 0 |
| 0 | 0 | 0 | 0 |
| 1 | 0 | 0 | 1 |
| 0 | 0 | 0 | 0 |
| 1 | 0 | 0 | 0 |
| 1 | 0 | 0 | 0 |
| 0 | 0 | 0 | 0 |
| 0 | 0 | 0 | 1 |

| | | | |
|---|---|---|---|
| 0 | 0 | 0 | 0 |
| 0 | 0 | 0 | 0 |
| 0 | 0 | 0 | 0 |
| 0 | 0 | 0 | 0 |
| 0 | 0 | 0 | 0 |
| 0 | 0 | 0 | 0 |
| 0 | 0 | 0 | 0 |
| 0 | 0 | 0 | 0 |
| 0 | 0 | 0 | 0 |
| 0 | 0 | 0 | 0 |
| 0 | 0 | 0 | 0 |
| 0 | 0 | 0 | 0 |
| 0 | 0 | 0 | 0 |
| 0 | 0 | 0 | 0 |
| 0 | 0 | 0 | 0 |
| 0 | 0 | 0 | 0 |
| 0 | 0 | 0 | 0 |
| 0 | 0 | 0 | 0 |
| 0 | 0 | 0 | 0 |
| 0 | 0 | 0 | 0 |
| 0 | 0 | 0 | 0 |
| 0 | 0 | 0 | 0 |
| 0 | 0 | 0 | 0 |
| 0 | 0 | 0 | 0 |
| 0 | 0 | 0 | 0 |
| 0 | 0 | 0 | 0 |
| 0 | 0 | 1 | 0 |
| 0 | 0 | 1 | 0 |
| 0 | 0 | 1 | 0 |
| 0 | 0 | 1 | 0 |
| 0 | 0 | 0 | 0 |
| 0 | 0 | 0 | 0 |
| 0 | 0 | 0 | 0 |
| 1 | 0 | 0 | 0 |
| 1 | 0 | 0 | 0 |
| 0 | 0 | 0 | 0 |
| 0 | 0 | 0 | 0 |
| 0 | 0 | 0 | 0 |
| 0 | 0 | 0 | 0 |
| 1 | 0 | 0 | 0 |
| 0 | 0 | 0 | 0 |
| 0 | 0 | 0 | 0 |
| 0 | 0 | 0 | 0 |
| 0 | 0 | 0 | 0 |
| 0 | 0 | 0 | 0 |
| 0 | 1 | 0 | 0 |
| 0 | 0 | 0 | 0 |
| 0 | 0 | 0 | 0 |
| 0 | 0 | 0 | 0 |
| 0 | 0 | 0 | 0 |
| 0 | 0 | 0 | 0 |
| 0 | 0 | 0 | 0 |
| 0 | 0 | 0 | 0 |
| 0 | 0 | 0 | 0 |

| | | | |
|---|---|---|---|
| 0 | 0 | 0 | 0 |
| 0 | 0 | 0 | 0 |
| 0 | 0 | 0 | 0 |
| 0 | 0 | 0 | 0 |
| 0 | 0 | 0 | 0 |
| 0 | 0 | 0 | 1 |
| 0 | 0 | 0 | 0 |
| 0 | 0 | 0 | 0 |
| 0 | 0 | 0 | 0 |
| 0 | 0 | 0 | 0 |
| 0 | 0 | 0 | 0 |
| 0 | 0 | 0 | 0 |
| 0 | 0 | 0 | 0 |
| 0 | 1 | 0 | 0 |
| 0 | 1 | 0 | 0 |
| 0 | 1 | 0 | 0 |
| 0 | 1 | 0 | 0 |
| 0 | 1 | 0 | 0 |
| 0 | 0 | 0 | 0 |
| 0 | 0 | 0 | 0 |
| 0 | 0 | 0 | 0 |
| 0 | 0 | 0 | 0 |
| 0 | 0 | 0 | 0 |
| 0 | 0 | 0 | 0 |
| 0 | 0 | 0 | 0 |
| 0 | 0 | 0 | 0 |
| 0 | 0 | 0 | 0 |
| 0 | 0 | 0 | 0 |
| 0 | 0 | 0 | 0 |
| 0 | 0 | 0 | 0 |
| 0 | 0 | 0 | 0 |
| 0 | 0 | 0 | 0 |
| 0 | 0 | 0 | 0 |
| 0 | 0 | 0 | 0 |
| 0 | 0 | 0 | 0 |
| 0 | 0 | 0 | 0 |
| 0 | 0 | 0 | 0 |
| 0 | 0 | 0 | 0 |
| 0 | 0 | 0 | 0 |
| 0 | 0 | 0 | 0 |
| 0 | 0 | 0 | 0 |
| 0 | 0 | 0 | 0 |
| 0 | 0 | 0 | 0 |
| 0 | 0 | 0 | 0 |
| 0 | 0 | 0 | 0 |
| 0 | 0 | 0 | 0 |
| 0 | 0 | 0 | 0 |
| 0 | 0 | 0 | 0 |
| 0 | 0 | 0 | 0 |
| 0 | 0 | 0 | 0 |
| 0 | 0 | 0 | 0 |
| 0 | 0 | 0 | 0 |
| 0 | 0 | 0 | 0 |
| 0 | 0 | 0 | 0 |

| | | | |
|---|---|---|---|
| 0 | 0 | 0 | 0 |
| 0 | 0 | 0 | 0 |
| 0 | 0 | 0 | 0 |
| 1 | 0 | 0 | 0 |
| 0 | 0 | 0 | 0 |
| 0 | 0 | 0 | 0 |
| 0 | 0 | 0 | 0 |
| 0 | 0 | 0 | 0 |
| 0 | 0 | 0 | 0 |
| 0 | 0 | 0 | 0 |
| 0 | 0 | 0 | 0 |
| 0 | 0 | 0 | 0 |
| 0 | 0 | 0 | 0 |
| 0 | 0 | 0 | 0 |
| 0 | 0 | 0 | 0 |
| 0 | 0 | 0 | 0 |
| 0 | 0 | 0 | 0 |
| 0 | 0 | 0 | 0 |
| 0 | 0 | 0 | 0 |
| 0 | 0 | 0 | 0 |
| 1 | 1 | 0 | 0 |
| 1 | 1 | 0 | 0 |
| 0 | 0 | 0 | 0 |
| 0 | 1 | 0 | 0 |
| 0 | 0 | 0 | 0 |
| 0 | 0 | 0 | 0 |
| 0 | 0 | 0 | 0 |
| 0 | 0 | 0 | 0 |
| 0 | 0 | 0 | 0 |
| 0 | 0 | 0 | 0 |
| 0 | 0 | 0 | 0 |
| 0 | 0 | 0 | 0 |
| 0 | 0 | 0 | 0 |
| 0 | 0 | 0 | 0 |
| 0 | 0 | 0 | 0 |
| 0 | 0 | 0 | 0 |
| 0 | 0 | 0 | 0 |
| 0 | 0 | 0 | 0 |
| 0 | 0 | 0 | 0 |
| 1 | 0 | 0 | 0 |
| 1 | 0 | 0 | 0 |
| 0 | 0 | 0 | 0 |
| 0 | 0 | 0 | 0 |
| 1 | 0 | 0 | 0 |
| 0 | 0 | 0 | 0 |
| 0 | 0 | 0 | 0 |
| 0 | 0 | 0 | 0 |
| 0 | 0 | 0 | 0 |
| 0 | 0 | 0 | 0 |
| 0 | 0 | 0 | 0 |
| 0 | 0 | 0 | 0 |
| 0 | 0 | 0 | 0 |
| 0 | 0 | 0 | 0 |
| 0 | 0 | 0 | 0 |

| | | | |
|---|---|---|---|
| 0 | 0 | 0 | 0 |
| 0 | 0 | 0 | 0 |
| 0 | 0 | 0 | 0 |
| 0 | 0 | 0 | 0 |
| 0 | 0 | 0 | 0 |
| 1 | 0 | 0 | 0 |
| 0 | 0 | 0 | 0 |
| 0 | 0 | 0 | 0 |
| 0 | 1 | 0 | 0 |
| 1 | 1 | 0 | 0 |
| 0 | 0 | 0 | 0 |
| 1 | 1 | 0 | 0 |
| 1 | 0 | 0 | 0 |
| 0 | 0 | 0 | 0 |
| 0 | 0 | 0 | 0 |
| 0 | 0 | 0 | 0 |
| 1 | 1 | 0 | 0 |
| 0 | 0 | 0 | 0 |
| 0 | 0 | 0 | 0 |
| 1 | 0 | 0 | 0 |
| 0 | 0 | 0 | 0 |
| 0 | 0 | 0 | 0 |
| 0 | 0 | 0 | 0 |
| 0 | 0 | 0 | 0 |
| 0 | 0 | 0 | 0 |
| 0 | 0 | 0 | 0 |
| 0 | 0 | 0 | 0 |
| 0 | 0 | 0 | 0 |
| 0 | 0 | 0 | 0 |
| 0 | 0 | 0 | 0 |
| 1 | 0 | 0 | 0 |
| 0 | 0 | 0 | 0 |
| 0 | 0 | 0 | 0 |
| 0 | 0 | 0 | 0 |
| 0 | 0 | 0 | 0 |
| 1 | 0 | 0 | 0 |
| 0 | 0 | 0 | 0 |
| 0 | 0 | 0 | 0 |
| 0 | 0 | 0 | 0 |
| 0 | 0 | 0 | 0 |
| 0 | 0 | 0 | 0 |
| 0 | 0 | 0 | 0 |
| 0 | 0 | 0 | 0 |
| 0 | 0 | 0 | 0 |
| 0 | 0 | 0 | 0 |
| 0 | 0 | 0 | 0 |
| 0 | 0 | 0 | 0 |
| 0 | 0 | 0 | 0 |
| 0 | 0 | 0 | 0 |
| 0 | 0 | 0 | 0 |
| 0 | 0 | 0 | 0 |
| 0 | 0 | 0 | 0 |
| 0 | 0 | 0 | 0 |
| 0 | 0 | 0 | 0 |

| | | | |
|---|---|---|---|
| 0 | 0 | 0 | 0 |
| 0 | 0 | 0 | 0 |
| 0 | 0 | 0 | 0 |
| 0 | 0 | 0 | 0 |
| 0 | 0 | 0 | 0 |
| 0 | 0 | 0 | 0 |
| 0 | 0 | 0 | 0 |
| 0 | 0 | 0 | 0 |
| 0 | 0 | 0 | 0 |
| 0 | 0 | 0 | 0 |
| 0 | 0 | 0 | 0 |
| 0 | 0 | 0 | 0 |
| 0 | 0 | 0 | 0 |
| 0 | 0 | 0 | 0 |
| 0 | 0 | 0 | 1 |
| 0 | 0 | 0 | 0 |
| 0 | 0 | 0 | 0 |
| 0 | 0 | 0 | 0 |
| 0 | 0 | 0 | 0 |
| 0 | 0 | 0 | 0 |
| 0 | 0 | 0 | 0 |
| 0 | 0 | 0 | 0 |
| 0 | 0 | 0 | 0 |
| 0 | 0 | 0 | 0 |
| 0 | 0 | 0 | 0 |
| 0 | 0 | 0 | 0 |
| 0 | 0 | 0 | 0 |
| 0 | 0 | 0 | 0 |
| 0 | 0 | 0 | 0 |
| 0 | 0 | 0 | 0 |
| 0 | 0 | 0 | 0 |
| 0 | 0 | 0 | 0 |
| 0 | 0 | 0 | 0 |
| 0 | 0 | 0 | 0 |
| 0 | 0 | 0 | 0 |
| 0 | 0 | 0 | 0 |
| 0 | 0 | 0 | 0 |
| 0 | 0 | 0 | 0 |
| 0 | 0 | 0 | 0 |
| 0 | 0 | 1 | 0 |
| 0 | 0 | 0 | 0 |
| 0 | 0 | 0 | 0 |
| 0 | 0 | 0 | 0 |
| 1 | 0 | 0 | 0 |
| 0 | 0 | 0 | 0 |
| 0 | 0 | 0 | 0 |
| 0 | 0 | 0 | 0 |
| 0 | 0 | 0 | 0 |
| 1 | 0 | 0 | 1 |
| 0 | 0 | 0 | 0 |
| 0 | 0 | 0 | 0 |
| 0 | 0 | 0 | 0 |
| 0 | 0 | 0 | 0 |
| 0 | 0 | 0 | 0 |

| | | | |
|---|---|---|---|
| 0 | 0 | 0 | 0 |
| 0 | 0 | 0 | 0 |
| 0 | 0 | 0 | 0 |
| 0 | 0 | 0 | 0 |
| 0 | 0 | 0 | 0 |
| 0 | 0 | 0 | 0 |
| 0 | 0 | 0 | 0 |
| 0 | 0 | 0 | 0 |
| 0 | 0 | 0 | 0 |
| 1 | 0 | 0 | 0 |
| 0 | 0 | 0 | 0 |
| 0 | 0 | 0 | 0 |
| 0 | 0 | 0 | 0 |
| 0 | 0 | 0 | 0 |
| 1 | 0 | 0 | 0 |
| 0 | 0 | 0 | 0 |
| 0 | 0 | 0 | 0 |
| 0 | 0 | 0 | 0 |
| 0 | 0 | 0 | 0 |
| 0 | 0 | 0 | 0 |
| 0 | 0 | 0 | 0 |
| 0 | 0 | 0 | 0 |
| 0 | 0 | 0 | 0 |
| 0 | 0 | 0 | 0 |
| 1 | 0 | 0 | 0 |
| 0 | 0 | 0 | 0 |
| 0 | 0 | 0 | 0 |
| 0 | 0 | 0 | 0 |
| 0 | 0 | 0 | 0 |
| 0 | 0 | 0 | 0 |
| 0 | 0 | 0 | 0 |
| 0 | 0 | 0 | 0 |
| 0 | 0 | 0 | 0 |
| 0 | 0 | 0 | 0 |
| 1 | 0 | 0 | 0 |
| 1 | 0 | 0 | 0 |
| 0 | 0 | 0 | 0 |
| 0 | 0 | 0 | 0 |
| 1 | 0 | 0 | 0 |
| 0 | 0 | 1 | 0 |
| 0 | 0 | 1 | 0 |
| 0 | 0 | 1 | 0 |
| 0 | 0 | 1 | 0 |
| 0 | 0 | 0 | 0 |
| 1 | 0 | 0 | 0 |
| 0 | 0 | 0 | 0 |
| 0 | 0 | 0 | 0 |
| 0 | 0 | 0 | 0 |
| 0 | 0 | 1 | 0 |
| 0 | 0 | 0 | 0 |
| 0 | 0 | 0 | 0 |
| 0 | 0 | 0 | 0 |
| 0 | 0 | 0 | 0 |
| 0 | 0 | 0 | 0 |

| | | | |
|---|---|---|---|
| 0 | 0 | 0 | 0 |
| 0 | 0 | 0 | 0 |
| 0 | 0 | 0 | 0 |
| 0 | 0 | 0 | 0 |
| 0 | 0 | 0 | 0 |
| 0 | 0 | 0 | 0 |
| 0 | 0 | 0 | 0 |
| 0 | 0 | 0 | 0 |
| 0 | 0 | 0 | 0 |
| 0 | 0 | 0 | 0 |
| 0 | 1 | 0 | 0 |
| 0 | 0 | 0 | 0 |
| 0 | 0 | 0 | 0 |
| 0 | 0 | 0 | 0 |
| 0 | 0 | 0 | 0 |
| 0 | 1 | 0 | 0 |
| 0 | 0 | 0 | 0 |
| 0 | 0 | 0 | 0 |
| 0 | 0 | 0 | 0 |
| 0 | 0 | 0 | 0 |
| 0 | 0 | 0 | 0 |
| 0 | 0 | 0 | 0 |
| 0 | 0 | 0 | 0 |
| 0 | 0 | 0 | 0 |
| 0 | 0 | 0 | 0 |
| 0 | 0 | 0 | 0 |
| 0 | 0 | 0 | 0 |
| 0 | 0 | 0 | 0 |
| 0 | 0 | 0 | 0 |
| 0 | 0 | 0 | 0 |
| 0 | 0 | 0 | 0 |
| 0 | 0 | 0 | 0 |
| 0 | 0 | 0 | 0 |
| 0 | 0 | 0 | 0 |
| 0 | 0 | 0 | 0 |
| 0 | 0 | 0 | 0 |
| 0 | 0 | 0 | 0 |
| 0 | 0 | 0 | 0 |
| 0 | 0 | 0 | 0 |
| 0 | 0 | 0 | 0 |
| 0 | 0 | 0 | 0 |
| 0 | 0 | 0 | 0 |
| 0 | 0 | 0 | 0 |
| 0 | 0 | 0 | 0 |
| 1 | 1 | 0 | 0 |
| 0 | 0 | 0 | 0 |
| 0 | 0 | 0 | 0 |
| 0 | 0 | 0 | 0 |
| 0 | 0 | 0 | 0 |
| 0 | 0 | 0 | 0 |
| 0 | 0 | 0 | 0 |
| 0 | 0 | 0 | 0 |
| 0 | 0 | 0 | 0 |
| 0 | 0 | 0 | 0 |

| | | | |
|---|---|---|---|
| 0 | 0 | 0 | 0 |
| 0 | 0 | 0 | 0 |
| 0 | 0 | 1 | 0 |
| 1 | 0 | 0 | 0 |
| 0 | 0 | 0 | 0 |
| 0 | 0 | 0 | 0 |
| 0 | 0 | 0 | 0 |
| 0 | 0 | 0 | 0 |
| 0 | 0 | 0 | 0 |
| 0 | 0 | 0 | 0 |
| 0 | 0 | 0 | 0 |
| 0 | 0 | 0 | 0 |
| 1 | 0 | 0 | 0 |
| 0 | 0 | 0 | 0 |
| 0 | 0 | 0 | 0 |
| 0 | 0 | 0 | 0 |
| 0 | 0 | 0 | 0 |
| 0 | 0 | 0 | 0 |
| 0 | 0 | 0 | 0 |
| 0 | 0 | 0 | 0 |
| 0 | 0 | 0 | 0 |
| 0 | 0 | 0 | 0 |
| 0 | 0 | 0 | 0 |
| 0 | 0 | 0 | 0 |
| 0 | 0 | 0 | 0 |
| 0 | 0 | 0 | 0 |
| 0 | 0 | 0 | 0 |
| 0 | 0 | 0 | 0 |
| 0 | 0 | 0 | 0 |
| 0 | 0 | 0 | 0 |
| 0 | 0 | 0 | 0 |
| 0 | 0 | 0 | 0 |
| 0 | 0 | 0 | 0 |
| 0 | 0 | 0 | 0 |
| 0 | 0 | 0 | 0 |
| 0 | 0 | 0 | 0 |
| 0 | 0 | 0 | 0 |
| 0 | 0 | 0 | 0 |
| 0 | 0 | 0 | 0 |
| 0 | 0 | 0 | 0 |
| 0 | 1 | 0 | 0 |
| 1 | 1 | 0 | 0 |
| 0 | 0 | 0 | 0 |
| 0 | 0 | 0 | 0 |
| 0 | 0 | 0 | 0 |
| 0 | 0 | 0 | 0 |
| 0 | 0 | 0 | 0 |
| 0 | 0 | 0 | 0 |
| 0 | 1 | 0 | 0 |
| 0 | 0 | 0 | 0 |
| 0 | 0 | 0 | 0 |
| 0 | 0 | 0 | 0 |
| 0 | 0 | 0 | 1 |
| 0 | 0 | 0 | 0 |

| | | | |
|---|---|---|---|
| 0 | 0 | 0 | 1 |
| 0 | 0 | 0 | 1 |
| 0 | 0 | 0 | 1 |
| 0 | 0 | 0 | 0 |
| 0 | 0 | 0 | 0 |
| 0 | 0 | 0 | 0 |
| 0 | 0 | 0 | 0 |
| 1 | 0 | 0 | 0 |
| 0 | 0 | 0 | 0 |
| 1 | 1 | 0 | 0 |
| 0 | 1 | 0 | 0 |
| 0 | 0 | 0 | 0 |
| 1 | 1 | 0 | 0 |
| 0 | 0 | 0 | 0 |
| 0 | 0 | 0 | 0 |
| 0 | 0 | 0 | 0 |
| 0 | 0 | 0 | 0 |
| 1 | 0 | 0 | 0 |
| 0 | 0 | 0 | 0 |
| 0 | 0 | 0 | 0 |
| 0 | 0 | 0 | 0 |
| 0 | 0 | 0 | 0 |
| 0 | 0 | 0 | 0 |
| 0 | 0 | 0 | 0 |
| 0 | 0 | 0 | 0 |
| 0 | 0 | 0 | 0 |
| 0 | 0 | 0 | 0 |
| 0 | 0 | 0 | 0 |
| 0 | 0 | 0 | 0 |
| 0 | 0 | 0 | 0 |
| 0 | 0 | 0 | 0 |
| 1 | 0 | 0 | 0 |
| 0 | 0 | 0 | 0 |
| 0 | 0 | 0 | 0 |
| 0 | 0 | 0 | 0 |
| 0 | 0 | 0 | 0 |
| 0 | 0 | 0 | 0 |
| 0 | 0 | 0 | 0 |
| 0 | 0 | 0 | 0 |
| 0 | 0 | 0 | 0 |
| 0 | 0 | 0 | 0 |
| 1 | 1 | 0 | 0 |
| 0 | 0 | 0 | 0 |
| 0 | 0 | 0 | 0 |
| 1 | 0 | 0 | 0 |
| 0 | 1 | 0 | 0 |
| 0 | 0 | 0 | 0 |
| 0 | 0 | 0 | 0 |
| 0 | 1 | 0 | 0 |
| 1 | 1 | 0 | 0 |
| 1 | 1 | 0 | 0 |
| 1 | 1 | 0 | 0 |
| 1 | 1 | 0 | 0 |
| 1 | 1 | 0 | 0 |

| | | | |
|---|---|---|---|
| 0 | 0 | 0 | 0 |
| 0 | 0 | 0 | 0 |
| 0 | 0 | 0 | 0 |
| 0 | 0 | 0 | 0 |
| 0 | 0 | 0 | 0 |
| 1 | 0 | 0 | 0 |
| 1 | 0 | 0 | 0 |
| 0 | 0 | 0 | 0 |
| 0 | 0 | 0 | 0 |
| 1 | 0 | 0 | 0 |
| 0 | 0 | 0 | 0 |
| 0 | 0 | 1 | 0 |
| 0 | 0 | 0 | 0 |
| 0 | 0 | 0 | 0 |
| 0 | 0 | 0 | 0 |
| 0 | 0 | 0 | 0 |
| 0 | 0 | 0 | 0 |
| 0 | 0 | 0 | 0 |
| 0 | 0 | 0 | 0 |
| 0 | 0 | 0 | 0 |
| 1 | 1 | 0 | 0 |
| 1 | 0 | 0 | 0 |
| 0 | 0 | 0 | 0 |
| 0 | 0 | 1 | 0 |
| 0 | 0 | 0 | 0 |
| 0 | 0 | 1 | 0 |
| 0 | 0 | 0 | 0 |
| 0 | 0 | 0 | 0 |
| 0 | 0 | 0 | 0 |
| 0 | 0 | 0 | 0 |
| 0 | 0 | 0 | 0 |
| 0 | 0 | 0 | 0 |
| 0 | 0 | 0 | 0 |
| 0 | 0 | 0 | 0 |
| 0 | 0 | 0 | 0 |
| 0 | 0 | 0 | 0 |
| 1 | 0 | 0 | 0 |
| 0 | 0 | 0 | 0 |
| 1 | 0 | 0 | 0 |
| 1 | 0 | 0 | 0 |
| 1 | 0 | 0 | 0 |
| 0 | 0 | 0 | 0 |
| 0 | 0 | 0 | 0 |
| 0 | 0 | 0 | 0 |
| 0 | 0 | 0 | 0 |
| 0 | 0 | 0 | 0 |
| 0 | 0 | 0 | 0 |
| 1 | 0 | 1 | 0 |
| 0 | 0 | 0 | 0 |
| 0 | 0 | 0 | 0 |
| 0 | 0 | 0 | 0 |
| 0 | 0 | 0 | 0 |
| 1 | 1 | 0 | 0 |
| 0 | 0 | 0 | 0 |

| | | | |
|---|---|---|---|
| 0 | 0 | 0 | 0 |
| 0 | 1 | 0 | 0 |
| 0 | 0 | 0 | 0 |
| 0 | 0 | 0 | 0 |
| 0 | 0 | 0 | 0 |
| 0 | 0 | 0 | 0 |
| 0 | 0 | 0 | 0 |
| 0 | 0 | 0 | 0 |
| 0 | 0 | 0 | 0 |
| 0 | 0 | 0 | 0 |
| 0 | 0 | 0 | 0 |
| 0 | 0 | 0 | 0 |
| 0 | 0 | 0 | 0 |
| 0 | 0 | 0 | 0 |
| 0 | 0 | 0 | 0 |
| 0 | 0 | 0 | 0 |
| 0 | 0 | 0 | 0 |
| 0 | 0 | 0 | 0 |
| 0 | 0 | 0 | 0 |
| 0 | 0 | 0 | 0 |
| 1 | 0 | 0 | 0 |
| 0 | 0 | 0 | 0 |
| 0 | 0 | 0 | 0 |
| 0 | 0 | 0 | 0 |
| 1 | 0 | 0 | 0 |
| 0 | 0 | 0 | 0 |
| 0 | 0 | 0 | 0 |
| 0 | 0 | 0 | 1 |
| 0 | 0 | 0 | 0 |
| 0 | 0 | 0 | 0 |
| 0 | 0 | 0 | 0 |
| 0 | 0 | 0 | 0 |
| 0 | 0 | 0 | 0 |
| 0 | 0 | 0 | 0 |
| 0 | 0 | 0 | 0 |
| 0 | 0 | 0 | 0 |
| 0 | 0 | 0 | 0 |
| 0 | 0 | 0 | 0 |
| 0 | 0 | 0 | 0 |
| 0 | 0 | 0 | 0 |
| 0 | 0 | 0 | 0 |
| 0 | 0 | 0 | 0 |
| 0 | 0 | 0 | 0 |
| 0 | 0 | 0 | 0 |
| 0 | 0 | 0 | 0 |
| 0 | 0 | 0 | 0 |
| 0 | 0 | 0 | 0 |
| 0 | 0 | 0 | 0 |
| 0 | 0 | 0 | 0 |
| 0 | 0 | 0 | 0 |
| 0 | 0 | 0 | 0 |
| 0 | 0 | 0 | 0 |
| 0 | 0 | 0 | 0 |
| 0 | 0 | 0 | 0 |

| | | | |
|---|---|---|---|
| 0 | 0 | 0 | 0 |
| 0 | 0 | 0 | 0 |
| 0 | 0 | 0 | 0 |
| 0 | 0 | 0 | 0 |
| 0 | 0 | 0 | 0 |
| 0 | 0 | 0 | 0 |
| 0 | 0 | 0 | 0 |
| 0 | 0 | 0 | 0 |
| 0 | 0 | 0 | 0 |
| 0 | 0 | 0 | 0 |
| 0 | 0 | 0 | 0 |
| 0 | 0 | 0 | 0 |
| 0 | 0 | 0 | 0 |
| 0 | 0 | 0 | 0 |
| 0 | 0 | 0 | 0 |
| 1 | 0 | 0 | 0 |
| 0 | 0 | 0 | 0 |
| 0 | 0 | 0 | 0 |
| 0 | 1 | 0 | 0 |
| 0 | 0 | 0 | 0 |
| 0 | 0 | 0 | 0 |
| 0 | 0 | 0 | 0 |
| 0 | 0 | 0 | 0 |
| 0 | 0 | 0 | 0 |
| 0 | 0 | 0 | 0 |
| 1 | 0 | 0 | 0 |
| 0 | 0 | 0 | 0 |
| 0 | 0 | 0 | 0 |
| 0 | 0 | 0 | 0 |
| 0 | 0 | 0 | 0 |
| 0 | 0 | 0 | 0 |
| 0 | 0 | 0 | 0 |
| 0 | 0 | 0 | 0 |
| 1 | 1 | 0 | 0 |
| 0 | 0 | 0 | 0 |
| 0 | 0 | 0 | 0 |
| 0 | 0 | 0 | 0 |
| 0 | 0 | 0 | 0 |
| 0 | 0 | 0 | 0 |
| 0 | 0 | 0 | 0 |
| 0 | 0 | 0 | 0 |
| 0 | 0 | 0 | 0 |
| 0 | 0 | 0 | 0 |
| 0 | 0 | 0 | 0 |
| 0 | 0 | 0 | 0 |
| 0 | 0 | 0 | 0 |
| 0 | 0 | 0 | 0 |
| 1 | 0 | 0 | 0 |
| 0 | 0 | 0 | 0 |
| 0 | 0 | 0 | 0 |
| 0 | 0 | 0 | 0 |
| 0 | 0 | 0 | 0 |
| 0 | 0 | 0 | 0 |
| 1 | 0 | 0 | 1 |

| | | | |
|---|---|---|---|
| 0 | 0 | 0 | 0 |
| 0 | 0 | 0 | 0 |
| 0 | 0 | 1 | 0 |
| 0 | 0 | 0 | 0 |
| 0 | 0 | 0 | 0 |
| 0 | 0 | 0 | 0 |
| 0 | 0 | 0 | 0 |
| 0 | 0 | 0 | 0 |
| 0 | 0 | 0 | 0 |
| 0 | 0 | 0 | 0 |
| 0 | 0 | 0 | 0 |
| 0 | 0 | 0 | 0 |
| 0 | 0 | 0 | 0 |
| 0 | 0 | 0 | 0 |
| 0 | 0 | 0 | 0 |
| 0 | 0 | 0 | 0 |
| 0 | 0 | 0 | 0 |
| 0 | 0 | 0 | 0 |
| 0 | 0 | 0 | 0 |
| 0 | 0 | 0 | 0 |
| 0 | 0 | 0 | 0 |
| 0 | 0 | 0 | 0 |
| 0 | 0 | 0 | 0 |
| 0 | 0 | 0 | 0 |
| 0 | 0 | 0 | 0 |
| 0 | 0 | 0 | 0 |
| 0 | 0 | 0 | 0 |
| 0 | 0 | 0 | 0 |
| 0 | 0 | 0 | 0 |
| 0 | 0 | 0 | 0 |
| 0 | 0 | 0 | 0 |
| 1 | 0 | 0 | 0 |
| 0 | 0 | 0 | 0 |
| 0 | 0 | 0 | 0 |
| 0 | 0 | 0 | 0 |
| 0 | 0 | 0 | 0 |
| 0 | 0 | 0 | 0 |
| 0 | 0 | 0 | 0 |
| 0 | 0 | 0 | 0 |
| 0 | 0 | 0 | 0 |
| 1 | 1 | 0 | 0 |
| 0 | 0 | 0 | 0 |
| 0 | 0 | 0 | 0 |
| 0 | 0 | 0 | 0 |
| 0 | 0 | 0 | 0 |
| 1 | 1 | 0 | 0 |
| 0 | 0 | 0 | 0 |
| 1 | 1 | 0 | 0 |
| 0 | 0 | 0 | 0 |
| 1 | 0 | 0 | 0 |
| 1 | 0 | 0 | 0 |
| 1 | 0 | 0 | 0 |
| 0 | 0 | 0 | 0 |
| 0 | 0 | 0 | 0 |

| | | | |
|---|---|---|---|
| 1 | 0 | 0 | 0 |
| 0 | 0 | 0 | 0 |
| 0 | 0 | 0 | 0 |
| 0 | 0 | 0 | 0 |
| 0 | 0 | 0 | 0 |
| 0 | 0 | 0 | 0 |
| 0 | 0 | 0 | 0 |
| 0 | 0 | 0 | 0 |
| 0 | 0 | 0 | 0 |
| 0 | 0 | 0 | 0 |
| 0 | 0 | 0 | 0 |
| 0 | 0 | 0 | 0 |
| 0 | 0 | 0 | 0 |
| 0 | 0 | 0 | 0 |
| 0 | 0 | 0 | 0 |
| 1 | 1 | 1 | 0 |
| 0 | 0 | 0 | 0 |
| 1 | 0 | 0 | 0 |
| 0 | 0 | 0 | 0 |
| 0 | 0 | 0 | 0 |
| 0 | 0 | 0 | 0 |
| 0 | 1 | 0 | 0 |
| 0 | 0 | 0 | 0 |
| 0 | 1 | 0 | 0 |
| 0 | 1 | 0 | 0 |
| 0 | 0 | 0 | 0 |
| 0 | 0 | 0 | 0 |
| 0 | 0 | 0 | 0 |
| 0 | 0 | 0 | 0 |
| 0 | 0 | 0 | 0 |
| 0 | 0 | 0 | 0 |
| 0 | 0 | 0 | 0 |
| 1 | 1 | 0 | 0 |
| 0 | 0 | 0 | 0 |
| 0 | 0 | 0 | 0 |
| 0 | 0 | 0 | 0 |
| 0 | 1 | 0 | 0 |
| 1 | 1 | 0 | 0 |
| 0 | 0 | 0 | 0 |
| 0 | 0 | 0 | 0 |
| 0 | 0 | 0 | 0 |
| 0 | 0 | 0 | 0 |
| 0 | 0 | 0 | 0 |
| 0 | 0 | 0 | 0 |
| 1 | 1 | 0 | 0 |
| 0 | 0 | 0 | 0 |
| 1 | 0 | 0 | 0 |
| 1 | 1 | 0 | 0 |
| 0 | 0 | 0 | 0 |
| 0 | 0 | 0 | 0 |
| 1 | 1 | 0 | 0 |
| 1 | 1 | 0 | 0 |
| 0 | 0 | 0 | 0 |
| 1 | 1 | 0 | 0 |

| | | | |
|---|---|---|---|
| 0 | 0 | 0 | 0 |
| 0 | 0 | 0 | 0 |
| 0 | 0 | 0 | 0 |
| 1 | 1 | 0 | 0 |
| 0 | 0 | 0 | 0 |
| 0 | 0 | 0 | 0 |
| 0 | 0 | 0 | 0 |
| 0 | 0 | 0 | 0 |
| 1 | 0 | 0 | 0 |
| 0 | 0 | 0 | 0 |
| 0 | 0 | 0 | 0 |
| 0 | 0 | 0 | 0 |
| 1 | 0 | 1 | 0 |
| 1 | 0 | 0 | 0 |
| 0 | 0 | 0 | 0 |
| 0 | 0 | 0 | 0 |
| 0 | 0 | 0 | 0 |
| 0 | 0 | 0 | 0 |
| 0 | 0 | 0 | 0 |
| 0 | 0 | 0 | 0 |
| 0 | 0 | 0 | 0 |
| 0 | 0 | 0 | 0 |
| 0 | 0 | 0 | 0 |
| 1 | 0 | 0 | 0 |
| 0 | 0 | 0 | 0 |
| 0 | 0 | 0 | 0 |
| 0 | 0 | 0 | 0 |
| 0 | 0 | 0 | 0 |
| 0 | 0 | 0 | 0 |
| 0 | 0 | 0 | 0 |
| 0 | 0 | 0 | 0 |
| 0 | 0 | 0 | 0 |
| 0 | 0 | 0 | 0 |
| 0 | 0 | 0 | 0 |
| 0 | 0 | 0 | 0 |
| 0 | 0 | 0 | 0 |
| 0 | 0 | 0 | 0 |
| 0 | 0 | 0 | 0 |
| 0 | 0 | 0 | 0 |
| 0 | 0 | 0 | 0 |
| 0 | 0 | 0 | 0 |
| 0 | 0 | 0 | 0 |
| 0 | 0 | 0 | 0 |
| 1 | 1 | 0 | 0 |
| 1 | 1 | 0 | 0 |
| 0 | 0 | 0 | 0 |
| 0 | 0 | 0 | 0 |
| 0 | 1 | 0 | 0 |
| 0 | 1 | 0 | 0 |
| 0 | 1 | 0 | 0 |
| 0 | 0 | 0 | 0 |
| 0 | 0 | 0 | 0 |
| 0 | 0 | 0 | 0 |
| 0 | 0 | 0 | 0 |

| | | | |
|---|---|---|---|
| 0 | 0 | 0 | 0 |
| 0 | 0 | 0 | 0 |
| 0 | 0 | 0 | 0 |
| 0 | 0 | 0 | 0 |
| 1 | 0 | 0 | 0 |
| 0 | 0 | 0 | 0 |
| 0 | 0 | 0 | 0 |
| 1 | 0 | 0 | 0 |
| 1 | 0 | 0 | 0 |
| 1 | 0 | 0 | 0 |
| 1 | 0 | 0 | 0 |
| 1 | 0 | 0 | 0 |
| 0 | 0 | 0 | 0 |
| 1 | 0 | 0 | 0 |
| 1 | 0 | 0 | 0 |
| 0 | 0 | 0 | 0 |
| 0 | 0 | 0 | 0 |
| 0 | 0 | 0 | 0 |
| 1 | 0 | 0 | 0 |
| 1 | 0 | 0 | 0 |
| 1 | 0 | 0 | 0 |
| 1 | 0 | 0 | 0 |
| 0 | 0 | 0 | 0 |
| 0 | 0 | 0 | 0 |
| 0 | 0 | 0 | 0 |
| 0 | 0 | 0 | 0 |
| 0 | 0 | 0 | 0 |
| 0 | 0 | 0 | 0 |
| 0 | 0 | 0 | 0 |
| 0 | 0 | 0 | 0 |
| 0 | 0 | 0 | 0 |
| 0 | 0 | 0 | 0 |
| 1 | 0 | 0 | 0 |
| 0 | 0 | 0 | 0 |
| 0 | 0 | 0 | 0 |
| 1 | 0 | 0 | 0 |
| 0 | 0 | 0 | 0 |
| 0 | 0 | 0 | 0 |
| 0 | 0 | 0 | 0 |
| 0 | 0 | 0 | 0 |
| 0 | 0 | 0 | 0 |
| 0 | 0 | 0 | 0 |
| 0 | 0 | 0 | 0 |
| 0 | 0 | 0 | 0 |
| 0 | 0 | 0 | 0 |
| 0 | 0 | 0 | 0 |
| 0 | 0 | 0 | 0 |
| 0 | 0 | 0 | 0 |
| 0 | 0 | 0 | 0 |
| 0 | 0 | 0 | 0 |
| 0 | 0 | 0 | 0 |
| 0 | 0 | 0 | 0 |
| 0 | 0 | 0 | 0 |
| 0 | 0 | 0 | 0 |

| | | | |
|---|---|---|---|
| 0 | 0 | 0 | 0 |
| 0 | 0 | 0 | 0 |
| 0 | 0 | 0 | 0 |
| 0 | 0 | 0 | 0 |
| 0 | 0 | 0 | 0 |
| 0 | 0 | 0 | 0 |
| 0 | 0 | 0 | 0 |
| 0 | 0 | 0 | 0 |
| 0 | 0 | 0 | 0 |
| 0 | 0 | 0 | 0 |
| 0 | 0 | 0 | 0 |
| 0 | 0 | 0 | 1 |
| 0 | 0 | 0 | 0 |
| 0 | 0 | 0 | 0 |
| 0 | 0 | 0 | 0 |
| 0 | 0 | 0 | 0 |
| 0 | 0 | 0 | 0 |
| 0 | 0 | 0 | 1 |
| 0 | 0 | 0 | 0 |
| 0 | 0 | 0 | 0 |
| 0 | 0 | 0 | 1 |
| 0 | 0 | 0 | 0 |
| 0 | 0 | 0 | 0 |
| 0 | 0 | 0 | 0 |
| 0 | 0 | 0 | 0 |
| 0 | 0 | 0 | 0 |
| 0 | 0 | 0 | 0 |
| 0 | 0 | 0 | 0 |
| 0 | 0 | 0 | 1 |
| 0 | 0 | 0 | 0 |
| 0 | 0 | 0 | 0 |
| 0 | 0 | 0 | 0 |
| 0 | 0 | 0 | 0 |
| 0 | 0 | 0 | 0 |
| 0 | 0 | 0 | 0 |
| 0 | 0 | 0 | 0 |
| 0 | 0 | 0 | 0 |
| 0 | 0 | 0 | 0 |
| 0 | 0 | 0 | 0 |
| 0 | 0 | 0 | 0 |
| 0 | 0 | 0 | 0 |
| 0 | 0 | 0 | 0 |
| 0 | 0 | 0 | 0 |
| 0 | 0 | 0 | 0 |
| 0 | 0 | 0 | 0 |
| 0 | 0 | 0 | 0 |
| 0 | 0 | 0 | 0 |
| 0 | 0 | 0 | 0 |
| 0 | 0 | 0 | 0 |
| 0 | 0 | 0 | 0 |
| 0 | 0 | 0 | 0 |
| 0 | 0 | 0 | 0 |
| 0 | 0 | 0 | 0 |
| 0 | 0 | 0 | 0 |

| | | | |
|---|---|---|---|
| 0 | 0 | 0 | 0 |
| 0 | 0 | 0 | 0 |
| 1 | 0 | 0 | 0 |
| 0 | 0 | 0 | 0 |
| 1 | 0 | 0 | 0 |
| 0 | 0 | 0 | 0 |
| 0 | 0 | 0 | 0 |
| 0 | 0 | 0 | 0 |
| 0 | 0 | 0 | 0 |
| 1 | 1 | 0 | 0 |
| 0 | 0 | 1 | 0 |
| 0 | 0 | 0 | 0 |
| 0 | 0 | 0 | 0 |
| 0 | 0 | 0 | 0 |
| 0 | 0 | 0 | 0 |
| 0 | 0 | 0 | 0 |
| 0 | 0 | 0 | 0 |
| 0 | 0 | 0 | 0 |
| 0 | 0 | 0 | 0 |
| 0 | 0 | 0 | 0 |
| 0 | 0 | 0 | 0 |
| 0 | 0 | 0 | 0 |
| 0 | 0 | 0 | 0 |
| 0 | 0 | 0 | 0 |
| 0 | 0 | 0 | 0 |
| 0 | 0 | 0 | 0 |
| 0 | 0 | 0 | 0 |
| 0 | 0 | 0 | 0 |
| 0 | 0 | 0 | 0 |
| 0 | 0 | 0 | 0 |
| 0 | 0 | 0 | 0 |
| 0 | 0 | 0 | 0 |
| 0 | 0 | 0 | 0 |
| 0 | 0 | 0 | 0 |
| 0 | 0 | 0 | 0 |
| 0 | 0 | 0 | 0 |
| 0 | 0 | 0 | 0 |
| 0 | 0 | 0 | 0 |
| 0 | 0 | 0 | 0 |
| 0 | 0 | 0 | 0 |
| 0 | 0 | 0 | 0 |
| 0 | 0 | 0 | 0 |
| 0 | 0 | 0 | 0 |
| 0 | 0 | 0 | 0 |
| 0 | 0 | 0 | 0 |
| 1 | 0 | 0 | 0 |
| 0 | 0 | 0 | 0 |
| 0 | 0 | 0 | 0 |
| 0 | 0 | 0 | 0 |
| 0 | 0 | 0 | 0 |
| 1 | 0 | 0 | 0 |
| 0 | 0 | 0 | 0 |
| 0 | 0 | 0 | 0 |
| 0 | 0 | 0 | 0 |

| | | | |
|---|---|---|---|
| 0 | 0 | 0 | 0 |
| 0 | 0 | 0 | 0 |
| 0 | 0 | 0 | 0 |
| 0 | 0 | 0 | 0 |
| 0 | 0 | 0 | 0 |
| 0 | 0 | 0 | 0 |
| 0 | 0 | 0 | 0 |
| 0 | 0 | 0 | 0 |
| 1 | 1 | 0 | 0 |
| 0 | 0 | 0 | 0 |
| 0 | 0 | 0 | 0 |
| 0 | 0 | 0 | 0 |
| 0 | 0 | 0 | 0 |
| 0 | 0 | 0 | 0 |
| 0 | 1 | 0 | 0 |
| 0 | 1 | 0 | 0 |
| 0 | 1 | 0 | 0 |
| 0 | 0 | 0 | 0 |
| 0 | 0 | 0 | 0 |
| 0 | 0 | 0 | 0 |
| 0 | 1 | 0 | 0 |
| 0 | 0 | 0 | 0 |
| 0 | 0 | 0 | 0 |
| 1 | 1 | 0 | 0 |
| 0 | 1 | 0 | 0 |
| 0 | 1 | 0 | 0 |
| 0 | 0 | 0 | 0 |
| 0 | 0 | 0 | 0 |
| 0 | 0 | 0 | 0 |
| 0 | 0 | 0 | 0 |
| 0 | 0 | 0 | 0 |
| 0 | 0 | 0 | 0 |
| 0 | 0 | 0 | 0 |
| 0 | 0 | 0 | 0 |
| 0 | 0 | 0 | 0 |
| 0 | 0 | 0 | 0 |
| 0 | 0 | 0 | 0 |
| 1 | 0 | 0 | 0 |
| 0 | 0 | 0 | 0 |
| 0 | 0 | 0 | 0 |
| 0 | 0 | 0 | 0 |
| 0 | 0 | 0 | 0 |
| 0 | 0 | 0 | 0 |
| 0 | 0 | 0 | 0 |
| 0 | 0 | 0 | 0 |
| 0 | 0 | 0 | 0 |
| 0 | 0 | 0 | 0 |
| 0 | 0 | 0 | 0 |
| 0 | 0 | 0 | 0 |
| 0 | 0 | 0 | 0 |
| 0 | 0 | 0 | 0 |
| 0 | 0 | 0 | 0 |
| 0 | 0 | 0 | 0 |
| 0 | 0 | 0 | 0 |

| | | | |
|---|---|---|---|
| 0 | 0 | 0 | 0 |
| 0 | 0 | 0 | 0 |
| 0 | 0 | 0 | 0 |
| 0 | 0 | 1 | 0 |
| 0 | 0 | 0 | 0 |
| 0 | 0 | 0 | 0 |
| 0 | 0 | 0 | 0 |
| 0 | 0 | 0 | 0 |
| 0 | 0 | 0 | 0 |
| 0 | 0 | 0 | 0 |
| 0 | 0 | 0 | 0 |
| 0 | 0 | 0 | 0 |
| 1 | 0 | 0 | 0 |
| 0 | 0 | 0 | 0 |
| 0 | 0 | 0 | 0 |
| 0 | 0 | 0 | 0 |
| 0 | 1 | 0 | 0 |
| 0 | 0 | 0 | 0 |
| 0 | 0 | 0 | 0 |
| 0 | 0 | 0 | 0 |
| 0 | 0 | 0 | 0 |
| 0 | 0 | 0 | 0 |
| 0 | 0 | 0 | 0 |
| 0 | 0 | 0 | 0 |
| 0 | 0 | 0 | 0 |
| 0 | 0 | 0 | 0 |
| 0 | 0 | 0 | 0 |
| 0 | 0 | 0 | 0 |
| 0 | 0 | 0 | 0 |
| 0 | 0 | 0 | 0 |
| 1 | 0 | 0 | 0 |
| 1 | 0 | 0 | 0 |
| 1 | 0 | 0 | 0 |
| 0 | 0 | 0 | 0 |
| 1 | 0 | 0 | 0 |
| 1 | 0 | 0 | 0 |
| 1 | 0 | 0 | 0 |
| 0 | 0 | 0 | 0 |
| 0 | 0 | 0 | 0 |
| 0 | 0 | 0 | 0 |
| 0 | 0 | 0 | 0 |
| 0 | 0 | 0 | 0 |
| 0 | 1 | 0 | 0 |
| 0 | 1 | 0 | 0 |
| 0 | 0 | 0 | 0 |
| 0 | 0 | 0 | 0 |
| 1 | 0 | 0 | 0 |
| 0 | 0 | 0 | 0 |
| 0 | 0 | 0 | 0 |
| 0 | 0 | 0 | 0 |
| 0 | 0 | 0 | 0 |
| 0 | 0 | 0 | 0 |
| 1 | 0 | 0 | 0 |
| 0 | 0 | 0 | 0 |

| | | | |
|---|---|---|---|
| 0 | 0 | 0 | 0 |
| 1 | 0 | 0 | 0 |
| 1 | 0 | 0 | 0 |
| 0 | 0 | 0 | 0 |
| 0 | 0 | 0 | 0 |
| 0 | 0 | 0 | 0 |
| 0 | 0 | 0 | 0 |
| 0 | 1 | 0 | 0 |
| 0 | 0 | 0 | 0 |
| 0 | 0 | 0 | 0 |
| 0 | 0 | 0 | 0 |
| 0 | 0 | 0 | 0 |
| 0 | 0 | 0 | 0 |
| 0 | 0 | 0 | 0 |
| 1 | 0 | 0 | 0 |
| 0 | 0 | 0 | 0 |
| 0 | 0 | 0 | 0 |
| 0 | 0 | 0 | 0 |
| 0 | 0 | 0 | 0 |
| 1 | 0 | 0 | 0 |
| 1 | 0 | 0 | 0 |
| 0 | 0 | 0 | 0 |
| 1 | 0 | 0 | 0 |
| 1 | 0 | 0 | 0 |
| 0 | 0 | 0 | 0 |
| 0 | 0 | 0 | 0 |
| 0 | 0 | 0 | 0 |
| 0 | 0 | 0 | 0 |
| 0 | 0 | 0 | 0 |
| 0 | 0 | 0 | 0 |
| 0 | 0 | 0 | 0 |
| 0 | 0 | 0 | 0 |
| 0 | 0 | 0 | 0 |
| 0 | 0 | 0 | 0 |
| 0 | 0 | 0 | 0 |
| 0 | 0 | 0 | 0 |
| 0 | 0 | 0 | 0 |
| 0 | 0 | 0 | 0 |
| 0 | 0 | 0 | 0 |
| 0 | 0 | 0 | 0 |
| 0 | 0 | 0 | 0 |
| 0 | 0 | 0 | 0 |
| 0 | 0 | 0 | 0 |
| 0 | 0 | 0 | 0 |
| 0 | 0 | 0 | 0 |
| 0 | 0 | 0 | 0 |
| 0 | 0 | 0 | 0 |
| 0 | 0 | 0 | 0 |
| 0 | 0 | 0 | 0 |
| 0 | 0 | 0 | 0 |
| 0 | 0 | 0 | 0 |
| 0 | 0 | 0 | 0 |
| 0 | 0 | 0 | 0 |
| 1 | 1 | 0 | 1 |

| | | | |
|---|---|---|---|
| 0 | 0 | 0 | 0 |
| 0 | 0 | 0 | 0 |
| 0 | 0 | 0 | 0 |
| 0 | 0 | 0 | 0 |
| 0 | 1 | 0 | 0 |
| 0 | 0 | 0 | 0 |
| 0 | 0 | 0 | 0 |
| 0 | 1 | 0 | 0 |
| 0 | 0 | 0 | 0 |
| 0 | 0 | 0 | 0 |
| 0 | 0 | 0 | 0 |
| 0 | 0 | 0 | 0 |
| 0 | 0 | 0 | 0 |
| 0 | 0 | 0 | 0 |
| 0 | 1 | 0 | 1 |
| 0 | 1 | 0 | 0 |
| 0 | 0 | 0 | 0 |
| 0 | 0 | 0 | 0 |
| 0 | 1 | 0 | 0 |
| 0 | 1 | 0 | 1 |
| 0 | 0 | 0 | 0 |
| 0 | 0 | 0 | 0 |
| 0 | 0 | 0 | 0 |
| 0 | 0 | 0 | 0 |
| 0 | 0 | 0 | 0 |
| 1 | 0 | 0 | 0 |
| 0 | 0 | 0 | 0 |
| 0 | 0 | 1 | 0 |
| 0 | 0 | 1 | 0 |
| 0 | 0 | 1 | 0 |
| 0 | 0 | 0 | 0 |
| 0 | 0 | 0 | 0 |
| 0 | 0 | 0 | 0 |
| 0 | 0 | 0 | 0 |
| 0 | 0 | 0 | 0 |
| 0 | 0 | 1 | 0 |
| 0 | 0 | 0 | 0 |
| 0 | 0 | 0 | 0 |
| 1 | 0 | 1 | 0 |
| 0 | 0 | 1 | 0 |
| 1 | 1 | 0 | 0 |
| 0 | 1 | 0 | 0 |
| 0 | 0 | 0 | 0 |
| 0 | 0 | 0 | 0 |
| 1 | 0 | 0 | 0 |
| 0 | 0 | 0 | 0 |
| 0 | 0 | 1 | 0 |
| 0 | 0 | 0 | 0 |
| 0 | 0 | 0 | 0 |
| 0 | 0 | 0 | 0 |
| 0 | 0 | 0 | 0 |
| 0 | 0 | 0 | 0 |
| 0 | 0 | 0 | 0 |
| 0 | 0 | 0 | 0 |

| | | | |
|---|---|---|---|
| 0 | 0 | 0 | 0 |
| 0 | 0 | 0 | 0 |
| 0 | 0 | 0 | 0 |
| 0 | 0 | 0 | 0 |
| 0 | 0 | 0 | 0 |
| 0 | 0 | 0 | 0 |
| 0 | 0 | 0 | 0 |
| 1 | 1 | 0 | 0 |
| 0 | 0 | 0 | 0 |
| 0 | 0 | 0 | 0 |
| 0 | 0 | 0 | 0 |
| 0 | 0 | 0 | 1 |
| 0 | 0 | 0 | 0 |
| 0 | 0 | 0 | 0 |
| 0 | 0 | 0 | 0 |
| 0 | 0 | 0 | 0 |
| 0 | 0 | 0 | 0 |
| 0 | 0 | 0 | 0 |
| 0 | 0 | 0 | 0 |
| 0 | 0 | 0 | 0 |
| 0 | 0 | 0 | 0 |
| 0 | 0 | 0 | 0 |
| 0 | 0 | 0 | 0 |
| 0 | 0 | 0 | 0 |
| 0 | 0 | 0 | 0 |
| 0 | 0 | 0 | 0 |
| 0 | 0 | 0 | 0 |
| 0 | 0 | 0 | 0 |
| 0 | 0 | 0 | 0 |
| 0 | 1 | 0 | 0 |
| 0 | 1 | 0 | 0 |
| 0 | 0 | 0 | 0 |
| 0 | 0 | 0 | 0 |
| 0 | 0 | 0 | 0 |
| 0 | 0 | 0 | 0 |
| 0 | 1 | 0 | 0 |
| 0 | 0 | 0 | 0 |
| 0 | 0 | 0 | 0 |
| 0 | 0 | 0 | 0 |
| 0 | 0 | 0 | 0 |
| 0 | 0 | 0 | 0 |
| 1 | 1 | 0 | 0 |
| 1 | 0 | 0 | 0 |
| 0 | 0 | 0 | 0 |
| 0 | 0 | 0 | 0 |
| 0 | 0 | 0 | 0 |
| 0 | 0 | 0 | 0 |
| 0 | 0 | 0 | 0 |
| 0 | 0 | 0 | 0 |
| 0 | 0 | 0 | 0 |
| 0 | 0 | 0 | 0 |
| 0 | 0 | 0 | 0 |
| 0 | 0 | 0 | 0 |
| 0 | 0 | 0 | 0 |

| | | | |
|---|---|---|---|
| 0 | 0 | 0 | 0 |
| 0 | 0 | 0 | 0 |
| 0 | 0 | 0 | 0 |
| 0 | 0 | 0 | 0 |
| 0 | 0 | 0 | 0 |
| 0 | 0 | 0 | 0 |
| 0 | 0 | 0 | 0 |
| 0 | 0 | 0 | 0 |
| 1 | 0 | 0 | 0 |
| 1 | 0 | 0 | 0 |
| 0 | 0 | 0 | 0 |
| 0 | 0 | 0 | 0 |
| 0 | 0 | 0 | 0 |
| 1 | 0 | 0 | 0 |
| 0 | 0 | 0 | 0 |
| 0 | 0 | 0 | 0 |
| 0 | 0 | 0 | 0 |
| 0 | 0 | 0 | 0 |
| 0 | 0 | 0 | 0 |
| 1 | 1 | 0 | 0 |
| 0 | 0 | 1 | 0 |
| 0 | 0 | 0 | 0 |
| 0 | 0 | 0 | 0 |
| 0 | 0 | 0 | 0 |
| 0 | 0 | 0 | 0 |
| 0 | 0 | 0 | 0 |
| 0 | 0 | 0 | 0 |
| 0 | 0 | 0 | 0 |
| 0 | 0 | 0 | 0 |
| 0 | 0 | 0 | 0 |
| 0 | 0 | 0 | 0 |
| 0 | 0 | 0 | 0 |
| 0 | 0 | 0 | 0 |
| 0 | 0 | 0 | 0 |
| 0 | 0 | 0 | 0 |
| 0 | 0 | 0 | 0 |
| 0 | 0 | 0 | 0 |
| 0 | 0 | 0 | 0 |
| 0 | 0 | 0 | 0 |
| 0 | 0 | 0 | 0 |
| 0 | 0 | 0 | 0 |
| 0 | 0 | 0 | 0 |
| 0 | 0 | 0 | 0 |
| 0 | 0 | 0 | 0 |
| 0 | 0 | 0 | 0 |
| 0 | 0 | 0 | 0 |
| 1 | 0 | 1 | 0 |
| 1 | 0 | 0 | 0 |
| 0 | 0 | 0 | 0 |
| 0 | 0 | 0 | 0 |
| 0 | 0 | 1 | 0 |
| 0 | 0 | 0 | 0 |
| 0 | 0 | 0 | 0 |
| 0 | 0 | 0 | 0 |

| | | | |
|---|---|---|---|
| 0 | 0 | 0 | 0 |
| 0 | 0 | 0 | 0 |
| 0 | 0 | 0 | 0 |
| 0 | 0 | 0 | 0 |
| 0 | 0 | 0 | 0 |
| 0 | 0 | 0 | 0 |
| 0 | 0 | 0 | 0 |
| 0 | 0 | 0 | 0 |
| 0 | 0 | 0 | 0 |
| 0 | 0 | 0 | 0 |
| 0 | 0 | 0 | 0 |
| 0 | 0 | 0 | 0 |
| 0 | 0 | 0 | 0 |
| 0 | 0 | 0 | 0 |
| 0 | 0 | 0 | 0 |
| 0 | 0 | 0 | 0 |
| 0 | 0 | 0 | 0 |
| 0 | 0 | 0 | 0 |
| 0 | 0 | 0 | 0 |
| 1 | 0 | 0 | 0 |
| 0 | 0 | 0 | 0 |
| 0 | 0 | 0 | 0 |
| 0 | 0 | 0 | 0 |
| 0 | 0 | 0 | 0 |
| 0 | 0 | 0 | 0 |
| 0 | 0 | 0 | 0 |
| 0 | 0 | 0 | 0 |
| 0 | 0 | 0 | 0 |
| 0 | 0 | 0 | 0 |
| 0 | 0 | 0 | 0 |
| 0 | 0 | 0 | 0 |
| 0 | 0 | 0 | 0 |
| 0 | 0 | 0 | 0 |
| 0 | 0 | 0 | 0 |
| 0 | 0 | 0 | 0 |
| 0 | 0 | 0 | 0 |
| 0 | 0 | 0 | 0 |
| 0 | 0 | 0 | 0 |
| 1 | 0 | 0 | 1 |
| 0 | 0 | 0 | 0 |
| 0 | 0 | 0 | 0 |
| 0 | 0 | 0 | 0 |
| 0 | 0 | 0 | 1 |
| 0 | 0 | 0 | 1 |
| 0 | 0 | 0 | 1 |
| 0 | 0 | 0 | 0 |
| 0 | 0 | 0 | 0 |
| 0 | 0 | 0 | 0 |
| 0 | 0 | 0 | 0 |
| 0 | 0 | 0 | 0 |
| 0 | 0 | 0 | 0 |
| 0 | 0 | 0 | 0 |
| 0 | 0 | 0 | 0 |
| 0 | 0 | 0 | 0 |

| | | | |
|---|---|---|---|
| 0 | 0 | 0 | 0 |
| 0 | 0 | 0 | 0 |
| 0 | 0 | 0 | 0 |
| 0 | 0 | 0 | 0 |
| 0 | 0 | 0 | 0 |
| 0 | 0 | 0 | 0 |
| 0 | 0 | 0 | 0 |
| 0 | 0 | 0 | 0 |
| 0 | 0 | 0 | 0 |
| 0 | 0 | 0 | 0 |
| 0 | 0 | 0 | 0 |
| 0 | 0 | 0 | 0 |
| 0 | 0 | 0 | 0 |
| 0 | 0 | 0 | 0 |
| 0 | 0 | 0 | 0 |
| 0 | 0 | 0 | 0 |
| 0 | 0 | 0 | 0 |
| 0 | 0 | 0 | 0 |
| 0 | 0 | 0 | 0 |
| 0 | 0 | 0 | 0 |
| 0 | 0 | 0 | 0 |
| 0 | 0 | 0 | 0 |
| 0 | 0 | 0 | 0 |
| 0 | 0 | 0 | 0 |
| 0 | 0 | 0 | 0 |
| 0 | 0 | 0 | 0 |
| 0 | 0 | 0 | 0 |
| 0 | 0 | 0 | 0 |
| 0 | 0 | 0 | 0 |
| 0 | 0 | 0 | 0 |
| 0 | 0 | 0 | 0 |
| 0 | 0 | 0 | 0 |
| 1 | 1 | 0 | 1 |
| 1 | 1 | 0 | 1 |
| 1 | 0 | 0 | 0 |
| 0 | 0 | 0 | 0 |
| 0 | 0 | 0 | 0 |
| 0 | 0 | 0 | 0 |
| 0 | 0 | 0 | 0 |
| 0 | 0 | 0 | 0 |
| 0 | 0 | 0 | 0 |
| 0 | 0 | 0 | 0 |
| 1 | 1 | 0 | 0 |
| 0 | 0 | 0 | 0 |
| 1 | 0 | 0 | 0 |
| 0 | 0 | 0 | 0 |
| 0 | 0 | 0 | 0 |
| 0 | 0 | 0 | 0 |
| 0 | 0 | 0 | 0 |
| 0 | 0 | 0 | 0 |
| 1 | 0 | 0 | 0 |
| 0 | 0 | 0 | 0 |
| 0 | 0 | 0 | 0 |
| 0 | 0 | 0 | 0 |

| | | | |
|---|---|---|---|
| 0 | 0 | 0 | 0 |
| 0 | 0 | 0 | 0 |
| 0 | 0 | 0 | 0 |
| 0 | 0 | 0 | 0 |
| 0 | 0 | 0 | 0 |
| 0 | 0 | 0 | 0 |
| 0 | 0 | 0 | 0 |
| 0 | 0 | 0 | 0 |
| 0 | 0 | 0 | 0 |
| 0 | 0 | 0 | 0 |
| 0 | 0 | 0 | 0 |
| 0 | 0 | 0 | 0 |
| 0 | 0 | 0 | 0 |
| 0 | 0 | 0 | 0 |
| 0 | 0 | 0 | 0 |
| 0 | 0 | 0 | 0 |
| 0 | 0 | 0 | 0 |
| 0 | 0 | 0 | 0 |
| 0 | 0 | 0 | 0 |
| 0 | 0 | 0 | 1 |
| 0 | 0 | 0 | 0 |
| 0 | 0 | 0 | 1 |
| 0 | 0 | 0 | 1 |
| 0 | 0 | 0 | 1 |
| 0 | 0 | 0 | 0 |
| 0 | 0 | 0 | 0 |
| 0 | 0 | 0 | 0 |
| 0 | 0 | 0 | 0 |
| 0 | 0 | 0 | 0 |
| 0 | 0 | 0 | 0 |
| 0 | 0 | 0 | 0 |
| 0 | 0 | 0 | 0 |
| 0 | 0 | 0 | 0 |
| 0 | 0 | 0 | 0 |
| 0 | 0 | 0 | 0 |
| 0 | 1 | 0 | 0 |
| 0 | 0 | 0 | 0 |
| 0 | 0 | 0 | 0 |
| 0 | 1 | 0 | 0 |
| 0 | 0 | 0 | 0 |
| 0 | 0 | 0 | 0 |
| 0 | 0 | 0 | 0 |
| 0 | 0 | 0 | 0 |
| 0 | 0 | 0 | 0 |
| 0 | 0 | 0 | 0 |
| 0 | 0 | 0 | 0 |
| 0 | 0 | 0 | 0 |
| 0 | 0 | 0 | 0 |
| 0 | 0 | 0 | 0 |
| 0 | 0 | 0 | 0 |
| 0 | 0 | 0 | 0 |
| 0 | 0 | 0 | 0 |
| 0 | 0 | 0 | 0 |
| 0 | 0 | 0 | 0 |

| | | | |
|---|---|---|---|
| 0 | 0 | 0 | 0 |
| 0 | 0 | 0 | 0 |
| 0 | 0 | 0 | 0 |
| 0 | 0 | 0 | 0 |
| 0 | 0 | 0 | 0 |
| 0 | 0 | 0 | 0 |
| 0 | 0 | 0 | 0 |
| 0 | 0 | 0 | 0 |
| 0 | 0 | 1 | 0 |
| 0 | 0 | 1 | 0 |
| 0 | 0 | 0 | 0 |
| 0 | 0 | 0 | 0 |
| 0 | 0 | 0 | 0 |
| 0 | 0 | 0 | 0 |
| 0 | 0 | 0 | 0 |
| 0 | 1 | 0 | 0 |
| 0 | 0 | 0 | 0 |
| 0 | 0 | 0 | 0 |
| 1 | 1 | 0 | 0 |
| 0 | 0 | 0 | 0 |
| 0 | 0 | 0 | 0 |
| 0 | 0 | 0 | 0 |
| 0 | 0 | 0 | 0 |
| 0 | 0 | 0 | 0 |
| 0 | 0 | 0 | 0 |
| 0 | 0 | 0 | 0 |
| 0 | 0 | 0 | 0 |
| 0 | 0 | 0 | 0 |
| 0 | 0 | 0 | 0 |
| 0 | 0 | 0 | 0 |
| 0 | 0 | 0 | 0 |
| 0 | 0 | 0 | 0 |
| 0 | 0 | 0 | 0 |
| 0 | 0 | 0 | 0 |
| 0 | 0 | 0 | 0 |
| 0 | 0 | 0 | 0 |
| 0 | 0 | 0 | 0 |
| 0 | 0 | 0 | 0 |
| 0 | 0 | 0 | 0 |
| 0 | 0 | 0 | 0 |
| 0 | 0 | 0 | 0 |
| 0 | 0 | 0 | 0 |
| 0 | 0 | 0 | 0 |
| 0 | 0 | 0 | 0 |
| 0 | 0 | 0 | 0 |
| 0 | 0 | 0 | 0 |
| 0 | 0 | 0 | 0 |
| 0 | 0 | 0 | 0 |
| 0 | 0 | 0 | 0 |
| 0 | 0 | 0 | 0 |
| 0 | 0 | 0 | 0 |
| 0 | 0 | 0 | 0 |
| 0 | 0 | 0 | 0 |
| 1 | 0 | 0 | 0 |

| | | | |
|---|---|---|---|
| 0 | 0 | 0 | 0 |
| 0 | 0 | 0 | 0 |
| 0 | 0 | 0 | 0 |
| 0 | 0 | 0 | 0 |
| 0 | 0 | 0 | 0 |
| 0 | 0 | 0 | 0 |
| 0 | 0 | 0 | 0 |
| 0 | 0 | 0 | 0 |
| 0 | 0 | 0 | 0 |
| 0 | 0 | 0 | 0 |
| 1 | 0 | 0 | 0 |
| 0 | 0 | 0 | 0 |
| 0 | 0 | 0 | 0 |
| 0 | 0 | 0 | 0 |
| 1 | 1 | 0 | 1 |
| 0 | 0 | 0 | 0 |
| 1 | 0 | 0 | 0 |
| 1 | 1 | 0 | 0 |
| 0 | 0 | 0 | 0 |
| 0 | 0 | 0 | 0 |
| 0 | 0 | 0 | 0 |
| 0 | 0 | 0 | 0 |
| 0 | 0 | 0 | 0 |
| 0 | 0 | 0 | 1 |
| 0 | 0 | 0 | 0 |
| 0 | 0 | 0 | 0 |
| 0 | 0 | 0 | 0 |
| 0 | 0 | 0 | 1 |
| 0 | 0 | 0 | 0 |
| 0 | 0 | 0 | 0 |
| 0 | 0 | 0 | 0 |
| 0 | 0 | 0 | 0 |
| 0 | 0 | 0 | 0 |
| 0 | 0 | 0 | 0 |
| 0 | 0 | 0 | 0 |
| 0 | 0 | 0 | 0 |
| 0 | 0 | 0 | 0 |
| 0 | 0 | 0 | 0 |
| 0 | 0 | 0 | 0 |
| 0 | 0 | 0 | 0 |
| 0 | 0 | 0 | 0 |
| 0 | 0 | 0 | 0 |
| 0 | 0 | 0 | 0 |
| 0 | 0 | 0 | 0 |
| 0 | 0 | 1 | 0 |
| 0 | 0 | 0 | 0 |
| 0 | 0 | 0 | 0 |
| 0 | 0 | 0 | 0 |
| 0 | 0 | 0 | 0 |
| 0 | 0 | 0 | 0 |
| 0 | 0 | 0 | 0 |
| 0 | 0 | 0 | 0 |
| 0 | 0 | 0 | 0 |
| 0 | 0 | 0 | 0 |

| | | | |
|---|---|---|---|
| 0 | 0 | 0 | 0 |
| 0 | 0 | 0 | 0 |
| 0 | 0 | 0 | 0 |
| 0 | 0 | 0 | 0 |
| 0 | 0 | 0 | 0 |
| 0 | 0 | 0 | 0 |
| 1 | 0 | 0 | 1 |
| 1 | 0 | 0 | 0 |
| 0 | 0 | 0 | 0 |
| 0 | 0 | 0 | 0 |
| 0 | 0 | 0 | 0 |
| 0 | 0 | 0 | 0 |
| 0 | 0 | 0 | 0 |
| 0 | 0 | 0 | 0 |
| 0 | 0 | 0 | 0 |
| 0 | 0 | 0 | 0 |
| 0 | 0 | 0 | 0 |
| 0 | 0 | 0 | 0 |
| 0 | 0 | 0 | 0 |
| 0 | 0 | 0 | 0 |
| 1 | 0 | 0 | 0 |
| 0 | 0 | 0 | 0 |
| 0 | 0 | 0 | 0 |
| 0 | 0 | 0 | 0 |
| 0 | 0 | 0 | 0 |
| 0 | 0 | 0 | 0 |
| 0 | 0 | 0 | 0 |
| 0 | 0 | 0 | 0 |
| 0 | 0 | 0 | 0 |
| 0 | 0 | 0 | 0 |
| 0 | 0 | 0 | 0 |
| 0 | 0 | 0 | 0 |
| 0 | 0 | 0 | 0 |
| 0 | 0 | 0 | 0 |
| 0 | 0 | 0 | 0 |
| 0 | 0 | 0 | 0 |
| 0 | 0 | 0 | 0 |
| 0 | 0 | 0 | 0 |
| 0 | 0 | 0 | 0 |
| 0 | 0 | 0 | 0 |
| 0 | 0 | 0 | 0 |
| 0 | 0 | 0 | 0 |
| 0 | 0 | 0 | 0 |
| 0 | 0 | 0 | 0 |
| 0 | 0 | 0 | 0 |
| 0 | 0 | 0 | 0 |
| 0 | 0 | 0 | 0 |
| 0 | 0 | 0 | 0 |
| 0 | 0 | 0 | 0 |
| 0 | 0 | 0 | 0 |
| 0 | 0 | 0 | 0 |
| 0 | 0 | 0 | 0 |
| 0 | 0 | 0 | 0 |
| 0 | 0 | 0 | 0 |

| | | | |
|---|---|---|---|
| 0 | 0 | 0 | 0 |
| 0 | 0 | 0 | 0 |
| 0 | 0 | 0 | 0 |
| 1 | 0 | 0 | 0 |
| 0 | 0 | 0 | 0 |
| 1 | 0 | 0 | 0 |
| 0 | 0 | 0 | 0 |
| 0 | 0 | 0 | 0 |
| 1 | 0 | 0 | 0 |
| 0 | 0 | 0 | 0 |
| 0 | 0 | 0 | 0 |
| 0 | 0 | 0 | 0 |
| 0 | 0 | 0 | 0 |
| 0 | 0 | 0 | 0 |
| 0 | 0 | 0 | 0 |
| 0 | 0 | 0 | 0 |
| 0 | 0 | 0 | 0 |
| 0 | 0 | 0 | 0 |
| 1 | 0 | 0 | 0 |
| 0 | 0 | 0 | 0 |
| 0 | 0 | 0 | 0 |
| 0 | 0 | 0 | 0 |
| 0 | 0 | 0 | 0 |
| 0 | 0 | 0 | 0 |
| 0 | 0 | 0 | 0 |
| 0 | 0 | 0 | 0 |
| 0 | 0 | 0 | 0 |
| 0 | 0 | 0 | 0 |
| 0 | 1 | 0 | 0 |
| 0 | 0 | 0 | 0 |
| 0 | 0 | 0 | 0 |
| 0 | 0 | 0 | 0 |
| 0 | 1 | 0 | 0 |
| 1 | 1 | 0 | 0 |
| 1 | 0 | 0 | 0 |
| 0 | 0 | 0 | 0 |
| 0 | 0 | 0 | 0 |
| 0 | 0 | 0 | 0 |
| 0 | 0 | 0 | 0 |
| 0 | 0 | 0 | 0 |
| 1 | 0 | 0 | 0 |
| 0 | 0 | 0 | 0 |
| 0 | 0 | 0 | 0 |
| 0 | 0 | 0 | 0 |
| 0 | 0 | 0 | 0 |
| 0 | 0 | 0 | 0 |
| 0 | 0 | 0 | 0 |
| 0 | 0 | 0 | 0 |
| 0 | 0 | 0 | 0 |
| 0 | 0 | 0 | 0 |
| 0 | 0 | 0 | 0 |
| 0 | 0 | 0 | 0 |
| 0 | 0 | 0 | 0 |
| 1 | 0 | 0 | 0 |

| | | | |
|---|---|---|---|
| 1 | 0 | 0 | 1 |
| 0 | 0 | 0 | 0 |
| 0 | 0 | 0 | 0 |
| 0 | 0 | 0 | 0 |
| 0 | 0 | 0 | 0 |
| 0 | 0 | 0 | 0 |
| 0 | 0 | 0 | 0 |
| 0 | 0 | 0 | 0 |
| 0 | 0 | 0 | 0 |
| 0 | 0 | 0 | 0 |
| 0 | 0 | 0 | 0 |
| 0 | 0 | 0 | 0 |
| 0 | 0 | 0 | 0 |
| 0 | 0 | 0 | 0 |
| 0 | 0 | 0 | 0 |
| 0 | 0 | 0 | 0 |
| 0 | 0 | 0 | 0 |
| 0 | 0 | 0 | 0 |
| 0 | 0 | 0 | 0 |
| 0 | 0 | 0 | 0 |
| 0 | 0 | 0 | 0 |
| 0 | 0 | 0 | 0 |
| 0 | 1 | 0 | 0 |
| 0 | 0 | 0 | 0 |
| 0 | 0 | 0 | 0 |
| 0 | 0 | 0 | 0 |
| 0 | 0 | 0 | 0 |
| 0 | 0 | 0 | 0 |
| 0 | 0 | 0 | 0 |
| 0 | 0 | 0 | 0 |
| 0 | 0 | 0 | 0 |
| 0 | 0 | 0 | 0 |
| 0 | 0 | 0 | 0 |
| 0 | 0 | 0 | 0 |
| 0 | 0 | 0 | 0 |
| 0 | 0 | 0 | 0 |
| 0 | 0 | 0 | 0 |
| 0 | 0 | 0 | 0 |
| 0 | 0 | 0 | 0 |
| 0 | 0 | 0 | 0 |
| 0 | 0 | 0 | 0 |
| 0 | 0 | 0 | 0 |
| 0 | 0 | 0 | 0 |
| 0 | 0 | 0 | 0 |
| 0 | 0 | 0 | 0 |
| 0 | 0 | 0 | 0 |
| 0 | 0 | 0 | 0 |
| 0 | 0 | 0 | 0 |
| 0 | 0 | 0 | 0 |
| 0 | 0 | 0 | 0 |
| 0 | 0 | 0 | 0 |
| 0 | 0 | 0 | 0 |
| 0 | 0 | 0 | 0 |
| 0 | 0 | 0 | 0 |

| | | | |
|---|---|---|---|
| 0 | 0 | 0 | 0 |
| 0 | 0 | 0 | 0 |
| 0 | 0 | 0 | 0 |
| 0 | 0 | 0 | 0 |
| 0 | 0 | 0 | 0 |
| 0 | 0 | 0 | 0 |
| 0 | 1 | 0 | 0 |
| 0 | 1 | 0 | 0 |
| 0 | 0 | 0 | 0 |
| 0 | 0 | 0 | 0 |
| 0 | 0 | 0 | 0 |
| 0 | 0 | 0 | 0 |
| 0 | 0 | 0 | 0 |
| 0 | 0 | 0 | 0 |
| 0 | 0 | 0 | 0 |
| 0 | 0 | 0 | 0 |
| 1 | 0 | 0 | 0 |
| 0 | 0 | 0 | 0 |
| 1 | 0 | 0 | 0 |
| 1 | 0 | 0 | 0 |
| 1 | 0 | 0 | 0 |
| 1 | 0 | 0 | 0 |
| 0 | 0 | 0 | 0 |
| 0 | 0 | 0 | 0 |
| 0 | 0 | 0 | 0 |
| 0 | 0 | 0 | 0 |
| 0 | 0 | 0 | 0 |
| 0 | 0 | 0 | 0 |
| 0 | 0 | 0 | 0 |
| 0 | 0 | 0 | 0 |
| 0 | 0 | 0 | 0 |
| 0 | 0 | 0 | 0 |
| 0 | 0 | 0 | 0 |
| 0 | 0 | 0 | 0 |
| 0 | 0 | 0 | 0 |
| 0 | 0 | 1 | 0 |
| 1 | 0 | 0 | 0 |
| 1 | 0 | 0 | 0 |
| 1 | 0 | 0 | 0 |
| 0 | 0 | 0 | 0 |
| 0 | 0 | 0 | 0 |
| 0 | 0 | 0 | 0 |
| 0 | 0 | 0 | 0 |
| 1 | 0 | 0 | 0 |
| 1 | 0 | 0 | 0 |
| 1 | 0 | 0 | 0 |
| 0 | 0 | 0 | 0 |
| 0 | 0 | 0 | 0 |
| 1 | 0 | 0 | 0 |
| 0 | 0 | 0 | 0 |
| 0 | 0 | 0 | 0 |
| 1 | 0 | 0 | 0 |
| 0 | 0 | 0 | 0 |
| 0 | 0 | 0 | 0 |

| | | | |
|---|---|---|---|
| 0 | 0 | 0 | 0 |
| 0 | 0 | 0 | 0 |
| 0 | 0 | 0 | 0 |
| 0 | 0 | 0 | 0 |
| 0 | 0 | 0 | 0 |
| 0 | 0 | 0 | 0 |
| 0 | 0 | 0 | 0 |
| 0 | 0 | 0 | 0 |
| 0 | 0 | 0 | 0 |
| 0 | 0 | 0 | 0 |
| 0 | 0 | 0 | 0 |
| 0 | 0 | 0 | 0 |
| 0 | 0 | 0 | 0 |
| 0 | 0 | 0 | 0 |
| 0 | 0 | 0 | 0 |
| 0 | 0 | 0 | 0 |
| 0 | 0 | 0 | 0 |
| 0 | 0 | 0 | 0 |
| 0 | 0 | 0 | 0 |
| 0 | 0 | 0 | 0 |
| 0 | 0 | 0 | 0 |
| 0 | 0 | 0 | 0 |
| 1 | 1 | 0 | 0 |
| 0 | 0 | 0 | 0 |
| 0 | 1 | 0 | 0 |
| 0 | 0 | 0 | 0 |
| 0 | 0 | 0 | 0 |
| 0 | 0 | 0 | 0 |
| 0 | 0 | 0 | 0 |
| 0 | 0 | 0 | 0 |
| 0 | 0 | 0 | 0 |
| 0 | 0 | 0 | 0 |
| 0 | 0 | 0 | 0 |
| 0 | 0 | 0 | 0 |
| 0 | 0 | 0 | 0 |
| 0 | 0 | 0 | 0 |
| 0 | 0 | 0 | 0 |
| 0 | 0 | 0 | 0 |
| 0 | 0 | 0 | 0 |
| 0 | 0 | 0 | 0 |
| 0 | 0 | 0 | 0 |
| 0 | 0 | 0 | 0 |
| 0 | 0 | 0 | 0 |
| 0 | 0 | 0 | 0 |
| 0 | 0 | 0 | 0 |
| 1 | 0 | 0 | 1 |
| 0 | 0 | 0 | 0 |
| 0 | 0 | 0 | 0 |
| 0 | 0 | 0 | 0 |
| 1 | 0 | 0 | 0 |
| 1 | 0 | 0 | 0 |
| 1 | 0 | 0 | 0 |
| 0 | 0 | 0 | 0 |
| 0 | 0 | 0 | 0 |

| | | | |
|---|---|---|---|
| 0 | 0 | 0 | 0 |
| 0 | 0 | 0 | 0 |
| 0 | 0 | 0 | 0 |
| 0 | 0 | 0 | 0 |
| 0 | 0 | 0 | 0 |
| 0 | 0 | 0 | 0 |
| 0 | 0 | 0 | 0 |
| 0 | 0 | 0 | 0 |
| 0 | 0 | 0 | 0 |
| 0 | 0 | 0 | 0 |
| 0 | 0 | 0 | 1 |
| 0 | 0 | 0 | 0 |
| 0 | 0 | 0 | 0 |
| 0 | 0 | 0 | 0 |
| 0 | 0 | 0 | 0 |
| 0 | 0 | 0 | 0 |
| 0 | 0 | 0 | 0 |
| 0 | 0 | 0 | 0 |
| 0 | 0 | 0 | 0 |
| 0 | 0 | 0 | 0 |
| 0 | 0 | 0 | 0 |
| 0 | 0 | 0 | 0 |
| 0 | 0 | 0 | 0 |
| 0 | 0 | 0 | 0 |
| 0 | 0 | 0 | 0 |
| 0 | 0 | 0 | 0 |
| 0 | 0 | 0 | 0 |
| 0 | 0 | 0 | 0 |
| 0 | 0 | 0 | 0 |
| 0 | 0 | 0 | 0 |
| 0 | 1 | 0 | 0 |
| 0 | 0 | 0 | 0 |
| 1 | 1 | 0 | 0 |
| 1 | 1 | 0 | 0 |
| 0 | 0 | 0 | 0 |
| 0 | 0 | 0 | 0 |
| 0 | 0 | 0 | 0 |
| 0 | 0 | 0 | 0 |
| 0 | 0 | 0 | 0 |
| 0 | 0 | 0 | 0 |
| 0 | 0 | 0 | 0 |
| 0 | 1 | 0 | 0 |
| 0 | 0 | 0 | 0 |
| 0 | 0 | 0 | 0 |
| 0 | 0 | 0 | 0 |
| 1 | 0 | 0 | 0 |
| 1 | 0 | 0 | 0 |
| 1 | 0 | 0 | 0 |
| 1 | 1 | 1 | 0 |
| 0 | 0 | 0 | 0 |
| 1 | 1 | 1 | 1 |
| 0 | 0 | 0 | 0 |
| 0 | 0 | 0 | 0 |
| 0 | 0 | 0 | 0 |

| | | | |
|---|---|---|---|
| 0 | 0 | 0 | 0 |
| 0 | 0 | 0 | 0 |
| 0 | 0 | 0 | 0 |
| 0 | 0 | 0 | 0 |
| 1 | 0 | 0 | 0 |
| 1 | 0 | 0 | 0 |
| 0 | 0 | 0 | 0 |
| 0 | 0 | 0 | 0 |
| 0 | 0 | 0 | 0 |
| 0 | 0 | 0 | 0 |
| 0 | 0 | 0 | 0 |
| 0 | 0 | 0 | 0 |
| 0 | 0 | 0 | 0 |
| 0 | 0 | 0 | 0 |
| 0 | 0 | 0 | 0 |
| 0 | 0 | 0 | 0 |
| 0 | 0 | 0 | 0 |
| 0 | 0 | 0 | 0 |
| 0 | 0 | 0 | 0 |
| 0 | 0 | 0 | 0 |
| 0 | 0 | 0 | 0 |
| 0 | 0 | 0 | 0 |
| 0 | 0 | 0 | 0 |
| 0 | 0 | 0 | 0 |
| 0 | 0 | 0 | 0 |
| 0 | 0 | 0 | 0 |
| 0 | 0 | 0 | 0 |
| 0 | 0 | 0 | 0 |
| 0 | 0 | 0 | 0 |
| 0 | 0 | 0 | 0 |
| 0 | 0 | 0 | 0 |
| 0 | 0 | 0 | 0 |
| 0 | 0 | 0 | 0 |
| 0 | 0 | 0 | 0 |
| 0 | 0 | 0 | 0 |
| 0 | 0 | 0 | 0 |
| 0 | 0 | 0 | 0 |
| 0 | 0 | 0 | 0 |
| 0 | 0 | 0 | 0 |
| 0 | 0 | 0 | 0 |
| 0 | 0 | 0 | 0 |
| 0 | 0 | 0 | 0 |
| 0 | 0 | 0 | 0 |
| 0 | 0 | 0 | 0 |
| 0 | 0 | 0 | 0 |
| 0 | 0 | 0 | 0 |
| 0 | 0 | 0 | 0 |
| 0 | 0 | 0 | 0 |
| 0 | 0 | 0 | 0 |
| 0 | 0 | 0 | 0 |
| 0 | 0 | 0 | 0 |
| 0 | 0 | 0 | 0 |
| 0 | 0 | 0 | 0 |
| 0 | 1 | 0 | 0 |

| | | | |
|---|---|---|---|
| 0 | 0 | 0 | 0 |
| 0 | 0 | 0 | 0 |
| 0 | 0 | 0 | 0 |
| 0 | 0 | 0 | 0 |
| 0 | 0 | 0 | 0 |
| 0 | 0 | 0 | 0 |
| 0 | 0 | 0 | 0 |
| 0 | 0 | 0 | 0 |
| 0 | 0 | 0 | 0 |
| 0 | 0 | 0 | 0 |
| 0 | 0 | 0 | 0 |
| 0 | 0 | 0 | 0 |
| 0 | 0 | 0 | 0 |
| 0 | 0 | 0 | 0 |
| 0 | 0 | 0 | 0 |
| 0 | 0 | 0 | 0 |
| 0 | 0 | 0 | 0 |
| 0 | 0 | 0 | 0 |
| 0 | 0 | 0 | 0 |
| 0 | 0 | 0 | 0 |
| 0 | 0 | 0 | 0 |
| 0 | 0 | 0 | 0 |
| 0 | 0 | 0 | 0 |
| 0 | 0 | 0 | 0 |
| 0 | 0 | 0 | 0 |
| 0 | 0 | 0 | 0 |
| 0 | 0 | 0 | 0 |
| 0 | 0 | 0 | 0 |
| 0 | 0 | 0 | 0 |
| 0 | 0 | 0 | 0 |
| 0 | 0 | 0 | 0 |
| 0 | 0 | 0 | 0 |
| 0 | 0 | 0 | 0 |
| 0 | 0 | 0 | 0 |
| 0 | 0 | 0 | 0 |
| 0 | 0 | 0 | 0 |
| 0 | 0 | 0 | 0 |
| 0 | 0 | 0 | 0 |
| 0 | 0 | 0 | 0 |
| 0 | 0 | 0 | 0 |
| 0 | 0 | 0 | 0 |
| 0 | 0 | 0 | 0 |
| 0 | 0 | 0 | 0 |
| 0 | 0 | 0 | 0 |
| 0 | 0 | 0 | 0 |
| 0 | 0 | 0 | 0 |
| 0 | 0 | 0 | 0 |
| 0 | 0 | 0 | 0 |
| 0 | 0 | 0 | 0 |
| 0 | 0 | 0 | 0 |
| 0 | 0 | 0 | 0 |
| 0 | 0 | 0 | 0 |
| 0 | 0 | 0 | 0 |
| 0 | 0 | 0 | 0 |

| | | | |
|---|---|---|---|
| 0 | 0 | 0 | 0 |
| 0 | 0 | 0 | 0 |
| 0 | 0 | 0 | 0 |
| 0 | 0 | 0 | 0 |
| 0 | 0 | 0 | 0 |
| 0 | 0 | 0 | 0 |
| 0 | 0 | 0 | 0 |
| 0 | 0 | 0 | 0 |
| 0 | 0 | 0 | 0 |
| 0 | 0 | 0 | 0 |
| 0 | 0 | 0 | 0 |
| 0 | 0 | 1 | 0 |
| 0 | 0 | 0 | 0 |
| 0 | 0 | 1 | 0 |
| 0 | 0 | 1 | 0 |
| 0 | 0 | 1 | 0 |
| 0 | 0 | 1 | 0 |
| 0 | 0 | 0 | 0 |
| 0 | 0 | 0 | 0 |
| 0 | 0 | 0 | 0 |
| 1 | 0 | 0 | 0 |

| glycan_binding_site | water_binding_site | water_only_binding_site | non_binding_site |
|---|---|---|---|
| 0 | 0 | 0 | 0 |
| 0 | 0 | 0 | 0 |
| 0 | 0 | 0 | 1 |
| 0 | 0 | 0 | 0 |
| 0 | 0 | 0 | 0 |
| 0 | 0 | 0 | 0 |
| 0 | 0 | 0 | 1 |
| 0 | 0 | 0 | 1 |
| 0 | 0 | 0 | 0 |
| 0 | 0 | 0 | 0 |
| 0 | 0 | 0 | 0 |
| 0 | 0 | 0 | 0 |
| 0 | 0 | 0 | 1 |
| 1 | 0 | 0 | 0 |
| 0 | 0 | 0 | 0 |
| 0 | 0 | 0 | 0 |
| 0 | 0 | 0 | 0 |
| 0 | 0 | 0 | 0 |
| 0 | 0 | 0 | 0 |
| 0 | 0 | 0 | 0 |
| 0 | 1 | 0 | 0 |
| 1 | 1 | 0 | 0 |
| 1 | 1 | 0 | 0 |
| 0 | 1 | 0 | 0 |
| 0 | 0 | 0 | 0 |
| 0 | 0 | 0 | 1 |
| 0 | 0 | 0 | 0 |
| 0 | 0 | 0 | 1 |
| 0 | 0 | 0 | 0 |
| 0 | 0 | 0 | 1 |
| 0 | 0 | 0 | 1 |
| 0 | 1 | 0 | 0 |
| 0 | 1 | 0 | 0 |
| 0 | 1 | 0 | 0 |
| 0 | 1 | 0 | 0 |
| 0 | 0 | 0 | 1 |
| 0 | 0 | 0 | 0 |
| 0 | 0 | 0 | 0 |
| 0 | 1 | 0 | 0 |
| 0 | 1 | 0 | 0 |
| 0 | 1 | 0 | 0 |
| 0 | 1 | 0 | 0 |
| 0 | 1 | 0 | 0 |
| 0 | 1 | 0 | 0 |
| 0 | 1 | 0 | 0 |
| 0 | 1 | 0 | 0 |
| 0 | 1 | 0 | 0 |
| 0 | 1 | 0 | 0 |
| 0 | 1 | 0 | 0 |
| 0 | 1 | 0 | 0 |
| 0 | 1 | 0 | 0 |
| 0 | 1 | 0 | 0 |
| 0 | 0 | 0 | 0 |

| | | | |
|---|---|---|---|
| 0 | 1 | 0 | 0 |
| 0 | 1 | 0 | 0 |
| 0 | 1 | 0 | 0 |
| 0 | 1 | 0 | 0 |
| 0 | 1 | 0 | 0 |
| 0 | 1 | 0 | 0 |
| 0 | 0 | 0 | 0 |
| 0 | 1 | 0 | 0 |
| 0 | 1 | 0 | 0 |
| 0 | 1 | 0 | 0 |
| 0 | 1 | 0 | 0 |
| 0 | 1 | 0 | 0 |
| 0 | 1 | 0 | 0 |
| 0 | 1 | 0 | 0 |
| 0 | 1 | 0 | 0 |
| 0 | 1 | 0 | 0 |
| 0 | 1 | 0 | 0 |
| 0 | 1 | 0 | 0 |
| 0 | 1 | 0 | 0 |
| 0 | 0 | 0 | 0 |
| 0 | 0 | 0 | 0 |
| 0 | 0 | 0 | 0 |
| 0 | 0 | 0 | 0 |
| 0 | 0 | 0 | 0 |
| 0 | 0 | 0 | 0 |
| 0 | 0 | 0 | 0 |
| 0 | 0 | 0 | 0 |
| 0 | 0 | 0 | 0 |
| 0 | 0 | 0 | 0 |
| 0 | 0 | 0 | 0 |
| 0 | 0 | 0 | 0 |
| 0 | 0 | 0 | 0 |
| 0 | 0 | 0 | 0 |
| 0 | 0 | 0 | 0 |
| 0 | 0 | 0 | 0 |
| 0 | 0 | 0 | 0 |
| 0 | 0 | 0 | 0 |
| 0 | 0 | 0 | 0 |
| 0 | 0 | 0 | 0 |
| 0 | 0 | 0 | 0 |
| 1 | 0 | 0 | 0 |
| 0 | 0 | 0 | 0 |
| 1 | 0 | 0 | 0 |
| 1 | 0 | 0 | 0 |
| 0 | 0 | 0 | 0 |
| 0 | 0 | 0 | 1 |
| 0 | 0 | 0 | 1 |
| 0 | 0 | 0 | 0 |
| 1 | 0 | 0 | 0 |
| 0 | 0 | 0 | 0 |
| 0 | 0 | 0 | 0 |
| 0 | 0 | 0 | 1 |
| 0 | 0 | 0 | 1 |
| 0 | 0 | 0 | 0 |

| | | | |
|---|---|---|---|
| 0 | 0 | 0 | 1 |
| 0 | 0 | 0 | 1 |
| 0 | 0 | 0 | 0 |
| 0 | 0 | 0 | 0 |
| 0 | 0 | 0 | 0 |
| 0 | 0 | 0 | 0 |
| 0 | 0 | 0 | 1 |
| 0 | 0 | 0 | 0 |
| 0 | 0 | 0 | 0 |
| 0 | 0 | 0 | 0 |
| 0 | 0 | 0 | 0 |
| 0 | 0 | 0 | 0 |
| 0 | 0 | 0 | 0 |
| 1 | 0 | 0 | 0 |
| 1 | 0 | 0 | 0 |
| 0 | 0 | 0 | 1 |
| 0 | 0 | 0 | 0 |
| 0 | 0 | 0 | 0 |
| 0 | 0 | 0 | 0 |
| 0 | 0 | 0 | 0 |
| 0 | 0 | 0 | 0 |
| 0 | 0 | 0 | 0 |
| 0 | 0 | 0 | 0 |
| 0 | 0 | 0 | 0 |
| 0 | 0 | 0 | 0 |
| 0 | 0 | 0 | 0 |
| 0 | 0 | 0 | 0 |
| 0 | 0 | 0 | 0 |
| 0 | 0 | 0 | 0 |
| 0 | 0 | 0 | 0 |
| 0 | 0 | 0 | 0 |
| 0 | 0 | 0 | 0 |
| 0 | 0 | 0 | 0 |
| 0 | 0 | 0 | 0 |
| 0 | 0 | 0 | 0 |
| 0 | 0 | 0 | 0 |
| 0 | 0 | 0 | 0 |
| 0 | 0 | 0 | 0 |
| 0 | 0 | 0 | 0 |
| 0 | 0 | 0 | 0 |
| 0 | 0 | 0 | 0 |
| 0 | 0 | 0 | 0 |
| 0 | 0 | 0 | 0 |
| 0 | 0 | 0 | 0 |
| 0 | 0 | 0 | 0 |
| 0 | 1 | 0 | 0 |
| 1 | 0 | 0 | 0 |
| 0 | 0 | 0 | 0 |
| 0 | 0 | 0 | 0 |
| 0 | 0 | 0 | 0 |
| 0 | 0 | 0 | 0 |
| 0 | 0 | 0 | 0 |
| 0 | 0 | 0 | 0 |
| 0 | 1 | 1 | 1 |

| | | | |
|---|---|---|---|
| 0 | 1 | 1 | 1 |
| 0 | 1 | 1 | 1 |
| 0 | 1 | 1 | 1 |
| 0 | 0 | 0 | 0 |
| 0 | 0 | 0 | 0 |
| 0 | 1 | 1 | 1 |
| 0 | 1 | 0 | 0 |
| 0 | 1 | 1 | 1 |
| 0 | 1 | 1 | 1 |
| 0 | 0 | 0 | 1 |
| 0 | 1 | 1 | 1 |
| 0 | 1 | 0 | 0 |
| 0 | 1 | 1 | 1 |
| 0 | 0 | 0 | 0 |
| 0 | 1 | 1 | 1 |
| 0 | 0 | 0 | 1 |
| 0 | 1 | 1 | 1 |
| 0 | 1 | 1 | 1 |
| 0 | 1 | 1 | 1 |
| 0 | 1 | 0 | 0 |
| 0 | 1 | 0 | 0 |
| 0 | 0 | 0 | 1 |
| 0 | 0 | 0 | 0 |
| 0 | 0 | 0 | 0 |
| 0 | 0 | 0 | 0 |
| 0 | 0 | 0 | 1 |
| 0 | 0 | 0 | 0 |
| 0 | 1 | 0 | 0 |
| 0 | 0 | 0 | 1 |
| 0 | 0 | 0 | 1 |
| 0 | 0 | 0 | 1 |
| 0 | 0 | 0 | 1 |
| 0 | 1 | 1 | 1 |
| 0 | 0 | 0 | 1 |
| 0 | 0 | 0 | 0 |
| 0 | 1 | 1 | 1 |
| 1 | 1 | 0 | 0 |
| 1 | 1 | 0 | 0 |
| 0 | 1 | 1 | 1 |
| 0 | 1 | 1 | 1 |
| 0 | 0 | 0 | 1 |
| 0 | 1 | 0 | 0 |
| 0 | 1 | 0 | 0 |
| 0 | 0 | 0 | 1 |
| 0 | 1 | 1 | 1 |
| 0 | 1 | 1 | 1 |
| 0 | 0 | 0 | 0 |
| 0 | 0 | 0 | 0 |
| 0 | 0 | 0 | 0 |
| 0 | 0 | 0 | 0 |
| 0 | 0 | 0 | 0 |
| 0 | 0 | 0 | 1 |
| 0 | 0 | 0 | 1 |
| 0 | 0 | 0 | 0 |

| | | | |
|---|---|---|---|
| 0 | 0 | 0 | 0 |
| 0 | 0 | 0 | 0 |
| 0 | 1 | 0 | 0 |
| 0 | 0 | 0 | 1 |
| 0 | 0 | 0 | 1 |
| 0 | 0 | 0 | 1 |
| 0 | 0 | 0 | 0 |
| 0 | 0 | 0 | 0 |
| 0 | 0 | 0 | 1 |
| 0 | 0 | 0 | 1 |
| 1 | 0 | 0 | 0 |
| 0 | 0 | 0 | 1 |
| 1 | 0 | 0 | 0 |
| 0 | 0 | 0 | 1 |
| 0 | 0 | 0 | 1 |
| 0 | 0 | 0 | 1 |
| 0 | 0 | 0 | 1 |
| 0 | 0 | 0 | 0 |
| 0 | 0 | 0 | 0 |
| 0 | 0 | 0 | 0 |
| 0 | 0 | 0 | 0 |
| 0 | 0 | 0 | 1 |
| 0 | 0 | 0 | 0 |
| 0 | 0 | 0 | 1 |
| 0 | 0 | 0 | 1 |
| 0 | 0 | 0 | 0 |
| 0 | 1 | 0 | 0 |
| 0 | 0 | 0 | 0 |
| 0 | 1 | 0 | 0 |
| 0 | 1 | 0 | 0 |
| 0 | 1 | 0 | 0 |
| 0 | 0 | 0 | 0 |
| 0 | 0 | 0 | 1 |
| 0 | 0 | 0 | 1 |
| 0 | 0 | 0 | 0 |
| 0 | 0 | 0 | 1 |
| 0 | 0 | 0 | 1 |
| 0 | 0 | 0 | 0 |
| 0 | 0 | 0 | 1 |
| 0 | 0 | 0 | 1 |
| 0 | 0 | 0 | 0 |
| 0 | 0 | 0 | 1 |
| 0 | 0 | 0 | 1 |
| 0 | 0 | 0 | 1 |
| 0 | 0 | 0 | 0 |
| 0 | 0 | 0 | 0 |
| 0 | 1 | 0 | 0 |
| 0 | 1 | 0 | 0 |
| 0 | 1 | 0 | 0 |
| 0 | 1 | 0 | 0 |
| 0 | 1 | 0 | 0 |
| 0 | 1 | 0 | 0 |
| 0 | 1 | 0 | 0 |
| 0 | 1 | 0 | 0 |

| | | | |
|---|---|---|---|
| 0 | 1 | 0 | 0 |
| 0 | 1 | 0 | 0 |
| 0 | 1 | 0 | 0 |
| 0 | 0 | 0 | 0 |
| 0 | 0 | 0 | 0 |
| 0 | 1 | 0 | 0 |
| 0 | 0 | 0 | 1 |
| 0 | 0 | 0 | 1 |
| 0 | 0 | 0 | 0 |
| 0 | 0 | 0 | 0 |
| 0 | 0 | 0 | 0 |
| 0 | 0 | 0 | 0 |
| 0 | 0 | 0 | 0 |
| 0 | 0 | 0 | 1 |
| 0 | 0 | 0 | 1 |
| 0 | 0 | 0 | 0 |
| 0 | 0 | 0 | 1 |
| 0 | 0 | 0 | 0 |
| 0 | 0 | 0 | 1 |
| 0 | 0 | 0 | 0 |
| 0 | 0 | 0 | 1 |
| 0 | 0 | 0 | 1 |
| 0 | 0 | 0 | 1 |
| 0 | 0 | 0 | 0 |
| 0 | 0 | 0 | 0 |
| 0 | 0 | 0 | 0 |
| 0 | 0 | 0 | 0 |
| 0 | 0 | 0 | 0 |
| 0 | 0 | 0 | 0 |
| 0 | 0 | 0 | 0 |
| 0 | 0 | 0 | 1 |
| 0 | 0 | 0 | 1 |
| 0 | 0 | 0 | 0 |
| 0 | 0 | 0 | 0 |
| 0 | 0 | 0 | 0 |
| 0 | 0 | 0 | 1 |
| 0 | 0 | 0 | 0 |
| 0 | 0 | 0 | 0 |
| 0 | 0 | 0 | 0 |
| 0 | 0 | 0 | 0 |
| 0 | 0 | 0 | 1 |
| 0 | 0 | 0 | 1 |
| 0 | 0 | 0 | 0 |
| 0 | 0 | 0 | 1 |
| 0 | 0 | 0 | 0 |
| 0 | 0 | 0 | 0 |
| 0 | 0 | 0 | 0 |
| 0 | 0 | 0 | 1 |
| 0 | 0 | 0 | 1 |
| 0 | 0 | 0 | 0 |
| 0 | 0 | 0 | 1 |
| 0 | 0 | 0 | 1 |
| 0 | 0 | 0 | 0 |
| 0 | 0 | 0 | 1 |

| | | | |
|---|---|---|---|
| 0 | 0 | 0 | 0 |
| 0 | 0 | 0 | 1 |
| 0 | 0 | 0 | 1 |
| 0 | 0 | 0 | 0 |
| 0 | 0 | 0 | 0 |
| 0 | 0 | 0 | 0 |
| 0 | 0 | 0 | 1 |
| 0 | 0 | 0 | 0 |
| 0 | 0 | 0 | 0 |
| 0 | 0 | 0 | 1 |
| 0 | 0 | 0 | 1 |
| 0 | 0 | 0 | 1 |
| 0 | 0 | 0 | 1 |
| 0 | 0 | 0 | 1 |
| 0 | 0 | 0 | 0 |
| 0 | 0 | 0 | 1 |
| 0 | 0 | 0 | 0 |
| 0 | 0 | 0 | 0 |
| 0 | 0 | 0 | 1 |
| 0 | 0 | 0 | 0 |
| 0 | 1 | 0 | 0 |
| 0 | 1 | 0 | 0 |
| 0 | 0 | 0 | 0 |
| 0 | 0 | 0 | 0 |
| 0 | 0 | 0 | 1 |
| 0 | 0 | 0 | 1 |
| 0 | 0 | 0 | 0 |
| 0 | 0 | 0 | 0 |
| 0 | 0 | 0 | 0 |
| 0 | 0 | 0 | 0 |
| 0 | 0 | 0 | 0 |
| 0 | 0 | 0 | 0 |
| 0 | 0 | 0 | 0 |
| 0 | 0 | 0 | 0 |
| 0 | 0 | 0 | 1 |
| 0 | 0 | 0 | 1 |
| 0 | 0 | 0 | 0 |
| 0 | 0 | 0 | 0 |
| 0 | 0 | 0 | 0 |
| 0 | 0 | 0 | 0 |
| 0 | 0 | 0 | 0 |
| 0 | 0 | 0 | 0 |
| 0 | 0 | 0 | 1 |
| 0 | 0 | 0 | 0 |
| 0 | 1 | 0 | 0 |
| 0 | 1 | 0 | 0 |
| 0 | 1 | 0 | 0 |
| 0 | 0 | 0 | 0 |
| 1 | 1 | 0 | 0 |
| 0 | 0 | 0 | 1 |
| 0 | 0 | 0 | 0 |
| 0 | 0 | 0 | 0 |
| 0 | 0 | 0 | 0 |
| 0 | 0 | 0 | 0 |

| | | | |
|---|---|---|---|
| 0 | 0 | 0 | 0 |
| 0 | 0 | 0 | 0 |
| 0 | 0 | 0 | 0 |
| 0 | 0 | 0 | 0 |
| 0 | 0 | 0 | 1 |
| 0 | 0 | 0 | 0 |
| 0 | 0 | 0 | 0 |
| 0 | 0 | 0 | 0 |
| 0 | 0 | 0 | 0 |
| 0 | 0 | 0 | 0 |
| 0 | 1 | 0 | 0 |
| 0 | 0 | 0 | 0 |
| 0 | 0 | 0 | 0 |
| 0 | 0 | 0 | 0 |
| 0 | 0 | 0 | 0 |
| 0 | 0 | 0 | 0 |
| 0 | 0 | 0 | 0 |
| 0 | 0 | 0 | 0 |
| 0 | 0 | 0 | 1 |
| 0 | 0 | 0 | 0 |
| 0 | 0 | 0 | 0 |
| 0 | 0 | 0 | 0 |
| 0 | 0 | 0 | 0 |
| 0 | 0 | 0 | 1 |
| 0 | 0 | 0 | 0 |
| 0 | 0 | 0 | 0 |
| 0 | 0 | 0 | 0 |
| 0 | 0 | 0 | 1 |
| 0 | 0 | 0 | 1 |
| 0 | 0 | 0 | 1 |
| 0 | 0 | 0 | 0 |
| 0 | 0 | 0 | 0 |
| 0 | 0 | 0 | 0 |
| 0 | 0 | 0 | 0 |
| 0 | 0 | 0 | 0 |
| 0 | 0 | 0 | 1 |
| 0 | 0 | 0 | 1 |
| 0 | 0 | 0 | 0 |
| 0 | 0 | 0 | 0 |
| 0 | 0 | 0 | 1 |
| 0 | 0 | 0 | 1 |
| 0 | 0 | 0 | 1 |
| 0 | 0 | 0 | 1 |
| 0 | 0 | 0 | 1 |
| 0 | 0 | 0 | 0 |
| 0 | 0 | 0 | 0 |
| 0 | 0 | 0 | 0 |
| 0 | 0 | 0 | 0 |
| 0 | 0 | 0 | 0 |
| 0 | 0 | 0 | 1 |
| 0 | 0 | 0 | 1 |
| 0 | 0 | 0 | 0 |
| 0 | 0 | 0 | 1 |
| 0 | 0 | 0 | 0 |

| | | | |
|---|---|---|---|
| 0 | 0 | 0 | 1 |
| 0 | 0 | 0 | 1 |
| 0 | 0 | 0 | 0 |
| 0 | 0 | 0 | 1 |
| 0 | 0 | 0 | 1 |
| 0 | 0 | 0 | 1 |
| 0 | 0 | 0 | 1 |
| 0 | 0 | 0 | 0 |
| 0 | 0 | 0 | 1 |
| 0 | 0 | 0 | 1 |
| 0 | 0 | 0 | 0 |
| 0 | 0 | 0 | 1 |
| 0 | 0 | 0 | 1 |
| 0 | 0 | 0 | 1 |
| 0 | 0 | 0 | 0 |
| 0 | 0 | 0 | 0 |
| 0 | 0 | 0 | 1 |
| 0 | 0 | 0 | 1 |
| 0 | 0 | 0 | 0 |
| 0 | 0 | 0 | 0 |
| 0 | 0 | 0 | 0 |
| 0 | 0 | 0 | 0 |
| 0 | 0 | 0 | 0 |
| 0 | 1 | 0 | 0 |
| 0 | 1 | 0 | 0 |
| 0 | 0 | 0 | 1 |
| 0 | 0 | 0 | 0 |
| 0 | 0 | 0 | 0 |
| 0 | 0 | 0 | 0 |
| 0 | 0 | 0 | 1 |
| 0 | 0 | 0 | 0 |
| 0 | 1 | 0 | 0 |
| 0 | 0 | 0 | 1 |
| 0 | 0 | 0 | 0 |
| 0 | 0 | 0 | 0 |
| 0 | 0 | 0 | 0 |
| 0 | 1 | 0 | 0 |
| 0 | 1 | 0 | 0 |
| 0 | 1 | 0 | 0 |
| 0 | 1 | 0 | 0 |
| 0 | 0 | 0 | 1 |
| 0 | 0 | 0 | 0 |
| 0 | 0 | 0 | 1 |
| 0 | 0 | 0 | 0 |
| 0 | 0 | 0 | 0 |
| 1 | 0 | 0 | 0 |
| 0 | 0 | 0 | 0 |
| 1 | 0 | 0 | 0 |
| 0 | 0 | 0 | 0 |
| 0 | 0 | 0 | 0 |
| 0 | 0 | 0 | 1 |
| 0 | 0 | 0 | 0 |
| 0 | 0 | 0 | 0 |
| 0 | 0 | 0 | 0 |

| | | | |
|---|---|---|---|
| 0 | 0 | 0 | 1 |
| 0 | 0 | 0 | 0 |
| 0 | 0 | 0 | 1 |
| 0 | 0 | 0 | 1 |
| 1 | 0 | 0 | 0 |
| 0 | 0 | 0 | 0 |
| 1 | 0 | 0 | 0 |
| 1 | 0 | 0 | 0 |
| 0 | 0 | 0 | 0 |
| 0 | 0 | 0 | 1 |
| 0 | 0 | 0 | 1 |
| 0 | 0 | 0 | 1 |
| 0 | 0 | 0 | 1 |
| 1 | 0 | 0 | 0 |
| 0 | 0 | 0 | 1 |
| 0 | 0 | 0 | 0 |
| 0 | 0 | 0 | 0 |
| 0 | 0 | 0 | 0 |
| 1 | 0 | 0 | 0 |
| 0 | 0 | 0 | 1 |
| 1 | 0 | 0 | 0 |
| 0 | 0 | 0 | 1 |
| 0 | 0 | 0 | 0 |
| 0 | 0 | 0 | 1 |
| 0 | 0 | 0 | 1 |
| 0 | 0 | 0 | 1 |
| 0 | 0 | 0 | 0 |
| 0 | 0 | 0 | 1 |
| 0 | 0 | 0 | 1 |
| 0 | 0 | 0 | 1 |
| 0 | 0 | 0 | 0 |
| 0 | 0 | 0 | 0 |
| 0 | 0 | 0 | 1 |
| 0 | 0 | 0 | 1 |
| 0 | 0 | 0 | 0 |
| 0 | 0 | 0 | 0 |
| 0 | 0 | 0 | 0 |
| 1 | 0 | 0 | 0 |
| 0 | 0 | 0 | 0 |
| 0 | 0 | 0 | 1 |
| 0 | 0 | 0 | 0 |
| 0 | 0 | 0 | 0 |
| 0 | 0 | 0 | 0 |
| 0 | 0 | 0 | 0 |
| 0 | 0 | 0 | 1 |
| 1 | 0 | 0 | 0 |
| 0 | 0 | 0 | 0 |
| 0 | 0 | 0 | 0 |
| 0 | 0 | 0 | 0 |
| 1 | 0 | 0 | 0 |
| 0 | 0 | 0 | 0 |
| 0 | 1 | 0 | 0 |
| 0 | 0 | 0 | 1 |
| 0 | 0 | 0 | 0 |

| | | | |
|---|---|---|---|
| 0 | 0 | 0 | 0 |
| 0 | 1 | 0 | 0 |
| 0 | 1 | 1 | 1 |
| 0 | 1 | 1 | 1 |
| 0 | 0 | 0 | 1 |
| 0 | 0 | 0 | 0 |
| 0 | 0 | 0 | 1 |
| 0 | 0 | 0 | 1 |
| 0 | 1 | 0 | 0 |
| 0 | 0 | 0 | 0 |
| 1 | 0 | 0 | 0 |
| 1 | 0 | 0 | 0 |
| 0 | 0 | 0 | 1 |
| 1 | 0 | 0 | 0 |
| 0 | 0 | 0 | 0 |
| 0 | 0 | 0 | 0 |
| 0 | 1 | 0 | 0 |
| 0 | 0 | 0 | 0 |
| 0 | 0 | 0 | 0 |
| 0 | 0 | 0 | 1 |
| 0 | 0 | 0 | 0 |
| 0 | 0 | 0 | 0 |
| 0 | 0 | 0 | 0 |
| 0 | 0 | 0 | 0 |
| 0 | 0 | 0 | 0 |
| 0 | 0 | 0 | 1 |
| 0 | 0 | 0 | 1 |
| 0 | 0 | 0 | 1 |
| 0 | 0 | 0 | 0 |
| 0 | 0 | 0 | 0 |
| 0 | 0 | 0 | 0 |
| 0 | 0 | 0 | 0 |
| 0 | 0 | 0 | 0 |
| 0 | 1 | 0 | 0 |
| 0 | 1 | 0 | 0 |
| 0 | 0 | 0 | 1 |
| 0 | 0 | 0 | 0 |
| 0 | 0 | 0 | 0 |
| 0 | 0 | 0 | 1 |
| 0 | 0 | 0 | 1 |
| 0 | 0 | 0 | 1 |
| 0 | 0 | 0 | 0 |
| 0 | 0 | 0 | 0 |
| 0 | 0 | 0 | 0 |
| 0 | 0 | 0 | 0 |
| 0 | 0 | 0 | 0 |
| 0 | 0 | 0 | 0 |
| 0 | 0 | 0 | 1 |
| 0 | 0 | 0 | 1 |
| 0 | 0 | 0 | 0 |
| 0 | 0 | 0 | 0 |
| 0 | 0 | 0 | 0 |
| 0 | 0 | 0 | 0 |
| 0 | 0 | 0 | 1 |

| | | | |
|---|---|---|---|
| 0 | 0 | 0 | 1 |
| 0 | 0 | 0 | 0 |
| 0 | 0 | 0 | 1 |
| 0 | 1 | 0 | 0 |
| 0 | 0 | 0 | 1 |
| 0 | 0 | 0 | 1 |
| 0 | 0 | 0 | 0 |
| 0 | 1 | 1 | 1 |
| 0 | 0 | 0 | 1 |
| 0 | 1 | 1 | 1 |
| 0 | 1 | 1 | 1 |
| 0 | 0 | 0 | 1 |
| 0 | 0 | 0 | 1 |
| 0 | 0 | 0 | 1 |
| 0 | 0 | 0 | 1 |
| 0 | 0 | 0 | 1 |
| 0 | 0 | 0 | 0 |
| 0 | 0 | 0 | 0 |
| 0 | 0 | 0 | 1 |
| 0 | 0 | 0 | 0 |
| 0 | 0 | 0 | 0 |
| 0 | 0 | 0 | 0 |
| 0 | 0 | 0 | 1 |
| 0 | 0 | 0 | 0 |
| 0 | 0 | 0 | 0 |
| 0 | 0 | 0 | 1 |
| 0 | 0 | 0 | 1 |
| 0 | 0 | 0 | 1 |
| 0 | 1 | 0 | 0 |
| 0 | 0 | 0 | 0 |
| 0 | 0 | 0 | 1 |
| 0 | 0 | 0 | 0 |
| 0 | 0 | 0 | 0 |
| 0 | 0 | 0 | 0 |
| 0 | 0 | 0 | 0 |
| 0 | 0 | 0 | 0 |
| 0 | 1 | 1 | 1 |
| 0 | 0 | 0 | 1 |
| 0 | 0 | 0 | 1 |
| 0 | 0 | 0 | 1 |
| 0 | 0 | 0 | 0 |
| 0 | 0 | 0 | 1 |
| 0 | 0 | 0 | 0 |
| 0 | 0 | 0 | 1 |
| 1 | 0 | 0 | 0 |
| 0 | 0 | 0 | 1 |
| 0 | 0 | 0 | 0 |
| 0 | 0 | 0 | 0 |
| 1 | 0 | 0 | 0 |
| 0 | 0 | 0 | 0 |
| 0 | 0 | 0 | 0 |
| 0 | 0 | 0 | 0 |
| 0 | 0 | 0 | 0 |
| 0 | 0 | 0 | 0 |

| | | | |
|---|---|---|---|
| 0 | 0 | 0 | 0 |
| 0 | 0 | 0 | 0 |
| 0 | 0 | 0 | 0 |
| 0 | 0 | 0 | 0 |
| 0 | 0 | 0 | 0 |
| 0 | 0 | 0 | 0 |
| 0 | 0 | 0 | 0 |
| 0 | 0 | 0 | 1 |
| 0 | 1 | 0 | 0 |
| 0 | 1 | 0 | 0 |
| 0 | 0 | 0 | 0 |
| 0 | 1 | 1 | 1 |
| 0 | 0 | 0 | 0 |
| 0 | 1 | 0 | 0 |
| 0 | 1 | 0 | 0 |
| 0 | 1 | 0 | 0 |
| 0 | 1 | 0 | 0 |
| 0 | 1 | 0 | 0 |
| 0 | 1 | 0 | 0 |
| 0 | 1 | 0 | 0 |
| 0 | 1 | 0 | 0 |
| 0 | 0 | 0 | 0 |
| 0 | 1 | 0 | 0 |
| 0 | 1 | 0 | 0 |
| 0 | 1 | 0 | 0 |
| 0 | 1 | 0 | 0 |
| 0 | 1 | 0 | 0 |
| 0 | 1 | 0 | 0 |
| 0 | 1 | 0 | 0 |
| 0 | 1 | 0 | 0 |
| 0 | 1 | 0 | 0 |
| 0 | 1 | 0 | 0 |
| 0 | 1 | 0 | 0 |
| 0 | 1 | 0 | 0 |
| 0 | 1 | 0 | 0 |
| 0 | 1 | 0 | 0 |
| 0 | 1 | 0 | 0 |
| 0 | 1 | 0 | 0 |
| 0 | 1 | 0 | 0 |
| 0 | 1 | 0 | 0 |
| 0 | 1 | 1 | 1 |
| 0 | 1 | 0 | 0 |
| 0 | 1 | 0 | 0 |
| 0 | 1 | 0 | 0 |
| 0 | 1 | 0 | 0 |
| 0 | 0 | 0 | 0 |
| 0 | 0 | 0 | 0 |
| 0 | 0 | 0 | 0 |
| 0 | 0 | 0 | 0 |
| 0 | 0 | 0 | 1 |
| 0 | 0 | 0 | 1 |
| 0 | 1 | 1 | 1 |
| 1 | 0 | 0 | 0 |
| 0 | 0 | 0 | 0 |

| | | | |
|---|---|---|---|
| 0 | 0 | 0 | 0 |
| 0 | 0 | 0 | 0 |
| 1 | 0 | 0 | 0 |
| 1 | 0 | 0 | 0 |
| 0 | 0 | 0 | 0 |
| 0 | 0 | 0 | 0 |
| 0 | 0 | 0 | 0 |
| 0 | 0 | 0 | 1 |
| 0 | 0 | 0 | 1 |
| 0 | 0 | 0 | 0 |
| 0 | 0 | 0 | 0 |
| 0 | 0 | 0 | 1 |
| 0 | 0 | 0 | 0 |
| 0 | 0 | 0 | 0 |
| 0 | 0 | 0 | 0 |
| 0 | 0 | 0 | 0 |
| 0 | 0 | 0 | 0 |
| 0 | 0 | 0 | 1 |
| 0 | 0 | 0 | 0 |
| 0 | 0 | 0 | 1 |
| 0 | 0 | 0 | 0 |
| 0 | 0 | 0 | 0 |
| 0 | 0 | 0 | 1 |
| 0 | 0 | 0 | 0 |
| 0 | 0 | 0 | 0 |
| 0 | 0 | 0 | 1 |
| 0 | 0 | 0 | 0 |
| 0 | 0 | 0 | 0 |
| 0 | 0 | 0 | 0 |
| 0 | 0 | 0 | 0 |
| 0 | 0 | 0 | 0 |
| 0 | 0 | 0 | 0 |
| 0 | 0 | 0 | 1 |
| 0 | 0 | 0 | 0 |
| 0 | 1 | 0 | 0 |
| 0 | 0 | 0 | 0 |
| 0 | 0 | 0 | 1 |
| 0 | 0 | 0 | 0 |
| 0 | 0 | 0 | 0 |
| 0 | 0 | 0 | 0 |
| 0 | 1 | 0 | 0 |
| 0 | 1 | 0 | 0 |
| 0 | 1 | 0 | 0 |
| 0 | 1 | 0 | 0 |
| 0 | 1 | 0 | 0 |
| 0 | 0 | 0 | 1 |
| 0 | 0 | 0 | 0 |
| 0 | 1 | 0 | 0 |
| 0 | 0 | 0 | 0 |
| 0 | 1 | 0 | 0 |
| 0 | 0 | 0 | 0 |
| 0 | 1 | 0 | 0 |
| 0 | 0 | 0 | 1 |
| 0 | 0 | 0 | 1 |

| | | | |
|---|---|---|---|
| 0 | 0 | 0 | 1 |
| 0 | 0 | 0 | 1 |
| 0 | 0 | 0 | 0 |
| 0 | 0 | 0 | 0 |
| 0 | 1 | 0 | 0 |
| 0 | 1 | 0 | 0 |
| 0 | 0 | 0 | 0 |
| 0 | 0 | 0 | 1 |
| 0 | 0 | 0 | 0 |
| 0 | 0 | 0 | 1 |
| 0 | 0 | 0 | 1 |
| 0 | 0 | 0 | 1 |
| 0 | 0 | 0 | 0 |
| 0 | 0 | 0 | 0 |
| 0 | 0 | 0 | 0 |
| 0 | 0 | 0 | 0 |
| 0 | 0 | 0 | 0 |
| 1 | 0 | 0 | 0 |
| 0 | 0 | 0 | 1 |
| 0 | 0 | 0 | 1 |
| 0 | 0 | 0 | 1 |
| 0 | 0 | 0 | 0 |
| 0 | 0 | 0 | 1 |
| 0 | 0 | 0 | 0 |
| 0 | 0 | 0 | 0 |
| 0 | 0 | 0 | 0 |
| 0 | 0 | 0 | 0 |
| 0 | 0 | 0 | 0 |
| 0 | 0 | 0 | 0 |
| 0 | 0 | 0 | 0 |
| 0 | 0 | 0 | 0 |
| 0 | 0 | 0 | 0 |
| 0 | 0 | 0 | 0 |
| 0 | 0 | 0 | 0 |
| 0 | 0 | 0 | 0 |
| 0 | 0 | 0 | 0 |
| 0 | 0 | 0 | 0 |
| 0 | 0 | 0 | 0 |
| 1 | 0 | 0 | 0 |
| 0 | 0 | 0 | 0 |
| 0 | 1 | 0 | 0 |
| 0 | 0 | 0 | 1 |
| 0 | 0 | 0 | 0 |
| 0 | 1 | 0 | 0 |
| 0 | 0 | 0 | 0 |
| 0 | 0 | 0 | 0 |
| 0 | 0 | 0 | 1 |
| 0 | 0 | 0 | 0 |
| 0 | 0 | 0 | 0 |
| 0 | 0 | 0 | 0 |
| 0 | 0 | 0 | 0 |
| 0 | 0 | 0 | 0 |
| 0 | 0 | 0 | 1 |
| 0 | 0 | 0 | 1 |

| | | | |
|---|---|---|---|
| 0 | 0 | 0 | 0 |
| 0 | 0 | 0 | 0 |
| 0 | 0 | 0 | 1 |
| 0 | 0 | 0 | 1 |
| 0 | 0 | 0 | 1 |
| 0 | 1 | 0 | 0 |
| 0 | 0 | 0 | 1 |
| 0 | 0 | 0 | 1 |
| 0 | 0 | 0 | 0 |
| 0 | 0 | 0 | 0 |
| 0 | 0 | 0 | 0 |
| 0 | 0 | 0 | 0 |
| 0 | 0 | 0 | 0 |
| 0 | 0 | 0 | 1 |
| 0 | 0 | 0 | 1 |
| 0 | 0 | 0 | 1 |
| 0 | 0 | 0 | 1 |
| 0 | 0 | 0 | 1 |
| 0 | 0 | 0 | 0 |
| 0 | 0 | 0 | 0 |
| 0 | 0 | 0 | 1 |
| 0 | 0 | 0 | 1 |
| 0 | 0 | 0 | 0 |
| 0 | 1 | 0 | 0 |
| 0 | 0 | 0 | 1 |
| 0 | 0 | 0 | 1 |
| 0 | 1 | 0 | 0 |
| 0 | 1 | 1 | 1 |
| 0 | 0 | 0 | 0 |
| 0 | 0 | 0 | 0 |
| 0 | 0 | 0 | 0 |
| 0 | 0 | 0 | 0 |
| 0 | 0 | 0 | 0 |
| 1 | 0 | 0 | 0 |
| 1 | 1 | 0 | 0 |
| 0 | 0 | 0 | 0 |
| 0 | 0 | 0 | 0 |
| 0 | 0 | 0 | 0 |
| 0 | 0 | 0 | 1 |
| 0 | 0 | 0 | 0 |
| 0 | 0 | 0 | 1 |
| 0 | 0 | 0 | 1 |
| 0 | 0 | 0 | 1 |
| 0 | 0 | 0 | 1 |
| 0 | 0 | 0 | 1 |
| 0 | 0 | 0 | 0 |
| 0 | 0 | 0 | 0 |
| 0 | 0 | 0 | 0 |
| 0 | 0 | 0 | 0 |
| 0 | 0 | 0 | 0 |
| 0 | 0 | 0 | 0 |
| 0 | 0 | 0 | 0 |
| 0 | 0 | 0 | 0 |
| 0 | 0 | 0 | 1 |

| | | | |
|---|---|---|---|
| 0 | 0 | 0 | 1 |
| 0 | 0 | 0 | 0 |
| 0 | 0 | 0 | 0 |
| 0 | 0 | 0 | 0 |
| 0 | 0 | 0 | 1 |
| 0 | 0 | 0 | 0 |
| 0 | 0 | 0 | 0 |
| 0 | 0 | 0 | 0 |
| 0 | 0 | 0 | 1 |
| 0 | 0 | 0 | 1 |
| 0 | 0 | 0 | 0 |
| 0 | 0 | 0 | 0 |
| 0 | 0 | 0 | 0 |
| 0 | 0 | 0 | 1 |
| 0 | 0 | 0 | 0 |
| 0 | 0 | 0 | 0 |
| 0 | 0 | 0 | 0 |
| 0 | 1 | 0 | 0 |
| 0 | 0 | 0 | 0 |
| 0 | 0 | 0 | 0 |
| 0 | 0 | 0 | 0 |
| 0 | 0 | 0 | 1 |
| 0 | 0 | 0 | 0 |
| 0 | 0 | 0 | 1 |
| 0 | 0 | 0 | 0 |
| 0 | 0 | 0 | 0 |
| 0 | 0 | 0 | 1 |
| 0 | 0 | 0 | 0 |
| 0 | 0 | 0 | 0 |
| 0 | 0 | 0 | 0 |
| 0 | 0 | 0 | 0 |
| 0 | 0 | 0 | 0 |
| 0 | 0 | 0 | 0 |
| 0 | 0 | 0 | 0 |
| 0 | 0 | 0 | 1 |
| 0 | 1 | 0 | 0 |
| 0 | 0 | 0 | 0 |
| 0 | 0 | 0 | 0 |
| 0 | 0 | 0 | 0 |
| 0 | 0 | 0 | 0 |
| 0 | 0 | 0 | 0 |
| 0 | 0 | 0 | 0 |
| 0 | 0 | 0 | 0 |
| 0 | 0 | 0 | 0 |
| 0 | 0 | 0 | 0 |
| 0 | 0 | 0 | 0 |
| 0 | 0 | 0 | 0 |
| 0 | 0 | 0 | 1 |
| 0 | 0 | 0 | 0 |
| 0 | 0 | 0 | 1 |
| 0 | 0 | 0 | 0 |
| 0 | 0 | 0 | 0 |
| 0 | 0 | 0 | 0 |
| 0 | 0 | 0 | 0 |

| | | | |
|---|---|---|---|
| 0 | 0 | 0 | 0 |
| 0 | 0 | 0 | 0 |
| 0 | 0 | 0 | 1 |
| 0 | 0 | 0 | 0 |
| 0 | 0 | 0 | 0 |
| 0 | 0 | 0 | 1 |
| 0 | 0 | 0 | 1 |
| 0 | 0 | 0 | 1 |
| 0 | 0 | 0 | 0 |
| 0 | 0 | 0 | 1 |
| 0 | 0 | 0 | 0 |
| 0 | 0 | 0 | 0 |
| 0 | 0 | 0 | 0 |
| 0 | 0 | 0 | 1 |
| 0 | 0 | 0 | 0 |
| 0 | 0 | 0 | 1 |
| 0 | 0 | 0 | 0 |
| 0 | 0 | 0 | 0 |
| 1 | 0 | 0 | 0 |
| 0 | 0 | 0 | 1 |
| 0 | 0 | 0 | 1 |
| 0 | 0 | 0 | 0 |
| 0 | 0 | 0 | 1 |
| 0 | 0 | 0 | 0 |
| 0 | 0 | 0 | 0 |
| 0 | 0 | 0 | 1 |
| 0 | 0 | 0 | 1 |
| 0 | 0 | 0 | 0 |
| 0 | 0 | 0 | 1 |
| 1 | 0 | 0 | 0 |
| 0 | 0 | 0 | 1 |
| 0 | 0 | 0 | 1 |
| 0 | 0 | 0 | 1 |
| 0 | 0 | 0 | 1 |
| 0 | 0 | 0 | 1 |
| 0 | 0 | 0 | 1 |
| 0 | 0 | 0 | 1 |
| 0 | 0 | 0 | 1 |
| 0 | 0 | 0 | 1 |
| 0 | 0 | 0 | 1 |
| 0 | 0 | 0 | 1 |
| 0 | 0 | 0 | 1 |
| 0 | 0 | 0 | 1 |
| 0 | 0 | 0 | 1 |
| 0 | 0 | 0 | 1 |
| 0 | 0 | 0 | 1 |
| 0 | 0 | 0 | 1 |
| 0 | 0 | 0 | 1 |
| 0 | 0 | 0 | 1 |
| 0 | 0 | 0 | 1 |
| 0 | 0 | 0 | 1 |
| 0 | 0 | 0 | 1 |
| 0 | 0 | 0 | 1 |
| 0 | 0 | 0 | 1 |

| | | | |
|---|---|---|---|
| 0 | 0 | 0 | 1 |
| 0 | 0 | 0 | 1 |
| 0 | 0 | 0 | 1 |
| 0 | 0 | 0 | 1 |
| 0 | 0 | 0 | 1 |
| 0 | 0 | 0 | 1 |
| 0 | 0 | 0 | 1 |
| 0 | 0 | 0 | 1 |
| 0 | 0 | 0 | 1 |
| 0 | 0 | 0 | 1 |
| 0 | 0 | 0 | 1 |
| 0 | 0 | 0 | 0 |
| 0 | 0 | 0 | 0 |
| 0 | 0 | 0 | 0 |
| 0 | 0 | 0 | 1 |
| 0 | 0 | 0 | 0 |
| 0 | 0 | 0 | 1 |
| 0 | 0 | 0 | 0 |
| 0 | 0 | 0 | 1 |
| 0 | 0 | 0 | 1 |
| 0 | 0 | 0 | 0 |
| 0 | 0 | 0 | 1 |
| 0 | 0 | 0 | 1 |
| 0 | 0 | 0 | 0 |
| 0 | 0 | 0 | 0 |
| 0 | 0 | 0 | 1 |
| 0 | 0 | 0 | 1 |
| 0 | 0 | 0 | 0 |
| 0 | 0 | 0 | 0 |
| 0 | 0 | 0 | 0 |
| 0 | 0 | 0 | 0 |
| 0 | 0 | 0 | 0 |
| 0 | 0 | 0 | 1 |
| 0 | 0 | 0 | 1 |
| 0 | 0 | 0 | 1 |
| 0 | 0 | 0 | 1 |
| 0 | 0 | 0 | 1 |
| 0 | 0 | 0 | 1 |
| 0 | 0 | 0 | 1 |
| 0 | 0 | 0 | 0 |
| 0 | 0 | 0 | 0 |
| 0 | 0 | 0 | 1 |
| 0 | 0 | 0 | 0 |
| 0 | 0 | 0 | 0 |
| 0 | 0 | 0 | 0 |
| 0 | 1 | 1 | 1 |
| 0 | 0 | 0 | 0 |
| 0 | 0 | 0 | 1 |
| 0 | 0 | 0 | 1 |
| 0 | 1 | 0 | 0 |
| 0 | 0 | 0 | 0 |
| 0 | 0 | 0 | 0 |
| 0 | 0 | 0 | 1 |
| 0 | 0 | 0 | 0 |

| | | | |
|---|---|---|---|
| 0 | 0 | 0 | 0 |
| 0 | 0 | 0 | 1 |
| 0 | 0 | 0 | 1 |
| 0 | 0 | 0 | 1 |
| 0 | 1 | 0 | 0 |
| 0 | 0 | 0 | 1 |
| 0 | 0 | 0 | 0 |
| 0 | 0 | 0 | 1 |
| 0 | 0 | 0 | 1 |
| 0 | 0 | 0 | 0 |
| 0 | 1 | 1 | 1 |
| 0 | 0 | 0 | 1 |
| 0 | 0 | 0 | 1 |
| 0 | 0 | 0 | 0 |
| 0 | 0 | 0 | 1 |
| 0 | 0 | 0 | 1 |
| 0 | 0 | 0 | 0 |
| 0 | 0 | 0 | 1 |
| 0 | 0 | 0 | 0 |
| 0 | 0 | 0 | 0 |
| 0 | 0 | 0 | 0 |
| 0 | 0 | 0 | 0 |
| 0 | 0 | 0 | 0 |
| 0 | 0 | 0 | 0 |
| 0 | 0 | 0 | 0 |
| 0 | 0 | 0 | 0 |
| 0 | 0 | 0 | 0 |
| 0 | 0 | 0 | 0 |
| 0 | 0 | 0 | 0 |
| 0 | 0 | 0 | 0 |
| 0 | 0 | 0 | 0 |
| 0 | 0 | 0 | 0 |
| 0 | 0 | 0 | 0 |
| 0 | 0 | 0 | 0 |
| 0 | 0 | 0 | 0 |
| 0 | 0 | 0 | 0 |
| 0 | 0 | 0 | 1 |
| 0 | 1 | 0 | 0 |
| 0 | 0 | 0 | 0 |
| 0 | 0 | 0 | 1 |
| 0 | 0 | 0 | 1 |
| 0 | 0 | 0 | 0 |
| 0 | 0 | 0 | 0 |
| 0 | 1 | 0 | 0 |
| 0 | 0 | 0 | 0 |
| 0 | 0 | 0 | 0 |
| 0 | 0 | 0 | 0 |
| 0 | 0 | 0 | 0 |
| 0 | 0 | 0 | 1 |
| 0 | 0 | 0 | 1 |
| 0 | 0 | 0 | 0 |
| 0 | 0 | 0 | 1 |
| 0 | 0 | 0 | 0 |
| 0 | 0 | 0 | 0 |

| | | | |
|---|---|---|---|
| 0 | 0 | 0 | 0 |
| 0 | 0 | 0 | 0 |
| 0 | 0 | 0 | 0 |
| 0 | 0 | 0 | 0 |
| 0 | 0 | 0 | 0 |
| 0 | 0 | 0 | 0 |
| 0 | 0 | 0 | 0 |
| 0 | 0 | 0 | 0 |
| 0 | 0 | 0 | 0 |
| 0 | 0 | 0 | 0 |
| 0 | 0 | 0 | 0 |
| 0 | 0 | 0 | 0 |
| 0 | 0 | 0 | 0 |
| 0 | 0 | 0 | 0 |
| 0 | 0 | 0 | 1 |
| 0 | 0 | 0 | 0 |
| 0 | 1 | 0 | 0 |
| 0 | 0 | 0 | 0 |
| 0 | 0 | 0 | 1 |
| 0 | 0 | 0 | 0 |
| 0 | 0 | 0 | 0 |
| 0 | 1 | 0 | 0 |
| 0 | 1 | 0 | 0 |
| 0 | 0 | 0 | 0 |
| 0 | 0 | 0 | 0 |
| 0 | 0 | 0 | 1 |
| 0 | 0 | 0 | 0 |
| 0 | 0 | 0 | 0 |
| 0 | 0 | 0 | 0 |
| 0 | 0 | 0 | 0 |
| 0 | 0 | 0 | 1 |
| 0 | 0 | 0 | 0 |
| 0 | 0 | 0 | 1 |
| 0 | 0 | 0 | 1 |
| 1 | 0 | 0 | 0 |
| 0 | 0 | 0 | 0 |
| 0 | 0 | 0 | 1 |
| 0 | 0 | 0 | 1 |
| 0 | 0 | 0 | 1 |
| 0 | 0 | 0 | 1 |
| 0 | 0 | 0 | 0 |
| 0 | 0 | 0 | 0 |
| 0 | 0 | 0 | 0 |
| 0 | 0 | 0 | 0 |
| 0 | 0 | 0 | 0 |
| 0 | 0 | 0 | 0 |
| 0 | 0 | 0 | 0 |
| 0 | 0 | 0 | 0 |
| 0 | 0 | 0 | 1 |
| 0 | 0 | 0 | 1 |
| 0 | 0 | 0 | 0 |
| 0 | 0 | 0 | 0 |
| 0 | 0 | 0 | 0 |
| 0 | 0 | 0 | 1 |

| | | | |
|---|---|---|---|
| 0 | 0 | 0 | 1 |
| 0 | 0 | 0 | 1 |
| 0 | 0 | 0 | 1 |
| 0 | 0 | 0 | 0 |
| 0 | 0 | 0 | 1 |
| 0 | 0 | 0 | 1 |
| 0 | 0 | 0 | 0 |
| 0 | 0 | 0 | 1 |
| 0 | 0 | 0 | 0 |
| 0 | 0 | 0 | 0 |
| 0 | 0 | 0 | 0 |
| 0 | 0 | 0 | 0 |
| 0 | 0 | 0 | 0 |
| 0 | 0 | 0 | 0 |
| 0 | 0 | 0 | 0 |
| 0 | 0 | 0 | 0 |
| 0 | 0 | 0 | 1 |
| 0 | 0 | 0 | 0 |
| 0 | 0 | 0 | 0 |
| 0 | 0 | 0 | 1 |
| 0 | 0 | 0 | 0 |
| 0 | 0 | 0 | 0 |
| 0 | 0 | 0 | 1 |
| 0 | 0 | 0 | 0 |
| 0 | 0 | 0 | 1 |
| 0 | 0 | 0 | 1 |
| 0 | 0 | 0 | 1 |
| 0 | 0 | 0 | 0 |
| 0 | 0 | 0 | 0 |
| 0 | 0 | 0 | 0 |
| 0 | 0 | 0 | 1 |
| 0 | 0 | 0 | 0 |
| 0 | 0 | 0 | 0 |
| 0 | 0 | 0 | 0 |
| 0 | 0 | 0 | 1 |
| 0 | 0 | 0 | 1 |
| 0 | 0 | 0 | 0 |
| 0 | 0 | 0 | 0 |
| 0 | 0 | 0 | 0 |
| 0 | 0 | 0 | 1 |
| 0 | 0 | 0 | 1 |
| 0 | 0 | 0 | 1 |
| 0 | 0 | 0 | 1 |
| 0 | 0 | 0 | 0 |
| 0 | 0 | 0 | 1 |
| 0 | 0 | 0 | 0 |
| 0 | 0 | 0 | 1 |
| 0 | 0 | 0 | 0 |
| 0 | 0 | 0 | 0 |
| 0 | 0 | 0 | 1 |
| 0 | 0 | 0 | 1 |
| 0 | 1 | 0 | 0 |
| 0 | 0 | 0 | 0 |
| 0 | 0 | 0 | 1 |

| | | | |
|---|---|---|---|
| 0 | 0 | 0 | 1 |
| 0 | 0 | 0 | 1 |
| 0 | 0 | 0 | 1 |
| 0 | 0 | 0 | 1 |
| 0 | 1 | 0 | 0 |
| 0 | 1 | 1 | 1 |
| 0 | 0 | 0 | 1 |
| 0 | 1 | 0 | 0 |
| 0 | 1 | 0 | 0 |
| 0 | 0 | 0 | 1 |
| 0 | 0 | 0 | 0 |
| 0 | 0 | 0 | 0 |
| 1 | 0 | 0 | 0 |
| 1 | 0 | 0 | 0 |
| 0 | 0 | 0 | 0 |
| 1 | 0 | 0 | 0 |
| 1 | 0 | 0 | 0 |
| 1 | 0 | 0 | 0 |
| 1 | 0 | 0 | 0 |
| 0 | 0 | 0 | 1 |
| 0 | 0 | 0 | 1 |
| 0 | 0 | 0 | 1 |
| 0 | 0 | 0 | 1 |
| 0 | 0 | 0 | 1 |
| 0 | 0 | 0 | 0 |
| 0 | 0 | 0 | 1 |
| 1 | 0 | 0 | 0 |
| 1 | 0 | 0 | 0 |
| 0 | 0 | 0 | 1 |
| 0 | 0 | 0 | 1 |
| 0 | 0 | 0 | 1 |
| 0 | 0 | 0 | 1 |
| 0 | 0 | 0 | 1 |
| 0 | 0 | 0 | 0 |
| 0 | 0 | 0 | 0 |
| 0 | 0 | 0 | 1 |
| 0 | 0 | 0 | 0 |
| 0 | 0 | 0 | 1 |
| 0 | 0 | 0 | 0 |
| 0 | 1 | 0 | 0 |
| 0 | 0 | 0 | 0 |
| 0 | 0 | 0 | 0 |
| 0 | 0 | 0 | 1 |
| 0 | 0 | 0 | 0 |
| 0 | 0 | 0 | 0 |
| 1 | 0 | 0 | 0 |
| 1 | 0 | 0 | 0 |
| 1 | 0 | 0 | 0 |
| 0 | 0 | 0 | 0 |
| 0 | 0 | 0 | 0 |
| 0 | 0 | 0 | 0 |
| 1 | 0 | 0 | 0 |
| 1 | 0 | 0 | 0 |
| 0 | 0 | 0 | 1 |

| | | | |
|---|---|---|---|
| 0 | 0 | 0 | 0 |
| 0 | 0 | 0 | 1 |
| 0 | 0 | 0 | 0 |
| 0 | 0 | 0 | 0 |
| 0 | 1 | 0 | 0 |
| 0 | 0 | 0 | 1 |
| 0 | 0 | 0 | 0 |
| 0 | 0 | 0 | 1 |
| 0 | 0 | 0 | 1 |
| 0 | 0 | 0 | 0 |
| 0 | 0 | 0 | 1 |
| 0 | 0 | 0 | 1 |
| 0 | 0 | 0 | 1 |
| 0 | 0 | 0 | 1 |
| 0 | 0 | 0 | 1 |
| 0 | 1 | 0 | 0 |
| 0 | 0 | 0 | 0 |
| 0 | 0 | 0 | 1 |
| 0 | 0 | 0 | 0 |
| 0 | 0 | 0 | 1 |
| 0 | 0 | 0 | 0 |
| 0 | 0 | 0 | 0 |
| 0 | 0 | 0 | 0 |
| 0 | 0 | 0 | 0 |
| 0 | 1 | 0 | 0 |
| 0 | 0 | 0 | 0 |
| 0 | 0 | 0 | 0 |
| 0 | 0 | 0 | 0 |
| 0 | 0 | 0 | 0 |
| 0 | 0 | 0 | 0 |
| 0 | 0 | 0 | 0 |
| 0 | 0 | 0 | 0 |
| 0 | 0 | 0 | 1 |
| 0 | 0 | 0 | 1 |
| 0 | 0 | 0 | 0 |
| 0 | 0 | 0 | 0 |
| 0 | 0 | 0 | 1 |
| 0 | 0 | 0 | 0 |
| 0 | 0 | 0 | 1 |
| 0 | 1 | 0 | 0 |
| 0 | 1 | 0 | 0 |
| 0 | 0 | 0 | 0 |
| 0 | 0 | 0 | 1 |
| 0 | 0 | 0 | 1 |
| 0 | 1 | 0 | 0 |
| 0 | 1 | 0 | 0 |
| 0 | 0 | 0 | 0 |
| 0 | 0 | 0 | 0 |
| 0 | 0 | 0 | 0 |
| 0 | 0 | 0 | 0 |
| 0 | 0 | 0 | 1 |
| 0 | 1 | 0 | 0 |
| 0 | 1 | 0 | 0 |
| 0 | 0 | 0 | 1 |

| | | | |
|---|---|---|---|
| 0 | 0 | 0 | 0 |
| 0 | 0 | 0 | 1 |
| 0 | 0 | 0 | 0 |
| 0 | 0 | 0 | 0 |
| 0 | 0 | 0 | 0 |
| 0 | 0 | 0 | 0 |
| 0 | 0 | 0 | 0 |
| 0 | 0 | 0 | 0 |
| 0 | 0 | 0 | 0 |
| 0 | 0 | 0 | 0 |
| 0 | 0 | 0 | 0 |
| 0 | 0 | 0 | 1 |
| 0 | 0 | 0 | 0 |
| 0 | 0 | 0 | 1 |
| 0 | 0 | 0 | 1 |
| 0 | 1 | 1 | 1 |
| 0 | 1 | 0 | 0 |
| 0 | 1 | 0 | 0 |
| 0 | 1 | 1 | 1 |
| 0 | 1 | 0 | 0 |
| 0 | 0 | 0 | 1 |
| 0 | 0 | 0 | 0 |
| 0 | 0 | 0 | 1 |
| 0 | 0 | 0 | 0 |
| 0 | 0 | 0 | 1 |
| 0 | 0 | 0 | 1 |
| 0 | 0 | 0 | 1 |
| 0 | 0 | 0 | 0 |
| 0 | 0 | 0 | 1 |
| 0 | 0 | 0 | 1 |
| 0 | 0 | 0 | 1 |
| 1 | 0 | 0 | 0 |
| 0 | 0 | 0 | 1 |
| 0 | 0 | 0 | 0 |
| 0 | 0 | 0 | 0 |
| 0 | 0 | 0 | 0 |
| 0 | 0 | 0 | 1 |
| 0 | 0 | 0 | 1 |
| 0 | 0 | 0 | 0 |
| 0 | 0 | 0 | 0 |
| 0 | 0 | 0 | 0 |
| 0 | 0 | 0 | 0 |
| 0 | 1 | 0 | 0 |
| 0 | 0 | 0 | 0 |
| 0 | 0 | 0 | 0 |
| 0 | 0 | 0 | 0 |
| 0 | 0 | 0 | 1 |
| 0 | 0 | 0 | 1 |
| 0 | 0 | 0 | 0 |
| 0 | 0 | 0 | 0 |
| 0 | 0 | 0 | 0 |
| 0 | 0 | 0 | 1 |
| 0 | 0 | 0 | 1 |
| 0 | 0 | 0 | 0 |

| | | | |
|---|---|---|---|
| 0 | 0 | 0 | 0 |
| 0 | 0 | 0 | 1 |
| 0 | 0 | 0 | 1 |
| 0 | 0 | 0 | 1 |
| 0 | 0 | 0 | 0 |
| 0 | 0 | 0 | 0 |
| 0 | 0 | 0 | 0 |
| 0 | 0 | 0 | 0 |
| 0 | 0 | 0 | 0 |
| 0 | 0 | 0 | 0 |
| 0 | 0 | 0 | 1 |
| 0 | 0 | 0 | 0 |
| 0 | 0 | 0 | 0 |
| 0 | 0 | 0 | 0 |
| 0 | 1 | 1 | 1 |
| 0 | 0 | 0 | 1 |
| 0 | 1 | 0 | 0 |
| 0 | 0 | 0 | 1 |
| 0 | 0 | 0 | 0 |
| 0 | 0 | 0 | 0 |
| 0 | 0 | 0 | 1 |
| 0 | 1 | 0 | 0 |
| 0 | 1 | 0 | 0 |
| 0 | 0 | 0 | 1 |
| 0 | 1 | 0 | 0 |
| 0 | 1 | 0 | 0 |
| 0 | 0 | 0 | 0 |
| 0 | 0 | 0 | 1 |
| 0 | 1 | 0 | 0 |
| 0 | 0 | 0 | 0 |
| 0 | 0 | 0 | 0 |
| 0 | 0 | 0 | 1 |
| 0 | 0 | 0 | 1 |
| 0 | 0 | 0 | 1 |
| 0 | 0 | 0 | 1 |
| 0 | 0 | 0 | 1 |
| 0 | 0 | 0 | 1 |
| 0 | 0 | 0 | 1 |
| 0 | 0 | 0 | 1 |
| 0 | 0 | 0 | 1 |
| 0 | 0 | 0 | 1 |
| 0 | 0 | 0 | 1 |
| 1 | 0 | 0 | 0 |
| 0 | 0 | 0 | 1 |
| 0 | 0 | 0 | 1 |
| 0 | 0 | 0 | 1 |
| 0 | 0 | 0 | 1 |
| 0 | 0 | 0 | 1 |
| 0 | 0 | 0 | 1 |
| 0 | 0 | 0 | 1 |
| 0 | 0 | 0 | 1 |
| 0 | 0 | 0 | 1 |
| 0 | 0 | 0 | 1 |
| 0 | 0 | 0 | 1 |

| | | | |
|---|---|---|---|
| 0 | 0 | 0 | 1 |
| 0 | 0 | 0 | 1 |
| 0 | 0 | 0 | 1 |
| 0 | 0 | 0 | 1 |
| 0 | 0 | 0 | 1 |
| 0 | 0 | 0 | 1 |
| 0 | 0 | 0 | 1 |
| 0 | 0 | 0 | 1 |
| 0 | 0 | 0 | 0 |
| 0 | 0 | 0 | 0 |
| 0 | 0 | 0 | 0 |
| 0 | 0 | 0 | 0 |
| 0 | 0 | 0 | 0 |
| 0 | 0 | 0 | 0 |
| 0 | 0 | 0 | 0 |
| 0 | 0 | 0 | 0 |
| 0 | 0 | 0 | 0 |
| 0 | 0 | 0 | 0 |
| 0 | 0 | 0 | 0 |
| 0 | 0 | 0 | 0 |
| 1 | 0 | 0 | 0 |
| 1 | 0 | 0 | 0 |
| 0 | 0 | 0 | 0 |
| 0 | 0 | 0 | 0 |
| 0 | 0 | 0 | 0 |
| 0 | 0 | 0 | 1 |
| 0 | 0 | 0 | 1 |
| 0 | 0 | 0 | 0 |
| 0 | 0 | 0 | 0 |
| 1 | 0 | 0 | 0 |
| 1 | 0 | 0 | 0 |
| 0 | 0 | 0 | 0 |
| 1 | 0 | 0 | 0 |
| 1 | 0 | 0 | 0 |
| 0 | 0 | 0 | 0 |
| 0 | 0 | 0 | 0 |
| 0 | 0 | 0 | 0 |
| 0 | 0 | 0 | 0 |
| 0 | 0 | 0 | 0 |
| 0 | 0 | 0 | 0 |
| 0 | 0 | 0 | 0 |
| 0 | 0 | 0 | 0 |
| 1 | 0 | 0 | 0 |
| 1 | 0 | 0 | 0 |
| 1 | 0 | 0 | 0 |
| 0 | 0 | 0 | 0 |
| 0 | 0 | 0 | 1 |
| 0 | 0 | 0 | 1 |
| 0 | 0 | 0 | 1 |
| 0 | 0 | 0 | 1 |
| 0 | 0 | 0 | 1 |
| 0 | 0 | 0 | 1 |
| 0 | 0 | 0 | 0 |
| 0 | 0 | 0 | 0 |

| | | | |
|---|---|---|---|
| 0 | 0 | 0 | 0 |
| 0 | 0 | 0 | 0 |
| 0 | 0 | 0 | 0 |
| 0 | 0 | 0 | 0 |
| 0 | 0 | 0 | 1 |
| 0 | 0 | 0 | 0 |
| 0 | 0 | 0 | 0 |
| 0 | 0 | 0 | 0 |
| 0 | 0 | 0 | 1 |
| 0 | 0 | 0 | 0 |
| 1 | 0 | 0 | 0 |
| 0 | 0 | 0 | 1 |
| 0 | 0 | 0 | 0 |
| 0 | 0 | 0 | 0 |
| 0 | 0 | 0 | 0 |
| 0 | 0 | 0 | 0 |
| 0 | 0 | 0 | 1 |
| 0 | 0 | 0 | 0 |
| 0 | 0 | 0 | 0 |
| 0 | 0 | 0 | 1 |
| 0 | 1 | 0 | 0 |
| 0 | 0 | 0 | 0 |
| 0 | 0 | 0 | 0 |
| 0 | 0 | 0 | 0 |
| 0 | 0 | 0 | 0 |
| 0 | 0 | 0 | 1 |
| 0 | 0 | 0 | 1 |
| 0 | 0 | 0 | 1 |
| 0 | 0 | 0 | 1 |
| 0 | 0 | 0 | 1 |
| 0 | 0 | 0 | 1 |
| 0 | 0 | 0 | 1 |
| 0 | 0 | 0 | 1 |
| 0 | 0 | 0 | 1 |
| 0 | 0 | 0 | 0 |
| 0 | 0 | 0 | 0 |
| 0 | 0 | 0 | 0 |
| 0 | 0 | 0 | 0 |
| 0 | 0 | 0 | 0 |
| 0 | 0 | 0 | 0 |
| 0 | 0 | 0 | 1 |
| 0 | 0 | 0 | 1 |
| 0 | 0 | 0 | 1 |
| 0 | 0 | 0 | 0 |
| 0 | 0 | 0 | 0 |
| 0 | 0 | 0 | 0 |
| 0 | 0 | 0 | 1 |
| 0 | 0 | 0 | 1 |
| 0 | 0 | 0 | 0 |
| 0 | 0 | 0 | 0 |
| 0 | 0 | 0 | 1 |
| 0 | 0 | 0 | 1 |
| 0 | 0 | 0 | 1 |
| 0 | 0 | 0 | 1 |

| | | | |
|---|---|---|---|
| 0 | 0 | 0 | 1 |
| 0 | 0 | 0 | 1 |
| 0 | 0 | 0 | 1 |
| 0 | 0 | 0 | 0 |
| 0 | 0 | 0 | 0 |
| 0 | 0 | 0 | 1 |
| 0 | 0 | 0 | 1 |
| 0 | 0 | 0 | 1 |
| 0 | 0 | 0 | 1 |
| 0 | 0 | 0 | 1 |
| 0 | 0 | 0 | 0 |
| 0 | 0 | 0 | 1 |
| 0 | 1 | 0 | 0 |
| 0 | 0 | 0 | 0 |
| 0 | 0 | 0 | 0 |
| 0 | 0 | 0 | 1 |
| 0 | 0 | 0 | 1 |
| 0 | 0 | 0 | 1 |
| 0 | 0 | 0 | 1 |
| 0 | 0 | 0 | 1 |
| 0 | 0 | 0 | 0 |
| 0 | 0 | 0 | 1 |
| 0 | 0 | 0 | 1 |
| 0 | 0 | 0 | 0 |
| 0 | 0 | 0 | 1 |
| 0 | 0 | 0 | 0 |
| 0 | 1 | 0 | 0 |
| 0 | 0 | 0 | 1 |
| 0 | 0 | 0 | 1 |
| 0 | 0 | 0 | 1 |
| 0 | 0 | 0 | 1 |
| 0 | 0 | 0 | 1 |
| 0 | 0 | 0 | 1 |
| 0 | 0 | 0 | 1 |
| 0 | 0 | 0 | 1 |
| 0 | 0 | 0 | 0 |
| 0 | 0 | 0 | 1 |
| 0 | 0 | 0 | 1 |
| 0 | 0 | 0 | 0 |
| 0 | 0 | 0 | 0 |
| 0 | 0 | 0 | 1 |
| 0 | 0 | 0 | 1 |
| 1 | 0 | 0 | 0 |
| 1 | 0 | 0 | 0 |
| 0 | 0 | 0 | 0 |
| 0 | 0 | 0 | 0 |
| 1 | 0 | 0 | 0 |
| 1 | 0 | 0 | 0 |
| 0 | 0 | 0 | 1 |
| 1 | 0 | 0 | 0 |
| 1 | 0 | 0 | 0 |
| 1 | 0 | 0 | 0 |
| 1 | 0 | 0 | 0 |
| 1 | 0 | 0 | 0 |

| | | | |
|---|---|---|---|
| 0 | 0 | 0 | 0 |
| 0 | 0 | 0 | 0 |
| 1 | 0 | 0 | 0 |
| 1 | 0 | 0 | 0 |
| 1 | 0 | 0 | 0 |
| 1 | 0 | 0 | 0 |
| 0 | 0 | 0 | 0 |
| 0 | 0 | 0 | 0 |
| 0 | 0 | 0 | 0 |
| 0 | 0 | 0 | 1 |
| 0 | 0 | 0 | 1 |
| 0 | 0 | 0 | 0 |
| 0 | 0 | 0 | 1 |
| 0 | 0 | 0 | 0 |
| 0 | 0 | 0 | 0 |
| 0 | 0 | 0 | 1 |
| 0 | 0 | 0 | 0 |
| 0 | 0 | 0 | 1 |
| 0 | 0 | 0 | 1 |
| 0 | 0 | 0 | 1 |
| 0 | 0 | 0 | 1 |
| 0 | 0 | 0 | 1 |
| 0 | 0 | 0 | 1 |
| 0 | 0 | 0 | 0 |
| 0 | 0 | 0 | 0 |
| 0 | 0 | 0 | 1 |
| 0 | 0 | 0 | 1 |
| 0 | 0 | 0 | 1 |
| 0 | 0 | 0 | 0 |
| 0 | 0 | 0 | 0 |
| 0 | 0 | 0 | 1 |
| 0 | 0 | 0 | 1 |
| 0 | 0 | 0 | 1 |
| 0 | 0 | 0 | 0 |
| 0 | 0 | 0 | 0 |
| 0 | 0 | 0 | 1 |
| 0 | 0 | 0 | 0 |
| 0 | 0 | 0 | 1 |
| 0 | 0 | 0 | 1 |
| 0 | 0 | 0 | 0 |
| 0 | 0 | 0 | 0 |
| 0 | 0 | 0 | 0 |
| 0 | 0 | 0 | 0 |
| 0 | 0 | 0 | 0 |
| 0 | 0 | 0 | 0 |
| 0 | 0 | 0 | 1 |
| 0 | 0 | 0 | 1 |
| 0 | 0 | 0 | 0 |
| 0 | 0 | 0 | 0 |
| 0 | 0 | 0 | 0 |
| 0 | 0 | 0 | 1 |
| 0 | 0 | 0 | 1 |
| 0 | 0 | 0 | 0 |
| 0 | 0 | 0 | 0 |

| | | | |
|---|---|---|---|
| 0 | 0 | 0 | 1 |
| 0 | 0 | 0 | 1 |
| 0 | 0 | 0 | 1 |
| 0 | 0 | 0 | 1 |
| 0 | 1 | 0 | 0 |
| 0 | 0 | 0 | 0 |
| 0 | 0 | 0 | 1 |
| 0 | 0 | 0 | 0 |
| 0 | 1 | 1 | 1 |
| 0 | 0 | 0 | 0 |
| 0 | 0 | 0 | 1 |
| 0 | 0 | 0 | 1 |
| 0 | 0 | 0 | 1 |
| 0 | 0 | 0 | 1 |
| 0 | 0 | 0 | 1 |
| 0 | 0 | 0 | 1 |
| 0 | 0 | 0 | 0 |
| 0 | 0 | 0 | 1 |
| 0 | 0 | 0 | 1 |
| 0 | 0 | 0 | 0 |
| 0 | 0 | 0 | 0 |
| 0 | 0 | 0 | 1 |
| 0 | 0 | 0 | 1 |
| 0 | 0 | 0 | 0 |
| 0 | 0 | 0 | 1 |
| 0 | 0 | 0 | 0 |
| 0 | 0 | 0 | 0 |
| 0 | 0 | 0 | 0 |
| 0 | 0 | 0 | 0 |
| 0 | 0 | 0 | 0 |
| 0 | 0 | 0 | 0 |
| 0 | 0 | 0 | 0 |
| 0 | 0 | 0 | 1 |
| 0 | 0 | 0 | 0 |
| 0 | 0 | 0 | 1 |
| 0 | 0 | 0 | 1 |
| 0 | 0 | 0 | 1 |
| 0 | 0 | 0 | 1 |
| 0 | 0 | 0 | 0 |
| 0 | 0 | 0 | 1 |
| 0 | 0 | 0 | 1 |
| 0 | 0 | 0 | 1 |
| 0 | 0 | 0 | 1 |
| 0 | 0 | 0 | 0 |
| 0 | 0 | 0 | 1 |
| 0 | 0 | 0 | 1 |
| 0 | 0 | 0 | 1 |
| 0 | 0 | 0 | 1 |
| 0 | 0 | 0 | 1 |
| 0 | 0 | 0 | 1 |
| 0 | 0 | 0 | 0 |
| 0 | 0 | 0 | 0 |
| 0 | 0 | 0 | 0 |
| 0 | 0 | 0 | 1 |

| | | | |
|---|---|---|---|
| 0 | 0 | 0 | 0 |
| 0 | 0 | 0 | 0 |
| 0 | 0 | 0 | 0 |
| 0 | 0 | 0 | 0 |
| 1 | 0 | 0 | 0 |
| 1 | 0 | 0 | 0 |
| 0 | 0 | 0 | 0 |
| 0 | 0 | 0 | 0 |
| 0 | 0 | 0 | 1 |
| 0 | 0 | 0 | 1 |
| 0 | 0 | 0 | 0 |
| 0 | 0 | 0 | 1 |
| 0 | 0 | 0 | 1 |
| 0 | 0 | 0 | 0 |
| 0 | 0 | 0 | 0 |
| 0 | 0 | 0 | 0 |
| 0 | 0 | 0 | 0 |
| 0 | 0 | 0 | 0 |
| 0 | 0 | 0 | 0 |
| 0 | 0 | 0 | 1 |
| 0 | 0 | 0 | 1 |
| 0 | 0 | 0 | 0 |
| 0 | 0 | 0 | 1 |
| 0 | 1 | 0 | 0 |
| 0 | 0 | 0 | 1 |
| 0 | 0 | 0 | 1 |
| 0 | 0 | 0 | 1 |
| 0 | 0 | 0 | 1 |
| 0 | 0 | 0 | 1 |
| 0 | 0 | 0 | 0 |
| 0 | 0 | 0 | 1 |
| 0 | 0 | 0 | 1 |
| 0 | 0 | 0 | 0 |
| 0 | 0 | 0 | 1 |
| 0 | 0 | 0 | 0 |
| 0 | 0 | 0 | 0 |
| 0 | 0 | 0 | 0 |
| 0 | 0 | 0 | 0 |
| 0 | 0 | 0 | 0 |
| 0 | 0 | 0 | 0 |
| 0 | 0 | 0 | 0 |
| 0 | 0 | 0 | 0 |
| 0 | 0 | 0 | 1 |
| 0 | 0 | 0 | 0 |
| 0 | 0 | 0 | 1 |
| 0 | 0 | 0 | 0 |
| 0 | 0 | 0 | 1 |
| 0 | 0 | 0 | 1 |
| 0 | 0 | 0 | 1 |
| 0 | 0 | 0 | 0 |
| 0 | 0 | 0 | 1 |
| 0 | 0 | 0 | 1 |
| 0 | 0 | 0 | 1 |
| 0 | 0 | 0 | 1 |

| | | | |
|---|---|---|---|
| 1 | 0 | 0 | 0 |
| 0 | 0 | 0 | 1 |
| 0 | 0 | 0 | 1 |
| 0 | 0 | 0 | 1 |
| 0 | 0 | 0 | 1 |
| 0 | 0 | 0 | 1 |
| 0 | 0 | 0 | 1 |
| 0 | 0 | 0 | 1 |
| 0 | 0 | 0 | 1 |
| 0 | 0 | 0 | 1 |
| 0 | 0 | 0 | 0 |
| 0 | 0 | 0 | 1 |
| 0 | 0 | 0 | 1 |
| 0 | 0 | 0 | 1 |
| 0 | 0 | 0 | 0 |
| 0 | 0 | 0 | 0 |
| 0 | 0 | 0 | 0 |
| 0 | 0 | 0 | 0 |
| 0 | 0 | 0 | 0 |
| 0 | 0 | 0 | 1 |
| 0 | 0 | 0 | 1 |
| 0 | 0 | 0 | 1 |
| 0 | 0 | 0 | 0 |
| 0 | 0 | 0 | 1 |
| 0 | 0 | 0 | 1 |
| 0 | 0 | 0 | 1 |
| 0 | 0 | 0 | 1 |
| 0 | 0 | 0 | 1 |
| 0 | 0 | 0 | 0 |
| 0 | 0 | 0 | 0 |
| 0 | 0 | 0 | 0 |
| 0 | 0 | 0 | 0 |
| 0 | 0 | 0 | 0 |
| 0 | 0 | 0 | 0 |
| 0 | 0 | 0 | 1 |
| 0 | 0 | 0 | 1 |
| 0 | 0 | 0 | 0 |
| 0 | 0 | 0 | 0 |
| 0 | 0 | 0 | 0 |
| 0 | 0 | 0 | 0 |
| 0 | 0 | 0 | 0 |
| 0 | 0 | 0 | 0 |
| 0 | 0 | 0 | 1 |
| 0 | 0 | 0 | 1 |
| 0 | 0 | 0 | 0 |
| 0 | 0 | 0 | 1 |
| 0 | 0 | 0 | 1 |
| 0 | 0 | 0 | 1 |
| 0 | 0 | 0 | 1 |
| 0 | 0 | 0 | 1 |
| 0 | 0 | 0 | 1 |
| 0 | 0 | 0 | 1 |
| 0 | 0 | 0 | 1 |
| 0 | 0 | 0 | 1 |

| | | | |
|---|---|---|---|
| 0 | 0 | 0 | 0 |
| 0 | 0 | 0 | 1 |
| 0 | 0 | 0 | 0 |
| 0 | 0 | 0 | 1 |
| 0 | 0 | 0 | 1 |
| 0 | 1 | 1 | 1 |
| 0 | 1 | 0 | 0 |
| 0 | 1 | 0 | 0 |
| 0 | 0 | 0 | 1 |
| 0 | 0 | 0 | 1 |
| 0 | 0 | 0 | 1 |
| 0 | 0 | 0 | 1 |
| 0 | 0 | 0 | 1 |
| 0 | 0 | 0 | 1 |
| 0 | 0 | 0 | 0 |
| 0 | 0 | 0 | 0 |
| 0 | 0 | 0 | 0 |
| 0 | 0 | 0 | 0 |
| 0 | 0 | 0 | 1 |
| 0 | 0 | 0 | 0 |
| 0 | 0 | 0 | 0 |
| 0 | 0 | 0 | 0 |
| 0 | 0 | 0 | 0 |
| 0 | 0 | 0 | 0 |
| 0 | 0 | 0 | 1 |
| 0 | 0 | 0 | 0 |
| 0 | 0 | 0 | 1 |
| 0 | 0 | 0 | 1 |
| 0 | 0 | 0 | 1 |
| 0 | 0 | 0 | 0 |
| 0 | 0 | 0 | 1 |
| 0 | 0 | 0 | 1 |
| 0 | 0 | 0 | 0 |
| 0 | 0 | 0 | 0 |
| 0 | 0 | 0 | 1 |
| 0 | 0 | 0 | 1 |
| 0 | 0 | 0 | 1 |
| 0 | 0 | 0 | 1 |
| 0 | 0 | 0 | 1 |
| 0 | 0 | 0 | 0 |
| 0 | 0 | 0 | 0 |
| 0 | 0 | 0 | 0 |
| 0 | 0 | 0 | 0 |
| 0 | 0 | 0 | 0 |
| 0 | 0 | 0 | 0 |
| 0 | 0 | 0 | 1 |
| 0 | 0 | 0 | 1 |
| 0 | 0 | 0 | 0 |
| 0 | 0 | 0 | 0 |
| 0 | 0 | 0 | 1 |
| 0 | 0 | 0 | 0 |
| 0 | 0 | 0 | 0 |
| 0 | 0 | 0 | 1 |
| 0 | 0 | 0 | 1 |

| | | | |
|---|---|---|---|
| 0 | 0 | 0 | 1 |
| 0 | 0 | 0 | 0 |
| 0 | 0 | 0 | 0 |
| 0 | 0 | 0 | 1 |
| 0 | 0 | 0 | 1 |
| 0 | 0 | 0 | 0 |
| 0 | 0 | 0 | 0 |
| 0 | 0 | 0 | 0 |
| 0 | 0 | 0 | 0 |
| 0 | 0 | 0 | 0 |
| 0 | 0 | 0 | 0 |
| 0 | 0 | 0 | 0 |
| 0 | 0 | 0 | 0 |
| 0 | 0 | 0 | 0 |
| 0 | 0 | 0 | 0 |
| 0 | 0 | 0 | 1 |
| 0 | 0 | 0 | 0 |
| 0 | 0 | 0 | 1 |
| 0 | 0 | 0 | 1 |
| 0 | 0 | 0 | 1 |
| 0 | 0 | 0 | 0 |
| 0 | 0 | 0 | 1 |
| 0 | 0 | 0 | 0 |
| 0 | 0 | 0 | 0 |
| 0 | 0 | 0 | 1 |
| 0 | 0 | 0 | 0 |
| 0 | 0 | 0 | 1 |
| 0 | 0 | 0 | 1 |
| 0 | 0 | 0 | 1 |
| 0 | 0 | 0 | 0 |
| 0 | 0 | 0 | 0 |
| 0 | 0 | 0 | 0 |
| 0 | 0 | 0 | 0 |
| 0 | 0 | 0 | 0 |
| 0 | 0 | 0 | 0 |
| 0 | 0 | 0 | 0 |
| 0 | 0 | 0 | 0 |
| 0 | 0 | 0 | 1 |
| 0 | 0 | 0 | 1 |
| 0 | 0 | 0 | 0 |
| 0 | 0 | 0 | 0 |
| 0 | 0 | 0 | 0 |
| 0 | 0 | 0 | 0 |
| 0 | 0 | 0 | 0 |
| 0 | 0 | 0 | 0 |
| 0 | 0 | 0 | 0 |
| 0 | 0 | 0 | 1 |
| 0 | 0 | 0 | 0 |
| 0 | 0 | 0 | 0 |
| 0 | 0 | 0 | 0 |
| 0 | 0 | 0 | 0 |
| 0 | 0 | 0 | 1 |
| 0 | 0 | 0 | 0 |
| 0 | 0 | 0 | 0 |

| | | | |
|---|---|---|---|
| 0 | 0 | 0 | 0 |
| 0 | 0 | 0 | 0 |
| 0 | 0 | 0 | 0 |
| 0 | 0 | 0 | 0 |
| 0 | 0 | 0 | 0 |
| 0 | 0 | 0 | 0 |
| 0 | 0 | 0 | 0 |
| 0 | 0 | 0 | 0 |
| 1 | 0 | 0 | 0 |
| 0 | 0 | 0 | 0 |
| 0 | 0 | 0 | 1 |
| 0 | 1 | 0 | 0 |
| 0 | 1 | 1 | 1 |
| 0 | 0 | 0 | 0 |
| 0 | 0 | 0 | 1 |
| 0 | 0 | 0 | 1 |
| 0 | 0 | 0 | 1 |
| 0 | 0 | 0 | 1 |
| 0 | 0 | 0 | 0 |
| 0 | 0 | 0 | 0 |
| 0 | 0 | 0 | 1 |
| 0 | 0 | 0 | 0 |
| 0 | 0 | 0 | 0 |
| 0 | 0 | 0 | 1 |
| 0 | 0 | 0 | 0 |
| 0 | 0 | 0 | 0 |
| 0 | 0 | 0 | 0 |
| 0 | 0 | 0 | 1 |
| 0 | 0 | 0 | 0 |
| 0 | 0 | 0 | 0 |
| 0 | 0 | 0 | 1 |
| 1 | 0 | 0 | 0 |
| 0 | 0 | 0 | 1 |
| 0 | 0 | 0 | 1 |
| 1 | 0 | 0 | 0 |
| 1 | 0 | 0 | 0 |
| 1 | 0 | 0 | 0 |
| 0 | 0 | 0 | 1 |
| 0 | 0 | 0 | 1 |
| 0 | 0 | 0 | 0 |
| 1 | 0 | 0 | 0 |
| 0 | 1 | 0 | 0 |
| 0 | 1 | 0 | 0 |
| 1 | 0 | 0 | 0 |
| 1 | 0 | 0 | 0 |
| 0 | 0 | 0 | 0 |
| 0 | 0 | 0 | 1 |
| 0 | 0 | 0 | 1 |
| 0 | 0 | 0 | 1 |
| 0 | 0 | 0 | 1 |
| 1 | 0 | 0 | 0 |
| 0 | 0 | 0 | 0 |
| 0 | 0 | 0 | 0 |
| 0 | 0 | 0 | 0 |

| | | | |
|---|---|---|---|
| 0 | 0 | 0 | 0 |
| 0 | 0 | 0 | 1 |
| 0 | 0 | 0 | 0 |
| 0 | 0 | 0 | 0 |
| 0 | 0 | 0 | 0 |
| 0 | 0 | 0 | 1 |
| 0 | 0 | 0 | 0 |
| 0 | 0 | 0 | 0 |
| 0 | 0 | 0 | 1 |
| 0 | 0 | 0 | 0 |
| 0 | 0 | 0 | 1 |
| 0 | 0 | 0 | 1 |
| 0 | 0 | 0 | 1 |
| 0 | 0 | 0 | 0 |
| 0 | 0 | 0 | 1 |
| 0 | 0 | 0 | 0 |
| 0 | 0 | 0 | 1 |
| 0 | 0 | 0 | 1 |
| 0 | 0 | 0 | 0 |
| 0 | 0 | 0 | 0 |
| 0 | 0 | 0 | 0 |
| 0 | 0 | 0 | 0 |
| 0 | 0 | 0 | 0 |
| 0 | 0 | 0 | 0 |
| 0 | 0 | 0 | 0 |
| 0 | 0 | 0 | 1 |
| 0 | 0 | 0 | 1 |
| 0 | 0 | 0 | 1 |
| 0 | 0 | 0 | 1 |
| 0 | 0 | 0 | 1 |
| 0 | 0 | 0 | 1 |
| 0 | 0 | 0 | 1 |
| 0 | 0 | 0 | 0 |
| 0 | 0 | 0 | 0 |
| 0 | 0 | 0 | 0 |
| 0 | 0 | 0 | 0 |
| 0 | 0 | 0 | 0 |
| 0 | 0 | 0 | 0 |
| 0 | 0 | 0 | 1 |
| 0 | 0 | 0 | 1 |
| 0 | 0 | 0 | 1 |
| 0 | 0 | 0 | 1 |
| 0 | 0 | 0 | 1 |
| 0 | 0 | 0 | 1 |
| 0 | 1 | 0 | 0 |
| 0 | 0 | 0 | 0 |
| 0 | 0 | 0 | 1 |
| 0 | 0 | 0 | 0 |
| 0 | 0 | 0 | 0 |
| 0 | 0 | 0 | 0 |
| 0 | 0 | 0 | 0 |
| 0 | 0 | 0 | 0 |
| 0 | 0 | 0 | 0 |
| 0 | 0 | 0 | 0 |

| | | | |
|---|---|---|---|
| 0 | 0 | 0 | 1 |
| 0 | 0 | 0 | 1 |
| 1 | 0 | 0 | 0 |
| 1 | 0 | 0 | 0 |
| 0 | 0 | 0 | 1 |
| 0 | 0 | 0 | 0 |
| 0 | 0 | 0 | 0 |
| 0 | 0 | 0 | 1 |
| 0 | 0 | 0 | 1 |
| 0 | 0 | 0 | 0 |
| 0 | 0 | 0 | 1 |
| 0 | 0 | 0 | 1 |
| 0 | 0 | 0 | 0 |
| 0 | 0 | 0 | 0 |
| 0 | 0 | 0 | 0 |
| 0 | 0 | 0 | 0 |
| 0 | 0 | 0 | 1 |
| 0 | 0 | 0 | 1 |
| 0 | 0 | 0 | 1 |
| 0 | 0 | 0 | 1 |
| 0 | 0 | 0 | 1 |
| 0 | 0 | 0 | 1 |
| 0 | 0 | 0 | 1 |
| 0 | 0 | 0 | 1 |
| 0 | 0 | 0 | 1 |
| 0 | 0 | 0 | 0 |
| 0 | 0 | 0 | 1 |
| 0 | 0 | 0 | 1 |
| 0 | 0 | 0 | 1 |
| 0 | 0 | 0 | 1 |
| 0 | 0 | 0 | 1 |
| 0 | 0 | 0 | 1 |
| 0 | 0 | 0 | 1 |
| 0 | 0 | 0 | 1 |
| 0 | 0 | 0 | 1 |
| 0 | 0 | 0 | 1 |
| 0 | 0 | 0 | 1 |
| 0 | 0 | 0 | 0 |
| 0 | 0 | 0 | 1 |
| 0 | 0 | 0 | 1 |
| 0 | 0 | 0 | 0 |
| 0 | 0 | 0 | 1 |
| 0 | 0 | 0 | 1 |
| 0 | 0 | 0 | 0 |
| 0 | 0 | 0 | 0 |
| 0 | 0 | 0 | 1 |
| 0 | 0 | 0 | 1 |
| 0 | 0 | 0 | 1 |
| 0 | 0 | 0 | 1 |
| 0 | 0 | 0 | 1 |
| 0 | 0 | 0 | 1 |
| 0 | 0 | 0 | 0 |
| 0 | 0 | 0 | 0 |
| 0 | 0 | 0 | 0 |

| | | | |
|---|---|---|---|
| 0 | 0 | 0 | 0 |
| 0 | 0 | 0 | 1 |
| 0 | 0 | 0 | 1 |
| 0 | 0 | 0 | 1 |
| 0 | 0 | 0 | 1 |
| 0 | 0 | 0 | 1 |
| 0 | 0 | 0 | 1 |
| 0 | 0 | 0 | 1 |
| 0 | 0 | 0 | 0 |
| 0 | 0 | 0 | 1 |
| 0 | 0 | 0 | 1 |
| 0 | 0 | 0 | 1 |
| 0 | 0 | 0 | 1 |
| 0 | 0 | 0 | 1 |
| 0 | 0 | 0 | 1 |
| 0 | 0 | 0 | 1 |
| 0 | 0 | 0 | 1 |
| 0 | 0 | 0 | 1 |
| 0 | 0 | 0 | 0 |
| 0 | 0 | 0 | 0 |
| 0 | 0 | 0 | 0 |
| 0 | 0 | 0 | 0 |
| 0 | 0 | 0 | 1 |
| 0 | 0 | 0 | 0 |
| 0 | 0 | 0 | 0 |
| 0 | 0 | 0 | 1 |
| 0 | 0 | 0 | 1 |
| 0 | 0 | 0 | 1 |
| 0 | 0 | 0 | 1 |
| 0 | 0 | 0 | 1 |
| 0 | 0 | 0 | 1 |
| 0 | 0 | 0 | 1 |
| 0 | 0 | 0 | 0 |
| 0 | 0 | 0 | 1 |
| 0 | 0 | 0 | 1 |
| 0 | 0 | 0 | 1 |
| 0 | 0 | 0 | 0 |
| 0 | 0 | 0 | 1 |
| 0 | 0 | 0 | 0 |
| 0 | 0 | 0 | 0 |
| 0 | 0 | 0 | 0 |
| 0 | 0 | 0 | 0 |
| 0 | 0 | 0 | 0 |
| 0 | 0 | 0 | 1 |
| 0 | 0 | 0 | 1 |
| 0 | 0 | 0 | 1 |
| 0 | 0 | 0 | 0 |
| 0 | 0 | 0 | 0 |
| 0 | 0 | 0 | 0 |
| 0 | 0 | 0 | 0 |
| 0 | 0 | 0 | 0 |
| 0 | 0 | 0 | 0 |
| 0 | 0 | 0 | 0 |
| 0 | 0 | 0 | 1 |

| | | | |
|---|---|---|---|
| 0 | 0 | 0 | 0 |
| 0 | 0 | 0 | 0 |
| 0 | 0 | 0 | 1 |
| 1 | 0 | 0 | 0 |
| 1 | 0 | 0 | 0 |
| 0 | 0 | 0 | 1 |
| 0 | 0 | 0 | 1 |
| 0 | 0 | 0 | 1 |
| 0 | 0 | 0 | 1 |
| 0 | 0 | 0 | 1 |
| 0 | 0 | 0 | 1 |
| 0 | 0 | 0 | 0 |
| 0 | 0 | 0 | 1 |
| 0 | 0 | 0 | 1 |
| 0 | 0 | 0 | 0 |
| 0 | 0 | 0 | 1 |
| 0 | 0 | 0 | 0 |
| 0 | 0 | 0 | 1 |
| 0 | 0 | 0 | 0 |
| 1 | 0 | 0 | 0 |
| 0 | 0 | 0 | 1 |
| 0 | 0 | 0 | 1 |
| 0 | 0 | 0 | 0 |
| 0 | 0 | 0 | 1 |
| 0 | 0 | 0 | 0 |
| 0 | 0 | 0 | 1 |
| 0 | 0 | 0 | 1 |
| 0 | 0 | 0 | 0 |
| 0 | 0 | 0 | 1 |
| 0 | 0 | 0 | 0 |
| 0 | 0 | 0 | 1 |
| 0 | 0 | 0 | 0 |
| 0 | 0 | 0 | 0 |
| 0 | 0 | 0 | 0 |
| 0 | 0 | 0 | 1 |
| 0 | 0 | 0 | 0 |
| 0 | 0 | 0 | 1 |
| 0 | 0 | 0 | 1 |
| 0 | 0 | 0 | 0 |
| 0 | 0 | 0 | 1 |
| 0 | 0 | 0 | 1 |
| 0 | 0 | 0 | 1 |
| 0 | 0 | 0 | 1 |
| 0 | 0 | 0 | 0 |
| 0 | 0 | 0 | 0 |
| 0 | 0 | 0 | 1 |
| 0 | 0 | 0 | 1 |
| 0 | 0 | 0 | 1 |
| 0 | 1 | 0 | 0 |
| 0 | 0 | 0 | 1 |
| 0 | 0 | 0 | 0 |
| 0 | 0 | 0 | 1 |
| 0 | 0 | 0 | 1 |
| 0 | 0 | 0 | 0 |

| | | | |
|---|---|---|---|
| 0 | 0 | 0 | 1 |
| 0 | 0 | 0 | 1 |
| 0 | 0 | 0 | 1 |
| 0 | 0 | 0 | 1 |
| 0 | 0 | 0 | 1 |
| 0 | 0 | 0 | 1 |
| 0 | 0 | 0 | 1 |
| 0 | 0 | 0 | 0 |
| 0 | 0 | 0 | 1 |
| 0 | 0 | 0 | 1 |
| 0 | 0 | 0 | 1 |
| 0 | 0 | 0 | 1 |
| 0 | 0 | 0 | 0 |
| 0 | 0 | 0 | 1 |
| 0 | 0 | 0 | 0 |
| 0 | 0 | 0 | 0 |
| 0 | 0 | 0 | 1 |
| 0 | 0 | 0 | 1 |
| 0 | 0 | 0 | 1 |
| 0 | 0 | 0 | 1 |
| 0 | 0 | 0 | 0 |
| 0 | 0 | 0 | 0 |
| 0 | 0 | 0 | 1 |
| 0 | 0 | 0 | 1 |
| 0 | 0 | 0 | 0 |
| 0 | 0 | 0 | 1 |
| 0 | 0 | 0 | 0 |
| 0 | 0 | 0 | 0 |
| 0 | 0 | 0 | 0 |
| 0 | 0 | 0 | 0 |
| 0 | 0 | 0 | 1 |
| 0 | 0 | 0 | 1 |
| 0 | 0 | 0 | 0 |
| 0 | 1 | 0 | 0 |
| 0 | 0 | 0 | 0 |
| 0 | 0 | 0 | 0 |
| 0 | 0 | 0 | 0 |
| 0 | 0 | 0 | 0 |
| 0 | 0 | 0 | 0 |
| 0 | 0 | 0 | 1 |
| 0 | 0 | 0 | 1 |
| 0 | 0 | 0 | 1 |
| 0 | 0 | 0 | 1 |
| 0 | 0 | 0 | 1 |
| 0 | 0 | 0 | 1 |
| 0 | 0 | 0 | 0 |
| 0 | 0 | 0 | 1 |
| 0 | 0 | 0 | 1 |
| 0 | 0 | 0 | 1 |
| 0 | 0 | 0 | 0 |
| 0 | 0 | 0 | 0 |
| 0 | 0 | 0 | 1 |
| 0 | 0 | 0 | 1 |
| 0 | 0 | 0 | 1 |

| | | | |
|---|---|---|---|
| 0 | 0 | 0 | 1 |
| 0 | 0 | 0 | 1 |
| 0 | 0 | 0 | 1 |
| 0 | 0 | 0 | 1 |
| 0 | 0 | 0 | 0 |
| 0 | 0 | 0 | 1 |
| 0 | 0 | 0 | 0 |
| 0 | 0 | 0 | 0 |
| 0 | 0 | 0 | 0 |
| 0 | 0 | 0 | 0 |
| 0 | 0 | 0 | 0 |
| 0 | 0 | 0 | 1 |
| 0 | 0 | 0 | 0 |
| 0 | 0 | 0 | 0 |
| 0 | 0 | 0 | 1 |
| 0 | 0 | 0 | 1 |
| 0 | 0 | 0 | 1 |
| 0 | 0 | 0 | 1 |
| 0 | 0 | 0 | 1 |
| 0 | 0 | 0 | 1 |
| 0 | 0 | 0 | 1 |
| 0 | 0 | 0 | 1 |
| 0 | 0 | 0 | 1 |
| 0 | 0 | 0 | 0 |
| 0 | 0 | 0 | 1 |
| 0 | 0 | 0 | 1 |
| 0 | 0 | 0 | 1 |
| 0 | 0 | 0 | 0 |
| 0 | 1 | 1 | 1 |
| 0 | 1 | 1 | 1 |
| 0 | 0 | 0 | 0 |
| 0 | 0 | 0 | 0 |
| 0 | 0 | 0 | 1 |
| 0 | 0 | 0 | 0 |
| 0 | 0 | 0 | 1 |
| 0 | 0 | 0 | 1 |
| 0 | 0 | 0 | 1 |
| 0 | 0 | 0 | 1 |
| 0 | 0 | 0 | 1 |
| 0 | 0 | 0 | 1 |
| 0 | 0 | 0 | 1 |
| 0 | 0 | 0 | 1 |
| 0 | 0 | 0 | 1 |
| 0 | 0 | 0 | 1 |
| 0 | 0 | 0 | 1 |
| 0 | 0 | 0 | 1 |
| 0 | 0 | 0 | 1 |
| 0 | 0 | 0 | 1 |
| 0 | 0 | 0 | 0 |
| 0 | 0 | 0 | 1 |
| 0 | 0 | 0 | 1 |
| 0 | 0 | 0 | 1 |
| 0 | 0 | 0 | 1 |
| 0 | 0 | 0 | 1 |

| | | | |
|---|---|---|---|
| 0 | 0 | 0 | 1 |
| 0 | 0 | 0 | 1 |
| 0 | 0 | 0 | 1 |
| 0 | 0 | 0 | 1 |
| 0 | 0 | 0 | 1 |
| 0 | 0 | 0 | 1 |
| 0 | 0 | 0 | 1 |
| 0 | 0 | 0 | 1 |
| 0 | 0 | 0 | 1 |
| 0 | 0 | 0 | 1 |
| 0 | 0 | 0 | 0 |
| 0 | 0 | 0 | 0 |
| 0 | 0 | 0 | 0 |
| 1 | 0 | 0 | 0 |
| 1 | 0 | 0 | 0 |
| 0 | 0 | 0 | 1 |
| 1 | 0 | 0 | 0 |
| 0 | 0 | 0 | 0 |
| 0 | 0 | 0 | 0 |
| 0 | 0 | 0 | 0 |
| 0 | 0 | 0 | 0 |
| 1 | 0 | 0 | 0 |
| 0 | 0 | 0 | 1 |
| 0 | 0 | 0 | 0 |
| 1 | 0 | 0 | 0 |
| 1 | 0 | 0 | 0 |
| 0 | 0 | 0 | 1 |
| 0 | 0 | 0 | 0 |
| 0 | 0 | 0 | 1 |
| 0 | 0 | 0 | 1 |
| 0 | 0 | 0 | 1 |
| 0 | 0 | 0 | 0 |
| 0 | 0 | 0 | 1 |
| 0 | 0 | 0 | 1 |
| 0 | 0 | 0 | 0 |
| 0 | 0 | 0 | 0 |
| 0 | 0 | 0 | 0 |
| 0 | 0 | 0 | 0 |
| 0 | 0 | 0 | 0 |
| 0 | 0 | 0 | 1 |
| 0 | 0 | 0 | 1 |
| 0 | 0 | 0 | 0 |
| 0 | 0 | 0 | 1 |
| 0 | 0 | 0 | 0 |
| 0 | 0 | 0 | 1 |
| 0 | 0 | 0 | 1 |
| 0 | 0 | 0 | 1 |
| 0 | 1 | 0 | 0 |
| 0 | 0 | 0 | 0 |
| 0 | 0 | 0 | 0 |
| 0 | 0 | 0 | 0 |
| 0 | 0 | 0 | 1 |
| 0 | 0 | 0 | 0 |
| 0 | 0 | 0 | 1 |

| | | | |
|---|---|---|---|
| 0 | 0 | 0 | 1 |
| 0 | 0 | 0 | 1 |
| 0 | 0 | 0 | 1 |
| 0 | 0 | 0 | 1 |
| 0 | 0 | 0 | 1 |
| 0 | 0 | 0 | 0 |
| 0 | 0 | 0 | 0 |
| 0 | 0 | 0 | 0 |
| 0 | 0 | 0 | 0 |
| 0 | 0 | 0 | 1 |
| 1 | 0 | 0 | 0 |
| 1 | 0 | 0 | 0 |
| 0 | 0 | 0 | 0 |
| 0 | 0 | 0 | 0 |
| 0 | 0 | 0 | 0 |
| 0 | 0 | 0 | 0 |
| 0 | 0 | 0 | 1 |
| 0 | 0 | 0 | 1 |
| 0 | 0 | 0 | 1 |
| 0 | 0 | 0 | 1 |
| 0 | 0 | 0 | 1 |
| 0 | 0 | 0 | 1 |
| 0 | 0 | 0 | 1 |
| 0 | 0 | 0 | 1 |
| 0 | 0 | 0 | 1 |
| 0 | 0 | 0 | 1 |
| 0 | 0 | 0 | 1 |
| 0 | 0 | 0 | 1 |
| 0 | 1 | 0 | 0 |
| 0 | 0 | 0 | 1 |
| 0 | 0 | 0 | 1 |
| 0 | 1 | 1 | 1 |
| 0 | 1 | 1 | 1 |
| 0 | 0 | 0 | 0 |
| 0 | 0 | 0 | 1 |
| 0 | 0 | 0 | 0 |
| 0 | 0 | 0 | 1 |
| 0 | 0 | 0 | 0 |
| 0 | 0 | 0 | 1 |
| 0 | 0 | 0 | 0 |
| 0 | 0 | 0 | 1 |
| 0 | 0 | 0 | 1 |
| 0 | 0 | 0 | 1 |
| 0 | 0 | 0 | 0 |
| 0 | 0 | 0 | 1 |
| 0 | 0 | 0 | 1 |
| 0 | 0 | 0 | 1 |
| 0 | 0 | 0 | 1 |
| 0 | 0 | 0 | 1 |
| 0 | 0 | 0 | 1 |
| 0 | 0 | 0 | 1 |
| 0 | 0 | 0 | 1 |
| 0 | 0 | 0 | 0 |
| 0 | 0 | 0 | 0 |

| | | | |
|---|---|---|---|
| 0 | 0 | 0 | 1 |
| 0 | 0 | 0 | 0 |
| 0 | 0 | 0 | 1 |
| 0 | 0 | 0 | 1 |
| 0 | 0 | 0 | 0 |
| 0 | 0 | 0 | 0 |
| 0 | 0 | 0 | 0 |
| 0 | 0 | 0 | 1 |
| 0 | 0 | 0 | 1 |
| 0 | 0 | 0 | 0 |
| 0 | 0 | 0 | 0 |
| 0 | 0 | 0 | 0 |
| 0 | 0 | 0 | 0 |
| 0 | 0 | 0 | 0 |
| 0 | 0 | 0 | 1 |
| 0 | 0 | 0 | 0 |
| 0 | 0 | 0 | 0 |
| 0 | 0 | 0 | 0 |
| 0 | 0 | 0 | 1 |
| 0 | 0 | 0 | 1 |
| 0 | 0 | 0 | 1 |
| 0 | 0 | 0 | 1 |
| 0 | 0 | 0 | 1 |
| 0 | 0 | 0 | 0 |
| 0 | 0 | 0 | 1 |
| 0 | 0 | 0 | 1 |
| 0 | 0 | 0 | 0 |
| 0 | 0 | 0 | 0 |
| 0 | 0 | 0 | 1 |
| 0 | 0 | 0 | 1 |
| 0 | 0 | 0 | 0 |
| 0 | 0 | 0 | 1 |
| 0 | 0 | 0 | 1 |
| 0 | 0 | 0 | 1 |
| 0 | 0 | 0 | 1 |
| 0 | 0 | 0 | 0 |
| 0 | 0 | 0 | 1 |
| 0 | 0 | 0 | 0 |
| 0 | 0 | 0 | 0 |
| 0 | 0 | 0 | 1 |
| 0 | 0 | 0 | 1 |
| 0 | 0 | 0 | 0 |
| 0 | 0 | 0 | 0 |
| 0 | 0 | 0 | 1 |
| 0 | 0 | 0 | 0 |
| 0 | 0 | 0 | 0 |
| 0 | 0 | 0 | 0 |
| 0 | 0 | 0 | 1 |
| 0 | 0 | 0 | 0 |
| 0 | 0 | 0 | 0 |
| 0 | 0 | 0 | 0 |
| 0 | 0 | 0 | 1 |
| 0 | 0 | 0 | 0 |
| 0 | 0 | 0 | 1 |

| | | | |
|---|---|---|---|
| 0 | 0 | 0 | 1 |
| 0 | 0 | 0 | 1 |
| 0 | 0 | 0 | 1 |
| 0 | 0 | 0 | 1 |
| 0 | 0 | 0 | 0 |
| 0 | 0 | 0 | 0 |
| 0 | 0 | 0 | 1 |
| 0 | 0 | 0 | 1 |
| 0 | 0 | 0 | 1 |
| 0 | 0 | 0 | 0 |
| 0 | 0 | 0 | 1 |
| 0 | 0 | 0 | 1 |
| 0 | 1 | 0 | 0 |
| 0 | 0 | 0 | 1 |
| 0 | 0 | 0 | 1 |
| 0 | 0 | 0 | 1 |
| 0 | 0 | 0 | 1 |
| 0 | 0 | 0 | 1 |
| 0 | 0 | 0 | 1 |
| 0 | 0 | 0 | 1 |
| 0 | 0 | 0 | 1 |
| 0 | 0 | 0 | 1 |
| 0 | 0 | 0 | 1 |
| 0 | 0 | 0 | 0 |
| 0 | 0 | 0 | 1 |
| 1 | 0 | 0 | 0 |
| 0 | 0 | 0 | 1 |
| 0 | 0 | 0 | 0 |
| 0 | 0 | 0 | 1 |
| 0 | 0 | 0 | 0 |
| 0 | 0 | 0 | 0 |
| 0 | 0 | 0 | 0 |
| 0 | 0 | 0 | 0 |
| 0 | 0 | 0 | 0 |
| 0 | 0 | 0 | 1 |
| 0 | 0 | 0 | 0 |
| 0 | 0 | 0 | 1 |
| 0 | 0 | 0 | 0 |
| 0 | 0 | 0 | 1 |
| 0 | 0 | 0 | 1 |
| 0 | 0 | 0 | 0 |
| 0 | 0 | 0 | 0 |
| 0 | 0 | 0 | 0 |
| 0 | 0 | 0 | 0 |
| 0 | 0 | 0 | 0 |
| 0 | 0 | 0 | 0 |
| 0 | 0 | 0 | 0 |
| 0 | 0 | 0 | 0 |
| 0 | 0 | 0 | 1 |
| 0 | 0 | 0 | 0 |
| 0 | 0 | 0 | 1 |
| 0 | 0 | 0 | 1 |
| 0 | 0 | 0 | 1 |
| 0 | 0 | 0 | 1 |

| | | | |
|---|---|---|---|
| 0 | 0 | 0 | 1 |
| 0 | 0 | 0 | 1 |
| 0 | 0 | 0 | 1 |
| 0 | 0 | 0 | 1 |
| 0 | 0 | 0 | 1 |
| 0 | 0 | 0 | 1 |
| 0 | 0 | 0 | 0 |
| 0 | 0 | 0 | 0 |
| 0 | 0 | 0 | 0 |
| 0 | 0 | 0 | 0 |
| 0 | 0 | 0 | 0 |
| 0 | 0 | 0 | 0 |
| 0 | 0 | 0 | 0 |
| 0 | 0 | 0 | 0 |
| 0 | 0 | 0 | 0 |
| 0 | 0 | 0 | 0 |
| 1 | 0 | 0 | 0 |
| 0 | 0 | 0 | 0 |
| 0 | 0 | 0 | 1 |
| 0 | 0 | 0 | 0 |
| 0 | 0 | 0 | 1 |
| 0 | 0 | 0 | 1 |
| 0 | 0 | 0 | 1 |
| 0 | 0 | 0 | 1 |
| 0 | 0 | 0 | 1 |
| 0 | 0 | 0 | 0 |
| 0 | 0 | 0 | 0 |
| 0 | 0 | 0 | 0 |
| 0 | 0 | 0 | 0 |
| 0 | 0 | 0 | 0 |
| 0 | 0 | 0 | 0 |
| 1 | 0 | 0 | 0 |
| 0 | 0 | 0 | 0 |
| 1 | 0 | 0 | 0 |
| 0 | 0 | 0 | 1 |
| 1 | 0 | 0 | 0 |
| 1 | 0 | 0 | 0 |
| 0 | 0 | 0 | 0 |
| 1 | 0 | 0 | 0 |
| 0 | 0 | 0 | 1 |
| 0 | 0 | 0 | 0 |
| 0 | 0 | 0 | 0 |
| 0 | 0 | 0 | 1 |
| 0 | 0 | 0 | 1 |
| 0 | 0 | 0 | 1 |
| 0 | 0 | 0 | 1 |
| 0 | 0 | 0 | 1 |
| 0 | 0 | 0 | 0 |
| 0 | 0 | 0 | 0 |
| 0 | 0 | 0 | 0 |
| 0 | 0 | 0 | 0 |
| 0 | 0 | 0 | 1 |
| 0 | 0 | 0 | 0 |
| 0 | 0 | 0 | 1 |

| | | | |
|---|---|---|---|
| 0 | 0 | 0 | 1 |
| 0 | 0 | 0 | 1 |
| 0 | 0 | 0 | 1 |
| 0 | 0 | 0 | 1 |
| 0 | 0 | 0 | 1 |
| 0 | 0 | 0 | 0 |
| 0 | 0 | 0 | 1 |
| 0 | 0 | 0 | 1 |
| 0 | 0 | 0 | 1 |
| 0 | 0 | 0 | 1 |
| 0 | 0 | 0 | 0 |
| 0 | 1 | 0 | 0 |
| 0 | 1 | 0 | 0 |
| 0 | 0 | 0 | 1 |
| 0 | 0 | 0 | 1 |
| 0 | 0 | 0 | 1 |
| 0 | 0 | 0 | 1 |
| 0 | 0 | 0 | 0 |
| 0 | 0 | 0 | 1 |
| 0 | 0 | 0 | 1 |
| 0 | 0 | 0 | 1 |
| 0 | 0 | 0 | 1 |
| 0 | 0 | 0 | 0 |
| 0 | 0 | 0 | 1 |
| 0 | 0 | 0 | 0 |
| 0 | 0 | 0 | 0 |
| 0 | 0 | 0 | 0 |
| 0 | 0 | 0 | 0 |
| 0 | 0 | 0 | 1 |
| 0 | 0 | 0 | 0 |
| 0 | 0 | 0 | 1 |
| 0 | 0 | 0 | 1 |
| 0 | 0 | 0 | 1 |
| 0 | 0 | 0 | 1 |
| 0 | 0 | 0 | 1 |
| 0 | 0 | 0 | 1 |
| 0 | 0 | 0 | 1 |
| 0 | 0 | 0 | 0 |
| 0 | 0 | 0 | 0 |
| 0 | 0 | 0 | 0 |
| 0 | 1 | 0 | 0 |
| 0 | 0 | 0 | 0 |
| 0 | 0 | 0 | 1 |
| 0 | 0 | 0 | 0 |
| 0 | 0 | 0 | 0 |
| 0 | 0 | 0 | 1 |
| 0 | 0 | 0 | 0 |
| 0 | 1 | 0 | 0 |
| 0 | 0 | 0 | 0 |
| 1 | 0 | 0 | 0 |
| 0 | 0 | 0 | 1 |
| 0 | 0 | 0 | 1 |
| 0 | 0 | 0 | 1 |
| 0 | 0 | 0 | 0 |

| | | | |
|---|---|---|---|
| 0 | 0 | 0 | 0 |
| 0 | 0 | 0 | 1 |
| 0 | 0 | 0 | 0 |
| 0 | 0 | 0 | 1 |
| 0 | 0 | 0 | 1 |
| 0 | 0 | 0 | 1 |
| 0 | 0 | 0 | 1 |
| 0 | 0 | 0 | 1 |
| 0 | 1 | 0 | 0 |
| 0 | 0 | 0 | 1 |
| 0 | 0 | 0 | 0 |
| 0 | 0 | 0 | 0 |
| 0 | 0 | 0 | 0 |
| 0 | 0 | 0 | 0 |
| 0 | 0 | 0 | 1 |
| 0 | 0 | 0 | 1 |
| 0 | 0 | 0 | 0 |
| 0 | 0 | 0 | 0 |
| 0 | 0 | 0 | 1 |
| 0 | 0 | 0 | 1 |
| 0 | 0 | 0 | 0 |
| 0 | 0 | 0 | 1 |
| 0 | 0 | 0 | 1 |
| 0 | 0 | 0 | 1 |
| 0 | 0 | 0 | 0 |
| 0 | 0 | 0 | 1 |
| 0 | 1 | 0 | 0 |
| 0 | 1 | 0 | 0 |
| 0 | 1 | 0 | 0 |
| 0 | 0 | 0 | 1 |
| 0 | 0 | 0 | 0 |
| 0 | 0 | 0 | 0 |
| 0 | 0 | 0 | 0 |
| 0 | 0 | 0 | 1 |
| 0 | 0 | 0 | 0 |
| 0 | 0 | 0 | 1 |
| 0 | 0 | 0 | 0 |
| 0 | 0 | 0 | 0 |
| 0 | 1 | 0 | 0 |
| 0 | 0 | 0 | 0 |
| 0 | 0 | 0 | 0 |
| 0 | 0 | 0 | 1 |
| 0 | 0 | 0 | 0 |
| 0 | 1 | 0 | 0 |
| 0 | 1 | 0 | 0 |
| 1 | 0 | 0 | 0 |
| 0 | 0 | 0 | 0 |
| 1 | 0 | 0 | 0 |
| 1 | 0 | 0 | 0 |
| 1 | 0 | 0 | 0 |
| 0 | 0 | 0 | 0 |
| 0 | 0 | 0 | 0 |
| 0 | 0 | 0 | 1 |
| 0 | 0 | 0 | 0 |

| | | | |
|---|---|---|---|
| 0 | 0 | 0 | 1 |
| 0 | 0 | 0 | 1 |
| 0 | 0 | 0 | 1 |
| 0 | 0 | 0 | 1 |
| 0 | 0 | 0 | 1 |
| 0 | 0 | 0 | 1 |
| 0 | 0 | 0 | 1 |
| 0 | 0 | 0 | 1 |
| 0 | 0 | 0 | 1 |
| 0 | 0 | 0 | 0 |
| 0 | 0 | 0 | 0 |
| 0 | 0 | 0 | 1 |
| 0 | 0 | 0 | 1 |
| 0 | 0 | 0 | 1 |
| 0 | 0 | 0 | 0 |
| 0 | 0 | 0 | 1 |
| 0 | 0 | 0 | 1 |
| 0 | 0 | 0 | 0 |
| 0 | 0 | 0 | 1 |
| 0 | 0 | 0 | 1 |
| 0 | 0 | 0 | 1 |
| 0 | 0 | 0 | 0 |
| 0 | 0 | 0 | 0 |
| 0 | 0 | 0 | 0 |
| 0 | 0 | 0 | 0 |
| 0 | 0 | 0 | 0 |
| 0 | 0 | 0 | 0 |
| 0 | 0 | 0 | 0 |
| 0 | 0 | 0 | 1 |
| 0 | 0 | 0 | 0 |
| 1 | 0 | 0 | 0 |
| 0 | 0 | 0 | 1 |
| 0 | 0 | 0 | 1 |
| 0 | 0 | 0 | 1 |
| 0 | 0 | 0 | 1 |
| 0 | 0 | 0 | 1 |
| 0 | 0 | 0 | 1 |
| 0 | 0 | 0 | 1 |
| 0 | 0 | 0 | 1 |
| 0 | 0 | 0 | 1 |
| 0 | 0 | 0 | 1 |
| 0 | 0 | 0 | 1 |
| 0 | 0 | 0 | 1 |
| 0 | 0 | 0 | 0 |
| 0 | 0 | 0 | 1 |
| 0 | 0 | 0 | 0 |
| 0 | 0 | 0 | 1 |
| 0 | 0 | 0 | 1 |
| 0 | 0 | 0 | 1 |
| 0 | 0 | 0 | 1 |
| 0 | 0 | 0 | 0 |
| 0 | 0 | 0 | 1 |
| 0 | 0 | 0 | 1 |
| 0 | 0 | 0 | 0 |

| | | | |
|---|---|---|---|
| 0 | 0 | 0 | 0 |
| 0 | 0 | 0 | 0 |
| 0 | 0 | 0 | 0 |
| 0 | 0 | 0 | 0 |
| 0 | 0 | 0 | 1 |
| 0 | 0 | 0 | 1 |
| 0 | 0 | 0 | 1 |
| 0 | 0 | 0 | 0 |
| 0 | 0 | 0 | 0 |
| 0 | 0 | 0 | 0 |
| 0 | 0 | 0 | 0 |
| 0 | 0 | 0 | 1 |
| 0 | 0 | 0 | 1 |
| 0 | 0 | 0 | 1 |
| 0 | 0 | 0 | 1 |
| 0 | 0 | 0 | 0 |
| 0 | 0 | 0 | 0 |
| 0 | 0 | 0 | 1 |
| 0 | 0 | 0 | 1 |
| 0 | 0 | 0 | 1 |
| 0 | 0 | 0 | 1 |
| 0 | 0 | 0 | 0 |
| 0 | 0 | 0 | 1 |
| 0 | 0 | 0 | 1 |
| 0 | 0 | 0 | 0 |
| 0 | 0 | 0 | 0 |
| 0 | 0 | 0 | 0 |
| 0 | 0 | 0 | 0 |
| 0 | 0 | 0 | 1 |
| 0 | 0 | 0 | 1 |
| 0 | 0 | 0 | 1 |
| 0 | 0 | 0 | 0 |
| 0 | 0 | 0 | 1 |
| 0 | 0 | 0 | 1 |
| 0 | 0 | 0 | 1 |
| 0 | 0 | 0 | 1 |
| 0 | 0 | 0 | 1 |
| 0 | 0 | 0 | 1 |
| 0 | 0 | 0 | 1 |
| 0 | 0 | 0 | 1 |
| 0 | 0 | 0 | 1 |
| 0 | 0 | 0 | 1 |
| 0 | 0 | 0 | 0 |
| 0 | 0 | 0 | 1 |
| 0 | 0 | 0 | 1 |
| 0 | 0 | 0 | 1 |
| 0 | 0 | 0 | 1 |
| 0 | 0 | 0 | 1 |
| 0 | 0 | 0 | 1 |
| 0 | 0 | 0 | 0 |
| 0 | 0 | 0 | 1 |
| 0 | 0 | 0 | 1 |
| 0 | 0 | 0 | 1 |
| 0 | 0 | 0 | 1 |

| | | | |
|---|---|---|---|
| 0 | 0 | 0 | 0 |
| 0 | 0 | 0 | 1 |
| 0 | 0 | 0 | 1 |
| 0 | 0 | 0 | 1 |
| 0 | 0 | 0 | 1 |
| 0 | 0 | 0 | 1 |
| 0 | 0 | 0 | 1 |
| 0 | 1 | 0 | 0 |
| 0 | 0 | 0 | 1 |
| 0 | 0 | 0 | 1 |
| 0 | 0 | 0 | 0 |
| 0 | 0 | 0 | 0 |
| 0 | 0 | 0 | 0 |
| 0 | 0 | 0 | 1 |
| 0 | 0 | 0 | 0 |
| 0 | 0 | 0 | 0 |
| 0 | 0 | 0 | 0 |
| 0 | 0 | 0 | 0 |
| 0 | 0 | 0 | 1 |
| 0 | 0 | 0 | 0 |
| 0 | 0 | 0 | 0 |
| 0 | 0 | 0 | 0 |
| 0 | 0 | 0 | 0 |
| 0 | 0 | 0 | 0 |
| 0 | 0 | 0 | 0 |
| 0 | 0 | 0 | 0 |
| 0 | 0 | 0 | 0 |
| 0 | 0 | 0 | 0 |
| 0 | 0 | 0 | 0 |
| 0 | 0 | 0 | 0 |
| 0 | 0 | 0 | 0 |
| 0 | 0 | 0 | 0 |
| 0 | 0 | 0 | 1 |
| 0 | 0 | 0 | 0 |
| 0 | 0 | 0 | 1 |
| 0 | 0 | 0 | 0 |
| 0 | 0 | 0 | 0 |
| 0 | 0 | 0 | 1 |
| 0 | 0 | 0 | 1 |
| 0 | 0 | 0 | 1 |
| 0 | 0 | 0 | 1 |
| 0 | 0 | 0 | 0 |
| 0 | 0 | 0 | 0 |
| 0 | 0 | 0 | 0 |
| 0 | 0 | 0 | 1 |
| 0 | 0 | 0 | 0 |
| 0 | 0 | 0 | 1 |
| 0 | 0 | 0 | 0 |
| 0 | 0 | 0 | 1 |
| 0 | 0 | 0 | 1 |
| 0 | 0 | 0 | 1 |
| 0 | 0 | 0 | 1 |
| 0 | 0 | 0 | 1 |
| 0 | 0 | 0 | 1 |

| | | | |
|---|---|---|---|
| 0 | 0 | 0 | 1 |
| 0 | 0 | 0 | 1 |
| 0 | 0 | 0 | 1 |
| 0 | 0 | 0 | 1 |
| 0 | 0 | 0 | 1 |
| 0 | 0 | 0 | 0 |
| 0 | 0 | 0 | 0 |
| 0 | 0 | 0 | 1 |
| 0 | 0 | 0 | 0 |
| 0 | 0 | 0 | 0 |
| 0 | 0 | 0 | 1 |
| 0 | 0 | 0 | 1 |
| 0 | 0 | 0 | 1 |
| 0 | 0 | 0 | 1 |
| 0 | 0 | 0 | 1 |
| 0 | 0 | 0 | 1 |
| 0 | 0 | 0 | 1 |
| 0 | 0 | 0 | 1 |
| 0 | 0 | 0 | 0 |
| 0 | 0 | 0 | 0 |
| 0 | 0 | 0 | 1 |
| 0 | 0 | 0 | 1 |
| 0 | 0 | 0 | 1 |
| 0 | 0 | 0 | 0 |
| 0 | 0 | 0 | 1 |
| 0 | 0 | 0 | 0 |
| 0 | 0 | 0 | 0 |
| 0 | 0 | 0 | 1 |
| 0 | 0 | 0 | 1 |
| 0 | 0 | 0 | 1 |
| 0 | 0 | 0 | 0 |
| 0 | 0 | 0 | 1 |
| 0 | 0 | 0 | 0 |
| 0 | 0 | 0 | 0 |
| 0 | 0 | 0 | 0 |
| 0 | 0 | 0 | 0 |
| 0 | 0 | 0 | 0 |
| 0 | 0 | 0 | 1 |
| 0 | 0 | 0 | 1 |
| 0 | 0 | 0 | 1 |
| 0 | 0 | 0 | 0 |
| 0 | 0 | 0 | 0 |
| 0 | 0 | 0 | 0 |
| 0 | 0 | 0 | 0 |
| 0 | 0 | 0 | 1 |
| 0 | 0 | 0 | 1 |
| 0 | 0 | 0 | 1 |
| 0 | 0 | 0 | 0 |
| 0 | 0 | 0 | 0 |
| 0 | 0 | 0 | 0 |
| 0 | 0 | 0 | 1 |
| 0 | 0 | 0 | 1 |
| 0 | 0 | 0 | 1 |
| 0 | 0 | 0 | 1 |

| | | | |
|---|---|---|---|
| 0 | 0 | 0 | 1 |
| 0 | 0 | 0 | 1 |
| 0 | 0 | 0 | 1 |
| 0 | 0 | 0 | 0 |
| 0 | 0 | 0 | 0 |
| 0 | 0 | 0 | 1 |
| 0 | 0 | 0 | 0 |
| 0 | 0 | 0 | 1 |
| 0 | 0 | 0 | 1 |
| 0 | 0 | 0 | 1 |
| 0 | 0 | 0 | 1 |
| 0 | 0 | 0 | 0 |
| 0 | 0 | 0 | 1 |
| 0 | 0 | 0 | 0 |
| 0 | 0 | 0 | 1 |
| 0 | 0 | 0 | 0 |
| 0 | 0 | 0 | 1 |
| 0 | 0 | 0 | 1 |
| 0 | 0 | 0 | 1 |
| 0 | 0 | 0 | 0 |
| 0 | 0 | 0 | 0 |
| 0 | 0 | 0 | 0 |
| 0 | 0 | 0 | 0 |
| 0 | 0 | 0 | 0 |
| 0 | 0 | 0 | 0 |
| 0 | 0 | 0 | 1 |
| 0 | 0 | 0 | 0 |
| 0 | 0 | 0 | 1 |
| 0 | 0 | 0 | 1 |
| 0 | 0 | 0 | 1 |
| 0 | 0 | 0 | 1 |
| 0 | 0 | 0 | 1 |
| 0 | 0 | 0 | 0 |
| 0 | 1 | 0 | 0 |
| 0 | 0 | 0 | 1 |
| 0 | 0 | 0 | 1 |
| 0 | 1 | 0 | 0 |
| 0 | 1 | 0 | 0 |
| 0 | 0 | 0 | 0 |
| 0 | 1 | 1 | 1 |
| 0 | 0 | 0 | 0 |
| 0 | 0 | 0 | 0 |
| 0 | 0 | 0 | 1 |
| 0 | 0 | 0 | 0 |
| 0 | 0 | 0 | 1 |
| 0 | 0 | 0 | 1 |
| 0 | 0 | 0 | 1 |
| 0 | 0 | 0 | 1 |
| 0 | 0 | 0 | 1 |
| 1 | 0 | 0 | 0 |
| 0 | 0 | 0 | 1 |
| 0 | 0 | 0 | 1 |
| 0 | 0 | 0 | 1 |
| 0 | 0 | 0 | 1 |

| | | | |
|---|---|---|---|
| 0 | 0 | 0 | 1 |
| 0 | 0 | 0 | 1 |
| 0 | 0 | 0 | 1 |
| 0 | 0 | 0 | 0 |
| 0 | 0 | 0 | 1 |
| 0 | 0 | 0 | 1 |
| 0 | 0 | 0 | 1 |
| 0 | 0 | 0 | 0 |
| 0 | 0 | 0 | 1 |
| 0 | 0 | 0 | 1 |
| 0 | 0 | 0 | 1 |
| 0 | 0 | 0 | 1 |
| 0 | 0 | 0 | 1 |
| 0 | 0 | 0 | 1 |
| 0 | 0 | 0 | 1 |
| 0 | 0 | 0 | 1 |
| 0 | 0 | 0 | 0 |
| 0 | 0 | 0 | 0 |
| 0 | 0 | 0 | 1 |
| 0 | 0 | 0 | 1 |
| 0 | 0 | 0 | 1 |
| 0 | 0 | 0 | 1 |
| 0 | 0 | 0 | 1 |
| 0 | 0 | 0 | 1 |
| 0 | 0 | 0 | 1 |
| 0 | 0 | 0 | 1 |
| 0 | 0 | 0 | 1 |
| 0 | 0 | 0 | 0 |
| 0 | 0 | 0 | 0 |
| 0 | 0 | 0 | 1 |
| 0 | 0 | 0 | 1 |
| 0 | 0 | 0 | 0 |
| 0 | 0 | 0 | 1 |
| 0 | 0 | 0 | 0 |
| 1 | 0 | 0 | 0 |
| 1 | 0 | 0 | 0 |
| 1 | 0 | 0 | 0 |
| 1 | 0 | 0 | 0 |
| 0 | 0 | 0 | 1 |
| 0 | 0 | 0 | 1 |
| 0 | 0 | 0 | 1 |
| 0 | 0 | 0 | 1 |
| 0 | 0 | 0 | 1 |
| 0 | 0 | 0 | 1 |
| 0 | 0 | 0 | 0 |
| 0 | 0 | 0 | 0 |
| 0 | 0 | 0 | 0 |
| 0 | 0 | 0 | 1 |
| 0 | 0 | 0 | 1 |
| 0 | 0 | 0 | 1 |
| 0 | 0 | 0 | 1 |
| 0 | 0 | 0 | 0 |
| 0 | 0 | 0 | 0 |
| 0 | 0 | 0 | 0 |

| | | | |
|---|---|---|---|
| 0 | 0 | 0 | 0 |
| 0 | 0 | 0 | 0 |
| 0 | 0 | 0 | 1 |
| 0 | 0 | 0 | 0 |
| 0 | 0 | 0 | 0 |
| 0 | 0 | 0 | 0 |
| 0 | 0 | 0 | 0 |
| 0 | 0 | 0 | 1 |
| 0 | 0 | 0 | 1 |
| 0 | 0 | 0 | 1 |
| 0 | 0 | 0 | 1 |
| 0 | 0 | 0 | 1 |
| 0 | 0 | 0 | 1 |
| 0 | 0 | 0 | 1 |
| 0 | 0 | 0 | 1 |
| 0 | 0 | 0 | 1 |
| 0 | 0 | 0 | 1 |
| 0 | 0 | 0 | 1 |
| 0 | 0 | 0 | 1 |
| 0 | 0 | 0 | 1 |
| 0 | 0 | 0 | 0 |
| 0 | 0 | 0 | 0 |
| 0 | 0 | 0 | 0 |
| 0 | 0 | 0 | 0 |
| 0 | 0 | 0 | 0 |
| 0 | 0 | 0 | 1 |
| 0 | 0 | 0 | 1 |
| 0 | 0 | 0 | 0 |
| 0 | 0 | 0 | 1 |
| 0 | 0 | 0 | 1 |
| 0 | 0 | 0 | 0 |
| 0 | 0 | 0 | 0 |
| 0 | 0 | 0 | 0 |
| 0 | 0 | 0 | 1 |
| 0 | 0 | 0 | 0 |
| 0 | 0 | 0 | 1 |
| 0 | 0 | 0 | 1 |
| 0 | 0 | 0 | 0 |
| 0 | 0 | 0 | 1 |
| 0 | 0 | 0 | 1 |
| 0 | 0 | 0 | 1 |
| 0 | 0 | 0 | 1 |
| 0 | 0 | 0 | 1 |
| 0 | 0 | 0 | 1 |
| 0 | 0 | 0 | 1 |
| 0 | 0 | 0 | 1 |
| 0 | 0 | 0 | 1 |
| 0 | 0 | 0 | 1 |
| 0 | 0 | 0 | 0 |
| 0 | 0 | 0 | 1 |
| 1 | 0 | 0 | 0 |
| 1 | 0 | 0 | 0 |
| 0 | 0 | 0 | 1 |
| 0 | 0 | 0 | 1 |

| | | | |
|---|---|---|---|
| 1 | 0 | 0 | 0 |
| 0 | 0 | 0 | 1 |
| 0 | 0 | 0 | 1 |
| 0 | 0 | 0 | 1 |
| 0 | 0 | 0 | 1 |
| 1 | 0 | 0 | 0 |
| 1 | 0 | 0 | 0 |
| 0 | 0 | 0 | 1 |
| 0 | 0 | 0 | 1 |
| 0 | 0 | 0 | 1 |
| 0 | 0 | 0 | 1 |
| 0 | 0 | 0 | 0 |
| 0 | 0 | 0 | 1 |
| 0 | 0 | 0 | 1 |
| 0 | 0 | 0 | 1 |
| 0 | 0 | 0 | 1 |
| 0 | 0 | 0 | 0 |
| 0 | 0 | 0 | 0 |
| 0 | 0 | 0 | 1 |
| 0 | 0 | 0 | 1 |
| 0 | 0 | 0 | 1 |
| 0 | 0 | 0 | 1 |
| 0 | 0 | 0 | 1 |
| 0 | 0 | 0 | 1 |
| 0 | 0 | 0 | 0 |
| 0 | 0 | 0 | 1 |
| 0 | 0 | 0 | 1 |
| 0 | 0 | 0 | 1 |
| 0 | 0 | 0 | 1 |
| 0 | 0 | 0 | 1 |
| 0 | 0 | 0 | 1 |
| 0 | 0 | 0 | 0 |
| 0 | 0 | 0 | 0 |
| 0 | 0 | 0 | 1 |
| 0 | 0 | 0 | 1 |
| 0 | 0 | 0 | 0 |
| 1 | 0 | 0 | 0 |
| 0 | 0 | 0 | 0 |
| 1 | 0 | 0 | 0 |
| 0 | 0 | 0 | 0 |
| 0 | 0 | 0 | 0 |
| 0 | 0 | 0 | 0 |
| 0 | 0 | 0 | 0 |
| 0 | 0 | 0 | 0 |
| 0 | 0 | 0 | 0 |
| 0 | 0 | 0 | 0 |
| 0 | 0 | 0 | 0 |
| 1 | 0 | 0 | 0 |
| 0 | 0 | 0 | 1 |
| 0 | 0 | 0 | 1 |
| 0 | 0 | 0 | 1 |
| 0 | 0 | 0 | 1 |
| 0 | 0 | 0 | 0 |
| 0 | 0 | 0 | 1 |

| | | | |
|---|---|---|---|
| 0 | 0 | 0 | 1 |
| 0 | 0 | 0 | 1 |
| 0 | 0 | 0 | 1 |
| 0 | 0 | 0 | 1 |
| 0 | 0 | 0 | 1 |
| 0 | 0 | 0 | 1 |
| 0 | 0 | 0 | 1 |
| 0 | 0 | 0 | 0 |
| 0 | 0 | 0 | 0 |
| 0 | 0 | 0 | 1 |
| 0 | 0 | 0 | 1 |
| 0 | 0 | 0 | 1 |
| 0 | 0 | 0 | 1 |
| 0 | 0 | 0 | 0 |
| 0 | 0 | 0 | 1 |
| 0 | 0 | 0 | 1 |
| 0 | 1 | 0 | 0 |
| 0 | 0 | 0 | 1 |
| 1 | 0 | 0 | 0 |
| 0 | 0 | 0 | 1 |
| 0 | 0 | 0 | 1 |
| 0 | 0 | 0 | 1 |
| 0 | 0 | 0 | 1 |
| 0 | 0 | 0 | 0 |
| 0 | 0 | 0 | 1 |
| 0 | 0 | 0 | 1 |
| 0 | 0 | 0 | 1 |
| 0 | 0 | 0 | 1 |
| 0 | 0 | 0 | 1 |
| 0 | 0 | 0 | 1 |
| 0 | 0 | 0 | 1 |
| 1 | 0 | 0 | 0 |
| 0 | 0 | 0 | 1 |
| 0 | 0 | 0 | 1 |
| 0 | 0 | 0 | 1 |
| 0 | 0 | 0 | 1 |
| 0 | 0 | 0 | 1 |
| 0 | 0 | 0 | 1 |
| 1 | 0 | 0 | 0 |
| 0 | 0 | 0 | 1 |
| 0 | 0 | 0 | 1 |
| 0 | 0 | 0 | 1 |
| 0 | 0 | 0 | 1 |
| 0 | 0 | 0 | 1 |
| 0 | 0 | 0 | 1 |
| 0 | 0 | 0 | 0 |
| 0 | 0 | 0 | 1 |
| 0 | 0 | 0 | 1 |
| 0 | 0 | 0 | 1 |
| 0 | 0 | 0 | 1 |
| 0 | 0 | 0 | 1 |
| 0 | 0 | 0 | 1 |
| 0 | 0 | 0 | 1 |
| 0 | 0 | 0 | 1 |

| | | | |
|---|---|---|---|
| 0 | 0 | 0 | 1 |
| 0 | 0 | 0 | 1 |
| 0 | 0 | 0 | 1 |
| 0 | 0 | 0 | 1 |
| 0 | 0 | 0 | 1 |
| 0 | 0 | 0 | 1 |
| 0 | 0 | 0 | 1 |
| 0 | 0 | 0 | 1 |
| 0 | 0 | 0 | 1 |
| 0 | 0 | 0 | 1 |
| 0 | 0 | 0 | 1 |
| 0 | 0 | 0 | 1 |
| 0 | 0 | 0 | 1 |
| 0 | 0 | 0 | 1 |
| 0 | 0 | 0 | 1 |
| 1 | 0 | 0 | 0 |
| 0 | 0 | 0 | 1 |
| 0 | 0 | 0 | 0 |
| 0 | 0 | 0 | 1 |
| 0 | 0 | 0 | 1 |
| 1 | 0 | 0 | 0 |
| 0 | 0 | 0 | 1 |
| 0 | 0 | 0 | 1 |
| 0 | 0 | 0 | 1 |
| 0 | 0 | 0 | 0 |
| 0 | 0 | 0 | 1 |
| 0 | 0 | 0 | 0 |
| 0 | 0 | 0 | 1 |
| 0 | 0 | 0 | 1 |
| 0 | 0 | 0 | 1 |
| 0 | 0 | 0 | 1 |
| 0 | 0 | 0 | 1 |
| 0 | 0 | 0 | 1 |
| 0 | 0 | 0 | 1 |
| 0 | 0 | 0 | 1 |
| 0 | 0 | 0 | 1 |
| 0 | 0 | 0 | 1 |
| 0 | 0 | 0 | 1 |
| 0 | 0 | 0 | 1 |
| 0 | 0 | 0 | 1 |
| 0 | 0 | 0 | 1 |
| 0 | 0 | 0 | 1 |
| 0 | 0 | 0 | 1 |
| 0 | 0 | 0 | 0 |
| 0 | 0 | 0 | 1 |
| 0 | 0 | 0 | 1 |
| 0 | 0 | 0 | 0 |
| 0 | 0 | 0 | 0 |
| 0 | 0 | 0 | 1 |
| 0 | 0 | 0 | 0 |
| 0 | 0 | 0 | 0 |
| 0 | 0 | 0 | 0 |
| 0 | 0 | 0 | 1 |
| 0 | 0 | 0 | 1 |

| | | | |
|---|---|---|---|
| 0 | 0 | 0 | 1 |
| 0 | 0 | 0 | 1 |
| 0 | 0 | 0 | 1 |
| 0 | 0 | 0 | 1 |
| 0 | 0 | 0 | 1 |
| 0 | 0 | 0 | 1 |
| 0 | 0 | 0 | 1 |
| 1 | 0 | 0 | 0 |
| 0 | 0 | 0 | 1 |
| 0 | 0 | 0 | 1 |
| 0 | 0 | 0 | 1 |
| 0 | 0 | 0 | 1 |
| 0 | 0 | 0 | 1 |
| 0 | 0 | 0 | 1 |
| 0 | 0 | 0 | 0 |
| 0 | 0 | 0 | 0 |
| 0 | 0 | 0 | 1 |
| 0 | 0 | 0 | 0 |
| 0 | 0 | 0 | 1 |
| 0 | 0 | 0 | 1 |
| 0 | 0 | 0 | 0 |
| 0 | 0 | 0 | 1 |
| 0 | 0 | 0 | 1 |
| 0 | 0 | 0 | 1 |
| 0 | 0 | 0 | 0 |
| 0 | 0 | 0 | 1 |
| 0 | 0 | 0 | 1 |
| 1 | 0 | 0 | 0 |
| 0 | 0 | 0 | 0 |
| 1 | 0 | 0 | 0 |
| 1 | 0 | 0 | 0 |
| 0 | 0 | 0 | 1 |
| 0 | 0 | 0 | 1 |
| 0 | 0 | 0 | 1 |
| 0 | 0 | 0 | 1 |
| 0 | 0 | 0 | 1 |
| 0 | 0 | 0 | 0 |
| 0 | 0 | 0 | 0 |
| 0 | 0 | 0 | 1 |
| 0 | 0 | 0 | 1 |
| 0 | 0 | 0 | 1 |
| 0 | 0 | 0 | 1 |
| 0 | 0 | 0 | 0 |
| 0 | 0 | 0 | 1 |
| 0 | 0 | 0 | 1 |
| 0 | 0 | 0 | 1 |
| 0 | 0 | 0 | 1 |
| 0 | 0 | 0 | 1 |
| 0 | 0 | 0 | 0 |
| 0 | 0 | 0 | 0 |
| 0 | 0 | 0 | 0 |
| 0 | 0 | 0 | 0 |
| 0 | 0 | 0 | 0 |
| 0 | 0 | 0 | 1 |

| | | | |
|---|---|---|---|
| 0 | 0 | 0 | 0 |
| 0 | 0 | 0 | 0 |
| 0 | 0 | 0 | 1 |
| 1 | 0 | 0 | 0 |
| 0 | 0 | 0 | 1 |
| 0 | 0 | 0 | 1 |
| 0 | 0 | 0 | 1 |
| 0 | 0 | 0 | 1 |
| 0 | 0 | 0 | 1 |
| 1 | 0 | 0 | 0 |
| 0 | 0 | 0 | 1 |
| 0 | 0 | 0 | 1 |
| 0 | 0 | 0 | 1 |
| 0 | 0 | 0 | 1 |
| 0 | 0 | 0 | 1 |
| 0 | 0 | 0 | 1 |
| 0 | 0 | 0 | 1 |
| 0 | 0 | 0 | 1 |
| 0 | 0 | 0 | 1 |
| 0 | 0 | 0 | 1 |
| 0 | 0 | 0 | 1 |
| 0 | 0 | 0 | 1 |
| 0 | 0 | 0 | 1 |
| 0 | 0 | 0 | 1 |
| 0 | 0 | 0 | 1 |
| 0 | 0 | 0 | 1 |
| 0 | 0 | 0 | 1 |
| 0 | 0 | 0 | 1 |
| 0 | 0 | 0 | 1 |
| 0 | 0 | 0 | 1 |
| 0 | 0 | 0 | 1 |
| 0 | 0 | 0 | 1 |
| 1 | 0 | 0 | 0 |
| 0 | 0 | 0 | 1 |
| 0 | 0 | 0 | 1 |
| 0 | 0 | 0 | 1 |
| 0 | 0 | 0 | 1 |
| 0 | 0 | 0 | 1 |
| 0 | 0 | 0 | 1 |
| 0 | 0 | 0 | 1 |
| 0 | 0 | 0 | 1 |
| 0 | 0 | 0 | 1 |
| 0 | 0 | 0 | 1 |
| 0 | 0 | 0 | 1 |
| 0 | 0 | 0 | 1 |
| 0 | 0 | 0 | 1 |
| 0 | 0 | 0 | 1 |
| 0 | 0 | 0 | 0 |
| 0 | 1 | 0 | 0 |
| 0 | 0 | 0 | 0 |
| 0 | 0 | 0 | 0 |
| 0 | 1 | 0 | 0 |
| 0 | 0 | 0 | 1 |
| 0 | 0 | 0 | 0 |

| | | | |
|---|---|---|---|
| 0 | 0 | 0 | 1 |
| 0 | 0 | 0 | 0 |
| 0 | 0 | 0 | 1 |
| 0 | 0 | 0 | 1 |
| 0 | 0 | 0 | 1 |
| 1 | 0 | 0 | 0 |
| 0 | 0 | 0 | 0 |
| 0 | 0 | 0 | 0 |
| 1 | 0 | 0 | 0 |
| 0 | 0 | 0 | 0 |
| 0 | 0 | 0 | 1 |
| 0 | 0 | 0 | 0 |
| 0 | 0 | 0 | 1 |
| 0 | 0 | 0 | 1 |
| 0 | 0 | 0 | 1 |
| 1 | 0 | 0 | 0 |
| 0 | 0 | 0 | 1 |
| 0 | 0 | 0 | 1 |
| 0 | 0 | 0 | 0 |
| 0 | 0 | 0 | 0 |
| 1 | 0 | 0 | 0 |
| 1 | 0 | 0 | 0 |
| 0 | 0 | 0 | 0 |
| 0 | 0 | 0 | 0 |
| 0 | 0 | 0 | 1 |
| 0 | 0 | 0 | 0 |
| 0 | 0 | 0 | 0 |
| 0 | 0 | 0 | 0 |
| 0 | 0 | 0 | 0 |
| 0 | 0 | 0 | 0 |
| 0 | 0 | 0 | 0 |
| 0 | 0 | 0 | 1 |
| 0 | 0 | 0 | 1 |
| 0 | 0 | 0 | 0 |
| 0 | 0 | 0 | 0 |
| 0 | 0 | 0 | 1 |
| 0 | 0 | 0 | 0 |
| 0 | 0 | 0 | 0 |
| 0 | 0 | 0 | 0 |
| 0 | 0 | 0 | 0 |
| 0 | 0 | 0 | 0 |
| 0 | 0 | 0 | 1 |
| 0 | 0 | 0 | 1 |
| 0 | 0 | 0 | 1 |
| 0 | 0 | 0 | 0 |
| 0 | 0 | 0 | 0 |
| 0 | 0 | 0 | 0 |
| 0 | 0 | 0 | 1 |
| 0 | 0 | 0 | 1 |
| 0 | 0 | 0 | 1 |
| 0 | 0 | 0 | 1 |
| 0 | 0 | 0 | 1 |
| 0 | 0 | 0 | 1 |
| 0 | 0 | 0 | 0 |

| | | | |
|---|---|---|---|
| 0 | 0 | 0 | 1 |
| 0 | 0 | 0 | 1 |
| 0 | 0 | 0 | 1 |
| 0 | 0 | 0 | 1 |
| 1 | 0 | 0 | 0 |
| 0 | 0 | 0 | 0 |
| 0 | 0 | 0 | 1 |
| 0 | 0 | 0 | 1 |
| 0 | 0 | 0 | 1 |
| 0 | 0 | 0 | 1 |
| 0 | 0 | 0 | 1 |
| 0 | 0 | 0 | 0 |
| 0 | 0 | 0 | 1 |
| 0 | 0 | 0 | 0 |
| 0 | 0 | 0 | 0 |
| 0 | 0 | 0 | 0 |
| 0 | 0 | 0 | 0 |
| 0 | 0 | 0 | 0 |
| 0 | 1 | 0 | 0 |
| 0 | 1 | 0 | 0 |
| 0 | 0 | 0 | 0 |
| 0 | 0 | 0 | 0 |
| 0 | 0 | 0 | 0 |
| 0 | 0 | 0 | 1 |
| 0 | 0 | 0 | 1 |
| 0 | 0 | 0 | 1 |
| 0 | 0 | 0 | 1 |
| 0 | 0 | 0 | 1 |
| 0 | 0 | 0 | 1 |
| 0 | 0 | 0 | 1 |
| 0 | 0 | 0 | 1 |
| 0 | 0 | 0 | 0 |
| 0 | 0 | 0 | 1 |
| 0 | 0 | 0 | 0 |
| 0 | 0 | 0 | 0 |
| 0 | 0 | 0 | 0 |
| 0 | 0 | 0 | 0 |
| 0 | 0 | 0 | 1 |
| 0 | 0 | 0 | 1 |
| 0 | 0 | 0 | 1 |
| 0 | 0 | 0 | 1 |
| 0 | 0 | 0 | 1 |
| 0 | 0 | 0 | 1 |
| 0 | 0 | 0 | 1 |
| 0 | 0 | 0 | 1 |
| 1 | 0 | 0 | 0 |
| 0 | 0 | 0 | 1 |
| 0 | 0 | 0 | 1 |
| 0 | 0 | 0 | 1 |
| 1 | 0 | 0 | 0 |
| 1 | 0 | 0 | 0 |
| 0 | 0 | 0 | 0 |
| 0 | 0 | 0 | 1 |
| 0 | 0 | 0 | 1 |

| | | | |
|---|---|---|---|
| 0 | 0 | 0 | 1 |
| 0 | 0 | 0 | 1 |
| 0 | 0 | 0 | 1 |
| 0 | 0 | 0 | 0 |
| 0 | 0 | 0 | 1 |
| 0 | 0 | 0 | 1 |
| 0 | 0 | 0 | 0 |
| 0 | 0 | 0 | 1 |
| 0 | 0 | 0 | 1 |
| 0 | 0 | 0 | 1 |
| 0 | 0 | 0 | 0 |
| 0 | 0 | 0 | 0 |
| 0 | 0 | 0 | 1 |
| 0 | 0 | 0 | 1 |
| 0 | 0 | 0 | 1 |
| 0 | 0 | 0 | 1 |
| 0 | 0 | 0 | 0 |
| 0 | 0 | 0 | 0 |
| 0 | 0 | 0 | 1 |
| 0 | 0 | 0 | 0 |
| 0 | 0 | 0 | 0 |
| 0 | 0 | 0 | 0 |
| 0 | 0 | 0 | 1 |
| 0 | 0 | 0 | 0 |
| 0 | 0 | 0 | 0 |
| 0 | 0 | 0 | 1 |
| 0 | 0 | 0 | 0 |
| 0 | 0 | 0 | 1 |
| 0 | 0 | 0 | 1 |
| 0 | 0 | 0 | 0 |
| 0 | 0 | 0 | 1 |
| 0 | 0 | 0 | 1 |
| 0 | 0 | 0 | 0 |
| 0 | 0 | 0 | 1 |
| 0 | 0 | 0 | 1 |
| 0 | 0 | 0 | 0 |
| 0 | 0 | 0 | 1 |
| 0 | 0 | 0 | 0 |
| 0 | 0 | 0 | 1 |
| 0 | 0 | 0 | 0 |
| 0 | 0 | 0 | 0 |
| 0 | 0 | 0 | 1 |
| 0 | 0 | 0 | 0 |
| 0 | 0 | 0 | 0 |
| 0 | 0 | 0 | 1 |
| 0 | 0 | 0 | 1 |
| 0 | 0 | 0 | 1 |
| 0 | 0 | 0 | 1 |
| 0 | 0 | 0 | 1 |
| 0 | 0 | 0 | 1 |
| 0 | 0 | 0 | 1 |
| 0 | 0 | 0 | 1 |
| 0 | 0 | 0 | 1 |
| 0 | 0 | 0 | 1 |

| | | | |
|---|---|---|---|
| 0 | 0 | 0 | 1 |
| 0 | 0 | 0 | 1 |
| 0 | 0 | 0 | 1 |
| 0 | 0 | 0 | 0 |
| 0 | 0 | 0 | 1 |
| 0 | 0 | 0 | 0 |
| 0 | 0 | 0 | 1 |
| 0 | 0 | 0 | 1 |
| 0 | 0 | 0 | 0 |
| 0 | 0 | 0 | 0 |
| 0 | 0 | 0 | 1 |
| 0 | 0 | 0 | 0 |
| 0 | 1 | 0 | 0 |
| 0 | 0 | 0 | 0 |
| 0 | 0 | 0 | 1 |
| 0 | 0 | 0 | 0 |
| 0 | 0 | 0 | 0 |
| 0 | 0 | 0 | 1 |
| 1 | 0 | 0 | 0 |
| 1 | 0 | 0 | 0 |
| 1 | 0 | 0 | 0 |
| 1 | 0 | 0 | 0 |
| 0 | 0 | 0 | 1 |
| 0 | 0 | 0 | 0 |
| 0 | 0 | 0 | 0 |
| 0 | 0 | 0 | 0 |
| 0 | 0 | 0 | 0 |
| 0 | 0 | 0 | 0 |
| 0 | 0 | 0 | 0 |
| 0 | 0 | 0 | 0 |
| 0 | 0 | 0 | 0 |
| 0 | 0 | 0 | 0 |
| 0 | 0 | 0 | 0 |
| 0 | 0 | 0 | 1 |
| 0 | 0 | 0 | 1 |
| 0 | 0 | 0 | 0 |
| 0 | 0 | 0 | 1 |
| 0 | 0 | 0 | 0 |
| 0 | 0 | 0 | 0 |
| 0 | 0 | 0 | 0 |
| 0 | 0 | 0 | 0 |
| 0 | 0 | 0 | 1 |
| 0 | 0 | 0 | 0 |
| 0 | 0 | 0 | 0 |
| 0 | 0 | 0 | 1 |
| 0 | 0 | 0 | 1 |
| 0 | 0 | 0 | 1 |
| 0 | 0 | 0 | 1 |
| 0 | 0 | 0 | 1 |
| 0 | 0 | 0 | 1 |
| 0 | 0 | 0 | 1 |
| 0 | 0 | 0 | 1 |
| 0 | 0 | 0 | 1 |
| 0 | 0 | 0 | 1 |

| | | | |
|---|---|---|---|
| 0 | 0 | 0 | 1 |
| 0 | 0 | 0 | 0 |
| 0 | 0 | 0 | 1 |
| 0 | 0 | 0 | 1 |
| 0 | 0 | 0 | 1 |
| 0 | 0 | 0 | 1 |
| 0 | 0 | 0 | 1 |
| 0 | 0 | 0 | 1 |
| 0 | 0 | 0 | 1 |
| 0 | 0 | 0 | 1 |
| 0 | 0 | 0 | 1 |
| 0 | 0 | 0 | 0 |
| 0 | 0 | 0 | 1 |
| 0 | 0 | 0 | 1 |
| 0 | 0 | 0 | 0 |
| 0 | 0 | 0 | 1 |
| 0 | 0 | 0 | 1 |
| 0 | 0 | 0 | 1 |
| 0 | 0 | 0 | 1 |
| 0 | 0 | 0 | 1 |
| 1 | 0 | 0 | 0 |
| 0 | 0 | 0 | 1 |
| 1 | 0 | 0 | 0 |
| 1 | 0 | 0 | 0 |
| 0 | 0 | 0 | 1 |
| 0 | 0 | 0 | 0 |
| 0 | 0 | 0 | 1 |
| 0 | 0 | 0 | 1 |
| 0 | 0 | 0 | 1 |
| 0 | 0 | 0 | 1 |
| 0 | 0 | 0 | 1 |
| 0 | 0 | 0 | 0 |
| 0 | 0 | 0 | 1 |
| 0 | 0 | 0 | 0 |
| 0 | 0 | 0 | 1 |
| 0 | 0 | 0 | 1 |
| 0 | 0 | 0 | 1 |
| 0 | 0 | 0 | 1 |
| 0 | 0 | 0 | 1 |
| 0 | 0 | 0 | 0 |
| 0 | 0 | 0 | 1 |
| 0 | 0 | 0 | 1 |
| 0 | 0 | 0 | 1 |
| 0 | 0 | 0 | 0 |
| 0 | 0 | 0 | 1 |
| 0 | 0 | 0 | 1 |
| 0 | 0 | 0 | 1 |
| 1 | 0 | 0 | 0 |
| 0 | 0 | 0 | 0 |
| 0 | 0 | 0 | 1 |
| 0 | 0 | 0 | 1 |
| 0 | 0 | 0 | 0 |
| 1 | 0 | 0 | 0 |
| 0 | 0 | 0 | 0 |

| | | | |
|---|---|---|---|
| 1 | 0 | 0 | 0 |
| 0 | 0 | 0 | 1 |
| 0 | 0 | 0 | 0 |
| 0 | 0 | 0 | 0 |
| 0 | 0 | 0 | 0 |
| 0 | 0 | 0 | 1 |
| 0 | 0 | 0 | 1 |
| 0 | 0 | 0 | 1 |
| 0 | 0 | 0 | 1 |
| 0 | 0 | 0 | 0 |
| 0 | 0 | 0 | 1 |
| 0 | 0 | 0 | 1 |
| 0 | 0 | 0 | 1 |
| 0 | 0 | 0 | 1 |
| 0 | 0 | 0 | 0 |
| 0 | 0 | 0 | 1 |
| 0 | 0 | 0 | 1 |
| 0 | 0 | 0 | 0 |
| 0 | 0 | 0 | 1 |
| 0 | 0 | 0 | 0 |
| 0 | 0 | 0 | 0 |
| 0 | 0 | 0 | 1 |
| 0 | 0 | 0 | 0 |
| 0 | 0 | 0 | 1 |
| 0 | 0 | 0 | 0 |
| 0 | 0 | 0 | 0 |
| 0 | 0 | 0 | 1 |
| 0 | 0 | 0 | 1 |
| 1 | 0 | 0 | 0 |
| 0 | 0 | 0 | 1 |
| 0 | 0 | 0 | 1 |
| 0 | 0 | 0 | 1 |
| 0 | 0 | 0 | 1 |
| 0 | 0 | 0 | 1 |
| 0 | 0 | 0 | 0 |
| 0 | 0 | 0 | 0 |
| 0 | 0 | 0 | 1 |
| 0 | 0 | 0 | 0 |
| 0 | 0 | 0 | 0 |
| 0 | 0 | 0 | 0 |
| 0 | 0 | 0 | 0 |
| 0 | 0 | 0 | 0 |
| 0 | 0 | 0 | 0 |
| 0 | 0 | 0 | 1 |
| 0 | 0 | 0 | 0 |
| 0 | 0 | 0 | 1 |
| 0 | 0 | 0 | 1 |
| 0 | 0 | 0 | 0 |
| 0 | 0 | 0 | 0 |
| 0 | 0 | 0 | 0 |
| 0 | 0 | 0 | 0 |
| 0 | 0 | 0 | 0 |
| 0 | 0 | 0 | 1 |
| 0 | 0 | 0 | 1 |

| | | | |
|---|---|---|---|
| 0 | 0 | 0 | 1 |
| 0 | 0 | 0 | 0 |
| 0 | 0 | 0 | 0 |
| 0 | 0 | 0 | 0 |
| 0 | 0 | 0 | 0 |
| 0 | 0 | 0 | 0 |
| 0 | 0 | 0 | 0 |
| 0 | 0 | 0 | 1 |
| 0 | 0 | 0 | 1 |
| 0 | 0 | 0 | 1 |
| 0 | 0 | 0 | 0 |
| 0 | 0 | 0 | 0 |
| 0 | 0 | 0 | 1 |
| 0 | 0 | 0 | 0 |
| 0 | 0 | 0 | 1 |
| 0 | 0 | 0 | 0 |
| 0 | 0 | 0 | 1 |
| 0 | 0 | 0 | 1 |
| 0 | 0 | 0 | 1 |
| 0 | 0 | 0 | 1 |
| 0 | 0 | 0 | 1 |
| 0 | 0 | 0 | 1 |
| 0 | 0 | 0 | 1 |
| 1 | 0 | 0 | 0 |
| 0 | 0 | 0 | 0 |
| 0 | 0 | 0 | 1 |
| 0 | 0 | 0 | 1 |
| 0 | 0 | 0 | 1 |
| 0 | 0 | 0 | 1 |
| 0 | 0 | 0 | 0 |
| 0 | 0 | 0 | 0 |
| 0 | 0 | 0 | 0 |
| 0 | 0 | 0 | 0 |
| 0 | 0 | 0 | 1 |
| 0 | 0 | 0 | 0 |
| 0 | 0 | 0 | 0 |
| 0 | 0 | 0 | 0 |
| 0 | 0 | 0 | 0 |
| 0 | 0 | 0 | 0 |
| 0 | 0 | 0 | 1 |
| 0 | 0 | 0 | 1 |
| 0 | 0 | 0 | 0 |
| 0 | 0 | 0 | 1 |
| 0 | 0 | 0 | 1 |
| 0 | 0 | 0 | 0 |
| 0 | 0 | 0 | 1 |
| 0 | 0 | 0 | 1 |
| 0 | 0 | 0 | 1 |
| 0 | 0 | 0 | 1 |
| 0 | 0 | 0 | 1 |
| 0 | 0 | 0 | 0 |
| 0 | 0 | 0 | 1 |
| 0 | 0 | 0 | 0 |
| 0 | 0 | 0 | 0 |

| | | | |
|---|---|---|---|
| 0 | 0 | 0 | 1 |
| 0 | 0 | 0 | 1 |
| 0 | 0 | 0 | 0 |
| 0 | 0 | 0 | 0 |
| 0 | 0 | 0 | 1 |
| 0 | 0 | 0 | 1 |
| 0 | 0 | 0 | 1 |
| 0 | 0 | 0 | 1 |
| 0 | 0 | 0 | 1 |
| 0 | 0 | 0 | 1 |
| 0 | 0 | 0 | 1 |
| 0 | 0 | 0 | 1 |
| 0 | 0 | 0 | 0 |
| 0 | 0 | 0 | 1 |
| 0 | 0 | 0 | 1 |
| 0 | 0 | 0 | 1 |
| 0 | 0 | 0 | 1 |
| 0 | 0 | 0 | 1 |
| 0 | 0 | 0 | 1 |
| 0 | 0 | 0 | 1 |
| 0 | 0 | 0 | 1 |
| 0 | 0 | 0 | 1 |
| 0 | 0 | 0 | 1 |
| 0 | 0 | 0 | 1 |
| 0 | 0 | 0 | 1 |
| 0 | 0 | 0 | 1 |
| 0 | 0 | 0 | 1 |
| 0 | 0 | 0 | 1 |
| 0 | 0 | 0 | 1 |
| 0 | 0 | 0 | 1 |
| 0 | 0 | 0 | 1 |
| 0 | 0 | 0 | 0 |
| 0 | 0 | 0 | 1 |
| 0 | 0 | 0 | 1 |
| 0 | 0 | 0 | 1 |
| 0 | 0 | 0 | 1 |
| 0 | 0 | 0 | 1 |
| 0 | 0 | 0 | 1 |
| 0 | 0 | 0 | 1 |
| 0 | 0 | 0 | 1 |
| 0 | 0 | 0 | 0 |
| 0 | 0 | 0 | 0 |
| 0 | 0 | 0 | 1 |
| 0 | 0 | 0 | 1 |
| 0 | 0 | 0 | 1 |
| 0 | 0 | 0 | 1 |
| 0 | 0 | 0 | 1 |
| 0 | 0 | 0 | 1 |
| 0 | 0 | 0 | 0 |
| 0 | 0 | 0 | 0 |
| 0 | 0 | 0 | 1 |
| 0 | 0 | 0 | 1 |
| 0 | 0 | 0 | 0 |
| 0 | 0 | 0 | 1 |

| | | | |
|---|---|---|---|
| 0 | 0 | 0 | 0 |
| 0 | 0 | 0 | 0 |
| 0 | 0 | 0 | 0 |
| 0 | 0 | 0 | 1 |
| 0 | 0 | 0 | 1 |
| 0 | 0 | 0 | 1 |
| 0 | 0 | 0 | 1 |
| 0 | 0 | 0 | 0 |
| 0 | 0 | 0 | 0 |
| 0 | 0 | 0 | 0 |
| 0 | 0 | 0 | 0 |
| 0 | 0 | 0 | 1 |
| 0 | 0 | 0 | 0 |
| 0 | 0 | 0 | 1 |
| 0 | 0 | 0 | 1 |
| 0 | 0 | 0 | 1 |
| 0 | 0 | 0 | 1 |
| 0 | 0 | 0 | 0 |
| 0 | 0 | 0 | 1 |
| 0 | 0 | 0 | 1 |
| 0 | 0 | 0 | 1 |
| 0 | 0 | 0 | 1 |
| 0 | 0 | 0 | 1 |
| 0 | 0 | 0 | 1 |
| 0 | 0 | 0 | 1 |
| 0 | 0 | 0 | 1 |
| 0 | 0 | 0 | 1 |
| 0 | 0 | 0 | 1 |
| 0 | 0 | 0 | 1 |
| 0 | 0 | 0 | 1 |
| 0 | 0 | 0 | 0 |
| 0 | 0 | 0 | 0 |
| 0 | 0 | 0 | 1 |
| 0 | 0 | 0 | 1 |
| 0 | 0 | 0 | 1 |
| 0 | 0 | 0 | 1 |
| 0 | 0 | 0 | 1 |
| 0 | 0 | 0 | 1 |
| 0 | 0 | 0 | 1 |
| 0 | 0 | 0 | 1 |
| 0 | 0 | 0 | 1 |
| 0 | 0 | 0 | 0 |
| 0 | 0 | 0 | 1 |
| 0 | 0 | 0 | 1 |
| 0 | 0 | 0 | 0 |
| 0 | 0 | 0 | 0 |
| 0 | 0 | 0 | 1 |
| 0 | 0 | 0 | 1 |
| 0 | 0 | 0 | 0 |
| 0 | 0 | 0 | 0 |
| 0 | 0 | 0 | 0 |
| 0 | 0 | 0 | 0 |
| 0 | 0 | 0 | 0 |
| 0 | 0 | 0 | 0 |

| | | | |
|---|---|---|---|
| 0 | 0 | 0 | 1 |
| 0 | 0 | 0 | 1 |
| 0 | 0 | 0 | 1 |
| 0 | 0 | 0 | 1 |
| 0 | 0 | 0 | 1 |
| 0 | 0 | 0 | 0 |
| 0 | 0 | 0 | 0 |
| 0 | 0 | 0 | 0 |
| 0 | 0 | 0 | 0 |
| 0 | 0 | 0 | 0 |
| 0 | 0 | 0 | 1 |
| 0 | 0 | 0 | 0 |
| 0 | 0 | 0 | 0 |
| 0 | 0 | 0 | 0 |
| 0 | 0 | 0 | 1 |
| 0 | 0 | 0 | 0 |
| 0 | 0 | 0 | 1 |
| 0 | 0 | 0 | 1 |
| 1 | 0 | 0 | 0 |
| 1 | 0 | 0 | 0 |
| 0 | 0 | 0 | 0 |
| 0 | 0 | 0 | 0 |
| 0 | 0 | 0 | 1 |
| 0 | 0 | 0 | 0 |
| 0 | 0 | 0 | 0 |
| 0 | 0 | 0 | 0 |
| 0 | 0 | 0 | 0 |
| 0 | 0 | 0 | 1 |
| 0 | 0 | 0 | 1 |
| 0 | 0 | 0 | 1 |
| 0 | 0 | 0 | 1 |
| 0 | 0 | 0 | 1 |
| 0 | 0 | 0 | 0 |
| 0 | 0 | 0 | 1 |
| 0 | 0 | 0 | 0 |
| 0 | 0 | 0 | 0 |
| 0 | 0 | 0 | 0 |
| 0 | 0 | 0 | 0 |
| 0 | 0 | 0 | 0 |
| 0 | 0 | 0 | 0 |
| 0 | 0 | 0 | 0 |
| 1 | 0 | 0 | 0 |
| 0 | 0 | 0 | 0 |
| 0 | 0 | 0 | 1 |
| 0 | 0 | 0 | 1 |
| 0 | 0 | 0 | 1 |
| 0 | 0 | 0 | 1 |
| 0 | 0 | 0 | 0 |
| 0 | 0 | 0 | 0 |
| 0 | 0 | 0 | 0 |
| 0 | 0 | 0 | 1 |
| 0 | 0 | 0 | 1 |
| 0 | 0 | 0 | 0 |
| 0 | 0 | 0 | 1 |

| | | | |
|---|---|---|---|
| 0 | 0 | 0 | 1 |
| 0 | 0 | 0 | 0 |
| 0 | 0 | 0 | 1 |
| 0 | 0 | 0 | 1 |
| 0 | 0 | 0 | 1 |
| 0 | 0 | 0 | 1 |
| 0 | 0 | 0 | 1 |
| 0 | 0 | 0 | 1 |
| 0 | 0 | 0 | 1 |
| 0 | 0 | 0 | 1 |
| 0 | 0 | 0 | 1 |
| 0 | 0 | 0 | 1 |
| 0 | 0 | 0 | 0 |
| 0 | 0 | 0 | 0 |
| 0 | 0 | 0 | 0 |
| 0 | 0 | 0 | 0 |
| 0 | 0 | 0 | 0 |
| 0 | 0 | 0 | 0 |
| 0 | 0 | 0 | 0 |
| 0 | 0 | 0 | 1 |
| 1 | 1 | 0 | 0 |
| 0 | 0 | 0 | 0 |
| 0 | 0 | 0 | 0 |
| 0 | 0 | 0 | 0 |
| 0 | 0 | 0 | 0 |
| 0 | 0 | 0 | 1 |
| 0 | 0 | 0 | 1 |
| 0 | 0 | 0 | 0 |
| 0 | 0 | 0 | 1 |
| 0 | 0 | 0 | 0 |
| 0 | 0 | 0 | 0 |
| 0 | 0 | 0 | 1 |
| 0 | 0 | 0 | 0 |
| 0 | 0 | 0 | 1 |
| 0 | 0 | 0 | 1 |
| 0 | 0 | 0 | 1 |
| 0 | 0 | 0 | 1 |
| 0 | 0 | 0 | 1 |
| 0 | 0 | 0 | 1 |
| 0 | 0 | 0 | 1 |
| 0 | 0 | 0 | 0 |
| 0 | 0 | 0 | 1 |
| 0 | 0 | 0 | 1 |
| 0 | 0 | 0 | 0 |
| 0 | 0 | 0 | 0 |
| 0 | 0 | 0 | 1 |
| 0 | 0 | 0 | 0 |
| 0 | 0 | 0 | 0 |
| 0 | 0 | 0 | 1 |
| 0 | 0 | 0 | 1 |
| 0 | 0 | 0 | 0 |
| 0 | 0 | 0 | 1 |
| 0 | 0 | 0 | 1 |
| 0 | 0 | 0 | 1 |

| | | | |
|---|---|---|---|
| 0 | 0 | 0 | 1 |
| 0 | 0 | 0 | 1 |
| 0 | 0 | 0 | 1 |
| 0 | 0 | 0 | 1 |
| 0 | 0 | 0 | 0 |
| 0 | 0 | 0 | 0 |
| 0 | 0 | 0 | 1 |
| 0 | 0 | 0 | 1 |
| 0 | 0 | 0 | 1 |
| 0 | 0 | 0 | 1 |
| 0 | 0 | 0 | 1 |
| 0 | 0 | 0 | 0 |
| 0 | 0 | 0 | 0 |
| 0 | 0 | 0 | 1 |
| 0 | 0 | 0 | 1 |
| 0 | 0 | 0 | 0 |
| 0 | 0 | 0 | 1 |
| 0 | 0 | 0 | 1 |
| 0 | 0 | 0 | 0 |
| 0 | 0 | 0 | 1 |
| 0 | 0 | 0 | 1 |
| 0 | 0 | 0 | 1 |
| 0 | 0 | 0 | 1 |
| 0 | 0 | 0 | 0 |
| 0 | 0 | 0 | 1 |
| 0 | 0 | 0 | 0 |
| 0 | 0 | 0 | 1 |
| 0 | 0 | 0 | 1 |
| 1 | 0 | 0 | 0 |
| 1 | 0 | 0 | 0 |
| 0 | 0 | 0 | 0 |
| 0 | 0 | 0 | 0 |
| 0 | 0 | 0 | 0 |
| 0 | 0 | 0 | 0 |
| 0 | 0 | 0 | 1 |
| 0 | 0 | 0 | 1 |
| 0 | 0 | 0 | 1 |
| 1 | 0 | 0 | 0 |
| 0 | 0 | 0 | 1 |
| 0 | 0 | 0 | 1 |
| 0 | 0 | 0 | 0 |
| 0 | 0 | 0 | 0 |
| 0 | 0 | 0 | 0 |
| 0 | 0 | 0 | 1 |
| 0 | 0 | 0 | 0 |
| 0 | 0 | 0 | 1 |
| 0 | 0 | 0 | 1 |
| 0 | 0 | 0 | 0 |
| 0 | 0 | 0 | 1 |
| 0 | 0 | 0 | 1 |
| 0 | 0 | 0 | 1 |
| 0 | 1 | 0 | 0 |
| 0 | 0 | 0 | 1 |
| 0 | 0 | 0 | 0 |

| | | | |
|---|---|---|---|
| 0 | 0 | 0 | 1 |
| 0 | 0 | 0 | 0 |
| 0 | 0 | 0 | 0 |
| 0 | 0 | 0 | 1 |
| 0 | 0 | 0 | 1 |
| 0 | 0 | 0 | 1 |
| 0 | 0 | 0 | 0 |
| 0 | 0 | 0 | 1 |
| 0 | 0 | 0 | 1 |
| 0 | 0 | 0 | 1 |
| 0 | 0 | 0 | 1 |
| 0 | 0 | 0 | 1 |
| 0 | 0 | 0 | 1 |
| 0 | 0 | 0 | 1 |
| 0 | 0 | 0 | 1 |
| 0 | 0 | 0 | 1 |
| 0 | 0 | 0 | 1 |
| 0 | 0 | 0 | 1 |
| 0 | 0 | 0 | 1 |
| 0 | 0 | 0 | 1 |
| 0 | 0 | 0 | 1 |
| 0 | 0 | 0 | 0 |
| 0 | 0 | 0 | 1 |
| 0 | 0 | 0 | 1 |
| 0 | 0 | 0 | 1 |
| 0 | 0 | 0 | 1 |
| 0 | 0 | 0 | 0 |
| 0 | 0 | 0 | 1 |
| 0 | 0 | 0 | 0 |
| 0 | 0 | 0 | 1 |
| 0 | 0 | 0 | 1 |
| 0 | 0 | 0 | 0 |
| 0 | 0 | 0 | 1 |
| 0 | 1 | 1 | 1 |
| 0 | 0 | 0 | 1 |
| 0 | 0 | 0 | 1 |
| 0 | 0 | 0 | 1 |
| 0 | 0 | 0 | 1 |
| 0 | 0 | 0 | 0 |
| 0 | 0 | 0 | 0 |
| 0 | 0 | 0 | 0 |
| 0 | 0 | 0 | 1 |
| 0 | 0 | 0 | 1 |
| 0 | 0 | 0 | 1 |
| 0 | 0 | 0 | 1 |
| 0 | 1 | 0 | 0 |
| 0 | 0 | 0 | 1 |
| 0 | 0 | 0 | 0 |
| 0 | 0 | 0 | 1 |
| 0 | 0 | 0 | 0 |
| 0 | 0 | 0 | 0 |
| 0 | 0 | 0 | 0 |
| 0 | 0 | 0 | 1 |
| 0 | 0 | 0 | 1 |

| | | | |
|---|---|---|---|
| 0 | 0 | 0 | 0 |
| 0 | 0 | 0 | 0 |
| 0 | 0 | 0 | 1 |
| 0 | 0 | 0 | 1 |
| 0 | 0 | 0 | 1 |
| 0 | 0 | 0 | 1 |
| 0 | 0 | 0 | 1 |
| 0 | 0 | 0 | 1 |
| 0 | 0 | 0 | 1 |
| 0 | 0 | 0 | 1 |
| 0 | 0 | 0 | 1 |
| 0 | 0 | 0 | 1 |
| 0 | 0 | 0 | 1 |
| 0 | 0 | 0 | 1 |
| 0 | 0 | 0 | 1 |
| 0 | 0 | 0 | 0 |
| 0 | 0 | 0 | 0 |
| 0 | 0 | 0 | 0 |
| 0 | 0 | 0 | 1 |
| 0 | 0 | 0 | 1 |
| 0 | 0 | 0 | 1 |
| 0 | 0 | 0 | 0 |
| 0 | 0 | 0 | 0 |
| 0 | 0 | 0 | 0 |
| 0 | 0 | 0 | 0 |
| 0 | 0 | 0 | 1 |
| 0 | 0 | 0 | 1 |
| 0 | 0 | 0 | 1 |
| 0 | 0 | 0 | 1 |
| 0 | 0 | 0 | 1 |
| 0 | 0 | 0 | 0 |
| 0 | 0 | 0 | 1 |
| 0 | 0 | 0 | 0 |
| 0 | 0 | 0 | 1 |
| 0 | 1 | 1 | 1 |
| 0 | 1 | 1 | 1 |
| 0 | 0 | 0 | 0 |
| 0 | 0 | 0 | 0 |
| 0 | 0 | 0 | 1 |
| 0 | 0 | 0 | 1 |
| 0 | 0 | 0 | 1 |
| 0 | 0 | 0 | 0 |
| 0 | 0 | 0 | 1 |
| 0 | 0 | 0 | 1 |
| 0 | 0 | 0 | 0 |
| 0 | 0 | 0 | 1 |
| 0 | 0 | 0 | 0 |
| 0 | 0 | 0 | 0 |
| 0 | 0 | 0 | 1 |
| 0 | 0 | 0 | 1 |
| 0 | 0 | 0 | 0 |
| 0 | 0 | 0 | 0 |
| 0 | 0 | 0 | 1 |
| 0 | 0 | 0 | 0 |

| | | | |
|---|---|---|---|
| 0 | 0 | 0 | 1 |
| 0 | 0 | 0 | 0 |
| 0 | 0 | 0 | 1 |
| 0 | 0 | 0 | 0 |
| 0 | 0 | 0 | 1 |
| 0 | 0 | 0 | 1 |
| 0 | 0 | 0 | 1 |
| 0 | 0 | 0 | 1 |
| 0 | 0 | 0 | 0 |
| 0 | 0 | 0 | 1 |
| 0 | 0 | 0 | 1 |
| 0 | 0 | 0 | 1 |
| 0 | 0 | 0 | 0 |
| 0 | 0 | 0 | 0 |
| 0 | 0 | 0 | 0 |
| 0 | 0 | 0 | 1 |
| 0 | 0 | 0 | 0 |
| 0 | 0 | 0 | 1 |
| 0 | 1 | 0 | 0 |
| 0 | 0 | 0 | 0 |
| 0 | 0 | 0 | 0 |
| 0 | 1 | 0 | 0 |
| 0 | 0 | 0 | 0 |
| 1 | 0 | 0 | 0 |
| 0 | 0 | 0 | 1 |
| 0 | 0 | 0 | 1 |
| 0 | 0 | 0 | 0 |
| 0 | 0 | 0 | 0 |
| 0 | 0 | 0 | 0 |
| 0 | 0 | 0 | 0 |
| 0 | 0 | 0 | 1 |
| 0 | 0 | 0 | 1 |
| 0 | 0 | 0 | 1 |
| 0 | 0 | 0 | 1 |
| 0 | 0 | 0 | 1 |
| 1 | 0 | 0 | 0 |
| 1 | 0 | 0 | 0 |
| 0 | 0 | 0 | 1 |
| 0 | 0 | 0 | 1 |
| 0 | 0 | 0 | 1 |
| 0 | 0 | 0 | 1 |
| 0 | 0 | 0 | 1 |
| 0 | 0 | 0 | 1 |
| 0 | 0 | 0 | 0 |
| 0 | 0 | 0 | 0 |
| 0 | 0 | 0 | 0 |
| 0 | 0 | 0 | 1 |
| 0 | 0 | 0 | 0 |
| 0 | 0 | 0 | 0 |
| 0 | 0 | 0 | 0 |
| 0 | 0 | 0 | 1 |
| 0 | 0 | 0 | 1 |
| 0 | 0 | 0 | 1 |
| 0 | 0 | 0 | 1 |

| | | | |
|---|---|---|---|
| 0 | 0 | 0 | 1 |
| 0 | 0 | 0 | 1 |
| 0 | 0 | 0 | 1 |
| 0 | 0 | 0 | 1 |
| 0 | 0 | 0 | 0 |
| 0 | 0 | 0 | 1 |
| 0 | 0 | 0 | 0 |
| 0 | 0 | 0 | 0 |
| 0 | 0 | 0 | 0 |
| 0 | 0 | 0 | 0 |
| 0 | 0 | 0 | 0 |
| 0 | 0 | 0 | 0 |
| 0 | 0 | 0 | 1 |
| 0 | 0 | 0 | 0 |
| 0 | 0 | 0 | 0 |
| 0 | 0 | 0 | 1 |
| 0 | 0 | 0 | 1 |
| 0 | 0 | 0 | 1 |
| 0 | 0 | 0 | 0 |
| 0 | 0 | 0 | 0 |
| 0 | 0 | 0 | 0 |
| 0 | 0 | 0 | 0 |
| 0 | 0 | 0 | 1 |
| 0 | 0 | 0 | 1 |
| 0 | 0 | 0 | 1 |
| 0 | 0 | 0 | 1 |
| 0 | 0 | 0 | 1 |
| 0 | 0 | 0 | 0 |
| 0 | 0 | 0 | 0 |
| 0 | 0 | 0 | 0 |
| 0 | 0 | 0 | 0 |
| 0 | 0 | 0 | 0 |
| 0 | 0 | 0 | 0 |
| 0 | 0 | 0 | 1 |
| 0 | 0 | 0 | 1 |
| 0 | 0 | 0 | 0 |
| 0 | 0 | 0 | 1 |
| 0 | 0 | 0 | 1 |
| 0 | 0 | 0 | 1 |
| 0 | 0 | 0 | 1 |
| 0 | 0 | 0 | 0 |
| 0 | 0 | 0 | 1 |
| 0 | 0 | 0 | 0 |
| 0 | 0 | 0 | 1 |
| 0 | 0 | 0 | 1 |
| 0 | 0 | 0 | 1 |
| 0 | 0 | 0 | 1 |
| 0 | 0 | 0 | 0 |
| 0 | 0 | 0 | 1 |
| 0 | 0 | 0 | 1 |
| 0 | 0 | 0 | 1 |
| 0 | 0 | 0 | 1 |
| 0 | 0 | 0 | 1 |
| 0 | 0 | 0 | 1 |

| | | | |
|---|---|---|---|
| 0 | 0 | 0 | 1 |
| 0 | 0 | 0 | 1 |
| 0 | 0 | 0 | 1 |
| 0 | 0 | 0 | 1 |
| 0 | 0 | 0 | 1 |
| 0 | 0 | 0 | 1 |
| 0 | 0 | 0 | 1 |
| 0 | 0 | 0 | 1 |
| 0 | 0 | 0 | 1 |
| 0 | 0 | 0 | 1 |
| 0 | 0 | 0 | 1 |
| 0 | 0 | 0 | 0 |
| 0 | 0 | 0 | 1 |
| 0 | 0 | 0 | 1 |
| 0 | 0 | 0 | 1 |
| 0 | 0 | 0 | 1 |
| 0 | 0 | 0 | 1 |
| 0 | 0 | 0 | 0 |
| 0 | 0 | 0 | 0 |
| 0 | 0 | 0 | 0 |
| 0 | 0 | 0 | 0 |
| 0 | 0 | 0 | 1 |
| 1 | 0 | 0 | 0 |
| 0 | 0 | 0 | 1 |
| 0 | 0 | 0 | 1 |
| 0 | 0 | 0 | 1 |
| 0 | 0 | 0 | 1 |
| 0 | 0 | 0 | 1 |
| 0 | 0 | 0 | 0 |
| 0 | 0 | 0 | 0 |
| 0 | 0 | 0 | 1 |
| 0 | 0 | 0 | 1 |
| 1 | 0 | 0 | 0 |
| 0 | 0 | 0 | 1 |
| 0 | 0 | 0 | 0 |
| 1 | 0 | 0 | 0 |
| 1 | 0 | 0 | 0 |
| 0 | 0 | 0 | 1 |
| 1 | 0 | 0 | 0 |
| 0 | 0 | 0 | 1 |
| 1 | 0 | 0 | 0 |
| 0 | 0 | 0 | 1 |
| 0 | 0 | 0 | 0 |
| 0 | 0 | 0 | 1 |
| 0 | 0 | 0 | 1 |
| 0 | 0 | 0 | 1 |
| 0 | 0 | 0 | 1 |
| 0 | 0 | 0 | 1 |
| 0 | 0 | 0 | 1 |
| 0 | 0 | 0 | 0 |
| 0 | 0 | 0 | 1 |
| 1 | 0 | 0 | 0 |
| 0 | 0 | 0 | 1 |
| 0 | 0 | 0 | 1 |

| | | | |
|---|---|---|---|
| 0 | 0 | 0 | 0 |
| 0 | 0 | 0 | 0 |
| 0 | 0 | 0 | 0 |
| 0 | 0 | 0 | 0 |
| 0 | 0 | 0 | 0 |
| 0 | 0 | 0 | 1 |
| 0 | 0 | 0 | 0 |
| 0 | 0 | 0 | 1 |
| 0 | 0 | 0 | 1 |
| 0 | 0 | 0 | 0 |
| 0 | 0 | 0 | 0 |
| 0 | 0 | 0 | 0 |
| 0 | 0 | 0 | 1 |
| 0 | 0 | 0 | 1 |
| 0 | 0 | 0 | 1 |
| 0 | 0 | 0 | 1 |
| 0 | 0 | 0 | 1 |
| 0 | 0 | 0 | 1 |
| 0 | 0 | 0 | 1 |
| 0 | 0 | 0 | 1 |
| 0 | 0 | 0 | 1 |
| 0 | 0 | 0 | 1 |
| 0 | 0 | 0 | 1 |
| 0 | 0 | 0 | 1 |
| 0 | 0 | 0 | 1 |
| 0 | 0 | 0 | 1 |
| 0 | 0 | 0 | 1 |
| 0 | 0 | 0 | 1 |
| 0 | 0 | 0 | 1 |
| 0 | 0 | 0 | 1 |
| 0 | 0 | 0 | 1 |
| 0 | 0 | 0 | 1 |
| 0 | 0 | 0 | 1 |
| 0 | 0 | 0 | 1 |
| 0 | 0 | 0 | 1 |
| 0 | 0 | 0 | 1 |
| 0 | 0 | 0 | 1 |
| 0 | 0 | 0 | 1 |
| 0 | 0 | 0 | 1 |
| 0 | 0 | 0 | 1 |
| 0 | 0 | 0 | 1 |
| 0 | 0 | 0 | 1 |
| 0 | 0 | 0 | 1 |
| 0 | 0 | 0 | 1 |
| 0 | 0 | 0 | 1 |
| 0 | 0 | 0 | 0 |
| 0 | 0 | 0 | 1 |
| 0 | 0 | 0 | 1 |
| 0 | 0 | 0 | 1 |
| 0 | 0 | 0 | 1 |
| 0 | 0 | 0 | 0 |
| 0 | 0 | 0 | 1 |
| 0 | 0 | 0 | 1 |
| 0 | 0 | 0 | 1 |

| | | | |
|---|---|---|---|
| 0 | 0 | 0 | 1 |
| 0 | 0 | 0 | 1 |
| 0 | 0 | 0 | 1 |
| 0 | 0 | 0 | 1 |
| 0 | 0 | 0 | 1 |
| 0 | 0 | 0 | 1 |
| 0 | 0 | 0 | 1 |
| 0 | 0 | 0 | 0 |
| 0 | 0 | 0 | 0 |
| 0 | 0 | 0 | 1 |
| 0 | 0 | 0 | 1 |
| 0 | 0 | 0 | 1 |
| 0 | 0 | 0 | 1 |
| 0 | 0 | 0 | 1 |
| 0 | 0 | 0 | 0 |
| 0 | 0 | 0 | 0 |
| 0 | 0 | 0 | 0 |
| 0 | 0 | 0 | 1 |
| 0 | 0 | 0 | 1 |
| 0 | 0 | 0 | 1 |
| 0 | 0 | 0 | 0 |
| 0 | 0 | 0 | 0 |
| 0 | 0 | 0 | 0 |
| 0 | 0 | 0 | 0 |
| 0 | 0 | 0 | 0 |
| 0 | 0 | 0 | 0 |
| 0 | 0 | 0 | 1 |
| 0 | 0 | 0 | 1 |
| 0 | 0 | 0 | 0 |
| 0 | 0 | 0 | 1 |
| 0 | 0 | 0 | 1 |
| 0 | 0 | 0 | 1 |
| 0 | 0 | 0 | 0 |
| 0 | 0 | 0 | 1 |
| 0 | 0 | 0 | 1 |
| 0 | 0 | 0 | 1 |
| 0 | 0 | 0 | 0 |
| 0 | 0 | 0 | 0 |
| 0 | 0 | 0 | 0 |
| 0 | 0 | 0 | 1 |
| 0 | 0 | 0 | 1 |
| 0 | 0 | 0 | 1 |
| 0 | 0 | 0 | 1 |
| 0 | 0 | 0 | 1 |
| 0 | 0 | 0 | 1 |
| 0 | 0 | 0 | 0 |
| 0 | 0 | 0 | 1 |
| 0 | 0 | 0 | 1 |
| 0 | 0 | 0 | 1 |
| 0 | 0 | 0 | 1 |
| 0 | 0 | 0 | 1 |
| 0 | 0 | 0 | 1 |
| 0 | 0 | 0 | 1 |
| 0 | 0 | 0 | 1 |

| | | | |
|---|---|---|---|
| 0 | 0 | 0 | 1 |
| 0 | 0 | 0 | 1 |
| 0 | 0 | 0 | 1 |
| 0 | 0 | 0 | 0 |
| 0 | 0 | 0 | 1 |
| 0 | 0 | 0 | 1 |
| 0 | 0 | 0 | 1 |
| 0 | 0 | 0 | 0 |
| 0 | 0 | 0 | 0 |
| 0 | 0 | 0 | 1 |
| 0 | 0 | 0 | 1 |
| 0 | 0 | 0 | 1 |
| 0 | 0 | 0 | 0 |
| 0 | 0 | 0 | 1 |
| 0 | 0 | 0 | 1 |
| 0 | 0 | 0 | 1 |
| 0 | 0 | 0 | 0 |
| 0 | 0 | 0 | 1 |
| 0 | 0 | 0 | 1 |
| 0 | 0 | 0 | 1 |
| 0 | 0 | 0 | 1 |
| 0 | 0 | 0 | 1 |
| 0 | 0 | 0 | 1 |
| 0 | 0 | 0 | 1 |
| 0 | 0 | 0 | 1 |
| 0 | 0 | 0 | 1 |
| 0 | 0 | 0 | 1 |
| 0 | 0 | 0 | 0 |
| 0 | 0 | 0 | 0 |
| 0 | 0 | 0 | 0 |
| 0 | 0 | 0 | 0 |
| 0 | 0 | 0 | 0 |
| 0 | 0 | 0 | 0 |
| 0 | 0 | 0 | 1 |
| 0 | 0 | 0 | 0 |
| 0 | 0 | 0 | 0 |
| 0 | 0 | 0 | 0 |
| 0 | 0 | 0 | 1 |
| 0 | 0 | 0 | 1 |
| 0 | 0 | 0 | 1 |
| 0 | 0 | 0 | 1 |
| 0 | 0 | 0 | 1 |
| 0 | 0 | 0 | 0 |
| 0 | 0 | 0 | 0 |
| 0 | 0 | 0 | 1 |
| 0 | 0 | 0 | 1 |
| 0 | 0 | 0 | 0 |
| 0 | 0 | 0 | 1 |
| 0 | 0 | 0 | 1 |
| 0 | 0 | 0 | 1 |
| 0 | 0 | 0 | 1 |
| 0 | 0 | 0 | 1 |
| 0 | 0 | 0 | 0 |
| 0 | 0 | 0 | 0 |

| | | | |
|---|---|---|---|
| 0 | 0 | 0 | 0 |
| 0 | 0 | 0 | 0 |
| 0 | 0 | 0 | 0 |
| 0 | 0 | 0 | 0 |
| 0 | 0 | 0 | 0 |
| 0 | 0 | 0 | 0 |
| 0 | 0 | 0 | 0 |
| 0 | 0 | 0 | 0 |
| 0 | 0 | 0 | 1 |
| 0 | 0 | 0 | 0 |
| 0 | 0 | 0 | 1 |
| 0 | 0 | 0 | 1 |
| 0 | 0 | 0 | 1 |
| 0 | 0 | 0 | 1 |
| 0 | 0 | 0 | 0 |
| 0 | 0 | 0 | 1 |
| 0 | 0 | 0 | 1 |
| 0 | 0 | 0 | 1 |
| 0 | 0 | 0 | 1 |
| 0 | 0 | 0 | 0 |
| 0 | 0 | 0 | 0 |
| 0 | 0 | 0 | 1 |
| 0 | 0 | 0 | 0 |
| 0 | 0 | 0 | 0 |
| 0 | 0 | 0 | 0 |
| 0 | 0 | 0 | 0 |
| 0 | 0 | 0 | 0 |
| 0 | 0 | 0 | 1 |
| 0 | 0 | 0 | 1 |
| 0 | 0 | 0 | 1 |
| 0 | 0 | 0 | 0 |
| 0 | 0 | 0 | 1 |
| 0 | 0 | 0 | 1 |
| 0 | 0 | 0 | 1 |
| 0 | 0 | 0 | 1 |
| 0 | 0 | 0 | 1 |
| 0 | 0 | 0 | 1 |
| 0 | 0 | 0 | 1 |
| 0 | 0 | 0 | 1 |
| 0 | 0 | 0 | 1 |
| 0 | 0 | 0 | 1 |
| 0 | 0 | 0 | 0 |
| 0 | 0 | 0 | 1 |
| 0 | 0 | 0 | 1 |
| 0 | 0 | 0 | 1 |
| 0 | 0 | 0 | 1 |
| 0 | 0 | 0 | 1 |
| 0 | 0 | 0 | 1 |
| 0 | 0 | 0 | 0 |
| 0 | 0 | 0 | 1 |
| 0 | 0 | 0 | 1 |
| 0 | 0 | 0 | 1 |
| 0 | 0 | 0 | 0 |
| 0 | 0 | 0 | 0 |

| | | | |
|---|---|---|---|
| 0 | 0 | 0 | 1 |
| 0 | 0 | 0 | 1 |
| 0 | 0 | 0 | 1 |
| 0 | 0 | 0 | 1 |
| 0 | 0 | 0 | 0 |
| 1 | 0 | 0 | 0 |
| 0 | 0 | 0 | 1 |
| 0 | 0 | 0 | 0 |
| 0 | 0 | 0 | 1 |
| 0 | 0 | 0 | 1 |
| 0 | 0 | 0 | 0 |
| 0 | 0 | 0 | 1 |
| 0 | 0 | 0 | 0 |
| 0 | 0 | 0 | 0 |
| 0 | 0 | 0 | 0 |
| 0 | 0 | 0 | 0 |
| 0 | 0 | 0 | 1 |
| 0 | 0 | 0 | 1 |
| 0 | 0 | 0 | 0 |
| 0 | 0 | 0 | 0 |
| 0 | 0 | 0 | 0 |
| 0 | 0 | 0 | 0 |
| 0 | 0 | 0 | 0 |
| 0 | 0 | 0 | 1 |
| 0 | 0 | 0 | 1 |
| 0 | 0 | 0 | 0 |
| 0 | 0 | 0 | 1 |
| 0 | 0 | 0 | 0 |
| 0 | 0 | 0 | 0 |
| 0 | 0 | 0 | 0 |
| 0 | 0 | 0 | 1 |
| 1 | 0 | 0 | 0 |
| 1 | 0 | 0 | 0 |
| 0 | 0 | 0 | 1 |
| 0 | 0 | 0 | 1 |
| 0 | 0 | 0 | 0 |
| 0 | 0 | 0 | 0 |
| 0 | 0 | 0 | 1 |
| 0 | 0 | 0 | 0 |
| 0 | 0 | 0 | 0 |
| 0 | 1 | 0 | 0 |
| 0 | 0 | 0 | 0 |
| 0 | 0 | 0 | 1 |
| 0 | 0 | 0 | 1 |
| 0 | 0 | 0 | 0 |
| 0 | 0 | 0 | 1 |
| 0 | 0 | 0 | 0 |
| 0 | 0 | 0 | 1 |
| 0 | 0 | 0 | 1 |
| 0 | 0 | 0 | 1 |
| 0 | 0 | 0 | 1 |
| 0 | 0 | 0 | 1 |
| 0 | 0 | 0 | 1 |
| 0 | 0 | 0 | 1 |

| | | | |
|---|---|---|---|
| 0 | 0 | 0 | 1 |
| 0 | 0 | 0 | 1 |
| 0 | 0 | 0 | 1 |
| 0 | 0 | 0 | 1 |
| 0 | 0 | 0 | 1 |
| 0 | 0 | 0 | 1 |
| 0 | 0 | 0 | 1 |
| 0 | 0 | 0 | 0 |
| 0 | 0 | 0 | 1 |
| 1 | 0 | 0 | 0 |
| 0 | 0 | 0 | 0 |
| 0 | 0 | 0 | 0 |
| 0 | 0 | 0 | 1 |
| 0 | 0 | 0 | 1 |
| 0 | 0 | 0 | 0 |
| 0 | 0 | 0 | 1 |
| 0 | 0 | 0 | 1 |
| 0 | 0 | 0 | 1 |
| 0 | 0 | 0 | 1 |
| 0 | 0 | 0 | 1 |
| 0 | 0 | 0 | 1 |
| 0 | 0 | 0 | 1 |
| 0 | 0 | 0 | 1 |
| 0 | 0 | 0 | 1 |
| 0 | 0 | 0 | 1 |
| 0 | 0 | 0 | 1 |
| 0 | 0 | 0 | 1 |
| 0 | 0 | 0 | 1 |
| 0 | 0 | 0 | 1 |
| 0 | 0 | 0 | 0 |
| 0 | 0 | 0 | 0 |
| 0 | 0 | 0 | 1 |
| 0 | 0 | 0 | 1 |
| 0 | 0 | 0 | 1 |
| 0 | 0 | 0 | 1 |
| 0 | 0 | 0 | 0 |
| 0 | 0 | 0 | 1 |
| 0 | 0 | 0 | 1 |
| 0 | 0 | 0 | 1 |
| 0 | 0 | 0 | 1 |
| 0 | 0 | 0 | 1 |
| 0 | 0 | 0 | 0 |
| 0 | 0 | 0 | 0 |
| 0 | 0 | 0 | 0 |
| 0 | 0 | 0 | 1 |
| 0 | 0 | 0 | 1 |
| 0 | 0 | 0 | 1 |
| 0 | 0 | 0 | 1 |
| 0 | 0 | 0 | 1 |
| 0 | 0 | 0 | 1 |
| 0 | 0 | 0 | 1 |
| 0 | 0 | 0 | 1 |
| 0 | 0 | 0 | 1 |
| 0 | 0 | 0 | 0 |

| | | | |
|---|---|---|---|
| 0 | 0 | 0 | 1 |
| 0 | 0 | 0 | 1 |
| 0 | 0 | 0 | 1 |
| 0 | 0 | 0 | 1 |
| 0 | 0 | 0 | 1 |
| 0 | 0 | 0 | 1 |
| 0 | 0 | 0 | 1 |
| 0 | 0 | 0 | 1 |
| 0 | 0 | 0 | 0 |
| 0 | 0 | 0 | 0 |
| 0 | 0 | 0 | 1 |
| 0 | 0 | 0 | 1 |
| 0 | 0 | 0 | 1 |
| 0 | 0 | 0 | 0 |
| 0 | 0 | 0 | 1 |
| 0 | 0 | 0 | 1 |
| 0 | 0 | 0 | 1 |
| 1 | 0 | 0 | 0 |
| 1 | 0 | 0 | 0 |
| 0 | 0 | 0 | 0 |
| 0 | 0 | 0 | 0 |
| 0 | 0 | 0 | 1 |
| 0 | 0 | 0 | 1 |
| 0 | 0 | 0 | 1 |
| 0 | 0 | 0 | 1 |
| 1 | 0 | 0 | 0 |
| 0 | 0 | 0 | 1 |
| 0 | 0 | 0 | 1 |
| 0 | 0 | 0 | 1 |
| 0 | 0 | 0 | 1 |
| 0 | 0 | 0 | 1 |
| 0 | 0 | 0 | 1 |
| 0 | 0 | 0 | 1 |
| 0 | 0 | 0 | 1 |
| 0 | 0 | 0 | 1 |
| 0 | 0 | 0 | 1 |
| 0 | 0 | 0 | 1 |
| 0 | 0 | 0 | 1 |
| 0 | 0 | 0 | 1 |
| 0 | 0 | 0 | 0 |
| 0 | 0 | 0 | 1 |
| 0 | 0 | 0 | 1 |
| 0 | 0 | 0 | 0 |
| 0 | 0 | 0 | 1 |
| 0 | 0 | 0 | 1 |
| 0 | 0 | 0 | 1 |
| 0 | 0 | 0 | 0 |
| 0 | 0 | 0 | 0 |
| 0 | 0 | 0 | 1 |
| 0 | 0 | 0 | 1 |
| 0 | 0 | 0 | 0 |
| 0 | 0 | 0 | 1 |
| 0 | 0 | 0 | 1 |
| 0 | 0 | 0 | 1 |

| | | | |
|---|---|---|---|
| 0 | 0 | 0 | 1 |
| 0 | 0 | 0 | 1 |
| 0 | 0 | 0 | 1 |
| 0 | 0 | 0 | 1 |
| 0 | 0 | 0 | 1 |
| 0 | 0 | 0 | 1 |
| 0 | 0 | 0 | 1 |
| 0 | 0 | 0 | 1 |
| 0 | 0 | 0 | 1 |
| 0 | 0 | 0 | 1 |
| 0 | 0 | 0 | 1 |
| 0 | 0 | 0 | 1 |
| 0 | 0 | 0 | 1 |
| 0 | 0 | 0 | 0 |
| 0 | 0 | 0 | 0 |
| 0 | 0 | 0 | 0 |
| 0 | 0 | 0 | 0 |
| 0 | 0 | 0 | 1 |
| 0 | 0 | 0 | 1 |
| 0 | 0 | 0 | 0 |
| 0 | 0 | 0 | 1 |
| 0 | 0 | 0 | 1 |
| 0 | 0 | 0 | 1 |
| 0 | 0 | 0 | 1 |
| 0 | 0 | 0 | 1 |
| 0 | 0 | 0 | 0 |
| 0 | 0 | 0 | 1 |
| 0 | 0 | 0 | 1 |
| 0 | 0 | 0 | 1 |
| 0 | 0 | 0 | 0 |
| 0 | 0 | 0 | 0 |
| 0 | 0 | 0 | 1 |
| 0 | 0 | 0 | 1 |
| 0 | 0 | 0 | 1 |
| 0 | 0 | 0 | 1 |
| 0 | 0 | 0 | 1 |
| 0 | 0 | 0 | 1 |
| 0 | 0 | 0 | 1 |
| 0 | 0 | 0 | 0 |
| 0 | 0 | 0 | 1 |
| 0 | 0 | 0 | 1 |
| 0 | 0 | 0 | 0 |
| 0 | 0 | 0 | 0 |
| 0 | 0 | 0 | 0 |
| 0 | 0 | 0 | 0 |
| 0 | 0 | 0 | 1 |
| 0 | 0 | 0 | 0 |
| 0 | 0 | 0 | 1 |
| 0 | 0 | 0 | 1 |
| 0 | 0 | 0 | 1 |
| 0 | 0 | 0 | 1 |
| 0 | 0 | 0 | 1 |
| 0 | 0 | 0 | 1 |
| 0 | 0 | 0 | 1 |

| | | | |
|---|---|---|---|
| 0 | 0 | 0 | 1 |
| 0 | 0 | 0 | 1 |
| 0 | 0 | 0 | 0 |
| 0 | 0 | 0 | 1 |
| 0 | 0 | 0 | 1 |
| 0 | 0 | 0 | 1 |
| 0 | 0 | 0 | 1 |
| 0 | 0 | 0 | 1 |
| 0 | 0 | 0 | 1 |
| 0 | 0 | 0 | 1 |
| 0 | 0 | 0 | 1 |
| 0 | 0 | 0 | 1 |
| 0 | 0 | 0 | 1 |
| 0 | 0 | 0 | 1 |
| 0 | 0 | 0 | 1 |
| 0 | 0 | 0 | 1 |
| 0 | 0 | 0 | 1 |
| 0 | 0 | 0 | 1 |
| 0 | 0 | 0 | 1 |
| 0 | 0 | 0 | 1 |
| 0 | 0 | 0 | 1 |
| 0 | 0 | 0 | 1 |
| 0 | 0 | 0 | 1 |
| 0 | 0 | 0 | 1 |
| 0 | 0 | 0 | 1 |
| 0 | 0 | 0 | 1 |
| 0 | 0 | 0 | 1 |
| 0 | 0 | 0 | 1 |
| 0 | 0 | 0 | 1 |
| 0 | 0 | 0 | 1 |
| 1 | 0 | 0 | 0 |
| 0 | 0 | 0 | 1 |
| 0 | 0 | 0 | 0 |
| 0 | 0 | 0 | 0 |
| 0 | 0 | 0 | 0 |
| 0 | 0 | 0 | 0 |
| 0 | 0 | 0 | 1 |
| 0 | 0 | 0 | 0 |
| 0 | 0 | 0 | 1 |
| 0 | 0 | 0 | 1 |
| 1 | 0 | 0 | 0 |
| 1 | 0 | 0 | 0 |
| 0 | 0 | 0 | 0 |
| 0 | 0 | 0 | 1 |
| 0 | 0 | 0 | 0 |
| 0 | 0 | 0 | 0 |
| 0 | 0 | 0 | 1 |
| 0 | 0 | 0 | 1 |
| 0 | 0 | 0 | 1 |
| 0 | 0 | 0 | 1 |
| 0 | 0 | 0 | 0 |
| 0 | 0 | 0 | 1 |
| 0 | 0 | 0 | 1 |
| 0 | 0 | 0 | 1 |

| | | | |
|---|---|---|---|
| 0 | 0 | 0 | 1 |
| 0 | 0 | 0 | 1 |
| 0 | 0 | 0 | 1 |
| 0 | 0 | 0 | 1 |
| 0 | 0 | 0 | 1 |
| 1 | 0 | 0 | 0 |
| 0 | 0 | 0 | 1 |
| 0 | 0 | 0 | 0 |
| 0 | 0 | 0 | 0 |
| 0 | 0 | 0 | 0 |
| 0 | 0 | 0 | 1 |
| 0 | 0 | 0 | 0 |
| 0 | 0 | 0 | 1 |
| 0 | 0 | 0 | 1 |
| 0 | 0 | 0 | 1 |
| 0 | 1 | 1 | 1 |
| 0 | 0 | 0 | 1 |
| 0 | 0 | 0 | 0 |
| 0 | 0 | 0 | 1 |
| 0 | 0 | 0 | 0 |
| 0 | 0 | 0 | 1 |
| 0 | 0 | 0 | 0 |
| 0 | 0 | 0 | 0 |
| 0 | 0 | 0 | 0 |
| 0 | 0 | 0 | 1 |
| 0 | 0 | 0 | 1 |
| 0 | 0 | 0 | 1 |
| 0 | 0 | 0 | 1 |
| 0 | 0 | 0 | 0 |
| 0 | 0 | 0 | 1 |
| 0 | 0 | 0 | 1 |
| 0 | 0 | 0 | 1 |
| 0 | 0 | 0 | 1 |
| 0 | 0 | 0 | 1 |
| 0 | 0 | 0 | 0 |
| 0 | 0 | 0 | 0 |
| 0 | 0 | 0 | 1 |
| 0 | 0 | 0 | 0 |
| 0 | 0 | 0 | 0 |
| 0 | 0 | 0 | 1 |
| 0 | 0 | 0 | 0 |
| 0 | 0 | 0 | 0 |
| 0 | 0 | 0 | 1 |
| 0 | 0 | 0 | 1 |
| 0 | 0 | 0 | 1 |
| 0 | 0 | 0 | 1 |
| 0 | 0 | 0 | 1 |
| 0 | 0 | 0 | 1 |
| 0 | 0 | 0 | 1 |
| 0 | 0 | 0 | 1 |
| 0 | 0 | 0 | 1 |
| 0 | 0 | 0 | 1 |
| 0 | 0 | 0 | 1 |
| 0 | 0 | 0 | 1 |

| | | | |
|---|---|---|---|
| 0 | 0 | 0 | 1 |
| 0 | 0 | 0 | 0 |
| 0 | 0 | 0 | 1 |
| 0 | 0 | 0 | 1 |
| 0 | 0 | 0 | 0 |
| 0 | 0 | 0 | 1 |
| 0 | 0 | 0 | 1 |
| 0 | 0 | 0 | 1 |
| 0 | 0 | 0 | 0 |
| 0 | 0 | 0 | 0 |
| 0 | 0 | 0 | 1 |
| 0 | 0 | 0 | 0 |
| 0 | 0 | 0 | 0 |
| 0 | 0 | 0 | 1 |
| 0 | 0 | 0 | 1 |
| 0 | 0 | 0 | 0 |
| 0 | 0 | 0 | 1 |
| 0 | 0 | 0 | 0 |
| 0 | 0 | 0 | 0 |
| 0 | 0 | 0 | 1 |
| 0 | 0 | 0 | 0 |
| 0 | 0 | 0 | 0 |
| 1 | 0 | 0 | 0 |
| 0 | 0 | 0 | 1 |
| 0 | 0 | 0 | 1 |
| 0 | 0 | 0 | 0 |
| 0 | 0 | 0 | 1 |
| 0 | 0 | 0 | 1 |
| 0 | 0 | 0 | 1 |
| 0 | 0 | 0 | 1 |
| 0 | 0 | 0 | 0 |
| 0 | 0 | 0 | 1 |
| 0 | 0 | 0 | 0 |
| 0 | 0 | 0 | 1 |
| 0 | 0 | 0 | 0 |
| 0 | 0 | 0 | 0 |
| 0 | 0 | 0 | 1 |
| 0 | 0 | 0 | 1 |
| 0 | 0 | 0 | 1 |
| 0 | 0 | 0 | 1 |
| 0 | 0 | 0 | 1 |
| 0 | 0 | 0 | 1 |
| 0 | 0 | 0 | 1 |
| 0 | 0 | 0 | 1 |
| 0 | 0 | 0 | 1 |
| 0 | 0 | 0 | 1 |
| 0 | 1 | 1 | 1 |
| 0 | 0 | 0 | 1 |
| 0 | 0 | 0 | 1 |
| 0 | 0 | 0 | 1 |
| 0 | 0 | 0 | 1 |
| 0 | 0 | 0 | 1 |
| 0 | 0 | 0 | 1 |
| 0 | 0 | 0 | 0 |

| | | | |
|---|---|---|---|
| 0 | 0 | 0 | 1 |
| 0 | 0 | 0 | 1 |
| 0 | 0 | 0 | 1 |
| 0 | 0 | 0 | 1 |
| 0 | 0 | 0 | 0 |
| 0 | 0 | 0 | 1 |
| 0 | 0 | 0 | 1 |
| 0 | 0 | 0 | 1 |
| 0 | 0 | 0 | 1 |
| 0 | 0 | 0 | 1 |
| 0 | 0 | 0 | 0 |
| 0 | 0 | 0 | 1 |
| 0 | 0 | 0 | 1 |
| 0 | 0 | 0 | 1 |
| 0 | 0 | 0 | 0 |
| 0 | 0 | 0 | 0 |
| 0 | 0 | 0 | 0 |
| 0 | 0 | 0 | 0 |
| 0 | 0 | 0 | 1 |
| 0 | 0 | 0 | 0 |
| 0 | 0 | 0 | 0 |
| 1 | 0 | 0 | 0 |
| 0 | 0 | 0 | 1 |
| 0 | 0 | 0 | 0 |
| 1 | 0 | 0 | 0 |
| 0 | 0 | 0 | 1 |
| 1 | 0 | 0 | 0 |
| 0 | 0 | 0 | 0 |
| 1 | 0 | 0 | 0 |
| 1 | 0 | 0 | 0 |
| 1 | 0 | 0 | 0 |
| 0 | 0 | 0 | 1 |
| 0 | 0 | 0 | 1 |
| 0 | 0 | 0 | 1 |
| 0 | 0 | 0 | 1 |
| 0 | 0 | 0 | 1 |
| 0 | 0 | 0 | 1 |
| 0 | 1 | 0 | 0 |
| 0 | 0 | 0 | 0 |
| 0 | 0 | 0 | 1 |
| 0 | 0 | 0 | 1 |
| 0 | 0 | 0 | 1 |
| 0 | 0 | 0 | 0 |
| 0 | 0 | 0 | 1 |
| 0 | 0 | 0 | 0 |
| 0 | 0 | 0 | 1 |
| 0 | 0 | 0 | 1 |
| 0 | 0 | 0 | 1 |
| 0 | 1 | 0 | 0 |
| 0 | 1 | 0 | 0 |
| 0 | 1 | 0 | 0 |
| 0 | 0 | 0 | 1 |
| 0 | 0 | 0 | 1 |
| 0 | 0 | 0 | 0 |

| | | | |
|---|---|---|---|
| 0 | 0 | 0 | 1 |
| 0 | 0 | 0 | 0 |
| 0 | 0 | 0 | 1 |
| 0 | 1 | 0 | 0 |
| 0 | 1 | 0 | 0 |
| 0 | 0 | 0 | 1 |
| 0 | 1 | 0 | 0 |
| 0 | 1 | 0 | 0 |
| 0 | 0 | 0 | 1 |
| 0 | 0 | 0 | 1 |
| 0 | 0 | 0 | 1 |
| 0 | 0 | 0 | 1 |
| 0 | 0 | 0 | 1 |
| 0 | 0 | 0 | 1 |
| 0 | 0 | 0 | 1 |
| 0 | 0 | 0 | 1 |
| 0 | 0 | 0 | 1 |
| 0 | 0 | 0 | 1 |
| 0 | 0 | 0 | 1 |
| 0 | 0 | 0 | 1 |
| 1 | 1 | 0 | 0 |
| 0 | 0 | 0 | 1 |
| 0 | 0 | 0 | 1 |
| 0 | 0 | 0 | 1 |
| 0 | 0 | 0 | 1 |
| 0 | 0 | 0 | 0 |
| 0 | 0 | 0 | 0 |
| 0 | 0 | 0 | 1 |
| 0 | 0 | 0 | 1 |
| 0 | 0 | 0 | 1 |
| 0 | 0 | 0 | 1 |
| 0 | 0 | 0 | 1 |
| 0 | 0 | 0 | 1 |
| 0 | 0 | 0 | 1 |
| 0 | 0 | 0 | 1 |
| 0 | 0 | 0 | 1 |
| 0 | 0 | 0 | 1 |
| 0 | 0 | 0 | 1 |
| 0 | 0 | 0 | 1 |
| 0 | 0 | 0 | 1 |
| 0 | 0 | 0 | 1 |
| 0 | 0 | 0 | 1 |
| 0 | 0 | 0 | 1 |
| 0 | 0 | 0 | 0 |
| 0 | 0 | 0 | 1 |
| 0 | 0 | 0 | 0 |
| 0 | 0 | 0 | 0 |
| 0 | 0 | 0 | 0 |
| 0 | 0 | 0 | 0 |
| 0 | 0 | 0 | 0 |
| 0 | 0 | 0 | 0 |
| 0 | 0 | 0 | 1 |
| 0 | 0 | 0 | 1 |
| 0 | 0 | 0 | 1 |

| | | | |
|---|---|---|---|
| 0 | 0 | 0 | 1 |
| 0 | 0 | 0 | 1 |
| 0 | 0 | 0 | 1 |
| 0 | 0 | 0 | 0 |
| 0 | 0 | 0 | 1 |
| 0 | 0 | 0 | 0 |
| 0 | 0 | 0 | 1 |
| 0 | 0 | 0 | 1 |
| 0 | 0 | 0 | 0 |
| 0 | 0 | 0 | 1 |
| 0 | 0 | 0 | 1 |
| 0 | 0 | 0 | 1 |
| 0 | 0 | 0 | 1 |
| 0 | 0 | 0 | 1 |
| 0 | 0 | 0 | 1 |
| 0 | 0 | 0 | 0 |
| 0 | 0 | 0 | 0 |
| 0 | 0 | 0 | 0 |
| 0 | 0 | 0 | 0 |
| 0 | 0 | 0 | 0 |
| 0 | 0 | 0 | 1 |
| 0 | 0 | 0 | 1 |
| 0 | 0 | 0 | 1 |
| 0 | 0 | 0 | 1 |
| 0 | 0 | 0 | 0 |
| 0 | 0 | 0 | 1 |
| 0 | 0 | 0 | 1 |
| 0 | 0 | 0 | 1 |
| 0 | 0 | 0 | 0 |
| 0 | 0 | 0 | 1 |
| 0 | 0 | 0 | 0 |
| 0 | 0 | 0 | 0 |
| 0 | 0 | 0 | 0 |
| 0 | 1 | 0 | 0 |
| 0 | 1 | 0 | 0 |
| 0 | 0 | 0 | 1 |
| 0 | 0 | 0 | 0 |
| 0 | 0 | 0 | 0 |
| 0 | 1 | 0 | 0 |
| 0 | 1 | 0 | 0 |
| 0 | 1 | 0 | 0 |
| 0 | 0 | 0 | 0 |
| 0 | 0 | 0 | 0 |
| 0 | 0 | 0 | 0 |
| 0 | 0 | 0 | 0 |
| 0 | 0 | 0 | 0 |
| 0 | 0 | 0 | 0 |
| 0 | 0 | 0 | 0 |
| 0 | 0 | 0 | 1 |
| 0 | 0 | 0 | 0 |
| 0 | 0 | 0 | 1 |
| 0 | 0 | 0 | 1 |
| 0 | 0 | 0 | 1 |
| 0 | 0 | 0 | 0 |

| | | | |
|---|---|---|---|
| 0 | 0 | 0 | 0 |
| 0 | 0 | 0 | 1 |
| 0 | 0 | 0 | 1 |
| 0 | 0 | 0 | 1 |
| 0 | 0 | 0 | 1 |
| 0 | 0 | 0 | 0 |
| 0 | 0 | 0 | 1 |
| 0 | 0 | 0 | 1 |
| 0 | 0 | 0 | 1 |
| 0 | 0 | 0 | 1 |
| 0 | 0 | 0 | 1 |
| 0 | 0 | 0 | 1 |
| 0 | 0 | 0 | 1 |
| 1 | 0 | 0 | 0 |
| 0 | 0 | 0 | 1 |
| 0 | 0 | 0 | 1 |
| 0 | 0 | 0 | 1 |
| 0 | 0 | 0 | 0 |
| 0 | 0 | 0 | 1 |
| 0 | 0 | 0 | 1 |
| 0 | 0 | 0 | 1 |
| 0 | 0 | 0 | 1 |
| 0 | 0 | 0 | 0 |
| 0 | 0 | 0 | 1 |
| 0 | 0 | 0 | 1 |
| 0 | 0 | 0 | 0 |
| 0 | 0 | 0 | 0 |
| 0 | 0 | 0 | 0 |
| 0 | 0 | 0 | 1 |
| 0 | 0 | 0 | 1 |
| 0 | 0 | 0 | 1 |
| 0 | 0 | 0 | 1 |
| 0 | 0 | 0 | 1 |
| 0 | 0 | 0 | 1 |
| 0 | 0 | 0 | 1 |
| 0 | 0 | 0 | 1 |
| 0 | 0 | 0 | 1 |
| 0 | 0 | 0 | 1 |
| 0 | 0 | 0 | 1 |
| 0 | 0 | 0 | 1 |
| 0 | 0 | 0 | 1 |
| 0 | 0 | 0 | 1 |
| 0 | 0 | 0 | 1 |
| 0 | 0 | 0 | 1 |
| 0 | 0 | 0 | 1 |
| 0 | 0 | 0 | 1 |
| 0 | 0 | 0 | 1 |
| 0 | 0 | 0 | 1 |
| 0 | 0 | 0 | 1 |
| 0 | 0 | 0 | 1 |
| 0 | 0 | 0 | 1 |
| 0 | 0 | 0 | 1 |
| 0 | 0 | 0 | 1 |

| | | | |
|---|---|---|---|
| 0 | 0 | 0 | 1 |
| 0 | 0 | 0 | 0 |
| 0 | 0 | 0 | 1 |
| 0 | 0 | 0 | 1 |
| 0 | 0 | 0 | 1 |
| 0 | 0 | 0 | 0 |
| 0 | 0 | 0 | 0 |
| 0 | 0 | 0 | 0 |
| 0 | 0 | 0 | 1 |
| 0 | 0 | 0 | 1 |
| 0 | 0 | 0 | 1 |
| 0 | 0 | 0 | 1 |
| 0 | 0 | 0 | 1 |
| 0 | 0 | 0 | 1 |
| 0 | 0 | 0 | 0 |
| 0 | 0 | 0 | 0 |
| 0 | 0 | 0 | 0 |
| 0 | 0 | 0 | 1 |
| 0 | 0 | 0 | 0 |
| 0 | 0 | 0 | 0 |
| 0 | 0 | 0 | 0 |
| 0 | 0 | 0 | 0 |
| 0 | 0 | 0 | 0 |
| 0 | 0 | 0 | 0 |
| 0 | 0 | 0 | 1 |
| 0 | 0 | 0 | 1 |
| 0 | 0 | 0 | 1 |
| 0 | 0 | 0 | 1 |
| 0 | 0 | 0 | 1 |
| 0 | 0 | 0 | 1 |
| 0 | 0 | 0 | 1 |
| 0 | 0 | 0 | 1 |
| 0 | 0 | 0 | 0 |
| 0 | 0 | 0 | 1 |
| 0 | 0 | 0 | 1 |
| 0 | 0 | 0 | 0 |
| 0 | 0 | 0 | 0 |
| 0 | 0 | 0 | 0 |
| 0 | 0 | 0 | 0 |
| 0 | 0 | 0 | 1 |
| 0 | 0 | 0 | 1 |
| 0 | 0 | 0 | 1 |
| 0 | 0 | 0 | 1 |
| 0 | 0 | 0 | 0 |
| 0 | 0 | 0 | 0 |
| 0 | 0 | 0 | 0 |
| 0 | 0 | 0 | 1 |
| 0 | 0 | 0 | 1 |
| 0 | 0 | 0 | 0 |
| 0 | 0 | 0 | 1 |
| 0 | 0 | 0 | 1 |
| 0 | 0 | 0 | 0 |
| 0 | 0 | 0 | 1 |
| 0 | 0 | 0 | 1 |

| | | | |
|---|---|---|---|
| 0 | 0 | 0 | 1 |
| 0 | 0 | 0 | 1 |
| 0 | 0 | 0 | 1 |
| 0 | 0 | 0 | 1 |
| 0 | 0 | 0 | 1 |
| 0 | 0 | 0 | 1 |
| 0 | 0 | 0 | 1 |
| 0 | 0 | 0 | 1 |
| 0 | 0 | 0 | 0 |
| 0 | 0 | 0 | 0 |
| 0 | 0 | 0 | 1 |
| 0 | 0 | 0 | 0 |
| 0 | 0 | 0 | 0 |
| 0 | 0 | 0 | 0 |
| 0 | 0 | 0 | 1 |
| 0 | 0 | 0 | 1 |
| 0 | 0 | 0 | 0 |
| 0 | 0 | 0 | 0 |
| 0 | 0 | 0 | 0 |
| 0 | 0 | 0 | 1 |
| 0 | 0 | 0 | 1 |
| 0 | 0 | 0 | 1 |
| 0 | 0 | 0 | 0 |
| 0 | 0 | 0 | 1 |
| 0 | 0 | 0 | 0 |
| 0 | 0 | 0 | 1 |
| 0 | 0 | 0 | 1 |
| 0 | 0 | 0 | 1 |
| 0 | 0 | 0 | 1 |
| 0 | 0 | 0 | 0 |
| 0 | 0 | 0 | 0 |
| 0 | 0 | 0 | 0 |
| 0 | 0 | 0 | 1 |
| 0 | 0 | 0 | 1 |
| 0 | 0 | 0 | 1 |
| 0 | 0 | 0 | 1 |
| 0 | 0 | 0 | 1 |
| 0 | 0 | 0 | 1 |
| 0 | 0 | 0 | 1 |
| 0 | 0 | 0 | 1 |
| 0 | 0 | 0 | 0 |
| 0 | 0 | 0 | 1 |
| 0 | 0 | 0 | 1 |
| 0 | 0 | 0 | 1 |
| 0 | 0 | 0 | 1 |
| 0 | 0 | 0 | 0 |
| 0 | 0 | 0 | 1 |
| 0 | 0 | 0 | 0 |
| 0 | 0 | 0 | 1 |
| 0 | 0 | 0 | 0 |
| 0 | 0 | 0 | 0 |
| 0 | 0 | 0 | 0 |
| 0 | 0 | 0 | 1 |
| 0 | 0 | 0 | 1 |

| | | | |
|---|---|---|---|
| 0 | 0 | 0 | 1 |
| 0 | 0 | 0 | 1 |
| 0 | 0 | 0 | 1 |
| 0 | 0 | 0 | 1 |
| 0 | 0 | 0 | 1 |
| 0 | 0 | 0 | 1 |
| 0 | 0 | 0 | 1 |
| 0 | 0 | 0 | 0 |
| 0 | 0 | 0 | 0 |
| 0 | 0 | 0 | 1 |
| 0 | 0 | 0 | 0 |
| 0 | 0 | 0 | 1 |
| 0 | 0 | 0 | 1 |
| 0 | 0 | 0 | 1 |
| 0 | 0 | 0 | 1 |
| 0 | 0 | 0 | 1 |
| 0 | 0 | 0 | 1 |
| 0 | 0 | 0 | 1 |
| 0 | 0 | 0 | 1 |
| 0 | 0 | 0 | 1 |
| 0 | 0 | 0 | 1 |
| 0 | 0 | 0 | 1 |
| 0 | 0 | 0 | 1 |
| 0 | 0 | 0 | 1 |
| 0 | 0 | 0 | 1 |
| 0 | 0 | 0 | 1 |
| 0 | 0 | 0 | 1 |
| 0 | 0 | 0 | 1 |
| 0 | 0 | 0 | 0 |
| 0 | 0 | 0 | 1 |
| 0 | 0 | 0 | 0 |
| 0 | 0 | 0 | 1 |
| 0 | 0 | 0 | 0 |
| 0 | 0 | 0 | 0 |
| 0 | 0 | 0 | 1 |
| 0 | 0 | 0 | 0 |
| 0 | 0 | 0 | 0 |
| 0 | 0 | 0 | 1 |
| 0 | 0 | 0 | 1 |
| 0 | 0 | 0 | 1 |
| 0 | 0 | 0 | 1 |
| 0 | 0 | 0 | 0 |
| 0 | 0 | 0 | 0 |
| 0 | 0 | 0 | 1 |
| 0 | 0 | 0 | 1 |
| 0 | 0 | 0 | 0 |
| 0 | 0 | 0 | 0 |
| 0 | 0 | 0 | 0 |
| 0 | 0 | 0 | 0 |
| 0 | 0 | 0 | 1 |
| 0 | 0 | 0 | 0 |
| 0 | 0 | 0 | 0 |
| 0 | 0 | 0 | 1 |
| 0 | 0 | 0 | 1 |

| | | | |
|---|---|---|---|
| 0 | 0 | 0 | 1 |
| 0 | 0 | 0 | 1 |
| 0 | 0 | 0 | 1 |
| 0 | 0 | 0 | 1 |
| 0 | 0 | 0 | 0 |
| 0 | 0 | 0 | 0 |
| 0 | 0 | 0 | 0 |
| 0 | 0 | 0 | 0 |
| 0 | 0 | 0 | 1 |
| 0 | 0 | 0 | 0 |
| 0 | 0 | 0 | 0 |
| 0 | 0 | 0 | 0 |
| 0 | 0 | 0 | 0 |
| 0 | 0 | 0 | 1 |
| 0 | 0 | 0 | 1 |
| 0 | 0 | 0 | 1 |
| 0 | 0 | 0 | 1 |
| 0 | 0 | 0 | 1 |
| 0 | 0 | 0 | 1 |
| 0 | 0 | 0 | 1 |
| 0 | 0 | 0 | 1 |
| 0 | 0 | 0 | 1 |
| 0 | 0 | 0 | 1 |
| 0 | 0 | 0 | 1 |
| 0 | 0 | 0 | 1 |
| 0 | 0 | 0 | 0 |
| 0 | 0 | 0 | 0 |
| 0 | 0 | 0 | 0 |
| 0 | 0 | 0 | 0 |
| 0 | 0 | 0 | 1 |
| 0 | 0 | 0 | 1 |
| 0 | 0 | 0 | 1 |
| 0 | 0 | 0 | 1 |
| 0 | 0 | 0 | 1 |
| 0 | 0 | 0 | 1 |
| 0 | 0 | 0 | 1 |
| 0 | 0 | 0 | 1 |
| 0 | 0 | 0 | 1 |
| 0 | 0 | 0 | 1 |
| 0 | 0 | 0 | 1 |
| 0 | 0 | 0 | 1 |
| 0 | 0 | 0 | 1 |
| 0 | 0 | 0 | 1 |
| 0 | 0 | 0 | 1 |
| 0 | 0 | 0 | 1 |
| 0 | 0 | 0 | 1 |
| 0 | 0 | 0 | 1 |
| 0 | 0 | 0 | 1 |
| 0 | 0 | 0 | 1 |
| 0 | 0 | 0 | 1 |
| 0 | 0 | 0 | 1 |
| 0 | 0 | 0 | 1 |
| 0 | 0 | 0 | 1 |
| 0 | 0 | 0 | 0 |

| | | | |
|---|---|---|---|
| 0 | 0 | 0 | 1 |
| 0 | 0 | 0 | 1 |
| 0 | 0 | 0 | 1 |
| 0 | 0 | 0 | 1 |
| 0 | 0 | 0 | 1 |
| 0 | 0 | 0 | 1 |
| 0 | 0 | 0 | 1 |
| 0 | 0 | 0 | 1 |
| 0 | 0 | 0 | 1 |
| 0 | 0 | 0 | 1 |
| 0 | 0 | 0 | 1 |
| 0 | 0 | 0 | 1 |
| 0 | 0 | 0 | 1 |
| 0 | 0 | 0 | 1 |
| 0 | 0 | 0 | 1 |
| 0 | 0 | 0 | 1 |
| 0 | 0 | 0 | 1 |
| 0 | 0 | 0 | 1 |
| 0 | 0 | 0 | 1 |
| 0 | 0 | 0 | 1 |
| 0 | 0 | 0 | 1 |
| 0 | 0 | 0 | 1 |
| 0 | 0 | 0 | 1 |
| 0 | 0 | 0 | 1 |
| 0 | 0 | 0 | 1 |
| 0 | 0 | 0 | 1 |
| 0 | 0 | 0 | 1 |
| 0 | 0 | 0 | 1 |
| 0 | 0 | 0 | 1 |
| 0 | 0 | 0 | 1 |
| 0 | 0 | 0 | 1 |
| 0 | 0 | 0 | 1 |
| 0 | 0 | 0 | 1 |
| 0 | 0 | 0 | 1 |
| 0 | 0 | 0 | 1 |
| 0 | 0 | 0 | 1 |
| 0 | 0 | 0 | 1 |
| 0 | 0 | 0 | 1 |
| 0 | 0 | 0 | 1 |
| 0 | 0 | 0 | 1 |
| 0 | 0 | 0 | 1 |
| 0 | 0 | 0 | 1 |
| 0 | 0 | 0 | 1 |
| 0 | 0 | 0 | 1 |
| 0 | 0 | 0 | 1 |
| 0 | 1 | 1 | 1 |
| 0 | 0 | 0 | 0 |
| 0 | 0 | 0 | 1 |
| 0 | 0 | 0 | 1 |
| 0 | 0 | 0 | 1 |
| 0 | 0 | 0 | 1 |
| 0 | 0 | 0 | 1 |
| 0 | 0 | 0 | 1 |
| 0 | 0 | 0 | 1 |

| | | | |
|---|---|---|---|
| 0 | 0 | 0 | 1 |
| 0 | 0 | 0 | 1 |
| 0 | 0 | 0 | 0 |
| 0 | 0 | 0 | 1 |
| 0 | 0 | 0 | 1 |
| 0 | 0 | 0 | 1 |
| 0 | 0 | 0 | 0 |
| 0 | 0 | 0 | 1 |
| 0 | 0 | 0 | 1 |
| 0 | 0 | 0 | 1 |
| 0 | 0 | 0 | 1 |
| 0 | 0 | 0 | 0 |
| 0 | 0 | 0 | 1 |
| 0 | 0 | 0 | 0 |
| 0 | 0 | 0 | 0 |
| 0 | 0 | 0 | 0 |
| 0 | 0 | 0 | 0 |
| 0 | 0 | 0 | 0 |
| 0 | 0 | 0 | 0 |
| 0 | 0 | 0 | 0 |
| 0 | 0 | 0 | 0 |

| surface_non_binding_site | non_binding_site_single_water | surface_non_binding_site_single_water |
|---|---|---|
| 0 | 0 | 0 |
| 0 | 0 | 0 |
| 0 | 0 | 0 |
| 0 | 0 | 0 |
| 0 | 0 | 0 |
| 0 | 0 | 0 |
| 1 | 1 | 1 |
| 1 | 1 | 1 |
| 0 | 0 | 0 |
| 0 | 0 | 0 |
| 0 | 0 | 0 |
| 0 | 0 | 0 |
| 0 | 0 | 0 |
| 0 | 0 | 0 |
| 0 | 0 | 0 |
| 0 | 0 | 0 |
| 0 | 0 | 0 |
| 0 | 0 | 0 |
| 0 | 0 | 0 |
| 0 | 0 | 0 |
| 0 | 0 | 0 |
| 0 | 0 | 0 |
| 0 | 0 | 0 |
| 0 | 0 | 0 |
| 0 | 0 | 0 |
| 1 | 0 | 0 |
| 0 | 0 | 0 |
| 1 | 1 | 1 |
| 0 | 0 | 0 |
| 0 | 1 | 0 |
| 0 | 1 | 0 |
| 0 | 0 | 0 |
| 0 | 0 | 0 |
| 0 | 0 | 0 |
| 0 | 0 | 0 |
| 1 | 1 | 1 |
| 0 | 0 | 0 |
| 0 | 0 | 0 |
| 0 | 0 | 0 |
| 0 | 0 | 0 |
| 0 | 0 | 0 |
| 0 | 0 | 0 |
| 0 | 0 | 0 |
| 0 | 0 | 0 |
| 0 | 0 | 0 |
| 0 | 0 | 0 |
| 0 | 0 | 0 |
| 0 | 0 | 0 |
| 0 | 0 | 0 |
| 0 | 0 | 0 |
| 0 | 0 | 0 |
| 0 | 0 | 0 |
| 0 | 0 | 0 |

| | | |
|---|---|---|
| 0 | 0 | 0 |
| 0 | 0 | 0 |
| 0 | 0 | 0 |
| 0 | 0 | 0 |
| 0 | 0 | 0 |
| 0 | 0 | 0 |
| 0 | 0 | 0 |
| 0 | 0 | 0 |
| 0 | 0 | 0 |
| 0 | 0 | 0 |
| 0 | 0 | 0 |
| 0 | 0 | 0 |
| 0 | 0 | 0 |
| 0 | 0 | 0 |
| 0 | 0 | 0 |
| 0 | 0 | 0 |
| 0 | 0 | 0 |
| 0 | 0 | 0 |
| 0 | 0 | 0 |
| 0 | 0 | 0 |
| 0 | 0 | 0 |
| 0 | 0 | 0 |
| 0 | 0 | 0 |
| 0 | 0 | 0 |
| 0 | 0 | 0 |
| 0 | 0 | 0 |
| 0 | 0 | 0 |
| 0 | 0 | 0 |
| 0 | 0 | 0 |
| 0 | 0 | 0 |
| 0 | 0 | 0 |
| 0 | 0 | 0 |
| 0 | 0 | 0 |
| 0 | 0 | 0 |
| 0 | 0 | 0 |
| 0 | 0 | 0 |
| 0 | 0 | 0 |
| 0 | 0 | 0 |
| 0 | 0 | 0 |
| 0 | 0 | 0 |
| 0 | 0 | 0 |
| 0 | 0 | 0 |
| 0 | 0 | 0 |
| 0 | 0 | 0 |
| 0 | 0 | 0 |
| 1 | 1 | 1 |
| 0 | 1 | 0 |
| 0 | 0 | 0 |
| 0 | 0 | 0 |
| 0 | 0 | 0 |
| 0 | 0 | 0 |
| 1 | 1 | 1 |
| 1 | 1 | 1 |
| 0 | 0 | 0 |

| | | |
|---|---|---|
| 0 | 1 | 0 |
| 0 | 1 | 0 |
| 0 | 0 | 0 |
| 0 | 0 | 0 |
| 0 | 0 | 0 |
| 0 | 0 | 0 |
| 0 | 1 | 0 |
| 0 | 0 | 0 |
| 0 | 0 | 0 |
| 0 | 0 | 0 |
| 0 | 0 | 0 |
| 0 | 0 | 0 |
| 0 | 0 | 0 |
| 0 | 0 | 0 |
| 0 | 0 | 0 |
| 1 | 1 | 1 |
| 0 | 0 | 0 |
| 0 | 0 | 0 |
| 0 | 0 | 0 |
| 0 | 0 | 0 |
| 0 | 0 | 0 |
| 0 | 0 | 0 |
| 0 | 0 | 0 |
| 0 | 0 | 0 |
| 0 | 0 | 0 |
| 0 | 0 | 0 |
| 0 | 0 | 0 |
| 0 | 0 | 0 |
| 0 | 0 | 0 |
| 0 | 0 | 0 |
| 0 | 0 | 0 |
| 0 | 0 | 0 |
| 0 | 0 | 0 |
| 0 | 0 | 0 |
| 0 | 0 | 0 |
| 0 | 0 | 0 |
| 0 | 0 | 0 |
| 0 | 0 | 0 |
| 0 | 0 | 0 |
| 0 | 0 | 0 |
| 0 | 0 | 0 |
| 0 | 0 | 0 |
| 0 | 0 | 0 |
| 0 | 0 | 0 |
| 0 | 0 | 0 |
| 0 | 0 | 0 |
| 0 | 0 | 0 |
| 0 | 0 | 0 |
| 0 | 0 | 0 |
| 0 | 0 | 0 |
| 0 | 0 | 0 |
| 0 | 0 | 0 |
| 0 | 0 | 0 |
| 0 | 0 | 0 |

| | | |
|---|---|---|
| 1 | 0 | 0 |
| 1 | 0 | 0 |
| 1 | 0 | 0 |
| 0 | 0 | 0 |
| 0 | 0 | 0 |
| 0 | 0 | 0 |
| 0 | 0 | 0 |
| 1 | 0 | 0 |
| 0 | 0 | 0 |
| 1 | 0 | 0 |
| 1 | 0 | 0 |
| 0 | 0 | 0 |
| 1 | 0 | 0 |
| 0 | 0 | 0 |
| 0 | 0 | 0 |
| 1 | 0 | 0 |
| 1 | 0 | 0 |
| 1 | 0 | 0 |
| 1 | 0 | 0 |
| 0 | 0 | 0 |
| 0 | 0 | 0 |
| 0 | 1 | 0 |
| 0 | 0 | 0 |
| 0 | 0 | 0 |
| 0 | 0 | 0 |
| 1 | 0 | 0 |
| 0 | 0 | 0 |
| 0 | 0 | 0 |
| 1 | 0 | 0 |
| 1 | 0 | 0 |
| 1 | 1 | 1 |
| 0 | 0 | 0 |
| 0 | 0 | 0 |
| 0 | 1 | 0 |
| 0 | 0 | 0 |
| 0 | 0 | 0 |
| 0 | 0 | 0 |
| 0 | 0 | 0 |
| 0 | 0 | 0 |
| 1 | 0 | 0 |
| 1 | 0 | 0 |
| 0 | 0 | 0 |
| 0 | 0 | 0 |
| 1 | 1 | 1 |
| 1 | 0 | 0 |
| 1 | 0 | 0 |
| 0 | 0 | 0 |
| 0 | 0 | 0 |
| 0 | 0 | 0 |
| 0 | 0 | 0 |
| 0 | 0 | 0 |
| 0 | 1 | 0 |
| 1 | 0 | 0 |
| 0 | 0 | 0 |

| | | |
|---|---|---|
| 0 | 0 | 0 |
| 0 | 0 | 0 |
| 0 | 0 | 0 |
| 1 | 1 | 1 |
| 1 | 1 | 1 |
| 1 | 0 | 0 |
| 0 | 0 | 0 |
| 0 | 0 | 0 |
| 1 | 0 | 0 |
| 1 | 0 | 0 |
| 0 | 0 | 0 |
| 0 | 1 | 0 |
| 0 | 0 | 0 |
| 1 | 0 | 0 |
| 0 | 1 | 0 |
| 1 | 1 | 1 |
| 1 | 1 | 1 |
| 0 | 0 | 0 |
| 0 | 0 | 0 |
| 0 | 0 | 0 |
| 0 | 0 | 0 |
| 1 | 0 | 0 |
| 0 | 0 | 0 |
| 0 | 1 | 0 |
| 1 | 1 | 1 |
| 0 | 0 | 0 |
| 0 | 0 | 0 |
| 0 | 0 | 0 |
| 0 | 0 | 0 |
| 0 | 0 | 0 |
| 0 | 0 | 0 |
| 0 | 0 | 0 |
| 1 | 0 | 0 |
| 1 | 0 | 0 |
| 0 | 0 | 0 |
| 1 | 0 | 0 |
| 0 | 0 | 0 |
| 0 | 0 | 0 |
| 1 | 0 | 0 |
| 0 | 1 | 0 |
| 0 | 0 | 0 |
| 1 | 0 | 0 |
| 1 | 0 | 0 |
| 1 | 0 | 0 |
| 0 | 0 | 0 |
| 0 | 0 | 0 |
| 0 | 0 | 0 |
| 0 | 0 | 0 |
| 0 | 0 | 0 |
| 0 | 0 | 0 |
| 0 | 0 | 0 |
| 0 | 0 | 0 |
| 0 | 0 | 0 |
| 0 | 0 | 0 |

| | | |
|---|---|---|
| 0 | 0 | 0 |
| 0 | 0 | 0 |
| 0 | 0 | 0 |
| 0 | 0 | 0 |
| 0 | 0 | 0 |
| 0 | 0 | 0 |
| 0 | 1 | 0 |
| 0 | 1 | 0 |
| 0 | 0 | 0 |
| 0 | 0 | 0 |
| 0 | 0 | 0 |
| 0 | 0 | 0 |
| 0 | 0 | 0 |
| 0 | 1 | 0 |
| 0 | 1 | 0 |
| 0 | 0 | 0 |
| 0 | 1 | 0 |
| 0 | 0 | 0 |
| 1 | 1 | 1 |
| 0 | 0 | 0 |
| 1 | 1 | 1 |
| 1 | 0 | 0 |
| 1 | 1 | 1 |
| 0 | 0 | 0 |
| 0 | 0 | 0 |
| 0 | 0 | 0 |
| 0 | 0 | 0 |
| 0 | 0 | 0 |
| 0 | 0 | 0 |
| 0 | 0 | 0 |
| 0 | 0 | 0 |
| 1 | 0 | 0 |
| 0 | 0 | 0 |
| 0 | 0 | 0 |
| 0 | 0 | 0 |
| 0 | 0 | 0 |
| 0 | 0 | 0 |
| 0 | 0 | 0 |
| 0 | 0 | 0 |
| 0 | 0 | 0 |
| 0 | 1 | 0 |
| 0 | 1 | 0 |
| 0 | 0 | 0 |
| 0 | 1 | 0 |
| 0 | 0 | 0 |
| 0 | 0 | 0 |
| 0 | 0 | 0 |
| 1 | 1 | 1 |
| 1 | 1 | 1 |
| 0 | 0 | 0 |
| 0 | 1 | 0 |
| 1 | 1 | 1 |
| 0 | 0 | 0 |
| 0 | 1 | 0 |

| | | |
|---|---|---|
| 0 | 0 | 0 |
| 1 | 1 | 1 |
| 0 | 0 | 0 |
| 0 | 0 | 0 |
| 0 | 0 | 0 |
| 0 | 0 | 0 |
| 0 | 1 | 0 |
| 0 | 0 | 0 |
| 0 | 0 | 0 |
| 1 | 1 | 1 |
| 1 | 1 | 1 |
| 1 | 1 | 1 |
| 1 | 1 | 1 |
| 1 | 1 | 1 |
| 0 | 0 | 0 |
| 1 | 1 | 1 |
| 0 | 0 | 0 |
| 0 | 0 | 0 |
| 1 | 1 | 1 |
| 0 | 0 | 0 |
| 0 | 0 | 0 |
| 0 | 0 | 0 |
| 0 | 0 | 0 |
| 0 | 0 | 0 |
| 1 | 1 | 1 |
| 1 | 0 | 0 |
| 0 | 0 | 0 |
| 0 | 0 | 0 |
| 0 | 0 | 0 |
| 0 | 0 | 0 |
| 0 | 0 | 0 |
| 0 | 0 | 0 |
| 0 | 0 | 0 |
| 0 | 0 | 0 |
| 1 | 1 | 1 |
| 1 | 1 | 1 |
| 0 | 0 | 0 |
| 0 | 0 | 0 |
| 0 | 0 | 0 |
| 0 | 0 | 0 |
| 0 | 0 | 0 |
| 0 | 0 | 0 |
| 0 | 1 | 0 |
| 0 | 0 | 0 |
| 0 | 0 | 0 |
| 0 | 0 | 0 |
| 0 | 0 | 0 |
| 0 | 0 | 0 |
| 0 | 0 | 0 |
| 1 | 1 | 1 |
| 0 | 0 | 0 |
| 0 | 0 | 0 |
| 0 | 0 | 0 |
| 0 | 0 | 0 |

| | | |
|---|---|---|
| 0 | 0 | 0 |
| 0 | 0 | 0 |
| 0 | 0 | 0 |
| 0 | 0 | 0 |
| 1 | 1 | 1 |
| 0 | 0 | 0 |
| 0 | 0 | 0 |
| 0 | 0 | 0 |
| 0 | 0 | 0 |
| 0 | 0 | 0 |
| 0 | 0 | 0 |
| 0 | 0 | 0 |
| 0 | 0 | 0 |
| 0 | 0 | 0 |
| 0 | 0 | 0 |
| 0 | 0 | 0 |
| 0 | 0 | 0 |
| 0 | 0 | 0 |
| 1 | 1 | 1 |
| 0 | 0 | 0 |
| 0 | 0 | 0 |
| 0 | 0 | 0 |
| 0 | 0 | 0 |
| 1 | 0 | 0 |
| 0 | 0 | 0 |
| 0 | 0 | 0 |
| 0 | 0 | 0 |
| 1 | 0 | 0 |
| 1 | 0 | 0 |
| 1 | 0 | 0 |
| 0 | 0 | 0 |
| 0 | 0 | 0 |
| 0 | 0 | 0 |
| 0 | 0 | 0 |
| 0 | 0 | 0 |
| 1 | 0 | 0 |
| 1 | 0 | 0 |
| 0 | 0 | 0 |
| 0 | 0 | 0 |
| 1 | 0 | 0 |
| 1 | 0 | 0 |
| 1 | 0 | 0 |
| 0 | 1 | 0 |
| 1 | 1 | 1 |
| 0 | 0 | 0 |
| 0 | 0 | 0 |
| 0 | 0 | 0 |
| 0 | 0 | 0 |
| 0 | 0 | 0 |
| 0 | 1 | 0 |
| 1 | 1 | 1 |
| 0 | 0 | 0 |
| 1 | 1 | 1 |
| 0 | 0 | 0 |

| | | |
|---|---|---|
| 1 | 1 | 1 |
| 1 | 1 | 1 |
| 0 | 0 | 0 |
| 1 | 1 | 1 |
| 1 | 1 | 1 |
| 1 | 1 | 1 |
| 1 | 1 | 1 |
| 0 | 0 | 0 |
| 1 | 0 | 0 |
| 1 | 1 | 1 |
| 0 | 0 | 0 |
| 1 | 1 | 1 |
| 1 | 1 | 1 |
| 0 | 1 | 0 |
| 0 | 0 | 0 |
| 0 | 0 | 0 |
| 1 | 1 | 1 |
| 1 | 1 | 1 |
| 0 | 0 | 0 |
| 0 | 0 | 0 |
| 0 | 0 | 0 |
| 0 | 0 | 0 |
| 0 | 0 | 0 |
| 0 | 0 | 0 |
| 0 | 0 | 0 |
| 1 | 1 | 1 |
| 0 | 0 | 0 |
| 0 | 0 | 0 |
| 0 | 0 | 0 |
| 1 | 0 | 0 |
| 0 | 0 | 0 |
| 0 | 0 | 0 |
| 0 | 0 | 0 |
| 0 | 0 | 0 |
| 0 | 0 | 0 |
| 0 | 0 | 0 |
| 0 | 0 | 0 |
| 0 | 0 | 0 |
| 0 | 0 | 0 |
| 0 | 0 | 0 |
| 0 | 1 | 0 |
| 0 | 0 | 0 |
| 1 | 1 | 1 |
| 0 | 0 | 0 |
| 0 | 0 | 0 |
| 0 | 0 | 0 |
| 0 | 0 | 0 |
| 0 | 0 | 0 |
| 0 | 0 | 0 |
| 0 | 0 | 0 |
| 0 | 1 | 0 |
| 0 | 0 | 0 |
| 0 | 0 | 0 |
| 0 | 0 | 0 |

| | | |
|---|---|---|
| 1 | 1 | 1 |
| 0 | 0 | 0 |
| 0 | 1 | 0 |
| 0 | 1 | 0 |
| 0 | 0 | 0 |
| 0 | 0 | 0 |
| 0 | 0 | 0 |
| 0 | 0 | 0 |
| 0 | 0 | 0 |
| 0 | 1 | 0 |
| 1 | 1 | 1 |
| 1 | 1 | 1 |
| 1 | 1 | 1 |
| 0 | 0 | 0 |
| 1 | 1 | 1 |
| 0 | 0 | 0 |
| 0 | 0 | 0 |
| 0 | 0 | 0 |
| 0 | 0 | 0 |
| 1 | 0 | 0 |
| 0 | 0 | 0 |
| 1 | 1 | 1 |
| 0 | 0 | 0 |
| 0 | 1 | 0 |
| 1 | 1 | 1 |
| 1 | 1 | 1 |
| 0 | 0 | 0 |
| 1 | 1 | 1 |
| 1 | 1 | 1 |
| 1 | 1 | 1 |
| 0 | 0 | 0 |
| 0 | 0 | 0 |
| 1 | 1 | 1 |
| 1 | 1 | 1 |
| 0 | 0 | 0 |
| 0 | 0 | 0 |
| 0 | 0 | 0 |
| 0 | 0 | 0 |
| 0 | 0 | 0 |
| 1 | 1 | 1 |
| 0 | 0 | 0 |
| 0 | 0 | 0 |
| 0 | 0 | 0 |
| 0 | 0 | 0 |
| 1 | 0 | 0 |
| 0 | 0 | 0 |
| 0 | 0 | 0 |
| 0 | 0 | 0 |
| 0 | 0 | 0 |
| 0 | 0 | 0 |
| 0 | 0 | 0 |
| 0 | 0 | 0 |
| 1 | 0 | 0 |
| 0 | 0 | 0 |

| | | |
|---|---|---|
| 0 | 0 | 0 |
| 0 | 0 | 0 |
| 0 | 0 | 0 |
| 0 | 0 | 0 |
| 0 | 0 | 0 |
| 0 | 0 | 0 |
| 0 | 0 | 0 |
| 1 | 0 | 0 |
| 0 | 0 | 0 |
| 0 | 0 | 0 |
| 0 | 0 | 0 |
| 0 | 0 | 0 |
| 1 | 1 | 1 |
| 0 | 0 | 0 |
| 0 | 0 | 0 |
| 0 | 0 | 0 |
| 0 | 0 | 0 |
| 0 | 0 | 0 |
| 0 | 0 | 0 |
| 1 | 1 | 1 |
| 0 | 0 | 0 |
| 0 | 0 | 0 |
| 0 | 0 | 0 |
| 0 | 0 | 0 |
| 0 | 0 | 0 |
| 1 | 0 | 0 |
| 1 | 1 | 1 |
| 0 | 1 | 0 |
| 0 | 0 | 0 |
| 0 | 0 | 0 |
| 0 | 0 | 0 |
| 0 | 0 | 0 |
| 0 | 0 | 0 |
| 0 | 0 | 0 |
| 0 | 0 | 0 |
| 1 | 0 | 0 |
| 0 | 0 | 0 |
| 0 | 0 | 0 |
| 1 | 1 | 1 |
| 1 | 0 | 0 |
| 0 | 1 | 0 |
| 0 | 0 | 0 |
| 0 | 0 | 0 |
| 0 | 0 | 0 |
| 0 | 0 | 0 |
| 0 | 0 | 0 |
| 0 | 0 | 0 |
| 1 | 1 | 1 |
| 1 | 1 | 1 |
| 0 | 0 | 0 |
| 0 | 0 | 0 |
| 0 | 0 | 0 |
| 0 | 0 | 0 |
| 1 | 1 | 1 |

| | | |
|---|---|---|
| 1 | 1 | 1 |
| 0 | 0 | 0 |
| 0 | 1 | 0 |
| 0 | 0 | 0 |
| 1 | 0 | 0 |
| 1 | 0 | 0 |
| 0 | 0 | 0 |
| 1 | 0 | 0 |
| 1 | 0 | 0 |
| 1 | 0 | 0 |
| 1 | 0 | 0 |
| 1 | 0 | 0 |
| 1 | 1 | 1 |
| 1 | 1 | 1 |
| 1 | 1 | 1 |
| 1 | 1 | 1 |
| 0 | 0 | 0 |
| 0 | 0 | 0 |
| 1 | 0 | 0 |
| 0 | 0 | 0 |
| 0 | 0 | 0 |
| 0 | 0 | 0 |
| 1 | 0 | 0 |
| 0 | 0 | 0 |
| 0 | 0 | 0 |
| 1 | 1 | 1 |
| 1 | 0 | 0 |
| 1 | 0 | 0 |
| 0 | 0 | 0 |
| 0 | 0 | 0 |
| 0 | 0 | 0 |
| 0 | 0 | 0 |
| 0 | 0 | 0 |
| 0 | 0 | 0 |
| 0 | 0 | 0 |
| 0 | 0 | 0 |
| 1 | 0 | 0 |
| 1 | 1 | 1 |
| 1 | 1 | 1 |
| 1 | 1 | 1 |
| 0 | 0 | 0 |
| 1 | 1 | 1 |
| 0 | 0 | 0 |
| 1 | 1 | 1 |
| 0 | 0 | 0 |
| 0 | 1 | 0 |
| 0 | 0 | 0 |
| 0 | 0 | 0 |
| 0 | 0 | 0 |
| 0 | 0 | 0 |
| 0 | 0 | 0 |
| 0 | 0 | 0 |
| 0 | 0 | 0 |
| 0 | 0 | 0 |

| | | |
|---|---|---|
| 0 | 0 | 0 |
| 0 | 0 | 0 |
| 0 | 0 | 0 |
| 0 | 0 | 0 |
| 0 | 0 | 0 |
| 0 | 0 | 0 |
| 0 | 0 | 0 |
| 1 | 1 | 1 |
| 0 | 0 | 0 |
| 0 | 0 | 0 |
| 0 | 0 | 0 |
| 1 | 0 | 0 |
| 0 | 0 | 0 |
| 0 | 0 | 0 |
| 0 | 0 | 0 |
| 0 | 0 | 0 |
| 0 | 0 | 0 |
| 0 | 0 | 0 |
| 0 | 0 | 0 |
| 0 | 0 | 0 |
| 0 | 0 | 0 |
| 0 | 0 | 0 |
| 0 | 0 | 0 |
| 0 | 0 | 0 |
| 0 | 0 | 0 |
| 0 | 0 | 0 |
| 0 | 0 | 0 |
| 0 | 0 | 0 |
| 0 | 0 | 0 |
| 0 | 0 | 0 |
| 0 | 0 | 0 |
| 0 | 0 | 0 |
| 0 | 0 | 0 |
| 0 | 0 | 0 |
| 0 | 0 | 0 |
| 0 | 0 | 0 |
| 0 | 0 | 0 |
| 0 | 0 | 0 |
| 0 | 0 | 0 |
| 0 | 0 | 0 |
| 1 | 0 | 0 |
| 0 | 0 | 0 |
| 0 | 0 | 0 |
| 0 | 0 | 0 |
| 0 | 0 | 0 |
| 0 | 0 | 0 |
| 0 | 0 | 0 |
| 0 | 0 | 0 |
| 0 | 0 | 0 |
| 0 | 1 | 0 |
| 0 | 1 | 0 |
| 1 | 0 | 0 |
| 0 | 0 | 0 |
| 0 | 0 | 0 |

| | | |
|---|---|---|
| 0 | 0 | 0 |
| 0 | 0 | 0 |
| 0 | 0 | 0 |
| 0 | 0 | 0 |
| 0 | 0 | 0 |
| 0 | 0 | 0 |
| 0 | 0 | 0 |
| 1 | 0 | 0 |
| 1 | 0 | 0 |
| 0 | 0 | 0 |
| 0 | 0 | 0 |
| 1 | 0 | 0 |
| 0 | 0 | 0 |
| 0 | 0 | 0 |
| 0 | 0 | 0 |
| 0 | 0 | 0 |
| 0 | 0 | 0 |
| 1 | 0 | 0 |
| 0 | 0 | 0 |
| 1 | 0 | 0 |
| 0 | 0 | 0 |
| 0 | 0 | 0 |
| 1 | 1 | 1 |
| 0 | 0 | 0 |
| 0 | 0 | 0 |
| 1 | 0 | 0 |
| 0 | 0 | 0 |
| 0 | 0 | 0 |
| 0 | 0 | 0 |
| 0 | 0 | 0 |
| 0 | 0 | 0 |
| 0 | 0 | 0 |
| 1 | 0 | 0 |
| 0 | 0 | 0 |
| 0 | 0 | 0 |
| 0 | 0 | 0 |
| 1 | 1 | 1 |
| 0 | 0 | 0 |
| 0 | 0 | 0 |
| 0 | 0 | 0 |
| 0 | 0 | 0 |
| 0 | 0 | 0 |
| 0 | 0 | 0 |
| 0 | 0 | 0 |
| 0 | 0 | 0 |
| 0 | 0 | 0 |
| 0 | 0 | 0 |
| 0 | 0 | 0 |
| 0 | 0 | 0 |
| 0 | 0 | 0 |
| 0 | 0 | 0 |
| 0 | 0 | 0 |
| 0 | 0 | 0 |
| 0 | 0 | 0 |

| | | |
|---|---|---|
| 1 | 0 | 0 |
| 0 | 1 | 0 |
| 0 | 0 | 0 |
| 0 | 0 | 0 |
| 0 | 0 | 0 |
| 0 | 0 | 0 |
| 0 | 0 | 0 |
| 0 | 0 | 0 |
| 0 | 0 | 0 |
| 1 | 0 | 0 |
| 0 | 1 | 0 |
| 1 | 1 | 1 |
| 0 | 0 | 0 |
| 0 | 0 | 0 |
| 0 | 0 | 0 |
| 0 | 0 | 0 |
| 0 | 0 | 0 |
| 0 | 0 | 0 |
| 1 | 1 | 1 |
| 1 | 0 | 0 |
| 1 | 0 | 0 |
| 0 | 0 | 0 |
| 0 | 1 | 0 |
| 0 | 0 | 0 |
| 0 | 0 | 0 |
| 0 | 0 | 0 |
| 0 | 0 | 0 |
| 0 | 0 | 0 |
| 0 | 0 | 0 |
| 0 | 0 | 0 |
| 0 | 0 | 0 |
| 0 | 0 | 0 |
| 0 | 0 | 0 |
| 0 | 0 | 0 |
| 0 | 0 | 0 |
| 0 | 0 | 0 |
| 0 | 0 | 0 |
| 0 | 0 | 0 |
| 0 | 0 | 0 |
| 0 | 0 | 0 |
| 0 | 0 | 0 |
| 1 | 0 | 0 |
| 0 | 0 | 0 |
| 0 | 0 | 0 |
| 0 | 0 | 0 |
| 0 | 0 | 0 |
| 1 | 0 | 0 |
| 0 | 0 | 0 |
| 0 | 0 | 0 |
| 0 | 0 | 0 |
| 0 | 0 | 0 |
| 0 | 0 | 0 |
| 1 | 0 | 0 |
| 0 | 1 | 0 |

| | | |
|---|---|---|
| 0 | 0 | 0 |
| 0 | 0 | 0 |
| 1 | 0 | 0 |
| 1 | 0 | 0 |
| 1 | 0 | 0 |
| 0 | 0 | 0 |
| 1 | 0 | 0 |
| 0 | 0 | 0 |
| 0 | 0 | 0 |
| 0 | 0 | 0 |
| 0 | 0 | 0 |
| 0 | 0 | 0 |
| 0 | 0 | 0 |
| 1 | 0 | 0 |
| 1 | 0 | 0 |
| 1 | 0 | 0 |
| 0 | 0 | 0 |
| 1 | 1 | 1 |
| 0 | 0 | 0 |
| 0 | 0 | 0 |
| 1 | 0 | 0 |
| 1 | 0 | 0 |
| 0 | 0 | 0 |
| 0 | 0 | 0 |
| 1 | 0 | 0 |
| 0 | 0 | 0 |
| 0 | 0 | 0 |
| 1 | 0 | 0 |
| 0 | 0 | 0 |
| 0 | 0 | 0 |
| 0 | 0 | 0 |
| 0 | 0 | 0 |
| 0 | 0 | 0 |
| 0 | 0 | 0 |
| 0 | 0 | 0 |
| 0 | 0 | 0 |
| 0 | 0 | 0 |
| 0 | 0 | 0 |
| 0 | 0 | 0 |
| 0 | 0 | 0 |
| 1 | 0 | 0 |
| 1 | 0 | 0 |
| 1 | 0 | 0 |
| 1 | 1 | 1 |
| 1 | 1 | 1 |
| 0 | 0 | 0 |
| 0 | 0 | 0 |
| 0 | 0 | 0 |
| 0 | 0 | 0 |
| 0 | 0 | 0 |
| 0 | 0 | 0 |
| 0 | 0 | 0 |
| 0 | 0 | 0 |
| 1 | 0 | 0 |

| | | |
|---|---|---|
| 1 | 0 | 0 |
| 0 | 0 | 0 |
| 0 | 0 | 0 |
| 0 | 0 | 0 |
| 1 | 0 | 0 |
| 0 | 0 | 0 |
| 0 | 0 | 0 |
| 0 | 0 | 0 |
| 1 | 0 | 0 |
| 1 | 0 | 0 |
| 0 | 0 | 0 |
| 0 | 0 | 0 |
| 0 | 0 | 0 |
| 1 | 0 | 0 |
| 0 | 0 | 0 |
| 0 | 0 | 0 |
| 0 | 0 | 0 |
| 0 | 0 | 0 |
| 0 | 0 | 0 |
| 0 | 0 | 0 |
| 0 | 0 | 0 |
| 1 | 1 | 1 |
| 0 | 0 | 0 |
| 0 | 1 | 0 |
| 0 | 0 | 0 |
| 0 | 0 | 0 |
| 0 | 0 | 0 |
| 0 | 0 | 0 |
| 0 | 0 | 0 |
| 0 | 0 | 0 |
| 0 | 0 | 0 |
| 0 | 0 | 0 |
| 0 | 0 | 0 |
| 0 | 0 | 0 |
| 0 | 1 | 0 |
| 0 | 0 | 0 |
| 0 | 0 | 0 |
| 0 | 0 | 0 |
| 0 | 0 | 0 |
| 0 | 0 | 0 |
| 0 | 0 | 0 |
| 0 | 0 | 0 |
| 0 | 0 | 0 |
| 0 | 0 | 0 |
| 0 | 0 | 0 |
| 0 | 0 | 0 |
| 0 | 0 | 0 |
| 1 | 1 | 1 |
| 0 | 0 | 0 |
| 1 | 1 | 1 |
| 0 | 0 | 0 |
| 0 | 0 | 0 |
| 0 | 0 | 0 |
| 0 | 0 | 0 |

| | | |
|---|---|---|
| 0 | 0 | 0 |
| 0 | 0 | 0 |
| 1 | 1 | 1 |
| 0 | 0 | 0 |
| 0 | 0 | 0 |
| 0 | 1 | 0 |
| 0 | 1 | 0 |
| 0 | 1 | 0 |
| 0 | 0 | 0 |
| 0 | 1 | 0 |
| 0 | 0 | 0 |
| 0 | 0 | 0 |
| 0 | 0 | 0 |
| 1 | 1 | 1 |
| 0 | 0 | 0 |
| 0 | 1 | 0 |
| 0 | 0 | 0 |
| 0 | 0 | 0 |
| 0 | 0 | 0 |
| 1 | 1 | 1 |
| 1 | 1 | 1 |
| 0 | 0 | 0 |
| 1 | 1 | 1 |
| 0 | 0 | 0 |
| 0 | 0 | 0 |
| 1 | 1 | 1 |
| 1 | 1 | 1 |
| 0 | 0 | 0 |
| 1 | 1 | 1 |
| 0 | 0 | 0 |
| 0 | 1 | 0 |
| 0 | 1 | 0 |
| 1 | 1 | 1 |
| 0 | 1 | 0 |
| 1 | 1 | 1 |
| 1 | 1 | 1 |
| 1 | 1 | 1 |
| 1 | 1 | 1 |
| 1 | 1 | 1 |
| 1 | 1 | 1 |
| 1 | 1 | 1 |
| 1 | 1 | 1 |
| 0 | 1 | 0 |
| 0 | 1 | 0 |
| 0 | 1 | 0 |
| 0 | 1 | 0 |
| 0 | 1 | 0 |
| 0 | 1 | 0 |
| 0 | 1 | 0 |
| 0 | 1 | 0 |
| 0 | 1 | 0 |
| 0 | 1 | 0 |
| 0 | 1 | 0 |
| 0 | 1 | 0 |

| | | |
|---|---|---|
| 0 | 1 | 0 |
| 1 | 1 | 1 |
| 1 | 1 | 1 |
| 1 | 1 | 1 |
| 1 | 1 | 1 |
| 1 | 1 | 1 |
| 1 | 1 | 1 |
| 0 | 1 | 0 |
| 0 | 1 | 0 |
| 0 | 1 | 0 |
| 0 | 1 | 0 |
| 0 | 0 | 0 |
| 0 | 0 | 0 |
| 0 | 0 | 0 |
| 1 | 0 | 0 |
| 0 | 0 | 0 |
| 1 | 0 | 0 |
| 0 | 0 | 0 |
| 1 | 0 | 0 |
| 1 | 1 | 1 |
| 0 | 0 | 0 |
| 1 | 1 | 1 |
| 1 | 1 | 1 |
| 0 | 0 | 0 |
| 0 | 0 | 0 |
| 1 | 1 | 1 |
| 1 | 1 | 1 |
| 0 | 0 | 0 |
| 0 | 0 | 0 |
| 0 | 0 | 0 |
| 0 | 0 | 0 |
| 0 | 0 | 0 |
| 0 | 1 | 0 |
| 0 | 1 | 0 |
| 1 | 1 | 1 |
| 1 | 1 | 1 |
| 1 | 1 | 1 |
| 1 | 1 | 1 |
| 1 | 1 | 1 |
| 0 | 0 | 0 |
| 0 | 0 | 0 |
| 0 | 1 | 0 |
| 0 | 0 | 0 |
| 0 | 0 | 0 |
| 0 | 0 | 0 |
| 1 | 0 | 0 |
| 0 | 0 | 0 |
| 1 | 0 | 0 |
| 1 | 0 | 0 |
| 0 | 0 | 0 |
| 0 | 0 | 0 |
| 0 | 0 | 0 |
| 0 | 0 | 0 |
| 0 | 0 | 0 |

| | | |
|---|---|---|
| 0 | 0 | 0 |
| 0 | 1 | 0 |
| 1 | 1 | 1 |
| 1 | 1 | 1 |
| 0 | 0 | 0 |
| 1 | 1 | 1 |
| 0 | 0 | 0 |
| 1 | 1 | 1 |
| 1 | 1 | 1 |
| 0 | 0 | 0 |
| 1 | 0 | 0 |
| 1 | 1 | 1 |
| 1 | 1 | 1 |
| 0 | 0 | 0 |
| 1 | 0 | 0 |
| 1 | 0 | 0 |
| 0 | 0 | 0 |
| 1 | 0 | 0 |
| 0 | 0 | 0 |
| 0 | 0 | 0 |
| 0 | 0 | 0 |
| 0 | 0 | 0 |
| 0 | 0 | 0 |
| 0 | 0 | 0 |
| 0 | 0 | 0 |
| 0 | 0 | 0 |
| 0 | 0 | 0 |
| 0 | 0 | 0 |
| 0 | 0 | 0 |
| 0 | 0 | 0 |
| 0 | 0 | 0 |
| 0 | 0 | 0 |
| 0 | 0 | 0 |
| 0 | 0 | 0 |
| 0 | 0 | 0 |
| 0 | 0 | 0 |
| 1 | 0 | 0 |
| 0 | 0 | 0 |
| 0 | 0 | 0 |
| 0 | 0 | 0 |
| 0 | 0 | 0 |
| 0 | 0 | 0 |
| 0 | 0 | 0 |
| 0 | 0 | 0 |
| 0 | 0 | 0 |
| 0 | 0 | 0 |
| 0 | 0 | 0 |
| 0 | 0 | 0 |
| 1 | 0 | 0 |
| 1 | 0 | 0 |
| 0 | 0 | 0 |
| 1 | 1 | 1 |
| 0 | 0 | 0 |
| 0 | 0 | 0 |

| | | |
|---|---|---|
| 0 | 0 | 0 |
| 0 | 0 | 0 |
| 0 | 0 | 0 |
| 0 | 0 | 0 |
| 0 | 0 | 0 |
| 0 | 0 | 0 |
| 0 | 0 | 0 |
| 0 | 0 | 0 |
| 0 | 0 | 0 |
| 0 | 0 | 0 |
| 0 | 0 | 0 |
| 0 | 0 | 0 |
| 0 | 0 | 0 |
| 0 | 0 | 0 |
| 0 | 1 | 0 |
| 0 | 0 | 0 |
| 0 | 0 | 0 |
| 0 | 0 | 0 |
| 1 | 0 | 0 |
| 0 | 0 | 0 |
| 0 | 0 | 0 |
| 0 | 0 | 0 |
| 0 | 0 | 0 |
| 0 | 0 | 0 |
| 0 | 0 | 0 |
| 1 | 0 | 0 |
| 0 | 0 | 0 |
| 0 | 0 | 0 |
| 0 | 0 | 0 |
| 0 | 0 | 0 |
| 1 | 1 | 1 |
| 0 | 0 | 0 |
| 1 | 1 | 1 |
| 1 | 0 | 0 |
| 0 | 0 | 0 |
| 0 | 0 | 0 |
| 1 | 1 | 1 |
| 1 | 1 | 1 |
| 1 | 1 | 1 |
| 1 | 0 | 0 |
| 0 | 0 | 0 |
| 0 | 0 | 0 |
| 0 | 0 | 0 |
| 0 | 0 | 0 |
| 0 | 0 | 0 |
| 0 | 0 | 0 |
| 0 | 0 | 0 |
| 0 | 0 | 0 |
| 1 | 1 | 1 |
| 1 | 1 | 1 |
| 0 | 0 | 0 |
| 0 | 0 | 0 |
| 0 | 0 | 0 |
| 0 | 1 | 0 |

| | | |
|---|---|---|
| 1 | 1 | 1 |
| 1 | 1 | 1 |
| 1 | 0 | 0 |
| 0 | 0 | 0 |
| 1 | 0 | 0 |
| 1 | 1 | 1 |
| 0 | 0 | 0 |
| 1 | 1 | 1 |
| 0 | 0 | 0 |
| 0 | 0 | 0 |
| 0 | 0 | 0 |
| 0 | 0 | 0 |
| 0 | 0 | 0 |
| 0 | 0 | 0 |
| 0 | 0 | 0 |
| 0 | 0 | 0 |
| 0 | 1 | 0 |
| 0 | 0 | 0 |
| 0 | 0 | 0 |
| 1 | 0 | 0 |
| 0 | 0 | 0 |
| 0 | 0 | 0 |
| 1 | 1 | 1 |
| 0 | 0 | 0 |
| 1 | 1 | 1 |
| 1 | 1 | 1 |
| 0 | 1 | 0 |
| 0 | 0 | 0 |
| 0 | 0 | 0 |
| 0 | 0 | 0 |
| 1 | 1 | 1 |
| 0 | 0 | 0 |
| 0 | 0 | 0 |
| 0 | 0 | 0 |
| 0 | 1 | 0 |
| 0 | 1 | 0 |
| 0 | 0 | 0 |
| 0 | 0 | 0 |
| 0 | 0 | 0 |
| 1 | 1 | 1 |
| 1 | 1 | 1 |
| 0 | 1 | 0 |
| 0 | 1 | 0 |
| 0 | 0 | 0 |
| 1 | 1 | 1 |
| 0 | 0 | 0 |
| 1 | 1 | 1 |
| 0 | 0 | 0 |
| 0 | 0 | 0 |
| 1 | 1 | 1 |
| 0 | 1 | 0 |
| 0 | 0 | 0 |
| 0 | 0 | 0 |
| 1 | 1 | 1 |

| | | |
|---|---|---|
| 1 | 1 | 1 |
| 1 | 1 | 1 |
| 0 | 1 | 0 |
| 0 | 0 | 0 |
| 0 | 0 | 0 |
| 1 | 0 | 0 |
| 1 | 0 | 0 |
| 0 | 0 | 0 |
| 0 | 0 | 0 |
| 1 | 1 | 1 |
| 0 | 0 | 0 |
| 0 | 0 | 0 |
| 0 | 0 | 0 |
| 0 | 0 | 0 |
| 0 | 0 | 0 |
| 0 | 0 | 0 |
| 0 | 0 | 0 |
| 0 | 0 | 0 |
| 0 | 0 | 0 |
| 0 | 1 | 0 |
| 1 | 1 | 1 |
| 1 | 1 | 1 |
| 1 | 1 | 1 |
| 1 | 1 | 1 |
| 0 | 0 | 0 |
| 0 | 1 | 0 |
| 0 | 0 | 0 |
| 0 | 0 | 0 |
| 1 | 1 | 1 |
| 1 | 1 | 1 |
| 1 | 1 | 1 |
| 1 | 1 | 1 |
| 1 | 1 | 1 |
| 0 | 0 | 0 |
| 0 | 0 | 0 |
| 1 | 1 | 1 |
| 0 | 0 | 0 |
| 0 | 1 | 0 |
| 0 | 0 | 0 |
| 0 | 0 | 0 |
| 0 | 0 | 0 |
| 0 | 0 | 0 |
| 0 | 1 | 0 |
| 0 | 0 | 0 |
| 0 | 0 | 0 |
| 0 | 0 | 0 |
| 0 | 0 | 0 |
| 0 | 0 | 0 |
| 0 | 0 | 0 |
| 0 | 0 | 0 |
| 0 | 0 | 0 |
| 0 | 0 | 0 |
| 0 | 0 | 0 |
| 0 | 1 | 0 |

| | | |
|---|---|---|
| 0 | 0 | 0 |
| 0 | 1 | 0 |
| 0 | 0 | 0 |
| 0 | 0 | 0 |
| 0 | 0 | 0 |
| 1 | 1 | 1 |
| 0 | 0 | 0 |
| 1 | 1 | 1 |
| 0 | 1 | 0 |
| 0 | 0 | 0 |
| 0 | 1 | 0 |
| 0 | 1 | 0 |
| 0 | 1 | 0 |
| 0 | 1 | 0 |
| 0 | 1 | 0 |
| 0 | 0 | 0 |
| 0 | 0 | 0 |
| 0 | 1 | 0 |
| 0 | 0 | 0 |
| 1 | 1 | 1 |
| 0 | 0 | 0 |
| 0 | 0 | 0 |
| 0 | 0 | 0 |
| 0 | 0 | 0 |
| 0 | 0 | 0 |
| 0 | 0 | 0 |
| 0 | 0 | 0 |
| 0 | 0 | 0 |
| 0 | 0 | 0 |
| 0 | 0 | 0 |
| 0 | 0 | 0 |
| 0 | 0 | 0 |
| 1 | 0 | 0 |
| 1 | 0 | 0 |
| 0 | 0 | 0 |
| 0 | 0 | 0 |
| 0 | 0 | 0 |
| 0 | 0 | 0 |
| 1 | 0 | 0 |
| 0 | 0 | 0 |
| 0 | 0 | 0 |
| 0 | 0 | 0 |
| 1 | 0 | 0 |
| 0 | 1 | 0 |
| 0 | 0 | 0 |
| 0 | 0 | 0 |
| 0 | 0 | 0 |
| 0 | 0 | 0 |
| 0 | 0 | 0 |
| 0 | 0 | 0 |
| 1 | 0 | 0 |
| 0 | 0 | 0 |
| 0 | 0 | 0 |
| 1 | 0 | 0 |

| | | |
|---|---|---|
| 0 | 0 | 0 |
| 1 | 0 | 0 |
| 0 | 0 | 0 |
| 0 | 0 | 0 |
| 0 | 0 | 0 |
| 0 | 0 | 0 |
| 0 | 0 | 0 |
| 0 | 0 | 0 |
| 0 | 0 | 0 |
| 0 | 0 | 0 |
| 0 | 0 | 0 |
| 1 | 1 | 1 |
| 0 | 0 | 0 |
| 1 | 1 | 1 |
| 1 | 0 | 0 |
| 1 | 0 | 0 |
| 0 | 0 | 0 |
| 0 | 0 | 0 |
| 0 | 0 | 0 |
| 0 | 0 | 0 |
| 1 | 0 | 0 |
| 0 | 0 | 0 |
| 0 | 1 | 0 |
| 0 | 0 | 0 |
| 1 | 1 | 1 |
| 1 | 1 | 1 |
| 1 | 1 | 1 |
| 0 | 0 | 0 |
| 1 | 1 | 1 |
| 1 | 1 | 1 |
| 1 | 1 | 1 |
| 0 | 0 | 0 |
| 1 | 1 | 1 |
| 0 | 0 | 0 |
| 0 | 0 | 0 |
| 0 | 0 | 0 |
| 0 | 1 | 0 |
| 1 | 1 | 1 |
| 0 | 0 | 0 |
| 0 | 0 | 0 |
| 0 | 0 | 0 |
| 0 | 0 | 0 |
| 0 | 0 | 0 |
| 0 | 0 | 0 |
| 0 | 0 | 0 |
| 0 | 0 | 0 |
| 1 | 1 | 1 |
| 1 | 1 | 1 |
| 0 | 0 | 0 |
| 0 | 0 | 0 |
| 0 | 0 | 0 |
| 1 | 1 | 1 |
| 1 | 1 | 1 |
| 0 | 0 | 0 |

| | | |
|---|---|---|
| 0 | 0 | 0 |
| 0 | 1 | 0 |
| 0 | 1 | 0 |
| 1 | 1 | 1 |
| 0 | 0 | 0 |
| 0 | 0 | 0 |
| 0 | 0 | 0 |
| 0 | 0 | 0 |
| 0 | 0 | 0 |
| 0 | 0 | 0 |
| 1 | 1 | 1 |
| 0 | 0 | 0 |
| 0 | 0 | 0 |
| 0 | 0 | 0 |
| 1 | 0 | 0 |
| 1 | 1 | 1 |
| 0 | 0 | 0 |
| 1 | 0 | 0 |
| 0 | 0 | 0 |
| 0 | 0 | 0 |
| 1 | 0 | 0 |
| 0 | 0 | 0 |
| 0 | 0 | 0 |
| 1 | 0 | 0 |
| 0 | 0 | 0 |
| 0 | 0 | 0 |
| 0 | 0 | 0 |
| 1 | 0 | 0 |
| 0 | 0 | 0 |
| 0 | 0 | 0 |
| 0 | 0 | 0 |
| 1 | 1 | 1 |
| 1 | 1 | 1 |
| 1 | 1 | 1 |
| 0 | 1 | 0 |
| 0 | 1 | 0 |
| 0 | 1 | 0 |
| 1 | 1 | 1 |
| 1 | 1 | 1 |
| 1 | 1 | 1 |
| 0 | 1 | 0 |
| 1 | 1 | 1 |
| 0 | 0 | 0 |
| 0 | 1 | 0 |
| 1 | 1 | 1 |
| 1 | 1 | 1 |
| 0 | 1 | 0 |
| 1 | 1 | 1 |
| 1 | 1 | 1 |
| 1 | 1 | 1 |
| 1 | 1 | 1 |
| 1 | 1 | 1 |
| 1 | 1 | 1 |
| 1 | 1 | 1 |

| | | |
|---|---|---|
| 0 | 1 | 0 |
| 0 | 1 | 0 |
| 1 | 1 | 1 |
| 1 | 1 | 1 |
| 1 | 1 | 1 |
| 1 | 1 | 1 |
| 1 | 1 | 1 |
| 1 | 1 | 1 |
| 0 | 0 | 0 |
| 0 | 0 | 0 |
| 0 | 0 | 0 |
| 0 | 0 | 0 |
| 0 | 0 | 0 |
| 0 | 0 | 0 |
| 0 | 0 | 0 |
| 0 | 0 | 0 |
| 0 | 0 | 0 |
| 0 | 0 | 0 |
| 0 | 0 | 0 |
| 0 | 0 | 0 |
| 0 | 0 | 0 |
| 0 | 0 | 0 |
| 0 | 0 | 0 |
| 0 | 0 | 0 |
| 0 | 0 | 0 |
| 1 | 1 | 1 |
| 1 | 1 | 1 |
| 0 | 0 | 0 |
| 0 | 0 | 0 |
| 0 | 0 | 0 |
| 0 | 0 | 0 |
| 0 | 0 | 0 |
| 0 | 0 | 0 |
| 0 | 0 | 0 |
| 0 | 0 | 0 |
| 0 | 0 | 0 |
| 0 | 0 | 0 |
| 0 | 0 | 0 |
| 0 | 0 | 0 |
| 0 | 0 | 0 |
| 0 | 0 | 0 |
| 0 | 0 | 0 |
| 0 | 0 | 0 |
| 0 | 0 | 0 |
| 0 | 0 | 0 |
| 0 | 0 | 0 |
| 1 | 1 | 1 |
| 0 | 1 | 0 |
| 1 | 1 | 1 |
| 1 | 1 | 1 |
| 1 | 1 | 1 |
| 1 | 1 | 1 |
| 0 | 0 | 0 |
| 0 | 0 | 0 |

| | | |
|---|---|---|
| 0 | 0 | 0 |
| 0 | 0 | 0 |
| 0 | 0 | 0 |
| 0 | 0 | 0 |
| 0 | 1 | 0 |
| 0 | 0 | 0 |
| 0 | 0 | 0 |
| 0 | 0 | 0 |
| 1 | 1 | 1 |
| 0 | 0 | 0 |
| 0 | 0 | 0 |
| 0 | 1 | 0 |
| 0 | 0 | 0 |
| 0 | 0 | 0 |
| 0 | 0 | 0 |
| 0 | 0 | 0 |
| 1 | 1 | 1 |
| 0 | 0 | 0 |
| 0 | 0 | 0 |
| 1 | 1 | 1 |
| 0 | 0 | 0 |
| 0 | 0 | 0 |
| 0 | 0 | 0 |
| 0 | 0 | 0 |
| 0 | 0 | 0 |
| 0 | 1 | 0 |
| 0 | 1 | 0 |
| 1 | 1 | 1 |
| 1 | 1 | 1 |
| 0 | 1 | 0 |
| 1 | 1 | 1 |
| 1 | 1 | 1 |
| 1 | 0 | 0 |
| 1 | 0 | 0 |
| 0 | 0 | 0 |
| 0 | 0 | 0 |
| 0 | 0 | 0 |
| 0 | 0 | 0 |
| 0 | 0 | 0 |
| 0 | 0 | 0 |
| 1 | 1 | 1 |
| 1 | 1 | 1 |
| 1 | 1 | 1 |
| 0 | 0 | 0 |
| 0 | 0 | 0 |
| 0 | 0 | 0 |
| 1 | 1 | 1 |
| 0 | 1 | 0 |
| 0 | 0 | 0 |
| 0 | 0 | 0 |
| 1 | 1 | 1 |
| 1 | 1 | 1 |
| 1 | 1 | 1 |
| 1 | 1 | 1 |

| | | |
|---|---|---|
| 0 | 1 | 0 |
| 1 | 1 | 1 |
| 0 | 1 | 0 |
| 0 | 0 | 0 |
| 0 | 0 | 0 |
| 1 | 1 | 1 |
| 1 | 1 | 1 |
| 1 | 1 | 1 |
| 1 | 1 | 1 |
| 1 | 1 | 1 |
| 0 | 0 | 0 |
| 0 | 1 | 0 |
| 0 | 0 | 0 |
| 0 | 0 | 0 |
| 0 | 0 | 0 |
| 1 | 1 | 1 |
| 1 | 1 | 1 |
| 0 | 1 | 0 |
| 1 | 1 | 1 |
| 1 | 1 | 1 |
| 0 | 0 | 0 |
| 1 | 1 | 1 |
| 1 | 1 | 1 |
| 0 | 0 | 0 |
| 1 | 0 | 0 |
| 0 | 0 | 0 |
| 0 | 0 | 0 |
| 1 | 1 | 1 |
| 1 | 1 | 1 |
| 1 | 1 | 1 |
| 1 | 1 | 1 |
| 1 | 1 | 1 |
| 1 | 1 | 1 |
| 1 | 1 | 1 |
| 1 | 1 | 1 |
| 0 | 0 | 0 |
| 1 | 1 | 1 |
| 1 | 1 | 1 |
| 0 | 0 | 0 |
| 0 | 0 | 0 |
| 1 | 1 | 1 |
| 0 | 1 | 0 |
| 0 | 0 | 0 |
| 0 | 0 | 0 |
| 0 | 0 | 0 |
| 0 | 0 | 0 |
| 0 | 0 | 0 |
| 0 | 0 | 0 |
| 1 | 1 | 1 |
| 0 | 0 | 0 |
| 0 | 0 | 0 |
| 0 | 0 | 0 |
| 0 | 0 | 0 |
| 0 | 0 | 0 |

| | | |
|---|---|---|
| 0 | 0 | 0 |
| 0 | 0 | 0 |
| 0 | 0 | 0 |
| 0 | 0 | 0 |
| 0 | 0 | 0 |
| 0 | 0 | 0 |
| 0 | 0 | 0 |
| 0 | 0 | 0 |
| 0 | 0 | 0 |
| 0 | 1 | 0 |
| 0 | 1 | 0 |
| 0 | 0 | 0 |
| 1 | 1 | 1 |
| 0 | 0 | 0 |
| 0 | 0 | 0 |
| 0 | 1 | 0 |
| 0 | 0 | 0 |
| 1 | 0 | 0 |
| 1 | 1 | 1 |
| 1 | 1 | 1 |
| 0 | 0 | 0 |
| 0 | 1 | 0 |
| 0 | 1 | 0 |
| 0 | 0 | 0 |
| 0 | 0 | 0 |
| 0 | 1 | 0 |
| 0 | 1 | 0 |
| 1 | 1 | 1 |
| 0 | 0 | 0 |
| 0 | 0 | 0 |
| 1 | 1 | 1 |
| 1 | 1 | 1 |
| 0 | 0 | 0 |
| 0 | 0 | 0 |
| 0 | 0 | 0 |
| 0 | 1 | 0 |
| 0 | 0 | 0 |
| 0 | 1 | 0 |
| 0 | 1 | 0 |
| 0 | 0 | 0 |
| 0 | 0 | 0 |
| 0 | 0 | 0 |
| 0 | 0 | 0 |
| 0 | 0 | 0 |
| 0 | 0 | 0 |
| 1 | 1 | 1 |
| 0 | 1 | 0 |
| 0 | 0 | 0 |
| 0 | 0 | 0 |
| 0 | 0 | 0 |
| 1 | 1 | 1 |
| 1 | 1 | 1 |
| 0 | 0 | 0 |
| 0 | 0 | 0 |

| | | |
|---|---|---|
| 1 | 1 | 1 |
| 1 | 1 | 1 |
| 1 | 1 | 1 |
| 1 | 1 | 1 |
| 0 | 0 | 0 |
| 0 | 0 | 0 |
| 1 | 0 | 0 |
| 0 | 0 | 0 |
| 1 | 0 | 0 |
| 0 | 0 | 0 |
| 1 | 1 | 1 |
| 1 | 1 | 1 |
| 1 | 1 | 1 |
| 1 | 1 | 1 |
| 1 | 1 | 1 |
| 1 | 1 | 1 |
| 0 | 0 | 0 |
| 1 | 1 | 1 |
| 1 | 1 | 1 |
| 0 | 0 | 0 |
| 0 | 0 | 0 |
| 0 | 1 | 0 |
| 0 | 1 | 0 |
| 0 | 0 | 0 |
| 0 | 1 | 0 |
| 0 | 0 | 0 |
| 0 | 0 | 0 |
| 0 | 0 | 0 |
| 0 | 0 | 0 |
| 0 | 0 | 0 |
| 0 | 0 | 0 |
| 0 | 0 | 0 |
| 0 | 1 | 0 |
| 0 | 0 | 0 |
| 1 | 1 | 1 |
| 1 | 1 | 1 |
| 1 | 1 | 1 |
| 1 | 1 | 1 |
| 0 | 0 | 0 |
| 1 | 1 | 1 |
| 1 | 1 | 1 |
| 1 | 1 | 1 |
| 1 | 1 | 1 |
| 0 | 0 | 0 |
| 1 | 1 | 1 |
| 1 | 1 | 1 |
| 1 | 1 | 1 |
| 1 | 1 | 1 |
| 1 | 1 | 1 |
| 1 | 1 | 1 |
| 0 | 0 | 0 |
| 0 | 0 | 0 |
| 0 | 0 | 0 |
| 0 | 1 | 0 |

| | | |
|---|---|---|
| 0 | 0 | 0 |
| 0 | 0 | 0 |
| 0 | 0 | 0 |
| 0 | 0 | 0 |
| 0 | 0 | 0 |
| 0 | 0 | 0 |
| 0 | 0 | 0 |
| 0 | 0 | 0 |
| 1 | 1 | 1 |
| 1 | 1 | 1 |
| 0 | 0 | 0 |
| 1 | 1 | 1 |
| 1 | 1 | 1 |
| 0 | 0 | 0 |
| 0 | 0 | 0 |
| 0 | 0 | 0 |
| 0 | 0 | 0 |
| 0 | 0 | 0 |
| 0 | 0 | 0 |
| 1 | 1 | 1 |
| 1 | 1 | 1 |
| 0 | 0 | 0 |
| 1 | 1 | 1 |
| 0 | 0 | 0 |
| 1 | 1 | 1 |
| 1 | 1 | 1 |
| 1 | 1 | 1 |
| 0 | 1 | 0 |
| 0 | 1 | 0 |
| 0 | 0 | 0 |
| 0 | 1 | 0 |
| 1 | 1 | 1 |
| 0 | 0 | 0 |
| 1 | 1 | 1 |
| 0 | 0 | 0 |
| 0 | 0 | 0 |
| 0 | 0 | 0 |
| 0 | 0 | 0 |
| 0 | 0 | 0 |
| 0 | 0 | 0 |
| 0 | 0 | 0 |
| 0 | 0 | 0 |
| 1 | 1 | 1 |
| 0 | 0 | 0 |
| 1 | 1 | 1 |
| 0 | 0 | 0 |
| 1 | 0 | 0 |
| 1 | 1 | 1 |
| 1 | 1 | 1 |
| 0 | 0 | 0 |
| 1 | 1 | 1 |
| 1 | 1 | 1 |
| 0 | 1 | 0 |
| 1 | 1 | 1 |

| | | |
|---|---|---|
| 0 | 0 | 0 |
| 0 | 1 | 0 |
| 1 | 1 | 1 |
| 0 | 1 | 0 |
| 0 | 1 | 0 |
| 0 | 1 | 0 |
| 0 | 1 | 0 |
| 1 | 1 | 1 |
| 1 | 1 | 1 |
| 1 | 1 | 1 |
| 0 | 0 | 0 |
| 1 | 1 | 1 |
| 1 | 1 | 1 |
| 0 | 1 | 0 |
| 0 | 0 | 0 |
| 0 | 0 | 0 |
| 0 | 0 | 0 |
| 0 | 0 | 0 |
| 0 | 0 | 0 |
| 0 | 1 | 0 |
| 0 | 1 | 0 |
| 1 | 1 | 1 |
| 0 | 0 | 0 |
| 0 | 1 | 0 |
| 0 | 1 | 0 |
| 0 | 1 | 0 |
| 1 | 1 | 1 |
| 1 | 1 | 1 |
| 0 | 0 | 0 |
| 0 | 0 | 0 |
| 0 | 0 | 0 |
| 0 | 0 | 0 |
| 0 | 0 | 0 |
| 0 | 0 | 0 |
| 1 | 1 | 1 |
| 1 | 1 | 1 |
| 0 | 0 | 0 |
| 0 | 0 | 0 |
| 0 | 0 | 0 |
| 0 | 0 | 0 |
| 0 | 0 | 0 |
| 0 | 0 | 0 |
| 1 | 1 | 1 |
| 1 | 1 | 1 |
| 0 | 0 | 0 |
| 1 | 1 | 1 |
| 1 | 1 | 1 |
| 1 | 1 | 1 |
| 1 | 1 | 1 |
| 1 | 1 | 1 |
| 1 | 0 | 0 |
| 0 | 1 | 0 |
| 0 | 1 | 0 |
| 1 | 0 | 0 |

| | | |
|---|---|---|
| 0 | 0 | 0 |
| 1 | 1 | 1 |
| 0 | 0 | 0 |
| 0 | 1 | 0 |
| 1 | 0 | 0 |
| 0 | 0 | 0 |
| 0 | 0 | 0 |
| 0 | 0 | 0 |
| 1 | 0 | 0 |
| 1 | 0 | 0 |
| 0 | 0 | 0 |
| 1 | 1 | 1 |
| 1 | 0 | 0 |
| 1 | 0 | 0 |
| 0 | 0 | 0 |
| 0 | 0 | 0 |
| 0 | 0 | 0 |
| 0 | 0 | 0 |
| 1 | 1 | 1 |
| 0 | 0 | 0 |
| 0 | 0 | 0 |
| 0 | 0 | 0 |
| 0 | 0 | 0 |
| 0 | 0 | 0 |
| 1 | 1 | 1 |
| 0 | 0 | 0 |
| 1 | 1 | 1 |
| 0 | 1 | 0 |
| 1 | 1 | 1 |
| 0 | 0 | 0 |
| 1 | 1 | 1 |
| 1 | 1 | 1 |
| 0 | 0 | 0 |
| 0 | 0 | 0 |
| 1 | 1 | 1 |
| 1 | 1 | 1 |
| 1 | 1 | 1 |
| 0 | 1 | 0 |
| 1 | 0 | 0 |
| 0 | 0 | 0 |
| 0 | 0 | 0 |
| 0 | 0 | 0 |
| 0 | 0 | 0 |
| 0 | 0 | 0 |
| 0 | 0 | 0 |
| 0 | 1 | 0 |
| 0 | 1 | 0 |
| 0 | 0 | 0 |
| 0 | 0 | 0 |
| 1 | 1 | 1 |
| 0 | 0 | 0 |
| 0 | 0 | 0 |
| 0 | 1 | 0 |
| 1 | 1 | 1 |

| | | |
|---|---|---|
| 1 | 1 | 1 |
| 0 | 0 | 0 |
| 0 | 0 | 0 |
| 1 | 0 | 0 |
| 1 | 1 | 1 |
| 0 | 0 | 0 |
| 0 | 0 | 0 |
| 0 | 0 | 0 |
| 0 | 0 | 0 |
| 0 | 0 | 0 |
| 0 | 0 | 0 |
| 0 | 0 | 0 |
| 0 | 0 | 0 |
| 0 | 0 | 0 |
| 0 | 0 | 0 |
| 0 | 1 | 0 |
| 0 | 0 | 0 |
| 1 | 1 | 1 |
| 1 | 0 | 0 |
| 1 | 1 | 1 |
| 0 | 0 | 0 |
| 1 | 1 | 1 |
| 0 | 0 | 0 |
| 0 | 0 | 0 |
| 1 | 1 | 1 |
| 0 | 0 | 0 |
| 0 | 1 | 0 |
| 0 | 1 | 0 |
| 0 | 1 | 0 |
| 0 | 0 | 0 |
| 0 | 0 | 0 |
| 0 | 0 | 0 |
| 0 | 0 | 0 |
| 0 | 0 | 0 |
| 0 | 0 | 0 |
| 0 | 0 | 0 |
| 0 | 0 | 0 |
| 1 | 1 | 1 |
| 1 | 0 | 0 |
| 0 | 0 | 0 |
| 0 | 0 | 0 |
| 0 | 0 | 0 |
| 0 | 0 | 0 |
| 0 | 0 | 0 |
| 0 | 0 | 0 |
| 0 | 0 | 0 |
| 1 | 1 | 1 |
| 0 | 0 | 0 |
| 0 | 0 | 0 |
| 0 | 0 | 0 |
| 0 | 0 | 0 |
| 0 | 1 | 0 |
| 0 | 0 | 0 |
| 0 | 0 | 0 |

| | | |
|---|---|---|
| 0 | 0 | 0 |
| 0 | 0 | 0 |
| 0 | 0 | 0 |
| 0 | 0 | 0 |
| 0 | 0 | 0 |
| 0 | 0 | 0 |
| 0 | 0 | 0 |
| 0 | 0 | 0 |
| 0 | 0 | 0 |
| 0 | 0 | 0 |
| 1 | 1 | 1 |
| 0 | 0 | 0 |
| 1 | 0 | 0 |
| 0 | 0 | 0 |
| 0 | 1 | 0 |
| 1 | 1 | 1 |
| 1 | 1 | 1 |
| 1 | 1 | 1 |
| 0 | 0 | 0 |
| 0 | 0 | 0 |
| 1 | 1 | 1 |
| 0 | 0 | 0 |
| 0 | 0 | 0 |
| 1 | 1 | 1 |
| 0 | 0 | 0 |
| 0 | 0 | 0 |
| 0 | 0 | 0 |
| 1 | 0 | 0 |
| 0 | 0 | 0 |
| 0 | 0 | 0 |
| 1 | 0 | 0 |
| 0 | 0 | 0 |
| 1 | 0 | 0 |
| 1 | 0 | 0 |
| 0 | 0 | 0 |
| 0 | 0 | 0 |
| 0 | 0 | 0 |
| 1 | 0 | 0 |
| 1 | 0 | 0 |
| 0 | 0 | 0 |
| 0 | 0 | 0 |
| 0 | 0 | 0 |
| 0 | 0 | 0 |
| 0 | 0 | 0 |
| 0 | 0 | 0 |
| 0 | 0 | 0 |
| 0 | 0 | 0 |
| 0 | 0 | 0 |
| 0 | 0 | 0 |
| 0 | 0 | 0 |
| 0 | 0 | 0 |
| 0 | 0 | 0 |
| 0 | 0 | 0 |
| 0 | 0 | 0 |

| | | |
|---|---|---|
| 0 | 0 | 0 |
| 1 | 0 | 0 |
| 0 | 0 | 0 |
| 0 | 0 | 0 |
| 0 | 0 | 0 |
| 1 | 0 | 0 |
| 0 | 0 | 0 |
| 0 | 0 | 0 |
| 1 | 1 | 1 |
| 0 | 0 | 0 |
| 0 | 1 | 0 |
| 1 | 1 | 1 |
| 1 | 1 | 1 |
| 0 | 0 | 0 |
| 1 | 1 | 1 |
| 0 | 0 | 0 |
| 1 | 1 | 1 |
| 1 | 1 | 1 |
| 0 | 0 | 0 |
| 0 | 0 | 0 |
| 0 | 0 | 0 |
| 0 | 0 | 0 |
| 0 | 0 | 0 |
| 0 | 0 | 0 |
| 0 | 0 | 0 |
| 0 | 1 | 0 |
| 1 | 1 | 1 |
| 1 | 1 | 1 |
| 1 | 0 | 0 |
| 1 | 1 | 1 |
| 1 | 1 | 1 |
| 0 | 1 | 0 |
| 0 | 0 | 0 |
| 0 | 0 | 0 |
| 0 | 0 | 0 |
| 0 | 0 | 0 |
| 0 | 0 | 0 |
| 0 | 0 | 0 |
| 0 | 1 | 0 |
| 0 | 1 | 0 |
| 0 | 1 | 0 |
| 0 | 1 | 0 |
| 0 | 1 | 0 |
| 0 | 1 | 0 |
| 0 | 0 | 0 |
| 0 | 0 | 0 |
| 0 | 1 | 0 |
| 0 | 0 | 0 |
| 0 | 0 | 0 |
| 0 | 0 | 0 |
| 0 | 0 | 0 |
| 0 | 0 | 0 |
| 0 | 0 | 0 |
| 0 | 0 | 0 |

| | | |
|---|---|---|
| 1 | 1 | 1 |
| 1 | 1 | 1 |
| 0 | 0 | 0 |
| 0 | 0 | 0 |
| 1 | 1 | 1 |
| 0 | 0 | 0 |
| 0 | 0 | 0 |
| 1 | 1 | 1 |
| 1 | 1 | 1 |
| 0 | 0 | 0 |
| 1 | 1 | 1 |
| 1 | 1 | 1 |
| 0 | 0 | 0 |
| 0 | 0 | 0 |
| 0 | 0 | 0 |
| 0 | 0 | 0 |
| 0 | 1 | 0 |
| 0 | 1 | 0 |
| 1 | 1 | 1 |
| 1 | 1 | 1 |
| 1 | 1 | 1 |
| 0 | 1 | 0 |
| 1 | 1 | 1 |
| 1 | 1 | 1 |
| 1 | 1 | 1 |
| 0 | 0 | 0 |
| 1 | 1 | 1 |
| 1 | 1 | 1 |
| 1 | 1 | 1 |
| 0 | 1 | 0 |
| 0 | 1 | 0 |
| 1 | 1 | 1 |
| 1 | 1 | 1 |
| 1 | 1 | 1 |
| 1 | 1 | 1 |
| 1 | 1 | 1 |
| 1 | 1 | 1 |
| 0 | 0 | 0 |
| 0 | 0 | 0 |
| 1 | 0 | 0 |
| 0 | 0 | 0 |
| 1 | 1 | 1 |
| 1 | 1 | 1 |
| 0 | 0 | 0 |
| 0 | 0 | 0 |
| 0 | 1 | 0 |
| 0 | 1 | 0 |
| 0 | 1 | 0 |
| 1 | 1 | 1 |
| 1 | 1 | 1 |
| 1 | 1 | 1 |
| 0 | 0 | 0 |
| 0 | 0 | 0 |
| 0 | 0 | 0 |

| | | |
|---|---|---|
| 0 | 0 | 0 |
| 1 | 1 | 1 |
| 1 | 1 | 1 |
| 1 | 1 | 1 |
| 1 | 1 | 1 |
| 1 | 1 | 1 |
| 1 | 1 | 1 |
| 1 | 1 | 1 |
| 0 | 0 | 0 |
| 1 | 1 | 1 |
| 1 | 1 | 1 |
| 1 | 1 | 1 |
| 0 | 1 | 0 |
| 0 | 1 | 0 |
| 1 | 1 | 1 |
| 1 | 1 | 1 |
| 0 | 1 | 0 |
| 0 | 1 | 0 |
| 0 | 0 | 0 |
| 0 | 0 | 0 |
| 0 | 0 | 0 |
| 0 | 0 | 0 |
| 0 | 1 | 0 |
| 0 | 0 | 0 |
| 0 | 0 | 0 |
| 1 | 1 | 1 |
| 1 | 1 | 1 |
| 1 | 1 | 1 |
| 1 | 1 | 1 |
| 1 | 1 | 1 |
| 1 | 1 | 1 |
| 1 | 1 | 1 |
| 0 | 0 | 0 |
| 1 | 1 | 1 |
| 1 | 1 | 1 |
| 0 | 1 | 0 |
| 0 | 0 | 0 |
| 1 | 1 | 1 |
| 0 | 0 | 0 |
| 0 | 0 | 0 |
| 0 | 0 | 0 |
| 0 | 0 | 0 |
| 0 | 0 | 0 |
| 1 | 1 | 1 |
| 1 | 1 | 1 |
| 1 | 1 | 1 |
| 0 | 0 | 0 |
| 0 | 0 | 0 |
| 0 | 0 | 0 |
| 0 | 0 | 0 |
| 0 | 0 | 0 |
| 0 | 0 | 0 |
| 0 | 0 | 0 |
| 0 | 1 | 0 |

| | | |
|---|---|---|
| 0 | 0 | 0 |
| 0 | 0 | 0 |
| 0 | 1 | 0 |
| 0 | 0 | 0 |
| 0 | 0 | 0 |
| 1 | 1 | 1 |
| 1 | 1 | 1 |
| 1 | 0 | 0 |
| 1 | 1 | 1 |
| 1 | 1 | 1 |
| 1 | 1 | 1 |
| 0 | 0 | 0 |
| 1 | 1 | 1 |
| 1 | 1 | 1 |
| 0 | 0 | 0 |
| 1 | 1 | 1 |
| 0 | 0 | 0 |
| 1 | 1 | 1 |
| 0 | 0 | 0 |
| 0 | 0 | 0 |
| 1 | 1 | 1 |
| 1 | 1 | 1 |
| 0 | 0 | 0 |
| 1 | 1 | 1 |
| 0 | 0 | 0 |
| 1 | 1 | 1 |
| 1 | 1 | 1 |
| 0 | 0 | 0 |
| 1 | 1 | 1 |
| 0 | 0 | 0 |
| 0 | 1 | 0 |
| 0 | 0 | 0 |
| 0 | 0 | 0 |
| 0 | 0 | 0 |
| 1 | 1 | 1 |
| 0 | 0 | 0 |
| 1 | 1 | 1 |
| 1 | 1 | 1 |
| 0 | 0 | 0 |
| 1 | 1 | 1 |
| 1 | 1 | 1 |
| 1 | 1 | 1 |
| 1 | 1 | 1 |
| 0 | 0 | 0 |
| 0 | 0 | 0 |
| 1 | 1 | 1 |
| 1 | 1 | 1 |
| 1 | 0 | 0 |
| 0 | 0 | 0 |
| 1 | 0 | 0 |
| 0 | 0 | 0 |
| 1 | 1 | 1 |
| 1 | 1 | 1 |
| 0 | 0 | 0 |

| | | |
|---|---|---|
| 1 | 1 | 1 |
| 1 | 1 | 1 |
| 1 | 1 | 1 |
| 1 | 1 | 1 |
| 0 | 1 | 0 |
| 1 | 1 | 1 |
| 1 | 1 | 1 |
| 0 | 0 | 0 |
| 1 | 1 | 1 |
| 1 | 1 | 1 |
| 1 | 1 | 1 |
| 1 | 1 | 1 |
| 0 | 0 | 0 |
| 1 | 1 | 1 |
| 0 | 0 | 0 |
| 0 | 0 | 0 |
| 1 | 1 | 1 |
| 1 | 1 | 1 |
| 1 | 1 | 1 |
| 1 | 1 | 1 |
| 0 | 0 | 0 |
| 0 | 0 | 0 |
| 1 | 1 | 1 |
| 1 | 1 | 1 |
| 0 | 0 | 0 |
| 1 | 1 | 1 |
| 0 | 0 | 0 |
| 0 | 0 | 0 |
| 0 | 0 | 0 |
| 0 | 0 | 0 |
| 0 | 1 | 0 |
| 0 | 1 | 0 |
| 0 | 0 | 0 |
| 0 | 0 | 0 |
| 0 | 0 | 0 |
| 0 | 0 | 0 |
| 0 | 0 | 0 |
| 0 | 0 | 0 |
| 0 | 0 | 0 |
| 1 | 0 | 0 |
| 1 | 1 | 1 |
| 0 | 1 | 0 |
| 0 | 1 | 0 |
| 0 | 1 | 0 |
| 0 | 1 | 0 |
| 0 | 0 | 0 |
| 0 | 1 | 0 |
| 0 | 1 | 0 |
| 1 | 1 | 1 |
| 0 | 0 | 0 |
| 0 | 0 | 0 |
| 1 | 1 | 1 |
| 1 | 1 | 1 |
| 1 | 1 | 1 |

| | | |
|---|---|---|
| 1 | 1 | 1 |
| 1 | 1 | 1 |
| 1 | 1 | 1 |
| 1 | 1 | 1 |
| 0 | 0 | 0 |
| 1 | 1 | 1 |
| 0 | 0 | 0 |
| 0 | 0 | 0 |
| 0 | 0 | 0 |
| 0 | 0 | 0 |
| 0 | 0 | 0 |
| 1 | 1 | 1 |
| 0 | 0 | 0 |
| 0 | 0 | 0 |
| 1 | 1 | 1 |
| 1 | 1 | 1 |
| 1 | 1 | 1 |
| 1 | 1 | 1 |
| 1 | 1 | 1 |
| 1 | 1 | 1 |
| 1 | 1 | 1 |
| 1 | 1 | 1 |
| 1 | 1 | 1 |
| 0 | 0 | 0 |
| 1 | 1 | 1 |
| 1 | 1 | 1 |
| 1 | 1 | 1 |
| 0 | 0 | 0 |
| 1 | 0 | 0 |
| 1 | 0 | 0 |
| 0 | 0 | 0 |
| 0 | 0 | 0 |
| 1 | 1 | 1 |
| 0 | 0 | 0 |
| 0 | 1 | 0 |
| 1 | 1 | 1 |
| 1 | 1 | 1 |
| 1 | 1 | 1 |
| 1 | 1 | 1 |
| 1 | 1 | 1 |
| 1 | 1 | 1 |
| 1 | 1 | 1 |
| 1 | 1 | 1 |
| 1 | 1 | 1 |
| 1 | 1 | 1 |
| 1 | 1 | 1 |
| 1 | 1 | 1 |
| 1 | 1 | 1 |
| 0 | 0 | 0 |
| 0 | 1 | 0 |
| 1 | 1 | 1 |
| 1 | 0 | 0 |
| 1 | 0 | 0 |
| 1 | 1 | 1 |

| | | |
|---|---|---|
| 1 | 1 | 1 |
| 0 | 1 | 0 |
| 1 | 1 | 1 |
| 0 | 1 | 0 |
| 1 | 0 | 0 |
| 1 | 1 | 1 |
| 1 | 1 | 1 |
| 1 | 1 | 1 |
| 1 | 1 | 1 |
| 1 | 1 | 1 |
| 0 | 0 | 0 |
| 0 | 0 | 0 |
| 0 | 0 | 0 |
| 0 | 0 | 0 |
| 0 | 0 | 0 |
| 1 | 0 | 0 |
| 0 | 0 | 0 |
| 0 | 0 | 0 |
| 0 | 0 | 0 |
| 0 | 0 | 0 |
| 0 | 0 | 0 |
| 0 | 0 | 0 |
| 0 | 1 | 0 |
| 0 | 0 | 0 |
| 0 | 0 | 0 |
| 0 | 0 | 0 |
| 1 | 1 | 1 |
| 0 | 0 | 0 |
| 0 | 1 | 0 |
| 0 | 1 | 0 |
| 0 | 1 | 0 |
| 0 | 0 | 0 |
| 1 | 1 | 1 |
| 1 | 1 | 1 |
| 0 | 0 | 0 |
| 0 | 0 | 0 |
| 0 | 0 | 0 |
| 0 | 0 | 0 |
| 0 | 0 | 0 |
| 1 | 0 | 0 |
| 1 | 0 | 0 |
| 0 | 0 | 0 |
| 1 | 0 | 0 |
| 0 | 0 | 0 |
| 1 | 1 | 1 |
| 1 | 0 | 0 |
| 1 | 1 | 1 |
| 0 | 0 | 0 |
| 0 | 0 | 0 |
| 0 | 0 | 0 |
| 0 | 0 | 0 |
| 1 | 1 | 1 |
| 0 | 0 | 0 |
| 1 | 1 | 1 |

| | | |
|---|---|---|
| 1 | 1 | 1 |
| 1 | 1 | 1 |
| 1 | 1 | 1 |
| 1 | 1 | 1 |
| 1 | 1 | 1 |
| 0 | 0 | 0 |
| 0 | 0 | 0 |
| 0 | 0 | 0 |
| 0 | 0 | 0 |
| 0 | 1 | 0 |
| 0 | 0 | 0 |
| 0 | 0 | 0 |
| 0 | 0 | 0 |
| 0 | 0 | 0 |
| 0 | 0 | 0 |
| 0 | 0 | 0 |
| 1 | 1 | 1 |
| 0 | 1 | 0 |
| 1 | 1 | 1 |
| 1 | 1 | 1 |
| 1 | 1 | 1 |
| 1 | 1 | 1 |
| 1 | 1 | 1 |
| 1 | 1 | 1 |
| 1 | 1 | 1 |
| 1 | 1 | 1 |
| 1 | 1 | 1 |
| 1 | 1 | 1 |
| 0 | 0 | 0 |
| 0 | 1 | 0 |
| 0 | 1 | 0 |
| 1 | 0 | 0 |
| 1 | 0 | 0 |
| 0 | 0 | 0 |
| 1 | 1 | 1 |
| 0 | 0 | 0 |
| 1 | 0 | 0 |
| 0 | 0 | 0 |
| 1 | 1 | 1 |
| 0 | 0 | 0 |
| 1 | 1 | 1 |
| 1 | 1 | 1 |
| 0 | 1 | 0 |
| 0 | 0 | 0 |
| 1 | 1 | 1 |
| 1 | 1 | 1 |
| 1 | 1 | 1 |
| 1 | 1 | 1 |
| 1 | 1 | 1 |
| 1 | 1 | 1 |
| 1 | 1 | 1 |
| 1 | 0 | 0 |
| 0 | 0 | 0 |
| 0 | 0 | 0 |

| | | |
|---|---|---|
| 1 | 1 | 1 |
| 0 | 0 | 0 |
| 1 | 1 | 1 |
| 1 | 1 | 1 |
| 0 | 0 | 0 |
| 0 | 0 | 0 |
| 0 | 0 | 0 |
| 1 | 1 | 1 |
| 1 | 1 | 1 |
| 0 | 0 | 0 |
| 0 | 0 | 0 |
| 0 | 0 | 0 |
| 0 | 0 | 0 |
| 0 | 0 | 0 |
| 1 | 1 | 1 |
| 0 | 0 | 0 |
| 0 | 0 | 0 |
| 0 | 0 | 0 |
| 1 | 1 | 1 |
| 1 | 1 | 1 |
| 1 | 1 | 1 |
| 1 | 1 | 1 |
| 1 | 1 | 1 |
| 0 | 0 | 0 |
| 0 | 1 | 0 |
| 0 | 1 | 0 |
| 0 | 0 | 0 |
| 0 | 0 | 0 |
| 1 | 1 | 1 |
| 1 | 1 | 1 |
| 0 | 0 | 0 |
| 1 | 1 | 1 |
| 1 | 1 | 1 |
| 0 | 1 | 0 |
| 1 | 1 | 1 |
| 0 | 0 | 0 |
| 0 | 1 | 0 |
| 0 | 0 | 0 |
| 0 | 0 | 0 |
| 0 | 1 | 0 |
| 0 | 1 | 0 |
| 0 | 0 | 0 |
| 0 | 0 | 0 |
| 0 | 1 | 0 |
| 0 | 0 | 0 |
| 0 | 0 | 0 |
| 0 | 0 | 0 |
| 0 | 1 | 0 |
| 0 | 0 | 0 |
| 0 | 0 | 0 |
| 0 | 0 | 0 |
| 1 | 1 | 1 |
| 0 | 0 | 0 |
| 1 | 1 | 1 |

| | | |
|---|---|---|
| 1 | 1 | 1 |
| 1 | 1 | 1 |
| 1 | 1 | 1 |
| 1 | 1 | 1 |
| 0 | 0 | 0 |
| 0 | 0 | 0 |
| 1 | 1 | 1 |
| 1 | 0 | 0 |
| 1 | 1 | 1 |
| 0 | 0 | 0 |
| 1 | 0 | 0 |
| 1 | 0 | 0 |
| 0 | 0 | 0 |
| 1 | 1 | 1 |
| 1 | 1 | 1 |
| 1 | 1 | 1 |
| 1 | 1 | 1 |
| 1 | 1 | 1 |
| 1 | 1 | 1 |
| 1 | 1 | 1 |
| 1 | 1 | 1 |
| 0 | 1 | 0 |
| 1 | 1 | 1 |
| 0 | 0 | 0 |
| 1 | 1 | 1 |
| 0 | 0 | 0 |
| 1 | 1 | 1 |
| 0 | 0 | 0 |
| 1 | 1 | 1 |
| 0 | 0 | 0 |
| 0 | 0 | 0 |
| 0 | 0 | 0 |
| 0 | 0 | 0 |
| 0 | 0 | 0 |
| 0 | 1 | 0 |
| 0 | 0 | 0 |
| 1 | 1 | 1 |
| 0 | 0 | 0 |
| 1 | 1 | 1 |
| 1 | 1 | 1 |
| 0 | 0 | 0 |
| 0 | 0 | 0 |
| 0 | 0 | 0 |
| 0 | 0 | 0 |
| 0 | 0 | 0 |
| 0 | 0 | 0 |
| 0 | 0 | 0 |
| 0 | 0 | 0 |
| 1 | 1 | 1 |
| 0 | 0 | 0 |
| 1 | 0 | 0 |
| 1 | 0 | 0 |
| 1 | 1 | 1 |
| 1 | 1 | 1 |

| | | |
|---|---|---|
| 1 | 1 | 1 |
| 1 | 1 | 1 |
| 1 | 1 | 1 |
| 0 | 1 | 0 |
| 0 | 1 | 0 |
| 0 | 1 | 0 |
| 0 | 0 | 0 |
| 0 | 0 | 0 |
| 0 | 0 | 0 |
| 0 | 0 | 0 |
| 0 | 0 | 0 |
| 0 | 0 | 0 |
| 0 | 0 | 0 |
| 0 | 0 | 0 |
| 0 | 0 | 0 |
| 0 | 0 | 0 |
| 0 | 0 | 0 |
| 0 | 0 | 0 |
| 1 | 1 | 1 |
| 0 | 0 | 0 |
| 0 | 1 | 0 |
| 0 | 1 | 0 |
| 0 | 1 | 0 |
| 1 | 1 | 1 |
| 0 | 1 | 0 |
| 0 | 0 | 0 |
| 0 | 0 | 0 |
| 0 | 0 | 0 |
| 0 | 0 | 0 |
| 0 | 0 | 0 |
| 0 | 0 | 0 |
| 0 | 0 | 0 |
| 0 | 0 | 0 |
| 0 | 0 | 0 |
| 1 | 1 | 1 |
| 0 | 0 | 0 |
| 0 | 0 | 0 |
| 0 | 0 | 0 |
| 0 | 0 | 0 |
| 0 | 1 | 0 |
| 0 | 0 | 0 |
| 0 | 0 | 0 |
| 1 | 1 | 1 |
| 0 | 1 | 0 |
| 1 | 1 | 1 |
| 0 | 1 | 0 |
| 0 | 1 | 0 |
| 0 | 0 | 0 |
| 0 | 0 | 0 |
| 0 | 0 | 0 |
| 0 | 0 | 0 |
| 0 | 1 | 0 |
| 0 | 0 | 0 |
| 1 | 1 | 1 |

| | | |
|---|---|---|
| 0 | 1 | 0 |
| 1 | 1 | 1 |
| 1 | 1 | 1 |
| 1 | 1 | 1 |
| 1 | 1 | 1 |
| 0 | 0 | 0 |
| 0 | 1 | 0 |
| 1 | 1 | 1 |
| 1 | 1 | 1 |
| 1 | 1 | 1 |
| 0 | 0 | 0 |
| 0 | 0 | 0 |
| 0 | 0 | 0 |
| 0 | 0 | 0 |
| 0 | 0 | 0 |
| 0 | 1 | 0 |
| 1 | 0 | 0 |
| 0 | 0 | 0 |
| 1 | 1 | 1 |
| 1 | 1 | 1 |
| 1 | 1 | 1 |
| 1 | 1 | 1 |
| 0 | 0 | 0 |
| 1 | 1 | 1 |
| 0 | 0 | 0 |
| 0 | 0 | 0 |
| 0 | 0 | 0 |
| 0 | 0 | 0 |
| 1 | 1 | 1 |
| 0 | 0 | 0 |
| 1 | 1 | 1 |
| 0 | 1 | 0 |
| 0 | 1 | 0 |
| 0 | 1 | 0 |
| 1 | 1 | 1 |
| 1 | 1 | 1 |
| 0 | 1 | 0 |
| 0 | 0 | 0 |
| 0 | 0 | 0 |
| 0 | 0 | 0 |
| 0 | 0 | 0 |
| 0 | 0 | 0 |
| 1 | 1 | 1 |
| 0 | 0 | 0 |
| 0 | 0 | 0 |
| 1 | 1 | 1 |
| 0 | 0 | 0 |
| 0 | 0 | 0 |
| 0 | 0 | 0 |
| 0 | 0 | 0 |
| 1 | 1 | 1 |
| 1 | 1 | 1 |
| 1 | 1 | 1 |
| 0 | 0 | 0 |

| | | |
|---|---|---|
| 0 | 0 | 0 |
| 1 | 1 | 1 |
| 0 | 0 | 0 |
| 0 | 1 | 0 |
| 1 | 0 | 0 |
| 1 | 0 | 0 |
| 1 | 0 | 0 |
| 1 | 1 | 1 |
| 0 | 0 | 0 |
| 1 | 1 | 1 |
| 0 | 0 | 0 |
| 0 | 0 | 0 |
| 0 | 0 | 0 |
| 0 | 0 | 0 |
| 1 | 1 | 1 |
| 1 | 1 | 1 |
| 0 | 0 | 0 |
| 0 | 0 | 0 |
| 0 | 1 | 0 |
| 1 | 1 | 1 |
| 0 | 0 | 0 |
| 1 | 0 | 0 |
| 1 | 0 | 0 |
| 1 | 1 | 1 |
| 0 | 0 | 0 |
| 1 | 0 | 0 |
| 0 | 0 | 0 |
| 0 | 0 | 0 |
| 0 | 0 | 0 |
| 1 | 1 | 1 |
| 0 | 0 | 0 |
| 0 | 0 | 0 |
| 0 | 0 | 0 |
| 1 | 1 | 1 |
| 0 | 0 | 0 |
| 1 | 1 | 1 |
| 0 | 0 | 0 |
| 0 | 0 | 0 |
| 0 | 0 | 0 |
| 0 | 0 | 0 |
| 0 | 0 | 0 |
| 1 | 1 | 1 |
| 0 | 0 | 0 |
| 0 | 0 | 0 |
| 0 | 0 | 0 |
| 0 | 0 | 0 |
| 0 | 0 | 0 |
| 0 | 0 | 0 |
| 0 | 0 | 0 |
| 0 | 0 | 0 |
| 0 | 0 | 0 |
| 0 | 0 | 0 |
| 1 | 1 | 1 |
| 0 | 0 | 0 |

| | | |
|---|---|---|
| 0 | 1 | 0 |
| 1 | 1 | 1 |
| 1 | 1 | 1 |
| 1 | 1 | 1 |
| 1 | 1 | 1 |
| 1 | 1 | 1 |
| 1 | 1 | 1 |
| 1 | 1 | 1 |
| 1 | 1 | 1 |
| 0 | 0 | 0 |
| 0 | 0 | 0 |
| 1 | 1 | 1 |
| 1 | 1 | 1 |
| 1 | 1 | 1 |
| 0 | 0 | 0 |
| 1 | 1 | 1 |
| 1 | 1 | 1 |
| 0 | 0 | 0 |
| 0 | 1 | 0 |
| 0 | 1 | 0 |
| 0 | 1 | 0 |
| 0 | 0 | 0 |
| 0 | 0 | 0 |
| 0 | 0 | 0 |
| 0 | 0 | 0 |
| 0 | 0 | 0 |
| 0 | 0 | 0 |
| 0 | 0 | 0 |
| 1 | 1 | 1 |
| 0 | 0 | 0 |
| 0 | 0 | 0 |
| 1 | 1 | 1 |
| 1 | 1 | 1 |
| 1 | 1 | 1 |
| 1 | 1 | 1 |
| 1 | 1 | 1 |
| 1 | 1 | 1 |
| 1 | 1 | 1 |
| 1 | 1 | 1 |
| 1 | 1 | 1 |
| 1 | 1 | 1 |
| 1 | 1 | 1 |
| 1 | 1 | 1 |
| 0 | 0 | 0 |
| 1 | 1 | 1 |
| 0 | 0 | 0 |
| 1 | 1 | 1 |
| 1 | 1 | 1 |
| 1 | 1 | 1 |
| 1 | 1 | 1 |
| 0 | 0 | 0 |
| 1 | 1 | 1 |
| 1 | 1 | 1 |
| 0 | 0 | 0 |

| | | |
|---|---|---|
| 0 | 0 | 0 |
| 0 | 0 | 0 |
| 0 | 0 | 0 |
| 0 | 0 | 0 |
| 1 | 1 | 1 |
| 0 | 1 | 0 |
| 1 | 1 | 1 |
| 0 | 0 | 0 |
| 0 | 0 | 0 |
| 0 | 0 | 0 |
| 0 | 0 | 0 |
| 1 | 1 | 1 |
| 1 | 1 | 1 |
| 1 | 1 | 1 |
| 1 | 1 | 1 |
| 0 | 0 | 0 |
| 0 | 0 | 0 |
| 0 | 1 | 0 |
| 0 | 1 | 0 |
| 1 | 1 | 1 |
| 1 | 1 | 1 |
| 0 | 0 | 0 |
| 1 | 1 | 1 |
| 0 | 1 | 0 |
| 0 | 0 | 0 |
| 0 | 0 | 0 |
| 0 | 0 | 0 |
| 0 | 0 | 0 |
| 1 | 1 | 1 |
| 1 | 1 | 1 |
| 1 | 1 | 1 |
| 0 | 0 | 0 |
| 1 | 1 | 1 |
| 1 | 1 | 1 |
| 1 | 1 | 1 |
| 1 | 1 | 1 |
| 1 | 1 | 1 |
| 1 | 1 | 1 |
| 1 | 1 | 1 |
| 1 | 1 | 1 |
| 1 | 1 | 1 |
| 1 | 1 | 1 |
| 0 | 0 | 0 |
| 0 | 1 | 0 |
| 0 | 1 | 0 |
| 1 | 1 | 1 |
| 0 | 1 | 0 |
| 1 | 1 | 1 |
| 1 | 1 | 1 |
| 0 | 0 | 0 |
| 1 | 1 | 1 |
| 1 | 1 | 1 |
| 1 | 1 | 1 |
| 1 | 1 | 1 |

| | | |
|---|---|---|
| 0 | 0 | 0 |
| 1 | 1 | 1 |
| 0 | 0 | 0 |
| 0 | 1 | 0 |
| 1 | 1 | 1 |
| 1 | 1 | 1 |
| 1 | 1 | 1 |
| 0 | 0 | 0 |
| 1 | 1 | 1 |
| 1 | 1 | 1 |
| 0 | 0 | 0 |
| 0 | 0 | 0 |
| 0 | 0 | 0 |
| 1 | 1 | 1 |
| 0 | 0 | 0 |
| 0 | 0 | 0 |
| 0 | 0 | 0 |
| 0 | 0 | 0 |
| 0 | 1 | 0 |
| 0 | 0 | 0 |
| 0 | 0 | 0 |
| 0 | 0 | 0 |
| 0 | 0 | 0 |
| 0 | 0 | 0 |
| 0 | 0 | 0 |
| 0 | 0 | 0 |
| 0 | 0 | 0 |
| 0 | 0 | 0 |
| 0 | 0 | 0 |
| 0 | 0 | 0 |
| 0 | 0 | 0 |
| 0 | 0 | 0 |
| 1 | 1 | 1 |
| 0 | 0 | 0 |
| 1 | 1 | 1 |
| 0 | 0 | 0 |
| 0 | 0 | 0 |
| 1 | 1 | 1 |
| 1 | 1 | 1 |
| 0 | 1 | 0 |
| 0 | 1 | 0 |
| 0 | 0 | 0 |
| 0 | 0 | 0 |
| 0 | 0 | 0 |
| 1 | 1 | 1 |
| 0 | 0 | 0 |
| 1 | 1 | 1 |
| 0 | 0 | 0 |
| 0 | 1 | 0 |
| 0 | 1 | 0 |
| 0 | 1 | 0 |
| 0 | 1 | 0 |
| 0 | 1 | 0 |
| 1 | 1 | 1 |

| | | |
|---|---|---|
| 1 | 1 | 1 |
| 1 | 1 | 1 |
| 0 | 1 | 0 |
| 1 | 1 | 1 |
| 1 | 1 | 1 |
| 0 | 0 | 0 |
| 0 | 0 | 0 |
| 1 | 1 | 1 |
| 0 | 0 | 0 |
| 0 | 0 | 0 |
| 1 | 1 | 1 |
| 0 | 1 | 0 |
| 1 | 1 | 1 |
| 1 | 1 | 1 |
| 1 | 1 | 1 |
| 0 | 0 | 0 |
| 1 | 1 | 1 |
| 1 | 1 | 1 |
| 0 | 0 | 0 |
| 0 | 0 | 0 |
| 0 | 0 | 0 |
| 1 | 1 | 1 |
| 1 | 1 | 1 |
| 0 | 0 | 0 |
| 1 | 1 | 1 |
| 0 | 0 | 0 |
| 0 | 0 | 0 |
| 1 | 1 | 1 |
| 1 | 1 | 1 |
| 1 | 1 | 1 |
| 0 | 0 | 0 |
| 1 | 0 | 0 |
| 0 | 0 | 0 |
| 0 | 0 | 0 |
| 0 | 0 | 0 |
| 0 | 0 | 0 |
| 0 | 0 | 0 |
| 1 | 1 | 1 |
| 0 | 0 | 0 |
| 0 | 1 | 0 |
| 0 | 0 | 0 |
| 0 | 0 | 0 |
| 0 | 0 | 0 |
| 0 | 0 | 0 |
| 1 | 1 | 1 |
| 0 | 1 | 0 |
| 0 | 1 | 0 |
| 0 | 0 | 0 |
| 0 | 0 | 0 |
| 0 | 0 | 0 |
| 1 | 1 | 1 |
| 0 | 1 | 0 |
| 1 | 1 | 1 |
| 1 | 1 | 1 |

| | | |
|---|---|---|
| 1 | 1 | 1 |
| 0 | 1 | 0 |
| 0 | 1 | 0 |
| 0 | 0 | 0 |
| 0 | 0 | 0 |
| 1 | 1 | 1 |
| 0 | 0 | 0 |
| 1 | 1 | 1 |
| 1 | 1 | 1 |
| 1 | 1 | 1 |
| 1 | 1 | 1 |
| 0 | 0 | 0 |
| 1 | 0 | 0 |
| 0 | 0 | 0 |
| 1 | 1 | 1 |
| 0 | 0 | 0 |
| 0 | 0 | 0 |
| 0 | 0 | 0 |
| 1 | 1 | 1 |
| 0 | 0 | 0 |
| 0 | 0 | 0 |
| 0 | 0 | 0 |
| 0 | 0 | 0 |
| 0 | 0 | 0 |
| 0 | 0 | 0 |
| 1 | 1 | 1 |
| 0 | 0 | 0 |
| 0 | 1 | 0 |
| 0 | 1 | 0 |
| 0 | 1 | 0 |
| 0 | 1 | 0 |
| 0 | 1 | 0 |
| 0 | 0 | 0 |
| 0 | 0 | 0 |
| 1 | 1 | 1 |
| 1 | 1 | 1 |
| 0 | 0 | 0 |
| 0 | 0 | 0 |
| 0 | 0 | 0 |
| 1 | 0 | 0 |
| 0 | 0 | 0 |
| 0 | 0 | 0 |
| 1 | 1 | 1 |
| 0 | 0 | 0 |
| 1 | 1 | 1 |
| 1 | 1 | 1 |
| 1 | 1 | 1 |
| 1 | 1 | 1 |
| 1 | 1 | 1 |
| 0 | 0 | 0 |
| 1 | 1 | 1 |
| 1 | 1 | 1 |
| 1 | 1 | 1 |
| 0 | 1 | 0 |

| | | |
|---|---|---|
| 1 | 1 | 1 |
| 1 | 1 | 1 |
| 1 | 1 | 1 |
| 0 | 0 | 0 |
| 1 | 1 | 1 |
| 1 | 1 | 1 |
| 0 | 1 | 0 |
| 0 | 0 | 0 |
| 1 | 1 | 1 |
| 1 | 1 | 1 |
| 1 | 1 | 1 |
| 1 | 1 | 1 |
| 1 | 1 | 1 |
| 0 | 1 | 0 |
| 0 | 1 | 0 |
| 0 | 1 | 0 |
| 0 | 0 | 0 |
| 0 | 0 | 0 |
| 0 | 1 | 0 |
| 1 | 1 | 1 |
| 0 | 1 | 0 |
| 1 | 1 | 1 |
| 0 | 1 | 0 |
| 1 | 1 | 1 |
| 1 | 1 | 1 |
| 1 | 1 | 1 |
| 1 | 1 | 1 |
| 0 | 0 | 0 |
| 0 | 0 | 0 |
| 1 | 1 | 1 |
| 1 | 1 | 1 |
| 0 | 0 | 0 |
| 0 | 1 | 0 |
| 0 | 0 | 0 |
| 0 | 0 | 0 |
| 0 | 0 | 0 |
| 0 | 0 | 0 |
| 0 | 0 | 0 |
| 1 | 1 | 1 |
| 1 | 1 | 1 |
| 1 | 1 | 1 |
| 0 | 1 | 0 |
| 1 | 1 | 1 |
| 1 | 1 | 1 |
| 0 | 0 | 0 |
| 0 | 0 | 0 |
| 0 | 0 | 0 |
| 1 | 1 | 1 |
| 1 | 1 | 1 |
| 1 | 1 | 1 |
| 1 | 1 | 1 |
| 0 | 0 | 0 |
| 0 | 0 | 0 |
| 0 | 0 | 0 |

| | | |
|---|---|---|
| 0 | 0 | 0 |
| 0 | 0 | 0 |
| 1 | 1 | 1 |
| 0 | 0 | 0 |
| 0 | 0 | 0 |
| 0 | 0 | 0 |
| 0 | 0 | 0 |
| 1 | 1 | 1 |
| 1 | 1 | 1 |
| 1 | 1 | 1 |
| 1 | 1 | 1 |
| 1 | 1 | 1 |
| 0 | 1 | 0 |
| 0 | 1 | 0 |
| 1 | 1 | 1 |
| 1 | 1 | 1 |
| 1 | 1 | 1 |
| 1 | 1 | 1 |
| 1 | 1 | 1 |
| 1 | 1 | 1 |
| 0 | 0 | 0 |
| 0 | 0 | 0 |
| 0 | 0 | 0 |
| 0 | 0 | 0 |
| 0 | 0 | 0 |
| 1 | 1 | 1 |
| 1 | 1 | 1 |
| 0 | 0 | 0 |
| 0 | 1 | 0 |
| 0 | 1 | 0 |
| 0 | 0 | 0 |
| 0 | 0 | 0 |
| 0 | 0 | 0 |
| 1 | 1 | 1 |
| 0 | 0 | 0 |
| 1 | 1 | 1 |
| 0 | 1 | 0 |
| 0 | 0 | 0 |
| 1 | 1 | 1 |
| 1 | 1 | 1 |
| 0 | 1 | 0 |
| 0 | 1 | 0 |
| 1 | 1 | 1 |
| 1 | 1 | 1 |
| 1 | 1 | 1 |
| 1 | 1 | 1 |
| 1 | 1 | 1 |
| 1 | 1 | 1 |
| 0 | 0 | 0 |
| 1 | 1 | 1 |
| 0 | 0 | 0 |
| 0 | 0 | 0 |
| 1 | 1 | 1 |
| 0 | 1 | 0 |

| | | |
|---|---|---|
| 0 | 0 | 0 |
| 1 | 1 | 1 |
| 1 | 1 | 1 |
| 1 | 1 | 1 |
| 1 | 1 | 1 |
| 0 | 0 | 0 |
| 0 | 0 | 0 |
| 1 | 1 | 1 |
| 0 | 1 | 0 |
| 1 | 1 | 1 |
| 1 | 1 | 1 |
| 0 | 0 | 0 |
| 0 | 1 | 0 |
| 1 | 1 | 1 |
| 0 | 1 | 0 |
| 1 | 1 | 1 |
| 0 | 0 | 0 |
| 0 | 0 | 0 |
| 1 | 1 | 1 |
| 1 | 1 | 1 |
| 1 | 1 | 1 |
| 1 | 1 | 1 |
| 1 | 1 | 1 |
| 1 | 1 | 1 |
| 0 | 0 | 0 |
| 1 | 1 | 1 |
| 1 | 1 | 1 |
| 1 | 1 | 1 |
| 1 | 1 | 1 |
| 1 | 1 | 1 |
| 0 | 1 | 0 |
| 0 | 0 | 0 |
| 0 | 0 | 0 |
| 0 | 1 | 0 |
| 1 | 1 | 1 |
| 0 | 0 | 0 |
| 0 | 0 | 0 |
| 0 | 0 | 0 |
| 0 | 0 | 0 |
| 0 | 0 | 0 |
| 0 | 0 | 0 |
| 0 | 0 | 0 |
| 0 | 0 | 0 |
| 0 | 0 | 0 |
| 0 | 0 | 0 |
| 0 | 0 | 0 |
| 0 | 0 | 0 |
| 0 | 0 | 0 |
| 1 | 0 | 0 |
| 1 | 1 | 1 |
| 1 | 0 | 0 |
| 1 | 1 | 1 |
| 0 | 0 | 0 |
| 1 | 1 | 1 |

| | | |
|---|---|---|
| 1 | 1 | 1 |
| 1 | 1 | 1 |
| 1 | 1 | 1 |
| 1 | 1 | 1 |
| 1 | 1 | 1 |
| 1 | 1 | 1 |
| 1 | 1 | 1 |
| 0 | 0 | 0 |
| 0 | 0 | 0 |
| 1 | 0 | 0 |
| 1 | 1 | 1 |
| 1 | 1 | 1 |
| 1 | 1 | 1 |
| 0 | 0 | 0 |
| 1 | 1 | 1 |
| 1 | 1 | 1 |
| 0 | 0 | 0 |
| 1 | 1 | 1 |
| 0 | 0 | 0 |
| 0 | 1 | 0 |
| 1 | 1 | 1 |
| 1 | 1 | 1 |
| 1 | 1 | 1 |
| 0 | 0 | 0 |
| 1 | 1 | 1 |
| 1 | 1 | 1 |
| 0 | 1 | 0 |
| 0 | 1 | 0 |
| 0 | 1 | 0 |
| 0 | 1 | 0 |
| 1 | 1 | 1 |
| 0 | 0 | 0 |
| 1 | 1 | 1 |
| 0 | 1 | 0 |
| 1 | 1 | 1 |
| 1 | 1 | 1 |
| 1 | 1 | 1 |
| 1 | 1 | 1 |
| 0 | 0 | 0 |
| 1 | 1 | 1 |
| 1 | 1 | 1 |
| 1 | 1 | 1 |
| 0 | 1 | 0 |
| 1 | 1 | 1 |
| 1 | 1 | 1 |
| 0 | 0 | 0 |
| 0 | 1 | 0 |
| 0 | 1 | 0 |
| 1 | 0 | 0 |
| 1 | 0 | 0 |
| 0 | 1 | 0 |
| 1 | 0 | 0 |
| 0 | 0 | 0 |
| 1 | 0 | 0 |

| | | |
|---|---|---|
| 1 | 0 | 0 |
| 1 | 0 | 0 |
| 1 | 0 | 0 |
| 1 | 0 | 0 |
| 0 | 0 | 0 |
| 0 | 0 | 0 |
| 0 | 0 | 0 |
| 0 | 0 | 0 |
| 1 | 0 | 0 |
| 1 | 0 | 0 |
| 1 | 0 | 0 |
| 1 | 0 | 0 |
| 1 | 0 | 0 |
| 0 | 0 | 0 |
| 1 | 0 | 0 |
| 0 | 0 | 0 |
| 1 | 0 | 0 |
| 0 | 0 | 0 |
| 1 | 0 | 0 |
| 0 | 0 | 0 |
| 0 | 0 | 0 |
| 1 | 0 | 0 |
| 1 | 0 | 0 |
| 0 | 0 | 0 |
| 0 | 0 | 0 |
| 1 | 0 | 0 |
| 0 | 0 | 0 |
| 0 | 0 | 0 |
| 0 | 0 | 0 |
| 1 | 0 | 0 |
| 1 | 0 | 0 |
| 0 | 0 | 0 |
| 1 | 0 | 0 |
| 1 | 0 | 0 |
| 1 | 0 | 0 |
| 1 | 0 | 0 |
| 1 | 0 | 0 |
| 1 | 0 | 0 |
| 1 | 0 | 0 |
| 1 | 0 | 0 |
| 1 | 0 | 0 |
| 1 | 0 | 0 |
| 1 | 0 | 0 |
| 0 | 0 | 0 |
| 1 | 1 | 1 |
| 1 | 1 | 1 |
| 0 | 0 | 0 |
| 0 | 0 | 0 |
| 0 | 1 | 0 |
| 0 | 0 | 0 |
| 0 | 0 | 0 |
| 0 | 0 | 0 |
| 1 | 1 | 1 |
| 1 | 1 | 1 |

| | | |
|---|---|---|
| 1 | 1 | 1 |
| 1 | 1 | 1 |
| 1 | 1 | 1 |
| 1 | 1 | 1 |
| 1 | 1 | 1 |
| 1 | 1 | 1 |
| 1 | 1 | 1 |
| 0 | 0 | 0 |
| 1 | 1 | 1 |
| 1 | 1 | 1 |
| 0 | 1 | 0 |
| 0 | 1 | 0 |
| 1 | 0 | 0 |
| 1 | 1 | 1 |
| 0 | 0 | 0 |
| 0 | 0 | 0 |
| 0 | 1 | 0 |
| 0 | 0 | 0 |
| 1 | 1 | 1 |
| 1 | 1 | 1 |
| 0 | 0 | 0 |
| 0 | 1 | 0 |
| 0 | 1 | 0 |
| 1 | 1 | 1 |
| 0 | 0 | 0 |
| 1 | 1 | 1 |
| 0 | 1 | 0 |
| 0 | 0 | 0 |
| 0 | 0 | 0 |
| 0 | 0 | 0 |
| 0 | 0 | 0 |
| 1 | 1 | 1 |
| 1 | 1 | 1 |
| 0 | 1 | 0 |
| 1 | 1 | 1 |
| 1 | 1 | 1 |
| 0 | 0 | 0 |
| 0 | 0 | 0 |
| 1 | 1 | 1 |
| 1 | 1 | 1 |
| 1 | 1 | 1 |
| 1 | 1 | 1 |
| 0 | 0 | 0 |
| 0 | 1 | 0 |
| 1 | 1 | 1 |
| 1 | 1 | 1 |
| 1 | 1 | 1 |
| 1 | 1 | 1 |
| 0 | 0 | 0 |
| 0 | 0 | 0 |
| 0 | 0 | 0 |
| 0 | 0 | 0 |
| 0 | 0 | 0 |
| 1 | 1 | 1 |

| | | |
|---|---|---|
| 0 | 0 | 0 |
| 0 | 0 | 0 |
| 0 | 1 | 0 |
| 0 | 0 | 0 |
| 1 | 1 | 1 |
| 0 | 1 | 0 |
| 0 | 1 | 0 |
| 1 | 1 | 1 |
| 1 | 1 | 1 |
| 0 | 0 | 0 |
| 1 | 1 | 1 |
| 0 | 1 | 0 |
| 1 | 1 | 1 |
| 0 | 1 | 0 |
| 0 | 1 | 0 |
| 0 | 1 | 0 |
| 0 | 1 | 0 |
| 1 | 1 | 1 |
| 1 | 1 | 1 |
| 0 | 1 | 0 |
| 0 | 1 | 0 |
| 0 | 1 | 0 |
| 0 | 1 | 0 |
| 0 | 1 | 0 |
| 0 | 1 | 0 |
| 0 | 1 | 0 |
| 0 | 1 | 0 |
| 0 | 1 | 0 |
| 0 | 1 | 0 |
| 0 | 1 | 0 |
| 1 | 1 | 1 |
| 1 | 1 | 1 |
| 0 | 0 | 0 |
| 1 | 1 | 1 |
| 1 | 1 | 1 |
| 1 | 1 | 1 |
| 1 | 1 | 1 |
| 0 | 1 | 0 |
| 1 | 1 | 1 |
| 1 | 1 | 1 |
| 1 | 1 | 1 |
| 1 | 1 | 1 |
| 1 | 1 | 1 |
| 0 | 1 | 0 |
| 0 | 1 | 0 |
| 1 | 1 | 1 |
| 1 | 1 | 1 |
| 0 | 0 | 0 |
| 0 | 0 | 0 |
| 0 | 0 | 0 |
| 0 | 0 | 0 |
| 0 | 0 | 0 |
| 1 | 1 | 1 |
| 0 | 0 | 0 |

| | | |
|---|---|---|
| 1 | 1 | 1 |
| 0 | 0 | 0 |
| 1 | 1 | 1 |
| 0 | 1 | 0 |
| 0 | 1 | 0 |
| 0 | 0 | 0 |
| 0 | 0 | 0 |
| 0 | 0 | 0 |
| 0 | 0 | 0 |
| 0 | 0 | 0 |
| 0 | 1 | 0 |
| 0 | 0 | 0 |
| 0 | 1 | 0 |
| 1 | 1 | 1 |
| 0 | 1 | 0 |
| 0 | 0 | 0 |
| 1 | 1 | 1 |
| 1 | 1 | 1 |
| 0 | 0 | 0 |
| 0 | 0 | 0 |
| 0 | 0 | 0 |
| 0 | 0 | 0 |
| 0 | 0 | 0 |
| 0 | 0 | 0 |
| 0 | 1 | 0 |
| 0 | 0 | 0 |
| 0 | 0 | 0 |
| 0 | 0 | 0 |
| 0 | 0 | 0 |
| 0 | 0 | 0 |
| 0 | 0 | 0 |
| 1 | 1 | 1 |
| 1 | 1 | 1 |
| 0 | 0 | 0 |
| 0 | 0 | 0 |
| 1 | 0 | 0 |
| 0 | 0 | 0 |
| 0 | 0 | 0 |
| 0 | 0 | 0 |
| 0 | 0 | 0 |
| 0 | 0 | 0 |
| 1 | 0 | 0 |
| 1 | 1 | 1 |
| 1 | 1 | 1 |
| 0 | 0 | 0 |
| 0 | 0 | 0 |
| 0 | 0 | 0 |
| 1 | 1 | 1 |
| 1 | 1 | 1 |
| 1 | 1 | 1 |
| 1 | 1 | 1 |
| 1 | 1 | 1 |
| 1 | 1 | 1 |
| 0 | 0 | 0 |

| | | |
|---|---|---|
| 1 | 1 | 1 |
| 1 | 1 | 1 |
| 1 | 1 | 1 |
| 1 | 1 | 1 |
| 0 | 0 | 0 |
| 0 | 0 | 0 |
| 1 | 0 | 0 |
| 1 | 0 | 0 |
| 0 | 1 | 0 |
| 1 | 1 | 1 |
| 1 | 1 | 1 |
| 0 | 0 | 0 |
| 1 | 1 | 1 |
| 0 | 0 | 0 |
| 0 | 0 | 0 |
| 0 | 0 | 0 |
| 0 | 0 | 0 |
| 0 | 0 | 0 |
| 0 | 0 | 0 |
| 0 | 0 | 0 |
| 0 | 0 | 0 |
| 0 | 0 | 0 |
| 0 | 0 | 0 |
| 0 | 1 | 0 |
| 0 | 1 | 0 |
| 0 | 1 | 0 |
| 1 | 1 | 1 |
| 1 | 1 | 1 |
| 1 | 1 | 1 |
| 1 | 1 | 1 |
| 1 | 1 | 1 |
| 0 | 0 | 0 |
| 1 | 1 | 1 |
| 0 | 0 | 0 |
| 0 | 0 | 0 |
| 0 | 0 | 0 |
| 0 | 0 | 0 |
| 1 | 1 | 1 |
| 1 | 1 | 1 |
| 0 | 1 | 0 |
| 1 | 1 | 1 |
| 1 | 1 | 1 |
| 1 | 1 | 1 |
| 0 | 1 | 0 |
| 1 | 1 | 1 |
| 0 | 0 | 0 |
| 1 | 1 | 1 |
| 1 | 1 | 1 |
| 1 | 1 | 1 |
| 0 | 0 | 0 |
| 0 | 0 | 0 |
| 0 | 0 | 0 |
| 0 | 1 | 0 |
| 0 | 1 | 0 |

| | | |
|---|---|---|
| 0 | 1 | 0 |
| 0 | 1 | 0 |
| 0 | 1 | 0 |
| 0 | 0 | 0 |
| 0 | 1 | 0 |
| 1 | 1 | 1 |
| 0 | 0 | 0 |
| 0 | 1 | 0 |
| 1 | 1 | 1 |
| 1 | 1 | 1 |
| 0 | 0 | 0 |
| 0 | 0 | 0 |
| 1 | 1 | 1 |
| 1 | 1 | 1 |
| 0 | 1 | 0 |
| 0 | 1 | 0 |
| 0 | 0 | 0 |
| 0 | 0 | 0 |
| 1 | 1 | 1 |
| 0 | 0 | 0 |
| 0 | 0 | 0 |
| 0 | 0 | 0 |
| 0 | 1 | 0 |
| 0 | 0 | 0 |
| 0 | 0 | 0 |
| 1 | 1 | 1 |
| 0 | 0 | 0 |
| 0 | 1 | 0 |
| 1 | 1 | 1 |
| 0 | 0 | 0 |
| 0 | 1 | 0 |
| 1 | 1 | 1 |
| 0 | 0 | 0 |
| 0 | 1 | 0 |
| 0 | 1 | 0 |
| 0 | 0 | 0 |
| 1 | 1 | 1 |
| 0 | 0 | 0 |
| 1 | 1 | 1 |
| 0 | 0 | 0 |
| 0 | 0 | 0 |
| 1 | 0 | 0 |
| 0 | 0 | 0 |
| 0 | 0 | 0 |
| 1 | 1 | 1 |
| 1 | 0 | 0 |
| 0 | 1 | 0 |
| 1 | 1 | 1 |
| 1 | 1 | 1 |
| 1 | 1 | 1 |
| 1 | 1 | 1 |
| 0 | 1 | 0 |
| 0 | 1 | 0 |
| 0 | 1 | 0 |

| | | |
|---|---|---|
| 0 | 1 | 0 |
| 0 | 1 | 0 |
| 0 | 1 | 0 |
| 0 | 0 | 0 |
| 1 | 0 | 0 |
| 0 | 0 | 0 |
| 1 | 1 | 1 |
| 1 | 0 | 0 |
| 0 | 0 | 0 |
| 0 | 0 | 0 |
| 1 | 1 | 1 |
| 0 | 0 | 0 |
| 0 | 0 | 0 |
| 0 | 0 | 0 |
| 1 | 0 | 0 |
| 0 | 0 | 0 |
| 0 | 0 | 0 |
| 0 | 1 | 0 |
| 0 | 0 | 0 |
| 0 | 0 | 0 |
| 0 | 0 | 0 |
| 0 | 0 | 0 |
| 1 | 1 | 1 |
| 0 | 0 | 0 |
| 0 | 0 | 0 |
| 0 | 0 | 0 |
| 0 | 0 | 0 |
| 0 | 0 | 0 |
| 0 | 0 | 0 |
| 0 | 0 | 0 |
| 0 | 0 | 0 |
| 0 | 0 | 0 |
| 0 | 0 | 0 |
| 0 | 1 | 0 |
| 0 | 1 | 0 |
| 0 | 0 | 0 |
| 1 | 1 | 1 |
| 0 | 0 | 0 |
| 0 | 0 | 0 |
| 0 | 0 | 0 |
| 0 | 0 | 0 |
| 1 | 1 | 1 |
| 0 | 0 | 0 |
| 0 | 0 | 0 |
| 1 | 1 | 1 |
| 0 | 1 | 0 |
| 1 | 1 | 1 |
| 1 | 1 | 1 |
| 1 | 1 | 1 |
| 0 | 1 | 0 |
| 1 | 1 | 1 |
| 1 | 1 | 1 |
| 1 | 1 | 1 |
| 1 | 1 | 1 |

| | | |
|---|---|---|
| 1 | 1 | 1 |
| 0 | 0 | 0 |
| 1 | 1 | 1 |
| 1 | 1 | 1 |
| 1 | 1 | 1 |
| 1 | 1 | 1 |
| 1 | 1 | 1 |
| 1 | 1 | 1 |
| 1 | 1 | 1 |
| 0 | 1 | 0 |
| 1 | 1 | 1 |
| 0 | 0 | 0 |
| 1 | 0 | 0 |
| 0 | 1 | 0 |
| 0 | 0 | 0 |
| 0 | 1 | 0 |
| 0 | 1 | 0 |
| 1 | 1 | 1 |
| 1 | 1 | 1 |
| 1 | 1 | 1 |
| 0 | 0 | 0 |
| 0 | 1 | 0 |
| 0 | 0 | 0 |
| 0 | 0 | 0 |
| 1 | 1 | 1 |
| 0 | 0 | 0 |
| 1 | 1 | 1 |
| 1 | 1 | 1 |
| 1 | 1 | 1 |
| 1 | 1 | 1 |
| 1 | 1 | 1 |
| 0 | 0 | 0 |
| 1 | 1 | 1 |
| 0 | 0 | 0 |
| 1 | 1 | 1 |
| 0 | 1 | 0 |
| 0 | 1 | 0 |
| 0 | 1 | 0 |
| 1 | 1 | 1 |
| 0 | 0 | 0 |
| 1 | 1 | 1 |
| 1 | 1 | 1 |
| 0 | 1 | 0 |
| 0 | 0 | 0 |
| 1 | 1 | 1 |
| 1 | 1 | 1 |
| 1 | 1 | 1 |
| 0 | 0 | 0 |
| 0 | 0 | 0 |
| 1 | 1 | 1 |
| 1 | 1 | 1 |
| 0 | 0 | 0 |
| 0 | 0 | 0 |
| 0 | 0 | 0 |

| | | |
|---|---|---|
| 0 | 0 | 0 |
| 1 | 1 | 1 |
| 0 | 0 | 0 |
| 0 | 0 | 0 |
| 0 | 0 | 0 |
| 1 | 1 | 1 |
| 1 | 1 | 1 |
| 0 | 1 | 0 |
| 1 | 1 | 1 |
| 0 | 0 | 0 |
| 1 | 1 | 1 |
| 0 | 1 | 0 |
| 1 | 1 | 1 |
| 1 | 1 | 1 |
| 0 | 0 | 0 |
| 0 | 1 | 0 |
| 1 | 1 | 1 |
| 0 | 0 | 0 |
| 1 | 1 | 1 |
| 0 | 0 | 0 |
| 0 | 0 | 0 |
| 1 | 1 | 1 |
| 0 | 0 | 0 |
| 1 | 1 | 1 |
| 0 | 0 | 0 |
| 0 | 0 | 0 |
| 1 | 1 | 1 |
| 0 | 1 | 0 |
| 0 | 0 | 0 |
| 1 | 1 | 1 |
| 1 | 1 | 1 |
| 1 | 1 | 1 |
| 1 | 1 | 1 |
| 0 | 1 | 0 |
| 0 | 0 | 0 |
| 0 | 0 | 0 |
| 1 | 1 | 1 |
| 0 | 0 | 0 |
| 0 | 0 | 0 |
| 0 | 0 | 0 |
| 0 | 0 | 0 |
| 0 | 0 | 0 |
| 0 | 0 | 0 |
| 0 | 1 | 0 |
| 0 | 0 | 0 |
| 1 | 1 | 1 |
| 0 | 1 | 0 |
| 0 | 0 | 0 |
| 0 | 0 | 0 |
| 0 | 0 | 0 |
| 0 | 0 | 0 |
| 0 | 0 | 0 |
| 1 | 1 | 1 |
| 1 | 1 | 1 |

| | | |
|---|---|---|
| 1 | 1 | 1 |
| 0 | 0 | 0 |
| 0 | 0 | 0 |
| 0 | 0 | 0 |
| 0 | 0 | 0 |
| 0 | 0 | 0 |
| 0 | 0 | 0 |
| 1 | 0 | 0 |
| 1 | 0 | 0 |
| 1 | 1 | 1 |
| 0 | 0 | 0 |
| 0 | 0 | 0 |
| 1 | 1 | 1 |
| 0 | 0 | 0 |
| 1 | 1 | 1 |
| 0 | 0 | 0 |
| 1 | 1 | 1 |
| 1 | 1 | 1 |
| 0 | 1 | 0 |
| 0 | 1 | 0 |
| 0 | 1 | 0 |
| 1 | 1 | 1 |
| 0 | 1 | 0 |
| 0 | 0 | 0 |
| 0 | 0 | 0 |
| 1 | 1 | 1 |
| 1 | 1 | 1 |
| 1 | 1 | 1 |
| 1 | 1 | 1 |
| 0 | 0 | 0 |
| 0 | 0 | 0 |
| 0 | 0 | 0 |
| 0 | 0 | 0 |
| 1 | 1 | 1 |
| 0 | 0 | 0 |
| 0 | 0 | 0 |
| 0 | 0 | 0 |
| 0 | 0 | 0 |
| 0 | 0 | 0 |
| 1 | 1 | 1 |
| 1 | 1 | 1 |
| 0 | 0 | 0 |
| 1 | 1 | 1 |
| 1 | 1 | 1 |
| 0 | 0 | 0 |
| 1 | 0 | 0 |
| 1 | 1 | 1 |
| 0 | 1 | 0 |
| 1 | 0 | 0 |
| 0 | 1 | 0 |
| 0 | 0 | 0 |
| 1 | 1 | 1 |
| 0 | 0 | 0 |
| 0 | 0 | 0 |

| | | |
|---|---|---|
| 0 | 1 | 0 |
| 1 | 1 | 1 |
| 0 | 0 | 0 |
| 0 | 0 | 0 |
| 1 | 1 | 1 |
| 1 | 1 | 1 |
| 1 | 1 | 1 |
| 1 | 1 | 1 |
| 1 | 1 | 1 |
| 1 | 1 | 1 |
| 1 | 1 | 1 |
| 1 | 1 | 1 |
| 0 | 0 | 0 |
| 1 | 1 | 1 |
| 1 | 1 | 1 |
| 1 | 1 | 1 |
| 1 | 1 | 1 |
| 1 | 1 | 1 |
| 1 | 1 | 1 |
| 1 | 1 | 1 |
| 1 | 1 | 1 |
| 1 | 1 | 1 |
| 1 | 1 | 1 |
| 1 | 1 | 1 |
| 1 | 1 | 1 |
| 1 | 1 | 1 |
| 0 | 1 | 0 |
| 1 | 1 | 1 |
| 1 | 1 | 1 |
| 1 | 1 | 1 |
| 1 | 1 | 1 |
| 0 | 0 | 0 |
| 1 | 1 | 1 |
| 0 | 1 | 0 |
| 1 | 1 | 1 |
| 1 | 1 | 1 |
| 0 | 1 | 0 |
| 1 | 1 | 1 |
| 1 | 1 | 1 |
| 1 | 1 | 1 |
| 0 | 0 | 0 |
| 0 | 0 | 0 |
| 1 | 1 | 1 |
| 1 | 1 | 1 |
| 1 | 1 | 1 |
| 1 | 0 | 0 |
| 1 | 0 | 0 |
| 1 | 1 | 1 |
| 0 | 0 | 0 |
| 0 | 0 | 0 |
| 1 | 1 | 1 |
| 0 | 1 | 0 |
| 0 | 0 | 0 |
| 1 | 1 | 1 |

| | | |
|---|---|---|
| 0 | 0 | 0 |
| 0 | 0 | 0 |
| 0 | 0 | 0 |
| 1 | 1 | 1 |
| 1 | 1 | 1 |
| 1 | 1 | 1 |
| 1 | 1 | 1 |
| 0 | 0 | 0 |
| 0 | 0 | 0 |
| 0 | 0 | 0 |
| 0 | 0 | 0 |
| 1 | 1 | 1 |
| 0 | 0 | 0 |
| 1 | 1 | 1 |
| 1 | 1 | 1 |
| 1 | 1 | 1 |
| 1 | 1 | 1 |
| 0 | 0 | 0 |
| 1 | 1 | 1 |
| 1 | 1 | 1 |
| 1 | 1 | 1 |
| 1 | 1 | 1 |
| 0 | 1 | 0 |
| 1 | 1 | 1 |
| 1 | 1 | 1 |
| 0 | 1 | 0 |
| 1 | 1 | 1 |
| 1 | 1 | 1 |
| 1 | 1 | 1 |
| 1 | 1 | 1 |
| 0 | 0 | 0 |
| 0 | 0 | 0 |
| 1 | 1 | 1 |
| 1 | 1 | 1 |
| 1 | 1 | 1 |
| 1 | 1 | 1 |
| 1 | 1 | 1 |
| 1 | 1 | 1 |
| 1 | 1 | 1 |
| 1 | 1 | 1 |
| 1 | 1 | 1 |
| 0 | 0 | 0 |
| 1 | 1 | 1 |
| 1 | 1 | 1 |
| 0 | 0 | 0 |
| 0 | 0 | 0 |
| 1 | 1 | 1 |
| 1 | 1 | 1 |
| 0 | 0 | 0 |
| 0 | 0 | 0 |
| 0 | 0 | 0 |
| 0 | 0 | 0 |
| 0 | 0 | 0 |
| 0 | 0 | 0 |

| | | |
|---|---|---|
| 1 | 1 | 1 |
| 1 | 1 | 1 |
| 1 | 1 | 1 |
| 1 | 1 | 1 |
| 1 | 1 | 1 |
| 0 | 0 | 0 |
| 0 | 0 | 0 |
| 0 | 0 | 0 |
| 0 | 0 | 0 |
| 0 | 0 | 0 |
| 0 | 1 | 0 |
| 0 | 0 | 0 |
| 0 | 0 | 0 |
| 0 | 0 | 0 |
| 0 | 1 | 0 |
| 0 | 0 | 0 |
| 0 | 1 | 0 |
| 0 | 1 | 0 |
| 0 | 0 | 0 |
| 0 | 0 | 0 |
| 0 | 0 | 0 |
| 0 | 0 | 0 |
| 1 | 0 | 0 |
| 0 | 0 | 0 |
| 0 | 0 | 0 |
| 0 | 0 | 0 |
| 0 | 0 | 0 |
| 1 | 1 | 1 |
| 1 | 1 | 1 |
| 1 | 1 | 1 |
| 1 | 1 | 1 |
| 1 | 1 | 1 |
| 0 | 0 | 0 |
| 0 | 1 | 0 |
| 0 | 0 | 0 |
| 0 | 0 | 0 |
| 0 | 0 | 0 |
| 0 | 0 | 0 |
| 0 | 0 | 0 |
| 0 | 0 | 0 |
| 0 | 0 | 0 |
| 0 | 0 | 0 |
| 0 | 0 | 0 |
| 1 | 1 | 1 |
| 1 | 1 | 1 |
| 1 | 1 | 1 |
| 1 | 1 | 1 |
| 0 | 0 | 0 |
| 0 | 0 | 0 |
| 0 | 0 | 0 |
| 1 | 1 | 1 |
| 1 | 1 | 1 |
| 0 | 0 | 0 |
| 1 | 1 | 1 |

| | | |
|---|---|---|
| 1 | 1 | 1 |
| 0 | 0 | 0 |
| 0 | 1 | 0 |
| 0 | 1 | 0 |
| 1 | 1 | 1 |
| 1 | 1 | 1 |
| 0 | 1 | 0 |
| 1 | 1 | 1 |
| 0 | 1 | 0 |
| 1 | 1 | 1 |
| 1 | 1 | 1 |
| 1 | 1 | 1 |
| 0 | 0 | 0 |
| 0 | 0 | 0 |
| 0 | 0 | 0 |
| 0 | 0 | 0 |
| 0 | 0 | 0 |
| 0 | 0 | 0 |
| 0 | 0 | 0 |
| 0 | 1 | 0 |
| 0 | 0 | 0 |
| 0 | 0 | 0 |
| 0 | 0 | 0 |
| 0 | 0 | 0 |
| 0 | 0 | 0 |
| 1 | 1 | 1 |
| 1 | 1 | 1 |
| 0 | 0 | 0 |
| 0 | 1 | 0 |
| 0 | 0 | 0 |
| 0 | 0 | 0 |
| 1 | 1 | 1 |
| 0 | 0 | 0 |
| 1 | 1 | 1 |
| 1 | 1 | 1 |
| 1 | 1 | 1 |
| 1 | 0 | 0 |
| 1 | 1 | 1 |
| 1 | 1 | 1 |
| 1 | 1 | 1 |
| 0 | 0 | 0 |
| 1 | 1 | 1 |
| 1 | 1 | 1 |
| 0 | 0 | 0 |
| 0 | 0 | 0 |
| 1 | 1 | 1 |
| 0 | 0 | 0 |
| 0 | 0 | 0 |
| 1 | 1 | 1 |
| 1 | 1 | 1 |
| 0 | 0 | 0 |
| 1 | 1 | 1 |
| 1 | 1 | 1 |
| 1 | 1 | 1 |

| | | |
|---|---|---|
| 1 | 1 | 1 |
| 1 | 1 | 1 |
| 1 | 1 | 1 |
| 1 | 1 | 1 |
| 0 | 0 | 0 |
| 0 | 0 | 0 |
| 1 | 1 | 1 |
| 1 | 1 | 1 |
| 1 | 1 | 1 |
| 0 | 1 | 0 |
| 1 | 1 | 1 |
| 0 | 0 | 0 |
| 0 | 0 | 0 |
| 1 | 1 | 1 |
| 1 | 1 | 1 |
| 0 | 0 | 0 |
| 0 | 1 | 0 |
| 1 | 1 | 1 |
| 0 | 0 | 0 |
| 1 | 1 | 1 |
| 1 | 1 | 1 |
| 1 | 1 | 1 |
| 1 | 1 | 1 |
| 0 | 0 | 0 |
| 1 | 1 | 1 |
| 0 | 0 | 0 |
| 1 | 1 | 1 |
| 1 | 1 | 1 |
| 0 | 0 | 0 |
| 0 | 0 | 0 |
| 0 | 0 | 0 |
| 0 | 0 | 0 |
| 0 | 0 | 0 |
| 0 | 0 | 0 |
| 1 | 1 | 1 |
| 1 | 1 | 1 |
| 1 | 1 | 1 |
| 0 | 0 | 0 |
| 1 | 1 | 1 |
| 1 | 1 | 1 |
| 0 | 0 | 0 |
| 0 | 0 | 0 |
| 0 | 0 | 0 |
| 1 | 1 | 1 |
| 0 | 0 | 0 |
| 0 | 1 | 0 |
| 0 | 1 | 0 |
| 0 | 0 | 0 |
| 1 | 1 | 1 |
| 1 | 1 | 1 |
| 1 | 1 | 1 |
| 0 | 0 | 0 |
| 1 | 1 | 1 |
| 0 | 0 | 0 |

| | | |
|---|---|---|
| 1 | 1 | 1 |
| 0 | 0 | 0 |
| 0 | 0 | 0 |
| 0 | 1 | 0 |
| 1 | 1 | 1 |
| 1 | 1 | 1 |
| 0 | 0 | 0 |
| 1 | 1 | 1 |
| 0 | 1 | 0 |
| 0 | 1 | 0 |
| 1 | 1 | 1 |
| 1 | 1 | 1 |
| 1 | 1 | 1 |
| 1 | 1 | 1 |
| 1 | 1 | 1 |
| 1 | 1 | 1 |
| 1 | 1 | 1 |
| 1 | 1 | 1 |
| 1 | 1 | 1 |
| 1 | 1 | 1 |
| 1 | 1 | 1 |
| 0 | 0 | 0 |
| 0 | 1 | 0 |
| 0 | 1 | 0 |
| 1 | 1 | 1 |
| 1 | 1 | 1 |
| 0 | 0 | 0 |
| 1 | 1 | 1 |
| 0 | 0 | 0 |
| 0 | 1 | 0 |
| 1 | 1 | 1 |
| 0 | 0 | 0 |
| 0 | 0 | 0 |
| 1 | 0 | 0 |
| 1 | 1 | 1 |
| 1 | 1 | 1 |
| 1 | 1 | 1 |
| 1 | 1 | 1 |
| 0 | 0 | 0 |
| 0 | 0 | 0 |
| 0 | 0 | 0 |
| 1 | 1 | 1 |
| 1 | 1 | 1 |
| 0 | 1 | 0 |
| 0 | 1 | 0 |
| 0 | 0 | 0 |
| 1 | 1 | 1 |
| 0 | 0 | 0 |
| 1 | 1 | 1 |
| 0 | 0 | 0 |
| 0 | 0 | 0 |
| 0 | 0 | 0 |
| 1 | 1 | 1 |
| 1 | 1 | 1 |

| | | |
|---|---|---|
| 0 | 0 | 0 |
| 0 | 0 | 0 |
| 1 | 1 | 1 |
| 1 | 1 | 1 |
| 1 | 1 | 1 |
| 1 | 1 | 1 |
| 1 | 1 | 1 |
| 1 | 1 | 1 |
| 1 | 1 | 1 |
| 1 | 1 | 1 |
| 1 | 1 | 1 |
| 1 | 1 | 1 |
| 1 | 1 | 1 |
| 1 | 1 | 1 |
| 1 | 1 | 1 |
| 0 | 0 | 0 |
| 0 | 0 | 0 |
| 0 | 0 | 0 |
| 1 | 1 | 1 |
| 1 | 1 | 1 |
| 0 | 1 | 0 |
| 0 | 0 | 0 |
| 0 | 0 | 0 |
| 0 | 0 | 0 |
| 0 | 0 | 0 |
| 1 | 1 | 1 |
| 1 | 1 | 1 |
| 1 | 1 | 1 |
| 1 | 1 | 1 |
| 1 | 1 | 1 |
| 0 | 0 | 0 |
| 0 | 1 | 0 |
| 0 | 0 | 0 |
| 1 | 1 | 1 |
| 1 | 0 | 0 |
| 1 | 0 | 0 |
| 0 | 0 | 0 |
| 0 | 0 | 0 |
| 0 | 1 | 0 |
| 1 | 1 | 1 |
| 1 | 0 | 0 |
| 0 | 0 | 0 |
| 1 | 1 | 1 |
| 0 | 1 | 0 |
| 0 | 0 | 0 |
| 1 | 0 | 0 |
| 0 | 0 | 0 |
| 0 | 0 | 0 |
| 1 | 1 | 1 |
| 0 | 1 | 0 |
| 0 | 0 | 0 |
| 0 | 0 | 0 |
| 1 | 1 | 1 |
| 0 | 0 | 0 |

| | | |
|---|---|---|
| 0 | 1 | 0 |
| 0 | 0 | 0 |
| 1 | 0 | 0 |
| 0 | 0 | 0 |
| 1 | 1 | 1 |
| 1 | 1 | 1 |
| 1 | 1 | 1 |
| 0 | 1 | 0 |
| 0 | 0 | 0 |
| 1 | 1 | 1 |
| 0 | 1 | 0 |
| 0 | 1 | 0 |
| 0 | 0 | 0 |
| 0 | 0 | 0 |
| 0 | 0 | 0 |
| 1 | 1 | 1 |
| 0 | 0 | 0 |
| 0 | 1 | 0 |
| 0 | 0 | 0 |
| 0 | 0 | 0 |
| 0 | 0 | 0 |
| 0 | 0 | 0 |
| 0 | 0 | 0 |
| 0 | 0 | 0 |
| 0 | 1 | 0 |
| 0 | 1 | 0 |
| 0 | 0 | 0 |
| 0 | 0 | 0 |
| 0 | 0 | 0 |
| 0 | 0 | 0 |
| 1 | 1 | 1 |
| 1 | 1 | 1 |
| 1 | 1 | 1 |
| 1 | 1 | 1 |
| 1 | 1 | 1 |
| 0 | 0 | 0 |
| 0 | 0 | 0 |
| 0 | 1 | 0 |
| 1 | 1 | 1 |
| 1 | 1 | 1 |
| 0 | 1 | 0 |
| 1 | 1 | 1 |
| 1 | 1 | 1 |
| 0 | 0 | 0 |
| 0 | 0 | 0 |
| 0 | 0 | 0 |
| 1 | 1 | 1 |
| 0 | 0 | 0 |
| 0 | 0 | 0 |
| 0 | 0 | 0 |
| 1 | 1 | 1 |
| 1 | 1 | 1 |
| 0 | 1 | 0 |
| 0 | 1 | 0 |

| | | |
|---|---|---|
| 0 | 1 | 0 |
| 1 | 1 | 1 |
| 1 | 0 | 0 |
| 1 | 1 | 1 |
| 0 | 0 | 0 |
| 1 | 0 | 0 |
| 0 | 0 | 0 |
| 0 | 0 | 0 |
| 0 | 0 | 0 |
| 0 | 0 | 0 |
| 0 | 0 | 0 |
| 0 | 0 | 0 |
| 0 | 1 | 0 |
| 0 | 0 | 0 |
| 0 | 0 | 0 |
| 1 | 1 | 1 |
| 1 | 1 | 1 |
| 1 | 1 | 1 |
| 0 | 0 | 0 |
| 0 | 0 | 0 |
| 0 | 0 | 0 |
| 0 | 0 | 0 |
| 0 | 1 | 0 |
| 0 | 1 | 0 |
| 1 | 1 | 1 |
| 1 | 1 | 1 |
| 1 | 1 | 1 |
| 0 | 0 | 0 |
| 0 | 0 | 0 |
| 0 | 0 | 0 |
| 0 | 0 | 0 |
| 0 | 0 | 0 |
| 0 | 0 | 0 |
| 0 | 1 | 0 |
| 1 | 1 | 1 |
| 0 | 0 | 0 |
| 1 | 1 | 1 |
| 1 | 1 | 1 |
| 1 | 1 | 1 |
| 1 | 1 | 1 |
| 0 | 0 | 0 |
| 1 | 1 | 1 |
| 0 | 0 | 0 |
| 0 | 1 | 0 |
| 1 | 1 | 1 |
| 1 | 1 | 1 |
| 1 | 1 | 1 |
| 0 | 0 | 0 |
| 1 | 1 | 1 |
| 1 | 1 | 1 |
| 1 | 1 | 1 |
| 1 | 1 | 1 |
| 1 | 1 | 1 |
| 0 | 1 | 0 |

| | | |
|---|---|---|
| 0 | 1 | 0 |
| 1 | 1 | 1 |
| 1 | 1 | 1 |
| 0 | 1 | 0 |
| 1 | 1 | 1 |
| 0 | 1 | 0 |
| 0 | 1 | 0 |
| 0 | 1 | 0 |
| 0 | 1 | 0 |
| 0 | 1 | 0 |
| 0 | 1 | 0 |
| 0 | 0 | 0 |
| 0 | 1 | 0 |
| 0 | 1 | 0 |
| 0 | 1 | 0 |
| 0 | 1 | 0 |
| 0 | 1 | 0 |
| 0 | 0 | 0 |
| 0 | 0 | 0 |
| 0 | 0 | 0 |
| 0 | 0 | 0 |
| 0 | 1 | 0 |
| 0 | 0 | 0 |
| 1 | 1 | 1 |
| 1 | 1 | 1 |
| 1 | 1 | 1 |
| 1 | 1 | 1 |
| 0 | 1 | 0 |
| 0 | 0 | 0 |
| 0 | 0 | 0 |
| 1 | 1 | 1 |
| 1 | 1 | 1 |
| 0 | 0 | 0 |
| 1 | 1 | 1 |
| 0 | 0 | 0 |
| 0 | 0 | 0 |
| 0 | 0 | 0 |
| 1 | 1 | 1 |
| 0 | 0 | 0 |
| 1 | 1 | 1 |
| 0 | 0 | 0 |
| 1 | 1 | 1 |
| 0 | 0 | 0 |
| 1 | 0 | 0 |
| 1 | 0 | 0 |
| 1 | 0 | 0 |
| 1 | 0 | 0 |
| 0 | 1 | 0 |
| 0 | 1 | 0 |
| 0 | 0 | 0 |
| 1 | 0 | 0 |
| 0 | 0 | 0 |
| 1 | 0 | 0 |
| 1 | 1 | 1 |

| | | |
|---|---|---|
| 0 | 0 | 0 |
| 0 | 0 | 0 |
| 0 | 0 | 0 |
| 0 | 0 | 0 |
| 0 | 0 | 0 |
| 1 | 1 | 1 |
| 0 | 0 | 0 |
| 0 | 0 | 0 |
| 1 | 1 | 1 |
| 0 | 0 | 0 |
| 0 | 0 | 0 |
| 0 | 0 | 0 |
| 0 | 1 | 0 |
| 0 | 1 | 0 |
| 0 | 1 | 0 |
| 0 | 1 | 0 |
| 0 | 1 | 0 |
| 0 | 1 | 0 |
| 0 | 1 | 0 |
| 0 | 1 | 0 |
| 0 | 1 | 0 |
| 0 | 1 | 0 |
| 0 | 1 | 0 |
| 0 | 1 | 0 |
| 0 | 1 | 0 |
| 0 | 1 | 0 |
| 0 | 1 | 0 |
| 0 | 1 | 0 |
| 0 | 1 | 0 |
| 0 | 1 | 0 |
| 0 | 1 | 0 |
| 0 | 1 | 0 |
| 0 | 1 | 0 |
| 0 | 1 | 0 |
| 0 | 1 | 0 |
| 0 | 1 | 0 |
| 0 | 1 | 0 |
| 0 | 1 | 0 |
| 0 | 1 | 0 |
| 0 | 1 | 0 |
| 0 | 1 | 0 |
| 0 | 1 | 0 |
| 0 | 1 | 0 |
| 0 | 1 | 0 |
| 0 | 0 | 0 |
| 1 | 1 | 1 |
| 1 | 1 | 1 |
| 1 | 1 | 1 |
| 1 | 1 | 1 |
| 0 | 0 | 0 |
| 1 | 1 | 1 |
| 1 | 1 | 1 |
| 0 | 1 | 0 |

| | | |
|---|---|---|
| 1 | 1 | 1 |
| 1 | 1 | 1 |
| 1 | 1 | 1 |
| 1 | 1 | 1 |
| 0 | 1 | 0 |
| 0 | 1 | 0 |
| 0 | 1 | 0 |
| 0 | 0 | 0 |
| 0 | 0 | 0 |
| 1 | 1 | 1 |
| 0 | 1 | 0 |
| 1 | 1 | 1 |
| 1 | 1 | 1 |
| 1 | 1 | 1 |
| 0 | 0 | 0 |
| 0 | 0 | 0 |
| 0 | 0 | 0 |
| 1 | 1 | 1 |
| 1 | 0 | 0 |
| 1 | 1 | 1 |
| 0 | 0 | 0 |
| 0 | 0 | 0 |
| 0 | 0 | 0 |
| 0 | 0 | 0 |
| 0 | 0 | 0 |
| 0 | 0 | 0 |
| 0 | 1 | 0 |
| 1 | 1 | 1 |
| 0 | 0 | 0 |
| 1 | 1 | 1 |
| 1 | 1 | 1 |
| 1 | 1 | 1 |
| 0 | 0 | 0 |
| 1 | 1 | 1 |
| 1 | 1 | 1 |
| 1 | 1 | 1 |
| 0 | 0 | 0 |
| 0 | 0 | 0 |
| 0 | 0 | 0 |
| 1 | 1 | 1 |
| 1 | 1 | 1 |
| 1 | 1 | 1 |
| 1 | 1 | 1 |
| 1 | 1 | 1 |
| 0 | 1 | 0 |
| 0 | 0 | 0 |
| 1 | 1 | 1 |
| 0 | 1 | 0 |
| 1 | 1 | 1 |
| 0 | 1 | 0 |
| 1 | 1 | 1 |
| 1 | 1 | 1 |
| 0 | 1 | 0 |
| 1 | 1 | 1 |

| | | |
|---|---|---|
| 1 | 1 | 1 |
| 0 | 1 | 0 |
| 1 | 1 | 1 |
| 0 | 0 | 0 |
| 1 | 1 | 1 |
| 1 | 1 | 1 |
| 1 | 1 | 1 |
| 0 | 0 | 0 |
| 0 | 0 | 0 |
| 1 | 1 | 1 |
| 1 | 1 | 1 |
| 1 | 1 | 1 |
| 0 | 0 | 0 |
| 0 | 1 | 0 |
| 0 | 1 | 0 |
| 0 | 1 | 0 |
| 0 | 0 | 0 |
| 1 | 1 | 1 |
| 1 | 1 | 1 |
| 1 | 1 | 1 |
| 1 | 1 | 1 |
| 1 | 1 | 1 |
| 1 | 1 | 1 |
| 1 | 1 | 1 |
| 0 | 1 | 0 |
| 1 | 1 | 1 |
| 0 | 1 | 0 |
| 0 | 0 | 0 |
| 0 | 0 | 0 |
| 0 | 0 | 0 |
| 0 | 0 | 0 |
| 0 | 0 | 0 |
| 0 | 0 | 0 |
| 0 | 1 | 0 |
| 0 | 0 | 0 |
| 0 | 0 | 0 |
| 0 | 0 | 0 |
| 1 | 1 | 1 |
| 1 | 1 | 1 |
| 1 | 1 | 1 |
| 1 | 1 | 1 |
| 0 | 1 | 0 |
| 0 | 0 | 0 |
| 0 | 0 | 0 |
| 1 | 1 | 1 |
| 1 | 1 | 1 |
| 0 | 0 | 0 |
| 1 | 1 | 1 |
| 1 | 1 | 1 |
| 1 | 1 | 1 |
| 1 | 1 | 1 |
| 1 | 1 | 1 |
| 0 | 0 | 0 |
| 0 | 0 | 0 |

| | | |
|---|---|---|
| 0 | 0 | 0 |
| 0 | 0 | 0 |
| 0 | 0 | 0 |
| 0 | 0 | 0 |
| 0 | 0 | 0 |
| 0 | 0 | 0 |
| 0 | 0 | 0 |
| 0 | 0 | 0 |
| 1 | 1 | 1 |
| 0 | 0 | 0 |
| 1 | 1 | 1 |
| 1 | 1 | 1 |
| 1 | 1 | 1 |
| 1 | 1 | 1 |
| 0 | 0 | 0 |
| 1 | 1 | 1 |
| 1 | 1 | 1 |
| 1 | 1 | 1 |
| 1 | 1 | 1 |
| 0 | 0 | 0 |
| 0 | 0 | 0 |
| 1 | 1 | 1 |
| 0 | 0 | 0 |
| 0 | 0 | 0 |
| 0 | 0 | 0 |
| 0 | 0 | 0 |
| 0 | 0 | 0 |
| 0 | 1 | 0 |
| 0 | 1 | 0 |
| 0 | 1 | 0 |
| 0 | 0 | 0 |
| 1 | 1 | 1 |
| 1 | 1 | 1 |
| 1 | 1 | 1 |
| 1 | 1 | 1 |
| 1 | 1 | 1 |
| 1 | 1 | 1 |
| 1 | 1 | 1 |
| 1 | 1 | 1 |
| 1 | 1 | 1 |
| 0 | 1 | 0 |
| 0 | 0 | 0 |
| 1 | 1 | 1 |
| 1 | 1 | 1 |
| 1 | 1 | 1 |
| 0 | 1 | 0 |
| 1 | 1 | 1 |
| 1 | 1 | 1 |
| 0 | 0 | 0 |
| 0 | 1 | 0 |
| 1 | 1 | 1 |
| 1 | 1 | 1 |
| 0 | 0 | 0 |
| 0 | 0 | 0 |

| | | |
|---|---|---|
| 1 | 1 | 1 |
| 1 | 1 | 1 |
| 0 | 1 | 0 |
| 1 | 0 | 0 |
| 0 | 0 | 0 |
| 0 | 0 | 0 |
| 0 | 1 | 0 |
| 0 | 0 | 0 |
| 1 | 1 | 1 |
| 1 | 1 | 1 |
| 0 | 0 | 0 |
| 1 | 0 | 0 |
| 0 | 0 | 0 |
| 0 | 0 | 0 |
| 0 | 0 | 0 |
| 0 | 0 | 0 |
| 1 | 1 | 1 |
| 1 | 1 | 1 |
| 0 | 0 | 0 |
| 0 | 0 | 0 |
| 0 | 0 | 0 |
| 0 | 0 | 0 |
| 0 | 0 | 0 |
| 1 | 1 | 1 |
| 1 | 1 | 1 |
| 0 | 0 | 0 |
| 1 | 1 | 1 |
| 0 | 0 | 0 |
| 0 | 0 | 0 |
| 0 | 0 | 0 |
| 0 | 1 | 0 |
| 0 | 0 | 0 |
| 0 | 0 | 0 |
| 1 | 1 | 1 |
| 1 | 1 | 1 |
| 0 | 0 | 0 |
| 0 | 0 | 0 |
| 1 | 0 | 0 |
| 0 | 0 | 0 |
| 0 | 0 | 0 |
| 0 | 0 | 0 |
| 0 | 0 | 0 |
| 0 | 1 | 0 |
| 1 | 1 | 1 |
| 0 | 0 | 0 |
| 0 | 1 | 0 |
| 0 | 0 | 0 |
| 1 | 1 | 1 |
| 1 | 1 | 1 |
| 1 | 1 | 1 |
| 1 | 1 | 1 |
| 1 | 1 | 1 |
| 0 | 1 | 0 |
| 1 | 1 | 1 |

| | | |
|---|---|---|
| 1 | 1 | 1 |
| 1 | 1 | 1 |
| 1 | 1 | 1 |
| 0 | 1 | 0 |
| 1 | 1 | 1 |
| 1 | 1 | 1 |
| 0 | 1 | 0 |
| 0 | 0 | 0 |
| 1 | 1 | 1 |
| 0 | 0 | 0 |
| 0 | 0 | 0 |
| 0 | 0 | 0 |
| 0 | 1 | 0 |
| 1 | 1 | 1 |
| 0 | 0 | 0 |
| 0 | 1 | 0 |
| 0 | 1 | 0 |
| 0 | 1 | 0 |
| 0 | 1 | 0 |
| 0 | 1 | 0 |
| 1 | 1 | 1 |
| 1 | 1 | 1 |
| 1 | 1 | 1 |
| 0 | 1 | 0 |
| 1 | 1 | 1 |
| 1 | 1 | 1 |
| 1 | 1 | 1 |
| 1 | 0 | 0 |
| 1 | 1 | 1 |
| 0 | 0 | 0 |
| 0 | 0 | 0 |
| 1 | 1 | 1 |
| 1 | 1 | 1 |
| 0 | 1 | 0 |
| 0 | 1 | 0 |
| 0 | 0 | 0 |
| 1 | 1 | 1 |
| 1 | 1 | 1 |
| 1 | 1 | 1 |
| 1 | 1 | 1 |
| 1 | 1 | 1 |
| 0 | 0 | 0 |
| 0 | 0 | 0 |
| 0 | 0 | 0 |
| 1 | 1 | 1 |
| 1 | 1 | 1 |
| 1 | 1 | 1 |
| 1 | 1 | 1 |
| 1 | 1 | 1 |
| 1 | 1 | 1 |
| 1 | 1 | 1 |
| 1 | 1 | 1 |
| 1 | 1 | 1 |
| 0 | 0 | 0 |

| | | |
|---|---|---|
| 1 | 1 | 1 |
| 1 | 1 | 1 |
| 1 | 1 | 1 |
| 1 | 1 | 1 |
| 1 | 1 | 1 |
| 1 | 1 | 1 |
| 1 | 1 | 1 |
| 0 | 1 | 0 |
| 0 | 0 | 0 |
| 0 | 0 | 0 |
| 1 | 1 | 1 |
| 0 | 1 | 0 |
| 0 | 1 | 0 |
| 0 | 0 | 0 |
| 1 | 1 | 1 |
| 1 | 1 | 1 |
| 1 | 1 | 1 |
| 0 | 0 | 0 |
| 0 | 0 | 0 |
| 0 | 0 | 0 |
| 0 | 0 | 0 |
| 0 | 1 | 0 |
| 1 | 1 | 1 |
| 1 | 1 | 1 |
| 1 | 1 | 1 |
| 0 | 0 | 0 |
| 1 | 1 | 1 |
| 0 | 1 | 0 |
| 1 | 1 | 1 |
| 1 | 1 | 1 |
| 1 | 1 | 1 |
| 1 | 1 | 1 |
| 1 | 1 | 1 |
| 1 | 1 | 1 |
| 1 | 1 | 1 |
| 1 | 1 | 1 |
| 1 | 1 | 1 |
| 1 | 1 | 1 |
| 1 | 1 | 1 |
| 0 | 0 | 0 |
| 1 | 1 | 1 |
| 1 | 1 | 1 |
| 0 | 0 | 0 |
| 1 | 1 | 1 |
| 1 | 1 | 1 |
| 1 | 1 | 1 |
| 0 | 0 | 0 |
| 0 | 0 | 0 |
| 1 | 1 | 1 |
| 1 | 1 | 1 |
| 0 | 0 | 0 |
| 1 | 1 | 1 |
| 0 | 1 | 0 |
| 1 | 1 | 1 |

| | | |
|---|---|---|
| 1 | 1 | 1 |
| 1 | 1 | 1 |
| 1 | 1 | 1 |
| 1 | 1 | 1 |
| 1 | 1 | 1 |
| 1 | 1 | 1 |
| 1 | 1 | 1 |
| 0 | 1 | 0 |
| 1 | 1 | 1 |
| 1 | 1 | 1 |
| 1 | 1 | 1 |
| 1 | 1 | 1 |
| 1 | 1 | 1 |
| 0 | 0 | 0 |
| 0 | 0 | 0 |
| 0 | 0 | 0 |
| 0 | 0 | 0 |
| 0 | 1 | 0 |
| 1 | 1 | 1 |
| 0 | 0 | 0 |
| 1 | 1 | 1 |
| 1 | 0 | 0 |
| 1 | 1 | 1 |
| 1 | 1 | 1 |
| 1 | 1 | 1 |
| 0 | 0 | 0 |
| 0 | 1 | 0 |
| 1 | 1 | 1 |
| 1 | 1 | 1 |
| 0 | 0 | 0 |
| 0 | 0 | 0 |
| 1 | 1 | 1 |
| 1 | 1 | 1 |
| 1 | 1 | 1 |
| 1 | 1 | 1 |
| 1 | 1 | 1 |
| 1 | 1 | 1 |
| 1 | 1 | 1 |
| 0 | 0 | 0 |
| 1 | 1 | 1 |
| 1 | 1 | 1 |
| 0 | 0 | 0 |
| 0 | 0 | 0 |
| 0 | 0 | 0 |
| 0 | 0 | 0 |
| 1 | 1 | 1 |
| 0 | 0 | 0 |
| 1 | 1 | 1 |
| 1 | 1 | 1 |
| 1 | 1 | 1 |
| 0 | 1 | 0 |
| 1 | 0 | 0 |
| 1 | 0 | 0 |
| 1 | 0 | 0 |

| | | |
|---|---|---|
| 1 | 1 | 1 |
| 1 | 1 | 1 |
| 0 | 0 | 0 |
| 1 | 1 | 1 |
| 1 | 1 | 1 |
| 1 | 1 | 1 |
| 1 | 1 | 1 |
| 0 | 1 | 0 |
| 1 | 1 | 1 |
| 1 | 1 | 1 |
| 1 | 1 | 1 |
| 0 | 1 | 0 |
| 0 | 1 | 0 |
| 0 | 1 | 0 |
| 0 | 1 | 0 |
| 0 | 1 | 0 |
| 0 | 1 | 0 |
| 0 | 1 | 0 |
| 0 | 1 | 0 |
| 0 | 1 | 0 |
| 0 | 1 | 0 |
| 0 | 1 | 0 |
| 0 | 1 | 0 |
| 0 | 1 | 0 |
| 0 | 1 | 0 |
| 0 | 1 | 0 |
| 0 | 1 | 0 |
| 0 | 1 | 0 |
| 1 | 1 | 1 |
| 1 | 1 | 1 |
| 0 | 0 | 0 |
| 1 | 1 | 1 |
| 0 | 0 | 0 |
| 0 | 0 | 0 |
| 0 | 0 | 0 |
| 0 | 0 | 0 |
| 1 | 1 | 1 |
| 0 | 0 | 0 |
| 1 | 1 | 1 |
| 1 | 1 | 1 |
| 0 | 0 | 0 |
| 0 | 0 | 0 |
| 0 | 0 | 0 |
| 1 | 1 | 1 |
| 0 | 0 | 0 |
| 0 | 0 | 0 |
| 1 | 1 | 1 |
| 1 | 1 | 1 |
| 1 | 1 | 1 |
| 1 | 1 | 1 |
| 0 | 0 | 0 |
| 1 | 1 | 1 |
| 1 | 1 | 1 |
| 1 | 1 | 1 |

| | | |
|---|---|---|
| 1 | 1 | 1 |
| 1 | 1 | 1 |
| 1 | 1 | 1 |
| 1 | 1 | 1 |
| 1 | 1 | 1 |
| 0 | 0 | 0 |
| 1 | 1 | 1 |
| 0 | 0 | 0 |
| 0 | 0 | 0 |
| 0 | 0 | 0 |
| 1 | 1 | 1 |
| 0 | 0 | 0 |
| 1 | 1 | 1 |
| 1 | 1 | 1 |
| 1 | 1 | 1 |
| 1 | 0 | 0 |
| 1 | 1 | 1 |
| 0 | 0 | 0 |
| 1 | 1 | 1 |
| 0 | 0 | 0 |
| 1 | 1 | 1 |
| 0 | 0 | 0 |
| 0 | 0 | 0 |
| 0 | 0 | 0 |
| 1 | 1 | 1 |
| 0 | 1 | 0 |
| 1 | 1 | 1 |
| 1 | 1 | 1 |
| 0 | 0 | 0 |
| 1 | 1 | 1 |
| 1 | 1 | 1 |
| 1 | 1 | 1 |
| 1 | 1 | 1 |
| 1 | 1 | 1 |
| 0 | 0 | 0 |
| 0 | 0 | 0 |
| 1 | 1 | 1 |
| 0 | 0 | 0 |
| 0 | 0 | 0 |
| 0 | 1 | 0 |
| 0 | 0 | 0 |
| 0 | 0 | 0 |
| 0 | 1 | 0 |
| 0 | 1 | 0 |
| 1 | 1 | 1 |
| 0 | 1 | 0 |
| 1 | 1 | 1 |
| 1 | 1 | 1 |
| 1 | 1 | 1 |
| 1 | 1 | 1 |
| 1 | 1 | 1 |
| 1 | 1 | 1 |
| 1 | 1 | 1 |
| 1 | 1 | 1 |

| | | |
|---|---|---|
| 1 | 1 | 1 |
| 0 | 0 | 0 |
| 0 | 1 | 0 |
| 1 | 1 | 1 |
| 0 | 0 | 0 |
| 1 | 1 | 1 |
| 1 | 1 | 1 |
| 1 | 1 | 1 |
| 0 | 0 | 0 |
| 0 | 0 | 0 |
| 1 | 1 | 1 |
| 0 | 0 | 0 |
| 0 | 0 | 0 |
| 1 | 1 | 1 |
| 1 | 1 | 1 |
| 0 | 0 | 0 |
| 0 | 1 | 0 |
| 0 | 0 | 0 |
| 0 | 0 | 0 |
| 1 | 1 | 1 |
| 0 | 0 | 0 |
| 0 | 0 | 0 |
| 0 | 0 | 0 |
| 1 | 1 | 1 |
| 0 | 1 | 0 |
| 0 | 0 | 0 |
| 1 | 1 | 1 |
| 1 | 1 | 1 |
| 1 | 1 | 1 |
| 1 | 1 | 1 |
| 0 | 0 | 0 |
| 1 | 1 | 1 |
| 0 | 0 | 0 |
| 1 | 1 | 1 |
| 0 | 0 | 0 |
| 0 | 0 | 0 |
| 1 | 1 | 1 |
| 1 | 1 | 1 |
| 1 | 1 | 1 |
| 1 | 1 | 1 |
| 1 | 1 | 1 |
| 1 | 1 | 1 |
| 1 | 1 | 1 |
| 1 | 1 | 1 |
| 1 | 1 | 1 |
| 1 | 1 | 1 |
| 1 | 0 | 0 |
| 1 | 1 | 1 |
| 1 | 1 | 1 |
| 1 | 1 | 1 |
| 1 | 1 | 1 |
| 1 | 1 | 1 |
| 1 | 1 | 1 |
| 0 | 0 | 0 |

| | | |
|---|---|---|
| 1 | 1 | 1 |
| 1 | 1 | 1 |
| 1 | 1 | 1 |
| 1 | 1 | 1 |
| 0 | 0 | 0 |
| 1 | 1 | 1 |
| 0 | 1 | 0 |
| 0 | 1 | 0 |
| 0 | 1 | 0 |
| 1 | 1 | 1 |
| 0 | 0 | 0 |
| 1 | 1 | 1 |
| 1 | 1 | 1 |
| 1 | 1 | 1 |
| 0 | 0 | 0 |
| 0 | 0 | 0 |
| 0 | 0 | 0 |
| 0 | 0 | 0 |
| 0 | 1 | 0 |
| 0 | 0 | 0 |
| 0 | 0 | 0 |
| 0 | 0 | 0 |
| 1 | 1 | 1 |
| 0 | 0 | 0 |
| 0 | 0 | 0 |
| 1 | 1 | 1 |
| 0 | 0 | 0 |
| 0 | 0 | 0 |
| 0 | 0 | 0 |
| 0 | 0 | 0 |
| 0 | 0 | 0 |
| 1 | 1 | 1 |
| 1 | 1 | 1 |
| 0 | 1 | 0 |
| 1 | 1 | 1 |
| 1 | 1 | 1 |
| 1 | 1 | 1 |
| 0 | 0 | 0 |
| 0 | 0 | 0 |
| 0 | 1 | 0 |
| 1 | 1 | 1 |
| 0 | 0 | 0 |
| 0 | 0 | 0 |
| 0 | 1 | 0 |
| 0 | 0 | 0 |
| 1 | 1 | 1 |
| 1 | 1 | 1 |
| 1 | 1 | 1 |
| 0 | 0 | 0 |
| 0 | 0 | 0 |
| 0 | 0 | 0 |
| 1 | 1 | 1 |
| 0 | 1 | 0 |
| 0 | 0 | 0 |

| | | |
|---|---|---|
| 1 | 1 | 1 |
| 0 | 0 | 0 |
| 1 | 1 | 1 |
| 0 | 0 | 0 |
| 0 | 0 | 0 |
| 0 | 1 | 0 |
| 0 | 0 | 0 |
| 0 | 0 | 0 |
| 1 | 1 | 1 |
| 1 | 1 | 1 |
| 1 | 1 | 1 |
| 1 | 1 | 1 |
| 1 | 1 | 1 |
| 1 | 1 | 1 |
| 1 | 1 | 1 |
| 1 | 1 | 1 |
| 1 | 1 | 1 |
| 0 | 1 | 0 |
| 1 | 1 | 1 |
| 1 | 1 | 1 |
| 0 | 0 | 0 |
| 0 | 1 | 0 |
| 0 | 1 | 0 |
| 0 | 1 | 0 |
| 0 | 1 | 0 |
| 0 | 0 | 0 |
| 0 | 0 | 0 |
| 1 | 1 | 1 |
| 1 | 1 | 1 |
| 1 | 1 | 1 |
| 1 | 1 | 1 |
| 1 | 1 | 1 |
| 1 | 1 | 1 |
| 0 | 1 | 0 |
| 1 | 1 | 1 |
| 1 | 1 | 1 |
| 1 | 1 | 1 |
| 1 | 1 | 1 |
| 0 | 1 | 0 |
| 1 | 1 | 1 |
| 1 | 1 | 1 |
| 0 | 1 | 0 |
| 1 | 1 | 1 |
| 0 | 0 | 0 |
| 1 | 1 | 1 |
| 0 | 0 | 0 |
| 0 | 0 | 0 |
| 0 | 0 | 0 |
| 0 | 0 | 0 |
| 0 | 0 | 0 |
| 0 | 0 | 0 |
| 0 | 1 | 0 |
| 1 | 1 | 1 |
| 1 | 1 | 1 |

| | | |
|---|---|---|
| 1 | 1 | 1 |
| 1 | 1 | 1 |
| 1 | 1 | 1 |
| 0 | 0 | 0 |
| 1 | 1 | 1 |
| 0 | 0 | 0 |
| 1 | 1 | 1 |
| 1 | 1 | 1 |
| 0 | 0 | 0 |
| 1 | 1 | 1 |
| 1 | 1 | 1 |
| 0 | 1 | 0 |
| 1 | 1 | 1 |
| 1 | 1 | 1 |
| 1 | 1 | 1 |
| 0 | 0 | 0 |
| 0 | 0 | 0 |
| 0 | 0 | 0 |
| 0 | 0 | 0 |
| 0 | 0 | 0 |
| 1 | 1 | 1 |
| 1 | 1 | 1 |
| 1 | 1 | 1 |
| 1 | 1 | 1 |
| 0 | 0 | 0 |
| 1 | 1 | 1 |
| 1 | 1 | 1 |
| 1 | 1 | 1 |
| 0 | 0 | 0 |
| 1 | 1 | 1 |
| 0 | 0 | 0 |
| 0 | 0 | 0 |
| 0 | 0 | 0 |
| 0 | 0 | 0 |
| 0 | 0 | 0 |
| 1 | 1 | 1 |
| 0 | 0 | 0 |
| 0 | 0 | 0 |
| 0 | 0 | 0 |
| 0 | 0 | 0 |
| 0 | 0 | 0 |
| 0 | 0 | 0 |
| 0 | 0 | 0 |
| 0 | 0 | 0 |
| 0 | 0 | 0 |
| 0 | 0 | 0 |
| 0 | 0 | 0 |
| 0 | 0 | 0 |
| 0 | 0 | 0 |
| 1 | 1 | 1 |
| 0 | 0 | 0 |
| 1 | 1 | 1 |
| 1 | 1 | 1 |
| 0 | 1 | 0 |
| 0 | 0 | 0 |

| | | |
|---|---|---|
| 0 | 0 | 0 |
| 0 | 1 | 0 |
| 1 | 1 | 1 |
| 1 | 1 | 1 |
| 1 | 1 | 1 |
| 0 | 0 | 0 |
| 1 | 1 | 1 |
| 0 | 1 | 0 |
| 1 | 1 | 1 |
| 0 | 1 | 0 |
| 0 | 1 | 0 |
| 0 | 1 | 0 |
| 1 | 1 | 1 |
| 0 | 0 | 0 |
| 0 | 1 | 0 |
| 1 | 1 | 1 |
| 1 | 1 | 1 |
| 0 | 0 | 0 |
| 1 | 1 | 1 |
| 1 | 1 | 1 |
| 1 | 1 | 1 |
| 1 | 1 | 1 |
| 0 | 0 | 0 |
| 1 | 1 | 1 |
| 1 | 1 | 1 |
| 0 | 0 | 0 |
| 0 | 0 | 0 |
| 0 | 0 | 0 |
| 1 | 1 | 1 |
| 0 | 1 | 0 |
| 1 | 1 | 1 |
| 1 | 1 | 1 |
| 1 | 1 | 1 |
| 1 | 1 | 1 |
| 1 | 1 | 1 |
| 1 | 1 | 1 |
| 1 | 1 | 1 |
| 1 | 1 | 1 |
| 1 | 1 | 1 |
| 1 | 1 | 1 |
| 1 | 1 | 1 |
| 1 | 1 | 1 |
| 1 | 1 | 1 |
| 1 | 1 | 1 |
| 1 | 1 | 1 |
| 1 | 1 | 1 |
| 1 | 1 | 1 |
| 1 | 1 | 1 |
| 1 | 0 | 0 |
| 1 | 1 | 1 |
| 1 | 1 | 1 |
| 1 | 1 | 1 |
| 1 | 1 | 1 |
| 1 | 1 | 1 |

| | | |
|---|---|---|
| 1 | 1 | 1 |
| 0 | 0 | 0 |
| 1 | 1 | 1 |
| 1 | 1 | 1 |
| 1 | 1 | 1 |
| 0 | 0 | 0 |
| 0 | 0 | 0 |
| 0 | 0 | 0 |
| 1 | 1 | 1 |
| 1 | 1 | 1 |
| 0 | 1 | 0 |
| 1 | 1 | 1 |
| 1 | 1 | 1 |
| 1 | 1 | 1 |
| 0 | 0 | 0 |
| 0 | 0 | 0 |
| 0 | 0 | 0 |
| 1 | 1 | 1 |
| 0 | 0 | 0 |
| 0 | 0 | 0 |
| 0 | 0 | 0 |
| 0 | 0 | 0 |
| 0 | 0 | 0 |
| 0 | 0 | 0 |
| 0 | 1 | 0 |
| 1 | 1 | 1 |
| 1 | 1 | 1 |
| 0 | 1 | 0 |
| 0 | 1 | 0 |
| 1 | 1 | 1 |
| 1 | 1 | 1 |
| 1 | 1 | 1 |
| 0 | 0 | 0 |
| 1 | 1 | 1 |
| 0 | 1 | 0 |
| 0 | 0 | 0 |
| 0 | 0 | 0 |
| 0 | 0 | 0 |
| 0 | 0 | 0 |
| 1 | 1 | 1 |
| 0 | 1 | 0 |
| 1 | 1 | 1 |
| 1 | 1 | 1 |
| 0 | 0 | 0 |
| 0 | 0 | 0 |
| 0 | 0 | 0 |
| 0 | 1 | 0 |
| 1 | 1 | 1 |
| 0 | 0 | 0 |
| 1 | 1 | 1 |
| 1 | 1 | 1 |
| 0 | 0 | 0 |
| 1 | 1 | 1 |
| 0 | 1 | 0 |

| | | |
|---|---|---|
| 1 | 1 | 1 |
| 0 | 1 | 0 |
| 1 | 1 | 1 |
| 1 | 1 | 1 |
| 1 | 1 | 1 |
| 1 | 1 | 1 |
| 0 | 1 | 0 |
| 1 | 1 | 1 |
| 0 | 0 | 0 |
| 0 | 0 | 0 |
| 1 | 1 | 1 |
| 0 | 0 | 0 |
| 0 | 0 | 0 |
| 0 | 0 | 0 |
| 1 | 1 | 1 |
| 1 | 1 | 1 |
| 0 | 0 | 0 |
| 0 | 0 | 0 |
| 0 | 0 | 0 |
| 1 | 1 | 1 |
| 1 | 1 | 1 |
| 1 | 1 | 1 |
| 0 | 0 | 0 |
| 1 | 1 | 1 |
| 0 | 0 | 0 |
| 1 | 1 | 1 |
| 1 | 1 | 1 |
| 1 | 1 | 1 |
| 1 | 1 | 1 |
| 0 | 0 | 0 |
| 0 | 0 | 0 |
| 0 | 0 | 0 |
| 1 | 1 | 1 |
| 0 | 1 | 0 |
| 1 | 1 | 1 |
| 1 | 1 | 1 |
| 1 | 1 | 1 |
| 0 | 1 | 0 |
| 0 | 1 | 0 |
| 0 | 1 | 0 |
| 0 | 0 | 0 |
| 1 | 1 | 1 |
| 1 | 1 | 1 |
| 1 | 1 | 1 |
| 1 | 1 | 1 |
| 0 | 0 | 0 |
| 1 | 0 | 0 |
| 0 | 0 | 0 |
| 1 | 1 | 1 |
| 0 | 0 | 0 |
| 0 | 0 | 0 |
| 0 | 0 | 0 |
| 0 | 1 | 0 |
| 1 | 0 | 0 |

| | | |
|---|---|---|
| 1 | 0 | 0 |
| 0 | 0 | 0 |
| 0 | 1 | 0 |
| 1 | 1 | 1 |
| 0 | 1 | 0 |
| 0 | 1 | 0 |
| 0 | 1 | 0 |
| 0 | 0 | 0 |
| 0 | 0 | 0 |
| 0 | 1 | 0 |
| 0 | 0 | 0 |
| 1 | 1 | 1 |
| 1 | 1 | 1 |
| 1 | 1 | 1 |
| 0 | 1 | 0 |
| 1 | 1 | 1 |
| 1 | 1 | 1 |
| 1 | 1 | 1 |
| 1 | 1 | 1 |
| 0 | 1 | 0 |
| 1 | 1 | 1 |
| 1 | 1 | 1 |
| 1 | 1 | 1 |
| 1 | 1 | 1 |
| 0 | 1 | 0 |
| 0 | 1 | 0 |
| 0 | 1 | 0 |
| 0 | 1 | 0 |
| 0 | 0 | 0 |
| 0 | 1 | 0 |
| 0 | 0 | 0 |
| 1 | 1 | 1 |
| 0 | 0 | 0 |
| 0 | 0 | 0 |
| 0 | 1 | 0 |
| 0 | 0 | 0 |
| 0 | 0 | 0 |
| 1 | 1 | 1 |
| 0 | 1 | 0 |
| 0 | 1 | 0 |
| 1 | 1 | 1 |
| 0 | 0 | 0 |
| 0 | 0 | 0 |
| 1 | 1 | 1 |
| 1 | 1 | 1 |
| 0 | 0 | 0 |
| 0 | 0 | 0 |
| 0 | 0 | 0 |
| 0 | 0 | 0 |
| 0 | 1 | 0 |
| 0 | 0 | 0 |
| 0 | 0 | 0 |
| 1 | 1 | 1 |
| 1 | 1 | 1 |

| | | |
|---|---|---|
| 1 | 1 | 1 |
| 1 | 1 | 1 |
| 1 | 1 | 1 |
| 1 | 1 | 1 |
| 0 | 0 | 0 |
| 0 | 0 | 0 |
| 0 | 0 | 0 |
| 0 | 0 | 0 |
| 1 | 1 | 1 |
| 0 | 0 | 0 |
| 0 | 0 | 0 |
| 0 | 0 | 0 |
| 0 | 0 | 0 |
| 1 | 1 | 1 |
| 1 | 0 | 0 |
| 0 | 1 | 0 |
| 1 | 1 | 1 |
| 1 | 1 | 1 |
| 1 | 1 | 1 |
| 0 | 1 | 0 |
| 1 | 1 | 1 |
| 1 | 1 | 1 |
| 1 | 1 | 1 |
| 1 | 1 | 1 |
| 1 | 1 | 1 |
| 0 | 0 | 0 |
| 0 | 0 | 0 |
| 0 | 0 | 0 |
| 0 | 0 | 0 |
| 0 | 1 | 0 |
| 0 | 1 | 0 |
| 0 | 1 | 0 |
| 1 | 1 | 1 |
| 0 | 1 | 0 |
| 0 | 1 | 0 |
| 1 | 1 | 1 |
| 1 | 1 | 1 |
| 1 | 1 | 1 |
| 1 | 1 | 1 |
| 1 | 1 | 1 |
| 1 | 1 | 1 |
| 1 | 1 | 1 |
| 1 | 1 | 1 |
| 1 | 1 | 1 |
| 1 | 1 | 1 |
| 1 | 1 | 1 |
| 1 | 1 | 1 |
| 1 | 1 | 1 |
| 1 | 1 | 1 |
| 1 | 1 | 1 |
| 1 | 1 | 1 |
| 1 | 1 | 1 |
| 1 | 1 | 1 |
| 0 | 0 | 0 |

| | | |
|---|---|---|
| 1 | 1 | 1 |
| 1 | 1 | 1 |
| 1 | 1 | 1 |
| 0 | 1 | 0 |
| 0 | 1 | 0 |
| 0 | 1 | 0 |
| 0 | 1 | 0 |
| 0 | 1 | 0 |
| 1 | 1 | 1 |
| 1 | 1 | 1 |
| 1 | 1 | 1 |
| 1 | 1 | 1 |
| 1 | 1 | 1 |
| 1 | 1 | 1 |
| 1 | 1 | 1 |
| 1 | 1 | 1 |
| 1 | 1 | 1 |
| 1 | 1 | 1 |
| 1 | 1 | 1 |
| 1 | 1 | 1 |
| 1 | 1 | 1 |
| 1 | 1 | 1 |
| 1 | 1 | 1 |
| 1 | 1 | 1 |
| 1 | 1 | 1 |
| 1 | 1 | 1 |
| 1 | 1 | 1 |
| 1 | 1 | 1 |
| 1 | 1 | 1 |
| 1 | 1 | 1 |
| 1 | 1 | 1 |
| 1 | 1 | 1 |
| 1 | 1 | 1 |
| 0 | 1 | 0 |
| 1 | 1 | 1 |
| 1 | 1 | 1 |
| 1 | 1 | 1 |
| 1 | 1 | 1 |
| 1 | 1 | 1 |
| 1 | 1 | 1 |
| 0 | 1 | 0 |
| 1 | 1 | 1 |
| 1 | 1 | 1 |
| 1 | 1 | 1 |
| 0 | 1 | 0 |
| 1 | 0 | 0 |
| 0 | 0 | 0 |
| 1 | 1 | 1 |
| 0 | 1 | 0 |
| 1 | 1 | 1 |
| 1 | 1 | 1 |
| 0 | 1 | 0 |
| 0 | 1 | 0 |
| 0 | 1 | 0 |

| | | |
|---|---|---|
| 0 | 1 | 0 |
| 1 | 1 | 1 |
| 0 | 0 | 0 |
| 1 | 1 | 1 |
| 1 | 1 | 1 |
| 1 | 1 | 1 |
| 0 | 0 | 0 |
| 1 | 1 | 1 |
| 1 | 1 | 1 |
| 0 | 1 | 0 |
| 0 | 1 | 0 |
| 0 | 0 | 0 |
| 1 | 1 | 1 |
| 0 | 0 | 0 |
| 0 | 0 | 0 |
| 0 | 0 | 0 |
| 0 | 0 | 0 |
| 0 | 0 | 0 |
| 0 | 0 | 0 |
| 0 | 0 | 0 |
| 0 | 0 | 0 |

| organism_scientific | evo_cons | cluster_id | pdb_id | chain_id | resi | resn | change | uniprot_id |
|---|---|---|---|---|---|---|---|---|
| HOMO SAPIENS | 5 | 87 | 3uzw | A | 106 | LEU | p.Leu106Phe | P51857 |
| HOMO SAPIENS | 3 | 87 | 3uzw | A | 133 | PRO | p.Pro133Arg | P51857 |
| HOMO SAPIENS | 0 | 87 | 3uzw | A | 198 | PRO | p.Pro198Leu | P51857 |
| HOMO SAPIENS | 7 | 87 | 3uzw | A | 261 | ARG | p.Arg261Cys | P51857 |
| HOMO SAPIENS | 0 | 4693 | 3mk1 | A | 3 | PRO | p.Pro25Leu | P05187 |
| HOMO SAPIENS | 8 | 4830 | 2h64 | B | 101 | CYS | p.Cys124Arg | P36894 |
| HOMO SAPIENS | 2 | 4830 | 2h64 | B | 116 | PRO | p.Pro139Ser | P36894 |
| HOMO SAPIENS | 0 | 6937 | 3dar | A | 186 | MET | p.Met186Thr | P21802 |
| HOMO SAPIENS | 3 | 5 | 1aut | C | 30 | GLN | p.Gln226His | P04070 |
| HOMO SAPIENS | 1 | 5 | 1aut | C | 68 | LEU | p.Leu265Phe | P04070 |
| HOMO SAPIENS | 0 | 5 | 1aut | C | 75 | ARG | p.Arg272Cys | P04070 |
| HOMO SAPIENS | 6 | 5 | 1aut | C | 92 | PRO | p.Pro289Leu | P04070 |
| HOMO SAPIENS | 0 | 5 | 1aut | C | 104 | ALA | p.Ala301Val | P04070 |
| HOMO SAPIENS | 1 | 5 | 1aut | C | 112 | ALA | p.Ala309Thr | P04070 |
| HOMO SAPIENS | 2 | 5 | 1aut | C | 115 | SER | p.Ser312Leu | P04070 |
| HOMO SAPIENS | 1 | 5 | 1aut | C | 133 | GLY | p.Gly334Ser | P04070 |
| HOMO SAPIENS | 0 | 5 | 1aut | C | 147 | ARG | p.Arg348Ter | P04070 |
| HOMO SAPIENS | 8 | 5 | 1aut | C | 194 | ASP | p.Asp401Asn | P04070 |
| HOMO SAPIENS | 7 | 5 | 1aut | C | 237 | TRP | p.Trp444Cys | P04070 |
| HOMO SAPIENS | 0 | 5 | 1aut | C | 238 | ILE | p.Ile445Met | P04070 |
| HOMO SAPIENS | 8 | 43 | 1aiy | F | 10 | HIS | p.His34Asp | P01308 |
| HOMO SAPIENS | 2 | 43 | 1aiy | F | 22 | ARG | p.Arg46Gln | P01308 |
| HOMO SAPIENS | 4 | 43 | 1aiy | F | 23 | GLY | p.Gly47Val | P01308 |
| HOMO SAPIENS | 7 | 43 | 1aiy | F | 24 | PHE | p.Phe48Ser | P01308 |
| HOMO SAPIENS | 5 | 24 | 2fg8 | A | 33 | THR | p.Thr30Ile | P02792 |
| HOMO SAPIENS | 4 | 24 | 2fg8 | A | 99 | ALA | p.Ala96Thr | P02792 |
| HOMO SAPIENS | 5 | 24 | 2fg8 | A | 107 | GLU | p.Glu104Ter | P02792 |
| HOMO SAPIENS | 3 | 5409 | 5bnq | A | 199 | MET | p.Met199Lys | O14788 |
| HOMO SAPIENS | -1 | 839 | 3swc | A | 959 | GLN | p.Gln990His | Q9H9B1 |
| HOMO SAPIENS | -1 | 839 | 3swc | A | 968 | ALA | p.Ala999Val | Q9H9B1 |
| HOMO SAPIENS | -1 | 839 | 3swc | A | 973 | SER | p.Ser1004Ile | Q9H9B1 |
| HOMO SAPIENS | 8 | 17 | 1eou | A | 18 | LYS | p.Lys18Glu | P00918 |
| HOMO SAPIENS | 2 | 17 | 1eou | A | 107 | HIS | p.His107Tyr | P00918 |
| HOMO SAPIENS | 1 | 17 | 1eou | A | 236 | PRO | p.Pro236His | P00918 |
| HOMO SAPIENS | 6 | 17 | 1eou | A | 252 | ASN | p.Asn252Asp | P00918 |
| HOMO SAPIENS | 2 | 9140 | 5c4v | A | 445 | ARG | p.Arg445Ter | Q13485 |
| HOMO SAPIENS | 7 | 9140 | 5c4v | A | 496 | ARG | p.Arg496Cys | Q13485 |
| HOMO SAPIENS | -1 | 42 | 4pwi | A | 6 | GLY | p.Gly26Ser | P02766 |
| HOMO SAPIENS | 6 | 42 | 4pwi | A | 10 | CYS | p.Cys30Arg | P02766 |
| HOMO SAPIENS | 8 | 42 | 4pwi | A | 12 | LEU | p.Leu32Pro | P02766 |
| HOMO SAPIENS | 9 | 42 | 4pwi | A | 18 | ASP | p.Asp38Gly | P02766 |
| HOMO SAPIENS | 8 | 42 | 4pwi | A | 20 | VAL | p.Val40Ile | P02766 |
| HOMO SAPIENS | 4 | 42 | 4pwi | A | 30 | MET | p.Val50Ala | P02766 |
| HOMO SAPIENS | 4 | 42 | 4pwi | A | 30 | MET | p.Val50Leu | P02766 |
| HOMO SAPIENS | 6 | 42 | 4pwi | A | 33 | PHE | p.Phe53Ile | P02766 |
| HOMO SAPIENS | 7 | 42 | 4pwi | A | 36 | ALA | p.Ala56Pro | P02766 |
| HOMO SAPIENS | 7 | 42 | 4pwi | A | 42 | GLU | p.Glu62Gly | P02766 |
| HOMO SAPIENS | 8 | 42 | 4pwi | A | 44 | PHE | p.Phe64Ser | P02766 |
| HOMO SAPIENS | 4 | 42 | 4pwi | A | 47 | GLY | p.Gly67Ala | P02766 |
| HOMO SAPIENS | 6 | 42 | 4pwi | A | 49 | THR | p.Thr69Ala | P02766 |
| HOMO SAPIENS | 6 | 42 | 4pwi | A | 50 | SER | p.Ser70Ile | P02766 |
| HOMO SAPIENS | 6 | 42 | 4pwi | A | 50 | SER | p.Ser70Arg | P02766 |
| HOMO SAPIENS | 0 | 42 | 4pwi | A | 53 | GLY | p.Gly73Glu | P02766 |

| | | | | | | | | |
|---|---|---|---|---|---|---|---|---|
| HOMO SAPIENS | 7 | 42 | 4pwi | A | 55 | LEU | p.Leu75Pro | P02766 |
| HOMO SAPIENS | 6 | 42 | 4pwi | A | 58 | LEU | p.Leu78His | P02766 |
| HOMO SAPIENS | 7 | 42 | 4pwi | A | 61 | GLU | p.Glu81Lys | P02766 |
| HOMO SAPIENS | 8 | 42 | 4pwi | A | 64 | PHE | p.Phe84Leu | P02766 |
| HOMO SAPIENS | 8 | 42 | 4pwi | A | 68 | ILE | p.Ile88Leu | P02766 |
| HOMO SAPIENS | 8 | 42 | 4pwi | A | 69 | TYR | p.Tyr89His | P02766 |
| HOMO SAPIENS | 4 | 42 | 4pwi | A | 71 | VAL | p.Val91Ala | P02766 |
| HOMO SAPIENS | 6 | 42 | 4pwi | A | 77 | SER | p.Ser97Tyr | P02766 |
| HOMO SAPIENS | 4 | 42 | 4pwi | A | 84 | ILE | p.Ile104Ser | P02766 |
| HOMO SAPIENS | 8 | 42 | 4pwi | A | 89 | GLU | p.Glu109Gln | P02766 |
| HOMO SAPIENS | 8 | 42 | 4pwi | A | 97 | ALA | p.Ala117Gly | P02766 |
| HOMO SAPIENS | 7 | 42 | 4pwi | A | 99 | ASP | p.Asp119Asn | P02766 |
| HOMO SAPIENS | 9 | 42 | 4pwi | A | 107 | ILE | p.Ile127Val | P02766 |
| HOMO SAPIENS | 9 | 42 | 4pwi | A | 111 | LEU | p.Leu131Met | P02766 |
| HOMO SAPIENS | 7 | 42 | 4pwi | A | 114 | TYR | p.Tyr134Cys | P02766 |
| HOMO SAPIENS | 7 | 42 | 4pwi | A | 114 | TYR | p.Tyr134His | P02766 |
| HOMO SAPIENS | 7 | 42 | 4pwi | A | 122 | VAL | p.Val142Ile | P02766 |
| HOMO SAPIENS | -1 | 42 | 4pwi | A | 126 | LYS | p.Lys146Arg | P02766 |
| HOMO SAPIENS | -1 | 42 | 4pwi | A | 127 | GLU | p.Glu147Ter | P02766 |
| HOMO SAPIENS | -1 | 1312 | 4dce | A | 1275 | ARG | p.Arg1275Gln | Q9UM73 |
| HOMO SAPIENS | -1 | 1312 | 4dce | A | 1280 | ALA | p.Ala1280Val | Q9UM73 |
| HOMO SAPIENS | 1 | 20 | 1y0x | X | 234 | ALA | p.Ala234Thr | P10828 |
| HOMO SAPIENS | 0 | 20 | 1y0x | X | 243 | ARG | p.Arg243Trp | P10828 |
| HOMO SAPIENS | 3 | 20 | 1y0x | X | 316 | ARG | p.Arg316His | P10828 |
| HOMO SAPIENS | 1 | 20 | 1y0x | X | 317 | ALA | p.Ala317Thr | P10828 |
| HOMO SAPIENS | 2 | 20 | 1y0x | X | 320 | ARG | p.Arg320Cys | P10828 |
| HOMO SAPIENS | 2 | 20 | 1y0x | X | 320 | ARG | p.Arg320His | P10828 |
| HOMO SAPIENS | 0 | 20 | 1y0x | X | 332 | GLY | p.Gly332Arg | P10828 |
| HOMO SAPIENS | 1 | 20 | 1y0x | X | 338 | ARG | p.Arg338Trp | P10828 |
| HOMO SAPIENS | 0 | 20 | 1y0x | X | 340 | GLN | p.Gln340His | P10828 |
| HOMO SAPIENS | 0 | 20 | 1y0x | X | 345 | GLY | p.Gly345Val | P10828 |
| HOMO SAPIENS | 0 | 20 | 1y0x | X | 345 | GLY | p.Gly345Ser | P10828 |
| HOMO SAPIENS | 0 | 20 | 1y0x | X | 347 | GLY | p.Gly347Glu | P10828 |
| HOMO SAPIENS | 1 | 20 | 1y0x | X | 438 | ARG | p.Arg438His | P10828 |
| HOMO SAPIENS | 0 | 20 | 1y0x | X | 442 | MET | p.Met442Val | P10828 |
| HOMO SAPIENS | 0 | 20 | 1y0x | X | 443 | LYS | p.Lys443Glu | P10828 |
| HOMO SAPIENS | 0 | 20 | 1y0x | X | 446 | CYS | p.Cys446Arg | P10828 |
| HOMO SAPIENS | 2 | 20 | 1y0x | X | 453 | PRO | p.Pro453His | P10828 |
| HOMO SAPIENS | 2 | 20 | 1y0x | X | 453 | PRO | p.Pro453Thr | P10828 |
| HOMO SAPIENS | 3 | 20 | 1y0x | X | 459 | PHE | p.Phe459Cys | P10828 |
| HOMO SAPIENS | 0 | 4409 | 3qb7 | D | 87 | GLU | p.Glu109Lys | P31785 |
| HOMO SAPIENS | 1 | 4409 | 3qb7 | D | 92 | GLY | p.Gly114Asp | P31785 |
| HOMO SAPIENS | 1 | 4409 | 3qb7 | D | 93 | CYS | p.Cys115Arg | P31785 |
| HOMO SAPIENS | 0 | 4409 | 3qb7 | D | 131 | ILE | p.Ile153Asn | P31785 |
| HOMO SAPIENS | 9 | 4409 | 3qb7 | D | 200 | ARG | p.Arg222Cys | P31785 |
| HOMO SAPIENS | 0 | 3891 | 3ije | B | 33 | LEU | p.Leu59Pro | P05106 |
| HOMO SAPIENS | 0 | 3891 | 3ije | B | 117 | LEU | p.Leu143Trp | P05106 |
| HOMO SAPIENS | 3 | 3891 | 3ije | B | 119 | ASP | p.Asp145Tyr | P05106 |
| HOMO SAPIENS | 2 | 3891 | 3ije | B | 143 | ARG | p.Arg169Gln | P05106 |
| HOMO SAPIENS | 2 | 3891 | 3ije | B | 214 | ARG | p.Arg240Gln | P05106 |
| HOMO SAPIENS | 2 | 3891 | 3ije | B | 214 | ARG | p.Arg240Trp | P05106 |
| HOMO SAPIENS | 2 | 3891 | 3ije | B | 374 | CYS | p.Cys400Tyr | P05106 |
| HOMO SAPIENS | 1 | 3891 | 3ije | B | 407 | PRO | p.Pro433Ala | P05106 |
| HOMO SAPIENS | 0 | 3891 | 3ije | B | 489 | ARG | p.Arg515Gln | P05106 |

| | | | | | | | | |
|---|---|---|---|---|---|---|---|---|
| HOMO SAPIENS | -1 | 9613 | 5mke | B | 28 | ARG | p.Arg28Pro | Q13563 |
| HOMO SAPIENS | -1 | 9613 | 5mke | B | 190 | ALA | p.Ala190Thr | Q13563 |
| HOMO SAPIENS | 4 | 9613 | 5mke | B | 262 | VAL | p.Val262Met | Q13563 |
| HOMO SAPIENS | 0 | 9613 | 5mke | B | 482 | PHE | p.Phe482Cys | Q13563 |
| HOMO SAPIENS | 4 | 9613 | 5mke | B | 511 | ASP | p.Asp511Val | Q13563 |
| HOMO SAPIENS | 5 | 9613 | 5mke | B | 516 | VAL | p.Val516Leu | Q13563 |
| HOMO SAPIENS | -1 | 9613 | 5mke | B | 807 | ARG | p.Arg807Gln | Q13563 |
| HOMO SAPIENS | 4 | 2428 | 1olh | C | 337 | ARG | p.Arg337Cys | P04637 |
| HOMO SAPIENS | 4 | 2428 | 1olh | C | 337 | ARG | p.Arg337His | P04637 |
| HOMO SAPIENS | 8 | 2428 | 1olh | C | 338 | PHE | p.Phe338Leu | P04637 |
| HOMO SAPIENS | 8 | 2428 | 1olh | C | 339 | GLU | p.Glu339Lys | P04637 |
| HOMO SAPIENS | 7 | 2428 | 1olh | C | 344 | LEU | p.Leu344Pro | P04637 |
| HOMO SAPIENS | 0 | 2428 | 1olh | C | 360 | GLY | p.Gly360Ala | P04637 |
| HOMO SAPIENS | 7 | 4867 | 1mox | B | 404 | GLY | p.Gly428Asp | P00533 |
| HOMO SAPIENS | 5 | 4867 | 1mox | B | 497 | ARG | p.Arg521Lys | P00533 |
| HOMO SAPIENS | 1 | 4630 | 1fu1 | A | 43 | TRP | p.Trp43Arg | Q13426 |
| HOMO SAPIENS | 2 | 4630 | 1fu1 | A | 161 | ARG | p.Arg161Ter | Q13426 |
| HOMO SAPIENS | 2 | 15 | 1lfy | B | 2 | HIS | p.His3Gln | P68871 |
| HOMO SAPIENS | 2 | 15 | 1lfy | B | 6 | GLU | p.Glu7Ala | P68871 |
| HOMO SAPIENS | 2 | 15 | 1lfy | B | 6 | GLU | p.Glu7Lys | P68871 |
| HOMO SAPIENS | 4 | 15 | 1lfy | B | 9 | SER | p.Ser10Cys | P68871 |
| HOMO SAPIENS | 6 | 15 | 1lfy | B | 15 | TRP | p.Trp16Ter | P68871 |
| HOMO SAPIENS | 3 | 15 | 1lfy | B | 17 | LYS | p.Lys18Glu | P68871 |
| HOMO SAPIENS | 6 | 15 | 1lfy | B | 19 | ASN | p.Asn20Ser | P68871 |
| HOMO SAPIENS | 6 | 15 | 1lfy | B | 22 | GLU | p.Glu23Ala | P68871 |
| HOMO SAPIENS | 6 | 15 | 1lfy | B | 22 | GLU | p.Glu23Lys | P68871 |
| HOMO SAPIENS | 8 | 15 | 1lfy | B | 26 | GLU | p.Glu27Lys | P68871 |
| HOMO SAPIENS | 8 | 15 | 1lfy | B | 28 | LEU | p.Leu29Pro | P68871 |
| HOMO SAPIENS | 8 | 15 | 1lfy | B | 30 | ARG | p.Arg31Ser | P68871 |
| HOMO SAPIENS | 8 | 15 | 1lfy | B | 30 | ARG | p.Arg31Lys | P68871 |
| HOMO SAPIENS | 6 | 15 | 1lfy | B | 39 | GLN | p.Gln40Ter | P68871 |
| HOMO SAPIENS | 2 | 15 | 1lfy | B | 43 | GLU | p.Glu44Gln | P68871 |
| HOMO SAPIENS | 6 | 15 | 1lfy | B | 57 | ASN | p.Asn58Lys | P68871 |
| HOMO SAPIENS | 6 | 15 | 1lfy | B | 59 | LYS | p.Lys60Asn | P68871 |
| HOMO SAPIENS | 8 | 15 | 1lfy | B | 61 | LYS | p.Lys62Glu | P68871 |
| HOMO SAPIENS | 4 | 15 | 1lfy | B | 73 | ASP | p.Asp74Tyr | P68871 |
| HOMO SAPIENS | 4 | 15 | 1lfy | B | 87 | THR | p.Thr88Ile | P68871 |
| HOMO SAPIENS | 4 | 15 | 1lfy | B | 87 | THR | p.Thr88Pro | P68871 |
| HOMO SAPIENS | 5 | 15 | 1lfy | B | 90 | GLU | p.Glu91Lys | P68871 |
| HOMO SAPIENS | 6 | 15 | 1lfy | B | 91 | LEU | p.Leu92Pro | P68871 |
| HOMO SAPIENS | 5 | 15 | 1lfy | B | 97 | HIS | p.His98Gln | P68871 |
| HOMO SAPIENS | 6 | 15 | 1lfy | B | 110 | LEU | p.Leu111Pro | P68871 |
| HOMO SAPIENS | 6 | 15 | 1lfy | B | 113 | VAL | p.Val114Glu | P68871 |
| HOMO SAPIENS | 2 | 15 | 1lfy | B | 119 | GLY | p.Gly120Ala | P68871 |
| HOMO SAPIENS | 4 | 15 | 1lfy | B | 121 | GLU | p.Glu122Gln | P68871 |
| HOMO SAPIENS | 5 | 15 | 1lfy | B | 134 | VAL | p.Val135Glu | P68871 |
| HOMO SAPIENS | 3 | 15 | 1lfy | B | 143 | HIS | p.His144Gln | P68871 |
| HOMO SAPIENS | 4 | 91 | 5d9k | A | 75 | GLY | p.Gly75Val | P51812 |
| HOMO SAPIENS | 5 | 91 | 5d9k | A | 82 | VAL | p.Val82Phe | P51812 |
| HOMO SAPIENS | 0 | 91 | 5d9k | A | 114 | ARG | p.Arg114Trp | P51812 |
| HOMO SAPIENS | 4 | 91 | 5d9k | A | 189 | ILE | p.Ile189Lys | P51812 |
| HOMO SAPIENS | -1 | 91 | 5d9k | A | 227 | SER | p.Ser227Ala | P51812 |
| HOMO SAPIENS | 1 | 91 | 5d9k | A | 268 | PHE | p.Phe268Ser | P51812 |
| HOMO SAPIENS | 0 | 954 | 2vt0 | A | 15 | VAL | p.Val54Leu | P04062 |

| | | | | | | | | |
|---|---|---|---|---|---|---|---|---|
| HOMO SAPIENS | 5 | 954 | 2vt0 | A | 46 | GLY | p.Gly85Glu | P04062 |
| HOMO SAPIENS | 5 | 954 | 2vt0 | A | 48 | ARG | p.Arg87Trp | P04062 |
| HOMO SAPIENS | 6 | 954 | 2vt0 | A | 79 | LYS | p.Lys118Asn | P04062 |
| HOMO SAPIENS | 0 | 954 | 2vt0 | A | 120 | ARG | p.Arg159Gln | P04062 |
| HOMO SAPIENS | 0 | 954 | 2vt0 | A | 120 | ARG | p.Arg159Trp | P04062 |
| HOMO SAPIENS | 0 | 954 | 2vt0 | A | 122 | PRO | p.Pro161Ser | P04062 |
| HOMO SAPIENS | 5 | 954 | 2vt0 | A | 131 | ARG | p.Arg170Leu | P04062 |
| HOMO SAPIENS | 5 | 954 | 2vt0 | A | 135 | TYR | p.Tyr174Cys | P04062 |
| HOMO SAPIENS | 0 | 954 | 2vt0 | A | 157 | LYS | p.Lys196Gln | P04062 |
| HOMO SAPIENS | 5 | 954 | 2vt0 | A | 163 | ARG | p.Arg202Gln | P04062 |
| HOMO SAPIENS | 5 | 954 | 2vt0 | A | 170 | ARG | p.Arg209Cys | P04062 |
| HOMO SAPIENS | 5 | 954 | 2vt0 | A | 188 | ASN | p.Asn227Ser | P04062 |
| HOMO SAPIENS | 3 | 954 | 2vt0 | A | 196 | SER | p.Ser235Pro | P04062 |
| HOMO SAPIENS | 5 | 954 | 2vt0 | A | 202 | GLY | p.Gly241Arg | P04062 |
| HOMO SAPIENS | 0 | 954 | 2vt0 | A | 212 | TYR | p.Tyr251His | P04062 |
| HOMO SAPIENS | 0 | 954 | 2vt0 | A | 213 | PHE | p.Phe252Ile | P04062 |
| HOMO SAPIENS | 5 | 954 | 2vt0 | A | 216 | PHE | p.Phe255Tyr | P04062 |
| HOMO SAPIENS | 6 | 954 | 2vt0 | A | 251 | PHE | p.Phe290Leu | P04062 |
| HOMO SAPIENS | 5 | 954 | 2vt0 | A | 255 | HIS | p.His294Gln | P04062 |
| HOMO SAPIENS | 6 | 954 | 2vt0 | A | 257 | ARG | p.Arg296Gln | P04062 |
| HOMO SAPIENS | 5 | 954 | 2vt0 | A | 289 | PRO | p.Pro328Leu | P04062 |
| HOMO SAPIENS | 0 | 954 | 2vt0 | A | 309 | ALA | p.Ala348Val | P04062 |
| HOMO SAPIENS | 0 | 954 | 2vt0 | A | 311 | HIS | p.His350Arg | P04062 |
| HOMO SAPIENS | 6 | 954 | 2vt0 | A | 312 | TRP | p.Trp351Cys | P04062 |
| HOMO SAPIENS | 5 | 954 | 2vt0 | A | 323 | THR | p.Thr362Ile | P04062 |
| HOMO SAPIENS | 5 | 954 | 2vt0 | A | 325 | GLY | p.Gly364Arg | P04062 |
| HOMO SAPIENS | 6 | 954 | 2vt0 | A | 342 | CYS | p.Cys381Gly | P04062 |
| HOMO SAPIENS | 5 | 954 | 2vt0 | A | 353 | ARG | p.Arg392Gly | P04062 |
| HOMO SAPIENS | 3 | 954 | 2vt0 | A | 364 | SER | p.Ser403Thr | P04062 |
| HOMO SAPIENS | 5 | 954 | 2vt0 | A | 369 | THR | p.Thr408Met | P04062 |
| HOMO SAPIENS | 3 | 954 | 2vt0 | A | 370 | ASN | p.Asn409Ser | P04062 |
| HOMO SAPIENS | 0 | 954 | 2vt0 | A | 371 | LEU | p.Leu410Val | P04062 |
| HOMO SAPIENS | 0 | 954 | 2vt0 | A | 375 | VAL | p.Val414Leu | P04062 |
| HOMO SAPIENS | 0 | 954 | 2vt0 | A | 377 | GLY | p.Gly416Ser | P04062 |
| HOMO SAPIENS | 8 | 954 | 2vt0 | A | 398 | VAL | p.Val437Phe | P04062 |
| HOMO SAPIENS | 0 | 954 | 2vt0 | A | 401 | PRO | p.Pro440Leu | P04062 |
| HOMO SAPIENS | 3 | 954 | 2vt0 | A | 409 | ASP | p.Asp448His | P04062 |
| HOMO SAPIENS | 3 | 954 | 2vt0 | A | 409 | ASP | p.Asp448Val | P04062 |
| HOMO SAPIENS | 0 | 954 | 2vt0 | A | 415 | PRO | p.Pro454Arg | P04062 |
| HOMO SAPIENS | 5 | 954 | 2vt0 | A | 444 | LEU | p.Leu483Arg | P04062 |
| HOMO SAPIENS | 5 | 954 | 2vt0 | A | 456 | ALA | p.Ala495Pro | P04062 |
| HOMO SAPIENS | 5 | 954 | 2vt0 | A | 463 | ARG | p.Arg502Cys | P04062 |
| HOMO SAPIENS | 5 | 954 | 2vt0 | A | 463 | ARG | p.Arg502His | P04062 |
| HOMO SAPIENS | 5 | 954 | 2vt0 | A | 478 | GLY | p.Gly517Ser | P04062 |
| HOMO SAPIENS | 5 | 954 | 2vt0 | A | 496 | ARG | p.Arg535Cys | P04062 |
| HOMO SAPIENS | 5 | 954 | 2vt0 | A | 496 | ARG | p.Arg535His | P04062 |
| HOMO SAPIENS | 1 | 9037 | 2rqb | A | 946 | ALA | p.Ala946Thr | Q9BYX4 |
| HOMO SAPIENS | 0 | 253 | 5E+092 | A | 308 | LEU | p.Leu308Pro | P37173 |
| HOMO SAPIENS | 0 | 253 | 5E+092 | A | 315 | THR | p.Thr315Met | P37173 |
| HOMO SAPIENS | 0 | 253 | 5E+092 | A | 328 | HIS | p.His328Gln | P37173 |
| HOMO SAPIENS | 0 | 253 | 5E+092 | A | 336 | TYR | p.Tyr336Asn | P37173 |
| HOMO SAPIENS | 0 | 253 | 5E+092 | A | 348 | ARG | p.Arg348Cys | P37173 |
| HOMO SAPIENS | 0 | 253 | 5E+092 | A | 355 | ALA | p.Ala355Pro | P37173 |
| HOMO SAPIENS | 1 | 253 | 5E+092 | A | 356 | ARG | p.Arg356Gly | P37173 |

| | | | | | | | | |
|---|---|---|---|---|---|---|---|---|
| HOMO SAPIENS | 0 | 253 | 5E+092 | A | 357 | GLY | p.Gly357Trp | P37173 |
| HOMO SAPIENS | 1 | 253 | 5E+092 | A | 403 | ARG | p.Arg403His | P37173 |
| HOMO SAPIENS | 5 | 253 | 5E+092 | A | 449 | SER | p.Ser449Phe | P37173 |
| HOMO SAPIENS | 0 | 253 | 5E+092 | A | 460 | ARG | p.Arg460Cys | P37173 |
| HOMO SAPIENS | 0 | 253 | 5E+092 | A | 460 | ARG | p.Arg460His | P37173 |
| HOMO SAPIENS | 2 | 253 | 5E+092 | A | 526 | GLU | p.Glu526Gln | P37173 |
| HOMO SAPIENS | 5 | 253 | 5E+092 | A | 528 | ARG | p.Arg528Cys | P37173 |
| HOMO SAPIENS | 5 | 253 | 5E+092 | A | 528 | ARG | p.Arg528His | P37173 |
| HOMO SAPIENS | 6 | 158 | 5amc | A | 231 | ARG | p.Arg260Leu | P12821 |
| HOMO SAPIENS | 7 | 158 | 5amc | A | 237 | TYR | p.Tyr266Ter | P12821 |
| HOMO SAPIENS | 6 | 158 | 5amc | A | 325 | GLY | p.Gly354Arg | P12821 |
| HOMO SAPIENS | 0 | 158 | 5amc | A | 467 | ARG | p.Arg496Ter | P12821 |
| HOMO SAPIENS | 4 | 158 | 5amc | A | 479 | ARG | p.Arg508Ter | P12821 |
| HOMO SAPIENS | 0 | 158 | 5amc | A | 563 | ASP | p.Asp592Gly | P12821 |
| HOMO SAPIENS | -1 | 33062 | 1gr3 | A | 545 | GLY | p.Gly545Arg | Q03692 |
| HOMO SAPIENS | 5 | 33062 | 1gr3 | A | 591 | CYS | p.Cys591Arg | Q03692 |
| HOMO SAPIENS | 2 | 33062 | 1gr3 | A | 595 | GLY | p.Gly595Glu | Q03692 |
| HOMO SAPIENS | 0 | 33062 | 1gr3 | A | 597 | TYR | p.Tyr597Cys | Q03692 |
| HOMO SAPIENS | 5 | 33062 | 1gr3 | A | 598 | TYR | p.Tyr598Asp | Q03692 |
| HOMO SAPIENS | 7 | 33062 | 1gr3 | A | 600 | SER | p.Ser600Pro | Q03692 |
| HOMO SAPIENS | 1 | 33062 | 1gr3 | A | 614 | LEU | p.Leu614Pro | Q03692 |
| HOMO SAPIENS | 6 | 33062 | 1gr3 | A | 651 | TRP | p.Trp651Arg | Q03692 |
| HOMO SAPIENS | 7 | 33062 | 1gr3 | A | 671 | SER | p.Ser671Pro | Q03692 |
| HOMO SAPIENS | -1 | 38257 | 2l2i | B | 87 | PRO | p.Pro87Ala | Q13351 |
| HOMO SAPIENS | 2 | 2036 | 1ghq | A | 231 | GLY | p.Gly1224Asp | P01024 |
| HOMO SAPIENS | 5 | 3594 | 4cde | F | 388 | TYR | p.Tyr412Cys | P53634 |
| HOMO SAPIENS | 9 | 3594 | 4cde | F | 405 | TRP | p.Trp429Cys | P53634 |
| HOMO SAPIENS | 0 | 3594 | 4cde | F | 429 | ILE | p.Ile453Val | P53634 |
| HOMO SAPIENS | 7 | 18 | 1jkb | A | 34 | TRP | p.Trp52Ter | P61626 |
| HOMO SAPIENS | 9 | 18 | 1jkb | A | 70 | THR | p.Thr88Asn | P61626 |
| HOMO SAPIENS | 8 | 18 | 1jkb | A | 121 | VAL | p.Val139Ile | P61626 |
| HOMO SAPIENS | 2 | 414 | 3fj6 | A | 136 | ARG | p.Arg135Cys | Q02127 |
| HOMO SAPIENS | 1 | 414 | 3fj6 | A | 153 | GLY | p.Gly152Arg | Q02127 |
| HOMO SAPIENS | 0 | 414 | 3fj6 | A | 200 | ARG | p.Arg199Cys | Q02127 |
| HOMO SAPIENS | 0 | 414 | 3fj6 | A | 203 | GLY | p.Gly202Ala | Q02127 |
| HOMO SAPIENS | 0 | 414 | 3fj6 | A | 245 | ARG | p.Arg244Trp | Q02127 |
| HOMO SAPIENS | 0 | 414 | 3fj6 | A | 342 | ALA | p.Ala341Val | Q02127 |
| HOMO SAPIENS | 1 | 414 | 3fj6 | A | 347 | ARG | p.Arg346Trp | Q02127 |
| HOMO SAPIENS | 6 | 103 | 1u3w | A | 77 | GLY | p.Gly78Ter | P00326 |
| HOMO SAPIENS | -1 | 8369 | 4ux8 | F | 16 | ARG | p.Arg93Trp | P39905 |
| HOMO SAPIENS | 0 | 366 | 2zjc | A | 32 | GLY | p.Arg108Trp | P01375 |
| HOMO SAPIENS | 0 | 1457 | 3v9i | A | 352 | LEU | p.Ser352Leu | P30038 |
| HOMO SAPIENS | 0 | 1457 | 3v9i | A | 470 | VAL | p.Val470Ile | P30038 |
| HOMO SAPIENS | 0 | 1457 | 3v9i | A | 473 | THR | p.Thr473Ala | P30038 |
| HOMO SAPIENS | 1 | 2 | 4wo5 | A | 462 | ARG | p.Arg462Ile | P15056 |
| HOMO SAPIENS | 3 | 2 | 4wo5 | A | 463 | ILE | p.Ile463Ser | P15056 |
| HOMO SAPIENS | 1 | 2 | 4wo5 | A | 466 | GLY | p.Gly466Val | P15056 |
| HOMO SAPIENS | 1 | 2 | 4wo5 | A | 468 | PHE | p.Phe468Ser | P15056 |
| HOMO SAPIENS | 2 | 2 | 4wo5 | A | 469 | GLY | p.Gly469Ala | P15056 |
| HOMO SAPIENS | 2 | 2 | 4wo5 | A | 469 | GLY | p.Gly469Arg | P15056 |
| HOMO SAPIENS | 2 | 2 | 4wo5 | A | 485 | LEU | p.Leu485Phe | P15056 |
| HOMO SAPIENS | 0 | 2 | 4wo5 | A | 501 | GLU | p.Glu501Gly | P15056 |
| HOMO SAPIENS | 0 | 2 | 4wo5 | A | 501 | GLU | p.Glu501Lys | P15056 |
| HOMO SAPIENS | 7 | 2 | 4wo5 | A | 581 | ASN | p.Asn581Asp | P15056 |

| | | | | | | | | |
|---|---|---|---|---|---|---|---|---|
| HOMO SAPIENS | 8 | 2 | 4wo5 | A | 594 | ASP | p.Asp594Gly | P15056 |
| HOMO SAPIENS | 0 | 2 | 4wo5 | A | 597 | LEU | p.Leu597Arg | P15056 |
| HOMO SAPIENS | 0 | 2 | 4wo5 | A | 597 | LEU | p.Leu597Val | P15056 |
| HOMO SAPIENS | 0 | 2 | 4wo5 | A | 600 | VAL | p.Val600Glu | P15056 |
| HOMO SAPIENS | 2 | 2 | 4wo5 | A | 601 | LYS | p.Lys601Glu | P15056 |
| HOMO SAPIENS | 8 | 2 | 4wo5 | A | 638 | ASP | p.Asp638Glu | P15056 |
| HOMO SAPIENS | -1 | 30366 | 5bqd | B | 19 | ALA | p.Ala19Ser | Q99593 |
| HOMO SAPIENS | -1 | 30366 | 5bqd | B | 23 | PRO | p.Pro23His | Q99593 |
| HOMO SAPIENS | 1 | 30366 | 5bqd | B | 49 | GLN | p.Gln49Lys | Q99593 |
| HOMO SAPIENS | 2 | 30366 | 5bqd | B | 54 | ILE | p.Ile54Thr | Q99593 |
| HOMO SAPIENS | 8 | 30366 | 5bqd | B | 80 | GLY | p.Gly80Arg | Q99593 |
| HOMO SAPIENS | 6 | 30366 | 5bqd | B | 106 | ILE | p.Ile106Val | Q99593 |
| HOMO SAPIENS | 7 | 30366 | 5bqd | B | 111 | ASP | p.Asp111Tyr | Q99593 |
| HOMO SAPIENS | -1 | 30366 | 5bqd | B | 237 | ARG | p.Arg237Gln | Q99593 |
| HOMO SAPIENS | -1 | 30366 | 5bqd | B | 237 | ARG | p.Arg237Trp | Q99593 |
| HOMO SAPIENS | 5 | 4789 | 1n4m | A | 567 | SER | p.Ser567Leu | P06400 |
| HOMO SAPIENS | 0 | 4789 | 1n4m | A | 658 | ALA | p.Ala658Asp | P06400 |
| HOMO SAPIENS | 7 | 4789 | 1n4m | A | 661 | ARG | p.Arg661Trp | P06400 |
| HOMO SAPIENS | 6 | 4789 | 1n4m | A | 668 | ARG | p.Arg668His | P06400 |
| HOMO SAPIENS | 8 | 4789 | 1n4m | A | 697 | ASP | p.Asp697Glu | P06400 |
| HOMO SAPIENS | 6 | 4789 | 1n4m | A | 712 | CYS | p.Cys712Arg | P06400 |
| HOMO SAPIENS | 1 | 1798 | 2r5g | A | 101 | HIS | p.His101Gln | O15247 |
| HOMO SAPIENS | 0 | 1798 | 2r5g | A | 160 | PRO | p.Pro160Ala | O15247 |
| HOMO SAPIENS | 2 | 22 | 4x0t | A | 82 | ARG | p.Arg110Ter | P49419 |
| HOMO SAPIENS | 0 | 22 | 4x0t | A | 149 | ALA | p.Ala177Pro | P49419 |
| HOMO SAPIENS | 4 | 22 | 4x0t | A | 164 | THR | p.Thr192Met | P49419 |
| HOMO SAPIENS | 9 | 22 | 4x0t | A | 167 | ASN | p.Asn195Ser | P49419 |
| HOMO SAPIENS | 0 | 22 | 4x0t | A | 175 | TRP | p.Trp203Gly | P49419 |
| HOMO SAPIENS | 0 | 22 | 4x0t | A | 194 | THR | p.Thr222Ala | P49419 |
| HOMO SAPIENS | 0 | 22 | 4x0t | A | 273 | ASN | p.Asn301Ile | P49419 |
| HOMO SAPIENS | 0 | 22 | 4x0t | A | 307 | ARG | p.Arg335Gln | P49419 |
| HOMO SAPIENS | 3 | 22 | 4x0t | A | 313 | SER | p.Ser341Asn | P49419 |
| HOMO SAPIENS | 0 | 22 | 4x0t | A | 329 | ILE | p.Ile357Val | P49419 |
| HOMO SAPIENS | 1 | 22 | 4x0t | A | 380 | TYR | p.Tyr408Ter | P49419 |
| HOMO SAPIENS | 9 | 22 | 4x0t | A | 399 | GLU | p.Glu427Gln | P49419 |
| HOMO SAPIENS | 0 | 22 | 4x0t | A | 406 | TYR | p.Tyr434Cys | P49419 |
| HOMO SAPIENS | 0 | 22 | 4x0t | A | 431 | ILE | p.Ile459Phe | P49419 |
| HOMO SAPIENS | 4 | 22 | 4x0t | A | 477 | GLY | p.Gly505Arg | P49419 |
| HOMO SAPIENS | 0 | 22 | 4x0t | A | 495 | THR | p.Thr523Ala | P49419 |
| HOMO SAPIENS | 8 | 11042 | 1y6b | A | 1145 | PRO | p.Pro1147Ser | P35968 |
| HOMO SAPIENS | -1 | 3867 | 2xij | A | 18 | GLN | p.Gln18Ter | P22033 |
| HOMO SAPIENS | -1 | 3867 | 2xij | A | 31 | ARG | p.Arg31Ter | P22033 |
| HOMO SAPIENS | 2 | 3867 | 2xij | A | 93 | ARG | p.Arg93His | P22033 |
| HOMO SAPIENS | 0 | 3867 | 2xij | A | 94 | GLY | p.Gly94Arg | P22033 |
| HOMO SAPIENS | 2 | 3867 | 2xij | A | 95 | PRO | p.Pro95Leu | P22033 |
| HOMO SAPIENS | 2 | 3867 | 2xij | A | 105 | TRP | p.Trp105Arg | P22033 |
| HOMO SAPIENS | 2 | 3867 | 2xij | A | 108 | ARG | p.Arg108Cys | P22033 |
| HOMO SAPIENS | 3 | 3867 | 2xij | A | 154 | ARG | p.Arg154His | P22033 |
| HOMO SAPIENS | 3 | 3867 | 2xij | A | 156 | ASP | p.Asp156Val | P22033 |
| HOMO SAPIENS | 0 | 3867 | 2xij | A | 189 | ASN | p.Asn189Ile | P22033 |
| HOMO SAPIENS | 0 | 3867 | 2xij | A | 191 | ALA | p.Ala191Glu | P22033 |
| HOMO SAPIENS | 0 | 3867 | 2xij | A | 203 | GLY | p.Gly203Arg | P22033 |
| HOMO SAPIENS | 2 | 3867 | 2xij | A | 215 | GLY | p.Gly215Ser | P22033 |
| HOMO SAPIENS | 0 | 3867 | 2xij | A | 219 | ASN | p.Asn219Tyr | P22033 |

| | | | | | | | | |
|---|---|---|---|---|---|---|---|---|
| HOMO SAPIENS | 3 | 3867 | 2xij | A | 228 | ARG | p.Arg228Ter | P22033 |
| HOMO SAPIENS | 1 | 3867 | 2xij | A | 297 | THR | p.Thr297Ile | P22033 |
| HOMO SAPIENS | 0 | 3867 | 2xij | A | 326 | ARG | p.Arg326Lys | P22033 |
| HOMO SAPIENS | 3 | 3867 | 2xij | A | 369 | ARG | p.Arg369His | P22033 |
| HOMO SAPIENS | 3 | 3867 | 2xij | A | 369 | ARG | p.Arg369Cys | P22033 |
| HOMO SAPIENS | 0 | 3867 | 2xij | A | 370 | THR | p.Thr370Pro | P22033 |
| HOMO SAPIENS | 0 | 3867 | 2xij | A | 377 | ALA | p.Ala377Glu | P22033 |
| HOMO SAPIENS | 2 | 3867 | 2xij | A | 426 | GLY | p.Gly426Ala | P22033 |
| HOMO SAPIENS | 2 | 3867 | 2xij | A | 427 | GLY | p.Gly427Asp | P22033 |
| HOMO SAPIENS | 1 | 3867 | 2xij | A | 499 | THR | p.Ala499Thr | P22033 |
| HOMO SAPIENS | 2 | 3867 | 2xij | A | 532 | ARG | p.Arg532His | P22033 |
| HOMO SAPIENS | 3 | 3867 | 2xij | A | 555 | ALA | p.Ala555Thr | P22033 |
| HOMO SAPIENS | 0 | 3867 | 2xij | A | 573 | PHE | p.Phe573Ser | P22033 |
| HOMO SAPIENS | 3 | 3867 | 2xij | A | 623 | GLY | p.Gly623Arg | P22033 |
| HOMO SAPIENS | 5 | 3867 | 2xij | A | 671 | VAL | p.Ile671Val | P22033 |
| HOMO SAPIENS | 4 | 3867 | 2xij | A | 694 | ARG | p.Arg694Trp | P22033 |
| HOMO SAPIENS | 1 | 3867 | 2xij | A | 700 | MET | p.Met700Lys | P22033 |
| HOMO SAPIENS | 8 | 3867 | 2xij | A | 703 | GLY | p.Gly703Arg | P22033 |
| HOMO SAPIENS | 0 | 3867 | 2xij | A | 717 | GLY | p.Gly717Val | P22033 |
| HOMO SAPIENS | 1 | 3867 | 2xij | A | 727 | ARG | p.Arg727Ter | P22033 |
| HOMO SAPIENS | -1 | 52 | 3k91 | B | 72 | GLY | p.Gly73Ser | P00441 |
| HOMO SAPIENS | 6 | 52 | 3k91 | B | 96 | ASP | p.Asp97Asn | P00441 |
| HOMO SAPIENS | 5 | 52 | 3k91 | B | 112 | ILE | p.Ile113Thr | P00441 |
| HOMO SAPIENS | -1 | 52 | 3k91 | B | 134 | SER | p.Ser135Asn | P00441 |
| HOMO SAPIENS | 0 | 3950 | 4fom | A | 296 | ASN | p.Asn296His | Q9NQS3 |
| HOMO SAPIENS | 1 | 10 | 1de4 | A | 31 | VAL | p.Val53Met | Q30201 |
| HOMO SAPIENS | 0 | 10 | 1de4 | A | 37 | VAL | p.Val59Met | Q30201 |
| HOMO SAPIENS | 0 | 10 | 1de4 | A | 71 | GLY | p.Gly93Arg | Q30201 |
| HOMO SAPIENS | 0 | 10 | 1de4 | A | 83 | ILE | p.Ile105Thr | Q30201 |
| HOMO SAPIENS | 0 | 10 | 1de4 | A | 105 | GLN | p.Gln127His | Q30201 |
| HOMO SAPIENS | 1 | 10 | 1de4 | A | 146 | GLU | p.Glu168Ter | Q30201 |
| HOMO SAPIENS | 2 | 10 | 1de4 | A | 261 | GLN | p.Gln283Pro | Q30201 |
| HOMO SAPIENS | 0 | 8038 | 4x7n | A | 586 | ILE | p.Ile586Leu | Q9NZJ5 |
| HOMO SAPIENS | 0 | 8038 | 4x7n | A | 588 | ARG | p.Arg588Gln | Q9NZJ5 |
| HOMO SAPIENS | 0 | 35175 | 5ugw | A | 1062 | ALA | p.Ala1062Thr | O14746 |
| HOMO SAPIENS | 2 | 35175 | 5ugw | A | 1090 | VAL | p.Val1090Met | O14746 |
| HOMO SAPIENS | 0 | 1722 | 4z3k | C | 147 | ARG | p.Arg150Gly | P35270 |
| HOMO SAPIENS | 0 | 1722 | 4z3k | C | 160 | PRO | p.Pro163Leu | P35270 |
| HOMO SAPIENS | 0 | 1722 | 4z3k | C | 207 | ARG | p.Arg210Trp | P35270 |
| HOMO SAPIENS | 0 | 1722 | 4z3k | C | 216 | ARG | p.Arg219Ter | P35270 |
| HOMO SAPIENS | 0 | 1722 | 4z3k | C | 248 | LYS | p.Lys251Ter | P35270 |
| HOMO SAPIENS | 1 | 2832 | 4x8v | L | 115 | HIS | p.His175Gln | P08709 |
| HOMO SAPIENS | 0 | 825 | 1emh | A | 242 | PHE | p.Phe251Ser | P13051 |
| HOMO SAPIENS | 0 | 122 | 5uwk | A | 109 | ARG | p.Arg109Ter | P45452 |
| HOMO SAPIENS | 1 | 122 | 5uwk | A | 170 | LYS | p.Lys170Thr | P45452 |
| HOMO SAPIENS | 9 | 122 | 5uwk | A | 232 | HIS | p.His232Asn | P45452 |
| HOMO SAPIENS | 5 | 122 | 5uwk | A | 257 | ASP | p.Asp257Val | P45452 |
| HOMO SAPIENS | 9 | 1074 | 3p5i | A | 264 | TRP | p.Trp264Arg | Q9UJ71 |
| HOMO SAPIENS | 0 | 146 | 3r3g | A | 14C | GLU | p.Glu352Lys | P00734 |
| HOMO SAPIENS | 3 | 9522 | 1wyq | A | 83 | LEU | p.Leu253Pro | O15020 |
| HOMO SAPIENS | 8 | 67 | 2vgb | A | 130 | SER | p.Ser130Tyr | P30613 |
| HOMO SAPIENS | 8 | 67 | 2vgb | A | 163 | ARG | p.Arg163Cys | P30613 |
| HOMO SAPIENS | 4 | 67 | 2vgb | A | 241 | GLU | p.Glu241Ter | P30613 |
| HOMO SAPIENS | 9 | 67 | 2vgb | A | 384 | THR | p.Thr384Met | P30613 |

| | | | | | | | | |
|---|---|---|---|---|---|---|---|---|
| HOMO SAPIENS | 0 | 67 | 2vgb | A | 421 | GLN | p.Gln421Lys | P30613 |
| HOMO SAPIENS | 6 | 67 | 2vgb | A | 479 | ARG | p.Arg479His | P30613 |
| HOMO SAPIENS | 8 | 67 | 2vgb | A | 510 | ARG | p.Arg510Gln | P30613 |
| HOMO SAPIENS | 4 | 67 | 2vgb | A | 569 | ARG | p.Arg569Trp | P30613 |
| HOMO SAPIENS | 0 | 9088 | 5v44 | A | 232 | ASN | p.Asn232Lys | Q9NZJ4 |
| HOMO SAPIENS | 4 | 16 | 3lbn | A | 12 | GLY | p.Gly12Val | P01112 |
| HOMO SAPIENS | 4 | 16 | 3lbn | A | 12 | GLY | p.Gly12Ser | P01112 |
| HOMO SAPIENS | 4 | 16 | 3lbn | A | 13 | GLY | p.Gly13Cys | P01112 |
| HOMO SAPIENS | 4 | 16 | 3lbn | A | 13 | GLY | p.Gly13Asp | P01112 |
| HOMO SAPIENS | 5 | 16 | 3lbn | A | 22 | GLN | p.Gln22Lys | P01112 |
| HOMO SAPIENS | 5 | 16 | 3lbn | A | 58 | THR | p.Thr58Ile | P01112 |
| HOMO SAPIENS | 3 | 16 | 3lbn | A | 61 | GLN | p.Gln61Lys | P01112 |
| HOMO SAPIENS | 1 | 16 | 3lbn | A | 63 | GLU | p.Glu63Lys | P01112 |
| HOMO SAPIENS | 9 | 16 | 3lbn | A | 117 | LYS | p.Lys117Arg | P01112 |
| HOMO SAPIENS | 6 | 16 | 3lbn | A | 123 | ARG | p.Arg123Cys | P01112 |
| HOMO SAPIENS | 1 | 16 | 3lbn | A | 138 | GLY | p.Gly138Ser | P01112 |
| HOMO SAPIENS | 9 | 16 | 3lbn | A | 146 | ALA | p.Ala146Thr | P01112 |
| HOMO SAPIENS | 9 | 16 | 3lbn | A | 146 | ALA | p.Ala146Val | P01112 |
| HOMO SAPIENS | 1 | 359 | 5nvy | C | 63 | LEU | p.Leu63Pro | P40337 |
| HOMO SAPIENS | 1 | 359 | 5nvy | C | 64 | ARG | p.Arg64Pro | P40337 |
| HOMO SAPIENS | 1 | 359 | 5nvy | C | 70 | GLU | p.Glu70Lys | P40337 |
| HOMO SAPIENS | 0 | 359 | 5nvy | C | 84 | VAL | p.Val84Leu | P40337 |
| HOMO SAPIENS | 1 | 359 | 5nvy | C | 88 | TRP | p.Trp88Ser | P40337 |
| HOMO SAPIENS | 1 | 359 | 5nvy | C | 93 | GLY | p.Gly93Cys | P40337 |
| HOMO SAPIENS | 1 | 359 | 5nvy | C | 98 | TYR | p.Tyr98His | P40337 |
| HOMO SAPIENS | 1 | 359 | 5nvy | C | 111 | SER | p.Ser111Arg | P40337 |
| HOMO SAPIENS | 1 | 359 | 5nvy | C | 112 | TYR | p.Tyr112His | P40337 |
| HOMO SAPIENS | 1 | 359 | 5nvy | C | 126 | ASP | p.Asp126Tyr | P40337 |
| HOMO SAPIENS | 1 | 359 | 5nvy | C | 130 | VAL | p.Val130Leu | P40337 |
| HOMO SAPIENS | 1 | 359 | 5nvy | C | 135 | LEU | p.Leu135Phe | P40337 |
| HOMO SAPIENS | 9 | 359 | 5nvy | C | 161 | ARG | p.Arg161Gly | P40337 |
| HOMO SAPIENS | 7 | 359 | 5nvy | C | 162 | CYS | p.Cys162Phe | P40337 |
| HOMO SAPIENS | 4 | 359 | 5nvy | C | 163 | LEU | p.Leu163Pro | P40337 |
| HOMO SAPIENS | 3 | 359 | 5nvy | C | 164 | GLN | p.Gln164Arg | P40337 |
| HOMO SAPIENS | 9 | 359 | 5nvy | C | 166 | VAL | p.Val166Phe | P40337 |
| HOMO SAPIENS | 7 | 359 | 5nvy | C | 167 | ARG | p.Arg167Gly | P40337 |
| HOMO SAPIENS | 7 | 359 | 5nvy | C | 167 | ARG | p.Arg167Gln | P40337 |
| HOMO SAPIENS | 1 | 359 | 5nvy | C | 183 | SER | p.Ser183Ter | P40337 |
| HOMO SAPIENS | 2 | 359 | 5nvy | C | 188 | LEU | p.Leu188Val | P40337 |
| HOMO SAPIENS | 1 | 359 | 5nvy | C | 191 | HIS | p.His191Asp | P40337 |
| HOMO SAPIENS | 1 | 359 | 5nvy | C | 192 | PRO | p.Pro192Ser | P40337 |
| HOMO SAPIENS | 1 | 359 | 5nvy | C | 200 | ARG | p.Arg200Trp | P40337 |
| HOMO SAPIENS | -1 | 359 | 5nvy | C | 211 | MET | p.Met211Leu | P40337 |
| HOMO SAPIENS | 0 | 14099 | 4btm | A | 282 | ARG | p.Arg258Gln | Q5TCY1 |
| HOMO SAPIENS | 0 | 696 | 5h1b | A | 150 | ARG | p.Arg150Gln | Q06609 |
| HOMO SAPIENS | -1 | 208 | 2bxl | A | 1 | ASP | p.Asp25Val | P02768 |
| HOMO SAPIENS | 0 | 208 | 2bxl | A | 269 | ASP | p.Asp293Gly | P02768 |
| HOMO SAPIENS | 0 | 208 | 2bxl | A | 570 | GLU | p.Glu594Lys | P02768 |
| HOMO SAPIENS | 0 | 208 | 2bxl | A | 573 | LYS | p.Lys597Glu | P02768 |
| HOMO SAPIENS | -1 | 12813 | 2uzx | B | 35 | MET | p.Met35Leu | P08581 |
| HOMO SAPIENS | 1 | 12813 | 2uzx | B | 156 | SER | p.Ser156Leu | P08581 |
| HOMO SAPIENS | 0 | 12813 | 2uzx | B | 168 | GLU | p.Glu168Asp | P08581 |
| HOMO SAPIENS | 2 | 12813 | 2uzx | B | 211 | LEU | p.Leu211Trp | P08581 |
| HOMO SAPIENS | 2 | 12813 | 2uzx | B | 218 | ARG | p.Arg218Ser | P08581 |

| | | | | | | | | |
|---|---|---|---|---|---|---|---|---|
| HOMO SAPIENS | 1 | 12813 | 2uzx | B | 238 | LEU | p.Leu238Ser | P08581 |
| HOMO SAPIENS | 4 | 12813 | 2uzx | B | 320 | ALA | p.Ala320Val | P08581 |
| HOMO SAPIENS | 1 | 12813 | 2uzx | B | 375 | ASN | p.Asn375Ser | P08581 |
| HOMO SAPIENS | 0 | 12813 | 2uzx | B | 575 | LEU | p.Leu575Ile | P08581 |
| HOMO SAPIENS | 1 | 10780 | 4ejx | A | 348 | ARG | p.Arg367Cys | P00450 |
| HOMO SAPIENS | 2 | 10780 | 4ejx | A | 525 | ASP | p.Asp544Glu | P00450 |
| HOMO SAPIENS | 1 | 1407 | 5jn5 | A | 88 | ILE | p.Ile88Val | P36871 |
| HOMO SAPIENS | 7 | 1407 | 5jn5 | A | 115 | THR | p.Thr115Ala | P36871 |
| HOMO SAPIENS | 3 | 1407 | 5jn5 | A | 121 | GLY | p.Gly121Arg | P36871 |
| HOMO SAPIENS | 2 | 1407 | 5jn5 | A | 221 | ARG | p.Arg221Cys | P36871 |
| HOMO SAPIENS | 6 | 1407 | 5jn5 | A | 263 | TYR | p.Asp263Tyr | P36871 |
| HOMO SAPIENS | 3 | 1407 | 5jn5 | A | 420 | TYR | p.Tyr420His | P36871 |
| HOMO SAPIENS | 0 | 1407 | 5jn5 | A | 501 | VAL | p.Val501Ile | P36871 |
| HOMO SAPIENS | -1 | 1878 | 1yow | A | 255 | ARG | p.Arg255Leu | Q13285 |
| HOMO SAPIENS | 6 | 1878 | 1yow | A | 293 | ASP | p.Asp293Asn | Q13285 |
| HOMO SAPIENS | 1 | 1878 | 1yow | A | 437 | LEU | p.Leu437Gln | Q13285 |
| HOMO SAPIENS | 0 | 3071 | 2wpm | E | 145 | ARG | p.Arg191His | P00740 |
| HOMO SAPIENS | 0 | 3071 | 2wpm | E | 145 | ARG | p.Arg191Cys | P00740 |
| HOMO SAPIENS | 3 | 589 | 3ggc | B | 15 | GLY | p.Gly16Ser | P00492 |
| HOMO SAPIENS | 4 | 589 | 3ggc | B | 40 | LEU | p.Leu41Pro | P00492 |
| HOMO SAPIENS | 2 | 589 | 3ggc | B | 50 | ARG | p.Arg51Gly | P00492 |
| HOMO SAPIENS | 5 | 589 | 3ggc | B | 51 | ASP | p.Asp52Gly | P00492 |
| HOMO SAPIENS | 0 | 589 | 3ggc | B | 57 | GLY | p.Gly58Arg | P00492 |
| HOMO SAPIENS | 6 | 589 | 3ggc | B | 69 | GLY | p.Gly70Glu | P00492 |
| HOMO SAPIENS | 1 | 589 | 3ggc | B | 73 | PHE | p.Phe74Leu | P00492 |
| HOMO SAPIENS | 0 | 589 | 3ggc | B | 77 | LEU | p.Leu78Val | P00492 |
| HOMO SAPIENS | 3 | 589 | 3ggc | B | 79 | ASP | p.Asp80Val | P00492 |
| HOMO SAPIENS | 0 | 589 | 3ggc | B | 103 | SER | p.Ser104Arg | P00492 |
| HOMO SAPIENS | 0 | 589 | 3ggc | B | 109 | SER | p.Ser110Leu | P00492 |
| HOMO SAPIENS | 1 | 589 | 3ggc | B | 129 | VAL | p.Val130Asp | P00492 |
| HOMO SAPIENS | 4 | 589 | 3ggc | B | 131 | ILE | p.Ile132Met | P00492 |
| HOMO SAPIENS | 7 | 589 | 3ggc | B | 142 | MET | p.Met143Lys | P00492 |
| HOMO SAPIENS | 0 | 589 | 3ggc | B | 160 | ALA | p.Ala161Ser | P00492 |
| HOMO SAPIENS | 6 | 589 | 3ggc | B | 167 | THR | p.Thr168Ile | P00492 |
| HOMO SAPIENS | 0 | 589 | 3ggc | B | 169 | ARG | p.Arg170Ter | P00492 |
| HOMO SAPIENS | 5 | 589 | 3ggc | B | 175 | PRO | p.Pro176Leu | P00492 |
| HOMO SAPIENS | 8 | 589 | 3ggc | B | 193 | ASP | p.Asp194Glu | P00492 |
| HOMO SAPIENS | 8 | 589 | 3ggc | B | 193 | ASP | p.Asp194Asn | P00492 |
| HOMO SAPIENS | 8 | 589 | 3ggc | B | 198 | PHE | p.Phe199Val | P00492 |
| HOMO SAPIENS | 8 | 589 | 3ggc | B | 200 | ASP | p.Asp201Gly | P00492 |
| HOMO SAPIENS | 0 | 4269 | 2pla | B | 174 | GLU | p.Glu174Lys | Q8N335 |
| HOMO SAPIENS | 1 | 4269 | 2pla | B | 220 | ARG | p.Arg220His | Q8N335 |
| HOMO SAPIENS | 4 | 4269 | 2pla | B | 280 | ALA | p.Ala280Val | Q8N335 |
| HOMO SAPIENS | 0 | 2712 | 1ii4 | E | 252 | TRP | p.Ser252Leu | P21802 |
| HOMO SAPIENS | 0 | 2712 | 1ii4 | E | 253 | PRO | p.Pro253Arg | P21802 |
| HOMO SAPIENS | 5 | 2712 | 1ii4 | E | 267 | SER | p.Ser267Pro | P21802 |
| HOMO SAPIENS | 0 | 2712 | 1ii4 | E | 278 | CYS | p.Cys278Phe | P21802 |
| HOMO SAPIENS | 0 | 2712 | 1ii4 | E | 289 | GLN | p.Gln289Pro | P21802 |
| HOMO SAPIENS | 0 | 2712 | 1ii4 | E | 290 | TRP | p.Trp290Cys | P21802 |
| HOMO SAPIENS | 0 | 2712 | 1ii4 | E | 290 | TRP | p.Trp290Arg | P21802 |
| HOMO SAPIENS | 0 | 2712 | 1ii4 | E | 292 | LYS | p.Lys292Glu | P21802 |
| HOMO SAPIENS | 0 | 2712 | 1ii4 | E | 315 | ALA | p.Ala315Ser | P21802 |
| HOMO SAPIENS | 0 | 2712 | 1ii4 | E | 321 | ASP | p.Asp321Ala | P21802 |
| HOMO SAPIENS | 1 | 2712 | 1ii4 | E | 328 | TYR | p.Tyr328Cys | P21802 |

| | | | | | | | | |
|---|---|---|---|---|---|---|---|---|
| HOMO SAPIENS | 9 | 2712 | 1ii4 | E | 340 | TYR | p.Tyr340His | P21802 |
| HOMO SAPIENS | 1 | 2712 | 1ii4 | E | 341 | THR | p.Thr341Pro | P21802 |
| HOMO SAPIENS | 0 | 2712 | 1ii4 | E | 342 | CYS | p.Cys342Arg | P21802 |
| HOMO SAPIENS | 0 | 2712 | 1ii4 | E | 342 | CYS | p.Cys342Trp | P21802 |
| HOMO SAPIENS | 0 | 2712 | 1ii4 | E | 344 | ALA | p.Ala344Gly | P21802 |
| HOMO SAPIENS | 0 | 2712 | 1ii4 | E | 347 | SER | p.Ser347Cys | P21802 |
| HOMO SAPIENS | 0 | 2712 | 1ii4 | E | 351 | SER | p.Ser351Cys | P21802 |
| HOMO SAPIENS | 1 | 2712 | 1ii4 | E | 354 | SER | p.Ser354Cys | P21802 |
| HOMO SAPIENS | 2 | 53 | 4pj1 | H | 5 | ASP | p.Asp29Gly | P10809 |
| HOMO SAPIENS | 0 | 53 | 4pj1 | H | 74 | VAL | p.Val98Ile | P10809 |
| HOMO SAPIENS | 0 | 29968 | 2fdb | M | 71 | LYS | p.Lys89Glu | P55075 |
| HOMO SAPIENS | 2 | 29968 | 2fdb | M | 98 | ARG | p.Arg116Gly | P55075 |
| HOMO SAPIENS | 2 | 29968 | 2fdb | M | 122 | GLY | p.Gly140Ser | P55075 |
| HOMO SAPIENS | 8 | 5061 | 5f85 | B | 60 | GLY | p.Gly106Ser | P00740 |
| HOMO SAPIENS | 0 | 35944 | 1h6h | A | 85 | THR | p.Thr85Asn | Q15080 |
| HOMO SAPIENS | 8 | 35944 | 1h6h | A | 105 | ARG | p.Arg105Gln | Q15080 |
| HOMO SAPIENS | 7 | 12016 | 1hj7 | A | 295 | ASN | p.Asn316Ser | P01130 |
| HOMO SAPIENS | 5 | 12016 | 1hj7 | A | 302 | GLY | p.Gly323Ser | P01130 |
| HOMO SAPIENS | 6 | 12016 | 1hj7 | A | 319 | CYS | p.Cys340Trp | P01130 |
| HOMO SAPIENS | 7 | 12016 | 1hj7 | A | 329 | ARG | p.Arg350Ter | P01130 |
| HOMO SAPIENS | 5 | 12016 | 1hj7 | A | 331 | CYS | p.Cys352Ter | P01130 |
| HOMO SAPIENS | 2 | 12016 | 1hj7 | A | 345 | GLN | p.Gln366Arg | P01130 |
| HOMO SAPIENS | 2 | 1481 | 2i07 | A | 239 | TYR | p.Tyr261Ter | P01024 |
| HOMO SAPIENS | -1 | 1481 | 2i07 | A | 292 | LEU | p.Pro314Leu | P01024 |
| HOMO SAPIENS | 2 | 1481 | 2i07 | A | 326 | ILE | p.Ile348Val | P01024 |
| HOMO SAPIENS | 2 | 1481 | 2i07 | A | 413 | GLU | p.Glu435Lys | P01024 |
| HOMO SAPIENS | 4 | 1481 | 2i07 | A | 447 | GLU | p.Glu469Asp | P01024 |
| HOMO SAPIENS | 2 | 1481 | 2i07 | A | 597 | VAL | p.Val619Met | P01024 |
| HOMO SAPIENS | 5 | 1481 | 2i07 | A | 603 | ILE | p.Ile625Phe | P01024 |
| HOMO SAPIENS | 0 | 1481 | 2i07 | A | 611 | LYS | p.Lys633Arg | P01024 |
| HOMO SAPIENS | 0 | 11123 | 3c5n | A | 315 | GLY | p.Gly368Trp | O00294 |
| HOMO SAPIENS | 8 | 11123 | 3c5n | A | 329 | PHE | p.Phe382Ser | O00294 |
| HOMO SAPIENS | 5 | 11123 | 3c5n | A | 429 | ARG | p.Arg482Trp | O00294 |
| HOMO SAPIENS | 8 | 11123 | 3c5n | A | 438 | PHE | p.Phe491Leu | O00294 |
| HOMO SAPIENS | 3 | 4398 | 3fx0 | B | 1288 | ALA | p.Ala288Gly | Q9Y6K9 |
| HOMO SAPIENS | 7 | 4398 | 3fx0 | B | 1315 | GLU | p.Glu315Ala | Q9Y6K9 |
| HOMO SAPIENS | 8 | 4398 | 3fx0 | B | 1319 | ARG | p.Arg319Gln | Q9Y6K9 |
| HOMO SAPIENS | 6 | 3989 | 3qd6 | S | 83 | CYS | p.Cys83Arg | P25942 |
| HOMO SAPIENS | 1 | 3989 | 3qd6 | S | 124 | SER | p.Ser124Leu | P25942 |
| HOMO SAPIENS | 2 | 24003 | 2m0v | A | 225 | GLU | p.Glu225Lys | O14745 |
| HOMO SAPIENS | 2 | 46 | 4aqd | A | 70 | ASP | p.Asp98Gly | P06276 |
| HOMO SAPIENS | 4 | 46 | 4aqd | A | 115 | GLY | p.Gly143Asp | P06276 |
| HOMO SAPIENS | 3 | 46 | 4aqd | A | 184 | ALA | p.Ala212Val | P06276 |
| HOMO SAPIENS | 0 | 46 | 4aqd | A | 225 | GLY | p.Gly253Ter | P06276 |
| HOMO SAPIENS | 2 | 46 | 4aqd | A | 243 | THR | p.Thr271Met | P06276 |
| HOMO SAPIENS | 2 | 46 | 4aqd | A | 255 | GLU | p.Glu283Asp | P06276 |
| HOMO SAPIENS | 5 | 46 | 4aqd | A | 267 | LYS | p.Lys295Arg | P06276 |
| HOMO SAPIENS | 3 | 46 | 4aqd | A | 271 | GLU | p.Glu299Ter | P06276 |
| HOMO SAPIENS | 2 | 46 | 4aqd | A | 307 | LEU | p.Leu335Pro | P06276 |
| HOMO SAPIENS | 1 | 46 | 4aqd | A | 330 | LEU | p.Leu358Ile | P06276 |
| HOMO SAPIENS | 0 | 46 | 4aqd | A | 497 | GLU | p.Glu525Val | P06276 |
| HOMO SAPIENS | 3 | 786 | 1ols | B | 51 | GLY | p.Gly101Asp | P21953 |
| HOMO SAPIENS | 2 | 786 | 1ols | B | 87 | ALA | p.Ala137Val | P21953 |
| HOMO SAPIENS | 0 | 786 | 1ols | B | 110 | ILE | p.Ile160Ser | P21953 |

| | | | | | | | | |
|---|---|---|---|---|---|---|---|---|
| HOMO SAPIENS | 4 | 786 | 1ols | B | 120 | ARG | p.Arg170Cys | P21953 |
| HOMO SAPIENS | 2 | 786 | 1ols | B | 126 | ASN | p.Asn176Tyr | P21953 |
| HOMO SAPIENS | 0 | 786 | 1ols | B | 133 | ARG | p.Arg183Gly | P21953 |
| HOMO SAPIENS | 0 | 786 | 1ols | B | 133 | ARG | p.Arg183Pro | P21953 |
| HOMO SAPIENS | 0 | 786 | 1ols | B | 135 | PRO | p.Pro185Leu | P21953 |
| HOMO SAPIENS | 3 | 786 | 1ols | B | 156 | HIS | p.His206Tyr | P21953 |
| HOMO SAPIENS | 0 | 786 | 1ols | B | 228 | GLY | p.Gly278Ser | P21953 |
| HOMO SAPIENS | 1 | 786 | 1ols | B | 235 | ARG | p.Arg285Ter | P21953 |
| HOMO SAPIENS | 6 | 786 | 1ols | B | 274 | ARG | p.Arg324Ter | P21953 |
| HOMO SAPIENS | 1 | 831 | 4zt1 | B | 48 | VAL | p.Val202Ile | P12830 |
| HOMO SAPIENS | 6 | 831 | 4zt1 | B | 70 | ARG | p.Arg224Cys | P12830 |
| HOMO SAPIENS | 5 | 831 | 4zt1 | B | 79 | HIS | p.His233Gln | P12830 |
| HOMO SAPIENS | 4 | 831 | 4zt1 | B | 120 | GLY | p.Gly274Ser | P12830 |
| HOMO SAPIENS | 7 | 831 | 4zt1 | B | 187 | TYR | p.Tyr341Ter | P12830 |
| HOMO SAPIENS | 0 | 8497 | 3sla | A | 287 | ASN | p.Asn287Ser | P35222 |
| HOMO SAPIENS | 7 | 23986 | 2ke1 | A | 301 | VAL | p.Val301Met | O43918 |
| HOMO SAPIENS | 0 | 4967 | 1rj7 | B | 334 | ARG | p.Arg334His | Q92838 |
| HOMO SAPIENS | 0 | 4967 | 1rj7 | B | 338 | THR | p.Thr338Met | Q92838 |
| HOMO SAPIENS | 3 | 4967 | 1rj7 | B | 349 | ALA | p.Ala349Thr | Q92838 |
| HOMO SAPIENS | 2 | 4967 | 1rj7 | B | 358 | GLN | p.Gln358Glu | Q92838 |
| HOMO SAPIENS | 1 | 4967 | 1rj7 | B | 375 | LYS | p.Lys375Glu | Q92838 |
| HOMO SAPIENS | 9 | 7623 | 1d4t | A | 32 | ARG | p.Arg32Thr | O60880 |
| HOMO SAPIENS | 6 | 7623 | 1d4t | A | 55 | ARG | p.Arg55Leu | O60880 |
| HOMO SAPIENS | 0 | 7623 | 1d4t | A | 68 | THR | p.Thr68Ile | O60880 |
| HOMO SAPIENS | 6 | 705 | 3idh | A | 59 | PRO | p.Pro59Ser | P35557 |
| HOMO SAPIENS | 5 | 705 | 3idh | A | 124 | ASP | p.Asp124Asn | P35557 |
| HOMO SAPIENS | 4 | 705 | 3idh | A | 131 | SER | p.Ser131Pro | P35557 |
| HOMO SAPIENS | 0 | 705 | 3idh | A | 147 | GLY | p.Gly147Asp | P35557 |
| HOMO SAPIENS | 5 | 705 | 3idh | A | 186 | ARG | p.Arg186Ter | P35557 |
| HOMO SAPIENS | 0 | 705 | 3idh | A | 202 | MET | p.Met202Thr | P35557 |
| HOMO SAPIENS | 2 | 705 | 3idh | A | 210 | MET | p.Met210Lys | P35557 |
| HOMO SAPIENS | 1 | 705 | 3idh | A | 215 | TYR | p.Tyr215Ter | P35557 |
| HOMO SAPIENS | 0 | 705 | 3idh | A | 226 | VAL | p.Val226Met | P35557 |
| HOMO SAPIENS | 9 | 705 | 3idh | A | 228 | THR | p.Thr228Met | P35557 |
| HOMO SAPIENS | 9 | 705 | 3idh | A | 256 | GLU | p.Glu256Lys | P35557 |
| HOMO SAPIENS | 2 | 705 | 3idh | A | 261 | GLY | p.Gly261Arg | P35557 |
| HOMO SAPIENS | 6 | 705 | 3idh | A | 279 | GLU | p.Glu279Gln | P35557 |
| HOMO SAPIENS | 7 | 705 | 3idh | A | 299 | GLY | p.Gly299Arg | P35557 |
| HOMO SAPIENS | 2 | 705 | 3idh | A | 318 | GLY | p.Gly318Trp | P35557 |
| HOMO SAPIENS | 4 | 705 | 3idh | A | 378 | ALA | p.Ala378Thr | P35557 |
| HOMO SAPIENS | 0 | 705 | 3idh | A | 387 | ALA | p.Ala387Glu | P35557 |
| HOMO SAPIENS | 5 | 705 | 3idh | A | 414 | LYS | p.Lys414Glu | P35557 |
| HOMO SAPIENS | 0 | 705 | 3idh | A | 455 | VAL | p.Val455Met | P35557 |
| HOMO SAPIENS | -1 | 705 | 3idh | A | 462 | MET | p.Met462Ile | P35557 |
| HOMO SAPIENS | 1 | 509 | 4nef | A | 22 | LEU | p.Leu22Val | P41181 |
| HOMO SAPIENS | 4 | 509 | 4nef | A | 57 | GLN | p.Gln57Pro | P41181 |
| HOMO SAPIENS | 8 | 509 | 4nef | A | 64 | GLY | p.Gly64Arg | P41181 |
| HOMO SAPIENS | 7 | 509 | 4nef | A | 68 | ASN | p.Asn68Ser | P41181 |
| HOMO SAPIENS | 1 | 509 | 4nef | A | 100 | GLY | p.Gly100Val | P41181 |
| HOMO SAPIENS | 0 | 509 | 4nef | A | 125 | THR | p.Thr125Met | P41181 |
| HOMO SAPIENS | 0 | 509 | 4nef | A | 126 | THR | p.Thr126Met | P41181 |
| HOMO SAPIENS | 4 | 509 | 4nef | A | 146 | PHE | p.Phe146Leu | P41181 |
| HOMO SAPIENS | 0 | 509 | 4nef | A | 175 | GLY | p.Gly175Arg | P41181 |
| HOMO SAPIENS | 5 | 509 | 4nef | A | 181 | CYS | p.Cys181Trp | P41181 |

| | | | | | | | | | |
|---|---|---|---|---|---|---|---|---|---|
| HOMO SAPIENS | 9 | 509 | 4nef | A | | 187 | ARG | p.Arg187Cys | P41181 |
| HOMO SAPIENS | 0 | 509 | 4nef | A | | 190 | ALA | p.Ala190Thr | P41181 |
| HOMO SAPIENS | 0 | 509 | 4nef | A | | 216 | SER | p.Ser216Pro | P41181 |
| HOMO SAPIENS | 4 | 2242 | 1z70 | X | | 1155 | SER | p.Ser155Pro | Q8NBK3 |
| HOMO SAPIENS | 3 | 2242 | 1z70 | X | | 1218 | CYS | p.Cys218Tyr | Q8NBK3 |
| HOMO SAPIENS | 0 | 2242 | 1z70 | X | | 1279 | ALA | p.Ala279Val | Q8NBK3 |
| HOMO SAPIENS | -1 | 2242 | 1z70 | X | | 1336 | OCS | p.Cys336Arg | Q8NBK3 |
| HOMO SAPIENS | 4 | 2242 | 1z70 | X | | 1345 | ARG | p.Arg345Cys | Q8NBK3 |
| HOMO SAPIENS | 2 | 2242 | 1z70 | X | | 1348 | ALA | p.Ala348Pro | Q8NBK3 |
| HOMO SAPIENS | 5 | 2242 | 1z70 | X | | 1349 | ARG | p.Arg349Gln | Q8NBK3 |
| HOMO SAPIENS | 5 | 2242 | 1z70 | X | | 1349 | ARG | p.Arg349Trp | Q8NBK3 |
| HOMO SAPIENS | 3 | 2242 | 1z70 | X | | 1359 | SER | p.Ser359Ter | Q8NBK3 |
| HOMO SAPIENS | 5 | 8269 | 5wrr | B | | 136 | ARG | p.Arg136Ter | Q96MK3 |
| HOMO SAPIENS | 1 | 8269 | 5wrr | B | | 332 | LYS | p.Asn332Lys | Q96MK3 |
| HOMO SAPIENS | 3 | 8269 | 5wrr | B | | 478 | ARG | p.Arg478Ter | Q96MK3 |
| HOMO SAPIENS | 5 | 8269 | 5wrr | B | | 504 | THR | p.Thr504Ile | Q96MK3 |
| HOMO SAPIENS | 5 | 1411 | 4h1s | B | | 221 | SER | p.Ser221Ter | P21589 |
| HOMO SAPIENS | 1 | 1411 | 4h1s | B | | 358 | CYS | p.Cys358Tyr | P21589 |
| HOMO SAPIENS | 2 | 23 | 5vvd | B | | 298 | GLU | p.Glu298Asp | P29474 |
| HOMO SAPIENS | 0 | 19 | 5jlh | E | | 88 | THR | p.Thr89Ile | P63261 |
| HOMO SAPIENS | 6 | 19 | 5jlh | E | | 117 | LYS | p.Lys118Met | P63261 |
| HOMO SAPIENS | 5 | 19 | 5jlh | E | | 263 | PRO | p.Pro264Leu | P63261 |
| HOMO SAPIENS | 1 | 19 | 5jlh | E | | 277 | THR | p.Thr278Ile | P63261 |
| HOMO SAPIENS | 6 | 19 | 5jlh | E | | 331 | PRO | p.Pro332Ala | P63261 |
| HOMO SAPIENS | 5 | 19 | 5jlh | E | | 369 | VAL | p.Val370Ala | P63261 |
| HOMO SAPIENS | 0 | 14306 | 4w4o | C | | 92 | ARG | p.Arg92Ter | P12314 |
| HOMO SAPIENS | 3 | 603 | 1nuh | A | | 4 | THR | p.Thr5Ile | P06744 |
| HOMO SAPIENS | 2 | 603 | 1nuh | A | | 19 | HIS | p.His20Pro | P06744 |
| HOMO SAPIENS | 8 | 603 | 1nuh | A | | 158 | GLY | p.Gly159Ser | P06744 |
| HOMO SAPIENS | 6 | 603 | 1nuh | A | | 223 | THR | p.Thr224Met | P06744 |
| HOMO SAPIENS | 0 | 603 | 1nuh | A | | 338 | LEU | p.Leu339Pro | P06744 |
| HOMO SAPIENS | 6 | 603 | 1nuh | A | | 342 | GLN | p.Gln343Arg | P06744 |
| HOMO SAPIENS | 6 | 603 | 1nuh | A | | 374 | THR | p.Thr375Arg | P06744 |
| HOMO SAPIENS | 4 | 603 | 1nuh | A | | 524 | ILE | p.Ile525Thr | P06744 |
| HOMO SAPIENS | 4 | 603 | 1nuh | A | | 538 | ASP | p.Asp539Asn | P06744 |
| HOMO SAPIENS | 1 | 3107 | 3kzj | A | | 1197 | VAL | p.Val1197Ala | P08603 |
| HOMO SAPIENS | 9 | 1748 | 2noh | A | | 46 | ARG | p.Arg46Gln | O15527 |
| HOMO SAPIENS | 2 | 11750 | 3s4y | A | | 33 | ARG | p.Arg33His | Q9H3S4 |
| HOMO SAPIENS | 2 | 11750 | 3s4y | A | | 40 | LEU | p.Leu40Pro | Q9H3S4 |
| HOMO SAPIENS | 4 | 11750 | 3s4y | A | | 142 | LEU | p.Leu142Phe | Q9H3S4 |
| HOMO SAPIENS | 5 | 11750 | 3s4y | A | | 219 | ASN | p.Asn219Ser | Q9H3S4 |
| HOMO SAPIENS | 1 | 2591 | 5vnj | A | | 382 | PHE | p.Phe382Leu | Q15436 |
| HOMO SAPIENS | 2 | 2591 | 5vnj | A | | 702 | MET | p.Met702Val | Q15436 |
| HOMO SAPIENS | 1 | 2426 | 5m8t | A | | 87 | ARG | p.Arg87Trp | P17643 |
| HOMO SAPIENS | 1 | 2426 | 5m8t | A | | 166 | SER | p.Ser166Ter | P17643 |
| HOMO SAPIENS | 0 | 2426 | 5m8t | A | | 356 | ARG | p.Arg356Gln | P17643 |
| HOMO SAPIENS | 2 | 2426 | 5m8t | A | | 374 | SER | p.Arg374Ter | P17643 |
| HOMO SAPIENS | 6 | 2426 | 5m8t | A | | 378 | ASN | p.Asn378Ser | P17643 |
| HOMO SAPIENS | 6 | 2426 | 5m8t | A | | 382 | LEU | p.Leu382Pro | P17643 |
| HOMO SAPIENS | 7 | 10097 | 4n6v | | 1 | 46 | GLN | p.Gln46Ter | P04080 |
| HOMO SAPIENS | 3 | 10097 | 4n6v | | 1 | 65 | VAL | p.Val65Ile | P04080 |
| HOMO SAPIENS | 8 | 10097 | 4n6v | | 1 | 68 | ARG | p.Arg68Ter | P04080 |
| HOMO SAPIENS | 2 | 859 | 4z9a | A | | 105 | GLN | p.Gln106Arg | P24666 |
| HOMO SAPIENS | 8 | 1420 | 5nan | F | | 95 | SER | p.Ser95Leu | Q96PD4 |

| | | | | | | | | |
|---|---|---|---|---|---|---|---|---|
| HOMO SAPIENS | 6 | 1420 | 5nan | F | 126 | GLU | p.Glu126Gly | Q96PD4 |
| HOMO SAPIENS | 8 | 1420 | 5nan | F | 155 | VAL | p.Val155Ile | Q96PD4 |
| HOMO SAPIENS | 6 | 1420 | 5nan | F | 157 | PRO | p.Pro157Ser | Q96PD4 |
| HOMO SAPIENS | -1 | 1420 | 5nan | F | 161 | HIS | p.His161Arg | Q96PD4 |
| HOMO SAPIENS | 3 | 288 | 5i96 | B | 140 | GLN | p.Arg140Gly | P48735 |
| HOMO SAPIENS | 3 | 288 | 5i96 | B | 140 | GLN | p.Arg140Gln | P48735 |
| HOMO SAPIENS | 2 | 288 | 5i96 | B | 261 | ARG | p.Arg261His | P48735 |
| HOMO SAPIENS | 2 | 288 | 5i96 | B | 435 | THR | p.Thr435Met | P48735 |
| HOMO SAPIENS | 9 | 120 | 1tup | B | 107 | TYR | p.Tyr107His | P04637 |
| HOMO SAPIENS | 8 | 120 | 1tup | B | 110 | ARG | p.Arg110Leu | P04637 |
| HOMO SAPIENS | 5 | 120 | 1tup | B | 120 | LYS | p.Lys120Glu | P04637 |
| HOMO SAPIENS | 4 | 120 | 1tup | B | 133 | MET | p.Met133Thr | P04637 |
| HOMO SAPIENS | 3 | 120 | 1tup | B | 134 | PHE | p.Phe134Cys | P04637 |
| HOMO SAPIENS | 8 | 120 | 1tup | B | 136 | GLN | p.Gln136His | P04637 |
| HOMO SAPIENS | 9 | 120 | 1tup | B | 138 | ALA | p.Ala138Pro | P04637 |
| HOMO SAPIENS | 9 | 120 | 1tup | B | 151 | PRO | p.Pro151Ser | P04637 |
| HOMO SAPIENS | 9 | 120 | 1tup | B | 152 | PRO | p.Pro152Leu | P04637 |
| HOMO SAPIENS | 9 | 120 | 1tup | B | 158 | ARG | p.Arg158Cys | P04637 |
| HOMO SAPIENS | 9 | 120 | 1tup | B | 158 | ARG | p.Arg158His | P04637 |
| HOMO SAPIENS | 9 | 120 | 1tup | B | 175 | ARG | p.Arg175Cys | P04637 |
| HOMO SAPIENS | 9 | 120 | 1tup | B | 175 | ARG | p.Arg175Leu | P04637 |
| HOMO SAPIENS | 8 | 120 | 1tup | B | 181 | ARG | p.Arg181Leu | P04637 |
| HOMO SAPIENS | 9 | 120 | 1tup | B | 189 | ALA | p.Ala189Val | P04637 |
| HOMO SAPIENS | 9 | 120 | 1tup | B | 196 | ARG | p.Arg196Gly | P04637 |
| HOMO SAPIENS | 9 | 120 | 1tup | B | 213 | ARG | p.Arg213Gln | P04637 |
| HOMO SAPIENS | 9 | 120 | 1tup | B | 213 | ARG | p.Arg213Ter | P04637 |
| HOMO SAPIENS | 9 | 120 | 1tup | B | 214 | HIS | p.His214Gln | P04637 |
| HOMO SAPIENS | 9 | 120 | 1tup | B | 220 | TYR | p.Tyr220Cys | P04637 |
| HOMO SAPIENS | 0 | 120 | 1tup | B | 234 | TYR | p.Tyr234Cys | P04637 |
| HOMO SAPIENS | 9 | 120 | 1tup | B | 237 | MET | p.Met237Ile | P04637 |
| HOMO SAPIENS | 0 | 120 | 1tup | B | 238 | CYS | p.Cys238Tyr | P04637 |
| HOMO SAPIENS | 9 | 120 | 1tup | B | 241 | SER | p.Ser241Phe | P04637 |
| HOMO SAPIENS | 9 | 120 | 1tup | B | 242 | CYS | p.Cys242Ser | P04637 |
| HOMO SAPIENS | 9 | 120 | 1tup | B | 245 | GLY | p.Gly245Cys | P04637 |
| HOMO SAPIENS | 9 | 120 | 1tup | B | 245 | GLY | p.Gly245Asp | P04637 |
| HOMO SAPIENS | 9 | 120 | 1tup | B | 248 | ARG | p.Arg248Leu | P04637 |
| HOMO SAPIENS | 9 | 120 | 1tup | B | 248 | ARG | p.Arg248Trp | P04637 |
| HOMO SAPIENS | 9 | 120 | 1tup | B | 251 | ILE | p.Ile251Ser | P04637 |
| HOMO SAPIENS | 9 | 120 | 1tup | B | 252 | LEU | p.Leu252Pro | P04637 |
| HOMO SAPIENS | 9 | 120 | 1tup | B | 258 | GLU | p.Glu258Lys | P04637 |
| HOMO SAPIENS | 8 | 120 | 1tup | B | 272 | VAL | p.Val272Leu | P04637 |
| HOMO SAPIENS | 9 | 120 | 1tup | B | 273 | ARG | p.Arg273Cys | P04637 |
| HOMO SAPIENS | 9 | 120 | 1tup | B | 273 | ARG | p.Arg273His | P04637 |
| HOMO SAPIENS | 9 | 120 | 1tup | B | 277 | CYS | p.Cys277Phe | P04637 |
| HOMO SAPIENS | 9 | 120 | 1tup | B | 280 | ARG | p.Arg280Ile | P04637 |
| HOMO SAPIENS | 9 | 120 | 1tup | B | 282 | ARG | p.Arg282Trp | P04637 |
| HOMO SAPIENS | 9 | 120 | 1tup | B | 283 | ARG | p.Arg283His | P04637 |
| HOMO SAPIENS | 9 | 120 | 1tup | B | 285 | GLU | p.Glu285Val | P04637 |
| HOMO SAPIENS | -1 | 120 | 1tup | B | 292 | LYS | p.Lys292Ile | P04637 |
| HOMO SAPIENS | -1 | 120 | 1tup | B | 298 | GLU | p.Glu298Lys | P04637 |
| HOMO SAPIENS | -1 | 2791 | 4ygb | A | 39 | VAL | p.Val39Ala | P49257 |
| HOMO SAPIENS | 2 | 106 | 4x6j | A | 32 | GLY | p.Gly146Arg | P43235 |
| HOMO SAPIENS | 0 | 106 | 4x6j | A | 76 | GLN | p.Gln190Ter | P43235 |
| HOMO SAPIENS | 7 | 106 | 4x6j | A | 198 | ARG | p.Arg312Ter | P43235 |

| | | | | | | | | |
|---|---|---|---|---|---|---|---|---|
| HOMO SAPIENS | 5 | 490 | 2hlo | B | 172 | LEU | p.Leu202Gln | P02675 |
| HOMO SAPIENS | 0 | 490 | 2hlo | B | 353 | LEU | p.Leu383Arg | P02675 |
| HOMO SAPIENS | 4 | 490 | 2hlo | B | 400 | GLY | p.Gly430Asp | P02675 |
| HOMO SAPIENS | 7 | 490 | 2hlo | B | 448 | ARG | p.Arg478Lys | P02675 |
| HOMO SAPIENS | 1 | 289 | 4p13 | B | 18 | GLU | p.Glu43Lys | P11310 |
| HOMO SAPIENS | 2 | 289 | 4p13 | B | 20 | GLN | p.Gln45Arg | P11310 |
| HOMO SAPIENS | 5 | 289 | 4p13 | B | 28 | ARG | p.Arg53Cys | P11310 |
| HOMO SAPIENS | 1 | 289 | 4p13 | B | 42 | TYR | p.Tyr67His | P11310 |
| HOMO SAPIENS | 1 | 289 | 4p13 | B | 53 | ILE | p.Ile78Thr | P11310 |
| HOMO SAPIENS | 1 | 289 | 4p13 | B | 59 | LEU | p.Leu84Phe | P11310 |
| HOMO SAPIENS | 2 | 289 | 4p13 | B | 96 | THR | p.Thr121Ile | P11310 |
| HOMO SAPIENS | 0 | 289 | 4p13 | B | 124 | MET | p.Met149Ile | P11310 |
| HOMO SAPIENS | 5 | 289 | 4p13 | B | 143 | ASP | p.Asp168Ala | P11310 |
| HOMO SAPIENS | 7 | 289 | 4p13 | B | 168 | THR | p.Thr193Ala | P11310 |
| HOMO SAPIENS | 1 | 289 | 4p13 | B | 170 | GLY | p.Gly195Arg | P11310 |
| HOMO SAPIENS | 2 | 289 | 4p13 | B | 181 | ARG | p.Arg206Cys | P11310 |
| HOMO SAPIENS | 2 | 289 | 4p13 | B | 181 | ARG | p.Arg206His | P11310 |
| HOMO SAPIENS | 8 | 289 | 4p13 | B | 203 | THR | p.Thr228Ile | P11310 |
| HOMO SAPIENS | 2 | 289 | 4p13 | B | 208 | ILE | p.Ile233Asn | P11310 |
| HOMO SAPIENS | 1 | 289 | 4p13 | B | 219 | CYS | p.Cys244Arg | P11310 |
| HOMO SAPIENS | 6 | 289 | 4p13 | B | 220 | SER | p.Ser245Leu | P11310 |
| HOMO SAPIENS | 5 | 289 | 4p13 | B | 228 | GLU | p.Glu253Lys | P11310 |
| HOMO SAPIENS | 2 | 289 | 4p13 | B | 241 | ASP | p.Asp266Gly | P11310 |
| HOMO SAPIENS | 1 | 289 | 4p13 | B | 242 | GLY | p.Gly267Arg | P11310 |
| HOMO SAPIENS | 7 | 289 | 4p13 | B | 256 | ARG | p.Arg281Thr | P11310 |
| HOMO SAPIENS | 4 | 289 | 4p13 | B | 269 | ARG | p.Arg294Thr | P11310 |
| HOMO SAPIENS | 4 | 289 | 4p13 | B | 304 | GLU | p.Lys329Glu | P11310 |
| HOMO SAPIENS | 1 | 289 | 4p13 | B | 324 | ARG | p.Arg349Ter | P11310 |
| HOMO SAPIENS | 3 | 289 | 4p13 | B | 326 | THR | p.Thr351Ile | P11310 |
| HOMO SAPIENS | 7 | 289 | 4p13 | B | 350 | ILE | p.Ile375Thr | P11310 |
| HOMO SAPIENS | 3 | 289 | 4p13 | B | 359 | GLU | p.Glu384Ter | P11310 |
| HOMO SAPIENS | 1 | 4563 | 3i2b | A | 16 | ARG | p.Arg16Cys | Q03393 |
| HOMO SAPIENS | 5 | 4563 | 3i2b | A | 25 | ARG | p.Arg25Gln | Q03393 |
| HOMO SAPIENS | 1 | 4563 | 3i2b | A | 47 | ASN | p.Asn47Asp | Q03393 |
| HOMO SAPIENS | 8 | 4563 | 3i2b | A | 52 | ASN | p.Asn52Ser | Q03393 |
| HOMO SAPIENS | 1 | 4563 | 3i2b | A | 56 | VAL | p.Val56Met | Q03393 |
| HOMO SAPIENS | 1 | 4563 | 3i2b | A | 87 | PRO | p.Pro87Ser | Q03393 |
| HOMO SAPIENS | 1 | 4563 | 3i2b | A | 96 | ASP | p.Asp96Asn | Q03393 |
| HOMO SAPIENS | 0 | 4563 | 3i2b | A | 99 | TYR | p.Tyr99Ter | Q03393 |
| HOMO SAPIENS | 2 | 4563 | 3i2b | A | 116 | ASP | p.Asp116Gly | Q03393 |
| HOMO SAPIENS | 2 | 41 | 3exx | A | 1 | GLY | p.Gly90Cys | P01308 |
| HOMO SAPIENS | 5 | 41 | 3exx | A | 3 | VAL | p.Val92Leu | P01308 |
| HOMO SAPIENS | 4 | 41 | 3exx | A | 7 | CYS | p.Cys96Ser | P01308 |
| HOMO SAPIENS | 9 | 41 | 3exx | A | 19 | TYR | p.Tyr108Cys | P01308 |
| HOMO SAPIENS | 9 | 131 | 4mjo | D | 118 | ASP | p.Asp119Asn | P09467 |
| HOMO SAPIENS | 0 | 131 | 4mjo | D | 163 | GLY | p.Gly164Ser | P09467 |
| HOMO SAPIENS | 0 | 131 | 4mjo | D | 176 | ALA | p.Ala177Asp | P09467 |
| HOMO SAPIENS | 5 | 131 | 4mjo | D | 193 | PHE | p.Phe194Ser | P09467 |
| HOMO SAPIENS | 4 | 131 | 4mjo | D | 217 | ARG | p.Arg218Lys | P09467 |
| HOMO SAPIENS | 4 | 131 | 4mjo | D | 223 | VAL | p.Val224Ile | P09467 |
| HOMO SAPIENS | 4 | 131 | 4mjo | D | 232 | PHE | p.Phe233Ile | P09467 |
| HOMO SAPIENS | 5 | 131 | 4mjo | D | 243 | ARG | p.Arg244Trp | P09467 |
| HOMO SAPIENS | 0 | 131 | 4mjo | D | 283 | PRO | p.Pro284Arg | P09467 |
| HOMO SAPIENS | 0 | 131 | 4mjo | D | 288 | MET | p.Met289Leu | P09467 |

| | | | | | | | | |
|---|---|---|---|---|---|---|---|---|
| HOMO SAPIENS | 0 | 131 | 4mjo | D | 323 | ASP | p.Asp324Asn | P09467 |
| HOMO SAPIENS | -1 | 13122 | 3hfc | A | 583 | ARG | p.Arg583Cys | P51787 |
| HOMO SAPIENS | 1 | 13122 | 3hfc | A | 587 | THR | p.Thr587Met | P51787 |
| HOMO SAPIENS | 0 | 13122 | 3hfc | A | 589 | GLY | p.Gly589Asp | P51787 |
| HOMO SAPIENS | 4 | 3002 | 3nbq | B | 94 | ARG | p.Arg94Cys | Q16831 |
| HOMO SAPIENS | 1 | 3616 | 3dbp | A | 97 | ILE | p.Ile286Asn | P11172 |
| HOMO SAPIENS | 0 | 3616 | 3dbp | A | 240 | GLY | p.Gly429Arg | P11172 |
| HOMO SAPIENS | 0 | 3616 | 3dbp | A | 257 | ILE | p.Ile446Val | P11172 |
| HOMO SAPIENS | 1 | 584 | 1drf | A | 79 | LEU | p.Leu80Phe | P00374 |
| HOMO SAPIENS | 3 | 1609 | 3qa2 | B | 49 | VAL | p.Val49Ile | P50053 |
| HOMO SAPIENS | 0 | 20560 | 5mug | A | 101 | HIS | p.His101Gln | P49638 |
| HOMO SAPIENS | 2 | 20560 | 5mug | A | 120 | ALA | p.Ala120Thr | P49638 |
| HOMO SAPIENS | 3 | 20560 | 5mug | A | 192 | ARG | p.Arg192His | P49638 |
| HOMO SAPIENS | 3 | 20560 | 5mug | A | 221 | ARG | p.Arg221Trp | P49638 |
| HOMO SAPIENS | 4 | 256 | 3vtv | A | 65 | THR | p.Glu50Lys | Q9GZQ8 |
| HOMO SAPIENS | 4 | 256 | 3vtv | A | 113 | VAL | p.Met98Lys | Q9GZQ8 |
| HOMO SAPIENS | 4 | 5397 | 1m61 | A | 192 | ARG | p.Arg192Trp | P43403 |
| HOMO SAPIENS | 5 | 39881 | 1aut | L | 107 | HIS | p.His149Pro | P04070 |
| HOMO SAPIENS | 5 | 4010 | 4uf8 | I | 33 | ALA | p.Ala33Val | Q13501 |
| HOMO SAPIENS | 3 | 313 | 5w49 | A | 98 | PRO | p.Pro98Leu | P23526 |
| HOMO SAPIENS | 2 | 313 | 5w49 | A | 143 | TYR | p.Tyr143Cys | P23526 |
| HOMO SAPIENS | -1 | 1643 | 2m8t | A | 102 | PRO | p.Pro102Leu | P04156 |
| HOMO SAPIENS | -1 | 1643 | 2m8t | A | 105 | PRO | p.Pro105Leu | P04156 |
| HOMO SAPIENS | -1 | 1643 | 2m8t | A | 117 | ALA | p.Ala117Val | P04156 |
| HOMO SAPIENS | 5 | 1643 | 2m8t | A | 129 | MET | p.Met129Val | P04156 |
| HOMO SAPIENS | 5 | 1643 | 2m8t | A | 131 | GLY | p.Gly131Val | P04156 |
| HOMO SAPIENS | 2 | 1643 | 2m8t | A | 142 | GLY | p.Gly142Ser | P04156 |
| HOMO SAPIENS | 3 | 1643 | 2m8t | A | 171 | ASN | p.Asn171Ser | P04156 |
| HOMO SAPIENS | 9 | 1643 | 2m8t | A | 178 | ASP | p.Asp178Asn | P04156 |
| HOMO SAPIENS | 9 | 1643 | 2m8t | A | 180 | VAL | p.Val180Ile | P04156 |
| HOMO SAPIENS | 9 | 1643 | 2m8t | A | 183 | THR | p.Thr183Ala | P04156 |
| HOMO SAPIENS | 9 | 1643 | 2m8t | A | 187 | HIS | p.His187Arg | P04156 |
| HOMO SAPIENS | 8 | 1643 | 2m8t | A | 198 | PHE | p.Phe198Ser | P04156 |
| HOMO SAPIENS | 9 | 1643 | 2m8t | A | 200 | GLU | p.Glu200Lys | P04156 |
| HOMO SAPIENS | 9 | 1643 | 2m8t | A | 208 | ARG | p.Arg208His | P04156 |
| HOMO SAPIENS | 3 | 1643 | 2m8t | A | 210 | VAL | p.Val210Ile | P04156 |
| HOMO SAPIENS | 6 | 1643 | 2m8t | A | 217 | GLN | p.Gln217Arg | P04156 |
| HOMO SAPIENS | 1 | 29 | 5kos | A | 151 | GLY | p.Gly217Arg | P00797 |
| HOMO SAPIENS | 1 | 29 | 5kos | A | 164 | ARG | p.Arg230Lys | P00797 |
| HOMO SAPIENS | 3 | 504 | 4zkv | B | 37 | ARG | p.Arg37Pro | P49773 |
| HOMO SAPIENS | 2 | 504 | 4zkv | B | 62 | GLN | p.Gln62Ter | P49773 |
| HOMO SAPIENS | 2 | 504 | 4zkv | B | 84 | CYS | p.Cys84Arg | P49773 |
| HOMO SAPIENS | 4 | 504 | 4zkv | B | 93 | GLY | p.Gly93Asp | P49773 |
| HOMO SAPIENS | 9 | 504 | 4zkv | B | 112 | HIS | p.His112Asn | P49773 |
| HOMO SAPIENS | 0 | 74 | 5vwo | C | 54 | ASN | p.Asn54Lys | P04181 |
| HOMO SAPIENS | 0 | 74 | 5vwo | C | 55 | TYR | p.Tyr55His | P04181 |
| HOMO SAPIENS | 2 | 74 | 5vwo | C | 90 | GLN | p.Gln90Glu | P04181 |
| HOMO SAPIENS | 6 | 74 | 5vwo | C | 91 | GLY | p.Gly91Glu | P04181 |
| HOMO SAPIENS | 2 | 74 | 5vwo | C | 93 | CYS | p.Cys93Phe | P04181 |
| HOMO SAPIENS | 1 | 74 | 5vwo | C | 154 | ARG | p.Arg154Leu | P04181 |
| HOMO SAPIENS | 0 | 74 | 5vwo | C | 180 | ARG | p.Arg180Thr | P04181 |
| HOMO SAPIENS | 0 | 74 | 5vwo | C | 199 | PRO | p.Pro199Gln | P04181 |
| HOMO SAPIENS | 4 | 74 | 5vwo | C | 209 | TYR | p.Tyr209Ter | P04181 |
| HOMO SAPIENS | 0 | 74 | 5vwo | C | 226 | ALA | p.Ala226Val | P04181 |

| | | | | | | | | |
|---|---|---|---|---|---|---|---|---|
| HOMO SAPIENS | 5 | 74 | 5vwo | C | 233 | GLN | p.Gln233Arg | P04181 |
| HOMO SAPIENS | 5 | 74 | 5vwo | C | 241 | PRO | p.Pro241Leu | P04181 |
| HOMO SAPIENS | 2 | 74 | 5vwo | C | 245 | TYR | p.Tyr245Cys | P04181 |
| HOMO SAPIENS | 6 | 74 | 5vwo | C | 250 | ARG | p.Arg250Pro | P04181 |
| HOMO SAPIENS | 6 | 74 | 5vwo | C | 250 | ARG | p.Arg250Ter | P04181 |
| HOMO SAPIENS | 8 | 74 | 5vwo | C | 267 | THR | p.Thr267Ile | P04181 |
| HOMO SAPIENS | 0 | 74 | 5vwo | C | 270 | ALA | p.Ala270Pro | P04181 |
| HOMO SAPIENS | 0 | 74 | 5vwo | C | 271 | ARG | p.Arg271Lys | P04181 |
| HOMO SAPIENS | 1 | 74 | 5vwo | C | 299 | TYR | p.Tyr299Ter | P04181 |
| HOMO SAPIENS | 0 | 74 | 5vwo | C | 318 | GLU | p.Glu318Lys | P04181 |
| HOMO SAPIENS | 1 | 74 | 5vwo | C | 319 | HIS | p.His319Tyr | P04181 |
| HOMO SAPIENS | 1 | 74 | 5vwo | C | 331 | ARG | p.Arg331Ter | P04181 |
| HOMO SAPIENS | 0 | 74 | 5vwo | C | 332 | VAL | p.Val332Met | P04181 |
| HOMO SAPIENS | 1 | 74 | 5vwo | C | 353 | GLY | p.Gly353Asp | P04181 |
| HOMO SAPIENS | 2 | 74 | 5vwo | C | 375 | GLY | p.Gly375Ala | P04181 |
| HOMO SAPIENS | 0 | 74 | 5vwo | C | 394 | CYS | p.Cys394Arg | P04181 |
| HOMO SAPIENS | 0 | 74 | 5vwo | C | 402 | LEU | p.Leu402Pro | P04181 |
| HOMO SAPIENS | 0 | 74 | 5vwo | C | 436 | ILE | p.Ile436Asn | P04181 |
| HOMO SAPIENS | 7 | 24356 | 4az0 | A | 104 | VAL | p.Val132Met | P10619 |
| HOMO SAPIENS | 4 | 24356 | 4az0 | A | 221 | TYR | p.Tyr249Asn | P10619 |
| HOMO SAPIENS | 3 | 245 | 1rsz | A | 24 | ARG | p.Arg24Ter | P00491 |
| HOMO SAPIENS | 3 | 245 | 1rsz | A | 51 | SER | p.Gly51Ser | P00491 |
| HOMO SAPIENS | 1 | 245 | 1rsz | A | 58 | ARG | p.Arg58Ter | P00491 |
| HOMO SAPIENS | 4 | 245 | 1rsz | A | 89 | GLU | p.Glu89Lys | P00491 |
| HOMO SAPIENS | 6 | 245 | 1rsz | A | 128 | ASP | p.Asp128Gly | P00491 |
| HOMO SAPIENS | 0 | 245 | 1rsz | A | 174 | ALA | p.Ala174Pro | P00491 |
| HOMO SAPIENS | 8 | 245 | 1rsz | A | 192 | TYR | p.Tyr192Cys | P00491 |
| HOMO SAPIENS | 8 | 245 | 1rsz | A | 234 | ARG | p.Arg234Pro | P00491 |
| HOMO SAPIENS | 0 | 37 | 5tby | E | 22 | GLU | p.Glu22Lys | P10916 |
| HOMO SAPIENS | 1 | 37 | 5tby | E | 86 | PHE | p.Phe86Ser | P10916 |
| HOMO SAPIENS | 9 | 71 | 5m9j | A | 12 | GLN | p.Gln36Leu | P03950 |
| HOMO SAPIENS | 3 | 71 | 5m9j | A | 17 | LYS | p.Lys41Glu | P03950 |
| HOMO SAPIENS | 3 | 71 | 5m9j | A | 17 | LYS | p.Lys41Ile | P03950 |
| HOMO SAPIENS | 4 | 71 | 5m9j | A | 28 | SER | p.Ser52Asn | P03950 |
| HOMO SAPIENS | 7 | 71 | 5m9j | A | 31 | ARG | p.Arg55Lys | P03950 |
| HOMO SAPIENS | 9 | 71 | 5m9j | A | 39 | CYS | p.Cys63Trp | P03950 |
| HOMO SAPIENS | 9 | 71 | 5m9j | A | 40 | LYS | p.Lys64Ile | P03950 |
| HOMO SAPIENS | 2 | 71 | 5m9j | A | 60 | GLU | p.Lys84Glu | P03950 |
| HOMO SAPIENS | 0 | 71 | 5m9j | A | 112 | PRO | p.Pro136Leu | P03950 |
| HOMO SAPIENS | 9 | 71 | 5m9j | A | 113 | VAL | p.Val137Ile | P03950 |
| HOMO SAPIENS | 3 | 326 | 5dqi | A | 209 | GLY | p.Gly209Val | Q9Y253 |
| HOMO SAPIENS | 1 | 326 | 5dqi | A | 242 | SER | p.Ser242Ter | Q9Y253 |
| HOMO SAPIENS | 1 | 326 | 5dqi | A | 303 | ARG | p.Arg303Ter | Q9Y253 |
| HOMO SAPIENS | 1 | 8203 | 3iu3 | K | 132 | VAL | p.Val153Ile | P01589 |
| HOMO SAPIENS | 0 | 4995 | 5ffo | H | 234 | THR | p.Thr263Ile | P01137 |
| HOMO SAPIENS | 5 | 38609 | 2ruk | A | 49 | ASP | p.Asp49His | P04637 |
| HOMO SAPIENS | 5 | 38609 | 2ruk | A | 58 | PRO | p.Pro58Arg | P04637 |
| HOMO SAPIENS | 0 | 1607 | 6pah | A | 117 | THR | p.Thr117Ile | P00439 |
| HOMO SAPIENS | 3 | 1607 | 6pah | A | 145 | ASP | p.Asp145Val | P00439 |
| HOMO SAPIENS | 5 | 1607 | 6pah | A | 158 | ARG | p.Arg158Gln | P00439 |
| HOMO SAPIENS | 5 | 1607 | 6pah | A | 158 | ARG | p.Arg158Trp | P00439 |
| HOMO SAPIENS | 5 | 1607 | 6pah | A | 169 | ARG | p.Arg169His | P00439 |
| HOMO SAPIENS | 5 | 1607 | 6pah | A | 169 | ARG | p.Arg169Cys | P00439 |
| HOMO SAPIENS | 3 | 1607 | 6pah | A | 176 | ARG | p.Arg176Leu | P00439 |

| | | | | | | | | |
|---|---|---|---|---|---|---|---|---|
| HOMO SAPIENS | 3 | 1607 | 6pah | A | 177 | VAL | p.Val177Leu | P00439 |
| HOMO SAPIENS | 1 | 1607 | 6pah | A | 194 | LEU | p.Leu194Pro | P00439 |
| HOMO SAPIENS | 1 | 1607 | 6pah | A | 204 | TYR | p.Tyr204Cys | P00439 |
| HOMO SAPIENS | 1 | 1607 | 6pah | A | 204 | TYR | p.Tyr204Ter | P00439 |
| HOMO SAPIENS | 2 | 1607 | 6pah | A | 230 | VAL | p.Val230Ile | P00439 |
| HOMO SAPIENS | 4 | 1607 | 6pah | A | 232 | GLN | p.Gln232Ter | P00439 |
| HOMO SAPIENS | 4 | 1607 | 6pah | A | 241 | ARG | p.Arg241Cys | P00439 |
| HOMO SAPIENS | 4 | 1607 | 6pah | A | 241 | ARG | p.Arg241His | P00439 |
| HOMO SAPIENS | 5 | 1607 | 6pah | A | 243 | ARG | p.Arg243Gln | P00439 |
| HOMO SAPIENS | 5 | 1607 | 6pah | A | 243 | ARG | p.Arg243Ter | P00439 |
| HOMO SAPIENS | 7 | 1607 | 6pah | A | 249 | LEU | p.Leu249Phe | P00439 |
| HOMO SAPIENS | 5 | 1607 | 6pah | A | 252 | ARG | p.Arg252Gly | P00439 |
| HOMO SAPIENS | 5 | 1607 | 6pah | A | 261 | ARG | p.Arg261Ter | P00439 |
| HOMO SAPIENS | 4 | 1607 | 6pah | A | 267 | GLN | p.Gln267Leu | P00439 |
| HOMO SAPIENS | 1 | 1607 | 6pah | A | 272 | GLY | p.Gly272Ter | P00439 |
| HOMO SAPIENS | 2 | 1607 | 6pah | A | 277 | TYR | p.Tyr277Asp | P00439 |
| HOMO SAPIENS | 7 | 1607 | 6pah | A | 280 | GLU | p.Glu280Lys | P00439 |
| HOMO SAPIENS | 8 | 1607 | 6pah | A | 281 | PRO | p.Pro281Leu | P00439 |
| HOMO SAPIENS | 0 | 1607 | 6pah | A | 299 | PHE | p.Phe299Cys | P00439 |
| HOMO SAPIENS | 5 | 1607 | 6pah | A | 300 | ALA | p.Ala300Ser | P00439 |
| HOMO SAPIENS | 5 | 1607 | 6pah | A | 309 | ALA | p.Ala309Asp | P00439 |
| HOMO SAPIENS | 4 | 1607 | 6pah | A | 320 | LYS | p.Lys320Asn | P00439 |
| HOMO SAPIENS | 0 | 1607 | 6pah | A | 345 | ALA | p.Ala345Ser | P00439 |
| HOMO SAPIENS | 0 | 1607 | 6pah | A | 348 | LEU | p.Leu348Val | P00439 |
| HOMO SAPIENS | 6 | 1607 | 6pah | A | 356 | TYR | p.Tyr356Ter | P00439 |
| HOMO SAPIENS | 2 | 1607 | 6pah | A | 380 | THR | p.Thr380Met | P00439 |
| HOMO SAPIENS | 0 | 1607 | 6pah | A | 386 | TYR | p.Tyr386Cys | P00439 |
| HOMO SAPIENS | 5 | 1607 | 6pah | A | 390 | GLU | p.Glu390Gly | P00439 |
| HOMO SAPIENS | 5 | 1607 | 6pah | A | 403 | ALA | p.Ala403Val | P00439 |
| HOMO SAPIENS | 4 | 1607 | 6pah | A | 408 | ARG | p.Arg408Trp | P00439 |
| HOMO SAPIENS | 3 | 1607 | 6pah | A | 413 | ARG | p.Arg413Pro | P00439 |
| HOMO SAPIENS | 4 | 1607 | 6pah | A | 414 | TYR | p.Tyr414Cys | P00439 |
| HOMO SAPIENS | 4 | 1607 | 6pah | A | 414 | TYR | p.Tyr414His | P00439 |
| HOMO SAPIENS | 4 | 1607 | 6pah | A | 415 | ASP | p.Asp415Asn | P00439 |
| HOMO SAPIENS | -1 | 676 | 5dc5 | A | 7 | PRO | p.Pro7Leu | Q9BY41 |
| HOMO SAPIENS | 0 | 676 | 5dc5 | A | 311 | THR | p.Thr311Met | Q9BY41 |
| HOMO SAPIENS | 1 | 124 | 5ftk | E | 95 | ARG | p.Arg95Gly | P55072 |
| HOMO SAPIENS | 1 | 124 | 5ftk | E | 155 | ARG | p.Arg155Cys | P55072 |
| HOMO SAPIENS | 1 | 124 | 5ftk | E | 155 | ARG | p.Arg155His | P55072 |
| HOMO SAPIENS | 1 | 124 | 5ftk | E | 159 | ARG | p.Arg159His | P55072 |
| HOMO SAPIENS | 1 | 124 | 5ftk | E | 159 | ARG | p.Arg159Gly | P55072 |
| HOMO SAPIENS | 0 | 124 | 5ftk | E | 191 | ARG | p.Arg191Gln | P55072 |
| HOMO SAPIENS | 0 | 124 | 5ftk | E | 232 | ALA | p.Ala232Glu | P55072 |
| HOMO SAPIENS | 5 | 5442 | 1fgg | B | 223 | GLY | p.Gly223Ser | O94766 |
| HOMO SAPIENS | 0 | 5442 | 1fgg | B | 277 | ARG | p.Arg277Gln | O94766 |
| HOMO SAPIENS | 0 | 9072 | 2gys | B | 225 | GLU | p.Glu249Gln | P32927 |
| HOMO SAPIENS | 0 | 9072 | 2gys | B | 247 | SER | p.Ser271Ter | P32927 |
| HOMO SAPIENS | 0 | 9072 | 2gys | B | 294 | ASP | p.Asp312Asn | P32927 |
| HOMO SAPIENS | 0 | 1484 | 4zrj | A | 62 | PHE | p.Phe62Ser | P35240 |
| HOMO SAPIENS | 3 | 1484 | 4zrj | A | 205 | MET | p.Met205Val | P35240 |
| HOMO SAPIENS | 3 | 1484 | 4zrj | A | 301 | ILE | p.Ile301Met | P35240 |
| HOMO SAPIENS | 4 | 40923 | 2enq | A | 68 | ILE | p.Ile391Met | P42336 |
| HOMO SAPIENS | 1 | 6179 | 5ofv | A | 125 | THR | p.Thr125Met | O43175 |
| HOMO SAPIENS | 0 | 6179 | 5ofv | A | 135 | ARG | p.Arg135Trp | O43175 |

| | | | | | | | | |
|---|---|---|---|---|---|---|---|---|
| HOMO SAPIENS | 1 | 6179 | 5ofv | A | 163 | ARG | p.Arg163Gln | O43175 |
| HOMO SAPIENS | 9 | 6179 | 5ofv | A | 261 | VAL | p.Val261Met | O43175 |
| HOMO SAPIENS | 7 | 34118 | 2i5o | A | 24 | MET | p.Met647Leu | Q9Y253 |
| HOMO SAPIENS | 8 | 3234 | 5c84 | A | 311 | LYS | p.Lys311Asn | P98170 |
| HOMO SAPIENS | 3 | 3234 | 5c84 | A | 321 | ALA | p.Ala321Gly | P98170 |
| HOMO SAPIENS | -1 | 15129 | 1zy8 | K | 1 | GLY | p.Gly54Ser | O00330 |
| HOMO SAPIENS | -1 | 15129 | 1zy8 | K | 48 | THR | p.Thr101Ala | O00330 |
| HOMO SAPIENS | -1 | 15129 | 1zy8 | K | 105 | GLU | p.Glu158Asp | O00330 |
| HOMO SAPIENS | 8 | 15129 | 1zy8 | K | 136 | ARG | p.Arg189His | O00330 |
| HOMO SAPIENS | -1 | 15129 | 1zy8 | K | 195 | GLN | p.Gln248Ter | O00330 |
| HOMO SAPIENS | 8 | 202 | 4bzb | A | 123 | HIS | p.His123Pro | Q9Y3Z3 |
| HOMO SAPIENS | 4 | 202 | 4bzb | A | 145 | ARG | p.Arg145Ter | Q9Y3Z3 |
| HOMO SAPIENS | 6 | 202 | 4bzb | A | 164 | ARG | p.Arg164Ter | Q9Y3Z3 |
| HOMO SAPIENS | 3 | 202 | 4bzb | A | 201 | ILE | p.Ile201Asn | Q9Y3Z3 |
| HOMO SAPIENS | 0 | 202 | 4bzb | A | 209 | GLY | p.Gly209Ser | Q9Y3Z3 |
| HOMO SAPIENS | 0 | 202 | 4bzb | A | 254 | MET | p.Met254Val | Q9Y3Z3 |
| HOMO SAPIENS | 5 | 202 | 4bzb | A | 531 | ARG | p.Arg531Ser | Q9Y3Z3 |
| HOMO SAPIENS | 6 | 202 | 4bzb | A | 548 | GLN | p.Gln548Ter | Q9Y3Z3 |
| HOMO SAPIENS | 3 | 30947 | 5lgd | A | 90 | PRO | p.Pro90Ser | P16671 |
| HOMO SAPIENS | 6 | 30947 | 5lgd | A | 254 | PHE | p.Phe254Leu | P16671 |
| HOMO SAPIENS | 0 | 30947 | 5lgd | A | 413 | ILE | p.Ile413Leu | P16671 |
| HOMO SAPIENS | 2 | 3906 | 5jtc | A | 735 | ARG | p.Arg735Trp | Q12797 |
| HOMO SAPIENS | 1 | 45 | 5uyx | C | 146 | GLU | p.Glu146Val | P48643 |
| HOMO SAPIENS | 1 | 45 | 5uyx | C | 147 | HIS | p.His147Arg | P48643 |
| HOMO SAPIENS | 0 | 45 | 5uyx | C | 362 | ILE | p.Ile362Met | P48643 |
| HOMO SAPIENS | 8 | 39596 | 1mhn | A | 95 | GLY | p.Gly95Arg | Q16637 |
| HOMO SAPIENS | 4 | 39596 | 1mhn | A | 111 | ALA | p.Ala111Gly | Q16637 |
| HOMO SAPIENS | 9 | 39596 | 1mhn | A | 116 | ILE | p.Ile116Phe | Q16637 |
| HOMO SAPIENS | 1 | 39596 | 1mhn | A | 136 | GLN | p.Gln136Glu | Q16637 |
| HOMO SAPIENS | 1 | 6408 | 5xjy | A | 85 | PRO | p.Pro85Leu | O95477 |
| HOMO SAPIENS | 0 | 6408 | 5xjy | A | 219 | ARG | p.Arg219Lys | O95477 |
| HOMO SAPIENS | 0 | 6408 | 5xjy | A | 230 | ARG | p.Arg230Cys | O95477 |
| HOMO SAPIENS | 1 | 6408 | 5xjy | A | 281 | MET | p.Met281Thr | O95477 |
| HOMO SAPIENS | -1 | 6408 | 5xjy | A | 343 | ALA | p.Ala343Gly | O95477 |
| HOMO SAPIENS | 1 | 6408 | 5xjy | A | 399 | VAL | p.Val399Ala | O95477 |
| HOMO SAPIENS | 1 | 6408 | 5xjy | A | 496 | ARG | p.Arg496Trp | O95477 |
| HOMO SAPIENS | 1 | 6408 | 5xjy | A | 590 | TRP | p.Trp590Ser | O95477 |
| HOMO SAPIENS | 1 | 6408 | 5xjy | A | 638 | ARG | p.Arg638Gln | O95477 |
| HOMO SAPIENS | 4 | 6408 | 5xjy | A | 697 | ALA | p.Ala697Thr | O95477 |
| HOMO SAPIENS | 1 | 6408 | 5xjy | A | 771 | VAL | p.Val771Met | O95477 |
| HOMO SAPIENS | 1 | 6408 | 5xjy | A | 774 | THR | p.Thr774Pro | O95477 |
| HOMO SAPIENS | 1 | 6408 | 5xjy | A | 776 | LYS | p.Lys776Asn | O95477 |
| HOMO SAPIENS | 0 | 6408 | 5xjy | A | 825 | VAL | p.Val825Ile | O95477 |
| HOMO SAPIENS | -1 | 6408 | 5xjy | A | 868 | GLU | p.Glu868Lys | O95477 |
| HOMO SAPIENS | -1 | 6408 | 5xjy | A | 883 | ILE | p.Ile883Met | O95477 |
| HOMO SAPIENS | 0 | 6408 | 5xjy | A | 935 | ASN | p.Asn935Ser | O95477 |
| HOMO SAPIENS | 0 | 6408 | 5xjy | A | 935 | ASN | p.Asn935His | O95477 |
| HOMO SAPIENS | 0 | 6408 | 5xjy | A | 937 | ALA | p.Ala937Val | O95477 |
| HOMO SAPIENS | 0 | 6408 | 5xjy | A | 1018 | ASP | p.Asp1018Gly | O95477 |
| HOMO SAPIENS | 0 | 6408 | 5xjy | A | 1096 | VAL | p.Val1096Ile | O95477 |
| HOMO SAPIENS | 0 | 6408 | 5xjy | A | 1099 | ASP | p.Asp1099Tyr | O95477 |
| HOMO SAPIENS | 0 | 6408 | 5xjy | A | 1172 | GLU | p.Glu1172Asp | O95477 |
| HOMO SAPIENS | 0 | 6408 | 5xjy | A | 1181 | SER | p.Ser1181Phe | O95477 |
| HOMO SAPIENS | 0 | 6408 | 5xjy | A | 1182 | ALA | p.Ala1182Thr | O95477 |

| | | | | | | | | |
|---|---|---|---|---|---|---|---|---|
| HOMO SAPIENS | -1 | 6408 | 5xjy | A | 1289 | ASP | p.Asp1289Asn | O95477 |
| HOMO SAPIENS | 0 | 6408 | 5xjy | A | 1399 | THR | p.Thr1399Ser | O95477 |
| HOMO SAPIENS | 1 | 6408 | 5xjy | A | 1408 | LEU | p.Leu1408Phe | O95477 |
| HOMO SAPIENS | 0 | 6408 | 5xjy | A | 1587 | LYS | p.Lys1587Arg | O95477 |
| HOMO SAPIENS | 1 | 6408 | 5xjy | A | 1674 | VAL | p.Val1674Ile | O95477 |
| HOMO SAPIENS | 1 | 6408 | 5xjy | A | 1680 | ARG | p.Arg1680Trp | O95477 |
| HOMO SAPIENS | 1 | 6408 | 5xjy | A | 1680 | ARG | p.Arg1680Gln | O95477 |
| HOMO SAPIENS | 0 | 6408 | 5xjy | A | 1800 | ASN | p.Asn1800His | O95477 |
| HOMO SAPIENS | -1 | 6408 | 5xjy | A | 1925 | ARG | p.Arg1925Gln | O95477 |
| HOMO SAPIENS | 0 | 6408 | 5xjy | A | 2009 | PHE | p.Phe2009Ser | O95477 |
| HOMO SAPIENS | -1 | 6408 | 5xjy | A | 2243 | ASP | p.Asp2243Glu | O95477 |
| HOMO SAPIENS | 4 | 23760 | 2mut | A | 231 | LEU | p.Phe231Leu | P07992 |
| HOMO SAPIENS | 6 | 1460 | 2hre | B | 96 | ARG | p.Arg96Gln | P22830 |
| HOMO SAPIENS | 8 | 1460 | 2hre | B | 129 | GLY | p.Gly129Arg | P22830 |
| HOMO SAPIENS | 9 | 1460 | 2hre | B | 172 | GLU | p.Glu172Lys | P22830 |
| HOMO SAPIENS | 4 | 1460 | 2hre | B | 285 | GLN | p.Gln285Arg | P22830 |
| HOMO SAPIENS | 5 | 1460 | 2hre | B | 362 | VAL | p.Val362Gly | P22830 |
| HOMO SAPIENS | 6 | 1460 | 2hre | B | 411 | CYS | p.Cys411Gly | P22830 |
| HOMO SAPIENS | 6 | 1460 | 2hre | B | 417 | PHE | p.Phe417Ser | P22830 |
| HOMO SAPIENS | 0 | 2210 | 5bug | A | 19 | ASP | p.Asp19Asn | P60484 |
| HOMO SAPIENS | 0 | 2210 | 5bug | A | 35 | MET | p.Met35Arg | P60484 |
| HOMO SAPIENS | 1 | 2210 | 5bug | A | 61 | HIS | p.His61Asp | P60484 |
| HOMO SAPIENS | 0 | 2210 | 5bug | A | 68 | TYR | p.Tyr68His | P60484 |
| HOMO SAPIENS | 8 | 2210 | 5bug | A | 70 | LEU | p.Leu70Pro | P60484 |
| HOMO SAPIENS | 6 | 2210 | 5bug | A | 93 | HIS | p.His93Arg | P60484 |
| HOMO SAPIENS | 1 | 2210 | 5bug | A | 112 | LEU | p.Leu112Pro | P60484 |
| HOMO SAPIENS | 0 | 2210 | 5bug | A | 121 | ALA | p.Ala121Gly | P60484 |
| HOMO SAPIENS | 3 | 2210 | 5bug | A | 123 | HIS | p.His123Arg | P60484 |
| HOMO SAPIENS | 7 | 2210 | 5bug | A | 124 | CYS | p.Cys124Arg | P60484 |
| HOMO SAPIENS | 9 | 2210 | 5bug | A | 129 | GLY | p.Gly129Glu | P60484 |
| HOMO SAPIENS | 9 | 2210 | 5bug | A | 130 | ARG | p.Arg130Gln | P60484 |
| HOMO SAPIENS | 9 | 2210 | 5bug | A | 130 | ARG | p.Arg130Gly | P60484 |
| HOMO SAPIENS | 0 | 2210 | 5bug | A | 132 | GLY | p.Gly132Val | P60484 |
| HOMO SAPIENS | 0 | 2210 | 5bug | A | 170 | SER | p.Ser170Arg | P60484 |
| HOMO SAPIENS | 1 | 2210 | 5bug | A | 173 | ARG | p.Arg173His | P60484 |
| HOMO SAPIENS | 0 | 2210 | 5bug | A | 217 | VAL | p.Val217Ile | P60484 |
| HOMO SAPIENS | 0 | 2210 | 5bug | A | 241 | PHE | p.Phe241Ser | P60484 |
| HOMO SAPIENS | 0 | 2210 | 5bug | A | 250 | CYS | p.Cys250Ter | P60484 |
| HOMO SAPIENS | 1 | 2210 | 5bug | A | 252 | ASP | p.Asp252Gly | P60484 |
| HOMO SAPIENS | 3 | 123 | 2ibu | A | 93 | ASN | p.Asn93Ser | P24752 |
| HOMO SAPIENS | 0 | 123 | 2ibu | A | 145 | GLN | p.Gln145Glu | P24752 |
| HOMO SAPIENS | 0 | 123 | 2ibu | A | 152 | GLY | p.Gly152Ala | P24752 |
| HOMO SAPIENS | 2 | 123 | 2ibu | A | 158 | ASN | p.Asn158Asp | P24752 |
| HOMO SAPIENS | 2 | 123 | 2ibu | A | 158 | ASN | p.Asn158Ser | P24752 |
| HOMO SAPIENS | 5 | 123 | 2ibu | A | 183 | GLY | p.Gly183Arg | P24752 |
| HOMO SAPIENS | 3 | 123 | 2ibu | A | 272 | GLN | p.Gln272Ter | P24752 |
| HOMO SAPIENS | 3 | 123 | 2ibu | A | 312 | ILE | p.Ile312Thr | P24752 |
| HOMO SAPIENS | 1 | 123 | 2ibu | A | 333 | ALA | p.Ala333Pro | P24752 |
| HOMO SAPIENS | 5 | 123 | 2ibu | A | 379 | GLY | p.Gly379Val | P24752 |
| HOMO SAPIENS | 0 | 123 | 2ibu | A | 380 | ALA | p.Ala380Thr | P24752 |
| HOMO SAPIENS | 2 | 369 | 5nrw | A | 1175 | TYR | p.Tyr1175Cys | Q92793 |
| HOMO SAPIENS | 2 | 830 | 2vc2 | A | 116 | LEU | p.Leu147Val | P08514 |
| HOMO SAPIENS | 0 | 830 | 2vc2 | A | 183 | LEU | p.Leu214Pro | P08514 |
| HOMO SAPIENS | 4 | 830 | 2vc2 | A | 242 | GLY | p.Gly273Asp | P08514 |

| | | | | | | | | |
|---|---|---|---|---|---|---|---|---|
| HOMO SAPIENS | 6 | 830 | 2vc2 | A | 324 | GLU | p.Glu355Lys | P08514 |
| HOMO SAPIENS | -1 | 10353 | 5ejc | E | 989 | ALA | p.Ala989Ser | Q92574 |
| HOMO SAPIENS | 0 | 15448 | 3lgg | A | 21 | GLY | p.Gly47Arg | Q9NZK5 |
| HOMO SAPIENS | 0 | 15448 | 3lgg | A | 21 | GLY | p.Gly47Val | Q9NZK5 |
| HOMO SAPIENS | 9 | 15448 | 3lgg | A | 86 | HIS | p.His112Gln | Q9NZK5 |
| HOMO SAPIENS | 0 | 15448 | 3lgg | A | 143 | ARG | p.Arg169Gln | Q9NZK5 |
| HOMO SAPIENS | 3 | 15448 | 3lgg | A | 221 | ALA | p.Ala247Val | Q9NZK5 |
| HOMO SAPIENS | 0 | 15448 | 3lgg | A | 225 | PRO | p.Pro251Leu | Q9NZK5 |
| HOMO SAPIENS | 0 | 15448 | 3lgg | A | 283 | MET | p.Met309Ile | Q9NZK5 |
| HOMO SAPIENS | 2 | 15448 | 3lgg | A | 334 | THR | p.Thr360Ala | Q9NZK5 |
| HOMO SAPIENS | 7 | 15448 | 3lgg | A | 357 | GLY | p.Gly383Ser | Q9NZK5 |
| HOMO SAPIENS | 0 | 15448 | 3lgg | A | 427 | TYR | p.Tyr453Cys | Q9NZK5 |
| HOMO SAPIENS | 0 | 872 | 2y3i | A | 87 | LEU | p.Leu88Pro | P00558 |
| HOMO SAPIENS | 1 | 872 | 2y3i | A | 163 | ASP | p.Asp164Val | P00558 |
| HOMO SAPIENS | 4 | 872 | 2y3i | A | 205 | ARG | p.Arg206Pro | P00558 |
| HOMO SAPIENS | 4 | 872 | 2y3i | A | 267 | ASP | p.Asp268Asn | P00558 |
| HOMO SAPIENS | 8 | 872 | 2y3i | A | 284 | ASP | p.Asp285Val | P00558 |
| HOMO SAPIENS | 4 | 8 | 5jco | B | 188 | ILE | p.Ile188Leu | Q71U36 |
| HOMO SAPIENS | 7 | 8 | 5jco | B | 263 | PRO | p.Pro263Thr | Q71U36 |
| HOMO SAPIENS | 6 | 8 | 5jco | B | 264 | ARG | p.Arg264Cys | Q71U36 |
| HOMO SAPIENS | 7 | 8 | 5jco | B | 402 | ARG | p.Arg402His | Q71U36 |
| HOMO SAPIENS | 7 | 8 | 5jco | B | 419 | SER | p.Ser419Leu | Q71U36 |
| HOMO SAPIENS | -1 | 880 | 4nwf | A | 2 | THR | p.Thr2Ile | Q06124 |
| HOMO SAPIENS | 2 | 880 | 4nwf | A | 58 | ASN | p.Asn58Ser | Q06124 |
| HOMO SAPIENS | 4 | 880 | 4nwf | A | 61 | ASP | p.Asp61Gly | Q06124 |
| HOMO SAPIENS | 2 | 880 | 4nwf | A | 62 | TYR | p.Tyr62Asp | Q06124 |
| HOMO SAPIENS | 3 | 880 | 4nwf | A | 63 | TYR | p.Tyr63Cys | Q06124 |
| HOMO SAPIENS | 2 | 880 | 4nwf | A | 69 | GLU | p.Glu69Gln | Q06124 |
| HOMO SAPIENS | 2 | 880 | 4nwf | A | 72 | ALA | p.Ala72Gly | Q06124 |
| HOMO SAPIENS | 2 | 880 | 4nwf | A | 72 | ALA | p.Ala72Ser | Q06124 |
| HOMO SAPIENS | 3 | 880 | 4nwf | A | 73 | THR | p.Thr73Ile | Q06124 |
| HOMO SAPIENS | 5 | 880 | 4nwf | A | 76 | GLU | p.Glu76Ala | Q06124 |
| HOMO SAPIENS | 5 | 880 | 4nwf | A | 76 | GLU | p.Glu76Lys | Q06124 |
| HOMO SAPIENS | 3 | 880 | 4nwf | A | 79 | GLN | p.Gln79Arg | Q06124 |
| HOMO SAPIENS | 5 | 880 | 4nwf | A | 152 | ARG | p.Arg152His | Q06124 |
| HOMO SAPIENS | 2 | 880 | 4nwf | A | 268 | GLY | p.Gly268Cys | Q06124 |
| HOMO SAPIENS | 9 | 880 | 4nwf | A | 276 | LYS | p.Lys276Arg | Q06124 |
| HOMO SAPIENS | 9 | 880 | 4nwf | A | 279 | TYR | p.Tyr279Cys | Q06124 |
| HOMO SAPIENS | 5 | 880 | 4nwf | A | 285 | PHE | p.Phe285Ser | Q06124 |
| HOMO SAPIENS | 0 | 880 | 4nwf | A | 308 | ASP | p.Asn308Ser | Q06124 |
| HOMO SAPIENS | 0 | 880 | 4nwf | A | 308 | ASP | p.Asn308Asp | Q06124 |
| HOMO SAPIENS | 5 | 880 | 4nwf | A | 392 | ALA | p.Ala392Thr | Q06124 |
| HOMO SAPIENS | 1 | 880 | 4nwf | A | 411 | THR | p.Thr415Met | Q06124 |
| HOMO SAPIENS | 0 | 880 | 4nwf | A | 461 | ALA | p.Ala465Thr | Q06124 |
| HOMO SAPIENS | 0 | 880 | 4nwf | A | 464 | GLY | p.Gly468Ala | Q06124 |
| HOMO SAPIENS | 1 | 880 | 4nwf | A | 491 | PRO | p.Pro495Ser | Q06124 |
| HOMO SAPIENS | 1 | 880 | 4nwf | A | 502 | SER | p.Ser506Thr | Q06124 |
| HOMO SAPIENS | 8 | 880 | 4nwf | A | 510 | GLN | p.Gln514Pro | Q06124 |
| HOMO SAPIENS | 4 | 1872 | 3wf1 | D | 109 | ARG | p.Arg109Trp | P16278 |
| HOMO SAPIENS | 0 | 1872 | 3wf1 | D | 521 | ARG | p.Arg521Cys | P16278 |
| HOMO SAPIENS | 5 | 910 | 5xte | A | 45 | SER | p.Ser45Phe | O14949 |
| HOMO SAPIENS | 2 | 9017 | 5f0k | A | 382 | LYS | p.Lys382Arg | Q96QK1 |
| HOMO SAPIENS | 1 | 710 | 4lsj | A | 559 | ILE | p.Ile559Asn | P04150 |
| HOMO SAPIENS | 5 | 710 | 4lsj | A | 571 | VAL | p.Val571Ala | P04150 |

| | | | | | | | | |
|---|---|---|---|---|---|---|---|---|
| HOMO SAPIENS | 0 | 710 | 4lsj | A | 641 | ASP | p.Asp641Val | P04150 |
| HOMO SAPIENS | 1 | 710 | 4lsj | A | 678 | ASP | p.Asp678Glu | P04150 |
| HOMO SAPIENS | 2 | 710 | 4lsj | A | 679 | GLY | p.Gly679Ser | P04150 |
| HOMO SAPIENS | 1 | 710 | 4lsj | A | 737 | PHE | p.Phe737Leu | P04150 |
| HOMO SAPIENS | 1 | 710 | 4lsj | A | 747 | ILE | p.Ile747Met | P04150 |
| HOMO SAPIENS | 1 | 710 | 4lsj | A | 753 | LEU | p.Leu753Phe | P04150 |
| HOMO SAPIENS | 1 | 710 | 4lsj | A | 773 | LEU | p.Leu773Pro | P04150 |
| HOMO SAPIENS | 1 | 2145 | 1av1 | B | 71 | PHE | p.Phe95Tyr | P02647 |
| HOMO SAPIENS | 1 | 2145 | 1av1 | B | 128 | GLU | p.Glu152Lys | P02647 |
| HOMO SAPIENS | 1 | 2145 | 1av1 | B | 136 | GLU | p.Glu160Lys | P02647 |
| HOMO SAPIENS | 2 | 2145 | 1av1 | B | 143 | PRO | p.Pro167Arg | P02647 |
| HOMO SAPIENS | 6 | 2145 | 1av1 | B | 164 | ALA | p.Ala188Ser | P02647 |
| HOMO SAPIENS | 7 | 2145 | 1av1 | B | 173 | ARG | p.Arg197Cys | P02647 |
| HOMO SAPIENS | 2 | 2145 | 1av1 | B | 198 | GLU | p.Glu222Lys | P02647 |
| HOMO SAPIENS | -1 | 3432 | 2beu | A | 1 | SER | p.Ser46Ter | P12694 |
| HOMO SAPIENS | 5 | 3432 | 2beu | A | 79 | ARG | p.Arg124Trp | P12694 |
| HOMO SAPIENS | 7 | 3432 | 2beu | A | 114 | ARG | p.Arg159Gln | P12694 |
| HOMO SAPIENS | 6 | 3432 | 2beu | A | 166 | THR | p.Thr211Met | P12694 |
| HOMO SAPIENS | 4 | 3432 | 2beu | A | 175 | ALA | p.Ala220Val | P12694 |
| HOMO SAPIENS | 5 | 3432 | 2beu | A | 204 | GLY | p.Gly249Ser | P12694 |
| HOMO SAPIENS | 4 | 3432 | 2beu | A | 209 | ALA | p.Ala254Asp | P12694 |
| HOMO SAPIENS | 6 | 3432 | 2beu | A | 220 | ARG | p.Arg265Trp | P12694 |
| HOMO SAPIENS | 6 | 3432 | 2beu | A | 234 | TYR | p.Tyr279Cys | P12694 |
| HOMO SAPIENS | 5 | 3432 | 2beu | A | 240 | ALA | p.Ala285Pro | P12694 |
| HOMO SAPIENS | 4 | 3432 | 2beu | A | 245 | GLY | p.Gly290Arg | P12694 |
| HOMO SAPIENS | 5 | 3432 | 2beu | A | 269 | ARG | p.Arg314Ter | P12694 |
| HOMO SAPIENS | 3 | 3432 | 2beu | A | 301 | ARG | p.Arg346Cys | P12694 |
| HOMO SAPIENS | 5 | 3432 | 2beu | A | 359 | ASN | p.Asn404Ser | P12694 |
| HOMO SAPIENS | 4 | 3432 | 2beu | A | 393 | TYR | p.Tyr438Asn | P12694 |
| HOMO SAPIENS | 8 | 4541 | 1rr8 | C | 533 | ASP | p.Asp533Gly | P11387 |
| HOMO SAPIENS | 2 | 3247 | 1tzn | d | 1105 | GLY | p.Gly105Asp | P58335 |
| HOMO SAPIENS | 3 | 3247 | 1tzn | d | 1111 | ARG | p.Arg111Cys | P58335 |
| HOMO SAPIENS | 2 | 3247 | 1tzn | d | 1189 | ILE | p.Ile189Thr | P58335 |
| HOMO SAPIENS | 9 | 467 | 1suv | C | 58 | ASP | p.Asp77Asn | P02787 |
| HOMO SAPIENS | 2 | 467 | 1suv | C | 277 | ASP | p.Asp296Gly | P02787 |
| HOMO SAPIENS | 9 | 6485 | 1xwd | B | 51 | CYS | p.Cys69Gly | P01225 |
| HOMO SAPIENS | 1 | 694 | 1wuu | A | 28 | PRO | p.Pro28Thr | P51570 |
| HOMO SAPIENS | 0 | 694 | 1wuu | A | 32 | VAL | p.Val32Met | P51570 |
| HOMO SAPIENS | 0 | 694 | 1wuu | A | 198 | ALA | p.Ala198Val | P51570 |
| HOMO SAPIENS | 0 | 143 | 4nkx | A | 93 | PHE | p.Phe93Cys | P05093 |
| HOMO SAPIENS | 3 | 143 | 4nkx | A | 96 | ARG | p.Arg96Trp | P05093 |
| HOMO SAPIENS | 3 | 143 | 4nkx | A | 96 | ARG | p.Arg96Gln | P05093 |
| HOMO SAPIENS | 0 | 143 | 4nkx | A | 106 | SER | p.Ser106Pro | P05093 |
| HOMO SAPIENS | 1 | 143 | 4nkx | A | 114 | PHE | p.Phe114Val | P05093 |
| HOMO SAPIENS | 1 | 143 | 4nkx | A | 116 | ASP | p.Asp116Val | P05093 |
| HOMO SAPIENS | 3 | 143 | 4nkx | A | 125 | ARG | p.Arg125Gln | P05093 |
| HOMO SAPIENS | 0 | 143 | 4nkx | A | 201 | TYR | p.Tyr201Asp | P05093 |
| HOMO SAPIENS | 0 | 143 | 4nkx | A | 239 | ARG | p.Arg239Ter | P05093 |
| HOMO SAPIENS | 0 | 143 | 4nkx | A | 329 | TYR | p.Tyr329Asp | P05093 |
| HOMO SAPIENS | 1 | 143 | 4nkx | A | 342 | PRO | p.Pro342Thr | P05093 |
| HOMO SAPIENS | 2 | 143 | 4nkx | A | 347 | ARG | p.Arg347His | P05093 |
| HOMO SAPIENS | 2 | 143 | 4nkx | A | 347 | ARG | p.Arg347Cys | P05093 |
| HOMO SAPIENS | 1 | 143 | 4nkx | A | 358 | ARG | p.Arg358Gln | P05093 |
| HOMO SAPIENS | 0 | 143 | 4nkx | A | 362 | ARG | p.Arg362Cys | P05093 |

| | | | | | | | | |
|---|---|---|---|---|---|---|---|---|
| HOMO SAPIENS | 1 | 143 | 4nkx | A | 406 | TRP | p.Trp406Arg | P05093 |
| HOMO SAPIENS | 0 | 143 | 4nkx | A | 428 | PRO | p.Pro428Leu | P05093 |
| HOMO SAPIENS | 6 | 1441 | 1uuh | A | 46 | ARG | p.Arg46Gly | P16070 |
| HOMO SAPIENS | 3 | 82 | 4wlv | D | 37 | GLY | p.Gly37Arg | P40926 |
| HOMO SAPIENS | 0 | 82 | 4wlv | D | 133 | PRO | p.Pro133Leu | P40926 |
| HOMO SAPIENS | 2 | 392 | 5xtd | A | 56 | SER | p.Ser56Pro | P49821 |
| HOMO SAPIENS | 2 | 392 | 5xtd | A | 59 | ARG | p.Arg59Ter | P49821 |
| HOMO SAPIENS | 2 | 392 | 5xtd | A | 74 | ASP | p.Asp74Asn | P49821 |
| HOMO SAPIENS | 4 | 392 | 5xtd | A | 116 | ASN | p.Asn116Lys | P49821 |
| HOMO SAPIENS | 4 | 392 | 5xtd | A | 117 | ALA | p.Ala117Thr | P49821 |
| HOMO SAPIENS | 5 | 392 | 5xtd | A | 122 | PRO | p.Pro122Leu | P49821 |
| HOMO SAPIENS | 3 | 392 | 5xtd | A | 160 | GLY | p.Gly160Glu | P49821 |
| HOMO SAPIENS | 4 | 392 | 5xtd | A | 214 | GLU | p.Glu214Lys | P49821 |
| HOMO SAPIENS | 2 | 392 | 5xtd | A | 357 | MET | p.Met357Thr | P49821 |
| HOMO SAPIENS | 4 | 392 | 5xtd | A | 423 | THR | p.Thr423Met | P49821 |
| HOMO SAPIENS | 0 | 22593 | 3pdf | A | 8 | TYR | p.Tyr32Ter | P53634 |
| HOMO SAPIENS | 0 | 22593 | 3pdf | A | 15 | TRP | p.Trp39Ser | P53634 |
| HOMO SAPIENS | 0 | 22593 | 3pdf | A | 103 | HIS | p.His127Pro | P53634 |
| HOMO SAPIENS | 9 | 22593 | 3pdf | A | 228 | GLN | p.Gln252Leu | P53634 |
| HOMO SAPIENS | 3 | 22593 | 3pdf | A | 262 | GLN | p.Gln286Arg | P53634 |
| HOMO SAPIENS | 4 | 22593 | 3pdf | A | 277 | GLY | p.Gly301Ser | P53634 |
| HOMO SAPIENS | 0 | 22593 | 3pdf | A | 323 | TYR | p.Tyr347Cys | P53634 |
| HOMO SAPIENS | 5 | 3461 | 4c2a | A | 1306 | GLN | p.Arg1306Trp | P04275 |
| HOMO SAPIENS | 5 | 3461 | 4c2a | A | 1374 | ARG | p.Arg1374Cys | P04275 |
| HOMO SAPIENS | 5 | 3461 | 4c2a | A | 1379 | ARG | p.Arg1379Cys | P04275 |
| HOMO SAPIENS | 4 | 3461 | 4c2a | A | 1380 | ILE | p.Ile1380Val | P04275 |
| HOMO SAPIENS | 0 | 3461 | 4c2a | A | 1381 | ALA | p.Thr1381Ala | P04275 |
| HOMO SAPIENS | 5 | 3461 | 4c2a | A | 1399 | ARG | p.Arg1399His | P04275 |
| HOMO SAPIENS | 5 | 3461 | 4c2a | A | 1435 | ASN | p.Asn1435Ser | P04275 |
| HOMO SAPIENS | 0 | 4533 | 5uj7 | C | 174 | TYR | p.Tyr174Cys | O43929 |
| HOMO SAPIENS | 0 | 4533 | 5uj7 | C | 202 | LEU | p.Leu202Val | O43929 |
| HOMO SAPIENS | 0 | 1374 | 3j07 | O | 39 | PRO | p.Pro39Leu | P02511 |
| HOMO SAPIENS | 0 | 1374 | 3j07 | O | 107 | ARG | p.Arg107His | P02511 |
| HOMO SAPIENS | 0 | 1374 | 3j07 | O | 157 | ARG | p.Arg157His | P02511 |
| HOMO SAPIENS | -1 | 942 | 1s9i | B | 57 | PHE | p.Phe57Cys | P36507 |
| HOMO SAPIENS | -1 | 942 | 1s9i | B | 57 | PHE | p.Phe57Val | P36507 |
| HOMO SAPIENS | 2 | 942 | 1s9i | B | 128 | PRO | p.Pro128Gln | P36507 |
| HOMO SAPIENS | 1 | 942 | 1s9i | B | 134 | TYR | p.Tyr134His | P36507 |
| HOMO SAPIENS | 1 | 942 | 1s9i | B | 188 | HIS | p.His188Gln | P36507 |
| HOMO SAPIENS | 0 | 942 | 1s9i | B | 262 | VAL | p.Val262Phe | P36507 |
| HOMO SAPIENS | 0 | 942 | 1s9i | B | 263 | GLY | p.Gly263Arg | P36507 |
| HOMO SAPIENS | -1 | 942 | 1s9i | B | 298 | PRO | p.Pro298Leu | P36507 |
| HOMO SAPIENS | -1 | 942 | 1s9i | B | 305 | VAL | p.Val305Ile | P36507 |
| HOMO SAPIENS | 0 | 3245 | 1fbv | A | 367 | GLN | p.Gln367Pro | P22681 |
| HOMO SAPIENS | 0 | 3245 | 1fbv | A | 371 | TYR | p.Tyr371His | P22681 |
| HOMO SAPIENS | 1 | 3245 | 1fbv | A | 382 | LYS | p.Lys382Glu | P22681 |
| HOMO SAPIENS | 2 | 3245 | 1fbv | A | 390 | ASP | p.Asp390Tyr | P22681 |
| HOMO SAPIENS | 9 | 3245 | 1fbv | A | 396 | CYS | p.Cys396Arg | P22681 |
| HOMO SAPIENS | 7 | 3245 | 1fbv | A | 420 | ARG | p.Arg420Gln | P22681 |
| HOMO SAPIENS | 1 | 3245 | 1fbv | A | 429 | ILE | p.Ile429Met | P22681 |
| HOMO SAPIENS | 0 | 2604 | 2pbd | P | 111 | LEU | p.Leu112Met | P07737 |
| HOMO SAPIENS | 3 | 2604 | 2pbd | P | 116 | GLU | p.Glu117Gly | P07737 |
| HOMO SAPIENS | 4 | 2013 | 3b7w | A | 472 | ARG | p.Arg472Gly | Q08AH3 |
| HOMO SAPIENS | 0 | 24346 | 3sh5 | A | 85 | ARG | p.Arg4281His | P98160 |

| | | | | | | | | |
|---|---|---|---|---|---|---|---|---|
| HOMO SAPIENS | 0 | 24346 | 3sh5 | A | 122 | ARG | p.Arg4318Trp | P98160 |
| HOMO SAPIENS | 0 | 24346 | 3sh5 | A | 135 | SER | p.Ser4331Asn | P98160 |
| HOMO SAPIENS | 0 | 24346 | 3sh5 | A | 136 | VAL | p.Val4332Ile | P98160 |
| HOMO SAPIENS | 0 | 707 | 4p2g | A | 26 | MET | p.Met26Ile | Q99497 |
| HOMO SAPIENS | 5 | 707 | 4p2g | A | 39 | ALA | p.Ala39Ser | Q99497 |
| HOMO SAPIENS | 5 | 707 | 4p2g | A | 64 | GLU | p.Glu64Asp | Q99497 |
| HOMO SAPIENS | 4 | 707 | 4p2g | A | 149 | ASP | p.Asp149Ala | Q99497 |
| HOMO SAPIENS | 6 | 707 | 4p2g | A | 163 | GLU | p.Glu163Lys | Q99497 |
| HOMO SAPIENS | 6 | 707 | 4p2g | A | 166 | LEU | p.Leu166Pro | Q99497 |
| HOMO SAPIENS | 3 | 5121 | 4xss | E | 31 | GLY | p.Gly58Arg | P06213 |
| HOMO SAPIENS | 2 | 5121 | 4xss | E | 119 | ILE | p.Ile146Met | P06213 |
| HOMO SAPIENS | 8 | 5121 | 4xss | E | 209 | HIS | p.His236Arg | P06213 |
| HOMO SAPIENS | 2 | 5121 | 4xss | E | 234 | ASP | p.Asp261Glu | P06213 |
| HOMO SAPIENS | 1 | 5121 | 4xss | E | 276 | GLN | p.Gln303His | P06213 |
| HOMO SAPIENS | 1 | 27790 | 5erp | B | 140 | THR | p.Thr275Met | Q02487 |
| HOMO SAPIENS | 0 | 27790 | 5erp | B | 150 | ILE | p.Ile285Thr | Q02487 |
| HOMO SAPIENS | 1 | 27790 | 5erp | B | 205 | THR | p.Thr340Ala | Q02487 |
| HOMO SAPIENS | 1 | 27790 | 5erp | B | 223 | THR | p.Thr358Ile | Q02487 |
| HOMO SAPIENS | 0 | 27790 | 5erp | B | 301 | GLY | p.Gly436Ala | Q02487 |
| HOMO SAPIENS | -1 | 27790 | 5erp | B | 315 | ARG | p.Arg450Ser | Q02487 |
| HOMO SAPIENS | 0 | 27790 | 5erp | B | 369 | SER | p.Ser504Gly | Q02487 |
| HOMO SAPIENS | 0 | 27790 | 5erp | B | 385 | ILE | p.Ile520Thr | Q02487 |
| HOMO SAPIENS | 0 | 27790 | 5erp | B | 461 | ALA | p.Ala596Val | Q02487 |
| HOMO SAPIENS | 0 | 27790 | 5erp | B | 503 | GLN | p.Gln638His | Q02487 |
| HOMO SAPIENS | 5 | 6599 | 4nsc | A | 129 | ARG | p.Arg129Leu | Q9BPX6 |
| HOMO SAPIENS | -1 | 6599 | 4nsc | A | 183 | GLN | p.Gln183Ter | Q9BPX6 |
| HOMO SAPIENS | 0 | 20420 | 3ghm | A | 398 | ARG | p.Arg398His | Q76LX8 |
| HOMO SAPIENS | 0 | 20420 | 3ghm | A | 448 | GLN | p.Gln448Glu | Q76LX8 |
| HOMO SAPIENS | 1 | 20420 | 3ghm | A | 449 | GLN | p.Gln449Ter | Q76LX8 |
| HOMO SAPIENS | 0 | 20420 | 3ghm | A | 456 | GLN | p.Gln456His | Q76LX8 |
| HOMO SAPIENS | 1 | 20420 | 3ghm | A | 528 | ARG | p.Arg528Gly | Q76LX8 |
| HOMO SAPIENS | 0 | 20420 | 3ghm | A | 618 | PRO | p.Pro618Ala | Q76LX8 |
| HOMO SAPIENS | 0 | 20420 | 3ghm | A | 625 | ARG | p.Arg625His | Q76LX8 |
| HOMO SAPIENS | 0 | 275 | 4rmz | A | 293 | GLN | p.Gln293Ter | Q9NWZ3 |
| HOMO SAPIENS | 0 | 275 | 4rmz | A | 428 | ALA | p.Ala428Thr | Q9NWZ3 |
| HOMO SAPIENS | 2 | 10458 | 3puf | B | 138 | LEU | p.Leu138Phe | Q5TBB1 |
| HOMO SAPIENS | 6 | 10458 | 3puf | B | 177 | ALA | p.Ala177Thr | Q5TBB1 |
| HOMO SAPIENS | -1 | 10458 | 3puf | B | 185 | VAL | p.Val185Gly | Q5TBB1 |
| HOMO SAPIENS | 0 | 26648 | 2n90 | A | 36 | ARG | p.Arg444Gln | P04629 |
| HOMO SAPIENS | 2 | 977 | 1dnw | D | 166 | ALA | p.Ala332Val | P05164 |
| HOMO SAPIENS | 4 | 977 | 1dnw | D | 333 | ARG | p.Arg499Cys | P05164 |
| HOMO SAPIENS | 1 | 977 | 1dnw | D | 403 | ARG | p.Arg569Trp | P05164 |
| HOMO SAPIENS | 0 | 25704 | 3ajb | A | 82 | GLN | p.Gln82Arg | P56589 |
| HOMO SAPIENS | 3 | 255 | 3t1p | A | 38 | TYR | p.Tyr62Ter | P01009 |
| HOMO SAPIENS | 3 | 255 | 3t1p | A | 39 | ARG | p.Arg63Cys | P01009 |
| HOMO SAPIENS | 0 | 255 | 3t1p | A | 101 | ARG | p.Arg125His | P01009 |
| HOMO SAPIENS | 0 | 255 | 3t1p | A | 212 | GLN | p.Gln236Arg | P01009 |
| HOMO SAPIENS | 0 | 255 | 3t1p | A | 213 | VAL | p.Val237Ala | P01009 |
| HOMO SAPIENS | 1 | 255 | 3t1p | A | 269 | HIS | p.His293Gln | P01009 |
| HOMO SAPIENS | 8 | 255 | 3t1p | A | 341 | ASP | p.Asp365Asn | P01009 |
| HOMO SAPIENS | 9 | 255 | 3t1p | A | 342 | GLU | p.Glu366Lys | P01009 |
| HOMO SAPIENS | 0 | 255 | 3t1p | A | 369 | PRO | p.Pro393Ser | P01009 |
| HOMO SAPIENS | 0 | 255 | 3t1p | A | 376 | GLU | p.Glu400Asp | P01009 |
| HOMO SAPIENS | -1 | 5175 | 3zxi | A | 35 | ALA | p.Ala35Asp | Q9Y2Z4 |

| | | | | | | | | | |
|---|---|---|---|---|---|---|---|---|---|
| HOMO SAPIENS | 0 | 5175 | 3zxi | A | | 52 | PHE | p.Phe52Leu | Q9Y2Z4 |
| HOMO SAPIENS | -1 | 5175 | 3zxi | A | | 68 | GLY | p.Gly68Ser | Q9Y2Z4 |
| HOMO SAPIENS | 0 | 5175 | 3zxi | A | | 179 | LYS | p.Lys179Gln | Q9Y2Z4 |
| HOMO SAPIENS | 0 | 5175 | 3zxi | A | | 191 | GLY | p.Gly191Val | Q9Y2Z4 |
| HOMO SAPIENS | 0 | 5175 | 3zxi | A | | 251 | ILE | p.Ile251Val | Q9Y2Z4 |
| HOMO SAPIENS | 2 | 5175 | 3zxi | A | | 295 | ASP | p.Asp295Asn | Q9Y2Z4 |
| HOMO SAPIENS | 0 | 5175 | 3zxi | A | | 311 | ASP | p.Asp311Glu | Q9Y2Z4 |
| HOMO SAPIENS | 0 | 5175 | 3zxi | A | | 360 | ARG | p.Arg360Ter | Q9Y2Z4 |
| HOMO SAPIENS | 1 | 3016 | 3exg | | 3 | 4 | ASP | p.Asp33His | P08559 |
| HOMO SAPIENS | 1 | 3016 | 3exg | | 3 | 176 | PHE | p.Phe205Leu | P08559 |
| HOMO SAPIENS | 1 | 3016 | 3exg | | 3 | 214 | TYR | p.Tyr243Asn | P08559 |
| HOMO SAPIENS | 1 | 3016 | 3exg | | 3 | 229 | ASP | p.Asp258Ala | P08559 |
| HOMO SAPIENS | 0 | 3016 | 3exg | | 3 | 233 | VAL | p.Val262Ile | P08559 |
| HOMO SAPIENS | 1 | 3016 | 3exg | | 3 | 234 | ARG | p.Arg263Gly | P08559 |
| HOMO SAPIENS | 0 | 3016 | 3exg | | 3 | 253 | MET | p.Met282Leu | P08559 |
| HOMO SAPIENS | 7 | 3016 | 3exg | | 3 | 259 | ARG | p.Arg288His | P08559 |
| HOMO SAPIENS | -1 | 3016 | 3exg | | 3 | 273 | ARG | p.Arg302Cys | P08559 |
| HOMO SAPIENS | 2 | 3016 | 3exg | | 3 | 286 | ASP | p.Asp315Asn | P08559 |
| HOMO SAPIENS | 1 | 3016 | 3exg | | 3 | 349 | ARG | p.Arg378His | P08559 |
| HOMO SAPIENS | 5 | 6729 | 4a0l | E | | 572 | ARG | p.Arg572Cys | Q13620 |
| HOMO SAPIENS | 0 | 1067 | 4kyo | A | | 9 | THR | p.Thr9Asn | P21549 |
| HOMO SAPIENS | 0 | 1067 | 4kyo | A | | 11 | PRO | p.Pro11Leu | P21549 |
| HOMO SAPIENS | 0 | 1067 | 4kyo | A | | 22 | ASN | p.Asn22Ser | P21549 |
| HOMO SAPIENS | 4 | 1067 | 4kyo | A | | 25 | LEU | p.Leu25Arg | P21549 |
| HOMO SAPIENS | 7 | 1067 | 4kyo | A | | 36 | ARG | p.Arg36His | P21549 |
| HOMO SAPIENS | 3 | 1067 | 4kyo | A | | 41 | GLY | p.Gly41Arg | P21549 |
| HOMO SAPIENS | 0 | 1067 | 4kyo | A | | 42 | GLY | p.Gly42Glu | P21549 |
| HOMO SAPIENS | 3 | 1067 | 4kyo | A | | 56 | ILE | p.Ile56Asn | P21549 |
| HOMO SAPIENS | 3 | 1067 | 4kyo | A | | 59 | GLU | p.Glu59Lys | P21549 |
| HOMO SAPIENS | 3 | 1067 | 4kyo | A | | 66 | TYR | p.Tyr66Ter | P21549 |
| HOMO SAPIENS | 3 | 1067 | 4kyo | A | | 111 | ARG | p.Arg111Leu | P21549 |
| HOMO SAPIENS | 7 | 1067 | 4kyo | A | | 118 | ARG | p.Arg118His | P21549 |
| HOMO SAPIENS | 3 | 1067 | 4kyo | A | | 122 | ARG | p.Arg122Ter | P21549 |
| HOMO SAPIENS | 3 | 1067 | 4kyo | A | | 124 | HIS | p.His124Pro | P21549 |
| HOMO SAPIENS | 0 | 1067 | 4kyo | A | | 152 | PHE | p.Phe152Ile | P21549 |
| HOMO SAPIENS | 1 | 1067 | 4kyo | A | | 156 | GLY | p.Gly156Arg | P21549 |
| HOMO SAPIENS | 0 | 1067 | 4kyo | A | | 161 | GLY | p.Gly161Arg | P21549 |
| HOMO SAPIENS | 3 | 1067 | 4kyo | A | | 166 | LEU | p.Leu166Pro | P21549 |
| HOMO SAPIENS | 5 | 1067 | 4kyo | A | | 170 | GLY | p.Gly170Arg | P21549 |
| HOMO SAPIENS | 5 | 1067 | 4kyo | A | | 186 | ALA | p.Ala186Val | P21549 |
| HOMO SAPIENS | 3 | 1067 | 4kyo | A | | 187 | SER | p.Ser187Phe | P21549 |
| HOMO SAPIENS | 3 | 1067 | 4kyo | A | | 197 | ARG | p.Arg197Pro | P21549 |
| HOMO SAPIENS | 5 | 1067 | 4kyo | A | | 201 | ASP | p.Asp201Glu | P21549 |
| HOMO SAPIENS | 0 | 1067 | 4kyo | A | | 202 | ILE | p.Ile202Asn | P21549 |
| HOMO SAPIENS | 1 | 1067 | 4kyo | A | | 205 | SER | p.Ser205Pro | P21549 |
| HOMO SAPIENS | 1 | 1067 | 4kyo | A | | 205 | SER | p.Ser205Ter | P21549 |
| HOMO SAPIENS | 2 | 1067 | 4kyo | A | | 216 | GLY | p.Gly216Arg | P21549 |
| HOMO SAPIENS | 5 | 1067 | 4kyo | A | | 233 | ARG | p.Arg233Cys | P21549 |
| HOMO SAPIENS | 5 | 1067 | 4kyo | A | | 244 | ILE | p.Ile244Thr | P21549 |
| HOMO SAPIENS | 5 | 1067 | 4kyo | A | | 244 | ILE | p.Ile244Met | P21549 |
| HOMO SAPIENS | 4 | 1067 | 4kyo | A | | 253 | CYS | p.Cys253Arg | P21549 |
| HOMO SAPIENS | 7 | 1067 | 4kyo | A | | 274 | GLU | p.Glu274Asp | P21549 |
| HOMO SAPIENS | 7 | 1067 | 4kyo | A | | 279 | ILE | p.Ile279Thr | P21549 |
| HOMO SAPIENS | 6 | 1067 | 4kyo | A | | 295 | ALA | p.Ala295Thr | P21549 |

| | | | | | | | | |
|---|---|---|---|---|---|---|---|---|
| HOMO SAPIENS | 5 | 1067 | 4kyo | A | 340 | ILE | p.Ile340Met | P21549 |
| HOMO SAPIENS | 1 | 790 | 2ric | C | 270 | SER | p.Ser290Thr | P35247 |
| HOMO SAPIENS | 0 | 790 | 2ric | C | 289 | GLU | p.Glu309Lys | P35247 |
| HOMO SAPIENS | 1 | 212 | 5sxe | A | 545 | GLU | p.Glu545Lys | P42336 |
| HOMO SAPIENS | 3 | 212 | 5sxe | A | 546 | GLN | p.Gln546Glu | P42336 |
| HOMO SAPIENS | 2 | 212 | 5sxe | A | 1047 | HIS | p.His1047Leu | P42336 |
| HOMO SAPIENS | 7 | 152 | 2zqq | F | 197 | ARG | p.Arg197Ter | Q13825 |
| HOMO SAPIENS | 0 | 152 | 2zqq | F | 248 | ALA | p.Ala248Thr | Q13825 |
| HOMO SAPIENS | 0 | 152 | 2zqq | F | 275 | ALA | p.Ala275Val | Q13825 |
| HOMO SAPIENS | 5 | 152 | 2zqq | F | 331 | LYS | p.Lys331Ter | Q13825 |
| HOMO SAPIENS | 6 | 17801 | 2j8l | A | 283 | PHE | p.Phe301Leu | P03951 |
| HOMO SAPIENS | 3 | 17801 | 2j8l | A | 308 | ARG | p.Arg326Cys | P03951 |
| HOMO SAPIENS | 0 | 17801 | 2j8l | A | 321 | CYS | p.Cys339Phe | P03951 |
| HOMO SAPIENS | 3 | 17801 | 2j8l | A | 351 | TYR | p.Tyr369Ter | P03951 |
| HOMO SAPIENS | 7 | 1802 | 5db0 | A | 22 | LEU | p.Leu22Arg | O00255 |
| HOMO SAPIENS | 7 | 1802 | 5db0 | A | 26 | GLU | p.Glu26Lys | O00255 |
| HOMO SAPIENS | 7 | 1802 | 5db0 | A | 139 | HIS | p.His139Asp | O00255 |
| HOMO SAPIENS | 0 | 1802 | 5db0 | A | 171 | ARG | p.Arg176Gln | O00255 |
| HOMO SAPIENS | 0 | 1802 | 5db0 | A | 184 | VAL | p.Val189Glu | O00255 |
| HOMO SAPIENS | 7 | 1802 | 5db0 | A | 255 | GLU | p.Glu260Lys | O00255 |
| HOMO SAPIENS | 4 | 1802 | 5db0 | A | 418 | ASP | p.Asp423Asn | O00255 |
| HOMO SAPIENS | 0 | 1802 | 5db0 | A | 436 | TRP | p.Trp441Arg | O00255 |
| HOMO SAPIENS | -1 | 1802 | 5db0 | A | 541 | ALA | p.Thr546Ala | O00255 |
| HOMO SAPIENS | 4 | 1802 | 5db0 | A | 552 | THR | p.Thr557Ser | O00255 |
| HOMO SAPIENS | 0 | 3450 | 2yuz | A | 36 | PRO | p.Pro461Gln | Q00872 |
| HOMO SAPIENS | 1 | 3450 | 2yuz | A | 56 | GLN | p.His481Gln | Q00872 |
| HOMO SAPIENS | 2 | 3450 | 2yuz | A | 68 | ALA | p.Ala493Val | Q00872 |
| HOMO SAPIENS | 0 | 3211 | 3rqg | A | 72 | VAL | p.Val72Leu | Q9BUL8 |
| HOMO SAPIENS | 2 | 4168 | 4q4f | A | 121 | ARG | p.Arg121Ter | Q14108 |
| HOMO SAPIENS | 2 | 4168 | 4q4f | A | 121 | ARG | p.Arg121Gln | Q14108 |
| HOMO SAPIENS | 2 | 4168 | 4q4f | A | 159 | MET | p.Met159Val | Q14108 |
| HOMO SAPIENS | 4 | 4168 | 4q4f | A | 363 | HIS | p.His363Asn | Q14108 |
| HOMO SAPIENS | 2 | 4168 | 4q4f | A | 396 | VAL | p.Val396Ile | Q14108 |
| HOMO SAPIENS | 0 | 1448 | 1y97 | A | 87 | ARG | p.Arg130Ter | Q9BQ50 |
| HOMO SAPIENS | 0 | 3444 | 4iwp | A | 151 | SER | p.Ser151Phe | Q9UHD2 |
| HOMO SAPIENS | 0 | 3444 | 4iwp | A | 401 | LYS | p.Lys401Glu | Q9UHD2 |
| HOMO SAPIENS | 0 | 116 | 1zy8 | B | 194 | GLY | p.Gly229Cys | P09622 |
| HOMO SAPIENS | 0 | 116 | 1zy8 | B | 255 | ASP | p.Asp290Gly | P09622 |
| HOMO SAPIENS | 2 | 116 | 1zy8 | B | 326 | MET | p.Met361Val | P09622 |
| HOMO SAPIENS | 0 | 116 | 1zy8 | B | 340 | GLU | p.Glu375Lys | P09622 |
| HOMO SAPIENS | 2 | 116 | 1zy8 | B | 358 | ILE | p.Ile393Thr | P09622 |
| HOMO SAPIENS | 1 | 116 | 1zy8 | B | 453 | PRO | p.Pro488Leu | P09622 |
| HOMO SAPIENS | 7 | 127 | 4qo7 | B | 160 | SER | p.Ser161Arg | P00338 |
| HOMO SAPIENS | 0 | 2025 | 1wrl | A | 8 | ALA | p.Ala8Val | P63316 |
| HOMO SAPIENS | 8 | 2025 | 1wrl | A | 24 | PHE | p.Phe24Leu | P63316 |
| HOMO SAPIENS | 0 | 2025 | 1wrl | A | 84 | SER | p.Cys84Tyr | P63316 |
| HOMO SAPIENS | 8 | 3791 | 1mhd | B | 93 | ARG | p.Arg93Ter | P84022 |
| HOMO SAPIENS | 0 | 1575 | 4phc | A | 205 | GLY | p.Gly205Asp | P12081 |
| HOMO SAPIENS | 0 | 1575 | 4phc | A | 454 | TYR | p.Tyr454Ser | P12081 |
| HOMO SAPIENS | 0 | 39620 | 1ozs | B | 135 | ARG | p.Arg136Gln | P19429 |
| HOMO SAPIENS | 5 | 39620 | 1ozs | B | 140 | ARG | p.Arg141Trp | P19429 |
| HOMO SAPIENS | 5 | 39620 | 1ozs | B | 142 | THR | p.Thr143Asn | P19429 |
| HOMO SAPIENS | 5 | 39620 | 1ozs | B | 143 | LEU | p.Leu144Gln | P19429 |
| HOMO SAPIENS | 5 | 39620 | 1ozs | B | 144 | ARG | p.Arg145Gly | P19429 |

| | | | | | | | | |
|---|---|---|---|---|---|---|---|---|
| HOMO SAPIENS | 5 | 39620 | 1ozs | B | 144 | ARG | p.Arg145Gln | P19429 |
| HOMO SAPIENS | -1 | 20934 | 4nuv | D | 42 | GLY | p.Gly42Asp | Q16570 |
| HOMO SAPIENS | -1 | 40565 | 5lcw | S | 4 | VAL | p.Val4Met | O60566 |
| HOMO SAPIENS | 0 | 40565 | 5lcw | S | 40 | THR | p.Thr40Met | O60566 |
| HOMO SAPIENS | 4 | 40565 | 5lcw | S | 198 | ALA | p.Glu390Asp | O60566 |
| HOMO SAPIENS | 4 | 758 | 4oiu | A | 726 | ARG | p.Arg727Leu | P10275 |
| HOMO SAPIENS | 4 | 758 | 4oiu | A | 730 | VAL | p.Val731Met | P10275 |
| HOMO SAPIENS | 2 | 758 | 4oiu | A | 798 | GLN | p.Gln799Glu | P10275 |
| HOMO SAPIENS | 3 | 758 | 4oiu | A | 855 | ARG | p.Arg856His | P10275 |
| HOMO SAPIENS | 1 | 758 | 4oiu | A | 886 | MET | p.Met887Val | P10275 |
| HOMO SAPIENS | 0 | 3303 | 4ie0 | A | 134 | ALA | p.Ala134Thr | Q9C0B1 |
| HOMO SAPIENS | 9 | 3303 | 4ie0 | A | 316 | ARG | p.Arg316Gln | Q9C0B1 |
| HOMO SAPIENS | 1 | 41502 | 3fgh | A | 136 | PRO | p.Pro178Leu | Q00059 |
| HOMO SAPIENS | 5 | 175 | 3ceh | B | 58 | THR | p.Thr59Lys | P06737 |
| HOMO SAPIENS | 8 | 175 | 3ceh | B | 93 | ARG | p.Arg94Ter | P06737 |
| HOMO SAPIENS | 5 | 175 | 3ceh | B | 221 | VAL | p.Val222Ile | P06737 |
| HOMO SAPIENS | 0 | 175 | 3ceh | B | 338 | ASN | p.Asn339Ser | P06737 |
| HOMO SAPIENS | 1 | 175 | 3ceh | B | 372 | PHE | p.Phe373Leu | P06737 |
| HOMO SAPIENS | 9 | 175 | 3ceh | B | 376 | ASN | p.Asn377Lys | P06737 |
| HOMO SAPIENS | 8 | 175 | 3ceh | B | 398 | ARG | p.Arg399Ter | P06737 |
| HOMO SAPIENS | 5 | 175 | 3ceh | B | 424 | ARG | p.Arg425Pro | P06737 |
| HOMO SAPIENS | 9 | 175 | 3ceh | B | 455 | VAL | p.Val456Met | P06737 |
| HOMO SAPIENS | 8 | 175 | 3ceh | B | 490 | ARG | p.Arg491Cys | P06737 |
| HOMO SAPIENS | 8 | 175 | 3ceh | B | 631 | ASN | p.Asn632Ile | P06737 |
| HOMO SAPIENS | 9 | 175 | 3ceh | B | 648 | TYR | p.Tyr649Ter | P06737 |
| HOMO SAPIENS | 9 | 175 | 3ceh | B | 672 | GLU | p.Glu673Lys | P06737 |
| HOMO SAPIENS | 9 | 175 | 3ceh | B | 674 | SER | p.Ser675Leu | P06737 |
| HOMO SAPIENS | 0 | 175 | 3ceh | B | 805 | ILE | p.Ile806Leu | P06737 |
| HOMO SAPIENS | 0 | 175 | 3ceh | B | 820 | TYR | p.Tyr821His | P06737 |
| HOMO SAPIENS | 1 | 6750 | 4fdj | A | 47 | GLY | p.Gly47Arg | P34059 |
| HOMO SAPIENS | 9 | 6750 | 4fdj | A | 60 | ASP | p.Asp60Asn | P34059 |
| HOMO SAPIENS | 5 | 6750 | 4fdj | A | 67 | LEU | p.Leu67Met | P34059 |
| HOMO SAPIENS | 0 | 6750 | 4fdj | A | 69 | PHE | p.Phe69Val | P34059 |
| HOMO SAPIENS | 3 | 6750 | 4fdj | A | 94 | ARG | p.Arg94Gly | P34059 |
| HOMO SAPIENS | 0 | 6750 | 4fdj | A | 113 | ILE | p.Ile113Phe | P34059 |
| HOMO SAPIENS | 0 | 6750 | 4fdj | A | 138 | VAL | p.Val138Ala | P34059 |
| HOMO SAPIENS | 0 | 6750 | 4fdj | A | 139 | GLY | p.Gly139Ser | P34059 |
| HOMO SAPIENS | 0 | 6750 | 4fdj | A | 151 | PRO | p.Pro151Thr | P34059 |
| HOMO SAPIENS | 0 | 6750 | 4fdj | A | 155 | GLY | p.Gly155Arg | P34059 |
| HOMO SAPIENS | 0 | 6750 | 4fdj | A | 159 | TRP | p.Trp159Ter | P34059 |
| HOMO SAPIENS | 0 | 6750 | 4fdj | A | 162 | SER | p.Ser162Phe | P34059 |
| HOMO SAPIENS | 0 | 6750 | 4fdj | A | 200 | THR | p.Thr200Met | P34059 |
| HOMO SAPIENS | 0 | 6750 | 4fdj | A | 204 | ASN | p.Asn204Lys | P34059 |
| HOMO SAPIENS | 0 | 6750 | 4fdj | A | 231 | ALA | p.Ala231Gly | P34059 |
| HOMO SAPIENS | 2 | 6750 | 4fdj | A | 235 | THR | p.Thr235Lys | P34059 |
| HOMO SAPIENS | 1 | 6750 | 4fdj | A | 259 | ARG | p.Arg259Gln | P34059 |
| HOMO SAPIENS | 0 | 6750 | 4fdj | A | 287 | SER | p.Ser287Leu | P34059 |
| HOMO SAPIENS | 0 | 6750 | 4fdj | A | 291 | ALA | p.Ala291Thr | P34059 |
| HOMO SAPIENS | 3 | 6750 | 4fdj | A | 301 | GLY | p.Gly301Cys | P34059 |
| HOMO SAPIENS | 0 | 6750 | 4fdj | A | 312 | THR | p.Thr312Ser | P34059 |
| HOMO SAPIENS | 0 | 6750 | 4fdj | A | 318 | MET | p.Met318Arg | P34059 |
| HOMO SAPIENS | 0 | 6750 | 4fdj | A | 340 | GLY | p.Gly340Asp | P34059 |
| HOMO SAPIENS | 0 | 6750 | 4fdj | A | 386 | ARG | p.Arg386Cys | P34059 |
| HOMO SAPIENS | 0 | 6750 | 4fdj | A | 391 | MET | p.Met391Val | P34059 |

| | | | | | | | | |
|---|---|---|---|---|---|---|---|---|
| HOMO SAPIENS | 0 | 6750 | 4fdj | A | 392 | ALA | p.Ala392Val | P34059 |
| HOMO SAPIENS | 0 | 6750 | 4fdj | A | 393 | ALA | p.Ala393Ser | P34059 |
| HOMO SAPIENS | 0 | 6750 | 4fdj | A | 459 | ALA | p.Ala459Val | P34059 |
| HOMO SAPIENS | 0 | 6750 | 4fdj | A | 473 | GLN | p.Gln473Ter | P34059 |
| HOMO SAPIENS | 2 | 6750 | 4fdj | A | 480 | VAL | p.Val480Phe | P34059 |
| HOMO SAPIENS | 0 | 6750 | 4fdj | A | 487 | ASN | p.Asn487Ser | P34059 |
| HOMO SAPIENS | 0 | 6750 | 4fdj | A | 488 | VAL | p.Val488Met | P34059 |
| HOMO SAPIENS | 0 | 6750 | 4fdj | A | 520 | TRP | p.Trp520Ter | P34059 |
| HOMO SAPIENS | 4 | 1088 | 2p3u | A | 102 | GLU | p.Glu142Lys | P00742 |
| HOMO SAPIENS | 6 | 370 | 1nr1 | D | 73 | ASP | p.Asp126His | P00367 |
| HOMO SAPIENS | 3 | 370 | 1nr1 | D | 221 | ARG | p.Arg274Cys | P00367 |
| HOMO SAPIENS | 2 | 370 | 1nr1 | D | 265 | ARG | p.Arg318Lys | P00367 |
| HOMO SAPIENS | 1 | 370 | 1nr1 | D | 269 | ARG | p.Arg322His | P00367 |
| HOMO SAPIENS | 1 | 370 | 1nr1 | D | 296 | GLU | p.Glu349Ala | P00367 |
| HOMO SAPIENS | 1 | 370 | 1nr1 | D | 445 | SER | p.Ser498Leu | P00367 |
| HOMO SAPIENS | 1 | 370 | 1nr1 | D | 446 | GLY | p.Gly499Asp | P00367 |
| HOMO SAPIENS | 1 | 370 | 1nr1 | D | 446 | GLY | p.Gly499Ser | P00367 |
| HOMO SAPIENS | 1 | 370 | 1nr1 | D | 448 | SER | p.Ser501Pro | P00367 |
| HOMO SAPIENS | 2 | 370 | 1nr1 | D | 454 | HIS | p.His507Tyr | P00367 |
| HOMO SAPIENS | 1 | 7720 | 2he0 | A | 147 | HIS | p.His2018Gln | P46531 |
| HOMO SAPIENS | 2 | 11844 | 5l0v | A | 218 | ARG | p.Arg218Ter | Q8NBL1 |
| HOMO SAPIENS | 2 | 11844 | 5l0v | A | 279 | ARG | p.Arg279Trp | Q8NBL1 |
| HOMO SAPIENS | 9 | 1485 | 5va0 | A | 477 | ARG | p.Arg477His | P04150 |
| HOMO SAPIENS | 0 | 235 | 2fun | B | 2248 | ARG | p.Arg248Trp | Q14790 |
| HOMO SAPIENS | 4 | 235 | 2fun | B | 2285 | HIS | p.Asp285His | Q14790 |
| HOMO SAPIENS | 0 | 4822 | 5y3n | A | 307 | ARG | p.Arg307Gly | Q12931 |
| HOMO SAPIENS | 8 | 32928 | 4m6w | A | 1886 | HIS | p.His1886Tyr | Q8IYD8 |
| HOMO SAPIENS | 0 | 2711 | 4y5v | F | 22 | GLY | p.Gly46Glu | P19235 |
| HOMO SAPIENS | -1 | 2711 | 4y5v | F | 166 | ALA | p.Ala190Ser | P19235 |
| HOMO SAPIENS | 4 | 5161 | 5bpu | A | 42 | HIS | p.His42Arg | Q00604 |
| HOMO SAPIENS | 0 | 5161 | 5bpu | A | 44 | TYR | p.Tyr44Cys | Q00604 |
| HOMO SAPIENS | 0 | 5161 | 5bpu | A | 45 | VAL | p.Val45Glu | Q00604 |
| HOMO SAPIENS | 0 | 5161 | 5bpu | A | 60 | VAL | p.Val60Glu | Q00604 |
| HOMO SAPIENS | 0 | 5161 | 5bpu | A | 61 | LEU | p.Leu61Phe | Q00604 |
| HOMO SAPIENS | 7 | 5161 | 5bpu | A | 69 | CYS | p.Cys69Ser | Q00604 |
| HOMO SAPIENS | 0 | 5161 | 5bpu | A | 75 | SER | p.Ser75Cys | Q00604 |
| HOMO SAPIENS | 0 | 5161 | 5bpu | A | 90 | ARG | p.Arg90Pro | Q00604 |
| HOMO SAPIENS | 9 | 5161 | 5bpu | A | 96 | CYS | p.Cys96Tyr | Q00604 |
| HOMO SAPIENS | 9 | 5161 | 5bpu | A | 96 | CYS | p.Cys96Trp | Q00604 |
| HOMO SAPIENS | 0 | 5161 | 5bpu | A | 101 | SER | p.Ser101Phe | Q00604 |
| HOMO SAPIENS | 0 | 5161 | 5bpu | A | 105 | ALA | p.Ala105Thr | Q00604 |
| HOMO SAPIENS | 0 | 5161 | 5bpu | A | 110 | CYS | p.Cys110Gly | Q00604 |
| HOMO SAPIENS | 0 | 5161 | 5bpu | A | 121 | ARG | p.Arg121Trp | Q00604 |
| HOMO SAPIENS | 0 | 5161 | 5bpu | A | 121 | ARG | p.Arg121Leu | Q00604 |
| HOMO SAPIENS | 0 | 5161 | 5bpu | A | 124 | LEU | p.Leu124Phe | Q00604 |
| HOMO SAPIENS | 8 | 5161 | 5bpu | A | 128 | CYS | p.Cys128Ter | Q00604 |
| HOMO SAPIENS | 1 | 13944 | 5xqh | A | 238 | ARG | p.Arg238His | Q9GZN7 |
| HOMO SAPIENS | -1 | 7508 | 5jl6 | A | 39 | TRP | p.Trp39Arg | P11511 |
| HOMO SAPIENS | 0 | 7508 | 5jl6 | A | 201 | THR | p.Thr201Met | P11511 |
| HOMO SAPIENS | 0 | 7508 | 5jl6 | A | 210 | GLU | p.Glu210Lys | P11511 |
| HOMO SAPIENS | 1 | 7508 | 5jl6 | A | 264 | ARG | p.Arg264Cys | P11511 |
| HOMO SAPIENS | 2 | 7508 | 5jl6 | A | 365 | ARG | p.Arg365Gln | P11511 |
| HOMO SAPIENS | 7 | 7508 | 5jl6 | A | 375 | ARG | p.Arg375Cys | P11511 |
| HOMO SAPIENS | 8 | 7508 | 5jl6 | A | 435 | ARG | p.Arg435Cys | P11511 |

| | | | | | | | | |
|---|---|---|---|---|---|---|---|---|
| HOMO SAPIENS | 9 | 7508 | 5jl6 | A | 437 | CYS | p.Cys437Tyr | P11511 |
| HOMO SAPIENS | -1 | 30206 | 5l95 | B | 246 | ARG | p.Arg246Ter | Q9GZZ9 |
| HOMO SAPIENS | 4 | 875 | 3efh | B | 43 | GLU | p.Glu43Asp | P60891 |
| HOMO SAPIENS | 5 | 875 | 3efh | B | 65 | ASP | p.Asp65Asn | P60891 |
| HOMO SAPIENS | 0 | 875 | 3efh | B | 87 | ALA | p.Ala87Thr | P60891 |
| HOMO SAPIENS | 2 | 875 | 3efh | B | 115 | MET | p.Met115Thr | P60891 |
| HOMO SAPIENS | 8 | 875 | 3efh | B | 133 | GLN | p.Gln133Pro | P60891 |
| HOMO SAPIENS | 8 | 875 | 3efh | B | 152 | LEU | p.Leu152Pro | P60891 |
| HOMO SAPIENS | 7 | 875 | 3efh | B | 290 | ILE | p.Ile290Thr | P60891 |
| HOMO SAPIENS | 6 | 875 | 3efh | B | 306 | GLY | p.Gly306Arg | P60891 |
| HOMO SAPIENS | 0 | 7978 | 5edm | A | 122 | MET | p.Thr165Met | P00734 |
| HOMO SAPIENS | -1 | 7978 | 5edm | A | 257 | ARG | p.Arg314Cys | P00734 |
| HOMO SAPIENS | 0 | 1716 | 1b98 | M | 126 | ARG | p.Arg206Gln | P34130 |
| HOMO SAPIENS | 0 | 1716 | 1b98 | M | 126 | ARG | p.Arg206Trp | P34130 |
| HOMO SAPIENS | 9 | 3960 | 2qyp | B | 28 | ILE | p.Ile338Val | P07602 |
| HOMO SAPIENS | 1 | 3960 | 2qyp | B | 39 | LEU | p.Leu349Pro | P07602 |
| HOMO SAPIENS | 0 | 31414 | 5u1d | A | 333 | ILE | p.Ile393Val | Q03518 |
| HOMO SAPIENS | 0 | 31414 | 5u1d | A | 637 | ASP | p.Asp697Gly | Q03518 |
| HOMO SAPIENS | 0 | 31414 | 5u1d | A | 659 | ARG | p.Arg719Gln | Q03518 |
| HOMO SAPIENS | 3 | 119 | 2pho | A | 11 | ILE | p.Ile11Thr | P05089 |
| HOMO SAPIENS | 8 | 119 | 2pho | A | 21 | ARG | p.Arg21Ter | P05089 |
| HOMO SAPIENS | 0 | 119 | 2pho | A | 122 | TRP | p.Trp122Ter | P05089 |
| HOMO SAPIENS | 6 | 119 | 2pho | A | 138 | GLY | p.Gly138Val | P05089 |
| HOMO SAPIENS | 5 | 119 | 2pho | A | 290 | THR | p.Thr290Ser | P05089 |
| HOMO SAPIENS | 5 | 119 | 2pho | A | 291 | ARG | p.Arg291Ter | P05089 |
| HOMO SAPIENS | 0 | 3732 | 2i99 | A | 247 | TYR | p.Tyr247Ter | Q14894 |
| HOMO SAPIENS | -1 | 1382 | 4ffx | A | 2 | ALA | p.Ala2Val | P30566 |
| HOMO SAPIENS | 1 | 1382 | 4ffx | A | 72 | ILE | p.Ile72Met | P30566 |
| HOMO SAPIENS | 1 | 1382 | 4ffx | A | 100 | PRO | p.Pro100Ala | P30566 |
| HOMO SAPIENS | 0 | 1382 | 4ffx | A | 114 | TYR | p.Tyr114His | P30566 |
| HOMO SAPIENS | 1 | 1382 | 4ffx | A | 141 | ARG | p.Arg141Trp | P30566 |
| HOMO SAPIENS | 0 | 1382 | 4ffx | A | 147 | LYS | p.Lys147Met | P30566 |
| HOMO SAPIENS | 3 | 1382 | 4ffx | A | 190 | ARG | p.Arg190Gln | P30566 |
| HOMO SAPIENS | 3 | 1382 | 4ffx | A | 190 | ARG | p.Arg190Ter | P30566 |
| HOMO SAPIENS | 0 | 1382 | 4ffx | A | 217 | HIS | p.His217Asp | P30566 |
| HOMO SAPIENS | 2 | 1382 | 4ffx | A | 246 | LYS | p.Lys246Glu | P30566 |
| HOMO SAPIENS | 5 | 1382 | 4ffx | A | 303 | ARG | p.Arg303Cys | P30566 |
| HOMO SAPIENS | 1 | 1382 | 4ffx | A | 318 | PRO | p.Pro318Leu | P30566 |
| HOMO SAPIENS | 6 | 1382 | 4ffx | A | 332 | ASP | p.Asp332His | P30566 |
| HOMO SAPIENS | 6 | 1382 | 4ffx | A | 337 | ARG | p.Arg337Ter | P30566 |
| HOMO SAPIENS | 1 | 1382 | 4ffx | A | 396 | ARG | p.Arg396His | P30566 |
| HOMO SAPIENS | 1 | 1382 | 4ffx | A | 426 | ARG | p.Arg426His | P30566 |
| HOMO SAPIENS | 0 | 1382 | 4ffx | A | 438 | SER | p.Ser438Pro | P30566 |
| HOMO SAPIENS | 1 | 1382 | 4ffx | A | 447 | SER | p.Ser447Pro | P30566 |
| HOMO SAPIENS | 2 | 1382 | 4ffx | A | 448 | SER | p.Ser448Pro | P30566 |
| HOMO SAPIENS | 2 | 1382 | 4ffx | A | 450 | THR | p.Thr450Ser | P30566 |
| HOMO SAPIENS | 0 | 6212 | 1ao3 | A | 46 | SER | p.Ser1731Thr | P04275 |
| HOMO SAPIENS | 0 | 1711 | 3f9c | A | 279 | VAL | p.Val279Phe | Q13093 |
| HOMO SAPIENS | 0 | 6112 | 4fvl | A | 312 | HIS | p.His312Arg | P45452 |
| HOMO SAPIENS | 0 | 6112 | 4fvl | A | 323 | THR | p.Thr323Met | P45452 |
| HOMO SAPIENS | 0 | 6112 | 4fvl | A | 390 | ASP | p.Asp390Gly | P45452 |
| HOMO SAPIENS | 0 | 6112 | 4fvl | A | 458 | ARG | p.Arg458Cys | P45452 |
| HOMO SAPIENS | 2 | 6180 | 5ggn | A | 97 | ARG | p.Arg97Trp | Q8WZA1 |
| HOMO SAPIENS | 2 | 6180 | 5ggn | A | 215 | ARG | p.Arg215Ter | Q8WZA1 |

| | | | | | | | | |
|---|---|---|---|---|---|---|---|---|
| HOMO SAPIENS | -1 | 5526 | 4ak3 | A | 11 | ASP | p.Asp1232Asn | P02461 |
| HOMO SAPIENS | 6 | 5526 | 4ak3 | A | 38 | ALA | p.Ala1259Thr | P02461 |
| HOMO SAPIENS | 0 | 5526 | 4ak3 | A | 92 | LYS | p.Lys1313Arg | P02461 |
| HOMO SAPIENS | 0 | 5526 | 4ak3 | A | 132 | GLN | p.His1353Gln | P02461 |
| HOMO SAPIENS | 4 | 16616 | 1hci | B | 433 | SER | p.Ser433Leu | P35609 |
| HOMO SAPIENS | 1 | 16616 | 1hci | B | 475 | ASP | p.Asp475Asn | P35609 |
| HOMO SAPIENS | 1 | 16616 | 1hci | B | 476 | ALA | p.Ala476Ser | P35609 |
| HOMO SAPIENS | 2 | 16616 | 1hci | B | 507 | MET | p.Met507Val | P35609 |
| HOMO SAPIENS | 1 | 16616 | 1hci | B | 604 | MET | p.Met604Val | P35609 |
| HOMO SAPIENS | 2 | 16616 | 1hci | B | 640 | ARG | p.Arg640His | P35609 |
| HOMO SAPIENS | 1 | 16616 | 1hci | B | 721 | ARG | p.Arg721Cys | P35609 |
| HOMO SAPIENS | -1 | 3402 | 5iaa | B | 57 | MET | p.Met57Val | Q9GZZ9 |
| HOMO SAPIENS | 7 | 3250 | 3irq | B | 193 | PRO | p.Pro193Ala | P55265 |
| HOMO SAPIENS | 2 | 15200 | 2wns | A | 30 | SER | p.Ser30Gly | P11172 |
| HOMO SAPIENS | 0 | 15200 | 2wns | A | 96 | ARG | p.Arg96Gly | P11172 |
| HOMO SAPIENS | 0 | 15200 | 2wns | A | 109 | VAL | p.Val109Gly | P11172 |
| HOMO SAPIENS | 2 | 15759 | 4x4w | B | 140 | LEU | p.Leu166Ser | Q96Q11 |
| HOMO SAPIENS | 0 | 15759 | 4x4w | B | 197 | ILE | p.Ile223Thr | Q96Q11 |
| HOMO SAPIENS | 0 | 3156 | 3w03 | B | 14 | ALA | p.Ala14Thr | Q9H9Q4 |
| HOMO SAPIENS | 0 | 3156 | 3w03 | B | 57 | ARG | p.Arg57Gly | Q9H9Q4 |
| HOMO SAPIENS | 0 | 3156 | 3w03 | B | 123 | CYS | p.Cys123Arg | Q9H9Q4 |
| HOMO SAPIENS | 0 | 11693 | 3ojy | A | 63 | GLN | p.Gln93Lys | P07357 |
| HOMO SAPIENS | 0 | 1421 | 2qc8 | E | 275 | GLU | p.Glu275Asp | P15104 |
| HOMO SAPIENS | 9 | 1421 | 2qc8 | E | 324 | ARG | p.Arg324Cys | P15104 |
| HOMO SAPIENS | 6 | 1421 | 2qc8 | E | 341 | ARG | p.Arg341Cys | P15104 |
| HOMO SAPIENS | 6 | 2850 | 3qvb | A | 83 | THR | p.Thr83Met | P46976 |
| HOMO SAPIENS | 8 | 2850 | 3qvb | A | 102 | ASP | p.Asp102His | P46976 |
| HOMO SAPIENS | 1 | 24314 | 4m9p | A | 480 | PRO | p.Pro480Leu | P21333 |
| HOMO SAPIENS | 0 | 24314 | 4m9p | A | 528 | VAL | p.Val528Met | P21333 |
| HOMO SAPIENS | 0 | 24314 | 4m9p | A | 590 | VAL | p.Val590Ile | P21333 |
| HOMO SAPIENS | 0 | 24314 | 4m9p | A | 605 | ASP | p.Asp605Asn | P21333 |
| HOMO SAPIENS | 1 | 24314 | 4m9p | A | 637 | PRO | p.Pro637Gln | P21333 |
| HOMO SAPIENS | 0 | 24314 | 4m9p | A | 656 | LEU | p.Leu656Phe | P21333 |
| HOMO SAPIENS | 2 | 24314 | 4m9p | A | 711 | VAL | p.Val711Asp | P21333 |
| HOMO SAPIENS | 0 | 4951 | 5joe | A | 40 | THR | p.Thr9621Ser | Q8WZ42 |
| HOMO SAPIENS | 1 | 4951 | 5joe | A | 41 | GLU | p.Glu9622Gln | Q8WZ42 |
| HOMO SAPIENS | 0 | 4951 | 5joe | A | 82 | ALA | p.Ala9663Ser | Q8WZ42 |
| HOMO SAPIENS | 4 | 4951 | 5joe | A | 89 | ILE | p.Ile9670Asn | Q8WZ42 |
| HOMO SAPIENS | 0 | 9664 | 3c6m | A | 56 | GLY | p.Gly56Ser | P52788 |
| HOMO SAPIENS | 0 | 9664 | 3c6m | A | 132 | VAL | p.Val132Gly | P52788 |
| HOMO SAPIENS | 1 | 9664 | 3c6m | A | 190 | THR | p.Thr190Ile | P52788 |
| HOMO SAPIENS | -1 | 12118 | 5hcd | B | 19 | GLN | p.Gln19Ter | P01031 |
| HOMO SAPIENS | 0 | 1583 | 2b4x | L | 7 | ILE | p.Ile39Asn | P01008 |
| HOMO SAPIENS | 0 | 1583 | 2b4x | L | 24 | ARG | p.Arg56Cys | P01008 |
| HOMO SAPIENS | 0 | 1583 | 2b4x | L | 47 | ARG | p.Arg79Cys | P01008 |
| HOMO SAPIENS | 0 | 1583 | 2b4x | L | 47 | ARG | p.Arg79His | P01008 |
| HOMO SAPIENS | 1 | 1583 | 2b4x | L | 95 | CYS | p.Cys127Arg | P01008 |
| HOMO SAPIENS | 1 | 1583 | 2b4x | L | 99 | LEU | p.Leu131Phe | P01008 |
| HOMO SAPIENS | 0 | 1583 | 2b4x | L | 116 | SER | p.Ser148Pro | P01008 |
| HOMO SAPIENS | 0 | 1583 | 2b4x | L | 129 | ARG | p.Arg161Gln | P01008 |
| HOMO SAPIENS | 0 | 1583 | 2b4x | L | 135 | ASN | p.Asn167Thr | P01008 |
| HOMO SAPIENS | 2 | 1583 | 2b4x | L | 187 | ASN | p.Asn219Asp | P01008 |
| HOMO SAPIENS | 0 | 1583 | 2b4x | L | 191 | SER | p.Ser223Pro | P01008 |
| HOMO SAPIENS | 2 | 1583 | 2b4x | L | 349 | SER | p.Ser381Pro | P01008 |

| | | | | | | | | |
|---|---|---|---|---|---|---|---|---|
| HOMO SAPIENS | 0 | 1583 | 2b4x | L | 382 | ALA | p.Ala414Thr | P01008 |
| HOMO SAPIENS | 0 | 1583 | 2b4x | L | 387 | ALA | p.Ala419Val | P01008 |
| HOMO SAPIENS | 0 | 1583 | 2b4x | L | 392 | GLY | p.Gly424Asp | P01008 |
| HOMO SAPIENS | 0 | 1583 | 2b4x | L | 393 | ARG | p.Arg425His | P01008 |
| HOMO SAPIENS | 0 | 1583 | 2b4x | L | 393 | ARG | p.Arg425Cys | P01008 |
| HOMO SAPIENS | 0 | 1583 | 2b4x | L | 394 | SER | p.Ser426Leu | P01008 |
| HOMO SAPIENS | 0 | 1583 | 2b4x | L | 404 | ALA | p.Ala436Thr | P01008 |
| HOMO SAPIENS | 2 | 1583 | 2b4x | L | 407 | PRO | p.Pro439Leu | P01008 |
| HOMO SAPIENS | 2 | 1583 | 2b4x | L | 429 | PRO | p.Pro461Leu | P01008 |
| HOMO SAPIENS | 0 | 11647 | 2a1t | S | 78 | ALA | p.Ala78Ser | P38117 |
| HOMO SAPIENS | 0 | 11647 | 2a1t | S | 98 | ARG | p.Arg98Cys | P38117 |
| HOMO SAPIENS | 8 | 11647 | 2a1t | S | 128 | ASP | p.Asp128Asn | P38117 |
| HOMO SAPIENS | 5 | 11647 | 2a1t | S | 154 | THR | p.Thr154Met | P38117 |
| HOMO SAPIENS | 6 | 11647 | 2a1t | S | 164 | ARG | p.Arg164Gln | P38117 |
| HOMO SAPIENS | 5 | 11647 | 2a1t | S | 174 | ARG | p.Arg174His | P38117 |
| HOMO SAPIENS | -1 | 11200 | 4tx5 | A | 4 | ILE | p.Ile59Val | Q9NR28 |
| HOMO SAPIENS | 0 | 11200 | 4tx5 | A | 169 | GLY | p.Gly224Arg | Q9NR28 |
| HOMO SAPIENS | 3 | 417 | 2vr2 | A | 6 | ARG | p.Arg6Gln | Q14117 |
| HOMO SAPIENS | 1 | 417 | 2vr2 | A | 334 | GLN | p.Gln334Arg | Q14117 |
| HOMO SAPIENS | 2 | 417 | 2vr2 | A | 360 | TRP | p.Trp360Arg | Q14117 |
| HOMO SAPIENS | 0 | 417 | 2vr2 | A | 379 | SER | p.Ser379Arg | Q14117 |
| HOMO SAPIENS | 0 | 417 | 2vr2 | A | 435 | GLY | p.Gly435Arg | Q14117 |
| HOMO SAPIENS | 0 | 12498 | 3eo3 | B | 423 | ILE | p.Ile423Val | Q9Y223 |
| HOMO SAPIENS | 5 | 12498 | 3eo3 | B | 572 | VAL | p.Val572Leu | Q9Y223 |
| HOMO SAPIENS | 0 | 12498 | 3eo3 | B | 576 | GLY | p.Gly576Glu | Q9Y223 |
| HOMO SAPIENS | 0 | 12498 | 3eo3 | B | 587 | ILE | p.Ile587Thr | Q9Y223 |
| HOMO SAPIENS | 0 | 12498 | 3eo3 | B | 615 | SER | p.Ser615Ter | Q9Y223 |
| HOMO SAPIENS | 1 | 12498 | 3eo3 | B | 631 | ALA | p.Ala631Val | Q9Y223 |
| HOMO SAPIENS | 1 | 12498 | 3eo3 | B | 669 | GLY | p.Gly669Arg | Q9Y223 |
| HOMO SAPIENS | 1 | 12498 | 3eo3 | B | 696 | VAL | p.Val696Met | Q9Y223 |
| HOMO SAPIENS | 2 | 12498 | 3eo3 | B | 712 | MET | p.Met712Thr | Q9Y223 |
| HOMO SAPIENS | 0 | 1568 | 1r74 | A | 49 | LEU | p.Leu50Pro | Q14749 |
| HOMO SAPIENS | 0 | 1568 | 1r74 | A | 176 | HIS | p.His177Asn | Q14749 |
| HOMO SAPIENS | 3 | 147 | 5eig | A | 67 | THR | p.Thr67Ile | P32929 |
| HOMO SAPIENS | 3 | 147 | 5eig | A | 240 | GLN | p.Gln240Glu | P32929 |
| HOMO SAPIENS | -1 | 147 | 5eig | A | 403 | SER | p.Ser403Ile | P32929 |
| HOMO SAPIENS | 8 | 20036 | 2lsr | A | 31 | ARG | p.Arg31Ter | Q9Y6N9 |
| HOMO SAPIENS | -1 | 41382 | 4uis | B | 431 | UNK | p.Ala431Glu | P49768 |
| HOMO SAPIENS | -1 | 8536 | 3i6w | B | 85 | PRO | p.Pro85Leu | O96017 |
| HOMO SAPIENS | 0 | 8536 | 3i6w | B | 93 | TRP | p.Trp93Ter | O96017 |
| HOMO SAPIENS | 0 | 8536 | 3i6w | B | 95 | ARG | p.Arg95Ter | O96017 |
| HOMO SAPIENS | 0 | 8536 | 3i6w | B | 117 | ARG | p.Arg117Gly | O96017 |
| HOMO SAPIENS | 0 | 8536 | 3i6w | B | 119 | LYS | p.Lys119Arg | O96017 |
| HOMO SAPIENS | 0 | 8536 | 3i6w | B | 137 | ARG | p.Arg137Gln | O96017 |
| HOMO SAPIENS | 0 | 8536 | 3i6w | B | 137 | ARG | p.Arg137Ter | O96017 |
| HOMO SAPIENS | 0 | 8536 | 3i6w | B | 145 | ARG | p.Arg145Trp | O96017 |
| HOMO SAPIENS | 0 | 8536 | 3i6w | B | 160 | ILE | p.Ile160Met | O96017 |
| HOMO SAPIENS | 0 | 41322 | 4bxl | C | 46 | GLU | p.Glu46Lys | P37840 |
| HOMO SAPIENS | 5 | 41322 | 4bxl | C | 53 | ALA | p.Ala53Thr | P37840 |
| HOMO SAPIENS | 2 | 24966 | 3hve | A | 15 | ARG | p.Arg15Ser | Q9H2C0 |
| HOMO SAPIENS | 1 | 24966 | 3hve | A | 138 | ARG | p.Arg138His | Q9H2C0 |
| HOMO SAPIENS | 0 | 24966 | 3hve | A | 201 | ARG | p.Arg201Ter | Q9H2C0 |
| HOMO SAPIENS | 0 | 11680 | 5an9 | J | 8 | ASN | p.Asn8Lys | Q9Y3A5 |
| HOMO SAPIENS | 1 | 11680 | 5an9 | J | 33 | LYS | p.Lys33Arg | Q9Y3A5 |

| | | | | | | | | |
|---|---|---|---|---|---|---|---|---|
| HOMO SAPIENS | 1 | 11680 | 5an9 | J | 126 | ARG | p.Arg126Thr | Q9Y3A5 |
| HOMO SAPIENS | 1 | 11680 | 5an9 | J | 130 | VAL | p.Val130Met | Q9Y3A5 |
| HOMO SAPIENS | 1 | 11680 | 5an9 | J | 169 | ARG | p.Arg169Cys | Q9Y3A5 |
| HOMO SAPIENS | 1 | 11680 | 5an9 | J | 218 | ARG | p.Arg218Ter | Q9Y3A5 |
| HOMO SAPIENS | 4 | 978 | 3zs0 | B | 7 | TYR | p.Tyr173Cys | P05164 |
| HOMO SAPIENS | 1 | 978 | 3zs0 | B | 85 | MET | p.Met251Thr | P05164 |
| HOMO SAPIENS | 4 | 5140 | 2l4r | A | 52 | CYS | p.Cys52Ter | Q12809 |
| HOMO SAPIENS | 4 | 5140 | 2l4r | A | 63 | PRO | p.Pro63His | Q12809 |
| HOMO SAPIENS | 0 | 5140 | 2l4r | A | 100 | ARG | p.Arg100Gly | Q12809 |
| HOMO SAPIENS | 4 | 5140 | 2l4r | A | 129 | PHE | p.Phe129Leu | Q12809 |
| HOMO SAPIENS | 0 | 25836 | 2gy7 | B | 103 | GLU | p.Glu103Asp | Q02763 |
| HOMO SAPIENS | 0 | 25836 | 2gy7 | B | 148 | ILE | p.Ile148Thr | Q02763 |
| HOMO SAPIENS | 0 | 25836 | 2gy7 | B | 226 | ALA | p.Ala226Val | Q02763 |
| HOMO SAPIENS | 0 | 25836 | 2gy7 | B | 391 | THR | p.Thr391Ile | Q02763 |
| HOMO SAPIENS | 4 | 12563 | 5kwy | D | 20 | GLU | p.Glu20Ter | P61916 |
| HOMO SAPIENS | 2 | 12563 | 5kwy | D | 30 | VAL | p.Val30Met | P61916 |
| HOMO SAPIENS | 4 | 12563 | 5kwy | D | 39 | VAL | p.Val39Met | P61916 |
| HOMO SAPIENS | 8 | 12563 | 5kwy | D | 67 | SER | p.Ser67Pro | P61916 |
| HOMO SAPIENS | 8 | 12563 | 5kwy | D | 99 | CYS | p.Cys99Arg | P61916 |
| HOMO SAPIENS | 2 | 12563 | 5kwy | D | 118 | GLU | p.Glu118Ter | P61916 |
| HOMO SAPIENS | 2 | 12563 | 5kwy | D | 120 | PRO | p.Pro120Ser | P61916 |
| HOMO SAPIENS | -1 | 1679 | 1pmk | A | -2 | VAL | p.Val374Phe | P00747 |
| HOMO SAPIENS | -1 | 5315 | 2w2n | P | 53 | ALA | p.Ala53Val | Q8NBP7 |
| HOMO SAPIENS | 8 | 13496 | 2yad | A | 138 | THR | p.Asn138Thr | P11686 |
| HOMO SAPIENS | -1 | 13496 | 2yad | A | 173 | ARG | p.Arg167Gln | P11686 |
| HOMO SAPIENS | 2 | 13496 | 2yad | A | 186 | SER | p.Asn186Ser | P11686 |
| HOMO SAPIENS | 0 | 4426 | 3qd6 | C | 123 | ALA | p.Ala123Glu | P29965 |
| HOMO SAPIENS | 1 | 4426 | 3qd6 | C | 140 | TRP | p.Trp140Gly | P29965 |
| HOMO SAPIENS | 0 | 4426 | 3qd6 | C | 155 | LEU | p.Leu155Pro | P29965 |
| HOMO SAPIENS | 6 | 4426 | 3qd6 | C | 227 | GLY | p.Gly227Val | P29965 |
| HOMO SAPIENS | 0 | 4426 | 3qd6 | C | 235 | ALA | p.Ala235Pro | P29965 |
| HOMO SAPIENS | 0 | 10734 | 3dyd | B | 45 | PRO | p.Pro45Ser | P17735 |
| HOMO SAPIENS | -1 | 10734 | 3dyd | B | 57 | ARG | p.Arg57Ter | P17735 |
| HOMO SAPIENS | 8 | 10734 | 3dyd | B | 417 | ARG | p.Arg417Ter | P17735 |
| HOMO SAPIENS | 0 | 6956 | 4pzs | A | 1600 | LEU | p.Leu1600Phe | Q09472 |
| HOMO SAPIENS | 0 | 9370 | 3hfv | A | 21 | SER | p.Ser57Cys | O95363 |
| HOMO SAPIENS | 1 | 9370 | 3hfv | A | 25 | ASP | p.Asp61Glu | O95363 |
| HOMO SAPIENS | 1 | 9370 | 3hfv | A | 130 | ALA | p.Ala166Thr | O95363 |
| HOMO SAPIENS | 5 | 9370 | 3hfv | A | 148 | ASP | p.Asp184Asn | O95363 |
| HOMO SAPIENS | 0 | 9370 | 3hfv | A | 187 | ARG | p.Arg223Cys | O95363 |
| HOMO SAPIENS | 0 | 9370 | 3hfv | A | 210 | THR | p.Thr246Met | O95363 |
| HOMO SAPIENS | 1 | 9370 | 3hfv | A | 244 | ASN | p.Asn280Ser | O95363 |
| HOMO SAPIENS | 2 | 9370 | 3hfv | A | 288 | TYR | p.Tyr324Cys | O95363 |
| HOMO SAPIENS | 5 | 9370 | 3hfv | A | 293 | ILE | p.Ile329Thr | O95363 |
| HOMO SAPIENS | 1 | 9370 | 3hfv | A | 355 | ASP | p.Asp391Val | O95363 |
| HOMO SAPIENS | 1 | 9370 | 3hfv | A | 371 | THR | p.Thr407Met | O95363 |
| HOMO SAPIENS | 1 | 9370 | 3hfv | A | 383 | ARG | p.Arg419Cys | O95363 |
| HOMO SAPIENS | 1 | 9370 | 3hfv | A | 387 | ARG | p.Arg423Gln | O95363 |
| HOMO SAPIENS | 0 | 14401 | 1q3x | A | 371 | ASP | p.Asp371Tyr | O00187 |
| HOMO SAPIENS | 0 | 14401 | 1q3x | A | 377 | VAL | p.Val377Ala | O00187 |
| HOMO SAPIENS | 0 | 14401 | 1q3x | A | 439 | ARG | p.Arg439His | O00187 |
| HOMO SAPIENS | 0 | 14401 | 1q3x | A | 577 | GLN | p.Gln577His | O00187 |
| HOMO SAPIENS | 2 | 18270 | 3teq | A | 429 | ARG | p.Arg429Cys | Q13586 |
| HOMO SAPIENS | -1 | 16081 | 4mqt | A | 231 | VAL | p.Val231Ile | P08172 |

| | | | | | | | | |
|---|---|---|---|---|---|---|---|---|
| HOMO SAPIENS | 9 | 6607 | 3whu | B | 81 | ASP | p.Asp81His | Q8NI22 |
| HOMO SAPIENS | 9 | 6607 | 3whu | B | 129 | ASP | p.Asp129Glu | Q8NI22 |
| HOMO SAPIENS | 9 | 6607 | 3whu | B | 136 | ILE | p.Ile136Thr | Q8NI22 |
| HOMO SAPIENS | 0 | 7208 | 2e9w | D | 11 | ASN | p.Asn36Ser | P21583 |
| HOMO SAPIENS | 0 | 24566 | 4bdx | A | 10 | LEU | p.Leu140Val | P00748 |
| HOMO SAPIENS | 5 | 24566 | 4bdx | A | 77 | PRO | p.Ala207Pro | P00748 |
| HOMO SAPIENS | 7 | 606 | 3qnf | A | 319 | MET | p.Met319Val | Q9NZ08 |
| HOMO SAPIENS | 8 | 4097 | 1ydl | A | 21 | LEU | p.Leu21Pro | Q6ZYL4 |
| HOMO SAPIENS | 6 | 4097 | 1ydl | A | 56 | ARG | p.Arg56Ter | Q6ZYL4 |
| HOMO SAPIENS | 8 | 36691 | 1dqb | A | 42 | ARG | p.Arg403Lys | P07204 |
| HOMO SAPIENS | 0 | 5642 | 3j7y | X | 59 | ARG | p.Arg59Pro | Q13084 |
| HOMO SAPIENS | 1 | 5642 | 3j7y | X | 204 | VAL | p.Val204Met | Q13084 |
| HOMO SAPIENS | 1 | 1119 | 3nwu | B | 252 | ALA | p.Ala252Thr | Q92743 |
| HOMO SAPIENS | 3 | 1119 | 3nwu | B | 274 | ARG | p.Arg274Gln | Q92743 |
| HOMO SAPIENS | 9 | 1119 | 3nwu | B | 297 | VAL | p.Val297Met | Q92743 |
| HOMO SAPIENS | -1 | 1119 | 3nwu | B | 302 | ARG | p.Arg302Ter | Q92743 |
| HOMO SAPIENS | 9 | 1119 | 3nwu | B | 321 | ALA | p.Ala321Thr | Q92743 |
| HOMO SAPIENS | -1 | 1119 | 3nwu | B | 370 | ARG | p.Arg370Ter | Q92743 |
| HOMO SAPIENS | 0 | 416 | 4ycu | B | 105 | LEU | p.Leu105His | Q15046 |
| HOMO SAPIENS | 0 | 416 | 4ycu | B | 145 | TYR | p.Tyr145His | Q15046 |
| HOMO SAPIENS | 0 | 416 | 4ycu | B | 200 | PRO | p.Pro200Leu | Q15046 |
| HOMO SAPIENS | 0 | 416 | 4ycu | B | 201 | TYR | p.Tyr201His | Q15046 |
| HOMO SAPIENS | 0 | 416 | 4ycu | B | 420 | ARG | p.Arg420Cys | Q15046 |
| HOMO SAPIENS | 9 | 3297 | 4fpb | A | 680 | THR | p.Thr680Ser | P19367 |
| HOMO SAPIENS | 0 | 6771 | 2zt6 | A | 201 | ALA | p.Ala255Val | P41250 |
| HOMO SAPIENS | 0 | 6771 | 2zt6 | A | 202 | ASP | p.Asp256His | P41250 |
| HOMO SAPIENS | 0 | 6771 | 2zt6 | A | 334 | ARG | p.Arg388Gln | P41250 |
| HOMO SAPIENS | -1 | 6771 | 2zt6 | A | 439 | ASN | p.Asn493Ser | P41250 |
| HOMO SAPIENS | -1 | 6771 | 2zt6 | A | 500 | ASP | p.Asp554Asn | P41250 |
| HOMO SAPIENS | 1 | 6771 | 2zt6 | A | 581 | SER | p.Ser635Leu | P41250 |
| HOMO SAPIENS | -1 | 6771 | 2zt6 | A | 684 | GLU | p.Glu738Lys | P41250 |
| HOMO SAPIENS | 0 | 2687 | 4grw | B | 276 | VAL | p.Val298Phe | P29460 |
| HOMO SAPIENS | 6 | 913 | 1m75 | B | 45 | ASP | p.Asp57Glu | Q16836 |
| HOMO SAPIENS | 2 | 913 | 1m75 | B | 74 | PRO | p.Leu86Pro | Q16836 |
| HOMO SAPIENS | 3 | 913 | 1m75 | B | 214 | TYR | p.Tyr226His | Q16836 |
| HOMO SAPIENS | 3 | 913 | 1m75 | B | 224 | ARG | p.Arg236Ter | Q16836 |
| HOMO SAPIENS | 4 | 913 | 1m75 | B | 246 | PRO | p.Pro258Leu | Q16836 |
| HOMO SAPIENS | 2 | 4713 | 4djn | B | 40 | ARG | p.Arg40Cys | Q7LG56 |
| HOMO SAPIENS | 1 | 4713 | 4djn | B | 41 | ARG | p.Arg41Pro | Q7LG56 |
| HOMO SAPIENS | 2 | 4713 | 4djn | B | 110 | ARG | p.Arg110His | Q7LG56 |
| HOMO SAPIENS | 2 | 4713 | 4djn | B | 121 | ARG | p.Arg121His | Q7LG56 |
| HOMO SAPIENS | 9 | 4713 | 4djn | B | 131 | GLU | p.Glu131Lys | Q7LG56 |
| HOMO SAPIENS | 2 | 4713 | 4djn | B | 182 | THR | p.Thr182Ala | Q7LG56 |
| HOMO SAPIENS | 6 | 4713 | 4djn | B | 194 | GLU | p.Glu194Lys | Q7LG56 |
| HOMO SAPIENS | 2 | 4713 | 4djn | B | 211 | ARG | p.Arg211Lys | Q7LG56 |
| HOMO SAPIENS | 0 | 4713 | 4djn | B | 224 | ILE | p.Ile224Ser | Q7LG56 |
| HOMO SAPIENS | 3 | 4713 | 4djn | B | 229 | GLY | p.Gly229Val | Q7LG56 |
| HOMO SAPIENS | 2 | 4713 | 4djn | B | 236 | CYS | p.Cys236Phe | Q7LG56 |
| HOMO SAPIENS | 0 | 4713 | 4djn | B | 273 | GLY | p.Gly273Ser | Q7LG56 |
| HOMO SAPIENS | -1 | 4713 | 4djn | B | 282 | MSE | p.Met282Ile | Q7LG56 |
| HOMO SAPIENS | 3 | 4713 | 4djn | B | 284 | GLN | p.Gln284Ter | Q7LG56 |
| HOMO SAPIENS | 3 | 4713 | 4djn | B | 305 | ALA | p.Ala305Ser | Q7LG56 |
| HOMO SAPIENS | -1 | 8974 | 5n7i | A | 500 | ARG | p.Arg500Ter | Q8N4S9 |
| HOMO SAPIENS | 0 | 17072 | 3fxi | B | 299 | ASP | p.Asp299Gly | O00206 |

| | | | | | | | | |
|---|---|---|---|---|---|---|---|---|
| HOMO SAPIENS | 1 | 14355 | 2wwk | T | 55 | ILE | p.Ile34306Asn | Q8WZ42 |
| HOMO SAPIENS | 0 | 14355 | 2wwk | T | 64 | LEU | p.Leu34315Pro | Q8WZ42 |
| HOMO SAPIENS | 2 | 17797 | 1xu1 | S | 72 | ARG | p.Arg72His | O14836 |
| HOMO SAPIENS | -1 | 10836 | 4ztu | A | 39 | GLN | p.Gln39Arg | P54098 |
| HOMO SAPIENS | -1 | 10836 | 4ztu | A | 42 | ARG | p.Arg42Gln | P54098 |
| HOMO SAPIENS | -1 | 10836 | 4ztu | A | 45 | GLN | p.Gln45Arg | P54098 |
| HOMO SAPIENS | -1 | 10836 | 4ztu | A | 52 | GLN | p.Gln52Glu | P54098 |
| HOMO SAPIENS | 0 | 10836 | 4ztu | A | 88 | PHE | p.Phe88Leu | P54098 |
| HOMO SAPIENS | 1 | 10836 | 4ztu | A | 194 | ALA | p.Ala194Val | P54098 |
| HOMO SAPIENS | 1 | 10836 | 4ztu | A | 227 | ARG | p.Arg227Trp | P54098 |
| HOMO SAPIENS | 0 | 10836 | 4ztu | A | 232 | ARG | p.Arg232His | P54098 |
| HOMO SAPIENS | 1 | 10836 | 4ztu | A | 285 | GLN | p.Gln285His | P54098 |
| HOMO SAPIENS | 1 | 10836 | 4ztu | A | 434 | TYR | p.Tyr434His | P54098 |
| HOMO SAPIENS | -1 | 10836 | 4ztu | A | 511 | SER | p.Ser511Asn | P54098 |
| HOMO SAPIENS | 0 | 10836 | 4ztu | A | 533 | CYS | p.Cys533Trp | P54098 |
| HOMO SAPIENS | 1 | 10836 | 4ztu | A | 546 | ARG | p.Arg546Cys | P54098 |
| HOMO SAPIENS | 1 | 10836 | 4ztu | A | 587 | PRO | p.Pro587Leu | P54098 |
| HOMO SAPIENS | 0 | 10836 | 4ztu | A | 603 | MET | p.Met603Thr | P54098 |
| HOMO SAPIENS | 0 | 10836 | 4ztu | A | 623 | LEU | p.Leu623Trp | P54098 |
| HOMO SAPIENS | -1 | 10836 | 4ztu | A | 627 | ARG | p.Arg627Trp | P54098 |
| HOMO SAPIENS | -1 | 10836 | 4ztu | A | 628 | ARG | p.Arg628Pro | P54098 |
| HOMO SAPIENS | 0 | 10836 | 4ztu | A | 635 | PRO | p.Pro635Leu | P54098 |
| HOMO SAPIENS | 0 | 10836 | 4ztu | A | 662 | GLU | p.Glu662Lys | P54098 |
| HOMO SAPIENS | -1 | 10836 | 4ztu | A | 674 | GLY | p.Gly674Asp | P54098 |
| HOMO SAPIENS | -1 | 10836 | 4ztu | A | 722 | ARG | p.Arg722His | P54098 |
| HOMO SAPIENS | -1 | 10836 | 4ztu | A | 737 | GLY | p.Gly737Arg | P54098 |
| HOMO SAPIENS | 0 | 10836 | 4ztu | A | 748 | TRP | p.Trp748Ser | P54098 |
| HOMO SAPIENS | 0 | 10836 | 4ztu | A | 749 | PHE | p.Phe749Ser | P54098 |
| HOMO SAPIENS | 1 | 10836 | 4ztu | A | 807 | ARG | p.Arg807Cys | P54098 |
| HOMO SAPIENS | 1 | 10836 | 4ztu | A | 831 | TYR | p.Tyr831Cys | P54098 |
| HOMO SAPIENS | 8 | 10836 | 4ztu | A | 848 | GLY | p.Gly848Ser | P54098 |
| HOMO SAPIENS | 1 | 10836 | 4ztu | A | 852 | ARG | p.Arg852Cys | P54098 |
| HOMO SAPIENS | 9 | 10836 | 4ztu | A | 853 | ARG | p.Arg853Trp | P54098 |
| HOMO SAPIENS | 1 | 10836 | 4ztu | A | 864 | ASN | p.Asn864Ser | P54098 |
| HOMO SAPIENS | 1 | 10836 | 4ztu | A | 879 | GLN | p.Gln879Arg | P54098 |
| HOMO SAPIENS | 1 | 10836 | 4ztu | A | 914 | THR | p.Thr914Pro | P54098 |
| HOMO SAPIENS | 6 | 10836 | 4ztu | A | 932 | HIS | p.His932Tyr | P54098 |
| HOMO SAPIENS | 2 | 10836 | 4ztu | A | 944 | GLU | p.Glu944Lys | P54098 |
| HOMO SAPIENS | 6 | 10836 | 4ztu | A | 955 | TYR | p.Tyr955Cys | P54098 |
| HOMO SAPIENS | 1 | 10836 | 4ztu | A | 957 | ALA | p.Ala957Ser | P54098 |
| HOMO SAPIENS | 1 | 10836 | 4ztu | A | 957 | ALA | p.Ala957Val | P54098 |
| HOMO SAPIENS | 1 | 10836 | 4ztu | A | 964 | ARG | p.Arg964Cys | P54098 |
| HOMO SAPIENS | 1 | 10836 | 4ztu | A | 966 | LEU | p.Leu966Arg | P54098 |
| HOMO SAPIENS | 0 | 10836 | 4ztu | A | 1051 | GLY | p.Gly1051Arg | P54098 |
| HOMO SAPIENS | 1 | 10836 | 4ztu | A | 1073 | PRO | p.Pro1073Leu | P54098 |
| HOMO SAPIENS | 1 | 10836 | 4ztu | A | 1096 | ARG | p.Arg1096Cys | P54098 |
| HOMO SAPIENS | 1 | 10836 | 4ztu | A | 1096 | ARG | p.Arg1096His | P54098 |
| HOMO SAPIENS | 0 | 10836 | 4ztu | A | 1116 | MET | p.Met1116Thr | P54098 |
| HOMO SAPIENS | 1 | 10836 | 4ztu | A | 1142 | ARG | p.Arg1142Trp | P54098 |
| HOMO SAPIENS | 1 | 10836 | 4ztu | A | 1146 | ARG | p.Arg1146Cys | P54098 |
| HOMO SAPIENS | 1 | 10836 | 4ztu | A | 1157 | ASN | p.Asn1157Ser | P54098 |
| HOMO SAPIENS | -1 | 10836 | 4ztu | A | 1234 | ARG | p.Arg1234Gln | P54098 |
| HOMO SAPIENS | -1 | 10836 | 4ztu | A | 1236 | GLN | p.Gln1236His | P54098 |
| HOMO SAPIENS | 5 | 2666 | 1jph | A | 62 | PRO | p.Pro62Leu | P06132 |

| | | | | | | | | |
|---|---|---|---|---|---|---|---|---|
| HOMO SAPIENS | 5 | 2666 | 1jph | A | 116 | GLN | p.Gln116Ter | P06132 |
| HOMO SAPIENS | 0 | 2666 | 1jph | A | 165 | MET | p.Met165Arg | P06132 |
| HOMO SAPIENS | 6 | 2666 | 1jph | A | 167 | GLU | p.Glu167Lys | P06132 |
| HOMO SAPIENS | 0 | 2666 | 1jph | A | 195 | LEU | p.Leu195Phe | P06132 |
| HOMO SAPIENS | 7 | 2666 | 1jph | A | 253 | LEU | p.Leu253Gln | P06132 |
| HOMO SAPIENS | 0 | 2666 | 1jph | A | 281 | GLY | p.Gly281Glu | P06132 |
| HOMO SAPIENS | 9 | 2666 | 1jph | A | 304 | ASN | p.Asn304Lys | P06132 |
| HOMO SAPIENS | 8 | 2666 | 1jph | A | 311 | TYR | p.Tyr311Cys | P06132 |
| HOMO SAPIENS | 5 | 2666 | 1jph | A | 332 | ARG | p.Arg332His | P06132 |
| HOMO SAPIENS | 1 | 889 | 1m51 | A | 184 | LEU | p.Val184Leu | P35558 |
| HOMO SAPIENS | 0 | 889 | 1m51 | A | 267 | VAL | p.Ile267Val | P35558 |
| HOMO SAPIENS | 2 | 889 | 1m51 | A | 276 | GLU | p.Glu276Lys | P35558 |
| HOMO SAPIENS | 6 | 889 | 1m51 | A | 368 | VAL | p.Val368Ile | P35558 |
| HOMO SAPIENS | 0 | 889 | 1m51 | A | 579 | SER | p.Ser579Arg | P35558 |
| HOMO SAPIENS | 1 | 4084 | 2gjx | A | 28 | GLN | p.Gln28Ter | P06865 |
| HOMO SAPIENS | 0 | 4084 | 2gjx | A | 39 | LEU | p.Leu39Arg | P06865 |
| HOMO SAPIENS | 0 | 4084 | 2gjx | A | 127 | LEU | p.Leu127Arg | P06865 |
| HOMO SAPIENS | 6 | 4084 | 2gjx | A | 137 | ARG | p.Arg137Ter | P06865 |
| HOMO SAPIENS | 6 | 4084 | 2gjx | A | 170 | ARG | p.Arg170Gln | P06865 |
| HOMO SAPIENS | 9 | 4084 | 2gjx | A | 178 | ARG | p.Arg178Cys | P06865 |
| HOMO SAPIENS | 9 | 4084 | 2gjx | A | 178 | ARG | p.Arg178His | P06865 |
| HOMO SAPIENS | 5 | 4084 | 2gjx | A | 180 | TYR | p.Tyr180His | P06865 |
| HOMO SAPIENS | 5 | 4084 | 2gjx | A | 180 | TYR | p.Tyr180Ter | P06865 |
| HOMO SAPIENS | 1 | 4084 | 2gjx | A | 197 | LYS | p.Lys197Thr | P06865 |
| HOMO SAPIENS | 0 | 4084 | 2gjx | A | 200 | VAL | p.Val200Met | P06865 |
| HOMO SAPIENS | 9 | 4084 | 2gjx | A | 204 | HIS | p.His204Arg | P06865 |
| HOMO SAPIENS | 5 | 4084 | 2gjx | A | 210 | SER | p.Ser210Phe | P06865 |
| HOMO SAPIENS | 0 | 4084 | 2gjx | A | 211 | PHE | p.Phe211Ser | P06865 |
| HOMO SAPIENS | 2 | 4084 | 2gjx | A | 250 | GLY | p.Gly250Asp | P06865 |
| HOMO SAPIENS | 9 | 4084 | 2gjx | A | 258 | ASP | p.Asp258His | P06865 |
| HOMO SAPIENS | 0 | 4084 | 2gjx | A | 269 | GLY | p.Gly269Ser | P06865 |
| HOMO SAPIENS | 0 | 4084 | 2gjx | A | 301 | MET | p.Met301Arg | P06865 |
| HOMO SAPIENS | 3 | 4084 | 2gjx | A | 393 | ARG | p.Arg393Ter | P06865 |
| HOMO SAPIENS | 3 | 4084 | 2gjx | A | 393 | ARG | p.Arg393Pro | P06865 |
| HOMO SAPIENS | 0 | 4084 | 2gjx | A | 399 | ASN | p.Asn399Asp | P06865 |
| HOMO SAPIENS | 2 | 4084 | 2gjx | A | 420 | TRP | p.Trp420Cys | P06865 |
| HOMO SAPIENS | 1 | 4084 | 2gjx | A | 434 | PHE | p.Phe434Leu | P06865 |
| HOMO SAPIENS | 0 | 4084 | 2gjx | A | 454 | GLY | p.Gly454Ser | P06865 |
| HOMO SAPIENS | 1 | 4084 | 2gjx | A | 474 | TRP | p.Trp474Cys | P06865 |
| HOMO SAPIENS | 0 | 4084 | 2gjx | A | 479 | ALA | p.Ala479Thr | P06865 |
| HOMO SAPIENS | 5 | 4084 | 2gjx | A | 485 | TRP | p.Trp485Arg | P06865 |
| HOMO SAPIENS | 4 | 4084 | 2gjx | A | 499 | ARG | p.Arg499Cys | P06865 |
| HOMO SAPIENS | 4 | 4084 | 2gjx | A | 499 | ARG | p.Arg499His | P06865 |
| HOMO SAPIENS | 2 | 4084 | 2gjx | A | 504 | ARG | p.Arg504Cys | P06865 |
| HOMO SAPIENS | 2 | 4084 | 2gjx | A | 504 | ARG | p.Arg504His | P06865 |
| HOMO SAPIENS | -1 | 1943 | 1xmj | A | 417 | ASN | p.Asn417Lys | P13569 |
| HOMO SAPIENS | -1 | 1943 | 1xmj | A | 422 | SER | p.Ser422Phe | P13569 |
| HOMO SAPIENS | 1 | 1943 | 1xmj | A | 438 | THR | p.Thr438Ala | P13569 |
| HOMO SAPIENS | 0 | 1943 | 1xmj | A | 455 | ALA | p.Ala455Glu | P13569 |
| HOMO SAPIENS | 4 | 1943 | 1xmj | A | 456 | VAL | p.Val456Ala | P13569 |
| HOMO SAPIENS | 9 | 1943 | 1xmj | A | 458 | GLY | p.Gly458Val | P13569 |
| HOMO SAPIENS | 8 | 1943 | 1xmj | A | 466 | SER | p.Ser466Ter | P13569 |
| HOMO SAPIENS | 0 | 1943 | 1xmj | A | 467 | LEU | p.Leu467Pro | P13569 |
| HOMO SAPIENS | 1 | 1943 | 1xmj | A | 470 | MET | p.Val470Met | P13569 |

| | | | | | | | | |
|---|---|---|---|---|---|---|---|---|
| HOMO SAPIENS | 5 | 1943 | 1xmj | A | 480 | GLY | p.Gly480Cys | P13569 |
| HOMO SAPIENS | 3 | 1943 | 1xmj | A | 492 | SER | p.Ser492Phe | P13569 |
| HOMO SAPIENS | 8 | 1943 | 1xmj | A | 493 | GLN | p.Gln493Ter | P13569 |
| HOMO SAPIENS | 5 | 1943 | 1xmj | A | 506 | ILE | p.Ile506Met | P13569 |
| HOMO SAPIENS | 3 | 1943 | 1xmj | A | 520 | VAL | p.Val520Phe | P13569 |
| HOMO SAPIENS | 1 | 1943 | 1xmj | A | 528 | GLU | p.Glu528Asp | P13569 |
| HOMO SAPIENS | 1 | 1943 | 1xmj | A | 534 | ALA | p.Ala534Glu | P13569 |
| HOMO SAPIENS | 4 | 1943 | 1xmj | A | 542 | GLY | p.Gly542Ter | P13569 |
| HOMO SAPIENS | 6 | 1943 | 1xmj | A | 549 | SER | p.Ser549Asn | P13569 |
| HOMO SAPIENS | 6 | 1943 | 1xmj | A | 549 | SER | p.Ser549Arg | P13569 |
| HOMO SAPIENS | 6 | 1943 | 1xmj | A | 549 | SER | p.Ser549Arg | P13569 |
| HOMO SAPIENS | 6 | 1943 | 1xmj | A | 551 | GLY | p.Gly551Asp | P13569 |
| HOMO SAPIENS | 6 | 1943 | 1xmj | A | 551 | GLY | p.Gly551Ser | P13569 |
| HOMO SAPIENS | 6 | 1943 | 1xmj | A | 553 | GLN | p.Arg553Gln | P13569 |
| HOMO SAPIENS | 6 | 1943 | 1xmj | A | 553 | GLN | p.Arg553Ter | P13569 |
| HOMO SAPIENS | 0 | 1943 | 1xmj | A | 559 | ALA | p.Ala559Thr | P13569 |
| HOMO SAPIENS | 7 | 1943 | 1xmj | A | 560 | ARG | p.Arg560Lys | P13569 |
| HOMO SAPIENS | 4 | 1943 | 1xmj | A | 562 | VAL | p.Val562Ile | P13569 |
| HOMO SAPIENS | 3 | 1943 | 1xmj | A | 563 | TYR | p.Tyr563Asn | P13569 |
| HOMO SAPIENS | 3 | 1943 | 1xmj | A | 569 | TYR | p.Tyr569Asp | P13569 |
| HOMO SAPIENS | 3 | 1943 | 1xmj | A | 585 | GLU | p.Glu585Ter | P13569 |
| HOMO SAPIENS | 1 | 1943 | 1xmj | A | 618 | ILE | p.Ile618Thr | P13569 |
| HOMO SAPIENS | 9 | 1943 | 1xmj | A | 622 | GLY | p.Gly622Asp | P13569 |
| HOMO SAPIENS | -1 | 1943 | 1xmj | A | 648 | ASP | p.Asp648Val | P13569 |
| HOMO SAPIENS | 0 | 1943 | 1xmj | A | 673 | GLY | p.Gly673Ter | P13569 |
| HOMO SAPIENS | 5 | 5111 | 4rec | A | 749 | ARG | p.Arg749Ter | Q9Y2M0 |
| HOMO SAPIENS | -1 | 4580 | 4l0d | B | 2 | GLY | p.Pro2Leu | P35520 |
| HOMO SAPIENS | -1 | 4580 | 4l0d | B | 12 | PRO | p.Pro12Ser | P35520 |
| HOMO SAPIENS | -1 | 4580 | 4l0d | B | 18 | ARG | p.Arg18Cys | P35520 |
| HOMO SAPIENS | 1 | 4580 | 4l0d | B | 49 | PRO | p.Pro49Leu | P35520 |
| HOMO SAPIENS | 2 | 4580 | 4l0d | B | 70 | PRO | p.Pro70Leu | P35520 |
| HOMO SAPIENS | 1 | 4580 | 4l0d | B | 102 | LYS | p.Lys102Gln | P35520 |
| HOMO SAPIENS | 7 | 4580 | 4l0d | B | 114 | ALA | p.Ala114Val | P35520 |
| HOMO SAPIENS | 5 | 4580 | 4l0d | B | 116 | GLY | p.Gly116Arg | P35520 |
| HOMO SAPIENS | 8 | 4580 | 4l0d | B | 121 | ARG | p.Arg121Cys | P35520 |
| HOMO SAPIENS | 8 | 4580 | 4l0d | B | 121 | ARG | p.Arg121His | P35520 |
| HOMO SAPIENS | 5 | 4580 | 4l0d | B | 125 | ARG | p.Arg125Gln | P35520 |
| HOMO SAPIENS | 4 | 4580 | 4l0d | B | 134 | GLY | p.Gly134Arg | P35520 |
| HOMO SAPIENS | 4 | 4580 | 4l0d | B | 139 | GLY | p.Gly139Arg | P35520 |
| HOMO SAPIENS | 0 | 4580 | 4l0d | B | 145 | PRO | p.Pro145Leu | P35520 |
| HOMO SAPIENS | 0 | 4580 | 4l0d | B | 148 | GLY | p.Gly148Arg | P35520 |
| HOMO SAPIENS | 0 | 4580 | 4l0d | B | 168 | VAL | p.Val168Met | P35520 |
| HOMO SAPIENS | 2 | 4580 | 4l0d | B | 191 | THR | p.Thr191Met | P35520 |
| HOMO SAPIENS | 5 | 4580 | 4l0d | B | 234 | ASP | p.Asp234Asn | P35520 |
| HOMO SAPIENS | 0 | 4580 | 4l0d | B | 257 | THR | p.Thr257Met | P35520 |
| HOMO SAPIENS | 4 | 4580 | 4l0d | B | 262 | THR | p.Thr262Met | P35520 |
| HOMO SAPIENS | 0 | 4580 | 4l0d | B | 278 | ILE | p.Ile278Thr | P35520 |
| HOMO SAPIENS | 2 | 4580 | 4l0d | B | 307 | GLY | p.Gly307Ser | P35520 |
| HOMO SAPIENS | 4 | 4580 | 4l0d | B | 320 | VAL | p.Val320Ala | P35520 |
| HOMO SAPIENS | 2 | 4580 | 4l0d | B | 336 | ARG | p.Arg336Cys | P35520 |
| HOMO SAPIENS | 2 | 4580 | 4l0d | B | 336 | ARG | p.Arg336His | P35520 |
| HOMO SAPIENS | 0 | 4580 | 4l0d | B | 351 | GLY | p.Gly351Arg | P35520 |
| HOMO SAPIENS | 0 | 4580 | 4l0d | B | 353 | THR | p.Thr353Met | P35520 |
| HOMO SAPIENS | 0 | 4580 | 4l0d | B | 371 | VAL | p.Val371Met | P35520 |

| | | | | | | | | |
|---|---|---|---|---|---|---|---|---|
| HOMO SAPIENS | 2 | 4580 | 4l0d | B | 379 | ARG | p.Arg379Gln | P35520 |
| HOMO SAPIENS | 2 | 4580 | 4l0d | B | 379 | ARG | p.Arg379Trp | P35520 |
| HOMO SAPIENS | 2 | 4580 | 4l0d | B | 384 | LYS | p.Lys384Glu | P35520 |
| HOMO SAPIENS | 0 | 4580 | 4l0d | B | 422 | PRO | p.Pro422Leu | P35520 |
| HOMO SAPIENS | 0 | 4580 | 4l0d | B | 444 | ASP | p.Asp444Asn | P35520 |
| HOMO SAPIENS | 0 | 4580 | 4l0d | B | 466 | SER | p.Ser466Leu | P35520 |
| HOMO SAPIENS | 0 | 4580 | 4l0d | B | 539 | LEU | p.Leu539Ser | P35520 |
| HOMO SAPIENS | 0 | 4580 | 4l0d | B | 548 | ARG | p.Arg548Gln | P35520 |
| HOMO SAPIENS | 1 | 6536 | 3ulu | A | 412 | LEU | p.Leu412Phe | O15455 |
| HOMO SAPIENS | 1 | 28493 | 3ii6 | X | 814 | ARG | p.Arg814Ter | P49917 |
| HOMO SAPIENS | 1 | 28493 | 3ii6 | X | 857 | ALA | p.Ala857Thr | P49917 |
| HOMO SAPIENS | 9 | 3543 | 2q51 | A | 24 | GLU | p.Glu24Gly | P45381 |
| HOMO SAPIENS | 1 | 3543 | 2q51 | A | 27 | GLY | p.Gly27Arg | P45381 |
| HOMO SAPIENS | 9 | 3543 | 2q51 | A | 71 | ARG | p.Arg71His | P45381 |
| HOMO SAPIENS | -1 | 3543 | 2q51 | A | 82 | MSE | p.Met82Thr | P45381 |
| HOMO SAPIENS | 0 | 3543 | 2q51 | A | 143 | ILE | p.Ile143Phe | P45381 |
| HOMO SAPIENS | 1 | 3543 | 2q51 | A | 152 | CYS | p.Cys152Arg | P45381 |
| HOMO SAPIENS | 2 | 3543 | 2q51 | A | 218 | CYS | p.Cys218Ter | P45381 |
| HOMO SAPIENS | 1 | 3543 | 2q51 | A | 231 | TYR | p.Tyr231Cys | P45381 |
| HOMO SAPIENS | 1 | 3543 | 2q51 | A | 231 | TYR | p.Tyr231Ter | P45381 |
| HOMO SAPIENS | 2 | 3543 | 2q51 | A | 274 | GLY | p.Gly274Arg | P45381 |
| HOMO SAPIENS | 6 | 3543 | 2q51 | A | 285 | GLU | p.Glu285Ala | P45381 |
| HOMO SAPIENS | 5 | 3543 | 2q51 | A | 287 | ALA | p.Ala287Thr | P45381 |
| HOMO SAPIENS | 2 | 3543 | 2q51 | A | 305 | ALA | p.Ala305Glu | P45381 |
| HOMO SAPIENS | 2 | 24687 | 1hfi | A | 25 | SER | p.Ser890Ile | P08603 |
| HOMO SAPIENS | 0 | 1508 | 3wn5 | C | 45 | LEU | p.Leu66Arg | P08637 |
| HOMO SAPIENS | 0 | 1508 | 3wn5 | C | 155 | VAL | p.Phe176Val | P08637 |
| HOMO SAPIENS | 0 | 750 | 5tpa | A | 54 | VAL | p.Val452Met | Q12879 |
| HOMO SAPIENS | 9 | 750 | 5tpa | A | 120 | ARG | p.Arg518Cys | Q12879 |
| HOMO SAPIENS | 0 | 750 | 5tpa | A | 199 | ALA | p.Ala716Thr | Q12879 |
| HOMO SAPIENS | 4 | 1363 | 3hg2 | B | 40 | PRO | p.Pro40Ser | P06280 |
| HOMO SAPIENS | 3 | 1363 | 3hg2 | B | 65 | SER | p.Ser65Thr | P06280 |
| HOMO SAPIENS | 4 | 1363 | 3hg2 | B | 66 | GLU | p.Glu66Gln | P06280 |
| HOMO SAPIENS | 2 | 1363 | 3hg2 | B | 80 | GLY | p.Gly80Asp | P06280 |
| HOMO SAPIENS | 3 | 1363 | 3hg2 | B | 112 | ARG | p.Arg112Cys | P06280 |
| HOMO SAPIENS | 3 | 1363 | 3hg2 | B | 112 | ARG | p.Arg112His | P06280 |
| HOMO SAPIENS | 0 | 1363 | 3hg2 | B | 139 | ASN | p.Asn139Ser | P06280 |
| HOMO SAPIENS | 7 | 1363 | 3hg2 | B | 156 | ALA | p.Ala156Thr | P06280 |
| HOMO SAPIENS | 5 | 1363 | 3hg2 | B | 162 | TRP | p.Trp162Arg | P06280 |
| HOMO SAPIENS | 5 | 1363 | 3hg2 | B | 207 | TYR | p.Tyr207His | P06280 |
| HOMO SAPIENS | 0 | 1363 | 3hg2 | B | 215 | ASN | p.Asn215Ser | P06280 |
| HOMO SAPIENS | 0 | 1363 | 3hg2 | B | 264 | ASP | p.Asp264Val | P06280 |
| HOMO SAPIENS | 5 | 1363 | 3hg2 | B | 266 | ASP | p.Asp266Val | P06280 |
| HOMO SAPIENS | 2 | 1363 | 3hg2 | B | 269 | VAL | p.Val269Ala | P06280 |
| HOMO SAPIENS | 3 | 1363 | 3hg2 | B | 272 | ASN | p.Asn272Ser | P06280 |
| HOMO SAPIENS | 3 | 1363 | 3hg2 | B | 279 | GLN | p.Gln279Glu | P06280 |
| HOMO SAPIENS | 0 | 1363 | 3hg2 | B | 289 | ILE | p.Ile289Val | P06280 |
| HOMO SAPIENS | 0 | 1363 | 3hg2 | B | 296 | MET | p.Met296Val | P06280 |
| HOMO SAPIENS | 0 | 1363 | 3hg2 | B | 296 | MET | p.Met296Ile | P06280 |
| HOMO SAPIENS | 0 | 1363 | 3hg2 | B | 297 | SER | p.Ser297Phe | P06280 |
| HOMO SAPIENS | 2 | 1363 | 3hg2 | B | 301 | ARG | p.Arg301Gln | P06280 |
| HOMO SAPIENS | 3 | 1363 | 3hg2 | B | 327 | GLN | p.Gln327Lys | P06280 |
| HOMO SAPIENS | 0 | 1363 | 3hg2 | B | 328 | GLY | p.Gly328Arg | P06280 |
| HOMO SAPIENS | 0 | 1363 | 3hg2 | B | 328 | GLY | p.Gly328Ala | P06280 |

| | | | | | | | | |
|---|---|---|---|---|---|---|---|---|
| HOMO SAPIENS | 2 | 1363 | 3hg2 | B | 331 | LEU | p.Leu331Phe | P06280 |
| HOMO SAPIENS | 0 | 1363 | 3hg2 | B | 342 | ARG | p.Arg342Gln | P06280 |
| HOMO SAPIENS | 1 | 1363 | 3hg2 | B | 361 | GLY | p.Gly361Arg | P06280 |
| HOMO SAPIENS | 1 | 1363 | 3hg2 | B | 363 | ARG | p.Arg363His | P06280 |
| HOMO SAPIENS | 1 | 1363 | 3hg2 | B | 385 | THR | p.Thr385Ala | P06280 |
| HOMO SAPIENS | 1 | 1363 | 3hg2 | B | 399 | TRP | p.Trp399Ser | P06280 |
| HOMO SAPIENS | 1 | 1363 | 3hg2 | B | 410 | THR | p.Thr410Ala | P06280 |
| HOMO SAPIENS | 2 | 3904 | 3caa | B | 385 | MET | p.Met414Val | P01011 |
| HOMO SAPIENS | 0 | 558 | 5eub | A | 56 | VAL | p.Val56Ile | Q13126 |
| HOMO SAPIENS | 0 | 558 | 5eub | A | 189 | TRP | p.Trp189Leu | Q13126 |
| HOMO SAPIENS | 0 | 558 | 5eub | A | 218 | ALA | p.Ala218Gly | Q13126 |
| HOMO SAPIENS | 1 | 3865 | 1cb5 | B | 443 | ILE | p.Ile443Val | Q13867 |
| HOMO SAPIENS | 2 | 14329 | 4m92 | A | 24 | ILE | p.Ile65Val | Q13454 |
| HOMO SAPIENS | 1 | 7381 | 5kz5 | k | 50 | GLY | p.Gly50Glu | Q9H1K1 |
| HOMO SAPIENS | 0 | 1102 | 4fdh | E | 86 | PRO | p.Pro86Ala | P19099 |
| HOMO SAPIENS | 0 | 1102 | 4fdh | E | 118 | ALA | p.Ala118Thr | P19099 |
| HOMO SAPIENS | 0 | 1102 | 4fdh | E | 159 | PRO | p.Pro159Leu | P19099 |
| HOMO SAPIENS | 0 | 1102 | 4fdh | E | 173 | LYS | p.Lys173Arg | P19099 |
| HOMO SAPIENS | 0 | 1102 | 4fdh | E | 181 | ARG | p.Arg181Trp | P19099 |
| HOMO SAPIENS | 0 | 1102 | 4fdh | E | 185 | THR | p.Thr185Ile | P19099 |
| HOMO SAPIENS | 0 | 1102 | 4fdh | E | 198 | GLU | p.Glu198Asp | P19099 |
| HOMO SAPIENS | 0 | 1102 | 4fdh | E | 214 | HIS | p.His214Asp | P19099 |
| HOMO SAPIENS | 0 | 1102 | 4fdh | E | 251 | LYS | p.Lys251Arg | P19099 |
| HOMO SAPIENS | 0 | 1102 | 4fdh | E | 255 | GLU | p.Glu255Ter | P19099 |
| HOMO SAPIENS | 0 | 1102 | 4fdh | E | 281 | ASN | p.Asn281Ser | P19099 |
| HOMO SAPIENS | 0 | 1102 | 4fdh | E | 320 | ALA | p.Ala320Val | P19099 |
| HOMO SAPIENS | 0 | 1102 | 4fdh | E | 339 | ILE | p.Ile339Thr | P19099 |
| HOMO SAPIENS | 0 | 1102 | 4fdh | E | 341 | ARG | p.Arg341Cys | P19099 |
| HOMO SAPIENS | 2 | 1102 | 4fdh | E | 374 | ARG | p.Arg374Trp | P19099 |
| HOMO SAPIENS | 0 | 1102 | 4fdh | E | 386 | VAL | p.Val386Ala | P19099 |
| HOMO SAPIENS | 0 | 1102 | 4fdh | E | 390 | LEU | p.Leu390Phe | P19099 |
| HOMO SAPIENS | -1 | 1102 | 4fdh | E | 435 | GLY | p.Gly435Ser | P19099 |
| HOMO SAPIENS | 0 | 1102 | 4fdh | E | 461 | LEU | p.Leu461Pro | P19099 |
| HOMO SAPIENS | 0 | 1102 | 4fdh | E | 498 | THR | p.Thr498Ala | P19099 |
| HOMO SAPIENS | 6 | 10292 | 2of5 | F | 128 | GLY | p.Gly128Arg | P78560 |
| HOMO SAPIENS | 6 | 10292 | 2of5 | F | 170 | ARG | p.Arg170Cys | P78560 |
| HOMO SAPIENS | -1 | 18965 | 3ghg | A | 16 | ARG | p.Arg35Cys | P02671 |
| HOMO SAPIENS | -1 | 18965 | 3ghg | A | 16 | ARG | p.Arg35His | P02671 |
| HOMO SAPIENS | -1 | 18965 | 3ghg | A | 283 | PRO | p.Pro302Ala | P02671 |
| HOMO SAPIENS | -1 | 18965 | 3ghg | A | 312 | THR | p.Thr331Ala | P02671 |
| HOMO SAPIENS | -1 | 18965 | 3ghg | A | 381 | SER | p.Ser400Phe | P02671 |
| HOMO SAPIENS | -1 | 18965 | 3ghg | A | 463 | VAL | p.Val482Met | P02671 |
| HOMO SAPIENS | -1 | 18965 | 3ghg | A | 526 | GLU | p.Glu545Val | P02671 |
| HOMO SAPIENS | -1 | 18965 | 3ghg | A | 554 | ARG | p.Arg573Leu | P02671 |
| HOMO SAPIENS | 8 | 3023 | 4ttb | A | 101 | ARG | p.Arg101Trp | Q6PHW0 |
| HOMO SAPIENS | 2 | 3023 | 4ttb | A | 116 | ILE | p.Ile116Thr | Q6PHW0 |
| HOMO SAPIENS | 0 | 3023 | 4ttb | A | 220 | ALA | p.Ala220Thr | Q6PHW0 |
| HOMO SAPIENS | 2 | 3023 | 4ttb | A | 271 | GLU | p.Glu271Lys | Q6PHW0 |
| HOMO SAPIENS | 3 | 33226 | 1ya5 | T | 11 | SER | p.Ser11Ter | O15273 |
| HOMO SAPIENS | 6 | 33226 | 1ya5 | T | 64 | SER | p.Ser64Leu | O15273 |
| HOMO SAPIENS | 6 | 33226 | 1ya5 | T | 70 | ARG | p.Arg70Gln | O15273 |
| HOMO SAPIENS | 3 | 33226 | 1ya5 | T | 75 | GLY | p.Gly75Ser | O15273 |
| HOMO SAPIENS | 0 | 33226 | 1ya5 | T | 87 | ARG | p.Arg87Trp | O15273 |
| HOMO SAPIENS | 2 | 5667 | 3dax | A | 398 | PRO | p.Pro398Ala | P22680 |

| | | | | | | | | |
|---|---|---|---|---|---|---|---|---|
| HOMO SAPIENS | 0 | 12861 | 5mya | A | 464 | ASN | p.Asn464His | Q02763 |
| HOMO SAPIENS | 2 | 12861 | 5mya | A | 486 | VAL | p.Val486Ile | Q02763 |
| HOMO SAPIENS | 0 | 12861 | 5mya | A | 522 | ARG | p.Arg522His | Q02763 |
| HOMO SAPIENS | 2 | 12861 | 5mya | A | 600 | VAL | p.Val600Leu | Q02763 |
| HOMO SAPIENS | 0 | 12861 | 5mya | A | 634 | LEU | p.Leu634Phe | Q02763 |
| HOMO SAPIENS | 0 | 12861 | 5mya | A | 684 | HIS | p.His684Gln | Q02763 |
| HOMO SAPIENS | 0 | 12861 | 5mya | A | 724 | ALA | p.Ala724Thr | Q02763 |
| HOMO SAPIENS | 0 | 7743 | 3qzw | G | 90 | GLY | p.Gly111Ser | P01732 |
| HOMO SAPIENS | 2 | 9498 | 3kci | A | 4267 | ASP | p.Asp4267Glu | O95714 |
| HOMO SAPIENS | 5 | 201 | 3ips | B | 415 | ARG | p.Arg415Gln | Q13133 |
| HOMO SAPIENS | 3 | 229 | 2q50 | A | 68 | LEU | p.Leu68Pro | Q9UBQ7 |
| HOMO SAPIENS | 2 | 229 | 2q50 | A | 99 | ARG | p.Arg99Ter | Q9UBQ7 |
| HOMO SAPIENS | 3 | 229 | 2q50 | A | 160 | GLY | p.Gly160Arg | Q9UBQ7 |
| HOMO SAPIENS | 5 | 229 | 2q50 | A | 165 | GLY | p.Gly165Asp | Q9UBQ7 |
| HOMO SAPIENS | 4 | 229 | 2q50 | A | 302 | ARG | p.Arg302Cys | Q9UBQ7 |
| HOMO SAPIENS | 4 | 229 | 2q50 | A | 302 | ARG | p.Arg302His | Q9UBQ7 |
| HOMO SAPIENS | -1 | 229 | 2q50 | A | 322 | MSE | p.Met322Arg | Q9UBQ7 |
| HOMO SAPIENS | -1 | 4519 | 5clw | A | 40 | PRO | p.Pro40Thr | Q04446 |
| HOMO SAPIENS | 1 | 4519 | 5clw | A | 59 | ILE | p.Ile59Thr | Q04446 |
| HOMO SAPIENS | 0 | 4519 | 5clw | A | 190 | ARG | p.Arg190Gly | Q04446 |
| HOMO SAPIENS | 1 | 4519 | 5clw | A | 224 | LEU | p.Leu224Pro | Q04446 |
| HOMO SAPIENS | 0 | 4519 | 5clw | A | 236 | GLN | p.Gln236His | Q04446 |
| HOMO SAPIENS | 1 | 4519 | 5clw | A | 254 | THR | p.Thr254Ala | Q04446 |
| HOMO SAPIENS | 0 | 4519 | 5clw | A | 257 | PHE | p.Phe257Leu | Q04446 |
| HOMO SAPIENS | 1 | 4519 | 5clw | A | 262 | ARG | p.Arg262Cys | Q04446 |
| HOMO SAPIENS | 3 | 4519 | 5clw | A | 329 | TYR | p.Tyr329Ser | Q04446 |
| HOMO SAPIENS | 0 | 4519 | 5clw | A | 507 | THR | p.Thr507Ala | Q04446 |
| HOMO SAPIENS | 0 | 4519 | 5clw | A | 515 | ARG | p.Arg515Cys | Q04446 |
| HOMO SAPIENS | 0 | 4519 | 5clw | A | 515 | ARG | p.Arg515His | Q04446 |
| HOMO SAPIENS | 0 | 4519 | 5clw | A | 524 | ARG | p.Arg524Gln | Q04446 |
| HOMO SAPIENS | 0 | 4519 | 5clw | A | 524 | ARG | p.Arg524Ter | Q04446 |
| HOMO SAPIENS | 0 | 4519 | 5clw | A | 545 | HIS | p.His545Arg | Q04446 |
| HOMO SAPIENS | 2 | 4519 | 5clw | A | 548 | TRP | p.Trp548Ter | Q04446 |
| HOMO SAPIENS | 0 | 4519 | 5clw | A | 592 | GLU | p.Glu592Ter | Q04446 |
| HOMO SAPIENS | 0 | 4519 | 5clw | A | 628 | HIS | p.His628Arg | Q04446 |
| HOMO SAPIENS | 0 | 4519 | 5clw | A | 637 | ARG | p.Arg637Ter | Q04446 |
| HOMO SAPIENS | 0 | 4519 | 5clw | A | 673 | ALA | p.Ala673Thr | Q04446 |
| HOMO SAPIENS | 1 | 10405 | 3bg3 | D | 507 | VAL | p.Val507Ile | P11498 |
| HOMO SAPIENS | 0 | 10405 | 3bg3 | D | 583 | ARG | p.Arg583Leu | P11498 |
| HOMO SAPIENS | 1 | 10405 | 3bg3 | D | 589 | LYS | p.Lys589Asn | P11498 |
| HOMO SAPIENS | 6 | 10405 | 3bg3 | D | 610 | ALA | p.Ala610Thr | P11498 |
| HOMO SAPIENS | 2 | 10405 | 3bg3 | D | 631 | ARG | p.Arg631Gln | P11498 |
| HOMO SAPIENS | 1 | 10405 | 3bg3 | D | 732 | ARG | p.Arg732Gln | P11498 |
| HOMO SAPIENS | 6 | 10405 | 3bg3 | D | 743 | MET | p.Met743Ile | P11498 |
| HOMO SAPIENS | 0 | 10405 | 3bg3 | D | 750 | THR | p.Thr750Met | P11498 |
| HOMO SAPIENS | -1 | 13751 | 2o8d | B | 350 | ALA | p.Ala350Val | P52701 |
| HOMO SAPIENS | -1 | 13751 | 2o8d | B | 352 | VAL | p.Val352Ile | P52701 |
| HOMO SAPIENS | -1 | 13751 | 2o8d | B | 355 | GLY | p.Gly355Ser | P52701 |
| HOMO SAPIENS | 0 | 13751 | 2o8d | B | 385 | ARG | p.Arg385Trp | P52701 |
| HOMO SAPIENS | 0 | 13751 | 2o8d | B | 390 | ASP | p.Asp390Glu | P52701 |
| HOMO SAPIENS | 0 | 13751 | 2o8d | B | 396 | LEU | p.Leu396Val | P52701 |
| HOMO SAPIENS | 0 | 13751 | 2o8d | B | 449 | LEU | p.Leu449Pro | P52701 |
| HOMO SAPIENS | 0 | 13751 | 2o8d | B | 468 | ARG | p.Arg468His | P52701 |
| HOMO SAPIENS | 0 | 13751 | 2o8d | B | 503 | SER | p.Ser503Cys | P52701 |

| | | | | | | | | |
|---|---|---|---|---|---|---|---|---|
| HOMO SAPIENS | 1 | 13751 | 2o8d | B | 509 | VAL | p.Val509Ala | P52701 |
| HOMO SAPIENS | 0 | 13751 | 2o8d | B | 675 | GLU | p.Glu675Asp | P52701 |
| HOMO SAPIENS | 0 | 13751 | 2o8d | B | 795 | ILE | p.Ile795Thr | P52701 |
| HOMO SAPIENS | 0 | 13751 | 2o8d | B | 850 | TYR | p.Tyr850Ter | P52701 |
| HOMO SAPIENS | 0 | 13751 | 2o8d | B | 878 | VAL | p.Val878Ala | P52701 |
| HOMO SAPIENS | 0 | 13751 | 2o8d | B | 911 | ARG | p.Arg911Ter | P52701 |
| HOMO SAPIENS | 0 | 13751 | 2o8d | B | 922 | ARG | p.Arg922Ter | P52701 |
| HOMO SAPIENS | 0 | 13751 | 2o8d | B | 969 | TYR | p.Tyr969Cys | P52701 |
| HOMO SAPIENS | 0 | 13751 | 2o8d | B | 1005 | ARG | p.Arg1005Ter | P52701 |
| HOMO SAPIENS | 0 | 13751 | 2o8d | B | 1035 | ARG | p.Arg1035Ter | P52701 |
| HOMO SAPIENS | 1 | 13751 | 2o8d | B | 1048 | GLN | p.Gln1048Ter | P52701 |
| HOMO SAPIENS | 1 | 13751 | 2o8d | B | 1051 | VAL | p.Val1051Ile | P52701 |
| HOMO SAPIENS | 0 | 13751 | 2o8d | B | 1054 | ILE | p.Ile1054Phe | P52701 |
| HOMO SAPIENS | 0 | 13751 | 2o8d | B | 1054 | ILE | p.Ile1054Met | P52701 |
| HOMO SAPIENS | 0 | 13751 | 2o8d | B | 1163 | GLU | p.Glu1163Val | P52701 |
| HOMO SAPIENS | 0 | 13751 | 2o8d | B | 1237 | GLU | p.Glu1237Asp | P52701 |
| HOMO SAPIENS | 0 | 13751 | 2o8d | B | 1242 | ARG | p.Arg1242His | P52701 |
| HOMO SAPIENS | -1 | 13751 | 2o8d | B | 1278 | PRO | p.Pro1278Thr | P52701 |
| HOMO SAPIENS | 0 | 13751 | 2o8d | B | 1304 | ARG | p.Arg1304Lys | P52701 |
| HOMO SAPIENS | 0 | 13751 | 2o8d | B | 1317 | HIS | p.His1317Gln | P52701 |
| HOMO SAPIENS | 0 | 13751 | 2o8d | B | 1329 | SER | p.Ser1329Leu | P52701 |
| HOMO SAPIENS | 3 | 13273 | 4g8k | A | 265 | GLU | p.Glu265Ter | Q05823 |
| HOMO SAPIENS | -1 | 11088 | 5uj7 | A | 666 | ARG | p.Arg666Trp | Q13415 |
| HOMO SAPIENS | 4 | 11088 | 5uj7 | A | 720 | ARG | p.Arg720Gln | Q13415 |
| HOMO SAPIENS | 0 | 11088 | 5uj7 | A | 816 | MET | p.Met816Thr | Q13415 |
| HOMO SAPIENS | 1 | 4352 | 5va1 | A | 421 | THR | p.Thr421Met | Q12809 |
| HOMO SAPIENS | 0 | 4352 | 5va1 | A | 451 | PRO | p.Pro451Leu | Q12809 |
| HOMO SAPIENS | 0 | 4352 | 5va1 | A | 470 | ASN | p.Asn470Asp | Q12809 |
| HOMO SAPIENS | 0 | 4352 | 5va1 | A | 490 | ALA | p.Ala490Thr | Q12809 |
| HOMO SAPIENS | 0 | 4352 | 5va1 | A | 491 | VAL | p.Val491Ile | Q12809 |
| HOMO SAPIENS | 1 | 4352 | 5va1 | A | 533 | VAL | p.Val533Gly | Q12809 |
| HOMO SAPIENS | 0 | 4352 | 5va1 | A | 552 | LEU | p.Leu552Ser | Q12809 |
| HOMO SAPIENS | 0 | 4352 | 5va1 | A | 558 | ALA | p.Ala558Pro | Q12809 |
| HOMO SAPIENS | 1 | 4352 | 5va1 | A | 561 | ALA | p.Ala561Val | Q12809 |
| HOMO SAPIENS | 0 | 4352 | 5va1 | A | 572 | GLY | p.Gly572Arg | Q12809 |
| HOMO SAPIENS | -1 | 4352 | 5va1 | A | 582 | ARG | p.Arg582Cys | Q12809 |
| HOMO SAPIENS | 0 | 4352 | 5va1 | A | 584 | GLY | p.Gly584Ser | Q12809 |
| HOMO SAPIENS | 0 | 4352 | 5va1 | A | 588 | ASN | p.Asn588Lys | Q12809 |
| HOMO SAPIENS | 0 | 4352 | 5va1 | A | 593 | ILE | p.Ile593Arg | Q12809 |
| HOMO SAPIENS | 1 | 4352 | 5va1 | A | 627 | PHE | p.Phe627Leu | Q12809 |
| HOMO SAPIENS | 0 | 4352 | 5va1 | A | 630 | VAL | p.Val630Leu | Q12809 |
| HOMO SAPIENS | 0 | 4352 | 5va1 | A | 652 | TYR | p.Tyr652Ter | Q12809 |
| HOMO SAPIENS | 0 | 4352 | 5va1 | A | 711 | ILE | p.Ile711Val | Q12809 |
| HOMO SAPIENS | 0 | 4352 | 5va1 | A | 735 | SER | p.Ser735Leu | Q12809 |
| HOMO SAPIENS | 0 | 4352 | 5va1 | A | 744 | ARG | p.Arg744Ter | Q12809 |
| HOMO SAPIENS | 0 | 4352 | 5va1 | A | 784 | ARG | p.Arg784Trp | Q12809 |
| HOMO SAPIENS | 1 | 4352 | 5va1 | A | 820 | GLY | p.Gly820Arg | Q12809 |
| HOMO SAPIENS | 5 | 4352 | 5va1 | A | 822 | VAL | p.Val822Met | Q12809 |
| HOMO SAPIENS | 1 | 4352 | 5va1 | A | 863 | ARG | p.Arg863Ter | Q12809 |
| HOMO SAPIENS | 5 | 303 | 2obv | A | 55 | ALA | p.Ala55Asp | Q00266 |
| HOMO SAPIENS | 7 | 303 | 2obv | A | 259 | ALA | p.Ala259Gly | Q00266 |
| HOMO SAPIENS | 9 | 303 | 2obv | A | 264 | ARG | p.Arg264His | Q00266 |
| HOMO SAPIENS | 9 | 303 | 2obv | A | 264 | ARG | p.Arg264Cys | Q00266 |
| HOMO SAPIENS | 0 | 303 | 2obv | A | 305 | LEU | p.Leu305Pro | Q00266 |

| | | | | | | | | |
|---|---|---|---|---|---|---|---|---|
| HOMO SAPIENS | 7 | 303 | 2obv | A | 322 | ILE | p.Ile322Met | Q00266 |
| HOMO SAPIENS | 0 | 303 | 2obv | A | 336 | GLY | p.Gly336Arg | Q00266 |
| HOMO SAPIENS | 0 | 303 | 2obv | A | 357 | PRO | p.Pro357Leu | Q00266 |
| HOMO SAPIENS | 0 | 2077 | 4zxb | E | 462 | ASN | p.Asn489Ser | P06213 |
| HOMO SAPIENS | -1 | 2077 | 4zxb | E | 519 | ASP | p.Asp546Glu | P06213 |
| HOMO SAPIENS | 2 | 2077 | 4zxb | E | 819 | THR | p.Thr858Ala | P06213 |
| HOMO SAPIENS | 0 | 2077 | 4zxb | E | 907 | ASP | p.Asp946Glu | P06213 |
| HOMO SAPIENS | 7 | 3274 | 1wsv | B | 14 | HIS | p.His42Arg | P48728 |
| HOMO SAPIENS | 0 | 3274 | 1wsv | B | 19 | GLY | p.Gly47Arg | P48728 |
| HOMO SAPIENS | 4 | 3274 | 1wsv | B | 45 | ARG | p.Arg73Cys | P48728 |
| HOMO SAPIENS | 1 | 3274 | 1wsv | B | 49 | SER | p.Ser77Leu | P48728 |
| HOMO SAPIENS | 3 | 3274 | 1wsv | B | 241 | GLY | p.Gly269Asp | P48728 |
| HOMO SAPIENS | 7 | 3274 | 1wsv | B | 248 | ASP | p.Asp276His | P48728 |
| HOMO SAPIENS | 0 | 3274 | 1wsv | B | 272 | MET | p.Met300Val | P48728 |
| HOMO SAPIENS | 4 | 3274 | 1wsv | B | 292 | ARG | p.Arg320His | P48728 |
| HOMO SAPIENS | 0 | 3274 | 1wsv | B | 354 | ARG | p.Arg382Gln | P48728 |
| HOMO SAPIENS | -1 | 435 | 3uyt | A | 44 | THR | p.Thr44Ala | P48730 |
| HOMO SAPIENS | 1 | 4698 | 5ife | C | 666 | LYS | p.Lys666Asn | O75533 |
| HOMO SAPIENS | 1 | 16917 | 1apy | C | 37 | GLY | p.Gly60Asp | P20933 |
| HOMO SAPIENS | 0 | 16917 | 1apy | C | 49 | SER | p.Ser72Pro | P20933 |
| HOMO SAPIENS | 0 | 16917 | 1apy | C | 84 | ARG | p.Arg107Ter | P20933 |
| HOMO SAPIENS | 0 | 16917 | 1apy | C | 126 | SER | p.Thr149Ser | P20933 |
| HOMO SAPIENS | 0 | 16917 | 1apy | C | 135 | TRP | p.Trp158Ter | P20933 |
| HOMO SAPIENS | 0 | 16917 | 1apy | C | 140 | CYS | p.Cys163Ser | P20933 |
| HOMO SAPIENS | 0 | 16917 | 1apy | C | 145 | TRP | p.Trp168Ter | P20933 |
| HOMO SAPIENS | 0 | 10413 | 4frw | A | 185 | THR | p.Thr185Met | Q96NY8 |
| HOMO SAPIENS | 0 | 5217 | 5n6w | G | 96 | TRP | p.Trp96Arg | O15537 |
| HOMO SAPIENS | 1 | 5217 | 5n6w | G | 102 | ARG | p.Arg102Trp | O15537 |
| HOMO SAPIENS | 0 | 5217 | 5n6w | G | 109 | GLY | p.Gly109Arg | O15537 |
| HOMO SAPIENS | -1 | 5217 | 5n6w | G | 223 | CYS | p.Cys223Arg | O15537 |
| HOMO SAPIENS | 0 | 9485 | 4z6y | E | 138 | ALA | p.Ala138Asp | Q9P0N9 |
| HOMO SAPIENS | 3 | 4504 | 4mwt | B | 411 | GLY | p.Gly439Ser | P10619 |
| HOMO SAPIENS | 3 | 4504 | 4mwt | B | 412 | PHE | p.Phe440Val | P10619 |
| HOMO SAPIENS | 6 | 6391 | 4kxy | A | 318 | ARG | p.Arg318Cys | P29401 |
| HOMO SAPIENS | 6 | 19546 | 2bx6 | A | 87 | THR | p.Thr87Ile | O75695 |
| HOMO SAPIENS | 1 | 19546 | 2bx6 | A | 118 | ARG | p.Arg118His | O75695 |
| HOMO SAPIENS | 1 | 19546 | 2bx6 | A | 201 | ILE | p.Ile201Val | O75695 |
| HOMO SAPIENS | 1 | 19546 | 2bx6 | A | 282 | ARG | p.Arg282Trp | O75695 |
| HOMO SAPIENS | 1 | 5265 | 5usr | H | 68 | ARG | p.Arg68Leu | Q9HD34 |
| HOMO SAPIENS | 2 | 1012 | 4nyi | S | 569 | LEU | p.Leu569Val | Q07889 |
| HOMO SAPIENS | 2 | 1012 | 4nyi | S | 655 | PRO | p.Pro655Leu | Q07889 |
| HOMO SAPIENS | 4 | 1012 | 4nyi | S | 663 | ILE | p.Ile663Met | Q07889 |
| HOMO SAPIENS | 2 | 1012 | 4nyi | S | 702 | TYR | p.Tyr702Cys | Q07889 |
| HOMO SAPIENS | 2 | 1012 | 4nyi | S | 708 | ALA | p.Ala708Thr | Q07889 |
| HOMO SAPIENS | 4 | 1012 | 4nyi | S | 733 | ILE | p.Ile733Phe | Q07889 |
| HOMO SAPIENS | 1 | 1848 | 3g1n | A | 4013 | ARG | p.Arg4013Trp | Q7Z6Z7 |
| HOMO SAPIENS | 1 | 1848 | 3g1n | A | 4187 | ARG | p.Arg4187Cys | Q7Z6Z7 |
| HOMO SAPIENS | 1 | 77 | 4pod | A | 41 | CYS | p.Cys79Tyr | P60174 |
| HOMO SAPIENS | 9 | 77 | 4pod | A | 104 | GLU | p.Glu142Asp | P60174 |
| HOMO SAPIENS | 1 | 77 | 4pod | A | 122 | GLY | p.Gly160Arg | P60174 |
| HOMO SAPIENS | -1 | 8672 | 3thz | A | 8 | THR | p.Thr8Met | P43246 |
| HOMO SAPIENS | 0 | 8672 | 3thz | A | 19 | PHE | p.Phe19Leu | P43246 |
| HOMO SAPIENS | 0 | 8672 | 3thz | A | 23 | PHE | p.Phe23Leu | P43246 |
| HOMO SAPIENS | 0 | 8672 | 3thz | A | 96 | ARG | p.Arg96His | P43246 |

| | | | | | | | | |
|---|---|---|---|---|---|---|---|---|
| HOMO SAPIENS | 0 | 8672 | 3thz | A | 102 | VAL | p.Val102Ile | P43246 |
| HOMO SAPIENS | 0 | 8672 | 3thz | A | 106 | ARG | p.Arg106Lys | P43246 |
| HOMO SAPIENS | 0 | 8672 | 3thz | A | 127 | ASN | p.Asn127Ser | P43246 |
| HOMO SAPIENS | 0 | 8672 | 3thz | A | 128 | LEU | p.Leu128Val | P43246 |
| HOMO SAPIENS | 0 | 8672 | 3thz | A | 163 | VAL | p.Val163Gly | P43246 |
| HOMO SAPIENS | 0 | 8672 | 3thz | A | 169 | ILE | p.Ile169Val | P43246 |
| HOMO SAPIENS | 0 | 8672 | 3thz | A | 315 | GLY | p.Gly315Asp | P43246 |
| HOMO SAPIENS | -1 | 8672 | 3thz | A | 322 | GLY | p.Gly322Asp | P43246 |
| HOMO SAPIENS | 0 | 8672 | 3thz | A | 331 | ASN | p.Asn331Asp | P43246 |
| HOMO SAPIENS | 1 | 8672 | 3thz | A | 390 | LEU | p.Leu390Phe | P43246 |
| HOMO SAPIENS | 0 | 8672 | 3thz | A | 419 | GLN | p.Gln419Lys | P43246 |
| HOMO SAPIENS | 0 | 8672 | 3thz | A | 447 | PHE | p.Phe447Leu | P43246 |
| HOMO SAPIENS | 0 | 8672 | 3thz | A | 524 | ARG | p.Arg524Pro | P43246 |
| HOMO SAPIENS | 0 | 8672 | 3thz | A | 539 | PHE | p.Phe539Leu | P43246 |
| HOMO SAPIENS | -1 | 8672 | 3thz | A | 546 | LYS | p.Lys546Asn | P43246 |
| HOMO SAPIENS | 0 | 8672 | 3thz | A | 564 | THR | p.Thr564Ala | P43246 |
| HOMO SAPIENS | 0 | 8672 | 3thz | A | 577 | ILE | p.Ile577Thr | P43246 |
| HOMO SAPIENS | 0 | 8672 | 3thz | A | 588 | TYR | p.Tyr588Ter | P43246 |
| HOMO SAPIENS | 0 | 8672 | 3thz | A | 622 | PRO | p.Pro622Leu | P43246 |
| HOMO SAPIENS | 0 | 8672 | 3thz | A | 629 | GLN | p.Gln629Arg | P43246 |
| HOMO SAPIENS | 0 | 8672 | 3thz | A | 636 | ALA | p.Ala636Pro | P43246 |
| HOMO SAPIENS | 2 | 8672 | 3thz | A | 642 | VAL | p.Val642Ile | P43246 |
| HOMO SAPIENS | 1 | 8672 | 3thz | A | 654 | ASP | p.Asp654Glu | P43246 |
| HOMO SAPIENS | 0 | 8672 | 3thz | A | 662 | GLN | p.Gln662His | P43246 |
| HOMO SAPIENS | 2 | 8672 | 3thz | A | 681 | GLN | p.Gln681His | P43246 |
| HOMO SAPIENS | 0 | 8672 | 3thz | A | 707 | CYS | p.Cys707Ser | P43246 |
| HOMO SAPIENS | 0 | 8672 | 3thz | A | 809 | GLU | p.Glu809Lys | P43246 |
| HOMO SAPIENS | 0 | 8672 | 3thz | A | 813 | MET | p.Met813Val | P43246 |
| HOMO SAPIENS | 1 | 8672 | 3thz | A | 834 | ALA | p.Ala834Thr | P43246 |
| HOMO SAPIENS | 0 | 8672 | 3thz | A | 849 | LEU | p.Leu849Val | P43246 |
| HOMO SAPIENS | -1 | 8672 | 3thz | A | 860 | SER | p.Ser860Leu | P43246 |
| HOMO SAPIENS | -1 | 8672 | 3thz | A | 934 | THR | p.Thr934Lys | P43246 |
| HOMO SAPIENS | 2 | 292 | 1i3k | A | 34 | ASN | p.Asn34Ser | Q14376 |
| HOMO SAPIENS | 7 | 292 | 1i3k | A | 90 | GLY | p.Gly90Glu | Q14376 |
| HOMO SAPIENS | 2 | 292 | 1i3k | A | 103 | ASP | p.Asp103Gly | Q14376 |
| HOMO SAPIENS | 2 | 292 | 1i3k | A | 136 | TYR | p.Tyr136Ter | Q14376 |
| HOMO SAPIENS | 0 | 292 | 1i3k | A | 169 | ARG | p.Arg169Trp | Q14376 |
| HOMO SAPIENS | 0 | 292 | 1i3k | A | 183 | LEU | p.Leu183Pro | Q14376 |
| HOMO SAPIENS | 0 | 292 | 1i3k | A | 302 | GLY | p.Gly302Asp | Q14376 |
| HOMO SAPIENS | 0 | 292 | 1i3k | A | 313 | LEU | p.Leu313Met | Q14376 |
| HOMO SAPIENS | 1 | 2084 | 5vc8 | A | 295 | GLN | p.Gln295Leu | O96028 |
| HOMO SAPIENS | -1 | 10157 | 5wrw | A | 7 | GLY | p.Gly7Ala | O76094 |
| HOMO SAPIENS | -1 | 10157 | 5wrw | A | 7 | GLY | p.Gly7Arg | O76094 |
| HOMO SAPIENS | -1 | 10157 | 5wrw | A | 8 | GLY | p.Gly8Ala | O76094 |
| HOMO SAPIENS | 0 | 3113 | 3in5 | A | 29 | GLU | p.Glu29Lys | Q9UBT6 |
| HOMO SAPIENS | 5 | 4546 | 5in3 | B | 44 | VAL | p.Val44Leu | P07902 |
| HOMO SAPIENS | -1 | 4546 | 5in3 | B | 51 | ARG | p.Arg51Leu | P07902 |
| HOMO SAPIENS | -1 | 4546 | 5in3 | B | 62 | LEU | p.Leu62Met | P07902 |
| HOMO SAPIENS | 4 | 4546 | 5in3 | B | 66 | PRO | p.Pro66His | P07902 |
| HOMO SAPIENS | 2 | 4546 | 5in3 | B | 74 | LEU | p.Leu74Pro | P07902 |
| HOMO SAPIENS | 5 | 4546 | 5in3 | B | 97 | ASN | p.Asn97Ser | P07902 |
| HOMO SAPIENS | 1 | 4546 | 5in3 | B | 135 | SER | p.Ser135Leu | P07902 |
| HOMO SAPIENS | 5 | 4546 | 5in3 | B | 138 | THR | p.Thr138Met | P07902 |
| HOMO SAPIENS | 7 | 4546 | 5in3 | B | 139 | LEU | p.Leu139Pro | P07902 |

| | | | | | | | | |
|---|---|---|---|---|---|---|---|---|
| HOMO SAPIENS | 2 | 4546 | 5in3 | B | 142 | MET | p.Met142Lys | P07902 |
| HOMO SAPIENS | 1 | 4546 | 5in3 | B | 143 | SER | p.Ser143Leu | P07902 |
| HOMO SAPIENS | 2 | 4546 | 5in3 | B | 148 | ARG | p.Arg148Gly | P07902 |
| HOMO SAPIENS | 2 | 4546 | 5in3 | B | 148 | ARG | p.Arg148Gln | P07902 |
| HOMO SAPIENS | 5 | 4546 | 5in3 | B | 183 | PRO | p.Pro183Thr | P07902 |
| HOMO SAPIENS | 5 | 4546 | 5in3 | B | 185 | PRO | p.Pro185His | P07902 |
| HOMO SAPIENS | 5 | 4546 | 5in3 | B | 186 | HIS | p.His186Tyr | P07902 |
| HOMO SAPIENS | 7 | 4546 | 5in3 | B | 188 | GLN | p.Gln188Arg | P07902 |
| HOMO SAPIENS | 5 | 4546 | 5in3 | B | 195 | LEU | p.Leu195Pro | P07902 |
| HOMO SAPIENS | 4 | 4546 | 5in3 | B | 201 | ARG | p.Arg201His | P07902 |
| HOMO SAPIENS | 4 | 4546 | 5in3 | B | 201 | ARG | p.Arg201Cys | P07902 |
| HOMO SAPIENS | 4 | 4546 | 5in3 | B | 203 | GLU | p.Glu203Lys | P07902 |
| HOMO SAPIENS | 2 | 4546 | 5in3 | B | 209 | TYR | p.Tyr209Cys | P07902 |
| HOMO SAPIENS | 5 | 4546 | 5in3 | B | 223 | ARG | p.Arg223Ser | P07902 |
| HOMO SAPIENS | 5 | 4546 | 5in3 | B | 226 | LEU | p.Leu226Pro | P07902 |
| HOMO SAPIENS | 5 | 4546 | 5in3 | B | 226 | LEU | p.Leu226Val | P07902 |
| HOMO SAPIENS | 2 | 4546 | 5in3 | B | 227 | LEU | p.Leu227Pro | P07902 |
| HOMO SAPIENS | 8 | 4546 | 5in3 | B | 258 | ARG | p.Arg258Cys | P07902 |
| HOMO SAPIENS | 1 | 4546 | 5in3 | B | 265 | PRO | p.Pro265Ala | P07902 |
| HOMO SAPIENS | 8 | 4546 | 5in3 | B | 268 | THR | p.Thr268Asn | P07902 |
| HOMO SAPIENS | 7 | 4546 | 5in3 | B | 278 | ILE | p.Ile278Asn | P07902 |
| HOMO SAPIENS | 5 | 4546 | 5in3 | B | 285 | LYS | p.Lys285Asn | P07902 |
| HOMO SAPIENS | 1 | 4546 | 5in3 | B | 319 | HIS | p.His319Gln | P07902 |
| HOMO SAPIENS | 4 | 4546 | 5in3 | B | 321 | HIS | p.His321Tyr | P07902 |
| HOMO SAPIENS | 4 | 4546 | 5in3 | B | 325 | PRO | p.Pro325Leu | P07902 |
| HOMO SAPIENS | 2 | 4546 | 5in3 | B | 328 | ARG | p.Arg328His | P07902 |
| HOMO SAPIENS | 2 | 4546 | 5in3 | B | 329 | SER | p.Ser329Phe | P07902 |
| HOMO SAPIENS | 5 | 4546 | 5in3 | B | 334 | LYS | p.Lys334Arg | P07902 |
| HOMO SAPIENS | 6 | 1290 | 5uav | C | 119 | ARG | p.Arg119Gly | P32322 |
| HOMO SAPIENS | 6 | 1290 | 5uav | C | 119 | ARG | p.Arg119His | P32322 |
| HOMO SAPIENS | 3 | 1290 | 5uav | C | 251 | ARG | p.Arg251His | P32322 |
| HOMO SAPIENS | 7 | 1290 | 5uav | C | 266 | ARG | p.Arg266Gln | P32322 |
| HOMO SAPIENS | 0 | 9618 | 2r7e | A | 22 | GLY | p.Gly41Cys | P00451 |
| HOMO SAPIENS | 2 | 9618 | 2r7e | A | 113 | GLU | p.Glu132Asp | P00451 |
| HOMO SAPIENS | -1 | 9618 | 2r7e | A | 336 | ARG | p.Arg355Ter | P00451 |
| HOMO SAPIENS | 0 | 9618 | 2r7e | A | 372 | ARG | p.Arg391His | P00451 |
| HOMO SAPIENS | 0 | 9618 | 2r7e | A | 373 | SER | p.Ser392Leu | P00451 |
| HOMO SAPIENS | 0 | 9618 | 2r7e | A | 373 | SER | p.Ser392Pro | P00451 |
| HOMO SAPIENS | 3 | 9618 | 2r7e | A | 386 | ILE | p.Ile405Ser | P00451 |
| HOMO SAPIENS | 6 | 9618 | 2r7e | A | 390 | GLU | p.Glu409Gly | P00451 |
| HOMO SAPIENS | 2 | 9618 | 2r7e | A | 412 | LEU | p.Leu431Phe | P00451 |
| HOMO SAPIENS | 5 | 9618 | 2r7e | A | 425 | LYS | p.Lys444Arg | P00451 |
| HOMO SAPIENS | 0 | 9618 | 2r7e | A | 484 | ARG | p.Arg503His | P00451 |
| HOMO SAPIENS | 2 | 9618 | 2r7e | A | 531 | ARG | p.Arg550His | P00451 |
| HOMO SAPIENS | 3 | 9618 | 2r7e | A | 535 | SER | p.Ser554Gly | P00451 |
| HOMO SAPIENS | 2 | 9618 | 2r7e | A | 593 | ARG | p.Arg612Cys | P00451 |
| HOMO SAPIENS | 0 | 9618 | 2r7e | A | 704 | ALA | p.Ala723Thr | P00451 |
| HOMO SAPIENS | 0 | 9618 | 2r7e | A | 720 | GLU | p.Glu739Lys | P00451 |
| HOMO SAPIENS | 1 | 13085 | 3ud1 | A | 1075 | ILE | p.Thr1075Ile | P16157 |
| HOMO SAPIENS | -1 | 3289 | 2fy4 | A | 2 | ALA | p.Ala120Thr | P28329 |
| HOMO SAPIENS | 7 | 3289 | 2fy4 | A | 18 | VAL | p.Val136Met | P28329 |
| HOMO SAPIENS | 5 | 3289 | 2fy4 | A | 92 | LEU | p.Leu210Pro | P28329 |
| HOMO SAPIENS | 2 | 3289 | 2fy4 | A | 93 | PRO | p.Pro211Ala | P28329 |
| HOMO SAPIENS | 4 | 3289 | 2fy4 | A | 131 | LEU | p.Leu249Val | P28329 |

| | | | | | | | | |
|---|---|---|---|---|---|---|---|---|
| HOMO SAPIENS | 2 | 3289 | 2fy4 | A | 181 | PRO | p.Pro299Leu | P28329 |
| HOMO SAPIENS | 0 | 3289 | 2fy4 | A | 218 | ILE | p.Ile336Thr | P28329 |
| HOMO SAPIENS | 2 | 3289 | 2fy4 | A | 261 | ASP | p.Asp379His | P28329 |
| HOMO SAPIENS | 0 | 3289 | 2fy4 | A | 302 | ARG | p.Arg420Cys | P28329 |
| HOMO SAPIENS | 5 | 3289 | 2fy4 | A | 323 | GLU | p.Glu441Lys | P28329 |
| HOMO SAPIENS | 0 | 3289 | 2fy4 | A | 343 | MET | p.Val461Met | P28329 |
| HOMO SAPIENS | 2 | 3289 | 2fy4 | A | 364 | ARG | p.Arg482Gly | P28329 |
| HOMO SAPIENS | 3 | 3289 | 2fy4 | A | 380 | SER | p.Ser498Leu | P28329 |
| HOMO SAPIENS | 6 | 3289 | 2fy4 | A | 388 | VAL | p.Val506Leu | P28329 |
| HOMO SAPIENS | 7 | 3289 | 2fy4 | A | 429 | HIS | p.His547Gln | P28329 |
| HOMO SAPIENS | 8 | 3289 | 2fy4 | A | 442 | ARG | p.Arg560His | P28329 |
| HOMO SAPIENS | 3 | 3289 | 2fy4 | A | 443 | ARG | p.Arg561Gln | P28329 |
| HOMO SAPIENS | 0 | 4137 | 3hhr | B | 38 | CYS | p.Cys56Ter | P10912 |
| HOMO SAPIENS | 0 | 4137 | 3hhr | B | 43 | ARG | p.Arg61Ter | P10912 |
| HOMO SAPIENS | 0 | 4137 | 3hhr | B | 44 | GLU | p.Glu62Lys | P10912 |
| HOMO SAPIENS | 0 | 4137 | 3hhr | B | 96 | PHE | p.Phe114Ser | P10912 |
| HOMO SAPIENS | 5 | 4137 | 3hhr | B | 131 | PRO | p.Pro149Gln | P10912 |
| HOMO SAPIENS | 0 | 4137 | 3hhr | B | 144 | VAL | p.Val162Ile | P10912 |
| HOMO SAPIENS | 0 | 4137 | 3hhr | B | 150 | HIS | p.His168Gln | P10912 |
| HOMO SAPIENS | 0 | 4137 | 3hhr | B | 152 | ASP | p.Asp170His | P10912 |
| HOMO SAPIENS | 0 | 4137 | 3hhr | B | 153 | ILE | p.Ile171Thr | P10912 |
| HOMO SAPIENS | 0 | 4137 | 3hhr | B | 154 | GLN | p.Gln172Pro | P10912 |
| HOMO SAPIENS | 0 | 4137 | 3hhr | B | 155 | VAL | p.Val173Gly | P10912 |
| HOMO SAPIENS | 0 | 4137 | 3hhr | B | 217 | ARG | p.Arg235Ter | P10912 |
| HOMO SAPIENS | 5 | 4137 | 3hhr | B | 224 | GLU | p.Glu242Asp | P10912 |
| HOMO SAPIENS | 5 | 4137 | 3hhr | B | 224 | GLU | p.Glu242Ter | P10912 |
| HOMO SAPIENS | 6 | 2493 | 5bpb | A | 69 | HIS | p.His69Tyr | Q9ULV1 |
| HOMO SAPIENS | 6 | 2493 | 5bpb | A | 105 | MET | p.Met105Val | Q9ULV1 |
| HOMO SAPIENS | -1 | 2493 | 5bpb | A | 168 | PRO | p.Pro168Ser | Q9ULV1 |
| HOMO SAPIENS | 0 | 32123 | 4ccz | A | 314 | ASP | p.Asp314Asn | Q99707 |
| HOMO SAPIENS | 0 | 32123 | 4ccz | A | 345 | VAL | p.Val345Phe | Q99707 |
| HOMO SAPIENS | 1 | 32123 | 4ccz | A | 585 | ARG | p.Arg585Ter | Q99707 |
| HOMO SAPIENS | 4 | 4618 | 2ag4 | A | 26 | PRO | p.Pro55Leu | P17900 |
| HOMO SAPIENS | 4 | 4618 | 2ag4 | A | 30 | VAL | p.Ile59Val | P17900 |
| HOMO SAPIENS | 2 | 4618 | 2ag4 | A | 40 | VAL | p.Met69Val | P17900 |
| HOMO SAPIENS | 2 | 4618 | 2ag4 | A | 124 | VAL | p.Val153Ala | P17900 |
| HOMO SAPIENS | 8 | 4618 | 2ag4 | A | 140 | ARG | p.Arg169Pro | P17900 |
| HOMO SAPIENS | 5 | 1082 | 1hk0 | X | 14 | ARG | p.Arg15Cys | P07320 |
| HOMO SAPIENS | 7 | 1082 | 1hk0 | X | 23 | PRO | p.Pro24Thr | P07320 |
| HOMO SAPIENS | 6 | 1082 | 1hk0 | X | 36 | ARG | p.Arg37Ser | P07320 |
| HOMO SAPIENS | 5 | 1082 | 1hk0 | X | 55 | TYR | p.Tyr56Ter | P07320 |
| HOMO SAPIENS | 3 | 1082 | 1hk0 | X | 58 | ARG | p.Arg59His | P07320 |
| HOMO SAPIENS | 7 | 1082 | 1hk0 | X | 88 | HIS | p.His88Gln | P07320 |
| HOMO SAPIENS | 6 | 1082 | 1hk0 | X | 95 | ARG | p.Arg95Ser | P07320 |
| HOMO SAPIENS | 5 | 1082 | 1hk0 | X | 126 | VAL | p.Val126Met | P07320 |
| HOMO SAPIENS | 8 | 1082 | 1hk0 | X | 157 | TRP | p.Trp157Ter | P07320 |
| HOMO SAPIENS | 0 | 22879 | 3j0a | B | 644 | ILE | p.Ile644Phe | O60602 |
| HOMO SAPIENS | 8 | 6 | 2o23 | B | 122 | LEU | p.Leu122Val | Q99714 |
| HOMO SAPIENS | 1 | 6 | 2o23 | B | 130 | ARG | p.Arg130Cys | Q99714 |
| HOMO SAPIENS | 1 | 6 | 2o23 | B | 247 | ASN | p.Asn247Ser | Q99714 |
| HOMO SAPIENS | 0 | 6 | 2o23 | B | 249 | GLU | p.Glu249Gln | Q99714 |
| HOMO SAPIENS | 1 | 12910 | 4hl4 | A | 67 | ARG | p.Arg67Ter | Q96BZ9 |
| HOMO SAPIENS | 8 | 16105 | 1lmj | A | 7 | GLU | p.Glu1073Lys | P35555 |
| HOMO SAPIENS | 8 | 16105 | 1lmj | A | 8 | CYS | p.Cys1074Arg | P35555 |

| | | | | | | | | |
|---|---|---|---|---|---|---|---|---|
| HOMO SAPIENS | 9 | 16105 | 1lmj | A | 51 | CYS | p.Cys1117Tyr | P35555 |
| HOMO SAPIENS | 9 | 16105 | 1lmj | A | 63 | CYS | p.Cys1129Tyr | P35555 |
| HOMO SAPIENS | 9 | 16105 | 1lmj | A | 65 | ASN | p.Asn1131Tyr | P35555 |
| HOMO SAPIENS | 4 | 16105 | 1lmj | A | 71 | ARG | p.Arg1137Pro | P35555 |
| HOMO SAPIENS | 1 | 16105 | 1lmj | A | 82 | PRO | p.Pro1148Ala | P35555 |
| HOMO SAPIENS | 0 | 4795 | 3s5f | A | 106 | LEU | p.Leu106Ser | Q16595 |
| HOMO SAPIENS | 0 | 4795 | 3s5f | A | 130 | GLY | p.Gly130Val | Q16595 |
| HOMO SAPIENS | 0 | 4795 | 3s5f | A | 154 | ILE | p.Ile154Phe | Q16595 |
| HOMO SAPIENS | -1 | 4795 | 3s5f | A | 209 | ASP | p.Asp209Gly | Q16595 |
| HOMO SAPIENS | -1 | 2884 | 4lli | A | 1655 | ASP | p.Asp1655Glu | Q9Y4I1 |
| HOMO SAPIENS | 0 | 1786 | 7pck | C | 56P | TYR | p.Tyr71Ter | P43235 |
| HOMO SAPIENS | 0 | 1786 | 7pck | C | 64P | GLY | p.Gly79Glu | P43235 |
| HOMO SAPIENS | 7 | 38286 | 2l53 | B | 1914 | ARG | p.Arg1914Gly | Q14524 |
| HOMO SAPIENS | 1 | 38286 | 2l53 | B | 1924 | ALA | p.Ala1924Thr | Q14524 |
| HOMO SAPIENS | 0 | 31747 | 3ksy | A | 25 | PRO | p.Pro25Ser | Q07889 |
| HOMO SAPIENS | 3 | 31747 | 3ksy | A | 37 | THR | p.Thr37Ala | Q07889 |
| HOMO SAPIENS | 0 | 31747 | 3ksy | A | 54 | GLN | p.Gln54His | Q07889 |
| HOMO SAPIENS | -1 | 31747 | 3ksy | A | 190 | ASP | p.Asp190Glu | Q07889 |
| HOMO SAPIENS | 2 | 31747 | 3ksy | A | 266 | THR | p.Thr266Lys | Q07889 |
| HOMO SAPIENS | 1 | 31747 | 3ksy | A | 269 | MET | p.Met269Arg | Q07889 |
| HOMO SAPIENS | 1 | 31747 | 3ksy | A | 340 | PRO | p.Pro340Ser | Q07889 |
| HOMO SAPIENS | 1 | 31747 | 3ksy | A | 423 | ASN | p.Asn423Lys | Q07889 |
| HOMO SAPIENS | 1 | 31747 | 3ksy | A | 424 | GLU | p.Glu424Lys | Q07889 |
| HOMO SAPIENS | 1 | 31747 | 3ksy | A | 432 | TRP | p.Trp432Arg | Q07889 |
| HOMO SAPIENS | 1 | 31747 | 3ksy | A | 552 | ARG | p.Arg552Gly | Q07889 |
| HOMO SAPIENS | 1 | 31747 | 3ksy | A | 552 | ARG | p.Arg552Ser | Q07889 |
| HOMO SAPIENS | 3 | 7529 | 6jdw | A | 93 | ILE | p.Ile93Met | P50440 |
| HOMO SAPIENS | 2 | 7529 | 6jdw | A | 110 | GLN | p.Gln110His | P50440 |
| HOMO SAPIENS | 5 | 7529 | 6jdw | A | 149 | TRP | p.Trp149Ter | P50440 |
| HOMO SAPIENS | 0 | 7529 | 6jdw | A | 169 | ARG | p.Arg169Ter | P50440 |
| HOMO SAPIENS | 0 | 7529 | 6jdw | A | 260 | ARG | p.Arg260Ter | P50440 |
| HOMO SAPIENS | 3 | 7529 | 6jdw | A | 344 | ILE | p.Ile344Val | P50440 |
| HOMO SAPIENS | 5 | 7529 | 6jdw | A | 418 | LEU | p.Leu418Ile | P50440 |
| HOMO SAPIENS | 1 | 14788 | 1eak | A | 101 | ARG | p.Arg101His | P08253 |
| HOMO SAPIENS | 0 | 14788 | 1eak | A | 383 | ASP | p.Asp383Glu | P08253 |
| HOMO SAPIENS | 7 | 14788 | 1eak | A | 404 | GLN | p.Glu404Lys | P08253 |
| HOMO SAPIENS | 2 | 14788 | 1eak | A | 447 | ALA | p.Ala447Val | P08253 |
| HOMO SAPIENS | 0 | 1255 | 5xtc | p | 8 | PRO | p.Pro8His | Q9Y6M9 |
| HOMO SAPIENS | 0 | 1255 | 5xtc | p | 146 | PRO | p.Pro146Ser | Q9Y6M9 |
| HOMO SAPIENS | 3 | 6721 | 1a22 | A | 25 | PHE | p.Phe51Tyr | P01241 |
| HOMO SAPIENS | 2 | 6721 | 1a22 | A | 95 | SER | p.Ser121Arg | P01241 |
| HOMO SAPIENS | 2 | 6721 | 1a22 | A | 110 | VAL | p.Val136Ile | P01241 |
| HOMO SAPIENS | 0 | 1380 | 5i6z | A | 605 | LYS | p.Lys605Asn | P31645 |
| HOMO SAPIENS | 0 | 941 | 1qki | D | 48 | ILE | p.Ile48Thr | P11413 |
| HOMO SAPIENS | 3 | 941 | 1qki | D | 68 | VAL | p.Val68Met | P11413 |
| HOMO SAPIENS | 2 | 941 | 1qki | D | 126 | ASN | p.Asn126Asp | P11413 |
| HOMO SAPIENS | 8 | 941 | 1qki | D | 163 | GLY | p.Gly163Ser | P11413 |
| HOMO SAPIENS | 6 | 941 | 1qki | D | 165 | ASN | p.Asn165Asp | P11413 |
| HOMO SAPIENS | 4 | 941 | 1qki | D | 181 | ASP | p.Asp181Val | P11413 |
| HOMO SAPIENS | 4 | 941 | 1qki | D | 182 | ARG | p.Arg182Trp | P11413 |
| HOMO SAPIENS | 5 | 941 | 1qki | D | 188 | SER | p.Ser188Phe | P11413 |
| HOMO SAPIENS | 9 | 941 | 1qki | D | 198 | ARG | p.Arg198Cys | P11413 |
| HOMO SAPIENS | 8 | 941 | 1qki | D | 213 | VAL | p.Val213Leu | P11413 |
| HOMO SAPIENS | 5 | 941 | 1qki | D | 282 | ASP | p.Asp282His | P11413 |

| | | | | | | | | |
|---|---|---|---|---|---|---|---|---|
| HOMO SAPIENS | 7 | 941 | 1qki | D | 285 | ARG | p.Arg285His | P11413 |
| HOMO SAPIENS | 5 | 941 | 1qki | D | 291 | VAL | p.Val291Met | P11413 |
| HOMO SAPIENS | 8 | 941 | 1qki | D | 323 | LEU | p.Leu323Pro | P11413 |
| HOMO SAPIENS | 3 | 941 | 1qki | D | 335 | ALA | p.Ala335Thr | P11413 |
| HOMO SAPIENS | 6 | 941 | 1qki | D | 350 | ASP | p.Asp350His | P11413 |
| HOMO SAPIENS | 5 | 941 | 1qki | D | 393 | ARG | p.Arg393His | P11413 |
| HOMO SAPIENS | 5 | 941 | 1qki | D | 454 | ARG | p.Arg454Cys | P11413 |
| HOMO SAPIENS | 2 | 941 | 1qki | D | 459 | LEU | p.Arg459Leu | P11413 |
| HOMO SAPIENS | 4 | 941 | 1qki | D | 463 | ARG | p.Arg463His | P11413 |
| HOMO SAPIENS | 4 | 2483 | 1tij | A | 68 | LEU | p.Leu94Gln | P01034 |
| HOMO SAPIENS | 2 | 20632 | 3v5b | A | 331 | ARG | p.Arg331Pro | P02545 |
| HOMO SAPIENS | 2 | 20632 | 3v5b | A | 374 | HIS | p.His374Gln | P02545 |
| HOMO SAPIENS | 4 | 13929 | 5xtd | J | 75 | ARG | p.Arg75His | Q16795 |
| HOMO SAPIENS | 3 | 13929 | 5xtd | J | 243 | VAL | p.Val243Ala | Q16795 |
| HOMO SAPIENS | 1 | 13929 | 5xtd | J | 294 | PRO | p.Pro294Leu | Q16795 |
| HOMO SAPIENS | 2 | 13929 | 5xtd | J | 321 | ARG | p.Arg321Pro | Q16795 |
| HOMO SAPIENS | 0 | 9411 | 5thl | A | 41 | ARG | p.Gly41Arg | P54577 |
| HOMO SAPIENS | 0 | 9411 | 5thl | A | 196 | GLU | p.Glu196Lys | P54577 |
| HOMO SAPIENS | 0 | 9411 | 5thl | A | 316 | ALA | p.Ala316Ser | P54577 |
| HOMO SAPIENS | 0 | 9411 | 5thl | A | 339 | ALA | p.Ala339Thr | P54577 |
| HOMO SAPIENS | 4 | 16273 | 2e8n | A | 46 | THR | p.Thr940Ile | P29317 |
| HOMO SAPIENS | 7 | 16273 | 2e8n | A | 54 | GLY | p.Gly948Trp | P29317 |
| HOMO SAPIENS | 6 | 16273 | 2e8n | A | 65 | ALA | p.Ala959Thr | P29317 |
| HOMO SAPIENS | 7 | 30387 | 2k9y | A | 541 | VAL | p.Val541Met | P29317 |
| HOMO SAPIENS | 0 | 6961 | 4nha | A | 843 | ILE | p.Ile853Val | Q9UPY3 |
| HOMO SAPIENS | -1 | 6961 | 4nha | A | 862 | ALA | p.Ala872Thr | Q9UPY3 |
| HOMO SAPIENS | 3 | 15708 | 4pzg | A | 16 | ARG | p.Arg16Ter | Q9Y5X0 |
| HOMO SAPIENS | 4 | 15708 | 4pzg | A | 51 | ARG | p.Arg51Gln | Q9Y5X0 |
| HOMO SAPIENS | 3 | 30166 | 4o00 | A | 44 | ARG | p.Arg14257Ter | Q8WZ42 |
| HOMO SAPIENS | 2 | 30166 | 4o00 | A | 54 | ARG | p.Arg14267His | Q8WZ42 |
| HOMO SAPIENS | 8 | 11158 | 4hca | A | 275 | TRP | p.Trp274Arg | P23771 |
| HOMO SAPIENS | 2 | 4328 | 4e0s | A | 1476 | ARG | p.Arg1476Ter | P01031 |
| HOMO SAPIENS | 0 | 11462 | 1dx5 | I | 455 | ALA | p.Ala473Val | P07204 |
| HOMO SAPIENS | -1 | 11150 | 2fic | A | 45 | LYS | p.Lys35Asn | O00499 |
| HOMO SAPIENS | 6 | 11150 | 2fic | A | 161 | ASP | p.Asp151Asn | O00499 |
| HOMO SAPIENS | 8 | 27382 | 4mqv | B | 17 | ARG | p.Arg17Ter | Q9NZC9 |
| HOMO SAPIENS | -1 | 37991 | 1q0p | A | 231 | GLN | p.Gln256Ter | P00751 |
| HOMO SAPIENS | 1 | 443 | 5xte | U | 215 | LEU | p.Leu215Phe | P08574 |
| HOMO SAPIENS | 8 | 21471 | 5y4e | A | 1461 | ASN | p.Thr692Ala | Q01484 |
| HOMO SAPIENS | -1 | 35443 | 3um3 | B | 129 | ASP | p.Asp129Val | Q9H444 |
| HOMO SAPIENS | -1 | 35443 | 3um3 | B | 161 | GLU | p.Glu161Lys | Q9H444 |
| HOMO SAPIENS | 6 | 28781 | 2n2a | B | 111 | ILE | p.Ile654Val | P04626 |
| HOMO SAPIENS | 7 | 28781 | 2n2a | B | 112 | ILE | p.Ile655Val | P04626 |
| HOMO SAPIENS | -1 | 3346 | 4ztz | C | 61 | GLY | p.Gly61Arg | Q9UHN1 |
| HOMO SAPIENS | 4 | 3346 | 4ztz | C | 103 | GLY | p.Gly103Ser | Q9UHN1 |
| HOMO SAPIENS | 4 | 3346 | 4ztz | C | 133 | HIS | p.His133Tyr | Q9UHN1 |
| HOMO SAPIENS | -1 | 3346 | 4ztz | C | 153 | LEU | p.Leu153Val | Q9UHN1 |
| HOMO SAPIENS | -1 | 3346 | 4ztz | C | 169 | ALA | p.Ala169Thr | Q9UHN1 |
| HOMO SAPIENS | 7 | 3346 | 4ztz | C | 369 | ARG | p.Arg369Gly | Q9UHN1 |
| HOMO SAPIENS | 4 | 3346 | 4ztz | C | 386 | ASP | p.Asp386Glu | Q9UHN1 |
| HOMO SAPIENS | 4 | 3346 | 4ztz | C | 451 | GLY | p.Gly451Glu | Q9UHN1 |
| HOMO SAPIENS | 4 | 2131 | 5fo9 | E | 1549 | LEU | p.Leu1549Met | P01024 |
| HOMO SAPIENS | 2 | 2131 | 5fo9 | E | 1619 | SER | p.Ser1619Arg | P01024 |
| HOMO SAPIENS | 3 | 12662 | 4jhp | C | 75 | ILE | p.Ile75Val | Q92834 |

| | | | | | | | | |
|---|---|---|---|---|---|---|---|---|
| HOMO SAPIENS | 1 | 12662 | 4jhp | C | 173 | GLY | p.Gly173Arg | Q92834 |
| HOMO SAPIENS | 0 | 12662 | 4jhp | C | 184 | GLN | p.Gln184His | Q92834 |
| HOMO SAPIENS | 3 | 12662 | 4jhp | C | 262 | ALA | p.Ala262Gly | Q92834 |
| HOMO SAPIENS | 3 | 12662 | 4jhp | C | 302 | CYS | p.Cys302Tyr | Q92834 |
| HOMO SAPIENS | 5 | 1093 | 2jg9 | A | 186 | GLN | p.Gln208Ter | P02745 |
| HOMO SAPIENS | 0 | 13181 | 4r4l | A | 12 | MET | p.Met13Ile | Q13976 |
| HOMO SAPIENS | 8 | 25939 | 4iyp | A | 106 | ARG | p.Arg106Gln | P78318 |
| HOMO SAPIENS | 1 | 217 | 2xsx | A | 71 | SER | p.Asn71Ser | P13929 |
| HOMO SAPIENS | 0 | 217 | 2xsx | A | 76 | ALA | p.Ala76Pro | P13929 |
| HOMO SAPIENS | 1 | 217 | 2xsx | A | 85 | ALA | p.Val85Ala | P13929 |
| HOMO SAPIENS | 8 | 217 | 2xsx | A | 156 | GLY | p.Gly156Asp | P13929 |
| HOMO SAPIENS | 5 | 217 | 2xsx | A | 332 | VAL | p.Val332Ile | P13929 |
| HOMO SAPIENS | 7 | 217 | 2xsx | A | 374 | GLY | p.Gly374Glu | P13929 |
| HOMO SAPIENS | 1 | 5508 | 3szm | A | 682 | THR | p.Thr682Asn | Q8NEM0 |
| HOMO SAPIENS | 0 | 5508 | 3szm | A | 715 | TRP | p.Trp715Ter | Q8NEM0 |
| HOMO SAPIENS | 1 | 5508 | 3szm | A | 761 | VAL | p.Ala761Val | Q8NEM0 |
| HOMO SAPIENS | 1 | 5508 | 3szm | A | 828 | PRO | p.Pro828Ser | Q8NEM0 |
| HOMO SAPIENS | 4 | 5508 | 3szm | A | 829 | GLU | p.Glu829Asp | Q8NEM0 |
| HOMO SAPIENS | 2 | 16353 | 2q0z | X | 171 | GLY | p.Gly1978Ala | O75643 |
| HOMO SAPIENS | 2 | 16353 | 2q0z | X | 188 | ALA | p.Ala1995Thr | O75643 |
| HOMO SAPIENS | 0 | 4127 | 3swr | A | 860 | GLY | p.Gly860Arg | P26358 |
| HOMO SAPIENS | -1 | 4127 | 3swr | A | 1122 | SER | p.Ser1122Cys | P26358 |
| HOMO SAPIENS | 0 | 4127 | 3swr | A | 1216 | ASP | p.Asp1216Glu | P26358 |
| HOMO SAPIENS | 0 | 6382 | 3o96 | A | 17 | GLU | p.Glu17Lys | P31749 |
| HOMO SAPIENS | 6 | 14314 | 5xtc | Q | 71 | PRO | p.Pro71Leu | O75306 |
| HOMO SAPIENS | 2 | 5886 | 4fao | E | 22 | ARG | p.Arg40His | Q13705 |
| HOMO SAPIENS | 1 | 2534 | 3mp4 | A | 37 | GLU | p.Glu37Ter | P35914 |
| HOMO SAPIENS | 3 | 2534 | 3mp4 | A | 41 | MET | p.Arg41Gln | P35914 |
| HOMO SAPIENS | 1 | 2534 | 3mp4 | A | 70 | VAL | p.Val70Leu | P35914 |
| HOMO SAPIENS | 2 | 2196 | 3ifq | A | 142 | ARG | p.Arg142His | P14923 |
| HOMO SAPIENS | 2 | 2196 | 3ifq | A | 153 | ASP | p.Asp153Glu | P14923 |
| HOMO SAPIENS | 3 | 2196 | 3ifq | A | 176 | ARG | p.Arg176Gln | P14923 |
| HOMO SAPIENS | 3 | 2196 | 3ifq | A | 265 | ARG | p.Arg265His | P14923 |
| HOMO SAPIENS | 2 | 2196 | 3ifq | A | 301 | GLU | p.Glu301Gly | P14923 |
| HOMO SAPIENS | 0 | 2196 | 3ifq | A | 456 | VAL | p.Val456Ile | P14923 |
| HOMO SAPIENS | 3 | 2196 | 3ifq | A | 603 | VAL | p.Val603Leu | P14923 |
| HOMO SAPIENS | 0 | 2196 | 3ifq | A | 648 | VAL | p.Val648Ile | P14923 |
| HOMO SAPIENS | 1 | 8746 | 1cgj | I | 32 | PRO | p.Pro55Ser | P00995 |
| HOMO SAPIENS | 1 | 8746 | 1cgj | I | 44 | ARG | p.Arg67His | P00995 |
| HOMO SAPIENS | 1 | 8746 | 1cgj | I | 44 | ARG | p.Arg67Cys | P00995 |
| HOMO SAPIENS | 3 | 3212 | 1syq | A | 105 | ARG | p.Arg105Ter | P18206 |
| HOMO SAPIENS | 3 | 3212 | 1syq | A | 126 | PHE | p.Phe126Leu | P18206 |
| HOMO SAPIENS | 4 | 10028 | 3eto | A | 1609 | ARG | p.Arg1608His | P46531 |
| HOMO SAPIENS | 1 | 10028 | 3eto | A | 1672 | VAL | p.Val1671Ile | P46531 |
| HOMO SAPIENS | 5 | 10028 | 3eto | A | 1692 | GLN | p.Gln1691His | P46531 |
| HOMO SAPIENS | 3 | 3749 | 5xtc | Z | 70 | GLY | p.Gly70Ter | O43676 |
| HOMO SAPIENS | 2 | 17039 | 3brv | B | 57 | GLU | p.Glu57Lys | Q9Y6K9 |
| HOMO SAPIENS | 8 | 3589 | 2fh3 | A | 433 | VAL | p.Val460Leu | P06396 |
| HOMO SAPIENS | 1 | 3589 | 2fh3 | A | 454 | ARG | p.Arg481Cys | P06396 |
| HOMO SAPIENS | 4 | 3589 | 2fh3 | A | 498 | ARG | p.Arg525His | P06396 |
| HOMO SAPIENS | 2 | 3589 | 2fh3 | A | 520 | TYR | p.Tyr547Cys | P06396 |
| HOMO SAPIENS | 1 | 3589 | 2fh3 | A | 529 | GLY | p.Gly556Arg | P06396 |
| HOMO SAPIENS | 2 | 3589 | 2fh3 | A | 536 | THR | p.Thr563Ser | P06396 |
| HOMO SAPIENS | 2 | 3589 | 2fh3 | A | 579 | VAL | p.Val606Met | P06396 |

| | | | | | | | | |
|---|---|---|---|---|---|---|---|---|
| HOMO SAPIENS | 1 | 3589 | 2fh3 | A | 589 | THR | p.Thr616Met | P06396 |
| HOMO SAPIENS | 5 | 3589 | 2fh3 | A | 668 | THR | p.Thr695Met | P06396 |
| HOMO SAPIENS | 0 | 148 | 1lcl | A | 130 | ILE | p.Ile130Ser | Q05315 |
| HOMO SAPIENS | -1 | 24727 | 4fj3 | P | 237 | ALA | p.Ala237Thr | P04049 |
| HOMO SAPIENS | 0 | 24727 | 4fj3 | P | 257 | SER | p.Ser257Leu | P04049 |
| HOMO SAPIENS | 0 | 24727 | 4fj3 | P | 261 | PRO | p.Pro261Ala | P04049 |
| HOMO SAPIENS | 0 | 2949 | 5xtb | B | 77 | ARG | p.Arg77Gly | O00217 |
| HOMO SAPIENS | 0 | 2949 | 5xtb | B | 79 | PRO | p.Pro79Leu | O00217 |
| HOMO SAPIENS | 0 | 2949 | 5xtb | B | 100 | ALA | p.Ala100Val | O00217 |
| HOMO SAPIENS | 1 | 2949 | 5xtb | B | 115 | LYS | p.Lys115Glu | O00217 |
| HOMO SAPIENS | 0 | 2949 | 5xtb | B | 138 | ARG | p.Arg138His | O00217 |
| HOMO SAPIENS | 1 | 3959 | 5k5s | B | 25 | ARG | p.Arg25Ter | P41180 |
| HOMO SAPIENS | 4 | 3959 | 5k5s | B | 39 | PRO | p.Pro39Ala | P41180 |
| HOMO SAPIENS | 0 | 3959 | 5k5s | B | 47 | LYS | p.Lys47Asn | P41180 |
| HOMO SAPIENS | 2 | 3959 | 5k5s | B | 62 | ARG | p.Arg62Met | P41180 |
| HOMO SAPIENS | 2 | 3959 | 5k5s | B | 66 | ARG | p.Arg66Cys | P41180 |
| HOMO SAPIENS | 4 | 3959 | 5k5s | B | 90 | ASN | p.Asn90Thr | P41180 |
| HOMO SAPIENS | 1 | 3959 | 5k5s | B | 116 | ALA | p.Ala116Thr | P41180 |
| HOMO SAPIENS | 1 | 3959 | 5k5s | B | 118 | ASN | p.Asn118Lys | P41180 |
| HOMO SAPIENS | -1 | 3959 | 5k5s | B | 125 | LEU | p.Leu125Pro | P41180 |
| HOMO SAPIENS | -1 | 3959 | 5k5s | B | 127 | GLU | p.Glu127Ala | P41180 |
| HOMO SAPIENS | -1 | 3959 | 5k5s | B | 128 | PHE | p.Phe128Leu | P41180 |
| HOMO SAPIENS | -1 | 3959 | 5k5s | B | 131 | CYS | p.Cys131Trp | P41180 |
| HOMO SAPIENS | 1 | 3959 | 5k5s | B | 138 | THR | p.Thr138Met | P41180 |
| HOMO SAPIENS | 0 | 3959 | 5k5s | B | 143 | GLY | p.Gly143Glu | P41180 |
| HOMO SAPIENS | 5 | 3959 | 5k5s | B | 151 | THR | p.Thr151Met | P41180 |
| HOMO SAPIENS | 4 | 3959 | 5k5s | B | 180 | PHE | p.Phe180Cys | P41180 |
| HOMO SAPIENS | 0 | 3959 | 5k5s | B | 185 | ARG | p.Arg185Gln | P41180 |
| HOMO SAPIENS | 0 | 3959 | 5k5s | B | 185 | ARG | p.Arg185Ter | P41180 |
| HOMO SAPIENS | 2 | 3959 | 5k5s | B | 191 | GLU | p.Glu191Lys | P41180 |
| HOMO SAPIENS | 2 | 3959 | 5k5s | B | 227 | ARG | p.Arg227Leu | P41180 |
| HOMO SAPIENS | 2 | 3959 | 5k5s | B | 250 | GLU | p.Glu250Lys | P41180 |
| HOMO SAPIENS | 6 | 3959 | 5k5s | B | 297 | GLU | p.Glu297Lys | P41180 |
| HOMO SAPIENS | 1 | 3959 | 5k5s | B | 336 | LYS | p.Lys336Asn | P41180 |
| HOMO SAPIENS | 6 | 3959 | 5k5s | B | 429 | HIS | p.His429Tyr | P41180 |
| HOMO SAPIENS | 0 | 3959 | 5k5s | B | 445 | THR | p.Thr445Ala | P41180 |
| HOMO SAPIENS | 3 | 3959 | 5k5s | B | 509 | GLY | p.Gly509Arg | P41180 |
| HOMO SAPIENS | 1 | 3959 | 5k5s | B | 553 | GLY | p.Gly553Arg | P41180 |
| HOMO SAPIENS | 3 | 3959 | 5k5s | B | 592 | ASN | p.Asn592Ser | P41180 |
| HOMO SAPIENS | -1 | 3959 | 5k5s | B | 604 | GLU | p.Glu604Lys | P41180 |
| HOMO SAPIENS | 4 | 2564 | 5l0c | C | 975 | ARG | p.Arg975Trp | P18206 |
| HOMO SAPIENS | 4 | 2564 | 5l0c | C | 975 | ARG | p.Arg975Leu | P18206 |
| HOMO SAPIENS | 1 | 5695 | 5i8r | A | 294 | ARG | p.Arg294Gln | P17405 |
| HOMO SAPIENS | 0 | 5695 | 5i8r | A | 485 | ALA | p.Ala485Val | P17405 |
| HOMO SAPIENS | 1 | 5695 | 5i8r | A | 506 | GLY | p.Gly506Arg | P17405 |
| HOMO SAPIENS | -1 | 1978 | 3avs | A | 1060 | HIS | p.His1060Leu | O15550 |
| HOMO SAPIENS | 0 | 11173 | 4v0c | B | 362 | LYS | p.Lys362Arg | P51787 |
| HOMO SAPIENS | 3 | 11173 | 4v0c | B | 381 | CYS | p.Cys381Tyr | P51787 |
| HOMO SAPIENS | 7 | 11173 | 4v0c | B | 518 | ARG | p.Arg518Gly | P51787 |
| HOMO SAPIENS | 2 | 11173 | 4v0c | B | 525 | ALA | p.Ala525Thr | P51787 |
| HOMO SAPIENS | 0 | 11173 | 4v0c | B | 530 | GLN | p.Gln530Ter | P51787 |
| HOMO SAPIENS | -1 | 11173 | 4v0c | B | 536 | TYR | p.Tyr536Ter | P51787 |
| HOMO SAPIENS | 2 | 1113 | 5xtd | N | 60 | ARG | p.Arg60Ter | Q9UI09 |
| HOMO SAPIENS | 3 | 6061 | 2mmz | A | 148 | LEU | p.Leu148Ser | Q86SX6 |

| | | | | | | | | |
|---|---|---|---|---|---|---|---|---|
| HOMO SAPIENS | -1 | 26081 | 3eu7 | A | 837 | GLU | p.Glu837Ter | Q86YC2 |
| HOMO SAPIENS | 0 | 26081 | 3eu7 | A | 864 | PRO | p.Pro864Ser | Q86YC2 |
| HOMO SAPIENS | 5 | 26081 | 3eu7 | A | 891 | CYS | p.Cys891Trp | Q86YC2 |
| HOMO SAPIENS | 0 | 26081 | 3eu7 | A | 932 | VAL | p.Val932Met | Q86YC2 |
| HOMO SAPIENS | 5 | 26081 | 3eu7 | A | 961 | LEU | p.Leu961Met | Q86YC2 |
| HOMO SAPIENS | 5 | 26081 | 3eu7 | A | 1038 | TRP | p.Trp1038Ter | Q86YC2 |
| HOMO SAPIENS | -1 | 26081 | 3eu7 | A | 1086 | ARG | p.Arg1086Ter | Q86YC2 |
| HOMO SAPIENS | 0 | 26081 | 3eu7 | A | 1122 | ASP | p.Asp1122Glu | Q86YC2 |
| HOMO SAPIENS | 5 | 26081 | 3eu7 | A | 1158 | HIS | p.His1158Gln | Q86YC2 |
| HOMO SAPIENS | 5 | 26081 | 3eu7 | A | 1183 | TYR | p.Tyr1183Ter | Q86YC2 |
| HOMO SAPIENS | 4 | 199 | 1qo5 | C | 45 | ARG | p.Arg46Trp | P05062 |
| HOMO SAPIENS | 5 | 199 | 1qo5 | C | 147 | TRP | p.Trp148Arg | P05062 |
| HOMO SAPIENS | 0 | 199 | 1qo5 | C | 149 | ALA | p.Ala150Pro | P05062 |
| HOMO SAPIENS | 0 | 199 | 1qo5 | C | 174 | ALA | p.Ala175Asp | P05062 |
| HOMO SAPIENS | 0 | 199 | 1qo5 | C | 334 | ASN | p.Asn335Lys | P05062 |
| HOMO SAPIENS | 0 | 199 | 1qo5 | C | 337 | ALA | p.Ala338Val | P05062 |
| HOMO SAPIENS | 5 | 199 | 1qo5 | C | 342 | TYR | p.Tyr343His | P05062 |
| HOMO SAPIENS | 0 | 6910 | 4zht | B | 59 | ARG | p.Arg59Ter | Q9Y223 |
| HOMO SAPIENS | 4 | 6910 | 4zht | B | 129 | ARG | p.Arg129Ter | Q9Y223 |
| HOMO SAPIENS | 4 | 6910 | 4zht | B | 129 | ARG | p.Arg129Gln | Q9Y223 |
| HOMO SAPIENS | 1 | 6910 | 4zht | B | 162 | ARG | p.Arg162Cys | Q9Y223 |
| HOMO SAPIENS | 0 | 6910 | 4zht | B | 176 | ASP | p.Asp176Val | Q9Y223 |
| HOMO SAPIENS | 0 | 6910 | 4zht | B | 216 | VAL | p.Val216Ala | Q9Y223 |
| HOMO SAPIENS | 0 | 6910 | 4zht | B | 246 | ARG | p.Arg246Trp | Q9Y223 |
| HOMO SAPIENS | 0 | 6910 | 4zht | B | 263 | ARG | p.Arg263Leu | Q9Y223 |
| HOMO SAPIENS | 0 | 6910 | 4zht | B | 266 | ARG | p.Arg266Gln | Q9Y223 |
| HOMO SAPIENS | 0 | 6910 | 4zht | B | 266 | ARG | p.Arg266Trp | Q9Y223 |
| HOMO SAPIENS | 0 | 6910 | 4zht | B | 277 | ARG | p.Arg277Cys | Q9Y223 |
| HOMO SAPIENS | 0 | 6910 | 4zht | B | 378 | ASP | p.Asp378Tyr | Q9Y223 |
| HOMO SAPIENS | 3 | 211 | 5kxi | C | 262 | VAL | p.Val287Leu | P17787 |
| HOMO SAPIENS | 0 | 211 | 5kxi | C | 369 | PHE | p.Phe478Leu | P17787 |
| HOMO SAPIENS | 0 | 2454 | 5iy9 | V | null | PHE | p.Phe99Ser | P19447 |
| HOMO SAPIENS | 0 | 2454 | 5iy9 | V | null | ARG | p.Arg109Ter | P19447 |
| HOMO SAPIENS | 0 | 2454 | 5iy9 | V | null | THR | p.Thr119Pro | P19447 |
| HOMO SAPIENS | 0 | 2454 | 5iy9 | V | 425 | ARG | p.Arg425Ter | P19447 |
| HOMO SAPIENS | 1 | 2454 | 5iy9 | V | 545 | GLN | p.Gln545Ter | P19447 |
| HOMO SAPIENS | -1 | 3805 | 3rbq | A | 57 | LYS | p.Lys57Ter | Q13432 |
| HOMO SAPIENS | 2 | 2649 | 4ido | A | 43 | ASP | p.Asp43Glu | Q8WXF7 |
| HOMO SAPIENS | 2 | 2649 | 4ido | A | 157 | LEU | p.Leu157Trp | Q8WXF7 |
| HOMO SAPIENS | 2 | 2649 | 4ido | A | 193 | PHE | p.Phe193Cys | Q8WXF7 |
| HOMO SAPIENS | 7 | 2649 | 4ido | A | 217 | ARG | p.Arg217Gln | Q8WXF7 |
| HOMO SAPIENS | 4 | 2649 | 4ido | A | 239 | ARG | p.Arg239Cys | Q8WXF7 |
| HOMO SAPIENS | 3 | 2649 | 4ido | A | 258 | HIS | p.His258Arg | Q8WXF7 |
| HOMO SAPIENS | 4 | 2649 | 4ido | A | 259 | SER | p.Ser259Tyr | Q8WXF7 |
| HOMO SAPIENS | 4 | 2649 | 4ido | A | 408 | MET | p.Met408Val | Q8WXF7 |
| HOMO SAPIENS | 2 | 2649 | 4ido | A | 415 | ARG | p.Arg415Trp | Q8WXF7 |
| HOMO SAPIENS | 3 | 2649 | 4ido | A | 416 | ARG | p.Arg416Cys | Q8WXF7 |
| HOMO SAPIENS | 5 | 2203 | 3wff | A | 810 | LEU | p.Ser810Leu | P08235 |
| HOMO SAPIENS | 4 | 12648 | 5ep6 | B | 696 | GLU | p.Glu696Lys | Q9UHD2 |
| HOMO SAPIENS | 0 | 30622 | 2pe4 | A | 90 | GLU | p.Glu90Asp | Q12794 |
| HOMO SAPIENS | 0 | 30622 | 2pe4 | A | 256 | GLY | p.Gly256Arg | Q12794 |
| HOMO SAPIENS | 1 | 30622 | 2pe4 | A | 268 | GLU | p.Glu268Lys | Q12794 |
| HOMO SAPIENS | 0 | 30622 | 2pe4 | A | 392 | THR | p.Thr392Met | Q12794 |
| HOMO SAPIENS | 5 | 5411 | 1lya | A | 93 | SER | p.Ser157Leu | P07339 |

| | | | | | | | | |
|---|---|---|---|---|---|---|---|---|
| HOMO SAPIENS | 3 | 5743 | 2uzp | A | 189 | ASN | p.Asn189Lys | P05129 |
| HOMO SAPIENS | 1 | 12529 | 3kfd | J | 21 | ASN | p.Asn45Ser | P36897 |
| HOMO SAPIENS | 1 | 1063 | 2e1q | B | 149 | ARG | p.Arg149Cys | P47989 |
| HOMO SAPIENS | -1 | 1063 | 2e1q | B | 172 | GLY | p.Gly172Arg | P47989 |
| HOMO SAPIENS | 4 | 1063 | 2e1q | B | 228 | ARG | p.Arg228Ter | P47989 |
| HOMO SAPIENS | 5 | 1063 | 2e1q | B | 395 | LYS | p.Lys395Met | P47989 |
| HOMO SAPIENS | 2 | 1063 | 2e1q | B | 646 | ILE | p.Ile646Val | P47989 |
| HOMO SAPIENS | 4 | 1063 | 2e1q | B | 703 | ILE | p.Ile703Val | P47989 |
| HOMO SAPIENS | 8 | 2525 | 1kqn | F | 18 | ASN | p.Asn18Ser | Q9HAN9 |
| HOMO SAPIENS | 0 | 2525 | 1kqn | F | 151 | VAL | p.Val151Phe | Q9HAN9 |
| HOMO SAPIENS | 0 | 2525 | 1kqn | F | 153 | LEU | p.Leu153Val | Q9HAN9 |
| HOMO SAPIENS | 0 | 2525 | 1kqn | F | 169 | TRP | p.Trp169Ter | Q9HAN9 |
| HOMO SAPIENS | 0 | 2525 | 1kqn | F | 207 | ARG | p.Arg207Trp | Q9HAN9 |
| HOMO SAPIENS | 0 | 2525 | 1kqn | F | 273 | ASN | p.Asn273Asp | Q9HAN9 |
| HOMO SAPIENS | 0 | 12299 | 5duq | A | 432 | VAL | p.Val454Glu | P05155 |
| HOMO SAPIENS | 0 | 12299 | 5duq | A | 436 | ALA | p.Ala458Thr | P05155 |
| HOMO SAPIENS | 0 | 12299 | 5duq | A | 443 | ALA | p.Ala465Val | P05155 |
| HOMO SAPIENS | 0 | 12299 | 5duq | A | 444 | ARG | p.Arg466His | P05155 |
| HOMO SAPIENS | 0 | 12299 | 5duq | A | 458 | MET | p.Val480Met | P05155 |
| HOMO SAPIENS | 0 | 327 | 1uhp | A | 36 | ASP | p.Asp274Asn | Q9UPQ7 |
| HOMO SAPIENS | 0 | 64 | 2mo5 | A | 54 | ALA | p.Ala55Thr | P12104 |
| HOMO SAPIENS | 2 | 35969 | 1jrh | I | 60 | CYS | p.Cys77Tyr | P15260 |
| HOMO SAPIENS | 4 | 35969 | 1jrh | I | 70 | ILE | p.Ile87Thr | P15260 |
| HOMO SAPIENS | 2 | 1068 | 1mu7 | A | 304 | ARG | p.Arg304Gln | Q9NUW8 |
| HOMO SAPIENS | 8 | 1068 | 1mu7 | A | 493 | HIS | p.His493Arg | Q9NUW8 |
| HOMO SAPIENS | 0 | 32360 | 2kqp | A | 31 | ARG | p.Arg55Cys | P01308 |
| HOMO SAPIENS | 0 | 32360 | 2kqp | A | 65 | ARG | p.Arg89His | P01308 |
| HOMO SAPIENS | 0 | 32360 | 2kqp | A | 65 | ARG | p.Arg89Cys | P01308 |
| HOMO SAPIENS | 6 | 10805 | 3nks | A | 12 | ILE | p.Ile12Thr | P50336 |
| HOMO SAPIENS | 0 | 10805 | 3nks | A | 20 | HIS | p.His20Pro | P50336 |
| HOMO SAPIENS | 0 | 10805 | 3nks | A | 59 | ARG | p.Arg59Trp | P50336 |
| HOMO SAPIENS | 0 | 10805 | 3nks | A | 168 | ARG | p.Arg168Cys | P50336 |
| HOMO SAPIENS | 0 | 10805 | 3nks | A | 168 | ARG | p.Arg168His | P50336 |
| HOMO SAPIENS | 2 | 10805 | 3nks | A | 232 | GLY | p.Gly232Arg | P50336 |
| HOMO SAPIENS | 3 | 10805 | 3nks | A | 256 | PRO | p.Pro256Arg | P50336 |
| HOMO SAPIENS | 0 | 10805 | 3nks | A | 304 | ARG | p.Arg304His | P50336 |
| HOMO SAPIENS | 0 | 10805 | 3nks | A | 349 | ASP | p.Asp349Ala | P50336 |
| HOMO SAPIENS | 9 | 7008 | 3u2l | A | 125 | GLN | p.Gln125Ter | P55789 |
| HOMO SAPIENS | 4 | 7008 | 3u2l | A | 172 | HIS | p.His172Asn | P55789 |
| HOMO SAPIENS | 4 | 7008 | 3u2l | A | 194 | ARG | p.Arg194His | P55789 |
| HOMO SAPIENS | 3 | 7008 | 3u2l | A | 198 | GLY | p.Gly198Ser | P55789 |
| HOMO SAPIENS | 0 | 20821 | 3e0g | A | 65 | HIS | p.His116Tyr | P42702 |
| HOMO SAPIENS | 3 | 1151 | 5l9v | A | 278 | ASP | p.Asp278Asn | Q9GZT9 |
| HOMO SAPIENS | 4 | 1151 | 5l9v | A | 317 | CYS | p.Pro317Arg | Q9GZT9 |
| HOMO SAPIENS | 5 | 1151 | 5l9v | A | 371 | ARG | p.Arg371His | Q9GZT9 |
| HOMO SAPIENS | 2 | 3935 | 1gwb | A | 36 | ARG | p.Arg36Gly | P07359 |
| HOMO SAPIENS | 1 | 3935 | 1gwb | A | 73 | LEU | p.Leu73Phe | P07359 |
| HOMO SAPIENS | 1 | 3935 | 1gwb | A | 172 | ALA | p.Ala172Val | P07359 |
| HOMO SAPIENS | 0 | 3935 | 1gwb | A | 249 | GLY | p.Gly249Val | P07359 |
| HOMO SAPIENS | 0 | 3935 | 1gwb | A | 255 | MET | p.Met255Val | P07359 |
| HOMO SAPIENS | 2 | 14541 | 5cuz | A | 138 | GLU | p.Glu138Lys | Q9H3L0 |
| HOMO SAPIENS | 2 | 14541 | 5cuz | A | 143 | SER | p.Ser143Ile | Q9H3L0 |
| HOMO SAPIENS | 4 | 14541 | 5cuz | A | 182 | THR | p.Thr182Asn | Q9H3L0 |
| HOMO SAPIENS | 6 | 14541 | 5cuz | A | 206 | ASN | p.Asn206Ile | Q9H3L0 |

| | | | | | | | | |
|---|---|---|---|---|---|---|---|---|
| HOMO SAPIENS | 0 | 14541 | 5cuz | A | 249 | TYR | p.Tyr249Cys | Q9H3L0 |
| HOMO SAPIENS | 4 | 14541 | 5cuz | A | 250 | ARG | p.Arg250Ter | Q9H3L0 |
| HOMO SAPIENS | 2 | 14541 | 5cuz | A | 259 | LEU | p.Leu259Pro | Q9H3L0 |
| HOMO SAPIENS | 0 | 4206 | 4fao | D | 30 | PRO | p.Pro30Ser | P37023 |
| HOMO SAPIENS | 1 | 4206 | 4fao | D | 50 | TRP | p.Trp50Cys | P37023 |
| HOMO SAPIENS | 0 | 16553 | 3v30 | A | 251 | GLN | p.Gln251Glu | O14593 |
| HOMO SAPIENS | 0 | 5012 | 5hbf | A | 442 | ALA | p.Ala442Gly | Q13231 |
| HOMO SAPIENS | 5 | 4844 | 5tdc | A | 136 | HIS | p.His136Arg | Q8IWV7 |
| HOMO SAPIENS | 0 | 15435 | 4cgz | A | 643 | ARG | p.Arg643His | P54132 |
| HOMO SAPIENS | 1 | 15435 | 4cgz | A | 672 | GLN | p.Gln672Arg | P54132 |
| HOMO SAPIENS | 1 | 15435 | 4cgz | A | 700 | GLN | p.Gln700Ter | P54132 |
| HOMO SAPIENS | 0 | 15435 | 4cgz | A | 707 | PRO | p.Pro707Ser | P54132 |
| HOMO SAPIENS | 0 | 15435 | 4cgz | A | 755 | LYS | p.Lys755Glu | P54132 |
| HOMO SAPIENS | 0 | 15435 | 4cgz | A | 881 | TRP | p.Trp881Ter | P54132 |
| HOMO SAPIENS | 2 | 15435 | 4cgz | A | 899 | ARG | p.Arg899Ter | P54132 |
| HOMO SAPIENS | 0 | 15435 | 4cgz | A | 1036 | CYS | p.Cys1036Phe | P54132 |
| HOMO SAPIENS | 0 | 15435 | 4cgz | A | 1055 | CYS | p.Cys1055Ser | P54132 |
| HOMO SAPIENS | 0 | 15435 | 4cgz | A | 1139 | ARG | p.Arg1139Ter | P54132 |
| HOMO SAPIENS | -1 | 15435 | 4cgz | A | 1198 | VAL | p.Val1198Met | P54132 |
| HOMO SAPIENS | -1 | 15435 | 4cgz | A | 1205 | VAL | p.Val1205Ile | P54132 |
| HOMO SAPIENS | 0 | 15435 | 4cgz | A | 1251 | GLU | p.Glu1251Gln | P54132 |
| HOMO SAPIENS | 0 | 11185 | 1kb4 | A | 30 | ARG | p.Arg30Ter | P11473 |
| HOMO SAPIENS | 5 | 11185 | 1kb4 | A | 33 | GLY | p.Gly33Asp | P11473 |
| HOMO SAPIENS | 5 | 11185 | 1kb4 | A | 46 | GLY | p.Gly46Asp | P11473 |
| HOMO SAPIENS | 6 | 11185 | 1kb4 | A | 50 | ARG | p.Arg50Gln | P11473 |
| HOMO SAPIENS | 9 | 11185 | 1kb4 | A | 73 | ARG | p.Arg73Gln | P11473 |
| HOMO SAPIENS | 9 | 11185 | 1kb4 | A | 80 | ARG | p.Arg80Gln | P11473 |
| HOMO SAPIENS | 0 | 5382 | 3fl7 | A | 41 | LEU | p.Leu41Val | P29317 |
| HOMO SAPIENS | 2 | 5382 | 3fl7 | A | 175 | ARG | p.Arg175Cys | P29317 |
| HOMO SAPIENS | 1 | 5382 | 3fl7 | A | 195 | ARG | p.Arg195Cys | P29317 |
| HOMO SAPIENS | 0 | 5382 | 3fl7 | A | 232 | ASP | p.Asp232Gly | P29317 |
| HOMO SAPIENS | 0 | 5382 | 3fl7 | A | 277 | SER | p.Ser277Leu | P29317 |
| HOMO SAPIENS | 0 | 5382 | 3fl7 | A | 315 | ARG | p.Arg315Gln | P29317 |
| HOMO SAPIENS | 7 | 5382 | 3fl7 | A | 350 | PRO | p.Pro350Thr | P29317 |
| HOMO SAPIENS | 0 | 5382 | 3fl7 | A | 391 | GLY | p.Gly391Arg | P29317 |
| HOMO SAPIENS | 0 | 5382 | 3fl7 | A | 418 | VAL | p.Val418Ile | P29317 |
| HOMO SAPIENS | 0 | 5382 | 3fl7 | A | 511 | THR | p.Thr511Met | P29317 |
| HOMO SAPIENS | 2 | 29168 | 4p22 | B | 144 | VAL | p.Val144Ile | P22314 |
| HOMO SAPIENS | 5 | 4348 | 1bf5 | A | 241 | ARG | p.Arg241Gln | P42224 |
| HOMO SAPIENS | 2 | 4348 | 1bf5 | A | 266 | VAL | p.Val266Ile | P42224 |
| HOMO SAPIENS | 4 | 4348 | 1bf5 | A | 320 | GLU | p.Glu320Gln | P42224 |
| HOMO SAPIENS | 1 | 4348 | 1bf5 | A | 372 | ASN | p.Asn372Lys | P42224 |
| HOMO SAPIENS | 1 | 4348 | 1bf5 | A | 447 | ASP | p.Asp447Glu | P42224 |
| HOMO SAPIENS | 2 | 4348 | 1bf5 | A | 463 | GLN | p.Gln463His | P42224 |
| HOMO SAPIENS | 4 | 4348 | 1bf5 | A | 600 | LEU | p.Leu600Pro | P42224 |
| HOMO SAPIENS | 0 | 4348 | 1bf5 | A | 706 | LEU | p.Leu706Ser | P42224 |
| HOMO SAPIENS | 4 | 4991 | 4bej | B | 141 | ASN | p.Asn141Ser | O00429 |
| HOMO SAPIENS | 0 | 4991 | 4bej | B | 395 | ALA | p.Ala395Asp | O00429 |
| HOMO SAPIENS | 2 | 11792 | 1fvo | A | 40 | ARG | p.Arg40Cys | P00480 |
| HOMO SAPIENS | 2 | 11792 | 1fvo | A | 40 | ARG | p.Arg40His | P00480 |
| HOMO SAPIENS | 0 | 11792 | 1fvo | A | 45 | LEU | p.Leu45Pro | P00480 |
| HOMO SAPIENS | 1 | 11792 | 1fvo | A | 47 | ASN | p.Asn47Ile | P00480 |
| HOMO SAPIENS | 0 | 11792 | 1fvo | A | 50 | GLY | p.Gly50Arg | P00480 |
| HOMO SAPIENS | 5 | 11792 | 1fvo | A | 87 | GLU | p.Glu87Lys | P00480 |

| | | | | | | | | |
|---|---|---|---|---|---|---|---|---|
| HOMO SAPIENS | 3 | 11792 | 1fvo | A | 111 | LEU | p.Leu111Pro | P00480 |
| HOMO SAPIENS | 1 | 11792 | 1fvo | A | 125 | THR | p.Thr125Met | P00480 |
| HOMO SAPIENS | 3 | 11792 | 1fvo | A | 129 | ARG | p.Arg129His | P00480 |
| HOMO SAPIENS | 9 | 11792 | 1fvo | A | 141 | ARG | p.Arg141Pro | P00480 |
| HOMO SAPIENS | 0 | 11792 | 1fvo | A | 206 | MET | p.Met206Arg | P00480 |
| HOMO SAPIENS | 0 | 11792 | 1fvo | A | 207 | SER | p.Ser207Arg | P00480 |
| HOMO SAPIENS | 0 | 11792 | 1fvo | A | 216 | GLN | p.Gln216Glu | P00480 |
| HOMO SAPIENS | 1 | 11792 | 1fvo | A | 225 | PRO | p.Pro225Leu | P00480 |
| HOMO SAPIENS | 0 | 11792 | 1fvo | A | 270 | ARG | p.Gln270Arg | P00480 |
| HOMO SAPIENS | 0 | 11792 | 1fvo | A | 270 | ARG | p.Gln270Ter | P00480 |
| HOMO SAPIENS | 2 | 11792 | 1fvo | A | 277 | ARG | p.Arg277Trp | P00480 |
| HOMO SAPIENS | 0 | 3721 | 4r3z | B | 2 | ASP | p.Asp2Gly | P54136 |
| HOMO SAPIENS | 0 | 3721 | 4r3z | B | 456 | SER | p.Ser456Leu | P54136 |
| HOMO SAPIENS | 5 | 3721 | 4r3z | B | 512 | ARG | p.Arg512Gln | P54136 |
| HOMO SAPIENS | 3 | 5749 | 3hl2 | A | 239 | ALA | p.Ala239Thr | Q9HD40 |
| HOMO SAPIENS | 1 | 5749 | 3hl2 | A | 334 | TYR | p.Tyr334Cys | Q9HD40 |
| HOMO SAPIENS | 1 | 5749 | 3hl2 | A | 452 | LYS | p.Lys452Asn | Q9HD40 |
| HOMO SAPIENS | -1 | 5749 | 3hl2 | A | 465 | ARG | p.Arg465Leu | Q9HD40 |
| HOMO SAPIENS | -1 | 5749 | 3hl2 | A | 489 | ASP | p.Asp489Gly | Q9HD40 |
| HOMO SAPIENS | 5 | 1721 | 3k15 | A | 1652 | MET | p.Met1652Ile | P38398 |
| HOMO SAPIENS | 5 | 1721 | 3k15 | A | 1652 | MET | p.Met1652Lys | P38398 |
| HOMO SAPIENS | 5 | 1721 | 3k15 | A | 1662 | PHE | p.Phe1662Cys | P38398 |
| HOMO SAPIENS | 5 | 1721 | 3k15 | A | 1664 | LEU | p.Leu1664Pro | P38398 |
| HOMO SAPIENS | 0 | 1721 | 3k15 | A | 1665 | VAL | p.Val1665Met | P38398 |
| HOMO SAPIENS | 5 | 1721 | 3k15 | A | 1690 | LYS | p.Lys1690Gln | P38398 |
| HOMO SAPIENS | 5 | 1721 | 3k15 | A | 1692 | ASP | p.Asp1692Asn | P38398 |
| HOMO SAPIENS | 5 | 1721 | 3k15 | A | 1699 | ARG | p.Arg1699Gln | P38398 |
| HOMO SAPIENS | 5 | 1721 | 3k15 | A | 1699 | ARG | p.Arg1699Trp | P38398 |
| HOMO SAPIENS | 3 | 1721 | 3k15 | A | 1706 | GLY | p.Gly1706Glu | P38398 |
| HOMO SAPIENS | 5 | 1721 | 3k15 | A | 1720 | THR | p.Thr1720Ala | P38398 |
| HOMO SAPIENS | 5 | 1721 | 3k15 | A | 1751 | ARG | p.Arg1751Pro | P38398 |
| HOMO SAPIENS | 4 | 1721 | 3k15 | A | 1775 | MET | p.Met1775Arg | P38398 |
| HOMO SAPIENS | 2 | 1721 | 3k15 | A | 1788 | GLY | p.Gly1788Val | P38398 |
| HOMO SAPIENS | 5 | 1721 | 3k15 | A | 1794 | GLU | p.Glu1794Asp | P38398 |
| HOMO SAPIENS | 5 | 1721 | 3k15 | A | 1804 | VAL | p.Val1804Asp | P38398 |
| HOMO SAPIENS | 3 | 1721 | 3k15 | A | 1815 | TRP | p.Trp1815Ter | P38398 |
| HOMO SAPIENS | 1 | 1721 | 3k15 | A | 1819 | ASN | p.Asn1819Ser | P38398 |
| HOMO SAPIENS | 5 | 1721 | 3k15 | A | 1856 | PRO | p.Pro1856Ser | P38398 |
| HOMO SAPIENS | 5 | 1721 | 3k15 | A | 1858 | ILE | p.Ile1858Leu | P38398 |
| HOMO SAPIENS | 5 | 1721 | 3k15 | A | 1858 | ILE | p.Ile1858Thr | P38398 |
| HOMO SAPIENS | 5 | 1721 | 3k15 | A | 1859 | PRO | p.Pro1859Arg | P38398 |
| HOMO SAPIENS | 2 | 22086 | 3kq4 | B | 1297 | PRO | p.Pro1297Leu | O60494 |
| HOMO SAPIENS | 1 | 12135 | 2k9j | B | 724 | ARG | p.Arg750Ter | P05106 |
| HOMO SAPIENS | -1 | 24027 | 2qtl | A | 175 | LEU | p.Ser202Leu | Q9UBK8 |
| HOMO SAPIENS | 0 | 24027 | 2qtl | A | 257 | SER | p.Ser284Thr | Q9UBK8 |
| HOMO SAPIENS | 0 | 24027 | 2qtl | A | 333 | LEU | p.Leu360Val | Q9UBK8 |
| HOMO SAPIENS | 0 | 24027 | 2qtl | A | 350 | LYS | p.Lys377Arg | Q9UBK8 |
| HOMO SAPIENS | 0 | 24027 | 2qtl | A | 415 | ARG | p.Arg442Cys | Q9UBK8 |
| HOMO SAPIENS | 2 | 24027 | 2qtl | A | 450 | PRO | p.Pro477Arg | Q9UBK8 |
| HOMO SAPIENS | 4 | 24027 | 2qtl | A | 487 | GLY | p.Gly514Arg | Q9UBK8 |
| HOMO SAPIENS | 1 | 24027 | 2qtl | A | 595 | HIS | p.His622Tyr | Q9UBK8 |
| HOMO SAPIENS | 0 | 5758 | 3di2 | B | 46 | THR | p.Thr66Ile | P16871 |
| HOMO SAPIENS | 1 | 5758 | 3di2 | B | 118 | VAL | p.Ile138Val | P16871 |
| HOMO SAPIENS | 4 | 848 | 5oa3 | Y | 79 | LEU | p.Leu79Val | P62847 |

| | | | | | | | | |
|---|---|---|---|---|---|---|---|---|
| HOMO SAPIENS | 2 | 848 | 5oa3 | Y | 124 | ASN | p.Asn124Ser | P62847 |
| HOMO SAPIENS | 0 | 81 | 3s5o | A | 36 | ALA | p.Ala36Glu | Q86XE5 |
| HOMO SAPIENS | 0 | 81 | 3s5o | A | 39 | TYR | p.Tyr39Ter | Q86XE5 |
| HOMO SAPIENS | 0 | 81 | 3s5o | A | 45 | PRO | p.Pro45Leu | Q86XE5 |
| HOMO SAPIENS | 0 | 81 | 3s5o | A | 70 | ARG | p.Arg70Ter | Q86XE5 |
| HOMO SAPIENS | 0 | 81 | 3s5o | A | 70 | ARG | p.Arg70Pro | Q86XE5 |
| HOMO SAPIENS | 1 | 81 | 3s5o | A | 97 | ARG | p.Arg97Cys | Q86XE5 |
| HOMO SAPIENS | 4 | 81 | 3s5o | A | 113 | GLU | p.Glu113Lys | Q86XE5 |
| HOMO SAPIENS | 1 | 81 | 3s5o | A | 190 | PRO | p.Pro190Leu | Q86XE5 |
| HOMO SAPIENS | 0 | 81 | 3s5o | A | 245 | VAL | p.Val245Phe | Q86XE5 |
| HOMO SAPIENS | 0 | 81 | 3s5o | A | 257 | CYS | p.Cys257Gly | Q86XE5 |
| HOMO SAPIENS | 0 | 81 | 3s5o | A | 287 | GLY | p.Gly287Val | Q86XE5 |
| HOMO SAPIENS | 7 | 81 | 3s5o | A | 303 | ARG | p.Arg303Cys | Q86XE5 |
| HOMO SAPIENS | 0 | 81 | 3s5o | A | 325 | GLY | p.Gly325Ser | Q86XE5 |
| HOMO SAPIENS | -1 | 622 | 5era | A | 12 | GLY | p.Gly12Arg | P29033 |
| HOMO SAPIENS | -1 | 622 | 5era | A | 12 | GLY | p.Gly12Asp | P29033 |
| HOMO SAPIENS | -1 | 622 | 5era | A | 17 | SER | p.Ser17Phe | P29033 |
| HOMO SAPIENS | 5 | 622 | 5era | A | 24 | TRP | p.Trp24Ter | P29033 |
| HOMO SAPIENS | 0 | 622 | 5era | A | 32 | ARG | p.Arg32Cys | P29033 |
| HOMO SAPIENS | 5 | 622 | 5era | A | 34 | MET | p.Met34Thr | P29033 |
| HOMO SAPIENS | 5 | 622 | 5era | A | 36 | LEU | p.Leu36Pro | P29033 |
| HOMO SAPIENS | 5 | 622 | 5era | A | 37 | VAL | p.Val37Ile | P29033 |
| HOMO SAPIENS | 5 | 622 | 5era | A | 40 | ALA | p.Ala40Glu | P29033 |
| HOMO SAPIENS | 5 | 622 | 5era | A | 44 | TRP | p.Trp44Cys | P29033 |
| HOMO SAPIENS | 5 | 622 | 5era | A | 45 | GLY | p.Gly45Glu | P29033 |
| HOMO SAPIENS | 5 | 622 | 5era | A | 47 | GLU | p.Glu47Ter | P29033 |
| HOMO SAPIENS | 5 | 622 | 5era | A | 50 | ASP | p.Asp50Asn | P29033 |
| HOMO SAPIENS | 5 | 622 | 5era | A | 54 | ASN | p.Asn54Lys | P29033 |
| HOMO SAPIENS | 5 | 622 | 5era | A | 57 | GLN | p.Gln57Ter | P29033 |
| HOMO SAPIENS | 5 | 622 | 5era | A | 59 | GLY | p.Gly59Ala | P29033 |
| HOMO SAPIENS | 5 | 622 | 5era | A | 59 | GLY | p.Gly59Ser | P29033 |
| HOMO SAPIENS | 5 | 622 | 5era | A | 66 | ASP | p.Asp66His | P29033 |
| HOMO SAPIENS | 5 | 622 | 5era | A | 68 | TYR | p.Tyr68Cys | P29033 |
| HOMO SAPIENS | 5 | 622 | 5era | A | 70 | PRO | p.Pro70Ala | P29033 |
| HOMO SAPIENS | 5 | 622 | 5era | A | 73 | HIS | p.His73Arg | P29033 |
| HOMO SAPIENS | 5 | 622 | 5era | A | 75 | ARG | p.Arg75Trp | P29033 |
| HOMO SAPIENS | 5 | 622 | 5era | A | 75 | ARG | p.Arg75Gln | P29033 |
| HOMO SAPIENS | 2 | 622 | 5era | A | 77 | TRP | p.Trp77Arg | P29033 |
| HOMO SAPIENS | 2 | 622 | 5era | A | 77 | TRP | p.Trp77Ter | P29033 |
| HOMO SAPIENS | 0 | 622 | 5era | A | 80 | GLN | p.Gln80Ter | P29033 |
| HOMO SAPIENS | 0 | 622 | 5era | A | 80 | GLN | p.Gln80Pro | P29033 |
| HOMO SAPIENS | 5 | 622 | 5era | A | 81 | LEU | p.Leu81Val | P29033 |
| HOMO SAPIENS | 5 | 622 | 5era | A | 82 | ILE | p.Ile82Met | P29033 |
| HOMO SAPIENS | 5 | 622 | 5era | A | 83 | PHE | p.Phe83Leu | P29033 |
| HOMO SAPIENS | 5 | 622 | 5era | A | 84 | VAL | p.Val84Leu | P29033 |
| HOMO SAPIENS | 5 | 622 | 5era | A | 90 | LEU | p.Leu90Pro | P29033 |
| HOMO SAPIENS | 5 | 622 | 5era | A | 95 | VAL | p.Val95Met | P29033 |
| HOMO SAPIENS | -1 | 622 | 5era | A | 100 | HIS | p.His100Tyr | P29033 |
| HOMO SAPIENS | -1 | 622 | 5era | A | 113 | SER | p.Ser113Arg | P29033 |
| HOMO SAPIENS | -1 | 622 | 5era | A | 119 | GLU | p.Glu119Lys | P29033 |
| HOMO SAPIENS | -1 | 622 | 5era | A | 122 | LYS | p.Lys122Ile | P29033 |
| HOMO SAPIENS | -1 | 622 | 5era | A | 123 | THR | p.Thr123Asn | P29033 |
| HOMO SAPIENS | -1 | 622 | 5era | A | 124 | GLN | p.Gln124Ter | P29033 |
| HOMO SAPIENS | 5 | 622 | 5era | A | 139 | SER | p.Ser139Asn | P29033 |

| | | | | | | | | |
|---|---|---|---|---|---|---|---|---|
| HOMO SAPIENS | 5 | 622 | 5era | A | 143 | ARG | p.Arg143Trp | P29033 |
| HOMO SAPIENS | 5 | 622 | 5era | A | 143 | ARG | p.Arg143Gln | P29033 |
| HOMO SAPIENS | 0 | 622 | 5era | A | 147 | GLU | p.Glu147Lys | P29033 |
| HOMO SAPIENS | 7 | 622 | 5era | A | 155 | TYR | p.Tyr155Ter | P29033 |
| HOMO SAPIENS | 2 | 622 | 5era | A | 159 | ASP | p.Asp159Val | P29033 |
| HOMO SAPIENS | 5 | 622 | 5era | A | 163 | MET | p.Met163Val | P29033 |
| HOMO SAPIENS | 5 | 622 | 5era | A | 165 | ARG | p.Arg165Trp | P29033 |
| HOMO SAPIENS | 5 | 622 | 5era | A | 167 | VAL | p.Val167Met | P29033 |
| HOMO SAPIENS | 5 | 622 | 5era | A | 169 | CYS | p.Cys169Tyr | P29033 |
| HOMO SAPIENS | 5 | 622 | 5era | A | 179 | ASP | p.Asp179Asn | P29033 |
| HOMO SAPIENS | 0 | 622 | 5era | A | 199 | SER | p.Ser199Phe | P29033 |
| HOMO SAPIENS | 0 | 622 | 5era | A | 202 | CYS | p.Cys202Phe | P29033 |
| HOMO SAPIENS | 5 | 622 | 5era | A | 203 | ILE | p.Ile203Thr | P29033 |
| HOMO SAPIENS | 5 | 622 | 5era | A | 206 | ASN | p.Asn206Ser | P29033 |
| HOMO SAPIENS | 0 | 8103 | 5dot | A | 163 | GLY | p.Gly163Ter | P31327 |
| HOMO SAPIENS | 0 | 8103 | 5dot | A | 337 | HIS | p.His337Arg | P31327 |
| HOMO SAPIENS | 3 | 8103 | 5dot | A | 344 | THR | p.Thr344Ala | P31327 |
| HOMO SAPIENS | 1 | 8103 | 5dot | A | 405 | THR | p.Thr405Asn | P31327 |
| HOMO SAPIENS | 5 | 8103 | 5dot | A | 544 | THR | p.Thr544Met | P31327 |
| HOMO SAPIENS | 5 | 8103 | 5dot | A | 716 | ASN | p.Asn716Lys | P31327 |
| HOMO SAPIENS | -1 | 8103 | 5dot | A | 780 | ARG | p.Arg780His | P31327 |
| HOMO SAPIENS | 0 | 8103 | 5dot | A | 982 | GLY | p.Gly982Asp | P31327 |
| HOMO SAPIENS | 3 | 8103 | 5dot | A | 1119 | ALA | p.Ala1119Ser | P31327 |
| HOMO SAPIENS | 1 | 8103 | 5dot | A | 1376 | GLY | p.Gly1376Ser | P31327 |
| HOMO SAPIENS | 4 | 6833 | 1l8l | B | 27 | ARG | p.Arg27Ser | P78330 |
| HOMO SAPIENS | 3 | 6833 | 1l8l | B | 32 | ASP | p.Asp32Gly | P78330 |
| HOMO SAPIENS | 3 | 6833 | 1l8l | B | 32 | ASP | p.Asp32Asn | P78330 |
| HOMO SAPIENS | 1 | 6833 | 1l8l | B | 49 | ARG | p.Arg49Trp | P78330 |
| HOMO SAPIENS | 0 | 6833 | 1l8l | B | 52 | MET | p.Met52Thr | P78330 |
| HOMO SAPIENS | 0 | 6833 | 1l8l | B | 83 | GLN | p.Gln83His | P78330 |
| HOMO SAPIENS | 6 | 810 | 2iw2 | A | 185 | ARG | p.Arg184Gln | P12955 |
| HOMO SAPIENS | 0 | 810 | 2iw2 | A | 213 | ALA | p.Ala212Pro | P12955 |
| HOMO SAPIENS | 1 | 810 | 2iw2 | A | 266 | ARG | p.Arg265Ter | P12955 |
| HOMO SAPIENS | 9 | 810 | 2iw2 | A | 277 | ASP | p.Asp276Asn | P12955 |
| HOMO SAPIENS | 0 | 810 | 2iw2 | A | 279 | GLY | p.Gly278Asp | P12955 |
| HOMO SAPIENS | 9 | 810 | 2iw2 | A | 378 | HIS | p.His377Gln | P12955 |
| HOMO SAPIENS | 0 | 810 | 2iw2 | A | 389 | ARG | p.Arg388His | P12955 |
| HOMO SAPIENS | 0 | 810 | 2iw2 | A | 436 | PHE | p.Leu435Phe | P12955 |
| HOMO SAPIENS | 0 | 810 | 2iw2 | A | 449 | GLY | p.Gly448Arg | P12955 |
| HOMO SAPIENS | -1 | 1530 | 5fwj | A | 388 | ALA | p.Ala388Pro | P41229 |
| HOMO SAPIENS | 4 | 1530 | 5fwj | A | 451 | SER | p.Ser451Arg | P41229 |
| HOMO SAPIENS | 3 | 1530 | 5fwj | A | 731 | LEU | p.Leu731Phe | P41229 |
| HOMO SAPIENS | 3 | 578 | 5net | B | 123 | ALA | p.Ala143Thr | P18564 |
| HOMO SAPIENS | 6 | 578 | 5net | B | 176 | PRO | p.Pro196Thr | P18564 |
| HOMO SAPIENS | 3 | 578 | 5net | B | 280 | GLU | p.Glu300Lys | P18564 |
| HOMO SAPIENS | -1 | 5247 | 4j15 | A | 274 | ALA | p.Ala274Val | P14868 |
| HOMO SAPIENS | -1 | 5247 | 4j15 | A | 280 | HIS | p.His280Leu | P14868 |
| HOMO SAPIENS | 2 | 5247 | 4j15 | A | 460 | ARG | p.Arg460His | P14868 |
| HOMO SAPIENS | 5 | 5247 | 4j15 | A | 494 | ARG | p.Arg494Cys | P14868 |
| HOMO SAPIENS | 7 | 2566 | 1b55 | A | 28 | ARG | p.Arg28His | Q06187 |
| HOMO SAPIENS | 5 | 2566 | 1b55 | A | 33 | THR | p.Thr33Pro | Q06187 |
| HOMO SAPIENS | 0 | 2566 | 1b55 | A | 113 | VAL | p.Val113Asp | Q06187 |
| HOMO SAPIENS | 0 | 356 | 5a2s | A | 786 | ALA | p.Ala786Thr | P56524 |
| HOMO SAPIENS | 5 | 2960 | 4cak | A | 553 | ARG | p.Arg584Ter | P08514 |

| | | | | | | | | |
|---|---|---|---|---|---|---|---|---|
| HOMO SAPIENS | 5 | 2960 | 4cak | A | 565 | ILE | p.Ile596Thr | P08514 |
| HOMO SAPIENS | -1 | 2960 | 4cak | A | 841 | LEU | p.Leu872Met | P08514 |
| HOMO SAPIENS | -1 | 2960 | 4cak | A | 843 | ILE | p.Ile874Ser | P08514 |
| HOMO SAPIENS | 0 | 5054 | 4k8r | B | 216 | PHE | p.Phe216Leu | Q8NBP7 |
| HOMO SAPIENS | 1 | 5054 | 4k8r | B | 417 | HIS | p.His417Gln | Q8NBP7 |
| HOMO SAPIENS | 1 | 5054 | 4k8r | B | 425 | ASN | p.Asn425Ser | Q8NBP7 |
| HOMO SAPIENS | 1 | 5054 | 4k8r | B | 443 | ALA | p.Ala443Thr | Q8NBP7 |
| HOMO SAPIENS | 0 | 27283 | 3n5n | X | 71 | HIS | p.His82Arg | Q9UIF7 |
| HOMO SAPIENS | 1 | 27283 | 3n5n | X | 83 | ARG | p.Arg94Ter | Q9UIF7 |
| HOMO SAPIENS | 8 | 27283 | 3n5n | X | 89 | TRP | p.Trp100Ter | Q9UIF7 |
| HOMO SAPIENS | 8 | 27283 | 3n5n | X | 90 | TYR | p.Tyr101Ter | Q9UIF7 |
| HOMO SAPIENS | 7 | 27283 | 3n5n | X | 95 | ARG | p.Arg106Trp | Q9UIF7 |
| HOMO SAPIENS | 7 | 27283 | 3n5n | X | 117 | TRP | p.Trp128Arg | Q9UIF7 |
| HOMO SAPIENS | 0 | 27283 | 3n5n | X | 137 | GLY | p.Gly148Arg | Q9UIF7 |
| HOMO SAPIENS | 5 | 27283 | 3n5n | X | 142 | TRP | p.Trp153Ter | Q9UIF7 |
| HOMO SAPIENS | 0 | 27283 | 3n5n | X | 165 | TYR | p.Tyr176Cys | Q9UIF7 |
| HOMO SAPIENS | 0 | 27283 | 3n5n | X | 168 | ARG | p.Arg179His | Q9UIF7 |
| HOMO SAPIENS | 0 | 27283 | 3n5n | X | 168 | ARG | p.Arg179Cys | Q9UIF7 |
| HOMO SAPIENS | 0 | 27283 | 3n5n | X | 196 | GLN | p.Gln207Ter | Q9UIF7 |
| HOMO SAPIENS | 8 | 27283 | 3n5n | X | 201 | VAL | p.Val212Met | Q9UIF7 |
| HOMO SAPIENS | 9 | 27283 | 3n5n | X | 227 | ARG | p.Arg238Trp | Q9UIF7 |
| HOMO SAPIENS | 9 | 27283 | 3n5n | X | 231 | ARG | p.Arg242Cys | Q9UIF7 |
| HOMO SAPIENS | 9 | 27283 | 3n5n | X | 231 | ARG | p.Arg242His | Q9UIF7 |
| HOMO SAPIENS | 3 | 27283 | 3n5n | X | 232 | VAL | p.Val243Phe | Q9UIF7 |
| HOMO SAPIENS | 3 | 27283 | 3n5n | X | 246 | GLN | p.Gln257Ter | Q9UIF7 |
| HOMO SAPIENS | 9 | 27283 | 3n5n | X | 272 | GLY | p.Gly283Glu | Q9UIF7 |
| HOMO SAPIENS | 9 | 27283 | 3n5n | X | 281 | PRO | p.Pro292Leu | Q9UIF7 |
| HOMO SAPIENS | -1 | 27283 | 3n5n | X | 314 | ASP | p.Asp325Glu | Q9UIF7 |
| HOMO SAPIENS | 0 | 27283 | 3n5n | X | 324 | GLN | p.Gln335His | Q9UIF7 |
| HOMO SAPIENS | 0 | 27283 | 3n5n | X | 324 | GLN | p.Gln335Arg | Q9UIF7 |
| HOMO SAPIENS | 0 | 27283 | 3n5n | X | 324 | GLN | p.Gln335Ter | Q9UIF7 |
| HOMO SAPIENS | -1 | 20852 | 4p60 | B | 243 | ARG | p.Arg243Lys | O75746 |
| HOMO SAPIENS | -1 | 1875 | 5dd2 | G | 152 | VAL | p.Val179Leu | P06396 |
| HOMO SAPIENS | 9 | 2126 | 3n1p | B | 46 | PRO | p.Pro46Leu | Q14623 |
| HOMO SAPIENS | 8 | 2126 | 3n1p | B | 76 | GLU | p.Glu76Ala | Q14623 |
| HOMO SAPIENS | 9 | 2126 | 3n1p | B | 77 | ARG | p.Arg77Ser | Q14623 |
| HOMO SAPIENS | 6 | 2126 | 3n1p | B | 95 | GLU | p.Glu95Lys | Q14623 |
| HOMO SAPIENS | 9 | 2126 | 3n1p | B | 100 | ASP | p.Asp100Glu | Q14623 |
| HOMO SAPIENS | 9 | 2126 | 3n1p | B | 131 | GLU | p.Glu131Lys | Q14623 |
| HOMO SAPIENS | 0 | 2126 | 3n1p | B | 190 | VAL | p.Val190Ala | Q14623 |
| HOMO SAPIENS | 5 | 40410 | 3ioz | B | 70 | GLN | p.Gln70Leu | P20963 |
| HOMO SAPIENS | 1 | 5754 | 5lkz | A | 1108 | THR | p.Tyr1191Ter | Q09472 |
| HOMO SAPIENS | 3 | 8159 | 1tiu | A | 3 | GLU | p.Lys12679Glu | Q8WZ42 |
| HOMO SAPIENS | 4 | 3275 | 4ldq | A | 210 | ARG | p.Arg210Gln | Q9BV20 |
| HOMO SAPIENS | 0 | 8143 | 4miv | B | 66 | SER | p.Ser66Trp | P51688 |
| HOMO SAPIENS | 1 | 8143 | 4miv | B | 74 | ARG | p.Arg74Cys | P51688 |
| HOMO SAPIENS | 0 | 8143 | 4miv | B | 128 | PRO | p.Pro128Leu | P51688 |
| HOMO SAPIENS | 1 | 8143 | 4miv | B | 150 | ARG | p.Arg150Gln | P51688 |
| HOMO SAPIENS | 0 | 8143 | 4miv | B | 191 | GLY | p.Gly191Arg | P51688 |
| HOMO SAPIENS | 0 | 8143 | 4miv | B | 206 | ARG | p.Arg206Pro | P51688 |
| HOMO SAPIENS | 0 | 8143 | 4miv | B | 225 | PHE | p.Phe225Leu | P51688 |
| HOMO SAPIENS | 0 | 8143 | 4miv | B | 233 | ARG | p.Arg233Ter | P51688 |
| HOMO SAPIENS | 0 | 8143 | 4miv | B | 234 | ALA | p.Ala234Gly | P51688 |
| HOMO SAPIENS | 0 | 8143 | 4miv | B | 245 | ARG | p.Arg245His | P51688 |

| | | | | | | | | |
|---|---|---|---|---|---|---|---|---|
| HOMO SAPIENS | 0 | 8143 | 4miv | B | 293 | PRO | p.Pro293Ser | P51688 |
| HOMO SAPIENS | 1 | 8143 | 4miv | B | 298 | SER | p.Ser298Pro | P51688 |
| HOMO SAPIENS | 0 | 8143 | 4miv | B | 361 | VAL | p.Val361Ile | P51688 |
| HOMO SAPIENS | 0 | 8143 | 4miv | B | 369 | GLU | p.Glu369Lys | P51688 |
| HOMO SAPIENS | 0 | 8143 | 4miv | B | 372 | MET | p.Met372Ile | P51688 |
| HOMO SAPIENS | 1 | 8143 | 4miv | B | 380 | GLN | p.Gln380Arg | P51688 |
| HOMO SAPIENS | 0 | 8143 | 4miv | B | 387 | VAL | p.Val387Met | P51688 |
| HOMO SAPIENS | 0 | 8143 | 4miv | B | 394 | MET | p.Met394Ile | P51688 |
| HOMO SAPIENS | 0 | 8143 | 4miv | B | 433 | ARG | p.Arg433Gln | P51688 |
| HOMO SAPIENS | 1 | 8143 | 4miv | B | 446 | HIS | p.His446Arg | P51688 |
| HOMO SAPIENS | 5 | 8143 | 4miv | B | 447 | GLU | p.Glu447Lys | P51688 |
| HOMO SAPIENS | 2 | 8143 | 4miv | B | 456 | ARG | p.Arg456His | P51688 |
| HOMO SAPIENS | 1 | 21880 | 2oo9 | B | 857 | LEU | p.Leu857Phe | P22681 |
| HOMO SAPIENS | 0 | 9467 | 1s1d | A | 154 | LEU | p.Leu224Pro | Q8WVQ1 |
| HOMO SAPIENS | 0 | 9467 | 1s1d | A | 156 | VAL | p.Val226Met | Q8WVQ1 |
| HOMO SAPIENS | 0 | 9467 | 1s1d | A | 229 | PRO | p.Pro299Leu | Q8WVQ1 |
| HOMO SAPIENS | 4 | 9467 | 1s1d | A | 230 | ARG | p.Arg300Cys | Q8WVQ1 |
| HOMO SAPIENS | 4 | 9467 | 1s1d | A | 230 | ARG | p.Arg300His | Q8WVQ1 |
| HOMO SAPIENS | 0 | 9467 | 1s1d | A | 253 | ALA | p.Ala323Thr | Q8WVQ1 |
| HOMO SAPIENS | 2 | 9467 | 1s1d | A | 305 | MET | p.Met375Leu | Q8WVQ1 |
| HOMO SAPIENS | -1 | 2361 | 5llm | A | 5 | ASN | p.Asn5Lys | P43003 |
| HOMO SAPIENS | 0 | 2361 | 5llm | A | 71 | TYR | p.Tyr71Cys | P43003 |
| HOMO SAPIENS | -1 | 2361 | 5llm | A | 211 | LYS | p.Glu219Asp | Q15758 |
| HOMO SAPIENS | 8 | 2361 | 5llm | A | 270 | PRO | p.Pro290Arg | P43003 |
| HOMO SAPIENS | 1 | 2361 | 5llm | A | 321 | THR | p.Ala329Thr | Q15758 |
| HOMO SAPIENS | 1 | 2361 | 5llm | A | 365 | ARG | p.Arg385His | P43003 |
| HOMO SAPIENS | 1 | 2361 | 5llm | A | 479 | ARG | p.Arg499Gln | P43003 |
| HOMO SAPIENS | 3 | 5412 | 4obz | D | 165 | PHE | p.Phe229Ile | P07339 |
| HOMO SAPIENS | 2 | 5412 | 4obz | D | 187 | ASP | p.Asp251Asn | P07339 |
| HOMO SAPIENS | 0 | 5412 | 4obz | D | 218 | GLY | p.Gly282Arg | P07339 |
| HOMO SAPIENS | 0 | 5412 | 4obz | D | 245 | ARG | p.Arg309His | P07339 |
| HOMO SAPIENS | 2 | 5412 | 4obz | D | 319 | TRP | p.Trp383Cys | P07339 |
| HOMO SAPIENS | 0 | 3544 | 5h64 | A | 277 | MET | p.Met277Ile | P42345 |
| HOMO SAPIENS | 0 | 14230 | 2kbx | A | 145 | ARG | p.Arg145Ser | Q13418 |
| HOMO SAPIENS | 2 | 34970 | 4xhz | A | 834 | ARG | p.Arg855Trp | Q96QU1 |
| HOMO SAPIENS | 4 | 34970 | 4xhz | A | 854 | SER | p.Ser875Ter | Q96QU1 |
| HOMO SAPIENS | 1 | 34970 | 4xhz | A | 908 | ARG | p.Arg929Gln | Q96QU1 |
| HOMO SAPIENS | 1 | 34970 | 4xhz | A | 941 | ARG | p.Arg962Cys | Q96QU1 |
| HOMO SAPIENS | 0 | 34970 | 4xhz | A | 970 | ARG | p.Arg991Ter | Q96QU1 |
| HOMO SAPIENS | 0 | 34970 | 4xhz | A | 976 | GLU | p.Glu997Gly | Q96QU1 |
| HOMO SAPIENS | 0 | 25734 | 4asc | A | 345 | SER | p.Asn345Ser | Q2TBA0 |
| HOMO SAPIENS | 0 | 25734 | 4asc | A | 514 | VAL | p.Val514Met | Q2TBA0 |
| HOMO SAPIENS | 6 | 25734 | 4asc | A | 528 | GLU | p.Glu528Lys | Q2TBA0 |
| HOMO SAPIENS | 0 | 25734 | 4asc | A | 580 | LEU | p.Leu580Ile | Q2TBA0 |
| HOMO SAPIENS | 8 | 2356 | 3kse | D | 22 | LYS | p.Lys22Ter | P01040 |
| HOMO SAPIENS | 1 | 2356 | 3kse | D | 58 | ARG | p.Arg58Ter | P01040 |
| HOMO SAPIENS | 2 | 6000 | 1jqd | A | 60 | GLY | p.Gly60Asp | P50135 |
| HOMO SAPIENS | 9 | 6000 | 1jqd | A | 208 | LEU | p.Leu208Pro | P50135 |
| HOMO SAPIENS | 0 | 14741 | 5eh1 | A | 58 | THR | p.Thr58Arg | P38484 |
| HOMO SAPIENS | 0 | 14741 | 5eh1 | A | 168 | THR | p.Thr168Asn | P38484 |
| HOMO SAPIENS | 1 | 4814 | 4kxk | D | 427 | ARG | p.Arg427Ter | P50542 |
| HOMO SAPIENS | 0 | 188 | 2xt3 | A | 49 | ASP | p.Asp52Asn | Q2M1P5 |
| HOMO SAPIENS | 3 | 188 | 2xt3 | A | 142 | TYR | p.Tyr145Ser | Q2M1P5 |
| HOMO SAPIENS | 9 | 8681 | 5tby | D | 57 | ALA | p.Ala57Asp | P08590 |

| | | | | | | | | |
|---|---|---|---|---|---|---|---|---|
| HOMO SAPIENS | 3 | 8681 | 5tby | D | 62 | ASP | p.Asp62Asn | P08590 |
| HOMO SAPIENS | 0 | 8681 | 5tby | D | 65 | PRO | p.Pro65Arg | P08590 |
| HOMO SAPIENS | 7 | 8681 | 5tby | D | 79 | VAL | p.Val79Ile | P08590 |
| HOMO SAPIENS | 0 | 8681 | 5tby | D | 143 | GLU | p.Glu143Lys | P08590 |
| HOMO SAPIENS | 0 | 8681 | 5tby | D | 149 | MET | p.Met149Val | P08590 |
| HOMO SAPIENS | 3 | 8681 | 5tby | D | 154 | ARG | p.Arg154His | P08590 |
| HOMO SAPIENS | 6 | 30984 | 4xwh | A | 140 | TYR | p.Tyr140Cys | P54802 |
| HOMO SAPIENS | 4 | 30984 | 4xwh | A | 141 | SER | p.Ser141Thr | P54802 |
| HOMO SAPIENS | 2 | 30984 | 4xwh | A | 234 | ARG | p.Arg234Cys | P54802 |
| HOMO SAPIENS | 2 | 30984 | 4xwh | A | 336 | GLU | p.Glu336Ter | P54802 |
| HOMO SAPIENS | 6 | 30984 | 4xwh | A | 482 | ARG | p.Arg482Trp | P54802 |
| HOMO SAPIENS | 2 | 30984 | 4xwh | A | 521 | PRO | p.Pro521Leu | P54802 |
| HOMO SAPIENS | 0 | 30984 | 4xwh | A | 565 | ARG | p.Arg565Gln | P54802 |
| HOMO SAPIENS | 0 | 30984 | 4xwh | A | 565 | ARG | p.Arg565Trp | P54802 |
| HOMO SAPIENS | 6 | 30984 | 4xwh | A | 626 | ARG | p.Arg626Ter | P54802 |
| HOMO SAPIENS | 6 | 30984 | 4xwh | A | 643 | ARG | p.Arg643His | P54802 |
| HOMO SAPIENS | 6 | 30984 | 4xwh | A | 643 | ARG | p.Arg643Cys | P54802 |
| HOMO SAPIENS | 2 | 30984 | 4xwh | A | 674 | ARG | p.Arg674His | P54802 |
| HOMO SAPIENS | 2 | 30984 | 4xwh | A | 737 | ARG | p.Arg737Gly | P54802 |
| HOMO SAPIENS | 5 | 10545 | 1oqe | N | 6 | PRO | p.Pro21Arg | Q96RJ3 |
| HOMO SAPIENS | 1 | 16995 | 5d3a | B | 12 | MET | p.Met947Arg | Q7Z4S6 |
| HOMO SAPIENS | 1 | 16995 | 5d3a | B | 12 | MET | p.Met947Val | Q7Z4S6 |
| HOMO SAPIENS | 2 | 16995 | 5d3a | B | 19 | ARG | p.Arg954Gln | Q7Z4S6 |
| HOMO SAPIENS | 2 | 16995 | 5d3a | B | 19 | ARG | p.Arg954Trp | Q7Z4S6 |
| HOMO SAPIENS | 0 | 16995 | 5d3a | B | 75 | ILE | p.Ile1010Thr | Q7Z4S6 |
| HOMO SAPIENS | 0 | 17091 | 2yin | B | 1324 | GLN | p.Gln1324Ter | Q92608 |
| HOMO SAPIENS | 5 | 27606 | 1lm7 | B | 2366 | ARG | p.Arg2366Cys | P15924 |
| HOMO SAPIENS | 2 | 15731 | 1bi7 | B | 24 | ARG | p.Arg24Pro | P42771 |
| HOMO SAPIENS | 0 | 15731 | 1bi7 | B | 28 | VAL | p.Val28Gly | P42771 |
| HOMO SAPIENS | 0 | 15731 | 1bi7 | B | 49 | ILE | p.Ile49Thr | P42771 |
| HOMO SAPIENS | 0 | 15731 | 1bi7 | B | 53 | MET | p.Met53Ile | P42771 |
| HOMO SAPIENS | 2 | 15731 | 1bi7 | B | 56 | SER | p.Ser56Ile | P42771 |
| HOMO SAPIENS | 1 | 15731 | 1bi7 | B | 57 | ALA | p.Ala57Val | P42771 |
| HOMO SAPIENS | 4 | 15731 | 1bi7 | B | 59 | VAL | p.Val59Gly | P42771 |
| HOMO SAPIENS | 9 | 15731 | 1bi7 | B | 89 | GLY | p.Gly89Asp | P42771 |
| HOMO SAPIENS | 0 | 15731 | 1bi7 | B | 101 | GLY | p.Gly101Trp | P42771 |
| HOMO SAPIENS | 0 | 15731 | 1bi7 | B | 114 | PRO | p.Pro114Ser | P42771 |
| HOMO SAPIENS | 9 | 15731 | 1bi7 | B | 123 | HIS | p.His123Gln | P42771 |
| HOMO SAPIENS | 8 | 15731 | 1bi7 | B | 126 | VAL | p.Val126Asp | P42771 |
| HOMO SAPIENS | 7 | 15731 | 1bi7 | B | 127 | ALA | p.Ala127Ser | P42771 |
| HOMO SAPIENS | -1 | 15731 | 1bi7 | B | 148 | ALA | p.Ala148Thr | P42771 |
| HOMO SAPIENS | -1 | 19858 | 3jbk | M | 866 | ALA | p.Ala934Val | P18206 |
| HOMO SAPIENS | 0 | 19484 | 3ee6 | B | 62 | SER | p.Ser62Leu | O14773 |
| HOMO SAPIENS | 1 | 19484 | 3ee6 | B | 66 | GLN | p.Gln66Ter | O14773 |
| HOMO SAPIENS | 0 | 19484 | 3ee6 | B | 100 | GLN | p.Gln100Arg | O14773 |
| HOMO SAPIENS | 0 | 19484 | 3ee6 | B | 104 | LEU | p.Leu104Ter | O14773 |
| HOMO SAPIENS | 0 | 19484 | 3ee6 | B | 127 | ARG | p.Arg127Gln | O14773 |
| HOMO SAPIENS | 0 | 19484 | 3ee6 | B | 127 | ARG | p.Arg127Ter | O14773 |
| HOMO SAPIENS | -1 | 19484 | 3ee6 | B | 185 | ARG | p.Arg185Cys | O14773 |
| HOMO SAPIENS | 0 | 19484 | 3ee6 | B | 206 | ARG | p.Arg206Cys | O14773 |
| HOMO SAPIENS | 0 | 19484 | 3ee6 | B | 208 | ARG | p.Arg208Ter | O14773 |
| HOMO SAPIENS | 0 | 19484 | 3ee6 | B | 214 | GLN | p.Gln214Ter | O14773 |
| HOMO SAPIENS | 1 | 19484 | 3ee6 | B | 276 | ASP | p.Asp276Val | O14773 |
| HOMO SAPIENS | 0 | 19484 | 3ee6 | B | 284 | GLY | p.Gly284Val | O14773 |

| | | | | | | | | |
|---|---|---|---|---|---|---|---|---|
| HOMO SAPIENS | 0 | 19484 | 3ee6 | B | 286 | ASN | p.Asn286Ser | O14773 |
| HOMO SAPIENS | 0 | 19484 | 3ee6 | B | 339 | ARG | p.Arg339Gly | O14773 |
| HOMO SAPIENS | 2 | 19484 | 3ee6 | B | 345 | MET | p.Met345Leu | O14773 |
| HOMO SAPIENS | 0 | 19484 | 3ee6 | B | 365 | CYS | p.Cys365Arg | O14773 |
| HOMO SAPIENS | 0 | 19484 | 3ee6 | B | 365 | CYS | p.Cys365Tyr | O14773 |
| HOMO SAPIENS | 0 | 19484 | 3ee6 | B | 373 | GLN | p.Gln373Glu | O14773 |
| HOMO SAPIENS | 6 | 19484 | 3ee6 | B | 418 | ARG | p.Arg418Gln | O14773 |
| HOMO SAPIENS | 7 | 19484 | 3ee6 | B | 422 | GLN | p.Gln422His | O14773 |
| HOMO SAPIENS | 0 | 19484 | 3ee6 | B | 447 | ARG | p.Arg447His | O14773 |
| HOMO SAPIENS | 0 | 19484 | 3ee6 | B | 466 | VAL | p.Val466Met | O14773 |
| HOMO SAPIENS | 0 | 4654 | 5t5w | A | 47 | LYS | p.Lys47Glu | Q08334 |
| HOMO SAPIENS | 0 | 4654 | 5t5w | A | 141 | GLU | p.Glu141Ter | Q08334 |
| HOMO SAPIENS | -1 | 18815 | 4wj7 | A | 53 | VAL | p.Val53Ile | Q9BSQ5 |
| HOMO SAPIENS | 0 | 18815 | 4wj7 | A | 120 | VAL | p.Val120Ile | Q9BSQ5 |
| HOMO SAPIENS | 0 | 18815 | 4wj7 | A | 198 | LEU | p.Leu198Arg | Q9BSQ5 |
| HOMO SAPIENS | 0 | 4602 | 1mjd | A | 47 | SER | p.Ser47Arg | O43602 |
| HOMO SAPIENS | 6 | 4602 | 1mjd | A | 59 | ARG | p.Arg59Leu | O43602 |
| HOMO SAPIENS | 7 | 4602 | 1mjd | A | 62 | ASP | p.Asp62Asn | O43602 |
| HOMO SAPIENS | 7 | 4602 | 1mjd | A | 71 | ALA | p.Ala71Ser | O43602 |
| HOMO SAPIENS | 7 | 4602 | 1mjd | A | 78 | ARG | p.Arg78Leu | O43602 |
| HOMO SAPIENS | 7 | 4602 | 1mjd | A | 89 | ARG | p.Arg89Gly | O43602 |
| HOMO SAPIENS | 8 | 4602 | 1mjd | A | 104 | ILE | p.Ile104Phe | O43602 |
| HOMO SAPIENS | 7 | 4602 | 1mjd | A | 125 | TYR | p.Tyr125His | O43602 |
| HOMO SAPIENS | 5 | 34467 | 1n0w | B | 1529 | GLY | p.Gly1529Arg | P51587 |
| HOMO SAPIENS | 8 | 5738 | 1u31 | A | 130 | ASN | p.Asn1009Lys | Q13423 |
| HOMO SAPIENS | 0 | 5760 | 2i6s | A | 298 | GLU | p.Glu318Asp | P06681 |
| HOMO SAPIENS | 0 | 5760 | 2i6s | A | 573 | ARG | p.Arg593Gln | P06681 |
| HOMO SAPIENS | 0 | 5760 | 2i6s | A | 592 | SER | p.Ser612Asn | P06681 |
| HOMO SAPIENS | 0 | 5760 | 2i6s | A | 621 | VAL | p.Val641Ala | P06681 |
| HOMO SAPIENS | 1 | 10977 | 2fby | A | 114 | VAL | p.Val114Ile | Q14191 |
| HOMO SAPIENS | 0 | 10977 | 2fby | A | 135 | LYS | p.Lys135Glu | Q14191 |
| HOMO SAPIENS | -1 | 1175 | 1qrk | A | 34 | VAL | p.Val35Leu | P00488 |
| HOMO SAPIENS | 4 | 1175 | 1qrk | A | 564 | PRO | p.Pro565Leu | P00488 |
| HOMO SAPIENS | 0 | 1175 | 1qrk | A | 650 | VAL | p.Val651Ile | P00488 |
| HOMO SAPIENS | 0 | 1175 | 1qrk | A | 651 | GLU | p.Gln652Glu | P00488 |
| HOMO SAPIENS | 3 | 909 | 4h0w | A | 375 | GLU | p.Glu394Lys | P02787 |
| HOMO SAPIENS | 0 | 909 | 4h0w | A | 458 | ALA | p.Ala477Pro | P02787 |
| HOMO SAPIENS | 6 | 909 | 4h0w | A | 515 | TYR | p.Tyr534Ter | P02787 |
| HOMO SAPIENS | 7 | 909 | 4h0w | A | 570 | PRO | p.Pro589Ser | P02787 |
| HOMO SAPIENS | 3 | 909 | 4h0w | A | 652 | GLY | p.Gly671Glu | P02787 |
| HOMO SAPIENS | 5 | 1964 | 2car | A | 151 | TRP | p.Trp151Ter | Q9BY32 |
| HOMO SAPIENS | 7 | 1964 | 2car | A | 178 | ARG | p.Arg178Cys | Q9BY32 |
| HOMO SAPIENS | 0 | 144 | 2n0a | J | 30 | ALA | p.Ala30Pro | P37840 |
| HOMO SAPIENS | 1 | 12288 | 4q58 | D | 1225 | ARG | p.Arg1225His | P16144 |
| HOMO SAPIENS | 2 | 12288 | 4q58 | D | 1281 | ARG | p.Arg1281Trp | P16144 |
| HOMO SAPIENS | 2 | 12288 | 4q58 | D | 1287 | ASN | p.Asn1287Lys | P16144 |
| HOMO SAPIENS | 0 | 5085 | 3s94 | A | 360 | ARG | p.Arg360His | O75581 |
| HOMO SAPIENS | 0 | 5085 | 3s94 | A | 433 | ASN | p.Asn433Ser | O75581 |
| HOMO SAPIENS | 2 | 5085 | 3s94 | A | 611 | ARG | p.Arg611Cys | O75581 |
| HOMO SAPIENS | 0 | 15452 | 4dur | A | 19 | LYS | p.Lys38Glu | P00747 |
| HOMO SAPIENS | 2 | 15452 | 4dur | A | 216 | ARG | p.Arg235His | P00747 |
| HOMO SAPIENS | 7 | 8406 | 4wo1 | D | 28 | VAL | p.Val55Leu | O43914 |
| HOMO SAPIENS | 2 | 895 | 5xte | K | 183 | ARG | p.Arg183Trp | P22695 |
| HOMO SAPIENS | 2 | 4191 | 4wnd | A | 120 | ARG | p.Arg127Ter | P81274 |

| | | | | | | | | |
|---|---|---|---|---|---|---|---|---|
| HOMO SAPIENS | 2 | 4191 | 4wnd | A | 120 | ARG | p.Arg127Gln | P81274 |
| HOMO SAPIENS | 1 | 4191 | 4wnd | A | 349 | GLY | p.Gly356Arg | P81274 |
| HOMO SAPIENS | 0 | 41734 | 2kbi | A | 1777 | VAL | p.Val1777Met | Q14524 |
| HOMO SAPIENS | 1 | 41734 | 2kbi | A | 1784 | GLU | p.Glu1784Lys | Q14524 |
| HOMO SAPIENS | 5 | 41734 | 2kbi | A | 1795 | TYR | p.Tyr1795Cys | Q14524 |
| HOMO SAPIENS | 5 | 41734 | 2kbi | A | 1795 | TYR | p.Tyr1795His | Q14524 |
| HOMO SAPIENS | 7 | 41734 | 2kbi | A | 1826 | ARG | p.Arg1826Cys | Q14524 |
| HOMO SAPIENS | 3 | 2119 | 1pl0 | A | 426 | LYS | p.Lys426Arg | P31939 |
| HOMO SAPIENS | 5 | 3475 | 5u62 | C | 50 | LEU | p.Leu50Ser | Q15910 |
| HOMO SAPIENS | 3 | 379 | 4uez | A | 200 | TYR | p.Tyr200Ter | P15085 |
| HOMO SAPIENS | 0 | 8195 | 5xaw | B | 67 | ARG | p.Arg67Trp | Q8IWT1 |
| HOMO SAPIENS | 0 | 8195 | 5xaw | B | 94 | THR | p.Thr94Met | Q8IWT1 |
| HOMO SAPIENS | 1 | 5711 | 1fo3 | A | 265 | ARG | p.Arg265His | Q9UKM7 |
| HOMO SAPIENS | 4 | 5711 | 1fo3 | A | 334 | ARG | p.Arg334Cys | Q9UKM7 |
| HOMO SAPIENS | 1 | 5711 | 1fo3 | A | 478 | LYS | p.Lys478Arg | Q9UKM7 |
| HOMO SAPIENS | 1 | 5711 | 1fo3 | A | 682 | ASP | p.Asp682Glu | Q9UKM7 |
| HOMO SAPIENS | 0 | 170 | 2n3y | A | 41 | GLY | p.Gly42Ser | P99999 |
| HOMO SAPIENS | 0 | 4122 | 3w81 | A | 33 | HIS | p.His33Gln | P35475 |
| HOMO SAPIENS | 0 | 4122 | 3w81 | A | 64 | TYR | p.Tyr64Ter | P35475 |
| HOMO SAPIENS | 0 | 4122 | 3w81 | A | 70 | GLN | p.Gln70Ter | P35475 |
| HOMO SAPIENS | 0 | 4122 | 3w81 | A | 75 | ALA | p.Ala75Thr | P35475 |
| HOMO SAPIENS | 0 | 4122 | 3w81 | A | 79 | ALA | p.Ala79Val | P35475 |
| HOMO SAPIENS | 0 | 4122 | 3w81 | A | 79 | ALA | p.Ala79Thr | P35475 |
| HOMO SAPIENS | 0 | 4122 | 3w81 | A | 89 | ARG | p.Arg89Gln | P35475 |
| HOMO SAPIENS | 0 | 4122 | 3w81 | A | 99 | THR | p.Thr99Ile | P35475 |
| HOMO SAPIENS | 0 | 4122 | 3w81 | A | 105 | ARG | p.Arg105Gln | P35475 |
| HOMO SAPIENS | 0 | 4122 | 3w81 | A | 238 | LEU | p.Leu238Gln | P35475 |
| HOMO SAPIENS | 0 | 4122 | 3w81 | A | 300 | ALA | p.Ala300Thr | P35475 |
| HOMO SAPIENS | 0 | 4122 | 3w81 | A | 322 | VAL | p.Val322Glu | P35475 |
| HOMO SAPIENS | 0 | 4122 | 3w81 | A | 327 | ALA | p.Ala327Pro | P35475 |
| HOMO SAPIENS | 0 | 4122 | 3w81 | A | 346 | LEU | p.Leu346Arg | P35475 |
| HOMO SAPIENS | 0 | 4122 | 3w81 | A | 349 | ASP | p.Asp349Asn | P35475 |
| HOMO SAPIENS | 0 | 4122 | 3w81 | A | 361 | ALA | p.Ala361Thr | P35475 |
| HOMO SAPIENS | 0 | 4122 | 3w81 | A | 366 | THR | p.Thr366Pro | P35475 |
| HOMO SAPIENS | 0 | 4122 | 3w81 | A | 402 | TRP | p.Trp402Ter | P35475 |
| HOMO SAPIENS | 0 | 4122 | 3w81 | A | 409 | GLY | p.Gly409Arg | P35475 |
| HOMO SAPIENS | 0 | 4122 | 3w81 | A | 454 | VAL | p.Val454Ile | P35475 |
| HOMO SAPIENS | 1 | 4122 | 3w81 | A | 490 | LEU | p.Leu490Pro | P35475 |
| HOMO SAPIENS | 1 | 4122 | 3w81 | A | 492 | ARG | p.Arg492Pro | P35475 |
| HOMO SAPIENS | 0 | 4122 | 3w81 | A | 533 | PRO | p.Pro533Arg | P35475 |
| HOMO SAPIENS | 0 | 4122 | 3w81 | A | 591 | ALA | p.Ala591Thr | P35475 |
| HOMO SAPIENS | 7 | 4122 | 3w81 | A | 619 | ARG | p.Arg619Gly | P35475 |
| HOMO SAPIENS | 0 | 15325 | 3wvz | B | 54 | VAL | p.Val54Leu | Q53FT3 |
| HOMO SAPIENS | 3 | 3228 | 5iz8 | A | 421 | TRP | p.Trp421Ter | P25054 |
| HOMO SAPIENS | 2 | 3228 | 5iz8 | A | 437 | PRO | p.Pro437Arg | P25054 |
| HOMO SAPIENS | 3 | 3228 | 5iz8 | A | 499 | ARG | p.Arg499Ter | P25054 |
| HOMO SAPIENS | -1 | 29229 | 5buo | A | 539 | SER | p.Ser614Gly | P05067 |
| HOMO SAPIENS | -1 | 34931 | 4n7v | C | 54 | SER | p.Ser54Leu | O94986 |
| HOMO SAPIENS | 0 | 2450 | 5kzx | A | 123 | GLY | p.Gly123Glu | P10253 |
| HOMO SAPIENS | 1 | 2450 | 5kzx | A | 190 | ARG | p.Arg190His | P10253 |
| HOMO SAPIENS | -1 | 2450 | 5kzx | A | 199 | HIS | p.His199Arg | P10253 |
| HOMO SAPIENS | 0 | 2450 | 5kzx | A | 219 | GLY | p.Gly219Arg | P10253 |
| HOMO SAPIENS | 0 | 2450 | 5kzx | A | 220 | VAL | p.Val220Leu | P10253 |
| HOMO SAPIENS | 0 | 2450 | 5kzx | A | 222 | VAL | p.Val222Met | P10253 |

| | | | | | | | | |
|---|---|---|---|---|---|---|---|---|
| HOMO SAPIENS | 0 | 2450 | 5kzx | A | 237 | ALA | p.Ala237Val | P10253 |
| HOMO SAPIENS | 0 | 2450 | 5kzx | A | 251 | SER | p.Ser251Leu | P10253 |
| HOMO SAPIENS | 0 | 2450 | 5kzx | A | 254 | SER | p.Ser254Leu | P10253 |
| HOMO SAPIENS | 0 | 2450 | 5kzx | A | 262 | GLU | p.Glu262Lys | P10253 |
| HOMO SAPIENS | 0 | 2450 | 5kzx | A | 293 | GLY | p.Gly293Arg | P10253 |
| HOMO SAPIENS | 0 | 2450 | 5kzx | A | 299 | LEU | p.Leu299Arg | P10253 |
| HOMO SAPIENS | 0 | 2450 | 5kzx | A | 309 | GLY | p.Gly309Arg | P10253 |
| HOMO SAPIENS | 0 | 2450 | 5kzx | A | 318 | MET | p.Met318Thr | P10253 |
| HOMO SAPIENS | 0 | 2450 | 5kzx | A | 355 | LEU | p.Leu355Pro | P10253 |
| HOMO SAPIENS | 1 | 2450 | 5kzx | A | 361 | PRO | p.Pro361Leu | P10253 |
| HOMO SAPIENS | 4 | 2450 | 5kzx | A | 408 | MET | p.Met408Val | P10253 |
| HOMO SAPIENS | 1 | 2450 | 5kzx | A | 429 | GLN | p.Gln429Glu | P10253 |
| HOMO SAPIENS | 4 | 2450 | 5kzx | A | 437 | ARG | p.Arg437Cys | P10253 |
| HOMO SAPIENS | 0 | 2450 | 5kzx | A | 439 | MET | p.Met439Lys | P10253 |
| HOMO SAPIENS | 1 | 2450 | 5kzx | A | 448 | SER | p.Ser448Thr | P10253 |
| HOMO SAPIENS | 0 | 2450 | 5kzx | A | 451 | PRO | p.Pro451Arg | P10253 |
| HOMO SAPIENS | 1 | 2450 | 5kzx | A | 459 | ASP | p.Asp459His | P10253 |
| HOMO SAPIENS | 8 | 2450 | 5kzx | A | 481 | TRP | p.Trp481Arg | P10253 |
| HOMO SAPIENS | 1 | 2450 | 5kzx | A | 489 | ASP | p.Asp489Asn | P10253 |
| HOMO SAPIENS | 0 | 2450 | 5kzx | A | 521 | GLU | p.Glu521Lys | P10253 |
| HOMO SAPIENS | 0 | 2450 | 5kzx | A | 529 | SER | p.Ser529Val | P10253 |
| HOMO SAPIENS | 0 | 2450 | 5kzx | A | 545 | PRO | p.Pro545Leu | P10253 |
| HOMO SAPIENS | 0 | 2450 | 5kzx | A | 552 | LEU | p.Leu552Pro | P10253 |
| HOMO SAPIENS | 0 | 2450 | 5kzx | A | 576 | GLY | p.Gly576Ser | P10253 |
| HOMO SAPIENS | 9 | 2450 | 5kzx | A | 600 | ARG | p.Arg600His | P10253 |
| HOMO SAPIENS | 8 | 2450 | 5kzx | A | 615 | GLY | p.Gly615Arg | P10253 |
| HOMO SAPIENS | 3 | 2450 | 5kzx | A | 619 | SER | p.Ser619Asn | P10253 |
| HOMO SAPIENS | 0 | 2450 | 5kzx | A | 638 | GLY | p.Gly638Trp | P10253 |
| HOMO SAPIENS | 0 | 2450 | 5kzx | A | 643 | GLY | p.Gly643Arg | P10253 |
| HOMO SAPIENS | 9 | 2450 | 5kzx | A | 645 | ASP | p.Asp645Glu | P10253 |
| HOMO SAPIENS | 9 | 2450 | 5kzx | A | 645 | ASP | p.Asp645His | P10253 |
| HOMO SAPIENS | 0 | 2450 | 5kzx | A | 648 | GLY | p.Gly648Ser | P10253 |
| HOMO SAPIENS | 5 | 2450 | 5kzx | A | 660 | ARG | p.Arg660His | P10253 |
| HOMO SAPIENS | 8 | 2450 | 5kzx | A | 672 | ARG | p.Arg672Gln | P10253 |
| HOMO SAPIENS | 0 | 2450 | 5kzx | A | 702 | ARG | p.Arg702Leu | P10253 |
| HOMO SAPIENS | 3 | 2450 | 5kzx | A | 725 | ARG | p.Arg725Trp | P10253 |
| HOMO SAPIENS | 0 | 2450 | 5kzx | A | 746 | TRP | p.Trp746Cys | P10253 |
| HOMO SAPIENS | 0 | 2450 | 5kzx | A | 780 | ILE | p.Val780Ile | P10253 |
| HOMO SAPIENS | 1 | 2450 | 5kzx | A | 816 | VAL | p.Val816Ile | P10253 |
| HOMO SAPIENS | 0 | 2450 | 5kzx | A | 838 | GLN | p.Gln838Ter | P10253 |
| HOMO SAPIENS | 2 | 2450 | 5kzx | A | 854 | ARG | p.Arg854Ter | P10253 |
| HOMO SAPIENS | 3 | 2450 | 5kzx | A | 870 | ARG | p.Arg870Ter | P10253 |
| HOMO SAPIENS | 0 | 2450 | 5kzx | A | 890 | VAL | p.Val890Leu | P10253 |
| HOMO SAPIENS | 6 | 1882 | 4km6 | A | 175 | CYS | p.Cys175Ter | P15328 |
| HOMO SAPIENS | 3 | 14767 | 1lm5 | A | 2628 | LEU | p.Leu2628Pro | P15924 |
| HOMO SAPIENS | 5 | 14767 | 1lm5 | A | 2655 | ALA | p.Ala2655Gly | P15924 |
| HOMO SAPIENS | 3 | 14767 | 1lm5 | A | 2731 | TYR | p.Tyr2731His | P15924 |
| HOMO SAPIENS | 1 | 14767 | 1lm5 | A | 2767 | THR | p.Thr2767Asn | P15924 |
| HOMO SAPIENS | -1 | 14767 | 1lm5 | A | 2819 | MET | p.Met2819Leu | P15924 |
| HOMO SAPIENS | -1 | 11246 | 4y5o | A | 289 | SER | p.Ser289Asn | Q9BSQ5 |
| HOMO SAPIENS | 0 | 7380 | 5kz5 | I | 60 | ARG | p.Arg60His | Q16595 |
| HOMO SAPIENS | 0 | 7380 | 5kz5 | I | 76 | MET | p.Met76Val | Q16595 |
| HOMO SAPIENS | 2 | 22187 | 3fby | A | 328 | CYS | p.Cys328Arg | P49747 |
| HOMO SAPIENS | 2 | 22187 | 3fby | A | 342 | ASP | p.Asp342Tyr | P49747 |

| | | | | | | | | |
|---|---|---|---|---|---|---|---|---|
| HOMO SAPIENS | 2 | 22187 | 3fby | A | 348 | CYS | p.Cys348Arg | P49747 |
| HOMO SAPIENS | 2 | 22187 | 3fby | A | 386 | ASN | p.Asn386Asp | P49747 |
| HOMO SAPIENS | 2 | 22187 | 3fby | A | 453 | ASN | p.Asn453Ser | P49747 |
| HOMO SAPIENS | 2 | 22187 | 3fby | A | 468 | CYS | p.Cys468Tyr | P49747 |
| HOMO SAPIENS | 3 | 22187 | 3fby | A | 472 | ASP | p.Asp472Tyr | P49747 |
| HOMO SAPIENS | 3 | 22187 | 3fby | A | 473 | ASP | p.Asp473Gly | P49747 |
| HOMO SAPIENS | 2 | 22187 | 3fby | A | 523 | ASN | p.Asn523Lys | P49747 |
| HOMO SAPIENS | 1 | 22187 | 3fby | A | 718 | ARG | p.Arg718Trp | P49747 |
| HOMO SAPIENS | 1 | 22187 | 3fby | A | 719 | GLY | p.Gly719Asp | P49747 |
| HOMO SAPIENS | 0 | 22187 | 3fby | A | 756 | GLN | p.Gln756Arg | P49747 |
| HOMO SAPIENS | -1 | 2784 | 5kvh | A | 88 | MET | p.Met88Val | O95831 |
| HOMO SAPIENS | -1 | 2784 | 5kvh | A | 96 | ASN | p.Asn96Ser | O95831 |
| HOMO SAPIENS | 0 | 2784 | 5kvh | A | 282 | THR | p.Thr282Met | O95831 |
| HOMO SAPIENS | 1 | 2784 | 5kvh | A | 366 | ASN | p.Asn366Ser | O95831 |
| HOMO SAPIENS | 3 | 2784 | 5kvh | A | 463 | ARG | p.Arg463Ile | O95831 |
| HOMO SAPIENS | 0 | 2784 | 5kvh | A | 498 | VAL | p.Val498Met | O95831 |
| HOMO SAPIENS | -1 | 2784 | 5kvh | A | 549 | ALA | p.Ala549Val | O95831 |
| HOMO SAPIENS | 0 | 38316 | 2lnk | C | 1933 | ARG | p.Arg1933Ter | P35579 |
| HOMO SAPIENS | 0 | 33808 | 2miu | A | 29 | VAL | p.Val29Met | Q14192 |
| HOMO SAPIENS | 1 | 33808 | 2miu | A | 37 | ALA | p.Ala37Ser | Q14192 |
| HOMO SAPIENS | 2 | 20994 | 2hey | R | 65 | ARG | p.Arg65Cys | P43489 |
| HOMO SAPIENS | -1 | 5687 | 2xwj | I | 7 | ARG | p.Arg32Gln | P00751 |
| HOMO SAPIENS | -1 | 5687 | 2xwj | I | 7 | ARG | p.Arg32Trp | P00751 |
| HOMO SAPIENS | 0 | 5687 | 2xwj | I | 49 | ARG | p.Arg74His | P00751 |
| HOMO SAPIENS | 0 | 5687 | 2xwj | I | 177 | ARG | p.Arg202Gly | P00751 |
| HOMO SAPIENS | 0 | 5687 | 2xwj | I | 217 | ILE | p.Ile242Leu | P00751 |
| HOMO SAPIENS | -1 | 5687 | 2xwj | I | 227 | GLY | p.Gly252Ser | P00751 |
| HOMO SAPIENS | 0 | 5687 | 2xwj | I | 483 | HIS | p.His508Gln | P00751 |
| HOMO SAPIENS | 0 | 5687 | 2xwj | I | 508 | LYS | p.Lys533Arg | P00751 |
| HOMO SAPIENS | 0 | 5687 | 2xwj | I | 540 | LYS | p.Lys565Glu | P00751 |
| HOMO SAPIENS | 0 | 5687 | 2xwj | I | 541 | GLU | p.Glu566Ala | P00751 |
| HOMO SAPIENS | 2 | 953 | 3d4j | A | 249 | PHE | p.Phe249Ser | P53602 |
| HOMO SAPIENS | 1 | 953 | 3d4j | A | 292 | ASN | p.Asn292Ser | P53602 |
| HOMO SAPIENS | 0 | 3300 | 1s9c | A | 140 | ASN | p.Asn457Tyr | P51659 |
| HOMO SAPIENS | 2 | 3300 | 1s9c | A | 167 | ILE | p.Ile484Val | P51659 |
| HOMO SAPIENS | 0 | 3300 | 1s9c | A | 174 | ALA | p.Ala491Thr | P51659 |
| HOMO SAPIENS | 8 | 3300 | 1s9c | A | 193 | ASP | p.Asp510Asn | P51659 |
| HOMO SAPIENS | 3 | 3300 | 1s9c | A | 194 | TRP | p.Trp511Arg | P51659 |
| HOMO SAPIENS | 0 | 3300 | 1s9c | A | 242 | VAL | p.Ile559Val | P51659 |
| HOMO SAPIENS | 0 | 10810 | 5i25 | A | 5 | GLN | p.Gln23Ter | P03951 |
| HOMO SAPIENS | 0 | 10810 | 5i25 | A | 55 | TRP | p.Trp73Ter | P03951 |
| HOMO SAPIENS | 0 | 10810 | 5i25 | A | 91 | ALA | p.Ala109Thr | P03951 |
| HOMO SAPIENS | 0 | 10810 | 5i25 | A | 117 | GLU | p.Glu135Ter | P03951 |
| HOMO SAPIENS | 0 | 10810 | 5i25 | A | 118 | CYS | p.Cys136Ter | P03951 |
| HOMO SAPIENS | 0 | 10810 | 5i25 | A | 128 | CYS | p.Cys146Ter | P03951 |
| HOMO SAPIENS | 0 | 10810 | 5i25 | A | 210 | ARG | p.Arg228Ter | P03951 |
| HOMO SAPIENS | 0 | 10810 | 5i25 | A | 226 | GLN | p.Gln244Ter | P03951 |
| HOMO SAPIENS | 0 | 10810 | 5i25 | A | 233 | GLN | p.Gln251Ter | P03951 |
| HOMO SAPIENS | 0 | 16668 | 3lbx | A | 28 | ARG | p.Arg28Leu | P02549 |
| HOMO SAPIENS | 0 | 16668 | 3lbx | A | 28 | ARG | p.Arg28Cys | P02549 |
| HOMO SAPIENS | 1 | 16668 | 3lbx | A | 41 | ARG | p.Arg41Trp | P02549 |
| HOMO SAPIENS | 1 | 16668 | 3lbx | A | 45 | ARG | p.Arg45Ser | P02549 |
| HOMO SAPIENS | 2 | 16668 | 3lbx | A | 48 | LYS | p.Lys48Arg | P02549 |
| HOMO SAPIENS | 1 | 16668 | 3lbx | A | 152 | ASP | p.Asp152Asn | P02549 |

| | | | | | | | | |
|---|---|---|---|---|---|---|---|---|
| HOMO SAPIENS | 0 | 226 | 2ble | A | 256 | ILE | p.Phe256Ile | P36959 |
| HOMO SAPIENS | 2 | 36808 | 4c7n | A | 3 | SER | p.Ser357Pro | O75030 |
| HOMO SAPIENS | 0 | 4092 | 5jnx | A | 113 | CYS | p.Cys113Arg | O15118 |
| HOMO SAPIENS | 4 | 4092 | 5jnx | A | 215 | HIS | p.His215Arg | O15118 |
| HOMO SAPIENS | 6 | 4092 | 5jnx | A | 237 | PRO | p.Pro237Ser | O15118 |
| HOMO SAPIENS | 0 | 4092 | 5jnx | A | 363 | ALA | p.Ala363Val | O15118 |
| HOMO SAPIENS | 0 | 4092 | 5jnx | A | 378 | VAL | p.Val378Ala | O15118 |
| HOMO SAPIENS | 8 | 4092 | 5jnx | A | 404 | ARG | p.Arg404Gln | O15118 |
| HOMO SAPIENS | 8 | 4092 | 5jnx | A | 411 | ARG | p.Arg411Gln | O15118 |
| HOMO SAPIENS | 2 | 4092 | 5jnx | A | 474 | PRO | p.Pro474Leu | O15118 |
| HOMO SAPIENS | 8 | 4092 | 5jnx | A | 511 | THR | p.Thr511Met | O15118 |
| HOMO SAPIENS | 0 | 4092 | 5jnx | A | 642 | MET | p.Met642Ile | O15118 |
| HOMO SAPIENS | 2 | 4092 | 5jnx | A | 664 | VAL | p.Val664Met | O15118 |
| HOMO SAPIENS | 0 | 4092 | 5jnx | A | 734 | SER | p.Ser734Ile | O15118 |
| HOMO SAPIENS | -1 | 4092 | 5jnx | A | 775 | GLN | p.Gln775Pro | O15118 |
| HOMO SAPIENS | 0 | 4092 | 5jnx | A | 858 | ILE | p.Ile858Val | O15118 |
| HOMO SAPIENS | 0 | 4092 | 5jnx | A | 889 | VAL | p.Val889Met | O15118 |
| HOMO SAPIENS | 0 | 4092 | 5jnx | A | 910 | GLY | p.Gly910Ser | O15118 |
| HOMO SAPIENS | 0 | 4092 | 5jnx | A | 911 | GLY | p.Gly911Ser | O15118 |
| HOMO SAPIENS | 0 | 4092 | 5jnx | A | 928 | GLN | p.Gln928Pro | O15118 |
| HOMO SAPIENS | 0 | 4092 | 5jnx | A | 931 | ASN | p.Asn931Lys | O15118 |
| HOMO SAPIENS | 0 | 4092 | 5jnx | A | 940 | SER | p.Ser940Leu | O15118 |
| HOMO SAPIENS | 0 | 4092 | 5jnx | A | 950 | VAL | p.Val950Met | O15118 |
| HOMO SAPIENS | 0 | 4092 | 5jnx | A | 954 | SER | p.Ser954Leu | O15118 |
| HOMO SAPIENS | 0 | 4092 | 5jnx | A | 958 | ARG | p.Arg958Gln | O15118 |
| HOMO SAPIENS | 0 | 4092 | 5jnx | A | 958 | ARG | p.Arg958Ter | O15118 |
| HOMO SAPIENS | -1 | 4092 | 5jnx | A | 978 | ARG | p.Arg978Cys | O15118 |
| HOMO SAPIENS | 0 | 4092 | 5jnx | A | 992 | GLY | p.Gly992Trp | O15118 |
| HOMO SAPIENS | 0 | 4092 | 5jnx | A | 1007 | PRO | p.Pro1007Ala | O15118 |
| HOMO SAPIENS | 0 | 4092 | 5jnx | A | 1019 | TYR | p.Tyr1019Cys | O15118 |
| HOMO SAPIENS | 2 | 4092 | 5jnx | A | 1035 | ALA | p.Ala1035Val | O15118 |
| HOMO SAPIENS | 2 | 4092 | 5jnx | A | 1054 | ALA | p.Ala1054Thr | O15118 |
| HOMO SAPIENS | 2 | 4092 | 5jnx | A | 1061 | ILE | p.Ile1061Thr | O15118 |
| HOMO SAPIENS | 0 | 4092 | 5jnx | A | 1073 | GLY | p.Gly1073Ser | O15118 |
| HOMO SAPIENS | 2 | 4092 | 5jnx | A | 1088 | TYR | p.Tyr1088Cys | O15118 |
| HOMO SAPIENS | 2 | 4092 | 5jnx | A | 1142 | MET | p.Met1142Thr | O15118 |
| HOMO SAPIENS | 2 | 4092 | 5jnx | A | 1156 | ASN | p.Asn1156Ser | O15118 |
| HOMO SAPIENS | 0 | 4092 | 5jnx | A | 1179 | MET | p.Met1179Val | O15118 |
| HOMO SAPIENS | 0 | 4092 | 5jnx | A | 1186 | ARG | p.Arg1186His | O15118 |
| HOMO SAPIENS | 0 | 4092 | 5jnx | A | 1187 | ALA | p.Ala1187Gly | O15118 |
| HOMO SAPIENS | 2 | 4092 | 5jnx | A | 1189 | GLU | p.Glu1189Gly | O15118 |
| HOMO SAPIENS | 2 | 4092 | 5jnx | A | 1200 | SER | p.Ser1200Gly | O15118 |
| HOMO SAPIENS | 2 | 4092 | 5jnx | A | 1205 | THR | p.Thr1205Lys | O15118 |
| HOMO SAPIENS | 0 | 4092 | 5jnx | A | 1213 | LEU | p.Leu1213Phe | O15118 |
| HOMO SAPIENS | -1 | 4092 | 5jnx | A | 1266 | ARG | p.Arg1266Gln | O15118 |
| HOMO SAPIENS | 9 | 16498 | 4i6o | A | 64 | ARG | p.Arg735Trp | P01024 |
| HOMO SAPIENS | 1 | 21902 | 3un9 | C | 751 | ARG | p.Arg751His | Q86UT6 |
| HOMO SAPIENS | 7 | 20391 | 2h32 | B | 105 | PRO | p.Pro142Leu | P15814 |
| HOMO SAPIENS | 6 | 1635 | 4x45 | A | 65 | ASP | p.Asp65Val | P07741 |
| HOMO SAPIENS | 1 | 1635 | 4x45 | A | 98 | TRP | p.Trp98Ter | P07741 |
| HOMO SAPIENS | 0 | 1635 | 4x45 | A | 110 | LEU | p.Leu110Pro | P07741 |
| HOMO SAPIENS | 7 | 1635 | 4x45 | A | 136 | MET | p.Met136Thr | P07741 |
| HOMO SAPIENS | 1 | 1635 | 4x45 | A | 150 | VAL | p.Val150Phe | P07741 |
| HOMO SAPIENS | 4 | 34810 | 2rpc | A | 16 | CYS | p.Cys253Ser | O60481 |

| | | | | | | | | |
|---|---|---|---|---|---|---|---|---|
| HOMO SAPIENS | 1 | 34810 | 2rpc | A | 18 | TRP | p.Trp255Gly | O60481 |
| HOMO SAPIENS | 1 | 14359 | 4hxi | A | 78 | MET | p.Met78Val | Q9UH77 |
| HOMO SAPIENS | 0 | 5342 | 4d9t | A | 628 | GLU | p.Glu628Asp | P51812 |
| HOMO SAPIENS | -1 | 5342 | 4d9t | A | 723 | ARG | p.Arg723His | P51812 |
| HOMO SAPIENS | -1 | 5342 | 4d9t | A | 729 | ARG | p.Arg729Gln | P51812 |
| HOMO SAPIENS | 1 | 9619 | 3j2s | B | 1781 | ARG | p.Arg1800Cys | P00451 |
| HOMO SAPIENS | 0 | 9619 | 3j2s | B | 1823 | MET | p.Met1842Ile | P00451 |
| HOMO SAPIENS | 1 | 9619 | 3j2s | B | 1825 | PRO | p.Pro1844Ser | P00451 |
| HOMO SAPIENS | 0 | 9619 | 3j2s | B | 1826 | THR | p.Thr1845Pro | P00451 |
| HOMO SAPIENS | 0 | 9619 | 3j2s | B | 1834 | ALA | p.Ala1853Val | P00451 |
| HOMO SAPIENS | 0 | 9619 | 3j2s | B | 1846 | ASP | p.Asp1865Asn | P00451 |
| HOMO SAPIENS | 1 | 9619 | 3j2s | B | 1848 | HIS | p.His1867Arg | P00451 |
| HOMO SAPIENS | 0 | 9619 | 3j2s | B | 1854 | PRO | p.Pro1873Arg | P00451 |
| HOMO SAPIENS | 1 | 9619 | 3j2s | B | 1885 | GLU | p.Glu1904Lys | P00451 |
| HOMO SAPIENS | 0 | 9619 | 3j2s | B | 1922 | ASN | p.Asn1941Ser | P00451 |
| HOMO SAPIENS | 1 | 9619 | 3j2s | B | 1941 | ARG | p.Arg1960Gln | P00451 |
| HOMO SAPIENS | 4 | 9619 | 3j2s | B | 2105 | TYR | p.Tyr2124Cys | P00451 |
| HOMO SAPIENS | 6 | 9619 | 3j2s | B | 2150 | ARG | p.Arg2169His | P00451 |
| HOMO SAPIENS | 0 | 9619 | 3j2s | B | 2159 | ARG | p.Arg2178Cys | P00451 |
| HOMO SAPIENS | 0 | 9619 | 3j2s | B | 2159 | ARG | p.Arg2178His | P00451 |
| HOMO SAPIENS | 2 | 9619 | 3j2s | B | 2238 | MET | p.Met2257Val | P00451 |
| HOMO SAPIENS | 2 | 9619 | 3j2s | B | 2300 | PRO | p.Pro2319Ser | P00451 |
| HOMO SAPIENS | 1 | 667 | 1z2m | A | 127 | GLU | p.Glu127Ter | P05161 |
| HOMO SAPIENS | 0 | 15717 | 2nn6 | G | 30 | GLY | p.Gly31Ala | Q9NQT5 |
| HOMO SAPIENS | -1 | 15717 | 2nn6 | G | 55 | ASN | p.Asn56His | Q9NQT5 |
| HOMO SAPIENS | 7 | 15717 | 2nn6 | G | 131 | ASP | p.Asp132Ala | Q9NQT5 |
| HOMO SAPIENS | 7 | 784 | 3rkq | A | 161 | ARG | p.Arg161Pro | P52952 |
| HOMO SAPIENS | 6 | 784 | 3rkq | A | 178 | THR | p.Thr178Met | P52952 |
| HOMO SAPIENS | 7 | 784 | 3rkq | A | 183 | LYS | p.Lys183Glu | P52952 |
| HOMO SAPIENS | 8 | 784 | 3rkq | A | 190 | ARG | p.Arg190Cys | P52952 |
| HOMO SAPIENS | 6 | 385 | 30000 | C | 70 | ALA | p.Ala70Thr | P07954 |
| HOMO SAPIENS | 2 | 385 | 30000 | C | 101 | ARG | p.Arg101Pro | P07954 |
| HOMO SAPIENS | 2 | 385 | 30000 | C | 174 | PRO | p.Pro174Arg | P07954 |
| HOMO SAPIENS | 6 | 385 | 30000 | C | 187 | SER | p.Ser187Ter | P07954 |
| HOMO SAPIENS | 9 | 385 | 30000 | C | 233 | ARG | p.Arg233His | P07954 |
| HOMO SAPIENS | 3 | 385 | 30000 | C | 273 | ALA | p.Ala273Thr | P07954 |
| HOMO SAPIENS | 4 | 385 | 30000 | C | 334 | SER | p.Ser334Arg | P07954 |
| HOMO SAPIENS | 6 | 385 | 30000 | C | 340 | ASN | p.Asn340Lys | P07954 |
| HOMO SAPIENS | 8 | 385 | 30000 | C | 343 | ARG | p.Arg343Ter | P07954 |
| HOMO SAPIENS | 3 | 385 | 30000 | C | 350 | ARG | p.Arg350Trp | P07954 |
| HOMO SAPIENS | 4 | 385 | 30000 | C | 376 | GLN | p.Gln376Pro | P07954 |
| HOMO SAPIENS | 6 | 385 | 30000 | C | 434 | CYS | p.Cys434Ter | P07954 |
| HOMO SAPIENS | 0 | 3316 | 4f91 | B | 934 | THR | p.Thr934Ala | O75643 |
| HOMO SAPIENS | 0 | 3316 | 4f91 | B | 1002 | ASN | p.Asn1002Ser | O75643 |
| HOMO SAPIENS | 0 | 3316 | 4f91 | B | 1087 | SER | p.Ser1087Leu | O75643 |
| HOMO SAPIENS | 4 | 3316 | 4f91 | B | 1389 | VAL | p.Val1389Ile | O75643 |
| HOMO SAPIENS | 0 | 619 | 5hbt | B | 195 | ASP | p.Asp240His | P02708 |
| HOMO SAPIENS | 1 | 619 | 5hbt | B | 209 | ARG | p.Arg254Leu | P02708 |
| HOMO SAPIENS | 0 | 18171 | 2y25 | B | 1408 | ASP | p.Asp1408Asn | P52179 |
| HOMO SAPIENS | 0 | 18171 | 2y25 | B | 1453 | MET | p.Met1453Thr | P52179 |
| HOMO SAPIENS | 0 | 18171 | 2y25 | B | 1459 | ILE | p.Ile1459Lys | P52179 |
| HOMO SAPIENS | 0 | 18171 | 2y25 | B | 1558 | PHE | p.Phe1558Leu | P52179 |
| HOMO SAPIENS | 0 | 18171 | 2y25 | B | 1573 | ARG | p.Arg1573Gln | P52179 |
| HOMO SAPIENS | 7 | 18171 | 2y25 | B | 1663 | VAL | p.Val1663Met | P52179 |

| | | | | | | | | |
|---|---|---|---|---|---|---|---|---|
| HOMO SAPIENS | 8 | 14262 | 2fcw | B | 117 | SER | p.Ser138Ter | P01130 |
| HOMO SAPIENS | 8 | 14262 | 2fcw | B | 118 | ASP | p.Asp139Glu | P01130 |
| HOMO SAPIENS | 9 | 14262 | 2fcw | B | 119 | GLU | p.Glu140Lys | P01130 |
| HOMO SAPIENS | 7 | 14262 | 2fcw | B | 122 | CYS | p.Cys143Ter | P01130 |
| HOMO SAPIENS | 6 | 14262 | 2fcw | B | 134 | CYS | p.Cys155Ter | P01130 |
| HOMO SAPIENS | 7 | 14262 | 2fcw | B | 147 | ASP | p.Asp168Glu | P01130 |
| HOMO SAPIENS | 4 | 2624 | 3b7b | A | 855 | VAL | p.Val886Met | Q9H9B1 |
| HOMO SAPIENS | 7 | 2624 | 3b7b | A | 868 | ILE | p.Ile899Val | Q9H9B1 |
| HOMO SAPIENS | 0 | 2624 | 3b7b | A | 892 | GLU | p.Glu923Lys | Q9H9B1 |
| HOMO SAPIENS | 0 | 4791 | 5ctd | B | 68 | LEU | p.Pro549Ala | Q14055 |
| HOMO SAPIENS | 5 | 460 | 4ui1 | D | 80 | CYS | p.Cys80Arg | Q6ZVN8 |
| HOMO SAPIENS | 3 | 460 | 4ui1 | D | 101 | LEU | p.Leu101Pro | Q6ZVN8 |
| HOMO SAPIENS | 0 | 40782 | 1sy6 | A | 47 | LYS | p.Lys69Ter | P09693 |
| HOMO SAPIENS | -1 | 1092 | 2idx | A | 62 | THR | p.Thr62Met | Q96EY8 |
| HOMO SAPIENS | 6 | 1092 | 2idx | A | 183 | HIS | p.His183Leu | Q96EY8 |
| HOMO SAPIENS | 7 | 1092 | 2idx | A | 186 | ARG | p.Arg186Trp | Q96EY8 |
| HOMO SAPIENS | 6 | 1092 | 2idx | A | 190 | ARG | p.Arg190Cys | Q96EY8 |
| HOMO SAPIENS | 6 | 1092 | 2idx | A | 190 | ARG | p.Arg190His | Q96EY8 |
| HOMO SAPIENS | 7 | 1092 | 2idx | A | 191 | ARG | p.Arg191Trp | Q96EY8 |
| HOMO SAPIENS | 7 | 1092 | 2idx | A | 191 | ARG | p.Arg191Gln | Q96EY8 |
| HOMO SAPIENS | 6 | 1092 | 2idx | A | 195 | ARG | p.Arg195His | Q96EY8 |
| HOMO SAPIENS | 1 | 1092 | 2idx | A | 234 | GLN | p.Gln234Ter | Q96EY8 |
| HOMO SAPIENS | 0 | 1092 | 2idx | A | 239 | MET | p.Met239Lys | Q96EY8 |
| HOMO SAPIENS | -1 | 6418 | 5fur | G | 11 | THR | p.Thr11Ile | P21675 |
| HOMO SAPIENS | -1 | 6418 | 5fur | G | 290 | LEU | p.Leu269Val | P21675 |
| HOMO SAPIENS | 0 | 6100 | 1wiq | A | 240 | ARG | p.Arg265Trp | P01730 |
| HOMO SAPIENS | 0 | 13757 | 4ye6 | A | 45 | GLY | p.Gly45Val | P47897 |
| HOMO SAPIENS | 1 | 13757 | 4ye6 | A | 57 | TYR | p.Tyr57His | P47897 |
| HOMO SAPIENS | 8 | 13757 | 4ye6 | A | 403 | ARG | p.Arg403Trp | P47897 |
| HOMO SAPIENS | 7 | 13757 | 4ye6 | A | 524 | ARG | p.Arg524Trp | P47897 |
| HOMO SAPIENS | 6 | 13757 | 4ye6 | A | 690 | ARG | p.Arg690Cys | P47897 |
| HOMO SAPIENS | 0 | 26958 | 2lzl | A | 370 | GLY | p.Gly370Cys | P22607 |
| HOMO SAPIENS | 0 | 26958 | 2lzl | A | 371 | SER | p.Ser371Cys | P22607 |
| HOMO SAPIENS | 0 | 26958 | 2lzl | A | 373 | TYR | p.Tyr373Cys | P22607 |
| HOMO SAPIENS | 0 | 26958 | 2lzl | A | 380 | GLY | p.Gly380Arg | P22607 |
| HOMO SAPIENS | 0 | 26958 | 2lzl | A | 384 | PHE | p.Phe384Leu | P22607 |
| HOMO SAPIENS | 5 | 26958 | 2lzl | A | 391 | ALA | p.Ala391Glu | P22607 |
| HOMO SAPIENS | 6 | 11325 | 2wa6 | A | 121 | TYR | p.Tyr121Phe | O75369 |
| HOMO SAPIENS | 2 | 11325 | 2wa6 | A | 161 | PHE | p.Phe161Cys | O75369 |
| HOMO SAPIENS | 0 | 11325 | 2wa6 | A | 171 | LEU | p.Leu171Arg | O75369 |
| HOMO SAPIENS | 2 | 11325 | 2wa6 | A | 173 | ALA | p.Ala173Val | O75369 |
| HOMO SAPIENS | 0 | 11325 | 2wa6 | A | 202 | MET | p.Met202Val | O75369 |
| HOMO SAPIENS | 3 | 11325 | 2wa6 | A | 227 | GLU | p.Glu227Lys | O75369 |
| HOMO SAPIENS | 0 | 11325 | 2wa6 | A | 234 | LEU | p.Leu234Val | O75369 |
| HOMO SAPIENS | 2 | 11325 | 2wa6 | A | 235 | SER | p.Ser235Pro | O75369 |
| HOMO SAPIENS | 9 | 7240 | 3tgx | A | 137 | TYR | p.Tyr156Cys | Q9HBE5 |
| HOMO SAPIENS | 0 | 35479 | 4elj | A | 307 | THR | p.Thr307Ile | P06400 |
| HOMO SAPIENS | 0 | 35479 | 4elj | A | 310 | GLY | p.Gly310Glu | P06400 |
| HOMO SAPIENS | 0 | 35479 | 4elj | A | 377 | THR | p.Thr377Ser | P06400 |
| HOMO SAPIENS | 9 | 36706 | 1he7 | A | 359 | TYR | p.Tyr359Cys | P04629 |
| HOMO SAPIENS | 4 | 6538 | 4d0e | A | 504 | ARG | p.Arg504His | P46531 |
| HOMO SAPIENS | 1 | 26518 | 5fm5 | M | 600 | GLU | p.Glu600Val | P52179 |
| HOMO SAPIENS | -1 | 26518 | 5fm5 | M | 651 | ARG | p.Arg651Gln | P52179 |
| HOMO SAPIENS | -1 | 26518 | 5fm5 | M | 660 | PRO | p.Pro660Ala | P52179 |

| | | | | | | | | |
|---|---|---|---|---|---|---|---|---|
| HOMO SAPIENS | -1 | 26518 | 5fm5 | M | 688 | THR | p.Thr688Ser | P52179 |
| HOMO SAPIENS | -1 | 26518 | 5fm5 | M | 704 | GLU | p.Glu704Lys | P52179 |
| HOMO SAPIENS | -1 | 26518 | 5fm5 | M | 727 | THR | p.Thr727Ala | P52179 |
| HOMO SAPIENS | 2 | 27054 | 4rhp | B | 209 | LEU | p.Leu209Val | O75208 |
| HOMO SAPIENS | -1 | 27054 | 4rhp | B | 288 | LYS | p.Lys288Asn | O75208 |
| HOMO SAPIENS | 5 | 2384 | 5fqd | B | 419 | ARG | p.Arg419Ter | Q96SW2 |
| HOMO SAPIENS | 4 | 5154 | 2hl3 | A | 56 | LYS | p.Lys56Arg | Q14203 |
| HOMO SAPIENS | 0 | 5154 | 2hl3 | A | 59 | GLY | p.Gly59Ser | Q14203 |
| HOMO SAPIENS | 0 | 5154 | 2hl3 | A | 71 | GLY | p.Gly71Ala | Q14203 |
| HOMO SAPIENS | 0 | 5154 | 2hl3 | A | 71 | GLY | p.Gly71Arg | Q14203 |
| HOMO SAPIENS | 7 | 5154 | 2hl3 | A | 72 | THR | p.Thr72Pro | Q14203 |
| HOMO SAPIENS | 2 | 5154 | 2hl3 | A | 74 | GLN | p.Gln74Pro | Q14203 |
| HOMO SAPIENS | 2 | 20039 | 2l1g | A | 9 | GLY | p.Gly9Cys | Q9NVV9 |
| HOMO SAPIENS | 5 | 20039 | 2l1g | A | 81 | PHE | p.Phe81Leu | Q9NVV9 |
| HOMO SAPIENS | -1 | 97 | 5uz7 | A | 201 | ARG | p.Arg201His | P63092 |
| HOMO SAPIENS | -1 | 97 | 5uz7 | A | 201 | ARG | p.Arg201Cys | P63092 |
| HOMO SAPIENS | 0 | 3257 | 4b7w | B | 278 | GLU | p.Glu278Asp | P41235 |
| HOMO SAPIENS | 1 | 3257 | 4b7w | B | 331 | ARG | p.Arg331Cys | P41235 |
| HOMO SAPIENS | 0 | 2814 | 4qgk | B | 10 | GLN | p.Gln10Ter | P51648 |
| HOMO SAPIENS | 0 | 2814 | 4qgk | B | 177 | ARG | p.Arg177Ter | P51648 |
| HOMO SAPIENS | 9 | 2814 | 4qgk | B | 184 | THR | p.Thr184Met | P51648 |
| HOMO SAPIENS | 1 | 2814 | 4qgk | B | 188 | ALA | p.Ala188Val | P51648 |
| HOMO SAPIENS | 0 | 2814 | 4qgk | B | 214 | CYS | p.Cys214Tyr | P51648 |
| HOMO SAPIENS | 0 | 2814 | 4qgk | B | 245 | ASP | p.Asp245Asn | P51648 |
| HOMO SAPIENS | 0 | 2814 | 4qgk | B | 266 | LYS | p.Lys266Asn | P51648 |
| HOMO SAPIENS | 0 | 2814 | 4qgk | B | 315 | PRO | p.Pro315Ser | P51648 |
| HOMO SAPIENS | 0 | 2814 | 4qgk | B | 386 | ASN | p.Asn386Ser | P51648 |
| HOMO SAPIENS | -1 | 5486 | 5u6o | B | 47 | GLY | p.Gly47Val | O60741 |
| HOMO SAPIENS | 1 | 21475 | 3pdy | A | 679 | THR | p.Thr789Ile | Q15149 |
| HOMO SAPIENS | 0 | 32890 | 4djc | B | 1501 | LEU | p.Leu1501Val | Q14524 |
| HOMO SAPIENS | -1 | 32890 | 4djc | B | 1512 | ARG | p.Arg1512Trp | Q14524 |
| HOMO SAPIENS | 0 | 26683 | 2xze | B | 38 | ARG | p.Arg38Cys | O95630 |
| HOMO SAPIENS | 0 | 26683 | 2xze | B | 100 | PHE | p.Phe100Tyr | O95630 |
| HOMO SAPIENS | -1 | 23175 | 5dka | A | 163 | SER | p.Ser163Leu | Q96EQ8 |
| HOMO SAPIENS | -1 | 23175 | 5dka | A | 174 | ARG | p.Arg174Cys | Q96EQ8 |
| HOMO SAPIENS | 6 | 21177 | 3k1r | A | 155 | ARG | p.Arg155Ter | Q9Y6N9 |
| HOMO SAPIENS | 0 | 1237 | 2wwm | O | 40 | PRO | p.Pro40Arg | O75147 |
| HOMO SAPIENS | 1 | 2543 | 5ewl | B | 64 | VAL | p.Val64Leu | Q13224 |
| HOMO SAPIENS | 2 | 2543 | 5ewl | B | 271 | ALA | p.Ala271Glu | Q13224 |
| HOMO SAPIENS | 3 | 34262 | 2mng | A | 613 | PHE | p.Phe613Leu | Q9Y3Q4 |
| HOMO SAPIENS | 7 | 34262 | 2mng | A | 672 | SER | p.Ser672Arg | Q9Y3Q4 |
| HOMO SAPIENS | 4 | 1667 | 1f05 | A | 321 | LYS | p.Lys321Met | P37837 |
| HOMO SAPIENS | 0 | 25179 | 2knc | B | 752 | SER | p.Ser778Pro | P05106 |
| HOMO SAPIENS | 0 | 2185 | 2ig7 | A | 50 | TYR | p.Tyr50Cys | Q9Y259 |
| HOMO SAPIENS | 0 | 2185 | 2ig7 | A | 224 | ASN | p.Asn224His | Q9Y259 |
| HOMO SAPIENS | 0 | 2185 | 2ig7 | A | 270 | TYR | p.Tyr270Ter | Q9Y259 |
| HOMO SAPIENS | 0 | 2185 | 2ig7 | A | 328 | GLN | p.Gln328Arg | Q9Y259 |
| HOMO SAPIENS | 3 | 2201 | 2r0o | A | 179 | PRO | p.Pro179Leu | O43707 |
| HOMO SAPIENS | 3 | 2201 | 2r0o | A | 255 | GLU | p.Lys255Glu | O43707 |
| HOMO SAPIENS | 3 | 2201 | 2r0o | A | 259 | THR | p.Thr259Ile | O43707 |
| HOMO SAPIENS | 4 | 2201 | 2r0o | A | 262 | SER | p.Ser262Pro | O43707 |
| HOMO SAPIENS | -1 | 31221 | 2wvr | C | 44 | THR | p.Thr44Pro | Q9H211 |
| HOMO SAPIENS | -1 | 31221 | 2wvr | C | 135 | ALA | p.Ala135Val | Q9H211 |
| HOMO SAPIENS | -1 | 31221 | 2wvr | C | 456 | GLU | p.Glu456Ala | Q9H211 |

| | | | | | | | | |
|---|---|---|---|---|---|---|---|---|
| HOMO SAPIENS | -1 | 31221 | 2wvr | C | 462 | ARG | p.Arg462Gln | Q9H211 |
| HOMO SAPIENS | -1 | 31221 | 2wvr | C | 468 | GLU | p.Glu468Lys | Q9H211 |
| HOMO SAPIENS | -1 | 31221 | 2wvr | C | 537 | ALA | p.Ala537Val | Q9H211 |
| HOMO SAPIENS | 5 | 2576 | 5o3l | C | 363 | VAL | p.Val680Ile | P10636 |
| HOMO SAPIENS | 5 | 1345 | 1czt | A | 37 | ARG | p.Arg2102Cys | P12259 |
| HOMO SAPIENS | 1 | 1345 | 1czt | A | 83 | MET | p.Met2148Thr | P12259 |
| HOMO SAPIENS | 1 | 1345 | 1czt | A | 157 | ASP | p.Asp2222Gly | P12259 |
| HOMO SAPIENS | 8 | 35985 | 1lo1 | A | 110 | ALA | p.Ala110Val | O95718 |
| HOMO SAPIENS | 1 | 617 | 2ocp | B | 44 | GLU | p.Glu44Lys | Q16854 |
| HOMO SAPIENS | 5 | 617 | 2ocp | B | 46 | ASN | p.Asn46Ser | Q16854 |
| HOMO SAPIENS | 0 | 617 | 2ocp | B | 59 | THR | p.Thr59Met | Q16854 |
| HOMO SAPIENS | 7 | 617 | 2ocp | B | 142 | ARG | p.Arg142Lys | Q16854 |
| HOMO SAPIENS | 3 | 617 | 2ocp | B | 154 | ASN | p.Asn154Lys | Q16854 |
| HOMO SAPIENS | 3 | 617 | 2ocp | B | 170 | GLN | p.Gln170Arg | Q16854 |
| HOMO SAPIENS | 3 | 617 | 2ocp | B | 227 | GLU | p.Glu227Lys | Q16854 |
| HOMO SAPIENS | 3 | 617 | 2ocp | B | 233 | LYS | p.Lys233Glu | Q16854 |
| HOMO SAPIENS | 3 | 617 | 2ocp | B | 255 | ASP | p.Asp255Tyr | Q16854 |
| HOMO SAPIENS | -1 | 38975 | 5itz | D | 421 | GLN | p.Gln421His | Q9HC77 |
| HOMO SAPIENS | 2 | 6235 | 2f8v | A | 15 | VAL | p.Val15Ile | Q8WZ42 |
| HOMO SAPIENS | 0 | 6235 | 2f8v | A | 54 | VAL | p.Val54Met | Q8WZ42 |
| HOMO SAPIENS | 0 | 6235 | 2f8v | A | 59 | SER | p.Ser59Arg | Q8WZ42 |
| HOMO SAPIENS | 0 | 6235 | 2f8v | A | 60 | ASP | p.Asp60Tyr | Q8WZ42 |
| HOMO SAPIENS | 0 | 12630 | 1ext | B | 12 | ASP | p.Asp41Glu | P19438 |
| HOMO SAPIENS | 3 | 12630 | 1ext | B | 30 | CYS | p.Cys59Arg | P19438 |
| HOMO SAPIENS | 3 | 12630 | 1ext | B | 30 | CYS | p.Cys59Ser | P19438 |
| HOMO SAPIENS | 5 | 12630 | 1ext | B | 33 | CYS | p.Cys62Tyr | P19438 |
| HOMO SAPIENS | 5 | 12630 | 1ext | B | 33 | CYS | p.Cys62Gly | P19438 |
| HOMO SAPIENS | 1 | 12630 | 1ext | B | 46 | PRO | p.Pro75Leu | P19438 |
| HOMO SAPIENS | 3 | 12630 | 1ext | B | 50 | THR | p.Thr79Met | P19438 |
| HOMO SAPIENS | 6 | 12630 | 1ext | B | 52 | CYS | p.Cys81Phe | P19438 |
| HOMO SAPIENS | 6 | 12630 | 1ext | B | 70 | CYS | p.Cys99Ser | P19438 |
| HOMO SAPIENS | 5 | 12630 | 1ext | B | 88 | CYS | p.Cys117Arg | P19438 |
| HOMO SAPIENS | 5 | 12630 | 1ext | B | 88 | CYS | p.Cys117Tyr | P19438 |
| HOMO SAPIENS | -1 | 12630 | 1ext | B | 170 | ILE | p.Ile199Asn | P19438 |
| HOMO SAPIENS | 0 | 994 | 1aos | B | 95 | ARG | p.Arg95Cys | P04424 |
| HOMO SAPIENS | 0 | 994 | 1aos | B | 111 | ARG | p.Arg111Gln | P04424 |
| HOMO SAPIENS | 0 | 994 | 1aos | B | 131 | THR | p.Thr131Met | P04424 |
| HOMO SAPIENS | 6 | 994 | 1aos | B | 178 | VAL | p.Val178Met | P04424 |
| HOMO SAPIENS | 9 | 994 | 1aos | B | 182 | ARG | p.Arg182Gln | P04424 |
| HOMO SAPIENS | 9 | 994 | 1aos | B | 186 | ARG | p.Arg186Trp | P04424 |
| HOMO SAPIENS | 8 | 994 | 1aos | B | 193 | ARG | p.Arg193Gln | P04424 |
| HOMO SAPIENS | 5 | 994 | 1aos | B | 217 | ARG | p.Arg217Ter | P04424 |
| HOMO SAPIENS | 0 | 994 | 1aos | B | 286 | GLN | p.Gln286Arg | P04424 |
| HOMO SAPIENS | 5 | 994 | 1aos | B | 379 | ARG | p.Arg379Cys | P04424 |
| HOMO SAPIENS | 4 | 16066 | 5mv9 | A | 1719 | TYR | p.Tyr1719Cys | Q13402 |
| HOMO SAPIENS | -1 | 16066 | 5mv9 | A | 1739 | ARG | p.Arg1739Ter | Q13402 |
| HOMO SAPIENS | 4 | 16066 | 5mv9 | A | 1775 | ILE | p.Ile1775Thr | Q13402 |
| HOMO SAPIENS | 2 | 16066 | 5mv9 | A | 1786 | SER | p.Ser1786Thr | Q13402 |
| HOMO SAPIENS | 4 | 16066 | 5mv9 | A | 1798 | GLN | p.Gln1798Ter | Q13402 |
| HOMO SAPIENS | 0 | 16066 | 5mv9 | A | 1858 | LEU | p.Leu1858Pro | Q13402 |
| HOMO SAPIENS | 6 | 16066 | 5mv9 | A | 1873 | ARG | p.Arg1873Trp | Q13402 |
| HOMO SAPIENS | 6 | 16066 | 5mv9 | A | 1873 | ARG | p.Arg1873Leu | Q13402 |
| HOMO SAPIENS | 6 | 16066 | 5mv9 | A | 1883 | ARG | p.Arg1883Gln | Q13402 |
| HOMO SAPIENS | 4 | 16066 | 5mv9 | A | 1887 | PRO | p.Pro1887Leu | Q13402 |

| | | | | | | | | |
|---|---|---|---|---|---|---|---|---|
| HOMO SAPIENS | 5 | 16066 | 5mv9 | A | 1942 | GLY | p.Gly1942Ter | Q13402 |
| HOMO SAPIENS | 5 | 16066 | 5mv9 | A | 1954 | LEU | p.Leu1954Ile | Q13402 |
| HOMO SAPIENS | 5 | 16066 | 5mv9 | A | 1956 | VAL | p.Val1956Ile | Q13402 |
| HOMO SAPIENS | 5 | 16066 | 5mv9 | A | 1967 | ARG | p.Arg1967Ter | Q13402 |
| HOMO SAPIENS | 5 | 16066 | 5mv9 | A | 1990 | GLN | p.Gln1990Ter | Q13402 |
| HOMO SAPIENS | 2 | 16066 | 5mv9 | A | 2005 | LYS | p.Lys2005Glu | Q13402 |
| HOMO SAPIENS | 1 | 16066 | 5mv9 | A | 2072 | VAL | p.Val2072Ile | Q13402 |
| HOMO SAPIENS | 4 | 16066 | 5mv9 | A | 2109 | THR | p.Thr2109Ile | Q13402 |
| HOMO SAPIENS | 4 | 16066 | 5mv9 | A | 2142 | ASP | p.Asp2142Asn | Q13402 |
| HOMO SAPIENS | -1 | 16066 | 5mv9 | A | 2214 | GLY | p.Gly2214Ser | Q13402 |
| HOMO SAPIENS | 7 | 27942 | 1deb | A | 24 | ARG | p.Arg24Ter | P25054 |
| HOMO SAPIENS | 0 | 876 | 5hnr | B | 12 | PHE | p.Phe12Leu | P13716 |
| HOMO SAPIENS | 0 | 876 | 5hnr | B | 59 | LYS | p.Lys59Asn | P13716 |
| HOMO SAPIENS | -1 | 876 | 5hnr | B | 133 | GLY | p.Gly133Arg | P13716 |
| HOMO SAPIENS | 2 | 876 | 5hnr | B | 240 | ARG | p.Arg240Trp | P13716 |
| HOMO SAPIENS | 0 | 876 | 5hnr | B | 274 | ALA | p.Ala274Thr | P13716 |
| HOMO SAPIENS | 0 | 876 | 5hnr | B | 275 | VAL | p.Val275Met | P13716 |
| HOMO SAPIENS | 0 | 1142 | 3qe2 | A | 503 | VAL | p.Ala500Val | P16435 |
| HOMO SAPIENS | 8 | 14358 | 4dkd | C | 32 | VAL | p.Val32Gly | P07333 |
| HOMO SAPIENS | 0 | 14358 | 4dkd | C | 60 | LEU | p.Leu60Val | P07333 |
| HOMO SAPIENS | 7 | 14358 | 4dkd | C | 75 | THR | p.Thr75Ile | P07333 |
| HOMO SAPIENS | -1 | 14358 | 4dkd | C | 90 | PRO | p.Pro90Thr | P07333 |
| HOMO SAPIENS | 0 | 14358 | 4dkd | C | 106 | ARG | p.Arg106Trp | P07333 |
| HOMO SAPIENS | 0 | 14358 | 4dkd | C | 241 | ASN | p.Asn241Asp | P07333 |
| HOMO SAPIENS | 0 | 14358 | 4dkd | C | 245 | ALA | p.Ala245Ser | P07333 |
| HOMO SAPIENS | 0 | 14358 | 4dkd | C | 255 | ASN | p.Asn255Ile | P07333 |
| HOMO SAPIENS | 0 | 14358 | 4dkd | C | 268 | VAL | p.Val268Ile | P07333 |
| HOMO SAPIENS | 0 | 14358 | 4dkd | C | 279 | VAL | p.Val279Met | P07333 |
| HOMO SAPIENS | 0 | 14358 | 4dkd | C | 294 | ARG | p.Arg294Gln | P07333 |
| HOMO SAPIENS | 5 | 8968 | 3ppy | A | 1565 | VAL | p.Val1565Leu | P04275 |
| HOMO SAPIENS | 4 | 8968 | 3ppy | A | 1584 | TYR | p.Tyr1584Cys | P04275 |
| HOMO SAPIENS | 5 | 5341 | 3k3n | A | 279 | PHE | p.Phe315Ser | Q9UPP1 |
| HOMO SAPIENS | 0 | 29463 | 1q7l | A | 197 | ARG | p.Arg197Trp | Q03154 |
| HOMO SAPIENS | 0 | 33761 | 2leh | B | 30 | ASP | p.Asp30Asn | Q16637 |
| HOMO SAPIENS | 6 | 33761 | 2leh | B | 44 | ASP | p.Asp44Val | Q16637 |
| HOMO SAPIENS | -1 | 12249 | 2pnr | C | 224 | GLU | p.Glu310Lys | P10515 |
| HOMO SAPIENS | -1 | 12249 | 2pnr | C | 232 | VAL | p.Val318Ala | P10515 |
| HOMO SAPIENS | -1 | 7 | 5whk | B | -9 | ALA | p.Ala11Pro | P61769 |
| HOMO SAPIENS | 8 | 20298 | 4euw | A | 154 | PHE | p.Phe154Leu | P48436 |
| HOMO SAPIENS | 9 | 20298 | 4euw | A | 158 | ALA | p.Ala158Thr | P48436 |
| HOMO SAPIENS | 5 | 20298 | 4euw | A | 165 | HIS | p.His165Tyr | P48436 |
| HOMO SAPIENS | 3 | 20298 | 4euw | A | 169 | HIS | p.His169Gln | P48436 |
| HOMO SAPIENS | 0 | 20298 | 4euw | A | 173 | LYS | p.Lys173Glu | P48436 |
| HOMO SAPIENS | 1 | 16465 | 2jgx | A | 402 | TYR | p.Tyr402His | P08603 |
| HOMO SAPIENS | 0 | 8993 | 3v8o | A | 659 | LEU | p.Leu659Arg | Q14315 |
| HOMO SAPIENS | 6 | 8993 | 3v8o | A | 693 | ASP | p.Asp693Ala | Q14315 |
| HOMO SAPIENS | 0 | 8993 | 3v8o | A | 709 | ALA | p.Ala709Thr | Q14315 |
| HOMO SAPIENS | 2 | 3852 | 5es4 | B | 106 | ASP | p.Asp128Asn | P05107 |
| HOMO SAPIENS | 2 | 3852 | 5es4 | B | 116 | SER | p.Ser138Pro | P05107 |
| HOMO SAPIENS | 1 | 3852 | 5es4 | B | 127 | LEU | p.Leu149Pro | P05107 |
| HOMO SAPIENS | 0 | 3852 | 5es4 | B | 147 | GLY | p.Gly169Arg | P05107 |
| HOMO SAPIENS | 3 | 3852 | 5es4 | B | 156 | PRO | p.Pro178Leu | P05107 |
| HOMO SAPIENS | 0 | 3852 | 5es4 | B | 174 | LYS | p.Lys196Thr | P05107 |
| HOMO SAPIENS | 2 | 3852 | 5es4 | B | 251 | GLY | p.Gly273Arg | P05107 |

| | | | | | | | | |
|---|---|---|---|---|---|---|---|---|
| HOMO SAPIENS | 2 | 3852 | 5es4 | B | 329 | ASN | p.Asn351Ser | P05107 |
| HOMO SAPIENS | 2 | 3852 | 5es4 | B | 571 | ARG | p.Arg593Cys | P05107 |
| HOMO SAPIENS | 7 | 5962 | 5fn2 | B | 146 | MET | p.Met146Leu | P49768 |
| HOMO SAPIENS | 3 | 5962 | 5fn2 | B | 166 | LEU | p.Leu166Pro | P49768 |
| HOMO SAPIENS | 1 | 5962 | 5fn2 | B | 233 | MET | p.Met233Leu | P49768 |
| HOMO SAPIENS | 1 | 5962 | 5fn2 | B | 261 | VAL | p.Val261Gly | P49768 |
| HOMO SAPIENS | 1 | 5962 | 5fn2 | B | 266 | GLY | p.Gly266Ser | P49768 |
| HOMO SAPIENS | -1 | 5962 | 5fn2 | B | 333 | ASP | p.Asp333Gly | P49768 |
| HOMO SAPIENS | 6 | 1918 | 3dlx | A | 58 | THR | p.Thr58Met | P55809 |
| HOMO SAPIENS | 5 | 1918 | 3dlx | A | 133 | VAL | p.Val133Glu | P55809 |
| HOMO SAPIENS | 5 | 1918 | 3dlx | A | 219 | GLY | p.Gly219Glu | P55809 |
| HOMO SAPIENS | 0 | 1918 | 3dlx | A | 221 | VAL | p.Val221Met | P55809 |
| HOMO SAPIENS | 5 | 1918 | 3dlx | A | 283 | SER | p.Ser283Ter | P55809 |
| HOMO SAPIENS | 4 | 1918 | 3dlx | A | 324 | GLY | p.Gly324Glu | P55809 |
| HOMO SAPIENS | 0 | 1918 | 3dlx | A | 456 | CYS | p.Cys456Phe | P55809 |
| HOMO SAPIENS | 0 | 9010 | 2wfr | A | 162 | LEU | p.Leu162Pro | O43323 |
| HOMO SAPIENS | 5 | 3770 | 5n38 | A | 56 | VAL | p.Val56Glu | O60260 |
| HOMO SAPIENS | 1 | 3770 | 5n38 | A | 161 | LYS | p.Lys161Asn | O60260 |
| HOMO SAPIENS | 1 | 3770 | 5n38 | A | 167 | SER | p.Ser167Asn | O60260 |
| HOMO SAPIENS | 1 | 3770 | 5n38 | A | 211 | LYS | p.Lys211Asn | O60260 |
| HOMO SAPIENS | 1 | 3770 | 5n38 | A | 212 | CYS | p.Cys212Tyr | O60260 |
| HOMO SAPIENS | 1 | 3770 | 5n38 | A | 275 | ARG | p.Arg275Trp | O60260 |
| HOMO SAPIENS | 2 | 3770 | 5n38 | A | 334 | ARG | p.Arg334Cys | O60260 |
| HOMO SAPIENS | 1 | 3770 | 5n38 | A | 366 | ARG | p.Arg366Trp | O60260 |
| HOMO SAPIENS | -1 | 3770 | 5n38 | A | 380 | VAL | p.Val380Leu | O60260 |
| HOMO SAPIENS | 1 | 3770 | 5n38 | A | 394 | ASP | p.Asp394Asn | O60260 |
| HOMO SAPIENS | 1 | 3770 | 5n38 | A | 430 | GLY | p.Gly430Asp | O60260 |
| HOMO SAPIENS | -1 | 2467 | 4xx1 | A | 10 | PRO | p.Pro34Leu | P04180 |
| HOMO SAPIENS | 4 | 2467 | 4xx1 | A | 83 | TYR | p.Tyr107Ter | P04180 |
| HOMO SAPIENS | 4 | 2467 | 4xx1 | A | 93 | ALA | p.Ala117Thr | P04180 |
| HOMO SAPIENS | 4 | 2467 | 4xx1 | A | 123 | THR | p.Thr147Ile | P04180 |
| HOMO SAPIENS | 4 | 2467 | 4xx1 | A | 135 | ARG | p.Arg159Trp | P04180 |
| HOMO SAPIENS | 3 | 2467 | 4xx1 | A | 158 | ARG | p.Arg182Cys | P04180 |
| HOMO SAPIENS | 0 | 2467 | 4xx1 | A | 209 | LEU | p.Leu233Pro | P04180 |
| HOMO SAPIENS | 1 | 2467 | 4xx1 | A | 228 | ASN | p.Asn252Lys | P04180 |
| HOMO SAPIENS | 4 | 2467 | 4xx1 | A | 252 | MET | p.Met276Lys | P04180 |
| HOMO SAPIENS | 3 | 2467 | 4xx1 | A | 293 | MET | p.Met317Ile | P04180 |
| HOMO SAPIENS | 5 | 2467 | 4xx1 | A | 321 | THR | p.Thr345Met | P04180 |
| HOMO SAPIENS | 9 | 2467 | 4xx1 | A | 347 | THR | p.Thr371Met | P04180 |
| HOMO SAPIENS | 1 | 5834 | 3qrf | G | 371 | PHE | p.Phe371Cys | Q9BZS1 |
| HOMO SAPIENS | 1 | 5834 | 3qrf | G | 373 | PHE | p.Phe373Ala | Q9BZS1 |
| HOMO SAPIENS | 9 | 5834 | 3qrf | G | 384 | ALA | p.Ala384Thr | Q9BZS1 |
| HOMO SAPIENS | 6 | 5834 | 3qrf | G | 397 | ARG | p.Arg397Trp | Q9BZS1 |
| HOMO SAPIENS | -1 | 11869 | 4km8 | A | 15 | PRO | p.Pro15Leu | Q9UMX1 |
| HOMO SAPIENS | 5 | 262 | 4kbo | A | 126 | ARG | p.Arg126Trp | P38646 |
| HOMO SAPIENS | 3 | 262 | 4kbo | A | 128 | TYR | p.Tyr128Cys | P38646 |
| HOMO SAPIENS | -1 | 11347 | 4uf9 | D | 117 | ALA | p.Ala117Val | Q13501 |
| HOMO SAPIENS | 0 | 2173 | 4ah6 | A | 45 | SER | p.Ser45Gly | Q6PI48 |
| HOMO SAPIENS | 2 | 2173 | 4ah6 | A | 152 | CYS | p.Cys152Phe | Q6PI48 |
| HOMO SAPIENS | 4 | 2173 | 4ah6 | A | 179 | ARG | p.Arg179His | Q6PI48 |
| HOMO SAPIENS | 0 | 2173 | 4ah6 | A | 196 | LYS | p.Lys196Arg | Q6PI48 |
| HOMO SAPIENS | 8 | 2173 | 4ah6 | A | 263 | ARG | p.Arg263Gln | Q6PI48 |
| HOMO SAPIENS | 1 | 2173 | 4ah6 | A | 338 | GLY | p.Gly338Glu | Q6PI48 |
| HOMO SAPIENS | 0 | 2173 | 4ah6 | A | 425 | GLU | p.Glu425Ter | Q6PI48 |

| | | | | | | | | |
|---|---|---|---|---|---|---|---|---|
| HOMO SAPIENS | 1 | 2173 | 4ah6 | A | 548 | LEU | p.Leu548Met | Q6PI48 |
| HOMO SAPIENS | 0 | 2173 | 4ah6 | A | 609 | ARG | p.Arg609Trp | Q6PI48 |
| HOMO SAPIENS | 2 | 2173 | 4ah6 | A | 613 | LEU | p.Leu613Phe | Q6PI48 |
| HOMO SAPIENS | 0 | 2173 | 4ah6 | A | 626 | LEU | p.Leu626Gln | Q6PI48 |
| HOMO SAPIENS | 0 | 2173 | 4ah6 | A | 626 | LEU | p.Leu626Val | Q6PI48 |
| HOMO SAPIENS | 7 | 3203 | 2cn1 | A | 87 | ASP | p.Asp137Val | Q9H0P0 |
| HOMO SAPIENS | 8 | 3203 | 2cn1 | A | 166 | GLN | p.Gln216Ter | Q9H0P0 |
| HOMO SAPIENS | 8 | 3203 | 2cn1 | A | 179 | ASN | p.Asn229Ser | Q9H0P0 |
| HOMO SAPIENS | 7 | 3203 | 2cn1 | A | 230 | GLY | p.Gly280Arg | Q9H0P0 |
| HOMO SAPIENS | 2 | 3424 | 2a1t | R | 116 | GLY | p.Gly116Arg | P13804 |
| HOMO SAPIENS | 2 | 3424 | 2a1t | R | 157 | VAL | p.Val157Gly | P13804 |
| HOMO SAPIENS | 2 | 3424 | 2a1t | R | 171 | THR | p.Thr171Ile | P13804 |
| HOMO SAPIENS | 0 | 3424 | 2a1t | R | 178 | THR | p.Thr178Arg | P13804 |
| HOMO SAPIENS | 0 | 3424 | 2a1t | R | 194 | VAL | p.Val194Met | P13804 |
| HOMO SAPIENS | 1 | 3424 | 2a1t | R | 209 | ARG | p.Arg209Ter | P13804 |
| HOMO SAPIENS | 6 | 3424 | 2a1t | R | 266 | THR | p.Thr266Met | P13804 |
| HOMO SAPIENS | 0 | 27598 | 1g1c | A | 26 | ARG | p.Arg2098Gly | Q8WZ42 |
| HOMO SAPIENS | 1 | 27598 | 1g1c | A | 46 | ILE | p.Ile2118Thr | Q8WZ42 |
| HOMO SAPIENS | 0 | 32885 | 4bsk | A | 149 | ASN | p.Asn149Asp | P35916 |
| HOMO SAPIENS | -1 | 8074 | 3ghg | B | 17 | ARG | p.Arg47Ter | P02675 |
| HOMO SAPIENS | -1 | 447 | 5g4p | A | 190 | ALA | p.Ala190Thr | P84074 |
| HOMO SAPIENS | 0 | 10761 | 5u1d | B | 379 | VAL | p.Val379Ile | Q03519 |
| HOMO SAPIENS | 0 | 10761 | 5u1d | B | 665 | THR | p.Thr665Ala | Q03519 |
| HOMO SAPIENS | 5 | 29251 | 5fn7 | A | 79 | ILE | p.Ile294Leu | P08575 |
| HOMO SAPIENS | -1 | 27727 | 4ky9 | A | 56 | LYS | p.Lys56Glu | P02730 |
| HOMO SAPIENS | 2 | 27727 | 4ky9 | A | 72 | GLU | p.Glu72Asp | P02730 |
| HOMO SAPIENS | 2 | 27727 | 4ky9 | A | 90 | GLU | p.Glu90Lys | P02730 |
| HOMO SAPIENS | 1 | 27727 | 4ky9 | A | 130 | GLY | p.Gly130Arg | P02730 |
| HOMO SAPIENS | 2 | 27727 | 4ky9 | A | 150 | ARG | p.Arg150Ter | P02730 |
| HOMO SAPIENS | 0 | 27727 | 4ky9 | A | 327 | PRO | p.Pro327Arg | P02730 |
| HOMO SAPIENS | 2 | 13202 | 2fxo | B | 870 | ARG | p.Arg870His | P12883 |
| HOMO SAPIENS | 2 | 13202 | 2fxo | B | 870 | ARG | p.Arg870Cys | P12883 |
| HOMO SAPIENS | 4 | 13202 | 2fxo | B | 904 | ARG | p.Arg904Cys | P12883 |
| HOMO SAPIENS | 4 | 13202 | 2fxo | B | 906 | ASP | p.Asp906Gly | P12883 |
| HOMO SAPIENS | 2 | 13202 | 2fxo | B | 908 | LEU | p.Leu908Val | P12883 |
| HOMO SAPIENS | 2 | 13202 | 2fxo | B | 923 | ASN | p.Asn923Lys | P12883 |
| HOMO SAPIENS | 7 | 13202 | 2fxo | B | 935 | GLU | p.Glu935Lys | P12883 |
| HOMO SAPIENS | 7 | 13202 | 2fxo | B | 935 | GLU | p.Glu935Val | P12883 |
| HOMO SAPIENS | 8 | 13202 | 2fxo | B | 949 | GLU | p.Glu949Lys | P12883 |
| HOMO SAPIENS | 0 | 13200 | 2a1s | A | 349 | ARG | p.Arg349Trp | O95453 |
| HOMO SAPIENS | 0 | 13200 | 2a1s | A | 421 | LYS | p.Lys421Arg | O95453 |
| HOMO SAPIENS | 6 | 3196 | 5xtb | T | 115 | CYS | p.Cys115Tyr | O75380 |
| HOMO SAPIENS | 0 | 1108 | 5ukf | A | 89 | ARG | p.Arg89Gln | Q99986 |
| HOMO SAPIENS | 0 | 1108 | 5ukf | A | 119 | HIS | p.His119Arg | Q99986 |
| HOMO SAPIENS | 0 | 1108 | 5ukf | A | 133 | ARG | p.Arg133Cys | Q99986 |
| HOMO SAPIENS | 0 | 1108 | 5ukf | A | 236 | VAL | p.Val236Leu | Q99986 |
| HOMO SAPIENS | 0 | 1108 | 5ukf | A | 321 | ARG | p.Arg321Cys | Q99986 |
| HOMO SAPIENS | -1 | 1108 | 5ukf | A | 358 | ARG | p.Arg358Ter | Q99986 |
| HOMO SAPIENS | 0 | 5842 | 4yli | C | 204 | GLY | p.Gly204Ser | Q9BWP8 |
| HOMO SAPIENS | -1 | 31362 | 4wqo | A | 25 | PRO | p.Pro25Leu | P40337 |
| HOMO SAPIENS | 2 | 10107 | 2j8h | A | 34 | VAL | p.Val31886Ile | Q8WZ42 |
| HOMO SAPIENS | 8 | 20190 | 1wm5 | A | 77 | ARG | p.Arg77Gln | P19878 |
| HOMO SAPIENS | 1 | 20190 | 1wm5 | A | 100 | GLN | p.Gln100Ter | P19878 |
| HOMO SAPIENS | 0 | 20190 | 1wm5 | A | 102 | ARG | p.Arg102Ter | P19878 |

| | | | | | | | | | |
|---|---|---|---|---|---|---|---|---|---|
| HOMO SAPIENS | 0 | 20190 | 1wm5 | A | | 128 | ALA | p.Ala128Val | P19878 |
| HOMO SAPIENS | 0 | 20190 | 1wm5 | A | | 181 | LYS | p.Lys181Arg | P19878 |
| HOMO SAPIENS | 7 | 1278 | 5a2q | a | | 33 | ASP | p.Asp33Asn | P62854 |
| HOMO SAPIENS | 3 | 19515 | 2obd | A | | 11 | ILE | p.Ile28Met | P11597 |
| HOMO SAPIENS | 3 | 19515 | 2obd | A | | 89 | VAL | p.Val106Met | P11597 |
| HOMO SAPIENS | 3 | 19515 | 2obd | A | | 137 | ARG | p.Arg154Trp | P11597 |
| HOMO SAPIENS | 3 | 19515 | 2obd | A | | 139 | ASP | p.Asp156Asn | P11597 |
| HOMO SAPIENS | 3 | 19515 | 2obd | A | | 204 | SER | p.Ser221Arg | P11597 |
| HOMO SAPIENS | 0 | 19515 | 2obd | A | | 297 | GLU | p.Glu314Lys | P11597 |
| HOMO SAPIENS | 0 | 19515 | 2obd | A | | 314 | GLY | p.Gly331Ser | P11597 |
| HOMO SAPIENS | 0 | 19515 | 2obd | A | | 332 | SER | p.Ser349Tyr | P11597 |
| HOMO SAPIENS | 0 | 19515 | 2obd | A | | 368 | VAL | p.Val385Met | P11597 |
| HOMO SAPIENS | 0 | 19515 | 2obd | A | | 373 | ALA | p.Ala390Pro | P11597 |
| HOMO SAPIENS | 3 | 19515 | 2obd | A | | 405 | ILE | p.Val422Ile | P11597 |
| HOMO SAPIENS | 0 | 19515 | 2obd | A | | 451 | ARG | p.Arg468Gln | P11597 |
| HOMO SAPIENS | 0 | 19515 | 2obd | A | | 451 | ARG | p.Arg468Ter | P11597 |
| HOMO SAPIENS | 0 | 6626 | 3hop | A | | 39 | ALA | p.Ala39Gly | P21333 |
| HOMO SAPIENS | 1 | 6626 | 3hop | A | | 82 | GLU | p.Glu82Val | P21333 |
| HOMO SAPIENS | 1 | 6626 | 3hop | A | | 128 | ALA | p.Ala128Val | P21333 |
| HOMO SAPIENS | 2 | 6626 | 3hop | A | | 203 | ASP | p.Asp203Tyr | P21333 |
| HOMO SAPIENS | 1 | 6626 | 3hop | A | | 207 | PRO | p.Pro207Leu | P21333 |
| HOMO SAPIENS | 2 | 6626 | 3hop | A | | 210 | CYS | p.Cys210Phe | P21333 |
| HOMO SAPIENS | 7 | 6626 | 3hop | A | | 254 | GLU | p.Glu254Lys | P21333 |
| HOMO SAPIENS | 4 | 4210 | 2avg | A | | 8 | VAL | p.Val158Met | Q14896 |
| HOMO SAPIENS | 6 | 4210 | 2avg | A | | 15 | GLU | p.Glu165Asp | Q14896 |
| HOMO SAPIENS | 4 | 4210 | 2avg | A | | 27 | ARG | p.Arg177His | Q14896 |
| HOMO SAPIENS | 0 | 4210 | 2avg | A | | 28 | VAL | p.Val178Met | Q14896 |
| HOMO SAPIENS | 6 | 4210 | 2avg | A | | 39 | VAL | p.Val189Ile | Q14896 |
| HOMO SAPIENS | 2 | 4210 | 2avg | A | | 52 | LYS | p.Lys202Gln | Q14896 |
| HOMO SAPIENS | 4 | 4210 | 2avg | A | | 67 | SER | p.Ser217Gly | Q14896 |
| HOMO SAPIENS | 6 | 4210 | 2avg | A | | 69 | VAL | p.Val219Leu | Q14896 |
| HOMO SAPIENS | 2 | 4210 | 2avg | A | | 86 | GLY | p.Ser236Gly | Q14896 |
| HOMO SAPIENS | 4 | 4210 | 2avg | A | | 108 | GLU | p.Glu258Lys | Q14896 |
| HOMO SAPIENS | 0 | 22237 | 1w6k | A | | 588 | GLY | p.Gly588Ser | P48449 |
| HOMO SAPIENS | 0 | 10765 | 4a5w | B | | 98 | GLU | p.Glu98Ala | P13671 |
| HOMO SAPIENS | 0 | 10765 | 4a5w | B | | 596 | ARG | p.Arg596Ter | P13671 |
| HOMO SAPIENS | -1 | 15515 | 3hx3 | A | | 11 | MSE | p.Met10Lys | P12271 |
| HOMO SAPIENS | 8 | 15515 | 3hx3 | A | | 64 | ARG | p.Arg64Gln | P12271 |
| HOMO SAPIENS | 0 | 15515 | 3hx3 | A | | 182 | PHE | p.Phe182Cys | P12271 |
| HOMO SAPIENS | 0 | 15515 | 3hx3 | A | | 216 | ARG | p.Arg216Gln | P12271 |
| HOMO SAPIENS | -1 | 15515 | 3hx3 | A | | 226 | MSE | p.Met226Lys | P12271 |
| HOMO SAPIENS | 0 | 15515 | 3hx3 | A | | 234 | TRP | p.Arg234Trp | P12271 |
| HOMO SAPIENS | 0 | 15515 | 3hx3 | A | | 273 | LEU | p.Leu273Phe | P12271 |
| HOMO SAPIENS | 0 | 15515 | 3hx3 | A | | 292 | THR | p.Thr292Met | P12271 |
| HOMO SAPIENS | 0 | 5152 | 2www | D | | 95 | GLN | p.Gln95Ter | Q8IVH4 |
| HOMO SAPIENS | 0 | 5152 | 2www | D | | 133 | GLN | p.Gln133Ter | Q8IVH4 |
| HOMO SAPIENS | 0 | 5152 | 2www | D | | 145 | ARG | p.Arg145Ter | Q8IVH4 |
| HOMO SAPIENS | 0 | 5152 | 2www | D | | 207 | TYR | p.Tyr207Cys | Q8IVH4 |
| HOMO SAPIENS | 8 | 5152 | 2www | D | | 330 | ARG | p.Arg330Ter | Q8IVH4 |
| HOMO SAPIENS | 0 | 5152 | 2www | D | | 363 | GLN | p.Gln363His | Q8IVH4 |
| HOMO SAPIENS | 1 | 1626 | 1ilr | | 1 | 105 | SER | p.Ser130Arg | P18510 |
| HOMO SAPIENS | -1 | 929 | 5loh | A | | 12 | GLY | p.Gly12Arg | Q96GX5 |
| HOMO SAPIENS | 9 | 3206 | 1x63 | A | | 21 | CYS | p.Cys104Arg | Q13642 |
| HOMO SAPIENS | 8 | 3206 | 1x63 | A | | 39 | TRP | p.Trp122Ser | Q13642 |

| | | | | | | | | |
|---|---|---|---|---|---|---|---|---|
| HOMO SAPIENS | 9 | 3206 | 1x63 | A | 40 | HIS | p.His123Tyr | Q13642 |
| HOMO SAPIENS | 9 | 3206 | 1x63 | A | 40 | HIS | p.His123Leu | Q13642 |
| HOMO SAPIENS | 9 | 3206 | 1x63 | A | 40 | HIS | p.His123Gln | Q13642 |
| HOMO SAPIENS | 3 | 3206 | 1x63 | A | 49 | CYS | p.Cys132Phe | Q13642 |
| HOMO SAPIENS | 3 | 3206 | 1x63 | A | 67 | CYS | p.Cys150Tyr | Q13642 |
| HOMO SAPIENS | 3 | 3206 | 1x63 | A | 70 | CYS | p.Cys153Arg | Q13642 |
| HOMO SAPIENS | 3 | 3206 | 1x63 | A | 70 | CYS | p.Cys153Tyr | Q13642 |
| HOMO SAPIENS | 5 | 1052 | 3ecs | E | 208 | ASN | p.Asn208Tyr | Q14232 |
| HOMO SAPIENS | 2 | 1052 | 3ecs | E | 275 | TYR | p.Tyr275Cys | Q14232 |
| HOMO SAPIENS | 0 | 24253 | 5i35 | A | 271 | ARG | p.Arg271Cys | Q8NI60 |
| HOMO SAPIENS | 0 | 24253 | 5i35 | A | 272 | GLY | p.Gly272Asp | Q8NI60 |
| HOMO SAPIENS | 0 | 24253 | 5i35 | A | 299 | ARG | p.Arg299Trp | Q8NI60 |
| HOMO SAPIENS | 1 | 24253 | 5i35 | A | 339 | ALA | p.Ala339Thr | Q8NI60 |
| HOMO SAPIENS | 0 | 24253 | 5i35 | A | 348 | ARG | p.Arg348Ter | Q8NI60 |
| HOMO SAPIENS | 0 | 24253 | 5i35 | A | 395 | ILE | p.Ile395Met | Q8NI60 |
| HOMO SAPIENS | 0 | 24253 | 5i35 | A | 396 | ASP | p.Asp396Glu | Q8NI60 |
| HOMO SAPIENS | 0 | 24253 | 5i35 | A | 480 | PHE | p.Phe480Leu | Q8NI60 |
| HOMO SAPIENS | 2 | 24253 | 5i35 | A | 511 | THR | p.Thr511Met | Q8NI60 |
| HOMO SAPIENS | 0 | 24253 | 5i35 | A | 514 | TYR | p.Tyr514Cys | Q8NI60 |
| HOMO SAPIENS | 0 | 24253 | 5i35 | A | 538 | ARG | p.Arg538Gln | Q8NI60 |
| HOMO SAPIENS | 0 | 24253 | 5i35 | A | 549 | GLY | p.Gly549Ser | Q8NI60 |
| HOMO SAPIENS | 0 | 24253 | 5i35 | A | 551 | GLU | p.Glu551Lys | Q8NI60 |
| HOMO SAPIENS | -1 | 24253 | 5i35 | A | 555 | MSE | p.Met555Ile | Q8NI60 |
| HOMO SAPIENS | 0 | 24253 | 5i35 | A | 568 | GLU | p.Glu568Ter | Q8NI60 |
| HOMO SAPIENS | 0 | 24253 | 5i35 | A | 602 | PRO | p.Pro602Arg | Q8NI60 |
| HOMO SAPIENS | -1 | 15505 | 4goj | C | 22 | GLY | p.Gly22Val | Q13432 |
| HOMO SAPIENS | 5 | 23902 | 2lrr | A | 765 | HIS | p.His765Gln | P38935 |
| HOMO SAPIENS | 1 | 29650 | 2vj2 | A | 234 | CYS | p.Cys234Tyr | P78504 |
| HOMO SAPIENS | 0 | 29650 | 2vj2 | A | 255 | TYR | p.Tyr255Ter | P78504 |
| HOMO SAPIENS | 3 | 29650 | 2vj2 | A | 274 | GLY | p.Gly274Asp | P78504 |
| HOMO SAPIENS | 1 | 105 | 5he0 | B | 77 | GLY | p.Gly77Ser | P62873 |
| HOMO SAPIENS | 2 | 105 | 5he0 | B | 80 | ILE | p.Ile80Thr | P62873 |
| HOMO SAPIENS | 3 | 4587 | 3rgh | A | 159 | ASP | p.Asp1159Ala | P21333 |
| HOMO SAPIENS | 3 | 4587 | 3rgh | A | 184 | ASP | p.Asp1184Glu | P21333 |
| HOMO SAPIENS | 2 | 4587 | 3rgh | A | 186 | SER | p.Ser1186Leu | P21333 |
| HOMO SAPIENS | 4 | 4587 | 3rgh | A | 188 | ALA | p.Ala1188Thr | P21333 |
| HOMO SAPIENS | 2 | 4587 | 3rgh | A | 199 | SER | p.Ser1199Leu | P21333 |
| HOMO SAPIENS | 0 | 20283 | 4n3w | A | 1278 | GLU | p.Glu1278Lys | Q92793 |
| HOMO SAPIENS | 1 | 14294 | 2d9q | B | 208 | MET | p.Met231Thr | Q99062 |
| HOMO SAPIENS | 6 | 14294 | 2d9q | B | 297 | ASP | p.Asp320Asn | Q99062 |
| HOMO SAPIENS | 0 | 11642 | 3bpt | A | 66 | ARG | p.Arg66Trp | Q6NVY1 |
| HOMO SAPIENS | 0 | 11642 | 3bpt | A | 122 | TYR | p.Tyr122Cys | Q6NVY1 |
| HOMO SAPIENS | 0 | 11642 | 3bpt | A | 345 | GLY | p.Gly345Ser | Q6NVY1 |
| HOMO SAPIENS | 1 | 3942 | 3sby | A | 60 | ASP | p.Asp60His | Q9Y4U1 |
| HOMO SAPIENS | 0 | 3942 | 3sby | A | 111 | ARG | p.Arg111Ter | Q9Y4U1 |
| HOMO SAPIENS | 1 | 3942 | 3sby | A | 116 | LEU | p.Leu116Pro | Q9Y4U1 |
| HOMO SAPIENS | 1 | 3942 | 3sby | A | 130 | TYR | p.Tyr130Cys | Q9Y4U1 |
| HOMO SAPIENS | 1 | 3942 | 3sby | A | 132 | ARG | p.Arg132Ter | Q9Y4U1 |
| HOMO SAPIENS | 1 | 3942 | 3sby | A | 147 | GLY | p.Gly147Ala | Q9Y4U1 |
| HOMO SAPIENS | 1 | 3942 | 3sby | A | 161 | ARG | p.Arg161Gln | Q9Y4U1 |
| HOMO SAPIENS | 1 | 3942 | 3sby | A | 161 | ARG | p.Arg161Ter | Q9Y4U1 |
| HOMO SAPIENS | 1 | 3942 | 3sby | A | 203 | TRP | p.Trp203Ter | Q9Y4U1 |
| HOMO SAPIENS | 1 | 3942 | 3sby | A | 203 | TRP | p.Trp203Ter | Q9Y4U1 |
| HOMO SAPIENS | 1 | 3942 | 3sby | A | 205 | TYR | p.Tyr205Ter | Q9Y4U1 |

| | | | | | | | | | |
|---|---|---|---|---|---|---|---|---|---|
| HOMO SAPIENS | 0 | 38340 | 2mbh | B | | 39 | MET | p.Met39Leu | Q13351 |
| HOMO SAPIENS | 2 | 7360 | 4k30 | A | | 430 | LEU | p.Leu430Pro | Q8N159 |
| HOMO SAPIENS | 2 | 7360 | 4k30 | A | | 433 | GLU | p.Glu433Asp | Q8N159 |
| HOMO SAPIENS | 7 | 7360 | 4k30 | A | | 484 | TRP | p.Trp484Arg | Q8N159 |
| HOMO SAPIENS | 1 | 41903 | 2xs6 | A | | 151 | PRO | p.Pro151Ser | O00459 |
| HOMO SAPIENS | 0 | 172 | 5glh | A | | 183 | ALA | p.Ala183Gly | P24530 |
| HOMO SAPIENS | 0 | 172 | 5glh | A | | 201 | ARG | p.Arg201Ter | P24530 |
| HOMO SAPIENS | 0 | 172 | 5glh | A | | 244 | THR | p.Thr244Met | P24530 |
| HOMO SAPIENS | 0 | 172 | 5glh | A | | 253 | ARG | p.Arg253Ter | P24530 |
| HOMO SAPIENS | 9 | 172 | 5glh | A | | 1017 | ILE | p.Leu17Phe | P00720 |
| HOMO SAPIENS | -1 | 3342 | 1ljm | A | | 1058 | HIS | p.His58Asn | Q01196 |
| HOMO SAPIENS | 4 | 3342 | 1ljm | A | | 1174 | ARG | p.Arg174Gln | Q01196 |
| HOMO SAPIENS | 5 | 25607 | 1p22 | C | | 32 | ASP | p.Asp32Ala | P35222 |
| HOMO SAPIENS | 5 | 25607 | 1p22 | C | | 32 | ASP | p.Asp32Tyr | P35222 |
| HOMO SAPIENS | -1 | 25607 | 1p22 | C | | 33 | SEP | p.Ser33Phe | P35222 |
| HOMO SAPIENS | 5 | 25607 | 1p22 | C | | 34 | GLY | p.Gly34Glu | P35222 |
| HOMO SAPIENS | -1 | 25607 | 1p22 | C | | 37 | SEP | p.Ser37Cys | P35222 |
| HOMO SAPIENS | -1 | 25607 | 1p22 | C | | 41 | THR | p.Thr41Ile | P35222 |
| HOMO SAPIENS | 1 | 48 | 4uow | | 9 | 60 | GLN | p.Arg34111His | Q8WZ42 |
| HOMO SAPIENS | 0 | 48 | 4uow | | 9 | 64 | LEU | p.Val34115Ala | Q8WZ42 |
| HOMO SAPIENS | 0 | 1127 | 5c01 | A | | 638 | ARG | p.Arg638Ter | P29597 |
| HOMO SAPIENS | 0 | 1127 | 5c01 | A | | 684 | ILE | p.Ile684Ser | P29597 |
| HOMO SAPIENS | 0 | 1127 | 5c01 | A | | 703 | ARG | p.Arg703Trp | P29597 |
| HOMO SAPIENS | 5 | 16389 | 2aq0 | B | | 128 | ALA | p.Ala860Asp | Q92889 |
| HOMO SAPIENS | 2 | 16389 | 2aq0 | B | | 141 | ILE | p.Ile873Val | Q92889 |
| HOMO SAPIENS | 5 | 16389 | 2aq0 | B | | 143 | GLU | p.Glu875Gly | Q92889 |
| HOMO SAPIENS | 0 | 16389 | 2aq0 | B | | 180 | GLY | p.Gly912Arg | Q92889 |
| HOMO SAPIENS | 0 | 15816 | 2xrc | A | | 89 | THR | p.Thr107Ala | P05156 |
| HOMO SAPIENS | 0 | 15816 | 2xrc | A | | 133 | ASN | p.Asn151Ser | P05156 |
| HOMO SAPIENS | 0 | 15816 | 2xrc | A | | 185 | THR | p.Thr203Ile | P05156 |
| HOMO SAPIENS | 0 | 15816 | 2xrc | A | | 225 | GLY | p.Gly243Asp | P05156 |
| HOMO SAPIENS | 0 | 15816 | 2xrc | A | | 243 | GLY | p.Gly261Ala | P05156 |
| HOMO SAPIENS | 0 | 15816 | 2xrc | A | | 269 | GLY | p.Gly287Arg | P05156 |
| HOMO SAPIENS | -1 | 15816 | 2xrc | A | | 288 | ILE | p.Ile306Val | P05156 |
| HOMO SAPIENS | -1 | 15816 | 2xrc | A | | 388 | ARG | p.Arg406His | P05156 |
| HOMO SAPIENS | 7 | 15816 | 2xrc | A | | 398 | ILE | p.Ile416Leu | P05156 |
| HOMO SAPIENS | 9 | 15816 | 2xrc | A | | 400 | HIS | p.His418Leu | P05156 |
| HOMO SAPIENS | 0 | 15816 | 2xrc | A | | 423 | LYS | p.Lys441Arg | P05156 |
| HOMO SAPIENS | -1 | 15816 | 2xrc | A | | 530 | GLU | p.Glu548Gln | P05156 |
| HOMO SAPIENS | 0 | 15816 | 2xrc | A | | 535 | PRO | p.Pro553Ala | P05156 |
| HOMO SAPIENS | 0 | 15816 | 2xrc | A | | 536 | GLU | p.Glu554Gly | P05156 |
| HOMO SAPIENS | 1 | 12385 | 4zlp | B | | 1450 | ALA | p.Ala1450Pro | Q9UM47 |
| HOMO SAPIENS | 7 | 12385 | 4zlp | B | | 1482 | GLY | p.Gly1482Arg | Q9UM47 |
| HOMO SAPIENS | 2 | 12385 | 4zlp | B | | 1518 | LEU | p.Leu1518Met | Q9UM47 |
| HOMO SAPIENS | 0 | 12385 | 4zlp | B | | 1560 | ARG | p.Arg1560Pro | Q9UM47 |
| HOMO SAPIENS | 0 | 1227 | 4a04 | A | | 148 | PHE | p.Phe148Tyr | O43435 |
| HOMO SAPIENS | 4 | 1227 | 4a04 | A | | 194 | HIS | p.His194Gln | O43435 |
| HOMO SAPIENS | 2 | 8017 | 3ei4 | B | | 244 | LYS | p.Lys244Glu | Q92466 |
| HOMO SAPIENS | 1 | 8017 | 3ei4 | B | | 273 | ARG | p.Arg273His | Q92466 |
| HOMO SAPIENS | 3 | 36112 | 2k8f | B | | 9 | SER | p.Ser9Arg | P04637 |
| HOMO SAPIENS | 2 | 12880 | 5ggf | A | | 265 | ARG | p.Arg265His | Q8WZA1 |
| HOMO SAPIENS | 2 | 12880 | 5ggf | A | | 287 | ILE | p.Ile287Ser | Q8WZA1 |
| HOMO SAPIENS | 2 | 12880 | 5ggf | A | | 311 | ARG | p.Arg311Gln | Q8WZA1 |
| HOMO SAPIENS | 2 | 12880 | 5ggf | A | | 425 | TRP | p.Trp425Ser | Q8WZA1 |

| | | | | | | | | |
|---|---|---|---|---|---|---|---|---|
| HOMO SAPIENS | 2 | 12880 | 5ggf | A | 442 | ARG | p.Arg442Cys | Q8WZA1 |
| HOMO SAPIENS | 2 | 12880 | 5ggf | A | 448 | GLY | p.Gly448Arg | Q8WZA1 |
| HOMO SAPIENS | 0 | 12880 | 5ggf | A | 490 | CYS | p.Cys490Tyr | Q8WZA1 |
| HOMO SAPIENS | 0 | 12880 | 5ggf | A | 493 | PRO | p.Pro493Arg | Q8WZA1 |
| HOMO SAPIENS | 3 | 12880 | 5ggf | A | 550 | SER | p.Ser550Asn | Q8WZA1 |
| HOMO SAPIENS | 1 | 12880 | 5ggf | A | 556 | ASP | p.Asp556Asn | Q8WZA1 |
| HOMO SAPIENS | 0 | 12880 | 5ggf | A | 580 | ARG | p.Arg580Ter | Q8WZA1 |
| HOMO SAPIENS | 2 | 12880 | 5ggf | A | 605 | ARG | p.Arg605Pro | Q8WZA1 |
| HOMO SAPIENS | 0 | 12880 | 5ggf | A | 623 | VAL | p.Met623Val | Q8WZA1 |
| HOMO SAPIENS | 1 | 10187 | 4qw2 | A | 145 | ALA | p.Ala145Ser | Q06787 |
| HOMO SAPIENS | 1 | 10187 | 4qw2 | A | 174 | ASN | p.Asn174Lys | Q06787 |
| HOMO SAPIENS | 0 | 24002 | 2m0t | A | 110 | LEU | p.Leu110Val | O14745 |
| HOMO SAPIENS | 9 | 3126 | 4mjh | A | 99 | LEU | p.Leu99Met | P04792 |
| HOMO SAPIENS | 1 | 3126 | 4mjh | A | 127 | ARG | p.Arg127Trp | P04792 |
| HOMO SAPIENS | 1 | 3126 | 4mjh | A | 127 | ARG | p.Arg127Leu | P04792 |
| HOMO SAPIENS | 1 | 3126 | 4mjh | A | 135 | SER | p.Ser135Phe | P04792 |
| HOMO SAPIENS | 2 | 3126 | 4mjh | A | 136 | ARG | p.Arg136Trp | P04792 |
| HOMO SAPIENS | 1 | 3126 | 4mjh | A | 140 | ARG | p.Arg140Gly | P04792 |
| HOMO SAPIENS | 0 | 3126 | 4mjh | A | 151 | THR | p.Thr151Ile | P04792 |
| HOMO SAPIENS | -1 | 15780 | 2jlp | A | 213 | ARG | p.Arg231Gly | P08294 |
| HOMO SAPIENS | 8 | 3578 | 1apz | D | 203 | GLY | p.Gly226Asp | P20933 |
| HOMO SAPIENS | 1 | 3578 | 1apz | D | 279 | GLY | p.Gly302Arg | P20933 |
| HOMO SAPIENS | 0 | 3578 | 1apz | D | 283 | CYS | p.Cys306Arg | P20933 |
| HOMO SAPIENS | 5 | 1335 | 5xtd | C | 115 | ARG | p.Arg115Cys | O75251 |
| HOMO SAPIENS | 0 | 1335 | 5xtd | C | 122 | VAL | p.Val122Met | O75251 |
| HOMO SAPIENS | 2 | 5063 | 2le4 | A | 37 | ARG | p.Arg74Pro | P48431 |
| HOMO SAPIENS | 1 | 25870 | 2o10 | A | 44 | LEU | p.Leu44Pro | P50461 |
| HOMO SAPIENS | 4 | 25870 | 2o10 | A | 58 | CYS | p.Cys58Gly | P50461 |
| HOMO SAPIENS | 0 | 18748 | 5fii | C | 39 | PHE | p.Phe39Leu | P00439 |
| HOMO SAPIENS | 6 | 18748 | 5fii | C | 48 | LEU | p.Leu48Ser | P00439 |
| HOMO SAPIENS | 2 | 18748 | 5fii | C | 65 | ILE | p.Ile65Asn | P00439 |
| HOMO SAPIENS | 3 | 18748 | 5fii | C | 95 | ILE | p.Ile95Phe | P00439 |
| HOMO SAPIENS | 1 | 14261 | 2vay | B | 1539 | ARG | p.Arg1539Cys | Q13698 |
| HOMO SAPIENS | 0 | 2494 | 4dx1 | A | 183 | GLU | p.Glu183Asp | Q9HBA0 |
| HOMO SAPIENS | 0 | 2494 | 4dx1 | A | 217 | ALA | p.Ala217Ser | Q9HBA0 |
| HOMO SAPIENS | 0 | 2494 | 4dx1 | A | 217 | ALA | p.Ala217Val | Q9HBA0 |
| HOMO SAPIENS | 4 | 2494 | 4dx1 | A | 224 | ARG | p.Arg224Trp | Q9HBA0 |
| HOMO SAPIENS | 4 | 2494 | 4dx1 | A | 254 | VAL | p.Val254Leu | Q9HBA0 |
| HOMO SAPIENS | 0 | 2494 | 4dx1 | A | 269 | ARG | p.Arg269Cys | Q9HBA0 |
| HOMO SAPIENS | 0 | 2494 | 4dx1 | A | 269 | ARG | p.Arg269His | Q9HBA0 |
| HOMO SAPIENS | 0 | 2494 | 4dx1 | A | 278 | GLU | p.Glu278Lys | Q9HBA0 |
| HOMO SAPIENS | 0 | 2494 | 4dx1 | A | 315 | ARG | p.Arg315Trp | Q9HBA0 |
| HOMO SAPIENS | 0 | 2494 | 4dx1 | A | 316 | ARG | p.Arg316Cys | Q9HBA0 |
| HOMO SAPIENS | 0 | 2494 | 4dx1 | A | 331 | ILE | p.Ile331Phe | Q9HBA0 |
| HOMO SAPIENS | 0 | 2494 | 4dx1 | A | 333 | ASP | p.Asp333Gly | Q9HBA0 |
| HOMO SAPIENS | 1 | 2494 | 4dx1 | A | 365 | VAL | p.Val365Met | Q9HBA0 |
| HOMO SAPIENS | 0 | 23998 | 2l1r | B | 161 | ARG | p.Arg162Trp | P19429 |
| HOMO SAPIENS | 0 | 8354 | 4f0x | D | 187 | SER | p.Ser187Ter | O95822 |
| HOMO SAPIENS | 2 | 8354 | 4f0x | D | 259 | GLY | p.Gly259Ala | O95822 |
| HOMO SAPIENS | 2 | 8354 | 4f0x | D | 284 | THR | p.Thr284Ala | O95822 |
| HOMO SAPIENS | 3 | 8354 | 4f0x | D | 309 | LYS | p.Lys309Gln | O95822 |
| HOMO SAPIENS | 0 | 21182 | 4apo | A | 228 | GLN | p.Gln228Lys | O00170 |
| HOMO SAPIENS | 1 | 21182 | 4apo | A | 307 | GLN | p.Gln307Arg | O00170 |
| HOMO SAPIENS | 0 | 2287 | 5of4 | B | 112 | ARG | p.Arg112His | P18074 |

| | | | | | | | | |
|---|---|---|---|---|---|---|---|---|
| HOMO SAPIENS | 0 | 2287 | 5of4 | B | 312 | ASP | p.Asp312Asn | P18074 |
| HOMO SAPIENS | 3 | 2287 | 5of4 | B | 461 | LEU | p.Leu461Val | P18074 |
| HOMO SAPIENS | 0 | 2287 | 5of4 | B | 485 | LEU | p.Leu485Pro | P18074 |
| HOMO SAPIENS | 1 | 2287 | 5of4 | B | 541 | SER | p.Ser541Arg | P18074 |
| HOMO SAPIENS | 0 | 2287 | 5of4 | B | 616 | ARG | p.Arg616Pro | P18074 |
| HOMO SAPIENS | 0 | 2287 | 5of4 | B | 616 | ARG | p.Arg616Trp | P18074 |
| HOMO SAPIENS | 1 | 2287 | 5of4 | B | 658 | ARG | p.Arg658Cys | P18074 |
| HOMO SAPIENS | 1 | 2287 | 5of4 | B | 681 | ASP | p.Asp681Asn | P18074 |
| HOMO SAPIENS | 1 | 2287 | 5of4 | B | 683 | ARG | p.Arg683Trp | P18074 |
| HOMO SAPIENS | 0 | 2287 | 5of4 | B | 713 | GLY | p.Gly713Arg | P18074 |
| HOMO SAPIENS | 0 | 2287 | 5of4 | B | 717 | ALA | p.Ala717Gly | P18074 |
| HOMO SAPIENS | 0 | 2287 | 5of4 | B | 722 | ARG | p.Arg722Trp | P18074 |
| HOMO SAPIENS | 0 | 2287 | 5of4 | B | 725 | ALA | p.Ala725Pro | P18074 |
| HOMO SAPIENS | 4 | 12188 | 4its | A | 122 | ARG | p.Arg122Gly | Q9Y606 |
| HOMO SAPIENS | 4 | 12188 | 4its | A | 133 | ASP | p.Asp133Asn | Q9Y606 |
| HOMO SAPIENS | 4 | 12188 | 4its | A | 144 | ARG | p.Arg144Trp | Q9Y606 |
| HOMO SAPIENS | 0 | 31529 | 1kgd | A | 761 | ARG | p.Arg766Gln | O14936 |
| HOMO SAPIENS | 1 | 31529 | 1kgd | A | 824 | GLN | p.Gln829Arg | O14936 |
| HOMO SAPIENS | 0 | 10906 | 2h63 | C | 18 | ARG | p.Arg18Ter | P53004 |
| HOMO SAPIENS | 9 | 27767 | 4wyq | A | 280 | LEU | p.Leu278Phe | Q9UPY3 |
| HOMO SAPIENS | 0 | 27767 | 4wyq | A | 297 | SER | p.Ser295Cys | Q9UPY3 |
| HOMO SAPIENS | 1 | 27767 | 4wyq | A | 334 | TYR | p.Tyr332Ter | Q9UPY3 |
| HOMO SAPIENS | 7 | 7775 | 5sv7 | C | 136 | SER | p.Ser136Cys | Q9NZJ5 |
| HOMO SAPIENS | 2 | 7775 | 5sv7 | C | 166 | GLN | p.Gln166Arg | Q9NZJ5 |
| HOMO SAPIENS | 2 | 7775 | 5sv7 | C | 332 | GLU | p.Glu332Ter | Q9NZJ5 |
| HOMO SAPIENS | 0 | 21449 | 2e9w | B | 84 | THR | p.Thr84Met | P10721 |
| HOMO SAPIENS | 0 | 21449 | 2e9w | B | 168 | ALA | p.Ala168Glu | P10721 |
| HOMO SAPIENS | 0 | 21449 | 2e9w | B | 178 | ALA | p.Ala178Thr | P10721 |
| HOMO SAPIENS | 0 | 21449 | 2e9w | B | 293 | ASN | p.Asn293Ser | P10721 |
| HOMO SAPIENS | -1 | 21449 | 2e9w | B | 318 | MET | p.Met318Leu | P10721 |
| HOMO SAPIENS | 0 | 21449 | 2e9w | B | 374 | VAL | p.Val374Ile | P10721 |
| HOMO SAPIENS | 6 | 3010 | 3flg | A | 66 | SER | p.Ser270Pro | Q9UBC3 |
| HOMO SAPIENS | 1 | 33918 | 2xzz | A | 760 | ARG | p.Arg760Ter | P22735 |
| HOMO SAPIENS | 2 | 1254 | 5xtc | w | 65 | ASN | p.Asn65Ser | O95299 |
| HOMO SAPIENS | 1 | 1254 | 5xtc | w | 238 | GLU | p.Glu238Lys | O95299 |
| HOMO SAPIENS | 3 | 10245 | 3siv | B | 194 | ALA | p.Ala194Glu | Q8WWY3 |
| HOMO SAPIENS | 4 | 10245 | 3siv | B | 216 | ALA | p.Ala216Pro | Q8WWY3 |
| HOMO SAPIENS | 0 | 2670 | 3dwb | A | 341 | THR | p.Thr341Ile | P42892 |
| HOMO SAPIENS | 1 | 2670 | 3dwb | A | 462 | SER | p.Ser462Asn | P42892 |
| HOMO SAPIENS | 1 | 2670 | 3dwb | A | 754 | ARG | p.Arg754Cys | P42892 |
| HOMO SAPIENS | 1 | 2961 | 5xtb | O | 40 | VAL | p.Val40Met | P19404 |
| HOMO SAPIENS | 5 | 2961 | 5xtb | O | 172 | ILE | p.Ile172Thr | P19404 |
| HOMO SAPIENS | 5 | 2961 | 5xtb | O | 190 | ASP | p.Asp190Gly | P19404 |
| HOMO SAPIENS | 5 | 29655 | 2xdg | A | 57 | ALA | p.Ala57Thr | Q02643 |
| HOMO SAPIENS | 4 | 29655 | 2xdg | A | 72 | GLU | p.Glu72Ter | Q02643 |
| HOMO SAPIENS | 0 | 29655 | 2xdg | A | 121 | GLU | p.Glu121Asp | Q02643 |
| HOMO SAPIENS | -1 | 17935 | 1hyn | P | 38 | ASP | p.Asp38Ala | P02730 |
| HOMO SAPIENS | 7 | 12825 | 1g82 | D | 99 | SER | p.Ser99Asn | P31371 |
| HOMO SAPIENS | 6 | 24927 | 1jm7 | B | 85 | VAL | p.Val85Leu | Q99728 |
| HOMO SAPIENS | 1 | 24927 | 1jm7 | B | 112 | ARG | p.Arg112Ter | Q99728 |
| HOMO SAPIENS | 0 | 24927 | 1jm7 | B | 116 | HIS | p.His116Tyr | Q99728 |
| HOMO SAPIENS | 7 | 92 | 5hde | A | 61 | LYS | p.Lys61Arg | Q05209 |
| HOMO SAPIENS | 0 | 17246 | 1szb | A | 84 | ARG | p.Arg99Gln | O00187 |
| HOMO SAPIENS | 1 | 17246 | 1szb | A | 105 | ASP | p.Asp120Gly | O00187 |

| | | | | | | | | |
|---|---|---|---|---|---|---|---|---|
| HOMO SAPIENS | 0 | 17246 | 1szb | A | 111 | PRO | p.Pro126Leu | O00187 |
| HOMO SAPIENS | 9 | 17246 | 1szb | A | 141 | CYS | p.Cys156Tyr | O00187 |
| HOMO SAPIENS | 0 | 32473 | 1b72 | A | 207 | ARG | p.Arg207Cys | P14653 |
| HOMO SAPIENS | 0 | 1920 | 4yzf | B | 488 | VAL | p.Val488Met | P02730 |
| HOMO SAPIENS | 0 | 1920 | 4yzf | B | 589 | ARG | p.Arg589Cys | P02730 |
| HOMO SAPIENS | 0 | 1920 | 4yzf | B | 589 | ARG | p.Arg589His | P02730 |
| HOMO SAPIENS | 0 | 1920 | 4yzf | B | 602 | ARG | p.Arg602Pro | P02730 |
| HOMO SAPIENS | 0 | 1920 | 4yzf | B | 613 | SER | p.Ser613Phe | P02730 |
| HOMO SAPIENS | -1 | 1920 | 4yzf | B | 646 | ARG | p.Arg646Gln | P02730 |
| HOMO SAPIENS | -1 | 1920 | 4yzf | B | 646 | ARG | p.Arg646Trp | P02730 |
| HOMO SAPIENS | 0 | 1920 | 4yzf | B | 701 | GLY | p.Gly701Asp | P02730 |
| HOMO SAPIENS | 0 | 1920 | 4yzf | B | 760 | ARG | p.Arg760Gln | P02730 |
| HOMO SAPIENS | 0 | 1920 | 4yzf | B | 771 | GLY | p.Gly771Asp | P02730 |
| HOMO SAPIENS | 0 | 1920 | 4yzf | B | 773 | SER | p.Ser773Pro | P02730 |
| HOMO SAPIENS | 0 | 1920 | 4yzf | B | 837 | THR | p.Thr837Ala | P02730 |
| HOMO SAPIENS | 0 | 1920 | 4yzf | B | 854 | PRO | p.Pro854Leu | P02730 |
| HOMO SAPIENS | 0 | 1920 | 4yzf | B | 858 | ALA | p.Ala858Asp | P02730 |
| HOMO SAPIENS | 0 | 1920 | 4yzf | B | 862 | VAL | p.Val862Ile | P02730 |
| HOMO SAPIENS | 0 | 1920 | 4yzf | B | 868 | PRO | p.Pro868Leu | P02730 |
| HOMO SAPIENS | 0 | 1920 | 4yzf | B | 870 | ARG | p.Arg870Trp | P02730 |
| HOMO SAPIENS | 0 | 9863 | 2juj | A | 54 | VAL | p.Val904Ile | P22681 |
| HOMO SAPIENS | 0 | 21934 | 2na7 | A | 184 | ILE | p.Ile184Val | P25445 |
| HOMO SAPIENS | -1 | 6746 | 1lw3 | A | 3 | THR | p.Lys3Thr | Q13614 |
| HOMO SAPIENS | -1 | 6746 | 1lw3 | A | 44 | VAL | p.Val44Ile | Q13614 |
| HOMO SAPIENS | 3 | 6746 | 1lw3 | A | 183 | ALA | p.Ala183Ser | Q13614 |
| HOMO SAPIENS | 4 | 6746 | 1lw3 | A | 482 | GLN | p.Gln482Ter | Q13614 |
| HOMO SAPIENS | 4 | 6746 | 1lw3 | A | 502 | GLU | p.Glu502Gln | Q13614 |
| HOMO SAPIENS | 3 | 6746 | 1lw3 | A | 545 | ASN | p.Asn545Ser | Q13614 |
| HOMO SAPIENS | 0 | 6746 | 1lw3 | A | 586 | ARG | p.Arg586Trp | Q13614 |
| HOMO SAPIENS | -1 | 6746 | 1lw3 | A | 602 | ALA | p.Ala602Gly | Q13614 |
| HOMO SAPIENS | 1 | 319 | 4ch9 | B | 336 | ARG | p.Arg336Ile | Q9UH77 |
| HOMO SAPIENS | 3 | 319 | 4ch9 | B | 340 | ALA | p.Ala340Val | Q9UH77 |
| HOMO SAPIENS | 0 | 319 | 4ch9 | B | 384 | ARG | p.Arg384Gln | Q9UH77 |
| HOMO SAPIENS | 5 | 319 | 4ch9 | B | 426 | PRO | p.Pro426Leu | Q9UH77 |
| HOMO SAPIENS | 3 | 319 | 4ch9 | B | 431 | ARG | p.Arg431Gln | Q9UH77 |
| HOMO SAPIENS | 2 | 319 | 4ch9 | B | 575 | ARG | p.Arg575Trp | Q9UH77 |
| HOMO SAPIENS | 0 | 574 | 2qkk | A | 142 | VAL | p.Val142Ile | O60930 |
| HOMO SAPIENS | 0 | 24021 | 2n8y | A | 752 | ARG | p.Arg752Gln | P12814 |
| HOMO SAPIENS | 0 | 18693 | 2y7j | B | 44 | ARG | p.Arg44Ter | P15735 |
| HOMO SAPIENS | 0 | 18693 | 2y7j | B | 106 | VAL | p.Val106Glu | P15735 |
| HOMO SAPIENS | 3 | 18693 | 2y7j | B | 189 | GLY | p.Gly189Glu | P15735 |
| HOMO SAPIENS | 0 | 17087 | 2ifg | A | 237 | THR | p.Thr237Met | P04629 |
| HOMO SAPIENS | 0 | 7980 | 1qub | A | 247 | VAL | p.Val266Leu | P02749 |
| HOMO SAPIENS | -1 | 2205 | 1h7u | A | 18 | ILE | p.Ile18Val | P54278 |
| HOMO SAPIENS | -1 | 2205 | 1h7u | A | 20 | ARG | p.Arg20Gln | P54278 |
| HOMO SAPIENS | 9 | 2205 | 1h7u | A | 46 | SER | p.Ser46Ile | P54278 |
| HOMO SAPIENS | 0 | 2205 | 1h7u | A | 60 | ASP | p.Asp60Glu | P54278 |
| HOMO SAPIENS | 0 | 2205 | 1h7u | A | 80 | PHE | p.Phe80Leu | P54278 |
| HOMO SAPIENS | 0 | 2205 | 1h7u | A | 118 | SER | p.Ser118Arg | P54278 |
| HOMO SAPIENS | 0 | 2205 | 1h7u | A | 126 | HIS | p.His126Gln | P54278 |
| HOMO SAPIENS | 0 | 2205 | 1h7u | A | 128 | SER | p.Ser128Leu | P54278 |
| HOMO SAPIENS | 1 | 2205 | 1h7u | A | 134 | ARG | p.Arg134Ter | P54278 |
| HOMO SAPIENS | 0 | 2205 | 1h7u | A | 166 | LEU | p.Leu166Pro | P54278 |
| HOMO SAPIENS | 1 | 2205 | 1h7u | A | 191 | TYR | p.Tyr191Cys | P54278 |

| | | | | | | | | |
|---|---|---|---|---|---|---|---|---|
| HOMO SAPIENS | 0 | 2205 | 1h7u | A | 211 | ARG | p.Arg211Ter | P54278 |
| HOMO SAPIENS | 0 | 2205 | 1h7u | A | 233 | GLN | p.Gln233Ter | P54278 |
| HOMO SAPIENS | 0 | 2205 | 1h7u | A | 255 | TYR | p.Tyr255Ter | P54278 |
| HOMO SAPIENS | 0 | 2205 | 1h7u | A | 277 | THR | p.Thr277Lys | P54278 |
| HOMO SAPIENS | 0 | 2205 | 1h7u | A | 301 | LYS | p.Lys301Asn | P54278 |
| HOMO SAPIENS | 0 | 2205 | 1h7u | A | 312 | MET | p.Met312Ile | P54278 |
| HOMO SAPIENS | 0 | 2205 | 1h7u | A | 312 | MET | p.Met312Thr | P54278 |
| HOMO SAPIENS | 0 | 2205 | 1h7u | A | 315 | ARG | p.Arg315Ter | P54278 |
| HOMO SAPIENS | 0 | 2205 | 1h7u | A | 318 | TYR | p.Tyr318Cys | P54278 |
| HOMO SAPIENS | 0 | 2205 | 1h7u | A | 335 | ASN | p.Asn335Ile | P54278 |
| HOMO SAPIENS | 0 | 2205 | 1h7u | A | 360 | ILE | p.Ile360Met | P54278 |
| HOMO SAPIENS | 6 | 20754 | 5mpo | B | 7 | VAL | p.Val7Phe | O96033 |
| HOMO SAPIENS | 4 | 20754 | 5mpo | B | 64 | VAL | p.Val64Ile | O96033 |
| HOMO SAPIENS | 5 | 34944 | 4qam | B | 1114 | ASP | p.Asp1114Gly | Q96KN7 |
| HOMO SAPIENS | 5 | 34944 | 4qam | B | 1240 | GLY | p.Gly1240Glu | Q96KN7 |
| HOMO SAPIENS | 9 | 40496 | 3vnn | A | 278 | ARG | p.Arg278His | P49917 |
| HOMO SAPIENS | 6 | 30131 | 4dt7 | E | 40 | PRO | p.Pro210Leu | P04070 |
| HOMO SAPIENS | -1 | 30131 | 4dt7 | E | 50 | ARG | p.Arg220Trp | P04070 |
| HOMO SAPIENS | 7 | 15311 | 2pqt | A | 64 | ARG | p.Arg64Trp | P18440 |
| HOMO SAPIENS | 2 | 15311 | 2pqt | A | 149 | VAL | p.Val149Ile | P18440 |
| HOMO SAPIENS | -1 | 19581 | 3hy8 | A | 33 | ASP | p.Asp33Val | Q9NVS9 |
| HOMO SAPIENS | 5 | 19581 | 3hy8 | A | 116 | ARG | p.Arg116Gln | Q9NVS9 |
| HOMO SAPIENS | 4 | 19581 | 3hy8 | A | 225 | ARG | p.Arg225His | Q9NVS9 |
| HOMO SAPIENS | 2 | 19581 | 3hy8 | A | 229 | TRP | p.Arg229Trp | Q9NVS9 |
| HOMO SAPIENS | 2 | 19581 | 3hy8 | A | 229 | TRP | p.Arg229Gln | Q9NVS9 |
| HOMO SAPIENS | 0 | 10266 | 3t1i | A | 41 | ASP | p.Asp41His | P49959 |
| HOMO SAPIENS | 1 | 10266 | 3t1i | A | 305 | ARG | p.Arg305Trp | P49959 |
| HOMO SAPIENS | 0 | 10266 | 3t1i | A | 308 | PHE | p.Phe308Leu | P49959 |
| HOMO SAPIENS | 0 | 10266 | 3t1i | A | 334 | SER | p.Ser334Arg | P49959 |
| HOMO SAPIENS | 1 | 1600 | 3rbf | A | 61 | GLU | p.Glu61Asp | P20711 |
| HOMO SAPIENS | 0 | 1600 | 3rbf | A | 91 | ALA | p.Ala91Val | P20711 |
| HOMO SAPIENS | -1 | 1600 | 3rbf | A | 102 | GLY | p.Gly102Ser | P20711 |
| HOMO SAPIENS | 3 | 1600 | 3rbf | A | 147 | SER | p.Ser147Arg | P20711 |
| HOMO SAPIENS | 1 | 1600 | 3rbf | A | 210 | PRO | p.Pro210Leu | P20711 |
| HOMO SAPIENS | 0 | 1600 | 3rbf | A | 217 | MET | p.Met217Val | P20711 |
| HOMO SAPIENS | 4 | 1600 | 3rbf | A | 250 | SER | p.Ser250Phe | P20711 |
| HOMO SAPIENS | 2 | 1600 | 3rbf | A | 275 | ALA | p.Ala275Thr | P20711 |
| HOMO SAPIENS | 1 | 1600 | 3rbf | A | 309 | PHE | p.Phe309Leu | P20711 |
| HOMO SAPIENS | -1 | 1600 | 3rbf | A | 347 | ARG | p.Arg347Gln | P20711 |
| HOMO SAPIENS | 0 | 1600 | 3rbf | A | 462 | ARG | p.Arg462Gln | P20711 |
| HOMO SAPIENS | 6 | 20830 | 5y4f | A | 714 | THR | p.Thr715Ile | Q01484 |
| HOMO SAPIENS | 7 | 20830 | 5y4f | A | 719 | ASP | p.Asp720Glu | Q01484 |
| HOMO SAPIENS | 0 | 20830 | 5y4f | A | 814 | VAL | p.Val815Ile | Q01484 |
| HOMO SAPIENS | -1 | 302 | 2c9y | A | 9 | GLU | p.Glu9Ter | P54819 |
| HOMO SAPIENS | 0 | 302 | 2c9y | A | 103 | ARG | p.Arg103Trp | P54819 |
| HOMO SAPIENS | 3 | 302 | 2c9y | A | 165 | ASP | p.Asp165Gly | P54819 |
| HOMO SAPIENS | 0 | 302 | 2c9y | A | 182 | ALA | p.Ala182Asp | P54819 |
| HOMO SAPIENS | 2 | 302 | 2c9y | A | 186 | ARG | p.Arg186Cys | P54819 |
| HOMO SAPIENS | 0 | 302 | 2c9y | A | 209 | ALA | p.Ala209Thr | P54819 |
| HOMO SAPIENS | -1 | 16920 | 1jlj | B | 10 | ASN | p.Asn10Tyr | Q9NQX3 |
| HOMO SAPIENS | 9 | 8462 | 1zbq | B | 16 | GLY | p.Gly16Ser | P51659 |
| HOMO SAPIENS | 5 | 8462 | 1zbq | B | 19 | ALA | p.Ala19Gly | P51659 |
| HOMO SAPIENS | 1 | 8462 | 1zbq | B | 34 | ALA | p.Ala34Val | P51659 |
| HOMO SAPIENS | 1 | 8462 | 1zbq | B | 106 | ARG | p.Arg106His | P51659 |

| | | | | | | | | |
|---|---|---|---|---|---|---|---|---|
| HOMO SAPIENS | 0 | 8462 | 1zbq | B | 140 | LYS | p.Lys140Asn | P51659 |
| HOMO SAPIENS | 0 | 8462 | 1zbq | B | 292 | THR | p.Thr292Ser | P51659 |
| HOMO SAPIENS | 0 | 4375 | 4ndi | A | 199 | ARG | p.Arg213His | Q7Z2E3 |
| HOMO SAPIENS | 3 | 4375 | 4ndi | A | 201 | HIS | p.His215Arg | Q7Z2E3 |
| HOMO SAPIENS | 0 | 4375 | 4ndi | A | 206 | PRO | p.Pro220Leu | Q7Z2E3 |
| HOMO SAPIENS | 2 | 4375 | 4ndi | A | 223 | LEU | p.Leu237Phe | Q7Z2E3 |
| HOMO SAPIENS | 0 | 4375 | 4ndi | A | 245 | ARG | p.Arg259His | Q7Z2E3 |
| HOMO SAPIENS | 0 | 4375 | 4ndi | A | 263 | VAL | p.Val277Gly | Q7Z2E3 |
| HOMO SAPIENS | 0 | 4375 | 4ndi | A | 279 | TRP | p.Trp293Ter | Q7Z2E3 |
| HOMO SAPIENS | 0 | 27004 | 4ccg | Y | 321 | ASP | p.Asp321Glu | Q9NW38 |
| HOMO SAPIENS | -1 | 27004 | 4ccg | Y | 372 | GLY | p.Gly372Ala | Q9NW38 |
| HOMO SAPIENS | -1 | 6383 | 5fai | A | 34 | ALA | p.Ala34Gly | Q92979 |
| HOMO SAPIENS | 8 | 6383 | 5fai | A | 86 | ASP | p.Asp86Gly | Q92979 |
| HOMO SAPIENS | 2 | 24585 | 4mi5 | A | 684 | ARG | p.Arg679Cys | Q15910 |
| HOMO SAPIENS | 0 | 15967 | 3nhb | A | 723 | ARG | p.Arg723Gln | Q9NP58 |
| HOMO SAPIENS | 5 | 18152 | 1oey | A | 389 | HIS | p.His389Gln | P19878 |
| HOMO SAPIENS | 5 | 18152 | 1oey | A | 395 | ARG | p.Arg395Trp | P19878 |
| HOMO SAPIENS | -1 | 1044 | 1li1 | F | 2 | VAL | p.Val1486Ile | P08572 |
| HOMO SAPIENS | 6 | 1044 | 1li1 | F | 37 | GLU | p.Glu1521Lys | P08572 |
| HOMO SAPIENS | 5 | 1044 | 1li1 | F | 157 | SER | p.Ser1639Leu | P08572 |
| HOMO SAPIENS | 5 | 1044 | 1li1 | F | 179 | ARG | p.Arg1662His | P08572 |
| HOMO SAPIENS | 5 | 1044 | 1li1 | F | 180 | GLY | p.Gly1663Arg | P08572 |
| HOMO SAPIENS | 6 | 20076 | 4uyi | A | 671 | LEU | p.Leu671Ser | Q8IY92 |
| HOMO SAPIENS | 0 | 19588 | 1jmj | A | 189 | ARG | p.Arg208His | P05546 |
| HOMO SAPIENS | -1 | 38023 | 1ry7 | B | 65 | GLY | p.Gly65Arg | P22607 |
| HOMO SAPIENS | -1 | 38023 | 1ry7 | B | 139 | ASP | p.Asp139Glu | P22607 |
| HOMO SAPIENS | 0 | 38023 | 1ry7 | B | 248 | ARG | p.Arg248Cys | P22607 |
| HOMO SAPIENS | 0 | 38023 | 1ry7 | B | 249 | SER | p.Ser249Cys | P22607 |
| HOMO SAPIENS | 0 | 38023 | 1ry7 | B | 250 | PRO | p.Pro250Arg | P22607 |
| HOMO SAPIENS | 0 | 38023 | 1ry7 | B | 322 | GLU | p.Glu322Lys | P22607 |
| HOMO SAPIENS | 5 | 16884 | 5eli | A | 33 | GLN | p.Gln33Ter | Q9NZC2 |
| HOMO SAPIENS | 0 | 16884 | 5eli | A | 44 | TRP | p.Trp44Ter | Q9NZC2 |
| HOMO SAPIENS | 0 | 16884 | 5eli | A | 47 | ARG | p.Arg47His | Q9NZC2 |
| HOMO SAPIENS | 0 | 16884 | 5eli | A | 66 | THR | p.Thr66Met | Q9NZC2 |
| HOMO SAPIENS | 0 | 16884 | 5eli | A | 78 | TRP | p.Trp78Ter | Q9NZC2 |
| HOMO SAPIENS | 1 | 16884 | 5eli | A | 96 | THR | p.Thr96Lys | Q9NZC2 |
| HOMO SAPIENS | 9 | 16884 | 5eli | A | 126 | VAL | p.Val126Gly | Q9NZC2 |
| HOMO SAPIENS | -1 | 12267 | 3w5o | A | 3 | ALA | p.Ala3Val | P49917 |
| HOMO SAPIENS | 1 | 12267 | 3w5o | A | 9 | THR | p.Thr9Ile | P49917 |
| HOMO SAPIENS | 2 | 12267 | 3w5o | A | 188 | ARG | p.Arg188Gln | P49917 |
| HOMO SAPIENS | 0 | 12267 | 3w5o | A | 469 | GLY | p.Gly469Glu | P49917 |
| HOMO SAPIENS | 1 | 12267 | 3w5o | A | 580 | ARG | p.Arg580Ter | P49917 |
| HOMO SAPIENS | 0 | 12267 | 3w5o | A | 600 | GLU | p.Glu600Lys | P49917 |
| HOMO SAPIENS | 9 | 2839 | 2hw5 | B | 54 | ARG | p.Arg54Cys | P30084 |
| HOMO SAPIENS | 9 | 2839 | 2hw5 | B | 59 | ASN | p.Asn59Ser | P30084 |
| HOMO SAPIENS | 0 | 2839 | 2hw5 | B | 273 | LYS | p.Lys273Glu | P30084 |
| HOMO SAPIENS | 1 | 12266 | 2o2k | B | 939 | GLY | p.Gly939Cys | Q99707 |
| HOMO SAPIENS | 5 | 12266 | 2o2k | B | 1164 | ARG | p.Arg1164His | Q99707 |
| HOMO SAPIENS | 1 | 12266 | 2o2k | B | 1166 | LEU | p.Leu1166Met | Q99707 |
| HOMO SAPIENS | 5 | 12266 | 2o2k | B | 1173 | PRO | p.Pro1173Leu | Q99707 |
| HOMO SAPIENS | 5 | 12266 | 2o2k | B | 1222 | ASN | p.Asn1222Ser | Q99707 |
| HOMO SAPIENS | 0 | 30891 | 3ch4 | B | 138 | ARG | p.Arg138Ter | Q15126 |
| HOMO SAPIENS | -1 | 4433 | 1qlz | A | 54 | GLY | p.Gly54Ser | P04156 |
| HOMO SAPIENS | 0 | 22568 | 3cxl | A | 20 | LEU | p.Leu20Phe | P15882 |

| | | | | | | | | |
|---|---|---|---|---|---|---|---|---|
| HOMO SAPIENS | 0 | 22568 | 3cxl | A | 126 | ILE | p.Ile126Met | P15882 |
| HOMO SAPIENS | 0 | 22568 | 3cxl | A | 141 | PRO | p.Pro141Leu | P15882 |
| HOMO SAPIENS | 0 | 22568 | 3cxl | A | 143 | TYR | p.Tyr143His | P15882 |
| HOMO SAPIENS | 0 | 22568 | 3cxl | A | 223 | ALA | p.Ala223Val | P15882 |
| HOMO SAPIENS | 0 | 22568 | 3cxl | A | 228 | GLY | p.Gly228Ser | P15882 |
| HOMO SAPIENS | 0 | 22568 | 3cxl | A | 252 | PRO | p.Pro252Gln | P15882 |
| HOMO SAPIENS | 0 | 22568 | 3cxl | A | 313 | GLU | p.Glu313Lys | P15882 |
| HOMO SAPIENS | 3 | 2668 | 5eqg | A | 34 | ASN | p.Asn34Ile | P11166 |
| HOMO SAPIENS | 3 | 2668 | 5eqg | A | 77 | MET | p.Met77Val | P11166 |
| HOMO SAPIENS | 6 | 2668 | 5eqg | A | 91 | GLY | p.Gly91Asp | P11166 |
| HOMO SAPIENS | 6 | 2668 | 5eqg | A | 92 | ARG | p.Arg92Trp | P11166 |
| HOMO SAPIENS | 6 | 2668 | 5eqg | A | 93 | ARG | p.Arg93Trp | P11166 |
| HOMO SAPIENS | 0 | 2668 | 5eqg | A | 95 | SER | p.Ser95Ile | P11166 |
| HOMO SAPIENS | 3 | 2668 | 5eqg | A | 126 | ARG | p.Arg126His | P11166 |
| HOMO SAPIENS | 3 | 2668 | 5eqg | A | 126 | ARG | p.Arg126Cys | P11166 |
| HOMO SAPIENS | 4 | 2668 | 5eqg | A | 232 | ARG | p.Arg232Cys | P11166 |
| HOMO SAPIENS | 3 | 2668 | 5eqg | A | 249 | ARG | p.Arg249Gln | P11166 |
| HOMO SAPIENS | 3 | 2668 | 5eqg | A | 255 | LYS | p.Lys255Arg | P11166 |
| HOMO SAPIENS | 0 | 2668 | 5eqg | A | 275 | ALA | p.Ala275Thr | P11166 |
| HOMO SAPIENS | 4 | 2668 | 5eqg | A | 299 | GLU | p.Glu299Lys | P11166 |
| HOMO SAPIENS | 6 | 2668 | 5eqg | A | 314 | GLY | p.Gly314Ser | P11166 |
| HOMO SAPIENS | 7 | 2668 | 5eqg | A | 315 | ILE | p.Ile315Met | P11166 |
| HOMO SAPIENS | -1 | 2668 | 5eqg | A | 458 | ARG | p.Arg458Trp | P11166 |
| HOMO SAPIENS | -1 | 2668 | 5eqg | A | 468 | ARG | p.Arg468Trp | P11166 |
| HOMO SAPIENS | 3 | 2291 | 2amy | A | 31 | PHE | p.Phe31Leu | O15305 |
| HOMO SAPIENS | 7 | 2291 | 2amy | A | 32 | LEU | p.Leu32Arg | O15305 |
| HOMO SAPIENS | 7 | 2291 | 2amy | A | 65 | ASP | p.Asp65Tyr | O15305 |
| HOMO SAPIENS | 1 | 2291 | 2amy | A | 108 | ALA | p.Ala108Val | O15305 |
| HOMO SAPIENS | 3 | 2291 | 2amy | A | 113 | PRO | p.Pro113Leu | O15305 |
| HOMO SAPIENS | 3 | 2291 | 2amy | A | 113 | PRO | p.Pro113Ser | O15305 |
| HOMO SAPIENS | 6 | 2291 | 2amy | A | 117 | GLY | p.Gly117Arg | O15305 |
| HOMO SAPIENS | 6 | 2291 | 2amy | A | 119 | PHE | p.Phe119Leu | O15305 |
| HOMO SAPIENS | 6 | 2291 | 2amy | A | 123 | ARG | p.Arg123Gln | O15305 |
| HOMO SAPIENS | 0 | 2291 | 2amy | A | 129 | VAL | p.Val129Met | O15305 |
| HOMO SAPIENS | 7 | 2291 | 2amy | A | 139 | GLU | p.Glu139Lys | O15305 |
| HOMO SAPIENS | 5 | 2291 | 2amy | A | 141 | ARG | p.Arg141His | O15305 |
| HOMO SAPIENS | 7 | 2291 | 2amy | A | 144 | PHE | p.Phe144Leu | O15305 |
| HOMO SAPIENS | 6 | 2291 | 2amy | A | 148 | ASP | p.Asp148Asn | O15305 |
| HOMO SAPIENS | 6 | 2291 | 2amy | A | 153 | ILE | p.Ile153Thr | O15305 |
| HOMO SAPIENS | 2 | 2291 | 2amy | A | 157 | PHE | p.Phe157Ser | O15305 |
| HOMO SAPIENS | 1 | 2291 | 2amy | A | 162 | ARG | p.Arg162Trp | O15305 |
| HOMO SAPIENS | 6 | 2291 | 2amy | A | 188 | ASP | p.Asp188Gly | O15305 |
| HOMO SAPIENS | 5 | 2291 | 2amy | A | 194 | ARG | p.Arg194Ter | O15305 |
| HOMO SAPIENS | 0 | 2291 | 2amy | A | 207 | PHE | p.Phe207Ser | O15305 |
| HOMO SAPIENS | 5 | 2291 | 2amy | A | 208 | GLY | p.Gly208Ala | O15305 |
| HOMO SAPIENS | 1 | 2291 | 2amy | A | 216 | ASN | p.Asn216Ile | O15305 |
| HOMO SAPIENS | 5 | 2291 | 2amy | A | 218 | HIS | p.His218Leu | O15305 |
| HOMO SAPIENS | 6 | 2291 | 2amy | A | 223 | ASP | p.Asp223Glu | O15305 |
| HOMO SAPIENS | 6 | 2291 | 2amy | A | 226 | THR | p.Thr226Ser | O15305 |
| HOMO SAPIENS | 5 | 2291 | 2amy | A | 231 | VAL | p.Val231Met | O15305 |
| HOMO SAPIENS | 1 | 2291 | 2amy | A | 237 | THR | p.Thr237Met | O15305 |
| HOMO SAPIENS | 1 | 2291 | 2amy | A | 241 | CYS | p.Cys241Ser | O15305 |
| HOMO SAPIENS | 5 | 35907 | 1d6g | A | 21 | GLY | p.Gly21Arg | P32238 |
| HOMO SAPIENS | -1 | 33209 | 1x9d | A | 197 | PRO | p.Pro197Leu | Q9UKM7 |

| | | | | | | | | |
|---|---|---|---|---|---|---|---|---|
| HOMO SAPIENS | -1 | 33209 | 1x9d | A | 201 | PRO | p.Pro201Leu | Q9UKM7 |
| HOMO SAPIENS | 6 | 21305 | 4yep | B | 1326 | ARG | p.Arg1326Ter | P24043 |
| HOMO SAPIENS | 6 | 21305 | 4yep | B | 1350 | ARG | p.Arg1350Ter | P24043 |
| HOMO SAPIENS | 0 | 21553 | 3qyb | A | 1120 | VAL | p.Val1119Ala | O60343 |
| HOMO SAPIENS | 2 | 21553 | 3qyb | A | 1148 | THR | p.Thr1147Met | O60343 |
| HOMO SAPIENS | -1 | 30645 | 3m0c | C | -17 | TRP | p.Trp4Ter | P01130 |
| HOMO SAPIENS | -1 | 30645 | 3m0c | C | -17 | TRP | p.Trp4Ter | P01130 |
| HOMO SAPIENS | -1 | 30645 | 3m0c | C | 12 | GLN | p.Gln33Ter | P01130 |
| HOMO SAPIENS | -1 | 30645 | 3m0c | C | 15 | ASP | p.Asp36Glu | P01130 |
| HOMO SAPIENS | -1 | 30645 | 3m0c | C | 28 | SER | p.Ser49Arg | P01130 |
| HOMO SAPIENS | -1 | 30645 | 3m0c | C | 41 | THR | p.Thr62Met | P01130 |
| HOMO SAPIENS | -1 | 30645 | 3m0c | C | 69 | ASP | p.Asp90Asn | P01130 |
| HOMO SAPIENS | -1 | 30645 | 3m0c | C | 76 | ASN | p.Asn97Lys | P01130 |
| HOMO SAPIENS | -1 | 30645 | 3m0c | C | 78 | SER | p.Ser99Ter | P01130 |
| HOMO SAPIENS | -1 | 30645 | 3m0c | C | 81 | GLN | p.Gln102Ter | P01130 |
| HOMO SAPIENS | -1 | 30645 | 3m0c | C | 84 | PRO | p.Pro105Ser | P01130 |
| HOMO SAPIENS | -1 | 30645 | 3m0c | C | 194 | ARG | p.Arg215Cys | P01130 |
| HOMO SAPIENS | -1 | 30645 | 3m0c | C | 198 | GLY | p.Gly219Asp | P01130 |
| HOMO SAPIENS | -1 | 30645 | 3m0c | C | 201 | CYS | p.Cys222Tyr | P01130 |
| HOMO SAPIENS | -1 | 30645 | 3m0c | C | 201 | CYS | p.Cys222Ter | P01130 |
| HOMO SAPIENS | -1 | 30645 | 3m0c | C | 206 | ASP | p.Asp227Glu | P01130 |
| HOMO SAPIENS | -1 | 30645 | 3m0c | C | 249 | CYS | p.Cys270Ter | P01130 |
| HOMO SAPIENS | -1 | 30645 | 3m0c | C | 250 | VAL | p.Val271Ile | P01130 |
| HOMO SAPIENS | 0 | 30645 | 3m0c | C | 255 | CYS | p.Cys276Tyr | P01130 |
| HOMO SAPIENS | 0 | 30645 | 3m0c | C | 255 | CYS | p.Cys276Ter | P01130 |
| HOMO SAPIENS | 0 | 30645 | 3m0c | C | 256 | GLU | p.Glu277Lys | P01130 |
| HOMO SAPIENS | 0 | 30645 | 3m0c | C | 265 | SER | p.Ser286Arg | P01130 |
| HOMO SAPIENS | 0 | 30645 | 3m0c | C | 284 | TRP | p.Trp305Ter | P01130 |
| HOMO SAPIENS | 0 | 30645 | 3m0c | C | 292 | CYS | p.Cys313Ter | P01130 |
| HOMO SAPIENS | 1 | 30645 | 3m0c | C | 385 | ARG | p.Arg406Trp | P01130 |
| HOMO SAPIENS | 0 | 30645 | 3m0c | C | 395 | ARG | p.Arg416Gln | P01130 |
| HOMO SAPIENS | 0 | 30645 | 3m0c | C | 398 | TYR | p.Tyr419Ter | P01130 |
| HOMO SAPIENS | 1 | 30645 | 3m0c | C | 415 | VAL | p.Val436Ala | P01130 |
| HOMO SAPIENS | 1 | 30645 | 3m0c | C | 424 | ASP | p.Asp445Glu | P01130 |
| HOMO SAPIENS | 1 | 30645 | 3m0c | C | 425 | LEU | p.Leu446Val | P01130 |
| HOMO SAPIENS | -1 | 30645 | 3m0c | C | 440 | GLY | p.Gly461Cys | P01130 |
| HOMO SAPIENS | -1 | 30645 | 3m0c | C | 441 | VAL | p.Val462Ile | P01130 |
| HOMO SAPIENS | 1 | 30645 | 3m0c | C | 447 | VAL | p.Val468Ile | P01130 |
| HOMO SAPIENS | 1 | 30645 | 3m0c | C | 489 | THR | p.Thr510Met | P01130 |
| HOMO SAPIENS | 0 | 30645 | 3m0c | C | 524 | GLY | p.Gly545Glu | P01130 |
| HOMO SAPIENS | 1 | 30645 | 3m0c | C | 615 | GLY | p.Gly636Ser | P01130 |
| HOMO SAPIENS | 1 | 30645 | 3m0c | C | 625 | LEU | p.Leu646Ile | P01130 |
| HOMO SAPIENS | 1 | 30645 | 3m0c | C | 630 | ASP | p.Asp651Gly | P01130 |
| HOMO SAPIENS | 3 | 30645 | 3m0c | C | 646 | CYS | p.Cys667Tyr | P01130 |
| HOMO SAPIENS | 4 | 30645 | 3m0c | C | 654 | GLY | p.Gly675Ser | P01130 |
| HOMO SAPIENS | 6 | 30645 | 3m0c | C | 660 | CYS | p.Cys681Ter | P01130 |
| HOMO SAPIENS | 5 | 30645 | 3m0c | C | 676 | ALA | p.Ala697Pro | P01130 |
| HOMO SAPIENS | -1 | 30645 | 3m0c | C | 749 | GLN | p.Gln770Ter | P01130 |
| HOMO SAPIENS | -1 | 41323 | 4bxo | A | 1812 | PRO | p.Pro1812Ala | Q8IYD8 |
| HOMO SAPIENS | 5 | 8335 | 3h0t | C | 12 | GLY | p.Gly71Asp | P81172 |
| HOMO SAPIENS | 6 | 609 | 1bhg | B | 148 | PRO | p.Pro148Ser | P08236 |
| HOMO SAPIENS | 1 | 609 | 1bhg | B | 176 | LEU | p.Leu176Phe | P08236 |
| HOMO SAPIENS | 6 | 609 | 1bhg | B | 216 | ARG | p.Arg216Trp | P08236 |
| HOMO SAPIENS | 0 | 609 | 1bhg | B | 350 | LYS | p.Lys350Asn | P08236 |

| | | | | | | | | |
|---|---|---|---|---|---|---|---|---|
| HOMO SAPIENS | 1 | 609 | 1bhg | B | 354 | ALA | p.Ala354Val | P08236 |
| HOMO SAPIENS | 1 | 609 | 1bhg | B | 357 | ARG | p.Arg357Ter | P08236 |
| HOMO SAPIENS | 0 | 609 | 1bhg | B | 382 | ARG | p.Arg382Cys | P08236 |
| HOMO SAPIENS | 0 | 609 | 1bhg | B | 495 | TYR | p.Tyr495Cys | P08236 |
| HOMO SAPIENS | 2 | 609 | 1bhg | B | 507 | TRP | p.Trp507Ter | P08236 |
| HOMO SAPIENS | 0 | 609 | 1bhg | B | 577 | ARG | p.Arg577Leu | P08236 |
| HOMO SAPIENS | 1 | 609 | 1bhg | B | 611 | ARG | p.Arg611Trp | P08236 |
| HOMO SAPIENS | 0 | 609 | 1bhg | B | 619 | ALA | p.Ala619Val | P08236 |
| HOMO SAPIENS | 0 | 609 | 1bhg | B | 627 | TRP | p.Trp627Cys | P08236 |
| HOMO SAPIENS | 5 | 3422 | 1x9n | A | 566 | GLU | p.Glu566Lys | P18858 |
| HOMO SAPIENS | 2 | 3422 | 1x9n | A | 771 | ARG | p.Arg771Trp | P18858 |
| HOMO SAPIENS | 0 | 3279 | 5tby | B | 797 | ALA | p.Ala797Thr | P12883 |
| HOMO SAPIENS | -1 | 3279 | 5tby | B | 982 | MET | p.Met982Thr | P12883 |
| HOMO SAPIENS | -1 | 3279 | 5tby | B | 992 | LEU | p.Leu992Met | P12883 |
| HOMO SAPIENS | -1 | 3279 | 5tby | B | 1004 | GLN | p.Gln1004Glu | P12883 |
| HOMO SAPIENS | -1 | 3279 | 5tby | B | 1013 | GLU | p.Glu1013Lys | P12883 |
| HOMO SAPIENS | -1 | 3279 | 5tby | B | 1032 | ASP | p.Asp1032Asn | P12883 |
| HOMO SAPIENS | -1 | 3279 | 5tby | B | 1053 | ARG | p.Arg1053Gln | P12883 |
| HOMO SAPIENS | -1 | 3279 | 5tby | B | 1117 | GLU | p.Glu1117Ter | P12883 |
| HOMO SAPIENS | -1 | 3279 | 5tby | B | 1193 | ARG | p.Arg1193His | P12883 |
| HOMO SAPIENS | -1 | 3279 | 5tby | B | 1205 | GLU | p.Glu1205Lys | P12883 |
| HOMO SAPIENS | -1 | 3279 | 5tby | B | 1207 | ILE | p.Ile1207Met | P12883 |
| HOMO SAPIENS | -1 | 3279 | 5tby | B | 1250 | ARG | p.Arg1250Pro | P12883 |
| HOMO SAPIENS | -1 | 3279 | 5tby | B | 1259 | HIS | p.His1259Gln | P12883 |
| HOMO SAPIENS | -1 | 3279 | 5tby | B | 1272 | ASP | p.Asp1272Asn | P12883 |
| HOMO SAPIENS | -1 | 3279 | 5tby | B | 1295 | GLU | p.Glu1295Lys | P12883 |
| HOMO SAPIENS | -1 | 3279 | 5tby | B | 1335 | SER | p.Ser1335Leu | P12883 |
| HOMO SAPIENS | -1 | 3279 | 5tby | B | 1399 | GLU | p.Glu1399Asp | P12883 |
| HOMO SAPIENS | -1 | 3279 | 5tby | B | 1413 | SER | p.Ser1413Ter | P12883 |
| HOMO SAPIENS | -1 | 3279 | 5tby | B | 1459 | LYS | p.Lys1459Asn | P12883 |
| HOMO SAPIENS | -1 | 3279 | 5tby | B | 1467 | LEU | p.Leu1467Val | P12883 |
| HOMO SAPIENS | -1 | 3279 | 5tby | B | 1491 | SER | p.Ser1491Cys | P12883 |
| HOMO SAPIENS | -1 | 3279 | 5tby | B | 1496 | GLU | p.Glu1496Ala | P12883 |
| HOMO SAPIENS | -1 | 3279 | 5tby | B | 1500 | ARG | p.Arg1500Pro | P12883 |
| HOMO SAPIENS | -1 | 3279 | 5tby | B | 1606 | ARG | p.Arg1606His | P12883 |
| HOMO SAPIENS | -1 | 3279 | 5tby | B | 1664 | ASN | p.Asn1664Lys | P12883 |
| HOMO SAPIENS | -1 | 3279 | 5tby | B | 1712 | ARG | p.Arg1712Trp | P12883 |
| HOMO SAPIENS | -1 | 3279 | 5tby | B | 1769 | LEU | p.Leu1769Met | P12883 |
| HOMO SAPIENS | -1 | 3279 | 5tby | B | 1808 | GLY | p.Gly1808Ser | P12883 |
| HOMO SAPIENS | -1 | 3279 | 5tby | B | 1818 | ARG | p.Arg1818Trp | P12883 |
| HOMO SAPIENS | -1 | 3279 | 5tby | B | 1820 | ARG | p.Arg1820Trp | P12883 |
| HOMO SAPIENS | -1 | 3279 | 5tby | B | 1844 | GLU | p.Glu1844Lys | P12883 |
| HOMO SAPIENS | -1 | 3279 | 5tby | B | 1845 | ARG | p.Arg1845Trp | P12883 |
| HOMO SAPIENS | -1 | 3279 | 5tby | B | 1854 | THR | p.Thr1854Met | P12883 |
| HOMO SAPIENS | -1 | 3279 | 5tby | B | 1909 | ARG | p.Arg1909Trp | P12883 |
| HOMO SAPIENS | 2 | 2260 | 4kl1 | A | 553 | ASP | p.Asp553Asn | Q9Y3Q4 |
| HOMO SAPIENS | 0 | 40570 | 5lst | A | 522 | ARG | p.Arg522Cys | O94761 |
| HOMO SAPIENS | 0 | 40570 | 5lst | A | 522 | ARG | p.Arg522His | O94761 |
| HOMO SAPIENS | 0 | 40570 | 5lst | A | 591 | PRO | p.Pro591Leu | O94761 |
| HOMO SAPIENS | 0 | 40570 | 5lst | A | 793 | PRO | p.Pro793Ser | O94761 |
| HOMO SAPIENS | 4 | 40570 | 5lst | A | 799 | VAL | p.Val799Met | O94761 |
| HOMO SAPIENS | 0 | 40570 | 5lst | A | 1004 | ARG | p.Arg1004Trp | O94761 |
| HOMO SAPIENS | 0 | 40570 | 5lst | A | 1005 | ARG | p.Arg1005Gln | O94761 |
| HOMO SAPIENS | 0 | 40570 | 5lst | A | 1021 | ARG | p.Arg1021Gln | O94761 |

| | | | | | | | | |
|---|---|---|---|---|---|---|---|---|
| HOMO SAPIENS | 0 | 40570 | 5lst | A | 1043 | LEU | p.Leu1043Pro | O94761 |
| HOMO SAPIENS | 0 | 40570 | 5lst | A | 1045 | ALA | p.Ala1045Thr | O94761 |
| HOMO SAPIENS | 0 | 40570 | 5lst | A | 1105 | GLY | p.Gly1105Asp | O94761 |
| HOMO SAPIENS | 0 | 40570 | 5lst | A | 1105 | GLY | p.Gly1105Ser | O94761 |
| HOMO SAPIENS | -1 | 40570 | 5lst | A | 1113 | GLY | p.Gly1113Arg | O94761 |
| HOMO SAPIENS | -1 | 18865 | 1gxc | A | 64 | GLU | p.Glu64Lys | O96017 |
| HOMO SAPIENS | -1 | 11673 | 3dzu | D | 85 | PRO | p.Pro113Gln | P37231 |
| HOMO SAPIENS | 0 | 19383 | 3b96 | A | 63 | LYS | p.Lys103Arg | P49748 |
| HOMO SAPIENS | 4 | 19383 | 3b96 | A | 140 | ALA | p.Ala180Thr | P49748 |
| HOMO SAPIENS | 1 | 19383 | 3b96 | A | 145 | GLY | p.Gly185Ser | P49748 |
| HOMO SAPIENS | 0 | 19383 | 3b96 | A | 220 | THR | p.Thr260Met | P49748 |
| HOMO SAPIENS | 5 | 19383 | 3b96 | A | 243 | VAL | p.Val283Ala | P49748 |
| HOMO SAPIENS | 0 | 19383 | 3b96 | A | 249 | GLY | p.Gly289Arg | P49748 |
| HOMO SAPIENS | 0 | 19383 | 3b96 | A | 254 | GLY | p.Gly294Glu | P49748 |
| HOMO SAPIENS | 7 | 19383 | 3b96 | A | 326 | ARG | p.Arg366Cys | P49748 |
| HOMO SAPIENS | 7 | 19383 | 3b96 | A | 326 | ARG | p.Arg366His | P49748 |
| HOMO SAPIENS | 5 | 19383 | 3b96 | A | 342 | LYS | p.Lys382Gln | P49748 |
| HOMO SAPIENS | 0 | 19383 | 3b96 | A | 369 | THR | p.Thr409Met | P49748 |
| HOMO SAPIENS | 5 | 19383 | 3b96 | A | 376 | ALA | p.Ala416Ser | P49748 |
| HOMO SAPIENS | 0 | 19383 | 3b96 | A | 385 | ALA | p.Ala425Thr | P49748 |
| HOMO SAPIENS | 5 | 19383 | 3b96 | A | 399 | GLY | p.Gly439Asp | P49748 |
| HOMO SAPIENS | 7 | 19383 | 3b96 | A | 401 | GLY | p.Gly441Asp | P49748 |
| HOMO SAPIENS | 9 | 19383 | 3b96 | A | 413 | ARG | p.Arg453Gln | P49748 |
| HOMO SAPIENS | 2 | 19383 | 3b96 | A | 418 | PHE | p.Phe458Leu | P49748 |
| HOMO SAPIENS | 6 | 19383 | 3b96 | A | 419 | ARG | p.Arg459Trp | P49748 |
| HOMO SAPIENS | 6 | 19383 | 3b96 | A | 419 | ARG | p.Arg459Gln | P49748 |
| HOMO SAPIENS | -1 | 19383 | 3b96 | A | 471 | ARG | p.Arg511Gln | P49748 |
| HOMO SAPIENS | 0 | 19383 | 3b96 | A | 573 | ARG | p.Arg613Trp | P49748 |
| HOMO SAPIENS | 0 | 19383 | 3b96 | A | 575 | ARG | p.Arg615Gln | P49748 |
| HOMO SAPIENS | 0 | 22286 | 3k2s | A | 3 | PRO | p.Pro27His | P02647 |
| HOMO SAPIENS | 0 | 22286 | 3k2s | A | 4 | PRO | p.Pro28Arg | P02647 |
| HOMO SAPIENS | 0 | 22286 | 3k2s | A | 10 | ARG | p.Arg34Leu | P02647 |
| HOMO SAPIENS | 0 | 22286 | 3k2s | A | 26 | GLY | p.Gly50Arg | P02647 |
| HOMO SAPIENS | 0 | 22286 | 3k2s | A | 37 | ALA | p.Ala61Thr | P02647 |
| HOMO SAPIENS | 1 | 1622 | 2qrv | D | 631 | ARG | p.Arg635Trp | Q9Y6K1 |
| HOMO SAPIENS | 1 | 1622 | 2qrv | D | 900 | PRO | p.Pro904Leu | Q9Y6K1 |
| HOMO SAPIENS | 6 | 40796 | 1v18 | B | 1509 | LEU | p.Leu1509Val | P25054 |
| HOMO SAPIENS | 1 | 22375 | 4pki | G | 1214 | LYS | p.Asp214Asn | P28289 |
| HOMO SAPIENS | 1 | 16307 | 5mv8 | B | 483 | ALA | p.Ala483Ser | Q9Y6N9 |
| HOMO SAPIENS | 6 | 16307 | 5mv8 | B | 506 | ASN | p.Asn506Lys | Q9Y6N9 |
| HOMO SAPIENS | 4 | 16307 | 5mv8 | B | 519 | ASP | p.Glu519Asp | Q9Y6N9 |
| HOMO SAPIENS | 0 | 16307 | 5mv8 | B | 530 | GLY | p.Gly530Arg | Q9Y6N9 |
| HOMO SAPIENS | 4 | 36221 | 4cvo | A | 127 | VAL | p.Val127Ile | Q03468 |
| HOMO SAPIENS | 7 | 36221 | 4cvo | A | 134 | ARG | p.Arg134Trp | Q03468 |
| HOMO SAPIENS | 0 | 36221 | 4cvo | A | 156 | GLN | p.Gln156Ter | Q03468 |
| HOMO SAPIENS | 0 | 10829 | 4a11 | B | 13 | GLU | p.Glu13Ter | Q13216 |
| HOMO SAPIENS | 1 | 10829 | 4a11 | B | 100 | TYR | p.Tyr100Ter | Q13216 |
| HOMO SAPIENS | 0 | 10829 | 4a11 | B | 160 | ALA | p.Ala160Val | Q13216 |
| HOMO SAPIENS | 1 | 6759 | 5lev | A | 29 | ILE | p.Ile29Phe | Q9H3H5 |
| HOMO SAPIENS | 0 | 6759 | 5lev | A | 108 | MET | p.Met108Ile | Q9H3H5 |
| HOMO SAPIENS | 2 | 6759 | 5lev | A | 117 | VAL | p.Val117Ile | Q9H3H5 |
| HOMO SAPIENS | 1 | 6759 | 5lev | A | 170 | TYR | p.Tyr170Cys | Q9H3H5 |
| HOMO SAPIENS | 0 | 6759 | 5lev | A | 264 | GLY | p.Val264Gly | Q9H3H5 |
| HOMO SAPIENS | 0 | 6759 | 5lev | A | 332 | PHE | p.Phe332Val | Q9H3H5 |

| | | | | | | | | |
|---|---|---|---|---|---|---|---|---|
| HOMO SAPIENS | 7 | 6759 | 5lev | A | 393 | ILE | p.Ile393Val | Q9H3H5 |
| HOMO SAPIENS | -1 | 33473 | 5m5e | C | 9 | ASN | p.Asn31Ser | P31785 |
| HOMO SAPIENS | 0 | 26155 | 1ic8 | B | 98 | ALA | p.Ala98Val | P20823 |
| HOMO SAPIENS | 0 | 15329 | 4iqr | B | 127 | ARG | p.Arg136Trp | P41235 |
| HOMO SAPIENS | 0 | 20725 | 4bsj | A | 350 | GLU | p.Glu350Val | P35916 |
| HOMO SAPIENS | 0 | 20725 | 4bsj | A | 494 | THR | p.Thr494Ala | P35916 |
| HOMO SAPIENS | 0 | 20725 | 4bsj | A | 527 | ASN | p.Asn527Ser | P35916 |
| HOMO SAPIENS | 6 | 29959 | 1xiw | B | 47 | ARG | p.Arg68Ter | P04234 |
| HOMO SAPIENS | 2 | 29959 | 1xiw | B | 72 | CYS | p.Cys93Ter | P04234 |
| HOMO SAPIENS | 3 | 20242 | 5tjc | A | 102 | ARG | p.Arg102Ter | Q9GZU1 |
| HOMO SAPIENS | 3 | 20242 | 5tjc | A | 232 | THR | p.Thr232Pro | Q9GZU1 |
| HOMO SAPIENS | 7 | 9590 | 4ap8 | A | 50 | THR | p.Thr50Ala | O96007 |
| HOMO SAPIENS | 4 | 9590 | 4ap8 | A | 168 | GLU | p.Glu168Lys | O96007 |
| HOMO SAPIENS | -1 | 4297 | 5do7 | B | 19 | ASP | p.Asp19His | Q9H221 |
| HOMO SAPIENS | -1 | 4297 | 5do7 | B | 54 | TYR | p.Tyr54Cys | Q9H221 |
| HOMO SAPIENS | -1 | 4297 | 5do7 | B | 80 | CYS | p.Cys80Tyr | Q9H221 |
| HOMO SAPIENS | 5 | 4297 | 5do7 | B | 107 | SER | p.Ser107Ter | Q9H221 |
| HOMO SAPIENS | 1 | 4297 | 5do7 | B | 164 | ARG | p.Arg164Pro | Q9H221 |
| HOMO SAPIENS | 0 | 4297 | 5do7 | B | 210 | VAL | p.Val210Met | Q9H221 |
| HOMO SAPIENS | 0 | 4297 | 5do7 | B | 231 | PRO | p.Pro231Thr | Q9H221 |
| HOMO SAPIENS | 0 | 4297 | 5do7 | B | 263 | ARG | p.Arg263Gln | Q9H221 |
| HOMO SAPIENS | 0 | 4297 | 5do7 | B | 361 | TRP | p.Trp361Ter | Q9H221 |
| HOMO SAPIENS | 0 | 4297 | 5do7 | B | 400 | THR | p.Thr400Lys | Q9H221 |
| HOMO SAPIENS | 0 | 4297 | 5do7 | B | 549 | ALA | p.Ala549Pro | Q9H221 |
| HOMO SAPIENS | 0 | 4297 | 5do7 | B | 596 | LEU | p.Leu596Arg | Q9H221 |
| HOMO SAPIENS | 0 | 4297 | 5do7 | B | 632 | VAL | p.Val632Ala | Q9H221 |
| HOMO SAPIENS | 0 | 4297 | 5do7 | B | 658 | TYR | p.Tyr658Ter | Q9H221 |
| HOMO SAPIENS | 2 | 9356 | 4igg | B | 206 | GLN | p.Gln206His | P35221 |
| HOMO SAPIENS | 6 | 2179 | 2wk6 | A | 44 | ARG | p.Arg44Gln | P19971 |
| HOMO SAPIENS | 6 | 2179 | 2wk6 | A | 81 | ARG | p.Arg81Gln | P19971 |
| HOMO SAPIENS | 6 | 2179 | 2wk6 | A | 145 | GLY | p.Gly145Arg | P19971 |
| HOMO SAPIENS | 7 | 2179 | 2wk6 | A | 146 | ARG | p.Arg146His | P19971 |
| HOMO SAPIENS | 7 | 2179 | 2wk6 | A | 153 | GLY | p.Gly153Ser | P19971 |
| HOMO SAPIENS | 0 | 2179 | 2wk6 | A | 173 | MET | p.Met173Ile | P19971 |
| HOMO SAPIENS | 6 | 2179 | 2wk6 | A | 202 | ARG | p.Arg202Lys | P19971 |
| HOMO SAPIENS | 6 | 2179 | 2wk6 | A | 208 | VAL | p.Val208Met | P19971 |
| HOMO SAPIENS | 5 | 2179 | 2wk6 | A | 222 | LYS | p.Lys222Arg | P19971 |
| HOMO SAPIENS | 5 | 2179 | 2wk6 | A | 289 | GLU | p.Glu289Ala | P19971 |
| HOMO SAPIENS | 2 | 2179 | 2wk6 | A | 311 | GLY | p.Gly311Arg | P19971 |
| HOMO SAPIENS | 2 | 2179 | 2wk6 | A | 376 | ARG | p.Arg376Pro | P19971 |
| HOMO SAPIENS | 2 | 2179 | 2wk6 | A | 465 | ALA | p.Ala465Thr | P19971 |
| HOMO SAPIENS | -1 | 11856 | 3orh | A | 3 | ALA | p.Ala3Ser | Q14353 |
| HOMO SAPIENS | 1 | 11856 | 3orh | A | 20 | TRP | p.Trp20Ser | Q14353 |
| HOMO SAPIENS | 1 | 11856 | 3orh | A | 50 | MET | p.Met50Leu | Q14353 |
| HOMO SAPIENS | 1 | 11856 | 3orh | A | 110 | VAL | p.Val110Phe | Q14353 |
| HOMO SAPIENS | 8 | 11856 | 3orh | A | 164 | GLY | p.Gly164Asp | Q14353 |
| HOMO SAPIENS | 1 | 11856 | 3orh | A | 169 | CYS | p.Cys169Tyr | Q14353 |
| HOMO SAPIENS | 1 | 11856 | 3orh | A | 174 | TRP | p.Trp174Ter | Q14353 |
| HOMO SAPIENS | 2 | 11856 | 3orh | A | 196 | ALA | p.Ala196Val | Q14353 |
| HOMO SAPIENS | 1 | 11856 | 3orh | A | 209 | THR | p.Thr209Met | Q14353 |
| HOMO SAPIENS | 1 | 11856 | 3orh | A | 219 | ASP | p.Asp219Asn | Q14353 |
| HOMO SAPIENS | 1 | 11856 | 3orh | A | 224 | ALA | p.Ala224Thr | Q14353 |
| HOMO SAPIENS | 1 | 13244 | 2m6i | D | 218 | ARG | p.Arg246Leu | P23415 |
| HOMO SAPIENS | 1 | 13244 | 2m6i | D | 244 | ILE | p.Ile272Asn | P23415 |

| | | | | | | | | |
|---|---|---|---|---|---|---|---|---|
| HOMO SAPIENS | 5 | 13244 | 2m6i | D | 250 | PRO | p.Pro278Thr | P23415 |
| HOMO SAPIENS | 6 | 13244 | 2m6i | D | 266 | GLN | p.Gln294His | P23415 |
| HOMO SAPIENS | 1 | 13244 | 2m6i | D | 271 | ARG | p.Arg299Leu | P23415 |
| HOMO SAPIENS | 1 | 13244 | 2m6i | D | 276 | LYS | p.Lys304Glu | P23415 |
| HOMO SAPIENS | 1 | 13244 | 2m6i | D | 279 | TYR | p.Tyr307Cys | P23415 |
| HOMO SAPIENS | 0 | 13244 | 2m6i | D | 316 | GLY | p.Gly378Ser | P23415 |
| HOMO SAPIENS | 1 | 13244 | 2m6i | D | 400 | ARG | p.Arg428His | P23415 |
| HOMO SAPIENS | 3 | 1967 | 2i6b | A | 301 | ALA | p.Ala318Glu | P55263 |
| HOMO SAPIENS | 4 | 11096 | 1h8b | A | 61 | ALA | p.Ala882Thr | P35609 |
| HOMO SAPIENS | 0 | 15423 | 5ngz | A | 2 | GLN | p.Gln2Glu | Q9NPD8 |
| HOMO SAPIENS | 3 | 30475 | 3lpw | A | 21 | SER | p.Ser22894Tyr | Q8WZ42 |
| HOMO SAPIENS | 6 | 32687 | 2arf | A | 1041 | ARG | p.Arg1041Trp | P35670 |
| HOMO SAPIENS | 8 | 32687 | 2arf | A | 1061 | GLY | p.Gly1061Glu | P35670 |
| HOMO SAPIENS | 6 | 32687 | 2arf | A | 1064 | GLU | p.Glu1064Ala | P35670 |
| HOMO SAPIENS | 2 | 32687 | 2arf | A | 1069 | HIS | p.His1069Gln | P35670 |
| HOMO SAPIENS | 2 | 32687 | 2arf | A | 1088 | LEU | p.Leu1088Ter | P35670 |
| HOMO SAPIENS | 0 | 32687 | 2arf | A | 1099 | GLY | p.Gly1099Ser | P35670 |
| HOMO SAPIENS | 4 | 32687 | 2arf | A | 1106 | VAL | p.Val1106Asp | P35670 |
| HOMO SAPIENS | 2 | 32687 | 2arf | A | 1142 | GLN | p.Gln1142His | P35670 |
| HOMO SAPIENS | 8 | 32687 | 2arf | A | 1148 | ILE | p.Ile1148Thr | P35670 |
| HOMO SAPIENS | 6 | 32687 | 2arf | A | 1151 | ARG | p.Arg1151Cys | P35670 |
| HOMO SAPIENS | 4 | 32687 | 2arf | A | 1169 | MET | p.Met1169Val | P35670 |
| HOMO SAPIENS | 4 | 32687 | 2arf | A | 1173 | GLU | p.Glu1173Lys | P35670 |
| HOMO SAPIENS | 2 | 32687 | 2arf | A | 1176 | GLY | p.Gly1176Arg | P35670 |
| HOMO SAPIENS | 3 | 15544 | 5uq3 | C | 575 | PRO | p.Pro69Leu | P46527 |
| HOMO SAPIENS | 6 | 15544 | 5uq3 | C | 592 | GLU | p.Glu86Asp | P46527 |
| HOMO SAPIENS | 6 | 15544 | 5uq3 | C | 599 | ARG | p.Arg93Trp | P46527 |
| HOMO SAPIENS | -1 | 15544 | 5uq3 | C | 615 | VAL | p.Val109Gly | P46527 |
| HOMO SAPIENS | -1 | 15544 | 5uq3 | C | 631 | SER | p.Ser125Pro | P46527 |
| HOMO SAPIENS | -1 | 15544 | 5uq3 | C | 639 | PRO | p.Pro133Ser | P46527 |
| HOMO SAPIENS | 2 | 2703 | 4wxq | A | 245 | CYS | p.Cys245Tyr | Q99972 |
| HOMO SAPIENS | 0 | 2703 | 4wxq | A | 252 | GLY | p.Gly252Arg | Q99972 |
| HOMO SAPIENS | 1 | 2703 | 4wxq | A | 337 | GLN | p.Gln337Arg | Q99972 |
| HOMO SAPIENS | 5 | 2703 | 4wxq | A | 347 | TYR | p.Tyr347Ter | Q99972 |
| HOMO SAPIENS | 3 | 2703 | 4wxq | A | 364 | GLY | p.Gly364Val | Q99972 |
| HOMO SAPIENS | 2 | 2703 | 4wxq | A | 367 | GLY | p.Gly367Arg | Q99972 |
| HOMO SAPIENS | 2 | 2703 | 4wxq | A | 368 | GLN | p.Gln368Ter | Q99972 |
| HOMO SAPIENS | 6 | 2703 | 4wxq | A | 370 | PRO | p.Pro370Leu | Q99972 |
| HOMO SAPIENS | 1 | 2703 | 4wxq | A | 380 | ASP | p.Asp380His | Q99972 |
| HOMO SAPIENS | 8 | 2703 | 4wxq | A | 399 | GLY | p.Gly399Val | Q99972 |
| HOMO SAPIENS | 0 | 2703 | 4wxq | A | 423 | LYS | p.Lys423Glu | Q99972 |
| HOMO SAPIENS | 3 | 2703 | 4wxq | A | 433 | CYS | p.Cys433Arg | Q99972 |
| HOMO SAPIENS | 3 | 2703 | 4wxq | A | 437 | TYR | p.Tyr437His | Q99972 |
| HOMO SAPIENS | 2 | 2703 | 4wxq | A | 477 | ILE | p.Ile477Ser | Q99972 |
| HOMO SAPIENS | 3 | 2703 | 4wxq | A | 480 | ASN | p.Asn480Lys | Q99972 |
| HOMO SAPIENS | -1 | 34100 | 2fot | C | 1171 | GLN | p.Gln1173Pro | Q13813 |
| HOMO SAPIENS | 0 | 8773 | 1kw2 | A | 432 | ASP | p.Asp432Glu | P02774 |
| HOMO SAPIENS | 0 | 8773 | 1kw2 | A | 436 | LYS | p.Lys436Thr | P02774 |
| HOMO SAPIENS | 0 | 8402 | 4dzo | B | 649 | TYR | p.Tyr649Ter | Q9Y6D9 |
| HOMO SAPIENS | 0 | 39962 | 1nst | A | 611 | GLY | p.Gly611Ser | P52848 |
| HOMO SAPIENS | 2 | 22578 | 3dof | B | 45 | MET | p.Met45Arg | Q9Y2Y0 |
| HOMO SAPIENS | 1 | 23290 | 5j1s | A | 292 | ILE | p.Ile292Thr | O14656 |
| HOMO SAPIENS | 4 | 9505 | 4j1y | A | 534 | ARG | p.Arg534Trp | P09871 |
| HOMO SAPIENS | 9 | 260 | 2h57 | A | 31 | THR | p.Thr31Met | Q9H0F7 |

| | | | | | | | | |
|---|---|---|---|---|---|---|---|---|
| HOMO SAPIENS | 0 | 260 | 2h57 | A | 91 | ILE | p.Ile91Thr | Q9H0F7 |
| HOMO SAPIENS | 0 | 260 | 2h57 | A | 121 | ARG | p.Arg121His | Q9H0F7 |
| HOMO SAPIENS | 0 | 260 | 2h57 | A | 122 | ARG | p.Arg122Ter | Q9H0F7 |
| HOMO SAPIENS | 2 | 260 | 2h57 | A | 169 | GLY | p.Gly169Ala | Q9H0F7 |
| HOMO SAPIENS | 4 | 260 | 2h57 | A | 170 | LEU | p.Leu170Trp | Q9H0F7 |
| HOMO SAPIENS | 4 | 34023 | 2cr3 | A | 18 | GLY | p.Gly48Ser | P11362 |
| HOMO SAPIENS | 0 | 37667 | 3ii7 | A | 420 | ARG | p.Arg420Cys | Q8IXQ5 |
| HOMO SAPIENS | 0 | 41720 | 2k9u | A | 68 | ARG | p.Arg2364His | Q14315 |
| HOMO SAPIENS | 1 | 29250 | 5fmv | A | 568 | HIS | p.His568Gln | P08575 |
| HOMO SAPIENS | 1 | 1280 | 1fb1 | A | 69 | PRO | p.Pro69Leu | P30793 |
| HOMO SAPIENS | 3 | 1280 | 1fb1 | A | 88 | ARG | p.Arg88Trp | P30793 |
| HOMO SAPIENS | 2 | 1280 | 1fb1 | A | 108 | GLY | p.Gly108Asp | P30793 |
| HOMO SAPIENS | 4 | 1280 | 1fb1 | A | 134 | ASP | p.Asp134Val | P30793 |
| HOMO SAPIENS | 6 | 1280 | 1fb1 | A | 135 | ILE | p.Ile135Lys | P30793 |
| HOMO SAPIENS | 9 | 1280 | 1fb1 | A | 144 | HIS | p.His144Pro | P30793 |
| HOMO SAPIENS | 9 | 1280 | 1fb1 | A | 184 | ARG | p.Arg184His | P30793 |
| HOMO SAPIENS | 1 | 1280 | 1fb1 | A | 196 | ALA | p.Ala196Ser | P30793 |
| HOMO SAPIENS | 3 | 1280 | 1fb1 | A | 201 | GLY | p.Gly201Glu | P30793 |
| HOMO SAPIENS | 2 | 1280 | 1fb1 | A | 204 | VAL | p.Val204Ile | P30793 |
| HOMO SAPIENS | 2 | 1280 | 1fb1 | A | 211 | MET | p.Met211Ile | P30793 |
| HOMO SAPIENS | 0 | 1280 | 1fb1 | A | 221 | MET | p.Met221Thr | P30793 |
| HOMO SAPIENS | 3 | 1280 | 1fb1 | A | 224 | LYS | p.Lys224Arg | P30793 |
| HOMO SAPIENS | 2 | 1280 | 1fb1 | A | 249 | ARG | p.Arg249Ser | P30793 |
| HOMO SAPIENS | 8 | 41352 | 4k12 | A | 47 | ILE | p.Ile551Thr | P08603 |
| HOMO SAPIENS | 9 | 20703 | 5nir | A | 57 | CYS | p.Cys57Tyr | P02458 |
| HOMO SAPIENS | 9 | 20703 | 5nir | A | 64 | CYS | p.Cys64Ter | P02458 |
| HOMO SAPIENS | 0 | 20087 | 4cmn | A | 299 | MET | p.Met299Ile | Q01968 |
| HOMO SAPIENS | 1 | 20819 | 3aaf | A | 979 | ILE | p.Ile979Met | Q14191 |
| HOMO SAPIENS | 1 | 20819 | 3aaf | A | 984 | LEU | p.Leu984Ile | Q14191 |
| HOMO SAPIENS | 3 | 20819 | 3aaf | A | 987 | ARG | p.Arg987Ter | Q14191 |
| HOMO SAPIENS | -1 | 20819 | 3aaf | A | 1074 | PHE | p.Leu1074Phe | Q14191 |
| HOMO SAPIENS | 0 | 3619 | 3dem | A | 206 | ASN | p.Asn225Ser | P48740 |
| HOMO SAPIENS | 1 | 3619 | 3dem | A | 269 | ARG | p.Arg288Leu | P48740 |
| HOMO SAPIENS | 0 | 3105 | 3ooi | A | 164 | TYR | p.Tyr2015His | Q96L73 |
| HOMO SAPIENS | 6 | 15828 | 2lkx | A | 16 | LEU | p.Leu100Gln | Q99697 |
| HOMO SAPIENS | 8 | 15828 | 2lkx | A | 24 | ARG | p.Arg108His | Q99697 |
| HOMO SAPIENS | 2 | 15828 | 2lkx | A | 30 | THR | p.Thr114Pro | Q99697 |
| HOMO SAPIENS | 8 | 15828 | 2lkx | A | 45 | VAL | p.Val129Leu | Q99697 |
| HOMO SAPIENS | 8 | 15828 | 2lkx | A | 46 | ARG | p.Arg130Trp | Q99697 |
| HOMO SAPIENS | 9 | 15828 | 2lkx | A | 53 | ARG | p.Arg137Pro | Q99697 |
| HOMO SAPIENS | 9 | 13017 | 4irq | A | 141 | ARG | p.Arg141Trp | Q9UBV7 |
| HOMO SAPIENS | 0 | 13017 | 4irq | A | 186 | ALA | p.Ala186Asp | Q9UBV7 |
| HOMO SAPIENS | 0 | 13017 | 4irq | A | 206 | LEU | p.Leu206Pro | Q9UBV7 |
| HOMO SAPIENS | 1 | 13017 | 4irq | A | 214 | CYS | p.Cys214Tyr | Q9UBV7 |
| HOMO SAPIENS | 0 | 13017 | 4irq | A | 270 | ARG | p.Arg270Cys | Q9UBV7 |
| HOMO SAPIENS | -1 | 13053 | 4mqw | X | 307 | ALA | p.Ala307Thr | P23945 |
| HOMO SAPIENS | 0 | 19406 | 4b3g | A | 43 | ARG | p.Arg43Ter | P38935 |
| HOMO SAPIENS | 0 | 19406 | 4b3g | A | 46 | CYS | p.Cys46Ter | P38935 |
| HOMO SAPIENS | 0 | 19406 | 4b3g | A | 75 | ALA | p.Ala75Thr | P38935 |
| HOMO SAPIENS | 0 | 19406 | 4b3g | A | 201 | LEU | p.Leu201Ser | P38935 |
| HOMO SAPIENS | 3 | 19406 | 4b3g | A | 213 | HIS | p.His213Arg | P38935 |
| HOMO SAPIENS | 0 | 19406 | 4b3g | A | 275 | ILE | p.Ile275Val | P38935 |
| HOMO SAPIENS | -1 | 19406 | 4b3g | A | 320 | ARG | p.Arg320Ter | P38935 |
| HOMO SAPIENS | 0 | 19406 | 4b3g | A | 361 | LEU | p.Leu361Pro | P38935 |

| | | | | | | | | |
|---|---|---|---|---|---|---|---|---|
| HOMO SAPIENS | 0 | 19406 | 4b3g | A | 369 | PHE | p.Phe369Leu | P38935 |
| HOMO SAPIENS | 0 | 19406 | 4b3g | A | 493 | THR | p.Thr493Ile | P38935 |
| HOMO SAPIENS | 0 | 19406 | 4b3g | A | 496 | CYS | p.Cys496Ter | P38935 |
| HOMO SAPIENS | 0 | 19406 | 4b3g | A | 514 | GLU | p.Glu514Lys | P38935 |
| HOMO SAPIENS | 0 | 19406 | 4b3g | A | 528 | ALA | p.Ala528Thr | P38935 |
| HOMO SAPIENS | 0 | 19406 | 4b3g | A | 531 | PRO | p.Pro531Thr | P38935 |
| HOMO SAPIENS | 0 | 19406 | 4b3g | A | 580 | VAL | p.Val580Ile | P38935 |
| HOMO SAPIENS | 6 | 19406 | 4b3g | A | 603 | ARG | p.Arg603His | P38935 |
| HOMO SAPIENS | 0 | 21501 | 5jde | A | 40 | SER | p.Ser2831Asn | Q8WZ42 |
| HOMO SAPIENS | 5 | 21501 | 5jde | A | 189 | ALA | p.Ala2980Thr | Q8WZ42 |
| HOMO SAPIENS | 6 | 15008 | 3mop | H | 12 | ARG | p.Arg12Cys | Q9NWZ3 |
| HOMO SAPIENS | 8 | 17455 | 3jtc | C | 15 | ARG | p.Arg57Trp | P04070 |
| HOMO SAPIENS | -1 | 17455 | 3jtc | C | 20 | CGU | p.Glu62Ala | P04070 |
| HOMO SAPIENS | 0 | 27795 | 5gox | B | 625 | TYR | p.Tyr625Ter | Q92878 |
| HOMO SAPIENS | 0 | 27795 | 5gox | B | 653 | SER | p.Ser653Ter | Q92878 |
| HOMO SAPIENS | -1 | 6558 | 5nug | A | 137 | VAL | p.Val137Ile | Q14204 |
| HOMO SAPIENS | -1 | 6558 | 5nug | A | 567 | ARG | p.Arg567His | Q14204 |
| HOMO SAPIENS | -1 | 6558 | 5nug | A | 1140 | THR | p.Thr1140Met | Q14204 |
| HOMO SAPIENS | -1 | 6558 | 5nug | A | 1991 | ASP | p.Asp1991Asn | Q14204 |
| HOMO SAPIENS | -1 | 6558 | 5nug | A | 2401 | LYS | p.Lys2401Asn | Q14204 |
| HOMO SAPIENS | 0 | 6558 | 5nug | A | 2488 | ARG | p.Arg2488His | Q14204 |
| HOMO SAPIENS | 0 | 6558 | 5nug | A | 3907 | HIS | p.His3907Gln | Q14204 |
| HOMO SAPIENS | 0 | 6558 | 5nug | A | 3981 | THR | p.Thr3981Arg | Q14204 |
| HOMO SAPIENS | 0 | 6558 | 5nug | A | 4394 | THR | p.Thr4394Met | Q14204 |
| HOMO SAPIENS | 0 | 6558 | 5nug | A | 4573 | ASN | p.Asn4573Ser | Q14204 |
| HOMO SAPIENS | 0 | 6558 | 5nug | A | 4603 | SER | p.Ser4603Ile | Q14204 |
| HOMO SAPIENS | 1 | 4244 | 1umk | A | 57 | ARG | p.Arg58Gln | P00387 |
| HOMO SAPIENS | 1 | 4244 | 1umk | A | 72 | LEU | p.Leu73Pro | P00387 |
| HOMO SAPIENS | 1 | 4244 | 1umk | A | 105 | VAL | p.Val106Met | P00387 |
| HOMO SAPIENS | 0 | 4244 | 1umk | A | 116 | THR | p.Thr117Ser | P00387 |
| HOMO SAPIENS | 8 | 4244 | 1umk | A | 127 | SER | p.Ser128Pro | P00387 |
| HOMO SAPIENS | 0 | 4244 | 1umk | A | 148 | LEU | p.Leu149Pro | P00387 |
| HOMO SAPIENS | 0 | 4244 | 1umk | A | 159 | ARG | p.Arg160Ter | P00387 |
| HOMO SAPIENS | 0 | 4244 | 1umk | A | 203 | CYS | p.Cys204Arg | P00387 |
| HOMO SAPIENS | 0 | 4244 | 1umk | A | 203 | CYS | p.Cys204Tyr | P00387 |
| HOMO SAPIENS | 2 | 4244 | 1umk | A | 239 | ASP | p.Asp240Gly | P00387 |
| HOMO SAPIENS | 0 | 4244 | 1umk | A | 291 | GLY | p.Gly292Asp | P00387 |
| HOMO SAPIENS | 4 | 34495 | 1s35 | A | 1151 | ASP | p.Asn1151Asp | P11277 |
| HOMO SAPIENS | 3 | 30110 | 3zqs | A | 224 | THR | p.Thr224Ala | Q9NW38 |
| HOMO SAPIENS | 3 | 17588 | 5dse | B | 53 | LEU | p.Leu53Pro | Q9BYI3 |
| HOMO SAPIENS | 0 | 1564 | 4yqf | B | 72 | HIS | p.His347Gln | Q9UHD8 |
| HOMO SAPIENS | 0 | 17673 | 4jnw | A | 32 | ARG | p.Arg32186His | Q8WZ42 |
| HOMO SAPIENS | 3 | 17673 | 4jnw | A | 140 | ILE | p.Ile32294Val | Q8WZ42 |
| HOMO SAPIENS | 5 | 17673 | 4jnw | A | 269 | ALA | p.Ala32423Thr | Q8WZ42 |
| HOMO SAPIENS | 3 | 12858 | 3s97 | C | 302 | LEU | p.Leu302Pro | Q12860 |
| HOMO SAPIENS | 0 | 6147 | 4nn0 | A | 114 | ARG | p.Arg114Gln | Q9BXJ0 |
| HOMO SAPIENS | 0 | 7549 | 1tr2 | A | 277 | LEU | p.Leu277Met | P18206 |
| HOMO SAPIENS | 1 | 7549 | 1tr2 | A | 413 | ALA | p.Ala413Thr | P18206 |
| HOMO SAPIENS | 1 | 7549 | 1tr2 | A | 519 | ILE | p.Ile519Leu | P18206 |
| HOMO SAPIENS | 5 | 7549 | 1tr2 | A | 547 | ARG | p.Arg547Ter | P18206 |
| HOMO SAPIENS | 2 | 7549 | 1tr2 | A | 636 | HIS | p.His636Arg | P18206 |
| HOMO SAPIENS | 2 | 7549 | 1tr2 | A | 824 | ILE | p.Ile824Met | P18206 |
| HOMO SAPIENS | 2 | 13900 | 3gro | A | 45 | CYS | p.Cys45Tyr | P50897 |
| HOMO SAPIENS | 2 | 13900 | 3gro | A | 75 | THR | p.Thr75Pro | P50897 |

| | | | | | | | | |
|---|---|---|---|---|---|---|---|---|
| HOMO SAPIENS | 2 | 13900 | 3gro | A | 79 | ASP | p.Asp79Gly | P50897 |
| HOMO SAPIENS | 5 | 13900 | 3gro | A | 91 | GLN | p.Gln91Pro | P50897 |
| HOMO SAPIENS | 5 | 13900 | 3gro | A | 96 | CYS | p.Cys96Tyr | P50897 |
| HOMO SAPIENS | 0 | 13900 | 3gro | A | 104 | LYS | p.Lys104Ter | P50897 |
| HOMO SAPIENS | 2 | 13900 | 3gro | A | 108 | GLY | p.Gly108Arg | P50897 |
| HOMO SAPIENS | 2 | 13900 | 3gro | A | 109 | TYR | p.Tyr109Asp | P50897 |
| HOMO SAPIENS | 0 | 13900 | 3gro | A | 122 | ARG | p.Arg122Trp | P50897 |
| HOMO SAPIENS | 0 | 13900 | 3gro | A | 138 | SER | p.Ser138Leu | P50897 |
| HOMO SAPIENS | 5 | 13900 | 3gro | A | 151 | ARG | p.Arg151Ter | P50897 |
| HOMO SAPIENS | 7 | 13900 | 3gro | A | 164 | ARG | p.Arg164Ter | P50897 |
| HOMO SAPIENS | 0 | 13900 | 3gro | A | 177 | GLN | p.Gln177Glu | P50897 |
| HOMO SAPIENS | 0 | 13900 | 3gro | A | 181 | VAL | p.Val181Leu | P50897 |
| HOMO SAPIENS | 0 | 13900 | 3gro | A | 184 | GLU | p.Glu184Lys | P50897 |
| HOMO SAPIENS | 5 | 13900 | 3gro | A | 186 | TRP | p.Trp186Ter | P50897 |
| HOMO SAPIENS | 0 | 13900 | 3gro | A | 219 | LEU | p.Leu219Gln | P50897 |
| HOMO SAPIENS | 0 | 13900 | 3gro | A | 222 | LEU | p.Leu222Pro | P50897 |
| HOMO SAPIENS | 5 | 13900 | 3gro | A | 296 | TRP | p.Trp296Ter | P50897 |
| HOMO SAPIENS | 6 | 35028 | 5mwb | A | 444 | CYS | p.Cys444Tyr | Q04721 |
| HOMO SAPIENS | 0 | 35028 | 5mwb | A | 514 | ASN | p.Asn514Ser | Q04721 |
| HOMO SAPIENS | 5 | 1483 | 4gl2 | A | 822 | ARG | p.Arg822Gln | Q9BYX4 |
| HOMO SAPIENS | 0 | 1483 | 4gl2 | A | 843 | ARG | p.His843Arg | Q9BYX4 |
| HOMO SAPIENS | -1 | 28838 | 3cl3 | D | 153 | LEU | p.Leu153Arg | Q9Y6K9 |
| HOMO SAPIENS | 2 | 22563 | 4zel | B | 79 | ARG | p.Arg79Trp | P09172 |
| HOMO SAPIENS | 2 | 22563 | 4zel | B | 101 | VAL | p.Val101Met | P09172 |
| HOMO SAPIENS | 2 | 22563 | 4zel | B | 114 | ASP | p.Asp114Glu | P09172 |
| HOMO SAPIENS | 0 | 22563 | 4zel | B | 201 | ASN | p.Asn201Ser | P09172 |
| HOMO SAPIENS | 2 | 22563 | 4zel | B | 211 | ALA | p.Ala211Thr | P09172 |
| HOMO SAPIENS | -1 | 22563 | 4zel | B | 290 | ASP | p.Asp290Asn | P09172 |
| HOMO SAPIENS | 2 | 22563 | 4zel | B | 318 | ALA | p.Ala318Ser | P09172 |
| HOMO SAPIENS | 2 | 22563 | 4zel | B | 345 | ASP | p.Asp345Asn | P09172 |
| HOMO SAPIENS | 2 | 22563 | 4zel | B | 362 | ALA | p.Ala362Glu | P09172 |
| HOMO SAPIENS | 2 | 22563 | 4zel | B | 549 | ARG | p.Arg549Cys | P09172 |
| HOMO SAPIENS | 9 | 9780 | 1y6k | R | 36 | TYR | p.Tyr57Cys | Q13651 |
| HOMO SAPIENS | 1 | 9780 | 1y6k | R | 40 | LEU | p.Leu61Val | Q13651 |
| HOMO SAPIENS | 1 | 9780 | 1y6k | R | 63 | THR | p.Thr84Ile | Q13651 |
| HOMO SAPIENS | 9 | 9780 | 1y6k | R | 80 | ARG | p.Arg101Trp | Q13651 |
| HOMO SAPIENS | 0 | 9780 | 1y6k | R | 120 | GLY | p.Gly141Arg | Q13651 |
| HOMO SAPIENS | 0 | 9780 | 1y6k | R | 138 | SER | p.Ser159Gly | Q13651 |
| HOMO SAPIENS | 0 | 9780 | 1y6k | R | 203 | ILE | p.Ile224Val | Q13651 |
| HOMO SAPIENS | 0 | 16019 | 3rep | A | 211 | ARG | p.Arg211Cys | Q13418 |
| HOMO SAPIENS | 3 | 7879 | 5ec3 | A | 33 | ALA | p.Ala33Thr | P32754 |
| HOMO SAPIENS | 4 | 7879 | 5ec3 | A | 113 | ARG | p.Arg113Gln | P32754 |
| HOMO SAPIENS | 3 | 7879 | 5ec3 | A | 340 | VAL | p.Val340Leu | P32754 |
| HOMO SAPIENS | 0 | 8904 | 1qk9 | A | 6 | LYS | p.Lys82Arg | P51608 |
| HOMO SAPIENS | 4 | 8904 | 1qk9 | A | 24 | LEU | p.Leu100Val | P51608 |
| HOMO SAPIENS | 7 | 8904 | 1qk9 | A | 30 | ARG | p.Arg106Trp | P51608 |
| HOMO SAPIENS | 6 | 8904 | 1qk9 | A | 57 | ARG | p.Arg133Cys | P51608 |
| HOMO SAPIENS | 3 | 8904 | 1qk9 | A | 61 | GLU | p.Glu137Gly | P51608 |
| HOMO SAPIENS | 2 | 8904 | 1qk9 | A | 64 | ALA | p.Ala140Val | P51608 |
| HOMO SAPIENS | 5 | 8904 | 1qk9 | A | 79 | PHE | p.Phe155Ser | P51608 |
| HOMO SAPIENS | 3 | 8904 | 1qk9 | A | 82 | THR | p.Thr158Met | P51608 |
| HOMO SAPIENS | 1 | 23121 | 3e9k | A | 198 | THR | p.Thr198Ala | Q16719 |
| HOMO SAPIENS | -1 | 26090 | 3tje | F | 19 | LEU | p.Leu35Met | P25445 |
| HOMO SAPIENS | 8 | 26090 | 3tje | F | 106 | THR | p.Thr122Ile | P25445 |

| | | | | | | | | |
|---|---|---|---|---|---|---|---|---|
| HOMO SAPIENS | 2 | 36404 | 5cqr | A | 765 | GLY | p.Gly765Glu | O95163 |
| HOMO SAPIENS | 2 | 36404 | 5cqr | A | 816 | ILE | p.Ile816Leu | O95163 |
| HOMO SAPIENS | 2 | 36404 | 5cqr | A | 830 | ILE | p.Ile830Met | O95163 |
| HOMO SAPIENS | 5 | 36404 | 5cqr | A | 939 | TYR | p.Tyr939Ter | O95163 |
| HOMO SAPIENS | 2 | 36404 | 5cqr | A | 952 | LYS | p.Lys952Ile | O95163 |
| HOMO SAPIENS | 5 | 36404 | 5cqr | A | 1013 | GLY | p.Gly1013Ser | O95163 |
| HOMO SAPIENS | 5 | 36404 | 5cqr | A | 1072 | CYS | p.Cys1072Ser | O95163 |
| HOMO SAPIENS | -1 | 36404 | 5cqr | A | 1158 | PRO | p.Pro1158Leu | O95163 |
| HOMO SAPIENS | 1 | 34149 | 2k40 | A | 18 | ASN | p.Asn125Ser | Q9UBX0 |
| HOMO SAPIENS | 0 | 34149 | 2k40 | A | 42 | LEU | p.Glu149Lys | Q9UBX0 |
| HOMO SAPIENS | 9 | 34149 | 2k40 | A | 53 | ARG | p.Arg160Cys | Q9UBX0 |
| HOMO SAPIENS | 0 | 34149 | 2k40 | A | 63 | SER | p.Ser170Leu | Q9UBX0 |
| HOMO SAPIENS | 2 | 2468 | 3vp6 | A | 484 | LEU | p.Leu484Met | Q99259 |
| HOMO SAPIENS | 0 | 2468 | 3vp6 | A | 532 | ARG | p.Arg532Gln | Q99259 |
| HOMO SAPIENS | 5 | 1253 | 5xtd | S | 8 | GLY | p.Gly8Arg | O15239 |
| HOMO SAPIENS | 7 | 1253 | 5xtd | S | 32 | GLY | p.Gly32Arg | O15239 |
| HOMO SAPIENS | 7 | 1253 | 5xtd | S | 37 | ARG | p.Arg37Ser | O15239 |
| HOMO SAPIENS | -1 | 13292 | 3u1k | A | 121 | VAL | p.Ile121Val | Q8TCS8 |
| HOMO SAPIENS | 2 | 13292 | 3u1k | A | 165 | PRO | p.Pro165Ser | Q8TCS8 |
| HOMO SAPIENS | 2 | 13292 | 3u1k | A | 192 | ARG | p.Arg192Gln | Q8TCS8 |
| HOMO SAPIENS | 1 | 13292 | 3u1k | A | 315 | LYS | p.Lys315Arg | Q8TCS8 |
| HOMO SAPIENS | 4 | 13292 | 3u1k | A | 509 | VAL | p.Val509Ile | Q8TCS8 |
| HOMO SAPIENS | 7 | 13292 | 3u1k | A | 531 | THR | p.Thr531Arg | Q8TCS8 |
| HOMO SAPIENS | 0 | 13292 | 3u1k | A | 590 | ASN | p.Asn590Asp | Q8TCS8 |
| HOMO SAPIENS | 0 | 27048 | 4p5w | B | 184 | ARG | p.Arg184Ter | Q9UJS0 |
| HOMO SAPIENS | 0 | 27048 | 4p5w | B | 225 | SER | p.Ser225Ter | Q9UJS0 |
| HOMO SAPIENS | -1 | 27048 | 4p5w | B | 631 | ALA | p.Arg360Ter | Q9UJS0 |
| HOMO SAPIENS | 0 | 24240 | 4mdr | A | 318 | ARG | p.Arg318Ter | O00189 |
| HOMO SAPIENS | 0 | 24240 | 4mdr | A | 338 | ARG | p.Arg338Ter | O00189 |
| HOMO SAPIENS | 0 | 13429 | 4cg4 | A | 479 | PHE | p.Phe479Leu | O15553 |
| HOMO SAPIENS | 0 | 13429 | 4cg4 | A | 487 | VAL | p.Val487Leu | O15553 |
| HOMO SAPIENS | 3 | 13429 | 4cg4 | A | 653 | ARG | p.Arg653His | O15553 |
| HOMO SAPIENS | 1 | 13429 | 4cg4 | A | 680 | MET | p.Met680Ile | O15553 |
| HOMO SAPIENS | 1 | 13429 | 4cg4 | A | 688 | TYR | p.Tyr688Ter | O15553 |
| HOMO SAPIENS | 0 | 13429 | 4cg4 | A | 694 | MET | p.Met694Ile | O15553 |
| HOMO SAPIENS | 0 | 13429 | 4cg4 | A | 694 | MET | p.Met694Val | O15553 |
| HOMO SAPIENS | 1 | 13429 | 4cg4 | A | 726 | VAL | p.Val726Ala | O15553 |
| HOMO SAPIENS | 0 | 13429 | 4cg4 | A | 761 | ARG | p.Arg761His | O15553 |
| HOMO SAPIENS | 0 | 2079 | 4okh | A | 691 | GLY | p.Gly691Arg | P20807 |
| HOMO SAPIENS | 2 | 2079 | 4okh | A | 748 | ARG | p.Arg748Ter | P20807 |
| HOMO SAPIENS | 0 | 2079 | 4okh | A | 763 | TYR | p.Tyr763Cys | P20807 |
| HOMO SAPIENS | 1 | 2079 | 4okh | A | 769 | ARG | p.Arg769Gln | P20807 |
| HOMO SAPIENS | 9 | 2079 | 4okh | A | 777 | ILE | p.Ile777Val | P20807 |
| HOMO SAPIENS | 8 | 2079 | 4okh | A | 778 | ASP | p.Asp778His | P20807 |
| HOMO SAPIENS | 5 | 2079 | 4okh | A | 780 | ASP | p.Asp780His | P20807 |
| HOMO SAPIENS | 1 | 1200 | 2gi7 | A | 142 | ARG | p.Arg162Trp | Q9HCN6 |
| HOMO SAPIENS | 4 | 9199 | 3ezq | C | 260 | ASP | p.Asp260Val | P25445 |
| HOMO SAPIENS | 0 | 36522 | 5ms9 | A | 122 | ARG | p.Arg122Cys | P35555 |
| HOMO SAPIENS | 0 | 36522 | 5ms9 | A | 240 | ARG | p.Arg240Cys | P35555 |
| HOMO SAPIENS | 0 | 17448 | 3g2f | A | 225 | ARG | p.Arg225His | Q13873 |
| HOMO SAPIENS | 0 | 17448 | 3g2f | A | 347 | CYS | p.Cys347Tyr | Q13873 |
| HOMO SAPIENS | 3 | 17448 | 3g2f | A | 485 | ASP | p.Asp485Gly | Q13873 |
| HOMO SAPIENS | 7 | 17448 | 3g2f | A | 491 | ARG | p.Arg491Gln | Q13873 |
| HOMO SAPIENS | 7 | 17448 | 3g2f | A | 491 | ARG | p.Arg491Trp | Q13873 |

| | | | | | | | | |
|---|---|---|---|---|---|---|---|---|
| HOMO SAPIENS | 0 | 4861 | 4ux8 | A | 56 | LEU | p.Leu56Met | P07949 |
| HOMO SAPIENS | 1 | 4861 | 4ux8 | A | 88 | ILE | p.Ile88Leu | P07949 |
| HOMO SAPIENS | 0 | 4861 | 4ux8 | A | 114 | ARG | p.Arg114His | P07949 |
| HOMO SAPIENS | 5 | 4861 | 4ux8 | A | 180 | ARG | p.Arg180Gln | P07949 |
| HOMO SAPIENS | 0 | 4861 | 4ux8 | A | 208 | GLU | p.Glu208Asp | P07949 |
| HOMO SAPIENS | 0 | 4861 | 4ux8 | A | 373 | ALA | p.Ala373Val | P07949 |
| HOMO SAPIENS | 0 | 4861 | 4ux8 | A | 375 | LEU | p.Leu375Gln | P07949 |
| HOMO SAPIENS | -1 | 4861 | 4ux8 | A | 386 | ALA | p.Ala386Val | P07949 |
| HOMO SAPIENS | 0 | 4861 | 4ux8 | A | 446 | GLY | p.Gly446Arg | P07949 |
| HOMO SAPIENS | 0 | 4861 | 4ux8 | A | 480 | GLU | p.Glu480Lys | P07949 |
| HOMO SAPIENS | 0 | 4861 | 4ux8 | A | 489 | ASP | p.Asp489Asn | P07949 |
| HOMO SAPIENS | -1 | 4861 | 4ux8 | A | 510 | ALA | p.Ala510Val | P07949 |
| HOMO SAPIENS | -1 | 4861 | 4ux8 | A | 533 | GLY | p.Gly533Cys | P07949 |
| HOMO SAPIENS | -1 | 4861 | 4ux8 | A | 567 | ASP | p.Asp567Asn | P07949 |
| HOMO SAPIENS | -1 | 4861 | 4ux8 | A | 609 | CYS | p.Cys609Tyr | P07949 |
| HOMO SAPIENS | -1 | 4861 | 4ux8 | A | 609 | CYS | p.Cys609Gly | P07949 |
| HOMO SAPIENS | -1 | 4861 | 4ux8 | A | 611 | CYS | p.Cys611Trp | P07949 |
| HOMO SAPIENS | -1 | 4861 | 4ux8 | A | 618 | CYS | p.Cys618Gly | P07949 |
| HOMO SAPIENS | -1 | 4861 | 4ux8 | A | 618 | CYS | p.Cys618Phe | P07949 |
| HOMO SAPIENS | -1 | 4861 | 4ux8 | A | 620 | CYS | p.Cys620Gly | P07949 |
| HOMO SAPIENS | -1 | 4861 | 4ux8 | A | 620 | CYS | p.Cys620Ser | P07949 |
| HOMO SAPIENS | -1 | 4861 | 4ux8 | A | 620 | CYS | p.Cys620Trp | P07949 |
| HOMO SAPIENS | -1 | 4861 | 4ux8 | A | 623 | GLU | p.Glu623Lys | P07949 |
| HOMO SAPIENS | -1 | 4861 | 4ux8 | A | 631 | ASP | p.Asp631Gly | P07949 |
| HOMO SAPIENS | -1 | 4861 | 4ux8 | A | 631 | ASP | p.Asp631Glu | P07949 |
| HOMO SAPIENS | -1 | 4861 | 4ux8 | A | 634 | CYS | p.Cys634Gly | P07949 |
| HOMO SAPIENS | -1 | 4861 | 4ux8 | A | 634 | CYS | p.Cys634Phe | P07949 |
| HOMO SAPIENS | -1 | 4861 | 4ux8 | A | 634 | CYS | p.Cys634Trp | P07949 |
| HOMO SAPIENS | 0 | 36308 | 4nkq | B | 123 | PRO | p.Pro142Leu | P15509 |
| HOMO SAPIENS | 0 | 36308 | 4nkq | B | 145 | ARG | p.Arg164Gln | P15509 |
| HOMO SAPIENS | -1 | 36308 | 4nkq | B | 177 | GLY | p.Gly196Arg | P15509 |
| HOMO SAPIENS | 0 | 36006 | 1oqa | A | 107 | HIS | p.His1860Pro | P38398 |
| HOMO SAPIENS | 7 | 15394 | 2ozo | A | 465 | ARG | p.Arg465His | P43403 |
| HOMO SAPIENS | 7 | 15394 | 2ozo | A | 465 | ARG | p.Arg465Cys | P43403 |
| HOMO SAPIENS | 4 | 15394 | 2ozo | A | 518 | SER | p.Ser518Arg | P43403 |
| HOMO SAPIENS | 1 | 20813 | 3o5z | A | 19 | ARG | p.Arg19Trp | O00459 |
| HOMO SAPIENS | 2 | 4329 | 2bb5 | A | 5 | ILE | p.Ile23Val | P20062 |
| HOMO SAPIENS | 4 | 4329 | 2bb5 | A | 59 | LYS | p.Lys77Met | P20062 |
| HOMO SAPIENS | 4 | 4329 | 2bb5 | A | 197 | ARG | p.Arg215Trp | P20062 |
| HOMO SAPIENS | 1 | 4329 | 2bb5 | A | 241 | PRO | p.Arg259Pro | P20062 |
| HOMO SAPIENS | 2 | 4329 | 2bb5 | A | 315 | THR | p.Thr333Lys | P20062 |
| HOMO SAPIENS | 5 | 4329 | 2bb5 | A | 330 | SER | p.Ser348Phe | P20062 |
| HOMO SAPIENS | 4 | 4329 | 2bb5 | A | 358 | LEU | p.Leu376Ser | P20062 |
| HOMO SAPIENS | 5 | 4329 | 2bb5 | A | 381 | ARG | p.Arg399Gln | P20062 |
| HOMO SAPIENS | 4 | 18294 | 4r30 | A | 163 | ASN | p.Asn163Asp | O95278 |
| HOMO SAPIENS | 6 | 18294 | 4r30 | A | 227 | ALA | p.Ala227Val | O95278 |
| HOMO SAPIENS | 4 | 18294 | 4r30 | A | 241 | ARG | p.Arg241Ter | O95278 |
| HOMO SAPIENS | 4 | 18294 | 4r30 | A | 325 | ARG | p.Arg325His | O95278 |
| HOMO SAPIENS | 0 | 2632 | 4xem | A | 22 | GLU | p.Glu22Lys | P49588 |
| HOMO SAPIENS | 0 | 2632 | 4xem | A | 275 | GLY | p.Gly275Asp | P49588 |
| HOMO SAPIENS | 1 | 2632 | 4xem | A | 329 | ARG | p.Arg329His | P49588 |
| HOMO SAPIENS | 0 | 11431 | 5t4n | A | 284 | GLN | p.Gln305Glu | Q96QU1 |
| HOMO SAPIENS | 0 | 11431 | 5t4n | A | 315 | ARG | p.Arg336Ter | Q96QU1 |
| HOMO SAPIENS | 0 | 11431 | 5t4n | A | 322 | ARG | p.Arg343Lys | Q96QU1 |

| | | | | | | | | |
|---|---|---|---|---|---|---|---|---|
| HOMO SAPIENS | 1 | 11431 | 5t4n | A | 326 | LEU | p.Leu347Phe | Q96QU1 |
| HOMO SAPIENS | 0 | 11431 | 5t4n | A | 373 | TYR | p.Tyr394Cys | Q96QU1 |
| HOMO SAPIENS | 0 | 11431 | 5t4n | A | 398 | ASN | p.Asn419Ser | Q96QU1 |
| HOMO SAPIENS | 0 | 11431 | 5t4n | A | 414 | ALA | p.Asp435Ala | Q96QU1 |
| HOMO SAPIENS | 0 | 11431 | 5t4n | A | 433 | VAL | p.Val454Phe | Q96QU1 |
| HOMO SAPIENS | 1 | 11431 | 5t4n | A | 507 | VAL | p.Val528Asp | Q96QU1 |
| HOMO SAPIENS | 0 | 11431 | 5t4n | A | 547 | ALA | p.Ala568Ser | Q96QU1 |
| HOMO SAPIENS | 3 | 3194 | 5xtd | L | 84 | ARG | p.Arg84His | O43181 |
| HOMO SAPIENS | 1 | 3194 | 5xtd | L | 130 | THR | p.Thr130Ser | O43181 |
| HOMO SAPIENS | 8 | 3194 | 5xtd | L | 136 | SER | p.Ser136Ala | O43181 |
| HOMO SAPIENS | 0 | 40749 | 1nun | A | 106 | CYS | p.Cys106Phe | O15520 |
| HOMO SAPIENS | 9 | 40749 | 1nun | A | 156 | ILE | p.Ile156Arg | O15520 |
| HOMO SAPIENS | 1 | 40749 | 1nun | A | 193 | ARG | p.Arg193Ter | O15520 |
| HOMO SAPIENS | 0 | 40749 | 1nun | A | 207 | HIS | p.His207Pro | O15520 |
| HOMO SAPIENS | 0 | 7026 | 3n1f | D | 875 | SER | p.Ser875Gly | Q4KMG0 |
| HOMO SAPIENS | 6 | 23802 | 3rzv | A | 424 | ARG | p.Arg424Ter | O95630 |
| HOMO SAPIENS | 0 | 36720 | 1jm7 | A | 12 | GLN | p.Gln12Ter | P38398 |
| HOMO SAPIENS | 3 | 36720 | 1jm7 | A | 23 | GLU | p.Glu23Asp | P38398 |
| HOMO SAPIENS | 1 | 36720 | 1jm7 | A | 45 | LYS | p.Lys45Gln | P38398 |
| HOMO SAPIENS | 8 | 36720 | 1jm7 | A | 47 | CYS | p.Cys47Ter | P38398 |
| HOMO SAPIENS | 0 | 36720 | 1jm7 | A | 59 | SER | p.Ser59Ter | P38398 |
| HOMO SAPIENS | 8 | 36720 | 1jm7 | A | 61 | CYS | p.Cys61Gly | P38398 |
| HOMO SAPIENS | 8 | 36720 | 1jm7 | A | 63 | LEU | p.Leu63Ter | P38398 |
| HOMO SAPIENS | 7 | 36720 | 1jm7 | A | 64 | CYS | p.Cys64Gly | P38398 |
| HOMO SAPIENS | 0 | 36720 | 1jm7 | A | 101 | TYR | p.Tyr101Ter | P38398 |
| HOMO SAPIENS | 0 | 36720 | 1jm7 | A | null | TYR | p.Tyr105Cys | P38398 |
| HOMO SAPIENS | 0 | 28500 | 3kuz | A | 1217 | CYS | p.Cys1217Arg | O60486 |
| HOMO SAPIENS | 5 | 8097 | 3p56 | C | 69 | ARG | p.Arg69Trp | Q8TDP1 |
| HOMO SAPIENS | 5 | 8097 | 3p56 | C | 143 | LYS | p.Lys143Ile | Q8TDP1 |
| HOMO SAPIENS | 2 | 30073 | 3nmz | A | 380 | ARG | p.Arg380Gln | P25054 |
| HOMO SAPIENS | 1 | 22573 | 1cjy | A | 111 | SER | p.Ser111Pro | P47712 |
| HOMO SAPIENS | 0 | 22573 | 1cjy | A | 485 | ARG | p.Arg485His | P47712 |
| HOMO SAPIENS | 2 | 965 | 3E+077 | A | 99 | ALA | p.Ala99Val | Q9Y617 |
| HOMO SAPIENS | 2 | 965 | 3E+077 | A | 100 | ASP | p.Asp100Ala | Q9Y617 |
| HOMO SAPIENS | 4 | 1503 | 3kxc | A | 62 | ARG | p.Arg62Trp | O43617 |
| HOMO SAPIENS | 6 | 9418 | 1hdr | A | 22 | GLY | p.Gly23Asp | P09417 |
| HOMO SAPIENS | 2 | 9418 | 1hdr | A | 35 | TRP | p.Trp36Arg | P09417 |
| HOMO SAPIENS | 2 | 9418 | 1hdr | A | 107 | TRP | p.Trp108Gly | P09417 |
| HOMO SAPIENS | 6 | 9418 | 1hdr | A | 149 | TYR | p.Tyr150Cys | P09417 |
| HOMO SAPIENS | -1 | 12079 | 3tnu | B | 373 | GLY | p.Gly373Ala | P13647 |
| HOMO SAPIENS | 1 | 12079 | 3tnu | B | 463 | LEU | p.Leu463Pro | P13647 |
| HOMO SAPIENS | 1 | 12079 | 3tnu | B | 471 | ARG | p.Arg471Cys | P13647 |
| HOMO SAPIENS | 0 | 8135 | 3rbn | B | 89 | SER | p.Ser556Ile | P40692 |
| HOMO SAPIENS | 0 | 8135 | 3rbn | B | 111 | GLU | p.Glu578Gly | P40692 |
| HOMO SAPIENS | 2 | 8135 | 3rbn | B | 114 | PRO | p.Pro581Leu | P40692 |
| HOMO SAPIENS | 0 | 8135 | 3rbn | B | 133 | GLU | p.Glu600Gly | P40692 |
| HOMO SAPIENS | 2 | 8135 | 3rbn | B | 136 | PRO | p.Pro603Arg | P40692 |
| HOMO SAPIENS | 2 | 8135 | 3rbn | B | 136 | PRO | p.Pro603Leu | P40692 |
| HOMO SAPIENS | 7 | 8135 | 3rbn | B | 140 | LEU | p.Leu607His | P40692 |
| HOMO SAPIENS | 2 | 8135 | 3rbn | B | 151 | LYS | p.Lys618Ala | P40692 |
| HOMO SAPIENS | 2 | 8135 | 3rbn | B | 151 | LYS | p.Lys618Thr | P40692 |
| HOMO SAPIENS | 5 | 8135 | 3rbn | B | 181 | PRO | p.Pro648Ser | P40692 |
| HOMO SAPIENS | 2 | 8135 | 3rbn | B | 188 | ILE | p.Ile655Val | P40692 |
| HOMO SAPIENS | 2 | 8135 | 3rbn | B | 214 | ALA | p.Ala681Thr | P40692 |

| | | | | | | | | |
|---|---|---|---|---|---|---|---|---|
| HOMO SAPIENS | 2 | 8135 | 3rbn | B | 220 | ARG | p.Arg687Trp | P40692 |
| HOMO SAPIENS | 2 | 8135 | 3rbn | B | 222 | GLN | p.Gln689Arg | P40692 |
| HOMO SAPIENS | -1 | 8135 | 3rbn | B | 236 | GLN | p.Gln701Ter | P40692 |
| HOMO SAPIENS | 7 | 8135 | 3rbn | B | 251 | HIS | p.His718Tyr | P40692 |
| HOMO SAPIENS | 0 | 8135 | 3rbn | B | 284 | LYS | p.Lys751Arg | P40692 |
| HOMO SAPIENS | 1 | 21998 | 5d68 | A | 355 | LEU | p.Leu355Phe | O00522 |
| HOMO SAPIENS | 3 | 21998 | 5d68 | A | 380 | THR | p.Thr380Met | O00522 |
| HOMO SAPIENS | 0 | 21998 | 5d68 | A | 415 | ILE | p.Ile415Met | O00522 |
| HOMO SAPIENS | 0 | 33789 | 2m0c | A | 13 | ARG | p.Arg218Gln | Q9H161 |
| HOMO SAPIENS | 7 | 33789 | 2m0c | A | 52 | ARG | p.Arg257Thr | Q9H161 |
| HOMO SAPIENS | 0 | 33789 | 2m0c | A | 67 | ARG | p.Arg272Pro | Q9H161 |
| HOMO SAPIENS | 6 | 8263 | 2osg | A | 51 | GLY | p.Gly355Arg | Q9UDY2 |
| HOMO SAPIENS | 5 | 8263 | 2osg | A | 60 | THR | p.Thr364Met | Q9UDY2 |
| HOMO SAPIENS | 1 | 2475 | 5E+026 | B | 234 | THR | p.Thr234Ala | Q9BZ23 |
| HOMO SAPIENS | 2 | 2475 | 5E+026 | B | 264 | ARG | p.Arg264Trp | Q9BZ23 |
| HOMO SAPIENS | 4 | 2475 | 5E+026 | B | 286 | ARG | p.Arg286Cys | Q9BZ23 |
| HOMO SAPIENS | 0 | 2475 | 5E+026 | B | 357 | ARG | p.Arg357Trp | Q9BZ23 |
| HOMO SAPIENS | 2 | 2475 | 5E+026 | B | 481 | ARG | p.Arg481Ter | Q9BZ23 |
| HOMO SAPIENS | 8 | 2475 | 5E+026 | B | 521 | GLY | p.Gly521Arg | Q9BZ23 |
| HOMO SAPIENS | 2 | 2475 | 5E+026 | B | 528 | THR | p.Thr528Met | Q9BZ23 |
| HOMO SAPIENS | 1 | 3742 | 5xtd | E | 108 | HIS | p.His134Tyr | P56556 |
| HOMO SAPIENS | 1 | 16725 | 3g04 | C | 36 | ASP | p.Asp36His | P16473 |
| HOMO SAPIENS | 3 | 16725 | 3g04 | C | 52 | PRO | p.Pro52Thr | P16473 |
| HOMO SAPIENS | 1 | 16725 | 3g04 | C | 162 | PRO | p.Pro162Ala | P16473 |
| HOMO SAPIENS | -1 | 12366 | 5d2d | C | 750 | PRO | p.Pro750Leu | P13569 |
| HOMO SAPIENS | 0 | 12366 | 5d2d | C | 764 | ARG | p.Arg764Ter | P13569 |
| HOMO SAPIENS | 0 | 34845 | 2y23 | A | 1151 | ASP | p.Asp1151Glu | P52179 |
| HOMO SAPIENS | 0 | 34845 | 2y23 | A | 1180 | ASP | p.Asp1180Ala | P52179 |
| HOMO SAPIENS | 6 | 36740 | 3lbx | B | 2053 | ALA | p.Ala2053Pro | P11277 |
| HOMO SAPIENS | 0 | 19289 | 3ihj | A | 404 | ARG | p.Arg404Ter | Q8TD30 |
| HOMO SAPIENS | 5 | 1034 | 5xtd | Q | 228 | ARG | p.Arg228Gln | O75306 |
| HOMO SAPIENS | 5 | 1034 | 5xtd | Q | 229 | PRO | p.Pro229Gln | O75306 |
| HOMO SAPIENS | 4 | 1034 | 5xtd | Q | 323 | ARG | p.Arg323Gln | O75306 |
| HOMO SAPIENS | 4 | 1034 | 5xtd | Q | 352 | PRO | p.Pro352Ala | O75306 |
| HOMO SAPIENS | 5 | 1034 | 5xtd | Q | 413 | SER | p.Ser413Pro | O75306 |
| HOMO SAPIENS | 1 | 2772 | 2vaf | A | 1047 | THR | p.Thr66Ala | O14958 |
| HOMO SAPIENS | 5 | 2772 | 2vaf | A | 1057 | VAL | p.Val76Met | O14958 |
| HOMO SAPIENS | 0 | 2772 | 2vaf | A | 1148 | LEU | p.Leu167His | O14958 |
| HOMO SAPIENS | 1 | 2772 | 2vaf | A | 1149 | ILE | p.Ile168Thr | O14958 |
| HOMO SAPIENS | 8 | 2772 | 2vaf | A | 1225 | HIS | p.His244Arg | O14958 |
| HOMO SAPIENS | 8 | 2772 | 2vaf | A | 1225 | HIS | p.His244Tyr | O14958 |
| HOMO SAPIENS | 7 | 2772 | 2vaf | A | 1251 | ILE | p.Ile270Thr | O14958 |
| HOMO SAPIENS | 3 | 2772 | 2vaf | A | 1288 | ASP | p.Asp307His | O14958 |
| HOMO SAPIENS | 5 | 2772 | 2vaf | A | 1289 | PRO | p.Pro308Leu | O14958 |
| HOMO SAPIENS | 1 | 31885 | 2aex | A | 132 | ALA | p.Ala132Val | P36551 |
| HOMO SAPIENS | 1 | 31885 | 2aex | A | 272 | ASN | p.Asn272His | P36551 |
| HOMO SAPIENS | 8 | 31885 | 2aex | A | 279 | GLY | p.Gly279Arg | P36551 |
| HOMO SAPIENS | 5 | 31885 | 2aex | A | 294 | VAL | p.Val294Ile | P36551 |
| HOMO SAPIENS | 5 | 31885 | 2aex | A | 295 | HIS | p.His295Asp | P36551 |
| HOMO SAPIENS | 1 | 31885 | 2aex | A | 331 | ARG | p.Arg331Trp | P36551 |
| HOMO SAPIENS | 3 | 31885 | 2aex | A | 352 | ARG | p.Arg352Cys | P36551 |
| HOMO SAPIENS | 1 | 31885 | 2aex | A | 404 | LYS | p.Lys404Glu | P36551 |
| HOMO SAPIENS | 2 | 31885 | 2aex | A | 447 | ARG | p.Arg447Cys | P36551 |
| HOMO SAPIENS | 5 | 11721 | 2r3v | A | 20 | HIS | p.His20Pro | Q03426 |

| | | | | | | | | |
|---|---|---|---|---|---|---|---|---|
| HOMO SAPIENS | 1 | 11721 | 2r3v | A | 116 | TYR | p.Tyr116His | Q03426 |
| HOMO SAPIENS | 1 | 11721 | 2r3v | A | 148 | ALA | p.Ala148Thr | Q03426 |
| HOMO SAPIENS | 1 | 11721 | 2r3v | A | 167 | PRO | p.Pro167Leu | Q03426 |
| HOMO SAPIENS | 6 | 11721 | 2r3v | A | 203 | VAL | p.Val203Ala | Q03426 |
| HOMO SAPIENS | 2 | 11721 | 2r3v | A | 215 | ARG | p.Arg215Ter | Q03426 |
| HOMO SAPIENS | 2 | 11721 | 2r3v | A | 237 | THR | p.Thr237Ser | Q03426 |
| HOMO SAPIENS | 0 | 11721 | 2r3v | A | 268 | ILE | p.Ile268Thr | Q03426 |
| HOMO SAPIENS | 0 | 11721 | 2r3v | A | 268 | ILE | p.Ile268Val | Q03426 |
| HOMO SAPIENS | 0 | 11721 | 2r3v | A | 301 | ASN | p.Asn301Ser | Q03426 |
| HOMO SAPIENS | 6 | 11721 | 2r3v | A | 310 | VAL | p.Val310Met | Q03426 |
| HOMO SAPIENS | 5 | 11721 | 2r3v | A | 334 | ALA | p.Ala334Thr | Q03426 |
| HOMO SAPIENS | 2 | 11721 | 2r3v | A | 377 | VAL | p.Val377Ile | Q03426 |
| HOMO SAPIENS | 1 | 11721 | 2r3v | A | 386 | ASP | p.Asp386Asn | Q03426 |
| HOMO SAPIENS | 1 | 11721 | 2r3v | A | 388 | ARG | p.Arg388Ter | Q03426 |
| HOMO SAPIENS | 8 | 5010 | 2hgs | A | 125 | ARG | p.Arg125Cys | P48637 |
| HOMO SAPIENS | 1 | 5010 | 2hgs | A | 164 | ARG | p.Arg164Gln | P48637 |
| HOMO SAPIENS | 0 | 5010 | 2hgs | A | 219 | ASP | p.Asp219Gly | P48637 |
| HOMO SAPIENS | 7 | 5010 | 2hgs | A | 267 | ARG | p.Arg267Trp | P48637 |
| HOMO SAPIENS | 0 | 5010 | 2hgs | A | 283 | ARG | p.Arg283Cys | P48637 |
| HOMO SAPIENS | 0 | 6892 | 5erd | B | 7 | VAL | p.Val56Met | Q14126 |
| HOMO SAPIENS | 1 | 6892 | 5erd | B | 25 | HIS | p.His74Arg | Q14126 |
| HOMO SAPIENS | 0 | 6892 | 5erd | B | 40 | TYR | p.Tyr89Cys | Q14126 |
| HOMO SAPIENS | 0 | 6892 | 5erd | B | 97 | ARG | p.Arg146Leu | Q14126 |
| HOMO SAPIENS | 7 | 6892 | 5erd | B | 217 | ASN | p.Asn266Ser | Q14126 |
| HOMO SAPIENS | 0 | 6892 | 5erd | B | 243 | ARG | p.Arg292His | Q14126 |
| HOMO SAPIENS | 0 | 6892 | 5erd | B | 244 | ILE | p.Ile293Val | Q14126 |
| HOMO SAPIENS | 0 | 6892 | 5erd | B | 343 | VAL | p.Val392Ile | Q14126 |
| HOMO SAPIENS | 4 | 6892 | 5erd | B | 350 | GLU | p.Glu399Lys | Q14126 |
| HOMO SAPIENS | 0 | 6892 | 5erd | B | 410 | TYR | p.Tyr459Cys | Q14126 |
| HOMO SAPIENS | 0 | 6892 | 5erd | B | 466 | VAL | p.Val515Ile | Q14126 |
| HOMO SAPIENS | 0 | 6892 | 5erd | B | 482 | PHE | p.Phe531Cys | Q14126 |
| HOMO SAPIENS | 0 | 6892 | 5erd | B | 484 | VAL | p.Val533Ile | Q14126 |
| HOMO SAPIENS | 0 | 6892 | 5erd | B | 545 | LEU | p.Leu594Pro | Q14126 |
| HOMO SAPIENS | 0 | 1338 | 2ax4 | A | 58 | THR | p.Thr48Arg | O95340 |
| HOMO SAPIENS | 7 | 8675 | 2zkm | X | 385 | LYS | p.Lys385Arg | Q00722 |
| HOMO SAPIENS | 1 | 11769 | 2wya | B | 54 | VAL | p.Val54Met | P54868 |
| HOMO SAPIENS | 1 | 11769 | 2wya | B | 116 | ARG | p.Arg116Cys | P54868 |
| HOMO SAPIENS | 5 | 11769 | 2wya | B | 167 | TYR | p.Tyr167Cys | P54868 |
| HOMO SAPIENS | 1 | 11769 | 2wya | B | 174 | PHE | p.Phe174Leu | P54868 |
| HOMO SAPIENS | 0 | 11769 | 2wya | B | 212 | GLY | p.Gly212Arg | P54868 |
| HOMO SAPIENS | 1 | 11769 | 2wya | B | 288 | ARG | p.Arg288Ter | P54868 |
| HOMO SAPIENS | 0 | 11769 | 2wya | B | 375 | SER | p.Ser375Cys | P54868 |
| HOMO SAPIENS | 4 | 11769 | 2wya | B | 500 | ARG | p.Arg500His | P54868 |
| HOMO SAPIENS | 0 | 3102 | 2iwo | A | 1880 | ARG | p.Arg1908Lys | O75970 |
| HOMO SAPIENS | 6 | 14296 | 2m9w | A | 38 | GLY | p.Gly296Ser | P43694 |
| HOMO SAPIENS | 0 | 14296 | 2m9w | A | 58 | GLN | p.Gln316Glu | P43694 |
| HOMO SAPIENS | 2 | 704 | 5ekd | B | 239 | ARG | p.Arg239Ter | Q9UGM6 |
| HOMO SAPIENS | 2 | 704 | 5ekd | B | 264 | TYR | p.Tyr264Cys | Q9UGM6 |
| HOMO SAPIENS | 6 | 2228 | 1ey2 | A | 47 | SER | p.Ser47Ter | Q93099 |
| HOMO SAPIENS | 6 | 2228 | 1ey2 | A | 48 | ALA | p.Ala48Ser | Q93099 |
| HOMO SAPIENS | 6 | 2228 | 1ey2 | A | 53 | ARG | p.Arg53Gln | Q93099 |
| HOMO SAPIENS | 3 | 2228 | 1ey2 | A | 80 | HIS | p.Gln80His | Q93099 |
| HOMO SAPIENS | 0 | 2228 | 1ey2 | A | 120 | CYS | p.Cys120Trp | Q93099 |
| HOMO SAPIENS | 3 | 2228 | 1ey2 | A | 122 | ALA | p.Ala122Val | Q93099 |

| | | | | | | | | |
|---|---|---|---|---|---|---|---|---|
| HOMO SAPIENS | 3 | 2228 | 1ey2 | A | 161 | GLY | p.Gly161Arg | Q93099 |
| HOMO SAPIENS | 6 | 2228 | 1ey2 | A | 225 | ARG | p.Arg225His | Q93099 |
| HOMO SAPIENS | 0 | 2228 | 1ey2 | A | 230 | PRO | p.Pro230Ser | Q93099 |
| HOMO SAPIENS | 6 | 2228 | 1ey2 | A | 270 | GLY | p.Gly270Arg | Q93099 |
| HOMO SAPIENS | 6 | 2228 | 1ey2 | A | 330 | ARG | p.Arg330Ser | Q93099 |
| HOMO SAPIENS | -1 | 2228 | 1ey2 | A | 368 | MSE | p.Met368Val | Q93099 |
| HOMO SAPIENS | 6 | 2228 | 1ey2 | A | 371 | HIS | p.His371Arg | Q93099 |
| HOMO SAPIENS | 6 | 2228 | 1ey2 | A | 401 | GLU | p.Glu401Gln | Q93099 |
| HOMO SAPIENS | 5 | 34648 | 2k27 | A | 31 | ARG | p.Arg31His | Q06710 |
| HOMO SAPIENS | 7 | 34648 | 2k27 | A | 40 | GLN | p.Gln40Pro | Q06710 |
| HOMO SAPIENS | 5 | 34648 | 2k27 | A | 57 | CYS | p.Cys57Tyr | Q06710 |
| HOMO SAPIENS | 5 | 34648 | 2k27 | A | 62 | LEU | p.Leu62Arg | Q06710 |
| HOMO SAPIENS | 1 | 21399 | 3edu | A | 1819 | GLU | p.Glu1819Ter | P11277 |
| HOMO SAPIENS | 1 | 18153 | 1oey | J | 245 | TYR | p.Tyr245Ter | Q15080 |
| HOMO SAPIENS | 2 | 41567 | 3sw0 | X | 1050 | ASN | p.Asn1050Tyr | P08603 |
| HOMO SAPIENS | 5 | 41567 | 3sw0 | X | 1059 | ILE | p.Ile1059Thr | P08603 |
| HOMO SAPIENS | 0 | 36362 | 4wrm | A | 362 | HIS | p.His362Arg | P07333 |
| HOMO SAPIENS | -1 | 36362 | 4wrm | A | 413 | GLY | p.Gly413Ser | P07333 |
| HOMO SAPIENS | 0 | 31255 | 2w2j | A | 100 | SER | p.Ser100Pro | P35219 |
| HOMO SAPIENS | 0 | 31255 | 2w2j | A | 100 | SER | p.Ser100Leu | P35219 |
| HOMO SAPIENS | 5 | 31255 | 2w2j | A | 237 | ARG | p.Arg237Gln | P35219 |
| HOMO SAPIENS | -1 | 26450 | 1th1 | C | 1395 | SER | p.Ser1395Cys | P25054 |
| HOMO SAPIENS | -1 | 26450 | 1th1 | C | 1413 | MET | p.Met1413Val | P25054 |
| HOMO SAPIENS | -1 | 26450 | 1th1 | C | 1446 | ALA | p.Ala1446Thr | P25054 |
| HOMO SAPIENS | -1 | 26450 | 1th1 | C | 1454 | LYS | p.Lys1454Glu | P25054 |
| HOMO SAPIENS | 0 | 26450 | 1th1 | C | 1474 | ALA | p.Ala1474Thr | P25054 |
| HOMO SAPIENS | 1 | 981 | 5xtb | F | 14 | LEU | p.Leu14Val | O43678 |
| HOMO SAPIENS | 1 | 11069 | 3kq4 | A | 46 | SER | p.Ser46Leu | P27352 |
| HOMO SAPIENS | 1 | 11069 | 3kq4 | A | 97 | MET | p.Met97Thr | P27352 |
| HOMO SAPIENS | 1 | 11069 | 3kq4 | A | 255 | ASN | p.Asn255Ser | P27352 |
| HOMO SAPIENS | 1 | 11069 | 3kq4 | A | 330 | ASN | p.Asn330Lys | P27352 |
| HOMO SAPIENS | 0 | 41489 | 3erb | A | 53 | PRO | p.Pro73Leu | P06681 |
| HOMO SAPIENS | 6 | 8715 | 2bq8 | X | 23 | THR | p.Thr44Met | P13686 |
| HOMO SAPIENS | 0 | 8715 | 2bq8 | X | 88 | GLY | p.Gly109Arg | P13686 |
| HOMO SAPIENS | 6 | 8715 | 2bq8 | X | 127 | VAL | p.Val148Met | P13686 |
| HOMO SAPIENS | 2 | 8715 | 2bq8 | X | 179 | VAL | p.Val200Met | P13686 |
| HOMO SAPIENS | 6 | 8715 | 2bq8 | X | 251 | ARG | p.Arg272Cys | P13686 |
| HOMO SAPIENS | 7 | 8112 | 3p56 | A | 37 | GLY | p.Gly37Ser | O75792 |
| HOMO SAPIENS | -1 | 8112 | 3p56 | A | 202 | LEU | p.Leu202Ser | O75792 |
| HOMO SAPIENS | 1 | 8112 | 3p56 | A | 205 | ASP | p.Asp205Glu | O75792 |
| HOMO SAPIENS | 4 | 8112 | 3p56 | A | 212 | ASN | p.Asn212Ile | O75792 |
| HOMO SAPIENS | 0 | 8112 | 3p56 | A | 235 | ARG | p.Arg235Gln | O75792 |
| HOMO SAPIENS | 4 | 8112 | 3p56 | A | 240 | THR | p.Thr240Met | O75792 |
| HOMO SAPIENS | 0 | 8112 | 3p56 | A | 291 | ARG | p.Arg291His | O75792 |
| HOMO SAPIENS | 0 | 37603 | 2yqg | A | 10 | ARG | p.Arg49His | Q14126 |
| HOMO SAPIENS | 0 | 21178 | 3k1r | B | 458 | ASP | p.Asp458Val | Q495M9 |
| HOMO SAPIENS | -1 | 36828 | 4fq3 | B | 507 | GLY | p.Gly507Asp | P35637 |
| HOMO SAPIENS | 5 | 36828 | 4fq3 | B | 517 | HIS | p.His517Gln | P35637 |
| HOMO SAPIENS | 8 | 36828 | 4fq3 | B | 518 | ARG | p.Arg518Lys | P35637 |
| HOMO SAPIENS | 6 | 36828 | 4fq3 | B | 521 | ARG | p.Arg521Cys | P35637 |
| HOMO SAPIENS | 6 | 36828 | 4fq3 | B | 521 | ARG | p.Arg521His | P35637 |
| HOMO SAPIENS | 0 | 6363 | 3j7y | c | null | HIS | p.His13Arg | Q9H9J2 |
| HOMO SAPIENS | 2 | 6363 | 3j7y | c | 105 | ARG | p.Arg105Cys | Q9H9J2 |
| HOMO SAPIENS | 2 | 6363 | 3j7y | c | 138 | THR | p.Thr138Ile | Q9H9J2 |

| | | | | | | | | |
|---|---|---|---|---|---|---|---|---|
| HOMO SAPIENS | 1 | 6363 | 3j7y | c | 156 | LEU | p.Leu156Arg | Q9H9J2 |
| HOMO SAPIENS | 1 | 6782 | 5l46 | A | 109 | HIS | p.His109Arg | Q9UI17 |
| HOMO SAPIENS | 0 | 23951 | 5t76 | A | 175 | GLY | p.Gly931Ser | P49588 |
| HOMO SAPIENS | 8 | 12604 | 3p0l | A | 182 | ARG | p.Arg182Leu | P49675 |
| HOMO SAPIENS | 2 | 12604 | 3p0l | A | 187 | VAL | p.Val187Met | P49675 |
| HOMO SAPIENS | 2 | 12604 | 3p0l | A | 188 | ARG | p.Arg188Cys | P49675 |
| HOMO SAPIENS | 0 | 12604 | 3p0l | A | 193 | ARG | p.Arg193Ter | P49675 |
| HOMO SAPIENS | 9 | 12604 | 3p0l | A | 217 | ARG | p.Arg217Thr | P49675 |
| HOMO SAPIENS | 4 | 12604 | 3p0l | A | 218 | ALA | p.Ala218Val | P49675 |
| HOMO SAPIENS | 0 | 12604 | 3p0l | A | 250 | TRP | p.Trp250Ter | P49675 |
| HOMO SAPIENS | 1 | 12604 | 3p0l | A | 258 | GLN | p.Gln258Ter | P49675 |
| HOMO SAPIENS | -1 | 27765 | 4wkr | A | 4 | GLU | p.Glu4Lys | Q4G0J3 |
| HOMO SAPIENS | 2 | 3587 | 4kit | C | 2301 | PRO | p.Pro2301Thr | Q6P2Q9 |
| HOMO SAPIENS | 3 | 3587 | 4kit | C | 2304 | PHE | p.Phe2304Leu | Q6P2Q9 |
| HOMO SAPIENS | 3 | 3587 | 4kit | C | 2309 | HIS | p.His2309Pro | Q6P2Q9 |
| HOMO SAPIENS | 3 | 3587 | 4kit | C | 2310 | ARG | p.Arg2310Lys | Q6P2Q9 |
| HOMO SAPIENS | 0 | 9073 | 1n69 | B | 21 | ASN | p.Asn215His | P07602 |
| HOMO SAPIENS | 0 | 9073 | 1n69 | B | 23 | THR | p.Thr217Ile | P07602 |
| HOMO SAPIENS | 0 | 9073 | 1n69 | B | 47 | CYS | p.Cys241Ser | P07602 |
| HOMO SAPIENS | 0 | 475 | 2vd5 | A | 225 | THR | p.Thr225Met | Q09013 |
| HOMO SAPIENS | 1 | 9066 | 4mzv | A | 31 | ASN | p.Asn31Lys | P16422 |
| HOMO SAPIENS | 1 | 9066 | 4mzv | A | 47 | GLN | p.Gln47His | P16422 |
| HOMO SAPIENS | 3 | 9066 | 4mzv | A | 66 | CYS | p.Cys66Tyr | P16422 |
| HOMO SAPIENS | 1 | 9066 | 4mzv | A | 102 | SER | p.Ser102Gly | P16422 |
| HOMO SAPIENS | 5 | 9066 | 4mzv | A | 115 | THR | p.Met115Thr | P16422 |
| HOMO SAPIENS | 1 | 9066 | 4mzv | A | 138 | ARG | p.Arg138Ter | P16422 |
| HOMO SAPIENS | 1 | 9066 | 4mzv | A | 153 | ARG | p.Arg153Thr | P16422 |
| HOMO SAPIENS | 1 | 9066 | 4mzv | A | 165 | ALA | p.Ala165Thr | P16422 |
| HOMO SAPIENS | 8 | 1827 | 2nz2 | A | 14 | GLY | p.Gly14Ser | P00966 |
| HOMO SAPIENS | 5 | 1827 | 2nz2 | A | 18 | SER | p.Ser18Leu | P00966 |
| HOMO SAPIENS | 3 | 1827 | 2nz2 | A | 86 | ARG | p.Arg86Cys | P00966 |
| HOMO SAPIENS | 3 | 1827 | 2nz2 | A | 86 | ARG | p.Arg86His | P00966 |
| HOMO SAPIENS | 1 | 1827 | 2nz2 | A | 179 | TRP | p.Trp179Arg | P00966 |
| HOMO SAPIENS | 7 | 1827 | 2nz2 | A | 180 | SER | p.Ser180Asn | P00966 |
| HOMO SAPIENS | 7 | 1827 | 2nz2 | A | 191 | GLU | p.Glu191Lys | P00966 |
| HOMO SAPIENS | 4 | 1827 | 2nz2 | A | 256 | GLU | p.Glu256Lys | P00966 |
| HOMO SAPIENS | 4 | 1827 | 2nz2 | A | 263 | VAL | p.Val263Met | P00966 |
| HOMO SAPIENS | 5 | 1827 | 2nz2 | A | 265 | ARG | p.Arg265Cys | P00966 |
| HOMO SAPIENS | 5 | 1827 | 2nz2 | A | 268 | ILE | p.Ile268Met | P00966 |
| HOMO SAPIENS | 6 | 1827 | 2nz2 | A | 269 | VAL | p.Val269Met | P00966 |
| HOMO SAPIENS | 7 | 1827 | 2nz2 | A | 272 | ARG | p.Arg272Cys | P00966 |
| HOMO SAPIENS | 3 | 1827 | 2nz2 | A | 304 | ARG | p.Arg304Trp | P00966 |
| HOMO SAPIENS | 3 | 1827 | 2nz2 | A | 307 | ARG | p.Arg307Cys | P00966 |
| HOMO SAPIENS | 3 | 1827 | 2nz2 | A | 310 | LYS | p.Lys310Gln | P00966 |
| HOMO SAPIENS | 2 | 1827 | 2nz2 | A | 362 | GLY | p.Gly362Val | P00966 |
| HOMO SAPIENS | 2 | 1827 | 2nz2 | A | 363 | ARG | p.Arg363Trp | P00966 |
| HOMO SAPIENS | 2 | 1827 | 2nz2 | A | 363 | ARG | p.Arg363Gln | P00966 |
| HOMO SAPIENS | 1 | 1827 | 2nz2 | A | 390 | GLY | p.Gly390Arg | P00966 |
| HOMO SAPIENS | 0 | 29524 | 5vbn | B | 2165 | ARG | p.Arg2165His | Q07864 |
| HOMO SAPIENS | 2 | 29524 | 5vbn | B | 2255 | ILE | p.Ile2255Phe | Q07864 |
| HOMO SAPIENS | 2 | 29524 | 5vbn | B | 2274 | LEU | p.Leu2274Val | Q07864 |
| HOMO SAPIENS | 0 | 31897 | 3t0o | A | 184 | CYS | p.Cys184Arg | O00584 |
| HOMO SAPIENS | -1 | 31897 | 3t0o | A | 215 | GLU | p.Glu215Lys | O00584 |
| HOMO SAPIENS | 0 | 31897 | 3t0o | A | 236 | ARG | p.Arg236Trp | O00584 |

| | | | | | | | | |
|---|---|---|---|---|---|---|---|---|
| HOMO SAPIENS | 7 | 18490 | 4pcw | B | 31 | GLU | p.Glu32Ter | P20930 |
| HOMO SAPIENS | -1 | 39865 | 2h9e | L | 152 | GLY | p.Gly192Arg | P00742 |
| HOMO SAPIENS | 2 | 4309 | 2x0b | F | 174 | THR | p.Thr207Met | P01019 |
| HOMO SAPIENS | 3 | 4309 | 2x0b | F | 342 | ARG | p.Arg375Gln | P01019 |
| HOMO SAPIENS | 2 | 3094 | 1zll | B | 9 | ARG | p.Arg9Cys | P26678 |
| HOMO SAPIENS | 9 | 3094 | 1zll | B | 39 | LEU | p.Leu39Ter | P26678 |
| HOMO SAPIENS | 5 | 3094 | 1zll | B | 49 | VAL | p.Val49Met | P26678 |
| HOMO SAPIENS | -1 | 41498 | 3ffd | P | 8 | LEU | p.Leu44Pro | P12272 |
| HOMO SAPIENS | 1 | 20697 | 1igr | A | 407 | ARG | p.Arg437His | P08069 |
| HOMO SAPIENS | 0 | 20697 | 1igr | A | 416 | MET | p.Met446Val | P08069 |
| HOMO SAPIENS | -1 | 20697 | 1igr | A | 462 | VAL | p.Val492Ile | P08069 |
| HOMO SAPIENS | 0 | 3973 | 4cbz | A | 37 | LEU | p.Leu37Ser | P78504 |
| HOMO SAPIENS | 0 | 3973 | 4cbz | A | 45 | VAL | p.Val45Leu | P78504 |
| HOMO SAPIENS | 0 | 3973 | 4cbz | A | 134 | LEU | p.Leu134Met | P78504 |
| HOMO SAPIENS | -1 | 3973 | 4cbz | A | 146 | VAL | p.Val146Ile | P78504 |
| HOMO SAPIENS | 0 | 3973 | 4cbz | A | 176 | VAL | p.Val176Ile | P78504 |
| HOMO SAPIENS | 0 | 3973 | 4cbz | A | 184 | ARG | p.Arg184Cys | P78504 |
| HOMO SAPIENS | 6 | 1697 | 4hmy | S | 33 | ARG | p.Arg33Trp | Q96PC3 |
| HOMO SAPIENS | 2 | 23752 | 5j4o | A | 13 | ARG | p.Arg1611Ser | P02549 |
| HOMO SAPIENS | 0 | 23752 | 5j4o | A | 42 | ALA | p.Ala1640Gly | P02549 |
| HOMO SAPIENS | 0 | 23752 | 5j4o | A | 108 | HIS | p.His1706Gln | P02549 |
| HOMO SAPIENS | 6 | 23752 | 5j4o | A | 206 | LEU | p.Leu1804Phe | P02549 |
| HOMO SAPIENS | 8 | 531 | 2qz4 | A | 349 | GLY | p.Gly349Ser | Q9UQ90 |
| HOMO SAPIENS | 0 | 531 | 2qz4 | A | 486 | ARG | p.Arg486Gln | Q9UQ90 |
| HOMO SAPIENS | 1 | 531 | 2qz4 | A | 510 | ALA | p.Ala510Val | Q9UQ90 |
| HOMO SAPIENS | 0 | 531 | 2qz4 | A | 529 | ALA | p.Ala529Val | Q9UQ90 |
| HOMO SAPIENS | 0 | 531 | 2qz4 | A | 558 | LYS | p.Lys558Ter | Q9UQ90 |
| HOMO SAPIENS | 1 | 531 | 2qz4 | A | 559 | LYS | p.Lys559Ter | Q9UQ90 |
| HOMO SAPIENS | 2 | 7736 | 1fnh | A | 114 | TRP | p.Trp1834Arg | P02751 |
| HOMO SAPIENS | 0 | 7736 | 1fnh | A | 163 | LEU | p.Leu1883Arg | P02751 |
| HOMO SAPIENS | 3 | 20337 | 5wy5 | B | 264 | LEU | p.Leu264Phe | Q96MG7 |
| HOMO SAPIENS | 8 | 36096 | 2jku | A | 643 | GLY | p.Gly668Arg | P05165 |
| HOMO SAPIENS | 6 | 36096 | 2jku | A | 663 | CYS | p.Cys688Arg | P05165 |
| HOMO SAPIENS | 0 | 9850 | 3kbt | A | 1606 | TYR | p.Tyr1606Ter | P11277 |
| HOMO SAPIENS | 4 | 6876 | 4jkq | A | 217 | ILE | p.Ile217Val | Q92736 |
| HOMO SAPIENS | 5 | 6876 | 4jkq | A | 420 | ARG | p.Arg420Trp | Q92736 |
| HOMO SAPIENS | 2 | 27603 | 1itv | B | 62 | PRO | p.Arg574Pro | P14780 |
| HOMO SAPIENS | 4 | 27603 | 1itv | B | 156 | ARG | p.Arg668Gln | P14780 |
| HOMO SAPIENS | 2 | 22954 | 4f52 | E | 48 | ILE | p.Ile48Met | Q92990 |
| HOMO SAPIENS | 0 | 22954 | 4f52 | E | 91 | ASP | p.Asp91Asn | Q92990 |
| HOMO SAPIENS | 0 | 22954 | 4f52 | E | 146 | ALA | p.Ala146Thr | Q92990 |
| HOMO SAPIENS | 0 | 22954 | 4f52 | E | 181 | ILE | p.Ile181Lys | Q92990 |
| HOMO SAPIENS | 0 | 22954 | 4f52 | E | 194 | ASN | p.Asn194Ser | Q92990 |
| HOMO SAPIENS | 2 | 22954 | 4f52 | E | 254 | PRO | p.Pro254Arg | Q92990 |
| HOMO SAPIENS | 2 | 22954 | 4f52 | E | 300 | ILE | p.Ile300Thr | Q92990 |
| HOMO SAPIENS | 2 | 22954 | 4f52 | E | 336 | LEU | p.Leu336Ser | Q92990 |
| HOMO SAPIENS | 0 | 22954 | 4f52 | E | 353 | TYR | p.Tyr353His | Q92990 |
| HOMO SAPIENS | 0 | 22954 | 4f52 | E | 387 | MET | p.Met387Val | Q92990 |
| HOMO SAPIENS | 4 | 2946 | 5gl7 | A | 344 | TYR | p.Tyr344Cys | P56192 |
| HOMO SAPIENS | 0 | 2946 | 5gl7 | A | 370 | PHE | p.Phe370Leu | P56192 |
| HOMO SAPIENS | 0 | 2946 | 5gl7 | A | 393 | ALA | p.Ala393Thr | P56192 |
| HOMO SAPIENS | 0 | 2946 | 5gl7 | A | 523 | ILE | p.Ile523Thr | P56192 |
| HOMO SAPIENS | 0 | 2946 | 5gl7 | A | 567 | SER | p.Ser567Leu | P56192 |
| HOMO SAPIENS | 0 | 2946 | 5gl7 | A | 800 | PRO | p.Pro800Thr | P56192 |

| | | | | | | | | |
|---|---|---|---|---|---|---|---|---|
| HOMO SAPIENS | 3 | 34961 | 4wy4 | C | 44 | TYR | p.Tyr44His | O95721 |
| HOMO SAPIENS | 8 | 34961 | 4wy4 | C | 89 | GLU | p.Glu89Lys | O95721 |
| HOMO SAPIENS | 0 | 9427 | 4chl | B | 21 | ALA | p.Ala21Ser | O95571 |
| HOMO SAPIENS | 2 | 9427 | 4chl | B | 62 | ALA | p.Ala62Thr | O95571 |
| HOMO SAPIENS | 0 | 9427 | 4chl | B | 114 | ILE | p.Ile114Phe | O95571 |
| HOMO SAPIENS | 2 | 9427 | 4chl | B | 159 | ARG | p.Arg159His | O95571 |
| HOMO SAPIENS | 8 | 9427 | 4chl | B | 163 | ARG | p.Arg163Trp | O95571 |
| HOMO SAPIENS | 8 | 9427 | 4chl | B | 163 | ARG | p.Arg163Gln | O95571 |
| HOMO SAPIENS | 2 | 9427 | 4chl | B | 185 | LEU | p.Leu185Arg | O95571 |
| HOMO SAPIENS | 0 | 40917 | 2e8o | A | 11 | PRO | p.Pro26Leu | Q9Y3Z3 |
| HOMO SAPIENS | -1 | 39190 | 4gow | A | 556 | TYR | p.Tyr556Ter | P56696 |
| HOMO SAPIENS | 0 | 2399 | 2mkv | A | 8 | GLY | p.Gly10Ser | P54710 |
| HOMO SAPIENS | 1 | 2399 | 2mkv | A | 24 | VAL | p.Val26Ile | P54710 |
| HOMO SAPIENS | 4 | 2399 | 2mkv | A | 39 | GLY | p.Gly41Arg | P54710 |
| HOMO SAPIENS | 0 | 20801 | 2zej | B | 1441 | ARG | p.Arg1441Cys | Q5S007 |
| HOMO SAPIENS | 0 | 20801 | 2zej | B | 1441 | ARG | p.Arg1441His | Q5S007 |
| HOMO SAPIENS | 5 | 1019 | 5xtb | M | 62 | ARG | p.Arg62Gln | P28331 |
| HOMO SAPIENS | 1 | 1019 | 5xtb | M | 85 | ALA | p.Ala85Thr | P28331 |
| HOMO SAPIENS | 5 | 1019 | 5xtb | M | 231 | LEU | p.Leu231Val | P28331 |
| HOMO SAPIENS | 2 | 1019 | 5xtb | M | 241 | ARG | p.Arg241Trp | P28331 |
| HOMO SAPIENS | 1 | 1019 | 5xtb | M | 252 | ASP | p.Asp252Gly | P28331 |
| HOMO SAPIENS | 1 | 1019 | 5xtb | M | 408 | ARG | p.Arg408Cys | P28331 |
| HOMO SAPIENS | 2 | 1019 | 5xtb | M | 417 | ARG | p.Arg417Ser | P28331 |
| HOMO SAPIENS | 1 | 1019 | 5xtb | M | 557 | ARG | p.Arg557Ter | P28331 |
| HOMO SAPIENS | 0 | 1019 | 5xtb | M | 567 | ILE | p.Ile567Val | P28331 |
| HOMO SAPIENS | 1 | 1019 | 5xtb | M | 595 | THR | p.Thr595Ala | P28331 |
| HOMO SAPIENS | 1 | 1019 | 5xtb | M | 703 | ALA | p.Ala703Thr | P28331 |
| HOMO SAPIENS | -1 | 3437 | 5hyn | A | 185 | ASP | p.Asp185His | Q15910 |
| HOMO SAPIENS | -1 | 3437 | 5hyn | A | 482 | ALA | p.Ala482Thr | Q15910 |
| HOMO SAPIENS | 0 | 8201 | 2wfd | A | 373 | TYR | p.Tyr373Cys | Q9P2J5 |
| HOMO SAPIENS | 1 | 3590 | 3edi | A | 90 | VAL | p.Val238Ala | O43897 |
| HOMO SAPIENS | 6 | 4555 | 1ytq | A | 64 | ALA | p.Ala65Ser | P43320 |
| HOMO SAPIENS | 0 | 4555 | 1ytq | A | 194 | TRP | p.Trp195Gly | P43320 |
| HOMO SAPIENS | 0 | 1013 | 2nn6 | C | 272 | SER | p.Ser272Thr | Q96B26 |
| HOMO SAPIENS | -1 | 29026 | 5m73 | D | 530 | ILE | p.Ile530Asn | O76094 |
| HOMO SAPIENS | 8 | 2252 | 1d5v | A | 57 | SER | p.Ser125Leu | Q99958 |
| HOMO SAPIENS | 1 | 4337 | 3ecr | A | 29 | ARG | p.Arg26His | P08397 |
| HOMO SAPIENS | 2 | 4337 | 3ecr | A | 34 | ALA | p.Ala31Thr | P08397 |
| HOMO SAPIENS | 2 | 4337 | 3ecr | A | 37 | GLN | p.Gln34Lys | P08397 |
| HOMO SAPIENS | 1 | 4337 | 3ecr | A | 58 | ALA | p.Ala55Ser | P08397 |
| HOMO SAPIENS | 0 | 4337 | 3ecr | A | 84 | LEU | p.Leu81Pro | P08397 |
| HOMO SAPIENS | 0 | 4337 | 3ecr | A | 114 | GLY | p.Gly111Arg | P08397 |
| HOMO SAPIENS | 1 | 4337 | 3ecr | A | 119 | ARG | p.Arg116Trp | P08397 |
| HOMO SAPIENS | 1 | 4337 | 3ecr | A | 152 | ARG | p.Arg149Gln | P08397 |
| HOMO SAPIENS | 1 | 4337 | 3ecr | A | 170 | ARG | p.Arg167Trp | P08397 |
| HOMO SAPIENS | 1 | 4337 | 3ecr | A | 176 | ARG | p.Arg173Gln | P08397 |
| HOMO SAPIENS | 0 | 4337 | 3ecr | A | 180 | LEU | p.Leu177Arg | P08397 |
| HOMO SAPIENS | 0 | 4337 | 3ecr | A | 181 | ASP | p.Asp178Asn | P08397 |
| HOMO SAPIENS | 1 | 4337 | 3ecr | A | 198 | ARG | p.Arg195Cys | P08397 |
| HOMO SAPIENS | 0 | 4337 | 3ecr | A | 201 | TRP | p.Trp198Ter | P08397 |
| HOMO SAPIENS | 1 | 4337 | 3ecr | A | 204 | ARG | p.Arg201Trp | P08397 |
| HOMO SAPIENS | 1 | 4337 | 3ecr | A | 219 | GLY | p.Gly216Asp | P08397 |
| HOMO SAPIENS | 1 | 4337 | 3ecr | A | 226 | GLU | p.Glu223Lys | P08397 |
| HOMO SAPIENS | 0 | 4337 | 3ecr | A | 248 | LEU | p.Leu245Arg | P08397 |

| | | | | | | | | |
|---|---|---|---|---|---|---|---|---|
| HOMO SAPIENS | 1 | 4337 | 3ecr | A | 250 | CYS | p.Cys247Arg | P08397 |
| HOMO SAPIENS | 0 | 4337 | 3ecr | A | 253 | GLU | p.Glu250Lys | P08397 |
| HOMO SAPIENS | 0 | 4337 | 3ecr | A | 255 | ALA | p.Ala252Thr | P08397 |
| HOMO SAPIENS | 0 | 4337 | 3ecr | A | 255 | ALA | p.Ala252Val | P08397 |
| HOMO SAPIENS | 0 | 4337 | 3ecr | A | 259 | HIS | p.His256Asn | P08397 |
| HOMO SAPIENS | 0 | 4337 | 3ecr | A | 324 | ARG | p.Arg321His | P08397 |
| HOMO SAPIENS | -1 | 4337 | 3ecr | A | 362 | ASP | p.Asp359Asn | P08397 |
| HOMO SAPIENS | 1 | 1204 | 5lwv | A | 854 | ILE | p.Ala864Thr | O15294 |
| HOMO SAPIENS | 1 | 25016 | 1m4u | A | 35 | PRO | p.Pro35Arg | Q13253 |
| HOMO SAPIENS | 1 | 25016 | 1m4u | A | 35 | PRO | p.Pro35Ser | Q13253 |
| HOMO SAPIENS | 1 | 25016 | 1m4u | A | 167 | ARG | p.Arg167Gly | Q13253 |
| HOMO SAPIENS | 1 | 25016 | 1m4u | A | 184 | CYS | p.Cys184Tyr | Q13253 |
| HOMO SAPIENS | 0 | 25016 | 1m4u | A | 189 | GLY | p.Gly189Cys | Q13253 |
| HOMO SAPIENS | 1 | 25016 | 1m4u | A | 204 | ARG | p.Arg204Leu | Q13253 |
| HOMO SAPIENS | 1 | 25016 | 1m4u | A | 205 | TRP | p.Trp205Cys | Q13253 |
| HOMO SAPIENS | 1 | 25016 | 1m4u | A | 217 | TRP | p.Trp217Gly | Q13253 |
| HOMO SAPIENS | 0 | 25016 | 1m4u | A | 222 | TYR | p.Tyr222Cys | Q13253 |
| HOMO SAPIENS | 0 | 25016 | 1m4u | A | 222 | TYR | p.Tyr222Asp | Q13253 |
| HOMO SAPIENS | 1 | 25016 | 1m4u | A | 223 | PRO | p.Pro223Leu | Q13253 |
| HOMO SAPIENS | 0 | 1394 | 1um7 | A | 95 | ALA | p.Ala463Val | Q92796 |
| HOMO SAPIENS | 0 | 21294 | 4jpz | B | 1788 | GLU | p.Glu1788Asp | Q99250 |
| HOMO SAPIENS | 9 | 24068 | 2db6 | A | 47 | ARG | p.Arg119Cys | Q96MF2 |
| HOMO SAPIENS | 5 | 1869 | 4omt | A | 39 | ARG | p.Arg39Leu | P08237 |
| HOMO SAPIENS | 2 | 1869 | 4omt | A | 100 | ARG | p.Arg100Gln | P08237 |
| HOMO SAPIENS | 7 | 1869 | 4omt | A | 543 | ASP | p.Asp543Ala | P08237 |
| HOMO SAPIENS | 1 | 1869 | 4omt | A | 686 | TRP | p.Trp686Cys | P08237 |
| HOMO SAPIENS | 1 | 1869 | 4omt | A | 696 | ARG | p.Arg696His | P08237 |
| HOMO SAPIENS | 0 | 12163 | 4wxx | A | 495 | TYR | p.Tyr495Cys | P26358 |
| HOMO SAPIENS | 0 | 12163 | 4wxx | A | 528 | ILE | p.Ile528Met | P26358 |
| HOMO SAPIENS | 0 | 12391 | 1j1d | C | 36 | LYS | p.Lys36Gln | P19429 |
| HOMO SAPIENS | 0 | 12391 | 1j1d | C | 82 | PRO | p.Pro82Ser | P19429 |
| HOMO SAPIENS | 7 | 12391 | 1j1d | C | 102 | ALA | p.Ala102Thr | P19429 |
| HOMO SAPIENS | 1 | 5000 | 4cca | A | 17 | GLY | p.Gly17Arg | Q15833 |
| HOMO SAPIENS | 0 | 5000 | 4cca | A | 205 | VAL | p.Val205Ile | Q15833 |
| HOMO SAPIENS | 0 | 5000 | 4cca | A | 209 | LEU | p.Leu209Pro | Q15833 |
| HOMO SAPIENS | 1 | 5000 | 4cca | A | 459 | ARG | p.Arg459Trp | Q15833 |
| HOMO SAPIENS | 7 | 5000 | 4cca | A | 477 | PRO | p.Pro477Leu | Q15833 |
| HOMO SAPIENS | -1 | 5000 | 4cca | A | 526 | ILE | p.Ile526Val | Q15833 |
| HOMO SAPIENS | 0 | 5000 | 4cca | A | 541 | GLY | p.Gly541Ser | Q15833 |
| HOMO SAPIENS | 0 | 5000 | 4cca | A | 555 | ARG | p.Arg555Gly | Q15833 |
| HOMO SAPIENS | 5 | 28551 | 3vcm | P | 26 | ARG | p.Arg49Ter | P00797 |
| HOMO SAPIENS | 6 | 36453 | 5dzz | A | 2070 | ASP | p.Asp2070His | P15924 |
| HOMO SAPIENS | 0 | 36453 | 5dzz | A | 2166 | ARG | p.Arg2166Ter | P15924 |
| HOMO SAPIENS | 0 | 20268 | 2da6 | A | 96 | ASN | p.Asn321Ser | P35680 |
| HOMO SAPIENS | 1 | 1405 | 3iar | A | 8 | ASP | p.Asp8Asn | P00813 |
| HOMO SAPIENS | 8 | 1405 | 3iar | A | 15 | HIS | p.His15Asp | P00813 |
| HOMO SAPIENS | 6 | 1405 | 3iar | A | 74 | GLY | p.Gly74Val | P00813 |
| HOMO SAPIENS | 6 | 1405 | 3iar | A | 80 | LYS | p.Lys80Arg | P00813 |
| HOMO SAPIENS | 5 | 1405 | 3iar | A | 97 | TYR | p.Tyr97Cys | P00813 |
| HOMO SAPIENS | 5 | 1405 | 3iar | A | 101 | ARG | p.Arg101Leu | P00813 |
| HOMO SAPIENS | 5 | 1405 | 3iar | A | 101 | ARG | p.Arg101Trp | P00813 |
| HOMO SAPIENS | 5 | 1405 | 3iar | A | 106 | LEU | p.Leu106Val | P00813 |
| HOMO SAPIENS | 0 | 1405 | 3iar | A | 107 | LEU | p.Leu107Pro | P00813 |
| HOMO SAPIENS | 9 | 1405 | 3iar | A | 156 | ARG | p.Arg156Cys | P00813 |

| | | | | | | | | |
|---|---|---|---|---|---|---|---|---|
| HOMO SAPIENS | 6 | 1405 | 3iar | A | 211 | ARG | p.Arg211His | P00813 |
| HOMO SAPIENS | 0 | 1405 | 3iar | A | 216 | GLY | p.Gly216Arg | P00813 |
| HOMO SAPIENS | 0 | 1405 | 3iar | A | 291 | SER | p.Ser291Leu | P00813 |
| HOMO SAPIENS | 6 | 1405 | 3iar | A | 304 | LEU | p.Leu304Arg | P00813 |
| HOMO SAPIENS | 0 | 1405 | 3iar | A | 329 | ALA | p.Ala329Val | P00813 |
| HOMO SAPIENS | -1 | 31340 | 4atu | I | 186 | ARG | p.Arg186His | O43602 |
| HOMO SAPIENS | -1 | 31340 | 4atu | I | 192 | ARG | p.Arg192Trp | O43602 |
| HOMO SAPIENS | -1 | 31340 | 4atu | I | 196 | ARG | p.Arg196His | O43602 |
| HOMO SAPIENS | -1 | 31340 | 4atu | I | 203 | THR | p.Thr203Arg | O43602 |
| HOMO SAPIENS | -1 | 31340 | 4atu | I | 286 | ALA | p.Ala286Ser | O43602 |
| HOMO SAPIENS | 4 | 30734 | 1fsu | A | 95 | ARG | p.Arg95Gln | P15848 |
| HOMO SAPIENS | 0 | 30734 | 1fsu | A | 117 | CYS | p.Cys117Arg | P15848 |
| HOMO SAPIENS | 0 | 30734 | 1fsu | A | 137 | GLY | p.Gly137Val | P15848 |
| HOMO SAPIENS | 0 | 30734 | 1fsu | A | 210 | TYR | p.Tyr210Cys | P15848 |
| HOMO SAPIENS | 0 | 30734 | 1fsu | A | 236 | LEU | p.Leu236Pro | P15848 |
| HOMO SAPIENS | 1 | 30734 | 1fsu | A | 315 | ARG | p.Arg315Gln | P15848 |
| HOMO SAPIENS | 0 | 30734 | 1fsu | A | 358 | VAL | p.Val358Met | P15848 |
| HOMO SAPIENS | 2 | 30734 | 1fsu | A | 376 | VAL | p.Val376Met | P15848 |
| HOMO SAPIENS | 0 | 30734 | 1fsu | A | 384 | SER | p.Ser384Asn | P15848 |
| HOMO SAPIENS | 0 | 30734 | 1fsu | A | 393 | HIS | p.His393Pro | P15848 |
| HOMO SAPIENS | 0 | 30734 | 1fsu | A | 405 | CYS | p.Cys405Tyr | P15848 |
| HOMO SAPIENS | 0 | 30734 | 1fsu | A | 458 | ASN | p.Asn458Ser | P15848 |
| HOMO SAPIENS | 1 | 30734 | 1fsu | A | 484 | ARG | p.Arg484Gly | P15848 |
| HOMO SAPIENS | 0 | 38072 | 1wmv | A | 51 | PRO | p.Pro98Leu | Q9NZC7 |
| HOMO SAPIENS | -1 | 5674 | 1j1e | C | 185 | ASN | p.Asn185Lys | P19429 |
| HOMO SAPIENS | -1 | 5674 | 1j1e | C | 192 | ARG | p.Arg192His | P19429 |
| HOMO SAPIENS | -1 | 5674 | 1j1e | C | 196 | ASP | p.Asp196Asn | P19429 |
| HOMO SAPIENS | -1 | 5674 | 1j1e | C | 206 | LYS | p.Lys206Gln | P19429 |
| HOMO SAPIENS | 0 | 38067 | 1wlp | A | 13 | PRO | p.Pro156Gln | P13498 |
| HOMO SAPIENS | -1 | 20441 | 3vfd | A | 289 | HIS | p.His289Tyr | Q9UBP0 |
| HOMO SAPIENS | 2 | 20441 | 3vfd | A | 344 | ILE | p.Ile344Lys | Q9UBP0 |
| HOMO SAPIENS | 0 | 20441 | 3vfd | A | 362 | SER | p.Ser362Cys | Q9UBP0 |
| HOMO SAPIENS | 1 | 20441 | 3vfd | A | 386 | ASN | p.Asn386Ser | Q9UBP0 |
| HOMO SAPIENS | 9 | 20441 | 3vfd | A | 441 | ASP | p.Asp441Gly | Q9UBP0 |
| HOMO SAPIENS | -1 | 20441 | 3vfd | A | 448 | CYS | p.Cys448Tyr | Q9UBP0 |
| HOMO SAPIENS | 2 | 20441 | 3vfd | A | 500 | ARG | p.Arg499Cys | Q9UBP0 |
| HOMO SAPIENS | 1 | 20441 | 3vfd | A | 563 | ARG | p.Arg562Gly | Q9UBP0 |
| HOMO SAPIENS | 3 | 18471 | 4dx8 | J | 26 | ARG | p.Arg26Gln | O00522 |
| HOMO SAPIENS | -1 | 18471 | 4dx8 | J | 101 | GLY | p.Gly101Val | O00522 |
| HOMO SAPIENS | -1 | 34735 | 2ly4 | B | 83 | ALA | p.Ala83Glu | P04637 |
| HOMO SAPIENS | 0 | 31128 | 4cvh | A | 116 | ARG | p.Arg116Cys | A4D126 |
| HOMO SAPIENS | 8 | 31128 | 4cvh | A | 215 | GLN | p.Gln215Ter | A4D126 |
| HOMO SAPIENS | 0 | 31128 | 4cvh | A | 268 | ARG | p.Arg268Ter | A4D126 |
| HOMO SAPIENS | 0 | 31128 | 4cvh | A | 316 | THR | p.Thr316Lys | A4D126 |
| HOMO SAPIENS | 1 | 7164 | 3mt5 | A | 369 | ASP | p.Asp434Gly | Q12791 |
| HOMO SAPIENS | 2 | 26812 | 2wtk | F | 49 | TYR | p.Tyr49Asp | Q15831 |
| HOMO SAPIENS | 0 | 26812 | 2wtk | F | 67 | LEU | p.Leu67Pro | Q15831 |
| HOMO SAPIENS | 4 | 26812 | 2wtk | F | 104 | ARG | p.Arg104Ser | Q15831 |
| HOMO SAPIENS | 0 | 26812 | 2wtk | F | 135 | GLY | p.Gly135Arg | Q15831 |
| HOMO SAPIENS | 0 | 26812 | 2wtk | F | 163 | GLY | p.Gly163Asp | Q15831 |
| HOMO SAPIENS | 2 | 26812 | 2wtk | F | 194 | ALA | p.Asp194Asn | Q15831 |
| HOMO SAPIENS | 2 | 1357 | 5xtd | W | 57 | ARG | p.Arg57His | Q9P0J0 |
| HOMO SAPIENS | 0 | 28086 | 4nn2 | A | 229 | HIS | p.His229Arg | Q8IWS0 |
| HOMO SAPIENS | 0 | 28086 | 4nn2 | A | 234 | LYS | p.Lys234Glu | Q8IWS0 |

| | | | | | | | | |
|---|---|---|---|---|---|---|---|---|
| HOMO SAPIENS | 0 | 28086 | 4nn2 | A | 257 | ARG | p.Arg257Gly | Q8IWS0 |
| HOMO SAPIENS | 5 | 31182 | 2a1x | A | 176 | GLN | p.Gln176Lys | O14832 |
| HOMO SAPIENS | 1 | 31182 | 2a1x | A | 204 | GLY | p.Gly204Ser | O14832 |
| HOMO SAPIENS | 0 | 31182 | 2a1x | A | 269 | ASN | p.Asn269His | O14832 |
| HOMO SAPIENS | 3 | 31182 | 2a1x | A | 275 | ARG | p.Arg275Gln | O14832 |
| HOMO SAPIENS | 3 | 31182 | 2a1x | A | 275 | ARG | p.Arg275Trp | O14832 |
| HOMO SAPIENS | 4 | 2823 | 5xtb | P | 145 | THR | p.Thr145Ile | O75489 |
| HOMO SAPIENS | 5 | 2823 | 5xtb | P | 199 | ARG | p.Arg199Trp | O75489 |
| HOMO SAPIENS | 0 | 8645 | 2h5g | A | 372 | SER | p.Ser372Tyr | P54886 |
| HOMO SAPIENS | 3 | 8645 | 2h5g | A | 715 | ASP | p.Asp715His | P54886 |
| HOMO SAPIENS | 0 | 8645 | 2h5g | A | 765 | ARG | p.Arg765Gln | P54886 |
| HOMO SAPIENS | 0 | 8645 | 2h5g | A | 782 | TYR | p.Tyr782Cys | P54886 |
| HOMO SAPIENS | 0 | 8645 | 2h5g | A | 784 | HIS | p.His784Tyr | P54886 |
| HOMO SAPIENS | 1 | 13992 | 2cue | A | 39 | ARG | p.Arg242Thr | P26367 |
| HOMO SAPIENS | 9 | 13992 | 2cue | A | 55 | PHE | p.Phe258Ser | P26367 |
| HOMO SAPIENS | 0 | 30418 | 2xsw | A | 303 | VAL | p.Val303Met | Q9NRR6 |
| HOMO SAPIENS | 0 | 30418 | 2xsw | A | 315 | PRO | p.Pro315Leu | Q9NRR6 |
| HOMO SAPIENS | 0 | 30418 | 2xsw | A | 341 | GLY | p.Gly341Ser | Q9NRR6 |
| HOMO SAPIENS | 0 | 30418 | 2xsw | A | 378 | ARG | p.Arg378Cys | Q9NRR6 |
| HOMO SAPIENS | 0 | 30418 | 2xsw | A | 435 | ARG | p.Arg435Gln | Q9NRR6 |
| HOMO SAPIENS | 0 | 30418 | 2xsw | A | 515 | ARG | p.Arg515Trp | Q9NRR6 |
| HOMO SAPIENS | 1 | 30418 | 2xsw | A | 526 | PRO | p.Pro526Leu | Q9NRR6 |
| HOMO SAPIENS | 0 | 30418 | 2xsw | A | 563 | ARG | p.Arg563His | Q9NRR6 |
| HOMO SAPIENS | 2 | 30418 | 2xsw | A | 585 | ARG | p.Arg585His | Q9NRR6 |
| HOMO SAPIENS | 2 | 11167 | 4y99 | B | 205 | ARG | p.Arg215Leu | P45379 |
| HOMO SAPIENS | 5 | 11167 | 4y99 | B | 244 | GLU | p.Glu254Asp | P45379 |
| HOMO SAPIENS | 1 | 11167 | 4y99 | B | 253 | LYS | p.Lys263Arg | P45379 |
| HOMO SAPIENS | 8 | 11167 | 4y99 | B | 265 | ARG | p.Arg275Gln | P45379 |
| HOMO SAPIENS | 3 | 11167 | 4y99 | B | 270 | ASP | p.Asp280Asn | P45379 |
| HOMO SAPIENS | -1 | 11167 | 4y99 | B | 278 | ARG | p.Arg288Pro | P45379 |
| HOMO SAPIENS | -1 | 11167 | 4y99 | B | 286 | ARG | p.Arg296Cys | P45379 |
| HOMO SAPIENS | -1 | 11167 | 4y99 | B | 286 | ARG | p.Arg296His | P45379 |
| HOMO SAPIENS | 0 | 15993 | 4po6 | A | 197 | ARG | p.Arg197His | P29597 |
| HOMO SAPIENS | -1 | 15993 | 4po6 | A | 362 | VAL | p.Val362Phe | P29597 |
| HOMO SAPIENS | -1 | 15993 | 4po6 | A | 363 | GLY | p.Gly363Ser | P29597 |
| HOMO SAPIENS | 0 | 5646 | 5uak | A | 7 | GLU | p.Glu7Ter | P13569 |
| HOMO SAPIENS | 0 | 5646 | 5uak | A | 27 | GLY | p.Gly27Ter | P13569 |
| HOMO SAPIENS | 0 | 5646 | 5uak | A | 46 | ALA | p.Ala46Asp | P13569 |
| HOMO SAPIENS | 0 | 5646 | 5uak | A | 60 | GLU | p.Glu60Ter | P13569 |
| HOMO SAPIENS | 0 | 5646 | 5uak | A | 67 | PRO | p.Pro67Leu | P13569 |
| HOMO SAPIENS | 0 | 5646 | 5uak | A | 74 | ARG | p.Arg74Trp | P13569 |
| HOMO SAPIENS | 0 | 5646 | 5uak | A | 75 | ARG | p.Arg75Ter | P13569 |
| HOMO SAPIENS | 0 | 5646 | 5uak | A | 85 | GLY | p.Gly85Glu | P13569 |
| HOMO SAPIENS | 0 | 5646 | 5uak | A | 91 | GLY | p.Gly91Arg | P13569 |
| HOMO SAPIENS | 0 | 5646 | 5uak | A | 92 | GLU | p.Glu92Lys | P13569 |
| HOMO SAPIENS | 0 | 5646 | 5uak | A | 98 | GLN | p.Gln98Arg | P13569 |
| HOMO SAPIENS | 0 | 5646 | 5uak | A | 98 | GLN | p.Gln98Ter | P13569 |
| HOMO SAPIENS | 0 | 5646 | 5uak | A | 109 | TYR | p.Tyr109Cys | P13569 |
| HOMO SAPIENS | 0 | 5646 | 5uak | A | 110 | ASP | p.Asp110His | P13569 |
| HOMO SAPIENS | 0 | 5646 | 5uak | A | 117 | ARG | p.Arg117Cys | P13569 |
| HOMO SAPIENS | 0 | 5646 | 5uak | A | 117 | ARG | p.Arg117His | P13569 |
| HOMO SAPIENS | 0 | 5646 | 5uak | A | 125 | ILE | p.Ile125Thr | P13569 |
| HOMO SAPIENS | 0 | 5646 | 5uak | A | 148 | ILE | p.Ile148Thr | P13569 |
| HOMO SAPIENS | 0 | 5646 | 5uak | A | 170 | ARG | p.Arg170His | P13569 |

| | | | | | | | | |
|---|---|---|---|---|---|---|---|---|
| HOMO SAPIENS | 0 | 5646 | 5uak | A | 178 | GLY | p.Gly178Arg | P13569 |
| HOMO SAPIENS | 0 | 5646 | 5uak | A | 206 | LEU | p.Leu206Trp | P13569 |
| HOMO SAPIENS | 0 | 5646 | 5uak | A | 373 | ASP | p.Asp373Asn | P13569 |
| HOMO SAPIENS | -1 | 5646 | 5uak | A | 709 | ARG | p.Arg709Ter | P13569 |
| HOMO SAPIENS | -1 | 5646 | 5uak | A | 732 | LEU | p.Leu732Ter | P13569 |
| HOMO SAPIENS | -1 | 5646 | 5uak | A | 785 | ARG | p.Arg785Ter | P13569 |
| HOMO SAPIENS | -1 | 5646 | 5uak | A | 831 | GLU | p.Glu831Ter | P13569 |
| HOMO SAPIENS | -1 | 5646 | 5uak | A | 836 | ASP | p.Asp836Tyr | P13569 |
| HOMO SAPIENS | 0 | 5646 | 5uak | A | 851 | ARG | p.Arg851Ter | P13569 |
| HOMO SAPIENS | 0 | 5646 | 5uak | A | 913 | TYR | p.Tyr913Cys | P13569 |
| HOMO SAPIENS | 0 | 5646 | 5uak | A | 913 | TYR | p.Tyr913Ter | P13569 |
| HOMO SAPIENS | 0 | 5646 | 5uak | A | 945 | SER | p.Ser945Leu | P13569 |
| HOMO SAPIENS | 0 | 5646 | 5uak | A | 949 | HIS | p.His949Tyr | P13569 |
| HOMO SAPIENS | 0 | 5646 | 5uak | A | 970 | GLY | p.Gly970Asp | P13569 |
| HOMO SAPIENS | 0 | 5646 | 5uak | A | 1027 | ILE | p.Ile1027Thr | P13569 |
| HOMO SAPIENS | 0 | 5646 | 5uak | A | 1032 | TYR | p.Tyr1032Cys | P13569 |
| HOMO SAPIENS | 0 | 5646 | 5uak | A | 1061 | GLY | p.Gly1061Arg | P13569 |
| HOMO SAPIENS | 1 | 5646 | 5uak | A | 1065 | LEU | p.Leu1065Pro | P13569 |
| HOMO SAPIENS | 1 | 5646 | 5uak | A | 1066 | ARG | p.Arg1066Cys | P13569 |
| HOMO SAPIENS | 1 | 5646 | 5uak | A | 1066 | ARG | p.Arg1066His | P13569 |
| HOMO SAPIENS | 1 | 5646 | 5uak | A | 1067 | ALA | p.Ala1067Thr | P13569 |
| HOMO SAPIENS | 0 | 5646 | 5uak | A | 1069 | GLY | p.Gly1069Arg | P13569 |
| HOMO SAPIENS | 0 | 5646 | 5uak | A | 1070 | ARG | p.Arg1070Trp | P13569 |
| HOMO SAPIENS | 1 | 5646 | 5uak | A | 1071 | GLN | p.Gln1071Pro | P13569 |
| HOMO SAPIENS | 0 | 5646 | 5uak | A | 1074 | PHE | p.Phe1074Leu | P13569 |
| HOMO SAPIENS | 0 | 5646 | 5uak | A | 1077 | LEU | p.Leu1077Pro | P13569 |
| HOMO SAPIENS | 0 | 5646 | 5uak | A | 1089 | TRP | p.Trp1089Ter | P13569 |
| HOMO SAPIENS | 0 | 5646 | 5uak | A | 1092 | TYR | p.Tyr1092Ter | P13569 |
| HOMO SAPIENS | 0 | 5646 | 5uak | A | 1101 | MET | p.Met1101Lys | P13569 |
| HOMO SAPIENS | 0 | 5646 | 5uak | A | 1156 | LEU | p.Leu1156Phe | P13569 |
| HOMO SAPIENS | 0 | 5646 | 5uak | A | 1158 | ARG | p.Arg1158Ter | P13569 |
| HOMO SAPIENS | 0 | 5646 | 5uak | A | 1162 | ARG | p.Arg1162Leu | P13569 |
| HOMO SAPIENS | 0 | 5646 | 5uak | A | 1162 | ARG | p.Arg1162Ter | P13569 |
| HOMO SAPIENS | -1 | 5646 | 5uak | A | 1476 | GLN | p.Gln1476Ter | P13569 |
| HOMO SAPIENS | 0 | 22763 | 1imv | A | 52 | MET | p.Thr72Met | P36955 |
| HOMO SAPIENS | 0 | 22763 | 1imv | A | 301 | TYR | p.Tyr321Ter | P36955 |
| HOMO SAPIENS | 0 | 23179 | 5fql | A | 160 | PRO | p.Pro160Arg | P22304 |
| HOMO SAPIENS | 0 | 23179 | 5fql | A | 172 | ARG | p.Arg172Ter | P22304 |
| HOMO SAPIENS | 0 | 23179 | 5fql | A | 214 | THR | p.Thr214Met | P22304 |
| HOMO SAPIENS | 0 | 23179 | 5fql | A | 309 | THR | p.Thr309Ala | P22304 |
| HOMO SAPIENS | 0 | 23179 | 5fql | A | 333 | SER | p.Ser333Leu | P22304 |
| HOMO SAPIENS | 0 | 23179 | 5fql | A | 443 | ARG | p.Arg443Ter | P22304 |
| HOMO SAPIENS | 0 | 23179 | 5fql | A | 468 | ARG | p.Arg468Leu | P22304 |
| HOMO SAPIENS | 0 | 23179 | 5fql | A | 468 | ARG | p.Arg468Trp | P22304 |
| HOMO SAPIENS | 0 | 23179 | 5fql | A | 502 | TRP | p.Trp502Ser | P22304 |
| HOMO SAPIENS | 9 | 20989 | 1nfi | F | 185 | THR | p.Thr185Met | P25963 |
| HOMO SAPIENS | 4 | 20989 | 1nfi | F | 194 | GLY | p.Gly194Ala | P25963 |
| HOMO SAPIENS | 0 | 11538 | 3eab | A | 162 | VAL | p.Val162Ile | Q9UBP0 |
| HOMO SAPIENS | 0 | 33399 | 2p39 | A | 71 | SER | p.Ser71Gly | Q9GZV9 |
| HOMO SAPIENS | 3 | 33399 | 2p39 | A | 96 | MET | p.Met96Thr | Q9GZV9 |
| HOMO SAPIENS | 0 | 33399 | 2p39 | A | 129 | SER | p.Ser129Phe | Q9GZV9 |
| HOMO SAPIENS | -1 | 33399 | 2p39 | A | 172 | PRO | p.Pro172Leu | Q9GZV9 |
| HOMO SAPIENS | -1 | 33399 | 2p39 | A | 176 | ARG | p.Arg176Gln | Q9GZV9 |
| HOMO SAPIENS | -1 | 33399 | 2p39 | A | 179 | ARG | p.Arg179Trp | Q9GZV9 |

| | | | | | | | | |
|---|---|---|---|---|---|---|---|---|
| HOMO SAPIENS | -1 | 33399 | 2p39 | A | 179 | ARG | p.Arg179Gln | Q9GZV9 |
| HOMO SAPIENS | 0 | 16217 | 2dne | A | 79 | ILE | p.Ile163Leu | P10515 |
| HOMO SAPIENS | 8 | 7666 | 4kbq | A | 65 | ASN | p.Asn65Ser | Q9UNE7 |
| HOMO SAPIENS | 1 | 6407 | 1ikt | A | 71 | THR | p.Thr687Ile | P51659 |
| HOMO SAPIENS | 0 | 6407 | 1ikt | A | 112 | MET | p.Met728Val | P51659 |
| HOMO SAPIENS | 0 | 5849 | 4ihb | A | 67 | VAL | p.Val67Asp | O75923 |
| HOMO SAPIENS | 6 | 17703 | 5fcn | A | 20 | HIS | p.His99Gln | Q66GS9 |
| HOMO SAPIENS | 8 | 17703 | 5fcn | A | 33 | ARG | p.Arg112His | Q66GS9 |
| HOMO SAPIENS | 0 | 34095 | 2E+073 | A | 101 | HIS | p.His101Tyr | P05129 |
| HOMO SAPIENS | -1 | 41358 | 4liq | E | 507 | THR | p.Thr507Met | P07333 |
| HOMO SAPIENS | -1 | 21290 | 1fch | A | 235 | MET | p.Met272Arg | P50542 |
| HOMO SAPIENS | 5 | 23431 | 6pax | A | 23 | ARG | p.Arg26Gly | P26367 |
| HOMO SAPIENS | 1 | 23431 | 6pax | A | 61 | GLY | p.Gly64Val | P26367 |
| HOMO SAPIENS | 1 | 23431 | 6pax | A | 65 | PRO | p.Pro68Ser | P26367 |
| HOMO SAPIENS | 1 | 23431 | 6pax | A | 116 | SER | p.Ser119Arg | P26367 |
| HOMO SAPIENS | 9 | 23431 | 6pax | A | 123 | VAL | p.Val126Asp | P26367 |
| HOMO SAPIENS | 9 | 23431 | 6pax | A | 125 | ARG | p.Arg128Cys | P26367 |
| HOMO SAPIENS | 9 | 33003 | 4zrj | B | 535 | LEU | p.Leu535Pro | P35240 |
| HOMO SAPIENS | 7 | 33003 | 4zrj | B | 538 | GLN | p.Gln538Pro | P35240 |
| HOMO SAPIENS | 0 | 33003 | 4zrj | B | 585 | TRP | p.Ala585Thr | P35240 |
| HOMO SAPIENS | 0 | 3800 | 2nzj | A | 193 | VAL | p.Val193Met | O75628 |

| gene_name | max_cons | max_bs_id | max_pdb_id | max_chain_id |
|---|---|---|---|---|
| AKR1D1 | None | None | None | None |
| AKR1D1 | None | None | None | None |
| AKR1D1 | None | None | None | None |
| AKR1D1 | None | None | None | None |
| ALPP | None | None | None | None |
| BMPR1A | None | None | None | None |
| BMPR1A | None | None | None | None |
| FGFR2 | None | None | None | None |
| PROC | None | None | None | None |
| PROC | None | None | None | None |
| PROC | None | None | None | None |
| PROC | None | None | None | None |
| PROC | None | None | None | None |
| PROC | None | None | None | None |
| PROC | None | None | None | None |
| PROC | None | None | None | None |
| PROC | None | None | None | None |
| PROC | None | None | None | None |
| PROC | None | None | None | None |
| PROC | None | None | None | None |
| INS | 1 | 2 | 3zs2 | J |
| INS | 0.85 | 1 | 4ajx | D |
| INS | 1 | 1 | 3zs2 | J |
| INS | 1 | 1 | 3zs2 | J |
| FTL | None | None | None | None |
| FTL | None | None | None | None |
| FTL | None | None | None | None |
| TNFSF11 | None | None | None | None |
| EHMT1 | None | None | None | None |
| EHMT1 | None | None | None | None |
| EHMT1 | None | None | None | None |
| CA2 | 0.939 | 1 | 4idr | X |
| CA2 | 0.939 | 1 | 4idr | X |
| CA2 | 0.902 | 5 | 4idr | X |
| CA2 | 0.902 | 5 | 4idr | X |
| SMAD4 | None | None | None | None |
| SMAD4 | None | None | None | None |
| TTR | None | None | None | None |
| TTR | 0.961 | 7 | 3grb | A |
| TTR | 0.891 | 4 | 3dk2 | B |
| TTR | 0.947 | 3 | 1tta | A |
| TTR | 0.923 | 1 | 3grb | B |
| TTR | 0.891 | 1 | 4y9b | B |
| TTR | 0.891 | 1 | 4y9b | B |
| TTR | 0.882 | 15 | 3grb | A |
| TTR | 0.909 | 5 | 4tl4 | A |
| TTR | 0.836 | 7 | 1z7j | B |
| TTR | 0.882 | 15 | 3grb | A |
| TTR | 0.891 | 5 | 3i9i | B |
| TTR | 0.902 | 9 | 3m1o | B |
| TTR | 0.902 | 9 | 3m1o | B |
| TTR | 0.902 | 9 | 3m1o | B |
| TTR | None | None | None | None |

| | | | | |
|---|---|---|---|---|
| TTR | 0.902 | 9 | 3m1o | B |
| TTR | 0.836 | 7 | 1bze | A |
| TTR | 0.846 | 16 | 3gps | A |
| TTR | 0.846 | 16 | 3gps | A |
| TTR | 0.909 | 5 | 4tl4 | A |
| TTR | 0.909 | 5 | 4tl4 | A |
| TTR | None | None | None | None |
| TTR | 0.945 | 4 | 4deu | A |
| TTR | 0.941 | 3 | 3did | B |
| TTR | 0.941 | 3 | 3did | B |
| TTR | 0.868 | 6 | 4l1t | A |
| TTR | 0.882 | 19 | 3gps | B |
| TTR | 0.891 | 4 | 3dk2 | B |
| TTR | 0.941 | 3 | 3did | B |
| TTR | 0.923 | 1 | 4hiq | A |
| TTR | 0.923 | 1 | 4hiq | A |
| TTR | 0.961 | 7 | 3grb | A |
| TTR | 0.839 | 7 | 1tsh | B |
| TTR | 0.891 | 6 | 1ttb | B |
| ALK | None | None | None | None |
| ALK | None | None | None | None |
| THRB | None | None | None | None |
| THRB | None | None | None | None |
| THRB | None | None | None | None |
| THRB | None | None | None | None |
| THRB | None | None | None | None |
| THRB | None | None | None | None |
| THRB | None | None | None | None |
| THRB | None | None | None | None |
| THRB | None | None | None | None |
| THRB | None | None | None | None |
| THRB | None | None | None | None |
| THRB | None | None | None | None |
| THRB | None | None | None | None |
| THRB | None | None | None | None |
| THRB | None | None | None | None |
| THRB | None | None | None | None |
| THRB | None | None | None | None |
| THRB | None | None | None | None |
| THRB | None | None | None | None |
| IL2RG | None | None | None | None |
| IL2RG | None | None | None | None |
| IL2RG | None | None | None | None |
| IL2RG | None | None | None | None |
| IL2RG | None | None | None | None |
| ITGB3 | None | None | None | None |
| ITGB3 | None | None | None | None |
| ITGB3 | None | None | None | None |
| ITGB3 | None | None | None | None |
| ITGB3 | None | None | None | None |
| ITGB3 | None | None | None | None |
| ITGB3 | None | None | None | None |
| ITGB3 | None | None | None | None |
| ITGB3 | None | None | None | None |

| | | | | |
|---|---|---|---|---|
| PKD2 | None | None | None | None |
| PKD2 | None | None | None | None |
| PKD2 | None | None | None | None |
| PKD2 | None | None | None | None |
| PKD2 | None | None | None | None |
| PKD2 | None | None | None | None |
| PKD2 | None | None | None | None |
| TP53 | None | None | None | None |
| TP53 | None | None | None | None |
| TP53 | None | None | None | None |
| TP53 | None | None | None | None |
| TP53 | None | None | None | None |
| TP53 | None | None | None | None |
| EGFR | None | None | None | None |
| EGFR | None | None | None | None |
| XRCC4 | None | None | None | None |
| XRCC4 | None | None | None | None |
| HBB | None | None | None | None |
| HBB | None | None | None | None |
| HBB | None | None | None | None |
| HBB | None | None | None | None |
| HBB | None | None | None | None |
| HBB | None | None | None | None |
| HBB | None | None | None | None |
| HBB | None | None | None | None |
| HBB | None | None | None | None |
| HBB | None | None | None | None |
| HBB | None | None | None | None |
| HBB | None | None | None | None |
| HBB | None | None | None | None |
| HBB | None | None | None | None |
| HBB | None | None | None | None |
| HBB | None | None | None | None |
| HBB | None | None | None | None |
| HBB | None | None | None | None |
| HBB | None | None | None | None |
| HBB | None | None | None | None |
| HBB | None | None | None | None |
| HBB | None | None | None | None |
| HBB | None | None | None | None |
| HBB | None | None | None | None |
| HBB | None | None | None | None |
| HBB | None | None | None | None |
| HBB | None | None | None | None |
| HBB | None | None | None | None |
| HBB | 0.733 | 1 | 1yff | D |
| HBB | None | None | None | None |
| RPS6KA3 | None | None | None | None |
| RPS6KA3 | None | None | None | None |
| RPS6KA3 | None | None | None | None |
| RPS6KA3 | None | None | None | None |
| RPS6KA3 | None | None | None | None |
| RPS6KA3 | None | None | None | None |
| GBA | 1 | 15 | 3gxd | B |

| | | | | | |
|---|---|---|---|---|---|
| GBA | | 1 | 55 | 3gxd | B |
| GBA | | 1 | 15 | 3gxd | B |
| GBA | | 0.75 | 45 | 3ril | A |
| GBA | None | None | | None | None |
| GBA | None | None | | None | None |
| GBA | | 1 | 12 | 1ogs | A |
| GBA | | 0.75 | 122 | 3gxi | A |
| GBA | | 1 | 2 | 1ogs | A |
| GBA | | 1 | 11 | 1ogs | A |
| GBA | None | None | | None | None |
| GBA | | 0.75 | 45 | 3ril | A |
| GBA | | 1 | 18 | 1ogs | A |
| GBA | | 1 | 3 | 1ogs | A |
| GBA | None | None | | None | None |
| GBA | | 1 | 2 | 1ogs | A |
| GBA | None | None | | None | None |
| GBA | | 0.75 | 76 | 3gxm | C |
| GBA | | 1 | 3 | 1ogs | A |
| GBA | | 1 | 3 | 1ogs | A |
| GBA | | 0.75 | 62 | 3ril | A |
| GBA | | 1 | 28 | 1ogs | A |
| GBA | None | None | | None | None |
| GBA | None | None | | None | None |
| GBA | None | None | | None | None |
| GBA | None | None | | None | None |
| GBA | None | None | | None | None |
| GBA | None | None | | None | None |
| GBA | | 0.75 | 49 | 3ril | A |
| GBA | None | None | | None | None |
| GBA | None | None | | None | None |
| GBA | None | None | | None | None |
| GBA | None | None | | None | None |
| GBA | | 0.75 | 21 | 3gxi | A |
| GBA | None | None | | None | None |
| GBA | None | None | | None | None |
| GBA | | 0.75 | 25 | 3keh | B |
| GBA | | 0.75 | 70 | 3ril | A |
| GBA | | 0.75 | 70 | 3ril | A |
| GBA | | 1 | 15 | 3gxd | B |
| GBA | | 1 | 44 | 3gxd | B |
| GBA | None | None | | None | None |
| GBA | | 1 | 44 | 3gxd | B |
| GBA | | 1 | 44 | 3gxd | B |
| GBA | None | None | | None | None |
| GBA | | 0.75 | 94 | 3gxi | A |
| GBA | | 0.75 | 94 | 3gxi | A |
| IFIH1 | None | None | | None | None |
| TGFBR2 | None | None | | None | None |
| TGFBR2 | None | None | | None | None |
| TGFBR2 | None | None | | None | None |
| TGFBR2 | None | None | | None | None |
| TGFBR2 | None | None | | None | None |
| TGFBR2 | None | None | | None | None |
| TGFBR2 | None | None | | None | None |

| | | | | |
|---|---|---|---|---|
| TGFBR2 | None | None | None | None |
| TGFBR2 | None | None | None | None |
| TGFBR2 | 0.742 | 1 | 5E+091 | A |
| TGFBR2 | None | None | None | None |
| TGFBR2 | None | None | None | None |
| TGFBR2 | None | None | None | None |
| TGFBR2 | None | None | None | None |
| TGFBR2 | None | None | None | None |
| ACE | None | None | None | None |
| ACE | None | None | None | None |
| ACE | None | None | None | None |
| ACE | None | None | None | None |
| ACE | None | None | None | None |
| ACE | None | None | None | None |
| COL10A1 | None | None | None | None |
| COL10A1 | None | None | None | None |
| COL10A1 | None | None | None | None |
| COL10A1 | None | None | None | None |
| COL10A1 | None | None | None | None |
| COL10A1 | None | None | None | None |
| COL10A1 | None | None | None | None |
| COL10A1 | None | None | None | None |
| COL10A1 | None | None | None | None |
| KLF1 | None | None | None | None |
| C3 | None | None | None | None |
| CTSC | None | None | None | None |
| CTSC | 0.934 | 1 | 2djg | C |
| CTSC | None | None | None | None |
| LYZ | 0.877 | 3 | 1bb5 | A |
| LYZ | 0.923 | 1 | 1lz4 | A |
| LYZ | 0.852 | 2 | 1ubz | A |
| DHODH | None | None | None | None |
| DHODH | None | None | None | None |
| DHODH | None | None | None | None |
| DHODH | None | None | None | None |
| DHODH | None | None | None | None |
| DHODH | None | None | None | None |
| DHODH | None | None | None | None |
| ADH1C | None | None | None | None |
| GDNF | None | None | None | None |
| TNF | None | None | None | None |
| ALDH4A1 | None | None | None | None |
| ALDH4A1 | None | None | None | None |
| ALDH4A1 | None | None | None | None |
| BRAF | None | None | None | None |
| BRAF | None | None | None | None |
| BRAF | 0.752 | 2 | 3psb | A |
| BRAF | 0.752 | 2 | 3psb | A |
| BRAF | 0.737 | 1 | 3pri | A |
| BRAF | 0.737 | 1 | 3pri | A |
| BRAF | 0.737 | 1 | 3pri | A |
| BRAF | 0.737 | 1 | 3prf | A |
| BRAF | 0.737 | 1 | 3prf | A |
| BRAF | 0.752 | 2 | 3psb | A |

| | | | | |
|---|---|---|---|---|
| BRAF | 0.752 | 2 | 3psb | A |
| BRAF | 0.752 | 2 | 3psb | A |
| BRAF | 0.752 | 2 | 3psb | A |
| BRAF | None | None | None | None |
| BRAF | None | None | None | None |
| BRAF | 0.734 | 1 | 5csx | A |
| TBX5 | None | None | None | None |
| TBX5 | None | None | None | None |
| TBX5 | None | None | None | None |
| TBX5 | None | None | None | None |
| TBX5 | None | None | None | None |
| TBX5 | None | None | None | None |
| TBX5 | None | None | None | None |
| TBX5 | None | None | None | None |
| TBX5 | None | None | None | None |
| RB1 | None | None | None | None |
| RB1 | None | None | None | None |
| RB1 | None | None | None | None |
| RB1 | None | None | None | None |
| RB1 | None | None | None | None |
| RB1 | None | None | None | None |
| CLIC2 | None | None | None | None |
| CLIC2 | None | None | None | None |
| ALDH7A1 | None | None | None | None |
| ALDH7A1 | None | None | None | None |
| ALDH7A1 | None | None | None | None |
| ALDH7A1 | None | None | None | None |
| ALDH7A1 | None | None | None | None |
| ALDH7A1 | None | None | None | None |
| ALDH7A1 | None | None | None | None |
| ALDH7A1 | None | None | None | None |
| ALDH7A1 | None | None | None | None |
| ALDH7A1 | None | None | None | None |
| ALDH7A1 | None | None | None | None |
| ALDH7A1 | None | None | None | None |
| ALDH7A1 | None | None | None | None |
| ALDH7A1 | None | None | None | None |
| ALDH7A1 | None | None | None | None |
| ALDH7A1 | None | None | None | None |
| KDR | None | None | None | None |
| MUT | None | None | None | None |
| MUT | None | None | None | None |
| MUT | None | None | None | None |
| MUT | None | None | None | None |
| MUT | None | None | None | None |
| MUT | None | None | None | None |
| MUT | None | None | None | None |
| MUT | None | None | None | None |
| MUT | None | None | None | None |
| MUT | None | None | None | None |
| MUT | None | None | None | None |
| MUT | None | None | None | None |
| MUT | None | None | None | None |
| MUT | None | None | None | None |

| | | | | |
|---|---|---|---|---|
| MUT | None | None | None | None |
| MUT | None | None | None | None |
| MUT | None | None | None | None |
| MUT | None | None | None | None |
| MUT | None | None | None | None |
| MUT | None | None | None | None |
| MUT | None | None | None | None |
| MUT | None | None | None | None |
| MUT | None | None | None | None |
| MUT | None | None | None | None |
| MUT | None | None | None | None |
| MUT | None | None | None | None |
| MUT | None | None | None | None |
| MUT | None | None | None | None |
| MUT | None | None | None | None |
| MUT | None | None | None | None |
| MUT | None | None | None | None |
| MUT | None | None | None | None |
| MUT | None | None | None | None |
| MUT | None | None | None | None |
| SOD1 | 0.778 | 1 | 2gbu | A |
| SOD1 | 0.727 | 4 | 4xcr | A |
| SOD1 | None | None | None | None |
| SOD1 | None | None | None | None |
| NECTIN3 | None | None | None | None |
| HFE | None | None | None | None |
| HFE | None | None | None | None |
| HFE | None | None | None | None |
| HFE | None | None | None | None |
| HFE | None | None | None | None |
| HFE | None | None | None | None |
| HFE | None | None | None | None |
| EIF2AK3 | None | None | None | None |
| EIF2AK3 | None | None | None | None |
| TERT | None | None | None | None |
| TERT | None | None | None | None |
| SPR | None | None | None | None |
| SPR | None | None | None | None |
| SPR | None | None | None | None |
| SPR | None | None | None | None |
| SPR | None | None | None | None |
| F7 | None | None | None | None |
| UNG | None | None | None | None |
| MMP13 | None | None | None | None |
| MMP13 | 0.786 | 1 | 3tvc | A |
| MMP13 | 0.766 | 1 | 3zxh | B |
| MMP13 | 0.766 | 1 | 3zxh | B |
| CD207 | None | None | None | None |
| F2 | 1 | 2 | 1nrq | L |
| SPTBN2 | None | None | None | None |
| PKLR | None | None | None | None |
| PKLR | None | None | None | None |
| PKLR | None | None | None | None |
| PKLR | None | None | None | None |

| | | | | |
|---|---|---|---|---|
| PKLR | None | None | None | None |
| PKLR | None | None | None | None |
| PKLR | None | None | None | None |
| PKLR | None | None | None | None |
| SACS | None | None | None | None |
| HRAS | None | None | None | None |
| HRAS | None | None | None | None |
| HRAS | None | None | None | None |
| HRAS | None | None | None | None |
| HRAS | None | None | None | None |
| HRAS | 0.742 | 1 | 1ctq | A |
| HRAS | None | None | None | None |
| HRAS | None | None | None | None |
| HRAS | None | None | None | None |
| HRAS | None | None | None | None |
| HRAS | None | None | None | None |
| HRAS | None | None | None | None |
| HRAS | None | None | None | None |
| VHL | None | None | None | None |
| VHL | None | None | None | None |
| VHL | None | None | None | None |
| VHL | None | None | None | None |
| VHL | None | None | None | None |
| VHL | None | None | None | None |
| VHL | None | None | None | None |
| VHL | None | None | None | None |
| VHL | None | None | None | None |
| VHL | None | None | None | None |
| VHL | None | None | None | None |
| VHL | None | None | None | None |
| VHL | None | None | None | None |
| VHL | None | None | None | None |
| VHL | None | None | None | None |
| VHL | None | None | None | None |
| VHL | None | None | None | None |
| VHL | None | None | None | None |
| VHL | None | None | None | None |
| VHL | None | None | None | None |
| VHL | None | None | None | None |
| VHL | None | None | None | None |
| VHL | None | None | None | None |
| VHL | None | None | None | None |
| VHL | None | None | None | None |
| TTBK1 | None | None | None | None |
| RAD51 | None | None | None | None |
| ALB | None | None | None | None |
| ALB | None | None | None | None |
| ALB | None | None | None | None |
| ALB | None | None | None | None |
| MET | None | None | None | None |
| MET | None | None | None | None |
| MET | None | None | None | None |
| MET | None | None | None | None |
| MET | None | None | None | None |

| | | | | |
|---|---|---|---|---|
| MET | None | None | None | None |
| MET | None | None | None | None |
| MET | None | None | None | None |
| MET | None | None | None | None |
| CP | None | None | None | None |
| CP | None | None | None | None |
| PGM1 | None | None | None | None |
| PGM1 | None | None | None | None |
| PGM1 | None | None | None | None |
| PGM1 | None | None | None | None |
| PGM1 | None | None | None | None |
| PGM1 | None | None | None | None |
| PGM1 | None | None | None | None |
| NR5A1 | None | None | None | None |
| NR5A1 | None | None | None | None |
| NR5A1 | None | None | None | None |
| F9 | None | None | None | None |
| F9 | None | None | None | None |
| HPRT1 | None | None | None | None |
| HPRT1 | None | None | None | None |
| HPRT1 | None | None | None | None |
| HPRT1 | None | None | None | None |
| HPRT1 | None | None | None | None |
| HPRT1 | 0.871 | 3 | 4rac | B |
| HPRT1 | 0.806 | 2 | 4ran | D |
| HPRT1 | None | None | None | None |
| HPRT1 | None | None | None | None |
| HPRT1 | None | None | None | None |
| HPRT1 | None | None | None | None |
| HPRT1 | None | None | None | None |
| HPRT1 | None | None | None | None |
| HPRT1 | 0.703 | 4 | 5w8v | A |
| HPRT1 | None | None | None | None |
| HPRT1 | None | None | None | None |
| HPRT1 | None | None | None | None |
| HPRT1 | None | None | None | None |
| HPRT1 | 0.871 | 3 | 4rac | B |
| HPRT1 | 0.871 | 3 | 4rac | B |
| HPRT1 | 0.765 | 2 | 4rao | D |
| HPRT1 | 0.774 | 2 | 4rac | B |
| GPD1L | None | None | None | None |
| GPD1L | None | None | None | None |
| GPD1L | None | None | None | None |
| FGFR2 | None | None | None | None |
| FGFR2 | None | None | None | None |
| FGFR2 | None | None | None | None |
| FGFR2 | None | None | None | None |
| FGFR2 | None | None | None | None |
| FGFR2 | None | None | None | None |
| FGFR2 | None | None | None | None |
| FGFR2 | None | None | None | None |
| FGFR2 | None | None | None | None |
| FGFR2 | None | None | None | None |
| FGFR2 | None | None | None | None |

| | | | | |
|---|---|---|---|---|
| FGFR2 | None | None | None | None |
| FGFR2 | None | None | None | None |
| FGFR2 | None | None | None | None |
| FGFR2 | None | None | None | None |
| FGFR2 | None | None | None | None |
| FGFR2 | None | None | None | None |
| FGFR2 | None | None | None | None |
| FGFR2 | None | None | None | None |
| HSPD1 | None | None | None | None |
| HSPD1 | None | None | None | None |
| FGF8 | None | None | None | None |
| FGF8 | None | None | None | None |
| FGF8 | None | None | None | None |
| F9 | None | None | None | None |
| NCF4 | None | None | None | None |
| NCF4 | None | None | None | None |
| LDLR | None | None | None | None |
| LDLR | None | None | None | None |
| LDLR | None | None | None | None |
| LDLR | None | None | None | None |
| LDLR | None | None | None | None |
| LDLR | None | None | None | None |
| C3 | None | None | None | None |
| C3 | None | None | None | None |
| C3 | None | None | None | None |
| C3 | None | None | None | None |
| C3 | None | None | None | None |
| C3 | None | None | None | None |
| C3 | None | None | None | None |
| C3 | None | None | None | None |
| TULP1 | None | None | None | None |
| TULP1 | None | None | None | None |
| TULP1 | None | None | None | None |
| TULP1 | None | None | None | None |
| IKBKG | None | None | None | None |
| IKBKG | None | None | None | None |
| IKBKG | None | None | None | None |
| CD40 | None | None | None | None |
| CD40 | None | None | None | None |
| SLC9A3R1 | None | None | None | None |
| BCHE | None | None | None | None |
| BCHE | None | None | None | None |
| BCHE | None | None | None | None |
| BCHE | None | None | None | None |
| BCHE | None | None | None | None |
| BCHE | None | None | None | None |
| BCHE | None | None | None | None |
| BCHE | None | None | None | None |
| BCHE | None | None | None | None |
| BCHE | None | None | None | None |
| BCHE | None | None | None | None |
| BCKDHB | 0.833 | 1 | 1v11 | B |
| BCKDHB | None | None | None | None |
| BCKDHB | None | None | None | None |

| | | | | | |
|---|---|---|---|---|---|
| BCKDHB | None | None | None | | None |
| BCKDHB | 0.833 | | 10 | 1olx | B |
| BCKDHB | 0.833 | | 1 | 1v11 | B |
| BCKDHB | 0.833 | | 1 | 1v11 | B |
| BCKDHB | None | None | None | | None |
| BCKDHB | None | None | None | | None |
| BCKDHB | None | None | None | | None |
| BCKDHB | None | None | None | | None |
| BCKDHB | 0.857 | | 1 | 2bff | B |
| CDH1 | None | None | None | | None |
| CDH1 | None | None | None | | None |
| CDH1 | None | None | None | | None |
| CDH1 | None | None | None | | None |
| CDH1 | None | None | None | | None |
| CTNNB1 | None | None | None | | None |
| AIRE | None | None | None | | None |
| EDA | 1 | | 1 | 1rj8 | A |
| EDA | None | None | None | | None |
| EDA | None | None | None | | None |
| EDA | None | None | None | | None |
| EDA | None | None | None | | None |
| SH2D1A | None | None | None | | None |
| SH2D1A | None | None | None | | None |
| SH2D1A | None | None | None | | None |
| GCK | None | None | None | | None |
| GCK | None | None | None | | None |
| GCK | None | None | None | | None |
| GCK | None | None | None | | None |
| GCK | None | None | None | | None |
| GCK | None | None | None | | None |
| GCK | None | None | None | | None |
| GCK | None | None | None | | None |
| GCK | None | None | None | | None |
| GCK | 0.778 | | 5 | 3id8 | A |
| GCK | 0.833 | | 1 | 4dhy | A |
| GCK | None | None | None | | None |
| GCK | None | None | None | | None |
| GCK | None | None | None | | None |
| GCK | None | None | None | | None |
| GCK | None | None | None | | None |
| GCK | None | None | None | | None |
| GCK | None | None | None | | None |
| GCK | None | None | None | | None |
| GCK | None | None | None | | None |
| AQP2 | None | None | None | | None |
| AQP2 | None | None | None | | None |
| AQP2 | None | None | None | | None |
| AQP2 | None | None | None | | None |
| AQP2 | None | None | None | | None |
| AQP2 | None | None | None | | None |
| AQP2 | None | None | None | | None |
| AQP2 | None | None | None | | None |
| AQP2 | None | None | None | | None |
| AQP2 | None | None | None | | None |

| | | | | |
|---|---|---|---|---|
| AQP2 | None | None | None | None |
| AQP2 | None | None | None | None |
| AQP2 | None | None | None | None |
| SUMF1 | 0.75 | 6 | 1z70 | X |
| SUMF1 | None | None | None | None |
| SUMF1 | None | None | None | None |
| SUMF1 | None | None | None | None |
| SUMF1 | 0.875 | 35 | 2afy | X |
| SUMF1 | None | None | None | None |
| SUMF1 | 0.75 | 17 | 1y1j | X |
| SUMF1 | 0.75 | 17 | 1y1j | X |
| SUMF1 | None | None | None | None |
| FAM20A | None | None | None | None |
| FAM20A | None | None | None | None |
| FAM20A | None | None | None | None |
| FAM20A | None | None | None | None |
| NT5E | None | None | None | None |
| NT5E | None | None | None | None |
| NOS3 | None | None | None | None |
| ACTG1 | None | None | None | None |
| ACTG1 | None | None | None | None |
| ACTG1 | None | None | None | None |
| ACTG1 | None | None | None | None |
| ACTG1 | None | None | None | None |
| ACTG1 | None | None | None | None |
| FCGR1A | None | None | None | None |
| GPI | None | None | None | None |
| GPI | None | None | None | None |
| GPI | 1 | 3 | 1jiq | A |
| GPI | None | None | None | None |
| GPI | None | None | None | None |
| GPI | None | None | None | None |
| GPI | None | None | None | None |
| GPI | None | None | None | None |
| GPI | None | None | None | None |
| CFH | None | None | None | None |
| OGG1 | 0.895 | 13 | 1m3h | A |
| TPK1 | None | None | None | None |
| TPK1 | None | None | None | None |
| TPK1 | None | None | None | None |
| TPK1 | None | None | None | None |
| SEC23A | None | None | None | None |
| SEC23A | None | None | None | None |
| TYRP1 | None | None | None | None |
| TYRP1 | None | None | None | None |
| TYRP1 | None | None | None | None |
| TYRP1 | None | None | None | None |
| TYRP1 | None | None | None | None |
| TYRP1 | None | None | None | None |
| CSTB | None | None | None | None |
| CSTB | None | None | None | None |
| CSTB | None | None | None | None |
| ACP1 | None | None | None | None |
| IL17F | None | None | None | None |

| | | | | |
|---|---|---|---|---|
| IL17F | None | None | None | None |
| IL17F | None | None | None | None |
| IL17F | None | None | None | None |
| IL17F | None | None | None | None |
| IDH2 | None | None | None | None |
| IDH2 | None | None | None | None |
| IDH2 | None | None | None | None |
| IDH2 | None | None | None | None |
| TP53 | 0.925 | 11 | 2ocj | A |
| TP53 | 0.8 | 16 | 2ahi | D |
| TP53 | None | None | None | None |
| TP53 | 0.725 | 27 | 5g4o | A |
| TP53 | None | None | None | None |
| TP53 | 0.9 | 1 | 4agp | A |
| TP53 | 0.9 | 1 | 4agp | A |
| TP53 | 0.925 | 10 | 4hje | A |
| TP53 | 0.7 | 29 | 3d05 | A |
| TP53 | 0.875 | 7 | 3kmd | B |
| TP53 | 0.875 | 7 | 3kmd | B |
| TP53 | 0.925 | 6 | 3kz8 | B |
| TP53 | 0.925 | 6 | 3kz8 | B |
| TP53 | None | None | None | None |
| TP53 | 0.925 | 5 | 3kmd | B |
| TP53 | 0.925 | 5 | 3kmd | B |
| TP53 | 0.925 | 8 | 3kz8 | B |
| TP53 | 0.925 | 8 | 3kz8 | B |
| TP53 | 0.925 | 1 | 3kmd | B |
| TP53 | 0.9 | 11 | 2pcx | A |
| TP53 | 0.9 | 1 | 5aol | A |
| TP53 | 0.85 | 5 | 5lgy | A |
| TP53 | 0.9 | 1 | 3q06 | B |
| TP53 | 0.725 | 19 | 5lgy | A |
| TP53 | 0.775 | 14 | 4mzi | A |
| TP53 | 0.825 | 8 | 1kzy | A |
| TP53 | 0.825 | 8 | 1kzy | A |
| TP53 | 0.8 | 11 | 4ibt | A |
| TP53 | 0.8 | 11 | 4ibt | A |
| TP53 | 0.725 | 15 | 5lgy | A |
| TP53 | 0.725 | 29 | 2h1l | O |
| TP53 | 0.875 | 7 | 3kmd | B |
| TP53 | 0.725 | 15 | 5lgy | A |
| TP53 | 0.8 | 10 | 4agp | A |
| TP53 | 0.8 | 10 | 4agp | A |
| TP53 | 0.8 | 10 | 4agp | A |
| TP53 | 0.8 | 10 | 4agp | A |
| TP53 | None | None | None | None |
| TP53 | None | None | None | None |
| TP53 | None | None | None | None |
| TP53 | None | None | None | None |
| TP53 | None | None | None | None |
| LMAN1 | None | None | None | None |
| CTSK | 0.75 | 2 | 4n79 | A |
| CTSK | None | None | None | None |
| CTSK | None | None | None | None |

| | | | | |
|---|---|---|---|---|
| FGB | None | None | None | None |
| FGB | None | None | None | None |
| FGB | None | None | None | None |
| FGB | None | None | None | None |
| ACADM | None | None | None | None |
| ACADM | None | None | None | None |
| ACADM | None | None | None | None |
| ACADM | None | None | None | None |
| ACADM | None | None | None | None |
| ACADM | None | None | None | None |
| ACADM | None | None | None | None |
| ACADM | None | None | None | None |
| ACADM | None | None | None | None |
| ACADM | None | None | None | None |
| ACADM | None | None | None | None |
| ACADM | None | None | None | None |
| ACADM | None | None | None | None |
| ACADM | None | None | None | None |
| ACADM | None | None | None | None |
| ACADM | None | None | None | None |
| ACADM | None | None | None | None |
| ACADM | None | None | None | None |
| ACADM | None | None | None | None |
| ACADM | None | None | None | None |
| ACADM | None | None | None | None |
| ACADM | None | None | None | None |
| ACADM | None | None | None | None |
| ACADM | None | None | None | None |
| ACADM | None | None | None | None |
| ACADM | None | None | None | None |
| ACADM | None | None | None | None |
| PTS | None | None | None | None |
| PTS | None | None | None | None |
| PTS | None | None | None | None |
| PTS | 0.778 | 1 | 3i2b | D |
| PTS | None | None | None | None |
| PTS | None | None | None | None |
| PTS | None | None | None | None |
| PTS | None | None | None | None |
| PTS | None | None | None | None |
| INS | 0.75 | 2 | 3zqr | E |
| INS | 1 | 5 | 2l1z | A |
| INS | 1 | 5 | 2l1z | A |
| INS | 1 | 1 | 1his | A |
| FBP1 | 0.71 | 26 | 2y5l | C |
| FBP1 | None | None | None | None |
| FBP1 | None | None | None | None |
| FBP1 | 0.7 | 10 | 2y5k | A |
| FBP1 | None | None | None | None |
| FBP1 | 0.733 | 10 | 3kc1 | A |
| FBP1 | None | None | None | None |
| FBP1 | 0.8 | 11 | 2y5k | A |
| FBP1 | None | None | None | None |
| FBP1 | None | None | None | None |

| | | | | |
|---|---|---|---|---|
| FBP1 | None | None | None | None |
| KCNQ1 | None | None | None | None |
| KCNQ1 | None | None | None | None |
| KCNQ1 | None | None | None | None |
| UPP1 | 0.706 | 4 | 3euf | A |
| UMPS | 1 | 3 | 2qcl | A |
| UMPS | None | None | None | None |
| UMPS | None | None | None | None |
| DHFR | None | None | None | None |
| KHK | None | None | None | None |
| TTPA | None | None | None | None |
| TTPA | None | None | None | None |
| TTPA | None | None | None | None |
| TTPA | None | None | None | None |
| MAP1LC3B | None | None | None | None |
| MAP1LC3B | None | None | None | None |
| ZAP70 | None | None | None | None |
| PROC | None | None | None | None |
| SQSTM1 | None | None | None | None |
| AHCY | None | None | None | None |
| AHCY | None | None | None | None |
| PRNP | None | None | None | None |
| PRNP | None | None | None | None |
| PRNP | None | None | None | None |
| PRNP | None | None | None | None |
| PRNP | None | None | None | None |
| PRNP | None | None | None | None |
| PRNP | None | None | None | None |
| PRNP | None | None | None | None |
| PRNP | None | None | None | None |
| PRNP | None | None | None | None |
| PRNP | None | None | None | None |
| PRNP | None | None | None | None |
| PRNP | None | None | None | None |
| PRNP | None | None | None | None |
| PRNP | None | None | None | None |
| PRNP | None | None | None | None |
| REN | None | None | None | None |
| REN | None | None | None | None |
| HINT1 | None | None | None | None |
| HINT1 | 0.897 | 1 | 5km4 | A |
| HINT1 | None | None | None | None |
| HINT1 | None | None | None | None |
| HINT1 | 0.923 | 1 | 5ipc | A |
| OAT | None | None | None | None |
| OAT | None | None | None | None |
| OAT | None | None | None | None |
| OAT | None | None | None | None |
| OAT | None | None | None | None |
| OAT | None | None | None | None |
| OAT | None | None | None | None |
| OAT | None | None | None | None |
| OAT | None | None | None | None |
| OAT | None | None | None | None |

| | | | | |
|---|---|---|---|---|
| OAT | None | None | None | None |
| OAT | None | None | None | None |
| OAT | None | None | None | None |
| OAT | None | None | None | None |
| OAT | None | None | None | None |
| OAT | 0.758 | 1 | 2byl | A |
| OAT | None | None | None | None |
| OAT | None | None | None | None |
| OAT | None | None | None | None |
| OAT | None | None | None | None |
| OAT | None | None | None | None |
| OAT | None | None | None | None |
| OAT | None | None | None | None |
| OAT | None | None | None | None |
| OAT | None | None | None | None |
| OAT | None | None | None | None |
| OAT | None | None | None | None |
| OAT | None | None | None | None |
| CTSA | None | None | None | None |
| CTSA | None | None | None | None |
| PNP | None | None | None | None |
| PNP | None | None | None | None |
| PNP | None | None | None | None |
| PNP | 0.706 | 5 | 4ear | C |
| PNP | None | None | None | None |
| PNP | None | None | None | None |
| PNP | 0.933 | 1 | 3ggs | C |
| PNP | 0.833 | 1 | 3d1v | A |
| MYL2 | None | None | None | None |
| MYL2 | None | None | None | None |
| ANG | None | None | None | None |
| ANG | None | None | None | None |
| ANG | None | None | None | None |
| ANG | None | None | None | None |
| ANG | 0.761 | 1 | 5eop | A |
| ANG | None | None | None | None |
| ANG | None | None | None | None |
| ANG | None | None | None | None |
| ANG | None | None | None | None |
| ANG | None | None | None | None |
| POLH | None | None | None | None |
| POLH | None | None | None | None |
| POLH | None | None | None | None |
| IL2RA | None | None | None | None |
| TGFB1 | None | None | None | None |
| TP53 | None | None | None | None |
| TP53 | None | None | None | None |
| PAH | None | None | None | None |
| PAH | None | None | None | None |
| PAH | None | None | None | None |
| PAH | None | None | None | None |
| PAH | None | None | None | None |
| PAH | None | None | None | None |
| PAH | None | None | None | None |

| | | | | |
|---|---|---|---|---|
| PAH | None | None | None | None |
| PAH | None | None | None | None |
| PAH | None | None | None | None |
| PAH | None | None | None | None |
| PAH | None | None | None | None |
| PAH | None | None | None | None |
| PAH | None | None | None | None |
| PAH | None | None | None | None |
| PAH | None | None | None | None |
| PAH | None | None | None | None |
| PAH | None | None | None | None |
| PAH | None | None | None | None |
| PAH | None | None | None | None |
| PAH | None | None | None | None |
| PAH | None | None | None | None |
| PAH | None | None | None | None |
| PAH | None | None | None | None |
| PAH | 0.76 | 1 | 1tdw | A |
| PAH | None | None | None | None |
| PAH | None | None | None | None |
| PAH | None | None | None | None |
| PAH | None | None | None | None |
| PAH | None | None | None | None |
| PAH | None | None | None | None |
| PAH | None | None | None | None |
| PAH | None | None | None | None |
| PAH | None | None | None | None |
| PAH | None | None | None | None |
| PAH | None | None | None | None |
| PAH | None | None | None | None |
| PAH | None | None | None | None |
| PAH | None | None | None | None |
| PAH | None | None | None | None |
| PAH | None | None | None | None |
| HDAC8 | None | None | None | None |
| HDAC8 | 0.732 | 3 | 3rqd | A |
| VCP | None | None | None | None |
| VCP | None | None | None | None |
| VCP | None | None | None | None |
| VCP | None | None | None | None |
| VCP | None | None | None | None |
| VCP | None | None | None | None |
| VCP | None | None | None | None |
| B3GAT3 | None | None | None | None |
| B3GAT3 | None | None | None | None |
| CSF2RB | None | None | None | None |
| CSF2RB | None | None | None | None |
| CSF2RB | None | None | None | None |
| NF2 | None | None | None | None |
| NF2 | None | None | None | None |
| NF2 | None | None | None | None |
| PIK3CA | None | None | None | None |
| PHGDH | None | None | None | None |
| PHGDH | None | None | None | None |

| | | | | |
|---|---|---|---|---|
| PHGDH | None | None | None | None |
| PHGDH | None | None | None | None |
| POLH | None | None | None | None |
| XIAP | None | None | None | None |
| XIAP | None | None | None | None |
| PDHX | None | None | None | None |
| PDHX | None | None | None | None |
| PDHX | None | None | None | None |
| PDHX | None | None | None | None |
| PDHX | None | None | None | None |
| SAMHD1 | None | None | None | None |
| SAMHD1 | None | None | None | None |
| SAMHD1 | None | None | None | None |
| SAMHD1 | None | None | None | None |
| SAMHD1 | None | None | None | None |
| SAMHD1 | None | None | None | None |
| SAMHD1 | None | None | None | None |
| SAMHD1 | None | None | None | None |
| CD36 | None | None | None | None |
| CD36 | None | None | None | None |
| CD36 | None | None | None | None |
| ASPH | None | None | None | None |
| CCT5 | None | None | None | None |
| CCT5 | None | None | None | None |
| CCT5 | None | None | None | None |
| SMN1 | None | None | None | None |
| SMN1 | None | None | None | None |
| SMN1 | None | None | None | None |
| SMN1 | None | None | None | None |
| ABCA1 | None | None | None | None |
| ABCA1 | None | None | None | None |
| ABCA1 | None | None | None | None |
| ABCA1 | None | None | None | None |
| ABCA1 | None | None | None | None |
| ABCA1 | None | None | None | None |
| ABCA1 | None | None | None | None |
| ABCA1 | None | None | None | None |
| ABCA1 | None | None | None | None |
| ABCA1 | None | None | None | None |
| ABCA1 | None | None | None | None |
| ABCA1 | None | None | None | None |
| ABCA1 | None | None | None | None |
| ABCA1 | None | None | None | None |
| ABCA1 | None | None | None | None |
| ABCA1 | None | None | None | None |
| ABCA1 | None | None | None | None |
| ABCA1 | None | None | None | None |
| ABCA1 | None | None | None | None |
| ABCA1 | None | None | None | None |
| ABCA1 | None | None | None | None |
| ABCA1 | None | None | None | None |
| ABCA1 | None | None | None | None |
| ABCA1 | None | None | None | None |

| | | | | |
|---|---|---|---|---|
| ABCA1 | None | None | None | None |
| ABCA1 | None | None | None | None |
| ABCA1 | None | None | None | None |
| ABCA1 | None | None | None | None |
| ABCA1 | None | None | None | None |
| ABCA1 | None | None | None | None |
| ABCA1 | None | None | None | None |
| ABCA1 | None | None | None | None |
| ABCA1 | None | None | None | None |
| ABCA1 | None | None | None | None |
| ABCA1 | None | None | None | None |
| ERCC1 | None | None | None | None |
| FECH | None | None | None | None |
| FECH | None | None | None | None |
| FECH | None | None | None | None |
| FECH | None | None | None | None |
| FECH | None | None | None | None |
| FECH | None | None | None | None |
| FECH | None | None | None | None |
| PTEN | None | None | None | None |
| PTEN | None | None | None | None |
| PTEN | None | None | None | None |
| PTEN | None | None | None | None |
| PTEN | None | None | None | None |
| PTEN | None | None | None | None |
| PTEN | None | None | None | None |
| PTEN | None | None | None | None |
| PTEN | None | None | None | None |
| PTEN | None | None | None | None |
| PTEN | None | None | None | None |
| PTEN | None | None | None | None |
| PTEN | None | None | None | None |
| PTEN | None | None | None | None |
| PTEN | None | None | None | None |
| PTEN | None | None | None | None |
| PTEN | None | None | None | None |
| PTEN | None | None | None | None |
| PTEN | None | None | None | None |
| PTEN | None | None | None | None |
| ACAT1 | None | None | None | None |
| ACAT1 | None | None | None | None |
| ACAT1 | None | None | None | None |
| ACAT1 | None | None | None | None |
| ACAT1 | None | None | None | None |
| ACAT1 | None | None | None | None |
| ACAT1 | 0.833 | 6 | 2ib7 | C |
| ACAT1 | None | None | None | None |
| ACAT1 | None | None | None | None |
| ACAT1 | None | None | None | None |
| ACAT1 | 0.703 | 16 | 2ibu | A |
| CREBBP | None | None | None | None |
| ITGA2B | None | None | None | None |
| ITGA2B | None | None | None | None |
| ITGA2B | None | None | None | None |

| | | | | |
|---|---|---|---|---|
| ITGA2B | None | None | None | None |
| TSC1 | None | None | None | None |
| ADA2 | None | None | None | None |
| ADA2 | None | None | None | None |
| ADA2 | 0.75 | 1 | 3lgd | A |
| ADA2 | None | None | None | None |
| ADA2 | None | None | None | None |
| ADA2 | None | None | None | None |
| ADA2 | None | None | None | None |
| ADA2 | None | None | None | None |
| ADA2 | 1 | 1 | 3lgd | B |
| ADA2 | None | None | None | None |
| PGK1 | None | None | None | None |
| PGK1 | None | None | None | None |
| PGK1 | None | None | None | None |
| PGK1 | None | None | None | None |
| PGK1 | None | None | None | None |
| TUBA1A | None | None | None | None |
| TUBA1A | None | None | None | None |
| TUBA1A | None | None | None | None |
| TUBA1A | None | None | None | None |
| TUBA1A | None | None | None | None |
| PTPN11 | None | None | None | None |
| PTPN11 | None | None | None | None |
| PTPN11 | None | None | None | None |
| PTPN11 | None | None | None | None |
| PTPN11 | None | None | None | None |
| PTPN11 | None | None | None | None |
| PTPN11 | None | None | None | None |
| PTPN11 | None | None | None | None |
| PTPN11 | None | None | None | None |
| PTPN11 | None | None | None | None |
| PTPN11 | None | None | None | None |
| PTPN11 | None | None | None | None |
| PTPN11 | None | None | None | None |
| PTPN11 | None | None | None | None |
| PTPN11 | None | None | None | None |
| PTPN11 | 0.776 | 1 | 3zm1 | A |
| PTPN11 | None | None | None | None |
| PTPN11 | None | None | None | None |
| PTPN11 | None | None | None | None |
| PTPN11 | None | None | None | None |
| PTPN11 | None | None | None | None |
| PTPN11 | 0.776 | 1 | 3zm1 | A |
| PTPN11 | None | None | None | None |
| PTPN11 | None | None | None | None |
| PTPN11 | None | None | None | None |
| PTPN11 | None | None | None | None |
| GLB1 | None | None | None | None |
| GLB1 | None | None | None | None |
| UQCRQ | None | None | None | None |
| VPS35 | None | None | None | None |
| NR3C1 | None | None | None | None |
| NR3C1 | None | None | None | None |

| | | | | |
|---|---|---|---|---|
| NR3C1 | None | None | None | None |
| NR3C1 | None | None | None | None |
| NR3C1 | None | None | None | None |
| NR3C1 | None | None | None | None |
| NR3C1 | None | None | None | None |
| NR3C1 | None | None | None | None |
| NR3C1 | None | None | None | None |
| APOA1 | None | None | None | None |
| APOA1 | None | None | None | None |
| APOA1 | None | None | None | None |
| APOA1 | None | None | None | None |
| APOA1 | None | None | None | None |
| APOA1 | None | None | None | None |
| APOA1 | None | None | None | None |
| BCKDHA | None | None | None | None |
| BCKDHA | None | None | None | None |
| BCKDHA | 0.963 | 1 | 1ols | A |
| BCKDHA | None | None | None | None |
| BCKDHA | None | None | None | None |
| BCKDHA | None | None | None | None |
| BCKDHA | None | None | None | None |
| BCKDHA | 1 | 1 | 2beu | A |
| BCKDHA | 0.84 | 2 | 2j9f | A |
| BCKDHA | None | None | None | None |
| BCKDHA | None | None | None | None |
| BCKDHA | None | None | None | None |
| BCKDHA | None | None | None | None |
| BCKDHA | None | None | None | None |
| BCKDHA | None | None | None | None |
| TOP1 | None | None | None | None |
| ANTXR2 | None | None | None | None |
| ANTXR2 | None | None | None | None |
| ANTXR2 | None | None | None | None |
| TF | None | None | None | None |
| TF | None | None | None | None |
| FSHB | None | None | None | None |
| GALK1 | None | None | None | None |
| GALK1 | None | None | None | None |
| GALK1 | None | None | None | None |
| CYP17A1 | None | None | None | None |
| CYP17A1 | None | None | None | None |
| CYP17A1 | None | None | None | None |
| CYP17A1 | None | None | None | None |
| CYP17A1 | None | None | None | None |
| CYP17A1 | None | None | None | None |
| CYP17A1 | None | None | None | None |
| CYP17A1 | None | None | None | None |
| CYP17A1 | None | None | None | None |
| CYP17A1 | None | None | None | None |
| CYP17A1 | None | None | None | None |
| CYP17A1 | None | None | None | None |
| CYP17A1 | None | None | None | None |
| CYP17A1 | None | None | None | None |
| CYP17A1 | None | None | None | None |

| | | | | |
|---|---|---|---|---|
| CYP17A1 | None | None | None | None |
| CYP17A1 | None | None | None | None |
| CD44 | None | None | None | None |
| MDH2 | None | None | None | None |
| MDH2 | None | None | None | None |
| NDUFV1 | None | None | None | None |
| NDUFV1 | None | None | None | None |
| NDUFV1 | None | None | None | None |
| NDUFV1 | None | None | None | None |
| NDUFV1 | None | None | None | None |
| NDUFV1 | None | None | None | None |
| NDUFV1 | None | None | None | None |
| NDUFV1 | None | None | None | None |
| NDUFV1 | None | None | None | None |
| NDUFV1 | None | None | None | None |
| CTSC | None | None | None | None |
| CTSC | None | None | None | None |
| CTSC | None | None | None | None |
| CTSC | None | None | None | None |
| CTSC | None | None | None | None |
| CTSC | None | None | None | None |
| CTSC | None | None | None | None |
| VWF | None | None | None | None |
| VWF | None | None | None | None |
| VWF | None | None | None | None |
| VWF | None | None | None | None |
| VWF | None | None | None | None |
| VWF | None | None | None | None |
| VWF | None | None | None | None |
| ORC4 | None | None | None | None |
| ORC4 | None | None | None | None |
| CRYAB | None | None | None | None |
| CRYAB | None | None | None | None |
| CRYAB | None | None | None | None |
| MAP2K2 | None | None | None | None |
| MAP2K2 | None | None | None | None |
| MAP2K2 | None | None | None | None |
| MAP2K2 | None | None | None | None |
| MAP2K2 | None | None | None | None |
| MAP2K2 | None | None | None | None |
| MAP2K2 | None | None | None | None |
| MAP2K2 | None | None | None | None |
| MAP2K2 | None | None | None | None |
| CBL | None | None | None | None |
| CBL | None | None | None | None |
| CBL | None | None | None | None |
| CBL | None | None | None | None |
| CBL | None | None | None | None |
| CBL | None | None | None | None |
| CBL | None | None | None | None |
| PFN1 | None | None | None | None |
| PFN1 | 0.75 | 2 | 1fik | A |
| ACSM2A | None | None | None | None |
| HSPG2 | None | None | None | None |

| | | | | |
|---|---|---|---|---|
| HSPG2 | None | None | None | None |
| HSPG2 | None | None | None | None |
| HSPG2 | None | None | None | None |
| PARK7 | None | None | None | None |
| PARK7 | 0.769 | 4 | 1pe0 | A |
| PARK7 | 0.808 | 2 | 1pe0 | A |
| PARK7 | None | None | None | None |
| PARK7 | 0.889 | 1 | 3cz9 | A |
| PARK7 | 0.75 | 2 | 1pdw | B |
| INSR | None | None | None | None |
| INSR | None | None | None | None |
| INSR | None | None | None | None |
| INSR | None | None | None | None |
| INSR | None | None | None | None |
| DSC2 | None | None | None | None |
| DSC2 | None | None | None | None |
| DSC2 | None | None | None | None |
| DSC2 | None | None | None | None |
| DSC2 | None | None | None | None |
| DSC2 | None | None | None | None |
| DSC2 | None | None | None | None |
| DSC2 | None | None | None | None |
| DSC2 | None | None | None | None |
| DSC2 | None | None | None | None |
| MICU1 | None | None | None | None |
| MICU1 | None | None | None | None |
| ADAMTS13 | None | None | None | None |
| ADAMTS13 | None | None | None | None |
| ADAMTS13 | None | None | None | None |
| ADAMTS13 | None | None | None | None |
| ADAMTS13 | None | None | None | None |
| ADAMTS13 | None | None | None | None |
| ADAMTS13 | None | None | None | None |
| IRAK4 | None | None | None | None |
| IRAK4 | None | None | None | None |
| RNASEH2B | None | None | None | None |
| RNASEH2B | None | None | None | None |
| RNASEH2B | None | None | None | None |
| NTRK1 | None | None | None | None |
| MPO | 0.75 | 3 | 1d5l | C |
| MPO | None | None | None | None |
| MPO | None | None | None | None |
| PEX3 | None | None | None | None |
| SERPINA1 | None | None | None | None |
| SERPINA1 | None | None | None | None |
| SERPINA1 | None | None | None | None |
| SERPINA1 | None | None | None | None |
| SERPINA1 | None | None | None | None |
| SERPINA1 | None | None | None | None |
| SERPINA1 | None | None | None | None |
| SERPINA1 | None | None | None | None |
| SERPINA1 | None | None | None | None |
| SERPINA1 | None | None | None | None |
| YARS2 | None | None | None | None |

| | | | | |
|---|---|---|---|---|
| YARS2 | None | None | None | None |
| YARS2 | None | None | None | None |
| YARS2 | None | None | None | None |
| YARS2 | None | None | None | None |
| YARS2 | 0.781 | 1 | 2pid | B |
| YARS2 | None | None | None | None |
| YARS2 | None | None | None | None |
| YARS2 | None | None | None | None |
| PDHA1 | None | None | None | None |
| PDHA1 | None | None | None | None |
| PDHA1 | None | None | None | None |
| PDHA1 | None | None | None | None |
| PDHA1 | None | None | None | None |
| PDHA1 | None | None | None | None |
| PDHA1 | None | None | None | None |
| PDHA1 | 0.96 | 1 | 3exh | C |
| PDHA1 | None | None | None | None |
| PDHA1 | None | None | None | None |
| PDHA1 | None | None | None | None |
| CUL4B | None | None | None | None |
| AGXT | None | None | None | None |
| AGXT | None | None | None | None |
| AGXT | None | None | None | None |
| AGXT | None | None | None | None |
| AGXT | 0.786 | 7 | 4kxk | C |
| AGXT | None | None | None | None |
| AGXT | None | None | None | None |
| AGXT | None | None | None | None |
| AGXT | None | None | None | None |
| AGXT | None | None | None | None |
| AGXT | None | None | None | None |
| AGXT | None | None | None | None |
| AGXT | None | None | None | None |
| AGXT | None | None | None | None |
| AGXT | None | None | None | None |
| AGXT | None | None | None | None |
| AGXT | None | None | None | None |
| AGXT | None | None | None | None |
| AGXT | None | None | None | None |
| AGXT | 0.75 | 2 | 5luc | B |
| AGXT | 0.722 | 3 | 5hhy | B |
| AGXT | None | None | None | None |
| AGXT | None | None | None | None |
| AGXT | None | None | None | None |
| AGXT | 0.778 | 1 | 5hhy | B |
| AGXT | 0.778 | 1 | 5hhy | B |
| AGXT | None | None | None | None |
| AGXT | None | None | None | None |
| AGXT | None | None | None | None |
| AGXT | None | None | None | None |
| AGXT | None | None | None | None |
| AGXT | 0.786 | 7 | 4kxk | C |
| AGXT | 0.846 | 1 | 4kxk | A |
| AGXT | None | None | None | None |

| | | | | |
|---|---|---|---|---|
| AGXT | None | None | None | None |
| SFTPD | None | None | None | None |
| SFTPD | None | None | None | None |
| PIK3CA | None | None | None | None |
| PIK3CA | None | None | None | None |
| PIK3CA | None | None | None | None |
| AUH | None | None | None | None |
| AUH | None | None | None | None |
| AUH | None | None | None | None |
| AUH | None | None | None | None |
| F11 | None | None | None | None |
| F11 | None | None | None | None |
| F11 | None | None | None | None |
| F11 | None | None | None | None |
| MEN1 | None | None | None | None |
| MEN1 | 0.8 | 54 | 4i80 | A |
| MEN1 | 0.8 | 84 | 5db0 | A |
| MEN1 | 0.8 | 30 | 5ddb | A |
| MEN1 | 1 | 2 | 4i80 | A |
| MEN1 | 0.8 | 17 | 5db0 | A |
| MEN1 | None | None | None | None |
| MEN1 | None | None | None | None |
| MEN1 | None | None | None | None |
| MEN1 | None | None | None | None |
| MYBPC1 | None | None | None | None |
| MYBPC1 | None | None | None | None |
| MYBPC1 | None | None | None | None |
| PDCD10 | None | None | None | None |
| SCARB2 | None | None | None | None |
| SCARB2 | None | None | None | None |
| SCARB2 | None | None | None | None |
| SCARB2 | None | None | None | None |
| SCARB2 | None | None | None | None |
| TREX2 | None | None | None | None |
| TBK1 | None | None | None | None |
| TBK1 | None | None | None | None |
| DLD | None | None | None | None |
| DLD | None | None | None | None |
| DLD | None | None | None | None |
| DLD | None | None | None | None |
| DLD | None | None | None | None |
| DLD | None | None | None | None |
| LDHA | 0.802 | 1 | 4qt0 | A |
| TNNC1 | None | None | None | None |
| TNNC1 | None | None | None | None |
| TNNC1 | None | None | None | None |
| SMAD3 | None | None | None | None |
| HARS | None | None | None | None |
| HARS | None | None | None | None |
| TNNI3 | None | None | None | None |
| TNNI3 | None | None | None | None |
| TNNI3 | None | None | None | None |
| TNNI3 | None | None | None | None |
| TNNI3 | None | None | None | None |

| | | | | |
|---|---|---|---|---|
| TNNI3 | None | None | None | None |
| ACKR1 | None | None | None | None |
| BUB1B | None | None | None | None |
| BUB1B | None | None | None | None |
| BUB1B | None | None | None | None |
| AR | None | None | None | None |
| AR | None | None | None | None |
| AR | None | None | None | None |
| AR | None | None | None | None |
| AR | None | None | None | None |
| FTO | None | None | None | None |
| FTO | None | None | None | None |
| TFAM | None | None | None | None |
| PYGL | None | None | None | None |
| PYGL | 0.778 | 3 | 1fa9 | A |
| PYGL | None | None | None | None |
| PYGL | 0.722 | 2 | 1fa9 | A |
| PYGL | None | None | None | None |
| PYGL | None | None | None | None |
| PYGL | None | None | None | None |
| PYGL | None | None | None | None |
| PYGL | 0.722 | 2 | 1fc0 | B |
| PYGL | 0.778 | 3 | 1fa9 | A |
| PYGL | None | None | None | None |
| PYGL | 0.722 | 12 | 2ati | A |
| PYGL | 0.722 | 1 | 3ceh | B |
| PYGL | None | None | None | None |
| PYGL | None | None | None | None |
| PYGL | 0.722 | 2 | 1fc0 | B |
| GALNS | None | None | None | None |
| GALNS | None | None | None | None |
| GALNS | None | None | None | None |
| GALNS | None | None | None | None |
| GALNS | None | None | None | None |
| GALNS | None | None | None | None |
| GALNS | None | None | None | None |
| GALNS | None | None | None | None |
| GALNS | None | None | None | None |
| GALNS | None | None | None | None |
| GALNS | None | None | None | None |
| GALNS | None | None | None | None |
| GALNS | None | None | None | None |
| GALNS | None | None | None | None |
| GALNS | None | None | None | None |
| GALNS | None | None | None | None |
| GALNS | None | None | None | None |
| GALNS | None | None | None | None |
| GALNS | None | None | None | None |
| GALNS | None | None | None | None |
| GALNS | None | None | None | None |
| GALNS | None | None | None | None |
| GALNS | None | None | None | None |
| GALNS | None | None | None | None |
| GALNS | None | None | None | None |
| GALNS | None | None | None | None |

| | | | | |
|---|---|---|---|---|
| GALNS | None | None | None | None |
| GALNS | None | None | None | None |
| GALNS | None | None | None | None |
| GALNS | None | None | None | None |
| GALNS | None | None | None | None |
| GALNS | None | None | None | None |
| GALNS | None | None | None | None |
| GALNS | None | None | None | None |
| F10 | None | None | None | None |
| GLUD1 | None | None | None | None |
| GLUD1 | None | None | None | None |
| GLUD1 | None | None | None | None |
| GLUD1 | None | None | None | None |
| GLUD1 | None | None | None | None |
| GLUD1 | None | None | None | None |
| GLUD1 | None | None | None | None |
| GLUD1 | None | None | None | None |
| GLUD1 | None | None | None | None |
| GLUD1 | None | None | None | None |
| NOTCH1 | None | None | None | None |
| POGLUT1 | None | None | None | None |
| POGLUT1 | None | None | None | None |
| NR3C1 | None | None | None | None |
| CASP8 | None | None | None | None |
| CASP8 | None | None | None | None |
| TRAP1 | None | None | None | None |
| FANCM | None | None | None | None |
| EPOR | None | None | None | None |
| EPOR | None | None | None | None |
| NDP | None | None | None | None |
| NDP | None | None | None | None |
| NDP | None | None | None | None |
| NDP | None | None | None | None |
| NDP | None | None | None | None |
| NDP | None | None | None | None |
| NDP | None | None | None | None |
| NDP | None | None | None | None |
| NDP | None | None | None | None |
| NDP | None | None | None | None |
| NDP | None | None | None | None |
| NDP | None | None | None | None |
| NDP | None | None | None | None |
| NDP | None | None | None | None |
| NDP | None | None | None | None |
| NDP | None | None | None | None |
| NDP | None | None | None | None |
| ROGDI | None | None | None | None |
| CYP19A1 | None | None | None | None |
| CYP19A1 | None | None | None | None |
| CYP19A1 | None | None | None | None |
| CYP19A1 | None | None | None | None |
| CYP19A1 | None | None | None | None |
| CYP19A1 | None | None | None | None |
| CYP19A1 | None | None | None | None |

| | | | | |
|---|---|---|---|---|
| CYP19A1 | None | None | None | None |
| UBA5 | None | None | None | None |
| PRPS1 | None | None | None | None |
| PRPS1 | None | None | None | None |
| PRPS1 | None | None | None | None |
| PRPS1 | None | None | None | None |
| PRPS1 | None | None | None | None |
| PRPS1 | None | None | None | None |
| PRPS1 | None | None | None | None |
| PRPS1 | None | None | None | None |
| F2 | None | None | None | None |
| F2 | None | None | None | None |
| NTF4 | None | None | None | None |
| NTF4 | None | None | None | None |
| PSAP | None | None | None | None |
| PSAP | None | None | None | None |
| TAP1 | None | None | None | None |
| TAP1 | None | None | None | None |
| TAP1 | None | None | None | None |
| ARG1 | None | None | None | None |
| ARG1 | 0.714 | 1 | 2pha | A |
| ARG1 | None | None | None | None |
| ARG1 | None | None | None | None |
| ARG1 | None | None | None | None |
| ARG1 | None | None | None | None |
| CRYM | None | None | None | None |
| ADSL | None | None | None | None |
| ADSL | None | None | None | None |
| ADSL | None | None | None | None |
| ADSL | None | None | None | None |
| ADSL | None | None | None | None |
| ADSL | None | None | None | None |
| ADSL | None | None | None | None |
| ADSL | None | None | None | None |
| ADSL | None | None | None | None |
| ADSL | None | None | None | None |
| ADSL | None | None | None | None |
| ADSL | None | None | None | None |
| ADSL | None | None | None | None |
| ADSL | None | None | None | None |
| ADSL | None | None | None | None |
| ADSL | None | None | None | None |
| ADSL | None | None | None | None |
| ADSL | None | None | None | None |
| ADSL | None | None | None | None |
| ADSL | None | None | None | None |
| VWF | None | None | None | None |
| PLA2G7 | None | None | None | None |
| MMP13 | None | None | None | None |
| MMP13 | None | None | None | None |
| MMP13 | None | None | None | None |
| MMP13 | None | None | None | None |
| POMGNT1 | None | None | None | None |
| POMGNT1 | None | None | None | None |

| | | | | |
|---|---|---|---|---|
| COL3A1 | None | None | None | None |
| COL3A1 | None | None | None | None |
| COL3A1 | None | None | None | None |
| COL3A1 | None | None | None | None |
| ACTN2 | None | None | None | None |
| ACTN2 | None | None | None | None |
| ACTN2 | None | None | None | None |
| ACTN2 | None | None | None | None |
| ACTN2 | None | None | None | None |
| ACTN2 | None | None | None | None |
| ACTN2 | None | None | None | None |
| UBA5 | None | None | None | None |
| ADAR | 0.75 | 1 | 1qbj | B |
| UMPS | None | None | None | None |
| UMPS | None | None | None | None |
| UMPS | None | None | None | None |
| TRNT1 | None | None | None | None |
| TRNT1 | None | None | None | None |
| NHEJ1 | None | None | None | None |
| NHEJ1 | None | None | None | None |
| NHEJ1 | None | None | None | None |
| C8A | None | None | None | None |
| GLUL | None | None | None | None |
| GLUL | None | None | None | None |
| GLUL | None | None | None | None |
| GYG1 | None | None | None | None |
| GYG1 | 0.923 | 1 | 3u2u | B |
| FLNA | None | None | None | None |
| FLNA | None | None | None | None |
| FLNA | None | None | None | None |
| FLNA | None | None | None | None |
| FLNA | None | None | None | None |
| FLNA | None | None | None | None |
| FLNA | None | None | None | None |
| TTN | None | None | None | None |
| TTN | None | None | None | None |
| TTN | None | None | None | None |
| TTN | None | None | None | None |
| SMS | None | None | None | None |
| SMS | None | None | None | None |
| SMS | None | None | None | None |
| C5 | None | None | None | None |
| SERPINC1 | None | None | None | None |
| SERPINC1 | None | None | None | None |
| SERPINC1 | None | None | None | None |
| SERPINC1 | None | None | None | None |
| SERPINC1 | None | None | None | None |
| SERPINC1 | None | None | None | None |
| SERPINC1 | None | None | None | None |
| SERPINC1 | None | None | None | None |
| SERPINC1 | None | None | None | None |
| SERPINC1 | None | None | None | None |
| SERPINC1 | None | None | None | None |
| SERPINC1 | None | None | None | None |

| | | | | |
|---|---|---|---|---|
| SERPINC1 | None | None | None | None |
| SERPINC1 | None | None | None | None |
| SERPINC1 | None | None | None | None |
| SERPINC1 | None | None | None | None |
| SERPINC1 | None | None | None | None |
| SERPINC1 | None | None | None | None |
| SERPINC1 | None | None | None | None |
| SERPINC1 | None | None | None | None |
| SERPINC1 | None | None | None | None |
| ETFB | None | None | None | None |
| ETFB | None | None | None | None |
| ETFB | None | None | None | None |
| ETFB | None | None | None | None |
| ETFB | None | None | None | None |
| ETFB | None | None | None | None |
| DIABLO | None | None | None | None |
| DIABLO | None | None | None | None |
| DPYS | None | None | None | None |
| DPYS | None | None | None | None |
| DPYS | None | None | None | None |
| DPYS | None | None | None | None |
| DPYS | None | None | None | None |
| GNE | None | None | None | None |
| GNE | None | None | None | None |
| GNE | None | None | None | None |
| GNE | None | None | None | None |
| GNE | None | None | None | None |
| GNE | None | None | None | None |
| GNE | None | None | None | None |
| GNE | None | None | None | None |
| GNE | None | None | None | None |
| GNMT | None | None | None | None |
| GNMT | None | None | None | None |
| CTH | None | None | None | None |
| CTH | None | None | None | None |
| CTH | None | None | None | None |
| USH1C | None | None | None | None |
| PSEN1 | None | None | None | None |
| CHEK2 | None | None | None | None |
| CHEK2 | None | None | None | None |
| CHEK2 | None | None | None | None |
| CHEK2 | None | None | None | None |
| CHEK2 | None | None | None | None |
| CHEK2 | None | None | None | None |
| CHEK2 | None | None | None | None |
| CHEK2 | None | None | None | None |
| CHEK2 | None | None | None | None |
| SNCA | None | None | None | None |
| SNCA | None | None | None | None |
| GAN | None | None | None | None |
| GAN | None | None | None | None |
| GAN | None | None | None | None |
| SBDS | None | None | None | None |
| SBDS | None | None | None | None |

| | | | | |
|---|---|---|---|---|
| SBDS | None | None | None | None |
| SBDS | None | None | None | None |
| SBDS | None | None | None | None |
| SBDS | None | None | None | None |
| MPO | 1 | 3 | 3f9p | A |
| MPO | None | None | None | None |
| KCNH2 | None | None | None | None |
| KCNH2 | None | None | None | None |
| KCNH2 | 0.75 | 1 | 4hp9 | A |
| KCNH2 | None | None | None | None |
| TEK | None | None | None | None |
| TEK | None | None | None | None |
| TEK | None | None | None | None |
| TEK | None | None | None | None |
| NPC2 | None | None | None | None |
| NPC2 | None | None | None | None |
| NPC2 | None | None | None | None |
| NPC2 | None | None | None | None |
| NPC2 | None | None | None | None |
| NPC2 | None | None | None | None |
| NPC2 | None | None | None | None |
| PLG | None | None | None | None |
| PCSK9 | None | None | None | None |
| SFTPC | None | None | None | None |
| SFTPC | None | None | None | None |
| SFTPC | None | None | None | None |
| CD40LG | None | None | None | None |
| CD40LG | None | None | None | None |
| CD40LG | None | None | None | None |
| CD40LG | None | None | None | None |
| CD40LG | None | None | None | None |
| TAT | None | None | None | None |
| TAT | None | None | None | None |
| TAT | None | None | None | None |
| EP300 | None | None | None | None |
| FARS2 | None | None | None | None |
| FARS2 | None | None | None | None |
| FARS2 | None | None | None | None |
| FARS2 | None | None | None | None |
| FARS2 | None | None | None | None |
| FARS2 | None | None | None | None |
| FARS2 | None | None | None | None |
| FARS2 | None | None | None | None |
| FARS2 | None | None | None | None |
| FARS2 | None | None | None | None |
| FARS2 | None | None | None | None |
| FARS2 | None | None | None | None |
| FARS2 | None | None | None | None |
| MASP2 | None | None | None | None |
| MASP2 | None | None | None | None |
| MASP2 | None | None | None | None |
| MASP2 | None | None | None | None |
| STIM1 | None | None | None | None |
| CHRM2 | None | None | None | None |

| | | | | |
|---|---|---|---|---|
| MCFD2 | None | None | None | None |
| MCFD2 | None | None | None | None |
| MCFD2 | None | None | None | None |
| KITLG | None | None | None | None |
| F12 | None | None | None | None |
| F12 | None | None | None | None |
| ERAP1 | None | None | None | None |
| GTF2H5 | None | None | None | None |
| GTF2H5 | None | None | None | None |
| THBD | None | None | None | None |
| MRPL28 | None | None | None | None |
| MRPL28 | None | None | None | None |
| HTRA1 | None | None | None | None |
| HTRA1 | None | None | None | None |
| HTRA1 | None | None | None | None |
| HTRA1 | None | None | None | None |
| HTRA1 | None | None | None | None |
| HTRA1 | None | None | None | None |
| KARS | None | None | None | None |
| KARS | None | None | None | None |
| KARS | None | None | None | None |
| KARS | None | None | None | None |
| KARS | None | None | None | None |
| HK1 | 0.722 | 11 | 4fpb | B |
| GARS | None | None | None | None |
| GARS | None | None | None | None |
| GARS | None | None | None | None |
| GARS | None | None | None | None |
| GARS | None | None | None | None |
| GARS | None | None | None | None |
| GARS | None | None | None | None |
| IL12B | None | None | None | None |
| HADH | None | None | None | None |
| HADH | None | None | None | None |
| HADH | None | None | None | None |
| HADH | None | None | None | None |
| HADH | None | None | None | None |
| RRM2B | None | None | None | None |
| RRM2B | None | None | None | None |
| RRM2B | None | None | None | None |
| RRM2B | None | None | None | None |
| RRM2B | None | None | None | None |
| RRM2B | None | None | None | None |
| RRM2B | None | None | None | None |
| RRM2B | None | None | None | None |
| RRM2B | None | None | None | None |
| RRM2B | None | None | None | None |
| RRM2B | None | None | None | None |
| RRM2B | None | None | None | None |
| RRM2B | None | None | None | None |
| RRM2B | None | None | None | None |
| RRM2B | None | None | None | None |
| MARVELD2 | None | None | None | None |
| TLR4 | None | None | None | None |

| | | | | |
|---|---|---|---|---|
| TTN | None | None | None | None |
| TTN | None | None | None | None |
| TNFRSF13B | None | None | None | None |
| POLG | None | None | None | None |
| POLG | None | None | None | None |
| POLG | None | None | None | None |
| POLG | None | None | None | None |
| POLG | None | None | None | None |
| POLG | None | None | None | None |
| POLG | None | None | None | None |
| POLG | None | None | None | None |
| POLG | None | None | None | None |
| POLG | None | None | None | None |
| POLG | None | None | None | None |
| POLG | None | None | None | None |
| POLG | None | None | None | None |
| POLG | None | None | None | None |
| POLG | None | None | None | None |
| POLG | None | None | None | None |
| POLG | None | None | None | None |
| POLG | None | None | None | None |
| POLG | None | None | None | None |
| POLG | None | None | None | None |
| POLG | None | None | None | None |
| POLG | None | None | None | None |
| POLG | None | None | None | None |
| POLG | None | None | None | None |
| POLG | None | None | None | None |
| POLG | None | None | None | None |
| POLG | None | None | None | None |
| POLG | None | None | None | None |
| POLG | None | None | None | None |
| POLG | None | None | None | None |
| POLG | None | None | None | None |
| POLG | None | None | None | None |
| POLG | None | None | None | None |
| POLG | None | None | None | None |
| POLG | None | None | None | None |
| POLG | None | None | None | None |
| POLG | None | None | None | None |
| POLG | None | None | None | None |
| POLG | None | None | None | None |
| POLG | None | None | None | None |
| POLG | None | None | None | None |
| POLG | None | None | None | None |
| POLG | None | None | None | None |
| POLG | None | None | None | None |
| POLG | None | None | None | None |
| POLG | None | None | None | None |
| POLG | None | None | None | None |
| POLG | None | None | None | None |
| POLG | None | None | None | None |
| POLG | None | None | None | None |
| UROD | None | None | None | None |

| | | | | |
|---|---|---|---|---|
| UROD | None | None | None | None |
| UROD | None | None | None | None |
| UROD | None | None | None | None |
| UROD | None | None | None | None |
| UROD | None | None | None | None |
| UROD | 0.789 | 6 | 3gvw | A |
| UROD | 0.842 | 2 | 3gvv | A |
| UROD | 0.737 | 31 | 1r3q | A |
| UROD | None | None | None | None |
| PCK1 | None | None | None | None |
| PCK1 | None | None | None | None |
| PCK1 | None | None | None | None |
| PCK1 | None | None | None | None |
| PCK1 | None | None | None | None |
| HEXA | None | None | None | None |
| HEXA | None | None | None | None |
| HEXA | None | None | None | None |
| HEXA | None | None | None | None |
| HEXA | None | None | None | None |
| HEXA | None | None | None | None |
| HEXA | None | None | None | None |
| HEXA | None | None | None | None |
| HEXA | None | None | None | None |
| HEXA | None | None | None | None |
| HEXA | None | None | None | None |
| HEXA | None | None | None | None |
| HEXA | None | None | None | None |
| HEXA | None | None | None | None |
| HEXA | None | None | None | None |
| HEXA | None | None | None | None |
| HEXA | None | None | None | None |
| HEXA | None | None | None | None |
| HEXA | None | None | None | None |
| HEXA | None | None | None | None |
| HEXA | None | None | None | None |
| HEXA | None | None | None | None |
| HEXA | None | None | None | None |
| HEXA | None | None | None | None |
| HEXA | None | None | None | None |
| HEXA | None | None | None | None |
| HEXA | None | None | None | None |
| HEXA | None | None | None | None |
| HEXA | None | None | None | None |
| HEXA | None | None | None | None |
| CFTR | None | None | None | None |
| CFTR | None | None | None | None |
| CFTR | None | None | None | None |
| CFTR | None | None | None | None |
| CFTR | None | None | None | None |
| CFTR | None | None | None | None |
| CFTR | None | None | None | None |
| CFTR | None | None | None | None |
| CFTR | None | None | None | None |

| | | | | |
|---|---|---|---|---|
| CFTR | None | None | None | None |
| CFTR | None | None | None | None |
| CFTR | None | None | None | None |
| CFTR | None | None | None | None |
| CFTR | None | None | None | None |
| CFTR | None | None | None | None |
| CFTR | None | None | None | None |
| CFTR | None | None | None | None |
| CFTR | None | None | None | None |
| CFTR | None | None | None | None |
| CFTR | None | None | None | None |
| CFTR | None | None | None | None |
| CFTR | None | None | None | None |
| CFTR | None | None | None | None |
| CFTR | None | None | None | None |
| CFTR | None | None | None | None |
| CFTR | None | None | None | None |
| CFTR | None | None | None | None |
| CFTR | None | None | None | None |
| CFTR | None | None | None | None |
| CFTR | None | None | None | None |
| CFTR | None | None | None | None |
| CFTR | None | None | None | None |
| CFTR | None | None | None | None |
| CFTR | None | None | None | None |
| FAN1 | None | None | None | None |
| CBS | None | None | None | None |
| CBS | None | None | None | None |
| CBS | None | None | None | None |
| CBS | None | None | None | None |
| CBS | None | None | None | None |
| CBS | None | None | None | None |
| CBS | None | None | None | None |
| CBS | None | None | None | None |
| CBS | None | None | None | None |
| CBS | None | None | None | None |
| CBS | None | None | None | None |
| CBS | None | None | None | None |
| CBS | None | None | None | None |
| CBS | None | None | None | None |
| CBS | None | None | None | None |
| CBS | None | None | None | None |
| CBS | None | None | None | None |
| CBS | None | None | None | None |
| CBS | None | None | None | None |
| CBS | None | None | None | None |
| CBS | None | None | None | None |
| CBS | None | None | None | None |
| CBS | None | None | None | None |
| CBS | None | None | None | None |
| CBS | None | None | None | None |
| CBS | None | None | None | None |
| CBS | None | None | None | None |
| CBS | None | None | None | None |

| | | | | |
|---|---|---|---|---|
| CBS | None | None | None | None |
| CBS | None | None | None | None |
| CBS | None | None | None | None |
| CBS | None | None | None | None |
| CBS | None | None | None | None |
| CBS | None | None | None | None |
| CBS | None | None | None | None |
| CBS | None | None | None | None |
| TLR3 | None | None | None | None |
| LIG4 | None | None | None | None |
| LIG4 | None | None | None | None |
| ASPA | 0.769 | 1 | 2i3c | B |
| ASPA | 0.769 | 1 | 2i3c | B |
| ASPA | None | None | None | None |
| ASPA | None | None | None | None |
| ASPA | None | None | None | None |
| ASPA | None | None | None | None |
| ASPA | None | None | None | None |
| ASPA | None | None | None | None |
| ASPA | None | None | None | None |
| ASPA | None | None | None | None |
| ASPA | None | None | None | None |
| ASPA | None | None | None | None |
| ASPA | None | None | None | None |
| CFH | None | None | None | None |
| FCGR3A | None | None | None | None |
| FCGR3A | None | None | None | None |
| GRIN2A | None | None | None | None |
| GRIN2A | None | None | None | None |
| GRIN2A | None | None | None | None |
| GLA | None | None | None | None |
| GLA | None | None | None | None |
| GLA | None | None | None | None |
| GLA | None | None | None | None |
| GLA | None | None | None | None |
| GLA | None | None | None | None |
| GLA | None | None | None | None |
| GLA | None | None | None | None |
| GLA | None | None | None | None |
| GLA | None | None | None | None |
| GLA | None | None | None | None |
| GLA | 0.75 | 3 | 3gxn | B |
| GLA | 0.75 | 3 | 3gxn | B |
| GLA | None | None | None | None |
| GLA | None | None | None | None |
| GLA | None | None | None | None |
| GLA | None | None | None | None |
| GLA | None | None | None | None |
| GLA | None | None | None | None |
| GLA | None | None | None | None |
| GLA | None | None | None | None |
| GLA | None | None | None | None |
| GLA | None | None | None | None |
| GLA | None | None | None | None |

| | | | | |
|---|---|---|---|---|
| GLA | None | None | None | None |
| GLA | None | None | None | None |
| GLA | None | None | None | None |
| GLA | None | None | None | None |
| GLA | None | None | None | None |
| GLA | None | None | None | None |
| GLA | None | None | None | None |
| SERPINA3 | None | None | None | None |
| MTAP | None | None | None | None |
| MTAP | None | None | None | None |
| MTAP | None | None | None | None |
| BLMH | None | None | None | None |
| TUSC3 | None | None | None | None |
| ISCU | None | None | None | None |
| CYP11B2 | None | None | None | None |
| CYP11B2 | None | None | None | None |
| CYP11B2 | None | None | None | None |
| CYP11B2 | None | None | None | None |
| CYP11B2 | None | None | None | None |
| CYP11B2 | None | None | None | None |
| CYP11B2 | None | None | None | None |
| CYP11B2 | None | None | None | None |
| CYP11B2 | None | None | None | None |
| CYP11B2 | None | None | None | None |
| CYP11B2 | None | None | None | None |
| CYP11B2 | None | None | None | None |
| CYP11B2 | None | None | None | None |
| CYP11B2 | None | None | None | None |
| CYP11B2 | None | None | None | None |
| CYP11B2 | None | None | None | None |
| CYP11B2 | None | None | None | None |
| CYP11B2 | None | None | None | None |
| CYP11B2 | None | None | None | None |
| CYP11B2 | None | None | None | None |
| CRADD | None | None | None | None |
| CRADD | None | None | None | None |
| FGA | None | None | None | None |
| FGA | None | None | None | None |
| FGA | None | None | None | None |
| FGA | None | None | None | None |
| FGA | None | None | None | None |
| FGA | None | None | None | None |
| FGA | None | None | None | None |
| FGA | None | None | None | None |
| IYD | 1 | 1 | 4ttb | A |
| IYD | None | None | None | None |
| IYD | None | None | None | None |
| IYD | None | None | None | None |
| TCAP | None | None | None | None |
| TCAP | None | None | None | None |
| TCAP | None | None | None | None |
| TCAP | None | None | None | None |
| TCAP | None | None | None | None |
| CYP7A1 | None | None | None | None |

| | | | | |
|---|---|---|---|---|
| TEK | None | None | None | None |
| TEK | None | None | None | None |
| TEK | None | None | None | None |
| TEK | None | None | None | None |
| TEK | None | None | None | None |
| TEK | None | None | None | None |
| TEK | None | None | None | None |
| CD8A | None | None | None | None |
| HERC2 | None | None | None | None |
| NR1H3 | None | None | None | None |
| GRHPR | None | None | None | None |
| GRHPR | None | None | None | None |
| GRHPR | None | None | None | None |
| GRHPR | None | None | None | None |
| GRHPR | None | None | None | None |
| GRHPR | None | None | None | None |
| GRHPR | None | None | None | None |
| GBE1 | None | None | None | None |
| GBE1 | None | None | None | None |
| GBE1 | None | None | None | None |
| GBE1 | None | None | None | None |
| GBE1 | None | None | None | None |
| GBE1 | None | None | None | None |
| GBE1 | None | None | None | None |
| GBE1 | None | None | None | None |
| GBE1 | None | None | None | None |
| GBE1 | None | None | None | None |
| GBE1 | None | None | None | None |
| GBE1 | None | None | None | None |
| GBE1 | None | None | None | None |
| GBE1 | None | None | None | None |
| GBE1 | None | None | None | None |
| GBE1 | None | None | None | None |
| GBE1 | None | None | None | None |
| GBE1 | None | None | None | None |
| GBE1 | None | None | None | None |
| PC | None | None | None | None |
| PC | None | None | None | None |
| PC | None | None | None | None |
| PC | None | None | None | None |
| PC | None | None | None | None |
| PC | None | None | None | None |
| PC | None | None | None | None |
| PC | None | None | None | None |
| MSH6 | None | None | None | None |
| MSH6 | None | None | None | None |
| MSH6 | None | None | None | None |
| MSH6 | None | None | None | None |
| MSH6 | None | None | None | None |
| MSH6 | None | None | None | None |
| MSH6 | None | None | None | None |
| MSH6 | None | None | None | None |
| MSH6 | None | None | None | None |

| | | | | |
|---|---|---|---|---|
| MSH6 | None | None | None | None |
| MSH6 | None | None | None | None |
| MSH6 | None | None | None | None |
| MSH6 | None | None | None | None |
| MSH6 | None | None | None | None |
| MSH6 | None | None | None | None |
| MSH6 | None | None | None | None |
| MSH6 | None | None | None | None |
| MSH6 | None | None | None | None |
| MSH6 | None | None | None | None |
| MSH6 | None | None | None | None |
| MSH6 | None | None | None | None |
| MSH6 | None | None | None | None |
| MSH6 | None | None | None | None |
| MSH6 | None | None | None | None |
| MSH6 | None | None | None | None |
| MSH6 | None | None | None | None |
| MSH6 | None | None | None | None |
| MSH6 | None | None | None | None |
| MSH6 | None | None | None | None |
| MSH6 | None | None | None | None |
| RNASEL | None | None | None | None |
| ORC1 | None | None | None | None |
| ORC1 | None | None | None | None |
| ORC1 | None | None | None | None |
| KCNH2 | None | None | None | None |
| KCNH2 | None | None | None | None |
| KCNH2 | None | None | None | None |
| KCNH2 | None | None | None | None |
| KCNH2 | None | None | None | None |
| KCNH2 | None | None | None | None |
| KCNH2 | None | None | None | None |
| KCNH2 | None | None | None | None |
| KCNH2 | None | None | None | None |
| KCNH2 | None | None | None | None |
| KCNH2 | None | None | None | None |
| KCNH2 | None | None | None | None |
| KCNH2 | None | None | None | None |
| KCNH2 | None | None | None | None |
| KCNH2 | None | None | None | None |
| KCNH2 | None | None | None | None |
| KCNH2 | None | None | None | None |
| KCNH2 | None | None | None | None |
| KCNH2 | None | None | None | None |
| KCNH2 | None | None | None | None |
| KCNH2 | None | None | None | None |
| KCNH2 | None | None | None | None |
| KCNH2 | None | None | None | None |
| KCNH2 | None | None | None | None |
| MAT1A | None | None | None | None |
| MAT1A | None | None | None | None |
| MAT1A | None | None | None | None |
| MAT1A | None | None | None | None |
| MAT1A | None | None | None | None |

| | | | | |
|---|---|---|---|---|
| MAT1A | None | None | None | None |
| MAT1A | None | None | None | None |
| MAT1A | None | None | None | None |
| INSR | None | None | None | None |
| INSR | None | None | None | None |
| INSR | None | None | None | None |
| INSR | None | None | None | None |
| AMT | None | None | None | None |
| AMT | None | None | None | None |
| AMT | None | None | None | None |
| AMT | None | None | None | None |
| AMT | None | None | None | None |
| AMT | None | None | None | None |
| AMT | None | None | None | None |
| AMT | None | None | None | None |
| AMT | None | None | None | None |
| CSNK1D | None | None | None | None |
| SF3B1 | None | None | None | None |
| AGA | None | None | None | None |
| AGA | None | None | None | None |
| AGA | None | None | None | None |
| AGA | None | None | None | None |
| AGA | None | None | None | None |
| AGA | None | None | None | None |
| AGA | None | None | None | None |
| NECTIN4 | None | None | None | None |
| RS1 | None | None | None | None |
| RS1 | None | None | None | None |
| RS1 | None | None | None | None |
| RS1 | None | None | None | None |
| TBC1D7 | None | None | None | None |
| CTSA | None | None | None | None |
| CTSA | None | None | None | None |
| TKT | None | None | None | None |
| RP2 | None | None | None | None |
| RP2 | None | None | None | None |
| RP2 | None | None | None | None |
| RP2 | None | None | None | None |
| LYRM4 | None | None | None | None |
| SOS1 | None | None | None | None |
| SOS1 | None | None | None | None |
| SOS1 | None | None | None | None |
| SOS1 | None | None | None | None |
| SOS1 | None | None | None | None |
| SOS1 | None | None | None | None |
| HUWE1 | None | None | None | None |
| HUWE1 | None | None | None | None |
| TPI1 | None | None | None | None |
| TPI1 | 0.886 | 1 | 4unk | B |
| TPI1 | None | None | None | None |
| MSH2 | None | None | None | None |
| MSH2 | None | None | None | None |
| MSH2 | None | None | None | None |
| MSH2 | None | None | None | None |

| | | | | |
|---|---|---|---|---|
| MSH2 | None | None | None | None |
| MSH2 | None | None | None | None |
| MSH2 | None | None | None | None |
| MSH2 | None | None | None | None |
| MSH2 | None | None | None | None |
| MSH2 | None | None | None | None |
| MSH2 | None | None | None | None |
| MSH2 | None | None | None | None |
| MSH2 | None | None | None | None |
| MSH2 | None | None | None | None |
| MSH2 | None | None | None | None |
| MSH2 | None | None | None | None |
| MSH2 | None | None | None | None |
| MSH2 | None | None | None | None |
| MSH2 | None | None | None | None |
| MSH2 | None | None | None | None |
| MSH2 | None | None | None | None |
| MSH2 | None | None | None | None |
| MSH2 | None | None | None | None |
| MSH2 | None | None | None | None |
| MSH2 | None | None | None | None |
| MSH2 | None | None | None | None |
| MSH2 | None | None | None | None |
| MSH2 | None | None | None | None |
| MSH2 | None | None | None | None |
| MSH2 | None | None | None | None |
| MSH2 | None | None | None | None |
| MSH2 | None | None | None | None |
| MSH2 | None | None | None | None |
| MSH2 | None | None | None | None |
| MSH2 | None | None | None | None |
| MSH2 | None | None | None | None |
| GALE | None | None | None | None |
| GALE | 0.78 | 1 | 1i3n | A |
| GALE | None | None | None | None |
| GALE | None | None | None | None |
| GALE | None | None | None | None |
| GALE | None | None | None | None |
| GALE | None | None | None | None |
| GALE | None | None | None | None |
| NSD2 | None | None | None | None |
| SRP72 | None | None | None | None |
| SRP72 | None | None | None | None |
| SRP72 | None | None | None | None |
| POLK | None | None | None | None |
| GALT | None | None | None | None |
| GALT | None | None | None | None |
| GALT | None | None | None | None |
| GALT | None | None | None | None |
| GALT | None | None | None | None |
| GALT | None | None | None | None |
| GALT | None | None | None | None |
| GALT | None | None | None | None |
| GALT | None | None | None | None |

| | | | | |
|---|---|---|---|---|
| GALT | None | None | None | None |
| GALT | None | None | None | None |
| GALT | None | None | None | None |
| GALT | None | None | None | None |
| GALT | None | None | None | None |
| GALT | None | None | None | None |
| GALT | None | None | None | None |
| GALT | None | None | None | None |
| GALT | None | None | None | None |
| GALT | None | None | None | None |
| GALT | None | None | None | None |
| GALT | None | None | None | None |
| GALT | None | None | None | None |
| GALT | None | None | None | None |
| GALT | None | None | None | None |
| GALT | None | None | None | None |
| GALT | None | None | None | None |
| GALT | None | None | None | None |
| GALT | None | None | None | None |
| GALT | None | None | None | None |
| GALT | None | None | None | None |
| GALT | None | None | None | None |
| GALT | None | None | None | None |
| GALT | None | None | None | None |
| GALT | None | None | None | None |
| GALT | None | None | None | None |
| GALT | None | None | None | None |
| GALT | None | None | None | None |
| PYCR1 | 0.75 | 11 | 5uav | A |
| PYCR1 | 0.75 | 11 | 5uav | A |
| PYCR1 | None | None | None | None |
| PYCR1 | None | None | None | None |
| F8 | None | None | None | None |
| F8 | None | None | None | None |
| F8 | None | None | None | None |
| F8 | None | None | None | None |
| F8 | None | None | None | None |
| F8 | None | None | None | None |
| F8 | None | None | None | None |
| F8 | None | None | None | None |
| F8 | None | None | None | None |
| F8 | None | None | None | None |
| F8 | None | None | None | None |
| F8 | None | None | None | None |
| F8 | None | None | None | None |
| F8 | None | None | None | None |
| F8 | None | None | None | None |
| F8 | None | None | None | None |
| ANK1 | None | None | None | None |
| CHAT | None | None | None | None |
| CHAT | None | None | None | None |
| CHAT | None | None | None | None |
| CHAT | None | None | None | None |
| CHAT | None | None | None | None |

| | | | | |
|---|---|---|---|---|
| CHAT | None | None | None | None |
| CHAT | None | None | None | None |
| CHAT | None | None | None | None |
| CHAT | None | None | None | None |
| CHAT | None | None | None | None |
| CHAT | None | None | None | None |
| CHAT | None | None | None | None |
| CHAT | None | None | None | None |
| CHAT | None | None | None | None |
| CHAT | None | None | None | None |
| CHAT | None | None | None | None |
| CHAT | None | None | None | None |
| GHR | None | None | None | None |
| GHR | None | None | None | None |
| GHR | None | None | None | None |
| GHR | None | None | None | None |
| GHR | None | None | None | None |
| GHR | None | None | None | None |
| GHR | None | None | None | None |
| GHR | None | None | None | None |
| GHR | None | None | None | None |
| GHR | None | None | None | None |
| GHR | None | None | None | None |
| GHR | None | None | None | None |
| GHR | None | None | None | None |
| GHR | None | None | None | None |
| FZD4 | None | None | None | None |
| FZD4 | None | None | None | None |
| FZD4 | None | None | None | None |
| MTR | None | None | None | None |
| MTR | None | None | None | None |
| MTR | None | None | None | None |
| GM2A | None | None | None | None |
| GM2A | None | None | None | None |
| GM2A | None | None | None | None |
| GM2A | None | None | None | None |
| GM2A | None | None | None | None |
| CRYGD | None | None | None | None |
| CRYGD | None | None | None | None |
| CRYGD | None | None | None | None |
| CRYGD | None | None | None | None |
| CRYGD | None | None | None | None |
| CRYGD | None | None | None | None |
| CRYGD | None | None | None | None |
| CRYGD | None | None | None | None |
| CRYGD | None | None | None | None |
| TLR5 | None | None | None | None |
| HSD17B10 | 0.855 | 1 | 1u7t | B |
| HSD17B10 | None | None | None | None |
| HSD17B10 | None | None | None | None |
| HSD17B10 | None | None | None | None |
| TBC1D20 | None | None | None | None |
| FBN1 | None | None | None | None |
| FBN1 | None | None | None | None |

| | | | | |
|---|---|---|---|---|
| FBN1 | None | None | None | None |
| FBN1 | None | None | None | None |
| FBN1 | None | None | None | None |
| FBN1 | None | None | None | None |
| FBN1 | None | None | None | None |
| FXN | None | None | None | None |
| FXN | None | None | None | None |
| FXN | None | None | None | None |
| FXN | None | None | None | None |
| MYO5A | None | None | None | None |
| CTSK | None | None | None | None |
| CTSK | None | None | None | None |
| SCN5A | None | None | None | None |
| SCN5A | None | None | None | None |
| SOS1 | None | None | None | None |
| SOS1 | None | None | None | None |
| SOS1 | None | None | None | None |
| SOS1 | None | None | None | None |
| SOS1 | None | None | None | None |
| SOS1 | None | None | None | None |
| SOS1 | None | None | None | None |
| SOS1 | None | None | None | None |
| SOS1 | None | None | None | None |
| SOS1 | None | None | None | None |
| SOS1 | None | None | None | None |
| SOS1 | None | None | None | None |
| GATM | None | None | None | None |
| GATM | None | None | None | None |
| GATM | 0.75 | 4 | 4jdw | A |
| GATM | None | None | None | None |
| GATM | None | None | None | None |
| GATM | 0.75 | 6 | 6jdw | A |
| GATM | 0.75 | 11 | 6jdw | A |
| MMP2 | None | None | None | None |
| MMP2 | None | None | None | None |
| MMP2 | None | None | None | None |
| MMP2 | None | None | None | None |
| NDUFB9 | None | None | None | None |
| NDUFB9 | None | None | None | None |
| GH1 | None | None | None | None |
| GH1 | None | None | None | None |
| GH1 | None | None | None | None |
| SLC6A4 | None | None | None | None |
| G6PD | None | None | None | None |
| G6PD | None | None | None | None |
| G6PD | None | None | None | None |
| G6PD | None | None | None | None |
| G6PD | None | None | None | None |
| G6PD | None | None | None | None |
| G6PD | None | None | None | None |
| G6PD | None | None | None | None |
| G6PD | None | None | None | None |
| G6PD | None | None | None | None |
| G6PD | None | None | None | None |

| | | | | |
|---|---|---|---|---|
| G6PD | None | None | None | None |
| G6PD | None | None | None | None |
| G6PD | None | None | None | None |
| G6PD | None | None | None | None |
| G6PD | None | None | None | None |
| G6PD | None | None | None | None |
| G6PD | None | None | None | None |
| G6PD | None | None | None | None |
| G6PD | None | None | None | None |
| CST3 | None | None | None | None |
| LMNA | None | None | None | None |
| LMNA | None | None | None | None |
| NDUFA9 | None | None | None | None |
| NDUFA9 | None | None | None | None |
| NDUFA9 | None | None | None | None |
| NDUFA9 | None | None | None | None |
| YARS | None | None | None | None |
| YARS | None | None | None | None |
| YARS | None | None | None | None |
| YARS | None | None | None | None |
| EPHA2 | None | None | None | None |
| EPHA2 | None | None | None | None |
| EPHA2 | None | None | None | None |
| EPHA2 | None | None | None | None |
| DICER1 | None | None | None | None |
| DICER1 | None | None | None | None |
| SNX10 | None | None | None | None |
| SNX10 | None | None | None | None |
| TTN | None | None | None | None |
| TTN | None | None | None | None |
| GATA3 | None | None | None | None |
| C5 | None | None | None | None |
| THBD | None | None | None | None |
| BIN1 | None | None | None | None |
| BIN1 | None | None | None | None |
| SMARCAL1 | None | None | None | None |
| CFB | None | None | None | None |
| CYC1 | None | None | None | None |
| ANK2 | None | None | None | None |
| CHMP4B | None | None | None | None |
| CHMP4B | None | None | None | None |
| ERBB2 | None | None | None | None |
| ERBB2 | None | None | None | None |
| POLG2 | None | None | None | None |
| POLG2 | None | None | None | None |
| POLG2 | None | None | None | None |
| POLG2 | None | None | None | None |
| POLG2 | None | None | None | None |
| POLG2 | None | None | None | None |
| POLG2 | None | None | None | None |
| POLG2 | None | None | None | None |
| C3 | None | None | None | None |
| C3 | None | None | None | None |
| RPGR | None | None | None | None |

| | | | | |
|---|---|---|---|---|
| RPGR | None | None | None | None |
| RPGR | None | None | None | None |
| RPGR | None | None | None | None |
| RPGR | None | None | None | None |
| C1QA | None | None | None | None |
| PRKG1 | None | None | None | None |
| IGBP1 | None | None | None | None |
| ENO3 | None | None | None | None |
| ENO3 | None | None | None | None |
| ENO3 | None | None | None | None |
| ENO3 | None | None | None | None |
| ENO3 | None | None | None | None |
| ENO3 | 0.897 | 2 | 2xsx | B |
| MCPH1 | None | None | None | None |
| MCPH1 | None | None | None | None |
| MCPH1 | None | None | None | None |
| MCPH1 | None | None | None | None |
| MCPH1 | None | None | None | None |
| SNRNP200 | None | None | None | None |
| SNRNP200 | None | None | None | None |
| DNMT1 | None | None | None | None |
| DNMT1 | None | None | None | None |
| DNMT1 | None | None | None | None |
| AKT1 | None | None | None | None |
| NDUFS2 | None | None | None | None |
| ACVR2B | None | None | None | None |
| HMGCL | None | None | None | None |
| HMGCL | None | None | None | None |
| HMGCL | None | None | None | None |
| JUP | None | None | None | None |
| JUP | None | None | None | None |
| JUP | None | None | None | None |
| JUP | None | None | None | None |
| JUP | None | None | None | None |
| JUP | None | None | None | None |
| JUP | None | None | None | None |
| JUP | None | None | None | None |
| SPINK1 | None | None | None | None |
| SPINK1 | None | None | None | None |
| SPINK1 | None | None | None | None |
| VCL | None | None | None | None |
| VCL | None | None | None | None |
| NOTCH1 | None | None | None | None |
| NOTCH1 | None | None | None | None |
| NOTCH1 | None | None | None | None |
| NDUFB3 | None | None | None | None |
| IKBKG | None | None | None | None |
| GSN | None | None | None | None |
| GSN | None | None | None | None |
| GSN | None | None | None | None |
| GSN | None | None | None | None |
| GSN | None | None | None | None |
| GSN | None | None | None | None |
| GSN | None | None | None | None |

| | | | | |
|---|---|---|---|---|
| GSN | None | None | None | None |
| GSN | None | None | None | None |
| CLC | None | None | None | None |
| RAF1 | None | None | None | None |
| RAF1 | None | None | None | None |
| RAF1 | None | None | None | None |
| NDUFS8 | None | None | None | None |
| NDUFS8 | None | None | None | None |
| NDUFS8 | None | None | None | None |
| NDUFS8 | None | None | None | None |
| NDUFS8 | None | None | None | None |
| CASR | None | None | None | None |
| CASR | None | None | None | None |
| CASR | None | None | None | None |
| CASR | None | None | None | None |
| CASR | None | None | None | None |
| CASR | None | None | None | None |
| CASR | None | None | None | None |
| CASR | None | None | None | None |
| CASR | None | None | None | None |
| CASR | None | None | None | None |
| CASR | None | None | None | None |
| CASR | None | None | None | None |
| CASR | None | None | None | None |
| CASR | None | None | None | None |
| CASR | None | None | None | None |
| CASR | None | None | None | None |
| CASR | None | None | None | None |
| CASR | None | None | None | None |
| CASR | None | None | None | None |
| CASR | None | None | None | None |
| CASR | None | None | None | None |
| CASR | None | None | None | None |
| CASR | None | None | None | None |
| CASR | None | None | None | None |
| CASR | None | None | None | None |
| CASR | None | None | None | None |
| CASR | None | None | None | None |
| CASR | None | None | None | None |
| CASR | None | None | None | None |
| VCL | None | None | None | None |
| VCL | None | None | None | None |
| SMPD1 | None | None | None | None |
| SMPD1 | None | None | None | None |
| SMPD1 | None | None | None | None |
| KDM6A | None | None | None | None |
| KCNQ1 | None | None | None | None |
| KCNQ1 | None | None | None | None |
| KCNQ1 | None | None | None | None |
| KCNQ1 | None | None | None | None |
| KCNQ1 | None | None | None | None |
| KCNQ1 | None | None | None | None |
| NDUFA12 | None | None | None | None |
| GLRX5 | None | None | None | None |

| | | | | |
|---|---|---|---|---|
| PALB2 | None | None | None | None |
| PALB2 | None | None | None | None |
| PALB2 | None | None | None | None |
| PALB2 | None | None | None | None |
| PALB2 | None | None | None | None |
| PALB2 | None | None | None | None |
| PALB2 | None | None | None | None |
| PALB2 | None | None | None | None |
| PALB2 | None | None | None | None |
| PALB2 | None | None | None | None |
| ALDOB | None | None | None | None |
| ALDOB | 0.731 | 2 | 1xdm | A |
| ALDOB | 0.731 | 1 | 1qo5 | B |
| ALDOB | None | None | None | None |
| ALDOB | None | None | None | None |
| ALDOB | None | None | None | None |
| ALDOB | None | None | None | None |
| GNE | None | None | None | None |
| GNE | None | None | None | None |
| GNE | None | None | None | None |
| GNE | None | None | None | None |
| GNE | None | None | None | None |
| GNE | None | None | None | None |
| GNE | None | None | None | None |
| GNE | None | None | None | None |
| GNE | None | None | None | None |
| GNE | None | None | None | None |
| GNE | None | None | None | None |
| GNE | None | None | None | None |
| CHRNB2 | None | None | None | None |
| CHRNB2 | None | None | None | None |
| ERCC3 | None | None | None | None |
| ERCC3 | None | None | None | None |
| ERCC3 | None | None | None | None |
| ERCC3 | None | None | None | None |
| ERCC3 | None | None | None | None |
| UNC119 | None | None | None | None |
| ATL1 | None | None | None | None |
| ATL1 | None | None | None | None |
| ATL1 | None | None | None | None |
| ATL1 | 0.8 | 1 | 3q5d | A |
| ATL1 | None | None | None | None |
| ATL1 | None | None | None | None |
| ATL1 | None | None | None | None |
| ATL1 | None | None | None | None |
| ATL1 | None | None | None | None |
| ATL1 | None | None | None | None |
| NR3C2 | 0.701 | 1 | 2oax | A |
| TBK1 | None | None | None | None |
| HYAL1 | None | None | None | None |
| HYAL1 | None | None | None | None |
| HYAL1 | None | None | None | None |
| HYAL1 | None | None | None | None |
| CTSD | None | None | None | None |

| | | | | | |
|---|---|---|---|---|---|
| PRKCG | None | None | | None | None |
| TGFBR1 | None | None | | None | None |
| XDH | None | None | | None | None |
| XDH | None | None | | None | None |
| XDH | None | None | | None | None |
| XDH | None | None | | None | None |
| XDH | None | None | | None | None |
| XDH | None | None | | None | None |
| NMNAT1 | 0.833 | | 1 | 1kqn | E |
| NMNAT1 | None | None | | None | None |
| NMNAT1 | None | None | | None | None |
| NMNAT1 | None | None | | None | None |
| NMNAT1 | None | None | | None | None |
| NMNAT1 | None | None | | None | None |
| SERPING1 | None | None | | None | None |
| SERPING1 | None | None | | None | None |
| SERPING1 | None | None | | None | None |
| SERPING1 | None | None | | None | None |
| SERPING1 | None | None | | None | None |
| PDZRN3 | None | None | | None | None |
| FABP2 | None | None | | None | None |
| IFNGR1 | None | None | | None | None |
| IFNGR1 | None | None | | None | None |
| TDP1 | None | None | | None | None |
| TDP1 | None | None | | None | None |
| INS | None | None | | None | None |
| INS | 0.769 | | 1 | 2r34 | C |
| INS | 0.769 | | 1 | 2r34 | C |
| PPOX | 0.857 | | 1 | 4ivm | B |
| PPOX | None | None | | None | None |
| PPOX | None | None | | None | None |
| PPOX | None | None | | None | None |
| PPOX | None | None | | None | None |
| PPOX | None | None | | None | None |
| PPOX | None | None | | None | None |
| PPOX | None | None | | None | None |
| PPOX | None | None | | None | None |
| GFER | None | None | | None | None |
| GFER | 0.769 | | 4 | 3tk0 | A |
| GFER | None | None | | None | None |
| GFER | None | None | | None | None |
| LIFR | None | None | | None | None |
| EGLN1 | None | None | | None | None |
| EGLN1 | 0.889 | | 6 | 5v18 | A |
| EGLN1 | 0.944 | | 2 | 5a3u | C |
| GP1BA | None | None | | None | None |
| GP1BA | None | None | | None | None |
| GP1BA | None | None | | None | None |
| GP1BA | None | None | | None | None |
| GP1BA | None | None | | None | None |
| MMADHC | None | None | | None | None |
| MMADHC | None | None | | None | None |
| MMADHC | None | None | | None | None |
| MMADHC | None | None | | None | None |

| | | | | |
|---|---|---|---|---|
| MMADHC | None | None | None | None |
| MMADHC | None | None | None | None |
| MMADHC | None | None | None | None |
| ACVRL1 | None | None | None | None |
| ACVRL1 | None | None | None | None |
| RFXANK | None | None | None | None |
| CHIT1 | None | None | None | None |
| UBR1 | None | None | None | None |
| BLM | None | None | None | None |
| BLM | None | None | None | None |
| BLM | None | None | None | None |
| BLM | None | None | None | None |
| BLM | None | None | None | None |
| BLM | None | None | None | None |
| BLM | None | None | None | None |
| BLM | None | None | None | None |
| BLM | None | None | None | None |
| BLM | None | None | None | None |
| BLM | None | None | None | None |
| BLM | None | None | None | None |
| BLM | None | None | None | None |
| VDR | None | None | None | None |
| VDR | None | None | None | None |
| VDR | None | None | None | None |
| VDR | None | None | None | None |
| VDR | None | None | None | None |
| VDR | None | None | None | None |
| EPHA2 | None | None | None | None |
| EPHA2 | None | None | None | None |
| EPHA2 | None | None | None | None |
| EPHA2 | None | None | None | None |
| EPHA2 | None | None | None | None |
| EPHA2 | None | None | None | None |
| EPHA2 | None | None | None | None |
| EPHA2 | None | None | None | None |
| EPHA2 | None | None | None | None |
| EPHA2 | None | None | None | None |
| UBA1 | None | None | None | None |
| STAT1 | None | None | None | None |
| STAT1 | None | None | None | None |
| STAT1 | None | None | None | None |
| STAT1 | None | None | None | None |
| STAT1 | None | None | None | None |
| STAT1 | None | None | None | None |
| STAT1 | None | None | None | None |
| STAT1 | None | None | None | None |
| DNM1L | None | None | None | None |
| DNM1L | None | None | None | None |
| OTC | None | None | None | None |
| OTC | None | None | None | None |
| OTC | None | None | None | None |
| OTC | None | None | None | None |
| OTC | None | None | None | None |
| OTC | None | None | None | None |

| | | | | |
|---|---|---|---|---|
| OTC | None | None | None | None |
| OTC | None | None | None | None |
| OTC | None | None | None | None |
| OTC | None | None | None | None |
| OTC | None | None | None | None |
| OTC | None | None | None | None |
| OTC | None | None | None | None |
| OTC | None | None | None | None |
| OTC | None | None | None | None |
| OTC | None | None | None | None |
| OTC | None | None | None | None |
| RARS | None | None | None | None |
| RARS | None | None | None | None |
| RARS | None | None | None | None |
| SEPSECS | None | None | None | None |
| SEPSECS | None | None | None | None |
| SEPSECS | None | None | None | None |
| SEPSECS | None | None | None | None |
| SEPSECS | None | None | None | None |
| BRCA1 | None | None | None | None |
| BRCA1 | None | None | None | None |
| BRCA1 | None | None | None | None |
| BRCA1 | None | None | None | None |
| BRCA1 | None | None | None | None |
| BRCA1 | None | None | None | None |
| BRCA1 | None | None | None | None |
| BRCA1 | None | None | None | None |
| BRCA1 | None | None | None | None |
| BRCA1 | None | None | None | None |
| BRCA1 | None | None | None | None |
| BRCA1 | None | None | None | None |
| BRCA1 | None | None | None | None |
| BRCA1 | None | None | None | None |
| BRCA1 | None | None | None | None |
| BRCA1 | None | None | None | None |
| BRCA1 | None | None | None | None |
| BRCA1 | None | None | None | None |
| BRCA1 | None | None | None | None |
| BRCA1 | None | None | None | None |
| BRCA1 | None | None | None | None |
| BRCA1 | None | None | None | None |
| CUBN | None | None | None | None |
| ITGB3 | None | None | None | None |
| MTRR | None | None | None | None |
| MTRR | None | None | None | None |
| MTRR | None | None | None | None |
| MTRR | None | None | None | None |
| MTRR | None | None | None | None |
| MTRR | None | None | None | None |
| MTRR | None | None | None | None |
| MTRR | None | None | None | None |
| IL7R | None | None | None | None |
| IL7R | None | None | None | None |
| RPS24 | None | None | None | None |

| | | | | |
|---|---|---|---|---|
| RPS24 | None | None | None | None |
| HOGA1 | None | None | None | None |
| HOGA1 | None | None | None | None |
| HOGA1 | None | None | None | None |
| HOGA1 | None | None | None | None |
| HOGA1 | None | None | None | None |
| HOGA1 | None | None | None | None |
| HOGA1 | 0.791 | 1 | 3s5n | A |
| HOGA1 | None | None | None | None |
| HOGA1 | None | None | None | None |
| HOGA1 | None | None | None | None |
| HOGA1 | None | None | None | None |
| HOGA1 | None | None | None | None |
| HOGA1 | None | None | None | None |
| GJB2 | None | None | None | None |
| GJB2 | None | None | None | None |
| GJB2 | None | None | None | None |
| GJB2 | None | None | None | None |
| GJB2 | None | None | None | None |
| GJB2 | None | None | None | None |
| GJB2 | None | None | None | None |
| GJB2 | None | None | None | None |
| GJB2 | None | None | None | None |
| GJB2 | None | None | None | None |
| GJB2 | None | None | None | None |
| GJB2 | None | None | None | None |
| GJB2 | None | None | None | None |
| GJB2 | None | None | None | None |
| GJB2 | None | None | None | None |
| GJB2 | None | None | None | None |
| GJB2 | None | None | None | None |
| GJB2 | None | None | None | None |
| GJB2 | None | None | None | None |
| GJB2 | None | None | None | None |
| GJB2 | None | None | None | None |
| GJB2 | None | None | None | None |
| GJB2 | None | None | None | None |
| GJB2 | None | None | None | None |
| GJB2 | None | None | None | None |
| GJB2 | None | None | None | None |
| GJB2 | None | None | None | None |
| GJB2 | None | None | None | None |
| GJB2 | None | None | None | None |
| GJB2 | None | None | None | None |
| GJB2 | None | None | None | None |
| GJB2 | None | None | None | None |
| GJB2 | None | None | None | None |
| GJB2 | None | None | None | None |
| GJB2 | None | None | None | None |
| GJB2 | None | None | None | None |
| GJB2 | None | None | None | None |
| GJB2 | None | None | None | None |
| GJB2 | None | None | None | None |
| GJB2 | None | None | None | None |

| | | | | |
|---|---|---|---|---|
| GJB2 | None | None | None | None |
| GJB2 | None | None | None | None |
| GJB2 | None | None | None | None |
| GJB2 | None | None | None | None |
| GJB2 | None | None | None | None |
| GJB2 | None | None | None | None |
| GJB2 | None | None | None | None |
| GJB2 | None | None | None | None |
| GJB2 | None | None | None | None |
| GJB2 | None | None | None | None |
| GJB2 | None | None | None | None |
| GJB2 | None | None | None | None |
| GJB2 | None | None | None | None |
| GJB2 | None | None | None | None |
| CPS1 | None | None | None | None |
| CPS1 | None | None | None | None |
| CPS1 | None | None | None | None |
| CPS1 | None | None | None | None |
| CPS1 | None | None | None | None |
| CPS1 | None | None | None | None |
| CPS1 | None | None | None | None |
| CPS1 | None | None | None | None |
| CPS1 | None | None | None | None |
| CPS1 | None | None | None | None |
| PSPH | None | None | None | None |
| PSPH | None | None | None | None |
| PSPH | None | None | None | None |
| PSPH | None | None | None | None |
| PSPH | None | None | None | None |
| PSPH | None | None | None | None |
| PEPD | None | None | None | None |
| PEPD | None | None | None | None |
| PEPD | None | None | None | None |
| PEPD | None | None | None | None |
| PEPD | None | None | None | None |
| PEPD | None | None | None | None |
| PEPD | None | None | None | None |
| PEPD | None | None | None | None |
| PEPD | None | None | None | None |
| KDM5C | None | None | None | None |
| KDM5C | None | None | None | None |
| KDM5C | None | None | None | None |
| ITGB6 | None | None | None | None |
| ITGB6 | None | None | None | None |
| ITGB6 | None | None | None | None |
| DARS | None | None | None | None |
| DARS | None | None | None | None |
| DARS | None | None | None | None |
| DARS | None | None | None | None |
| BTK | None | None | None | None |
| BTK | None | None | None | None |
| BTK | None | None | None | None |
| HDAC4 | None | None | None | None |
| ITGA2B | None | None | None | None |

| | | | | |
|---|---|---|---|---|
| ITGA2B | None | None | None | None |
| ITGA2B | None | None | None | None |
| ITGA2B | None | None | None | None |
| PCSK9 | None | None | None | None |
| PCSK9 | None | None | None | None |
| PCSK9 | None | None | None | None |
| PCSK9 | None | None | None | None |
| MUTYH | None | None | None | None |
| MUTYH | None | None | None | None |
| MUTYH | None | None | None | None |
| MUTYH | None | None | None | None |
| MUTYH | None | None | None | None |
| MUTYH | None | None | None | None |
| MUTYH | None | None | None | None |
| MUTYH | None | None | None | None |
| MUTYH | None | None | None | None |
| MUTYH | None | None | None | None |
| MUTYH | None | None | None | None |
| MUTYH | None | None | None | None |
| MUTYH | None | None | None | None |
| MUTYH | None | None | None | None |
| MUTYH | None | None | None | None |
| MUTYH | None | None | None | None |
| MUTYH | None | None | None | None |
| MUTYH | None | None | None | None |
| MUTYH | None | None | None | None |
| MUTYH | None | None | None | None |
| MUTYH | None | None | None | None |
| MUTYH | None | None | None | None |
| MUTYH | None | None | None | None |
| MUTYH | None | None | None | None |
| SLC25A12 | None | None | None | None |
| GSN | None | None | None | None |
| IHH | 0.909 | 1 | 3n1p | B |
| IHH | None | None | None | None |
| IHH | None | None | None | None |
| IHH | 0.818 | 2 | 3n1p | B |
| IHH | 0.727 | 12 | 3n1p | B |
| IHH | None | None | None | None |
| IHH | 0.818 | 7 | 3n1p | B |
| CD247 | None | None | None | None |
| EP300 | None | None | None | None |
| TTN | None | None | None | None |
| MRI1 | None | None | None | None |
| SGSH | None | None | None | None |
| SGSH | None | None | None | None |
| SGSH | None | None | None | None |
| SGSH | None | None | None | None |
| SGSH | None | None | None | None |
| SGSH | None | None | None | None |
| SGSH | None | None | None | None |
| SGSH | None | None | None | None |
| SGSH | None | None | None | None |
| SGSH | None | None | None | None |

| | | | | |
|---|---|---|---|---|
| SGSH | None | None | None | None |
| SGSH | None | None | None | None |
| SGSH | None | None | None | None |
| SGSH | None | None | None | None |
| SGSH | None | None | None | None |
| SGSH | None | None | None | None |
| SGSH | None | None | None | None |
| SGSH | None | None | None | None |
| SGSH | None | None | None | None |
| SGSH | None | None | None | None |
| SGSH | None | None | None | None |
| SGSH | None | None | None | None |
| CBL | None | None | None | None |
| CANT1 | None | None | None | None |
| CANT1 | None | None | None | None |
| CANT1 | None | None | None | None |
| CANT1 | None | None | None | None |
| CANT1 | None | None | None | None |
| CANT1 | None | None | None | None |
| CANT1 | None | None | None | None |
| SLC1A3 | None | None | None | None |
| SLC1A3 | None | None | None | None |
| SLC1A5 | None | None | None | None |
| SLC1A3 | None | None | None | None |
| SLC1A5 | None | None | None | None |
| SLC1A3 | None | None | None | None |
| SLC1A3 | None | None | None | None |
| CTSD | None | None | None | None |
| CTSD | None | None | None | None |
| CTSD | None | None | None | None |
| CTSD | None | None | None | None |
| CTSD | None | None | None | None |
| MTOR | None | None | None | None |
| ILK | None | None | None | None |
| PCDH15 | None | None | None | None |
| PCDH15 | None | None | None | None |
| PCDH15 | None | None | None | None |
| PCDH15 | None | None | None | None |
| PCDH15 | None | None | None | None |
| PCDH15 | None | None | None | None |
| KLHL40 | None | None | None | None |
| KLHL40 | None | None | None | None |
| KLHL40 | None | None | None | None |
| KLHL40 | None | None | None | None |
| CSTA | None | None | None | None |
| CSTA | None | None | None | None |
| HNMT | None | None | None | None |
| HNMT | None | None | None | None |
| IFNGR2 | None | None | None | None |
| IFNGR2 | None | None | None | None |
| PEX5 | None | None | None | None |
| KIF7 | None | None | None | None |
| KIF7 | None | None | None | None |
| MYL3 | None | None | None | None |

| | | | | |
|---|---|---|---|---|
| MYL3 | None | None | None | None |
| MYL3 | None | None | None | None |
| MYL3 | None | None | None | None |
| MYL3 | None | None | None | None |
| MYL3 | None | None | None | None |
| MYL3 | None | None | None | None |
| NAGLU | None | None | None | None |
| NAGLU | None | None | None | None |
| NAGLU | None | None | None | None |
| NAGLU | None | None | None | None |
| NAGLU | None | None | None | None |
| NAGLU | None | None | None | None |
| NAGLU | None | None | None | None |
| NAGLU | None | None | None | None |
| NAGLU | None | None | None | None |
| NAGLU | None | None | None | None |
| NAGLU | None | None | None | None |
| NAGLU | None | None | None | None |
| NAGLU | None | None | None | None |
| TNFRSF13C | None | None | None | None |
| KIF21A | None | None | None | None |
| KIF21A | None | None | None | None |
| KIF21A | None | None | None | None |
| KIF21A | None | None | None | None |
| KIF21A | None | None | None | None |
| DOCK2 | None | None | None | None |
| DSP | None | None | None | None |
| CDKN2A | None | None | None | None |
| CDKN2A | None | None | None | None |
| CDKN2A | None | None | None | None |
| CDKN2A | None | None | None | None |
| CDKN2A | None | None | None | None |
| CDKN2A | None | None | None | None |
| CDKN2A | None | None | None | None |
| CDKN2A | None | None | None | None |
| CDKN2A | None | None | None | None |
| CDKN2A | None | None | None | None |
| CDKN2A | None | None | None | None |
| CDKN2A | None | None | None | None |
| CDKN2A | None | None | None | None |
| CDKN2A | None | None | None | None |
| VCL | None | None | None | None |
| TPP1 | None | None | None | None |
| TPP1 | None | None | None | None |
| TPP1 | None | None | None | None |
| TPP1 | None | None | None | None |
| TPP1 | None | None | None | None |
| TPP1 | None | None | None | None |
| TPP1 | None | None | None | None |
| TPP1 | None | None | None | None |
| TPP1 | None | None | None | None |
| TPP1 | None | None | None | None |
| TPP1 | None | None | None | None |
| TPP1 | None | None | None | None |

| | | | | |
|---|---|---|---|---|
| TPP1 | None | None | None | None |
| TPP1 | None | None | None | None |
| TPP1 | None | None | None | None |
| TPP1 | None | None | None | None |
| TPP1 | None | None | None | None |
| TPP1 | None | None | None | None |
| TPP1 | None | None | None | None |
| TPP1 | None | None | None | None |
| TPP1 | None | None | None | None |
| TPP1 | None | None | None | None |
| IL10RB | None | None | None | None |
| IL10RB | None | None | None | None |
| CCM2 | None | None | None | None |
| CCM2 | None | None | None | None |
| CCM2 | None | None | None | None |
| DCX | None | None | None | None |
| DCX | None | None | None | None |
| DCX | None | None | None | None |
| DCX | None | None | None | None |
| DCX | None | None | None | None |
| DCX | None | None | None | None |
| DCX | None | None | None | None |
| DCX | None | None | None | None |
| BRCA2 | None | None | None | None |
| NNT | None | None | None | None |
| C2 | None | None | None | None |
| C2 | None | None | None | None |
| C2 | None | None | None | None |
| C2 | None | None | None | None |
| WRN | None | None | None | None |
| WRN | None | None | None | None |
| F13A1 | None | None | None | None |
| F13A1 | None | None | None | None |
| F13A1 | None | None | None | None |
| F13A1 | None | None | None | None |
| TF | None | None | None | None |
| TF | None | None | None | None |
| TF | None | None | None | None |
| TF | None | None | None | None |
| TF | None | None | None | None |
| ITPA | None | None | None | None |
| ITPA | None | None | None | None |
| SNCA | None | None | None | None |
| ITGB4 | None | None | None | None |
| ITGB4 | None | None | None | None |
| ITGB4 | None | None | None | None |
| LRP6 | None | None | None | None |
| LRP6 | None | None | None | None |
| LRP6 | None | None | None | None |
| PLG | None | None | None | None |
| PLG | None | None | None | None |
| TYROBP | None | None | None | None |
| UQCRC2 | None | None | None | None |
| GPSM2 | None | None | None | None |

| | | | | |
|---|---|---|---|---|
| GPSM2 | None | None | None | None |
| GPSM2 | None | None | None | None |
| SCN5A | None | None | None | None |
| SCN5A | None | None | None | None |
| SCN5A | None | None | None | None |
| SCN5A | None | None | None | None |
| SCN5A | None | None | None | None |
| ATIC | None | None | None | None |
| EZH2 | None | None | None | None |
| CPA1 | None | None | None | None |
| SCN4B | None | None | None | None |
| SCN4B | None | None | None | None |
| MAN1B1 | None | None | None | None |
| MAN1B1 | None | None | None | None |
| MAN1B1 | None | None | None | None |
| MAN1B1 | None | None | None | None |
| CYCS | 0.74 | 1 | 2n9i | A |
| IDUA | None | None | None | None |
| IDUA | None | None | None | None |
| IDUA | None | None | None | None |
| IDUA | None | None | None | None |
| IDUA | None | None | None | None |
| IDUA | None | None | None | None |
| IDUA | None | None | None | None |
| IDUA | None | None | None | None |
| IDUA | None | None | None | None |
| IDUA | None | None | None | None |
| IDUA | None | None | None | None |
| IDUA | None | None | None | None |
| IDUA | None | None | None | None |
| IDUA | None | None | None | None |
| IDUA | None | None | None | None |
| IDUA | None | None | None | None |
| IDUA | None | None | None | None |
| IDUA | None | None | None | None |
| IDUA | None | None | None | None |
| IDUA | None | None | None | None |
| IDUA | None | None | None | None |
| IDUA | None | None | None | None |
| IDUA | None | None | None | None |
| IDUA | None | None | None | None |
| IDUA | None | None | None | None |
| HIKESHI | None | None | None | None |
| APC | None | None | None | None |
| APC | None | None | None | None |
| APC | None | None | None | None |
| APP | None | None | None | None |
| CEP152 | None | None | None | None |
| GAA | None | None | None | None |
| GAA | None | None | None | None |
| GAA | None | None | None | None |
| GAA | None | None | None | None |
| GAA | None | None | None | None |
| GAA | None | None | None | None |

| | | | | |
|---|---|---|---|---|
| GAA | None | None | None | None |
| GAA | None | None | None | None |
| GAA | None | None | None | None |
| GAA | None | None | None | None |
| GAA | None | None | None | None |
| GAA | None | None | None | None |
| GAA | None | None | None | None |
| GAA | None | None | None | None |
| GAA | None | None | None | None |
| GAA | None | None | None | None |
| GAA | None | None | None | None |
| GAA | None | None | None | None |
| GAA | None | None | None | None |
| GAA | None | None | None | None |
| GAA | None | None | None | None |
| GAA | None | None | None | None |
| GAA | None | None | None | None |
| GAA | None | None | None | None |
| GAA | None | None | None | None |
| GAA | None | None | None | None |
| GAA | None | None | None | None |
| GAA | None | None | None | None |
| GAA | None | None | None | None |
| GAA | None | None | None | None |
| GAA | None | None | None | None |
| GAA | None | None | None | None |
| GAA | None | None | None | None |
| GAA | None | None | None | None |
| GAA | None | None | None | None |
| GAA | None | None | None | None |
| GAA | None | None | None | None |
| GAA | None | None | None | None |
| GAA | None | None | None | None |
| GAA | None | None | None | None |
| GAA | None | None | None | None |
| GAA | None | None | None | None |
| GAA | None | None | None | None |
| GAA | None | None | None | None |
| GAA | None | None | None | None |
| GAA | None | None | None | None |
| GAA | None | None | None | None |
| GAA | None | None | None | None |
| GAA | None | None | None | None |
| FOLR1 | None | None | None | None |
| DSP | None | None | None | None |
| DSP | None | None | None | None |
| DSP | None | None | None | None |
| DSP | None | None | None | None |
| DSP | None | None | None | None |
| CCM2 | None | None | None | None |
| FXN | None | None | None | None |
| FXN | None | None | None | None |
| COMP | None | None | None | None |
| COMP | None | None | None | None |

| | | | | |
|---|---|---|---|---|
| COMP | None | None | None | None |
| COMP | None | None | None | None |
| COMP | None | None | None | None |
| COMP | None | None | None | None |
| COMP | None | None | None | None |
| COMP | None | None | None | None |
| COMP | None | None | None | None |
| COMP | None | None | None | None |
| COMP | None | None | None | None |
| COMP | None | None | None | None |
| AIFM1 | None | None | None | None |
| AIFM1 | None | None | None | None |
| AIFM1 | None | None | None | None |
| AIFM1 | None | None | None | None |
| AIFM1 | None | None | None | None |
| AIFM1 | None | None | None | None |
| AIFM1 | None | None | None | None |
| MYH9 | None | None | None | None |
| FHL2 | None | None | None | None |
| FHL2 | None | None | None | None |
| TNFRSF4 | None | None | None | None |
| CFB | None | None | None | None |
| CFB | None | None | None | None |
| CFB | None | None | None | None |
| CFB | None | None | None | None |
| CFB | None | None | None | None |
| CFB | None | None | None | None |
| CFB | None | None | None | None |
| CFB | None | None | None | None |
| CFB | None | None | None | None |
| CFB | None | None | None | None |
| MVD | None | None | None | None |
| MVD | None | None | None | None |
| HSD17B4 | None | None | None | None |
| HSD17B4 | None | None | None | None |
| HSD17B4 | None | None | None | None |
| HSD17B4 | None | None | None | None |
| HSD17B4 | None | None | None | None |
| HSD17B4 | None | None | None | None |
| F11 | None | None | None | None |
| F11 | None | None | None | None |
| F11 | None | None | None | None |
| F11 | None | None | None | None |
| F11 | None | None | None | None |
| F11 | None | None | None | None |
| F11 | None | None | None | None |
| F11 | None | None | None | None |
| F11 | None | None | None | None |
| SPTA1 | None | None | None | None |
| SPTA1 | None | None | None | None |
| SPTA1 | None | None | None | None |
| SPTA1 | None | None | None | None |
| SPTA1 | None | None | None | None |
| SPTA1 | None | None | None | None |

| | | | | |
|---|---|---|---|---|
| GMPR | None | None | None | None |
| MITF | None | None | None | None |
| NPC1 | None | None | None | None |
| NPC1 | None | None | None | None |
| NPC1 | None | None | None | None |
| NPC1 | None | None | None | None |
| NPC1 | None | None | None | None |
| NPC1 | None | None | None | None |
| NPC1 | None | None | None | None |
| NPC1 | None | None | None | None |
| NPC1 | None | None | None | None |
| NPC1 | None | None | None | None |
| NPC1 | None | None | None | None |
| NPC1 | None | None | None | None |
| NPC1 | None | None | None | None |
| NPC1 | None | None | None | None |
| NPC1 | None | None | None | None |
| NPC1 | None | None | None | None |
| NPC1 | None | None | None | None |
| NPC1 | None | None | None | None |
| NPC1 | None | None | None | None |
| NPC1 | None | None | None | None |
| NPC1 | None | None | None | None |
| NPC1 | None | None | None | None |
| NPC1 | None | None | None | None |
| NPC1 | None | None | None | None |
| NPC1 | None | None | None | None |
| NPC1 | None | None | None | None |
| NPC1 | None | None | None | None |
| NPC1 | None | None | None | None |
| NPC1 | None | None | None | None |
| NPC1 | None | None | None | None |
| NPC1 | None | None | None | None |
| NPC1 | None | None | None | None |
| NPC1 | None | None | None | None |
| NPC1 | None | None | None | None |
| NPC1 | None | None | None | None |
| NPC1 | None | None | None | None |
| NPC1 | None | None | None | None |
| NPC1 | None | None | None | None |
| NPC1 | None | None | None | None |
| NPC1 | None | None | None | None |
| NPC1 | None | None | None | None |
| NPC1 | None | None | None | None |
| NPC1 | None | None | None | None |
| C3 | None | None | None | None |
| NLRX1 | None | None | None | None |
| IGLL1 | None | None | None | None |
| APRT | 0.769 | 4 | 1zn9 | B |
| APRT | None | None | None | None |
| APRT | None | None | None | None |
| APRT | 0.923 | 1 | 1zn8 | A |
| APRT | None | None | None | None |
| ZIC3 | None | None | None | None |

| | | | | |
|---|---|---|---|---|
| ZIC3 | None | None | None | None |
| KLHL3 | None | None | None | None |
| RPS6KA3 | None | None | None | None |
| RPS6KA3 | None | None | None | None |
| RPS6KA3 | None | None | None | None |
| F8 | None | None | None | None |
| F8 | None | None | None | None |
| F8 | None | None | None | None |
| F8 | None | None | None | None |
| F8 | None | None | None | None |
| F8 | None | None | None | None |
| F8 | None | None | None | None |
| F8 | None | None | None | None |
| F8 | None | None | None | None |
| F8 | None | None | None | None |
| F8 | None | None | None | None |
| F8 | None | None | None | None |
| F8 | None | None | None | None |
| F8 | None | None | None | None |
| F8 | None | None | None | None |
| F8 | None | None | None | None |
| F8 | None | None | None | None |
| ISG15 | None | None | None | None |
| EXOSC3 | None | None | None | None |
| EXOSC3 | None | None | None | None |
| EXOSC3 | None | None | None | None |
| NKX2-5 | None | None | None | None |
| NKX2-5 | None | None | None | None |
| NKX2-5 | None | None | None | None |
| NKX2-5 | None | None | None | None |
| FH | None | None | None | None |
| FH | None | None | None | None |
| FH | None | None | None | None |
| FH | None | None | None | None |
| FH | None | None | None | None |
| FH | None | None | None | None |
| FH | None | None | None | None |
| FH | None | None | None | None |
| FH | None | None | None | None |
| FH | None | None | None | None |
| FH | None | None | None | None |
| FH | None | None | None | None |
| SNRNP200 | None | None | None | None |
| SNRNP200 | None | None | None | None |
| SNRNP200 | None | None | None | None |
| SNRNP200 | None | None | None | None |
| CHRNA1 | None | None | None | None |
| CHRNA1 | None | None | None | None |
| MYOM1 | None | None | None | None |
| MYOM1 | None | None | None | None |
| MYOM1 | None | None | None | None |
| MYOM1 | None | None | None | None |
| MYOM1 | None | None | None | None |
| MYOM1 | None | None | None | None |

| | | | | |
|---|---|---|---|---|
| LDLR | None | None | None | None |
| LDLR | None | None | None | None |
| LDLR | None | None | None | None |
| LDLR | None | None | None | None |
| LDLR | None | None | None | None |
| LDLR | None | None | None | None |
| EHMT1 | None | None | None | None |
| EHMT1 | None | None | None | None |
| EHMT1 | None | None | None | None |
| COL9A2 | None | None | None | None |
| HFE2 | None | None | None | None |
| HFE2 | None | None | None | None |
| CD3G | None | None | None | None |
| MMAB | None | None | None | None |
| MMAB | None | None | None | None |
| MMAB | None | None | None | None |
| MMAB | None | None | None | None |
| MMAB | None | None | None | None |
| MMAB | 0.7 | 2 | 2idx | B |
| MMAB | 0.7 | 2 | 2idx | B |
| MMAB | None | None | None | None |
| MMAB | None | None | None | None |
| MMAB | None | None | None | None |
| TAF1 | None | None | None | None |
| TAF1 | None | None | None | None |
| CD4 | None | None | None | None |
| QARS | None | None | None | None |
| QARS | None | None | None | None |
| QARS | None | None | None | None |
| QARS | None | None | None | None |
| QARS | None | None | None | None |
| FGFR3 | None | None | None | None |
| FGFR3 | None | None | None | None |
| FGFR3 | None | None | None | None |
| FGFR3 | None | None | None | None |
| FGFR3 | None | None | None | None |
| FGFR3 | None | None | None | None |
| FLNB | None | None | None | None |
| FLNB | None | None | None | None |
| FLNB | None | None | None | None |
| FLNB | None | None | None | None |
| FLNB | None | None | None | None |
| FLNB | None | None | None | None |
| FLNB | None | None | None | None |
| FLNB | None | None | None | None |
| IL21R | None | None | None | None |
| RB1 | None | None | None | None |
| RB1 | None | None | None | None |
| RB1 | None | None | None | None |
| NTRK1 | None | None | None | None |
| NOTCH1 | None | None | None | None |
| MYOM1 | None | None | None | None |
| MYOM1 | None | None | None | None |
| MYOM1 | None | None | None | None |

| | | | | |
|---|---|---|---|---|
| MYOM1 | None | None | None | None |
| MYOM1 | None | None | None | None |
| MYOM1 | None | None | None | None |
| COQ9 | None | None | None | None |
| COQ9 | None | None | None | None |
| CRBN | None | None | None | None |
| DCTN1 | None | None | None | None |
| DCTN1 | None | None | None | None |
| DCTN1 | None | None | None | None |
| DCTN1 | None | None | None | None |
| DCTN1 | None | None | None | None |
| DCTN1 | None | None | None | None |
| THAP1 | None | None | None | None |
| THAP1 | None | None | None | None |
| GNAS | None | None | None | None |
| GNAS | None | None | None | None |
| HNF4A | None | None | None | None |
| HNF4A | None | None | None | None |
| ALDH3A2 | None | None | None | None |
| ALDH3A2 | None | None | None | None |
| ALDH3A2 | None | None | None | None |
| ALDH3A2 | None | None | None | None |
| ALDH3A2 | None | None | None | None |
| ALDH3A2 | None | None | None | None |
| ALDH3A2 | None | None | None | None |
| ALDH3A2 | None | None | None | None |
| ALDH3A2 | None | None | None | None |
| HCN1 | None | None | None | None |
| PLEC | None | None | None | None |
| SCN5A | None | None | None | None |
| SCN5A | None | None | None | None |
| STAMBP | None | None | None | None |
| STAMBP | None | None | None | None |
| RNF125 | None | None | None | None |
| RNF125 | None | None | None | None |
| USH1C | None | None | None | None |
| OBSL1 | None | None | None | None |
| GRIN2B | None | None | None | None |
| GRIN2B | None | None | None | None |
| HCN4 | None | None | None | None |
| HCN4 | None | None | None | None |
| TALDO1 | None | None | None | None |
| ITGB3 | None | None | None | None |
| CHKB | None | None | None | None |
| CHKB | None | None | None | None |
| CHKB | None | None | None | None |
| CHKB | None | None | None | None |
| ACTN4 | None | None | None | None |
| ACTN4 | None | None | None | None |
| ACTN4 | None | None | None | None |
| ACTN4 | None | None | None | None |
| CDT1 | None | None | None | None |
| CDT1 | None | None | None | None |
| CDT1 | None | None | None | None |

| | | | | |
|---|---|---|---|---|
| CDT1 | None | None | None | None |
| CDT1 | None | None | None | None |
| CDT1 | None | None | None | None |
| MAPT | None | None | None | None |
| F5 | None | None | None | None |
| F5 | None | None | None | None |
| F5 | None | None | None | None |
| ESRRB | None | None | None | None |
| DGUOK | None | None | None | None |
| DGUOK | None | None | None | None |
| DGUOK | None | None | None | None |
| DGUOK | None | None | None | None |
| DGUOK | 0.765 | 1 | 2ocp | A |
| DGUOK | None | None | None | None |
| DGUOK | None | None | None | None |
| DGUOK | None | None | None | None |
| DGUOK | None | None | None | None |
| CENPJ | None | None | None | None |
| TTN | None | None | None | None |
| TTN | None | None | None | None |
| TTN | None | None | None | None |
| TTN | None | None | None | None |
| TNFRSF1A | None | None | None | None |
| TNFRSF1A | None | None | None | None |
| TNFRSF1A | None | None | None | None |
| TNFRSF1A | None | None | None | None |
| TNFRSF1A | None | None | None | None |
| TNFRSF1A | None | None | None | None |
| TNFRSF1A | None | None | None | None |
| TNFRSF1A | None | None | None | None |
| TNFRSF1A | None | None | None | None |
| TNFRSF1A | None | None | None | None |
| TNFRSF1A | None | None | None | None |
| TNFRSF1A | None | None | None | None |
| ASL | None | None | None | None |
| ASL | None | None | None | None |
| ASL | None | None | None | None |
| ASL | None | None | None | None |
| ASL | None | None | None | None |
| ASL | None | None | None | None |
| ASL | None | None | None | None |
| ASL | None | None | None | None |
| ASL | None | None | None | None |
| ASL | None | None | None | None |
| MYO7A | None | None | None | None |
| MYO7A | None | None | None | None |
| MYO7A | None | None | None | None |
| MYO7A | None | None | None | None |
| MYO7A | None | None | None | None |
| MYO7A | None | None | None | None |
| MYO7A | None | None | None | None |
| MYO7A | None | None | None | None |
| MYO7A | None | None | None | None |
| MYO7A | None | None | None | None |

| | | | | |
|---|---|---|---|---|
| MYO7A | None | None | None | None |
| MYO7A | None | None | None | None |
| MYO7A | None | None | None | None |
| MYO7A | None | None | None | None |
| MYO7A | None | None | None | None |
| MYO7A | None | None | None | None |
| MYO7A | None | None | None | None |
| MYO7A | None | None | None | None |
| MYO7A | None | None | None | None |
| MYO7A | None | None | None | None |
| APC | None | None | None | None |
| ALAD | None | None | None | None |
| ALAD | None | None | None | None |
| ALAD | None | None | None | None |
| ALAD | None | None | None | None |
| ALAD | None | None | None | None |
| ALAD | None | None | None | None |
| POR | None | None | None | None |
| CSF1R | None | None | None | None |
| CSF1R | None | None | None | None |
| CSF1R | None | None | None | None |
| CSF1R | None | None | None | None |
| CSF1R | None | None | None | None |
| CSF1R | None | None | None | None |
| CSF1R | None | None | None | None |
| CSF1R | None | None | None | None |
| CSF1R | None | None | None | None |
| CSF1R | None | None | None | None |
| CSF1R | None | None | None | None |
| VWF | None | None | None | None |
| VWF | None | None | None | None |
| PHF8 | None | None | None | None |
| ACY1 | None | None | None | None |
| SMN1 | None | None | None | None |
| SMN1 | None | None | None | None |
| DLAT | None | None | None | None |
| DLAT | None | None | None | None |
| B2M | None | None | None | None |
| SOX9 | None | None | None | None |
| SOX9 | None | None | None | None |
| SOX9 | None | None | None | None |
| SOX9 | None | None | None | None |
| SOX9 | None | None | None | None |
| CFH | None | None | None | None |
| FLNC | None | None | None | None |
| FLNC | None | None | None | None |
| FLNC | None | None | None | None |
| ITGB2 | None | None | None | None |
| ITGB2 | None | None | None | None |
| ITGB2 | None | None | None | None |
| ITGB2 | None | None | None | None |
| ITGB2 | None | None | None | None |
| ITGB2 | None | None | None | None |
| ITGB2 | None | None | None | None |

| | | | | |
|---|---|---|---|---|
| ITGB2 | None | None | None | None |
| ITGB2 | None | None | None | None |
| PSEN1 | None | None | None | None |
| PSEN1 | None | None | None | None |
| PSEN1 | None | None | None | None |
| PSEN1 | None | None | None | None |
| PSEN1 | None | None | None | None |
| PSEN1 | None | None | None | None |
| OXCT1 | None | None | None | None |
| OXCT1 | None | None | None | None |
| OXCT1 | None | None | None | None |
| OXCT1 | None | None | None | None |
| OXCT1 | None | None | None | None |
| OXCT1 | None | None | None | None |
| OXCT1 | None | None | None | None |
| DHH | None | None | None | None |
| PRKN | None | None | None | None |
| PRKN | None | None | None | None |
| PRKN | None | None | None | None |
| PRKN | None | None | None | None |
| PRKN | None | None | None | None |
| PRKN | None | None | None | None |
| PRKN | None | None | None | None |
| PRKN | None | None | None | None |
| PRKN | None | None | None | None |
| PRKN | None | None | None | None |
| PRKN | None | None | None | None |
| LCAT | None | None | None | None |
| LCAT | None | None | None | None |
| LCAT | None | None | None | None |
| LCAT | None | None | None | None |
| LCAT | None | None | None | None |
| LCAT | None | None | None | None |
| LCAT | None | None | None | None |
| LCAT | None | None | None | None |
| LCAT | None | None | None | None |
| LCAT | None | None | None | None |
| LCAT | None | None | None | None |
| LCAT | None | None | None | None |
| FOXP3 | None | None | None | None |
| FOXP3 | None | None | None | None |
| FOXP3 | None | None | None | None |
| FOXP3 | None | None | None | None |
| SUFU | None | None | None | None |
| HSPA9 | None | None | None | None |
| HSPA9 | None | None | None | None |
| SQSTM1 | None | None | None | None |
| DARS2 | None | None | None | None |
| DARS2 | None | None | None | None |
| DARS2 | None | None | None | None |
| DARS2 | None | None | None | None |
| DARS2 | None | None | None | None |
| DARS2 | None | None | None | None |
| DARS2 | None | None | None | None |

| | | | | |
|---|---|---|---|---|
| DARS2 | None | None | None | None |
| DARS2 | None | None | None | None |
| DARS2 | None | None | None | None |
| DARS2 | None | None | None | None |
| DARS2 | None | None | None | None |
| NT5C3A | None | None | None | None |
| NT5C3A | None | None | None | None |
| NT5C3A | None | None | None | None |
| NT5C3A | None | None | None | None |
| ETFA | None | None | None | None |
| ETFA | None | None | None | None |
| ETFA | None | None | None | None |
| ETFA | None | None | None | None |
| ETFA | None | None | None | None |
| ETFA | None | None | None | None |
| ETFA | None | None | None | None |
| TTN | None | None | None | None |
| TTN | None | None | None | None |
| FLT4 | None | None | None | None |
| FGB | None | None | None | None |
| HPCA | None | None | None | None |
| TAP2 | None | None | None | None |
| TAP2 | None | None | None | None |
| PTPRC | None | None | None | None |
| SLC4A1 | None | None | None | None |
| SLC4A1 | None | None | None | None |
| SLC4A1 | None | None | None | None |
| SLC4A1 | None | None | None | None |
| SLC4A1 | None | None | None | None |
| SLC4A1 | None | None | None | None |
| MYH7 | None | None | None | None |
| MYH7 | None | None | None | None |
| MYH7 | None | None | None | None |
| MYH7 | None | None | None | None |
| MYH7 | None | None | None | None |
| MYH7 | None | None | None | None |
| MYH7 | None | None | None | None |
| MYH7 | None | None | None | None |
| MYH7 | None | None | None | None |
| PARN | None | None | None | None |
| PARN | None | None | None | None |
| NDUFS6 | None | None | None | None |
| VRK1 | None | None | None | None |
| VRK1 | None | None | None | None |
| VRK1 | None | None | None | None |
| VRK1 | None | None | None | None |
| VRK1 | None | None | None | None |
| VRK1 | None | None | None | None |
| COLEC11 | None | None | None | None |
| VHL | None | None | None | None |
| TTN | None | None | None | None |
| NCF2 | None | None | None | None |
| NCF2 | None | None | None | None |
| NCF2 | None | None | None | None |

| | | | | |
|---|---|---|---|---|
| NCF2 | None | None | None | None |
| NCF2 | None | None | None | None |
| RPS26 | None | None | None | None |
| CETP | None | None | None | None |
| CETP | None | None | None | None |
| CETP | None | None | None | None |
| CETP | None | None | None | None |
| CETP | None | None | None | None |
| CETP | None | None | None | None |
| CETP | None | None | None | None |
| CETP | None | None | None | None |
| CETP | None | None | None | None |
| CETP | None | None | None | None |
| CETP | None | None | None | None |
| CETP | None | None | None | None |
| CETP | None | None | None | None |
| FLNA | None | None | None | None |
| FLNA | None | None | None | None |
| FLNA | None | None | None | None |
| FLNA | None | None | None | None |
| FLNA | None | None | None | None |
| FLNA | None | None | None | None |
| FLNA | None | None | None | None |
| MYBPC3 | None | None | None | None |
| MYBPC3 | None | None | None | None |
| MYBPC3 | None | None | None | None |
| MYBPC3 | None | None | None | None |
| MYBPC3 | None | None | None | None |
| MYBPC3 | None | None | None | None |
| MYBPC3 | None | None | None | None |
| MYBPC3 | None | None | None | None |
| MYBPC3 | None | None | None | None |
| MYBPC3 | None | None | None | None |
| LSS | None | None | None | None |
| C6 | None | None | None | None |
| C6 | None | None | None | None |
| RLBP1 | None | None | None | None |
| RLBP1 | None | None | None | None |
| RLBP1 | None | None | None | None |
| RLBP1 | None | None | None | None |
| RLBP1 | None | None | None | None |
| RLBP1 | None | None | None | None |
| RLBP1 | None | None | None | None |
| RLBP1 | None | None | None | None |
| MMAA | None | None | None | None |
| MMAA | None | None | None | None |
| MMAA | None | None | None | None |
| MMAA | None | None | None | None |
| MMAA | None | None | None | None |
| MMAA | None | None | None | None |
| IL1RN | None | None | None | None |
| MASTL | None | None | None | None |
| FHL1 | None | None | None | None |
| FHL1 | None | None | None | None |

| | | | | |
|---|---|---|---|---|
| FHL1 | None | None | None | None |
| FHL1 | None | None | None | None |
| FHL1 | None | None | None | None |
| FHL1 | None | None | None | None |
| FHL1 | None | None | None | None |
| FHL1 | None | None | None | None |
| FHL1 | None | None | None | None |
| EIF2B1 | None | None | None | None |
| EIF2B1 | None | None | None | None |
| COQ8A | None | None | None | None |
| COQ8A | None | None | None | None |
| COQ8A | None | None | None | None |
| COQ8A | None | None | None | None |
| COQ8A | None | None | None | None |
| COQ8A | None | None | None | None |
| COQ8A | None | None | None | None |
| COQ8A | None | None | None | None |
| COQ8A | None | None | None | None |
| COQ8A | None | None | None | None |
| COQ8A | None | None | None | None |
| COQ8A | None | None | None | None |
| COQ8A | None | None | None | None |
| COQ8A | None | None | None | None |
| COQ8A | None | None | None | None |
| COQ8A | None | None | None | None |
| UNC119 | None | None | None | None |
| IGHMBP2 | None | None | None | None |
| JAG1 | None | None | None | None |
| JAG1 | None | None | None | None |
| JAG1 | None | None | None | None |
| GNB1 | None | None | None | None |
| GNB1 | None | None | None | None |
| FLNA | None | None | None | None |
| FLNA | None | None | None | None |
| FLNA | None | None | None | None |
| FLNA | None | None | None | None |
| FLNA | None | None | None | None |
| CREBBP | None | None | None | None |
| CSF3R | None | None | None | None |
| CSF3R | None | None | None | None |
| HIBCH | None | None | None | None |
| HIBCH | None | None | None | None |
| HIBCH | None | None | None | None |
| MMACHC | None | None | None | None |
| MMACHC | None | None | None | None |
| MMACHC | None | None | None | None |
| MMACHC | None | None | None | None |
| MMACHC | None | None | None | None |
| MMACHC | None | None | None | None |
| MMACHC | None | None | None | None |
| MMACHC | None | None | None | None |
| MMACHC | None | None | None | None |
| MMACHC | None | None | None | None |
| MMACHC | None | None | None | None |

| | | | | |
|---|---|---|---|---|
| KLF1 | None | None | None | None |
| NAGS | None | None | None | None |
| NAGS | None | None | None | None |
| NAGS | None | None | None | None |
| PIK3R2 | None | None | None | None |
| EDNRB | None | None | None | None |
| EDNRB | None | None | None | None |
| EDNRB | None | None | None | None |
| EDNRB | None | None | None | None |
| E | None | None | None | None |
| RUNX1 | None | None | None | None |
| RUNX1 | None | None | None | None |
| CTNNB1 | None | None | None | None |
| CTNNB1 | None | None | None | None |
| CTNNB1 | None | None | None | None |
| CTNNB1 | None | None | None | None |
| CTNNB1 | None | None | None | None |
| CTNNB1 | None | None | None | None |
| TTN | None | None | None | None |
| TTN | None | None | None | None |
| TYK2 | None | None | None | None |
| TYK2 | None | None | None | None |
| TYK2 | None | None | None | None |
| ERCC4 | None | None | None | None |
| ERCC4 | None | None | None | None |
| ERCC4 | None | None | None | None |
| ERCC4 | None | None | None | None |
| CFI | None | None | None | None |
| CFI | None | None | None | None |
| CFI | None | None | None | None |
| CFI | None | None | None | None |
| CFI | None | None | None | None |
| CFI | None | None | None | None |
| CFI | None | None | None | None |
| CFI | None | None | None | None |
| CFI | None | None | None | None |
| CFI | None | None | None | None |
| CFI | None | None | None | None |
| CFI | None | None | None | None |
| CFI | None | None | None | None |
| CFI | None | None | None | None |
| NOTCH3 | None | None | None | None |
| NOTCH3 | None | None | None | None |
| NOTCH3 | None | None | None | None |
| NOTCH3 | None | None | None | None |
| TBX1 | None | None | None | None |
| TBX1 | None | None | None | None |
| DDB2 | None | None | None | None |
| DDB2 | None | None | None | None |
| TP53 | None | None | None | None |
| POMGNT1 | None | None | None | None |
| POMGNT1 | None | None | None | None |
| POMGNT1 | None | None | None | None |
| POMGNT1 | None | None | None | None |

| | | | | |
|---|---|---|---|---|
| POMGNT1 | None | None | None | None |
| POMGNT1 | None | None | None | None |
| POMGNT1 | None | None | None | None |
| POMGNT1 | None | None | None | None |
| POMGNT1 | None | None | None | None |
| POMGNT1 | None | None | None | None |
| POMGNT1 | None | None | None | None |
| POMGNT1 | None | None | None | None |
| POMGNT1 | None | None | None | None |
| FMR1 | None | None | None | None |
| FMR1 | None | None | None | None |
| SLC9A3R1 | None | None | None | None |
| HSPB1 | None | None | None | None |
| HSPB1 | None | None | None | None |
| HSPB1 | None | None | None | None |
| HSPB1 | None | None | None | None |
| HSPB1 | None | None | None | None |
| HSPB1 | None | None | None | None |
| HSPB1 | None | None | None | None |
| SOD3 | None | None | None | None |
| AGA | 1 | 2 | 1apz | D |
| AGA | None | None | None | None |
| AGA | None | None | None | None |
| NDUFS7 | None | None | None | None |
| NDUFS7 | None | None | None | None |
| SOX2 | None | None | None | None |
| CSRP3 | None | None | None | None |
| CSRP3 | None | None | None | None |
| PAH | None | None | None | None |
| PAH | None | None | None | None |
| PAH | None | None | None | None |
| PAH | None | None | None | None |
| CACNA1S | None | None | None | None |
| TRPV4 | None | None | None | None |
| TRPV4 | None | None | None | None |
| TRPV4 | None | None | None | None |
| TRPV4 | None | None | None | None |
| TRPV4 | None | None | None | None |
| TRPV4 | None | None | None | None |
| TRPV4 | None | None | None | None |
| TRPV4 | None | None | None | None |
| TRPV4 | None | None | None | None |
| TRPV4 | None | None | None | None |
| TRPV4 | None | None | None | None |
| TRPV4 | None | None | None | None |
| TRPV4 | None | None | None | None |
| TNNI3 | None | None | None | None |
| MLYCD | None | None | None | None |
| MLYCD | None | None | None | None |
| MLYCD | None | None | None | None |
| MLYCD | None | None | None | None |
| AIP | None | None | None | None |
| AIP | None | None | None | None |
| ERCC2 | None | None | None | None |

| | | | | |
|---|---|---|---|---|
| ERCC2 | None | None | None | None |
| ERCC2 | None | None | None | None |
| ERCC2 | None | None | None | None |
| ERCC2 | None | None | None | None |
| ERCC2 | None | None | None | None |
| ERCC2 | None | None | None | None |
| ERCC2 | None | None | None | None |
| ERCC2 | None | None | None | None |
| ERCC2 | None | None | None | None |
| ERCC2 | None | None | None | None |
| ERCC2 | None | None | None | None |
| ERCC2 | None | None | None | None |
| ERCC2 | None | None | None | None |
| PUS1 | None | None | None | None |
| PUS1 | None | None | None | None |
| PUS1 | None | None | None | None |
| CASK | None | None | None | None |
| CASK | None | None | None | None |
| BLVRA | None | None | None | None |
| DICER1 | None | None | None | None |
| DICER1 | None | None | None | None |
| DICER1 | None | None | None | None |
| EIF2AK3 | None | None | None | None |
| EIF2AK3 | None | None | None | None |
| EIF2AK3 | None | None | None | None |
| KIT | None | None | None | None |
| KIT | None | None | None | None |
| KIT | None | None | None | None |
| KIT | None | None | None | None |
| KIT | None | None | None | None |
| KIT | None | None | None | None |
| DNMT3B | None | None | None | None |
| TGM1 | None | None | None | None |
| NDUFA10 | None | None | None | None |
| NDUFA10 | None | None | None | None |
| PRPF31 | None | None | None | None |
| PRPF31 | None | None | None | None |
| ECE1 | None | None | None | None |
| ECE1 | None | None | None | None |
| ECE1 | None | None | None | None |
| NDUFV2 | None | None | None | None |
| NDUFV2 | None | None | None | None |
| NDUFV2 | None | None | None | None |
| GHRHR | None | None | None | None |
| GHRHR | None | None | None | None |
| GHRHR | None | None | None | None |
| SLC4A1 | None | None | None | None |
| FGF9 | None | None | None | None |
| BARD1 | None | None | None | None |
| BARD1 | None | None | None | None |
| BARD1 | None | None | None | None |
| PTPN12 | 0.741 | 2 | 5j8r | D |
| MASP2 | None | None | None | None |
| MASP2 | None | None | None | None |

| | | | | |
|---|---|---|---|---|
| MASP2 | None | None | None | None |
| MASP2 | None | None | None | None |
| HOXB1 | None | None | None | None |
| SLC4A1 | None | None | None | None |
| SLC4A1 | None | None | None | None |
| SLC4A1 | None | None | None | None |
| SLC4A1 | None | None | None | None |
| SLC4A1 | None | None | None | None |
| SLC4A1 | None | None | None | None |
| SLC4A1 | None | None | None | None |
| SLC4A1 | None | None | None | None |
| SLC4A1 | None | None | None | None |
| SLC4A1 | None | None | None | None |
| SLC4A1 | None | None | None | None |
| SLC4A1 | None | None | None | None |
| SLC4A1 | None | None | None | None |
| SLC4A1 | None | None | None | None |
| SLC4A1 | None | None | None | None |
| SLC4A1 | None | None | None | None |
| SLC4A1 | None | None | None | None |
| CBL | None | None | None | None |
| FAS | None | None | None | None |
| MTMR2 | None | None | None | None |
| MTMR2 | None | None | None | None |
| MTMR2 | None | None | None | None |
| MTMR2 | None | None | None | None |
| MTMR2 | None | None | None | None |
| MTMR2 | None | None | None | None |
| MTMR2 | None | None | None | None |
| MTMR2 | None | None | None | None |
| KLHL3 | None | None | None | None |
| KLHL3 | None | None | None | None |
| KLHL3 | None | None | None | None |
| KLHL3 | 0.733 | 1 | 5nkp | A |
| KLHL3 | None | None | None | None |
| KLHL3 | None | None | None | None |
| RNASEH1 | None | None | None | None |
| ACTN1 | None | None | None | None |
| PHKG2 | None | None | None | None |
| PHKG2 | None | None | None | None |
| PHKG2 | None | None | None | None |
| NTRK1 | None | None | None | None |
| APOH | None | None | None | None |
| PMS2 | None | None | None | None |
| PMS2 | None | None | None | None |
| PMS2 | 0.797 | 1 | 1ea6 | A |
| PMS2 | None | None | None | None |
| PMS2 | None | None | None | None |
| PMS2 | None | None | None | None |
| PMS2 | None | None | None | None |
| PMS2 | None | None | None | None |
| PMS2 | None | None | None | None |
| PMS2 | None | None | None | None |
| PMS2 | None | None | None | None |

| | | | | |
|---|---|---|---|---|
| PMS2 | None | None | None | None |
| PMS2 | None | None | None | None |
| PMS2 | None | None | None | None |
| PMS2 | None | None | None | None |
| PMS2 | None | None | None | None |
| PMS2 | None | None | None | None |
| PMS2 | None | None | None | None |
| PMS2 | None | None | None | None |
| PMS2 | None | None | None | None |
| PMS2 | None | None | None | None |
| PMS2 | None | None | None | None |
| MOCS2 | None | None | None | None |
| MOCS2 | None | None | None | None |
| RPGRIP1 | None | None | None | None |
| RPGRIP1 | None | None | None | None |
| LIG4 | None | None | None | None |
| PROC | None | None | None | None |
| PROC | None | None | None | None |
| NAT1 | None | None | None | None |
| NAT1 | None | None | None | None |
| PNPO | None | None | None | None |
| PNPO | None | None | None | None |
| PNPO | None | None | None | None |
| PNPO | None | None | None | None |
| PNPO | None | None | None | None |
| MRE11 | None | None | None | None |
| MRE11 | None | None | None | None |
| MRE11 | None | None | None | None |
| MRE11 | None | None | None | None |
| DDC | None | None | None | None |
| DDC | None | None | None | None |
| DDC | None | None | None | None |
| DDC | None | None | None | None |
| DDC | None | None | None | None |
| DDC | 0.909 | 7 | 3rch | B |
| DDC | 0.909 | 7 | 3rch | B |
| DDC | None | None | None | None |
| DDC | None | None | None | None |
| DDC | None | None | None | None |
| DDC | None | None | None | None |
| ANK2 | None | None | None | None |
| ANK2 | None | None | None | None |
| ANK2 | None | None | None | None |
| AK2 | None | None | None | None |
| AK2 | None | None | None | None |
| AK2 | None | None | None | None |
| AK2 | None | None | None | None |
| AK2 | None | None | None | None |
| AK2 | None | None | None | None |
| GPHN | None | None | None | None |
| HSD17B4 | None | None | None | None |
| HSD17B4 | None | None | None | None |
| HSD17B4 | None | None | None | None |
| HSD17B4 | None | None | None | None |

| | | | | |
|---|---|---|---|---|
| HSD17B4 | None | None | None | None |
| HSD17B4 | None | None | None | None |
| APTX | None | None | None | None |
| APTX | None | None | None | None |
| APTX | None | None | None | None |
| APTX | None | None | None | None |
| APTX | None | None | None | None |
| APTX | None | None | None | None |
| APTX | None | None | None | None |
| FANCL | None | None | None | None |
| FANCL | None | None | None | None |
| EMG1 | None | None | None | None |
| EMG1 | None | None | None | None |
| EZH2 | None | None | None | None |
| ABCB6 | None | None | None | None |
| NCF2 | None | None | None | None |
| NCF2 | None | None | None | None |
| COL4A2 | None | None | None | None |
| COL4A2 | 1 | 1 | 1li1 | F |
| COL4A2 | None | None | None | None |
| COL4A2 | None | None | None | None |
| COL4A2 | 0.75 | 16 | 1li1 | C |
| SLX4 | None | None | None | None |
| SERPIND1 | None | None | None | None |
| FGFR3 | None | None | None | None |
| FGFR3 | None | None | None | None |
| FGFR3 | None | None | None | None |
| FGFR3 | None | None | None | None |
| FGFR3 | None | None | None | None |
| FGFR3 | None | None | None | None |
| TREM2 | None | None | None | None |
| TREM2 | None | None | None | None |
| TREM2 | None | None | None | None |
| TREM2 | None | None | None | None |
| TREM2 | None | None | None | None |
| TREM2 | None | None | None | None |
| TREM2 | None | None | None | None |
| LIG4 | None | None | None | None |
| LIG4 | None | None | None | None |
| LIG4 | None | None | None | None |
| LIG4 | None | None | None | None |
| LIG4 | None | None | None | None |
| LIG4 | None | None | None | None |
| ECHS1 | None | None | None | None |
| ECHS1 | None | None | None | None |
| ECHS1 | None | None | None | None |
| MTR | None | None | None | None |
| MTR | None | None | None | None |
| MTR | None | None | None | None |
| MTR | None | None | None | None |
| MTR | None | None | None | None |
| PMVK | None | None | None | None |
| PRNP | None | None | None | None |
| CHN1 | None | None | None | None |

| | | | | |
|---|---|---|---|---|
| CHN1 | None | None | None | None |
| CHN1 | None | None | None | None |
| CHN1 | None | None | None | None |
| CHN1 | None | None | None | None |
| CHN1 | None | None | None | None |
| CHN1 | None | None | None | None |
| CHN1 | None | None | None | None |
| SLC2A1 | None | None | None | None |
| SLC2A1 | None | None | None | None |
| SLC2A1 | None | None | None | None |
| SLC2A1 | None | None | None | None |
| SLC2A1 | None | None | None | None |
| SLC2A1 | None | None | None | None |
| SLC2A1 | None | None | None | None |
| SLC2A1 | None | None | None | None |
| SLC2A1 | None | None | None | None |
| SLC2A1 | None | None | None | None |
| SLC2A1 | None | None | None | None |
| SLC2A1 | None | None | None | None |
| SLC2A1 | None | None | None | None |
| SLC2A1 | None | None | None | None |
| SLC2A1 | None | None | None | None |
| SLC2A1 | None | None | None | None |
| SLC2A1 | None | None | None | None |
| PMM2 | None | None | None | None |
| PMM2 | None | None | None | None |
| PMM2 | None | None | None | None |
| PMM2 | None | None | None | None |
| PMM2 | None | None | None | None |
| PMM2 | None | None | None | None |
| PMM2 | None | None | None | None |
| PMM2 | None | None | None | None |
| PMM2 | None | None | None | None |
| PMM2 | None | None | None | None |
| PMM2 | None | None | None | None |
| PMM2 | None | None | None | None |
| PMM2 | None | None | None | None |
| PMM2 | None | None | None | None |
| PMM2 | None | None | None | None |
| PMM2 | None | None | None | None |
| PMM2 | None | None | None | None |
| PMM2 | None | None | None | None |
| PMM2 | None | None | None | None |
| PMM2 | None | None | None | None |
| PMM2 | None | None | None | None |
| PMM2 | None | None | None | None |
| PMM2 | None | None | None | None |
| PMM2 | None | None | None | None |
| PMM2 | None | None | None | None |
| PMM2 | None | None | None | None |
| PMM2 | None | None | None | None |
| PMM2 | None | None | None | None |
| CCKAR | None | None | None | None |
| MAN1B1 | None | None | None | None |

| | | | | |
|---|---|---|---|---|
| MAN1B1 | None | None | None | None |
| LAMA2 | None | None | None | None |
| LAMA2 | None | None | None | None |
| TBC1D4 | None | None | None | None |
| TBC1D4 | None | None | None | None |
| LDLR | None | None | None | None |
| LDLR | None | None | None | None |
| LDLR | None | None | None | None |
| LDLR | None | None | None | None |
| LDLR | None | None | None | None |
| LDLR | None | None | None | None |
| LDLR | None | None | None | None |
| LDLR | None | None | None | None |
| LDLR | None | None | None | None |
| LDLR | None | None | None | None |
| LDLR | None | None | None | None |
| LDLR | None | None | None | None |
| LDLR | None | None | None | None |
| LDLR | None | None | None | None |
| LDLR | None | None | None | None |
| LDLR | None | None | None | None |
| LDLR | None | None | None | None |
| LDLR | None | None | None | None |
| LDLR | None | None | None | None |
| LDLR | None | None | None | None |
| LDLR | None | None | None | None |
| LDLR | None | None | None | None |
| LDLR | None | None | None | None |
| LDLR | None | None | None | None |
| LDLR | None | None | None | None |
| LDLR | None | None | None | None |
| LDLR | None | None | None | None |
| LDLR | None | None | None | None |
| LDLR | None | None | None | None |
| LDLR | None | None | None | None |
| LDLR | None | None | None | None |
| LDLR | None | None | None | None |
| LDLR | None | None | None | None |
| LDLR | None | None | None | None |
| LDLR | None | None | None | None |
| LDLR | None | None | None | None |
| LDLR | None | None | None | None |
| LDLR | None | None | None | None |
| LDLR | None | None | None | None |
| LDLR | None | None | None | None |
| LDLR | None | None | None | None |
| LDLR | None | None | None | None |
| LDLR | None | None | None | None |
| FANCM | None | None | None | None |
| HAMP | None | None | None | None |
| GUSB | None | None | None | None |
| GUSB | None | None | None | None |
| GUSB | None | None | None | None |
| GUSB | None | None | None | None |

| | | | | |
|---|---|---|---|---|
| GUSB | None | None | None | None |
| GUSB | None | None | None | None |
| GUSB | None | None | None | None |
| GUSB | None | None | None | None |
| GUSB | None | None | None | None |
| GUSB | None | None | None | None |
| GUSB | None | None | None | None |
| GUSB | None | None | None | None |
| GUSB | None | None | None | None |
| LIG1 | None | None | None | None |
| LIG1 | None | None | None | None |
| MYH7 | None | None | None | None |
| MYH7 | None | None | None | None |
| MYH7 | None | None | None | None |
| MYH7 | None | None | None | None |
| MYH7 | None | None | None | None |
| MYH7 | None | None | None | None |
| MYH7 | None | None | None | None |
| MYH7 | None | None | None | None |
| MYH7 | None | None | None | None |
| MYH7 | None | None | None | None |
| MYH7 | None | None | None | None |
| MYH7 | None | None | None | None |
| MYH7 | None | None | None | None |
| MYH7 | None | None | None | None |
| MYH7 | None | None | None | None |
| MYH7 | None | None | None | None |
| MYH7 | None | None | None | None |
| MYH7 | None | None | None | None |
| MYH7 | None | None | None | None |
| MYH7 | None | None | None | None |
| MYH7 | None | None | None | None |
| MYH7 | None | None | None | None |
| MYH7 | None | None | None | None |
| MYH7 | None | None | None | None |
| MYH7 | None | None | None | None |
| MYH7 | None | None | None | None |
| MYH7 | None | None | None | None |
| MYH7 | None | None | None | None |
| MYH7 | None | None | None | None |
| MYH7 | None | None | None | None |
| MYH7 | None | None | None | None |
| MYH7 | None | None | None | None |
| MYH7 | None | None | None | None |
| MYH7 | None | None | None | None |
| HCN4 | None | None | None | None |
| RECQL4 | None | None | None | None |
| RECQL4 | None | None | None | None |
| RECQL4 | None | None | None | None |
| RECQL4 | None | None | None | None |
| RECQL4 | None | None | None | None |
| RECQL4 | None | None | None | None |
| RECQL4 | None | None | None | None |
| RECQL4 | None | None | None | None |

| | | | | |
|---|---|---|---|---|
| RECQL4 | None | None | None | None |
| RECQL4 | None | None | None | None |
| RECQL4 | None | None | None | None |
| RECQL4 | None | None | None | None |
| RECQL4 | None | None | None | None |
| CHEK2 | None | None | None | None |
| PPARG | None | None | None | None |
| ACADVL | None | None | None | None |
| ACADVL | None | None | None | None |
| ACADVL | None | None | None | None |
| ACADVL | None | None | None | None |
| ACADVL | None | None | None | None |
| ACADVL | None | None | None | None |
| ACADVL | None | None | None | None |
| ACADVL | None | None | None | None |
| ACADVL | None | None | None | None |
| ACADVL | None | None | None | None |
| ACADVL | None | None | None | None |
| ACADVL | None | None | None | None |
| ACADVL | None | None | None | None |
| ACADVL | None | None | None | None |
| ACADVL | None | None | None | None |
| ACADVL | None | None | None | None |
| ACADVL | None | None | None | None |
| ACADVL | None | None | None | None |
| ACADVL | None | None | None | None |
| ACADVL | None | None | None | None |
| ACADVL | None | None | None | None |
| APOA1 | None | None | None | None |
| APOA1 | None | None | None | None |
| APOA1 | None | None | None | None |
| APOA1 | None | None | None | None |
| APOA1 | None | None | None | None |
| DNMT3A | None | None | None | None |
| DNMT3A | None | None | None | None |
| APC | None | None | None | None |
| TMOD1 | None | None | None | None |
| USH1C | None | None | None | None |
| USH1C | None | None | None | None |
| USH1C | None | None | None | None |
| USH1C | None | None | None | None |
| ERCC6 | None | None | None | None |
| ERCC6 | None | None | None | None |
| ERCC6 | None | None | None | None |
| ERCC8 | None | None | None | None |
| ERCC8 | None | None | None | None |
| ERCC8 | None | None | None | None |
| DPAGT1 | None | None | None | None |
| DPAGT1 | None | None | None | None |
| DPAGT1 | None | None | None | None |
| DPAGT1 | None | None | None | None |
| DPAGT1 | None | None | None | None |
| DPAGT1 | None | None | None | None |

| | | | | |
|---|---|---|---|---|
| DPAGT1 | None | None | None | None |
| IL2RG | None | None | None | None |
| HNF1A | None | None | None | None |
| HNF4A | None | None | None | None |
| FLT4 | None | None | None | None |
| FLT4 | None | None | None | None |
| FLT4 | None | None | None | None |
| CD3D | None | None | None | None |
| CD3D | None | None | None | None |
| MCOLN1 | None | None | None | None |
| MCOLN1 | None | None | None | None |
| MOCS2 | None | None | None | None |
| MOCS2 | None | None | None | None |
| ABCG8 | None | None | None | None |
| ABCG8 | None | None | None | None |
| ABCG8 | None | None | None | None |
| ABCG8 | None | None | None | None |
| ABCG8 | None | None | None | None |
| ABCG8 | None | None | None | None |
| ABCG8 | None | None | None | None |
| ABCG8 | None | None | None | None |
| ABCG8 | None | None | None | None |
| ABCG8 | None | None | None | None |
| ABCG8 | None | None | None | None |
| ABCG8 | None | None | None | None |
| ABCG8 | None | None | None | None |
| ABCG8 | None | None | None | None |
| CTNNA1 | None | None | None | None |
| TYMP | None | None | None | None |
| TYMP | None | None | None | None |
| TYMP | None | None | None | None |
| TYMP | None | None | None | None |
| TYMP | None | None | None | None |
| TYMP | None | None | None | None |
| TYMP | None | None | None | None |
| TYMP | None | None | None | None |
| TYMP | None | None | None | None |
| TYMP | None | None | None | None |
| TYMP | None | None | None | None |
| TYMP | None | None | None | None |
| TYMP | None | None | None | None |
| GAMT | None | None | None | None |
| GAMT | None | None | None | None |
| GAMT | None | None | None | None |
| GAMT | None | None | None | None |
| GAMT | None | None | None | None |
| GAMT | None | None | None | None |
| GAMT | None | None | None | None |
| GAMT | None | None | None | None |
| GAMT | None | None | None | None |
| GAMT | None | None | None | None |
| GAMT | None | None | None | None |
| GLRA1 | None | None | None | None |
| GLRA1 | None | None | None | None |

| | | | | |
|---|---|---|---|---|
| GLRA1 | None | None | None | None |
| GLRA1 | None | None | None | None |
| GLRA1 | None | None | None | None |
| GLRA1 | None | None | None | None |
| GLRA1 | None | None | None | None |
| GLRA1 | None | None | None | None |
| GLRA1 | None | None | None | None |
| ADK | None | None | None | None |
| ACTN2 | None | None | None | None |
| UBE2T | None | None | None | None |
| TTN | None | None | None | None |
| ATP7B | None | None | None | None |
| ATP7B | None | None | None | None |
| ATP7B | None | None | None | None |
| ATP7B | None | None | None | None |
| ATP7B | None | None | None | None |
| ATP7B | None | None | None | None |
| ATP7B | None | None | None | None |
| ATP7B | None | None | None | None |
| ATP7B | None | None | None | None |
| ATP7B | None | None | None | None |
| ATP7B | None | None | None | None |
| ATP7B | None | None | None | None |
| ATP7B | None | None | None | None |
| CDKN1B | None | None | None | None |
| CDKN1B | None | None | None | None |
| CDKN1B | None | None | None | None |
| CDKN1B | None | None | None | None |
| CDKN1B | None | None | None | None |
| CDKN1B | None | None | None | None |
| MYOC | None | None | None | None |
| MYOC | None | None | None | None |
| MYOC | None | None | None | None |
| MYOC | None | None | None | None |
| MYOC | None | None | None | None |
| MYOC | None | None | None | None |
| MYOC | None | None | None | None |
| MYOC | None | None | None | None |
| MYOC | None | None | None | None |
| MYOC | None | None | None | None |
| MYOC | None | None | None | None |
| MYOC | None | None | None | None |
| MYOC | None | None | None | None |
| MYOC | None | None | None | None |
| MYOC | None | None | None | None |
| SPTAN1 | None | None | None | None |
| GC | None | None | None | None |
| GC | None | None | None | None |
| MAD1L1 | None | None | None | None |
| NDST1 | None | None | None | None |
| ARL2BP | None | None | None | None |
| TOR1A | None | None | None | None |
| C1S | None | None | None | None |
| ARL6 | None | None | None | None |

| | | | | |
|---|---|---|---|---|
| ARL6 | None | None | None | None |
| ARL6 | None | None | None | None |
| ARL6 | None | None | None | None |
| ARL6 | None | None | None | None |
| ARL6 | None | None | None | None |
| FGFR1 | None | None | None | None |
| KLHL7 | None | None | None | None |
| FLNC | None | None | None | None |
| PTPRC | None | None | None | None |
| GCH1 | None | None | None | None |
| GCH1 | None | None | None | None |
| GCH1 | None | None | None | None |
| GCH1 | None | None | None | None |
| GCH1 | None | None | None | None |
| GCH1 | None | None | None | None |
| GCH1 | None | None | None | None |
| GCH1 | None | None | None | None |
| GCH1 | None | None | None | None |
| GCH1 | None | None | None | None |
| GCH1 | None | None | None | None |
| GCH1 | None | None | None | None |
| GCH1 | None | None | None | None |
| GCH1 | None | None | None | None |
| CFH | None | None | None | None |
| COL2A1 | None | None | None | None |
| COL2A1 | None | None | None | None |
| OCRL | None | None | None | None |
| WRN | None | None | None | None |
| WRN | None | None | None | None |
| WRN | None | None | None | None |
| WRN | None | None | None | None |
| MASP1 | None | None | None | None |
| MASP1 | None | None | None | None |
| NSD1 | None | None | None | None |
| PITX2 | None | None | None | None |
| PITX2 | None | None | None | None |
| PITX2 | None | None | None | None |
| PITX2 | None | None | None | None |
| PITX2 | None | None | None | None |
| PITX2 | None | None | None | None |
| B4GALT7 | 0.917 | 1 | 4irq | A |
| B4GALT7 | None | None | None | None |
| B4GALT7 | None | None | None | None |
| B4GALT7 | None | None | None | None |
| B4GALT7 | None | None | None | None |
| FSHR | None | None | None | None |
| IGHMBP2 | None | None | None | None |
| IGHMBP2 | None | None | None | None |
| IGHMBP2 | None | None | None | None |
| IGHMBP2 | None | None | None | None |
| IGHMBP2 | None | None | None | None |
| IGHMBP2 | None | None | None | None |
| IGHMBP2 | None | None | None | None |
| IGHMBP2 | None | None | None | None |

| | | | | |
|---|---|---|---|---|
| IGHMBP2 | None | None | None | None |
| IGHMBP2 | None | None | None | None |
| IGHMBP2 | None | None | None | None |
| IGHMBP2 | None | None | None | None |
| IGHMBP2 | None | None | None | None |
| IGHMBP2 | None | None | None | None |
| IGHMBP2 | None | None | None | None |
| IGHMBP2 | None | None | None | None |
| TTN | None | None | None | None |
| TTN | None | None | None | None |
| IRAK4 | None | None | None | None |
| PROC | None | None | None | None |
| PROC | None | None | None | None |
| RAD50 | None | None | None | None |
| RAD50 | None | None | None | None |
| DYNC1H1 | None | None | None | None |
| DYNC1H1 | None | None | None | None |
| DYNC1H1 | None | None | None | None |
| DYNC1H1 | None | None | None | None |
| DYNC1H1 | None | None | None | None |
| DYNC1H1 | None | None | None | None |
| DYNC1H1 | None | None | None | None |
| DYNC1H1 | None | None | None | None |
| DYNC1H1 | None | None | None | None |
| DYNC1H1 | None | None | None | None |
| DYNC1H1 | None | None | None | None |
| CYB5R3 | None | None | None | None |
| CYB5R3 | None | None | None | None |
| CYB5R3 | None | None | None | None |
| CYB5R3 | None | None | None | None |
| CYB5R3 | None | None | None | None |
| CYB5R3 | None | None | None | None |
| CYB5R3 | None | None | None | None |
| CYB5R3 | None | None | None | None |
| CYB5R3 | None | None | None | None |
| CYB5R3 | None | None | None | None |
| CYB5R3 | None | None | None | None |
| SPTB | None | None | None | None |
| FANCL | None | None | None | None |
| FAM126A | None | None | None | None |
| SEPT9 | None | None | None | None |
| TTN | None | None | None | None |
| TTN | None | None | None | None |
| TTN | None | None | None | None |
| CNTN1 | None | None | None | None |
| C1QTNF5 | None | None | None | None |
| VCL | None | None | None | None |
| VCL | None | None | None | None |
| VCL | None | None | None | None |
| VCL | None | None | None | None |
| VCL | None | None | None | None |
| VCL | None | None | None | None |
| PPT1 | None | None | None | None |
| PPT1 | None | None | None | None |

| | | | | |
|---|---|---|---|---|
| PPT1 | None | None | None | None |
| PPT1 | None | None | None | None |
| PPT1 | None | None | None | None |
| PPT1 | None | None | None | None |
| PPT1 | None | None | None | None |
| PPT1 | None | None | None | None |
| PPT1 | None | None | None | None |
| PPT1 | None | None | None | None |
| PPT1 | None | None | None | None |
| PPT1 | None | None | None | None |
| PPT1 | None | None | None | None |
| PPT1 | None | None | None | None |
| PPT1 | None | None | None | None |
| PPT1 | None | None | None | None |
| PPT1 | None | None | None | None |
| PPT1 | None | None | None | None |
| PPT1 | None | None | None | None |
| NOTCH2 | None | None | None | None |
| NOTCH2 | None | None | None | None |
| IFIH1 | None | None | None | None |
| IFIH1 | None | None | None | None |
| IKBKG | None | None | None | None |
| DBH | None | None | None | None |
| DBH | None | None | None | None |
| DBH | None | None | None | None |
| DBH | None | None | None | None |
| DBH | None | None | None | None |
| DBH | None | None | None | None |
| DBH | None | None | None | None |
| DBH | None | None | None | None |
| DBH | None | None | None | None |
| DBH | None | None | None | None |
| IL10RA | None | None | None | None |
| IL10RA | None | None | None | None |
| IL10RA | None | None | None | None |
| IL10RA | None | None | None | None |
| IL10RA | None | None | None | None |
| IL10RA | None | None | None | None |
| IL10RA | None | None | None | None |
| ILK | None | None | None | None |
| HPD | None | None | None | None |
| HPD | None | None | None | None |
| HPD | None | None | None | None |
| MECP2 | None | None | None | None |
| MECP2 | None | None | None | None |
| MECP2 | None | None | None | None |
| MECP2 | None | None | None | None |
| MECP2 | None | None | None | None |
| MECP2 | None | None | None | None |
| MECP2 | None | None | None | None |
| MECP2 | None | None | None | None |
| KYNU | None | None | None | None |
| FAS | None | None | None | None |
| FAS | None | None | None | None |

| | | | | |
|---|---|---|---|---|
| ELP1 | None | None | None | None |
| ELP1 | None | None | None | None |
| ELP1 | None | None | None | None |
| ELP1 | None | None | None | None |
| ELP1 | None | None | None | None |
| ELP1 | None | None | None | None |
| ELP1 | None | None | None | None |
| ELP1 | None | None | None | None |
| HESX1 | None | None | None | None |
| HESX1 | None | None | None | None |
| HESX1 | None | None | None | None |
| HESX1 | None | None | None | None |
| GAD1 | None | None | None | None |
| GAD1 | None | None | None | None |
| NDUFA1 | None | None | None | None |
| NDUFA1 | None | None | None | None |
| NDUFA1 | None | None | None | None |
| PNPT1 | None | None | None | None |
| PNPT1 | None | None | None | None |
| PNPT1 | None | None | None | None |
| PNPT1 | None | None | None | None |
| PNPT1 | None | None | None | None |
| PNPT1 | None | None | None | None |
| PNPT1 | None | None | None | None |
| SLC25A13 | None | None | None | None |
| SLC25A13 | None | None | None | None |
| SLC25A13 | None | None | None | None |
| AP4M1 | None | None | None | None |
| AP4M1 | None | None | None | None |
| MEFV | None | None | None | None |
| MEFV | None | None | None | None |
| MEFV | None | None | None | None |
| MEFV | None | None | None | None |
| MEFV | None | None | None | None |
| MEFV | None | None | None | None |
| MEFV | None | None | None | None |
| MEFV | None | None | None | None |
| MEFV | None | None | None | None |
| CAPN3 | None | None | None | None |
| CAPN3 | None | None | None | None |
| CAPN3 | None | None | None | None |
| CAPN3 | None | None | None | None |
| CAPN3 | None | None | None | None |
| CAPN3 | None | None | None | None |
| CAPN3 | None | None | None | None |
| GP6 | None | None | None | None |
| FAS | None | None | None | None |
| FBN1 | None | None | None | None |
| FBN1 | None | None | None | None |
| BMPR2 | None | None | None | None |
| BMPR2 | None | None | None | None |
| BMPR2 | None | None | None | None |
| BMPR2 | None | None | None | None |
| BMPR2 | None | None | None | None |

| | | | | |
|---|---|---|---|---|
| RET | None | None | None | None |
| RET | None | None | None | None |
| RET | None | None | None | None |
| RET | None | None | None | None |
| RET | None | None | None | None |
| RET | None | None | None | None |
| RET | None | None | None | None |
| RET | None | None | None | None |
| RET | None | None | None | None |
| RET | None | None | None | None |
| RET | None | None | None | None |
| RET | None | None | None | None |
| RET | None | None | None | None |
| RET | None | None | None | None |
| RET | None | None | None | None |
| RET | None | None | None | None |
| RET | None | None | None | None |
| RET | None | None | None | None |
| RET | None | None | None | None |
| RET | None | None | None | None |
| RET | None | None | None | None |
| RET | None | None | None | None |
| RET | None | None | None | None |
| RET | None | None | None | None |
| RET | None | None | None | None |
| RET | None | None | None | None |
| RET | None | None | None | None |
| RET | None | None | None | None |
| CSF2RA | None | None | None | None |
| CSF2RA | None | None | None | None |
| CSF2RA | None | None | None | None |
| BRCA1 | None | None | None | None |
| ZAP70 | None | None | None | None |
| ZAP70 | None | None | None | None |
| ZAP70 | None | None | None | None |
| PIK3R2 | None | None | None | None |
| TCN2 | None | None | None | None |
| TCN2 | None | None | None | None |
| TCN2 | None | None | None | None |
| TCN2 | None | None | None | None |
| TCN2 | None | None | None | None |
| TCN2 | None | None | None | None |
| TCN2 | None | None | None | None |
| TCN2 | None | None | None | None |
| EPM2A | None | None | None | None |
| EPM2A | None | None | None | None |
| EPM2A | None | None | None | None |
| EPM2A | None | None | None | None |
| AARS | None | None | None | None |
| AARS | None | None | None | None |
| AARS | None | None | None | None |
| PCDH15 | None | None | None | None |
| PCDH15 | None | None | None | None |
| PCDH15 | None | None | None | None |

| | | | | |
|---|---|---|---|---|
| PCDH15 | None | None | None | None |
| PCDH15 | None | None | None | None |
| PCDH15 | None | None | None | None |
| PCDH15 | None | None | None | None |
| PCDH15 | None | None | None | None |
| PCDH15 | None | None | None | None |
| PCDH15 | None | None | None | None |
| NDUFS4 | None | None | None | None |
| NDUFS4 | None | None | None | None |
| NDUFS4 | None | None | None | None |
| FGF10 | None | None | None | None |
| FGF10 | None | None | None | None |
| FGF10 | None | None | None | None |
| FGF10 | None | None | None | None |
| CDON | None | None | None | None |
| STAMBP | 0.909 | 2 | 3rzv | A |
| BRCA1 | None | None | None | None |
| BRCA1 | None | None | None | None |
| BRCA1 | None | None | None | None |
| BRCA1 | None | None | None | None |
| BRCA1 | None | None | None | None |
| BRCA1 | None | None | None | None |
| BRCA1 | None | None | None | None |
| BRCA1 | None | None | None | None |
| BRCA1 | None | None | None | None |
| BRCA1 | None | None | None | None |
| PLXNC1 | None | None | None | None |
| RNASEH2C | None | None | None | None |
| RNASEH2C | None | None | None | None |
| APC | None | None | None | None |
| PLA2G4A | None | None | None | None |
| PLA2G4A | None | None | None | None |
| PSAT1 | None | None | None | None |
| PSAT1 | None | None | None | None |
| TRAPPC3 | None | None | None | None |
| QDPR | None | None | None | None |
| QDPR | None | None | None | None |
| QDPR | None | None | None | None |
| QDPR | None | None | None | None |
| KRT5 | None | None | None | None |
| KRT5 | None | None | None | None |
| KRT5 | None | None | None | None |
| MLH1 | None | None | None | None |
| MLH1 | None | None | None | None |
| MLH1 | None | None | None | None |
| MLH1 | None | None | None | None |
| MLH1 | None | None | None | None |
| MLH1 | None | None | None | None |
| MLH1 | None | None | None | None |
| MLH1 | None | None | None | None |
| MLH1 | None | None | None | None |
| MLH1 | None | None | None | None |
| MLH1 | None | None | None | None |
| MLH1 | None | None | None | None |

| | | | | |
|---|---|---|---|---|
| MLH1 | None | None | None | None |
| MLH1 | None | None | None | None |
| MLH1 | None | None | None | None |
| MLH1 | None | None | None | None |
| MLH1 | None | None | None | None |
| KRIT1 | None | None | None | None |
| KRIT1 | None | None | None | None |
| KRIT1 | None | None | None | None |
| ALX4 | None | None | None | None |
| ALX4 | None | None | None | None |
| ALX4 | None | None | None | None |
| TJP2 | None | None | None | None |
| TJP2 | None | None | None | None |
| PANK2 | None | None | None | None |
| PANK2 | None | None | None | None |
| PANK2 | None | None | None | None |
| PANK2 | None | None | None | None |
| PANK2 | None | None | None | None |
| PANK2 | None | None | None | None |
| PANK2 | None | None | None | None |
| NDUFA6 | None | None | None | None |
| TSHR | None | None | None | None |
| TSHR | None | None | None | None |
| TSHR | None | None | None | None |
| CFTR | None | None | None | None |
| CFTR | None | None | None | None |
| MYOM1 | None | None | None | None |
| MYOM1 | None | None | None | None |
| SPTB | None | None | None | None |
| GPT2 | None | None | None | None |
| NDUFS2 | None | None | None | None |
| NDUFS2 | None | None | None | None |
| NDUFS2 | None | None | None | None |
| NDUFS2 | None | None | None | None |
| NDUFS2 | None | None | None | None |
| CASQ2 | None | None | None | None |
| CASQ2 | None | None | None | None |
| CASQ2 | None | None | None | None |
| CASQ2 | None | None | None | None |
| CASQ2 | None | None | None | None |
| CASQ2 | None | None | None | None |
| CASQ2 | None | None | None | None |
| CASQ2 | None | None | None | None |
| CASQ2 | None | None | None | None |
| CPOX | None | None | None | None |
| CPOX | None | None | None | None |
| CPOX | 0.857 | 2 | 2aex | A |
| CPOX | None | None | None | None |
| CPOX | None | None | None | None |
| CPOX | None | None | None | None |
| CPOX | None | None | None | None |
| CPOX | None | None | None | None |
| CPOX | None | None | None | None |
| MVK | None | None | None | None |

| | | | | |
|---|---|---|---|---|
| MVK | None | None | None | None |
| MVK | None | None | None | None |
| MVK | None | None | None | None |
| MVK | None | None | None | None |
| MVK | None | None | None | None |
| MVK | None | None | None | None |
| MVK | None | None | None | None |
| MVK | None | None | None | None |
| MVK | None | None | None | None |
| MVK | None | None | None | None |
| MVK | None | None | None | None |
| MVK | None | None | None | None |
| MVK | None | None | None | None |
| MVK | None | None | None | None |
| GSS | None | None | None | None |
| GSS | None | None | None | None |
| GSS | None | None | None | None |
| GSS | None | None | None | None |
| GSS | None | None | None | None |
| DSG2 | None | None | None | None |
| DSG2 | None | None | None | None |
| DSG2 | None | None | None | None |
| DSG2 | None | None | None | None |
| DSG2 | None | None | None | None |
| DSG2 | None | None | None | None |
| DSG2 | None | None | None | None |
| DSG2 | None | None | None | None |
| DSG2 | None | None | None | None |
| DSG2 | None | None | None | None |
| DSG2 | None | None | None | None |
| DSG2 | None | None | None | None |
| DSG2 | None | None | None | None |
| DSG2 | None | None | None | None |
| PAPSS2 | None | None | None | None |
| PLCB2 | None | None | None | None |
| HMGCS2 | None | None | None | None |
| HMGCS2 | None | None | None | None |
| HMGCS2 | 0.938 | 1 | 2wya | C |
| HMGCS2 | None | None | None | None |
| HMGCS2 | None | None | None | None |
| HMGCS2 | None | None | None | None |
| HMGCS2 | None | None | None | None |
| HMGCS2 | None | None | None | None |
| MPDZ | None | None | None | None |
| GATA4 | None | None | None | None |
| GATA4 | None | None | None | None |
| WARS2 | None | None | None | None |
| WARS2 | None | None | None | None |
| HGD | 1 | 47 | 1eyb | A |
| HGD | 1 | 47 | 1eyb | A |
| HGD | 1 | 70 | 1eyb | A |
| HGD | None | None | None | None |
| HGD | None | None | None | None |
| HGD | None | None | None | None |

| | | | | |
|---|---|---|---|---|
| HGD | None | None | None | None |
| HGD | None | None | None | None |
| HGD | None | None | None | None |
| HGD | 1 | 1 | 1ey2 | A |
| HGD | 1 | 9 | 1ey2 | A |
| HGD | None | None | None | None |
| HGD | 1 | 10 | 1eyb | A |
| HGD | 1 | 7 | 1ey2 | A |
| PAX8 | None | None | None | None |
| PAX8 | None | None | None | None |
| PAX8 | None | None | None | None |
| PAX8 | None | None | None | None |
| SPTB | None | None | None | None |
| NCF4 | None | None | None | None |
| CFH | None | None | None | None |
| CFH | None | None | None | None |
| CSF1R | None | None | None | None |
| CSF1R | None | None | None | None |
| CA8 | None | None | None | None |
| CA8 | None | None | None | None |
| CA8 | 0.748 | 7 | 2w2j | A |
| APC | None | None | None | None |
| APC | None | None | None | None |
| APC | None | None | None | None |
| APC | None | None | None | None |
| APC | None | None | None | None |
| NDUFA2 | None | None | None | None |
| GIF | None | None | None | None |
| GIF | None | None | None | None |
| GIF | None | None | None | None |
| GIF | None | None | None | None |
| C2 | None | None | None | None |
| ACP5 | None | None | None | None |
| ACP5 | None | None | None | None |
| ACP5 | None | None | None | None |
| ACP5 | None | None | None | None |
| ACP5 | None | None | None | None |
| RNASEH2A | None | None | None | None |
| RNASEH2A | None | None | None | None |
| RNASEH2A | None | None | None | None |
| RNASEH2A | None | None | None | None |
| RNASEH2A | None | None | None | None |
| RNASEH2A | None | None | None | None |
| RNASEH2A | None | None | None | None |
| DSG2 | None | None | None | None |
| USH1G | None | None | None | None |
| FUS | None | None | None | None |
| FUS | None | None | None | None |
| FUS | None | None | None | None |
| FUS | None | None | None | None |
| FUS | None | None | None | None |
| MRPL44 | None | None | None | None |
| MRPL44 | None | None | None | None |
| MRPL44 | None | None | None | None |

| | | | | |
|---|---|---|---|---|
| MRPL44 | None | None | None | None |
| DMGDH | None | None | None | None |
| AARS | None | None | None | None |
| STAR | None | None | None | None |
| STAR | None | None | None | None |
| STAR | None | None | None | None |
| STAR | None | None | None | None |
| STAR | None | None | None | None |
| STAR | None | None | None | None |
| STAR | None | None | None | None |
| STAR | None | None | None | None |
| LARP7 | None | None | None | None |
| PRPF8 | None | None | None | None |
| PRPF8 | None | None | None | None |
| PRPF8 | None | None | None | None |
| PRPF8 | None | None | None | None |
| PSAP | None | None | None | None |
| PSAP | None | None | None | None |
| PSAP | None | None | None | None |
| DMPK | None | None | None | None |
| EPCAM | None | None | None | None |
| EPCAM | None | None | None | None |
| EPCAM | None | None | None | None |
| EPCAM | None | None | None | None |
| EPCAM | None | None | None | None |
| EPCAM | None | None | None | None |
| EPCAM | None | None | None | None |
| EPCAM | None | None | None | None |
| ASS1 | None | None | None | None |
| ASS1 | None | None | None | None |
| ASS1 | None | None | None | None |
| ASS1 | None | None | None | None |
| ASS1 | None | None | None | None |
| ASS1 | 0.857 | 20 | 2nz2 | A |
| ASS1 | 0.857 | 9 | 2nz2 | A |
| ASS1 | None | None | None | None |
| ASS1 | None | None | None | None |
| ASS1 | None | None | None | None |
| ASS1 | 0.857 | 9 | 2nz2 | A |
| ASS1 | 0.857 | 9 | 2nz2 | A |
| ASS1 | 0.857 | 9 | 2nz2 | A |
| ASS1 | None | None | None | None |
| ASS1 | None | None | None | None |
| ASS1 | None | None | None | None |
| ASS1 | None | None | None | None |
| ASS1 | None | None | None | None |
| ASS1 | None | None | None | None |
| ASS1 | None | None | None | None |
| POLE | None | None | None | None |
| POLE | None | None | None | None |
| POLE | None | None | None | None |
| RNASET2 | None | None | None | None |
| RNASET2 | None | None | None | None |
| RNASET2 | None | None | None | None |

| | | | | |
|---|---|---|---|---|
| FLG | None | None | None | None |
| F10 | None | None | None | None |
| AGT | None | None | None | None |
| AGT | None | None | None | None |
| PLN | None | None | None | None |
| PLN | None | None | None | None |
| PLN | None | None | None | None |
| PTHLH | None | None | None | None |
| IGF1R | None | None | None | None |
| IGF1R | None | None | None | None |
| IGF1R | None | None | None | None |
| JAG1 | None | None | None | None |
| JAG1 | None | None | None | None |
| JAG1 | None | None | None | None |
| JAG1 | None | None | None | None |
| JAG1 | None | None | None | None |
| JAG1 | None | None | None | None |
| AP1S3 | None | None | None | None |
| SPTA1 | None | None | None | None |
| SPTA1 | None | None | None | None |
| SPTA1 | None | None | None | None |
| SPTA1 | None | None | None | None |
| SPG7 | None | None | None | None |
| SPG7 | None | None | None | None |
| SPG7 | None | None | None | None |
| SPG7 | None | None | None | None |
| SPG7 | None | None | None | None |
| SPG7 | None | None | None | None |
| FN1 | None | None | None | None |
| FN1 | None | None | None | None |
| NSMCE3 | None | None | None | None |
| PCCA | None | None | None | None |
| PCCA | None | None | None | None |
| SPTB | None | None | None | None |
| RYR2 | None | None | None | None |
| RYR2 | None | None | None | None |
| MMP9 | None | None | None | None |
| MMP9 | None | None | None | None |
| GLMN | None | None | None | None |
| GLMN | None | None | None | None |
| GLMN | None | None | None | None |
| GLMN | None | None | None | None |
| GLMN | None | None | None | None |
| GLMN | None | None | None | None |
| GLMN | None | None | None | None |
| GLMN | None | None | None | None |
| GLMN | None | None | None | None |
| GLMN | None | None | None | None |
| MARS | None | None | None | None |
| MARS | None | None | None | None |
| MARS | None | None | None | None |
| MARS | None | None | None | None |
| MARS | None | None | None | None |
| MARS | None | None | None | None |

| | | | | |
|---|---|---|---|---|
| SNAP29 | None | None | None | None |
| SNAP29 | None | None | None | None |
| ETHE1 | None | None | None | None |
| ETHE1 | None | None | None | None |
| ETHE1 | None | None | None | None |
| ETHE1 | None | None | None | None |
| ETHE1 | None | None | None | None |
| ETHE1 | None | None | None | None |
| ETHE1 | None | None | None | None |
| SAMHD1 | None | None | None | None |
| KCNQ4 | None | None | None | None |
| FXYD2 | None | None | None | None |
| FXYD2 | None | None | None | None |
| FXYD2 | None | None | None | None |
| LRRK2 | None | None | None | None |
| LRRK2 | None | None | None | None |
| NDUFS1 | None | None | None | None |
| NDUFS1 | None | None | None | None |
| NDUFS1 | None | None | None | None |
| NDUFS1 | None | None | None | None |
| NDUFS1 | None | None | None | None |
| NDUFS1 | None | None | None | None |
| NDUFS1 | None | None | None | None |
| NDUFS1 | None | None | None | None |
| NDUFS1 | None | None | None | None |
| NDUFS1 | None | None | None | None |
| NDUFS1 | None | None | None | None |
| EZH2 | None | None | None | None |
| EZH2 | None | None | None | None |
| LARS | None | None | None | None |
| TLL1 | None | None | None | None |
| CRYBB2 | None | None | None | None |
| CRYBB2 | None | None | None | None |
| EXOSC8 | None | None | None | None |
| SRP72 | None | None | None | None |
| FOXC2 | None | None | None | None |
| HMBS | None | None | None | None |
| HMBS | None | None | None | None |
| HMBS | None | None | None | None |
| HMBS | None | None | None | None |
| HMBS | None | None | None | None |
| HMBS | None | None | None | None |
| HMBS | None | None | None | None |
| HMBS | None | None | None | None |
| HMBS | None | None | None | None |
| HMBS | None | None | None | None |
| HMBS | None | None | None | None |
| HMBS | None | None | None | None |
| HMBS | None | None | None | None |
| HMBS | None | None | None | None |
| HMBS | None | None | None | None |
| HMBS | None | None | None | None |
| HMBS | None | None | None | None |
| HMBS | None | None | None | None |

| | | | | |
|---|---|---|---|---|
| HMBS | None | None | None | None |
| HMBS | None | None | None | None |
| HMBS | None | None | None | None |
| HMBS | None | None | None | None |
| HMBS | None | None | None | None |
| HMBS | None | None | None | None |
| HMBS | None | None | None | None |
| OGT | None | None | None | None |
| NOG | None | None | None | None |
| NOG | None | None | None | None |
| NOG | None | None | None | None |
| NOG | None | None | None | None |
| NOG | None | None | None | None |
| NOG | None | None | None | None |
| NOG | None | None | None | None |
| NOG | None | None | None | None |
| NOG | None | None | None | None |
| NOG | None | None | None | None |
| NOG | None | None | None | None |
| DLG3 | None | None | None | None |
| SCN2A | None | None | None | None |
| STAC3 | None | None | None | None |
| PFKM | None | None | None | None |
| PFKM | None | None | None | None |
| PFKM | None | None | None | None |
| PFKM | None | None | None | None |
| PFKM | None | None | None | None |
| DNMT1 | None | None | None | None |
| DNMT1 | None | None | None | None |
| TNNI3 | None | None | None | None |
| TNNI3 | None | None | None | None |
| TNNI3 | None | None | None | None |
| STXBP2 | None | None | None | None |
| STXBP2 | None | None | None | None |
| STXBP2 | None | None | None | None |
| STXBP2 | None | None | None | None |
| STXBP2 | None | None | None | None |
| STXBP2 | None | None | None | None |
| STXBP2 | None | None | None | None |
| STXBP2 | None | None | None | None |
| REN | None | None | None | None |
| DSP | None | None | None | None |
| DSP | None | None | None | None |
| HNF1B | None | None | None | None |
| ADA | None | None | None | None |
| ADA | None | None | None | None |
| ADA | None | None | None | None |
| ADA | None | None | None | None |
| ADA | None | None | None | None |
| ADA | None | None | None | None |
| ADA | None | None | None | None |
| ADA | None | None | None | None |
| ADA | None | None | None | None |
| ADA | None | None | None | None |

| | | | | |
|---|---|---|---|---|
| ADA | None | None | None | None |
| ADA | None | None | None | None |
| ADA | None | None | None | None |
| ADA | None | None | None | None |
| ADA | None | None | None | None |
| DCX | None | None | None | None |
| DCX | None | None | None | None |
| DCX | None | None | None | None |
| DCX | None | None | None | None |
| DCX | None | None | None | None |
| ARSB | None | None | None | None |
| ARSB | None | None | None | None |
| ARSB | None | None | None | None |
| ARSB | None | None | None | None |
| ARSB | None | None | None | None |
| ARSB | None | None | None | None |
| ARSB | None | None | None | None |
| ARSB | None | None | None | None |
| ARSB | None | None | None | None |
| ARSB | None | None | None | None |
| ARSB | None | None | None | None |
| ARSB | None | None | None | None |
| ARSB | None | None | None | None |
| WWOX | None | None | None | None |
| TNNI3 | None | None | None | None |
| TNNI3 | None | None | None | None |
| TNNI3 | None | None | None | None |
| TNNI3 | None | None | None | None |
| CYBA | None | None | None | None |
| SPAST | None | None | None | None |
| SPAST | None | None | None | None |
| SPAST | None | None | None | None |
| SPAST | None | None | None | None |
| SPAST | None | None | None | None |
| SPAST | None | None | None | None |
| SPAST | None | None | None | None |
| SPAST | None | None | None | None |
| KRIT1 | None | None | None | None |
| KRIT1 | None | None | None | None |
| TP53 | None | None | None | None |
| ISPD | None | None | None | None |
| ISPD | None | None | None | None |
| ISPD | None | None | None | None |
| ISPD | None | None | None | None |
| KCNMA1 | None | None | None | None |
| STK11 | None | None | None | None |
| STK11 | None | None | None | None |
| STK11 | None | None | None | None |
| STK11 | None | None | None | None |
| STK11 | None | None | None | None |
| STK11 | None | None | None | None |
| NDUFA13 | None | None | None | None |
| PHF6 | None | None | None | None |
| PHF6 | None | None | None | None |

| | | | | |
|---|---|---|---|---|
| PHF6 | None | None | None | None |
| PHYH | None | None | None | None |
| PHYH | None | None | None | None |
| PHYH | None | None | None | None |
| PHYH | None | None | None | None |
| PHYH | None | None | None | None |
| NDUFS3 | None | None | None | None |
| NDUFS3 | None | None | None | None |
| ALDH18A1 | None | None | None | None |
| ALDH18A1 | None | None | None | None |
| ALDH18A1 | None | None | None | None |
| ALDH18A1 | None | None | None | None |
| ALDH18A1 | None | None | None | None |
| PAX6 | None | None | None | None |
| PAX6 | None | None | None | None |
| INPP5E | None | None | None | None |
| INPP5E | None | None | None | None |
| INPP5E | None | None | None | None |
| INPP5E | None | None | None | None |
| INPP5E | None | None | None | None |
| INPP5E | None | None | None | None |
| INPP5E | None | None | None | None |
| INPP5E | None | None | None | None |
| INPP5E | None | None | None | None |
| TNNT2 | None | None | None | None |
| TNNT2 | None | None | None | None |
| TNNT2 | None | None | None | None |
| TNNT2 | None | None | None | None |
| TNNT2 | None | None | None | None |
| TNNT2 | None | None | None | None |
| TNNT2 | None | None | None | None |
| TNNT2 | None | None | None | None |
| TYK2 | None | None | None | None |
| TYK2 | None | None | None | None |
| TYK2 | None | None | None | None |
| CFTR | None | None | None | None |
| CFTR | None | None | None | None |
| CFTR | None | None | None | None |
| CFTR | None | None | None | None |
| CFTR | None | None | None | None |
| CFTR | None | None | None | None |
| CFTR | None | None | None | None |
| CFTR | None | None | None | None |
| CFTR | None | None | None | None |
| CFTR | None | None | None | None |
| CFTR | None | None | None | None |
| CFTR | None | None | None | None |
| CFTR | None | None | None | None |
| CFTR | None | None | None | None |
| CFTR | None | None | None | None |
| CFTR | None | None | None | None |
| CFTR | None | None | None | None |
| CFTR | None | None | None | None |
| CFTR | None | None | None | None |

| | | | | |
|---|---|---|---|---|
| CFTR | None | None | None | None |
| CFTR | None | None | None | None |
| CFTR | None | None | None | None |
| CFTR | None | None | None | None |
| CFTR | None | None | None | None |
| CFTR | None | None | None | None |
| CFTR | None | None | None | None |
| CFTR | None | None | None | None |
| CFTR | None | None | None | None |
| CFTR | None | None | None | None |
| CFTR | None | None | None | None |
| CFTR | None | None | None | None |
| CFTR | None | None | None | None |
| CFTR | None | None | None | None |
| CFTR | None | None | None | None |
| CFTR | None | None | None | None |
| CFTR | None | None | None | None |
| CFTR | None | None | None | None |
| CFTR | None | None | None | None |
| CFTR | None | None | None | None |
| CFTR | None | None | None | None |
| CFTR | None | None | None | None |
| CFTR | None | None | None | None |
| CFTR | None | None | None | None |
| CFTR | None | None | None | None |
| CFTR | None | None | None | None |
| CFTR | None | None | None | None |
| CFTR | None | None | None | None |
| CFTR | None | None | None | None |
| CFTR | None | None | None | None |
| CFTR | None | None | None | None |
| CFTR | None | None | None | None |
| CFTR | None | None | None | None |
| CFTR | None | None | None | None |
| SERPINF1 | None | None | None | None |
| SERPINF1 | None | None | None | None |
| IDS | None | None | None | None |
| IDS | None | None | None | None |
| IDS | None | None | None | None |
| IDS | None | None | None | None |
| IDS | None | None | None | None |
| IDS | None | None | None | None |
| IDS | None | None | None | None |
| IDS | None | None | None | None |
| IDS | None | None | None | None |
| NFKBIA | 0.7 | 1 | 1nfi | F |
| NFKBIA | None | None | None | None |
| SPAST | None | None | None | None |
| FGF23 | None | None | None | None |
| FGF23 | None | None | None | None |
| FGF23 | None | None | None | None |
| FGF23 | None | None | None | None |
| FGF23 | None | None | None | None |
| FGF23 | None | None | None | None |

| | | | | |
|---|---|---|---|---|
| FGF23 | None | None | None | None |
| DLAT | None | None | None | None |
| STUB1 | None | None | None | None |
| HSD17B4 | None | None | None | None |
| HSD17B4 | None | None | None | None |
| DYSF | None | None | None | None |
| CEP135 | None | None | None | None |
| CEP135 | None | None | None | None |
| PRKCG | None | None | None | None |
| CSF1R | None | None | None | None |
| PEX5 | None | None | None | None |
| PAX6 | None | None | None | None |
| PAX6 | None | None | None | None |
| PAX6 | None | None | None | None |
| PAX6 | None | None | None | None |
| PAX6 | None | None | None | None |
| PAX6 | None | None | None | None |
| NF2 | None | None | None | None |
| NF2 | None | None | None | None |
| NF2 | None | None | None | None |
| REM1 | None | None | None | None |

| source_id | classification | binding_site | binding_site_single_water | protein_binding_site |
|---|---|---|---|---|
| rs121918343 | pathogenic | 1 | 1 | 0 |
| rs267606649 | pathogenic | 1 | 1 | 1 |
| rs121918342 | pathogenic | 0 | 1 | 0 |
| rs267606650 | pathogenic | 1 | 1 | 1 |
| rs1130335 | benign | 1 | 1 | 1 |
| rs199476087 | pathogenic | 1 | 1 | 1 |
| rs772163112 | benign | 0 | 0 | 0 |
| rs755793 | benign | 0 | 0 | 0 |
| rs121918155 | pathogenic | 1 | 1 | 0 |
| rs121918156 | pathogenic | 1 | 1 | 0 |
| rs121918154 | pathogenic | 1 | 1 | 1 |
| rs121918151 | pathogenic | 1 | 1 | 1 |
| rs121918144 | pathogenic | 0 | 1 | 0 |
| rs121918146 | pathogenic | 1 | 1 | 1 |
| rs121918160 | pathogenic | 1 | 1 | 1 |
| rs121918150 | pathogenic | 1 | 1 | 1 |
| rs121918141 | pathogenic | 1 | 1 | 1 |
| rs142742242 | pathogenic | 1 | 1 | 1 |
| rs121918142 | pathogenic | 1 | 1 | 1 |
| rs121918157 | pathogenic | 1 | 1 | 1 |
| rs121918101 | pathogenic | 1 | 1 | 1 |
| rs121908260 | pathogenic | 1 | 1 | 1 |
| rs80356667 | pathogenic | 1 | 1 | 1 |
| rs80356668 | pathogenic | 1 | 1 | 1 |
| rs397514540 | pathogenic | 1 | 1 | 1 |
| rs104894685 | pathogenic | 0 | 1 | 0 |
| rs199869995 | pathogenic | 1 | 1 | 1 |
| rs121909072 | pathogenic | 0 | 0 | 0 |
| rs143669310 | benign | 1 | 1 | 1 |
| rs747347498 | benign | 0 | 0 | 0 |
| rs138283222 | benign | 0 | 0 | 0 |
| rs118203931 | pathogenic | 1 | 1 | 1 |
| rs118203933 | pathogenic | 1 | 1 | 0 |
| rs118203932 | pathogenic | 1 | 1 | 0 |
| rs2228063 | benign | 1 | 1 | 1 |
| rs377767360 | pathogenic | 0 | 0 | 0 |
| rs397518413 | pathogenic | 1 | 1 | 1 |
| rs1800458 | benign | 1 | 1 | 0 |
| rs121918083 | pathogenic | 1 | 1 | 1 |
| rs121918094 | pathogenic | 1 | 1 | 1 |
| rs121918098 | pathogenic | 1 | 1 | 1 |
| rs121918093 | pathogenic | 1 | 1 | 1 |
| rs79977247 | pathogenic | 1 | 1 | 1 |
| rs28933979 | pathogenic | 1 | 1 | 1 |
| rs121918068 | pathogenic | 1 | 1 | 1 |
| rs121918077 | pathogenic | 1 | 1 | 1 |
| rs11541796 | pathogenic | 1 | 1 | 1 |
| rs104894665 | pathogenic | 1 | 1 | 1 |
| rs121918090 | pathogenic | 1 | 1 | 1 |
| rs121918081 | pathogenic | 1 | 1 | 1 |
| rs121918080 | pathogenic | 1 | 1 | 1 |
| rs121918076 | pathogenic | 1 | 1 | 1 |
| rs121918097 | pathogenic | 1 | 1 | 1 |

| | | | | |
|---|---|---|---|---|
| rs121918079 | pathogenic | 1 | 1 | 1 |
| rs121918069 | pathogenic | 1 | 1 | 1 |
| rs121918086 | pathogenic | 1 | 1 | 1 |
| rs121918091 | pathogenic | 1 | 1 | 1 |
| rs121918085 | pathogenic | 1 | 1 | 1 |
| rs121918100 | pathogenic | 1 | 1 | 1 |
| rs121918084 | pathogenic | 1 | 1 | 1 |
| rs121918071 | pathogenic | 1 | 1 | 1 |
| rs121918072 | pathogenic | 1 | 1 | 1 |
| rs121918082 | pathogenic | 1 | 1 | 1 |
| rs121918087 | pathogenic | 1 | 1 | 1 |
| rs76410435 | benign | 1 | 1 | 1 |
| rs121918089 | pathogenic | 1 | 1 | 1 |
| rs121918073 | pathogenic | 1 | 1 | 1 |
| rs121918075 | pathogenic | 1 | 1 | 1 |
| rs121918088 | pathogenic | 1 | 1 | 1 |
| rs76992529 | pathogenic | 1 | 1 | 1 |
| rs536294863 | benign | 1 | 1 | 1 |
| rs730881162 | benign | 1 | 1 | 1 |
| rs113994087 | pathogenic | 1 | 1 | 1 |
| rs74716434 | benign | 1 | 1 | 1 |
| rs121918694 | pathogenic | 1 | 1 | 1 |
| rs121918707 | pathogenic | 1 | 1 | 1 |
| rs121918695 | pathogenic | 1 | 1 | 0 |
| rs121918690 | pathogenic | 1 | 1 | 0 |
| rs121918696 | pathogenic | 1 | 1 | 0 |
| rs121918693 | pathogenic | 1 | 1 | 0 |
| rs28999969 | pathogenic | 1 | 1 | 1 |
| rs121918697 | pathogenic | 1 | 1 | 1 |
| rs121918688 | pathogenic | 1 | 1 | 0 |
| rs28999970 | pathogenic | 1 | 1 | 0 |
| rs121918686 | pathogenic | 1 | 1 | 0 |
| rs28999971 | pathogenic | 1 | 1 | 1 |
| rs121918698 | pathogenic | 1 | 1 | 1 |
| rs121918691 | pathogenic | 1 | 1 | 0 |
| rs121918692 | pathogenic | 1 | 1 | 1 |
| rs121918703 | pathogenic | 1 | 1 | 1 |
| rs121918687 | pathogenic | 1 | 1 | 1 |
| rs28933408 | pathogenic | 1 | 1 | 1 |
| rs121918702 | pathogenic | 1 | 1 | 1 |
| rs17875899 | benign | 1 | 1 | 1 |
| rs111033620 | pathogenic | 1 | 1 | 1 |
| rs111033622 | pathogenic | 1 | 1 | 1 |
| rs111033621 | pathogenic | 1 | 1 | 1 |
| rs111033618 | pathogenic | 1 | 1 | 1 |
| rs5918 | benign | 0 | 0 | 0 |
| rs121918452 | pathogenic | 0 | 0 | 0 |
| rs121918445 | pathogenic | 1 | 1 | 1 |
| rs5917 | benign | 1 | 1 | 1 |
| rs121918444 | pathogenic | 1 | 1 | 1 |
| rs121918446 | pathogenic | 1 | 1 | 1 |
| rs121918449 | pathogenic | 0 | 0 | 0 |
| rs121918448 | benign | 0 | 0 | 0 |
| rs13306487 | benign | 1 | 1 | 1 |

| | | | | |
|---|---|---|---|---|
| rs1805044 | benign | 0 | 0 | 0 |
| rs117078377 | benign | 0 | 0 | 0 |
| rs138132026 | benign | 1 | 1 | 1 |
| rs75762896 | benign | 1 | 1 | 0 |
| rs121918043 | pathogenic | 1 | 1 | 0 |
| rs143581690 | benign | 1 | 1 | 0 |
| rs147654263 | benign | 0 | 0 | 0 |
| rs587782529 | pathogenic | 1 | 1 | 1 |
| rs121912664 | pathogenic | 1 | 1 | 1 |
| rs150293825 | benign | 1 | 1 | 1 |
| rs17882252 | benign | 1 | 1 | 1 |
| rs121912662 | pathogenic | 1 | 1 | 1 |
| rs35993958 | benign | 1 | 1 | 1 |
| rs606231253 | pathogenic | 1 | 1 | 1 |
| rs2227983 | benign | 1 | 1 | 1 |
| rs587779351 | pathogenic | 0 | 0 | 0 |
| rs779773463 | pathogenic | 1 | 1 | 1 |
| rs713040 | benign | 1 | 1 | 1 |
| rs334 | pathogenic | 1 | 1 | 1 |
| rs33930165 | pathogenic | 1 | 1 | 1 |
| rs33918131 | benign | 1 | 1 | 1 |
| rs63750783 | pathogenic | 1 | 1 | 1 |
| rs33986703 | pathogenic | 1 | 1 | 1 |
| rs33972047 | pathogenic | 1 | 1 | 1 |
| rs33936254 | pathogenic | 1 | 1 | 1 |
| rs33959855 | benign | 1 | 1 | 1 |
| rs33950507 | pathogenic | 1 | 1 | 1 |
| rs33916412 | pathogenic | 1 | 1 | 0 |
| rs1135071 | pathogenic | 1 | 1 | 1 |
| rs33960103 | pathogenic | 1 | 1 | 1 |
| rs11549407 | pathogenic | 1 | 1 | 1 |
| rs33922842 | pathogenic | 1 | 1 | 1 |
| rs35278874 | benign | 1 | 1 | 1 |
| rs34621955 | benign | 1 | 1 | 1 |
| rs33995148 | pathogenic | 1 | 1 | 1 |
| rs33945705 | benign | 1 | 1 | 1 |
| rs33993568 | benign | 1 | 1 | 1 |
| rs35553496 | benign | 1 | 1 | 1 |
| rs33913712 | pathogenic | 1 | 1 | 1 |
| rs33917785 | pathogenic | 1 | 1 | 1 |
| rs34515413 | benign | 1 | 1 | 1 |
| rs35256489 | pathogenic | 1 | 1 | 0 |
| rs34484056 | pathogenic | 1 | 1 | 1 |
| rs33947020 | benign | 1 | 1 | 1 |
| rs33946267 | pathogenic | 1 | 1 | 1 |
| rs33966761 | pathogenic | 1 | 1 | 1 |
| rs36020563 | benign | 1 | 1 | 1 |
| rs122454124 | pathogenic | 1 | 1 | 1 |
| rs122454126 | pathogenic | 1 | 1 | 0 |
| rs122454127 | pathogenic | 1 | 1 | 1 |
| rs122454130 | pathogenic | 1 | 1 | 1 |
| rs122454125 | pathogenic | 1 | 1 | 1 |
| rs122454131 | pathogenic | 1 | 1 | 0 |
| rs121908302 | pathogenic | 0 | 1 | 0 |

| | | | | |
|---|---|---|---|---|
| rs77829017 | pathogenic | 0 | 1 | 0 |
| rs1141814 | pathogenic | 0 | 1 | 0 |
| rs121908312 | pathogenic | 0 | 1 | 0 |
| rs79653797 | pathogenic | 1 | 1 | 0 |
| rs439898 | pathogenic | 1 | 1 | 0 |
| rs121908299 | pathogenic | 0 | 1 | 0 |
| rs80356763 | pathogenic | 1 | 1 | 1 |
| rs781152868 | pathogenic | 0 | 1 | 0 |
| rs121908297 | pathogenic | 0 | 1 | 0 |
| rs398123531 | benign | 0 | 1 | 0 |
| rs398123532 | pathogenic | 0 | 1 | 0 |
| rs364897 | pathogenic | 1 | 1 | 1 |
| rs1064644 | pathogenic | 0 | 1 | 0 |
| rs409652 | pathogenic | 1 | 1 | 1 |
| rs121908300 | pathogenic | 0 | 1 | 0 |
| rs381737 | pathogenic | 0 | 1 | 0 |
| rs74500255 | pathogenic | 0 | 1 | 0 |
| rs121908313 | pathogenic | 0 | 1 | 0 |
| rs367968666 | pathogenic | 0 | 1 | 0 |
| rs78973108 | pathogenic | 1 | 1 | 1 |
| rs121908298 | pathogenic | 1 | 1 | 1 |
| rs78396650 | pathogenic | 0 | 0 | 0 |
| rs78198234 | pathogenic | 1 | 1 | 0 |
| rs121908304 | pathogenic | 1 | 1 | 1 |
| rs76539814 | pathogenic | 1 | 1 | 1 |
| rs121908305 | pathogenic | 0 | 1 | 0 |
| rs121908306 | pathogenic | 1 | 1 | 0 |
| rs121908308 | pathogenic | 1 | 1 | 1 |
| rs121908307 | pathogenic | 0 | 1 | 0 |
| rs75548401 | benign | 0 | 1 | 0 |
| rs76763715 | pathogenic | 0 | 0 | 0 |
| rs121908314 | pathogenic | 0 | 1 | 0 |
| rs398123528 | pathogenic | 0 | 1 | 0 |
| rs121908311 | pathogenic | 0 | 0 | 0 |
| rs121908310 | pathogenic | 1 | 1 | 0 |
| rs74598136 | pathogenic | 0 | 1 | 0 |
| rs1064651 | pathogenic | 1 | 1 | 1 |
| rs77369218 | pathogenic | 1 | 1 | 1 |
| rs121908295 | pathogenic | 0 | 1 | 0 |
| rs421016 | pathogenic | 0 | 1 | 0 |
| rs368060 | pathogenic | 0 | 1 | 0 |
| rs80356771 | pathogenic | 1 | 1 | 1 |
| rs80356772 | pathogenic | 1 | 1 | 1 |
| rs121908301 | pathogenic | 0 | 0 | 0 |
| rs747506979 | pathogenic | 0 | 1 | 0 |
| rs75822236 | pathogenic | 0 | 1 | 0 |
| rs1990760 | benign | 1 | 1 | 0 |
| rs28934568 | pathogenic | 1 | 1 | 0 |
| rs34833812 | benign | 1 | 1 | 0 |
| rs193922666 | benign | 1 | 1 | 0 |
| rs104893812 | pathogenic | 1 | 1 | 0 |
| rs144701411 | benign | 0 | 0 | 0 |
| rs104893813 | pathogenic | 0 | 1 | 0 |
| rs199660234 | pathogenic | 1 | 1 | 0 |

| | | | | |
|---|---|---|---|---|
| rs104893814 | pathogenic | 1 | 1 | 0 |
| rs143095746 | benign | 1 | 1 | 1 |
| rs104893807 | pathogenic | 1 | 1 | 0 |
| rs104893811 | pathogenic | 0 | 0 | 0 |
| rs104893816 | pathogenic | 0 | 0 | 0 |
| rs121918714 | pathogenic | 0 | 1 | 0 |
| rs104893810 | pathogenic | 1 | 1 | 0 |
| rs104893815 | pathogenic | 1 | 1 | 0 |
| rs4303 | benign | 0 | 1 | 0 |
| rs121912704 | pathogenic | 0 | 1 | 0 |
| rs56394458 | benign | 1 | 1 | 0 |
| rs397514688 | pathogenic | 0 | 0 | 0 |
| rs367797185 | pathogenic | 1 | 1 | 0 |
| rs12709426 | benign | 0 | 1 | 0 |
| rs2228547 | benign | 0 | 0 | 0 |
| rs111033546 | pathogenic | 0 | 0 | 0 |
| rs111033553 | pathogenic | 0 | 0 | 0 |
| rs111033554 | pathogenic | 1 | 1 | 1 |
| rs111033544 | pathogenic | 1 | 1 | 1 |
| rs111033555 | pathogenic | 1 | 1 | 1 |
| rs111033545 | pathogenic | 1 | 1 | 1 |
| rs111033549 | pathogenic | 0 | 1 | 0 |
| rs111033552 | pathogenic | 1 | 1 | 1 |
| rs752204035 | benign | 0 | 0 | 0 |
| rs11569534 | benign | 0 | 0 | 0 |
| rs28937571 | pathogenic | 1 | 1 | 1 |
| rs104894215 | pathogenic | 1 | 1 | 1 |
| rs3888798 | benign | 1 | 1 | 1 |
| rs764263034 | benign | 1 | 1 | 1 |
| rs1800973 | benign | 1 | 1 | 1 |
| rs367623154 | benign | 1 | 1 | 1 |
| rs201230446 | pathogenic | 1 | 1 | 1 |
| rs267606766 | pathogenic | 0 | 1 | 0 |
| rs267606769 | pathogenic | 0 | 1 | 0 |
| rs267606767 | pathogenic | 1 | 1 | 0 |
| rs267606768 | pathogenic | 0 | 1 | 0 |
| rs61733129 | benign | 0 | 1 | 0 |
| rs201947120 | pathogenic | 1 | 1 | 1 |
| rs283413 | benign | 0 | 1 | 0 |
| rs36119840 | benign | 0 | 0 | 0 |
| rs281865419 | pathogenic | 1 | 1 | 1 |
| rs137852937 | pathogenic | 0 | 1 | 0 |
| rs2230709 | benign | 0 | 1 | 0 |
| rs6695033 | benign | 0 | 1 | 0 |
| rs180177032 | pathogenic | 1 | 1 | 1 |
| rs180177033 | pathogenic | 1 | 1 | 1 |
| rs121913351 | pathogenic | 1 | 1 | 1 |
| rs397507473 | pathogenic | 1 | 1 | 1 |
| rs121913355 | pathogenic | 1 | 1 | 1 |
| rs121913357 | pathogenic | 1 | 1 | 1 |
| rs180177036 | pathogenic | 1 | 1 | 1 |
| rs180177039 | pathogenic | 1 | 1 | 1 |
| rs180177038 | pathogenic | 1 | 1 | 1 |
| rs180177040 | pathogenic | 1 | 1 | 1 |

| | | | | |
|---|---|---|---|---|
| rs121913338 | pathogenic | 1 | 1 | 1 |
| rs121913366 | pathogenic | 1 | 1 | 1 |
| rs121913369 | pathogenic | 1 | 1 | 1 |
| rs113488022 | pathogenic | 1 | 1 | 1 |
| rs121913364 | pathogenic | 1 | 1 | 1 |
| rs180177042 | pathogenic | 1 | 1 | 1 |
| rs200461617 | benign | 0 | 0 | 0 |
| rs141609745 | benign | 0 | 0 | 0 |
| rs104894383 | pathogenic | 1 | 1 | 1 |
| rs104894384 | pathogenic | 1 | 1 | 1 |
| rs104894381 | pathogenic | 1 | 1 | 0 |
| rs147710408 | benign | 1 | 1 | 1 |
| rs77357563 | benign | 1 | 1 | 1 |
| rs104894378 | pathogenic | 0 | 0 | 0 |
| rs104894382 | pathogenic | 0 | 0 | 0 |
| rs137853292 | pathogenic | 1 | 1 | 1 |
| rs587778834 | pathogenic | 0 | 0 | 0 |
| rs137853294 | pathogenic | 1 | 1 | 1 |
| rs551747882 | benign | 0 | 0 | 0 |
| rs3092903 | benign | 1 | 1 | 1 |
| rs137853296 | pathogenic | 0 | 0 | 0 |
| rs398122917 | pathogenic | 0 | 1 | 0 |
| rs41304992 | benign | 0 | 0 | 0 |
| rs121912708 | pathogenic | 1 | 1 | 1 |
| rs140102105 | pathogenic | 1 | 1 | 1 |
| rs376917645 | pathogenic | 1 | 1 | 0 |
| rs372660425 | pathogenic | 1 | 1 | 0 |
| rs555896752 | pathogenic | 1 | 1 | 1 |
| rs777829351 | pathogenic | 1 | 1 | 0 |
| rs121912711 | pathogenic | 1 | 1 | 0 |
| rs754449549 | pathogenic | 0 | 1 | 0 |
| rs141701364 | benign | 0 | 1 | 0 |
| rs761011399 | benign | 1 | 1 | 1 |
| rs121912710 | benign | 1 | 1 | 0 |
| rs121912707 | pathogenic | 1 | 1 | 0 |
| rs747597620 | pathogenic | 0 | 1 | 0 |
| rs186558364 | pathogenic | 1 | 1 | 1 |
| rs556400964 | pathogenic | 1 | 1 | 1 |
| rs61757684 | benign | 1 | 1 | 1 |
| rs121917766 | pathogenic | 1 | 1 | 1 |
| rs121918248 | pathogenic | 0 | 0 | 0 |
| rs398123278 | pathogenic | 0 | 0 | 0 |
| rs121918251 | pathogenic | 1 | 1 | 1 |
| rs727504022 | pathogenic | 0 | 0 | 0 |
| rs190834116 | pathogenic | 1 | 1 | 1 |
| rs121918249 | pathogenic | 1 | 1 | 0 |
| rs121918257 | pathogenic | 1 | 1 | 0 |
| rs142068898 | benign | 0 | 0 | 0 |
| rs757000253 | pathogenic | 0 | 0 | 0 |
| rs200908035 | pathogenic | 1 | 1 | 0 |
| rs760782399 | pathogenic | 0 | 0 | 0 |
| rs778702777 | pathogenic | 0 | 0 | 0 |
| rs121918258 | pathogenic | 1 | 1 | 0 |
| rs121918256 | pathogenic | 0 | 0 | 0 |

| | | | | |
|---|---|---|---|---|
| rs200596762 | pathogenic | 1 | 1 | 0 |
| rs547709692 | pathogenic | 0 | 0 | 0 |
| rs758577372 | pathogenic | 0 | 1 | 0 |
| rs564069299 | pathogenic | 1 | 1 | 1 |
| rs772552898 | pathogenic | 1 | 1 | 1 |
| rs368790885 | pathogenic | 1 | 1 | 1 |
| rs121918250 | pathogenic | 0 | 0 | 0 |
| rs533755473 | pathogenic | 1 | 1 | 1 |
| rs753288303 | pathogenic | 1 | 1 | 1 |
| rs2229385 | benign | 0 | 0 | 0 |
| rs1141321 | benign | 0 | 0 | 0 |
| rs753564352 | pathogenic | 0 | 0 | 0 |
| rs775593146 | pathogenic | 0 | 0 | 0 |
| rs121918254 | pathogenic | 0 | 0 | 0 |
| rs8589 | benign | 1 | 1 | 0 |
| rs777758903 | pathogenic | 0 | 0 | 0 |
| rs140600746 | pathogenic | 1 | 1 | 0 |
| rs121918255 | pathogenic | 1 | 1 | 0 |
| rs121918252 | pathogenic | 0 | 0 | 0 |
| rs779990936 | pathogenic | 1 | 1 | 0 |
| rs121912455 | pathogenic | 1 | 1 | 1 |
| rs121912459 | pathogenic | 1 | 1 | 1 |
| rs74315452 | pathogenic | 1 | 1 | 1 |
| rs121912451 | pathogenic | 1 | 1 | 1 |
| rs79006549 | pathogenic | 0 | 0 | 0 |
| rs28934889 | benign | 0 | 1 | 0 |
| rs111033557 | benign | 1 | 1 | 1 |
| rs28934597 | pathogenic | 1 | 1 | 1 |
| rs28934596 | pathogenic | 1 | 1 | 1 |
| rs28934595 | pathogenic | 1 | 1 | 1 |
| rs146519482 | pathogenic | 1 | 1 | 1 |
| rs111033563 | pathogenic | 1 | 1 | 1 |
| rs75385605 | benign | 1 | 1 | 1 |
| rs121908569 | pathogenic | 1 | 1 | 1 |
| rs35719940 | benign | 0 | 0 | 0 |
| rs121918664 | pathogenic | 0 | 0 | 0 |
| rs104893665 | pathogenic | 1 | 1 | 1 |
| rs104893666 | pathogenic | 1 | 1 | 1 |
| rs200845457 | benign | 1 | 1 | 1 |
| rs779204655 | pathogenic | 1 | 1 | 1 |
| rs121917747 | pathogenic | 1 | 1 | 1 |
| rs6042 | benign | 1 | 1 | 1 |
| rs104894380 | pathogenic | 0 | 0 | 0 |
| rs369083541 | pathogenic | 1 | 1 | 1 |
| rs145243532 | benign | 1 | 1 | 1 |
| rs121909500 | pathogenic | 1 | 1 | 1 |
| rs140993310 | benign | 1 | 1 | 1 |
| rs200837270 | pathogenic | 1 | 1 | 1 |
| rs121918484 | pathogenic | 1 | 1 | 1 |
| rs121918306 | pathogenic | 0 | 0 | 0 |
| rs118204089 | pathogenic | 1 | 1 | 0 |
| rs118204083 | pathogenic | 1 | 1 | 0 |
| rs201953584 | pathogenic | 1 | 1 | 1 |
| rs74315362 | pathogenic | 1 | 1 | 1 |

| | | | | |
|---|---|---|---|---|
| rs118204084 | pathogenic | 1 | 1 | 0 |
| rs118204085 | pathogenic | 1 | 1 | 1 |
| rs113403872 | pathogenic | 1 | 1 | 0 |
| rs1052176 | benign | 1 | 1 | 1 |
| rs2031640 | benign | 0 | 0 | 0 |
| rs104894230 | pathogenic | 1 | 1 | 1 |
| rs104894229 | pathogenic | 1 | 1 | 1 |
| rs104894228 | pathogenic | 1 | 1 | 1 |
| rs104894226 | pathogenic | 1 | 1 | 1 |
| rs121917757 | pathogenic | 1 | 1 | 1 |
| rs121917758 | pathogenic | 1 | 1 | 1 |
| rs28933406 | pathogenic | 1 | 1 | 1 |
| rs121917756 | pathogenic | 1 | 1 | 1 |
| rs104894227 | pathogenic | 1 | 1 | 1 |
| rs369106578 | pathogenic | 1 | 1 | 1 |
| rs397517142 | benign | 1 | 1 | 1 |
| rs104894231 | pathogenic | 1 | 1 | 1 |
| rs121917759 | pathogenic | 1 | 1 | 1 |
| rs104893827 | pathogenic | 0 | 0 | 0 |
| rs104893826 | pathogenic | 1 | 1 | 1 |
| rs5030802 | pathogenic | 1 | 1 | 1 |
| rs5030827 | pathogenic | 1 | 1 | 0 |
| rs119103277 | pathogenic | 1 | 1 | 1 |
| rs5030808 | pathogenic | 0 | 1 | 0 |
| rs5030809 | pathogenic | 1 | 1 | 1 |
| rs765978945 | pathogenic | 1 | 1 | 1 |
| rs104893824 | pathogenic | 1 | 1 | 1 |
| rs104893831 | pathogenic | 0 | 1 | 0 |
| rs104893830 | pathogenic | 0 | 1 | 0 |
| rs119103278 | pathogenic | 0 | 1 | 0 |
| rs5030818 | pathogenic | 1 | 1 | 1 |
| rs397516444 | pathogenic | 1 | 1 | 1 |
| rs28940297 | pathogenic | 1 | 1 | 1 |
| rs267607170 | pathogenic | 1 | 1 | 1 |
| rs104893825 | pathogenic | 1 | 1 | 1 |
| rs5030820 | pathogenic | 0 | 1 | 0 |
| rs5030821 | pathogenic | 0 | 1 | 0 |
| rs5030823 | pathogenic | 1 | 1 | 1 |
| rs5030824 | pathogenic | 1 | 1 | 1 |
| rs28940301 | pathogenic | 0 | 1 | 0 |
| rs28940300 | pathogenic | 0 | 1 | 0 |
| rs28940298 | pathogenic | 0 | 1 | 0 |
| rs200019083 | benign | 0 | 0 | 0 |
| rs759748655 | pathogenic | 0 | 0 | 0 |
| rs121917739 | pathogenic | 1 | 1 | 1 |
| rs75353611 | pathogenic | 1 | 1 | 1 |
| rs79744198 | benign | 1 | 1 | 1 |
| rs79228041 | pathogenic | 1 | 1 | 1 |
| rs80106970 | benign | 1 | 1 | 1 |
| rs375353223 | benign | 0 | 0 | 0 |
| rs56311081 | benign | 0 | 0 | 0 |
| rs55985569 | benign | 1 | 1 | 1 |
| rs45483396 | benign | 0 | 0 | 0 |
| rs35284565 | benign | 1 | 1 | 1 |

| | | | | |
|---|---|---|---|---|
| rs34349517 | benign | 0 | 0 | 0 |
| rs35776110 | benign | 0 | 0 | 0 |
| rs33917957 | benign | 1 | 1 | 1 |
| rs587778445 | benign | 0 | 0 | 0 |
| rs34624984 | benign | 0 | 0 | 0 |
| rs701753 | benign | 0 | 0 | 0 |
| rs855314 | benign | 0 | 0 | 0 |
| rs121918371 | pathogenic | 1 | 1 | 0 |
| rs398122912 | pathogenic | 0 | 1 | 0 |
| rs1126728 | benign | 0 | 0 | 0 |
| rs587777404 | pathogenic | 1 | 1 | 0 |
| rs11208257 | benign | 0 | 0 | 0 |
| rs6676290 | benign | 0 | 0 | 0 |
| rs104894118 | pathogenic | 0 | 0 | 0 |
| rs121918655 | pathogenic | 1 | 1 | 1 |
| rs104894120 | pathogenic | 1 | 1 | 1 |
| rs137852238 | pathogenic | 0 | 0 | 0 |
| rs137852237 | pathogenic | 0 | 0 | 0 |
| rs137852499 | pathogenic | 1 | 1 | 1 |
| rs137852480 | pathogenic | 1 | 1 | 1 |
| rs137852494 | pathogenic | 1 | 1 | 1 |
| rs137852502 | pathogenic | 1 | 1 | 1 |
| rs137852500 | pathogenic | 1 | 1 | 1 |
| rs137852487 | pathogenic | 1 | 1 | 1 |
| rs137852481 | pathogenic | 1 | 1 | 1 |
| rs137852501 | pathogenic | 0 | 0 | 0 |
| rs137852478 | pathogenic | 1 | 1 | 1 |
| rs137852485 | pathogenic | 1 | 1 | 1 |
| rs137852482 | pathogenic | 1 | 1 | 0 |
| rs137852483 | pathogenic | 0 | 1 | 0 |
| rs137852477 | pathogenic | 1 | 1 | 0 |
| rs137852496 | pathogenic | 1 | 1 | 0 |
| rs137852484 | pathogenic | 0 | 1 | 0 |
| rs137852498 | pathogenic | 1 | 1 | 0 |
| rs137852497 | pathogenic | 1 | 1 | 0 |
| rs137852493 | pathogenic | 1 | 1 | 0 |
| rs137852504 | pathogenic | 1 | 1 | 1 |
| rs267606863 | pathogenic | 1 | 1 | 1 |
| rs137852486 | pathogenic | 1 | 1 | 1 |
| rs137852479 | pathogenic | 1 | 1 | 1 |
| rs112122950 | benign | 0 | 0 | 0 |
| rs72546648 | benign | 1 | 1 | 1 |
| rs72552291 | pathogenic | 0 | 0 | 0 |
| rs79184941 | pathogenic | 1 | 1 | 1 |
| rs77543610 | pathogenic | 1 | 1 | 1 |
| rs121918505 | pathogenic | 1 | 1 | 1 |
| rs776587763 | pathogenic | 1 | 1 | 1 |
| rs121918497 | pathogenic | 1 | 1 | 1 |
| rs121918499 | pathogenic | 1 | 1 | 1 |
| rs121918501 | pathogenic | 1 | 1 | 1 |
| rs121918500 | pathogenic | 0 | 0 | 0 |
| rs121918504 | pathogenic | 1 | 1 | 1 |
| rs121918510 | pathogenic | 1 | 1 | 1 |
| rs121918493 | pathogenic | 1 | 1 | 1 |

| | | | | |
|---|---|---|---|---|
| rs121918489 | pathogenic | 0 | 0 | 0 |
| rs121918495 | pathogenic | 1 | 1 | 1 |
| rs121918488 | pathogenic | 0 | 0 | 0 |
| rs121918496 | pathogenic | 0 | 0 | 0 |
| rs121918492 | pathogenic | 1 | 1 | 1 |
| rs121918494 | pathogenic | 1 | 1 | 1 |
| rs121918502 | pathogenic | 1 | 1 | 1 |
| rs121918490 | pathogenic | 1 | 1 | 1 |
| rs72466451 | pathogenic | 1 | 1 | 1 |
| rs66468541 | pathogenic | 0 | 0 | 0 |
| rs137852662 | pathogenic | 0 | 0 | 0 |
| rs137852663 | pathogenic | 0 | 0 | 0 |
| rs606231407 | pathogenic | 0 | 0 | 0 |
| rs137852233 | pathogenic | 1 | 1 | 1 |
| rs112306225 | benign | 0 | 0 | 0 |
| rs387906808 | pathogenic | 1 | 1 | 0 |
| rs730882094 | pathogenic | 1 | 1 | 1 |
| rs373869746 | pathogenic | 1 | 1 | 1 |
| rs777524402 | pathogenic | 1 | 1 | 1 |
| rs769737896 | pathogenic | 0 | 1 | 0 |
| rs13306515 | benign | 1 | 1 | 0 |
| rs746982741 | pathogenic | 0 | 0 | 0 |
| rs2230200 | benign | 1 | 1 | 1 |
| rs1047286 | benign | 0 | 0 | 0 |
| rs141737564 | benign | 0 | 0 | 0 |
| rs774826179 | benign | 0 | 0 | 0 |
| rs11569422 | benign | 1 | 1 | 1 |
| rs146613648 | benign | 0 | 0 | 0 |
| rs144432231 | benign | 0 | 0 | 0 |
| rs140655115 | benign | 0 | 0 | 0 |
| rs387906837 | pathogenic | 1 | 1 | 0 |
| rs121909076 | pathogenic | 1 | 1 | 0 |
| rs121909077 | pathogenic | 0 | 0 | 0 |
| rs121909074 | pathogenic | 0 | 0 | 0 |
| rs137853330 | pathogenic | 1 | 1 | 1 |
| rs137853331 | pathogenic | 1 | 1 | 1 |
| rs137853332 | pathogenic | 1 | 1 | 1 |
| rs28931586 | pathogenic | 1 | 1 | 0 |
| rs11569321 | benign | 1 | 1 | 1 |
| rs119486097 | benign | 0 | 0 | 0 |
| rs1799807 | pathogenic | 1 | 1 | 1 |
| rs201820739 | pathogenic | 1 | 1 | 0 |
| rs114706984 | pathogenic | 1 | 1 | 0 |
| rs140080572 | pathogenic | 1 | 1 | 0 |
| rs28933389 | pathogenic | 0 | 1 | 0 |
| rs16849700 | benign | 1 | 1 | 0 |
| rs115624085 | pathogenic | 1 | 1 | 1 |
| rs747196387 | pathogenic | 1 | 1 | 1 |
| rs104893684 | pathogenic | 1 | 1 | 0 |
| rs121918557 | pathogenic | 1 | 1 | 0 |
| rs121918556 | pathogenic | 1 | 1 | 1 |
| rs398124571 | pathogenic | 1 | 1 | 1 |
| rs776631396 | pathogenic | 0 | 1 | 0 |
| rs398124576 | pathogenic | 1 | 1 | 1 |

| | | | | |
|---|---|---|---|---|
| rs398124581 | pathogenic | 1 | 1 | 1 |
| rs398124582 | pathogenic | 1 | 1 | 1 |
| rs149766077 | pathogenic | 0 | 1 | 0 |
| rs79761867 | pathogenic | 0 | 1 | 0 |
| rs148905512 | pathogenic | 0 | 1 | 0 |
| rs121965004 | pathogenic | 1 | 1 | 1 |
| rs386834233 | pathogenic | 0 | 1 | 0 |
| rs398124598 | pathogenic | 0 | 1 | 0 |
| rs398124603 | pathogenic | 1 | 1 | 1 |
| rs546716073 | benign | 1 | 1 | 1 |
| rs200310662 | benign | 1 | 1 | 1 |
| rs115494727 | benign | 1 | 1 | 1 |
| rs781513008 | benign | 0 | 0 | 0 |
| rs587776398 | benign | 1 | 1 | 0 |
| rs35288908 | benign | 1 | 1 | 1 |
| rs150634562 | pathogenic | 1 | 1 | 0 |
| rs142948132 | pathogenic | 1 | 1 | 1 |
| rs132630321 | pathogenic | 1 | 1 | 1 |
| rs132630317 | pathogenic | 1 | 1 | 1 |
| rs132630320 | pathogenic | 0 | 0 | 0 |
| rs781394318 | pathogenic | 1 | 1 | 1 |
| rs111033624 | pathogenic | 1 | 1 | 1 |
| rs111033630 | pathogenic | 1 | 1 | 1 |
| rs111033627 | pathogenic | 1 | 1 | 1 |
| rs193922287 | pathogenic | 1 | 1 | 1 |
| rs759072800 | pathogenic | 0 | 1 | 0 |
| rs104894010 | pathogenic | 0 | 0 | 0 |
| rs193922296 | pathogenic | 0 | 0 | 0 |
| rs104894006 | pathogenic | 1 | 1 | 1 |
| rs193922311 | pathogenic | 1 | 1 | 0 |
| rs80356654 | pathogenic | 1 | 1 | 0 |
| rs144723656 | benign | 1 | 1 | 1 |
| rs148311934 | pathogenic | 1 | 1 | 0 |
| rs80356655 | pathogenic | 1 | 1 | 1 |
| rs769268803 | pathogenic | 1 | 1 | 0 |
| rs104894008 | pathogenic | 0 | 1 | 0 |
| rs104894005 | pathogenic | 1 | 1 | 1 |
| rs104894009 | pathogenic | 1 | 1 | 0 |
| rs193922340 | pathogenic | 0 | 0 | 0 |
| rs104894016 | pathogenic | 0 | 1 | 0 |
| rs193921338 | pathogenic | 0 | 0 | 0 |
| rs193922272 | pathogenic | 1 | 1 | 1 |
| rs104894012 | pathogenic | 1 | 1 | 0 |
| rs193922285 | pathogenic | 1 | 1 | 1 |
| rs104894336 | pathogenic | 1 | 1 | 1 |
| rs28931580 | pathogenic | 1 | 1 | 1 |
| rs104894326 | pathogenic | 1 | 1 | 1 |
| rs104894331 | pathogenic | 0 | 0 | 0 |
| rs104894338 | pathogenic | 0 | 0 | 0 |
| rs104894333 | pathogenic | 1 | 1 | 1 |
| rs104894330 | pathogenic | 1 | 1 | 1 |
| rs143886391 | benign | 1 | 1 | 1 |
| rs104894335 | pathogenic | 1 | 1 | 1 |
| rs104894337 | pathogenic | 0 | 0 | 0 |

| | | | | |
|---|---|---|---|---|
| rs104894328 | pathogenic | 0 | 0 | 0 |
| rs104894341 | pathogenic | 1 | 1 | 0 |
| rs104894329 | pathogenic | 0 | 0 | 0 |
| rs137852850 | pathogenic | 1 | 1 | 1 |
| rs137852854 | pathogenic | 0 | 1 | 0 |
| rs137852849 | pathogenic | 0 | 1 | 0 |
| rs137852848 | pathogenic | 1 | 1 | 1 |
| rs137852852 | pathogenic | 0 | 1 | 0 |
| rs137852853 | pathogenic | 0 | 1 | 0 |
| rs137852847 | pathogenic | 0 | 1 | 0 |
| rs137852846 | pathogenic | 0 | 1 | 0 |
| rs137852844 | pathogenic | 0 | 1 | 0 |
| rs144411158 | pathogenic | 0 | 0 | 0 |
| rs2302234 | benign | 0 | 0 | 0 |
| rs139620139 | pathogenic | 0 | 0 | 0 |
| rs146689929 | benign | 0 | 0 | 0 |
| rs373328681 | pathogenic | 1 | 1 | 0 |
| rs387906620 | pathogenic | 1 | 1 | 1 |
| rs1799983 | benign | 0 | 1 | 0 |
| rs28999111 | pathogenic | 1 | 1 | 1 |
| rs104894544 | pathogenic | 1 | 1 | 1 |
| rs104894546 | pathogenic | 1 | 1 | 1 |
| rs28999112 | pathogenic | 0 | 1 | 0 |
| rs104894545 | pathogenic | 1 | 1 | 1 |
| rs104894547 | pathogenic | 1 | 1 | 1 |
| rs74315310 | pathogenic | 0 | 0 | 0 |
| rs267606852 | pathogenic | 0 | 1 | 0 |
| rs137853586 | pathogenic | 0 | 1 | 0 |
| rs137853582 | pathogenic | 1 | 1 | 1 |
| rs61754634 | pathogenic | 1 | 1 | 1 |
| rs137853587 | pathogenic | 0 | 1 | 0 |
| rs267606851 | pathogenic | 1 | 1 | 1 |
| rs267606853 | pathogenic | 1 | 1 | 1 |
| rs137853584 | pathogenic | 1 | 1 | 1 |
| rs137853585 | pathogenic | 1 | 1 | 1 |
| rs460184 | pathogenic | 1 | 1 | 1 |
| rs104893751 | pathogenic | 0 | 1 | 0 |
| rs77358162 | benign | 0 | 0 | 0 |
| rs387906936 | pathogenic | 0 | 0 | 0 |
| rs769525399 | pathogenic | 0 | 0 | 0 |
| rs371271054 | pathogenic | 1 | 1 | 0 |
| rs118204000 | pathogenic | 0 | 0 | 0 |
| rs138568622 | pathogenic | 1 | 1 | 1 |
| rs34509359 | benign | 0 | 0 | 0 |
| rs104894130 | pathogenic | 1 | 1 | 0 |
| rs281865424 | pathogenic | 0 | 0 | 0 |
| rs121912778 | pathogenic | 1 | 1 | 0 |
| rs771055145 | pathogenic | 1 | 1 | 0 |
| rs776174514 | pathogenic | 1 | 1 | 0 |
| rs545986367 | pathogenic | 1 | 1 | 1 |
| rs570768038 | benign | 1 | 1 | 1 |
| rs74315442 | pathogenic | 1 | 1 | 1 |
| rs79716074 | benign | 1 | 1 | 1 |
| rs748486078 | pathogenic | 1 | 1 | 1 |

| | | | | |
|---|---|---|---|---|
| rs2397084 | benign | 1 | 1 | 1 |
| rs11465553 | benign | 1 | 1 | 1 |
| rs144576902 | benign | 1 | 1 | 1 |
| rs763780 | benign | 1 | 1 | 1 |
| rs267606870 | pathogenic | 1 | 1 | 0 |
| rs121913502 | pathogenic | 1 | 1 | 0 |
| rs118101777 | benign | 1 | 1 | 1 |
| rs118053940 | benign | 0 | 0 | 0 |
| rs368771578 | benign | 1 | 1 | 1 |
| rs11540654 | pathogenic | 1 | 1 | 1 |
| rs121912658 | pathogenic | 1 | 1 | 1 |
| rs28934873 | pathogenic | 0 | 1 | 0 |
| rs780442292 | pathogenic | 1 | 1 | 1 |
| rs758781593 | benign | 1 | 1 | 1 |
| rs28934875 | pathogenic | 1 | 1 | 1 |
| rs28934874 | pathogenic | 1 | 1 | 0 |
| rs587782705 | pathogenic | 1 | 1 | 1 |
| rs587780068 | pathogenic | 1 | 1 | 1 |
| rs587782144 | pathogenic | 1 | 1 | 1 |
| rs138729528 | pathogenic | 1 | 1 | 1 |
| rs28934578 | pathogenic | 1 | 1 | 1 |
| rs397514495 | pathogenic | 1 | 1 | 1 |
| rs121912665 | pathogenic | 1 | 1 | 1 |
| rs397516435 | pathogenic | 1 | 1 | 1 |
| rs587778720 | pathogenic | 1 | 1 | 1 |
| rs397516436 | pathogenic | 1 | 1 | 1 |
| rs587781386 | benign | 1 | 1 | 1 |
| rs121912666 | pathogenic | 1 | 1 | 0 |
| rs587780073 | pathogenic | 1 | 1 | 1 |
| rs587782664 | pathogenic | 1 | 1 | 1 |
| rs730882005 | pathogenic | 1 | 1 | 1 |
| rs28934573 | pathogenic | 1 | 1 | 1 |
| rs121912655 | pathogenic | 1 | 1 | 1 |
| rs28934575 | pathogenic | 1 | 1 | 1 |
| rs121912656 | pathogenic | 1 | 1 | 1 |
| rs11540652 | pathogenic | 1 | 1 | 1 |
| rs121912651 | pathogenic | 1 | 1 | 1 |
| rs730882027 | pathogenic | 1 | 1 | 1 |
| rs121912653 | pathogenic | 1 | 1 | 1 |
| rs121912652 | pathogenic | 1 | 1 | 1 |
| rs121912657 | pathogenic | 0 | 1 | 0 |
| rs121913343 | pathogenic | 1 | 1 | 1 |
| rs28934576 | pathogenic | 1 | 1 | 1 |
| rs763098116 | pathogenic | 1 | 1 | 1 |
| rs121912660 | pathogenic | 1 | 1 | 1 |
| rs28934574 | pathogenic | 1 | 1 | 1 |
| rs371409680 | pathogenic | 1 | 1 | 1 |
| rs121912667 | pathogenic | 1 | 1 | 1 |
| rs121912663 | pathogenic | 1 | 1 | 1 |
| rs201744589 | pathogenic | 0 | 0 | 0 |
| rs33926449 | benign | 0 | 0 | 0 |
| rs74315302 | pathogenic | 0 | 1 | 0 |
| rs202040269 | pathogenic | 1 | 1 | 1 |
| rs375958814 | pathogenic | 1 | 1 | 1 |

| | | | | |
|---|---|---|---|---|
| rs121909624 | pathogenic | 1 | 1 | 1 |
| rs121909621 | pathogenic | 1 | 1 | 1 |
| rs121909622 | pathogenic | 1 | 1 | 0 |
| rs4220 | benign | 1 | 1 | 1 |
| rs147559466 | pathogenic | 1 | 1 | 1 |
| rs757434857 | pathogenic | 1 | 1 | 1 |
| rs398123072 | pathogenic | 1 | 1 | 1 |
| rs121434280 | pathogenic | 0 | 1 | 0 |
| rs398123074 | pathogenic | 0 | 1 | 0 |
| rs762114560 | pathogenic | 1 | 1 | 1 |
| rs121434283 | pathogenic | 1 | 1 | 0 |
| rs121434277 | pathogenic | 0 | 1 | 0 |
| rs745844469 | pathogenic | 1 | 1 | 1 |
| rs121434279 | pathogenic | 1 | 1 | 0 |
| rs121434278 | pathogenic | 1 | 1 | 1 |
| rs373715782 | pathogenic | 1 | 1 | 1 |
| rs200724875 | pathogenic | 1 | 1 | 1 |
| rs149678400 | pathogenic | 0 | 1 | 0 |
| rs770273135 | pathogenic | 1 | 1 | 1 |
| rs121434276 | pathogenic | 0 | 1 | 0 |
| rs121434281 | pathogenic | 1 | 1 | 0 |
| rs768884003 | pathogenic | 1 | 1 | 1 |
| rs201375579 | pathogenic | 0 | 0 | 0 |
| rs121434274 | pathogenic | 1 | 1 | 0 |
| rs121434282 | pathogenic | 1 | 1 | 0 |
| rs779759347 | pathogenic | 0 | 1 | 0 |
| rs77931234 | pathogenic | 1 | 1 | 1 |
| rs148207467 | pathogenic | 1 | 1 | 1 |
| rs766140986 | pathogenic | 1 | 1 | 1 |
| rs121434275 | pathogenic | 1 | 1 | 1 |
| rs748110745 | pathogenic | 1 | 1 | 1 |
| rs104894274 | pathogenic | 1 | 1 | 1 |
| rs104894273 | pathogenic | 0 | 1 | 0 |
| rs104894278 | pathogenic | 1 | 1 | 1 |
| rs104894275 | pathogenic | 1 | 1 | 1 |
| rs104894277 | pathogenic | 1 | 1 | 1 |
| rs104894276 | pathogenic | 0 | 0 | 0 |
| rs104894280 | pathogenic | 1 | 1 | 1 |
| rs145882709 | pathogenic | 1 | 1 | 1 |
| rs104894279 | pathogenic | 1 | 1 | 1 |
| rs80356670 | pathogenic | 1 | 1 | 1 |
| rs121918102 | pathogenic | 1 | 1 | 1 |
| rs80356671 | pathogenic | 1 | 1 | 1 |
| rs80356672 | pathogenic | 1 | 1 | 1 |
| rs758609113 | pathogenic | 1 | 1 | 0 |
| rs121918188 | pathogenic | 0 | 1 | 0 |
| rs121918189 | pathogenic | 1 | 1 | 0 |
| rs121918191 | pathogenic | 1 | 1 | 1 |
| rs1769259 | benign | 1 | 1 | 1 |
| rs140222720 | benign | 1 | 1 | 1 |
| rs2297085 | benign | 1 | 1 | 1 |
| rs200679026 | pathogenic | 1 | 1 | 1 |
| rs121918192 | pathogenic | 0 | 1 | 0 |
| rs145771944 | benign | 0 | 1 | 0 |

| | | | | |
|---|---|---|---|---|
| rs201508880 | pathogenic | 0 | 1 | 0 |
| rs17221854 | pathogenic | 0 | 0 | 0 |
| rs120074189 | pathogenic | 1 | 1 | 1 |
| rs120074190 | pathogenic | 1 | 1 | 1 |
| rs758600283 | benign | 1 | 1 | 1 |
| rs200305064 | pathogenic | 1 | 1 | 1 |
| rs121917891 | pathogenic | 1 | 1 | 1 |
| rs3772809 | benign | 0 | 1 | 0 |
| rs387906619 | pathogenic | 1 | 1 | 0 |
| rs2304681 | benign | 0 | 1 | 0 |
| rs121917849 | pathogenic | 0 | 0 | 0 |
| rs143010236 | pathogenic | 0 | 0 | 0 |
| rs121917850 | pathogenic | 1 | 1 | 0 |
| rs35916840 | pathogenic | 1 | 1 | 0 |
| rs28939688 | pathogenic | 1 | 1 | 1 |
| rs11258194 | benign | 1 | 1 | 1 |
| rs199840952 | pathogenic | 1 | 1 | 1 |
| rs121918159 | pathogenic | 1 | 1 | 1 |
| rs200396166 | pathogenic | 0 | 0 | 0 |
| rs757966746 | pathogenic | 0 | 1 | 0 |
| rs121918608 | pathogenic | 0 | 1 | 0 |
| rs74315401 | pathogenic | 1 | 1 | 1 |
| rs11538758 | pathogenic | 0 | 0 | 0 |
| rs74315402 | pathogenic | 1 | 1 | 1 |
| rs1799990 | pathogenic | 1 | 1 | 1 |
| rs74315410 | pathogenic | 1 | 1 | 1 |
| rs150351644 | benign | 1 | 1 | 1 |
| rs16990018 | benign | 1 | 1 | 1 |
| rs74315403 | pathogenic | 1 | 1 | 1 |
| rs74315408 | pathogenic | 1 | 1 | 1 |
| rs74315411 | pathogenic | 1 | 1 | 1 |
| rs74315413 | pathogenic | 1 | 1 | 1 |
| rs74315405 | pathogenic | 1 | 1 | 1 |
| rs28933385 | pathogenic | 1 | 1 | 1 |
| rs74315412 | pathogenic | 1 | 1 | 1 |
| rs74315407 | pathogenic | 1 | 1 | 1 |
| rs74315406 | pathogenic | 1 | 1 | 1 |
| rs11571117 | benign | 1 | 1 | 1 |
| rs121917742 | pathogenic | 1 | 1 | 1 |
| rs149782619 | pathogenic | 1 | 1 | 1 |
| rs397514492 | pathogenic | 1 | 1 | 1 |
| rs397514489 | pathogenic | 0 | 1 | 0 |
| rs397514493 | pathogenic | 1 | 1 | 1 |
| rs373849532 | pathogenic | 1 | 1 | 1 |
| rs121965048 | pathogenic | 1 | 1 | 1 |
| rs121965037 | pathogenic | 1 | 1 | 1 |
| rs121965060 | pathogenic | 0 | 1 | 0 |
| rs386833603 | pathogenic | 1 | 1 | 1 |
| rs121965038 | pathogenic | 1 | 1 | 1 |
| rs121965039 | pathogenic | 1 | 1 | 1 |
| rs121965040 | pathogenic | 1 | 1 | 1 |
| rs267606925 | pathogenic | 1 | 1 | 1 |
| rs121965056 | pathogenic | 0 | 1 | 0 |
| rs121965059 | pathogenic | 0 | 0 | 0 |

| | | | | |
|---|---|---|---|---|
| rs386833615 | pathogenic | 1 | 1 | 0 |
| rs121965051 | pathogenic | 1 | 1 | 0 |
| rs121965046 | pathogenic | 0 | 1 | 0 |
| rs121965052 | pathogenic | 0 | 1 | 0 |
| rs386833617 | pathogenic | 0 | 1 | 0 |
| rs386833618 | pathogenic | 1 | 1 | 0 |
| rs121965041 | pathogenic | 0 | 1 | 0 |
| rs121965042 | pathogenic | 0 | 1 | 0 |
| rs121965057 | pathogenic | 1 | 1 | 1 |
| rs386833621 | pathogenic | 1 | 1 | 1 |
| rs121965049 | pathogenic | 1 | 1 | 1 |
| rs386833623 | pathogenic | 1 | 1 | 1 |
| rs121965047 | pathogenic | 1 | 1 | 1 |
| rs121965053 | pathogenic | 0 | 1 | 0 |
| rs121965045 | pathogenic | 0 | 1 | 0 |
| rs121965054 | pathogenic | 0 | 1 | 0 |
| rs121965043 | pathogenic | 0 | 1 | 0 |
| rs386833598 | pathogenic | 0 | 0 | 0 |
| rs137854545 | pathogenic | 1 | 1 | 1 |
| rs137854544 | pathogenic | 1 | 1 | 1 |
| rs104894455 | pathogenic | 0 | 1 | 0 |
| rs1049564 | benign | 0 | 1 | 0 |
| rs104894460 | pathogenic | 1 | 1 | 1 |
| rs104894453 | pathogenic | 1 | 1 | 1 |
| rs104894450 | pathogenic | 0 | 1 | 0 |
| rs104894454 | pathogenic | 0 | 1 | 0 |
| rs104894452 | pathogenic | 1 | 1 | 1 |
| rs104894451 | pathogenic | 0 | 1 | 0 |
| rs104894368 | pathogenic | 1 | 1 | 1 |
| rs730880950 | pathogenic | 1 | 1 | 1 |
| rs121909535 | pathogenic | 1 | 1 | 1 |
| rs121909537 | pathogenic | 1 | 1 | 1 |
| rs121909536 | pathogenic | 1 | 1 | 1 |
| rs121909542 | pathogenic | 1 | 1 | 1 |
| rs121909538 | pathogenic | 1 | 1 | 1 |
| rs121909539 | pathogenic | 1 | 1 | 1 |
| rs121909540 | pathogenic | 1 | 1 | 1 |
| rs17560 | benign | 1 | 1 | 1 |
| rs121909543 | pathogenic | 0 | 1 | 0 |
| rs121909544 | pathogenic | 1 | 1 | 1 |
| rs2307456 | benign | 0 | 1 | 0 |
| rs745778317 | pathogenic | 0 | 1 | 0 |
| rs759607901 | pathogenic | 0 | 1 | 0 |
| rs201188114 | benign | 0 | 0 | 0 |
| rs1800472 | benign | 0 | 0 | 0 |
| rs587780728 | benign | 1 | 1 | 1 |
| rs144386518 | benign | 1 | 1 | 1 |
| rs281865439 | pathogenic | 1 | 1 | 1 |
| rs140175796 | pathogenic | 1 | 1 | 1 |
| rs5030843 | pathogenic | 1 | 1 | 0 |
| rs75166491 | pathogenic | 1 | 1 | 0 |
| rs199475679 | pathogenic | 1 | 1 | 1 |
| rs281865440 | pathogenic | 1 | 1 | 1 |
| rs74486803 | pathogenic | 0 | 1 | 0 |

| | | | | |
|---|---|---|---|---|
| rs199475602 | pathogenic | 0 | 1 | 0 |
| rs5030844 | pathogenic | 1 | 1 | 0 |
| rs62514927 | pathogenic | 1 | 1 | 1 |
| rs62514928 | pathogenic | 1 | 1 | 1 |
| rs62516152 | pathogenic | 0 | 1 | 0 |
| rs62507348 | pathogenic | 1 | 1 | 1 |
| rs76687508 | pathogenic | 1 | 1 | 1 |
| rs62508730 | pathogenic | 1 | 1 | 1 |
| rs62508588 | pathogenic | 0 | 1 | 0 |
| rs5030846 | pathogenic | 0 | 1 | 0 |
| rs74503222 | pathogenic | 1 | 1 | 0 |
| rs5030847 | pathogenic | 1 | 1 | 0 |
| rs5030850 | pathogenic | 1 | 1 | 1 |
| rs778154939 | pathogenic | 0 | 1 | 0 |
| rs62514952 | pathogenic | 1 | 1 | 1 |
| rs78655458 | pathogenic | 1 | 1 | 1 |
| rs62508698 | pathogenic | 1 | 1 | 0 |
| rs5030851 | pathogenic | 1 | 1 | 0 |
| rs62642933 | pathogenic | 1 | 1 | 1 |
| rs5030853 | pathogenic | 1 | 1 | 1 |
| rs62642935 | pathogenic | 1 | 1 | 1 |
| rs199475615 | pathogenic | 0 | 0 | 0 |
| rs62516062 | pathogenic | 1 | 1 | 0 |
| rs62516092 | pathogenic | 0 | 0 | 0 |
| rs62516095 | pathogenic | 1 | 1 | 1 |
| rs62642937 | pathogenic | 1 | 1 | 1 |
| rs62516141 | pathogenic | 0 | 1 | 0 |
| rs5030856 | pathogenic | 1 | 1 | 1 |
| rs5030857 | pathogenic | 1 | 1 | 1 |
| rs5030858 | pathogenic | 1 | 1 | 1 |
| rs79931499 | pathogenic | 1 | 1 | 1 |
| rs5030860 | pathogenic | 1 | 1 | 1 |
| rs281865437 | pathogenic | 1 | 1 | 1 |
| rs62644499 | pathogenic | 1 | 1 | 1 |
| rs147689487 | benign | 0 | 0 | 0 |
| rs397515417 | pathogenic | 1 | 1 | 1 |
| rs121909332 | pathogenic | 1 | 1 | 1 |
| rs121909330 | pathogenic | 1 | 1 | 0 |
| rs121909329 | pathogenic | 1 | 1 | 0 |
| rs121909335 | pathogenic | 1 | 1 | 1 |
| rs387906789 | pathogenic | 1 | 1 | 1 |
| rs121909334 | pathogenic | 1 | 1 | 1 |
| rs121909331 | pathogenic | 1 | 1 | 1 |
| rs372487178 | pathogenic | 1 | 1 | 1 |
| rs387906937 | pathogenic | 1 | 1 | 0 |
| rs16845 | benign | 1 | 1 | 1 |
| rs672601313 | pathogenic | 1 | 1 | 1 |
| rs117805308 | benign | 0 | 0 | 0 |
| rs121434261 | pathogenic | 1 | 1 | 1 |
| rs141629512 | benign | 0 | 0 | 0 |
| rs199957176 | benign | 1 | 1 | 1 |
| rs2230461 | benign | 1 | 1 | 1 |
| rs764618040 | pathogenic | 1 | 1 | 1 |
| rs267606949 | pathogenic | 1 | 1 | 1 |

| | | | | |
|---|---|---|---|---|
| rs587777483 | pathogenic | 1 | 1 | 1 |
| rs267606947 | pathogenic | 1 | 1 | 1 |
| rs6941583 | benign | 0 | 0 | 0 |
| rs749597959 | benign | 1 | 1 | 1 |
| rs182340753 | benign | 1 | 1 | 1 |
| rs146678221 | benign | 0 | 0 | 0 |
| rs11539202 | benign | 0 | 0 | 0 |
| rs148723565 | benign | 0 | 0 | 0 |
| rs61752925 | benign | 1 | 1 | 1 |
| rs113309941 | pathogenic | 0 | 0 | 0 |
| rs121434520 | pathogenic | 1 | 1 | 1 |
| rs121434517 | pathogenic | 1 | 1 | 1 |
| rs267607027 | pathogenic | 1 | 1 | 1 |
| rs138603088 | pathogenic | 0 | 0 | 0 |
| rs121434516 | pathogenic | 1 | 1 | 0 |
| rs121434521 | pathogenic | 0 | 0 | 0 |
| rs145735112 | benign | 1 | 1 | 1 |
| rs121434519 | pathogenic | 1 | 1 | 1 |
| rs75326924 | pathogenic | 1 | 1 | 0 |
| rs142186404 | pathogenic | 0 | 0 | 0 |
| rs121918035 | pathogenic | 0 | 0 | 0 |
| rs374385878 | pathogenic | 1 | 1 | 0 |
| rs11557652 | benign | 0 | 0 | 0 |
| rs118203986 | pathogenic | 1 | 1 | 1 |
| rs141675330 | benign | 1 | 1 | 1 |
| rs104893927 | pathogenic | 0 | 0 | 0 |
| rs104893935 | pathogenic | 0 | 0 | 0 |
| rs104893933 | pathogenic | 1 | 1 | 0 |
| rs104893934 | pathogenic | 0 | 0 | 0 |
| rs145183203 | benign | 1 | 1 | 0 |
| rs2230806 | benign | 0 | 0 | 0 |
| rs9282541 | benign | 0 | 0 | 0 |
| rs559110055 | benign | 0 | 0 | 0 |
| rs200030513 | benign | 0 | 0 | 0 |
| rs9282543 | benign | 0 | 0 | 0 |
| rs147675550 | benign | 0 | 0 | 0 |
| rs137854496 | pathogenic | 0 | 0 | 0 |
| rs374190304 | benign | 0 | 0 | 0 |
| rs114620717 | benign | 0 | 0 | 0 |
| rs2066718 | benign | 0 | 0 | 0 |
| rs35819696 | benign | 0 | 0 | 0 |
| rs138880920 | benign | 0 | 0 | 0 |
| rs2066715 | benign | 0 | 0 | 0 |
| rs35207495 | benign | 0 | 0 | 0 |
| rs2066714 | benign | 0 | 0 | 0 |
| rs28937313 | pathogenic | 0 | 0 | 0 |
| rs28937314 | pathogenic | 0 | 0 | 0 |
| rs137854495 | pathogenic | 0 | 0 | 0 |
| rs140365800 | benign | 0 | 0 | 0 |
| rs13306073 | benign | 0 | 0 | 0 |
| rs28933692 | pathogenic | 0 | 0 | 0 |
| rs33918808 | benign | 0 | 0 | 0 |
| rs76881554 | benign | 0 | 0 | 0 |
| rs143180998 | benign | 0 | 0 | 0 |

| | | | | |
|---|---|---|---|---|
| rs137854500 | pathogenic | 0 | 0 | 0 |
| rs538614702 | benign | 0 | 0 | 0 |
| rs201879964 | benign | 0 | 0 | 0 |
| rs2230808 | benign | 0 | 0 | 0 |
| rs138422574 | benign | 0 | 0 | 0 |
| rs137854498 | pathogenic | 0 | 0 | 0 |
| rs150125857 | benign | 0 | 0 | 0 |
| rs146292819 | pathogenic | 0 | 0 | 0 |
| rs142688906 | benign | 0 | 0 | 0 |
| rs137854499 | pathogenic | 0 | 0 | 0 |
| rs34879708 | benign | 0 | 0 | 0 |
| rs121913028 | pathogenic | 1 | 1 | 1 |
| rs1041951 | benign | 1 | 1 | 0 |
| rs138714094 | benign | 1 | 1 | 0 |
| rs200650502 | benign | 0 | 1 | 0 |
| rs370708663 | pathogenic | 1 | 1 | 1 |
| rs118204040 | pathogenic | 0 | 1 | 0 |
| rs146899669 | pathogenic | 1 | 1 | 0 |
| rs118204039 | pathogenic | 0 | 1 | 0 |
| rs121909233 | pathogenic | 0 | 0 | 0 |
| rs121909225 | pathogenic | 1 | 1 | 0 |
| rs121909236 | pathogenic | 0 | 0 | 0 |
| rs398123317 | pathogenic | 0 | 0 | 0 |
| rs121909226 | pathogenic | 1 | 1 | 0 |
| rs121909238 | pathogenic | 1 | 1 | 0 |
| rs121909230 | pathogenic | 0 | 0 | 0 |
| rs121909237 | pathogenic | 0 | 0 | 0 |
| rs121909222 | pathogenic | 1 | 1 | 0 |
| rs121909223 | pathogenic | 1 | 1 | 0 |
| rs121909218 | pathogenic | 1 | 1 | 0 |
| rs121909229 | pathogenic | 1 | 1 | 0 |
| rs121909224 | pathogenic | 1 | 1 | 0 |
| rs121909241 | pathogenic | 0 | 0 | 0 |
| rs121909221 | pathogenic | 0 | 0 | 0 |
| rs121913294 | pathogenic | 0 | 0 | 0 |
| rs121909234 | pathogenic | 0 | 0 | 0 |
| rs121909240 | pathogenic | 0 | 0 | 0 |
| rs767623493 | benign | 0 | 0 | 0 |
| rs121909239 | pathogenic | 0 | 0 | 0 |
| rs120074145 | pathogenic | 1 | 1 | 1 |
| rs120074148 | pathogenic | 1 | 1 | 1 |
| rs762991875 | pathogenic | 0 | 0 | 0 |
| rs148639841 | pathogenic | 1 | 1 | 1 |
| rs199524907 | pathogenic | 1 | 1 | 1 |
| rs120074141 | pathogenic | 1 | 1 | 1 |
| rs120074144 | pathogenic | 0 | 1 | 0 |
| rs120074146 | pathogenic | 1 | 1 | 1 |
| rs120074147 | pathogenic | 0 | 1 | 0 |
| rs120074143 | pathogenic | 0 | 1 | 0 |
| rs120074140 | pathogenic | 1 | 1 | 0 |
| rs28937315 | pathogenic | 1 | 1 | 1 |
| rs76066357 | benign | 1 | 1 | 1 |
| rs137852911 | pathogenic | 0 | 1 | 0 |
| rs137852907 | pathogenic | 1 | 1 | 0 |

| | | | | |
|---|---|---|---|---|
| rs137852910 | pathogenic | 1 | 1 | 1 |
| rs537585211 | benign | 0 | 0 | 0 |
| rs202134424 | pathogenic | 0 | 0 | 0 |
| rs200930463 | pathogenic | 0 | 0 | 0 |
| rs587777241 | pathogenic | 1 | 1 | 0 |
| rs77563738 | pathogenic | 0 | 0 | 0 |
| rs750868279 | pathogenic | 1 | 1 | 0 |
| rs148936893 | pathogenic | 0 | 0 | 0 |
| rs146597836 | pathogenic | 0 | 0 | 0 |
| rs775440641 | pathogenic | 1 | 1 | 1 |
| rs770689762 | pathogenic | 0 | 1 | 0 |
| rs376785840 | pathogenic | 0 | 0 | 0 |
| rs137852531 | pathogenic | 0 | 0 | 0 |
| rs137852538 | pathogenic | 1 | 1 | 0 |
| rs137852529 | pathogenic | 0 | 1 | 0 |
| rs137852528 | pathogenic | 0 | 1 | 0 |
| rs137852535 | pathogenic | 1 | 1 | 0 |
| rs137853045 | pathogenic | 0 | 1 | 0 |
| rs137853046 | pathogenic | 1 | 1 | 1 |
| rs137853043 | pathogenic | 1 | 1 | 1 |
| rs137853044 | pathogenic | 1 | 1 | 1 |
| rs137853047 | pathogenic | 1 | 1 | 1 |
| rs267606990 | pathogenic | 1 | 1 | 1 |
| rs751437780 | pathogenic | 1 | 1 | 0 |
| rs121918461 | pathogenic | 1 | 1 | 1 |
| rs121918460 | pathogenic | 1 | 1 | 1 |
| rs121918459 | pathogenic | 1 | 1 | 0 |
| rs397507511 | pathogenic | 1 | 1 | 1 |
| rs121918454 | pathogenic | 1 | 1 | 1 |
| rs121918453 | pathogenic | 1 | 1 | 1 |
| rs121918462 | pathogenic | 1 | 1 | 0 |
| rs121918465 | pathogenic | 1 | 1 | 0 |
| rs121918464 | pathogenic | 1 | 1 | 0 |
| rs121918466 | pathogenic | 1 | 1 | 1 |
| rs397507521 | pathogenic | 1 | 1 | 1 |
| rs397507527 | pathogenic | 1 | 1 | 1 |
| rs777603059 | pathogenic | 0 | 1 | 0 |
| rs121918456 | pathogenic | 1 | 1 | 1 |
| rs121918463 | pathogenic | 1 | 1 | 1 |
| rs121918455 | pathogenic | 0 | 1 | 0 |
| rs28933386 | pathogenic | 0 | 1 | 0 |
| rs774356443 | benign | 1 | 1 | 1 |
| rs121918467 | pathogenic | 1 | 1 | 1 |
| rs121918468 | pathogenic | 1 | 1 | 1 |
| rs121918469 | pathogenic | 1 | 1 | 1 |
| rs397507539 | pathogenic | 1 | 1 | 0 |
| rs121918458 | pathogenic | 1 | 1 | 1 |
| rs121918470 | pathogenic | 1 | 1 | 1 |
| rs35289681 | benign | 0 | 1 | 0 |
| rs4302331 | benign | 0 | 1 | 0 |
| rs11544803 | pathogenic | 1 | 1 | 1 |
| rs116254156 | benign | 0 | 0 | 0 |
| rs104893909 | pathogenic | 1 | 1 | 1 |
| rs104893911 | pathogenic | 1 | 1 | 1 |

| | | | | |
|---|---|---|---|---|
| rs104893908 | pathogenic | 1 | 1 | 0 |
| rs258751 | benign | 1 | 1 | 1 |
| rs104893914 | pathogenic | 1 | 1 | 0 |
| rs121909727 | pathogenic | 1 | 1 | 1 |
| rs104893910 | pathogenic | 1 | 1 | 1 |
| rs121909726 | pathogenic | 1 | 1 | 1 |
| rs104893912 | pathogenic | 1 | 1 | 1 |
| rs138407155 | benign | 1 | 1 | 1 |
| rs574061789 | benign | 1 | 1 | 1 |
| rs121912718 | pathogenic | 1 | 1 | 1 |
| rs121912719 | pathogenic | 1 | 1 | 1 |
| rs140770089 | benign | 1 | 1 | 1 |
| rs28931573 | pathogenic | 1 | 1 | 1 |
| rs121912717 | pathogenic | 1 | 1 | 1 |
| rs376456598 | pathogenic | 0 | 0 | 0 |
| rs398123499 | pathogenic | 1 | 1 | 1 |
| rs773048903 | pathogenic | 1 | 1 | 0 |
| rs398123503 | pathogenic | 1 | 1 | 1 |
| rs375785084 | pathogenic | 0 | 1 | 0 |
| rs137852874 | pathogenic | 1 | 1 | 1 |
| rs373713279 | pathogenic | 1 | 1 | 1 |
| rs137852873 | pathogenic | 1 | 1 | 0 |
| rs367823977 | pathogenic | 1 | 1 | 1 |
| rs398123508 | pathogenic | 1 | 1 | 1 |
| rs137852871 | pathogenic | 1 | 1 | 1 |
| rs753698250 | pathogenic | 0 | 1 | 0 |
| rs182923857 | pathogenic | 1 | 1 | 1 |
| rs148090804 | benign | 1 | 1 | 1 |
| rs137852870 | pathogenic | 1 | 1 | 1 |
| rs267607131 | pathogenic | 1 | 1 | 0 |
| rs137852902 | pathogenic | 0 | 0 | 0 |
| rs61741646 | benign | 1 | 1 | 1 |
| rs137852905 | pathogenic | 0 | 0 | 0 |
| rs121918681 | pathogenic | 0 | 1 | 0 |
| rs8177238 | benign | 1 | 1 | 0 |
| rs5030776 | pathogenic | 1 | 1 | 1 |
| rs104894572 | pathogenic | 0 | 0 | 0 |
| rs104894576 | pathogenic | 0 | 0 | 0 |
| rs80084721 | pathogenic | 0 | 0 | 0 |
| rs104894146 | pathogenic | 0 | 1 | 0 |
| rs104894138 | pathogenic | 1 | 1 | 0 |
| rs104894153 | pathogenic | 1 | 1 | 0 |
| rs104894135 | pathogenic | 1 | 1 | 0 |
| rs104894147 | pathogenic | 1 | 1 | 0 |
| rs104894148 | pathogenic | 1 | 1 | 0 |
| rs104894154 | pathogenic | 1 | 1 | 0 |
| rs104894150 | pathogenic | 1 | 1 | 0 |
| rs104894136 | pathogenic | 1 | 1 | 0 |
| rs104894144 | pathogenic | 0 | 0 | 0 |
| rs104894137 | pathogenic | 0 | 0 | 0 |
| rs61754278 | pathogenic | 1 | 1 | 0 |
| rs104894149 | pathogenic | 1 | 1 | 0 |
| rs104894139 | pathogenic | 1 | 1 | 0 |
| rs104894142 | pathogenic | 0 | 0 | 0 |

| | | | | |
|---|---|---|---|---|
| rs104894143 | pathogenic | 0 | 0 | 0 |
| rs104894145 | pathogenic | 0 | 0 | 0 |
| rs121909545 | benign | 0 | 1 | 0 |
| rs782308462 | pathogenic | 1 | 1 | 1 |
| rs375002796 | pathogenic | 0 | 1 | 0 |
| rs201727685 | pathogenic | 0 | 0 | 0 |
| rs768050261 | pathogenic | 1 | 1 | 1 |
| rs780851340 | benign | 0 | 0 | 0 |
| rs139824241 | benign | 1 | 1 | 0 |
| rs757486575 | pathogenic | 1 | 1 | 0 |
| rs750831299 | pathogenic | 1 | 1 | 1 |
| rs767602961 | pathogenic | 1 | 1 | 1 |
| rs121913661 | pathogenic | 1 | 1 | 1 |
| rs777692271 | pathogenic | 1 | 1 | 1 |
| rs121913659 | pathogenic | 1 | 1 | 1 |
| rs587777533 | pathogenic | 1 | 1 | 1 |
| rs104894210 | pathogenic | 0 | 0 | 0 |
| rs104894216 | pathogenic | 1 | 1 | 1 |
| rs104894207 | pathogenic | 1 | 1 | 1 |
| rs104894208 | pathogenic | 0 | 1 | 0 |
| rs104894214 | pathogenic | 1 | 1 | 1 |
| rs104894211 | pathogenic | 1 | 1 | 1 |
| rs61749384 | pathogenic | 0 | 0 | 0 |
| rs61750071 | pathogenic | 1 | 1 | 1 |
| rs61750074 | pathogenic | 0 | 0 | 0 |
| rs11063988 | benign | 0 | 0 | 0 |
| rs216311 | benign | 0 | 0 | 0 |
| rs1800382 | benign | 1 | 1 | 1 |
| rs11063987 | benign | 1 | 1 | 1 |
| rs387906847 | pathogenic | 1 | 1 | 1 |
| rs73003466 | benign | 0 | 0 | 0 |
| rs149787233 | benign | 1 | 1 | 1 |
| rs144451841 | pathogenic | 1 | 1 | 1 |
| rs141638421 | pathogenic | 1 | 1 | 1 |
| rs121434497 | pathogenic | 0 | 0 | 0 |
| rs121434498 | pathogenic | 0 | 0 | 0 |
| rs267607230 | pathogenic | 1 | 1 | 0 |
| rs121434499 | pathogenic | 1 | 1 | 0 |
| rs546749336 | benign | 1 | 1 | 1 |
| rs138873805 | benign | 0 | 0 | 0 |
| rs730880522 | pathogenic | 0 | 0 | 0 |
| rs200371894 | benign | 0 | 0 | 0 |
| rs730880509 | benign | 0 | 0 | 0 |
| rs267606704 | pathogenic | 1 | 1 | 1 |
| rs267606706 | pathogenic | 0 | 0 | 0 |
| rs267606705 | pathogenic | 1 | 1 | 1 |
| rs267606707 | pathogenic | 0 | 0 | 0 |
| rs387906665 | pathogenic | 1 | 1 | 1 |
| rs267606708 | pathogenic | 1 | 1 | 1 |
| rs148368481 | benign | 0 | 0 | 0 |
| rs13204 | benign | 0 | 0 | 0 |
| rs140547520 | pathogenic | 1 | 1 | 1 |
| rs755234990 | benign | 1 | 1 | 0 |
| rs151184052 | benign | 0 | 0 | 0 |

| | | | | |
|---|---|---|---|---|
| rs3736358 | benign | 0 | 0 | 0 |
| rs3736360 | benign | 0 | 0 | 0 |
| rs1138469 | benign | 0 | 0 | 0 |
| rs74315351 | pathogenic | 0 | 1 | 0 |
| rs137853051 | pathogenic | 1 | 1 | 0 |
| rs74315353 | pathogenic | 0 | 1 | 0 |
| rs74315352 | pathogenic | 0 | 1 | 0 |
| rs74315354 | pathogenic | 1 | 1 | 1 |
| rs28938172 | pathogenic | 1 | 1 | 1 |
| rs52836744 | pathogenic | 0 | 0 | 0 |
| rs121913159 | pathogenic | 1 | 1 | 1 |
| rs121913145 | pathogenic | 1 | 1 | 1 |
| rs891087 | benign | 1 | 1 | 1 |
| rs9282757 | benign | 1 | 1 | 1 |
| rs397517404 | pathogenic | 1 | 1 | 1 |
| rs199918720 | benign | 1 | 1 | 0 |
| rs368299411 | benign | 1 | 1 | 0 |
| rs139399951 | benign | 1 | 1 | 0 |
| rs763981974 | benign | 1 | 1 | 0 |
| rs144242114 | benign | 0 | 0 | 0 |
| rs750965801 | benign | 0 | 0 | 0 |
| rs561310777 | benign | 0 | 0 | 0 |
| rs148185335 | benign | 0 | 0 | 0 |
| rs147742157 | benign | 0 | 0 | 0 |
| rs375664373 | pathogenic | 1 | 1 | 1 |
| rs777327250 | pathogenic | 0 | 0 | 0 |
| rs121908471 | pathogenic | 1 | 1 | 0 |
| rs2301612 | pathogenic | 1 | 1 | 0 |
| rs121908476 | pathogenic | 0 | 0 | 0 |
| rs36220239 | benign | 0 | 0 | 0 |
| rs121908473 | pathogenic | 0 | 0 | 0 |
| rs28647808 | benign | 0 | 0 | 0 |
| rs36090624 | benign | 0 | 0 | 0 |
| rs121908002 | pathogenic | 1 | 1 | 1 |
| rs4251545 | benign | 1 | 1 | 1 |
| rs78705382 | pathogenic | 0 | 0 | 0 |
| rs75184679 | pathogenic | 1 | 1 | 1 |
| rs74555752 | pathogenic | 0 | 0 | 0 |
| rs56320207 | benign | 1 | 1 | 1 |
| rs28730837 | pathogenic | 1 | 1 | 1 |
| rs119469014 | pathogenic | 1 | 1 | 1 |
| rs119468010 | pathogenic | 1 | 1 | 1 |
| rs35220041 | benign | 0 | 0 | 0 |
| rs762321137 | pathogenic | 1 | 1 | 1 |
| rs28931570 | pathogenic | 1 | 1 | 1 |
| rs709932 | benign | 1 | 1 | 1 |
| rs746531546 | benign | 1 | 1 | 1 |
| rs6647 | benign | 1 | 1 | 1 |
| rs141095970 | benign | 1 | 1 | 1 |
| rs143370956 | benign | 1 | 1 | 1 |
| rs28929474 | pathogenic | 1 | 1 | 1 |
| rs61761869 | pathogenic | 1 | 1 | 1 |
| rs1303 | benign | 1 | 1 | 1 |
| rs149447502 | benign | 0 | 0 | 0 |

| | | | | |
|---|---|---|---|---|
| rs267607180 | pathogenic | 1 | 1 | 1 |
| rs150304121 | benign | 0 | 0 | 0 |
| rs147630375 | benign | 1 | 1 | 1 |
| rs11539445 | benign | 1 | 1 | 1 |
| rs372098364 | pathogenic | 1 | 1 | 0 |
| rs760072286 | benign | 0 | 0 | 0 |
| rs745595833 | pathogenic | 1 | 1 | 1 |
| rs587777214 | pathogenic | 0 | 0 | 0 |
| rs150318528 | benign | 0 | 0 | 0 |
| rs137853254 | pathogenic | 1 | 1 | 1 |
| rs137853255 | pathogenic | 0 | 0 | 0 |
| rs137853253 | pathogenic | 0 | 0 | 0 |
| rs202166915 | benign | 0 | 0 | 0 |
| rs137853259 | pathogenic | 0 | 0 | 0 |
| rs2229137 | benign | 0 | 0 | 0 |
| rs137853258 | pathogenic | 1 | 1 | 0 |
| rs137853252 | pathogenic | 1 | 1 | 1 |
| rs137853256 | pathogenic | 0 | 0 | 0 |
| rs137853250 | pathogenic | 1 | 1 | 1 |
| rs121434615 | pathogenic | 0 | 0 | 0 |
| rs115014558 | benign | 1 | 1 | 1 |
| rs34116584 | benign | 1 | 1 | 1 |
| rs34885252 | benign | 1 | 1 | 1 |
| rs180177262 | pathogenic | 1 | 1 | 1 |
| rs180177162 | pathogenic | 1 | 1 | 1 |
| rs121908523 | pathogenic | 1 | 1 | 1 |
| rs180177170 | pathogenic | 1 | 1 | 1 |
| rs180177180 | pathogenic | 1 | 1 | 1 |
| rs767586362 | pathogenic | 1 | 1 | 1 |
| rs121908521 | pathogenic | 1 | 1 | 1 |
| rs180177203 | pathogenic | 1 | 1 | 1 |
| rs138025751 | pathogenic | 1 | 1 | 1 |
| rs180177210 | pathogenic | 0 | 1 | 0 |
| rs180177211 | pathogenic | 0 | 1 | 0 |
| rs121908524 | pathogenic | 1 | 1 | 0 |
| rs121908530 | pathogenic | 1 | 1 | 0 |
| rs180177227 | pathogenic | 0 | 1 | 0 |
| rs180177230 | pathogenic | 1 | 1 | 1 |
| rs121908529 | pathogenic | 0 | 1 | 0 |
| rs117195882 | benign | 1 | 1 | 0 |
| rs180177238 | pathogenic | 1 | 1 | 0 |
| rs34664134 | benign | 1 | 1 | 1 |
| rs180177246 | pathogenic | 0 | 1 | 0 |
| rs536352238 | pathogenic | 0 | 0 | 0 |
| rs121908520 | pathogenic | 1 | 1 | 0 |
| rs180177248 | pathogenic | 1 | 1 | 0 |
| rs180177252 | pathogenic | 1 | 1 | 1 |
| rs121908526 | pathogenic | 1 | 1 | 1 |
| rs121908525 | pathogenic | 1 | 1 | 1 |
| rs147106773 | benign | 1 | 1 | 1 |
| rs180177264 | pathogenic | 0 | 1 | 0 |
| rs146525143 | pathogenic | 1 | 1 | 1 |
| rs140992177 | benign | 1 | 1 | 1 |
| rs13408961 | benign | 0 | 1 | 0 |

| | | | | |
|---|---|---|---|---|
| rs4426527 | benign | 1 | 1 | 1 |
| rs3088308 | benign | 0 | 1 | 0 |
| rs4469829 | benign | 1 | 1 | 0 |
| rs104886003 | pathogenic | 1 | 1 | 1 |
| rs121913286 | pathogenic | 1 | 1 | 1 |
| rs121913279 | pathogenic | 1 | 1 | 1 |
| rs121434636 | pathogenic | 1 | 1 | 1 |
| rs755299132 | pathogenic | 1 | 1 | 1 |
| rs748318386 | pathogenic | 1 | 1 | 1 |
| rs387906757 | pathogenic | 1 | 1 | 1 |
| rs121965064 | pathogenic | 1 | 1 | 1 |
| rs28934608 | pathogenic | 0 | 0 | 0 |
| rs5967 | benign | 1 | 1 | 1 |
| rs773905328 | pathogenic | 0 | 0 | 0 |
| rs104894256 | pathogenic | 0 | 1 | 0 |
| rs28931612 | pathogenic | 0 | 1 | 0 |
| rs104894263 | pathogenic | 1 | 1 | 1 |
| rs607969 | benign | 1 | 1 | 0 |
| rs104894262 | pathogenic | 0 | 1 | 0 |
| rs104894268 | pathogenic | 1 | 1 | 1 |
| rs104894264 | pathogenic | 0 | 1 | 0 |
| rs104894259 | pathogenic | 1 | 1 | 0 |
| rs2959656 | benign | 0 | 0 | 0 |
| rs121913035 | pathogenic | 1 | 1 | 0 |
| rs139605474 | benign | 0 | 0 | 0 |
| rs3817552 | benign | 0 | 0 | 0 |
| rs150541115 | benign | 0 | 0 | 0 |
| rs562763010 | benign | 1 | 1 | 1 |
| rs200053119 | pathogenic | 0 | 0 | 0 |
| rs73826386 | benign | 0 | 0 | 0 |
| rs143655258 | benign | 0 | 0 | 0 |
| rs758857853 | pathogenic | 1 | 1 | 0 |
| rs2228380 | benign | 0 | 0 | 0 |
| rs141078733 | benign | 1 | 1 | 1 |
| rs55824172 | pathogenic | 1 | 1 | 1 |
| rs756751089 | pathogenic | 1 | 1 | 1 |
| rs121964990 | pathogenic | 0 | 0 | 0 |
| rs143555186 | benign | 0 | 0 | 0 |
| rs121964993 | pathogenic | 1 | 1 | 0 |
| rs121964992 | pathogenic | 1 | 1 | 1 |
| rs121964991 | pathogenic | 1 | 1 | 0 |
| rs121964988 | pathogenic | 1 | 1 | 1 |
| rs4687 | benign | 1 | 1 | 1 |
| rs267607125 | pathogenic | 1 | 1 | 1 |
| rs751682878 | benign | 1 | 1 | 1 |
| rs267607126 | pathogenic | 1 | 1 | 1 |
| rs768713596 | pathogenic | 0 | 0 | 0 |
| rs147288996 | benign | 0 | 0 | 0 |
| rs387906639 | pathogenic | 1 | 1 | 1 |
| rs730881069 | pathogenic | 1 | 1 | 1 |
| rs730881071 | pathogenic | 1 | 1 | 1 |
| rs397516348 | pathogenic | 0 | 0 | 0 |
| rs121917760 | pathogenic | 0 | 0 | 0 |
| rs104894724 | pathogenic | 1 | 1 | 1 |

| | | | | |
|---|---|---|---|---|
| rs397516349 | pathogenic | 1 | 1 | 1 |
| rs12075 | benign | 0 | 0 | 0 |
| rs147549987 | benign | 0 | 0 | 0 |
| rs56079734 | benign | 0 | 0 | 0 |
| rs1017842 | benign | 1 | 1 | 1 |
| rs137852593 | pathogenic | 1 | 1 | 1 |
| rs137852571 | pathogenic | 1 | 1 | 1 |
| rs137852591 | pathogenic | 1 | 1 | 1 |
| rs9332971 | pathogenic | 1 | 1 | 1 |
| rs755226547 | pathogenic | 1 | 1 | 1 |
| rs79206939 | benign | 0 | 0 | 0 |
| rs121918214 | pathogenic | 1 | 1 | 0 |
| rs757075712 | pathogenic | 1 | 1 | 0 |
| rs150483902 | benign | 1 | 1 | 1 |
| rs113993973 | pathogenic | 0 | 1 | 0 |
| rs946616 | benign | 0 | 0 | 0 |
| rs113993976 | pathogenic | 1 | 1 | 0 |
| rs151022264 | benign | 0 | 1 | 0 |
| rs113993977 | pathogenic | 1 | 1 | 0 |
| rs113993978 | pathogenic | 1 | 1 | 0 |
| rs2228499 | benign | 0 | 1 | 0 |
| rs113993979 | pathogenic | 1 | 1 | 0 |
| rs113993980 | pathogenic | 1 | 1 | 0 |
| rs113993983 | pathogenic | 0 | 1 | 0 |
| rs150547274 | pathogenic | 1 | 1 | 0 |
| rs113993984 | pathogenic | 1 | 1 | 0 |
| rs113993986 | pathogenic | 1 | 1 | 0 |
| rs34313873 | benign | 0 | 1 | 0 |
| rs113993988 | pathogenic | 1 | 1 | 0 |
| rs199638097 | pathogenic | 1 | 1 | 1 |
| rs118204447 | pathogenic | 1 | 1 | 1 |
| rs11862754 | benign | 0 | 0 | 0 |
| rs118204445 | pathogenic | 0 | 0 | 0 |
| rs118204441 | pathogenic | 0 | 0 | 0 |
| rs118204438 | pathogenic | 0 | 0 | 0 |
| rs118204436 | pathogenic | 0 | 0 | 0 |
| rs146093755 | pathogenic | 0 | 0 | 0 |
| rs781439830 | pathogenic | 0 | 0 | 0 |
| rs398123438 | pathogenic | 0 | 0 | 0 |
| rs398123439 | pathogenic | 0 | 0 | 0 |
| rs118204444 | pathogenic | 0 | 0 | 0 |
| rs7187889 | benign | 0 | 0 | 0 |
| rs118204435 | pathogenic | 1 | 1 | 0 |
| rs34745339 | benign | 0 | 0 | 0 |
| rs398123440 | pathogenic | 0 | 0 | 0 |
| rs118204442 | pathogenic | 0 | 0 | 0 |
| rs770053354 | pathogenic | 0 | 0 | 0 |
| rs118204448 | pathogenic | 0 | 0 | 0 |
| rs118204443 | pathogenic | 0 | 0 | 0 |
| rs118204446 | pathogenic | 0 | 0 | 0 |
| rs746756997 | pathogenic | 0 | 0 | 0 |
| rs267606838 | pathogenic | 0 | 0 | 0 |
| rs118204437 | pathogenic | 0 | 0 | 0 |
| rs398123429 | pathogenic | 0 | 0 | 0 |

| | | | | |
|---|---|---|---|---|
| rs398123430 | pathogenic | 0 | 0 | 0 |
| rs2303269 | benign | 0 | 0 | 0 |
| rs114703967 | benign | 0 | 0 | 0 |
| rs118204439 | pathogenic | 0 | 0 | 0 |
| rs151296605 | benign | 0 | 0 | 0 |
| rs118204440 | pathogenic | 0 | 0 | 0 |
| rs78127134 | benign | 0 | 0 | 0 |
| rs372893383 | pathogenic | 0 | 0 | 0 |
| rs61753266 | pathogenic | 1 | 1 | 1 |
| rs139579928 | benign | 1 | 1 | 1 |
| rs56275071 | pathogenic | 1 | 1 | 1 |
| rs121909736 | pathogenic | 1 | 1 | 0 |
| rs121909737 | pathogenic | 1 | 1 | 0 |
| rs121909735 | pathogenic | 1 | 1 | 0 |
| rs121909731 | pathogenic | 1 | 1 | 1 |
| rs121909734 | pathogenic | 1 | 1 | 1 |
| rs121909733 | pathogenic | 1 | 1 | 1 |
| rs121909732 | pathogenic | 1 | 1 | 1 |
| rs121909730 | pathogenic | 1 | 1 | 1 |
| rs202198360 | benign | 1 | 1 | 1 |
| rs587777294 | pathogenic | 1 | 1 | 0 |
| rs587777297 | pathogenic | 1 | 1 | 0 |
| rs104893913 | pathogenic | 1 | 1 | 0 |
| rs17860424 | pathogenic | 1 | 1 | 1 |
| rs1045485 | benign | 1 | 1 | 1 |
| rs13926 | benign | 0 | 0 | 0 |
| rs79343837 | benign | 0 | 0 | 0 |
| rs45516306 | benign | 1 | 1 | 1 |
| rs139849355 | benign | 1 | 1 | 1 |
| rs104894874 | pathogenic | 1 | 1 | 1 |
| rs104894870 | pathogenic | 1 | 1 | 1 |
| rs137852221 | pathogenic | 1 | 1 | 1 |
| rs104894869 | pathogenic | 1 | 1 | 1 |
| rs104894880 | pathogenic | 1 | 1 | 1 |
| rs104894872 | pathogenic | 1 | 1 | 1 |
| rs104894868 | pathogenic | 1 | 1 | 1 |
| rs104894867 | pathogenic | 1 | 1 | 1 |
| rs104894871 | pathogenic | 1 | 1 | 1 |
| rs104894877 | pathogenic | 1 | 1 | 1 |
| rs104894883 | pathogenic | 1 | 1 | 1 |
| rs104894875 | pathogenic | 1 | 1 | 1 |
| rs104894876 | pathogenic | 1 | 1 | 1 |
| rs104894878 | pathogenic | 1 | 1 | 1 |
| rs137852220 | pathogenic | 1 | 1 | 1 |
| rs28933684 | pathogenic | 1 | 1 | 1 |
| rs104894873 | pathogenic | 1 | 1 | 1 |
| rs138409264 | benign | 0 | 0 | 0 |
| rs2236722 | benign | 0 | 0 | 0 |
| rs28757184 | benign | 0 | 0 | 0 |
| rs121434538 | pathogenic | 0 | 0 | 0 |
| rs700519 | benign | 0 | 0 | 0 |
| rs80051519 | pathogenic | 0 | 0 | 0 |
| rs121434536 | pathogenic | 1 | 1 | 0 |
| rs121434534 | pathogenic | 1 | 1 | 0 |

| | | | | |
|---|---|---|---|---|
| rs78310315 | pathogenic | 1 | 1 | 0 |
| rs540839115 | pathogenic | 1 | 1 | 1 |
| rs80338731 | pathogenic | 1 | 1 | 1 |
| rs180177151 | pathogenic | 1 | 1 | 1 |
| rs180177152 | pathogenic | 0 | 0 | 0 |
| rs80338732 | pathogenic | 1 | 1 | 1 |
| rs80338675 | pathogenic | 1 | 1 | 1 |
| rs80338676 | pathogenic | 1 | 1 | 1 |
| rs180177153 | pathogenic | 0 | 0 | 0 |
| rs180177154 | pathogenic | 1 | 1 | 1 |
| rs5896 | benign | 1 | 1 | 1 |
| rs121918477 | pathogenic | 0 | 0 | 0 |
| rs121918428 | pathogenic | 1 | 1 | 1 |
| rs121918427 | pathogenic | 1 | 1 | 1 |
| rs544300820 | benign | 1 | 1 | 1 |
| rs121918110 | pathogenic | 1 | 1 | 1 |
| rs1057141 | benign | 0 | 0 | 0 |
| rs1135216 | benign | 1 | 1 | 1 |
| rs121917702 | pathogenic | 1 | 1 | 1 |
| rs28941474 | pathogenic | 0 | 0 | 0 |
| rs104893944 | pathogenic | 1 | 1 | 1 |
| rs104893947 | pathogenic | 1 | 1 | 0 |
| rs104893943 | pathogenic | 1 | 1 | 1 |
| rs104893942 | pathogenic | 1 | 1 | 1 |
| rs104893940 | pathogenic | 1 | 1 | 1 |
| rs34045013 | benign | 0 | 0 | 0 |
| rs143083947 | pathogenic | 0 | 0 | 0 |
| rs145786986 | benign | 0 | 0 | 0 |
| rs119450942 | pathogenic | 0 | 0 | 0 |
| rs374259530 | pathogenic | 0 | 0 | 0 |
| rs756210458 | pathogenic | 0 | 0 | 0 |
| rs11089991 | benign | 0 | 0 | 0 |
| rs28941471 | pathogenic | 0 | 1 | 0 |
| rs750614500 | pathogenic | 0 | 1 | 0 |
| rs199761158 | benign | 1 | 1 | 1 |
| rs119450944 | pathogenic | 1 | 1 | 1 |
| rs373458753 | pathogenic | 1 | 1 | 1 |
| rs202064195 | pathogenic | 1 | 1 | 1 |
| rs776496275 | pathogenic | 1 | 1 | 1 |
| rs761493155 | pathogenic | 1 | 1 | 1 |
| rs763542069 | pathogenic | 0 | 0 | 0 |
| rs119450941 | pathogenic | 0 | 0 | 0 |
| rs119450940 | pathogenic | 0 | 0 | 0 |
| rs777821034 | pathogenic | 1 | 1 | 1 |
| rs771121666 | pathogenic | 1 | 1 | 1 |
| rs372895468 | pathogenic | 1 | 1 | 1 |
| rs61750603 | pathogenic | 0 | 0 | 0 |
| rs76863441 | pathogenic | 0 | 0 | 0 |
| rs142601143 | benign | 1 | 1 | 1 |
| rs185832993 | benign | 1 | 1 | 0 |
| rs17860568 | benign | 0 | 0 | 0 |
| rs142064825 | benign | 0 | 0 | 0 |
| rs564076183 | benign | 0 | 0 | 0 |
| rs386834034 | pathogenic | 0 | 0 | 0 |

| | | | | |
|---|---|---|---|---|
| rs147663769 | benign | 0 | 0 | 0 |
| rs776478974 | benign | 0 | 0 | 0 |
| rs111840783 | benign | 0 | 0 | 0 |
| rs1516446 | benign | 1 | 1 | 1 |
| rs143749154 | benign | 1 | 1 | 1 |
| rs80257412 | benign | 0 | 0 | 0 |
| rs142943120 | benign | 0 | 0 | 0 |
| rs144553482 | benign | 0 | 0 | 0 |
| rs35997569 | benign | 0 | 0 | 0 |
| rs772909106 | pathogenic | 0 | 0 | 0 |
| rs149433837 | benign | 1 | 1 | 1 |
| rs532178791 | pathogenic | 0 | 0 | 0 |
| rs145588689 | pathogenic | 1 | 1 | 0 |
| rs17843776 | benign | 1 | 1 | 0 |
| rs121917890 | pathogenic | 1 | 1 | 1 |
| rs121917892 | pathogenic | 0 | 0 | 0 |
| rs606231289 | pathogenic | 0 | 0 | 0 |
| rs370011798 | pathogenic | 0 | 0 | 0 |
| rs34689457 | benign | 0 | 0 | 0 |
| rs118204451 | pathogenic | 0 | 0 | 0 |
| rs118204452 | pathogenic | 1 | 1 | 1 |
| rs652785 | benign | 0 | 0 | 0 |
| rs35378126 | benign | 0 | 0 | 0 |
| rs80358214 | pathogenic | 1 | 1 | 1 |
| rs80358215 | pathogenic | 0 | 1 | 0 |
| rs267606858 | pathogenic | 1 | 1 | 1 |
| rs143137713 | pathogenic | 1 | 1 | 1 |
| rs782168634 | benign | 0 | 0 | 0 |
| rs143873938 | benign | 0 | 0 | 0 |
| rs782776511 | benign | 0 | 0 | 0 |
| rs201093148 | benign | 0 | 0 | 0 |
| rs267606815 | pathogenic | 0 | 0 | 0 |
| rs137853311 | pathogenic | 0 | 0 | 0 |
| rs267606817 | pathogenic | 0 | 0 | 0 |
| rs72650006 | benign | 0 | 0 | 0 |
| rs727503638 | benign | 1 | 1 | 0 |
| rs189286381 | benign | 0 | 0 | 0 |
| rs376644553 | benign | 0 | 0 | 0 |
| rs121434610 | pathogenic | 1 | 1 | 1 |
| rs267607076 | pathogenic | 1 | 1 | 1 |
| rs201275730 | benign | 0 | 0 | 0 |
| rs121909587 | pathogenic | 0 | 0 | 0 |
| rs121909558 | pathogenic | 1 | 1 | 0 |
| rs28929469 | pathogenic | 1 | 1 | 0 |
| rs121909547 | pathogenic | 1 | 1 | 0 |
| rs121909552 | pathogenic | 1 | 1 | 0 |
| rs121909573 | pathogenic | 1 | 1 | 1 |
| rs121909567 | pathogenic | 1 | 1 | 0 |
| rs121909569 | pathogenic | 0 | 0 | 0 |
| rs121909563 | pathogenic | 1 | 1 | 0 |
| rs121909570 | pathogenic | 1 | 1 | 1 |
| rs121909571 | pathogenic | 1 | 1 | 0 |
| rs121909572 | pathogenic | 1 | 1 | 1 |
| rs121909565 | pathogenic | 1 | 1 | 1 |

| | | | | |
|---|---|---|---|---|
| rs121909557 | pathogenic | 1 | 1 | 1 |
| rs121909568 | pathogenic | 1 | 1 | 1 |
| rs121909566 | pathogenic | 1 | 1 | 1 |
| rs121909549 | pathogenic | 1 | 1 | 1 |
| rs121909554 | pathogenic | 1 | 1 | 1 |
| rs121909550 | pathogenic | 1 | 1 | 1 |
| rs121909546 | pathogenic | 1 | 1 | 0 |
| rs121909555 | pathogenic | 1 | 1 | 1 |
| rs121909564 | pathogenic | 1 | 1 | 1 |
| rs548046212 | pathogenic | 0 | 0 | 0 |
| rs147353781 | benign | 0 | 0 | 0 |
| rs104894678 | pathogenic | 1 | 1 | 1 |
| rs1130426 | benign | 0 | 0 | 0 |
| rs104894677 | pathogenic | 1 | 1 | 1 |
| rs141917423 | benign | 1 | 1 | 1 |
| rs574777883 | benign | 0 | 0 | 0 |
| rs150199226 | benign | 1 | 1 | 1 |
| rs199618701 | benign | 0 | 1 | 0 |
| rs121964923 | pathogenic | 0 | 0 | 0 |
| rs121964924 | pathogenic | 0 | 0 | 0 |
| rs201258823 | pathogenic | 0 | 1 | 0 |
| rs267606773 | pathogenic | 0 | 0 | 0 |
| rs35638832 | benign | 0 | 0 | 0 |
| rs121908632 | pathogenic | 1 | 1 | 1 |
| rs121908625 | pathogenic | 1 | 1 | 1 |
| rs748949603 | pathogenic | 0 | 0 | 0 |
| rs757523840 | pathogenic | 0 | 0 | 0 |
| rs62541771 | pathogenic | 0 | 0 | 0 |
| rs776384541 | pathogenic | 1 | 1 | 0 |
| rs121908627 | pathogenic | 1 | 1 | 0 |
| rs28937594 | pathogenic | 0 | 0 | 0 |
| rs121907888 | pathogenic | 0 | 0 | 0 |
| rs121907889 | pathogenic | 0 | 1 | 0 |
| rs28941785 | pathogenic | 1 | 1 | 1 |
| rs28941786 | pathogenic | 1 | 1 | 1 |
| rs1021737 | benign | 0 | 0 | 0 |
| rs121908370 | pathogenic | 1 | 1 | 1 |
| rs63750083 | pathogenic | 0 | 0 | 0 |
| rs17883862 | benign | 0 | 0 | 0 |
| rs587782070 | pathogenic | 1 | 1 | 1 |
| rs587781269 | pathogenic | 1 | 1 | 1 |
| rs28909982 | pathogenic | 1 | 1 | 1 |
| rs373648967 | benign | 1 | 1 | 1 |
| rs368570187 | benign | 1 | 1 | 1 |
| rs730881701 | pathogenic | 1 | 1 | 1 |
| rs137853007 | pathogenic | 0 | 0 | 0 |
| rs575910805 | benign | 0 | 0 | 0 |
| rs104893875 | pathogenic | 1 | 1 | 1 |
| rs104893877 | pathogenic | 1 | 1 | 1 |
| rs119485093 | pathogenic | 1 | 1 | 1 |
| rs119485092 | pathogenic | 0 | 0 | 0 |
| rs119485090 | pathogenic | 0 | 0 | 0 |
| rs28942099 | pathogenic | 1 | 1 | 1 |
| rs373730800 | pathogenic | 1 | 1 | 1 |

| | | | | |
|---|---|---|---|---|
| rs113993995 | pathogenic | 0 | 0 | 0 |
| rs201070132 | pathogenic | 0 | 0 | 0 |
| rs113993996 | pathogenic | 0 | 0 | 0 |
| rs113993998 | pathogenic | 0 | 0 | 0 |
| rs78950939 | pathogenic | 1 | 1 | 1 |
| rs56378716 | pathogenic | 1 | 1 | 1 |
| rs754921704 | benign | 0 | 1 | 0 |
| rs766379103 | pathogenic | 1 | 1 | 0 |
| rs121912515 | pathogenic | 0 | 1 | 0 |
| rs764831888 | benign | 1 | 1 | 0 |
| rs572527340 | benign | 0 | 0 | 0 |
| rs35969327 | benign | 0 | 0 | 0 |
| rs35814893 | benign | 0 | 0 | 0 |
| rs34032300 | benign | 0 | 0 | 0 |
| rs80358260 | pathogenic | 0 | 0 | 0 |
| rs151220873 | benign | 0 | 0 | 0 |
| rs80358261 | pathogenic | 1 | 1 | 0 |
| rs11694 | pathogenic | 0 | 0 | 0 |
| rs80358264 | pathogenic | 0 | 0 | 0 |
| rs80358266 | pathogenic | 1 | 1 | 0 |
| rs104894458 | pathogenic | 1 | 1 | 1 |
| rs121918028 | pathogenic | 0 | 0 | 0 |
| rs11583680 | pathogenic | 0 | 0 | 0 |
| rs4715 | benign | 1 | 1 | 1 |
| rs34957318 | benign | 0 | 0 | 0 |
| rs1124 | benign | 1 | 1 | 1 |
| rs104894778 | pathogenic | 1 | 1 | 1 |
| rs104894777 | pathogenic | 1 | 1 | 1 |
| rs104894769 | pathogenic | 1 | 1 | 1 |
| rs104894768 | pathogenic | 1 | 1 | 1 |
| rs104894771 | pathogenic | 1 | 1 | 1 |
| rs575372249 | benign | 1 | 1 | 1 |
| rs118203914 | pathogenic | 0 | 0 | 0 |
| rs118203916 | pathogenic | 1 | 1 | 0 |
| rs140154690 | benign | 0 | 0 | 0 |
| rs34382405 | benign | 0 | 0 | 0 |
| rs73718082 | benign | 0 | 0 | 0 |
| rs764932724 | benign | 0 | 0 | 0 |
| rs554931092 | pathogenic | 1 | 1 | 0 |
| rs202060864 | pathogenic | 0 | 0 | 0 |
| rs116567033 | benign | 0 | 0 | 0 |
| rs11243011 | benign | 0 | 0 | 0 |
| rs142073519 | pathogenic | 0 | 0 | 0 |
| rs397514611 | pathogenic | 1 | 1 | 0 |
| rs397514612 | pathogenic | 0 | 0 | 0 |
| rs372054960 | pathogenic | 0 | 0 | 0 |
| rs775690041 | pathogenic | 0 | 0 | 0 |
| rs148921184 | pathogenic | 0 | 0 | 0 |
| rs12711521 | benign | 0 | 0 | 0 |
| rs2273346 | benign | 0 | 0 | 0 |
| rs12085877 | benign | 1 | 1 | 1 |
| rs144471433 | benign | 1 | 1 | 1 |
| rs397514671 | pathogenic | 1 | 1 | 1 |
| rs76394680 | benign | 0 | 0 | 0 |

| | | | | |
|---|---|---|---|---|
| rs78289603 | pathogenic | 1 | 1 | 1 |
| rs137852913 | pathogenic | 1 | 1 | 1 |
| rs137852914 | pathogenic | 1 | 1 | 1 |
| rs121918653 | pathogenic | 1 | 1 | 1 |
| rs35515200 | benign | 1 | 1 | 0 |
| rs17876030 | benign | 1 | 1 | 1 |
| rs556623443 | benign | 1 | 1 | 1 |
| rs121434365 | pathogenic | 1 | 1 | 1 |
| rs121434364 | pathogenic | 0 | 0 | 0 |
| rs41400249 | benign | 0 | 0 | 0 |
| rs149440376 | benign | 1 | 1 | 1 |
| rs181590179 | benign | 0 | 0 | 0 |
| rs113993968 | pathogenic | 0 | 0 | 0 |
| rs587776445 | pathogenic | 1 | 1 | 1 |
| rs113993969 | pathogenic | 1 | 1 | 1 |
| rs113993970 | pathogenic | 1 | 1 | 1 |
| rs587776449 | pathogenic | 1 | 1 | 1 |
| rs113993971 | pathogenic | 1 | 1 | 1 |
| rs267607194 | pathogenic | 1 | 1 | 1 |
| rs397514745 | pathogenic | 0 | 0 | 0 |
| rs201650281 | pathogenic | 0 | 0 | 0 |
| rs150529876 | benign | 1 | 1 | 1 |
| rs77573084 | benign | 0 | 0 | 0 |
| rs398122379 | pathogenic | 1 | 1 | 0 |
| rs765478968 | benign | 0 | 0 | 0 |
| rs201399681 | benign | 0 | 0 | 0 |
| rs17159287 | benign | 0 | 0 | 0 |
| rs538571144 | benign | 0 | 0 | 0 |
| rs137852647 | pathogenic | 0 | 0 | 0 |
| rs201358272 | pathogenic | 1 | 1 | 1 |
| rs181251337 | benign | 0 | 0 | 0 |
| rs3213119 | benign | 0 | 0 | 0 |
| rs137853102 | pathogenic | 1 | 1 | 0 |
| rs4956145 | benign | 0 | 0 | 0 |
| rs146036912 | pathogenic | 1 | 1 | 1 |
| rs375717077 | pathogenic | 1 | 1 | 1 |
| rs137853103 | pathogenic | 1 | 1 | 1 |
| rs776184830 | pathogenic | 1 | 1 | 1 |
| rs200273673 | pathogenic | 1 | 1 | 1 |
| rs267607025 | pathogenic | 1 | 1 | 1 |
| rs267607024 | pathogenic | 1 | 1 | 1 |
| rs515726188 | pathogenic | 1 | 1 | 1 |
| rs147315735 | benign | 0 | 0 | 0 |
| rs121918308 | pathogenic | 1 | 1 | 0 |
| rs515726195 | pathogenic | 0 | 0 | 0 |
| rs515726196 | pathogenic | 1 | 1 | 0 |
| rs121918311 | pathogenic | 0 | 1 | 0 |
| rs121918309 | pathogenic | 0 | 0 | 0 |
| rs387906891 | pathogenic | 0 | 0 | 0 |
| rs182614164 | pathogenic | 1 | 1 | 0 |
| rs121918307 | pathogenic | 0 | 0 | 0 |
| rs201440849 | benign | 0 | 0 | 0 |
| rs118203957 | pathogenic | 0 | 0 | 0 |
| rs4986790 | benign | 0 | 0 | 0 |

| | | | | |
|---|---|---|---|---|
| rs281864928 | pathogenic | 1 | 1 | 0 |
| rs267607156 | pathogenic | 0 | 0 | 0 |
| rs55916807 | benign | 0 | 0 | 0 |
| rs749750052 | benign | 0 | 0 | 0 |
| rs74382477 | benign | 0 | 0 | 0 |
| rs201016638 | benign | 0 | 0 | 0 |
| rs376683989 | benign | 0 | 0 | 0 |
| rs144439703 | pathogenic | 0 | 0 | 0 |
| rs779122022 | benign | 0 | 0 | 0 |
| rs121918056 | pathogenic | 0 | 0 | 0 |
| rs113994093 | pathogenic | 1 | 1 | 1 |
| rs141367015 | pathogenic | 0 | 0 | 0 |
| rs775538075 | pathogenic | 0 | 0 | 0 |
| rs121918055 | pathogenic | 0 | 0 | 0 |
| rs754246294 | benign | 1 | 1 | 1 |
| rs2307447 | benign | 1 | 1 | 1 |
| rs113994096 | pathogenic | 1 | 1 | 1 |
| rs367610201 | pathogenic | 1 | 1 | 1 |
| rs758438414 | pathogenic | 1 | 1 | 0 |
| rs121918046 | pathogenic | 0 | 0 | 0 |
| rs201871736 | benign | 0 | 0 | 0 |
| rs773994204 | benign | 0 | 0 | 0 |
| rs2307450 | benign | 1 | 1 | 1 |
| rs200257554 | benign | 0 | 0 | 0 |
| rs185645212 | benign | 0 | 0 | 0 |
| rs121918054 | pathogenic | 0 | 0 | 0 |
| rs113994097 | pathogenic | 0 | 0 | 0 |
| rs202037973 | pathogenic | 0 | 0 | 0 |
| rs769827124 | pathogenic | 1 | 1 | 0 |
| rs41549716 | benign | 1 | 1 | 1 |
| rs113994098 | pathogenic | 1 | 1 | 0 |
| rs144500145 | pathogenic | 1 | 1 | 0 |
| rs121918053 | pathogenic | 1 | 1 | 0 |
| rs121918050 | pathogenic | 1 | 1 | 0 |
| rs368587966 | pathogenic | 0 | 0 | 0 |
| rs139590686 | pathogenic | 0 | 0 | 0 |
| rs121918048 | pathogenic | 1 | 1 | 0 |
| rs768653086 | pathogenic | 1 | 1 | 0 |
| rs113994099 | pathogenic | 1 | 1 | 0 |
| rs121918051 | pathogenic | 1 | 1 | 0 |
| rs753160398 | pathogenic | 1 | 1 | 0 |
| rs201477273 | pathogenic | 1 | 1 | 0 |
| rs142347031 | pathogenic | 0 | 0 | 0 |
| rs121918049 | pathogenic | 0 | 0 | 0 |
| rs267606959 | pathogenic | 1 | 1 | 0 |
| rs201732356 | pathogenic | 0 | 0 | 0 |
| rs368435864 | pathogenic | 0 | 0 | 0 |
| rs764036283 | pathogenic | 0 | 0 | 0 |
| rs2307442 | benign | 0 | 0 | 0 |
| rs2307440 | benign | 0 | 0 | 0 |
| rs548076633 | pathogenic | 0 | 1 | 0 |
| rs750792237 | benign | 0 | 0 | 0 |
| rs3087374 | benign | 0 | 0 | 0 |
| rs121918060 | pathogenic | 0 | 1 | 0 |

| | | | | |
|---|---|---|---|---|
| rs397514765 | pathogenic | 1 | 1 | 1 |
| rs121918063 | pathogenic | 0 | 0 | 0 |
| rs121918058 | pathogenic | 1 | 1 | 1 |
| rs121918064 | pathogenic | 0 | 0 | 0 |
| rs36033115 | benign | 0 | 1 | 0 |
| rs121918057 | pathogenic | 0 | 1 | 0 |
| rs121918065 | pathogenic | 1 | 1 | 1 |
| rs121918061 | pathogenic | 1 | 1 | 1 |
| rs121918066 | pathogenic | 0 | 1 | 0 |
| rs707555 | benign | 0 | 1 | 0 |
| rs8192708 | benign | 0 | 1 | 0 |
| rs11552145 | benign | 0 | 0 | 0 |
| rs1804160 | benign | 0 | 1 | 0 |
| rs34103538 | benign | 0 | 1 | 0 |
| rs751393950 | pathogenic | 1 | 1 | 1 |
| rs121907979 | pathogenic | 1 | 1 | 1 |
| rs121907975 | pathogenic | 1 | 1 | 1 |
| rs121907962 | pathogenic | 1 | 1 | 1 |
| rs121907957 | pathogenic | 0 | 0 | 0 |
| rs121907953 | pathogenic | 1 | 1 | 1 |
| rs28941770 | pathogenic | 1 | 1 | 1 |
| rs28941771 | pathogenic | 1 | 1 | 1 |
| rs121907969 | pathogenic | 1 | 1 | 1 |
| rs121907973 | pathogenic | 1 | 1 | 1 |
| rs1800429 | pathogenic | 0 | 0 | 0 |
| rs121907976 | pathogenic | 1 | 1 | 0 |
| rs121907961 | pathogenic | 0 | 0 | 0 |
| rs121907974 | pathogenic | 0 | 0 | 0 |
| rs121907959 | pathogenic | 0 | 0 | 0 |
| rs121907971 | pathogenic | 1 | 1 | 0 |
| rs121907954 | pathogenic | 0 | 0 | 0 |
| rs121907977 | pathogenic | 0 | 0 | 0 |
| rs121907963 | pathogenic | 1 | 1 | 0 |
| rs370266293 | pathogenic | 1 | 1 | 0 |
| rs1800430 | benign | 0 | 0 | 0 |
| rs121907958 | pathogenic | 0 | 0 | 0 |
| rs202173526 | pathogenic | 0 | 0 | 0 |
| rs121907978 | pathogenic | 0 | 0 | 0 |
| rs121907981 | pathogenic | 0 | 1 | 0 |
| rs145012038 | benign | 1 | 1 | 1 |
| rs121907968 | pathogenic | 1 | 1 | 1 |
| rs121907966 | pathogenic | 1 | 1 | 1 |
| rs121907956 | pathogenic | 1 | 1 | 1 |
| rs28942071 | pathogenic | 1 | 1 | 1 |
| rs121907955 | pathogenic | 1 | 1 | 1 |
| rs4727853 | benign | 0 | 0 | 0 |
| rs201880593 | benign | 0 | 0 | 0 |
| rs201434579 | benign | 1 | 1 | 1 |
| rs74551128 | pathogenic | 1 | 1 | 0 |
| rs193922500 | pathogenic | 0 | 0 | 0 |
| rs121909009 | pathogenic | 1 | 1 | 0 |
| rs121908805 | pathogenic | 1 | 1 | 0 |
| rs139573311 | pathogenic | 0 | 0 | 0 |
| rs213950 | benign | 0 | 0 | 0 |

| | | | | |
|---|---|---|---|---|
| rs79282516 | pathogenic | 0 | 0 | 0 |
| rs121909017 | pathogenic | 1 | 1 | 1 |
| rs77101217 | pathogenic | 1 | 1 | 1 |
| rs1800092 | pathogenic | 0 | 1 | 0 |
| rs77646904 | pathogenic | 0 | 0 | 0 |
| rs1800095 | benign | 1 | 1 | 1 |
| rs387906368 | pathogenic | 1 | 1 | 1 |
| rs113993959 | pathogenic | 1 | 1 | 1 |
| rs121908755 | pathogenic | 1 | 1 | 1 |
| rs121909005 | pathogenic | 1 | 1 | 1 |
| rs121908757 | pathogenic | 1 | 1 | 1 |
| rs75527207 | pathogenic | 1 | 1 | 0 |
| rs121909013 | pathogenic | 1 | 1 | 0 |
| rs121909044 | pathogenic | 1 | 1 | 0 |
| rs74597325 | pathogenic | 1 | 1 | 0 |
| rs75549581 | pathogenic | 0 | 0 | 0 |
| rs80055610 | pathogenic | 1 | 1 | 1 |
| rs1800097 | pathogenic | 0 | 0 | 0 |
| rs121909006 | pathogenic | 0 | 1 | 0 |
| rs397508276 | pathogenic | 0 | 0 | 0 |
| rs397508296 | pathogenic | 1 | 1 | 1 |
| rs139468767 | pathogenic | 0 | 0 | 0 |
| rs121908759 | pathogenic | 1 | 1 | 0 |
| rs121909033 | pathogenic | 1 | 1 | 0 |
| rs397508331 | pathogenic | 0 | 0 | 0 |
| rs387907279 | pathogenic | 1 | 1 | 1 |
| rs546530618 | benign | 0 | 0 | 0 |
| rs558259739 | benign | 0 | 0 | 0 |
| rs201827340 | benign | 0 | 0 | 0 |
| rs148865119 | pathogenic | 1 | 1 | 0 |
| rs2229413 | benign | 1 | 1 | 1 |
| rs34040148 | benign | 1 | 1 | 1 |
| rs121964964 | pathogenic | 1 | 1 | 1 |
| rs760214620 | pathogenic | 1 | 1 | 1 |
| rs775992753 | pathogenic | 1 | 1 | 1 |
| rs770095972 | pathogenic | 1 | 1 | 1 |
| rs781444670 | pathogenic | 1 | 1 | 1 |
| rs147474549 | pathogenic | 0 | 0 | 0 |
| rs121964965 | pathogenic | 0 | 1 | 0 |
| rs121964963 | pathogenic | 1 | 1 | 1 |
| rs755952006 | pathogenic | 1 | 1 | 1 |
| rs121964970 | pathogenic | 1 | 1 | 1 |
| rs121964973 | pathogenic | 1 | 1 | 1 |
| rs773734233 | pathogenic | 1 | 1 | 1 |
| rs758236584 | pathogenic | 1 | 1 | 1 |
| rs149119723 | pathogenic | 1 | 1 | 0 |
| rs5742905 | pathogenic | 0 | 0 | 0 |
| rs121964962 | pathogenic | 1 | 1 | 1 |
| rs781567152 | pathogenic | 1 | 1 | 0 |
| rs398123151 | pathogenic | 1 | 1 | 1 |
| rs760417941 | pathogenic | 1 | 1 | 1 |
| rs774926464 | pathogenic | 0 | 0 | 0 |
| rs121964972 | pathogenic | 1 | 1 | 0 |
| rs372010465 | pathogenic | 1 | 1 | 1 |

| | | | | |
|---|---|---|---|---|
| rs763036586 | pathogenic | 1 | 1 | 1 |
| rs769080151 | pathogenic | 1 | 1 | 1 |
| rs121964967 | pathogenic | 1 | 1 | 1 |
| rs28934892 | pathogenic | 1 | 1 | 0 |
| rs28934891 | pathogenic | 1 | 1 | 1 |
| rs121964971 | pathogenic | 1 | 1 | 1 |
| rs121964968 | pathogenic | 1 | 1 | 1 |
| rs150828989 | benign | 1 | 1 | 1 |
| rs3775291 | benign | 1 | 1 | 0 |
| rs104894419 | pathogenic | 1 | 1 | 1 |
| rs2232642 | benign | 0 | 0 | 0 |
| rs104894551 | pathogenic | 1 | 1 | 0 |
| rs766328537 | pathogenic | 0 | 1 | 0 |
| rs104894553 | pathogenic | 1 | 1 | 0 |
| rs80099330 | benign | 0 | 0 | 0 |
| rs199565861 | pathogenic | 0 | 0 | 0 |
| rs104894548 | pathogenic | 0 | 0 | 0 |
| rs104894549 | pathogenic | 0 | 0 | 0 |
| rs104894550 | pathogenic | 1 | 1 | 1 |
| rs12948217 | pathogenic | 1 | 1 | 1 |
| rs761064915 | pathogenic | 0 | 0 | 0 |
| rs28940279 | pathogenic | 1 | 1 | 1 |
| rs774323189 | pathogenic | 1 | 1 | 1 |
| rs28940574 | pathogenic | 0 | 0 | 0 |
| rs515299 | benign | 1 | 1 | 1 |
| rs10127939 | benign | 1 | 1 | 1 |
| rs396991 | benign | 1 | 1 | 1 |
| rs145956175 | benign | 0 | 1 | 0 |
| rs747838255 | pathogenic | 1 | 1 | 1 |
| rs762659685 | pathogenic | 1 | 1 | 0 |
| rs104894831 | pathogenic | 0 | 1 | 0 |
| rs104894848 | pathogenic | 1 | 1 | 0 |
| rs104894833 | pathogenic | 0 | 1 | 0 |
| rs781838005 | pathogenic | 0 | 1 | 0 |
| rs104894834 | pathogenic | 1 | 1 | 0 |
| rs372966991 | pathogenic | 1 | 1 | 0 |
| rs138886989 | benign | 1 | 1 | 1 |
| rs28935195 | pathogenic | 0 | 1 | 0 |
| rs28935196 | pathogenic | 0 | 1 | 0 |
| rs372416832 | pathogenic | 1 | 1 | 1 |
| rs28935197 | pathogenic | 1 | 1 | 0 |
| rs28935486 | pathogenic | 1 | 1 | 0 |
| rs28935487 | pathogenic | 1 | 1 | 0 |
| rs28935488 | pathogenic | 1 | 1 | 1 |
| rs28935495 | pathogenic | 1 | 1 | 1 |
| rs28935485 | pathogenic | 1 | 1 | 1 |
| rs140329381 | pathogenic | 0 | 1 | 0 |
| rs104894830 | pathogenic | 0 | 1 | 0 |
| rs104894846 | pathogenic | 0 | 1 | 0 |
| rs28935489 | pathogenic | 0 | 1 | 0 |
| rs104894828 | pathogenic | 1 | 1 | 0 |
| rs28935491 | pathogenic | 1 | 1 | 0 |
| rs104894832 | pathogenic | 1 | 1 | 0 |
| rs28935492 | pathogenic | 1 | 1 | 0 |

| | | | | |
|---|---|---|---|---|
| rs730880437 | benign | 1 | 1 | 0 |
| rs28935493 | pathogenic | 0 | 1 | 0 |
| rs28935494 | pathogenic | 1 | 1 | 1 |
| rs111422676 | pathogenic | 1 | 1 | 1 |
| rs397515869 | benign | 1 | 1 | 0 |
| rs782449839 | pathogenic | 0 | 1 | 0 |
| rs104894852 | pathogenic | 1 | 1 | 1 |
| rs116929575 | pathogenic | 1 | 1 | 1 |
| rs7023954 | benign | 0 | 0 | 0 |
| rs184520335 | benign | 1 | 1 | 1 |
| rs143080527 | benign | 0 | 0 | 0 |
| rs1050565 | benign | 0 | 0 | 0 |
| rs11545035 | benign | 0 | 0 | 0 |
| rs267607190 | pathogenic | 1 | 1 | 1 |
| rs200283987 | benign | 0 | 0 | 0 |
| rs372556807 | benign | 1 | 1 | 0 |
| rs563073392 | benign | 0 | 0 | 0 |
| rs4539 | benign | 0 | 0 | 0 |
| rs28931609 | pathogenic | 1 | 1 | 1 |
| rs121912978 | pathogenic | 1 | 1 | 1 |
| rs104894072 | pathogenic | 1 | 1 | 1 |
| rs551933154 | benign | 1 | 1 | 1 |
| rs752962897 | benign | 1 | 1 | 0 |
| rs121912977 | pathogenic | 1 | 1 | 0 |
| rs4537 | benign | 1 | 1 | 1 |
| rs201830462 | benign | 0 | 0 | 0 |
| rs4544 | benign | 0 | 0 | 0 |
| rs562670189 | benign | 0 | 0 | 0 |
| rs4538 | benign | 0 | 1 | 0 |
| rs61757294 | pathogenic | 0 | 0 | 0 |
| rs5313 | benign | 0 | 0 | 0 |
| rs4545 | benign | 0 | 0 | 0 |
| rs72554627 | pathogenic | 1 | 1 | 1 |
| rs72554626 | pathogenic | 1 | 1 | 1 |
| rs387906861 | pathogenic | 1 | 1 | 1 |
| rs749655461 | pathogenic | 1 | 1 | 1 |
| rs121909606 | pathogenic | 1 | 1 | 1 |
| rs121909607 | pathogenic | 1 | 1 | 1 |
| rs200203992 | benign | 0 | 0 | 0 |
| rs6050 | benign | 0 | 0 | 0 |
| rs184635235 | benign | 0 | 0 | 0 |
| rs139146037 | benign | 0 | 0 | 0 |
| rs121909612 | pathogenic | 0 | 0 | 0 |
| rs78506343 | pathogenic | 0 | 0 | 0 |
| rs121918138 | pathogenic | 1 | 1 | 1 |
| rs121918139 | pathogenic | 1 | 1 | 1 |
| rs121918140 | pathogenic | 0 | 0 | 0 |
| rs36063028 | benign | 1 | 1 | 1 |
| rs45495192 | benign | 1 | 1 | 1 |
| rs45458802 | benign | 1 | 1 | 1 |
| rs552865793 | pathogenic | 1 | 1 | 1 |
| rs753744791 | pathogenic | 1 | 1 | 1 |
| rs777518512 | benign | 1 | 1 | 1 |
| rs142708991 | benign | 1 | 1 | 0 |

| | | | | |
|---|---|---|---|---|
| rs746069957 | benign | 0 | 0 | 0 |
| rs1334811 | benign | 0 | 0 | 0 |
| rs138894008 | benign | 1 | 1 | 0 |
| rs35030851 | benign | 1 | 1 | 0 |
| rs35378598 | benign | 0 | 0 | 0 |
| rs35818764 | benign | 1 | 1 | 1 |
| rs4631561 | benign | 1 | 1 | 1 |
| rs121918660 | pathogenic | 0 | 0 | 0 |
| rs373175587 | benign | 0 | 0 | 0 |
| rs61731956 | pathogenic | 1 | 1 | 1 |
| rs180177305 | pathogenic | 0 | 0 | 0 |
| rs119490108 | pathogenic | 0 | 0 | 0 |
| rs180177312 | pathogenic | 1 | 1 | 0 |
| rs180177314 | pathogenic | 1 | 1 | 0 |
| rs180177322 | pathogenic | 1 | 1 | 1 |
| rs180177323 | pathogenic | 1 | 1 | 1 |
| rs180177325 | pathogenic | 0 | 0 | 0 |
| rs35196441 | benign | 0 | 0 | 0 |
| rs28763904 | benign | 0 | 0 | 0 |
| rs2229519 | benign | 0 | 0 | 0 |
| rs137852886 | pathogenic | 0 | 0 | 0 |
| rs137852892 | pathogenic | 0 | 0 | 0 |
| rs770427750 | pathogenic | 0 | 0 | 0 |
| rs137852887 | pathogenic | 0 | 0 | 0 |
| rs137852893 | pathogenic | 0 | 0 | 0 |
| rs80338671 | pathogenic | 1 | 1 | 0 |
| rs2228389 | benign | 0 | 0 | 0 |
| rs80338672 | pathogenic | 0 | 0 | 0 |
| rs201958741 | pathogenic | 0 | 0 | 0 |
| rs80338673 | pathogenic | 0 | 0 | 0 |
| rs137852888 | pathogenic | 0 | 0 | 0 |
| rs137852889 | pathogenic | 0 | 0 | 0 |
| rs137852894 | pathogenic | 0 | 0 | 0 |
| rs137852890 | pathogenic | 0 | 0 | 0 |
| rs137852891 | pathogenic | 0 | 0 | 0 |
| rs766935302 | pathogenic | 0 | 0 | 0 |
| rs193074572 | benign | 0 | 0 | 0 |
| rs555210692 | benign | 1 | 1 | 0 |
| rs119103242 | pathogenic | 0 | 1 | 0 |
| rs143725101 | benign | 0 | 1 | 0 |
| rs28940589 | pathogenic | 1 | 1 | 1 |
| rs113994145 | pathogenic | 0 | 0 | 0 |
| rs773872994 | benign | 0 | 0 | 0 |
| rs28940590 | pathogenic | 1 | 1 | 0 |
| rs145400751 | benign | 1 | 1 | 1 |
| rs587782331 | benign | 0 | 0 | 0 |
| rs730881787 | benign | 0 | 0 | 0 |
| rs587778531 | benign | 0 | 0 | 0 |
| rs781652274 | benign | 0 | 0 | 0 |
| rs55882234 | benign | 0 | 0 | 0 |
| rs2020908 | benign | 0 | 0 | 0 |
| rs63750741 | pathogenic | 1 | 1 | 1 |
| rs41295268 | benign | 1 | 1 | 1 |
| rs63750897 | benign | 1 | 1 | 0 |

| | | | | |
|---|---|---|---|---|
| rs63751005 | benign | 1 | 1 | 0 |
| rs587779223 | benign | 0 | 0 | 0 |
| rs202127474 | benign | 0 | 0 | 0 |
| rs374230313 | benign | 0 | 0 | 0 |
| rs2020912 | benign | 0 | 0 | 0 |
| rs63751017 | pathogenic | 0 | 0 | 0 |
| rs587779246 | pathogenic | 0 | 0 | 0 |
| rs63749919 | pathogenic | 0 | 0 | 0 |
| rs63750563 | pathogenic | 1 | 1 | 1 |
| rs63749999 | pathogenic | 0 | 0 | 0 |
| rs200492211 | pathogenic | 0 | 0 | 0 |
| rs576269342 | benign | 0 | 0 | 0 |
| rs267608075 | benign | 0 | 0 | 0 |
| rs149605979 | benign | 0 | 0 | 0 |
| rs63750252 | benign | 0 | 0 | 0 |
| rs754289472 | benign | 0 | 0 | 0 |
| rs63750119 | pathogenic | 0 | 0 | 0 |
| rs587782109 | benign | 0 | 0 | 0 |
| rs34625968 | benign | 1 | 1 | 1 |
| rs764786814 | benign | 1 | 1 | 1 |
| rs199594809 | benign | 1 | 1 | 1 |
| rs74315364 | pathogenic | 1 | 1 | 1 |
| rs201253919 | pathogenic | 0 | 0 | 0 |
| rs387906828 | pathogenic | 1 | 1 | 1 |
| rs34521609 | benign | 1 | 1 | 1 |
| rs199472894 | pathogenic | 0 | 0 | 0 |
| rs199472902 | pathogenic | 0 | 0 | 0 |
| rs121912505 | pathogenic | 0 | 0 | 0 |
| rs28928905 | pathogenic | 0 | 0 | 0 |
| rs374376640 | benign | 0 | 0 | 0 |
| rs202102276 | benign | 0 | 0 | 0 |
| rs199472918 | pathogenic | 0 | 0 | 0 |
| rs121912516 | pathogenic | 1 | 1 | 1 |
| rs121912504 | pathogenic | 0 | 0 | 0 |
| rs9333649 | pathogenic | 0 | 0 | 0 |
| rs121912508 | pathogenic | 0 | 0 | 0 |
| rs199473428 | pathogenic | 1 | 1 | 1 |
| rs104894021 | pathogenic | 0 | 0 | 0 |
| rs28928904 | pathogenic | 1 | 1 | 1 |
| rs199473039 | benign | 1 | 1 | 1 |
| rs199472958 | pathogenic | 1 | 1 | 1 |
| rs1137617 | pathogenic | 1 | 1 | 1 |
| rs199473532 | pathogenic | 1 | 1 | 1 |
| rs199472988 | pathogenic | 0 | 0 | 0 |
| rs189014161 | pathogenic | 0 | 0 | 0 |
| rs12720441 | pathogenic | 0 | 0 | 0 |
| rs199473001 | pathogenic | 1 | 1 | 0 |
| rs121912506 | pathogenic | 1 | 1 | 0 |
| rs773724817 | pathogenic | 1 | 1 | 1 |
| rs118204002 | pathogenic | 1 | 1 | 1 |
| rs138556525 | pathogenic | 1 | 1 | 1 |
| rs72558181 | pathogenic | 1 | 1 | 1 |
| rs118204005 | pathogenic | 1 | 1 | 1 |
| rs118204004 | pathogenic | 0 | 0 | 0 |

| | | | | |
|---|---|---|---|---|
| rs118204001 | pathogenic | 1 | 1 | 1 |
| rs118204006 | pathogenic | 1 | 1 | 1 |
| rs118204003 | pathogenic | 0 | 0 | 0 |
| rs121913147 | pathogenic | 1 | 1 | 0 |
| rs2229429 | benign | 1 | 1 | 0 |
| rs182552223 | benign | 0 | 0 | 0 |
| rs146588336 | benign | 0 | 0 | 0 |
| rs121964983 | pathogenic | 0 | 1 | 0 |
| rs121964982 | pathogenic | 0 | 0 | 0 |
| rs386833679 | pathogenic | 0 | 0 | 0 |
| rs386833680 | pathogenic | 0 | 0 | 0 |
| rs121964981 | pathogenic | 1 | 1 | 0 |
| rs121964984 | pathogenic | 0 | 0 | 0 |
| rs144971200 | benign | 0 | 0 | 0 |
| rs121964985 | pathogenic | 1 | 1 | 0 |
| rs141246107 | benign | 0 | 0 | 0 |
| rs104894561 | pathogenic | 1 | 1 | 0 |
| rs377023736 | pathogenic | 0 | 0 | 0 |
| rs121964907 | pathogenic | 1 | 1 | 1 |
| rs121964909 | pathogenic | 1 | 1 | 1 |
| rs765070743 | pathogenic | 0 | 0 | 0 |
| rs2228119 | benign | 0 | 0 | 0 |
| rs745976989 | pathogenic | 1 | 1 | 1 |
| rs121964904 | pathogenic | 0 | 0 | 0 |
| rs386833430 | pathogenic | 1 | 1 | 1 |
| rs267606992 | pathogenic | 0 | 0 | 0 |
| rs61752063 | pathogenic | 0 | 0 | 0 |
| rs61752067 | pathogenic | 1 | 1 | 1 |
| rs104894934 | pathogenic | 0 | 0 | 0 |
| rs104894929 | pathogenic | 1 | 1 | 1 |
| rs80189640 | benign | 0 | 0 | 0 |
| rs137854547 | pathogenic | 1 | 1 | 1 |
| rs137854540 | pathogenic | 1 | 1 | 1 |
| rs782092363 | pathogenic | 1 | 1 | 1 |
| rs147783915 | benign | 0 | 0 | 0 |
| rs28933687 | pathogenic | 1 | 1 | 1 |
| rs149503319 | benign | 0 | 0 | 0 |
| rs1805147 | benign | 0 | 0 | 0 |
| rs587777218 | pathogenic | 1 | 1 | 1 |
| rs200786705 | benign | 0 | 0 | 0 |
| rs56219475 | benign | 0 | 0 | 0 |
| rs587781172 | benign | 0 | 0 | 0 |
| rs757094189 | pathogenic | 0 | 0 | 0 |
| rs140811086 | benign | 1 | 1 | 1 |
| rs574088829 | pathogenic | 1 | 1 | 1 |
| rs121918525 | pathogenic | 0 | 0 | 0 |
| rs121918527 | pathogenic | 0 | 0 | 0 |
| rs121964848 | pathogenic | 0 | 1 | 0 |
| rs121964845 | pathogenic | 1 | 1 | 1 |
| rs121964846 | pathogenic | 0 | 1 | 0 |
| rs17217716 | benign | 1 | 1 | 1 |
| rs141711342 | benign | 0 | 0 | 0 |
| rs372619120 | benign | 0 | 0 | 0 |
| rs63750002 | benign | 1 | 1 | 1 |

| | | | | |
|---|---|---|---|---|
| rs193922373 | benign | 0 | 0 | 0 |
| rs41295286 | benign | 0 | 0 | 0 |
| rs17217772 | benign | 0 | 0 | 0 |
| rs145649774 | benign | 0 | 0 | 0 |
| rs63750214 | pathogenic | 0 | 0 | 0 |
| rs63750716 | benign | 0 | 0 | 0 |
| rs202026056 | benign | 0 | 0 | 0 |
| rs4987188 | benign | 1 | 1 | 0 |
| rs267607938 | benign | 0 | 0 | 0 |
| rs17224367 | benign | 0 | 0 | 0 |
| rs63750006 | benign | 0 | 0 | 0 |
| rs587781373 | benign | 0 | 0 | 0 |
| rs63751207 | pathogenic | 1 | 1 | 0 |
| rs730881759 | benign | 0 | 0 | 0 |
| rs372350768 | benign | 1 | 1 | 1 |
| rs55778204 | benign | 1 | 1 | 0 |
| rs63749910 | benign | 0 | 0 | 0 |
| rs63750844 | benign | 0 | 0 | 0 |
| rs28929483 | pathogenic | 0 | 0 | 0 |
| rs61756468 | benign | 0 | 0 | 0 |
| rs63750875 | pathogenic | 1 | 1 | 0 |
| rs776528054 | benign | 1 | 1 | 0 |
| rs751939698 | benign | 0 | 0 | 0 |
| rs587780685 | benign | 0 | 0 | 0 |
| rs730881763 | benign | 1 | 1 | 0 |
| rs373226409 | benign | 0 | 0 | 0 |
| rs202145681 | benign | 1 | 1 | 1 |
| rs63749841 | benign | 1 | 1 | 1 |
| rs63750757 | benign | 1 | 1 | 1 |
| rs587778527 | benign | 1 | 1 | 1 |
| rs63750849 | benign | 0 | 0 | 0 |
| rs587779969 | benign | 0 | 0 | 0 |
| rs121908046 | pathogenic | 1 | 1 | 1 |
| rs28940882 | pathogenic | 1 | 1 | 0 |
| rs28940883 | pathogenic | 1 | 1 | 1 |
| rs727503943 | pathogenic | 1 | 1 | 1 |
| rs137853859 | pathogenic | 1 | 1 | 1 |
| rs121908045 | pathogenic | 1 | 1 | 0 |
| rs137853861 | pathogenic | 1 | 1 | 0 |
| rs3180383 | pathogenic | 0 | 1 | 0 |
| rs144431814 | benign | 0 | 0 | 0 |
| rs139502866 | benign | 0 | 0 | 0 |
| rs17524437 | benign | 0 | 0 | 0 |
| rs760960779 | benign | 0 | 0 | 0 |
| rs148960463 | pathogenic | 0 | 0 | 0 |
| rs111033647 | pathogenic | 1 | 1 | 1 |
| rs111033648 | pathogenic | 0 | 0 | 0 |
| rs1800461 | benign | 0 | 0 | 0 |
| rs111033656 | pathogenic | 0 | 0 | 0 |
| rs111033663 | pathogenic | 1 | 1 | 0 |
| rs111033669 | pathogenic | 1 | 1 | 1 |
| rs111033690 | pathogenic | 0 | 0 | 0 |
| rs111033686 | pathogenic | 0 | 0 | 0 |
| rs111033687 | pathogenic | 0 | 0 | 0 |

| | | | | |
|---|---|---|---|---|
| rs111033695 | pathogenic | 0 | 0 | 0 |
| rs111033697 | pathogenic | 0 | 0 | 0 |
| rs111033693 | pathogenic | 0 | 0 | 0 |
| rs111033694 | pathogenic | 0 | 0 | 0 |
| rs111033721 | pathogenic | 1 | 1 | 1 |
| rs111033722 | pathogenic | 0 | 0 | 0 |
| rs111033725 | pathogenic | 1 | 1 | 0 |
| rs75391579 | pathogenic | 1 | 1 | 1 |
| rs111033728 | pathogenic | 1 | 1 | 1 |
| rs111033735 | pathogenic | 1 | 1 | 1 |
| rs111033739 | pathogenic | 1 | 1 | 1 |
| rs111033736 | pathogenic | 0 | 0 | 0 |
| rs111033744 | pathogenic | 1 | 1 | 1 |
| rs111033750 | pathogenic | 1 | 1 | 1 |
| rs111033752 | pathogenic | 0 | 0 | 0 |
| rs111033751 | pathogenic | 0 | 0 | 0 |
| rs111033846 | pathogenic | 0 | 0 | 0 |
| rs368166217 | pathogenic | 0 | 0 | 0 |
| rs111033764 | pathogenic | 0 | 0 | 0 |
| rs111033847 | benign | 0 | 0 | 0 |
| rs111033778 | pathogenic | 0 | 0 | 0 |
| rs111033773 | pathogenic | 0 | 0 | 0 |
| rs111033792 | pathogenic | 0 | 0 | 0 |
| rs367543266 | pathogenic | 1 | 1 | 1 |
| rs111033794 | pathogenic | 0 | 0 | 0 |
| rs111033802 | pathogenic | 0 | 0 | 0 |
| rs111033803 | pathogenic | 0 | 0 | 0 |
| rs111033809 | pathogenic | 1 | 1 | 0 |
| rs121918376 | pathogenic | 0 | 1 | 0 |
| rs121918377 | pathogenic | 0 | 1 | 0 |
| rs121918378 | pathogenic | 1 | 1 | 1 |
| rs121918374 | pathogenic | 1 | 1 | 1 |
| rs137852379 | pathogenic | 0 | 0 | 0 |
| rs137852388 | pathogenic | 1 | 1 | 1 |
| rs137852368 | pathogenic | 0 | 0 | 0 |
| rs28935499 | pathogenic | 0 | 0 | 0 |
| rs28933668 | pathogenic | 0 | 0 | 0 |
| rs28933669 | pathogenic | 0 | 0 | 0 |
| rs28933670 | pathogenic | 0 | 0 | 0 |
| rs28933671 | pathogenic | 0 | 0 | 0 |
| rs28933672 | pathogenic | 0 | 0 | 0 |
| rs28937272 | pathogenic | 0 | 0 | 0 |
| rs35383156 | benign | 0 | 0 | 0 |
| rs137852418 | pathogenic | 0 | 0 | 0 |
| rs137852419 | pathogenic | 0 | 0 | 0 |
| rs137852428 | pathogenic | 0 | 0 | 0 |
| rs137852436 | pathogenic | 0 | 0 | 0 |
| rs28937285 | pathogenic | 0 | 0 | 0 |
| rs35213384 | benign | 1 | 1 | 1 |
| rs3810950 | benign | 0 | 0 | 0 |
| rs201479289 | pathogenic | 0 | 0 | 0 |
| rs121912820 | pathogenic | 0 | 1 | 0 |
| rs121912815 | pathogenic | 0 | 1 | 0 |
| rs115510708 | benign | 0 | 0 | 0 |

| | | | | |
|---|---|---|---|---|
| rs868749 | benign | 0 | 0 | 0 |
| rs121912823 | pathogenic | 0 | 0 | 0 |
| rs115212829 | benign | 0 | 0 | 0 |
| rs121912822 | pathogenic | 0 | 0 | 0 |
| rs121912816 | pathogenic | 0 | 1 | 0 |
| rs4838544 | benign | 0 | 0 | 0 |
| rs121912818 | pathogenic | 0 | 0 | 0 |
| rs121912821 | pathogenic | 0 | 0 | 0 |
| rs121912817 | pathogenic | 0 | 0 | 0 |
| rs8178992 | benign | 0 | 0 | 0 |
| rs121912819 | pathogenic | 1 | 1 | 0 |
| rs80097077 | benign | 1 | 1 | 0 |
| rs121909359 | pathogenic | 1 | 1 | 0 |
| rs121909358 | pathogenic | 1 | 1 | 1 |
| rs121909361 | pathogenic | 1 | 1 | 1 |
| rs121909357 | pathogenic | 0 | 1 | 0 |
| rs121909365 | pathogenic | 1 | 1 | 1 |
| rs6413484 | pathogenic | 1 | 1 | 1 |
| rs121909373 | pathogenic | 1 | 1 | 1 |
| rs121909366 | pathogenic | 1 | 1 | 1 |
| rs121909367 | pathogenic | 1 | 1 | 1 |
| rs121909368 | pathogenic | 1 | 1 | 1 |
| rs121909369 | pathogenic | 0 | 0 | 0 |
| rs121909363 | pathogenic | 1 | 1 | 1 |
| rs45588036 | pathogenic | 1 | 1 | 1 |
| rs121909364 | pathogenic | 1 | 1 | 1 |
| rs80358282 | benign | 0 | 0 | 0 |
| rs80358284 | pathogenic | 1 | 1 | 1 |
| rs61735303 | benign | 0 | 0 | 0 |
| rs2229274 | benign | 0 | 0 | 0 |
| rs145006491 | benign | 0 | 0 | 0 |
| rs121913580 | pathogenic | 1 | 1 | 0 |
| rs730882196 | pathogenic | 0 | 0 | 0 |
| rs153477 | benign | 0 | 0 | 0 |
| rs153478 | benign | 1 | 1 | 0 |
| rs61740602 | benign | 1 | 1 | 0 |
| rs104893892 | pathogenic | 1 | 1 | 0 |
| rs121909595 | pathogenic | 1 | 1 | 1 |
| rs28931605 | pathogenic | 1 | 1 | 1 |
| rs121909597 | pathogenic | 0 | 1 | 0 |
| rs202233735 | pathogenic | 0 | 1 | 0 |
| rs121909596 | pathogenic | 1 | 1 | 1 |
| rs75156162 | benign | 0 | 1 | 0 |
| rs2305430 | benign | 1 | 1 | 1 |
| rs150318966 | benign | 0 | 0 | 0 |
| rs121909598 | pathogenic | 0 | 1 | 0 |
| rs5744175 | benign | 0 | 0 | 0 |
| rs28935476 | pathogenic | 1 | 1 | 1 |
| rs28935475 | pathogenic | 1 | 1 | 1 |
| rs122461163 | pathogenic | 1 | 1 | 1 |
| rs62626305 | pathogenic | 1 | 1 | 1 |
| rs587777157 | pathogenic | 0 | 0 | 0 |
| rs137854478 | pathogenic | 1 | 1 | 0 |
| rs137854465 | pathogenic | 0 | 0 | 0 |

| | | | | |
|---|---|---|---|---|
| rs137854470 | pathogenic | 0 | 0 | 0 |
| rs137854482 | pathogenic | 0 | 0 | 0 |
| rs137854473 | pathogenic | 0 | 0 | 0 |
| rs137854456 | pathogenic | 0 | 0 | 0 |
| rs140598 | benign | 0 | 0 | 0 |
| rs104894105 | pathogenic | 1 | 1 | 1 |
| rs104894107 | pathogenic | 1 | 1 | 1 |
| rs104894106 | pathogenic | 1 | 1 | 1 |
| rs74621026 | benign | 1 | 1 | 1 |
| rs2290332 | benign | 0 | 0 | 0 |
| rs780202604 | pathogenic | 1 | 1 | 1 |
| rs74315305 | pathogenic | 1 | 1 | 0 |
| rs766751878 | pathogenic | 1 | 1 | 1 |
| rs137854603 | pathogenic | 1 | 1 | 1 |
| rs139592595 | benign | 1 | 1 | 0 |
| rs150565592 | benign | 1 | 1 | 1 |
| rs763520126 | benign | 0 | 0 | 0 |
| rs55980502 | benign | 0 | 0 | 0 |
| rs137852812 | pathogenic | 0 | 0 | 0 |
| rs137852813 | pathogenic | 0 | 0 | 0 |
| rs190222208 | benign | 0 | 0 | 0 |
| rs138459502 | pathogenic | 0 | 0 | 0 |
| rs730881041 | pathogenic | 0 | 0 | 0 |
| rs267607080 | pathogenic | 0 | 0 | 0 |
| rs137852814 | pathogenic | 0 | 0 | 0 |
| rs267607079 | pathogenic | 0 | 0 | 0 |
| rs192378417 | benign | 0 | 0 | 0 |
| rs1288775 | benign | 0 | 0 | 0 |
| rs80338737 | pathogenic | 1 | 1 | 1 |
| rs397514708 | pathogenic | 0 | 0 | 0 |
| rs775933965 | pathogenic | 0 | 0 | 0 |
| rs747608698 | benign | 0 | 1 | 0 |
| rs1145086 | benign | 0 | 1 | 0 |
| rs121912953 | pathogenic | 1 | 1 | 0 |
| rs243849 | benign | 0 | 0 | 0 |
| rs121912955 | pathogenic | 1 | 1 | 1 |
| rs17859943 | benign | 0 | 1 | 0 |
| rs755557988 | benign | 1 | 1 | 1 |
| rs10195 | benign | 0 | 0 | 0 |
| rs61735357 | benign | 1 | 1 | 1 |
| rs137882374 | benign | 0 | 0 | 0 |
| rs5388 | benign | 0 | 0 | 0 |
| rs6352 | benign | 0 | 0 | 0 |
| rs76645461 | pathogenic | 1 | 1 | 0 |
| rs1050828 | pathogenic | 0 | 0 | 0 |
| rs1050829 | pathogenic | 0 | 0 | 0 |
| rs137852314 | pathogenic | 0 | 0 | 0 |
| rs137852331 | pathogenic | 0 | 0 | 0 |
| rs5030872 | pathogenic | 0 | 0 | 0 |
| rs267606836 | pathogenic | 0 | 0 | 0 |
| rs5030868 | pathogenic | 0 | 0 | 0 |
| rs137852330 | pathogenic | 0 | 1 | 0 |
| rs137852326 | pathogenic | 1 | 1 | 1 |
| rs137852318 | pathogenic | 1 | 1 | 1 |

| | | | | |
|---|---|---|---|---|
| rs74575103 | benign | 0 | 0 | 0 |
| rs137852327 | pathogenic | 1 | 1 | 1 |
| rs76723693 | pathogenic | 0 | 0 | 0 |
| rs5030869 | pathogenic | 0 | 0 | 0 |
| rs34193178 | pathogenic | 1 | 1 | 1 |
| rs137852316 | pathogenic | 1 | 1 | 0 |
| rs398123546 | pathogenic | 1 | 1 | 1 |
| rs72554665 | pathogenic | 0 | 0 | 0 |
| rs72554664 | pathogenic | 0 | 0 | 0 |
| rs28939068 | pathogenic | 1 | 1 | 1 |
| rs59301204 | pathogenic | 1 | 1 | 1 |
| rs143715750 | benign | 1 | 1 | 1 |
| rs35263902 | pathogenic | 1 | 1 | 1 |
| rs202214518 | benign | 1 | 1 | 1 |
| rs34076756 | benign | 0 | 0 | 0 |
| rs199592341 | pathogenic | 1 | 1 | 0 |
| rs121908833 | pathogenic | 1 | 1 | 1 |
| rs121908834 | pathogenic | 1 | 1 | 0 |
| rs138454151 | benign | 0 | 0 | 0 |
| rs141323599 | benign | 0 | 0 | 0 |
| rs137853200 | pathogenic | 0 | 0 | 0 |
| rs137853199 | pathogenic | 0 | 0 | 0 |
| rs139787163 | benign | 0 | 0 | 0 |
| rs61731097 | benign | 1 | 1 | 1 |
| rs61729795 | benign | 0 | 0 | 0 |
| rs149242330 | benign | 0 | 0 | 0 |
| rs587777490 | pathogenic | 1 | 1 | 1 |
| rs398123011 | pathogenic | 1 | 1 | 0 |
| rs775186117 | pathogenic | 0 | 0 | 0 |
| rs72677237 | benign | 0 | 0 | 0 |
| rs104894163 | pathogenic | 1 | 1 | 1 |
| rs121909588 | pathogenic | 0 | 0 | 0 |
| rs1042579 | benign | 0 | 0 | 0 |
| rs121909273 | pathogenic | 0 | 0 | 0 |
| rs121909274 | pathogenic | 0 | 0 | 0 |
| rs119473034 | pathogenic | 1 | 1 | 1 |
| rs398123065 | pathogenic | 0 | 0 | 0 |
| rs587777042 | pathogenic | 1 | 1 | 0 |
| rs761413864 | pathogenic | 1 | 1 | 1 |
| rs118203965 | pathogenic | 0 | 0 | 0 |
| rs118203966 | pathogenic | 0 | 0 | 0 |
| rs1801201 | benign | 1 | 1 | 0 |
| rs1136201 | benign | 1 | 1 | 1 |
| rs144148008 | benign | 0 | 0 | 0 |
| rs200118378 | benign | 1 | 1 | 1 |
| rs112984118 | benign | 1 | 1 | 1 |
| rs149446102 | benign | 1 | 1 | 1 |
| rs1427463 | benign | 0 | 0 | 0 |
| rs201936720 | benign | 1 | 1 | 1 |
| rs61751983 | benign | 1 | 1 | 1 |
| rs104894632 | pathogenic | 1 | 1 | 1 |
| rs149202905 | benign | 0 | 0 | 0 |
| rs2230210 | benign | 1 | 1 | 1 |
| rs111631988 | benign | 0 | 0 | 0 |

| | | | | |
|---|---|---|---|---|
| rs137852550 | pathogenic | 0 | 0 | 0 |
| rs5963403 | benign | 0 | 0 | 0 |
| rs138018739 | benign | 0 | 0 | 0 |
| rs62640590 | benign | 0 | 0 | 0 |
| rs121909581 | pathogenic | 1 | 1 | 1 |
| rs202017913 | benign | 1 | 1 | 1 |
| rs143163513 | benign | 0 | 0 | 0 |
| rs238238 | benign | 0 | 1 | 0 |
| rs143945974 | benign | 0 | 0 | 0 |
| rs238239 | benign | 1 | 1 | 1 |
| rs121918403 | pathogenic | 0 | 1 | 0 |
| rs61735456 | benign | 0 | 1 | 0 |
| rs121918404 | pathogenic | 1 | 1 | 0 |
| rs12674488 | benign | 0 | 0 | 0 |
| rs201599657 | pathogenic | 0 | 0 | 0 |
| rs1057090 | benign | 0 | 0 | 0 |
| rs1057091 | benign | 0 | 0 | 0 |
| rs587783740 | benign | 0 | 0 | 0 |
| rs75956769 | benign | 0 | 0 | 0 |
| rs201691299 | benign | 0 | 0 | 0 |
| rs62621087 | benign | 0 | 0 | 0 |
| rs201308454 | benign | 0 | 0 | 0 |
| rs145657360 | benign | 0 | 0 | 0 |
| rs121434592 | pathogenic | 1 | 1 | 0 |
| rs375650413 | benign | 0 | 0 | 0 |
| rs121434437 | pathogenic | 1 | 1 | 0 |
| rs763494292 | pathogenic | 0 | 0 | 0 |
| rs121964997 | pathogenic | 1 | 1 | 0 |
| rs121964996 | pathogenic | 0 | 0 | 0 |
| rs41283425 | benign | 1 | 1 | 1 |
| rs376289528 | benign | 1 | 1 | 1 |
| rs144171604 | benign | 1 | 1 | 1 |
| rs782440692 | pathogenic | 1 | 1 | 1 |
| rs782058451 | pathogenic | 1 | 1 | 1 |
| rs78437817 | benign | 0 | 0 | 0 |
| rs200327969 | benign | 1 | 1 | 1 |
| rs143043662 | benign | 0 | 0 | 0 |
| rs111966833 | benign | 1 | 1 | 1 |
| rs35523678 | benign | 0 | 0 | 0 |
| rs515726208 | pathogenic | 0 | 0 | 0 |
| rs397517239 | pathogenic | 1 | 1 | 1 |
| rs148966602 | benign | 1 | 1 | 1 |
| rs76371972 | benign | 1 | 1 | 1 |
| rs2229968 | benign | 1 | 1 | 1 |
| rs61751538 | benign | 1 | 1 | 1 |
| rs200800978 | pathogenic | 1 | 1 | 1 |
| rs148695964 | benign | 1 | 1 | 1 |
| rs140042418 | benign | 1 | 1 | 1 |
| rs116185403 | benign | 0 | 0 | 0 |
| rs142828669 | benign | 1 | 1 | 1 |
| rs139832048 | benign | 0 | 1 | 0 |
| rs58750568 | benign | 0 | 1 | 0 |
| rs77681311 | benign | 0 | 0 | 0 |
| rs151208452 | benign | 0 | 0 | 0 |

| | | | | |
|---|---|---|---|---|
| rs76463933 | benign | 0 | 0 | 0 |
| rs144434647 | benign | 0 | 0 | 0 |
| rs117361485 | benign | 0 | 0 | 0 |
| rs587777588 | pathogenic | 0 | 0 | 0 |
| rs80338796 | pathogenic | 0 | 0 | 0 |
| rs121434594 | pathogenic | 0 | 0 | 0 |
| rs146766138 | pathogenic | 1 | 1 | 1 |
| rs28939679 | pathogenic | 1 | 1 | 1 |
| rs748754134 | pathogenic | 1 | 1 | 0 |
| rs764276946 | pathogenic | 1 | 1 | 1 |
| rs111033588 | pathogenic | 1 | 1 | 1 |
| rs201633414 | pathogenic | 1 | 1 | 1 |
| rs121909262 | pathogenic | 1 | 1 | 1 |
| rs104893702 | pathogenic | 1 | 1 | 1 |
| rs121909265 | pathogenic | 1 | 1 | 1 |
| rs121909266 | pathogenic | 1 | 1 | 1 |
| rs193922439 | pathogenic | 1 | 1 | 0 |
| rs104893691 | pathogenic | 1 | 1 | 1 |
| rs104893695 | pathogenic | 0 | 0 | 0 |
| rs104893708 | pathogenic | 1 | 1 | 1 |
| rs121909260 | pathogenic | 0 | 0 | 0 |
| rs104893696 | pathogenic | 0 | 0 | 0 |
| rs121909267 | pathogenic | 0 | 0 | 0 |
| rs121909263 | pathogenic | 0 | 0 | 0 |
| rs121909264 | pathogenic | 0 | 0 | 0 |
| rs104893694 | pathogenic | 1 | 1 | 1 |
| rs121909268 | pathogenic | 1 | 1 | 1 |
| rs104893689 | pathogenic | 1 | 1 | 0 |
| rs104893707 | pathogenic | 1 | 1 | 0 |
| rs104893697 | pathogenic | 1 | 1 | 1 |
| rs28936684 | pathogenic | 1 | 1 | 1 |
| rs62269092 | benign | 1 | 1 | 0 |
| rs121909259 | pathogenic | 1 | 1 | 0 |
| rs548403340 | benign | 1 | 1 | 0 |
| rs142818334 | benign | 0 | 0 | 0 |
| rs12493789 | benign | 1 | 1 | 1 |
| rs193922423 | pathogenic | 1 | 1 | 0 |
| rs104893719 | pathogenic | 1 | 1 | 1 |
| rs117375173 | benign | 1 | 1 | 0 |
| rs104893712 | pathogenic | 0 | 0 | 0 |
| rs121917776 | pathogenic | 1 | 1 | 1 |
| rs767325003 | benign | 1 | 1 | 1 |
| rs35824453 | benign | 0 | 0 | 0 |
| rs141641266 | benign | 0 | 0 | 0 |
| rs1050239 | benign | 0 | 0 | 0 |
| rs141303384 | benign | 0 | 0 | 0 |
| rs12720458 | pathogenic | 0 | 0 | 0 |
| rs368507376 | pathogenic | 1 | 1 | 1 |
| rs17215500 | pathogenic | 1 | 1 | 1 |
| rs120074188 | pathogenic | 1 | 1 | 1 |
| rs397508097 | pathogenic | 1 | 1 | 1 |
| rs138551008 | pathogenic | 0 | 0 | 0 |
| rs387907139 | pathogenic | 1 | 1 | 1 |
| rs765487627 | pathogenic | 0 | 0 | 0 |

| | | | | |
|---|---|---|---|---|
| rs587778587 | benign | 0 | 0 | 0 |
| rs45568339 | benign | 0 | 0 | 0 |
| rs78179744 | benign | 0 | 0 | 0 |
| rs45624036 | benign | 0 | 0 | 0 |
| rs61755166 | benign | 0 | 0 | 0 |
| rs180177132 | pathogenic | 1 | 1 | 1 |
| rs587776527 | pathogenic | 0 | 0 | 0 |
| rs373783514 | benign | 0 | 0 | 0 |
| rs730881896 | benign | 0 | 0 | 0 |
| rs118203998 | pathogenic | 0 | 0 | 0 |
| rs41281039 | pathogenic | 1 | 1 | 1 |
| rs118204430 | pathogenic | 1 | 1 | 1 |
| rs1800546 | pathogenic | 1 | 1 | 0 |
| rs76917243 | pathogenic | 0 | 1 | 0 |
| rs78340951 | pathogenic | 0 | 1 | 0 |
| rs77718928 | pathogenic | 0 | 0 | 0 |
| rs369586696 | pathogenic | 0 | 1 | 0 |
| rs745517517 | pathogenic | 1 | 1 | 0 |
| rs372872777 | pathogenic | 0 | 0 | 0 |
| rs748704459 | pathogenic | 0 | 0 | 0 |
| rs769215411 | pathogenic | 0 | 0 | 0 |
| rs139425890 | pathogenic | 0 | 0 | 0 |
| rs779694939 | pathogenic | 1 | 1 | 0 |
| rs773729410 | pathogenic | 0 | 0 | 0 |
| rs121908623 | pathogenic | 1 | 1 | 1 |
| rs121908622 | pathogenic | 1 | 1 | 1 |
| rs121908621 | pathogenic | 1 | 1 | 1 |
| rs762106720 | pathogenic | 1 | 1 | 0 |
| rs199877522 | pathogenic | 0 | 0 | 0 |
| rs74315291 | pathogenic | 1 | 1 | 1 |
| rs79137415 | benign | 0 | 0 | 0 |
| rs121913045 | pathogenic | 0 | 0 | 0 |
| rs34295337 | pathogenic | 0 | 0 | 0 |
| rs121913046 | pathogenic | 0 | 0 | 0 |
| rs121913047 | pathogenic | 0 | 0 | 0 |
| rs121913048 | pathogenic | 0 | 0 | 0 |
| rs267607166 | pathogenic | 0 | 0 | 0 |
| rs17850684 | benign | 1 | 1 | 1 |
| rs119476051 | pathogenic | 1 | 1 | 1 |
| rs17850683 | benign | 1 | 1 | 1 |
| rs119476049 | pathogenic | 1 | 1 | 1 |
| rs119476046 | pathogenic | 1 | 1 | 1 |
| rs119476048 | pathogenic | 0 | 0 | 0 |
| rs119476047 | pathogenic | 1 | 1 | 1 |
| rs28939094 | pathogenic | 1 | 1 | 1 |
| rs119476050 | pathogenic | 0 | 0 | 0 |
| rs387906941 | pathogenic | 1 | 1 | 1 |
| rs41511344 | pathogenic | 1 | 1 | 0 |
| rs748112833 | pathogenic | 1 | 1 | 1 |
| rs74342080 | benign | 1 | 1 | 0 |
| rs116482870 | benign | 0 | 0 | 0 |
| rs104893743 | pathogenic | 0 | 0 | 0 |
| rs117179004 | benign | 0 | 0 | 0 |
| rs587779409 | pathogenic | 1 | 1 | 1 |

| | | | | |
|---|---|---|---|---|
| rs3745406 | benign | 1 | 1 | 0 |
| rs387906696 | pathogenic | 0 | 0 | 0 |
| rs72549369 | pathogenic | 1 | 1 | 0 |
| rs45523133 | benign | 0 | 0 | 0 |
| rs119460972 | pathogenic | 0 | 1 | 0 |
| rs34929837 | benign | 0 | 1 | 0 |
| rs17323225 | benign | 0 | 1 | 0 |
| rs17011368 | benign | 0 | 0 | 0 |
| rs748902766 | pathogenic | 1 | 1 | 1 |
| rs387907292 | pathogenic | 0 | 0 | 0 |
| rs387907293 | pathogenic | 1 | 1 | 0 |
| rs371526758 | pathogenic | 1 | 1 | 1 |
| rs142968179 | pathogenic | 1 | 1 | 1 |
| rs387907291 | pathogenic | 1 | 1 | 0 |
| rs121907949 | pathogenic | 0 | 0 | 0 |
| rs121907947 | pathogenic | 0 | 0 | 0 |
| rs121907950 | pathogenic | 1 | 1 | 1 |
| rs121907948 | pathogenic | 1 | 1 | 1 |
| rs4926 | benign | 0 | 0 | 0 |
| rs748809996 | pathogenic | 0 | 0 | 0 |
| rs1799883 | benign | 1 | 1 | 0 |
| rs104893974 | pathogenic | 0 | 1 | 0 |
| rs104893973 | pathogenic | 0 | 1 | 0 |
| rs34452707 | benign | 0 | 0 | 0 |
| rs119467003 | pathogenic | 1 | 1 | 1 |
| rs121908261 | pathogenic | 0 | 1 | 0 |
| rs28933985 | pathogenic | 1 | 1 | 1 |
| rs80356669 | pathogenic | 1 | 1 | 1 |
| rs28936677 | pathogenic | 1 | 1 | 0 |
| rs121918326 | pathogenic | 0 | 0 | 0 |
| rs121918324 | pathogenic | 1 | 1 | 0 |
| rs121918325 | pathogenic | 1 | 1 | 0 |
| rs41270025 | pathogenic | 1 | 1 | 0 |
| rs121918323 | pathogenic | 0 | 0 | 0 |
| rs12735723 | pathogenic | 1 | 1 | 0 |
| rs36013429 | benign | 0 | 0 | 0 |
| rs28936676 | pathogenic | 1 | 1 | 0 |
| rs771809901 | pathogenic | 1 | 1 | 1 |
| rs758350837 | benign | 1 | 1 | 0 |
| rs121908192 | pathogenic | 1 | 1 | 1 |
| rs147568500 | pathogenic | 1 | 1 | 1 |
| rs3729734 | benign | 0 | 0 | 0 |
| rs371536347 | benign | 1 | 1 | 1 |
| rs80358193 | pathogenic | 1 | 1 | 1 |
| rs119476044 | pathogenic | 1 | 1 | 1 |
| rs111292798 | benign | 1 | 1 | 1 |
| rs121908063 | pathogenic | 1 | 1 | 0 |
| rs121908065 | pathogenic | 1 | 1 | 1 |
| rs121908062 | pathogenic | 1 | 1 | 1 |
| rs121908064 | pathogenic | 1 | 1 | 1 |
| rs61746421 | benign | 1 | 1 | 1 |
| rs34886916 | benign | 1 | 1 | 1 |
| rs118204045 | pathogenic | 0 | 0 | 0 |
| rs138607412 | benign | 1 | 1 | 1 |

| | | | | |
|---|---|---|---|---|
| rs118204046 | pathogenic | 0 | 0 | 0 |
| rs118204048 | pathogenic | 0 | 0 | 0 |
| rs118204044 | pathogenic | 0 | 0 | 0 |
| rs149664056 | benign | 0 | 0 | 0 |
| rs121909285 | pathogenic | 0 | 0 | 0 |
| rs1802498 | benign | 0 | 0 | 0 |
| rs1065761 | benign | 0 | 0 | 0 |
| rs119477054 | pathogenic | 0 | 0 | 0 |
| rs12720097 | benign | 0 | 0 | 0 |
| rs747281324 | pathogenic | 1 | 1 | 0 |
| rs367543028 | pathogenic | 1 | 1 | 0 |
| rs146077918 | benign | 0 | 0 | 0 |
| rs142551229 | benign | 0 | 0 | 0 |
| rs367543039 | pathogenic | 0 | 0 | 0 |
| rs587779884 | pathogenic | 1 | 1 | 0 |
| rs137853153 | pathogenic | 0 | 0 | 0 |
| rs367543029 | pathogenic | 0 | 0 | 0 |
| rs587783037 | pathogenic | 1 | 1 | 0 |
| rs142928725 | benign | 0 | 0 | 0 |
| rs28385141 | benign | 0 | 0 | 0 |
| rs587779888 | pathogenic | 0 | 0 | 0 |
| rs121909801 | pathogenic | 1 | 1 | 1 |
| rs121909790 | pathogenic | 1 | 1 | 0 |
| rs121909797 | pathogenic | 1 | 1 | 0 |
| rs121909794 | pathogenic | 1 | 1 | 1 |
| rs121909791 | pathogenic | 1 | 1 | 1 |
| rs121909793 | pathogenic | 1 | 1 | 1 |
| rs147977279 | benign | 1 | 1 | 1 |
| rs147352564 | benign | 0 | 0 | 0 |
| rs368758045 | benign | 1 | 1 | 1 |
| rs114498261 | benign | 1 | 1 | 0 |
| rs35484156 | benign | 0 | 0 | 0 |
| rs139176878 | benign | 0 | 0 | 0 |
| rs11543934 | benign | 0 | 0 | 0 |
| rs34192549 | benign | 0 | 0 | 0 |
| rs568736295 | benign | 0 | 0 | 0 |
| rs55747232 | benign | 0 | 0 | 0 |
| rs781887272 | benign | 0 | 0 | 0 |
| rs146273341 | benign | 0 | 0 | 0 |
| rs41473544 | benign | 0 | 0 | 0 |
| rs137852680 | pathogenic | 0 | 0 | 0 |
| rs55891000 | benign | 0 | 0 | 0 |
| rs140351189 | benign | 0 | 0 | 0 |
| rs137852679 | pathogenic | 1 | 1 | 0 |
| rs137852678 | pathogenic | 0 | 0 | 0 |
| rs137852677 | pathogenic | 1 | 1 | 1 |
| rs138620818 | benign | 0 | 0 | 0 |
| rs121908531 | pathogenic | 0 | 0 | 0 |
| rs72554307 | pathogenic | 0 | 0 | 0 |
| rs72554308 | pathogenic | 0 | 0 | 0 |
| rs72554312 | pathogenic | 1 | 1 | 0 |
| rs67939655 | pathogenic | 0 | 0 | 0 |
| rs67486158 | pathogenic | 0 | 0 | 0 |
| rs72554338 | pathogenic | 1 | 1 | 1 |

| | | | | |
|---|---|---|---|---|
| rs1800324 | pathogenic | 1 | 1 | 1 |
| rs72554356 | pathogenic | 1 | 1 | 1 |
| rs66656800 | pathogenic | 1 | 1 | 1 |
| rs68026851 | pathogenic | 1 | 1 | 1 |
| rs72558412 | pathogenic | 0 | 0 | 0 |
| rs72558415 | pathogenic | 0 | 0 | 0 |
| rs72558423 | pathogenic | 0 | 0 | 0 |
| rs67120076 | pathogenic | 1 | 1 | 1 |
| rs1800328 | pathogenic | 1 | 1 | 1 |
| rs72558451 | pathogenic | 1 | 1 | 1 |
| rs72558454 | pathogenic | 1 | 1 | 1 |
| rs672601372 | pathogenic | 0 | 0 | 0 |
| rs139644798 | pathogenic | 0 | 0 | 0 |
| rs369398935 | pathogenic | 0 | 0 | 0 |
| rs267607035 | pathogenic | 0 | 0 | 0 |
| rs267607036 | pathogenic | 1 | 1 | 1 |
| rs2302566 | benign | 1 | 1 | 0 |
| rs115959591 | benign | 0 | 0 | 0 |
| rs773876739 | pathogenic | 0 | 0 | 0 |
| rs1799967 | benign | 0 | 0 | 0 |
| rs80356968 | benign | 0 | 0 | 0 |
| rs28897695 | benign | 1 | 1 | 1 |
| rs80357314 | benign | 0 | 0 | 0 |
| rs80357169 | benign | 0 | 0 | 0 |
| rs397507239 | benign | 1 | 1 | 1 |
| rs80187739 | pathogenic | 1 | 1 | 1 |
| rs41293459 | pathogenic | 1 | 1 | 1 |
| rs55770810 | pathogenic | 1 | 1 | 1 |
| rs80356860 | pathogenic | 0 | 0 | 0 |
| rs56195342 | benign | 0 | 0 | 0 |
| rs80357442 | benign | 0 | 0 | 0 |
| rs41293463 | pathogenic | 1 | 1 | 1 |
| rs80357069 | pathogenic | 0 | 0 | 0 |
| rs397509275 | benign | 0 | 0 | 0 |
| rs80356920 | benign | 0 | 0 | 0 |
| rs397509284 | pathogenic | 0 | 0 | 0 |
| rs80357286 | benign | 0 | 0 | 0 |
| rs80357274 | benign | 0 | 0 | 0 |
| rs765656957 | benign | 0 | 0 | 0 |
| rs755427809 | benign | 0 | 0 | 0 |
| rs80357322 | benign | 0 | 0 | 0 |
| rs121434430 | pathogenic | 0 | 0 | 0 |
| rs121918450 | pathogenic | 1 | 1 | 1 |
| rs1532268 | benign | 0 | 0 | 0 |
| rs2303080 | benign | 0 | 0 | 0 |
| rs10064631 | benign | 0 | 0 | 0 |
| rs162036 | benign | 0 | 0 | 0 |
| rs2287780 | benign | 0 | 0 | 0 |
| rs16879334 | benign | 0 | 0 | 0 |
| rs137853061 | pathogenic | 1 | 1 | 0 |
| rs10380 | benign | 0 | 0 | 0 |
| rs1494558 | benign | 0 | 0 | 0 |
| rs1494555 | benign | 0 | 0 | 0 |
| rs192994060 | benign | 0 | 0 | 0 |

| | | | | |
|---|---|---|---|---|
| rs147128920 | benign | 1 | 1 | 0 |
| rs201803986 | pathogenic | 0 | 0 | 0 |
| rs746419489 | pathogenic | 0 | 1 | 0 |
| rs764396564 | pathogenic | 0 | 0 | 0 |
| rs758304537 | pathogenic | 0 | 0 | 0 |
| rs267606763 | pathogenic | 0 | 0 | 0 |
| rs267606762 | pathogenic | 0 | 0 | 0 |
| rs150702945 | pathogenic | 1 | 1 | 1 |
| rs202047589 | pathogenic | 0 | 0 | 0 |
| rs755562733 | pathogenic | 0 | 0 | 0 |
| rs267606764 | pathogenic | 1 | 1 | 1 |
| rs138207257 | pathogenic | 1 | 1 | 0 |
| rs149150736 | pathogenic | 1 | 1 | 1 |
| rs777046879 | pathogenic | 0 | 0 | 0 |
| rs104894408 | pathogenic | 1 | 1 | 1 |
| rs1801002 | pathogenic | 1 | 1 | 1 |
| rs28929485 | pathogenic | 1 | 1 | 1 |
| rs104894396 | pathogenic | 1 | 1 | 1 |
| rs371024165 | pathogenic | 0 | 0 | 0 |
| rs35887622 | pathogenic | 1 | 1 | 1 |
| rs587783644 | pathogenic | 1 | 1 | 1 |
| rs72474224 | pathogenic | 1 | 1 | 1 |
| rs111033296 | pathogenic | 1 | 1 | 1 |
| rs104894407 | pathogenic | 1 | 1 | 1 |
| rs72561723 | pathogenic | 1 | 1 | 1 |
| rs104894398 | pathogenic | 1 | 1 | 1 |
| rs28931594 | pathogenic | 1 | 1 | 1 |
| rs104894412 | pathogenic | 1 | 1 | 1 |
| rs111033297 | pathogenic | 1 | 1 | 1 |
| rs104894404 | pathogenic | 1 | 1 | 1 |
| rs104894410 | pathogenic | 1 | 1 | 1 |
| rs104894403 | pathogenic | 1 | 1 | 1 |
| rs397516870 | benign | 0 | 0 | 0 |
| rs200023879 | pathogenic | 1 | 1 | 1 |
| rs121912968 | pathogenic | 0 | 0 | 0 |
| rs104894402 | pathogenic | 1 | 1 | 1 |
| rs28931593 | pathogenic | 1 | 1 | 1 |
| rs104894397 | pathogenic | 0 | 0 | 0 |
| rs80338944 | pathogenic | 0 | 0 | 0 |
| rs199883710 | pathogenic | 0 | 0 | 0 |
| rs727504302 | pathogenic | 0 | 0 | 0 |
| rs145216882 | benign | 1 | 1 | 1 |
| rs781534323 | pathogenic | 1 | 1 | 1 |
| rs111033218 | benign | 1 | 1 | 1 |
| rs104894409 | pathogenic | 0 | 0 | 0 |
| rs80338945 | pathogenic | 1 | 1 | 1 |
| rs111033299 | pathogenic | 0 | 0 | 0 |
| rs143343083 | pathogenic | 1 | 1 | 1 |
| rs80338946 | pathogenic | 0 | 0 | 0 |
| rs150529554 | pathogenic | 0 | 0 | 0 |
| rs111033295 | pathogenic | 0 | 0 | 0 |
| rs111033188 | benign | 0 | 0 | 0 |
| rs397516874 | pathogenic | 0 | 0 | 0 |
| rs76434661 | pathogenic | 0 | 0 | 0 |

| | | | | |
|---|---|---|---|---|
| rs80338948 | pathogenic | 0 | 0 | 0 |
| rs104894401 | pathogenic | 0 | 0 | 0 |
| rs767178508 | pathogenic | 0 | 0 | 0 |
| rs772264564 | pathogenic | 0 | 0 | 0 |
| rs28931592 | pathogenic | 0 | 0 | 0 |
| rs80338949 | pathogenic | 1 | 1 | 1 |
| rs376898963 | benign | 1 | 1 | 1 |
| rs111033360 | benign | 0 | 0 | 0 |
| rs774518779 | pathogenic | 1 | 1 | 1 |
| rs28931595 | pathogenic | 1 | 1 | 1 |
| rs771748289 | pathogenic | 0 | 0 | 0 |
| rs104894406 | pathogenic | 0 | 0 | 0 |
| rs76838169 | benign | 0 | 0 | 0 |
| rs111033294 | pathogenic | 0 | 0 | 0 |
| rs200214298 | pathogenic | 0 | 0 | 0 |
| rs28940283 | pathogenic | 0 | 1 | 0 |
| rs1047883 | benign | 0 | 0 | 0 |
| rs138254425 | benign | 0 | 0 | 0 |
| rs121912592 | pathogenic | 1 | 1 | 0 |
| rs369061090 | pathogenic | 1 | 1 | 0 |
| rs758724746 | pathogenic | 0 | 1 | 0 |
| rs121912595 | pathogenic | 0 | 0 | 0 |
| rs76340296 | benign | 0 | 0 | 0 |
| rs140578009 | benign | 1 | 1 | 1 |
| rs74445297 | benign | 0 | 0 | 0 |
| rs78599516 | benign | 1 | 1 | 0 |
| rs104894035 | pathogenic | 1 | 1 | 0 |
| rs79451216 | benign | 0 | 0 | 0 |
| rs104894036 | pathogenic | 0 | 0 | 0 |
| rs73343757 | benign | 0 | 0 | 0 |
| rs121917722 | pathogenic | 1 | 1 | 1 |
| rs747700126 | pathogenic | 0 | 1 | 0 |
| rs121917725 | pathogenic | 1 | 1 | 1 |
| rs121917721 | pathogenic | 1 | 1 | 1 |
| rs121917723 | pathogenic | 1 | 1 | 0 |
| rs17569 | benign | 1 | 1 | 1 |
| rs2230062 | benign | 1 | 1 | 1 |
| rs17570 | benign | 0 | 0 | 0 |
| rs121917724 | pathogenic | 0 | 1 | 0 |
| rs199422235 | pathogenic | 0 | 0 | 0 |
| rs199422237 | pathogenic | 1 | 1 | 1 |
| rs199422234 | pathogenic | 1 | 1 | 0 |
| rs140015315 | pathogenic | 1 | 1 | 1 |
| rs730880298 | pathogenic | 1 | 1 | 1 |
| rs779692470 | pathogenic | 0 | 0 | 0 |
| rs369152939 | pathogenic | 0 | 0 | 0 |
| rs527236040 | pathogenic | 0 | 0 | 0 |
| rs587776985 | pathogenic | 1 | 1 | 1 |
| rs147077598 | pathogenic | 1 | 1 | 1 |
| rs128620185 | pathogenic | 1 | 1 | 1 |
| rs128620189 | pathogenic | 0 | 0 | 0 |
| rs128621190 | pathogenic | 0 | 0 | 0 |
| rs61754648 | benign | 0 | 0 | 0 |
| rs137852906 | pathogenic | 0 | 0 | 0 |

| | | | | |
|---|---|---|---|---|
| rs76811038 | pathogenic | 0 | 0 | 0 |
| rs149468422 | benign | 0 | 0 | 0 |
| rs5911 | benign | 0 | 0 | 0 |
| rs28942112 | pathogenic | 1 | 1 | 1 |
| rs143275858 | pathogenic | 1 | 1 | 0 |
| rs28362261 | benign | 0 | 0 | 0 |
| rs28362263 | benign | 1 | 1 | 1 |
| rs558707786 | pathogenic | 0 | 0 | 0 |
| rs138775799 | pathogenic | 0 | 0 | 0 |
| rs748170941 | pathogenic | 0 | 0 | 0 |
| rs121908380 | pathogenic | 0 | 0 | 0 |
| rs765123255 | pathogenic | 1 | 1 | 1 |
| rs730881832 | pathogenic | 0 | 1 | 0 |
| rs587782165 | benign | 1 | 1 | 1 |
| rs762307622 | pathogenic | 0 | 0 | 0 |
| rs34612342 | pathogenic | 1 | 1 | 0 |
| rs143353451 | pathogenic | 0 | 1 | 0 |
| rs747993448 | pathogenic | 0 | 1 | 0 |
| rs376561094 | pathogenic | 0 | 0 | 0 |
| rs776487884 | pathogenic | 1 | 1 | 0 |
| rs34126013 | pathogenic | 1 | 1 | 0 |
| rs200495564 | pathogenic | 1 | 1 | 0 |
| rs140342925 | pathogenic | 1 | 1 | 0 |
| rs587780749 | pathogenic | 1 | 1 | 0 |
| rs773087549 | pathogenic | 1 | 1 | 0 |
| rs730881833 | pathogenic | 0 | 0 | 0 |
| rs374950566 | pathogenic | 1 | 1 | 0 |
| rs587780752 | benign | 0 | 0 | 0 |
| rs3219489 | benign | 0 | 0 | 0 |
| rs199742231 | benign | 0 | 0 | 0 |
| rs587780082 | pathogenic | 0 | 0 | 0 |
| rs35881803 | benign | 0 | 0 | 0 |
| rs41305623 | benign | 1 | 1 | 1 |
| rs121917856 | pathogenic | 1 | 1 | 1 |
| rs576284122 | benign | 0 | 0 | 0 |
| rs142036701 | benign | 0 | 0 | 0 |
| rs121917852 | pathogenic | 1 | 1 | 1 |
| rs121917854 | pathogenic | 1 | 1 | 1 |
| rs121917853 | pathogenic | 1 | 1 | 1 |
| rs121917857 | pathogenic | 0 | 1 | 0 |
| rs193922740 | pathogenic | 1 | 1 | 1 |
| rs565779970 | pathogenic | 1 | 1 | 1 |
| rs6723526 | benign | 0 | 0 | 0 |
| rs141094096 | pathogenic | 1 | 1 | 1 |
| rs104894637 | pathogenic | 0 | 0 | 0 |
| rs104894636 | pathogenic | 0 | 0 | 0 |
| rs104894642 | pathogenic | 0 | 0 | 0 |
| rs104894638 | pathogenic | 0 | 0 | 0 |
| rs753666460 | pathogenic | 0 | 0 | 0 |
| rs104894643 | pathogenic | 1 | 1 | 0 |
| rs34520362 | pathogenic | 0 | 0 | 0 |
| rs374621913 | pathogenic | 0 | 0 | 0 |
| rs113641837 | benign | 0 | 0 | 0 |
| rs104894635 | pathogenic | 0 | 0 | 0 |

| | | | | |
|---|---|---|---|---|
| rs143947056 | pathogenic | 0 | 0 | 0 |
| rs138504221 | pathogenic | 0 | 0 | 0 |
| rs9894254 | benign | 0 | 0 | 0 |
| rs104894640 | pathogenic | 1 | 1 | 1 |
| rs58786455 | benign | 0 | 0 | 0 |
| rs144143780 | pathogenic | 0 | 0 | 0 |
| rs62620232 | benign | 0 | 0 | 0 |
| rs34297805 | benign | 1 | 1 | 1 |
| rs104894641 | pathogenic | 0 | 0 | 0 |
| rs148876719 | benign | 0 | 0 | 0 |
| rs104894639 | pathogenic | 0 | 0 | 0 |
| rs7503034 | benign | 0 | 0 | 0 |
| rs201631570 | benign | 0 | 0 | 0 |
| rs150181226 | pathogenic | 0 | 0 | 0 |
| rs377546036 | pathogenic | 0 | 0 | 0 |
| rs267606700 | pathogenic | 0 | 0 | 0 |
| rs267606701 | pathogenic | 1 | 1 | 0 |
| rs267606699 | pathogenic | 1 | 1 | 0 |
| rs9903215 | benign | 0 | 0 | 0 |
| rs139486406 | benign | 0 | 0 | 0 |
| rs143791614 | benign | 0 | 0 | 0 |
| rs573483474 | benign | 0 | 0 | 0 |
| rs2032892 | benign | 0 | 0 | 0 |
| rs137852619 | pathogenic | 0 | 0 | 0 |
| rs200243548 | benign | 0 | 0 | 0 |
| rs115702388 | benign | 0 | 0 | 0 |
| rs138085358 | pathogenic | 0 | 0 | 0 |
| rs121912789 | pathogenic | 1 | 1 | 1 |
| rs763407972 | pathogenic | 1 | 1 | 1 |
| rs147278302 | benign | 0 | 0 | 0 |
| rs374540411 | benign | 0 | 0 | 0 |
| rs121912790 | pathogenic | 1 | 1 | 0 |
| rs752049917 | benign | 0 | 0 | 0 |
| rs139640491 | benign | 1 | 1 | 1 |
| rs138010738 | benign | 1 | 1 | 0 |
| rs201328768 | pathogenic | 1 | 1 | 0 |
| rs2135720 | benign | 1 | 1 | 0 |
| rs201816080 | benign | 1 | 1 | 0 |
| rs754391973 | pathogenic | 0 | 0 | 0 |
| rs139087859 | benign | 0 | 0 | 0 |
| rs6805421 | benign | 0 | 0 | 0 |
| rs146266900 | benign | 0 | 0 | 0 |
| rs397509419 | pathogenic | 0 | 0 | 0 |
| rs149980411 | benign | 0 | 0 | 0 |
| rs747711488 | pathogenic | 1 | 1 | 1 |
| rs149474339 | pathogenic | 1 | 1 | 1 |
| rs758252808 | pathogenic | 1 | 1 | 0 |
| rs745756308 | pathogenic | 0 | 0 | 0 |
| rs4986958 | benign | 0 | 0 | 0 |
| rs74315444 | pathogenic | 0 | 0 | 0 |
| rs61752137 | pathogenic | 0 | 0 | 0 |
| rs8179065 | benign | 1 | 1 | 1 |
| rs758361736 | pathogenic | 1 | 1 | 0 |
| rs139794067 | pathogenic | 1 | 1 | 1 |

| | | | | |
|---|---|---|---|---|
| rs730880954 | pathogenic | 1 | 1 | 1 |
| rs730880955 | pathogenic | 1 | 1 | 1 |
| rs150634297 | pathogenic | 0 | 0 | 0 |
| rs104893750 | pathogenic | 1 | 1 | 1 |
| rs104893748 | pathogenic | 1 | 1 | 1 |
| rs104893749 | pathogenic | 1 | 1 | 1 |
| rs753520553 | pathogenic | 1 | 1 | 0 |
| rs61737294 | benign | 0 | 0 | 0 |
| rs104894601 | pathogenic | 0 | 0 | 0 |
| rs376090795 | pathogenic | 0 | 0 | 0 |
| rs104894596 | pathogenic | 0 | 0 | 0 |
| rs104894595 | pathogenic | 0 | 0 | 0 |
| rs104894598 | pathogenic | 0 | 0 | 0 |
| rs104894597 | pathogenic | 0 | 0 | 0 |
| rs104894591 | pathogenic | 0 | 0 | 0 |
| rs104894593 | pathogenic | 0 | 0 | 0 |
| rs104894594 | pathogenic | 0 | 0 | 0 |
| rs104894590 | pathogenic | 0 | 0 | 0 |
| rs86312 | benign | 0 | 0 | 0 |
| rs77874543 | benign | 0 | 0 | 0 |
| rs121912590 | pathogenic | 1 | 1 | 1 |
| rs121912589 | pathogenic | 1 | 1 | 1 |
| rs121912586 | pathogenic | 1 | 1 | 1 |
| rs121912585 | pathogenic | 1 | 1 | 1 |
| rs121912587 | pathogenic | 1 | 1 | 1 |
| rs762909359 | pathogenic | 0 | 0 | 0 |
| rs28931610 | pathogenic | 0 | 0 | 0 |
| rs104894097 | pathogenic | 1 | 1 | 1 |
| rs775176191 | benign | 0 | 0 | 0 |
| rs199907548 | pathogenic | 0 | 0 | 0 |
| rs104894095 | pathogenic | 1 | 1 | 1 |
| rs104894109 | pathogenic | 1 | 1 | 1 |
| rs372266620 | benign | 1 | 1 | 1 |
| rs104894099 | pathogenic | 0 | 0 | 0 |
| rs137854599 | pathogenic | 1 | 1 | 1 |
| rs104894094 | pathogenic | 0 | 0 | 0 |
| rs104894104 | pathogenic | 0 | 0 | 0 |
| rs6413463 | benign | 1 | 1 | 1 |
| rs104894098 | pathogenic | 0 | 0 | 0 |
| rs6413464 | benign | 0 | 0 | 0 |
| rs3731249 | benign | 0 | 0 | 0 |
| rs16931179 | benign | 0 | 0 | 0 |
| rs2734715 | benign | 0 | 0 | 0 |
| rs759080581 | pathogenic | 0 | 0 | 0 |
| rs1800746 | benign | 0 | 0 | 0 |
| rs202189057 | pathogenic | 0 | 0 | 0 |
| rs121908204 | pathogenic | 0 | 0 | 0 |
| rs756564767 | pathogenic | 0 | 0 | 0 |
| rs34758634 | benign | 1 | 1 | 1 |
| rs28940573 | pathogenic | 0 | 0 | 0 |
| rs119455955 | pathogenic | 1 | 1 | 0 |
| rs752164603 | pathogenic | 1 | 1 | 0 |
| rs763162812 | pathogenic | 0 | 0 | 0 |
| rs119455957 | pathogenic | 0 | 0 | 0 |

| | | | | |
|---|---|---|---|---|
| rs119455958 | pathogenic | 1 | 1 | 0 |
| rs750428882 | pathogenic | 0 | 0 | 0 |
| rs141482368 | benign | 0 | 0 | 0 |
| rs119455953 | pathogenic | 0 | 0 | 0 |
| rs119455954 | pathogenic | 0 | 0 | 0 |
| rs112812685 | benign | 1 | 1 | 0 |
| rs138744051 | benign | 1 | 1 | 0 |
| rs121908200 | pathogenic | 0 | 0 | 0 |
| rs119455956 | pathogenic | 0 | 0 | 0 |
| rs78970155 | benign | 0 | 0 | 0 |
| rs2834167 | benign | 0 | 0 | 0 |
| rs387907326 | pathogenic | 1 | 1 | 1 |
| rs2107732 | benign | 0 | 0 | 0 |
| rs11552377 | benign | 0 | 0 | 0 |
| rs137852843 | pathogenic | 0 | 0 | 0 |
| rs104894783 | pathogenic | 0 | 0 | 0 |
| rs122457137 | pathogenic | 1 | 1 | 1 |
| rs104894779 | pathogenic | 1 | 1 | 1 |
| rs104894786 | pathogenic | 0 | 0 | 0 |
| rs104894784 | pathogenic | 0 | 0 | 0 |
| rs104894785 | pathogenic | 0 | 0 | 0 |
| rs201870761 | pathogenic | 0 | 0 | 0 |
| rs104894781 | pathogenic | 0 | 0 | 0 |
| rs28897728 | benign | 0 | 0 | 0 |
| rs370273690 | pathogenic | 1 | 1 | 0 |
| rs9332739 | benign | 0 | 0 | 0 |
| rs150878060 | benign | 0 | 0 | 0 |
| rs573509224 | benign | 0 | 0 | 0 |
| rs36221133 | benign | 0 | 0 | 0 |
| rs2230009 | benign | 0 | 0 | 0 |
| rs267607008 | pathogenic | 0 | 0 | 0 |
| rs5985 | benign | 1 | 1 | 1 |
| rs5982 | benign | 1 | 1 | 1 |
| rs5987 | benign | 0 | 0 | 0 |
| rs5988 | benign | 0 | 0 | 0 |
| rs121918680 | pathogenic | 1 | 1 | 0 |
| rs121918679 | pathogenic | 1 | 1 | 0 |
| rs7845 | benign | 1 | 1 | 0 |
| rs1049296 | benign | 1 | 1 | 1 |
| rs121918677 | pathogenic | 1 | 1 | 0 |
| rs200086262 | pathogenic | 1 | 1 | 0 |
| rs746930990 | pathogenic | 1 | 1 | 0 |
| rs104893878 | pathogenic | 1 | 1 | 1 |
| rs121912468 | pathogenic | 1 | 1 | 1 |
| rs121912467 | pathogenic | 1 | 1 | 1 |
| rs145351926 | benign | 1 | 1 | 1 |
| rs141212743 | pathogenic | 1 | 1 | 1 |
| rs397515473 | pathogenic | 1 | 1 | 0 |
| rs121918313 | pathogenic | 0 | 1 | 0 |
| rs73015965 | pathogenic | 0 | 0 | 0 |
| rs121918030 | pathogenic | 0 | 1 | 0 |
| rs77782321 | benign | 0 | 0 | 0 |
| rs374661051 | pathogenic | 1 | 1 | 1 |
| rs267606854 | pathogenic | 0 | 0 | 0 |

| | | | | |
|---|---|---|---|---|
| rs41279678 | benign | 0 | 0 | 0 |
| rs61754640 | benign | 0 | 0 | 0 |
| rs199473314 | pathogenic | 0 | 0 | 0 |
| rs137854601 | pathogenic | 0 | 0 | 0 |
| rs137854614 | pathogenic | 0 | 0 | 0 |
| rs137854615 | pathogenic | 0 | 0 | 0 |
| rs199473635 | pathogenic | 0 | 0 | 0 |
| rs121434478 | pathogenic | 1 | 1 | 1 |
| rs775407864 | pathogenic | 1 | 1 | 1 |
| rs12706927 | benign | 0 | 1 | 0 |
| rs775614261 | benign | 0 | 0 | 0 |
| rs768926261 | benign | 0 | 0 | 0 |
| rs114057640 | benign | 0 | 0 | 0 |
| rs387906886 | pathogenic | 1 | 1 | 0 |
| rs138658585 | benign | 0 | 0 | 0 |
| rs4880091 | benign | 0 | 0 | 0 |
| rs121918552 | pathogenic | 1 | 1 | 1 |
| rs10794537 | benign | 0 | 0 | 0 |
| rs121965022 | pathogenic | 1 | 1 | 0 |
| rs121965020 | pathogenic | 0 | 0 | 0 |
| rs758452450 | pathogenic | 0 | 0 | 0 |
| rs747981483 | benign | 0 | 0 | 0 |
| rs58037052 | benign | 0 | 0 | 0 |
| rs121965029 | pathogenic | 1 | 1 | 0 |
| rs147490060 | benign | 0 | 0 | 0 |
| rs3755955 | benign | 0 | 0 | 0 |
| rs148789453 | pathogenic | 0 | 0 | 0 |
| rs121965030 | pathogenic | 0 | 0 | 0 |
| rs76722191 | benign | 0 | 0 | 0 |
| rs199801029 | pathogenic | 0 | 0 | 0 |
| rs121965033 | pathogenic | 0 | 0 | 0 |
| rs368454909 | pathogenic | 1 | 1 | 0 |
| rs6831280 | benign | 0 | 0 | 0 |
| rs121965024 | pathogenic | 0 | 0 | 0 |
| rs121965019 | pathogenic | 0 | 0 | 0 |
| rs11934801 | benign | 0 | 0 | 0 |
| rs73066479 | benign | 0 | 0 | 0 |
| rs121965027 | pathogenic | 0 | 0 | 0 |
| rs121965026 | pathogenic | 1 | 1 | 0 |
| rs121965021 | pathogenic | 0 | 0 | 0 |
| rs398123257 | benign | 0 | 0 | 0 |
| rs121965031 | pathogenic | 0 | 0 | 0 |
| rs202003795 | pathogenic | 0 | 0 | 0 |
| rs559510809 | pathogenic | 0 | 0 | 0 |
| rs762936223 | benign | 0 | 0 | 0 |
| rs137854580 | pathogenic | 1 | 1 | 1 |
| rs112263157 | benign | 0 | 0 | 0 |
| rs2289181 | benign | 0 | 0 | 0 |
| rs138034915 | benign | 0 | 1 | 0 |
| rs528367092 | pathogenic | 0 | 1 | 0 |
| rs1042393 | benign | 0 | 0 | 0 |
| rs370950728 | pathogenic | 0 | 1 | 0 |
| rs530478036 | benign | 0 | 1 | 0 |
| rs374569672 | benign | 0 | 1 | 0 |

| | | | | |
|---|---|---|---|---|
| rs121907944 | pathogenic | 0 | 1 | 0 |
| rs200856561 | pathogenic | 0 | 1 | 0 |
| rs577915581 | pathogenic | 0 | 1 | 0 |
| rs201896815 | pathogenic | 0 | 1 | 0 |
| rs121907945 | pathogenic | 0 | 1 | 0 |
| rs121907940 | pathogenic | 0 | 1 | 0 |
| rs543300039 | pathogenic | 0 | 1 | 0 |
| rs121907936 | pathogenic | 0 | 1 | 0 |
| rs766074609 | pathogenic | 0 | 1 | 0 |
| rs755253527 | pathogenic | 0 | 1 | 0 |
| rs560575383 | pathogenic | 0 | 1 | 0 |
| rs528369909 | benign | 0 | 1 | 0 |
| rs770610356 | pathogenic | 0 | 1 | 0 |
| rs747610090 | pathogenic | 0 | 1 | 0 |
| rs145712232 | benign | 0 | 1 | 0 |
| rs7215458 | benign | 1 | 1 | 0 |
| rs535644999 | pathogenic | 0 | 1 | 0 |
| rs772883420 | pathogenic | 1 | 1 | 0 |
| rs398123169 | pathogenic | 0 | 1 | 0 |
| rs121907937 | pathogenic | 0 | 1 | 0 |
| rs121907941 | pathogenic | 1 | 1 | 0 |
| rs121907942 | pathogenic | 0 | 1 | 0 |
| rs779556619 | pathogenic | 0 | 1 | 0 |
| rs1800307 | benign | 0 | 1 | 0 |
| rs377544304 | pathogenic | 1 | 1 | 0 |
| rs549029029 | pathogenic | 0 | 1 | 0 |
| rs753269119 | pathogenic | 1 | 1 | 1 |
| rs757617999 | pathogenic | 0 | 1 | 0 |
| rs28937909 | pathogenic | 0 | 1 | 0 |
| rs28940868 | pathogenic | 0 | 1 | 0 |
| rs368438393 | pathogenic | 0 | 1 | 0 |
| rs536906561 | pathogenic | 0 | 1 | 0 |
| rs374143224 | pathogenic | 0 | 1 | 0 |
| rs778418246 | pathogenic | 0 | 1 | 0 |
| rs398123172 | pathogenic | 0 | 1 | 0 |
| rs121907938 | pathogenic | 0 | 1 | 0 |
| rs1800312 | pathogenic | 0 | 1 | 0 |
| rs1126690 | benign | 0 | 1 | 0 |
| rs1800314 | benign | 0 | 1 | 0 |
| rs369532274 | pathogenic | 0 | 1 | 0 |
| rs121907943 | pathogenic | 0 | 1 | 0 |
| rs780321415 | pathogenic | 0 | 1 | 0 |
| rs377286472 | benign | 0 | 1 | 0 |
| rs121918406 | pathogenic | 1 | 1 | 1 |
| rs147484870 | benign | 0 | 0 | 0 |
| rs193922671 | pathogenic | 0 | 0 | 0 |
| rs201397978 | benign | 1 | 1 | 1 |
| rs34884895 | benign | 1 | 1 | 1 |
| rs138329459 | benign | 0 | 0 | 0 |
| rs2289366 | benign | 1 | 1 | 1 |
| rs141858334 | benign | 1 | 1 | 1 |
| rs59907886 | benign | 1 | 1 | 1 |
| rs137852653 | pathogenic | 0 | 0 | 0 |
| rs137852652 | pathogenic | 0 | 0 | 0 |

| | | | | |
|---|---|---|---|---|
| rs137852656 | pathogenic | 0 | 0 | 0 |
| rs61739916 | benign | 0 | 0 | 0 |
| rs28936668 | pathogenic | 0 | 0 | 0 |
| rs137852651 | pathogenic | 0 | 0 | 0 |
| rs137852650 | pathogenic | 0 | 0 | 0 |
| rs28936669 | pathogenic | 0 | 0 | 0 |
| rs137852654 | pathogenic | 0 | 0 | 0 |
| rs28936368 | pathogenic | 1 | 1 | 0 |
| rs137852655 | pathogenic | 0 | 0 | 0 |
| rs61752496 | benign | 0 | 0 | 0 |
| rs750098055 | benign | 0 | 0 | 0 |
| rs764149793 | benign | 0 | 0 | 0 |
| rs724160017 | pathogenic | 0 | 1 | 0 |
| rs724160019 | pathogenic | 0 | 0 | 0 |
| rs202219398 | pathogenic | 1 | 1 | 0 |
| rs724160023 | pathogenic | 1 | 1 | 0 |
| rs761953453 | pathogenic | 0 | 0 | 0 |
| rs80338835 | pathogenic | 1 | 1 | 1 |
| rs56284940 | benign | 0 | 0 | 0 |
| rs74575749 | benign | 0 | 0 | 0 |
| rs587777075 | pathogenic | 1 | 1 | 1 |
| rs641153 | benign | 0 | 0 | 0 |
| rs12614 | benign | 0 | 0 | 0 |
| rs117314762 | benign | 0 | 0 | 0 |
| rs537478097 | benign | 1 | 1 | 1 |
| rs144812066 | benign | 0 | 0 | 0 |
| rs4151651 | benign | 0 | 0 | 0 |
| rs138207668 | benign | 1 | 1 | 1 |
| rs149101394 | benign | 1 | 1 | 1 |
| rs4151659 | benign | 1 | 1 | 1 |
| rs45484591 | benign | 1 | 1 | 1 |
| rs761991070 | pathogenic | 0 | 0 | 0 |
| rs755948940 | pathogenic | 0 | 0 | 0 |
| rs137853097 | pathogenic | 0 | 0 | 0 |
| rs775297352 | benign | 0 | 0 | 0 |
| rs28943591 | benign | 0 | 0 | 0 |
| rs191468413 | pathogenic | 1 | 1 | 0 |
| rs11539471 | benign | 1 | 1 | 1 |
| rs11205 | benign | 0 | 0 | 0 |
| rs768409400 | pathogenic | 0 | 0 | 0 |
| rs762013077 | pathogenic | 0 | 0 | 0 |
| rs768474112 | pathogenic | 0 | 0 | 0 |
| rs121965063 | pathogenic | 1 | 1 | 1 |
| rs143648758 | pathogenic | 0 | 0 | 0 |
| rs121965066 | pathogenic | 0 | 0 | 0 |
| rs752907087 | pathogenic | 0 | 0 | 0 |
| rs747702749 | pathogenic | 0 | 0 | 0 |
| rs754087775 | pathogenic | 0 | 0 | 0 |
| rs121918641 | pathogenic | 1 | 1 | 1 |
| rs121918642 | pathogenic | 1 | 1 | 1 |
| rs121918640 | pathogenic | 1 | 1 | 1 |
| rs121918637 | pathogenic | 1 | 1 | 1 |
| rs121918644 | pathogenic | 1 | 1 | 1 |
| rs16840544 | benign | 0 | 0 | 0 |

| | | | | |
|---|---|---|---|---|
| rs1042391 | benign | 1 | 1 | 1 |
| rs104893744 | pathogenic | 1 | 1 | 1 |
| rs120074136 | pathogenic | 0 | 0 | 0 |
| rs1805081 | benign | 1 | 1 | 0 |
| rs80358251 | benign | 0 | 0 | 0 |
| rs774588125 | benign | 0 | 0 | 0 |
| rs120074134 | pathogenic | 0 | 0 | 0 |
| rs139751448 | pathogenic | 0 | 0 | 0 |
| rs77080672 | benign | 0 | 0 | 0 |
| rs372445155 | pathogenic | 1 | 1 | 0 |
| rs13381670 | benign | 0 | 0 | 0 |
| rs1788799 | benign | 0 | 0 | 0 |
| rs376213990 | pathogenic | 0 | 0 | 0 |
| rs757475924 | pathogenic | 0 | 0 | 0 |
| rs80358253 | pathogenic | 0 | 0 | 0 |
| rs1805082 | benign | 0 | 0 | 0 |
| rs120074130 | pathogenic | 0 | 0 | 0 |
| rs768999208 | pathogenic | 0 | 0 | 0 |
| rs34302553 | benign | 0 | 0 | 0 |
| rs28940897 | pathogenic | 0 | 0 | 0 |
| rs1140458 | benign | 0 | 0 | 0 |
| rs143124972 | pathogenic | 0 | 0 | 0 |
| rs120074135 | pathogenic | 0 | 0 | 0 |
| rs543206298 | pathogenic | 0 | 0 | 0 |
| rs120074132 | pathogenic | 0 | 0 | 0 |
| rs759826138 | pathogenic | 0 | 0 | 0 |
| rs28942108 | pathogenic | 0 | 0 | 0 |
| rs80358254 | pathogenic | 0 | 0 | 0 |
| rs80358257 | pathogenic | 0 | 0 | 0 |
| rs781261962 | pathogenic | 0 | 0 | 0 |
| rs28942107 | pathogenic | 0 | 0 | 0 |
| rs80358258 | pathogenic | 0 | 0 | 0 |
| rs80358259 | pathogenic | 1 | 1 | 0 |
| rs141440861 | benign | 0 | 0 | 0 |
| rs28942106 | pathogenic | 0 | 0 | 0 |
| rs778878523 | pathogenic | 0 | 0 | 0 |
| rs28942105 | pathogenic | 0 | 0 | 0 |
| rs61731969 | benign | 0 | 0 | 0 |
| rs200444084 | pathogenic | 0 | 0 | 0 |
| rs113371321 | pathogenic | 0 | 0 | 0 |
| rs369098773 | pathogenic | 0 | 0 | 0 |
| rs35248744 | benign | 0 | 0 | 0 |
| rs758902805 | pathogenic | 0 | 0 | 0 |
| rs120074131 | pathogenic | 0 | 0 | 0 |
| rs1805084 | benign | 0 | 0 | 0 |
| rs117793540 | benign | 0 | 0 | 0 |
| rs530058063 | benign | 0 | 0 | 0 |
| rs1064422 | pathogenic | 1 | 1 | 1 |
| rs104894506 | pathogenic | 1 | 1 | 1 |
| rs104894507 | pathogenic | 1 | 1 | 1 |
| rs104894508 | pathogenic | 1 | 1 | 0 |
| rs28999113 | pathogenic | 1 | 1 | 0 |
| rs281860266 | pathogenic | 0 | 0 | 0 |
| rs104894961 | pathogenic | 1 | 1 | 0 |

| | | | | |
|---|---|---|---|---|
| rs122463168 | pathogenic | 0 | 0 | 0 |
| rs199469624 | pathogenic | 1 | 1 | 1 |
| rs150107747 | benign | 0 | 0 | 0 |
| rs144984628 | pathogenic | 0 | 0 | 0 |
| rs28935171 | pathogenic | 0 | 0 | 0 |
| rs137852443 | pathogenic | 1 | 1 | 0 |
| rs28933674 | pathogenic | 1 | 1 | 1 |
| rs28933675 | pathogenic | 1 | 1 | 1 |
| rs28933676 | pathogenic | 1 | 1 | 1 |
| rs28933677 | pathogenic | 1 | 1 | 1 |
| rs28933678 | pathogenic | 0 | 0 | 0 |
| rs28933679 | pathogenic | 1 | 1 | 1 |
| rs28933680 | pathogenic | 0 | 0 | 0 |
| rs28933681 | pathogenic | 0 | 0 | 0 |
| rs28933682 | pathogenic | 0 | 0 | 0 |
| rs28937294 | pathogenic | 1 | 1 | 0 |
| rs137852459 | pathogenic | 0 | 0 | 0 |
| rs137852461 | pathogenic | 0 | 0 | 0 |
| rs137852464 | pathogenic | 1 | 1 | 1 |
| rs137852465 | pathogenic | 1 | 1 | 1 |
| rs1800297 | benign | 1 | 1 | 0 |
| rs137852374 | pathogenic | 1 | 1 | 1 |
| rs672601312 | pathogenic | 1 | 1 | 1 |
| rs387907196 | pathogenic | 1 | 1 | 1 |
| rs148348866 | benign | 0 | 0 | 0 |
| rs141138948 | pathogenic | 1 | 1 | 1 |
| rs137852685 | pathogenic | 1 | 1 | 0 |
| rs104893900 | pathogenic | 1 | 1 | 0 |
| rs137852686 | pathogenic | 1 | 1 | 1 |
| rs104893906 | pathogenic | 1 | 1 | 0 |
| rs587782207 | pathogenic | 1 | 1 | 1 |
| rs75086406 | pathogenic | 0 | 0 | 0 |
| rs199822819 | pathogenic | 0 | 0 | 0 |
| rs398123166 | pathogenic | 1 | 1 | 1 |
| rs121913123 | pathogenic | 1 | 1 | 1 |
| rs772190176 | pathogenic | 0 | 1 | 0 |
| rs587782216 | pathogenic | 1 | 1 | 1 |
| rs398123159 | pathogenic | 1 | 1 | 1 |
| rs121913122 | pathogenic | 1 | 1 | 1 |
| rs755436052 | pathogenic | 1 | 1 | 1 |
| rs200796606 | pathogenic | 1 | 1 | 1 |
| rs2070080 | pathogenic | 0 | 1 | 0 |
| rs149616320 | benign | 0 | 0 | 0 |
| rs143529458 | benign | 0 | 0 | 0 |
| rs267607077 | pathogenic | 1 | 1 | 1 |
| rs143898031 | benign | 1 | 1 | 0 |
| rs148304857 | benign | 1 | 1 | 1 |
| rs137852809 | pathogenic | 0 | 0 | 0 |
| rs3765623 | benign | 0 | 0 | 0 |
| rs16944397 | benign | 0 | 0 | 0 |
| rs730880167 | benign | 0 | 0 | 0 |
| rs187108957 | benign | 0 | 0 | 0 |
| rs117342470 | benign | 0 | 0 | 0 |
| rs751200138 | pathogenic | 1 | 1 | 1 |

| | | | | |
|---|---|---|---|---|
| rs755799528 | pathogenic | 0 | 0 | 0 |
| rs537484504 | benign | 0 | 0 | 0 |
| rs748944640 | pathogenic | 0 | 0 | 0 |
| rs199774121 | pathogenic | 0 | 0 | 0 |
| rs766094434 | pathogenic | 1 | 1 | 0 |
| rs777321035 | pathogenic | 1 | 1 | 1 |
| rs146269774 | benign | 0 | 1 | 0 |
| rs144085805 | benign | 0 | 1 | 0 |
| rs367643028 | benign | 0 | 0 | 0 |
| rs42524 | benign | 0 | 0 | 0 |
| rs28940586 | pathogenic | 0 | 0 | 0 |
| rs74315327 | pathogenic | 1 | 1 | 1 |
| rs199676861 | pathogenic | 0 | 0 | 0 |
| rs115802744 | benign | 1 | 1 | 0 |
| rs752866643 | pathogenic | 1 | 1 | 1 |
| rs28941784 | pathogenic | 1 | 1 | 1 |
| rs398124434 | pathogenic | 1 | 1 | 1 |
| rs756414548 | pathogenic | 1 | 1 | 1 |
| rs376128990 | pathogenic | 1 | 1 | 1 |
| rs746219370 | pathogenic | 1 | 1 | 1 |
| rs756195708 | pathogenic | 1 | 1 | 1 |
| rs369296618 | pathogenic | 1 | 1 | 1 |
| rs9593 | benign | 1 | 1 | 1 |
| rs139510844 | benign | 0 | 0 | 0 |
| rs28382158 | benign | 0 | 0 | 0 |
| rs28919570 | pathogenic | 0 | 0 | 0 |
| rs587777331 | pathogenic | 0 | 0 | 0 |
| rs587777333 | pathogenic | 0 | 0 | 0 |
| rs587777332 | pathogenic | 0 | 0 | 0 |
| rs774980346 | pathogenic | 0 | 0 | 0 |
| rs370934093 | pathogenic | 0 | 0 | 0 |
| rs121913479 | pathogenic | 1 | 1 | 1 |
| rs121913484 | pathogenic | 0 | 0 | 0 |
| rs121913485 | pathogenic | 1 | 1 | 1 |
| rs28931614 | pathogenic | 1 | 1 | 1 |
| rs17881656 | benign | 1 | 1 | 1 |
| rs28931615 | pathogenic | 1 | 1 | 1 |
| rs147846832 | benign | 0 | 0 | 0 |
| rs80356506 | pathogenic | 0 | 0 | 0 |
| rs80356494 | pathogenic | 0 | 0 | 0 |
| rs121908894 | pathogenic | 0 | 0 | 0 |
| rs121908895 | pathogenic | 0 | 0 | 0 |
| rs80356508 | pathogenic | 0 | 0 | 0 |
| rs80356507 | pathogenic | 0 | 0 | 0 |
| rs121908896 | pathogenic | 0 | 0 | 0 |
| rs745372589 | pathogenic | 1 | 1 | 0 |
| rs183898408 | benign | 0 | 0 | 0 |
| rs200844292 | benign | 0 | 0 | 0 |
| rs146897002 | benign | 0 | 0 | 0 |
| rs121964869 | pathogenic | 1 | 1 | 1 |
| rs201768800 | benign | 1 | 1 | 1 |
| rs9807556 | benign | 1 | 1 | 1 |
| rs184721031 | benign | 0 | 0 | 0 |
| rs201104206 | benign | 0 | 0 | 0 |

| | | | | |
|---|---|---|---|---|
| rs188677538 | benign | 0 | 0 | 0 |
| rs149528866 | benign | 0 | 0 | 0 |
| rs115382168 | benign | 0 | 0 | 0 |
| rs78846023 | benign | 1 | 1 | 1 |
| rs61730662 | benign | 0 | 0 | 0 |
| rs121918368 | pathogenic | 0 | 0 | 0 |
| rs566433112 | pathogenic | 1 | 1 | 1 |
| rs121909342 | pathogenic | 0 | 0 | 0 |
| rs67586389 | pathogenic | 0 | 0 | 0 |
| rs72466485 | pathogenic | 0 | 0 | 0 |
| rs72466486 | pathogenic | 1 | 1 | 1 |
| rs72466487 | pathogenic | 1 | 1 | 1 |
| rs267607112 | pathogenic | 0 | 0 | 0 |
| rs118204013 | pathogenic | 0 | 0 | 0 |
| rs121913495 | pathogenic | 0 | 0 | 0 |
| rs11554273 | pathogenic | 0 | 0 | 0 |
| rs193922477 | pathogenic | 1 | 1 | 1 |
| rs193922479 | pathogenic | 1 | 1 | 1 |
| rs72547554 | pathogenic | 0 | 0 | 0 |
| rs72547561 | pathogenic | 1 | 1 | 1 |
| rs72547562 | pathogenic | 1 | 1 | 0 |
| rs115977487 | benign | 1 | 1 | 0 |
| rs72547564 | pathogenic | 0 | 0 | 0 |
| rs72547568 | pathogenic | 1 | 1 | 0 |
| rs72547569 | pathogenic | 1 | 1 | 1 |
| rs72547571 | pathogenic | 0 | 0 | 0 |
| rs72547575 | pathogenic | 1 | 1 | 1 |
| rs544994462 | benign | 0 | 0 | 0 |
| rs190470017 | benign | 0 | 0 | 0 |
| rs199473266 | pathogenic | 1 | 1 | 1 |
| rs137854602 | pathogenic | 0 | 0 | 0 |
| rs143739249 | pathogenic | 0 | 0 | 0 |
| rs397514697 | pathogenic | 1 | 1 | 1 |
| rs373764886 | pathogenic | 0 | 0 | 0 |
| rs370242930 | pathogenic | 0 | 0 | 0 |
| rs377145777 | pathogenic | 1 | 1 | 1 |
| rs191256186 | benign | 0 | 0 | 0 |
| rs150070901 | benign | 1 | 1 | 1 |
| rs138098032 | benign | 0 | 0 | 0 |
| rs117731813 | benign | 1 | 1 | 1 |
| rs104894488 | pathogenic | 1 | 1 | 1 |
| rs11302 | benign | 0 | 1 | 0 |
| rs121918447 | pathogenic | 1 | 1 | 1 |
| rs138205828 | benign | 1 | 1 | 0 |
| rs149858290 | benign | 0 | 0 | 0 |
| rs750764003 | pathogenic | 0 | 1 | 0 |
| rs141381896 | benign | 0 | 0 | 0 |
| rs149027682 | benign | 0 | 0 | 0 |
| rs121908415 | pathogenic | 0 | 0 | 0 |
| rs121908416 | pathogenic | 0 | 0 | 0 |
| rs121908417 | pathogenic | 0 | 0 | 0 |
| rs587780306 | benign | 0 | 0 | 0 |
| rs3218725 | benign | 0 | 0 | 0 |
| rs3218729 | benign | 0 | 0 | 0 |

| | | | | |
|---|---|---|---|---|
| rs387906917 | pathogenic | 0 | 0 | 0 |
| rs200652608 | pathogenic | 0 | 0 | 0 |
| rs3218721 | benign | 0 | 0 | 0 |
| rs63750869 | pathogenic | 1 | 1 | 1 |
| rs118203910 | pathogenic | 0 | 1 | 0 |
| rs9332701 | benign | 1 | 1 | 0 |
| rs6027 | benign | 0 | 0 | 0 |
| rs121909110 | pathogenic | 0 | 1 | 0 |
| rs762550967 | pathogenic | 1 | 1 | 0 |
| rs763615602 | pathogenic | 1 | 1 | 0 |
| rs144907403 | benign | 0 | 0 | 0 |
| rs104893631 | pathogenic | 1 | 1 | 0 |
| rs144181978 | pathogenic | 1 | 1 | 0 |
| rs74874677 | benign | 1 | 1 | 1 |
| rs104893632 | pathogenic | 0 | 1 | 0 |
| rs371342464 | benign | 1 | 1 | 1 |
| rs104893633 | pathogenic | 1 | 1 | 0 |
| rs201088712 | pathogenic | 0 | 0 | 0 |
| rs201857541 | benign | 1 | 1 | 1 |
| rs139517732 | pathogenic | 1 | 1 | 0 |
| rs191057824 | benign | 1 | 1 | 0 |
| rs35683768 | benign | 1 | 1 | 0 |
| rs104895271 | pathogenic | 0 | 0 | 0 |
| rs104895217 | pathogenic | 1 | 1 | 1 |
| rs104895223 | pathogenic | 1 | 1 | 1 |
| rs104895218 | pathogenic | 1 | 1 | 1 |
| rs104895225 | pathogenic | 1 | 1 | 1 |
| rs4149637 | benign | 1 | 1 | 1 |
| rs104895219 | pathogenic | 1 | 1 | 1 |
| rs104895220 | pathogenic | 1 | 1 | 1 |
| rs104895228 | pathogenic | 1 | 1 | 1 |
| rs104895221 | pathogenic | 1 | 1 | 1 |
| rs104895222 | pathogenic | 1 | 1 | 1 |
| rs104895247 | pathogenic | 0 | 0 | 0 |
| rs28940585 | pathogenic | 0 | 0 | 0 |
| rs561367199 | pathogenic | 1 | 1 | 1 |
| rs143793815 | benign | 0 | 0 | 0 |
| rs28941473 | pathogenic | 1 | 1 | 1 |
| rs751590073 | pathogenic | 1 | 1 | 1 |
| rs111407265 | benign | 1 | 1 | 1 |
| rs373697663 | pathogenic | 1 | 1 | 1 |
| rs369879957 | pathogenic | 0 | 0 | 0 |
| rs28941472 | pathogenic | 1 | 1 | 1 |
| rs28940287 | pathogenic | 1 | 1 | 1 |
| rs77625410 | benign | 0 | 0 | 0 |
| rs111033477 | benign | 0 | 0 | 0 |
| rs115123584 | benign | 0 | 0 | 0 |
| rs374576916 | benign | 0 | 0 | 0 |
| rs397516317 | pathogenic | 0 | 0 | 0 |
| rs368657015 | pathogenic | 0 | 0 | 0 |
| rs397516321 | pathogenic | 0 | 0 | 0 |
| rs397516322 | pathogenic | 0 | 0 | 0 |
| rs111033215 | pathogenic | 0 | 0 | 0 |
| rs199606180 | pathogenic | 0 | 0 | 0 |

| | | | | |
|---|---|---|---|---|
| rs111033192 | pathogenic | 0 | 0 | 0 |
| rs948962 | benign | 1 | 1 | 1 |
| rs142293185 | benign | 0 | 0 | 0 |
| rs376764423 | pathogenic | 0 | 0 | 0 |
| rs773844428 | pathogenic | 0 | 0 | 0 |
| rs186644871 | benign | 0 | 0 | 0 |
| rs200313391 | benign | 0 | 0 | 0 |
| rs377670513 | pathogenic | 0 | 0 | 0 |
| rs1132036 | benign | 0 | 0 | 0 |
| rs111033231 | benign | 0 | 0 | 0 |
| rs145945630 | pathogenic | 0 | 0 | 0 |
| rs121912984 | pathogenic | 1 | 1 | 1 |
| rs1800435 | benign | 0 | 1 | 0 |
| rs121912980 | pathogenic | 0 | 0 | 0 |
| rs121912982 | pathogenic | 1 | 1 | 1 |
| rs121912983 | pathogenic | 0 | 0 | 0 |
| rs121912981 | pathogenic | 0 | 0 | 0 |
| rs1057868 | benign | 0 | 0 | 0 |
| rs56048668 | benign | 0 | 0 | 0 |
| rs55865465 | benign | 0 | 0 | 0 |
| rs748096324 | benign | 1 | 1 | 0 |
| rs150475750 | benign | 0 | 0 | 0 |
| rs143025739 | benign | 1 | 1 | 0 |
| rs79702016 | benign | 1 | 1 | 0 |
| rs41338945 | benign | 0 | 0 | 0 |
| rs146406037 | benign | 1 | 1 | 1 |
| rs143274491 | benign | 0 | 0 | 0 |
| rs3829986 | benign | 0 | 0 | 0 |
| rs149168939 | benign | 0 | 0 | 0 |
| rs1800385 | benign | 0 | 0 | 0 |
| rs1800386 | pathogenic | 0 | 0 | 0 |
| rs121918524 | pathogenic | 1 | 1 | 1 |
| rs121912700 | pathogenic | 0 | 0 | 0 |
| rs104893930 | pathogenic | 1 | 1 | 1 |
| rs104893931 | pathogenic | 0 | 0 | 0 |
| rs116125936 | benign | 0 | 0 | 0 |
| rs627441 | benign | 0 | 0 | 0 |
| rs104894481 | pathogenic | 0 | 0 | 0 |
| rs137853129 | pathogenic | 0 | 0 | 0 |
| rs137853130 | pathogenic | 1 | 1 | 0 |
| rs28940282 | pathogenic | 0 | 0 | 0 |
| rs2229989 | benign | 0 | 0 | 0 |
| rs104894647 | pathogenic | 0 | 0 | 0 |
| rs1061170 | benign | 1 | 1 | 1 |
| rs576402053 | benign | 0 | 0 | 0 |
| rs34972246 | benign | 0 | 0 | 0 |
| rs192725607 | benign | 0 | 0 | 0 |
| rs137852615 | pathogenic | 1 | 1 | 0 |
| rs137852617 | pathogenic | 1 | 1 | 1 |
| rs137852611 | pathogenic | 0 | 0 | 0 |
| rs137852612 | pathogenic | 0 | 0 | 0 |
| rs137852614 | pathogenic | 1 | 1 | 1 |
| rs137852610 | pathogenic | 1 | 1 | 1 |
| rs137852618 | pathogenic | 1 | 1 | 1 |

| | | | | |
|---|---|---|---|---|
| rs137852613 | pathogenic | 1 | 1 | 0 |
| rs137852609 | pathogenic | 0 | 0 | 0 |
| rs63750306 | pathogenic | 1 | 1 | 1 |
| rs63750265 | pathogenic | 1 | 1 | 1 |
| rs63751287 | pathogenic | 1 | 1 | 1 |
| rs199723282 | benign | 0 | 0 | 0 |
| rs121917807 | pathogenic | 0 | 0 | 0 |
| rs121917809 | pathogenic | 0 | 0 | 0 |
| rs75134564 | pathogenic | 0 | 0 | 0 |
| rs267606930 | pathogenic | 1 | 1 | 0 |
| rs121909302 | pathogenic | 0 | 0 | 0 |
| rs121909303 | pathogenic | 0 | 0 | 0 |
| rs121909299 | pathogenic | 0 | 0 | 0 |
| rs121909301 | pathogenic | 0 | 0 | 0 |
| rs121909300 | pathogenic | 1 | 1 | 0 |
| rs111033589 | pathogenic | 0 | 0 | 0 |
| rs137853059 | pathogenic | 0 | 0 | 0 |
| rs137853057 | pathogenic | 1 | 1 | 1 |
| rs1801474 | benign | 0 | 0 | 0 |
| rs137853060 | pathogenic | 1 | 1 | 1 |
| rs137853058 | pathogenic | 1 | 1 | 1 |
| rs34424986 | pathogenic | 0 | 0 | 0 |
| rs199657839 | benign | 1 | 1 | 1 |
| rs56092260 | benign | 0 | 0 | 0 |
| rs1801582 | benign | 1 | 1 | 1 |
| rs1801334 | benign | 1 | 1 | 1 |
| rs191486604 | pathogenic | 0 | 0 | 0 |
| rs121908051 | pathogenic | 0 | 0 | 0 |
| rs121908055 | pathogenic | 1 | 1 | 0 |
| rs28940886 | pathogenic | 0 | 0 | 0 |
| rs121908050 | pathogenic | 0 | 0 | 0 |
| rs28940887 | pathogenic | 0 | 0 | 0 |
| rs387906300 | pathogenic | 0 | 0 | 0 |
| rs28942087 | pathogenic | 0 | 0 | 0 |
| rs121908049 | pathogenic | 1 | 1 | 0 |
| rs121908054 | pathogenic | 1 | 1 | 0 |
| rs121908048 | pathogenic | 0 | 0 | 0 |
| rs28940888 | pathogenic | 1 | 1 | 1 |
| rs121908053 | pathogenic | 1 | 1 | 0 |
| rs122467169 | pathogenic | 1 | 1 | 1 |
| rs122467172 | pathogenic | 1 | 1 | 1 |
| rs122467170 | pathogenic | 1 | 1 | 1 |
| rs28935477 | pathogenic | 1 | 1 | 0 |
| rs28942088 | pathogenic | 0 | 0 | 0 |
| rs751478142 | pathogenic | 1 | 1 | 0 |
| rs765368797 | pathogenic | 0 | 0 | 0 |
| rs147810437 | benign | 0 | 0 | 0 |
| rs121918209 | pathogenic | 1 | 1 | 1 |
| rs121918208 | pathogenic | 1 | 1 | 1 |
| rs121918210 | pathogenic | 1 | 1 | 1 |
| rs35515638 | benign | 1 | 1 | 1 |
| rs121918207 | pathogenic | 1 | 1 | 1 |
| rs141298312 | benign | 0 | 0 | 0 |
| rs121918211 | pathogenic | 0 | 0 | 0 |

| | | | | |
|---|---|---|---|---|
| rs149660059 | benign | 0 | 0 | 0 |
| rs200670286 | pathogenic | 1 | 1 | 1 |
| rs121918212 | pathogenic | 1 | 1 | 1 |
| rs121918213 | pathogenic | 1 | 1 | 1 |
| rs121918205 | pathogenic | 1 | 1 | 1 |
| rs104894025 | pathogenic | 1 | 1 | 1 |
| rs104894026 | pathogenic | 1 | 1 | 1 |
| rs104894028 | pathogenic | 0 | 1 | 0 |
| rs104894029 | pathogenic | 0 | 1 | 0 |
| rs119458971 | pathogenic | 0 | 0 | 0 |
| rs119458969 | pathogenic | 1 | 1 | 1 |
| rs1801591 | benign | 1 | 1 | 1 |
| rs140169311 | benign | 0 | 0 | 0 |
| rs753363278 | benign | 1 | 1 | 1 |
| rs199763682 | pathogenic | 0 | 0 | 0 |
| rs119458970 | pathogenic | 1 | 1 | 1 |
| rs763722868 | pathogenic | 0 | 0 | 0 |
| rs56404770 | benign | 0 | 0 | 0 |
| rs34221241 | benign | 0 | 0 | 0 |
| rs121909625 | pathogenic | 0 | 0 | 0 |
| rs550921485 | pathogenic | 0 | 0 | 0 |
| rs1800454 | benign | 0 | 0 | 0 |
| rs241447 | benign | 0 | 0 | 0 |
| rs2230606 | benign | 1 | 1 | 0 |
| rs5036 | pathogenic | 1 | 1 | 1 |
| rs13306788 | benign | 0 | 0 | 0 |
| rs28929480 | pathogenic | 0 | 0 | 0 |
| rs121912749 | pathogenic | 0 | 0 | 0 |
| rs56361140 | pathogenic | 0 | 0 | 0 |
| rs28931583 | pathogenic | 1 | 1 | 1 |
| rs36211715 | pathogenic | 1 | 1 | 1 |
| rs138049878 | pathogenic | 1 | 1 | 1 |
| rs727503253 | pathogenic | 1 | 1 | 1 |
| rs267606908 | pathogenic | 0 | 0 | 0 |
| rs121913631 | pathogenic | 1 | 1 | 1 |
| rs36211716 | benign | 1 | 1 | 1 |
| rs121913639 | pathogenic | 1 | 1 | 1 |
| rs730880761 | pathogenic | 1 | 1 | 1 |
| rs121913629 | pathogenic | 1 | 1 | 1 |
| rs754368658 | pathogenic | 0 | 0 | 0 |
| rs777090017 | pathogenic | 0 | 0 | 0 |
| rs267606913 | pathogenic | 1 | 1 | 1 |
| rs773138218 | pathogenic | 0 | 0 | 0 |
| rs371295780 | pathogenic | 0 | 0 | 0 |
| rs387906830 | pathogenic | 1 | 1 | 0 |
| rs771364038 | pathogenic | 0 | 1 | 0 |
| rs772731615 | pathogenic | 0 | 0 | 0 |
| rs137853063 | pathogenic | 0 | 0 | 0 |
| rs387907076 | pathogenic | 0 | 1 | 0 |
| rs35460768 | benign | 0 | 0 | 0 |
| rs2278196 | benign | 1 | 1 | 1 |
| rs119103275 | pathogenic | 0 | 0 | 0 |
| rs119103276 | pathogenic | 1 | 1 | 1 |
| rs374402066 | pathogenic | 1 | 1 | 1 |

| | | | | |
|---|---|---|---|---|
| rs119103274 | pathogenic | 0 | 0 | 0 |
| rs2274064 | benign | 0 | 0 | 0 |
| rs267607023 | pathogenic | 1 | 1 | 0 |
| rs142117489 | benign | 1 | 1 | 0 |
| rs371258270 | benign | 0 | 0 | 0 |
| rs34716057 | benign | 0 | 0 | 0 |
| rs201778606 | benign | 0 | 0 | 0 |
| rs201438792 | benign | 0 | 0 | 0 |
| rs140547417 | benign | 0 | 0 | 0 |
| rs5881 | benign | 0 | 0 | 0 |
| rs752298084 | benign | 0 | 0 | 0 |
| rs34855278 | benign | 0 | 0 | 0 |
| rs5880 | benign | 1 | 1 | 0 |
| rs5882 | benign | 0 | 0 | 0 |
| rs1800777 | benign | 0 | 0 | 0 |
| rs577301331 | benign | 0 | 0 | 0 |
| rs137853313 | pathogenic | 0 | 0 | 0 |
| rs28935169 | pathogenic | 0 | 0 | 0 |
| rs137853315 | pathogenic | 0 | 0 | 0 |
| rs137853314 | pathogenic | 0 | 0 | 0 |
| rs28935469 | pathogenic | 0 | 0 | 0 |
| rs137853318 | pathogenic | 0 | 0 | 0 |
| rs28935470 | pathogenic | 0 | 0 | 0 |
| rs3729986 | benign | 0 | 0 | 0 |
| rs730880619 | pathogenic | 0 | 0 | 0 |
| rs201012766 | benign | 0 | 0 | 0 |
| rs727503218 | pathogenic | 0 | 0 | 0 |
| rs11570052 | benign | 0 | 0 | 0 |
| rs730880623 | pathogenic | 0 | 0 | 0 |
| rs138753870 | benign | 0 | 0 | 0 |
| rs397516068 | pathogenic | 0 | 0 | 0 |
| rs3729989 | benign | 1 | 1 | 1 |
| rs397516074 | pathogenic | 0 | 0 | 0 |
| rs561449819 | pathogenic | 0 | 0 | 0 |
| rs121917779 | benign | 0 | 0 | 0 |
| rs142881576 | pathogenic | 0 | 0 | 0 |
| rs77384282 | benign | 0 | 0 | 0 |
| rs201865787 | benign | 0 | 0 | 0 |
| rs142244640 | benign | 0 | 0 | 0 |
| rs200488706 | benign | 0 | 0 | 0 |
| rs137853291 | pathogenic | 1 | 1 | 0 |
| rs28933990 | pathogenic | 1 | 1 | 0 |
| rs140570189 | benign | 0 | 0 | 0 |
| rs201588178 | benign | 0 | 0 | 0 |
| rs104893846 | pathogenic | 0 | 0 | 0 |
| rs754545360 | pathogenic | 0 | 1 | 0 |
| rs104893851 | pathogenic | 0 | 1 | 0 |
| rs104893849 | pathogenic | 0 | 0 | 0 |
| rs571038432 | pathogenic | 1 | 1 | 0 |
| rs2270655 | benign | 1 | 1 | 1 |
| rs315952 | benign | 0 | 0 | 0 |
| rs151176548 | benign | 0 | 0 | 0 |
| rs122459147 | pathogenic | 1 | 1 | 0 |
| rs122458140 | pathogenic | 0 | 0 | 0 |

| | | | | |
|---|---|---|---|---|
| rs122458142 | pathogenic | 1 | 1 | 0 |
| rs267606812 | pathogenic | 1 | 1 | 0 |
| rs267606813 | pathogenic | 1 | 1 | 0 |
| rs122458143 | pathogenic | 0 | 0 | 0 |
| rs122459146 | pathogenic | 0 | 0 | 0 |
| rs122458144 | pathogenic | 0 | 0 | 0 |
| rs122458145 | pathogenic | 0 | 0 | 0 |
| rs113994007 | pathogenic | 1 | 1 | 0 |
| rs758746181 | pathogenic | 1 | 1 | 1 |
| rs145034527 | pathogenic | 1 | 1 | 0 |
| rs119468006 | pathogenic | 1 | 1 | 0 |
| rs201908721 | pathogenic | 0 | 0 | 0 |
| rs755933881 | pathogenic | 1 | 1 | 0 |
| rs771578775 | pathogenic | 0 | 0 | 0 |
| rs17849927 | benign | 0 | 0 | 0 |
| rs139133094 | benign | 0 | 0 | 0 |
| rs12593 | benign | 0 | 0 | 0 |
| rs578189699 | pathogenic | 1 | 1 | 0 |
| rs119468008 | pathogenic | 0 | 0 | 0 |
| rs768524626 | benign | 0 | 0 | 0 |
| rs119468009 | pathogenic | 0 | 0 | 0 |
| rs119468004 | pathogenic | 0 | 0 | 0 |
| rs199874519 | pathogenic | 0 | 0 | 0 |
| rs376478331 | pathogenic | 0 | 0 | 0 |
| rs61995958 | pathogenic | 0 | 0 | 0 |
| rs199714731 | benign | 0 | 0 | 0 |
| rs149185954 | benign | 0 | 0 | 0 |
| rs121918353 | pathogenic | 0 | 0 | 0 |
| rs1131695 | benign | 0 | 0 | 0 |
| rs28939668 | pathogenic | 0 | 0 | 0 |
| rs758432471 | pathogenic | 1 | 1 | 1 |
| rs752746786 | pathogenic | 1 | 1 | 1 |
| rs28935471 | pathogenic | 0 | 0 | 0 |
| rs80338837 | pathogenic | 0 | 0 | 0 |
| rs137853312 | pathogenic | 0 | 0 | 0 |
| rs28935472 | pathogenic | 0 | 0 | 0 |
| rs28935473 | pathogenic | 0 | 0 | 0 |
| rs267606752 | pathogenic | 0 | 0 | 0 |
| rs3917973 | benign | 0 | 0 | 0 |
| rs3918018 | benign | 0 | 0 | 0 |
| rs757976755 | pathogenic | 0 | 0 | 0 |
| rs121918329 | pathogenic | 1 | 1 | 0 |
| rs770114459 | pathogenic | 0 | 0 | 0 |
| rs6662272 | benign | 0 | 0 | 0 |
| rs121918242 | pathogenic | 1 | 1 | 0 |
| rs121918240 | pathogenic | 1 | 1 | 0 |
| rs200094982 | pathogenic | 0 | 0 | 0 |
| rs121918241 | pathogenic | 0 | 0 | 0 |
| rs140522266 | pathogenic | 1 | 1 | 0 |
| rs121918243 | pathogenic | 1 | 1 | 0 |
| rs370596113 | pathogenic | 1 | 1 | 0 |
| rs398124295 | pathogenic | 1 | 1 | 0 |
| rs587776889 | pathogenic | 1 | 1 | 0 |
| rs747527726 | pathogenic | 1 | 1 | 0 |

| | | | | |
|---|---|---|---|---|
| rs112631212 | benign | 0 | 0 | 0 |
| rs104894605 | pathogenic | 0 | 0 | 0 |
| rs104894607 | pathogenic | 0 | 0 | 0 |
| rs104894606 | pathogenic | 0 | 0 | 0 |
| rs143873873 | benign | 0 | 0 | 0 |
| rs104894388 | pathogenic | 1 | 1 | 0 |
| rs104894391 | pathogenic | 1 | 1 | 0 |
| rs5350 | benign | 1 | 1 | 1 |
| rs104894390 | pathogenic | 1 | 1 | 1 |
| rs5346 | benign | 1 | 1 | 1 |
| rs121912500 | pathogenic | 0 | 0 | 0 |
| rs74315450 | pathogenic | 1 | 1 | 0 |
| rs121913396 | pathogenic | 1 | 1 | 1 |
| rs28931588 | pathogenic | 1 | 1 | 1 |
| rs121913400 | pathogenic | 0 | 0 | 0 |
| rs28931589 | pathogenic | 1 | 1 | 1 |
| rs121913403 | pathogenic | 0 | 0 | 0 |
| rs121913413 | pathogenic | 0 | 0 | 0 |
| rs760107623 | pathogenic | 1 | 1 | 0 |
| rs16866378 | benign | 1 | 1 | 0 |
| rs201025290 | pathogenic | 1 | 1 | 0 |
| rs12720356 | benign | 1 | 1 | 0 |
| rs55882956 | benign | 0 | 1 | 0 |
| rs4986933 | benign | 1 | 1 | 0 |
| rs2020957 | benign | 1 | 1 | 1 |
| rs1800124 | benign | 1 | 1 | 1 |
| rs150077735 | benign | 1 | 1 | 1 |
| rs201419000 | benign | 0 | 0 | 0 |
| rs772044176 | pathogenic | 0 | 0 | 0 |
| rs138346388 | benign | 0 | 0 | 0 |
| rs121964916 | pathogenic | 0 | 0 | 0 |
| rs112534524 | pathogenic | 0 | 0 | 0 |
| rs182078921 | benign | 1 | 1 | 1 |
| rs113273712 | benign | 0 | 0 | 0 |
| rs74817407 | benign | 1 | 1 | 1 |
| rs61733901 | benign | 1 | 1 | 1 |
| rs121964912 | pathogenic | 1 | 1 | 1 |
| rs41278047 | benign | 1 | 1 | 1 |
| rs7437875 | benign | 1 | 1 | 1 |
| rs113460688 | benign | 1 | 1 | 1 |
| rs754572081 | benign | 1 | 1 | 1 |
| rs201082692 | benign | 1 | 1 | 1 |
| rs202085803 | benign | 1 | 1 | 1 |
| rs141320511 | benign | 0 | 0 | 0 |
| rs78501403 | benign | 0 | 0 | 0 |
| rs28939675 | pathogenic | 0 | 0 | 0 |
| rs74315522 | pathogenic | 0 | 0 | 0 |
| rs121434639 | pathogenic | 1 | 1 | 0 |
| rs121434640 | pathogenic | 1 | 1 | 1 |
| rs757282628 | benign | 1 | 1 | 1 |
| rs386834010 | pathogenic | 0 | 0 | 0 |
| rs200863680 | pathogenic | 0 | 0 | 0 |
| rs193919336 | pathogenic | 1 | 1 | 0 |
| rs386834011 | pathogenic | 0 | 0 | 0 |

| | | | | |
|---|---|---|---|---|
| rs28940869 | pathogenic | 0 | 0 | 0 |
| rs386834014 | pathogenic | 1 | 1 | 0 |
| rs267606960 | pathogenic | 0 | 0 | 0 |
| rs28942068 | pathogenic | 0 | 0 | 0 |
| rs193919335 | pathogenic | 0 | 0 | 0 |
| rs74374973 | pathogenic | 0 | 0 | 0 |
| rs386834018 | pathogenic | 0 | 0 | 0 |
| rs267606962 | pathogenic | 0 | 0 | 0 |
| rs6659553 | benign | 0 | 0 | 0 |
| rs29281 | benign | 0 | 0 | 0 |
| rs782265090 | benign | 0 | 0 | 0 |
| rs35910969 | pathogenic | 0 | 0 | 0 |
| rs121909113 | pathogenic | 1 | 1 | 1 |
| rs29001571 | pathogenic | 1 | 1 | 1 |
| rs587781250 | pathogenic | 1 | 1 | 1 |
| rs28939680 | pathogenic | 1 | 1 | 1 |
| rs28939681 | pathogenic | 1 | 1 | 1 |
| rs121909112 | pathogenic | 1 | 1 | 1 |
| rs28937568 | pathogenic | 1 | 1 | 1 |
| rs1799895 | pathogenic | 0 | 0 | 0 |
| rs386833431 | pathogenic | 1 | 1 | 1 |
| rs121964905 | pathogenic | 1 | 1 | 1 |
| rs121964906 | pathogenic | 1 | 1 | 1 |
| rs11551664 | pathogenic | 1 | 1 | 1 |
| rs104894705 | pathogenic | 1 | 1 | 1 |
| rs104893805 | pathogenic | 0 | 0 | 0 |
| rs104894205 | pathogenic | 0 | 0 | 0 |
| rs104894204 | pathogenic | 1 | 1 | 0 |
| rs62642926 | pathogenic | 0 | 0 | 0 |
| rs5030841 | pathogenic | 1 | 1 | 1 |
| rs75193786 | pathogenic | 1 | 1 | 1 |
| rs62508682 | pathogenic | 0 | 0 | 0 |
| rs3850625 | benign | 1 | 1 | 1 |
| rs141908793 | benign | 0 | 0 | 0 |
| rs187864727 | benign | 0 | 0 | 0 |
| rs548909101 | benign | 0 | 0 | 0 |
| rs3825394 | benign | 0 | 1 | 0 |
| rs143548402 | benign | 0 | 0 | 0 |
| rs267607146 | pathogenic | 0 | 0 | 0 |
| rs267607144 | pathogenic | 0 | 0 | 0 |
| rs267607148 | pathogenic | 1 | 1 | 1 |
| rs267607143 | pathogenic | 0 | 0 | 0 |
| rs267607145 | pathogenic | 0 | 0 | 0 |
| rs121912636 | pathogenic | 1 | 1 | 1 |
| rs121912634 | pathogenic | 1 | 1 | 1 |
| rs570675468 | benign | 0 | 0 | 0 |
| rs368861241 | pathogenic | 1 | 1 | 1 |
| rs104894528 | pathogenic | 1 | 1 | 1 |
| rs77493891 | benign | 0 | 0 | 0 |
| rs62640903 | benign | 0 | 0 | 0 |
| rs149926091 | benign | 1 | 1 | 1 |
| rs641081 | benign | 0 | 0 | 0 |
| rs4930199 | benign | 0 | 0 | 0 |
| rs121913020 | pathogenic | 0 | 0 | 0 |

| | | | | |
|---|---|---|---|---|
| rs1799793 | benign | 0 | 0 | 0 |
| rs121913016 | pathogenic | 0 | 0 | 0 |
| rs121913025 | pathogenic | 0 | 0 | 0 |
| rs121913019 | pathogenic | 0 | 0 | 0 |
| rs376556895 | pathogenic | 1 | 1 | 1 |
| rs121913024 | pathogenic | 1 | 1 | 1 |
| rs121913021 | pathogenic | 0 | 0 | 0 |
| rs121913023 | pathogenic | 0 | 0 | 0 |
| rs41556519 | pathogenic | 0 | 0 | 0 |
| rs121913022 | pathogenic | 0 | 0 | 0 |
| rs144564120 | pathogenic | 0 | 0 | 0 |
| rs121913026 | pathogenic | 1 | 1 | 1 |
| rs121913018 | pathogenic | 1 | 1 | 1 |
| rs142954643 | benign | 0 | 0 | 0 |
| rs76655496 | benign | 0 | 0 | 0 |
| rs104894371 | pathogenic | 1 | 1 | 0 |
| rs137964936 | benign | 0 | 0 | 0 |
| rs759736132 | pathogenic | 0 | 0 | 0 |
| rs387906595 | pathogenic | 1 | 1 | 0 |
| rs768248216 | pathogenic | 0 | 0 | 0 |
| rs548231008 | benign | 0 | 0 | 0 |
| rs757094384 | benign | 0 | 0 | 0 |
| rs867529 | benign | 0 | 0 | 0 |
| rs13045 | benign | 1 | 1 | 1 |
| rs121908570 | pathogenic | 0 | 0 | 0 |
| rs201872586 | benign | 1 | 1 | 0 |
| rs149092990 | benign | 0 | 0 | 0 |
| rs115585711 | benign | 0 | 0 | 0 |
| rs137909416 | benign | 1 | 1 | 0 |
| rs143388949 | benign | 1 | 1 | 1 |
| rs73137716 | benign | 1 | 1 | 1 |
| rs121908947 | pathogenic | 1 | 1 | 1 |
| rs398122904 | pathogenic | 1 | 1 | 1 |
| rs35715497 | benign | 1 | 1 | 0 |
| rs35462421 | benign | 0 | 0 | 0 |
| rs119475043 | pathogenic | 0 | 0 | 0 |
| rs119475042 | pathogenic | 0 | 0 | 0 |
| rs1076669 | benign | 1 | 1 | 0 |
| rs141146885 | benign | 0 | 0 | 0 |
| rs3026906 | pathogenic | 0 | 0 | 0 |
| rs75362221 | benign | 1 | 1 | 1 |
| rs138472116 | benign | 1 | 1 | 1 |
| rs371040282 | pathogenic | 1 | 1 | 1 |
| rs4988496 | benign | 0 | 0 | 0 |
| rs121918117 | pathogenic | 1 | 1 | 1 |
| rs4988498 | benign | 0 | 0 | 0 |
| rs5035 | benign | 0 | 0 | 0 |
| rs121918322 | pathogenic | 1 | 1 | 1 |
| rs370359540 | benign | 0 | 0 | 0 |
| rs758972589 | pathogenic | 0 | 0 | 0 |
| rs144856889 | benign | 0 | 0 | 0 |
| rs121434623 | pathogenic | 1 | 1 | 1 |
| rs61735600 | benign | 0 | 0 | 0 |
| rs72550870 | benign | 1 | 1 | 1 |

| | | | | |
|---|---|---|---|---|
| rs56392418 | benign | 0 | 0 | 0 |
| rs41307788 | benign | 1 | 1 | 1 |
| rs387907239 | pathogenic | 1 | 1 | 0 |
| rs28931584 | pathogenic | 0 | 0 | 0 |
| rs121912745 | pathogenic | 0 | 0 | 0 |
| rs121912744 | pathogenic | 0 | 0 | 0 |
| rs121912754 | pathogenic | 1 | 1 | 1 |
| rs121912746 | pathogenic | 0 | 0 | 0 |
| rs121912757 | pathogenic | 0 | 0 | 0 |
| rs121912758 | pathogenic | 0 | 0 | 0 |
| rs121912748 | pathogenic | 0 | 0 | 0 |
| rs121912755 | pathogenic | 0 | 0 | 0 |
| rs121912741 | pathogenic | 0 | 0 | 0 |
| rs121912753 | pathogenic | 0 | 0 | 0 |
| rs121912750 | pathogenic | 0 | 0 | 0 |
| rs2285644 | pathogenic | 0 | 0 | 0 |
| rs121912751 | pathogenic | 0 | 0 | 0 |
| rs5026 | benign | 0 | 0 | 0 |
| rs121912759 | pathogenic | 0 | 0 | 0 |
| rs28931585 | pathogenic | 0 | 0 | 0 |
| rs17122769 | benign | 0 | 0 | 0 |
| rs28362322 | benign | 1 | 1 | 1 |
| rs3824874 | benign | 0 | 0 | 0 |
| rs146004831 | benign | 0 | 0 | 0 |
| rs142155860 | benign | 0 | 0 | 0 |
| rs121434404 | pathogenic | 0 | 0 | 0 |
| rs61735578 | benign | 1 | 1 | 1 |
| rs558018 | benign | 0 | 0 | 0 |
| rs61735577 | benign | 1 | 1 | 1 |
| rs76784113 | benign | 0 | 0 | 0 |
| rs199469640 | pathogenic | 0 | 0 | 0 |
| rs199469628 | pathogenic | 1 | 1 | 0 |
| rs199469629 | pathogenic | 0 | 1 | 0 |
| rs387907156 | pathogenic | 0 | 1 | 0 |
| rs199469643 | pathogenic | 0 | 0 | 0 |
| rs199469646 | pathogenic | 0 | 0 | 0 |
| rs766294940 | pathogenic | 0 | 0 | 0 |
| rs387907347 | pathogenic | 0 | 0 | 0 |
| rs137853590 | pathogenic | 1 | 1 | 1 |
| rs137853589 | pathogenic | 1 | 1 | 1 |
| rs137853588 | pathogenic | 1 | 1 | 1 |
| rs55909005 | benign | 0 | 0 | 0 |
| rs4581 | benign | 0 | 0 | 0 |
| rs63750123 | benign | 0 | 0 | 0 |
| rs10254120 | benign | 0 | 0 | 0 |
| rs121434629 | pathogenic | 1 | 1 | 0 |
| rs200313585 | benign | 0 | 0 | 0 |
| rs143162541 | benign | 1 | 1 | 0 |
| rs760615315 | benign | 0 | 0 | 0 |
| rs768488890 | benign | 1 | 1 | 1 |
| rs116373169 | benign | 1 | 1 | 0 |
| rs63750871 | pathogenic | 1 | 1 | 0 |
| rs116349687 | benign | 0 | 0 | 0 |
| rs375289386 | benign | 0 | 0 | 0 |

| | | | | |
|---|---|---|---|---|
| rs760228510 | pathogenic | 1 | 1 | 1 |
| rs587779343 | pathogenic | 1 | 1 | 1 |
| rs573125799 | benign | 0 | 0 | 0 |
| rs1805322 | benign | 0 | 0 | 0 |
| rs267608153 | pathogenic | 0 | 0 | 0 |
| rs139194813 | benign | 0 | 0 | 0 |
| rs530021751 | benign | 0 | 0 | 0 |
| rs200640585 | pathogenic | 0 | 0 | 0 |
| rs139438201 | benign | 0 | 0 | 0 |
| rs200513014 | benign | 0 | 0 | 0 |
| rs567102013 | benign | 0 | 0 | 0 |
| rs121908608 | pathogenic | 0 | 0 | 0 |
| rs121908606 | pathogenic | 0 | 0 | 0 |
| rs17103671 | benign | 0 | 0 | 0 |
| rs34725281 | benign | 0 | 0 | 0 |
| rs104894421 | pathogenic | 1 | 1 | 0 |
| rs121918145 | pathogenic | 1 | 1 | 1 |
| rs121918152 | pathogenic | 1 | 1 | 1 |
| rs56379106 | benign | 0 | 0 | 0 |
| rs4987076 | benign | 0 | 0 | 0 |
| rs370243877 | pathogenic | 0 | 0 | 0 |
| rs17679445 | benign | 1 | 1 | 1 |
| rs550423482 | pathogenic | 1 | 1 | 1 |
| rs104894629 | pathogenic | 1 | 1 | 1 |
| rs773450573 | pathogenic | 1 | 1 | 1 |
| rs116679717 | benign | 0 | 0 | 0 |
| rs372000848 | pathogenic | 0 | 0 | 0 |
| rs138595973 | benign | 0 | 0 | 0 |
| rs115244417 | benign | 0 | 0 | 0 |
| rs11575292 | benign | 0 | 0 | 0 |
| rs137853211 | pathogenic | 1 | 1 | 1 |
| rs137853207 | pathogenic | 0 | 0 | 0 |
| rs137853210 | pathogenic | 1 | 1 | 1 |
| rs6262 | benign | 0 | 0 | 0 |
| rs6263 | benign | 0 | 1 | 0 |
| rs137853208 | pathogenic | 0 | 1 | 0 |
| rs137853212 | pathogenic | 1 | 1 | 0 |
| rs137853209 | pathogenic | 1 | 1 | 1 |
| rs201951824 | pathogenic | 0 | 0 | 0 |
| rs11575542 | benign | 0 | 0 | 0 |
| rs199681788 | benign | 0 | 1 | 0 |
| rs748835902 | benign | 1 | 1 | 1 |
| rs753719949 | benign | 0 | 0 | 0 |
| rs267606647 | pathogenic | 0 | 0 | 0 |
| rs267606648 | pathogenic | 1 | 1 | 1 |
| rs267606643 | pathogenic | 0 | 1 | 0 |
| rs559947967 | pathogenic | 1 | 1 | 0 |
| rs267606645 | pathogenic | 1 | 1 | 1 |
| rs12116440 | benign | 0 | 0 | 0 |
| rs121908539 | pathogenic | 0 | 0 | 0 |
| rs137853096 | pathogenic | 1 | 1 | 0 |
| rs148363262 | benign | 1 | 1 | 0 |
| rs587777442 | pathogenic | 0 | 0 | 0 |
| rs25640 | benign | 1 | 1 | 1 |

| | | | | |
|---|---|---|---|---|
| rs28943589 | benign | 0 | 0 | 0 |
| rs1143650 | benign | 1 | 1 | 1 |
| rs150886026 | benign | 0 | 1 | 0 |
| rs121908133 | pathogenic | 1 | 1 | 0 |
| rs121908131 | pathogenic | 0 | 0 | 0 |
| rs754477154 | pathogenic | 0 | 0 | 0 |
| rs747334007 | benign | 0 | 0 | 0 |
| rs121908132 | pathogenic | 0 | 0 | 0 |
| rs104894103 | pathogenic | 1 | 1 | 0 |
| rs140088149 | benign | 0 | 0 | 0 |
| rs149803148 | benign | 0 | 0 | 0 |
| rs11064480 | benign | 0 | 0 | 0 |
| rs74435397 | pathogenic | 1 | 1 | 1 |
| rs587783626 | pathogenic | 1 | 1 | 1 |
| rs148211042 | pathogenic | 1 | 1 | 1 |
| rs17849502 | benign | 0 | 0 | 0 |
| rs13306575 | pathogenic | 1 | 1 | 1 |
| rs201426733 | benign | 0 | 0 | 0 |
| rs182278784 | benign | 1 | 1 | 1 |
| rs77786415 | benign | 1 | 1 | 1 |
| rs200192119 | benign | 1 | 1 | 1 |
| rs12877501 | benign | 1 | 1 | 1 |
| rs77985244 | benign | 1 | 1 | 1 |
| rs5907 | pathogenic | 1 | 1 | 0 |
| rs2305178 | benign | 0 | 0 | 0 |
| rs3135867 | benign | 0 | 0 | 0 |
| rs121913482 | pathogenic | 1 | 1 | 1 |
| rs121913483 | pathogenic | 1 | 1 | 1 |
| rs4647924 | pathogenic | 1 | 1 | 1 |
| rs121913111 | pathogenic | 1 | 1 | 1 |
| rs104894002 | pathogenic | 0 | 0 | 0 |
| rs104894001 | pathogenic | 0 | 0 | 0 |
| rs75932628 | benign | 0 | 0 | 0 |
| rs201258663 | pathogenic | 0 | 0 | 0 |
| rs104893998 | pathogenic | 0 | 0 | 0 |
| rs2234253 | benign | 1 | 1 | 0 |
| rs121908402 | pathogenic | 1 | 1 | 0 |
| rs1805389 | benign | 0 | 0 | 0 |
| rs1805388 | benign | 0 | 0 | 0 |
| rs748385144 | pathogenic | 0 | 0 | 0 |
| rs104894420 | pathogenic | 0 | 0 | 0 |
| rs104894418 | pathogenic | 0 | 0 | 0 |
| rs61731910 | benign | 0 | 0 | 0 |
| rs758723288 | pathogenic | 1 | 1 | 0 |
| rs201865375 | pathogenic | 1 | 1 | 0 |
| rs565090080 | pathogenic | 1 | 1 | 1 |
| rs113042166 | benign | 0 | 0 | 0 |
| rs61736326 | benign | 1 | 1 | 1 |
| rs12030699 | benign | 1 | 1 | 1 |
| rs121913578 | pathogenic | 1 | 1 | 1 |
| rs61739582 | benign | 0 | 0 | 0 |
| rs745983207 | pathogenic | 0 | 0 | 0 |
| rs763524380 | benign | 0 | 0 | 0 |
| rs121912792 | pathogenic | 0 | 0 | 0 |

| | | | | |
|---|---|---|---|---|
| rs121912793 | pathogenic | 0 | 0 | 0 |
| rs387906599 | pathogenic | 0 | 0 | 0 |
| rs121912794 | pathogenic | 0 | 1 | 0 |
| rs121912795 | pathogenic | 0 | 0 | 0 |
| rs121912796 | pathogenic | 1 | 1 | 0 |
| rs121912797 | pathogenic | 0 | 1 | 0 |
| rs121912798 | pathogenic | 1 | 1 | 1 |
| rs80359812 | pathogenic | 1 | 1 | 0 |
| rs776583130 | benign | 1 | 1 | 0 |
| rs80359814 | pathogenic | 1 | 1 | 0 |
| rs202060209 | pathogenic | 1 | 1 | 0 |
| rs267607061 | pathogenic | 1 | 1 | 0 |
| rs267607060 | pathogenic | 0 | 0 | 0 |
| rs80359816 | pathogenic | 1 | 1 | 0 |
| rs80359818 | pathogenic | 1 | 1 | 0 |
| rs387907313 | pathogenic | 0 | 0 | 0 |
| rs587784395 | benign | 0 | 0 | 0 |
| rs5811 | benign | 0 | 0 | 0 |
| rs121909740 | pathogenic | 1 | 1 | 0 |
| rs148518827 | benign | 1 | 1 | 0 |
| rs121909739 | pathogenic | 1 | 1 | 0 |
| rs780528770 | benign | 1 | 1 | 0 |
| rs13306758 | pathogenic | 0 | 0 | 0 |
| rs267607059 | pathogenic | 0 | 0 | 0 |
| rs61730638 | benign | 0 | 0 | 0 |
| rs104894533 | pathogenic | 0 | 0 | 0 |
| rs104894527 | pathogenic | 0 | 0 | 0 |
| rs200503569 | pathogenic | 1 | 1 | 1 |
| rs80338700 | pathogenic | 1 | 1 | 1 |
| rs765433263 | pathogenic | 1 | 1 | 1 |
| rs104894530 | pathogenic | 1 | 1 | 1 |
| rs80338701 | pathogenic | 1 | 1 | 1 |
| rs141498002 | pathogenic | 1 | 1 | 1 |
| rs104894525 | pathogenic | 0 | 0 | 0 |
| rs80338703 | pathogenic | 0 | 0 | 0 |
| rs28936415 | pathogenic | 1 | 1 | 0 |
| rs150719105 | pathogenic | 0 | 0 | 0 |
| rs148032587 | pathogenic | 0 | 0 | 0 |
| rs150577656 | pathogenic | 0 | 0 | 0 |
| rs190521996 | pathogenic | 0 | 0 | 0 |
| rs104894526 | pathogenic | 1 | 1 | 1 |
| rs80338704 | pathogenic | 0 | 0 | 0 |
| rs199562225 | pathogenic | 1 | 1 | 1 |
| rs532870929 | pathogenic | 0 | 0 | 0 |
| rs398123309 | pathogenic | 0 | 0 | 0 |
| rs78290141 | pathogenic | 0 | 0 | 0 |
| rs80338705 | pathogenic | 0 | 0 | 0 |
| rs104894531 | pathogenic | 1 | 1 | 1 |
| rs80338706 | pathogenic | 0 | 0 | 0 |
| rs80338707 | pathogenic | 0 | 0 | 0 |
| rs80338708 | pathogenic | 0 | 0 | 0 |
| rs80338709 | pathogenic | 0 | 0 | 0 |
| rs104893833 | benign | 0 | 0 | 0 |
| rs61744585 | benign | 0 | 0 | 0 |

| | | | | |
|---|---|---|---|---|
| rs73569515 | benign | 0 | 0 | 0 |
| rs398123373 | pathogenic | 0 | 0 | 0 |
| rs756854513 | pathogenic | 0 | 0 | 0 |
| rs58232698 | benign | 0 | 0 | 0 |
| rs9600455 | benign | 0 | 0 | 0 |
| rs201016593 | pathogenic | 0 | 0 | 0 |
| rs756039188 | pathogenic | 0 | 0 | 0 |
| rs121908024 | pathogenic | 0 | 0 | 0 |
| rs373144619 | benign | 0 | 0 | 0 |
| rs138078086 | benign | 0 | 0 | 0 |
| rs376207800 | pathogenic | 0 | 0 | 0 |
| rs749038326 | pathogenic | 1 | 1 | 0 |
| rs372845091 | benign | 0 | 0 | 0 |
| rs377271627 | pathogenic | 0 | 0 | 0 |
| rs563390335 | pathogenic | 0 | 0 | 0 |
| rs13306510 | benign | 0 | 0 | 0 |
| rs764042910 | pathogenic | 0 | 0 | 0 |
| rs201384282 | benign | 1 | 1 | 1 |
| rs730882086 | pathogenic | 1 | 1 | 1 |
| rs756613387 | pathogenic | 1 | 1 | 1 |
| rs121908028 | benign | 1 | 1 | 0 |
| rs773328511 | benign | 0 | 0 | 0 |
| rs749220643 | pathogenic | 1 | 1 | 0 |
| rs730882089 | pathogenic | 0 | 0 | 0 |
| rs146651743 | benign | 0 | 0 | 0 |
| rs148698650 | pathogenic | 0 | 0 | 0 |
| rs140241383 | benign | 0 | 0 | 0 |
| rs757252110 | pathogenic | 0 | 0 | 0 |
| rs13306512 | pathogenic | 1 | 1 | 1 |
| rs121908043 | pathogenic | 1 | 1 | 1 |
| rs773658037 | pathogenic | 0 | 0 | 0 |
| rs774439908 | pathogenic | 0 | 0 | 0 |
| rs779732323 | pathogenic | 1 | 1 | 0 |
| rs749780672 | pathogenic | 0 | 0 | 0 |
| rs375651668 | benign | 1 | 1 | 1 |
| rs193922568 | pathogenic | 1 | 1 | 0 |
| rs750363970 | benign | 1 | 1 | 0 |
| rs5932 | benign | 0 | 0 | 0 |
| rs755154048 | pathogenic | 1 | 1 | 0 |
| rs759876319 | pathogenic | 0 | 0 | 0 |
| rs764550980 | pathogenic | 1 | 1 | 0 |
| rs779940524 | pathogenic | 0 | 0 | 0 |
| rs563382937 | pathogenic | 1 | 1 | 1 |
| rs28942083 | pathogenic | 0 | 1 | 0 |
| rs770744861 | pathogenic | 0 | 1 | 0 |
| rs121908031 | pathogenic | 0 | 1 | 0 |
| rs776217028 | pathogenic | 0 | 1 | 0 |
| rs763625913 | pathogenic | 0 | 0 | 0 |
| rs3736772 | benign | 0 | 0 | 0 |
| rs104894696 | benign | 1 | 1 | 1 |
| rs121918177 | pathogenic | 0 | 1 | 0 |
| rs121918181 | pathogenic | 1 | 1 | 0 |
| rs121918174 | pathogenic | 0 | 1 | 0 |
| rs121918182 | pathogenic | 0 | 0 | 0 |

| | | | | |
|---|---|---|---|---|
| rs121918175 | pathogenic | 1 | 1 | 1 |
| rs121918185 | pathogenic | 1 | 1 | 1 |
| rs121918173 | pathogenic | 1 | 1 | 0 |
| rs121918178 | pathogenic | 1 | 1 | 1 |
| rs121918179 | pathogenic | 1 | 1 | 1 |
| rs121918183 | pathogenic | 0 | 0 | 0 |
| rs121918176 | pathogenic | 1 | 1 | 1 |
| rs121918172 | pathogenic | 0 | 1 | 0 |
| rs121918184 | pathogenic | 0 | 0 | 0 |
| rs121434560 | pathogenic | 1 | 1 | 0 |
| rs121434561 | pathogenic | 1 | 1 | 0 |
| rs3218716 | pathogenic | 1 | 1 | 1 |
| rs145532615 | benign | 0 | 0 | 0 |
| rs149840927 | pathogenic | 0 | 0 | 0 |
| rs730880762 | pathogenic | 0 | 0 | 0 |
| rs730880764 | pathogenic | 0 | 0 | 0 |
| rs730880765 | pathogenic | 0 | 0 | 0 |
| rs587782962 | pathogenic | 0 | 0 | 0 |
| rs141735183 | pathogenic | 0 | 0 | 0 |
| rs397516187 | pathogenic | 0 | 0 | 0 |
| rs727505026 | pathogenic | 0 | 0 | 0 |
| rs529700838 | pathogenic | 0 | 0 | 0 |
| rs540263945 | pathogenic | 0 | 0 | 0 |
| rs149103761 | benign | 0 | 0 | 0 |
| rs730880906 | pathogenic | 0 | 0 | 0 |
| rs730880785 | pathogenic | 0 | 0 | 0 |
| rs397516199 | pathogenic | 0 | 0 | 0 |
| rs730880725 | benign | 0 | 0 | 0 |
| rs730880796 | pathogenic | 0 | 0 | 0 |
| rs201307101 | benign | 0 | 0 | 0 |
| rs397516214 | pathogenic | 0 | 0 | 0 |
| rs3729823 | benign | 0 | 0 | 0 |
| rs397516218 | pathogenic | 0 | 0 | 0 |
| rs121913647 | pathogenic | 0 | 0 | 0 |
| rs373514686 | pathogenic | 0 | 0 | 0 |
| rs763538103 | pathogenic | 0 | 0 | 0 |
| rs121913650 | pathogenic | 0 | 0 | 0 |
| rs139222507 | pathogenic | 0 | 0 | 0 |
| rs369940645 | pathogenic | 0 | 0 | 0 |
| rs763073072 | pathogenic | 0 | 0 | 0 |
| rs145734640 | pathogenic | 0 | 0 | 0 |
| rs730880821 | pathogenic | 0 | 0 | 0 |
| rs28933098 | pathogenic | 0 | 0 | 0 |
| rs372381770 | pathogenic | 0 | 0 | 0 |
| rs146796870 | benign | 0 | 0 | 0 |
| rs104894485 | pathogenic | 1 | 1 | 1 |
| rs35407712 | benign | 0 | 0 | 0 |
| rs35842750 | benign | 0 | 0 | 0 |
| rs2721191 | benign | 0 | 0 | 0 |
| rs35098923 | benign | 0 | 0 | 0 |
| rs34293591 | benign | 1 | 1 | 0 |
| rs36023964 | benign | 0 | 0 | 0 |
| rs4251691 | benign | 0 | 0 | 0 |
| rs34666647 | benign | 0 | 0 | 0 |

| | | | | |
|---|---|---|---|---|
| rs4925828 | benign | 0 | 0 | 0 |
| rs35348691 | benign | 0 | 0 | 0 |
| rs36078464 | benign | 0 | 0 | 0 |
| rs34915097 | benign | 0 | 0 | 0 |
| rs35101495 | benign | 0 | 0 | 0 |
| rs141568342 | pathogenic | 0 | 0 | 0 |
| rs1800571 | pathogenic | 0 | 0 | 0 |
| rs140566084 | benign | 1 | 1 | 1 |
| rs727503791 | pathogenic | 1 | 1 | 0 |
| rs545215807 | pathogenic | 0 | 0 | 0 |
| rs113994168 | pathogenic | 0 | 0 | 0 |
| rs113994167 | pathogenic | 0 | 0 | 0 |
| rs200788251 | pathogenic | 0 | 0 | 0 |
| rs200573371 | pathogenic | 0 | 0 | 0 |
| rs771874163 | pathogenic | 1 | 1 | 1 |
| rs112406105 | pathogenic | 1 | 1 | 1 |
| rs118204015 | pathogenic | 1 | 1 | 1 |
| rs113994169 | pathogenic | 0 | 0 | 0 |
| rs118204018 | pathogenic | 0 | 1 | 0 |
| rs138834083 | pathogenic | 0 | 0 | 0 |
| rs533055438 | pathogenic | 1 | 1 | 1 |
| rs2309689 | pathogenic | 1 | 1 | 1 |
| rs138058572 | pathogenic | 1 | 1 | 1 |
| rs118204017 | pathogenic | 1 | 1 | 0 |
| rs766742117 | pathogenic | 1 | 1 | 1 |
| rs751995154 | pathogenic | 1 | 1 | 1 |
| rs200771970 | pathogenic | 0 | 0 | 0 |
| rs118204014 | pathogenic | 0 | 0 | 0 |
| rs148584617 | pathogenic | 1 | 1 | 1 |
| rs121912720 | pathogenic | 0 | 0 | 0 |
| rs121912721 | pathogenic | 0 | 0 | 0 |
| rs28929476 | pathogenic | 0 | 0 | 0 |
| rs28931574 | pathogenic | 1 | 1 | 1 |
| rs12718465 | benign | 0 | 0 | 0 |
| rs144689354 | pathogenic | 0 | 0 | 0 |
| rs149095705 | pathogenic | 0 | 0 | 0 |
| rs572839648 | benign | 1 | 1 | 1 |
| rs121909715 | pathogenic | 1 | 1 | 0 |
| rs34077456 | benign | 1 | 1 | 1 |
| rs397517876 | benign | 0 | 0 | 0 |
| rs1064074 | benign | 0 | 0 | 0 |
| rs142751309 | benign | 0 | 0 | 0 |
| rs116275562 | benign | 0 | 0 | 0 |
| rs148095899 | benign | 0 | 0 | 0 |
| rs751838040 | pathogenic | 0 | 0 | 0 |
| rs121434324 | pathogenic | 1 | 1 | 1 |
| rs143367518 | pathogenic | 0 | 0 | 0 |
| rs121434325 | pathogenic | 0 | 0 | 0 |
| rs397515328 | pathogenic | 0 | 0 | 0 |
| rs376039938 | pathogenic | 0 | 0 | 0 |
| rs387907243 | pathogenic | 0 | 0 | 0 |
| rs28934876 | pathogenic | 0 | 0 | 0 |
| rs387907245 | pathogenic | 0 | 0 | 0 |
| rs138544311 | benign | 0 | 0 | 0 |

| | | | | |
|---|---|---|---|---|
| rs643788 | benign | 0 | 0 | 0 |
| rs749977756 | benign | 0 | 0 | 0 |
| rs1800574 | benign | 0 | 0 | 0 |
| rs137853336 | pathogenic | 1 | 1 | 1 |
| rs111916034 | benign | 0 | 0 | 0 |
| rs307826 | benign | 0 | 0 | 0 |
| rs35874891 | benign | 0 | 0 | 0 |
| rs111033580 | pathogenic | 1 | 1 | 1 |
| rs111033581 | pathogenic | 1 | 1 | 1 |
| rs121908373 | pathogenic | 0 | 0 | 0 |
| rs767122713 | pathogenic | 0 | 0 | 0 |
| rs2233213 | benign | 0 | 0 | 0 |
| rs121908605 | pathogenic | 1 | 1 | 0 |
| rs11887534 | benign | 0 | 0 | 0 |
| rs4148211 | benign | 0 | 0 | 0 |
| rs80025980 | benign | 0 | 0 | 0 |
| rs137854891 | pathogenic | 1 | 1 | 0 |
| rs74358901 | benign | 0 | 0 | 0 |
| rs9282574 | benign | 0 | 0 | 0 |
| rs137852993 | pathogenic | 0 | 0 | 0 |
| rs137852990 | pathogenic | 0 | 0 | 0 |
| rs137852987 | pathogenic | 0 | 0 | 0 |
| rs4148217 | benign | 0 | 0 | 0 |
| rs140690030 | benign | 0 | 0 | 0 |
| rs137852992 | pathogenic | 0 | 0 | 0 |
| rs6544718 | benign | 0 | 0 | 0 |
| rs137852989 | pathogenic | 0 | 0 | 0 |
| rs150893072 | benign | 1 | 1 | 1 |
| rs28931613 | pathogenic | 1 | 1 | 1 |
| rs143789597 | benign | 1 | 1 | 1 |
| rs121913037 | pathogenic | 1 | 1 | 0 |
| rs188802138 | pathogenic | 1 | 1 | 0 |
| rs121913038 | pathogenic | 1 | 1 | 0 |
| rs146835206 | benign | 0 | 0 | 0 |
| rs121913041 | pathogenic | 1 | 1 | 1 |
| rs121913039 | pathogenic | 1 | 1 | 1 |
| rs149977726 | pathogenic | 1 | 1 | 0 |
| rs121913036 | pathogenic | 0 | 0 | 0 |
| rs121913040 | pathogenic | 0 | 0 | 0 |
| rs765007158 | benign | 0 | 0 | 0 |
| rs112723255 | benign | 0 | 0 | 0 |
| rs574164748 | benign | 0 | 0 | 0 |
| rs80338734 | pathogenic | 1 | 1 | 0 |
| rs104894694 | pathogenic | 1 | 1 | 0 |
| rs753198836 | pathogenic | 0 | 0 | 0 |
| rs760101382 | pathogenic | 0 | 0 | 0 |
| rs121909272 | pathogenic | 1 | 1 | 0 |
| rs370421531 | pathogenic | 0 | 0 | 0 |
| rs565109128 | benign | 0 | 0 | 0 |
| rs17851582 | benign | 0 | 0 | 0 |
| rs753228876 | benign | 0 | 0 | 0 |
| rs141471799 | benign | 0 | 0 | 0 |
| rs281864916 | pathogenic | 1 | 1 | 0 |
| rs121918409 | pathogenic | 1 | 1 | 1 |

| | | | | |
|---|---|---|---|---|
| rs121918413 | pathogenic | 1 | 1 | 1 |
| rs121918411 | pathogenic | 1 | 1 | 1 |
| rs121918408 | pathogenic | 1 | 1 | 1 |
| rs121918412 | pathogenic | 1 | 1 | 1 |
| rs121918410 | pathogenic | 1 | 1 | 1 |
| rs116474260 | benign | 1 | 1 | 1 |
| rs281864919 | pathogenic | 1 | 1 | 1 |
| rs397514452 | pathogenic | 1 | 1 | 0 |
| rs747843638 | benign | 0 | 0 | 0 |
| rs774357609 | pathogenic | 1 | 1 | 1 |
| rs201804005 | benign | 0 | 0 | 0 |
| rs746485916 | pathogenic | 0 | 0 | 0 |
| rs764131178 | pathogenic | 0 | 0 | 0 |
| rs374094065 | pathogenic | 0 | 0 | 0 |
| rs76151636 | pathogenic | 1 | 1 | 0 |
| rs753250853 | pathogenic | 0 | 0 | 0 |
| rs761632029 | pathogenic | 0 | 0 | 0 |
| rs775541743 | pathogenic | 0 | 0 | 0 |
| rs778749563 | pathogenic | 0 | 0 | 0 |
| rs60431989 | pathogenic | 1 | 1 | 0 |
| rs755554442 | pathogenic | 1 | 1 | 0 |
| rs749085322 | pathogenic | 0 | 0 | 0 |
| rs756029120 | pathogenic | 1 | 1 | 0 |
| rs137853279 | pathogenic | 1 | 1 | 0 |
| rs777354267 | pathogenic | 1 | 1 | 1 |
| rs142940486 | benign | 1 | 1 | 1 |
| rs140167393 | benign | 1 | 1 | 1 |
| rs2066827 | benign | 0 | 0 | 0 |
| rs765681672 | benign | 0 | 0 | 0 |
| rs137985549 | benign | 0 | 0 | 0 |
| rs74315340 | pathogenic | 1 | 1 | 1 |
| rs74315341 | pathogenic | 0 | 0 | 0 |
| rs74315335 | pathogenic | 0 | 0 | 0 |
| rs61730974 | benign | 0 | 0 | 0 |
| rs121909193 | pathogenic | 0 | 0 | 0 |
| rs74315334 | pathogenic | 0 | 0 | 0 |
| rs74315329 | pathogenic | 0 | 0 | 0 |
| rs74315330 | pathogenic | 0 | 0 | 0 |
| rs121909194 | pathogenic | 0 | 0 | 0 |
| rs28936694 | pathogenic | 0 | 0 | 0 |
| rs74315336 | pathogenic | 0 | 0 | 0 |
| rs74315338 | pathogenic | 1 | 1 | 1 |
| rs74315328 | pathogenic | 0 | 0 | 0 |
| rs74315331 | pathogenic | 0 | 0 | 0 |
| rs74315332 | pathogenic | 0 | 0 | 0 |
| rs141696651 | benign | 0 | 0 | 0 |
| rs7041 | benign | 0 | 0 | 0 |
| rs4588 | benign | 0 | 0 | 0 |
| rs121908981 | pathogenic | 1 | 1 | 1 |
| rs606231459 | pathogenic | 0 | 0 | 0 |
| rs398123053 | pathogenic | 0 | 0 | 0 |
| rs774259802 | benign | 0 | 0 | 0 |
| rs121909582 | pathogenic | 1 | 1 | 1 |
| rs104893680 | pathogenic | 1 | 1 | 1 |

| | | | | |
|---|---|---|---|---|
| rs137854907 | pathogenic | 0 | 0 | 0 |
| rs765715798 | pathogenic | 0 | 0 | 0 |
| rs104893678 | pathogenic | 0 | 0 | 0 |
| rs104893679 | pathogenic | 0 | 1 | 0 |
| rs104893681 | pathogenic | 1 | 1 | 0 |
| rs121909640 | pathogenic | 1 | 1 | 0 |
| rs780705654 | pathogenic | 0 | 0 | 0 |
| rs201672146 | benign | 1 | 1 | 0 |
| rs12136658 | benign | 0 | 0 | 0 |
| rs56127440 | benign | 0 | 0 | 0 |
| rs104894433 | pathogenic | 1 | 1 | 1 |
| rs104894435 | pathogenic | 0 | 1 | 0 |
| rs104894437 | pathogenic | 1 | 1 | 1 |
| rs104894441 | pathogenic | 1 | 1 | 1 |
| rs104894440 | pathogenic | 1 | 1 | 1 |
| rs104894445 | pathogenic | 1 | 1 | 1 |
| rs104894436 | pathogenic | 0 | 0 | 0 |
| rs104894438 | pathogenic | 0 | 0 | 0 |
| rs200891969 | pathogenic | 1 | 1 | 0 |
| rs104894443 | pathogenic | 1 | 1 | 1 |
| rs104894434 | pathogenic | 1 | 1 | 1 |
| rs41298442 | pathogenic | 1 | 1 | 1 |
| rs104894442 | pathogenic | 1 | 1 | 1 |
| rs35453854 | benign | 0 | 0 | 0 |
| rs121912898 | pathogenic | 0 | 0 | 0 |
| rs121912897 | pathogenic | 1 | 1 | 0 |
| rs138260625 | benign | 0 | 0 | 0 |
| rs147802438 | benign | 1 | 1 | 0 |
| rs146443310 | benign | 1 | 1 | 0 |
| rs747319628 | pathogenic | 1 | 1 | 0 |
| rs1801195 | benign | 0 | 0 | 0 |
| rs145045341 | benign | 1 | 1 | 0 |
| rs116001173 | benign | 1 | 1 | 0 |
| rs587784175 | pathogenic | 0 | 0 | 0 |
| rs104893857 | pathogenic | 0 | 0 | 0 |
| rs104893862 | pathogenic | 1 | 1 | 0 |
| rs104893858 | pathogenic | 1 | 1 | 1 |
| rs121909249 | pathogenic | 0 | 1 | 0 |
| rs121909248 | pathogenic | 1 | 1 | 1 |
| rs104893859 | pathogenic | 1 | 1 | 0 |
| rs187063864 | pathogenic | 1 | 1 | 0 |
| rs121917817 | pathogenic | 1 | 1 | 0 |
| rs121917818 | pathogenic | 0 | 0 | 0 |
| rs753594601 | pathogenic | 0 | 0 | 0 |
| rs28937869 | pathogenic | 1 | 1 | 0 |
| rs6165 | benign | 0 | 0 | 0 |
| rs200089714 | pathogenic | 1 | 1 | 0 |
| rs372000714 | pathogenic | 0 | 0 | 0 |
| rs2228206 | benign | 0 | 0 | 0 |
| rs560096 | benign | 0 | 0 | 0 |
| rs137852666 | pathogenic | 0 | 0 | 0 |
| rs10896380 | benign | 0 | 0 | 0 |
| rs773690764 | pathogenic | 0 | 0 | 0 |
| rs201060167 | pathogenic | 0 | 0 | 0 |

| | | | | |
|---|---|---|---|---|
| rs137852670 | pathogenic | 0 | 0 | 0 |
| rs780594709 | pathogenic | 0 | 0 | 0 |
| rs145226920 | pathogenic | 0 | 0 | 0 |
| rs137852665 | pathogenic | 0 | 0 | 0 |
| rs724159960 | pathogenic | 0 | 0 | 0 |
| rs756985703 | pathogenic | 0 | 0 | 0 |
| rs137852667 | pathogenic | 0 | 0 | 0 |
| rs151079750 | pathogenic | 1 | 1 | 0 |
| rs2306636 | benign | 0 | 0 | 0 |
| rs72647885 | benign | 1 | 1 | 0 |
| rs377584435 | pathogenic | 1 | 1 | 1 |
| rs757583846 | pathogenic | 1 | 1 | 0 |
| rs121918148 | pathogenic | 0 | 0 | 0 |
| rs149201802 | benign | 0 | 0 | 0 |
| rs587781904 | pathogenic | 1 | 1 | 1 |
| rs143421325 | benign | 0 | 0 | 0 |
| rs587781253 | pathogenic | 0 | 0 | 0 |
| rs587780331 | benign | 0 | 0 | 0 |
| rs151001016 | benign | 0 | 0 | 0 |
| rs150888094 | benign | 0 | 0 | 0 |
| rs149581331 | benign | 0 | 0 | 0 |
| rs774840535 | benign | 0 | 0 | 0 |
| rs138428684 | benign | 0 | 0 | 0 |
| rs149300055 | benign | 0 | 0 | 0 |
| rs571005578 | benign | 0 | 0 | 0 |
| rs200731839 | benign | 0 | 0 | 0 |
| rs121965007 | pathogenic | 0 | 0 | 0 |
| rs121965013 | pathogenic | 0 | 1 | 0 |
| rs121965009 | pathogenic | 0 | 0 | 0 |
| rs1800457 | benign | 1 | 1 | 0 |
| rs121965006 | pathogenic | 1 | 1 | 0 |
| rs121965008 | pathogenic | 0 | 0 | 0 |
| rs61732609 | pathogenic | 0 | 0 | 0 |
| rs121965011 | pathogenic | 0 | 0 | 0 |
| rs121965015 | pathogenic | 0 | 0 | 0 |
| rs121965018 | pathogenic | 1 | 1 | 0 |
| rs121965016 | pathogenic | 0 | 0 | 0 |
| rs77806 | benign | 0 | 0 | 0 |
| rs149731356 | benign | 0 | 0 | 0 |
| rs72549407 | pathogenic | 0 | 0 | 0 |
| rs200363929 | benign | 0 | 0 | 0 |
| rs376403708 | benign | 1 | 1 | 0 |
| rs56376197 | benign | 1 | 1 | 1 |
| rs200237973 | benign | 1 | 1 | 1 |
| rs548330075 | benign | 0 | 0 | 0 |
| rs148291350 | benign | 0 | 0 | 0 |
| rs71579353 | pathogenic | 0 | 0 | 0 |
| rs146278697 | benign | 0 | 0 | 0 |
| rs141033098 | benign | 0 | 0 | 0 |
| rs397517234 | pathogenic | 0 | 0 | 0 |
| rs71579374 | benign | 0 | 0 | 0 |
| rs140381835 | benign | 0 | 0 | 0 |
| rs137852702 | pathogenic | 0 | 0 | 0 |
| rs137852696 | pathogenic | 1 | 1 | 1 |

| | | | | |
|---|---|---|---|---|
| rs137852697 | pathogenic | 0 | 0 | 0 |
| rs386833639 | pathogenic | 0 | 0 | 0 |
| rs386833640 | pathogenic | 0 | 0 | 0 |
| rs386833641 | benign | 0 | 0 | 0 |
| rs137852701 | pathogenic | 0 | 0 | 0 |
| rs386833642 | pathogenic | 0 | 0 | 0 |
| rs137852695 | pathogenic | 0 | 0 | 0 |
| rs386833646 | pathogenic | 0 | 0 | 0 |
| rs137852700 | pathogenic | 1 | 1 | 0 |
| rs386833649 | pathogenic | 1 | 1 | 0 |
| rs386833650 | pathogenic | 0 | 0 | 0 |
| rs148412181 | pathogenic | 0 | 0 | 0 |
| rs386833655 | pathogenic | 0 | 0 | 0 |
| rs386833656 | pathogenic | 1 | 1 | 0 |
| rs137852698 | pathogenic | 0 | 0 | 0 |
| rs386833661 | pathogenic | 0 | 0 | 0 |
| rs386833670 | pathogenic | 0 | 0 | 0 |
| rs111033632 | pathogenic | 1 | 1 | 1 |
| rs34308573 | benign | 1 | 1 | 1 |
| rs376048533 | pathogenic | 1 | 1 | 0 |
| rs3747517 | benign | 1 | 1 | 0 |
| rs137853328 | pathogenic | 0 | 0 | 0 |
| rs77273740 | benign | 0 | 0 | 0 |
| rs267606760 | pathogenic | 0 | 0 | 0 |
| rs77576840 | pathogenic | 0 | 0 | 0 |
| rs45465204 | benign | 1 | 1 | 0 |
| rs5320 | benign | 0 | 0 | 0 |
| rs5324 | benign | 0 | 0 | 0 |
| rs4531 | benign | 0 | 0 | 0 |
| rs267606761 | pathogenic | 0 | 0 | 0 |
| rs75215331 | pathogenic | 0 | 0 | 0 |
| rs6271 | benign | 0 | 0 | 0 |
| rs201643277 | pathogenic | 0 | 0 | 0 |
| rs4252250 | benign | 0 | 0 | 0 |
| rs137853580 | pathogenic | 0 | 0 | 0 |
| rs368287711 | pathogenic | 0 | 0 | 0 |
| rs137853579 | pathogenic | 0 | 0 | 0 |
| rs3135932 | benign | 0 | 0 | 0 |
| rs2228055 | benign | 0 | 0 | 0 |
| rs140322345 | benign | 1 | 1 | 1 |
| rs1154510 | pathogenic | 0 | 0 | 0 |
| rs11833399 | benign | 0 | 0 | 0 |
| rs36023382 | benign | 1 | 1 | 1 |
| rs61754444 | benign | 0 | 0 | 0 |
| rs28935168 | pathogenic | 0 | 0 | 0 |
| rs28934907 | pathogenic | 0 | 0 | 0 |
| rs28934904 | pathogenic | 1 | 1 | 0 |
| rs61748392 | pathogenic | 1 | 1 | 0 |
| rs28934908 | pathogenic | 0 | 0 | 0 |
| rs28934905 | pathogenic | 0 | 0 | 0 |
| rs28934906 | pathogenic | 1 | 1 | 0 |
| rs606231307 | pathogenic | 0 | 0 | 0 |
| rs9333296 | benign | 0 | 0 | 0 |
| rs3218614 | benign | 0 | 0 | 0 |

| | | | | |
|---|---|---|---|---|
| rs2230792 | benign | 0 | 0 | 0 |
| rs2230793 | benign | 0 | 0 | 0 |
| rs2230794 | benign | 0 | 0 | 0 |
| rs749052963 | pathogenic | 0 | 0 | 0 |
| rs2230798 | benign | 0 | 0 | 0 |
| rs2230795 | benign | 0 | 0 | 0 |
| rs3204145 | benign | 0 | 0 | 0 |
| rs1538660 | benign | 0 | 0 | 0 |
| rs9878928 | benign | 0 | 0 | 0 |
| rs104893742 | pathogenic | 0 | 0 | 0 |
| rs28936702 | pathogenic | 0 | 0 | 0 |
| rs28936703 | pathogenic | 0 | 0 | 0 |
| rs769392 | benign | 0 | 0 | 0 |
| rs769402 | benign | 1 | 1 | 1 |
| rs104894884 | pathogenic | 1 | 1 | 1 |
| rs1801316 | benign | 1 | 1 | 1 |
| rs104894885 | pathogenic | 1 | 1 | 1 |
| rs782572 | benign | 0 | 0 | 0 |
| rs151166046 | pathogenic | 0 | 0 | 0 |
| rs772153760 | pathogenic | 1 | 1 | 0 |
| rs35405862 | benign | 0 | 0 | 0 |
| rs146571352 | pathogenic | 0 | 1 | 0 |
| rs374698153 | pathogenic | 0 | 0 | 0 |
| rs7594497 | benign | 0 | 0 | 0 |
| rs80338716 | pathogenic | 0 | 0 | 0 |
| rs80338719 | pathogenic | 1 | 1 | 1 |
| rs80338721 | pathogenic | 0 | 0 | 0 |
| rs730882249 | pathogenic | 0 | 0 | 0 |
| rs146262009 | pathogenic | 0 | 0 | 0 |
| rs104895083 | pathogenic | 1 | 1 | 1 |
| rs104895100 | benign | 1 | 1 | 1 |
| rs104895085 | pathogenic | 0 | 0 | 0 |
| rs28940580 | pathogenic | 0 | 0 | 0 |
| rs104895098 | pathogenic | 0 | 0 | 0 |
| rs28940578 | pathogenic | 0 | 0 | 0 |
| rs61752717 | pathogenic | 0 | 0 | 0 |
| rs28940579 | pathogenic | 0 | 0 | 0 |
| rs104895097 | pathogenic | 0 | 0 | 0 |
| rs140425651 | benign | 1 | 1 | 0 |
| rs768090444 | pathogenic | 0 | 0 | 0 |
| rs764459544 | pathogenic | 0 | 0 | 0 |
| rs80338802 | pathogenic | 1 | 1 | 1 |
| rs149969786 | pathogenic | 1 | 1 | 0 |
| rs115311625 | pathogenic | 1 | 1 | 1 |
| rs778768583 | pathogenic | 1 | 1 | 1 |
| rs892090 | benign | 0 | 0 | 0 |
| rs28929498 | pathogenic | 1 | 1 | 1 |
| rs137854467 | pathogenic | 0 | 0 | 0 |
| rs137854480 | pathogenic | 0 | 0 | 0 |
| rs148682262 | benign | 0 | 0 | 0 |
| rs137852744 | pathogenic | 0 | 0 | 0 |
| rs137852745 | pathogenic | 0 | 1 | 0 |
| rs137852749 | pathogenic | 0 | 1 | 0 |
| rs137852746 | pathogenic | 0 | 1 | 0 |

| | | | | |
|---|---|---|---|---|
| rs145633958 | benign | 0 | 0 | 0 |
| rs141679950 | benign | 0 | 0 | 0 |
| rs76397662 | benign | 1 | 1 | 1 |
| rs370736139 | benign | 0 | 0 | 0 |
| rs781750106 | benign | 0 | 0 | 0 |
| rs546866208 | benign | 0 | 0 | 0 |
| rs142338976 | benign | 0 | 0 | 0 |
| rs115272158 | benign | 0 | 0 | 0 |
| rs115423919 | benign | 0 | 0 | 0 |
| rs537874538 | benign | 0 | 0 | 0 |
| rs9282834 | benign | 0 | 0 | 0 |
| rs201745826 | benign | 0 | 0 | 0 |
| rs75873440 | pathogenic | 0 | 0 | 0 |
| rs147219360 | benign | 0 | 0 | 0 |
| rs77939446 | pathogenic | 0 | 0 | 0 |
| rs77558292 | pathogenic | 0 | 0 | 0 |
| rs80069458 | pathogenic | 0 | 0 | 0 |
| rs76262710 | pathogenic | 0 | 0 | 0 |
| rs79781594 | pathogenic | 0 | 0 | 0 |
| rs77316810 | pathogenic | 0 | 0 | 0 |
| rs77503355 | pathogenic | 0 | 0 | 0 |
| rs79890926 | pathogenic | 0 | 0 | 0 |
| rs377767402 | benign | 0 | 0 | 0 |
| rs121913308 | pathogenic | 0 | 0 | 0 |
| rs55846256 | benign | 0 | 0 | 0 |
| rs75076352 | pathogenic | 0 | 0 | 0 |
| rs75996173 | pathogenic | 0 | 0 | 0 |
| rs77709286 | pathogenic | 0 | 0 | 0 |
| rs151058706 | benign | 0 | 0 | 0 |
| rs138964358 | benign | 0 | 0 | 0 |
| rs137852353 | pathogenic | 1 | 1 | 0 |
| rs201196020 | benign | 0 | 0 | 0 |
| rs137853201 | pathogenic | 1 | 1 | 1 |
| rs113994174 | pathogenic | 1 | 1 | 1 |
| rs104893674 | pathogenic | 1 | 1 | 1 |
| rs779519249 | benign | 1 | 1 | 1 |
| rs9606756 | benign | 0 | 0 | 0 |
| rs75680863 | benign | 1 | 1 | 1 |
| rs35838082 | benign | 0 | 0 | 0 |
| rs1801198 | benign | 0 | 0 | 0 |
| rs117458738 | benign | 1 | 1 | 0 |
| rs9621049 | benign | 1 | 1 | 0 |
| rs1131603 | benign | 1 | 1 | 0 |
| rs4820889 | benign | 0 | 0 | 0 |
| rs777767978 | pathogenic | 1 | 1 | 0 |
| rs147399860 | benign | 1 | 1 | 1 |
| rs104893950 | pathogenic | 0 | 0 | 0 |
| rs777308612 | benign | 0 | 0 | 0 |
| rs151091410 | benign | 0 | 0 | 0 |
| rs11537667 | benign | 0 | 0 | 0 |
| rs267606621 | pathogenic | 1 | 1 | 0 |
| rs143058902 | benign | 0 | 0 | 0 |
| rs370261904 | pathogenic | 0 | 0 | 0 |
| rs141169746 | benign | 0 | 0 | 0 |

| | | | | |
|---|---|---|---|---|
| rs111033436 | benign | 0 | 0 | 0 |
| rs148533341 | benign | 0 | 0 | 0 |
| rs143827620 | benign | 0 | 0 | 0 |
| rs4935502 | benign | 0 | 0 | 0 |
| rs61735473 | benign | 0 | 0 | 0 |
| rs267606932 | pathogenic | 1 | 1 | 0 |
| rs61730754 | benign | 0 | 0 | 0 |
| rs374491359 | pathogenic | 1 | 1 | 1 |
| rs761800077 | benign | 1 | 1 | 1 |
| rs760391381 | benign | 1 | 1 | 1 |
| rs104893885 | pathogenic | 0 | 0 | 0 |
| rs104893886 | pathogenic | 1 | 1 | 1 |
| rs104893884 | pathogenic | 0 | 0 | 0 |
| rs147715509 | benign | 0 | 0 | 0 |
| rs115533243 | benign | 0 | 0 | 0 |
| rs397509390 | pathogenic | 0 | 1 | 0 |
| rs80357134 | pathogenic | 0 | 0 | 0 |
| rs766004110 | benign | 1 | 1 | 1 |
| rs769650474 | benign | 0 | 0 | 0 |
| rs398122635 | benign | 1 | 1 | 0 |
| rs199522616 | pathogenic | 0 | 0 | 0 |
| rs28897672 | pathogenic | 1 | 1 | 0 |
| rs80357086 | pathogenic | 1 | 1 | 1 |
| rs80357064 | pathogenic | 1 | 1 | 0 |
| rs80356936 | benign | 0 | 0 | 0 |
| rs28897673 | benign | 0 | 0 | 0 |
| rs146747324 | benign | 0 | 0 | 0 |
| rs78635798 | pathogenic | 0 | 0 | 0 |
| rs75146158 | pathogenic | 1 | 1 | 1 |
| rs587782886 | benign | 0 | 0 | 0 |
| rs121434634 | pathogenic | 0 | 0 | 0 |
| rs121434635 | pathogenic | 0 | 0 | 0 |
| rs587777778 | pathogenic | 0 | 0 | 0 |
| rs118203967 | pathogenic | 0 | 0 | 0 |
| rs751375244 | pathogenic | 1 | 1 | 1 |
| rs104893863 | pathogenic | 1 | 1 | 0 |
| rs104893865 | pathogenic | 0 | 0 | 0 |
| rs104893864 | pathogenic | 1 | 1 | 1 |
| rs104893866 | pathogenic | 1 | 1 | 0 |
| rs561738497 | benign | 0 | 0 | 0 |
| rs57599352 | pathogenic | 1 | 1 | 1 |
| rs267607448 | pathogenic | 1 | 1 | 1 |
| rs63751596 | pathogenic | 0 | 0 | 0 |
| rs63751612 | benign | 0 | 0 | 0 |
| rs63751684 | benign | 0 | 0 | 0 |
| rs267607861 | benign | 0 | 0 | 0 |
| rs35831931 | benign | 0 | 0 | 0 |
| rs63750876 | benign | 0 | 0 | 0 |
| rs41295284 | benign | 0 | 0 | 0 |
| rs35502531 | benign | 0 | 0 | 0 |
| rs63750449 | benign | 0 | 0 | 0 |
| rs63750899 | pathogenic | 0 | 0 | 0 |
| rs55907433 | benign | 0 | 0 | 0 |
| rs63750217 | pathogenic | 0 | 0 | 0 |

| | | | | |
|---|---|---|---|---|
| rs63751275 | pathogenic | 0 | 0 | 0 |
| rs63750702 | benign | 1 | 1 | 1 |
| rs63750114 | benign | 0 | 0 | 0 |
| rs2020873 | benign | 0 | 0 | 0 |
| rs140195825 | benign | 1 | 1 | 1 |
| rs182763805 | benign | 0 | 0 | 0 |
| rs533696377 | benign | 0 | 0 | 0 |
| rs41278788 | benign | 0 | 0 | 0 |
| rs104894193 | pathogenic | 1 | 1 | 0 |
| rs3824915 | benign | 1 | 1 | 1 |
| rs104894196 | pathogenic | 0 | 0 | 0 |
| rs78681604 | benign | 1 | 1 | 1 |
| rs77321498 | benign | 1 | 1 | 1 |
| rs137852965 | pathogenic | 0 | 0 | 0 |
| rs137852961 | pathogenic | 0 | 0 | 0 |
| rs137852962 | pathogenic | 1 | 1 | 0 |
| rs753376100 | pathogenic | 0 | 0 | 0 |
| rs137852968 | pathogenic | 1 | 1 | 1 |
| rs137852959 | pathogenic | 1 | 1 | 0 |
| rs137852967 | pathogenic | 0 | 0 | 0 |
| rs113437301 | benign | 1 | 1 | 1 |
| rs61747482 | benign | 1 | 1 | 1 |
| rs2234919 | benign | 1 | 1 | 0 |
| rs121908863 | pathogenic | 0 | 0 | 0 |
| rs140455771 | pathogenic | 0 | 0 | 0 |
| rs121908810 | pathogenic | 1 | 1 | 1 |
| rs143879853 | benign | 0 | 0 | 0 |
| rs188319622 | benign | 0 | 0 | 0 |
| rs121918645 | pathogenic | 0 | 0 | 0 |
| rs115352435 | pathogenic | 0 | 0 | 0 |
| rs121434427 | pathogenic | 1 | 1 | 1 |
| rs121434428 | pathogenic | 0 | 0 | 0 |
| rs35086265 | benign | 1 | 1 | 1 |
| rs11576415 | benign | 0 | 0 | 0 |
| rs121434429 | pathogenic | 1 | 1 | 1 |
| rs4074536 | benign | 1 | 1 | 1 |
| rs10801999 | benign | 0 | 0 | 0 |
| rs121434550 | pathogenic | 0 | 0 | 0 |
| rs148057999 | pathogenic | 0 | 0 | 0 |
| rs28730716 | benign | 0 | 0 | 0 |
| rs142036299 | benign | 0 | 0 | 0 |
| rs762381137 | pathogenic | 0 | 0 | 0 |
| rs121434549 | pathogenic | 0 | 0 | 0 |
| rs139228801 | pathogenic | 0 | 0 | 0 |
| rs147219463 | benign | 0 | 0 | 0 |
| rs1131857 | benign | 0 | 0 | 0 |
| rs121917874 | pathogenic | 0 | 1 | 0 |
| rs2228056 | benign | 0 | 0 | 0 |
| rs121917870 | pathogenic | 0 | 0 | 0 |
| rs121917866 | pathogenic | 0 | 0 | 0 |
| rs11921054 | benign | 0 | 0 | 0 |
| rs121917868 | pathogenic | 0 | 0 | 0 |
| rs28931603 | pathogenic | 0 | 0 | 0 |
| rs104895295 | pathogenic | 1 | 1 | 0 |

| | | | | |
|---|---|---|---|---|
| rs104895382 | pathogenic | 0 | 0 | 0 |
| rs104895298 | pathogenic | 0 | 0 | 0 |
| rs104895300 | pathogenic | 0 | 0 | 0 |
| rs104895332 | pathogenic | 0 | 0 | 0 |
| rs758026399 | pathogenic | 1 | 1 | 1 |
| rs104895366 | pathogenic | 0 | 0 | 0 |
| rs104895304 | pathogenic | 0 | 0 | 0 |
| rs759997079 | pathogenic | 0 | 0 | 0 |
| rs121917789 | pathogenic | 0 | 0 | 0 |
| rs104895319 | pathogenic | 0 | 0 | 0 |
| rs104895317 | pathogenic | 1 | 1 | 0 |
| rs28934897 | pathogenic | 0 | 0 | 0 |
| rs104895380 | benign | 0 | 0 | 0 |
| rs104895360 | pathogenic | 0 | 0 | 0 |
| rs28936396 | pathogenic | 1 | 1 | 0 |
| rs121909307 | pathogenic | 1 | 1 | 1 |
| rs28938472 | pathogenic | 1 | 1 | 0 |
| rs121909308 | pathogenic | 1 | 1 | 0 |
| rs121909309 | pathogenic | 0 | 0 | 0 |
| rs121913013 | benign | 1 | 1 | 1 |
| rs201855245 | benign | 1 | 1 | 1 |
| rs2230232 | benign | 1 | 1 | 0 |
| rs113451409 | benign | 0 | 0 | 0 |
| rs121913011 | pathogenic | 1 | 1 | 0 |
| rs185821167 | pathogenic | 1 | 1 | 0 |
| rs2230234 | benign | 0 | 0 | 0 |
| rs193922639 | benign | 1 | 1 | 0 |
| rs774863785 | pathogenic | 1 | 1 | 0 |
| rs576404380 | pathogenic | 1 | 1 | 0 |
| rs2230235 | benign | 1 | 1 | 0 |
| rs200484060 | pathogenic | 1 | 1 | 0 |
| rs199761749 | benign | 0 | 0 | 0 |
| rs199681901 | benign | 0 | 0 | 0 |
| rs121908951 | pathogenic | 0 | 0 | 0 |
| rs769251460 | benign | 0 | 0 | 0 |
| rs28937320 | pathogenic | 0 | 0 | 0 |
| rs200607527 | pathogenic | 0 | 0 | 0 |
| rs137852640 | pathogenic | 1 | 1 | 1 |
| rs137852636 | pathogenic | 1 | 1 | 1 |
| rs137852638 | pathogenic | 0 | 0 | 0 |
| rs142637231 | pathogenic | 0 | 0 | 0 |
| rs151328418 | benign | 0 | 1 | 0 |
| rs137852639 | pathogenic | 1 | 1 | 1 |
| rs34605667 | benign | 0 | 0 | 0 |
| rs104894073 | pathogenic | 1 | 1 | 0 |
| rs56298569 | pathogenic | 0 | 0 | 0 |
| rs757600616 | pathogenic | 0 | 0 | 0 |
| rs139194636 | pathogenic | 0 | 0 | 0 |
| rs369517993 | pathogenic | 1 | 1 | 1 |
| rs138846036 | benign | 1 | 1 | 1 |
| rs200808744 | pathogenic | 1 | 1 | 1 |
| rs2255543 | benign | 0 | 0 | 0 |
| rs149165166 | pathogenic | 0 | 0 | 0 |
| rs544956641 | pathogenic | 1 | 1 | 1 |

| | | | | |
|---|---|---|---|---|
| rs28941783 | pathogenic | 0 | 0 | 0 |
| rs562853291 | pathogenic | 1 | 1 | 1 |
| rs28942100 | pathogenic | 0 | 0 | 0 |
| rs120074174 | pathogenic | 1 | 1 | 1 |
| rs120074171 | pathogenic | 1 | 1 | 1 |
| rs120074173 | pathogenic | 0 | 0 | 0 |
| rs120074172 | pathogenic | 1 | 1 | 1 |
| rs767159114 | pathogenic | 1 | 1 | 1 |
| rs104893657 | pathogenic | 0 | 0 | 0 |
| rs104893656 | pathogenic | 0 | 0 | 0 |
| rs104893659 | pathogenic | 0 | 0 | 0 |
| rs104893658 | pathogenic | 0 | 0 | 0 |
| rs200386310 | pathogenic | 0 | 0 | 0 |
| rs2072712 | benign | 0 | 0 | 0 |
| rs35274867 | benign | 0 | 0 | 0 |
| rs35343172 | benign | 0 | 0 | 0 |
| rs10079250 | benign | 0 | 0 | 0 |
| rs34951517 | benign | 0 | 0 | 0 |
| rs267606695 | pathogenic | 0 | 0 | 0 |
| rs149391728 | benign | 0 | 0 | 0 |
| rs387906598 | pathogenic | 1 | 1 | 0 |
| rs137854578 | pathogenic | 0 | 0 | 0 |
| rs141519952 | benign | 0 | 0 | 0 |
| rs146572883 | benign | 0 | 0 | 0 |
| rs111866410 | benign | 0 | 0 | 0 |
| rs139387758 | benign | 1 | 1 | 1 |
| rs35672074 | benign | 1 | 1 | 1 |
| rs121434322 | pathogenic | 0 | 0 | 0 |
| rs150884181 | benign | 0 | 0 | 0 |
| rs35867471 | benign | 0 | 0 | 0 |
| rs2867802 | benign | 0 | 0 | 0 |
| rs137902889 | benign | 0 | 0 | 0 |
| rs369804864 | pathogenic | 0 | 0 | 0 |
| rs781050795 | pathogenic | 0 | 0 | 0 |
| rs2305799 | benign | 0 | 0 | 0 |
| rs2229531 | benign | 0 | 0 | 0 |
| rs147025508 | benign | 0 | 0 | 0 |
| rs76857106 | pathogenic | 0 | 0 | 0 |
| rs7247284 | benign | 0 | 0 | 0 |
| rs62619782 | benign | 0 | 0 | 0 |
| rs377244188 | pathogenic | 0 | 0 | 0 |
| rs75718910 | pathogenic | 0 | 0 | 0 |
| rs79843600 | pathogenic | 0 | 0 | 0 |
| rs75037667 | pathogenic | 1 | 1 | 1 |
| rs121913006 | pathogenic | 0 | 0 | 0 |
| rs397517925 | pathogenic | 1 | 1 | 1 |
| rs267606831 | pathogenic | 1 | 1 | 1 |
| rs121909667 | pathogenic | 1 | 1 | 1 |
| rs121909669 | pathogenic | 1 | 1 | 1 |
| rs121909668 | pathogenic | 1 | 1 | 1 |
| rs121909671 | pathogenic | 1 | 1 | 1 |
| rs534596036 | benign | 0 | 0 | 0 |
| rs144037629 | pathogenic | 0 | 0 | 0 |
| rs11546406 | benign | 0 | 0 | 0 |

| | | | | |
|---|---|---|---|---|
| rs143697995 | pathogenic | 0 | 0 | 0 |
| rs121908331 | pathogenic | 0 | 0 | 0 |
| rs149377346 | benign | 0 | 0 | 0 |
| rs104894086 | pathogenic | 1 | 1 | 0 |
| rs104894089 | pathogenic | 0 | 0 | 0 |
| rs104894090 | pathogenic | 1 | 1 | 0 |
| rs387907235 | pathogenic | 0 | 0 | 0 |
| rs137852689 | pathogenic | 0 | 0 | 0 |
| rs137852690 | pathogenic | 1 | 1 | 0 |
| rs104894087 | pathogenic | 0 | 0 | 0 |
| rs104894085 | pathogenic | 0 | 0 | 0 |
| rs79383654 | benign | 0 | 0 | 0 |
| rs121434239 | pathogenic | 0 | 0 | 0 |
| rs121434240 | pathogenic | 0 | 0 | 0 |
| rs121434236 | pathogenic | 0 | 0 | 0 |
| rs121434238 | pathogenic | 1 | 1 | 1 |
| rs121918107 | pathogenic | 1 | 1 | 1 |
| rs121918103 | pathogenic | 1 | 1 | 1 |
| rs121918104 | pathogenic | 1 | 1 | 1 |
| rs61747614 | benign | 1 | 1 | 1 |
| rs555329870 | benign | 0 | 0 | 0 |
| rs186094931 | benign | 0 | 0 | 0 |
| rs267606785 | pathogenic | 0 | 0 | 0 |
| rs34474955 | benign | 0 | 0 | 0 |
| rs1126497 | benign | 1 | 1 | 1 |
| rs397514661 | pathogenic | 0 | 0 | 0 |
| rs189732445 | benign | 0 | 0 | 0 |
| rs146685071 | benign | 0 | 0 | 0 |
| rs121908636 | pathogenic | 1 | 1 | 0 |
| rs121908643 | pathogenic | 0 | 0 | 0 |
| rs121908644 | pathogenic | 1 | 1 | 1 |
| rs575001023 | pathogenic | 1 | 1 | 1 |
| rs121908646 | pathogenic | 1 | 1 | 0 |
| rs121908638 | pathogenic | 1 | 1 | 1 |
| rs777828000 | pathogenic | 1 | 1 | 1 |
| rs74923032 | benign | 0 | 0 | 0 |
| rs192838388 | pathogenic | 1 | 1 | 1 |
| rs148918985 | pathogenic | 1 | 1 | 1 |
| rs773540764 | benign | 1 | 1 | 1 |
| rs370595480 | pathogenic | 1 | 1 | 1 |
| rs762387914 | pathogenic | 1 | 1 | 1 |
| rs121908642 | pathogenic | 1 | 1 | 1 |
| rs183276875 | pathogenic | 1 | 1 | 1 |
| rs121908648 | pathogenic | 1 | 1 | 1 |
| rs121908647 | pathogenic | 1 | 1 | 1 |
| rs121908640 | pathogenic | 1 | 1 | 1 |
| rs771937610 | pathogenic | 1 | 1 | 1 |
| rs121908641 | pathogenic | 1 | 1 | 1 |
| rs5745068 | benign | 0 | 0 | 0 |
| rs73155056 | benign | 1 | 1 | 1 |
| rs148788180 | benign | 0 | 0 | 0 |
| rs121918137 | pathogenic | 0 | 0 | 0 |
| rs41269593 | benign | 0 | 0 | 0 |
| rs11159 | benign | 1 | 1 | 0 |

| | | | | |
|---|---|---|---|---|
| rs114733570 | pathogenic | 1 | 1 | 0 |
| rs3211783 | benign | 0 | 0 | 0 |
| rs4762 | benign | 0 | 0 | 0 |
| rs74315283 | pathogenic | 0 | 0 | 0 |
| rs111033559 | pathogenic | 0 | 0 | 0 |
| rs111033560 | pathogenic | 1 | 1 | 1 |
| rs749962743 | benign | 0 | 0 | 0 |
| rs267606986 | pathogenic | 0 | 0 | 0 |
| rs34516635 | benign | 0 | 0 | 0 |
| rs117440569 | benign | 0 | 0 | 0 |
| rs61740877 | benign | 0 | 0 | 0 |
| rs121918352 | pathogenic | 0 | 0 | 0 |
| rs183974372 | benign | 0 | 0 | 0 |
| rs140511204 | benign | 1 | 1 | 0 |
| rs6040067 | benign | 0 | 0 | 0 |
| rs199674138 | benign | 0 | 0 | 0 |
| rs121918350 | pathogenic | 0 | 0 | 0 |
| rs138292988 | benign | 1 | 1 | 1 |
| rs145288947 | benign | 0 | 0 | 0 |
| rs369714529 | benign | 0 | 0 | 0 |
| rs200945419 | benign | 0 | 0 | 0 |
| rs116959874 | benign | 0 | 0 | 0 |
| rs141659620 | pathogenic | 1 | 1 | 0 |
| rs111475461 | benign | 0 | 0 | 0 |
| rs61755320 | pathogenic | 0 | 0 | 0 |
| rs748600162 | pathogenic | 1 | 1 | 1 |
| rs369227537 | pathogenic | 1 | 1 | 1 |
| rs372981030 | pathogenic | 1 | 1 | 1 |
| rs137854486 | pathogenic | 0 | 0 | 0 |
| rs137854487 | pathogenic | 0 | 0 | 0 |
| rs199905054 | pathogenic | 0 | 0 | 0 |
| rs771438170 | pathogenic | 0 | 0 | 0 |
| rs774949844 | pathogenic | 0 | 0 | 0 |
| rs4899145 | benign | 0 | 0 | 0 |
| rs200642525 | benign | 0 | 0 | 0 |
| rs190140598 | pathogenic | 0 | 0 | 0 |
| rs2250889 | benign | 0 | 0 | 0 |
| rs17577 | benign | 0 | 0 | 0 |
| rs142032681 | benign | 0 | 0 | 0 |
| rs144577963 | benign | 0 | 0 | 0 |
| rs61754623 | benign | 0 | 0 | 0 |
| rs139873200 | benign | 0 | 0 | 0 |
| rs532530906 | benign | 0 | 0 | 0 |
| rs145762716 | benign | 0 | 0 | 0 |
| rs34062020 | benign | 0 | 0 | 0 |
| rs35258161 | benign | 0 | 0 | 0 |
| rs149792649 | benign | 0 | 0 | 0 |
| rs78101774 | benign | 0 | 0 | 0 |
| rs766466297 | pathogenic | 0 | 1 | 0 |
| rs140467171 | pathogenic | 0 | 0 | 0 |
| rs141340466 | pathogenic | 0 | 0 | 0 |
| rs201555303 | pathogenic | 0 | 0 | 0 |
| rs143592405 | pathogenic | 0 | 0 | 0 |
| rs781249411 | pathogenic | 0 | 0 | 0 |

| | | | | |
|---|---|---|---|---|
| rs116644127 | benign | 0 | 0 | 0 |
| rs151146863 | benign | 1 | 1 | 1 |
| rs116440799 | benign | 0 | 0 | 0 |
| rs138958351 | benign | 0 | 0 | 0 |
| rs143319354 | benign | 0 | 0 | 0 |
| rs768669208 | pathogenic | 1 | 1 | 1 |
| rs28940289 | pathogenic | 1 | 1 | 0 |
| rs745656120 | pathogenic | 1 | 1 | 0 |
| rs387906987 | pathogenic | 0 | 0 | 0 |
| rs147240777 | benign | 0 | 0 | 0 |
| rs140945833 | benign | 0 | 0 | 0 |
| rs146614981 | benign | 0 | 0 | 0 |
| rs149878562 | benign | 0 | 0 | 0 |
| rs28938168 | pathogenic | 0 | 0 | 0 |
| rs33939927 | pathogenic | 1 | 1 | 1 |
| rs34995376 | pathogenic | 1 | 1 | 1 |
| rs201034481 | pathogenic | 1 | 1 | 1 |
| rs774069983 | benign | 0 | 0 | 0 |
| rs199422226 | pathogenic | 1 | 1 | 0 |
| rs199422225 | pathogenic | 1 | 1 | 1 |
| rs199422224 | pathogenic | 1 | 1 | 0 |
| rs149271416 | pathogenic | 1 | 1 | 0 |
| rs1801318 | benign | 1 | 1 | 1 |
| rs372691318 | pathogenic | 1 | 1 | 1 |
| rs147685849 | benign | 0 | 0 | 0 |
| rs387907199 | pathogenic | 0 | 0 | 0 |
| rs767122069 | pathogenic | 0 | 0 | 0 |
| rs2302427 | benign | 0 | 0 | 0 |
| rs201135441 | benign | 0 | 0 | 0 |
| rs201861847 | pathogenic | 0 | 0 | 0 |
| rs137852952 | pathogenic | 0 | 0 | 0 |
| rs16986560 | benign | 0 | 0 | 0 |
| rs147344332 | pathogenic | 1 | 1 | 1 |
| rs36027220 | pathogenic | 0 | 0 | 0 |
| rs192401229 | benign | 0 | 0 | 0 |
| rs121909106 | pathogenic | 1 | 1 | 0 |
| rs118204103 | pathogenic | 1 | 1 | 0 |
| rs118204104 | pathogenic | 1 | 1 | 0 |
| rs118204105 | pathogenic | 1 | 1 | 0 |
| rs118204106 | pathogenic | 0 | 0 | 0 |
| rs118204119 | pathogenic | 0 | 0 | 0 |
| rs118204107 | pathogenic | 0 | 0 | 0 |
| rs118204094 | pathogenic | 0 | 0 | 0 |
| rs118204098 | pathogenic | 1 | 1 | 0 |
| rs118204101 | pathogenic | 1 | 1 | 0 |
| rs118204096 | pathogenic | 1 | 1 | 0 |
| rs118204108 | pathogenic | 0 | 0 | 0 |
| rs536814318 | pathogenic | 0 | 0 | 0 |
| rs34413634 | pathogenic | 1 | 1 | 0 |
| rs118204100 | pathogenic | 0 | 0 | 0 |
| rs118204109 | pathogenic | 0 | 0 | 0 |
| rs118204116 | pathogenic | 1 | 1 | 0 |
| rs118204110 | pathogenic | 0 | 0 | 0 |
| rs118204099 | pathogenic | 0 | 0 | 0 |

| | | | | |
|---|---|---|---|---|
| rs118204111 | pathogenic | 0 | 0 | 0 |
| rs118204112 | pathogenic | 0 | 0 | 0 |
| rs118204113 | pathogenic | 0 | 0 | 0 |
| rs118204114 | pathogenic | 0 | 0 | 0 |
| rs118204115 | pathogenic | 0 | 0 | 0 |
| rs150428209 | benign | 0 | 0 | 0 |
| rs144949995 | benign | 0 | 0 | 0 |
| rs190023981 | benign | 0 | 0 | 0 |
| rs104894611 | pathogenic | 1 | 1 | 1 |
| rs28937580 | pathogenic | 1 | 1 | 1 |
| rs121908949 | pathogenic | 0 | 0 | 0 |
| rs104894612 | pathogenic | 1 | 1 | 1 |
| rs104894609 | pathogenic | 1 | 1 | 1 |
| rs104894610 | pathogenic | 1 | 1 | 1 |
| rs104894615 | pathogenic | 0 | 0 | 0 |
| rs104894603 | pathogenic | 0 | 0 | 0 |
| rs104894602 | pathogenic | 1 | 1 | 1 |
| rs121908948 | pathogenic | 1 | 1 | 1 |
| rs104894608 | pathogenic | 1 | 1 | 1 |
| rs371545673 | benign | 0 | 0 | 0 |
| rs199925238 | benign | 0 | 0 | 0 |
| rs146313451 | benign | 0 | 0 | 0 |
| rs121918193 | pathogenic | 1 | 1 | 0 |
| rs2228500 | benign | 0 | 0 | 0 |
| rs121918194 | pathogenic | 1 | 1 | 0 |
| rs121918196 | pathogenic | 0 | 0 | 0 |
| rs41291971 | benign | 0 | 0 | 0 |
| rs199473690 | pathogenic | 0 | 0 | 0 |
| rs2228613 | benign | 0 | 0 | 0 |
| rs267607130 | pathogenic | 1 | 1 | 1 |
| rs77615401 | benign | 1 | 1 | 1 |
| rs374618872 | benign | 1 | 1 | 1 |
| rs146165014 | benign | 0 | 0 | 0 |
| rs144586070 | benign | 0 | 0 | 0 |
| rs121918541 | pathogenic | 0 | 0 | 0 |
| rs142105943 | benign | 0 | 0 | 0 |
| rs121918540 | pathogenic | 0 | 0 | 0 |
| rs6791 | benign | 0 | 0 | 0 |
| rs61736587 | pathogenic | 0 | 0 | 0 |
| rs61736586 | benign | 0 | 0 | 0 |
| rs121917741 | pathogenic | 1 | 1 | 1 |
| rs41302885 | benign | 0 | 0 | 0 |
| rs141026028 | benign | 0 | 0 | 0 |
| rs193922493 | pathogenic | 0 | 0 | 0 |
| rs73598374 | benign | 0 | 0 | 0 |
| rs121908725 | pathogenic | 1 | 1 | 0 |
| rs199422328 | pathogenic | 0 | 1 | 0 |
| rs11555566 | benign | 1 | 1 | 1 |
| rs267606634 | pathogenic | 0 | 0 | 0 |
| rs121908714 | pathogenic | 1 | 1 | 0 |
| rs121908717 | pathogenic | 1 | 1 | 0 |
| rs267606635 | pathogenic | 1 | 1 | 0 |
| rs121908739 | pathogenic | 0 | 0 | 0 |
| rs121908735 | pathogenic | 0 | 1 | 0 |

| | | | | |
|---|---|---|---|---|
| rs121908716 | pathogenic | 0 | 1 | 0 |
| rs121908723 | pathogenic | 0 | 1 | 0 |
| rs121908721 | pathogenic | 0 | 0 | 0 |
| rs199422327 | pathogenic | 0 | 0 | 0 |
| rs121908715 | pathogenic | 0 | 0 | 0 |
| rs587783563 | pathogenic | 0 | 0 | 0 |
| rs104894780 | pathogenic | 0 | 0 | 0 |
| rs56030372 | pathogenic | 1 | 1 | 1 |
| rs104894782 | pathogenic | 1 | 1 | 1 |
| rs149495971 | benign | 0 | 0 | 0 |
| rs118203942 | pathogenic | 1 | 1 | 0 |
| rs118203939 | pathogenic | 0 | 0 | 0 |
| rs118203938 | pathogenic | 0 | 0 | 0 |
| rs118203943 | pathogenic | 0 | 0 | 0 |
| rs118203940 | pathogenic | 0 | 0 | 0 |
| rs727503809 | pathogenic | 0 | 0 | 0 |
| rs1065757 | benign | 0 | 0 | 0 |
| rs1071598 | benign | 0 | 0 | 0 |
| rs25414 | benign | 0 | 0 | 0 |
| rs118203944 | pathogenic | 0 | 0 | 0 |
| rs118203941 | pathogenic | 0 | 0 | 0 |
| rs144222167 | benign | 0 | 0 | 0 |
| rs201101343 | pathogenic | 0 | 0 | 0 |
| rs144601717 | benign | 0 | 0 | 0 |
| rs267607129 | pathogenic | 0 | 0 | 0 |
| rs104894729 | pathogenic | 0 | 0 | 0 |
| rs104894727 | pathogenic | 0 | 0 | 0 |
| rs104894725 | pathogenic | 0 | 0 | 0 |
| rs104894515 | pathogenic | 1 | 1 | 1 |
| rs74688377 | benign | 0 | 0 | 0 |
| rs121908513 | pathogenic | 1 | 1 | 0 |
| rs121908509 | pathogenic | 0 | 0 | 0 |
| rs121908514 | pathogenic | 1 | 1 | 0 |
| rs121908512 | pathogenic | 1 | 1 | 0 |
| rs121908510 | pathogenic | 0 | 0 | 0 |
| rs121908511 | pathogenic | 1 | 1 | 1 |
| rs121908518 | pathogenic | 1 | 1 | 1 |
| rs34358665 | benign | 0 | 0 | 0 |
| rs147568834 | benign | 0 | 0 | 0 |
| rs201717599 | benign | 0 | 0 | 0 |
| rs61744487 | benign | 0 | 0 | 0 |
| rs370627877 | pathogenic | 1 | 1 | 1 |
| rs368593151 | pathogenic | 1 | 1 | 1 |
| rs114363936 | benign | 0 | 0 | 0 |
| rs137853333 | pathogenic | 0 | 0 | 0 |
| rs137853080 | pathogenic | 1 | 1 | 1 |
| rs137853077 | pathogenic | 1 | 1 | 1 |
| rs780749732 | benign | 1 | 1 | 1 |
| rs137853081 | pathogenic | 1 | 1 | 0 |
| rs137853078 | pathogenic | 0 | 0 | 0 |
| rs121913315 | pathogenic | 1 | 1 | 1 |
| rs752513525 | pathogenic | 1 | 1 | 1 |
| rs104894918 | pathogenic | 0 | 0 | 0 |
| rs104894917 | pathogenic | 0 | 0 | 0 |

| | | | | |
|---|---|---|---|---|
| rs104894919 | pathogenic | 0 | 0 | 0 |
| rs28939672 | pathogenic | 0 | 0 | 0 |
| rs104894173 | pathogenic | 0 | 0 | 0 |
| rs104894179 | pathogenic | 0 | 0 | 0 |
| rs104894174 | pathogenic | 1 | 1 | 0 |
| rs104894178 | pathogenic | 1 | 1 | 0 |
| rs28939714 | pathogenic | 1 | 1 | 1 |
| rs104894270 | pathogenic | 1 | 1 | 1 |
| rs3765571 | benign | 0 | 0 | 0 |
| rs752669339 | pathogenic | 1 | 1 | 1 |
| rs537043237 | pathogenic | 1 | 1 | 1 |
| rs774047299 | pathogenic | 1 | 1 | 1 |
| rs121434583 | pathogenic | 1 | 1 | 1 |
| rs121907927 | pathogenic | 0 | 0 | 0 |
| rs121907925 | pathogenic | 0 | 1 | 0 |
| rs746212325 | pathogenic | 0 | 0 | 0 |
| rs754637179 | pathogenic | 0 | 0 | 0 |
| rs780882740 | pathogenic | 0 | 0 | 0 |
| rs121918130 | pathogenic | 0 | 0 | 0 |
| rs121918129 | pathogenic | 0 | 0 | 0 |
| rs13297509 | pathogenic | 0 | 0 | 0 |
| rs746867724 | pathogenic | 0 | 0 | 0 |
| rs121918128 | pathogenic | 0 | 0 | 0 |
| rs752300607 | pathogenic | 0 | 0 | 0 |
| rs121964860 | pathogenic | 0 | 0 | 0 |
| rs45466197 | pathogenic | 1 | 1 | 1 |
| rs3730238 | benign | 1 | 1 | 1 |
| rs730881125 | pathogenic | 1 | 1 | 1 |
| rs121964861 | pathogenic | 1 | 1 | 1 |
| rs397516484 | pathogenic | 0 | 0 | 0 |
| rs367785431 | pathogenic | 0 | 0 | 0 |
| rs141121678 | pathogenic | 0 | 0 | 0 |
| rs12720263 | benign | 0 | 0 | 0 |
| rs2304256 | benign | 0 | 0 | 0 |
| rs2304255 | benign | 0 | 0 | 0 |
| rs121909045 | pathogenic | 0 | 0 | 0 |
| rs397508796 | pathogenic | 0 | 0 | 0 |
| rs151020603 | pathogenic | 0 | 0 | 0 |
| rs77284892 | pathogenic | 0 | 0 | 0 |
| rs368505753 | pathogenic | 0 | 0 | 0 |
| rs115545701 | pathogenic | 0 | 0 | 0 |
| rs121908749 | pathogenic | 0 | 0 | 0 |
| rs75961395 | pathogenic | 0 | 0 | 0 |
| rs121908750 | pathogenic | 0 | 0 | 0 |
| rs121908751 | pathogenic | 0 | 0 | 0 |
| rs397508464 | pathogenic | 0 | 0 | 0 |
| rs397508461 | pathogenic | 0 | 0 | 0 |
| rs121909031 | pathogenic | 0 | 0 | 0 |
| rs113993958 | pathogenic | 0 | 0 | 0 |
| rs77834169 | pathogenic | 0 | 0 | 0 |
| rs78655421 | pathogenic | 0 | 0 | 0 |
| rs141723617 | benign | 0 | 0 | 0 |
| rs35516286 | benign | 0 | 0 | 0 |
| rs1800079 | pathogenic | 1 | 1 | 0 |

| | | | | |
|---|---|---|---|---|
| rs80282562 | pathogenic | 0 | 0 | 0 |
| rs121908752 | pathogenic | 0 | 0 | 0 |
| rs556880586 | pathogenic | 0 | 0 | 0 |
| rs121908760 | pathogenic | 0 | 0 | 0 |
| rs397508350 | pathogenic | 0 | 0 | 0 |
| rs374946172 | pathogenic | 0 | 0 | 0 |
| rs397508387 | pathogenic | 0 | 0 | 0 |
| rs201386642 | pathogenic | 0 | 0 | 0 |
| rs121909012 | pathogenic | 0 | 0 | 0 |
| rs121909008 | pathogenic | 0 | 0 | 0 |
| rs149790377 | pathogenic | 0 | 0 | 0 |
| rs397508442 | pathogenic | 0 | 0 | 0 |
| rs121909035 | pathogenic | 0 | 0 | 0 |
| rs386134230 | pathogenic | 0 | 0 | 0 |
| rs1800112 | benign | 0 | 0 | 0 |
| rs144055758 | pathogenic | 0 | 0 | 0 |
| rs142394380 | pathogenic | 0 | 0 | 0 |
| rs121909036 | pathogenic | 0 | 0 | 0 |
| rs78194216 | pathogenic | 0 | 0 | 0 |
| rs121909019 | pathogenic | 0 | 0 | 0 |
| rs121909020 | pathogenic | 0 | 0 | 0 |
| rs200321110 | pathogenic | 0 | 0 | 0 |
| rs202179988 | pathogenic | 0 | 0 | 0 |
| rs121909037 | pathogenic | 0 | 0 | 0 |
| rs186045772 | pathogenic | 0 | 0 | 0 |
| rs139304906 | pathogenic | 0 | 0 | 0 |
| rs78802634 | pathogenic | 0 | 0 | 0 |
| rs121908761 | pathogenic | 0 | 0 | 0 |
| rs36210737 | pathogenic | 0 | 0 | 0 |
| rs139729994 | pathogenic | 0 | 0 | 0 |
| rs79850223 | pathogenic | 0 | 0 | 0 |
| rs1800120 | benign | 0 | 0 | 0 |
| rs74767530 | pathogenic | 0 | 0 | 0 |
| rs374705585 | pathogenic | 0 | 0 | 0 |
| rs1136287 | benign | 0 | 0 | 0 |
| rs6828 | benign | 0 | 0 | 0 |
| rs104894856 | pathogenic | 0 | 0 | 0 |
| rs104894860 | pathogenic | 0 | 0 | 0 |
| rs61736892 | pathogenic | 0 | 0 | 0 |
| rs145807417 | benign | 0 | 0 | 0 |
| rs104894853 | pathogenic | 0 | 0 | 0 |
| rs199422227 | pathogenic | 0 | 0 | 0 |
| rs113993946 | pathogenic | 0 | 0 | 0 |
| rs199422231 | pathogenic | 0 | 0 | 0 |
| rs199422228 | pathogenic | 0 | 0 | 0 |
| rs142195196 | benign | 0 | 1 | 0 |
| rs148656104 | benign | 1 | 1 | 1 |
| rs141944844 | benign | 0 | 0 | 0 |
| rs104894342 | pathogenic | 0 | 0 | 0 |
| rs104894343 | pathogenic | 0 | 0 | 0 |
| rs104894344 | pathogenic | 0 | 0 | 0 |
| rs573322878 | benign | 0 | 0 | 0 |
| rs104894347 | pathogenic | 0 | 0 | 0 |
| rs28937882 | pathogenic | 0 | 0 | 0 |

| | | | | |
|---|---|---|---|---|
| rs193922702 | pathogenic | 0 | 0 | 0 |
| rs200786104 | benign | 0 | 0 | 0 |
| rs690016544 | pathogenic | 1 | 1 | 1 |
| rs28943592 | benign | 0 | 0 | 0 |
| rs28943594 | benign | 0 | 0 | 0 |
| rs121908957 | pathogenic | 0 | 0 | 0 |
| rs115408010 | benign | 1 | 1 | 1 |
| rs77591659 | benign | 0 | 0 | 0 |
| rs121918511 | pathogenic | 0 | 0 | 0 |
| rs538359948 | benign | 0 | 0 | 0 |
| rs76708142 | benign | 0 | 0 | 0 |
| rs121907913 | pathogenic | 1 | 1 | 0 |
| rs121907920 | pathogenic | 0 | 0 | 0 |
| rs121907923 | pathogenic | 1 | 1 | 0 |
| rs121907928 | pathogenic | 1 | 1 | 0 |
| rs121907919 | pathogenic | 1 | 1 | 0 |
| rs121907918 | pathogenic | 1 | 1 | 0 |
| rs74315493 | pathogenic | 1 | 1 | 1 |
| rs74315494 | pathogenic | 1 | 1 | 1 |
| rs145446060 | benign | 1 | 1 | 1 |
| rs550409143 | benign | 1 | 1 | 0 |

| compound_binding_site | cofactor_binding_site | nucleic_binding_site | ion_binding_site |
|---|---|---|---|
| 1 | 0 | 0 | 0 |
| 1 | 0 | 0 | 0 |
| 0 | 0 | 0 | 0 |
| 1 | 0 | 0 | 0 |
| 0 | 0 | 0 | 0 |
| 0 | 0 | 0 | 0 |
| 0 | 0 | 0 | 0 |
| 0 | 0 | 0 | 0 |
| 1 | 0 | 0 | 1 |
| 1 | 0 | 0 | 0 |
| 0 | 0 | 0 | 1 |
| 1 | 0 | 1 | 0 |
| 0 | 0 | 0 | 0 |
| 0 | 0 | 0 | 0 |
| 0 | 0 | 0 | 0 |
| 0 | 0 | 0 | 0 |
| 0 | 0 | 0 | 0 |
| 1 | 0 | 0 | 1 |
| 1 | 0 | 1 | 0 |
| 0 | 0 | 0 | 0 |
| 1 | 0 | 0 | 1 |
| 1 | 0 | 0 | 0 |
| 1 | 0 | 0 | 0 |
| 1 | 0 | 0 | 0 |
| 0 | 0 | 0 | 0 |
| 0 | 0 | 0 | 0 |
| 1 | 0 | 0 | 1 |
| 0 | 0 | 0 | 0 |
| 0 | 0 | 0 | 0 |
| 0 | 0 | 0 | 0 |
| 0 | 0 | 0 | 0 |
| 1 | 0 | 0 | 0 |
| 1 | 0 | 0 | 1 |
| 1 | 0 | 0 | 0 |
| 0 | 0 | 0 | 0 |
| 0 | 0 | 0 | 0 |
| 0 | 0 | 0 | 0 |
| 1 | 0 | 0 | 1 |
| 1 | 0 | 0 | 1 |
| 1 | 0 | 0 | 1 |
| 1 | 0 | 0 | 0 |
| 1 | 0 | 0 | 0 |
| 1 | 0 | 0 | 0 |
| 1 | 0 | 0 | 0 |
| 1 | 0 | 0 | 0 |
| 1 | 0 | 0 | 0 |
| 1 | 0 | 0 | 0 |
| 1 | 0 | 0 | 0 |
| 1 | 0 | 0 | 0 |
| 1 | 0 | 0 | 0 |
| 1 | 0 | 0 | 0 |
| 1 | 0 | 0 | 0 |
| 1 | 0 | 0 | 0 |

| | | | |
|---|---|---|---|
| 1 | 0 | 0 | 1 |
| 1 | 0 | 0 | 0 |
| 1 | 0 | 0 | 0 |
| 1 | 0 | 0 | 0 |
| 0 | 0 | 0 | 0 |
| 1 | 0 | 0 | 0 |
| 0 | 0 | 0 | 0 |
| 1 | 0 | 0 | 0 |
| 1 | 0 | 0 | 0 |
| 1 | 0 | 0 | 1 |
| 1 | 0 | 0 | 0 |
| 1 | 0 | 0 | 0 |
| 1 | 0 | 0 | 0 |
| 1 | 0 | 0 | 0 |
| 1 | 0 | 0 | 0 |
| 1 | 0 | 0 | 0 |
| 1 | 0 | 0 | 0 |
| 1 | 0 | 0 | 1 |
| 1 | 0 | 0 | 1 |
| 1 | 1 | 0 | 1 |
| 1 | 0 | 0 | 0 |
| 1 | 0 | 0 | 0 |
| 1 | 0 | 0 | 0 |
| 1 | 0 | 0 | 0 |
| 1 | 1 | 0 | 0 |
| 1 | 0 | 0 | 0 |
| 1 | 0 | 0 | 0 |
| 1 | 0 | 0 | 0 |
| 1 | 0 | 0 | 0 |
| 1 | 0 | 0 | 0 |
| 1 | 1 | 0 | 0 |
| 1 | 1 | 0 | 0 |
| 1 | 1 | 0 | 0 |
| 1 | 1 | 0 | 0 |
| 1 | 1 | 0 | 0 |
| 1 | 0 | 0 | 0 |
| 1 | 1 | 0 | 0 |
| 1 | 0 | 0 | 0 |
| 1 | 0 | 0 | 0 |
| 1 | 1 | 0 | 0 |
| 0 | 0 | 0 | 0 |
| 0 | 0 | 0 | 0 |
| 0 | 0 | 0 | 0 |
| 0 | 0 | 0 | 0 |
| 0 | 0 | 0 | 0 |
| 0 | 0 | 0 | 0 |
| 0 | 0 | 0 | 0 |
| 1 | 0 | 0 | 1 |
| 0 | 0 | 0 | 0 |
| 1 | 0 | 0 | 0 |
| 1 | 0 | 0 | 0 |
| 0 | 0 | 0 | 0 |
| 0 | 0 | 0 | 0 |
| 0 | 0 | 0 | 0 |

| | | | |
|---|---|---|---|
| 0 | 0 | 0 | 0 |
| 0 | 0 | 0 | 0 |
| 0 | 0 | 0 | 0 |
| 1 | 0 | 0 | 0 |
| 1 | 0 | 0 | 0 |
| 1 | 0 | 0 | 0 |
| 0 | 0 | 0 | 0 |
| 0 | 0 | 1 | 0 |
| 0 | 0 | 1 | 0 |
| 0 | 0 | 1 | 0 |
| 0 | 0 | 1 | 0 |
| 0 | 0 | 1 | 0 |
| 0 | 0 | 0 | 0 |
| 0 | 0 | 0 | 0 |
| 1 | 0 | 0 | 0 |
| 0 | 0 | 0 | 0 |
| 0 | 0 | 0 | 0 |
| 1 | 0 | 0 | 0 |
| 1 | 0 | 0 | 0 |
| 1 | 0 | 0 | 0 |
| 1 | 0 | 0 | 0 |
| 1 | 1 | 0 | 0 |
| 1 | 0 | 0 | 0 |
| 1 | 0 | 0 | 0 |
| 0 | 0 | 0 | 0 |
| 0 | 0 | 0 | 0 |
| 1 | 0 | 0 | 0 |
| 1 | 1 | 0 | 0 |
| 1 | 0 | 0 | 0 |
| 1 | 0 | 0 | 0 |
| 1 | 1 | 0 | 0 |
| 1 | 1 | 0 | 0 |
| 1 | 1 | 0 | 0 |
| 1 | 1 | 0 | 0 |
| 1 | 1 | 0 | 0 |
| 1 | 1 | 0 | 0 |
| 1 | 1 | 0 | 0 |
| 1 | 1 | 0 | 0 |
| 1 | 1 | 0 | 0 |
| 1 | 1 | 0 | 0 |
| 1 | 1 | 0 | 0 |
| 1 | 1 | 0 | 0 |
| 1 | 0 | 0 | 0 |
| 1 | 0 | 0 | 0 |
| 1 | 0 | 0 | 0 |
| 1 | 1 | 0 | 0 |
| 1 | 0 | 0 | 0 |
| 1 | 1 | 1 | 0 |
| 1 | 1 | 1 | 0 |
| 1 | 1 | 0 | 0 |
| 1 | 1 | 0 | 0 |
| 1 | 0 | 0 | 0 |
| 1 | 0 | 0 | 0 |
| 0 | 0 | 0 | 0 |

| | | | |
|---|---|---|---|
| 0 | 0 | 0 | 0 |
| 0 | 0 | 0 | 0 |
| 0 | 0 | 0 | 0 |
| 1 | 0 | 0 | 0 |
| 1 | 0 | 0 | 0 |
| 0 | 0 | 0 | 0 |
| 0 | 0 | 0 | 0 |
| 0 | 0 | 0 | 0 |
| 0 | 0 | 0 | 0 |
| 0 | 0 | 0 | 0 |
| 0 | 0 | 0 | 0 |
| 0 | 0 | 0 | 0 |
| 0 | 0 | 0 | 0 |
| 0 | 0 | 0 | 0 |
| 0 | 0 | 0 | 0 |
| 0 | 0 | 0 | 0 |
| 0 | 0 | 0 | 0 |
| 0 | 0 | 0 | 0 |
| 0 | 0 | 0 | 0 |
| 0 | 0 | 0 | 0 |
| 0 | 0 | 0 | 0 |
| 0 | 0 | 0 | 0 |
| 1 | 0 | 0 | 0 |
| 1 | 0 | 0 | 0 |
| 0 | 0 | 0 | 0 |
| 0 | 0 | 0 | 0 |
| 1 | 0 | 0 | 0 |
| 0 | 0 | 0 | 0 |
| 0 | 0 | 0 | 0 |
| 0 | 0 | 0 | 0 |
| 0 | 0 | 0 | 0 |
| 0 | 0 | 0 | 0 |
| 0 | 0 | 0 | 0 |
| 0 | 0 | 0 | 0 |
| 1 | 0 | 0 | 0 |
| 0 | 0 | 0 | 0 |
| 0 | 0 | 0 | 0 |
| 0 | 0 | 0 | 0 |
| 0 | 0 | 0 | 0 |
| 0 | 0 | 0 | 0 |
| 0 | 0 | 0 | 0 |
| 0 | 0 | 0 | 0 |
| 0 | 0 | 0 | 0 |
| 0 | 0 | 0 | 0 |
| 0 | 0 | 0 | 0 |
| 0 | 0 | 0 | 0 |
| 0 | 0 | 1 | 0 |
| 1 | 0 | 0 | 0 |
| 1 | 0 | 0 | 0 |
| 1 | 1 | 0 | 0 |
| 1 | 0 | 0 | 0 |
| 0 | 0 | 0 | 0 |
| 0 | 0 | 0 | 0 |
| 1 | 0 | 0 | 0 |

| | | | |
|---|---|---|---|
| 1 | 0 | 0 | 0 |
| 1 | 0 | 0 | 0 |
| 1 | 0 | 0 | 0 |
| 0 | 0 | 0 | 0 |
| 0 | 0 | 0 | 0 |
| 0 | 0 | 0 | 0 |
| 1 | 0 | 0 | 0 |
| 1 | 0 | 0 | 0 |
| 0 | 0 | 0 | 0 |
| 0 | 0 | 0 | 0 |
| 1 | 0 | 0 | 0 |
| 0 | 0 | 0 | 0 |
| 1 | 0 | 0 | 0 |
| 0 | 0 | 0 | 0 |
| 0 | 0 | 0 | 0 |
| 0 | 0 | 0 | 0 |
| 0 | 0 | 0 | 0 |
| 0 | 0 | 0 | 0 |
| 0 | 0 | 0 | 0 |
| 0 | 0 | 0 | 0 |
| 0 | 0 | 0 | 0 |
| 0 | 0 | 0 | 0 |
| 0 | 0 | 0 | 0 |
| 0 | 0 | 0 | 0 |
| 0 | 0 | 0 | 0 |
| 0 | 0 | 0 | 0 |
| 1 | 0 | 0 | 0 |
| 1 | 0 | 0 | 0 |
| 1 | 0 | 0 | 0 |
| 1 | 0 | 0 | 0 |
| 1 | 0 | 0 | 0 |
| 1 | 0 | 0 | 0 |
| 0 | 0 | 0 | 0 |
| 0 | 0 | 0 | 0 |
| 1 | 0 | 0 | 0 |
| 0 | 0 | 0 | 0 |
| 0 | 0 | 0 | 0 |
| 0 | 0 | 0 | 0 |
| 0 | 0 | 0 | 0 |
| 0 | 0 | 0 | 0 |
| 0 | 0 | 0 | 0 |
| 0 | 0 | 0 | 0 |
| 0 | 0 | 0 | 0 |
| 0 | 0 | 0 | 0 |
| 1 | 1 | 0 | 0 |
| 1 | 1 | 0 | 0 |
| 1 | 1 | 0 | 1 |
| 1 | 1 | 0 | 1 |
| 1 | 1 | 0 | 1 |
| 1 | 1 | 0 | 1 |
| 1 | 1 | 0 | 0 |
| 1 | 1 | 0 | 1 |
| 1 | 1 | 0 | 1 |
| 1 | 1 | 0 | 1 |

| | | | |
|---|---|---|---|
| 1 | 1 | 0 | 1 |
| 1 | 1 | 0 | 1 |
| 1 | 1 | 0 | 1 |
| 1 | 1 | 0 | 1 |
| 1 | 1 | 0 | 1 |
| 1 | 0 | 0 | 0 |
| 0 | 0 | 0 | 0 |
| 0 | 0 | 0 | 0 |
| 0 | 0 | 0 | 0 |
| 1 | 0 | 0 | 0 |
| 0 | 0 | 1 | 0 |
| 0 | 0 | 0 | 0 |
| 0 | 0 | 0 | 0 |
| 0 | 0 | 0 | 0 |
| 0 | 0 | 0 | 0 |
| 0 | 0 | 0 | 0 |
| 0 | 0 | 0 | 0 |
| 0 | 0 | 0 | 0 |
| 0 | 0 | 0 | 0 |
| 0 | 0 | 0 | 0 |
| 0 | 0 | 0 | 0 |
| 0 | 0 | 0 | 0 |
| 0 | 0 | 0 | 0 |
| 0 | 0 | 0 | 0 |
| 0 | 0 | 0 | 0 |
| 1 | 1 | 0 | 0 |
| 1 | 1 | 0 | 0 |
| 1 | 1 | 0 | 0 |
| 0 | 1 | 0 | 0 |
| 1 | 0 | 0 | 0 |
| 0 | 0 | 0 | 0 |
| 0 | 0 | 0 | 0 |
| 0 | 0 | 0 | 0 |
| 1 | 0 | 0 | 0 |
| 1 | 1 | 0 | 0 |
| 0 | 0 | 0 | 0 |
| 0 | 0 | 0 | 0 |
| 0 | 0 | 0 | 0 |
| 0 | 0 | 0 | 0 |
| 1 | 0 | 0 | 0 |
| 0 | 0 | 0 | 0 |
| 0 | 0 | 0 | 0 |
| 0 | 0 | 0 | 0 |
| 0 | 0 | 0 | 0 |
| 0 | 0 | 0 | 0 |
| 0 | 1 | 0 | 0 |
| 0 | 1 | 0 | 0 |
| 0 | 0 | 0 | 0 |
| 0 | 0 | 0 | 0 |
| 0 | 1 | 0 | 0 |
| 0 | 0 | 0 | 0 |
| 0 | 0 | 0 | 0 |
| 0 | 1 | 0 | 0 |
| 0 | 0 | 0 | 0 |

| | | | |
|---|---|---|---|
| 1 | 1 | 0 | 0 |
| 0 | 0 | 0 | 0 |
| 0 | 0 | 0 | 0 |
| 0 | 0 | 0 | 0 |
| 0 | 0 | 0 | 0 |
| 0 | 0 | 0 | 0 |
| 0 | 0 | 0 | 0 |
| 0 | 0 | 0 | 0 |
| 0 | 0 | 0 | 0 |
| 0 | 0 | 0 | 0 |
| 0 | 0 | 0 | 0 |
| 0 | 0 | 0 | 0 |
| 0 | 0 | 0 | 0 |
| 0 | 0 | 0 | 0 |
| 0 | 1 | 0 | 0 |
| 0 | 0 | 0 | 0 |
| 0 | 1 | 0 | 0 |
| 0 | 1 | 0 | 0 |
| 0 | 0 | 0 | 0 |
| 0 | 1 | 0 | 0 |
| 1 | 0 | 0 | 1 |
| 1 | 1 | 0 | 0 |
| 1 | 0 | 0 | 0 |
| 1 | 0 | 0 | 1 |
| 0 | 0 | 0 | 0 |
| 0 | 0 | 0 | 0 |
| 0 | 0 | 0 | 0 |
| 0 | 0 | 0 | 0 |
| 1 | 0 | 0 | 0 |
| 0 | 0 | 0 | 0 |
| 1 | 0 | 0 | 0 |
| 0 | 0 | 0 | 0 |
| 0 | 0 | 0 | 0 |
| 0 | 0 | 0 | 0 |
| 0 | 0 | 0 | 0 |
| 0 | 0 | 0 | 0 |
| 0 | 0 | 0 | 0 |
| 1 | 1 | 0 | 0 |
| 1 | 1 | 0 | 0 |
| 1 | 1 | 0 | 0 |
| 0 | 0 | 0 | 0 |
| 0 | 0 | 0 | 0 |
| 0 | 0 | 0 | 0 |
| 1 | 0 | 0 | 0 |
| 1 | 0 | 0 | 1 |
| 1 | 0 | 0 | 1 |
| 1 | 0 | 0 | 0 |
| 1 | 0 | 0 | 1 |
| 1 | 0 | 0 | 0 |
| 0 | 0 | 0 | 0 |
| 1 | 1 | 0 | 0 |
| 1 | 1 | 0 | 0 |
| 0 | 0 | 0 | 0 |
| 0 | 0 | 0 | 0 |

| | | | |
|---|---|---|---|
| 1 | 0 | 0 | 0 |
| 1 | 0 | 0 | 0 |
| 1 | 0 | 0 | 0 |
| 0 | 0 | 0 | 0 |
| 0 | 0 | 0 | 0 |
| 1 | 1 | 0 | 1 |
| 1 | 1 | 0 | 1 |
| 1 | 1 | 0 | 1 |
| 1 | 1 | 0 | 1 |
| 1 | 1 | 0 | 1 |
| 1 | 1 | 0 | 1 |
| 1 | 1 | 0 | 1 |
| 1 | 1 | 0 | 0 |
| 1 | 1 | 0 | 0 |
| 1 | 1 | 1 | 1 |
| 0 | 0 | 0 | 0 |
| 1 | 1 | 0 | 0 |
| 1 | 1 | 0 | 0 |
| 0 | 0 | 0 | 0 |
| 0 | 0 | 0 | 0 |
| 0 | 0 | 0 | 0 |
| 1 | 0 | 0 | 0 |
| 1 | 0 | 0 | 0 |
| 0 | 0 | 0 | 0 |
| 1 | 0 | 0 | 0 |
| 1 | 0 | 0 | 0 |
| 1 | 0 | 0 | 0 |
| 0 | 0 | 0 | 0 |
| 0 | 0 | 0 | 0 |
| 0 | 0 | 0 | 0 |
| 0 | 0 | 0 | 0 |
| 0 | 0 | 0 | 0 |
| 0 | 0 | 0 | 0 |
| 0 | 0 | 0 | 0 |
| 0 | 0 | 0 | 0 |
| 0 | 0 | 0 | 0 |
| 0 | 0 | 0 | 0 |
| 0 | 0 | 0 | 0 |
| 0 | 0 | 0 | 0 |
| 0 | 0 | 0 | 0 |
| 0 | 0 | 0 | 0 |
| 0 | 0 | 0 | 0 |
| 0 | 0 | 0 | 0 |
| 0 | 0 | 0 | 0 |
| 0 | 0 | 0 | 0 |
| 1 | 0 | 0 | 0 |
| 1 | 0 | 0 | 0 |
| 1 | 0 | 0 | 0 |
| 1 | 0 | 0 | 0 |
| 0 | 0 | 0 | 0 |
| 0 | 0 | 0 | 0 |
| 0 | 0 | 0 | 0 |
| 0 | 0 | 0 | 0 |
| 0 | 0 | 0 | 0 |

| | | | |
|---|---|---|---|
| 0 | 0 | 0 | 0 |
| 0 | 0 | 0 | 0 |
| 0 | 0 | 0 | 0 |
| 0 | 0 | 0 | 0 |
| 0 | 0 | 0 | 0 |
| 0 | 0 | 0 | 0 |
| 0 | 0 | 0 | 0 |
| 1 | 0 | 0 | 1 |
| 0 | 0 | 0 | 0 |
| 0 | 0 | 0 | 0 |
| 1 | 0 | 0 | 0 |
| 0 | 0 | 0 | 0 |
| 0 | 0 | 0 | 0 |
| 0 | 0 | 0 | 0 |
| 0 | 0 | 0 | 0 |
| 1 | 0 | 0 | 0 |
| 0 | 0 | 0 | 0 |
| 0 | 0 | 0 | 0 |
| 0 | 0 | 0 | 0 |
| 0 | 0 | 0 | 0 |
| 0 | 0 | 0 | 0 |
| 0 | 0 | 0 | 0 |
| 0 | 0 | 0 | 0 |
| 1 | 1 | 0 | 1 |
| 0 | 0 | 0 | 0 |
| 0 | 0 | 0 | 0 |
| 0 | 0 | 0 | 0 |
| 1 | 1 | 0 | 1 |
| 1 | 1 | 0 | 0 |
| 0 | 0 | 0 | 0 |
| 1 | 0 | 0 | 0 |
| 1 | 1 | 0 | 0 |
| 0 | 0 | 0 | 0 |
| 1 | 0 | 0 | 0 |
| 1 | 1 | 0 | 0 |
| 1 | 0 | 0 | 0 |
| 1 | 1 | 0 | 1 |
| 1 | 1 | 0 | 1 |
| 1 | 0 | 0 | 0 |
| 0 | 0 | 0 | 0 |
| 0 | 0 | 0 | 0 |
| 0 | 0 | 0 | 0 |
| 0 | 0 | 0 | 0 |
| 0 | 0 | 0 | 0 |
| 0 | 0 | 0 | 0 |
| 0 | 0 | 0 | 0 |
| 0 | 0 | 0 | 0 |
| 0 | 0 | 0 | 0 |
| 0 | 0 | 0 | 0 |
| 0 | 0 | 0 | 0 |
| 0 | 0 | 0 | 0 |
| 0 | 0 | 0 | 0 |
| 0 | 0 | 0 | 0 |
| 0 | 0 | 0 | 0 |

| | | | |
|---|---|---|---|
| 0 | 0 | 0 | 0 |
| 0 | 0 | 0 | 0 |
| 0 | 0 | 0 | 0 |
| 0 | 0 | 0 | 0 |
| 0 | 0 | 0 | 0 |
| 0 | 0 | 0 | 0 |
| 0 | 0 | 0 | 0 |
| 0 | 0 | 0 | 0 |
| 0 | 0 | 0 | 0 |
| 0 | 0 | 0 | 0 |
| 0 | 0 | 0 | 0 |
| 0 | 0 | 0 | 0 |
| 0 | 0 | 0 | 0 |
| 1 | 1 | 0 | 0 |
| 0 | 0 | 0 | 0 |
| 1 | 0 | 0 | 0 |
| 0 | 0 | 0 | 1 |
| 0 | 0 | 0 | 0 |
| 0 | 0 | 0 | 0 |
| 0 | 0 | 0 | 0 |
| 0 | 0 | 0 | 0 |
| 0 | 0 | 0 | 0 |
| 0 | 0 | 0 | 0 |
| 0 | 0 | 0 | 0 |
| 0 | 0 | 0 | 0 |
| 0 | 0 | 0 | 0 |
| 0 | 0 | 0 | 0 |
| 0 | 0 | 0 | 0 |
| 0 | 0 | 0 | 0 |
| 0 | 0 | 0 | 0 |
| 1 | 0 | 0 | 0 |
| 1 | 0 | 0 | 0 |
| 0 | 0 | 0 | 0 |
| 0 | 0 | 0 | 0 |
| 0 | 0 | 0 | 0 |
| 0 | 0 | 0 | 0 |
| 0 | 0 | 0 | 0 |
| 0 | 0 | 0 | 0 |
| 0 | 0 | 0 | 0 |
| 0 | 0 | 0 | 0 |
| 1 | 0 | 0 | 0 |
| 1 | 0 | 0 | 0 |
| 1 | 0 | 0 | 0 |
| 1 | 0 | 0 | 0 |
| 0 | 0 | 0 | 0 |
| 1 | 0 | 0 | 0 |
| 0 | 0 | 0 | 0 |
| 0 | 0 | 0 | 0 |
| 1 | 0 | 0 | 0 |
| 1 | 0 | 0 | 0 |
| 0 | 0 | 0 | 0 |
| 0 | 0 | 0 | 0 |
| 0 | 0 | 0 | 0 |
| 0 | 0 | 0 | 0 |

| | | | |
|---|---|---|---|
| 0 | 0 | 0 | 0 |
| 0 | 0 | 0 | 0 |
| 0 | 0 | 0 | 0 |
| 0 | 0 | 0 | 0 |
| 0 | 0 | 0 | 0 |
| 0 | 0 | 0 | 0 |
| 0 | 0 | 0 | 0 |
| 0 | 0 | 0 | 0 |
| 0 | 0 | 0 | 0 |
| 0 | 0 | 0 | 0 |
| 0 | 0 | 0 | 0 |
| 0 | 0 | 0 | 0 |
| 0 | 0 | 0 | 0 |
| 0 | 0 | 0 | 0 |
| 0 | 0 | 0 | 0 |
| 1 | 0 | 0 | 1 |
| 0 | 0 | 0 | 0 |
| 0 | 0 | 0 | 0 |
| 0 | 0 | 0 | 0 |
| 0 | 0 | 0 | 0 |
| 0 | 0 | 0 | 0 |
| 1 | 1 | 0 | 0 |
| 1 | 1 | 0 | 0 |
| 1 | 0 | 0 | 0 |
| 1 | 0 | 0 | 0 |
| 0 | 0 | 0 | 0 |
| 0 | 0 | 0 | 0 |
| 0 | 0 | 0 | 0 |
| 0 | 0 | 0 | 0 |
| 1 | 0 | 0 | 0 |
| 1 | 0 | 0 | 0 |
| 1 | 0 | 0 | 0 |
| 1 | 1 | 0 | 0 |
| 1 | 1 | 0 | 0 |
| 1 | 0 | 0 | 0 |
| 0 | 0 | 0 | 0 |
| 0 | 0 | 0 | 0 |
| 0 | 1 | 0 | 0 |
| 0 | 0 | 0 | 0 |
| 0 | 0 | 0 | 0 |
| 0 | 0 | 0 | 0 |
| 1 | 1 | 0 | 0 |
| 1 | 0 | 0 | 0 |
| 1 | 0 | 0 | 0 |
| 0 | 0 | 0 | 0 |
| 0 | 0 | 0 | 0 |
| 0 | 0 | 0 | 0 |
| 0 | 0 | 0 | 0 |
| 0 | 0 | 0 | 0 |
| 0 | 0 | 0 | 0 |
| 0 | 0 | 0 | 0 |
| 0 | 0 | 0 | 0 |
| 0 | 0 | 0 | 0 |
| 0 | 0 | 0 | 0 |

| | | | |
|---|---|---|---|
| 0 | 0 | 0 | 0 |
| 1 | 0 | 0 | 0 |
| 0 | 0 | 0 | 0 |
| 0 | 0 | 0 | 0 |
| 0 | 0 | 0 | 0 |
| 0 | 0 | 0 | 0 |
| 1 | 0 | 0 | 0 |
| 0 | 0 | 0 | 0 |
| 0 | 0 | 0 | 0 |
| 0 | 0 | 0 | 0 |
| 0 | 0 | 0 | 0 |
| 0 | 0 | 0 | 0 |
| 0 | 0 | 0 | 0 |
| 0 | 0 | 0 | 0 |
| 0 | 0 | 0 | 0 |
| 0 | 0 | 0 | 0 |
| 0 | 1 | 0 | 0 |
| 0 | 0 | 0 | 0 |
| 0 | 0 | 0 | 0 |
| 0 | 0 | 0 | 0 |
| 1 | 0 | 0 | 0 |
| 0 | 0 | 0 | 0 |
| 0 | 0 | 0 | 0 |
| 1 | 1 | 0 | 0 |
| 0 | 0 | 0 | 0 |
| 0 | 0 | 0 | 0 |
| 0 | 0 | 0 | 0 |
| 0 | 0 | 0 | 0 |
| 1 | 0 | 0 | 0 |
| 0 | 0 | 0 | 0 |
| 0 | 0 | 0 | 0 |
| 0 | 0 | 0 | 0 |
| 0 | 0 | 0 | 0 |
| 0 | 0 | 0 | 0 |
| 0 | 0 | 0 | 0 |
| 1 | 0 | 0 | 0 |
| 0 | 0 | 0 | 0 |
| 0 | 0 | 0 | 0 |
| 0 | 0 | 0 | 0 |
| 0 | 0 | 0 | 0 |
| 1 | 1 | 0 | 0 |
| 0 | 0 | 0 | 0 |
| 0 | 0 | 0 | 0 |
| 0 | 0 | 0 | 0 |
| 0 | 0 | 0 | 0 |
| 0 | 0 | 0 | 0 |
| 1 | 0 | 0 | 0 |
| 1 | 0 | 0 | 1 |
| 1 | 0 | 0 | 0 |
| 0 | 0 | 0 | 0 |
| 0 | 0 | 0 | 0 |
| 0 | 0 | 0 | 0 |
| 1 | 0 | 0 | 0 |
| 1 | 0 | 0 | 0 |

| | | | |
|---|---|---|---|
| 1 | 0 | 0 | 0 |
| 0 | 0 | 0 | 0 |
| 0 | 0 | 0 | 0 |
| 0 | 0 | 0 | 0 |
| 1 | 1 | 0 | 0 |
| 1 | 1 | 0 | 0 |
| 0 | 0 | 0 | 0 |
| 0 | 0 | 0 | 0 |
| 1 | 0 | 0 | 0 |
| 1 | 0 | 0 | 0 |
| 1 | 0 | 1 | 0 |
| 0 | 0 | 0 | 0 |
| 0 | 0 | 1 | 0 |
| 0 | 0 | 1 | 0 |
| 0 | 0 | 1 | 0 |
| 1 | 0 | 0 | 0 |
| 1 | 0 | 0 | 0 |
| 0 | 0 | 0 | 0 |
| 0 | 0 | 0 | 0 |
| 1 | 0 | 0 | 1 |
| 1 | 0 | 0 | 1 |
| 1 | 0 | 0 | 0 |
| 0 | 0 | 0 | 0 |
| 0 | 0 | 0 | 0 |
| 0 | 0 | 0 | 0 |
| 0 | 0 | 0 | 0 |
| 0 | 0 | 0 | 0 |
| 1 | 0 | 0 | 0 |
| 0 | 0 | 0 | 0 |
| 0 | 0 | 0 | 1 |
| 0 | 0 | 0 | 1 |
| 0 | 0 | 1 | 1 |
| 0 | 0 | 1 | 1 |
| 0 | 0 | 1 | 1 |
| 0 | 0 | 1 | 1 |
| 0 | 0 | 1 | 0 |
| 0 | 0 | 1 | 0 |
| 0 | 0 | 0 | 0 |
| 0 | 0 | 0 | 0 |
| 0 | 0 | 0 | 0 |
| 0 | 0 | 0 | 0 |
| 0 | 0 | 1 | 0 |
| 0 | 0 | 1 | 0 |
| 0 | 0 | 1 | 0 |
| 0 | 0 | 1 | 0 |
| 0 | 0 | 1 | 0 |
| 0 | 0 | 1 | 0 |
| 0 | 0 | 1 | 0 |
| 0 | 0 | 1 | 0 |
| 0 | 0 | 0 | 0 |
| 0 | 0 | 0 | 0 |
| 0 | 0 | 0 | 0 |
| 0 | 0 | 0 | 0 |
| 1 | 0 | 0 | 0 |

| | | | |
|---|---|---|---|
| 0 | 0 | 0 | 0 |
| 0 | 0 | 0 | 0 |
| 0 | 0 | 0 | 0 |
| 1 | 0 | 0 | 0 |
| 0 | 0 | 0 | 0 |
| 0 | 0 | 0 | 0 |
| 0 | 0 | 0 | 0 |
| 0 | 0 | 0 | 0 |
| 0 | 0 | 0 | 0 |
| 0 | 0 | 0 | 0 |
| 1 | 1 | 0 | 0 |
| 0 | 0 | 0 | 0 |
| 1 | 1 | 0 | 0 |
| 1 | 1 | 0 | 0 |
| 0 | 1 | 0 | 0 |
| 0 | 0 | 0 | 0 |
| 0 | 0 | 0 | 0 |
| 0 | 0 | 0 | 0 |
| 0 | 0 | 0 | 0 |
| 0 | 0 | 0 | 0 |
| 1 | 1 | 0 | 0 |
| 0 | 1 | 0 | 0 |
| 0 | 0 | 0 | 0 |
| 0 | 1 | 0 | 0 |
| 1 | 1 | 0 | 0 |
| 0 | 0 | 0 | 0 |
| 0 | 0 | 0 | 0 |
| 1 | 1 | 0 | 0 |
| 1 | 1 | 0 | 0 |
| 0 | 1 | 0 | 0 |
| 0 | 0 | 0 | 0 |
| 0 | 0 | 0 | 0 |
| 0 | 0 | 0 | 0 |
| 0 | 0 | 0 | 0 |
| 0 | 0 | 0 | 0 |
| 0 | 0 | 0 | 0 |
| 0 | 0 | 0 | 0 |
| 0 | 0 | 0 | 0 |
| 0 | 0 | 0 | 0 |
| 0 | 0 | 0 | 0 |
| 1 | 0 | 0 | 0 |
| 1 | 0 | 0 | 0 |
| 1 | 0 | 0 | 0 |
| 1 | 0 | 0 | 0 |
| 1 | 1 | 0 | 1 |
| 0 | 0 | 0 | 0 |
| 1 | 0 | 0 | 0 |
| 0 | 0 | 0 | 0 |
| 1 | 0 | 0 | 0 |
| 0 | 0 | 0 | 0 |
| 1 | 0 | 0 | 0 |
| 1 | 0 | 0 | 0 |
| 0 | 0 | 0 | 0 |
| 0 | 0 | 0 | 0 |

| | | | |
|---|---|---|---|
| 0 | 0 | 0 | 0 |
| 0 | 0 | 0 | 0 |
| 0 | 0 | 0 | 0 |
| 0 | 0 | 0 | 0 |
| 1 | 0 | 0 | 0 |
| 1 | 0 | 0 | 0 |
| 1 | 0 | 0 | 0 |
| 0 | 0 | 0 | 0 |
| 1 | 1 | 0 | 0 |
| 0 | 0 | 0 | 0 |
| 0 | 0 | 0 | 0 |
| 0 | 0 | 0 | 0 |
| 1 | 1 | 0 | 0 |
| 1 | 0 | 0 | 0 |
| 0 | 0 | 0 | 0 |
| 0 | 0 | 0 | 0 |
| 1 | 1 | 0 | 0 |
| 0 | 0 | 0 | 0 |
| 0 | 0 | 0 | 0 |
| 0 | 0 | 0 | 0 |
| 0 | 0 | 0 | 0 |
| 0 | 0 | 0 | 0 |
| 0 | 0 | 0 | 0 |
| 0 | 0 | 0 | 0 |
| 1 | 0 | 0 | 0 |
| 0 | 0 | 0 | 0 |
| 1 | 0 | 0 | 0 |
| 0 | 0 | 0 | 0 |
| 0 | 0 | 0 | 0 |
| 0 | 0 | 0 | 0 |
| 1 | 0 | 0 | 0 |
| 1 | 0 | 0 | 0 |
| 0 | 0 | 0 | 0 |
| 0 | 0 | 0 | 0 |
| 1 | 0 | 0 | 0 |
| 0 | 0 | 0 | 0 |
| 0 | 0 | 0 | 0 |
| 1 | 0 | 0 | 0 |
| 0 | 0 | 0 | 0 |
| 1 | 0 | 0 | 0 |
| 1 | 0 | 0 | 0 |
| 0 | 0 | 0 | 0 |
| 0 | 0 | 0 | 0 |
| 1 | 1 | 0 | 0 |
| 1 | 0 | 0 | 0 |
| 1 | 1 | 0 | 0 |
| 0 | 0 | 0 | 0 |
| 0 | 0 | 0 | 0 |
| 0 | 0 | 0 | 0 |
| 0 | 0 | 0 | 0 |
| 1 | 1 | 0 | 0 |
| 0 | 0 | 0 | 0 |
| 0 | 0 | 0 | 0 |
| 0 | 0 | 0 | 0 |

| | | | |
|---|---|---|---|
| 0 | 1 | 0 | 0 |
| 1 | 0 | 0 | 0 |
| 0 | 0 | 0 | 0 |
| 0 | 0 | 0 | 0 |
| 0 | 0 | 0 | 0 |
| 1 | 1 | 0 | 0 |
| 0 | 0 | 0 | 0 |
| 0 | 0 | 0 | 0 |
| 0 | 0 | 0 | 0 |
| 1 | 0 | 0 | 0 |
| 1 | 0 | 0 | 0 |
| 0 | 0 | 0 | 0 |
| 0 | 0 | 0 | 0 |
| 0 | 0 | 0 | 0 |
| 0 | 0 | 0 | 0 |
| 0 | 0 | 0 | 0 |
| 0 | 0 | 0 | 0 |
| 0 | 0 | 0 | 0 |
| 0 | 0 | 0 | 0 |
| 0 | 0 | 0 | 0 |
| 0 | 0 | 0 | 0 |
| 0 | 0 | 0 | 0 |
| 0 | 0 | 0 | 0 |
| 1 | 1 | 0 | 0 |
| 0 | 0 | 0 | 0 |
| 0 | 0 | 0 | 0 |
| 1 | 1 | 1 | 0 |
| 0 | 0 | 0 | 0 |
| 0 | 0 | 0 | 0 |
| 0 | 0 | 0 | 0 |
| 1 | 1 | 1 | 0 |
| 1 | 0 | 0 | 0 |
| 1 | 0 | 0 | 0 |
| 1 | 0 | 0 | 0 |
| 1 | 0 | 0 | 0 |
| 1 | 1 | 1 | 0 |
| 1 | 1 | 1 | 0 |
| 0 | 0 | 0 | 0 |
| 0 | 0 | 0 | 0 |
| 1 | 1 | 1 | 0 |
| 0 | 0 | 0 | 0 |
| 0 | 0 | 0 | 0 |
| 0 | 0 | 0 | 0 |
| 0 | 0 | 0 | 0 |
| 0 | 0 | 0 | 0 |
| 0 | 0 | 0 | 0 |
| 0 | 0 | 0 | 0 |
| 0 | 0 | 0 | 0 |
| 1 | 0 | 0 | 0 |
| 1 | 0 | 0 | 0 |
| 1 | 0 | 0 | 0 |
| 0 | 0 | 0 | 0 |
| 0 | 0 | 0 | 0 |
| 0 | 0 | 0 | 0 |

| | | | |
|---|---|---|---|
| 0 | 0 | 0 | 0 |
| 1 | 0 | 0 | 0 |
| 0 | 0 | 0 | 0 |
| 0 | 0 | 0 | 0 |
| 0 | 0 | 0 | 0 |
| 0 | 0 | 0 | 0 |
| 0 | 0 | 0 | 0 |
| 0 | 0 | 0 | 0 |
| 0 | 0 | 0 | 0 |
| 0 | 0 | 0 | 0 |
| 1 | 1 | 0 | 0 |
| 1 | 1 | 0 | 0 |
| 0 | 0 | 0 | 0 |
| 0 | 0 | 0 | 0 |
| 0 | 0 | 0 | 0 |
| 1 | 0 | 0 | 0 |
| 1 | 0 | 0 | 0 |
| 1 | 1 | 0 | 1 |
| 0 | 0 | 0 | 0 |
| 0 | 0 | 0 | 0 |
| 0 | 0 | 0 | 0 |
| 0 | 0 | 0 | 0 |
| 1 | 0 | 0 | 1 |
| 0 | 0 | 0 | 0 |
| 0 | 0 | 0 | 0 |
| 1 | 0 | 0 | 0 |
| 0 | 0 | 0 | 0 |
| 0 | 0 | 0 | 0 |
| 0 | 0 | 0 | 0 |
| 0 | 0 | 0 | 0 |
| 0 | 0 | 0 | 0 |
| 0 | 0 | 0 | 0 |
| 0 | 0 | 0 | 0 |
| 0 | 0 | 0 | 0 |
| 0 | 0 | 0 | 0 |
| 1 | 0 | 0 | 0 |
| 1 | 0 | 0 | 0 |
| 1 | 1 | 0 | 0 |
| 1 | 1 | 0 | 0 |
| 0 | 0 | 0 | 0 |
| 0 | 0 | 0 | 0 |
| 1 | 0 | 0 | 0 |
| 0 | 0 | 0 | 0 |
| 1 | 0 | 0 | 0 |
| 1 | 1 | 0 | 0 |
| 0 | 0 | 0 | 0 |
| 0 | 0 | 0 | 0 |
| 0 | 0 | 0 | 0 |
| 0 | 0 | 0 | 0 |
| 0 | 0 | 0 | 0 |
| 0 | 0 | 0 | 0 |
| 1 | 0 | 0 | 0 |
| 0 | 0 | 0 | 0 |
| 0 | 0 | 0 | 0 |

| | | | |
|---|---|---|---|
| 0 | 0 | 0 | 0 |
| 1 | 1 | 0 | 0 |
| 0 | 0 | 0 | 0 |
| 1 | 0 | 0 | 0 |
| 1 | 0 | 0 | 0 |
| 0 | 0 | 0 | 0 |
| 0 | 0 | 0 | 0 |
| 0 | 0 | 0 | 0 |
| 0 | 0 | 0 | 0 |
| 0 | 0 | 0 | 0 |
| 0 | 0 | 0 | 0 |
| 1 | 1 | 0 | 0 |
| 1 | 1 | 0 | 1 |
| 0 | 0 | 0 | 0 |
| 0 | 1 | 0 | 0 |
| 0 | 0 | 0 | 0 |
| 0 | 0 | 0 | 0 |
| 0 | 0 | 0 | 0 |
| 0 | 0 | 0 | 0 |
| 0 | 0 | 0 | 0 |
| 0 | 0 | 0 | 0 |
| 1 | 0 | 0 | 0 |
| 0 | 0 | 0 | 0 |
| 0 | 0 | 0 | 0 |
| 0 | 0 | 0 | 0 |
| 0 | 0 | 0 | 0 |
| 0 | 0 | 0 | 0 |
| 1 | 0 | 0 | 0 |
| 0 | 0 | 0 | 0 |
| 0 | 0 | 0 | 0 |
| 0 | 0 | 0 | 0 |
| 0 | 0 | 0 | 0 |
| 0 | 0 | 0 | 0 |
| 0 | 0 | 0 | 0 |
| 0 | 0 | 0 | 0 |
| 0 | 0 | 0 | 0 |
| 0 | 0 | 0 | 0 |
| 0 | 0 | 0 | 0 |
| 0 | 0 | 0 | 0 |
| 0 | 0 | 0 | 0 |
| 0 | 0 | 0 | 0 |
| 0 | 0 | 0 | 0 |
| 0 | 0 | 0 | 0 |
| 0 | 0 | 0 | 0 |
| 0 | 0 | 0 | 0 |
| 0 | 0 | 0 | 0 |
| 0 | 0 | 0 | 0 |
| 0 | 0 | 0 | 0 |
| 0 | 0 | 0 | 0 |
| 0 | 0 | 0 | 0 |
| 0 | 0 | 0 | 0 |
| 0 | 0 | 0 | 0 |
| 0 | 0 | 0 | 0 |
| 0 | 0 | 0 | 0 |

| | | | |
|---|---|---|---|
| 0 | 0 | 0 | 0 |
| 0 | 0 | 0 | 0 |
| 0 | 0 | 0 | 0 |
| 0 | 0 | 0 | 0 |
| 0 | 0 | 0 | 0 |
| 0 | 0 | 0 | 0 |
| 0 | 0 | 0 | 0 |
| 0 | 0 | 0 | 0 |
| 0 | 0 | 0 | 0 |
| 0 | 0 | 0 | 0 |
| 0 | 0 | 0 | 0 |
| 0 | 0 | 0 | 0 |
| 1 | 0 | 0 | 0 |
| 1 | 1 | 0 | 0 |
| 0 | 0 | 0 | 0 |
| 1 | 0 | 0 | 0 |
| 0 | 0 | 0 | 0 |
| 1 | 0 | 0 | 0 |
| 0 | 0 | 0 | 0 |
| 0 | 0 | 0 | 0 |
| 1 | 0 | 0 | 0 |
| 0 | 0 | 0 | 0 |
| 0 | 0 | 0 | 0 |
| 1 | 0 | 0 | 0 |
| 1 | 0 | 0 | 0 |
| 0 | 0 | 0 | 0 |
| 0 | 0 | 0 | 0 |
| 1 | 0 | 0 | 0 |
| 1 | 0 | 0 | 0 |
| 1 | 0 | 0 | 0 |
| 1 | 0 | 0 | 0 |
| 1 | 0 | 0 | 0 |
| 0 | 0 | 0 | 0 |
| 0 | 0 | 0 | 0 |
| 0 | 0 | 0 | 0 |
| 0 | 0 | 0 | 0 |
| 0 | 0 | 0 | 0 |
| 0 | 0 | 0 | 0 |
| 0 | 0 | 0 | 0 |
| 0 | 0 | 0 | 0 |
| 0 | 0 | 0 | 0 |
| 0 | 0 | 0 | 0 |
| 0 | 0 | 0 | 0 |
| 0 | 0 | 0 | 0 |
| 1 | 1 | 0 | 0 |
| 0 | 0 | 0 | 0 |
| 0 | 0 | 0 | 0 |
| 0 | 0 | 0 | 0 |
| 0 | 0 | 0 | 0 |
| 1 | 1 | 0 | 0 |
| 1 | 0 | 0 | 0 |
| 0 | 0 | 0 | 0 |
| 0 | 0 | 0 | 0 |
| 0 | 0 | 0 | 1 |

| | | | |
|---|---|---|---|
| 0 | 0 | 0 | 0 |
| 0 | 0 | 0 | 0 |
| 0 | 0 | 0 | 0 |
| 0 | 0 | 0 | 0 |
| 1 | 0 | 0 | 1 |
| 0 | 0 | 0 | 0 |
| 1 | 0 | 0 | 0 |
| 0 | 0 | 0 | 0 |
| 0 | 0 | 0 | 0 |
| 0 | 0 | 0 | 0 |
| 0 | 0 | 0 | 0 |
| 0 | 0 | 0 | 0 |
| 0 | 0 | 0 | 0 |
| 1 | 0 | 0 | 0 |
| 0 | 0 | 0 | 0 |
| 0 | 0 | 0 | 0 |
| 1 | 1 | 0 | 0 |
| 0 | 0 | 0 | 0 |
| 1 | 0 | 0 | 0 |
| 1 | 0 | 0 | 0 |
| 0 | 0 | 0 | 0 |
| 0 | 0 | 0 | 0 |
| 0 | 0 | 0 | 0 |
| 1 | 0 | 0 | 0 |
| 1 | 0 | 0 | 0 |
| 1 | 0 | 0 | 0 |
| 1 | 0 | 0 | 0 |
| 1 | 0 | 0 | 0 |
| 1 | 0 | 0 | 0 |
| 1 | 0 | 0 | 0 |
| 1 | 0 | 0 | 0 |
| 1 | 0 | 0 | 0 |
| 1 | 0 | 0 | 0 |
| 1 | 0 | 0 | 0 |
| 0 | 0 | 0 | 0 |
| 0 | 0 | 0 | 0 |
| 0 | 0 | 0 | 0 |
| 1 | 0 | 0 | 0 |
| 1 | 0 | 0 | 0 |
| 0 | 0 | 0 | 0 |
| 0 | 0 | 0 | 0 |
| 1 | 0 | 0 | 0 |
| 0 | 0 | 0 | 0 |
| 1 | 0 | 0 | 0 |
| 1 | 0 | 0 | 0 |
| 1 | 0 | 0 | 0 |
| 1 | 0 | 0 | 0 |
| 1 | 0 | 0 | 0 |
| 0 | 0 | 0 | 0 |
| 0 | 0 | 0 | 0 |
| 1 | 0 | 0 | 0 |
| 0 | 0 | 0 | 0 |
| 1 | 0 | 0 | 0 |
| 1 | 0 | 0 | 0 |

| | | | |
|---|---|---|---|
| 1 | 0 | 0 | 0 |
| 1 | 0 | 0 | 0 |
| 1 | 0 | 0 | 0 |
| 1 | 0 | 0 | 0 |
| 1 | 0 | 0 | 0 |
| 1 | 0 | 0 | 0 |
| 0 | 0 | 0 | 0 |
| 1 | 0 | 0 | 0 |
| 1 | 0 | 0 | 0 |
| 1 | 0 | 0 | 0 |
| 1 | 0 | 0 | 0 |
| 1 | 0 | 0 | 0 |
| 1 | 0 | 0 | 0 |
| 1 | 0 | 0 | 0 |
| 0 | 0 | 0 | 0 |
| 0 | 0 | 0 | 0 |
| 1 | 1 | 0 | 0 |
| 1 | 1 | 0 | 0 |
| 0 | 0 | 0 | 0 |
| 0 | 0 | 0 | 0 |
| 0 | 0 | 0 | 0 |
| 1 | 1 | 0 | 1 |
| 0 | 0 | 0 | 0 |
| 0 | 0 | 0 | 0 |
| 0 | 0 | 0 | 0 |
| 0 | 0 | 0 | 0 |
| 0 | 0 | 0 | 0 |
| 0 | 0 | 0 | 0 |
| 0 | 0 | 0 | 0 |
| 1 | 0 | 1 | 0 |
| 0 | 0 | 0 | 0 |
| 0 | 0 | 0 | 0 |
| 0 | 0 | 0 | 0 |
| 0 | 0 | 0 | 0 |
| 0 | 0 | 0 | 0 |
| 0 | 0 | 0 | 0 |
| 0 | 0 | 0 | 0 |
| 0 | 0 | 0 | 0 |
| 0 | 0 | 0 | 0 |
| 0 | 0 | 0 | 0 |
| 1 | 1 | 0 | 0 |
| 1 | 1 | 0 | 0 |
| 1 | 0 | 0 | 0 |
| 1 | 1 | 0 | 0 |
| 1 | 0 | 0 | 0 |
| 1 | 1 | 0 | 0 |
| 1 | 0 | 0 | 0 |
| 1 | 0 | 0 | 0 |
| 0 | 0 | 0 | 0 |
| 0 | 0 | 0 | 0 |
| 1 | 0 | 0 | 0 |
| 1 | 0 | 0 | 0 |
| 1 | 0 | 0 | 0 |
| 0 | 0 | 0 | 0 |

| | | | |
|---|---|---|---|
| 0 | 0 | 0 | 0 |
| 0 | 0 | 0 | 0 |
| 0 | 0 | 0 | 0 |
| 1 | 1 | 1 | 0 |
| 0 | 0 | 0 | 0 |
| 0 | 0 | 0 | 0 |
| 0 | 0 | 0 | 0 |
| 0 | 0 | 0 | 0 |
| 0 | 1 | 0 | 0 |
| 0 | 1 | 0 | 0 |
| 0 | 1 | 0 | 0 |
| 0 | 0 | 0 | 0 |
| 0 | 0 | 0 | 0 |
| 0 | 0 | 0 | 0 |
| 1 | 0 | 0 | 0 |
| 1 | 0 | 0 | 0 |
| 0 | 0 | 0 | 0 |
| 0 | 0 | 0 | 0 |
| 1 | 0 | 0 | 0 |
| 0 | 0 | 0 | 0 |
| 1 | 0 | 0 | 0 |
| 1 | 0 | 0 | 0 |
| 0 | 0 | 0 | 0 |
| 0 | 0 | 0 | 0 |
| 0 | 0 | 0 | 0 |
| 0 | 0 | 0 | 0 |
| 0 | 0 | 0 | 0 |
| 0 | 0 | 1 | 0 |
| 0 | 0 | 0 | 0 |
| 0 | 1 | 0 | 0 |
| 0 | 0 | 0 | 0 |
| 0 | 0 | 0 | 0 |
| 0 | 0 | 0 | 0 |
| 0 | 0 | 0 | 0 |
| 0 | 0 | 0 | 0 |
| 0 | 0 | 0 | 0 |
| 1 | 0 | 0 | 0 |
| 1 | 0 | 0 | 0 |
| 1 | 0 | 0 | 0 |
| 0 | 0 | 0 | 0 |
| 0 | 0 | 0 | 0 |
| 0 | 0 | 0 | 0 |
| 0 | 0 | 0 | 0 |
| 0 | 0 | 0 | 0 |
| 0 | 0 | 0 | 0 |
| 0 | 0 | 0 | 0 |
| 0 | 0 | 0 | 0 |
| 0 | 0 | 0 | 1 |
| 0 | 0 | 0 | 0 |
| 0 | 0 | 0 | 0 |
| 0 | 0 | 0 | 0 |
| 0 | 0 | 0 | 0 |
| 1 | 1 | 0 | 0 |
| 0 | 0 | 0 | 0 |

| | | | |
|---|---|---|---|
| 0 | 0 | 0 | 0 |
| 0 | 0 | 0 | 0 |
| 0 | 0 | 0 | 0 |
| 0 | 0 | 0 | 0 |
| 1 | 0 | 0 | 0 |
| 0 | 0 | 0 | 0 |
| 0 | 0 | 0 | 0 |
| 0 | 0 | 0 | 0 |
| 0 | 0 | 0 | 0 |
| 0 | 0 | 0 | 0 |
| 0 | 0 | 0 | 0 |
| 0 | 0 | 0 | 0 |
| 0 | 0 | 0 | 0 |
| 0 | 0 | 0 | 0 |
| 0 | 0 | 0 | 0 |
| 0 | 0 | 0 | 0 |
| 0 | 0 | 0 | 0 |
| 0 | 0 | 0 | 0 |
| 0 | 0 | 0 | 0 |
| 0 | 0 | 0 | 0 |
| 0 | 0 | 0 | 0 |
| 0 | 0 | 0 | 0 |
| 0 | 0 | 0 | 0 |
| 0 | 0 | 0 | 0 |
| 0 | 0 | 0 | 0 |
| 0 | 0 | 0 | 0 |
| 0 | 0 | 0 | 0 |
| 0 | 0 | 0 | 0 |
| 0 | 0 | 0 | 0 |
| 0 | 0 | 0 | 0 |
| 0 | 0 | 0 | 0 |
| 0 | 0 | 0 | 0 |
| 0 | 0 | 0 | 0 |
| 0 | 0 | 0 | 0 |
| 0 | 0 | 0 | 0 |
| 0 | 0 | 0 | 0 |
| 0 | 0 | 0 | 0 |
| 0 | 0 | 0 | 0 |
| 0 | 0 | 0 | 0 |
| 0 | 0 | 0 | 0 |
| 1 | 1 | 0 | 0 |
| 0 | 0 | 0 | 0 |
| 0 | 0 | 0 | 0 |
| 0 | 0 | 0 | 0 |
| 0 | 0 | 0 | 0 |
| 1 | 0 | 0 | 0 |
| 1 | 0 | 0 | 0 |
| 0 | 0 | 0 | 0 |
| 1 | 0 | 0 | 0 |
| 0 | 0 | 0 | 0 |
| 0 | 0 | 0 | 0 |
| 1 | 0 | 0 | 0 |
| 1 | 0 | 0 | 0 |
| 0 | 0 | 0 | 0 |

| | | | |
|---|---|---|---|
| 0 | 0 | 0 | 0 |
| 0 | 0 | 0 | 0 |
| 0 | 0 | 0 | 0 |
| 0 | 0 | 0 | 0 |
| 1 | 0 | 0 | 0 |
| 0 | 0 | 0 | 0 |
| 0 | 0 | 0 | 0 |
| 0 | 0 | 0 | 0 |
| 0 | 0 | 0 | 0 |
| 0 | 0 | 0 | 0 |
| 0 | 0 | 0 | 0 |
| 0 | 0 | 0 | 0 |
| 0 | 0 | 0 | 0 |
| 0 | 0 | 0 | 0 |
| 0 | 0 | 0 | 0 |
| 1 | 1 | 0 | 1 |
| 0 | 0 | 0 | 0 |
| 0 | 0 | 0 | 0 |
| 0 | 0 | 0 | 0 |
| 0 | 0 | 0 | 0 |
| 0 | 0 | 0 | 0 |
| 0 | 0 | 0 | 0 |
| 0 | 0 | 0 | 0 |
| 0 | 0 | 0 | 0 |
| 0 | 0 | 0 | 0 |
| 0 | 0 | 0 | 0 |
| 0 | 0 | 0 | 0 |
| 1 | 0 | 0 | 0 |
| 1 | 0 | 0 | 0 |
| 0 | 0 | 0 | 0 |
| 1 | 1 | 0 | 0 |
| 0 | 0 | 0 | 0 |
| 0 | 0 | 0 | 0 |
| 0 | 0 | 0 | 0 |
| 1 | 1 | 0 | 0 |
| 1 | 1 | 0 | 0 |
| 0 | 0 | 0 | 0 |
| 1 | 0 | 0 | 0 |
| 0 | 0 | 0 | 0 |
| 1 | 1 | 0 | 0 |
| 0 | 1 | 0 | 0 |
| 0 | 0 | 0 | 0 |
| 0 | 0 | 0 | 0 |
| 0 | 0 | 0 | 0 |
| 1 | 1 | 0 | 0 |
| 1 | 1 | 0 | 0 |
| 1 | 1 | 0 | 0 |
| 0 | 0 | 0 | 0 |
| 0 | 0 | 0 | 0 |
| 0 | 0 | 0 | 0 |
| 0 | 0 | 0 | 0 |
| 0 | 0 | 0 | 0 |
| 0 | 0 | 0 | 0 |
| 0 | 0 | 0 | 0 |

| | | | |
|---|---|---|---|
| 0 | 0 | 0 | 0 |
| 0 | 0 | 0 | 0 |
| 1 | 0 | 0 | 0 |
| 1 | 0 | 0 | 0 |
| 0 | 0 | 0 | 0 |
| 1 | 0 | 0 | 0 |
| 0 | 0 | 0 | 0 |
| 0 | 0 | 0 | 0 |
| 0 | 0 | 0 | 0 |
| 0 | 1 | 0 | 0 |
| 0 | 0 | 0 | 0 |
| 0 | 0 | 0 | 0 |
| 0 | 0 | 0 | 0 |
| 0 | 0 | 0 | 0 |
| 0 | 0 | 0 | 0 |
| 0 | 0 | 0 | 0 |
| 0 | 0 | 0 | 0 |
| 1 | 0 | 0 | 0 |
| 0 | 0 | 0 | 0 |
| 1 | 0 | 0 | 0 |
| 0 | 0 | 0 | 0 |
| 1 | 0 | 0 | 0 |
| 0 | 0 | 0 | 0 |
| 1 | 0 | 0 | 0 |
| 0 | 0 | 0 | 0 |
| 0 | 0 | 0 | 0 |
| 0 | 0 | 0 | 0 |
| 0 | 0 | 0 | 0 |
| 0 | 0 | 0 | 0 |
| 0 | 0 | 0 | 0 |
| 0 | 0 | 0 | 0 |
| 1 | 0 | 0 | 0 |
| 0 | 0 | 0 | 0 |
| 0 | 0 | 0 | 0 |
| 0 | 0 | 0 | 0 |
| 0 | 0 | 0 | 0 |
| 0 | 0 | 0 | 0 |
| 0 | 0 | 0 | 0 |
| 1 | 1 | 0 | 0 |
| 1 | 1 | 0 | 0 |
| 1 | 1 | 0 | 0 |
| 1 | 1 | 0 | 0 |
| 1 | 1 | 0 | 0 |
| 1 | 0 | 0 | 0 |
| 1 | 0 | 0 | 0 |
| 1 | 0 | 0 | 0 |
| 0 | 0 | 0 | 0 |
| 0 | 0 | 0 | 0 |
| 0 | 0 | 0 | 0 |
| 0 | 0 | 0 | 0 |
| 0 | 0 | 0 | 0 |
| 0 | 0 | 0 | 0 |
| 0 | 0 | 0 | 0 |
| 0 | 0 | 0 | 0 |

| | | | |
|---|---|---|---|
| 0 | 0 | 0 | 0 |
| 0 | 0 | 0 | 0 |
| 0 | 0 | 0 | 0 |
| 0 | 0 | 0 | 0 |
| 0 | 0 | 0 | 0 |
| 1 | 0 | 0 | 0 |
| 1 | 0 | 0 | 0 |
| 0 | 0 | 0 | 0 |
| 0 | 0 | 0 | 0 |
| 0 | 0 | 0 | 0 |
| 0 | 0 | 0 | 0 |
| 1 | 0 | 0 | 0 |
| 0 | 0 | 1 | 0 |
| 0 | 0 | 0 | 0 |
| 0 | 0 | 0 | 0 |
| 0 | 0 | 0 | 0 |
| 1 | 0 | 0 | 0 |
| 0 | 0 | 0 | 0 |
| 1 | 0 | 0 | 0 |
| 1 | 0 | 0 | 0 |
| 0 | 0 | 0 | 0 |
| 1 | 0 | 0 | 0 |
| 1 | 1 | 0 | 0 |
| 0 | 0 | 0 | 0 |
| 1 | 1 | 0 | 0 |
| 1 | 1 | 0 | 0 |
| 1 | 1 | 0 | 0 |
| 0 | 0 | 0 | 0 |
| 1 | 0 | 0 | 0 |
| 0 | 0 | 0 | 0 |
| 0 | 0 | 0 | 0 |
| 0 | 0 | 0 | 0 |
| 0 | 0 | 0 | 0 |
| 0 | 0 | 0 | 0 |
| 0 | 0 | 0 | 0 |
| 0 | 0 | 0 | 0 |
| 0 | 0 | 0 | 0 |
| 0 | 0 | 0 | 0 |
| 0 | 0 | 0 | 0 |
| 0 | 0 | 0 | 0 |
| 0 | 0 | 0 | 0 |
| 0 | 0 | 0 | 0 |
| 0 | 0 | 0 | 0 |
| 0 | 0 | 0 | 0 |
| 0 | 0 | 0 | 0 |
| 0 | 0 | 0 | 0 |
| 0 | 0 | 0 | 0 |
| 0 | 0 | 0 | 0 |
| 0 | 0 | 0 | 0 |
| 0 | 0 | 0 | 0 |
| 0 | 0 | 0 | 0 |
| 0 | 0 | 0 | 0 |
| 0 | 0 | 0 | 0 |
| 0 | 0 | 0 | 0 |

| | | | |
|---|---|---|---|
| 0 | 0 | 0 | 0 |
| 0 | 0 | 0 | 0 |
| 0 | 0 | 0 | 0 |
| 0 | 0 | 0 | 0 |
| 0 | 0 | 0 | 0 |
| 0 | 0 | 0 | 0 |
| 0 | 0 | 0 | 0 |
| 0 | 0 | 0 | 0 |
| 0 | 0 | 0 | 0 |
| 0 | 0 | 0 | 0 |
| 0 | 1 | 0 | 0 |
| 0 | 1 | 0 | 0 |
| 0 | 1 | 0 | 0 |
| 0 | 1 | 0 | 0 |
| 0 | 0 | 0 | 0 |
| 0 | 1 | 0 | 0 |
| 0 | 1 | 0 | 0 |
| 1 | 1 | 0 | 0 |
| 0 | 1 | 0 | 0 |
| 1 | 0 | 0 | 0 |
| 1 | 1 | 0 | 0 |
| 1 | 1 | 0 | 0 |
| 0 | 0 | 1 | 1 |
| 0 | 0 | 0 | 0 |
| 1 | 0 | 0 | 0 |
| 0 | 0 | 0 | 0 |
| 0 | 0 | 0 | 0 |
| 0 | 0 | 0 | 0 |
| 0 | 0 | 0 | 0 |
| 0 | 0 | 0 | 0 |
| 0 | 0 | 0 | 0 |
| 0 | 0 | 0 | 0 |
| 0 | 0 | 0 | 0 |
| 0 | 0 | 0 | 0 |
| 0 | 0 | 0 | 0 |
| 0 | 0 | 0 | 0 |
| 0 | 0 | 0 | 0 |
| 0 | 0 | 0 | 0 |
| 0 | 0 | 0 | 0 |
| 0 | 0 | 0 | 0 |
| 1 | 0 | 0 | 0 |
| 0 | 0 | 0 | 0 |
| 1 | 0 | 0 | 0 |
| 1 | 0 | 0 | 0 |
| 0 | 0 | 0 | 0 |
| 1 | 0 | 0 | 0 |
| 0 | 0 | 0 | 0 |
| 0 | 0 | 0 | 0 |
| 0 | 0 | 0 | 0 |
| 0 | 0 | 0 | 0 |
| 0 | 0 | 0 | 0 |
| 0 | 0 | 0 | 0 |
| 1 | 1 | 0 | 0 |
| 1 | 1 | 0 | 0 |

| | | | |
|---|---|---|---|
| 1 | 1 | 0 | 0 |
| 0 | 0 | 0 | 0 |
| 0 | 0 | 0 | 0 |
| 0 | 0 | 0 | 0 |
| 0 | 0 | 0 | 0 |
| 0 | 0 | 0 | 0 |
| 0 | 1 | 0 | 0 |
| 0 | 0 | 0 | 0 |
| 0 | 0 | 0 | 0 |
| 0 | 1 | 0 | 0 |
| 1 | 0 | 0 | 0 |
| 0 | 0 | 0 | 0 |
| 0 | 0 | 1 | 0 |
| 0 | 0 | 1 | 0 |
| 0 | 0 | 0 | 0 |
| 0 | 0 | 0 | 0 |
| 0 | 0 | 0 | 0 |
| 0 | 1 | 0 | 0 |
| 0 | 0 | 0 | 0 |
| 0 | 0 | 0 | 0 |
| 1 | 0 | 0 | 0 |
| 0 | 0 | 0 | 1 |
| 0 | 0 | 0 | 0 |
| 0 | 0 | 0 | 0 |
| 0 | 0 | 0 | 0 |
| 0 | 0 | 0 | 0 |
| 0 | 0 | 0 | 0 |
| 0 | 0 | 0 | 0 |
| 0 | 0 | 0 | 0 |
| 0 | 0 | 0 | 0 |
| 0 | 0 | 0 | 0 |
| 0 | 0 | 0 | 0 |
| 0 | 0 | 0 | 0 |
| 0 | 0 | 0 | 0 |
| 0 | 0 | 0 | 0 |
| 0 | 0 | 0 | 0 |
| 1 | 0 | 0 | 0 |
| 0 | 0 | 0 | 0 |
| 1 | 0 | 0 | 0 |
| 0 | 0 | 0 | 0 |
| 0 | 0 | 0 | 0 |
| 0 | 0 | 0 | 0 |
| 0 | 0 | 0 | 0 |
| 0 | 0 | 0 | 0 |
| 0 | 0 | 0 | 0 |
| 0 | 0 | 0 | 0 |
| 0 | 0 | 0 | 0 |
| 0 | 0 | 0 | 0 |
| 0 | 0 | 0 | 0 |
| 1 | 0 | 0 | 0 |
| 0 | 0 | 0 | 0 |
| 0 | 0 | 0 | 0 |
| 0 | 0 | 0 | 0 |
| 0 | 0 | 0 | 0 |

| | | | |
|---|---|---|---|
| 0 | 0 | 0 | 0 |
| 0 | 0 | 0 | 0 |
| 0 | 0 | 0 | 0 |
| 0 | 0 | 0 | 0 |
| 0 | 0 | 0 | 0 |
| 0 | 0 | 0 | 0 |
| 0 | 0 | 0 | 0 |
| 0 | 0 | 0 | 0 |
| 0 | 0 | 0 | 0 |
| 0 | 0 | 0 | 0 |
| 0 | 0 | 0 | 0 |
| 0 | 0 | 0 | 0 |
| 0 | 0 | 1 | 0 |
| 1 | 0 | 0 | 0 |
| 1 | 0 | 0 | 0 |
| 0 | 0 | 0 | 0 |
| 0 | 0 | 0 | 0 |
| 0 | 0 | 0 | 0 |
| 0 | 0 | 0 | 0 |
| 0 | 0 | 0 | 0 |
| 0 | 0 | 0 | 0 |
| 0 | 0 | 0 | 0 |
| 0 | 0 | 0 | 0 |
| 1 | 1 | 0 | 1 |
| 0 | 0 | 0 | 0 |
| 1 | 1 | 0 | 0 |
| 1 | 1 | 0 | 1 |
| 0 | 0 | 0 | 0 |
| 0 | 0 | 0 | 0 |
| 0 | 0 | 0 | 0 |
| 0 | 0 | 0 | 0 |
| 0 | 0 | 0 | 0 |
| 0 | 0 | 0 | 0 |
| 0 | 0 | 0 | 0 |
| 0 | 0 | 0 | 0 |
| 1 | 0 | 0 | 0 |
| 0 | 0 | 0 | 0 |
| 0 | 0 | 0 | 0 |
| 0 | 0 | 0 | 0 |
| 0 | 0 | 0 | 0 |
| 0 | 0 | 0 | 0 |
| 0 | 0 | 0 | 0 |
| 1 | 0 | 0 | 0 |
| 1 | 0 | 0 | 0 |
| 1 | 0 | 0 | 0 |
| 1 | 0 | 0 | 0 |
| 1 | 0 | 0 | 0 |
| 0 | 0 | 0 | 0 |
| 0 | 0 | 0 | 0 |
| 1 | 0 | 0 | 0 |
| 1 | 0 | 0 | 0 |
| 0 | 0 | 0 | 0 |
| 1 | 0 | 0 | 0 |
| 1 | 0 | 0 | 0 |

| | | | |
|---|---|---|---|
| 0 | 0 | 0 | 0 |
| 1 | 0 | 0 | 0 |
| 0 | 0 | 0 | 0 |
| 0 | 0 | 0 | 0 |
| 0 | 0 | 0 | 0 |
| 0 | 0 | 0 | 0 |
| 1 | 0 | 0 | 0 |
| 0 | 0 | 0 | 0 |
| 0 | 0 | 0 | 0 |
| 0 | 0 | 0 | 0 |
| 0 | 0 | 0 | 0 |
| 1 | 1 | 0 | 0 |
| 0 | 0 | 0 | 0 |
| 0 | 0 | 0 | 0 |
| 0 | 0 | 0 | 0 |
| 0 | 0 | 0 | 0 |
| 0 | 0 | 0 | 0 |
| 0 | 0 | 0 | 0 |
| 0 | 0 | 0 | 0 |
| 0 | 0 | 0 | 0 |
| 0 | 0 | 0 | 0 |
| 0 | 0 | 0 | 0 |
| 0 | 0 | 0 | 0 |
| 0 | 0 | 0 | 0 |
| 0 | 0 | 0 | 0 |
| 0 | 0 | 0 | 0 |
| 0 | 0 | 0 | 0 |
| 0 | 0 | 0 | 0 |
| 1 | 1 | 0 | 0 |
| 1 | 0 | 0 | 0 |
| 0 | 0 | 0 | 0 |
| 0 | 0 | 0 | 0 |
| 0 | 0 | 0 | 0 |
| 0 | 0 | 0 | 0 |
| 0 | 0 | 0 | 0 |
| 0 | 0 | 0 | 0 |
| 0 | 0 | 0 | 0 |
| 0 | 0 | 0 | 0 |
| 0 | 0 | 0 | 0 |
| 0 | 0 | 0 | 0 |
| 0 | 0 | 0 | 0 |
| 0 | 0 | 0 | 0 |
| 0 | 0 | 0 | 0 |
| 0 | 0 | 0 | 0 |
| 0 | 0 | 0 | 0 |
| 0 | 0 | 0 | 0 |
| 0 | 0 | 0 | 0 |
| 0 | 0 | 0 | 0 |
| 0 | 0 | 0 | 0 |
| 0 | 0 | 0 | 0 |
| 0 | 0 | 0 | 0 |
| 0 | 0 | 0 | 0 |
| 0 | 0 | 1 | 0 |
| 0 | 0 | 0 | 0 |

| | | | |
|---|---|---|---|
| 0 | 0 | 0 | 0 |
| 0 | 0 | 0 | 0 |
| 0 | 0 | 0 | 0 |
| 0 | 0 | 0 | 0 |
| 1 | 0 | 0 | 0 |
| 0 | 0 | 0 | 0 |
| 0 | 0 | 0 | 0 |
| 1 | 1 | 0 | 0 |
| 0 | 0 | 0 | 0 |
| 1 | 1 | 0 | 0 |
| 0 | 0 | 0 | 0 |
| 0 | 0 | 0 | 0 |
| 0 | 0 | 0 | 0 |
| 0 | 0 | 0 | 0 |
| 0 | 0 | 0 | 0 |
| 0 | 0 | 0 | 0 |
| 1 | 0 | 0 | 0 |
| 0 | 0 | 0 | 0 |
| 0 | 0 | 0 | 0 |
| 1 | 0 | 0 | 0 |
| 1 | 0 | 0 | 0 |
| 0 | 0 | 0 | 0 |
| 0 | 0 | 0 | 0 |
| 0 | 0 | 0 | 0 |
| 0 | 0 | 0 | 0 |
| 0 | 0 | 0 | 0 |
| 1 | 0 | 0 | 0 |
| 0 | 0 | 0 | 0 |
| 0 | 0 | 0 | 0 |
| 1 | 0 | 0 | 0 |
| 0 | 0 | 0 | 0 |
| 0 | 0 | 0 | 0 |
| 0 | 0 | 0 | 0 |
| 1 | 1 | 0 | 0 |
| 0 | 0 | 0 | 0 |
| 0 | 0 | 0 | 0 |
| 0 | 0 | 0 | 0 |
| 0 | 0 | 0 | 0 |
| 0 | 1 | 0 | 0 |
| 0 | 0 | 0 | 0 |
| 0 | 0 | 0 | 0 |
| 0 | 0 | 0 | 0 |
| 0 | 0 | 0 | 0 |
| 1 | 1 | 0 | 0 |
| 0 | 0 | 0 | 0 |
| 0 | 0 | 0 | 0 |
| 0 | 0 | 0 | 0 |
| 0 | 0 | 0 | 0 |
| 0 | 0 | 0 | 0 |
| 0 | 0 | 0 | 0 |
| 0 | 0 | 0 | 0 |
| 1 | 0 | 0 | 0 |
| 0 | 0 | 0 | 0 |
| 0 | 0 | 0 | 0 |

| | | | |
|---|---|---|---|
| 1 | 0 | 0 | 1 |
| 1 | 0 | 0 | 1 |
| 1 | 0 | 0 | 1 |
| 0 | 0 | 0 | 0 |
| 0 | 0 | 0 | 0 |
| 0 | 0 | 0 | 0 |
| 1 | 0 | 0 | 0 |
| 0 | 0 | 0 | 0 |
| 0 | 0 | 0 | 0 |
| 0 | 0 | 0 | 0 |
| 0 | 0 | 1 | 0 |
| 0 | 0 | 0 | 0 |
| 0 | 0 | 0 | 0 |
| 0 | 0 | 0 | 0 |
| 0 | 0 | 0 | 0 |
| 0 | 0 | 0 | 0 |
| 1 | 0 | 0 | 0 |
| 0 | 0 | 0 | 0 |
| 0 | 0 | 0 | 0 |
| 0 | 0 | 0 | 0 |
| 0 | 0 | 0 | 0 |
| 0 | 0 | 0 | 0 |
| 0 | 0 | 0 | 0 |
| 1 | 1 | 0 | 0 |
| 0 | 0 | 0 | 0 |
| 0 | 0 | 0 | 0 |
| 0 | 0 | 0 | 0 |
| 0 | 0 | 0 | 0 |
| 0 | 0 | 0 | 0 |
| 0 | 0 | 0 | 0 |
| 0 | 0 | 0 | 0 |
| 0 | 0 | 0 | 0 |
| 0 | 1 | 0 | 0 |
| 0 | 0 | 0 | 0 |
| 1 | 1 | 0 | 0 |
| 0 | 0 | 0 | 0 |
| 0 | 0 | 0 | 0 |
| 0 | 0 | 0 | 0 |
| 0 | 0 | 0 | 0 |
| 0 | 0 | 0 | 0 |
| 0 | 0 | 0 | 0 |
| 1 | 0 | 0 | 1 |
| 0 | 0 | 0 | 0 |
| 1 | 0 | 0 | 1 |
| 0 | 0 | 0 | 0 |
| 1 | 0 | 0 | 1 |
| 0 | 0 | 0 | 0 |
| 0 | 0 | 0 | 0 |
| 0 | 0 | 0 | 0 |
| 1 | 0 | 0 | 0 |
| 0 | 0 | 0 | 0 |
| 0 | 0 | 0 | 0 |
| 0 | 0 | 0 | 0 |
| 0 | 0 | 0 | 0 |

| | | | |
|---|---|---|---|
| 0 | 0 | 0 | 0 |
| 0 | 0 | 0 | 0 |
| 0 | 0 | 0 | 0 |
| 0 | 0 | 0 | 0 |
| 0 | 0 | 0 | 0 |
| 0 | 0 | 0 | 0 |
| 0 | 0 | 0 | 0 |
| 0 | 0 | 0 | 0 |
| 0 | 0 | 0 | 0 |
| 0 | 0 | 0 | 0 |
| 0 | 0 | 0 | 0 |
| 0 | 0 | 0 | 0 |
| 0 | 0 | 0 | 0 |
| 0 | 0 | 0 | 0 |
| 0 | 0 | 0 | 0 |
| 0 | 0 | 0 | 0 |
| 0 | 0 | 0 | 0 |
| 0 | 0 | 0 | 0 |
| 0 | 0 | 1 | 0 |
| 0 | 0 | 0 | 0 |
| 0 | 0 | 0 | 0 |
| 0 | 0 | 0 | 0 |
| 0 | 0 | 0 | 0 |
| 0 | 0 | 0 | 0 |
| 0 | 0 | 0 | 0 |
| 0 | 0 | 0 | 0 |
| 0 | 0 | 0 | 0 |
| 0 | 0 | 0 | 0 |
| 0 | 0 | 1 | 0 |
| 0 | 0 | 0 | 0 |
| 0 | 0 | 1 | 0 |
| 1 | 0 | 0 | 0 |
| 1 | 1 | 1 | 0 |
| 1 | 1 | 1 | 0 |
| 0 | 0 | 0 | 0 |
| 0 | 0 | 0 | 0 |
| 0 | 1 | 0 | 0 |
| 1 | 1 | 1 | 0 |
| 1 | 1 | 1 | 0 |
| 0 | 0 | 1 | 0 |
| 0 | 0 | 1 | 0 |
| 1 | 0 | 0 | 0 |
| 0 | 0 | 0 | 0 |
| 0 | 0 | 0 | 0 |
| 1 | 1 | 0 | 0 |
| 0 | 0 | 0 | 0 |
| 0 | 0 | 0 | 0 |
| 0 | 0 | 0 | 0 |
| 0 | 0 | 0 | 0 |
| 0 | 0 | 0 | 0 |
| 0 | 0 | 0 | 0 |
| 0 | 0 | 0 | 0 |
| 0 | 0 | 0 | 0 |
| 0 | 0 | 0 | 0 |

| | | | |
|---|---|---|---|
| 0 | 0 | 0 | 0 |
| 0 | 0 | 0 | 0 |
| 0 | 0 | 0 | 0 |
| 0 | 0 | 0 | 0 |
| 0 | 0 | 0 | 0 |
| 0 | 0 | 0 | 0 |
| 0 | 0 | 0 | 0 |
| 0 | 0 | 0 | 0 |
| 0 | 0 | 0 | 0 |
| 0 | 0 | 0 | 0 |
| 0 | 0 | 0 | 0 |
| 0 | 0 | 0 | 0 |
| 0 | 0 | 0 | 0 |
| 0 | 0 | 0 | 0 |
| 0 | 0 | 0 | 0 |
| 0 | 0 | 0 | 0 |
| 0 | 0 | 0 | 0 |
| 0 | 0 | 0 | 0 |
| 0 | 0 | 0 | 0 |
| 1 | 0 | 0 | 0 |
| 1 | 0 | 0 | 0 |
| 0 | 0 | 0 | 0 |
| 0 | 0 | 0 | 0 |
| 0 | 0 | 0 | 0 |
| 0 | 0 | 0 | 0 |
| 1 | 0 | 0 | 0 |
| 0 | 0 | 0 | 0 |
| 0 | 0 | 0 | 0 |
| 0 | 0 | 0 | 0 |
| 1 | 0 | 0 | 0 |
| 0 | 0 | 0 | 0 |
| 0 | 0 | 0 | 0 |
| 1 | 0 | 0 | 0 |
| 1 | 0 | 0 | 0 |
| 0 | 0 | 0 | 0 |
| 0 | 0 | 0 | 0 |
| 0 | 0 | 0 | 0 |
| 0 | 0 | 0 | 0 |
| 0 | 0 | 0 | 0 |
| 0 | 0 | 0 | 0 |
| 0 | 0 | 0 | 0 |
| 0 | 0 | 0 | 0 |
| 0 | 0 | 0 | 0 |
| 0 | 0 | 0 | 0 |
| 0 | 0 | 0 | 0 |
| 0 | 0 | 0 | 0 |
| 0 | 0 | 0 | 0 |
| 0 | 1 | 0 | 0 |
| 0 | 1 | 0 | 0 |
| 0 | 0 | 0 | 0 |
| 0 | 1 | 0 | 0 |
| 0 | 1 | 0 | 0 |
| 0 | 0 | 0 | 0 |
| 0 | 0 | 0 | 0 |

| | | | |
|---|---|---|---|
| 0 | 0 | 0 | 0 |
| 0 | 0 | 0 | 0 |
| 1 | 1 | 0 | 1 |
| 0 | 0 | 0 | 0 |
| 0 | 0 | 0 | 0 |
| 0 | 0 | 0 | 0 |
| 0 | 0 | 0 | 0 |
| 0 | 1 | 0 | 0 |
| 1 | 1 | 0 | 0 |
| 1 | 1 | 0 | 0 |
| 1 | 1 | 0 | 0 |
| 1 | 1 | 0 | 0 |
| 1 | 1 | 0 | 0 |
| 0 | 1 | 0 | 0 |
| 0 | 1 | 0 | 0 |
| 0 | 0 | 0 | 0 |
| 0 | 0 | 0 | 0 |
| 0 | 0 | 0 | 0 |
| 0 | 0 | 0 | 0 |
| 0 | 0 | 0 | 0 |
| 0 | 0 | 0 | 0 |
| 0 | 0 | 0 | 0 |
| 0 | 1 | 0 | 0 |
| 0 | 1 | 0 | 0 |
| 0 | 0 | 0 | 0 |
| 0 | 0 | 1 | 0 |
| 0 | 0 | 0 | 0 |
| 0 | 0 | 0 | 0 |
| 0 | 0 | 0 | 0 |
| 0 | 1 | 0 | 0 |
| 0 | 0 | 0 | 0 |
| 0 | 0 | 0 | 0 |
| 0 | 0 | 0 | 0 |
| 0 | 0 | 0 | 0 |
| 0 | 0 | 0 | 0 |
| 0 | 0 | 0 | 0 |
| 0 | 0 | 0 | 0 |
| 0 | 0 | 0 | 0 |
| 0 | 0 | 0 | 0 |
| 1 | 0 | 0 | 0 |
| 1 | 1 | 0 | 0 |
| 1 | 0 | 0 | 0 |
| 0 | 0 | 0 | 0 |
| 0 | 1 | 0 | 0 |
| 1 | 1 | 0 | 0 |
| 0 | 1 | 0 | 0 |
| 0 | 0 | 0 | 0 |
| 1 | 0 | 0 | 0 |
| 0 | 1 | 0 | 0 |
| 0 | 0 | 0 | 0 |
| 0 | 0 | 0 | 0 |
| 0 | 0 | 0 | 0 |
| 1 | 1 | 0 | 0 |
| 0 | 0 | 0 | 0 |

| | | | |
|---|---|---|---|
| 0 | 0 | 0 | 0 |
| 0 | 0 | 0 | 0 |
| 1 | 0 | 0 | 0 |
| 1 | 1 | 0 | 0 |
| 1 | 1 | 0 | 0 |
| 0 | 0 | 0 | 0 |
| 0 | 0 | 0 | 0 |
| 0 | 0 | 0 | 0 |
| 1 | 0 | 0 | 0 |
| 0 | 0 | 0 | 0 |
| 0 | 0 | 0 | 0 |
| 1 | 0 | 0 | 1 |
| 0 | 0 | 0 | 0 |
| 1 | 0 | 0 | 0 |
| 0 | 0 | 0 | 0 |
| 0 | 0 | 0 | 0 |
| 0 | 0 | 0 | 0 |
| 0 | 0 | 0 | 0 |
| 0 | 0 | 0 | 0 |
| 0 | 0 | 0 | 0 |
| 0 | 0 | 0 | 0 |
| 1 | 0 | 0 | 1 |
| 1 | 0 | 0 | 0 |
| 0 | 0 | 0 | 0 |
| 0 | 0 | 0 | 0 |
| 0 | 0 | 0 | 0 |
| 0 | 0 | 0 | 0 |
| 0 | 0 | 0 | 0 |
| 1 | 0 | 0 | 0 |
| 1 | 0 | 0 | 0 |
| 0 | 0 | 0 | 0 |
| 0 | 0 | 0 | 0 |
| 0 | 0 | 0 | 0 |
| 0 | 0 | 0 | 0 |
| 0 | 0 | 0 | 0 |
| 0 | 0 | 0 | 0 |
| 1 | 0 | 0 | 0 |
| 0 | 0 | 0 | 0 |
| 0 | 0 | 0 | 0 |
| 1 | 0 | 0 | 0 |
| 1 | 0 | 0 | 0 |
| 1 | 0 | 0 | 0 |
| 1 | 0 | 0 | 0 |
| 0 | 0 | 0 | 0 |
| 0 | 0 | 0 | 0 |
| 1 | 0 | 0 | 0 |
| 0 | 0 | 0 | 0 |
| 0 | 0 | 0 | 0 |
| 0 | 0 | 0 | 0 |
| 0 | 0 | 0 | 0 |
| 0 | 0 | 0 | 0 |
| 1 | 0 | 0 | 0 |
| 1 | 0 | 0 | 0 |
| 1 | 0 | 0 | 0 |

| | | | |
|---|---|---|---|
| 1 | 0 | 0 | 0 |
| 0 | 0 | 0 | 0 |
| 0 | 0 | 0 | 0 |
| 0 | 0 | 0 | 0 |
| 1 | 0 | 0 | 0 |
| 0 | 0 | 0 | 0 |
| 1 | 0 | 0 | 0 |
| 0 | 0 | 0 | 0 |
| 0 | 0 | 0 | 0 |
| 0 | 0 | 0 | 0 |
| 0 | 0 | 0 | 0 |
| 0 | 0 | 0 | 0 |
| 0 | 0 | 0 | 0 |
| 0 | 0 | 0 | 0 |
| 0 | 0 | 0 | 0 |
| 1 | 0 | 0 | 0 |
| 0 | 0 | 0 | 0 |
| 0 | 0 | 0 | 0 |
| 0 | 0 | 0 | 0 |
| 0 | 0 | 0 | 0 |
| 0 | 0 | 0 | 0 |
| 0 | 0 | 0 | 0 |
| 1 | 0 | 0 | 0 |
| 1 | 0 | 0 | 0 |
| 0 | 0 | 0 | 0 |
| 0 | 0 | 0 | 0 |
| 0 | 0 | 0 | 0 |
| 0 | 0 | 0 | 0 |
| 0 | 0 | 0 | 0 |
| 0 | 0 | 0 | 0 |
| 0 | 0 | 0 | 0 |
| 0 | 0 | 0 | 0 |
| 1 | 0 | 0 | 0 |
| 0 | 0 | 0 | 0 |
| 0 | 0 | 0 | 0 |
| 0 | 0 | 0 | 0 |
| 0 | 0 | 0 | 0 |
| 0 | 0 | 0 | 0 |
| 0 | 0 | 0 | 0 |
| 0 | 0 | 0 | 0 |
| 0 | 0 | 0 | 0 |
| 0 | 0 | 0 | 0 |
| 0 | 0 | 0 | 0 |
| 0 | 0 | 0 | 0 |
| 0 | 1 | 0 | 0 |
| 0 | 0 | 0 | 0 |
| 0 | 0 | 0 | 0 |
| 0 | 0 | 0 | 0 |
| 0 | 0 | 0 | 0 |
| 0 | 0 | 0 | 0 |
| 0 | 0 | 0 | 0 |
| 0 | 0 | 0 | 0 |
| 0 | 0 | 0 | 0 |
| 1 | 0 | 0 | 0 |

| | | | |
|---|---|---|---|
| 0 | 0 | 0 | 0 |
| 0 | 0 | 0 | 0 |
| 0 | 0 | 0 | 0 |
| 0 | 0 | 0 | 0 |
| 0 | 0 | 0 | 0 |
| 0 | 0 | 0 | 0 |
| 0 | 0 | 0 | 0 |
| 0 | 0 | 0 | 0 |
| 0 | 0 | 0 | 0 |
| 1 | 0 | 0 | 0 |
| 0 | 0 | 0 | 0 |
| 0 | 0 | 0 | 0 |
| 1 | 1 | 0 | 0 |
| 1 | 1 | 0 | 0 |
| 1 | 1 | 0 | 0 |
| 1 | 1 | 0 | 0 |
| 0 | 0 | 0 | 0 |
| 0 | 0 | 0 | 0 |
| 0 | 0 | 0 | 0 |
| 0 | 0 | 0 | 0 |
| 0 | 0 | 0 | 0 |
| 0 | 0 | 0 | 0 |
| 0 | 0 | 0 | 0 |
| 0 | 0 | 0 | 0 |
| 0 | 0 | 0 | 0 |
| 0 | 0 | 0 | 1 |
| 0 | 0 | 0 | 0 |
| 0 | 0 | 0 | 0 |
| 0 | 0 | 0 | 0 |
| 0 | 0 | 0 | 0 |
| 0 | 0 | 0 | 0 |
| 0 | 0 | 0 | 0 |
| 0 | 0 | 0 | 0 |
| 0 | 0 | 0 | 0 |
| 0 | 0 | 0 | 0 |
| 0 | 0 | 0 | 0 |
| 0 | 0 | 0 | 0 |
| 1 | 1 | 0 | 0 |
| 0 | 0 | 0 | 0 |
| 0 | 0 | 0 | 0 |
| 1 | 1 | 0 | 0 |
| 0 | 0 | 0 | 0 |
| 0 | 0 | 0 | 0 |
| 1 | 0 | 0 | 1 |
| 0 | 0 | 0 | 0 |
| 0 | 0 | 0 | 0 |
| 0 | 0 | 0 | 0 |
| 0 | 0 | 0 | 0 |
| 0 | 0 | 0 | 0 |
| 0 | 0 | 0 | 0 |
| 0 | 0 | 0 | 0 |
| 0 | 0 | 1 | 0 |
| 0 | 0 | 1 | 0 |
| 0 | 0 | 1 | 0 |

| | | | |
|---|---|---|---|
| 0 | 0 | 1 | 0 |
| 0 | 0 | 0 | 0 |
| 0 | 0 | 0 | 0 |
| 0 | 0 | 0 | 0 |
| 0 | 0 | 0 | 0 |
| 0 | 0 | 0 | 0 |
| 0 | 0 | 0 | 0 |
| 0 | 0 | 0 | 0 |
| 0 | 0 | 0 | 0 |
| 0 | 0 | 0 | 0 |
| 0 | 0 | 0 | 0 |
| 0 | 0 | 0 | 0 |
| 0 | 0 | 0 | 0 |
| 0 | 0 | 0 | 0 |
| 0 | 0 | 0 | 0 |
| 0 | 0 | 0 | 0 |
| 0 | 0 | 0 | 0 |
| 0 | 0 | 0 | 0 |
| 0 | 0 | 0 | 0 |
| 0 | 0 | 0 | 0 |
| 0 | 0 | 0 | 0 |
| 0 | 0 | 0 | 0 |
| 0 | 0 | 0 | 0 |
| 0 | 1 | 0 | 1 |
| 0 | 0 | 0 | 0 |
| 0 | 0 | 0 | 0 |
| 0 | 0 | 0 | 0 |
| 0 | 0 | 0 | 0 |
| 0 | 0 | 0 | 0 |
| 0 | 0 | 0 | 0 |
| 0 | 0 | 0 | 0 |
| 0 | 0 | 0 | 0 |
| 0 | 0 | 0 | 0 |
| 0 | 0 | 0 | 0 |
| 0 | 0 | 0 | 0 |
| 0 | 0 | 0 | 0 |
| 0 | 0 | 0 | 0 |
| 0 | 0 | 0 | 0 |
| 0 | 0 | 0 | 0 |
| 0 | 0 | 0 | 0 |
| 0 | 0 | 0 | 0 |
| 0 | 0 | 0 | 0 |
| 0 | 0 | 0 | 0 |
| 0 | 0 | 0 | 0 |
| 0 | 0 | 0 | 0 |
| 0 | 0 | 0 | 0 |
| 1 | 0 | 0 | 0 |
| 1 | 0 | 0 | 0 |
| 0 | 0 | 0 | 0 |
| 0 | 1 | 0 | 0 |
| 1 | 1 | 0 | 1 |
| 1 | 1 | 0 | 1 |
| 1 | 1 | 0 | 1 |
| 0 | 0 | 0 | 0 |

| | | | |
|---|---|---|---|
| 1 | 1 | 0 | 0 |
| 0 | 0 | 0 | 0 |
| 0 | 0 | 0 | 0 |
| 0 | 0 | 0 | 0 |
| 0 | 0 | 0 | 0 |
| 0 | 0 | 0 | 0 |
| 0 | 0 | 0 | 0 |
| 0 | 0 | 0 | 0 |
| 0 | 0 | 0 | 0 |
| 0 | 0 | 0 | 0 |
| 0 | 0 | 0 | 0 |
| 0 | 1 | 0 | 0 |
| 0 | 0 | 0 | 0 |
| 0 | 0 | 0 | 0 |
| 0 | 1 | 0 | 0 |
| 0 | 0 | 0 | 0 |
| 1 | 1 | 0 | 0 |
| 0 | 0 | 0 | 0 |
| 0 | 0 | 0 | 0 |
| 1 | 0 | 0 | 0 |
| 0 | 0 | 0 | 0 |
| 0 | 0 | 0 | 0 |
| 0 | 0 | 0 | 0 |
| 0 | 0 | 0 | 0 |
| 0 | 0 | 0 | 0 |
| 0 | 0 | 0 | 0 |
| 0 | 0 | 0 | 0 |
| 0 | 0 | 0 | 0 |
| 0 | 0 | 0 | 0 |
| 0 | 0 | 0 | 0 |
| 0 | 0 | 0 | 0 |
| 0 | 0 | 0 | 0 |
| 0 | 0 | 0 | 0 |
| 1 | 0 | 0 | 0 |
| 0 | 0 | 0 | 0 |
| 0 | 1 | 0 | 0 |
| 0 | 0 | 0 | 0 |
| 0 | 0 | 0 | 0 |
| 0 | 0 | 0 | 0 |
| 0 | 0 | 0 | 0 |
| 0 | 0 | 0 | 0 |
| 0 | 0 | 0 | 0 |
| 0 | 0 | 0 | 0 |
| 0 | 0 | 0 | 0 |
| 0 | 0 | 0 | 0 |
| 0 | 0 | 0 | 0 |
| 0 | 0 | 0 | 0 |
| 0 | 0 | 0 | 0 |
| 1 | 1 | 0 | 0 |
| 0 | 0 | 0 | 0 |
| 0 | 0 | 0 | 0 |
| 0 | 0 | 0 | 0 |
| 0 | 0 | 0 | 0 |
| 0 | 0 | 0 | 0 |

| | | | |
|---|---|---|---|
| 0 | 0 | 0 | 0 |
| 0 | 0 | 0 | 0 |
| 0 | 0 | 0 | 0 |
| 0 | 0 | 0 | 0 |
| 0 | 0 | 0 | 0 |
| 0 | 0 | 0 | 0 |
| 0 | 0 | 0 | 0 |
| 0 | 1 | 0 | 0 |
| 0 | 0 | 0 | 0 |
| 0 | 0 | 0 | 0 |
| 0 | 0 | 0 | 0 |
| 0 | 0 | 0 | 0 |
| 0 | 0 | 1 | 0 |
| 0 | 0 | 0 | 0 |
| 0 | 0 | 1 | 0 |
| 0 | 0 | 1 | 0 |
| 0 | 0 | 0 | 0 |
| 0 | 0 | 0 | 0 |
| 0 | 0 | 0 | 0 |
| 0 | 0 | 0 | 0 |
| 0 | 1 | 0 | 0 |
| 0 | 1 | 0 | 0 |
| 0 | 0 | 0 | 0 |
| 0 | 0 | 0 | 0 |
| 0 | 1 | 0 | 0 |
| 0 | 0 | 0 | 0 |
| 0 | 0 | 0 | 0 |
| 0 | 1 | 0 | 0 |
| 0 | 0 | 0 | 0 |
| 0 | 0 | 0 | 0 |
| 0 | 0 | 0 | 0 |
| 0 | 0 | 0 | 0 |
| 1 | 1 | 0 | 0 |
| 1 | 1 | 0 | 0 |
| 1 | 1 | 0 | 0 |
| 0 | 1 | 0 | 0 |
| 1 | 0 | 0 | 0 |
| 0 | 1 | 0 | 0 |
| 1 | 1 | 0 | 0 |
| 0 | 0 | 0 | 0 |
| 0 | 0 | 0 | 0 |
| 0 | 0 | 0 | 0 |
| 0 | 0 | 0 | 0 |
| 0 | 0 | 0 | 0 |
| 0 | 0 | 0 | 0 |
| 0 | 0 | 0 | 0 |
| 0 | 0 | 0 | 0 |
| 0 | 0 | 0 | 0 |
| 0 | 0 | 0 | 0 |
| 1 | 1 | 0 | 0 |
| 1 | 1 | 0 | 0 |
| 0 | 0 | 0 | 0 |
| 0 | 0 | 0 | 0 |
| 0 | 0 | 0 | 0 |

| | | | |
|---|---|---|---|
| 0 | 0 | 0 | 0 |
| 0 | 0 | 0 | 0 |
| 0 | 0 | 0 | 0 |
| 0 | 0 | 0 | 0 |
| 0 | 0 | 0 | 0 |
| 0 | 0 | 0 | 0 |
| 1 | 1 | 0 | 0 |
| 1 | 1 | 0 | 0 |
| 0 | 0 | 0 | 0 |
| 0 | 0 | 0 | 0 |
| 0 | 0 | 0 | 0 |
| 0 | 0 | 0 | 0 |
| 0 | 0 | 0 | 0 |
| 0 | 0 | 0 | 0 |
| 0 | 0 | 0 | 0 |
| 0 | 0 | 0 | 0 |
| 0 | 0 | 0 | 0 |
| 0 | 0 | 0 | 0 |
| 0 | 0 | 0 | 0 |
| 0 | 0 | 0 | 0 |
| 0 | 0 | 0 | 0 |
| 0 | 0 | 0 | 0 |
| 0 | 0 | 0 | 0 |
| 0 | 0 | 0 | 0 |
| 0 | 0 | 0 | 0 |
| 0 | 0 | 0 | 0 |
| 0 | 0 | 0 | 0 |
| 1 | 1 | 0 | 0 |
| 0 | 0 | 0 | 0 |
| 0 | 0 | 0 | 0 |
| 0 | 0 | 0 | 0 |
| 0 | 0 | 0 | 0 |
| 0 | 0 | 0 | 0 |
| 0 | 0 | 0 | 0 |
| 0 | 0 | 0 | 0 |
| 0 | 0 | 0 | 0 |
| 0 | 0 | 0 | 0 |
| 0 | 0 | 0 | 0 |
| 0 | 0 | 0 | 0 |
| 0 | 0 | 0 | 0 |
| 0 | 0 | 0 | 0 |
| 0 | 0 | 0 | 0 |
| 0 | 0 | 0 | 0 |
| 0 | 0 | 0 | 0 |
| 0 | 0 | 0 | 0 |
| 0 | 0 | 0 | 0 |
| 0 | 0 | 0 | 0 |
| 0 | 0 | 0 | 0 |
| 0 | 0 | 0 | 0 |
| 0 | 0 | 0 | 0 |
| 0 | 0 | 0 | 0 |
| 0 | 0 | 0 | 0 |
| 0 | 0 | 0 | 0 |
| 0 | 0 | 0 | 0 |

| | | | |
|---|---|---|---|
| 0 | 0 | 0 | 0 |
| 0 | 0 | 0 | 0 |
| 0 | 0 | 0 | 0 |
| 0 | 0 | 0 | 0 |
| 0 | 0 | 0 | 0 |
| 0 | 0 | 0 | 0 |
| 0 | 0 | 0 | 0 |
| 0 | 0 | 0 | 0 |
| 0 | 0 | 0 | 0 |
| 0 | 0 | 0 | 0 |
| 1 | 0 | 0 | 0 |
| 1 | 1 | 0 | 0 |
| 1 | 0 | 0 | 0 |
| 0 | 0 | 0 | 0 |
| 0 | 0 | 0 | 0 |
| 0 | 0 | 0 | 0 |
| 0 | 0 | 0 | 0 |
| 0 | 0 | 0 | 0 |
| 0 | 0 | 0 | 0 |
| 0 | 0 | 0 | 0 |
| 0 | 0 | 0 | 0 |
| 0 | 0 | 0 | 0 |
| 0 | 0 | 0 | 0 |
| 0 | 0 | 0 | 0 |
| 0 | 0 | 0 | 0 |
| 0 | 0 | 0 | 0 |
| 0 | 0 | 0 | 0 |
| 0 | 0 | 0 | 0 |
| 0 | 0 | 0 | 0 |
| 0 | 0 | 0 | 0 |
| 0 | 0 | 0 | 0 |
| 0 | 1 | 0 | 0 |
| 0 | 0 | 0 | 0 |
| 0 | 0 | 0 | 0 |
| 1 | 0 | 0 | 0 |
| 1 | 0 | 0 | 0 |
| 1 | 0 | 0 | 0 |
| 0 | 0 | 0 | 0 |
| 0 | 0 | 0 | 0 |
| 0 | 0 | 0 | 0 |
| 0 | 0 | 0 | 0 |
| 0 | 0 | 0 | 0 |
| 0 | 0 | 0 | 0 |
| 0 | 0 | 0 | 0 |
| 0 | 0 | 0 | 0 |
| 0 | 0 | 0 | 0 |
| 0 | 0 | 0 | 0 |
| 0 | 0 | 0 | 0 |
| 0 | 0 | 0 | 0 |
| 0 | 0 | 0 | 0 |
| 0 | 0 | 0 | 0 |
| 0 | 0 | 0 | 0 |
| 0 | 0 | 0 | 1 |
| 0 | 0 | 0 | 0 |

| | | | |
|---|---|---|---|
| 0 | 0 | 0 | 0 |
| 0 | 0 | 0 | 0 |
| 0 | 0 | 0 | 0 |
| 0 | 0 | 0 | 0 |
| 0 | 0 | 0 | 0 |
| 0 | 0 | 0 | 0 |
| 0 | 0 | 0 | 0 |
| 0 | 0 | 0 | 0 |
| 0 | 0 | 0 | 0 |
| 0 | 0 | 0 | 0 |
| 0 | 0 | 0 | 0 |
| 1 | 0 | 0 | 0 |
| 0 | 0 | 0 | 0 |
| 0 | 0 | 0 | 0 |
| 0 | 0 | 1 | 0 |
| 0 | 0 | 0 | 0 |
| 0 | 0 | 0 | 0 |
| 0 | 0 | 0 | 0 |
| 0 | 0 | 0 | 0 |
| 0 | 0 | 0 | 0 |
| 0 | 0 | 0 | 0 |
| 0 | 0 | 0 | 0 |
| 0 | 0 | 0 | 0 |
| 0 | 0 | 0 | 0 |
| 0 | 0 | 0 | 0 |
| 0 | 0 | 0 | 0 |
| 0 | 0 | 0 | 0 |
| 0 | 0 | 0 | 0 |
| 0 | 0 | 0 | 0 |
| 0 | 0 | 0 | 0 |
| 0 | 0 | 0 | 0 |
| 0 | 0 | 0 | 0 |
| 0 | 0 | 0 | 0 |
| 1 | 0 | 0 | 0 |
| 0 | 0 | 0 | 0 |
| 1 | 0 | 0 | 1 |
| 0 | 0 | 0 | 0 |
| 0 | 0 | 0 | 0 |
| 0 | 0 | 0 | 0 |
| 0 | 0 | 0 | 0 |
| 0 | 0 | 0 | 0 |
| 0 | 0 | 0 | 0 |
| 0 | 0 | 0 | 0 |
| 0 | 1 | 0 | 0 |
| 0 | 0 | 0 | 0 |
| 0 | 0 | 0 | 0 |
| 0 | 0 | 0 | 0 |
| 0 | 0 | 0 | 0 |
| 0 | 0 | 0 | 0 |
| 0 | 0 | 0 | 0 |
| 0 | 0 | 0 | 0 |
| 0 | 0 | 0 | 0 |
| 0 | 0 | 0 | 0 |
| 0 | 0 | 0 | 0 |

| | | | |
|---|---|---|---|
| 0 | 0 | 0 | 0 |
| 0 | 0 | 0 | 0 |
| 0 | 0 | 0 | 0 |
| 0 | 0 | 0 | 0 |
| 0 | 0 | 0 | 0 |
| 0 | 1 | 0 | 0 |
| 0 | 0 | 0 | 0 |
| 0 | 0 | 0 | 0 |
| 0 | 0 | 0 | 0 |
| 0 | 0 | 0 | 0 |
| 0 | 0 | 0 | 0 |
| 0 | 0 | 0 | 0 |
| 0 | 0 | 0 | 0 |
| 0 | 0 | 0 | 0 |
| 0 | 0 | 0 | 0 |
| 0 | 1 | 0 | 0 |
| 1 | 1 | 0 | 0 |
| 0 | 0 | 1 | 0 |
| 0 | 0 | 0 | 0 |
| 0 | 0 | 0 | 0 |
| 0 | 0 | 0 | 0 |
| 0 | 0 | 0 | 0 |
| 0 | 0 | 0 | 0 |
| 0 | 0 | 0 | 0 |
| 0 | 0 | 0 | 0 |
| 0 | 0 | 0 | 0 |
| 0 | 0 | 0 | 0 |
| 1 | 0 | 0 | 0 |
| 0 | 0 | 0 | 0 |
| 0 | 0 | 0 | 0 |
| 0 | 0 | 1 | 0 |
| 0 | 0 | 0 | 0 |
| 0 | 0 | 0 | 0 |
| 0 | 0 | 0 | 0 |
| 0 | 0 | 0 | 0 |
| 0 | 0 | 0 | 0 |
| 0 | 0 | 0 | 0 |
| 1 | 1 | 0 | 0 |
| 0 | 0 | 0 | 0 |
| 0 | 0 | 0 | 0 |
| 0 | 0 | 0 | 0 |
| 1 | 0 | 0 | 0 |
| 1 | 0 | 0 | 0 |
| 0 | 0 | 0 | 0 |
| 0 | 0 | 0 | 0 |
| 0 | 0 | 0 | 0 |
| 0 | 0 | 0 | 0 |
| 0 | 0 | 0 | 0 |
| 0 | 0 | 0 | 0 |
| 0 | 0 | 0 | 0 |
| 0 | 0 | 0 | 0 |
| 0 | 0 | 0 | 0 |
| 0 | 0 | 0 | 0 |
| 0 | 0 | 0 | 0 |

| | | | |
|---|---|---|---|
| 0 | 0 | 0 | 0 |
| 0 | 0 | 0 | 0 |
| 0 | 0 | 0 | 0 |
| 0 | 0 | 0 | 0 |
| 0 | 0 | 0 | 0 |
| 0 | 0 | 0 | 0 |
| 0 | 0 | 0 | 0 |
| 0 | 0 | 0 | 0 |
| 0 | 0 | 0 | 0 |
| 0 | 0 | 0 | 0 |
| 0 | 0 | 0 | 0 |
| 0 | 0 | 0 | 0 |
| 1 | 0 | 0 | 0 |
| 0 | 0 | 0 | 0 |
| 0 | 0 | 0 | 0 |
| 0 | 0 | 0 | 0 |
| 0 | 0 | 0 | 0 |
| 0 | 0 | 0 | 0 |
| 0 | 0 | 0 | 0 |
| 0 | 0 | 0 | 0 |
| 0 | 0 | 0 | 0 |
| 0 | 0 | 0 | 0 |
| 0 | 0 | 0 | 0 |
| 1 | 0 | 0 | 0 |
| 0 | 0 | 0 | 0 |
| 0 | 0 | 0 | 0 |
| 0 | 0 | 0 | 0 |
| 1 | 1 | 0 | 1 |
| 0 | 0 | 0 | 0 |
| 0 | 0 | 0 | 0 |
| 0 | 0 | 0 | 0 |
| 0 | 0 | 0 | 0 |
| 0 | 0 | 0 | 0 |
| 0 | 0 | 0 | 0 |
| 0 | 0 | 0 | 0 |
| 0 | 0 | 0 | 0 |
| 0 | 0 | 0 | 0 |
| 0 | 0 | 0 | 0 |
| 0 | 0 | 0 | 0 |
| 0 | 0 | 0 | 0 |
| 0 | 0 | 0 | 0 |
| 0 | 0 | 0 | 0 |
| 0 | 0 | 0 | 0 |
| 0 | 0 | 0 | 0 |
| 0 | 0 | 0 | 0 |
| 0 | 0 | 0 | 0 |
| 0 | 0 | 0 | 0 |
| 0 | 0 | 0 | 0 |
| 0 | 0 | 0 | 0 |
| 0 | 0 | 0 | 0 |
| 0 | 0 | 0 | 0 |
| 0 | 0 | 0 | 0 |
| 0 | 0 | 0 | 0 |
| 0 | 0 | 0 | 0 |

| | | | |
|---|---|---|---|
| 0 | 0 | 0 | 0 |
| 0 | 0 | 0 | 0 |
| 0 | 0 | 0 | 0 |
| 0 | 0 | 0 | 0 |
| 0 | 0 | 0 | 0 |
| 0 | 0 | 0 | 0 |
| 0 | 0 | 0 | 0 |
| 0 | 0 | 0 | 0 |
| 1 | 0 | 0 | 0 |
| 1 | 0 | 0 | 0 |
| 1 | 0 | 0 | 0 |
| 0 | 0 | 0 | 0 |
| 1 | 0 | 0 | 0 |
| 0 | 0 | 0 | 0 |
| 0 | 0 | 0 | 0 |
| 1 | 0 | 0 | 0 |
| 0 | 0 | 0 | 0 |
| 0 | 0 | 0 | 0 |
| 0 | 0 | 0 | 0 |
| 0 | 0 | 0 | 0 |
| 0 | 0 | 0 | 0 |
| 0 | 0 | 0 | 0 |
| 0 | 0 | 0 | 0 |
| 0 | 0 | 0 | 0 |
| 0 | 0 | 0 | 0 |
| 0 | 0 | 0 | 0 |
| 0 | 0 | 0 | 0 |
| 1 | 0 | 0 | 0 |
| 1 | 0 | 0 | 0 |
| 0 | 0 | 0 | 0 |
| 0 | 0 | 0 | 0 |
| 0 | 0 | 0 | 0 |
| 1 | 0 | 0 | 0 |
| 0 | 0 | 0 | 0 |
| 0 | 0 | 0 | 0 |
| 0 | 0 | 0 | 0 |
| 0 | 0 | 0 | 0 |
| 0 | 0 | 0 | 0 |
| 0 | 0 | 0 | 0 |
| 0 | 0 | 0 | 0 |
| 1 | 0 | 0 | 0 |
| 1 | 0 | 0 | 0 |
| 0 | 0 | 0 | 0 |
| 0 | 0 | 0 | 0 |
| 0 | 0 | 0 | 0 |
| 0 | 0 | 0 | 0 |
| 0 | 0 | 0 | 0 |
| 0 | 0 | 0 | 0 |
| 0 | 0 | 0 | 0 |
| 0 | 0 | 0 | 0 |
| 0 | 0 | 0 | 0 |
| 0 | 0 | 0 | 0 |
| 0 | 0 | 0 | 0 |

| | | | |
|---|---|---|---|
| 0 | 0 | 0 | 0 |
| 0 | 0 | 0 | 0 |
| 0 | 0 | 0 | 0 |
| 0 | 0 | 0 | 0 |
| 0 | 0 | 0 | 0 |
| 0 | 0 | 0 | 0 |
| 0 | 0 | 0 | 0 |
| 0 | 0 | 0 | 0 |
| 0 | 0 | 0 | 0 |
| 0 | 0 | 0 | 0 |
| 1 | 0 | 0 | 0 |
| 0 | 0 | 0 | 0 |
| 1 | 0 | 0 | 0 |
| 0 | 0 | 0 | 0 |
| 0 | 0 | 0 | 0 |
| 0 | 0 | 0 | 0 |
| 0 | 0 | 0 | 0 |
| 0 | 1 | 0 | 0 |
| 0 | 0 | 0 | 0 |
| 0 | 0 | 0 | 0 |
| 0 | 0 | 0 | 0 |
| 0 | 0 | 0 | 0 |
| 1 | 0 | 0 | 0 |
| 0 | 0 | 0 | 0 |
| 0 | 0 | 0 | 0 |
| 1 | 0 | 0 | 0 |
| 1 | 0 | 0 | 0 |
| 1 | 0 | 0 | 0 |
| 0 | 0 | 0 | 0 |
| 0 | 0 | 0 | 0 |
| 0 | 0 | 0 | 0 |
| 0 | 0 | 0 | 0 |
| 0 | 0 | 0 | 0 |
| 0 | 0 | 0 | 0 |
| 0 | 0 | 0 | 0 |
| 0 | 0 | 0 | 0 |
| 0 | 0 | 0 | 0 |
| 0 | 0 | 0 | 0 |
| 0 | 0 | 0 | 0 |
| 0 | 0 | 0 | 0 |
| 1 | 1 | 0 | 0 |
| 0 | 0 | 0 | 0 |
| 0 | 0 | 0 | 0 |
| 0 | 0 | 0 | 0 |
| 0 | 0 | 0 | 0 |
| 0 | 0 | 0 | 0 |
| 0 | 0 | 0 | 0 |
| 1 | 0 | 0 | 0 |
| 0 | 0 | 0 | 0 |
| 0 | 0 | 0 | 0 |
| 0 | 0 | 0 | 0 |
| 0 | 0 | 0 | 0 |
| 0 | 0 | 0 | 0 |
| 0 | 0 | 0 | 0 |

| | | | |
|---|---|---|---|
| 1 | 0 | 0 | 0 |
| 0 | 0 | 0 | 0 |
| 1 | 0 | 0 | 0 |
| 0 | 0 | 0 | 0 |
| 0 | 0 | 0 | 0 |
| 0 | 0 | 0 | 0 |
| 0 | 0 | 0 | 0 |
| 0 | 0 | 0 | 0 |
| 1 | 1 | 0 | 0 |
| 0 | 0 | 0 | 0 |
| 1 | 1 | 0 | 0 |
| 1 | 1 | 0 | 0 |
| 0 | 0 | 0 | 0 |
| 1 | 1 | 0 | 0 |
| 0 | 0 | 0 | 0 |
| 0 | 0 | 0 | 0 |
| 0 | 0 | 0 | 0 |
| 0 | 0 | 0 | 0 |
| 0 | 0 | 0 | 0 |
| 0 | 0 | 0 | 0 |
| 1 | 0 | 0 | 0 |
| 0 | 0 | 0 | 0 |
| 0 | 0 | 0 | 0 |
| 0 | 0 | 0 | 0 |
| 1 | 0 | 0 | 0 |
| 0 | 0 | 0 | 0 |
| 1 | 0 | 0 | 0 |
| 1 | 0 | 0 | 0 |
| 1 | 1 | 0 | 0 |
| 0 | 0 | 0 | 0 |
| 1 | 1 | 0 | 0 |
| 1 | 1 | 0 | 0 |
| 1 | 1 | 0 | 0 |
| 0 | 0 | 0 | 0 |
| 1 | 1 | 0 | 0 |
| 0 | 0 | 0 | 0 |
| 1 | 0 | 0 | 0 |
| 0 | 0 | 0 | 0 |
| 1 | 1 | 0 | 0 |
| 1 | 1 | 0 | 0 |
| 1 | 0 | 0 | 0 |
| 0 | 0 | 0 | 0 |
| 0 | 0 | 0 | 0 |
| 1 | 0 | 0 | 0 |
| 0 | 0 | 0 | 0 |
| 0 | 0 | 0 | 0 |
| 1 | 0 | 0 | 0 |
| 0 | 0 | 0 | 0 |
| 1 | 0 | 0 | 0 |
| 0 | 0 | 0 | 0 |
| 0 | 0 | 0 | 0 |
| 0 | 0 | 0 | 0 |
| 0 | 0 | 0 | 0 |
| 0 | 0 | 0 | 0 |

| | | | |
|---|---|---|---|
| 0 | 0 | 0 | 0 |
| 0 | 0 | 0 | 0 |
| 0 | 0 | 0 | 0 |
| 0 | 0 | 0 | 0 |
| 0 | 0 | 0 | 0 |
| 0 | 0 | 0 | 0 |
| 0 | 0 | 0 | 0 |
| 0 | 0 | 0 | 0 |
| 0 | 0 | 0 | 0 |
| 1 | 1 | 0 | 0 |
| 1 | 1 | 0 | 0 |
| 0 | 0 | 0 | 0 |
| 0 | 0 | 0 | 0 |
| 0 | 0 | 0 | 0 |
| 1 | 0 | 1 | 0 |
| 0 | 0 | 0 | 0 |
| 0 | 0 | 0 | 0 |
| 1 | 0 | 1 | 0 |
| 0 | 0 | 0 | 0 |
| 0 | 0 | 0 | 0 |
| 0 | 0 | 0 | 0 |
| 0 | 0 | 1 | 0 |
| 1 | 0 | 1 | 0 |
| 0 | 0 | 1 | 0 |
| 0 | 0 | 1 | 0 |
| 0 | 0 | 1 | 1 |
| 0 | 0 | 1 | 1 |
| 0 | 0 | 0 | 0 |
| 0 | 0 | 0 | 0 |
| 0 | 0 | 0 | 0 |
| 0 | 0 | 0 | 0 |
| 0 | 0 | 0 | 0 |
| 0 | 0 | 0 | 0 |
| 0 | 0 | 0 | 0 |
| 0 | 0 | 0 | 0 |
| 0 | 0 | 0 | 0 |
| 0 | 0 | 0 | 0 |
| 0 | 0 | 0 | 0 |
| 0 | 0 | 0 | 0 |
| 0 | 0 | 0 | 0 |
| 0 | 0 | 0 | 0 |
| 0 | 0 | 0 | 0 |
| 0 | 0 | 0 | 0 |
| 0 | 0 | 1 | 0 |
| 0 | 0 | 0 | 0 |
| 0 | 0 | 0 | 0 |
| 0 | 0 | 0 | 0 |
| 0 | 0 | 0 | 0 |
| 0 | 0 | 0 | 0 |
| 0 | 0 | 0 | 0 |
| 1 | 0 | 0 | 0 |
| 0 | 0 | 0 | 0 |
| 0 | 0 | 0 | 0 |
| 0 | 0 | 0 | 0 |

| | | | |
|---|---|---|---|
| 0 | 0 | 0 | 0 |
| 0 | 0 | 0 | 0 |
| 0 | 0 | 0 | 0 |
| 1 | 0 | 0 | 0 |
| 0 | 0 | 0 | 0 |
| 0 | 0 | 0 | 0 |
| 0 | 0 | 0 | 0 |
| 0 | 0 | 0 | 0 |
| 1 | 0 | 0 | 0 |
| 1 | 0 | 0 | 0 |
| 0 | 0 | 0 | 0 |
| 0 | 0 | 0 | 0 |
| 0 | 0 | 0 | 0 |
| 0 | 0 | 0 | 0 |
| 0 | 0 | 0 | 0 |
| 0 | 0 | 0 | 0 |
| 0 | 0 | 1 | 0 |
| 0 | 0 | 0 | 0 |
| 0 | 0 | 0 | 0 |
| 0 | 0 | 0 | 0 |
| 0 | 0 | 0 | 0 |
| 0 | 0 | 0 | 0 |
| 0 | 0 | 0 | 0 |
| 0 | 0 | 0 | 0 |
| 0 | 0 | 0 | 0 |
| 0 | 0 | 0 | 0 |
| 0 | 0 | 0 | 0 |
| 0 | 0 | 0 | 0 |
| 0 | 0 | 0 | 0 |
| 0 | 0 | 0 | 0 |
| 0 | 0 | 0 | 0 |
| 0 | 0 | 0 | 0 |
| 0 | 0 | 0 | 0 |
| 0 | 0 | 0 | 0 |
| 0 | 0 | 0 | 0 |
| 0 | 0 | 0 | 0 |
| 0 | 0 | 0 | 0 |
| 0 | 0 | 0 | 0 |
| 0 | 0 | 0 | 0 |
| 0 | 0 | 0 | 0 |
| 0 | 0 | 0 | 0 |
| 0 | 0 | 0 | 0 |
| 0 | 0 | 0 | 0 |
| 0 | 0 | 0 | 0 |
| 0 | 0 | 0 | 0 |
| 0 | 0 | 0 | 0 |
| 0 | 0 | 0 | 0 |
| 0 | 0 | 0 | 0 |
| 0 | 0 | 0 | 0 |
| 0 | 1 | 0 | 0 |
| 0 | 0 | 0 | 0 |
| 0 | 0 | 0 | 0 |
| 0 | 0 | 0 | 0 |
| 0 | 0 | 0 | 0 |

| | | | |
|---|---|---|---|
| 0 | 0 | 1 | 0 |
| 0 | 0 | 0 | 0 |
| 0 | 0 | 0 | 0 |
| 0 | 0 | 0 | 0 |
| 0 | 0 | 0 | 0 |
| 0 | 0 | 0 | 0 |
| 0 | 0 | 0 | 0 |
| 1 | 0 | 0 | 0 |
| 0 | 0 | 0 | 0 |
| 0 | 0 | 0 | 0 |
| 0 | 0 | 0 | 0 |
| 1 | 0 | 0 | 0 |
| 0 | 0 | 0 | 0 |
| 0 | 0 | 0 | 0 |
| 0 | 0 | 0 | 0 |
| 0 | 0 | 0 | 0 |
| 0 | 0 | 0 | 0 |
| 0 | 0 | 0 | 0 |
| 0 | 0 | 0 | 0 |
| 0 | 0 | 0 | 0 |
| 0 | 0 | 0 | 0 |
| 0 | 0 | 0 | 0 |
| 0 | 0 | 0 | 0 |
| 0 | 0 | 0 | 0 |
| 0 | 0 | 0 | 0 |
| 0 | 0 | 0 | 0 |
| 0 | 0 | 0 | 0 |
| 0 | 0 | 0 | 0 |
| 0 | 0 | 0 | 0 |
| 0 | 0 | 0 | 0 |
| 0 | 0 | 0 | 0 |
| 0 | 0 | 0 | 0 |
| 0 | 0 | 0 | 0 |
| 0 | 0 | 0 | 0 |
| 0 | 0 | 0 | 0 |
| 0 | 0 | 0 | 0 |
| 0 | 0 | 0 | 0 |
| 0 | 0 | 0 | 0 |
| 0 | 0 | 0 | 0 |
| 0 | 0 | 0 | 0 |
| 0 | 0 | 0 | 0 |
| 0 | 0 | 0 | 0 |
| 0 | 0 | 0 | 0 |
| 0 | 0 | 0 | 0 |
| 0 | 0 | 0 | 0 |
| 0 | 0 | 0 | 0 |
| 0 | 0 | 0 | 0 |
| 0 | 0 | 0 | 0 |
| 0 | 0 | 0 | 0 |
| 0 | 0 | 0 | 0 |
| 0 | 0 | 0 | 0 |
| 0 | 0 | 0 | 0 |
| 0 | 0 | 0 | 0 |
| 0 | 0 | 0 | 0 |

| | | | |
|---|---|---|---|
| 0 | 0 | 0 | 0 |
| 0 | 0 | 0 | 0 |
| 0 | 0 | 0 | 0 |
| 0 | 0 | 0 | 0 |
| 0 | 0 | 0 | 0 |
| 0 | 0 | 0 | 0 |
| 0 | 0 | 0 | 0 |
| 0 | 0 | 0 | 0 |
| 0 | 0 | 0 | 0 |
| 0 | 0 | 0 | 0 |
| 0 | 0 | 0 | 0 |
| 0 | 0 | 0 | 0 |
| 0 | 0 | 0 | 0 |
| 0 | 0 | 0 | 0 |
| 0 | 0 | 0 | 0 |
| 0 | 0 | 0 | 0 |
| 0 | 0 | 0 | 0 |
| 0 | 0 | 0 | 0 |
| 0 | 0 | 0 | 1 |
| 0 | 1 | 0 | 1 |
| 0 | 0 | 0 | 0 |
| 0 | 0 | 0 | 0 |
| 0 | 0 | 0 | 0 |
| 0 | 0 | 0 | 0 |
| 0 | 0 | 0 | 0 |
| 1 | 0 | 0 | 0 |
| 1 | 0 | 0 | 0 |
| 0 | 0 | 0 | 0 |
| 0 | 0 | 0 | 0 |
| 0 | 0 | 0 | 0 |
| 0 | 0 | 0 | 0 |
| 0 | 0 | 0 | 0 |
| 0 | 0 | 0 | 0 |
| 1 | 0 | 0 | 1 |
| 1 | 0 | 0 | 0 |
| 1 | 0 | 0 | 1 |
| 1 | 0 | 0 | 0 |
| 0 | 0 | 0 | 0 |
| 0 | 0 | 0 | 0 |
| 0 | 0 | 0 | 0 |
| 0 | 0 | 0 | 0 |
| 1 | 0 | 0 | 0 |
| 1 | 0 | 0 | 1 |
| 1 | 0 | 0 | 0 |
| 0 | 0 | 0 | 0 |
| 0 | 0 | 0 | 0 |
| 0 | 0 | 0 | 0 |
| 0 | 0 | 0 | 0 |
| 0 | 0 | 0 | 0 |
| 1 | 0 | 0 | 0 |
| 0 | 0 | 0 | 0 |
| 0 | 0 | 0 | 0 |
| 0 | 0 | 0 | 0 |
| 0 | 0 | 0 | 0 |

| | | | |
|---|---|---|---|
| 0 | 0 | 0 | 0 |
| 0 | 0 | 0 | 0 |
| 0 | 0 | 0 | 0 |
| 1 | 0 | 0 | 0 |
| 1 | 0 | 0 | 0 |
| 0 | 0 | 0 | 0 |
| 1 | 0 | 0 | 0 |
| 0 | 0 | 0 | 0 |
| 0 | 0 | 0 | 0 |
| 0 | 0 | 0 | 0 |
| 0 | 0 | 0 | 0 |
| 1 | 0 | 1 | 0 |
| 0 | 0 | 0 | 0 |
| 0 | 0 | 0 | 0 |
| 0 | 0 | 0 | 0 |
| 0 | 0 | 1 | 0 |
| 0 | 0 | 0 | 0 |
| 0 | 0 | 0 | 0 |
| 0 | 0 | 0 | 0 |
| 0 | 0 | 1 | 0 |
| 1 | 0 | 1 | 0 |
| 1 | 0 | 0 | 0 |
| 1 | 0 | 0 | 0 |
| 1 | 0 | 0 | 0 |
| 0 | 0 | 1 | 0 |
| 0 | 0 | 0 | 0 |
| 1 | 0 | 0 | 0 |
| 0 | 0 | 0 | 0 |
| 0 | 0 | 0 | 0 |
| 0 | 0 | 0 | 0 |
| 0 | 0 | 0 | 0 |
| 0 | 0 | 0 | 0 |
| 0 | 0 | 0 | 0 |
| 0 | 0 | 0 | 0 |
| 0 | 0 | 0 | 0 |
| 0 | 0 | 0 | 0 |
| 0 | 0 | 0 | 1 |
| 0 | 0 | 0 | 1 |
| 0 | 0 | 0 | 1 |
| 0 | 0 | 0 | 0 |
| 0 | 0 | 0 | 0 |
| 0 | 0 | 0 | 0 |
| 0 | 0 | 0 | 0 |
| 0 | 0 | 0 | 0 |
| 0 | 0 | 0 | 0 |
| 0 | 0 | 0 | 0 |
| 0 | 0 | 0 | 0 |
| 0 | 0 | 0 | 0 |
| 0 | 0 | 0 | 0 |
| 0 | 0 | 0 | 0 |
| 0 | 0 | 0 | 0 |
| 0 | 0 | 0 | 0 |
| 0 | 0 | 0 | 0 |
| 0 | 0 | 0 | 0 |

| | | | |
|---|---|---|---|
| 0 | 0 | 0 | 0 |
| 0 | 0 | 0 | 0 |
| 0 | 0 | 0 | 0 |
| 0 | 0 | 0 | 0 |
| 0 | 0 | 0 | 0 |
| 0 | 0 | 0 | 0 |
| 0 | 0 | 0 | 0 |
| 0 | 0 | 0 | 0 |
| 0 | 0 | 0 | 0 |
| 0 | 0 | 0 | 0 |
| 0 | 0 | 0 | 0 |
| 0 | 0 | 0 | 0 |
| 0 | 0 | 0 | 0 |
| 0 | 0 | 0 | 0 |
| 0 | 0 | 0 | 0 |
| 0 | 0 | 0 | 0 |
| 0 | 1 | 0 | 0 |
| 0 | 1 | 0 | 0 |
| 0 | 0 | 0 | 0 |
| 0 | 0 | 0 | 0 |
| 0 | 0 | 0 | 0 |
| 0 | 0 | 0 | 0 |
| 0 | 0 | 0 | 0 |
| 0 | 0 | 0 | 0 |
| 0 | 0 | 0 | 0 |
| 0 | 0 | 0 | 0 |
| 0 | 0 | 0 | 0 |
| 0 | 0 | 0 | 0 |
| 0 | 0 | 0 | 0 |
| 0 | 0 | 0 | 0 |
| 0 | 0 | 0 | 0 |
| 1 | 0 | 0 | 0 |
| 0 | 0 | 0 | 0 |
| 0 | 0 | 0 | 0 |
| 0 | 0 | 0 | 0 |
| 0 | 0 | 0 | 0 |
| 0 | 0 | 0 | 0 |
| 0 | 0 | 0 | 0 |
| 0 | 0 | 0 | 0 |
| 0 | 0 | 0 | 0 |
| 0 | 0 | 0 | 0 |
| 0 | 0 | 0 | 0 |
| 0 | 0 | 0 | 0 |
| 0 | 0 | 0 | 0 |
| 0 | 0 | 0 | 0 |
| 0 | 0 | 0 | 0 |
| 1 | 1 | 0 | 0 |
| 0 | 0 | 0 | 0 |
| 0 | 0 | 0 | 0 |
| 0 | 0 | 0 | 0 |
| 0 | 0 | 0 | 0 |
| 0 | 0 | 0 | 0 |
| 1 | 0 | 0 | 0 |
| 0 | 0 | 0 | 0 |

| | | | |
|---|---|---|---|
| 0 | 0 | 0 | 0 |
| 0 | 0 | 0 | 0 |
| 0 | 0 | 0 | 0 |
| 0 | 0 | 0 | 0 |
| 0 | 0 | 0 | 0 |
| 0 | 0 | 0 | 0 |
| 1 | 0 | 0 | 0 |
| 0 | 0 | 0 | 0 |
| 0 | 0 | 0 | 0 |
| 0 | 0 | 0 | 0 |
| 0 | 0 | 0 | 0 |
| 0 | 0 | 0 | 0 |
| 0 | 0 | 0 | 0 |
| 0 | 0 | 0 | 0 |
| 0 | 0 | 0 | 0 |
| 0 | 0 | 0 | 0 |
| 0 | 0 | 0 | 0 |
| 0 | 0 | 0 | 0 |
| 0 | 0 | 0 | 0 |
| 0 | 0 | 0 | 0 |
| 0 | 0 | 0 | 0 |
| 0 | 0 | 0 | 0 |
| 0 | 0 | 0 | 0 |
| 0 | 0 | 0 | 0 |
| 0 | 0 | 0 | 0 |
| 0 | 0 | 0 | 0 |
| 0 | 0 | 0 | 0 |
| 0 | 0 | 0 | 0 |
| 0 | 0 | 0 | 0 |
| 0 | 0 | 0 | 0 |
| 0 | 0 | 0 | 0 |
| 0 | 0 | 0 | 0 |
| 0 | 0 | 0 | 0 |
| 0 | 0 | 0 | 0 |
| 0 | 0 | 0 | 0 |
| 0 | 0 | 0 | 0 |
| 0 | 0 | 0 | 0 |
| 0 | 0 | 0 | 0 |
| 0 | 0 | 0 | 0 |
| 0 | 0 | 0 | 0 |
| 0 | 0 | 0 | 0 |
| 0 | 0 | 0 | 0 |
| 0 | 0 | 0 | 0 |
| 0 | 0 | 0 | 0 |
| 0 | 0 | 0 | 0 |
| 0 | 0 | 0 | 0 |
| 0 | 0 | 0 | 0 |
| 0 | 0 | 0 | 0 |
| 0 | 0 | 0 | 0 |
| 0 | 0 | 0 | 0 |
| 0 | 0 | 0 | 0 |
| 0 | 0 | 0 | 0 |
| 0 | 0 | 0 | 0 |
| 0 | 0 | 0 | 0 |

| | | | |
|---|---|---|---|
| 0 | 0 | 0 | 0 |
| 0 | 0 | 0 | 0 |
| 0 | 0 | 0 | 0 |
| 0 | 0 | 0 | 0 |
| 0 | 0 | 0 | 0 |
| 0 | 0 | 0 | 0 |
| 0 | 0 | 0 | 0 |
| 0 | 0 | 0 | 0 |
| 0 | 0 | 0 | 0 |
| 0 | 0 | 0 | 0 |
| 0 | 0 | 0 | 0 |
| 0 | 0 | 0 | 0 |
| 0 | 0 | 0 | 0 |
| 0 | 0 | 0 | 0 |
| 0 | 0 | 0 | 0 |
| 0 | 0 | 0 | 0 |
| 0 | 0 | 0 | 0 |
| 0 | 0 | 0 | 0 |
| 0 | 0 | 0 | 0 |
| 0 | 0 | 0 | 0 |
| 0 | 0 | 0 | 0 |
| 0 | 0 | 0 | 0 |
| 0 | 0 | 0 | 0 |
| 0 | 0 | 0 | 0 |
| 1 | 1 | 0 | 0 |
| 0 | 0 | 0 | 0 |
| 0 | 0 | 0 | 0 |
| 0 | 0 | 0 | 0 |
| 0 | 0 | 0 | 0 |
| 0 | 0 | 0 | 0 |
| 0 | 0 | 0 | 0 |
| 0 | 0 | 0 | 0 |
| 0 | 0 | 0 | 0 |
| 0 | 0 | 0 | 0 |
| 0 | 0 | 0 | 0 |
| 1 | 0 | 0 | 0 |
| 1 | 0 | 0 | 1 |
| 1 | 0 | 0 | 0 |
| 0 | 0 | 0 | 0 |
| 1 | 0 | 0 | 0 |
| 1 | 1 | 0 | 0 |
| 1 | 1 | 0 | 0 |
| 0 | 0 | 0 | 0 |
| 0 | 0 | 0 | 0 |
| 0 | 0 | 0 | 0 |
| 0 | 0 | 0 | 0 |
| 0 | 0 | 0 | 0 |
| 0 | 0 | 0 | 0 |
| 0 | 0 | 0 | 0 |
| 0 | 0 | 0 | 0 |
| 0 | 0 | 0 | 0 |
| 0 | 0 | 0 | 0 |
| 0 | 0 | 0 | 0 |
| 0 | 0 | 0 | 0 |

| | | | |
|---|---|---|---|
| 0 | 0 | 0 | 0 |
| 0 | 0 | 0 | 0 |
| 0 | 0 | 0 | 0 |
| 0 | 0 | 0 | 0 |
| 0 | 0 | 0 | 0 |
| 0 | 0 | 0 | 0 |
| 0 | 0 | 0 | 0 |
| 0 | 0 | 0 | 0 |
| 0 | 0 | 0 | 0 |
| 0 | 0 | 0 | 0 |
| 0 | 0 | 0 | 0 |
| 0 | 0 | 0 | 0 |
| 0 | 0 | 0 | 0 |
| 1 | 0 | 0 | 0 |
| 0 | 0 | 0 | 0 |
| 0 | 0 | 0 | 0 |
| 1 | 1 | 0 | 0 |
| 0 | 0 | 0 | 0 |
| 0 | 0 | 0 | 0 |
| 0 | 0 | 0 | 0 |
| 0 | 0 | 0 | 0 |
| 0 | 0 | 0 | 0 |
| 0 | 0 | 0 | 0 |
| 1 | 0 | 0 | 0 |
| 0 | 0 | 0 | 0 |
| 0 | 0 | 0 | 0 |
| 0 | 0 | 0 | 0 |
| 0 | 0 | 0 | 0 |
| 0 | 0 | 0 | 0 |
| 0 | 0 | 0 | 0 |
| 0 | 0 | 0 | 0 |
| 1 | 0 | 0 | 0 |
| 0 | 0 | 0 | 0 |
| 0 | 0 | 0 | 0 |
| 0 | 0 | 0 | 0 |
| 0 | 0 | 0 | 0 |
| 0 | 0 | 0 | 0 |
| 0 | 0 | 0 | 0 |
| 0 | 0 | 0 | 0 |
| 0 | 0 | 0 | 0 |
| 0 | 0 | 0 | 0 |
| 0 | 0 | 0 | 0 |
| 0 | 0 | 0 | 0 |
| 0 | 0 | 0 | 0 |
| 0 | 0 | 0 | 0 |
| 0 | 0 | 0 | 0 |
| 0 | 0 | 0 | 0 |
| 0 | 0 | 0 | 0 |
| 0 | 0 | 0 | 0 |
| 0 | 0 | 0 | 0 |
| 0 | 0 | 0 | 0 |
| 0 | 0 | 0 | 0 |
| 0 | 0 | 0 | 0 |
| 0 | 0 | 0 | 0 |

| | | | |
|---|---|---|---|
| 0 | 0 | 0 | 0 |
| 0 | 0 | 0 | 0 |
| 0 | 0 | 0 | 0 |
| 0 | 0 | 0 | 0 |
| 0 | 0 | 0 | 0 |
| 0 | 0 | 0 | 0 |
| 0 | 0 | 0 | 0 |
| 0 | 0 | 0 | 0 |
| 0 | 0 | 0 | 0 |
| 0 | 0 | 0 | 0 |
| 0 | 0 | 0 | 0 |
| 0 | 0 | 0 | 0 |
| 0 | 0 | 0 | 0 |
| 0 | 0 | 0 | 0 |
| 0 | 0 | 0 | 0 |
| 0 | 0 | 0 | 0 |
| 0 | 0 | 0 | 0 |
| 1 | 0 | 0 | 0 |
| 0 | 0 | 0 | 0 |
| 0 | 0 | 0 | 0 |
| 0 | 0 | 0 | 0 |
| 0 | 0 | 0 | 0 |
| 0 | 0 | 0 | 0 |
| 0 | 0 | 0 | 0 |
| 1 | 0 | 0 | 0 |
| 0 | 0 | 0 | 0 |
| 0 | 0 | 0 | 0 |
| 0 | 0 | 0 | 0 |
| 0 | 0 | 0 | 0 |
| 0 | 0 | 0 | 0 |
| 0 | 0 | 0 | 0 |
| 0 | 0 | 0 | 0 |
| 0 | 0 | 0 | 0 |
| 0 | 0 | 0 | 0 |
| 0 | 0 | 0 | 0 |
| 0 | 0 | 0 | 0 |
| 0 | 0 | 0 | 0 |
| 0 | 0 | 0 | 0 |
| 0 | 0 | 0 | 0 |
| 0 | 0 | 0 | 0 |
| 0 | 0 | 0 | 0 |
| 0 | 0 | 0 | 0 |
| 0 | 0 | 0 | 0 |
| 0 | 0 | 0 | 0 |
| 0 | 0 | 0 | 0 |
| 0 | 0 | 0 | 0 |
| 0 | 0 | 0 | 0 |
| 0 | 0 | 0 | 0 |
| 0 | 0 | 0 | 0 |
| 0 | 0 | 0 | 0 |
| 0 | 0 | 0 | 0 |
| 0 | 0 | 0 | 0 |
| 0 | 0 | 0 | 0 |
| 0 | 0 | 0 | 0 |

| | | | |
|---|---|---|---|
| 0 | 0 | 0 | 0 |
| 0 | 0 | 0 | 0 |
| 0 | 0 | 0 | 0 |
| 0 | 0 | 0 | 0 |
| 0 | 0 | 0 | 0 |
| 0 | 0 | 0 | 0 |
| 0 | 0 | 0 | 0 |
| 0 | 0 | 0 | 0 |
| 0 | 0 | 0 | 0 |
| 0 | 0 | 0 | 0 |
| 0 | 0 | 0 | 0 |
| 0 | 0 | 0 | 0 |
| 0 | 0 | 0 | 0 |
| 0 | 0 | 0 | 0 |
| 1 | 1 | 0 | 0 |
| 1 | 1 | 0 | 0 |
| 0 | 0 | 0 | 0 |
| 0 | 0 | 0 | 0 |
| 0 | 0 | 0 | 0 |
| 0 | 0 | 0 | 0 |
| 0 | 0 | 0 | 0 |
| 0 | 0 | 0 | 0 |
| 0 | 0 | 0 | 0 |
| 0 | 0 | 0 | 0 |
| 0 | 0 | 0 | 0 |
| 0 | 0 | 0 | 0 |
| 0 | 0 | 0 | 0 |
| 1 | 0 | 0 | 0 |
| 0 | 0 | 0 | 0 |
| 1 | 0 | 0 | 0 |
| 0 | 0 | 0 | 0 |
| 0 | 0 | 0 | 0 |
| 0 | 0 | 0 | 0 |
| 0 | 0 | 0 | 0 |
| 0 | 0 | 0 | 0 |
| 0 | 0 | 0 | 0 |
| 1 | 0 | 0 | 0 |
| 0 | 0 | 0 | 0 |
| 0 | 0 | 0 | 0 |
| 0 | 0 | 0 | 0 |
| 0 | 0 | 0 | 0 |
| 0 | 0 | 0 | 0 |
| 0 | 0 | 0 | 0 |
| 0 | 0 | 0 | 0 |
| 0 | 0 | 0 | 0 |
| 0 | 0 | 0 | 0 |
| 0 | 0 | 0 | 0 |
| 0 | 0 | 0 | 0 |
| 0 | 0 | 0 | 0 |
| 0 | 0 | 0 | 0 |
| 0 | 0 | 0 | 0 |
| 0 | 0 | 0 | 0 |
| 0 | 0 | 0 | 0 |
| 0 | 0 | 0 | 0 |

| | | | |
|---|---|---|---|
| 0 | 0 | 0 | 0 |
| 0 | 0 | 0 | 0 |
| 0 | 0 | 0 | 0 |
| 0 | 0 | 0 | 0 |
| 0 | 0 | 0 | 0 |
| 0 | 0 | 0 | 0 |
| 0 | 0 | 0 | 0 |
| 0 | 0 | 0 | 0 |
| 0 | 0 | 0 | 0 |
| 0 | 0 | 0 | 0 |
| 0 | 0 | 0 | 0 |
| 0 | 0 | 0 | 0 |
| 0 | 0 | 0 | 0 |
| 0 | 0 | 0 | 0 |
| 0 | 0 | 0 | 0 |
| 0 | 0 | 0 | 0 |
| 0 | 0 | 0 | 0 |
| 0 | 0 | 0 | 0 |
| 0 | 0 | 0 | 0 |
| 0 | 0 | 0 | 0 |
| 0 | 0 | 0 | 0 |
| 0 | 0 | 0 | 0 |
| 0 | 0 | 0 | 0 |
| 0 | 0 | 0 | 0 |
| 0 | 0 | 0 | 0 |
| 0 | 0 | 0 | 0 |
| 0 | 0 | 0 | 0 |
| 0 | 0 | 0 | 0 |
| 0 | 0 | 0 | 0 |
| 0 | 0 | 0 | 0 |
| 0 | 0 | 0 | 0 |
| 0 | 0 | 0 | 0 |
| 0 | 0 | 0 | 0 |
| 0 | 0 | 0 | 0 |
| 0 | 0 | 0 | 0 |
| 0 | 0 | 0 | 0 |
| 0 | 0 | 0 | 0 |
| 0 | 0 | 0 | 0 |
| 0 | 0 | 0 | 0 |
| 0 | 0 | 0 | 0 |
| 0 | 0 | 0 | 0 |
| 0 | 0 | 0 | 0 |
| 0 | 0 | 0 | 0 |
| 0 | 0 | 0 | 0 |
| 0 | 0 | 0 | 0 |
| 0 | 0 | 0 | 0 |
| 0 | 0 | 0 | 0 |
| 0 | 0 | 0 | 0 |
| 1 | 0 | 0 | 1 |
| 0 | 0 | 0 | 0 |
| 1 | 0 | 0 | 0 |
| 1 | 0 | 0 | 0 |
| 0 | 0 | 0 | 0 |
| 0 | 0 | 0 | 1 |

| | | | |
|---|---|---|---|
| 0 | 0 | 0 | 0 |
| 0 | 0 | 0 | 0 |
| 0 | 0 | 0 | 0 |
| 0 | 0 | 0 | 0 |
| 0 | 0 | 0 | 0 |
| 0 | 0 | 0 | 0 |
| 0 | 0 | 0 | 0 |
| 0 | 0 | 0 | 0 |
| 0 | 0 | 0 | 0 |
| 0 | 0 | 0 | 0 |
| 0 | 0 | 0 | 0 |
| 0 | 0 | 0 | 0 |
| 0 | 0 | 0 | 0 |
| 0 | 0 | 0 | 0 |
| 0 | 0 | 0 | 0 |
| 0 | 0 | 0 | 0 |
| 0 | 0 | 0 | 0 |
| 0 | 0 | 0 | 0 |
| 0 | 0 | 0 | 0 |
| 0 | 0 | 0 | 0 |
| 0 | 0 | 0 | 0 |
| 0 | 0 | 0 | 0 |
| 0 | 0 | 0 | 0 |
| 0 | 0 | 0 | 0 |
| 0 | 0 | 0 | 0 |
| 0 | 0 | 0 | 0 |
| 0 | 0 | 1 | 0 |
| 0 | 0 | 1 | 0 |
| 0 | 0 | 1 | 0 |
| 0 | 0 | 1 | 0 |
| 0 | 0 | 0 | 0 |
| 0 | 0 | 0 | 0 |
| 0 | 0 | 0 | 0 |
| 1 | 0 | 0 | 0 |
| 1 | 0 | 0 | 0 |
| 0 | 0 | 0 | 0 |
| 0 | 0 | 0 | 0 |
| 0 | 0 | 0 | 0 |
| 0 | 0 | 0 | 0 |
| 1 | 0 | 0 | 0 |
| 0 | 0 | 0 | 0 |
| 0 | 0 | 0 | 0 |
| 0 | 0 | 0 | 0 |
| 0 | 0 | 0 | 0 |
| 0 | 0 | 0 | 0 |
| 0 | 1 | 0 | 0 |
| 0 | 0 | 0 | 0 |
| 0 | 0 | 0 | 0 |
| 0 | 0 | 0 | 0 |
| 0 | 0 | 0 | 0 |
| 0 | 0 | 0 | 0 |
| 0 | 0 | 0 | 0 |
| 0 | 0 | 0 | 0 |
| 0 | 0 | 0 | 0 |

| | | | |
|---|---|---|---|
| 0 | 0 | 0 | 0 |
| 0 | 0 | 0 | 0 |
| 0 | 0 | 0 | 0 |
| 0 | 0 | 0 | 0 |
| 0 | 0 | 0 | 0 |
| 0 | 0 | 0 | 1 |
| 0 | 0 | 0 | 0 |
| 0 | 0 | 0 | 0 |
| 0 | 0 | 0 | 0 |
| 0 | 0 | 0 | 0 |
| 0 | 0 | 0 | 0 |
| 0 | 0 | 0 | 0 |
| 0 | 0 | 0 | 0 |
| 0 | 1 | 0 | 0 |
| 0 | 1 | 0 | 0 |
| 0 | 1 | 0 | 0 |
| 0 | 1 | 0 | 0 |
| 0 | 1 | 0 | 0 |
| 0 | 0 | 0 | 0 |
| 0 | 0 | 0 | 0 |
| 0 | 0 | 0 | 0 |
| 0 | 0 | 0 | 0 |
| 0 | 0 | 0 | 0 |
| 0 | 0 | 0 | 0 |
| 0 | 0 | 0 | 0 |
| 0 | 0 | 0 | 0 |
| 0 | 0 | 0 | 0 |
| 0 | 0 | 0 | 0 |
| 0 | 0 | 0 | 0 |
| 0 | 0 | 0 | 0 |
| 0 | 0 | 0 | 0 |
| 0 | 0 | 0 | 0 |
| 0 | 0 | 0 | 0 |
| 0 | 0 | 0 | 0 |
| 0 | 0 | 0 | 0 |
| 0 | 0 | 0 | 0 |
| 0 | 0 | 0 | 0 |
| 0 | 0 | 0 | 0 |
| 0 | 0 | 0 | 0 |
| 0 | 0 | 0 | 0 |
| 0 | 0 | 0 | 0 |
| 0 | 0 | 0 | 0 |
| 0 | 0 | 0 | 0 |
| 0 | 0 | 0 | 0 |
| 0 | 0 | 0 | 0 |
| 0 | 0 | 0 | 0 |
| 0 | 0 | 0 | 0 |
| 0 | 0 | 0 | 0 |
| 0 | 0 | 0 | 0 |
| 0 | 0 | 0 | 0 |
| 0 | 0 | 0 | 0 |
| 0 | 0 | 0 | 0 |
| 0 | 0 | 0 | 0 |
| 0 | 0 | 0 | 0 |

| | | | |
|---|---|---|---|
| 0 | 0 | 0 | 0 |
| 0 | 0 | 0 | 0 |
| 0 | 0 | 0 | 0 |
| 1 | 0 | 0 | 0 |
| 0 | 0 | 0 | 0 |
| 0 | 0 | 0 | 0 |
| 0 | 0 | 0 | 0 |
| 0 | 0 | 0 | 0 |
| 0 | 0 | 0 | 0 |
| 0 | 0 | 0 | 0 |
| 0 | 0 | 0 | 0 |
| 0 | 0 | 0 | 0 |
| 0 | 0 | 0 | 0 |
| 0 | 0 | 0 | 0 |
| 0 | 0 | 0 | 0 |
| 0 | 0 | 0 | 0 |
| 0 | 0 | 0 | 0 |
| 0 | 0 | 0 | 0 |
| 0 | 0 | 0 | 0 |
| 0 | 0 | 0 | 0 |
| 1 | 1 | 0 | 0 |
| 1 | 1 | 0 | 0 |
| 0 | 0 | 0 | 0 |
| 0 | 1 | 0 | 0 |
| 0 | 0 | 0 | 0 |
| 0 | 0 | 0 | 0 |
| 0 | 0 | 0 | 0 |
| 0 | 0 | 0 | 0 |
| 0 | 0 | 0 | 0 |
| 0 | 0 | 0 | 0 |
| 0 | 0 | 0 | 0 |
| 0 | 0 | 0 | 0 |
| 0 | 0 | 0 | 0 |
| 0 | 0 | 0 | 0 |
| 0 | 0 | 0 | 0 |
| 0 | 0 | 0 | 0 |
| 0 | 0 | 0 | 0 |
| 0 | 0 | 0 | 0 |
| 0 | 0 | 0 | 0 |
| 1 | 0 | 0 | 0 |
| 1 | 0 | 0 | 0 |
| 0 | 0 | 0 | 0 |
| 0 | 0 | 0 | 0 |
| 1 | 0 | 0 | 0 |
| 0 | 0 | 0 | 0 |
| 0 | 0 | 0 | 0 |
| 0 | 0 | 0 | 0 |
| 0 | 0 | 0 | 0 |
| 0 | 0 | 0 | 0 |
| 0 | 0 | 0 | 0 |
| 0 | 0 | 0 | 0 |
| 0 | 0 | 0 | 0 |
| 0 | 0 | 0 | 0 |
| 0 | 0 | 0 | 0 |

| | | | |
|---|---|---|---|
| 0 | 0 | 0 | 0 |
| 0 | 0 | 0 | 0 |
| 0 | 0 | 0 | 0 |
| 0 | 0 | 0 | 0 |
| 0 | 0 | 0 | 0 |
| 1 | 0 | 0 | 0 |
| 0 | 0 | 0 | 0 |
| 0 | 0 | 0 | 0 |
| 0 | 1 | 0 | 0 |
| 1 | 1 | 0 | 0 |
| 0 | 0 | 0 | 0 |
| 1 | 1 | 0 | 0 |
| 1 | 0 | 0 | 0 |
| 0 | 0 | 0 | 0 |
| 0 | 0 | 0 | 0 |
| 0 | 0 | 0 | 0 |
| 1 | 1 | 0 | 0 |
| 0 | 0 | 0 | 0 |
| 0 | 0 | 0 | 0 |
| 1 | 0 | 0 | 0 |
| 0 | 0 | 0 | 0 |
| 0 | 0 | 0 | 0 |
| 0 | 0 | 0 | 0 |
| 0 | 0 | 0 | 0 |
| 0 | 0 | 0 | 0 |
| 0 | 0 | 0 | 0 |
| 0 | 0 | 0 | 0 |
| 0 | 0 | 0 | 0 |
| 0 | 0 | 0 | 0 |
| 0 | 0 | 0 | 0 |
| 1 | 0 | 0 | 0 |
| 0 | 0 | 0 | 0 |
| 0 | 0 | 0 | 0 |
| 0 | 0 | 0 | 0 |
| 0 | 0 | 0 | 0 |
| 1 | 0 | 0 | 0 |
| 0 | 0 | 0 | 0 |
| 0 | 0 | 0 | 0 |
| 0 | 0 | 0 | 0 |
| 0 | 0 | 0 | 0 |
| 0 | 0 | 0 | 0 |
| 0 | 0 | 0 | 0 |
| 0 | 0 | 0 | 0 |
| 0 | 0 | 0 | 0 |
| 0 | 0 | 0 | 0 |
| 0 | 0 | 0 | 0 |
| 0 | 0 | 0 | 0 |
| 0 | 0 | 0 | 0 |
| 0 | 0 | 0 | 0 |
| 0 | 0 | 0 | 0 |
| 0 | 0 | 0 | 0 |
| 0 | 0 | 0 | 0 |
| 0 | 0 | 0 | 0 |
| 0 | 0 | 0 | 0 |

| | | | |
|---|---|---|---|
| 0 | 0 | 0 | 0 |
| 0 | 0 | 0 | 0 |
| 0 | 0 | 0 | 0 |
| 0 | 0 | 0 | 0 |
| 0 | 0 | 0 | 0 |
| 0 | 0 | 0 | 0 |
| 0 | 0 | 0 | 0 |
| 0 | 0 | 0 | 0 |
| 0 | 0 | 0 | 0 |
| 0 | 0 | 0 | 0 |
| 0 | 0 | 0 | 0 |
| 0 | 0 | 0 | 0 |
| 0 | 0 | 0 | 0 |
| 0 | 0 | 0 | 0 |
| 0 | 0 | 0 | 1 |
| 0 | 0 | 0 | 0 |
| 0 | 0 | 0 | 0 |
| 0 | 0 | 0 | 0 |
| 0 | 0 | 0 | 0 |
| 0 | 0 | 0 | 0 |
| 0 | 0 | 0 | 0 |
| 0 | 0 | 0 | 0 |
| 0 | 0 | 0 | 0 |
| 0 | 0 | 0 | 0 |
| 0 | 0 | 0 | 0 |
| 0 | 0 | 0 | 0 |
| 0 | 0 | 0 | 0 |
| 0 | 0 | 0 | 0 |
| 0 | 0 | 0 | 0 |
| 0 | 0 | 0 | 0 |
| 0 | 0 | 0 | 0 |
| 0 | 0 | 0 | 0 |
| 0 | 0 | 0 | 0 |
| 0 | 0 | 0 | 0 |
| 0 | 0 | 0 | 0 |
| 0 | 0 | 0 | 0 |
| 0 | 0 | 0 | 0 |
| 0 | 0 | 0 | 0 |
| 0 | 0 | 0 | 0 |
| 0 | 0 | 1 | 0 |
| 0 | 0 | 0 | 0 |
| 0 | 0 | 0 | 0 |
| 0 | 0 | 0 | 0 |
| 1 | 0 | 0 | 0 |
| 0 | 0 | 0 | 0 |
| 0 | 0 | 0 | 0 |
| 0 | 0 | 0 | 0 |
| 0 | 0 | 0 | 0 |
| 1 | 0 | 0 | 1 |
| 0 | 0 | 0 | 0 |
| 0 | 0 | 0 | 0 |
| 0 | 0 | 0 | 0 |
| 0 | 0 | 0 | 0 |
| 0 | 0 | 0 | 0 |

| | | | |
|---|---|---|---|
| 0 | 0 | 0 | 0 |
| 0 | 0 | 0 | 0 |
| 0 | 0 | 0 | 0 |
| 0 | 0 | 0 | 0 |
| 0 | 0 | 0 | 0 |
| 0 | 0 | 0 | 0 |
| 0 | 0 | 0 | 0 |
| 0 | 0 | 0 | 0 |
| 0 | 0 | 0 | 0 |
| 1 | 0 | 0 | 0 |
| 0 | 0 | 0 | 0 |
| 0 | 0 | 0 | 0 |
| 0 | 0 | 0 | 0 |
| 0 | 0 | 0 | 0 |
| 1 | 0 | 0 | 0 |
| 0 | 0 | 0 | 0 |
| 0 | 0 | 0 | 0 |
| 0 | 0 | 0 | 0 |
| 0 | 0 | 0 | 0 |
| 0 | 0 | 0 | 0 |
| 0 | 0 | 0 | 0 |
| 0 | 0 | 0 | 0 |
| 0 | 0 | 0 | 0 |
| 0 | 0 | 0 | 0 |
| 1 | 0 | 0 | 0 |
| 0 | 0 | 0 | 0 |
| 0 | 0 | 0 | 0 |
| 0 | 0 | 0 | 0 |
| 0 | 0 | 0 | 0 |
| 0 | 0 | 0 | 0 |
| 0 | 0 | 0 | 0 |
| 0 | 0 | 0 | 0 |
| 0 | 0 | 0 | 0 |
| 0 | 0 | 0 | 0 |
| 1 | 0 | 0 | 0 |
| 1 | 0 | 0 | 0 |
| 0 | 0 | 0 | 0 |
| 0 | 0 | 0 | 0 |
| 1 | 0 | 0 | 0 |
| 0 | 0 | 1 | 0 |
| 0 | 0 | 1 | 0 |
| 0 | 0 | 1 | 0 |
| 0 | 0 | 1 | 0 |
| 0 | 0 | 0 | 0 |
| 1 | 0 | 0 | 0 |
| 0 | 0 | 0 | 0 |
| 0 | 0 | 0 | 0 |
| 0 | 0 | 0 | 0 |
| 0 | 0 | 1 | 0 |
| 0 | 0 | 0 | 0 |
| 0 | 0 | 0 | 0 |
| 0 | 0 | 0 | 0 |
| 0 | 0 | 0 | 0 |
| 0 | 0 | 0 | 0 |

| | | | |
|---|---|---|---|
| 0 | 0 | 0 | 0 |
| 0 | 0 | 0 | 0 |
| 0 | 0 | 0 | 0 |
| 0 | 0 | 0 | 0 |
| 0 | 0 | 0 | 0 |
| 0 | 0 | 0 | 0 |
| 0 | 0 | 0 | 0 |
| 0 | 0 | 0 | 0 |
| 0 | 0 | 0 | 0 |
| 0 | 0 | 0 | 0 |
| 0 | 1 | 0 | 0 |
| 0 | 0 | 0 | 0 |
| 0 | 0 | 0 | 0 |
| 0 | 0 | 0 | 0 |
| 0 | 0 | 0 | 0 |
| 0 | 1 | 0 | 0 |
| 0 | 0 | 0 | 0 |
| 0 | 0 | 0 | 0 |
| 0 | 0 | 0 | 0 |
| 0 | 0 | 0 | 0 |
| 0 | 0 | 0 | 0 |
| 0 | 0 | 0 | 0 |
| 0 | 0 | 0 | 0 |
| 0 | 0 | 0 | 0 |
| 0 | 0 | 0 | 0 |
| 0 | 0 | 0 | 0 |
| 0 | 0 | 0 | 0 |
| 0 | 0 | 0 | 0 |
| 0 | 0 | 0 | 0 |
| 0 | 0 | 0 | 0 |
| 0 | 0 | 0 | 0 |
| 0 | 0 | 0 | 0 |
| 0 | 0 | 0 | 0 |
| 0 | 0 | 0 | 0 |
| 0 | 0 | 0 | 0 |
| 0 | 0 | 0 | 0 |
| 0 | 0 | 0 | 0 |
| 0 | 0 | 0 | 0 |
| 0 | 0 | 0 | 0 |
| 0 | 0 | 0 | 0 |
| 0 | 0 | 0 | 0 |
| 0 | 0 | 0 | 0 |
| 0 | 0 | 0 | 0 |
| 0 | 0 | 0 | 0 |
| 1 | 1 | 0 | 0 |
| 0 | 0 | 0 | 0 |
| 0 | 0 | 0 | 0 |
| 0 | 0 | 0 | 0 |
| 0 | 0 | 0 | 0 |
| 0 | 0 | 0 | 0 |
| 0 | 0 | 0 | 0 |
| 0 | 0 | 0 | 0 |
| 0 | 0 | 0 | 0 |
| 0 | 0 | 0 | 0 |

| | | | |
|---|---|---|---|
| 0 | 0 | 0 | 0 |
| 0 | 0 | 0 | 0 |
| 0 | 0 | 1 | 0 |
| 1 | 0 | 0 | 0 |
| 0 | 0 | 0 | 0 |
| 0 | 0 | 0 | 0 |
| 0 | 0 | 0 | 0 |
| 0 | 0 | 0 | 0 |
| 0 | 0 | 0 | 0 |
| 0 | 0 | 0 | 0 |
| 0 | 0 | 0 | 0 |
| 0 | 0 | 0 | 0 |
| 1 | 0 | 0 | 0 |
| 0 | 0 | 0 | 0 |
| 0 | 0 | 0 | 0 |
| 0 | 0 | 0 | 0 |
| 0 | 0 | 0 | 0 |
| 0 | 0 | 0 | 0 |
| 0 | 0 | 0 | 0 |
| 0 | 0 | 0 | 0 |
| 0 | 0 | 0 | 0 |
| 0 | 0 | 0 | 0 |
| 0 | 0 | 0 | 0 |
| 0 | 0 | 0 | 0 |
| 0 | 0 | 0 | 0 |
| 0 | 0 | 0 | 0 |
| 0 | 0 | 0 | 0 |
| 0 | 0 | 0 | 0 |
| 0 | 0 | 0 | 0 |
| 0 | 0 | 0 | 0 |
| 0 | 0 | 0 | 0 |
| 0 | 0 | 0 | 0 |
| 0 | 0 | 0 | 0 |
| 0 | 0 | 0 | 0 |
| 0 | 0 | 0 | 0 |
| 0 | 0 | 0 | 0 |
| 0 | 0 | 0 | 0 |
| 0 | 0 | 0 | 0 |
| 0 | 0 | 0 | 0 |
| 0 | 0 | 0 | 0 |
| 0 | 1 | 0 | 0 |
| 1 | 1 | 0 | 0 |
| 0 | 0 | 0 | 0 |
| 0 | 0 | 0 | 0 |
| 0 | 0 | 0 | 0 |
| 0 | 0 | 0 | 0 |
| 0 | 0 | 0 | 0 |
| 0 | 0 | 0 | 0 |
| 0 | 1 | 0 | 0 |
| 0 | 0 | 0 | 0 |
| 0 | 0 | 0 | 0 |
| 0 | 0 | 0 | 0 |
| 0 | 0 | 0 | 1 |
| 0 | 0 | 0 | 0 |

| | | | |
|---|---|---|---|
| 0 | 0 | 0 | 1 |
| 0 | 0 | 0 | 1 |
| 0 | 0 | 0 | 1 |
| 0 | 0 | 0 | 0 |
| 0 | 0 | 0 | 0 |
| 0 | 0 | 0 | 0 |
| 0 | 0 | 0 | 0 |
| 1 | 0 | 0 | 0 |
| 0 | 0 | 0 | 0 |
| 1 | 1 | 0 | 0 |
| 0 | 1 | 0 | 0 |
| 0 | 0 | 0 | 0 |
| 1 | 1 | 0 | 0 |
| 0 | 0 | 0 | 0 |
| 0 | 0 | 0 | 0 |
| 0 | 0 | 0 | 0 |
| 0 | 0 | 0 | 0 |
| 1 | 0 | 0 | 0 |
| 0 | 0 | 0 | 0 |
| 0 | 0 | 0 | 0 |
| 0 | 0 | 0 | 0 |
| 0 | 0 | 0 | 0 |
| 0 | 0 | 0 | 0 |
| 0 | 0 | 0 | 0 |
| 0 | 0 | 0 | 0 |
| 0 | 0 | 0 | 0 |
| 0 | 0 | 0 | 0 |
| 0 | 0 | 0 | 0 |
| 0 | 0 | 0 | 0 |
| 0 | 0 | 0 | 0 |
| 0 | 0 | 0 | 0 |
| 1 | 0 | 0 | 0 |
| 0 | 0 | 0 | 0 |
| 0 | 0 | 0 | 0 |
| 0 | 0 | 0 | 0 |
| 0 | 0 | 0 | 0 |
| 0 | 0 | 0 | 0 |
| 0 | 0 | 0 | 0 |
| 0 | 0 | 0 | 0 |
| 0 | 0 | 0 | 0 |
| 0 | 0 | 0 | 0 |
| 1 | 1 | 0 | 0 |
| 0 | 0 | 0 | 0 |
| 0 | 0 | 0 | 0 |
| 1 | 0 | 0 | 0 |
| 0 | 1 | 0 | 0 |
| 0 | 0 | 0 | 0 |
| 0 | 0 | 0 | 0 |
| 0 | 1 | 0 | 0 |
| 1 | 1 | 0 | 0 |
| 1 | 1 | 0 | 0 |
| 1 | 1 | 0 | 0 |
| 1 | 1 | 0 | 0 |
| 1 | 1 | 0 | 0 |

| | | | |
|---|---|---|---|
| 0 | 0 | 0 | 0 |
| 0 | 0 | 0 | 0 |
| 0 | 0 | 0 | 0 |
| 0 | 0 | 0 | 0 |
| 0 | 0 | 0 | 0 |
| 1 | 0 | 0 | 0 |
| 1 | 0 | 0 | 0 |
| 0 | 0 | 0 | 0 |
| 0 | 0 | 0 | 0 |
| 1 | 0 | 0 | 0 |
| 0 | 0 | 0 | 0 |
| 0 | 0 | 1 | 0 |
| 0 | 0 | 0 | 0 |
| 0 | 0 | 0 | 0 |
| 0 | 0 | 0 | 0 |
| 0 | 0 | 0 | 0 |
| 0 | 0 | 0 | 0 |
| 0 | 0 | 0 | 0 |
| 0 | 0 | 0 | 0 |
| 0 | 0 | 0 | 0 |
| 1 | 1 | 0 | 0 |
| 1 | 0 | 0 | 0 |
| 0 | 0 | 0 | 0 |
| 0 | 0 | 1 | 0 |
| 0 | 0 | 0 | 0 |
| 0 | 0 | 1 | 0 |
| 0 | 0 | 0 | 0 |
| 0 | 0 | 0 | 0 |
| 0 | 0 | 0 | 0 |
| 0 | 0 | 0 | 0 |
| 0 | 0 | 0 | 0 |
| 0 | 0 | 0 | 0 |
| 0 | 0 | 0 | 0 |
| 0 | 0 | 0 | 0 |
| 0 | 0 | 0 | 0 |
| 0 | 0 | 0 | 0 |
| 1 | 0 | 0 | 0 |
| 0 | 0 | 0 | 0 |
| 1 | 0 | 0 | 0 |
| 1 | 0 | 0 | 0 |
| 1 | 0 | 0 | 0 |
| 0 | 0 | 0 | 0 |
| 0 | 0 | 0 | 0 |
| 0 | 0 | 0 | 0 |
| 0 | 0 | 0 | 0 |
| 0 | 0 | 0 | 0 |
| 0 | 0 | 0 | 0 |
| 1 | 0 | 1 | 0 |
| 0 | 0 | 0 | 0 |
| 0 | 0 | 0 | 0 |
| 0 | 0 | 0 | 0 |
| 0 | 0 | 0 | 0 |
| 1 | 1 | 0 | 0 |
| 0 | 0 | 0 | 0 |

| | | | |
|---|---|---|---|
| 0 | 0 | 0 | 0 |
| 0 | 1 | 0 | 0 |
| 0 | 0 | 0 | 0 |
| 0 | 0 | 0 | 0 |
| 0 | 0 | 0 | 0 |
| 0 | 0 | 0 | 0 |
| 0 | 0 | 0 | 0 |
| 0 | 0 | 0 | 0 |
| 0 | 0 | 0 | 0 |
| 0 | 0 | 0 | 0 |
| 0 | 0 | 0 | 0 |
| 0 | 0 | 0 | 0 |
| 0 | 0 | 0 | 0 |
| 0 | 0 | 0 | 0 |
| 0 | 0 | 0 | 0 |
| 0 | 0 | 0 | 0 |
| 0 | 0 | 0 | 0 |
| 0 | 0 | 0 | 0 |
| 0 | 0 | 0 | 0 |
| 0 | 0 | 0 | 0 |
| 1 | 0 | 0 | 0 |
| 0 | 0 | 0 | 0 |
| 0 | 0 | 0 | 0 |
| 0 | 0 | 0 | 0 |
| 1 | 0 | 0 | 0 |
| 0 | 0 | 0 | 0 |
| 0 | 0 | 0 | 0 |
| 0 | 0 | 0 | 1 |
| 0 | 0 | 0 | 0 |
| 0 | 0 | 0 | 0 |
| 0 | 0 | 0 | 0 |
| 0 | 0 | 0 | 0 |
| 0 | 0 | 0 | 0 |
| 0 | 0 | 0 | 0 |
| 0 | 0 | 0 | 0 |
| 0 | 0 | 0 | 0 |
| 0 | 0 | 0 | 0 |
| 0 | 0 | 0 | 0 |
| 0 | 0 | 0 | 0 |
| 0 | 0 | 0 | 0 |
| 0 | 0 | 0 | 0 |
| 0 | 0 | 0 | 0 |
| 0 | 0 | 0 | 0 |
| 0 | 0 | 0 | 0 |
| 0 | 0 | 0 | 0 |
| 0 | 0 | 0 | 0 |
| 0 | 0 | 0 | 0 |
| 0 | 0 | 0 | 0 |
| 0 | 0 | 0 | 0 |
| 0 | 0 | 0 | 0 |
| 0 | 0 | 0 | 0 |
| 0 | 0 | 0 | 0 |
| 0 | 0 | 0 | 0 |
| 0 | 0 | 0 | 0 |

| | | | |
|---|---|---|---|
| 0 | 0 | 0 | 0 |
| 0 | 0 | 0 | 0 |
| 0 | 0 | 0 | 0 |
| 0 | 0 | 0 | 0 |
| 0 | 0 | 0 | 0 |
| 0 | 0 | 0 | 0 |
| 0 | 0 | 0 | 0 |
| 0 | 0 | 0 | 0 |
| 0 | 0 | 0 | 0 |
| 0 | 0 | 0 | 0 |
| 0 | 0 | 0 | 0 |
| 0 | 0 | 0 | 0 |
| 0 | 0 | 0 | 0 |
| 0 | 0 | 0 | 0 |
| 0 | 0 | 0 | 0 |
| 1 | 0 | 0 | 0 |
| 0 | 0 | 0 | 0 |
| 0 | 0 | 0 | 0 |
| 0 | 1 | 0 | 0 |
| 0 | 0 | 0 | 0 |
| 0 | 0 | 0 | 0 |
| 0 | 0 | 0 | 0 |
| 0 | 0 | 0 | 0 |
| 0 | 0 | 0 | 0 |
| 0 | 0 | 0 | 0 |
| 1 | 0 | 0 | 0 |
| 0 | 0 | 0 | 0 |
| 0 | 0 | 0 | 0 |
| 0 | 0 | 0 | 0 |
| 0 | 0 | 0 | 0 |
| 0 | 0 | 0 | 0 |
| 0 | 0 | 0 | 0 |
| 0 | 0 | 0 | 0 |
| 1 | 1 | 0 | 0 |
| 0 | 0 | 0 | 0 |
| 0 | 0 | 0 | 0 |
| 0 | 0 | 0 | 0 |
| 0 | 0 | 0 | 0 |
| 0 | 0 | 0 | 0 |
| 0 | 0 | 0 | 0 |
| 0 | 0 | 0 | 0 |
| 0 | 0 | 0 | 0 |
| 0 | 0 | 0 | 0 |
| 0 | 0 | 0 | 0 |
| 0 | 0 | 0 | 0 |
| 0 | 0 | 0 | 0 |
| 0 | 0 | 0 | 0 |
| 1 | 0 | 0 | 0 |
| 0 | 0 | 0 | 0 |
| 0 | 0 | 0 | 0 |
| 0 | 0 | 0 | 0 |
| 0 | 0 | 0 | 0 |
| 0 | 0 | 0 | 0 |
| 1 | 0 | 0 | 1 |

| | | | |
|---|---|---|---|
| 0 | 0 | 0 | 0 |
| 0 | 0 | 0 | 0 |
| 0 | 0 | 1 | 0 |
| 0 | 0 | 0 | 0 |
| 0 | 0 | 0 | 0 |
| 0 | 0 | 0 | 0 |
| 0 | 0 | 0 | 0 |
| 0 | 0 | 0 | 0 |
| 0 | 0 | 0 | 0 |
| 0 | 0 | 0 | 0 |
| 0 | 0 | 0 | 0 |
| 0 | 0 | 0 | 0 |
| 0 | 0 | 0 | 0 |
| 0 | 0 | 0 | 0 |
| 0 | 0 | 0 | 0 |
| 0 | 0 | 0 | 0 |
| 0 | 0 | 0 | 0 |
| 0 | 0 | 0 | 0 |
| 0 | 0 | 0 | 0 |
| 0 | 0 | 0 | 0 |
| 0 | 0 | 0 | 0 |
| 0 | 0 | 0 | 0 |
| 0 | 0 | 0 | 0 |
| 0 | 0 | 0 | 0 |
| 0 | 0 | 0 | 0 |
| 0 | 0 | 0 | 0 |
| 0 | 0 | 0 | 0 |
| 0 | 0 | 0 | 0 |
| 0 | 0 | 0 | 0 |
| 0 | 0 | 0 | 0 |
| 0 | 0 | 0 | 0 |
| 1 | 0 | 0 | 0 |
| 0 | 0 | 0 | 0 |
| 0 | 0 | 0 | 0 |
| 0 | 0 | 0 | 0 |
| 0 | 0 | 0 | 0 |
| 0 | 0 | 0 | 0 |
| 0 | 0 | 0 | 0 |
| 0 | 0 | 0 | 0 |
| 0 | 0 | 0 | 0 |
| 1 | 1 | 0 | 0 |
| 0 | 0 | 0 | 0 |
| 0 | 0 | 0 | 0 |
| 0 | 0 | 0 | 0 |
| 0 | 0 | 0 | 0 |
| 1 | 1 | 0 | 0 |
| 0 | 0 | 0 | 0 |
| 1 | 1 | 0 | 0 |
| 0 | 0 | 0 | 0 |
| 1 | 0 | 0 | 0 |
| 1 | 0 | 0 | 0 |
| 1 | 0 | 0 | 0 |
| 0 | 0 | 0 | 0 |
| 0 | 0 | 0 | 0 |

| | | | |
|---|---|---|---|
| 1 | 0 | 0 | 0 |
| 0 | 0 | 0 | 0 |
| 0 | 0 | 0 | 0 |
| 0 | 0 | 0 | 0 |
| 0 | 0 | 0 | 0 |
| 0 | 0 | 0 | 0 |
| 0 | 0 | 0 | 0 |
| 0 | 0 | 0 | 0 |
| 0 | 0 | 0 | 0 |
| 0 | 0 | 0 | 0 |
| 0 | 0 | 0 | 0 |
| 0 | 0 | 0 | 0 |
| 0 | 0 | 0 | 0 |
| 0 | 0 | 0 | 0 |
| 0 | 0 | 0 | 0 |
| 1 | 1 | 1 | 0 |
| 0 | 0 | 0 | 0 |
| 1 | 0 | 0 | 0 |
| 0 | 0 | 0 | 0 |
| 0 | 0 | 0 | 0 |
| 0 | 0 | 0 | 0 |
| 0 | 1 | 0 | 0 |
| 0 | 0 | 0 | 0 |
| 0 | 1 | 0 | 0 |
| 0 | 1 | 0 | 0 |
| 0 | 0 | 0 | 0 |
| 0 | 0 | 0 | 0 |
| 0 | 0 | 0 | 0 |
| 0 | 0 | 0 | 0 |
| 0 | 0 | 0 | 0 |
| 0 | 0 | 0 | 0 |
| 0 | 0 | 0 | 0 |
| 1 | 1 | 0 | 0 |
| 0 | 0 | 0 | 0 |
| 0 | 0 | 0 | 0 |
| 0 | 0 | 0 | 0 |
| 0 | 1 | 0 | 0 |
| 1 | 1 | 0 | 0 |
| 0 | 0 | 0 | 0 |
| 0 | 0 | 0 | 0 |
| 0 | 0 | 0 | 0 |
| 0 | 0 | 0 | 0 |
| 0 | 0 | 0 | 0 |
| 0 | 0 | 0 | 0 |
| 1 | 1 | 0 | 0 |
| 0 | 0 | 0 | 0 |
| 1 | 0 | 0 | 0 |
| 1 | 1 | 0 | 0 |
| 0 | 0 | 0 | 0 |
| 0 | 0 | 0 | 0 |
| 1 | 1 | 0 | 0 |
| 1 | 1 | 0 | 0 |
| 0 | 0 | 0 | 0 |
| 1 | 1 | 0 | 0 |

| | | | |
|---|---|---|---|
| 0 | 0 | 0 | 0 |
| 0 | 0 | 0 | 0 |
| 0 | 0 | 0 | 0 |
| 1 | 1 | 0 | 0 |
| 0 | 0 | 0 | 0 |
| 0 | 0 | 0 | 0 |
| 0 | 0 | 0 | 0 |
| 0 | 0 | 0 | 0 |
| 1 | 0 | 0 | 0 |
| 0 | 0 | 0 | 0 |
| 0 | 0 | 0 | 0 |
| 0 | 0 | 0 | 0 |
| 1 | 0 | 1 | 0 |
| 1 | 0 | 0 | 0 |
| 0 | 0 | 0 | 0 |
| 0 | 0 | 0 | 0 |
| 0 | 0 | 0 | 0 |
| 0 | 0 | 0 | 0 |
| 0 | 0 | 0 | 0 |
| 0 | 0 | 0 | 0 |
| 0 | 0 | 0 | 0 |
| 0 | 0 | 0 | 0 |
| 0 | 0 | 0 | 0 |
| 1 | 0 | 0 | 0 |
| 0 | 0 | 0 | 0 |
| 0 | 0 | 0 | 0 |
| 0 | 0 | 0 | 0 |
| 0 | 0 | 0 | 0 |
| 0 | 0 | 0 | 0 |
| 0 | 0 | 0 | 0 |
| 0 | 0 | 0 | 0 |
| 0 | 0 | 0 | 0 |
| 0 | 0 | 0 | 0 |
| 0 | 0 | 0 | 0 |
| 0 | 0 | 0 | 0 |
| 0 | 0 | 0 | 0 |
| 0 | 0 | 0 | 0 |
| 0 | 0 | 0 | 0 |
| 0 | 0 | 0 | 0 |
| 0 | 0 | 0 | 0 |
| 0 | 0 | 0 | 0 |
| 0 | 0 | 0 | 0 |
| 0 | 0 | 0 | 0 |
| 1 | 1 | 0 | 0 |
| 1 | 1 | 0 | 0 |
| 0 | 0 | 0 | 0 |
| 0 | 0 | 0 | 0 |
| 0 | 1 | 0 | 0 |
| 0 | 1 | 0 | 0 |
| 0 | 1 | 0 | 0 |
| 0 | 0 | 0 | 0 |
| 0 | 0 | 0 | 0 |
| 0 | 0 | 0 | 0 |
| 0 | 0 | 0 | 0 |

| | | | |
|---|---|---|---|
| 0 | 0 | 0 | 0 |
| 0 | 0 | 0 | 0 |
| 0 | 0 | 0 | 0 |
| 0 | 0 | 0 | 0 |
| 1 | 0 | 0 | 0 |
| 0 | 0 | 0 | 0 |
| 0 | 0 | 0 | 0 |
| 1 | 0 | 0 | 0 |
| 1 | 0 | 0 | 0 |
| 1 | 0 | 0 | 0 |
| 1 | 0 | 0 | 0 |
| 1 | 0 | 0 | 0 |
| 0 | 0 | 0 | 0 |
| 1 | 0 | 0 | 0 |
| 1 | 0 | 0 | 0 |
| 0 | 0 | 0 | 0 |
| 0 | 0 | 0 | 0 |
| 0 | 0 | 0 | 0 |
| 1 | 0 | 0 | 0 |
| 1 | 0 | 0 | 0 |
| 1 | 0 | 0 | 0 |
| 1 | 0 | 0 | 0 |
| 0 | 0 | 0 | 0 |
| 0 | 0 | 0 | 0 |
| 0 | 0 | 0 | 0 |
| 0 | 0 | 0 | 0 |
| 0 | 0 | 0 | 0 |
| 0 | 0 | 0 | 0 |
| 0 | 0 | 0 | 0 |
| 0 | 0 | 0 | 0 |
| 0 | 0 | 0 | 0 |
| 0 | 0 | 0 | 0 |
| 1 | 0 | 0 | 0 |
| 0 | 0 | 0 | 0 |
| 0 | 0 | 0 | 0 |
| 1 | 0 | 0 | 0 |
| 0 | 0 | 0 | 0 |
| 0 | 0 | 0 | 0 |
| 0 | 0 | 0 | 0 |
| 0 | 0 | 0 | 0 |
| 0 | 0 | 0 | 0 |
| 0 | 0 | 0 | 0 |
| 0 | 0 | 0 | 0 |
| 0 | 0 | 0 | 0 |
| 0 | 0 | 0 | 0 |
| 0 | 0 | 0 | 0 |
| 0 | 0 | 0 | 0 |
| 0 | 0 | 0 | 0 |
| 0 | 0 | 0 | 0 |
| 0 | 0 | 0 | 0 |
| 0 | 0 | 0 | 0 |
| 0 | 0 | 0 | 0 |
| 0 | 0 | 0 | 0 |
| 0 | 0 | 0 | 0 |

| | | | |
|---|---|---|---|
| 0 | 0 | 0 | 0 |
| 0 | 0 | 0 | 0 |
| 0 | 0 | 0 | 0 |
| 0 | 0 | 0 | 0 |
| 0 | 0 | 0 | 0 |
| 0 | 0 | 0 | 0 |
| 0 | 0 | 0 | 0 |
| 0 | 0 | 0 | 0 |
| 0 | 0 | 0 | 0 |
| 0 | 0 | 0 | 0 |
| 0 | 0 | 0 | 0 |
| 0 | 0 | 0 | 1 |
| 0 | 0 | 0 | 0 |
| 0 | 0 | 0 | 0 |
| 0 | 0 | 0 | 0 |
| 0 | 0 | 0 | 0 |
| 0 | 0 | 0 | 0 |
| 0 | 0 | 0 | 1 |
| 0 | 0 | 0 | 0 |
| 0 | 0 | 0 | 0 |
| 0 | 0 | 0 | 1 |
| 0 | 0 | 0 | 0 |
| 0 | 0 | 0 | 0 |
| 0 | 0 | 0 | 0 |
| 0 | 0 | 0 | 0 |
| 0 | 0 | 0 | 0 |
| 0 | 0 | 0 | 0 |
| 0 | 0 | 0 | 0 |
| 0 | 0 | 0 | 1 |
| 0 | 0 | 0 | 0 |
| 0 | 0 | 0 | 0 |
| 0 | 0 | 0 | 0 |
| 0 | 0 | 0 | 0 |
| 0 | 0 | 0 | 0 |
| 0 | 0 | 0 | 0 |
| 0 | 0 | 0 | 0 |
| 0 | 0 | 0 | 0 |
| 0 | 0 | 0 | 0 |
| 0 | 0 | 0 | 0 |
| 0 | 0 | 0 | 0 |
| 0 | 0 | 0 | 0 |
| 0 | 0 | 0 | 0 |
| 0 | 0 | 0 | 0 |
| 0 | 0 | 0 | 0 |
| 0 | 0 | 0 | 0 |
| 0 | 0 | 0 | 0 |
| 0 | 0 | 0 | 0 |
| 0 | 0 | 0 | 0 |
| 0 | 0 | 0 | 0 |
| 0 | 0 | 0 | 0 |
| 0 | 0 | 0 | 0 |
| 0 | 0 | 0 | 0 |
| 0 | 0 | 0 | 0 |
| 0 | 0 | 0 | 0 |

| | | | |
|---|---|---|---|
| 0 | 0 | 0 | 0 |
| 0 | 0 | 0 | 0 |
| 1 | 0 | 0 | 0 |
| 0 | 0 | 0 | 0 |
| 1 | 0 | 0 | 0 |
| 0 | 0 | 0 | 0 |
| 0 | 0 | 0 | 0 |
| 0 | 0 | 0 | 0 |
| 0 | 0 | 0 | 0 |
| 1 | 1 | 0 | 0 |
| 0 | 0 | 1 | 0 |
| 0 | 0 | 0 | 0 |
| 0 | 0 | 0 | 0 |
| 0 | 0 | 0 | 0 |
| 0 | 0 | 0 | 0 |
| 0 | 0 | 0 | 0 |
| 0 | 0 | 0 | 0 |
| 0 | 0 | 0 | 0 |
| 0 | 0 | 0 | 0 |
| 0 | 0 | 0 | 0 |
| 0 | 0 | 0 | 0 |
| 0 | 0 | 0 | 0 |
| 0 | 0 | 0 | 0 |
| 0 | 0 | 0 | 0 |
| 0 | 0 | 0 | 0 |
| 0 | 0 | 0 | 0 |
| 0 | 0 | 0 | 0 |
| 0 | 0 | 0 | 0 |
| 0 | 0 | 0 | 0 |
| 0 | 0 | 0 | 0 |
| 0 | 0 | 0 | 0 |
| 0 | 0 | 0 | 0 |
| 0 | 0 | 0 | 0 |
| 0 | 0 | 0 | 0 |
| 0 | 0 | 0 | 0 |
| 0 | 0 | 0 | 0 |
| 0 | 0 | 0 | 0 |
| 0 | 0 | 0 | 0 |
| 0 | 0 | 0 | 0 |
| 0 | 0 | 0 | 0 |
| 0 | 0 | 0 | 0 |
| 0 | 0 | 0 | 0 |
| 0 | 0 | 0 | 0 |
| 0 | 0 | 0 | 0 |
| 0 | 0 | 0 | 0 |
| 1 | 0 | 0 | 0 |
| 0 | 0 | 0 | 0 |
| 0 | 0 | 0 | 0 |
| 0 | 0 | 0 | 0 |
| 0 | 0 | 0 | 0 |
| 1 | 0 | 0 | 0 |
| 0 | 0 | 0 | 0 |
| 0 | 0 | 0 | 0 |
| 0 | 0 | 0 | 0 |

| | | | |
|---|---|---|---|
| 0 | 0 | 0 | 0 |
| 0 | 0 | 0 | 0 |
| 0 | 0 | 0 | 0 |
| 0 | 0 | 0 | 0 |
| 0 | 0 | 0 | 0 |
| 0 | 0 | 0 | 0 |
| 0 | 0 | 0 | 0 |
| 0 | 0 | 0 | 0 |
| 1 | 1 | 0 | 0 |
| 0 | 0 | 0 | 0 |
| 0 | 0 | 0 | 0 |
| 0 | 0 | 0 | 0 |
| 0 | 0 | 0 | 0 |
| 0 | 0 | 0 | 0 |
| 0 | 1 | 0 | 0 |
| 0 | 1 | 0 | 0 |
| 0 | 1 | 0 | 0 |
| 0 | 0 | 0 | 0 |
| 0 | 0 | 0 | 0 |
| 0 | 0 | 0 | 0 |
| 0 | 1 | 0 | 0 |
| 0 | 0 | 0 | 0 |
| 0 | 0 | 0 | 0 |
| 1 | 1 | 0 | 0 |
| 0 | 1 | 0 | 0 |
| 0 | 1 | 0 | 0 |
| 0 | 0 | 0 | 0 |
| 0 | 0 | 0 | 0 |
| 0 | 0 | 0 | 0 |
| 0 | 0 | 0 | 0 |
| 0 | 0 | 0 | 0 |
| 0 | 0 | 0 | 0 |
| 0 | 0 | 0 | 0 |
| 0 | 0 | 0 | 0 |
| 0 | 0 | 0 | 0 |
| 0 | 0 | 0 | 0 |
| 0 | 0 | 0 | 0 |
| 1 | 0 | 0 | 0 |
| 0 | 0 | 0 | 0 |
| 0 | 0 | 0 | 0 |
| 0 | 0 | 0 | 0 |
| 0 | 0 | 0 | 0 |
| 0 | 0 | 0 | 0 |
| 0 | 0 | 0 | 0 |
| 0 | 0 | 0 | 0 |
| 0 | 0 | 0 | 0 |
| 0 | 0 | 0 | 0 |
| 0 | 0 | 0 | 0 |
| 0 | 0 | 0 | 0 |
| 0 | 0 | 0 | 0 |
| 0 | 0 | 0 | 0 |
| 0 | 0 | 0 | 0 |
| 0 | 0 | 0 | 0 |
| 0 | 0 | 0 | 0 |

| | | | |
|---|---|---|---|
| 0 | 0 | 0 | 0 |
| 0 | 0 | 0 | 0 |
| 0 | 0 | 0 | 0 |
| 0 | 0 | 1 | 0 |
| 0 | 0 | 0 | 0 |
| 0 | 0 | 0 | 0 |
| 0 | 0 | 0 | 0 |
| 0 | 0 | 0 | 0 |
| 0 | 0 | 0 | 0 |
| 0 | 0 | 0 | 0 |
| 0 | 0 | 0 | 0 |
| 0 | 0 | 0 | 0 |
| 1 | 0 | 0 | 0 |
| 0 | 0 | 0 | 0 |
| 0 | 0 | 0 | 0 |
| 0 | 0 | 0 | 0 |
| 0 | 1 | 0 | 0 |
| 0 | 0 | 0 | 0 |
| 0 | 0 | 0 | 0 |
| 0 | 0 | 0 | 0 |
| 0 | 0 | 0 | 0 |
| 0 | 0 | 0 | 0 |
| 0 | 0 | 0 | 0 |
| 0 | 0 | 0 | 0 |
| 0 | 0 | 0 | 0 |
| 0 | 0 | 0 | 0 |
| 0 | 0 | 0 | 0 |
| 0 | 0 | 0 | 0 |
| 0 | 0 | 0 | 0 |
| 0 | 0 | 0 | 0 |
| 1 | 0 | 0 | 0 |
| 1 | 0 | 0 | 0 |
| 1 | 0 | 0 | 0 |
| 0 | 0 | 0 | 0 |
| 1 | 0 | 0 | 0 |
| 1 | 0 | 0 | 0 |
| 1 | 0 | 0 | 0 |
| 0 | 0 | 0 | 0 |
| 0 | 0 | 0 | 0 |
| 0 | 0 | 0 | 0 |
| 0 | 0 | 0 | 0 |
| 0 | 0 | 0 | 0 |
| 0 | 1 | 0 | 0 |
| 0 | 1 | 0 | 0 |
| 0 | 0 | 0 | 0 |
| 0 | 0 | 0 | 0 |
| 1 | 0 | 0 | 0 |
| 0 | 0 | 0 | 0 |
| 0 | 0 | 0 | 0 |
| 0 | 0 | 0 | 0 |
| 0 | 0 | 0 | 0 |
| 0 | 0 | 0 | 0 |
| 1 | 0 | 0 | 0 |
| 0 | 0 | 0 | 0 |

| | | | |
|---|---|---|---|
| 0 | 0 | 0 | 0 |
| 1 | 0 | 0 | 0 |
| 1 | 0 | 0 | 0 |
| 0 | 0 | 0 | 0 |
| 0 | 0 | 0 | 0 |
| 0 | 0 | 0 | 0 |
| 0 | 0 | 0 | 0 |
| 0 | 1 | 0 | 0 |
| 0 | 0 | 0 | 0 |
| 0 | 0 | 0 | 0 |
| 0 | 0 | 0 | 0 |
| 0 | 0 | 0 | 0 |
| 0 | 0 | 0 | 0 |
| 0 | 0 | 0 | 0 |
| 1 | 0 | 0 | 0 |
| 0 | 0 | 0 | 0 |
| 0 | 0 | 0 | 0 |
| 0 | 0 | 0 | 0 |
| 0 | 0 | 0 | 0 |
| 1 | 0 | 0 | 0 |
| 1 | 0 | 0 | 0 |
| 0 | 0 | 0 | 0 |
| 1 | 0 | 0 | 0 |
| 1 | 0 | 0 | 0 |
| 0 | 0 | 0 | 0 |
| 0 | 0 | 0 | 0 |
| 0 | 0 | 0 | 0 |
| 0 | 0 | 0 | 0 |
| 0 | 0 | 0 | 0 |
| 0 | 0 | 0 | 0 |
| 0 | 0 | 0 | 0 |
| 0 | 0 | 0 | 0 |
| 0 | 0 | 0 | 0 |
| 0 | 0 | 0 | 0 |
| 0 | 0 | 0 | 0 |
| 0 | 0 | 0 | 0 |
| 0 | 0 | 0 | 0 |
| 0 | 0 | 0 | 0 |
| 0 | 0 | 0 | 0 |
| 0 | 0 | 0 | 0 |
| 0 | 0 | 0 | 0 |
| 0 | 0 | 0 | 0 |
| 0 | 0 | 0 | 0 |
| 0 | 0 | 0 | 0 |
| 0 | 0 | 0 | 0 |
| 0 | 0 | 0 | 0 |
| 0 | 0 | 0 | 0 |
| 0 | 0 | 0 | 0 |
| 0 | 0 | 0 | 0 |
| 0 | 0 | 0 | 0 |
| 0 | 0 | 0 | 0 |
| 0 | 0 | 0 | 0 |
| 0 | 0 | 0 | 0 |
| 1 | 1 | 0 | 1 |

| | | | |
|---|---|---|---|
| 0 | 0 | 0 | 0 |
| 0 | 0 | 0 | 0 |
| 0 | 0 | 0 | 0 |
| 0 | 0 | 0 | 0 |
| 0 | 1 | 0 | 0 |
| 0 | 0 | 0 | 0 |
| 0 | 0 | 0 | 0 |
| 0 | 1 | 0 | 0 |
| 0 | 0 | 0 | 0 |
| 0 | 0 | 0 | 0 |
| 0 | 0 | 0 | 0 |
| 0 | 0 | 0 | 0 |
| 0 | 0 | 0 | 0 |
| 0 | 0 | 0 | 0 |
| 0 | 1 | 0 | 1 |
| 0 | 1 | 0 | 0 |
| 0 | 0 | 0 | 0 |
| 0 | 0 | 0 | 0 |
| 0 | 1 | 0 | 0 |
| 0 | 1 | 0 | 1 |
| 0 | 0 | 0 | 0 |
| 0 | 0 | 0 | 0 |
| 0 | 0 | 0 | 0 |
| 0 | 0 | 0 | 0 |
| 0 | 0 | 0 | 0 |
| 1 | 0 | 0 | 0 |
| 0 | 0 | 0 | 0 |
| 0 | 0 | 1 | 0 |
| 0 | 0 | 1 | 0 |
| 0 | 0 | 1 | 0 |
| 0 | 0 | 0 | 0 |
| 0 | 0 | 0 | 0 |
| 0 | 0 | 0 | 0 |
| 0 | 0 | 0 | 0 |
| 0 | 0 | 0 | 0 |
| 0 | 0 | 1 | 0 |
| 0 | 0 | 0 | 0 |
| 0 | 0 | 0 | 0 |
| 1 | 0 | 1 | 0 |
| 0 | 0 | 1 | 0 |
| 1 | 1 | 0 | 0 |
| 0 | 1 | 0 | 0 |
| 0 | 0 | 0 | 0 |
| 0 | 0 | 0 | 0 |
| 1 | 0 | 0 | 0 |
| 0 | 0 | 0 | 0 |
| 0 | 0 | 1 | 0 |
| 0 | 0 | 0 | 0 |
| 0 | 0 | 0 | 0 |
| 0 | 0 | 0 | 0 |
| 0 | 0 | 0 | 0 |
| 0 | 0 | 0 | 0 |
| 0 | 0 | 0 | 0 |
| 0 | 0 | 0 | 0 |

| | | | |
|---|---|---|---|
| 0 | 0 | 0 | 0 |
| 0 | 0 | 0 | 0 |
| 0 | 0 | 0 | 0 |
| 0 | 0 | 0 | 0 |
| 0 | 0 | 0 | 0 |
| 0 | 0 | 0 | 0 |
| 0 | 0 | 0 | 0 |
| 1 | 1 | 0 | 0 |
| 0 | 0 | 0 | 0 |
| 0 | 0 | 0 | 0 |
| 0 | 0 | 0 | 0 |
| 0 | 0 | 0 | 1 |
| 0 | 0 | 0 | 0 |
| 0 | 0 | 0 | 0 |
| 0 | 0 | 0 | 0 |
| 0 | 0 | 0 | 0 |
| 0 | 0 | 0 | 0 |
| 0 | 0 | 0 | 0 |
| 0 | 0 | 0 | 0 |
| 0 | 0 | 0 | 0 |
| 0 | 0 | 0 | 0 |
| 0 | 0 | 0 | 0 |
| 0 | 0 | 0 | 0 |
| 0 | 0 | 0 | 0 |
| 0 | 0 | 0 | 0 |
| 0 | 0 | 0 | 0 |
| 0 | 0 | 0 | 0 |
| 0 | 0 | 0 | 0 |
| 0 | 0 | 0 | 0 |
| 0 | 1 | 0 | 0 |
| 0 | 1 | 0 | 0 |
| 0 | 0 | 0 | 0 |
| 0 | 0 | 0 | 0 |
| 0 | 0 | 0 | 0 |
| 0 | 0 | 0 | 0 |
| 0 | 1 | 0 | 0 |
| 0 | 0 | 0 | 0 |
| 0 | 0 | 0 | 0 |
| 0 | 0 | 0 | 0 |
| 0 | 0 | 0 | 0 |
| 0 | 0 | 0 | 0 |
| 1 | 1 | 0 | 0 |
| 1 | 0 | 0 | 0 |
| 0 | 0 | 0 | 0 |
| 0 | 0 | 0 | 0 |
| 0 | 0 | 0 | 0 |
| 0 | 0 | 0 | 0 |
| 0 | 0 | 0 | 0 |
| 0 | 0 | 0 | 0 |
| 0 | 0 | 0 | 0 |
| 0 | 0 | 0 | 0 |
| 0 | 0 | 0 | 0 |
| 0 | 0 | 0 | 0 |
| 0 | 0 | 0 | 0 |

| | | | |
|---|---|---|---|
| 0 | 0 | 0 | 0 |
| 0 | 0 | 0 | 0 |
| 0 | 0 | 0 | 0 |
| 0 | 0 | 0 | 0 |
| 0 | 0 | 0 | 0 |
| 0 | 0 | 0 | 0 |
| 0 | 0 | 0 | 0 |
| 0 | 0 | 0 | 0 |
| 1 | 0 | 0 | 0 |
| 1 | 0 | 0 | 0 |
| 0 | 0 | 0 | 0 |
| 0 | 0 | 0 | 0 |
| 0 | 0 | 0 | 0 |
| 1 | 0 | 0 | 0 |
| 0 | 0 | 0 | 0 |
| 0 | 0 | 0 | 0 |
| 0 | 0 | 0 | 0 |
| 0 | 0 | 0 | 0 |
| 0 | 0 | 0 | 0 |
| 1 | 1 | 0 | 0 |
| 0 | 0 | 1 | 0 |
| 0 | 0 | 0 | 0 |
| 0 | 0 | 0 | 0 |
| 0 | 0 | 0 | 0 |
| 0 | 0 | 0 | 0 |
| 0 | 0 | 0 | 0 |
| 0 | 0 | 0 | 0 |
| 0 | 0 | 0 | 0 |
| 0 | 0 | 0 | 0 |
| 0 | 0 | 0 | 0 |
| 0 | 0 | 0 | 0 |
| 0 | 0 | 0 | 0 |
| 0 | 0 | 0 | 0 |
| 0 | 0 | 0 | 0 |
| 0 | 0 | 0 | 0 |
| 0 | 0 | 0 | 0 |
| 0 | 0 | 0 | 0 |
| 0 | 0 | 0 | 0 |
| 0 | 0 | 0 | 0 |
| 0 | 0 | 0 | 0 |
| 0 | 0 | 0 | 0 |
| 0 | 0 | 0 | 0 |
| 0 | 0 | 0 | 0 |
| 0 | 0 | 0 | 0 |
| 0 | 0 | 0 | 0 |
| 1 | 0 | 1 | 0 |
| 1 | 0 | 0 | 0 |
| 0 | 0 | 0 | 0 |
| 0 | 0 | 0 | 0 |
| 0 | 0 | 1 | 0 |
| 0 | 0 | 0 | 0 |
| 0 | 0 | 0 | 0 |
| 0 | 0 | 0 | 0 |

| | | | |
|---|---|---|---|
| 0 | 0 | 0 | 0 |
| 0 | 0 | 0 | 0 |
| 0 | 0 | 0 | 0 |
| 0 | 0 | 0 | 0 |
| 0 | 0 | 0 | 0 |
| 0 | 0 | 0 | 0 |
| 0 | 0 | 0 | 0 |
| 0 | 0 | 0 | 0 |
| 0 | 0 | 0 | 0 |
| 0 | 0 | 0 | 0 |
| 0 | 0 | 0 | 0 |
| 0 | 0 | 0 | 0 |
| 0 | 0 | 0 | 0 |
| 0 | 0 | 0 | 0 |
| 0 | 0 | 0 | 0 |
| 0 | 0 | 0 | 0 |
| 0 | 0 | 0 | 0 |
| 0 | 0 | 0 | 0 |
| 0 | 0 | 0 | 0 |
| 1 | 0 | 0 | 0 |
| 0 | 0 | 0 | 0 |
| 0 | 0 | 0 | 0 |
| 0 | 0 | 0 | 0 |
| 0 | 0 | 0 | 0 |
| 0 | 0 | 0 | 0 |
| 0 | 0 | 0 | 0 |
| 0 | 0 | 0 | 0 |
| 0 | 0 | 0 | 0 |
| 0 | 0 | 0 | 0 |
| 0 | 0 | 0 | 0 |
| 0 | 0 | 0 | 0 |
| 0 | 0 | 0 | 0 |
| 0 | 0 | 0 | 0 |
| 0 | 0 | 0 | 0 |
| 0 | 0 | 0 | 0 |
| 0 | 0 | 0 | 0 |
| 0 | 0 | 0 | 0 |
| 0 | 0 | 0 | 0 |
| 1 | 0 | 0 | 1 |
| 0 | 0 | 0 | 0 |
| 0 | 0 | 0 | 0 |
| 0 | 0 | 0 | 0 |
| 0 | 0 | 0 | 1 |
| 0 | 0 | 0 | 1 |
| 0 | 0 | 0 | 1 |
| 0 | 0 | 0 | 0 |
| 0 | 0 | 0 | 0 |
| 0 | 0 | 0 | 0 |
| 0 | 0 | 0 | 0 |
| 0 | 0 | 0 | 0 |
| 0 | 0 | 0 | 0 |
| 0 | 0 | 0 | 0 |
| 0 | 0 | 0 | 0 |
| 0 | 0 | 0 | 0 |

| | | | |
|---|---|---|---|
| 0 | 0 | 0 | 0 |
| 0 | 0 | 0 | 0 |
| 0 | 0 | 0 | 0 |
| 0 | 0 | 0 | 0 |
| 0 | 0 | 0 | 0 |
| 0 | 0 | 0 | 0 |
| 0 | 0 | 0 | 0 |
| 0 | 0 | 0 | 0 |
| 0 | 0 | 0 | 0 |
| 0 | 0 | 0 | 0 |
| 0 | 0 | 0 | 0 |
| 0 | 0 | 0 | 0 |
| 0 | 0 | 0 | 0 |
| 0 | 0 | 0 | 0 |
| 0 | 0 | 0 | 0 |
| 0 | 0 | 0 | 0 |
| 0 | 0 | 0 | 0 |
| 0 | 0 | 0 | 0 |
| 0 | 0 | 0 | 0 |
| 0 | 0 | 0 | 0 |
| 0 | 0 | 0 | 0 |
| 0 | 0 | 0 | 0 |
| 0 | 0 | 0 | 0 |
| 0 | 0 | 0 | 0 |
| 0 | 0 | 0 | 0 |
| 0 | 0 | 0 | 0 |
| 0 | 0 | 0 | 0 |
| 0 | 0 | 0 | 0 |
| 0 | 0 | 0 | 0 |
| 0 | 0 | 0 | 0 |
| 0 | 0 | 0 | 0 |
| 0 | 0 | 0 | 0 |
| 1 | 1 | 0 | 1 |
| 1 | 1 | 0 | 1 |
| 1 | 0 | 0 | 0 |
| 0 | 0 | 0 | 0 |
| 0 | 0 | 0 | 0 |
| 0 | 0 | 0 | 0 |
| 0 | 0 | 0 | 0 |
| 0 | 0 | 0 | 0 |
| 0 | 0 | 0 | 0 |
| 0 | 0 | 0 | 0 |
| 1 | 1 | 0 | 0 |
| 0 | 0 | 0 | 0 |
| 1 | 0 | 0 | 0 |
| 0 | 0 | 0 | 0 |
| 0 | 0 | 0 | 0 |
| 0 | 0 | 0 | 0 |
| 0 | 0 | 0 | 0 |
| 0 | 0 | 0 | 0 |
| 1 | 0 | 0 | 0 |
| 0 | 0 | 0 | 0 |
| 0 | 0 | 0 | 0 |
| 0 | 0 | 0 | 0 |

| | | | |
|---|---|---|---|
| 0 | 0 | 0 | 0 |
| 0 | 0 | 0 | 0 |
| 0 | 0 | 0 | 0 |
| 0 | 0 | 0 | 0 |
| 0 | 0 | 0 | 0 |
| 0 | 0 | 0 | 0 |
| 0 | 0 | 0 | 0 |
| 0 | 0 | 0 | 0 |
| 0 | 0 | 0 | 0 |
| 0 | 0 | 0 | 0 |
| 0 | 0 | 0 | 0 |
| 0 | 0 | 0 | 0 |
| 0 | 0 | 0 | 0 |
| 0 | 0 | 0 | 0 |
| 0 | 0 | 0 | 0 |
| 0 | 0 | 0 | 0 |
| 0 | 0 | 0 | 0 |
| 0 | 0 | 0 | 0 |
| 0 | 0 | 0 | 0 |
| 0 | 0 | 0 | 1 |
| 0 | 0 | 0 | 0 |
| 0 | 0 | 0 | 1 |
| 0 | 0 | 0 | 1 |
| 0 | 0 | 0 | 1 |
| 0 | 0 | 0 | 0 |
| 0 | 0 | 0 | 0 |
| 0 | 0 | 0 | 0 |
| 0 | 0 | 0 | 0 |
| 0 | 0 | 0 | 0 |
| 0 | 0 | 0 | 0 |
| 0 | 0 | 0 | 0 |
| 0 | 0 | 0 | 0 |
| 0 | 0 | 0 | 0 |
| 0 | 0 | 0 | 0 |
| 0 | 0 | 0 | 0 |
| 0 | 1 | 0 | 0 |
| 0 | 0 | 0 | 0 |
| 0 | 0 | 0 | 0 |
| 0 | 1 | 0 | 0 |
| 0 | 0 | 0 | 0 |
| 0 | 0 | 0 | 0 |
| 0 | 0 | 0 | 0 |
| 0 | 0 | 0 | 0 |
| 0 | 0 | 0 | 0 |
| 0 | 0 | 0 | 0 |
| 0 | 0 | 0 | 0 |
| 0 | 0 | 0 | 0 |
| 0 | 0 | 0 | 0 |
| 0 | 0 | 0 | 0 |
| 0 | 0 | 0 | 0 |
| 0 | 0 | 0 | 0 |
| 0 | 0 | 0 | 0 |
| 0 | 0 | 0 | 0 |
| 0 | 0 | 0 | 0 |

| | | | |
|---|---|---|---|
| 0 | 0 | 0 | 0 |
| 0 | 0 | 0 | 0 |
| 0 | 0 | 0 | 0 |
| 0 | 0 | 0 | 0 |
| 0 | 0 | 0 | 0 |
| 0 | 0 | 0 | 0 |
| 0 | 0 | 0 | 0 |
| 0 | 0 | 0 | 0 |
| 0 | 0 | 1 | 0 |
| 0 | 0 | 1 | 0 |
| 0 | 0 | 0 | 0 |
| 0 | 0 | 0 | 0 |
| 0 | 0 | 0 | 0 |
| 0 | 0 | 0 | 0 |
| 0 | 0 | 0 | 0 |
| 0 | 1 | 0 | 0 |
| 0 | 0 | 0 | 0 |
| 0 | 0 | 0 | 0 |
| 1 | 1 | 0 | 0 |
| 0 | 0 | 0 | 0 |
| 0 | 0 | 0 | 0 |
| 0 | 0 | 0 | 0 |
| 0 | 0 | 0 | 0 |
| 0 | 0 | 0 | 0 |
| 0 | 0 | 0 | 0 |
| 0 | 0 | 0 | 0 |
| 0 | 0 | 0 | 0 |
| 0 | 0 | 0 | 0 |
| 0 | 0 | 0 | 0 |
| 0 | 0 | 0 | 0 |
| 0 | 0 | 0 | 0 |
| 0 | 0 | 0 | 0 |
| 0 | 0 | 0 | 0 |
| 0 | 0 | 0 | 0 |
| 0 | 0 | 0 | 0 |
| 0 | 0 | 0 | 0 |
| 0 | 0 | 0 | 0 |
| 0 | 0 | 0 | 0 |
| 0 | 0 | 0 | 0 |
| 0 | 0 | 0 | 0 |
| 0 | 0 | 0 | 0 |
| 0 | 0 | 0 | 0 |
| 0 | 0 | 0 | 0 |
| 0 | 0 | 0 | 0 |
| 0 | 0 | 0 | 0 |
| 0 | 0 | 0 | 0 |
| 0 | 0 | 0 | 0 |
| 0 | 0 | 0 | 0 |
| 0 | 0 | 0 | 0 |
| 0 | 0 | 0 | 0 |
| 0 | 0 | 0 | 0 |
| 0 | 0 | 0 | 0 |
| 0 | 0 | 0 | 0 |
| 0 | 0 | 0 | 0 |
| 1 | 0 | 0 | 0 |

| | | | |
|---|---|---|---|
| 0 | 0 | 0 | 0 |
| 0 | 0 | 0 | 0 |
| 0 | 0 | 0 | 0 |
| 0 | 0 | 0 | 0 |
| 0 | 0 | 0 | 0 |
| 0 | 0 | 0 | 0 |
| 0 | 0 | 0 | 0 |
| 0 | 0 | 0 | 0 |
| 0 | 0 | 0 | 0 |
| 0 | 0 | 0 | 0 |
| 1 | 0 | 0 | 0 |
| 0 | 0 | 0 | 0 |
| 0 | 0 | 0 | 0 |
| 0 | 0 | 0 | 0 |
| 1 | 1 | 0 | 1 |
| 0 | 0 | 0 | 0 |
| 1 | 0 | 0 | 0 |
| 1 | 1 | 0 | 0 |
| 0 | 0 | 0 | 0 |
| 0 | 0 | 0 | 0 |
| 0 | 0 | 0 | 0 |
| 0 | 0 | 0 | 0 |
| 0 | 0 | 0 | 0 |
| 0 | 0 | 0 | 1 |
| 0 | 0 | 0 | 0 |
| 0 | 0 | 0 | 0 |
| 0 | 0 | 0 | 0 |
| 0 | 0 | 0 | 1 |
| 0 | 0 | 0 | 0 |
| 0 | 0 | 0 | 0 |
| 0 | 0 | 0 | 0 |
| 0 | 0 | 0 | 0 |
| 0 | 0 | 0 | 0 |
| 0 | 0 | 0 | 0 |
| 0 | 0 | 0 | 0 |
| 0 | 0 | 0 | 0 |
| 0 | 0 | 0 | 0 |
| 0 | 0 | 0 | 0 |
| 0 | 0 | 0 | 0 |
| 0 | 0 | 0 | 0 |
| 0 | 0 | 0 | 0 |
| 0 | 0 | 0 | 0 |
| 0 | 0 | 0 | 0 |
| 0 | 0 | 0 | 0 |
| 0 | 0 | 1 | 0 |
| 0 | 0 | 0 | 0 |
| 0 | 0 | 0 | 0 |
| 0 | 0 | 0 | 0 |
| 0 | 0 | 0 | 0 |
| 0 | 0 | 0 | 0 |
| 0 | 0 | 0 | 0 |
| 0 | 0 | 0 | 0 |
| 0 | 0 | 0 | 0 |
| 0 | 0 | 0 | 0 |

| | | | |
|---|---|---|---|
| 0 | 0 | 0 | 0 |
| 0 | 0 | 0 | 0 |
| 0 | 0 | 0 | 0 |
| 0 | 0 | 0 | 0 |
| 0 | 0 | 0 | 0 |
| 0 | 0 | 0 | 0 |
| 1 | 0 | 0 | 1 |
| 1 | 0 | 0 | 0 |
| 0 | 0 | 0 | 0 |
| 0 | 0 | 0 | 0 |
| 0 | 0 | 0 | 0 |
| 0 | 0 | 0 | 0 |
| 0 | 0 | 0 | 0 |
| 0 | 0 | 0 | 0 |
| 0 | 0 | 0 | 0 |
| 0 | 0 | 0 | 0 |
| 0 | 0 | 0 | 0 |
| 0 | 0 | 0 | 0 |
| 0 | 0 | 0 | 0 |
| 0 | 0 | 0 | 0 |
| 1 | 0 | 0 | 0 |
| 0 | 0 | 0 | 0 |
| 0 | 0 | 0 | 0 |
| 0 | 0 | 0 | 0 |
| 0 | 0 | 0 | 0 |
| 0 | 0 | 0 | 0 |
| 0 | 0 | 0 | 0 |
| 0 | 0 | 0 | 0 |
| 0 | 0 | 0 | 0 |
| 0 | 0 | 0 | 0 |
| 0 | 0 | 0 | 0 |
| 0 | 0 | 0 | 0 |
| 0 | 0 | 0 | 0 |
| 0 | 0 | 0 | 0 |
| 0 | 0 | 0 | 0 |
| 0 | 0 | 0 | 0 |
| 0 | 0 | 0 | 0 |
| 0 | 0 | 0 | 0 |
| 0 | 0 | 0 | 0 |
| 0 | 0 | 0 | 0 |
| 0 | 0 | 0 | 0 |
| 0 | 0 | 0 | 0 |
| 0 | 0 | 0 | 0 |
| 0 | 0 | 0 | 0 |
| 0 | 0 | 0 | 0 |
| 0 | 0 | 0 | 0 |
| 0 | 0 | 0 | 0 |
| 0 | 0 | 0 | 0 |
| 0 | 0 | 0 | 0 |
| 0 | 0 | 0 | 0 |
| 0 | 0 | 0 | 0 |
| 0 | 0 | 0 | 0 |
| 0 | 0 | 0 | 0 |
| 0 | 0 | 0 | 0 |

| | | | |
|---|---|---|---|
| 0 | 0 | 0 | 0 |
| 0 | 0 | 0 | 0 |
| 0 | 0 | 0 | 0 |
| 1 | 0 | 0 | 0 |
| 0 | 0 | 0 | 0 |
| 1 | 0 | 0 | 0 |
| 0 | 0 | 0 | 0 |
| 0 | 0 | 0 | 0 |
| 1 | 0 | 0 | 0 |
| 0 | 0 | 0 | 0 |
| 0 | 0 | 0 | 0 |
| 0 | 0 | 0 | 0 |
| 0 | 0 | 0 | 0 |
| 0 | 0 | 0 | 0 |
| 0 | 0 | 0 | 0 |
| 0 | 0 | 0 | 0 |
| 0 | 0 | 0 | 0 |
| 0 | 0 | 0 | 0 |
| 1 | 0 | 0 | 0 |
| 0 | 0 | 0 | 0 |
| 0 | 0 | 0 | 0 |
| 0 | 0 | 0 | 0 |
| 0 | 0 | 0 | 0 |
| 0 | 0 | 0 | 0 |
| 0 | 0 | 0 | 0 |
| 0 | 0 | 0 | 0 |
| 0 | 0 | 0 | 0 |
| 0 | 0 | 0 | 0 |
| 0 | 1 | 0 | 0 |
| 0 | 0 | 0 | 0 |
| 0 | 0 | 0 | 0 |
| 0 | 0 | 0 | 0 |
| 0 | 1 | 0 | 0 |
| 1 | 1 | 0 | 0 |
| 1 | 0 | 0 | 0 |
| 0 | 0 | 0 | 0 |
| 0 | 0 | 0 | 0 |
| 0 | 0 | 0 | 0 |
| 0 | 0 | 0 | 0 |
| 0 | 0 | 0 | 0 |
| 1 | 0 | 0 | 0 |
| 0 | 0 | 0 | 0 |
| 0 | 0 | 0 | 0 |
| 0 | 0 | 0 | 0 |
| 0 | 0 | 0 | 0 |
| 0 | 0 | 0 | 0 |
| 0 | 0 | 0 | 0 |
| 0 | 0 | 0 | 0 |
| 0 | 0 | 0 | 0 |
| 0 | 0 | 0 | 0 |
| 0 | 0 | 0 | 0 |
| 0 | 0 | 0 | 0 |
| 0 | 0 | 0 | 0 |
| 1 | 0 | 0 | 0 |

| | | | |
|---|---|---|---|
| 1 | 0 | 0 | 1 |
| 0 | 0 | 0 | 0 |
| 0 | 0 | 0 | 0 |
| 0 | 0 | 0 | 0 |
| 0 | 0 | 0 | 0 |
| 0 | 0 | 0 | 0 |
| 0 | 0 | 0 | 0 |
| 0 | 0 | 0 | 0 |
| 0 | 0 | 0 | 0 |
| 0 | 0 | 0 | 0 |
| 0 | 0 | 0 | 0 |
| 0 | 0 | 0 | 0 |
| 0 | 0 | 0 | 0 |
| 0 | 0 | 0 | 0 |
| 0 | 0 | 0 | 0 |
| 0 | 0 | 0 | 0 |
| 0 | 0 | 0 | 0 |
| 0 | 0 | 0 | 0 |
| 0 | 0 | 0 | 0 |
| 0 | 0 | 0 | 0 |
| 0 | 0 | 0 | 0 |
| 0 | 0 | 0 | 0 |
| 0 | 1 | 0 | 0 |
| 0 | 0 | 0 | 0 |
| 0 | 0 | 0 | 0 |
| 0 | 0 | 0 | 0 |
| 0 | 0 | 0 | 0 |
| 0 | 0 | 0 | 0 |
| 0 | 0 | 0 | 0 |
| 0 | 0 | 0 | 0 |
| 0 | 0 | 0 | 0 |
| 0 | 0 | 0 | 0 |
| 0 | 0 | 0 | 0 |
| 0 | 0 | 0 | 0 |
| 0 | 0 | 0 | 0 |
| 0 | 0 | 0 | 0 |
| 0 | 0 | 0 | 0 |
| 0 | 0 | 0 | 0 |
| 0 | 0 | 0 | 0 |
| 0 | 0 | 0 | 0 |
| 0 | 0 | 0 | 0 |
| 0 | 0 | 0 | 0 |
| 0 | 0 | 0 | 0 |
| 0 | 0 | 0 | 0 |
| 0 | 0 | 0 | 0 |
| 0 | 0 | 0 | 0 |
| 0 | 0 | 0 | 0 |
| 0 | 0 | 0 | 0 |
| 0 | 0 | 0 | 0 |
| 0 | 0 | 0 | 0 |
| 0 | 0 | 0 | 0 |
| 0 | 0 | 0 | 0 |
| 0 | 0 | 0 | 0 |
| 0 | 0 | 0 | 0 |

| | | | |
|---|---|---|---|
| 0 | 0 | 0 | 0 |
| 0 | 0 | 0 | 0 |
| 0 | 0 | 0 | 0 |
| 0 | 0 | 0 | 0 |
| 0 | 0 | 0 | 0 |
| 0 | 0 | 0 | 0 |
| 0 | 1 | 0 | 0 |
| 0 | 1 | 0 | 0 |
| 0 | 0 | 0 | 0 |
| 0 | 0 | 0 | 0 |
| 0 | 0 | 0 | 0 |
| 0 | 0 | 0 | 0 |
| 0 | 0 | 0 | 0 |
| 0 | 0 | 0 | 0 |
| 0 | 0 | 0 | 0 |
| 0 | 0 | 0 | 0 |
| 1 | 0 | 0 | 0 |
| 0 | 0 | 0 | 0 |
| 1 | 0 | 0 | 0 |
| 1 | 0 | 0 | 0 |
| 1 | 0 | 0 | 0 |
| 1 | 0 | 0 | 0 |
| 0 | 0 | 0 | 0 |
| 0 | 0 | 0 | 0 |
| 0 | 0 | 0 | 0 |
| 0 | 0 | 0 | 0 |
| 0 | 0 | 0 | 0 |
| 0 | 0 | 0 | 0 |
| 0 | 0 | 0 | 0 |
| 0 | 0 | 0 | 0 |
| 0 | 0 | 0 | 0 |
| 0 | 0 | 0 | 0 |
| 0 | 0 | 0 | 0 |
| 0 | 0 | 0 | 0 |
| 0 | 0 | 0 | 0 |
| 0 | 0 | 1 | 0 |
| 1 | 0 | 0 | 0 |
| 1 | 0 | 0 | 0 |
| 1 | 0 | 0 | 0 |
| 0 | 0 | 0 | 0 |
| 0 | 0 | 0 | 0 |
| 0 | 0 | 0 | 0 |
| 0 | 0 | 0 | 0 |
| 1 | 0 | 0 | 0 |
| 1 | 0 | 0 | 0 |
| 1 | 0 | 0 | 0 |
| 0 | 0 | 0 | 0 |
| 0 | 0 | 0 | 0 |
| 1 | 0 | 0 | 0 |
| 0 | 0 | 0 | 0 |
| 0 | 0 | 0 | 0 |
| 1 | 0 | 0 | 0 |
| 0 | 0 | 0 | 0 |
| 0 | 0 | 0 | 0 |

| | | | |
|---|---|---|---|
| 0 | 0 | 0 | 0 |
| 0 | 0 | 0 | 0 |
| 0 | 0 | 0 | 0 |
| 0 | 0 | 0 | 0 |
| 0 | 0 | 0 | 0 |
| 0 | 0 | 0 | 0 |
| 0 | 0 | 0 | 0 |
| 0 | 0 | 0 | 0 |
| 0 | 0 | 0 | 0 |
| 0 | 0 | 0 | 0 |
| 0 | 0 | 0 | 0 |
| 0 | 0 | 0 | 0 |
| 0 | 0 | 0 | 0 |
| 0 | 0 | 0 | 0 |
| 0 | 0 | 0 | 0 |
| 0 | 0 | 0 | 0 |
| 0 | 0 | 0 | 0 |
| 0 | 0 | 0 | 0 |
| 0 | 0 | 0 | 0 |
| 0 | 0 | 0 | 0 |
| 0 | 0 | 0 | 0 |
| 0 | 0 | 0 | 0 |
| 1 | 1 | 0 | 0 |
| 0 | 0 | 0 | 0 |
| 0 | 1 | 0 | 0 |
| 0 | 0 | 0 | 0 |
| 0 | 0 | 0 | 0 |
| 0 | 0 | 0 | 0 |
| 0 | 0 | 0 | 0 |
| 0 | 0 | 0 | 0 |
| 0 | 0 | 0 | 0 |
| 0 | 0 | 0 | 0 |
| 0 | 0 | 0 | 0 |
| 0 | 0 | 0 | 0 |
| 0 | 0 | 0 | 0 |
| 0 | 0 | 0 | 0 |
| 0 | 0 | 0 | 0 |
| 0 | 0 | 0 | 0 |
| 0 | 0 | 0 | 0 |
| 0 | 0 | 0 | 0 |
| 0 | 0 | 0 | 0 |
| 0 | 0 | 0 | 0 |
| 0 | 0 | 0 | 0 |
| 0 | 0 | 0 | 0 |
| 0 | 0 | 0 | 0 |
| 1 | 0 | 0 | 1 |
| 0 | 0 | 0 | 0 |
| 0 | 0 | 0 | 0 |
| 0 | 0 | 0 | 0 |
| 1 | 0 | 0 | 0 |
| 1 | 0 | 0 | 0 |
| 1 | 0 | 0 | 0 |
| 0 | 0 | 0 | 0 |
| 0 | 0 | 0 | 0 |

| | | | |
|---|---|---|---|
| 0 | 0 | 0 | 0 |
| 0 | 0 | 0 | 0 |
| 0 | 0 | 0 | 0 |
| 0 | 0 | 0 | 0 |
| 0 | 0 | 0 | 0 |
| 0 | 0 | 0 | 0 |
| 0 | 0 | 0 | 0 |
| 0 | 0 | 0 | 0 |
| 0 | 0 | 0 | 0 |
| 0 | 0 | 0 | 0 |
| 0 | 0 | 0 | 1 |
| 0 | 0 | 0 | 0 |
| 0 | 0 | 0 | 0 |
| 0 | 0 | 0 | 0 |
| 0 | 0 | 0 | 0 |
| 0 | 0 | 0 | 0 |
| 0 | 0 | 0 | 0 |
| 0 | 0 | 0 | 0 |
| 0 | 0 | 0 | 0 |
| 0 | 0 | 0 | 0 |
| 0 | 0 | 0 | 0 |
| 0 | 0 | 0 | 0 |
| 0 | 0 | 0 | 0 |
| 0 | 0 | 0 | 0 |
| 0 | 0 | 0 | 0 |
| 0 | 0 | 0 | 0 |
| 0 | 0 | 0 | 0 |
| 0 | 0 | 0 | 0 |
| 0 | 0 | 0 | 0 |
| 0 | 0 | 0 | 0 |
| 0 | 1 | 0 | 0 |
| 0 | 0 | 0 | 0 |
| 1 | 1 | 0 | 0 |
| 1 | 1 | 0 | 0 |
| 0 | 0 | 0 | 0 |
| 0 | 0 | 0 | 0 |
| 0 | 0 | 0 | 0 |
| 0 | 0 | 0 | 0 |
| 0 | 0 | 0 | 0 |
| 0 | 0 | 0 | 0 |
| 0 | 0 | 0 | 0 |
| 0 | 1 | 0 | 0 |
| 0 | 0 | 0 | 0 |
| 0 | 0 | 0 | 0 |
| 0 | 0 | 0 | 0 |
| 1 | 0 | 0 | 0 |
| 1 | 0 | 0 | 0 |
| 1 | 0 | 0 | 0 |
| 1 | 1 | 1 | 0 |
| 0 | 0 | 0 | 0 |
| 1 | 1 | 1 | 1 |
| 0 | 0 | 0 | 0 |
| 0 | 0 | 0 | 0 |
| 0 | 0 | 0 | 0 |

| | | | |
|---|---|---|---|
| 0 | 0 | 0 | 0 |
| 0 | 0 | 0 | 0 |
| 0 | 0 | 0 | 0 |
| 0 | 0 | 0 | 0 |
| 1 | 0 | 0 | 0 |
| 1 | 0 | 0 | 0 |
| 0 | 0 | 0 | 0 |
| 0 | 0 | 0 | 0 |
| 0 | 0 | 0 | 0 |
| 0 | 0 | 0 | 0 |
| 0 | 0 | 0 | 0 |
| 0 | 0 | 0 | 0 |
| 0 | 0 | 0 | 0 |
| 0 | 0 | 0 | 0 |
| 0 | 0 | 0 | 0 |
| 0 | 0 | 0 | 0 |
| 0 | 0 | 0 | 0 |
| 0 | 0 | 0 | 0 |
| 0 | 0 | 0 | 0 |
| 0 | 0 | 0 | 0 |
| 0 | 0 | 0 | 0 |
| 0 | 0 | 0 | 0 |
| 0 | 0 | 0 | 0 |
| 0 | 0 | 0 | 0 |
| 0 | 0 | 0 | 0 |
| 0 | 0 | 0 | 0 |
| 0 | 0 | 0 | 0 |
| 0 | 0 | 0 | 0 |
| 0 | 0 | 0 | 0 |
| 0 | 0 | 0 | 0 |
| 0 | 0 | 0 | 0 |
| 0 | 0 | 0 | 0 |
| 0 | 0 | 0 | 0 |
| 0 | 0 | 0 | 0 |
| 0 | 0 | 0 | 0 |
| 0 | 0 | 0 | 0 |
| 0 | 0 | 0 | 0 |
| 0 | 0 | 0 | 0 |
| 0 | 0 | 0 | 0 |
| 0 | 0 | 0 | 0 |
| 0 | 0 | 0 | 0 |
| 0 | 0 | 0 | 0 |
| 0 | 0 | 0 | 0 |
| 0 | 0 | 0 | 0 |
| 0 | 0 | 0 | 0 |
| 0 | 0 | 0 | 0 |
| 0 | 0 | 0 | 0 |
| 0 | 0 | 0 | 0 |
| 0 | 0 | 0 | 0 |
| 0 | 0 | 0 | 0 |
| 0 | 0 | 0 | 0 |
| 0 | 0 | 0 | 0 |
| 0 | 0 | 0 | 0 |
| 0 | 1 | 0 | 0 |

| | | | |
|---|---|---|---|
| 0 | 0 | 0 | 0 |
| 0 | 0 | 0 | 0 |
| 0 | 0 | 0 | 0 |
| 0 | 0 | 0 | 0 |
| 0 | 0 | 0 | 0 |
| 0 | 0 | 0 | 0 |
| 0 | 0 | 0 | 0 |
| 0 | 0 | 0 | 0 |
| 0 | 0 | 0 | 0 |
| 0 | 0 | 0 | 0 |
| 0 | 0 | 0 | 0 |
| 0 | 0 | 0 | 0 |
| 0 | 0 | 0 | 0 |
| 0 | 0 | 0 | 0 |
| 0 | 0 | 0 | 0 |
| 0 | 0 | 0 | 0 |
| 0 | 0 | 0 | 0 |
| 0 | 0 | 0 | 0 |
| 0 | 0 | 0 | 0 |
| 0 | 0 | 0 | 0 |
| 0 | 0 | 0 | 0 |
| 0 | 0 | 0 | 0 |
| 0 | 0 | 0 | 0 |
| 0 | 0 | 0 | 0 |
| 0 | 0 | 0 | 0 |
| 0 | 0 | 0 | 0 |
| 0 | 0 | 0 | 0 |
| 0 | 0 | 0 | 0 |
| 0 | 0 | 0 | 0 |
| 0 | 0 | 0 | 0 |
| 0 | 0 | 0 | 0 |
| 0 | 0 | 0 | 0 |
| 0 | 0 | 0 | 0 |
| 0 | 0 | 0 | 0 |
| 0 | 0 | 0 | 0 |
| 0 | 0 | 0 | 0 |
| 0 | 0 | 0 | 0 |
| 0 | 0 | 0 | 0 |
| 0 | 0 | 0 | 0 |
| 0 | 0 | 0 | 0 |
| 0 | 0 | 0 | 0 |
| 0 | 0 | 0 | 0 |
| 0 | 0 | 0 | 0 |
| 0 | 0 | 0 | 0 |
| 0 | 0 | 0 | 0 |
| 0 | 0 | 0 | 0 |
| 0 | 0 | 0 | 0 |
| 0 | 0 | 0 | 0 |
| 0 | 0 | 0 | 0 |
| 0 | 0 | 0 | 0 |
| 0 | 0 | 0 | 0 |
| 0 | 0 | 0 | 0 |
| 0 | 0 | 0 | 0 |
| 0 | 0 | 0 | 0 |

| | | | |
|---|---|---|---|
| 0 | 0 | 0 | 0 |
| 0 | 0 | 0 | 0 |
| 0 | 0 | 0 | 0 |
| 0 | 0 | 0 | 0 |
| 0 | 0 | 0 | 0 |
| 0 | 0 | 0 | 0 |
| 0 | 0 | 0 | 0 |
| 0 | 0 | 0 | 0 |
| 0 | 0 | 0 | 0 |
| 0 | 0 | 0 | 0 |
| 0 | 0 | 0 | 0 |
| 0 | 0 | 1 | 0 |
| 0 | 0 | 0 | 0 |
| 0 | 0 | 1 | 0 |
| 0 | 0 | 1 | 0 |
| 0 | 0 | 1 | 0 |
| 0 | 0 | 1 | 0 |
| 0 | 0 | 0 | 0 |
| 0 | 0 | 0 | 0 |
| 0 | 0 | 0 | 0 |
| 1 | 0 | 0 | 0 |

| glycan_binding_site | water_binding_site | water_only_binding_site | non_binding_site |
|---|---|---|---|
| 0 | 0 | 0 | 0 |
| 0 | 0 | 0 | 0 |
| 0 | 0 | 0 | 1 |
| 0 | 0 | 0 | 0 |
| 0 | 0 | 0 | 0 |
| 0 | 0 | 0 | 0 |
| 0 | 0 | 0 | 1 |
| 0 | 0 | 0 | 1 |
| 0 | 0 | 0 | 0 |
| 0 | 0 | 0 | 0 |
| 0 | 0 | 0 | 0 |
| 0 | 0 | 0 | 0 |
| 0 | 0 | 0 | 1 |
| 1 | 0 | 0 | 0 |
| 0 | 0 | 0 | 0 |
| 0 | 0 | 0 | 0 |
| 0 | 0 | 0 | 0 |
| 0 | 0 | 0 | 0 |
| 0 | 0 | 0 | 0 |
| 0 | 0 | 0 | 0 |
| 0 | 1 | 0 | 0 |
| 1 | 1 | 0 | 0 |
| 1 | 1 | 0 | 0 |
| 0 | 1 | 0 | 0 |
| 0 | 0 | 0 | 0 |
| 0 | 0 | 0 | 1 |
| 0 | 0 | 0 | 0 |
| 0 | 0 | 0 | 1 |
| 0 | 0 | 0 | 0 |
| 0 | 0 | 0 | 1 |
| 0 | 0 | 0 | 1 |
| 0 | 1 | 0 | 0 |
| 0 | 1 | 0 | 0 |
| 0 | 1 | 0 | 0 |
| 0 | 1 | 0 | 0 |
| 0 | 0 | 0 | 1 |
| 0 | 0 | 0 | 0 |
| 0 | 0 | 0 | 0 |
| 0 | 1 | 0 | 0 |
| 0 | 1 | 0 | 0 |
| 0 | 1 | 0 | 0 |
| 0 | 1 | 0 | 0 |
| 0 | 1 | 0 | 0 |
| 0 | 1 | 0 | 0 |
| 0 | 1 | 0 | 0 |
| 0 | 1 | 0 | 0 |
| 0 | 1 | 0 | 0 |
| 0 | 1 | 0 | 0 |
| 0 | 1 | 0 | 0 |
| 0 | 1 | 0 | 0 |
| 0 | 1 | 0 | 0 |
| 0 | 1 | 0 | 0 |
| 0 | 0 | 0 | 0 |

| | | | |
|---|---|---|---|
| 0 | 1 | 0 | 0 |
| 0 | 1 | 0 | 0 |
| 0 | 1 | 0 | 0 |
| 0 | 1 | 0 | 0 |
| 0 | 1 | 0 | 0 |
| 0 | 1 | 0 | 0 |
| 0 | 0 | 0 | 0 |
| 0 | 1 | 0 | 0 |
| 0 | 1 | 0 | 0 |
| 0 | 1 | 0 | 0 |
| 0 | 1 | 0 | 0 |
| 0 | 1 | 0 | 0 |
| 0 | 1 | 0 | 0 |
| 0 | 1 | 0 | 0 |
| 0 | 1 | 0 | 0 |
| 0 | 1 | 0 | 0 |
| 0 | 1 | 0 | 0 |
| 0 | 1 | 0 | 0 |
| 0 | 1 | 0 | 0 |
| 0 | 0 | 0 | 0 |
| 0 | 0 | 0 | 0 |
| 0 | 0 | 0 | 0 |
| 0 | 0 | 0 | 0 |
| 0 | 0 | 0 | 0 |
| 0 | 0 | 0 | 0 |
| 0 | 0 | 0 | 0 |
| 0 | 0 | 0 | 0 |
| 0 | 0 | 0 | 0 |
| 0 | 0 | 0 | 0 |
| 0 | 0 | 0 | 0 |
| 0 | 0 | 0 | 0 |
| 0 | 0 | 0 | 0 |
| 0 | 0 | 0 | 0 |
| 0 | 0 | 0 | 0 |
| 0 | 0 | 0 | 0 |
| 0 | 0 | 0 | 0 |
| 0 | 0 | 0 | 0 |
| 0 | 0 | 0 | 0 |
| 0 | 0 | 0 | 0 |
| 0 | 0 | 0 | 0 |
| 1 | 0 | 0 | 0 |
| 0 | 0 | 0 | 0 |
| 1 | 0 | 0 | 0 |
| 1 | 0 | 0 | 0 |
| 0 | 0 | 0 | 0 |
| 0 | 0 | 0 | 1 |
| 0 | 0 | 0 | 1 |
| 0 | 0 | 0 | 0 |
| 1 | 0 | 0 | 0 |
| 0 | 0 | 0 | 0 |
| 0 | 0 | 0 | 0 |
| 0 | 0 | 0 | 1 |
| 0 | 0 | 0 | 1 |
| 0 | 0 | 0 | 0 |

| | | | |
|---|---|---|---|
| 0 | 0 | 0 | 1 |
| 0 | 0 | 0 | 1 |
| 0 | 0 | 0 | 0 |
| 0 | 0 | 0 | 0 |
| 0 | 0 | 0 | 0 |
| 0 | 0 | 0 | 0 |
| 0 | 0 | 0 | 1 |
| 0 | 0 | 0 | 0 |
| 0 | 0 | 0 | 0 |
| 0 | 0 | 0 | 0 |
| 0 | 0 | 0 | 0 |
| 0 | 0 | 0 | 0 |
| 0 | 0 | 0 | 0 |
| 1 | 0 | 0 | 0 |
| 1 | 0 | 0 | 0 |
| 0 | 0 | 0 | 1 |
| 0 | 0 | 0 | 0 |
| 0 | 0 | 0 | 0 |
| 0 | 0 | 0 | 0 |
| 0 | 0 | 0 | 0 |
| 0 | 0 | 0 | 0 |
| 0 | 0 | 0 | 0 |
| 0 | 0 | 0 | 0 |
| 0 | 0 | 0 | 0 |
| 0 | 0 | 0 | 0 |
| 0 | 0 | 0 | 0 |
| 0 | 0 | 0 | 0 |
| 0 | 0 | 0 | 0 |
| 0 | 0 | 0 | 0 |
| 0 | 0 | 0 | 0 |
| 0 | 0 | 0 | 0 |
| 0 | 0 | 0 | 0 |
| 0 | 0 | 0 | 0 |
| 0 | 0 | 0 | 0 |
| 0 | 0 | 0 | 0 |
| 0 | 0 | 0 | 0 |
| 0 | 0 | 0 | 0 |
| 0 | 0 | 0 | 0 |
| 0 | 0 | 0 | 0 |
| 0 | 0 | 0 | 0 |
| 0 | 0 | 0 | 0 |
| 0 | 0 | 0 | 0 |
| 0 | 0 | 0 | 0 |
| 0 | 0 | 0 | 0 |
| 0 | 0 | 0 | 0 |
| 0 | 0 | 0 | 0 |
| 1 | 0 | 0 | 0 |
| 0 | 0 | 0 | 0 |
| 0 | 0 | 0 | 0 |
| 0 | 0 | 0 | 0 |
| 0 | 0 | 0 | 0 |
| 0 | 0 | 0 | 0 |
| 0 | 0 | 0 | 0 |
| 0 | 1 | 1 | 1 |

| | | | |
|---|---|---|---|
| 0 | 1 | 1 | 1 |
| 0 | 1 | 1 | 1 |
| 0 | 0 | 0 | 1 |
| 0 | 0 | 0 | 0 |
| 0 | 0 | 0 | 0 |
| 0 | 1 | 1 | 1 |
| 0 | 0 | 0 | 0 |
| 0 | 1 | 1 | 1 |
| 0 | 1 | 1 | 1 |
| 0 | 0 | 0 | 1 |
| 0 | 0 | 0 | 1 |
| 0 | 1 | 0 | 0 |
| 0 | 1 | 1 | 1 |
| 0 | 0 | 0 | 0 |
| 0 | 1 | 1 | 1 |
| 0 | 0 | 0 | 1 |
| 0 | 0 | 0 | 1 |
| 0 | 1 | 1 | 1 |
| 0 | 1 | 1 | 1 |
| 0 | 0 | 0 | 0 |
| 0 | 1 | 0 | 0 |
| 0 | 0 | 0 | 1 |
| 0 | 0 | 0 | 0 |
| 0 | 0 | 0 | 0 |
| 0 | 0 | 0 | 0 |
| 0 | 0 | 0 | 1 |
| 0 | 0 | 0 | 0 |
| 0 | 0 | 0 | 0 |
| 0 | 0 | 0 | 1 |
| 0 | 0 | 0 | 1 |
| 0 | 0 | 0 | 1 |
| 0 | 0 | 0 | 1 |
| 0 | 0 | 0 | 1 |
| 0 | 0 | 0 | 1 |
| 0 | 0 | 0 | 0 |
| 0 | 0 | 0 | 1 |
| 1 | 0 | 0 | 0 |
| 1 | 0 | 0 | 0 |
| 0 | 1 | 1 | 1 |
| 0 | 1 | 1 | 1 |
| 0 | 0 | 0 | 1 |
| 0 | 1 | 0 | 0 |
| 0 | 1 | 0 | 0 |
| 0 | 0 | 0 | 1 |
| 0 | 0 | 0 | 1 |
| 0 | 0 | 0 | 1 |
| 0 | 0 | 0 | 0 |
| 0 | 0 | 0 | 0 |
| 0 | 0 | 0 | 0 |
| 0 | 0 | 0 | 0 |
| 0 | 0 | 0 | 0 |
| 0 | 0 | 0 | 1 |
| 0 | 0 | 0 | 1 |
| 0 | 0 | 0 | 0 |

| | | | |
|---|---|---|---|
| 0 | 0 | 0 | 0 |
| 0 | 0 | 0 | 0 |
| 0 | 0 | 0 | 0 |
| 0 | 0 | 0 | 1 |
| 0 | 0 | 0 | 1 |
| 0 | 0 | 0 | 1 |
| 0 | 0 | 0 | 0 |
| 0 | 0 | 0 | 0 |
| 0 | 0 | 0 | 1 |
| 0 | 0 | 0 | 1 |
| 1 | 0 | 0 | 0 |
| 0 | 0 | 0 | 1 |
| 1 | 0 | 0 | 0 |
| 0 | 0 | 0 | 1 |
| 0 | 0 | 0 | 1 |
| 0 | 0 | 0 | 1 |
| 0 | 0 | 0 | 1 |
| 0 | 0 | 0 | 0 |
| 0 | 0 | 0 | 0 |
| 0 | 0 | 0 | 0 |
| 0 | 0 | 0 | 0 |
| 0 | 0 | 0 | 1 |
| 0 | 0 | 0 | 0 |
| 0 | 0 | 0 | 1 |
| 0 | 0 | 0 | 1 |
| 0 | 0 | 0 | 0 |
| 0 | 1 | 0 | 0 |
| 0 | 0 | 0 | 0 |
| 0 | 1 | 0 | 0 |
| 0 | 1 | 0 | 0 |
| 0 | 1 | 0 | 0 |
| 0 | 0 | 0 | 0 |
| 0 | 0 | 0 | 1 |
| 0 | 0 | 0 | 1 |
| 0 | 0 | 0 | 0 |
| 0 | 0 | 0 | 1 |
| 0 | 0 | 0 | 1 |
| 0 | 0 | 0 | 0 |
| 0 | 0 | 0 | 1 |
| 0 | 0 | 0 | 1 |
| 0 | 0 | 0 | 0 |
| 0 | 0 | 0 | 1 |
| 0 | 0 | 0 | 1 |
| 0 | 0 | 0 | 1 |
| 0 | 0 | 0 | 0 |
| 0 | 0 | 0 | 0 |
| 0 | 0 | 0 | 0 |
| 0 | 0 | 0 | 0 |
| 0 | 0 | 0 | 0 |
| 0 | 0 | 0 | 0 |
| 0 | 0 | 0 | 0 |
| 0 | 0 | 0 | 0 |
| 0 | 0 | 0 | 0 |
| 0 | 0 | 0 | 0 |

| | | | |
|---|---|---|---|
| 0 | 0 | 0 | 0 |
| 0 | 0 | 0 | 0 |
| 0 | 0 | 0 | 0 |
| 0 | 0 | 0 | 0 |
| 0 | 0 | 0 | 0 |
| 0 | 0 | 0 | 0 |
| 0 | 0 | 0 | 1 |
| 0 | 0 | 0 | 1 |
| 0 | 0 | 0 | 0 |
| 0 | 0 | 0 | 0 |
| 0 | 0 | 0 | 0 |
| 0 | 0 | 0 | 0 |
| 0 | 0 | 0 | 0 |
| 0 | 0 | 0 | 1 |
| 0 | 0 | 0 | 1 |
| 0 | 0 | 0 | 0 |
| 0 | 0 | 0 | 1 |
| 0 | 0 | 0 | 0 |
| 0 | 0 | 0 | 1 |
| 0 | 0 | 0 | 0 |
| 0 | 0 | 0 | 1 |
| 0 | 0 | 0 | 1 |
| 0 | 0 | 0 | 1 |
| 0 | 0 | 0 | 0 |
| 0 | 0 | 0 | 0 |
| 0 | 0 | 0 | 0 |
| 0 | 0 | 0 | 0 |
| 0 | 0 | 0 | 0 |
| 0 | 0 | 0 | 0 |
| 0 | 0 | 0 | 0 |
| 0 | 0 | 0 | 1 |
| 0 | 0 | 0 | 1 |
| 0 | 0 | 0 | 0 |
| 0 | 0 | 0 | 0 |
| 0 | 0 | 0 | 0 |
| 0 | 0 | 0 | 1 |
| 0 | 0 | 0 | 0 |
| 0 | 0 | 0 | 0 |
| 0 | 0 | 0 | 0 |
| 0 | 0 | 0 | 0 |
| 0 | 0 | 0 | 1 |
| 0 | 0 | 0 | 1 |
| 0 | 0 | 0 | 0 |
| 0 | 0 | 0 | 1 |
| 0 | 0 | 0 | 0 |
| 0 | 0 | 0 | 0 |
| 0 | 0 | 0 | 0 |
| 0 | 0 | 0 | 1 |
| 0 | 0 | 0 | 1 |
| 0 | 0 | 0 | 0 |
| 0 | 0 | 0 | 1 |
| 0 | 0 | 0 | 1 |
| 0 | 0 | 0 | 0 |
| 0 | 0 | 0 | 1 |

| | | | |
|---|---|---|---|
| 0 | 0 | 0 | 0 |
| 0 | 0 | 0 | 1 |
| 0 | 0 | 0 | 1 |
| 0 | 0 | 0 | 0 |
| 0 | 0 | 0 | 0 |
| 0 | 0 | 0 | 0 |
| 0 | 0 | 0 | 1 |
| 0 | 0 | 0 | 0 |
| 0 | 0 | 0 | 0 |
| 0 | 0 | 0 | 1 |
| 0 | 0 | 0 | 1 |
| 0 | 0 | 0 | 1 |
| 0 | 0 | 0 | 1 |
| 0 | 0 | 0 | 1 |
| 0 | 0 | 0 | 0 |
| 0 | 0 | 0 | 1 |
| 0 | 0 | 0 | 0 |
| 0 | 0 | 0 | 0 |
| 0 | 0 | 0 | 1 |
| 0 | 0 | 0 | 0 |
| 0 | 0 | 0 | 0 |
| 0 | 0 | 0 | 0 |
| 0 | 0 | 0 | 0 |
| 0 | 0 | 0 | 0 |
| 0 | 0 | 0 | 1 |
| 0 | 0 | 0 | 1 |
| 0 | 0 | 0 | 0 |
| 0 | 0 | 0 | 0 |
| 0 | 0 | 0 | 0 |
| 0 | 0 | 0 | 0 |
| 0 | 0 | 0 | 0 |
| 0 | 0 | 0 | 0 |
| 0 | 0 | 0 | 0 |
| 0 | 0 | 0 | 0 |
| 0 | 0 | 0 | 1 |
| 0 | 0 | 0 | 1 |
| 0 | 0 | 0 | 0 |
| 0 | 0 | 0 | 0 |
| 0 | 0 | 0 | 0 |
| 0 | 0 | 0 | 0 |
| 0 | 0 | 0 | 0 |
| 0 | 0 | 0 | 0 |
| 0 | 0 | 0 | 1 |
| 0 | 0 | 0 | 0 |
| 0 | 0 | 0 | 0 |
| 0 | 0 | 0 | 0 |
| 0 | 0 | 0 | 0 |
| 0 | 0 | 0 | 0 |
| 1 | 1 | 0 | 0 |
| 0 | 0 | 0 | 1 |
| 0 | 0 | 0 | 0 |
| 0 | 0 | 0 | 0 |
| 0 | 0 | 0 | 0 |
| 0 | 0 | 0 | 0 |

| | | | |
|---|---|---|---|
| 0 | 0 | 0 | 0 |
| 0 | 0 | 0 | 0 |
| 0 | 0 | 0 | 0 |
| 0 | 0 | 0 | 0 |
| 0 | 0 | 0 | 1 |
| 0 | 0 | 0 | 0 |
| 0 | 0 | 0 | 0 |
| 0 | 0 | 0 | 0 |
| 0 | 0 | 0 | 0 |
| 0 | 0 | 0 | 0 |
| 0 | 0 | 0 | 0 |
| 0 | 0 | 0 | 0 |
| 0 | 0 | 0 | 0 |
| 0 | 0 | 0 | 0 |
| 0 | 0 | 0 | 0 |
| 0 | 0 | 0 | 0 |
| 0 | 0 | 0 | 0 |
| 0 | 0 | 0 | 0 |
| 0 | 0 | 0 | 1 |
| 0 | 0 | 0 | 0 |
| 0 | 0 | 0 | 0 |
| 0 | 0 | 0 | 0 |
| 0 | 0 | 0 | 0 |
| 0 | 0 | 0 | 1 |
| 0 | 0 | 0 | 0 |
| 0 | 0 | 0 | 0 |
| 0 | 0 | 0 | 0 |
| 0 | 0 | 0 | 1 |
| 0 | 0 | 0 | 1 |
| 0 | 0 | 0 | 1 |
| 0 | 0 | 0 | 0 |
| 0 | 0 | 0 | 0 |
| 0 | 0 | 0 | 0 |
| 0 | 0 | 0 | 0 |
| 0 | 0 | 0 | 0 |
| 0 | 0 | 0 | 1 |
| 0 | 0 | 0 | 1 |
| 0 | 0 | 0 | 0 |
| 0 | 0 | 0 | 0 |
| 0 | 0 | 0 | 1 |
| 0 | 0 | 0 | 1 |
| 0 | 0 | 0 | 1 |
| 0 | 0 | 0 | 1 |
| 0 | 0 | 0 | 1 |
| 0 | 0 | 0 | 0 |
| 0 | 0 | 0 | 0 |
| 0 | 0 | 0 | 0 |
| 0 | 0 | 0 | 0 |
| 0 | 0 | 0 | 0 |
| 0 | 0 | 0 | 1 |
| 0 | 0 | 0 | 1 |
| 0 | 0 | 0 | 0 |
| 0 | 0 | 0 | 1 |
| 0 | 0 | 0 | 0 |

| | | | |
|---|---|---|---|
| 0 | 0 | 0 | 1 |
| 0 | 0 | 0 | 1 |
| 0 | 0 | 0 | 0 |
| 0 | 0 | 0 | 1 |
| 0 | 0 | 0 | 1 |
| 0 | 0 | 0 | 1 |
| 0 | 0 | 0 | 1 |
| 0 | 0 | 0 | 0 |
| 0 | 0 | 0 | 1 |
| 0 | 0 | 0 | 1 |
| 0 | 0 | 0 | 0 |
| 0 | 0 | 0 | 1 |
| 0 | 0 | 0 | 1 |
| 0 | 0 | 0 | 1 |
| 0 | 0 | 0 | 0 |
| 0 | 0 | 0 | 0 |
| 0 | 0 | 0 | 1 |
| 0 | 0 | 0 | 1 |
| 0 | 0 | 0 | 0 |
| 0 | 0 | 0 | 0 |
| 0 | 0 | 0 | 0 |
| 0 | 0 | 0 | 0 |
| 0 | 0 | 0 | 0 |
| 0 | 1 | 0 | 0 |
| 0 | 1 | 0 | 0 |
| 0 | 0 | 0 | 1 |
| 0 | 0 | 0 | 0 |
| 0 | 0 | 0 | 0 |
| 0 | 0 | 0 | 0 |
| 0 | 0 | 0 | 1 |
| 0 | 0 | 0 | 0 |
| 0 | 0 | 0 | 0 |
| 0 | 0 | 0 | 1 |
| 0 | 0 | 0 | 0 |
| 0 | 0 | 0 | 0 |
| 0 | 0 | 0 | 0 |
| 0 | 1 | 0 | 0 |
| 0 | 1 | 0 | 0 |
| 0 | 0 | 0 | 0 |
| 0 | 0 | 0 | 0 |
| 0 | 0 | 0 | 1 |
| 0 | 0 | 0 | 0 |
| 0 | 0 | 0 | 1 |
| 0 | 0 | 0 | 0 |
| 0 | 0 | 0 | 0 |
| 1 | 0 | 0 | 0 |
| 0 | 0 | 0 | 0 |
| 1 | 0 | 0 | 0 |
| 0 | 0 | 0 | 0 |
| 0 | 0 | 0 | 0 |
| 0 | 0 | 0 | 1 |
| 0 | 0 | 0 | 0 |
| 0 | 0 | 0 | 0 |
| 0 | 0 | 0 | 0 |

| | | | |
|---|---|---|---|
| 0 | 0 | 0 | 1 |
| 0 | 0 | 0 | 0 |
| 0 | 0 | 0 | 1 |
| 0 | 0 | 0 | 1 |
| 1 | 0 | 0 | 0 |
| 0 | 0 | 0 | 0 |
| 1 | 0 | 0 | 0 |
| 1 | 0 | 0 | 0 |
| 0 | 0 | 0 | 0 |
| 0 | 0 | 0 | 1 |
| 0 | 0 | 0 | 1 |
| 0 | 0 | 0 | 1 |
| 0 | 0 | 0 | 1 |
| 1 | 0 | 0 | 0 |
| 0 | 0 | 0 | 1 |
| 0 | 0 | 0 | 0 |
| 0 | 0 | 0 | 0 |
| 0 | 0 | 0 | 0 |
| 1 | 0 | 0 | 0 |
| 0 | 0 | 0 | 1 |
| 1 | 0 | 0 | 0 |
| 0 | 0 | 0 | 1 |
| 0 | 0 | 0 | 0 |
| 0 | 0 | 0 | 1 |
| 0 | 0 | 0 | 1 |
| 0 | 0 | 0 | 1 |
| 0 | 0 | 0 | 0 |
| 0 | 0 | 0 | 1 |
| 0 | 0 | 0 | 1 |
| 0 | 0 | 0 | 1 |
| 0 | 0 | 0 | 0 |
| 0 | 0 | 0 | 0 |
| 0 | 0 | 0 | 1 |
| 0 | 0 | 0 | 1 |
| 0 | 0 | 0 | 0 |
| 0 | 0 | 0 | 0 |
| 0 | 0 | 0 | 0 |
| 1 | 0 | 0 | 0 |
| 0 | 0 | 0 | 0 |
| 0 | 0 | 0 | 1 |
| 0 | 0 | 0 | 0 |
| 0 | 0 | 0 | 0 |
| 0 | 0 | 0 | 0 |
| 0 | 0 | 0 | 0 |
| 0 | 0 | 0 | 1 |
| 1 | 0 | 0 | 0 |
| 0 | 0 | 0 | 0 |
| 0 | 0 | 0 | 0 |
| 0 | 0 | 0 | 0 |
| 1 | 0 | 0 | 0 |
| 0 | 0 | 0 | 0 |
| 0 | 1 | 0 | 0 |
| 0 | 0 | 0 | 1 |
| 0 | 0 | 0 | 0 |

| | | | |
|---|---|---|---|
| 0 | 0 | 0 | 0 |
| 0 | 1 | 0 | 0 |
| 0 | 1 | 1 | 1 |
| 0 | 1 | 1 | 1 |
| 0 | 0 | 0 | 1 |
| 0 | 0 | 0 | 0 |
| 0 | 0 | 0 | 1 |
| 0 | 0 | 0 | 1 |
| 0 | 1 | 0 | 0 |
| 0 | 0 | 0 | 0 |
| 1 | 0 | 0 | 0 |
| 1 | 0 | 0 | 0 |
| 0 | 0 | 0 | 1 |
| 1 | 0 | 0 | 0 |
| 0 | 0 | 0 | 0 |
| 0 | 0 | 0 | 0 |
| 0 | 1 | 0 | 0 |
| 0 | 0 | 0 | 0 |
| 0 | 0 | 0 | 0 |
| 0 | 0 | 0 | 1 |
| 0 | 0 | 0 | 0 |
| 0 | 0 | 0 | 0 |
| 0 | 0 | 0 | 0 |
| 0 | 0 | 0 | 0 |
| 0 | 0 | 0 | 0 |
| 0 | 0 | 0 | 1 |
| 0 | 0 | 0 | 1 |
| 0 | 0 | 0 | 1 |
| 0 | 0 | 0 | 0 |
| 0 | 0 | 0 | 0 |
| 0 | 0 | 0 | 0 |
| 0 | 0 | 0 | 0 |
| 0 | 0 | 0 | 0 |
| 0 | 0 | 0 | 0 |
| 0 | 1 | 0 | 0 |
| 0 | 0 | 0 | 1 |
| 0 | 0 | 0 | 0 |
| 0 | 0 | 0 | 0 |
| 0 | 0 | 0 | 1 |
| 0 | 0 | 0 | 1 |
| 0 | 0 | 0 | 1 |
| 0 | 0 | 0 | 0 |
| 0 | 0 | 0 | 0 |
| 0 | 0 | 0 | 0 |
| 0 | 0 | 0 | 0 |
| 0 | 0 | 0 | 0 |
| 0 | 0 | 0 | 0 |
| 0 | 0 | 0 | 1 |
| 0 | 0 | 0 | 1 |
| 0 | 0 | 0 | 0 |
| 0 | 0 | 0 | 0 |
| 0 | 0 | 0 | 0 |
| 0 | 0 | 0 | 0 |
| 0 | 0 | 0 | 1 |

| | | | |
|---|---|---|---|
| 0 | 0 | 0 | 1 |
| 0 | 0 | 0 | 0 |
| 0 | 0 | 0 | 1 |
| 0 | 0 | 0 | 0 |
| 0 | 0 | 0 | 1 |
| 0 | 0 | 0 | 1 |
| 0 | 0 | 0 | 0 |
| 0 | 1 | 1 | 1 |
| 0 | 0 | 0 | 1 |
| 0 | 0 | 0 | 1 |
| 0 | 0 | 0 | 1 |
| 0 | 0 | 0 | 1 |
| 0 | 0 | 0 | 1 |
| 0 | 0 | 0 | 1 |
| 0 | 0 | 0 | 1 |
| 0 | 0 | 0 | 1 |
| 0 | 0 | 0 | 0 |
| 0 | 0 | 0 | 0 |
| 0 | 0 | 0 | 1 |
| 0 | 0 | 0 | 0 |
| 0 | 0 | 0 | 0 |
| 0 | 0 | 0 | 0 |
| 0 | 0 | 0 | 1 |
| 0 | 0 | 0 | 0 |
| 0 | 0 | 0 | 0 |
| 0 | 0 | 0 | 1 |
| 0 | 0 | 0 | 1 |
| 0 | 0 | 0 | 1 |
| 0 | 1 | 0 | 0 |
| 0 | 0 | 0 | 0 |
| 0 | 0 | 0 | 1 |
| 0 | 0 | 0 | 0 |
| 0 | 0 | 0 | 0 |
| 0 | 0 | 0 | 0 |
| 0 | 0 | 0 | 0 |
| 0 | 0 | 0 | 0 |
| 0 | 1 | 1 | 1 |
| 0 | 0 | 0 | 1 |
| 0 | 0 | 0 | 1 |
| 0 | 0 | 0 | 1 |
| 0 | 0 | 0 | 0 |
| 0 | 0 | 0 | 1 |
| 0 | 0 | 0 | 0 |
| 0 | 0 | 0 | 1 |
| 1 | 0 | 0 | 0 |
| 0 | 0 | 0 | 1 |
| 0 | 0 | 0 | 0 |
| 0 | 0 | 0 | 0 |
| 1 | 0 | 0 | 0 |
| 0 | 0 | 0 | 0 |
| 0 | 0 | 0 | 0 |
| 0 | 0 | 0 | 0 |
| 0 | 0 | 0 | 0 |
| 0 | 0 | 0 | 0 |

| | | | |
|---|---|---|---|
| 0 | 0 | 0 | 0 |
| 0 | 0 | 0 | 0 |
| 0 | 0 | 0 | 0 |
| 0 | 0 | 0 | 0 |
| 0 | 0 | 0 | 0 |
| 0 | 0 | 0 | 0 |
| 0 | 0 | 0 | 0 |
| 0 | 0 | 0 | 1 |
| 0 | 1 | 0 | 0 |
| 0 | 1 | 0 | 0 |
| 0 | 0 | 0 | 0 |
| 0 | 0 | 0 | 1 |
| 0 | 0 | 0 | 0 |
| 0 | 1 | 0 | 0 |
| 0 | 1 | 0 | 0 |
| 0 | 1 | 0 | 0 |
| 0 | 0 | 0 | 0 |
| 0 | 1 | 0 | 0 |
| 0 | 1 | 0 | 0 |
| 0 | 1 | 0 | 0 |
| 0 | 1 | 0 | 0 |
| 0 | 0 | 0 | 0 |
| 0 | 1 | 0 | 0 |
| 0 | 1 | 0 | 0 |
| 0 | 1 | 0 | 0 |
| 0 | 1 | 0 | 0 |
| 0 | 1 | 0 | 0 |
| 0 | 1 | 0 | 0 |
| 0 | 1 | 0 | 0 |
| 0 | 1 | 0 | 0 |
| 0 | 1 | 0 | 0 |
| 0 | 0 | 0 | 0 |
| 0 | 0 | 0 | 0 |
| 0 | 1 | 0 | 0 |
| 0 | 1 | 0 | 0 |
| 0 | 1 | 0 | 0 |
| 0 | 1 | 0 | 0 |
| 0 | 0 | 0 | 0 |
| 0 | 0 | 0 | 0 |
| 0 | 1 | 0 | 0 |
| 0 | 0 | 0 | 1 |
| 0 | 1 | 0 | 0 |
| 0 | 1 | 0 | 0 |
| 0 | 1 | 0 | 0 |
| 0 | 1 | 0 | 0 |
| 0 | 0 | 0 | 0 |
| 0 | 0 | 0 | 0 |
| 0 | 0 | 0 | 0 |
| 0 | 0 | 0 | 0 |
| 0 | 0 | 0 | 1 |
| 0 | 0 | 0 | 1 |
| 0 | 0 | 0 | 1 |
| 1 | 0 | 0 | 0 |
| 0 | 0 | 0 | 0 |

| | | | |
|---|---|---|---|
| 0 | 0 | 0 | 0 |
| 0 | 0 | 0 | 0 |
| 1 | 0 | 0 | 0 |
| 1 | 0 | 0 | 0 |
| 0 | 0 | 0 | 0 |
| 0 | 0 | 0 | 0 |
| 0 | 0 | 0 | 0 |
| 0 | 0 | 0 | 1 |
| 0 | 0 | 0 | 1 |
| 0 | 0 | 0 | 0 |
| 0 | 0 | 0 | 0 |
| 0 | 0 | 0 | 1 |
| 0 | 0 | 0 | 0 |
| 0 | 0 | 0 | 0 |
| 0 | 0 | 0 | 0 |
| 0 | 0 | 0 | 0 |
| 0 | 0 | 0 | 0 |
| 0 | 0 | 0 | 1 |
| 0 | 0 | 0 | 0 |
| 0 | 0 | 0 | 1 |
| 0 | 0 | 0 | 0 |
| 0 | 0 | 0 | 0 |
| 0 | 0 | 0 | 1 |
| 0 | 0 | 0 | 0 |
| 0 | 0 | 0 | 0 |
| 0 | 0 | 0 | 1 |
| 0 | 0 | 0 | 0 |
| 0 | 0 | 0 | 0 |
| 0 | 0 | 0 | 0 |
| 0 | 0 | 0 | 0 |
| 0 | 0 | 0 | 0 |
| 0 | 0 | 0 | 0 |
| 0 | 0 | 0 | 1 |
| 0 | 0 | 0 | 0 |
| 0 | 0 | 0 | 0 |
| 0 | 0 | 0 | 0 |
| 0 | 0 | 0 | 1 |
| 0 | 0 | 0 | 0 |
| 0 | 0 | 0 | 0 |
| 0 | 0 | 0 | 0 |
| 0 | 0 | 0 | 0 |
| 0 | 1 | 0 | 0 |
| 0 | 1 | 0 | 0 |
| 0 | 1 | 0 | 0 |
| 0 | 0 | 0 | 0 |
| 0 | 0 | 0 | 1 |
| 0 | 0 | 0 | 0 |
| 0 | 0 | 0 | 0 |
| 0 | 0 | 0 | 0 |
| 0 | 0 | 0 | 0 |
| 0 | 0 | 0 | 0 |
| 0 | 1 | 0 | 0 |
| 0 | 0 | 0 | 1 |
| 0 | 0 | 0 | 1 |

| | | | |
|---|---|---|---|
| 0 | 0 | 0 | 1 |
| 0 | 0 | 0 | 1 |
| 0 | 0 | 0 | 0 |
| 0 | 0 | 0 | 0 |
| 0 | 0 | 0 | 0 |
| 0 | 1 | 0 | 0 |
| 0 | 0 | 0 | 0 |
| 0 | 0 | 0 | 1 |
| 0 | 0 | 0 | 0 |
| 0 | 0 | 0 | 1 |
| 0 | 0 | 0 | 1 |
| 0 | 0 | 0 | 1 |
| 0 | 0 | 0 | 0 |
| 0 | 0 | 0 | 0 |
| 0 | 0 | 0 | 0 |
| 0 | 0 | 0 | 0 |
| 0 | 0 | 0 | 0 |
| 1 | 0 | 0 | 0 |
| 0 | 0 | 0 | 1 |
| 0 | 0 | 0 | 1 |
| 0 | 0 | 0 | 1 |
| 0 | 0 | 0 | 0 |
| 0 | 0 | 0 | 1 |
| 0 | 0 | 0 | 0 |
| 0 | 0 | 0 | 0 |
| 0 | 0 | 0 | 0 |
| 0 | 0 | 0 | 0 |
| 0 | 0 | 0 | 0 |
| 0 | 0 | 0 | 0 |
| 0 | 0 | 0 | 0 |
| 0 | 0 | 0 | 0 |
| 0 | 0 | 0 | 0 |
| 0 | 0 | 0 | 0 |
| 0 | 0 | 0 | 0 |
| 0 | 0 | 0 | 0 |
| 0 | 0 | 0 | 0 |
| 0 | 0 | 0 | 0 |
| 0 | 0 | 0 | 0 |
| 1 | 0 | 0 | 0 |
| 0 | 0 | 0 | 0 |
| 0 | 1 | 0 | 0 |
| 0 | 0 | 0 | 1 |
| 0 | 0 | 0 | 0 |
| 0 | 1 | 0 | 0 |
| 0 | 0 | 0 | 0 |
| 0 | 0 | 0 | 0 |
| 0 | 0 | 0 | 1 |
| 0 | 0 | 0 | 0 |
| 0 | 0 | 0 | 0 |
| 0 | 0 | 0 | 0 |
| 0 | 0 | 0 | 0 |
| 0 | 0 | 0 | 0 |
| 0 | 0 | 0 | 1 |
| 0 | 0 | 0 | 1 |

| | | | |
|---|---|---|---|
| 0 | 0 | 0 | 0 |
| 0 | 0 | 0 | 0 |
| 0 | 0 | 0 | 1 |
| 0 | 0 | 0 | 1 |
| 0 | 0 | 0 | 1 |
| 0 | 0 | 0 | 0 |
| 0 | 0 | 0 | 1 |
| 0 | 0 | 0 | 1 |
| 0 | 0 | 0 | 0 |
| 0 | 0 | 0 | 0 |
| 0 | 0 | 0 | 0 |
| 0 | 0 | 0 | 0 |
| 0 | 0 | 0 | 0 |
| 0 | 0 | 0 | 1 |
| 0 | 0 | 0 | 1 |
| 0 | 0 | 0 | 1 |
| 0 | 0 | 0 | 1 |
| 0 | 0 | 0 | 1 |
| 0 | 0 | 0 | 0 |
| 0 | 0 | 0 | 0 |
| 0 | 0 | 0 | 1 |
| 0 | 0 | 0 | 1 |
| 0 | 0 | 0 | 0 |
| 0 | 0 | 0 | 0 |
| 0 | 0 | 0 | 1 |
| 0 | 0 | 0 | 1 |
| 0 | 1 | 0 | 0 |
| 0 | 1 | 1 | 1 |
| 0 | 0 | 0 | 0 |
| 0 | 0 | 0 | 0 |
| 0 | 0 | 0 | 0 |
| 0 | 0 | 0 | 0 |
| 0 | 0 | 0 | 0 |
| 1 | 0 | 0 | 0 |
| 1 | 0 | 0 | 0 |
| 0 | 0 | 0 | 0 |
| 0 | 0 | 0 | 0 |
| 0 | 0 | 0 | 0 |
| 0 | 0 | 0 | 1 |
| 0 | 0 | 0 | 0 |
| 0 | 0 | 0 | 1 |
| 0 | 0 | 0 | 1 |
| 0 | 0 | 0 | 1 |
| 0 | 0 | 0 | 1 |
| 0 | 0 | 0 | 1 |
| 0 | 0 | 0 | 0 |
| 0 | 0 | 0 | 0 |
| 0 | 0 | 0 | 0 |
| 0 | 0 | 0 | 0 |
| 0 | 0 | 0 | 0 |
| 0 | 0 | 0 | 0 |
| 0 | 0 | 0 | 0 |
| 0 | 0 | 0 | 0 |
| 0 | 0 | 0 | 1 |

| | | | |
|---|---|---|---|
| 0 | 0 | 0 | 1 |
| 0 | 0 | 0 | 0 |
| 0 | 0 | 0 | 0 |
| 0 | 0 | 0 | 0 |
| 0 | 0 | 0 | 1 |
| 0 | 0 | 0 | 0 |
| 0 | 0 | 0 | 0 |
| 0 | 0 | 0 | 0 |
| 0 | 0 | 0 | 1 |
| 0 | 0 | 0 | 1 |
| 0 | 0 | 0 | 0 |
| 0 | 0 | 0 | 0 |
| 0 | 0 | 0 | 0 |
| 0 | 0 | 0 | 1 |
| 0 | 0 | 0 | 0 |
| 0 | 0 | 0 | 0 |
| 0 | 0 | 0 | 0 |
| 0 | 0 | 0 | 0 |
| 0 | 0 | 0 | 0 |
| 0 | 0 | 0 | 0 |
| 0 | 0 | 0 | 0 |
| 0 | 0 | 0 | 1 |
| 0 | 0 | 0 | 0 |
| 0 | 0 | 0 | 1 |
| 0 | 0 | 0 | 0 |
| 0 | 0 | 0 | 0 |
| 0 | 0 | 0 | 1 |
| 0 | 0 | 0 | 0 |
| 0 | 0 | 0 | 0 |
| 0 | 0 | 0 | 0 |
| 0 | 0 | 0 | 0 |
| 0 | 0 | 0 | 0 |
| 0 | 0 | 0 | 0 |
| 0 | 0 | 0 | 0 |
| 0 | 0 | 0 | 1 |
| 0 | 0 | 0 | 0 |
| 0 | 0 | 0 | 0 |
| 0 | 0 | 0 | 0 |
| 0 | 0 | 0 | 0 |
| 0 | 0 | 0 | 0 |
| 0 | 0 | 0 | 0 |
| 0 | 0 | 0 | 0 |
| 0 | 0 | 0 | 0 |
| 0 | 0 | 0 | 0 |
| 0 | 0 | 0 | 0 |
| 0 | 0 | 0 | 0 |
| 0 | 0 | 0 | 0 |
| 0 | 0 | 0 | 1 |
| 0 | 0 | 0 | 0 |
| 0 | 0 | 0 | 1 |
| 0 | 0 | 0 | 0 |
| 0 | 0 | 0 | 0 |
| 0 | 0 | 0 | 0 |
| 0 | 0 | 0 | 0 |

| | | | |
|---|---|---|---|
| 0 | 0 | 0 | 0 |
| 0 | 0 | 0 | 0 |
| 0 | 0 | 0 | 1 |
| 0 | 0 | 0 | 0 |
| 0 | 0 | 0 | 0 |
| 0 | 0 | 0 | 1 |
| 0 | 0 | 0 | 1 |
| 0 | 0 | 0 | 1 |
| 0 | 0 | 0 | 0 |
| 0 | 0 | 0 | 1 |
| 0 | 0 | 0 | 0 |
| 0 | 0 | 0 | 0 |
| 0 | 0 | 0 | 0 |
| 0 | 0 | 0 | 1 |
| 0 | 0 | 0 | 0 |
| 0 | 0 | 0 | 1 |
| 0 | 0 | 0 | 0 |
| 0 | 0 | 0 | 0 |
| 1 | 0 | 0 | 0 |
| 0 | 0 | 0 | 1 |
| 0 | 0 | 0 | 1 |
| 0 | 0 | 0 | 0 |
| 0 | 0 | 0 | 1 |
| 0 | 0 | 0 | 0 |
| 0 | 0 | 0 | 0 |
| 0 | 0 | 0 | 1 |
| 0 | 0 | 0 | 1 |
| 0 | 0 | 0 | 0 |
| 0 | 0 | 0 | 1 |
| 1 | 0 | 0 | 0 |
| 0 | 0 | 0 | 1 |
| 0 | 0 | 0 | 1 |
| 0 | 0 | 0 | 1 |
| 0 | 0 | 0 | 1 |
| 0 | 0 | 0 | 1 |
| 0 | 0 | 0 | 1 |
| 0 | 0 | 0 | 1 |
| 0 | 0 | 0 | 1 |
| 0 | 0 | 0 | 1 |
| 0 | 0 | 0 | 1 |
| 0 | 0 | 0 | 1 |
| 0 | 0 | 0 | 1 |
| 0 | 0 | 0 | 1 |
| 0 | 0 | 0 | 1 |
| 0 | 0 | 0 | 1 |
| 0 | 0 | 0 | 1 |
| 0 | 0 | 0 | 1 |
| 0 | 0 | 0 | 1 |
| 0 | 0 | 0 | 1 |
| 0 | 0 | 0 | 1 |
| 0 | 0 | 0 | 1 |
| 0 | 0 | 0 | 1 |
| 0 | 0 | 0 | 1 |
| 0 | 0 | 0 | 1 |

| | | | |
|---|---|---|---|
| 0 | 0 | 0 | 1 |
| 0 | 0 | 0 | 1 |
| 0 | 0 | 0 | 1 |
| 0 | 0 | 0 | 1 |
| 0 | 0 | 0 | 1 |
| 0 | 0 | 0 | 1 |
| 0 | 0 | 0 | 1 |
| 0 | 0 | 0 | 1 |
| 0 | 0 | 0 | 1 |
| 0 | 0 | 0 | 1 |
| 0 | 0 | 0 | 1 |
| 0 | 0 | 0 | 0 |
| 0 | 0 | 0 | 0 |
| 0 | 0 | 0 | 0 |
| 0 | 0 | 0 | 1 |
| 0 | 0 | 0 | 0 |
| 0 | 0 | 0 | 1 |
| 0 | 0 | 0 | 0 |
| 0 | 0 | 0 | 1 |
| 0 | 0 | 0 | 1 |
| 0 | 0 | 0 | 0 |
| 0 | 0 | 0 | 1 |
| 0 | 0 | 0 | 1 |
| 0 | 0 | 0 | 0 |
| 0 | 0 | 0 | 0 |
| 0 | 0 | 0 | 1 |
| 0 | 0 | 0 | 1 |
| 0 | 0 | 0 | 0 |
| 0 | 0 | 0 | 0 |
| 0 | 0 | 0 | 0 |
| 0 | 0 | 0 | 0 |
| 0 | 0 | 0 | 0 |
| 0 | 0 | 0 | 1 |
| 0 | 0 | 0 | 1 |
| 0 | 0 | 0 | 1 |
| 0 | 0 | 0 | 1 |
| 0 | 0 | 0 | 1 |
| 0 | 0 | 0 | 1 |
| 0 | 0 | 0 | 1 |
| 0 | 0 | 0 | 0 |
| 0 | 0 | 0 | 0 |
| 0 | 0 | 0 | 1 |
| 0 | 0 | 0 | 0 |
| 0 | 0 | 0 | 0 |
| 0 | 0 | 0 | 0 |
| 0 | 1 | 1 | 1 |
| 0 | 0 | 0 | 0 |
| 0 | 0 | 0 | 1 |
| 0 | 0 | 0 | 1 |
| 0 | 0 | 0 | 0 |
| 0 | 0 | 0 | 0 |
| 0 | 0 | 0 | 0 |
| 0 | 0 | 0 | 1 |
| 0 | 0 | 0 | 0 |

| | | | |
|---|---|---|---|
| 0 | 0 | 0 | 0 |
| 0 | 0 | 0 | 1 |
| 0 | 0 | 0 | 1 |
| 0 | 0 | 0 | 1 |
| 0 | 0 | 0 | 0 |
| 0 | 0 | 0 | 1 |
| 0 | 0 | 0 | 0 |
| 0 | 0 | 0 | 1 |
| 0 | 0 | 0 | 1 |
| 0 | 0 | 0 | 0 |
| 0 | 1 | 1 | 1 |
| 0 | 0 | 0 | 1 |
| 0 | 0 | 0 | 1 |
| 0 | 0 | 0 | 0 |
| 0 | 0 | 0 | 1 |
| 0 | 0 | 0 | 1 |
| 0 | 0 | 0 | 0 |
| 0 | 0 | 0 | 1 |
| 0 | 0 | 0 | 0 |
| 0 | 0 | 0 | 0 |
| 0 | 0 | 0 | 0 |
| 0 | 0 | 0 | 0 |
| 0 | 0 | 0 | 0 |
| 0 | 0 | 0 | 0 |
| 0 | 0 | 0 | 0 |
| 0 | 0 | 0 | 0 |
| 0 | 0 | 0 | 0 |
| 0 | 0 | 0 | 0 |
| 0 | 0 | 0 | 0 |
| 0 | 0 | 0 | 0 |
| 0 | 0 | 0 | 0 |
| 0 | 0 | 0 | 0 |
| 0 | 0 | 0 | 0 |
| 0 | 0 | 0 | 0 |
| 0 | 0 | 0 | 0 |
| 0 | 0 | 0 | 0 |
| 0 | 0 | 0 | 1 |
| 0 | 0 | 0 | 0 |
| 0 | 0 | 0 | 0 |
| 0 | 0 | 0 | 1 |
| 0 | 0 | 0 | 1 |
| 0 | 0 | 0 | 0 |
| 0 | 0 | 0 | 0 |
| 0 | 0 | 0 | 0 |
| 0 | 0 | 0 | 0 |
| 0 | 0 | 0 | 0 |
| 0 | 0 | 0 | 0 |
| 0 | 0 | 0 | 0 |
| 0 | 0 | 0 | 1 |
| 0 | 0 | 0 | 1 |
| 0 | 0 | 0 | 0 |
| 0 | 0 | 0 | 1 |
| 0 | 0 | 0 | 0 |
| 0 | 0 | 0 | 0 |

| | | | |
|---|---|---|---|
| 0 | 0 | 0 | 0 |
| 0 | 0 | 0 | 0 |
| 0 | 0 | 0 | 0 |
| 0 | 0 | 0 | 0 |
| 0 | 0 | 0 | 0 |
| 0 | 0 | 0 | 0 |
| 0 | 0 | 0 | 0 |
| 0 | 0 | 0 | 0 |
| 0 | 0 | 0 | 0 |
| 0 | 0 | 0 | 0 |
| 0 | 0 | 0 | 0 |
| 0 | 0 | 0 | 0 |
| 0 | 0 | 0 | 0 |
| 0 | 0 | 0 | 0 |
| 0 | 0 | 0 | 1 |
| 0 | 0 | 0 | 0 |
| 0 | 1 | 0 | 0 |
| 0 | 0 | 0 | 0 |
| 0 | 0 | 0 | 1 |
| 0 | 0 | 0 | 0 |
| 0 | 0 | 0 | 0 |
| 0 | 1 | 0 | 0 |
| 0 | 1 | 0 | 0 |
| 0 | 0 | 0 | 0 |
| 0 | 0 | 0 | 0 |
| 0 | 0 | 0 | 1 |
| 0 | 0 | 0 | 0 |
| 0 | 0 | 0 | 0 |
| 0 | 0 | 0 | 0 |
| 0 | 0 | 0 | 0 |
| 0 | 0 | 0 | 1 |
| 0 | 0 | 0 | 0 |
| 0 | 0 | 0 | 1 |
| 0 | 0 | 0 | 1 |
| 1 | 0 | 0 | 0 |
| 0 | 0 | 0 | 0 |
| 0 | 0 | 0 | 1 |
| 0 | 0 | 0 | 1 |
| 0 | 0 | 0 | 1 |
| 0 | 0 | 0 | 1 |
| 0 | 0 | 0 | 0 |
| 0 | 0 | 0 | 0 |
| 0 | 0 | 0 | 0 |
| 0 | 0 | 0 | 0 |
| 0 | 0 | 0 | 0 |
| 0 | 0 | 0 | 0 |
| 0 | 0 | 0 | 0 |
| 0 | 0 | 0 | 0 |
| 0 | 0 | 0 | 1 |
| 0 | 0 | 0 | 1 |
| 0 | 0 | 0 | 0 |
| 0 | 0 | 0 | 0 |
| 0 | 0 | 0 | 0 |
| 0 | 0 | 0 | 1 |

| | | | |
|---|---|---|---|
| 0 | 0 | 0 | 1 |
| 0 | 0 | 0 | 1 |
| 0 | 0 | 0 | 1 |
| 0 | 0 | 0 | 0 |
| 0 | 0 | 0 | 1 |
| 0 | 0 | 0 | 1 |
| 0 | 0 | 0 | 0 |
| 0 | 0 | 0 | 1 |
| 0 | 0 | 0 | 0 |
| 0 | 0 | 0 | 0 |
| 0 | 0 | 0 | 0 |
| 0 | 0 | 0 | 0 |
| 0 | 0 | 0 | 0 |
| 0 | 0 | 0 | 0 |
| 0 | 0 | 0 | 0 |
| 0 | 0 | 0 | 0 |
| 0 | 0 | 0 | 1 |
| 0 | 0 | 0 | 0 |
| 0 | 0 | 0 | 0 |
| 0 | 0 | 0 | 1 |
| 0 | 0 | 0 | 0 |
| 0 | 0 | 0 | 0 |
| 0 | 0 | 0 | 1 |
| 0 | 0 | 0 | 0 |
| 0 | 0 | 0 | 1 |
| 0 | 0 | 0 | 1 |
| 0 | 0 | 0 | 1 |
| 0 | 0 | 0 | 0 |
| 0 | 0 | 0 | 0 |
| 0 | 0 | 0 | 0 |
| 0 | 0 | 0 | 1 |
| 0 | 0 | 0 | 0 |
| 0 | 0 | 0 | 0 |
| 0 | 0 | 0 | 0 |
| 0 | 0 | 0 | 1 |
| 0 | 0 | 0 | 1 |
| 0 | 0 | 0 | 0 |
| 0 | 0 | 0 | 0 |
| 0 | 0 | 0 | 0 |
| 0 | 0 | 0 | 1 |
| 0 | 0 | 0 | 1 |
| 0 | 0 | 0 | 1 |
| 0 | 0 | 0 | 1 |
| 0 | 0 | 0 | 0 |
| 0 | 0 | 0 | 1 |
| 0 | 0 | 0 | 0 |
| 0 | 0 | 0 | 1 |
| 0 | 0 | 0 | 0 |
| 0 | 0 | 0 | 0 |
| 0 | 0 | 0 | 1 |
| 0 | 0 | 0 | 1 |
| 0 | 0 | 0 | 0 |
| 0 | 0 | 0 | 0 |
| 0 | 0 | 0 | 1 |

| | | | |
|---|---|---|---|
| 0 | 0 | 0 | 1 |
| 0 | 0 | 0 | 1 |
| 0 | 0 | 0 | 1 |
| 0 | 0 | 0 | 1 |
| 0 | 0 | 0 | 0 |
| 0 | 1 | 1 | 1 |
| 0 | 0 | 0 | 1 |
| 0 | 1 | 0 | 0 |
| 0 | 0 | 0 | 0 |
| 0 | 0 | 0 | 1 |
| 0 | 0 | 0 | 0 |
| 0 | 0 | 0 | 0 |
| 1 | 0 | 0 | 0 |
| 1 | 0 | 0 | 0 |
| 0 | 0 | 0 | 0 |
| 1 | 0 | 0 | 0 |
| 1 | 0 | 0 | 0 |
| 1 | 0 | 0 | 0 |
| 1 | 0 | 0 | 0 |
| 0 | 0 | 0 | 1 |
| 0 | 0 | 0 | 1 |
| 0 | 0 | 0 | 1 |
| 0 | 0 | 0 | 1 |
| 0 | 0 | 0 | 1 |
| 0 | 0 | 0 | 0 |
| 0 | 0 | 0 | 1 |
| 1 | 0 | 0 | 0 |
| 1 | 0 | 0 | 0 |
| 0 | 0 | 0 | 1 |
| 0 | 0 | 0 | 1 |
| 0 | 0 | 0 | 1 |
| 0 | 0 | 0 | 1 |
| 0 | 0 | 0 | 1 |
| 0 | 0 | 0 | 0 |
| 0 | 0 | 0 | 0 |
| 0 | 0 | 0 | 1 |
| 0 | 0 | 0 | 0 |
| 0 | 0 | 0 | 1 |
| 0 | 0 | 0 | 0 |
| 0 | 0 | 0 | 0 |
| 0 | 0 | 0 | 0 |
| 0 | 0 | 0 | 0 |
| 0 | 0 | 0 | 1 |
| 0 | 0 | 0 | 0 |
| 0 | 0 | 0 | 0 |
| 1 | 0 | 0 | 0 |
| 1 | 0 | 0 | 0 |
| 1 | 0 | 0 | 0 |
| 0 | 0 | 0 | 0 |
| 0 | 0 | 0 | 0 |
| 0 | 0 | 0 | 0 |
| 1 | 0 | 0 | 0 |
| 1 | 0 | 0 | 0 |
| 0 | 0 | 0 | 1 |

| | | | |
|---|---|---|---|
| 0 | 0 | 0 | 0 |
| 0 | 0 | 0 | 1 |
| 0 | 0 | 0 | 0 |
| 0 | 0 | 0 | 0 |
| 0 | 0 | 0 | 0 |
| 0 | 0 | 0 | 1 |
| 0 | 0 | 0 | 0 |
| 0 | 0 | 0 | 1 |
| 0 | 0 | 0 | 1 |
| 0 | 0 | 0 | 0 |
| 0 | 0 | 0 | 1 |
| 0 | 0 | 0 | 1 |
| 0 | 0 | 0 | 1 |
| 0 | 0 | 0 | 1 |
| 0 | 0 | 0 | 1 |
| 0 | 1 | 0 | 0 |
| 0 | 0 | 0 | 0 |
| 0 | 0 | 0 | 1 |
| 0 | 0 | 0 | 0 |
| 0 | 0 | 0 | 1 |
| 0 | 0 | 0 | 0 |
| 0 | 0 | 0 | 0 |
| 0 | 0 | 0 | 0 |
| 0 | 0 | 0 | 0 |
| 0 | 0 | 0 | 0 |
| 0 | 0 | 0 | 0 |
| 0 | 0 | 0 | 0 |
| 0 | 0 | 0 | 0 |
| 0 | 0 | 0 | 0 |
| 0 | 0 | 0 | 0 |
| 0 | 0 | 0 | 0 |
| 0 | 0 | 0 | 0 |
| 0 | 0 | 0 | 1 |
| 0 | 0 | 0 | 1 |
| 0 | 0 | 0 | 0 |
| 0 | 0 | 0 | 0 |
| 0 | 0 | 0 | 1 |
| 0 | 0 | 0 | 0 |
| 0 | 0 | 0 | 1 |
| 0 | 0 | 0 | 0 |
| 0 | 0 | 0 | 0 |
| 0 | 0 | 0 | 0 |
| 0 | 0 | 0 | 1 |
| 0 | 0 | 0 | 1 |
| 0 | 0 | 0 | 0 |
| 0 | 0 | 0 | 0 |
| 0 | 0 | 0 | 0 |
| 0 | 0 | 0 | 0 |
| 0 | 0 | 0 | 0 |
| 0 | 0 | 0 | 0 |
| 0 | 0 | 0 | 1 |
| 0 | 0 | 0 | 0 |
| 0 | 1 | 0 | 0 |
| 0 | 0 | 0 | 1 |

| | | | |
|---|---|---|---|
| 0 | 0 | 0 | 0 |
| 0 | 0 | 0 | 1 |
| 0 | 0 | 0 | 0 |
| 0 | 0 | 0 | 0 |
| 0 | 0 | 0 | 0 |
| 0 | 0 | 0 | 0 |
| 0 | 0 | 0 | 0 |
| 0 | 0 | 0 | 0 |
| 0 | 0 | 0 | 0 |
| 0 | 0 | 0 | 0 |
| 0 | 0 | 0 | 0 |
| 0 | 0 | 0 | 1 |
| 0 | 0 | 0 | 0 |
| 0 | 0 | 0 | 1 |
| 0 | 0 | 0 | 1 |
| 0 | 1 | 1 | 1 |
| 0 | 1 | 0 | 0 |
| 0 | 1 | 0 | 0 |
| 0 | 1 | 1 | 1 |
| 0 | 1 | 0 | 0 |
| 0 | 0 | 0 | 1 |
| 0 | 0 | 0 | 0 |
| 0 | 0 | 0 | 1 |
| 0 | 0 | 0 | 0 |
| 0 | 0 | 0 | 1 |
| 0 | 0 | 0 | 1 |
| 0 | 0 | 0 | 1 |
| 0 | 0 | 0 | 0 |
| 0 | 0 | 0 | 1 |
| 0 | 0 | 0 | 1 |
| 0 | 0 | 0 | 1 |
| 1 | 0 | 0 | 0 |
| 0 | 0 | 0 | 1 |
| 0 | 0 | 0 | 0 |
| 0 | 0 | 0 | 0 |
| 0 | 0 | 0 | 0 |
| 0 | 0 | 0 | 1 |
| 0 | 0 | 0 | 1 |
| 0 | 0 | 0 | 0 |
| 0 | 0 | 0 | 0 |
| 0 | 0 | 0 | 0 |
| 0 | 0 | 0 | 0 |
| 0 | 1 | 0 | 0 |
| 0 | 0 | 0 | 0 |
| 0 | 0 | 0 | 0 |
| 0 | 0 | 0 | 0 |
| 0 | 0 | 0 | 1 |
| 0 | 0 | 0 | 1 |
| 0 | 0 | 0 | 0 |
| 0 | 0 | 0 | 0 |
| 0 | 0 | 0 | 0 |
| 0 | 0 | 0 | 1 |
| 0 | 0 | 0 | 1 |
| 0 | 0 | 0 | 0 |

| | | | |
|---|---|---|---|
| 0 | 0 | 0 | 0 |
| 0 | 0 | 0 | 1 |
| 0 | 0 | 0 | 1 |
| 0 | 0 | 0 | 1 |
| 0 | 0 | 0 | 0 |
| 0 | 0 | 0 | 0 |
| 0 | 0 | 0 | 0 |
| 0 | 0 | 0 | 0 |
| 0 | 0 | 0 | 0 |
| 0 | 0 | 0 | 0 |
| 0 | 0 | 0 | 1 |
| 0 | 0 | 0 | 0 |
| 0 | 0 | 0 | 0 |
| 0 | 0 | 0 | 0 |
| 0 | 0 | 0 | 1 |
| 0 | 0 | 0 | 1 |
| 0 | 0 | 0 | 0 |
| 0 | 0 | 0 | 1 |
| 0 | 0 | 0 | 0 |
| 0 | 0 | 0 | 0 |
| 0 | 0 | 0 | 1 |
| 0 | 0 | 0 | 0 |
| 0 | 0 | 0 | 0 |
| 0 | 0 | 0 | 1 |
| 0 | 0 | 0 | 0 |
| 0 | 0 | 0 | 0 |
| 0 | 0 | 0 | 0 |
| 0 | 0 | 0 | 1 |
| 0 | 0 | 0 | 0 |
| 0 | 0 | 0 | 0 |
| 0 | 0 | 0 | 0 |
| 0 | 0 | 0 | 1 |
| 0 | 0 | 0 | 1 |
| 0 | 0 | 0 | 1 |
| 0 | 0 | 0 | 1 |
| 0 | 0 | 0 | 1 |
| 0 | 0 | 0 | 1 |
| 0 | 0 | 0 | 1 |
| 0 | 0 | 0 | 1 |
| 0 | 0 | 0 | 1 |
| 0 | 0 | 0 | 1 |
| 0 | 0 | 0 | 1 |
| 1 | 0 | 0 | 0 |
| 0 | 0 | 0 | 1 |
| 0 | 0 | 0 | 1 |
| 0 | 0 | 0 | 1 |
| 0 | 0 | 0 | 1 |
| 0 | 0 | 0 | 1 |
| 0 | 0 | 0 | 1 |
| 0 | 0 | 0 | 1 |
| 0 | 0 | 0 | 1 |
| 0 | 0 | 0 | 1 |
| 0 | 0 | 0 | 1 |
| 0 | 0 | 0 | 1 |

| | | | |
|---|---|---|---|
| 0 | 0 | 0 | 1 |
| 0 | 0 | 0 | 1 |
| 0 | 0 | 0 | 1 |
| 0 | 0 | 0 | 1 |
| 0 | 0 | 0 | 1 |
| 0 | 0 | 0 | 1 |
| 0 | 0 | 0 | 1 |
| 0 | 0 | 0 | 1 |
| 0 | 0 | 0 | 0 |
| 0 | 0 | 0 | 0 |
| 0 | 0 | 0 | 0 |
| 0 | 0 | 0 | 0 |
| 0 | 0 | 0 | 0 |
| 0 | 0 | 0 | 0 |
| 0 | 0 | 0 | 0 |
| 0 | 0 | 0 | 0 |
| 0 | 0 | 0 | 0 |
| 0 | 0 | 0 | 0 |
| 0 | 0 | 0 | 0 |
| 0 | 0 | 0 | 0 |
| 1 | 0 | 0 | 0 |
| 1 | 0 | 0 | 0 |
| 0 | 0 | 0 | 0 |
| 0 | 0 | 0 | 0 |
| 0 | 0 | 0 | 0 |
| 0 | 0 | 0 | 1 |
| 0 | 0 | 0 | 1 |
| 0 | 0 | 0 | 0 |
| 0 | 0 | 0 | 0 |
| 1 | 0 | 0 | 0 |
| 1 | 0 | 0 | 0 |
| 0 | 0 | 0 | 0 |
| 1 | 0 | 0 | 0 |
| 1 | 0 | 0 | 0 |
| 0 | 0 | 0 | 0 |
| 0 | 0 | 0 | 0 |
| 0 | 0 | 0 | 0 |
| 0 | 0 | 0 | 0 |
| 0 | 0 | 0 | 0 |
| 0 | 0 | 0 | 0 |
| 0 | 0 | 0 | 0 |
| 0 | 0 | 0 | 0 |
| 1 | 0 | 0 | 0 |
| 1 | 0 | 0 | 0 |
| 1 | 0 | 0 | 0 |
| 0 | 0 | 0 | 0 |
| 0 | 0 | 0 | 1 |
| 0 | 0 | 0 | 1 |
| 0 | 0 | 0 | 1 |
| 0 | 0 | 0 | 1 |
| 0 | 0 | 0 | 1 |
| 0 | 0 | 0 | 1 |
| 0 | 0 | 0 | 0 |
| 0 | 0 | 0 | 0 |

| | | | |
|---|---|---|---|
| 0 | 0 | 0 | 0 |
| 0 | 0 | 0 | 0 |
| 0 | 0 | 0 | 0 |
| 0 | 0 | 0 | 0 |
| 0 | 0 | 0 | 1 |
| 0 | 0 | 0 | 0 |
| 0 | 0 | 0 | 0 |
| 0 | 0 | 0 | 0 |
| 0 | 0 | 0 | 1 |
| 0 | 0 | 0 | 0 |
| 1 | 0 | 0 | 0 |
| 0 | 0 | 0 | 1 |
| 0 | 0 | 0 | 0 |
| 0 | 0 | 0 | 0 |
| 0 | 0 | 0 | 0 |
| 0 | 0 | 0 | 0 |
| 0 | 0 | 0 | 1 |
| 0 | 0 | 0 | 0 |
| 0 | 0 | 0 | 0 |
| 0 | 0 | 0 | 1 |
| 0 | 0 | 0 | 0 |
| 0 | 0 | 0 | 0 |
| 0 | 0 | 0 | 0 |
| 0 | 0 | 0 | 0 |
| 0 | 0 | 0 | 0 |
| 0 | 0 | 0 | 1 |
| 0 | 0 | 0 | 1 |
| 0 | 0 | 0 | 1 |
| 0 | 0 | 0 | 1 |
| 0 | 0 | 0 | 1 |
| 0 | 0 | 0 | 1 |
| 0 | 0 | 0 | 1 |
| 0 | 0 | 0 | 1 |
| 0 | 0 | 0 | 1 |
| 0 | 0 | 0 | 0 |
| 0 | 0 | 0 | 0 |
| 0 | 0 | 0 | 0 |
| 0 | 0 | 0 | 0 |
| 0 | 0 | 0 | 0 |
| 0 | 0 | 0 | 0 |
| 0 | 0 | 0 | 1 |
| 0 | 0 | 0 | 1 |
| 0 | 0 | 0 | 1 |
| 0 | 0 | 0 | 0 |
| 0 | 0 | 0 | 0 |
| 0 | 0 | 0 | 0 |
| 0 | 0 | 0 | 1 |
| 0 | 0 | 0 | 1 |
| 0 | 0 | 0 | 0 |
| 0 | 0 | 0 | 0 |
| 0 | 0 | 0 | 1 |
| 0 | 0 | 0 | 1 |
| 0 | 0 | 0 | 1 |
| 0 | 0 | 0 | 1 |

| | | | |
|---|---|---|---|
| 0 | 0 | 0 | 1 |
| 0 | 0 | 0 | 1 |
| 0 | 0 | 0 | 1 |
| 0 | 0 | 0 | 0 |
| 0 | 0 | 0 | 0 |
| 0 | 0 | 0 | 1 |
| 0 | 0 | 0 | 1 |
| 0 | 0 | 0 | 1 |
| 0 | 0 | 0 | 1 |
| 0 | 0 | 0 | 1 |
| 0 | 0 | 0 | 0 |
| 0 | 0 | 0 | 1 |
| 0 | 0 | 0 | 0 |
| 0 | 0 | 0 | 0 |
| 0 | 0 | 0 | 0 |
| 0 | 0 | 0 | 1 |
| 0 | 0 | 0 | 1 |
| 0 | 0 | 0 | 1 |
| 0 | 0 | 0 | 1 |
| 0 | 0 | 0 | 1 |
| 0 | 0 | 0 | 0 |
| 0 | 0 | 0 | 1 |
| 0 | 0 | 0 | 1 |
| 0 | 0 | 0 | 0 |
| 0 | 0 | 0 | 1 |
| 0 | 0 | 0 | 0 |
| 0 | 1 | 0 | 0 |
| 0 | 0 | 0 | 1 |
| 0 | 0 | 0 | 1 |
| 0 | 0 | 0 | 1 |
| 0 | 0 | 0 | 1 |
| 0 | 0 | 0 | 1 |
| 0 | 0 | 0 | 1 |
| 0 | 0 | 0 | 1 |
| 0 | 0 | 0 | 1 |
| 0 | 0 | 0 | 0 |
| 0 | 0 | 0 | 1 |
| 0 | 0 | 0 | 1 |
| 0 | 0 | 0 | 0 |
| 0 | 0 | 0 | 0 |
| 0 | 0 | 0 | 1 |
| 0 | 0 | 0 | 1 |
| 1 | 0 | 0 | 0 |
| 1 | 0 | 0 | 0 |
| 0 | 0 | 0 | 0 |
| 0 | 0 | 0 | 0 |
| 1 | 0 | 0 | 0 |
| 1 | 0 | 0 | 0 |
| 0 | 0 | 0 | 1 |
| 1 | 0 | 0 | 0 |
| 1 | 0 | 0 | 0 |
| 1 | 0 | 0 | 0 |
| 1 | 0 | 0 | 0 |
| 1 | 0 | 0 | 0 |

| | | | |
|---|---|---|---|
| 0 | 0 | 0 | 0 |
| 0 | 0 | 0 | 0 |
| 1 | 0 | 0 | 0 |
| 1 | 0 | 0 | 0 |
| 1 | 0 | 0 | 0 |
| 1 | 0 | 0 | 0 |
| 0 | 0 | 0 | 0 |
| 0 | 0 | 0 | 0 |
| 0 | 0 | 0 | 0 |
| 0 | 0 | 0 | 1 |
| 0 | 0 | 0 | 1 |
| 0 | 0 | 0 | 0 |
| 0 | 0 | 0 | 1 |
| 0 | 0 | 0 | 0 |
| 0 | 0 | 0 | 0 |
| 0 | 0 | 0 | 1 |
| 0 | 0 | 0 | 0 |
| 0 | 0 | 0 | 1 |
| 0 | 0 | 0 | 1 |
| 0 | 0 | 0 | 1 |
| 0 | 0 | 0 | 1 |
| 0 | 0 | 0 | 1 |
| 0 | 0 | 0 | 1 |
| 0 | 0 | 0 | 0 |
| 0 | 0 | 0 | 0 |
| 0 | 0 | 0 | 1 |
| 0 | 0 | 0 | 1 |
| 0 | 0 | 0 | 1 |
| 0 | 0 | 0 | 0 |
| 0 | 0 | 0 | 0 |
| 0 | 0 | 0 | 1 |
| 0 | 0 | 0 | 1 |
| 0 | 0 | 0 | 1 |
| 0 | 0 | 0 | 0 |
| 0 | 0 | 0 | 0 |
| 0 | 0 | 0 | 1 |
| 0 | 0 | 0 | 0 |
| 0 | 0 | 0 | 1 |
| 0 | 0 | 0 | 1 |
| 0 | 0 | 0 | 0 |
| 0 | 0 | 0 | 0 |
| 0 | 0 | 0 | 0 |
| 0 | 0 | 0 | 0 |
| 0 | 0 | 0 | 0 |
| 0 | 0 | 0 | 0 |
| 0 | 0 | 0 | 1 |
| 0 | 0 | 0 | 1 |
| 0 | 0 | 0 | 0 |
| 0 | 0 | 0 | 0 |
| 0 | 0 | 0 | 0 |
| 0 | 0 | 0 | 1 |
| 0 | 0 | 0 | 1 |
| 0 | 0 | 0 | 0 |
| 0 | 0 | 0 | 0 |

| | | | |
|---|---|---|---|
| 0 | 0 | 0 | 1 |
| 0 | 0 | 0 | 1 |
| 0 | 0 | 0 | 1 |
| 0 | 0 | 0 | 1 |
| 0 | 1 | 0 | 0 |
| 0 | 0 | 0 | 0 |
| 0 | 0 | 0 | 1 |
| 0 | 0 | 0 | 0 |
| 0 | 0 | 0 | 1 |
| 0 | 0 | 0 | 0 |
| 0 | 0 | 0 | 1 |
| 0 | 0 | 0 | 1 |
| 0 | 0 | 0 | 1 |
| 0 | 0 | 0 | 1 |
| 0 | 0 | 0 | 1 |
| 0 | 0 | 0 | 1 |
| 0 | 0 | 0 | 0 |
| 0 | 0 | 0 | 1 |
| 0 | 0 | 0 | 1 |
| 0 | 0 | 0 | 0 |
| 0 | 0 | 0 | 0 |
| 0 | 0 | 0 | 1 |
| 0 | 0 | 0 | 1 |
| 0 | 0 | 0 | 0 |
| 0 | 0 | 0 | 1 |
| 0 | 0 | 0 | 0 |
| 0 | 0 | 0 | 0 |
| 0 | 0 | 0 | 0 |
| 0 | 0 | 0 | 0 |
| 0 | 0 | 0 | 0 |
| 0 | 0 | 0 | 0 |
| 0 | 0 | 0 | 0 |
| 0 | 0 | 0 | 1 |
| 0 | 0 | 0 | 0 |
| 0 | 0 | 0 | 1 |
| 0 | 0 | 0 | 1 |
| 0 | 0 | 0 | 1 |
| 0 | 0 | 0 | 1 |
| 0 | 0 | 0 | 0 |
| 0 | 0 | 0 | 1 |
| 0 | 0 | 0 | 1 |
| 0 | 0 | 0 | 1 |
| 0 | 0 | 0 | 1 |
| 0 | 0 | 0 | 0 |
| 0 | 0 | 0 | 1 |
| 0 | 0 | 0 | 1 |
| 0 | 0 | 0 | 1 |
| 0 | 0 | 0 | 1 |
| 0 | 0 | 0 | 1 |
| 0 | 0 | 0 | 1 |
| 0 | 0 | 0 | 0 |
| 0 | 0 | 0 | 0 |
| 0 | 0 | 0 | 0 |
| 0 | 0 | 0 | 1 |

| | | | |
|---|---|---|---|
| 0 | 0 | 0 | 0 |
| 0 | 0 | 0 | 0 |
| 0 | 0 | 0 | 0 |
| 0 | 0 | 0 | 0 |
| 1 | 0 | 0 | 0 |
| 1 | 0 | 0 | 0 |
| 0 | 0 | 0 | 0 |
| 0 | 0 | 0 | 0 |
| 0 | 0 | 0 | 1 |
| 0 | 0 | 0 | 1 |
| 0 | 0 | 0 | 0 |
| 0 | 0 | 0 | 1 |
| 0 | 0 | 0 | 1 |
| 0 | 0 | 0 | 0 |
| 0 | 0 | 0 | 0 |
| 0 | 0 | 0 | 0 |
| 0 | 0 | 0 | 0 |
| 0 | 0 | 0 | 0 |
| 0 | 0 | 0 | 0 |
| 0 | 0 | 0 | 1 |
| 0 | 0 | 0 | 1 |
| 0 | 0 | 0 | 0 |
| 0 | 0 | 0 | 1 |
| 0 | 0 | 0 | 0 |
| 0 | 0 | 0 | 1 |
| 0 | 0 | 0 | 1 |
| 0 | 0 | 0 | 1 |
| 0 | 0 | 0 | 1 |
| 0 | 0 | 0 | 1 |
| 0 | 0 | 0 | 0 |
| 0 | 0 | 0 | 1 |
| 0 | 0 | 0 | 1 |
| 0 | 0 | 0 | 0 |
| 0 | 0 | 0 | 1 |
| 0 | 0 | 0 | 0 |
| 0 | 0 | 0 | 0 |
| 0 | 0 | 0 | 0 |
| 0 | 0 | 0 | 0 |
| 0 | 0 | 0 | 0 |
| 0 | 0 | 0 | 0 |
| 0 | 0 | 0 | 0 |
| 0 | 0 | 0 | 0 |
| 0 | 0 | 0 | 1 |
| 0 | 0 | 0 | 0 |
| 0 | 0 | 0 | 1 |
| 0 | 0 | 0 | 0 |
| 0 | 0 | 0 | 1 |
| 0 | 0 | 0 | 1 |
| 0 | 0 | 0 | 1 |
| 0 | 0 | 0 | 0 |
| 0 | 0 | 0 | 1 |
| 0 | 0 | 0 | 1 |
| 0 | 0 | 0 | 1 |
| 0 | 0 | 0 | 1 |

| | | | |
|---|---|---|---|
| 1 | 0 | 0 | 0 |
| 0 | 0 | 0 | 1 |
| 0 | 0 | 0 | 1 |
| 0 | 0 | 0 | 1 |
| 0 | 0 | 0 | 1 |
| 0 | 0 | 0 | 1 |
| 0 | 0 | 0 | 1 |
| 0 | 0 | 0 | 1 |
| 0 | 0 | 0 | 1 |
| 0 | 0 | 0 | 1 |
| 0 | 0 | 0 | 0 |
| 0 | 0 | 0 | 1 |
| 0 | 0 | 0 | 1 |
| 0 | 0 | 0 | 1 |
| 0 | 0 | 0 | 0 |
| 0 | 0 | 0 | 0 |
| 0 | 0 | 0 | 0 |
| 0 | 0 | 0 | 0 |
| 0 | 0 | 0 | 0 |
| 0 | 0 | 0 | 1 |
| 0 | 0 | 0 | 1 |
| 0 | 0 | 0 | 1 |
| 0 | 0 | 0 | 0 |
| 0 | 0 | 0 | 1 |
| 0 | 0 | 0 | 1 |
| 0 | 0 | 0 | 1 |
| 0 | 0 | 0 | 1 |
| 0 | 0 | 0 | 1 |
| 0 | 0 | 0 | 0 |
| 0 | 0 | 0 | 0 |
| 0 | 0 | 0 | 0 |
| 0 | 0 | 0 | 0 |
| 0 | 0 | 0 | 0 |
| 0 | 0 | 0 | 0 |
| 0 | 0 | 0 | 1 |
| 0 | 0 | 0 | 1 |
| 0 | 0 | 0 | 0 |
| 0 | 0 | 0 | 0 |
| 0 | 0 | 0 | 0 |
| 0 | 0 | 0 | 0 |
| 0 | 0 | 0 | 0 |
| 0 | 0 | 0 | 0 |
| 0 | 0 | 0 | 1 |
| 0 | 0 | 0 | 1 |
| 0 | 0 | 0 | 0 |
| 0 | 0 | 0 | 1 |
| 0 | 0 | 0 | 1 |
| 0 | 0 | 0 | 1 |
| 0 | 0 | 0 | 1 |
| 0 | 0 | 0 | 1 |
| 0 | 0 | 0 | 1 |
| 0 | 0 | 0 | 1 |
| 0 | 0 | 0 | 1 |
| 0 | 0 | 0 | 1 |

| | | | |
|---|---|---|---|
| 0 | 0 | 0 | 0 |
| 0 | 0 | 0 | 1 |
| 0 | 0 | 0 | 0 |
| 0 | 0 | 0 | 1 |
| 0 | 0 | 0 | 1 |
| 0 | 0 | 0 | 1 |
| 0 | 1 | 0 | 0 |
| 0 | 0 | 0 | 0 |
| 0 | 0 | 0 | 1 |
| 0 | 0 | 0 | 1 |
| 0 | 0 | 0 | 1 |
| 0 | 0 | 0 | 1 |
| 0 | 0 | 0 | 1 |
| 0 | 0 | 0 | 1 |
| 0 | 0 | 0 | 0 |
| 0 | 0 | 0 | 0 |
| 0 | 0 | 0 | 0 |
| 0 | 0 | 0 | 0 |
| 0 | 0 | 0 | 1 |
| 0 | 0 | 0 | 0 |
| 0 | 0 | 0 | 0 |
| 0 | 0 | 0 | 0 |
| 0 | 0 | 0 | 0 |
| 0 | 0 | 0 | 0 |
| 0 | 0 | 0 | 1 |
| 0 | 0 | 0 | 0 |
| 0 | 0 | 0 | 1 |
| 0 | 0 | 0 | 1 |
| 0 | 0 | 0 | 1 |
| 0 | 0 | 0 | 0 |
| 0 | 0 | 0 | 1 |
| 0 | 0 | 0 | 1 |
| 0 | 0 | 0 | 0 |
| 0 | 0 | 0 | 0 |
| 0 | 0 | 0 | 1 |
| 0 | 0 | 0 | 1 |
| 0 | 0 | 0 | 1 |
| 0 | 0 | 0 | 1 |
| 0 | 0 | 0 | 1 |
| 0 | 0 | 0 | 0 |
| 0 | 0 | 0 | 0 |
| 0 | 0 | 0 | 0 |
| 0 | 0 | 0 | 0 |
| 0 | 0 | 0 | 0 |
| 0 | 0 | 0 | 0 |
| 0 | 0 | 0 | 1 |
| 0 | 0 | 0 | 1 |
| 0 | 0 | 0 | 0 |
| 0 | 0 | 0 | 0 |
| 0 | 0 | 0 | 1 |
| 0 | 0 | 0 | 0 |
| 0 | 0 | 0 | 0 |
| 0 | 0 | 0 | 1 |
| 0 | 0 | 0 | 1 |

| | | | |
|---|---|---|---|
| 0 | 0 | 0 | 1 |
| 0 | 0 | 0 | 0 |
| 0 | 0 | 0 | 0 |
| 0 | 0 | 0 | 1 |
| 0 | 0 | 0 | 1 |
| 0 | 0 | 0 | 0 |
| 0 | 0 | 0 | 0 |
| 0 | 0 | 0 | 0 |
| 0 | 0 | 0 | 0 |
| 0 | 0 | 0 | 0 |
| 0 | 0 | 0 | 0 |
| 0 | 0 | 0 | 0 |
| 0 | 0 | 0 | 0 |
| 0 | 0 | 0 | 0 |
| 0 | 0 | 0 | 0 |
| 0 | 0 | 0 | 1 |
| 0 | 0 | 0 | 0 |
| 0 | 0 | 0 | 1 |
| 0 | 0 | 0 | 1 |
| 0 | 0 | 0 | 1 |
| 0 | 0 | 0 | 0 |
| 0 | 0 | 0 | 1 |
| 0 | 0 | 0 | 0 |
| 0 | 0 | 0 | 0 |
| 0 | 0 | 0 | 1 |
| 0 | 0 | 0 | 0 |
| 0 | 0 | 0 | 1 |
| 0 | 0 | 0 | 1 |
| 0 | 0 | 0 | 1 |
| 0 | 0 | 0 | 0 |
| 0 | 0 | 0 | 0 |
| 0 | 0 | 0 | 0 |
| 0 | 0 | 0 | 0 |
| 0 | 0 | 0 | 0 |
| 0 | 0 | 0 | 0 |
| 0 | 0 | 0 | 0 |
| 0 | 0 | 0 | 0 |
| 0 | 0 | 0 | 1 |
| 0 | 0 | 0 | 1 |
| 0 | 0 | 0 | 0 |
| 0 | 0 | 0 | 0 |
| 0 | 0 | 0 | 0 |
| 0 | 0 | 0 | 0 |
| 0 | 0 | 0 | 0 |
| 0 | 0 | 0 | 0 |
| 0 | 0 | 0 | 0 |
| 0 | 0 | 0 | 1 |
| 0 | 0 | 0 | 0 |
| 0 | 0 | 0 | 0 |
| 0 | 0 | 0 | 0 |
| 0 | 0 | 0 | 0 |
| 0 | 0 | 0 | 1 |
| 0 | 0 | 0 | 0 |
| 0 | 0 | 0 | 0 |

| | | | |
|---|---|---|---|
| 0 | 0 | 0 | 0 |
| 0 | 0 | 0 | 0 |
| 0 | 0 | 0 | 0 |
| 0 | 0 | 0 | 0 |
| 0 | 0 | 0 | 0 |
| 0 | 0 | 0 | 0 |
| 0 | 0 | 0 | 0 |
| 0 | 0 | 0 | 0 |
| 1 | 0 | 0 | 0 |
| 0 | 0 | 0 | 0 |
| 0 | 0 | 0 | 1 |
| 0 | 0 | 0 | 0 |
| 0 | 0 | 0 | 1 |
| 0 | 0 | 0 | 0 |
| 0 | 0 | 0 | 1 |
| 0 | 0 | 0 | 1 |
| 0 | 0 | 0 | 1 |
| 0 | 0 | 0 | 1 |
| 0 | 0 | 0 | 0 |
| 0 | 0 | 0 | 0 |
| 0 | 0 | 0 | 1 |
| 0 | 0 | 0 | 0 |
| 0 | 0 | 0 | 0 |
| 0 | 0 | 0 | 1 |
| 0 | 0 | 0 | 0 |
| 0 | 0 | 0 | 0 |
| 0 | 0 | 0 | 0 |
| 0 | 0 | 0 | 1 |
| 0 | 0 | 0 | 0 |
| 0 | 0 | 0 | 0 |
| 0 | 0 | 0 | 1 |
| 1 | 0 | 0 | 0 |
| 0 | 0 | 0 | 1 |
| 0 | 0 | 0 | 1 |
| 1 | 0 | 0 | 0 |
| 1 | 0 | 0 | 0 |
| 1 | 0 | 0 | 0 |
| 0 | 0 | 0 | 1 |
| 0 | 0 | 0 | 1 |
| 0 | 0 | 0 | 0 |
| 1 | 0 | 0 | 0 |
| 0 | 0 | 0 | 0 |
| 0 | 0 | 0 | 0 |
| 1 | 0 | 0 | 0 |
| 1 | 0 | 0 | 0 |
| 0 | 0 | 0 | 0 |
| 0 | 0 | 0 | 1 |
| 0 | 0 | 0 | 1 |
| 0 | 0 | 0 | 1 |
| 0 | 0 | 0 | 1 |
| 1 | 0 | 0 | 0 |
| 0 | 0 | 0 | 0 |
| 0 | 0 | 0 | 0 |
| 0 | 0 | 0 | 0 |

| | | | |
|---|---|---|---|
| 0 | 0 | 0 | 0 |
| 0 | 0 | 0 | 1 |
| 0 | 0 | 0 | 0 |
| 0 | 0 | 0 | 0 |
| 0 | 0 | 0 | 0 |
| 0 | 0 | 0 | 1 |
| 0 | 0 | 0 | 0 |
| 0 | 0 | 0 | 0 |
| 0 | 0 | 0 | 1 |
| 0 | 0 | 0 | 0 |
| 0 | 0 | 0 | 1 |
| 0 | 0 | 0 | 1 |
| 0 | 0 | 0 | 1 |
| 0 | 0 | 0 | 0 |
| 0 | 0 | 0 | 1 |
| 0 | 0 | 0 | 0 |
| 0 | 0 | 0 | 1 |
| 0 | 0 | 0 | 1 |
| 0 | 0 | 0 | 0 |
| 0 | 0 | 0 | 0 |
| 0 | 0 | 0 | 0 |
| 0 | 0 | 0 | 0 |
| 0 | 0 | 0 | 0 |
| 0 | 0 | 0 | 0 |
| 0 | 0 | 0 | 0 |
| 0 | 0 | 0 | 1 |
| 0 | 0 | 0 | 1 |
| 0 | 0 | 0 | 1 |
| 0 | 0 | 0 | 1 |
| 0 | 0 | 0 | 1 |
| 0 | 0 | 0 | 1 |
| 0 | 0 | 0 | 1 |
| 0 | 0 | 0 | 0 |
| 0 | 0 | 0 | 0 |
| 0 | 0 | 0 | 0 |
| 0 | 0 | 0 | 0 |
| 0 | 0 | 0 | 0 |
| 0 | 0 | 0 | 0 |
| 0 | 0 | 0 | 1 |
| 0 | 0 | 0 | 1 |
| 0 | 0 | 0 | 1 |
| 0 | 0 | 0 | 1 |
| 0 | 0 | 0 | 1 |
| 0 | 0 | 0 | 1 |
| 0 | 1 | 0 | 0 |
| 0 | 0 | 0 | 0 |
| 0 | 0 | 0 | 1 |
| 0 | 0 | 0 | 0 |
| 0 | 0 | 0 | 0 |
| 0 | 0 | 0 | 0 |
| 0 | 0 | 0 | 0 |
| 0 | 0 | 0 | 0 |
| 0 | 0 | 0 | 0 |
| 0 | 0 | 0 | 0 |

| | | | |
|---|---|---|---|
| 0 | 0 | 0 | 1 |
| 0 | 0 | 0 | 1 |
| 1 | 0 | 0 | 0 |
| 1 | 0 | 0 | 0 |
| 0 | 0 | 0 | 1 |
| 0 | 0 | 0 | 0 |
| 0 | 0 | 0 | 0 |
| 0 | 0 | 0 | 1 |
| 0 | 0 | 0 | 1 |
| 0 | 0 | 0 | 0 |
| 0 | 0 | 0 | 1 |
| 0 | 0 | 0 | 1 |
| 0 | 0 | 0 | 0 |
| 0 | 0 | 0 | 0 |
| 0 | 0 | 0 | 0 |
| 0 | 0 | 0 | 0 |
| 0 | 0 | 0 | 1 |
| 0 | 0 | 0 | 1 |
| 0 | 0 | 0 | 1 |
| 0 | 0 | 0 | 1 |
| 0 | 0 | 0 | 1 |
| 0 | 0 | 0 | 1 |
| 0 | 0 | 0 | 1 |
| 0 | 0 | 0 | 1 |
| 0 | 0 | 0 | 1 |
| 0 | 0 | 0 | 0 |
| 0 | 0 | 0 | 1 |
| 0 | 0 | 0 | 1 |
| 0 | 0 | 0 | 1 |
| 0 | 0 | 0 | 1 |
| 0 | 0 | 0 | 1 |
| 0 | 0 | 0 | 1 |
| 0 | 0 | 0 | 1 |
| 0 | 0 | 0 | 1 |
| 0 | 0 | 0 | 1 |
| 0 | 0 | 0 | 1 |
| 0 | 0 | 0 | 1 |
| 0 | 0 | 0 | 0 |
| 0 | 0 | 0 | 1 |
| 0 | 0 | 0 | 1 |
| 0 | 0 | 0 | 0 |
| 0 | 0 | 0 | 1 |
| 0 | 0 | 0 | 1 |
| 0 | 0 | 0 | 0 |
| 0 | 0 | 0 | 0 |
| 0 | 0 | 0 | 1 |
| 0 | 0 | 0 | 1 |
| 0 | 0 | 0 | 1 |
| 0 | 0 | 0 | 1 |
| 0 | 0 | 0 | 1 |
| 0 | 0 | 0 | 1 |
| 0 | 0 | 0 | 0 |
| 0 | 0 | 0 | 0 |
| 0 | 0 | 0 | 0 |

| | | | |
|---|---|---|---|
| 0 | 0 | 0 | 0 |
| 0 | 0 | 0 | 1 |
| 0 | 0 | 0 | 1 |
| 0 | 0 | 0 | 1 |
| 0 | 0 | 0 | 1 |
| 0 | 0 | 0 | 1 |
| 0 | 0 | 0 | 1 |
| 0 | 0 | 0 | 1 |
| 0 | 0 | 0 | 0 |
| 0 | 0 | 0 | 1 |
| 0 | 0 | 0 | 1 |
| 0 | 0 | 0 | 1 |
| 0 | 0 | 0 | 1 |
| 0 | 0 | 0 | 1 |
| 0 | 0 | 0 | 1 |
| 0 | 0 | 0 | 1 |
| 0 | 0 | 0 | 1 |
| 0 | 0 | 0 | 1 |
| 0 | 0 | 0 | 0 |
| 0 | 0 | 0 | 0 |
| 0 | 0 | 0 | 0 |
| 0 | 0 | 0 | 0 |
| 0 | 0 | 0 | 1 |
| 0 | 0 | 0 | 0 |
| 0 | 0 | 0 | 0 |
| 0 | 0 | 0 | 1 |
| 0 | 0 | 0 | 1 |
| 0 | 0 | 0 | 1 |
| 0 | 0 | 0 | 1 |
| 0 | 0 | 0 | 1 |
| 0 | 0 | 0 | 1 |
| 0 | 0 | 0 | 1 |
| 0 | 0 | 0 | 0 |
| 0 | 0 | 0 | 1 |
| 0 | 0 | 0 | 1 |
| 0 | 0 | 0 | 1 |
| 0 | 0 | 0 | 0 |
| 0 | 0 | 0 | 1 |
| 0 | 0 | 0 | 0 |
| 0 | 0 | 0 | 0 |
| 0 | 0 | 0 | 0 |
| 0 | 0 | 0 | 0 |
| 0 | 0 | 0 | 0 |
| 0 | 0 | 0 | 1 |
| 0 | 0 | 0 | 1 |
| 0 | 0 | 0 | 1 |
| 0 | 0 | 0 | 0 |
| 0 | 0 | 0 | 0 |
| 0 | 0 | 0 | 0 |
| 0 | 0 | 0 | 0 |
| 0 | 0 | 0 | 0 |
| 0 | 0 | 0 | 0 |
| 0 | 0 | 0 | 0 |
| 0 | 0 | 0 | 1 |

| | | | |
|---|---|---|---|
| 0 | 0 | 0 | 0 |
| 0 | 0 | 0 | 0 |
| 0 | 0 | 0 | 1 |
| 1 | 0 | 0 | 0 |
| 1 | 0 | 0 | 0 |
| 0 | 0 | 0 | 1 |
| 0 | 0 | 0 | 1 |
| 0 | 0 | 0 | 1 |
| 0 | 0 | 0 | 1 |
| 0 | 0 | 0 | 1 |
| 0 | 0 | 0 | 1 |
| 0 | 0 | 0 | 0 |
| 0 | 0 | 0 | 1 |
| 0 | 0 | 0 | 1 |
| 0 | 0 | 0 | 0 |
| 0 | 0 | 0 | 1 |
| 0 | 0 | 0 | 0 |
| 0 | 0 | 0 | 1 |
| 0 | 0 | 0 | 0 |
| 1 | 0 | 0 | 0 |
| 0 | 0 | 0 | 1 |
| 0 | 0 | 0 | 1 |
| 0 | 0 | 0 | 0 |
| 0 | 0 | 0 | 1 |
| 0 | 0 | 0 | 0 |
| 0 | 0 | 0 | 1 |
| 0 | 0 | 0 | 1 |
| 0 | 0 | 0 | 0 |
| 0 | 0 | 0 | 1 |
| 0 | 0 | 0 | 0 |
| 0 | 0 | 0 | 1 |
| 0 | 0 | 0 | 0 |
| 0 | 0 | 0 | 0 |
| 0 | 0 | 0 | 0 |
| 0 | 0 | 0 | 1 |
| 0 | 0 | 0 | 0 |
| 0 | 0 | 0 | 1 |
| 0 | 0 | 0 | 1 |
| 0 | 0 | 0 | 0 |
| 0 | 0 | 0 | 1 |
| 0 | 0 | 0 | 1 |
| 0 | 0 | 0 | 1 |
| 0 | 0 | 0 | 1 |
| 0 | 0 | 0 | 0 |
| 0 | 0 | 0 | 0 |
| 0 | 0 | 0 | 1 |
| 0 | 0 | 0 | 1 |
| 0 | 0 | 0 | 1 |
| 0 | 1 | 0 | 0 |
| 0 | 0 | 0 | 1 |
| 0 | 0 | 0 | 0 |
| 0 | 0 | 0 | 1 |
| 0 | 0 | 0 | 1 |
| 0 | 0 | 0 | 0 |

| | | | |
|---|---|---|---|
| 0 | 0 | 0 | 1 |
| 0 | 0 | 0 | 1 |
| 0 | 0 | 0 | 1 |
| 0 | 0 | 0 | 1 |
| 0 | 0 | 0 | 1 |
| 0 | 0 | 0 | 1 |
| 0 | 0 | 0 | 1 |
| 0 | 0 | 0 | 0 |
| 0 | 0 | 0 | 1 |
| 0 | 0 | 0 | 1 |
| 0 | 0 | 0 | 1 |
| 0 | 0 | 0 | 1 |
| 0 | 0 | 0 | 0 |
| 0 | 0 | 0 | 1 |
| 0 | 0 | 0 | 0 |
| 0 | 0 | 0 | 0 |
| 0 | 0 | 0 | 1 |
| 0 | 0 | 0 | 1 |
| 0 | 0 | 0 | 1 |
| 0 | 0 | 0 | 1 |
| 0 | 0 | 0 | 0 |
| 0 | 0 | 0 | 0 |
| 0 | 0 | 0 | 1 |
| 0 | 0 | 0 | 1 |
| 0 | 0 | 0 | 0 |
| 0 | 0 | 0 | 1 |
| 0 | 0 | 0 | 0 |
| 0 | 0 | 0 | 0 |
| 0 | 0 | 0 | 0 |
| 0 | 0 | 0 | 0 |
| 0 | 0 | 0 | 1 |
| 0 | 0 | 0 | 1 |
| 0 | 0 | 0 | 0 |
| 0 | 0 | 0 | 0 |
| 0 | 0 | 0 | 0 |
| 0 | 0 | 0 | 0 |
| 0 | 0 | 0 | 0 |
| 0 | 0 | 0 | 0 |
| 0 | 0 | 0 | 0 |
| 0 | 0 | 0 | 1 |
| 0 | 0 | 0 | 1 |
| 0 | 0 | 0 | 1 |
| 0 | 0 | 0 | 1 |
| 0 | 0 | 0 | 1 |
| 0 | 0 | 0 | 1 |
| 0 | 0 | 0 | 0 |
| 0 | 0 | 0 | 1 |
| 0 | 0 | 0 | 1 |
| 0 | 0 | 0 | 1 |
| 0 | 0 | 0 | 0 |
| 0 | 0 | 0 | 0 |
| 0 | 0 | 0 | 1 |
| 0 | 0 | 0 | 1 |
| 0 | 0 | 0 | 1 |

| | | | |
|---|---|---|---|
| 0 | 0 | 0 | 1 |
| 0 | 0 | 0 | 1 |
| 0 | 0 | 0 | 1 |
| 0 | 0 | 0 | 1 |
| 0 | 0 | 0 | 0 |
| 0 | 0 | 0 | 1 |
| 0 | 0 | 0 | 0 |
| 0 | 0 | 0 | 0 |
| 0 | 0 | 0 | 0 |
| 0 | 0 | 0 | 0 |
| 0 | 0 | 0 | 0 |
| 0 | 0 | 0 | 1 |
| 0 | 0 | 0 | 0 |
| 0 | 0 | 0 | 0 |
| 0 | 0 | 0 | 1 |
| 0 | 0 | 0 | 1 |
| 0 | 0 | 0 | 1 |
| 0 | 0 | 0 | 1 |
| 0 | 0 | 0 | 1 |
| 0 | 0 | 0 | 1 |
| 0 | 0 | 0 | 1 |
| 0 | 0 | 0 | 1 |
| 0 | 0 | 0 | 1 |
| 0 | 0 | 0 | 0 |
| 0 | 0 | 0 | 1 |
| 0 | 0 | 0 | 1 |
| 0 | 0 | 0 | 1 |
| 0 | 0 | 0 | 0 |
| 0 | 0 | 0 | 1 |
| 0 | 0 | 0 | 1 |
| 0 | 0 | 0 | 0 |
| 0 | 0 | 0 | 0 |
| 0 | 0 | 0 | 1 |
| 0 | 0 | 0 | 0 |
| 0 | 0 | 0 | 1 |
| 0 | 0 | 0 | 1 |
| 0 | 0 | 0 | 1 |
| 0 | 0 | 0 | 1 |
| 0 | 0 | 0 | 1 |
| 0 | 0 | 0 | 1 |
| 0 | 0 | 0 | 1 |
| 0 | 0 | 0 | 1 |
| 0 | 0 | 0 | 1 |
| 0 | 0 | 0 | 1 |
| 0 | 0 | 0 | 1 |
| 0 | 0 | 0 | 1 |
| 0 | 0 | 0 | 1 |
| 0 | 0 | 0 | 1 |
| 0 | 0 | 0 | 0 |
| 0 | 0 | 0 | 1 |
| 0 | 0 | 0 | 1 |
| 0 | 0 | 0 | 1 |
| 0 | 0 | 0 | 1 |
| 0 | 0 | 0 | 1 |

| | | | |
|---|---|---|---|
| 0 | 0 | 0 | 1 |
| 0 | 0 | 0 | 1 |
| 0 | 0 | 0 | 1 |
| 0 | 0 | 0 | 1 |
| 0 | 0 | 0 | 1 |
| 0 | 0 | 0 | 1 |
| 0 | 0 | 0 | 1 |
| 0 | 0 | 0 | 1 |
| 0 | 0 | 0 | 1 |
| 0 | 0 | 0 | 1 |
| 0 | 0 | 0 | 0 |
| 0 | 0 | 0 | 0 |
| 0 | 0 | 0 | 0 |
| 1 | 0 | 0 | 0 |
| 1 | 0 | 0 | 0 |
| 0 | 0 | 0 | 1 |
| 1 | 0 | 0 | 0 |
| 0 | 0 | 0 | 0 |
| 0 | 0 | 0 | 0 |
| 0 | 0 | 0 | 0 |
| 0 | 0 | 0 | 0 |
| 1 | 0 | 0 | 0 |
| 0 | 0 | 0 | 1 |
| 0 | 0 | 0 | 0 |
| 1 | 0 | 0 | 0 |
| 1 | 0 | 0 | 0 |
| 0 | 0 | 0 | 1 |
| 0 | 0 | 0 | 0 |
| 0 | 0 | 0 | 1 |
| 0 | 0 | 0 | 1 |
| 0 | 0 | 0 | 1 |
| 0 | 0 | 0 | 0 |
| 0 | 0 | 0 | 1 |
| 0 | 0 | 0 | 1 |
| 0 | 0 | 0 | 0 |
| 0 | 0 | 0 | 0 |
| 0 | 0 | 0 | 0 |
| 0 | 0 | 0 | 0 |
| 0 | 0 | 0 | 0 |
| 0 | 0 | 0 | 1 |
| 0 | 0 | 0 | 1 |
| 0 | 0 | 0 | 0 |
| 0 | 0 | 0 | 1 |
| 0 | 0 | 0 | 0 |
| 0 | 0 | 0 | 1 |
| 0 | 0 | 0 | 1 |
| 0 | 0 | 0 | 1 |
| 0 | 1 | 0 | 0 |
| 0 | 0 | 0 | 0 |
| 0 | 0 | 0 | 0 |
| 0 | 0 | 0 | 0 |
| 0 | 0 | 0 | 1 |
| 0 | 0 | 0 | 0 |
| 0 | 0 | 0 | 1 |

| | | | |
|---|---|---|---|
| 0 | 0 | 0 | 1 |
| 0 | 0 | 0 | 1 |
| 0 | 0 | 0 | 1 |
| 0 | 0 | 0 | 1 |
| 0 | 0 | 0 | 1 |
| 0 | 0 | 0 | 0 |
| 0 | 0 | 0 | 0 |
| 0 | 0 | 0 | 0 |
| 0 | 0 | 0 | 0 |
| 0 | 0 | 0 | 1 |
| 1 | 0 | 0 | 0 |
| 1 | 0 | 0 | 0 |
| 0 | 0 | 0 | 0 |
| 0 | 0 | 0 | 0 |
| 0 | 0 | 0 | 0 |
| 0 | 0 | 0 | 0 |
| 0 | 0 | 0 | 1 |
| 0 | 0 | 0 | 1 |
| 0 | 0 | 0 | 1 |
| 0 | 0 | 0 | 1 |
| 0 | 0 | 0 | 1 |
| 0 | 0 | 0 | 1 |
| 0 | 0 | 0 | 1 |
| 0 | 0 | 0 | 1 |
| 0 | 0 | 0 | 1 |
| 0 | 0 | 0 | 1 |
| 0 | 0 | 0 | 1 |
| 0 | 0 | 0 | 1 |
| 0 | 0 | 0 | 0 |
| 0 | 0 | 0 | 1 |
| 0 | 0 | 0 | 1 |
| 0 | 0 | 0 | 1 |
| 0 | 0 | 0 | 1 |
| 0 | 0 | 0 | 0 |
| 0 | 0 | 0 | 1 |
| 0 | 0 | 0 | 0 |
| 0 | 0 | 0 | 1 |
| 0 | 0 | 0 | 0 |
| 0 | 0 | 0 | 1 |
| 0 | 0 | 0 | 0 |
| 0 | 0 | 0 | 1 |
| 0 | 0 | 0 | 1 |
| 0 | 0 | 0 | 1 |
| 0 | 0 | 0 | 0 |
| 0 | 0 | 0 | 1 |
| 0 | 0 | 0 | 1 |
| 0 | 0 | 0 | 1 |
| 0 | 0 | 0 | 1 |
| 0 | 0 | 0 | 1 |
| 0 | 0 | 0 | 1 |
| 0 | 0 | 0 | 1 |
| 0 | 0 | 0 | 1 |
| 0 | 0 | 0 | 0 |
| 0 | 0 | 0 | 0 |

| | | | |
|---|---|---|---|
| 0 | 0 | 0 | 1 |
| 0 | 0 | 0 | 0 |
| 0 | 0 | 0 | 1 |
| 0 | 0 | 0 | 1 |
| 0 | 0 | 0 | 0 |
| 0 | 0 | 0 | 0 |
| 0 | 0 | 0 | 0 |
| 0 | 0 | 0 | 1 |
| 0 | 0 | 0 | 1 |
| 0 | 0 | 0 | 0 |
| 0 | 0 | 0 | 0 |
| 0 | 0 | 0 | 0 |
| 0 | 0 | 0 | 0 |
| 0 | 0 | 0 | 0 |
| 0 | 0 | 0 | 1 |
| 0 | 0 | 0 | 0 |
| 0 | 0 | 0 | 0 |
| 0 | 0 | 0 | 0 |
| 0 | 0 | 0 | 1 |
| 0 | 0 | 0 | 1 |
| 0 | 0 | 0 | 1 |
| 0 | 0 | 0 | 1 |
| 0 | 0 | 0 | 1 |
| 0 | 0 | 0 | 0 |
| 0 | 0 | 0 | 1 |
| 0 | 0 | 0 | 1 |
| 0 | 0 | 0 | 0 |
| 0 | 0 | 0 | 0 |
| 0 | 0 | 0 | 1 |
| 0 | 0 | 0 | 1 |
| 0 | 0 | 0 | 0 |
| 0 | 0 | 0 | 1 |
| 0 | 0 | 0 | 1 |
| 0 | 0 | 0 | 1 |
| 0 | 0 | 0 | 1 |
| 0 | 0 | 0 | 0 |
| 0 | 0 | 0 | 1 |
| 0 | 0 | 0 | 0 |
| 0 | 0 | 0 | 0 |
| 0 | 0 | 0 | 1 |
| 0 | 0 | 0 | 1 |
| 0 | 0 | 0 | 0 |
| 0 | 0 | 0 | 0 |
| 0 | 0 | 0 | 1 |
| 0 | 0 | 0 | 0 |
| 0 | 0 | 0 | 0 |
| 0 | 0 | 0 | 0 |
| 0 | 0 | 0 | 1 |
| 0 | 0 | 0 | 0 |
| 0 | 0 | 0 | 0 |
| 0 | 0 | 0 | 0 |
| 0 | 0 | 0 | 1 |
| 0 | 0 | 0 | 0 |
| 0 | 0 | 0 | 1 |

| | | | |
|---|---|---|---|
| 0 | 0 | 0 | 1 |
| 0 | 0 | 0 | 1 |
| 0 | 0 | 0 | 1 |
| 0 | 0 | 0 | 1 |
| 0 | 0 | 0 | 0 |
| 0 | 0 | 0 | 0 |
| 0 | 0 | 0 | 1 |
| 0 | 0 | 0 | 1 |
| 0 | 0 | 0 | 1 |
| 0 | 0 | 0 | 0 |
| 0 | 0 | 0 | 1 |
| 0 | 0 | 0 | 1 |
| 0 | 1 | 0 | 0 |
| 0 | 0 | 0 | 1 |
| 0 | 0 | 0 | 1 |
| 0 | 0 | 0 | 1 |
| 0 | 0 | 0 | 1 |
| 0 | 0 | 0 | 1 |
| 0 | 0 | 0 | 1 |
| 0 | 0 | 0 | 1 |
| 0 | 0 | 0 | 1 |
| 0 | 0 | 0 | 1 |
| 0 | 0 | 0 | 1 |
| 0 | 0 | 0 | 0 |
| 0 | 0 | 0 | 1 |
| 1 | 0 | 0 | 0 |
| 0 | 0 | 0 | 1 |
| 0 | 0 | 0 | 0 |
| 0 | 0 | 0 | 1 |
| 0 | 0 | 0 | 0 |
| 0 | 0 | 0 | 0 |
| 0 | 0 | 0 | 0 |
| 0 | 0 | 0 | 0 |
| 0 | 0 | 0 | 0 |
| 0 | 0 | 0 | 1 |
| 0 | 0 | 0 | 0 |
| 0 | 0 | 0 | 1 |
| 0 | 0 | 0 | 0 |
| 0 | 0 | 0 | 1 |
| 0 | 0 | 0 | 1 |
| 0 | 0 | 0 | 0 |
| 0 | 0 | 0 | 0 |
| 0 | 0 | 0 | 0 |
| 0 | 0 | 0 | 0 |
| 0 | 0 | 0 | 0 |
| 0 | 0 | 0 | 0 |
| 0 | 0 | 0 | 0 |
| 0 | 0 | 0 | 0 |
| 0 | 0 | 0 | 1 |
| 0 | 0 | 0 | 0 |
| 0 | 0 | 0 | 1 |
| 0 | 0 | 0 | 1 |
| 0 | 0 | 0 | 1 |
| 0 | 0 | 0 | 1 |

| | | | |
|---|---|---|---|
| 0 | 0 | 0 | 1 |
| 0 | 0 | 0 | 1 |
| 0 | 0 | 0 | 1 |
| 0 | 0 | 0 | 1 |
| 0 | 0 | 0 | 1 |
| 0 | 0 | 0 | 1 |
| 0 | 0 | 0 | 0 |
| 0 | 0 | 0 | 0 |
| 0 | 0 | 0 | 0 |
| 0 | 0 | 0 | 0 |
| 0 | 0 | 0 | 0 |
| 0 | 0 | 0 | 0 |
| 0 | 0 | 0 | 0 |
| 0 | 0 | 0 | 0 |
| 0 | 0 | 0 | 0 |
| 0 | 0 | 0 | 0 |
| 1 | 0 | 0 | 0 |
| 0 | 0 | 0 | 0 |
| 0 | 0 | 0 | 1 |
| 0 | 0 | 0 | 0 |
| 0 | 0 | 0 | 1 |
| 0 | 0 | 0 | 1 |
| 0 | 0 | 0 | 1 |
| 0 | 0 | 0 | 1 |
| 0 | 0 | 0 | 1 |
| 0 | 0 | 0 | 0 |
| 0 | 0 | 0 | 0 |
| 0 | 0 | 0 | 0 |
| 0 | 0 | 0 | 0 |
| 0 | 0 | 0 | 0 |
| 0 | 0 | 0 | 0 |
| 1 | 0 | 0 | 0 |
| 0 | 0 | 0 | 0 |
| 1 | 0 | 0 | 0 |
| 0 | 0 | 0 | 1 |
| 1 | 0 | 0 | 0 |
| 1 | 0 | 0 | 0 |
| 0 | 0 | 0 | 0 |
| 1 | 0 | 0 | 0 |
| 0 | 0 | 0 | 1 |
| 0 | 0 | 0 | 0 |
| 0 | 0 | 0 | 0 |
| 0 | 0 | 0 | 1 |
| 0 | 0 | 0 | 1 |
| 0 | 0 | 0 | 1 |
| 0 | 0 | 0 | 1 |
| 0 | 0 | 0 | 1 |
| 0 | 0 | 0 | 0 |
| 0 | 0 | 0 | 0 |
| 0 | 0 | 0 | 0 |
| 0 | 0 | 0 | 0 |
| 0 | 0 | 0 | 1 |
| 0 | 0 | 0 | 0 |
| 0 | 0 | 0 | 1 |

| | | | |
|---|---|---|---|
| 0 | 0 | 0 | 1 |
| 0 | 0 | 0 | 1 |
| 0 | 0 | 0 | 1 |
| 0 | 0 | 0 | 1 |
| 0 | 0 | 0 | 1 |
| 0 | 0 | 0 | 0 |
| 0 | 0 | 0 | 1 |
| 0 | 0 | 0 | 1 |
| 0 | 0 | 0 | 1 |
| 0 | 0 | 0 | 1 |
| 0 | 0 | 0 | 0 |
| 0 | 0 | 0 | 0 |
| 0 | 0 | 0 | 0 |
| 0 | 0 | 0 | 1 |
| 0 | 0 | 0 | 1 |
| 0 | 0 | 0 | 1 |
| 0 | 0 | 0 | 1 |
| 0 | 0 | 0 | 0 |
| 0 | 0 | 0 | 1 |
| 0 | 0 | 0 | 1 |
| 0 | 0 | 0 | 1 |
| 0 | 0 | 0 | 1 |
| 0 | 0 | 0 | 0 |
| 0 | 0 | 0 | 1 |
| 0 | 0 | 0 | 0 |
| 0 | 0 | 0 | 0 |
| 0 | 0 | 0 | 0 |
| 0 | 0 | 0 | 0 |
| 0 | 0 | 0 | 1 |
| 0 | 0 | 0 | 0 |
| 0 | 0 | 0 | 1 |
| 0 | 0 | 0 | 1 |
| 0 | 0 | 0 | 1 |
| 0 | 0 | 0 | 1 |
| 0 | 0 | 0 | 1 |
| 0 | 0 | 0 | 1 |
| 0 | 0 | 0 | 1 |
| 0 | 0 | 0 | 0 |
| 0 | 0 | 0 | 0 |
| 0 | 0 | 0 | 0 |
| 0 | 1 | 0 | 0 |
| 0 | 0 | 0 | 0 |
| 0 | 0 | 0 | 1 |
| 0 | 0 | 0 | 0 |
| 0 | 0 | 0 | 0 |
| 0 | 0 | 0 | 1 |
| 0 | 0 | 0 | 0 |
| 0 | 0 | 0 | 0 |
| 0 | 0 | 0 | 0 |
| 1 | 0 | 0 | 0 |
| 0 | 0 | 0 | 1 |
| 0 | 0 | 0 | 1 |
| 0 | 0 | 0 | 1 |
| 0 | 0 | 0 | 0 |

| | | | |
|---|---|---|---|
| 0 | 0 | 0 | 0 |
| 0 | 0 | 0 | 1 |
| 0 | 0 | 0 | 0 |
| 0 | 0 | 0 | 1 |
| 0 | 0 | 0 | 1 |
| 0 | 0 | 0 | 1 |
| 0 | 0 | 0 | 1 |
| 0 | 0 | 0 | 1 |
| 0 | 1 | 0 | 0 |
| 0 | 0 | 0 | 1 |
| 0 | 0 | 0 | 0 |
| 0 | 0 | 0 | 0 |
| 0 | 0 | 0 | 0 |
| 0 | 0 | 0 | 0 |
| 0 | 0 | 0 | 1 |
| 0 | 0 | 0 | 1 |
| 0 | 0 | 0 | 0 |
| 0 | 0 | 0 | 0 |
| 0 | 0 | 0 | 1 |
| 0 | 0 | 0 | 1 |
| 0 | 0 | 0 | 0 |
| 0 | 0 | 0 | 1 |
| 0 | 0 | 0 | 1 |
| 0 | 0 | 0 | 1 |
| 0 | 0 | 0 | 0 |
| 0 | 0 | 0 | 1 |
| 0 | 0 | 0 | 0 |
| 0 | 0 | 0 | 0 |
| 0 | 1 | 0 | 0 |
| 0 | 0 | 0 | 1 |
| 0 | 0 | 0 | 0 |
| 0 | 0 | 0 | 0 |
| 0 | 0 | 0 | 0 |
| 0 | 0 | 0 | 1 |
| 0 | 0 | 0 | 0 |
| 0 | 0 | 0 | 1 |
| 0 | 0 | 0 | 0 |
| 0 | 0 | 0 | 0 |
| 0 | 0 | 0 | 0 |
| 0 | 0 | 0 | 0 |
| 0 | 0 | 0 | 0 |
| 0 | 0 | 0 | 1 |
| 0 | 0 | 0 | 0 |
| 0 | 1 | 0 | 0 |
| 0 | 1 | 0 | 0 |
| 1 | 0 | 0 | 0 |
| 0 | 0 | 0 | 0 |
| 1 | 0 | 0 | 0 |
| 1 | 0 | 0 | 0 |
| 1 | 0 | 0 | 0 |
| 0 | 0 | 0 | 0 |
| 0 | 0 | 0 | 0 |
| 0 | 0 | 0 | 1 |
| 0 | 0 | 0 | 0 |

| | | | |
|---|---|---|---|
| 0 | 0 | 0 | 1 |
| 0 | 0 | 0 | 1 |
| 0 | 0 | 0 | 1 |
| 0 | 0 | 0 | 1 |
| 0 | 0 | 0 | 1 |
| 0 | 0 | 0 | 1 |
| 0 | 0 | 0 | 1 |
| 0 | 0 | 0 | 1 |
| 0 | 0 | 0 | 1 |
| 0 | 0 | 0 | 0 |
| 0 | 0 | 0 | 0 |
| 0 | 0 | 0 | 1 |
| 0 | 0 | 0 | 1 |
| 0 | 0 | 0 | 1 |
| 0 | 0 | 0 | 0 |
| 0 | 0 | 0 | 1 |
| 0 | 0 | 0 | 1 |
| 0 | 0 | 0 | 0 |
| 0 | 0 | 0 | 1 |
| 0 | 0 | 0 | 1 |
| 0 | 0 | 0 | 1 |
| 0 | 0 | 0 | 0 |
| 0 | 0 | 0 | 0 |
| 0 | 0 | 0 | 0 |
| 0 | 0 | 0 | 0 |
| 0 | 0 | 0 | 0 |
| 0 | 0 | 0 | 0 |
| 0 | 0 | 0 | 0 |
| 0 | 0 | 0 | 1 |
| 0 | 0 | 0 | 0 |
| 1 | 0 | 0 | 0 |
| 0 | 0 | 0 | 1 |
| 0 | 0 | 0 | 1 |
| 0 | 0 | 0 | 1 |
| 0 | 0 | 0 | 1 |
| 0 | 0 | 0 | 1 |
| 0 | 0 | 0 | 1 |
| 0 | 0 | 0 | 1 |
| 0 | 0 | 0 | 1 |
| 0 | 0 | 0 | 1 |
| 0 | 0 | 0 | 1 |
| 0 | 0 | 0 | 1 |
| 0 | 0 | 0 | 1 |
| 0 | 0 | 0 | 0 |
| 0 | 0 | 0 | 1 |
| 0 | 0 | 0 | 0 |
| 0 | 0 | 0 | 1 |
| 0 | 0 | 0 | 1 |
| 0 | 0 | 0 | 1 |
| 0 | 0 | 0 | 1 |
| 0 | 0 | 0 | 0 |
| 0 | 0 | 0 | 1 |
| 0 | 0 | 0 | 1 |
| 0 | 0 | 0 | 0 |

| | | | |
|---|---|---|---|
| 0 | 0 | 0 | 0 |
| 0 | 0 | 0 | 0 |
| 0 | 0 | 0 | 0 |
| 0 | 0 | 0 | 0 |
| 0 | 0 | 0 | 1 |
| 0 | 0 | 0 | 1 |
| 0 | 0 | 0 | 1 |
| 0 | 0 | 0 | 0 |
| 0 | 0 | 0 | 0 |
| 0 | 0 | 0 | 0 |
| 0 | 0 | 0 | 0 |
| 0 | 0 | 0 | 1 |
| 0 | 0 | 0 | 1 |
| 0 | 0 | 0 | 1 |
| 0 | 0 | 0 | 1 |
| 0 | 0 | 0 | 0 |
| 0 | 0 | 0 | 0 |
| 0 | 0 | 0 | 1 |
| 0 | 0 | 0 | 1 |
| 0 | 0 | 0 | 1 |
| 0 | 0 | 0 | 1 |
| 0 | 0 | 0 | 0 |
| 0 | 0 | 0 | 1 |
| 0 | 0 | 0 | 1 |
| 0 | 0 | 0 | 0 |
| 0 | 0 | 0 | 0 |
| 0 | 0 | 0 | 0 |
| 0 | 0 | 0 | 0 |
| 0 | 0 | 0 | 1 |
| 0 | 0 | 0 | 1 |
| 0 | 0 | 0 | 1 |
| 0 | 0 | 0 | 0 |
| 0 | 0 | 0 | 1 |
| 0 | 0 | 0 | 1 |
| 0 | 0 | 0 | 1 |
| 0 | 0 | 0 | 1 |
| 0 | 0 | 0 | 1 |
| 0 | 0 | 0 | 1 |
| 0 | 0 | 0 | 1 |
| 0 | 0 | 0 | 1 |
| 0 | 0 | 0 | 1 |
| 0 | 0 | 0 | 1 |
| 0 | 0 | 0 | 0 |
| 0 | 0 | 0 | 1 |
| 0 | 0 | 0 | 1 |
| 0 | 0 | 0 | 1 |
| 0 | 0 | 0 | 1 |
| 0 | 0 | 0 | 1 |
| 0 | 0 | 0 | 1 |
| 0 | 0 | 0 | 0 |
| 0 | 0 | 0 | 1 |
| 0 | 0 | 0 | 1 |
| 0 | 0 | 0 | 1 |
| 0 | 0 | 0 | 1 |

| | | | |
|---|---|---|---|
| 0 | 0 | 0 | 0 |
| 0 | 0 | 0 | 1 |
| 0 | 0 | 0 | 1 |
| 0 | 0 | 0 | 1 |
| 0 | 0 | 0 | 1 |
| 0 | 0 | 0 | 1 |
| 0 | 0 | 0 | 1 |
| 0 | 0 | 0 | 0 |
| 0 | 0 | 0 | 1 |
| 0 | 0 | 0 | 1 |
| 0 | 0 | 0 | 0 |
| 0 | 0 | 0 | 0 |
| 0 | 0 | 0 | 0 |
| 0 | 0 | 0 | 1 |
| 0 | 0 | 0 | 0 |
| 0 | 0 | 0 | 0 |
| 0 | 0 | 0 | 0 |
| 0 | 0 | 0 | 0 |
| 0 | 0 | 0 | 1 |
| 0 | 0 | 0 | 0 |
| 0 | 0 | 0 | 0 |
| 0 | 0 | 0 | 0 |
| 0 | 0 | 0 | 0 |
| 0 | 0 | 0 | 0 |
| 0 | 0 | 0 | 0 |
| 0 | 0 | 0 | 0 |
| 0 | 0 | 0 | 0 |
| 0 | 0 | 0 | 0 |
| 0 | 0 | 0 | 0 |
| 0 | 0 | 0 | 0 |
| 0 | 0 | 0 | 0 |
| 0 | 0 | 0 | 0 |
| 0 | 0 | 0 | 1 |
| 0 | 0 | 0 | 0 |
| 0 | 0 | 0 | 1 |
| 0 | 0 | 0 | 0 |
| 0 | 0 | 0 | 0 |
| 0 | 0 | 0 | 1 |
| 0 | 0 | 0 | 1 |
| 0 | 0 | 0 | 1 |
| 0 | 0 | 0 | 1 |
| 0 | 0 | 0 | 0 |
| 0 | 0 | 0 | 0 |
| 0 | 0 | 0 | 0 |
| 0 | 0 | 0 | 1 |
| 0 | 0 | 0 | 0 |
| 0 | 0 | 0 | 1 |
| 0 | 0 | 0 | 0 |
| 0 | 0 | 0 | 1 |
| 0 | 0 | 0 | 1 |
| 0 | 0 | 0 | 1 |
| 0 | 0 | 0 | 1 |
| 0 | 0 | 0 | 1 |
| 0 | 0 | 0 | 1 |

| | | | |
|---|---|---|---|
| 0 | 0 | 0 | 1 |
| 0 | 0 | 0 | 1 |
| 0 | 0 | 0 | 1 |
| 0 | 0 | 0 | 1 |
| 0 | 0 | 0 | 1 |
| 0 | 0 | 0 | 0 |
| 0 | 0 | 0 | 0 |
| 0 | 0 | 0 | 1 |
| 0 | 0 | 0 | 0 |
| 0 | 0 | 0 | 0 |
| 0 | 0 | 0 | 1 |
| 0 | 0 | 0 | 1 |
| 0 | 0 | 0 | 1 |
| 0 | 0 | 0 | 1 |
| 0 | 0 | 0 | 1 |
| 0 | 0 | 0 | 1 |
| 0 | 0 | 0 | 1 |
| 0 | 0 | 0 | 1 |
| 0 | 0 | 0 | 0 |
| 0 | 0 | 0 | 0 |
| 0 | 0 | 0 | 1 |
| 0 | 0 | 0 | 1 |
| 0 | 0 | 0 | 1 |
| 0 | 0 | 0 | 0 |
| 0 | 0 | 0 | 1 |
| 0 | 0 | 0 | 0 |
| 0 | 0 | 0 | 0 |
| 0 | 0 | 0 | 1 |
| 0 | 0 | 0 | 1 |
| 0 | 0 | 0 | 1 |
| 0 | 0 | 0 | 0 |
| 0 | 0 | 0 | 1 |
| 0 | 0 | 0 | 0 |
| 0 | 0 | 0 | 0 |
| 0 | 0 | 0 | 0 |
| 0 | 0 | 0 | 0 |
| 0 | 0 | 0 | 0 |
| 0 | 0 | 0 | 1 |
| 0 | 0 | 0 | 1 |
| 0 | 0 | 0 | 1 |
| 0 | 0 | 0 | 0 |
| 0 | 0 | 0 | 0 |
| 0 | 0 | 0 | 0 |
| 0 | 0 | 0 | 0 |
| 0 | 0 | 0 | 1 |
| 0 | 0 | 0 | 1 |
| 0 | 0 | 0 | 1 |
| 0 | 0 | 0 | 0 |
| 0 | 0 | 0 | 0 |
| 0 | 0 | 0 | 0 |
| 0 | 0 | 0 | 1 |
| 0 | 0 | 0 | 1 |
| 0 | 0 | 0 | 1 |
| 0 | 0 | 0 | 1 |

| | | | |
|---|---|---|---|
| 0 | 0 | 0 | 1 |
| 0 | 0 | 0 | 1 |
| 0 | 0 | 0 | 1 |
| 0 | 0 | 0 | 0 |
| 0 | 0 | 0 | 0 |
| 0 | 0 | 0 | 1 |
| 0 | 0 | 0 | 0 |
| 0 | 0 | 0 | 1 |
| 0 | 0 | 0 | 1 |
| 0 | 0 | 0 | 1 |
| 0 | 0 | 0 | 1 |
| 0 | 0 | 0 | 0 |
| 0 | 0 | 0 | 1 |
| 0 | 0 | 0 | 0 |
| 0 | 0 | 0 | 1 |
| 0 | 0 | 0 | 0 |
| 0 | 0 | 0 | 1 |
| 0 | 0 | 0 | 1 |
| 0 | 0 | 0 | 1 |
| 0 | 0 | 0 | 0 |
| 0 | 0 | 0 | 0 |
| 0 | 0 | 0 | 0 |
| 0 | 0 | 0 | 0 |
| 0 | 0 | 0 | 0 |
| 0 | 0 | 0 | 0 |
| 0 | 0 | 0 | 1 |
| 0 | 0 | 0 | 0 |
| 0 | 0 | 0 | 1 |
| 0 | 0 | 0 | 1 |
| 0 | 0 | 0 | 1 |
| 0 | 0 | 0 | 1 |
| 0 | 0 | 0 | 1 |
| 0 | 0 | 0 | 0 |
| 0 | 1 | 0 | 0 |
| 0 | 0 | 0 | 1 |
| 0 | 0 | 0 | 1 |
| 0 | 1 | 0 | 0 |
| 0 | 0 | 0 | 0 |
| 0 | 0 | 0 | 0 |
| 0 | 1 | 1 | 1 |
| 0 | 0 | 0 | 0 |
| 0 | 0 | 0 | 0 |
| 0 | 0 | 0 | 1 |
| 0 | 0 | 0 | 0 |
| 0 | 0 | 0 | 1 |
| 0 | 0 | 0 | 1 |
| 0 | 0 | 0 | 1 |
| 0 | 0 | 0 | 1 |
| 0 | 0 | 0 | 1 |
| 1 | 0 | 0 | 0 |
| 0 | 0 | 0 | 1 |
| 0 | 0 | 0 | 1 |
| 0 | 0 | 0 | 1 |
| 0 | 0 | 0 | 1 |

| | | | |
|---|---|---|---|
| 0 | 0 | 0 | 1 |
| 0 | 0 | 0 | 1 |
| 0 | 0 | 0 | 1 |
| 0 | 0 | 0 | 0 |
| 0 | 0 | 0 | 1 |
| 0 | 0 | 0 | 1 |
| 0 | 0 | 0 | 1 |
| 0 | 0 | 0 | 0 |
| 0 | 0 | 0 | 1 |
| 0 | 0 | 0 | 1 |
| 0 | 0 | 0 | 1 |
| 0 | 0 | 0 | 1 |
| 0 | 0 | 0 | 1 |
| 0 | 0 | 0 | 1 |
| 0 | 0 | 0 | 1 |
| 0 | 0 | 0 | 1 |
| 0 | 0 | 0 | 0 |
| 0 | 0 | 0 | 0 |
| 0 | 0 | 0 | 1 |
| 0 | 0 | 0 | 1 |
| 0 | 0 | 0 | 1 |
| 0 | 0 | 0 | 1 |
| 0 | 0 | 0 | 1 |
| 0 | 0 | 0 | 1 |
| 0 | 0 | 0 | 1 |
| 0 | 0 | 0 | 1 |
| 0 | 0 | 0 | 1 |
| 0 | 0 | 0 | 0 |
| 0 | 0 | 0 | 0 |
| 0 | 0 | 0 | 1 |
| 0 | 0 | 0 | 1 |
| 0 | 0 | 0 | 0 |
| 0 | 0 | 0 | 1 |
| 0 | 0 | 0 | 0 |
| 1 | 0 | 0 | 0 |
| 1 | 0 | 0 | 0 |
| 1 | 0 | 0 | 0 |
| 1 | 0 | 0 | 0 |
| 0 | 0 | 0 | 1 |
| 0 | 0 | 0 | 1 |
| 0 | 0 | 0 | 1 |
| 0 | 0 | 0 | 1 |
| 0 | 0 | 0 | 1 |
| 0 | 0 | 0 | 1 |
| 0 | 0 | 0 | 0 |
| 0 | 0 | 0 | 0 |
| 0 | 0 | 0 | 0 |
| 0 | 0 | 0 | 1 |
| 0 | 0 | 0 | 1 |
| 0 | 0 | 0 | 1 |
| 0 | 0 | 0 | 1 |
| 0 | 0 | 0 | 0 |
| 0 | 0 | 0 | 0 |
| 0 | 0 | 0 | 0 |

| | | | |
|---|---|---|---|
| 0 | 0 | 0 | 0 |
| 0 | 0 | 0 | 0 |
| 0 | 0 | 0 | 1 |
| 0 | 0 | 0 | 0 |
| 0 | 0 | 0 | 0 |
| 0 | 0 | 0 | 0 |
| 0 | 0 | 0 | 0 |
| 0 | 0 | 0 | 1 |
| 0 | 0 | 0 | 1 |
| 0 | 0 | 0 | 1 |
| 0 | 0 | 0 | 1 |
| 0 | 0 | 0 | 1 |
| 0 | 0 | 0 | 1 |
| 0 | 0 | 0 | 1 |
| 0 | 0 | 0 | 1 |
| 0 | 0 | 0 | 1 |
| 0 | 0 | 0 | 1 |
| 0 | 0 | 0 | 1 |
| 0 | 0 | 0 | 1 |
| 0 | 0 | 0 | 1 |
| 0 | 0 | 0 | 0 |
| 0 | 0 | 0 | 0 |
| 0 | 0 | 0 | 0 |
| 0 | 0 | 0 | 0 |
| 0 | 0 | 0 | 0 |
| 0 | 0 | 0 | 1 |
| 0 | 0 | 0 | 1 |
| 0 | 0 | 0 | 0 |
| 0 | 0 | 0 | 1 |
| 0 | 0 | 0 | 1 |
| 0 | 0 | 0 | 0 |
| 0 | 0 | 0 | 0 |
| 0 | 0 | 0 | 0 |
| 0 | 0 | 0 | 1 |
| 0 | 0 | 0 | 0 |
| 0 | 0 | 0 | 1 |
| 0 | 0 | 0 | 1 |
| 0 | 0 | 0 | 0 |
| 0 | 0 | 0 | 1 |
| 0 | 0 | 0 | 1 |
| 0 | 0 | 0 | 1 |
| 0 | 0 | 0 | 1 |
| 0 | 0 | 0 | 1 |
| 0 | 0 | 0 | 1 |
| 0 | 0 | 0 | 1 |
| 0 | 0 | 0 | 1 |
| 0 | 0 | 0 | 1 |
| 0 | 0 | 0 | 1 |
| 0 | 0 | 0 | 0 |
| 0 | 0 | 0 | 1 |
| 1 | 0 | 0 | 0 |
| 1 | 0 | 0 | 0 |
| 0 | 0 | 0 | 1 |
| 0 | 0 | 0 | 1 |

| | | | |
|---|---|---|---|
| 1 | 0 | 0 | 0 |
| 0 | 0 | 0 | 1 |
| 0 | 0 | 0 | 1 |
| 0 | 0 | 0 | 1 |
| 0 | 0 | 0 | 1 |
| 1 | 0 | 0 | 0 |
| 1 | 0 | 0 | 0 |
| 0 | 0 | 0 | 1 |
| 0 | 0 | 0 | 1 |
| 0 | 0 | 0 | 1 |
| 0 | 0 | 0 | 1 |
| 0 | 0 | 0 | 0 |
| 0 | 0 | 0 | 1 |
| 0 | 0 | 0 | 1 |
| 0 | 0 | 0 | 1 |
| 0 | 0 | 0 | 1 |
| 0 | 0 | 0 | 0 |
| 0 | 0 | 0 | 0 |
| 0 | 0 | 0 | 1 |
| 0 | 0 | 0 | 1 |
| 0 | 0 | 0 | 1 |
| 0 | 0 | 0 | 1 |
| 0 | 0 | 0 | 1 |
| 0 | 0 | 0 | 1 |
| 0 | 0 | 0 | 0 |
| 0 | 0 | 0 | 1 |
| 0 | 0 | 0 | 1 |
| 0 | 0 | 0 | 1 |
| 0 | 0 | 0 | 1 |
| 0 | 0 | 0 | 1 |
| 0 | 0 | 0 | 1 |
| 0 | 0 | 0 | 0 |
| 0 | 0 | 0 | 0 |
| 0 | 0 | 0 | 1 |
| 0 | 0 | 0 | 1 |
| 0 | 0 | 0 | 0 |
| 1 | 0 | 0 | 0 |
| 0 | 0 | 0 | 0 |
| 1 | 0 | 0 | 0 |
| 0 | 0 | 0 | 0 |
| 0 | 0 | 0 | 0 |
| 0 | 0 | 0 | 0 |
| 0 | 0 | 0 | 0 |
| 0 | 0 | 0 | 0 |
| 0 | 0 | 0 | 0 |
| 0 | 0 | 0 | 0 |
| 0 | 0 | 0 | 0 |
| 1 | 0 | 0 | 0 |
| 0 | 0 | 0 | 1 |
| 0 | 0 | 0 | 1 |
| 0 | 0 | 0 | 1 |
| 0 | 0 | 0 | 1 |
| 0 | 0 | 0 | 0 |
| 0 | 0 | 0 | 1 |

| | | | |
|---|---|---|---|
| 0 | 0 | 0 | 1 |
| 0 | 0 | 0 | 1 |
| 0 | 0 | 0 | 1 |
| 0 | 0 | 0 | 1 |
| 0 | 0 | 0 | 1 |
| 0 | 0 | 0 | 1 |
| 0 | 0 | 0 | 1 |
| 0 | 0 | 0 | 0 |
| 0 | 0 | 0 | 0 |
| 0 | 0 | 0 | 1 |
| 0 | 0 | 0 | 1 |
| 0 | 0 | 0 | 1 |
| 0 | 0 | 0 | 1 |
| 0 | 0 | 0 | 0 |
| 0 | 0 | 0 | 1 |
| 0 | 0 | 0 | 1 |
| 0 | 0 | 0 | 0 |
| 0 | 0 | 0 | 1 |
| 1 | 0 | 0 | 0 |
| 0 | 0 | 0 | 1 |
| 0 | 0 | 0 | 1 |
| 0 | 0 | 0 | 1 |
| 0 | 0 | 0 | 1 |
| 0 | 0 | 0 | 0 |
| 0 | 0 | 0 | 1 |
| 0 | 0 | 0 | 1 |
| 0 | 0 | 0 | 1 |
| 0 | 0 | 0 | 1 |
| 0 | 0 | 0 | 1 |
| 0 | 0 | 0 | 1 |
| 0 | 0 | 0 | 1 |
| 1 | 0 | 0 | 0 |
| 0 | 0 | 0 | 1 |
| 0 | 0 | 0 | 1 |
| 0 | 0 | 0 | 1 |
| 0 | 0 | 0 | 1 |
| 0 | 0 | 0 | 1 |
| 0 | 0 | 0 | 1 |
| 1 | 0 | 0 | 0 |
| 0 | 0 | 0 | 1 |
| 0 | 0 | 0 | 1 |
| 0 | 0 | 0 | 1 |
| 0 | 0 | 0 | 1 |
| 0 | 0 | 0 | 1 |
| 0 | 0 | 0 | 1 |
| 0 | 0 | 0 | 0 |
| 0 | 0 | 0 | 1 |
| 0 | 0 | 0 | 1 |
| 0 | 0 | 0 | 1 |
| 0 | 0 | 0 | 1 |
| 0 | 0 | 0 | 1 |
| 0 | 0 | 0 | 1 |
| 0 | 0 | 0 | 1 |
| 0 | 0 | 0 | 1 |

| | | | |
|---|---|---|---|
| 0 | 0 | 0 | 1 |
| 0 | 0 | 0 | 1 |
| 0 | 0 | 0 | 1 |
| 0 | 0 | 0 | 1 |
| 0 | 0 | 0 | 1 |
| 0 | 0 | 0 | 1 |
| 0 | 0 | 0 | 1 |
| 0 | 0 | 0 | 1 |
| 0 | 0 | 0 | 1 |
| 0 | 0 | 0 | 1 |
| 0 | 0 | 0 | 1 |
| 0 | 0 | 0 | 1 |
| 0 | 0 | 0 | 1 |
| 0 | 0 | 0 | 1 |
| 0 | 0 | 0 | 1 |
| 1 | 0 | 0 | 0 |
| 0 | 0 | 0 | 1 |
| 0 | 0 | 0 | 0 |
| 0 | 0 | 0 | 1 |
| 0 | 0 | 0 | 1 |
| 1 | 0 | 0 | 0 |
| 0 | 0 | 0 | 1 |
| 0 | 0 | 0 | 1 |
| 0 | 0 | 0 | 1 |
| 0 | 0 | 0 | 0 |
| 0 | 0 | 0 | 1 |
| 0 | 0 | 0 | 0 |
| 0 | 0 | 0 | 1 |
| 0 | 0 | 0 | 1 |
| 0 | 0 | 0 | 1 |
| 0 | 0 | 0 | 1 |
| 0 | 0 | 0 | 1 |
| 0 | 0 | 0 | 1 |
| 0 | 0 | 0 | 1 |
| 0 | 0 | 0 | 1 |
| 0 | 0 | 0 | 1 |
| 0 | 0 | 0 | 1 |
| 0 | 0 | 0 | 1 |
| 0 | 0 | 0 | 1 |
| 0 | 0 | 0 | 1 |
| 0 | 0 | 0 | 1 |
| 0 | 0 | 0 | 1 |
| 0 | 0 | 0 | 1 |
| 0 | 0 | 0 | 0 |
| 0 | 0 | 0 | 1 |
| 0 | 0 | 0 | 1 |
| 0 | 0 | 0 | 0 |
| 0 | 0 | 0 | 0 |
| 0 | 0 | 0 | 1 |
| 0 | 0 | 0 | 0 |
| 0 | 0 | 0 | 0 |
| 0 | 0 | 0 | 0 |
| 0 | 0 | 0 | 1 |
| 0 | 0 | 0 | 1 |

| | | | |
|---|---|---|---|
| 0 | 0 | 0 | 1 |
| 0 | 0 | 0 | 1 |
| 0 | 0 | 0 | 1 |
| 0 | 0 | 0 | 1 |
| 0 | 0 | 0 | 1 |
| 0 | 0 | 0 | 1 |
| 0 | 0 | 0 | 1 |
| 1 | 0 | 0 | 0 |
| 0 | 0 | 0 | 1 |
| 0 | 0 | 0 | 1 |
| 0 | 0 | 0 | 1 |
| 0 | 0 | 0 | 1 |
| 0 | 0 | 0 | 1 |
| 0 | 0 | 0 | 1 |
| 0 | 0 | 0 | 0 |
| 0 | 0 | 0 | 0 |
| 0 | 0 | 0 | 1 |
| 0 | 0 | 0 | 0 |
| 0 | 0 | 0 | 1 |
| 0 | 0 | 0 | 1 |
| 0 | 0 | 0 | 0 |
| 0 | 0 | 0 | 1 |
| 0 | 0 | 0 | 1 |
| 0 | 0 | 0 | 1 |
| 0 | 0 | 0 | 0 |
| 0 | 0 | 0 | 1 |
| 0 | 0 | 0 | 1 |
| 1 | 0 | 0 | 0 |
| 0 | 0 | 0 | 0 |
| 1 | 0 | 0 | 0 |
| 1 | 0 | 0 | 0 |
| 0 | 0 | 0 | 1 |
| 0 | 0 | 0 | 1 |
| 0 | 0 | 0 | 1 |
| 0 | 0 | 0 | 1 |
| 0 | 0 | 0 | 1 |
| 0 | 0 | 0 | 0 |
| 0 | 0 | 0 | 0 |
| 0 | 0 | 0 | 1 |
| 0 | 0 | 0 | 1 |
| 0 | 0 | 0 | 1 |
| 0 | 0 | 0 | 1 |
| 0 | 0 | 0 | 0 |
| 0 | 0 | 0 | 1 |
| 0 | 0 | 0 | 1 |
| 0 | 0 | 0 | 1 |
| 0 | 0 | 0 | 1 |
| 0 | 0 | 0 | 1 |
| 0 | 0 | 0 | 0 |
| 0 | 0 | 0 | 0 |
| 0 | 0 | 0 | 0 |
| 0 | 0 | 0 | 0 |
| 0 | 0 | 0 | 0 |
| 0 | 0 | 0 | 1 |

| | | | |
|---|---|---|---|
| 0 | 0 | 0 | 0 |
| 0 | 0 | 0 | 0 |
| 0 | 0 | 0 | 1 |
| 1 | 0 | 0 | 0 |
| 0 | 0 | 0 | 1 |
| 0 | 0 | 0 | 1 |
| 0 | 0 | 0 | 1 |
| 0 | 0 | 0 | 1 |
| 0 | 0 | 0 | 1 |
| 1 | 0 | 0 | 0 |
| 0 | 0 | 0 | 1 |
| 0 | 0 | 0 | 1 |
| 0 | 0 | 0 | 1 |
| 0 | 0 | 0 | 1 |
| 0 | 0 | 0 | 1 |
| 0 | 0 | 0 | 1 |
| 0 | 0 | 0 | 1 |
| 0 | 0 | 0 | 1 |
| 0 | 0 | 0 | 1 |
| 0 | 0 | 0 | 1 |
| 0 | 0 | 0 | 1 |
| 0 | 0 | 0 | 1 |
| 0 | 0 | 0 | 1 |
| 0 | 0 | 0 | 1 |
| 0 | 0 | 0 | 1 |
| 0 | 0 | 0 | 1 |
| 0 | 0 | 0 | 1 |
| 0 | 0 | 0 | 1 |
| 0 | 0 | 0 | 1 |
| 0 | 0 | 0 | 1 |
| 0 | 0 | 0 | 1 |
| 0 | 0 | 0 | 1 |
| 1 | 0 | 0 | 0 |
| 0 | 0 | 0 | 1 |
| 0 | 0 | 0 | 1 |
| 0 | 0 | 0 | 1 |
| 0 | 0 | 0 | 1 |
| 0 | 0 | 0 | 1 |
| 0 | 0 | 0 | 1 |
| 0 | 0 | 0 | 1 |
| 0 | 0 | 0 | 1 |
| 0 | 0 | 0 | 1 |
| 0 | 0 | 0 | 1 |
| 0 | 0 | 0 | 1 |
| 0 | 0 | 0 | 1 |
| 0 | 0 | 0 | 1 |
| 0 | 0 | 0 | 1 |
| 0 | 0 | 0 | 0 |
| 0 | 0 | 0 | 0 |
| 0 | 0 | 0 | 0 |
| 0 | 0 | 0 | 0 |
| 0 | 1 | 0 | 0 |
| 0 | 0 | 0 | 1 |
| 0 | 0 | 0 | 0 |

| | | | |
|---|---|---|---|
| 0 | 0 | 0 | 1 |
| 0 | 0 | 0 | 0 |
| 0 | 0 | 0 | 1 |
| 0 | 0 | 0 | 1 |
| 0 | 0 | 0 | 1 |
| 1 | 0 | 0 | 0 |
| 0 | 0 | 0 | 0 |
| 0 | 0 | 0 | 0 |
| 1 | 0 | 0 | 0 |
| 0 | 0 | 0 | 0 |
| 0 | 0 | 0 | 1 |
| 0 | 0 | 0 | 0 |
| 0 | 0 | 0 | 1 |
| 0 | 0 | 0 | 1 |
| 0 | 0 | 0 | 1 |
| 1 | 0 | 0 | 0 |
| 0 | 0 | 0 | 1 |
| 0 | 0 | 0 | 1 |
| 0 | 0 | 0 | 0 |
| 0 | 0 | 0 | 0 |
| 1 | 0 | 0 | 0 |
| 1 | 0 | 0 | 0 |
| 0 | 0 | 0 | 0 |
| 0 | 0 | 0 | 0 |
| 0 | 0 | 0 | 1 |
| 0 | 0 | 0 | 0 |
| 0 | 0 | 0 | 0 |
| 0 | 0 | 0 | 0 |
| 0 | 0 | 0 | 0 |
| 0 | 0 | 0 | 0 |
| 0 | 0 | 0 | 0 |
| 0 | 0 | 0 | 1 |
| 0 | 0 | 0 | 1 |
| 0 | 0 | 0 | 0 |
| 0 | 0 | 0 | 0 |
| 0 | 0 | 0 | 1 |
| 0 | 0 | 0 | 0 |
| 0 | 0 | 0 | 0 |
| 0 | 0 | 0 | 0 |
| 0 | 0 | 0 | 0 |
| 0 | 0 | 0 | 0 |
| 0 | 0 | 0 | 1 |
| 0 | 0 | 0 | 1 |
| 0 | 0 | 0 | 1 |
| 0 | 0 | 0 | 0 |
| 0 | 0 | 0 | 0 |
| 0 | 0 | 0 | 0 |
| 0 | 0 | 0 | 1 |
| 0 | 0 | 0 | 1 |
| 0 | 0 | 0 | 1 |
| 0 | 0 | 0 | 1 |
| 0 | 0 | 0 | 1 |
| 0 | 0 | 0 | 1 |
| 0 | 0 | 0 | 0 |

| | | | |
|---|---|---|---|
| 0 | 0 | 0 | 1 |
| 0 | 0 | 0 | 1 |
| 0 | 0 | 0 | 1 |
| 0 | 0 | 0 | 1 |
| 1 | 0 | 0 | 0 |
| 0 | 0 | 0 | 0 |
| 0 | 0 | 0 | 1 |
| 0 | 0 | 0 | 1 |
| 0 | 0 | 0 | 1 |
| 0 | 0 | 0 | 1 |
| 0 | 0 | 0 | 1 |
| 0 | 0 | 0 | 0 |
| 0 | 0 | 0 | 1 |
| 0 | 0 | 0 | 0 |
| 0 | 0 | 0 | 0 |
| 0 | 0 | 0 | 0 |
| 0 | 0 | 0 | 0 |
| 0 | 0 | 0 | 0 |
| 0 | 0 | 0 | 0 |
| 0 | 0 | 0 | 0 |
| 0 | 0 | 0 | 0 |
| 0 | 0 | 0 | 0 |
| 0 | 0 | 0 | 0 |
| 0 | 0 | 0 | 1 |
| 0 | 0 | 0 | 1 |
| 0 | 0 | 0 | 1 |
| 0 | 0 | 0 | 1 |
| 0 | 0 | 0 | 1 |
| 0 | 0 | 0 | 1 |
| 0 | 0 | 0 | 1 |
| 0 | 0 | 0 | 1 |
| 0 | 0 | 0 | 0 |
| 0 | 0 | 0 | 1 |
| 0 | 0 | 0 | 0 |
| 0 | 0 | 0 | 0 |
| 0 | 0 | 0 | 0 |
| 0 | 0 | 0 | 0 |
| 0 | 0 | 0 | 1 |
| 0 | 0 | 0 | 1 |
| 0 | 0 | 0 | 1 |
| 0 | 0 | 0 | 1 |
| 0 | 0 | 0 | 1 |
| 0 | 0 | 0 | 1 |
| 0 | 0 | 0 | 1 |
| 0 | 0 | 0 | 1 |
| 1 | 0 | 0 | 0 |
| 0 | 0 | 0 | 1 |
| 0 | 0 | 0 | 1 |
| 0 | 0 | 0 | 1 |
| 1 | 0 | 0 | 0 |
| 1 | 0 | 0 | 0 |
| 0 | 0 | 0 | 0 |
| 0 | 0 | 0 | 1 |
| 0 | 0 | 0 | 1 |

| | | | |
|---|---|---|---|
| 0 | 0 | 0 | 1 |
| 0 | 0 | 0 | 1 |
| 0 | 0 | 0 | 1 |
| 0 | 0 | 0 | 0 |
| 0 | 0 | 0 | 1 |
| 0 | 0 | 0 | 1 |
| 0 | 0 | 0 | 0 |
| 0 | 0 | 0 | 1 |
| 0 | 0 | 0 | 1 |
| 0 | 0 | 0 | 1 |
| 0 | 0 | 0 | 0 |
| 0 | 0 | 0 | 0 |
| 0 | 0 | 0 | 1 |
| 0 | 0 | 0 | 1 |
| 0 | 0 | 0 | 1 |
| 0 | 0 | 0 | 1 |
| 0 | 0 | 0 | 0 |
| 0 | 0 | 0 | 0 |
| 0 | 0 | 0 | 1 |
| 0 | 0 | 0 | 0 |
| 0 | 0 | 0 | 0 |
| 0 | 0 | 0 | 0 |
| 0 | 0 | 0 | 1 |
| 0 | 0 | 0 | 0 |
| 0 | 0 | 0 | 0 |
| 0 | 0 | 0 | 1 |
| 0 | 0 | 0 | 0 |
| 0 | 0 | 0 | 1 |
| 0 | 0 | 0 | 1 |
| 0 | 0 | 0 | 0 |
| 0 | 0 | 0 | 1 |
| 0 | 0 | 0 | 1 |
| 0 | 0 | 0 | 0 |
| 0 | 0 | 0 | 1 |
| 0 | 0 | 0 | 1 |
| 0 | 0 | 0 | 0 |
| 0 | 0 | 0 | 1 |
| 0 | 0 | 0 | 0 |
| 0 | 0 | 0 | 1 |
| 0 | 0 | 0 | 0 |
| 0 | 0 | 0 | 0 |
| 0 | 0 | 0 | 1 |
| 0 | 0 | 0 | 0 |
| 0 | 0 | 0 | 0 |
| 0 | 0 | 0 | 1 |
| 0 | 0 | 0 | 1 |
| 0 | 0 | 0 | 1 |
| 0 | 0 | 0 | 1 |
| 0 | 0 | 0 | 1 |
| 0 | 0 | 0 | 1 |
| 0 | 0 | 0 | 1 |
| 0 | 0 | 0 | 1 |
| 0 | 0 | 0 | 1 |
| 0 | 0 | 0 | 1 |

| | | | |
|---|---|---|---|
| 0 | 0 | 0 | 1 |
| 0 | 0 | 0 | 1 |
| 0 | 0 | 0 | 1 |
| 0 | 0 | 0 | 0 |
| 0 | 0 | 0 | 1 |
| 0 | 0 | 0 | 0 |
| 0 | 0 | 0 | 1 |
| 0 | 0 | 0 | 1 |
| 0 | 0 | 0 | 0 |
| 0 | 0 | 0 | 0 |
| 0 | 0 | 0 | 1 |
| 0 | 0 | 0 | 0 |
| 0 | 0 | 0 | 0 |
| 0 | 0 | 0 | 0 |
| 0 | 0 | 0 | 1 |
| 0 | 0 | 0 | 0 |
| 0 | 0 | 0 | 0 |
| 0 | 0 | 0 | 1 |
| 1 | 0 | 0 | 0 |
| 1 | 0 | 0 | 0 |
| 1 | 0 | 0 | 0 |
| 1 | 0 | 0 | 0 |
| 0 | 0 | 0 | 1 |
| 0 | 0 | 0 | 0 |
| 0 | 0 | 0 | 0 |
| 0 | 0 | 0 | 0 |
| 0 | 0 | 0 | 0 |
| 0 | 0 | 0 | 0 |
| 0 | 0 | 0 | 0 |
| 0 | 0 | 0 | 0 |
| 0 | 0 | 0 | 0 |
| 0 | 0 | 0 | 0 |
| 0 | 0 | 0 | 0 |
| 0 | 0 | 0 | 1 |
| 0 | 0 | 0 | 1 |
| 0 | 0 | 0 | 0 |
| 0 | 0 | 0 | 1 |
| 0 | 0 | 0 | 0 |
| 0 | 0 | 0 | 0 |
| 0 | 0 | 0 | 0 |
| 0 | 0 | 0 | 0 |
| 0 | 0 | 0 | 1 |
| 0 | 0 | 0 | 0 |
| 0 | 0 | 0 | 0 |
| 0 | 0 | 0 | 1 |
| 0 | 0 | 0 | 1 |
| 0 | 0 | 0 | 1 |
| 0 | 0 | 0 | 1 |
| 0 | 0 | 0 | 1 |
| 0 | 0 | 0 | 1 |
| 0 | 0 | 0 | 1 |
| 0 | 0 | 0 | 1 |
| 0 | 0 | 0 | 1 |
| 0 | 0 | 0 | 1 |

| | | | |
|---|---|---|---|
| 0 | 0 | 0 | 1 |
| 0 | 0 | 0 | 0 |
| 0 | 0 | 0 | 1 |
| 0 | 0 | 0 | 1 |
| 0 | 0 | 0 | 1 |
| 0 | 0 | 0 | 1 |
| 0 | 0 | 0 | 1 |
| 0 | 0 | 0 | 1 |
| 0 | 0 | 0 | 1 |
| 0 | 0 | 0 | 1 |
| 0 | 0 | 0 | 1 |
| 0 | 0 | 0 | 0 |
| 0 | 0 | 0 | 1 |
| 0 | 0 | 0 | 1 |
| 0 | 0 | 0 | 0 |
| 0 | 0 | 0 | 1 |
| 0 | 0 | 0 | 1 |
| 0 | 0 | 0 | 1 |
| 0 | 0 | 0 | 1 |
| 0 | 0 | 0 | 1 |
| 1 | 0 | 0 | 0 |
| 0 | 0 | 0 | 1 |
| 1 | 0 | 0 | 0 |
| 1 | 0 | 0 | 0 |
| 0 | 0 | 0 | 1 |
| 0 | 0 | 0 | 0 |
| 0 | 0 | 0 | 1 |
| 0 | 0 | 0 | 1 |
| 0 | 0 | 0 | 1 |
| 0 | 0 | 0 | 1 |
| 0 | 0 | 0 | 1 |
| 0 | 0 | 0 | 0 |
| 0 | 0 | 0 | 1 |
| 0 | 0 | 0 | 0 |
| 0 | 0 | 0 | 1 |
| 0 | 0 | 0 | 1 |
| 0 | 0 | 0 | 1 |
| 0 | 0 | 0 | 1 |
| 0 | 0 | 0 | 1 |
| 0 | 0 | 0 | 0 |
| 0 | 0 | 0 | 1 |
| 0 | 0 | 0 | 1 |
| 0 | 0 | 0 | 1 |
| 0 | 0 | 0 | 0 |
| 0 | 0 | 0 | 1 |
| 0 | 0 | 0 | 1 |
| 0 | 0 | 0 | 1 |
| 1 | 0 | 0 | 0 |
| 0 | 0 | 0 | 0 |
| 0 | 0 | 0 | 1 |
| 0 | 0 | 0 | 1 |
| 0 | 0 | 0 | 0 |
| 1 | 0 | 0 | 0 |
| 0 | 0 | 0 | 0 |

| | | | |
|---|---|---|---|
| 1 | 0 | 0 | 0 |
| 0 | 0 | 0 | 1 |
| 0 | 0 | 0 | 0 |
| 0 | 0 | 0 | 0 |
| 0 | 0 | 0 | 0 |
| 0 | 0 | 0 | 1 |
| 0 | 0 | 0 | 1 |
| 0 | 0 | 0 | 1 |
| 0 | 0 | 0 | 1 |
| 0 | 0 | 0 | 0 |
| 0 | 0 | 0 | 1 |
| 0 | 0 | 0 | 1 |
| 0 | 0 | 0 | 1 |
| 0 | 0 | 0 | 1 |
| 0 | 0 | 0 | 0 |
| 0 | 0 | 0 | 1 |
| 0 | 0 | 0 | 1 |
| 0 | 0 | 0 | 0 |
| 0 | 0 | 0 | 1 |
| 0 | 0 | 0 | 0 |
| 0 | 0 | 0 | 0 |
| 0 | 0 | 0 | 1 |
| 0 | 0 | 0 | 0 |
| 0 | 0 | 0 | 1 |
| 0 | 0 | 0 | 0 |
| 0 | 0 | 0 | 0 |
| 0 | 0 | 0 | 1 |
| 0 | 0 | 0 | 1 |
| 1 | 0 | 0 | 0 |
| 0 | 0 | 0 | 1 |
| 0 | 0 | 0 | 1 |
| 0 | 0 | 0 | 1 |
| 0 | 0 | 0 | 1 |
| 0 | 0 | 0 | 1 |
| 0 | 0 | 0 | 0 |
| 0 | 0 | 0 | 0 |
| 0 | 0 | 0 | 1 |
| 0 | 0 | 0 | 0 |
| 0 | 0 | 0 | 0 |
| 0 | 0 | 0 | 0 |
| 0 | 0 | 0 | 0 |
| 0 | 0 | 0 | 0 |
| 0 | 0 | 0 | 0 |
| 0 | 0 | 0 | 1 |
| 0 | 0 | 0 | 0 |
| 0 | 0 | 0 | 1 |
| 0 | 0 | 0 | 1 |
| 0 | 0 | 0 | 0 |
| 0 | 0 | 0 | 0 |
| 0 | 0 | 0 | 0 |
| 0 | 0 | 0 | 0 |
| 0 | 0 | 0 | 0 |
| 0 | 0 | 0 | 1 |
| 0 | 0 | 0 | 1 |

| | | | |
|---|---|---|---|
| 0 | 0 | 0 | 1 |
| 0 | 0 | 0 | 0 |
| 0 | 0 | 0 | 0 |
| 0 | 0 | 0 | 0 |
| 0 | 0 | 0 | 0 |
| 0 | 0 | 0 | 0 |
| 0 | 0 | 0 | 0 |
| 0 | 0 | 0 | 1 |
| 0 | 0 | 0 | 1 |
| 0 | 0 | 0 | 1 |
| 0 | 0 | 0 | 0 |
| 0 | 0 | 0 | 0 |
| 0 | 0 | 0 | 1 |
| 0 | 0 | 0 | 0 |
| 0 | 0 | 0 | 1 |
| 0 | 0 | 0 | 0 |
| 0 | 0 | 0 | 1 |
| 0 | 0 | 0 | 1 |
| 0 | 0 | 0 | 1 |
| 0 | 0 | 0 | 1 |
| 0 | 0 | 0 | 1 |
| 0 | 0 | 0 | 1 |
| 0 | 0 | 0 | 1 |
| 1 | 0 | 0 | 0 |
| 0 | 0 | 0 | 0 |
| 0 | 0 | 0 | 1 |
| 0 | 0 | 0 | 1 |
| 0 | 0 | 0 | 1 |
| 0 | 0 | 0 | 1 |
| 0 | 0 | 0 | 0 |
| 0 | 0 | 0 | 0 |
| 0 | 0 | 0 | 0 |
| 0 | 0 | 0 | 0 |
| 0 | 0 | 0 | 1 |
| 0 | 0 | 0 | 0 |
| 0 | 0 | 0 | 0 |
| 0 | 0 | 0 | 0 |
| 0 | 0 | 0 | 0 |
| 0 | 0 | 0 | 0 |
| 0 | 0 | 0 | 1 |
| 0 | 0 | 0 | 1 |
| 0 | 0 | 0 | 0 |
| 0 | 0 | 0 | 1 |
| 0 | 0 | 0 | 1 |
| 0 | 0 | 0 | 0 |
| 0 | 0 | 0 | 1 |
| 0 | 0 | 0 | 1 |
| 0 | 0 | 0 | 1 |
| 0 | 0 | 0 | 1 |
| 0 | 0 | 0 | 1 |
| 0 | 0 | 0 | 0 |
| 0 | 0 | 0 | 1 |
| 0 | 0 | 0 | 0 |
| 0 | 0 | 0 | 0 |

| | | | |
|---|---|---|---|
| 0 | 0 | 0 | 1 |
| 0 | 0 | 0 | 1 |
| 0 | 0 | 0 | 0 |
| 0 | 0 | 0 | 0 |
| 0 | 0 | 0 | 1 |
| 0 | 0 | 0 | 1 |
| 0 | 0 | 0 | 1 |
| 0 | 0 | 0 | 1 |
| 0 | 0 | 0 | 1 |
| 0 | 0 | 0 | 1 |
| 0 | 0 | 0 | 1 |
| 0 | 0 | 0 | 1 |
| 0 | 0 | 0 | 0 |
| 0 | 0 | 0 | 1 |
| 0 | 0 | 0 | 1 |
| 0 | 0 | 0 | 1 |
| 0 | 0 | 0 | 1 |
| 0 | 0 | 0 | 1 |
| 0 | 0 | 0 | 1 |
| 0 | 0 | 0 | 1 |
| 0 | 0 | 0 | 1 |
| 0 | 0 | 0 | 1 |
| 0 | 0 | 0 | 1 |
| 0 | 0 | 0 | 1 |
| 0 | 0 | 0 | 1 |
| 0 | 0 | 0 | 1 |
| 0 | 0 | 0 | 1 |
| 0 | 0 | 0 | 1 |
| 0 | 0 | 0 | 1 |
| 0 | 0 | 0 | 1 |
| 0 | 0 | 0 | 1 |
| 0 | 0 | 0 | 0 |
| 0 | 0 | 0 | 1 |
| 0 | 0 | 0 | 1 |
| 0 | 0 | 0 | 1 |
| 0 | 0 | 0 | 1 |
| 0 | 0 | 0 | 1 |
| 0 | 0 | 0 | 1 |
| 0 | 0 | 0 | 1 |
| 0 | 0 | 0 | 1 |
| 0 | 0 | 0 | 0 |
| 0 | 0 | 0 | 0 |
| 0 | 0 | 0 | 1 |
| 0 | 0 | 0 | 1 |
| 0 | 0 | 0 | 1 |
| 0 | 0 | 0 | 1 |
| 0 | 0 | 0 | 1 |
| 0 | 0 | 0 | 1 |
| 0 | 0 | 0 | 0 |
| 0 | 0 | 0 | 0 |
| 0 | 0 | 0 | 1 |
| 0 | 0 | 0 | 1 |
| 0 | 0 | 0 | 0 |
| 0 | 0 | 0 | 1 |

| | | | |
|---|---|---|---|
| 0 | 0 | 0 | 0 |
| 0 | 0 | 0 | 0 |
| 0 | 0 | 0 | 0 |
| 0 | 0 | 0 | 1 |
| 0 | 0 | 0 | 1 |
| 0 | 0 | 0 | 1 |
| 0 | 0 | 0 | 1 |
| 0 | 0 | 0 | 0 |
| 0 | 0 | 0 | 0 |
| 0 | 0 | 0 | 0 |
| 0 | 0 | 0 | 0 |
| 0 | 0 | 0 | 1 |
| 0 | 0 | 0 | 0 |
| 0 | 0 | 0 | 1 |
| 0 | 0 | 0 | 1 |
| 0 | 0 | 0 | 1 |
| 0 | 0 | 0 | 1 |
| 0 | 0 | 0 | 0 |
| 0 | 0 | 0 | 1 |
| 0 | 0 | 0 | 1 |
| 0 | 0 | 0 | 1 |
| 0 | 0 | 0 | 1 |
| 0 | 0 | 0 | 1 |
| 0 | 0 | 0 | 1 |
| 0 | 0 | 0 | 1 |
| 0 | 0 | 0 | 1 |
| 0 | 0 | 0 | 1 |
| 0 | 0 | 0 | 1 |
| 0 | 0 | 0 | 1 |
| 0 | 0 | 0 | 1 |
| 0 | 0 | 0 | 0 |
| 0 | 0 | 0 | 0 |
| 0 | 0 | 0 | 1 |
| 0 | 0 | 0 | 1 |
| 0 | 0 | 0 | 1 |
| 0 | 0 | 0 | 1 |
| 0 | 0 | 0 | 1 |
| 0 | 0 | 0 | 1 |
| 0 | 0 | 0 | 1 |
| 0 | 0 | 0 | 1 |
| 0 | 0 | 0 | 1 |
| 0 | 0 | 0 | 0 |
| 0 | 0 | 0 | 1 |
| 0 | 0 | 0 | 1 |
| 0 | 0 | 0 | 0 |
| 0 | 0 | 0 | 0 |
| 0 | 0 | 0 | 1 |
| 0 | 0 | 0 | 1 |
| 0 | 0 | 0 | 0 |
| 0 | 0 | 0 | 0 |
| 0 | 0 | 0 | 0 |
| 0 | 0 | 0 | 0 |
| 0 | 0 | 0 | 0 |
| 0 | 0 | 0 | 0 |

| | | | |
|---|---|---|---|
| 0 | 0 | 0 | 1 |
| 0 | 0 | 0 | 1 |
| 0 | 0 | 0 | 1 |
| 0 | 0 | 0 | 1 |
| 0 | 0 | 0 | 1 |
| 0 | 0 | 0 | 0 |
| 0 | 0 | 0 | 0 |
| 0 | 0 | 0 | 0 |
| 0 | 0 | 0 | 0 |
| 0 | 0 | 0 | 0 |
| 0 | 0 | 0 | 1 |
| 0 | 0 | 0 | 0 |
| 0 | 0 | 0 | 0 |
| 0 | 0 | 0 | 0 |
| 0 | 0 | 0 | 1 |
| 0 | 0 | 0 | 0 |
| 0 | 0 | 0 | 1 |
| 0 | 0 | 0 | 1 |
| 1 | 0 | 0 | 0 |
| 1 | 0 | 0 | 0 |
| 0 | 0 | 0 | 0 |
| 0 | 0 | 0 | 0 |
| 0 | 0 | 0 | 1 |
| 0 | 0 | 0 | 0 |
| 0 | 0 | 0 | 0 |
| 0 | 0 | 0 | 0 |
| 0 | 0 | 0 | 0 |
| 0 | 0 | 0 | 1 |
| 0 | 0 | 0 | 1 |
| 0 | 0 | 0 | 1 |
| 0 | 0 | 0 | 1 |
| 0 | 0 | 0 | 1 |
| 0 | 0 | 0 | 0 |
| 0 | 0 | 0 | 1 |
| 0 | 0 | 0 | 0 |
| 0 | 0 | 0 | 0 |
| 0 | 0 | 0 | 0 |
| 0 | 0 | 0 | 0 |
| 0 | 0 | 0 | 0 |
| 0 | 0 | 0 | 0 |
| 0 | 0 | 0 | 0 |
| 1 | 0 | 0 | 0 |
| 0 | 0 | 0 | 0 |
| 0 | 0 | 0 | 1 |
| 0 | 0 | 0 | 1 |
| 0 | 0 | 0 | 1 |
| 0 | 0 | 0 | 1 |
| 0 | 0 | 0 | 0 |
| 0 | 0 | 0 | 0 |
| 0 | 0 | 0 | 0 |
| 0 | 0 | 0 | 1 |
| 0 | 0 | 0 | 1 |
| 0 | 0 | 0 | 0 |
| 0 | 0 | 0 | 1 |

| | | | |
|---|---|---|---|
| 0 | 0 | 0 | 1 |
| 0 | 0 | 0 | 0 |
| 0 | 0 | 0 | 1 |
| 0 | 0 | 0 | 1 |
| 0 | 0 | 0 | 1 |
| 0 | 0 | 0 | 1 |
| 0 | 0 | 0 | 1 |
| 0 | 0 | 0 | 1 |
| 0 | 0 | 0 | 1 |
| 0 | 0 | 0 | 1 |
| 0 | 0 | 0 | 1 |
| 0 | 0 | 0 | 1 |
| 0 | 0 | 0 | 0 |
| 0 | 0 | 0 | 0 |
| 0 | 0 | 0 | 0 |
| 0 | 0 | 0 | 0 |
| 0 | 0 | 0 | 0 |
| 0 | 0 | 0 | 0 |
| 0 | 0 | 0 | 0 |
| 0 | 0 | 0 | 1 |
| 1 | 1 | 0 | 0 |
| 0 | 0 | 0 | 0 |
| 0 | 0 | 0 | 0 |
| 0 | 0 | 0 | 0 |
| 0 | 0 | 0 | 0 |
| 0 | 0 | 0 | 1 |
| 0 | 0 | 0 | 1 |
| 0 | 0 | 0 | 0 |
| 0 | 0 | 0 | 1 |
| 0 | 0 | 0 | 0 |
| 0 | 0 | 0 | 0 |
| 0 | 0 | 0 | 1 |
| 0 | 0 | 0 | 0 |
| 0 | 0 | 0 | 1 |
| 0 | 0 | 0 | 1 |
| 0 | 0 | 0 | 1 |
| 0 | 0 | 0 | 1 |
| 0 | 0 | 0 | 1 |
| 0 | 0 | 0 | 1 |
| 0 | 0 | 0 | 1 |
| 0 | 0 | 0 | 0 |
| 0 | 0 | 0 | 1 |
| 0 | 0 | 0 | 1 |
| 0 | 0 | 0 | 0 |
| 0 | 0 | 0 | 0 |
| 0 | 0 | 0 | 1 |
| 0 | 0 | 0 | 0 |
| 0 | 0 | 0 | 0 |
| 0 | 0 | 0 | 1 |
| 0 | 0 | 0 | 1 |
| 0 | 0 | 0 | 0 |
| 0 | 0 | 0 | 1 |
| 0 | 0 | 0 | 1 |
| 0 | 0 | 0 | 1 |

| | | | |
|---|---|---|---|
| 0 | 0 | 0 | 1 |
| 0 | 0 | 0 | 1 |
| 0 | 0 | 0 | 1 |
| 0 | 0 | 0 | 1 |
| 0 | 0 | 0 | 0 |
| 0 | 0 | 0 | 0 |
| 0 | 0 | 0 | 1 |
| 0 | 0 | 0 | 1 |
| 0 | 0 | 0 | 1 |
| 0 | 0 | 0 | 1 |
| 0 | 0 | 0 | 1 |
| 0 | 0 | 0 | 0 |
| 0 | 0 | 0 | 0 |
| 0 | 0 | 0 | 1 |
| 0 | 0 | 0 | 1 |
| 0 | 0 | 0 | 0 |
| 0 | 0 | 0 | 1 |
| 0 | 0 | 0 | 1 |
| 0 | 0 | 0 | 0 |
| 0 | 0 | 0 | 1 |
| 0 | 0 | 0 | 1 |
| 0 | 0 | 0 | 1 |
| 0 | 0 | 0 | 1 |
| 0 | 0 | 0 | 0 |
| 0 | 0 | 0 | 1 |
| 0 | 0 | 0 | 0 |
| 0 | 0 | 0 | 1 |
| 0 | 0 | 0 | 1 |
| 1 | 0 | 0 | 0 |
| 1 | 0 | 0 | 0 |
| 0 | 0 | 0 | 0 |
| 0 | 0 | 0 | 0 |
| 0 | 0 | 0 | 0 |
| 0 | 0 | 0 | 0 |
| 0 | 0 | 0 | 1 |
| 0 | 0 | 0 | 1 |
| 0 | 0 | 0 | 1 |
| 1 | 0 | 0 | 0 |
| 0 | 0 | 0 | 1 |
| 0 | 0 | 0 | 1 |
| 0 | 0 | 0 | 0 |
| 0 | 0 | 0 | 0 |
| 0 | 0 | 0 | 0 |
| 0 | 0 | 0 | 1 |
| 0 | 0 | 0 | 0 |
| 0 | 0 | 0 | 1 |
| 0 | 0 | 0 | 1 |
| 0 | 0 | 0 | 0 |
| 0 | 0 | 0 | 1 |
| 0 | 0 | 0 | 1 |
| 0 | 0 | 0 | 1 |
| 0 | 0 | 0 | 0 |
| 0 | 0 | 0 | 1 |
| 0 | 0 | 0 | 0 |

| | | | |
|---|---|---|---|
| 0 | 0 | 0 | 1 |
| 0 | 0 | 0 | 0 |
| 0 | 0 | 0 | 0 |
| 0 | 0 | 0 | 1 |
| 0 | 0 | 0 | 1 |
| 0 | 0 | 0 | 1 |
| 0 | 0 | 0 | 0 |
| 0 | 0 | 0 | 1 |
| 0 | 0 | 0 | 1 |
| 0 | 0 | 0 | 1 |
| 0 | 0 | 0 | 1 |
| 0 | 0 | 0 | 1 |
| 0 | 0 | 0 | 1 |
| 0 | 0 | 0 | 1 |
| 0 | 0 | 0 | 1 |
| 0 | 0 | 0 | 1 |
| 0 | 0 | 0 | 1 |
| 0 | 0 | 0 | 1 |
| 0 | 0 | 0 | 1 |
| 0 | 0 | 0 | 1 |
| 0 | 0 | 0 | 1 |
| 0 | 0 | 0 | 0 |
| 0 | 0 | 0 | 1 |
| 0 | 0 | 0 | 1 |
| 0 | 0 | 0 | 1 |
| 0 | 0 | 0 | 1 |
| 0 | 0 | 0 | 0 |
| 0 | 0 | 0 | 1 |
| 0 | 0 | 0 | 0 |
| 0 | 0 | 0 | 1 |
| 0 | 0 | 0 | 1 |
| 0 | 0 | 0 | 0 |
| 0 | 0 | 0 | 1 |
| 0 | 0 | 0 | 1 |
| 0 | 0 | 0 | 1 |
| 0 | 0 | 0 | 1 |
| 0 | 0 | 0 | 1 |
| 0 | 0 | 0 | 1 |
| 0 | 0 | 0 | 0 |
| 0 | 0 | 0 | 0 |
| 0 | 0 | 0 | 0 |
| 0 | 0 | 0 | 1 |
| 0 | 0 | 0 | 1 |
| 0 | 0 | 0 | 1 |
| 0 | 0 | 0 | 1 |
| 0 | 0 | 0 | 0 |
| 0 | 0 | 0 | 1 |
| 0 | 0 | 0 | 0 |
| 0 | 0 | 0 | 1 |
| 0 | 0 | 0 | 0 |
| 0 | 0 | 0 | 0 |
| 0 | 0 | 0 | 0 |
| 0 | 0 | 0 | 1 |
| 0 | 0 | 0 | 1 |

| | | | |
|---|---|---|---|
| 0 | 0 | 0 | 0 |
| 0 | 0 | 0 | 0 |
| 0 | 0 | 0 | 1 |
| 0 | 0 | 0 | 1 |
| 0 | 0 | 0 | 1 |
| 0 | 0 | 0 | 1 |
| 0 | 0 | 0 | 1 |
| 0 | 0 | 0 | 1 |
| 0 | 0 | 0 | 1 |
| 0 | 0 | 0 | 1 |
| 0 | 0 | 0 | 1 |
| 0 | 0 | 0 | 1 |
| 0 | 0 | 0 | 1 |
| 0 | 0 | 0 | 1 |
| 0 | 0 | 0 | 1 |
| 0 | 0 | 0 | 0 |
| 0 | 0 | 0 | 0 |
| 0 | 0 | 0 | 0 |
| 0 | 0 | 0 | 1 |
| 0 | 0 | 0 | 1 |
| 0 | 0 | 0 | 1 |
| 0 | 0 | 0 | 0 |
| 0 | 0 | 0 | 0 |
| 0 | 0 | 0 | 0 |
| 0 | 0 | 0 | 0 |
| 0 | 0 | 0 | 1 |
| 0 | 0 | 0 | 1 |
| 0 | 0 | 0 | 1 |
| 0 | 0 | 0 | 1 |
| 0 | 0 | 0 | 1 |
| 0 | 0 | 0 | 0 |
| 0 | 0 | 0 | 1 |
| 0 | 0 | 0 | 0 |
| 0 | 0 | 0 | 1 |
| 0 | 1 | 1 | 1 |
| 0 | 1 | 1 | 1 |
| 0 | 0 | 0 | 0 |
| 0 | 0 | 0 | 0 |
| 0 | 0 | 0 | 1 |
| 0 | 0 | 0 | 1 |
| 0 | 0 | 0 | 1 |
| 0 | 0 | 0 | 0 |
| 0 | 0 | 0 | 1 |
| 0 | 0 | 0 | 1 |
| 0 | 0 | 0 | 0 |
| 0 | 0 | 0 | 1 |
| 0 | 0 | 0 | 0 |
| 0 | 0 | 0 | 0 |
| 0 | 0 | 0 | 1 |
| 0 | 0 | 0 | 1 |
| 0 | 0 | 0 | 0 |
| 0 | 0 | 0 | 0 |
| 0 | 0 | 0 | 1 |
| 0 | 0 | 0 | 0 |

| | | | |
|---|---|---|---|
| 0 | 0 | 0 | 1 |
| 0 | 0 | 0 | 0 |
| 0 | 0 | 0 | 1 |
| 0 | 0 | 0 | 0 |
| 0 | 0 | 0 | 1 |
| 0 | 0 | 0 | 1 |
| 0 | 0 | 0 | 1 |
| 0 | 0 | 0 | 1 |
| 0 | 0 | 0 | 0 |
| 0 | 0 | 0 | 1 |
| 0 | 0 | 0 | 1 |
| 0 | 0 | 0 | 1 |
| 0 | 0 | 0 | 0 |
| 0 | 0 | 0 | 0 |
| 0 | 0 | 0 | 0 |
| 0 | 0 | 0 | 1 |
| 0 | 0 | 0 | 0 |
| 0 | 0 | 0 | 1 |
| 0 | 1 | 0 | 0 |
| 0 | 0 | 0 | 0 |
| 0 | 0 | 0 | 0 |
| 0 | 0 | 0 | 0 |
| 0 | 0 | 0 | 0 |
| 1 | 0 | 0 | 0 |
| 0 | 0 | 0 | 1 |
| 0 | 0 | 0 | 1 |
| 0 | 0 | 0 | 0 |
| 0 | 0 | 0 | 0 |
| 0 | 0 | 0 | 0 |
| 0 | 0 | 0 | 0 |
| 0 | 0 | 0 | 1 |
| 0 | 0 | 0 | 1 |
| 0 | 0 | 0 | 1 |
| 0 | 0 | 0 | 1 |
| 0 | 0 | 0 | 1 |
| 1 | 0 | 0 | 0 |
| 1 | 0 | 0 | 0 |
| 0 | 0 | 0 | 1 |
| 0 | 0 | 0 | 1 |
| 0 | 0 | 0 | 1 |
| 0 | 0 | 0 | 1 |
| 0 | 0 | 0 | 1 |
| 0 | 0 | 0 | 1 |
| 0 | 0 | 0 | 0 |
| 0 | 0 | 0 | 0 |
| 0 | 0 | 0 | 0 |
| 0 | 0 | 0 | 1 |
| 0 | 0 | 0 | 0 |
| 0 | 0 | 0 | 0 |
| 0 | 0 | 0 | 0 |
| 0 | 0 | 0 | 1 |
| 0 | 0 | 0 | 1 |
| 0 | 0 | 0 | 1 |
| 0 | 0 | 0 | 1 |

| | | | |
|---|---|---|---|
| 0 | 0 | 0 | 1 |
| 0 | 0 | 0 | 1 |
| 0 | 0 | 0 | 1 |
| 0 | 0 | 0 | 1 |
| 0 | 0 | 0 | 0 |
| 0 | 0 | 0 | 1 |
| 0 | 0 | 0 | 0 |
| 0 | 0 | 0 | 0 |
| 0 | 0 | 0 | 0 |
| 0 | 0 | 0 | 0 |
| 0 | 0 | 0 | 0 |
| 0 | 0 | 0 | 0 |
| 0 | 0 | 0 | 1 |
| 0 | 0 | 0 | 0 |
| 0 | 0 | 0 | 0 |
| 0 | 0 | 0 | 1 |
| 0 | 0 | 0 | 1 |
| 0 | 0 | 0 | 1 |
| 0 | 0 | 0 | 0 |
| 0 | 0 | 0 | 0 |
| 0 | 0 | 0 | 0 |
| 0 | 0 | 0 | 0 |
| 0 | 0 | 0 | 1 |
| 0 | 0 | 0 | 1 |
| 0 | 0 | 0 | 1 |
| 0 | 0 | 0 | 1 |
| 0 | 0 | 0 | 1 |
| 0 | 0 | 0 | 0 |
| 0 | 0 | 0 | 0 |
| 0 | 0 | 0 | 0 |
| 0 | 0 | 0 | 0 |
| 0 | 0 | 0 | 0 |
| 0 | 0 | 0 | 0 |
| 0 | 0 | 0 | 1 |
| 0 | 0 | 0 | 1 |
| 0 | 0 | 0 | 0 |
| 0 | 0 | 0 | 1 |
| 0 | 0 | 0 | 1 |
| 0 | 0 | 0 | 1 |
| 0 | 0 | 0 | 1 |
| 0 | 0 | 0 | 0 |
| 0 | 0 | 0 | 1 |
| 0 | 0 | 0 | 0 |
| 0 | 0 | 0 | 1 |
| 0 | 0 | 0 | 1 |
| 0 | 0 | 0 | 1 |
| 0 | 0 | 0 | 1 |
| 0 | 0 | 0 | 0 |
| 0 | 0 | 0 | 1 |
| 0 | 0 | 0 | 1 |
| 0 | 0 | 0 | 1 |
| 0 | 0 | 0 | 1 |
| 0 | 0 | 0 | 1 |
| 0 | 0 | 0 | 1 |

| | | | |
|---|---|---|---|
| 0 | 0 | 0 | 1 |
| 0 | 0 | 0 | 1 |
| 0 | 0 | 0 | 1 |
| 0 | 0 | 0 | 1 |
| 0 | 0 | 0 | 1 |
| 0 | 0 | 0 | 1 |
| 0 | 0 | 0 | 1 |
| 0 | 0 | 0 | 1 |
| 0 | 0 | 0 | 1 |
| 0 | 0 | 0 | 1 |
| 0 | 0 | 0 | 1 |
| 0 | 0 | 0 | 0 |
| 0 | 0 | 0 | 1 |
| 0 | 0 | 0 | 1 |
| 0 | 0 | 0 | 1 |
| 0 | 0 | 0 | 1 |
| 0 | 0 | 0 | 1 |
| 0 | 0 | 0 | 0 |
| 0 | 0 | 0 | 0 |
| 0 | 0 | 0 | 0 |
| 0 | 0 | 0 | 0 |
| 0 | 0 | 0 | 1 |
| 1 | 0 | 0 | 0 |
| 0 | 0 | 0 | 1 |
| 0 | 0 | 0 | 1 |
| 0 | 0 | 0 | 1 |
| 0 | 0 | 0 | 1 |
| 0 | 0 | 0 | 1 |
| 0 | 0 | 0 | 0 |
| 0 | 0 | 0 | 0 |
| 0 | 0 | 0 | 1 |
| 0 | 0 | 0 | 1 |
| 1 | 0 | 0 | 0 |
| 0 | 0 | 0 | 1 |
| 0 | 0 | 0 | 0 |
| 1 | 0 | 0 | 0 |
| 1 | 0 | 0 | 0 |
| 0 | 0 | 0 | 1 |
| 1 | 0 | 0 | 0 |
| 0 | 0 | 0 | 1 |
| 1 | 0 | 0 | 0 |
| 0 | 0 | 0 | 1 |
| 0 | 0 | 0 | 0 |
| 0 | 0 | 0 | 1 |
| 0 | 0 | 0 | 1 |
| 0 | 0 | 0 | 1 |
| 0 | 0 | 0 | 1 |
| 0 | 0 | 0 | 1 |
| 0 | 0 | 0 | 1 |
| 0 | 0 | 0 | 0 |
| 0 | 0 | 0 | 1 |
| 1 | 0 | 0 | 0 |
| 0 | 0 | 0 | 1 |
| 0 | 0 | 0 | 1 |

| | | | |
|---|---|---|---|
| 0 | 0 | 0 | 0 |
| 0 | 0 | 0 | 0 |
| 0 | 0 | 0 | 0 |
| 0 | 0 | 0 | 0 |
| 0 | 0 | 0 | 0 |
| 0 | 0 | 0 | 1 |
| 0 | 0 | 0 | 0 |
| 0 | 0 | 0 | 1 |
| 0 | 0 | 0 | 1 |
| 0 | 0 | 0 | 0 |
| 0 | 0 | 0 | 0 |
| 0 | 0 | 0 | 0 |
| 0 | 0 | 0 | 1 |
| 0 | 0 | 0 | 1 |
| 0 | 0 | 0 | 1 |
| 0 | 0 | 0 | 1 |
| 0 | 0 | 0 | 1 |
| 0 | 0 | 0 | 1 |
| 0 | 0 | 0 | 1 |
| 0 | 0 | 0 | 1 |
| 0 | 0 | 0 | 1 |
| 0 | 0 | 0 | 1 |
| 0 | 0 | 0 | 1 |
| 0 | 0 | 0 | 1 |
| 0 | 0 | 0 | 1 |
| 0 | 0 | 0 | 1 |
| 0 | 0 | 0 | 1 |
| 0 | 0 | 0 | 1 |
| 0 | 0 | 0 | 1 |
| 0 | 0 | 0 | 1 |
| 0 | 0 | 0 | 1 |
| 0 | 0 | 0 | 1 |
| 0 | 0 | 0 | 1 |
| 0 | 0 | 0 | 1 |
| 0 | 0 | 0 | 1 |
| 0 | 0 | 0 | 1 |
| 0 | 0 | 0 | 1 |
| 0 | 0 | 0 | 1 |
| 0 | 0 | 0 | 1 |
| 0 | 0 | 0 | 1 |
| 0 | 0 | 0 | 1 |
| 0 | 0 | 0 | 1 |
| 0 | 0 | 0 | 1 |
| 0 | 0 | 0 | 1 |
| 0 | 0 | 0 | 1 |
| 0 | 0 | 0 | 0 |
| 0 | 0 | 0 | 1 |
| 0 | 0 | 0 | 1 |
| 0 | 0 | 0 | 1 |
| 0 | 0 | 0 | 1 |
| 0 | 0 | 0 | 0 |
| 0 | 0 | 0 | 1 |
| 0 | 0 | 0 | 1 |
| 0 | 0 | 0 | 1 |

| | | | |
|---|---|---|---|
| 0 | 0 | 0 | 1 |
| 0 | 0 | 0 | 1 |
| 0 | 0 | 0 | 1 |
| 0 | 0 | 0 | 1 |
| 0 | 0 | 0 | 1 |
| 0 | 0 | 0 | 1 |
| 0 | 0 | 0 | 1 |
| 0 | 0 | 0 | 0 |
| 0 | 0 | 0 | 0 |
| 0 | 0 | 0 | 1 |
| 0 | 0 | 0 | 1 |
| 0 | 0 | 0 | 1 |
| 0 | 0 | 0 | 1 |
| 0 | 0 | 0 | 1 |
| 0 | 0 | 0 | 0 |
| 0 | 0 | 0 | 0 |
| 0 | 0 | 0 | 0 |
| 0 | 0 | 0 | 1 |
| 0 | 0 | 0 | 1 |
| 0 | 0 | 0 | 1 |
| 0 | 0 | 0 | 0 |
| 0 | 0 | 0 | 0 |
| 0 | 0 | 0 | 0 |
| 0 | 0 | 0 | 0 |
| 0 | 0 | 0 | 0 |
| 0 | 0 | 0 | 0 |
| 0 | 0 | 0 | 1 |
| 0 | 0 | 0 | 1 |
| 0 | 0 | 0 | 0 |
| 0 | 0 | 0 | 1 |
| 0 | 0 | 0 | 1 |
| 0 | 0 | 0 | 1 |
| 0 | 0 | 0 | 0 |
| 0 | 0 | 0 | 1 |
| 0 | 0 | 0 | 1 |
| 0 | 0 | 0 | 1 |
| 0 | 0 | 0 | 0 |
| 0 | 0 | 0 | 0 |
| 0 | 0 | 0 | 0 |
| 0 | 0 | 0 | 1 |
| 0 | 0 | 0 | 1 |
| 0 | 0 | 0 | 1 |
| 0 | 0 | 0 | 1 |
| 0 | 0 | 0 | 1 |
| 0 | 0 | 0 | 1 |
| 0 | 0 | 0 | 0 |
| 0 | 0 | 0 | 1 |
| 0 | 0 | 0 | 1 |
| 0 | 0 | 0 | 1 |
| 0 | 0 | 0 | 1 |
| 0 | 0 | 0 | 1 |
| 0 | 0 | 0 | 1 |
| 0 | 0 | 0 | 1 |
| 0 | 0 | 0 | 1 |

| | | | |
|---|---|---|---|
| 0 | 0 | 0 | 1 |
| 0 | 0 | 0 | 1 |
| 0 | 0 | 0 | 1 |
| 0 | 0 | 0 | 0 |
| 0 | 0 | 0 | 1 |
| 0 | 0 | 0 | 1 |
| 0 | 0 | 0 | 1 |
| 0 | 0 | 0 | 0 |
| 0 | 0 | 0 | 0 |
| 0 | 0 | 0 | 1 |
| 0 | 0 | 0 | 1 |
| 0 | 0 | 0 | 1 |
| 0 | 0 | 0 | 0 |
| 0 | 0 | 0 | 1 |
| 0 | 0 | 0 | 1 |
| 0 | 0 | 0 | 1 |
| 0 | 0 | 0 | 0 |
| 0 | 0 | 0 | 1 |
| 0 | 0 | 0 | 1 |
| 0 | 0 | 0 | 1 |
| 0 | 0 | 0 | 1 |
| 0 | 0 | 0 | 1 |
| 0 | 0 | 0 | 1 |
| 0 | 0 | 0 | 1 |
| 0 | 0 | 0 | 1 |
| 0 | 0 | 0 | 1 |
| 0 | 0 | 0 | 1 |
| 0 | 0 | 0 | 0 |
| 0 | 0 | 0 | 0 |
| 0 | 0 | 0 | 0 |
| 0 | 0 | 0 | 0 |
| 0 | 0 | 0 | 0 |
| 0 | 0 | 0 | 0 |
| 0 | 0 | 0 | 1 |
| 0 | 0 | 0 | 0 |
| 0 | 0 | 0 | 0 |
| 0 | 0 | 0 | 0 |
| 0 | 0 | 0 | 1 |
| 0 | 0 | 0 | 1 |
| 0 | 0 | 0 | 1 |
| 0 | 0 | 0 | 1 |
| 0 | 0 | 0 | 1 |
| 0 | 0 | 0 | 0 |
| 0 | 0 | 0 | 0 |
| 0 | 0 | 0 | 1 |
| 0 | 0 | 0 | 1 |
| 0 | 0 | 0 | 0 |
| 0 | 0 | 0 | 1 |
| 0 | 0 | 0 | 1 |
| 0 | 0 | 0 | 1 |
| 0 | 0 | 0 | 1 |
| 0 | 0 | 0 | 1 |
| 0 | 0 | 0 | 0 |
| 0 | 0 | 0 | 0 |

| | | | |
|---|---|---|---|
| 0 | 0 | 0 | 0 |
| 0 | 0 | 0 | 0 |
| 0 | 0 | 0 | 0 |
| 0 | 0 | 0 | 0 |
| 0 | 0 | 0 | 0 |
| 0 | 0 | 0 | 0 |
| 0 | 0 | 0 | 0 |
| 0 | 0 | 0 | 0 |
| 0 | 0 | 0 | 1 |
| 0 | 0 | 0 | 0 |
| 0 | 0 | 0 | 1 |
| 0 | 0 | 0 | 1 |
| 0 | 0 | 0 | 1 |
| 0 | 0 | 0 | 1 |
| 0 | 0 | 0 | 0 |
| 0 | 0 | 0 | 1 |
| 0 | 0 | 0 | 1 |
| 0 | 0 | 0 | 1 |
| 0 | 0 | 0 | 1 |
| 0 | 0 | 0 | 0 |
| 0 | 0 | 0 | 0 |
| 0 | 0 | 0 | 1 |
| 0 | 0 | 0 | 0 |
| 0 | 0 | 0 | 0 |
| 0 | 0 | 0 | 0 |
| 0 | 0 | 0 | 0 |
| 0 | 0 | 0 | 0 |
| 0 | 0 | 0 | 1 |
| 0 | 0 | 0 | 1 |
| 0 | 0 | 0 | 1 |
| 0 | 0 | 0 | 0 |
| 0 | 0 | 0 | 1 |
| 0 | 0 | 0 | 1 |
| 0 | 0 | 0 | 1 |
| 0 | 0 | 0 | 1 |
| 0 | 0 | 0 | 1 |
| 0 | 0 | 0 | 1 |
| 0 | 0 | 0 | 1 |
| 0 | 0 | 0 | 1 |
| 0 | 0 | 0 | 1 |
| 0 | 0 | 0 | 1 |
| 0 | 0 | 0 | 0 |
| 0 | 0 | 0 | 1 |
| 0 | 0 | 0 | 1 |
| 0 | 0 | 0 | 1 |
| 0 | 0 | 0 | 1 |
| 0 | 0 | 0 | 1 |
| 0 | 0 | 0 | 1 |
| 0 | 0 | 0 | 0 |
| 0 | 0 | 0 | 1 |
| 0 | 0 | 0 | 1 |
| 0 | 0 | 0 | 1 |
| 0 | 0 | 0 | 0 |
| 0 | 0 | 0 | 0 |

| | | | |
|---|---|---|---|
| 0 | 0 | 0 | 1 |
| 0 | 0 | 0 | 1 |
| 0 | 0 | 0 | 1 |
| 0 | 0 | 0 | 1 |
| 0 | 0 | 0 | 0 |
| 1 | 0 | 0 | 0 |
| 0 | 0 | 0 | 1 |
| 0 | 0 | 0 | 0 |
| 0 | 0 | 0 | 1 |
| 0 | 0 | 0 | 1 |
| 0 | 0 | 0 | 0 |
| 0 | 0 | 0 | 1 |
| 0 | 0 | 0 | 0 |
| 0 | 0 | 0 | 0 |
| 0 | 0 | 0 | 0 |
| 0 | 0 | 0 | 0 |
| 0 | 0 | 0 | 1 |
| 0 | 0 | 0 | 1 |
| 0 | 0 | 0 | 0 |
| 0 | 0 | 0 | 0 |
| 0 | 0 | 0 | 0 |
| 0 | 0 | 0 | 0 |
| 0 | 0 | 0 | 0 |
| 0 | 0 | 0 | 1 |
| 0 | 0 | 0 | 1 |
| 0 | 0 | 0 | 0 |
| 0 | 0 | 0 | 1 |
| 0 | 0 | 0 | 0 |
| 0 | 0 | 0 | 0 |
| 0 | 0 | 0 | 0 |
| 0 | 0 | 0 | 1 |
| 1 | 0 | 0 | 0 |
| 1 | 0 | 0 | 0 |
| 0 | 0 | 0 | 1 |
| 0 | 0 | 0 | 1 |
| 0 | 0 | 0 | 0 |
| 0 | 0 | 0 | 0 |
| 0 | 0 | 0 | 1 |
| 0 | 0 | 0 | 0 |
| 0 | 0 | 0 | 0 |
| 0 | 1 | 0 | 0 |
| 0 | 0 | 0 | 0 |
| 0 | 0 | 0 | 1 |
| 0 | 0 | 0 | 1 |
| 0 | 0 | 0 | 0 |
| 0 | 0 | 0 | 1 |
| 0 | 0 | 0 | 0 |
| 0 | 0 | 0 | 1 |
| 0 | 0 | 0 | 1 |
| 0 | 0 | 0 | 1 |
| 0 | 0 | 0 | 1 |
| 0 | 0 | 0 | 1 |
| 0 | 0 | 0 | 1 |
| 0 | 0 | 0 | 1 |

| | | | |
|---|---|---|---|
| 0 | 0 | 0 | 1 |
| 0 | 0 | 0 | 1 |
| 0 | 0 | 0 | 1 |
| 0 | 0 | 0 | 1 |
| 0 | 0 | 0 | 1 |
| 0 | 0 | 0 | 1 |
| 0 | 0 | 0 | 1 |
| 0 | 0 | 0 | 0 |
| 0 | 0 | 0 | 1 |
| 1 | 0 | 0 | 0 |
| 0 | 0 | 0 | 0 |
| 0 | 0 | 0 | 0 |
| 0 | 0 | 0 | 1 |
| 0 | 0 | 0 | 1 |
| 0 | 0 | 0 | 0 |
| 0 | 0 | 0 | 1 |
| 0 | 0 | 0 | 1 |
| 0 | 0 | 0 | 1 |
| 0 | 0 | 0 | 1 |
| 0 | 0 | 0 | 1 |
| 0 | 0 | 0 | 1 |
| 0 | 0 | 0 | 1 |
| 0 | 0 | 0 | 1 |
| 0 | 0 | 0 | 1 |
| 0 | 0 | 0 | 1 |
| 0 | 0 | 0 | 1 |
| 0 | 0 | 0 | 1 |
| 0 | 0 | 0 | 1 |
| 0 | 0 | 0 | 1 |
| 0 | 0 | 0 | 0 |
| 0 | 0 | 0 | 0 |
| 0 | 0 | 0 | 1 |
| 0 | 0 | 0 | 1 |
| 0 | 0 | 0 | 1 |
| 0 | 0 | 0 | 1 |
| 0 | 0 | 0 | 0 |
| 0 | 0 | 0 | 1 |
| 0 | 0 | 0 | 1 |
| 0 | 0 | 0 | 1 |
| 0 | 0 | 0 | 1 |
| 0 | 0 | 0 | 1 |
| 0 | 0 | 0 | 0 |
| 0 | 0 | 0 | 0 |
| 0 | 0 | 0 | 0 |
| 0 | 0 | 0 | 1 |
| 0 | 0 | 0 | 1 |
| 0 | 0 | 0 | 1 |
| 0 | 0 | 0 | 1 |
| 0 | 0 | 0 | 1 |
| 0 | 0 | 0 | 1 |
| 0 | 0 | 0 | 1 |
| 0 | 0 | 0 | 1 |
| 0 | 0 | 0 | 1 |
| 0 | 0 | 0 | 0 |

| | | | |
|---|---|---|---|
| 0 | 0 | 0 | 1 |
| 0 | 0 | 0 | 1 |
| 0 | 0 | 0 | 1 |
| 0 | 0 | 0 | 1 |
| 0 | 0 | 0 | 1 |
| 0 | 0 | 0 | 1 |
| 0 | 0 | 0 | 1 |
| 0 | 0 | 0 | 1 |
| 0 | 0 | 0 | 0 |
| 0 | 0 | 0 | 0 |
| 0 | 0 | 0 | 1 |
| 0 | 0 | 0 | 1 |
| 0 | 0 | 0 | 1 |
| 0 | 0 | 0 | 0 |
| 0 | 0 | 0 | 1 |
| 0 | 0 | 0 | 1 |
| 0 | 0 | 0 | 1 |
| 1 | 0 | 0 | 0 |
| 1 | 0 | 0 | 0 |
| 0 | 0 | 0 | 0 |
| 0 | 0 | 0 | 0 |
| 0 | 0 | 0 | 1 |
| 0 | 0 | 0 | 1 |
| 0 | 0 | 0 | 1 |
| 0 | 0 | 0 | 1 |
| 1 | 0 | 0 | 0 |
| 0 | 0 | 0 | 1 |
| 0 | 0 | 0 | 1 |
| 0 | 0 | 0 | 1 |
| 0 | 0 | 0 | 1 |
| 0 | 0 | 0 | 1 |
| 0 | 0 | 0 | 1 |
| 0 | 0 | 0 | 1 |
| 0 | 0 | 0 | 1 |
| 0 | 0 | 0 | 1 |
| 0 | 0 | 0 | 1 |
| 0 | 0 | 0 | 1 |
| 0 | 0 | 0 | 1 |
| 0 | 0 | 0 | 1 |
| 0 | 0 | 0 | 0 |
| 0 | 0 | 0 | 1 |
| 0 | 0 | 0 | 1 |
| 0 | 0 | 0 | 0 |
| 0 | 0 | 0 | 1 |
| 0 | 0 | 0 | 1 |
| 0 | 0 | 0 | 1 |
| 0 | 0 | 0 | 0 |
| 0 | 0 | 0 | 0 |
| 0 | 0 | 0 | 1 |
| 0 | 0 | 0 | 1 |
| 0 | 0 | 0 | 0 |
| 0 | 0 | 0 | 1 |
| 0 | 0 | 0 | 1 |
| 0 | 0 | 0 | 1 |

| | | | |
|---|---|---|---|
| 0 | 0 | 0 | 1 |
| 0 | 0 | 0 | 1 |
| 0 | 0 | 0 | 1 |
| 0 | 0 | 0 | 1 |
| 0 | 0 | 0 | 1 |
| 0 | 0 | 0 | 1 |
| 0 | 0 | 0 | 1 |
| 0 | 0 | 0 | 1 |
| 0 | 0 | 0 | 1 |
| 0 | 0 | 0 | 1 |
| 0 | 0 | 0 | 1 |
| 0 | 0 | 0 | 1 |
| 0 | 0 | 0 | 1 |
| 0 | 0 | 0 | 0 |
| 0 | 0 | 0 | 0 |
| 0 | 0 | 0 | 0 |
| 0 | 0 | 0 | 0 |
| 0 | 0 | 0 | 1 |
| 0 | 0 | 0 | 1 |
| 0 | 0 | 0 | 0 |
| 0 | 0 | 0 | 1 |
| 0 | 0 | 0 | 1 |
| 0 | 0 | 0 | 1 |
| 0 | 0 | 0 | 1 |
| 0 | 0 | 0 | 1 |
| 0 | 0 | 0 | 0 |
| 0 | 0 | 0 | 1 |
| 0 | 0 | 0 | 1 |
| 0 | 0 | 0 | 1 |
| 0 | 0 | 0 | 0 |
| 0 | 0 | 0 | 0 |
| 0 | 0 | 0 | 1 |
| 0 | 0 | 0 | 1 |
| 0 | 0 | 0 | 1 |
| 0 | 0 | 0 | 1 |
| 0 | 0 | 0 | 1 |
| 0 | 0 | 0 | 1 |
| 0 | 0 | 0 | 1 |
| 0 | 0 | 0 | 0 |
| 0 | 0 | 0 | 1 |
| 0 | 0 | 0 | 1 |
| 0 | 0 | 0 | 0 |
| 0 | 0 | 0 | 0 |
| 0 | 0 | 0 | 0 |
| 0 | 0 | 0 | 0 |
| 0 | 0 | 0 | 1 |
| 0 | 0 | 0 | 0 |
| 0 | 0 | 0 | 1 |
| 0 | 0 | 0 | 1 |
| 0 | 0 | 0 | 1 |
| 0 | 0 | 0 | 1 |
| 0 | 0 | 0 | 1 |
| 0 | 0 | 0 | 1 |
| 0 | 0 | 0 | 1 |

| | | | |
|---|---|---|---|
| 0 | 0 | 0 | 1 |
| 0 | 0 | 0 | 1 |
| 0 | 0 | 0 | 0 |
| 0 | 0 | 0 | 1 |
| 0 | 0 | 0 | 1 |
| 0 | 0 | 0 | 1 |
| 0 | 0 | 0 | 1 |
| 0 | 0 | 0 | 1 |
| 0 | 0 | 0 | 1 |
| 0 | 0 | 0 | 1 |
| 0 | 0 | 0 | 1 |
| 0 | 0 | 0 | 1 |
| 0 | 0 | 0 | 1 |
| 0 | 0 | 0 | 1 |
| 0 | 0 | 0 | 1 |
| 0 | 0 | 0 | 1 |
| 0 | 0 | 0 | 1 |
| 0 | 0 | 0 | 1 |
| 0 | 0 | 0 | 1 |
| 0 | 0 | 0 | 1 |
| 0 | 0 | 0 | 1 |
| 0 | 0 | 0 | 1 |
| 0 | 0 | 0 | 1 |
| 0 | 0 | 0 | 1 |
| 0 | 0 | 0 | 1 |
| 0 | 0 | 0 | 1 |
| 0 | 0 | 0 | 1 |
| 0 | 0 | 0 | 1 |
| 0 | 0 | 0 | 1 |
| 0 | 0 | 0 | 1 |
| 1 | 0 | 0 | 0 |
| 0 | 0 | 0 | 1 |
| 0 | 0 | 0 | 0 |
| 0 | 0 | 0 | 0 |
| 0 | 0 | 0 | 0 |
| 0 | 0 | 0 | 0 |
| 0 | 0 | 0 | 1 |
| 0 | 0 | 0 | 0 |
| 0 | 0 | 0 | 1 |
| 0 | 0 | 0 | 1 |
| 1 | 0 | 0 | 0 |
| 1 | 0 | 0 | 0 |
| 0 | 0 | 0 | 0 |
| 0 | 0 | 0 | 1 |
| 0 | 0 | 0 | 0 |
| 0 | 0 | 0 | 0 |
| 0 | 0 | 0 | 1 |
| 0 | 0 | 0 | 1 |
| 0 | 0 | 0 | 1 |
| 0 | 0 | 0 | 1 |
| 0 | 0 | 0 | 0 |
| 0 | 0 | 0 | 1 |
| 0 | 0 | 0 | 1 |
| 0 | 0 | 0 | 1 |

| | | | |
|---|---|---|---|
| 0 | 0 | 0 | 1 |
| 0 | 0 | 0 | 1 |
| 0 | 0 | 0 | 1 |
| 0 | 0 | 0 | 1 |
| 0 | 0 | 0 | 1 |
| 1 | 0 | 0 | 0 |
| 0 | 0 | 0 | 1 |
| 0 | 0 | 0 | 0 |
| 0 | 0 | 0 | 0 |
| 0 | 0 | 0 | 0 |
| 0 | 0 | 0 | 1 |
| 0 | 0 | 0 | 0 |
| 0 | 0 | 0 | 1 |
| 0 | 0 | 0 | 1 |
| 0 | 0 | 0 | 1 |
| 0 | 1 | 1 | 1 |
| 0 | 0 | 0 | 1 |
| 0 | 0 | 0 | 0 |
| 0 | 0 | 0 | 1 |
| 0 | 0 | 0 | 0 |
| 0 | 0 | 0 | 1 |
| 0 | 0 | 0 | 0 |
| 0 | 0 | 0 | 0 |
| 0 | 0 | 0 | 0 |
| 0 | 0 | 0 | 1 |
| 0 | 0 | 0 | 1 |
| 0 | 0 | 0 | 1 |
| 0 | 0 | 0 | 1 |
| 0 | 0 | 0 | 0 |
| 0 | 0 | 0 | 1 |
| 0 | 0 | 0 | 1 |
| 0 | 0 | 0 | 1 |
| 0 | 0 | 0 | 1 |
| 0 | 0 | 0 | 1 |
| 0 | 0 | 0 | 0 |
| 0 | 0 | 0 | 0 |
| 0 | 0 | 0 | 1 |
| 0 | 0 | 0 | 0 |
| 0 | 0 | 0 | 0 |
| 0 | 0 | 0 | 1 |
| 0 | 0 | 0 | 0 |
| 0 | 0 | 0 | 0 |
| 0 | 0 | 0 | 1 |
| 0 | 0 | 0 | 1 |
| 0 | 0 | 0 | 1 |
| 0 | 0 | 0 | 1 |
| 0 | 0 | 0 | 1 |
| 0 | 0 | 0 | 1 |
| 0 | 0 | 0 | 1 |
| 0 | 0 | 0 | 1 |
| 0 | 0 | 0 | 1 |
| 0 | 0 | 0 | 1 |
| 0 | 0 | 0 | 1 |
| 0 | 0 | 0 | 1 |

| | | | |
|---|---|---|---|
| 0 | 0 | 0 | 1 |
| 0 | 0 | 0 | 0 |
| 0 | 0 | 0 | 1 |
| 0 | 0 | 0 | 1 |
| 0 | 0 | 0 | 0 |
| 0 | 0 | 0 | 1 |
| 0 | 0 | 0 | 1 |
| 0 | 0 | 0 | 1 |
| 0 | 0 | 0 | 0 |
| 0 | 0 | 0 | 0 |
| 0 | 0 | 0 | 1 |
| 0 | 0 | 0 | 0 |
| 0 | 0 | 0 | 0 |
| 0 | 0 | 0 | 1 |
| 0 | 0 | 0 | 1 |
| 0 | 0 | 0 | 0 |
| 0 | 0 | 0 | 1 |
| 0 | 0 | 0 | 0 |
| 0 | 0 | 0 | 0 |
| 0 | 0 | 0 | 1 |
| 0 | 0 | 0 | 0 |
| 0 | 0 | 0 | 0 |
| 1 | 0 | 0 | 0 |
| 0 | 0 | 0 | 1 |
| 0 | 0 | 0 | 1 |
| 0 | 0 | 0 | 0 |
| 0 | 0 | 0 | 1 |
| 0 | 0 | 0 | 1 |
| 0 | 0 | 0 | 1 |
| 0 | 0 | 0 | 1 |
| 0 | 0 | 0 | 0 |
| 0 | 0 | 0 | 1 |
| 0 | 0 | 0 | 0 |
| 0 | 0 | 0 | 1 |
| 0 | 0 | 0 | 0 |
| 0 | 0 | 0 | 0 |
| 0 | 0 | 0 | 1 |
| 0 | 0 | 0 | 1 |
| 0 | 0 | 0 | 1 |
| 0 | 0 | 0 | 1 |
| 0 | 0 | 0 | 1 |
| 0 | 0 | 0 | 1 |
| 0 | 0 | 0 | 1 |
| 0 | 0 | 0 | 1 |
| 0 | 0 | 0 | 1 |
| 0 | 0 | 0 | 1 |
| 0 | 1 | 1 | 1 |
| 0 | 0 | 0 | 1 |
| 0 | 0 | 0 | 1 |
| 0 | 0 | 0 | 1 |
| 0 | 0 | 0 | 1 |
| 0 | 0 | 0 | 1 |
| 0 | 0 | 0 | 1 |
| 0 | 0 | 0 | 0 |

| | | | |
|---|---|---|---|
| 0 | 0 | 0 | 1 |
| 0 | 0 | 0 | 1 |
| 0 | 0 | 0 | 1 |
| 0 | 0 | 0 | 1 |
| 0 | 0 | 0 | 0 |
| 0 | 0 | 0 | 1 |
| 0 | 0 | 0 | 1 |
| 0 | 0 | 0 | 1 |
| 0 | 0 | 0 | 1 |
| 0 | 0 | 0 | 1 |
| 0 | 0 | 0 | 0 |
| 0 | 0 | 0 | 1 |
| 0 | 0 | 0 | 1 |
| 0 | 0 | 0 | 1 |
| 0 | 0 | 0 | 0 |
| 0 | 0 | 0 | 0 |
| 0 | 0 | 0 | 0 |
| 0 | 0 | 0 | 0 |
| 0 | 0 | 0 | 1 |
| 0 | 0 | 0 | 0 |
| 0 | 0 | 0 | 0 |
| 1 | 0 | 0 | 0 |
| 0 | 0 | 0 | 1 |
| 0 | 0 | 0 | 0 |
| 1 | 0 | 0 | 0 |
| 0 | 0 | 0 | 1 |
| 1 | 0 | 0 | 0 |
| 0 | 0 | 0 | 0 |
| 1 | 0 | 0 | 0 |
| 1 | 0 | 0 | 0 |
| 1 | 0 | 0 | 0 |
| 0 | 0 | 0 | 1 |
| 0 | 0 | 0 | 1 |
| 0 | 0 | 0 | 1 |
| 0 | 0 | 0 | 1 |
| 0 | 0 | 0 | 1 |
| 0 | 0 | 0 | 1 |
| 0 | 1 | 0 | 0 |
| 0 | 0 | 0 | 0 |
| 0 | 0 | 0 | 1 |
| 0 | 0 | 0 | 1 |
| 0 | 0 | 0 | 1 |
| 0 | 0 | 0 | 0 |
| 0 | 0 | 0 | 1 |
| 0 | 0 | 0 | 0 |
| 0 | 0 | 0 | 1 |
| 0 | 0 | 0 | 1 |
| 0 | 0 | 0 | 1 |
| 0 | 1 | 0 | 0 |
| 0 | 1 | 0 | 0 |
| 0 | 1 | 0 | 0 |
| 0 | 0 | 0 | 1 |
| 0 | 0 | 0 | 1 |
| 0 | 0 | 0 | 0 |

| | | | |
|---|---|---|---|
| 0 | 0 | 0 | 1 |
| 0 | 0 | 0 | 0 |
| 0 | 0 | 0 | 1 |
| 0 | 1 | 0 | 0 |
| 0 | 1 | 0 | 0 |
| 0 | 0 | 0 | 1 |
| 0 | 1 | 0 | 0 |
| 0 | 1 | 0 | 0 |
| 0 | 0 | 0 | 1 |
| 0 | 0 | 0 | 1 |
| 0 | 0 | 0 | 1 |
| 0 | 0 | 0 | 1 |
| 0 | 0 | 0 | 1 |
| 0 | 0 | 0 | 1 |
| 0 | 0 | 0 | 1 |
| 0 | 0 | 0 | 1 |
| 0 | 0 | 0 | 1 |
| 0 | 0 | 0 | 1 |
| 0 | 0 | 0 | 1 |
| 0 | 0 | 0 | 1 |
| 1 | 0 | 0 | 0 |
| 0 | 0 | 0 | 1 |
| 0 | 0 | 0 | 1 |
| 0 | 0 | 0 | 1 |
| 0 | 0 | 0 | 1 |
| 0 | 0 | 0 | 0 |
| 0 | 0 | 0 | 0 |
| 0 | 0 | 0 | 1 |
| 0 | 0 | 0 | 1 |
| 0 | 0 | 0 | 1 |
| 0 | 0 | 0 | 1 |
| 0 | 0 | 0 | 1 |
| 0 | 0 | 0 | 1 |
| 0 | 0 | 0 | 1 |
| 0 | 0 | 0 | 1 |
| 0 | 0 | 0 | 1 |
| 0 | 0 | 0 | 1 |
| 0 | 0 | 0 | 1 |
| 0 | 0 | 0 | 1 |
| 0 | 0 | 0 | 1 |
| 0 | 0 | 0 | 1 |
| 0 | 0 | 0 | 1 |
| 0 | 0 | 0 | 1 |
| 0 | 0 | 0 | 0 |
| 0 | 0 | 0 | 1 |
| 0 | 0 | 0 | 0 |
| 0 | 0 | 0 | 0 |
| 0 | 0 | 0 | 0 |
| 0 | 0 | 0 | 0 |
| 0 | 0 | 0 | 0 |
| 0 | 0 | 0 | 0 |
| 0 | 0 | 0 | 1 |
| 0 | 0 | 0 | 1 |
| 0 | 0 | 0 | 1 |

| | | | |
|---|---|---|---|
| 0 | 0 | 0 | 1 |
| 0 | 0 | 0 | 1 |
| 0 | 0 | 0 | 1 |
| 0 | 0 | 0 | 0 |
| 0 | 0 | 0 | 1 |
| 0 | 0 | 0 | 0 |
| 0 | 0 | 0 | 1 |
| 0 | 0 | 0 | 1 |
| 0 | 0 | 0 | 0 |
| 0 | 0 | 0 | 1 |
| 0 | 0 | 0 | 1 |
| 0 | 0 | 0 | 1 |
| 0 | 0 | 0 | 1 |
| 0 | 0 | 0 | 1 |
| 0 | 0 | 0 | 1 |
| 0 | 0 | 0 | 0 |
| 0 | 0 | 0 | 0 |
| 0 | 0 | 0 | 0 |
| 0 | 0 | 0 | 0 |
| 0 | 0 | 0 | 0 |
| 0 | 0 | 0 | 1 |
| 0 | 0 | 0 | 1 |
| 0 | 0 | 0 | 1 |
| 0 | 0 | 0 | 1 |
| 0 | 0 | 0 | 0 |
| 0 | 0 | 0 | 1 |
| 0 | 0 | 0 | 1 |
| 0 | 0 | 0 | 1 |
| 0 | 0 | 0 | 0 |
| 0 | 0 | 0 | 1 |
| 0 | 0 | 0 | 0 |
| 0 | 0 | 0 | 0 |
| 0 | 0 | 0 | 0 |
| 0 | 1 | 0 | 0 |
| 0 | 1 | 0 | 0 |
| 0 | 0 | 0 | 1 |
| 0 | 0 | 0 | 0 |
| 0 | 0 | 0 | 0 |
| 0 | 1 | 0 | 0 |
| 0 | 1 | 0 | 0 |
| 0 | 1 | 0 | 0 |
| 0 | 0 | 0 | 0 |
| 0 | 0 | 0 | 0 |
| 0 | 0 | 0 | 0 |
| 0 | 0 | 0 | 0 |
| 0 | 0 | 0 | 0 |
| 0 | 0 | 0 | 0 |
| 0 | 0 | 0 | 0 |
| 0 | 0 | 0 | 1 |
| 0 | 0 | 0 | 0 |
| 0 | 0 | 0 | 1 |
| 0 | 0 | 0 | 1 |
| 0 | 0 | 0 | 1 |
| 0 | 0 | 0 | 0 |

| | | | |
|---|---|---|---|
| 0 | 0 | 0 | 0 |
| 0 | 0 | 0 | 1 |
| 0 | 0 | 0 | 1 |
| 0 | 0 | 0 | 1 |
| 0 | 0 | 0 | 1 |
| 0 | 0 | 0 | 0 |
| 0 | 0 | 0 | 1 |
| 0 | 0 | 0 | 1 |
| 0 | 0 | 0 | 1 |
| 0 | 0 | 0 | 1 |
| 0 | 0 | 0 | 1 |
| 0 | 0 | 0 | 1 |
| 0 | 0 | 0 | 1 |
| 1 | 0 | 0 | 0 |
| 0 | 0 | 0 | 1 |
| 0 | 0 | 0 | 1 |
| 0 | 0 | 0 | 1 |
| 0 | 0 | 0 | 0 |
| 0 | 0 | 0 | 1 |
| 0 | 0 | 0 | 1 |
| 0 | 0 | 0 | 1 |
| 0 | 0 | 0 | 1 |
| 0 | 0 | 0 | 0 |
| 0 | 0 | 0 | 1 |
| 0 | 0 | 0 | 1 |
| 0 | 0 | 0 | 0 |
| 0 | 0 | 0 | 0 |
| 0 | 0 | 0 | 0 |
| 0 | 0 | 0 | 1 |
| 0 | 0 | 0 | 1 |
| 0 | 0 | 0 | 1 |
| 0 | 0 | 0 | 1 |
| 0 | 0 | 0 | 1 |
| 0 | 0 | 0 | 1 |
| 0 | 0 | 0 | 1 |
| 0 | 0 | 0 | 1 |
| 0 | 0 | 0 | 1 |
| 0 | 0 | 0 | 1 |
| 0 | 0 | 0 | 1 |
| 0 | 0 | 0 | 1 |
| 0 | 0 | 0 | 1 |
| 0 | 0 | 0 | 1 |
| 0 | 0 | 0 | 1 |
| 0 | 0 | 0 | 1 |
| 0 | 0 | 0 | 1 |
| 0 | 0 | 0 | 1 |
| 0 | 0 | 0 | 1 |
| 0 | 0 | 0 | 1 |
| 0 | 0 | 0 | 1 |
| 0 | 0 | 0 | 1 |
| 0 | 0 | 0 | 1 |
| 0 | 0 | 0 | 1 |
| 0 | 0 | 0 | 1 |

| | | | |
|---|---|---|---|
| 0 | 0 | 0 | 1 |
| 0 | 0 | 0 | 0 |
| 0 | 0 | 0 | 1 |
| 0 | 0 | 0 | 1 |
| 0 | 0 | 0 | 1 |
| 0 | 0 | 0 | 0 |
| 0 | 0 | 0 | 0 |
| 0 | 0 | 0 | 0 |
| 0 | 0 | 0 | 1 |
| 0 | 0 | 0 | 1 |
| 0 | 0 | 0 | 1 |
| 0 | 0 | 0 | 1 |
| 0 | 0 | 0 | 1 |
| 0 | 0 | 0 | 1 |
| 0 | 0 | 0 | 0 |
| 0 | 0 | 0 | 0 |
| 0 | 0 | 0 | 0 |
| 0 | 0 | 0 | 1 |
| 0 | 0 | 0 | 0 |
| 0 | 0 | 0 | 0 |
| 0 | 0 | 0 | 0 |
| 0 | 0 | 0 | 0 |
| 0 | 0 | 0 | 0 |
| 0 | 0 | 0 | 0 |
| 0 | 0 | 0 | 1 |
| 0 | 0 | 0 | 1 |
| 0 | 0 | 0 | 1 |
| 0 | 0 | 0 | 1 |
| 0 | 0 | 0 | 1 |
| 0 | 0 | 0 | 1 |
| 0 | 0 | 0 | 1 |
| 0 | 0 | 0 | 1 |
| 0 | 0 | 0 | 0 |
| 0 | 0 | 0 | 1 |
| 0 | 0 | 0 | 1 |
| 0 | 0 | 0 | 0 |
| 0 | 0 | 0 | 0 |
| 0 | 0 | 0 | 0 |
| 0 | 0 | 0 | 0 |
| 0 | 0 | 0 | 1 |
| 0 | 0 | 0 | 1 |
| 0 | 0 | 0 | 1 |
| 0 | 0 | 0 | 1 |
| 0 | 0 | 0 | 0 |
| 0 | 0 | 0 | 0 |
| 0 | 0 | 0 | 0 |
| 0 | 0 | 0 | 1 |
| 0 | 0 | 0 | 1 |
| 0 | 0 | 0 | 0 |
| 0 | 0 | 0 | 1 |
| 0 | 0 | 0 | 1 |
| 0 | 0 | 0 | 0 |
| 0 | 0 | 0 | 1 |
| 0 | 0 | 0 | 1 |

| | | | |
|---|---|---|---|
| 0 | 0 | 0 | 1 |
| 0 | 0 | 0 | 1 |
| 0 | 0 | 0 | 1 |
| 0 | 0 | 0 | 1 |
| 0 | 0 | 0 | 1 |
| 0 | 0 | 0 | 1 |
| 0 | 0 | 0 | 1 |
| 0 | 0 | 0 | 1 |
| 0 | 0 | 0 | 0 |
| 0 | 0 | 0 | 0 |
| 0 | 0 | 0 | 1 |
| 0 | 0 | 0 | 0 |
| 0 | 0 | 0 | 0 |
| 0 | 0 | 0 | 0 |
| 0 | 0 | 0 | 1 |
| 0 | 0 | 0 | 1 |
| 0 | 0 | 0 | 0 |
| 0 | 0 | 0 | 0 |
| 0 | 0 | 0 | 0 |
| 0 | 0 | 0 | 1 |
| 0 | 0 | 0 | 1 |
| 0 | 0 | 0 | 1 |
| 0 | 0 | 0 | 0 |
| 0 | 0 | 0 | 1 |
| 0 | 0 | 0 | 0 |
| 0 | 0 | 0 | 1 |
| 0 | 0 | 0 | 1 |
| 0 | 0 | 0 | 1 |
| 0 | 0 | 0 | 1 |
| 0 | 0 | 0 | 0 |
| 0 | 0 | 0 | 0 |
| 0 | 0 | 0 | 0 |
| 0 | 0 | 0 | 1 |
| 0 | 0 | 0 | 1 |
| 0 | 0 | 0 | 1 |
| 0 | 0 | 0 | 1 |
| 0 | 0 | 0 | 1 |
| 0 | 0 | 0 | 1 |
| 0 | 0 | 0 | 1 |
| 0 | 0 | 0 | 1 |
| 0 | 0 | 0 | 0 |
| 0 | 0 | 0 | 1 |
| 0 | 0 | 0 | 1 |
| 0 | 0 | 0 | 1 |
| 0 | 0 | 0 | 1 |
| 0 | 0 | 0 | 0 |
| 0 | 0 | 0 | 1 |
| 0 | 0 | 0 | 0 |
| 0 | 0 | 0 | 1 |
| 0 | 0 | 0 | 0 |
| 0 | 0 | 0 | 0 |
| 0 | 0 | 0 | 0 |
| 0 | 0 | 0 | 1 |
| 0 | 0 | 0 | 1 |

| | | | |
|---|---|---|---|
| 0 | 0 | 0 | 1 |
| 0 | 0 | 0 | 1 |
| 0 | 0 | 0 | 1 |
| 0 | 0 | 0 | 1 |
| 0 | 0 | 0 | 1 |
| 0 | 0 | 0 | 1 |
| 0 | 0 | 0 | 1 |
| 0 | 0 | 0 | 0 |
| 0 | 0 | 0 | 0 |
| 0 | 0 | 0 | 1 |
| 0 | 0 | 0 | 0 |
| 0 | 0 | 0 | 1 |
| 0 | 0 | 0 | 1 |
| 0 | 0 | 0 | 1 |
| 0 | 0 | 0 | 1 |
| 0 | 0 | 0 | 1 |
| 0 | 0 | 0 | 1 |
| 0 | 0 | 0 | 1 |
| 0 | 0 | 0 | 1 |
| 0 | 0 | 0 | 1 |
| 0 | 0 | 0 | 1 |
| 0 | 0 | 0 | 1 |
| 0 | 0 | 0 | 1 |
| 0 | 0 | 0 | 1 |
| 0 | 0 | 0 | 1 |
| 0 | 0 | 0 | 1 |
| 0 | 0 | 0 | 1 |
| 0 | 0 | 0 | 1 |
| 0 | 0 | 0 | 0 |
| 0 | 0 | 0 | 1 |
| 0 | 0 | 0 | 0 |
| 0 | 0 | 0 | 1 |
| 0 | 0 | 0 | 0 |
| 0 | 0 | 0 | 0 |
| 0 | 0 | 0 | 1 |
| 0 | 0 | 0 | 0 |
| 0 | 0 | 0 | 0 |
| 0 | 0 | 0 | 1 |
| 0 | 0 | 0 | 1 |
| 0 | 0 | 0 | 1 |
| 0 | 0 | 0 | 1 |
| 0 | 0 | 0 | 0 |
| 0 | 0 | 0 | 0 |
| 0 | 0 | 0 | 1 |
| 0 | 0 | 0 | 1 |
| 0 | 0 | 0 | 0 |
| 0 | 0 | 0 | 0 |
| 0 | 0 | 0 | 0 |
| 0 | 0 | 0 | 0 |
| 0 | 0 | 0 | 1 |
| 0 | 0 | 0 | 0 |
| 0 | 0 | 0 | 0 |
| 0 | 0 | 0 | 1 |
| 0 | 0 | 0 | 1 |

| | | | |
|---|---|---|---|
| 0 | 0 | 0 | 1 |
| 0 | 0 | 0 | 1 |
| 0 | 0 | 0 | 1 |
| 0 | 0 | 0 | 1 |
| 0 | 0 | 0 | 0 |
| 0 | 0 | 0 | 0 |
| 0 | 0 | 0 | 0 |
| 0 | 0 | 0 | 0 |
| 0 | 0 | 0 | 1 |
| 0 | 0 | 0 | 0 |
| 0 | 0 | 0 | 0 |
| 0 | 0 | 0 | 0 |
| 0 | 0 | 0 | 0 |
| 0 | 0 | 0 | 1 |
| 0 | 0 | 0 | 1 |
| 0 | 0 | 0 | 1 |
| 0 | 0 | 0 | 1 |
| 0 | 0 | 0 | 1 |
| 0 | 0 | 0 | 1 |
| 0 | 0 | 0 | 1 |
| 0 | 0 | 0 | 1 |
| 0 | 0 | 0 | 1 |
| 0 | 0 | 0 | 1 |
| 0 | 0 | 0 | 1 |
| 0 | 0 | 0 | 1 |
| 0 | 0 | 0 | 0 |
| 0 | 0 | 0 | 0 |
| 0 | 0 | 0 | 0 |
| 0 | 0 | 0 | 0 |
| 0 | 0 | 0 | 1 |
| 0 | 0 | 0 | 1 |
| 0 | 0 | 0 | 1 |
| 0 | 0 | 0 | 1 |
| 0 | 0 | 0 | 1 |
| 0 | 0 | 0 | 1 |
| 0 | 0 | 0 | 1 |
| 0 | 0 | 0 | 1 |
| 0 | 0 | 0 | 1 |
| 0 | 0 | 0 | 1 |
| 0 | 0 | 0 | 1 |
| 0 | 0 | 0 | 1 |
| 0 | 0 | 0 | 1 |
| 0 | 0 | 0 | 1 |
| 0 | 0 | 0 | 1 |
| 0 | 0 | 0 | 1 |
| 0 | 0 | 0 | 1 |
| 0 | 0 | 0 | 1 |
| 0 | 0 | 0 | 1 |
| 0 | 0 | 0 | 1 |
| 0 | 0 | 0 | 1 |
| 0 | 0 | 0 | 1 |
| 0 | 0 | 0 | 1 |
| 0 | 0 | 0 | 1 |
| 0 | 0 | 0 | 0 |

| | | | |
|---|---|---|---|
| 0 | 0 | 0 | 1 |
| 0 | 0 | 0 | 1 |
| 0 | 0 | 0 | 1 |
| 0 | 0 | 0 | 1 |
| 0 | 0 | 0 | 1 |
| 0 | 0 | 0 | 1 |
| 0 | 0 | 0 | 1 |
| 0 | 0 | 0 | 1 |
| 0 | 0 | 0 | 1 |
| 0 | 0 | 0 | 1 |
| 0 | 0 | 0 | 1 |
| 0 | 0 | 0 | 1 |
| 0 | 0 | 0 | 1 |
| 0 | 0 | 0 | 1 |
| 0 | 0 | 0 | 1 |
| 0 | 0 | 0 | 1 |
| 0 | 0 | 0 | 1 |
| 0 | 0 | 0 | 1 |
| 0 | 0 | 0 | 1 |
| 0 | 0 | 0 | 1 |
| 0 | 0 | 0 | 1 |
| 0 | 0 | 0 | 1 |
| 0 | 0 | 0 | 1 |
| 0 | 0 | 0 | 1 |
| 0 | 0 | 0 | 1 |
| 0 | 0 | 0 | 1 |
| 0 | 0 | 0 | 1 |
| 0 | 0 | 0 | 1 |
| 0 | 0 | 0 | 1 |
| 0 | 0 | 0 | 1 |
| 0 | 0 | 0 | 1 |
| 0 | 0 | 0 | 1 |
| 0 | 0 | 0 | 1 |
| 0 | 0 | 0 | 1 |
| 0 | 0 | 0 | 1 |
| 0 | 0 | 0 | 1 |
| 0 | 0 | 0 | 1 |
| 0 | 0 | 0 | 1 |
| 0 | 0 | 0 | 1 |
| 0 | 0 | 0 | 1 |
| 0 | 0 | 0 | 1 |
| 0 | 0 | 0 | 1 |
| 0 | 0 | 0 | 1 |
| 0 | 0 | 0 | 1 |
| 0 | 0 | 0 | 1 |
| 0 | 0 | 0 | 1 |
| 0 | 0 | 0 | 0 |
| 0 | 0 | 0 | 1 |
| 0 | 0 | 0 | 1 |
| 0 | 0 | 0 | 1 |
| 0 | 0 | 0 | 1 |
| 0 | 0 | 0 | 1 |
| 0 | 0 | 0 | 1 |
| 0 | 0 | 0 | 1 |

| | | | |
|---|---|---|---|
| 0 | 0 | 0 | 1 |
| 0 | 0 | 0 | 1 |
| 0 | 0 | 0 | 0 |
| 0 | 0 | 0 | 1 |
| 0 | 0 | 0 | 1 |
| 0 | 0 | 0 | 1 |
| 0 | 0 | 0 | 0 |
| 0 | 0 | 0 | 1 |
| 0 | 0 | 0 | 1 |
| 0 | 0 | 0 | 1 |
| 0 | 0 | 0 | 1 |
| 0 | 0 | 0 | 0 |
| 0 | 0 | 0 | 1 |
| 0 | 0 | 0 | 0 |
| 0 | 0 | 0 | 0 |
| 0 | 0 | 0 | 0 |
| 0 | 0 | 0 | 0 |
| 0 | 0 | 0 | 0 |
| 0 | 0 | 0 | 0 |
| 0 | 0 | 0 | 0 |
| 0 | 0 | 0 | 0 |

| surface_non_binding_site | non_binding_site_single_water | surface_non_binding_site_single_water |
|---|---|---|
| 0 | 0 | 0 |
| 0 | 0 | 0 |
| 0 | 0 | 0 |
| 0 | 0 | 0 |
| 0 | 0 | 0 |
| 0 | 0 | 0 |
| 1 | 1 | 1 |
| 1 | 1 | 1 |
| 0 | 0 | 0 |
| 0 | 0 | 0 |
| 0 | 0 | 0 |
| 0 | 0 | 0 |
| 0 | 0 | 0 |
| 0 | 0 | 0 |
| 0 | 0 | 0 |
| 0 | 0 | 0 |
| 0 | 0 | 0 |
| 0 | 0 | 0 |
| 0 | 0 | 0 |
| 0 | 0 | 0 |
| 0 | 0 | 0 |
| 0 | 0 | 0 |
| 0 | 0 | 0 |
| 0 | 0 | 0 |
| 0 | 0 | 0 |
| 1 | 0 | 0 |
| 0 | 0 | 0 |
| 1 | 1 | 1 |
| 0 | 0 | 0 |
| 0 | 1 | 0 |
| 0 | 1 | 0 |
| 0 | 0 | 0 |
| 0 | 0 | 0 |
| 0 | 0 | 0 |
| 0 | 0 | 0 |
| 1 | 1 | 1 |
| 0 | 0 | 0 |
| 0 | 0 | 0 |
| 0 | 0 | 0 |
| 0 | 0 | 0 |
| 0 | 0 | 0 |
| 0 | 0 | 0 |
| 0 | 0 | 0 |
| 0 | 0 | 0 |
| 0 | 0 | 0 |
| 0 | 0 | 0 |
| 0 | 0 | 0 |
| 0 | 0 | 0 |
| 0 | 0 | 0 |
| 0 | 0 | 0 |
| 0 | 0 | 0 |
| 0 | 0 | 0 |
| 0 | 0 | 0 |

| | | |
|---|---|---|
| 0 | 0 | 0 |
| 0 | 0 | 0 |
| 0 | 0 | 0 |
| 0 | 0 | 0 |
| 0 | 0 | 0 |
| 0 | 0 | 0 |
| 0 | 0 | 0 |
| 0 | 0 | 0 |
| 0 | 0 | 0 |
| 0 | 0 | 0 |
| 0 | 0 | 0 |
| 0 | 0 | 0 |
| 0 | 0 | 0 |
| 0 | 0 | 0 |
| 0 | 0 | 0 |
| 0 | 0 | 0 |
| 0 | 0 | 0 |
| 0 | 0 | 0 |
| 0 | 0 | 0 |
| 0 | 0 | 0 |
| 0 | 0 | 0 |
| 0 | 0 | 0 |
| 0 | 0 | 0 |
| 0 | 0 | 0 |
| 0 | 0 | 0 |
| 0 | 0 | 0 |
| 0 | 0 | 0 |
| 0 | 0 | 0 |
| 0 | 0 | 0 |
| 0 | 0 | 0 |
| 0 | 0 | 0 |
| 0 | 0 | 0 |
| 0 | 0 | 0 |
| 0 | 0 | 0 |
| 0 | 0 | 0 |
| 0 | 0 | 0 |
| 0 | 0 | 0 |
| 0 | 0 | 0 |
| 0 | 0 | 0 |
| 0 | 0 | 0 |
| 0 | 0 | 0 |
| 0 | 0 | 0 |
| 0 | 0 | 0 |
| 0 | 0 | 0 |
| 0 | 0 | 0 |
| 1 | 1 | 1 |
| 0 | 1 | 0 |
| 0 | 0 | 0 |
| 0 | 0 | 0 |
| 0 | 0 | 0 |
| 0 | 0 | 0 |
| 1 | 1 | 1 |
| 1 | 1 | 1 |
| 0 | 0 | 0 |

| | | |
|---|---|---|
| 0 | 1 | 0 |
| 0 | 1 | 0 |
| 0 | 0 | 0 |
| 0 | 0 | 0 |
| 0 | 0 | 0 |
| 0 | 0 | 0 |
| 0 | 1 | 0 |
| 0 | 0 | 0 |
| 0 | 0 | 0 |
| 0 | 0 | 0 |
| 0 | 0 | 0 |
| 0 | 0 | 0 |
| 0 | 0 | 0 |
| 0 | 0 | 0 |
| 0 | 0 | 0 |
| 1 | 1 | 1 |
| 0 | 0 | 0 |
| 0 | 0 | 0 |
| 0 | 0 | 0 |
| 0 | 0 | 0 |
| 0 | 0 | 0 |
| 0 | 0 | 0 |
| 0 | 0 | 0 |
| 0 | 0 | 0 |
| 0 | 0 | 0 |
| 0 | 0 | 0 |
| 0 | 0 | 0 |
| 0 | 0 | 0 |
| 0 | 0 | 0 |
| 0 | 0 | 0 |
| 0 | 0 | 0 |
| 0 | 0 | 0 |
| 0 | 0 | 0 |
| 0 | 0 | 0 |
| 0 | 0 | 0 |
| 0 | 0 | 0 |
| 0 | 0 | 0 |
| 0 | 0 | 0 |
| 0 | 0 | 0 |
| 0 | 0 | 0 |
| 0 | 0 | 0 |
| 0 | 0 | 0 |
| 0 | 0 | 0 |
| 0 | 0 | 0 |
| 0 | 0 | 0 |
| 0 | 0 | 0 |
| 0 | 0 | 0 |
| 0 | 0 | 0 |
| 0 | 0 | 0 |
| 0 | 0 | 0 |
| 0 | 0 | 0 |
| 0 | 0 | 0 |
| 0 | 0 | 0 |
| 0 | 0 | 0 |

| | | |
|---|---|---|
| 1 | 0 | 0 |
| 1 | 0 | 0 |
| 1 | 0 | 0 |
| 0 | 0 | 0 |
| 0 | 0 | 0 |
| 0 | 0 | 0 |
| 0 | 0 | 0 |
| 1 | 0 | 0 |
| 0 | 0 | 0 |
| 1 | 0 | 0 |
| 1 | 0 | 0 |
| 0 | 0 | 0 |
| 1 | 0 | 0 |
| 0 | 0 | 0 |
| 0 | 0 | 0 |
| 1 | 0 | 0 |
| 1 | 0 | 0 |
| 1 | 0 | 0 |
| 1 | 0 | 0 |
| 0 | 0 | 0 |
| 0 | 0 | 0 |
| 0 | 1 | 0 |
| 0 | 0 | 0 |
| 0 | 0 | 0 |
| 0 | 0 | 0 |
| 1 | 0 | 0 |
| 0 | 0 | 0 |
| 0 | 0 | 0 |
| 1 | 0 | 0 |
| 1 | 0 | 0 |
| 1 | 1 | 1 |
| 0 | 0 | 0 |
| 0 | 0 | 0 |
| 0 | 1 | 0 |
| 0 | 0 | 0 |
| 0 | 0 | 0 |
| 0 | 0 | 0 |
| 0 | 0 | 0 |
| 0 | 0 | 0 |
| 1 | 0 | 0 |
| 1 | 0 | 0 |
| 0 | 0 | 0 |
| 0 | 0 | 0 |
| 1 | 1 | 1 |
| 1 | 0 | 0 |
| 1 | 0 | 0 |
| 0 | 0 | 0 |
| 0 | 0 | 0 |
| 0 | 0 | 0 |
| 0 | 0 | 0 |
| 0 | 0 | 0 |
| 0 | 1 | 0 |
| 1 | 0 | 0 |
| 0 | 0 | 0 |

| | | |
|---|---|---|
| 0 | 0 | 0 |
| 0 | 0 | 0 |
| 0 | 0 | 0 |
| 1 | 1 | 1 |
| 1 | 1 | 1 |
| 1 | 0 | 0 |
| 0 | 0 | 0 |
| 0 | 0 | 0 |
| 1 | 0 | 0 |
| 1 | 0 | 0 |
| 0 | 0 | 0 |
| 0 | 1 | 0 |
| 0 | 0 | 0 |
| 1 | 0 | 0 |
| 0 | 1 | 0 |
| 1 | 1 | 1 |
| 1 | 1 | 1 |
| 0 | 0 | 0 |
| 0 | 0 | 0 |
| 0 | 0 | 0 |
| 0 | 0 | 0 |
| 1 | 0 | 0 |
| 0 | 0 | 0 |
| 0 | 1 | 0 |
| 1 | 1 | 1 |
| 0 | 0 | 0 |
| 0 | 0 | 0 |
| 0 | 0 | 0 |
| 0 | 0 | 0 |
| 0 | 0 | 0 |
| 0 | 0 | 0 |
| 0 | 0 | 0 |
| 1 | 0 | 0 |
| 1 | 0 | 0 |
| 0 | 0 | 0 |
| 1 | 0 | 0 |
| 0 | 0 | 0 |
| 0 | 0 | 0 |
| 1 | 0 | 0 |
| 0 | 1 | 0 |
| 0 | 0 | 0 |
| 1 | 0 | 0 |
| 1 | 0 | 0 |
| 1 | 0 | 0 |
| 0 | 0 | 0 |
| 0 | 0 | 0 |
| 0 | 0 | 0 |
| 0 | 0 | 0 |
| 0 | 0 | 0 |
| 0 | 0 | 0 |
| 0 | 0 | 0 |
| 0 | 0 | 0 |
| 0 | 0 | 0 |
| 0 | 0 | 0 |

| | | |
|---|---|---|
| 0 | 0 | 0 |
| 0 | 0 | 0 |
| 0 | 0 | 0 |
| 0 | 0 | 0 |
| 0 | 0 | 0 |
| 0 | 0 | 0 |
| 0 | 1 | 0 |
| 0 | 1 | 0 |
| 0 | 0 | 0 |
| 0 | 0 | 0 |
| 0 | 0 | 0 |
| 0 | 0 | 0 |
| 0 | 0 | 0 |
| 0 | 1 | 0 |
| 0 | 1 | 0 |
| 0 | 0 | 0 |
| 0 | 1 | 0 |
| 0 | 0 | 0 |
| 1 | 1 | 1 |
| 0 | 0 | 0 |
| 1 | 1 | 1 |
| 1 | 0 | 0 |
| 1 | 1 | 1 |
| 0 | 0 | 0 |
| 0 | 0 | 0 |
| 0 | 0 | 0 |
| 0 | 0 | 0 |
| 0 | 0 | 0 |
| 0 | 0 | 0 |
| 0 | 0 | 0 |
| 0 | 0 | 0 |
| 1 | 0 | 0 |
| 0 | 0 | 0 |
| 0 | 0 | 0 |
| 0 | 0 | 0 |
| 0 | 0 | 0 |
| 0 | 0 | 0 |
| 0 | 0 | 0 |
| 0 | 0 | 0 |
| 0 | 0 | 0 |
| 0 | 1 | 0 |
| 0 | 1 | 0 |
| 0 | 0 | 0 |
| 0 | 1 | 0 |
| 0 | 0 | 0 |
| 0 | 0 | 0 |
| 0 | 0 | 0 |
| 1 | 1 | 1 |
| 1 | 1 | 1 |
| 0 | 0 | 0 |
| 0 | 1 | 0 |
| 1 | 1 | 1 |
| 0 | 0 | 0 |
| 0 | 1 | 0 |

| | | |
|---|---|---|
| 0 | 0 | 0 |
| 1 | 1 | 1 |
| 0 | 0 | 0 |
| 0 | 0 | 0 |
| 0 | 0 | 0 |
| 0 | 0 | 0 |
| 0 | 1 | 0 |
| 0 | 0 | 0 |
| 0 | 0 | 0 |
| 1 | 1 | 1 |
| 1 | 1 | 1 |
| 1 | 1 | 1 |
| 1 | 1 | 1 |
| 1 | 1 | 1 |
| 0 | 0 | 0 |
| 1 | 1 | 1 |
| 0 | 0 | 0 |
| 0 | 0 | 0 |
| 1 | 1 | 1 |
| 0 | 0 | 0 |
| 0 | 0 | 0 |
| 0 | 0 | 0 |
| 0 | 0 | 0 |
| 0 | 0 | 0 |
| 1 | 1 | 1 |
| 1 | 0 | 0 |
| 0 | 0 | 0 |
| 0 | 0 | 0 |
| 0 | 0 | 0 |
| 0 | 0 | 0 |
| 0 | 0 | 0 |
| 0 | 0 | 0 |
| 0 | 0 | 0 |
| 0 | 0 | 0 |
| 1 | 1 | 1 |
| 1 | 1 | 1 |
| 0 | 0 | 0 |
| 0 | 0 | 0 |
| 0 | 0 | 0 |
| 0 | 0 | 0 |
| 0 | 0 | 0 |
| 0 | 0 | 0 |
| 0 | 1 | 0 |
| 0 | 0 | 0 |
| 0 | 0 | 0 |
| 0 | 0 | 0 |
| 0 | 0 | 0 |
| 0 | 0 | 0 |
| 0 | 0 | 0 |
| 1 | 1 | 1 |
| 0 | 0 | 0 |
| 0 | 0 | 0 |
| 0 | 0 | 0 |
| 0 | 0 | 0 |

| | | |
|---|---|---|
| 0 | 0 | 0 |
| 0 | 0 | 0 |
| 0 | 0 | 0 |
| 0 | 0 | 0 |
| 1 | 1 | 1 |
| 0 | 0 | 0 |
| 0 | 0 | 0 |
| 0 | 0 | 0 |
| 0 | 0 | 0 |
| 0 | 0 | 0 |
| 0 | 0 | 0 |
| 0 | 0 | 0 |
| 0 | 0 | 0 |
| 0 | 0 | 0 |
| 0 | 0 | 0 |
| 0 | 0 | 0 |
| 0 | 0 | 0 |
| 0 | 0 | 0 |
| 1 | 1 | 1 |
| 0 | 0 | 0 |
| 0 | 0 | 0 |
| 0 | 0 | 0 |
| 0 | 0 | 0 |
| 1 | 0 | 0 |
| 0 | 0 | 0 |
| 0 | 0 | 0 |
| 0 | 0 | 0 |
| 1 | 0 | 0 |
| 1 | 0 | 0 |
| 1 | 0 | 0 |
| 0 | 0 | 0 |
| 0 | 0 | 0 |
| 0 | 0 | 0 |
| 0 | 0 | 0 |
| 0 | 0 | 0 |
| 1 | 0 | 0 |
| 1 | 0 | 0 |
| 0 | 0 | 0 |
| 0 | 0 | 0 |
| 1 | 0 | 0 |
| 1 | 0 | 0 |
| 1 | 0 | 0 |
| 0 | 1 | 0 |
| 1 | 1 | 1 |
| 0 | 0 | 0 |
| 0 | 0 | 0 |
| 0 | 0 | 0 |
| 0 | 0 | 0 |
| 0 | 0 | 0 |
| 0 | 1 | 0 |
| 1 | 1 | 1 |
| 0 | 0 | 0 |
| 1 | 1 | 1 |
| 0 | 0 | 0 |

| | | |
|---|---|---|
| 1 | 1 | 1 |
| 1 | 1 | 1 |
| 0 | 0 | 0 |
| 1 | 1 | 1 |
| 1 | 1 | 1 |
| 1 | 1 | 1 |
| 1 | 1 | 1 |
| 0 | 0 | 0 |
| 1 | 0 | 0 |
| 1 | 1 | 1 |
| 0 | 0 | 0 |
| 1 | 1 | 1 |
| 1 | 1 | 1 |
| 0 | 1 | 0 |
| 0 | 0 | 0 |
| 0 | 0 | 0 |
| 1 | 1 | 1 |
| 1 | 1 | 1 |
| 0 | 0 | 0 |
| 0 | 0 | 0 |
| 0 | 0 | 0 |
| 0 | 0 | 0 |
| 0 | 0 | 0 |
| 0 | 0 | 0 |
| 0 | 0 | 0 |
| 1 | 1 | 1 |
| 0 | 0 | 0 |
| 0 | 0 | 0 |
| 0 | 0 | 0 |
| 1 | 0 | 0 |
| 0 | 0 | 0 |
| 0 | 0 | 0 |
| 0 | 0 | 0 |
| 0 | 0 | 0 |
| 0 | 0 | 0 |
| 0 | 0 | 0 |
| 0 | 0 | 0 |
| 0 | 0 | 0 |
| 0 | 0 | 0 |
| 0 | 0 | 0 |
| 0 | 1 | 0 |
| 0 | 0 | 0 |
| 1 | 1 | 1 |
| 0 | 0 | 0 |
| 0 | 0 | 0 |
| 0 | 0 | 0 |
| 0 | 0 | 0 |
| 0 | 0 | 0 |
| 0 | 0 | 0 |
| 0 | 0 | 0 |
| 0 | 1 | 0 |
| 0 | 0 | 0 |
| 0 | 0 | 0 |
| 0 | 0 | 0 |

| | | |
|---|---|---|
| 1 | 1 | 1 |
| 0 | 0 | 0 |
| 0 | 1 | 0 |
| 0 | 1 | 0 |
| 0 | 0 | 0 |
| 0 | 0 | 0 |
| 0 | 0 | 0 |
| 0 | 0 | 0 |
| 0 | 0 | 0 |
| 0 | 1 | 0 |
| 1 | 1 | 1 |
| 1 | 1 | 1 |
| 1 | 1 | 1 |
| 0 | 0 | 0 |
| 1 | 1 | 1 |
| 0 | 0 | 0 |
| 0 | 0 | 0 |
| 0 | 0 | 0 |
| 0 | 0 | 0 |
| 1 | 0 | 0 |
| 0 | 0 | 0 |
| 1 | 1 | 1 |
| 0 | 0 | 0 |
| 0 | 1 | 0 |
| 1 | 1 | 1 |
| 1 | 1 | 1 |
| 0 | 0 | 0 |
| 1 | 1 | 1 |
| 1 | 1 | 1 |
| 1 | 1 | 1 |
| 0 | 0 | 0 |
| 0 | 0 | 0 |
| 1 | 1 | 1 |
| 1 | 1 | 1 |
| 0 | 0 | 0 |
| 0 | 0 | 0 |
| 0 | 0 | 0 |
| 0 | 0 | 0 |
| 0 | 0 | 0 |
| 1 | 1 | 1 |
| 0 | 0 | 0 |
| 0 | 0 | 0 |
| 0 | 0 | 0 |
| 0 | 0 | 0 |
| 1 | 0 | 0 |
| 0 | 0 | 0 |
| 0 | 0 | 0 |
| 0 | 0 | 0 |
| 0 | 0 | 0 |
| 0 | 0 | 0 |
| 0 | 0 | 0 |
| 0 | 0 | 0 |
| 1 | 0 | 0 |
| 0 | 0 | 0 |

| | | |
|---|---|---|
| 0 | 0 | 0 |
| 0 | 0 | 0 |
| 0 | 0 | 0 |
| 0 | 0 | 0 |
| 0 | 0 | 0 |
| 0 | 0 | 0 |
| 0 | 0 | 0 |
| 1 | 0 | 0 |
| 0 | 0 | 0 |
| 0 | 0 | 0 |
| 0 | 0 | 0 |
| 0 | 0 | 0 |
| 1 | 1 | 1 |
| 0 | 0 | 0 |
| 0 | 0 | 0 |
| 0 | 0 | 0 |
| 0 | 0 | 0 |
| 0 | 0 | 0 |
| 0 | 0 | 0 |
| 1 | 1 | 1 |
| 0 | 0 | 0 |
| 0 | 0 | 0 |
| 0 | 0 | 0 |
| 0 | 0 | 0 |
| 0 | 0 | 0 |
| 1 | 0 | 0 |
| 1 | 1 | 1 |
| 0 | 1 | 0 |
| 0 | 0 | 0 |
| 0 | 0 | 0 |
| 0 | 0 | 0 |
| 0 | 0 | 0 |
| 0 | 0 | 0 |
| 0 | 0 | 0 |
| 0 | 0 | 0 |
| 1 | 0 | 0 |
| 0 | 0 | 0 |
| 0 | 0 | 0 |
| 1 | 1 | 1 |
| 1 | 0 | 0 |
| 0 | 1 | 0 |
| 0 | 0 | 0 |
| 0 | 0 | 0 |
| 0 | 0 | 0 |
| 0 | 0 | 0 |
| 0 | 0 | 0 |
| 0 | 0 | 0 |
| 1 | 1 | 1 |
| 1 | 1 | 1 |
| 0 | 0 | 0 |
| 0 | 0 | 0 |
| 0 | 0 | 0 |
| 0 | 0 | 0 |
| 1 | 1 | 1 |

| | | |
|---|---|---|
| 1 | 1 | 1 |
| 0 | 0 | 0 |
| 0 | 1 | 0 |
| 0 | 0 | 0 |
| 1 | 0 | 0 |
| 1 | 0 | 0 |
| 0 | 0 | 0 |
| 1 | 0 | 0 |
| 1 | 0 | 0 |
| 1 | 0 | 0 |
| 1 | 0 | 0 |
| 1 | 0 | 0 |
| 1 | 1 | 1 |
| 1 | 1 | 1 |
| 1 | 1 | 1 |
| 1 | 1 | 1 |
| 0 | 0 | 0 |
| 0 | 0 | 0 |
| 1 | 0 | 0 |
| 0 | 0 | 0 |
| 0 | 0 | 0 |
| 0 | 0 | 0 |
| 1 | 0 | 0 |
| 0 | 0 | 0 |
| 0 | 0 | 0 |
| 1 | 1 | 1 |
| 1 | 0 | 0 |
| 1 | 0 | 0 |
| 0 | 0 | 0 |
| 0 | 0 | 0 |
| 0 | 0 | 0 |
| 0 | 0 | 0 |
| 0 | 0 | 0 |
| 0 | 0 | 0 |
| 0 | 0 | 0 |
| 0 | 0 | 0 |
| 1 | 0 | 0 |
| 1 | 1 | 1 |
| 1 | 1 | 1 |
| 1 | 1 | 1 |
| 0 | 0 | 0 |
| 1 | 1 | 1 |
| 0 | 0 | 0 |
| 1 | 1 | 1 |
| 0 | 0 | 0 |
| 0 | 1 | 0 |
| 0 | 0 | 0 |
| 0 | 0 | 0 |
| 0 | 0 | 0 |
| 0 | 0 | 0 |
| 0 | 0 | 0 |
| 0 | 0 | 0 |
| 0 | 0 | 0 |
| 0 | 0 | 0 |

| | | |
|---|---|---|
| 0 | 0 | 0 |
| 0 | 0 | 0 |
| 0 | 0 | 0 |
| 0 | 0 | 0 |
| 0 | 0 | 0 |
| 0 | 0 | 0 |
| 0 | 0 | 0 |
| 1 | 1 | 1 |
| 0 | 0 | 0 |
| 0 | 0 | 0 |
| 0 | 0 | 0 |
| 1 | 0 | 0 |
| 0 | 0 | 0 |
| 0 | 0 | 0 |
| 0 | 0 | 0 |
| 0 | 0 | 0 |
| 0 | 0 | 0 |
| 0 | 0 | 0 |
| 0 | 0 | 0 |
| 0 | 0 | 0 |
| 0 | 0 | 0 |
| 0 | 0 | 0 |
| 0 | 0 | 0 |
| 0 | 0 | 0 |
| 0 | 0 | 0 |
| 0 | 0 | 0 |
| 0 | 0 | 0 |
| 0 | 0 | 0 |
| 0 | 0 | 0 |
| 0 | 0 | 0 |
| 0 | 0 | 0 |
| 0 | 0 | 0 |
| 0 | 0 | 0 |
| 0 | 0 | 0 |
| 0 | 0 | 0 |
| 0 | 0 | 0 |
| 0 | 0 | 0 |
| 0 | 0 | 0 |
| 0 | 0 | 0 |
| 0 | 0 | 0 |
| 1 | 0 | 0 |
| 0 | 0 | 0 |
| 0 | 0 | 0 |
| 0 | 0 | 0 |
| 0 | 0 | 0 |
| 0 | 0 | 0 |
| 0 | 0 | 0 |
| 0 | 0 | 0 |
| 0 | 0 | 0 |
| 0 | 1 | 0 |
| 0 | 1 | 0 |
| 1 | 0 | 0 |
| 0 | 0 | 0 |
| 0 | 0 | 0 |

| | | |
|---|---|---|
| 0 | 0 | 0 |
| 0 | 0 | 0 |
| 0 | 0 | 0 |
| 0 | 0 | 0 |
| 0 | 0 | 0 |
| 0 | 0 | 0 |
| 0 | 0 | 0 |
| 1 | 0 | 0 |
| 1 | 0 | 0 |
| 0 | 0 | 0 |
| 0 | 0 | 0 |
| 1 | 0 | 0 |
| 0 | 0 | 0 |
| 0 | 0 | 0 |
| 0 | 0 | 0 |
| 0 | 0 | 0 |
| 0 | 0 | 0 |
| 1 | 0 | 0 |
| 0 | 0 | 0 |
| 1 | 0 | 0 |
| 0 | 0 | 0 |
| 0 | 0 | 0 |
| 1 | 1 | 1 |
| 0 | 0 | 0 |
| 0 | 0 | 0 |
| 1 | 0 | 0 |
| 0 | 0 | 0 |
| 0 | 0 | 0 |
| 0 | 0 | 0 |
| 0 | 0 | 0 |
| 0 | 0 | 0 |
| 0 | 0 | 0 |
| 1 | 0 | 0 |
| 0 | 0 | 0 |
| 0 | 0 | 0 |
| 0 | 0 | 0 |
| 1 | 1 | 1 |
| 0 | 0 | 0 |
| 0 | 0 | 0 |
| 0 | 0 | 0 |
| 0 | 0 | 0 |
| 0 | 0 | 0 |
| 0 | 0 | 0 |
| 0 | 0 | 0 |
| 0 | 0 | 0 |
| 0 | 0 | 0 |
| 0 | 0 | 0 |
| 0 | 0 | 0 |
| 0 | 0 | 0 |
| 0 | 0 | 0 |
| 0 | 0 | 0 |
| 0 | 0 | 0 |
| 0 | 0 | 0 |
| 0 | 0 | 0 |

| | | |
|---|---|---|
| 1 | 0 | 0 |
| 0 | 1 | 0 |
| 0 | 0 | 0 |
| 0 | 0 | 0 |
| 0 | 0 | 0 |
| 0 | 0 | 0 |
| 0 | 0 | 0 |
| 0 | 0 | 0 |
| 0 | 0 | 0 |
| 1 | 0 | 0 |
| 0 | 1 | 0 |
| 1 | 1 | 1 |
| 0 | 0 | 0 |
| 0 | 0 | 0 |
| 0 | 0 | 0 |
| 0 | 0 | 0 |
| 0 | 0 | 0 |
| 0 | 0 | 0 |
| 1 | 1 | 1 |
| 1 | 0 | 0 |
| 1 | 0 | 0 |
| 0 | 0 | 0 |
| 0 | 1 | 0 |
| 0 | 0 | 0 |
| 0 | 0 | 0 |
| 0 | 0 | 0 |
| 0 | 0 | 0 |
| 0 | 0 | 0 |
| 0 | 0 | 0 |
| 0 | 0 | 0 |
| 0 | 0 | 0 |
| 0 | 0 | 0 |
| 0 | 0 | 0 |
| 0 | 0 | 0 |
| 0 | 0 | 0 |
| 0 | 0 | 0 |
| 0 | 0 | 0 |
| 0 | 0 | 0 |
| 0 | 0 | 0 |
| 0 | 0 | 0 |
| 0 | 0 | 0 |
| 1 | 0 | 0 |
| 0 | 0 | 0 |
| 0 | 0 | 0 |
| 0 | 0 | 0 |
| 0 | 0 | 0 |
| 1 | 0 | 0 |
| 0 | 0 | 0 |
| 0 | 0 | 0 |
| 0 | 0 | 0 |
| 0 | 0 | 0 |
| 0 | 0 | 0 |
| 1 | 0 | 0 |
| 0 | 1 | 0 |

| | | |
|---|---|---|
| 0 | 0 | 0 |
| 0 | 0 | 0 |
| 1 | 0 | 0 |
| 1 | 0 | 0 |
| 1 | 0 | 0 |
| 0 | 0 | 0 |
| 1 | 0 | 0 |
| 0 | 0 | 0 |
| 0 | 0 | 0 |
| 0 | 0 | 0 |
| 0 | 0 | 0 |
| 0 | 0 | 0 |
| 0 | 0 | 0 |
| 1 | 0 | 0 |
| 1 | 0 | 0 |
| 1 | 0 | 0 |
| 0 | 0 | 0 |
| 1 | 1 | 1 |
| 0 | 0 | 0 |
| 0 | 0 | 0 |
| 1 | 0 | 0 |
| 1 | 0 | 0 |
| 0 | 0 | 0 |
| 0 | 0 | 0 |
| 1 | 0 | 0 |
| 0 | 0 | 0 |
| 0 | 0 | 0 |
| 1 | 0 | 0 |
| 0 | 0 | 0 |
| 0 | 0 | 0 |
| 0 | 0 | 0 |
| 0 | 0 | 0 |
| 0 | 0 | 0 |
| 0 | 0 | 0 |
| 0 | 0 | 0 |
| 0 | 0 | 0 |
| 0 | 0 | 0 |
| 0 | 0 | 0 |
| 0 | 0 | 0 |
| 0 | 0 | 0 |
| 1 | 0 | 0 |
| 1 | 0 | 0 |
| 1 | 0 | 0 |
| 1 | 1 | 1 |
| 1 | 1 | 1 |
| 0 | 0 | 0 |
| 0 | 0 | 0 |
| 0 | 0 | 0 |
| 0 | 0 | 0 |
| 0 | 0 | 0 |
| 0 | 0 | 0 |
| 0 | 0 | 0 |
| 0 | 0 | 0 |
| 1 | 0 | 0 |

| | | |
|---|---|---|
| 1 | 0 | 0 |
| 0 | 0 | 0 |
| 0 | 0 | 0 |
| 0 | 0 | 0 |
| 1 | 0 | 0 |
| 0 | 0 | 0 |
| 0 | 0 | 0 |
| 0 | 0 | 0 |
| 1 | 0 | 0 |
| 1 | 0 | 0 |
| 0 | 0 | 0 |
| 0 | 0 | 0 |
| 0 | 0 | 0 |
| 1 | 0 | 0 |
| 0 | 0 | 0 |
| 0 | 0 | 0 |
| 0 | 0 | 0 |
| 0 | 0 | 0 |
| 0 | 0 | 0 |
| 0 | 0 | 0 |
| 0 | 0 | 0 |
| 1 | 1 | 1 |
| 0 | 0 | 0 |
| 0 | 1 | 0 |
| 0 | 0 | 0 |
| 0 | 0 | 0 |
| 0 | 0 | 0 |
| 0 | 0 | 0 |
| 0 | 0 | 0 |
| 0 | 0 | 0 |
| 0 | 0 | 0 |
| 0 | 0 | 0 |
| 0 | 0 | 0 |
| 0 | 0 | 0 |
| 0 | 1 | 0 |
| 0 | 0 | 0 |
| 0 | 0 | 0 |
| 0 | 0 | 0 |
| 0 | 0 | 0 |
| 0 | 0 | 0 |
| 0 | 0 | 0 |
| 0 | 0 | 0 |
| 0 | 0 | 0 |
| 0 | 0 | 0 |
| 0 | 0 | 0 |
| 0 | 0 | 0 |
| 0 | 0 | 0 |
| 1 | 1 | 1 |
| 0 | 0 | 0 |
| 1 | 1 | 1 |
| 0 | 0 | 0 |
| 0 | 0 | 0 |
| 0 | 0 | 0 |
| 0 | 0 | 0 |

| | | |
|---|---|---|
| 0 | 0 | 0 |
| 0 | 0 | 0 |
| 1 | 1 | 1 |
| 0 | 0 | 0 |
| 0 | 0 | 0 |
| 0 | 1 | 0 |
| 0 | 1 | 0 |
| 0 | 1 | 0 |
| 0 | 0 | 0 |
| 0 | 1 | 0 |
| 0 | 0 | 0 |
| 0 | 0 | 0 |
| 0 | 0 | 0 |
| 1 | 1 | 1 |
| 0 | 0 | 0 |
| 0 | 1 | 0 |
| 0 | 0 | 0 |
| 0 | 0 | 0 |
| 0 | 0 | 0 |
| 1 | 1 | 1 |
| 1 | 1 | 1 |
| 0 | 0 | 0 |
| 1 | 1 | 1 |
| 0 | 0 | 0 |
| 0 | 0 | 0 |
| 1 | 1 | 1 |
| 1 | 1 | 1 |
| 0 | 0 | 0 |
| 1 | 1 | 1 |
| 0 | 0 | 0 |
| 0 | 1 | 0 |
| 0 | 1 | 0 |
| 1 | 1 | 1 |
| 0 | 1 | 0 |
| 1 | 1 | 1 |
| 1 | 1 | 1 |
| 1 | 1 | 1 |
| 1 | 1 | 1 |
| 1 | 1 | 1 |
| 1 | 1 | 1 |
| 1 | 1 | 1 |
| 1 | 1 | 1 |
| 0 | 1 | 0 |
| 0 | 1 | 0 |
| 0 | 1 | 0 |
| 0 | 1 | 0 |
| 0 | 1 | 0 |
| 0 | 1 | 0 |
| 0 | 1 | 0 |
| 0 | 1 | 0 |
| 0 | 1 | 0 |
| 0 | 1 | 0 |
| 0 | 1 | 0 |
| 0 | 1 | 0 |

| | | |
|---|---|---|
| 0 | 1 | 0 |
| 1 | 1 | 1 |
| 1 | 1 | 1 |
| 1 | 1 | 1 |
| 1 | 1 | 1 |
| 1 | 1 | 1 |
| 1 | 1 | 1 |
| 0 | 1 | 0 |
| 0 | 1 | 0 |
| 0 | 1 | 0 |
| 0 | 1 | 0 |
| 0 | 0 | 0 |
| 0 | 0 | 0 |
| 0 | 0 | 0 |
| 1 | 0 | 0 |
| 0 | 0 | 0 |
| 1 | 0 | 0 |
| 0 | 0 | 0 |
| 1 | 0 | 0 |
| 1 | 1 | 1 |
| 0 | 0 | 0 |
| 1 | 1 | 1 |
| 1 | 1 | 1 |
| 0 | 0 | 0 |
| 0 | 0 | 0 |
| 1 | 1 | 1 |
| 1 | 1 | 1 |
| 0 | 0 | 0 |
| 0 | 0 | 0 |
| 0 | 0 | 0 |
| 0 | 0 | 0 |
| 0 | 0 | 0 |
| 0 | 1 | 0 |
| 0 | 1 | 0 |
| 1 | 1 | 1 |
| 1 | 1 | 1 |
| 1 | 1 | 1 |
| 1 | 1 | 1 |
| 1 | 1 | 1 |
| 0 | 0 | 0 |
| 0 | 0 | 0 |
| 0 | 1 | 0 |
| 0 | 0 | 0 |
| 0 | 0 | 0 |
| 0 | 0 | 0 |
| 1 | 0 | 0 |
| 0 | 0 | 0 |
| 1 | 0 | 0 |
| 1 | 0 | 0 |
| 0 | 0 | 0 |
| 0 | 0 | 0 |
| 0 | 0 | 0 |
| 0 | 0 | 0 |
| 0 | 0 | 0 |

| | | |
|---|---|---|
| 0 | 0 | 0 |
| 0 | 1 | 0 |
| 1 | 1 | 1 |
| 1 | 1 | 1 |
| 0 | 0 | 0 |
| 1 | 1 | 1 |
| 0 | 0 | 0 |
| 1 | 1 | 1 |
| 1 | 1 | 1 |
| 0 | 0 | 0 |
| 1 | 0 | 0 |
| 1 | 1 | 1 |
| 1 | 1 | 1 |
| 0 | 0 | 0 |
| 1 | 0 | 0 |
| 1 | 0 | 0 |
| 0 | 0 | 0 |
| 1 | 0 | 0 |
| 0 | 0 | 0 |
| 0 | 0 | 0 |
| 0 | 0 | 0 |
| 0 | 0 | 0 |
| 0 | 0 | 0 |
| 0 | 0 | 0 |
| 0 | 0 | 0 |
| 0 | 0 | 0 |
| 0 | 0 | 0 |
| 0 | 0 | 0 |
| 0 | 0 | 0 |
| 0 | 0 | 0 |
| 0 | 0 | 0 |
| 0 | 0 | 0 |
| 0 | 0 | 0 |
| 0 | 0 | 0 |
| 0 | 0 | 0 |
| 0 | 0 | 0 |
| 1 | 0 | 0 |
| 0 | 0 | 0 |
| 0 | 0 | 0 |
| 0 | 0 | 0 |
| 0 | 0 | 0 |
| 0 | 0 | 0 |
| 0 | 0 | 0 |
| 0 | 0 | 0 |
| 0 | 0 | 0 |
| 0 | 0 | 0 |
| 0 | 0 | 0 |
| 0 | 0 | 0 |
| 1 | 0 | 0 |
| 1 | 0 | 0 |
| 0 | 0 | 0 |
| 1 | 1 | 1 |
| 0 | 0 | 0 |
| 0 | 0 | 0 |

| | | |
|---|---|---|
| 0 | 0 | 0 |
| 0 | 0 | 0 |
| 0 | 0 | 0 |
| 0 | 0 | 0 |
| 0 | 0 | 0 |
| 0 | 0 | 0 |
| 0 | 0 | 0 |
| 0 | 0 | 0 |
| 0 | 0 | 0 |
| 0 | 0 | 0 |
| 0 | 0 | 0 |
| 0 | 0 | 0 |
| 0 | 0 | 0 |
| 0 | 0 | 0 |
| 0 | 1 | 0 |
| 0 | 0 | 0 |
| 0 | 0 | 0 |
| 0 | 0 | 0 |
| 1 | 0 | 0 |
| 0 | 0 | 0 |
| 0 | 0 | 0 |
| 0 | 0 | 0 |
| 0 | 0 | 0 |
| 0 | 0 | 0 |
| 0 | 0 | 0 |
| 1 | 0 | 0 |
| 0 | 0 | 0 |
| 0 | 0 | 0 |
| 0 | 0 | 0 |
| 0 | 0 | 0 |
| 1 | 1 | 1 |
| 0 | 0 | 0 |
| 1 | 1 | 1 |
| 1 | 0 | 0 |
| 0 | 0 | 0 |
| 0 | 0 | 0 |
| 1 | 1 | 1 |
| 1 | 1 | 1 |
| 1 | 1 | 1 |
| 1 | 0 | 0 |
| 0 | 0 | 0 |
| 0 | 0 | 0 |
| 0 | 0 | 0 |
| 0 | 0 | 0 |
| 0 | 0 | 0 |
| 0 | 0 | 0 |
| 0 | 0 | 0 |
| 0 | 0 | 0 |
| 1 | 1 | 1 |
| 1 | 1 | 1 |
| 0 | 0 | 0 |
| 0 | 0 | 0 |
| 0 | 0 | 0 |
| 0 | 1 | 0 |

| | | |
|---|---|---|
| 1 | 1 | 1 |
| 1 | 1 | 1 |
| 1 | 0 | 0 |
| 0 | 0 | 0 |
| 1 | 0 | 0 |
| 1 | 1 | 1 |
| 0 | 0 | 0 |
| 1 | 1 | 1 |
| 0 | 0 | 0 |
| 0 | 0 | 0 |
| 0 | 0 | 0 |
| 0 | 0 | 0 |
| 0 | 0 | 0 |
| 0 | 0 | 0 |
| 0 | 0 | 0 |
| 0 | 0 | 0 |
| 0 | 1 | 0 |
| 0 | 0 | 0 |
| 0 | 0 | 0 |
| 1 | 0 | 0 |
| 0 | 0 | 0 |
| 0 | 0 | 0 |
| 1 | 1 | 1 |
| 0 | 0 | 0 |
| 1 | 1 | 1 |
| 1 | 1 | 1 |
| 0 | 1 | 0 |
| 0 | 0 | 0 |
| 0 | 0 | 0 |
| 0 | 0 | 0 |
| 1 | 1 | 1 |
| 0 | 0 | 0 |
| 0 | 0 | 0 |
| 0 | 0 | 0 |
| 0 | 1 | 0 |
| 0 | 1 | 0 |
| 0 | 0 | 0 |
| 0 | 0 | 0 |
| 0 | 0 | 0 |
| 1 | 1 | 1 |
| 1 | 1 | 1 |
| 0 | 1 | 0 |
| 0 | 1 | 0 |
| 0 | 0 | 0 |
| 1 | 1 | 1 |
| 0 | 0 | 0 |
| 1 | 1 | 1 |
| 0 | 0 | 0 |
| 0 | 0 | 0 |
| 1 | 1 | 1 |
| 0 | 1 | 0 |
| 0 | 0 | 0 |
| 0 | 0 | 0 |
| 1 | 1 | 1 |

| | | |
|---|---|---|
| 1 | 1 | 1 |
| 1 | 1 | 1 |
| 0 | 1 | 0 |
| 0 | 0 | 0 |
| 0 | 0 | 0 |
| 1 | 0 | 0 |
| 1 | 0 | 0 |
| 0 | 0 | 0 |
| 0 | 0 | 0 |
| 1 | 1 | 1 |
| 0 | 0 | 0 |
| 0 | 0 | 0 |
| 0 | 0 | 0 |
| 0 | 0 | 0 |
| 0 | 0 | 0 |
| 0 | 0 | 0 |
| 0 | 0 | 0 |
| 0 | 0 | 0 |
| 0 | 0 | 0 |
| 0 | 1 | 0 |
| 1 | 1 | 1 |
| 1 | 1 | 1 |
| 1 | 1 | 1 |
| 1 | 1 | 1 |
| 0 | 0 | 0 |
| 0 | 1 | 0 |
| 0 | 0 | 0 |
| 0 | 0 | 0 |
| 1 | 1 | 1 |
| 1 | 1 | 1 |
| 1 | 1 | 1 |
| 1 | 1 | 1 |
| 1 | 1 | 1 |
| 0 | 0 | 0 |
| 0 | 0 | 0 |
| 1 | 1 | 1 |
| 0 | 0 | 0 |
| 0 | 1 | 0 |
| 0 | 0 | 0 |
| 0 | 0 | 0 |
| 0 | 0 | 0 |
| 0 | 0 | 0 |
| 0 | 1 | 0 |
| 0 | 0 | 0 |
| 0 | 0 | 0 |
| 0 | 0 | 0 |
| 0 | 0 | 0 |
| 0 | 0 | 0 |
| 0 | 0 | 0 |
| 0 | 0 | 0 |
| 0 | 0 | 0 |
| 0 | 0 | 0 |
| 0 | 0 | 0 |
| 0 | 1 | 0 |

| | | |
|---|---|---|
| 0 | 0 | 0 |
| 0 | 1 | 0 |
| 0 | 0 | 0 |
| 0 | 0 | 0 |
| 0 | 0 | 0 |
| 1 | 1 | 1 |
| 0 | 0 | 0 |
| 1 | 1 | 1 |
| 0 | 1 | 0 |
| 0 | 0 | 0 |
| 0 | 1 | 0 |
| 0 | 1 | 0 |
| 0 | 1 | 0 |
| 0 | 1 | 0 |
| 0 | 1 | 0 |
| 0 | 0 | 0 |
| 0 | 0 | 0 |
| 0 | 1 | 0 |
| 0 | 0 | 0 |
| 1 | 1 | 1 |
| 0 | 0 | 0 |
| 0 | 0 | 0 |
| 0 | 0 | 0 |
| 0 | 0 | 0 |
| 0 | 0 | 0 |
| 0 | 0 | 0 |
| 0 | 0 | 0 |
| 0 | 0 | 0 |
| 0 | 0 | 0 |
| 0 | 0 | 0 |
| 0 | 0 | 0 |
| 0 | 0 | 0 |
| 1 | 0 | 0 |
| 1 | 0 | 0 |
| 0 | 0 | 0 |
| 0 | 0 | 0 |
| 0 | 0 | 0 |
| 0 | 0 | 0 |
| 1 | 0 | 0 |
| 0 | 0 | 0 |
| 0 | 0 | 0 |
| 0 | 0 | 0 |
| 1 | 0 | 0 |
| 0 | 1 | 0 |
| 0 | 0 | 0 |
| 0 | 0 | 0 |
| 0 | 0 | 0 |
| 0 | 0 | 0 |
| 0 | 0 | 0 |
| 0 | 0 | 0 |
| 1 | 0 | 0 |
| 0 | 0 | 0 |
| 0 | 0 | 0 |
| 1 | 0 | 0 |

| | | |
|---|---|---|
| 0 | 0 | 0 |
| 1 | 0 | 0 |
| 0 | 0 | 0 |
| 0 | 0 | 0 |
| 0 | 0 | 0 |
| 0 | 0 | 0 |
| 0 | 0 | 0 |
| 0 | 0 | 0 |
| 0 | 0 | 0 |
| 0 | 0 | 0 |
| 0 | 0 | 0 |
| 1 | 1 | 1 |
| 0 | 0 | 0 |
| 1 | 1 | 1 |
| 1 | 0 | 0 |
| 1 | 0 | 0 |
| 0 | 0 | 0 |
| 0 | 0 | 0 |
| 0 | 0 | 0 |
| 0 | 0 | 0 |
| 1 | 0 | 0 |
| 0 | 0 | 0 |
| 0 | 1 | 0 |
| 0 | 0 | 0 |
| 1 | 1 | 1 |
| 1 | 1 | 1 |
| 1 | 1 | 1 |
| 0 | 0 | 0 |
| 1 | 1 | 1 |
| 1 | 1 | 1 |
| 1 | 1 | 1 |
| 0 | 0 | 0 |
| 1 | 1 | 1 |
| 0 | 0 | 0 |
| 0 | 0 | 0 |
| 0 | 0 | 0 |
| 0 | 1 | 0 |
| 1 | 1 | 1 |
| 0 | 0 | 0 |
| 0 | 0 | 0 |
| 0 | 0 | 0 |
| 0 | 0 | 0 |
| 0 | 0 | 0 |
| 0 | 0 | 0 |
| 0 | 0 | 0 |
| 0 | 0 | 0 |
| 1 | 1 | 1 |
| 1 | 1 | 1 |
| 0 | 0 | 0 |
| 0 | 0 | 0 |
| 0 | 0 | 0 |
| 1 | 1 | 1 |
| 1 | 1 | 1 |
| 0 | 0 | 0 |

| | | |
|---|---|---|
| 0 | 0 | 0 |
| 0 | 1 | 0 |
| 0 | 1 | 0 |
| 1 | 1 | 1 |
| 0 | 0 | 0 |
| 0 | 0 | 0 |
| 0 | 0 | 0 |
| 0 | 0 | 0 |
| 0 | 0 | 0 |
| 0 | 0 | 0 |
| 1 | 1 | 1 |
| 0 | 0 | 0 |
| 0 | 0 | 0 |
| 0 | 0 | 0 |
| 1 | 0 | 0 |
| 1 | 1 | 1 |
| 0 | 0 | 0 |
| 1 | 0 | 0 |
| 0 | 0 | 0 |
| 0 | 0 | 0 |
| 1 | 0 | 0 |
| 0 | 0 | 0 |
| 0 | 0 | 0 |
| 1 | 0 | 0 |
| 0 | 0 | 0 |
| 0 | 0 | 0 |
| 0 | 0 | 0 |
| 1 | 0 | 0 |
| 0 | 0 | 0 |
| 0 | 0 | 0 |
| 0 | 0 | 0 |
| 1 | 1 | 1 |
| 1 | 1 | 1 |
| 1 | 1 | 1 |
| 0 | 1 | 0 |
| 0 | 1 | 0 |
| 0 | 1 | 0 |
| 1 | 1 | 1 |
| 1 | 1 | 1 |
| 1 | 1 | 1 |
| 0 | 1 | 0 |
| 1 | 1 | 1 |
| 0 | 0 | 0 |
| 0 | 1 | 0 |
| 1 | 1 | 1 |
| 1 | 1 | 1 |
| 0 | 1 | 0 |
| 1 | 1 | 1 |
| 1 | 1 | 1 |
| 1 | 1 | 1 |
| 1 | 1 | 1 |
| 1 | 1 | 1 |
| 1 | 1 | 1 |
| 1 | 1 | 1 |

| | | |
|---|---|---|
| 0 | 1 | 0 |
| 0 | 1 | 0 |
| 1 | 1 | 1 |
| 1 | 1 | 1 |
| 1 | 1 | 1 |
| 1 | 1 | 1 |
| 1 | 1 | 1 |
| 1 | 1 | 1 |
| 0 | 0 | 0 |
| 0 | 0 | 0 |
| 0 | 0 | 0 |
| 0 | 0 | 0 |
| 0 | 0 | 0 |
| 0 | 0 | 0 |
| 0 | 0 | 0 |
| 0 | 0 | 0 |
| 0 | 0 | 0 |
| 0 | 0 | 0 |
| 0 | 0 | 0 |
| 0 | 0 | 0 |
| 0 | 0 | 0 |
| 0 | 0 | 0 |
| 0 | 0 | 0 |
| 0 | 0 | 0 |
| 0 | 0 | 0 |
| 1 | 1 | 1 |
| 1 | 1 | 1 |
| 0 | 0 | 0 |
| 0 | 0 | 0 |
| 0 | 0 | 0 |
| 0 | 0 | 0 |
| 0 | 0 | 0 |
| 0 | 0 | 0 |
| 0 | 0 | 0 |
| 0 | 0 | 0 |
| 0 | 0 | 0 |
| 0 | 0 | 0 |
| 0 | 0 | 0 |
| 0 | 0 | 0 |
| 0 | 0 | 0 |
| 0 | 0 | 0 |
| 0 | 0 | 0 |
| 0 | 0 | 0 |
| 0 | 0 | 0 |
| 0 | 0 | 0 |
| 0 | 0 | 0 |
| 1 | 1 | 1 |
| 0 | 1 | 0 |
| 1 | 1 | 1 |
| 1 | 1 | 1 |
| 1 | 1 | 1 |
| 1 | 1 | 1 |
| 0 | 0 | 0 |
| 0 | 0 | 0 |

| | | |
|---|---|---|
| 0 | 0 | 0 |
| 0 | 0 | 0 |
| 0 | 0 | 0 |
| 0 | 0 | 0 |
| 0 | 1 | 0 |
| 0 | 0 | 0 |
| 0 | 0 | 0 |
| 0 | 0 | 0 |
| 1 | 1 | 1 |
| 0 | 0 | 0 |
| 0 | 0 | 0 |
| 0 | 1 | 0 |
| 0 | 0 | 0 |
| 0 | 0 | 0 |
| 0 | 0 | 0 |
| 0 | 0 | 0 |
| 1 | 1 | 1 |
| 0 | 0 | 0 |
| 0 | 0 | 0 |
| 1 | 1 | 1 |
| 0 | 0 | 0 |
| 0 | 0 | 0 |
| 0 | 0 | 0 |
| 0 | 0 | 0 |
| 0 | 0 | 0 |
| 0 | 1 | 0 |
| 0 | 1 | 0 |
| 1 | 1 | 1 |
| 1 | 1 | 1 |
| 0 | 1 | 0 |
| 1 | 1 | 1 |
| 1 | 1 | 1 |
| 1 | 0 | 0 |
| 1 | 0 | 0 |
| 0 | 0 | 0 |
| 0 | 0 | 0 |
| 0 | 0 | 0 |
| 0 | 0 | 0 |
| 0 | 0 | 0 |
| 0 | 0 | 0 |
| 1 | 1 | 1 |
| 1 | 1 | 1 |
| 1 | 1 | 1 |
| 0 | 0 | 0 |
| 0 | 0 | 0 |
| 0 | 0 | 0 |
| 1 | 1 | 1 |
| 0 | 1 | 0 |
| 0 | 0 | 0 |
| 0 | 0 | 0 |
| 1 | 1 | 1 |
| 1 | 1 | 1 |
| 1 | 1 | 1 |
| 1 | 1 | 1 |

| | | |
|---|---|---|
| 0 | 1 | 0 |
| 1 | 1 | 1 |
| 0 | 1 | 0 |
| 0 | 0 | 0 |
| 0 | 0 | 0 |
| 1 | 1 | 1 |
| 1 | 1 | 1 |
| 1 | 1 | 1 |
| 1 | 1 | 1 |
| 1 | 1 | 1 |
| 0 | 0 | 0 |
| 0 | 1 | 0 |
| 0 | 0 | 0 |
| 0 | 0 | 0 |
| 0 | 0 | 0 |
| 1 | 1 | 1 |
| 1 | 1 | 1 |
| 0 | 1 | 0 |
| 1 | 1 | 1 |
| 1 | 1 | 1 |
| 0 | 0 | 0 |
| 1 | 1 | 1 |
| 1 | 1 | 1 |
| 0 | 0 | 0 |
| 1 | 0 | 0 |
| 0 | 0 | 0 |
| 0 | 0 | 0 |
| 1 | 1 | 1 |
| 1 | 1 | 1 |
| 1 | 1 | 1 |
| 1 | 1 | 1 |
| 1 | 1 | 1 |
| 1 | 1 | 1 |
| 1 | 1 | 1 |
| 1 | 1 | 1 |
| 0 | 0 | 0 |
| 1 | 1 | 1 |
| 1 | 1 | 1 |
| 0 | 0 | 0 |
| 0 | 0 | 0 |
| 1 | 1 | 1 |
| 0 | 1 | 0 |
| 0 | 0 | 0 |
| 0 | 0 | 0 |
| 0 | 0 | 0 |
| 0 | 0 | 0 |
| 0 | 0 | 0 |
| 0 | 0 | 0 |
| 1 | 1 | 1 |
| 0 | 0 | 0 |
| 0 | 0 | 0 |
| 0 | 0 | 0 |
| 0 | 0 | 0 |
| 0 | 0 | 0 |

| | | |
|---|---|---|
| 0 | 0 | 0 |
| 0 | 0 | 0 |
| 0 | 0 | 0 |
| 0 | 0 | 0 |
| 0 | 0 | 0 |
| 0 | 0 | 0 |
| 0 | 0 | 0 |
| 0 | 0 | 0 |
| 0 | 0 | 0 |
| 0 | 1 | 0 |
| 0 | 1 | 0 |
| 0 | 0 | 0 |
| 1 | 1 | 1 |
| 0 | 0 | 0 |
| 0 | 0 | 0 |
| 0 | 1 | 0 |
| 0 | 0 | 0 |
| 1 | 0 | 0 |
| 1 | 1 | 1 |
| 1 | 1 | 1 |
| 0 | 0 | 0 |
| 0 | 1 | 0 |
| 0 | 1 | 0 |
| 0 | 0 | 0 |
| 0 | 0 | 0 |
| 0 | 1 | 0 |
| 0 | 1 | 0 |
| 1 | 1 | 1 |
| 0 | 0 | 0 |
| 0 | 0 | 0 |
| 1 | 1 | 1 |
| 1 | 1 | 1 |
| 0 | 0 | 0 |
| 0 | 0 | 0 |
| 0 | 0 | 0 |
| 0 | 1 | 0 |
| 0 | 0 | 0 |
| 0 | 1 | 0 |
| 0 | 1 | 0 |
| 0 | 0 | 0 |
| 0 | 0 | 0 |
| 0 | 0 | 0 |
| 0 | 0 | 0 |
| 0 | 0 | 0 |
| 0 | 0 | 0 |
| 1 | 1 | 1 |
| 0 | 1 | 0 |
| 0 | 0 | 0 |
| 0 | 0 | 0 |
| 0 | 0 | 0 |
| 1 | 1 | 1 |
| 1 | 1 | 1 |
| 0 | 0 | 0 |
| 0 | 0 | 0 |

| | | |
|---|---|---|
| 1 | 1 | 1 |
| 1 | 1 | 1 |
| 1 | 1 | 1 |
| 1 | 1 | 1 |
| 0 | 0 | 0 |
| 0 | 0 | 0 |
| 1 | 0 | 0 |
| 0 | 0 | 0 |
| 1 | 0 | 0 |
| 0 | 0 | 0 |
| 1 | 1 | 1 |
| 1 | 1 | 1 |
| 1 | 1 | 1 |
| 1 | 1 | 1 |
| 1 | 1 | 1 |
| 1 | 1 | 1 |
| 0 | 0 | 0 |
| 1 | 1 | 1 |
| 1 | 1 | 1 |
| 0 | 0 | 0 |
| 0 | 0 | 0 |
| 0 | 1 | 0 |
| 0 | 1 | 0 |
| 0 | 0 | 0 |
| 0 | 1 | 0 |
| 0 | 0 | 0 |
| 0 | 0 | 0 |
| 0 | 0 | 0 |
| 0 | 0 | 0 |
| 0 | 0 | 0 |
| 0 | 0 | 0 |
| 0 | 0 | 0 |
| 0 | 1 | 0 |
| 0 | 0 | 0 |
| 1 | 1 | 1 |
| 1 | 1 | 1 |
| 1 | 1 | 1 |
| 1 | 1 | 1 |
| 0 | 0 | 0 |
| 1 | 1 | 1 |
| 1 | 1 | 1 |
| 1 | 1 | 1 |
| 1 | 1 | 1 |
| 0 | 0 | 0 |
| 1 | 1 | 1 |
| 1 | 1 | 1 |
| 1 | 1 | 1 |
| 1 | 1 | 1 |
| 1 | 1 | 1 |
| 1 | 1 | 1 |
| 0 | 0 | 0 |
| 0 | 0 | 0 |
| 0 | 0 | 0 |
| 0 | 1 | 0 |

| | | |
|---|---|---|
| 0 | 0 | 0 |
| 0 | 0 | 0 |
| 0 | 0 | 0 |
| 0 | 0 | 0 |
| 0 | 0 | 0 |
| 0 | 0 | 0 |
| 0 | 0 | 0 |
| 0 | 0 | 0 |
| 1 | 1 | 1 |
| 1 | 1 | 1 |
| 0 | 0 | 0 |
| 1 | 1 | 1 |
| 1 | 1 | 1 |
| 0 | 0 | 0 |
| 0 | 0 | 0 |
| 0 | 0 | 0 |
| 0 | 0 | 0 |
| 0 | 0 | 0 |
| 0 | 0 | 0 |
| 1 | 1 | 1 |
| 1 | 1 | 1 |
| 0 | 0 | 0 |
| 1 | 1 | 1 |
| 0 | 0 | 0 |
| 1 | 1 | 1 |
| 1 | 1 | 1 |
| 1 | 1 | 1 |
| 0 | 1 | 0 |
| 0 | 1 | 0 |
| 0 | 0 | 0 |
| 0 | 1 | 0 |
| 1 | 1 | 1 |
| 0 | 0 | 0 |
| 1 | 1 | 1 |
| 0 | 0 | 0 |
| 0 | 0 | 0 |
| 0 | 0 | 0 |
| 0 | 0 | 0 |
| 0 | 0 | 0 |
| 0 | 0 | 0 |
| 0 | 0 | 0 |
| 0 | 0 | 0 |
| 1 | 1 | 1 |
| 0 | 0 | 0 |
| 1 | 1 | 1 |
| 0 | 0 | 0 |
| 1 | 0 | 0 |
| 1 | 1 | 1 |
| 1 | 1 | 1 |
| 0 | 0 | 0 |
| 1 | 1 | 1 |
| 1 | 1 | 1 |
| 0 | 1 | 0 |
| 1 | 1 | 1 |

| | | |
|---|---|---|
| 0 | 0 | 0 |
| 0 | 1 | 0 |
| 1 | 1 | 1 |
| 0 | 1 | 0 |
| 0 | 1 | 0 |
| 0 | 1 | 0 |
| 0 | 1 | 0 |
| 1 | 1 | 1 |
| 1 | 1 | 1 |
| 1 | 1 | 1 |
| 0 | 0 | 0 |
| 1 | 1 | 1 |
| 1 | 1 | 1 |
| 0 | 1 | 0 |
| 0 | 0 | 0 |
| 0 | 0 | 0 |
| 0 | 0 | 0 |
| 0 | 0 | 0 |
| 0 | 0 | 0 |
| 0 | 1 | 0 |
| 0 | 1 | 0 |
| 1 | 1 | 1 |
| 0 | 0 | 0 |
| 0 | 1 | 0 |
| 0 | 1 | 0 |
| 0 | 1 | 0 |
| 1 | 1 | 1 |
| 1 | 1 | 1 |
| 0 | 0 | 0 |
| 0 | 0 | 0 |
| 0 | 0 | 0 |
| 0 | 0 | 0 |
| 0 | 0 | 0 |
| 0 | 0 | 0 |
| 1 | 1 | 1 |
| 1 | 1 | 1 |
| 0 | 0 | 0 |
| 0 | 0 | 0 |
| 0 | 0 | 0 |
| 0 | 0 | 0 |
| 0 | 0 | 0 |
| 0 | 0 | 0 |
| 1 | 1 | 1 |
| 1 | 1 | 1 |
| 0 | 0 | 0 |
| 1 | 1 | 1 |
| 1 | 1 | 1 |
| 1 | 1 | 1 |
| 1 | 1 | 1 |
| 1 | 1 | 1 |
| 1 | 0 | 0 |
| 0 | 1 | 0 |
| 0 | 1 | 0 |
| 1 | 0 | 0 |

| | | |
|---|---|---|
| 0 | 0 | 0 |
| 1 | 1 | 1 |
| 0 | 0 | 0 |
| 0 | 1 | 0 |
| 1 | 0 | 0 |
| 0 | 0 | 0 |
| 0 | 0 | 0 |
| 0 | 0 | 0 |
| 1 | 0 | 0 |
| 1 | 0 | 0 |
| 0 | 0 | 0 |
| 1 | 1 | 1 |
| 1 | 0 | 0 |
| 1 | 0 | 0 |
| 0 | 0 | 0 |
| 0 | 0 | 0 |
| 0 | 0 | 0 |
| 0 | 0 | 0 |
| 1 | 1 | 1 |
| 0 | 0 | 0 |
| 0 | 0 | 0 |
| 0 | 0 | 0 |
| 0 | 0 | 0 |
| 0 | 0 | 0 |
| 1 | 1 | 1 |
| 0 | 0 | 0 |
| 1 | 1 | 1 |
| 0 | 1 | 0 |
| 1 | 1 | 1 |
| 0 | 0 | 0 |
| 1 | 1 | 1 |
| 1 | 1 | 1 |
| 0 | 0 | 0 |
| 0 | 0 | 0 |
| 1 | 1 | 1 |
| 1 | 1 | 1 |
| 1 | 1 | 1 |
| 0 | 1 | 0 |
| 1 | 0 | 0 |
| 0 | 0 | 0 |
| 0 | 0 | 0 |
| 0 | 0 | 0 |
| 0 | 0 | 0 |
| 0 | 0 | 0 |
| 0 | 0 | 0 |
| 0 | 1 | 0 |
| 0 | 1 | 0 |
| 0 | 0 | 0 |
| 0 | 0 | 0 |
| 1 | 1 | 1 |
| 0 | 0 | 0 |
| 0 | 0 | 0 |
| 0 | 1 | 0 |
| 1 | 1 | 1 |

| | | |
|---|---|---|
| 1 | 1 | 1 |
| 0 | 0 | 0 |
| 0 | 0 | 0 |
| 1 | 0 | 0 |
| 1 | 1 | 1 |
| 0 | 0 | 0 |
| 0 | 0 | 0 |
| 0 | 0 | 0 |
| 0 | 0 | 0 |
| 0 | 0 | 0 |
| 0 | 0 | 0 |
| 0 | 0 | 0 |
| 0 | 0 | 0 |
| 0 | 0 | 0 |
| 0 | 0 | 0 |
| 0 | 1 | 0 |
| 0 | 0 | 0 |
| 1 | 1 | 1 |
| 1 | 0 | 0 |
| 1 | 1 | 1 |
| 0 | 0 | 0 |
| 1 | 1 | 1 |
| 0 | 0 | 0 |
| 0 | 0 | 0 |
| 1 | 1 | 1 |
| 0 | 0 | 0 |
| 0 | 1 | 0 |
| 0 | 1 | 0 |
| 0 | 1 | 0 |
| 0 | 0 | 0 |
| 0 | 0 | 0 |
| 0 | 0 | 0 |
| 0 | 0 | 0 |
| 0 | 0 | 0 |
| 0 | 0 | 0 |
| 0 | 0 | 0 |
| 0 | 0 | 0 |
| 1 | 1 | 1 |
| 1 | 0 | 0 |
| 0 | 0 | 0 |
| 0 | 0 | 0 |
| 0 | 0 | 0 |
| 0 | 0 | 0 |
| 0 | 0 | 0 |
| 0 | 0 | 0 |
| 0 | 0 | 0 |
| 1 | 1 | 1 |
| 0 | 0 | 0 |
| 0 | 0 | 0 |
| 0 | 0 | 0 |
| 0 | 0 | 0 |
| 0 | 1 | 0 |
| 0 | 0 | 0 |
| 0 | 0 | 0 |

| | | |
|---|---|---|
| 0 | 0 | 0 |
| 0 | 0 | 0 |
| 0 | 0 | 0 |
| 0 | 0 | 0 |
| 0 | 0 | 0 |
| 0 | 0 | 0 |
| 0 | 0 | 0 |
| 0 | 0 | 0 |
| 0 | 0 | 0 |
| 0 | 0 | 0 |
| 1 | 1 | 1 |
| 0 | 0 | 0 |
| 1 | 0 | 0 |
| 0 | 0 | 0 |
| 0 | 1 | 0 |
| 1 | 1 | 1 |
| 1 | 1 | 1 |
| 1 | 1 | 1 |
| 0 | 0 | 0 |
| 0 | 0 | 0 |
| 1 | 1 | 1 |
| 0 | 0 | 0 |
| 0 | 0 | 0 |
| 1 | 1 | 1 |
| 0 | 0 | 0 |
| 0 | 0 | 0 |
| 0 | 0 | 0 |
| 1 | 0 | 0 |
| 0 | 0 | 0 |
| 0 | 0 | 0 |
| 1 | 0 | 0 |
| 0 | 0 | 0 |
| 1 | 0 | 0 |
| 1 | 0 | 0 |
| 0 | 0 | 0 |
| 0 | 0 | 0 |
| 0 | 0 | 0 |
| 1 | 0 | 0 |
| 1 | 0 | 0 |
| 0 | 0 | 0 |
| 0 | 0 | 0 |
| 0 | 0 | 0 |
| 0 | 0 | 0 |
| 0 | 0 | 0 |
| 0 | 0 | 0 |
| 0 | 0 | 0 |
| 0 | 0 | 0 |
| 0 | 0 | 0 |
| 0 | 0 | 0 |
| 0 | 0 | 0 |
| 0 | 0 | 0 |
| 0 | 0 | 0 |
| 0 | 0 | 0 |
| 0 | 0 | 0 |

| | | |
|---|---|---|
| 0 | 0 | 0 |
| 1 | 0 | 0 |
| 0 | 0 | 0 |
| 0 | 0 | 0 |
| 0 | 0 | 0 |
| 1 | 0 | 0 |
| 0 | 0 | 0 |
| 0 | 0 | 0 |
| 1 | 1 | 1 |
| 0 | 0 | 0 |
| 0 | 1 | 0 |
| 1 | 1 | 1 |
| 1 | 1 | 1 |
| 0 | 0 | 0 |
| 1 | 1 | 1 |
| 0 | 0 | 0 |
| 1 | 1 | 1 |
| 1 | 1 | 1 |
| 0 | 0 | 0 |
| 0 | 0 | 0 |
| 0 | 0 | 0 |
| 0 | 0 | 0 |
| 0 | 0 | 0 |
| 0 | 0 | 0 |
| 0 | 0 | 0 |
| 0 | 1 | 0 |
| 1 | 1 | 1 |
| 1 | 1 | 1 |
| 1 | 0 | 0 |
| 1 | 1 | 1 |
| 1 | 1 | 1 |
| 0 | 1 | 0 |
| 0 | 0 | 0 |
| 0 | 0 | 0 |
| 0 | 0 | 0 |
| 0 | 0 | 0 |
| 0 | 0 | 0 |
| 0 | 0 | 0 |
| 0 | 1 | 0 |
| 0 | 1 | 0 |
| 0 | 1 | 0 |
| 0 | 1 | 0 |
| 0 | 1 | 0 |
| 0 | 1 | 0 |
| 0 | 0 | 0 |
| 0 | 0 | 0 |
| 0 | 1 | 0 |
| 0 | 0 | 0 |
| 0 | 0 | 0 |
| 0 | 0 | 0 |
| 0 | 0 | 0 |
| 0 | 0 | 0 |
| 0 | 0 | 0 |
| 0 | 0 | 0 |

| | | |
|---|---|---|
| 1 | 1 | 1 |
| 1 | 1 | 1 |
| 0 | 0 | 0 |
| 0 | 0 | 0 |
| 1 | 1 | 1 |
| 0 | 0 | 0 |
| 0 | 0 | 0 |
| 1 | 1 | 1 |
| 1 | 1 | 1 |
| 0 | 0 | 0 |
| 1 | 1 | 1 |
| 1 | 1 | 1 |
| 0 | 0 | 0 |
| 0 | 0 | 0 |
| 0 | 0 | 0 |
| 0 | 0 | 0 |
| 0 | 1 | 0 |
| 0 | 1 | 0 |
| 1 | 1 | 1 |
| 1 | 1 | 1 |
| 1 | 1 | 1 |
| 0 | 1 | 0 |
| 1 | 1 | 1 |
| 1 | 1 | 1 |
| 1 | 1 | 1 |
| 0 | 0 | 0 |
| 1 | 1 | 1 |
| 1 | 1 | 1 |
| 1 | 1 | 1 |
| 0 | 1 | 0 |
| 0 | 1 | 0 |
| 1 | 1 | 1 |
| 1 | 1 | 1 |
| 1 | 1 | 1 |
| 1 | 1 | 1 |
| 1 | 1 | 1 |
| 1 | 1 | 1 |
| 0 | 0 | 0 |
| 0 | 0 | 0 |
| 1 | 0 | 0 |
| 0 | 0 | 0 |
| 1 | 1 | 1 |
| 1 | 1 | 1 |
| 0 | 0 | 0 |
| 0 | 0 | 0 |
| 0 | 1 | 0 |
| 0 | 1 | 0 |
| 0 | 1 | 0 |
| 1 | 1 | 1 |
| 1 | 1 | 1 |
| 1 | 1 | 1 |
| 0 | 0 | 0 |
| 0 | 0 | 0 |
| 0 | 0 | 0 |

| | | |
|---|---|---|
| 0 | 0 | 0 |
| 1 | 1 | 1 |
| 1 | 1 | 1 |
| 1 | 1 | 1 |
| 1 | 1 | 1 |
| 1 | 1 | 1 |
| 1 | 1 | 1 |
| 1 | 1 | 1 |
| 0 | 0 | 0 |
| 1 | 1 | 1 |
| 1 | 1 | 1 |
| 1 | 1 | 1 |
| 0 | 1 | 0 |
| 0 | 1 | 0 |
| 1 | 1 | 1 |
| 1 | 1 | 1 |
| 0 | 1 | 0 |
| 0 | 1 | 0 |
| 0 | 0 | 0 |
| 0 | 0 | 0 |
| 0 | 0 | 0 |
| 0 | 0 | 0 |
| 0 | 1 | 0 |
| 0 | 0 | 0 |
| 0 | 0 | 0 |
| 1 | 1 | 1 |
| 1 | 1 | 1 |
| 1 | 1 | 1 |
| 1 | 1 | 1 |
| 1 | 1 | 1 |
| 1 | 1 | 1 |
| 1 | 1 | 1 |
| 0 | 0 | 0 |
| 1 | 1 | 1 |
| 1 | 1 | 1 |
| 0 | 1 | 0 |
| 0 | 0 | 0 |
| 1 | 1 | 1 |
| 0 | 0 | 0 |
| 0 | 0 | 0 |
| 0 | 0 | 0 |
| 0 | 0 | 0 |
| 0 | 0 | 0 |
| 1 | 1 | 1 |
| 1 | 1 | 1 |
| 1 | 1 | 1 |
| 0 | 0 | 0 |
| 0 | 0 | 0 |
| 0 | 0 | 0 |
| 0 | 0 | 0 |
| 0 | 0 | 0 |
| 0 | 0 | 0 |
| 0 | 0 | 0 |
| 0 | 1 | 0 |

| | | |
|---|---|---|
| 0 | 0 | 0 |
| 0 | 0 | 0 |
| 0 | 1 | 0 |
| 0 | 0 | 0 |
| 0 | 0 | 0 |
| 1 | 1 | 1 |
| 1 | 1 | 1 |
| 1 | 0 | 0 |
| 1 | 1 | 1 |
| 1 | 1 | 1 |
| 1 | 1 | 1 |
| 0 | 0 | 0 |
| 1 | 1 | 1 |
| 1 | 1 | 1 |
| 0 | 0 | 0 |
| 1 | 1 | 1 |
| 0 | 0 | 0 |
| 1 | 1 | 1 |
| 0 | 0 | 0 |
| 0 | 0 | 0 |
| 1 | 1 | 1 |
| 1 | 1 | 1 |
| 0 | 0 | 0 |
| 1 | 1 | 1 |
| 0 | 0 | 0 |
| 1 | 1 | 1 |
| 1 | 1 | 1 |
| 0 | 0 | 0 |
| 1 | 1 | 1 |
| 0 | 0 | 0 |
| 0 | 1 | 0 |
| 0 | 0 | 0 |
| 0 | 0 | 0 |
| 0 | 0 | 0 |
| 1 | 1 | 1 |
| 0 | 0 | 0 |
| 1 | 1 | 1 |
| 1 | 1 | 1 |
| 0 | 0 | 0 |
| 1 | 1 | 1 |
| 1 | 1 | 1 |
| 1 | 1 | 1 |
| 1 | 1 | 1 |
| 0 | 0 | 0 |
| 0 | 0 | 0 |
| 1 | 1 | 1 |
| 1 | 1 | 1 |
| 1 | 0 | 0 |
| 0 | 0 | 0 |
| 1 | 0 | 0 |
| 0 | 0 | 0 |
| 1 | 1 | 1 |
| 1 | 1 | 1 |
| 0 | 0 | 0 |

| | | |
|---|---|---|
| 1 | 1 | 1 |
| 1 | 1 | 1 |
| 1 | 1 | 1 |
| 1 | 1 | 1 |
| 0 | 1 | 0 |
| 1 | 1 | 1 |
| 1 | 1 | 1 |
| 0 | 0 | 0 |
| 1 | 1 | 1 |
| 1 | 1 | 1 |
| 1 | 1 | 1 |
| 1 | 1 | 1 |
| 0 | 0 | 0 |
| 1 | 1 | 1 |
| 0 | 0 | 0 |
| 0 | 0 | 0 |
| 1 | 1 | 1 |
| 1 | 1 | 1 |
| 1 | 1 | 1 |
| 1 | 1 | 1 |
| 0 | 0 | 0 |
| 0 | 0 | 0 |
| 1 | 1 | 1 |
| 1 | 1 | 1 |
| 0 | 0 | 0 |
| 1 | 1 | 1 |
| 0 | 0 | 0 |
| 0 | 0 | 0 |
| 0 | 0 | 0 |
| 0 | 0 | 0 |
| 0 | 1 | 0 |
| 0 | 1 | 0 |
| 0 | 0 | 0 |
| 0 | 0 | 0 |
| 0 | 0 | 0 |
| 0 | 0 | 0 |
| 0 | 0 | 0 |
| 0 | 0 | 0 |
| 0 | 0 | 0 |
| 1 | 0 | 0 |
| 1 | 1 | 1 |
| 0 | 1 | 0 |
| 0 | 1 | 0 |
| 0 | 1 | 0 |
| 0 | 1 | 0 |
| 0 | 0 | 0 |
| 0 | 1 | 0 |
| 0 | 1 | 0 |
| 1 | 1 | 1 |
| 0 | 0 | 0 |
| 0 | 0 | 0 |
| 1 | 1 | 1 |
| 1 | 1 | 1 |
| 1 | 1 | 1 |

| | | |
|---|---|---|
| 1 | 1 | 1 |
| 1 | 1 | 1 |
| 1 | 1 | 1 |
| 1 | 1 | 1 |
| 0 | 0 | 0 |
| 1 | 1 | 1 |
| 0 | 0 | 0 |
| 0 | 0 | 0 |
| 0 | 0 | 0 |
| 0 | 0 | 0 |
| 0 | 0 | 0 |
| 1 | 1 | 1 |
| 0 | 0 | 0 |
| 0 | 0 | 0 |
| 1 | 1 | 1 |
| 1 | 1 | 1 |
| 1 | 1 | 1 |
| 1 | 1 | 1 |
| 1 | 1 | 1 |
| 1 | 1 | 1 |
| 1 | 1 | 1 |
| 1 | 1 | 1 |
| 1 | 1 | 1 |
| 0 | 0 | 0 |
| 1 | 1 | 1 |
| 1 | 1 | 1 |
| 1 | 1 | 1 |
| 0 | 0 | 0 |
| 1 | 0 | 0 |
| 1 | 0 | 0 |
| 0 | 0 | 0 |
| 0 | 0 | 0 |
| 1 | 1 | 1 |
| 0 | 0 | 0 |
| 0 | 1 | 0 |
| 1 | 1 | 1 |
| 1 | 1 | 1 |
| 1 | 1 | 1 |
| 1 | 1 | 1 |
| 1 | 1 | 1 |
| 1 | 1 | 1 |
| 1 | 1 | 1 |
| 1 | 1 | 1 |
| 1 | 1 | 1 |
| 1 | 1 | 1 |
| 1 | 1 | 1 |
| 1 | 1 | 1 |
| 1 | 1 | 1 |
| 0 | 0 | 0 |
| 0 | 1 | 0 |
| 1 | 1 | 1 |
| 1 | 0 | 0 |
| 1 | 0 | 0 |
| 1 | 1 | 1 |

| | | |
|---|---|---|
| 1 | 1 | 1 |
| 0 | 1 | 0 |
| 1 | 1 | 1 |
| 0 | 1 | 0 |
| 1 | 0 | 0 |
| 1 | 1 | 1 |
| 1 | 1 | 1 |
| 1 | 1 | 1 |
| 1 | 1 | 1 |
| 1 | 1 | 1 |
| 0 | 0 | 0 |
| 0 | 0 | 0 |
| 0 | 0 | 0 |
| 0 | 0 | 0 |
| 0 | 0 | 0 |
| 1 | 0 | 0 |
| 0 | 0 | 0 |
| 0 | 0 | 0 |
| 0 | 0 | 0 |
| 0 | 0 | 0 |
| 0 | 0 | 0 |
| 0 | 0 | 0 |
| 0 | 1 | 0 |
| 0 | 0 | 0 |
| 0 | 0 | 0 |
| 0 | 0 | 0 |
| 1 | 1 | 1 |
| 0 | 0 | 0 |
| 0 | 1 | 0 |
| 0 | 1 | 0 |
| 0 | 1 | 0 |
| 0 | 0 | 0 |
| 1 | 1 | 1 |
| 1 | 1 | 1 |
| 0 | 0 | 0 |
| 0 | 0 | 0 |
| 0 | 0 | 0 |
| 0 | 0 | 0 |
| 0 | 0 | 0 |
| 1 | 0 | 0 |
| 1 | 0 | 0 |
| 0 | 0 | 0 |
| 1 | 0 | 0 |
| 0 | 0 | 0 |
| 1 | 1 | 1 |
| 1 | 0 | 0 |
| 1 | 1 | 1 |
| 0 | 0 | 0 |
| 0 | 0 | 0 |
| 0 | 0 | 0 |
| 0 | 0 | 0 |
| 1 | 1 | 1 |
| 0 | 0 | 0 |
| 1 | 1 | 1 |

| | | |
|---|---|---|
| 1 | 1 | 1 |
| 1 | 1 | 1 |
| 1 | 1 | 1 |
| 1 | 1 | 1 |
| 1 | 1 | 1 |
| 0 | 0 | 0 |
| 0 | 0 | 0 |
| 0 | 0 | 0 |
| 0 | 0 | 0 |
| 0 | 1 | 0 |
| 0 | 0 | 0 |
| 0 | 0 | 0 |
| 0 | 0 | 0 |
| 0 | 0 | 0 |
| 0 | 0 | 0 |
| 0 | 0 | 0 |
| 1 | 1 | 1 |
| 0 | 1 | 0 |
| 1 | 1 | 1 |
| 1 | 1 | 1 |
| 1 | 1 | 1 |
| 1 | 1 | 1 |
| 1 | 1 | 1 |
| 1 | 1 | 1 |
| 1 | 1 | 1 |
| 1 | 1 | 1 |
| 1 | 1 | 1 |
| 1 | 1 | 1 |
| 0 | 0 | 0 |
| 0 | 1 | 0 |
| 0 | 1 | 0 |
| 1 | 0 | 0 |
| 1 | 0 | 0 |
| 0 | 0 | 0 |
| 1 | 1 | 1 |
| 0 | 0 | 0 |
| 1 | 0 | 0 |
| 0 | 0 | 0 |
| 1 | 1 | 1 |
| 0 | 0 | 0 |
| 1 | 1 | 1 |
| 1 | 1 | 1 |
| 0 | 1 | 0 |
| 0 | 0 | 0 |
| 1 | 1 | 1 |
| 1 | 1 | 1 |
| 1 | 1 | 1 |
| 1 | 1 | 1 |
| 1 | 1 | 1 |
| 1 | 1 | 1 |
| 1 | 1 | 1 |
| 1 | 0 | 0 |
| 0 | 0 | 0 |
| 0 | 0 | 0 |

| | | |
|---|---|---|
| 1 | 1 | 1 |
| 0 | 0 | 0 |
| 1 | 1 | 1 |
| 1 | 1 | 1 |
| 0 | 0 | 0 |
| 0 | 0 | 0 |
| 0 | 0 | 0 |
| 1 | 1 | 1 |
| 1 | 1 | 1 |
| 0 | 0 | 0 |
| 0 | 0 | 0 |
| 0 | 0 | 0 |
| 0 | 0 | 0 |
| 0 | 0 | 0 |
| 1 | 1 | 1 |
| 0 | 0 | 0 |
| 0 | 0 | 0 |
| 0 | 0 | 0 |
| 1 | 1 | 1 |
| 1 | 1 | 1 |
| 1 | 1 | 1 |
| 1 | 1 | 1 |
| 1 | 1 | 1 |
| 0 | 0 | 0 |
| 0 | 1 | 0 |
| 0 | 1 | 0 |
| 0 | 0 | 0 |
| 0 | 0 | 0 |
| 1 | 1 | 1 |
| 1 | 1 | 1 |
| 0 | 0 | 0 |
| 1 | 1 | 1 |
| 1 | 1 | 1 |
| 0 | 1 | 0 |
| 1 | 1 | 1 |
| 0 | 0 | 0 |
| 0 | 1 | 0 |
| 0 | 0 | 0 |
| 0 | 0 | 0 |
| 0 | 1 | 0 |
| 0 | 1 | 0 |
| 0 | 0 | 0 |
| 0 | 0 | 0 |
| 0 | 1 | 0 |
| 0 | 0 | 0 |
| 0 | 0 | 0 |
| 0 | 0 | 0 |
| 0 | 1 | 0 |
| 0 | 0 | 0 |
| 0 | 0 | 0 |
| 0 | 0 | 0 |
| 1 | 1 | 1 |
| 0 | 0 | 0 |
| 1 | 1 | 1 |

| | | |
|---|---|---|
| 1 | 1 | 1 |
| 1 | 1 | 1 |
| 1 | 1 | 1 |
| 1 | 1 | 1 |
| 0 | 0 | 0 |
| 0 | 0 | 0 |
| 1 | 1 | 1 |
| 1 | 0 | 0 |
| 1 | 1 | 1 |
| 0 | 0 | 0 |
| 1 | 0 | 0 |
| 1 | 0 | 0 |
| 0 | 0 | 0 |
| 1 | 1 | 1 |
| 1 | 1 | 1 |
| 1 | 1 | 1 |
| 1 | 1 | 1 |
| 1 | 1 | 1 |
| 1 | 1 | 1 |
| 1 | 1 | 1 |
| 1 | 1 | 1 |
| 0 | 1 | 0 |
| 1 | 1 | 1 |
| 0 | 0 | 0 |
| 1 | 1 | 1 |
| 0 | 0 | 0 |
| 1 | 1 | 1 |
| 0 | 0 | 0 |
| 1 | 1 | 1 |
| 0 | 0 | 0 |
| 0 | 0 | 0 |
| 0 | 0 | 0 |
| 0 | 0 | 0 |
| 0 | 0 | 0 |
| 0 | 1 | 0 |
| 0 | 0 | 0 |
| 1 | 1 | 1 |
| 0 | 0 | 0 |
| 1 | 1 | 1 |
| 1 | 1 | 1 |
| 0 | 0 | 0 |
| 0 | 0 | 0 |
| 0 | 0 | 0 |
| 0 | 0 | 0 |
| 0 | 0 | 0 |
| 0 | 0 | 0 |
| 0 | 0 | 0 |
| 0 | 0 | 0 |
| 1 | 1 | 1 |
| 0 | 0 | 0 |
| 1 | 0 | 0 |
| 1 | 0 | 0 |
| 1 | 1 | 1 |
| 1 | 1 | 1 |

| | | |
|---|---|---|
| 1 | 1 | 1 |
| 1 | 1 | 1 |
| 1 | 1 | 1 |
| 0 | 1 | 0 |
| 0 | 1 | 0 |
| 0 | 1 | 0 |
| 0 | 0 | 0 |
| 0 | 0 | 0 |
| 0 | 0 | 0 |
| 0 | 0 | 0 |
| 0 | 0 | 0 |
| 0 | 0 | 0 |
| 0 | 0 | 0 |
| 0 | 0 | 0 |
| 0 | 0 | 0 |
| 0 | 0 | 0 |
| 0 | 0 | 0 |
| 0 | 0 | 0 |
| 1 | 1 | 1 |
| 0 | 0 | 0 |
| 0 | 1 | 0 |
| 0 | 1 | 0 |
| 0 | 1 | 0 |
| 1 | 1 | 1 |
| 0 | 1 | 0 |
| 0 | 0 | 0 |
| 0 | 0 | 0 |
| 0 | 0 | 0 |
| 0 | 0 | 0 |
| 0 | 0 | 0 |
| 0 | 0 | 0 |
| 0 | 0 | 0 |
| 0 | 0 | 0 |
| 0 | 0 | 0 |
| 1 | 1 | 1 |
| 0 | 0 | 0 |
| 0 | 0 | 0 |
| 0 | 0 | 0 |
| 0 | 0 | 0 |
| 0 | 1 | 0 |
| 0 | 0 | 0 |
| 0 | 0 | 0 |
| 1 | 1 | 1 |
| 0 | 1 | 0 |
| 1 | 1 | 1 |
| 0 | 1 | 0 |
| 0 | 1 | 0 |
| 0 | 0 | 0 |
| 0 | 0 | 0 |
| 0 | 0 | 0 |
| 0 | 0 | 0 |
| 0 | 1 | 0 |
| 0 | 0 | 0 |
| 1 | 1 | 1 |

| | | |
|---|---|---|
| 0 | 1 | 0 |
| 1 | 1 | 1 |
| 1 | 1 | 1 |
| 1 | 1 | 1 |
| 1 | 1 | 1 |
| 0 | 0 | 0 |
| 0 | 1 | 0 |
| 1 | 1 | 1 |
| 1 | 1 | 1 |
| 1 | 1 | 1 |
| 0 | 0 | 0 |
| 0 | 0 | 0 |
| 0 | 0 | 0 |
| 0 | 0 | 0 |
| 0 | 0 | 0 |
| 0 | 1 | 0 |
| 1 | 0 | 0 |
| 0 | 0 | 0 |
| 1 | 1 | 1 |
| 1 | 1 | 1 |
| 1 | 1 | 1 |
| 1 | 1 | 1 |
| 0 | 0 | 0 |
| 1 | 1 | 1 |
| 0 | 0 | 0 |
| 0 | 0 | 0 |
| 0 | 0 | 0 |
| 0 | 0 | 0 |
| 1 | 1 | 1 |
| 0 | 0 | 0 |
| 1 | 1 | 1 |
| 0 | 1 | 0 |
| 0 | 1 | 0 |
| 0 | 1 | 0 |
| 1 | 1 | 1 |
| 1 | 1 | 1 |
| 0 | 1 | 0 |
| 0 | 0 | 0 |
| 0 | 0 | 0 |
| 0 | 0 | 0 |
| 0 | 0 | 0 |
| 0 | 0 | 0 |
| 1 | 1 | 1 |
| 0 | 0 | 0 |
| 0 | 0 | 0 |
| 1 | 1 | 1 |
| 0 | 0 | 0 |
| 0 | 0 | 0 |
| 0 | 0 | 0 |
| 0 | 0 | 0 |
| 1 | 1 | 1 |
| 1 | 1 | 1 |
| 1 | 1 | 1 |
| 0 | 0 | 0 |

| | | |
|---|---|---|
| 0 | 0 | 0 |
| 1 | 1 | 1 |
| 0 | 0 | 0 |
| 0 | 1 | 0 |
| 1 | 0 | 0 |
| 1 | 0 | 0 |
| 1 | 0 | 0 |
| 1 | 1 | 1 |
| 0 | 0 | 0 |
| 1 | 1 | 1 |
| 0 | 0 | 0 |
| 0 | 0 | 0 |
| 0 | 0 | 0 |
| 0 | 0 | 0 |
| 1 | 1 | 1 |
| 1 | 1 | 1 |
| 0 | 0 | 0 |
| 0 | 0 | 0 |
| 0 | 1 | 0 |
| 1 | 1 | 1 |
| 0 | 0 | 0 |
| 1 | 0 | 0 |
| 1 | 0 | 0 |
| 1 | 1 | 1 |
| 0 | 0 | 0 |
| 1 | 0 | 0 |
| 0 | 0 | 0 |
| 0 | 0 | 0 |
| 0 | 0 | 0 |
| 1 | 1 | 1 |
| 0 | 0 | 0 |
| 0 | 0 | 0 |
| 0 | 0 | 0 |
| 1 | 1 | 1 |
| 0 | 0 | 0 |
| 1 | 1 | 1 |
| 0 | 0 | 0 |
| 0 | 0 | 0 |
| 0 | 0 | 0 |
| 0 | 0 | 0 |
| 0 | 0 | 0 |
| 1 | 1 | 1 |
| 0 | 0 | 0 |
| 0 | 0 | 0 |
| 0 | 0 | 0 |
| 0 | 0 | 0 |
| 0 | 0 | 0 |
| 0 | 0 | 0 |
| 0 | 0 | 0 |
| 0 | 0 | 0 |
| 0 | 0 | 0 |
| 0 | 0 | 0 |
| 1 | 1 | 1 |
| 0 | 0 | 0 |

| | | |
|---|---|---|
| 0 | 1 | 0 |
| 1 | 1 | 1 |
| 1 | 1 | 1 |
| 1 | 1 | 1 |
| 1 | 1 | 1 |
| 1 | 1 | 1 |
| 1 | 1 | 1 |
| 1 | 1 | 1 |
| 1 | 1 | 1 |
| 0 | 0 | 0 |
| 0 | 0 | 0 |
| 1 | 1 | 1 |
| 1 | 1 | 1 |
| 1 | 1 | 1 |
| 0 | 0 | 0 |
| 1 | 1 | 1 |
| 1 | 1 | 1 |
| 0 | 0 | 0 |
| 0 | 1 | 0 |
| 0 | 1 | 0 |
| 0 | 1 | 0 |
| 0 | 0 | 0 |
| 0 | 0 | 0 |
| 0 | 0 | 0 |
| 0 | 0 | 0 |
| 0 | 0 | 0 |
| 0 | 0 | 0 |
| 0 | 0 | 0 |
| 1 | 1 | 1 |
| 0 | 0 | 0 |
| 0 | 0 | 0 |
| 1 | 1 | 1 |
| 1 | 1 | 1 |
| 1 | 1 | 1 |
| 1 | 1 | 1 |
| 1 | 1 | 1 |
| 1 | 1 | 1 |
| 1 | 1 | 1 |
| 1 | 1 | 1 |
| 1 | 1 | 1 |
| 1 | 1 | 1 |
| 1 | 1 | 1 |
| 1 | 1 | 1 |
| 0 | 0 | 0 |
| 1 | 1 | 1 |
| 0 | 0 | 0 |
| 1 | 1 | 1 |
| 1 | 1 | 1 |
| 1 | 1 | 1 |
| 1 | 1 | 1 |
| 0 | 0 | 0 |
| 1 | 1 | 1 |
| 1 | 1 | 1 |
| 0 | 0 | 0 |

| | | |
|---|---|---|
| 0 | 0 | 0 |
| 0 | 0 | 0 |
| 0 | 0 | 0 |
| 0 | 0 | 0 |
| 1 | 1 | 1 |
| 0 | 1 | 0 |
| 1 | 1 | 1 |
| 0 | 0 | 0 |
| 0 | 0 | 0 |
| 0 | 0 | 0 |
| 0 | 0 | 0 |
| 1 | 1 | 1 |
| 1 | 1 | 1 |
| 1 | 1 | 1 |
| 1 | 1 | 1 |
| 0 | 0 | 0 |
| 0 | 0 | 0 |
| 0 | 1 | 0 |
| 0 | 1 | 0 |
| 1 | 1 | 1 |
| 1 | 1 | 1 |
| 0 | 0 | 0 |
| 1 | 1 | 1 |
| 0 | 1 | 0 |
| 0 | 0 | 0 |
| 0 | 0 | 0 |
| 0 | 0 | 0 |
| 0 | 0 | 0 |
| 1 | 1 | 1 |
| 1 | 1 | 1 |
| 1 | 1 | 1 |
| 0 | 0 | 0 |
| 1 | 1 | 1 |
| 1 | 1 | 1 |
| 1 | 1 | 1 |
| 1 | 1 | 1 |
| 1 | 1 | 1 |
| 1 | 1 | 1 |
| 1 | 1 | 1 |
| 1 | 1 | 1 |
| 1 | 1 | 1 |
| 1 | 1 | 1 |
| 0 | 0 | 0 |
| 0 | 1 | 0 |
| 0 | 1 | 0 |
| 1 | 1 | 1 |
| 0 | 1 | 0 |
| 1 | 1 | 1 |
| 1 | 1 | 1 |
| 0 | 0 | 0 |
| 1 | 1 | 1 |
| 1 | 1 | 1 |
| 1 | 1 | 1 |
| 1 | 1 | 1 |

| | | |
|---|---|---|
| 0 | 0 | 0 |
| 1 | 1 | 1 |
| 0 | 0 | 0 |
| 0 | 1 | 0 |
| 1 | 1 | 1 |
| 1 | 1 | 1 |
| 1 | 1 | 1 |
| 0 | 0 | 0 |
| 1 | 1 | 1 |
| 1 | 1 | 1 |
| 0 | 0 | 0 |
| 0 | 0 | 0 |
| 0 | 0 | 0 |
| 1 | 1 | 1 |
| 0 | 0 | 0 |
| 0 | 0 | 0 |
| 0 | 0 | 0 |
| 0 | 0 | 0 |
| 0 | 1 | 0 |
| 0 | 0 | 0 |
| 0 | 0 | 0 |
| 0 | 0 | 0 |
| 0 | 0 | 0 |
| 0 | 0 | 0 |
| 0 | 0 | 0 |
| 0 | 0 | 0 |
| 0 | 0 | 0 |
| 0 | 0 | 0 |
| 0 | 0 | 0 |
| 0 | 0 | 0 |
| 0 | 0 | 0 |
| 0 | 0 | 0 |
| 1 | 1 | 1 |
| 0 | 0 | 0 |
| 1 | 1 | 1 |
| 0 | 0 | 0 |
| 0 | 0 | 0 |
| 1 | 1 | 1 |
| 1 | 1 | 1 |
| 0 | 1 | 0 |
| 0 | 1 | 0 |
| 0 | 0 | 0 |
| 0 | 0 | 0 |
| 0 | 0 | 0 |
| 1 | 1 | 1 |
| 0 | 0 | 0 |
| 1 | 1 | 1 |
| 0 | 0 | 0 |
| 0 | 1 | 0 |
| 0 | 1 | 0 |
| 0 | 1 | 0 |
| 0 | 1 | 0 |
| 0 | 1 | 0 |
| 1 | 1 | 1 |

| | | |
|---|---|---|
| 1 | 1 | 1 |
| 1 | 1 | 1 |
| 0 | 1 | 0 |
| 1 | 1 | 1 |
| 1 | 1 | 1 |
| 0 | 0 | 0 |
| 0 | 0 | 0 |
| 1 | 1 | 1 |
| 0 | 0 | 0 |
| 0 | 0 | 0 |
| 1 | 1 | 1 |
| 0 | 1 | 0 |
| 1 | 1 | 1 |
| 1 | 1 | 1 |
| 1 | 1 | 1 |
| 0 | 0 | 0 |
| 1 | 1 | 1 |
| 1 | 1 | 1 |
| 0 | 0 | 0 |
| 0 | 0 | 0 |
| 0 | 0 | 0 |
| 1 | 1 | 1 |
| 1 | 1 | 1 |
| 0 | 0 | 0 |
| 1 | 1 | 1 |
| 0 | 0 | 0 |
| 0 | 0 | 0 |
| 1 | 1 | 1 |
| 1 | 1 | 1 |
| 1 | 1 | 1 |
| 0 | 0 | 0 |
| 1 | 0 | 0 |
| 0 | 0 | 0 |
| 0 | 0 | 0 |
| 0 | 0 | 0 |
| 0 | 0 | 0 |
| 0 | 0 | 0 |
| 1 | 1 | 1 |
| 0 | 0 | 0 |
| 0 | 1 | 0 |
| 0 | 0 | 0 |
| 0 | 0 | 0 |
| 0 | 0 | 0 |
| 0 | 0 | 0 |
| 1 | 1 | 1 |
| 0 | 1 | 0 |
| 0 | 1 | 0 |
| 0 | 0 | 0 |
| 0 | 0 | 0 |
| 0 | 0 | 0 |
| 1 | 1 | 1 |
| 0 | 1 | 0 |
| 1 | 1 | 1 |
| 1 | 1 | 1 |

| | | |
|---|---|---|
| 1 | 1 | 1 |
| 0 | 1 | 0 |
| 0 | 1 | 0 |
| 0 | 0 | 0 |
| 0 | 0 | 0 |
| 1 | 1 | 1 |
| 0 | 0 | 0 |
| 1 | 1 | 1 |
| 1 | 1 | 1 |
| 1 | 1 | 1 |
| 1 | 1 | 1 |
| 0 | 0 | 0 |
| 1 | 0 | 0 |
| 0 | 0 | 0 |
| 1 | 1 | 1 |
| 0 | 0 | 0 |
| 0 | 0 | 0 |
| 0 | 0 | 0 |
| 1 | 1 | 1 |
| 0 | 0 | 0 |
| 0 | 0 | 0 |
| 0 | 0 | 0 |
| 0 | 0 | 0 |
| 0 | 0 | 0 |
| 0 | 0 | 0 |
| 1 | 1 | 1 |
| 0 | 0 | 0 |
| 0 | 1 | 0 |
| 0 | 1 | 0 |
| 0 | 1 | 0 |
| 0 | 1 | 0 |
| 0 | 1 | 0 |
| 0 | 0 | 0 |
| 0 | 0 | 0 |
| 1 | 1 | 1 |
| 1 | 1 | 1 |
| 0 | 0 | 0 |
| 0 | 0 | 0 |
| 0 | 0 | 0 |
| 1 | 0 | 0 |
| 0 | 0 | 0 |
| 0 | 0 | 0 |
| 1 | 1 | 1 |
| 0 | 0 | 0 |
| 1 | 1 | 1 |
| 1 | 1 | 1 |
| 1 | 1 | 1 |
| 1 | 1 | 1 |
| 1 | 1 | 1 |
| 0 | 0 | 0 |
| 1 | 1 | 1 |
| 1 | 1 | 1 |
| 1 | 1 | 1 |
| 0 | 1 | 0 |

| | | |
|---|---|---|
| 1 | 1 | 1 |
| 1 | 1 | 1 |
| 1 | 1 | 1 |
| 0 | 0 | 0 |
| 1 | 1 | 1 |
| 1 | 1 | 1 |
| 0 | 1 | 0 |
| 0 | 0 | 0 |
| 1 | 1 | 1 |
| 1 | 1 | 1 |
| 1 | 1 | 1 |
| 1 | 1 | 1 |
| 1 | 1 | 1 |
| 0 | 1 | 0 |
| 0 | 1 | 0 |
| 0 | 1 | 0 |
| 0 | 0 | 0 |
| 0 | 0 | 0 |
| 0 | 1 | 0 |
| 1 | 1 | 1 |
| 0 | 1 | 0 |
| 1 | 1 | 1 |
| 0 | 1 | 0 |
| 1 | 1 | 1 |
| 1 | 1 | 1 |
| 1 | 1 | 1 |
| 1 | 1 | 1 |
| 0 | 0 | 0 |
| 0 | 0 | 0 |
| 1 | 1 | 1 |
| 1 | 1 | 1 |
| 0 | 0 | 0 |
| 0 | 1 | 0 |
| 0 | 0 | 0 |
| 0 | 0 | 0 |
| 0 | 0 | 0 |
| 0 | 0 | 0 |
| 0 | 0 | 0 |
| 1 | 1 | 1 |
| 1 | 1 | 1 |
| 1 | 1 | 1 |
| 0 | 1 | 0 |
| 1 | 1 | 1 |
| 1 | 1 | 1 |
| 0 | 0 | 0 |
| 0 | 0 | 0 |
| 0 | 0 | 0 |
| 1 | 1 | 1 |
| 1 | 1 | 1 |
| 1 | 1 | 1 |
| 1 | 1 | 1 |
| 0 | 0 | 0 |
| 0 | 0 | 0 |
| 0 | 0 | 0 |

| | | |
|---|---|---|
| 0 | 0 | 0 |
| 0 | 0 | 0 |
| 1 | 1 | 1 |
| 0 | 0 | 0 |
| 0 | 0 | 0 |
| 0 | 0 | 0 |
| 0 | 0 | 0 |
| 1 | 1 | 1 |
| 1 | 1 | 1 |
| 1 | 1 | 1 |
| 1 | 1 | 1 |
| 1 | 1 | 1 |
| 0 | 1 | 0 |
| 0 | 1 | 0 |
| 1 | 1 | 1 |
| 1 | 1 | 1 |
| 1 | 1 | 1 |
| 1 | 1 | 1 |
| 1 | 1 | 1 |
| 1 | 1 | 1 |
| 0 | 0 | 0 |
| 0 | 0 | 0 |
| 0 | 0 | 0 |
| 0 | 0 | 0 |
| 0 | 0 | 0 |
| 1 | 1 | 1 |
| 1 | 1 | 1 |
| 0 | 0 | 0 |
| 0 | 1 | 0 |
| 0 | 1 | 0 |
| 0 | 0 | 0 |
| 0 | 0 | 0 |
| 0 | 0 | 0 |
| 1 | 1 | 1 |
| 0 | 0 | 0 |
| 1 | 1 | 1 |
| 0 | 1 | 0 |
| 0 | 0 | 0 |
| 1 | 1 | 1 |
| 1 | 1 | 1 |
| 0 | 1 | 0 |
| 0 | 1 | 0 |
| 1 | 1 | 1 |
| 1 | 1 | 1 |
| 1 | 1 | 1 |
| 1 | 1 | 1 |
| 1 | 1 | 1 |
| 1 | 1 | 1 |
| 0 | 0 | 0 |
| 1 | 1 | 1 |
| 0 | 0 | 0 |
| 0 | 0 | 0 |
| 1 | 1 | 1 |
| 0 | 1 | 0 |

| | | |
|---|---|---|
| 0 | 0 | 0 |
| 1 | 1 | 1 |
| 1 | 1 | 1 |
| 1 | 1 | 1 |
| 1 | 1 | 1 |
| 0 | 0 | 0 |
| 0 | 0 | 0 |
| 1 | 1 | 1 |
| 0 | 1 | 0 |
| 1 | 1 | 1 |
| 1 | 1 | 1 |
| 0 | 0 | 0 |
| 0 | 1 | 0 |
| 1 | 1 | 1 |
| 0 | 1 | 0 |
| 1 | 1 | 1 |
| 0 | 0 | 0 |
| 0 | 0 | 0 |
| 1 | 1 | 1 |
| 1 | 1 | 1 |
| 1 | 1 | 1 |
| 1 | 1 | 1 |
| 1 | 1 | 1 |
| 1 | 1 | 1 |
| 0 | 0 | 0 |
| 1 | 1 | 1 |
| 1 | 1 | 1 |
| 1 | 1 | 1 |
| 1 | 1 | 1 |
| 1 | 1 | 1 |
| 0 | 1 | 0 |
| 0 | 0 | 0 |
| 0 | 0 | 0 |
| 0 | 1 | 0 |
| 1 | 1 | 1 |
| 0 | 0 | 0 |
| 0 | 0 | 0 |
| 0 | 0 | 0 |
| 0 | 0 | 0 |
| 0 | 0 | 0 |
| 0 | 0 | 0 |
| 0 | 0 | 0 |
| 0 | 0 | 0 |
| 0 | 0 | 0 |
| 0 | 0 | 0 |
| 0 | 0 | 0 |
| 0 | 0 | 0 |
| 0 | 0 | 0 |
| 1 | 0 | 0 |
| 1 | 1 | 1 |
| 1 | 0 | 0 |
| 1 | 1 | 1 |
| 0 | 0 | 0 |
| 1 | 1 | 1 |

| | | |
|---|---|---|
| 1 | 1 | 1 |
| 1 | 1 | 1 |
| 1 | 1 | 1 |
| 1 | 1 | 1 |
| 1 | 1 | 1 |
| 1 | 1 | 1 |
| 1 | 1 | 1 |
| 0 | 0 | 0 |
| 0 | 0 | 0 |
| 1 | 0 | 0 |
| 1 | 1 | 1 |
| 1 | 1 | 1 |
| 1 | 1 | 1 |
| 0 | 0 | 0 |
| 1 | 1 | 1 |
| 1 | 1 | 1 |
| 0 | 0 | 0 |
| 1 | 1 | 1 |
| 0 | 0 | 0 |
| 0 | 1 | 0 |
| 1 | 1 | 1 |
| 1 | 1 | 1 |
| 1 | 1 | 1 |
| 0 | 0 | 0 |
| 1 | 1 | 1 |
| 1 | 1 | 1 |
| 0 | 1 | 0 |
| 0 | 1 | 0 |
| 0 | 1 | 0 |
| 0 | 1 | 0 |
| 1 | 1 | 1 |
| 0 | 0 | 0 |
| 1 | 1 | 1 |
| 0 | 1 | 0 |
| 1 | 1 | 1 |
| 1 | 1 | 1 |
| 1 | 1 | 1 |
| 1 | 1 | 1 |
| 0 | 0 | 0 |
| 1 | 1 | 1 |
| 1 | 1 | 1 |
| 1 | 1 | 1 |
| 0 | 1 | 0 |
| 1 | 1 | 1 |
| 1 | 1 | 1 |
| 0 | 0 | 0 |
| 0 | 1 | 0 |
| 0 | 1 | 0 |
| 1 | 0 | 0 |
| 1 | 0 | 0 |
| 0 | 1 | 0 |
| 1 | 0 | 0 |
| 0 | 0 | 0 |
| 1 | 0 | 0 |

| | | |
|---|---|---|
| 1 | 0 | 0 |
| 1 | 0 | 0 |
| 1 | 0 | 0 |
| 1 | 0 | 0 |
| 0 | 0 | 0 |
| 0 | 0 | 0 |
| 0 | 0 | 0 |
| 0 | 0 | 0 |
| 1 | 0 | 0 |
| 1 | 0 | 0 |
| 1 | 0 | 0 |
| 1 | 0 | 0 |
| 1 | 0 | 0 |
| 0 | 0 | 0 |
| 1 | 0 | 0 |
| 0 | 0 | 0 |
| 1 | 0 | 0 |
| 0 | 0 | 0 |
| 1 | 0 | 0 |
| 0 | 0 | 0 |
| 0 | 0 | 0 |
| 1 | 0 | 0 |
| 1 | 0 | 0 |
| 0 | 0 | 0 |
| 0 | 0 | 0 |
| 1 | 0 | 0 |
| 0 | 0 | 0 |
| 0 | 0 | 0 |
| 0 | 0 | 0 |
| 1 | 0 | 0 |
| 1 | 0 | 0 |
| 0 | 0 | 0 |
| 1 | 0 | 0 |
| 1 | 0 | 0 |
| 1 | 0 | 0 |
| 1 | 0 | 0 |
| 1 | 0 | 0 |
| 1 | 0 | 0 |
| 1 | 0 | 0 |
| 1 | 0 | 0 |
| 1 | 0 | 0 |
| 1 | 0 | 0 |
| 1 | 0 | 0 |
| 0 | 0 | 0 |
| 1 | 1 | 1 |
| 1 | 1 | 1 |
| 0 | 0 | 0 |
| 0 | 0 | 0 |
| 0 | 1 | 0 |
| 0 | 0 | 0 |
| 0 | 0 | 0 |
| 0 | 0 | 0 |
| 1 | 1 | 1 |
| 1 | 1 | 1 |

| | | |
|---|---|---|
| 1 | 1 | 1 |
| 1 | 1 | 1 |
| 1 | 1 | 1 |
| 1 | 1 | 1 |
| 1 | 1 | 1 |
| 1 | 1 | 1 |
| 1 | 1 | 1 |
| 0 | 0 | 0 |
| 1 | 1 | 1 |
| 1 | 1 | 1 |
| 0 | 1 | 0 |
| 0 | 1 | 0 |
| 1 | 0 | 0 |
| 1 | 1 | 1 |
| 0 | 0 | 0 |
| 0 | 0 | 0 |
| 0 | 1 | 0 |
| 0 | 0 | 0 |
| 1 | 1 | 1 |
| 1 | 1 | 1 |
| 0 | 0 | 0 |
| 0 | 1 | 0 |
| 0 | 1 | 0 |
| 1 | 1 | 1 |
| 0 | 0 | 0 |
| 1 | 1 | 1 |
| 0 | 1 | 0 |
| 0 | 0 | 0 |
| 0 | 0 | 0 |
| 0 | 0 | 0 |
| 0 | 0 | 0 |
| 1 | 1 | 1 |
| 1 | 1 | 1 |
| 0 | 1 | 0 |
| 1 | 1 | 1 |
| 1 | 1 | 1 |
| 0 | 0 | 0 |
| 0 | 0 | 0 |
| 1 | 1 | 1 |
| 1 | 1 | 1 |
| 1 | 1 | 1 |
| 1 | 1 | 1 |
| 0 | 0 | 0 |
| 0 | 1 | 0 |
| 1 | 1 | 1 |
| 1 | 1 | 1 |
| 1 | 1 | 1 |
| 1 | 1 | 1 |
| 0 | 0 | 0 |
| 0 | 0 | 0 |
| 0 | 0 | 0 |
| 0 | 0 | 0 |
| 0 | 0 | 0 |
| 1 | 1 | 1 |

| | | |
|---|---|---|
| 0 | 0 | 0 |
| 0 | 0 | 0 |
| 0 | 1 | 0 |
| 0 | 0 | 0 |
| 1 | 1 | 1 |
| 0 | 1 | 0 |
| 0 | 1 | 0 |
| 1 | 1 | 1 |
| 1 | 1 | 1 |
| 0 | 0 | 0 |
| 1 | 1 | 1 |
| 0 | 1 | 0 |
| 1 | 1 | 1 |
| 0 | 1 | 0 |
| 0 | 1 | 0 |
| 0 | 1 | 0 |
| 0 | 1 | 0 |
| 1 | 1 | 1 |
| 1 | 1 | 1 |
| 0 | 1 | 0 |
| 0 | 1 | 0 |
| 0 | 1 | 0 |
| 0 | 1 | 0 |
| 0 | 1 | 0 |
| 0 | 1 | 0 |
| 0 | 1 | 0 |
| 0 | 1 | 0 |
| 0 | 1 | 0 |
| 0 | 1 | 0 |
| 0 | 1 | 0 |
| 1 | 1 | 1 |
| 1 | 1 | 1 |
| 0 | 0 | 0 |
| 1 | 1 | 1 |
| 1 | 1 | 1 |
| 1 | 1 | 1 |
| 1 | 1 | 1 |
| 0 | 1 | 0 |
| 1 | 1 | 1 |
| 1 | 1 | 1 |
| 1 | 1 | 1 |
| 1 | 1 | 1 |
| 1 | 1 | 1 |
| 0 | 1 | 0 |
| 0 | 1 | 0 |
| 1 | 1 | 1 |
| 1 | 1 | 1 |
| 0 | 0 | 0 |
| 0 | 0 | 0 |
| 0 | 0 | 0 |
| 0 | 0 | 0 |
| 0 | 0 | 0 |
| 1 | 1 | 1 |
| 0 | 0 | 0 |

| | | |
|---|---|---|
| 1 | 1 | 1 |
| 0 | 0 | 0 |
| 1 | 1 | 1 |
| 0 | 1 | 0 |
| 0 | 1 | 0 |
| 0 | 0 | 0 |
| 0 | 0 | 0 |
| 0 | 0 | 0 |
| 0 | 0 | 0 |
| 0 | 0 | 0 |
| 0 | 1 | 0 |
| 0 | 0 | 0 |
| 0 | 1 | 0 |
| 1 | 1 | 1 |
| 0 | 1 | 0 |
| 0 | 0 | 0 |
| 1 | 1 | 1 |
| 1 | 1 | 1 |
| 0 | 0 | 0 |
| 0 | 0 | 0 |
| 0 | 0 | 0 |
| 0 | 0 | 0 |
| 0 | 0 | 0 |
| 0 | 0 | 0 |
| 0 | 1 | 0 |
| 0 | 0 | 0 |
| 0 | 0 | 0 |
| 0 | 0 | 0 |
| 0 | 0 | 0 |
| 0 | 0 | 0 |
| 0 | 0 | 0 |
| 1 | 1 | 1 |
| 1 | 1 | 1 |
| 0 | 0 | 0 |
| 0 | 0 | 0 |
| 1 | 0 | 0 |
| 0 | 0 | 0 |
| 0 | 0 | 0 |
| 0 | 0 | 0 |
| 0 | 0 | 0 |
| 0 | 0 | 0 |
| 1 | 0 | 0 |
| 1 | 1 | 1 |
| 1 | 1 | 1 |
| 0 | 0 | 0 |
| 0 | 0 | 0 |
| 0 | 0 | 0 |
| 1 | 1 | 1 |
| 1 | 1 | 1 |
| 1 | 1 | 1 |
| 1 | 1 | 1 |
| 1 | 1 | 1 |
| 1 | 1 | 1 |
| 0 | 0 | 0 |

| | | |
|---|---|---|
| 1 | 1 | 1 |
| 1 | 1 | 1 |
| 1 | 1 | 1 |
| 1 | 1 | 1 |
| 0 | 0 | 0 |
| 0 | 0 | 0 |
| 1 | 0 | 0 |
| 1 | 0 | 0 |
| 0 | 1 | 0 |
| 1 | 1 | 1 |
| 1 | 1 | 1 |
| 0 | 0 | 0 |
| 1 | 1 | 1 |
| 0 | 0 | 0 |
| 0 | 0 | 0 |
| 0 | 0 | 0 |
| 0 | 0 | 0 |
| 0 | 0 | 0 |
| 0 | 0 | 0 |
| 0 | 0 | 0 |
| 0 | 0 | 0 |
| 0 | 0 | 0 |
| 0 | 0 | 0 |
| 0 | 1 | 0 |
| 0 | 1 | 0 |
| 0 | 1 | 0 |
| 1 | 1 | 1 |
| 1 | 1 | 1 |
| 1 | 1 | 1 |
| 1 | 1 | 1 |
| 1 | 1 | 1 |
| 0 | 0 | 0 |
| 1 | 1 | 1 |
| 0 | 0 | 0 |
| 0 | 0 | 0 |
| 0 | 0 | 0 |
| 0 | 0 | 0 |
| 1 | 1 | 1 |
| 1 | 1 | 1 |
| 0 | 1 | 0 |
| 1 | 1 | 1 |
| 1 | 1 | 1 |
| 1 | 1 | 1 |
| 0 | 1 | 0 |
| 1 | 1 | 1 |
| 0 | 0 | 0 |
| 1 | 1 | 1 |
| 1 | 1 | 1 |
| 1 | 1 | 1 |
| 0 | 0 | 0 |
| 0 | 0 | 0 |
| 0 | 0 | 0 |
| 0 | 1 | 0 |
| 0 | 1 | 0 |

| | | |
|---|---|---|
| 0 | 1 | 0 |
| 0 | 1 | 0 |
| 0 | 1 | 0 |
| 0 | 0 | 0 |
| 0 | 1 | 0 |
| 1 | 1 | 1 |
| 0 | 0 | 0 |
| 0 | 1 | 0 |
| 1 | 1 | 1 |
| 1 | 1 | 1 |
| 0 | 0 | 0 |
| 0 | 0 | 0 |
| 1 | 1 | 1 |
| 1 | 1 | 1 |
| 0 | 1 | 0 |
| 0 | 1 | 0 |
| 0 | 0 | 0 |
| 0 | 0 | 0 |
| 1 | 1 | 1 |
| 0 | 0 | 0 |
| 0 | 0 | 0 |
| 0 | 0 | 0 |
| 0 | 1 | 0 |
| 0 | 0 | 0 |
| 0 | 0 | 0 |
| 1 | 1 | 1 |
| 0 | 0 | 0 |
| 0 | 1 | 0 |
| 1 | 1 | 1 |
| 0 | 0 | 0 |
| 0 | 1 | 0 |
| 1 | 1 | 1 |
| 0 | 0 | 0 |
| 0 | 1 | 0 |
| 0 | 1 | 0 |
| 0 | 0 | 0 |
| 1 | 1 | 1 |
| 0 | 0 | 0 |
| 1 | 1 | 1 |
| 0 | 0 | 0 |
| 0 | 0 | 0 |
| 1 | 0 | 0 |
| 0 | 0 | 0 |
| 0 | 0 | 0 |
| 1 | 1 | 1 |
| 1 | 0 | 0 |
| 0 | 1 | 0 |
| 1 | 1 | 1 |
| 1 | 1 | 1 |
| 1 | 1 | 1 |
| 1 | 1 | 1 |
| 0 | 1 | 0 |
| 0 | 1 | 0 |
| 0 | 1 | 0 |

| | | |
|---|---|---|
| 0 | 1 | 0 |
| 0 | 1 | 0 |
| 0 | 1 | 0 |
| 0 | 0 | 0 |
| 1 | 0 | 0 |
| 0 | 0 | 0 |
| 1 | 1 | 1 |
| 1 | 0 | 0 |
| 0 | 0 | 0 |
| 0 | 0 | 0 |
| 1 | 1 | 1 |
| 0 | 0 | 0 |
| 0 | 0 | 0 |
| 0 | 0 | 0 |
| 1 | 0 | 0 |
| 0 | 0 | 0 |
| 0 | 0 | 0 |
| 0 | 1 | 0 |
| 0 | 0 | 0 |
| 0 | 0 | 0 |
| 0 | 0 | 0 |
| 0 | 0 | 0 |
| 1 | 1 | 1 |
| 0 | 0 | 0 |
| 0 | 0 | 0 |
| 0 | 0 | 0 |
| 0 | 0 | 0 |
| 0 | 0 | 0 |
| 0 | 0 | 0 |
| 0 | 0 | 0 |
| 0 | 0 | 0 |
| 0 | 0 | 0 |
| 0 | 0 | 0 |
| 0 | 1 | 0 |
| 0 | 1 | 0 |
| 0 | 0 | 0 |
| 1 | 1 | 1 |
| 0 | 0 | 0 |
| 0 | 0 | 0 |
| 0 | 0 | 0 |
| 0 | 0 | 0 |
| 1 | 1 | 1 |
| 0 | 0 | 0 |
| 0 | 0 | 0 |
| 1 | 1 | 1 |
| 0 | 1 | 0 |
| 1 | 1 | 1 |
| 1 | 1 | 1 |
| 1 | 1 | 1 |
| 0 | 1 | 0 |
| 1 | 1 | 1 |
| 1 | 1 | 1 |
| 1 | 1 | 1 |
| 1 | 1 | 1 |

| | | |
|---|---|---|
| 1 | 1 | 1 |
| 0 | 0 | 0 |
| 1 | 1 | 1 |
| 1 | 1 | 1 |
| 1 | 1 | 1 |
| 1 | 1 | 1 |
| 1 | 1 | 1 |
| 1 | 1 | 1 |
| 1 | 1 | 1 |
| 0 | 1 | 0 |
| 1 | 1 | 1 |
| 0 | 0 | 0 |
| 1 | 0 | 0 |
| 0 | 1 | 0 |
| 0 | 0 | 0 |
| 0 | 1 | 0 |
| 0 | 1 | 0 |
| 1 | 1 | 1 |
| 1 | 1 | 1 |
| 1 | 1 | 1 |
| 0 | 0 | 0 |
| 0 | 1 | 0 |
| 0 | 0 | 0 |
| 0 | 0 | 0 |
| 1 | 1 | 1 |
| 0 | 0 | 0 |
| 1 | 1 | 1 |
| 1 | 1 | 1 |
| 1 | 1 | 1 |
| 1 | 1 | 1 |
| 1 | 1 | 1 |
| 0 | 0 | 0 |
| 1 | 1 | 1 |
| 0 | 0 | 0 |
| 1 | 1 | 1 |
| 0 | 1 | 0 |
| 0 | 1 | 0 |
| 0 | 1 | 0 |
| 1 | 1 | 1 |
| 0 | 0 | 0 |
| 1 | 1 | 1 |
| 1 | 1 | 1 |
| 0 | 1 | 0 |
| 0 | 0 | 0 |
| 1 | 1 | 1 |
| 1 | 1 | 1 |
| 1 | 1 | 1 |
| 0 | 0 | 0 |
| 0 | 0 | 0 |
| 1 | 1 | 1 |
| 1 | 1 | 1 |
| 0 | 0 | 0 |
| 0 | 0 | 0 |
| 0 | 0 | 0 |

| | | |
|---|---|---|
| 0 | 0 | 0 |
| 1 | 1 | 1 |
| 0 | 0 | 0 |
| 0 | 0 | 0 |
| 0 | 0 | 0 |
| 1 | 1 | 1 |
| 1 | 1 | 1 |
| 0 | 1 | 0 |
| 1 | 1 | 1 |
| 0 | 0 | 0 |
| 1 | 1 | 1 |
| 0 | 1 | 0 |
| 1 | 1 | 1 |
| 1 | 1 | 1 |
| 0 | 0 | 0 |
| 0 | 1 | 0 |
| 1 | 1 | 1 |
| 0 | 0 | 0 |
| 1 | 1 | 1 |
| 0 | 0 | 0 |
| 0 | 0 | 0 |
| 1 | 1 | 1 |
| 0 | 0 | 0 |
| 1 | 1 | 1 |
| 0 | 0 | 0 |
| 0 | 0 | 0 |
| 1 | 1 | 1 |
| 0 | 1 | 0 |
| 0 | 0 | 0 |
| 1 | 1 | 1 |
| 1 | 1 | 1 |
| 1 | 1 | 1 |
| 1 | 1 | 1 |
| 0 | 1 | 0 |
| 0 | 0 | 0 |
| 0 | 0 | 0 |
| 1 | 1 | 1 |
| 0 | 0 | 0 |
| 0 | 0 | 0 |
| 0 | 0 | 0 |
| 0 | 0 | 0 |
| 0 | 0 | 0 |
| 0 | 0 | 0 |
| 0 | 1 | 0 |
| 0 | 0 | 0 |
| 1 | 1 | 1 |
| 0 | 1 | 0 |
| 0 | 0 | 0 |
| 0 | 0 | 0 |
| 0 | 0 | 0 |
| 0 | 0 | 0 |
| 0 | 0 | 0 |
| 1 | 1 | 1 |
| 1 | 1 | 1 |

| | | |
|---|---|---|
| 1 | 1 | 1 |
| 0 | 0 | 0 |
| 0 | 0 | 0 |
| 0 | 0 | 0 |
| 0 | 0 | 0 |
| 0 | 0 | 0 |
| 0 | 0 | 0 |
| 1 | 0 | 0 |
| 1 | 0 | 0 |
| 1 | 1 | 1 |
| 0 | 0 | 0 |
| 0 | 0 | 0 |
| 1 | 1 | 1 |
| 0 | 0 | 0 |
| 1 | 1 | 1 |
| 0 | 0 | 0 |
| 1 | 1 | 1 |
| 1 | 1 | 1 |
| 0 | 1 | 0 |
| 0 | 1 | 0 |
| 0 | 1 | 0 |
| 1 | 1 | 1 |
| 0 | 1 | 0 |
| 0 | 0 | 0 |
| 0 | 0 | 0 |
| 1 | 1 | 1 |
| 1 | 1 | 1 |
| 1 | 1 | 1 |
| 1 | 1 | 1 |
| 0 | 0 | 0 |
| 0 | 0 | 0 |
| 0 | 0 | 0 |
| 0 | 0 | 0 |
| 1 | 1 | 1 |
| 0 | 0 | 0 |
| 0 | 0 | 0 |
| 0 | 0 | 0 |
| 0 | 0 | 0 |
| 0 | 0 | 0 |
| 1 | 1 | 1 |
| 1 | 1 | 1 |
| 0 | 0 | 0 |
| 1 | 1 | 1 |
| 1 | 1 | 1 |
| 0 | 0 | 0 |
| 1 | 0 | 0 |
| 1 | 1 | 1 |
| 0 | 1 | 0 |
| 1 | 0 | 0 |
| 0 | 1 | 0 |
| 0 | 0 | 0 |
| 1 | 1 | 1 |
| 0 | 0 | 0 |
| 0 | 0 | 0 |

| | | |
|---|---|---|
| 0 | 1 | 0 |
| 1 | 1 | 1 |
| 0 | 0 | 0 |
| 0 | 0 | 0 |
| 1 | 1 | 1 |
| 1 | 1 | 1 |
| 1 | 1 | 1 |
| 1 | 1 | 1 |
| 1 | 1 | 1 |
| 1 | 1 | 1 |
| 1 | 1 | 1 |
| 1 | 1 | 1 |
| 0 | 0 | 0 |
| 1 | 1 | 1 |
| 1 | 1 | 1 |
| 1 | 1 | 1 |
| 1 | 1 | 1 |
| 1 | 1 | 1 |
| 1 | 1 | 1 |
| 1 | 1 | 1 |
| 1 | 1 | 1 |
| 1 | 1 | 1 |
| 1 | 1 | 1 |
| 1 | 1 | 1 |
| 1 | 1 | 1 |
| 1 | 1 | 1 |
| 0 | 1 | 0 |
| 1 | 1 | 1 |
| 1 | 1 | 1 |
| 1 | 1 | 1 |
| 1 | 1 | 1 |
| 0 | 0 | 0 |
| 1 | 1 | 1 |
| 0 | 1 | 0 |
| 1 | 1 | 1 |
| 1 | 1 | 1 |
| 0 | 1 | 0 |
| 1 | 1 | 1 |
| 1 | 1 | 1 |
| 1 | 1 | 1 |
| 0 | 0 | 0 |
| 0 | 0 | 0 |
| 1 | 1 | 1 |
| 1 | 1 | 1 |
| 1 | 1 | 1 |
| 1 | 0 | 0 |
| 1 | 0 | 0 |
| 1 | 1 | 1 |
| 0 | 0 | 0 |
| 0 | 0 | 0 |
| 1 | 1 | 1 |
| 0 | 1 | 0 |
| 0 | 0 | 0 |
| 1 | 1 | 1 |

| | | |
|---|---|---|
| 0 | 0 | 0 |
| 0 | 0 | 0 |
| 0 | 0 | 0 |
| 1 | 1 | 1 |
| 1 | 1 | 1 |
| 1 | 1 | 1 |
| 1 | 1 | 1 |
| 0 | 0 | 0 |
| 0 | 0 | 0 |
| 0 | 0 | 0 |
| 0 | 0 | 0 |
| 1 | 1 | 1 |
| 0 | 0 | 0 |
| 1 | 1 | 1 |
| 1 | 1 | 1 |
| 1 | 1 | 1 |
| 1 | 1 | 1 |
| 0 | 0 | 0 |
| 1 | 1 | 1 |
| 1 | 1 | 1 |
| 1 | 1 | 1 |
| 1 | 1 | 1 |
| 0 | 1 | 0 |
| 1 | 1 | 1 |
| 1 | 1 | 1 |
| 0 | 1 | 0 |
| 1 | 1 | 1 |
| 1 | 1 | 1 |
| 1 | 1 | 1 |
| 1 | 1 | 1 |
| 0 | 0 | 0 |
| 0 | 0 | 0 |
| 1 | 1 | 1 |
| 1 | 1 | 1 |
| 1 | 1 | 1 |
| 1 | 1 | 1 |
| 1 | 1 | 1 |
| 1 | 1 | 1 |
| 1 | 1 | 1 |
| 1 | 1 | 1 |
| 1 | 1 | 1 |
| 0 | 0 | 0 |
| 1 | 1 | 1 |
| 1 | 1 | 1 |
| 0 | 0 | 0 |
| 0 | 0 | 0 |
| 1 | 1 | 1 |
| 1 | 1 | 1 |
| 0 | 0 | 0 |
| 0 | 0 | 0 |
| 0 | 0 | 0 |
| 0 | 0 | 0 |
| 0 | 0 | 0 |
| 0 | 0 | 0 |

| | | |
|---|---|---|
| 1 | 1 | 1 |
| 1 | 1 | 1 |
| 1 | 1 | 1 |
| 1 | 1 | 1 |
| 1 | 1 | 1 |
| 0 | 0 | 0 |
| 0 | 0 | 0 |
| 0 | 0 | 0 |
| 0 | 0 | 0 |
| 0 | 0 | 0 |
| 0 | 1 | 0 |
| 0 | 0 | 0 |
| 0 | 0 | 0 |
| 0 | 0 | 0 |
| 0 | 1 | 0 |
| 0 | 0 | 0 |
| 0 | 1 | 0 |
| 0 | 1 | 0 |
| 0 | 0 | 0 |
| 0 | 0 | 0 |
| 0 | 0 | 0 |
| 0 | 0 | 0 |
| 1 | 0 | 0 |
| 0 | 0 | 0 |
| 0 | 0 | 0 |
| 0 | 0 | 0 |
| 0 | 0 | 0 |
| 1 | 1 | 1 |
| 1 | 1 | 1 |
| 1 | 1 | 1 |
| 1 | 1 | 1 |
| 1 | 1 | 1 |
| 0 | 0 | 0 |
| 0 | 1 | 0 |
| 0 | 0 | 0 |
| 0 | 0 | 0 |
| 0 | 0 | 0 |
| 0 | 0 | 0 |
| 0 | 0 | 0 |
| 0 | 0 | 0 |
| 0 | 0 | 0 |
| 0 | 0 | 0 |
| 0 | 0 | 0 |
| 1 | 1 | 1 |
| 1 | 1 | 1 |
| 1 | 1 | 1 |
| 1 | 1 | 1 |
| 0 | 0 | 0 |
| 0 | 0 | 0 |
| 0 | 0 | 0 |
| 1 | 1 | 1 |
| 1 | 1 | 1 |
| 0 | 0 | 0 |
| 1 | 1 | 1 |

| | | |
|---|---|---|
| 1 | 1 | 1 |
| 0 | 0 | 0 |
| 0 | 1 | 0 |
| 0 | 1 | 0 |
| 1 | 1 | 1 |
| 1 | 1 | 1 |
| 0 | 1 | 0 |
| 1 | 1 | 1 |
| 0 | 1 | 0 |
| 1 | 1 | 1 |
| 1 | 1 | 1 |
| 1 | 1 | 1 |
| 0 | 0 | 0 |
| 0 | 0 | 0 |
| 0 | 0 | 0 |
| 0 | 0 | 0 |
| 0 | 0 | 0 |
| 0 | 0 | 0 |
| 0 | 0 | 0 |
| 0 | 1 | 0 |
| 0 | 0 | 0 |
| 0 | 0 | 0 |
| 0 | 0 | 0 |
| 0 | 0 | 0 |
| 0 | 0 | 0 |
| 1 | 1 | 1 |
| 1 | 1 | 1 |
| 0 | 0 | 0 |
| 0 | 1 | 0 |
| 0 | 0 | 0 |
| 0 | 0 | 0 |
| 1 | 1 | 1 |
| 0 | 0 | 0 |
| 1 | 1 | 1 |
| 1 | 1 | 1 |
| 1 | 1 | 1 |
| 1 | 0 | 0 |
| 1 | 1 | 1 |
| 1 | 1 | 1 |
| 1 | 1 | 1 |
| 0 | 0 | 0 |
| 1 | 1 | 1 |
| 1 | 1 | 1 |
| 0 | 0 | 0 |
| 0 | 0 | 0 |
| 1 | 1 | 1 |
| 0 | 0 | 0 |
| 0 | 0 | 0 |
| 1 | 1 | 1 |
| 1 | 1 | 1 |
| 0 | 0 | 0 |
| 1 | 1 | 1 |
| 1 | 1 | 1 |
| 1 | 1 | 1 |

| | | |
|---|---|---|
| 1 | 1 | 1 |
| 1 | 1 | 1 |
| 1 | 1 | 1 |
| 1 | 1 | 1 |
| 0 | 0 | 0 |
| 0 | 0 | 0 |
| 1 | 1 | 1 |
| 1 | 1 | 1 |
| 1 | 1 | 1 |
| 0 | 1 | 0 |
| 1 | 1 | 1 |
| 0 | 0 | 0 |
| 0 | 0 | 0 |
| 1 | 1 | 1 |
| 1 | 1 | 1 |
| 0 | 0 | 0 |
| 0 | 1 | 0 |
| 1 | 1 | 1 |
| 0 | 0 | 0 |
| 1 | 1 | 1 |
| 1 | 1 | 1 |
| 1 | 1 | 1 |
| 1 | 1 | 1 |
| 0 | 0 | 0 |
| 1 | 1 | 1 |
| 0 | 0 | 0 |
| 1 | 1 | 1 |
| 1 | 1 | 1 |
| 0 | 0 | 0 |
| 0 | 0 | 0 |
| 0 | 0 | 0 |
| 0 | 0 | 0 |
| 0 | 0 | 0 |
| 0 | 0 | 0 |
| 1 | 1 | 1 |
| 1 | 1 | 1 |
| 1 | 1 | 1 |
| 0 | 0 | 0 |
| 1 | 1 | 1 |
| 1 | 1 | 1 |
| 0 | 0 | 0 |
| 0 | 0 | 0 |
| 0 | 0 | 0 |
| 1 | 1 | 1 |
| 0 | 0 | 0 |
| 0 | 1 | 0 |
| 0 | 1 | 0 |
| 0 | 0 | 0 |
| 1 | 1 | 1 |
| 1 | 1 | 1 |
| 1 | 1 | 1 |
| 0 | 0 | 0 |
| 1 | 1 | 1 |
| 0 | 0 | 0 |

| | | |
|---|---|---|
| 1 | 1 | 1 |
| 0 | 0 | 0 |
| 0 | 0 | 0 |
| 0 | 1 | 0 |
| 1 | 1 | 1 |
| 1 | 1 | 1 |
| 0 | 0 | 0 |
| 1 | 1 | 1 |
| 0 | 1 | 0 |
| 0 | 1 | 0 |
| 1 | 1 | 1 |
| 1 | 1 | 1 |
| 1 | 1 | 1 |
| 1 | 1 | 1 |
| 1 | 1 | 1 |
| 1 | 1 | 1 |
| 1 | 1 | 1 |
| 1 | 1 | 1 |
| 1 | 1 | 1 |
| 1 | 1 | 1 |
| 1 | 1 | 1 |
| 0 | 0 | 0 |
| 0 | 1 | 0 |
| 0 | 1 | 0 |
| 1 | 1 | 1 |
| 1 | 1 | 1 |
| 0 | 0 | 0 |
| 1 | 1 | 1 |
| 0 | 0 | 0 |
| 0 | 1 | 0 |
| 1 | 1 | 1 |
| 0 | 0 | 0 |
| 0 | 0 | 0 |
| 1 | 0 | 0 |
| 1 | 1 | 1 |
| 1 | 1 | 1 |
| 1 | 1 | 1 |
| 1 | 1 | 1 |
| 0 | 0 | 0 |
| 0 | 0 | 0 |
| 0 | 0 | 0 |
| 1 | 1 | 1 |
| 1 | 1 | 1 |
| 0 | 1 | 0 |
| 0 | 1 | 0 |
| 0 | 0 | 0 |
| 1 | 1 | 1 |
| 0 | 0 | 0 |
| 1 | 1 | 1 |
| 0 | 0 | 0 |
| 0 | 0 | 0 |
| 0 | 0 | 0 |
| 1 | 1 | 1 |
| 1 | 1 | 1 |

| | | |
|---|---|---|
| 0 | 0 | 0 |
| 0 | 0 | 0 |
| 1 | 1 | 1 |
| 1 | 1 | 1 |
| 1 | 1 | 1 |
| 1 | 1 | 1 |
| 1 | 1 | 1 |
| 1 | 1 | 1 |
| 1 | 1 | 1 |
| 1 | 1 | 1 |
| 1 | 1 | 1 |
| 1 | 1 | 1 |
| 1 | 1 | 1 |
| 1 | 1 | 1 |
| 1 | 1 | 1 |
| 0 | 0 | 0 |
| 0 | 0 | 0 |
| 0 | 0 | 0 |
| 1 | 1 | 1 |
| 1 | 1 | 1 |
| 0 | 1 | 0 |
| 0 | 0 | 0 |
| 0 | 0 | 0 |
| 0 | 0 | 0 |
| 0 | 0 | 0 |
| 1 | 1 | 1 |
| 1 | 1 | 1 |
| 1 | 1 | 1 |
| 1 | 1 | 1 |
| 1 | 1 | 1 |
| 0 | 0 | 0 |
| 0 | 1 | 0 |
| 0 | 0 | 0 |
| 1 | 1 | 1 |
| 1 | 0 | 0 |
| 1 | 0 | 0 |
| 0 | 0 | 0 |
| 0 | 0 | 0 |
| 0 | 1 | 0 |
| 1 | 1 | 1 |
| 1 | 0 | 0 |
| 0 | 0 | 0 |
| 1 | 1 | 1 |
| 0 | 1 | 0 |
| 0 | 0 | 0 |
| 1 | 0 | 0 |
| 0 | 0 | 0 |
| 0 | 0 | 0 |
| 1 | 1 | 1 |
| 0 | 1 | 0 |
| 0 | 0 | 0 |
| 0 | 0 | 0 |
| 1 | 1 | 1 |
| 0 | 0 | 0 |

| | | |
|---|---|---|
| 0 | 1 | 0 |
| 0 | 0 | 0 |
| 1 | 0 | 0 |
| 0 | 0 | 0 |
| 1 | 1 | 1 |
| 1 | 1 | 1 |
| 1 | 1 | 1 |
| 0 | 1 | 0 |
| 0 | 0 | 0 |
| 1 | 1 | 1 |
| 0 | 1 | 0 |
| 0 | 1 | 0 |
| 0 | 0 | 0 |
| 0 | 0 | 0 |
| 0 | 0 | 0 |
| 1 | 1 | 1 |
| 0 | 0 | 0 |
| 0 | 1 | 0 |
| 0 | 0 | 0 |
| 0 | 0 | 0 |
| 0 | 0 | 0 |
| 0 | 0 | 0 |
| 0 | 0 | 0 |
| 0 | 0 | 0 |
| 0 | 1 | 0 |
| 0 | 1 | 0 |
| 0 | 0 | 0 |
| 0 | 0 | 0 |
| 0 | 0 | 0 |
| 0 | 0 | 0 |
| 1 | 1 | 1 |
| 1 | 1 | 1 |
| 1 | 1 | 1 |
| 1 | 1 | 1 |
| 1 | 1 | 1 |
| 0 | 0 | 0 |
| 0 | 0 | 0 |
| 0 | 1 | 0 |
| 1 | 1 | 1 |
| 1 | 1 | 1 |
| 0 | 1 | 0 |
| 1 | 1 | 1 |
| 1 | 1 | 1 |
| 0 | 0 | 0 |
| 0 | 0 | 0 |
| 0 | 0 | 0 |
| 1 | 1 | 1 |
| 0 | 0 | 0 |
| 0 | 0 | 0 |
| 0 | 0 | 0 |
| 1 | 1 | 1 |
| 1 | 1 | 1 |
| 0 | 1 | 0 |
| 0 | 1 | 0 |

| | | |
|---|---|---|
| 0 | 1 | 0 |
| 1 | 1 | 1 |
| 1 | 0 | 0 |
| 1 | 1 | 1 |
| 0 | 0 | 0 |
| 1 | 0 | 0 |
| 0 | 0 | 0 |
| 0 | 0 | 0 |
| 0 | 0 | 0 |
| 0 | 0 | 0 |
| 0 | 0 | 0 |
| 0 | 0 | 0 |
| 0 | 1 | 0 |
| 0 | 0 | 0 |
| 0 | 0 | 0 |
| 1 | 1 | 1 |
| 1 | 1 | 1 |
| 1 | 1 | 1 |
| 0 | 0 | 0 |
| 0 | 0 | 0 |
| 0 | 0 | 0 |
| 0 | 0 | 0 |
| 0 | 1 | 0 |
| 0 | 1 | 0 |
| 1 | 1 | 1 |
| 1 | 1 | 1 |
| 1 | 1 | 1 |
| 0 | 0 | 0 |
| 0 | 0 | 0 |
| 0 | 0 | 0 |
| 0 | 0 | 0 |
| 0 | 0 | 0 |
| 0 | 0 | 0 |
| 0 | 1 | 0 |
| 1 | 1 | 1 |
| 0 | 0 | 0 |
| 1 | 1 | 1 |
| 1 | 1 | 1 |
| 1 | 1 | 1 |
| 1 | 1 | 1 |
| 0 | 0 | 0 |
| 1 | 1 | 1 |
| 0 | 0 | 0 |
| 0 | 1 | 0 |
| 1 | 1 | 1 |
| 1 | 1 | 1 |
| 1 | 1 | 1 |
| 0 | 0 | 0 |
| 1 | 1 | 1 |
| 1 | 1 | 1 |
| 1 | 1 | 1 |
| 1 | 1 | 1 |
| 1 | 1 | 1 |
| 0 | 1 | 0 |

| | | |
|---|---|---|
| 0 | 1 | 0 |
| 1 | 1 | 1 |
| 1 | 1 | 1 |
| 0 | 1 | 0 |
| 1 | 1 | 1 |
| 0 | 1 | 0 |
| 0 | 1 | 0 |
| 0 | 1 | 0 |
| 0 | 1 | 0 |
| 0 | 1 | 0 |
| 0 | 1 | 0 |
| 0 | 0 | 0 |
| 0 | 1 | 0 |
| 0 | 1 | 0 |
| 0 | 1 | 0 |
| 0 | 1 | 0 |
| 0 | 1 | 0 |
| 0 | 0 | 0 |
| 0 | 0 | 0 |
| 0 | 0 | 0 |
| 0 | 0 | 0 |
| 0 | 1 | 0 |
| 0 | 0 | 0 |
| 1 | 1 | 1 |
| 1 | 1 | 1 |
| 1 | 1 | 1 |
| 1 | 1 | 1 |
| 0 | 1 | 0 |
| 0 | 0 | 0 |
| 0 | 0 | 0 |
| 1 | 1 | 1 |
| 1 | 1 | 1 |
| 0 | 0 | 0 |
| 1 | 1 | 1 |
| 0 | 0 | 0 |
| 0 | 0 | 0 |
| 0 | 0 | 0 |
| 1 | 1 | 1 |
| 0 | 0 | 0 |
| 1 | 1 | 1 |
| 0 | 0 | 0 |
| 1 | 1 | 1 |
| 0 | 0 | 0 |
| 1 | 0 | 0 |
| 1 | 0 | 0 |
| 1 | 0 | 0 |
| 1 | 0 | 0 |
| 0 | 1 | 0 |
| 0 | 1 | 0 |
| 0 | 0 | 0 |
| 1 | 0 | 0 |
| 0 | 0 | 0 |
| 1 | 0 | 0 |
| 1 | 1 | 1 |

| | | |
|---|---|---|
| 0 | 0 | 0 |
| 0 | 0 | 0 |
| 0 | 0 | 0 |
| 0 | 0 | 0 |
| 0 | 0 | 0 |
| 1 | 1 | 1 |
| 0 | 0 | 0 |
| 0 | 0 | 0 |
| 1 | 1 | 1 |
| 0 | 0 | 0 |
| 0 | 0 | 0 |
| 0 | 0 | 0 |
| 0 | 1 | 0 |
| 0 | 1 | 0 |
| 0 | 1 | 0 |
| 0 | 1 | 0 |
| 0 | 1 | 0 |
| 0 | 1 | 0 |
| 0 | 1 | 0 |
| 0 | 1 | 0 |
| 0 | 1 | 0 |
| 0 | 1 | 0 |
| 0 | 1 | 0 |
| 0 | 1 | 0 |
| 0 | 1 | 0 |
| 0 | 1 | 0 |
| 0 | 1 | 0 |
| 0 | 1 | 0 |
| 0 | 1 | 0 |
| 0 | 1 | 0 |
| 0 | 1 | 0 |
| 0 | 1 | 0 |
| 0 | 1 | 0 |
| 0 | 1 | 0 |
| 0 | 1 | 0 |
| 0 | 1 | 0 |
| 0 | 1 | 0 |
| 0 | 1 | 0 |
| 0 | 1 | 0 |
| 0 | 1 | 0 |
| 0 | 1 | 0 |
| 0 | 1 | 0 |
| 0 | 1 | 0 |
| 0 | 1 | 0 |
| 0 | 1 | 0 |
| 0 | 0 | 0 |
| 1 | 1 | 1 |
| 1 | 1 | 1 |
| 1 | 1 | 1 |
| 1 | 1 | 1 |
| 0 | 0 | 0 |
| 1 | 1 | 1 |
| 1 | 1 | 1 |
| 0 | 1 | 0 |

| | | |
|---|---|---|
| 1 | 1 | 1 |
| 1 | 1 | 1 |
| 1 | 1 | 1 |
| 1 | 1 | 1 |
| 0 | 1 | 0 |
| 0 | 1 | 0 |
| 0 | 1 | 0 |
| 0 | 0 | 0 |
| 0 | 0 | 0 |
| 1 | 1 | 1 |
| 0 | 1 | 0 |
| 1 | 1 | 1 |
| 1 | 1 | 1 |
| 1 | 1 | 1 |
| 0 | 0 | 0 |
| 0 | 0 | 0 |
| 0 | 0 | 0 |
| 1 | 1 | 1 |
| 1 | 0 | 0 |
| 1 | 1 | 1 |
| 0 | 0 | 0 |
| 0 | 0 | 0 |
| 0 | 0 | 0 |
| 0 | 0 | 0 |
| 0 | 0 | 0 |
| 0 | 0 | 0 |
| 0 | 1 | 0 |
| 1 | 1 | 1 |
| 0 | 0 | 0 |
| 1 | 1 | 1 |
| 1 | 1 | 1 |
| 1 | 1 | 1 |
| 0 | 0 | 0 |
| 1 | 1 | 1 |
| 1 | 1 | 1 |
| 1 | 1 | 1 |
| 0 | 0 | 0 |
| 0 | 0 | 0 |
| 0 | 0 | 0 |
| 1 | 1 | 1 |
| 1 | 1 | 1 |
| 1 | 1 | 1 |
| 1 | 1 | 1 |
| 1 | 1 | 1 |
| 0 | 1 | 0 |
| 0 | 0 | 0 |
| 1 | 1 | 1 |
| 0 | 1 | 0 |
| 1 | 1 | 1 |
| 0 | 1 | 0 |
| 1 | 1 | 1 |
| 1 | 1 | 1 |
| 0 | 1 | 0 |
| 1 | 1 | 1 |

| | | |
|---|---|---|
| 1 | 1 | 1 |
| 0 | 1 | 0 |
| 1 | 1 | 1 |
| 0 | 0 | 0 |
| 1 | 1 | 1 |
| 1 | 1 | 1 |
| 1 | 1 | 1 |
| 0 | 0 | 0 |
| 0 | 0 | 0 |
| 1 | 1 | 1 |
| 1 | 1 | 1 |
| 1 | 1 | 1 |
| 0 | 0 | 0 |
| 0 | 1 | 0 |
| 0 | 1 | 0 |
| 0 | 1 | 0 |
| 0 | 0 | 0 |
| 1 | 1 | 1 |
| 1 | 1 | 1 |
| 1 | 1 | 1 |
| 1 | 1 | 1 |
| 1 | 1 | 1 |
| 1 | 1 | 1 |
| 1 | 1 | 1 |
| 0 | 1 | 0 |
| 1 | 1 | 1 |
| 0 | 1 | 0 |
| 0 | 0 | 0 |
| 0 | 0 | 0 |
| 0 | 0 | 0 |
| 0 | 0 | 0 |
| 0 | 0 | 0 |
| 0 | 0 | 0 |
| 0 | 1 | 0 |
| 0 | 0 | 0 |
| 0 | 0 | 0 |
| 0 | 0 | 0 |
| 1 | 1 | 1 |
| 1 | 1 | 1 |
| 1 | 1 | 1 |
| 1 | 1 | 1 |
| 0 | 1 | 0 |
| 0 | 0 | 0 |
| 0 | 0 | 0 |
| 1 | 1 | 1 |
| 1 | 1 | 1 |
| 0 | 0 | 0 |
| 1 | 1 | 1 |
| 1 | 1 | 1 |
| 1 | 1 | 1 |
| 1 | 1 | 1 |
| 1 | 1 | 1 |
| 0 | 0 | 0 |
| 0 | 0 | 0 |

| | | |
|---|---|---|
| 0 | 0 | 0 |
| 0 | 0 | 0 |
| 0 | 0 | 0 |
| 0 | 0 | 0 |
| 0 | 0 | 0 |
| 0 | 0 | 0 |
| 0 | 0 | 0 |
| 0 | 0 | 0 |
| 1 | 1 | 1 |
| 0 | 0 | 0 |
| 1 | 1 | 1 |
| 1 | 1 | 1 |
| 1 | 1 | 1 |
| 1 | 1 | 1 |
| 0 | 0 | 0 |
| 1 | 1 | 1 |
| 1 | 1 | 1 |
| 1 | 1 | 1 |
| 1 | 1 | 1 |
| 0 | 0 | 0 |
| 0 | 0 | 0 |
| 1 | 1 | 1 |
| 0 | 0 | 0 |
| 0 | 0 | 0 |
| 0 | 0 | 0 |
| 0 | 0 | 0 |
| 0 | 0 | 0 |
| 0 | 1 | 0 |
| 0 | 1 | 0 |
| 0 | 1 | 0 |
| 0 | 0 | 0 |
| 1 | 1 | 1 |
| 1 | 1 | 1 |
| 1 | 1 | 1 |
| 1 | 1 | 1 |
| 1 | 1 | 1 |
| 1 | 1 | 1 |
| 1 | 1 | 1 |
| 1 | 1 | 1 |
| 1 | 1 | 1 |
| 0 | 1 | 0 |
| 0 | 0 | 0 |
| 1 | 1 | 1 |
| 1 | 1 | 1 |
| 1 | 1 | 1 |
| 0 | 1 | 0 |
| 1 | 1 | 1 |
| 1 | 1 | 1 |
| 0 | 0 | 0 |
| 0 | 1 | 0 |
| 1 | 1 | 1 |
| 1 | 1 | 1 |
| 0 | 0 | 0 |
| 0 | 0 | 0 |

| | | |
|---|---|---|
| 1 | 1 | 1 |
| 1 | 1 | 1 |
| 0 | 1 | 0 |
| 1 | 0 | 0 |
| 0 | 0 | 0 |
| 0 | 0 | 0 |
| 0 | 1 | 0 |
| 0 | 0 | 0 |
| 1 | 1 | 1 |
| 1 | 1 | 1 |
| 0 | 0 | 0 |
| 1 | 0 | 0 |
| 0 | 0 | 0 |
| 0 | 0 | 0 |
| 0 | 0 | 0 |
| 0 | 0 | 0 |
| 1 | 1 | 1 |
| 1 | 1 | 1 |
| 0 | 0 | 0 |
| 0 | 0 | 0 |
| 0 | 0 | 0 |
| 0 | 0 | 0 |
| 0 | 0 | 0 |
| 1 | 1 | 1 |
| 1 | 1 | 1 |
| 0 | 0 | 0 |
| 1 | 1 | 1 |
| 0 | 0 | 0 |
| 0 | 0 | 0 |
| 0 | 0 | 0 |
| 0 | 1 | 0 |
| 0 | 0 | 0 |
| 0 | 0 | 0 |
| 1 | 1 | 1 |
| 1 | 1 | 1 |
| 0 | 0 | 0 |
| 0 | 0 | 0 |
| 1 | 0 | 0 |
| 0 | 0 | 0 |
| 0 | 0 | 0 |
| 0 | 0 | 0 |
| 0 | 0 | 0 |
| 0 | 1 | 0 |
| 1 | 1 | 1 |
| 0 | 0 | 0 |
| 0 | 1 | 0 |
| 0 | 0 | 0 |
| 1 | 1 | 1 |
| 1 | 1 | 1 |
| 1 | 1 | 1 |
| 1 | 1 | 1 |
| 1 | 1 | 1 |
| 0 | 1 | 0 |
| 1 | 1 | 1 |

| | | |
|---|---|---|
| 1 | 1 | 1 |
| 1 | 1 | 1 |
| 1 | 1 | 1 |
| 0 | 1 | 0 |
| 1 | 1 | 1 |
| 1 | 1 | 1 |
| 0 | 1 | 0 |
| 0 | 0 | 0 |
| 1 | 1 | 1 |
| 0 | 0 | 0 |
| 0 | 0 | 0 |
| 0 | 0 | 0 |
| 0 | 1 | 0 |
| 1 | 1 | 1 |
| 0 | 0 | 0 |
| 0 | 1 | 0 |
| 0 | 1 | 0 |
| 0 | 1 | 0 |
| 0 | 1 | 0 |
| 0 | 1 | 0 |
| 1 | 1 | 1 |
| 1 | 1 | 1 |
| 1 | 1 | 1 |
| 0 | 1 | 0 |
| 1 | 1 | 1 |
| 1 | 1 | 1 |
| 1 | 1 | 1 |
| 1 | 0 | 0 |
| 1 | 1 | 1 |
| 0 | 0 | 0 |
| 0 | 0 | 0 |
| 1 | 1 | 1 |
| 1 | 1 | 1 |
| 0 | 1 | 0 |
| 0 | 1 | 0 |
| 0 | 0 | 0 |
| 1 | 1 | 1 |
| 1 | 1 | 1 |
| 1 | 1 | 1 |
| 1 | 1 | 1 |
| 1 | 1 | 1 |
| 0 | 0 | 0 |
| 0 | 0 | 0 |
| 0 | 0 | 0 |
| 1 | 1 | 1 |
| 1 | 1 | 1 |
| 1 | 1 | 1 |
| 1 | 1 | 1 |
| 1 | 1 | 1 |
| 1 | 1 | 1 |
| 1 | 1 | 1 |
| 1 | 1 | 1 |
| 1 | 1 | 1 |
| 0 | 0 | 0 |

| | | |
|---|---|---|
| 1 | 1 | 1 |
| 1 | 1 | 1 |
| 1 | 1 | 1 |
| 1 | 1 | 1 |
| 1 | 1 | 1 |
| 1 | 1 | 1 |
| 1 | 1 | 1 |
| 0 | 1 | 0 |
| 0 | 0 | 0 |
| 0 | 0 | 0 |
| 1 | 1 | 1 |
| 0 | 1 | 0 |
| 0 | 1 | 0 |
| 0 | 0 | 0 |
| 1 | 1 | 1 |
| 1 | 1 | 1 |
| 1 | 1 | 1 |
| 0 | 0 | 0 |
| 0 | 0 | 0 |
| 0 | 0 | 0 |
| 0 | 0 | 0 |
| 0 | 1 | 0 |
| 1 | 1 | 1 |
| 1 | 1 | 1 |
| 1 | 1 | 1 |
| 0 | 0 | 0 |
| 1 | 1 | 1 |
| 0 | 1 | 0 |
| 1 | 1 | 1 |
| 1 | 1 | 1 |
| 1 | 1 | 1 |
| 1 | 1 | 1 |
| 1 | 1 | 1 |
| 1 | 1 | 1 |
| 1 | 1 | 1 |
| 1 | 1 | 1 |
| 1 | 1 | 1 |
| 1 | 1 | 1 |
| 1 | 1 | 1 |
| 0 | 0 | 0 |
| 1 | 1 | 1 |
| 1 | 1 | 1 |
| 0 | 0 | 0 |
| 1 | 1 | 1 |
| 1 | 1 | 1 |
| 1 | 1 | 1 |
| 0 | 0 | 0 |
| 0 | 0 | 0 |
| 1 | 1 | 1 |
| 1 | 1 | 1 |
| 0 | 0 | 0 |
| 1 | 1 | 1 |
| 0 | 1 | 0 |
| 1 | 1 | 1 |

| | | |
|---|---|---|
| 1 | 1 | 1 |
| 1 | 1 | 1 |
| 1 | 1 | 1 |
| 1 | 1 | 1 |
| 1 | 1 | 1 |
| 1 | 1 | 1 |
| 1 | 1 | 1 |
| 0 | 1 | 0 |
| 1 | 1 | 1 |
| 1 | 1 | 1 |
| 1 | 1 | 1 |
| 1 | 1 | 1 |
| 1 | 1 | 1 |
| 0 | 0 | 0 |
| 0 | 0 | 0 |
| 0 | 0 | 0 |
| 0 | 0 | 0 |
| 0 | 1 | 0 |
| 1 | 1 | 1 |
| 0 | 0 | 0 |
| 1 | 1 | 1 |
| 1 | 0 | 0 |
| 1 | 1 | 1 |
| 1 | 1 | 1 |
| 1 | 1 | 1 |
| 0 | 0 | 0 |
| 0 | 1 | 0 |
| 1 | 1 | 1 |
| 1 | 1 | 1 |
| 0 | 0 | 0 |
| 0 | 0 | 0 |
| 1 | 1 | 1 |
| 1 | 1 | 1 |
| 1 | 1 | 1 |
| 1 | 1 | 1 |
| 1 | 1 | 1 |
| 1 | 1 | 1 |
| 1 | 1 | 1 |
| 0 | 0 | 0 |
| 1 | 1 | 1 |
| 1 | 1 | 1 |
| 0 | 0 | 0 |
| 0 | 0 | 0 |
| 0 | 0 | 0 |
| 0 | 0 | 0 |
| 1 | 1 | 1 |
| 0 | 0 | 0 |
| 1 | 1 | 1 |
| 1 | 1 | 1 |
| 1 | 1 | 1 |
| 0 | 1 | 0 |
| 1 | 0 | 0 |
| 1 | 0 | 0 |
| 1 | 0 | 0 |

| | | |
|---|---|---|
| 1 | 1 | 1 |
| 1 | 1 | 1 |
| 0 | 0 | 0 |
| 1 | 1 | 1 |
| 1 | 1 | 1 |
| 1 | 1 | 1 |
| 1 | 1 | 1 |
| 0 | 1 | 0 |
| 1 | 1 | 1 |
| 1 | 1 | 1 |
| 1 | 1 | 1 |
| 0 | 1 | 0 |
| 0 | 1 | 0 |
| 0 | 1 | 0 |
| 0 | 1 | 0 |
| 0 | 1 | 0 |
| 0 | 1 | 0 |
| 0 | 1 | 0 |
| 0 | 1 | 0 |
| 0 | 1 | 0 |
| 0 | 1 | 0 |
| 0 | 1 | 0 |
| 0 | 1 | 0 |
| 0 | 1 | 0 |
| 0 | 1 | 0 |
| 0 | 1 | 0 |
| 0 | 1 | 0 |
| 0 | 1 | 0 |
| 1 | 1 | 1 |
| 1 | 1 | 1 |
| 0 | 0 | 0 |
| 1 | 1 | 1 |
| 0 | 0 | 0 |
| 0 | 0 | 0 |
| 0 | 0 | 0 |
| 0 | 0 | 0 |
| 1 | 1 | 1 |
| 0 | 0 | 0 |
| 1 | 1 | 1 |
| 1 | 1 | 1 |
| 0 | 0 | 0 |
| 0 | 0 | 0 |
| 0 | 0 | 0 |
| 1 | 1 | 1 |
| 0 | 0 | 0 |
| 0 | 0 | 0 |
| 1 | 1 | 1 |
| 1 | 1 | 1 |
| 1 | 1 | 1 |
| 1 | 1 | 1 |
| 0 | 0 | 0 |
| 1 | 1 | 1 |
| 1 | 1 | 1 |
| 1 | 1 | 1 |

| | | |
|---|---|---|
| 1 | 1 | 1 |
| 1 | 1 | 1 |
| 1 | 1 | 1 |
| 1 | 1 | 1 |
| 1 | 1 | 1 |
| 0 | 0 | 0 |
| 1 | 1 | 1 |
| 0 | 0 | 0 |
| 0 | 0 | 0 |
| 0 | 0 | 0 |
| 1 | 1 | 1 |
| 0 | 0 | 0 |
| 1 | 1 | 1 |
| 1 | 1 | 1 |
| 1 | 1 | 1 |
| 1 | 0 | 0 |
| 1 | 1 | 1 |
| 0 | 0 | 0 |
| 1 | 1 | 1 |
| 0 | 0 | 0 |
| 1 | 1 | 1 |
| 0 | 0 | 0 |
| 0 | 0 | 0 |
| 0 | 0 | 0 |
| 1 | 1 | 1 |
| 0 | 1 | 0 |
| 1 | 1 | 1 |
| 1 | 1 | 1 |
| 0 | 0 | 0 |
| 1 | 1 | 1 |
| 1 | 1 | 1 |
| 1 | 1 | 1 |
| 1 | 1 | 1 |
| 1 | 1 | 1 |
| 0 | 0 | 0 |
| 0 | 0 | 0 |
| 1 | 1 | 1 |
| 0 | 0 | 0 |
| 0 | 0 | 0 |
| 0 | 1 | 0 |
| 0 | 0 | 0 |
| 0 | 0 | 0 |
| 0 | 1 | 0 |
| 0 | 1 | 0 |
| 1 | 1 | 1 |
| 0 | 1 | 0 |
| 1 | 1 | 1 |
| 1 | 1 | 1 |
| 1 | 1 | 1 |
| 1 | 1 | 1 |
| 1 | 1 | 1 |
| 1 | 1 | 1 |
| 1 | 1 | 1 |
| 1 | 1 | 1 |

| | | |
|---|---|---|
| 1 | 1 | 1 |
| 0 | 0 | 0 |
| 0 | 1 | 0 |
| 1 | 1 | 1 |
| 0 | 0 | 0 |
| 1 | 1 | 1 |
| 1 | 1 | 1 |
| 1 | 1 | 1 |
| 0 | 0 | 0 |
| 0 | 0 | 0 |
| 1 | 1 | 1 |
| 0 | 0 | 0 |
| 0 | 0 | 0 |
| 1 | 1 | 1 |
| 1 | 1 | 1 |
| 0 | 0 | 0 |
| 0 | 1 | 0 |
| 0 | 0 | 0 |
| 0 | 0 | 0 |
| 1 | 1 | 1 |
| 0 | 0 | 0 |
| 0 | 0 | 0 |
| 0 | 0 | 0 |
| 1 | 1 | 1 |
| 0 | 1 | 0 |
| 0 | 0 | 0 |
| 1 | 1 | 1 |
| 1 | 1 | 1 |
| 1 | 1 | 1 |
| 1 | 1 | 1 |
| 0 | 0 | 0 |
| 1 | 1 | 1 |
| 0 | 0 | 0 |
| 1 | 1 | 1 |
| 0 | 0 | 0 |
| 0 | 0 | 0 |
| 1 | 1 | 1 |
| 1 | 1 | 1 |
| 1 | 1 | 1 |
| 1 | 1 | 1 |
| 1 | 1 | 1 |
| 1 | 1 | 1 |
| 1 | 1 | 1 |
| 1 | 1 | 1 |
| 1 | 1 | 1 |
| 1 | 1 | 1 |
| 1 | 0 | 0 |
| 1 | 1 | 1 |
| 1 | 1 | 1 |
| 1 | 1 | 1 |
| 1 | 1 | 1 |
| 1 | 1 | 1 |
| 1 | 1 | 1 |
| 0 | 0 | 0 |

| | | |
|---|---|---|
| 1 | 1 | 1 |
| 1 | 1 | 1 |
| 1 | 1 | 1 |
| 1 | 1 | 1 |
| 0 | 0 | 0 |
| 1 | 1 | 1 |
| 0 | 1 | 0 |
| 0 | 1 | 0 |
| 0 | 1 | 0 |
| 1 | 1 | 1 |
| 0 | 0 | 0 |
| 1 | 1 | 1 |
| 1 | 1 | 1 |
| 1 | 1 | 1 |
| 0 | 0 | 0 |
| 0 | 0 | 0 |
| 0 | 0 | 0 |
| 0 | 0 | 0 |
| 0 | 1 | 0 |
| 0 | 0 | 0 |
| 0 | 0 | 0 |
| 0 | 0 | 0 |
| 1 | 1 | 1 |
| 0 | 0 | 0 |
| 0 | 0 | 0 |
| 1 | 1 | 1 |
| 0 | 0 | 0 |
| 0 | 0 | 0 |
| 0 | 0 | 0 |
| 0 | 0 | 0 |
| 0 | 0 | 0 |
| 1 | 1 | 1 |
| 1 | 1 | 1 |
| 0 | 1 | 0 |
| 1 | 1 | 1 |
| 1 | 1 | 1 |
| 1 | 1 | 1 |
| 0 | 0 | 0 |
| 0 | 0 | 0 |
| 0 | 1 | 0 |
| 1 | 1 | 1 |
| 0 | 0 | 0 |
| 0 | 0 | 0 |
| 0 | 1 | 0 |
| 0 | 0 | 0 |
| 1 | 1 | 1 |
| 1 | 1 | 1 |
| 1 | 1 | 1 |
| 0 | 0 | 0 |
| 0 | 0 | 0 |
| 0 | 0 | 0 |
| 1 | 1 | 1 |
| 0 | 1 | 0 |
| 0 | 0 | 0 |

| | | |
|---|---|---|
| 1 | 1 | 1 |
| 0 | 0 | 0 |
| 1 | 1 | 1 |
| 0 | 0 | 0 |
| 0 | 0 | 0 |
| 0 | 1 | 0 |
| 0 | 0 | 0 |
| 0 | 0 | 0 |
| 1 | 1 | 1 |
| 1 | 1 | 1 |
| 1 | 1 | 1 |
| 1 | 1 | 1 |
| 1 | 1 | 1 |
| 1 | 1 | 1 |
| 1 | 1 | 1 |
| 1 | 1 | 1 |
| 1 | 1 | 1 |
| 0 | 1 | 0 |
| 1 | 1 | 1 |
| 1 | 1 | 1 |
| 0 | 0 | 0 |
| 0 | 1 | 0 |
| 0 | 1 | 0 |
| 0 | 1 | 0 |
| 0 | 1 | 0 |
| 0 | 0 | 0 |
| 0 | 0 | 0 |
| 1 | 1 | 1 |
| 1 | 1 | 1 |
| 1 | 1 | 1 |
| 1 | 1 | 1 |
| 1 | 1 | 1 |
| 1 | 1 | 1 |
| 0 | 1 | 0 |
| 1 | 1 | 1 |
| 1 | 1 | 1 |
| 1 | 1 | 1 |
| 1 | 1 | 1 |
| 0 | 1 | 0 |
| 1 | 1 | 1 |
| 1 | 1 | 1 |
| 0 | 1 | 0 |
| 1 | 1 | 1 |
| 0 | 0 | 0 |
| 1 | 1 | 1 |
| 0 | 0 | 0 |
| 0 | 0 | 0 |
| 0 | 0 | 0 |
| 0 | 0 | 0 |
| 0 | 0 | 0 |
| 0 | 0 | 0 |
| 0 | 1 | 0 |
| 1 | 1 | 1 |
| 1 | 1 | 1 |

| | | |
|---|---|---|
| 1 | 1 | 1 |
| 1 | 1 | 1 |
| 1 | 1 | 1 |
| 0 | 0 | 0 |
| 1 | 1 | 1 |
| 0 | 0 | 0 |
| 1 | 1 | 1 |
| 1 | 1 | 1 |
| 0 | 0 | 0 |
| 1 | 1 | 1 |
| 1 | 1 | 1 |
| 0 | 1 | 0 |
| 1 | 1 | 1 |
| 1 | 1 | 1 |
| 1 | 1 | 1 |
| 0 | 0 | 0 |
| 0 | 0 | 0 |
| 0 | 0 | 0 |
| 0 | 0 | 0 |
| 0 | 0 | 0 |
| 1 | 1 | 1 |
| 1 | 1 | 1 |
| 1 | 1 | 1 |
| 1 | 1 | 1 |
| 0 | 0 | 0 |
| 1 | 1 | 1 |
| 1 | 1 | 1 |
| 1 | 1 | 1 |
| 0 | 0 | 0 |
| 1 | 1 | 1 |
| 0 | 0 | 0 |
| 0 | 0 | 0 |
| 0 | 0 | 0 |
| 0 | 0 | 0 |
| 0 | 0 | 0 |
| 1 | 1 | 1 |
| 0 | 0 | 0 |
| 0 | 0 | 0 |
| 0 | 0 | 0 |
| 0 | 0 | 0 |
| 0 | 0 | 0 |
| 0 | 0 | 0 |
| 0 | 0 | 0 |
| 0 | 0 | 0 |
| 0 | 0 | 0 |
| 0 | 0 | 0 |
| 0 | 0 | 0 |
| 0 | 0 | 0 |
| 1 | 1 | 1 |
| 0 | 0 | 0 |
| 1 | 1 | 1 |
| 1 | 1 | 1 |
| 0 | 1 | 0 |
| 0 | 0 | 0 |

| | | |
|---|---|---|
| 0 | 0 | 0 |
| 0 | 1 | 0 |
| 1 | 1 | 1 |
| 1 | 1 | 1 |
| 1 | 1 | 1 |
| 0 | 0 | 0 |
| 1 | 1 | 1 |
| 0 | 1 | 0 |
| 1 | 1 | 1 |
| 0 | 1 | 0 |
| 0 | 1 | 0 |
| 0 | 1 | 0 |
| 1 | 1 | 1 |
| 0 | 0 | 0 |
| 0 | 1 | 0 |
| 1 | 1 | 1 |
| 1 | 1 | 1 |
| 0 | 0 | 0 |
| 1 | 1 | 1 |
| 1 | 1 | 1 |
| 1 | 1 | 1 |
| 1 | 1 | 1 |
| 0 | 0 | 0 |
| 1 | 1 | 1 |
| 1 | 1 | 1 |
| 0 | 0 | 0 |
| 0 | 0 | 0 |
| 0 | 0 | 0 |
| 1 | 1 | 1 |
| 0 | 1 | 0 |
| 1 | 1 | 1 |
| 1 | 1 | 1 |
| 1 | 1 | 1 |
| 1 | 1 | 1 |
| 1 | 1 | 1 |
| 1 | 1 | 1 |
| 1 | 1 | 1 |
| 1 | 1 | 1 |
| 1 | 1 | 1 |
| 1 | 1 | 1 |
| 1 | 1 | 1 |
| 1 | 1 | 1 |
| 1 | 1 | 1 |
| 1 | 1 | 1 |
| 1 | 1 | 1 |
| 1 | 1 | 1 |
| 1 | 1 | 1 |
| 1 | 1 | 1 |
| 1 | 0 | 0 |
| 1 | 1 | 1 |
| 1 | 1 | 1 |
| 1 | 1 | 1 |
| 1 | 1 | 1 |
| 1 | 1 | 1 |

| | | |
|---|---|---|
| 1 | 1 | 1 |
| 0 | 0 | 0 |
| 1 | 1 | 1 |
| 1 | 1 | 1 |
| 1 | 1 | 1 |
| 0 | 0 | 0 |
| 0 | 0 | 0 |
| 0 | 0 | 0 |
| 1 | 1 | 1 |
| 1 | 1 | 1 |
| 0 | 1 | 0 |
| 1 | 1 | 1 |
| 1 | 1 | 1 |
| 1 | 1 | 1 |
| 0 | 0 | 0 |
| 0 | 0 | 0 |
| 0 | 0 | 0 |
| 1 | 1 | 1 |
| 0 | 0 | 0 |
| 0 | 0 | 0 |
| 0 | 0 | 0 |
| 0 | 0 | 0 |
| 0 | 0 | 0 |
| 0 | 0 | 0 |
| 0 | 1 | 0 |
| 1 | 1 | 1 |
| 1 | 1 | 1 |
| 0 | 1 | 0 |
| 0 | 1 | 0 |
| 1 | 1 | 1 |
| 1 | 1 | 1 |
| 1 | 1 | 1 |
| 0 | 0 | 0 |
| 1 | 1 | 1 |
| 0 | 1 | 0 |
| 0 | 0 | 0 |
| 0 | 0 | 0 |
| 0 | 0 | 0 |
| 0 | 0 | 0 |
| 1 | 1 | 1 |
| 0 | 1 | 0 |
| 1 | 1 | 1 |
| 1 | 1 | 1 |
| 0 | 0 | 0 |
| 0 | 0 | 0 |
| 0 | 0 | 0 |
| 0 | 1 | 0 |
| 1 | 1 | 1 |
| 0 | 0 | 0 |
| 1 | 1 | 1 |
| 1 | 1 | 1 |
| 0 | 0 | 0 |
| 1 | 1 | 1 |
| 0 | 1 | 0 |

| | | |
|---|---|---|
| 1 | 1 | 1 |
| 0 | 1 | 0 |
| 1 | 1 | 1 |
| 1 | 1 | 1 |
| 1 | 1 | 1 |
| 1 | 1 | 1 |
| 0 | 1 | 0 |
| 1 | 1 | 1 |
| 0 | 0 | 0 |
| 0 | 0 | 0 |
| 1 | 1 | 1 |
| 0 | 0 | 0 |
| 0 | 0 | 0 |
| 0 | 0 | 0 |
| 1 | 1 | 1 |
| 1 | 1 | 1 |
| 0 | 0 | 0 |
| 0 | 0 | 0 |
| 0 | 0 | 0 |
| 1 | 1 | 1 |
| 1 | 1 | 1 |
| 1 | 1 | 1 |
| 0 | 0 | 0 |
| 1 | 1 | 1 |
| 0 | 0 | 0 |
| 1 | 1 | 1 |
| 1 | 1 | 1 |
| 1 | 1 | 1 |
| 1 | 1 | 1 |
| 0 | 0 | 0 |
| 0 | 0 | 0 |
| 0 | 0 | 0 |
| 1 | 1 | 1 |
| 0 | 1 | 0 |
| 1 | 1 | 1 |
| 1 | 1 | 1 |
| 1 | 1 | 1 |
| 0 | 1 | 0 |
| 0 | 1 | 0 |
| 0 | 1 | 0 |
| 0 | 0 | 0 |
| 1 | 1 | 1 |
| 1 | 1 | 1 |
| 1 | 1 | 1 |
| 1 | 1 | 1 |
| 0 | 0 | 0 |
| 1 | 0 | 0 |
| 0 | 0 | 0 |
| 1 | 1 | 1 |
| 0 | 0 | 0 |
| 0 | 0 | 0 |
| 0 | 0 | 0 |
| 0 | 1 | 0 |
| 1 | 0 | 0 |

| | | |
|---|---|---|
| 1 | 0 | 0 |
| 0 | 0 | 0 |
| 0 | 1 | 0 |
| 1 | 1 | 1 |
| 0 | 1 | 0 |
| 0 | 1 | 0 |
| 0 | 1 | 0 |
| 0 | 0 | 0 |
| 0 | 0 | 0 |
| 0 | 1 | 0 |
| 0 | 0 | 0 |
| 1 | 1 | 1 |
| 1 | 1 | 1 |
| 1 | 1 | 1 |
| 0 | 1 | 0 |
| 1 | 1 | 1 |
| 1 | 1 | 1 |
| 1 | 1 | 1 |
| 1 | 1 | 1 |
| 0 | 1 | 0 |
| 1 | 1 | 1 |
| 1 | 1 | 1 |
| 1 | 1 | 1 |
| 1 | 1 | 1 |
| 0 | 1 | 0 |
| 0 | 1 | 0 |
| 0 | 1 | 0 |
| 0 | 1 | 0 |
| 0 | 0 | 0 |
| 0 | 1 | 0 |
| 0 | 0 | 0 |
| 1 | 1 | 1 |
| 0 | 0 | 0 |
| 0 | 0 | 0 |
| 0 | 1 | 0 |
| 0 | 0 | 0 |
| 0 | 0 | 0 |
| 1 | 1 | 1 |
| 0 | 1 | 0 |
| 0 | 1 | 0 |
| 1 | 1 | 1 |
| 0 | 0 | 0 |
| 0 | 0 | 0 |
| 1 | 1 | 1 |
| 1 | 1 | 1 |
| 0 | 0 | 0 |
| 0 | 0 | 0 |
| 0 | 0 | 0 |
| 0 | 0 | 0 |
| 0 | 1 | 0 |
| 0 | 0 | 0 |
| 0 | 0 | 0 |
| 1 | 1 | 1 |
| 1 | 1 | 1 |

| | | |
|---|---|---|
| 1 | 1 | 1 |
| 1 | 1 | 1 |
| 1 | 1 | 1 |
| 1 | 1 | 1 |
| 0 | 0 | 0 |
| 0 | 0 | 0 |
| 0 | 0 | 0 |
| 0 | 0 | 0 |
| 1 | 1 | 1 |
| 0 | 0 | 0 |
| 0 | 0 | 0 |
| 0 | 0 | 0 |
| 0 | 0 | 0 |
| 1 | 1 | 1 |
| 1 | 0 | 0 |
| 0 | 1 | 0 |
| 1 | 1 | 1 |
| 1 | 1 | 1 |
| 1 | 1 | 1 |
| 0 | 1 | 0 |
| 1 | 1 | 1 |
| 1 | 1 | 1 |
| 1 | 1 | 1 |
| 1 | 1 | 1 |
| 1 | 1 | 1 |
| 0 | 0 | 0 |
| 0 | 0 | 0 |
| 0 | 0 | 0 |
| 0 | 0 | 0 |
| 0 | 1 | 0 |
| 0 | 1 | 0 |
| 0 | 1 | 0 |
| 1 | 1 | 1 |
| 0 | 1 | 0 |
| 0 | 1 | 0 |
| 1 | 1 | 1 |
| 1 | 1 | 1 |
| 1 | 1 | 1 |
| 1 | 1 | 1 |
| 1 | 1 | 1 |
| 1 | 1 | 1 |
| 1 | 1 | 1 |
| 1 | 1 | 1 |
| 1 | 1 | 1 |
| 1 | 1 | 1 |
| 1 | 1 | 1 |
| 1 | 1 | 1 |
| 1 | 1 | 1 |
| 1 | 1 | 1 |
| 1 | 1 | 1 |
| 1 | 1 | 1 |
| 1 | 1 | 1 |
| 1 | 1 | 1 |
| 0 | 0 | 0 |

| | | |
|---|---|---|
| 1 | 1 | 1 |
| 1 | 1 | 1 |
| 1 | 1 | 1 |
| 0 | 1 | 0 |
| 0 | 1 | 0 |
| 0 | 1 | 0 |
| 0 | 1 | 0 |
| 0 | 1 | 0 |
| 1 | 1 | 1 |
| 1 | 1 | 1 |
| 1 | 1 | 1 |
| 1 | 1 | 1 |
| 1 | 1 | 1 |
| 1 | 1 | 1 |
| 1 | 1 | 1 |
| 1 | 1 | 1 |
| 1 | 1 | 1 |
| 1 | 1 | 1 |
| 1 | 1 | 1 |
| 1 | 1 | 1 |
| 1 | 1 | 1 |
| 1 | 1 | 1 |
| 1 | 1 | 1 |
| 1 | 1 | 1 |
| 1 | 1 | 1 |
| 1 | 1 | 1 |
| 1 | 1 | 1 |
| 1 | 1 | 1 |
| 1 | 1 | 1 |
| 1 | 1 | 1 |
| 1 | 1 | 1 |
| 1 | 1 | 1 |
| 1 | 1 | 1 |
| 1 | 1 | 1 |
| 0 | 1 | 0 |
| 1 | 1 | 1 |
| 1 | 1 | 1 |
| 1 | 1 | 1 |
| 1 | 1 | 1 |
| 1 | 1 | 1 |
| 1 | 1 | 1 |
| 0 | 1 | 0 |
| 1 | 1 | 1 |
| 1 | 1 | 1 |
| 1 | 1 | 1 |
| 0 | 1 | 0 |
| 1 | 0 | 0 |
| 0 | 0 | 0 |
| 1 | 1 | 1 |
| 0 | 1 | 0 |
| 1 | 1 | 1 |
| 1 | 1 | 1 |
| 0 | 1 | 0 |
| 0 | 1 | 0 |
| 0 | 1 | 0 |

| | | |
|---|---|---|
| 0 | 1 | 0 |
| 1 | 1 | 1 |
| 0 | 0 | 0 |
| 1 | 1 | 1 |
| 1 | 1 | 1 |
| 1 | 1 | 1 |
| 0 | 0 | 0 |
| 1 | 1 | 1 |
| 1 | 1 | 1 |
| 0 | 1 | 0 |
| 0 | 1 | 0 |
| 0 | 0 | 0 |
| 1 | 1 | 1 |
| 0 | 0 | 0 |
| 0 | 0 | 0 |
| 0 | 0 | 0 |
| 0 | 0 | 0 |
| 0 | 0 | 0 |
| 0 | 0 | 0 |
| 0 | 0 | 0 |
| 0 | 0 | 0 |

| organism_scientific | evo_cons | cluster_id | pdb_id | chain_id | resi | resn | change | uniprot_id |
|---|---|---|---|---|---|---|---|---|
| HOMO SAPIENS | 5 | 87 | 3uzw | A | 106 | LEU | p.Leu106Phe | P51857 |
| HOMO SAPIENS | 3 | 87 | 3uzw | A | 133 | PRO | p.Pro133Arg | P51857 |
| HOMO SAPIENS | 0 | 87 | 3uzw | A | 198 | PRO | p.Pro198Leu | P51857 |
| HOMO SAPIENS | 7 | 87 | 3uzw | A | 261 | ARG | p.Arg261Cys | P51857 |
| HOMO SAPIENS | 0 | 4693 | 3mk1 | A | 3 | PRO | p.Pro25Leu | P05187 |
| HOMO SAPIENS | 8 | 4830 | 2h64 | B | 101 | CYS | p.Cys124Arg | P36894 |
| HOMO SAPIENS | 2 | 4830 | 2h64 | B | 116 | PRO | p.Pro139Ser | P36894 |
| HOMO SAPIENS | 0 | 6937 | 3dar | A | 186 | MET | p.Met186Thr | P21802 |
| HOMO SAPIENS | 3 | 5 | 1aut | C | 30 | GLN | p.Gln226His | P04070 |
| HOMO SAPIENS | 1 | 5 | 1aut | C | 68 | LEU | p.Leu265Phe | P04070 |
| HOMO SAPIENS | 0 | 5 | 1aut | C | 75 | ARG | p.Arg272Cys | P04070 |
| HOMO SAPIENS | 6 | 5 | 1aut | C | 92 | PRO | p.Pro289Leu | P04070 |
| HOMO SAPIENS | 0 | 5 | 1aut | C | 104 | ALA | p.Ala301Val | P04070 |
| HOMO SAPIENS | 1 | 5 | 1aut | C | 112 | ALA | p.Ala309Thr | P04070 |
| HOMO SAPIENS | 2 | 5 | 1aut | C | 115 | SER | p.Ser312Leu | P04070 |
| HOMO SAPIENS | 1 | 5 | 1aut | C | 133 | GLY | p.Gly334Ser | P04070 |
| HOMO SAPIENS | 0 | 5 | 1aut | C | 147 | ARG | p.Arg348Ter | P04070 |
| HOMO SAPIENS | 8 | 5 | 1aut | C | 194 | ASP | p.Asp401Asn | P04070 |
| HOMO SAPIENS | 7 | 5 | 1aut | C | 237 | TRP | p.Trp444Cys | P04070 |
| HOMO SAPIENS | 0 | 5 | 1aut | C | 238 | ILE | p.Ile445Met | P04070 |
| HOMO SAPIENS | 8 | 43 | 1aiy | F | 10 | HIS | p.His34Asp | P01308 |
| HOMO SAPIENS | 2 | 43 | 1aiy | F | 22 | ARG | p.Arg46Gln | P01308 |
| HOMO SAPIENS | 4 | 43 | 1aiy | F | 23 | GLY | p.Gly47Val | P01308 |
| HOMO SAPIENS | 7 | 43 | 1aiy | F | 24 | PHE | p.Phe48Ser | P01308 |
| HOMO SAPIENS | 5 | 24 | 2fg8 | A | 33 | THR | p.Thr30Ile | P02792 |
| HOMO SAPIENS | 4 | 24 | 2fg8 | A | 99 | ALA | p.Ala96Thr | P02792 |
| HOMO SAPIENS | 5 | 24 | 2fg8 | A | 107 | GLU | p.Glu104Ter | P02792 |
| HOMO SAPIENS | 3 | 5409 | 5bnq | A | 199 | MET | p.Met199Lys | O14788 |
| HOMO SAPIENS | -1 | 839 | 3swc | A | 959 | GLN | p.Gln990His | Q9H9B1 |
| HOMO SAPIENS | -1 | 839 | 3swc | A | 968 | ALA | p.Ala999Val | Q9H9B1 |
| HOMO SAPIENS | -1 | 839 | 3swc | A | 973 | SER | p.Ser1004Ile | Q9H9B1 |
| HOMO SAPIENS | 8 | 17 | 1eou | A | 18 | LYS | p.Lys18Glu | P00918 |
| HOMO SAPIENS | 2 | 17 | 1eou | A | 107 | HIS | p.His107Tyr | P00918 |
| HOMO SAPIENS | 1 | 17 | 1eou | A | 236 | PRO | p.Pro236His | P00918 |
| HOMO SAPIENS | 6 | 17 | 1eou | A | 252 | ASN | p.Asn252Asp | P00918 |
| HOMO SAPIENS | 2 | 9140 | 5c4v | A | 445 | ARG | p.Arg445Ter | Q13485 |
| HOMO SAPIENS | 7 | 9140 | 5c4v | A | 496 | ARG | p.Arg496Cys | Q13485 |
| HOMO SAPIENS | -1 | 42 | 4pwi | A | 6 | GLY | p.Gly26Ser | P02766 |
| HOMO SAPIENS | 6 | 42 | 4pwi | A | 10 | CYS | p.Cys30Arg | P02766 |
| HOMO SAPIENS | 8 | 42 | 4pwi | A | 12 | LEU | p.Leu32Pro | P02766 |
| HOMO SAPIENS | 9 | 42 | 4pwi | A | 18 | ASP | p.Asp38Gly | P02766 |
| HOMO SAPIENS | 8 | 42 | 4pwi | A | 20 | VAL | p.Val40Ile | P02766 |
| HOMO SAPIENS | 4 | 42 | 4pwi | A | 30 | MET | p.Val50Ala | P02766 |
| HOMO SAPIENS | 4 | 42 | 4pwi | A | 30 | MET | p.Val50Leu | P02766 |
| HOMO SAPIENS | 6 | 42 | 4pwi | A | 33 | PHE | p.Phe53Ile | P02766 |
| HOMO SAPIENS | 7 | 42 | 4pwi | A | 36 | ALA | p.Ala56Pro | P02766 |
| HOMO SAPIENS | 7 | 42 | 4pwi | A | 42 | GLU | p.Glu62Gly | P02766 |
| HOMO SAPIENS | 8 | 42 | 4pwi | A | 44 | PHE | p.Phe64Ser | P02766 |
| HOMO SAPIENS | 4 | 42 | 4pwi | A | 47 | GLY | p.Gly67Ala | P02766 |
| HOMO SAPIENS | 6 | 42 | 4pwi | A | 49 | THR | p.Thr69Ala | P02766 |
| HOMO SAPIENS | 6 | 42 | 4pwi | A | 50 | SER | p.Ser70Ile | P02766 |
| HOMO SAPIENS | 6 | 42 | 4pwi | A | 50 | SER | p.Ser70Arg | P02766 |
| HOMO SAPIENS | 0 | 42 | 4pwi | A | 53 | GLY | p.Gly73Glu | P02766 |

| | | | | | | | | |
|---|---|---|---|---|---|---|---|---|
| HOMO SAPIENS | 7 | 42 | 4pwi | A | 55 | LEU | p.Leu75Pro | P02766 |
| HOMO SAPIENS | 6 | 42 | 4pwi | A | 58 | LEU | p.Leu78His | P02766 |
| HOMO SAPIENS | 7 | 42 | 4pwi | A | 61 | GLU | p.Glu81Lys | P02766 |
| HOMO SAPIENS | 8 | 42 | 4pwi | A | 64 | PHE | p.Phe84Leu | P02766 |
| HOMO SAPIENS | 8 | 42 | 4pwi | A | 68 | ILE | p.Ile88Leu | P02766 |
| HOMO SAPIENS | 8 | 42 | 4pwi | A | 69 | TYR | p.Tyr89His | P02766 |
| HOMO SAPIENS | 4 | 42 | 4pwi | A | 71 | VAL | p.Val91Ala | P02766 |
| HOMO SAPIENS | 6 | 42 | 4pwi | A | 77 | SER | p.Ser97Tyr | P02766 |
| HOMO SAPIENS | 4 | 42 | 4pwi | A | 84 | ILE | p.Ile104Ser | P02766 |
| HOMO SAPIENS | 8 | 42 | 4pwi | A | 89 | GLU | p.Glu109Gln | P02766 |
| HOMO SAPIENS | 8 | 42 | 4pwi | A | 97 | ALA | p.Ala117Gly | P02766 |
| HOMO SAPIENS | 7 | 42 | 4pwi | A | 99 | ASP | p.Asp119Asn | P02766 |
| HOMO SAPIENS | 9 | 42 | 4pwi | A | 107 | ILE | p.Ile127Val | P02766 |
| HOMO SAPIENS | 9 | 42 | 4pwi | A | 111 | LEU | p.Leu131Met | P02766 |
| HOMO SAPIENS | 7 | 42 | 4pwi | A | 114 | TYR | p.Tyr134Cys | P02766 |
| HOMO SAPIENS | 7 | 42 | 4pwi | A | 114 | TYR | p.Tyr134His | P02766 |
| HOMO SAPIENS | 7 | 42 | 4pwi | A | 122 | VAL | p.Val142Ile | P02766 |
| HOMO SAPIENS | -1 | 42 | 4pwi | A | 126 | LYS | p.Lys146Arg | P02766 |
| HOMO SAPIENS | -1 | 42 | 4pwi | A | 127 | GLU | p.Glu147Ter | P02766 |
| HOMO SAPIENS | -1 | 1312 | 4dce | A | 1275 | ARG | p.Arg1275Gln | Q9UM73 |
| HOMO SAPIENS | -1 | 1312 | 4dce | A | 1280 | ALA | p.Ala1280Val | Q9UM73 |
| HOMO SAPIENS | 1 | 20 | 1y0x | X | 234 | ALA | p.Ala234Thr | P10828 |
| HOMO SAPIENS | 0 | 20 | 1y0x | X | 243 | ARG | p.Arg243Trp | P10828 |
| HOMO SAPIENS | 3 | 20 | 1y0x | X | 316 | ARG | p.Arg316His | P10828 |
| HOMO SAPIENS | 1 | 20 | 1y0x | X | 317 | ALA | p.Ala317Thr | P10828 |
| HOMO SAPIENS | 2 | 20 | 1y0x | X | 320 | ARG | p.Arg320Cys | P10828 |
| HOMO SAPIENS | 2 | 20 | 1y0x | X | 320 | ARG | p.Arg320His | P10828 |
| HOMO SAPIENS | 0 | 20 | 1y0x | X | 332 | GLY | p.Gly332Arg | P10828 |
| HOMO SAPIENS | 1 | 20 | 1y0x | X | 338 | ARG | p.Arg338Trp | P10828 |
| HOMO SAPIENS | 0 | 20 | 1y0x | X | 340 | GLN | p.Gln340His | P10828 |
| HOMO SAPIENS | 0 | 20 | 1y0x | X | 345 | GLY | p.Gly345Val | P10828 |
| HOMO SAPIENS | 0 | 20 | 1y0x | X | 345 | GLY | p.Gly345Ser | P10828 |
| HOMO SAPIENS | 0 | 20 | 1y0x | X | 347 | GLY | p.Gly347Glu | P10828 |
| HOMO SAPIENS | 1 | 20 | 1y0x | X | 438 | ARG | p.Arg438His | P10828 |
| HOMO SAPIENS | 0 | 20 | 1y0x | X | 442 | MET | p.Met442Val | P10828 |
| HOMO SAPIENS | 0 | 20 | 1y0x | X | 443 | LYS | p.Lys443Glu | P10828 |
| HOMO SAPIENS | 0 | 20 | 1y0x | X | 446 | CYS | p.Cys446Arg | P10828 |
| HOMO SAPIENS | 2 | 20 | 1y0x | X | 453 | PRO | p.Pro453His | P10828 |
| HOMO SAPIENS | 2 | 20 | 1y0x | X | 453 | PRO | p.Pro453Thr | P10828 |
| HOMO SAPIENS | 3 | 20 | 1y0x | X | 459 | PHE | p.Phe459Cys | P10828 |
| HOMO SAPIENS | 0 | 4409 | 3qb7 | D | 87 | GLU | p.Glu109Lys | P31785 |
| HOMO SAPIENS | 1 | 4409 | 3qb7 | D | 92 | GLY | p.Gly114Asp | P31785 |
| HOMO SAPIENS | 1 | 4409 | 3qb7 | D | 93 | CYS | p.Cys115Arg | P31785 |
| HOMO SAPIENS | 0 | 4409 | 3qb7 | D | 131 | ILE | p.Ile153Asn | P31785 |
| HOMO SAPIENS | 9 | 4409 | 3qb7 | D | 200 | ARG | p.Arg222Cys | P31785 |
| HOMO SAPIENS | 0 | 3891 | 3ije | B | 33 | LEU | p.Leu59Pro | P05106 |
| HOMO SAPIENS | 0 | 3891 | 3ije | B | 117 | LEU | p.Leu143Trp | P05106 |
| HOMO SAPIENS | 3 | 3891 | 3ije | B | 119 | ASP | p.Asp145Tyr | P05106 |
| HOMO SAPIENS | 2 | 3891 | 3ije | B | 143 | ARG | p.Arg169Gln | P05106 |
| HOMO SAPIENS | 2 | 3891 | 3ije | B | 214 | ARG | p.Arg240Gln | P05106 |
| HOMO SAPIENS | 2 | 3891 | 3ije | B | 214 | ARG | p.Arg240Trp | P05106 |
| HOMO SAPIENS | 2 | 3891 | 3ije | B | 374 | CYS | p.Cys400Tyr | P05106 |
| HOMO SAPIENS | 1 | 3891 | 3ije | B | 407 | PRO | p.Pro433Ala | P05106 |
| HOMO SAPIENS | 0 | 3891 | 3ije | B | 489 | ARG | p.Arg515Gln | P05106 |

| | | | | | | | | |
|---|---|---|---|---|---|---|---|---|
| HOMO SAPIENS | -1 | 9613 | 5mke | B | 28 | ARG | p.Arg28Pro | Q13563 |
| HOMO SAPIENS | -1 | 9613 | 5mke | B | 190 | ALA | p.Ala190Thr | Q13563 |
| HOMO SAPIENS | 4 | 9613 | 5mke | B | 262 | VAL | p.Val262Met | Q13563 |
| HOMO SAPIENS | 0 | 9613 | 5mke | B | 482 | PHE | p.Phe482Cys | Q13563 |
| HOMO SAPIENS | 4 | 9613 | 5mke | B | 511 | ASP | p.Asp511Val | Q13563 |
| HOMO SAPIENS | 5 | 9613 | 5mke | B | 516 | VAL | p.Val516Leu | Q13563 |
| HOMO SAPIENS | -1 | 9613 | 5mke | B | 807 | ARG | p.Arg807Gln | Q13563 |
| HOMO SAPIENS | 4 | 2428 | 1olh | C | 337 | ARG | p.Arg337Cys | P04637 |
| HOMO SAPIENS | 4 | 2428 | 1olh | C | 337 | ARG | p.Arg337His | P04637 |
| HOMO SAPIENS | 8 | 2428 | 1olh | C | 338 | PHE | p.Phe338Leu | P04637 |
| HOMO SAPIENS | 8 | 2428 | 1olh | C | 339 | GLU | p.Glu339Lys | P04637 |
| HOMO SAPIENS | 7 | 2428 | 1olh | C | 344 | LEU | p.Leu344Pro | P04637 |
| HOMO SAPIENS | 0 | 2428 | 1olh | C | 360 | GLY | p.Gly360Ala | P04637 |
| HOMO SAPIENS | 7 | 4867 | 1mox | B | 404 | GLY | p.Gly428Asp | P00533 |
| HOMO SAPIENS | 5 | 4867 | 1mox | B | 497 | ARG | p.Arg521Lys | P00533 |
| HOMO SAPIENS | 1 | 4630 | 1fu1 | A | 43 | TRP | p.Trp43Arg | Q13426 |
| HOMO SAPIENS | 2 | 4630 | 1fu1 | A | 161 | ARG | p.Arg161Ter | Q13426 |
| HOMO SAPIENS | 2 | 15 | 1lfy | B | 2 | HIS | p.His3Gln | P68871 |
| HOMO SAPIENS | 2 | 15 | 1lfy | B | 6 | GLU | p.Glu7Ala | P68871 |
| HOMO SAPIENS | 2 | 15 | 1lfy | B | 6 | GLU | p.Glu7Lys | P68871 |
| HOMO SAPIENS | 4 | 15 | 1lfy | B | 9 | SER | p.Ser10Cys | P68871 |
| HOMO SAPIENS | 6 | 15 | 1lfy | B | 15 | TRP | p.Trp16Ter | P68871 |
| HOMO SAPIENS | 3 | 15 | 1lfy | B | 17 | LYS | p.Lys18Glu | P68871 |
| HOMO SAPIENS | 6 | 15 | 1lfy | B | 19 | ASN | p.Asn20Ser | P68871 |
| HOMO SAPIENS | 6 | 15 | 1lfy | B | 22 | GLU | p.Glu23Ala | P68871 |
| HOMO SAPIENS | 6 | 15 | 1lfy | B | 22 | GLU | p.Glu23Lys | P68871 |
| HOMO SAPIENS | 8 | 15 | 1lfy | B | 26 | GLU | p.Glu27Lys | P68871 |
| HOMO SAPIENS | 8 | 15 | 1lfy | B | 28 | LEU | p.Leu29Pro | P68871 |
| HOMO SAPIENS | 8 | 15 | 1lfy | B | 30 | ARG | p.Arg31Ser | P68871 |
| HOMO SAPIENS | 8 | 15 | 1lfy | B | 30 | ARG | p.Arg31Lys | P68871 |
| HOMO SAPIENS | 6 | 15 | 1lfy | B | 39 | GLN | p.Gln40Ter | P68871 |
| HOMO SAPIENS | 2 | 15 | 1lfy | B | 43 | GLU | p.Glu44Gln | P68871 |
| HOMO SAPIENS | 6 | 15 | 1lfy | B | 57 | ASN | p.Asn58Lys | P68871 |
| HOMO SAPIENS | 6 | 15 | 1lfy | B | 59 | LYS | p.Lys60Asn | P68871 |
| HOMO SAPIENS | 8 | 15 | 1lfy | B | 61 | LYS | p.Lys62Glu | P68871 |
| HOMO SAPIENS | 4 | 15 | 1lfy | B | 73 | ASP | p.Asp74Tyr | P68871 |
| HOMO SAPIENS | 4 | 15 | 1lfy | B | 87 | THR | p.Thr88Ile | P68871 |
| HOMO SAPIENS | 4 | 15 | 1lfy | B | 87 | THR | p.Thr88Pro | P68871 |
| HOMO SAPIENS | 5 | 15 | 1lfy | B | 90 | GLU | p.Glu91Lys | P68871 |
| HOMO SAPIENS | 6 | 15 | 1lfy | B | 91 | LEU | p.Leu92Pro | P68871 |
| HOMO SAPIENS | 5 | 15 | 1lfy | B | 97 | HIS | p.His98Gln | P68871 |
| HOMO SAPIENS | 6 | 15 | 1lfy | B | 110 | LEU | p.Leu111Pro | P68871 |
| HOMO SAPIENS | 6 | 15 | 1lfy | B | 113 | VAL | p.Val114Glu | P68871 |
| HOMO SAPIENS | 2 | 15 | 1lfy | B | 119 | GLY | p.Gly120Ala | P68871 |
| HOMO SAPIENS | 4 | 15 | 1lfy | B | 121 | GLU | p.Glu122Gln | P68871 |
| HOMO SAPIENS | 5 | 15 | 1lfy | B | 134 | VAL | p.Val135Glu | P68871 |
| HOMO SAPIENS | 3 | 15 | 1lfy | B | 143 | HIS | p.His144Gln | P68871 |
| HOMO SAPIENS | 4 | 91 | 5d9k | A | 75 | GLY | p.Gly75Val | P51812 |
| HOMO SAPIENS | 5 | 91 | 5d9k | A | 82 | VAL | p.Val82Phe | P51812 |
| HOMO SAPIENS | 0 | 91 | 5d9k | A | 114 | ARG | p.Arg114Trp | P51812 |
| HOMO SAPIENS | 4 | 91 | 5d9k | A | 189 | ILE | p.Ile189Lys | P51812 |
| HOMO SAPIENS | -1 | 91 | 5d9k | A | 227 | SER | p.Ser227Ala | P51812 |
| HOMO SAPIENS | 1 | 91 | 5d9k | A | 268 | PHE | p.Phe268Ser | P51812 |
| HOMO SAPIENS | 0 | 954 | 2vt0 | A | 15 | VAL | p.Val54Leu | P04062 |

| | | | | | | | | |
|---|---|---|---|---|---|---|---|---|
| HOMO SAPIENS | 5 | 954 | 2vt0 | A | 46 | GLY | p.Gly85Glu | P04062 |
| HOMO SAPIENS | 5 | 954 | 2vt0 | A | 48 | ARG | p.Arg87Trp | P04062 |
| HOMO SAPIENS | 6 | 954 | 2vt0 | A | 79 | LYS | p.Lys118Asn | P04062 |
| HOMO SAPIENS | 0 | 954 | 2vt0 | A | 120 | ARG | p.Arg159Gln | P04062 |
| HOMO SAPIENS | 0 | 954 | 2vt0 | A | 120 | ARG | p.Arg159Trp | P04062 |
| HOMO SAPIENS | 0 | 954 | 2vt0 | A | 122 | PRO | p.Pro161Ser | P04062 |
| HOMO SAPIENS | 5 | 954 | 2vt0 | A | 131 | ARG | p.Arg170Leu | P04062 |
| HOMO SAPIENS | 5 | 954 | 2vt0 | A | 135 | TYR | p.Tyr174Cys | P04062 |
| HOMO SAPIENS | 0 | 954 | 2vt0 | A | 157 | LYS | p.Lys196Gln | P04062 |
| HOMO SAPIENS | 5 | 954 | 2vt0 | A | 163 | ARG | p.Arg202Gln | P04062 |
| HOMO SAPIENS | 5 | 954 | 2vt0 | A | 170 | ARG | p.Arg209Cys | P04062 |
| HOMO SAPIENS | 5 | 954 | 2vt0 | A | 188 | ASN | p.Asn227Ser | P04062 |
| HOMO SAPIENS | 3 | 954 | 2vt0 | A | 196 | SER | p.Ser235Pro | P04062 |
| HOMO SAPIENS | 5 | 954 | 2vt0 | A | 202 | GLY | p.Gly241Arg | P04062 |
| HOMO SAPIENS | 0 | 954 | 2vt0 | A | 212 | TYR | p.Tyr251His | P04062 |
| HOMO SAPIENS | 0 | 954 | 2vt0 | A | 213 | PHE | p.Phe252Ile | P04062 |
| HOMO SAPIENS | 5 | 954 | 2vt0 | A | 216 | PHE | p.Phe255Tyr | P04062 |
| HOMO SAPIENS | 6 | 954 | 2vt0 | A | 251 | PHE | p.Phe290Leu | P04062 |
| HOMO SAPIENS | 5 | 954 | 2vt0 | A | 255 | HIS | p.His294Gln | P04062 |
| HOMO SAPIENS | 6 | 954 | 2vt0 | A | 257 | ARG | p.Arg296Gln | P04062 |
| HOMO SAPIENS | 5 | 954 | 2vt0 | A | 289 | PRO | p.Pro328Leu | P04062 |
| HOMO SAPIENS | 0 | 954 | 2vt0 | A | 309 | ALA | p.Ala348Val | P04062 |
| HOMO SAPIENS | 0 | 954 | 2vt0 | A | 311 | HIS | p.His350Arg | P04062 |
| HOMO SAPIENS | 6 | 954 | 2vt0 | A | 312 | TRP | p.Trp351Cys | P04062 |
| HOMO SAPIENS | 5 | 954 | 2vt0 | A | 323 | THR | p.Thr362Ile | P04062 |
| HOMO SAPIENS | 5 | 954 | 2vt0 | A | 325 | GLY | p.Gly364Arg | P04062 |
| HOMO SAPIENS | 6 | 954 | 2vt0 | A | 342 | CYS | p.Cys381Gly | P04062 |
| HOMO SAPIENS | 5 | 954 | 2vt0 | A | 353 | ARG | p.Arg392Gly | P04062 |
| HOMO SAPIENS | 3 | 954 | 2vt0 | A | 364 | SER | p.Ser403Thr | P04062 |
| HOMO SAPIENS | 5 | 954 | 2vt0 | A | 369 | THR | p.Thr408Met | P04062 |
| HOMO SAPIENS | 3 | 954 | 2vt0 | A | 370 | ASN | p.Asn409Ser | P04062 |
| HOMO SAPIENS | 0 | 954 | 2vt0 | A | 371 | LEU | p.Leu410Val | P04062 |
| HOMO SAPIENS | 0 | 954 | 2vt0 | A | 375 | VAL | p.Val414Leu | P04062 |
| HOMO SAPIENS | 0 | 954 | 2vt0 | A | 377 | GLY | p.Gly416Ser | P04062 |
| HOMO SAPIENS | 8 | 954 | 2vt0 | A | 398 | VAL | p.Val437Phe | P04062 |
| HOMO SAPIENS | 0 | 954 | 2vt0 | A | 401 | PRO | p.Pro440Leu | P04062 |
| HOMO SAPIENS | 3 | 954 | 2vt0 | A | 409 | ASP | p.Asp448His | P04062 |
| HOMO SAPIENS | 3 | 954 | 2vt0 | A | 409 | ASP | p.Asp448Val | P04062 |
| HOMO SAPIENS | 0 | 954 | 2vt0 | A | 415 | PRO | p.Pro454Arg | P04062 |
| HOMO SAPIENS | 5 | 954 | 2vt0 | A | 444 | LEU | p.Leu483Arg | P04062 |
| HOMO SAPIENS | 5 | 954 | 2vt0 | A | 456 | ALA | p.Ala495Pro | P04062 |
| HOMO SAPIENS | 5 | 954 | 2vt0 | A | 463 | ARG | p.Arg502Cys | P04062 |
| HOMO SAPIENS | 5 | 954 | 2vt0 | A | 463 | ARG | p.Arg502His | P04062 |
| HOMO SAPIENS | 5 | 954 | 2vt0 | A | 478 | GLY | p.Gly517Ser | P04062 |
| HOMO SAPIENS | 5 | 954 | 2vt0 | A | 496 | ARG | p.Arg535Cys | P04062 |
| HOMO SAPIENS | 5 | 954 | 2vt0 | A | 496 | ARG | p.Arg535His | P04062 |
| HOMO SAPIENS | 1 | 9037 | 2rqb | A | 946 | ALA | p.Ala946Thr | Q9BYX4 |
| HOMO SAPIENS | 0 | 253 | 5E+092 | A | 308 | LEU | p.Leu308Pro | P37173 |
| HOMO SAPIENS | 0 | 253 | 5E+092 | A | 315 | THR | p.Thr315Met | P37173 |
| HOMO SAPIENS | 0 | 253 | 5E+092 | A | 328 | HIS | p.His328Gln | P37173 |
| HOMO SAPIENS | 0 | 253 | 5E+092 | A | 336 | TYR | p.Tyr336Asn | P37173 |
| HOMO SAPIENS | 0 | 253 | 5E+092 | A | 348 | ARG | p.Arg348Cys | P37173 |
| HOMO SAPIENS | 0 | 253 | 5E+092 | A | 355 | ALA | p.Ala355Pro | P37173 |
| HOMO SAPIENS | 1 | 253 | 5E+092 | A | 356 | ARG | p.Arg356Gly | P37173 |

| | | | | | | | | |
|---|---|---|---|---|---|---|---|---|
| HOMO SAPIENS | 0 | 253 | 5E+092 | A | 357 | GLY | p.Gly357Trp | P37173 |
| HOMO SAPIENS | 1 | 253 | 5E+092 | A | 403 | ARG | p.Arg403His | P37173 |
| HOMO SAPIENS | 5 | 253 | 5E+092 | A | 449 | SER | p.Ser449Phe | P37173 |
| HOMO SAPIENS | 0 | 253 | 5E+092 | A | 460 | ARG | p.Arg460Cys | P37173 |
| HOMO SAPIENS | 0 | 253 | 5E+092 | A | 460 | ARG | p.Arg460His | P37173 |
| HOMO SAPIENS | 2 | 253 | 5E+092 | A | 526 | GLU | p.Glu526Gln | P37173 |
| HOMO SAPIENS | 5 | 253 | 5E+092 | A | 528 | ARG | p.Arg528Cys | P37173 |
| HOMO SAPIENS | 5 | 253 | 5E+092 | A | 528 | ARG | p.Arg528His | P37173 |
| HOMO SAPIENS | 6 | 158 | 5amc | A | 231 | ARG | p.Arg260Leu | P12821 |
| HOMO SAPIENS | 7 | 158 | 5amc | A | 237 | TYR | p.Tyr266Ter | P12821 |
| HOMO SAPIENS | 6 | 158 | 5amc | A | 325 | GLY | p.Gly354Arg | P12821 |
| HOMO SAPIENS | 0 | 158 | 5amc | A | 467 | ARG | p.Arg496Ter | P12821 |
| HOMO SAPIENS | 4 | 158 | 5amc | A | 479 | ARG | p.Arg508Ter | P12821 |
| HOMO SAPIENS | 0 | 158 | 5amc | A | 563 | ASP | p.Asp592Gly | P12821 |
| HOMO SAPIENS | -1 | 33062 | 1gr3 | A | 545 | GLY | p.Gly545Arg | Q03692 |
| HOMO SAPIENS | 5 | 33062 | 1gr3 | A | 591 | CYS | p.Cys591Arg | Q03692 |
| HOMO SAPIENS | 2 | 33062 | 1gr3 | A | 595 | GLY | p.Gly595Glu | Q03692 |
| HOMO SAPIENS | 0 | 33062 | 1gr3 | A | 597 | TYR | p.Tyr597Cys | Q03692 |
| HOMO SAPIENS | 5 | 33062 | 1gr3 | A | 598 | TYR | p.Tyr598Asp | Q03692 |
| HOMO SAPIENS | 7 | 33062 | 1gr3 | A | 600 | SER | p.Ser600Pro | Q03692 |
| HOMO SAPIENS | 1 | 33062 | 1gr3 | A | 614 | LEU | p.Leu614Pro | Q03692 |
| HOMO SAPIENS | 6 | 33062 | 1gr3 | A | 651 | TRP | p.Trp651Arg | Q03692 |
| HOMO SAPIENS | 7 | 33062 | 1gr3 | A | 671 | SER | p.Ser671Pro | Q03692 |
| HOMO SAPIENS | -1 | 38257 | 2l2i | B | 87 | PRO | p.Pro87Ala | Q13351 |
| HOMO SAPIENS | 2 | 2036 | 1ghq | A | 231 | GLY | p.Gly1224Asp | P01024 |
| HOMO SAPIENS | 5 | 3594 | 4cde | F | 388 | TYR | p.Tyr412Cys | P53634 |
| HOMO SAPIENS | 9 | 3594 | 4cde | F | 405 | TRP | p.Trp429Cys | P53634 |
| HOMO SAPIENS | 0 | 3594 | 4cde | F | 429 | ILE | p.Ile453Val | P53634 |
| HOMO SAPIENS | 7 | 18 | 1jkb | A | 34 | TRP | p.Trp52Ter | P61626 |
| HOMO SAPIENS | 9 | 18 | 1jkb | A | 70 | THR | p.Thr88Asn | P61626 |
| HOMO SAPIENS | 8 | 18 | 1jkb | A | 121 | VAL | p.Val139Ile | P61626 |
| HOMO SAPIENS | 2 | 414 | 3fj6 | A | 136 | ARG | p.Arg135Cys | Q02127 |
| HOMO SAPIENS | 1 | 414 | 3fj6 | A | 153 | GLY | p.Gly152Arg | Q02127 |
| HOMO SAPIENS | 0 | 414 | 3fj6 | A | 200 | ARG | p.Arg199Cys | Q02127 |
| HOMO SAPIENS | 0 | 414 | 3fj6 | A | 203 | GLY | p.Gly202Ala | Q02127 |
| HOMO SAPIENS | 0 | 414 | 3fj6 | A | 245 | ARG | p.Arg244Trp | Q02127 |
| HOMO SAPIENS | 0 | 414 | 3fj6 | A | 342 | ALA | p.Ala341Val | Q02127 |
| HOMO SAPIENS | 1 | 414 | 3fj6 | A | 347 | ARG | p.Arg346Trp | Q02127 |
| HOMO SAPIENS | 6 | 103 | 1u3w | A | 77 | GLY | p.Gly78Ter | P00326 |
| HOMO SAPIENS | -1 | 8369 | 4ux8 | F | 16 | ARG | p.Arg93Trp | P39905 |
| HOMO SAPIENS | 0 | 366 | 2zjc | A | 32 | GLY | p.Arg108Trp | P01375 |
| HOMO SAPIENS | 0 | 1457 | 3v9i | A | 352 | LEU | p.Ser352Leu | P30038 |
| HOMO SAPIENS | 0 | 1457 | 3v9i | A | 470 | VAL | p.Val470Ile | P30038 |
| HOMO SAPIENS | 0 | 1457 | 3v9i | A | 473 | THR | p.Thr473Ala | P30038 |
| HOMO SAPIENS | 1 | 2 | 4wo5 | A | 462 | ARG | p.Arg462Ile | P15056 |
| HOMO SAPIENS | 3 | 2 | 4wo5 | A | 463 | ILE | p.Ile463Ser | P15056 |
| HOMO SAPIENS | 1 | 2 | 4wo5 | A | 466 | GLY | p.Gly466Val | P15056 |
| HOMO SAPIENS | 1 | 2 | 4wo5 | A | 468 | PHE | p.Phe468Ser | P15056 |
| HOMO SAPIENS | 2 | 2 | 4wo5 | A | 469 | GLY | p.Gly469Ala | P15056 |
| HOMO SAPIENS | 2 | 2 | 4wo5 | A | 469 | GLY | p.Gly469Arg | P15056 |
| HOMO SAPIENS | 2 | 2 | 4wo5 | A | 485 | LEU | p.Leu485Phe | P15056 |
| HOMO SAPIENS | 0 | 2 | 4wo5 | A | 501 | GLU | p.Glu501Gly | P15056 |
| HOMO SAPIENS | 0 | 2 | 4wo5 | A | 501 | GLU | p.Glu501Lys | P15056 |
| HOMO SAPIENS | 7 | 2 | 4wo5 | A | 581 | ASN | p.Asn581Asp | P15056 |

| | | | | | | | | |
|---|---|---|---|---|---|---|---|---|
| HOMO SAPIENS | 8 | 2 | 4wo5 | A | 594 | ASP | p.Asp594Gly | P15056 |
| HOMO SAPIENS | 0 | 2 | 4wo5 | A | 597 | LEU | p.Leu597Arg | P15056 |
| HOMO SAPIENS | 0 | 2 | 4wo5 | A | 597 | LEU | p.Leu597Val | P15056 |
| HOMO SAPIENS | 0 | 2 | 4wo5 | A | 600 | VAL | p.Val600Glu | P15056 |
| HOMO SAPIENS | 2 | 2 | 4wo5 | A | 601 | LYS | p.Lys601Glu | P15056 |
| HOMO SAPIENS | 8 | 2 | 4wo5 | A | 638 | ASP | p.Asp638Glu | P15056 |
| HOMO SAPIENS | -1 | 30366 | 5bqd | B | 19 | ALA | p.Ala19Ser | Q99593 |
| HOMO SAPIENS | -1 | 30366 | 5bqd | B | 23 | PRO | p.Pro23His | Q99593 |
| HOMO SAPIENS | 1 | 30366 | 5bqd | B | 49 | GLN | p.Gln49Lys | Q99593 |
| HOMO SAPIENS | 2 | 30366 | 5bqd | B | 54 | ILE | p.Ile54Thr | Q99593 |
| HOMO SAPIENS | 8 | 30366 | 5bqd | B | 80 | GLY | p.Gly80Arg | Q99593 |
| HOMO SAPIENS | 6 | 30366 | 5bqd | B | 106 | ILE | p.Ile106Val | Q99593 |
| HOMO SAPIENS | 7 | 30366 | 5bqd | B | 111 | ASP | p.Asp111Tyr | Q99593 |
| HOMO SAPIENS | -1 | 30366 | 5bqd | B | 237 | ARG | p.Arg237Gln | Q99593 |
| HOMO SAPIENS | -1 | 30366 | 5bqd | B | 237 | ARG | p.Arg237Trp | Q99593 |
| HOMO SAPIENS | 5 | 4789 | 1n4m | A | 567 | SER | p.Ser567Leu | P06400 |
| HOMO SAPIENS | 0 | 4789 | 1n4m | A | 658 | ALA | p.Ala658Asp | P06400 |
| HOMO SAPIENS | 7 | 4789 | 1n4m | A | 661 | ARG | p.Arg661Trp | P06400 |
| HOMO SAPIENS | 6 | 4789 | 1n4m | A | 668 | ARG | p.Arg668His | P06400 |
| HOMO SAPIENS | 8 | 4789 | 1n4m | A | 697 | ASP | p.Asp697Glu | P06400 |
| HOMO SAPIENS | 6 | 4789 | 1n4m | A | 712 | CYS | p.Cys712Arg | P06400 |
| HOMO SAPIENS | 1 | 1798 | 2r5g | A | 101 | HIS | p.His101Gln | O15247 |
| HOMO SAPIENS | 0 | 1798 | 2r5g | A | 160 | PRO | p.Pro160Ala | O15247 |
| HOMO SAPIENS | 2 | 22 | 4x0t | A | 82 | ARG | p.Arg110Ter | P49419 |
| HOMO SAPIENS | 0 | 22 | 4x0t | A | 149 | ALA | p.Ala177Pro | P49419 |
| HOMO SAPIENS | 4 | 22 | 4x0t | A | 164 | THR | p.Thr192Met | P49419 |
| HOMO SAPIENS | 9 | 22 | 4x0t | A | 167 | ASN | p.Asn195Ser | P49419 |
| HOMO SAPIENS | 0 | 22 | 4x0t | A | 175 | TRP | p.Trp203Gly | P49419 |
| HOMO SAPIENS | 0 | 22 | 4x0t | A | 194 | THR | p.Thr222Ala | P49419 |
| HOMO SAPIENS | 0 | 22 | 4x0t | A | 273 | ASN | p.Asn301Ile | P49419 |
| HOMO SAPIENS | 0 | 22 | 4x0t | A | 307 | ARG | p.Arg335Gln | P49419 |
| HOMO SAPIENS | 3 | 22 | 4x0t | A | 313 | SER | p.Ser341Asn | P49419 |
| HOMO SAPIENS | 0 | 22 | 4x0t | A | 329 | ILE | p.Ile357Val | P49419 |
| HOMO SAPIENS | 1 | 22 | 4x0t | A | 380 | TYR | p.Tyr408Ter | P49419 |
| HOMO SAPIENS | 9 | 22 | 4x0t | A | 399 | GLU | p.Glu427Gln | P49419 |
| HOMO SAPIENS | 0 | 22 | 4x0t | A | 406 | TYR | p.Tyr434Cys | P49419 |
| HOMO SAPIENS | 0 | 22 | 4x0t | A | 431 | ILE | p.Ile459Phe | P49419 |
| HOMO SAPIENS | 4 | 22 | 4x0t | A | 477 | GLY | p.Gly505Arg | P49419 |
| HOMO SAPIENS | 0 | 22 | 4x0t | A | 495 | THR | p.Thr523Ala | P49419 |
| HOMO SAPIENS | 8 | 11042 | 1y6b | A | 1145 | PRO | p.Pro1147Ser | P35968 |
| HOMO SAPIENS | -1 | 3867 | 2xij | A | 18 | GLN | p.Gln18Ter | P22033 |
| HOMO SAPIENS | -1 | 3867 | 2xij | A | 31 | ARG | p.Arg31Ter | P22033 |
| HOMO SAPIENS | 2 | 3867 | 2xij | A | 93 | ARG | p.Arg93His | P22033 |
| HOMO SAPIENS | 0 | 3867 | 2xij | A | 94 | GLY | p.Gly94Arg | P22033 |
| HOMO SAPIENS | 2 | 3867 | 2xij | A | 95 | PRO | p.Pro95Leu | P22033 |
| HOMO SAPIENS | 2 | 3867 | 2xij | A | 105 | TRP | p.Trp105Arg | P22033 |
| HOMO SAPIENS | 2 | 3867 | 2xij | A | 108 | ARG | p.Arg108Cys | P22033 |
| HOMO SAPIENS | 3 | 3867 | 2xij | A | 154 | ARG | p.Arg154His | P22033 |
| HOMO SAPIENS | 3 | 3867 | 2xij | A | 156 | ASP | p.Asp156Val | P22033 |
| HOMO SAPIENS | 0 | 3867 | 2xij | A | 189 | ASN | p.Asn189Ile | P22033 |
| HOMO SAPIENS | 0 | 3867 | 2xij | A | 191 | ALA | p.Ala191Glu | P22033 |
| HOMO SAPIENS | 0 | 3867 | 2xij | A | 203 | GLY | p.Gly203Arg | P22033 |
| HOMO SAPIENS | 2 | 3867 | 2xij | A | 215 | GLY | p.Gly215Ser | P22033 |
| HOMO SAPIENS | 0 | 3867 | 2xij | A | 219 | ASN | p.Asn219Tyr | P22033 |

| | | | | | | | | |
|---|---|---|---|---|---|---|---|---|
| HOMO SAPIENS | 3 | 3867 | 2xij | A | 228 | ARG | p.Arg228Ter | P22033 |
| HOMO SAPIENS | 1 | 3867 | 2xij | A | 297 | THR | p.Thr297Ile | P22033 |
| HOMO SAPIENS | 0 | 3867 | 2xij | A | 326 | ARG | p.Arg326Lys | P22033 |
| HOMO SAPIENS | 3 | 3867 | 2xij | A | 369 | ARG | p.Arg369His | P22033 |
| HOMO SAPIENS | 3 | 3867 | 2xij | A | 369 | ARG | p.Arg369Cys | P22033 |
| HOMO SAPIENS | 0 | 3867 | 2xij | A | 370 | THR | p.Thr370Pro | P22033 |
| HOMO SAPIENS | 0 | 3867 | 2xij | A | 377 | ALA | p.Ala377Glu | P22033 |
| HOMO SAPIENS | 2 | 3867 | 2xij | A | 426 | GLY | p.Gly426Ala | P22033 |
| HOMO SAPIENS | 2 | 3867 | 2xij | A | 427 | GLY | p.Gly427Asp | P22033 |
| HOMO SAPIENS | 1 | 3867 | 2xij | A | 499 | THR | p.Ala499Thr | P22033 |
| HOMO SAPIENS | 2 | 3867 | 2xij | A | 532 | ARG | p.Arg532His | P22033 |
| HOMO SAPIENS | 3 | 3867 | 2xij | A | 555 | ALA | p.Ala555Thr | P22033 |
| HOMO SAPIENS | 0 | 3867 | 2xij | A | 573 | PHE | p.Phe573Ser | P22033 |
| HOMO SAPIENS | 3 | 3867 | 2xij | A | 623 | GLY | p.Gly623Arg | P22033 |
| HOMO SAPIENS | 5 | 3867 | 2xij | A | 671 | VAL | p.Ile671Val | P22033 |
| HOMO SAPIENS | 4 | 3867 | 2xij | A | 694 | ARG | p.Arg694Trp | P22033 |
| HOMO SAPIENS | 1 | 3867 | 2xij | A | 700 | MET | p.Met700Lys | P22033 |
| HOMO SAPIENS | 8 | 3867 | 2xij | A | 703 | GLY | p.Gly703Arg | P22033 |
| HOMO SAPIENS | 0 | 3867 | 2xij | A | 717 | GLY | p.Gly717Val | P22033 |
| HOMO SAPIENS | 1 | 3867 | 2xij | A | 727 | ARG | p.Arg727Ter | P22033 |
| HOMO SAPIENS | -1 | 52 | 3k91 | B | 72 | GLY | p.Gly73Ser | P00441 |
| HOMO SAPIENS | 6 | 52 | 3k91 | B | 96 | ASP | p.Asp97Asn | P00441 |
| HOMO SAPIENS | 5 | 52 | 3k91 | B | 112 | ILE | p.Ile113Thr | P00441 |
| HOMO SAPIENS | -1 | 52 | 3k91 | B | 134 | SER | p.Ser135Asn | P00441 |
| HOMO SAPIENS | 0 | 3950 | 4fom | A | 296 | ASN | p.Asn296His | Q9NQS3 |
| HOMO SAPIENS | 1 | 10 | 1de4 | A | 31 | VAL | p.Val53Met | Q30201 |
| HOMO SAPIENS | 0 | 10 | 1de4 | A | 37 | VAL | p.Val59Met | Q30201 |
| HOMO SAPIENS | 0 | 10 | 1de4 | A | 71 | GLY | p.Gly93Arg | Q30201 |
| HOMO SAPIENS | 0 | 10 | 1de4 | A | 83 | ILE | p.Ile105Thr | Q30201 |
| HOMO SAPIENS | 0 | 10 | 1de4 | A | 105 | GLN | p.Gln127His | Q30201 |
| HOMO SAPIENS | 1 | 10 | 1de4 | A | 146 | GLU | p.Glu168Ter | Q30201 |
| HOMO SAPIENS | 2 | 10 | 1de4 | A | 261 | GLN | p.Gln283Pro | Q30201 |
| HOMO SAPIENS | 0 | 8038 | 4x7n | A | 586 | ILE | p.Ile586Leu | Q9NZJ5 |
| HOMO SAPIENS | 0 | 8038 | 4x7n | A | 588 | ARG | p.Arg588Gln | Q9NZJ5 |
| HOMO SAPIENS | 0 | 35175 | 5ugw | A | 1062 | ALA | p.Ala1062Thr | O14746 |
| HOMO SAPIENS | 2 | 35175 | 5ugw | A | 1090 | VAL | p.Val1090Met | O14746 |
| HOMO SAPIENS | 0 | 1722 | 4z3k | C | 147 | ARG | p.Arg150Gly | P35270 |
| HOMO SAPIENS | 0 | 1722 | 4z3k | C | 160 | PRO | p.Pro163Leu | P35270 |
| HOMO SAPIENS | 0 | 1722 | 4z3k | C | 207 | ARG | p.Arg210Trp | P35270 |
| HOMO SAPIENS | 0 | 1722 | 4z3k | C | 216 | ARG | p.Arg219Ter | P35270 |
| HOMO SAPIENS | 0 | 1722 | 4z3k | C | 248 | LYS | p.Lys251Ter | P35270 |
| HOMO SAPIENS | 1 | 2832 | 4x8v | L | 115 | HIS | p.His175Gln | P08709 |
| HOMO SAPIENS | 0 | 825 | 1emh | A | 242 | PHE | p.Phe251Ser | P13051 |
| HOMO SAPIENS | 0 | 122 | 5uwk | A | 109 | ARG | p.Arg109Ter | P45452 |
| HOMO SAPIENS | 1 | 122 | 5uwk | A | 170 | LYS | p.Lys170Thr | P45452 |
| HOMO SAPIENS | 9 | 122 | 5uwk | A | 232 | HIS | p.His232Asn | P45452 |
| HOMO SAPIENS | 5 | 122 | 5uwk | A | 257 | ASP | p.Asp257Val | P45452 |
| HOMO SAPIENS | 9 | 1074 | 3p5i | A | 264 | TRP | p.Trp264Arg | Q9UJ71 |
| HOMO SAPIENS | 0 | 146 | 3r3g | A | 14C | GLU | p.Glu352Lys | P00734 |
| HOMO SAPIENS | 3 | 9522 | 1wyq | A | 83 | LEU | p.Leu253Pro | O15020 |
| HOMO SAPIENS | 8 | 67 | 2vgb | A | 130 | SER | p.Ser130Tyr | P30613 |
| HOMO SAPIENS | 8 | 67 | 2vgb | A | 163 | ARG | p.Arg163Cys | P30613 |
| HOMO SAPIENS | 4 | 67 | 2vgb | A | 241 | GLU | p.Glu241Ter | P30613 |
| HOMO SAPIENS | 9 | 67 | 2vgb | A | 384 | THR | p.Thr384Met | P30613 |

| | | | | | | | | |
|---|---|---|---|---|---|---|---|---|
| HOMO SAPIENS | 0 | 67 | 2vgb | A | 421 | GLN | p.Gln421Lys | P30613 |
| HOMO SAPIENS | 6 | 67 | 2vgb | A | 479 | ARG | p.Arg479His | P30613 |
| HOMO SAPIENS | 8 | 67 | 2vgb | A | 510 | ARG | p.Arg510Gln | P30613 |
| HOMO SAPIENS | 4 | 67 | 2vgb | A | 569 | ARG | p.Arg569Trp | P30613 |
| HOMO SAPIENS | 0 | 9088 | 5v44 | A | 232 | ASN | p.Asn232Lys | Q9NZJ4 |
| HOMO SAPIENS | 4 | 16 | 3lbn | A | 12 | GLY | p.Gly12Val | P01112 |
| HOMO SAPIENS | 4 | 16 | 3lbn | A | 12 | GLY | p.Gly12Ser | P01112 |
| HOMO SAPIENS | 4 | 16 | 3lbn | A | 13 | GLY | p.Gly13Cys | P01112 |
| HOMO SAPIENS | 4 | 16 | 3lbn | A | 13 | GLY | p.Gly13Asp | P01112 |
| HOMO SAPIENS | 5 | 16 | 3lbn | A | 22 | GLN | p.Gln22Lys | P01112 |
| HOMO SAPIENS | 5 | 16 | 3lbn | A | 58 | THR | p.Thr58Ile | P01112 |
| HOMO SAPIENS | 3 | 16 | 3lbn | A | 61 | GLN | p.Gln61Lys | P01112 |
| HOMO SAPIENS | 1 | 16 | 3lbn | A | 63 | GLU | p.Glu63Lys | P01112 |
| HOMO SAPIENS | 9 | 16 | 3lbn | A | 117 | LYS | p.Lys117Arg | P01112 |
| HOMO SAPIENS | 6 | 16 | 3lbn | A | 123 | ARG | p.Arg123Cys | P01112 |
| HOMO SAPIENS | 1 | 16 | 3lbn | A | 138 | GLY | p.Gly138Ser | P01112 |
| HOMO SAPIENS | 9 | 16 | 3lbn | A | 146 | ALA | p.Ala146Thr | P01112 |
| HOMO SAPIENS | 9 | 16 | 3lbn | A | 146 | ALA | p.Ala146Val | P01112 |
| HOMO SAPIENS | 1 | 359 | 5nvy | C | 63 | LEU | p.Leu63Pro | P40337 |
| HOMO SAPIENS | 1 | 359 | 5nvy | C | 64 | ARG | p.Arg64Pro | P40337 |
| HOMO SAPIENS | 1 | 359 | 5nvy | C | 70 | GLU | p.Glu70Lys | P40337 |
| HOMO SAPIENS | 0 | 359 | 5nvy | C | 84 | VAL | p.Val84Leu | P40337 |
| HOMO SAPIENS | 1 | 359 | 5nvy | C | 88 | TRP | p.Trp88Ser | P40337 |
| HOMO SAPIENS | 1 | 359 | 5nvy | C | 93 | GLY | p.Gly93Cys | P40337 |
| HOMO SAPIENS | 1 | 359 | 5nvy | C | 98 | TYR | p.Tyr98His | P40337 |
| HOMO SAPIENS | 1 | 359 | 5nvy | C | 111 | SER | p.Ser111Arg | P40337 |
| HOMO SAPIENS | 1 | 359 | 5nvy | C | 112 | TYR | p.Tyr112His | P40337 |
| HOMO SAPIENS | 1 | 359 | 5nvy | C | 126 | ASP | p.Asp126Tyr | P40337 |
| HOMO SAPIENS | 1 | 359 | 5nvy | C | 130 | VAL | p.Val130Leu | P40337 |
| HOMO SAPIENS | 1 | 359 | 5nvy | C | 135 | LEU | p.Leu135Phe | P40337 |
| HOMO SAPIENS | 9 | 359 | 5nvy | C | 161 | ARG | p.Arg161Gly | P40337 |
| HOMO SAPIENS | 7 | 359 | 5nvy | C | 162 | CYS | p.Cys162Phe | P40337 |
| HOMO SAPIENS | 4 | 359 | 5nvy | C | 163 | LEU | p.Leu163Pro | P40337 |
| HOMO SAPIENS | 3 | 359 | 5nvy | C | 164 | GLN | p.Gln164Arg | P40337 |
| HOMO SAPIENS | 9 | 359 | 5nvy | C | 166 | VAL | p.Val166Phe | P40337 |
| HOMO SAPIENS | 7 | 359 | 5nvy | C | 167 | ARG | p.Arg167Gly | P40337 |
| HOMO SAPIENS | 7 | 359 | 5nvy | C | 167 | ARG | p.Arg167Gln | P40337 |
| HOMO SAPIENS | 1 | 359 | 5nvy | C | 183 | SER | p.Ser183Ter | P40337 |
| HOMO SAPIENS | 2 | 359 | 5nvy | C | 188 | LEU | p.Leu188Val | P40337 |
| HOMO SAPIENS | 1 | 359 | 5nvy | C | 191 | HIS | p.His191Asp | P40337 |
| HOMO SAPIENS | 1 | 359 | 5nvy | C | 192 | PRO | p.Pro192Ser | P40337 |
| HOMO SAPIENS | 1 | 359 | 5nvy | C | 200 | ARG | p.Arg200Trp | P40337 |
| HOMO SAPIENS | -1 | 359 | 5nvy | C | 211 | MET | p.Met211Leu | P40337 |
| HOMO SAPIENS | 0 | 14099 | 4btm | A | 282 | ARG | p.Arg258Gln | Q5TCY1 |
| HOMO SAPIENS | 0 | 696 | 5h1b | A | 150 | ARG | p.Arg150Gln | Q06609 |
| HOMO SAPIENS | -1 | 208 | 2bxl | A | 1 | ASP | p.Asp25Val | P02768 |
| HOMO SAPIENS | 0 | 208 | 2bxl | A | 269 | ASP | p.Asp293Gly | P02768 |
| HOMO SAPIENS | 0 | 208 | 2bxl | A | 570 | GLU | p.Glu594Lys | P02768 |
| HOMO SAPIENS | 0 | 208 | 2bxl | A | 573 | LYS | p.Lys597Glu | P02768 |
| HOMO SAPIENS | -1 | 12813 | 2uzx | B | 35 | MET | p.Met35Leu | P08581 |
| HOMO SAPIENS | 1 | 12813 | 2uzx | B | 156 | SER | p.Ser156Leu | P08581 |
| HOMO SAPIENS | 0 | 12813 | 2uzx | B | 168 | GLU | p.Glu168Asp | P08581 |
| HOMO SAPIENS | 2 | 12813 | 2uzx | B | 211 | LEU | p.Leu211Trp | P08581 |
| HOMO SAPIENS | 2 | 12813 | 2uzx | B | 218 | ARG | p.Arg218Ser | P08581 |

| | | | | | | | | |
|---|---|---|---|---|---|---|---|---|
| HOMO SAPIENS | 1 | 12813 | 2uzx | B | 238 | LEU | p.Leu238Ser | P08581 |
| HOMO SAPIENS | 4 | 12813 | 2uzx | B | 320 | ALA | p.Ala320Val | P08581 |
| HOMO SAPIENS | 1 | 12813 | 2uzx | B | 375 | ASN | p.Asn375Ser | P08581 |
| HOMO SAPIENS | 0 | 12813 | 2uzx | B | 575 | LEU | p.Leu575Ile | P08581 |
| HOMO SAPIENS | 1 | 10780 | 4ejx | A | 348 | ARG | p.Arg367Cys | P00450 |
| HOMO SAPIENS | 2 | 10780 | 4ejx | A | 525 | ASP | p.Asp544Glu | P00450 |
| HOMO SAPIENS | 1 | 1407 | 5jn5 | A | 88 | ILE | p.Ile88Val | P36871 |
| HOMO SAPIENS | 7 | 1407 | 5jn5 | A | 115 | THR | p.Thr115Ala | P36871 |
| HOMO SAPIENS | 3 | 1407 | 5jn5 | A | 121 | GLY | p.Gly121Arg | P36871 |
| HOMO SAPIENS | 2 | 1407 | 5jn5 | A | 221 | ARG | p.Arg221Cys | P36871 |
| HOMO SAPIENS | 6 | 1407 | 5jn5 | A | 263 | TYR | p.Asp263Tyr | P36871 |
| HOMO SAPIENS | 3 | 1407 | 5jn5 | A | 420 | TYR | p.Tyr420His | P36871 |
| HOMO SAPIENS | 0 | 1407 | 5jn5 | A | 501 | VAL | p.Val501Ile | P36871 |
| HOMO SAPIENS | -1 | 1878 | 1yow | A | 255 | ARG | p.Arg255Leu | Q13285 |
| HOMO SAPIENS | 6 | 1878 | 1yow | A | 293 | ASP | p.Asp293Asn | Q13285 |
| HOMO SAPIENS | 1 | 1878 | 1yow | A | 437 | LEU | p.Leu437Gln | Q13285 |
| HOMO SAPIENS | 0 | 3071 | 2wpm | E | 145 | ARG | p.Arg191His | P00740 |
| HOMO SAPIENS | 0 | 3071 | 2wpm | E | 145 | ARG | p.Arg191Cys | P00740 |
| HOMO SAPIENS | 3 | 589 | 3ggc | B | 15 | GLY | p.Gly16Ser | P00492 |
| HOMO SAPIENS | 4 | 589 | 3ggc | B | 40 | LEU | p.Leu41Pro | P00492 |
| HOMO SAPIENS | 2 | 589 | 3ggc | B | 50 | ARG | p.Arg51Gly | P00492 |
| HOMO SAPIENS | 5 | 589 | 3ggc | B | 51 | ASP | p.Asp52Gly | P00492 |
| HOMO SAPIENS | 0 | 589 | 3ggc | B | 57 | GLY | p.Gly58Arg | P00492 |
| HOMO SAPIENS | 6 | 589 | 3ggc | B | 69 | GLY | p.Gly70Glu | P00492 |
| HOMO SAPIENS | 1 | 589 | 3ggc | B | 73 | PHE | p.Phe74Leu | P00492 |
| HOMO SAPIENS | 0 | 589 | 3ggc | B | 77 | LEU | p.Leu78Val | P00492 |
| HOMO SAPIENS | 3 | 589 | 3ggc | B | 79 | ASP | p.Asp80Val | P00492 |
| HOMO SAPIENS | 0 | 589 | 3ggc | B | 103 | SER | p.Ser104Arg | P00492 |
| HOMO SAPIENS | 0 | 589 | 3ggc | B | 109 | SER | p.Ser110Leu | P00492 |
| HOMO SAPIENS | 1 | 589 | 3ggc | B | 129 | VAL | p.Val130Asp | P00492 |
| HOMO SAPIENS | 4 | 589 | 3ggc | B | 131 | ILE | p.Ile132Met | P00492 |
| HOMO SAPIENS | 7 | 589 | 3ggc | B | 142 | MET | p.Met143Lys | P00492 |
| HOMO SAPIENS | 0 | 589 | 3ggc | B | 160 | ALA | p.Ala161Ser | P00492 |
| HOMO SAPIENS | 6 | 589 | 3ggc | B | 167 | THR | p.Thr168Ile | P00492 |
| HOMO SAPIENS | 0 | 589 | 3ggc | B | 169 | ARG | p.Arg170Ter | P00492 |
| HOMO SAPIENS | 5 | 589 | 3ggc | B | 175 | PRO | p.Pro176Leu | P00492 |
| HOMO SAPIENS | 8 | 589 | 3ggc | B | 193 | ASP | p.Asp194Glu | P00492 |
| HOMO SAPIENS | 8 | 589 | 3ggc | B | 193 | ASP | p.Asp194Asn | P00492 |
| HOMO SAPIENS | 8 | 589 | 3ggc | B | 198 | PHE | p.Phe199Val | P00492 |
| HOMO SAPIENS | 8 | 589 | 3ggc | B | 200 | ASP | p.Asp201Gly | P00492 |
| HOMO SAPIENS | 0 | 4269 | 2pla | B | 174 | GLU | p.Glu174Lys | Q8N335 |
| HOMO SAPIENS | 1 | 4269 | 2pla | B | 220 | ARG | p.Arg220His | Q8N335 |
| HOMO SAPIENS | 4 | 4269 | 2pla | B | 280 | ALA | p.Ala280Val | Q8N335 |
| HOMO SAPIENS | 0 | 2712 | 1ii4 | E | 252 | TRP | p.Ser252Leu | P21802 |
| HOMO SAPIENS | 0 | 2712 | 1ii4 | E | 253 | PRO | p.Pro253Arg | P21802 |
| HOMO SAPIENS | 5 | 2712 | 1ii4 | E | 267 | SER | p.Ser267Pro | P21802 |
| HOMO SAPIENS | 0 | 2712 | 1ii4 | E | 278 | CYS | p.Cys278Phe | P21802 |
| HOMO SAPIENS | 0 | 2712 | 1ii4 | E | 289 | GLN | p.Gln289Pro | P21802 |
| HOMO SAPIENS | 0 | 2712 | 1ii4 | E | 290 | TRP | p.Trp290Cys | P21802 |
| HOMO SAPIENS | 0 | 2712 | 1ii4 | E | 290 | TRP | p.Trp290Arg | P21802 |
| HOMO SAPIENS | 0 | 2712 | 1ii4 | E | 292 | LYS | p.Lys292Glu | P21802 |
| HOMO SAPIENS | 0 | 2712 | 1ii4 | E | 315 | ALA | p.Ala315Ser | P21802 |
| HOMO SAPIENS | 0 | 2712 | 1ii4 | E | 321 | ASP | p.Asp321Ala | P21802 |
| HOMO SAPIENS | 1 | 2712 | 1ii4 | E | 328 | TYR | p.Tyr328Cys | P21802 |

| | | | | | | | | |
|---|---|---|---|---|---|---|---|---|
| HOMO SAPIENS | 9 | 2712 | 1ii4 | E | 340 | TYR | p.Tyr340His | P21802 |
| HOMO SAPIENS | 1 | 2712 | 1ii4 | E | 341 | THR | p.Thr341Pro | P21802 |
| HOMO SAPIENS | 0 | 2712 | 1ii4 | E | 342 | CYS | p.Cys342Arg | P21802 |
| HOMO SAPIENS | 0 | 2712 | 1ii4 | E | 342 | CYS | p.Cys342Trp | P21802 |
| HOMO SAPIENS | 0 | 2712 | 1ii4 | E | 344 | ALA | p.Ala344Gly | P21802 |
| HOMO SAPIENS | 0 | 2712 | 1ii4 | E | 347 | SER | p.Ser347Cys | P21802 |
| HOMO SAPIENS | 0 | 2712 | 1ii4 | E | 351 | SER | p.Ser351Cys | P21802 |
| HOMO SAPIENS | 1 | 2712 | 1ii4 | E | 354 | SER | p.Ser354Cys | P21802 |
| HOMO SAPIENS | 2 | 53 | 4pj1 | H | 5 | ASP | p.Asp29Gly | P10809 |
| HOMO SAPIENS | 0 | 53 | 4pj1 | H | 74 | VAL | p.Val98Ile | P10809 |
| HOMO SAPIENS | 0 | 29968 | 2fdb | M | 71 | LYS | p.Lys89Glu | P55075 |
| HOMO SAPIENS | 2 | 29968 | 2fdb | M | 98 | ARG | p.Arg116Gly | P55075 |
| HOMO SAPIENS | 2 | 29968 | 2fdb | M | 122 | GLY | p.Gly140Ser | P55075 |
| HOMO SAPIENS | 8 | 5061 | 5f85 | B | 60 | GLY | p.Gly106Ser | P00740 |
| HOMO SAPIENS | 0 | 35944 | 1h6h | A | 85 | THR | p.Thr85Asn | Q15080 |
| HOMO SAPIENS | 8 | 35944 | 1h6h | A | 105 | ARG | p.Arg105Gln | Q15080 |
| HOMO SAPIENS | 7 | 12016 | 1hj7 | A | 295 | ASN | p.Asn316Ser | P01130 |
| HOMO SAPIENS | 5 | 12016 | 1hj7 | A | 302 | GLY | p.Gly323Ser | P01130 |
| HOMO SAPIENS | 6 | 12016 | 1hj7 | A | 319 | CYS | p.Cys340Trp | P01130 |
| HOMO SAPIENS | 7 | 12016 | 1hj7 | A | 329 | ARG | p.Arg350Ter | P01130 |
| HOMO SAPIENS | 5 | 12016 | 1hj7 | A | 331 | CYS | p.Cys352Ter | P01130 |
| HOMO SAPIENS | 2 | 12016 | 1hj7 | A | 345 | GLN | p.Gln366Arg | P01130 |
| HOMO SAPIENS | 2 | 1481 | 2i07 | A | 239 | TYR | p.Tyr261Ter | P01024 |
| HOMO SAPIENS | -1 | 1481 | 2i07 | A | 292 | LEU | p.Pro314Leu | P01024 |
| HOMO SAPIENS | 2 | 1481 | 2i07 | A | 326 | ILE | p.Ile348Val | P01024 |
| HOMO SAPIENS | 2 | 1481 | 2i07 | A | 413 | GLU | p.Glu435Lys | P01024 |
| HOMO SAPIENS | 4 | 1481 | 2i07 | A | 447 | GLU | p.Glu469Asp | P01024 |
| HOMO SAPIENS | 2 | 1481 | 2i07 | A | 597 | VAL | p.Val619Met | P01024 |
| HOMO SAPIENS | 5 | 1481 | 2i07 | A | 603 | ILE | p.Ile625Phe | P01024 |
| HOMO SAPIENS | 0 | 1481 | 2i07 | A | 611 | LYS | p.Lys633Arg | P01024 |
| HOMO SAPIENS | 0 | 11123 | 3c5n | A | 315 | GLY | p.Gly368Trp | O00294 |
| HOMO SAPIENS | 8 | 11123 | 3c5n | A | 329 | PHE | p.Phe382Ser | O00294 |
| HOMO SAPIENS | 5 | 11123 | 3c5n | A | 429 | ARG | p.Arg482Trp | O00294 |
| HOMO SAPIENS | 8 | 11123 | 3c5n | A | 438 | PHE | p.Phe491Leu | O00294 |
| HOMO SAPIENS | 3 | 4398 | 3fx0 | B | 1288 | ALA | p.Ala288Gly | Q9Y6K9 |
| HOMO SAPIENS | 7 | 4398 | 3fx0 | B | 1315 | GLU | p.Glu315Ala | Q9Y6K9 |
| HOMO SAPIENS | 8 | 4398 | 3fx0 | B | 1319 | ARG | p.Arg319Gln | Q9Y6K9 |
| HOMO SAPIENS | 6 | 3989 | 3qd6 | S | 83 | CYS | p.Cys83Arg | P25942 |
| HOMO SAPIENS | 1 | 3989 | 3qd6 | S | 124 | SER | p.Ser124Leu | P25942 |
| HOMO SAPIENS | 2 | 24003 | 2m0v | A | 225 | GLU | p.Glu225Lys | O14745 |
| HOMO SAPIENS | 2 | 46 | 4aqd | A | 70 | ASP | p.Asp98Gly | P06276 |
| HOMO SAPIENS | 4 | 46 | 4aqd | A | 115 | GLY | p.Gly143Asp | P06276 |
| HOMO SAPIENS | 3 | 46 | 4aqd | A | 184 | ALA | p.Ala212Val | P06276 |
| HOMO SAPIENS | 0 | 46 | 4aqd | A | 225 | GLY | p.Gly253Ter | P06276 |
| HOMO SAPIENS | 2 | 46 | 4aqd | A | 243 | THR | p.Thr271Met | P06276 |
| HOMO SAPIENS | 2 | 46 | 4aqd | A | 255 | GLU | p.Glu283Asp | P06276 |
| HOMO SAPIENS | 5 | 46 | 4aqd | A | 267 | LYS | p.Lys295Arg | P06276 |
| HOMO SAPIENS | 3 | 46 | 4aqd | A | 271 | GLU | p.Glu299Ter | P06276 |
| HOMO SAPIENS | 2 | 46 | 4aqd | A | 307 | LEU | p.Leu335Pro | P06276 |
| HOMO SAPIENS | 1 | 46 | 4aqd | A | 330 | LEU | p.Leu358Ile | P06276 |
| HOMO SAPIENS | 0 | 46 | 4aqd | A | 497 | GLU | p.Glu525Val | P06276 |
| HOMO SAPIENS | 3 | 786 | 1ols | B | 51 | GLY | p.Gly101Asp | P21953 |
| HOMO SAPIENS | 2 | 786 | 1ols | B | 87 | ALA | p.Ala137Val | P21953 |
| HOMO SAPIENS | 0 | 786 | 1ols | B | 110 | ILE | p.Ile160Ser | P21953 |

| | | | | | | | | |
|---|---|---|---|---|---|---|---|---|
| HOMO SAPIENS | 4 | 786 | 1ols | B | 120 | ARG | p.Arg170Cys | P21953 |
| HOMO SAPIENS | 2 | 786 | 1ols | B | 126 | ASN | p.Asn176Tyr | P21953 |
| HOMO SAPIENS | 0 | 786 | 1ols | B | 133 | ARG | p.Arg183Gly | P21953 |
| HOMO SAPIENS | 0 | 786 | 1ols | B | 133 | ARG | p.Arg183Pro | P21953 |
| HOMO SAPIENS | 0 | 786 | 1ols | B | 135 | PRO | p.Pro185Leu | P21953 |
| HOMO SAPIENS | 3 | 786 | 1ols | B | 156 | HIS | p.His206Tyr | P21953 |
| HOMO SAPIENS | 0 | 786 | 1ols | B | 228 | GLY | p.Gly278Ser | P21953 |
| HOMO SAPIENS | 1 | 786 | 1ols | B | 235 | ARG | p.Arg285Ter | P21953 |
| HOMO SAPIENS | 6 | 786 | 1ols | B | 274 | ARG | p.Arg324Ter | P21953 |
| HOMO SAPIENS | 1 | 831 | 4zt1 | B | 48 | VAL | p.Val202Ile | P12830 |
| HOMO SAPIENS | 6 | 831 | 4zt1 | B | 70 | ARG | p.Arg224Cys | P12830 |
| HOMO SAPIENS | 5 | 831 | 4zt1 | B | 79 | HIS | p.His233Gln | P12830 |
| HOMO SAPIENS | 4 | 831 | 4zt1 | B | 120 | GLY | p.Gly274Ser | P12830 |
| HOMO SAPIENS | 7 | 831 | 4zt1 | B | 187 | TYR | p.Tyr341Ter | P12830 |
| HOMO SAPIENS | 0 | 8497 | 3sla | A | 287 | ASN | p.Asn287Ser | P35222 |
| HOMO SAPIENS | 7 | 23986 | 2ke1 | A | 301 | VAL | p.Val301Met | O43918 |
| HOMO SAPIENS | 0 | 4967 | 1rj7 | B | 334 | ARG | p.Arg334His | Q92838 |
| HOMO SAPIENS | 0 | 4967 | 1rj7 | B | 338 | THR | p.Thr338Met | Q92838 |
| HOMO SAPIENS | 3 | 4967 | 1rj7 | B | 349 | ALA | p.Ala349Thr | Q92838 |
| HOMO SAPIENS | 2 | 4967 | 1rj7 | B | 358 | GLN | p.Gln358Glu | Q92838 |
| HOMO SAPIENS | 1 | 4967 | 1rj7 | B | 375 | LYS | p.Lys375Glu | Q92838 |
| HOMO SAPIENS | 9 | 7623 | 1d4t | A | 32 | ARG | p.Arg32Thr | O60880 |
| HOMO SAPIENS | 6 | 7623 | 1d4t | A | 55 | ARG | p.Arg55Leu | O60880 |
| HOMO SAPIENS | 0 | 7623 | 1d4t | A | 68 | THR | p.Thr68Ile | O60880 |
| HOMO SAPIENS | 6 | 705 | 3idh | A | 59 | PRO | p.Pro59Ser | P35557 |
| HOMO SAPIENS | 5 | 705 | 3idh | A | 124 | ASP | p.Asp124Asn | P35557 |
| HOMO SAPIENS | 4 | 705 | 3idh | A | 131 | SER | p.Ser131Pro | P35557 |
| HOMO SAPIENS | 0 | 705 | 3idh | A | 147 | GLY | p.Gly147Asp | P35557 |
| HOMO SAPIENS | 5 | 705 | 3idh | A | 186 | ARG | p.Arg186Ter | P35557 |
| HOMO SAPIENS | 0 | 705 | 3idh | A | 202 | MET | p.Met202Thr | P35557 |
| HOMO SAPIENS | 2 | 705 | 3idh | A | 210 | MET | p.Met210Lys | P35557 |
| HOMO SAPIENS | 1 | 705 | 3idh | A | 215 | TYR | p.Tyr215Ter | P35557 |
| HOMO SAPIENS | 0 | 705 | 3idh | A | 226 | VAL | p.Val226Met | P35557 |
| HOMO SAPIENS | 9 | 705 | 3idh | A | 228 | THR | p.Thr228Met | P35557 |
| HOMO SAPIENS | 9 | 705 | 3idh | A | 256 | GLU | p.Glu256Lys | P35557 |
| HOMO SAPIENS | 2 | 705 | 3idh | A | 261 | GLY | p.Gly261Arg | P35557 |
| HOMO SAPIENS | 6 | 705 | 3idh | A | 279 | GLU | p.Glu279Gln | P35557 |
| HOMO SAPIENS | 7 | 705 | 3idh | A | 299 | GLY | p.Gly299Arg | P35557 |
| HOMO SAPIENS | 2 | 705 | 3idh | A | 318 | GLY | p.Gly318Trp | P35557 |
| HOMO SAPIENS | 4 | 705 | 3idh | A | 378 | ALA | p.Ala378Thr | P35557 |
| HOMO SAPIENS | 0 | 705 | 3idh | A | 387 | ALA | p.Ala387Glu | P35557 |
| HOMO SAPIENS | 5 | 705 | 3idh | A | 414 | LYS | p.Lys414Glu | P35557 |
| HOMO SAPIENS | 0 | 705 | 3idh | A | 455 | VAL | p.Val455Met | P35557 |
| HOMO SAPIENS | -1 | 705 | 3idh | A | 462 | MET | p.Met462Ile | P35557 |
| HOMO SAPIENS | 1 | 509 | 4nef | A | 22 | LEU | p.Leu22Val | P41181 |
| HOMO SAPIENS | 4 | 509 | 4nef | A | 57 | GLN | p.Gln57Pro | P41181 |
| HOMO SAPIENS | 8 | 509 | 4nef | A | 64 | GLY | p.Gly64Arg | P41181 |
| HOMO SAPIENS | 7 | 509 | 4nef | A | 68 | ASN | p.Asn68Ser | P41181 |
| HOMO SAPIENS | 1 | 509 | 4nef | A | 100 | GLY | p.Gly100Val | P41181 |
| HOMO SAPIENS | 0 | 509 | 4nef | A | 125 | THR | p.Thr125Met | P41181 |
| HOMO SAPIENS | 0 | 509 | 4nef | A | 126 | THR | p.Thr126Met | P41181 |
| HOMO SAPIENS | 4 | 509 | 4nef | A | 146 | PHE | p.Phe146Leu | P41181 |
| HOMO SAPIENS | 0 | 509 | 4nef | A | 175 | GLY | p.Gly175Arg | P41181 |
| HOMO SAPIENS | 5 | 509 | 4nef | A | 181 | CYS | p.Cys181Trp | P41181 |

| | | | | | | | | | |
|---|---|---|---|---|---|---|---|---|---|
| HOMO SAPIENS | 9 | 509 | 4nef | A | | 187 | ARG | p.Arg187Cys | P41181 |
| HOMO SAPIENS | 0 | 509 | 4nef | A | | 190 | ALA | p.Ala190Thr | P41181 |
| HOMO SAPIENS | 0 | 509 | 4nef | A | | 216 | SER | p.Ser216Pro | P41181 |
| HOMO SAPIENS | 4 | 2242 | 1z70 | X | | 1155 | SER | p.Ser155Pro | Q8NBK3 |
| HOMO SAPIENS | 3 | 2242 | 1z70 | X | | 1218 | CYS | p.Cys218Tyr | Q8NBK3 |
| HOMO SAPIENS | 0 | 2242 | 1z70 | X | | 1279 | ALA | p.Ala279Val | Q8NBK3 |
| HOMO SAPIENS | -1 | 2242 | 1z70 | X | | 1336 | OCS | p.Cys336Arg | Q8NBK3 |
| HOMO SAPIENS | 4 | 2242 | 1z70 | X | | 1345 | ARG | p.Arg345Cys | Q8NBK3 |
| HOMO SAPIENS | 2 | 2242 | 1z70 | X | | 1348 | ALA | p.Ala348Pro | Q8NBK3 |
| HOMO SAPIENS | 5 | 2242 | 1z70 | X | | 1349 | ARG | p.Arg349Gln | Q8NBK3 |
| HOMO SAPIENS | 5 | 2242 | 1z70 | X | | 1349 | ARG | p.Arg349Trp | Q8NBK3 |
| HOMO SAPIENS | 3 | 2242 | 1z70 | X | | 1359 | SER | p.Ser359Ter | Q8NBK3 |
| HOMO SAPIENS | 5 | 8269 | 5wrr | B | | 136 | ARG | p.Arg136Ter | Q96MK3 |
| HOMO SAPIENS | 1 | 8269 | 5wrr | B | | 332 | LYS | p.Asn332Lys | Q96MK3 |
| HOMO SAPIENS | 3 | 8269 | 5wrr | B | | 478 | ARG | p.Arg478Ter | Q96MK3 |
| HOMO SAPIENS | 5 | 8269 | 5wrr | B | | 504 | THR | p.Thr504Ile | Q96MK3 |
| HOMO SAPIENS | 5 | 1411 | 4h1s | B | | 221 | SER | p.Ser221Ter | P21589 |
| HOMO SAPIENS | 1 | 1411 | 4h1s | B | | 358 | CYS | p.Cys358Tyr | P21589 |
| HOMO SAPIENS | 2 | 23 | 5vvd | B | | 298 | GLU | p.Glu298Asp | P29474 |
| HOMO SAPIENS | 0 | 19 | 5jlh | E | | 88 | THR | p.Thr89Ile | P63261 |
| HOMO SAPIENS | 6 | 19 | 5jlh | E | | 117 | LYS | p.Lys118Met | P63261 |
| HOMO SAPIENS | 5 | 19 | 5jlh | E | | 263 | PRO | p.Pro264Leu | P63261 |
| HOMO SAPIENS | 1 | 19 | 5jlh | E | | 277 | THR | p.Thr278Ile | P63261 |
| HOMO SAPIENS | 6 | 19 | 5jlh | E | | 331 | PRO | p.Pro332Ala | P63261 |
| HOMO SAPIENS | 5 | 19 | 5jlh | E | | 369 | VAL | p.Val370Ala | P63261 |
| HOMO SAPIENS | 0 | 14306 | 4w4o | C | | 92 | ARG | p.Arg92Ter | P12314 |
| HOMO SAPIENS | 3 | 603 | 1nuh | A | | 4 | THR | p.Thr5Ile | P06744 |
| HOMO SAPIENS | 2 | 603 | 1nuh | A | | 19 | HIS | p.His20Pro | P06744 |
| HOMO SAPIENS | 8 | 603 | 1nuh | A | | 158 | GLY | p.Gly159Ser | P06744 |
| HOMO SAPIENS | 6 | 603 | 1nuh | A | | 223 | THR | p.Thr224Met | P06744 |
| HOMO SAPIENS | 0 | 603 | 1nuh | A | | 338 | LEU | p.Leu339Pro | P06744 |
| HOMO SAPIENS | 6 | 603 | 1nuh | A | | 342 | GLN | p.Gln343Arg | P06744 |
| HOMO SAPIENS | 6 | 603 | 1nuh | A | | 374 | THR | p.Thr375Arg | P06744 |
| HOMO SAPIENS | 4 | 603 | 1nuh | A | | 524 | ILE | p.Ile525Thr | P06744 |
| HOMO SAPIENS | 4 | 603 | 1nuh | A | | 538 | ASP | p.Asp539Asn | P06744 |
| HOMO SAPIENS | 1 | 3107 | 3kzj | A | | 1197 | VAL | p.Val1197Ala | P08603 |
| HOMO SAPIENS | 9 | 1748 | 2noh | A | | 46 | ARG | p.Arg46Gln | O15527 |
| HOMO SAPIENS | 2 | 11750 | 3s4y | A | | 33 | ARG | p.Arg33His | Q9H3S4 |
| HOMO SAPIENS | 2 | 11750 | 3s4y | A | | 40 | LEU | p.Leu40Pro | Q9H3S4 |
| HOMO SAPIENS | 4 | 11750 | 3s4y | A | | 142 | LEU | p.Leu142Phe | Q9H3S4 |
| HOMO SAPIENS | 5 | 11750 | 3s4y | A | | 219 | ASN | p.Asn219Ser | Q9H3S4 |
| HOMO SAPIENS | 1 | 2591 | 5vnj | A | | 382 | PHE | p.Phe382Leu | Q15436 |
| HOMO SAPIENS | 2 | 2591 | 5vnj | A | | 702 | MET | p.Met702Val | Q15436 |
| HOMO SAPIENS | 1 | 2426 | 5m8t | A | | 87 | ARG | p.Arg87Trp | P17643 |
| HOMO SAPIENS | 1 | 2426 | 5m8t | A | | 166 | SER | p.Ser166Ter | P17643 |
| HOMO SAPIENS | 0 | 2426 | 5m8t | A | | 356 | ARG | p.Arg356Gln | P17643 |
| HOMO SAPIENS | 2 | 2426 | 5m8t | A | | 374 | SER | p.Arg374Ter | P17643 |
| HOMO SAPIENS | 6 | 2426 | 5m8t | A | | 378 | ASN | p.Asn378Ser | P17643 |
| HOMO SAPIENS | 6 | 2426 | 5m8t | A | | 382 | LEU | p.Leu382Pro | P17643 |
| HOMO SAPIENS | 7 | 10097 | 4n6v | | 1 | 46 | GLN | p.Gln46Ter | P04080 |
| HOMO SAPIENS | 3 | 10097 | 4n6v | | 1 | 65 | VAL | p.Val65Ile | P04080 |
| HOMO SAPIENS | 8 | 10097 | 4n6v | | 1 | 68 | ARG | p.Arg68Ter | P04080 |
| HOMO SAPIENS | 2 | 859 | 4z9a | A | | 105 | GLN | p.Gln106Arg | P24666 |
| HOMO SAPIENS | 8 | 1420 | 5nan | F | | 95 | SER | p.Ser95Leu | Q96PD4 |

| | | | | | | | | |
|---|---|---|---|---|---|---|---|---|
| HOMO SAPIENS | 6 | 1420 | 5nan | F | 126 | GLU | p.Glu126Gly | Q96PD4 |
| HOMO SAPIENS | 8 | 1420 | 5nan | F | 155 | VAL | p.Val155Ile | Q96PD4 |
| HOMO SAPIENS | 6 | 1420 | 5nan | F | 157 | PRO | p.Pro157Ser | Q96PD4 |
| HOMO SAPIENS | -1 | 1420 | 5nan | F | 161 | HIS | p.His161Arg | Q96PD4 |
| HOMO SAPIENS | 3 | 288 | 5i96 | B | 140 | GLN | p.Arg140Gly | P48735 |
| HOMO SAPIENS | 3 | 288 | 5i96 | B | 140 | GLN | p.Arg140Gln | P48735 |
| HOMO SAPIENS | 2 | 288 | 5i96 | B | 261 | ARG | p.Arg261His | P48735 |
| HOMO SAPIENS | 2 | 288 | 5i96 | B | 435 | THR | p.Thr435Met | P48735 |
| HOMO SAPIENS | 9 | 120 | 1tup | B | 107 | TYR | p.Tyr107His | P04637 |
| HOMO SAPIENS | 8 | 120 | 1tup | B | 110 | ARG | p.Arg110Leu | P04637 |
| HOMO SAPIENS | 5 | 120 | 1tup | B | 120 | LYS | p.Lys120Glu | P04637 |
| HOMO SAPIENS | 4 | 120 | 1tup | B | 133 | MET | p.Met133Thr | P04637 |
| HOMO SAPIENS | 3 | 120 | 1tup | B | 134 | PHE | p.Phe134Cys | P04637 |
| HOMO SAPIENS | 8 | 120 | 1tup | B | 136 | GLN | p.Gln136His | P04637 |
| HOMO SAPIENS | 9 | 120 | 1tup | B | 138 | ALA | p.Ala138Pro | P04637 |
| HOMO SAPIENS | 9 | 120 | 1tup | B | 151 | PRO | p.Pro151Ser | P04637 |
| HOMO SAPIENS | 9 | 120 | 1tup | B | 152 | PRO | p.Pro152Leu | P04637 |
| HOMO SAPIENS | 9 | 120 | 1tup | B | 158 | ARG | p.Arg158Cys | P04637 |
| HOMO SAPIENS | 9 | 120 | 1tup | B | 158 | ARG | p.Arg158His | P04637 |
| HOMO SAPIENS | 9 | 120 | 1tup | B | 175 | ARG | p.Arg175Cys | P04637 |
| HOMO SAPIENS | 9 | 120 | 1tup | B | 175 | ARG | p.Arg175Leu | P04637 |
| HOMO SAPIENS | 8 | 120 | 1tup | B | 181 | ARG | p.Arg181Leu | P04637 |
| HOMO SAPIENS | 9 | 120 | 1tup | B | 189 | ALA | p.Ala189Val | P04637 |
| HOMO SAPIENS | 9 | 120 | 1tup | B | 196 | ARG | p.Arg196Gly | P04637 |
| HOMO SAPIENS | 9 | 120 | 1tup | B | 213 | ARG | p.Arg213Gln | P04637 |
| HOMO SAPIENS | 9 | 120 | 1tup | B | 213 | ARG | p.Arg213Ter | P04637 |
| HOMO SAPIENS | 9 | 120 | 1tup | B | 214 | HIS | p.His214Gln | P04637 |
| HOMO SAPIENS | 9 | 120 | 1tup | B | 220 | TYR | p.Tyr220Cys | P04637 |
| HOMO SAPIENS | 0 | 120 | 1tup | B | 234 | TYR | p.Tyr234Cys | P04637 |
| HOMO SAPIENS | 9 | 120 | 1tup | B | 237 | MET | p.Met237Ile | P04637 |
| HOMO SAPIENS | 0 | 120 | 1tup | B | 238 | CYS | p.Cys238Tyr | P04637 |
| HOMO SAPIENS | 9 | 120 | 1tup | B | 241 | SER | p.Ser241Phe | P04637 |
| HOMO SAPIENS | 9 | 120 | 1tup | B | 242 | CYS | p.Cys242Ser | P04637 |
| HOMO SAPIENS | 9 | 120 | 1tup | B | 245 | GLY | p.Gly245Cys | P04637 |
| HOMO SAPIENS | 9 | 120 | 1tup | B | 245 | GLY | p.Gly245Asp | P04637 |
| HOMO SAPIENS | 9 | 120 | 1tup | B | 248 | ARG | p.Arg248Leu | P04637 |
| HOMO SAPIENS | 9 | 120 | 1tup | B | 248 | ARG | p.Arg248Trp | P04637 |
| HOMO SAPIENS | 9 | 120 | 1tup | B | 251 | ILE | p.Ile251Ser | P04637 |
| HOMO SAPIENS | 9 | 120 | 1tup | B | 252 | LEU | p.Leu252Pro | P04637 |
| HOMO SAPIENS | 9 | 120 | 1tup | B | 258 | GLU | p.Glu258Lys | P04637 |
| HOMO SAPIENS | 8 | 120 | 1tup | B | 272 | VAL | p.Val272Leu | P04637 |
| HOMO SAPIENS | 9 | 120 | 1tup | B | 273 | ARG | p.Arg273Cys | P04637 |
| HOMO SAPIENS | 9 | 120 | 1tup | B | 273 | ARG | p.Arg273His | P04637 |
| HOMO SAPIENS | 9 | 120 | 1tup | B | 277 | CYS | p.Cys277Phe | P04637 |
| HOMO SAPIENS | 9 | 120 | 1tup | B | 280 | ARG | p.Arg280Ile | P04637 |
| HOMO SAPIENS | 9 | 120 | 1tup | B | 282 | ARG | p.Arg282Trp | P04637 |
| HOMO SAPIENS | 9 | 120 | 1tup | B | 283 | ARG | p.Arg283His | P04637 |
| HOMO SAPIENS | 9 | 120 | 1tup | B | 285 | GLU | p.Glu285Val | P04637 |
| HOMO SAPIENS | -1 | 120 | 1tup | B | 292 | LYS | p.Lys292Ile | P04637 |
| HOMO SAPIENS | -1 | 120 | 1tup | B | 298 | GLU | p.Glu298Lys | P04637 |
| HOMO SAPIENS | -1 | 2791 | 4ygb | A | 39 | VAL | p.Val39Ala | P49257 |
| HOMO SAPIENS | 2 | 106 | 4x6j | A | 32 | GLY | p.Gly146Arg | P43235 |
| HOMO SAPIENS | 0 | 106 | 4x6j | A | 76 | GLN | p.Gln190Ter | P43235 |
| HOMO SAPIENS | 7 | 106 | 4x6j | A | 198 | ARG | p.Arg312Ter | P43235 |

| | | | | | | | | |
|---|---|---|---|---|---|---|---|---|
| HOMO SAPIENS | 5 | 490 | 2hlo | B | 172 | LEU | p.Leu202Gln | P02675 |
| HOMO SAPIENS | 0 | 490 | 2hlo | B | 353 | LEU | p.Leu383Arg | P02675 |
| HOMO SAPIENS | 4 | 490 | 2hlo | B | 400 | GLY | p.Gly430Asp | P02675 |
| HOMO SAPIENS | 7 | 490 | 2hlo | B | 448 | ARG | p.Arg478Lys | P02675 |
| HOMO SAPIENS | 1 | 289 | 4p13 | B | 18 | GLU | p.Glu43Lys | P11310 |
| HOMO SAPIENS | 2 | 289 | 4p13 | B | 20 | GLN | p.Gln45Arg | P11310 |
| HOMO SAPIENS | 5 | 289 | 4p13 | B | 28 | ARG | p.Arg53Cys | P11310 |
| HOMO SAPIENS | 1 | 289 | 4p13 | B | 42 | TYR | p.Tyr67His | P11310 |
| HOMO SAPIENS | 1 | 289 | 4p13 | B | 53 | ILE | p.Ile78Thr | P11310 |
| HOMO SAPIENS | 1 | 289 | 4p13 | B | 59 | LEU | p.Leu84Phe | P11310 |
| HOMO SAPIENS | 2 | 289 | 4p13 | B | 96 | THR | p.Thr121Ile | P11310 |
| HOMO SAPIENS | 0 | 289 | 4p13 | B | 124 | MET | p.Met149Ile | P11310 |
| HOMO SAPIENS | 5 | 289 | 4p13 | B | 143 | ASP | p.Asp168Ala | P11310 |
| HOMO SAPIENS | 7 | 289 | 4p13 | B | 168 | THR | p.Thr193Ala | P11310 |
| HOMO SAPIENS | 1 | 289 | 4p13 | B | 170 | GLY | p.Gly195Arg | P11310 |
| HOMO SAPIENS | 2 | 289 | 4p13 | B | 181 | ARG | p.Arg206Cys | P11310 |
| HOMO SAPIENS | 2 | 289 | 4p13 | B | 181 | ARG | p.Arg206His | P11310 |
| HOMO SAPIENS | 8 | 289 | 4p13 | B | 203 | THR | p.Thr228Ile | P11310 |
| HOMO SAPIENS | 2 | 289 | 4p13 | B | 208 | ILE | p.Ile233Asn | P11310 |
| HOMO SAPIENS | 1 | 289 | 4p13 | B | 219 | CYS | p.Cys244Arg | P11310 |
| HOMO SAPIENS | 6 | 289 | 4p13 | B | 220 | SER | p.Ser245Leu | P11310 |
| HOMO SAPIENS | 5 | 289 | 4p13 | B | 228 | GLU | p.Glu253Lys | P11310 |
| HOMO SAPIENS | 2 | 289 | 4p13 | B | 241 | ASP | p.Asp266Gly | P11310 |
| HOMO SAPIENS | 1 | 289 | 4p13 | B | 242 | GLY | p.Gly267Arg | P11310 |
| HOMO SAPIENS | 7 | 289 | 4p13 | B | 256 | ARG | p.Arg281Thr | P11310 |
| HOMO SAPIENS | 4 | 289 | 4p13 | B | 269 | ARG | p.Arg294Thr | P11310 |
| HOMO SAPIENS | 4 | 289 | 4p13 | B | 304 | GLU | p.Lys329Glu | P11310 |
| HOMO SAPIENS | 1 | 289 | 4p13 | B | 324 | ARG | p.Arg349Ter | P11310 |
| HOMO SAPIENS | 3 | 289 | 4p13 | B | 326 | THR | p.Thr351Ile | P11310 |
| HOMO SAPIENS | 7 | 289 | 4p13 | B | 350 | ILE | p.Ile375Thr | P11310 |
| HOMO SAPIENS | 3 | 289 | 4p13 | B | 359 | GLU | p.Glu384Ter | P11310 |
| HOMO SAPIENS | 1 | 4563 | 3i2b | A | 16 | ARG | p.Arg16Cys | Q03393 |
| HOMO SAPIENS | 5 | 4563 | 3i2b | A | 25 | ARG | p.Arg25Gln | Q03393 |
| HOMO SAPIENS | 1 | 4563 | 3i2b | A | 47 | ASN | p.Asn47Asp | Q03393 |
| HOMO SAPIENS | 8 | 4563 | 3i2b | A | 52 | ASN | p.Asn52Ser | Q03393 |
| HOMO SAPIENS | 1 | 4563 | 3i2b | A | 56 | VAL | p.Val56Met | Q03393 |
| HOMO SAPIENS | 1 | 4563 | 3i2b | A | 87 | PRO | p.Pro87Ser | Q03393 |
| HOMO SAPIENS | 1 | 4563 | 3i2b | A | 96 | ASP | p.Asp96Asn | Q03393 |
| HOMO SAPIENS | 0 | 4563 | 3i2b | A | 99 | TYR | p.Tyr99Ter | Q03393 |
| HOMO SAPIENS | 2 | 4563 | 3i2b | A | 116 | ASP | p.Asp116Gly | Q03393 |
| HOMO SAPIENS | 2 | 41 | 3exx | A | 1 | GLY | p.Gly90Cys | P01308 |
| HOMO SAPIENS | 5 | 41 | 3exx | A | 3 | VAL | p.Val92Leu | P01308 |
| HOMO SAPIENS | 4 | 41 | 3exx | A | 7 | CYS | p.Cys96Ser | P01308 |
| HOMO SAPIENS | 9 | 41 | 3exx | A | 19 | TYR | p.Tyr108Cys | P01308 |
| HOMO SAPIENS | 9 | 131 | 4mjo | D | 118 | ASP | p.Asp119Asn | P09467 |
| HOMO SAPIENS | 0 | 131 | 4mjo | D | 163 | GLY | p.Gly164Ser | P09467 |
| HOMO SAPIENS | 0 | 131 | 4mjo | D | 176 | ALA | p.Ala177Asp | P09467 |
| HOMO SAPIENS | 5 | 131 | 4mjo | D | 193 | PHE | p.Phe194Ser | P09467 |
| HOMO SAPIENS | 4 | 131 | 4mjo | D | 217 | ARG | p.Arg218Lys | P09467 |
| HOMO SAPIENS | 4 | 131 | 4mjo | D | 223 | VAL | p.Val224Ile | P09467 |
| HOMO SAPIENS | 4 | 131 | 4mjo | D | 232 | PHE | p.Phe233Ile | P09467 |
| HOMO SAPIENS | 5 | 131 | 4mjo | D | 243 | ARG | p.Arg244Trp | P09467 |
| HOMO SAPIENS | 0 | 131 | 4mjo | D | 283 | PRO | p.Pro284Arg | P09467 |
| HOMO SAPIENS | 0 | 131 | 4mjo | D | 288 | MET | p.Met289Leu | P09467 |

| | | | | | | | | |
|---|---|---|---|---|---|---|---|---|
| HOMO SAPIENS | 0 | 131 | 4mjo | D | 323 | ASP | p.Asp324Asn | P09467 |
| HOMO SAPIENS | -1 | 13122 | 3hfc | A | 583 | ARG | p.Arg583Cys | P51787 |
| HOMO SAPIENS | 1 | 13122 | 3hfc | A | 587 | THR | p.Thr587Met | P51787 |
| HOMO SAPIENS | 0 | 13122 | 3hfc | A | 589 | GLY | p.Gly589Asp | P51787 |
| HOMO SAPIENS | 4 | 3002 | 3nbq | B | 94 | ARG | p.Arg94Cys | Q16831 |
| HOMO SAPIENS | 1 | 3616 | 3dbp | A | 97 | ILE | p.Ile286Asn | P11172 |
| HOMO SAPIENS | 0 | 3616 | 3dbp | A | 240 | GLY | p.Gly429Arg | P11172 |
| HOMO SAPIENS | 0 | 3616 | 3dbp | A | 257 | ILE | p.Ile446Val | P11172 |
| HOMO SAPIENS | 1 | 584 | 1drf | A | 79 | LEU | p.Leu80Phe | P00374 |
| HOMO SAPIENS | 3 | 1609 | 3qa2 | B | 49 | VAL | p.Val49Ile | P50053 |
| HOMO SAPIENS | 0 | 20560 | 5mug | A | 101 | HIS | p.His101Gln | P49638 |
| HOMO SAPIENS | 2 | 20560 | 5mug | A | 120 | ALA | p.Ala120Thr | P49638 |
| HOMO SAPIENS | 3 | 20560 | 5mug | A | 192 | ARG | p.Arg192His | P49638 |
| HOMO SAPIENS | 3 | 20560 | 5mug | A | 221 | ARG | p.Arg221Trp | P49638 |
| HOMO SAPIENS | 4 | 256 | 3vtv | A | 65 | THR | p.Glu50Lys | Q9GZQ8 |
| HOMO SAPIENS | 4 | 256 | 3vtv | A | 113 | VAL | p.Met98Lys | Q9GZQ8 |
| HOMO SAPIENS | 4 | 5397 | 1m61 | A | 192 | ARG | p.Arg192Trp | P43403 |
| HOMO SAPIENS | 5 | 39881 | 1aut | L | 107 | HIS | p.His149Pro | P04070 |
| HOMO SAPIENS | 5 | 4010 | 4uf8 | I | 33 | ALA | p.Ala33Val | Q13501 |
| HOMO SAPIENS | 3 | 313 | 5w49 | A | 98 | PRO | p.Pro98Leu | P23526 |
| HOMO SAPIENS | 2 | 313 | 5w49 | A | 143 | TYR | p.Tyr143Cys | P23526 |
| HOMO SAPIENS | -1 | 1643 | 2m8t | A | 102 | PRO | p.Pro102Leu | P04156 |
| HOMO SAPIENS | -1 | 1643 | 2m8t | A | 105 | PRO | p.Pro105Leu | P04156 |
| HOMO SAPIENS | -1 | 1643 | 2m8t | A | 117 | ALA | p.Ala117Val | P04156 |
| HOMO SAPIENS | 5 | 1643 | 2m8t | A | 129 | MET | p.Met129Val | P04156 |
| HOMO SAPIENS | 5 | 1643 | 2m8t | A | 131 | GLY | p.Gly131Val | P04156 |
| HOMO SAPIENS | 2 | 1643 | 2m8t | A | 142 | GLY | p.Gly142Ser | P04156 |
| HOMO SAPIENS | 3 | 1643 | 2m8t | A | 171 | ASN | p.Asn171Ser | P04156 |
| HOMO SAPIENS | 9 | 1643 | 2m8t | A | 178 | ASP | p.Asp178Asn | P04156 |
| HOMO SAPIENS | 9 | 1643 | 2m8t | A | 180 | VAL | p.Val180Ile | P04156 |
| HOMO SAPIENS | 9 | 1643 | 2m8t | A | 183 | THR | p.Thr183Ala | P04156 |
| HOMO SAPIENS | 9 | 1643 | 2m8t | A | 187 | HIS | p.His187Arg | P04156 |
| HOMO SAPIENS | 8 | 1643 | 2m8t | A | 198 | PHE | p.Phe198Ser | P04156 |
| HOMO SAPIENS | 9 | 1643 | 2m8t | A | 200 | GLU | p.Glu200Lys | P04156 |
| HOMO SAPIENS | 9 | 1643 | 2m8t | A | 208 | ARG | p.Arg208His | P04156 |
| HOMO SAPIENS | 3 | 1643 | 2m8t | A | 210 | VAL | p.Val210Ile | P04156 |
| HOMO SAPIENS | 6 | 1643 | 2m8t | A | 217 | GLN | p.Gln217Arg | P04156 |
| HOMO SAPIENS | 1 | 29 | 5kos | A | 151 | GLY | p.Gly217Arg | P00797 |
| HOMO SAPIENS | 1 | 29 | 5kos | A | 164 | ARG | p.Arg230Lys | P00797 |
| HOMO SAPIENS | 3 | 504 | 4zkv | B | 37 | ARG | p.Arg37Pro | P49773 |
| HOMO SAPIENS | 2 | 504 | 4zkv | B | 62 | GLN | p.Gln62Ter | P49773 |
| HOMO SAPIENS | 2 | 504 | 4zkv | B | 84 | CYS | p.Cys84Arg | P49773 |
| HOMO SAPIENS | 4 | 504 | 4zkv | B | 93 | GLY | p.Gly93Asp | P49773 |
| HOMO SAPIENS | 9 | 504 | 4zkv | B | 112 | HIS | p.His112Asn | P49773 |
| HOMO SAPIENS | 0 | 74 | 5vwo | C | 54 | ASN | p.Asn54Lys | P04181 |
| HOMO SAPIENS | 0 | 74 | 5vwo | C | 55 | TYR | p.Tyr55His | P04181 |
| HOMO SAPIENS | 2 | 74 | 5vwo | C | 90 | GLN | p.Gln90Glu | P04181 |
| HOMO SAPIENS | 6 | 74 | 5vwo | C | 91 | GLY | p.Gly91Glu | P04181 |
| HOMO SAPIENS | 2 | 74 | 5vwo | C | 93 | CYS | p.Cys93Phe | P04181 |
| HOMO SAPIENS | 1 | 74 | 5vwo | C | 154 | ARG | p.Arg154Leu | P04181 |
| HOMO SAPIENS | 0 | 74 | 5vwo | C | 180 | ARG | p.Arg180Thr | P04181 |
| HOMO SAPIENS | 0 | 74 | 5vwo | C | 199 | PRO | p.Pro199Gln | P04181 |
| HOMO SAPIENS | 4 | 74 | 5vwo | C | 209 | TYR | p.Tyr209Ter | P04181 |
| HOMO SAPIENS | 0 | 74 | 5vwo | C | 226 | ALA | p.Ala226Val | P04181 |

| | | | | | | | | |
|---|---|---|---|---|---|---|---|---|
| HOMO SAPIENS | 5 | 74 | 5vwo | C | 233 | GLN | p.Gln233Arg | P04181 |
| HOMO SAPIENS | 5 | 74 | 5vwo | C | 241 | PRO | p.Pro241Leu | P04181 |
| HOMO SAPIENS | 2 | 74 | 5vwo | C | 245 | TYR | p.Tyr245Cys | P04181 |
| HOMO SAPIENS | 6 | 74 | 5vwo | C | 250 | ARG | p.Arg250Pro | P04181 |
| HOMO SAPIENS | 6 | 74 | 5vwo | C | 250 | ARG | p.Arg250Ter | P04181 |
| HOMO SAPIENS | 8 | 74 | 5vwo | C | 267 | THR | p.Thr267Ile | P04181 |
| HOMO SAPIENS | 0 | 74 | 5vwo | C | 270 | ALA | p.Ala270Pro | P04181 |
| HOMO SAPIENS | 0 | 74 | 5vwo | C | 271 | ARG | p.Arg271Lys | P04181 |
| HOMO SAPIENS | 1 | 74 | 5vwo | C | 299 | TYR | p.Tyr299Ter | P04181 |
| HOMO SAPIENS | 0 | 74 | 5vwo | C | 318 | GLU | p.Glu318Lys | P04181 |
| HOMO SAPIENS | 1 | 74 | 5vwo | C | 319 | HIS | p.His319Tyr | P04181 |
| HOMO SAPIENS | 1 | 74 | 5vwo | C | 331 | ARG | p.Arg331Ter | P04181 |
| HOMO SAPIENS | 0 | 74 | 5vwo | C | 332 | VAL | p.Val332Met | P04181 |
| HOMO SAPIENS | 1 | 74 | 5vwo | C | 353 | GLY | p.Gly353Asp | P04181 |
| HOMO SAPIENS | 2 | 74 | 5vwo | C | 375 | GLY | p.Gly375Ala | P04181 |
| HOMO SAPIENS | 0 | 74 | 5vwo | C | 394 | CYS | p.Cys394Arg | P04181 |
| HOMO SAPIENS | 0 | 74 | 5vwo | C | 402 | LEU | p.Leu402Pro | P04181 |
| HOMO SAPIENS | 0 | 74 | 5vwo | C | 436 | ILE | p.Ile436Asn | P04181 |
| HOMO SAPIENS | 7 | 24356 | 4az0 | A | 104 | VAL | p.Val132Met | P10619 |
| HOMO SAPIENS | 4 | 24356 | 4az0 | A | 221 | TYR | p.Tyr249Asn | P10619 |
| HOMO SAPIENS | 3 | 245 | 1rsz | A | 24 | ARG | p.Arg24Ter | P00491 |
| HOMO SAPIENS | 3 | 245 | 1rsz | A | 51 | SER | p.Gly51Ser | P00491 |
| HOMO SAPIENS | 1 | 245 | 1rsz | A | 58 | ARG | p.Arg58Ter | P00491 |
| HOMO SAPIENS | 4 | 245 | 1rsz | A | 89 | GLU | p.Glu89Lys | P00491 |
| HOMO SAPIENS | 6 | 245 | 1rsz | A | 128 | ASP | p.Asp128Gly | P00491 |
| HOMO SAPIENS | 0 | 245 | 1rsz | A | 174 | ALA | p.Ala174Pro | P00491 |
| HOMO SAPIENS | 8 | 245 | 1rsz | A | 192 | TYR | p.Tyr192Cys | P00491 |
| HOMO SAPIENS | 8 | 245 | 1rsz | A | 234 | ARG | p.Arg234Pro | P00491 |
| HOMO SAPIENS | 0 | 37 | 5tby | E | 22 | GLU | p.Glu22Lys | P10916 |
| HOMO SAPIENS | 1 | 37 | 5tby | E | 86 | PHE | p.Phe86Ser | P10916 |
| HOMO SAPIENS | 9 | 71 | 5m9j | A | 12 | GLN | p.Gln36Leu | P03950 |
| HOMO SAPIENS | 3 | 71 | 5m9j | A | 17 | LYS | p.Lys41Glu | P03950 |
| HOMO SAPIENS | 3 | 71 | 5m9j | A | 17 | LYS | p.Lys41Ile | P03950 |
| HOMO SAPIENS | 4 | 71 | 5m9j | A | 28 | SER | p.Ser52Asn | P03950 |
| HOMO SAPIENS | 7 | 71 | 5m9j | A | 31 | ARG | p.Arg55Lys | P03950 |
| HOMO SAPIENS | 9 | 71 | 5m9j | A | 39 | CYS | p.Cys63Trp | P03950 |
| HOMO SAPIENS | 9 | 71 | 5m9j | A | 40 | LYS | p.Lys64Ile | P03950 |
| HOMO SAPIENS | 2 | 71 | 5m9j | A | 60 | GLU | p.Lys84Glu | P03950 |
| HOMO SAPIENS | 0 | 71 | 5m9j | A | 112 | PRO | p.Pro136Leu | P03950 |
| HOMO SAPIENS | 9 | 71 | 5m9j | A | 113 | VAL | p.Val137Ile | P03950 |
| HOMO SAPIENS | 3 | 326 | 5dqi | A | 209 | GLY | p.Gly209Val | Q9Y253 |
| HOMO SAPIENS | 1 | 326 | 5dqi | A | 242 | SER | p.Ser242Ter | Q9Y253 |
| HOMO SAPIENS | 1 | 326 | 5dqi | A | 303 | ARG | p.Arg303Ter | Q9Y253 |
| HOMO SAPIENS | 1 | 8203 | 3iu3 | K | 132 | VAL | p.Val153Ile | P01589 |
| HOMO SAPIENS | 0 | 4995 | 5ffo | H | 234 | THR | p.Thr263Ile | P01137 |
| HOMO SAPIENS | 5 | 38609 | 2ruk | A | 49 | ASP | p.Asp49His | P04637 |
| HOMO SAPIENS | 5 | 38609 | 2ruk | A | 58 | PRO | p.Pro58Arg | P04637 |
| HOMO SAPIENS | 0 | 1607 | 6pah | A | 117 | THR | p.Thr117Ile | P00439 |
| HOMO SAPIENS | 3 | 1607 | 6pah | A | 145 | ASP | p.Asp145Val | P00439 |
| HOMO SAPIENS | 5 | 1607 | 6pah | A | 158 | ARG | p.Arg158Gln | P00439 |
| HOMO SAPIENS | 5 | 1607 | 6pah | A | 158 | ARG | p.Arg158Trp | P00439 |
| HOMO SAPIENS | 5 | 1607 | 6pah | A | 169 | ARG | p.Arg169His | P00439 |
| HOMO SAPIENS | 5 | 1607 | 6pah | A | 169 | ARG | p.Arg169Cys | P00439 |
| HOMO SAPIENS | 3 | 1607 | 6pah | A | 176 | ARG | p.Arg176Leu | P00439 |

| | | | | | | | | |
|---|---|---|---|---|---|---|---|---|
| HOMO SAPIENS | 3 | 1607 | 6pah | A | 177 | VAL | p.Val177Leu | P00439 |
| HOMO SAPIENS | 1 | 1607 | 6pah | A | 194 | LEU | p.Leu194Pro | P00439 |
| HOMO SAPIENS | 1 | 1607 | 6pah | A | 204 | TYR | p.Tyr204Cys | P00439 |
| HOMO SAPIENS | 1 | 1607 | 6pah | A | 204 | TYR | p.Tyr204Ter | P00439 |
| HOMO SAPIENS | 2 | 1607 | 6pah | A | 230 | VAL | p.Val230Ile | P00439 |
| HOMO SAPIENS | 4 | 1607 | 6pah | A | 232 | GLN | p.Gln232Ter | P00439 |
| HOMO SAPIENS | 4 | 1607 | 6pah | A | 241 | ARG | p.Arg241Cys | P00439 |
| HOMO SAPIENS | 4 | 1607 | 6pah | A | 241 | ARG | p.Arg241His | P00439 |
| HOMO SAPIENS | 5 | 1607 | 6pah | A | 243 | ARG | p.Arg243Gln | P00439 |
| HOMO SAPIENS | 5 | 1607 | 6pah | A | 243 | ARG | p.Arg243Ter | P00439 |
| HOMO SAPIENS | 7 | 1607 | 6pah | A | 249 | LEU | p.Leu249Phe | P00439 |
| HOMO SAPIENS | 5 | 1607 | 6pah | A | 252 | ARG | p.Arg252Gly | P00439 |
| HOMO SAPIENS | 5 | 1607 | 6pah | A | 261 | ARG | p.Arg261Ter | P00439 |
| HOMO SAPIENS | 4 | 1607 | 6pah | A | 267 | GLN | p.Gln267Leu | P00439 |
| HOMO SAPIENS | 1 | 1607 | 6pah | A | 272 | GLY | p.Gly272Ter | P00439 |
| HOMO SAPIENS | 2 | 1607 | 6pah | A | 277 | TYR | p.Tyr277Asp | P00439 |
| HOMO SAPIENS | 7 | 1607 | 6pah | A | 280 | GLU | p.Glu280Lys | P00439 |
| HOMO SAPIENS | 8 | 1607 | 6pah | A | 281 | PRO | p.Pro281Leu | P00439 |
| HOMO SAPIENS | 0 | 1607 | 6pah | A | 299 | PHE | p.Phe299Cys | P00439 |
| HOMO SAPIENS | 5 | 1607 | 6pah | A | 300 | ALA | p.Ala300Ser | P00439 |
| HOMO SAPIENS | 5 | 1607 | 6pah | A | 309 | ALA | p.Ala309Asp | P00439 |
| HOMO SAPIENS | 4 | 1607 | 6pah | A | 320 | LYS | p.Lys320Asn | P00439 |
| HOMO SAPIENS | 0 | 1607 | 6pah | A | 345 | ALA | p.Ala345Ser | P00439 |
| HOMO SAPIENS | 0 | 1607 | 6pah | A | 348 | LEU | p.Leu348Val | P00439 |
| HOMO SAPIENS | 6 | 1607 | 6pah | A | 356 | TYR | p.Tyr356Ter | P00439 |
| HOMO SAPIENS | 2 | 1607 | 6pah | A | 380 | THR | p.Thr380Met | P00439 |
| HOMO SAPIENS | 0 | 1607 | 6pah | A | 386 | TYR | p.Tyr386Cys | P00439 |
| HOMO SAPIENS | 5 | 1607 | 6pah | A | 390 | GLU | p.Glu390Gly | P00439 |
| HOMO SAPIENS | 5 | 1607 | 6pah | A | 403 | ALA | p.Ala403Val | P00439 |
| HOMO SAPIENS | 4 | 1607 | 6pah | A | 408 | ARG | p.Arg408Trp | P00439 |
| HOMO SAPIENS | 3 | 1607 | 6pah | A | 413 | ARG | p.Arg413Pro | P00439 |
| HOMO SAPIENS | 4 | 1607 | 6pah | A | 414 | TYR | p.Tyr414Cys | P00439 |
| HOMO SAPIENS | 4 | 1607 | 6pah | A | 414 | TYR | p.Tyr414His | P00439 |
| HOMO SAPIENS | 4 | 1607 | 6pah | A | 415 | ASP | p.Asp415Asn | P00439 |
| HOMO SAPIENS | -1 | 676 | 5dc5 | A | 7 | PRO | p.Pro7Leu | Q9BY41 |
| HOMO SAPIENS | 0 | 676 | 5dc5 | A | 311 | THR | p.Thr311Met | Q9BY41 |
| HOMO SAPIENS | 1 | 124 | 5ftk | E | 95 | ARG | p.Arg95Gly | P55072 |
| HOMO SAPIENS | 1 | 124 | 5ftk | E | 155 | ARG | p.Arg155Cys | P55072 |
| HOMO SAPIENS | 1 | 124 | 5ftk | E | 155 | ARG | p.Arg155His | P55072 |
| HOMO SAPIENS | 1 | 124 | 5ftk | E | 159 | ARG | p.Arg159His | P55072 |
| HOMO SAPIENS | 1 | 124 | 5ftk | E | 159 | ARG | p.Arg159Gly | P55072 |
| HOMO SAPIENS | 0 | 124 | 5ftk | E | 191 | ARG | p.Arg191Gln | P55072 |
| HOMO SAPIENS | 0 | 124 | 5ftk | E | 232 | ALA | p.Ala232Glu | P55072 |
| HOMO SAPIENS | 5 | 5442 | 1fgg | B | 223 | GLY | p.Gly223Ser | O94766 |
| HOMO SAPIENS | 0 | 5442 | 1fgg | B | 277 | ARG | p.Arg277Gln | O94766 |
| HOMO SAPIENS | 0 | 9072 | 2gys | B | 225 | GLU | p.Glu249Gln | P32927 |
| HOMO SAPIENS | 0 | 9072 | 2gys | B | 247 | SER | p.Ser271Ter | P32927 |
| HOMO SAPIENS | 0 | 9072 | 2gys | B | 294 | ASP | p.Asp312Asn | P32927 |
| HOMO SAPIENS | 0 | 1484 | 4zrj | A | 62 | PHE | p.Phe62Ser | P35240 |
| HOMO SAPIENS | 3 | 1484 | 4zrj | A | 205 | MET | p.Met205Val | P35240 |
| HOMO SAPIENS | 3 | 1484 | 4zrj | A | 301 | ILE | p.Ile301Met | P35240 |
| HOMO SAPIENS | 4 | 40923 | 2enq | A | 68 | ILE | p.Ile391Met | P42336 |
| HOMO SAPIENS | 1 | 6179 | 5ofv | A | 125 | THR | p.Thr125Met | O43175 |
| HOMO SAPIENS | 0 | 6179 | 5ofv | A | 135 | ARG | p.Arg135Trp | O43175 |

| | | | | | | | | |
|---|---|---|---|---|---|---|---|---|
| HOMO SAPIENS | 1 | 6179 | 5ofv | A | 163 | ARG | p.Arg163Gln | O43175 |
| HOMO SAPIENS | 9 | 6179 | 5ofv | A | 261 | VAL | p.Val261Met | O43175 |
| HOMO SAPIENS | 7 | 34118 | 2i5o | A | 24 | MET | p.Met647Leu | Q9Y253 |
| HOMO SAPIENS | 8 | 3234 | 5c84 | A | 311 | LYS | p.Lys311Asn | P98170 |
| HOMO SAPIENS | 3 | 3234 | 5c84 | A | 321 | ALA | p.Ala321Gly | P98170 |
| HOMO SAPIENS | -1 | 15129 | 1zy8 | K | 1 | GLY | p.Gly54Ser | O00330 |
| HOMO SAPIENS | -1 | 15129 | 1zy8 | K | 48 | THR | p.Thr101Ala | O00330 |
| HOMO SAPIENS | -1 | 15129 | 1zy8 | K | 105 | GLU | p.Glu158Asp | O00330 |
| HOMO SAPIENS | 8 | 15129 | 1zy8 | K | 136 | ARG | p.Arg189His | O00330 |
| HOMO SAPIENS | -1 | 15129 | 1zy8 | K | 195 | GLN | p.Gln248Ter | O00330 |
| HOMO SAPIENS | 8 | 202 | 4bzb | A | 123 | HIS | p.His123Pro | Q9Y3Z3 |
| HOMO SAPIENS | 4 | 202 | 4bzb | A | 145 | ARG | p.Arg145Ter | Q9Y3Z3 |
| HOMO SAPIENS | 6 | 202 | 4bzb | A | 164 | ARG | p.Arg164Ter | Q9Y3Z3 |
| HOMO SAPIENS | 3 | 202 | 4bzb | A | 201 | ILE | p.Ile201Asn | Q9Y3Z3 |
| HOMO SAPIENS | 0 | 202 | 4bzb | A | 209 | GLY | p.Gly209Ser | Q9Y3Z3 |
| HOMO SAPIENS | 0 | 202 | 4bzb | A | 254 | MET | p.Met254Val | Q9Y3Z3 |
| HOMO SAPIENS | 5 | 202 | 4bzb | A | 531 | ARG | p.Arg531Ser | Q9Y3Z3 |
| HOMO SAPIENS | 6 | 202 | 4bzb | A | 548 | GLN | p.Gln548Ter | Q9Y3Z3 |
| HOMO SAPIENS | 3 | 30947 | 5lgd | A | 90 | PRO | p.Pro90Ser | P16671 |
| HOMO SAPIENS | 6 | 30947 | 5lgd | A | 254 | PHE | p.Phe254Leu | P16671 |
| HOMO SAPIENS | 0 | 30947 | 5lgd | A | 413 | ILE | p.Ile413Leu | P16671 |
| HOMO SAPIENS | 2 | 3906 | 5jtc | A | 735 | ARG | p.Arg735Trp | Q12797 |
| HOMO SAPIENS | 1 | 45 | 5uyx | C | 146 | GLU | p.Glu146Val | P48643 |
| HOMO SAPIENS | 1 | 45 | 5uyx | C | 147 | HIS | p.His147Arg | P48643 |
| HOMO SAPIENS | 0 | 45 | 5uyx | C | 362 | ILE | p.Ile362Met | P48643 |
| HOMO SAPIENS | 8 | 39596 | 1mhn | A | 95 | GLY | p.Gly95Arg | Q16637 |
| HOMO SAPIENS | 4 | 39596 | 1mhn | A | 111 | ALA | p.Ala111Gly | Q16637 |
| HOMO SAPIENS | 9 | 39596 | 1mhn | A | 116 | ILE | p.Ile116Phe | Q16637 |
| HOMO SAPIENS | 1 | 39596 | 1mhn | A | 136 | GLN | p.Gln136Glu | Q16637 |
| HOMO SAPIENS | 1 | 6408 | 5xjy | A | 85 | PRO | p.Pro85Leu | O95477 |
| HOMO SAPIENS | 0 | 6408 | 5xjy | A | 219 | ARG | p.Arg219Lys | O95477 |
| HOMO SAPIENS | 0 | 6408 | 5xjy | A | 230 | ARG | p.Arg230Cys | O95477 |
| HOMO SAPIENS | 1 | 6408 | 5xjy | A | 281 | MET | p.Met281Thr | O95477 |
| HOMO SAPIENS | -1 | 6408 | 5xjy | A | 343 | ALA | p.Ala343Gly | O95477 |
| HOMO SAPIENS | 1 | 6408 | 5xjy | A | 399 | VAL | p.Val399Ala | O95477 |
| HOMO SAPIENS | 1 | 6408 | 5xjy | A | 496 | ARG | p.Arg496Trp | O95477 |
| HOMO SAPIENS | 1 | 6408 | 5xjy | A | 590 | TRP | p.Trp590Ser | O95477 |
| HOMO SAPIENS | 1 | 6408 | 5xjy | A | 638 | ARG | p.Arg638Gln | O95477 |
| HOMO SAPIENS | 4 | 6408 | 5xjy | A | 697 | ALA | p.Ala697Thr | O95477 |
| HOMO SAPIENS | 1 | 6408 | 5xjy | A | 771 | VAL | p.Val771Met | O95477 |
| HOMO SAPIENS | 1 | 6408 | 5xjy | A | 774 | THR | p.Thr774Pro | O95477 |
| HOMO SAPIENS | 1 | 6408 | 5xjy | A | 776 | LYS | p.Lys776Asn | O95477 |
| HOMO SAPIENS | 0 | 6408 | 5xjy | A | 825 | VAL | p.Val825Ile | O95477 |
| HOMO SAPIENS | -1 | 6408 | 5xjy | A | 868 | GLU | p.Glu868Lys | O95477 |
| HOMO SAPIENS | -1 | 6408 | 5xjy | A | 883 | ILE | p.Ile883Met | O95477 |
| HOMO SAPIENS | 0 | 6408 | 5xjy | A | 935 | ASN | p.Asn935Ser | O95477 |
| HOMO SAPIENS | 0 | 6408 | 5xjy | A | 935 | ASN | p.Asn935His | O95477 |
| HOMO SAPIENS | 0 | 6408 | 5xjy | A | 937 | ALA | p.Ala937Val | O95477 |
| HOMO SAPIENS | 0 | 6408 | 5xjy | A | 1018 | ASP | p.Asp1018Gly | O95477 |
| HOMO SAPIENS | 0 | 6408 | 5xjy | A | 1096 | VAL | p.Val1096Ile | O95477 |
| HOMO SAPIENS | 0 | 6408 | 5xjy | A | 1099 | ASP | p.Asp1099Tyr | O95477 |
| HOMO SAPIENS | 0 | 6408 | 5xjy | A | 1172 | GLU | p.Glu1172Asp | O95477 |
| HOMO SAPIENS | 0 | 6408 | 5xjy | A | 1181 | SER | p.Ser1181Phe | O95477 |
| HOMO SAPIENS | 0 | 6408 | 5xjy | A | 1182 | ALA | p.Ala1182Thr | O95477 |

| | | | | | | | | |
|---|---|---|---|---|---|---|---|---|
| HOMO SAPIENS | -1 | 6408 | 5xjy | A | 1289 | ASP | p.Asp1289Asn | O95477 |
| HOMO SAPIENS | 0 | 6408 | 5xjy | A | 1399 | THR | p.Thr1399Ser | O95477 |
| HOMO SAPIENS | 1 | 6408 | 5xjy | A | 1408 | LEU | p.Leu1408Phe | O95477 |
| HOMO SAPIENS | 0 | 6408 | 5xjy | A | 1587 | LYS | p.Lys1587Arg | O95477 |
| HOMO SAPIENS | 1 | 6408 | 5xjy | A | 1674 | VAL | p.Val1674Ile | O95477 |
| HOMO SAPIENS | 1 | 6408 | 5xjy | A | 1680 | ARG | p.Arg1680Trp | O95477 |
| HOMO SAPIENS | 1 | 6408 | 5xjy | A | 1680 | ARG | p.Arg1680Gln | O95477 |
| HOMO SAPIENS | 0 | 6408 | 5xjy | A | 1800 | ASN | p.Asn1800His | O95477 |
| HOMO SAPIENS | -1 | 6408 | 5xjy | A | 1925 | ARG | p.Arg1925Gln | O95477 |
| HOMO SAPIENS | 0 | 6408 | 5xjy | A | 2009 | PHE | p.Phe2009Ser | O95477 |
| HOMO SAPIENS | -1 | 6408 | 5xjy | A | 2243 | ASP | p.Asp2243Glu | O95477 |
| HOMO SAPIENS | 4 | 23760 | 2mut | A | 231 | LEU | p.Phe231Leu | P07992 |
| HOMO SAPIENS | 6 | 1460 | 2hre | B | 96 | ARG | p.Arg96Gln | P22830 |
| HOMO SAPIENS | 8 | 1460 | 2hre | B | 129 | GLY | p.Gly129Arg | P22830 |
| HOMO SAPIENS | 9 | 1460 | 2hre | B | 172 | GLU | p.Glu172Lys | P22830 |
| HOMO SAPIENS | 4 | 1460 | 2hre | B | 285 | GLN | p.Gln285Arg | P22830 |
| HOMO SAPIENS | 5 | 1460 | 2hre | B | 362 | VAL | p.Val362Gly | P22830 |
| HOMO SAPIENS | 6 | 1460 | 2hre | B | 411 | CYS | p.Cys411Gly | P22830 |
| HOMO SAPIENS | 6 | 1460 | 2hre | B | 417 | PHE | p.Phe417Ser | P22830 |
| HOMO SAPIENS | 0 | 2210 | 5bug | A | 19 | ASP | p.Asp19Asn | P60484 |
| HOMO SAPIENS | 0 | 2210 | 5bug | A | 35 | MET | p.Met35Arg | P60484 |
| HOMO SAPIENS | 1 | 2210 | 5bug | A | 61 | HIS | p.His61Asp | P60484 |
| HOMO SAPIENS | 0 | 2210 | 5bug | A | 68 | TYR | p.Tyr68His | P60484 |
| HOMO SAPIENS | 8 | 2210 | 5bug | A | 70 | LEU | p.Leu70Pro | P60484 |
| HOMO SAPIENS | 6 | 2210 | 5bug | A | 93 | HIS | p.His93Arg | P60484 |
| HOMO SAPIENS | 1 | 2210 | 5bug | A | 112 | LEU | p.Leu112Pro | P60484 |
| HOMO SAPIENS | 0 | 2210 | 5bug | A | 121 | ALA | p.Ala121Gly | P60484 |
| HOMO SAPIENS | 3 | 2210 | 5bug | A | 123 | HIS | p.His123Arg | P60484 |
| HOMO SAPIENS | 7 | 2210 | 5bug | A | 124 | CYS | p.Cys124Arg | P60484 |
| HOMO SAPIENS | 9 | 2210 | 5bug | A | 129 | GLY | p.Gly129Glu | P60484 |
| HOMO SAPIENS | 9 | 2210 | 5bug | A | 130 | ARG | p.Arg130Gln | P60484 |
| HOMO SAPIENS | 9 | 2210 | 5bug | A | 130 | ARG | p.Arg130Gly | P60484 |
| HOMO SAPIENS | 0 | 2210 | 5bug | A | 132 | GLY | p.Gly132Val | P60484 |
| HOMO SAPIENS | 0 | 2210 | 5bug | A | 170 | SER | p.Ser170Arg | P60484 |
| HOMO SAPIENS | 1 | 2210 | 5bug | A | 173 | ARG | p.Arg173His | P60484 |
| HOMO SAPIENS | 0 | 2210 | 5bug | A | 217 | VAL | p.Val217Ile | P60484 |
| HOMO SAPIENS | 0 | 2210 | 5bug | A | 241 | PHE | p.Phe241Ser | P60484 |
| HOMO SAPIENS | 0 | 2210 | 5bug | A | 250 | CYS | p.Cys250Ter | P60484 |
| HOMO SAPIENS | 1 | 2210 | 5bug | A | 252 | ASP | p.Asp252Gly | P60484 |
| HOMO SAPIENS | 3 | 123 | 2ibu | A | 93 | ASN | p.Asn93Ser | P24752 |
| HOMO SAPIENS | 0 | 123 | 2ibu | A | 145 | GLN | p.Gln145Glu | P24752 |
| HOMO SAPIENS | 0 | 123 | 2ibu | A | 152 | GLY | p.Gly152Ala | P24752 |
| HOMO SAPIENS | 2 | 123 | 2ibu | A | 158 | ASN | p.Asn158Asp | P24752 |
| HOMO SAPIENS | 2 | 123 | 2ibu | A | 158 | ASN | p.Asn158Ser | P24752 |
| HOMO SAPIENS | 5 | 123 | 2ibu | A | 183 | GLY | p.Gly183Arg | P24752 |
| HOMO SAPIENS | 3 | 123 | 2ibu | A | 272 | GLN | p.Gln272Ter | P24752 |
| HOMO SAPIENS | 3 | 123 | 2ibu | A | 312 | ILE | p.Ile312Thr | P24752 |
| HOMO SAPIENS | 1 | 123 | 2ibu | A | 333 | ALA | p.Ala333Pro | P24752 |
| HOMO SAPIENS | 5 | 123 | 2ibu | A | 379 | GLY | p.Gly379Val | P24752 |
| HOMO SAPIENS | 0 | 123 | 2ibu | A | 380 | ALA | p.Ala380Thr | P24752 |
| HOMO SAPIENS | 2 | 369 | 5nrw | A | 1175 | TYR | p.Tyr1175Cys | Q92793 |
| HOMO SAPIENS | 2 | 830 | 2vc2 | A | 116 | LEU | p.Leu147Val | P08514 |
| HOMO SAPIENS | 0 | 830 | 2vc2 | A | 183 | LEU | p.Leu214Pro | P08514 |
| HOMO SAPIENS | 4 | 830 | 2vc2 | A | 242 | GLY | p.Gly273Asp | P08514 |

| | | | | | | | | |
|---|---|---|---|---|---|---|---|---|
| HOMO SAPIENS | 6 | 830 | 2vc2 | A | 324 | GLU | p.Glu355Lys | P08514 |
| HOMO SAPIENS | -1 | 10353 | 5ejc | E | 989 | ALA | p.Ala989Ser | Q92574 |
| HOMO SAPIENS | 0 | 15448 | 3lgg | A | 21 | GLY | p.Gly47Arg | Q9NZK5 |
| HOMO SAPIENS | 0 | 15448 | 3lgg | A | 21 | GLY | p.Gly47Val | Q9NZK5 |
| HOMO SAPIENS | 9 | 15448 | 3lgg | A | 86 | HIS | p.His112Gln | Q9NZK5 |
| HOMO SAPIENS | 0 | 15448 | 3lgg | A | 143 | ARG | p.Arg169Gln | Q9NZK5 |
| HOMO SAPIENS | 3 | 15448 | 3lgg | A | 221 | ALA | p.Ala247Val | Q9NZK5 |
| HOMO SAPIENS | 0 | 15448 | 3lgg | A | 225 | PRO | p.Pro251Leu | Q9NZK5 |
| HOMO SAPIENS | 0 | 15448 | 3lgg | A | 283 | MET | p.Met309Ile | Q9NZK5 |
| HOMO SAPIENS | 2 | 15448 | 3lgg | A | 334 | THR | p.Thr360Ala | Q9NZK5 |
| HOMO SAPIENS | 7 | 15448 | 3lgg | A | 357 | GLY | p.Gly383Ser | Q9NZK5 |
| HOMO SAPIENS | 0 | 15448 | 3lgg | A | 427 | TYR | p.Tyr453Cys | Q9NZK5 |
| HOMO SAPIENS | 0 | 872 | 2y3i | A | 87 | LEU | p.Leu88Pro | P00558 |
| HOMO SAPIENS | 1 | 872 | 2y3i | A | 163 | ASP | p.Asp164Val | P00558 |
| HOMO SAPIENS | 4 | 872 | 2y3i | A | 205 | ARG | p.Arg206Pro | P00558 |
| HOMO SAPIENS | 4 | 872 | 2y3i | A | 267 | ASP | p.Asp268Asn | P00558 |
| HOMO SAPIENS | 8 | 872 | 2y3i | A | 284 | ASP | p.Asp285Val | P00558 |
| HOMO SAPIENS | 4 | 8 | 5jco | B | 188 | ILE | p.Ile188Leu | Q71U36 |
| HOMO SAPIENS | 7 | 8 | 5jco | B | 263 | PRO | p.Pro263Thr | Q71U36 |
| HOMO SAPIENS | 6 | 8 | 5jco | B | 264 | ARG | p.Arg264Cys | Q71U36 |
| HOMO SAPIENS | 7 | 8 | 5jco | B | 402 | ARG | p.Arg402His | Q71U36 |
| HOMO SAPIENS | 7 | 8 | 5jco | B | 419 | SER | p.Ser419Leu | Q71U36 |
| HOMO SAPIENS | -1 | 880 | 4nwf | A | 2 | THR | p.Thr2Ile | Q06124 |
| HOMO SAPIENS | 2 | 880 | 4nwf | A | 58 | ASN | p.Asn58Ser | Q06124 |
| HOMO SAPIENS | 4 | 880 | 4nwf | A | 61 | ASP | p.Asp61Gly | Q06124 |
| HOMO SAPIENS | 2 | 880 | 4nwf | A | 62 | TYR | p.Tyr62Asp | Q06124 |
| HOMO SAPIENS | 3 | 880 | 4nwf | A | 63 | TYR | p.Tyr63Cys | Q06124 |
| HOMO SAPIENS | 2 | 880 | 4nwf | A | 69 | GLU | p.Glu69Gln | Q06124 |
| HOMO SAPIENS | 2 | 880 | 4nwf | A | 72 | ALA | p.Ala72Gly | Q06124 |
| HOMO SAPIENS | 2 | 880 | 4nwf | A | 72 | ALA | p.Ala72Ser | Q06124 |
| HOMO SAPIENS | 3 | 880 | 4nwf | A | 73 | THR | p.Thr73Ile | Q06124 |
| HOMO SAPIENS | 5 | 880 | 4nwf | A | 76 | GLU | p.Glu76Ala | Q06124 |
| HOMO SAPIENS | 5 | 880 | 4nwf | A | 76 | GLU | p.Glu76Lys | Q06124 |
| HOMO SAPIENS | 3 | 880 | 4nwf | A | 79 | GLN | p.Gln79Arg | Q06124 |
| HOMO SAPIENS | 5 | 880 | 4nwf | A | 152 | ARG | p.Arg152His | Q06124 |
| HOMO SAPIENS | 2 | 880 | 4nwf | A | 268 | GLY | p.Gly268Cys | Q06124 |
| HOMO SAPIENS | 9 | 880 | 4nwf | A | 276 | LYS | p.Lys276Arg | Q06124 |
| HOMO SAPIENS | 9 | 880 | 4nwf | A | 279 | TYR | p.Tyr279Cys | Q06124 |
| HOMO SAPIENS | 5 | 880 | 4nwf | A | 285 | PHE | p.Phe285Ser | Q06124 |
| HOMO SAPIENS | 0 | 880 | 4nwf | A | 308 | ASP | p.Asn308Ser | Q06124 |
| HOMO SAPIENS | 0 | 880 | 4nwf | A | 308 | ASP | p.Asn308Asp | Q06124 |
| HOMO SAPIENS | 5 | 880 | 4nwf | A | 392 | ALA | p.Ala392Thr | Q06124 |
| HOMO SAPIENS | 1 | 880 | 4nwf | A | 411 | THR | p.Thr415Met | Q06124 |
| HOMO SAPIENS | 0 | 880 | 4nwf | A | 461 | ALA | p.Ala465Thr | Q06124 |
| HOMO SAPIENS | 0 | 880 | 4nwf | A | 464 | GLY | p.Gly468Ala | Q06124 |
| HOMO SAPIENS | 1 | 880 | 4nwf | A | 491 | PRO | p.Pro495Ser | Q06124 |
| HOMO SAPIENS | 1 | 880 | 4nwf | A | 502 | SER | p.Ser506Thr | Q06124 |
| HOMO SAPIENS | 8 | 880 | 4nwf | A | 510 | GLN | p.Gln514Pro | Q06124 |
| HOMO SAPIENS | 4 | 1872 | 3wf1 | D | 109 | ARG | p.Arg109Trp | P16278 |
| HOMO SAPIENS | 0 | 1872 | 3wf1 | D | 521 | ARG | p.Arg521Cys | P16278 |
| HOMO SAPIENS | 5 | 910 | 5xte | A | 45 | SER | p.Ser45Phe | O14949 |
| HOMO SAPIENS | 2 | 9017 | 5f0k | A | 382 | LYS | p.Lys382Arg | Q96QK1 |
| HOMO SAPIENS | 1 | 710 | 4lsj | A | 559 | ILE | p.Ile559Asn | P04150 |
| HOMO SAPIENS | 5 | 710 | 4lsj | A | 571 | VAL | p.Val571Ala | P04150 |

| | | | | | | | | |
|---|---|---|---|---|---|---|---|---|
| HOMO SAPIENS | 0 | 710 | 4lsj | A | 641 | ASP | p.Asp641Val | P04150 |
| HOMO SAPIENS | 1 | 710 | 4lsj | A | 678 | ASP | p.Asp678Glu | P04150 |
| HOMO SAPIENS | 2 | 710 | 4lsj | A | 679 | GLY | p.Gly679Ser | P04150 |
| HOMO SAPIENS | 1 | 710 | 4lsj | A | 737 | PHE | p.Phe737Leu | P04150 |
| HOMO SAPIENS | 1 | 710 | 4lsj | A | 747 | ILE | p.Ile747Met | P04150 |
| HOMO SAPIENS | 1 | 710 | 4lsj | A | 753 | LEU | p.Leu753Phe | P04150 |
| HOMO SAPIENS | 1 | 710 | 4lsj | A | 773 | LEU | p.Leu773Pro | P04150 |
| HOMO SAPIENS | 1 | 2145 | 1av1 | B | 71 | PHE | p.Phe95Tyr | P02647 |
| HOMO SAPIENS | 1 | 2145 | 1av1 | B | 128 | GLU | p.Glu152Lys | P02647 |
| HOMO SAPIENS | 1 | 2145 | 1av1 | B | 136 | GLU | p.Glu160Lys | P02647 |
| HOMO SAPIENS | 2 | 2145 | 1av1 | B | 143 | PRO | p.Pro167Arg | P02647 |
| HOMO SAPIENS | 6 | 2145 | 1av1 | B | 164 | ALA | p.Ala188Ser | P02647 |
| HOMO SAPIENS | 7 | 2145 | 1av1 | B | 173 | ARG | p.Arg197Cys | P02647 |
| HOMO SAPIENS | 2 | 2145 | 1av1 | B | 198 | GLU | p.Glu222Lys | P02647 |
| HOMO SAPIENS | -1 | 3432 | 2beu | A | 1 | SER | p.Ser46Ter | P12694 |
| HOMO SAPIENS | 5 | 3432 | 2beu | A | 79 | ARG | p.Arg124Trp | P12694 |
| HOMO SAPIENS | 7 | 3432 | 2beu | A | 114 | ARG | p.Arg159Gln | P12694 |
| HOMO SAPIENS | 6 | 3432 | 2beu | A | 166 | THR | p.Thr211Met | P12694 |
| HOMO SAPIENS | 4 | 3432 | 2beu | A | 175 | ALA | p.Ala220Val | P12694 |
| HOMO SAPIENS | 5 | 3432 | 2beu | A | 204 | GLY | p.Gly249Ser | P12694 |
| HOMO SAPIENS | 4 | 3432 | 2beu | A | 209 | ALA | p.Ala254Asp | P12694 |
| HOMO SAPIENS | 6 | 3432 | 2beu | A | 220 | ARG | p.Arg265Trp | P12694 |
| HOMO SAPIENS | 6 | 3432 | 2beu | A | 234 | TYR | p.Tyr279Cys | P12694 |
| HOMO SAPIENS | 5 | 3432 | 2beu | A | 240 | ALA | p.Ala285Pro | P12694 |
| HOMO SAPIENS | 4 | 3432 | 2beu | A | 245 | GLY | p.Gly290Arg | P12694 |
| HOMO SAPIENS | 5 | 3432 | 2beu | A | 269 | ARG | p.Arg314Ter | P12694 |
| HOMO SAPIENS | 3 | 3432 | 2beu | A | 301 | ARG | p.Arg346Cys | P12694 |
| HOMO SAPIENS | 5 | 3432 | 2beu | A | 359 | ASN | p.Asn404Ser | P12694 |
| HOMO SAPIENS | 4 | 3432 | 2beu | A | 393 | TYR | p.Tyr438Asn | P12694 |
| HOMO SAPIENS | 8 | 4541 | 1rr8 | C | 533 | ASP | p.Asp533Gly | P11387 |
| HOMO SAPIENS | 2 | 3247 | 1tzn | d | 1105 | GLY | p.Gly105Asp | P58335 |
| HOMO SAPIENS | 3 | 3247 | 1tzn | d | 1111 | ARG | p.Arg111Cys | P58335 |
| HOMO SAPIENS | 2 | 3247 | 1tzn | d | 1189 | ILE | p.Ile189Thr | P58335 |
| HOMO SAPIENS | 9 | 467 | 1suv | C | 58 | ASP | p.Asp77Asn | P02787 |
| HOMO SAPIENS | 2 | 467 | 1suv | C | 277 | ASP | p.Asp296Gly | P02787 |
| HOMO SAPIENS | 9 | 6485 | 1xwd | B | 51 | CYS | p.Cys69Gly | P01225 |
| HOMO SAPIENS | 1 | 694 | 1wuu | A | 28 | PRO | p.Pro28Thr | P51570 |
| HOMO SAPIENS | 0 | 694 | 1wuu | A | 32 | VAL | p.Val32Met | P51570 |
| HOMO SAPIENS | 0 | 694 | 1wuu | A | 198 | ALA | p.Ala198Val | P51570 |
| HOMO SAPIENS | 0 | 143 | 4nkx | A | 93 | PHE | p.Phe93Cys | P05093 |
| HOMO SAPIENS | 3 | 143 | 4nkx | A | 96 | ARG | p.Arg96Trp | P05093 |
| HOMO SAPIENS | 3 | 143 | 4nkx | A | 96 | ARG | p.Arg96Gln | P05093 |
| HOMO SAPIENS | 0 | 143 | 4nkx | A | 106 | SER | p.Ser106Pro | P05093 |
| HOMO SAPIENS | 1 | 143 | 4nkx | A | 114 | PHE | p.Phe114Val | P05093 |
| HOMO SAPIENS | 1 | 143 | 4nkx | A | 116 | ASP | p.Asp116Val | P05093 |
| HOMO SAPIENS | 3 | 143 | 4nkx | A | 125 | ARG | p.Arg125Gln | P05093 |
| HOMO SAPIENS | 0 | 143 | 4nkx | A | 201 | TYR | p.Tyr201Asp | P05093 |
| HOMO SAPIENS | 0 | 143 | 4nkx | A | 239 | ARG | p.Arg239Ter | P05093 |
| HOMO SAPIENS | 0 | 143 | 4nkx | A | 329 | TYR | p.Tyr329Asp | P05093 |
| HOMO SAPIENS | 1 | 143 | 4nkx | A | 342 | PRO | p.Pro342Thr | P05093 |
| HOMO SAPIENS | 2 | 143 | 4nkx | A | 347 | ARG | p.Arg347His | P05093 |
| HOMO SAPIENS | 2 | 143 | 4nkx | A | 347 | ARG | p.Arg347Cys | P05093 |
| HOMO SAPIENS | 1 | 143 | 4nkx | A | 358 | ARG | p.Arg358Gln | P05093 |
| HOMO SAPIENS | 0 | 143 | 4nkx | A | 362 | ARG | p.Arg362Cys | P05093 |

| | | | | | | | | |
|---|---|---|---|---|---|---|---|---|
| HOMO SAPIENS | 1 | 143 | 4nkx | A | 406 | TRP | p.Trp406Arg | P05093 |
| HOMO SAPIENS | 0 | 143 | 4nkx | A | 428 | PRO | p.Pro428Leu | P05093 |
| HOMO SAPIENS | 6 | 1441 | 1uuh | A | 46 | ARG | p.Arg46Gly | P16070 |
| HOMO SAPIENS | 3 | 82 | 4wlv | D | 37 | GLY | p.Gly37Arg | P40926 |
| HOMO SAPIENS | 0 | 82 | 4wlv | D | 133 | PRO | p.Pro133Leu | P40926 |
| HOMO SAPIENS | 2 | 392 | 5xtd | A | 56 | SER | p.Ser56Pro | P49821 |
| HOMO SAPIENS | 2 | 392 | 5xtd | A | 59 | ARG | p.Arg59Ter | P49821 |
| HOMO SAPIENS | 2 | 392 | 5xtd | A | 74 | ASP | p.Asp74Asn | P49821 |
| HOMO SAPIENS | 4 | 392 | 5xtd | A | 116 | ASN | p.Asn116Lys | P49821 |
| HOMO SAPIENS | 4 | 392 | 5xtd | A | 117 | ALA | p.Ala117Thr | P49821 |
| HOMO SAPIENS | 5 | 392 | 5xtd | A | 122 | PRO | p.Pro122Leu | P49821 |
| HOMO SAPIENS | 3 | 392 | 5xtd | A | 160 | GLY | p.Gly160Glu | P49821 |
| HOMO SAPIENS | 4 | 392 | 5xtd | A | 214 | GLU | p.Glu214Lys | P49821 |
| HOMO SAPIENS | 2 | 392 | 5xtd | A | 357 | MET | p.Met357Thr | P49821 |
| HOMO SAPIENS | 4 | 392 | 5xtd | A | 423 | THR | p.Thr423Met | P49821 |
| HOMO SAPIENS | 0 | 22593 | 3pdf | A | 8 | TYR | p.Tyr32Ter | P53634 |
| HOMO SAPIENS | 0 | 22593 | 3pdf | A | 15 | TRP | p.Trp39Ser | P53634 |
| HOMO SAPIENS | 0 | 22593 | 3pdf | A | 103 | HIS | p.His127Pro | P53634 |
| HOMO SAPIENS | 9 | 22593 | 3pdf | A | 228 | GLN | p.Gln252Leu | P53634 |
| HOMO SAPIENS | 3 | 22593 | 3pdf | A | 262 | GLN | p.Gln286Arg | P53634 |
| HOMO SAPIENS | 4 | 22593 | 3pdf | A | 277 | GLY | p.Gly301Ser | P53634 |
| HOMO SAPIENS | 0 | 22593 | 3pdf | A | 323 | TYR | p.Tyr347Cys | P53634 |
| HOMO SAPIENS | 5 | 3461 | 4c2a | A | 1306 | GLN | p.Arg1306Trp | P04275 |
| HOMO SAPIENS | 5 | 3461 | 4c2a | A | 1374 | ARG | p.Arg1374Cys | P04275 |
| HOMO SAPIENS | 5 | 3461 | 4c2a | A | 1379 | ARG | p.Arg1379Cys | P04275 |
| HOMO SAPIENS | 4 | 3461 | 4c2a | A | 1380 | ILE | p.Ile1380Val | P04275 |
| HOMO SAPIENS | 0 | 3461 | 4c2a | A | 1381 | ALA | p.Thr1381Ala | P04275 |
| HOMO SAPIENS | 5 | 3461 | 4c2a | A | 1399 | ARG | p.Arg1399His | P04275 |
| HOMO SAPIENS | 5 | 3461 | 4c2a | A | 1435 | ASN | p.Asn1435Ser | P04275 |
| HOMO SAPIENS | 0 | 4533 | 5uj7 | C | 174 | TYR | p.Tyr174Cys | O43929 |
| HOMO SAPIENS | 0 | 4533 | 5uj7 | C | 202 | LEU | p.Leu202Val | O43929 |
| HOMO SAPIENS | 0 | 1374 | 3j07 | O | 39 | PRO | p.Pro39Leu | P02511 |
| HOMO SAPIENS | 0 | 1374 | 3j07 | O | 107 | ARG | p.Arg107His | P02511 |
| HOMO SAPIENS | 0 | 1374 | 3j07 | O | 157 | ARG | p.Arg157His | P02511 |
| HOMO SAPIENS | -1 | 942 | 1s9i | B | 57 | PHE | p.Phe57Cys | P36507 |
| HOMO SAPIENS | -1 | 942 | 1s9i | B | 57 | PHE | p.Phe57Val | P36507 |
| HOMO SAPIENS | 2 | 942 | 1s9i | B | 128 | PRO | p.Pro128Gln | P36507 |
| HOMO SAPIENS | 1 | 942 | 1s9i | B | 134 | TYR | p.Tyr134His | P36507 |
| HOMO SAPIENS | 1 | 942 | 1s9i | B | 188 | HIS | p.His188Gln | P36507 |
| HOMO SAPIENS | 0 | 942 | 1s9i | B | 262 | VAL | p.Val262Phe | P36507 |
| HOMO SAPIENS | 0 | 942 | 1s9i | B | 263 | GLY | p.Gly263Arg | P36507 |
| HOMO SAPIENS | -1 | 942 | 1s9i | B | 298 | PRO | p.Pro298Leu | P36507 |
| HOMO SAPIENS | -1 | 942 | 1s9i | B | 305 | VAL | p.Val305Ile | P36507 |
| HOMO SAPIENS | 0 | 3245 | 1fbv | A | 367 | GLN | p.Gln367Pro | P22681 |
| HOMO SAPIENS | 0 | 3245 | 1fbv | A | 371 | TYR | p.Tyr371His | P22681 |
| HOMO SAPIENS | 1 | 3245 | 1fbv | A | 382 | LYS | p.Lys382Glu | P22681 |
| HOMO SAPIENS | 2 | 3245 | 1fbv | A | 390 | ASP | p.Asp390Tyr | P22681 |
| HOMO SAPIENS | 9 | 3245 | 1fbv | A | 396 | CYS | p.Cys396Arg | P22681 |
| HOMO SAPIENS | 7 | 3245 | 1fbv | A | 420 | ARG | p.Arg420Gln | P22681 |
| HOMO SAPIENS | 1 | 3245 | 1fbv | A | 429 | ILE | p.Ile429Met | P22681 |
| HOMO SAPIENS | 0 | 2604 | 2pbd | P | 111 | LEU | p.Leu112Met | P07737 |
| HOMO SAPIENS | 3 | 2604 | 2pbd | P | 116 | GLU | p.Glu117Gly | P07737 |
| HOMO SAPIENS | 4 | 2013 | 3b7w | A | 472 | ARG | p.Arg472Gly | Q08AH3 |
| HOMO SAPIENS | 0 | 24346 | 3sh5 | A | 85 | ARG | p.Arg4281His | P98160 |

| | | | | | | | | |
|---|---|---|---|---|---|---|---|---|
| HOMO SAPIENS | 0 | 24346 | 3sh5 | A | 122 | ARG | p.Arg4318Trp | P98160 |
| HOMO SAPIENS | 0 | 24346 | 3sh5 | A | 135 | SER | p.Ser4331Asn | P98160 |
| HOMO SAPIENS | 0 | 24346 | 3sh5 | A | 136 | VAL | p.Val4332Ile | P98160 |
| HOMO SAPIENS | 0 | 707 | 4p2g | A | 26 | MET | p.Met26Ile | Q99497 |
| HOMO SAPIENS | 5 | 707 | 4p2g | A | 39 | ALA | p.Ala39Ser | Q99497 |
| HOMO SAPIENS | 5 | 707 | 4p2g | A | 64 | GLU | p.Glu64Asp | Q99497 |
| HOMO SAPIENS | 4 | 707 | 4p2g | A | 149 | ASP | p.Asp149Ala | Q99497 |
| HOMO SAPIENS | 6 | 707 | 4p2g | A | 163 | GLU | p.Glu163Lys | Q99497 |
| HOMO SAPIENS | 6 | 707 | 4p2g | A | 166 | LEU | p.Leu166Pro | Q99497 |
| HOMO SAPIENS | 3 | 5121 | 4xss | E | 31 | GLY | p.Gly58Arg | P06213 |
| HOMO SAPIENS | 2 | 5121 | 4xss | E | 119 | ILE | p.Ile146Met | P06213 |
| HOMO SAPIENS | 8 | 5121 | 4xss | E | 209 | HIS | p.His236Arg | P06213 |
| HOMO SAPIENS | 2 | 5121 | 4xss | E | 234 | ASP | p.Asp261Glu | P06213 |
| HOMO SAPIENS | 1 | 5121 | 4xss | E | 276 | GLN | p.Gln303His | P06213 |
| HOMO SAPIENS | 1 | 27790 | 5erp | B | 140 | THR | p.Thr275Met | Q02487 |
| HOMO SAPIENS | 0 | 27790 | 5erp | B | 150 | ILE | p.Ile285Thr | Q02487 |
| HOMO SAPIENS | 1 | 27790 | 5erp | B | 205 | THR | p.Thr340Ala | Q02487 |
| HOMO SAPIENS | 1 | 27790 | 5erp | B | 223 | THR | p.Thr358Ile | Q02487 |
| HOMO SAPIENS | 0 | 27790 | 5erp | B | 301 | GLY | p.Gly436Ala | Q02487 |
| HOMO SAPIENS | -1 | 27790 | 5erp | B | 315 | ARG | p.Arg450Ser | Q02487 |
| HOMO SAPIENS | 0 | 27790 | 5erp | B | 369 | SER | p.Ser504Gly | Q02487 |
| HOMO SAPIENS | 0 | 27790 | 5erp | B | 385 | ILE | p.Ile520Thr | Q02487 |
| HOMO SAPIENS | 0 | 27790 | 5erp | B | 461 | ALA | p.Ala596Val | Q02487 |
| HOMO SAPIENS | 0 | 27790 | 5erp | B | 503 | GLN | p.Gln638His | Q02487 |
| HOMO SAPIENS | 5 | 6599 | 4nsc | A | 129 | ARG | p.Arg129Leu | Q9BPX6 |
| HOMO SAPIENS | -1 | 6599 | 4nsc | A | 183 | GLN | p.Gln183Ter | Q9BPX6 |
| HOMO SAPIENS | 0 | 20420 | 3ghm | A | 398 | ARG | p.Arg398His | Q76LX8 |
| HOMO SAPIENS | 0 | 20420 | 3ghm | A | 448 | GLN | p.Gln448Glu | Q76LX8 |
| HOMO SAPIENS | 1 | 20420 | 3ghm | A | 449 | GLN | p.Gln449Ter | Q76LX8 |
| HOMO SAPIENS | 0 | 20420 | 3ghm | A | 456 | GLN | p.Gln456His | Q76LX8 |
| HOMO SAPIENS | 1 | 20420 | 3ghm | A | 528 | ARG | p.Arg528Gly | Q76LX8 |
| HOMO SAPIENS | 0 | 20420 | 3ghm | A | 618 | PRO | p.Pro618Ala | Q76LX8 |
| HOMO SAPIENS | 0 | 20420 | 3ghm | A | 625 | ARG | p.Arg625His | Q76LX8 |
| HOMO SAPIENS | 0 | 275 | 4rmz | A | 293 | GLN | p.Gln293Ter | Q9NWZ3 |
| HOMO SAPIENS | 0 | 275 | 4rmz | A | 428 | ALA | p.Ala428Thr | Q9NWZ3 |
| HOMO SAPIENS | 2 | 10458 | 3puf | B | 138 | LEU | p.Leu138Phe | Q5TBB1 |
| HOMO SAPIENS | 6 | 10458 | 3puf | B | 177 | ALA | p.Ala177Thr | Q5TBB1 |
| HOMO SAPIENS | -1 | 10458 | 3puf | B | 185 | VAL | p.Val185Gly | Q5TBB1 |
| HOMO SAPIENS | 0 | 26648 | 2n90 | A | 36 | ARG | p.Arg444Gln | P04629 |
| HOMO SAPIENS | 2 | 977 | 1dnw | D | 166 | ALA | p.Ala332Val | P05164 |
| HOMO SAPIENS | 4 | 977 | 1dnw | D | 333 | ARG | p.Arg499Cys | P05164 |
| HOMO SAPIENS | 1 | 977 | 1dnw | D | 403 | ARG | p.Arg569Trp | P05164 |
| HOMO SAPIENS | 0 | 25704 | 3ajb | A | 82 | GLN | p.Gln82Arg | P56589 |
| HOMO SAPIENS | 3 | 255 | 3t1p | A | 38 | TYR | p.Tyr62Ter | P01009 |
| HOMO SAPIENS | 3 | 255 | 3t1p | A | 39 | ARG | p.Arg63Cys | P01009 |
| HOMO SAPIENS | 0 | 255 | 3t1p | A | 101 | ARG | p.Arg125His | P01009 |
| HOMO SAPIENS | 0 | 255 | 3t1p | A | 212 | GLN | p.Gln236Arg | P01009 |
| HOMO SAPIENS | 0 | 255 | 3t1p | A | 213 | VAL | p.Val237Ala | P01009 |
| HOMO SAPIENS | 1 | 255 | 3t1p | A | 269 | HIS | p.His293Gln | P01009 |
| HOMO SAPIENS | 8 | 255 | 3t1p | A | 341 | ASP | p.Asp365Asn | P01009 |
| HOMO SAPIENS | 9 | 255 | 3t1p | A | 342 | GLU | p.Glu366Lys | P01009 |
| HOMO SAPIENS | 0 | 255 | 3t1p | A | 369 | PRO | p.Pro393Ser | P01009 |
| HOMO SAPIENS | 0 | 255 | 3t1p | A | 376 | GLU | p.Glu400Asp | P01009 |
| HOMO SAPIENS | -1 | 5175 | 3zxi | A | 35 | ALA | p.Ala35Asp | Q9Y2Z4 |

| | | | | | | | | | |
|---|---|---|---|---|---|---|---|---|---|
| HOMO SAPIENS | 0 | 5175 | 3zxi | A | | 52 | PHE | p.Phe52Leu | Q9Y2Z4 |
| HOMO SAPIENS | -1 | 5175 | 3zxi | A | | 68 | GLY | p.Gly68Ser | Q9Y2Z4 |
| HOMO SAPIENS | 0 | 5175 | 3zxi | A | | 179 | LYS | p.Lys179Gln | Q9Y2Z4 |
| HOMO SAPIENS | 0 | 5175 | 3zxi | A | | 191 | GLY | p.Gly191Val | Q9Y2Z4 |
| HOMO SAPIENS | 0 | 5175 | 3zxi | A | | 251 | ILE | p.Ile251Val | Q9Y2Z4 |
| HOMO SAPIENS | 2 | 5175 | 3zxi | A | | 295 | ASP | p.Asp295Asn | Q9Y2Z4 |
| HOMO SAPIENS | 0 | 5175 | 3zxi | A | | 311 | ASP | p.Asp311Glu | Q9Y2Z4 |
| HOMO SAPIENS | 0 | 5175 | 3zxi | A | | 360 | ARG | p.Arg360Ter | Q9Y2Z4 |
| HOMO SAPIENS | 1 | 3016 | 3exg | | 3 | 4 | ASP | p.Asp33His | P08559 |
| HOMO SAPIENS | 1 | 3016 | 3exg | | 3 | 176 | PHE | p.Phe205Leu | P08559 |
| HOMO SAPIENS | 1 | 3016 | 3exg | | 3 | 214 | TYR | p.Tyr243Asn | P08559 |
| HOMO SAPIENS | 1 | 3016 | 3exg | | 3 | 229 | ASP | p.Asp258Ala | P08559 |
| HOMO SAPIENS | 0 | 3016 | 3exg | | 3 | 233 | VAL | p.Val262Ile | P08559 |
| HOMO SAPIENS | 1 | 3016 | 3exg | | 3 | 234 | ARG | p.Arg263Gly | P08559 |
| HOMO SAPIENS | 0 | 3016 | 3exg | | 3 | 253 | MET | p.Met282Leu | P08559 |
| HOMO SAPIENS | 7 | 3016 | 3exg | | 3 | 259 | ARG | p.Arg288His | P08559 |
| HOMO SAPIENS | -1 | 3016 | 3exg | | 3 | 273 | ARG | p.Arg302Cys | P08559 |
| HOMO SAPIENS | 2 | 3016 | 3exg | | 3 | 286 | ASP | p.Asp315Asn | P08559 |
| HOMO SAPIENS | 1 | 3016 | 3exg | | 3 | 349 | ARG | p.Arg378His | P08559 |
| HOMO SAPIENS | 5 | 6729 | 4a0l | E | | 572 | ARG | p.Arg572Cys | Q13620 |
| HOMO SAPIENS | 0 | 1067 | 4kyo | A | | 9 | THR | p.Thr9Asn | P21549 |
| HOMO SAPIENS | 0 | 1067 | 4kyo | A | | 11 | PRO | p.Pro11Leu | P21549 |
| HOMO SAPIENS | 0 | 1067 | 4kyo | A | | 22 | ASN | p.Asn22Ser | P21549 |
| HOMO SAPIENS | 4 | 1067 | 4kyo | A | | 25 | LEU | p.Leu25Arg | P21549 |
| HOMO SAPIENS | 7 | 1067 | 4kyo | A | | 36 | ARG | p.Arg36His | P21549 |
| HOMO SAPIENS | 3 | 1067 | 4kyo | A | | 41 | GLY | p.Gly41Arg | P21549 |
| HOMO SAPIENS | 0 | 1067 | 4kyo | A | | 42 | GLY | p.Gly42Glu | P21549 |
| HOMO SAPIENS | 3 | 1067 | 4kyo | A | | 56 | ILE | p.Ile56Asn | P21549 |
| HOMO SAPIENS | 3 | 1067 | 4kyo | A | | 59 | GLU | p.Glu59Lys | P21549 |
| HOMO SAPIENS | 3 | 1067 | 4kyo | A | | 66 | TYR | p.Tyr66Ter | P21549 |
| HOMO SAPIENS | 3 | 1067 | 4kyo | A | | 111 | ARG | p.Arg111Leu | P21549 |
| HOMO SAPIENS | 7 | 1067 | 4kyo | A | | 118 | ARG | p.Arg118His | P21549 |
| HOMO SAPIENS | 3 | 1067 | 4kyo | A | | 122 | ARG | p.Arg122Ter | P21549 |
| HOMO SAPIENS | 3 | 1067 | 4kyo | A | | 124 | HIS | p.His124Pro | P21549 |
| HOMO SAPIENS | 0 | 1067 | 4kyo | A | | 152 | PHE | p.Phe152Ile | P21549 |
| HOMO SAPIENS | 1 | 1067 | 4kyo | A | | 156 | GLY | p.Gly156Arg | P21549 |
| HOMO SAPIENS | 0 | 1067 | 4kyo | A | | 161 | GLY | p.Gly161Arg | P21549 |
| HOMO SAPIENS | 3 | 1067 | 4kyo | A | | 166 | LEU | p.Leu166Pro | P21549 |
| HOMO SAPIENS | 5 | 1067 | 4kyo | A | | 170 | GLY | p.Gly170Arg | P21549 |
| HOMO SAPIENS | 5 | 1067 | 4kyo | A | | 186 | ALA | p.Ala186Val | P21549 |
| HOMO SAPIENS | 3 | 1067 | 4kyo | A | | 187 | SER | p.Ser187Phe | P21549 |
| HOMO SAPIENS | 3 | 1067 | 4kyo | A | | 197 | ARG | p.Arg197Pro | P21549 |
| HOMO SAPIENS | 5 | 1067 | 4kyo | A | | 201 | ASP | p.Asp201Glu | P21549 |
| HOMO SAPIENS | 0 | 1067 | 4kyo | A | | 202 | ILE | p.Ile202Asn | P21549 |
| HOMO SAPIENS | 1 | 1067 | 4kyo | A | | 205 | SER | p.Ser205Pro | P21549 |
| HOMO SAPIENS | 1 | 1067 | 4kyo | A | | 205 | SER | p.Ser205Ter | P21549 |
| HOMO SAPIENS | 2 | 1067 | 4kyo | A | | 216 | GLY | p.Gly216Arg | P21549 |
| HOMO SAPIENS | 5 | 1067 | 4kyo | A | | 233 | ARG | p.Arg233Cys | P21549 |
| HOMO SAPIENS | 5 | 1067 | 4kyo | A | | 244 | ILE | p.Ile244Thr | P21549 |
| HOMO SAPIENS | 5 | 1067 | 4kyo | A | | 244 | ILE | p.Ile244Met | P21549 |
| HOMO SAPIENS | 4 | 1067 | 4kyo | A | | 253 | CYS | p.Cys253Arg | P21549 |
| HOMO SAPIENS | 7 | 1067 | 4kyo | A | | 274 | GLU | p.Glu274Asp | P21549 |
| HOMO SAPIENS | 7 | 1067 | 4kyo | A | | 279 | ILE | p.Ile279Thr | P21549 |
| HOMO SAPIENS | 6 | 1067 | 4kyo | A | | 295 | ALA | p.Ala295Thr | P21549 |

| | | | | | | | | |
|---|---|---|---|---|---|---|---|---|
| HOMO SAPIENS | 5 | 1067 | 4kyo | A | 340 | ILE | p.Ile340Met | P21549 |
| HOMO SAPIENS | 1 | 790 | 2ric | C | 270 | SER | p.Ser290Thr | P35247 |
| HOMO SAPIENS | 0 | 790 | 2ric | C | 289 | GLU | p.Glu309Lys | P35247 |
| HOMO SAPIENS | 1 | 212 | 5sxe | A | 545 | GLU | p.Glu545Lys | P42336 |
| HOMO SAPIENS | 3 | 212 | 5sxe | A | 546 | GLN | p.Gln546Glu | P42336 |
| HOMO SAPIENS | 2 | 212 | 5sxe | A | 1047 | HIS | p.His1047Leu | P42336 |
| HOMO SAPIENS | 7 | 152 | 2zqq | F | 197 | ARG | p.Arg197Ter | Q13825 |
| HOMO SAPIENS | 0 | 152 | 2zqq | F | 248 | ALA | p.Ala248Thr | Q13825 |
| HOMO SAPIENS | 0 | 152 | 2zqq | F | 275 | ALA | p.Ala275Val | Q13825 |
| HOMO SAPIENS | 5 | 152 | 2zqq | F | 331 | LYS | p.Lys331Ter | Q13825 |
| HOMO SAPIENS | 6 | 17801 | 2j8l | A | 283 | PHE | p.Phe301Leu | P03951 |
| HOMO SAPIENS | 3 | 17801 | 2j8l | A | 308 | ARG | p.Arg326Cys | P03951 |
| HOMO SAPIENS | 0 | 17801 | 2j8l | A | 321 | CYS | p.Cys339Phe | P03951 |
| HOMO SAPIENS | 3 | 17801 | 2j8l | A | 351 | TYR | p.Tyr369Ter | P03951 |
| HOMO SAPIENS | 7 | 1802 | 5db0 | A | 22 | LEU | p.Leu22Arg | O00255 |
| HOMO SAPIENS | 7 | 1802 | 5db0 | A | 26 | GLU | p.Glu26Lys | O00255 |
| HOMO SAPIENS | 7 | 1802 | 5db0 | A | 139 | HIS | p.His139Asp | O00255 |
| HOMO SAPIENS | 0 | 1802 | 5db0 | A | 171 | ARG | p.Arg176Gln | O00255 |
| HOMO SAPIENS | 0 | 1802 | 5db0 | A | 184 | VAL | p.Val189Glu | O00255 |
| HOMO SAPIENS | 7 | 1802 | 5db0 | A | 255 | GLU | p.Glu260Lys | O00255 |
| HOMO SAPIENS | 4 | 1802 | 5db0 | A | 418 | ASP | p.Asp423Asn | O00255 |
| HOMO SAPIENS | 0 | 1802 | 5db0 | A | 436 | TRP | p.Trp441Arg | O00255 |
| HOMO SAPIENS | -1 | 1802 | 5db0 | A | 541 | ALA | p.Thr546Ala | O00255 |
| HOMO SAPIENS | 4 | 1802 | 5db0 | A | 552 | THR | p.Thr557Ser | O00255 |
| HOMO SAPIENS | 0 | 3450 | 2yuz | A | 36 | PRO | p.Pro461Gln | Q00872 |
| HOMO SAPIENS | 1 | 3450 | 2yuz | A | 56 | GLN | p.His481Gln | Q00872 |
| HOMO SAPIENS | 2 | 3450 | 2yuz | A | 68 | ALA | p.Ala493Val | Q00872 |
| HOMO SAPIENS | 0 | 3211 | 3rqg | A | 72 | VAL | p.Val72Leu | Q9BUL8 |
| HOMO SAPIENS | 2 | 4168 | 4q4f | A | 121 | ARG | p.Arg121Ter | Q14108 |
| HOMO SAPIENS | 2 | 4168 | 4q4f | A | 121 | ARG | p.Arg121Gln | Q14108 |
| HOMO SAPIENS | 2 | 4168 | 4q4f | A | 159 | MET | p.Met159Val | Q14108 |
| HOMO SAPIENS | 4 | 4168 | 4q4f | A | 363 | HIS | p.His363Asn | Q14108 |
| HOMO SAPIENS | 2 | 4168 | 4q4f | A | 396 | VAL | p.Val396Ile | Q14108 |
| HOMO SAPIENS | 0 | 1448 | 1y97 | A | 87 | ARG | p.Arg130Ter | Q9BQ50 |
| HOMO SAPIENS | 0 | 3444 | 4iwp | A | 151 | SER | p.Ser151Phe | Q9UHD2 |
| HOMO SAPIENS | 0 | 3444 | 4iwp | A | 401 | LYS | p.Lys401Glu | Q9UHD2 |
| HOMO SAPIENS | 0 | 116 | 1zy8 | B | 194 | GLY | p.Gly229Cys | P09622 |
| HOMO SAPIENS | 0 | 116 | 1zy8 | B | 255 | ASP | p.Asp290Gly | P09622 |
| HOMO SAPIENS | 2 | 116 | 1zy8 | B | 326 | MET | p.Met361Val | P09622 |
| HOMO SAPIENS | 0 | 116 | 1zy8 | B | 340 | GLU | p.Glu375Lys | P09622 |
| HOMO SAPIENS | 2 | 116 | 1zy8 | B | 358 | ILE | p.Ile393Thr | P09622 |
| HOMO SAPIENS | 1 | 116 | 1zy8 | B | 453 | PRO | p.Pro488Leu | P09622 |
| HOMO SAPIENS | 7 | 127 | 4qo7 | B | 160 | SER | p.Ser161Arg | P00338 |
| HOMO SAPIENS | 0 | 2025 | 1wrl | A | 8 | ALA | p.Ala8Val | P63316 |
| HOMO SAPIENS | 8 | 2025 | 1wrl | A | 24 | PHE | p.Phe24Leu | P63316 |
| HOMO SAPIENS | 0 | 2025 | 1wrl | A | 84 | SER | p.Cys84Tyr | P63316 |
| HOMO SAPIENS | 8 | 3791 | 1mhd | B | 93 | ARG | p.Arg93Ter | P84022 |
| HOMO SAPIENS | 0 | 1575 | 4phc | A | 205 | GLY | p.Gly205Asp | P12081 |
| HOMO SAPIENS | 0 | 1575 | 4phc | A | 454 | TYR | p.Tyr454Ser | P12081 |
| HOMO SAPIENS | 0 | 39620 | 1ozs | B | 135 | ARG | p.Arg136Gln | P19429 |
| HOMO SAPIENS | 5 | 39620 | 1ozs | B | 140 | ARG | p.Arg141Trp | P19429 |
| HOMO SAPIENS | 5 | 39620 | 1ozs | B | 142 | THR | p.Thr143Asn | P19429 |
| HOMO SAPIENS | 5 | 39620 | 1ozs | B | 143 | LEU | p.Leu144Gln | P19429 |
| HOMO SAPIENS | 5 | 39620 | 1ozs | B | 144 | ARG | p.Arg145Gly | P19429 |

| | | | | | | | | |
|---|---|---|---|---|---|---|---|---|
| HOMO SAPIENS | 5 | 39620 | 1ozs | B | 144 | ARG | p.Arg145Gln | P19429 |
| HOMO SAPIENS | -1 | 20934 | 4nuv | D | 42 | GLY | p.Gly42Asp | Q16570 |
| HOMO SAPIENS | -1 | 40565 | 5lcw | S | 4 | VAL | p.Val4Met | O60566 |
| HOMO SAPIENS | 0 | 40565 | 5lcw | S | 40 | THR | p.Thr40Met | O60566 |
| HOMO SAPIENS | 4 | 40565 | 5lcw | S | 198 | ALA | p.Glu390Asp | O60566 |
| HOMO SAPIENS | 4 | 758 | 4oiu | A | 726 | ARG | p.Arg727Leu | P10275 |
| HOMO SAPIENS | 4 | 758 | 4oiu | A | 730 | VAL | p.Val731Met | P10275 |
| HOMO SAPIENS | 2 | 758 | 4oiu | A | 798 | GLN | p.Gln799Glu | P10275 |
| HOMO SAPIENS | 3 | 758 | 4oiu | A | 855 | ARG | p.Arg856His | P10275 |
| HOMO SAPIENS | 1 | 758 | 4oiu | A | 886 | MET | p.Met887Val | P10275 |
| HOMO SAPIENS | 0 | 3303 | 4ie0 | A | 134 | ALA | p.Ala134Thr | Q9C0B1 |
| HOMO SAPIENS | 9 | 3303 | 4ie0 | A | 316 | ARG | p.Arg316Gln | Q9C0B1 |
| HOMO SAPIENS | 1 | 41502 | 3fgh | A | 136 | PRO | p.Pro178Leu | Q00059 |
| HOMO SAPIENS | 5 | 175 | 3ceh | B | 58 | THR | p.Thr59Lys | P06737 |
| HOMO SAPIENS | 8 | 175 | 3ceh | B | 93 | ARG | p.Arg94Ter | P06737 |
| HOMO SAPIENS | 5 | 175 | 3ceh | B | 221 | VAL | p.Val222Ile | P06737 |
| HOMO SAPIENS | 0 | 175 | 3ceh | B | 338 | ASN | p.Asn339Ser | P06737 |
| HOMO SAPIENS | 1 | 175 | 3ceh | B | 372 | PHE | p.Phe373Leu | P06737 |
| HOMO SAPIENS | 9 | 175 | 3ceh | B | 376 | ASN | p.Asn377Lys | P06737 |
| HOMO SAPIENS | 8 | 175 | 3ceh | B | 398 | ARG | p.Arg399Ter | P06737 |
| HOMO SAPIENS | 5 | 175 | 3ceh | B | 424 | ARG | p.Arg425Pro | P06737 |
| HOMO SAPIENS | 9 | 175 | 3ceh | B | 455 | VAL | p.Val456Met | P06737 |
| HOMO SAPIENS | 8 | 175 | 3ceh | B | 490 | ARG | p.Arg491Cys | P06737 |
| HOMO SAPIENS | 8 | 175 | 3ceh | B | 631 | ASN | p.Asn632Ile | P06737 |
| HOMO SAPIENS | 9 | 175 | 3ceh | B | 648 | TYR | p.Tyr649Ter | P06737 |
| HOMO SAPIENS | 9 | 175 | 3ceh | B | 672 | GLU | p.Glu673Lys | P06737 |
| HOMO SAPIENS | 9 | 175 | 3ceh | B | 674 | SER | p.Ser675Leu | P06737 |
| HOMO SAPIENS | 0 | 175 | 3ceh | B | 805 | ILE | p.Ile806Leu | P06737 |
| HOMO SAPIENS | 0 | 175 | 3ceh | B | 820 | TYR | p.Tyr821His | P06737 |
| HOMO SAPIENS | 1 | 6750 | 4fdj | A | 47 | GLY | p.Gly47Arg | P34059 |
| HOMO SAPIENS | 9 | 6750 | 4fdj | A | 60 | ASP | p.Asp60Asn | P34059 |
| HOMO SAPIENS | 5 | 6750 | 4fdj | A | 67 | LEU | p.Leu67Met | P34059 |
| HOMO SAPIENS | 0 | 6750 | 4fdj | A | 69 | PHE | p.Phe69Val | P34059 |
| HOMO SAPIENS | 3 | 6750 | 4fdj | A | 94 | ARG | p.Arg94Gly | P34059 |
| HOMO SAPIENS | 0 | 6750 | 4fdj | A | 113 | ILE | p.Ile113Phe | P34059 |
| HOMO SAPIENS | 0 | 6750 | 4fdj | A | 138 | VAL | p.Val138Ala | P34059 |
| HOMO SAPIENS | 0 | 6750 | 4fdj | A | 139 | GLY | p.Gly139Ser | P34059 |
| HOMO SAPIENS | 0 | 6750 | 4fdj | A | 151 | PRO | p.Pro151Thr | P34059 |
| HOMO SAPIENS | 0 | 6750 | 4fdj | A | 155 | GLY | p.Gly155Arg | P34059 |
| HOMO SAPIENS | 0 | 6750 | 4fdj | A | 159 | TRP | p.Trp159Ter | P34059 |
| HOMO SAPIENS | 0 | 6750 | 4fdj | A | 162 | SER | p.Ser162Phe | P34059 |
| HOMO SAPIENS | 0 | 6750 | 4fdj | A | 200 | THR | p.Thr200Met | P34059 |
| HOMO SAPIENS | 0 | 6750 | 4fdj | A | 204 | ASN | p.Asn204Lys | P34059 |
| HOMO SAPIENS | 0 | 6750 | 4fdj | A | 231 | ALA | p.Ala231Gly | P34059 |
| HOMO SAPIENS | 2 | 6750 | 4fdj | A | 235 | THR | p.Thr235Lys | P34059 |
| HOMO SAPIENS | 1 | 6750 | 4fdj | A | 259 | ARG | p.Arg259Gln | P34059 |
| HOMO SAPIENS | 0 | 6750 | 4fdj | A | 287 | SER | p.Ser287Leu | P34059 |
| HOMO SAPIENS | 0 | 6750 | 4fdj | A | 291 | ALA | p.Ala291Thr | P34059 |
| HOMO SAPIENS | 3 | 6750 | 4fdj | A | 301 | GLY | p.Gly301Cys | P34059 |
| HOMO SAPIENS | 0 | 6750 | 4fdj | A | 312 | THR | p.Thr312Ser | P34059 |
| HOMO SAPIENS | 0 | 6750 | 4fdj | A | 318 | MET | p.Met318Arg | P34059 |
| HOMO SAPIENS | 0 | 6750 | 4fdj | A | 340 | GLY | p.Gly340Asp | P34059 |
| HOMO SAPIENS | 0 | 6750 | 4fdj | A | 386 | ARG | p.Arg386Cys | P34059 |
| HOMO SAPIENS | 0 | 6750 | 4fdj | A | 391 | MET | p.Met391Val | P34059 |

| | | | | | | | | |
|---|---|---|---|---|---|---|---|---|
| HOMO SAPIENS | 0 | 6750 | 4fdj | A | 392 | ALA | p.Ala392Val | P34059 |
| HOMO SAPIENS | 0 | 6750 | 4fdj | A | 393 | ALA | p.Ala393Ser | P34059 |
| HOMO SAPIENS | 0 | 6750 | 4fdj | A | 459 | ALA | p.Ala459Val | P34059 |
| HOMO SAPIENS | 0 | 6750 | 4fdj | A | 473 | GLN | p.Gln473Ter | P34059 |
| HOMO SAPIENS | 2 | 6750 | 4fdj | A | 480 | VAL | p.Val480Phe | P34059 |
| HOMO SAPIENS | 0 | 6750 | 4fdj | A | 487 | ASN | p.Asn487Ser | P34059 |
| HOMO SAPIENS | 0 | 6750 | 4fdj | A | 488 | VAL | p.Val488Met | P34059 |
| HOMO SAPIENS | 0 | 6750 | 4fdj | A | 520 | TRP | p.Trp520Ter | P34059 |
| HOMO SAPIENS | 4 | 1088 | 2p3u | A | 102 | GLU | p.Glu142Lys | P00742 |
| HOMO SAPIENS | 6 | 370 | 1nr1 | D | 73 | ASP | p.Asp126His | P00367 |
| HOMO SAPIENS | 3 | 370 | 1nr1 | D | 221 | ARG | p.Arg274Cys | P00367 |
| HOMO SAPIENS | 2 | 370 | 1nr1 | D | 265 | ARG | p.Arg318Lys | P00367 |
| HOMO SAPIENS | 1 | 370 | 1nr1 | D | 269 | ARG | p.Arg322His | P00367 |
| HOMO SAPIENS | 1 | 370 | 1nr1 | D | 296 | GLU | p.Glu349Ala | P00367 |
| HOMO SAPIENS | 1 | 370 | 1nr1 | D | 445 | SER | p.Ser498Leu | P00367 |
| HOMO SAPIENS | 1 | 370 | 1nr1 | D | 446 | GLY | p.Gly499Asp | P00367 |
| HOMO SAPIENS | 1 | 370 | 1nr1 | D | 446 | GLY | p.Gly499Ser | P00367 |
| HOMO SAPIENS | 1 | 370 | 1nr1 | D | 448 | SER | p.Ser501Pro | P00367 |
| HOMO SAPIENS | 2 | 370 | 1nr1 | D | 454 | HIS | p.His507Tyr | P00367 |
| HOMO SAPIENS | 1 | 7720 | 2he0 | A | 147 | HIS | p.His2018Gln | P46531 |
| HOMO SAPIENS | 2 | 11844 | 5l0v | A | 218 | ARG | p.Arg218Ter | Q8NBL1 |
| HOMO SAPIENS | 2 | 11844 | 5l0v | A | 279 | ARG | p.Arg279Trp | Q8NBL1 |
| HOMO SAPIENS | 9 | 1485 | 5va0 | A | 477 | ARG | p.Arg477His | P04150 |
| HOMO SAPIENS | 0 | 235 | 2fun | B | 2248 | ARG | p.Arg248Trp | Q14790 |
| HOMO SAPIENS | 4 | 235 | 2fun | B | 2285 | HIS | p.Asp285His | Q14790 |
| HOMO SAPIENS | 0 | 4822 | 5y3n | A | 307 | ARG | p.Arg307Gly | Q12931 |
| HOMO SAPIENS | 8 | 32928 | 4m6w | A | 1886 | HIS | p.His1886Tyr | Q8IYD8 |
| HOMO SAPIENS | 0 | 2711 | 4y5v | F | 22 | GLY | p.Gly46Glu | P19235 |
| HOMO SAPIENS | -1 | 2711 | 4y5v | F | 166 | ALA | p.Ala190Ser | P19235 |
| HOMO SAPIENS | 4 | 5161 | 5bpu | A | 42 | HIS | p.His42Arg | Q00604 |
| HOMO SAPIENS | 0 | 5161 | 5bpu | A | 44 | TYR | p.Tyr44Cys | Q00604 |
| HOMO SAPIENS | 0 | 5161 | 5bpu | A | 45 | VAL | p.Val45Glu | Q00604 |
| HOMO SAPIENS | 0 | 5161 | 5bpu | A | 60 | VAL | p.Val60Glu | Q00604 |
| HOMO SAPIENS | 0 | 5161 | 5bpu | A | 61 | LEU | p.Leu61Phe | Q00604 |
| HOMO SAPIENS | 7 | 5161 | 5bpu | A | 69 | CYS | p.Cys69Ser | Q00604 |
| HOMO SAPIENS | 0 | 5161 | 5bpu | A | 75 | SER | p.Ser75Cys | Q00604 |
| HOMO SAPIENS | 0 | 5161 | 5bpu | A | 90 | ARG | p.Arg90Pro | Q00604 |
| HOMO SAPIENS | 9 | 5161 | 5bpu | A | 96 | CYS | p.Cys96Tyr | Q00604 |
| HOMO SAPIENS | 9 | 5161 | 5bpu | A | 96 | CYS | p.Cys96Trp | Q00604 |
| HOMO SAPIENS | 0 | 5161 | 5bpu | A | 101 | SER | p.Ser101Phe | Q00604 |
| HOMO SAPIENS | 0 | 5161 | 5bpu | A | 105 | ALA | p.Ala105Thr | Q00604 |
| HOMO SAPIENS | 0 | 5161 | 5bpu | A | 110 | CYS | p.Cys110Gly | Q00604 |
| HOMO SAPIENS | 0 | 5161 | 5bpu | A | 121 | ARG | p.Arg121Trp | Q00604 |
| HOMO SAPIENS | 0 | 5161 | 5bpu | A | 121 | ARG | p.Arg121Leu | Q00604 |
| HOMO SAPIENS | 0 | 5161 | 5bpu | A | 124 | LEU | p.Leu124Phe | Q00604 |
| HOMO SAPIENS | 8 | 5161 | 5bpu | A | 128 | CYS | p.Cys128Ter | Q00604 |
| HOMO SAPIENS | 1 | 13944 | 5xqh | A | 238 | ARG | p.Arg238His | Q9GZN7 |
| HOMO SAPIENS | -1 | 7508 | 5jl6 | A | 39 | TRP | p.Trp39Arg | P11511 |
| HOMO SAPIENS | 0 | 7508 | 5jl6 | A | 201 | THR | p.Thr201Met | P11511 |
| HOMO SAPIENS | 0 | 7508 | 5jl6 | A | 210 | GLU | p.Glu210Lys | P11511 |
| HOMO SAPIENS | 1 | 7508 | 5jl6 | A | 264 | ARG | p.Arg264Cys | P11511 |
| HOMO SAPIENS | 2 | 7508 | 5jl6 | A | 365 | ARG | p.Arg365Gln | P11511 |
| HOMO SAPIENS | 7 | 7508 | 5jl6 | A | 375 | ARG | p.Arg375Cys | P11511 |
| HOMO SAPIENS | 8 | 7508 | 5jl6 | A | 435 | ARG | p.Arg435Cys | P11511 |

| | | | | | | | | |
|---|---|---|---|---|---|---|---|---|
| HOMO SAPIENS | 9 | 7508 | 5jl6 | A | 437 | CYS | p.Cys437Tyr | P11511 |
| HOMO SAPIENS | -1 | 30206 | 5l95 | B | 246 | ARG | p.Arg246Ter | Q9GZZ9 |
| HOMO SAPIENS | 4 | 875 | 3efh | B | 43 | GLU | p.Glu43Asp | P60891 |
| HOMO SAPIENS | 5 | 875 | 3efh | B | 65 | ASP | p.Asp65Asn | P60891 |
| HOMO SAPIENS | 0 | 875 | 3efh | B | 87 | ALA | p.Ala87Thr | P60891 |
| HOMO SAPIENS | 2 | 875 | 3efh | B | 115 | MET | p.Met115Thr | P60891 |
| HOMO SAPIENS | 8 | 875 | 3efh | B | 133 | GLN | p.Gln133Pro | P60891 |
| HOMO SAPIENS | 8 | 875 | 3efh | B | 152 | LEU | p.Leu152Pro | P60891 |
| HOMO SAPIENS | 7 | 875 | 3efh | B | 290 | ILE | p.Ile290Thr | P60891 |
| HOMO SAPIENS | 6 | 875 | 3efh | B | 306 | GLY | p.Gly306Arg | P60891 |
| HOMO SAPIENS | 0 | 7978 | 5edm | A | 122 | MET | p.Thr165Met | P00734 |
| HOMO SAPIENS | -1 | 7978 | 5edm | A | 257 | ARG | p.Arg314Cys | P00734 |
| HOMO SAPIENS | 0 | 1716 | 1b98 | M | 126 | ARG | p.Arg206Gln | P34130 |
| HOMO SAPIENS | 0 | 1716 | 1b98 | M | 126 | ARG | p.Arg206Trp | P34130 |
| HOMO SAPIENS | 9 | 3960 | 2qyp | B | 28 | ILE | p.Ile338Val | P07602 |
| HOMO SAPIENS | 1 | 3960 | 2qyp | B | 39 | LEU | p.Leu349Pro | P07602 |
| HOMO SAPIENS | 0 | 31414 | 5u1d | A | 333 | ILE | p.Ile393Val | Q03518 |
| HOMO SAPIENS | 0 | 31414 | 5u1d | A | 637 | ASP | p.Asp697Gly | Q03518 |
| HOMO SAPIENS | 0 | 31414 | 5u1d | A | 659 | ARG | p.Arg719Gln | Q03518 |
| HOMO SAPIENS | 3 | 119 | 2pho | A | 11 | ILE | p.Ile11Thr | P05089 |
| HOMO SAPIENS | 8 | 119 | 2pho | A | 21 | ARG | p.Arg21Ter | P05089 |
| HOMO SAPIENS | 0 | 119 | 2pho | A | 122 | TRP | p.Trp122Ter | P05089 |
| HOMO SAPIENS | 6 | 119 | 2pho | A | 138 | GLY | p.Gly138Val | P05089 |
| HOMO SAPIENS | 5 | 119 | 2pho | A | 290 | THR | p.Thr290Ser | P05089 |
| HOMO SAPIENS | 5 | 119 | 2pho | A | 291 | ARG | p.Arg291Ter | P05089 |
| HOMO SAPIENS | 0 | 3732 | 2i99 | A | 247 | TYR | p.Tyr247Ter | Q14894 |
| HOMO SAPIENS | -1 | 1382 | 4ffx | A | 2 | ALA | p.Ala2Val | P30566 |
| HOMO SAPIENS | 1 | 1382 | 4ffx | A | 72 | ILE | p.Ile72Met | P30566 |
| HOMO SAPIENS | 1 | 1382 | 4ffx | A | 100 | PRO | p.Pro100Ala | P30566 |
| HOMO SAPIENS | 0 | 1382 | 4ffx | A | 114 | TYR | p.Tyr114His | P30566 |
| HOMO SAPIENS | 1 | 1382 | 4ffx | A | 141 | ARG | p.Arg141Trp | P30566 |
| HOMO SAPIENS | 0 | 1382 | 4ffx | A | 147 | LYS | p.Lys147Met | P30566 |
| HOMO SAPIENS | 3 | 1382 | 4ffx | A | 190 | ARG | p.Arg190Gln | P30566 |
| HOMO SAPIENS | 3 | 1382 | 4ffx | A | 190 | ARG | p.Arg190Ter | P30566 |
| HOMO SAPIENS | 0 | 1382 | 4ffx | A | 217 | HIS | p.His217Asp | P30566 |
| HOMO SAPIENS | 2 | 1382 | 4ffx | A | 246 | LYS | p.Lys246Glu | P30566 |
| HOMO SAPIENS | 5 | 1382 | 4ffx | A | 303 | ARG | p.Arg303Cys | P30566 |
| HOMO SAPIENS | 1 | 1382 | 4ffx | A | 318 | PRO | p.Pro318Leu | P30566 |
| HOMO SAPIENS | 6 | 1382 | 4ffx | A | 332 | ASP | p.Asp332His | P30566 |
| HOMO SAPIENS | 6 | 1382 | 4ffx | A | 337 | ARG | p.Arg337Ter | P30566 |
| HOMO SAPIENS | 1 | 1382 | 4ffx | A | 396 | ARG | p.Arg396His | P30566 |
| HOMO SAPIENS | 1 | 1382 | 4ffx | A | 426 | ARG | p.Arg426His | P30566 |
| HOMO SAPIENS | 0 | 1382 | 4ffx | A | 438 | SER | p.Ser438Pro | P30566 |
| HOMO SAPIENS | 1 | 1382 | 4ffx | A | 447 | SER | p.Ser447Pro | P30566 |
| HOMO SAPIENS | 2 | 1382 | 4ffx | A | 448 | SER | p.Ser448Pro | P30566 |
| HOMO SAPIENS | 2 | 1382 | 4ffx | A | 450 | THR | p.Thr450Ser | P30566 |
| HOMO SAPIENS | 0 | 6212 | 1ao3 | A | 46 | SER | p.Ser1731Thr | P04275 |
| HOMO SAPIENS | 0 | 1711 | 3f9c | A | 279 | VAL | p.Val279Phe | Q13093 |
| HOMO SAPIENS | 0 | 6112 | 4fvl | A | 312 | HIS | p.His312Arg | P45452 |
| HOMO SAPIENS | 0 | 6112 | 4fvl | A | 323 | THR | p.Thr323Met | P45452 |
| HOMO SAPIENS | 0 | 6112 | 4fvl | A | 390 | ASP | p.Asp390Gly | P45452 |
| HOMO SAPIENS | 0 | 6112 | 4fvl | A | 458 | ARG | p.Arg458Cys | P45452 |
| HOMO SAPIENS | 2 | 6180 | 5ggn | A | 97 | ARG | p.Arg97Trp | Q8WZA1 |
| HOMO SAPIENS | 2 | 6180 | 5ggn | A | 215 | ARG | p.Arg215Ter | Q8WZA1 |

| | | | | | | | | |
|---|---|---|---|---|---|---|---|---|
| HOMO SAPIENS | -1 | 5526 | 4ak3 | A | 11 | ASP | p.Asp1232Asn | P02461 |
| HOMO SAPIENS | 6 | 5526 | 4ak3 | A | 38 | ALA | p.Ala1259Thr | P02461 |
| HOMO SAPIENS | 0 | 5526 | 4ak3 | A | 92 | LYS | p.Lys1313Arg | P02461 |
| HOMO SAPIENS | 0 | 5526 | 4ak3 | A | 132 | GLN | p.His1353Gln | P02461 |
| HOMO SAPIENS | 4 | 16616 | 1hci | B | 433 | SER | p.Ser433Leu | P35609 |
| HOMO SAPIENS | 1 | 16616 | 1hci | B | 475 | ASP | p.Asp475Asn | P35609 |
| HOMO SAPIENS | 1 | 16616 | 1hci | B | 476 | ALA | p.Ala476Ser | P35609 |
| HOMO SAPIENS | 2 | 16616 | 1hci | B | 507 | MET | p.Met507Val | P35609 |
| HOMO SAPIENS | 1 | 16616 | 1hci | B | 604 | MET | p.Met604Val | P35609 |
| HOMO SAPIENS | 2 | 16616 | 1hci | B | 640 | ARG | p.Arg640His | P35609 |
| HOMO SAPIENS | 1 | 16616 | 1hci | B | 721 | ARG | p.Arg721Cys | P35609 |
| HOMO SAPIENS | -1 | 3402 | 5iaa | B | 57 | MET | p.Met57Val | Q9GZZ9 |
| HOMO SAPIENS | 7 | 3250 | 3irq | B | 193 | PRO | p.Pro193Ala | P55265 |
| HOMO SAPIENS | 2 | 15200 | 2wns | A | 30 | SER | p.Ser30Gly | P11172 |
| HOMO SAPIENS | 0 | 15200 | 2wns | A | 96 | ARG | p.Arg96Gly | P11172 |
| HOMO SAPIENS | 0 | 15200 | 2wns | A | 109 | VAL | p.Val109Gly | P11172 |
| HOMO SAPIENS | 2 | 15759 | 4x4w | B | 140 | LEU | p.Leu166Ser | Q96Q11 |
| HOMO SAPIENS | 0 | 15759 | 4x4w | B | 197 | ILE | p.Ile223Thr | Q96Q11 |
| HOMO SAPIENS | 0 | 3156 | 3w03 | B | 14 | ALA | p.Ala14Thr | Q9H9Q4 |
| HOMO SAPIENS | 0 | 3156 | 3w03 | B | 57 | ARG | p.Arg57Gly | Q9H9Q4 |
| HOMO SAPIENS | 0 | 3156 | 3w03 | B | 123 | CYS | p.Cys123Arg | Q9H9Q4 |
| HOMO SAPIENS | 0 | 11693 | 3ojy | A | 63 | GLN | p.Gln93Lys | P07357 |
| HOMO SAPIENS | 0 | 1421 | 2qc8 | E | 275 | GLU | p.Glu275Asp | P15104 |
| HOMO SAPIENS | 9 | 1421 | 2qc8 | E | 324 | ARG | p.Arg324Cys | P15104 |
| HOMO SAPIENS | 6 | 1421 | 2qc8 | E | 341 | ARG | p.Arg341Cys | P15104 |
| HOMO SAPIENS | 6 | 2850 | 3qvb | A | 83 | THR | p.Thr83Met | P46976 |
| HOMO SAPIENS | 8 | 2850 | 3qvb | A | 102 | ASP | p.Asp102His | P46976 |
| HOMO SAPIENS | 1 | 24314 | 4m9p | A | 480 | PRO | p.Pro480Leu | P21333 |
| HOMO SAPIENS | 0 | 24314 | 4m9p | A | 528 | VAL | p.Val528Met | P21333 |
| HOMO SAPIENS | 0 | 24314 | 4m9p | A | 590 | VAL | p.Val590Ile | P21333 |
| HOMO SAPIENS | 0 | 24314 | 4m9p | A | 605 | ASP | p.Asp605Asn | P21333 |
| HOMO SAPIENS | 1 | 24314 | 4m9p | A | 637 | PRO | p.Pro637Gln | P21333 |
| HOMO SAPIENS | 0 | 24314 | 4m9p | A | 656 | LEU | p.Leu656Phe | P21333 |
| HOMO SAPIENS | 2 | 24314 | 4m9p | A | 711 | VAL | p.Val711Asp | P21333 |
| HOMO SAPIENS | 0 | 4951 | 5joe | A | 40 | THR | p.Thr9621Ser | Q8WZ42 |
| HOMO SAPIENS | 1 | 4951 | 5joe | A | 41 | GLU | p.Glu9622Gln | Q8WZ42 |
| HOMO SAPIENS | 0 | 4951 | 5joe | A | 82 | ALA | p.Ala9663Ser | Q8WZ42 |
| HOMO SAPIENS | 4 | 4951 | 5joe | A | 89 | ILE | p.Ile9670Asn | Q8WZ42 |
| HOMO SAPIENS | 0 | 9664 | 3c6m | A | 56 | GLY | p.Gly56Ser | P52788 |
| HOMO SAPIENS | 0 | 9664 | 3c6m | A | 132 | VAL | p.Val132Gly | P52788 |
| HOMO SAPIENS | 1 | 9664 | 3c6m | A | 190 | THR | p.Thr190Ile | P52788 |
| HOMO SAPIENS | -1 | 12118 | 5hcd | B | 19 | GLN | p.Gln19Ter | P01031 |
| HOMO SAPIENS | 0 | 1583 | 2b4x | L | 7 | ILE | p.Ile39Asn | P01008 |
| HOMO SAPIENS | 0 | 1583 | 2b4x | L | 24 | ARG | p.Arg56Cys | P01008 |
| HOMO SAPIENS | 0 | 1583 | 2b4x | L | 47 | ARG | p.Arg79Cys | P01008 |
| HOMO SAPIENS | 0 | 1583 | 2b4x | L | 47 | ARG | p.Arg79His | P01008 |
| HOMO SAPIENS | 1 | 1583 | 2b4x | L | 95 | CYS | p.Cys127Arg | P01008 |
| HOMO SAPIENS | 1 | 1583 | 2b4x | L | 99 | LEU | p.Leu131Phe | P01008 |
| HOMO SAPIENS | 0 | 1583 | 2b4x | L | 116 | SER | p.Ser148Pro | P01008 |
| HOMO SAPIENS | 0 | 1583 | 2b4x | L | 129 | ARG | p.Arg161Gln | P01008 |
| HOMO SAPIENS | 0 | 1583 | 2b4x | L | 135 | ASN | p.Asn167Thr | P01008 |
| HOMO SAPIENS | 2 | 1583 | 2b4x | L | 187 | ASN | p.Asn219Asp | P01008 |
| HOMO SAPIENS | 0 | 1583 | 2b4x | L | 191 | SER | p.Ser223Pro | P01008 |
| HOMO SAPIENS | 2 | 1583 | 2b4x | L | 349 | SER | p.Ser381Pro | P01008 |

| | | | | | | | | |
|---|---|---|---|---|---|---|---|---|
| HOMO SAPIENS | 0 | 1583 | 2b4x | L | 382 | ALA | p.Ala414Thr | P01008 |
| HOMO SAPIENS | 0 | 1583 | 2b4x | L | 387 | ALA | p.Ala419Val | P01008 |
| HOMO SAPIENS | 0 | 1583 | 2b4x | L | 392 | GLY | p.Gly424Asp | P01008 |
| HOMO SAPIENS | 0 | 1583 | 2b4x | L | 393 | ARG | p.Arg425His | P01008 |
| HOMO SAPIENS | 0 | 1583 | 2b4x | L | 393 | ARG | p.Arg425Cys | P01008 |
| HOMO SAPIENS | 0 | 1583 | 2b4x | L | 394 | SER | p.Ser426Leu | P01008 |
| HOMO SAPIENS | 0 | 1583 | 2b4x | L | 404 | ALA | p.Ala436Thr | P01008 |
| HOMO SAPIENS | 2 | 1583 | 2b4x | L | 407 | PRO | p.Pro439Leu | P01008 |
| HOMO SAPIENS | 2 | 1583 | 2b4x | L | 429 | PRO | p.Pro461Leu | P01008 |
| HOMO SAPIENS | 0 | 11647 | 2a1t | S | 78 | ALA | p.Ala78Ser | P38117 |
| HOMO SAPIENS | 0 | 11647 | 2a1t | S | 98 | ARG | p.Arg98Cys | P38117 |
| HOMO SAPIENS | 8 | 11647 | 2a1t | S | 128 | ASP | p.Asp128Asn | P38117 |
| HOMO SAPIENS | 5 | 11647 | 2a1t | S | 154 | THR | p.Thr154Met | P38117 |
| HOMO SAPIENS | 6 | 11647 | 2a1t | S | 164 | ARG | p.Arg164Gln | P38117 |
| HOMO SAPIENS | 5 | 11647 | 2a1t | S | 174 | ARG | p.Arg174His | P38117 |
| HOMO SAPIENS | -1 | 11200 | 4tx5 | A | 4 | ILE | p.Ile59Val | Q9NR28 |
| HOMO SAPIENS | 0 | 11200 | 4tx5 | A | 169 | GLY | p.Gly224Arg | Q9NR28 |
| HOMO SAPIENS | 3 | 417 | 2vr2 | A | 6 | ARG | p.Arg6Gln | Q14117 |
| HOMO SAPIENS | 1 | 417 | 2vr2 | A | 334 | GLN | p.Gln334Arg | Q14117 |
| HOMO SAPIENS | 2 | 417 | 2vr2 | A | 360 | TRP | p.Trp360Arg | Q14117 |
| HOMO SAPIENS | 0 | 417 | 2vr2 | A | 379 | SER | p.Ser379Arg | Q14117 |
| HOMO SAPIENS | 0 | 417 | 2vr2 | A | 435 | GLY | p.Gly435Arg | Q14117 |
| HOMO SAPIENS | 0 | 12498 | 3eo3 | B | 423 | ILE | p.Ile423Val | Q9Y223 |
| HOMO SAPIENS | 5 | 12498 | 3eo3 | B | 572 | VAL | p.Val572Leu | Q9Y223 |
| HOMO SAPIENS | 0 | 12498 | 3eo3 | B | 576 | GLY | p.Gly576Glu | Q9Y223 |
| HOMO SAPIENS | 0 | 12498 | 3eo3 | B | 587 | ILE | p.Ile587Thr | Q9Y223 |
| HOMO SAPIENS | 0 | 12498 | 3eo3 | B | 615 | SER | p.Ser615Ter | Q9Y223 |
| HOMO SAPIENS | 1 | 12498 | 3eo3 | B | 631 | ALA | p.Ala631Val | Q9Y223 |
| HOMO SAPIENS | 1 | 12498 | 3eo3 | B | 669 | GLY | p.Gly669Arg | Q9Y223 |
| HOMO SAPIENS | 1 | 12498 | 3eo3 | B | 696 | VAL | p.Val696Met | Q9Y223 |
| HOMO SAPIENS | 2 | 12498 | 3eo3 | B | 712 | MET | p.Met712Thr | Q9Y223 |
| HOMO SAPIENS | 0 | 1568 | 1r74 | A | 49 | LEU | p.Leu50Pro | Q14749 |
| HOMO SAPIENS | 0 | 1568 | 1r74 | A | 176 | HIS | p.His177Asn | Q14749 |
| HOMO SAPIENS | 3 | 147 | 5eig | A | 67 | THR | p.Thr67Ile | P32929 |
| HOMO SAPIENS | 3 | 147 | 5eig | A | 240 | GLN | p.Gln240Glu | P32929 |
| HOMO SAPIENS | -1 | 147 | 5eig | A | 403 | SER | p.Ser403Ile | P32929 |
| HOMO SAPIENS | 8 | 20036 | 2lsr | A | 31 | ARG | p.Arg31Ter | Q9Y6N9 |
| HOMO SAPIENS | -1 | 41382 | 4uis | B | 431 | UNK | p.Ala431Glu | P49768 |
| HOMO SAPIENS | -1 | 8536 | 3i6w | B | 85 | PRO | p.Pro85Leu | O96017 |
| HOMO SAPIENS | 0 | 8536 | 3i6w | B | 93 | TRP | p.Trp93Ter | O96017 |
| HOMO SAPIENS | 0 | 8536 | 3i6w | B | 95 | ARG | p.Arg95Ter | O96017 |
| HOMO SAPIENS | 0 | 8536 | 3i6w | B | 117 | ARG | p.Arg117Gly | O96017 |
| HOMO SAPIENS | 0 | 8536 | 3i6w | B | 119 | LYS | p.Lys119Arg | O96017 |
| HOMO SAPIENS | 0 | 8536 | 3i6w | B | 137 | ARG | p.Arg137Gln | O96017 |
| HOMO SAPIENS | 0 | 8536 | 3i6w | B | 137 | ARG | p.Arg137Ter | O96017 |
| HOMO SAPIENS | 0 | 8536 | 3i6w | B | 145 | ARG | p.Arg145Trp | O96017 |
| HOMO SAPIENS | 0 | 8536 | 3i6w | B | 160 | ILE | p.Ile160Met | O96017 |
| HOMO SAPIENS | 0 | 41322 | 4bxl | C | 46 | GLU | p.Glu46Lys | P37840 |
| HOMO SAPIENS | 5 | 41322 | 4bxl | C | 53 | ALA | p.Ala53Thr | P37840 |
| HOMO SAPIENS | 2 | 24966 | 3hve | A | 15 | ARG | p.Arg15Ser | Q9H2C0 |
| HOMO SAPIENS | 1 | 24966 | 3hve | A | 138 | ARG | p.Arg138His | Q9H2C0 |
| HOMO SAPIENS | 0 | 24966 | 3hve | A | 201 | ARG | p.Arg201Ter | Q9H2C0 |
| HOMO SAPIENS | 0 | 11680 | 5an9 | J | 8 | ASN | p.Asn8Lys | Q9Y3A5 |
| HOMO SAPIENS | 1 | 11680 | 5an9 | J | 33 | LYS | p.Lys33Arg | Q9Y3A5 |

| | | | | | | | | |
|---|---|---|---|---|---|---|---|---|
| HOMO SAPIENS | 1 | 11680 | 5an9 | J | 126 | ARG | p.Arg126Thr | Q9Y3A5 |
| HOMO SAPIENS | 1 | 11680 | 5an9 | J | 130 | VAL | p.Val130Met | Q9Y3A5 |
| HOMO SAPIENS | 1 | 11680 | 5an9 | J | 169 | ARG | p.Arg169Cys | Q9Y3A5 |
| HOMO SAPIENS | 1 | 11680 | 5an9 | J | 218 | ARG | p.Arg218Ter | Q9Y3A5 |
| HOMO SAPIENS | 4 | 978 | 3zs0 | B | 7 | TYR | p.Tyr173Cys | P05164 |
| HOMO SAPIENS | 1 | 978 | 3zs0 | B | 85 | MET | p.Met251Thr | P05164 |
| HOMO SAPIENS | 4 | 5140 | 2l4r | A | 52 | CYS | p.Cys52Ter | Q12809 |
| HOMO SAPIENS | 4 | 5140 | 2l4r | A | 63 | PRO | p.Pro63His | Q12809 |
| HOMO SAPIENS | 0 | 5140 | 2l4r | A | 100 | ARG | p.Arg100Gly | Q12809 |
| HOMO SAPIENS | 4 | 5140 | 2l4r | A | 129 | PHE | p.Phe129Leu | Q12809 |
| HOMO SAPIENS | 0 | 25836 | 2gy7 | B | 103 | GLU | p.Glu103Asp | Q02763 |
| HOMO SAPIENS | 0 | 25836 | 2gy7 | B | 148 | ILE | p.Ile148Thr | Q02763 |
| HOMO SAPIENS | 0 | 25836 | 2gy7 | B | 226 | ALA | p.Ala226Val | Q02763 |
| HOMO SAPIENS | 0 | 25836 | 2gy7 | B | 391 | THR | p.Thr391Ile | Q02763 |
| HOMO SAPIENS | 4 | 12563 | 5kwy | D | 20 | GLU | p.Glu20Ter | P61916 |
| HOMO SAPIENS | 2 | 12563 | 5kwy | D | 30 | VAL | p.Val30Met | P61916 |
| HOMO SAPIENS | 4 | 12563 | 5kwy | D | 39 | VAL | p.Val39Met | P61916 |
| HOMO SAPIENS | 8 | 12563 | 5kwy | D | 67 | SER | p.Ser67Pro | P61916 |
| HOMO SAPIENS | 8 | 12563 | 5kwy | D | 99 | CYS | p.Cys99Arg | P61916 |
| HOMO SAPIENS | 2 | 12563 | 5kwy | D | 118 | GLU | p.Glu118Ter | P61916 |
| HOMO SAPIENS | 2 | 12563 | 5kwy | D | 120 | PRO | p.Pro120Ser | P61916 |
| HOMO SAPIENS | -1 | 1679 | 1pmk | A | -2 | VAL | p.Val374Phe | P00747 |
| HOMO SAPIENS | -1 | 5315 | 2w2n | P | 53 | ALA | p.Ala53Val | Q8NBP7 |
| HOMO SAPIENS | 8 | 13496 | 2yad | A | 138 | THR | p.Asn138Thr | P11686 |
| HOMO SAPIENS | -1 | 13496 | 2yad | A | 173 | ARG | p.Arg167Gln | P11686 |
| HOMO SAPIENS | 2 | 13496 | 2yad | A | 186 | SER | p.Asn186Ser | P11686 |
| HOMO SAPIENS | 0 | 4426 | 3qd6 | C | 123 | ALA | p.Ala123Glu | P29965 |
| HOMO SAPIENS | 1 | 4426 | 3qd6 | C | 140 | TRP | p.Trp140Gly | P29965 |
| HOMO SAPIENS | 0 | 4426 | 3qd6 | C | 155 | LEU | p.Leu155Pro | P29965 |
| HOMO SAPIENS | 6 | 4426 | 3qd6 | C | 227 | GLY | p.Gly227Val | P29965 |
| HOMO SAPIENS | 0 | 4426 | 3qd6 | C | 235 | ALA | p.Ala235Pro | P29965 |
| HOMO SAPIENS | 0 | 10734 | 3dyd | B | 45 | PRO | p.Pro45Ser | P17735 |
| HOMO SAPIENS | -1 | 10734 | 3dyd | B | 57 | ARG | p.Arg57Ter | P17735 |
| HOMO SAPIENS | 8 | 10734 | 3dyd | B | 417 | ARG | p.Arg417Ter | P17735 |
| HOMO SAPIENS | 0 | 6956 | 4pzs | A | 1600 | LEU | p.Leu1600Phe | Q09472 |
| HOMO SAPIENS | 0 | 9370 | 3hfv | A | 21 | SER | p.Ser57Cys | O95363 |
| HOMO SAPIENS | 1 | 9370 | 3hfv | A | 25 | ASP | p.Asp61Glu | O95363 |
| HOMO SAPIENS | 1 | 9370 | 3hfv | A | 130 | ALA | p.Ala166Thr | O95363 |
| HOMO SAPIENS | 5 | 9370 | 3hfv | A | 148 | ASP | p.Asp184Asn | O95363 |
| HOMO SAPIENS | 0 | 9370 | 3hfv | A | 187 | ARG | p.Arg223Cys | O95363 |
| HOMO SAPIENS | 0 | 9370 | 3hfv | A | 210 | THR | p.Thr246Met | O95363 |
| HOMO SAPIENS | 1 | 9370 | 3hfv | A | 244 | ASN | p.Asn280Ser | O95363 |
| HOMO SAPIENS | 2 | 9370 | 3hfv | A | 288 | TYR | p.Tyr324Cys | O95363 |
| HOMO SAPIENS | 5 | 9370 | 3hfv | A | 293 | ILE | p.Ile329Thr | O95363 |
| HOMO SAPIENS | 1 | 9370 | 3hfv | A | 355 | ASP | p.Asp391Val | O95363 |
| HOMO SAPIENS | 1 | 9370 | 3hfv | A | 371 | THR | p.Thr407Met | O95363 |
| HOMO SAPIENS | 1 | 9370 | 3hfv | A | 383 | ARG | p.Arg419Cys | O95363 |
| HOMO SAPIENS | 1 | 9370 | 3hfv | A | 387 | ARG | p.Arg423Gln | O95363 |
| HOMO SAPIENS | 0 | 14401 | 1q3x | A | 371 | ASP | p.Asp371Tyr | O00187 |
| HOMO SAPIENS | 0 | 14401 | 1q3x | A | 377 | VAL | p.Val377Ala | O00187 |
| HOMO SAPIENS | 0 | 14401 | 1q3x | A | 439 | ARG | p.Arg439His | O00187 |
| HOMO SAPIENS | 0 | 14401 | 1q3x | A | 577 | GLN | p.Gln577His | O00187 |
| HOMO SAPIENS | 2 | 18270 | 3teq | A | 429 | ARG | p.Arg429Cys | Q13586 |
| HOMO SAPIENS | -1 | 16081 | 4mqt | A | 231 | VAL | p.Val231Ile | P08172 |

| | | | | | | | | |
|---|---|---|---|---|---|---|---|---|
| HOMO SAPIENS | 9 | 6607 | 3whu | B | 81 | ASP | p.Asp81His | Q8NI22 |
| HOMO SAPIENS | 9 | 6607 | 3whu | B | 129 | ASP | p.Asp129Glu | Q8NI22 |
| HOMO SAPIENS | 9 | 6607 | 3whu | B | 136 | ILE | p.Ile136Thr | Q8NI22 |
| HOMO SAPIENS | 0 | 7208 | 2e9w | D | 11 | ASN | p.Asn36Ser | P21583 |
| HOMO SAPIENS | 0 | 24566 | 4bdx | A | 10 | LEU | p.Leu140Val | P00748 |
| HOMO SAPIENS | 5 | 24566 | 4bdx | A | 77 | PRO | p.Ala207Pro | P00748 |
| HOMO SAPIENS | 7 | 606 | 3qnf | A | 319 | MET | p.Met319Val | Q9NZ08 |
| HOMO SAPIENS | 8 | 4097 | 1ydl | A | 21 | LEU | p.Leu21Pro | Q6ZYL4 |
| HOMO SAPIENS | 6 | 4097 | 1ydl | A | 56 | ARG | p.Arg56Ter | Q6ZYL4 |
| HOMO SAPIENS | 8 | 36691 | 1dqb | A | 42 | ARG | p.Arg403Lys | P07204 |
| HOMO SAPIENS | 0 | 5642 | 3j7y | X | 59 | ARG | p.Arg59Pro | Q13084 |
| HOMO SAPIENS | 1 | 5642 | 3j7y | X | 204 | VAL | p.Val204Met | Q13084 |
| HOMO SAPIENS | 1 | 1119 | 3nwu | B | 252 | ALA | p.Ala252Thr | Q92743 |
| HOMO SAPIENS | 3 | 1119 | 3nwu | B | 274 | ARG | p.Arg274Gln | Q92743 |
| HOMO SAPIENS | 9 | 1119 | 3nwu | B | 297 | VAL | p.Val297Met | Q92743 |
| HOMO SAPIENS | -1 | 1119 | 3nwu | B | 302 | ARG | p.Arg302Ter | Q92743 |
| HOMO SAPIENS | 9 | 1119 | 3nwu | B | 321 | ALA | p.Ala321Thr | Q92743 |
| HOMO SAPIENS | -1 | 1119 | 3nwu | B | 370 | ARG | p.Arg370Ter | Q92743 |
| HOMO SAPIENS | 0 | 416 | 4ycu | B | 105 | LEU | p.Leu105His | Q15046 |
| HOMO SAPIENS | 0 | 416 | 4ycu | B | 145 | TYR | p.Tyr145His | Q15046 |
| HOMO SAPIENS | 0 | 416 | 4ycu | B | 200 | PRO | p.Pro200Leu | Q15046 |
| HOMO SAPIENS | 0 | 416 | 4ycu | B | 201 | TYR | p.Tyr201His | Q15046 |
| HOMO SAPIENS | 0 | 416 | 4ycu | B | 420 | ARG | p.Arg420Cys | Q15046 |
| HOMO SAPIENS | 9 | 3297 | 4fpb | A | 680 | THR | p.Thr680Ser | P19367 |
| HOMO SAPIENS | 0 | 6771 | 2zt6 | A | 201 | ALA | p.Ala255Val | P41250 |
| HOMO SAPIENS | 0 | 6771 | 2zt6 | A | 202 | ASP | p.Asp256His | P41250 |
| HOMO SAPIENS | 0 | 6771 | 2zt6 | A | 334 | ARG | p.Arg388Gln | P41250 |
| HOMO SAPIENS | -1 | 6771 | 2zt6 | A | 439 | ASN | p.Asn493Ser | P41250 |
| HOMO SAPIENS | -1 | 6771 | 2zt6 | A | 500 | ASP | p.Asp554Asn | P41250 |
| HOMO SAPIENS | 1 | 6771 | 2zt6 | A | 581 | SER | p.Ser635Leu | P41250 |
| HOMO SAPIENS | -1 | 6771 | 2zt6 | A | 684 | GLU | p.Glu738Lys | P41250 |
| HOMO SAPIENS | 0 | 2687 | 4grw | B | 276 | VAL | p.Val298Phe | P29460 |
| HOMO SAPIENS | 6 | 913 | 1m75 | B | 45 | ASP | p.Asp57Glu | Q16836 |
| HOMO SAPIENS | 2 | 913 | 1m75 | B | 74 | PRO | p.Leu86Pro | Q16836 |
| HOMO SAPIENS | 3 | 913 | 1m75 | B | 214 | TYR | p.Tyr226His | Q16836 |
| HOMO SAPIENS | 3 | 913 | 1m75 | B | 224 | ARG | p.Arg236Ter | Q16836 |
| HOMO SAPIENS | 4 | 913 | 1m75 | B | 246 | PRO | p.Pro258Leu | Q16836 |
| HOMO SAPIENS | 2 | 4713 | 4djn | B | 40 | ARG | p.Arg40Cys | Q7LG56 |
| HOMO SAPIENS | 1 | 4713 | 4djn | B | 41 | ARG | p.Arg41Pro | Q7LG56 |
| HOMO SAPIENS | 2 | 4713 | 4djn | B | 110 | ARG | p.Arg110His | Q7LG56 |
| HOMO SAPIENS | 2 | 4713 | 4djn | B | 121 | ARG | p.Arg121His | Q7LG56 |
| HOMO SAPIENS | 9 | 4713 | 4djn | B | 131 | GLU | p.Glu131Lys | Q7LG56 |
| HOMO SAPIENS | 2 | 4713 | 4djn | B | 182 | THR | p.Thr182Ala | Q7LG56 |
| HOMO SAPIENS | 6 | 4713 | 4djn | B | 194 | GLU | p.Glu194Lys | Q7LG56 |
| HOMO SAPIENS | 2 | 4713 | 4djn | B | 211 | ARG | p.Arg211Lys | Q7LG56 |
| HOMO SAPIENS | 0 | 4713 | 4djn | B | 224 | ILE | p.Ile224Ser | Q7LG56 |
| HOMO SAPIENS | 3 | 4713 | 4djn | B | 229 | GLY | p.Gly229Val | Q7LG56 |
| HOMO SAPIENS | 2 | 4713 | 4djn | B | 236 | CYS | p.Cys236Phe | Q7LG56 |
| HOMO SAPIENS | 0 | 4713 | 4djn | B | 273 | GLY | p.Gly273Ser | Q7LG56 |
| HOMO SAPIENS | -1 | 4713 | 4djn | B | 282 | MSE | p.Met282Ile | Q7LG56 |
| HOMO SAPIENS | 3 | 4713 | 4djn | B | 284 | GLN | p.Gln284Ter | Q7LG56 |
| HOMO SAPIENS | 3 | 4713 | 4djn | B | 305 | ALA | p.Ala305Ser | Q7LG56 |
| HOMO SAPIENS | -1 | 8974 | 5n7i | A | 500 | ARG | p.Arg500Ter | Q8N4S9 |
| HOMO SAPIENS | 0 | 17072 | 3fxi | B | 299 | ASP | p.Asp299Gly | O00206 |

| | | | | | | | | |
|---|---|---|---|---|---|---|---|---|
| HOMO SAPIENS | 1 | 14355 | 2wwk | T | 55 | ILE | p.Ile34306Asn | Q8WZ42 |
| HOMO SAPIENS | 0 | 14355 | 2wwk | T | 64 | LEU | p.Leu34315Pro | Q8WZ42 |
| HOMO SAPIENS | 2 | 17797 | 1xu1 | S | 72 | ARG | p.Arg72His | O14836 |
| HOMO SAPIENS | -1 | 10836 | 4ztu | A | 39 | GLN | p.Gln39Arg | P54098 |
| HOMO SAPIENS | -1 | 10836 | 4ztu | A | 42 | ARG | p.Arg42Gln | P54098 |
| HOMO SAPIENS | -1 | 10836 | 4ztu | A | 45 | GLN | p.Gln45Arg | P54098 |
| HOMO SAPIENS | -1 | 10836 | 4ztu | A | 52 | GLN | p.Gln52Glu | P54098 |
| HOMO SAPIENS | 0 | 10836 | 4ztu | A | 88 | PHE | p.Phe88Leu | P54098 |
| HOMO SAPIENS | 1 | 10836 | 4ztu | A | 194 | ALA | p.Ala194Val | P54098 |
| HOMO SAPIENS | 1 | 10836 | 4ztu | A | 227 | ARG | p.Arg227Trp | P54098 |
| HOMO SAPIENS | 0 | 10836 | 4ztu | A | 232 | ARG | p.Arg232His | P54098 |
| HOMO SAPIENS | 1 | 10836 | 4ztu | A | 285 | GLN | p.Gln285His | P54098 |
| HOMO SAPIENS | 1 | 10836 | 4ztu | A | 434 | TYR | p.Tyr434His | P54098 |
| HOMO SAPIENS | -1 | 10836 | 4ztu | A | 511 | SER | p.Ser511Asn | P54098 |
| HOMO SAPIENS | 0 | 10836 | 4ztu | A | 533 | CYS | p.Cys533Trp | P54098 |
| HOMO SAPIENS | 1 | 10836 | 4ztu | A | 546 | ARG | p.Arg546Cys | P54098 |
| HOMO SAPIENS | 1 | 10836 | 4ztu | A | 587 | PRO | p.Pro587Leu | P54098 |
| HOMO SAPIENS | 0 | 10836 | 4ztu | A | 603 | MET | p.Met603Thr | P54098 |
| HOMO SAPIENS | 0 | 10836 | 4ztu | A | 623 | LEU | p.Leu623Trp | P54098 |
| HOMO SAPIENS | -1 | 10836 | 4ztu | A | 627 | ARG | p.Arg627Trp | P54098 |
| HOMO SAPIENS | -1 | 10836 | 4ztu | A | 628 | ARG | p.Arg628Pro | P54098 |
| HOMO SAPIENS | 0 | 10836 | 4ztu | A | 635 | PRO | p.Pro635Leu | P54098 |
| HOMO SAPIENS | 0 | 10836 | 4ztu | A | 662 | GLU | p.Glu662Lys | P54098 |
| HOMO SAPIENS | -1 | 10836 | 4ztu | A | 674 | GLY | p.Gly674Asp | P54098 |
| HOMO SAPIENS | -1 | 10836 | 4ztu | A | 722 | ARG | p.Arg722His | P54098 |
| HOMO SAPIENS | -1 | 10836 | 4ztu | A | 737 | GLY | p.Gly737Arg | P54098 |
| HOMO SAPIENS | 0 | 10836 | 4ztu | A | 748 | TRP | p.Trp748Ser | P54098 |
| HOMO SAPIENS | 0 | 10836 | 4ztu | A | 749 | PHE | p.Phe749Ser | P54098 |
| HOMO SAPIENS | 1 | 10836 | 4ztu | A | 807 | ARG | p.Arg807Cys | P54098 |
| HOMO SAPIENS | 1 | 10836 | 4ztu | A | 831 | TYR | p.Tyr831Cys | P54098 |
| HOMO SAPIENS | 8 | 10836 | 4ztu | A | 848 | GLY | p.Gly848Ser | P54098 |
| HOMO SAPIENS | 1 | 10836 | 4ztu | A | 852 | ARG | p.Arg852Cys | P54098 |
| HOMO SAPIENS | 9 | 10836 | 4ztu | A | 853 | ARG | p.Arg853Trp | P54098 |
| HOMO SAPIENS | 1 | 10836 | 4ztu | A | 864 | ASN | p.Asn864Ser | P54098 |
| HOMO SAPIENS | 1 | 10836 | 4ztu | A | 879 | GLN | p.Gln879Arg | P54098 |
| HOMO SAPIENS | 1 | 10836 | 4ztu | A | 914 | THR | p.Thr914Pro | P54098 |
| HOMO SAPIENS | 6 | 10836 | 4ztu | A | 932 | HIS | p.His932Tyr | P54098 |
| HOMO SAPIENS | 2 | 10836 | 4ztu | A | 944 | GLU | p.Glu944Lys | P54098 |
| HOMO SAPIENS | 6 | 10836 | 4ztu | A | 955 | TYR | p.Tyr955Cys | P54098 |
| HOMO SAPIENS | 1 | 10836 | 4ztu | A | 957 | ALA | p.Ala957Ser | P54098 |
| HOMO SAPIENS | 1 | 10836 | 4ztu | A | 957 | ALA | p.Ala957Val | P54098 |
| HOMO SAPIENS | 1 | 10836 | 4ztu | A | 964 | ARG | p.Arg964Cys | P54098 |
| HOMO SAPIENS | 1 | 10836 | 4ztu | A | 966 | LEU | p.Leu966Arg | P54098 |
| HOMO SAPIENS | 0 | 10836 | 4ztu | A | 1051 | GLY | p.Gly1051Arg | P54098 |
| HOMO SAPIENS | 1 | 10836 | 4ztu | A | 1073 | PRO | p.Pro1073Leu | P54098 |
| HOMO SAPIENS | 1 | 10836 | 4ztu | A | 1096 | ARG | p.Arg1096Cys | P54098 |
| HOMO SAPIENS | 1 | 10836 | 4ztu | A | 1096 | ARG | p.Arg1096His | P54098 |
| HOMO SAPIENS | 0 | 10836 | 4ztu | A | 1116 | MET | p.Met1116Thr | P54098 |
| HOMO SAPIENS | 1 | 10836 | 4ztu | A | 1142 | ARG | p.Arg1142Trp | P54098 |
| HOMO SAPIENS | 1 | 10836 | 4ztu | A | 1146 | ARG | p.Arg1146Cys | P54098 |
| HOMO SAPIENS | 1 | 10836 | 4ztu | A | 1157 | ASN | p.Asn1157Ser | P54098 |
| HOMO SAPIENS | -1 | 10836 | 4ztu | A | 1234 | ARG | p.Arg1234Gln | P54098 |
| HOMO SAPIENS | -1 | 10836 | 4ztu | A | 1236 | GLN | p.Gln1236His | P54098 |
| HOMO SAPIENS | 5 | 2666 | 1jph | A | 62 | PRO | p.Pro62Leu | P06132 |

| | | | | | | | | |
|---|---|---|---|---|---|---|---|---|
| HOMO SAPIENS | 5 | 2666 | 1jph | A | 116 | GLN | p.Gln116Ter | P06132 |
| HOMO SAPIENS | 0 | 2666 | 1jph | A | 165 | MET | p.Met165Arg | P06132 |
| HOMO SAPIENS | 6 | 2666 | 1jph | A | 167 | GLU | p.Glu167Lys | P06132 |
| HOMO SAPIENS | 0 | 2666 | 1jph | A | 195 | LEU | p.Leu195Phe | P06132 |
| HOMO SAPIENS | 7 | 2666 | 1jph | A | 253 | LEU | p.Leu253Gln | P06132 |
| HOMO SAPIENS | 0 | 2666 | 1jph | A | 281 | GLY | p.Gly281Glu | P06132 |
| HOMO SAPIENS | 9 | 2666 | 1jph | A | 304 | ASN | p.Asn304Lys | P06132 |
| HOMO SAPIENS | 8 | 2666 | 1jph | A | 311 | TYR | p.Tyr311Cys | P06132 |
| HOMO SAPIENS | 5 | 2666 | 1jph | A | 332 | ARG | p.Arg332His | P06132 |
| HOMO SAPIENS | 1 | 889 | 1m51 | A | 184 | LEU | p.Val184Leu | P35558 |
| HOMO SAPIENS | 0 | 889 | 1m51 | A | 267 | VAL | p.Ile267Val | P35558 |
| HOMO SAPIENS | 2 | 889 | 1m51 | A | 276 | GLU | p.Glu276Lys | P35558 |
| HOMO SAPIENS | 6 | 889 | 1m51 | A | 368 | VAL | p.Val368Ile | P35558 |
| HOMO SAPIENS | 0 | 889 | 1m51 | A | 579 | SER | p.Ser579Arg | P35558 |
| HOMO SAPIENS | 1 | 4084 | 2gjx | A | 28 | GLN | p.Gln28Ter | P06865 |
| HOMO SAPIENS | 0 | 4084 | 2gjx | A | 39 | LEU | p.Leu39Arg | P06865 |
| HOMO SAPIENS | 0 | 4084 | 2gjx | A | 127 | LEU | p.Leu127Arg | P06865 |
| HOMO SAPIENS | 6 | 4084 | 2gjx | A | 137 | ARG | p.Arg137Ter | P06865 |
| HOMO SAPIENS | 6 | 4084 | 2gjx | A | 170 | ARG | p.Arg170Gln | P06865 |
| HOMO SAPIENS | 9 | 4084 | 2gjx | A | 178 | ARG | p.Arg178Cys | P06865 |
| HOMO SAPIENS | 9 | 4084 | 2gjx | A | 178 | ARG | p.Arg178His | P06865 |
| HOMO SAPIENS | 5 | 4084 | 2gjx | A | 180 | TYR | p.Tyr180His | P06865 |
| HOMO SAPIENS | 5 | 4084 | 2gjx | A | 180 | TYR | p.Tyr180Ter | P06865 |
| HOMO SAPIENS | 1 | 4084 | 2gjx | A | 197 | LYS | p.Lys197Thr | P06865 |
| HOMO SAPIENS | 0 | 4084 | 2gjx | A | 200 | VAL | p.Val200Met | P06865 |
| HOMO SAPIENS | 9 | 4084 | 2gjx | A | 204 | HIS | p.His204Arg | P06865 |
| HOMO SAPIENS | 5 | 4084 | 2gjx | A | 210 | SER | p.Ser210Phe | P06865 |
| HOMO SAPIENS | 0 | 4084 | 2gjx | A | 211 | PHE | p.Phe211Ser | P06865 |
| HOMO SAPIENS | 2 | 4084 | 2gjx | A | 250 | GLY | p.Gly250Asp | P06865 |
| HOMO SAPIENS | 9 | 4084 | 2gjx | A | 258 | ASP | p.Asp258His | P06865 |
| HOMO SAPIENS | 0 | 4084 | 2gjx | A | 269 | GLY | p.Gly269Ser | P06865 |
| HOMO SAPIENS | 0 | 4084 | 2gjx | A | 301 | MET | p.Met301Arg | P06865 |
| HOMO SAPIENS | 3 | 4084 | 2gjx | A | 393 | ARG | p.Arg393Ter | P06865 |
| HOMO SAPIENS | 3 | 4084 | 2gjx | A | 393 | ARG | p.Arg393Pro | P06865 |
| HOMO SAPIENS | 0 | 4084 | 2gjx | A | 399 | ASN | p.Asn399Asp | P06865 |
| HOMO SAPIENS | 2 | 4084 | 2gjx | A | 420 | TRP | p.Trp420Cys | P06865 |
| HOMO SAPIENS | 1 | 4084 | 2gjx | A | 434 | PHE | p.Phe434Leu | P06865 |
| HOMO SAPIENS | 0 | 4084 | 2gjx | A | 454 | GLY | p.Gly454Ser | P06865 |
| HOMO SAPIENS | 1 | 4084 | 2gjx | A | 474 | TRP | p.Trp474Cys | P06865 |
| HOMO SAPIENS | 0 | 4084 | 2gjx | A | 479 | ALA | p.Ala479Thr | P06865 |
| HOMO SAPIENS | 5 | 4084 | 2gjx | A | 485 | TRP | p.Trp485Arg | P06865 |
| HOMO SAPIENS | 4 | 4084 | 2gjx | A | 499 | ARG | p.Arg499Cys | P06865 |
| HOMO SAPIENS | 4 | 4084 | 2gjx | A | 499 | ARG | p.Arg499His | P06865 |
| HOMO SAPIENS | 2 | 4084 | 2gjx | A | 504 | ARG | p.Arg504Cys | P06865 |
| HOMO SAPIENS | 2 | 4084 | 2gjx | A | 504 | ARG | p.Arg504His | P06865 |
| HOMO SAPIENS | -1 | 1943 | 1xmj | A | 417 | ASN | p.Asn417Lys | P13569 |
| HOMO SAPIENS | -1 | 1943 | 1xmj | A | 422 | SER | p.Ser422Phe | P13569 |
| HOMO SAPIENS | 1 | 1943 | 1xmj | A | 438 | THR | p.Thr438Ala | P13569 |
| HOMO SAPIENS | 0 | 1943 | 1xmj | A | 455 | ALA | p.Ala455Glu | P13569 |
| HOMO SAPIENS | 4 | 1943 | 1xmj | A | 456 | VAL | p.Val456Ala | P13569 |
| HOMO SAPIENS | 9 | 1943 | 1xmj | A | 458 | GLY | p.Gly458Val | P13569 |
| HOMO SAPIENS | 8 | 1943 | 1xmj | A | 466 | SER | p.Ser466Ter | P13569 |
| HOMO SAPIENS | 0 | 1943 | 1xmj | A | 467 | LEU | p.Leu467Pro | P13569 |
| HOMO SAPIENS | 1 | 1943 | 1xmj | A | 470 | MET | p.Val470Met | P13569 |

| | | | | | | | | |
|---|---|---|---|---|---|---|---|---|
| HOMO SAPIENS | 5 | 1943 | 1xmj | A | 480 | GLY | p.Gly480Cys | P13569 |
| HOMO SAPIENS | 3 | 1943 | 1xmj | A | 492 | SER | p.Ser492Phe | P13569 |
| HOMO SAPIENS | 8 | 1943 | 1xmj | A | 493 | GLN | p.Gln493Ter | P13569 |
| HOMO SAPIENS | 5 | 1943 | 1xmj | A | 506 | ILE | p.Ile506Met | P13569 |
| HOMO SAPIENS | 3 | 1943 | 1xmj | A | 520 | VAL | p.Val520Phe | P13569 |
| HOMO SAPIENS | 1 | 1943 | 1xmj | A | 528 | GLU | p.Glu528Asp | P13569 |
| HOMO SAPIENS | 1 | 1943 | 1xmj | A | 534 | ALA | p.Ala534Glu | P13569 |
| HOMO SAPIENS | 4 | 1943 | 1xmj | A | 542 | GLY | p.Gly542Ter | P13569 |
| HOMO SAPIENS | 6 | 1943 | 1xmj | A | 549 | SER | p.Ser549Asn | P13569 |
| HOMO SAPIENS | 6 | 1943 | 1xmj | A | 549 | SER | p.Ser549Arg | P13569 |
| HOMO SAPIENS | 6 | 1943 | 1xmj | A | 549 | SER | p.Ser549Arg | P13569 |
| HOMO SAPIENS | 6 | 1943 | 1xmj | A | 551 | GLY | p.Gly551Asp | P13569 |
| HOMO SAPIENS | 6 | 1943 | 1xmj | A | 551 | GLY | p.Gly551Ser | P13569 |
| HOMO SAPIENS | 6 | 1943 | 1xmj | A | 553 | GLN | p.Arg553Gln | P13569 |
| HOMO SAPIENS | 6 | 1943 | 1xmj | A | 553 | GLN | p.Arg553Ter | P13569 |
| HOMO SAPIENS | 0 | 1943 | 1xmj | A | 559 | ALA | p.Ala559Thr | P13569 |
| HOMO SAPIENS | 7 | 1943 | 1xmj | A | 560 | ARG | p.Arg560Lys | P13569 |
| HOMO SAPIENS | 4 | 1943 | 1xmj | A | 562 | VAL | p.Val562Ile | P13569 |
| HOMO SAPIENS | 3 | 1943 | 1xmj | A | 563 | TYR | p.Tyr563Asn | P13569 |
| HOMO SAPIENS | 3 | 1943 | 1xmj | A | 569 | TYR | p.Tyr569Asp | P13569 |
| HOMO SAPIENS | 3 | 1943 | 1xmj | A | 585 | GLU | p.Glu585Ter | P13569 |
| HOMO SAPIENS | 1 | 1943 | 1xmj | A | 618 | ILE | p.Ile618Thr | P13569 |
| HOMO SAPIENS | 9 | 1943 | 1xmj | A | 622 | GLY | p.Gly622Asp | P13569 |
| HOMO SAPIENS | -1 | 1943 | 1xmj | A | 648 | ASP | p.Asp648Val | P13569 |
| HOMO SAPIENS | 0 | 1943 | 1xmj | A | 673 | GLY | p.Gly673Ter | P13569 |
| HOMO SAPIENS | 5 | 5111 | 4rec | A | 749 | ARG | p.Arg749Ter | Q9Y2M0 |
| HOMO SAPIENS | -1 | 4580 | 4l0d | B | 2 | GLY | p.Pro2Leu | P35520 |
| HOMO SAPIENS | -1 | 4580 | 4l0d | B | 12 | PRO | p.Pro12Ser | P35520 |
| HOMO SAPIENS | -1 | 4580 | 4l0d | B | 18 | ARG | p.Arg18Cys | P35520 |
| HOMO SAPIENS | 1 | 4580 | 4l0d | B | 49 | PRO | p.Pro49Leu | P35520 |
| HOMO SAPIENS | 2 | 4580 | 4l0d | B | 70 | PRO | p.Pro70Leu | P35520 |
| HOMO SAPIENS | 1 | 4580 | 4l0d | B | 102 | LYS | p.Lys102Gln | P35520 |
| HOMO SAPIENS | 7 | 4580 | 4l0d | B | 114 | ALA | p.Ala114Val | P35520 |
| HOMO SAPIENS | 5 | 4580 | 4l0d | B | 116 | GLY | p.Gly116Arg | P35520 |
| HOMO SAPIENS | 8 | 4580 | 4l0d | B | 121 | ARG | p.Arg121Cys | P35520 |
| HOMO SAPIENS | 8 | 4580 | 4l0d | B | 121 | ARG | p.Arg121His | P35520 |
| HOMO SAPIENS | 5 | 4580 | 4l0d | B | 125 | ARG | p.Arg125Gln | P35520 |
| HOMO SAPIENS | 4 | 4580 | 4l0d | B | 134 | GLY | p.Gly134Arg | P35520 |
| HOMO SAPIENS | 4 | 4580 | 4l0d | B | 139 | GLY | p.Gly139Arg | P35520 |
| HOMO SAPIENS | 0 | 4580 | 4l0d | B | 145 | PRO | p.Pro145Leu | P35520 |
| HOMO SAPIENS | 0 | 4580 | 4l0d | B | 148 | GLY | p.Gly148Arg | P35520 |
| HOMO SAPIENS | 0 | 4580 | 4l0d | B | 168 | VAL | p.Val168Met | P35520 |
| HOMO SAPIENS | 2 | 4580 | 4l0d | B | 191 | THR | p.Thr191Met | P35520 |
| HOMO SAPIENS | 5 | 4580 | 4l0d | B | 234 | ASP | p.Asp234Asn | P35520 |
| HOMO SAPIENS | 0 | 4580 | 4l0d | B | 257 | THR | p.Thr257Met | P35520 |
| HOMO SAPIENS | 4 | 4580 | 4l0d | B | 262 | THR | p.Thr262Met | P35520 |
| HOMO SAPIENS | 0 | 4580 | 4l0d | B | 278 | ILE | p.Ile278Thr | P35520 |
| HOMO SAPIENS | 2 | 4580 | 4l0d | B | 307 | GLY | p.Gly307Ser | P35520 |
| HOMO SAPIENS | 4 | 4580 | 4l0d | B | 320 | VAL | p.Val320Ala | P35520 |
| HOMO SAPIENS | 2 | 4580 | 4l0d | B | 336 | ARG | p.Arg336Cys | P35520 |
| HOMO SAPIENS | 2 | 4580 | 4l0d | B | 336 | ARG | p.Arg336His | P35520 |
| HOMO SAPIENS | 0 | 4580 | 4l0d | B | 351 | GLY | p.Gly351Arg | P35520 |
| HOMO SAPIENS | 0 | 4580 | 4l0d | B | 353 | THR | p.Thr353Met | P35520 |
| HOMO SAPIENS | 0 | 4580 | 4l0d | B | 371 | VAL | p.Val371Met | P35520 |

| | | | | | | | | |
|---|---|---|---|---|---|---|---|---|
| HOMO SAPIENS | 2 | 4580 | 4l0d | B | 379 | ARG | p.Arg379Gln | P35520 |
| HOMO SAPIENS | 2 | 4580 | 4l0d | B | 379 | ARG | p.Arg379Trp | P35520 |
| HOMO SAPIENS | 2 | 4580 | 4l0d | B | 384 | LYS | p.Lys384Glu | P35520 |
| HOMO SAPIENS | 0 | 4580 | 4l0d | B | 422 | PRO | p.Pro422Leu | P35520 |
| HOMO SAPIENS | 0 | 4580 | 4l0d | B | 444 | ASP | p.Asp444Asn | P35520 |
| HOMO SAPIENS | 0 | 4580 | 4l0d | B | 466 | SER | p.Ser466Leu | P35520 |
| HOMO SAPIENS | 0 | 4580 | 4l0d | B | 539 | LEU | p.Leu539Ser | P35520 |
| HOMO SAPIENS | 0 | 4580 | 4l0d | B | 548 | ARG | p.Arg548Gln | P35520 |
| HOMO SAPIENS | 1 | 6536 | 3ulu | A | 412 | LEU | p.Leu412Phe | O15455 |
| HOMO SAPIENS | 1 | 28493 | 3ii6 | X | 814 | ARG | p.Arg814Ter | P49917 |
| HOMO SAPIENS | 1 | 28493 | 3ii6 | X | 857 | ALA | p.Ala857Thr | P49917 |
| HOMO SAPIENS | 9 | 3543 | 2q51 | A | 24 | GLU | p.Glu24Gly | P45381 |
| HOMO SAPIENS | 1 | 3543 | 2q51 | A | 27 | GLY | p.Gly27Arg | P45381 |
| HOMO SAPIENS | 9 | 3543 | 2q51 | A | 71 | ARG | p.Arg71His | P45381 |
| HOMO SAPIENS | -1 | 3543 | 2q51 | A | 82 | MSE | p.Met82Thr | P45381 |
| HOMO SAPIENS | 0 | 3543 | 2q51 | A | 143 | ILE | p.Ile143Phe | P45381 |
| HOMO SAPIENS | 1 | 3543 | 2q51 | A | 152 | CYS | p.Cys152Arg | P45381 |
| HOMO SAPIENS | 2 | 3543 | 2q51 | A | 218 | CYS | p.Cys218Ter | P45381 |
| HOMO SAPIENS | 1 | 3543 | 2q51 | A | 231 | TYR | p.Tyr231Cys | P45381 |
| HOMO SAPIENS | 1 | 3543 | 2q51 | A | 231 | TYR | p.Tyr231Ter | P45381 |
| HOMO SAPIENS | 2 | 3543 | 2q51 | A | 274 | GLY | p.Gly274Arg | P45381 |
| HOMO SAPIENS | 6 | 3543 | 2q51 | A | 285 | GLU | p.Glu285Ala | P45381 |
| HOMO SAPIENS | 5 | 3543 | 2q51 | A | 287 | ALA | p.Ala287Thr | P45381 |
| HOMO SAPIENS | 2 | 3543 | 2q51 | A | 305 | ALA | p.Ala305Glu | P45381 |
| HOMO SAPIENS | 2 | 24687 | 1hfi | A | 25 | SER | p.Ser890Ile | P08603 |
| HOMO SAPIENS | 0 | 1508 | 3wn5 | C | 45 | LEU | p.Leu66Arg | P08637 |
| HOMO SAPIENS | 0 | 1508 | 3wn5 | C | 155 | VAL | p.Phe176Val | P08637 |
| HOMO SAPIENS | 0 | 750 | 5tpa | A | 54 | VAL | p.Val452Met | Q12879 |
| HOMO SAPIENS | 9 | 750 | 5tpa | A | 120 | ARG | p.Arg518Cys | Q12879 |
| HOMO SAPIENS | 0 | 750 | 5tpa | A | 199 | ALA | p.Ala716Thr | Q12879 |
| HOMO SAPIENS | 4 | 1363 | 3hg2 | B | 40 | PRO | p.Pro40Ser | P06280 |
| HOMO SAPIENS | 3 | 1363 | 3hg2 | B | 65 | SER | p.Ser65Thr | P06280 |
| HOMO SAPIENS | 4 | 1363 | 3hg2 | B | 66 | GLU | p.Glu66Gln | P06280 |
| HOMO SAPIENS | 2 | 1363 | 3hg2 | B | 80 | GLY | p.Gly80Asp | P06280 |
| HOMO SAPIENS | 3 | 1363 | 3hg2 | B | 112 | ARG | p.Arg112Cys | P06280 |
| HOMO SAPIENS | 3 | 1363 | 3hg2 | B | 112 | ARG | p.Arg112His | P06280 |
| HOMO SAPIENS | 0 | 1363 | 3hg2 | B | 139 | ASN | p.Asn139Ser | P06280 |
| HOMO SAPIENS | 7 | 1363 | 3hg2 | B | 156 | ALA | p.Ala156Thr | P06280 |
| HOMO SAPIENS | 5 | 1363 | 3hg2 | B | 162 | TRP | p.Trp162Arg | P06280 |
| HOMO SAPIENS | 5 | 1363 | 3hg2 | B | 207 | TYR | p.Tyr207His | P06280 |
| HOMO SAPIENS | 0 | 1363 | 3hg2 | B | 215 | ASN | p.Asn215Ser | P06280 |
| HOMO SAPIENS | 0 | 1363 | 3hg2 | B | 264 | ASP | p.Asp264Val | P06280 |
| HOMO SAPIENS | 5 | 1363 | 3hg2 | B | 266 | ASP | p.Asp266Val | P06280 |
| HOMO SAPIENS | 2 | 1363 | 3hg2 | B | 269 | VAL | p.Val269Ala | P06280 |
| HOMO SAPIENS | 3 | 1363 | 3hg2 | B | 272 | ASN | p.Asn272Ser | P06280 |
| HOMO SAPIENS | 3 | 1363 | 3hg2 | B | 279 | GLN | p.Gln279Glu | P06280 |
| HOMO SAPIENS | 0 | 1363 | 3hg2 | B | 289 | ILE | p.Ile289Val | P06280 |
| HOMO SAPIENS | 0 | 1363 | 3hg2 | B | 296 | MET | p.Met296Val | P06280 |
| HOMO SAPIENS | 0 | 1363 | 3hg2 | B | 296 | MET | p.Met296Ile | P06280 |
| HOMO SAPIENS | 0 | 1363 | 3hg2 | B | 297 | SER | p.Ser297Phe | P06280 |
| HOMO SAPIENS | 2 | 1363 | 3hg2 | B | 301 | ARG | p.Arg301Gln | P06280 |
| HOMO SAPIENS | 3 | 1363 | 3hg2 | B | 327 | GLN | p.Gln327Lys | P06280 |
| HOMO SAPIENS | 0 | 1363 | 3hg2 | B | 328 | GLY | p.Gly328Arg | P06280 |
| HOMO SAPIENS | 0 | 1363 | 3hg2 | B | 328 | GLY | p.Gly328Ala | P06280 |

| | | | | | | | | |
|---|---|---|---|---|---|---|---|---|
| HOMO SAPIENS | 2 | 1363 | 3hg2 | B | 331 | LEU | p.Leu331Phe | P06280 |
| HOMO SAPIENS | 0 | 1363 | 3hg2 | B | 342 | ARG | p.Arg342Gln | P06280 |
| HOMO SAPIENS | 1 | 1363 | 3hg2 | B | 361 | GLY | p.Gly361Arg | P06280 |
| HOMO SAPIENS | 1 | 1363 | 3hg2 | B | 363 | ARG | p.Arg363His | P06280 |
| HOMO SAPIENS | 1 | 1363 | 3hg2 | B | 385 | THR | p.Thr385Ala | P06280 |
| HOMO SAPIENS | 1 | 1363 | 3hg2 | B | 399 | TRP | p.Trp399Ser | P06280 |
| HOMO SAPIENS | 1 | 1363 | 3hg2 | B | 410 | THR | p.Thr410Ala | P06280 |
| HOMO SAPIENS | 2 | 3904 | 3caa | B | 385 | MET | p.Met414Val | P01011 |
| HOMO SAPIENS | 0 | 558 | 5eub | A | 56 | VAL | p.Val56Ile | Q13126 |
| HOMO SAPIENS | 0 | 558 | 5eub | A | 189 | TRP | p.Trp189Leu | Q13126 |
| HOMO SAPIENS | 0 | 558 | 5eub | A | 218 | ALA | p.Ala218Gly | Q13126 |
| HOMO SAPIENS | 1 | 3865 | 1cb5 | B | 443 | ILE | p.Ile443Val | Q13867 |
| HOMO SAPIENS | 2 | 14329 | 4m92 | A | 24 | ILE | p.Ile65Val | Q13454 |
| HOMO SAPIENS | 1 | 7381 | 5kz5 | k | 50 | GLY | p.Gly50Glu | Q9H1K1 |
| HOMO SAPIENS | 0 | 1102 | 4fdh | E | 86 | PRO | p.Pro86Ala | P19099 |
| HOMO SAPIENS | 0 | 1102 | 4fdh | E | 118 | ALA | p.Ala118Thr | P19099 |
| HOMO SAPIENS | 0 | 1102 | 4fdh | E | 159 | PRO | p.Pro159Leu | P19099 |
| HOMO SAPIENS | 0 | 1102 | 4fdh | E | 173 | LYS | p.Lys173Arg | P19099 |
| HOMO SAPIENS | 0 | 1102 | 4fdh | E | 181 | ARG | p.Arg181Trp | P19099 |
| HOMO SAPIENS | 0 | 1102 | 4fdh | E | 185 | THR | p.Thr185Ile | P19099 |
| HOMO SAPIENS | 0 | 1102 | 4fdh | E | 198 | GLU | p.Glu198Asp | P19099 |
| HOMO SAPIENS | 0 | 1102 | 4fdh | E | 214 | HIS | p.His214Asp | P19099 |
| HOMO SAPIENS | 0 | 1102 | 4fdh | E | 251 | LYS | p.Lys251Arg | P19099 |
| HOMO SAPIENS | 0 | 1102 | 4fdh | E | 255 | GLU | p.Glu255Ter | P19099 |
| HOMO SAPIENS | 0 | 1102 | 4fdh | E | 281 | ASN | p.Asn281Ser | P19099 |
| HOMO SAPIENS | 0 | 1102 | 4fdh | E | 320 | ALA | p.Ala320Val | P19099 |
| HOMO SAPIENS | 0 | 1102 | 4fdh | E | 339 | ILE | p.Ile339Thr | P19099 |
| HOMO SAPIENS | 0 | 1102 | 4fdh | E | 341 | ARG | p.Arg341Cys | P19099 |
| HOMO SAPIENS | 2 | 1102 | 4fdh | E | 374 | ARG | p.Arg374Trp | P19099 |
| HOMO SAPIENS | 0 | 1102 | 4fdh | E | 386 | VAL | p.Val386Ala | P19099 |
| HOMO SAPIENS | 0 | 1102 | 4fdh | E | 390 | LEU | p.Leu390Phe | P19099 |
| HOMO SAPIENS | -1 | 1102 | 4fdh | E | 435 | GLY | p.Gly435Ser | P19099 |
| HOMO SAPIENS | 0 | 1102 | 4fdh | E | 461 | LEU | p.Leu461Pro | P19099 |
| HOMO SAPIENS | 0 | 1102 | 4fdh | E | 498 | THR | p.Thr498Ala | P19099 |
| HOMO SAPIENS | 6 | 10292 | 2of5 | F | 128 | GLY | p.Gly128Arg | P78560 |
| HOMO SAPIENS | 6 | 10292 | 2of5 | F | 170 | ARG | p.Arg170Cys | P78560 |
| HOMO SAPIENS | -1 | 18965 | 3ghg | A | 16 | ARG | p.Arg35Cys | P02671 |
| HOMO SAPIENS | -1 | 18965 | 3ghg | A | 16 | ARG | p.Arg35His | P02671 |
| HOMO SAPIENS | -1 | 18965 | 3ghg | A | 283 | PRO | p.Pro302Ala | P02671 |
| HOMO SAPIENS | -1 | 18965 | 3ghg | A | 312 | THR | p.Thr331Ala | P02671 |
| HOMO SAPIENS | -1 | 18965 | 3ghg | A | 381 | SER | p.Ser400Phe | P02671 |
| HOMO SAPIENS | -1 | 18965 | 3ghg | A | 463 | VAL | p.Val482Met | P02671 |
| HOMO SAPIENS | -1 | 18965 | 3ghg | A | 526 | GLU | p.Glu545Val | P02671 |
| HOMO SAPIENS | -1 | 18965 | 3ghg | A | 554 | ARG | p.Arg573Leu | P02671 |
| HOMO SAPIENS | 8 | 3023 | 4ttb | A | 101 | ARG | p.Arg101Trp | Q6PHW0 |
| HOMO SAPIENS | 2 | 3023 | 4ttb | A | 116 | ILE | p.Ile116Thr | Q6PHW0 |
| HOMO SAPIENS | 0 | 3023 | 4ttb | A | 220 | ALA | p.Ala220Thr | Q6PHW0 |
| HOMO SAPIENS | 2 | 3023 | 4ttb | A | 271 | GLU | p.Glu271Lys | Q6PHW0 |
| HOMO SAPIENS | 3 | 33226 | 1ya5 | T | 11 | SER | p.Ser11Ter | O15273 |
| HOMO SAPIENS | 6 | 33226 | 1ya5 | T | 64 | SER | p.Ser64Leu | O15273 |
| HOMO SAPIENS | 6 | 33226 | 1ya5 | T | 70 | ARG | p.Arg70Gln | O15273 |
| HOMO SAPIENS | 3 | 33226 | 1ya5 | T | 75 | GLY | p.Gly75Ser | O15273 |
| HOMO SAPIENS | 0 | 33226 | 1ya5 | T | 87 | ARG | p.Arg87Trp | O15273 |
| HOMO SAPIENS | 2 | 5667 | 3dax | A | 398 | PRO | p.Pro398Ala | P22680 |

| | | | | | | | | |
|---|---|---|---|---|---|---|---|---|
| HOMO SAPIENS | 0 | 12861 | 5mya | A | 464 | ASN | p.Asn464His | Q02763 |
| HOMO SAPIENS | 2 | 12861 | 5mya | A | 486 | VAL | p.Val486Ile | Q02763 |
| HOMO SAPIENS | 0 | 12861 | 5mya | A | 522 | ARG | p.Arg522His | Q02763 |
| HOMO SAPIENS | 2 | 12861 | 5mya | A | 600 | VAL | p.Val600Leu | Q02763 |
| HOMO SAPIENS | 0 | 12861 | 5mya | A | 634 | LEU | p.Leu634Phe | Q02763 |
| HOMO SAPIENS | 0 | 12861 | 5mya | A | 684 | HIS | p.His684Gln | Q02763 |
| HOMO SAPIENS | 0 | 12861 | 5mya | A | 724 | ALA | p.Ala724Thr | Q02763 |
| HOMO SAPIENS | 0 | 7743 | 3qzw | G | 90 | GLY | p.Gly111Ser | P01732 |
| HOMO SAPIENS | 2 | 9498 | 3kci | A | 4267 | ASP | p.Asp4267Glu | O95714 |
| HOMO SAPIENS | 5 | 201 | 3ips | B | 415 | ARG | p.Arg415Gln | Q13133 |
| HOMO SAPIENS | 3 | 229 | 2q50 | A | 68 | LEU | p.Leu68Pro | Q9UBQ7 |
| HOMO SAPIENS | 2 | 229 | 2q50 | A | 99 | ARG | p.Arg99Ter | Q9UBQ7 |
| HOMO SAPIENS | 3 | 229 | 2q50 | A | 160 | GLY | p.Gly160Arg | Q9UBQ7 |
| HOMO SAPIENS | 5 | 229 | 2q50 | A | 165 | GLY | p.Gly165Asp | Q9UBQ7 |
| HOMO SAPIENS | 4 | 229 | 2q50 | A | 302 | ARG | p.Arg302Cys | Q9UBQ7 |
| HOMO SAPIENS | 4 | 229 | 2q50 | A | 302 | ARG | p.Arg302His | Q9UBQ7 |
| HOMO SAPIENS | -1 | 229 | 2q50 | A | 322 | MSE | p.Met322Arg | Q9UBQ7 |
| HOMO SAPIENS | -1 | 4519 | 5clw | A | 40 | PRO | p.Pro40Thr | Q04446 |
| HOMO SAPIENS | 1 | 4519 | 5clw | A | 59 | ILE | p.Ile59Thr | Q04446 |
| HOMO SAPIENS | 0 | 4519 | 5clw | A | 190 | ARG | p.Arg190Gly | Q04446 |
| HOMO SAPIENS | 1 | 4519 | 5clw | A | 224 | LEU | p.Leu224Pro | Q04446 |
| HOMO SAPIENS | 0 | 4519 | 5clw | A | 236 | GLN | p.Gln236His | Q04446 |
| HOMO SAPIENS | 1 | 4519 | 5clw | A | 254 | THR | p.Thr254Ala | Q04446 |
| HOMO SAPIENS | 0 | 4519 | 5clw | A | 257 | PHE | p.Phe257Leu | Q04446 |
| HOMO SAPIENS | 1 | 4519 | 5clw | A | 262 | ARG | p.Arg262Cys | Q04446 |
| HOMO SAPIENS | 3 | 4519 | 5clw | A | 329 | TYR | p.Tyr329Ser | Q04446 |
| HOMO SAPIENS | 0 | 4519 | 5clw | A | 507 | THR | p.Thr507Ala | Q04446 |
| HOMO SAPIENS | 0 | 4519 | 5clw | A | 515 | ARG | p.Arg515Cys | Q04446 |
| HOMO SAPIENS | 0 | 4519 | 5clw | A | 515 | ARG | p.Arg515His | Q04446 |
| HOMO SAPIENS | 0 | 4519 | 5clw | A | 524 | ARG | p.Arg524Gln | Q04446 |
| HOMO SAPIENS | 0 | 4519 | 5clw | A | 524 | ARG | p.Arg524Ter | Q04446 |
| HOMO SAPIENS | 0 | 4519 | 5clw | A | 545 | HIS | p.His545Arg | Q04446 |
| HOMO SAPIENS | 2 | 4519 | 5clw | A | 548 | TRP | p.Trp548Ter | Q04446 |
| HOMO SAPIENS | 0 | 4519 | 5clw | A | 592 | GLU | p.Glu592Ter | Q04446 |
| HOMO SAPIENS | 0 | 4519 | 5clw | A | 628 | HIS | p.His628Arg | Q04446 |
| HOMO SAPIENS | 0 | 4519 | 5clw | A | 637 | ARG | p.Arg637Ter | Q04446 |
| HOMO SAPIENS | 0 | 4519 | 5clw | A | 673 | ALA | p.Ala673Thr | Q04446 |
| HOMO SAPIENS | 1 | 10405 | 3bg3 | D | 507 | VAL | p.Val507Ile | P11498 |
| HOMO SAPIENS | 0 | 10405 | 3bg3 | D | 583 | ARG | p.Arg583Leu | P11498 |
| HOMO SAPIENS | 1 | 10405 | 3bg3 | D | 589 | LYS | p.Lys589Asn | P11498 |
| HOMO SAPIENS | 6 | 10405 | 3bg3 | D | 610 | ALA | p.Ala610Thr | P11498 |
| HOMO SAPIENS | 2 | 10405 | 3bg3 | D | 631 | ARG | p.Arg631Gln | P11498 |
| HOMO SAPIENS | 1 | 10405 | 3bg3 | D | 732 | ARG | p.Arg732Gln | P11498 |
| HOMO SAPIENS | 6 | 10405 | 3bg3 | D | 743 | MET | p.Met743Ile | P11498 |
| HOMO SAPIENS | 0 | 10405 | 3bg3 | D | 750 | THR | p.Thr750Met | P11498 |
| HOMO SAPIENS | -1 | 13751 | 2o8d | B | 350 | ALA | p.Ala350Val | P52701 |
| HOMO SAPIENS | -1 | 13751 | 2o8d | B | 352 | VAL | p.Val352Ile | P52701 |
| HOMO SAPIENS | -1 | 13751 | 2o8d | B | 355 | GLY | p.Gly355Ser | P52701 |
| HOMO SAPIENS | 0 | 13751 | 2o8d | B | 385 | ARG | p.Arg385Trp | P52701 |
| HOMO SAPIENS | 0 | 13751 | 2o8d | B | 390 | ASP | p.Asp390Glu | P52701 |
| HOMO SAPIENS | 0 | 13751 | 2o8d | B | 396 | LEU | p.Leu396Val | P52701 |
| HOMO SAPIENS | 0 | 13751 | 2o8d | B | 449 | LEU | p.Leu449Pro | P52701 |
| HOMO SAPIENS | 0 | 13751 | 2o8d | B | 468 | ARG | p.Arg468His | P52701 |
| HOMO SAPIENS | 0 | 13751 | 2o8d | B | 503 | SER | p.Ser503Cys | P52701 |

| | | | | | | | | |
|---|---|---|---|---|---|---|---|---|
| HOMO SAPIENS | 1 | 13751 | 2o8d | B | 509 | VAL | p.Val509Ala | P52701 |
| HOMO SAPIENS | 0 | 13751 | 2o8d | B | 675 | GLU | p.Glu675Asp | P52701 |
| HOMO SAPIENS | 0 | 13751 | 2o8d | B | 795 | ILE | p.Ile795Thr | P52701 |
| HOMO SAPIENS | 0 | 13751 | 2o8d | B | 850 | TYR | p.Tyr850Ter | P52701 |
| HOMO SAPIENS | 0 | 13751 | 2o8d | B | 878 | VAL | p.Val878Ala | P52701 |
| HOMO SAPIENS | 0 | 13751 | 2o8d | B | 911 | ARG | p.Arg911Ter | P52701 |
| HOMO SAPIENS | 0 | 13751 | 2o8d | B | 922 | ARG | p.Arg922Ter | P52701 |
| HOMO SAPIENS | 0 | 13751 | 2o8d | B | 969 | TYR | p.Tyr969Cys | P52701 |
| HOMO SAPIENS | 0 | 13751 | 2o8d | B | 1005 | ARG | p.Arg1005Ter | P52701 |
| HOMO SAPIENS | 0 | 13751 | 2o8d | B | 1035 | ARG | p.Arg1035Ter | P52701 |
| HOMO SAPIENS | 1 | 13751 | 2o8d | B | 1048 | GLN | p.Gln1048Ter | P52701 |
| HOMO SAPIENS | 1 | 13751 | 2o8d | B | 1051 | VAL | p.Val1051Ile | P52701 |
| HOMO SAPIENS | 0 | 13751 | 2o8d | B | 1054 | ILE | p.Ile1054Phe | P52701 |
| HOMO SAPIENS | 0 | 13751 | 2o8d | B | 1054 | ILE | p.Ile1054Met | P52701 |
| HOMO SAPIENS | 0 | 13751 | 2o8d | B | 1163 | GLU | p.Glu1163Val | P52701 |
| HOMO SAPIENS | 0 | 13751 | 2o8d | B | 1237 | GLU | p.Glu1237Asp | P52701 |
| HOMO SAPIENS | 0 | 13751 | 2o8d | B | 1242 | ARG | p.Arg1242His | P52701 |
| HOMO SAPIENS | -1 | 13751 | 2o8d | B | 1278 | PRO | p.Pro1278Thr | P52701 |
| HOMO SAPIENS | 0 | 13751 | 2o8d | B | 1304 | ARG | p.Arg1304Lys | P52701 |
| HOMO SAPIENS | 0 | 13751 | 2o8d | B | 1317 | HIS | p.His1317Gln | P52701 |
| HOMO SAPIENS | 0 | 13751 | 2o8d | B | 1329 | SER | p.Ser1329Leu | P52701 |
| HOMO SAPIENS | 3 | 13273 | 4g8k | A | 265 | GLU | p.Glu265Ter | Q05823 |
| HOMO SAPIENS | -1 | 11088 | 5uj7 | A | 666 | ARG | p.Arg666Trp | Q13415 |
| HOMO SAPIENS | 4 | 11088 | 5uj7 | A | 720 | ARG | p.Arg720Gln | Q13415 |
| HOMO SAPIENS | 0 | 11088 | 5uj7 | A | 816 | MET | p.Met816Thr | Q13415 |
| HOMO SAPIENS | 1 | 4352 | 5va1 | A | 421 | THR | p.Thr421Met | Q12809 |
| HOMO SAPIENS | 0 | 4352 | 5va1 | A | 451 | PRO | p.Pro451Leu | Q12809 |
| HOMO SAPIENS | 0 | 4352 | 5va1 | A | 470 | ASN | p.Asn470Asp | Q12809 |
| HOMO SAPIENS | 0 | 4352 | 5va1 | A | 490 | ALA | p.Ala490Thr | Q12809 |
| HOMO SAPIENS | 0 | 4352 | 5va1 | A | 491 | VAL | p.Val491Ile | Q12809 |
| HOMO SAPIENS | 1 | 4352 | 5va1 | A | 533 | VAL | p.Val533Gly | Q12809 |
| HOMO SAPIENS | 0 | 4352 | 5va1 | A | 552 | LEU | p.Leu552Ser | Q12809 |
| HOMO SAPIENS | 0 | 4352 | 5va1 | A | 558 | ALA | p.Ala558Pro | Q12809 |
| HOMO SAPIENS | 1 | 4352 | 5va1 | A | 561 | ALA | p.Ala561Val | Q12809 |
| HOMO SAPIENS | 0 | 4352 | 5va1 | A | 572 | GLY | p.Gly572Arg | Q12809 |
| HOMO SAPIENS | -1 | 4352 | 5va1 | A | 582 | ARG | p.Arg582Cys | Q12809 |
| HOMO SAPIENS | 0 | 4352 | 5va1 | A | 584 | GLY | p.Gly584Ser | Q12809 |
| HOMO SAPIENS | 0 | 4352 | 5va1 | A | 588 | ASN | p.Asn588Lys | Q12809 |
| HOMO SAPIENS | 0 | 4352 | 5va1 | A | 593 | ILE | p.Ile593Arg | Q12809 |
| HOMO SAPIENS | 1 | 4352 | 5va1 | A | 627 | PHE | p.Phe627Leu | Q12809 |
| HOMO SAPIENS | 0 | 4352 | 5va1 | A | 630 | VAL | p.Val630Leu | Q12809 |
| HOMO SAPIENS | 0 | 4352 | 5va1 | A | 652 | TYR | p.Tyr652Ter | Q12809 |
| HOMO SAPIENS | 0 | 4352 | 5va1 | A | 711 | ILE | p.Ile711Val | Q12809 |
| HOMO SAPIENS | 0 | 4352 | 5va1 | A | 735 | SER | p.Ser735Leu | Q12809 |
| HOMO SAPIENS | 0 | 4352 | 5va1 | A | 744 | ARG | p.Arg744Ter | Q12809 |
| HOMO SAPIENS | 0 | 4352 | 5va1 | A | 784 | ARG | p.Arg784Trp | Q12809 |
| HOMO SAPIENS | 1 | 4352 | 5va1 | A | 820 | GLY | p.Gly820Arg | Q12809 |
| HOMO SAPIENS | 5 | 4352 | 5va1 | A | 822 | VAL | p.Val822Met | Q12809 |
| HOMO SAPIENS | 1 | 4352 | 5va1 | A | 863 | ARG | p.Arg863Ter | Q12809 |
| HOMO SAPIENS | 5 | 303 | 2obv | A | 55 | ALA | p.Ala55Asp | Q00266 |
| HOMO SAPIENS | 7 | 303 | 2obv | A | 259 | ALA | p.Ala259Gly | Q00266 |
| HOMO SAPIENS | 9 | 303 | 2obv | A | 264 | ARG | p.Arg264His | Q00266 |
| HOMO SAPIENS | 9 | 303 | 2obv | A | 264 | ARG | p.Arg264Cys | Q00266 |
| HOMO SAPIENS | 0 | 303 | 2obv | A | 305 | LEU | p.Leu305Pro | Q00266 |

| | | | | | | | | |
|---|---|---|---|---|---|---|---|---|
| HOMO SAPIENS | 7 | 303 | 2obv | A | 322 | ILE | p.Ile322Met | Q00266 |
| HOMO SAPIENS | 0 | 303 | 2obv | A | 336 | GLY | p.Gly336Arg | Q00266 |
| HOMO SAPIENS | 0 | 303 | 2obv | A | 357 | PRO | p.Pro357Leu | Q00266 |
| HOMO SAPIENS | 0 | 2077 | 4zxb | E | 462 | ASN | p.Asn489Ser | P06213 |
| HOMO SAPIENS | -1 | 2077 | 4zxb | E | 519 | ASP | p.Asp546Glu | P06213 |
| HOMO SAPIENS | 2 | 2077 | 4zxb | E | 819 | THR | p.Thr858Ala | P06213 |
| HOMO SAPIENS | 0 | 2077 | 4zxb | E | 907 | ASP | p.Asp946Glu | P06213 |
| HOMO SAPIENS | 7 | 3274 | 1wsv | B | 14 | HIS | p.His42Arg | P48728 |
| HOMO SAPIENS | 0 | 3274 | 1wsv | B | 19 | GLY | p.Gly47Arg | P48728 |
| HOMO SAPIENS | 4 | 3274 | 1wsv | B | 45 | ARG | p.Arg73Cys | P48728 |
| HOMO SAPIENS | 1 | 3274 | 1wsv | B | 49 | SER | p.Ser77Leu | P48728 |
| HOMO SAPIENS | 3 | 3274 | 1wsv | B | 241 | GLY | p.Gly269Asp | P48728 |
| HOMO SAPIENS | 7 | 3274 | 1wsv | B | 248 | ASP | p.Asp276His | P48728 |
| HOMO SAPIENS | 0 | 3274 | 1wsv | B | 272 | MET | p.Met300Val | P48728 |
| HOMO SAPIENS | 4 | 3274 | 1wsv | B | 292 | ARG | p.Arg320His | P48728 |
| HOMO SAPIENS | 0 | 3274 | 1wsv | B | 354 | ARG | p.Arg382Gln | P48728 |
| HOMO SAPIENS | -1 | 435 | 3uyt | A | 44 | THR | p.Thr44Ala | P48730 |
| HOMO SAPIENS | 1 | 4698 | 5ife | C | 666 | LYS | p.Lys666Asn | O75533 |
| HOMO SAPIENS | 1 | 16917 | 1apy | C | 37 | GLY | p.Gly60Asp | P20933 |
| HOMO SAPIENS | 0 | 16917 | 1apy | C | 49 | SER | p.Ser72Pro | P20933 |
| HOMO SAPIENS | 0 | 16917 | 1apy | C | 84 | ARG | p.Arg107Ter | P20933 |
| HOMO SAPIENS | 0 | 16917 | 1apy | C | 126 | SER | p.Thr149Ser | P20933 |
| HOMO SAPIENS | 0 | 16917 | 1apy | C | 135 | TRP | p.Trp158Ter | P20933 |
| HOMO SAPIENS | 0 | 16917 | 1apy | C | 140 | CYS | p.Cys163Ser | P20933 |
| HOMO SAPIENS | 0 | 16917 | 1apy | C | 145 | TRP | p.Trp168Ter | P20933 |
| HOMO SAPIENS | 0 | 10413 | 4frw | A | 185 | THR | p.Thr185Met | Q96NY8 |
| HOMO SAPIENS | 0 | 5217 | 5n6w | G | 96 | TRP | p.Trp96Arg | O15537 |
| HOMO SAPIENS | 1 | 5217 | 5n6w | G | 102 | ARG | p.Arg102Trp | O15537 |
| HOMO SAPIENS | 0 | 5217 | 5n6w | G | 109 | GLY | p.Gly109Arg | O15537 |
| HOMO SAPIENS | -1 | 5217 | 5n6w | G | 223 | CYS | p.Cys223Arg | O15537 |
| HOMO SAPIENS | 0 | 9485 | 4z6y | E | 138 | ALA | p.Ala138Asp | Q9P0N9 |
| HOMO SAPIENS | 3 | 4504 | 4mwt | B | 411 | GLY | p.Gly439Ser | P10619 |
| HOMO SAPIENS | 3 | 4504 | 4mwt | B | 412 | PHE | p.Phe440Val | P10619 |
| HOMO SAPIENS | 6 | 6391 | 4kxy | A | 318 | ARG | p.Arg318Cys | P29401 |
| HOMO SAPIENS | 6 | 19546 | 2bx6 | A | 87 | THR | p.Thr87Ile | O75695 |
| HOMO SAPIENS | 1 | 19546 | 2bx6 | A | 118 | ARG | p.Arg118His | O75695 |
| HOMO SAPIENS | 1 | 19546 | 2bx6 | A | 201 | ILE | p.Ile201Val | O75695 |
| HOMO SAPIENS | 1 | 19546 | 2bx6 | A | 282 | ARG | p.Arg282Trp | O75695 |
| HOMO SAPIENS | 1 | 5265 | 5usr | H | 68 | ARG | p.Arg68Leu | Q9HD34 |
| HOMO SAPIENS | 2 | 1012 | 4nyi | S | 569 | LEU | p.Leu569Val | Q07889 |
| HOMO SAPIENS | 2 | 1012 | 4nyi | S | 655 | PRO | p.Pro655Leu | Q07889 |
| HOMO SAPIENS | 4 | 1012 | 4nyi | S | 663 | ILE | p.Ile663Met | Q07889 |
| HOMO SAPIENS | 2 | 1012 | 4nyi | S | 702 | TYR | p.Tyr702Cys | Q07889 |
| HOMO SAPIENS | 2 | 1012 | 4nyi | S | 708 | ALA | p.Ala708Thr | Q07889 |
| HOMO SAPIENS | 4 | 1012 | 4nyi | S | 733 | ILE | p.Ile733Phe | Q07889 |
| HOMO SAPIENS | 1 | 1848 | 3g1n | A | 4013 | ARG | p.Arg4013Trp | Q7Z6Z7 |
| HOMO SAPIENS | 1 | 1848 | 3g1n | A | 4187 | ARG | p.Arg4187Cys | Q7Z6Z7 |
| HOMO SAPIENS | 1 | 77 | 4pod | A | 41 | CYS | p.Cys79Tyr | P60174 |
| HOMO SAPIENS | 9 | 77 | 4pod | A | 104 | GLU | p.Glu142Asp | P60174 |
| HOMO SAPIENS | 1 | 77 | 4pod | A | 122 | GLY | p.Gly160Arg | P60174 |
| HOMO SAPIENS | -1 | 8672 | 3thz | A | 8 | THR | p.Thr8Met | P43246 |
| HOMO SAPIENS | 0 | 8672 | 3thz | A | 19 | PHE | p.Phe19Leu | P43246 |
| HOMO SAPIENS | 0 | 8672 | 3thz | A | 23 | PHE | p.Phe23Leu | P43246 |
| HOMO SAPIENS | 0 | 8672 | 3thz | A | 96 | ARG | p.Arg96His | P43246 |

| | | | | | | | | |
|---|---|---|---|---|---|---|---|---|
| HOMO SAPIENS | 0 | 8672 | 3thz | A | 102 | VAL | p.Val102Ile | P43246 |
| HOMO SAPIENS | 0 | 8672 | 3thz | A | 106 | ARG | p.Arg106Lys | P43246 |
| HOMO SAPIENS | 0 | 8672 | 3thz | A | 127 | ASN | p.Asn127Ser | P43246 |
| HOMO SAPIENS | 0 | 8672 | 3thz | A | 128 | LEU | p.Leu128Val | P43246 |
| HOMO SAPIENS | 0 | 8672 | 3thz | A | 163 | VAL | p.Val163Gly | P43246 |
| HOMO SAPIENS | 0 | 8672 | 3thz | A | 169 | ILE | p.Ile169Val | P43246 |
| HOMO SAPIENS | 0 | 8672 | 3thz | A | 315 | GLY | p.Gly315Asp | P43246 |
| HOMO SAPIENS | -1 | 8672 | 3thz | A | 322 | GLY | p.Gly322Asp | P43246 |
| HOMO SAPIENS | 0 | 8672 | 3thz | A | 331 | ASN | p.Asn331Asp | P43246 |
| HOMO SAPIENS | 1 | 8672 | 3thz | A | 390 | LEU | p.Leu390Phe | P43246 |
| HOMO SAPIENS | 0 | 8672 | 3thz | A | 419 | GLN | p.Gln419Lys | P43246 |
| HOMO SAPIENS | 0 | 8672 | 3thz | A | 447 | PHE | p.Phe447Leu | P43246 |
| HOMO SAPIENS | 0 | 8672 | 3thz | A | 524 | ARG | p.Arg524Pro | P43246 |
| HOMO SAPIENS | 0 | 8672 | 3thz | A | 539 | PHE | p.Phe539Leu | P43246 |
| HOMO SAPIENS | -1 | 8672 | 3thz | A | 546 | LYS | p.Lys546Asn | P43246 |
| HOMO SAPIENS | 0 | 8672 | 3thz | A | 564 | THR | p.Thr564Ala | P43246 |
| HOMO SAPIENS | 0 | 8672 | 3thz | A | 577 | ILE | p.Ile577Thr | P43246 |
| HOMO SAPIENS | 0 | 8672 | 3thz | A | 588 | TYR | p.Tyr588Ter | P43246 |
| HOMO SAPIENS | 0 | 8672 | 3thz | A | 622 | PRO | p.Pro622Leu | P43246 |
| HOMO SAPIENS | 0 | 8672 | 3thz | A | 629 | GLN | p.Gln629Arg | P43246 |
| HOMO SAPIENS | 0 | 8672 | 3thz | A | 636 | ALA | p.Ala636Pro | P43246 |
| HOMO SAPIENS | 2 | 8672 | 3thz | A | 642 | VAL | p.Val642Ile | P43246 |
| HOMO SAPIENS | 1 | 8672 | 3thz | A | 654 | ASP | p.Asp654Glu | P43246 |
| HOMO SAPIENS | 0 | 8672 | 3thz | A | 662 | GLN | p.Gln662His | P43246 |
| HOMO SAPIENS | 2 | 8672 | 3thz | A | 681 | GLN | p.Gln681His | P43246 |
| HOMO SAPIENS | 0 | 8672 | 3thz | A | 707 | CYS | p.Cys707Ser | P43246 |
| HOMO SAPIENS | 0 | 8672 | 3thz | A | 809 | GLU | p.Glu809Lys | P43246 |
| HOMO SAPIENS | 0 | 8672 | 3thz | A | 813 | MET | p.Met813Val | P43246 |
| HOMO SAPIENS | 1 | 8672 | 3thz | A | 834 | ALA | p.Ala834Thr | P43246 |
| HOMO SAPIENS | 0 | 8672 | 3thz | A | 849 | LEU | p.Leu849Val | P43246 |
| HOMO SAPIENS | -1 | 8672 | 3thz | A | 860 | SER | p.Ser860Leu | P43246 |
| HOMO SAPIENS | -1 | 8672 | 3thz | A | 934 | THR | p.Thr934Lys | P43246 |
| HOMO SAPIENS | 2 | 292 | 1i3k | A | 34 | ASN | p.Asn34Ser | Q14376 |
| HOMO SAPIENS | 7 | 292 | 1i3k | A | 90 | GLY | p.Gly90Glu | Q14376 |
| HOMO SAPIENS | 2 | 292 | 1i3k | A | 103 | ASP | p.Asp103Gly | Q14376 |
| HOMO SAPIENS | 2 | 292 | 1i3k | A | 136 | TYR | p.Tyr136Ter | Q14376 |
| HOMO SAPIENS | 0 | 292 | 1i3k | A | 169 | ARG | p.Arg169Trp | Q14376 |
| HOMO SAPIENS | 0 | 292 | 1i3k | A | 183 | LEU | p.Leu183Pro | Q14376 |
| HOMO SAPIENS | 0 | 292 | 1i3k | A | 302 | GLY | p.Gly302Asp | Q14376 |
| HOMO SAPIENS | 0 | 292 | 1i3k | A | 313 | LEU | p.Leu313Met | Q14376 |
| HOMO SAPIENS | 1 | 2084 | 5vc8 | A | 295 | GLN | p.Gln295Leu | O96028 |
| HOMO SAPIENS | -1 | 10157 | 5wrw | A | 7 | GLY | p.Gly7Ala | O76094 |
| HOMO SAPIENS | -1 | 10157 | 5wrw | A | 7 | GLY | p.Gly7Arg | O76094 |
| HOMO SAPIENS | -1 | 10157 | 5wrw | A | 8 | GLY | p.Gly8Ala | O76094 |
| HOMO SAPIENS | 0 | 3113 | 3in5 | A | 29 | GLU | p.Glu29Lys | Q9UBT6 |
| HOMO SAPIENS | 5 | 4546 | 5in3 | B | 44 | VAL | p.Val44Leu | P07902 |
| HOMO SAPIENS | -1 | 4546 | 5in3 | B | 51 | ARG | p.Arg51Leu | P07902 |
| HOMO SAPIENS | -1 | 4546 | 5in3 | B | 62 | LEU | p.Leu62Met | P07902 |
| HOMO SAPIENS | 4 | 4546 | 5in3 | B | 66 | PRO | p.Pro66His | P07902 |
| HOMO SAPIENS | 2 | 4546 | 5in3 | B | 74 | LEU | p.Leu74Pro | P07902 |
| HOMO SAPIENS | 5 | 4546 | 5in3 | B | 97 | ASN | p.Asn97Ser | P07902 |
| HOMO SAPIENS | 1 | 4546 | 5in3 | B | 135 | SER | p.Ser135Leu | P07902 |
| HOMO SAPIENS | 5 | 4546 | 5in3 | B | 138 | THR | p.Thr138Met | P07902 |
| HOMO SAPIENS | 7 | 4546 | 5in3 | B | 139 | LEU | p.Leu139Pro | P07902 |

| | | | | | | | | |
|---|---|---|---|---|---|---|---|---|
| HOMO SAPIENS | 2 | 4546 | 5in3 | B | 142 | MET | p.Met142Lys | P07902 |
| HOMO SAPIENS | 1 | 4546 | 5in3 | B | 143 | SER | p.Ser143Leu | P07902 |
| HOMO SAPIENS | 2 | 4546 | 5in3 | B | 148 | ARG | p.Arg148Gly | P07902 |
| HOMO SAPIENS | 2 | 4546 | 5in3 | B | 148 | ARG | p.Arg148Gln | P07902 |
| HOMO SAPIENS | 5 | 4546 | 5in3 | B | 183 | PRO | p.Pro183Thr | P07902 |
| HOMO SAPIENS | 5 | 4546 | 5in3 | B | 185 | PRO | p.Pro185His | P07902 |
| HOMO SAPIENS | 5 | 4546 | 5in3 | B | 186 | HIS | p.His186Tyr | P07902 |
| HOMO SAPIENS | 7 | 4546 | 5in3 | B | 188 | GLN | p.Gln188Arg | P07902 |
| HOMO SAPIENS | 5 | 4546 | 5in3 | B | 195 | LEU | p.Leu195Pro | P07902 |
| HOMO SAPIENS | 4 | 4546 | 5in3 | B | 201 | ARG | p.Arg201His | P07902 |
| HOMO SAPIENS | 4 | 4546 | 5in3 | B | 201 | ARG | p.Arg201Cys | P07902 |
| HOMO SAPIENS | 4 | 4546 | 5in3 | B | 203 | GLU | p.Glu203Lys | P07902 |
| HOMO SAPIENS | 2 | 4546 | 5in3 | B | 209 | TYR | p.Tyr209Cys | P07902 |
| HOMO SAPIENS | 5 | 4546 | 5in3 | B | 223 | ARG | p.Arg223Ser | P07902 |
| HOMO SAPIENS | 5 | 4546 | 5in3 | B | 226 | LEU | p.Leu226Pro | P07902 |
| HOMO SAPIENS | 5 | 4546 | 5in3 | B | 226 | LEU | p.Leu226Val | P07902 |
| HOMO SAPIENS | 2 | 4546 | 5in3 | B | 227 | LEU | p.Leu227Pro | P07902 |
| HOMO SAPIENS | 8 | 4546 | 5in3 | B | 258 | ARG | p.Arg258Cys | P07902 |
| HOMO SAPIENS | 1 | 4546 | 5in3 | B | 265 | PRO | p.Pro265Ala | P07902 |
| HOMO SAPIENS | 8 | 4546 | 5in3 | B | 268 | THR | p.Thr268Asn | P07902 |
| HOMO SAPIENS | 7 | 4546 | 5in3 | B | 278 | ILE | p.Ile278Asn | P07902 |
| HOMO SAPIENS | 5 | 4546 | 5in3 | B | 285 | LYS | p.Lys285Asn | P07902 |
| HOMO SAPIENS | 1 | 4546 | 5in3 | B | 319 | HIS | p.His319Gln | P07902 |
| HOMO SAPIENS | 4 | 4546 | 5in3 | B | 321 | HIS | p.His321Tyr | P07902 |
| HOMO SAPIENS | 4 | 4546 | 5in3 | B | 325 | PRO | p.Pro325Leu | P07902 |
| HOMO SAPIENS | 2 | 4546 | 5in3 | B | 328 | ARG | p.Arg328His | P07902 |
| HOMO SAPIENS | 2 | 4546 | 5in3 | B | 329 | SER | p.Ser329Phe | P07902 |
| HOMO SAPIENS | 5 | 4546 | 5in3 | B | 334 | LYS | p.Lys334Arg | P07902 |
| HOMO SAPIENS | 6 | 1290 | 5uav | C | 119 | ARG | p.Arg119Gly | P32322 |
| HOMO SAPIENS | 6 | 1290 | 5uav | C | 119 | ARG | p.Arg119His | P32322 |
| HOMO SAPIENS | 3 | 1290 | 5uav | C | 251 | ARG | p.Arg251His | P32322 |
| HOMO SAPIENS | 7 | 1290 | 5uav | C | 266 | ARG | p.Arg266Gln | P32322 |
| HOMO SAPIENS | 0 | 9618 | 2r7e | A | 22 | GLY | p.Gly41Cys | P00451 |
| HOMO SAPIENS | 2 | 9618 | 2r7e | A | 113 | GLU | p.Glu132Asp | P00451 |
| HOMO SAPIENS | -1 | 9618 | 2r7e | A | 336 | ARG | p.Arg355Ter | P00451 |
| HOMO SAPIENS | 0 | 9618 | 2r7e | A | 372 | ARG | p.Arg391His | P00451 |
| HOMO SAPIENS | 0 | 9618 | 2r7e | A | 373 | SER | p.Ser392Leu | P00451 |
| HOMO SAPIENS | 0 | 9618 | 2r7e | A | 373 | SER | p.Ser392Pro | P00451 |
| HOMO SAPIENS | 3 | 9618 | 2r7e | A | 386 | ILE | p.Ile405Ser | P00451 |
| HOMO SAPIENS | 6 | 9618 | 2r7e | A | 390 | GLU | p.Glu409Gly | P00451 |
| HOMO SAPIENS | 2 | 9618 | 2r7e | A | 412 | LEU | p.Leu431Phe | P00451 |
| HOMO SAPIENS | 5 | 9618 | 2r7e | A | 425 | LYS | p.Lys444Arg | P00451 |
| HOMO SAPIENS | 0 | 9618 | 2r7e | A | 484 | ARG | p.Arg503His | P00451 |
| HOMO SAPIENS | 2 | 9618 | 2r7e | A | 531 | ARG | p.Arg550His | P00451 |
| HOMO SAPIENS | 3 | 9618 | 2r7e | A | 535 | SER | p.Ser554Gly | P00451 |
| HOMO SAPIENS | 2 | 9618 | 2r7e | A | 593 | ARG | p.Arg612Cys | P00451 |
| HOMO SAPIENS | 0 | 9618 | 2r7e | A | 704 | ALA | p.Ala723Thr | P00451 |
| HOMO SAPIENS | 0 | 9618 | 2r7e | A | 720 | GLU | p.Glu739Lys | P00451 |
| HOMO SAPIENS | 1 | 13085 | 3ud1 | A | 1075 | ILE | p.Thr1075Ile | P16157 |
| HOMO SAPIENS | -1 | 3289 | 2fy4 | A | 2 | ALA | p.Ala120Thr | P28329 |
| HOMO SAPIENS | 7 | 3289 | 2fy4 | A | 18 | VAL | p.Val136Met | P28329 |
| HOMO SAPIENS | 5 | 3289 | 2fy4 | A | 92 | LEU | p.Leu210Pro | P28329 |
| HOMO SAPIENS | 2 | 3289 | 2fy4 | A | 93 | PRO | p.Pro211Ala | P28329 |
| HOMO SAPIENS | 4 | 3289 | 2fy4 | A | 131 | LEU | p.Leu249Val | P28329 |

| | | | | | | | | |
|---|---|---|---|---|---|---|---|---|
| HOMO SAPIENS | 2 | 3289 | 2fy4 | A | 181 | PRO | p.Pro299Leu | P28329 |
| HOMO SAPIENS | 0 | 3289 | 2fy4 | A | 218 | ILE | p.Ile336Thr | P28329 |
| HOMO SAPIENS | 2 | 3289 | 2fy4 | A | 261 | ASP | p.Asp379His | P28329 |
| HOMO SAPIENS | 0 | 3289 | 2fy4 | A | 302 | ARG | p.Arg420Cys | P28329 |
| HOMO SAPIENS | 5 | 3289 | 2fy4 | A | 323 | GLU | p.Glu441Lys | P28329 |
| HOMO SAPIENS | 0 | 3289 | 2fy4 | A | 343 | MET | p.Val461Met | P28329 |
| HOMO SAPIENS | 2 | 3289 | 2fy4 | A | 364 | ARG | p.Arg482Gly | P28329 |
| HOMO SAPIENS | 3 | 3289 | 2fy4 | A | 380 | SER | p.Ser498Leu | P28329 |
| HOMO SAPIENS | 6 | 3289 | 2fy4 | A | 388 | VAL | p.Val506Leu | P28329 |
| HOMO SAPIENS | 7 | 3289 | 2fy4 | A | 429 | HIS | p.His547Gln | P28329 |
| HOMO SAPIENS | 8 | 3289 | 2fy4 | A | 442 | ARG | p.Arg560His | P28329 |
| HOMO SAPIENS | 3 | 3289 | 2fy4 | A | 443 | ARG | p.Arg561Gln | P28329 |
| HOMO SAPIENS | 0 | 4137 | 3hhr | B | 38 | CYS | p.Cys56Ter | P10912 |
| HOMO SAPIENS | 0 | 4137 | 3hhr | B | 43 | ARG | p.Arg61Ter | P10912 |
| HOMO SAPIENS | 0 | 4137 | 3hhr | B | 44 | GLU | p.Glu62Lys | P10912 |
| HOMO SAPIENS | 0 | 4137 | 3hhr | B | 96 | PHE | p.Phe114Ser | P10912 |
| HOMO SAPIENS | 5 | 4137 | 3hhr | B | 131 | PRO | p.Pro149Gln | P10912 |
| HOMO SAPIENS | 0 | 4137 | 3hhr | B | 144 | VAL | p.Val162Ile | P10912 |
| HOMO SAPIENS | 0 | 4137 | 3hhr | B | 150 | HIS | p.His168Gln | P10912 |
| HOMO SAPIENS | 0 | 4137 | 3hhr | B | 152 | ASP | p.Asp170His | P10912 |
| HOMO SAPIENS | 0 | 4137 | 3hhr | B | 153 | ILE | p.Ile171Thr | P10912 |
| HOMO SAPIENS | 0 | 4137 | 3hhr | B | 154 | GLN | p.Gln172Pro | P10912 |
| HOMO SAPIENS | 0 | 4137 | 3hhr | B | 155 | VAL | p.Val173Gly | P10912 |
| HOMO SAPIENS | 0 | 4137 | 3hhr | B | 217 | ARG | p.Arg235Ter | P10912 |
| HOMO SAPIENS | 5 | 4137 | 3hhr | B | 224 | GLU | p.Glu242Asp | P10912 |
| HOMO SAPIENS | 5 | 4137 | 3hhr | B | 224 | GLU | p.Glu242Ter | P10912 |
| HOMO SAPIENS | 6 | 2493 | 5bpb | A | 69 | HIS | p.His69Tyr | Q9ULV1 |
| HOMO SAPIENS | 6 | 2493 | 5bpb | A | 105 | MET | p.Met105Val | Q9ULV1 |
| HOMO SAPIENS | -1 | 2493 | 5bpb | A | 168 | PRO | p.Pro168Ser | Q9ULV1 |
| HOMO SAPIENS | 0 | 32123 | 4ccz | A | 314 | ASP | p.Asp314Asn | Q99707 |
| HOMO SAPIENS | 0 | 32123 | 4ccz | A | 345 | VAL | p.Val345Phe | Q99707 |
| HOMO SAPIENS | 1 | 32123 | 4ccz | A | 585 | ARG | p.Arg585Ter | Q99707 |
| HOMO SAPIENS | 4 | 4618 | 2ag4 | A | 26 | PRO | p.Pro55Leu | P17900 |
| HOMO SAPIENS | 4 | 4618 | 2ag4 | A | 30 | VAL | p.Ile59Val | P17900 |
| HOMO SAPIENS | 2 | 4618 | 2ag4 | A | 40 | VAL | p.Met69Val | P17900 |
| HOMO SAPIENS | 2 | 4618 | 2ag4 | A | 124 | VAL | p.Val153Ala | P17900 |
| HOMO SAPIENS | 8 | 4618 | 2ag4 | A | 140 | ARG | p.Arg169Pro | P17900 |
| HOMO SAPIENS | 5 | 1082 | 1hk0 | X | 14 | ARG | p.Arg15Cys | P07320 |
| HOMO SAPIENS | 7 | 1082 | 1hk0 | X | 23 | PRO | p.Pro24Thr | P07320 |
| HOMO SAPIENS | 6 | 1082 | 1hk0 | X | 36 | ARG | p.Arg37Ser | P07320 |
| HOMO SAPIENS | 5 | 1082 | 1hk0 | X | 55 | TYR | p.Tyr56Ter | P07320 |
| HOMO SAPIENS | 3 | 1082 | 1hk0 | X | 58 | ARG | p.Arg59His | P07320 |
| HOMO SAPIENS | 7 | 1082 | 1hk0 | X | 88 | HIS | p.His88Gln | P07320 |
| HOMO SAPIENS | 6 | 1082 | 1hk0 | X | 95 | ARG | p.Arg95Ser | P07320 |
| HOMO SAPIENS | 5 | 1082 | 1hk0 | X | 126 | VAL | p.Val126Met | P07320 |
| HOMO SAPIENS | 8 | 1082 | 1hk0 | X | 157 | TRP | p.Trp157Ter | P07320 |
| HOMO SAPIENS | 0 | 22879 | 3j0a | B | 644 | ILE | p.Ile644Phe | O60602 |
| HOMO SAPIENS | 8 | 6 | 2o23 | B | 122 | LEU | p.Leu122Val | Q99714 |
| HOMO SAPIENS | 1 | 6 | 2o23 | B | 130 | ARG | p.Arg130Cys | Q99714 |
| HOMO SAPIENS | 1 | 6 | 2o23 | B | 247 | ASN | p.Asn247Ser | Q99714 |
| HOMO SAPIENS | 0 | 6 | 2o23 | B | 249 | GLU | p.Glu249Gln | Q99714 |
| HOMO SAPIENS | 1 | 12910 | 4hl4 | A | 67 | ARG | p.Arg67Ter | Q96BZ9 |
| HOMO SAPIENS | 8 | 16105 | 1lmj | A | 7 | GLU | p.Glu1073Lys | P35555 |
| HOMO SAPIENS | 8 | 16105 | 1lmj | A | 8 | CYS | p.Cys1074Arg | P35555 |

| | | | | | | | | |
|---|---|---|---|---|---|---|---|---|
| HOMO SAPIENS | 9 | 16105 | 1lmj | A | 51 | CYS | p.Cys1117Tyr | P35555 |
| HOMO SAPIENS | 9 | 16105 | 1lmj | A | 63 | CYS | p.Cys1129Tyr | P35555 |
| HOMO SAPIENS | 9 | 16105 | 1lmj | A | 65 | ASN | p.Asn1131Tyr | P35555 |
| HOMO SAPIENS | 4 | 16105 | 1lmj | A | 71 | ARG | p.Arg1137Pro | P35555 |
| HOMO SAPIENS | 1 | 16105 | 1lmj | A | 82 | PRO | p.Pro1148Ala | P35555 |
| HOMO SAPIENS | 0 | 4795 | 3s5f | A | 106 | LEU | p.Leu106Ser | Q16595 |
| HOMO SAPIENS | 0 | 4795 | 3s5f | A | 130 | GLY | p.Gly130Val | Q16595 |
| HOMO SAPIENS | 0 | 4795 | 3s5f | A | 154 | ILE | p.Ile154Phe | Q16595 |
| HOMO SAPIENS | -1 | 4795 | 3s5f | A | 209 | ASP | p.Asp209Gly | Q16595 |
| HOMO SAPIENS | -1 | 2884 | 4lli | A | 1655 | ASP | p.Asp1655Glu | Q9Y4I1 |
| HOMO SAPIENS | 0 | 1786 | 7pck | C | 56P | TYR | p.Tyr71Ter | P43235 |
| HOMO SAPIENS | 0 | 1786 | 7pck | C | 64P | GLY | p.Gly79Glu | P43235 |
| HOMO SAPIENS | 7 | 38286 | 2l53 | B | 1914 | ARG | p.Arg1914Gly | Q14524 |
| HOMO SAPIENS | 1 | 38286 | 2l53 | B | 1924 | ALA | p.Ala1924Thr | Q14524 |
| HOMO SAPIENS | 0 | 31747 | 3ksy | A | 25 | PRO | p.Pro25Ser | Q07889 |
| HOMO SAPIENS | 3 | 31747 | 3ksy | A | 37 | THR | p.Thr37Ala | Q07889 |
| HOMO SAPIENS | 0 | 31747 | 3ksy | A | 54 | GLN | p.Gln54His | Q07889 |
| HOMO SAPIENS | -1 | 31747 | 3ksy | A | 190 | ASP | p.Asp190Glu | Q07889 |
| HOMO SAPIENS | 2 | 31747 | 3ksy | A | 266 | THR | p.Thr266Lys | Q07889 |
| HOMO SAPIENS | 1 | 31747 | 3ksy | A | 269 | MET | p.Met269Arg | Q07889 |
| HOMO SAPIENS | 1 | 31747 | 3ksy | A | 340 | PRO | p.Pro340Ser | Q07889 |
| HOMO SAPIENS | 1 | 31747 | 3ksy | A | 423 | ASN | p.Asn423Lys | Q07889 |
| HOMO SAPIENS | 1 | 31747 | 3ksy | A | 424 | GLU | p.Glu424Lys | Q07889 |
| HOMO SAPIENS | 1 | 31747 | 3ksy | A | 432 | TRP | p.Trp432Arg | Q07889 |
| HOMO SAPIENS | 1 | 31747 | 3ksy | A | 552 | ARG | p.Arg552Gly | Q07889 |
| HOMO SAPIENS | 1 | 31747 | 3ksy | A | 552 | ARG | p.Arg552Ser | Q07889 |
| HOMO SAPIENS | 3 | 7529 | 6jdw | A | 93 | ILE | p.Ile93Met | P50440 |
| HOMO SAPIENS | 2 | 7529 | 6jdw | A | 110 | GLN | p.Gln110His | P50440 |
| HOMO SAPIENS | 5 | 7529 | 6jdw | A | 149 | TRP | p.Trp149Ter | P50440 |
| HOMO SAPIENS | 0 | 7529 | 6jdw | A | 169 | ARG | p.Arg169Ter | P50440 |
| HOMO SAPIENS | 0 | 7529 | 6jdw | A | 260 | ARG | p.Arg260Ter | P50440 |
| HOMO SAPIENS | 3 | 7529 | 6jdw | A | 344 | ILE | p.Ile344Val | P50440 |
| HOMO SAPIENS | 5 | 7529 | 6jdw | A | 418 | LEU | p.Leu418Ile | P50440 |
| HOMO SAPIENS | 1 | 14788 | 1eak | A | 101 | ARG | p.Arg101His | P08253 |
| HOMO SAPIENS | 0 | 14788 | 1eak | A | 383 | ASP | p.Asp383Glu | P08253 |
| HOMO SAPIENS | 7 | 14788 | 1eak | A | 404 | GLN | p.Glu404Lys | P08253 |
| HOMO SAPIENS | 2 | 14788 | 1eak | A | 447 | ALA | p.Ala447Val | P08253 |
| HOMO SAPIENS | 0 | 1255 | 5xtc | p | 8 | PRO | p.Pro8His | Q9Y6M9 |
| HOMO SAPIENS | 0 | 1255 | 5xtc | p | 146 | PRO | p.Pro146Ser | Q9Y6M9 |
| HOMO SAPIENS | 3 | 6721 | 1a22 | A | 25 | PHE | p.Phe51Tyr | P01241 |
| HOMO SAPIENS | 2 | 6721 | 1a22 | A | 95 | SER | p.Ser121Arg | P01241 |
| HOMO SAPIENS | 2 | 6721 | 1a22 | A | 110 | VAL | p.Val136Ile | P01241 |
| HOMO SAPIENS | 0 | 1380 | 5i6z | A | 605 | LYS | p.Lys605Asn | P31645 |
| HOMO SAPIENS | 0 | 941 | 1qki | D | 48 | ILE | p.Ile48Thr | P11413 |
| HOMO SAPIENS | 3 | 941 | 1qki | D | 68 | VAL | p.Val68Met | P11413 |
| HOMO SAPIENS | 2 | 941 | 1qki | D | 126 | ASN | p.Asn126Asp | P11413 |
| HOMO SAPIENS | 8 | 941 | 1qki | D | 163 | GLY | p.Gly163Ser | P11413 |
| HOMO SAPIENS | 6 | 941 | 1qki | D | 165 | ASN | p.Asn165Asp | P11413 |
| HOMO SAPIENS | 4 | 941 | 1qki | D | 181 | ASP | p.Asp181Val | P11413 |
| HOMO SAPIENS | 4 | 941 | 1qki | D | 182 | ARG | p.Arg182Trp | P11413 |
| HOMO SAPIENS | 5 | 941 | 1qki | D | 188 | SER | p.Ser188Phe | P11413 |
| HOMO SAPIENS | 9 | 941 | 1qki | D | 198 | ARG | p.Arg198Cys | P11413 |
| HOMO SAPIENS | 8 | 941 | 1qki | D | 213 | VAL | p.Val213Leu | P11413 |
| HOMO SAPIENS | 5 | 941 | 1qki | D | 282 | ASP | p.Asp282His | P11413 |

| | | | | | | | | |
|---|---|---|---|---|---|---|---|---|
| HOMO SAPIENS | 7 | 941 | 1qki | D | 285 | ARG | p.Arg285His | P11413 |
| HOMO SAPIENS | 5 | 941 | 1qki | D | 291 | VAL | p.Val291Met | P11413 |
| HOMO SAPIENS | 8 | 941 | 1qki | D | 323 | LEU | p.Leu323Pro | P11413 |
| HOMO SAPIENS | 3 | 941 | 1qki | D | 335 | ALA | p.Ala335Thr | P11413 |
| HOMO SAPIENS | 6 | 941 | 1qki | D | 350 | ASP | p.Asp350His | P11413 |
| HOMO SAPIENS | 5 | 941 | 1qki | D | 393 | ARG | p.Arg393His | P11413 |
| HOMO SAPIENS | 5 | 941 | 1qki | D | 454 | ARG | p.Arg454Cys | P11413 |
| HOMO SAPIENS | 2 | 941 | 1qki | D | 459 | LEU | p.Arg459Leu | P11413 |
| HOMO SAPIENS | 4 | 941 | 1qki | D | 463 | ARG | p.Arg463His | P11413 |
| HOMO SAPIENS | 4 | 2483 | 1tij | A | 68 | LEU | p.Leu94Gln | P01034 |
| HOMO SAPIENS | 2 | 20632 | 3v5b | A | 331 | ARG | p.Arg331Pro | P02545 |
| HOMO SAPIENS | 2 | 20632 | 3v5b | A | 374 | HIS | p.His374Gln | P02545 |
| HOMO SAPIENS | 4 | 13929 | 5xtd | J | 75 | ARG | p.Arg75His | Q16795 |
| HOMO SAPIENS | 3 | 13929 | 5xtd | J | 243 | VAL | p.Val243Ala | Q16795 |
| HOMO SAPIENS | 1 | 13929 | 5xtd | J | 294 | PRO | p.Pro294Leu | Q16795 |
| HOMO SAPIENS | 2 | 13929 | 5xtd | J | 321 | ARG | p.Arg321Pro | Q16795 |
| HOMO SAPIENS | 0 | 9411 | 5thl | A | 41 | ARG | p.Gly41Arg | P54577 |
| HOMO SAPIENS | 0 | 9411 | 5thl | A | 196 | GLU | p.Glu196Lys | P54577 |
| HOMO SAPIENS | 0 | 9411 | 5thl | A | 316 | ALA | p.Ala316Ser | P54577 |
| HOMO SAPIENS | 0 | 9411 | 5thl | A | 339 | ALA | p.Ala339Thr | P54577 |
| HOMO SAPIENS | 4 | 16273 | 2e8n | A | 46 | THR | p.Thr940Ile | P29317 |
| HOMO SAPIENS | 7 | 16273 | 2e8n | A | 54 | GLY | p.Gly948Trp | P29317 |
| HOMO SAPIENS | 6 | 16273 | 2e8n | A | 65 | ALA | p.Ala959Thr | P29317 |
| HOMO SAPIENS | 7 | 30387 | 2k9y | A | 541 | VAL | p.Val541Met | P29317 |
| HOMO SAPIENS | 0 | 6961 | 4nha | A | 843 | ILE | p.Ile853Val | Q9UPY3 |
| HOMO SAPIENS | -1 | 6961 | 4nha | A | 862 | ALA | p.Ala872Thr | Q9UPY3 |
| HOMO SAPIENS | 3 | 15708 | 4pzg | A | 16 | ARG | p.Arg16Ter | Q9Y5X0 |
| HOMO SAPIENS | 4 | 15708 | 4pzg | A | 51 | ARG | p.Arg51Gln | Q9Y5X0 |
| HOMO SAPIENS | 3 | 30166 | 4o00 | A | 44 | ARG | p.Arg14257Ter | Q8WZ42 |
| HOMO SAPIENS | 2 | 30166 | 4o00 | A | 54 | ARG | p.Arg14267His | Q8WZ42 |
| HOMO SAPIENS | 8 | 11158 | 4hca | A | 275 | TRP | p.Trp274Arg | P23771 |
| HOMO SAPIENS | 2 | 4328 | 4e0s | A | 1476 | ARG | p.Arg1476Ter | P01031 |
| HOMO SAPIENS | 0 | 11462 | 1dx5 | I | 455 | ALA | p.Ala473Val | P07204 |
| HOMO SAPIENS | -1 | 11150 | 2fic | A | 45 | LYS | p.Lys35Asn | O00499 |
| HOMO SAPIENS | 6 | 11150 | 2fic | A | 161 | ASP | p.Asp151Asn | O00499 |
| HOMO SAPIENS | 8 | 27382 | 4mqv | B | 17 | ARG | p.Arg17Ter | Q9NZC9 |
| HOMO SAPIENS | -1 | 37991 | 1q0p | A | 231 | GLN | p.Gln256Ter | P00751 |
| HOMO SAPIENS | 1 | 443 | 5xte | U | 215 | LEU | p.Leu215Phe | P08574 |
| HOMO SAPIENS | 8 | 21471 | 5y4e | A | 1461 | ASN | p.Thr692Ala | Q01484 |
| HOMO SAPIENS | -1 | 35443 | 3um3 | B | 129 | ASP | p.Asp129Val | Q9H444 |
| HOMO SAPIENS | -1 | 35443 | 3um3 | B | 161 | GLU | p.Glu161Lys | Q9H444 |
| HOMO SAPIENS | 6 | 28781 | 2n2a | B | 111 | ILE | p.Ile654Val | P04626 |
| HOMO SAPIENS | 7 | 28781 | 2n2a | B | 112 | ILE | p.Ile655Val | P04626 |
| HOMO SAPIENS | -1 | 3346 | 4ztz | C | 61 | GLY | p.Gly61Arg | Q9UHN1 |
| HOMO SAPIENS | 4 | 3346 | 4ztz | C | 103 | GLY | p.Gly103Ser | Q9UHN1 |
| HOMO SAPIENS | 4 | 3346 | 4ztz | C | 133 | HIS | p.His133Tyr | Q9UHN1 |
| HOMO SAPIENS | -1 | 3346 | 4ztz | C | 153 | LEU | p.Leu153Val | Q9UHN1 |
| HOMO SAPIENS | -1 | 3346 | 4ztz | C | 169 | ALA | p.Ala169Thr | Q9UHN1 |
| HOMO SAPIENS | 7 | 3346 | 4ztz | C | 369 | ARG | p.Arg369Gly | Q9UHN1 |
| HOMO SAPIENS | 4 | 3346 | 4ztz | C | 386 | ASP | p.Asp386Glu | Q9UHN1 |
| HOMO SAPIENS | 4 | 3346 | 4ztz | C | 451 | GLY | p.Gly451Glu | Q9UHN1 |
| HOMO SAPIENS | 4 | 2131 | 5fo9 | E | 1549 | LEU | p.Leu1549Met | P01024 |
| HOMO SAPIENS | 2 | 2131 | 5fo9 | E | 1619 | SER | p.Ser1619Arg | P01024 |
| HOMO SAPIENS | 3 | 12662 | 4jhp | C | 75 | ILE | p.Ile75Val | Q92834 |

| | | | | | | | | |
|---|---|---|---|---|---|---|---|---|
| HOMO SAPIENS | 1 | 12662 | 4jhp | C | 173 | GLY | p.Gly173Arg | Q92834 |
| HOMO SAPIENS | 0 | 12662 | 4jhp | C | 184 | GLN | p.Gln184His | Q92834 |
| HOMO SAPIENS | 3 | 12662 | 4jhp | C | 262 | ALA | p.Ala262Gly | Q92834 |
| HOMO SAPIENS | 3 | 12662 | 4jhp | C | 302 | CYS | p.Cys302Tyr | Q92834 |
| HOMO SAPIENS | 5 | 1093 | 2jg9 | A | 186 | GLN | p.Gln208Ter | P02745 |
| HOMO SAPIENS | 0 | 13181 | 4r4l | A | 12 | MET | p.Met13Ile | Q13976 |
| HOMO SAPIENS | 8 | 25939 | 4iyp | A | 106 | ARG | p.Arg106Gln | P78318 |
| HOMO SAPIENS | 1 | 217 | 2xsx | A | 71 | SER | p.Asn71Ser | P13929 |
| HOMO SAPIENS | 0 | 217 | 2xsx | A | 76 | ALA | p.Ala76Pro | P13929 |
| HOMO SAPIENS | 1 | 217 | 2xsx | A | 85 | ALA | p.Val85Ala | P13929 |
| HOMO SAPIENS | 8 | 217 | 2xsx | A | 156 | GLY | p.Gly156Asp | P13929 |
| HOMO SAPIENS | 5 | 217 | 2xsx | A | 332 | VAL | p.Val332Ile | P13929 |
| HOMO SAPIENS | 7 | 217 | 2xsx | A | 374 | GLY | p.Gly374Glu | P13929 |
| HOMO SAPIENS | 1 | 5508 | 3szm | A | 682 | THR | p.Thr682Asn | Q8NEM0 |
| HOMO SAPIENS | 0 | 5508 | 3szm | A | 715 | TRP | p.Trp715Ter | Q8NEM0 |
| HOMO SAPIENS | 1 | 5508 | 3szm | A | 761 | VAL | p.Ala761Val | Q8NEM0 |
| HOMO SAPIENS | 1 | 5508 | 3szm | A | 828 | PRO | p.Pro828Ser | Q8NEM0 |
| HOMO SAPIENS | 4 | 5508 | 3szm | A | 829 | GLU | p.Glu829Asp | Q8NEM0 |
| HOMO SAPIENS | 2 | 16353 | 2q0z | X | 171 | GLY | p.Gly1978Ala | O75643 |
| HOMO SAPIENS | 2 | 16353 | 2q0z | X | 188 | ALA | p.Ala1995Thr | O75643 |
| HOMO SAPIENS | 0 | 4127 | 3swr | A | 860 | GLY | p.Gly860Arg | P26358 |
| HOMO SAPIENS | -1 | 4127 | 3swr | A | 1122 | SER | p.Ser1122Cys | P26358 |
| HOMO SAPIENS | 0 | 4127 | 3swr | A | 1216 | ASP | p.Asp1216Glu | P26358 |
| HOMO SAPIENS | 0 | 6382 | 3o96 | A | 17 | GLU | p.Glu17Lys | P31749 |
| HOMO SAPIENS | 6 | 14314 | 5xtc | Q | 71 | PRO | p.Pro71Leu | O75306 |
| HOMO SAPIENS | 2 | 5886 | 4fao | E | 22 | ARG | p.Arg40His | Q13705 |
| HOMO SAPIENS | 1 | 2534 | 3mp4 | A | 37 | GLU | p.Glu37Ter | P35914 |
| HOMO SAPIENS | 3 | 2534 | 3mp4 | A | 41 | MET | p.Arg41Gln | P35914 |
| HOMO SAPIENS | 1 | 2534 | 3mp4 | A | 70 | VAL | p.Val70Leu | P35914 |
| HOMO SAPIENS | 2 | 2196 | 3ifq | A | 142 | ARG | p.Arg142His | P14923 |
| HOMO SAPIENS | 2 | 2196 | 3ifq | A | 153 | ASP | p.Asp153Glu | P14923 |
| HOMO SAPIENS | 3 | 2196 | 3ifq | A | 176 | ARG | p.Arg176Gln | P14923 |
| HOMO SAPIENS | 3 | 2196 | 3ifq | A | 265 | ARG | p.Arg265His | P14923 |
| HOMO SAPIENS | 2 | 2196 | 3ifq | A | 301 | GLU | p.Glu301Gly | P14923 |
| HOMO SAPIENS | 0 | 2196 | 3ifq | A | 456 | VAL | p.Val456Ile | P14923 |
| HOMO SAPIENS | 3 | 2196 | 3ifq | A | 603 | VAL | p.Val603Leu | P14923 |
| HOMO SAPIENS | 0 | 2196 | 3ifq | A | 648 | VAL | p.Val648Ile | P14923 |
| HOMO SAPIENS | 1 | 8746 | 1cgj | I | 32 | PRO | p.Pro55Ser | P00995 |
| HOMO SAPIENS | 1 | 8746 | 1cgj | I | 44 | ARG | p.Arg67His | P00995 |
| HOMO SAPIENS | 1 | 8746 | 1cgj | I | 44 | ARG | p.Arg67Cys | P00995 |
| HOMO SAPIENS | 3 | 3212 | 1syq | A | 105 | ARG | p.Arg105Ter | P18206 |
| HOMO SAPIENS | 3 | 3212 | 1syq | A | 126 | PHE | p.Phe126Leu | P18206 |
| HOMO SAPIENS | 4 | 10028 | 3eto | A | 1609 | ARG | p.Arg1608His | P46531 |
| HOMO SAPIENS | 1 | 10028 | 3eto | A | 1672 | VAL | p.Val1671Ile | P46531 |
| HOMO SAPIENS | 5 | 10028 | 3eto | A | 1692 | GLN | p.Gln1691His | P46531 |
| HOMO SAPIENS | 3 | 3749 | 5xtc | Z | 70 | GLY | p.Gly70Ter | O43676 |
| HOMO SAPIENS | 2 | 17039 | 3brv | B | 57 | GLU | p.Glu57Lys | Q9Y6K9 |
| HOMO SAPIENS | 8 | 3589 | 2fh3 | A | 433 | VAL | p.Val460Leu | P06396 |
| HOMO SAPIENS | 1 | 3589 | 2fh3 | A | 454 | ARG | p.Arg481Cys | P06396 |
| HOMO SAPIENS | 4 | 3589 | 2fh3 | A | 498 | ARG | p.Arg525His | P06396 |
| HOMO SAPIENS | 2 | 3589 | 2fh3 | A | 520 | TYR | p.Tyr547Cys | P06396 |
| HOMO SAPIENS | 1 | 3589 | 2fh3 | A | 529 | GLY | p.Gly556Arg | P06396 |
| HOMO SAPIENS | 2 | 3589 | 2fh3 | A | 536 | THR | p.Thr563Ser | P06396 |
| HOMO SAPIENS | 2 | 3589 | 2fh3 | A | 579 | VAL | p.Val606Met | P06396 |

| | | | | | | | | |
|---|---|---|---|---|---|---|---|---|
| HOMO SAPIENS | 1 | 3589 | 2fh3 | A | 589 | THR | p.Thr616Met | P06396 |
| HOMO SAPIENS | 5 | 3589 | 2fh3 | A | 668 | THR | p.Thr695Met | P06396 |
| HOMO SAPIENS | 0 | 148 | 1lcl | A | 130 | ILE | p.Ile130Ser | Q05315 |
| HOMO SAPIENS | -1 | 24727 | 4fj3 | P | 237 | ALA | p.Ala237Thr | P04049 |
| HOMO SAPIENS | 0 | 24727 | 4fj3 | P | 257 | SER | p.Ser257Leu | P04049 |
| HOMO SAPIENS | 0 | 24727 | 4fj3 | P | 261 | PRO | p.Pro261Ala | P04049 |
| HOMO SAPIENS | 0 | 2949 | 5xtb | B | 77 | ARG | p.Arg77Gly | O00217 |
| HOMO SAPIENS | 0 | 2949 | 5xtb | B | 79 | PRO | p.Pro79Leu | O00217 |
| HOMO SAPIENS | 0 | 2949 | 5xtb | B | 100 | ALA | p.Ala100Val | O00217 |
| HOMO SAPIENS | 1 | 2949 | 5xtb | B | 115 | LYS | p.Lys115Glu | O00217 |
| HOMO SAPIENS | 0 | 2949 | 5xtb | B | 138 | ARG | p.Arg138His | O00217 |
| HOMO SAPIENS | 1 | 3959 | 5k5s | B | 25 | ARG | p.Arg25Ter | P41180 |
| HOMO SAPIENS | 4 | 3959 | 5k5s | B | 39 | PRO | p.Pro39Ala | P41180 |
| HOMO SAPIENS | 0 | 3959 | 5k5s | B | 47 | LYS | p.Lys47Asn | P41180 |
| HOMO SAPIENS | 2 | 3959 | 5k5s | B | 62 | ARG | p.Arg62Met | P41180 |
| HOMO SAPIENS | 2 | 3959 | 5k5s | B | 66 | ARG | p.Arg66Cys | P41180 |
| HOMO SAPIENS | 4 | 3959 | 5k5s | B | 90 | ASN | p.Asn90Thr | P41180 |
| HOMO SAPIENS | 1 | 3959 | 5k5s | B | 116 | ALA | p.Ala116Thr | P41180 |
| HOMO SAPIENS | 1 | 3959 | 5k5s | B | 118 | ASN | p.Asn118Lys | P41180 |
| HOMO SAPIENS | -1 | 3959 | 5k5s | B | 125 | LEU | p.Leu125Pro | P41180 |
| HOMO SAPIENS | -1 | 3959 | 5k5s | B | 127 | GLU | p.Glu127Ala | P41180 |
| HOMO SAPIENS | -1 | 3959 | 5k5s | B | 128 | PHE | p.Phe128Leu | P41180 |
| HOMO SAPIENS | -1 | 3959 | 5k5s | B | 131 | CYS | p.Cys131Trp | P41180 |
| HOMO SAPIENS | 1 | 3959 | 5k5s | B | 138 | THR | p.Thr138Met | P41180 |
| HOMO SAPIENS | 0 | 3959 | 5k5s | B | 143 | GLY | p.Gly143Glu | P41180 |
| HOMO SAPIENS | 5 | 3959 | 5k5s | B | 151 | THR | p.Thr151Met | P41180 |
| HOMO SAPIENS | 4 | 3959 | 5k5s | B | 180 | PHE | p.Phe180Cys | P41180 |
| HOMO SAPIENS | 0 | 3959 | 5k5s | B | 185 | ARG | p.Arg185Gln | P41180 |
| HOMO SAPIENS | 0 | 3959 | 5k5s | B | 185 | ARG | p.Arg185Ter | P41180 |
| HOMO SAPIENS | 2 | 3959 | 5k5s | B | 191 | GLU | p.Glu191Lys | P41180 |
| HOMO SAPIENS | 2 | 3959 | 5k5s | B | 227 | ARG | p.Arg227Leu | P41180 |
| HOMO SAPIENS | 2 | 3959 | 5k5s | B | 250 | GLU | p.Glu250Lys | P41180 |
| HOMO SAPIENS | 6 | 3959 | 5k5s | B | 297 | GLU | p.Glu297Lys | P41180 |
| HOMO SAPIENS | 1 | 3959 | 5k5s | B | 336 | LYS | p.Lys336Asn | P41180 |
| HOMO SAPIENS | 6 | 3959 | 5k5s | B | 429 | HIS | p.His429Tyr | P41180 |
| HOMO SAPIENS | 0 | 3959 | 5k5s | B | 445 | THR | p.Thr445Ala | P41180 |
| HOMO SAPIENS | 3 | 3959 | 5k5s | B | 509 | GLY | p.Gly509Arg | P41180 |
| HOMO SAPIENS | 1 | 3959 | 5k5s | B | 553 | GLY | p.Gly553Arg | P41180 |
| HOMO SAPIENS | 3 | 3959 | 5k5s | B | 592 | ASN | p.Asn592Ser | P41180 |
| HOMO SAPIENS | -1 | 3959 | 5k5s | B | 604 | GLU | p.Glu604Lys | P41180 |
| HOMO SAPIENS | 4 | 2564 | 5l0c | C | 975 | ARG | p.Arg975Trp | P18206 |
| HOMO SAPIENS | 4 | 2564 | 5l0c | C | 975 | ARG | p.Arg975Leu | P18206 |
| HOMO SAPIENS | 1 | 5695 | 5i8r | A | 294 | ARG | p.Arg294Gln | P17405 |
| HOMO SAPIENS | 0 | 5695 | 5i8r | A | 485 | ALA | p.Ala485Val | P17405 |
| HOMO SAPIENS | 1 | 5695 | 5i8r | A | 506 | GLY | p.Gly506Arg | P17405 |
| HOMO SAPIENS | -1 | 1978 | 3avs | A | 1060 | HIS | p.His1060Leu | O15550 |
| HOMO SAPIENS | 0 | 11173 | 4v0c | B | 362 | LYS | p.Lys362Arg | P51787 |
| HOMO SAPIENS | 3 | 11173 | 4v0c | B | 381 | CYS | p.Cys381Tyr | P51787 |
| HOMO SAPIENS | 7 | 11173 | 4v0c | B | 518 | ARG | p.Arg518Gly | P51787 |
| HOMO SAPIENS | 2 | 11173 | 4v0c | B | 525 | ALA | p.Ala525Thr | P51787 |
| HOMO SAPIENS | 0 | 11173 | 4v0c | B | 530 | GLN | p.Gln530Ter | P51787 |
| HOMO SAPIENS | -1 | 11173 | 4v0c | B | 536 | TYR | p.Tyr536Ter | P51787 |
| HOMO SAPIENS | 2 | 1113 | 5xtd | N | 60 | ARG | p.Arg60Ter | Q9UI09 |
| HOMO SAPIENS | 3 | 6061 | 2mmz | A | 148 | LEU | p.Leu148Ser | Q86SX6 |

| | | | | | | | | |
|---|---|---|---|---|---|---|---|---|
| HOMO SAPIENS | -1 | 26081 | 3eu7 | A | 837 | GLU | p.Glu837Ter | Q86YC2 |
| HOMO SAPIENS | 0 | 26081 | 3eu7 | A | 864 | PRO | p.Pro864Ser | Q86YC2 |
| HOMO SAPIENS | 5 | 26081 | 3eu7 | A | 891 | CYS | p.Cys891Trp | Q86YC2 |
| HOMO SAPIENS | 0 | 26081 | 3eu7 | A | 932 | VAL | p.Val932Met | Q86YC2 |
| HOMO SAPIENS | 5 | 26081 | 3eu7 | A | 961 | LEU | p.Leu961Met | Q86YC2 |
| HOMO SAPIENS | 5 | 26081 | 3eu7 | A | 1038 | TRP | p.Trp1038Ter | Q86YC2 |
| HOMO SAPIENS | -1 | 26081 | 3eu7 | A | 1086 | ARG | p.Arg1086Ter | Q86YC2 |
| HOMO SAPIENS | 0 | 26081 | 3eu7 | A | 1122 | ASP | p.Asp1122Glu | Q86YC2 |
| HOMO SAPIENS | 5 | 26081 | 3eu7 | A | 1158 | HIS | p.His1158Gln | Q86YC2 |
| HOMO SAPIENS | 5 | 26081 | 3eu7 | A | 1183 | TYR | p.Tyr1183Ter | Q86YC2 |
| HOMO SAPIENS | 4 | 199 | 1qo5 | C | 45 | ARG | p.Arg46Trp | P05062 |
| HOMO SAPIENS | 5 | 199 | 1qo5 | C | 147 | TRP | p.Trp148Arg | P05062 |
| HOMO SAPIENS | 0 | 199 | 1qo5 | C | 149 | ALA | p.Ala150Pro | P05062 |
| HOMO SAPIENS | 0 | 199 | 1qo5 | C | 174 | ALA | p.Ala175Asp | P05062 |
| HOMO SAPIENS | 0 | 199 | 1qo5 | C | 334 | ASN | p.Asn335Lys | P05062 |
| HOMO SAPIENS | 0 | 199 | 1qo5 | C | 337 | ALA | p.Ala338Val | P05062 |
| HOMO SAPIENS | 5 | 199 | 1qo5 | C | 342 | TYR | p.Tyr343His | P05062 |
| HOMO SAPIENS | 0 | 6910 | 4zht | B | 59 | ARG | p.Arg59Ter | Q9Y223 |
| HOMO SAPIENS | 4 | 6910 | 4zht | B | 129 | ARG | p.Arg129Ter | Q9Y223 |
| HOMO SAPIENS | 4 | 6910 | 4zht | B | 129 | ARG | p.Arg129Gln | Q9Y223 |
| HOMO SAPIENS | 1 | 6910 | 4zht | B | 162 | ARG | p.Arg162Cys | Q9Y223 |
| HOMO SAPIENS | 0 | 6910 | 4zht | B | 176 | ASP | p.Asp176Val | Q9Y223 |
| HOMO SAPIENS | 0 | 6910 | 4zht | B | 216 | VAL | p.Val216Ala | Q9Y223 |
| HOMO SAPIENS | 0 | 6910 | 4zht | B | 246 | ARG | p.Arg246Trp | Q9Y223 |
| HOMO SAPIENS | 0 | 6910 | 4zht | B | 263 | ARG | p.Arg263Leu | Q9Y223 |
| HOMO SAPIENS | 0 | 6910 | 4zht | B | 266 | ARG | p.Arg266Gln | Q9Y223 |
| HOMO SAPIENS | 0 | 6910 | 4zht | B | 266 | ARG | p.Arg266Trp | Q9Y223 |
| HOMO SAPIENS | 0 | 6910 | 4zht | B | 277 | ARG | p.Arg277Cys | Q9Y223 |
| HOMO SAPIENS | 0 | 6910 | 4zht | B | 378 | ASP | p.Asp378Tyr | Q9Y223 |
| HOMO SAPIENS | 3 | 211 | 5kxi | C | 262 | VAL | p.Val287Leu | P17787 |
| HOMO SAPIENS | 0 | 211 | 5kxi | C | 369 | PHE | p.Phe478Leu | P17787 |
| HOMO SAPIENS | 0 | 2454 | 5iy9 | V | null | PHE | p.Phe99Ser | P19447 |
| HOMO SAPIENS | 0 | 2454 | 5iy9 | V | null | ARG | p.Arg109Ter | P19447 |
| HOMO SAPIENS | 0 | 2454 | 5iy9 | V | null | THR | p.Thr119Pro | P19447 |
| HOMO SAPIENS | 0 | 2454 | 5iy9 | V | 425 | ARG | p.Arg425Ter | P19447 |
| HOMO SAPIENS | 1 | 2454 | 5iy9 | V | 545 | GLN | p.Gln545Ter | P19447 |
| HOMO SAPIENS | -1 | 3805 | 3rbq | A | 57 | LYS | p.Lys57Ter | Q13432 |
| HOMO SAPIENS | 2 | 2649 | 4ido | A | 43 | ASP | p.Asp43Glu | Q8WXF7 |
| HOMO SAPIENS | 2 | 2649 | 4ido | A | 157 | LEU | p.Leu157Trp | Q8WXF7 |
| HOMO SAPIENS | 2 | 2649 | 4ido | A | 193 | PHE | p.Phe193Cys | Q8WXF7 |
| HOMO SAPIENS | 7 | 2649 | 4ido | A | 217 | ARG | p.Arg217Gln | Q8WXF7 |
| HOMO SAPIENS | 4 | 2649 | 4ido | A | 239 | ARG | p.Arg239Cys | Q8WXF7 |
| HOMO SAPIENS | 3 | 2649 | 4ido | A | 258 | HIS | p.His258Arg | Q8WXF7 |
| HOMO SAPIENS | 4 | 2649 | 4ido | A | 259 | SER | p.Ser259Tyr | Q8WXF7 |
| HOMO SAPIENS | 4 | 2649 | 4ido | A | 408 | MET | p.Met408Val | Q8WXF7 |
| HOMO SAPIENS | 2 | 2649 | 4ido | A | 415 | ARG | p.Arg415Trp | Q8WXF7 |
| HOMO SAPIENS | 3 | 2649 | 4ido | A | 416 | ARG | p.Arg416Cys | Q8WXF7 |
| HOMO SAPIENS | 5 | 2203 | 3wff | A | 810 | LEU | p.Ser810Leu | P08235 |
| HOMO SAPIENS | 4 | 12648 | 5ep6 | B | 696 | GLU | p.Glu696Lys | Q9UHD2 |
| HOMO SAPIENS | 0 | 30622 | 2pe4 | A | 90 | GLU | p.Glu90Asp | Q12794 |
| HOMO SAPIENS | 0 | 30622 | 2pe4 | A | 256 | GLY | p.Gly256Arg | Q12794 |
| HOMO SAPIENS | 1 | 30622 | 2pe4 | A | 268 | GLU | p.Glu268Lys | Q12794 |
| HOMO SAPIENS | 0 | 30622 | 2pe4 | A | 392 | THR | p.Thr392Met | Q12794 |
| HOMO SAPIENS | 5 | 5411 | 1lya | A | 93 | SER | p.Ser157Leu | P07339 |

| | | | | | | | | |
|---|---|---|---|---|---|---|---|---|
| HOMO SAPIENS | 3 | 5743 | 2uzp | A | 189 | ASN | p.Asn189Lys | P05129 |
| HOMO SAPIENS | 1 | 12529 | 3kfd | J | 21 | ASN | p.Asn45Ser | P36897 |
| HOMO SAPIENS | 1 | 1063 | 2e1q | B | 149 | ARG | p.Arg149Cys | P47989 |
| HOMO SAPIENS | -1 | 1063 | 2e1q | B | 172 | GLY | p.Gly172Arg | P47989 |
| HOMO SAPIENS | 4 | 1063 | 2e1q | B | 228 | ARG | p.Arg228Ter | P47989 |
| HOMO SAPIENS | 5 | 1063 | 2e1q | B | 395 | LYS | p.Lys395Met | P47989 |
| HOMO SAPIENS | 2 | 1063 | 2e1q | B | 646 | ILE | p.Ile646Val | P47989 |
| HOMO SAPIENS | 4 | 1063 | 2e1q | B | 703 | ILE | p.Ile703Val | P47989 |
| HOMO SAPIENS | 8 | 2525 | 1kqn | F | 18 | ASN | p.Asn18Ser | Q9HAN9 |
| HOMO SAPIENS | 0 | 2525 | 1kqn | F | 151 | VAL | p.Val151Phe | Q9HAN9 |
| HOMO SAPIENS | 0 | 2525 | 1kqn | F | 153 | LEU | p.Leu153Val | Q9HAN9 |
| HOMO SAPIENS | 0 | 2525 | 1kqn | F | 169 | TRP | p.Trp169Ter | Q9HAN9 |
| HOMO SAPIENS | 0 | 2525 | 1kqn | F | 207 | ARG | p.Arg207Trp | Q9HAN9 |
| HOMO SAPIENS | 0 | 2525 | 1kqn | F | 273 | ASN | p.Asn273Asp | Q9HAN9 |
| HOMO SAPIENS | 0 | 12299 | 5duq | A | 432 | VAL | p.Val454Glu | P05155 |
| HOMO SAPIENS | 0 | 12299 | 5duq | A | 436 | ALA | p.Ala458Thr | P05155 |
| HOMO SAPIENS | 0 | 12299 | 5duq | A | 443 | ALA | p.Ala465Val | P05155 |
| HOMO SAPIENS | 0 | 12299 | 5duq | A | 444 | ARG | p.Arg466His | P05155 |
| HOMO SAPIENS | 0 | 12299 | 5duq | A | 458 | MET | p.Val480Met | P05155 |
| HOMO SAPIENS | 0 | 327 | 1uhp | A | 36 | ASP | p.Asp274Asn | Q9UPQ7 |
| HOMO SAPIENS | 0 | 64 | 2mo5 | A | 54 | ALA | p.Ala55Thr | P12104 |
| HOMO SAPIENS | 2 | 35969 | 1jrh | I | 60 | CYS | p.Cys77Tyr | P15260 |
| HOMO SAPIENS | 4 | 35969 | 1jrh | I | 70 | ILE | p.Ile87Thr | P15260 |
| HOMO SAPIENS | 2 | 1068 | 1mu7 | A | 304 | ARG | p.Arg304Gln | Q9NUW8 |
| HOMO SAPIENS | 8 | 1068 | 1mu7 | A | 493 | HIS | p.His493Arg | Q9NUW8 |
| HOMO SAPIENS | 0 | 32360 | 2kqp | A | 31 | ARG | p.Arg55Cys | P01308 |
| HOMO SAPIENS | 0 | 32360 | 2kqp | A | 65 | ARG | p.Arg89His | P01308 |
| HOMO SAPIENS | 0 | 32360 | 2kqp | A | 65 | ARG | p.Arg89Cys | P01308 |
| HOMO SAPIENS | 6 | 10805 | 3nks | A | 12 | ILE | p.Ile12Thr | P50336 |
| HOMO SAPIENS | 0 | 10805 | 3nks | A | 20 | HIS | p.His20Pro | P50336 |
| HOMO SAPIENS | 0 | 10805 | 3nks | A | 59 | ARG | p.Arg59Trp | P50336 |
| HOMO SAPIENS | 0 | 10805 | 3nks | A | 168 | ARG | p.Arg168Cys | P50336 |
| HOMO SAPIENS | 0 | 10805 | 3nks | A | 168 | ARG | p.Arg168His | P50336 |
| HOMO SAPIENS | 2 | 10805 | 3nks | A | 232 | GLY | p.Gly232Arg | P50336 |
| HOMO SAPIENS | 3 | 10805 | 3nks | A | 256 | PRO | p.Pro256Arg | P50336 |
| HOMO SAPIENS | 0 | 10805 | 3nks | A | 304 | ARG | p.Arg304His | P50336 |
| HOMO SAPIENS | 0 | 10805 | 3nks | A | 349 | ASP | p.Asp349Ala | P50336 |
| HOMO SAPIENS | 9 | 7008 | 3u2l | A | 125 | GLN | p.Gln125Ter | P55789 |
| HOMO SAPIENS | 4 | 7008 | 3u2l | A | 172 | HIS | p.His172Asn | P55789 |
| HOMO SAPIENS | 4 | 7008 | 3u2l | A | 194 | ARG | p.Arg194His | P55789 |
| HOMO SAPIENS | 3 | 7008 | 3u2l | A | 198 | GLY | p.Gly198Ser | P55789 |
| HOMO SAPIENS | 0 | 20821 | 3e0g | A | 65 | HIS | p.His116Tyr | P42702 |
| HOMO SAPIENS | 3 | 1151 | 5l9v | A | 278 | ASP | p.Asp278Asn | Q9GZT9 |
| HOMO SAPIENS | 4 | 1151 | 5l9v | A | 317 | CYS | p.Pro317Arg | Q9GZT9 |
| HOMO SAPIENS | 5 | 1151 | 5l9v | A | 371 | ARG | p.Arg371His | Q9GZT9 |
| HOMO SAPIENS | 2 | 3935 | 1gwb | A | 36 | ARG | p.Arg36Gly | P07359 |
| HOMO SAPIENS | 1 | 3935 | 1gwb | A | 73 | LEU | p.Leu73Phe | P07359 |
| HOMO SAPIENS | 1 | 3935 | 1gwb | A | 172 | ALA | p.Ala172Val | P07359 |
| HOMO SAPIENS | 0 | 3935 | 1gwb | A | 249 | GLY | p.Gly249Val | P07359 |
| HOMO SAPIENS | 0 | 3935 | 1gwb | A | 255 | MET | p.Met255Val | P07359 |
| HOMO SAPIENS | 2 | 14541 | 5cuz | A | 138 | GLU | p.Glu138Lys | Q9H3L0 |
| HOMO SAPIENS | 2 | 14541 | 5cuz | A | 143 | SER | p.Ser143Ile | Q9H3L0 |
| HOMO SAPIENS | 4 | 14541 | 5cuz | A | 182 | THR | p.Thr182Asn | Q9H3L0 |
| HOMO SAPIENS | 6 | 14541 | 5cuz | A | 206 | ASN | p.Asn206Ile | Q9H3L0 |

| | | | | | | | | |
|---|---|---|---|---|---|---|---|---|
| HOMO SAPIENS | 0 | 14541 | 5cuz | A | 249 | TYR | p.Tyr249Cys | Q9H3L0 |
| HOMO SAPIENS | 4 | 14541 | 5cuz | A | 250 | ARG | p.Arg250Ter | Q9H3L0 |
| HOMO SAPIENS | 2 | 14541 | 5cuz | A | 259 | LEU | p.Leu259Pro | Q9H3L0 |
| HOMO SAPIENS | 0 | 4206 | 4fao | D | 30 | PRO | p.Pro30Ser | P37023 |
| HOMO SAPIENS | 1 | 4206 | 4fao | D | 50 | TRP | p.Trp50Cys | P37023 |
| HOMO SAPIENS | 0 | 16553 | 3v30 | A | 251 | GLN | p.Gln251Glu | O14593 |
| HOMO SAPIENS | 0 | 5012 | 5hbf | A | 442 | ALA | p.Ala442Gly | Q13231 |
| HOMO SAPIENS | 5 | 4844 | 5tdc | A | 136 | HIS | p.His136Arg | Q8IWV7 |
| HOMO SAPIENS | 0 | 15435 | 4cgz | A | 643 | ARG | p.Arg643His | P54132 |
| HOMO SAPIENS | 1 | 15435 | 4cgz | A | 672 | GLN | p.Gln672Arg | P54132 |
| HOMO SAPIENS | 1 | 15435 | 4cgz | A | 700 | GLN | p.Gln700Ter | P54132 |
| HOMO SAPIENS | 0 | 15435 | 4cgz | A | 707 | PRO | p.Pro707Ser | P54132 |
| HOMO SAPIENS | 0 | 15435 | 4cgz | A | 755 | LYS | p.Lys755Glu | P54132 |
| HOMO SAPIENS | 0 | 15435 | 4cgz | A | 881 | TRP | p.Trp881Ter | P54132 |
| HOMO SAPIENS | 2 | 15435 | 4cgz | A | 899 | ARG | p.Arg899Ter | P54132 |
| HOMO SAPIENS | 0 | 15435 | 4cgz | A | 1036 | CYS | p.Cys1036Phe | P54132 |
| HOMO SAPIENS | 0 | 15435 | 4cgz | A | 1055 | CYS | p.Cys1055Ser | P54132 |
| HOMO SAPIENS | 0 | 15435 | 4cgz | A | 1139 | ARG | p.Arg1139Ter | P54132 |
| HOMO SAPIENS | -1 | 15435 | 4cgz | A | 1198 | VAL | p.Val1198Met | P54132 |
| HOMO SAPIENS | -1 | 15435 | 4cgz | A | 1205 | VAL | p.Val1205Ile | P54132 |
| HOMO SAPIENS | 0 | 15435 | 4cgz | A | 1251 | GLU | p.Glu1251Gln | P54132 |
| HOMO SAPIENS | 0 | 11185 | 1kb4 | A | 30 | ARG | p.Arg30Ter | P11473 |
| HOMO SAPIENS | 5 | 11185 | 1kb4 | A | 33 | GLY | p.Gly33Asp | P11473 |
| HOMO SAPIENS | 5 | 11185 | 1kb4 | A | 46 | GLY | p.Gly46Asp | P11473 |
| HOMO SAPIENS | 6 | 11185 | 1kb4 | A | 50 | ARG | p.Arg50Gln | P11473 |
| HOMO SAPIENS | 9 | 11185 | 1kb4 | A | 73 | ARG | p.Arg73Gln | P11473 |
| HOMO SAPIENS | 9 | 11185 | 1kb4 | A | 80 | ARG | p.Arg80Gln | P11473 |
| HOMO SAPIENS | 0 | 5382 | 3fl7 | A | 41 | LEU | p.Leu41Val | P29317 |
| HOMO SAPIENS | 2 | 5382 | 3fl7 | A | 175 | ARG | p.Arg175Cys | P29317 |
| HOMO SAPIENS | 1 | 5382 | 3fl7 | A | 195 | ARG | p.Arg195Cys | P29317 |
| HOMO SAPIENS | 0 | 5382 | 3fl7 | A | 232 | ASP | p.Asp232Gly | P29317 |
| HOMO SAPIENS | 0 | 5382 | 3fl7 | A | 277 | SER | p.Ser277Leu | P29317 |
| HOMO SAPIENS | 0 | 5382 | 3fl7 | A | 315 | ARG | p.Arg315Gln | P29317 |
| HOMO SAPIENS | 7 | 5382 | 3fl7 | A | 350 | PRO | p.Pro350Thr | P29317 |
| HOMO SAPIENS | 0 | 5382 | 3fl7 | A | 391 | GLY | p.Gly391Arg | P29317 |
| HOMO SAPIENS | 0 | 5382 | 3fl7 | A | 418 | VAL | p.Val418Ile | P29317 |
| HOMO SAPIENS | 0 | 5382 | 3fl7 | A | 511 | THR | p.Thr511Met | P29317 |
| HOMO SAPIENS | 2 | 29168 | 4p22 | B | 144 | VAL | p.Val144Ile | P22314 |
| HOMO SAPIENS | 5 | 4348 | 1bf5 | A | 241 | ARG | p.Arg241Gln | P42224 |
| HOMO SAPIENS | 2 | 4348 | 1bf5 | A | 266 | VAL | p.Val266Ile | P42224 |
| HOMO SAPIENS | 4 | 4348 | 1bf5 | A | 320 | GLU | p.Glu320Gln | P42224 |
| HOMO SAPIENS | 1 | 4348 | 1bf5 | A | 372 | ASN | p.Asn372Lys | P42224 |
| HOMO SAPIENS | 1 | 4348 | 1bf5 | A | 447 | ASP | p.Asp447Glu | P42224 |
| HOMO SAPIENS | 2 | 4348 | 1bf5 | A | 463 | GLN | p.Gln463His | P42224 |
| HOMO SAPIENS | 4 | 4348 | 1bf5 | A | 600 | LEU | p.Leu600Pro | P42224 |
| HOMO SAPIENS | 0 | 4348 | 1bf5 | A | 706 | LEU | p.Leu706Ser | P42224 |
| HOMO SAPIENS | 4 | 4991 | 4bej | B | 141 | ASN | p.Asn141Ser | O00429 |
| HOMO SAPIENS | 0 | 4991 | 4bej | B | 395 | ALA | p.Ala395Asp | O00429 |
| HOMO SAPIENS | 2 | 11792 | 1fvo | A | 40 | ARG | p.Arg40Cys | P00480 |
| HOMO SAPIENS | 2 | 11792 | 1fvo | A | 40 | ARG | p.Arg40His | P00480 |
| HOMO SAPIENS | 0 | 11792 | 1fvo | A | 45 | LEU | p.Leu45Pro | P00480 |
| HOMO SAPIENS | 1 | 11792 | 1fvo | A | 47 | ASN | p.Asn47Ile | P00480 |
| HOMO SAPIENS | 0 | 11792 | 1fvo | A | 50 | GLY | p.Gly50Arg | P00480 |
| HOMO SAPIENS | 5 | 11792 | 1fvo | A | 87 | GLU | p.Glu87Lys | P00480 |

| | | | | | | | | |
|---|---|---|---|---|---|---|---|---|
| HOMO SAPIENS | 3 | 11792 | 1fvo | A | 111 | LEU | p.Leu111Pro | P00480 |
| HOMO SAPIENS | 1 | 11792 | 1fvo | A | 125 | THR | p.Thr125Met | P00480 |
| HOMO SAPIENS | 3 | 11792 | 1fvo | A | 129 | ARG | p.Arg129His | P00480 |
| HOMO SAPIENS | 9 | 11792 | 1fvo | A | 141 | ARG | p.Arg141Pro | P00480 |
| HOMO SAPIENS | 0 | 11792 | 1fvo | A | 206 | MET | p.Met206Arg | P00480 |
| HOMO SAPIENS | 0 | 11792 | 1fvo | A | 207 | SER | p.Ser207Arg | P00480 |
| HOMO SAPIENS | 0 | 11792 | 1fvo | A | 216 | GLN | p.Gln216Glu | P00480 |
| HOMO SAPIENS | 1 | 11792 | 1fvo | A | 225 | PRO | p.Pro225Leu | P00480 |
| HOMO SAPIENS | 0 | 11792 | 1fvo | A | 270 | ARG | p.Gln270Arg | P00480 |
| HOMO SAPIENS | 0 | 11792 | 1fvo | A | 270 | ARG | p.Gln270Ter | P00480 |
| HOMO SAPIENS | 2 | 11792 | 1fvo | A | 277 | ARG | p.Arg277Trp | P00480 |
| HOMO SAPIENS | 0 | 3721 | 4r3z | B | 2 | ASP | p.Asp2Gly | P54136 |
| HOMO SAPIENS | 0 | 3721 | 4r3z | B | 456 | SER | p.Ser456Leu | P54136 |
| HOMO SAPIENS | 5 | 3721 | 4r3z | B | 512 | ARG | p.Arg512Gln | P54136 |
| HOMO SAPIENS | 3 | 5749 | 3hl2 | A | 239 | ALA | p.Ala239Thr | Q9HD40 |
| HOMO SAPIENS | 1 | 5749 | 3hl2 | A | 334 | TYR | p.Tyr334Cys | Q9HD40 |
| HOMO SAPIENS | 1 | 5749 | 3hl2 | A | 452 | LYS | p.Lys452Asn | Q9HD40 |
| HOMO SAPIENS | -1 | 5749 | 3hl2 | A | 465 | ARG | p.Arg465Leu | Q9HD40 |
| HOMO SAPIENS | -1 | 5749 | 3hl2 | A | 489 | ASP | p.Asp489Gly | Q9HD40 |
| HOMO SAPIENS | 5 | 1721 | 3k15 | A | 1652 | MET | p.Met1652Ile | P38398 |
| HOMO SAPIENS | 5 | 1721 | 3k15 | A | 1652 | MET | p.Met1652Lys | P38398 |
| HOMO SAPIENS | 5 | 1721 | 3k15 | A | 1662 | PHE | p.Phe1662Cys | P38398 |
| HOMO SAPIENS | 5 | 1721 | 3k15 | A | 1664 | LEU | p.Leu1664Pro | P38398 |
| HOMO SAPIENS | 0 | 1721 | 3k15 | A | 1665 | VAL | p.Val1665Met | P38398 |
| HOMO SAPIENS | 5 | 1721 | 3k15 | A | 1690 | LYS | p.Lys1690Gln | P38398 |
| HOMO SAPIENS | 5 | 1721 | 3k15 | A | 1692 | ASP | p.Asp1692Asn | P38398 |
| HOMO SAPIENS | 5 | 1721 | 3k15 | A | 1699 | ARG | p.Arg1699Gln | P38398 |
| HOMO SAPIENS | 5 | 1721 | 3k15 | A | 1699 | ARG | p.Arg1699Trp | P38398 |
| HOMO SAPIENS | 3 | 1721 | 3k15 | A | 1706 | GLY | p.Gly1706Glu | P38398 |
| HOMO SAPIENS | 5 | 1721 | 3k15 | A | 1720 | THR | p.Thr1720Ala | P38398 |
| HOMO SAPIENS | 5 | 1721 | 3k15 | A | 1751 | ARG | p.Arg1751Pro | P38398 |
| HOMO SAPIENS | 4 | 1721 | 3k15 | A | 1775 | MET | p.Met1775Arg | P38398 |
| HOMO SAPIENS | 2 | 1721 | 3k15 | A | 1788 | GLY | p.Gly1788Val | P38398 |
| HOMO SAPIENS | 5 | 1721 | 3k15 | A | 1794 | GLU | p.Glu1794Asp | P38398 |
| HOMO SAPIENS | 5 | 1721 | 3k15 | A | 1804 | VAL | p.Val1804Asp | P38398 |
| HOMO SAPIENS | 3 | 1721 | 3k15 | A | 1815 | TRP | p.Trp1815Ter | P38398 |
| HOMO SAPIENS | 1 | 1721 | 3k15 | A | 1819 | ASN | p.Asn1819Ser | P38398 |
| HOMO SAPIENS | 5 | 1721 | 3k15 | A | 1856 | PRO | p.Pro1856Ser | P38398 |
| HOMO SAPIENS | 5 | 1721 | 3k15 | A | 1858 | ILE | p.Ile1858Leu | P38398 |
| HOMO SAPIENS | 5 | 1721 | 3k15 | A | 1858 | ILE | p.Ile1858Thr | P38398 |
| HOMO SAPIENS | 5 | 1721 | 3k15 | A | 1859 | PRO | p.Pro1859Arg | P38398 |
| HOMO SAPIENS | 2 | 22086 | 3kq4 | B | 1297 | PRO | p.Pro1297Leu | O60494 |
| HOMO SAPIENS | 1 | 12135 | 2k9j | B | 724 | ARG | p.Arg750Ter | P05106 |
| HOMO SAPIENS | -1 | 24027 | 2qtl | A | 175 | LEU | p.Ser202Leu | Q9UBK8 |
| HOMO SAPIENS | 0 | 24027 | 2qtl | A | 257 | SER | p.Ser284Thr | Q9UBK8 |
| HOMO SAPIENS | 0 | 24027 | 2qtl | A | 333 | LEU | p.Leu360Val | Q9UBK8 |
| HOMO SAPIENS | 0 | 24027 | 2qtl | A | 350 | LYS | p.Lys377Arg | Q9UBK8 |
| HOMO SAPIENS | 0 | 24027 | 2qtl | A | 415 | ARG | p.Arg442Cys | Q9UBK8 |
| HOMO SAPIENS | 2 | 24027 | 2qtl | A | 450 | PRO | p.Pro477Arg | Q9UBK8 |
| HOMO SAPIENS | 4 | 24027 | 2qtl | A | 487 | GLY | p.Gly514Arg | Q9UBK8 |
| HOMO SAPIENS | 1 | 24027 | 2qtl | A | 595 | HIS | p.His622Tyr | Q9UBK8 |
| HOMO SAPIENS | 0 | 5758 | 3di2 | B | 46 | THR | p.Thr66Ile | P16871 |
| HOMO SAPIENS | 1 | 5758 | 3di2 | B | 118 | VAL | p.Ile138Val | P16871 |
| HOMO SAPIENS | 4 | 848 | 5oa3 | Y | 79 | LEU | p.Leu79Val | P62847 |

| | | | | | | | | |
|---|---|---|---|---|---|---|---|---|
| HOMO SAPIENS | 2 | 848 | 5oa3 | Y | 124 | ASN | p.Asn124Ser | P62847 |
| HOMO SAPIENS | 0 | 81 | 3s5o | A | 36 | ALA | p.Ala36Glu | Q86XE5 |
| HOMO SAPIENS | 0 | 81 | 3s5o | A | 39 | TYR | p.Tyr39Ter | Q86XE5 |
| HOMO SAPIENS | 0 | 81 | 3s5o | A | 45 | PRO | p.Pro45Leu | Q86XE5 |
| HOMO SAPIENS | 0 | 81 | 3s5o | A | 70 | ARG | p.Arg70Ter | Q86XE5 |
| HOMO SAPIENS | 0 | 81 | 3s5o | A | 70 | ARG | p.Arg70Pro | Q86XE5 |
| HOMO SAPIENS | 1 | 81 | 3s5o | A | 97 | ARG | p.Arg97Cys | Q86XE5 |
| HOMO SAPIENS | 4 | 81 | 3s5o | A | 113 | GLU | p.Glu113Lys | Q86XE5 |
| HOMO SAPIENS | 1 | 81 | 3s5o | A | 190 | PRO | p.Pro190Leu | Q86XE5 |
| HOMO SAPIENS | 0 | 81 | 3s5o | A | 245 | VAL | p.Val245Phe | Q86XE5 |
| HOMO SAPIENS | 0 | 81 | 3s5o | A | 257 | CYS | p.Cys257Gly | Q86XE5 |
| HOMO SAPIENS | 0 | 81 | 3s5o | A | 287 | GLY | p.Gly287Val | Q86XE5 |
| HOMO SAPIENS | 7 | 81 | 3s5o | A | 303 | ARG | p.Arg303Cys | Q86XE5 |
| HOMO SAPIENS | 0 | 81 | 3s5o | A | 325 | GLY | p.Gly325Ser | Q86XE5 |
| HOMO SAPIENS | -1 | 622 | 5era | A | 12 | GLY | p.Gly12Arg | P29033 |
| HOMO SAPIENS | -1 | 622 | 5era | A | 12 | GLY | p.Gly12Asp | P29033 |
| HOMO SAPIENS | -1 | 622 | 5era | A | 17 | SER | p.Ser17Phe | P29033 |
| HOMO SAPIENS | 5 | 622 | 5era | A | 24 | TRP | p.Trp24Ter | P29033 |
| HOMO SAPIENS | 0 | 622 | 5era | A | 32 | ARG | p.Arg32Cys | P29033 |
| HOMO SAPIENS | 5 | 622 | 5era | A | 34 | MET | p.Met34Thr | P29033 |
| HOMO SAPIENS | 5 | 622 | 5era | A | 36 | LEU | p.Leu36Pro | P29033 |
| HOMO SAPIENS | 5 | 622 | 5era | A | 37 | VAL | p.Val37Ile | P29033 |
| HOMO SAPIENS | 5 | 622 | 5era | A | 40 | ALA | p.Ala40Glu | P29033 |
| HOMO SAPIENS | 5 | 622 | 5era | A | 44 | TRP | p.Trp44Cys | P29033 |
| HOMO SAPIENS | 5 | 622 | 5era | A | 45 | GLY | p.Gly45Glu | P29033 |
| HOMO SAPIENS | 5 | 622 | 5era | A | 47 | GLU | p.Glu47Ter | P29033 |
| HOMO SAPIENS | 5 | 622 | 5era | A | 50 | ASP | p.Asp50Asn | P29033 |
| HOMO SAPIENS | 5 | 622 | 5era | A | 54 | ASN | p.Asn54Lys | P29033 |
| HOMO SAPIENS | 5 | 622 | 5era | A | 57 | GLN | p.Gln57Ter | P29033 |
| HOMO SAPIENS | 5 | 622 | 5era | A | 59 | GLY | p.Gly59Ala | P29033 |
| HOMO SAPIENS | 5 | 622 | 5era | A | 59 | GLY | p.Gly59Ser | P29033 |
| HOMO SAPIENS | 5 | 622 | 5era | A | 66 | ASP | p.Asp66His | P29033 |
| HOMO SAPIENS | 5 | 622 | 5era | A | 68 | TYR | p.Tyr68Cys | P29033 |
| HOMO SAPIENS | 5 | 622 | 5era | A | 70 | PRO | p.Pro70Ala | P29033 |
| HOMO SAPIENS | 5 | 622 | 5era | A | 73 | HIS | p.His73Arg | P29033 |
| HOMO SAPIENS | 5 | 622 | 5era | A | 75 | ARG | p.Arg75Trp | P29033 |
| HOMO SAPIENS | 5 | 622 | 5era | A | 75 | ARG | p.Arg75Gln | P29033 |
| HOMO SAPIENS | 2 | 622 | 5era | A | 77 | TRP | p.Trp77Arg | P29033 |
| HOMO SAPIENS | 2 | 622 | 5era | A | 77 | TRP | p.Trp77Ter | P29033 |
| HOMO SAPIENS | 0 | 622 | 5era | A | 80 | GLN | p.Gln80Ter | P29033 |
| HOMO SAPIENS | 0 | 622 | 5era | A | 80 | GLN | p.Gln80Pro | P29033 |
| HOMO SAPIENS | 5 | 622 | 5era | A | 81 | LEU | p.Leu81Val | P29033 |
| HOMO SAPIENS | 5 | 622 | 5era | A | 82 | ILE | p.Ile82Met | P29033 |
| HOMO SAPIENS | 5 | 622 | 5era | A | 83 | PHE | p.Phe83Leu | P29033 |
| HOMO SAPIENS | 5 | 622 | 5era | A | 84 | VAL | p.Val84Leu | P29033 |
| HOMO SAPIENS | 5 | 622 | 5era | A | 90 | LEU | p.Leu90Pro | P29033 |
| HOMO SAPIENS | 5 | 622 | 5era | A | 95 | VAL | p.Val95Met | P29033 |
| HOMO SAPIENS | -1 | 622 | 5era | A | 100 | HIS | p.His100Tyr | P29033 |
| HOMO SAPIENS | -1 | 622 | 5era | A | 113 | SER | p.Ser113Arg | P29033 |
| HOMO SAPIENS | -1 | 622 | 5era | A | 119 | GLU | p.Glu119Lys | P29033 |
| HOMO SAPIENS | -1 | 622 | 5era | A | 122 | LYS | p.Lys122Ile | P29033 |
| HOMO SAPIENS | -1 | 622 | 5era | A | 123 | THR | p.Thr123Asn | P29033 |
| HOMO SAPIENS | -1 | 622 | 5era | A | 124 | GLN | p.Gln124Ter | P29033 |
| HOMO SAPIENS | 5 | 622 | 5era | A | 139 | SER | p.Ser139Asn | P29033 |

| | | | | | | | | |
|---|---|---|---|---|---|---|---|---|
| HOMO SAPIENS | 5 | 622 | 5era | A | 143 | ARG | p.Arg143Trp | P29033 |
| HOMO SAPIENS | 5 | 622 | 5era | A | 143 | ARG | p.Arg143Gln | P29033 |
| HOMO SAPIENS | 0 | 622 | 5era | A | 147 | GLU | p.Glu147Lys | P29033 |
| HOMO SAPIENS | 7 | 622 | 5era | A | 155 | TYR | p.Tyr155Ter | P29033 |
| HOMO SAPIENS | 2 | 622 | 5era | A | 159 | ASP | p.Asp159Val | P29033 |
| HOMO SAPIENS | 5 | 622 | 5era | A | 163 | MET | p.Met163Val | P29033 |
| HOMO SAPIENS | 5 | 622 | 5era | A | 165 | ARG | p.Arg165Trp | P29033 |
| HOMO SAPIENS | 5 | 622 | 5era | A | 167 | VAL | p.Val167Met | P29033 |
| HOMO SAPIENS | 5 | 622 | 5era | A | 169 | CYS | p.Cys169Tyr | P29033 |
| HOMO SAPIENS | 5 | 622 | 5era | A | 179 | ASP | p.Asp179Asn | P29033 |
| HOMO SAPIENS | 0 | 622 | 5era | A | 199 | SER | p.Ser199Phe | P29033 |
| HOMO SAPIENS | 0 | 622 | 5era | A | 202 | CYS | p.Cys202Phe | P29033 |
| HOMO SAPIENS | 5 | 622 | 5era | A | 203 | ILE | p.Ile203Thr | P29033 |
| HOMO SAPIENS | 5 | 622 | 5era | A | 206 | ASN | p.Asn206Ser | P29033 |
| HOMO SAPIENS | 0 | 8103 | 5dot | A | 163 | GLY | p.Gly163Ter | P31327 |
| HOMO SAPIENS | 0 | 8103 | 5dot | A | 337 | HIS | p.His337Arg | P31327 |
| HOMO SAPIENS | 3 | 8103 | 5dot | A | 344 | THR | p.Thr344Ala | P31327 |
| HOMO SAPIENS | 1 | 8103 | 5dot | A | 405 | THR | p.Thr405Asn | P31327 |
| HOMO SAPIENS | 5 | 8103 | 5dot | A | 544 | THR | p.Thr544Met | P31327 |
| HOMO SAPIENS | 5 | 8103 | 5dot | A | 716 | ASN | p.Asn716Lys | P31327 |
| HOMO SAPIENS | -1 | 8103 | 5dot | A | 780 | ARG | p.Arg780His | P31327 |
| HOMO SAPIENS | 0 | 8103 | 5dot | A | 982 | GLY | p.Gly982Asp | P31327 |
| HOMO SAPIENS | 3 | 8103 | 5dot | A | 1119 | ALA | p.Ala1119Ser | P31327 |
| HOMO SAPIENS | 1 | 8103 | 5dot | A | 1376 | GLY | p.Gly1376Ser | P31327 |
| HOMO SAPIENS | 4 | 6833 | 1l8l | B | 27 | ARG | p.Arg27Ser | P78330 |
| HOMO SAPIENS | 3 | 6833 | 1l8l | B | 32 | ASP | p.Asp32Gly | P78330 |
| HOMO SAPIENS | 3 | 6833 | 1l8l | B | 32 | ASP | p.Asp32Asn | P78330 |
| HOMO SAPIENS | 1 | 6833 | 1l8l | B | 49 | ARG | p.Arg49Trp | P78330 |
| HOMO SAPIENS | 0 | 6833 | 1l8l | B | 52 | MET | p.Met52Thr | P78330 |
| HOMO SAPIENS | 0 | 6833 | 1l8l | B | 83 | GLN | p.Gln83His | P78330 |
| HOMO SAPIENS | 6 | 810 | 2iw2 | A | 185 | ARG | p.Arg184Gln | P12955 |
| HOMO SAPIENS | 0 | 810 | 2iw2 | A | 213 | ALA | p.Ala212Pro | P12955 |
| HOMO SAPIENS | 1 | 810 | 2iw2 | A | 266 | ARG | p.Arg265Ter | P12955 |
| HOMO SAPIENS | 9 | 810 | 2iw2 | A | 277 | ASP | p.Asp276Asn | P12955 |
| HOMO SAPIENS | 0 | 810 | 2iw2 | A | 279 | GLY | p.Gly278Asp | P12955 |
| HOMO SAPIENS | 9 | 810 | 2iw2 | A | 378 | HIS | p.His377Gln | P12955 |
| HOMO SAPIENS | 0 | 810 | 2iw2 | A | 389 | ARG | p.Arg388His | P12955 |
| HOMO SAPIENS | 0 | 810 | 2iw2 | A | 436 | PHE | p.Leu435Phe | P12955 |
| HOMO SAPIENS | 0 | 810 | 2iw2 | A | 449 | GLY | p.Gly448Arg | P12955 |
| HOMO SAPIENS | -1 | 1530 | 5fwj | A | 388 | ALA | p.Ala388Pro | P41229 |
| HOMO SAPIENS | 4 | 1530 | 5fwj | A | 451 | SER | p.Ser451Arg | P41229 |
| HOMO SAPIENS | 3 | 1530 | 5fwj | A | 731 | LEU | p.Leu731Phe | P41229 |
| HOMO SAPIENS | 3 | 578 | 5net | B | 123 | ALA | p.Ala143Thr | P18564 |
| HOMO SAPIENS | 6 | 578 | 5net | B | 176 | PRO | p.Pro196Thr | P18564 |
| HOMO SAPIENS | 3 | 578 | 5net | B | 280 | GLU | p.Glu300Lys | P18564 |
| HOMO SAPIENS | -1 | 5247 | 4j15 | A | 274 | ALA | p.Ala274Val | P14868 |
| HOMO SAPIENS | -1 | 5247 | 4j15 | A | 280 | HIS | p.His280Leu | P14868 |
| HOMO SAPIENS | 2 | 5247 | 4j15 | A | 460 | ARG | p.Arg460His | P14868 |
| HOMO SAPIENS | 5 | 5247 | 4j15 | A | 494 | ARG | p.Arg494Cys | P14868 |
| HOMO SAPIENS | 7 | 2566 | 1b55 | A | 28 | ARG | p.Arg28His | Q06187 |
| HOMO SAPIENS | 5 | 2566 | 1b55 | A | 33 | THR | p.Thr33Pro | Q06187 |
| HOMO SAPIENS | 0 | 2566 | 1b55 | A | 113 | VAL | p.Val113Asp | Q06187 |
| HOMO SAPIENS | 0 | 356 | 5a2s | A | 786 | ALA | p.Ala786Thr | P56524 |
| HOMO SAPIENS | 5 | 2960 | 4cak | A | 553 | ARG | p.Arg584Ter | P08514 |

| | | | | | | | | |
|---|---|---|---|---|---|---|---|---|
| HOMO SAPIENS | 5 | 2960 | 4cak | A | 565 | ILE | p.Ile596Thr | P08514 |
| HOMO SAPIENS | -1 | 2960 | 4cak | A | 841 | LEU | p.Leu872Met | P08514 |
| HOMO SAPIENS | -1 | 2960 | 4cak | A | 843 | ILE | p.Ile874Ser | P08514 |
| HOMO SAPIENS | 0 | 5054 | 4k8r | B | 216 | PHE | p.Phe216Leu | Q8NBP7 |
| HOMO SAPIENS | 1 | 5054 | 4k8r | B | 417 | HIS | p.His417Gln | Q8NBP7 |
| HOMO SAPIENS | 1 | 5054 | 4k8r | B | 425 | ASN | p.Asn425Ser | Q8NBP7 |
| HOMO SAPIENS | 1 | 5054 | 4k8r | B | 443 | ALA | p.Ala443Thr | Q8NBP7 |
| HOMO SAPIENS | 0 | 27283 | 3n5n | X | 71 | HIS | p.His82Arg | Q9UIF7 |
| HOMO SAPIENS | 1 | 27283 | 3n5n | X | 83 | ARG | p.Arg94Ter | Q9UIF7 |
| HOMO SAPIENS | 8 | 27283 | 3n5n | X | 89 | TRP | p.Trp100Ter | Q9UIF7 |
| HOMO SAPIENS | 8 | 27283 | 3n5n | X | 90 | TYR | p.Tyr101Ter | Q9UIF7 |
| HOMO SAPIENS | 7 | 27283 | 3n5n | X | 95 | ARG | p.Arg106Trp | Q9UIF7 |
| HOMO SAPIENS | 7 | 27283 | 3n5n | X | 117 | TRP | p.Trp128Arg | Q9UIF7 |
| HOMO SAPIENS | 0 | 27283 | 3n5n | X | 137 | GLY | p.Gly148Arg | Q9UIF7 |
| HOMO SAPIENS | 5 | 27283 | 3n5n | X | 142 | TRP | p.Trp153Ter | Q9UIF7 |
| HOMO SAPIENS | 0 | 27283 | 3n5n | X | 165 | TYR | p.Tyr176Cys | Q9UIF7 |
| HOMO SAPIENS | 0 | 27283 | 3n5n | X | 168 | ARG | p.Arg179His | Q9UIF7 |
| HOMO SAPIENS | 0 | 27283 | 3n5n | X | 168 | ARG | p.Arg179Cys | Q9UIF7 |
| HOMO SAPIENS | 0 | 27283 | 3n5n | X | 196 | GLN | p.Gln207Ter | Q9UIF7 |
| HOMO SAPIENS | 8 | 27283 | 3n5n | X | 201 | VAL | p.Val212Met | Q9UIF7 |
| HOMO SAPIENS | 9 | 27283 | 3n5n | X | 227 | ARG | p.Arg238Trp | Q9UIF7 |
| HOMO SAPIENS | 9 | 27283 | 3n5n | X | 231 | ARG | p.Arg242Cys | Q9UIF7 |
| HOMO SAPIENS | 9 | 27283 | 3n5n | X | 231 | ARG | p.Arg242His | Q9UIF7 |
| HOMO SAPIENS | 3 | 27283 | 3n5n | X | 232 | VAL | p.Val243Phe | Q9UIF7 |
| HOMO SAPIENS | 3 | 27283 | 3n5n | X | 246 | GLN | p.Gln257Ter | Q9UIF7 |
| HOMO SAPIENS | 9 | 27283 | 3n5n | X | 272 | GLY | p.Gly283Glu | Q9UIF7 |
| HOMO SAPIENS | 9 | 27283 | 3n5n | X | 281 | PRO | p.Pro292Leu | Q9UIF7 |
| HOMO SAPIENS | -1 | 27283 | 3n5n | X | 314 | ASP | p.Asp325Glu | Q9UIF7 |
| HOMO SAPIENS | 0 | 27283 | 3n5n | X | 324 | GLN | p.Gln335His | Q9UIF7 |
| HOMO SAPIENS | 0 | 27283 | 3n5n | X | 324 | GLN | p.Gln335Arg | Q9UIF7 |
| HOMO SAPIENS | 0 | 27283 | 3n5n | X | 324 | GLN | p.Gln335Ter | Q9UIF7 |
| HOMO SAPIENS | -1 | 20852 | 4p60 | B | 243 | ARG | p.Arg243Lys | O75746 |
| HOMO SAPIENS | -1 | 1875 | 5dd2 | G | 152 | VAL | p.Val179Leu | P06396 |
| HOMO SAPIENS | 9 | 2126 | 3n1p | B | 46 | PRO | p.Pro46Leu | Q14623 |
| HOMO SAPIENS | 8 | 2126 | 3n1p | B | 76 | GLU | p.Glu76Ala | Q14623 |
| HOMO SAPIENS | 9 | 2126 | 3n1p | B | 77 | ARG | p.Arg77Ser | Q14623 |
| HOMO SAPIENS | 6 | 2126 | 3n1p | B | 95 | GLU | p.Glu95Lys | Q14623 |
| HOMO SAPIENS | 9 | 2126 | 3n1p | B | 100 | ASP | p.Asp100Glu | Q14623 |
| HOMO SAPIENS | 9 | 2126 | 3n1p | B | 131 | GLU | p.Glu131Lys | Q14623 |
| HOMO SAPIENS | 0 | 2126 | 3n1p | B | 190 | VAL | p.Val190Ala | Q14623 |
| HOMO SAPIENS | 5 | 40410 | 3ioz | B | 70 | GLN | p.Gln70Leu | P20963 |
| HOMO SAPIENS | 1 | 5754 | 5lkz | A | 1108 | THR | p.Tyr1191Ter | Q09472 |
| HOMO SAPIENS | 3 | 8159 | 1tiu | A | 3 | GLU | p.Lys12679Glu | Q8WZ42 |
| HOMO SAPIENS | 4 | 3275 | 4ldq | A | 210 | ARG | p.Arg210Gln | Q9BV20 |
| HOMO SAPIENS | 0 | 8143 | 4miv | B | 66 | SER | p.Ser66Trp | P51688 |
| HOMO SAPIENS | 1 | 8143 | 4miv | B | 74 | ARG | p.Arg74Cys | P51688 |
| HOMO SAPIENS | 0 | 8143 | 4miv | B | 128 | PRO | p.Pro128Leu | P51688 |
| HOMO SAPIENS | 1 | 8143 | 4miv | B | 150 | ARG | p.Arg150Gln | P51688 |
| HOMO SAPIENS | 0 | 8143 | 4miv | B | 191 | GLY | p.Gly191Arg | P51688 |
| HOMO SAPIENS | 0 | 8143 | 4miv | B | 206 | ARG | p.Arg206Pro | P51688 |
| HOMO SAPIENS | 0 | 8143 | 4miv | B | 225 | PHE | p.Phe225Leu | P51688 |
| HOMO SAPIENS | 0 | 8143 | 4miv | B | 233 | ARG | p.Arg233Ter | P51688 |
| HOMO SAPIENS | 0 | 8143 | 4miv | B | 234 | ALA | p.Ala234Gly | P51688 |
| HOMO SAPIENS | 0 | 8143 | 4miv | B | 245 | ARG | p.Arg245His | P51688 |

| | | | | | | | | |
|---|---|---|---|---|---|---|---|---|
| HOMO SAPIENS | 0 | 8143 | 4miv | B | 293 | PRO | p.Pro293Ser | P51688 |
| HOMO SAPIENS | 1 | 8143 | 4miv | B | 298 | SER | p.Ser298Pro | P51688 |
| HOMO SAPIENS | 0 | 8143 | 4miv | B | 361 | VAL | p.Val361Ile | P51688 |
| HOMO SAPIENS | 0 | 8143 | 4miv | B | 369 | GLU | p.Glu369Lys | P51688 |
| HOMO SAPIENS | 0 | 8143 | 4miv | B | 372 | MET | p.Met372Ile | P51688 |
| HOMO SAPIENS | 1 | 8143 | 4miv | B | 380 | GLN | p.Gln380Arg | P51688 |
| HOMO SAPIENS | 0 | 8143 | 4miv | B | 387 | VAL | p.Val387Met | P51688 |
| HOMO SAPIENS | 0 | 8143 | 4miv | B | 394 | MET | p.Met394Ile | P51688 |
| HOMO SAPIENS | 0 | 8143 | 4miv | B | 433 | ARG | p.Arg433Gln | P51688 |
| HOMO SAPIENS | 1 | 8143 | 4miv | B | 446 | HIS | p.His446Arg | P51688 |
| HOMO SAPIENS | 5 | 8143 | 4miv | B | 447 | GLU | p.Glu447Lys | P51688 |
| HOMO SAPIENS | 2 | 8143 | 4miv | B | 456 | ARG | p.Arg456His | P51688 |
| HOMO SAPIENS | 1 | 21880 | 2oo9 | B | 857 | LEU | p.Leu857Phe | P22681 |
| HOMO SAPIENS | 0 | 9467 | 1s1d | A | 154 | LEU | p.Leu224Pro | Q8WVQ1 |
| HOMO SAPIENS | 0 | 9467 | 1s1d | A | 156 | VAL | p.Val226Met | Q8WVQ1 |
| HOMO SAPIENS | 0 | 9467 | 1s1d | A | 229 | PRO | p.Pro299Leu | Q8WVQ1 |
| HOMO SAPIENS | 4 | 9467 | 1s1d | A | 230 | ARG | p.Arg300Cys | Q8WVQ1 |
| HOMO SAPIENS | 4 | 9467 | 1s1d | A | 230 | ARG | p.Arg300His | Q8WVQ1 |
| HOMO SAPIENS | 0 | 9467 | 1s1d | A | 253 | ALA | p.Ala323Thr | Q8WVQ1 |
| HOMO SAPIENS | 2 | 9467 | 1s1d | A | 305 | MET | p.Met375Leu | Q8WVQ1 |
| HOMO SAPIENS | -1 | 2361 | 5llm | A | 5 | ASN | p.Asn5Lys | P43003 |
| HOMO SAPIENS | 0 | 2361 | 5llm | A | 71 | TYR | p.Tyr71Cys | P43003 |
| HOMO SAPIENS | -1 | 2361 | 5llm | A | 211 | LYS | p.Glu219Asp | Q15758 |
| HOMO SAPIENS | 8 | 2361 | 5llm | A | 270 | PRO | p.Pro290Arg | P43003 |
| HOMO SAPIENS | 1 | 2361 | 5llm | A | 321 | THR | p.Ala329Thr | Q15758 |
| HOMO SAPIENS | 1 | 2361 | 5llm | A | 365 | ARG | p.Arg385His | P43003 |
| HOMO SAPIENS | 1 | 2361 | 5llm | A | 479 | ARG | p.Arg499Gln | P43003 |
| HOMO SAPIENS | 3 | 5412 | 4obz | D | 165 | PHE | p.Phe229Ile | P07339 |
| HOMO SAPIENS | 2 | 5412 | 4obz | D | 187 | ASP | p.Asp251Asn | P07339 |
| HOMO SAPIENS | 0 | 5412 | 4obz | D | 218 | GLY | p.Gly282Arg | P07339 |
| HOMO SAPIENS | 0 | 5412 | 4obz | D | 245 | ARG | p.Arg309His | P07339 |
| HOMO SAPIENS | 2 | 5412 | 4obz | D | 319 | TRP | p.Trp383Cys | P07339 |
| HOMO SAPIENS | 0 | 3544 | 5h64 | A | 277 | MET | p.Met277Ile | P42345 |
| HOMO SAPIENS | 0 | 14230 | 2kbx | A | 145 | ARG | p.Arg145Ser | Q13418 |
| HOMO SAPIENS | 2 | 34970 | 4xhz | A | 834 | ARG | p.Arg855Trp | Q96QU1 |
| HOMO SAPIENS | 4 | 34970 | 4xhz | A | 854 | SER | p.Ser875Ter | Q96QU1 |
| HOMO SAPIENS | 1 | 34970 | 4xhz | A | 908 | ARG | p.Arg929Gln | Q96QU1 |
| HOMO SAPIENS | 1 | 34970 | 4xhz | A | 941 | ARG | p.Arg962Cys | Q96QU1 |
| HOMO SAPIENS | 0 | 34970 | 4xhz | A | 970 | ARG | p.Arg991Ter | Q96QU1 |
| HOMO SAPIENS | 0 | 34970 | 4xhz | A | 976 | GLU | p.Glu997Gly | Q96QU1 |
| HOMO SAPIENS | 0 | 25734 | 4asc | A | 345 | SER | p.Asn345Ser | Q2TBA0 |
| HOMO SAPIENS | 0 | 25734 | 4asc | A | 514 | VAL | p.Val514Met | Q2TBA0 |
| HOMO SAPIENS | 6 | 25734 | 4asc | A | 528 | GLU | p.Glu528Lys | Q2TBA0 |
| HOMO SAPIENS | 0 | 25734 | 4asc | A | 580 | LEU | p.Leu580Ile | Q2TBA0 |
| HOMO SAPIENS | 8 | 2356 | 3kse | D | 22 | LYS | p.Lys22Ter | P01040 |
| HOMO SAPIENS | 1 | 2356 | 3kse | D | 58 | ARG | p.Arg58Ter | P01040 |
| HOMO SAPIENS | 2 | 6000 | 1jqd | A | 60 | GLY | p.Gly60Asp | P50135 |
| HOMO SAPIENS | 9 | 6000 | 1jqd | A | 208 | LEU | p.Leu208Pro | P50135 |
| HOMO SAPIENS | 0 | 14741 | 5eh1 | A | 58 | THR | p.Thr58Arg | P38484 |
| HOMO SAPIENS | 0 | 14741 | 5eh1 | A | 168 | THR | p.Thr168Asn | P38484 |
| HOMO SAPIENS | 1 | 4814 | 4kxk | D | 427 | ARG | p.Arg427Ter | P50542 |
| HOMO SAPIENS | 0 | 188 | 2xt3 | A | 49 | ASP | p.Asp52Asn | Q2M1P5 |
| HOMO SAPIENS | 3 | 188 | 2xt3 | A | 142 | TYR | p.Tyr145Ser | Q2M1P5 |
| HOMO SAPIENS | 9 | 8681 | 5tby | D | 57 | ALA | p.Ala57Asp | P08590 |

| | | | | | | | | |
|---|---|---|---|---|---|---|---|---|
| HOMO SAPIENS | 3 | 8681 | 5tby | D | 62 | ASP | p.Asp62Asn | P08590 |
| HOMO SAPIENS | 0 | 8681 | 5tby | D | 65 | PRO | p.Pro65Arg | P08590 |
| HOMO SAPIENS | 7 | 8681 | 5tby | D | 79 | VAL | p.Val79Ile | P08590 |
| HOMO SAPIENS | 0 | 8681 | 5tby | D | 143 | GLU | p.Glu143Lys | P08590 |
| HOMO SAPIENS | 0 | 8681 | 5tby | D | 149 | MET | p.Met149Val | P08590 |
| HOMO SAPIENS | 3 | 8681 | 5tby | D | 154 | ARG | p.Arg154His | P08590 |
| HOMO SAPIENS | 6 | 30984 | 4xwh | A | 140 | TYR | p.Tyr140Cys | P54802 |
| HOMO SAPIENS | 4 | 30984 | 4xwh | A | 141 | SER | p.Ser141Thr | P54802 |
| HOMO SAPIENS | 2 | 30984 | 4xwh | A | 234 | ARG | p.Arg234Cys | P54802 |
| HOMO SAPIENS | 2 | 30984 | 4xwh | A | 336 | GLU | p.Glu336Ter | P54802 |
| HOMO SAPIENS | 6 | 30984 | 4xwh | A | 482 | ARG | p.Arg482Trp | P54802 |
| HOMO SAPIENS | 2 | 30984 | 4xwh | A | 521 | PRO | p.Pro521Leu | P54802 |
| HOMO SAPIENS | 0 | 30984 | 4xwh | A | 565 | ARG | p.Arg565Gln | P54802 |
| HOMO SAPIENS | 0 | 30984 | 4xwh | A | 565 | ARG | p.Arg565Trp | P54802 |
| HOMO SAPIENS | 6 | 30984 | 4xwh | A | 626 | ARG | p.Arg626Ter | P54802 |
| HOMO SAPIENS | 6 | 30984 | 4xwh | A | 643 | ARG | p.Arg643His | P54802 |
| HOMO SAPIENS | 6 | 30984 | 4xwh | A | 643 | ARG | p.Arg643Cys | P54802 |
| HOMO SAPIENS | 2 | 30984 | 4xwh | A | 674 | ARG | p.Arg674His | P54802 |
| HOMO SAPIENS | 2 | 30984 | 4xwh | A | 737 | ARG | p.Arg737Gly | P54802 |
| HOMO SAPIENS | 5 | 10545 | 1oqe | N | 6 | PRO | p.Pro21Arg | Q96RJ3 |
| HOMO SAPIENS | 1 | 16995 | 5d3a | B | 12 | MET | p.Met947Arg | Q7Z4S6 |
| HOMO SAPIENS | 1 | 16995 | 5d3a | B | 12 | MET | p.Met947Val | Q7Z4S6 |
| HOMO SAPIENS | 2 | 16995 | 5d3a | B | 19 | ARG | p.Arg954Gln | Q7Z4S6 |
| HOMO SAPIENS | 2 | 16995 | 5d3a | B | 19 | ARG | p.Arg954Trp | Q7Z4S6 |
| HOMO SAPIENS | 0 | 16995 | 5d3a | B | 75 | ILE | p.Ile1010Thr | Q7Z4S6 |
| HOMO SAPIENS | 0 | 17091 | 2yin | B | 1324 | GLN | p.Gln1324Ter | Q92608 |
| HOMO SAPIENS | 5 | 27606 | 1lm7 | B | 2366 | ARG | p.Arg2366Cys | P15924 |
| HOMO SAPIENS | 2 | 15731 | 1bi7 | B | 24 | ARG | p.Arg24Pro | P42771 |
| HOMO SAPIENS | 0 | 15731 | 1bi7 | B | 28 | VAL | p.Val28Gly | P42771 |
| HOMO SAPIENS | 0 | 15731 | 1bi7 | B | 49 | ILE | p.Ile49Thr | P42771 |
| HOMO SAPIENS | 0 | 15731 | 1bi7 | B | 53 | MET | p.Met53Ile | P42771 |
| HOMO SAPIENS | 2 | 15731 | 1bi7 | B | 56 | SER | p.Ser56Ile | P42771 |
| HOMO SAPIENS | 1 | 15731 | 1bi7 | B | 57 | ALA | p.Ala57Val | P42771 |
| HOMO SAPIENS | 4 | 15731 | 1bi7 | B | 59 | VAL | p.Val59Gly | P42771 |
| HOMO SAPIENS | 9 | 15731 | 1bi7 | B | 89 | GLY | p.Gly89Asp | P42771 |
| HOMO SAPIENS | 0 | 15731 | 1bi7 | B | 101 | GLY | p.Gly101Trp | P42771 |
| HOMO SAPIENS | 0 | 15731 | 1bi7 | B | 114 | PRO | p.Pro114Ser | P42771 |
| HOMO SAPIENS | 9 | 15731 | 1bi7 | B | 123 | HIS | p.His123Gln | P42771 |
| HOMO SAPIENS | 8 | 15731 | 1bi7 | B | 126 | VAL | p.Val126Asp | P42771 |
| HOMO SAPIENS | 7 | 15731 | 1bi7 | B | 127 | ALA | p.Ala127Ser | P42771 |
| HOMO SAPIENS | -1 | 15731 | 1bi7 | B | 148 | ALA | p.Ala148Thr | P42771 |
| HOMO SAPIENS | -1 | 19858 | 3jbk | M | 866 | ALA | p.Ala934Val | P18206 |
| HOMO SAPIENS | 0 | 19484 | 3ee6 | B | 62 | SER | p.Ser62Leu | O14773 |
| HOMO SAPIENS | 1 | 19484 | 3ee6 | B | 66 | GLN | p.Gln66Ter | O14773 |
| HOMO SAPIENS | 0 | 19484 | 3ee6 | B | 100 | GLN | p.Gln100Arg | O14773 |
| HOMO SAPIENS | 0 | 19484 | 3ee6 | B | 104 | LEU | p.Leu104Ter | O14773 |
| HOMO SAPIENS | 0 | 19484 | 3ee6 | B | 127 | ARG | p.Arg127Gln | O14773 |
| HOMO SAPIENS | 0 | 19484 | 3ee6 | B | 127 | ARG | p.Arg127Ter | O14773 |
| HOMO SAPIENS | -1 | 19484 | 3ee6 | B | 185 | ARG | p.Arg185Cys | O14773 |
| HOMO SAPIENS | 0 | 19484 | 3ee6 | B | 206 | ARG | p.Arg206Cys | O14773 |
| HOMO SAPIENS | 0 | 19484 | 3ee6 | B | 208 | ARG | p.Arg208Ter | O14773 |
| HOMO SAPIENS | 0 | 19484 | 3ee6 | B | 214 | GLN | p.Gln214Ter | O14773 |
| HOMO SAPIENS | 1 | 19484 | 3ee6 | B | 276 | ASP | p.Asp276Val | O14773 |
| HOMO SAPIENS | 0 | 19484 | 3ee6 | B | 284 | GLY | p.Gly284Val | O14773 |

| | | | | | | | | |
|---|---|---|---|---|---|---|---|---|
| HOMO SAPIENS | 0 | 19484 | 3ee6 | B | 286 | ASN | p.Asn286Ser | O14773 |
| HOMO SAPIENS | 0 | 19484 | 3ee6 | B | 339 | ARG | p.Arg339Gly | O14773 |
| HOMO SAPIENS | 2 | 19484 | 3ee6 | B | 345 | MET | p.Met345Leu | O14773 |
| HOMO SAPIENS | 0 | 19484 | 3ee6 | B | 365 | CYS | p.Cys365Arg | O14773 |
| HOMO SAPIENS | 0 | 19484 | 3ee6 | B | 365 | CYS | p.Cys365Tyr | O14773 |
| HOMO SAPIENS | 0 | 19484 | 3ee6 | B | 373 | GLN | p.Gln373Glu | O14773 |
| HOMO SAPIENS | 6 | 19484 | 3ee6 | B | 418 | ARG | p.Arg418Gln | O14773 |
| HOMO SAPIENS | 7 | 19484 | 3ee6 | B | 422 | GLN | p.Gln422His | O14773 |
| HOMO SAPIENS | 0 | 19484 | 3ee6 | B | 447 | ARG | p.Arg447His | O14773 |
| HOMO SAPIENS | 0 | 19484 | 3ee6 | B | 466 | VAL | p.Val466Met | O14773 |
| HOMO SAPIENS | 0 | 4654 | 5t5w | A | 47 | LYS | p.Lys47Glu | Q08334 |
| HOMO SAPIENS | 0 | 4654 | 5t5w | A | 141 | GLU | p.Glu141Ter | Q08334 |
| HOMO SAPIENS | -1 | 18815 | 4wj7 | A | 53 | VAL | p.Val53Ile | Q9BSQ5 |
| HOMO SAPIENS | 0 | 18815 | 4wj7 | A | 120 | VAL | p.Val120Ile | Q9BSQ5 |
| HOMO SAPIENS | 0 | 18815 | 4wj7 | A | 198 | LEU | p.Leu198Arg | Q9BSQ5 |
| HOMO SAPIENS | 0 | 4602 | 1mjd | A | 47 | SER | p.Ser47Arg | O43602 |
| HOMO SAPIENS | 6 | 4602 | 1mjd | A | 59 | ARG | p.Arg59Leu | O43602 |
| HOMO SAPIENS | 7 | 4602 | 1mjd | A | 62 | ASP | p.Asp62Asn | O43602 |
| HOMO SAPIENS | 7 | 4602 | 1mjd | A | 71 | ALA | p.Ala71Ser | O43602 |
| HOMO SAPIENS | 7 | 4602 | 1mjd | A | 78 | ARG | p.Arg78Leu | O43602 |
| HOMO SAPIENS | 7 | 4602 | 1mjd | A | 89 | ARG | p.Arg89Gly | O43602 |
| HOMO SAPIENS | 8 | 4602 | 1mjd | A | 104 | ILE | p.Ile104Phe | O43602 |
| HOMO SAPIENS | 7 | 4602 | 1mjd | A | 125 | TYR | p.Tyr125His | O43602 |
| HOMO SAPIENS | 5 | 34467 | 1n0w | B | 1529 | GLY | p.Gly1529Arg | P51587 |
| HOMO SAPIENS | 8 | 5738 | 1u31 | A | 130 | ASN | p.Asn1009Lys | Q13423 |
| HOMO SAPIENS | 0 | 5760 | 2i6s | A | 298 | GLU | p.Glu318Asp | P06681 |
| HOMO SAPIENS | 0 | 5760 | 2i6s | A | 573 | ARG | p.Arg593Gln | P06681 |
| HOMO SAPIENS | 0 | 5760 | 2i6s | A | 592 | SER | p.Ser612Asn | P06681 |
| HOMO SAPIENS | 0 | 5760 | 2i6s | A | 621 | VAL | p.Val641Ala | P06681 |
| HOMO SAPIENS | 1 | 10977 | 2fby | A | 114 | VAL | p.Val114Ile | Q14191 |
| HOMO SAPIENS | 0 | 10977 | 2fby | A | 135 | LYS | p.Lys135Glu | Q14191 |
| HOMO SAPIENS | -1 | 1175 | 1qrk | A | 34 | VAL | p.Val35Leu | P00488 |
| HOMO SAPIENS | 4 | 1175 | 1qrk | A | 564 | PRO | p.Pro565Leu | P00488 |
| HOMO SAPIENS | 0 | 1175 | 1qrk | A | 650 | VAL | p.Val651Ile | P00488 |
| HOMO SAPIENS | 0 | 1175 | 1qrk | A | 651 | GLU | p.Gln652Glu | P00488 |
| HOMO SAPIENS | 3 | 909 | 4h0w | A | 375 | GLU | p.Glu394Lys | P02787 |
| HOMO SAPIENS | 0 | 909 | 4h0w | A | 458 | ALA | p.Ala477Pro | P02787 |
| HOMO SAPIENS | 6 | 909 | 4h0w | A | 515 | TYR | p.Tyr534Ter | P02787 |
| HOMO SAPIENS | 7 | 909 | 4h0w | A | 570 | PRO | p.Pro589Ser | P02787 |
| HOMO SAPIENS | 3 | 909 | 4h0w | A | 652 | GLY | p.Gly671Glu | P02787 |
| HOMO SAPIENS | 5 | 1964 | 2car | A | 151 | TRP | p.Trp151Ter | Q9BY32 |
| HOMO SAPIENS | 7 | 1964 | 2car | A | 178 | ARG | p.Arg178Cys | Q9BY32 |
| HOMO SAPIENS | 0 | 144 | 2n0a | J | 30 | ALA | p.Ala30Pro | P37840 |
| HOMO SAPIENS | 1 | 12288 | 4q58 | D | 1225 | ARG | p.Arg1225His | P16144 |
| HOMO SAPIENS | 2 | 12288 | 4q58 | D | 1281 | ARG | p.Arg1281Trp | P16144 |
| HOMO SAPIENS | 2 | 12288 | 4q58 | D | 1287 | ASN | p.Asn1287Lys | P16144 |
| HOMO SAPIENS | 0 | 5085 | 3s94 | A | 360 | ARG | p.Arg360His | O75581 |
| HOMO SAPIENS | 0 | 5085 | 3s94 | A | 433 | ASN | p.Asn433Ser | O75581 |
| HOMO SAPIENS | 2 | 5085 | 3s94 | A | 611 | ARG | p.Arg611Cys | O75581 |
| HOMO SAPIENS | 0 | 15452 | 4dur | A | 19 | LYS | p.Lys38Glu | P00747 |
| HOMO SAPIENS | 2 | 15452 | 4dur | A | 216 | ARG | p.Arg235His | P00747 |
| HOMO SAPIENS | 7 | 8406 | 4wo1 | D | 28 | VAL | p.Val55Leu | O43914 |
| HOMO SAPIENS | 2 | 895 | 5xte | K | 183 | ARG | p.Arg183Trp | P22695 |
| HOMO SAPIENS | 2 | 4191 | 4wnd | A | 120 | ARG | p.Arg127Ter | P81274 |

| | | | | | | | | |
|---|---|---|---|---|---|---|---|---|
| HOMO SAPIENS | 2 | 4191 | 4wnd | A | 120 | ARG | p.Arg127Gln | P81274 |
| HOMO SAPIENS | 1 | 4191 | 4wnd | A | 349 | GLY | p.Gly356Arg | P81274 |
| HOMO SAPIENS | 0 | 41734 | 2kbi | A | 1777 | VAL | p.Val1777Met | Q14524 |
| HOMO SAPIENS | 1 | 41734 | 2kbi | A | 1784 | GLU | p.Glu1784Lys | Q14524 |
| HOMO SAPIENS | 5 | 41734 | 2kbi | A | 1795 | TYR | p.Tyr1795Cys | Q14524 |
| HOMO SAPIENS | 5 | 41734 | 2kbi | A | 1795 | TYR | p.Tyr1795His | Q14524 |
| HOMO SAPIENS | 7 | 41734 | 2kbi | A | 1826 | ARG | p.Arg1826Cys | Q14524 |
| HOMO SAPIENS | 3 | 2119 | 1pl0 | A | 426 | LYS | p.Lys426Arg | P31939 |
| HOMO SAPIENS | 5 | 3475 | 5u62 | C | 50 | LEU | p.Leu50Ser | Q15910 |
| HOMO SAPIENS | 3 | 379 | 4uez | A | 200 | TYR | p.Tyr200Ter | P15085 |
| HOMO SAPIENS | 0 | 8195 | 5xaw | B | 67 | ARG | p.Arg67Trp | Q8IWT1 |
| HOMO SAPIENS | 0 | 8195 | 5xaw | B | 94 | THR | p.Thr94Met | Q8IWT1 |
| HOMO SAPIENS | 1 | 5711 | 1fo3 | A | 265 | ARG | p.Arg265His | Q9UKM7 |
| HOMO SAPIENS | 4 | 5711 | 1fo3 | A | 334 | ARG | p.Arg334Cys | Q9UKM7 |
| HOMO SAPIENS | 1 | 5711 | 1fo3 | A | 478 | LYS | p.Lys478Arg | Q9UKM7 |
| HOMO SAPIENS | 1 | 5711 | 1fo3 | A | 682 | ASP | p.Asp682Glu | Q9UKM7 |
| HOMO SAPIENS | 0 | 170 | 2n3y | A | 41 | GLY | p.Gly42Ser | P99999 |
| HOMO SAPIENS | 0 | 4122 | 3w81 | A | 33 | HIS | p.His33Gln | P35475 |
| HOMO SAPIENS | 0 | 4122 | 3w81 | A | 64 | TYR | p.Tyr64Ter | P35475 |
| HOMO SAPIENS | 0 | 4122 | 3w81 | A | 70 | GLN | p.Gln70Ter | P35475 |
| HOMO SAPIENS | 0 | 4122 | 3w81 | A | 75 | ALA | p.Ala75Thr | P35475 |
| HOMO SAPIENS | 0 | 4122 | 3w81 | A | 79 | ALA | p.Ala79Val | P35475 |
| HOMO SAPIENS | 0 | 4122 | 3w81 | A | 79 | ALA | p.Ala79Thr | P35475 |
| HOMO SAPIENS | 0 | 4122 | 3w81 | A | 89 | ARG | p.Arg89Gln | P35475 |
| HOMO SAPIENS | 0 | 4122 | 3w81 | A | 99 | THR | p.Thr99Ile | P35475 |
| HOMO SAPIENS | 0 | 4122 | 3w81 | A | 105 | ARG | p.Arg105Gln | P35475 |
| HOMO SAPIENS | 0 | 4122 | 3w81 | A | 238 | LEU | p.Leu238Gln | P35475 |
| HOMO SAPIENS | 0 | 4122 | 3w81 | A | 300 | ALA | p.Ala300Thr | P35475 |
| HOMO SAPIENS | 0 | 4122 | 3w81 | A | 322 | VAL | p.Val322Glu | P35475 |
| HOMO SAPIENS | 0 | 4122 | 3w81 | A | 327 | ALA | p.Ala327Pro | P35475 |
| HOMO SAPIENS | 0 | 4122 | 3w81 | A | 346 | LEU | p.Leu346Arg | P35475 |
| HOMO SAPIENS | 0 | 4122 | 3w81 | A | 349 | ASP | p.Asp349Asn | P35475 |
| HOMO SAPIENS | 0 | 4122 | 3w81 | A | 361 | ALA | p.Ala361Thr | P35475 |
| HOMO SAPIENS | 0 | 4122 | 3w81 | A | 366 | THR | p.Thr366Pro | P35475 |
| HOMO SAPIENS | 0 | 4122 | 3w81 | A | 402 | TRP | p.Trp402Ter | P35475 |
| HOMO SAPIENS | 0 | 4122 | 3w81 | A | 409 | GLY | p.Gly409Arg | P35475 |
| HOMO SAPIENS | 0 | 4122 | 3w81 | A | 454 | VAL | p.Val454Ile | P35475 |
| HOMO SAPIENS | 1 | 4122 | 3w81 | A | 490 | LEU | p.Leu490Pro | P35475 |
| HOMO SAPIENS | 1 | 4122 | 3w81 | A | 492 | ARG | p.Arg492Pro | P35475 |
| HOMO SAPIENS | 0 | 4122 | 3w81 | A | 533 | PRO | p.Pro533Arg | P35475 |
| HOMO SAPIENS | 0 | 4122 | 3w81 | A | 591 | ALA | p.Ala591Thr | P35475 |
| HOMO SAPIENS | 7 | 4122 | 3w81 | A | 619 | ARG | p.Arg619Gly | P35475 |
| HOMO SAPIENS | 0 | 15325 | 3wvz | B | 54 | VAL | p.Val54Leu | Q53FT3 |
| HOMO SAPIENS | 3 | 3228 | 5iz8 | A | 421 | TRP | p.Trp421Ter | P25054 |
| HOMO SAPIENS | 2 | 3228 | 5iz8 | A | 437 | PRO | p.Pro437Arg | P25054 |
| HOMO SAPIENS | 3 | 3228 | 5iz8 | A | 499 | ARG | p.Arg499Ter | P25054 |
| HOMO SAPIENS | -1 | 29229 | 5buo | A | 539 | SER | p.Ser614Gly | P05067 |
| HOMO SAPIENS | -1 | 34931 | 4n7v | C | 54 | SER | p.Ser54Leu | O94986 |
| HOMO SAPIENS | 0 | 2450 | 5kzx | A | 123 | GLY | p.Gly123Glu | P10253 |
| HOMO SAPIENS | 1 | 2450 | 5kzx | A | 190 | ARG | p.Arg190His | P10253 |
| HOMO SAPIENS | -1 | 2450 | 5kzx | A | 199 | HIS | p.His199Arg | P10253 |
| HOMO SAPIENS | 0 | 2450 | 5kzx | A | 219 | GLY | p.Gly219Arg | P10253 |
| HOMO SAPIENS | 0 | 2450 | 5kzx | A | 220 | VAL | p.Val220Leu | P10253 |
| HOMO SAPIENS | 0 | 2450 | 5kzx | A | 222 | VAL | p.Val222Met | P10253 |

| | | | | | | | | |
|---|---|---|---|---|---|---|---|---|
| HOMO SAPIENS | 0 | 2450 | 5kzx | A | 237 | ALA | p.Ala237Val | P10253 |
| HOMO SAPIENS | 0 | 2450 | 5kzx | A | 251 | SER | p.Ser251Leu | P10253 |
| HOMO SAPIENS | 0 | 2450 | 5kzx | A | 254 | SER | p.Ser254Leu | P10253 |
| HOMO SAPIENS | 0 | 2450 | 5kzx | A | 262 | GLU | p.Glu262Lys | P10253 |
| HOMO SAPIENS | 0 | 2450 | 5kzx | A | 293 | GLY | p.Gly293Arg | P10253 |
| HOMO SAPIENS | 0 | 2450 | 5kzx | A | 299 | LEU | p.Leu299Arg | P10253 |
| HOMO SAPIENS | 0 | 2450 | 5kzx | A | 309 | GLY | p.Gly309Arg | P10253 |
| HOMO SAPIENS | 0 | 2450 | 5kzx | A | 318 | MET | p.Met318Thr | P10253 |
| HOMO SAPIENS | 0 | 2450 | 5kzx | A | 355 | LEU | p.Leu355Pro | P10253 |
| HOMO SAPIENS | 1 | 2450 | 5kzx | A | 361 | PRO | p.Pro361Leu | P10253 |
| HOMO SAPIENS | 4 | 2450 | 5kzx | A | 408 | MET | p.Met408Val | P10253 |
| HOMO SAPIENS | 1 | 2450 | 5kzx | A | 429 | GLN | p.Gln429Glu | P10253 |
| HOMO SAPIENS | 4 | 2450 | 5kzx | A | 437 | ARG | p.Arg437Cys | P10253 |
| HOMO SAPIENS | 0 | 2450 | 5kzx | A | 439 | MET | p.Met439Lys | P10253 |
| HOMO SAPIENS | 1 | 2450 | 5kzx | A | 448 | SER | p.Ser448Thr | P10253 |
| HOMO SAPIENS | 0 | 2450 | 5kzx | A | 451 | PRO | p.Pro451Arg | P10253 |
| HOMO SAPIENS | 1 | 2450 | 5kzx | A | 459 | ASP | p.Asp459His | P10253 |
| HOMO SAPIENS | 8 | 2450 | 5kzx | A | 481 | TRP | p.Trp481Arg | P10253 |
| HOMO SAPIENS | 1 | 2450 | 5kzx | A | 489 | ASP | p.Asp489Asn | P10253 |
| HOMO SAPIENS | 0 | 2450 | 5kzx | A | 521 | GLU | p.Glu521Lys | P10253 |
| HOMO SAPIENS | 0 | 2450 | 5kzx | A | 529 | SER | p.Ser529Val | P10253 |
| HOMO SAPIENS | 0 | 2450 | 5kzx | A | 545 | PRO | p.Pro545Leu | P10253 |
| HOMO SAPIENS | 0 | 2450 | 5kzx | A | 552 | LEU | p.Leu552Pro | P10253 |
| HOMO SAPIENS | 0 | 2450 | 5kzx | A | 576 | GLY | p.Gly576Ser | P10253 |
| HOMO SAPIENS | 9 | 2450 | 5kzx | A | 600 | ARG | p.Arg600His | P10253 |
| HOMO SAPIENS | 8 | 2450 | 5kzx | A | 615 | GLY | p.Gly615Arg | P10253 |
| HOMO SAPIENS | 3 | 2450 | 5kzx | A | 619 | SER | p.Ser619Asn | P10253 |
| HOMO SAPIENS | 0 | 2450 | 5kzx | A | 638 | GLY | p.Gly638Trp | P10253 |
| HOMO SAPIENS | 0 | 2450 | 5kzx | A | 643 | GLY | p.Gly643Arg | P10253 |
| HOMO SAPIENS | 9 | 2450 | 5kzx | A | 645 | ASP | p.Asp645Glu | P10253 |
| HOMO SAPIENS | 9 | 2450 | 5kzx | A | 645 | ASP | p.Asp645His | P10253 |
| HOMO SAPIENS | 0 | 2450 | 5kzx | A | 648 | GLY | p.Gly648Ser | P10253 |
| HOMO SAPIENS | 5 | 2450 | 5kzx | A | 660 | ARG | p.Arg660His | P10253 |
| HOMO SAPIENS | 8 | 2450 | 5kzx | A | 672 | ARG | p.Arg672Gln | P10253 |
| HOMO SAPIENS | 0 | 2450 | 5kzx | A | 702 | ARG | p.Arg702Leu | P10253 |
| HOMO SAPIENS | 3 | 2450 | 5kzx | A | 725 | ARG | p.Arg725Trp | P10253 |
| HOMO SAPIENS | 0 | 2450 | 5kzx | A | 746 | TRP | p.Trp746Cys | P10253 |
| HOMO SAPIENS | 0 | 2450 | 5kzx | A | 780 | ILE | p.Val780Ile | P10253 |
| HOMO SAPIENS | 1 | 2450 | 5kzx | A | 816 | VAL | p.Val816Ile | P10253 |
| HOMO SAPIENS | 0 | 2450 | 5kzx | A | 838 | GLN | p.Gln838Ter | P10253 |
| HOMO SAPIENS | 2 | 2450 | 5kzx | A | 854 | ARG | p.Arg854Ter | P10253 |
| HOMO SAPIENS | 3 | 2450 | 5kzx | A | 870 | ARG | p.Arg870Ter | P10253 |
| HOMO SAPIENS | 0 | 2450 | 5kzx | A | 890 | VAL | p.Val890Leu | P10253 |
| HOMO SAPIENS | 6 | 1882 | 4km6 | A | 175 | CYS | p.Cys175Ter | P15328 |
| HOMO SAPIENS | 3 | 14767 | 1lm5 | A | 2628 | LEU | p.Leu2628Pro | P15924 |
| HOMO SAPIENS | 5 | 14767 | 1lm5 | A | 2655 | ALA | p.Ala2655Gly | P15924 |
| HOMO SAPIENS | 3 | 14767 | 1lm5 | A | 2731 | TYR | p.Tyr2731His | P15924 |
| HOMO SAPIENS | 1 | 14767 | 1lm5 | A | 2767 | THR | p.Thr2767Asn | P15924 |
| HOMO SAPIENS | -1 | 14767 | 1lm5 | A | 2819 | MET | p.Met2819Leu | P15924 |
| HOMO SAPIENS | -1 | 11246 | 4y5o | A | 289 | SER | p.Ser289Asn | Q9BSQ5 |
| HOMO SAPIENS | 0 | 7380 | 5kz5 | I | 60 | ARG | p.Arg60His | Q16595 |
| HOMO SAPIENS | 0 | 7380 | 5kz5 | I | 76 | MET | p.Met76Val | Q16595 |
| HOMO SAPIENS | 2 | 22187 | 3fby | A | 328 | CYS | p.Cys328Arg | P49747 |
| HOMO SAPIENS | 2 | 22187 | 3fby | A | 342 | ASP | p.Asp342Tyr | P49747 |

| | | | | | | | | |
|---|---|---|---|---|---|---|---|---|
| HOMO SAPIENS | 2 | 22187 | 3fby | A | 348 | CYS | p.Cys348Arg | P49747 |
| HOMO SAPIENS | 2 | 22187 | 3fby | A | 386 | ASN | p.Asn386Asp | P49747 |
| HOMO SAPIENS | 2 | 22187 | 3fby | A | 453 | ASN | p.Asn453Ser | P49747 |
| HOMO SAPIENS | 2 | 22187 | 3fby | A | 468 | CYS | p.Cys468Tyr | P49747 |
| HOMO SAPIENS | 3 | 22187 | 3fby | A | 472 | ASP | p.Asp472Tyr | P49747 |
| HOMO SAPIENS | 3 | 22187 | 3fby | A | 473 | ASP | p.Asp473Gly | P49747 |
| HOMO SAPIENS | 2 | 22187 | 3fby | A | 523 | ASN | p.Asn523Lys | P49747 |
| HOMO SAPIENS | 1 | 22187 | 3fby | A | 718 | ARG | p.Arg718Trp | P49747 |
| HOMO SAPIENS | 1 | 22187 | 3fby | A | 719 | GLY | p.Gly719Asp | P49747 |
| HOMO SAPIENS | 0 | 22187 | 3fby | A | 756 | GLN | p.Gln756Arg | P49747 |
| HOMO SAPIENS | -1 | 2784 | 5kvh | A | 88 | MET | p.Met88Val | O95831 |
| HOMO SAPIENS | -1 | 2784 | 5kvh | A | 96 | ASN | p.Asn96Ser | O95831 |
| HOMO SAPIENS | 0 | 2784 | 5kvh | A | 282 | THR | p.Thr282Met | O95831 |
| HOMO SAPIENS | 1 | 2784 | 5kvh | A | 366 | ASN | p.Asn366Ser | O95831 |
| HOMO SAPIENS | 3 | 2784 | 5kvh | A | 463 | ARG | p.Arg463Ile | O95831 |
| HOMO SAPIENS | 0 | 2784 | 5kvh | A | 498 | VAL | p.Val498Met | O95831 |
| HOMO SAPIENS | -1 | 2784 | 5kvh | A | 549 | ALA | p.Ala549Val | O95831 |
| HOMO SAPIENS | 0 | 38316 | 2lnk | C | 1933 | ARG | p.Arg1933Ter | P35579 |
| HOMO SAPIENS | 0 | 33808 | 2miu | A | 29 | VAL | p.Val29Met | Q14192 |
| HOMO SAPIENS | 1 | 33808 | 2miu | A | 37 | ALA | p.Ala37Ser | Q14192 |
| HOMO SAPIENS | 2 | 20994 | 2hey | R | 65 | ARG | p.Arg65Cys | P43489 |
| HOMO SAPIENS | -1 | 5687 | 2xwj | I | 7 | ARG | p.Arg32Gln | P00751 |
| HOMO SAPIENS | -1 | 5687 | 2xwj | I | 7 | ARG | p.Arg32Trp | P00751 |
| HOMO SAPIENS | 0 | 5687 | 2xwj | I | 49 | ARG | p.Arg74His | P00751 |
| HOMO SAPIENS | 0 | 5687 | 2xwj | I | 177 | ARG | p.Arg202Gly | P00751 |
| HOMO SAPIENS | 0 | 5687 | 2xwj | I | 217 | ILE | p.Ile242Leu | P00751 |
| HOMO SAPIENS | -1 | 5687 | 2xwj | I | 227 | GLY | p.Gly252Ser | P00751 |
| HOMO SAPIENS | 0 | 5687 | 2xwj | I | 483 | HIS | p.His508Gln | P00751 |
| HOMO SAPIENS | 0 | 5687 | 2xwj | I | 508 | LYS | p.Lys533Arg | P00751 |
| HOMO SAPIENS | 0 | 5687 | 2xwj | I | 540 | LYS | p.Lys565Glu | P00751 |
| HOMO SAPIENS | 0 | 5687 | 2xwj | I | 541 | GLU | p.Glu566Ala | P00751 |
| HOMO SAPIENS | 2 | 953 | 3d4j | A | 249 | PHE | p.Phe249Ser | P53602 |
| HOMO SAPIENS | 1 | 953 | 3d4j | A | 292 | ASN | p.Asn292Ser | P53602 |
| HOMO SAPIENS | 0 | 3300 | 1s9c | A | 140 | ASN | p.Asn457Tyr | P51659 |
| HOMO SAPIENS | 2 | 3300 | 1s9c | A | 167 | ILE | p.Ile484Val | P51659 |
| HOMO SAPIENS | 0 | 3300 | 1s9c | A | 174 | ALA | p.Ala491Thr | P51659 |
| HOMO SAPIENS | 8 | 3300 | 1s9c | A | 193 | ASP | p.Asp510Asn | P51659 |
| HOMO SAPIENS | 3 | 3300 | 1s9c | A | 194 | TRP | p.Trp511Arg | P51659 |
| HOMO SAPIENS | 0 | 3300 | 1s9c | A | 242 | VAL | p.Ile559Val | P51659 |
| HOMO SAPIENS | 0 | 10810 | 5i25 | A | 5 | GLN | p.Gln23Ter | P03951 |
| HOMO SAPIENS | 0 | 10810 | 5i25 | A | 55 | TRP | p.Trp73Ter | P03951 |
| HOMO SAPIENS | 0 | 10810 | 5i25 | A | 91 | ALA | p.Ala109Thr | P03951 |
| HOMO SAPIENS | 0 | 10810 | 5i25 | A | 117 | GLU | p.Glu135Ter | P03951 |
| HOMO SAPIENS | 0 | 10810 | 5i25 | A | 118 | CYS | p.Cys136Ter | P03951 |
| HOMO SAPIENS | 0 | 10810 | 5i25 | A | 128 | CYS | p.Cys146Ter | P03951 |
| HOMO SAPIENS | 0 | 10810 | 5i25 | A | 210 | ARG | p.Arg228Ter | P03951 |
| HOMO SAPIENS | 0 | 10810 | 5i25 | A | 226 | GLN | p.Gln244Ter | P03951 |
| HOMO SAPIENS | 0 | 10810 | 5i25 | A | 233 | GLN | p.Gln251Ter | P03951 |
| HOMO SAPIENS | 0 | 16668 | 3lbx | A | 28 | ARG | p.Arg28Leu | P02549 |
| HOMO SAPIENS | 0 | 16668 | 3lbx | A | 28 | ARG | p.Arg28Cys | P02549 |
| HOMO SAPIENS | 1 | 16668 | 3lbx | A | 41 | ARG | p.Arg41Trp | P02549 |
| HOMO SAPIENS | 1 | 16668 | 3lbx | A | 45 | ARG | p.Arg45Ser | P02549 |
| HOMO SAPIENS | 2 | 16668 | 3lbx | A | 48 | LYS | p.Lys48Arg | P02549 |
| HOMO SAPIENS | 1 | 16668 | 3lbx | A | 152 | ASP | p.Asp152Asn | P02549 |

| | | | | | | | | |
|---|---|---|---|---|---|---|---|---|
| HOMO SAPIENS | 0 | 226 | 2ble | A | 256 | ILE | p.Phe256Ile | P36959 |
| HOMO SAPIENS | 2 | 36808 | 4c7n | A | 3 | SER | p.Ser357Pro | O75030 |
| HOMO SAPIENS | 0 | 4092 | 5jnx | A | 113 | CYS | p.Cys113Arg | O15118 |
| HOMO SAPIENS | 4 | 4092 | 5jnx | A | 215 | HIS | p.His215Arg | O15118 |
| HOMO SAPIENS | 6 | 4092 | 5jnx | A | 237 | PRO | p.Pro237Ser | O15118 |
| HOMO SAPIENS | 0 | 4092 | 5jnx | A | 363 | ALA | p.Ala363Val | O15118 |
| HOMO SAPIENS | 0 | 4092 | 5jnx | A | 378 | VAL | p.Val378Ala | O15118 |
| HOMO SAPIENS | 8 | 4092 | 5jnx | A | 404 | ARG | p.Arg404Gln | O15118 |
| HOMO SAPIENS | 8 | 4092 | 5jnx | A | 411 | ARG | p.Arg411Gln | O15118 |
| HOMO SAPIENS | 2 | 4092 | 5jnx | A | 474 | PRO | p.Pro474Leu | O15118 |
| HOMO SAPIENS | 8 | 4092 | 5jnx | A | 511 | THR | p.Thr511Met | O15118 |
| HOMO SAPIENS | 0 | 4092 | 5jnx | A | 642 | MET | p.Met642Ile | O15118 |
| HOMO SAPIENS | 2 | 4092 | 5jnx | A | 664 | VAL | p.Val664Met | O15118 |
| HOMO SAPIENS | 0 | 4092 | 5jnx | A | 734 | SER | p.Ser734Ile | O15118 |
| HOMO SAPIENS | -1 | 4092 | 5jnx | A | 775 | GLN | p.Gln775Pro | O15118 |
| HOMO SAPIENS | 0 | 4092 | 5jnx | A | 858 | ILE | p.Ile858Val | O15118 |
| HOMO SAPIENS | 0 | 4092 | 5jnx | A | 889 | VAL | p.Val889Met | O15118 |
| HOMO SAPIENS | 0 | 4092 | 5jnx | A | 910 | GLY | p.Gly910Ser | O15118 |
| HOMO SAPIENS | 0 | 4092 | 5jnx | A | 911 | GLY | p.Gly911Ser | O15118 |
| HOMO SAPIENS | 0 | 4092 | 5jnx | A | 928 | GLN | p.Gln928Pro | O15118 |
| HOMO SAPIENS | 0 | 4092 | 5jnx | A | 931 | ASN | p.Asn931Lys | O15118 |
| HOMO SAPIENS | 0 | 4092 | 5jnx | A | 940 | SER | p.Ser940Leu | O15118 |
| HOMO SAPIENS | 0 | 4092 | 5jnx | A | 950 | VAL | p.Val950Met | O15118 |
| HOMO SAPIENS | 0 | 4092 | 5jnx | A | 954 | SER | p.Ser954Leu | O15118 |
| HOMO SAPIENS | 0 | 4092 | 5jnx | A | 958 | ARG | p.Arg958Gln | O15118 |
| HOMO SAPIENS | 0 | 4092 | 5jnx | A | 958 | ARG | p.Arg958Ter | O15118 |
| HOMO SAPIENS | -1 | 4092 | 5jnx | A | 978 | ARG | p.Arg978Cys | O15118 |
| HOMO SAPIENS | 0 | 4092 | 5jnx | A | 992 | GLY | p.Gly992Trp | O15118 |
| HOMO SAPIENS | 0 | 4092 | 5jnx | A | 1007 | PRO | p.Pro1007Ala | O15118 |
| HOMO SAPIENS | 0 | 4092 | 5jnx | A | 1019 | TYR | p.Tyr1019Cys | O15118 |
| HOMO SAPIENS | 2 | 4092 | 5jnx | A | 1035 | ALA | p.Ala1035Val | O15118 |
| HOMO SAPIENS | 2 | 4092 | 5jnx | A | 1054 | ALA | p.Ala1054Thr | O15118 |
| HOMO SAPIENS | 2 | 4092 | 5jnx | A | 1061 | ILE | p.Ile1061Thr | O15118 |
| HOMO SAPIENS | 0 | 4092 | 5jnx | A | 1073 | GLY | p.Gly1073Ser | O15118 |
| HOMO SAPIENS | 2 | 4092 | 5jnx | A | 1088 | TYR | p.Tyr1088Cys | O15118 |
| HOMO SAPIENS | 2 | 4092 | 5jnx | A | 1142 | MET | p.Met1142Thr | O15118 |
| HOMO SAPIENS | 2 | 4092 | 5jnx | A | 1156 | ASN | p.Asn1156Ser | O15118 |
| HOMO SAPIENS | 0 | 4092 | 5jnx | A | 1179 | MET | p.Met1179Val | O15118 |
| HOMO SAPIENS | 0 | 4092 | 5jnx | A | 1186 | ARG | p.Arg1186His | O15118 |
| HOMO SAPIENS | 0 | 4092 | 5jnx | A | 1187 | ALA | p.Ala1187Gly | O15118 |
| HOMO SAPIENS | 2 | 4092 | 5jnx | A | 1189 | GLU | p.Glu1189Gly | O15118 |
| HOMO SAPIENS | 2 | 4092 | 5jnx | A | 1200 | SER | p.Ser1200Gly | O15118 |
| HOMO SAPIENS | 2 | 4092 | 5jnx | A | 1205 | THR | p.Thr1205Lys | O15118 |
| HOMO SAPIENS | 0 | 4092 | 5jnx | A | 1213 | LEU | p.Leu1213Phe | O15118 |
| HOMO SAPIENS | -1 | 4092 | 5jnx | A | 1266 | ARG | p.Arg1266Gln | O15118 |
| HOMO SAPIENS | 9 | 16498 | 4i6o | A | 64 | ARG | p.Arg735Trp | P01024 |
| HOMO SAPIENS | 1 | 21902 | 3un9 | C | 751 | ARG | p.Arg751His | Q86UT6 |
| HOMO SAPIENS | 7 | 20391 | 2h32 | B | 105 | PRO | p.Pro142Leu | P15814 |
| HOMO SAPIENS | 6 | 1635 | 4x45 | A | 65 | ASP | p.Asp65Val | P07741 |
| HOMO SAPIENS | 1 | 1635 | 4x45 | A | 98 | TRP | p.Trp98Ter | P07741 |
| HOMO SAPIENS | 0 | 1635 | 4x45 | A | 110 | LEU | p.Leu110Pro | P07741 |
| HOMO SAPIENS | 7 | 1635 | 4x45 | A | 136 | MET | p.Met136Thr | P07741 |
| HOMO SAPIENS | 1 | 1635 | 4x45 | A | 150 | VAL | p.Val150Phe | P07741 |
| HOMO SAPIENS | 4 | 34810 | 2rpc | A | 16 | CYS | p.Cys253Ser | O60481 |

| | | | | | | | | |
|---|---|---|---|---|---|---|---|---|
| HOMO SAPIENS | 1 | 34810 | 2rpc | A | 18 | TRP | p.Trp255Gly | O60481 |
| HOMO SAPIENS | 1 | 14359 | 4hxi | A | 78 | MET | p.Met78Val | Q9UH77 |
| HOMO SAPIENS | 0 | 5342 | 4d9t | A | 628 | GLU | p.Glu628Asp | P51812 |
| HOMO SAPIENS | -1 | 5342 | 4d9t | A | 723 | ARG | p.Arg723His | P51812 |
| HOMO SAPIENS | -1 | 5342 | 4d9t | A | 729 | ARG | p.Arg729Gln | P51812 |
| HOMO SAPIENS | 1 | 9619 | 3j2s | B | 1781 | ARG | p.Arg1800Cys | P00451 |
| HOMO SAPIENS | 0 | 9619 | 3j2s | B | 1823 | MET | p.Met1842Ile | P00451 |
| HOMO SAPIENS | 1 | 9619 | 3j2s | B | 1825 | PRO | p.Pro1844Ser | P00451 |
| HOMO SAPIENS | 0 | 9619 | 3j2s | B | 1826 | THR | p.Thr1845Pro | P00451 |
| HOMO SAPIENS | 0 | 9619 | 3j2s | B | 1834 | ALA | p.Ala1853Val | P00451 |
| HOMO SAPIENS | 0 | 9619 | 3j2s | B | 1846 | ASP | p.Asp1865Asn | P00451 |
| HOMO SAPIENS | 1 | 9619 | 3j2s | B | 1848 | HIS | p.His1867Arg | P00451 |
| HOMO SAPIENS | 0 | 9619 | 3j2s | B | 1854 | PRO | p.Pro1873Arg | P00451 |
| HOMO SAPIENS | 1 | 9619 | 3j2s | B | 1885 | GLU | p.Glu1904Lys | P00451 |
| HOMO SAPIENS | 0 | 9619 | 3j2s | B | 1922 | ASN | p.Asn1941Ser | P00451 |
| HOMO SAPIENS | 1 | 9619 | 3j2s | B | 1941 | ARG | p.Arg1960Gln | P00451 |
| HOMO SAPIENS | 4 | 9619 | 3j2s | B | 2105 | TYR | p.Tyr2124Cys | P00451 |
| HOMO SAPIENS | 6 | 9619 | 3j2s | B | 2150 | ARG | p.Arg2169His | P00451 |
| HOMO SAPIENS | 0 | 9619 | 3j2s | B | 2159 | ARG | p.Arg2178Cys | P00451 |
| HOMO SAPIENS | 0 | 9619 | 3j2s | B | 2159 | ARG | p.Arg2178His | P00451 |
| HOMO SAPIENS | 2 | 9619 | 3j2s | B | 2238 | MET | p.Met2257Val | P00451 |
| HOMO SAPIENS | 2 | 9619 | 3j2s | B | 2300 | PRO | p.Pro2319Ser | P00451 |
| HOMO SAPIENS | 1 | 667 | 1z2m | A | 127 | GLU | p.Glu127Ter | P05161 |
| HOMO SAPIENS | 0 | 15717 | 2nn6 | G | 30 | GLY | p.Gly31Ala | Q9NQT5 |
| HOMO SAPIENS | -1 | 15717 | 2nn6 | G | 55 | ASN | p.Asn56His | Q9NQT5 |
| HOMO SAPIENS | 7 | 15717 | 2nn6 | G | 131 | ASP | p.Asp132Ala | Q9NQT5 |
| HOMO SAPIENS | 7 | 784 | 3rkq | A | 161 | ARG | p.Arg161Pro | P52952 |
| HOMO SAPIENS | 6 | 784 | 3rkq | A | 178 | THR | p.Thr178Met | P52952 |
| HOMO SAPIENS | 7 | 784 | 3rkq | A | 183 | LYS | p.Lys183Glu | P52952 |
| HOMO SAPIENS | 8 | 784 | 3rkq | A | 190 | ARG | p.Arg190Cys | P52952 |
| HOMO SAPIENS | 6 | 385 | 30000 | C | 70 | ALA | p.Ala70Thr | P07954 |
| HOMO SAPIENS | 2 | 385 | 30000 | C | 101 | ARG | p.Arg101Pro | P07954 |
| HOMO SAPIENS | 2 | 385 | 30000 | C | 174 | PRO | p.Pro174Arg | P07954 |
| HOMO SAPIENS | 6 | 385 | 30000 | C | 187 | SER | p.Ser187Ter | P07954 |
| HOMO SAPIENS | 9 | 385 | 30000 | C | 233 | ARG | p.Arg233His | P07954 |
| HOMO SAPIENS | 3 | 385 | 30000 | C | 273 | ALA | p.Ala273Thr | P07954 |
| HOMO SAPIENS | 4 | 385 | 30000 | C | 334 | SER | p.Ser334Arg | P07954 |
| HOMO SAPIENS | 6 | 385 | 30000 | C | 340 | ASN | p.Asn340Lys | P07954 |
| HOMO SAPIENS | 8 | 385 | 30000 | C | 343 | ARG | p.Arg343Ter | P07954 |
| HOMO SAPIENS | 3 | 385 | 30000 | C | 350 | ARG | p.Arg350Trp | P07954 |
| HOMO SAPIENS | 4 | 385 | 30000 | C | 376 | GLN | p.Gln376Pro | P07954 |
| HOMO SAPIENS | 6 | 385 | 30000 | C | 434 | CYS | p.Cys434Ter | P07954 |
| HOMO SAPIENS | 0 | 3316 | 4f91 | B | 934 | THR | p.Thr934Ala | O75643 |
| HOMO SAPIENS | 0 | 3316 | 4f91 | B | 1002 | ASN | p.Asn1002Ser | O75643 |
| HOMO SAPIENS | 0 | 3316 | 4f91 | B | 1087 | SER | p.Ser1087Leu | O75643 |
| HOMO SAPIENS | 4 | 3316 | 4f91 | B | 1389 | VAL | p.Val1389Ile | O75643 |
| HOMO SAPIENS | 0 | 619 | 5hbt | B | 195 | ASP | p.Asp240His | P02708 |
| HOMO SAPIENS | 1 | 619 | 5hbt | B | 209 | ARG | p.Arg254Leu | P02708 |
| HOMO SAPIENS | 0 | 18171 | 2y25 | B | 1408 | ASP | p.Asp1408Asn | P52179 |
| HOMO SAPIENS | 0 | 18171 | 2y25 | B | 1453 | MET | p.Met1453Thr | P52179 |
| HOMO SAPIENS | 0 | 18171 | 2y25 | B | 1459 | ILE | p.Ile1459Lys | P52179 |
| HOMO SAPIENS | 0 | 18171 | 2y25 | B | 1558 | PHE | p.Phe1558Leu | P52179 |
| HOMO SAPIENS | 0 | 18171 | 2y25 | B | 1573 | ARG | p.Arg1573Gln | P52179 |
| HOMO SAPIENS | 7 | 18171 | 2y25 | B | 1663 | VAL | p.Val1663Met | P52179 |

| | | | | | | | | |
|---|---|---|---|---|---|---|---|---|
| HOMO SAPIENS | 8 | 14262 | 2fcw | B | 117 | SER | p.Ser138Ter | P01130 |
| HOMO SAPIENS | 8 | 14262 | 2fcw | B | 118 | ASP | p.Asp139Glu | P01130 |
| HOMO SAPIENS | 9 | 14262 | 2fcw | B | 119 | GLU | p.Glu140Lys | P01130 |
| HOMO SAPIENS | 7 | 14262 | 2fcw | B | 122 | CYS | p.Cys143Ter | P01130 |
| HOMO SAPIENS | 6 | 14262 | 2fcw | B | 134 | CYS | p.Cys155Ter | P01130 |
| HOMO SAPIENS | 7 | 14262 | 2fcw | B | 147 | ASP | p.Asp168Glu | P01130 |
| HOMO SAPIENS | 4 | 2624 | 3b7b | A | 855 | VAL | p.Val886Met | Q9H9B1 |
| HOMO SAPIENS | 7 | 2624 | 3b7b | A | 868 | ILE | p.Ile899Val | Q9H9B1 |
| HOMO SAPIENS | 0 | 2624 | 3b7b | A | 892 | GLU | p.Glu923Lys | Q9H9B1 |
| HOMO SAPIENS | 0 | 4791 | 5ctd | B | 68 | LEU | p.Pro549Ala | Q14055 |
| HOMO SAPIENS | 5 | 460 | 4ui1 | D | 80 | CYS | p.Cys80Arg | Q6ZVN8 |
| HOMO SAPIENS | 3 | 460 | 4ui1 | D | 101 | LEU | p.Leu101Pro | Q6ZVN8 |
| HOMO SAPIENS | 0 | 40782 | 1sy6 | A | 47 | LYS | p.Lys69Ter | P09693 |
| HOMO SAPIENS | -1 | 1092 | 2idx | A | 62 | THR | p.Thr62Met | Q96EY8 |
| HOMO SAPIENS | 6 | 1092 | 2idx | A | 183 | HIS | p.His183Leu | Q96EY8 |
| HOMO SAPIENS | 7 | 1092 | 2idx | A | 186 | ARG | p.Arg186Trp | Q96EY8 |
| HOMO SAPIENS | 6 | 1092 | 2idx | A | 190 | ARG | p.Arg190Cys | Q96EY8 |
| HOMO SAPIENS | 6 | 1092 | 2idx | A | 190 | ARG | p.Arg190His | Q96EY8 |
| HOMO SAPIENS | 7 | 1092 | 2idx | A | 191 | ARG | p.Arg191Trp | Q96EY8 |
| HOMO SAPIENS | 7 | 1092 | 2idx | A | 191 | ARG | p.Arg191Gln | Q96EY8 |
| HOMO SAPIENS | 6 | 1092 | 2idx | A | 195 | ARG | p.Arg195His | Q96EY8 |
| HOMO SAPIENS | 1 | 1092 | 2idx | A | 234 | GLN | p.Gln234Ter | Q96EY8 |
| HOMO SAPIENS | 0 | 1092 | 2idx | A | 239 | MET | p.Met239Lys | Q96EY8 |
| HOMO SAPIENS | -1 | 6418 | 5fur | G | 11 | THR | p.Thr11Ile | P21675 |
| HOMO SAPIENS | -1 | 6418 | 5fur | G | 290 | LEU | p.Leu269Val | P21675 |
| HOMO SAPIENS | 0 | 6100 | 1wiq | A | 240 | ARG | p.Arg265Trp | P01730 |
| HOMO SAPIENS | 0 | 13757 | 4ye6 | A | 45 | GLY | p.Gly45Val | P47897 |
| HOMO SAPIENS | 1 | 13757 | 4ye6 | A | 57 | TYR | p.Tyr57His | P47897 |
| HOMO SAPIENS | 8 | 13757 | 4ye6 | A | 403 | ARG | p.Arg403Trp | P47897 |
| HOMO SAPIENS | 7 | 13757 | 4ye6 | A | 524 | ARG | p.Arg524Trp | P47897 |
| HOMO SAPIENS | 6 | 13757 | 4ye6 | A | 690 | ARG | p.Arg690Cys | P47897 |
| HOMO SAPIENS | 0 | 26958 | 2lzl | A | 370 | GLY | p.Gly370Cys | P22607 |
| HOMO SAPIENS | 0 | 26958 | 2lzl | A | 371 | SER | p.Ser371Cys | P22607 |
| HOMO SAPIENS | 0 | 26958 | 2lzl | A | 373 | TYR | p.Tyr373Cys | P22607 |
| HOMO SAPIENS | 0 | 26958 | 2lzl | A | 380 | GLY | p.Gly380Arg | P22607 |
| HOMO SAPIENS | 0 | 26958 | 2lzl | A | 384 | PHE | p.Phe384Leu | P22607 |
| HOMO SAPIENS | 5 | 26958 | 2lzl | A | 391 | ALA | p.Ala391Glu | P22607 |
| HOMO SAPIENS | 6 | 11325 | 2wa6 | A | 121 | TYR | p.Tyr121Phe | O75369 |
| HOMO SAPIENS | 2 | 11325 | 2wa6 | A | 161 | PHE | p.Phe161Cys | O75369 |
| HOMO SAPIENS | 0 | 11325 | 2wa6 | A | 171 | LEU | p.Leu171Arg | O75369 |
| HOMO SAPIENS | 2 | 11325 | 2wa6 | A | 173 | ALA | p.Ala173Val | O75369 |
| HOMO SAPIENS | 0 | 11325 | 2wa6 | A | 202 | MET | p.Met202Val | O75369 |
| HOMO SAPIENS | 3 | 11325 | 2wa6 | A | 227 | GLU | p.Glu227Lys | O75369 |
| HOMO SAPIENS | 0 | 11325 | 2wa6 | A | 234 | LEU | p.Leu234Val | O75369 |
| HOMO SAPIENS | 2 | 11325 | 2wa6 | A | 235 | SER | p.Ser235Pro | O75369 |
| HOMO SAPIENS | 9 | 7240 | 3tgx | A | 137 | TYR | p.Tyr156Cys | Q9HBE5 |
| HOMO SAPIENS | 0 | 35479 | 4elj | A | 307 | THR | p.Thr307Ile | P06400 |
| HOMO SAPIENS | 0 | 35479 | 4elj | A | 310 | GLY | p.Gly310Glu | P06400 |
| HOMO SAPIENS | 0 | 35479 | 4elj | A | 377 | THR | p.Thr377Ser | P06400 |
| HOMO SAPIENS | 9 | 36706 | 1he7 | A | 359 | TYR | p.Tyr359Cys | P04629 |
| HOMO SAPIENS | 4 | 6538 | 4d0e | A | 504 | ARG | p.Arg504His | P46531 |
| HOMO SAPIENS | 1 | 26518 | 5fm5 | M | 600 | GLU | p.Glu600Val | P52179 |
| HOMO SAPIENS | -1 | 26518 | 5fm5 | M | 651 | ARG | p.Arg651Gln | P52179 |
| HOMO SAPIENS | -1 | 26518 | 5fm5 | M | 660 | PRO | p.Pro660Ala | P52179 |

| | | | | | | | | |
|---|---|---|---|---|---|---|---|---|
| HOMO SAPIENS | -1 | 26518 | 5fm5 | M | 688 | THR | p.Thr688Ser | P52179 |
| HOMO SAPIENS | -1 | 26518 | 5fm5 | M | 704 | GLU | p.Glu704Lys | P52179 |
| HOMO SAPIENS | -1 | 26518 | 5fm5 | M | 727 | THR | p.Thr727Ala | P52179 |
| HOMO SAPIENS | 2 | 27054 | 4rhp | B | 209 | LEU | p.Leu209Val | O75208 |
| HOMO SAPIENS | -1 | 27054 | 4rhp | B | 288 | LYS | p.Lys288Asn | O75208 |
| HOMO SAPIENS | 5 | 2384 | 5fqd | B | 419 | ARG | p.Arg419Ter | Q96SW2 |
| HOMO SAPIENS | 4 | 5154 | 2hl3 | A | 56 | LYS | p.Lys56Arg | Q14203 |
| HOMO SAPIENS | 0 | 5154 | 2hl3 | A | 59 | GLY | p.Gly59Ser | Q14203 |
| HOMO SAPIENS | 0 | 5154 | 2hl3 | A | 71 | GLY | p.Gly71Ala | Q14203 |
| HOMO SAPIENS | 0 | 5154 | 2hl3 | A | 71 | GLY | p.Gly71Arg | Q14203 |
| HOMO SAPIENS | 7 | 5154 | 2hl3 | A | 72 | THR | p.Thr72Pro | Q14203 |
| HOMO SAPIENS | 2 | 5154 | 2hl3 | A | 74 | GLN | p.Gln74Pro | Q14203 |
| HOMO SAPIENS | 2 | 20039 | 2l1g | A | 9 | GLY | p.Gly9Cys | Q9NVV9 |
| HOMO SAPIENS | 5 | 20039 | 2l1g | A | 81 | PHE | p.Phe81Leu | Q9NVV9 |
| HOMO SAPIENS | -1 | 97 | 5uz7 | A | 201 | ARG | p.Arg201His | P63092 |
| HOMO SAPIENS | -1 | 97 | 5uz7 | A | 201 | ARG | p.Arg201Cys | P63092 |
| HOMO SAPIENS | 0 | 3257 | 4b7w | B | 278 | GLU | p.Glu278Asp | P41235 |
| HOMO SAPIENS | 1 | 3257 | 4b7w | B | 331 | ARG | p.Arg331Cys | P41235 |
| HOMO SAPIENS | 0 | 2814 | 4qgk | B | 10 | GLN | p.Gln10Ter | P51648 |
| HOMO SAPIENS | 0 | 2814 | 4qgk | B | 177 | ARG | p.Arg177Ter | P51648 |
| HOMO SAPIENS | 9 | 2814 | 4qgk | B | 184 | THR | p.Thr184Met | P51648 |
| HOMO SAPIENS | 1 | 2814 | 4qgk | B | 188 | ALA | p.Ala188Val | P51648 |
| HOMO SAPIENS | 0 | 2814 | 4qgk | B | 214 | CYS | p.Cys214Tyr | P51648 |
| HOMO SAPIENS | 0 | 2814 | 4qgk | B | 245 | ASP | p.Asp245Asn | P51648 |
| HOMO SAPIENS | 0 | 2814 | 4qgk | B | 266 | LYS | p.Lys266Asn | P51648 |
| HOMO SAPIENS | 0 | 2814 | 4qgk | B | 315 | PRO | p.Pro315Ser | P51648 |
| HOMO SAPIENS | 0 | 2814 | 4qgk | B | 386 | ASN | p.Asn386Ser | P51648 |
| HOMO SAPIENS | -1 | 5486 | 5u6o | B | 47 | GLY | p.Gly47Val | O60741 |
| HOMO SAPIENS | 1 | 21475 | 3pdy | A | 679 | THR | p.Thr789Ile | Q15149 |
| HOMO SAPIENS | 0 | 32890 | 4djc | B | 1501 | LEU | p.Leu1501Val | Q14524 |
| HOMO SAPIENS | -1 | 32890 | 4djc | B | 1512 | ARG | p.Arg1512Trp | Q14524 |
| HOMO SAPIENS | 0 | 26683 | 2xze | B | 38 | ARG | p.Arg38Cys | O95630 |
| HOMO SAPIENS | 0 | 26683 | 2xze | B | 100 | PHE | p.Phe100Tyr | O95630 |
| HOMO SAPIENS | -1 | 23175 | 5dka | A | 163 | SER | p.Ser163Leu | Q96EQ8 |
| HOMO SAPIENS | -1 | 23175 | 5dka | A | 174 | ARG | p.Arg174Cys | Q96EQ8 |
| HOMO SAPIENS | 6 | 21177 | 3k1r | A | 155 | ARG | p.Arg155Ter | Q9Y6N9 |
| HOMO SAPIENS | 0 | 1237 | 2wwm | O | 40 | PRO | p.Pro40Arg | O75147 |
| HOMO SAPIENS | 1 | 2543 | 5ewl | B | 64 | VAL | p.Val64Leu | Q13224 |
| HOMO SAPIENS | 2 | 2543 | 5ewl | B | 271 | ALA | p.Ala271Glu | Q13224 |
| HOMO SAPIENS | 3 | 34262 | 2mng | A | 613 | PHE | p.Phe613Leu | Q9Y3Q4 |
| HOMO SAPIENS | 7 | 34262 | 2mng | A | 672 | SER | p.Ser672Arg | Q9Y3Q4 |
| HOMO SAPIENS | 4 | 1667 | 1f05 | A | 321 | LYS | p.Lys321Met | P37837 |
| HOMO SAPIENS | 0 | 25179 | 2knc | B | 752 | SER | p.Ser778Pro | P05106 |
| HOMO SAPIENS | 0 | 2185 | 2ig7 | A | 50 | TYR | p.Tyr50Cys | Q9Y259 |
| HOMO SAPIENS | 0 | 2185 | 2ig7 | A | 224 | ASN | p.Asn224His | Q9Y259 |
| HOMO SAPIENS | 0 | 2185 | 2ig7 | A | 270 | TYR | p.Tyr270Ter | Q9Y259 |
| HOMO SAPIENS | 0 | 2185 | 2ig7 | A | 328 | GLN | p.Gln328Arg | Q9Y259 |
| HOMO SAPIENS | 3 | 2201 | 2r0o | A | 179 | PRO | p.Pro179Leu | O43707 |
| HOMO SAPIENS | 3 | 2201 | 2r0o | A | 255 | GLU | p.Lys255Glu | O43707 |
| HOMO SAPIENS | 3 | 2201 | 2r0o | A | 259 | THR | p.Thr259Ile | O43707 |
| HOMO SAPIENS | 4 | 2201 | 2r0o | A | 262 | SER | p.Ser262Pro | O43707 |
| HOMO SAPIENS | -1 | 31221 | 2wvr | C | 44 | THR | p.Thr44Pro | Q9H211 |
| HOMO SAPIENS | -1 | 31221 | 2wvr | C | 135 | ALA | p.Ala135Val | Q9H211 |
| HOMO SAPIENS | -1 | 31221 | 2wvr | C | 456 | GLU | p.Glu456Ala | Q9H211 |

| | | | | | | | | |
|---|---|---|---|---|---|---|---|---|
| HOMO SAPIENS | -1 | 31221 | 2wvr | C | 462 | ARG | p.Arg462Gln | Q9H211 |
| HOMO SAPIENS | -1 | 31221 | 2wvr | C | 468 | GLU | p.Glu468Lys | Q9H211 |
| HOMO SAPIENS | -1 | 31221 | 2wvr | C | 537 | ALA | p.Ala537Val | Q9H211 |
| HOMO SAPIENS | 5 | 2576 | 5o3l | C | 363 | VAL | p.Val680Ile | P10636 |
| HOMO SAPIENS | 5 | 1345 | 1czt | A | 37 | ARG | p.Arg2102Cys | P12259 |
| HOMO SAPIENS | 1 | 1345 | 1czt | A | 83 | MET | p.Met2148Thr | P12259 |
| HOMO SAPIENS | 1 | 1345 | 1czt | A | 157 | ASP | p.Asp2222Gly | P12259 |
| HOMO SAPIENS | 8 | 35985 | 1lo1 | A | 110 | ALA | p.Ala110Val | O95718 |
| HOMO SAPIENS | 1 | 617 | 2ocp | B | 44 | GLU | p.Glu44Lys | Q16854 |
| HOMO SAPIENS | 5 | 617 | 2ocp | B | 46 | ASN | p.Asn46Ser | Q16854 |
| HOMO SAPIENS | 0 | 617 | 2ocp | B | 59 | THR | p.Thr59Met | Q16854 |
| HOMO SAPIENS | 7 | 617 | 2ocp | B | 142 | ARG | p.Arg142Lys | Q16854 |
| HOMO SAPIENS | 3 | 617 | 2ocp | B | 154 | ASN | p.Asn154Lys | Q16854 |
| HOMO SAPIENS | 3 | 617 | 2ocp | B | 170 | GLN | p.Gln170Arg | Q16854 |
| HOMO SAPIENS | 3 | 617 | 2ocp | B | 227 | GLU | p.Glu227Lys | Q16854 |
| HOMO SAPIENS | 3 | 617 | 2ocp | B | 233 | LYS | p.Lys233Glu | Q16854 |
| HOMO SAPIENS | 3 | 617 | 2ocp | B | 255 | ASP | p.Asp255Tyr | Q16854 |
| HOMO SAPIENS | -1 | 38975 | 5itz | D | 421 | GLN | p.Gln421His | Q9HC77 |
| HOMO SAPIENS | 2 | 6235 | 2f8v | A | 15 | VAL | p.Val15Ile | Q8WZ42 |
| HOMO SAPIENS | 0 | 6235 | 2f8v | A | 54 | VAL | p.Val54Met | Q8WZ42 |
| HOMO SAPIENS | 0 | 6235 | 2f8v | A | 59 | SER | p.Ser59Arg | Q8WZ42 |
| HOMO SAPIENS | 0 | 6235 | 2f8v | A | 60 | ASP | p.Asp60Tyr | Q8WZ42 |
| HOMO SAPIENS | 0 | 12630 | 1ext | B | 12 | ASP | p.Asp41Glu | P19438 |
| HOMO SAPIENS | 3 | 12630 | 1ext | B | 30 | CYS | p.Cys59Arg | P19438 |
| HOMO SAPIENS | 3 | 12630 | 1ext | B | 30 | CYS | p.Cys59Ser | P19438 |
| HOMO SAPIENS | 5 | 12630 | 1ext | B | 33 | CYS | p.Cys62Tyr | P19438 |
| HOMO SAPIENS | 5 | 12630 | 1ext | B | 33 | CYS | p.Cys62Gly | P19438 |
| HOMO SAPIENS | 1 | 12630 | 1ext | B | 46 | PRO | p.Pro75Leu | P19438 |
| HOMO SAPIENS | 3 | 12630 | 1ext | B | 50 | THR | p.Thr79Met | P19438 |
| HOMO SAPIENS | 6 | 12630 | 1ext | B | 52 | CYS | p.Cys81Phe | P19438 |
| HOMO SAPIENS | 6 | 12630 | 1ext | B | 70 | CYS | p.Cys99Ser | P19438 |
| HOMO SAPIENS | 5 | 12630 | 1ext | B | 88 | CYS | p.Cys117Arg | P19438 |
| HOMO SAPIENS | 5 | 12630 | 1ext | B | 88 | CYS | p.Cys117Tyr | P19438 |
| HOMO SAPIENS | -1 | 12630 | 1ext | B | 170 | ILE | p.Ile199Asn | P19438 |
| HOMO SAPIENS | 0 | 994 | 1aos | B | 95 | ARG | p.Arg95Cys | P04424 |
| HOMO SAPIENS | 0 | 994 | 1aos | B | 111 | ARG | p.Arg111Gln | P04424 |
| HOMO SAPIENS | 0 | 994 | 1aos | B | 131 | THR | p.Thr131Met | P04424 |
| HOMO SAPIENS | 6 | 994 | 1aos | B | 178 | VAL | p.Val178Met | P04424 |
| HOMO SAPIENS | 9 | 994 | 1aos | B | 182 | ARG | p.Arg182Gln | P04424 |
| HOMO SAPIENS | 9 | 994 | 1aos | B | 186 | ARG | p.Arg186Trp | P04424 |
| HOMO SAPIENS | 8 | 994 | 1aos | B | 193 | ARG | p.Arg193Gln | P04424 |
| HOMO SAPIENS | 5 | 994 | 1aos | B | 217 | ARG | p.Arg217Ter | P04424 |
| HOMO SAPIENS | 0 | 994 | 1aos | B | 286 | GLN | p.Gln286Arg | P04424 |
| HOMO SAPIENS | 5 | 994 | 1aos | B | 379 | ARG | p.Arg379Cys | P04424 |
| HOMO SAPIENS | 4 | 16066 | 5mv9 | A | 1719 | TYR | p.Tyr1719Cys | Q13402 |
| HOMO SAPIENS | -1 | 16066 | 5mv9 | A | 1739 | ARG | p.Arg1739Ter | Q13402 |
| HOMO SAPIENS | 4 | 16066 | 5mv9 | A | 1775 | ILE | p.Ile1775Thr | Q13402 |
| HOMO SAPIENS | 2 | 16066 | 5mv9 | A | 1786 | SER | p.Ser1786Thr | Q13402 |
| HOMO SAPIENS | 4 | 16066 | 5mv9 | A | 1798 | GLN | p.Gln1798Ter | Q13402 |
| HOMO SAPIENS | 0 | 16066 | 5mv9 | A | 1858 | LEU | p.Leu1858Pro | Q13402 |
| HOMO SAPIENS | 6 | 16066 | 5mv9 | A | 1873 | ARG | p.Arg1873Trp | Q13402 |
| HOMO SAPIENS | 6 | 16066 | 5mv9 | A | 1873 | ARG | p.Arg1873Leu | Q13402 |
| HOMO SAPIENS | 6 | 16066 | 5mv9 | A | 1883 | ARG | p.Arg1883Gln | Q13402 |
| HOMO SAPIENS | 4 | 16066 | 5mv9 | A | 1887 | PRO | p.Pro1887Leu | Q13402 |

| | | | | | | | | |
|---|---|---|---|---|---|---|---|---|
| HOMO SAPIENS | 5 | 16066 | 5mv9 | A | 1942 | GLY | p.Gly1942Ter | Q13402 |
| HOMO SAPIENS | 5 | 16066 | 5mv9 | A | 1954 | LEU | p.Leu1954Ile | Q13402 |
| HOMO SAPIENS | 5 | 16066 | 5mv9 | A | 1956 | VAL | p.Val1956Ile | Q13402 |
| HOMO SAPIENS | 5 | 16066 | 5mv9 | A | 1967 | ARG | p.Arg1967Ter | Q13402 |
| HOMO SAPIENS | 5 | 16066 | 5mv9 | A | 1990 | GLN | p.Gln1990Ter | Q13402 |
| HOMO SAPIENS | 2 | 16066 | 5mv9 | A | 2005 | LYS | p.Lys2005Glu | Q13402 |
| HOMO SAPIENS | 1 | 16066 | 5mv9 | A | 2072 | VAL | p.Val2072Ile | Q13402 |
| HOMO SAPIENS | 4 | 16066 | 5mv9 | A | 2109 | THR | p.Thr2109Ile | Q13402 |
| HOMO SAPIENS | 4 | 16066 | 5mv9 | A | 2142 | ASP | p.Asp2142Asn | Q13402 |
| HOMO SAPIENS | -1 | 16066 | 5mv9 | A | 2214 | GLY | p.Gly2214Ser | Q13402 |
| HOMO SAPIENS | 7 | 27942 | 1deb | A | 24 | ARG | p.Arg24Ter | P25054 |
| HOMO SAPIENS | 0 | 876 | 5hnr | B | 12 | PHE | p.Phe12Leu | P13716 |
| HOMO SAPIENS | 0 | 876 | 5hnr | B | 59 | LYS | p.Lys59Asn | P13716 |
| HOMO SAPIENS | -1 | 876 | 5hnr | B | 133 | GLY | p.Gly133Arg | P13716 |
| HOMO SAPIENS | 2 | 876 | 5hnr | B | 240 | ARG | p.Arg240Trp | P13716 |
| HOMO SAPIENS | 0 | 876 | 5hnr | B | 274 | ALA | p.Ala274Thr | P13716 |
| HOMO SAPIENS | 0 | 876 | 5hnr | B | 275 | VAL | p.Val275Met | P13716 |
| HOMO SAPIENS | 0 | 1142 | 3qe2 | A | 503 | VAL | p.Ala500Val | P16435 |
| HOMO SAPIENS | 8 | 14358 | 4dkd | C | 32 | VAL | p.Val32Gly | P07333 |
| HOMO SAPIENS | 0 | 14358 | 4dkd | C | 60 | LEU | p.Leu60Val | P07333 |
| HOMO SAPIENS | 7 | 14358 | 4dkd | C | 75 | THR | p.Thr75Ile | P07333 |
| HOMO SAPIENS | -1 | 14358 | 4dkd | C | 90 | PRO | p.Pro90Thr | P07333 |
| HOMO SAPIENS | 0 | 14358 | 4dkd | C | 106 | ARG | p.Arg106Trp | P07333 |
| HOMO SAPIENS | 0 | 14358 | 4dkd | C | 241 | ASN | p.Asn241Asp | P07333 |
| HOMO SAPIENS | 0 | 14358 | 4dkd | C | 245 | ALA | p.Ala245Ser | P07333 |
| HOMO SAPIENS | 0 | 14358 | 4dkd | C | 255 | ASN | p.Asn255Ile | P07333 |
| HOMO SAPIENS | 0 | 14358 | 4dkd | C | 268 | VAL | p.Val268Ile | P07333 |
| HOMO SAPIENS | 0 | 14358 | 4dkd | C | 279 | VAL | p.Val279Met | P07333 |
| HOMO SAPIENS | 0 | 14358 | 4dkd | C | 294 | ARG | p.Arg294Gln | P07333 |
| HOMO SAPIENS | 5 | 8968 | 3ppy | A | 1565 | VAL | p.Val1565Leu | P04275 |
| HOMO SAPIENS | 4 | 8968 | 3ppy | A | 1584 | TYR | p.Tyr1584Cys | P04275 |
| HOMO SAPIENS | 5 | 5341 | 3k3n | A | 279 | PHE | p.Phe315Ser | Q9UPP1 |
| HOMO SAPIENS | 0 | 29463 | 1q7l | A | 197 | ARG | p.Arg197Trp | Q03154 |
| HOMO SAPIENS | 0 | 33761 | 2leh | B | 30 | ASP | p.Asp30Asn | Q16637 |
| HOMO SAPIENS | 6 | 33761 | 2leh | B | 44 | ASP | p.Asp44Val | Q16637 |
| HOMO SAPIENS | -1 | 12249 | 2pnr | C | 224 | GLU | p.Glu310Lys | P10515 |
| HOMO SAPIENS | -1 | 12249 | 2pnr | C | 232 | VAL | p.Val318Ala | P10515 |
| HOMO SAPIENS | -1 | 7 | 5whk | B | -9 | ALA | p.Ala11Pro | P61769 |
| HOMO SAPIENS | 8 | 20298 | 4euw | A | 154 | PHE | p.Phe154Leu | P48436 |
| HOMO SAPIENS | 9 | 20298 | 4euw | A | 158 | ALA | p.Ala158Thr | P48436 |
| HOMO SAPIENS | 5 | 20298 | 4euw | A | 165 | HIS | p.His165Tyr | P48436 |
| HOMO SAPIENS | 3 | 20298 | 4euw | A | 169 | HIS | p.His169Gln | P48436 |
| HOMO SAPIENS | 0 | 20298 | 4euw | A | 173 | LYS | p.Lys173Glu | P48436 |
| HOMO SAPIENS | 1 | 16465 | 2jgx | A | 402 | TYR | p.Tyr402His | P08603 |
| HOMO SAPIENS | 0 | 8993 | 3v8o | A | 659 | LEU | p.Leu659Arg | Q14315 |
| HOMO SAPIENS | 6 | 8993 | 3v8o | A | 693 | ASP | p.Asp693Ala | Q14315 |
| HOMO SAPIENS | 0 | 8993 | 3v8o | A | 709 | ALA | p.Ala709Thr | Q14315 |
| HOMO SAPIENS | 2 | 3852 | 5es4 | B | 106 | ASP | p.Asp128Asn | P05107 |
| HOMO SAPIENS | 2 | 3852 | 5es4 | B | 116 | SER | p.Ser138Pro | P05107 |
| HOMO SAPIENS | 1 | 3852 | 5es4 | B | 127 | LEU | p.Leu149Pro | P05107 |
| HOMO SAPIENS | 0 | 3852 | 5es4 | B | 147 | GLY | p.Gly169Arg | P05107 |
| HOMO SAPIENS | 3 | 3852 | 5es4 | B | 156 | PRO | p.Pro178Leu | P05107 |
| HOMO SAPIENS | 0 | 3852 | 5es4 | B | 174 | LYS | p.Lys196Thr | P05107 |
| HOMO SAPIENS | 2 | 3852 | 5es4 | B | 251 | GLY | p.Gly273Arg | P05107 |

| | | | | | | | | |
|---|---|---|---|---|---|---|---|---|
| HOMO SAPIENS | 2 | 3852 | 5es4 | B | 329 | ASN | p.Asn351Ser | P05107 |
| HOMO SAPIENS | 2 | 3852 | 5es4 | B | 571 | ARG | p.Arg593Cys | P05107 |
| HOMO SAPIENS | 7 | 5962 | 5fn2 | B | 146 | MET | p.Met146Leu | P49768 |
| HOMO SAPIENS | 3 | 5962 | 5fn2 | B | 166 | LEU | p.Leu166Pro | P49768 |
| HOMO SAPIENS | 1 | 5962 | 5fn2 | B | 233 | MET | p.Met233Leu | P49768 |
| HOMO SAPIENS | 1 | 5962 | 5fn2 | B | 261 | VAL | p.Val261Gly | P49768 |
| HOMO SAPIENS | 1 | 5962 | 5fn2 | B | 266 | GLY | p.Gly266Ser | P49768 |
| HOMO SAPIENS | -1 | 5962 | 5fn2 | B | 333 | ASP | p.Asp333Gly | P49768 |
| HOMO SAPIENS | 6 | 1918 | 3dlx | A | 58 | THR | p.Thr58Met | P55809 |
| HOMO SAPIENS | 5 | 1918 | 3dlx | A | 133 | VAL | p.Val133Glu | P55809 |
| HOMO SAPIENS | 5 | 1918 | 3dlx | A | 219 | GLY | p.Gly219Glu | P55809 |
| HOMO SAPIENS | 0 | 1918 | 3dlx | A | 221 | VAL | p.Val221Met | P55809 |
| HOMO SAPIENS | 5 | 1918 | 3dlx | A | 283 | SER | p.Ser283Ter | P55809 |
| HOMO SAPIENS | 4 | 1918 | 3dlx | A | 324 | GLY | p.Gly324Glu | P55809 |
| HOMO SAPIENS | 0 | 1918 | 3dlx | A | 456 | CYS | p.Cys456Phe | P55809 |
| HOMO SAPIENS | 0 | 9010 | 2wfr | A | 162 | LEU | p.Leu162Pro | O43323 |
| HOMO SAPIENS | 5 | 3770 | 5n38 | A | 56 | VAL | p.Val56Glu | O60260 |
| HOMO SAPIENS | 1 | 3770 | 5n38 | A | 161 | LYS | p.Lys161Asn | O60260 |
| HOMO SAPIENS | 1 | 3770 | 5n38 | A | 167 | SER | p.Ser167Asn | O60260 |
| HOMO SAPIENS | 1 | 3770 | 5n38 | A | 211 | LYS | p.Lys211Asn | O60260 |
| HOMO SAPIENS | 1 | 3770 | 5n38 | A | 212 | CYS | p.Cys212Tyr | O60260 |
| HOMO SAPIENS | 1 | 3770 | 5n38 | A | 275 | ARG | p.Arg275Trp | O60260 |
| HOMO SAPIENS | 2 | 3770 | 5n38 | A | 334 | ARG | p.Arg334Cys | O60260 |
| HOMO SAPIENS | 1 | 3770 | 5n38 | A | 366 | ARG | p.Arg366Trp | O60260 |
| HOMO SAPIENS | -1 | 3770 | 5n38 | A | 380 | VAL | p.Val380Leu | O60260 |
| HOMO SAPIENS | 1 | 3770 | 5n38 | A | 394 | ASP | p.Asp394Asn | O60260 |
| HOMO SAPIENS | 1 | 3770 | 5n38 | A | 430 | GLY | p.Gly430Asp | O60260 |
| HOMO SAPIENS | -1 | 2467 | 4xx1 | A | 10 | PRO | p.Pro34Leu | P04180 |
| HOMO SAPIENS | 4 | 2467 | 4xx1 | A | 83 | TYR | p.Tyr107Ter | P04180 |
| HOMO SAPIENS | 4 | 2467 | 4xx1 | A | 93 | ALA | p.Ala117Thr | P04180 |
| HOMO SAPIENS | 4 | 2467 | 4xx1 | A | 123 | THR | p.Thr147Ile | P04180 |
| HOMO SAPIENS | 4 | 2467 | 4xx1 | A | 135 | ARG | p.Arg159Trp | P04180 |
| HOMO SAPIENS | 3 | 2467 | 4xx1 | A | 158 | ARG | p.Arg182Cys | P04180 |
| HOMO SAPIENS | 0 | 2467 | 4xx1 | A | 209 | LEU | p.Leu233Pro | P04180 |
| HOMO SAPIENS | 1 | 2467 | 4xx1 | A | 228 | ASN | p.Asn252Lys | P04180 |
| HOMO SAPIENS | 4 | 2467 | 4xx1 | A | 252 | MET | p.Met276Lys | P04180 |
| HOMO SAPIENS | 3 | 2467 | 4xx1 | A | 293 | MET | p.Met317Ile | P04180 |
| HOMO SAPIENS | 5 | 2467 | 4xx1 | A | 321 | THR | p.Thr345Met | P04180 |
| HOMO SAPIENS | 9 | 2467 | 4xx1 | A | 347 | THR | p.Thr371Met | P04180 |
| HOMO SAPIENS | 1 | 5834 | 3qrf | G | 371 | PHE | p.Phe371Cys | Q9BZS1 |
| HOMO SAPIENS | 1 | 5834 | 3qrf | G | 373 | PHE | p.Phe373Ala | Q9BZS1 |
| HOMO SAPIENS | 9 | 5834 | 3qrf | G | 384 | ALA | p.Ala384Thr | Q9BZS1 |
| HOMO SAPIENS | 6 | 5834 | 3qrf | G | 397 | ARG | p.Arg397Trp | Q9BZS1 |
| HOMO SAPIENS | -1 | 11869 | 4km8 | A | 15 | PRO | p.Pro15Leu | Q9UMX1 |
| HOMO SAPIENS | 5 | 262 | 4kbo | A | 126 | ARG | p.Arg126Trp | P38646 |
| HOMO SAPIENS | 3 | 262 | 4kbo | A | 128 | TYR | p.Tyr128Cys | P38646 |
| HOMO SAPIENS | -1 | 11347 | 4uf9 | D | 117 | ALA | p.Ala117Val | Q13501 |
| HOMO SAPIENS | 0 | 2173 | 4ah6 | A | 45 | SER | p.Ser45Gly | Q6PI48 |
| HOMO SAPIENS | 2 | 2173 | 4ah6 | A | 152 | CYS | p.Cys152Phe | Q6PI48 |
| HOMO SAPIENS | 4 | 2173 | 4ah6 | A | 179 | ARG | p.Arg179His | Q6PI48 |
| HOMO SAPIENS | 0 | 2173 | 4ah6 | A | 196 | LYS | p.Lys196Arg | Q6PI48 |
| HOMO SAPIENS | 8 | 2173 | 4ah6 | A | 263 | ARG | p.Arg263Gln | Q6PI48 |
| HOMO SAPIENS | 1 | 2173 | 4ah6 | A | 338 | GLY | p.Gly338Glu | Q6PI48 |
| HOMO SAPIENS | 0 | 2173 | 4ah6 | A | 425 | GLU | p.Glu425Ter | Q6PI48 |

| | | | | | | | | |
|---|---|---|---|---|---|---|---|---|
| HOMO SAPIENS | 1 | 2173 | 4ah6 | A | 548 | LEU | p.Leu548Met | Q6PI48 |
| HOMO SAPIENS | 0 | 2173 | 4ah6 | A | 609 | ARG | p.Arg609Trp | Q6PI48 |
| HOMO SAPIENS | 2 | 2173 | 4ah6 | A | 613 | LEU | p.Leu613Phe | Q6PI48 |
| HOMO SAPIENS | 0 | 2173 | 4ah6 | A | 626 | LEU | p.Leu626Gln | Q6PI48 |
| HOMO SAPIENS | 0 | 2173 | 4ah6 | A | 626 | LEU | p.Leu626Val | Q6PI48 |
| HOMO SAPIENS | 7 | 3203 | 2cn1 | A | 87 | ASP | p.Asp137Val | Q9H0P0 |
| HOMO SAPIENS | 8 | 3203 | 2cn1 | A | 166 | GLN | p.Gln216Ter | Q9H0P0 |
| HOMO SAPIENS | 8 | 3203 | 2cn1 | A | 179 | ASN | p.Asn229Ser | Q9H0P0 |
| HOMO SAPIENS | 7 | 3203 | 2cn1 | A | 230 | GLY | p.Gly280Arg | Q9H0P0 |
| HOMO SAPIENS | 2 | 3424 | 2a1t | R | 116 | GLY | p.Gly116Arg | P13804 |
| HOMO SAPIENS | 2 | 3424 | 2a1t | R | 157 | VAL | p.Val157Gly | P13804 |
| HOMO SAPIENS | 2 | 3424 | 2a1t | R | 171 | THR | p.Thr171Ile | P13804 |
| HOMO SAPIENS | 0 | 3424 | 2a1t | R | 178 | THR | p.Thr178Arg | P13804 |
| HOMO SAPIENS | 0 | 3424 | 2a1t | R | 194 | VAL | p.Val194Met | P13804 |
| HOMO SAPIENS | 1 | 3424 | 2a1t | R | 209 | ARG | p.Arg209Ter | P13804 |
| HOMO SAPIENS | 6 | 3424 | 2a1t | R | 266 | THR | p.Thr266Met | P13804 |
| HOMO SAPIENS | 0 | 27598 | 1g1c | A | 26 | ARG | p.Arg2098Gly | Q8WZ42 |
| HOMO SAPIENS | 1 | 27598 | 1g1c | A | 46 | ILE | p.Ile2118Thr | Q8WZ42 |
| HOMO SAPIENS | 0 | 32885 | 4bsk | A | 149 | ASN | p.Asn149Asp | P35916 |
| HOMO SAPIENS | -1 | 8074 | 3ghg | B | 17 | ARG | p.Arg47Ter | P02675 |
| HOMO SAPIENS | -1 | 447 | 5g4p | A | 190 | ALA | p.Ala190Thr | P84074 |
| HOMO SAPIENS | 0 | 10761 | 5u1d | B | 379 | VAL | p.Val379Ile | Q03519 |
| HOMO SAPIENS | 0 | 10761 | 5u1d | B | 665 | THR | p.Thr665Ala | Q03519 |
| HOMO SAPIENS | 5 | 29251 | 5fn7 | A | 79 | ILE | p.Ile294Leu | P08575 |
| HOMO SAPIENS | -1 | 27727 | 4ky9 | A | 56 | LYS | p.Lys56Glu | P02730 |
| HOMO SAPIENS | 2 | 27727 | 4ky9 | A | 72 | GLU | p.Glu72Asp | P02730 |
| HOMO SAPIENS | 2 | 27727 | 4ky9 | A | 90 | GLU | p.Glu90Lys | P02730 |
| HOMO SAPIENS | 1 | 27727 | 4ky9 | A | 130 | GLY | p.Gly130Arg | P02730 |
| HOMO SAPIENS | 2 | 27727 | 4ky9 | A | 150 | ARG | p.Arg150Ter | P02730 |
| HOMO SAPIENS | 0 | 27727 | 4ky9 | A | 327 | PRO | p.Pro327Arg | P02730 |
| HOMO SAPIENS | 2 | 13202 | 2fxo | B | 870 | ARG | p.Arg870His | P12883 |
| HOMO SAPIENS | 2 | 13202 | 2fxo | B | 870 | ARG | p.Arg870Cys | P12883 |
| HOMO SAPIENS | 4 | 13202 | 2fxo | B | 904 | ARG | p.Arg904Cys | P12883 |
| HOMO SAPIENS | 4 | 13202 | 2fxo | B | 906 | ASP | p.Asp906Gly | P12883 |
| HOMO SAPIENS | 2 | 13202 | 2fxo | B | 908 | LEU | p.Leu908Val | P12883 |
| HOMO SAPIENS | 2 | 13202 | 2fxo | B | 923 | ASN | p.Asn923Lys | P12883 |
| HOMO SAPIENS | 7 | 13202 | 2fxo | B | 935 | GLU | p.Glu935Lys | P12883 |
| HOMO SAPIENS | 7 | 13202 | 2fxo | B | 935 | GLU | p.Glu935Val | P12883 |
| HOMO SAPIENS | 8 | 13202 | 2fxo | B | 949 | GLU | p.Glu949Lys | P12883 |
| HOMO SAPIENS | 0 | 13200 | 2a1s | A | 349 | ARG | p.Arg349Trp | O95453 |
| HOMO SAPIENS | 0 | 13200 | 2a1s | A | 421 | LYS | p.Lys421Arg | O95453 |
| HOMO SAPIENS | 6 | 3196 | 5xtb | T | 115 | CYS | p.Cys115Tyr | O75380 |
| HOMO SAPIENS | 0 | 1108 | 5ukf | A | 89 | ARG | p.Arg89Gln | Q99986 |
| HOMO SAPIENS | 0 | 1108 | 5ukf | A | 119 | HIS | p.His119Arg | Q99986 |
| HOMO SAPIENS | 0 | 1108 | 5ukf | A | 133 | ARG | p.Arg133Cys | Q99986 |
| HOMO SAPIENS | 0 | 1108 | 5ukf | A | 236 | VAL | p.Val236Leu | Q99986 |
| HOMO SAPIENS | 0 | 1108 | 5ukf | A | 321 | ARG | p.Arg321Cys | Q99986 |
| HOMO SAPIENS | -1 | 1108 | 5ukf | A | 358 | ARG | p.Arg358Ter | Q99986 |
| HOMO SAPIENS | 0 | 5842 | 4yli | C | 204 | GLY | p.Gly204Ser | Q9BWP8 |
| HOMO SAPIENS | -1 | 31362 | 4wqo | A | 25 | PRO | p.Pro25Leu | P40337 |
| HOMO SAPIENS | 2 | 10107 | 2j8h | A | 34 | VAL | p.Val31886Ile | Q8WZ42 |
| HOMO SAPIENS | 8 | 20190 | 1wm5 | A | 77 | ARG | p.Arg77Gln | P19878 |
| HOMO SAPIENS | 1 | 20190 | 1wm5 | A | 100 | GLN | p.Gln100Ter | P19878 |
| HOMO SAPIENS | 0 | 20190 | 1wm5 | A | 102 | ARG | p.Arg102Ter | P19878 |

| | | | | | | | | | |
|---|---|---|---|---|---|---|---|---|---|
| HOMO SAPIENS | 0 | 20190 | 1wm5 | A | | 128 | ALA | p.Ala128Val | P19878 |
| HOMO SAPIENS | 0 | 20190 | 1wm5 | A | | 181 | LYS | p.Lys181Arg | P19878 |
| HOMO SAPIENS | 7 | 1278 | 5a2q | a | | 33 | ASP | p.Asp33Asn | P62854 |
| HOMO SAPIENS | 3 | 19515 | 2obd | A | | 11 | ILE | p.Ile28Met | P11597 |
| HOMO SAPIENS | 3 | 19515 | 2obd | A | | 89 | VAL | p.Val106Met | P11597 |
| HOMO SAPIENS | 3 | 19515 | 2obd | A | | 137 | ARG | p.Arg154Trp | P11597 |
| HOMO SAPIENS | 3 | 19515 | 2obd | A | | 139 | ASP | p.Asp156Asn | P11597 |
| HOMO SAPIENS | 3 | 19515 | 2obd | A | | 204 | SER | p.Ser221Arg | P11597 |
| HOMO SAPIENS | 0 | 19515 | 2obd | A | | 297 | GLU | p.Glu314Lys | P11597 |
| HOMO SAPIENS | 0 | 19515 | 2obd | A | | 314 | GLY | p.Gly331Ser | P11597 |
| HOMO SAPIENS | 0 | 19515 | 2obd | A | | 332 | SER | p.Ser349Tyr | P11597 |
| HOMO SAPIENS | 0 | 19515 | 2obd | A | | 368 | VAL | p.Val385Met | P11597 |
| HOMO SAPIENS | 0 | 19515 | 2obd | A | | 373 | ALA | p.Ala390Pro | P11597 |
| HOMO SAPIENS | 3 | 19515 | 2obd | A | | 405 | ILE | p.Val422Ile | P11597 |
| HOMO SAPIENS | 0 | 19515 | 2obd | A | | 451 | ARG | p.Arg468Gln | P11597 |
| HOMO SAPIENS | 0 | 19515 | 2obd | A | | 451 | ARG | p.Arg468Ter | P11597 |
| HOMO SAPIENS | 0 | 6626 | 3hop | A | | 39 | ALA | p.Ala39Gly | P21333 |
| HOMO SAPIENS | 1 | 6626 | 3hop | A | | 82 | GLU | p.Glu82Val | P21333 |
| HOMO SAPIENS | 1 | 6626 | 3hop | A | | 128 | ALA | p.Ala128Val | P21333 |
| HOMO SAPIENS | 2 | 6626 | 3hop | A | | 203 | ASP | p.Asp203Tyr | P21333 |
| HOMO SAPIENS | 1 | 6626 | 3hop | A | | 207 | PRO | p.Pro207Leu | P21333 |
| HOMO SAPIENS | 2 | 6626 | 3hop | A | | 210 | CYS | p.Cys210Phe | P21333 |
| HOMO SAPIENS | 7 | 6626 | 3hop | A | | 254 | GLU | p.Glu254Lys | P21333 |
| HOMO SAPIENS | 4 | 4210 | 2avg | A | | 8 | VAL | p.Val158Met | Q14896 |
| HOMO SAPIENS | 6 | 4210 | 2avg | A | | 15 | GLU | p.Glu165Asp | Q14896 |
| HOMO SAPIENS | 4 | 4210 | 2avg | A | | 27 | ARG | p.Arg177His | Q14896 |
| HOMO SAPIENS | 0 | 4210 | 2avg | A | | 28 | VAL | p.Val178Met | Q14896 |
| HOMO SAPIENS | 6 | 4210 | 2avg | A | | 39 | VAL | p.Val189Ile | Q14896 |
| HOMO SAPIENS | 2 | 4210 | 2avg | A | | 52 | LYS | p.Lys202Gln | Q14896 |
| HOMO SAPIENS | 4 | 4210 | 2avg | A | | 67 | SER | p.Ser217Gly | Q14896 |
| HOMO SAPIENS | 6 | 4210 | 2avg | A | | 69 | VAL | p.Val219Leu | Q14896 |
| HOMO SAPIENS | 2 | 4210 | 2avg | A | | 86 | GLY | p.Ser236Gly | Q14896 |
| HOMO SAPIENS | 4 | 4210 | 2avg | A | | 108 | GLU | p.Glu258Lys | Q14896 |
| HOMO SAPIENS | 0 | 22237 | 1w6k | A | | 588 | GLY | p.Gly588Ser | P48449 |
| HOMO SAPIENS | 0 | 10765 | 4a5w | B | | 98 | GLU | p.Glu98Ala | P13671 |
| HOMO SAPIENS | 0 | 10765 | 4a5w | B | | 596 | ARG | p.Arg596Ter | P13671 |
| HOMO SAPIENS | -1 | 15515 | 3hx3 | A | | 11 | MSE | p.Met10Lys | P12271 |
| HOMO SAPIENS | 8 | 15515 | 3hx3 | A | | 64 | ARG | p.Arg64Gln | P12271 |
| HOMO SAPIENS | 0 | 15515 | 3hx3 | A | | 182 | PHE | p.Phe182Cys | P12271 |
| HOMO SAPIENS | 0 | 15515 | 3hx3 | A | | 216 | ARG | p.Arg216Gln | P12271 |
| HOMO SAPIENS | -1 | 15515 | 3hx3 | A | | 226 | MSE | p.Met226Lys | P12271 |
| HOMO SAPIENS | 0 | 15515 | 3hx3 | A | | 234 | TRP | p.Arg234Trp | P12271 |
| HOMO SAPIENS | 0 | 15515 | 3hx3 | A | | 273 | LEU | p.Leu273Phe | P12271 |
| HOMO SAPIENS | 0 | 15515 | 3hx3 | A | | 292 | THR | p.Thr292Met | P12271 |
| HOMO SAPIENS | 0 | 5152 | 2www | D | | 95 | GLN | p.Gln95Ter | Q8IVH4 |
| HOMO SAPIENS | 0 | 5152 | 2www | D | | 133 | GLN | p.Gln133Ter | Q8IVH4 |
| HOMO SAPIENS | 0 | 5152 | 2www | D | | 145 | ARG | p.Arg145Ter | Q8IVH4 |
| HOMO SAPIENS | 0 | 5152 | 2www | D | | 207 | TYR | p.Tyr207Cys | Q8IVH4 |
| HOMO SAPIENS | 8 | 5152 | 2www | D | | 330 | ARG | p.Arg330Ter | Q8IVH4 |
| HOMO SAPIENS | 0 | 5152 | 2www | D | | 363 | GLN | p.Gln363His | Q8IVH4 |
| HOMO SAPIENS | 1 | 1626 | 1ilr | | 1 | 105 | SER | p.Ser130Arg | P18510 |
| HOMO SAPIENS | -1 | 929 | 5loh | A | | 12 | GLY | p.Gly12Arg | Q96GX5 |
| HOMO SAPIENS | 9 | 3206 | 1x63 | A | | 21 | CYS | p.Cys104Arg | Q13642 |
| HOMO SAPIENS | 8 | 3206 | 1x63 | A | | 39 | TRP | p.Trp122Ser | Q13642 |

| | | | | | | | | |
|---|---|---|---|---|---|---|---|---|
| HOMO SAPIENS | 9 | 3206 | 1x63 | A | 40 | HIS | p.His123Tyr | Q13642 |
| HOMO SAPIENS | 9 | 3206 | 1x63 | A | 40 | HIS | p.His123Leu | Q13642 |
| HOMO SAPIENS | 9 | 3206 | 1x63 | A | 40 | HIS | p.His123Gln | Q13642 |
| HOMO SAPIENS | 3 | 3206 | 1x63 | A | 49 | CYS | p.Cys132Phe | Q13642 |
| HOMO SAPIENS | 3 | 3206 | 1x63 | A | 67 | CYS | p.Cys150Tyr | Q13642 |
| HOMO SAPIENS | 3 | 3206 | 1x63 | A | 70 | CYS | p.Cys153Arg | Q13642 |
| HOMO SAPIENS | 3 | 3206 | 1x63 | A | 70 | CYS | p.Cys153Tyr | Q13642 |
| HOMO SAPIENS | 5 | 1052 | 3ecs | E | 208 | ASN | p.Asn208Tyr | Q14232 |
| HOMO SAPIENS | 2 | 1052 | 3ecs | E | 275 | TYR | p.Tyr275Cys | Q14232 |
| HOMO SAPIENS | 0 | 24253 | 5i35 | A | 271 | ARG | p.Arg271Cys | Q8NI60 |
| HOMO SAPIENS | 0 | 24253 | 5i35 | A | 272 | GLY | p.Gly272Asp | Q8NI60 |
| HOMO SAPIENS | 0 | 24253 | 5i35 | A | 299 | ARG | p.Arg299Trp | Q8NI60 |
| HOMO SAPIENS | 1 | 24253 | 5i35 | A | 339 | ALA | p.Ala339Thr | Q8NI60 |
| HOMO SAPIENS | 0 | 24253 | 5i35 | A | 348 | ARG | p.Arg348Ter | Q8NI60 |
| HOMO SAPIENS | 0 | 24253 | 5i35 | A | 395 | ILE | p.Ile395Met | Q8NI60 |
| HOMO SAPIENS | 0 | 24253 | 5i35 | A | 396 | ASP | p.Asp396Glu | Q8NI60 |
| HOMO SAPIENS | 0 | 24253 | 5i35 | A | 480 | PHE | p.Phe480Leu | Q8NI60 |
| HOMO SAPIENS | 2 | 24253 | 5i35 | A | 511 | THR | p.Thr511Met | Q8NI60 |
| HOMO SAPIENS | 0 | 24253 | 5i35 | A | 514 | TYR | p.Tyr514Cys | Q8NI60 |
| HOMO SAPIENS | 0 | 24253 | 5i35 | A | 538 | ARG | p.Arg538Gln | Q8NI60 |
| HOMO SAPIENS | 0 | 24253 | 5i35 | A | 549 | GLY | p.Gly549Ser | Q8NI60 |
| HOMO SAPIENS | 0 | 24253 | 5i35 | A | 551 | GLU | p.Glu551Lys | Q8NI60 |
| HOMO SAPIENS | -1 | 24253 | 5i35 | A | 555 | MSE | p.Met555Ile | Q8NI60 |
| HOMO SAPIENS | 0 | 24253 | 5i35 | A | 568 | GLU | p.Glu568Ter | Q8NI60 |
| HOMO SAPIENS | 0 | 24253 | 5i35 | A | 602 | PRO | p.Pro602Arg | Q8NI60 |
| HOMO SAPIENS | -1 | 15505 | 4goj | C | 22 | GLY | p.Gly22Val | Q13432 |
| HOMO SAPIENS | 5 | 23902 | 2lrr | A | 765 | HIS | p.His765Gln | P38935 |
| HOMO SAPIENS | 1 | 29650 | 2vj2 | A | 234 | CYS | p.Cys234Tyr | P78504 |
| HOMO SAPIENS | 0 | 29650 | 2vj2 | A | 255 | TYR | p.Tyr255Ter | P78504 |
| HOMO SAPIENS | 3 | 29650 | 2vj2 | A | 274 | GLY | p.Gly274Asp | P78504 |
| HOMO SAPIENS | 1 | 105 | 5he0 | B | 77 | GLY | p.Gly77Ser | P62873 |
| HOMO SAPIENS | 2 | 105 | 5he0 | B | 80 | ILE | p.Ile80Thr | P62873 |
| HOMO SAPIENS | 3 | 4587 | 3rgh | A | 159 | ASP | p.Asp1159Ala | P21333 |
| HOMO SAPIENS | 3 | 4587 | 3rgh | A | 184 | ASP | p.Asp1184Glu | P21333 |
| HOMO SAPIENS | 2 | 4587 | 3rgh | A | 186 | SER | p.Ser1186Leu | P21333 |
| HOMO SAPIENS | 4 | 4587 | 3rgh | A | 188 | ALA | p.Ala1188Thr | P21333 |
| HOMO SAPIENS | 2 | 4587 | 3rgh | A | 199 | SER | p.Ser1199Leu | P21333 |
| HOMO SAPIENS | 0 | 20283 | 4n3w | A | 1278 | GLU | p.Glu1278Lys | Q92793 |
| HOMO SAPIENS | 1 | 14294 | 2d9q | B | 208 | MET | p.Met231Thr | Q99062 |
| HOMO SAPIENS | 6 | 14294 | 2d9q | B | 297 | ASP | p.Asp320Asn | Q99062 |
| HOMO SAPIENS | 0 | 11642 | 3bpt | A | 66 | ARG | p.Arg66Trp | Q6NVY1 |
| HOMO SAPIENS | 0 | 11642 | 3bpt | A | 122 | TYR | p.Tyr122Cys | Q6NVY1 |
| HOMO SAPIENS | 0 | 11642 | 3bpt | A | 345 | GLY | p.Gly345Ser | Q6NVY1 |
| HOMO SAPIENS | 1 | 3942 | 3sby | A | 60 | ASP | p.Asp60His | Q9Y4U1 |
| HOMO SAPIENS | 0 | 3942 | 3sby | A | 111 | ARG | p.Arg111Ter | Q9Y4U1 |
| HOMO SAPIENS | 1 | 3942 | 3sby | A | 116 | LEU | p.Leu116Pro | Q9Y4U1 |
| HOMO SAPIENS | 1 | 3942 | 3sby | A | 130 | TYR | p.Tyr130Cys | Q9Y4U1 |
| HOMO SAPIENS | 1 | 3942 | 3sby | A | 132 | ARG | p.Arg132Ter | Q9Y4U1 |
| HOMO SAPIENS | 1 | 3942 | 3sby | A | 147 | GLY | p.Gly147Ala | Q9Y4U1 |
| HOMO SAPIENS | 1 | 3942 | 3sby | A | 161 | ARG | p.Arg161Gln | Q9Y4U1 |
| HOMO SAPIENS | 1 | 3942 | 3sby | A | 161 | ARG | p.Arg161Ter | Q9Y4U1 |
| HOMO SAPIENS | 1 | 3942 | 3sby | A | 203 | TRP | p.Trp203Ter | Q9Y4U1 |
| HOMO SAPIENS | 1 | 3942 | 3sby | A | 203 | TRP | p.Trp203Ter | Q9Y4U1 |
| HOMO SAPIENS | 1 | 3942 | 3sby | A | 205 | TYR | p.Tyr205Ter | Q9Y4U1 |

| | | | | | | | | |
|---|---|---|---|---|---|---|---|---|
| HOMO SAPIENS | 0 | 38340 | 2mbh | B | | 39 MET | p.Met39Leu | Q13351 |
| HOMO SAPIENS | 2 | 7360 | 4k30 | A | | 430 LEU | p.Leu430Pro | Q8N159 |
| HOMO SAPIENS | 2 | 7360 | 4k30 | A | | 433 GLU | p.Glu433Asp | Q8N159 |
| HOMO SAPIENS | 7 | 7360 | 4k30 | A | | 484 TRP | p.Trp484Arg | Q8N159 |
| HOMO SAPIENS | 1 | 41903 | 2xs6 | A | | 151 PRO | p.Pro151Ser | O00459 |
| HOMO SAPIENS | 0 | 172 | 5glh | A | | 183 ALA | p.Ala183Gly | P24530 |
| HOMO SAPIENS | 0 | 172 | 5glh | A | | 201 ARG | p.Arg201Ter | P24530 |
| HOMO SAPIENS | 0 | 172 | 5glh | A | | 244 THR | p.Thr244Met | P24530 |
| HOMO SAPIENS | 0 | 172 | 5glh | A | | 253 ARG | p.Arg253Ter | P24530 |
| HOMO SAPIENS | 9 | 172 | 5glh | A | | 1017 ILE | p.Leu17Phe | P00720 |
| HOMO SAPIENS | -1 | 3342 | 1ljm | A | | 1058 HIS | p.His58Asn | Q01196 |
| HOMO SAPIENS | 4 | 3342 | 1ljm | A | | 1174 ARG | p.Arg174Gln | Q01196 |
| HOMO SAPIENS | 5 | 25607 | 1p22 | C | | 32 ASP | p.Asp32Ala | P35222 |
| HOMO SAPIENS | 5 | 25607 | 1p22 | C | | 32 ASP | p.Asp32Tyr | P35222 |
| HOMO SAPIENS | -1 | 25607 | 1p22 | C | | 33 SEP | p.Ser33Phe | P35222 |
| HOMO SAPIENS | 5 | 25607 | 1p22 | C | | 34 GLY | p.Gly34Glu | P35222 |
| HOMO SAPIENS | -1 | 25607 | 1p22 | C | | 37 SEP | p.Ser37Cys | P35222 |
| HOMO SAPIENS | -1 | 25607 | 1p22 | C | | 41 THR | p.Thr41Ile | P35222 |
| HOMO SAPIENS | 1 | 48 | 4uow | | 9 | 60 GLN | p.Arg34111His | Q8WZ42 |
| HOMO SAPIENS | 0 | 48 | 4uow | | 9 | 64 LEU | p.Val34115Ala | Q8WZ42 |
| HOMO SAPIENS | 0 | 1127 | 5c01 | A | | 638 ARG | p.Arg638Ter | P29597 |
| HOMO SAPIENS | 0 | 1127 | 5c01 | A | | 684 ILE | p.Ile684Ser | P29597 |
| HOMO SAPIENS | 0 | 1127 | 5c01 | A | | 703 ARG | p.Arg703Trp | P29597 |
| HOMO SAPIENS | 5 | 16389 | 2aq0 | B | | 128 ALA | p.Ala860Asp | Q92889 |
| HOMO SAPIENS | 2 | 16389 | 2aq0 | B | | 141 ILE | p.Ile873Val | Q92889 |
| HOMO SAPIENS | 5 | 16389 | 2aq0 | B | | 143 GLU | p.Glu875Gly | Q92889 |
| HOMO SAPIENS | 0 | 16389 | 2aq0 | B | | 180 GLY | p.Gly912Arg | Q92889 |
| HOMO SAPIENS | 0 | 15816 | 2xrc | A | | 89 THR | p.Thr107Ala | P05156 |
| HOMO SAPIENS | 0 | 15816 | 2xrc | A | | 133 ASN | p.Asn151Ser | P05156 |
| HOMO SAPIENS | 0 | 15816 | 2xrc | A | | 185 THR | p.Thr203Ile | P05156 |
| HOMO SAPIENS | 0 | 15816 | 2xrc | A | | 225 GLY | p.Gly243Asp | P05156 |
| HOMO SAPIENS | 0 | 15816 | 2xrc | A | | 243 GLY | p.Gly261Ala | P05156 |
| HOMO SAPIENS | 0 | 15816 | 2xrc | A | | 269 GLY | p.Gly287Arg | P05156 |
| HOMO SAPIENS | -1 | 15816 | 2xrc | A | | 288 ILE | p.Ile306Val | P05156 |
| HOMO SAPIENS | -1 | 15816 | 2xrc | A | | 388 ARG | p.Arg406His | P05156 |
| HOMO SAPIENS | 7 | 15816 | 2xrc | A | | 398 ILE | p.Ile416Leu | P05156 |
| HOMO SAPIENS | 9 | 15816 | 2xrc | A | | 400 HIS | p.His418Leu | P05156 |
| HOMO SAPIENS | 0 | 15816 | 2xrc | A | | 423 LYS | p.Lys441Arg | P05156 |
| HOMO SAPIENS | -1 | 15816 | 2xrc | A | | 530 GLU | p.Glu548Gln | P05156 |
| HOMO SAPIENS | 0 | 15816 | 2xrc | A | | 535 PRO | p.Pro553Ala | P05156 |
| HOMO SAPIENS | 0 | 15816 | 2xrc | A | | 536 GLU | p.Glu554Gly | P05156 |
| HOMO SAPIENS | 1 | 12385 | 4zlp | B | | 1450 ALA | p.Ala1450Pro | Q9UM47 |
| HOMO SAPIENS | 7 | 12385 | 4zlp | B | | 1482 GLY | p.Gly1482Arg | Q9UM47 |
| HOMO SAPIENS | 2 | 12385 | 4zlp | B | | 1518 LEU | p.Leu1518Met | Q9UM47 |
| HOMO SAPIENS | 0 | 12385 | 4zlp | B | | 1560 ARG | p.Arg1560Pro | Q9UM47 |
| HOMO SAPIENS | 0 | 1227 | 4a04 | A | | 148 PHE | p.Phe148Tyr | O43435 |
| HOMO SAPIENS | 4 | 1227 | 4a04 | A | | 194 HIS | p.His194Gln | O43435 |
| HOMO SAPIENS | 2 | 8017 | 3ei4 | B | | 244 LYS | p.Lys244Glu | Q92466 |
| HOMO SAPIENS | 1 | 8017 | 3ei4 | B | | 273 ARG | p.Arg273His | Q92466 |
| HOMO SAPIENS | 3 | 36112 | 2k8f | B | | 9 SER | p.Ser9Arg | P04637 |
| HOMO SAPIENS | 2 | 12880 | 5ggf | A | | 265 ARG | p.Arg265His | Q8WZA1 |
| HOMO SAPIENS | 2 | 12880 | 5ggf | A | | 287 ILE | p.Ile287Ser | Q8WZA1 |
| HOMO SAPIENS | 2 | 12880 | 5ggf | A | | 311 ARG | p.Arg311Gln | Q8WZA1 |
| HOMO SAPIENS | 2 | 12880 | 5ggf | A | | 425 TRP | p.Trp425Ser | Q8WZA1 |

| | | | | | | | | |
|---|---|---|---|---|---|---|---|---|
| HOMO SAPIENS | 2 | 12880 | 5ggf | A | 442 | ARG | p.Arg442Cys | Q8WZA1 |
| HOMO SAPIENS | 2 | 12880 | 5ggf | A | 448 | GLY | p.Gly448Arg | Q8WZA1 |
| HOMO SAPIENS | 0 | 12880 | 5ggf | A | 490 | CYS | p.Cys490Tyr | Q8WZA1 |
| HOMO SAPIENS | 0 | 12880 | 5ggf | A | 493 | PRO | p.Pro493Arg | Q8WZA1 |
| HOMO SAPIENS | 3 | 12880 | 5ggf | A | 550 | SER | p.Ser550Asn | Q8WZA1 |
| HOMO SAPIENS | 1 | 12880 | 5ggf | A | 556 | ASP | p.Asp556Asn | Q8WZA1 |
| HOMO SAPIENS | 0 | 12880 | 5ggf | A | 580 | ARG | p.Arg580Ter | Q8WZA1 |
| HOMO SAPIENS | 2 | 12880 | 5ggf | A | 605 | ARG | p.Arg605Pro | Q8WZA1 |
| HOMO SAPIENS | 0 | 12880 | 5ggf | A | 623 | VAL | p.Met623Val | Q8WZA1 |
| HOMO SAPIENS | 1 | 10187 | 4qw2 | A | 145 | ALA | p.Ala145Ser | Q06787 |
| HOMO SAPIENS | 1 | 10187 | 4qw2 | A | 174 | ASN | p.Asn174Lys | Q06787 |
| HOMO SAPIENS | 0 | 24002 | 2m0t | A | 110 | LEU | p.Leu110Val | O14745 |
| HOMO SAPIENS | 9 | 3126 | 4mjh | A | 99 | LEU | p.Leu99Met | P04792 |
| HOMO SAPIENS | 1 | 3126 | 4mjh | A | 127 | ARG | p.Arg127Trp | P04792 |
| HOMO SAPIENS | 1 | 3126 | 4mjh | A | 127 | ARG | p.Arg127Leu | P04792 |
| HOMO SAPIENS | 1 | 3126 | 4mjh | A | 135 | SER | p.Ser135Phe | P04792 |
| HOMO SAPIENS | 2 | 3126 | 4mjh | A | 136 | ARG | p.Arg136Trp | P04792 |
| HOMO SAPIENS | 1 | 3126 | 4mjh | A | 140 | ARG | p.Arg140Gly | P04792 |
| HOMO SAPIENS | 0 | 3126 | 4mjh | A | 151 | THR | p.Thr151Ile | P04792 |
| HOMO SAPIENS | -1 | 15780 | 2jlp | A | 213 | ARG | p.Arg231Gly | P08294 |
| HOMO SAPIENS | 8 | 3578 | 1apz | D | 203 | GLY | p.Gly226Asp | P20933 |
| HOMO SAPIENS | 1 | 3578 | 1apz | D | 279 | GLY | p.Gly302Arg | P20933 |
| HOMO SAPIENS | 0 | 3578 | 1apz | D | 283 | CYS | p.Cys306Arg | P20933 |
| HOMO SAPIENS | 5 | 1335 | 5xtd | C | 115 | ARG | p.Arg115Cys | O75251 |
| HOMO SAPIENS | 0 | 1335 | 5xtd | C | 122 | VAL | p.Val122Met | O75251 |
| HOMO SAPIENS | 2 | 5063 | 2le4 | A | 37 | ARG | p.Arg74Pro | P48431 |
| HOMO SAPIENS | 1 | 25870 | 2o10 | A | 44 | LEU | p.Leu44Pro | P50461 |
| HOMO SAPIENS | 4 | 25870 | 2o10 | A | 58 | CYS | p.Cys58Gly | P50461 |
| HOMO SAPIENS | 0 | 18748 | 5fii | C | 39 | PHE | p.Phe39Leu | P00439 |
| HOMO SAPIENS | 6 | 18748 | 5fii | C | 48 | LEU | p.Leu48Ser | P00439 |
| HOMO SAPIENS | 2 | 18748 | 5fii | C | 65 | ILE | p.Ile65Asn | P00439 |
| HOMO SAPIENS | 3 | 18748 | 5fii | C | 95 | ILE | p.Ile95Phe | P00439 |
| HOMO SAPIENS | 1 | 14261 | 2vay | B | 1539 | ARG | p.Arg1539Cys | Q13698 |
| HOMO SAPIENS | 0 | 2494 | 4dx1 | A | 183 | GLU | p.Glu183Asp | Q9HBA0 |
| HOMO SAPIENS | 0 | 2494 | 4dx1 | A | 217 | ALA | p.Ala217Ser | Q9HBA0 |
| HOMO SAPIENS | 0 | 2494 | 4dx1 | A | 217 | ALA | p.Ala217Val | Q9HBA0 |
| HOMO SAPIENS | 4 | 2494 | 4dx1 | A | 224 | ARG | p.Arg224Trp | Q9HBA0 |
| HOMO SAPIENS | 4 | 2494 | 4dx1 | A | 254 | VAL | p.Val254Leu | Q9HBA0 |
| HOMO SAPIENS | 0 | 2494 | 4dx1 | A | 269 | ARG | p.Arg269Cys | Q9HBA0 |
| HOMO SAPIENS | 0 | 2494 | 4dx1 | A | 269 | ARG | p.Arg269His | Q9HBA0 |
| HOMO SAPIENS | 0 | 2494 | 4dx1 | A | 278 | GLU | p.Glu278Lys | Q9HBA0 |
| HOMO SAPIENS | 0 | 2494 | 4dx1 | A | 315 | ARG | p.Arg315Trp | Q9HBA0 |
| HOMO SAPIENS | 0 | 2494 | 4dx1 | A | 316 | ARG | p.Arg316Cys | Q9HBA0 |
| HOMO SAPIENS | 0 | 2494 | 4dx1 | A | 331 | ILE | p.Ile331Phe | Q9HBA0 |
| HOMO SAPIENS | 0 | 2494 | 4dx1 | A | 333 | ASP | p.Asp333Gly | Q9HBA0 |
| HOMO SAPIENS | 1 | 2494 | 4dx1 | A | 365 | VAL | p.Val365Met | Q9HBA0 |
| HOMO SAPIENS | 0 | 23998 | 2l1r | B | 161 | ARG | p.Arg162Trp | P19429 |
| HOMO SAPIENS | 0 | 8354 | 4f0x | D | 187 | SER | p.Ser187Ter | O95822 |
| HOMO SAPIENS | 2 | 8354 | 4f0x | D | 259 | GLY | p.Gly259Ala | O95822 |
| HOMO SAPIENS | 2 | 8354 | 4f0x | D | 284 | THR | p.Thr284Ala | O95822 |
| HOMO SAPIENS | 3 | 8354 | 4f0x | D | 309 | LYS | p.Lys309Gln | O95822 |
| HOMO SAPIENS | 0 | 21182 | 4apo | A | 228 | GLN | p.Gln228Lys | O00170 |
| HOMO SAPIENS | 1 | 21182 | 4apo | A | 307 | GLN | p.Gln307Arg | O00170 |
| HOMO SAPIENS | 0 | 2287 | 5of4 | B | 112 | ARG | p.Arg112His | P18074 |

| | | | | | | | | |
|---|---|---|---|---|---|---|---|---|
| HOMO SAPIENS | 0 | 2287 | 5of4 | B | 312 | ASP | p.Asp312Asn | P18074 |
| HOMO SAPIENS | 3 | 2287 | 5of4 | B | 461 | LEU | p.Leu461Val | P18074 |
| HOMO SAPIENS | 0 | 2287 | 5of4 | B | 485 | LEU | p.Leu485Pro | P18074 |
| HOMO SAPIENS | 1 | 2287 | 5of4 | B | 541 | SER | p.Ser541Arg | P18074 |
| HOMO SAPIENS | 0 | 2287 | 5of4 | B | 616 | ARG | p.Arg616Pro | P18074 |
| HOMO SAPIENS | 0 | 2287 | 5of4 | B | 616 | ARG | p.Arg616Trp | P18074 |
| HOMO SAPIENS | 1 | 2287 | 5of4 | B | 658 | ARG | p.Arg658Cys | P18074 |
| HOMO SAPIENS | 1 | 2287 | 5of4 | B | 681 | ASP | p.Asp681Asn | P18074 |
| HOMO SAPIENS | 1 | 2287 | 5of4 | B | 683 | ARG | p.Arg683Trp | P18074 |
| HOMO SAPIENS | 0 | 2287 | 5of4 | B | 713 | GLY | p.Gly713Arg | P18074 |
| HOMO SAPIENS | 0 | 2287 | 5of4 | B | 717 | ALA | p.Ala717Gly | P18074 |
| HOMO SAPIENS | 0 | 2287 | 5of4 | B | 722 | ARG | p.Arg722Trp | P18074 |
| HOMO SAPIENS | 0 | 2287 | 5of4 | B | 725 | ALA | p.Ala725Pro | P18074 |
| HOMO SAPIENS | 4 | 12188 | 4its | A | 122 | ARG | p.Arg122Gly | Q9Y606 |
| HOMO SAPIENS | 4 | 12188 | 4its | A | 133 | ASP | p.Asp133Asn | Q9Y606 |
| HOMO SAPIENS | 4 | 12188 | 4its | A | 144 | ARG | p.Arg144Trp | Q9Y606 |
| HOMO SAPIENS | 0 | 31529 | 1kgd | A | 761 | ARG | p.Arg766Gln | O14936 |
| HOMO SAPIENS | 1 | 31529 | 1kgd | A | 824 | GLN | p.Gln829Arg | O14936 |
| HOMO SAPIENS | 0 | 10906 | 2h63 | C | 18 | ARG | p.Arg18Ter | P53004 |
| HOMO SAPIENS | 9 | 27767 | 4wyq | A | 280 | LEU | p.Leu278Phe | Q9UPY3 |
| HOMO SAPIENS | 0 | 27767 | 4wyq | A | 297 | SER | p.Ser295Cys | Q9UPY3 |
| HOMO SAPIENS | 1 | 27767 | 4wyq | A | 334 | TYR | p.Tyr332Ter | Q9UPY3 |
| HOMO SAPIENS | 7 | 7775 | 5sv7 | C | 136 | SER | p.Ser136Cys | Q9NZJ5 |
| HOMO SAPIENS | 2 | 7775 | 5sv7 | C | 166 | GLN | p.Gln166Arg | Q9NZJ5 |
| HOMO SAPIENS | 2 | 7775 | 5sv7 | C | 332 | GLU | p.Glu332Ter | Q9NZJ5 |
| HOMO SAPIENS | 0 | 21449 | 2e9w | B | 84 | THR | p.Thr84Met | P10721 |
| HOMO SAPIENS | 0 | 21449 | 2e9w | B | 168 | ALA | p.Ala168Glu | P10721 |
| HOMO SAPIENS | 0 | 21449 | 2e9w | B | 178 | ALA | p.Ala178Thr | P10721 |
| HOMO SAPIENS | 0 | 21449 | 2e9w | B | 293 | ASN | p.Asn293Ser | P10721 |
| HOMO SAPIENS | -1 | 21449 | 2e9w | B | 318 | MET | p.Met318Leu | P10721 |
| HOMO SAPIENS | 0 | 21449 | 2e9w | B | 374 | VAL | p.Val374Ile | P10721 |
| HOMO SAPIENS | 6 | 3010 | 3flg | A | 66 | SER | p.Ser270Pro | Q9UBC3 |
| HOMO SAPIENS | 1 | 33918 | 2xzz | A | 760 | ARG | p.Arg760Ter | P22735 |
| HOMO SAPIENS | 2 | 1254 | 5xtc | w | 65 | ASN | p.Asn65Ser | O95299 |
| HOMO SAPIENS | 1 | 1254 | 5xtc | w | 238 | GLU | p.Glu238Lys | O95299 |
| HOMO SAPIENS | 3 | 10245 | 3siv | B | 194 | ALA | p.Ala194Glu | Q8WWY3 |
| HOMO SAPIENS | 4 | 10245 | 3siv | B | 216 | ALA | p.Ala216Pro | Q8WWY3 |
| HOMO SAPIENS | 0 | 2670 | 3dwb | A | 341 | THR | p.Thr341Ile | P42892 |
| HOMO SAPIENS | 1 | 2670 | 3dwb | A | 462 | SER | p.Ser462Asn | P42892 |
| HOMO SAPIENS | 1 | 2670 | 3dwb | A | 754 | ARG | p.Arg754Cys | P42892 |
| HOMO SAPIENS | 1 | 2961 | 5xtb | O | 40 | VAL | p.Val40Met | P19404 |
| HOMO SAPIENS | 5 | 2961 | 5xtb | O | 172 | ILE | p.Ile172Thr | P19404 |
| HOMO SAPIENS | 5 | 2961 | 5xtb | O | 190 | ASP | p.Asp190Gly | P19404 |
| HOMO SAPIENS | 5 | 29655 | 2xdg | A | 57 | ALA | p.Ala57Thr | Q02643 |
| HOMO SAPIENS | 4 | 29655 | 2xdg | A | 72 | GLU | p.Glu72Ter | Q02643 |
| HOMO SAPIENS | 0 | 29655 | 2xdg | A | 121 | GLU | p.Glu121Asp | Q02643 |
| HOMO SAPIENS | -1 | 17935 | 1hyn | P | 38 | ASP | p.Asp38Ala | P02730 |
| HOMO SAPIENS | 7 | 12825 | 1g82 | D | 99 | SER | p.Ser99Asn | P31371 |
| HOMO SAPIENS | 6 | 24927 | 1jm7 | B | 85 | VAL | p.Val85Leu | Q99728 |
| HOMO SAPIENS | 1 | 24927 | 1jm7 | B | 112 | ARG | p.Arg112Ter | Q99728 |
| HOMO SAPIENS | 0 | 24927 | 1jm7 | B | 116 | HIS | p.His116Tyr | Q99728 |
| HOMO SAPIENS | 7 | 92 | 5hde | A | 61 | LYS | p.Lys61Arg | Q05209 |
| HOMO SAPIENS | 0 | 17246 | 1szb | A | 84 | ARG | p.Arg99Gln | O00187 |
| HOMO SAPIENS | 1 | 17246 | 1szb | A | 105 | ASP | p.Asp120Gly | O00187 |

| | | | | | | | | |
|---|---|---|---|---|---|---|---|---|
| HOMO SAPIENS | 0 | 17246 | 1szb | A | 111 | PRO | p.Pro126Leu | O00187 |
| HOMO SAPIENS | 9 | 17246 | 1szb | A | 141 | CYS | p.Cys156Tyr | O00187 |
| HOMO SAPIENS | 0 | 32473 | 1b72 | A | 207 | ARG | p.Arg207Cys | P14653 |
| HOMO SAPIENS | 0 | 1920 | 4yzf | B | 488 | VAL | p.Val488Met | P02730 |
| HOMO SAPIENS | 0 | 1920 | 4yzf | B | 589 | ARG | p.Arg589Cys | P02730 |
| HOMO SAPIENS | 0 | 1920 | 4yzf | B | 589 | ARG | p.Arg589His | P02730 |
| HOMO SAPIENS | 0 | 1920 | 4yzf | B | 602 | ARG | p.Arg602Pro | P02730 |
| HOMO SAPIENS | 0 | 1920 | 4yzf | B | 613 | SER | p.Ser613Phe | P02730 |
| HOMO SAPIENS | -1 | 1920 | 4yzf | B | 646 | ARG | p.Arg646Gln | P02730 |
| HOMO SAPIENS | -1 | 1920 | 4yzf | B | 646 | ARG | p.Arg646Trp | P02730 |
| HOMO SAPIENS | 0 | 1920 | 4yzf | B | 701 | GLY | p.Gly701Asp | P02730 |
| HOMO SAPIENS | 0 | 1920 | 4yzf | B | 760 | ARG | p.Arg760Gln | P02730 |
| HOMO SAPIENS | 0 | 1920 | 4yzf | B | 771 | GLY | p.Gly771Asp | P02730 |
| HOMO SAPIENS | 0 | 1920 | 4yzf | B | 773 | SER | p.Ser773Pro | P02730 |
| HOMO SAPIENS | 0 | 1920 | 4yzf | B | 837 | THR | p.Thr837Ala | P02730 |
| HOMO SAPIENS | 0 | 1920 | 4yzf | B | 854 | PRO | p.Pro854Leu | P02730 |
| HOMO SAPIENS | 0 | 1920 | 4yzf | B | 858 | ALA | p.Ala858Asp | P02730 |
| HOMO SAPIENS | 0 | 1920 | 4yzf | B | 862 | VAL | p.Val862Ile | P02730 |
| HOMO SAPIENS | 0 | 1920 | 4yzf | B | 868 | PRO | p.Pro868Leu | P02730 |
| HOMO SAPIENS | 0 | 1920 | 4yzf | B | 870 | ARG | p.Arg870Trp | P02730 |
| HOMO SAPIENS | 0 | 9863 | 2juj | A | 54 | VAL | p.Val904Ile | P22681 |
| HOMO SAPIENS | 0 | 21934 | 2na7 | A | 184 | ILE | p.Ile184Val | P25445 |
| HOMO SAPIENS | -1 | 6746 | 1lw3 | A | 3 | THR | p.Lys3Thr | Q13614 |
| HOMO SAPIENS | -1 | 6746 | 1lw3 | A | 44 | VAL | p.Val44Ile | Q13614 |
| HOMO SAPIENS | 3 | 6746 | 1lw3 | A | 183 | ALA | p.Ala183Ser | Q13614 |
| HOMO SAPIENS | 4 | 6746 | 1lw3 | A | 482 | GLN | p.Gln482Ter | Q13614 |
| HOMO SAPIENS | 4 | 6746 | 1lw3 | A | 502 | GLU | p.Glu502Gln | Q13614 |
| HOMO SAPIENS | 3 | 6746 | 1lw3 | A | 545 | ASN | p.Asn545Ser | Q13614 |
| HOMO SAPIENS | 0 | 6746 | 1lw3 | A | 586 | ARG | p.Arg586Trp | Q13614 |
| HOMO SAPIENS | -1 | 6746 | 1lw3 | A | 602 | ALA | p.Ala602Gly | Q13614 |
| HOMO SAPIENS | 1 | 319 | 4ch9 | B | 336 | ARG | p.Arg336Ile | Q9UH77 |
| HOMO SAPIENS | 3 | 319 | 4ch9 | B | 340 | ALA | p.Ala340Val | Q9UH77 |
| HOMO SAPIENS | 0 | 319 | 4ch9 | B | 384 | ARG | p.Arg384Gln | Q9UH77 |
| HOMO SAPIENS | 5 | 319 | 4ch9 | B | 426 | PRO | p.Pro426Leu | Q9UH77 |
| HOMO SAPIENS | 3 | 319 | 4ch9 | B | 431 | ARG | p.Arg431Gln | Q9UH77 |
| HOMO SAPIENS | 2 | 319 | 4ch9 | B | 575 | ARG | p.Arg575Trp | Q9UH77 |
| HOMO SAPIENS | 0 | 574 | 2qkk | A | 142 | VAL | p.Val142Ile | O60930 |
| HOMO SAPIENS | 0 | 24021 | 2n8y | A | 752 | ARG | p.Arg752Gln | P12814 |
| HOMO SAPIENS | 0 | 18693 | 2y7j | B | 44 | ARG | p.Arg44Ter | P15735 |
| HOMO SAPIENS | 0 | 18693 | 2y7j | B | 106 | VAL | p.Val106Glu | P15735 |
| HOMO SAPIENS | 3 | 18693 | 2y7j | B | 189 | GLY | p.Gly189Glu | P15735 |
| HOMO SAPIENS | 0 | 17087 | 2ifg | A | 237 | THR | p.Thr237Met | P04629 |
| HOMO SAPIENS | 0 | 7980 | 1qub | A | 247 | VAL | p.Val266Leu | P02749 |
| HOMO SAPIENS | -1 | 2205 | 1h7u | A | 18 | ILE | p.Ile18Val | P54278 |
| HOMO SAPIENS | -1 | 2205 | 1h7u | A | 20 | ARG | p.Arg20Gln | P54278 |
| HOMO SAPIENS | 9 | 2205 | 1h7u | A | 46 | SER | p.Ser46Ile | P54278 |
| HOMO SAPIENS | 0 | 2205 | 1h7u | A | 60 | ASP | p.Asp60Glu | P54278 |
| HOMO SAPIENS | 0 | 2205 | 1h7u | A | 80 | PHE | p.Phe80Leu | P54278 |
| HOMO SAPIENS | 0 | 2205 | 1h7u | A | 118 | SER | p.Ser118Arg | P54278 |
| HOMO SAPIENS | 0 | 2205 | 1h7u | A | 126 | HIS | p.His126Gln | P54278 |
| HOMO SAPIENS | 0 | 2205 | 1h7u | A | 128 | SER | p.Ser128Leu | P54278 |
| HOMO SAPIENS | 1 | 2205 | 1h7u | A | 134 | ARG | p.Arg134Ter | P54278 |
| HOMO SAPIENS | 0 | 2205 | 1h7u | A | 166 | LEU | p.Leu166Pro | P54278 |
| HOMO SAPIENS | 1 | 2205 | 1h7u | A | 191 | TYR | p.Tyr191Cys | P54278 |

| | | | | | | | | |
|---|---|---|---|---|---|---|---|---|
| HOMO SAPIENS | 0 | 2205 | 1h7u | A | 211 | ARG | p.Arg211Ter | P54278 |
| HOMO SAPIENS | 0 | 2205 | 1h7u | A | 233 | GLN | p.Gln233Ter | P54278 |
| HOMO SAPIENS | 0 | 2205 | 1h7u | A | 255 | TYR | p.Tyr255Ter | P54278 |
| HOMO SAPIENS | 0 | 2205 | 1h7u | A | 277 | THR | p.Thr277Lys | P54278 |
| HOMO SAPIENS | 0 | 2205 | 1h7u | A | 301 | LYS | p.Lys301Asn | P54278 |
| HOMO SAPIENS | 0 | 2205 | 1h7u | A | 312 | MET | p.Met312Ile | P54278 |
| HOMO SAPIENS | 0 | 2205 | 1h7u | A | 312 | MET | p.Met312Thr | P54278 |
| HOMO SAPIENS | 0 | 2205 | 1h7u | A | 315 | ARG | p.Arg315Ter | P54278 |
| HOMO SAPIENS | 0 | 2205 | 1h7u | A | 318 | TYR | p.Tyr318Cys | P54278 |
| HOMO SAPIENS | 0 | 2205 | 1h7u | A | 335 | ASN | p.Asn335Ile | P54278 |
| HOMO SAPIENS | 0 | 2205 | 1h7u | A | 360 | ILE | p.Ile360Met | P54278 |
| HOMO SAPIENS | 6 | 20754 | 5mpo | B | 7 | VAL | p.Val7Phe | O96033 |
| HOMO SAPIENS | 4 | 20754 | 5mpo | B | 64 | VAL | p.Val64Ile | O96033 |
| HOMO SAPIENS | 5 | 34944 | 4qam | B | 1114 | ASP | p.Asp1114Gly | Q96KN7 |
| HOMO SAPIENS | 5 | 34944 | 4qam | B | 1240 | GLY | p.Gly1240Glu | Q96KN7 |
| HOMO SAPIENS | 9 | 40496 | 3vnn | A | 278 | ARG | p.Arg278His | P49917 |
| HOMO SAPIENS | 6 | 30131 | 4dt7 | E | 40 | PRO | p.Pro210Leu | P04070 |
| HOMO SAPIENS | -1 | 30131 | 4dt7 | E | 50 | ARG | p.Arg220Trp | P04070 |
| HOMO SAPIENS | 7 | 15311 | 2pqt | A | 64 | ARG | p.Arg64Trp | P18440 |
| HOMO SAPIENS | 2 | 15311 | 2pqt | A | 149 | VAL | p.Val149Ile | P18440 |
| HOMO SAPIENS | -1 | 19581 | 3hy8 | A | 33 | ASP | p.Asp33Val | Q9NVS9 |
| HOMO SAPIENS | 5 | 19581 | 3hy8 | A | 116 | ARG | p.Arg116Gln | Q9NVS9 |
| HOMO SAPIENS | 4 | 19581 | 3hy8 | A | 225 | ARG | p.Arg225His | Q9NVS9 |
| HOMO SAPIENS | 2 | 19581 | 3hy8 | A | 229 | TRP | p.Arg229Trp | Q9NVS9 |
| HOMO SAPIENS | 2 | 19581 | 3hy8 | A | 229 | TRP | p.Arg229Gln | Q9NVS9 |
| HOMO SAPIENS | 0 | 10266 | 3t1i | A | 41 | ASP | p.Asp41His | P49959 |
| HOMO SAPIENS | 1 | 10266 | 3t1i | A | 305 | ARG | p.Arg305Trp | P49959 |
| HOMO SAPIENS | 0 | 10266 | 3t1i | A | 308 | PHE | p.Phe308Leu | P49959 |
| HOMO SAPIENS | 0 | 10266 | 3t1i | A | 334 | SER | p.Ser334Arg | P49959 |
| HOMO SAPIENS | 1 | 1600 | 3rbf | A | 61 | GLU | p.Glu61Asp | P20711 |
| HOMO SAPIENS | 0 | 1600 | 3rbf | A | 91 | ALA | p.Ala91Val | P20711 |
| HOMO SAPIENS | -1 | 1600 | 3rbf | A | 102 | GLY | p.Gly102Ser | P20711 |
| HOMO SAPIENS | 3 | 1600 | 3rbf | A | 147 | SER | p.Ser147Arg | P20711 |
| HOMO SAPIENS | 1 | 1600 | 3rbf | A | 210 | PRO | p.Pro210Leu | P20711 |
| HOMO SAPIENS | 0 | 1600 | 3rbf | A | 217 | MET | p.Met217Val | P20711 |
| HOMO SAPIENS | 4 | 1600 | 3rbf | A | 250 | SER | p.Ser250Phe | P20711 |
| HOMO SAPIENS | 2 | 1600 | 3rbf | A | 275 | ALA | p.Ala275Thr | P20711 |
| HOMO SAPIENS | 1 | 1600 | 3rbf | A | 309 | PHE | p.Phe309Leu | P20711 |
| HOMO SAPIENS | -1 | 1600 | 3rbf | A | 347 | ARG | p.Arg347Gln | P20711 |
| HOMO SAPIENS | 0 | 1600 | 3rbf | A | 462 | ARG | p.Arg462Gln | P20711 |
| HOMO SAPIENS | 6 | 20830 | 5y4f | A | 714 | THR | p.Thr715Ile | Q01484 |
| HOMO SAPIENS | 7 | 20830 | 5y4f | A | 719 | ASP | p.Asp720Glu | Q01484 |
| HOMO SAPIENS | 0 | 20830 | 5y4f | A | 814 | VAL | p.Val815Ile | Q01484 |
| HOMO SAPIENS | -1 | 302 | 2c9y | A | 9 | GLU | p.Glu9Ter | P54819 |
| HOMO SAPIENS | 0 | 302 | 2c9y | A | 103 | ARG | p.Arg103Trp | P54819 |
| HOMO SAPIENS | 3 | 302 | 2c9y | A | 165 | ASP | p.Asp165Gly | P54819 |
| HOMO SAPIENS | 0 | 302 | 2c9y | A | 182 | ALA | p.Ala182Asp | P54819 |
| HOMO SAPIENS | 2 | 302 | 2c9y | A | 186 | ARG | p.Arg186Cys | P54819 |
| HOMO SAPIENS | 0 | 302 | 2c9y | A | 209 | ALA | p.Ala209Thr | P54819 |
| HOMO SAPIENS | -1 | 16920 | 1jlj | B | 10 | ASN | p.Asn10Tyr | Q9NQX3 |
| HOMO SAPIENS | 9 | 8462 | 1zbq | B | 16 | GLY | p.Gly16Ser | P51659 |
| HOMO SAPIENS | 5 | 8462 | 1zbq | B | 19 | ALA | p.Ala19Gly | P51659 |
| HOMO SAPIENS | 1 | 8462 | 1zbq | B | 34 | ALA | p.Ala34Val | P51659 |
| HOMO SAPIENS | 1 | 8462 | 1zbq | B | 106 | ARG | p.Arg106His | P51659 |

| | | | | | | | | |
|---|---|---|---|---|---|---|---|---|
| HOMO SAPIENS | 0 | 8462 | 1zbq | B | 140 | LYS | p.Lys140Asn | P51659 |
| HOMO SAPIENS | 0 | 8462 | 1zbq | B | 292 | THR | p.Thr292Ser | P51659 |
| HOMO SAPIENS | 0 | 4375 | 4ndi | A | 199 | ARG | p.Arg213His | Q7Z2E3 |
| HOMO SAPIENS | 3 | 4375 | 4ndi | A | 201 | HIS | p.His215Arg | Q7Z2E3 |
| HOMO SAPIENS | 0 | 4375 | 4ndi | A | 206 | PRO | p.Pro220Leu | Q7Z2E3 |
| HOMO SAPIENS | 2 | 4375 | 4ndi | A | 223 | LEU | p.Leu237Phe | Q7Z2E3 |
| HOMO SAPIENS | 0 | 4375 | 4ndi | A | 245 | ARG | p.Arg259His | Q7Z2E3 |
| HOMO SAPIENS | 0 | 4375 | 4ndi | A | 263 | VAL | p.Val277Gly | Q7Z2E3 |
| HOMO SAPIENS | 0 | 4375 | 4ndi | A | 279 | TRP | p.Trp293Ter | Q7Z2E3 |
| HOMO SAPIENS | 0 | 27004 | 4ccg | Y | 321 | ASP | p.Asp321Glu | Q9NW38 |
| HOMO SAPIENS | -1 | 27004 | 4ccg | Y | 372 | GLY | p.Gly372Ala | Q9NW38 |
| HOMO SAPIENS | -1 | 6383 | 5fai | A | 34 | ALA | p.Ala34Gly | Q92979 |
| HOMO SAPIENS | 8 | 6383 | 5fai | A | 86 | ASP | p.Asp86Gly | Q92979 |
| HOMO SAPIENS | 2 | 24585 | 4mi5 | A | 684 | ARG | p.Arg679Cys | Q15910 |
| HOMO SAPIENS | 0 | 15967 | 3nhb | A | 723 | ARG | p.Arg723Gln | Q9NP58 |
| HOMO SAPIENS | 5 | 18152 | 1oey | A | 389 | HIS | p.His389Gln | P19878 |
| HOMO SAPIENS | 5 | 18152 | 1oey | A | 395 | ARG | p.Arg395Trp | P19878 |
| HOMO SAPIENS | -1 | 1044 | 1li1 | F | 2 | VAL | p.Val1486Ile | P08572 |
| HOMO SAPIENS | 6 | 1044 | 1li1 | F | 37 | GLU | p.Glu1521Lys | P08572 |
| HOMO SAPIENS | 5 | 1044 | 1li1 | F | 157 | SER | p.Ser1639Leu | P08572 |
| HOMO SAPIENS | 5 | 1044 | 1li1 | F | 179 | ARG | p.Arg1662His | P08572 |
| HOMO SAPIENS | 5 | 1044 | 1li1 | F | 180 | GLY | p.Gly1663Arg | P08572 |
| HOMO SAPIENS | 6 | 20076 | 4uyi | A | 671 | LEU | p.Leu671Ser | Q8IY92 |
| HOMO SAPIENS | 0 | 19588 | 1jmj | A | 189 | ARG | p.Arg208His | P05546 |
| HOMO SAPIENS | -1 | 38023 | 1ry7 | B | 65 | GLY | p.Gly65Arg | P22607 |
| HOMO SAPIENS | -1 | 38023 | 1ry7 | B | 139 | ASP | p.Asp139Glu | P22607 |
| HOMO SAPIENS | 0 | 38023 | 1ry7 | B | 248 | ARG | p.Arg248Cys | P22607 |
| HOMO SAPIENS | 0 | 38023 | 1ry7 | B | 249 | SER | p.Ser249Cys | P22607 |
| HOMO SAPIENS | 0 | 38023 | 1ry7 | B | 250 | PRO | p.Pro250Arg | P22607 |
| HOMO SAPIENS | 0 | 38023 | 1ry7 | B | 322 | GLU | p.Glu322Lys | P22607 |
| HOMO SAPIENS | 5 | 16884 | 5eli | A | 33 | GLN | p.Gln33Ter | Q9NZC2 |
| HOMO SAPIENS | 0 | 16884 | 5eli | A | 44 | TRP | p.Trp44Ter | Q9NZC2 |
| HOMO SAPIENS | 0 | 16884 | 5eli | A | 47 | ARG | p.Arg47His | Q9NZC2 |
| HOMO SAPIENS | 0 | 16884 | 5eli | A | 66 | THR | p.Thr66Met | Q9NZC2 |
| HOMO SAPIENS | 0 | 16884 | 5eli | A | 78 | TRP | p.Trp78Ter | Q9NZC2 |
| HOMO SAPIENS | 1 | 16884 | 5eli | A | 96 | THR | p.Thr96Lys | Q9NZC2 |
| HOMO SAPIENS | 9 | 16884 | 5eli | A | 126 | VAL | p.Val126Gly | Q9NZC2 |
| HOMO SAPIENS | -1 | 12267 | 3w5o | A | 3 | ALA | p.Ala3Val | P49917 |
| HOMO SAPIENS | 1 | 12267 | 3w5o | A | 9 | THR | p.Thr9Ile | P49917 |
| HOMO SAPIENS | 2 | 12267 | 3w5o | A | 188 | ARG | p.Arg188Gln | P49917 |
| HOMO SAPIENS | 0 | 12267 | 3w5o | A | 469 | GLY | p.Gly469Glu | P49917 |
| HOMO SAPIENS | 1 | 12267 | 3w5o | A | 580 | ARG | p.Arg580Ter | P49917 |
| HOMO SAPIENS | 0 | 12267 | 3w5o | A | 600 | GLU | p.Glu600Lys | P49917 |
| HOMO SAPIENS | 9 | 2839 | 2hw5 | B | 54 | ARG | p.Arg54Cys | P30084 |
| HOMO SAPIENS | 9 | 2839 | 2hw5 | B | 59 | ASN | p.Asn59Ser | P30084 |
| HOMO SAPIENS | 0 | 2839 | 2hw5 | B | 273 | LYS | p.Lys273Glu | P30084 |
| HOMO SAPIENS | 1 | 12266 | 2o2k | B | 939 | GLY | p.Gly939Cys | Q99707 |
| HOMO SAPIENS | 5 | 12266 | 2o2k | B | 1164 | ARG | p.Arg1164His | Q99707 |
| HOMO SAPIENS | 1 | 12266 | 2o2k | B | 1166 | LEU | p.Leu1166Met | Q99707 |
| HOMO SAPIENS | 5 | 12266 | 2o2k | B | 1173 | PRO | p.Pro1173Leu | Q99707 |
| HOMO SAPIENS | 5 | 12266 | 2o2k | B | 1222 | ASN | p.Asn1222Ser | Q99707 |
| HOMO SAPIENS | 0 | 30891 | 3ch4 | B | 138 | ARG | p.Arg138Ter | Q15126 |
| HOMO SAPIENS | -1 | 4433 | 1qlz | A | 54 | GLY | p.Gly54Ser | P04156 |
| HOMO SAPIENS | 0 | 22568 | 3cxl | A | 20 | LEU | p.Leu20Phe | P15882 |

| | | | | | | | | |
|---|---|---|---|---|---|---|---|---|
| HOMO SAPIENS | 0 | 22568 | 3cxl | A | 126 | ILE | p.Ile126Met | P15882 |
| HOMO SAPIENS | 0 | 22568 | 3cxl | A | 141 | PRO | p.Pro141Leu | P15882 |
| HOMO SAPIENS | 0 | 22568 | 3cxl | A | 143 | TYR | p.Tyr143His | P15882 |
| HOMO SAPIENS | 0 | 22568 | 3cxl | A | 223 | ALA | p.Ala223Val | P15882 |
| HOMO SAPIENS | 0 | 22568 | 3cxl | A | 228 | GLY | p.Gly228Ser | P15882 |
| HOMO SAPIENS | 0 | 22568 | 3cxl | A | 252 | PRO | p.Pro252Gln | P15882 |
| HOMO SAPIENS | 0 | 22568 | 3cxl | A | 313 | GLU | p.Glu313Lys | P15882 |
| HOMO SAPIENS | 3 | 2668 | 5eqg | A | 34 | ASN | p.Asn34Ile | P11166 |
| HOMO SAPIENS | 3 | 2668 | 5eqg | A | 77 | MET | p.Met77Val | P11166 |
| HOMO SAPIENS | 6 | 2668 | 5eqg | A | 91 | GLY | p.Gly91Asp | P11166 |
| HOMO SAPIENS | 6 | 2668 | 5eqg | A | 92 | ARG | p.Arg92Trp | P11166 |
| HOMO SAPIENS | 6 | 2668 | 5eqg | A | 93 | ARG | p.Arg93Trp | P11166 |
| HOMO SAPIENS | 0 | 2668 | 5eqg | A | 95 | SER | p.Ser95Ile | P11166 |
| HOMO SAPIENS | 3 | 2668 | 5eqg | A | 126 | ARG | p.Arg126His | P11166 |
| HOMO SAPIENS | 3 | 2668 | 5eqg | A | 126 | ARG | p.Arg126Cys | P11166 |
| HOMO SAPIENS | 4 | 2668 | 5eqg | A | 232 | ARG | p.Arg232Cys | P11166 |
| HOMO SAPIENS | 3 | 2668 | 5eqg | A | 249 | ARG | p.Arg249Gln | P11166 |
| HOMO SAPIENS | 3 | 2668 | 5eqg | A | 255 | LYS | p.Lys255Arg | P11166 |
| HOMO SAPIENS | 0 | 2668 | 5eqg | A | 275 | ALA | p.Ala275Thr | P11166 |
| HOMO SAPIENS | 4 | 2668 | 5eqg | A | 299 | GLU | p.Glu299Lys | P11166 |
| HOMO SAPIENS | 6 | 2668 | 5eqg | A | 314 | GLY | p.Gly314Ser | P11166 |
| HOMO SAPIENS | 7 | 2668 | 5eqg | A | 315 | ILE | p.Ile315Met | P11166 |
| HOMO SAPIENS | -1 | 2668 | 5eqg | A | 458 | ARG | p.Arg458Trp | P11166 |
| HOMO SAPIENS | -1 | 2668 | 5eqg | A | 468 | ARG | p.Arg468Trp | P11166 |
| HOMO SAPIENS | 3 | 2291 | 2amy | A | 31 | PHE | p.Phe31Leu | O15305 |
| HOMO SAPIENS | 7 | 2291 | 2amy | A | 32 | LEU | p.Leu32Arg | O15305 |
| HOMO SAPIENS | 7 | 2291 | 2amy | A | 65 | ASP | p.Asp65Tyr | O15305 |
| HOMO SAPIENS | 1 | 2291 | 2amy | A | 108 | ALA | p.Ala108Val | O15305 |
| HOMO SAPIENS | 3 | 2291 | 2amy | A | 113 | PRO | p.Pro113Leu | O15305 |
| HOMO SAPIENS | 3 | 2291 | 2amy | A | 113 | PRO | p.Pro113Ser | O15305 |
| HOMO SAPIENS | 6 | 2291 | 2amy | A | 117 | GLY | p.Gly117Arg | O15305 |
| HOMO SAPIENS | 6 | 2291 | 2amy | A | 119 | PHE | p.Phe119Leu | O15305 |
| HOMO SAPIENS | 6 | 2291 | 2amy | A | 123 | ARG | p.Arg123Gln | O15305 |
| HOMO SAPIENS | 0 | 2291 | 2amy | A | 129 | VAL | p.Val129Met | O15305 |
| HOMO SAPIENS | 7 | 2291 | 2amy | A | 139 | GLU | p.Glu139Lys | O15305 |
| HOMO SAPIENS | 5 | 2291 | 2amy | A | 141 | ARG | p.Arg141His | O15305 |
| HOMO SAPIENS | 7 | 2291 | 2amy | A | 144 | PHE | p.Phe144Leu | O15305 |
| HOMO SAPIENS | 6 | 2291 | 2amy | A | 148 | ASP | p.Asp148Asn | O15305 |
| HOMO SAPIENS | 6 | 2291 | 2amy | A | 153 | ILE | p.Ile153Thr | O15305 |
| HOMO SAPIENS | 2 | 2291 | 2amy | A | 157 | PHE | p.Phe157Ser | O15305 |
| HOMO SAPIENS | 1 | 2291 | 2amy | A | 162 | ARG | p.Arg162Trp | O15305 |
| HOMO SAPIENS | 6 | 2291 | 2amy | A | 188 | ASP | p.Asp188Gly | O15305 |
| HOMO SAPIENS | 5 | 2291 | 2amy | A | 194 | ARG | p.Arg194Ter | O15305 |
| HOMO SAPIENS | 0 | 2291 | 2amy | A | 207 | PHE | p.Phe207Ser | O15305 |
| HOMO SAPIENS | 5 | 2291 | 2amy | A | 208 | GLY | p.Gly208Ala | O15305 |
| HOMO SAPIENS | 1 | 2291 | 2amy | A | 216 | ASN | p.Asn216Ile | O15305 |
| HOMO SAPIENS | 5 | 2291 | 2amy | A | 218 | HIS | p.His218Leu | O15305 |
| HOMO SAPIENS | 6 | 2291 | 2amy | A | 223 | ASP | p.Asp223Glu | O15305 |
| HOMO SAPIENS | 6 | 2291 | 2amy | A | 226 | THR | p.Thr226Ser | O15305 |
| HOMO SAPIENS | 5 | 2291 | 2amy | A | 231 | VAL | p.Val231Met | O15305 |
| HOMO SAPIENS | 1 | 2291 | 2amy | A | 237 | THR | p.Thr237Met | O15305 |
| HOMO SAPIENS | 1 | 2291 | 2amy | A | 241 | CYS | p.Cys241Ser | O15305 |
| HOMO SAPIENS | 5 | 35907 | 1d6g | A | 21 | GLY | p.Gly21Arg | P32238 |
| HOMO SAPIENS | -1 | 33209 | 1x9d | A | 197 | PRO | p.Pro197Leu | Q9UKM7 |

| | | | | | | | | |
|---|---|---|---|---|---|---|---|---|
| HOMO SAPIENS | -1 | 33209 | 1x9d | A | 201 | PRO | p.Pro201Leu | Q9UKM7 |
| HOMO SAPIENS | 6 | 21305 | 4yep | B | 1326 | ARG | p.Arg1326Ter | P24043 |
| HOMO SAPIENS | 6 | 21305 | 4yep | B | 1350 | ARG | p.Arg1350Ter | P24043 |
| HOMO SAPIENS | 0 | 21553 | 3qyb | A | 1120 | VAL | p.Val1119Ala | O60343 |
| HOMO SAPIENS | 2 | 21553 | 3qyb | A | 1148 | THR | p.Thr1147Met | O60343 |
| HOMO SAPIENS | -1 | 30645 | 3m0c | C | -17 | TRP | p.Trp4Ter | P01130 |
| HOMO SAPIENS | -1 | 30645 | 3m0c | C | -17 | TRP | p.Trp4Ter | P01130 |
| HOMO SAPIENS | -1 | 30645 | 3m0c | C | 12 | GLN | p.Gln33Ter | P01130 |
| HOMO SAPIENS | -1 | 30645 | 3m0c | C | 15 | ASP | p.Asp36Glu | P01130 |
| HOMO SAPIENS | -1 | 30645 | 3m0c | C | 28 | SER | p.Ser49Arg | P01130 |
| HOMO SAPIENS | -1 | 30645 | 3m0c | C | 41 | THR | p.Thr62Met | P01130 |
| HOMO SAPIENS | -1 | 30645 | 3m0c | C | 69 | ASP | p.Asp90Asn | P01130 |
| HOMO SAPIENS | -1 | 30645 | 3m0c | C | 76 | ASN | p.Asn97Lys | P01130 |
| HOMO SAPIENS | -1 | 30645 | 3m0c | C | 78 | SER | p.Ser99Ter | P01130 |
| HOMO SAPIENS | -1 | 30645 | 3m0c | C | 81 | GLN | p.Gln102Ter | P01130 |
| HOMO SAPIENS | -1 | 30645 | 3m0c | C | 84 | PRO | p.Pro105Ser | P01130 |
| HOMO SAPIENS | -1 | 30645 | 3m0c | C | 194 | ARG | p.Arg215Cys | P01130 |
| HOMO SAPIENS | -1 | 30645 | 3m0c | C | 198 | GLY | p.Gly219Asp | P01130 |
| HOMO SAPIENS | -1 | 30645 | 3m0c | C | 201 | CYS | p.Cys222Tyr | P01130 |
| HOMO SAPIENS | -1 | 30645 | 3m0c | C | 201 | CYS | p.Cys222Ter | P01130 |
| HOMO SAPIENS | -1 | 30645 | 3m0c | C | 206 | ASP | p.Asp227Glu | P01130 |
| HOMO SAPIENS | -1 | 30645 | 3m0c | C | 249 | CYS | p.Cys270Ter | P01130 |
| HOMO SAPIENS | -1 | 30645 | 3m0c | C | 250 | VAL | p.Val271Ile | P01130 |
| HOMO SAPIENS | 0 | 30645 | 3m0c | C | 255 | CYS | p.Cys276Tyr | P01130 |
| HOMO SAPIENS | 0 | 30645 | 3m0c | C | 255 | CYS | p.Cys276Ter | P01130 |
| HOMO SAPIENS | 0 | 30645 | 3m0c | C | 256 | GLU | p.Glu277Lys | P01130 |
| HOMO SAPIENS | 0 | 30645 | 3m0c | C | 265 | SER | p.Ser286Arg | P01130 |
| HOMO SAPIENS | 0 | 30645 | 3m0c | C | 284 | TRP | p.Trp305Ter | P01130 |
| HOMO SAPIENS | 0 | 30645 | 3m0c | C | 292 | CYS | p.Cys313Ter | P01130 |
| HOMO SAPIENS | 1 | 30645 | 3m0c | C | 385 | ARG | p.Arg406Trp | P01130 |
| HOMO SAPIENS | 0 | 30645 | 3m0c | C | 395 | ARG | p.Arg416Gln | P01130 |
| HOMO SAPIENS | 0 | 30645 | 3m0c | C | 398 | TYR | p.Tyr419Ter | P01130 |
| HOMO SAPIENS | 1 | 30645 | 3m0c | C | 415 | VAL | p.Val436Ala | P01130 |
| HOMO SAPIENS | 1 | 30645 | 3m0c | C | 424 | ASP | p.Asp445Glu | P01130 |
| HOMO SAPIENS | 1 | 30645 | 3m0c | C | 425 | LEU | p.Leu446Val | P01130 |
| HOMO SAPIENS | -1 | 30645 | 3m0c | C | 440 | GLY | p.Gly461Cys | P01130 |
| HOMO SAPIENS | -1 | 30645 | 3m0c | C | 441 | VAL | p.Val462Ile | P01130 |
| HOMO SAPIENS | 1 | 30645 | 3m0c | C | 447 | VAL | p.Val468Ile | P01130 |
| HOMO SAPIENS | 1 | 30645 | 3m0c | C | 489 | THR | p.Thr510Met | P01130 |
| HOMO SAPIENS | 0 | 30645 | 3m0c | C | 524 | GLY | p.Gly545Glu | P01130 |
| HOMO SAPIENS | 1 | 30645 | 3m0c | C | 615 | GLY | p.Gly636Ser | P01130 |
| HOMO SAPIENS | 1 | 30645 | 3m0c | C | 625 | LEU | p.Leu646Ile | P01130 |
| HOMO SAPIENS | 1 | 30645 | 3m0c | C | 630 | ASP | p.Asp651Gly | P01130 |
| HOMO SAPIENS | 3 | 30645 | 3m0c | C | 646 | CYS | p.Cys667Tyr | P01130 |
| HOMO SAPIENS | 4 | 30645 | 3m0c | C | 654 | GLY | p.Gly675Ser | P01130 |
| HOMO SAPIENS | 6 | 30645 | 3m0c | C | 660 | CYS | p.Cys681Ter | P01130 |
| HOMO SAPIENS | 5 | 30645 | 3m0c | C | 676 | ALA | p.Ala697Pro | P01130 |
| HOMO SAPIENS | -1 | 30645 | 3m0c | C | 749 | GLN | p.Gln770Ter | P01130 |
| HOMO SAPIENS | -1 | 41323 | 4bxo | A | 1812 | PRO | p.Pro1812Ala | Q8IYD8 |
| HOMO SAPIENS | 5 | 8335 | 3h0t | C | 12 | GLY | p.Gly71Asp | P81172 |
| HOMO SAPIENS | 6 | 609 | 1bhg | B | 148 | PRO | p.Pro148Ser | P08236 |
| HOMO SAPIENS | 1 | 609 | 1bhg | B | 176 | LEU | p.Leu176Phe | P08236 |
| HOMO SAPIENS | 6 | 609 | 1bhg | B | 216 | ARG | p.Arg216Trp | P08236 |
| HOMO SAPIENS | 0 | 609 | 1bhg | B | 350 | LYS | p.Lys350Asn | P08236 |

| | | | | | | | | |
|---|---|---|---|---|---|---|---|---|
| HOMO SAPIENS | 1 | 609 | 1bhg | B | 354 | ALA | p.Ala354Val | P08236 |
| HOMO SAPIENS | 1 | 609 | 1bhg | B | 357 | ARG | p.Arg357Ter | P08236 |
| HOMO SAPIENS | 0 | 609 | 1bhg | B | 382 | ARG | p.Arg382Cys | P08236 |
| HOMO SAPIENS | 0 | 609 | 1bhg | B | 495 | TYR | p.Tyr495Cys | P08236 |
| HOMO SAPIENS | 2 | 609 | 1bhg | B | 507 | TRP | p.Trp507Ter | P08236 |
| HOMO SAPIENS | 0 | 609 | 1bhg | B | 577 | ARG | p.Arg577Leu | P08236 |
| HOMO SAPIENS | 1 | 609 | 1bhg | B | 611 | ARG | p.Arg611Trp | P08236 |
| HOMO SAPIENS | 0 | 609 | 1bhg | B | 619 | ALA | p.Ala619Val | P08236 |
| HOMO SAPIENS | 0 | 609 | 1bhg | B | 627 | TRP | p.Trp627Cys | P08236 |
| HOMO SAPIENS | 5 | 3422 | 1x9n | A | 566 | GLU | p.Glu566Lys | P18858 |
| HOMO SAPIENS | 2 | 3422 | 1x9n | A | 771 | ARG | p.Arg771Trp | P18858 |
| HOMO SAPIENS | 0 | 3279 | 5tby | B | 797 | ALA | p.Ala797Thr | P12883 |
| HOMO SAPIENS | -1 | 3279 | 5tby | B | 982 | MET | p.Met982Thr | P12883 |
| HOMO SAPIENS | -1 | 3279 | 5tby | B | 992 | LEU | p.Leu992Met | P12883 |
| HOMO SAPIENS | -1 | 3279 | 5tby | B | 1004 | GLN | p.Gln1004Glu | P12883 |
| HOMO SAPIENS | -1 | 3279 | 5tby | B | 1013 | GLU | p.Glu1013Lys | P12883 |
| HOMO SAPIENS | -1 | 3279 | 5tby | B | 1032 | ASP | p.Asp1032Asn | P12883 |
| HOMO SAPIENS | -1 | 3279 | 5tby | B | 1053 | ARG | p.Arg1053Gln | P12883 |
| HOMO SAPIENS | -1 | 3279 | 5tby | B | 1117 | GLU | p.Glu1117Ter | P12883 |
| HOMO SAPIENS | -1 | 3279 | 5tby | B | 1193 | ARG | p.Arg1193His | P12883 |
| HOMO SAPIENS | -1 | 3279 | 5tby | B | 1205 | GLU | p.Glu1205Lys | P12883 |
| HOMO SAPIENS | -1 | 3279 | 5tby | B | 1207 | ILE | p.Ile1207Met | P12883 |
| HOMO SAPIENS | -1 | 3279 | 5tby | B | 1250 | ARG | p.Arg1250Pro | P12883 |
| HOMO SAPIENS | -1 | 3279 | 5tby | B | 1259 | HIS | p.His1259Gln | P12883 |
| HOMO SAPIENS | -1 | 3279 | 5tby | B | 1272 | ASP | p.Asp1272Asn | P12883 |
| HOMO SAPIENS | -1 | 3279 | 5tby | B | 1295 | GLU | p.Glu1295Lys | P12883 |
| HOMO SAPIENS | -1 | 3279 | 5tby | B | 1335 | SER | p.Ser1335Leu | P12883 |
| HOMO SAPIENS | -1 | 3279 | 5tby | B | 1399 | GLU | p.Glu1399Asp | P12883 |
| HOMO SAPIENS | -1 | 3279 | 5tby | B | 1413 | SER | p.Ser1413Ter | P12883 |
| HOMO SAPIENS | -1 | 3279 | 5tby | B | 1459 | LYS | p.Lys1459Asn | P12883 |
| HOMO SAPIENS | -1 | 3279 | 5tby | B | 1467 | LEU | p.Leu1467Val | P12883 |
| HOMO SAPIENS | -1 | 3279 | 5tby | B | 1491 | SER | p.Ser1491Cys | P12883 |
| HOMO SAPIENS | -1 | 3279 | 5tby | B | 1496 | GLU | p.Glu1496Ala | P12883 |
| HOMO SAPIENS | -1 | 3279 | 5tby | B | 1500 | ARG | p.Arg1500Pro | P12883 |
| HOMO SAPIENS | -1 | 3279 | 5tby | B | 1606 | ARG | p.Arg1606His | P12883 |
| HOMO SAPIENS | -1 | 3279 | 5tby | B | 1664 | ASN | p.Asn1664Lys | P12883 |
| HOMO SAPIENS | -1 | 3279 | 5tby | B | 1712 | ARG | p.Arg1712Trp | P12883 |
| HOMO SAPIENS | -1 | 3279 | 5tby | B | 1769 | LEU | p.Leu1769Met | P12883 |
| HOMO SAPIENS | -1 | 3279 | 5tby | B | 1808 | GLY | p.Gly1808Ser | P12883 |
| HOMO SAPIENS | -1 | 3279 | 5tby | B | 1818 | ARG | p.Arg1818Trp | P12883 |
| HOMO SAPIENS | -1 | 3279 | 5tby | B | 1820 | ARG | p.Arg1820Trp | P12883 |
| HOMO SAPIENS | -1 | 3279 | 5tby | B | 1844 | GLU | p.Glu1844Lys | P12883 |
| HOMO SAPIENS | -1 | 3279 | 5tby | B | 1845 | ARG | p.Arg1845Trp | P12883 |
| HOMO SAPIENS | -1 | 3279 | 5tby | B | 1854 | THR | p.Thr1854Met | P12883 |
| HOMO SAPIENS | -1 | 3279 | 5tby | B | 1909 | ARG | p.Arg1909Trp | P12883 |
| HOMO SAPIENS | 2 | 2260 | 4kl1 | A | 553 | ASP | p.Asp553Asn | Q9Y3Q4 |
| HOMO SAPIENS | 0 | 40570 | 5lst | A | 522 | ARG | p.Arg522Cys | O94761 |
| HOMO SAPIENS | 0 | 40570 | 5lst | A | 522 | ARG | p.Arg522His | O94761 |
| HOMO SAPIENS | 0 | 40570 | 5lst | A | 591 | PRO | p.Pro591Leu | O94761 |
| HOMO SAPIENS | 0 | 40570 | 5lst | A | 793 | PRO | p.Pro793Ser | O94761 |
| HOMO SAPIENS | 4 | 40570 | 5lst | A | 799 | VAL | p.Val799Met | O94761 |
| HOMO SAPIENS | 0 | 40570 | 5lst | A | 1004 | ARG | p.Arg1004Trp | O94761 |
| HOMO SAPIENS | 0 | 40570 | 5lst | A | 1005 | ARG | p.Arg1005Gln | O94761 |
| HOMO SAPIENS | 0 | 40570 | 5lst | A | 1021 | ARG | p.Arg1021Gln | O94761 |

| | | | | | | | | |
|---|---|---|---|---|---|---|---|---|
| HOMO SAPIENS | 0 | 40570 | 5lst | A | 1043 | LEU | p.Leu1043Pro | O94761 |
| HOMO SAPIENS | 0 | 40570 | 5lst | A | 1045 | ALA | p.Ala1045Thr | O94761 |
| HOMO SAPIENS | 0 | 40570 | 5lst | A | 1105 | GLY | p.Gly1105Asp | O94761 |
| HOMO SAPIENS | 0 | 40570 | 5lst | A | 1105 | GLY | p.Gly1105Ser | O94761 |
| HOMO SAPIENS | -1 | 40570 | 5lst | A | 1113 | GLY | p.Gly1113Arg | O94761 |
| HOMO SAPIENS | -1 | 18865 | 1gxc | A | 64 | GLU | p.Glu64Lys | O96017 |
| HOMO SAPIENS | -1 | 11673 | 3dzu | D | 85 | PRO | p.Pro113Gln | P37231 |
| HOMO SAPIENS | 0 | 19383 | 3b96 | A | 63 | LYS | p.Lys103Arg | P49748 |
| HOMO SAPIENS | 4 | 19383 | 3b96 | A | 140 | ALA | p.Ala180Thr | P49748 |
| HOMO SAPIENS | 1 | 19383 | 3b96 | A | 145 | GLY | p.Gly185Ser | P49748 |
| HOMO SAPIENS | 0 | 19383 | 3b96 | A | 220 | THR | p.Thr260Met | P49748 |
| HOMO SAPIENS | 5 | 19383 | 3b96 | A | 243 | VAL | p.Val283Ala | P49748 |
| HOMO SAPIENS | 0 | 19383 | 3b96 | A | 249 | GLY | p.Gly289Arg | P49748 |
| HOMO SAPIENS | 0 | 19383 | 3b96 | A | 254 | GLY | p.Gly294Glu | P49748 |
| HOMO SAPIENS | 7 | 19383 | 3b96 | A | 326 | ARG | p.Arg366Cys | P49748 |
| HOMO SAPIENS | 7 | 19383 | 3b96 | A | 326 | ARG | p.Arg366His | P49748 |
| HOMO SAPIENS | 5 | 19383 | 3b96 | A | 342 | LYS | p.Lys382Gln | P49748 |
| HOMO SAPIENS | 0 | 19383 | 3b96 | A | 369 | THR | p.Thr409Met | P49748 |
| HOMO SAPIENS | 5 | 19383 | 3b96 | A | 376 | ALA | p.Ala416Ser | P49748 |
| HOMO SAPIENS | 0 | 19383 | 3b96 | A | 385 | ALA | p.Ala425Thr | P49748 |
| HOMO SAPIENS | 5 | 19383 | 3b96 | A | 399 | GLY | p.Gly439Asp | P49748 |
| HOMO SAPIENS | 7 | 19383 | 3b96 | A | 401 | GLY | p.Gly441Asp | P49748 |
| HOMO SAPIENS | 9 | 19383 | 3b96 | A | 413 | ARG | p.Arg453Gln | P49748 |
| HOMO SAPIENS | 2 | 19383 | 3b96 | A | 418 | PHE | p.Phe458Leu | P49748 |
| HOMO SAPIENS | 6 | 19383 | 3b96 | A | 419 | ARG | p.Arg459Trp | P49748 |
| HOMO SAPIENS | 6 | 19383 | 3b96 | A | 419 | ARG | p.Arg459Gln | P49748 |
| HOMO SAPIENS | -1 | 19383 | 3b96 | A | 471 | ARG | p.Arg511Gln | P49748 |
| HOMO SAPIENS | 0 | 19383 | 3b96 | A | 573 | ARG | p.Arg613Trp | P49748 |
| HOMO SAPIENS | 0 | 19383 | 3b96 | A | 575 | ARG | p.Arg615Gln | P49748 |
| HOMO SAPIENS | 0 | 22286 | 3k2s | A | 3 | PRO | p.Pro27His | P02647 |
| HOMO SAPIENS | 0 | 22286 | 3k2s | A | 4 | PRO | p.Pro28Arg | P02647 |
| HOMO SAPIENS | 0 | 22286 | 3k2s | A | 10 | ARG | p.Arg34Leu | P02647 |
| HOMO SAPIENS | 0 | 22286 | 3k2s | A | 26 | GLY | p.Gly50Arg | P02647 |
| HOMO SAPIENS | 0 | 22286 | 3k2s | A | 37 | ALA | p.Ala61Thr | P02647 |
| HOMO SAPIENS | 1 | 1622 | 2qrv | D | 631 | ARG | p.Arg635Trp | Q9Y6K1 |
| HOMO SAPIENS | 1 | 1622 | 2qrv | D | 900 | PRO | p.Pro904Leu | Q9Y6K1 |
| HOMO SAPIENS | 6 | 40796 | 1v18 | B | 1509 | LEU | p.Leu1509Val | P25054 |
| HOMO SAPIENS | 1 | 22375 | 4pki | G | 1214 | LYS | p.Asp214Asn | P28289 |
| HOMO SAPIENS | 1 | 16307 | 5mv8 | B | 483 | ALA | p.Ala483Ser | Q9Y6N9 |
| HOMO SAPIENS | 6 | 16307 | 5mv8 | B | 506 | ASN | p.Asn506Lys | Q9Y6N9 |
| HOMO SAPIENS | 4 | 16307 | 5mv8 | B | 519 | ASP | p.Glu519Asp | Q9Y6N9 |
| HOMO SAPIENS | 0 | 16307 | 5mv8 | B | 530 | GLY | p.Gly530Arg | Q9Y6N9 |
| HOMO SAPIENS | 4 | 36221 | 4cvo | A | 127 | VAL | p.Val127Ile | Q03468 |
| HOMO SAPIENS | 7 | 36221 | 4cvo | A | 134 | ARG | p.Arg134Trp | Q03468 |
| HOMO SAPIENS | 0 | 36221 | 4cvo | A | 156 | GLN | p.Gln156Ter | Q03468 |
| HOMO SAPIENS | 0 | 10829 | 4a11 | B | 13 | GLU | p.Glu13Ter | Q13216 |
| HOMO SAPIENS | 1 | 10829 | 4a11 | B | 100 | TYR | p.Tyr100Ter | Q13216 |
| HOMO SAPIENS | 0 | 10829 | 4a11 | B | 160 | ALA | p.Ala160Val | Q13216 |
| HOMO SAPIENS | 1 | 6759 | 5lev | A | 29 | ILE | p.Ile29Phe | Q9H3H5 |
| HOMO SAPIENS | 0 | 6759 | 5lev | A | 108 | MET | p.Met108Ile | Q9H3H5 |
| HOMO SAPIENS | 2 | 6759 | 5lev | A | 117 | VAL | p.Val117Ile | Q9H3H5 |
| HOMO SAPIENS | 1 | 6759 | 5lev | A | 170 | TYR | p.Tyr170Cys | Q9H3H5 |
| HOMO SAPIENS | 0 | 6759 | 5lev | A | 264 | GLY | p.Val264Gly | Q9H3H5 |
| HOMO SAPIENS | 0 | 6759 | 5lev | A | 332 | PHE | p.Phe332Val | Q9H3H5 |

| | | | | | | | | |
|---|---|---|---|---|---|---|---|---|
| HOMO SAPIENS | 7 | 6759 | 5lev | A | 393 | ILE | p.Ile393Val | Q9H3H5 |
| HOMO SAPIENS | -1 | 33473 | 5m5e | C | 9 | ASN | p.Asn31Ser | P31785 |
| HOMO SAPIENS | 0 | 26155 | 1ic8 | B | 98 | ALA | p.Ala98Val | P20823 |
| HOMO SAPIENS | 0 | 15329 | 4iqr | B | 127 | ARG | p.Arg136Trp | P41235 |
| HOMO SAPIENS | 0 | 20725 | 4bsj | A | 350 | GLU | p.Glu350Val | P35916 |
| HOMO SAPIENS | 0 | 20725 | 4bsj | A | 494 | THR | p.Thr494Ala | P35916 |
| HOMO SAPIENS | 0 | 20725 | 4bsj | A | 527 | ASN | p.Asn527Ser | P35916 |
| HOMO SAPIENS | 6 | 29959 | 1xiw | B | 47 | ARG | p.Arg68Ter | P04234 |
| HOMO SAPIENS | 2 | 29959 | 1xiw | B | 72 | CYS | p.Cys93Ter | P04234 |
| HOMO SAPIENS | 3 | 20242 | 5tjc | A | 102 | ARG | p.Arg102Ter | Q9GZU1 |
| HOMO SAPIENS | 3 | 20242 | 5tjc | A | 232 | THR | p.Thr232Pro | Q9GZU1 |
| HOMO SAPIENS | 7 | 9590 | 4ap8 | A | 50 | THR | p.Thr50Ala | O96007 |
| HOMO SAPIENS | 4 | 9590 | 4ap8 | A | 168 | GLU | p.Glu168Lys | O96007 |
| HOMO SAPIENS | -1 | 4297 | 5do7 | B | 19 | ASP | p.Asp19His | Q9H221 |
| HOMO SAPIENS | -1 | 4297 | 5do7 | B | 54 | TYR | p.Tyr54Cys | Q9H221 |
| HOMO SAPIENS | -1 | 4297 | 5do7 | B | 80 | CYS | p.Cys80Tyr | Q9H221 |
| HOMO SAPIENS | 5 | 4297 | 5do7 | B | 107 | SER | p.Ser107Ter | Q9H221 |
| HOMO SAPIENS | 1 | 4297 | 5do7 | B | 164 | ARG | p.Arg164Pro | Q9H221 |
| HOMO SAPIENS | 0 | 4297 | 5do7 | B | 210 | VAL | p.Val210Met | Q9H221 |
| HOMO SAPIENS | 0 | 4297 | 5do7 | B | 231 | PRO | p.Pro231Thr | Q9H221 |
| HOMO SAPIENS | 0 | 4297 | 5do7 | B | 263 | ARG | p.Arg263Gln | Q9H221 |
| HOMO SAPIENS | 0 | 4297 | 5do7 | B | 361 | TRP | p.Trp361Ter | Q9H221 |
| HOMO SAPIENS | 0 | 4297 | 5do7 | B | 400 | THR | p.Thr400Lys | Q9H221 |
| HOMO SAPIENS | 0 | 4297 | 5do7 | B | 549 | ALA | p.Ala549Pro | Q9H221 |
| HOMO SAPIENS | 0 | 4297 | 5do7 | B | 596 | LEU | p.Leu596Arg | Q9H221 |
| HOMO SAPIENS | 0 | 4297 | 5do7 | B | 632 | VAL | p.Val632Ala | Q9H221 |
| HOMO SAPIENS | 0 | 4297 | 5do7 | B | 658 | TYR | p.Tyr658Ter | Q9H221 |
| HOMO SAPIENS | 2 | 9356 | 4igg | B | 206 | GLN | p.Gln206His | P35221 |
| HOMO SAPIENS | 6 | 2179 | 2wk6 | A | 44 | ARG | p.Arg44Gln | P19971 |
| HOMO SAPIENS | 6 | 2179 | 2wk6 | A | 81 | ARG | p.Arg81Gln | P19971 |
| HOMO SAPIENS | 6 | 2179 | 2wk6 | A | 145 | GLY | p.Gly145Arg | P19971 |
| HOMO SAPIENS | 7 | 2179 | 2wk6 | A | 146 | ARG | p.Arg146His | P19971 |
| HOMO SAPIENS | 7 | 2179 | 2wk6 | A | 153 | GLY | p.Gly153Ser | P19971 |
| HOMO SAPIENS | 0 | 2179 | 2wk6 | A | 173 | MET | p.Met173Ile | P19971 |
| HOMO SAPIENS | 6 | 2179 | 2wk6 | A | 202 | ARG | p.Arg202Lys | P19971 |
| HOMO SAPIENS | 6 | 2179 | 2wk6 | A | 208 | VAL | p.Val208Met | P19971 |
| HOMO SAPIENS | 5 | 2179 | 2wk6 | A | 222 | LYS | p.Lys222Arg | P19971 |
| HOMO SAPIENS | 5 | 2179 | 2wk6 | A | 289 | GLU | p.Glu289Ala | P19971 |
| HOMO SAPIENS | 2 | 2179 | 2wk6 | A | 311 | GLY | p.Gly311Arg | P19971 |
| HOMO SAPIENS | 2 | 2179 | 2wk6 | A | 376 | ARG | p.Arg376Pro | P19971 |
| HOMO SAPIENS | 2 | 2179 | 2wk6 | A | 465 | ALA | p.Ala465Thr | P19971 |
| HOMO SAPIENS | -1 | 11856 | 3orh | A | 3 | ALA | p.Ala3Ser | Q14353 |
| HOMO SAPIENS | 1 | 11856 | 3orh | A | 20 | TRP | p.Trp20Ser | Q14353 |
| HOMO SAPIENS | 1 | 11856 | 3orh | A | 50 | MET | p.Met50Leu | Q14353 |
| HOMO SAPIENS | 1 | 11856 | 3orh | A | 110 | VAL | p.Val110Phe | Q14353 |
| HOMO SAPIENS | 8 | 11856 | 3orh | A | 164 | GLY | p.Gly164Asp | Q14353 |
| HOMO SAPIENS | 1 | 11856 | 3orh | A | 169 | CYS | p.Cys169Tyr | Q14353 |
| HOMO SAPIENS | 1 | 11856 | 3orh | A | 174 | TRP | p.Trp174Ter | Q14353 |
| HOMO SAPIENS | 2 | 11856 | 3orh | A | 196 | ALA | p.Ala196Val | Q14353 |
| HOMO SAPIENS | 1 | 11856 | 3orh | A | 209 | THR | p.Thr209Met | Q14353 |
| HOMO SAPIENS | 1 | 11856 | 3orh | A | 219 | ASP | p.Asp219Asn | Q14353 |
| HOMO SAPIENS | 1 | 11856 | 3orh | A | 224 | ALA | p.Ala224Thr | Q14353 |
| HOMO SAPIENS | 1 | 13244 | 2m6i | D | 218 | ARG | p.Arg246Leu | P23415 |
| HOMO SAPIENS | 1 | 13244 | 2m6i | D | 244 | ILE | p.Ile272Asn | P23415 |

| | | | | | | | | |
|---|---|---|---|---|---|---|---|---|
| HOMO SAPIENS | 5 | 13244 | 2m6i | D | 250 | PRO | p.Pro278Thr | P23415 |
| HOMO SAPIENS | 6 | 13244 | 2m6i | D | 266 | GLN | p.Gln294His | P23415 |
| HOMO SAPIENS | 1 | 13244 | 2m6i | D | 271 | ARG | p.Arg299Leu | P23415 |
| HOMO SAPIENS | 1 | 13244 | 2m6i | D | 276 | LYS | p.Lys304Glu | P23415 |
| HOMO SAPIENS | 1 | 13244 | 2m6i | D | 279 | TYR | p.Tyr307Cys | P23415 |
| HOMO SAPIENS | 0 | 13244 | 2m6i | D | 316 | GLY | p.Gly378Ser | P23415 |
| HOMO SAPIENS | 1 | 13244 | 2m6i | D | 400 | ARG | p.Arg428His | P23415 |
| HOMO SAPIENS | 3 | 1967 | 2i6b | A | 301 | ALA | p.Ala318Glu | P55263 |
| HOMO SAPIENS | 4 | 11096 | 1h8b | A | 61 | ALA | p.Ala882Thr | P35609 |
| HOMO SAPIENS | 0 | 15423 | 5ngz | A | 2 | GLN | p.Gln2Glu | Q9NPD8 |
| HOMO SAPIENS | 3 | 30475 | 3lpw | A | 21 | SER | p.Ser22894Tyr | Q8WZ42 |
| HOMO SAPIENS | 6 | 32687 | 2arf | A | 1041 | ARG | p.Arg1041Trp | P35670 |
| HOMO SAPIENS | 8 | 32687 | 2arf | A | 1061 | GLY | p.Gly1061Glu | P35670 |
| HOMO SAPIENS | 6 | 32687 | 2arf | A | 1064 | GLU | p.Glu1064Ala | P35670 |
| HOMO SAPIENS | 2 | 32687 | 2arf | A | 1069 | HIS | p.His1069Gln | P35670 |
| HOMO SAPIENS | 2 | 32687 | 2arf | A | 1088 | LEU | p.Leu1088Ter | P35670 |
| HOMO SAPIENS | 0 | 32687 | 2arf | A | 1099 | GLY | p.Gly1099Ser | P35670 |
| HOMO SAPIENS | 4 | 32687 | 2arf | A | 1106 | VAL | p.Val1106Asp | P35670 |
| HOMO SAPIENS | 2 | 32687 | 2arf | A | 1142 | GLN | p.Gln1142His | P35670 |
| HOMO SAPIENS | 8 | 32687 | 2arf | A | 1148 | ILE | p.Ile1148Thr | P35670 |
| HOMO SAPIENS | 6 | 32687 | 2arf | A | 1151 | ARG | p.Arg1151Cys | P35670 |
| HOMO SAPIENS | 4 | 32687 | 2arf | A | 1169 | MET | p.Met1169Val | P35670 |
| HOMO SAPIENS | 4 | 32687 | 2arf | A | 1173 | GLU | p.Glu1173Lys | P35670 |
| HOMO SAPIENS | 2 | 32687 | 2arf | A | 1176 | GLY | p.Gly1176Arg | P35670 |
| HOMO SAPIENS | 3 | 15544 | 5uq3 | C | 575 | PRO | p.Pro69Leu | P46527 |
| HOMO SAPIENS | 6 | 15544 | 5uq3 | C | 592 | GLU | p.Glu86Asp | P46527 |
| HOMO SAPIENS | 6 | 15544 | 5uq3 | C | 599 | ARG | p.Arg93Trp | P46527 |
| HOMO SAPIENS | -1 | 15544 | 5uq3 | C | 615 | VAL | p.Val109Gly | P46527 |
| HOMO SAPIENS | -1 | 15544 | 5uq3 | C | 631 | SER | p.Ser125Pro | P46527 |
| HOMO SAPIENS | -1 | 15544 | 5uq3 | C | 639 | PRO | p.Pro133Ser | P46527 |
| HOMO SAPIENS | 2 | 2703 | 4wxq | A | 245 | CYS | p.Cys245Tyr | Q99972 |
| HOMO SAPIENS | 0 | 2703 | 4wxq | A | 252 | GLY | p.Gly252Arg | Q99972 |
| HOMO SAPIENS | 1 | 2703 | 4wxq | A | 337 | GLN | p.Gln337Arg | Q99972 |
| HOMO SAPIENS | 5 | 2703 | 4wxq | A | 347 | TYR | p.Tyr347Ter | Q99972 |
| HOMO SAPIENS | 3 | 2703 | 4wxq | A | 364 | GLY | p.Gly364Val | Q99972 |
| HOMO SAPIENS | 2 | 2703 | 4wxq | A | 367 | GLY | p.Gly367Arg | Q99972 |
| HOMO SAPIENS | 2 | 2703 | 4wxq | A | 368 | GLN | p.Gln368Ter | Q99972 |
| HOMO SAPIENS | 6 | 2703 | 4wxq | A | 370 | PRO | p.Pro370Leu | Q99972 |
| HOMO SAPIENS | 1 | 2703 | 4wxq | A | 380 | ASP | p.Asp380His | Q99972 |
| HOMO SAPIENS | 8 | 2703 | 4wxq | A | 399 | GLY | p.Gly399Val | Q99972 |
| HOMO SAPIENS | 0 | 2703 | 4wxq | A | 423 | LYS | p.Lys423Glu | Q99972 |
| HOMO SAPIENS | 3 | 2703 | 4wxq | A | 433 | CYS | p.Cys433Arg | Q99972 |
| HOMO SAPIENS | 3 | 2703 | 4wxq | A | 437 | TYR | p.Tyr437His | Q99972 |
| HOMO SAPIENS | 2 | 2703 | 4wxq | A | 477 | ILE | p.Ile477Ser | Q99972 |
| HOMO SAPIENS | 3 | 2703 | 4wxq | A | 480 | ASN | p.Asn480Lys | Q99972 |
| HOMO SAPIENS | -1 | 34100 | 2fot | C | 1171 | GLN | p.Gln1173Pro | Q13813 |
| HOMO SAPIENS | 0 | 8773 | 1kw2 | A | 432 | ASP | p.Asp432Glu | P02774 |
| HOMO SAPIENS | 0 | 8773 | 1kw2 | A | 436 | LYS | p.Lys436Thr | P02774 |
| HOMO SAPIENS | 0 | 8402 | 4dzo | B | 649 | TYR | p.Tyr649Ter | Q9Y6D9 |
| HOMO SAPIENS | 0 | 39962 | 1nst | A | 611 | GLY | p.Gly611Ser | P52848 |
| HOMO SAPIENS | 2 | 22578 | 3dof | B | 45 | MET | p.Met45Arg | Q9Y2Y0 |
| HOMO SAPIENS | 1 | 23290 | 5j1s | A | 292 | ILE | p.Ile292Thr | O14656 |
| HOMO SAPIENS | 4 | 9505 | 4j1y | A | 534 | ARG | p.Arg534Trp | P09871 |
| HOMO SAPIENS | 9 | 260 | 2h57 | A | 31 | THR | p.Thr31Met | Q9H0F7 |

| | | | | | | | | |
|---|---|---|---|---|---|---|---|---|
| HOMO SAPIENS | 0 | 260 | 2h57 | A | 91 | ILE | p.Ile91Thr | Q9H0F7 |
| HOMO SAPIENS | 0 | 260 | 2h57 | A | 121 | ARG | p.Arg121His | Q9H0F7 |
| HOMO SAPIENS | 0 | 260 | 2h57 | A | 122 | ARG | p.Arg122Ter | Q9H0F7 |
| HOMO SAPIENS | 2 | 260 | 2h57 | A | 169 | GLY | p.Gly169Ala | Q9H0F7 |
| HOMO SAPIENS | 4 | 260 | 2h57 | A | 170 | LEU | p.Leu170Trp | Q9H0F7 |
| HOMO SAPIENS | 4 | 34023 | 2cr3 | A | 18 | GLY | p.Gly48Ser | P11362 |
| HOMO SAPIENS | 0 | 37667 | 3ii7 | A | 420 | ARG | p.Arg420Cys | Q8IXQ5 |
| HOMO SAPIENS | 0 | 41720 | 2k9u | A | 68 | ARG | p.Arg2364His | Q14315 |
| HOMO SAPIENS | 1 | 29250 | 5fmv | A | 568 | HIS | p.His568Gln | P08575 |
| HOMO SAPIENS | 1 | 1280 | 1fb1 | A | 69 | PRO | p.Pro69Leu | P30793 |
| HOMO SAPIENS | 3 | 1280 | 1fb1 | A | 88 | ARG | p.Arg88Trp | P30793 |
| HOMO SAPIENS | 2 | 1280 | 1fb1 | A | 108 | GLY | p.Gly108Asp | P30793 |
| HOMO SAPIENS | 4 | 1280 | 1fb1 | A | 134 | ASP | p.Asp134Val | P30793 |
| HOMO SAPIENS | 6 | 1280 | 1fb1 | A | 135 | ILE | p.Ile135Lys | P30793 |
| HOMO SAPIENS | 9 | 1280 | 1fb1 | A | 144 | HIS | p.His144Pro | P30793 |
| HOMO SAPIENS | 9 | 1280 | 1fb1 | A | 184 | ARG | p.Arg184His | P30793 |
| HOMO SAPIENS | 1 | 1280 | 1fb1 | A | 196 | ALA | p.Ala196Ser | P30793 |
| HOMO SAPIENS | 3 | 1280 | 1fb1 | A | 201 | GLY | p.Gly201Glu | P30793 |
| HOMO SAPIENS | 2 | 1280 | 1fb1 | A | 204 | VAL | p.Val204Ile | P30793 |
| HOMO SAPIENS | 2 | 1280 | 1fb1 | A | 211 | MET | p.Met211Ile | P30793 |
| HOMO SAPIENS | 0 | 1280 | 1fb1 | A | 221 | MET | p.Met221Thr | P30793 |
| HOMO SAPIENS | 3 | 1280 | 1fb1 | A | 224 | LYS | p.Lys224Arg | P30793 |
| HOMO SAPIENS | 2 | 1280 | 1fb1 | A | 249 | ARG | p.Arg249Ser | P30793 |
| HOMO SAPIENS | 8 | 41352 | 4k12 | A | 47 | ILE | p.Ile551Thr | P08603 |
| HOMO SAPIENS | 9 | 20703 | 5nir | A | 57 | CYS | p.Cys57Tyr | P02458 |
| HOMO SAPIENS | 9 | 20703 | 5nir | A | 64 | CYS | p.Cys64Ter | P02458 |
| HOMO SAPIENS | 0 | 20087 | 4cmn | A | 299 | MET | p.Met299Ile | Q01968 |
| HOMO SAPIENS | 1 | 20819 | 3aaf | A | 979 | ILE | p.Ile979Met | Q14191 |
| HOMO SAPIENS | 1 | 20819 | 3aaf | A | 984 | LEU | p.Leu984Ile | Q14191 |
| HOMO SAPIENS | 3 | 20819 | 3aaf | A | 987 | ARG | p.Arg987Ter | Q14191 |
| HOMO SAPIENS | -1 | 20819 | 3aaf | A | 1074 | PHE | p.Leu1074Phe | Q14191 |
| HOMO SAPIENS | 0 | 3619 | 3dem | A | 206 | ASN | p.Asn225Ser | P48740 |
| HOMO SAPIENS | 1 | 3619 | 3dem | A | 269 | ARG | p.Arg288Leu | P48740 |
| HOMO SAPIENS | 0 | 3105 | 3ooi | A | 164 | TYR | p.Tyr2015His | Q96L73 |
| HOMO SAPIENS | 6 | 15828 | 2lkx | A | 16 | LEU | p.Leu100Gln | Q99697 |
| HOMO SAPIENS | 8 | 15828 | 2lkx | A | 24 | ARG | p.Arg108His | Q99697 |
| HOMO SAPIENS | 2 | 15828 | 2lkx | A | 30 | THR | p.Thr114Pro | Q99697 |
| HOMO SAPIENS | 8 | 15828 | 2lkx | A | 45 | VAL | p.Val129Leu | Q99697 |
| HOMO SAPIENS | 8 | 15828 | 2lkx | A | 46 | ARG | p.Arg130Trp | Q99697 |
| HOMO SAPIENS | 9 | 15828 | 2lkx | A | 53 | ARG | p.Arg137Pro | Q99697 |
| HOMO SAPIENS | 9 | 13017 | 4irq | A | 141 | ARG | p.Arg141Trp | Q9UBV7 |
| HOMO SAPIENS | 0 | 13017 | 4irq | A | 186 | ALA | p.Ala186Asp | Q9UBV7 |
| HOMO SAPIENS | 0 | 13017 | 4irq | A | 206 | LEU | p.Leu206Pro | Q9UBV7 |
| HOMO SAPIENS | 1 | 13017 | 4irq | A | 214 | CYS | p.Cys214Tyr | Q9UBV7 |
| HOMO SAPIENS | 0 | 13017 | 4irq | A | 270 | ARG | p.Arg270Cys | Q9UBV7 |
| HOMO SAPIENS | -1 | 13053 | 4mqw | X | 307 | ALA | p.Ala307Thr | P23945 |
| HOMO SAPIENS | 0 | 19406 | 4b3g | A | 43 | ARG | p.Arg43Ter | P38935 |
| HOMO SAPIENS | 0 | 19406 | 4b3g | A | 46 | CYS | p.Cys46Ter | P38935 |
| HOMO SAPIENS | 0 | 19406 | 4b3g | A | 75 | ALA | p.Ala75Thr | P38935 |
| HOMO SAPIENS | 0 | 19406 | 4b3g | A | 201 | LEU | p.Leu201Ser | P38935 |
| HOMO SAPIENS | 3 | 19406 | 4b3g | A | 213 | HIS | p.His213Arg | P38935 |
| HOMO SAPIENS | 0 | 19406 | 4b3g | A | 275 | ILE | p.Ile275Val | P38935 |
| HOMO SAPIENS | -1 | 19406 | 4b3g | A | 320 | ARG | p.Arg320Ter | P38935 |
| HOMO SAPIENS | 0 | 19406 | 4b3g | A | 361 | LEU | p.Leu361Pro | P38935 |

| | | | | | | | | |
|---|---|---|---|---|---|---|---|---|
| HOMO SAPIENS | 0 | 19406 | 4b3g | A | 369 | PHE | p.Phe369Leu | P38935 |
| HOMO SAPIENS | 0 | 19406 | 4b3g | A | 493 | THR | p.Thr493Ile | P38935 |
| HOMO SAPIENS | 0 | 19406 | 4b3g | A | 496 | CYS | p.Cys496Ter | P38935 |
| HOMO SAPIENS | 0 | 19406 | 4b3g | A | 514 | GLU | p.Glu514Lys | P38935 |
| HOMO SAPIENS | 0 | 19406 | 4b3g | A | 528 | ALA | p.Ala528Thr | P38935 |
| HOMO SAPIENS | 0 | 19406 | 4b3g | A | 531 | PRO | p.Pro531Thr | P38935 |
| HOMO SAPIENS | 0 | 19406 | 4b3g | A | 580 | VAL | p.Val580Ile | P38935 |
| HOMO SAPIENS | 6 | 19406 | 4b3g | A | 603 | ARG | p.Arg603His | P38935 |
| HOMO SAPIENS | 0 | 21501 | 5jde | A | 40 | SER | p.Ser2831Asn | Q8WZ42 |
| HOMO SAPIENS | 5 | 21501 | 5jde | A | 189 | ALA | p.Ala2980Thr | Q8WZ42 |
| HOMO SAPIENS | 6 | 15008 | 3mop | H | 12 | ARG | p.Arg12Cys | Q9NWZ3 |
| HOMO SAPIENS | 8 | 17455 | 3jtc | C | 15 | ARG | p.Arg57Trp | P04070 |
| HOMO SAPIENS | -1 | 17455 | 3jtc | C | 20 | CGU | p.Glu62Ala | P04070 |
| HOMO SAPIENS | 0 | 27795 | 5gox | B | 625 | TYR | p.Tyr625Ter | Q92878 |
| HOMO SAPIENS | 0 | 27795 | 5gox | B | 653 | SER | p.Ser653Ter | Q92878 |
| HOMO SAPIENS | -1 | 6558 | 5nug | A | 137 | VAL | p.Val137Ile | Q14204 |
| HOMO SAPIENS | -1 | 6558 | 5nug | A | 567 | ARG | p.Arg567His | Q14204 |
| HOMO SAPIENS | -1 | 6558 | 5nug | A | 1140 | THR | p.Thr1140Met | Q14204 |
| HOMO SAPIENS | -1 | 6558 | 5nug | A | 1991 | ASP | p.Asp1991Asn | Q14204 |
| HOMO SAPIENS | -1 | 6558 | 5nug | A | 2401 | LYS | p.Lys2401Asn | Q14204 |
| HOMO SAPIENS | 0 | 6558 | 5nug | A | 2488 | ARG | p.Arg2488His | Q14204 |
| HOMO SAPIENS | 0 | 6558 | 5nug | A | 3907 | HIS | p.His3907Gln | Q14204 |
| HOMO SAPIENS | 0 | 6558 | 5nug | A | 3981 | THR | p.Thr3981Arg | Q14204 |
| HOMO SAPIENS | 0 | 6558 | 5nug | A | 4394 | THR | p.Thr4394Met | Q14204 |
| HOMO SAPIENS | 0 | 6558 | 5nug | A | 4573 | ASN | p.Asn4573Ser | Q14204 |
| HOMO SAPIENS | 0 | 6558 | 5nug | A | 4603 | SER | p.Ser4603Ile | Q14204 |
| HOMO SAPIENS | 1 | 4244 | 1umk | A | 57 | ARG | p.Arg58Gln | P00387 |
| HOMO SAPIENS | 1 | 4244 | 1umk | A | 72 | LEU | p.Leu73Pro | P00387 |
| HOMO SAPIENS | 1 | 4244 | 1umk | A | 105 | VAL | p.Val106Met | P00387 |
| HOMO SAPIENS | 0 | 4244 | 1umk | A | 116 | THR | p.Thr117Ser | P00387 |
| HOMO SAPIENS | 8 | 4244 | 1umk | A | 127 | SER | p.Ser128Pro | P00387 |
| HOMO SAPIENS | 0 | 4244 | 1umk | A | 148 | LEU | p.Leu149Pro | P00387 |
| HOMO SAPIENS | 0 | 4244 | 1umk | A | 159 | ARG | p.Arg160Ter | P00387 |
| HOMO SAPIENS | 0 | 4244 | 1umk | A | 203 | CYS | p.Cys204Arg | P00387 |
| HOMO SAPIENS | 0 | 4244 | 1umk | A | 203 | CYS | p.Cys204Tyr | P00387 |
| HOMO SAPIENS | 2 | 4244 | 1umk | A | 239 | ASP | p.Asp240Gly | P00387 |
| HOMO SAPIENS | 0 | 4244 | 1umk | A | 291 | GLY | p.Gly292Asp | P00387 |
| HOMO SAPIENS | 4 | 34495 | 1s35 | A | 1151 | ASP | p.Asn1151Asp | P11277 |
| HOMO SAPIENS | 3 | 30110 | 3zqs | A | 224 | THR | p.Thr224Ala | Q9NW38 |
| HOMO SAPIENS | 3 | 17588 | 5dse | B | 53 | LEU | p.Leu53Pro | Q9BYI3 |
| HOMO SAPIENS | 0 | 1564 | 4yqf | B | 72 | HIS | p.His347Gln | Q9UHD8 |
| HOMO SAPIENS | 0 | 17673 | 4jnw | A | 32 | ARG | p.Arg32186His | Q8WZ42 |
| HOMO SAPIENS | 3 | 17673 | 4jnw | A | 140 | ILE | p.Ile32294Val | Q8WZ42 |
| HOMO SAPIENS | 5 | 17673 | 4jnw | A | 269 | ALA | p.Ala32423Thr | Q8WZ42 |
| HOMO SAPIENS | 3 | 12858 | 3s97 | C | 302 | LEU | p.Leu302Pro | Q12860 |
| HOMO SAPIENS | 0 | 6147 | 4nn0 | A | 114 | ARG | p.Arg114Gln | Q9BXJ0 |
| HOMO SAPIENS | 0 | 7549 | 1tr2 | A | 277 | LEU | p.Leu277Met | P18206 |
| HOMO SAPIENS | 1 | 7549 | 1tr2 | A | 413 | ALA | p.Ala413Thr | P18206 |
| HOMO SAPIENS | 1 | 7549 | 1tr2 | A | 519 | ILE | p.Ile519Leu | P18206 |
| HOMO SAPIENS | 5 | 7549 | 1tr2 | A | 547 | ARG | p.Arg547Ter | P18206 |
| HOMO SAPIENS | 2 | 7549 | 1tr2 | A | 636 | HIS | p.His636Arg | P18206 |
| HOMO SAPIENS | 2 | 7549 | 1tr2 | A | 824 | ILE | p.Ile824Met | P18206 |
| HOMO SAPIENS | 2 | 13900 | 3gro | A | 45 | CYS | p.Cys45Tyr | P50897 |
| HOMO SAPIENS | 2 | 13900 | 3gro | A | 75 | THR | p.Thr75Pro | P50897 |

| | | | | | | | | |
|---|---|---|---|---|---|---|---|---|
| HOMO SAPIENS | 2 | 13900 | 3gro | A | 79 | ASP | p.Asp79Gly | P50897 |
| HOMO SAPIENS | 5 | 13900 | 3gro | A | 91 | GLN | p.Gln91Pro | P50897 |
| HOMO SAPIENS | 5 | 13900 | 3gro | A | 96 | CYS | p.Cys96Tyr | P50897 |
| HOMO SAPIENS | 0 | 13900 | 3gro | A | 104 | LYS | p.Lys104Ter | P50897 |
| HOMO SAPIENS | 2 | 13900 | 3gro | A | 108 | GLY | p.Gly108Arg | P50897 |
| HOMO SAPIENS | 2 | 13900 | 3gro | A | 109 | TYR | p.Tyr109Asp | P50897 |
| HOMO SAPIENS | 0 | 13900 | 3gro | A | 122 | ARG | p.Arg122Trp | P50897 |
| HOMO SAPIENS | 0 | 13900 | 3gro | A | 138 | SER | p.Ser138Leu | P50897 |
| HOMO SAPIENS | 5 | 13900 | 3gro | A | 151 | ARG | p.Arg151Ter | P50897 |
| HOMO SAPIENS | 7 | 13900 | 3gro | A | 164 | ARG | p.Arg164Ter | P50897 |
| HOMO SAPIENS | 0 | 13900 | 3gro | A | 177 | GLN | p.Gln177Glu | P50897 |
| HOMO SAPIENS | 0 | 13900 | 3gro | A | 181 | VAL | p.Val181Leu | P50897 |
| HOMO SAPIENS | 0 | 13900 | 3gro | A | 184 | GLU | p.Glu184Lys | P50897 |
| HOMO SAPIENS | 5 | 13900 | 3gro | A | 186 | TRP | p.Trp186Ter | P50897 |
| HOMO SAPIENS | 0 | 13900 | 3gro | A | 219 | LEU | p.Leu219Gln | P50897 |
| HOMO SAPIENS | 0 | 13900 | 3gro | A | 222 | LEU | p.Leu222Pro | P50897 |
| HOMO SAPIENS | 5 | 13900 | 3gro | A | 296 | TRP | p.Trp296Ter | P50897 |
| HOMO SAPIENS | 6 | 35028 | 5mwb | A | 444 | CYS | p.Cys444Tyr | Q04721 |
| HOMO SAPIENS | 0 | 35028 | 5mwb | A | 514 | ASN | p.Asn514Ser | Q04721 |
| HOMO SAPIENS | 5 | 1483 | 4gl2 | A | 822 | ARG | p.Arg822Gln | Q9BYX4 |
| HOMO SAPIENS | 0 | 1483 | 4gl2 | A | 843 | ARG | p.His843Arg | Q9BYX4 |
| HOMO SAPIENS | -1 | 28838 | 3cl3 | D | 153 | LEU | p.Leu153Arg | Q9Y6K9 |
| HOMO SAPIENS | 2 | 22563 | 4zel | B | 79 | ARG | p.Arg79Trp | P09172 |
| HOMO SAPIENS | 2 | 22563 | 4zel | B | 101 | VAL | p.Val101Met | P09172 |
| HOMO SAPIENS | 2 | 22563 | 4zel | B | 114 | ASP | p.Asp114Glu | P09172 |
| HOMO SAPIENS | 0 | 22563 | 4zel | B | 201 | ASN | p.Asn201Ser | P09172 |
| HOMO SAPIENS | 2 | 22563 | 4zel | B | 211 | ALA | p.Ala211Thr | P09172 |
| HOMO SAPIENS | -1 | 22563 | 4zel | B | 290 | ASP | p.Asp290Asn | P09172 |
| HOMO SAPIENS | 2 | 22563 | 4zel | B | 318 | ALA | p.Ala318Ser | P09172 |
| HOMO SAPIENS | 2 | 22563 | 4zel | B | 345 | ASP | p.Asp345Asn | P09172 |
| HOMO SAPIENS | 2 | 22563 | 4zel | B | 362 | ALA | p.Ala362Glu | P09172 |
| HOMO SAPIENS | 2 | 22563 | 4zel | B | 549 | ARG | p.Arg549Cys | P09172 |
| HOMO SAPIENS | 9 | 9780 | 1y6k | R | 36 | TYR | p.Tyr57Cys | Q13651 |
| HOMO SAPIENS | 1 | 9780 | 1y6k | R | 40 | LEU | p.Leu61Val | Q13651 |
| HOMO SAPIENS | 1 | 9780 | 1y6k | R | 63 | THR | p.Thr84Ile | Q13651 |
| HOMO SAPIENS | 9 | 9780 | 1y6k | R | 80 | ARG | p.Arg101Trp | Q13651 |
| HOMO SAPIENS | 0 | 9780 | 1y6k | R | 120 | GLY | p.Gly141Arg | Q13651 |
| HOMO SAPIENS | 0 | 9780 | 1y6k | R | 138 | SER | p.Ser159Gly | Q13651 |
| HOMO SAPIENS | 0 | 9780 | 1y6k | R | 203 | ILE | p.Ile224Val | Q13651 |
| HOMO SAPIENS | 0 | 16019 | 3rep | A | 211 | ARG | p.Arg211Cys | Q13418 |
| HOMO SAPIENS | 3 | 7879 | 5ec3 | A | 33 | ALA | p.Ala33Thr | P32754 |
| HOMO SAPIENS | 4 | 7879 | 5ec3 | A | 113 | ARG | p.Arg113Gln | P32754 |
| HOMO SAPIENS | 3 | 7879 | 5ec3 | A | 340 | VAL | p.Val340Leu | P32754 |
| HOMO SAPIENS | 0 | 8904 | 1qk9 | A | 6 | LYS | p.Lys82Arg | P51608 |
| HOMO SAPIENS | 4 | 8904 | 1qk9 | A | 24 | LEU | p.Leu100Val | P51608 |
| HOMO SAPIENS | 7 | 8904 | 1qk9 | A | 30 | ARG | p.Arg106Trp | P51608 |
| HOMO SAPIENS | 6 | 8904 | 1qk9 | A | 57 | ARG | p.Arg133Cys | P51608 |
| HOMO SAPIENS | 3 | 8904 | 1qk9 | A | 61 | GLU | p.Glu137Gly | P51608 |
| HOMO SAPIENS | 2 | 8904 | 1qk9 | A | 64 | ALA | p.Ala140Val | P51608 |
| HOMO SAPIENS | 5 | 8904 | 1qk9 | A | 79 | PHE | p.Phe155Ser | P51608 |
| HOMO SAPIENS | 3 | 8904 | 1qk9 | A | 82 | THR | p.Thr158Met | P51608 |
| HOMO SAPIENS | 1 | 23121 | 3e9k | A | 198 | THR | p.Thr198Ala | Q16719 |
| HOMO SAPIENS | -1 | 26090 | 3tje | F | 19 | LEU | p.Leu35Met | P25445 |
| HOMO SAPIENS | 8 | 26090 | 3tje | F | 106 | THR | p.Thr122Ile | P25445 |

| | | | | | | | | |
|---|---|---|---|---|---|---|---|---|
| HOMO SAPIENS | 2 | 36404 | 5cqr | A | 765 | GLY | p.Gly765Glu | O95163 |
| HOMO SAPIENS | 2 | 36404 | 5cqr | A | 816 | ILE | p.Ile816Leu | O95163 |
| HOMO SAPIENS | 2 | 36404 | 5cqr | A | 830 | ILE | p.Ile830Met | O95163 |
| HOMO SAPIENS | 5 | 36404 | 5cqr | A | 939 | TYR | p.Tyr939Ter | O95163 |
| HOMO SAPIENS | 2 | 36404 | 5cqr | A | 952 | LYS | p.Lys952Ile | O95163 |
| HOMO SAPIENS | 5 | 36404 | 5cqr | A | 1013 | GLY | p.Gly1013Ser | O95163 |
| HOMO SAPIENS | 5 | 36404 | 5cqr | A | 1072 | CYS | p.Cys1072Ser | O95163 |
| HOMO SAPIENS | -1 | 36404 | 5cqr | A | 1158 | PRO | p.Pro1158Leu | O95163 |
| HOMO SAPIENS | 1 | 34149 | 2k40 | A | 18 | ASN | p.Asn125Ser | Q9UBX0 |
| HOMO SAPIENS | 0 | 34149 | 2k40 | A | 42 | LEU | p.Glu149Lys | Q9UBX0 |
| HOMO SAPIENS | 9 | 34149 | 2k40 | A | 53 | ARG | p.Arg160Cys | Q9UBX0 |
| HOMO SAPIENS | 0 | 34149 | 2k40 | A | 63 | SER | p.Ser170Leu | Q9UBX0 |
| HOMO SAPIENS | 2 | 2468 | 3vp6 | A | 484 | LEU | p.Leu484Met | Q99259 |
| HOMO SAPIENS | 0 | 2468 | 3vp6 | A | 532 | ARG | p.Arg532Gln | Q99259 |
| HOMO SAPIENS | 5 | 1253 | 5xtd | S | 8 | GLY | p.Gly8Arg | O15239 |
| HOMO SAPIENS | 7 | 1253 | 5xtd | S | 32 | GLY | p.Gly32Arg | O15239 |
| HOMO SAPIENS | 7 | 1253 | 5xtd | S | 37 | ARG | p.Arg37Ser | O15239 |
| HOMO SAPIENS | -1 | 13292 | 3u1k | A | 121 | VAL | p.Ile121Val | Q8TCS8 |
| HOMO SAPIENS | 2 | 13292 | 3u1k | A | 165 | PRO | p.Pro165Ser | Q8TCS8 |
| HOMO SAPIENS | 2 | 13292 | 3u1k | A | 192 | ARG | p.Arg192Gln | Q8TCS8 |
| HOMO SAPIENS | 1 | 13292 | 3u1k | A | 315 | LYS | p.Lys315Arg | Q8TCS8 |
| HOMO SAPIENS | 4 | 13292 | 3u1k | A | 509 | VAL | p.Val509Ile | Q8TCS8 |
| HOMO SAPIENS | 7 | 13292 | 3u1k | A | 531 | THR | p.Thr531Arg | Q8TCS8 |
| HOMO SAPIENS | 0 | 13292 | 3u1k | A | 590 | ASN | p.Asn590Asp | Q8TCS8 |
| HOMO SAPIENS | 0 | 27048 | 4p5w | B | 184 | ARG | p.Arg184Ter | Q9UJS0 |
| HOMO SAPIENS | 0 | 27048 | 4p5w | B | 225 | SER | p.Ser225Ter | Q9UJS0 |
| HOMO SAPIENS | -1 | 27048 | 4p5w | B | 631 | ALA | p.Arg360Ter | Q9UJS0 |
| HOMO SAPIENS | 0 | 24240 | 4mdr | A | 318 | ARG | p.Arg318Ter | O00189 |
| HOMO SAPIENS | 0 | 24240 | 4mdr | A | 338 | ARG | p.Arg338Ter | O00189 |
| HOMO SAPIENS | 0 | 13429 | 4cg4 | A | 479 | PHE | p.Phe479Leu | O15553 |
| HOMO SAPIENS | 0 | 13429 | 4cg4 | A | 487 | VAL | p.Val487Leu | O15553 |
| HOMO SAPIENS | 3 | 13429 | 4cg4 | A | 653 | ARG | p.Arg653His | O15553 |
| HOMO SAPIENS | 1 | 13429 | 4cg4 | A | 680 | MET | p.Met680Ile | O15553 |
| HOMO SAPIENS | 1 | 13429 | 4cg4 | A | 688 | TYR | p.Tyr688Ter | O15553 |
| HOMO SAPIENS | 0 | 13429 | 4cg4 | A | 694 | MET | p.Met694Ile | O15553 |
| HOMO SAPIENS | 0 | 13429 | 4cg4 | A | 694 | MET | p.Met694Val | O15553 |
| HOMO SAPIENS | 1 | 13429 | 4cg4 | A | 726 | VAL | p.Val726Ala | O15553 |
| HOMO SAPIENS | 0 | 13429 | 4cg4 | A | 761 | ARG | p.Arg761His | O15553 |
| HOMO SAPIENS | 0 | 2079 | 4okh | A | 691 | GLY | p.Gly691Arg | P20807 |
| HOMO SAPIENS | 2 | 2079 | 4okh | A | 748 | ARG | p.Arg748Ter | P20807 |
| HOMO SAPIENS | 0 | 2079 | 4okh | A | 763 | TYR | p.Tyr763Cys | P20807 |
| HOMO SAPIENS | 1 | 2079 | 4okh | A | 769 | ARG | p.Arg769Gln | P20807 |
| HOMO SAPIENS | 9 | 2079 | 4okh | A | 777 | ILE | p.Ile777Val | P20807 |
| HOMO SAPIENS | 8 | 2079 | 4okh | A | 778 | ASP | p.Asp778His | P20807 |
| HOMO SAPIENS | 5 | 2079 | 4okh | A | 780 | ASP | p.Asp780His | P20807 |
| HOMO SAPIENS | 1 | 1200 | 2gi7 | A | 142 | ARG | p.Arg162Trp | Q9HCN6 |
| HOMO SAPIENS | 4 | 9199 | 3ezq | C | 260 | ASP | p.Asp260Val | P25445 |
| HOMO SAPIENS | 0 | 36522 | 5ms9 | A | 122 | ARG | p.Arg122Cys | P35555 |
| HOMO SAPIENS | 0 | 36522 | 5ms9 | A | 240 | ARG | p.Arg240Cys | P35555 |
| HOMO SAPIENS | 0 | 17448 | 3g2f | A | 225 | ARG | p.Arg225His | Q13873 |
| HOMO SAPIENS | 0 | 17448 | 3g2f | A | 347 | CYS | p.Cys347Tyr | Q13873 |
| HOMO SAPIENS | 3 | 17448 | 3g2f | A | 485 | ASP | p.Asp485Gly | Q13873 |
| HOMO SAPIENS | 7 | 17448 | 3g2f | A | 491 | ARG | p.Arg491Gln | Q13873 |
| HOMO SAPIENS | 7 | 17448 | 3g2f | A | 491 | ARG | p.Arg491Trp | Q13873 |

| | | | | | | | | |
|---|---|---|---|---|---|---|---|---|
| HOMO SAPIENS | 0 | 4861 | 4ux8 | A | 56 | LEU | p.Leu56Met | P07949 |
| HOMO SAPIENS | 1 | 4861 | 4ux8 | A | 88 | ILE | p.Ile88Leu | P07949 |
| HOMO SAPIENS | 0 | 4861 | 4ux8 | A | 114 | ARG | p.Arg114His | P07949 |
| HOMO SAPIENS | 5 | 4861 | 4ux8 | A | 180 | ARG | p.Arg180Gln | P07949 |
| HOMO SAPIENS | 0 | 4861 | 4ux8 | A | 208 | GLU | p.Glu208Asp | P07949 |
| HOMO SAPIENS | 0 | 4861 | 4ux8 | A | 373 | ALA | p.Ala373Val | P07949 |
| HOMO SAPIENS | 0 | 4861 | 4ux8 | A | 375 | LEU | p.Leu375Gln | P07949 |
| HOMO SAPIENS | -1 | 4861 | 4ux8 | A | 386 | ALA | p.Ala386Val | P07949 |
| HOMO SAPIENS | 0 | 4861 | 4ux8 | A | 446 | GLY | p.Gly446Arg | P07949 |
| HOMO SAPIENS | 0 | 4861 | 4ux8 | A | 480 | GLU | p.Glu480Lys | P07949 |
| HOMO SAPIENS | 0 | 4861 | 4ux8 | A | 489 | ASP | p.Asp489Asn | P07949 |
| HOMO SAPIENS | -1 | 4861 | 4ux8 | A | 510 | ALA | p.Ala510Val | P07949 |
| HOMO SAPIENS | -1 | 4861 | 4ux8 | A | 533 | GLY | p.Gly533Cys | P07949 |
| HOMO SAPIENS | -1 | 4861 | 4ux8 | A | 567 | ASP | p.Asp567Asn | P07949 |
| HOMO SAPIENS | -1 | 4861 | 4ux8 | A | 609 | CYS | p.Cys609Tyr | P07949 |
| HOMO SAPIENS | -1 | 4861 | 4ux8 | A | 609 | CYS | p.Cys609Gly | P07949 |
| HOMO SAPIENS | -1 | 4861 | 4ux8 | A | 611 | CYS | p.Cys611Trp | P07949 |
| HOMO SAPIENS | -1 | 4861 | 4ux8 | A | 618 | CYS | p.Cys618Gly | P07949 |
| HOMO SAPIENS | -1 | 4861 | 4ux8 | A | 618 | CYS | p.Cys618Phe | P07949 |
| HOMO SAPIENS | -1 | 4861 | 4ux8 | A | 620 | CYS | p.Cys620Gly | P07949 |
| HOMO SAPIENS | -1 | 4861 | 4ux8 | A | 620 | CYS | p.Cys620Ser | P07949 |
| HOMO SAPIENS | -1 | 4861 | 4ux8 | A | 620 | CYS | p.Cys620Trp | P07949 |
| HOMO SAPIENS | -1 | 4861 | 4ux8 | A | 623 | GLU | p.Glu623Lys | P07949 |
| HOMO SAPIENS | -1 | 4861 | 4ux8 | A | 631 | ASP | p.Asp631Gly | P07949 |
| HOMO SAPIENS | -1 | 4861 | 4ux8 | A | 631 | ASP | p.Asp631Glu | P07949 |
| HOMO SAPIENS | -1 | 4861 | 4ux8 | A | 634 | CYS | p.Cys634Gly | P07949 |
| HOMO SAPIENS | -1 | 4861 | 4ux8 | A | 634 | CYS | p.Cys634Phe | P07949 |
| HOMO SAPIENS | -1 | 4861 | 4ux8 | A | 634 | CYS | p.Cys634Trp | P07949 |
| HOMO SAPIENS | 0 | 36308 | 4nkq | B | 123 | PRO | p.Pro142Leu | P15509 |
| HOMO SAPIENS | 0 | 36308 | 4nkq | B | 145 | ARG | p.Arg164Gln | P15509 |
| HOMO SAPIENS | -1 | 36308 | 4nkq | B | 177 | GLY | p.Gly196Arg | P15509 |
| HOMO SAPIENS | 0 | 36006 | 1oqa | A | 107 | HIS | p.His1860Pro | P38398 |
| HOMO SAPIENS | 7 | 15394 | 2ozo | A | 465 | ARG | p.Arg465His | P43403 |
| HOMO SAPIENS | 7 | 15394 | 2ozo | A | 465 | ARG | p.Arg465Cys | P43403 |
| HOMO SAPIENS | 4 | 15394 | 2ozo | A | 518 | SER | p.Ser518Arg | P43403 |
| HOMO SAPIENS | 1 | 20813 | 3o5z | A | 19 | ARG | p.Arg19Trp | O00459 |
| HOMO SAPIENS | 2 | 4329 | 2bb5 | A | 5 | ILE | p.Ile23Val | P20062 |
| HOMO SAPIENS | 4 | 4329 | 2bb5 | A | 59 | LYS | p.Lys77Met | P20062 |
| HOMO SAPIENS | 4 | 4329 | 2bb5 | A | 197 | ARG | p.Arg215Trp | P20062 |
| HOMO SAPIENS | 1 | 4329 | 2bb5 | A | 241 | PRO | p.Arg259Pro | P20062 |
| HOMO SAPIENS | 2 | 4329 | 2bb5 | A | 315 | THR | p.Thr333Lys | P20062 |
| HOMO SAPIENS | 5 | 4329 | 2bb5 | A | 330 | SER | p.Ser348Phe | P20062 |
| HOMO SAPIENS | 4 | 4329 | 2bb5 | A | 358 | LEU | p.Leu376Ser | P20062 |
| HOMO SAPIENS | 5 | 4329 | 2bb5 | A | 381 | ARG | p.Arg399Gln | P20062 |
| HOMO SAPIENS | 4 | 18294 | 4r30 | A | 163 | ASN | p.Asn163Asp | O95278 |
| HOMO SAPIENS | 6 | 18294 | 4r30 | A | 227 | ALA | p.Ala227Val | O95278 |
| HOMO SAPIENS | 4 | 18294 | 4r30 | A | 241 | ARG | p.Arg241Ter | O95278 |
| HOMO SAPIENS | 4 | 18294 | 4r30 | A | 325 | ARG | p.Arg325His | O95278 |
| HOMO SAPIENS | 0 | 2632 | 4xem | A | 22 | GLU | p.Glu22Lys | P49588 |
| HOMO SAPIENS | 0 | 2632 | 4xem | A | 275 | GLY | p.Gly275Asp | P49588 |
| HOMO SAPIENS | 1 | 2632 | 4xem | A | 329 | ARG | p.Arg329His | P49588 |
| HOMO SAPIENS | 0 | 11431 | 5t4n | A | 284 | GLN | p.Gln305Glu | Q96QU1 |
| HOMO SAPIENS | 0 | 11431 | 5t4n | A | 315 | ARG | p.Arg336Ter | Q96QU1 |
| HOMO SAPIENS | 0 | 11431 | 5t4n | A | 322 | ARG | p.Arg343Lys | Q96QU1 |

| | | | | | | | | |
|---|---|---|---|---|---|---|---|---|
| HOMO SAPIENS | 1 | 11431 | 5t4n | A | 326 | LEU | p.Leu347Phe | Q96QU1 |
| HOMO SAPIENS | 0 | 11431 | 5t4n | A | 373 | TYR | p.Tyr394Cys | Q96QU1 |
| HOMO SAPIENS | 0 | 11431 | 5t4n | A | 398 | ASN | p.Asn419Ser | Q96QU1 |
| HOMO SAPIENS | 0 | 11431 | 5t4n | A | 414 | ALA | p.Asp435Ala | Q96QU1 |
| HOMO SAPIENS | 0 | 11431 | 5t4n | A | 433 | VAL | p.Val454Phe | Q96QU1 |
| HOMO SAPIENS | 1 | 11431 | 5t4n | A | 507 | VAL | p.Val528Asp | Q96QU1 |
| HOMO SAPIENS | 0 | 11431 | 5t4n | A | 547 | ALA | p.Ala568Ser | Q96QU1 |
| HOMO SAPIENS | 3 | 3194 | 5xtd | L | 84 | ARG | p.Arg84His | O43181 |
| HOMO SAPIENS | 1 | 3194 | 5xtd | L | 130 | THR | p.Thr130Ser | O43181 |
| HOMO SAPIENS | 8 | 3194 | 5xtd | L | 136 | SER | p.Ser136Ala | O43181 |
| HOMO SAPIENS | 0 | 40749 | 1nun | A | 106 | CYS | p.Cys106Phe | O15520 |
| HOMO SAPIENS | 9 | 40749 | 1nun | A | 156 | ILE | p.Ile156Arg | O15520 |
| HOMO SAPIENS | 1 | 40749 | 1nun | A | 193 | ARG | p.Arg193Ter | O15520 |
| HOMO SAPIENS | 0 | 40749 | 1nun | A | 207 | HIS | p.His207Pro | O15520 |
| HOMO SAPIENS | 0 | 7026 | 3n1f | D | 875 | SER | p.Ser875Gly | Q4KMG0 |
| HOMO SAPIENS | 6 | 23802 | 3rzv | A | 424 | ARG | p.Arg424Ter | O95630 |
| HOMO SAPIENS | 0 | 36720 | 1jm7 | A | 12 | GLN | p.Gln12Ter | P38398 |
| HOMO SAPIENS | 3 | 36720 | 1jm7 | A | 23 | GLU | p.Glu23Asp | P38398 |
| HOMO SAPIENS | 1 | 36720 | 1jm7 | A | 45 | LYS | p.Lys45Gln | P38398 |
| HOMO SAPIENS | 8 | 36720 | 1jm7 | A | 47 | CYS | p.Cys47Ter | P38398 |
| HOMO SAPIENS | 0 | 36720 | 1jm7 | A | 59 | SER | p.Ser59Ter | P38398 |
| HOMO SAPIENS | 8 | 36720 | 1jm7 | A | 61 | CYS | p.Cys61Gly | P38398 |
| HOMO SAPIENS | 8 | 36720 | 1jm7 | A | 63 | LEU | p.Leu63Ter | P38398 |
| HOMO SAPIENS | 7 | 36720 | 1jm7 | A | 64 | CYS | p.Cys64Gly | P38398 |
| HOMO SAPIENS | 0 | 36720 | 1jm7 | A | 101 | TYR | p.Tyr101Ter | P38398 |
| HOMO SAPIENS | 0 | 36720 | 1jm7 | A | null | TYR | p.Tyr105Cys | P38398 |
| HOMO SAPIENS | 0 | 28500 | 3kuz | A | 1217 | CYS | p.Cys1217Arg | O60486 |
| HOMO SAPIENS | 5 | 8097 | 3p56 | C | 69 | ARG | p.Arg69Trp | Q8TDP1 |
| HOMO SAPIENS | 5 | 8097 | 3p56 | C | 143 | LYS | p.Lys143Ile | Q8TDP1 |
| HOMO SAPIENS | 2 | 30073 | 3nmz | A | 380 | ARG | p.Arg380Gln | P25054 |
| HOMO SAPIENS | 1 | 22573 | 1cjy | A | 111 | SER | p.Ser111Pro | P47712 |
| HOMO SAPIENS | 0 | 22573 | 1cjy | A | 485 | ARG | p.Arg485His | P47712 |
| HOMO SAPIENS | 2 | 965 | 3E+077 | A | 99 | ALA | p.Ala99Val | Q9Y617 |
| HOMO SAPIENS | 2 | 965 | 3E+077 | A | 100 | ASP | p.Asp100Ala | Q9Y617 |
| HOMO SAPIENS | 4 | 1503 | 3kxc | A | 62 | ARG | p.Arg62Trp | O43617 |
| HOMO SAPIENS | 6 | 9418 | 1hdr | A | 22 | GLY | p.Gly23Asp | P09417 |
| HOMO SAPIENS | 2 | 9418 | 1hdr | A | 35 | TRP | p.Trp36Arg | P09417 |
| HOMO SAPIENS | 2 | 9418 | 1hdr | A | 107 | TRP | p.Trp108Gly | P09417 |
| HOMO SAPIENS | 6 | 9418 | 1hdr | A | 149 | TYR | p.Tyr150Cys | P09417 |
| HOMO SAPIENS | -1 | 12079 | 3tnu | B | 373 | GLY | p.Gly373Ala | P13647 |
| HOMO SAPIENS | 1 | 12079 | 3tnu | B | 463 | LEU | p.Leu463Pro | P13647 |
| HOMO SAPIENS | 1 | 12079 | 3tnu | B | 471 | ARG | p.Arg471Cys | P13647 |
| HOMO SAPIENS | 0 | 8135 | 3rbn | B | 89 | SER | p.Ser556Ile | P40692 |
| HOMO SAPIENS | 0 | 8135 | 3rbn | B | 111 | GLU | p.Glu578Gly | P40692 |
| HOMO SAPIENS | 2 | 8135 | 3rbn | B | 114 | PRO | p.Pro581Leu | P40692 |
| HOMO SAPIENS | 0 | 8135 | 3rbn | B | 133 | GLU | p.Glu600Gly | P40692 |
| HOMO SAPIENS | 2 | 8135 | 3rbn | B | 136 | PRO | p.Pro603Arg | P40692 |
| HOMO SAPIENS | 2 | 8135 | 3rbn | B | 136 | PRO | p.Pro603Leu | P40692 |
| HOMO SAPIENS | 7 | 8135 | 3rbn | B | 140 | LEU | p.Leu607His | P40692 |
| HOMO SAPIENS | 2 | 8135 | 3rbn | B | 151 | LYS | p.Lys618Ala | P40692 |
| HOMO SAPIENS | 2 | 8135 | 3rbn | B | 151 | LYS | p.Lys618Thr | P40692 |
| HOMO SAPIENS | 5 | 8135 | 3rbn | B | 181 | PRO | p.Pro648Ser | P40692 |
| HOMO SAPIENS | 2 | 8135 | 3rbn | B | 188 | ILE | p.Ile655Val | P40692 |
| HOMO SAPIENS | 2 | 8135 | 3rbn | B | 214 | ALA | p.Ala681Thr | P40692 |

| | | | | | | | | |
|---|---|---|---|---|---|---|---|---|
| HOMO SAPIENS | 2 | 8135 | 3rbn | B | 220 | ARG | p.Arg687Trp | P40692 |
| HOMO SAPIENS | 2 | 8135 | 3rbn | B | 222 | GLN | p.Gln689Arg | P40692 |
| HOMO SAPIENS | -1 | 8135 | 3rbn | B | 236 | GLN | p.Gln701Ter | P40692 |
| HOMO SAPIENS | 7 | 8135 | 3rbn | B | 251 | HIS | p.His718Tyr | P40692 |
| HOMO SAPIENS | 0 | 8135 | 3rbn | B | 284 | LYS | p.Lys751Arg | P40692 |
| HOMO SAPIENS | 1 | 21998 | 5d68 | A | 355 | LEU | p.Leu355Phe | O00522 |
| HOMO SAPIENS | 3 | 21998 | 5d68 | A | 380 | THR | p.Thr380Met | O00522 |
| HOMO SAPIENS | 0 | 21998 | 5d68 | A | 415 | ILE | p.Ile415Met | O00522 |
| HOMO SAPIENS | 0 | 33789 | 2m0c | A | 13 | ARG | p.Arg218Gln | Q9H161 |
| HOMO SAPIENS | 7 | 33789 | 2m0c | A | 52 | ARG | p.Arg257Thr | Q9H161 |
| HOMO SAPIENS | 0 | 33789 | 2m0c | A | 67 | ARG | p.Arg272Pro | Q9H161 |
| HOMO SAPIENS | 6 | 8263 | 2osg | A | 51 | GLY | p.Gly355Arg | Q9UDY2 |
| HOMO SAPIENS | 5 | 8263 | 2osg | A | 60 | THR | p.Thr364Met | Q9UDY2 |
| HOMO SAPIENS | 1 | 2475 | 5E+026 | B | 234 | THR | p.Thr234Ala | Q9BZ23 |
| HOMO SAPIENS | 2 | 2475 | 5E+026 | B | 264 | ARG | p.Arg264Trp | Q9BZ23 |
| HOMO SAPIENS | 4 | 2475 | 5E+026 | B | 286 | ARG | p.Arg286Cys | Q9BZ23 |
| HOMO SAPIENS | 0 | 2475 | 5E+026 | B | 357 | ARG | p.Arg357Trp | Q9BZ23 |
| HOMO SAPIENS | 2 | 2475 | 5E+026 | B | 481 | ARG | p.Arg481Ter | Q9BZ23 |
| HOMO SAPIENS | 8 | 2475 | 5E+026 | B | 521 | GLY | p.Gly521Arg | Q9BZ23 |
| HOMO SAPIENS | 2 | 2475 | 5E+026 | B | 528 | THR | p.Thr528Met | Q9BZ23 |
| HOMO SAPIENS | 1 | 3742 | 5xtd | E | 108 | HIS | p.His134Tyr | P56556 |
| HOMO SAPIENS | 1 | 16725 | 3g04 | C | 36 | ASP | p.Asp36His | P16473 |
| HOMO SAPIENS | 3 | 16725 | 3g04 | C | 52 | PRO | p.Pro52Thr | P16473 |
| HOMO SAPIENS | 1 | 16725 | 3g04 | C | 162 | PRO | p.Pro162Ala | P16473 |
| HOMO SAPIENS | -1 | 12366 | 5d2d | C | 750 | PRO | p.Pro750Leu | P13569 |
| HOMO SAPIENS | 0 | 12366 | 5d2d | C | 764 | ARG | p.Arg764Ter | P13569 |
| HOMO SAPIENS | 0 | 34845 | 2y23 | A | 1151 | ASP | p.Asp1151Glu | P52179 |
| HOMO SAPIENS | 0 | 34845 | 2y23 | A | 1180 | ASP | p.Asp1180Ala | P52179 |
| HOMO SAPIENS | 6 | 36740 | 3lbx | B | 2053 | ALA | p.Ala2053Pro | P11277 |
| HOMO SAPIENS | 0 | 19289 | 3ihj | A | 404 | ARG | p.Arg404Ter | Q8TD30 |
| HOMO SAPIENS | 5 | 1034 | 5xtd | Q | 228 | ARG | p.Arg228Gln | O75306 |
| HOMO SAPIENS | 5 | 1034 | 5xtd | Q | 229 | PRO | p.Pro229Gln | O75306 |
| HOMO SAPIENS | 4 | 1034 | 5xtd | Q | 323 | ARG | p.Arg323Gln | O75306 |
| HOMO SAPIENS | 4 | 1034 | 5xtd | Q | 352 | PRO | p.Pro352Ala | O75306 |
| HOMO SAPIENS | 5 | 1034 | 5xtd | Q | 413 | SER | p.Ser413Pro | O75306 |
| HOMO SAPIENS | 1 | 2772 | 2vaf | A | 1047 | THR | p.Thr66Ala | O14958 |
| HOMO SAPIENS | 5 | 2772 | 2vaf | A | 1057 | VAL | p.Val76Met | O14958 |
| HOMO SAPIENS | 0 | 2772 | 2vaf | A | 1148 | LEU | p.Leu167His | O14958 |
| HOMO SAPIENS | 1 | 2772 | 2vaf | A | 1149 | ILE | p.Ile168Thr | O14958 |
| HOMO SAPIENS | 8 | 2772 | 2vaf | A | 1225 | HIS | p.His244Arg | O14958 |
| HOMO SAPIENS | 8 | 2772 | 2vaf | A | 1225 | HIS | p.His244Tyr | O14958 |
| HOMO SAPIENS | 7 | 2772 | 2vaf | A | 1251 | ILE | p.Ile270Thr | O14958 |
| HOMO SAPIENS | 3 | 2772 | 2vaf | A | 1288 | ASP | p.Asp307His | O14958 |
| HOMO SAPIENS | 5 | 2772 | 2vaf | A | 1289 | PRO | p.Pro308Leu | O14958 |
| HOMO SAPIENS | 1 | 31885 | 2aex | A | 132 | ALA | p.Ala132Val | P36551 |
| HOMO SAPIENS | 1 | 31885 | 2aex | A | 272 | ASN | p.Asn272His | P36551 |
| HOMO SAPIENS | 8 | 31885 | 2aex | A | 279 | GLY | p.Gly279Arg | P36551 |
| HOMO SAPIENS | 5 | 31885 | 2aex | A | 294 | VAL | p.Val294Ile | P36551 |
| HOMO SAPIENS | 5 | 31885 | 2aex | A | 295 | HIS | p.His295Asp | P36551 |
| HOMO SAPIENS | 1 | 31885 | 2aex | A | 331 | ARG | p.Arg331Trp | P36551 |
| HOMO SAPIENS | 3 | 31885 | 2aex | A | 352 | ARG | p.Arg352Cys | P36551 |
| HOMO SAPIENS | 1 | 31885 | 2aex | A | 404 | LYS | p.Lys404Glu | P36551 |
| HOMO SAPIENS | 2 | 31885 | 2aex | A | 447 | ARG | p.Arg447Cys | P36551 |
| HOMO SAPIENS | 5 | 11721 | 2r3v | A | 20 | HIS | p.His20Pro | Q03426 |

| | | | | | | | | |
|---|---|---|---|---|---|---|---|---|
| HOMO SAPIENS | 1 | 11721 | 2r3v | A | 116 | TYR | p.Tyr116His | Q03426 |
| HOMO SAPIENS | 1 | 11721 | 2r3v | A | 148 | ALA | p.Ala148Thr | Q03426 |
| HOMO SAPIENS | 1 | 11721 | 2r3v | A | 167 | PRO | p.Pro167Leu | Q03426 |
| HOMO SAPIENS | 6 | 11721 | 2r3v | A | 203 | VAL | p.Val203Ala | Q03426 |
| HOMO SAPIENS | 2 | 11721 | 2r3v | A | 215 | ARG | p.Arg215Ter | Q03426 |
| HOMO SAPIENS | 2 | 11721 | 2r3v | A | 237 | THR | p.Thr237Ser | Q03426 |
| HOMO SAPIENS | 0 | 11721 | 2r3v | A | 268 | ILE | p.Ile268Thr | Q03426 |
| HOMO SAPIENS | 0 | 11721 | 2r3v | A | 268 | ILE | p.Ile268Val | Q03426 |
| HOMO SAPIENS | 0 | 11721 | 2r3v | A | 301 | ASN | p.Asn301Ser | Q03426 |
| HOMO SAPIENS | 6 | 11721 | 2r3v | A | 310 | VAL | p.Val310Met | Q03426 |
| HOMO SAPIENS | 5 | 11721 | 2r3v | A | 334 | ALA | p.Ala334Thr | Q03426 |
| HOMO SAPIENS | 2 | 11721 | 2r3v | A | 377 | VAL | p.Val377Ile | Q03426 |
| HOMO SAPIENS | 1 | 11721 | 2r3v | A | 386 | ASP | p.Asp386Asn | Q03426 |
| HOMO SAPIENS | 1 | 11721 | 2r3v | A | 388 | ARG | p.Arg388Ter | Q03426 |
| HOMO SAPIENS | 8 | 5010 | 2hgs | A | 125 | ARG | p.Arg125Cys | P48637 |
| HOMO SAPIENS | 1 | 5010 | 2hgs | A | 164 | ARG | p.Arg164Gln | P48637 |
| HOMO SAPIENS | 0 | 5010 | 2hgs | A | 219 | ASP | p.Asp219Gly | P48637 |
| HOMO SAPIENS | 7 | 5010 | 2hgs | A | 267 | ARG | p.Arg267Trp | P48637 |
| HOMO SAPIENS | 0 | 5010 | 2hgs | A | 283 | ARG | p.Arg283Cys | P48637 |
| HOMO SAPIENS | 0 | 6892 | 5erd | B | 7 | VAL | p.Val56Met | Q14126 |
| HOMO SAPIENS | 1 | 6892 | 5erd | B | 25 | HIS | p.His74Arg | Q14126 |
| HOMO SAPIENS | 0 | 6892 | 5erd | B | 40 | TYR | p.Tyr89Cys | Q14126 |
| HOMO SAPIENS | 0 | 6892 | 5erd | B | 97 | ARG | p.Arg146Leu | Q14126 |
| HOMO SAPIENS | 7 | 6892 | 5erd | B | 217 | ASN | p.Asn266Ser | Q14126 |
| HOMO SAPIENS | 0 | 6892 | 5erd | B | 243 | ARG | p.Arg292His | Q14126 |
| HOMO SAPIENS | 0 | 6892 | 5erd | B | 244 | ILE | p.Ile293Val | Q14126 |
| HOMO SAPIENS | 0 | 6892 | 5erd | B | 343 | VAL | p.Val392Ile | Q14126 |
| HOMO SAPIENS | 4 | 6892 | 5erd | B | 350 | GLU | p.Glu399Lys | Q14126 |
| HOMO SAPIENS | 0 | 6892 | 5erd | B | 410 | TYR | p.Tyr459Cys | Q14126 |
| HOMO SAPIENS | 0 | 6892 | 5erd | B | 466 | VAL | p.Val515Ile | Q14126 |
| HOMO SAPIENS | 0 | 6892 | 5erd | B | 482 | PHE | p.Phe531Cys | Q14126 |
| HOMO SAPIENS | 0 | 6892 | 5erd | B | 484 | VAL | p.Val533Ile | Q14126 |
| HOMO SAPIENS | 0 | 6892 | 5erd | B | 545 | LEU | p.Leu594Pro | Q14126 |
| HOMO SAPIENS | 0 | 1338 | 2ax4 | A | 58 | THR | p.Thr48Arg | O95340 |
| HOMO SAPIENS | 7 | 8675 | 2zkm | X | 385 | LYS | p.Lys385Arg | Q00722 |
| HOMO SAPIENS | 1 | 11769 | 2wya | B | 54 | VAL | p.Val54Met | P54868 |
| HOMO SAPIENS | 1 | 11769 | 2wya | B | 116 | ARG | p.Arg116Cys | P54868 |
| HOMO SAPIENS | 5 | 11769 | 2wya | B | 167 | TYR | p.Tyr167Cys | P54868 |
| HOMO SAPIENS | 1 | 11769 | 2wya | B | 174 | PHE | p.Phe174Leu | P54868 |
| HOMO SAPIENS | 0 | 11769 | 2wya | B | 212 | GLY | p.Gly212Arg | P54868 |
| HOMO SAPIENS | 1 | 11769 | 2wya | B | 288 | ARG | p.Arg288Ter | P54868 |
| HOMO SAPIENS | 0 | 11769 | 2wya | B | 375 | SER | p.Ser375Cys | P54868 |
| HOMO SAPIENS | 4 | 11769 | 2wya | B | 500 | ARG | p.Arg500His | P54868 |
| HOMO SAPIENS | 0 | 3102 | 2iwo | A | 1880 | ARG | p.Arg1908Lys | O75970 |
| HOMO SAPIENS | 6 | 14296 | 2m9w | A | 38 | GLY | p.Gly296Ser | P43694 |
| HOMO SAPIENS | 0 | 14296 | 2m9w | A | 58 | GLN | p.Gln316Glu | P43694 |
| HOMO SAPIENS | 2 | 704 | 5ekd | B | 239 | ARG | p.Arg239Ter | Q9UGM6 |
| HOMO SAPIENS | 2 | 704 | 5ekd | B | 264 | TYR | p.Tyr264Cys | Q9UGM6 |
| HOMO SAPIENS | 6 | 2228 | 1ey2 | A | 47 | SER | p.Ser47Ter | Q93099 |
| HOMO SAPIENS | 6 | 2228 | 1ey2 | A | 48 | ALA | p.Ala48Ser | Q93099 |
| HOMO SAPIENS | 6 | 2228 | 1ey2 | A | 53 | ARG | p.Arg53Gln | Q93099 |
| HOMO SAPIENS | 3 | 2228 | 1ey2 | A | 80 | HIS | p.Gln80His | Q93099 |
| HOMO SAPIENS | 0 | 2228 | 1ey2 | A | 120 | CYS | p.Cys120Trp | Q93099 |
| HOMO SAPIENS | 3 | 2228 | 1ey2 | A | 122 | ALA | p.Ala122Val | Q93099 |

| | | | | | | | | |
|---|---|---|---|---|---|---|---|---|
| HOMO SAPIENS | 3 | 2228 | 1ey2 | A | 161 | GLY | p.Gly161Arg | Q93099 |
| HOMO SAPIENS | 6 | 2228 | 1ey2 | A | 225 | ARG | p.Arg225His | Q93099 |
| HOMO SAPIENS | 0 | 2228 | 1ey2 | A | 230 | PRO | p.Pro230Ser | Q93099 |
| HOMO SAPIENS | 6 | 2228 | 1ey2 | A | 270 | GLY | p.Gly270Arg | Q93099 |
| HOMO SAPIENS | 6 | 2228 | 1ey2 | A | 330 | ARG | p.Arg330Ser | Q93099 |
| HOMO SAPIENS | -1 | 2228 | 1ey2 | A | 368 | MSE | p.Met368Val | Q93099 |
| HOMO SAPIENS | 6 | 2228 | 1ey2 | A | 371 | HIS | p.His371Arg | Q93099 |
| HOMO SAPIENS | 6 | 2228 | 1ey2 | A | 401 | GLU | p.Glu401Gln | Q93099 |
| HOMO SAPIENS | 5 | 34648 | 2k27 | A | 31 | ARG | p.Arg31His | Q06710 |
| HOMO SAPIENS | 7 | 34648 | 2k27 | A | 40 | GLN | p.Gln40Pro | Q06710 |
| HOMO SAPIENS | 5 | 34648 | 2k27 | A | 57 | CYS | p.Cys57Tyr | Q06710 |
| HOMO SAPIENS | 5 | 34648 | 2k27 | A | 62 | LEU | p.Leu62Arg | Q06710 |
| HOMO SAPIENS | 1 | 21399 | 3edu | A | 1819 | GLU | p.Glu1819Ter | P11277 |
| HOMO SAPIENS | 1 | 18153 | 1oey | J | 245 | TYR | p.Tyr245Ter | Q15080 |
| HOMO SAPIENS | 2 | 41567 | 3sw0 | X | 1050 | ASN | p.Asn1050Tyr | P08603 |
| HOMO SAPIENS | 5 | 41567 | 3sw0 | X | 1059 | ILE | p.Ile1059Thr | P08603 |
| HOMO SAPIENS | 0 | 36362 | 4wrm | A | 362 | HIS | p.His362Arg | P07333 |
| HOMO SAPIENS | -1 | 36362 | 4wrm | A | 413 | GLY | p.Gly413Ser | P07333 |
| HOMO SAPIENS | 0 | 31255 | 2w2j | A | 100 | SER | p.Ser100Pro | P35219 |
| HOMO SAPIENS | 0 | 31255 | 2w2j | A | 100 | SER | p.Ser100Leu | P35219 |
| HOMO SAPIENS | 5 | 31255 | 2w2j | A | 237 | ARG | p.Arg237Gln | P35219 |
| HOMO SAPIENS | -1 | 26450 | 1th1 | C | 1395 | SER | p.Ser1395Cys | P25054 |
| HOMO SAPIENS | -1 | 26450 | 1th1 | C | 1413 | MET | p.Met1413Val | P25054 |
| HOMO SAPIENS | -1 | 26450 | 1th1 | C | 1446 | ALA | p.Ala1446Thr | P25054 |
| HOMO SAPIENS | -1 | 26450 | 1th1 | C | 1454 | LYS | p.Lys1454Glu | P25054 |
| HOMO SAPIENS | 0 | 26450 | 1th1 | C | 1474 | ALA | p.Ala1474Thr | P25054 |
| HOMO SAPIENS | 1 | 981 | 5xtb | F | 14 | LEU | p.Leu14Val | O43678 |
| HOMO SAPIENS | 1 | 11069 | 3kq4 | A | 46 | SER | p.Ser46Leu | P27352 |
| HOMO SAPIENS | 1 | 11069 | 3kq4 | A | 97 | MET | p.Met97Thr | P27352 |
| HOMO SAPIENS | 1 | 11069 | 3kq4 | A | 255 | ASN | p.Asn255Ser | P27352 |
| HOMO SAPIENS | 1 | 11069 | 3kq4 | A | 330 | ASN | p.Asn330Lys | P27352 |
| HOMO SAPIENS | 0 | 41489 | 3erb | A | 53 | PRO | p.Pro73Leu | P06681 |
| HOMO SAPIENS | 6 | 8715 | 2bq8 | X | 23 | THR | p.Thr44Met | P13686 |
| HOMO SAPIENS | 0 | 8715 | 2bq8 | X | 88 | GLY | p.Gly109Arg | P13686 |
| HOMO SAPIENS | 6 | 8715 | 2bq8 | X | 127 | VAL | p.Val148Met | P13686 |
| HOMO SAPIENS | 2 | 8715 | 2bq8 | X | 179 | VAL | p.Val200Met | P13686 |
| HOMO SAPIENS | 6 | 8715 | 2bq8 | X | 251 | ARG | p.Arg272Cys | P13686 |
| HOMO SAPIENS | 7 | 8112 | 3p56 | A | 37 | GLY | p.Gly37Ser | O75792 |
| HOMO SAPIENS | -1 | 8112 | 3p56 | A | 202 | LEU | p.Leu202Ser | O75792 |
| HOMO SAPIENS | 1 | 8112 | 3p56 | A | 205 | ASP | p.Asp205Glu | O75792 |
| HOMO SAPIENS | 4 | 8112 | 3p56 | A | 212 | ASN | p.Asn212Ile | O75792 |
| HOMO SAPIENS | 0 | 8112 | 3p56 | A | 235 | ARG | p.Arg235Gln | O75792 |
| HOMO SAPIENS | 4 | 8112 | 3p56 | A | 240 | THR | p.Thr240Met | O75792 |
| HOMO SAPIENS | 0 | 8112 | 3p56 | A | 291 | ARG | p.Arg291His | O75792 |
| HOMO SAPIENS | 0 | 37603 | 2yqg | A | 10 | ARG | p.Arg49His | Q14126 |
| HOMO SAPIENS | 0 | 21178 | 3k1r | B | 458 | ASP | p.Asp458Val | Q495M9 |
| HOMO SAPIENS | -1 | 36828 | 4fq3 | B | 507 | GLY | p.Gly507Asp | P35637 |
| HOMO SAPIENS | 5 | 36828 | 4fq3 | B | 517 | HIS | p.His517Gln | P35637 |
| HOMO SAPIENS | 8 | 36828 | 4fq3 | B | 518 | ARG | p.Arg518Lys | P35637 |
| HOMO SAPIENS | 6 | 36828 | 4fq3 | B | 521 | ARG | p.Arg521Cys | P35637 |
| HOMO SAPIENS | 6 | 36828 | 4fq3 | B | 521 | ARG | p.Arg521His | P35637 |
| HOMO SAPIENS | 0 | 6363 | 3j7y | c | null | HIS | p.His13Arg | Q9H9J2 |
| HOMO SAPIENS | 2 | 6363 | 3j7y | c | 105 | ARG | p.Arg105Cys | Q9H9J2 |
| HOMO SAPIENS | 2 | 6363 | 3j7y | c | 138 | THR | p.Thr138Ile | Q9H9J2 |

| | | | | | | | | |
|---|---|---|---|---|---|---|---|---|
| HOMO SAPIENS | 1 | 6363 | 3j7y | c | 156 | LEU | p.Leu156Arg | Q9H9J2 |
| HOMO SAPIENS | 1 | 6782 | 5l46 | A | 109 | HIS | p.His109Arg | Q9UI17 |
| HOMO SAPIENS | 0 | 23951 | 5t76 | A | 175 | GLY | p.Gly931Ser | P49588 |
| HOMO SAPIENS | 8 | 12604 | 3p0l | A | 182 | ARG | p.Arg182Leu | P49675 |
| HOMO SAPIENS | 2 | 12604 | 3p0l | A | 187 | VAL | p.Val187Met | P49675 |
| HOMO SAPIENS | 2 | 12604 | 3p0l | A | 188 | ARG | p.Arg188Cys | P49675 |
| HOMO SAPIENS | 0 | 12604 | 3p0l | A | 193 | ARG | p.Arg193Ter | P49675 |
| HOMO SAPIENS | 9 | 12604 | 3p0l | A | 217 | ARG | p.Arg217Thr | P49675 |
| HOMO SAPIENS | 4 | 12604 | 3p0l | A | 218 | ALA | p.Ala218Val | P49675 |
| HOMO SAPIENS | 0 | 12604 | 3p0l | A | 250 | TRP | p.Trp250Ter | P49675 |
| HOMO SAPIENS | 1 | 12604 | 3p0l | A | 258 | GLN | p.Gln258Ter | P49675 |
| HOMO SAPIENS | -1 | 27765 | 4wkr | A | 4 | GLU | p.Glu4Lys | Q4G0J3 |
| HOMO SAPIENS | 2 | 3587 | 4kit | C | 2301 | PRO | p.Pro2301Thr | Q6P2Q9 |
| HOMO SAPIENS | 3 | 3587 | 4kit | C | 2304 | PHE | p.Phe2304Leu | Q6P2Q9 |
| HOMO SAPIENS | 3 | 3587 | 4kit | C | 2309 | HIS | p.His2309Pro | Q6P2Q9 |
| HOMO SAPIENS | 3 | 3587 | 4kit | C | 2310 | ARG | p.Arg2310Lys | Q6P2Q9 |
| HOMO SAPIENS | 0 | 9073 | 1n69 | B | 21 | ASN | p.Asn215His | P07602 |
| HOMO SAPIENS | 0 | 9073 | 1n69 | B | 23 | THR | p.Thr217Ile | P07602 |
| HOMO SAPIENS | 0 | 9073 | 1n69 | B | 47 | CYS | p.Cys241Ser | P07602 |
| HOMO SAPIENS | 0 | 475 | 2vd5 | A | 225 | THR | p.Thr225Met | Q09013 |
| HOMO SAPIENS | 1 | 9066 | 4mzv | A | 31 | ASN | p.Asn31Lys | P16422 |
| HOMO SAPIENS | 1 | 9066 | 4mzv | A | 47 | GLN | p.Gln47His | P16422 |
| HOMO SAPIENS | 3 | 9066 | 4mzv | A | 66 | CYS | p.Cys66Tyr | P16422 |
| HOMO SAPIENS | 1 | 9066 | 4mzv | A | 102 | SER | p.Ser102Gly | P16422 |
| HOMO SAPIENS | 5 | 9066 | 4mzv | A | 115 | THR | p.Met115Thr | P16422 |
| HOMO SAPIENS | 1 | 9066 | 4mzv | A | 138 | ARG | p.Arg138Ter | P16422 |
| HOMO SAPIENS | 1 | 9066 | 4mzv | A | 153 | ARG | p.Arg153Thr | P16422 |
| HOMO SAPIENS | 1 | 9066 | 4mzv | A | 165 | ALA | p.Ala165Thr | P16422 |
| HOMO SAPIENS | 8 | 1827 | 2nz2 | A | 14 | GLY | p.Gly14Ser | P00966 |
| HOMO SAPIENS | 5 | 1827 | 2nz2 | A | 18 | SER | p.Ser18Leu | P00966 |
| HOMO SAPIENS | 3 | 1827 | 2nz2 | A | 86 | ARG | p.Arg86Cys | P00966 |
| HOMO SAPIENS | 3 | 1827 | 2nz2 | A | 86 | ARG | p.Arg86His | P00966 |
| HOMO SAPIENS | 1 | 1827 | 2nz2 | A | 179 | TRP | p.Trp179Arg | P00966 |
| HOMO SAPIENS | 7 | 1827 | 2nz2 | A | 180 | SER | p.Ser180Asn | P00966 |
| HOMO SAPIENS | 7 | 1827 | 2nz2 | A | 191 | GLU | p.Glu191Lys | P00966 |
| HOMO SAPIENS | 4 | 1827 | 2nz2 | A | 256 | GLU | p.Glu256Lys | P00966 |
| HOMO SAPIENS | 4 | 1827 | 2nz2 | A | 263 | VAL | p.Val263Met | P00966 |
| HOMO SAPIENS | 5 | 1827 | 2nz2 | A | 265 | ARG | p.Arg265Cys | P00966 |
| HOMO SAPIENS | 5 | 1827 | 2nz2 | A | 268 | ILE | p.Ile268Met | P00966 |
| HOMO SAPIENS | 6 | 1827 | 2nz2 | A | 269 | VAL | p.Val269Met | P00966 |
| HOMO SAPIENS | 7 | 1827 | 2nz2 | A | 272 | ARG | p.Arg272Cys | P00966 |
| HOMO SAPIENS | 3 | 1827 | 2nz2 | A | 304 | ARG | p.Arg304Trp | P00966 |
| HOMO SAPIENS | 3 | 1827 | 2nz2 | A | 307 | ARG | p.Arg307Cys | P00966 |
| HOMO SAPIENS | 3 | 1827 | 2nz2 | A | 310 | LYS | p.Lys310Gln | P00966 |
| HOMO SAPIENS | 2 | 1827 | 2nz2 | A | 362 | GLY | p.Gly362Val | P00966 |
| HOMO SAPIENS | 2 | 1827 | 2nz2 | A | 363 | ARG | p.Arg363Trp | P00966 |
| HOMO SAPIENS | 2 | 1827 | 2nz2 | A | 363 | ARG | p.Arg363Gln | P00966 |
| HOMO SAPIENS | 1 | 1827 | 2nz2 | A | 390 | GLY | p.Gly390Arg | P00966 |
| HOMO SAPIENS | 0 | 29524 | 5vbn | B | 2165 | ARG | p.Arg2165His | Q07864 |
| HOMO SAPIENS | 2 | 29524 | 5vbn | B | 2255 | ILE | p.Ile2255Phe | Q07864 |
| HOMO SAPIENS | 2 | 29524 | 5vbn | B | 2274 | LEU | p.Leu2274Val | Q07864 |
| HOMO SAPIENS | 0 | 31897 | 3t0o | A | 184 | CYS | p.Cys184Arg | O00584 |
| HOMO SAPIENS | -1 | 31897 | 3t0o | A | 215 | GLU | p.Glu215Lys | O00584 |
| HOMO SAPIENS | 0 | 31897 | 3t0o | A | 236 | ARG | p.Arg236Trp | O00584 |

| | | | | | | | | |
|---|---|---|---|---|---|---|---|---|
| HOMO SAPIENS | 7 | 18490 | 4pcw | B | 31 | GLU | p.Glu32Ter | P20930 |
| HOMO SAPIENS | -1 | 39865 | 2h9e | L | 152 | GLY | p.Gly192Arg | P00742 |
| HOMO SAPIENS | 2 | 4309 | 2x0b | F | 174 | THR | p.Thr207Met | P01019 |
| HOMO SAPIENS | 3 | 4309 | 2x0b | F | 342 | ARG | p.Arg375Gln | P01019 |
| HOMO SAPIENS | 2 | 3094 | 1zll | B | 9 | ARG | p.Arg9Cys | P26678 |
| HOMO SAPIENS | 9 | 3094 | 1zll | B | 39 | LEU | p.Leu39Ter | P26678 |
| HOMO SAPIENS | 5 | 3094 | 1zll | B | 49 | VAL | p.Val49Met | P26678 |
| HOMO SAPIENS | -1 | 41498 | 3ffd | P | 8 | LEU | p.Leu44Pro | P12272 |
| HOMO SAPIENS | 1 | 20697 | 1igr | A | 407 | ARG | p.Arg437His | P08069 |
| HOMO SAPIENS | 0 | 20697 | 1igr | A | 416 | MET | p.Met446Val | P08069 |
| HOMO SAPIENS | -1 | 20697 | 1igr | A | 462 | VAL | p.Val492Ile | P08069 |
| HOMO SAPIENS | 0 | 3973 | 4cbz | A | 37 | LEU | p.Leu37Ser | P78504 |
| HOMO SAPIENS | 0 | 3973 | 4cbz | A | 45 | VAL | p.Val45Leu | P78504 |
| HOMO SAPIENS | 0 | 3973 | 4cbz | A | 134 | LEU | p.Leu134Met | P78504 |
| HOMO SAPIENS | -1 | 3973 | 4cbz | A | 146 | VAL | p.Val146Ile | P78504 |
| HOMO SAPIENS | 0 | 3973 | 4cbz | A | 176 | VAL | p.Val176Ile | P78504 |
| HOMO SAPIENS | 0 | 3973 | 4cbz | A | 184 | ARG | p.Arg184Cys | P78504 |
| HOMO SAPIENS | 6 | 1697 | 4hmy | S | 33 | ARG | p.Arg33Trp | Q96PC3 |
| HOMO SAPIENS | 2 | 23752 | 5j4o | A | 13 | ARG | p.Arg1611Ser | P02549 |
| HOMO SAPIENS | 0 | 23752 | 5j4o | A | 42 | ALA | p.Ala1640Gly | P02549 |
| HOMO SAPIENS | 0 | 23752 | 5j4o | A | 108 | HIS | p.His1706Gln | P02549 |
| HOMO SAPIENS | 6 | 23752 | 5j4o | A | 206 | LEU | p.Leu1804Phe | P02549 |
| HOMO SAPIENS | 8 | 531 | 2qz4 | A | 349 | GLY | p.Gly349Ser | Q9UQ90 |
| HOMO SAPIENS | 0 | 531 | 2qz4 | A | 486 | ARG | p.Arg486Gln | Q9UQ90 |
| HOMO SAPIENS | 1 | 531 | 2qz4 | A | 510 | ALA | p.Ala510Val | Q9UQ90 |
| HOMO SAPIENS | 0 | 531 | 2qz4 | A | 529 | ALA | p.Ala529Val | Q9UQ90 |
| HOMO SAPIENS | 0 | 531 | 2qz4 | A | 558 | LYS | p.Lys558Ter | Q9UQ90 |
| HOMO SAPIENS | 1 | 531 | 2qz4 | A | 559 | LYS | p.Lys559Ter | Q9UQ90 |
| HOMO SAPIENS | 2 | 7736 | 1fnh | A | 114 | TRP | p.Trp1834Arg | P02751 |
| HOMO SAPIENS | 0 | 7736 | 1fnh | A | 163 | LEU | p.Leu1883Arg | P02751 |
| HOMO SAPIENS | 3 | 20337 | 5wy5 | B | 264 | LEU | p.Leu264Phe | Q96MG7 |
| HOMO SAPIENS | 8 | 36096 | 2jku | A | 643 | GLY | p.Gly668Arg | P05165 |
| HOMO SAPIENS | 6 | 36096 | 2jku | A | 663 | CYS | p.Cys688Arg | P05165 |
| HOMO SAPIENS | 0 | 9850 | 3kbt | A | 1606 | TYR | p.Tyr1606Ter | P11277 |
| HOMO SAPIENS | 4 | 6876 | 4jkq | A | 217 | ILE | p.Ile217Val | Q92736 |
| HOMO SAPIENS | 5 | 6876 | 4jkq | A | 420 | ARG | p.Arg420Trp | Q92736 |
| HOMO SAPIENS | 2 | 27603 | 1itv | B | 62 | PRO | p.Arg574Pro | P14780 |
| HOMO SAPIENS | 4 | 27603 | 1itv | B | 156 | ARG | p.Arg668Gln | P14780 |
| HOMO SAPIENS | 2 | 22954 | 4f52 | E | 48 | ILE | p.Ile48Met | Q92990 |
| HOMO SAPIENS | 0 | 22954 | 4f52 | E | 91 | ASP | p.Asp91Asn | Q92990 |
| HOMO SAPIENS | 0 | 22954 | 4f52 | E | 146 | ALA | p.Ala146Thr | Q92990 |
| HOMO SAPIENS | 0 | 22954 | 4f52 | E | 181 | ILE | p.Ile181Lys | Q92990 |
| HOMO SAPIENS | 0 | 22954 | 4f52 | E | 194 | ASN | p.Asn194Ser | Q92990 |
| HOMO SAPIENS | 2 | 22954 | 4f52 | E | 254 | PRO | p.Pro254Arg | Q92990 |
| HOMO SAPIENS | 2 | 22954 | 4f52 | E | 300 | ILE | p.Ile300Thr | Q92990 |
| HOMO SAPIENS | 2 | 22954 | 4f52 | E | 336 | LEU | p.Leu336Ser | Q92990 |
| HOMO SAPIENS | 0 | 22954 | 4f52 | E | 353 | TYR | p.Tyr353His | Q92990 |
| HOMO SAPIENS | 0 | 22954 | 4f52 | E | 387 | MET | p.Met387Val | Q92990 |
| HOMO SAPIENS | 4 | 2946 | 5gl7 | A | 344 | TYR | p.Tyr344Cys | P56192 |
| HOMO SAPIENS | 0 | 2946 | 5gl7 | A | 370 | PHE | p.Phe370Leu | P56192 |
| HOMO SAPIENS | 0 | 2946 | 5gl7 | A | 393 | ALA | p.Ala393Thr | P56192 |
| HOMO SAPIENS | 0 | 2946 | 5gl7 | A | 523 | ILE | p.Ile523Thr | P56192 |
| HOMO SAPIENS | 0 | 2946 | 5gl7 | A | 567 | SER | p.Ser567Leu | P56192 |
| HOMO SAPIENS | 0 | 2946 | 5gl7 | A | 800 | PRO | p.Pro800Thr | P56192 |

| | | | | | | | | |
|---|---|---|---|---|---|---|---|---|
| HOMO SAPIENS | 3 | 34961 | 4wy4 | C | 44 | TYR | p.Tyr44His | O95721 |
| HOMO SAPIENS | 8 | 34961 | 4wy4 | C | 89 | GLU | p.Glu89Lys | O95721 |
| HOMO SAPIENS | 0 | 9427 | 4chl | B | 21 | ALA | p.Ala21Ser | O95571 |
| HOMO SAPIENS | 2 | 9427 | 4chl | B | 62 | ALA | p.Ala62Thr | O95571 |
| HOMO SAPIENS | 0 | 9427 | 4chl | B | 114 | ILE | p.Ile114Phe | O95571 |
| HOMO SAPIENS | 2 | 9427 | 4chl | B | 159 | ARG | p.Arg159His | O95571 |
| HOMO SAPIENS | 8 | 9427 | 4chl | B | 163 | ARG | p.Arg163Trp | O95571 |
| HOMO SAPIENS | 8 | 9427 | 4chl | B | 163 | ARG | p.Arg163Gln | O95571 |
| HOMO SAPIENS | 2 | 9427 | 4chl | B | 185 | LEU | p.Leu185Arg | O95571 |
| HOMO SAPIENS | 0 | 40917 | 2e8o | A | 11 | PRO | p.Pro26Leu | Q9Y3Z3 |
| HOMO SAPIENS | -1 | 39190 | 4gow | A | 556 | TYR | p.Tyr556Ter | P56696 |
| HOMO SAPIENS | 0 | 2399 | 2mkv | A | 8 | GLY | p.Gly10Ser | P54710 |
| HOMO SAPIENS | 1 | 2399 | 2mkv | A | 24 | VAL | p.Val26Ile | P54710 |
| HOMO SAPIENS | 4 | 2399 | 2mkv | A | 39 | GLY | p.Gly41Arg | P54710 |
| HOMO SAPIENS | 0 | 20801 | 2zej | B | 1441 | ARG | p.Arg1441Cys | Q5S007 |
| HOMO SAPIENS | 0 | 20801 | 2zej | B | 1441 | ARG | p.Arg1441His | Q5S007 |
| HOMO SAPIENS | 5 | 1019 | 5xtb | M | 62 | ARG | p.Arg62Gln | P28331 |
| HOMO SAPIENS | 1 | 1019 | 5xtb | M | 85 | ALA | p.Ala85Thr | P28331 |
| HOMO SAPIENS | 5 | 1019 | 5xtb | M | 231 | LEU | p.Leu231Val | P28331 |
| HOMO SAPIENS | 2 | 1019 | 5xtb | M | 241 | ARG | p.Arg241Trp | P28331 |
| HOMO SAPIENS | 1 | 1019 | 5xtb | M | 252 | ASP | p.Asp252Gly | P28331 |
| HOMO SAPIENS | 1 | 1019 | 5xtb | M | 408 | ARG | p.Arg408Cys | P28331 |
| HOMO SAPIENS | 2 | 1019 | 5xtb | M | 417 | ARG | p.Arg417Ser | P28331 |
| HOMO SAPIENS | 1 | 1019 | 5xtb | M | 557 | ARG | p.Arg557Ter | P28331 |
| HOMO SAPIENS | 0 | 1019 | 5xtb | M | 567 | ILE | p.Ile567Val | P28331 |
| HOMO SAPIENS | 1 | 1019 | 5xtb | M | 595 | THR | p.Thr595Ala | P28331 |
| HOMO SAPIENS | 1 | 1019 | 5xtb | M | 703 | ALA | p.Ala703Thr | P28331 |
| HOMO SAPIENS | -1 | 3437 | 5hyn | A | 185 | ASP | p.Asp185His | Q15910 |
| HOMO SAPIENS | -1 | 3437 | 5hyn | A | 482 | ALA | p.Ala482Thr | Q15910 |
| HOMO SAPIENS | 0 | 8201 | 2wfd | A | 373 | TYR | p.Tyr373Cys | Q9P2J5 |
| HOMO SAPIENS | 1 | 3590 | 3edi | A | 90 | VAL | p.Val238Ala | O43897 |
| HOMO SAPIENS | 6 | 4555 | 1ytq | A | 64 | ALA | p.Ala65Ser | P43320 |
| HOMO SAPIENS | 0 | 4555 | 1ytq | A | 194 | TRP | p.Trp195Gly | P43320 |
| HOMO SAPIENS | 0 | 1013 | 2nn6 | C | 272 | SER | p.Ser272Thr | Q96B26 |
| HOMO SAPIENS | -1 | 29026 | 5m73 | D | 530 | ILE | p.Ile530Asn | O76094 |
| HOMO SAPIENS | 8 | 2252 | 1d5v | A | 57 | SER | p.Ser125Leu | Q99958 |
| HOMO SAPIENS | 1 | 4337 | 3ecr | A | 29 | ARG | p.Arg26His | P08397 |
| HOMO SAPIENS | 2 | 4337 | 3ecr | A | 34 | ALA | p.Ala31Thr | P08397 |
| HOMO SAPIENS | 2 | 4337 | 3ecr | A | 37 | GLN | p.Gln34Lys | P08397 |
| HOMO SAPIENS | 1 | 4337 | 3ecr | A | 58 | ALA | p.Ala55Ser | P08397 |
| HOMO SAPIENS | 0 | 4337 | 3ecr | A | 84 | LEU | p.Leu81Pro | P08397 |
| HOMO SAPIENS | 0 | 4337 | 3ecr | A | 114 | GLY | p.Gly111Arg | P08397 |
| HOMO SAPIENS | 1 | 4337 | 3ecr | A | 119 | ARG | p.Arg116Trp | P08397 |
| HOMO SAPIENS | 1 | 4337 | 3ecr | A | 152 | ARG | p.Arg149Gln | P08397 |
| HOMO SAPIENS | 1 | 4337 | 3ecr | A | 170 | ARG | p.Arg167Trp | P08397 |
| HOMO SAPIENS | 1 | 4337 | 3ecr | A | 176 | ARG | p.Arg173Gln | P08397 |
| HOMO SAPIENS | 0 | 4337 | 3ecr | A | 180 | LEU | p.Leu177Arg | P08397 |
| HOMO SAPIENS | 0 | 4337 | 3ecr | A | 181 | ASP | p.Asp178Asn | P08397 |
| HOMO SAPIENS | 1 | 4337 | 3ecr | A | 198 | ARG | p.Arg195Cys | P08397 |
| HOMO SAPIENS | 0 | 4337 | 3ecr | A | 201 | TRP | p.Trp198Ter | P08397 |
| HOMO SAPIENS | 1 | 4337 | 3ecr | A | 204 | ARG | p.Arg201Trp | P08397 |
| HOMO SAPIENS | 1 | 4337 | 3ecr | A | 219 | GLY | p.Gly216Asp | P08397 |
| HOMO SAPIENS | 1 | 4337 | 3ecr | A | 226 | GLU | p.Glu223Lys | P08397 |
| HOMO SAPIENS | 0 | 4337 | 3ecr | A | 248 | LEU | p.Leu245Arg | P08397 |

| | | | | | | | | |
|---|---|---|---|---|---|---|---|---|
| HOMO SAPIENS | 1 | 4337 | 3ecr | A | 250 | CYS | p.Cys247Arg | P08397 |
| HOMO SAPIENS | 0 | 4337 | 3ecr | A | 253 | GLU | p.Glu250Lys | P08397 |
| HOMO SAPIENS | 0 | 4337 | 3ecr | A | 255 | ALA | p.Ala252Thr | P08397 |
| HOMO SAPIENS | 0 | 4337 | 3ecr | A | 255 | ALA | p.Ala252Val | P08397 |
| HOMO SAPIENS | 0 | 4337 | 3ecr | A | 259 | HIS | p.His256Asn | P08397 |
| HOMO SAPIENS | 0 | 4337 | 3ecr | A | 324 | ARG | p.Arg321His | P08397 |
| HOMO SAPIENS | -1 | 4337 | 3ecr | A | 362 | ASP | p.Asp359Asn | P08397 |
| HOMO SAPIENS | 1 | 1204 | 5lwv | A | 854 | ILE | p.Ala864Thr | O15294 |
| HOMO SAPIENS | 1 | 25016 | 1m4u | A | 35 | PRO | p.Pro35Arg | Q13253 |
| HOMO SAPIENS | 1 | 25016 | 1m4u | A | 35 | PRO | p.Pro35Ser | Q13253 |
| HOMO SAPIENS | 1 | 25016 | 1m4u | A | 167 | ARG | p.Arg167Gly | Q13253 |
| HOMO SAPIENS | 1 | 25016 | 1m4u | A | 184 | CYS | p.Cys184Tyr | Q13253 |
| HOMO SAPIENS | 0 | 25016 | 1m4u | A | 189 | GLY | p.Gly189Cys | Q13253 |
| HOMO SAPIENS | 1 | 25016 | 1m4u | A | 204 | ARG | p.Arg204Leu | Q13253 |
| HOMO SAPIENS | 1 | 25016 | 1m4u | A | 205 | TRP | p.Trp205Cys | Q13253 |
| HOMO SAPIENS | 1 | 25016 | 1m4u | A | 217 | TRP | p.Trp217Gly | Q13253 |
| HOMO SAPIENS | 0 | 25016 | 1m4u | A | 222 | TYR | p.Tyr222Cys | Q13253 |
| HOMO SAPIENS | 0 | 25016 | 1m4u | A | 222 | TYR | p.Tyr222Asp | Q13253 |
| HOMO SAPIENS | 1 | 25016 | 1m4u | A | 223 | PRO | p.Pro223Leu | Q13253 |
| HOMO SAPIENS | 0 | 1394 | 1um7 | A | 95 | ALA | p.Ala463Val | Q92796 |
| HOMO SAPIENS | 0 | 21294 | 4jpz | B | 1788 | GLU | p.Glu1788Asp | Q99250 |
| HOMO SAPIENS | 9 | 24068 | 2db6 | A | 47 | ARG | p.Arg119Cys | Q96MF2 |
| HOMO SAPIENS | 5 | 1869 | 4omt | A | 39 | ARG | p.Arg39Leu | P08237 |
| HOMO SAPIENS | 2 | 1869 | 4omt | A | 100 | ARG | p.Arg100Gln | P08237 |
| HOMO SAPIENS | 7 | 1869 | 4omt | A | 543 | ASP | p.Asp543Ala | P08237 |
| HOMO SAPIENS | 1 | 1869 | 4omt | A | 686 | TRP | p.Trp686Cys | P08237 |
| HOMO SAPIENS | 1 | 1869 | 4omt | A | 696 | ARG | p.Arg696His | P08237 |
| HOMO SAPIENS | 0 | 12163 | 4wxx | A | 495 | TYR | p.Tyr495Cys | P26358 |
| HOMO SAPIENS | 0 | 12163 | 4wxx | A | 528 | ILE | p.Ile528Met | P26358 |
| HOMO SAPIENS | 0 | 12391 | 1j1d | C | 36 | LYS | p.Lys36Gln | P19429 |
| HOMO SAPIENS | 0 | 12391 | 1j1d | C | 82 | PRO | p.Pro82Ser | P19429 |
| HOMO SAPIENS | 7 | 12391 | 1j1d | C | 102 | ALA | p.Ala102Thr | P19429 |
| HOMO SAPIENS | 1 | 5000 | 4cca | A | 17 | GLY | p.Gly17Arg | Q15833 |
| HOMO SAPIENS | 0 | 5000 | 4cca | A | 205 | VAL | p.Val205Ile | Q15833 |
| HOMO SAPIENS | 0 | 5000 | 4cca | A | 209 | LEU | p.Leu209Pro | Q15833 |
| HOMO SAPIENS | 1 | 5000 | 4cca | A | 459 | ARG | p.Arg459Trp | Q15833 |
| HOMO SAPIENS | 7 | 5000 | 4cca | A | 477 | PRO | p.Pro477Leu | Q15833 |
| HOMO SAPIENS | -1 | 5000 | 4cca | A | 526 | ILE | p.Ile526Val | Q15833 |
| HOMO SAPIENS | 0 | 5000 | 4cca | A | 541 | GLY | p.Gly541Ser | Q15833 |
| HOMO SAPIENS | 0 | 5000 | 4cca | A | 555 | ARG | p.Arg555Gly | Q15833 |
| HOMO SAPIENS | 5 | 28551 | 3vcm | P | 26 | ARG | p.Arg49Ter | P00797 |
| HOMO SAPIENS | 6 | 36453 | 5dzz | A | 2070 | ASP | p.Asp2070His | P15924 |
| HOMO SAPIENS | 0 | 36453 | 5dzz | A | 2166 | ARG | p.Arg2166Ter | P15924 |
| HOMO SAPIENS | 0 | 20268 | 2da6 | A | 96 | ASN | p.Asn321Ser | P35680 |
| HOMO SAPIENS | 1 | 1405 | 3iar | A | 8 | ASP | p.Asp8Asn | P00813 |
| HOMO SAPIENS | 8 | 1405 | 3iar | A | 15 | HIS | p.His15Asp | P00813 |
| HOMO SAPIENS | 6 | 1405 | 3iar | A | 74 | GLY | p.Gly74Val | P00813 |
| HOMO SAPIENS | 6 | 1405 | 3iar | A | 80 | LYS | p.Lys80Arg | P00813 |
| HOMO SAPIENS | 5 | 1405 | 3iar | A | 97 | TYR | p.Tyr97Cys | P00813 |
| HOMO SAPIENS | 5 | 1405 | 3iar | A | 101 | ARG | p.Arg101Leu | P00813 |
| HOMO SAPIENS | 5 | 1405 | 3iar | A | 101 | ARG | p.Arg101Trp | P00813 |
| HOMO SAPIENS | 5 | 1405 | 3iar | A | 106 | LEU | p.Leu106Val | P00813 |
| HOMO SAPIENS | 0 | 1405 | 3iar | A | 107 | LEU | p.Leu107Pro | P00813 |
| HOMO SAPIENS | 9 | 1405 | 3iar | A | 156 | ARG | p.Arg156Cys | P00813 |

| | | | | | | | | |
|---|---|---|---|---|---|---|---|---|
| HOMO SAPIENS | 6 | 1405 | 3iar | A | 211 | ARG | p.Arg211His | P00813 |
| HOMO SAPIENS | 0 | 1405 | 3iar | A | 216 | GLY | p.Gly216Arg | P00813 |
| HOMO SAPIENS | 0 | 1405 | 3iar | A | 291 | SER | p.Ser291Leu | P00813 |
| HOMO SAPIENS | 6 | 1405 | 3iar | A | 304 | LEU | p.Leu304Arg | P00813 |
| HOMO SAPIENS | 0 | 1405 | 3iar | A | 329 | ALA | p.Ala329Val | P00813 |
| HOMO SAPIENS | -1 | 31340 | 4atu | I | 186 | ARG | p.Arg186His | O43602 |
| HOMO SAPIENS | -1 | 31340 | 4atu | I | 192 | ARG | p.Arg192Trp | O43602 |
| HOMO SAPIENS | -1 | 31340 | 4atu | I | 196 | ARG | p.Arg196His | O43602 |
| HOMO SAPIENS | -1 | 31340 | 4atu | I | 203 | THR | p.Thr203Arg | O43602 |
| HOMO SAPIENS | -1 | 31340 | 4atu | I | 286 | ALA | p.Ala286Ser | O43602 |
| HOMO SAPIENS | 4 | 30734 | 1fsu | A | 95 | ARG | p.Arg95Gln | P15848 |
| HOMO SAPIENS | 0 | 30734 | 1fsu | A | 117 | CYS | p.Cys117Arg | P15848 |
| HOMO SAPIENS | 0 | 30734 | 1fsu | A | 137 | GLY | p.Gly137Val | P15848 |
| HOMO SAPIENS | 0 | 30734 | 1fsu | A | 210 | TYR | p.Tyr210Cys | P15848 |
| HOMO SAPIENS | 0 | 30734 | 1fsu | A | 236 | LEU | p.Leu236Pro | P15848 |
| HOMO SAPIENS | 1 | 30734 | 1fsu | A | 315 | ARG | p.Arg315Gln | P15848 |
| HOMO SAPIENS | 0 | 30734 | 1fsu | A | 358 | VAL | p.Val358Met | P15848 |
| HOMO SAPIENS | 2 | 30734 | 1fsu | A | 376 | VAL | p.Val376Met | P15848 |
| HOMO SAPIENS | 0 | 30734 | 1fsu | A | 384 | SER | p.Ser384Asn | P15848 |
| HOMO SAPIENS | 0 | 30734 | 1fsu | A | 393 | HIS | p.His393Pro | P15848 |
| HOMO SAPIENS | 0 | 30734 | 1fsu | A | 405 | CYS | p.Cys405Tyr | P15848 |
| HOMO SAPIENS | 0 | 30734 | 1fsu | A | 458 | ASN | p.Asn458Ser | P15848 |
| HOMO SAPIENS | 1 | 30734 | 1fsu | A | 484 | ARG | p.Arg484Gly | P15848 |
| HOMO SAPIENS | 0 | 38072 | 1wmv | A | 51 | PRO | p.Pro98Leu | Q9NZC7 |
| HOMO SAPIENS | -1 | 5674 | 1j1e | C | 185 | ASN | p.Asn185Lys | P19429 |
| HOMO SAPIENS | -1 | 5674 | 1j1e | C | 192 | ARG | p.Arg192His | P19429 |
| HOMO SAPIENS | -1 | 5674 | 1j1e | C | 196 | ASP | p.Asp196Asn | P19429 |
| HOMO SAPIENS | -1 | 5674 | 1j1e | C | 206 | LYS | p.Lys206Gln | P19429 |
| HOMO SAPIENS | 0 | 38067 | 1wlp | A | 13 | PRO | p.Pro156Gln | P13498 |
| HOMO SAPIENS | -1 | 20441 | 3vfd | A | 289 | HIS | p.His289Tyr | Q9UBP0 |
| HOMO SAPIENS | 2 | 20441 | 3vfd | A | 344 | ILE | p.Ile344Lys | Q9UBP0 |
| HOMO SAPIENS | 0 | 20441 | 3vfd | A | 362 | SER | p.Ser362Cys | Q9UBP0 |
| HOMO SAPIENS | 1 | 20441 | 3vfd | A | 386 | ASN | p.Asn386Ser | Q9UBP0 |
| HOMO SAPIENS | 9 | 20441 | 3vfd | A | 441 | ASP | p.Asp441Gly | Q9UBP0 |
| HOMO SAPIENS | -1 | 20441 | 3vfd | A | 448 | CYS | p.Cys448Tyr | Q9UBP0 |
| HOMO SAPIENS | 2 | 20441 | 3vfd | A | 500 | ARG | p.Arg499Cys | Q9UBP0 |
| HOMO SAPIENS | 1 | 20441 | 3vfd | A | 563 | ARG | p.Arg562Gly | Q9UBP0 |
| HOMO SAPIENS | 3 | 18471 | 4dx8 | J | 26 | ARG | p.Arg26Gln | O00522 |
| HOMO SAPIENS | -1 | 18471 | 4dx8 | J | 101 | GLY | p.Gly101Val | O00522 |
| HOMO SAPIENS | -1 | 34735 | 2ly4 | B | 83 | ALA | p.Ala83Glu | P04637 |
| HOMO SAPIENS | 0 | 31128 | 4cvh | A | 116 | ARG | p.Arg116Cys | A4D126 |
| HOMO SAPIENS | 8 | 31128 | 4cvh | A | 215 | GLN | p.Gln215Ter | A4D126 |
| HOMO SAPIENS | 0 | 31128 | 4cvh | A | 268 | ARG | p.Arg268Ter | A4D126 |
| HOMO SAPIENS | 0 | 31128 | 4cvh | A | 316 | THR | p.Thr316Lys | A4D126 |
| HOMO SAPIENS | 1 | 7164 | 3mt5 | A | 369 | ASP | p.Asp434Gly | Q12791 |
| HOMO SAPIENS | 2 | 26812 | 2wtk | F | 49 | TYR | p.Tyr49Asp | Q15831 |
| HOMO SAPIENS | 0 | 26812 | 2wtk | F | 67 | LEU | p.Leu67Pro | Q15831 |
| HOMO SAPIENS | 4 | 26812 | 2wtk | F | 104 | ARG | p.Arg104Ser | Q15831 |
| HOMO SAPIENS | 0 | 26812 | 2wtk | F | 135 | GLY | p.Gly135Arg | Q15831 |
| HOMO SAPIENS | 0 | 26812 | 2wtk | F | 163 | GLY | p.Gly163Asp | Q15831 |
| HOMO SAPIENS | 2 | 26812 | 2wtk | F | 194 | ALA | p.Asp194Asn | Q15831 |
| HOMO SAPIENS | 2 | 1357 | 5xtd | W | 57 | ARG | p.Arg57His | Q9P0J0 |
| HOMO SAPIENS | 0 | 28086 | 4nn2 | A | 229 | HIS | p.His229Arg | Q8IWS0 |
| HOMO SAPIENS | 0 | 28086 | 4nn2 | A | 234 | LYS | p.Lys234Glu | Q8IWS0 |

| | | | | | | | | |
|---|---|---|---|---|---|---|---|---|
| HOMO SAPIENS | 0 | 28086 | 4nn2 | A | 257 | ARG | p.Arg257Gly | Q8IWS0 |
| HOMO SAPIENS | 5 | 31182 | 2a1x | A | 176 | GLN | p.Gln176Lys | O14832 |
| HOMO SAPIENS | 1 | 31182 | 2a1x | A | 204 | GLY | p.Gly204Ser | O14832 |
| HOMO SAPIENS | 0 | 31182 | 2a1x | A | 269 | ASN | p.Asn269His | O14832 |
| HOMO SAPIENS | 3 | 31182 | 2a1x | A | 275 | ARG | p.Arg275Gln | O14832 |
| HOMO SAPIENS | 3 | 31182 | 2a1x | A | 275 | ARG | p.Arg275Trp | O14832 |
| HOMO SAPIENS | 4 | 2823 | 5xtb | P | 145 | THR | p.Thr145Ile | O75489 |
| HOMO SAPIENS | 5 | 2823 | 5xtb | P | 199 | ARG | p.Arg199Trp | O75489 |
| HOMO SAPIENS | 0 | 8645 | 2h5g | A | 372 | SER | p.Ser372Tyr | P54886 |
| HOMO SAPIENS | 3 | 8645 | 2h5g | A | 715 | ASP | p.Asp715His | P54886 |
| HOMO SAPIENS | 0 | 8645 | 2h5g | A | 765 | ARG | p.Arg765Gln | P54886 |
| HOMO SAPIENS | 0 | 8645 | 2h5g | A | 782 | TYR | p.Tyr782Cys | P54886 |
| HOMO SAPIENS | 0 | 8645 | 2h5g | A | 784 | HIS | p.His784Tyr | P54886 |
| HOMO SAPIENS | 1 | 13992 | 2cue | A | 39 | ARG | p.Arg242Thr | P26367 |
| HOMO SAPIENS | 9 | 13992 | 2cue | A | 55 | PHE | p.Phe258Ser | P26367 |
| HOMO SAPIENS | 0 | 30418 | 2xsw | A | 303 | VAL | p.Val303Met | Q9NRR6 |
| HOMO SAPIENS | 0 | 30418 | 2xsw | A | 315 | PRO | p.Pro315Leu | Q9NRR6 |
| HOMO SAPIENS | 0 | 30418 | 2xsw | A | 341 | GLY | p.Gly341Ser | Q9NRR6 |
| HOMO SAPIENS | 0 | 30418 | 2xsw | A | 378 | ARG | p.Arg378Cys | Q9NRR6 |
| HOMO SAPIENS | 0 | 30418 | 2xsw | A | 435 | ARG | p.Arg435Gln | Q9NRR6 |
| HOMO SAPIENS | 0 | 30418 | 2xsw | A | 515 | ARG | p.Arg515Trp | Q9NRR6 |
| HOMO SAPIENS | 1 | 30418 | 2xsw | A | 526 | PRO | p.Pro526Leu | Q9NRR6 |
| HOMO SAPIENS | 0 | 30418 | 2xsw | A | 563 | ARG | p.Arg563His | Q9NRR6 |
| HOMO SAPIENS | 2 | 30418 | 2xsw | A | 585 | ARG | p.Arg585His | Q9NRR6 |
| HOMO SAPIENS | 2 | 11167 | 4y99 | B | 205 | ARG | p.Arg215Leu | P45379 |
| HOMO SAPIENS | 5 | 11167 | 4y99 | B | 244 | GLU | p.Glu254Asp | P45379 |
| HOMO SAPIENS | 1 | 11167 | 4y99 | B | 253 | LYS | p.Lys263Arg | P45379 |
| HOMO SAPIENS | 8 | 11167 | 4y99 | B | 265 | ARG | p.Arg275Gln | P45379 |
| HOMO SAPIENS | 3 | 11167 | 4y99 | B | 270 | ASP | p.Asp280Asn | P45379 |
| HOMO SAPIENS | -1 | 11167 | 4y99 | B | 278 | ARG | p.Arg288Pro | P45379 |
| HOMO SAPIENS | -1 | 11167 | 4y99 | B | 286 | ARG | p.Arg296Cys | P45379 |
| HOMO SAPIENS | -1 | 11167 | 4y99 | B | 286 | ARG | p.Arg296His | P45379 |
| HOMO SAPIENS | 0 | 15993 | 4po6 | A | 197 | ARG | p.Arg197His | P29597 |
| HOMO SAPIENS | -1 | 15993 | 4po6 | A | 362 | VAL | p.Val362Phe | P29597 |
| HOMO SAPIENS | -1 | 15993 | 4po6 | A | 363 | GLY | p.Gly363Ser | P29597 |
| HOMO SAPIENS | 0 | 5646 | 5uak | A | 7 | GLU | p.Glu7Ter | P13569 |
| HOMO SAPIENS | 0 | 5646 | 5uak | A | 27 | GLY | p.Gly27Ter | P13569 |
| HOMO SAPIENS | 0 | 5646 | 5uak | A | 46 | ALA | p.Ala46Asp | P13569 |
| HOMO SAPIENS | 0 | 5646 | 5uak | A | 60 | GLU | p.Glu60Ter | P13569 |
| HOMO SAPIENS | 0 | 5646 | 5uak | A | 67 | PRO | p.Pro67Leu | P13569 |
| HOMO SAPIENS | 0 | 5646 | 5uak | A | 74 | ARG | p.Arg74Trp | P13569 |
| HOMO SAPIENS | 0 | 5646 | 5uak | A | 75 | ARG | p.Arg75Ter | P13569 |
| HOMO SAPIENS | 0 | 5646 | 5uak | A | 85 | GLY | p.Gly85Glu | P13569 |
| HOMO SAPIENS | 0 | 5646 | 5uak | A | 91 | GLY | p.Gly91Arg | P13569 |
| HOMO SAPIENS | 0 | 5646 | 5uak | A | 92 | GLU | p.Glu92Lys | P13569 |
| HOMO SAPIENS | 0 | 5646 | 5uak | A | 98 | GLN | p.Gln98Arg | P13569 |
| HOMO SAPIENS | 0 | 5646 | 5uak | A | 98 | GLN | p.Gln98Ter | P13569 |
| HOMO SAPIENS | 0 | 5646 | 5uak | A | 109 | TYR | p.Tyr109Cys | P13569 |
| HOMO SAPIENS | 0 | 5646 | 5uak | A | 110 | ASP | p.Asp110His | P13569 |
| HOMO SAPIENS | 0 | 5646 | 5uak | A | 117 | ARG | p.Arg117Cys | P13569 |
| HOMO SAPIENS | 0 | 5646 | 5uak | A | 117 | ARG | p.Arg117His | P13569 |
| HOMO SAPIENS | 0 | 5646 | 5uak | A | 125 | ILE | p.Ile125Thr | P13569 |
| HOMO SAPIENS | 0 | 5646 | 5uak | A | 148 | ILE | p.Ile148Thr | P13569 |
| HOMO SAPIENS | 0 | 5646 | 5uak | A | 170 | ARG | p.Arg170His | P13569 |

| | | | | | | | | |
|---|---|---|---|---|---|---|---|---|
| HOMO SAPIENS | 0 | 5646 | 5uak | A | 178 | GLY | p.Gly178Arg | P13569 |
| HOMO SAPIENS | 0 | 5646 | 5uak | A | 206 | LEU | p.Leu206Trp | P13569 |
| HOMO SAPIENS | 0 | 5646 | 5uak | A | 373 | ASP | p.Asp373Asn | P13569 |
| HOMO SAPIENS | -1 | 5646 | 5uak | A | 709 | ARG | p.Arg709Ter | P13569 |
| HOMO SAPIENS | -1 | 5646 | 5uak | A | 732 | LEU | p.Leu732Ter | P13569 |
| HOMO SAPIENS | -1 | 5646 | 5uak | A | 785 | ARG | p.Arg785Ter | P13569 |
| HOMO SAPIENS | -1 | 5646 | 5uak | A | 831 | GLU | p.Glu831Ter | P13569 |
| HOMO SAPIENS | -1 | 5646 | 5uak | A | 836 | ASP | p.Asp836Tyr | P13569 |
| HOMO SAPIENS | 0 | 5646 | 5uak | A | 851 | ARG | p.Arg851Ter | P13569 |
| HOMO SAPIENS | 0 | 5646 | 5uak | A | 913 | TYR | p.Tyr913Cys | P13569 |
| HOMO SAPIENS | 0 | 5646 | 5uak | A | 913 | TYR | p.Tyr913Ter | P13569 |
| HOMO SAPIENS | 0 | 5646 | 5uak | A | 945 | SER | p.Ser945Leu | P13569 |
| HOMO SAPIENS | 0 | 5646 | 5uak | A | 949 | HIS | p.His949Tyr | P13569 |
| HOMO SAPIENS | 0 | 5646 | 5uak | A | 970 | GLY | p.Gly970Asp | P13569 |
| HOMO SAPIENS | 0 | 5646 | 5uak | A | 1027 | ILE | p.Ile1027Thr | P13569 |
| HOMO SAPIENS | 0 | 5646 | 5uak | A | 1032 | TYR | p.Tyr1032Cys | P13569 |
| HOMO SAPIENS | 0 | 5646 | 5uak | A | 1061 | GLY | p.Gly1061Arg | P13569 |
| HOMO SAPIENS | 1 | 5646 | 5uak | A | 1065 | LEU | p.Leu1065Pro | P13569 |
| HOMO SAPIENS | 1 | 5646 | 5uak | A | 1066 | ARG | p.Arg1066Cys | P13569 |
| HOMO SAPIENS | 1 | 5646 | 5uak | A | 1066 | ARG | p.Arg1066His | P13569 |
| HOMO SAPIENS | 1 | 5646 | 5uak | A | 1067 | ALA | p.Ala1067Thr | P13569 |
| HOMO SAPIENS | 0 | 5646 | 5uak | A | 1069 | GLY | p.Gly1069Arg | P13569 |
| HOMO SAPIENS | 0 | 5646 | 5uak | A | 1070 | ARG | p.Arg1070Trp | P13569 |
| HOMO SAPIENS | 1 | 5646 | 5uak | A | 1071 | GLN | p.Gln1071Pro | P13569 |
| HOMO SAPIENS | 0 | 5646 | 5uak | A | 1074 | PHE | p.Phe1074Leu | P13569 |
| HOMO SAPIENS | 0 | 5646 | 5uak | A | 1077 | LEU | p.Leu1077Pro | P13569 |
| HOMO SAPIENS | 0 | 5646 | 5uak | A | 1089 | TRP | p.Trp1089Ter | P13569 |
| HOMO SAPIENS | 0 | 5646 | 5uak | A | 1092 | TYR | p.Tyr1092Ter | P13569 |
| HOMO SAPIENS | 0 | 5646 | 5uak | A | 1101 | MET | p.Met1101Lys | P13569 |
| HOMO SAPIENS | 0 | 5646 | 5uak | A | 1156 | LEU | p.Leu1156Phe | P13569 |
| HOMO SAPIENS | 0 | 5646 | 5uak | A | 1158 | ARG | p.Arg1158Ter | P13569 |
| HOMO SAPIENS | 0 | 5646 | 5uak | A | 1162 | ARG | p.Arg1162Leu | P13569 |
| HOMO SAPIENS | 0 | 5646 | 5uak | A | 1162 | ARG | p.Arg1162Ter | P13569 |
| HOMO SAPIENS | -1 | 5646 | 5uak | A | 1476 | GLN | p.Gln1476Ter | P13569 |
| HOMO SAPIENS | 0 | 22763 | 1imv | A | 52 | MET | p.Thr72Met | P36955 |
| HOMO SAPIENS | 0 | 22763 | 1imv | A | 301 | TYR | p.Tyr321Ter | P36955 |
| HOMO SAPIENS | 0 | 23179 | 5fql | A | 160 | PRO | p.Pro160Arg | P22304 |
| HOMO SAPIENS | 0 | 23179 | 5fql | A | 172 | ARG | p.Arg172Ter | P22304 |
| HOMO SAPIENS | 0 | 23179 | 5fql | A | 214 | THR | p.Thr214Met | P22304 |
| HOMO SAPIENS | 0 | 23179 | 5fql | A | 309 | THR | p.Thr309Ala | P22304 |
| HOMO SAPIENS | 0 | 23179 | 5fql | A | 333 | SER | p.Ser333Leu | P22304 |
| HOMO SAPIENS | 0 | 23179 | 5fql | A | 443 | ARG | p.Arg443Ter | P22304 |
| HOMO SAPIENS | 0 | 23179 | 5fql | A | 468 | ARG | p.Arg468Leu | P22304 |
| HOMO SAPIENS | 0 | 23179 | 5fql | A | 468 | ARG | p.Arg468Trp | P22304 |
| HOMO SAPIENS | 0 | 23179 | 5fql | A | 502 | TRP | p.Trp502Ser | P22304 |
| HOMO SAPIENS | 9 | 20989 | 1nfi | F | 185 | THR | p.Thr185Met | P25963 |
| HOMO SAPIENS | 4 | 20989 | 1nfi | F | 194 | GLY | p.Gly194Ala | P25963 |
| HOMO SAPIENS | 0 | 11538 | 3eab | A | 162 | VAL | p.Val162Ile | Q9UBP0 |
| HOMO SAPIENS | 0 | 33399 | 2p39 | A | 71 | SER | p.Ser71Gly | Q9GZV9 |
| HOMO SAPIENS | 3 | 33399 | 2p39 | A | 96 | MET | p.Met96Thr | Q9GZV9 |
| HOMO SAPIENS | 0 | 33399 | 2p39 | A | 129 | SER | p.Ser129Phe | Q9GZV9 |
| HOMO SAPIENS | -1 | 33399 | 2p39 | A | 172 | PRO | p.Pro172Leu | Q9GZV9 |
| HOMO SAPIENS | -1 | 33399 | 2p39 | A | 176 | ARG | p.Arg176Gln | Q9GZV9 |
| HOMO SAPIENS | -1 | 33399 | 2p39 | A | 179 | ARG | p.Arg179Trp | Q9GZV9 |

| | | | | | | | | |
|---|---|---|---|---|---|---|---|---|
| HOMO SAPIENS | -1 | 33399 | 2p39 | A | 179 | ARG | p.Arg179Gln | Q9GZV9 |
| HOMO SAPIENS | 0 | 16217 | 2dne | A | 79 | ILE | p.Ile163Leu | P10515 |
| HOMO SAPIENS | 8 | 7666 | 4kbq | A | 65 | ASN | p.Asn65Ser | Q9UNE7 |
| HOMO SAPIENS | 1 | 6407 | 1ikt | A | 71 | THR | p.Thr687Ile | P51659 |
| HOMO SAPIENS | 0 | 6407 | 1ikt | A | 112 | MET | p.Met728Val | P51659 |
| HOMO SAPIENS | 0 | 5849 | 4ihb | A | 67 | VAL | p.Val67Asp | O75923 |
| HOMO SAPIENS | 6 | 17703 | 5fcn | A | 20 | HIS | p.His99Gln | Q66GS9 |
| HOMO SAPIENS | 8 | 17703 | 5fcn | A | 33 | ARG | p.Arg112His | Q66GS9 |
| HOMO SAPIENS | 0 | 34095 | 2E+073 | A | 101 | HIS | p.His101Tyr | P05129 |
| HOMO SAPIENS | -1 | 41358 | 4liq | E | 507 | THR | p.Thr507Met | P07333 |
| HOMO SAPIENS | -1 | 21290 | 1fch | A | 235 | MET | p.Met272Arg | P50542 |
| HOMO SAPIENS | 5 | 23431 | 6pax | A | 23 | ARG | p.Arg26Gly | P26367 |
| HOMO SAPIENS | 1 | 23431 | 6pax | A | 61 | GLY | p.Gly64Val | P26367 |
| HOMO SAPIENS | 1 | 23431 | 6pax | A | 65 | PRO | p.Pro68Ser | P26367 |
| HOMO SAPIENS | 1 | 23431 | 6pax | A | 116 | SER | p.Ser119Arg | P26367 |
| HOMO SAPIENS | 9 | 23431 | 6pax | A | 123 | VAL | p.Val126Asp | P26367 |
| HOMO SAPIENS | 9 | 23431 | 6pax | A | 125 | ARG | p.Arg128Cys | P26367 |
| HOMO SAPIENS | 9 | 33003 | 4zrj | B | 535 | LEU | p.Leu535Pro | P35240 |
| HOMO SAPIENS | 7 | 33003 | 4zrj | B | 538 | GLN | p.Gln538Pro | P35240 |
| HOMO SAPIENS | 0 | 33003 | 4zrj | B | 585 | TRP | p.Ala585Thr | P35240 |
| HOMO SAPIENS | 0 | 3800 | 2nzj | A | 193 | VAL | p.Val193Met | O75628 |

| gene_name | max_cons | max_bs_id | max_pdb_id | max_chain_id |
|---|---|---|---|---|
| AKR1D1 | None | None | None | None |
| AKR1D1 | None | None | None | None |
| AKR1D1 | None | None | None | None |
| AKR1D1 | None | None | None | None |
| ALPP | None | None | None | None |
| BMPR1A | None | None | None | None |
| BMPR1A | None | None | None | None |
| FGFR2 | None | None | None | None |
| PROC | None | None | None | None |
| PROC | None | None | None | None |
| PROC | None | None | None | None |
| PROC | None | None | None | None |
| PROC | None | None | None | None |
| PROC | None | None | None | None |
| PROC | None | None | None | None |
| PROC | None | None | None | None |
| PROC | None | None | None | None |
| PROC | None | None | None | None |
| PROC | None | None | None | None |
| PROC | None | None | None | None |
| INS | 1 | 2 | 3zs2 | J |
| INS | 0.85 | 1 | 4ajx | D |
| INS | 1 | 1 | 3zs2 | J |
| INS | 1 | 1 | 3zs2 | J |
| FTL | None | None | None | None |
| FTL | None | None | None | None |
| FTL | None | None | None | None |
| TNFSF11 | None | None | None | None |
| EHMT1 | None | None | None | None |
| EHMT1 | None | None | None | None |
| EHMT1 | None | None | None | None |
| CA2 | 0.939 | 1 | 4idr | X |
| CA2 | 0.939 | 1 | 4idr | X |
| CA2 | 0.902 | 5 | 4idr | X |
| CA2 | 0.902 | 5 | 4idr | X |
| SMAD4 | None | None | None | None |
| SMAD4 | None | None | None | None |
| TTR | None | None | None | None |
| TTR | 0.961 | 7 | 3grb | A |
| TTR | 0.891 | 4 | 3dk2 | B |
| TTR | 0.947 | 3 | 1tta | A |
| TTR | 0.923 | 1 | 3grb | B |
| TTR | 0.891 | 1 | 4y9b | B |
| TTR | 0.891 | 1 | 4y9b | B |
| TTR | 0.882 | 15 | 3grb | A |
| TTR | 0.909 | 5 | 4tl4 | A |
| TTR | 0.836 | 7 | 1z7j | B |
| TTR | 0.882 | 15 | 3grb | A |
| TTR | 0.891 | 5 | 3i9i | B |
| TTR | 0.902 | 9 | 3m1o | B |
| TTR | 0.902 | 9 | 3m1o | B |
| TTR | 0.902 | 9 | 3m1o | B |
| TTR | None | None | None | None |

| | | | | |
|---|---|---|---|---|
| TTR | 0.902 | 9 | 3m1o | B |
| TTR | 0.836 | 7 | 1bze | A |
| TTR | 0.846 | 16 | 3gps | A |
| TTR | 0.846 | 16 | 3gps | A |
| TTR | 0.909 | 5 | 4tl4 | A |
| TTR | 0.909 | 5 | 4tl4 | A |
| TTR | None | None | None | None |
| TTR | 0.945 | 4 | 4deu | A |
| TTR | 0.941 | 3 | 3did | B |
| TTR | 0.941 | 3 | 3did | B |
| TTR | 0.868 | 6 | 4l1t | A |
| TTR | 0.882 | 19 | 3gps | B |
| TTR | 0.891 | 4 | 3dk2 | B |
| TTR | 0.941 | 3 | 3did | B |
| TTR | 0.923 | 1 | 4hiq | A |
| TTR | 0.923 | 1 | 4hiq | A |
| TTR | 0.961 | 7 | 3grb | A |
| TTR | 0.839 | 7 | 1tsh | B |
| TTR | 0.891 | 6 | 1ttb | B |
| ALK | None | None | None | None |
| ALK | None | None | None | None |
| THRB | None | None | None | None |
| THRB | None | None | None | None |
| THRB | None | None | None | None |
| THRB | None | None | None | None |
| THRB | None | None | None | None |
| THRB | None | None | None | None |
| THRB | None | None | None | None |
| THRB | None | None | None | None |
| THRB | None | None | None | None |
| THRB | None | None | None | None |
| THRB | None | None | None | None |
| THRB | None | None | None | None |
| THRB | None | None | None | None |
| THRB | None | None | None | None |
| THRB | None | None | None | None |
| THRB | None | None | None | None |
| THRB | None | None | None | None |
| THRB | None | None | None | None |
| THRB | None | None | None | None |
| IL2RG | None | None | None | None |
| IL2RG | None | None | None | None |
| IL2RG | None | None | None | None |
| IL2RG | None | None | None | None |
| IL2RG | None | None | None | None |
| ITGB3 | None | None | None | None |
| ITGB3 | None | None | None | None |
| ITGB3 | None | None | None | None |
| ITGB3 | None | None | None | None |
| ITGB3 | None | None | None | None |
| ITGB3 | None | None | None | None |
| ITGB3 | None | None | None | None |
| ITGB3 | None | None | None | None |
| ITGB3 | None | None | None | None |

| | | | | |
|---|---|---|---|---|
| PKD2 | None | None | None | None |
| PKD2 | None | None | None | None |
| PKD2 | None | None | None | None |
| PKD2 | None | None | None | None |
| PKD2 | None | None | None | None |
| PKD2 | None | None | None | None |
| PKD2 | None | None | None | None |
| TP53 | None | None | None | None |
| TP53 | None | None | None | None |
| TP53 | None | None | None | None |
| TP53 | None | None | None | None |
| TP53 | None | None | None | None |
| TP53 | None | None | None | None |
| EGFR | None | None | None | None |
| EGFR | None | None | None | None |
| XRCC4 | None | None | None | None |
| XRCC4 | None | None | None | None |
| HBB | None | None | None | None |
| HBB | None | None | None | None |
| HBB | None | None | None | None |
| HBB | None | None | None | None |
| HBB | None | None | None | None |
| HBB | None | None | None | None |
| HBB | None | None | None | None |
| HBB | None | None | None | None |
| HBB | None | None | None | None |
| HBB | None | None | None | None |
| HBB | None | None | None | None |
| HBB | None | None | None | None |
| HBB | None | None | None | None |
| HBB | None | None | None | None |
| HBB | None | None | None | None |
| HBB | None | None | None | None |
| HBB | None | None | None | None |
| HBB | None | None | None | None |
| HBB | None | None | None | None |
| HBB | None | None | None | None |
| HBB | None | None | None | None |
| HBB | None | None | None | None |
| HBB | None | None | None | None |
| HBB | None | None | None | None |
| HBB | None | None | None | None |
| HBB | None | None | None | None |
| HBB | None | None | None | None |
| HBB | None | None | None | None |
| HBB | None | None | None | None |
| HBB | None | None | None | None |
| RPS6KA3 | None | None | None | None |
| RPS6KA3 | None | None | None | None |
| RPS6KA3 | None | None | None | None |
| RPS6KA3 | None | None | None | None |
| RPS6KA3 | None | None | None | None |
| RPS6KA3 | None | None | None | None |
| GBA | 1 | 15 | 3gxd | B |

| | | | | | |
|---|---|---|---|---|---|
| GBA | | 1 | 55 | 3gxd | B |
| GBA | | 1 | 15 | 3gxd | B |
| GBA | None | None | | None | None |
| GBA | None | None | | None | None |
| GBA | None | None | | None | None |
| GBA | | 1 | 12 | 1ogs | A |
| GBA | None | None | | None | None |
| GBA | | 1 | 2 | 1ogs | A |
| GBA | | 1 | 11 | 1ogs | A |
| GBA | None | None | | None | None |
| GBA | None | None | | None | None |
| GBA | | 1 | 18 | 1ogs | A |
| GBA | | 1 | 3 | 1ogs | A |
| GBA | None | None | | None | None |
| GBA | | 1 | 2 | 1ogs | A |
| GBA | None | None | | None | None |
| GBA | None | None | | None | None |
| GBA | | 1 | 3 | 1ogs | A |
| GBA | | 1 | 3 | 1ogs | A |
| GBA | None | None | | None | None |
| GBA | | 1 | 28 | 1ogs | A |
| GBA | None | None | | None | None |
| GBA | None | None | | None | None |
| GBA | None | None | | None | None |
| GBA | None | None | | None | None |
| GBA | None | None | | None | None |
| GBA | None | None | | None | None |
| GBA | None | None | | None | None |
| GBA | None | None | | None | None |
| GBA | None | None | | None | None |
| GBA | None | None | | None | None |
| GBA | None | None | | None | None |
| GBA | None | None | | None | None |
| GBA | None | None | | None | None |
| GBA | None | None | | None | None |
| GBA | None | None | | None | None |
| GBA | None | None | | None | None |
| GBA | None | None | | None | None |
| GBA | | 1 | 15 | 3gxd | B |
| GBA | | 1 | 44 | 3gxd | B |
| GBA | None | None | | None | None |
| GBA | | 1 | 44 | 3gxd | B |
| GBA | | 1 | 44 | 3gxd | B |
| GBA | None | None | | None | None |
| GBA | None | None | | None | None |
| GBA | None | None | | None | None |
| IFIH1 | None | None | | None | None |
| TGFBR2 | None | None | | None | None |
| TGFBR2 | None | None | | None | None |
| TGFBR2 | None | None | | None | None |
| TGFBR2 | None | None | | None | None |
| TGFBR2 | None | None | | None | None |
| TGFBR2 | None | None | | None | None |
| TGFBR2 | None | None | | None | None |

| | | | | |
|---|---|---|---|---|
| TGFBR2 | None | None | None | None |
| TGFBR2 | None | None | None | None |
| TGFBR2 | None | None | None | None |
| TGFBR2 | None | None | None | None |
| TGFBR2 | None | None | None | None |
| TGFBR2 | None | None | None | None |
| TGFBR2 | None | None | None | None |
| TGFBR2 | None | None | None | None |
| ACE | None | None | None | None |
| ACE | None | None | None | None |
| ACE | None | None | None | None |
| ACE | None | None | None | None |
| ACE | None | None | None | None |
| ACE | None | None | None | None |
| COL10A1 | None | None | None | None |
| COL10A1 | None | None | None | None |
| COL10A1 | None | None | None | None |
| COL10A1 | None | None | None | None |
| COL10A1 | None | None | None | None |
| COL10A1 | None | None | None | None |
| COL10A1 | None | None | None | None |
| COL10A1 | None | None | None | None |
| COL10A1 | None | None | None | None |
| KLF1 | None | None | None | None |
| C3 | None | None | None | None |
| CTSC | None | None | None | None |
| CTSC | 0.934 | 1 | 2djg | C |
| CTSC | None | None | None | None |
| LYZ | 0.877 | 3 | 1bb5 | A |
| LYZ | 0.923 | 1 | 1lz4 | A |
| LYZ | 0.852 | 2 | 1ubz | A |
| DHODH | None | None | None | None |
| DHODH | None | None | None | None |
| DHODH | None | None | None | None |
| DHODH | None | None | None | None |
| DHODH | None | None | None | None |
| DHODH | None | None | None | None |
| DHODH | None | None | None | None |
| ADH1C | None | None | None | None |
| GDNF | None | None | None | None |
| TNF | None | None | None | None |
| ALDH4A1 | None | None | None | None |
| ALDH4A1 | None | None | None | None |
| ALDH4A1 | None | None | None | None |
| BRAF | None | None | None | None |
| BRAF | None | None | None | None |
| BRAF | None | None | None | None |
| BRAF | None | None | None | None |
| BRAF | None | None | None | None |
| BRAF | None | None | None | None |
| BRAF | None | None | None | None |
| BRAF | None | None | None | None |
| BRAF | None | None | None | None |
| BRAF | None | None | None | None |

| | | | | |
|---|---|---|---|---|
| BRAF | None | None | None | None |
| BRAF | None | None | None | None |
| BRAF | None | None | None | None |
| BRAF | None | None | None | None |
| BRAF | None | None | None | None |
| BRAF | None | None | None | None |
| TBX5 | None | None | None | None |
| TBX5 | None | None | None | None |
| TBX5 | None | None | None | None |
| TBX5 | None | None | None | None |
| TBX5 | None | None | None | None |
| TBX5 | None | None | None | None |
| TBX5 | None | None | None | None |
| TBX5 | None | None | None | None |
| TBX5 | None | None | None | None |
| RB1 | None | None | None | None |
| RB1 | None | None | None | None |
| RB1 | None | None | None | None |
| RB1 | None | None | None | None |
| RB1 | None | None | None | None |
| RB1 | None | None | None | None |
| CLIC2 | None | None | None | None |
| CLIC2 | None | None | None | None |
| ALDH7A1 | None | None | None | None |
| ALDH7A1 | None | None | None | None |
| ALDH7A1 | None | None | None | None |
| ALDH7A1 | None | None | None | None |
| ALDH7A1 | None | None | None | None |
| ALDH7A1 | None | None | None | None |
| ALDH7A1 | None | None | None | None |
| ALDH7A1 | None | None | None | None |
| ALDH7A1 | None | None | None | None |
| ALDH7A1 | None | None | None | None |
| ALDH7A1 | None | None | None | None |
| ALDH7A1 | None | None | None | None |
| ALDH7A1 | None | None | None | None |
| ALDH7A1 | None | None | None | None |
| ALDH7A1 | None | None | None | None |
| ALDH7A1 | None | None | None | None |
| KDR | None | None | None | None |
| MUT | None | None | None | None |
| MUT | None | None | None | None |
| MUT | None | None | None | None |
| MUT | None | None | None | None |
| MUT | None | None | None | None |
| MUT | None | None | None | None |
| MUT | None | None | None | None |
| MUT | None | None | None | None |
| MUT | None | None | None | None |
| MUT | None | None | None | None |
| MUT | None | None | None | None |
| MUT | None | None | None | None |
| MUT | None | None | None | None |
| MUT | None | None | None | None |

| | | | | |
|---|---|---|---|---|
| MUT | None | None | None | None |
| MUT | None | None | None | None |
| MUT | None | None | None | None |
| MUT | None | None | None | None |
| MUT | None | None | None | None |
| MUT | None | None | None | None |
| MUT | None | None | None | None |
| MUT | None | None | None | None |
| MUT | None | None | None | None |
| MUT | None | None | None | None |
| MUT | None | None | None | None |
| MUT | None | None | None | None |
| MUT | None | None | None | None |
| MUT | None | None | None | None |
| MUT | None | None | None | None |
| MUT | None | None | None | None |
| MUT | None | None | None | None |
| MUT | None | None | None | None |
| MUT | None | None | None | None |
| MUT | None | None | None | None |
| SOD1 | None | None | None | None |
| SOD1 | None | None | None | None |
| SOD1 | None | None | None | None |
| SOD1 | None | None | None | None |
| NECTIN3 | None | None | None | None |
| HFE | None | None | None | None |
| HFE | None | None | None | None |
| HFE | None | None | None | None |
| HFE | None | None | None | None |
| HFE | None | None | None | None |
| HFE | None | None | None | None |
| HFE | None | None | None | None |
| EIF2AK3 | None | None | None | None |
| EIF2AK3 | None | None | None | None |
| TERT | None | None | None | None |
| TERT | None | None | None | None |
| SPR | None | None | None | None |
| SPR | None | None | None | None |
| SPR | None | None | None | None |
| SPR | None | None | None | None |
| SPR | None | None | None | None |
| F7 | None | None | None | None |
| UNG | None | None | None | None |
| MMP13 | None | None | None | None |
| MMP13 | None | None | None | None |
| MMP13 | None | None | None | None |
| MMP13 | None | None | None | None |
| CD207 | None | None | None | None |
| F2 | 1 | 2 | 1nrq | L |
| SPTBN2 | None | None | None | None |
| PKLR | None | None | None | None |
| PKLR | None | None | None | None |
| PKLR | None | None | None | None |
| PKLR | None | None | None | None |

| | | | | |
|---|---|---|---|---|
| PKLR | None | None | None | None |
| PKLR | None | None | None | None |
| PKLR | None | None | None | None |
| PKLR | None | None | None | None |
| SACS | None | None | None | None |
| HRAS | None | None | None | None |
| HRAS | None | None | None | None |
| HRAS | None | None | None | None |
| HRAS | None | None | None | None |
| HRAS | None | None | None | None |
| HRAS | None | None | None | None |
| HRAS | None | None | None | None |
| HRAS | None | None | None | None |
| HRAS | None | None | None | None |
| HRAS | None | None | None | None |
| HRAS | None | None | None | None |
| HRAS | None | None | None | None |
| HRAS | None | None | None | None |
| VHL | None | None | None | None |
| VHL | None | None | None | None |
| VHL | None | None | None | None |
| VHL | None | None | None | None |
| VHL | None | None | None | None |
| VHL | None | None | None | None |
| VHL | None | None | None | None |
| VHL | None | None | None | None |
| VHL | None | None | None | None |
| VHL | None | None | None | None |
| VHL | None | None | None | None |
| VHL | None | None | None | None |
| VHL | None | None | None | None |
| VHL | None | None | None | None |
| VHL | None | None | None | None |
| VHL | None | None | None | None |
| VHL | None | None | None | None |
| VHL | None | None | None | None |
| VHL | None | None | None | None |
| VHL | None | None | None | None |
| VHL | None | None | None | None |
| VHL | None | None | None | None |
| VHL | None | None | None | None |
| VHL | None | None | None | None |
| VHL | None | None | None | None |
| TTBK1 | None | None | None | None |
| RAD51 | None | None | None | None |
| ALB | None | None | None | None |
| ALB | None | None | None | None |
| ALB | None | None | None | None |
| ALB | None | None | None | None |
| MET | None | None | None | None |
| MET | None | None | None | None |
| MET | None | None | None | None |
| MET | None | None | None | None |
| MET | None | None | None | None |

| | | | | |
|---|---|---|---|---|
| MET | None | None | None | None |
| MET | None | None | None | None |
| MET | None | None | None | None |
| MET | None | None | None | None |
| CP | None | None | None | None |
| CP | None | None | None | None |
| PGM1 | None | None | None | None |
| PGM1 | None | None | None | None |
| PGM1 | None | None | None | None |
| PGM1 | None | None | None | None |
| PGM1 | None | None | None | None |
| PGM1 | None | None | None | None |
| PGM1 | None | None | None | None |
| NR5A1 | None | None | None | None |
| NR5A1 | None | None | None | None |
| NR5A1 | None | None | None | None |
| F9 | None | None | None | None |
| F9 | None | None | None | None |
| HPRT1 | None | None | None | None |
| HPRT1 | None | None | None | None |
| HPRT1 | None | None | None | None |
| HPRT1 | None | None | None | None |
| HPRT1 | None | None | None | None |
| HPRT1 | 0.871 | 3 | 4rac | B |
| HPRT1 | 0.806 | 2 | 4ran | D |
| HPRT1 | None | None | None | None |
| HPRT1 | None | None | None | None |
| HPRT1 | None | None | None | None |
| HPRT1 | None | None | None | None |
| HPRT1 | None | None | None | None |
| HPRT1 | None | None | None | None |
| HPRT1 | None | None | None | None |
| HPRT1 | None | None | None | None |
| HPRT1 | None | None | None | None |
| HPRT1 | None | None | None | None |
| HPRT1 | None | None | None | None |
| HPRT1 | 0.871 | 3 | 4rac | B |
| HPRT1 | 0.871 | 3 | 4rac | B |
| HPRT1 | None | None | None | None |
| HPRT1 | None | None | None | None |
| GPD1L | None | None | None | None |
| GPD1L | None | None | None | None |
| GPD1L | None | None | None | None |
| FGFR2 | None | None | None | None |
| FGFR2 | None | None | None | None |
| FGFR2 | None | None | None | None |
| FGFR2 | None | None | None | None |
| FGFR2 | None | None | None | None |
| FGFR2 | None | None | None | None |
| FGFR2 | None | None | None | None |
| FGFR2 | None | None | None | None |
| FGFR2 | None | None | None | None |
| FGFR2 | None | None | None | None |
| FGFR2 | None | None | None | None |

| | | | | |
|---|---|---|---|---|
| FGFR2 | None | None | None | None |
| FGFR2 | None | None | None | None |
| FGFR2 | None | None | None | None |
| FGFR2 | None | None | None | None |
| FGFR2 | None | None | None | None |
| FGFR2 | None | None | None | None |
| FGFR2 | None | None | None | None |
| FGFR2 | None | None | None | None |
| HSPD1 | None | None | None | None |
| HSPD1 | None | None | None | None |
| FGF8 | None | None | None | None |
| FGF8 | None | None | None | None |
| FGF8 | None | None | None | None |
| F9 | None | None | None | None |
| NCF4 | None | None | None | None |
| NCF4 | None | None | None | None |
| LDLR | None | None | None | None |
| LDLR | None | None | None | None |
| LDLR | None | None | None | None |
| LDLR | None | None | None | None |
| LDLR | None | None | None | None |
| LDLR | None | None | None | None |
| C3 | None | None | None | None |
| C3 | None | None | None | None |
| C3 | None | None | None | None |
| C3 | None | None | None | None |
| C3 | None | None | None | None |
| C3 | None | None | None | None |
| C3 | None | None | None | None |
| C3 | None | None | None | None |
| TULP1 | None | None | None | None |
| TULP1 | None | None | None | None |
| TULP1 | None | None | None | None |
| TULP1 | None | None | None | None |
| IKBKG | None | None | None | None |
| IKBKG | None | None | None | None |
| IKBKG | None | None | None | None |
| CD40 | None | None | None | None |
| CD40 | None | None | None | None |
| SLC9A3R1 | None | None | None | None |
| BCHE | None | None | None | None |
| BCHE | None | None | None | None |
| BCHE | None | None | None | None |
| BCHE | None | None | None | None |
| BCHE | None | None | None | None |
| BCHE | None | None | None | None |
| BCHE | None | None | None | None |
| BCHE | None | None | None | None |
| BCHE | None | None | None | None |
| BCHE | None | None | None | None |
| BCHE | None | None | None | None |
| BCKDHB | 0.833 | 1 | 1v11 | B |
| BCKDHB | None | None | None | None |
| BCKDHB | None | None | None | None |

| | | | | |
|---|---|---|---|---|
| BCKDHB | None | None | None | None |
| BCKDHB | 0.833 | 10 | 1olx | B |
| BCKDHB | 0.833 | 1 | 1v11 | B |
| BCKDHB | 0.833 | 1 | 1v11 | B |
| BCKDHB | None | None | None | None |
| BCKDHB | None | None | None | None |
| BCKDHB | None | None | None | None |
| BCKDHB | None | None | None | None |
| BCKDHB | 0.857 | 1 | 2bff | B |
| CDH1 | None | None | None | None |
| CDH1 | None | None | None | None |
| CDH1 | None | None | None | None |
| CDH1 | None | None | None | None |
| CDH1 | None | None | None | None |
| CTNNB1 | None | None | None | None |
| AIRE | None | None | None | None |
| EDA | 1 | 1 | 1rj8 | A |
| EDA | None | None | None | None |
| EDA | None | None | None | None |
| EDA | None | None | None | None |
| EDA | None | None | None | None |
| SH2D1A | None | None | None | None |
| SH2D1A | None | None | None | None |
| SH2D1A | None | None | None | None |
| GCK | None | None | None | None |
| GCK | None | None | None | None |
| GCK | None | None | None | None |
| GCK | None | None | None | None |
| GCK | None | None | None | None |
| GCK | None | None | None | None |
| GCK | None | None | None | None |
| GCK | None | None | None | None |
| GCK | None | None | None | None |
| GCK | None | None | None | None |
| GCK | 0.833 | 1 | 4dhy | A |
| GCK | None | None | None | None |
| GCK | None | None | None | None |
| GCK | None | None | None | None |
| GCK | None | None | None | None |
| GCK | None | None | None | None |
| GCK | None | None | None | None |
| GCK | None | None | None | None |
| GCK | None | None | None | None |
| GCK | None | None | None | None |
| AQP2 | None | None | None | None |
| AQP2 | None | None | None | None |
| AQP2 | None | None | None | None |
| AQP2 | None | None | None | None |
| AQP2 | None | None | None | None |
| AQP2 | None | None | None | None |
| AQP2 | None | None | None | None |
| AQP2 | None | None | None | None |
| AQP2 | None | None | None | None |
| AQP2 | None | None | None | None |

| | | | | |
|---|---|---|---|---|
| AQP2 | None | None | None | None |
| AQP2 | None | None | None | None |
| AQP2 | None | None | None | None |
| SUMF1 | None | None | None | None |
| SUMF1 | None | None | None | None |
| SUMF1 | None | None | None | None |
| SUMF1 | None | None | None | None |
| SUMF1 | 0.875 | 35 | 2afy | X |
| SUMF1 | None | None | None | None |
| SUMF1 | None | None | None | None |
| SUMF1 | None | None | None | None |
| SUMF1 | None | None | None | None |
| FAM20A | None | None | None | None |
| FAM20A | None | None | None | None |
| FAM20A | None | None | None | None |
| FAM20A | None | None | None | None |
| NT5E | None | None | None | None |
| NT5E | None | None | None | None |
| NOS3 | None | None | None | None |
| ACTG1 | None | None | None | None |
| ACTG1 | None | None | None | None |
| ACTG1 | None | None | None | None |
| ACTG1 | None | None | None | None |
| ACTG1 | None | None | None | None |
| ACTG1 | None | None | None | None |
| FCGR1A | None | None | None | None |
| GPI | None | None | None | None |
| GPI | None | None | None | None |
| GPI | 1 | 3 | 1jiq | A |
| GPI | None | None | None | None |
| GPI | None | None | None | None |
| GPI | None | None | None | None |
| GPI | None | None | None | None |
| GPI | None | None | None | None |
| GPI | None | None | None | None |
| CFH | None | None | None | None |
| OGG1 | 0.895 | 13 | 1m3h | A |
| TPK1 | None | None | None | None |
| TPK1 | None | None | None | None |
| TPK1 | None | None | None | None |
| TPK1 | None | None | None | None |
| SEC23A | None | None | None | None |
| SEC23A | None | None | None | None |
| TYRP1 | None | None | None | None |
| TYRP1 | None | None | None | None |
| TYRP1 | None | None | None | None |
| TYRP1 | None | None | None | None |
| TYRP1 | None | None | None | None |
| TYRP1 | None | None | None | None |
| CSTB | None | None | None | None |
| CSTB | None | None | None | None |
| CSTB | None | None | None | None |
| ACP1 | None | None | None | None |
| IL17F | None | None | None | None |

| | | | | |
|---|---|---|---|---|
| IL17F | None | None | None | None |
| IL17F | None | None | None | None |
| IL17F | None | None | None | None |
| IL17F | None | None | None | None |
| IDH2 | None | None | None | None |
| IDH2 | None | None | None | None |
| IDH2 | None | None | None | None |
| IDH2 | None | None | None | None |
| TP53 | 0.925 | 11 | 2ocj | A |
| TP53 | 0.8 | 16 | 2ahi | D |
| TP53 | None | None | None | None |
| TP53 | None | None | None | None |
| TP53 | None | None | None | None |
| TP53 | 0.9 | 1 | 4agp | A |
| TP53 | 0.9 | 1 | 4agp | A |
| TP53 | 0.925 | 10 | 4hje | A |
| TP53 | None | None | None | None |
| TP53 | 0.875 | 7 | 3kmd | B |
| TP53 | 0.875 | 7 | 3kmd | B |
| TP53 | 0.925 | 6 | 3kz8 | B |
| TP53 | 0.925 | 6 | 3kz8 | B |
| TP53 | None | None | None | None |
| TP53 | 0.925 | 5 | 3kmd | B |
| TP53 | 0.925 | 5 | 3kmd | B |
| TP53 | 0.925 | 8 | 3kz8 | B |
| TP53 | 0.925 | 8 | 3kz8 | B |
| TP53 | 0.925 | 1 | 3kmd | B |
| TP53 | 0.9 | 11 | 2pcx | A |
| TP53 | 0.9 | 1 | 5aol | A |
| TP53 | 0.85 | 5 | 5lgy | A |
| TP53 | 0.9 | 1 | 3q06 | B |
| TP53 | None | None | None | None |
| TP53 | None | None | None | None |
| TP53 | 0.825 | 8 | 1kzy | A |
| TP53 | 0.825 | 8 | 1kzy | A |
| TP53 | 0.8 | 11 | 4ibt | A |
| TP53 | 0.8 | 11 | 4ibt | A |
| TP53 | None | None | None | None |
| TP53 | None | None | None | None |
| TP53 | 0.875 | 7 | 3kmd | B |
| TP53 | None | None | None | None |
| TP53 | 0.8 | 10 | 4agp | A |
| TP53 | 0.8 | 10 | 4agp | A |
| TP53 | 0.8 | 10 | 4agp | A |
| TP53 | 0.8 | 10 | 4agp | A |
| TP53 | None | None | None | None |
| TP53 | None | None | None | None |
| TP53 | None | None | None | None |
| TP53 | None | None | None | None |
| TP53 | None | None | None | None |
| LMAN1 | None | None | None | None |
| CTSK | None | None | None | None |
| CTSK | None | None | None | None |
| CTSK | None | None | None | None |

| | | | | |
|---|---|---|---|---|
| FGB | None | None | None | None |
| FGB | None | None | None | None |
| FGB | None | None | None | None |
| FGB | None | None | None | None |
| ACADM | None | None | None | None |
| ACADM | None | None | None | None |
| ACADM | None | None | None | None |
| ACADM | None | None | None | None |
| ACADM | None | None | None | None |
| ACADM | None | None | None | None |
| ACADM | None | None | None | None |
| ACADM | None | None | None | None |
| ACADM | None | None | None | None |
| ACADM | None | None | None | None |
| ACADM | None | None | None | None |
| ACADM | None | None | None | None |
| ACADM | None | None | None | None |
| ACADM | None | None | None | None |
| ACADM | None | None | None | None |
| ACADM | None | None | None | None |
| ACADM | None | None | None | None |
| ACADM | None | None | None | None |
| ACADM | None | None | None | None |
| ACADM | None | None | None | None |
| ACADM | None | None | None | None |
| ACADM | None | None | None | None |
| ACADM | None | None | None | None |
| ACADM | None | None | None | None |
| ACADM | None | None | None | None |
| ACADM | None | None | None | None |
| ACADM | None | None | None | None |
| PTS | None | None | None | None |
| PTS | None | None | None | None |
| PTS | None | None | None | None |
| PTS | None | None | None | None |
| PTS | None | None | None | None |
| PTS | None | None | None | None |
| PTS | None | None | None | None |
| PTS | None | None | None | None |
| PTS | None | None | None | None |
| INS | None | None | None | None |
| INS | | 1 | 5 2l1z | A |
| INS | | 1 | 5 2l1z | A |
| INS | | 1 | 1 1his | A |
| FBP1 | None | None | None | None |
| FBP1 | None | None | None | None |
| FBP1 | None | None | None | None |
| FBP1 | None | None | None | None |
| FBP1 | None | None | None | None |
| FBP1 | None | None | None | None |
| FBP1 | None | None | None | None |
| FBP1 | | 0.8 | 11 2y5k | A |
| FBP1 | None | None | None | None |
| FBP1 | None | None | None | None |

| | | | | |
|---|---|---|---|---|
| FBP1 | None | None | None | None |
| KCNQ1 | None | None | None | None |
| KCNQ1 | None | None | None | None |
| KCNQ1 | None | None | None | None |
| UPP1 | None | None | None | None |
| UMPS | 1 | 3 | 2qcl | A |
| UMPS | None | None | None | None |
| UMPS | None | None | None | None |
| DHFR | None | None | None | None |
| KHK | None | None | None | None |
| TTPA | None | None | None | None |
| TTPA | None | None | None | None |
| TTPA | None | None | None | None |
| TTPA | None | None | None | None |
| MAP1LC3B | None | None | None | None |
| MAP1LC3B | None | None | None | None |
| ZAP70 | None | None | None | None |
| PROC | None | None | None | None |
| SQSTM1 | None | None | None | None |
| AHCY | None | None | None | None |
| AHCY | None | None | None | None |
| PRNP | None | None | None | None |
| PRNP | None | None | None | None |
| PRNP | None | None | None | None |
| PRNP | None | None | None | None |
| PRNP | None | None | None | None |
| PRNP | None | None | None | None |
| PRNP | None | None | None | None |
| PRNP | None | None | None | None |
| PRNP | None | None | None | None |
| PRNP | None | None | None | None |
| PRNP | None | None | None | None |
| PRNP | None | None | None | None |
| PRNP | None | None | None | None |
| PRNP | None | None | None | None |
| PRNP | None | None | None | None |
| PRNP | None | None | None | None |
| REN | None | None | None | None |
| REN | None | None | None | None |
| HINT1 | None | None | None | None |
| HINT1 | 0.897 | 1 | 5km4 | A |
| HINT1 | None | None | None | None |
| HINT1 | None | None | None | None |
| HINT1 | 0.923 | 1 | 5ipc | A |
| OAT | None | None | None | None |
| OAT | None | None | None | None |
| OAT | None | None | None | None |
| OAT | None | None | None | None |
| OAT | None | None | None | None |
| OAT | None | None | None | None |
| OAT | None | None | None | None |
| OAT | None | None | None | None |
| OAT | None | None | None | None |
| OAT | None | None | None | None |

| | | | | |
|---|---|---|---|---|
| OAT | None | None | None | None |
| OAT | None | None | None | None |
| OAT | None | None | None | None |
| OAT | None | None | None | None |
| OAT | None | None | None | None |
| OAT | None | None | None | None |
| OAT | None | None | None | None |
| OAT | None | None | None | None |
| OAT | None | None | None | None |
| OAT | None | None | None | None |
| OAT | None | None | None | None |
| OAT | None | None | None | None |
| OAT | None | None | None | None |
| OAT | None | None | None | None |
| OAT | None | None | None | None |
| OAT | None | None | None | None |
| OAT | None | None | None | None |
| OAT | None | None | None | None |
| CTSA | None | None | None | None |
| CTSA | None | None | None | None |
| PNP | None | None | None | None |
| PNP | None | None | None | None |
| PNP | None | None | None | None |
| PNP | None | None | None | None |
| PNP | None | None | None | None |
| PNP | None | None | None | None |
| PNP | 0.933 | 1 | 3ggs | C |
| PNP | 0.833 | 1 | 3d1v | A |
| MYL2 | None | None | None | None |
| MYL2 | None | None | None | None |
| ANG | None | None | None | None |
| ANG | None | None | None | None |
| ANG | None | None | None | None |
| ANG | None | None | None | None |
| ANG | None | None | None | None |
| ANG | None | None | None | None |
| ANG | None | None | None | None |
| ANG | None | None | None | None |
| ANG | None | None | None | None |
| ANG | None | None | None | None |
| POLH | None | None | None | None |
| POLH | None | None | None | None |
| POLH | None | None | None | None |
| IL2RA | None | None | None | None |
| TGFB1 | None | None | None | None |
| TP53 | None | None | None | None |
| TP53 | None | None | None | None |
| PAH | None | None | None | None |
| PAH | None | None | None | None |
| PAH | None | None | None | None |
| PAH | None | None | None | None |
| PAH | None | None | None | None |
| PAH | None | None | None | None |
| PAH | None | None | None | None |

| | | | | |
|---|---|---|---|---|
| PAH | None | None | None | None |
| PAH | None | None | None | None |
| PAH | None | None | None | None |
| PAH | None | None | None | None |
| PAH | None | None | None | None |
| PAH | None | None | None | None |
| PAH | None | None | None | None |
| PAH | None | None | None | None |
| PAH | None | None | None | None |
| PAH | None | None | None | None |
| PAH | None | None | None | None |
| PAH | None | None | None | None |
| PAH | None | None | None | None |
| PAH | None | None | None | None |
| PAH | None | None | None | None |
| PAH | None | None | None | None |
| PAH | None | None | None | None |
| PAH | None | None | None | None |
| PAH | None | None | None | None |
| PAH | None | None | None | None |
| PAH | None | None | None | None |
| PAH | None | None | None | None |
| PAH | None | None | None | None |
| PAH | None | None | None | None |
| PAH | None | None | None | None |
| PAH | None | None | None | None |
| PAH | None | None | None | None |
| PAH | None | None | None | None |
| PAH | None | None | None | None |
| PAH | None | None | None | None |
| PAH | None | None | None | None |
| PAH | None | None | None | None |
| PAH | None | None | None | None |
| PAH | None | None | None | None |
| HDAC8 | None | None | None | None |
| HDAC8 | None | None | None | None |
| VCP | None | None | None | None |
| VCP | None | None | None | None |
| VCP | None | None | None | None |
| VCP | None | None | None | None |
| VCP | None | None | None | None |
| VCP | None | None | None | None |
| VCP | None | None | None | None |
| B3GAT3 | None | None | None | None |
| B3GAT3 | None | None | None | None |
| CSF2RB | None | None | None | None |
| CSF2RB | None | None | None | None |
| CSF2RB | None | None | None | None |
| NF2 | None | None | None | None |
| NF2 | None | None | None | None |
| NF2 | None | None | None | None |
| PIK3CA | None | None | None | None |
| PHGDH | None | None | None | None |
| PHGDH | None | None | None | None |

| | | | | |
|---|---|---|---|---|
| PHGDH | None | None | None | None |
| PHGDH | None | None | None | None |
| POLH | None | None | None | None |
| XIAP | None | None | None | None |
| XIAP | None | None | None | None |
| PDHX | None | None | None | None |
| PDHX | None | None | None | None |
| PDHX | None | None | None | None |
| PDHX | None | None | None | None |
| PDHX | None | None | None | None |
| SAMHD1 | None | None | None | None |
| SAMHD1 | None | None | None | None |
| SAMHD1 | None | None | None | None |
| SAMHD1 | None | None | None | None |
| SAMHD1 | None | None | None | None |
| SAMHD1 | None | None | None | None |
| SAMHD1 | None | None | None | None |
| SAMHD1 | None | None | None | None |
| CD36 | None | None | None | None |
| CD36 | None | None | None | None |
| CD36 | None | None | None | None |
| ASPH | None | None | None | None |
| CCT5 | None | None | None | None |
| CCT5 | None | None | None | None |
| CCT5 | None | None | None | None |
| SMN1 | None | None | None | None |
| SMN1 | None | None | None | None |
| SMN1 | None | None | None | None |
| SMN1 | None | None | None | None |
| ABCA1 | None | None | None | None |
| ABCA1 | None | None | None | None |
| ABCA1 | None | None | None | None |
| ABCA1 | None | None | None | None |
| ABCA1 | None | None | None | None |
| ABCA1 | None | None | None | None |
| ABCA1 | None | None | None | None |
| ABCA1 | None | None | None | None |
| ABCA1 | None | None | None | None |
| ABCA1 | None | None | None | None |
| ABCA1 | None | None | None | None |
| ABCA1 | None | None | None | None |
| ABCA1 | None | None | None | None |
| ABCA1 | None | None | None | None |
| ABCA1 | None | None | None | None |
| ABCA1 | None | None | None | None |
| ABCA1 | None | None | None | None |
| ABCA1 | None | None | None | None |
| ABCA1 | None | None | None | None |
| ABCA1 | None | None | None | None |
| ABCA1 | None | None | None | None |
| ABCA1 | None | None | None | None |
| ABCA1 | None | None | None | None |
| ABCA1 | None | None | None | None |

| | | | | |
|---|---|---|---|---|
| ABCA1 | None | None | None | None |
| ABCA1 | None | None | None | None |
| ABCA1 | None | None | None | None |
| ABCA1 | None | None | None | None |
| ABCA1 | None | None | None | None |
| ABCA1 | None | None | None | None |
| ABCA1 | None | None | None | None |
| ABCA1 | None | None | None | None |
| ABCA1 | None | None | None | None |
| ABCA1 | None | None | None | None |
| ABCA1 | None | None | None | None |
| ERCC1 | None | None | None | None |
| FECH | None | None | None | None |
| FECH | None | None | None | None |
| FECH | None | None | None | None |
| FECH | None | None | None | None |
| FECH | None | None | None | None |
| FECH | None | None | None | None |
| FECH | None | None | None | None |
| PTEN | None | None | None | None |
| PTEN | None | None | None | None |
| PTEN | None | None | None | None |
| PTEN | None | None | None | None |
| PTEN | None | None | None | None |
| PTEN | None | None | None | None |
| PTEN | None | None | None | None |
| PTEN | None | None | None | None |
| PTEN | None | None | None | None |
| PTEN | None | None | None | None |
| PTEN | None | None | None | None |
| PTEN | None | None | None | None |
| PTEN | None | None | None | None |
| PTEN | None | None | None | None |
| PTEN | None | None | None | None |
| PTEN | None | None | None | None |
| PTEN | None | None | None | None |
| PTEN | None | None | None | None |
| PTEN | None | None | None | None |
| PTEN | None | None | None | None |
| ACAT1 | None | None | None | None |
| ACAT1 | None | None | None | None |
| ACAT1 | None | None | None | None |
| ACAT1 | None | None | None | None |
| ACAT1 | None | None | None | None |
| ACAT1 | None | None | None | None |
| ACAT1 | 0.833 | 6 | 2ib7 | C |
| ACAT1 | None | None | None | None |
| ACAT1 | None | None | None | None |
| ACAT1 | None | None | None | None |
| ACAT1 | None | None | None | None |
| CREBBP | None | None | None | None |
| ITGA2B | None | None | None | None |
| ITGA2B | None | None | None | None |
| ITGA2B | None | None | None | None |

| | | | | |
|---|---|---|---|---|
| ITGA2B | None | None | None | None |
| TSC1 | None | None | None | None |
| ADA2 | None | None | None | None |
| ADA2 | None | None | None | None |
| ADA2 | None | None | None | None |
| ADA2 | None | None | None | None |
| ADA2 | None | None | None | None |
| ADA2 | None | None | None | None |
| ADA2 | None | None | None | None |
| ADA2 | None | None | None | None |
| ADA2 | 1 | 1 | 3lgd | B |
| ADA2 | None | None | None | None |
| PGK1 | None | None | None | None |
| PGK1 | None | None | None | None |
| PGK1 | None | None | None | None |
| PGK1 | None | None | None | None |
| PGK1 | None | None | None | None |
| TUBA1A | None | None | None | None |
| TUBA1A | None | None | None | None |
| TUBA1A | None | None | None | None |
| TUBA1A | None | None | None | None |
| TUBA1A | None | None | None | None |
| PTPN11 | None | None | None | None |
| PTPN11 | None | None | None | None |
| PTPN11 | None | None | None | None |
| PTPN11 | None | None | None | None |
| PTPN11 | None | None | None | None |
| PTPN11 | None | None | None | None |
| PTPN11 | None | None | None | None |
| PTPN11 | None | None | None | None |
| PTPN11 | None | None | None | None |
| PTPN11 | None | None | None | None |
| PTPN11 | None | None | None | None |
| PTPN11 | None | None | None | None |
| PTPN11 | None | None | None | None |
| PTPN11 | None | None | None | None |
| PTPN11 | None | None | None | None |
| PTPN11 | None | None | None | None |
| PTPN11 | None | None | None | None |
| PTPN11 | None | None | None | None |
| PTPN11 | None | None | None | None |
| PTPN11 | None | None | None | None |
| PTPN11 | None | None | None | None |
| PTPN11 | None | None | None | None |
| PTPN11 | None | None | None | None |
| PTPN11 | None | None | None | None |
| PTPN11 | None | None | None | None |
| PTPN11 | None | None | None | None |
| GLB1 | None | None | None | None |
| GLB1 | None | None | None | None |
| UQCRQ | None | None | None | None |
| VPS35 | None | None | None | None |
| NR3C1 | None | None | None | None |
| NR3C1 | None | None | None | None |

| | | | | |
|---|---|---|---|---|
| NR3C1 | None | None | None | None |
| NR3C1 | None | None | None | None |
| NR3C1 | None | None | None | None |
| NR3C1 | None | None | None | None |
| NR3C1 | None | None | None | None |
| NR3C1 | None | None | None | None |
| NR3C1 | None | None | None | None |
| APOA1 | None | None | None | None |
| APOA1 | None | None | None | None |
| APOA1 | None | None | None | None |
| APOA1 | None | None | None | None |
| APOA1 | None | None | None | None |
| APOA1 | None | None | None | None |
| APOA1 | None | None | None | None |
| BCKDHA | None | None | None | None |
| BCKDHA | None | None | None | None |
| BCKDHA | 0.963 | 1 | 1ols | A |
| BCKDHA | None | None | None | None |
| BCKDHA | None | None | None | None |
| BCKDHA | None | None | None | None |
| BCKDHA | None | None | None | None |
| BCKDHA | 1 | 1 | 2beu | A |
| BCKDHA | 0.84 | 2 | 2j9f | A |
| BCKDHA | None | None | None | None |
| BCKDHA | None | None | None | None |
| BCKDHA | None | None | None | None |
| BCKDHA | None | None | None | None |
| BCKDHA | None | None | None | None |
| BCKDHA | None | None | None | None |
| TOP1 | None | None | None | None |
| ANTXR2 | None | None | None | None |
| ANTXR2 | None | None | None | None |
| ANTXR2 | None | None | None | None |
| TF | None | None | None | None |
| TF | None | None | None | None |
| FSHB | None | None | None | None |
| GALK1 | None | None | None | None |
| GALK1 | None | None | None | None |
| GALK1 | None | None | None | None |
| CYP17A1 | None | None | None | None |
| CYP17A1 | None | None | None | None |
| CYP17A1 | None | None | None | None |
| CYP17A1 | None | None | None | None |
| CYP17A1 | None | None | None | None |
| CYP17A1 | None | None | None | None |
| CYP17A1 | None | None | None | None |
| CYP17A1 | None | None | None | None |
| CYP17A1 | None | None | None | None |
| CYP17A1 | None | None | None | None |
| CYP17A1 | None | None | None | None |
| CYP17A1 | None | None | None | None |
| CYP17A1 | None | None | None | None |
| CYP17A1 | None | None | None | None |
| CYP17A1 | None | None | None | None |

| | | | | |
|---|---|---|---|---|
| CYP17A1 | None | None | None | None |
| CYP17A1 | None | None | None | None |
| CD44 | None | None | None | None |
| MDH2 | None | None | None | None |
| MDH2 | None | None | None | None |
| NDUFV1 | None | None | None | None |
| NDUFV1 | None | None | None | None |
| NDUFV1 | None | None | None | None |
| NDUFV1 | None | None | None | None |
| NDUFV1 | None | None | None | None |
| NDUFV1 | None | None | None | None |
| NDUFV1 | None | None | None | None |
| NDUFV1 | None | None | None | None |
| NDUFV1 | None | None | None | None |
| NDUFV1 | None | None | None | None |
| CTSC | None | None | None | None |
| CTSC | None | None | None | None |
| CTSC | None | None | None | None |
| CTSC | None | None | None | None |
| CTSC | None | None | None | None |
| CTSC | None | None | None | None |
| CTSC | None | None | None | None |
| VWF | None | None | None | None |
| VWF | None | None | None | None |
| VWF | None | None | None | None |
| VWF | None | None | None | None |
| VWF | None | None | None | None |
| VWF | None | None | None | None |
| VWF | None | None | None | None |
| ORC4 | None | None | None | None |
| ORC4 | None | None | None | None |
| CRYAB | None | None | None | None |
| CRYAB | None | None | None | None |
| CRYAB | None | None | None | None |
| MAP2K2 | None | None | None | None |
| MAP2K2 | None | None | None | None |
| MAP2K2 | None | None | None | None |
| MAP2K2 | None | None | None | None |
| MAP2K2 | None | None | None | None |
| MAP2K2 | None | None | None | None |
| MAP2K2 | None | None | None | None |
| MAP2K2 | None | None | None | None |
| MAP2K2 | None | None | None | None |
| CBL | None | None | None | None |
| CBL | None | None | None | None |
| CBL | None | None | None | None |
| CBL | None | None | None | None |
| CBL | None | None | None | None |
| CBL | None | None | None | None |
| CBL | None | None | None | None |
| PFN1 | None | None | None | None |
| PFN1 | None | None | None | None |
| ACSM2A | None | None | None | None |
| HSPG2 | None | None | None | None |

| | | | | |
|---|---|---|---|---|
| HSPG2 | None | None | None | None |
| HSPG2 | None | None | None | None |
| HSPG2 | None | None | None | None |
| PARK7 | None | None | None | None |
| PARK7 | None | None | None | None |
| PARK7 | 0.808 | 2 | 1pe0 | A |
| PARK7 | None | None | None | None |
| PARK7 | 0.889 | 1 | 3cz9 | A |
| PARK7 | None | None | None | None |
| INSR | None | None | None | None |
| INSR | None | None | None | None |
| INSR | None | None | None | None |
| INSR | None | None | None | None |
| INSR | None | None | None | None |
| DSC2 | None | None | None | None |
| DSC2 | None | None | None | None |
| DSC2 | None | None | None | None |
| DSC2 | None | None | None | None |
| DSC2 | None | None | None | None |
| DSC2 | None | None | None | None |
| DSC2 | None | None | None | None |
| DSC2 | None | None | None | None |
| DSC2 | None | None | None | None |
| DSC2 | None | None | None | None |
| MICU1 | None | None | None | None |
| MICU1 | None | None | None | None |
| ADAMTS13 | None | None | None | None |
| ADAMTS13 | None | None | None | None |
| ADAMTS13 | None | None | None | None |
| ADAMTS13 | None | None | None | None |
| ADAMTS13 | None | None | None | None |
| ADAMTS13 | None | None | None | None |
| ADAMTS13 | None | None | None | None |
| IRAK4 | None | None | None | None |
| IRAK4 | None | None | None | None |
| RNASEH2B | None | None | None | None |
| RNASEH2B | None | None | None | None |
| RNASEH2B | None | None | None | None |
| NTRK1 | None | None | None | None |
| MPO | None | None | None | None |
| MPO | None | None | None | None |
| MPO | None | None | None | None |
| PEX3 | None | None | None | None |
| SERPINA1 | None | None | None | None |
| SERPINA1 | None | None | None | None |
| SERPINA1 | None | None | None | None |
| SERPINA1 | None | None | None | None |
| SERPINA1 | None | None | None | None |
| SERPINA1 | None | None | None | None |
| SERPINA1 | None | None | None | None |
| SERPINA1 | None | None | None | None |
| SERPINA1 | None | None | None | None |
| SERPINA1 | None | None | None | None |
| YARS2 | None | None | None | None |

| | | | | |
|---|---|---|---|---|
| YARS2 | None | None | None | None |
| YARS2 | None | None | None | None |
| YARS2 | None | None | None | None |
| YARS2 | None | None | None | None |
| YARS2 | None | None | None | None |
| YARS2 | None | None | None | None |
| YARS2 | None | None | None | None |
| YARS2 | None | None | None | None |
| PDHA1 | None | None | None | None |
| PDHA1 | None | None | None | None |
| PDHA1 | None | None | None | None |
| PDHA1 | None | None | None | None |
| PDHA1 | None | None | None | None |
| PDHA1 | None | None | None | None |
| PDHA1 | None | None | None | None |
| PDHA1 | 0.96 | 1 | 3exh | C |
| PDHA1 | None | None | None | None |
| PDHA1 | None | None | None | None |
| PDHA1 | None | None | None | None |
| CUL4B | None | None | None | None |
| AGXT | None | None | None | None |
| AGXT | None | None | None | None |
| AGXT | None | None | None | None |
| AGXT | None | None | None | None |
| AGXT | None | None | None | None |
| AGXT | None | None | None | None |
| AGXT | None | None | None | None |
| AGXT | None | None | None | None |
| AGXT | None | None | None | None |
| AGXT | None | None | None | None |
| AGXT | None | None | None | None |
| AGXT | None | None | None | None |
| AGXT | None | None | None | None |
| AGXT | None | None | None | None |
| AGXT | None | None | None | None |
| AGXT | None | None | None | None |
| AGXT | None | None | None | None |
| AGXT | None | None | None | None |
| AGXT | None | None | None | None |
| AGXT | None | None | None | None |
| AGXT | None | None | None | None |
| AGXT | None | None | None | None |
| AGXT | None | None | None | None |
| AGXT | None | None | None | None |
| AGXT | None | None | None | None |
| AGXT | None | None | None | None |
| AGXT | None | None | None | None |
| AGXT | None | None | None | None |
| AGXT | None | None | None | None |
| AGXT | None | None | None | None |
| AGXT | None | None | None | None |
| AGXT | None | None | None | None |
| AGXT | 0.846 | 1 | 4kxk | A |
| AGXT | None | None | None | None |

| | | | | |
|---|---|---|---|---|
| AGXT | None | None | None | None |
| SFTPD | None | None | None | None |
| SFTPD | None | None | None | None |
| PIK3CA | None | None | None | None |
| PIK3CA | None | None | None | None |
| PIK3CA | None | None | None | None |
| AUH | None | None | None | None |
| AUH | None | None | None | None |
| AUH | None | None | None | None |
| AUH | None | None | None | None |
| F11 | None | None | None | None |
| F11 | None | None | None | None |
| F11 | None | None | None | None |
| F11 | None | None | None | None |
| MEN1 | None | None | None | None |
| MEN1 | 0.8 | 54 | 4i80 | A |
| MEN1 | 0.8 | 84 | 5db0 | A |
| MEN1 | 0.8 | 30 | 5ddb | A |
| MEN1 | 1 | 2 | 4i80 | A |
| MEN1 | 0.8 | 17 | 5db0 | A |
| MEN1 | None | None | None | None |
| MEN1 | None | None | None | None |
| MEN1 | None | None | None | None |
| MEN1 | None | None | None | None |
| MYBPC1 | None | None | None | None |
| MYBPC1 | None | None | None | None |
| MYBPC1 | None | None | None | None |
| PDCD10 | None | None | None | None |
| SCARB2 | None | None | None | None |
| SCARB2 | None | None | None | None |
| SCARB2 | None | None | None | None |
| SCARB2 | None | None | None | None |
| SCARB2 | None | None | None | None |
| TREX2 | None | None | None | None |
| TBK1 | None | None | None | None |
| TBK1 | None | None | None | None |
| DLD | None | None | None | None |
| DLD | None | None | None | None |
| DLD | None | None | None | None |
| DLD | None | None | None | None |
| DLD | None | None | None | None |
| DLD | None | None | None | None |
| LDHA | 0.802 | 1 | 4qt0 | A |
| TNNC1 | None | None | None | None |
| TNNC1 | None | None | None | None |
| TNNC1 | None | None | None | None |
| SMAD3 | None | None | None | None |
| HARS | None | None | None | None |
| HARS | None | None | None | None |
| TNNI3 | None | None | None | None |
| TNNI3 | None | None | None | None |
| TNNI3 | None | None | None | None |
| TNNI3 | None | None | None | None |
| TNNI3 | None | None | None | None |

| | | | | |
|---|---|---|---|---|
| TNNI3 | None | None | None | None |
| ACKR1 | None | None | None | None |
| BUB1B | None | None | None | None |
| BUB1B | None | None | None | None |
| BUB1B | None | None | None | None |
| AR | None | None | None | None |
| AR | None | None | None | None |
| AR | None | None | None | None |
| AR | None | None | None | None |
| AR | None | None | None | None |
| FTO | None | None | None | None |
| FTO | None | None | None | None |
| TFAM | None | None | None | None |
| PYGL | None | None | None | None |
| PYGL | None | None | None | None |
| PYGL | None | None | None | None |
| PYGL | None | None | None | None |
| PYGL | None | None | None | None |
| PYGL | None | None | None | None |
| PYGL | None | None | None | None |
| PYGL | None | None | None | None |
| PYGL | None | None | None | None |
| PYGL | None | None | None | None |
| PYGL | None | None | None | None |
| PYGL | None | None | None | None |
| PYGL | None | None | None | None |
| PYGL | None | None | None | None |
| PYGL | None | None | None | None |
| PYGL | None | None | None | None |
| GALNS | None | None | None | None |
| GALNS | None | None | None | None |
| GALNS | None | None | None | None |
| GALNS | None | None | None | None |
| GALNS | None | None | None | None |
| GALNS | None | None | None | None |
| GALNS | None | None | None | None |
| GALNS | None | None | None | None |
| GALNS | None | None | None | None |
| GALNS | None | None | None | None |
| GALNS | None | None | None | None |
| GALNS | None | None | None | None |
| GALNS | None | None | None | None |
| GALNS | None | None | None | None |
| GALNS | None | None | None | None |
| GALNS | None | None | None | None |
| GALNS | None | None | None | None |
| GALNS | None | None | None | None |
| GALNS | None | None | None | None |
| GALNS | None | None | None | None |
| GALNS | None | None | None | None |
| GALNS | None | None | None | None |
| GALNS | None | None | None | None |
| GALNS | None | None | None | None |
| GALNS | None | None | None | None |

| | | | | |
|---|---|---|---|---|
| GALNS | None | None | None | None |
| GALNS | None | None | None | None |
| GALNS | None | None | None | None |
| GALNS | None | None | None | None |
| GALNS | None | None | None | None |
| GALNS | None | None | None | None |
| GALNS | None | None | None | None |
| GALNS | None | None | None | None |
| F10 | None | None | None | None |
| GLUD1 | None | None | None | None |
| GLUD1 | None | None | None | None |
| GLUD1 | None | None | None | None |
| GLUD1 | None | None | None | None |
| GLUD1 | None | None | None | None |
| GLUD1 | None | None | None | None |
| GLUD1 | None | None | None | None |
| GLUD1 | None | None | None | None |
| GLUD1 | None | None | None | None |
| GLUD1 | None | None | None | None |
| NOTCH1 | None | None | None | None |
| POGLUT1 | None | None | None | None |
| POGLUT1 | None | None | None | None |
| NR3C1 | None | None | None | None |
| CASP8 | None | None | None | None |
| CASP8 | None | None | None | None |
| TRAP1 | None | None | None | None |
| FANCM | None | None | None | None |
| EPOR | None | None | None | None |
| EPOR | None | None | None | None |
| NDP | None | None | None | None |
| NDP | None | None | None | None |
| NDP | None | None | None | None |
| NDP | None | None | None | None |
| NDP | None | None | None | None |
| NDP | None | None | None | None |
| NDP | None | None | None | None |
| NDP | None | None | None | None |
| NDP | None | None | None | None |
| NDP | None | None | None | None |
| NDP | None | None | None | None |
| NDP | None | None | None | None |
| NDP | None | None | None | None |
| NDP | None | None | None | None |
| NDP | None | None | None | None |
| NDP | None | None | None | None |
| NDP | None | None | None | None |
| ROGDI | None | None | None | None |
| CYP19A1 | None | None | None | None |
| CYP19A1 | None | None | None | None |
| CYP19A1 | None | None | None | None |
| CYP19A1 | None | None | None | None |
| CYP19A1 | None | None | None | None |
| CYP19A1 | None | None | None | None |
| CYP19A1 | None | None | None | None |

| | | | | |
|---|---|---|---|---|
| CYP19A1 | None | None | None | None |
| UBA5 | None | None | None | None |
| PRPS1 | None | None | None | None |
| PRPS1 | None | None | None | None |
| PRPS1 | None | None | None | None |
| PRPS1 | None | None | None | None |
| PRPS1 | None | None | None | None |
| PRPS1 | None | None | None | None |
| PRPS1 | None | None | None | None |
| PRPS1 | None | None | None | None |
| F2 | None | None | None | None |
| F2 | None | None | None | None |
| NTF4 | None | None | None | None |
| NTF4 | None | None | None | None |
| PSAP | None | None | None | None |
| PSAP | None | None | None | None |
| TAP1 | None | None | None | None |
| TAP1 | None | None | None | None |
| TAP1 | None | None | None | None |
| ARG1 | None | None | None | None |
| ARG1 | None | None | None | None |
| ARG1 | None | None | None | None |
| ARG1 | None | None | None | None |
| ARG1 | None | None | None | None |
| ARG1 | None | None | None | None |
| CRYM | None | None | None | None |
| ADSL | None | None | None | None |
| ADSL | None | None | None | None |
| ADSL | None | None | None | None |
| ADSL | None | None | None | None |
| ADSL | None | None | None | None |
| ADSL | None | None | None | None |
| ADSL | None | None | None | None |
| ADSL | None | None | None | None |
| ADSL | None | None | None | None |
| ADSL | None | None | None | None |
| ADSL | None | None | None | None |
| ADSL | None | None | None | None |
| ADSL | None | None | None | None |
| ADSL | None | None | None | None |
| ADSL | None | None | None | None |
| ADSL | None | None | None | None |
| ADSL | None | None | None | None |
| ADSL | None | None | None | None |
| ADSL | None | None | None | None |
| ADSL | None | None | None | None |
| VWF | None | None | None | None |
| PLA2G7 | None | None | None | None |
| MMP13 | None | None | None | None |
| MMP13 | None | None | None | None |
| MMP13 | None | None | None | None |
| MMP13 | None | None | None | None |
| POMGNT1 | None | None | None | None |
| POMGNT1 | None | None | None | None |

| | | | | |
|---|---|---|---|---|
| COL3A1 | None | None | None | None |
| COL3A1 | None | None | None | None |
| COL3A1 | None | None | None | None |
| COL3A1 | None | None | None | None |
| ACTN2 | None | None | None | None |
| ACTN2 | None | None | None | None |
| ACTN2 | None | None | None | None |
| ACTN2 | None | None | None | None |
| ACTN2 | None | None | None | None |
| ACTN2 | None | None | None | None |
| ACTN2 | None | None | None | None |
| UBA5 | None | None | None | None |
| ADAR | None | None | None | None |
| UMPS | None | None | None | None |
| UMPS | None | None | None | None |
| UMPS | None | None | None | None |
| TRNT1 | None | None | None | None |
| TRNT1 | None | None | None | None |
| NHEJ1 | None | None | None | None |
| NHEJ1 | None | None | None | None |
| NHEJ1 | None | None | None | None |
| C8A | None | None | None | None |
| GLUL | None | None | None | None |
| GLUL | None | None | None | None |
| GLUL | None | None | None | None |
| GYG1 | None | None | None | None |
| GYG1 | 0.923 | 1 | 3u2u | B |
| FLNA | None | None | None | None |
| FLNA | None | None | None | None |
| FLNA | None | None | None | None |
| FLNA | None | None | None | None |
| FLNA | None | None | None | None |
| FLNA | None | None | None | None |
| FLNA | None | None | None | None |
| TTN | None | None | None | None |
| TTN | None | None | None | None |
| TTN | None | None | None | None |
| TTN | None | None | None | None |
| SMS | None | None | None | None |
| SMS | None | None | None | None |
| SMS | None | None | None | None |
| C5 | None | None | None | None |
| SERPINC1 | None | None | None | None |
| SERPINC1 | None | None | None | None |
| SERPINC1 | None | None | None | None |
| SERPINC1 | None | None | None | None |
| SERPINC1 | None | None | None | None |
| SERPINC1 | None | None | None | None |
| SERPINC1 | None | None | None | None |
| SERPINC1 | None | None | None | None |
| SERPINC1 | None | None | None | None |
| SERPINC1 | None | None | None | None |
| SERPINC1 | None | None | None | None |
| SERPINC1 | None | None | None | None |

| | | | | |
|---|---|---|---|---|
| SERPINC1 | None | None | None | None |
| SERPINC1 | None | None | None | None |
| SERPINC1 | None | None | None | None |
| SERPINC1 | None | None | None | None |
| SERPINC1 | None | None | None | None |
| SERPINC1 | None | None | None | None |
| SERPINC1 | None | None | None | None |
| SERPINC1 | None | None | None | None |
| SERPINC1 | None | None | None | None |
| ETFB | None | None | None | None |
| ETFB | None | None | None | None |
| ETFB | None | None | None | None |
| ETFB | None | None | None | None |
| ETFB | None | None | None | None |
| ETFB | None | None | None | None |
| DIABLO | None | None | None | None |
| DIABLO | None | None | None | None |
| DPYS | None | None | None | None |
| DPYS | None | None | None | None |
| DPYS | None | None | None | None |
| DPYS | None | None | None | None |
| DPYS | None | None | None | None |
| GNE | None | None | None | None |
| GNE | None | None | None | None |
| GNE | None | None | None | None |
| GNE | None | None | None | None |
| GNE | None | None | None | None |
| GNE | None | None | None | None |
| GNE | None | None | None | None |
| GNE | None | None | None | None |
| GNE | None | None | None | None |
| GNMT | None | None | None | None |
| GNMT | None | None | None | None |
| CTH | None | None | None | None |
| CTH | None | None | None | None |
| CTH | None | None | None | None |
| USH1C | None | None | None | None |
| PSEN1 | None | None | None | None |
| CHEK2 | None | None | None | None |
| CHEK2 | None | None | None | None |
| CHEK2 | None | None | None | None |
| CHEK2 | None | None | None | None |
| CHEK2 | None | None | None | None |
| CHEK2 | None | None | None | None |
| CHEK2 | None | None | None | None |
| CHEK2 | None | None | None | None |
| CHEK2 | None | None | None | None |
| SNCA | None | None | None | None |
| SNCA | None | None | None | None |
| GAN | None | None | None | None |
| GAN | None | None | None | None |
| GAN | None | None | None | None |
| SBDS | None | None | None | None |
| SBDS | None | None | None | None |

| | | | | |
|---|---|---|---|---|
| SBDS | None | None | None | None |
| SBDS | None | None | None | None |
| SBDS | None | None | None | None |
| SBDS | None | None | None | None |
| MPO | | 1 | 3 | 3f9p | A |
| MPO | None | None | None | None |
| KCNH2 | None | None | None | None |
| KCNH2 | None | None | None | None |
| KCNH2 | None | None | None | None |
| KCNH2 | None | None | None | None |
| TEK | None | None | None | None |
| TEK | None | None | None | None |
| TEK | None | None | None | None |
| TEK | None | None | None | None |
| NPC2 | None | None | None | None |
| NPC2 | None | None | None | None |
| NPC2 | None | None | None | None |
| NPC2 | None | None | None | None |
| NPC2 | None | None | None | None |
| NPC2 | None | None | None | None |
| NPC2 | None | None | None | None |
| PLG | None | None | None | None |
| PCSK9 | None | None | None | None |
| SFTPC | None | None | None | None |
| SFTPC | None | None | None | None |
| SFTPC | None | None | None | None |
| CD40LG | None | None | None | None |
| CD40LG | None | None | None | None |
| CD40LG | None | None | None | None |
| CD40LG | None | None | None | None |
| CD40LG | None | None | None | None |
| TAT | None | None | None | None |
| TAT | None | None | None | None |
| TAT | None | None | None | None |
| EP300 | None | None | None | None |
| FARS2 | None | None | None | None |
| FARS2 | None | None | None | None |
| FARS2 | None | None | None | None |
| FARS2 | None | None | None | None |
| FARS2 | None | None | None | None |
| FARS2 | None | None | None | None |
| FARS2 | None | None | None | None |
| FARS2 | None | None | None | None |
| FARS2 | None | None | None | None |
| FARS2 | None | None | None | None |
| FARS2 | None | None | None | None |
| FARS2 | None | None | None | None |
| FARS2 | None | None | None | None |
| MASP2 | None | None | None | None |
| MASP2 | None | None | None | None |
| MASP2 | None | None | None | None |
| MASP2 | None | None | None | None |
| STIM1 | None | None | None | None |
| CHRM2 | None | None | None | None |

| | | | | |
|---|---|---|---|---|
| MCFD2 | None | None | None | None |
| MCFD2 | None | None | None | None |
| MCFD2 | None | None | None | None |
| KITLG | None | None | None | None |
| F12 | None | None | None | None |
| F12 | None | None | None | None |
| ERAP1 | None | None | None | None |
| GTF2H5 | None | None | None | None |
| GTF2H5 | None | None | None | None |
| THBD | None | None | None | None |
| MRPL28 | None | None | None | None |
| MRPL28 | None | None | None | None |
| HTRA1 | None | None | None | None |
| HTRA1 | None | None | None | None |
| HTRA1 | None | None | None | None |
| HTRA1 | None | None | None | None |
| HTRA1 | None | None | None | None |
| HTRA1 | None | None | None | None |
| KARS | None | None | None | None |
| KARS | None | None | None | None |
| KARS | None | None | None | None |
| KARS | None | None | None | None |
| KARS | None | None | None | None |
| HK1 | None | None | None | None |
| GARS | None | None | None | None |
| GARS | None | None | None | None |
| GARS | None | None | None | None |
| GARS | None | None | None | None |
| GARS | None | None | None | None |
| GARS | None | None | None | None |
| GARS | None | None | None | None |
| IL12B | None | None | None | None |
| HADH | None | None | None | None |
| HADH | None | None | None | None |
| HADH | None | None | None | None |
| HADH | None | None | None | None |
| HADH | None | None | None | None |
| RRM2B | None | None | None | None |
| RRM2B | None | None | None | None |
| RRM2B | None | None | None | None |
| RRM2B | None | None | None | None |
| RRM2B | None | None | None | None |
| RRM2B | None | None | None | None |
| RRM2B | None | None | None | None |
| RRM2B | None | None | None | None |
| RRM2B | None | None | None | None |
| RRM2B | None | None | None | None |
| RRM2B | None | None | None | None |
| RRM2B | None | None | None | None |
| RRM2B | None | None | None | None |
| RRM2B | None | None | None | None |
| RRM2B | None | None | None | None |
| MARVELD2 | None | None | None | None |
| TLR4 | None | None | None | None |

| | | | | |
|---|---|---|---|---|
| TTN | None | None | None | None |
| TTN | None | None | None | None |
| TNFRSF13B | None | None | None | None |
| POLG | None | None | None | None |
| POLG | None | None | None | None |
| POLG | None | None | None | None |
| POLG | None | None | None | None |
| POLG | None | None | None | None |
| POLG | None | None | None | None |
| POLG | None | None | None | None |
| POLG | None | None | None | None |
| POLG | None | None | None | None |
| POLG | None | None | None | None |
| POLG | None | None | None | None |
| POLG | None | None | None | None |
| POLG | None | None | None | None |
| POLG | None | None | None | None |
| POLG | None | None | None | None |
| POLG | None | None | None | None |
| POLG | None | None | None | None |
| POLG | None | None | None | None |
| POLG | None | None | None | None |
| POLG | None | None | None | None |
| POLG | None | None | None | None |
| POLG | None | None | None | None |
| POLG | None | None | None | None |
| POLG | None | None | None | None |
| POLG | None | None | None | None |
| POLG | None | None | None | None |
| POLG | None | None | None | None |
| POLG | None | None | None | None |
| POLG | None | None | None | None |
| POLG | None | None | None | None |
| POLG | None | None | None | None |
| POLG | None | None | None | None |
| POLG | None | None | None | None |
| POLG | None | None | None | None |
| POLG | None | None | None | None |
| POLG | None | None | None | None |
| POLG | None | None | None | None |
| POLG | None | None | None | None |
| POLG | None | None | None | None |
| POLG | None | None | None | None |
| POLG | None | None | None | None |
| POLG | None | None | None | None |
| POLG | None | None | None | None |
| POLG | None | None | None | None |
| POLG | None | None | None | None |
| POLG | None | None | None | None |
| POLG | None | None | None | None |
| POLG | None | None | None | None |
| POLG | None | None | None | None |
| POLG | None | None | None | None |
| POLG | None | None | None | None |
| UROD | None | None | None | None |

| | | | | |
|---|---|---|---|---|
| UROD | None | None | None | None |
| UROD | None | None | None | None |
| UROD | None | None | None | None |
| UROD | None | None | None | None |
| UROD | None | None | None | None |
| UROD | None | None | None | None |
| UROD | 0.842 | 2 | 3gvv | A |
| UROD | None | None | None | None |
| UROD | None | None | None | None |
| PCK1 | None | None | None | None |
| PCK1 | None | None | None | None |
| PCK1 | None | None | None | None |
| PCK1 | None | None | None | None |
| PCK1 | None | None | None | None |
| HEXA | None | None | None | None |
| HEXA | None | None | None | None |
| HEXA | None | None | None | None |
| HEXA | None | None | None | None |
| HEXA | None | None | None | None |
| HEXA | None | None | None | None |
| HEXA | None | None | None | None |
| HEXA | None | None | None | None |
| HEXA | None | None | None | None |
| HEXA | None | None | None | None |
| HEXA | None | None | None | None |
| HEXA | None | None | None | None |
| HEXA | None | None | None | None |
| HEXA | None | None | None | None |
| HEXA | None | None | None | None |
| HEXA | None | None | None | None |
| HEXA | None | None | None | None |
| HEXA | None | None | None | None |
| HEXA | None | None | None | None |
| HEXA | None | None | None | None |
| HEXA | None | None | None | None |
| HEXA | None | None | None | None |
| HEXA | None | None | None | None |
| HEXA | None | None | None | None |
| HEXA | None | None | None | None |
| HEXA | None | None | None | None |
| HEXA | None | None | None | None |
| HEXA | None | None | None | None |
| HEXA | None | None | None | None |
| HEXA | None | None | None | None |
| CFTR | None | None | None | None |
| CFTR | None | None | None | None |
| CFTR | None | None | None | None |
| CFTR | None | None | None | None |
| CFTR | None | None | None | None |
| CFTR | None | None | None | None |
| CFTR | None | None | None | None |
| CFTR | None | None | None | None |
| CFTR | None | None | None | None |

| | | | | |
|---|---|---|---|---|
| CFTR | None | None | None | None |
| CFTR | None | None | None | None |
| CFTR | None | None | None | None |
| CFTR | None | None | None | None |
| CFTR | None | None | None | None |
| CFTR | None | None | None | None |
| CFTR | None | None | None | None |
| CFTR | None | None | None | None |
| CFTR | None | None | None | None |
| CFTR | None | None | None | None |
| CFTR | None | None | None | None |
| CFTR | None | None | None | None |
| CFTR | None | None | None | None |
| CFTR | None | None | None | None |
| CFTR | None | None | None | None |
| CFTR | None | None | None | None |
| CFTR | None | None | None | None |
| CFTR | None | None | None | None |
| CFTR | None | None | None | None |
| CFTR | None | None | None | None |
| CFTR | None | None | None | None |
| CFTR | None | None | None | None |
| CFTR | None | None | None | None |
| CFTR | None | None | None | None |
| CFTR | None | None | None | None |
| FAN1 | None | None | None | None |
| CBS | None | None | None | None |
| CBS | None | None | None | None |
| CBS | None | None | None | None |
| CBS | None | None | None | None |
| CBS | None | None | None | None |
| CBS | None | None | None | None |
| CBS | None | None | None | None |
| CBS | None | None | None | None |
| CBS | None | None | None | None |
| CBS | None | None | None | None |
| CBS | None | None | None | None |
| CBS | None | None | None | None |
| CBS | None | None | None | None |
| CBS | None | None | None | None |
| CBS | None | None | None | None |
| CBS | None | None | None | None |
| CBS | None | None | None | None |
| CBS | None | None | None | None |
| CBS | None | None | None | None |
| CBS | None | None | None | None |
| CBS | None | None | None | None |
| CBS | None | None | None | None |
| CBS | None | None | None | None |
| CBS | None | None | None | None |
| CBS | None | None | None | None |
| CBS | None | None | None | None |
| CBS | None | None | None | None |
| CBS | None | None | None | None |

| | | | | |
|---|---|---|---|---|
| CBS | None | None | None | None |
| CBS | None | None | None | None |
| CBS | None | None | None | None |
| CBS | None | None | None | None |
| CBS | None | None | None | None |
| CBS | None | None | None | None |
| CBS | None | None | None | None |
| CBS | None | None | None | None |
| TLR3 | None | None | None | None |
| LIG4 | None | None | None | None |
| LIG4 | None | None | None | None |
| ASPA | None | None | None | None |
| ASPA | None | None | None | None |
| ASPA | None | None | None | None |
| ASPA | None | None | None | None |
| ASPA | None | None | None | None |
| ASPA | None | None | None | None |
| ASPA | None | None | None | None |
| ASPA | None | None | None | None |
| ASPA | None | None | None | None |
| ASPA | None | None | None | None |
| ASPA | None | None | None | None |
| ASPA | None | None | None | None |
| ASPA | None | None | None | None |
| CFH | None | None | None | None |
| FCGR3A | None | None | None | None |
| FCGR3A | None | None | None | None |
| GRIN2A | None | None | None | None |
| GRIN2A | None | None | None | None |
| GRIN2A | None | None | None | None |
| GLA | None | None | None | None |
| GLA | None | None | None | None |
| GLA | None | None | None | None |
| GLA | None | None | None | None |
| GLA | None | None | None | None |
| GLA | None | None | None | None |
| GLA | None | None | None | None |
| GLA | None | None | None | None |
| GLA | None | None | None | None |
| GLA | None | None | None | None |
| GLA | None | None | None | None |
| GLA | None | None | None | None |
| GLA | None | None | None | None |
| GLA | None | None | None | None |
| GLA | None | None | None | None |
| GLA | None | None | None | None |
| GLA | None | None | None | None |
| GLA | None | None | None | None |
| GLA | None | None | None | None |
| GLA | None | None | None | None |
| GLA | None | None | None | None |
| GLA | None | None | None | None |
| GLA | None | None | None | None |
| GLA | None | None | None | None |
| GLA | None | None | None | None |

| | | | | |
|---|---|---|---|---|
| GLA | None | None | None | None |
| GLA | None | None | None | None |
| GLA | None | None | None | None |
| GLA | None | None | None | None |
| GLA | None | None | None | None |
| GLA | None | None | None | None |
| GLA | None | None | None | None |
| SERPINA3 | None | None | None | None |
| MTAP | None | None | None | None |
| MTAP | None | None | None | None |
| MTAP | None | None | None | None |
| BLMH | None | None | None | None |
| TUSC3 | None | None | None | None |
| ISCU | None | None | None | None |
| CYP11B2 | None | None | None | None |
| CYP11B2 | None | None | None | None |
| CYP11B2 | None | None | None | None |
| CYP11B2 | None | None | None | None |
| CYP11B2 | None | None | None | None |
| CYP11B2 | None | None | None | None |
| CYP11B2 | None | None | None | None |
| CYP11B2 | None | None | None | None |
| CYP11B2 | None | None | None | None |
| CYP11B2 | None | None | None | None |
| CYP11B2 | None | None | None | None |
| CYP11B2 | None | None | None | None |
| CYP11B2 | None | None | None | None |
| CYP11B2 | None | None | None | None |
| CYP11B2 | None | None | None | None |
| CYP11B2 | None | None | None | None |
| CYP11B2 | None | None | None | None |
| CYP11B2 | None | None | None | None |
| CYP11B2 | None | None | None | None |
| CYP11B2 | None | None | None | None |
| CRADD | None | None | None | None |
| CRADD | None | None | None | None |
| FGA | None | None | None | None |
| FGA | None | None | None | None |
| FGA | None | None | None | None |
| FGA | None | None | None | None |
| FGA | None | None | None | None |
| FGA | None | None | None | None |
| FGA | None | None | None | None |
| FGA | None | None | None | None |
| IYD | 1 | 1 | 4ttb | A |
| IYD | None | None | None | None |
| IYD | None | None | None | None |
| IYD | None | None | None | None |
| TCAP | None | None | None | None |
| TCAP | None | None | None | None |
| TCAP | None | None | None | None |
| TCAP | None | None | None | None |
| TCAP | None | None | None | None |
| CYP7A1 | None | None | None | None |

| | | | | |
|---|---|---|---|---|
| TEK | None | None | None | None |
| TEK | None | None | None | None |
| TEK | None | None | None | None |
| TEK | None | None | None | None |
| TEK | None | None | None | None |
| TEK | None | None | None | None |
| TEK | None | None | None | None |
| CD8A | None | None | None | None |
| HERC2 | None | None | None | None |
| NR1H3 | None | None | None | None |
| GRHPR | None | None | None | None |
| GRHPR | None | None | None | None |
| GRHPR | None | None | None | None |
| GRHPR | None | None | None | None |
| GRHPR | None | None | None | None |
| GRHPR | None | None | None | None |
| GRHPR | None | None | None | None |
| GBE1 | None | None | None | None |
| GBE1 | None | None | None | None |
| GBE1 | None | None | None | None |
| GBE1 | None | None | None | None |
| GBE1 | None | None | None | None |
| GBE1 | None | None | None | None |
| GBE1 | None | None | None | None |
| GBE1 | None | None | None | None |
| GBE1 | None | None | None | None |
| GBE1 | None | None | None | None |
| GBE1 | None | None | None | None |
| GBE1 | None | None | None | None |
| GBE1 | None | None | None | None |
| GBE1 | None | None | None | None |
| GBE1 | None | None | None | None |
| GBE1 | None | None | None | None |
| GBE1 | None | None | None | None |
| GBE1 | None | None | None | None |
| GBE1 | None | None | None | None |
| GBE1 | None | None | None | None |
| PC | None | None | None | None |
| PC | None | None | None | None |
| PC | None | None | None | None |
| PC | None | None | None | None |
| PC | None | None | None | None |
| PC | None | None | None | None |
| PC | None | None | None | None |
| PC | None | None | None | None |
| MSH6 | None | None | None | None |
| MSH6 | None | None | None | None |
| MSH6 | None | None | None | None |
| MSH6 | None | None | None | None |
| MSH6 | None | None | None | None |
| MSH6 | None | None | None | None |
| MSH6 | None | None | None | None |
| MSH6 | None | None | None | None |
| MSH6 | None | None | None | None |

| | | | | |
|---|---|---|---|---|
| MSH6 | None | None | None | None |
| MSH6 | None | None | None | None |
| MSH6 | None | None | None | None |
| MSH6 | None | None | None | None |
| MSH6 | None | None | None | None |
| MSH6 | None | None | None | None |
| MSH6 | None | None | None | None |
| MSH6 | None | None | None | None |
| MSH6 | None | None | None | None |
| MSH6 | None | None | None | None |
| MSH6 | None | None | None | None |
| MSH6 | None | None | None | None |
| MSH6 | None | None | None | None |
| MSH6 | None | None | None | None |
| MSH6 | None | None | None | None |
| MSH6 | None | None | None | None |
| MSH6 | None | None | None | None |
| MSH6 | None | None | None | None |
| MSH6 | None | None | None | None |
| MSH6 | None | None | None | None |
| MSH6 | None | None | None | None |
| RNASEL | None | None | None | None |
| ORC1 | None | None | None | None |
| ORC1 | None | None | None | None |
| ORC1 | None | None | None | None |
| KCNH2 | None | None | None | None |
| KCNH2 | None | None | None | None |
| KCNH2 | None | None | None | None |
| KCNH2 | None | None | None | None |
| KCNH2 | None | None | None | None |
| KCNH2 | None | None | None | None |
| KCNH2 | None | None | None | None |
| KCNH2 | None | None | None | None |
| KCNH2 | None | None | None | None |
| KCNH2 | None | None | None | None |
| KCNH2 | None | None | None | None |
| KCNH2 | None | None | None | None |
| KCNH2 | None | None | None | None |
| KCNH2 | None | None | None | None |
| KCNH2 | None | None | None | None |
| KCNH2 | None | None | None | None |
| KCNH2 | None | None | None | None |
| KCNH2 | None | None | None | None |
| KCNH2 | None | None | None | None |
| KCNH2 | None | None | None | None |
| KCNH2 | None | None | None | None |
| KCNH2 | None | None | None | None |
| KCNH2 | None | None | None | None |
| KCNH2 | None | None | None | None |
| MAT1A | None | None | None | None |
| MAT1A | None | None | None | None |
| MAT1A | None | None | None | None |
| MAT1A | None | None | None | None |
| MAT1A | None | None | None | None |

| | | | | |
|---|---|---|---|---|
| MAT1A | None | None | None | None |
| MAT1A | None | None | None | None |
| MAT1A | None | None | None | None |
| INSR | None | None | None | None |
| INSR | None | None | None | None |
| INSR | None | None | None | None |
| INSR | None | None | None | None |
| AMT | None | None | None | None |
| AMT | None | None | None | None |
| AMT | None | None | None | None |
| AMT | None | None | None | None |
| AMT | None | None | None | None |
| AMT | None | None | None | None |
| AMT | None | None | None | None |
| AMT | None | None | None | None |
| AMT | None | None | None | None |
| CSNK1D | None | None | None | None |
| SF3B1 | None | None | None | None |
| AGA | None | None | None | None |
| AGA | None | None | None | None |
| AGA | None | None | None | None |
| AGA | None | None | None | None |
| AGA | None | None | None | None |
| AGA | None | None | None | None |
| AGA | None | None | None | None |
| NECTIN4 | None | None | None | None |
| RS1 | None | None | None | None |
| RS1 | None | None | None | None |
| RS1 | None | None | None | None |
| RS1 | None | None | None | None |
| TBC1D7 | None | None | None | None |
| CTSA | None | None | None | None |
| CTSA | None | None | None | None |
| TKT | None | None | None | None |
| RP2 | None | None | None | None |
| RP2 | None | None | None | None |
| RP2 | None | None | None | None |
| RP2 | None | None | None | None |
| LYRM4 | None | None | None | None |
| SOS1 | None | None | None | None |
| SOS1 | None | None | None | None |
| SOS1 | None | None | None | None |
| SOS1 | None | None | None | None |
| SOS1 | None | None | None | None |
| SOS1 | None | None | None | None |
| HUWE1 | None | None | None | None |
| HUWE1 | None | None | None | None |
| TPI1 | None | None | None | None |
| TPI1 | 0.886 | 1 | 4unk | B |
| TPI1 | None | None | None | None |
| MSH2 | None | None | None | None |
| MSH2 | None | None | None | None |
| MSH2 | None | None | None | None |
| MSH2 | None | None | None | None |

| | | | | |
|---|---|---|---|---|
| MSH2 | None | None | None | None |
| MSH2 | None | None | None | None |
| MSH2 | None | None | None | None |
| MSH2 | None | None | None | None |
| MSH2 | None | None | None | None |
| MSH2 | None | None | None | None |
| MSH2 | None | None | None | None |
| MSH2 | None | None | None | None |
| MSH2 | None | None | None | None |
| MSH2 | None | None | None | None |
| MSH2 | None | None | None | None |
| MSH2 | None | None | None | None |
| MSH2 | None | None | None | None |
| MSH2 | None | None | None | None |
| MSH2 | None | None | None | None |
| MSH2 | None | None | None | None |
| MSH2 | None | None | None | None |
| MSH2 | None | None | None | None |
| MSH2 | None | None | None | None |
| MSH2 | None | None | None | None |
| MSH2 | None | None | None | None |
| MSH2 | None | None | None | None |
| MSH2 | None | None | None | None |
| MSH2 | None | None | None | None |
| MSH2 | None | None | None | None |
| MSH2 | None | None | None | None |
| MSH2 | None | None | None | None |
| MSH2 | None | None | None | None |
| MSH2 | None | None | None | None |
| MSH2 | None | None | None | None |
| MSH2 | None | None | None | None |
| MSH2 | None | None | None | None |
| GALE | None | None | None | None |
| GALE | None | None | None | None |
| GALE | None | None | None | None |
| GALE | None | None | None | None |
| GALE | None | None | None | None |
| GALE | None | None | None | None |
| GALE | None | None | None | None |
| GALE | None | None | None | None |
| NSD2 | None | None | None | None |
| SRP72 | None | None | None | None |
| SRP72 | None | None | None | None |
| SRP72 | None | None | None | None |
| POLK | None | None | None | None |
| GALT | None | None | None | None |
| GALT | None | None | None | None |
| GALT | None | None | None | None |
| GALT | None | None | None | None |
| GALT | None | None | None | None |
| GALT | None | None | None | None |
| GALT | None | None | None | None |
| GALT | None | None | None | None |
| GALT | None | None | None | None |

| | | | | |
|---|---|---|---|---|
| GALT | None | None | None | None |
| GALT | None | None | None | None |
| GALT | None | None | None | None |
| GALT | None | None | None | None |
| GALT | None | None | None | None |
| GALT | None | None | None | None |
| GALT | None | None | None | None |
| GALT | None | None | None | None |
| GALT | None | None | None | None |
| GALT | None | None | None | None |
| GALT | None | None | None | None |
| GALT | None | None | None | None |
| GALT | None | None | None | None |
| GALT | None | None | None | None |
| GALT | None | None | None | None |
| GALT | None | None | None | None |
| GALT | None | None | None | None |
| GALT | None | None | None | None |
| GALT | None | None | None | None |
| GALT | None | None | None | None |
| GALT | None | None | None | None |
| GALT | None | None | None | None |
| GALT | None | None | None | None |
| GALT | None | None | None | None |
| GALT | None | None | None | None |
| GALT | None | None | None | None |
| GALT | None | None | None | None |
| GALT | None | None | None | None |
| PYCR1 | None | None | None | None |
| PYCR1 | None | None | None | None |
| PYCR1 | None | None | None | None |
| PYCR1 | None | None | None | None |
| F8 | None | None | None | None |
| F8 | None | None | None | None |
| F8 | None | None | None | None |
| F8 | None | None | None | None |
| F8 | None | None | None | None |
| F8 | None | None | None | None |
| F8 | None | None | None | None |
| F8 | None | None | None | None |
| F8 | None | None | None | None |
| F8 | None | None | None | None |
| F8 | None | None | None | None |
| F8 | None | None | None | None |
| F8 | None | None | None | None |
| F8 | None | None | None | None |
| F8 | None | None | None | None |
| F8 | None | None | None | None |
| ANK1 | None | None | None | None |
| CHAT | None | None | None | None |
| CHAT | None | None | None | None |
| CHAT | None | None | None | None |
| CHAT | None | None | None | None |
| CHAT | None | None | None | None |

| | | | | |
|---|---|---|---|---|
| CHAT | None | None | None | None |
| CHAT | None | None | None | None |
| CHAT | None | None | None | None |
| CHAT | None | None | None | None |
| CHAT | None | None | None | None |
| CHAT | None | None | None | None |
| CHAT | None | None | None | None |
| CHAT | None | None | None | None |
| CHAT | None | None | None | None |
| CHAT | None | None | None | None |
| CHAT | None | None | None | None |
| CHAT | None | None | None | None |
| GHR | None | None | None | None |
| GHR | None | None | None | None |
| GHR | None | None | None | None |
| GHR | None | None | None | None |
| GHR | None | None | None | None |
| GHR | None | None | None | None |
| GHR | None | None | None | None |
| GHR | None | None | None | None |
| GHR | None | None | None | None |
| GHR | None | None | None | None |
| GHR | None | None | None | None |
| GHR | None | None | None | None |
| GHR | None | None | None | None |
| GHR | None | None | None | None |
| FZD4 | None | None | None | None |
| FZD4 | None | None | None | None |
| FZD4 | None | None | None | None |
| MTR | None | None | None | None |
| MTR | None | None | None | None |
| MTR | None | None | None | None |
| GM2A | None | None | None | None |
| GM2A | None | None | None | None |
| GM2A | None | None | None | None |
| GM2A | None | None | None | None |
| GM2A | None | None | None | None |
| CRYGD | None | None | None | None |
| CRYGD | None | None | None | None |
| CRYGD | None | None | None | None |
| CRYGD | None | None | None | None |
| CRYGD | None | None | None | None |
| CRYGD | None | None | None | None |
| CRYGD | None | None | None | None |
| CRYGD | None | None | None | None |
| CRYGD | None | None | None | None |
| TLR5 | None | None | None | None |
| HSD17B10 | 0.855 | 1 | 1u7t | B |
| HSD17B10 | None | None | None | None |
| HSD17B10 | None | None | None | None |
| HSD17B10 | None | None | None | None |
| TBC1D20 | None | None | None | None |
| FBN1 | None | None | None | None |
| FBN1 | None | None | None | None |

| | | | | |
|---|---|---|---|---|
| FBN1 | None | None | None | None |
| FBN1 | None | None | None | None |
| FBN1 | None | None | None | None |
| FBN1 | None | None | None | None |
| FBN1 | None | None | None | None |
| FXN | None | None | None | None |
| FXN | None | None | None | None |
| FXN | None | None | None | None |
| FXN | None | None | None | None |
| MYO5A | None | None | None | None |
| CTSK | None | None | None | None |
| CTSK | None | None | None | None |
| SCN5A | None | None | None | None |
| SCN5A | None | None | None | None |
| SOS1 | None | None | None | None |
| SOS1 | None | None | None | None |
| SOS1 | None | None | None | None |
| SOS1 | None | None | None | None |
| SOS1 | None | None | None | None |
| SOS1 | None | None | None | None |
| SOS1 | None | None | None | None |
| SOS1 | None | None | None | None |
| SOS1 | None | None | None | None |
| SOS1 | None | None | None | None |
| SOS1 | None | None | None | None |
| SOS1 | None | None | None | None |
| GATM | None | None | None | None |
| GATM | None | None | None | None |
| GATM | None | None | None | None |
| GATM | None | None | None | None |
| GATM | None | None | None | None |
| GATM | None | None | None | None |
| GATM | None | None | None | None |
| MMP2 | None | None | None | None |
| MMP2 | None | None | None | None |
| MMP2 | None | None | None | None |
| MMP2 | None | None | None | None |
| NDUFB9 | None | None | None | None |
| NDUFB9 | None | None | None | None |
| GH1 | None | None | None | None |
| GH1 | None | None | None | None |
| GH1 | None | None | None | None |
| SLC6A4 | None | None | None | None |
| G6PD | None | None | None | None |
| G6PD | None | None | None | None |
| G6PD | None | None | None | None |
| G6PD | None | None | None | None |
| G6PD | None | None | None | None |
| G6PD | None | None | None | None |
| G6PD | None | None | None | None |
| G6PD | None | None | None | None |
| G6PD | None | None | None | None |
| G6PD | None | None | None | None |
| G6PD | None | None | None | None |

| | | | | |
|---|---|---|---|---|
| G6PD | None | None | None | None |
| G6PD | None | None | None | None |
| G6PD | None | None | None | None |
| G6PD | None | None | None | None |
| G6PD | None | None | None | None |
| G6PD | None | None | None | None |
| G6PD | None | None | None | None |
| G6PD | None | None | None | None |
| G6PD | None | None | None | None |
| CST3 | None | None | None | None |
| LMNA | None | None | None | None |
| LMNA | None | None | None | None |
| NDUFA9 | None | None | None | None |
| NDUFA9 | None | None | None | None |
| NDUFA9 | None | None | None | None |
| NDUFA9 | None | None | None | None |
| YARS | None | None | None | None |
| YARS | None | None | None | None |
| YARS | None | None | None | None |
| YARS | None | None | None | None |
| EPHA2 | None | None | None | None |
| EPHA2 | None | None | None | None |
| EPHA2 | None | None | None | None |
| EPHA2 | None | None | None | None |
| DICER1 | None | None | None | None |
| DICER1 | None | None | None | None |
| SNX10 | None | None | None | None |
| SNX10 | None | None | None | None |
| TTN | None | None | None | None |
| TTN | None | None | None | None |
| GATA3 | None | None | None | None |
| C5 | None | None | None | None |
| THBD | None | None | None | None |
| BIN1 | None | None | None | None |
| BIN1 | None | None | None | None |
| SMARCAL1 | None | None | None | None |
| CFB | None | None | None | None |
| CYC1 | None | None | None | None |
| ANK2 | None | None | None | None |
| CHMP4B | None | None | None | None |
| CHMP4B | None | None | None | None |
| ERBB2 | None | None | None | None |
| ERBB2 | None | None | None | None |
| POLG2 | None | None | None | None |
| POLG2 | None | None | None | None |
| POLG2 | None | None | None | None |
| POLG2 | None | None | None | None |
| POLG2 | None | None | None | None |
| POLG2 | None | None | None | None |
| POLG2 | None | None | None | None |
| POLG2 | None | None | None | None |
| C3 | None | None | None | None |
| C3 | None | None | None | None |
| RPGR | None | None | None | None |

| | | | | |
|---|---|---|---|---|
| RPGR | None | None | None | None |
| RPGR | None | None | None | None |
| RPGR | None | None | None | None |
| RPGR | None | None | None | None |
| C1QA | None | None | None | None |
| PRKG1 | None | None | None | None |
| IGBP1 | None | None | None | None |
| ENO3 | None | None | None | None |
| ENO3 | None | None | None | None |
| ENO3 | None | None | None | None |
| ENO3 | None | None | None | None |
| ENO3 | None | None | None | None |
| ENO3 | 0.897 | 2 | 2xsx | B |
| MCPH1 | None | None | None | None |
| MCPH1 | None | None | None | None |
| MCPH1 | None | None | None | None |
| MCPH1 | None | None | None | None |
| MCPH1 | None | None | None | None |
| SNRNP200 | None | None | None | None |
| SNRNP200 | None | None | None | None |
| DNMT1 | None | None | None | None |
| DNMT1 | None | None | None | None |
| DNMT1 | None | None | None | None |
| AKT1 | None | None | None | None |
| NDUFS2 | None | None | None | None |
| ACVR2B | None | None | None | None |
| HMGCL | None | None | None | None |
| HMGCL | None | None | None | None |
| HMGCL | None | None | None | None |
| JUP | None | None | None | None |
| JUP | None | None | None | None |
| JUP | None | None | None | None |
| JUP | None | None | None | None |
| JUP | None | None | None | None |
| JUP | None | None | None | None |
| JUP | None | None | None | None |
| JUP | None | None | None | None |
| SPINK1 | None | None | None | None |
| SPINK1 | None | None | None | None |
| SPINK1 | None | None | None | None |
| VCL | None | None | None | None |
| VCL | None | None | None | None |
| NOTCH1 | None | None | None | None |
| NOTCH1 | None | None | None | None |
| NOTCH1 | None | None | None | None |
| NDUFB3 | None | None | None | None |
| IKBKG | None | None | None | None |
| GSN | None | None | None | None |
| GSN | None | None | None | None |
| GSN | None | None | None | None |
| GSN | None | None | None | None |
| GSN | None | None | None | None |
| GSN | None | None | None | None |
| GSN | None | None | None | None |

| | | | | |
|---|---|---|---|---|
| GSN | None | None | None | None |
| GSN | None | None | None | None |
| CLC | None | None | None | None |
| RAF1 | None | None | None | None |
| RAF1 | None | None | None | None |
| RAF1 | None | None | None | None |
| NDUFS8 | None | None | None | None |
| NDUFS8 | None | None | None | None |
| NDUFS8 | None | None | None | None |
| NDUFS8 | None | None | None | None |
| NDUFS8 | None | None | None | None |
| CASR | None | None | None | None |
| CASR | None | None | None | None |
| CASR | None | None | None | None |
| CASR | None | None | None | None |
| CASR | None | None | None | None |
| CASR | None | None | None | None |
| CASR | None | None | None | None |
| CASR | None | None | None | None |
| CASR | None | None | None | None |
| CASR | None | None | None | None |
| CASR | None | None | None | None |
| CASR | None | None | None | None |
| CASR | None | None | None | None |
| CASR | None | None | None | None |
| CASR | None | None | None | None |
| CASR | None | None | None | None |
| CASR | None | None | None | None |
| CASR | None | None | None | None |
| CASR | None | None | None | None |
| CASR | None | None | None | None |
| CASR | None | None | None | None |
| CASR | None | None | None | None |
| CASR | None | None | None | None |
| CASR | None | None | None | None |
| CASR | None | None | None | None |
| CASR | None | None | None | None |
| CASR | None | None | None | None |
| CASR | None | None | None | None |
| CASR | None | None | None | None |
| CASR | None | None | None | None |
| VCL | None | None | None | None |
| VCL | None | None | None | None |
| SMPD1 | None | None | None | None |
| SMPD1 | None | None | None | None |
| SMPD1 | None | None | None | None |
| KDM6A | None | None | None | None |
| KCNQ1 | None | None | None | None |
| KCNQ1 | None | None | None | None |
| KCNQ1 | None | None | None | None |
| KCNQ1 | None | None | None | None |
| KCNQ1 | None | None | None | None |
| KCNQ1 | None | None | None | None |
| NDUFA12 | None | None | None | None |
| GLRX5 | None | None | None | None |

| | | | | |
|---|---|---|---|---|
| PALB2 | None | None | None | None |
| PALB2 | None | None | None | None |
| PALB2 | None | None | None | None |
| PALB2 | None | None | None | None |
| PALB2 | None | None | None | None |
| PALB2 | None | None | None | None |
| PALB2 | None | None | None | None |
| PALB2 | None | None | None | None |
| PALB2 | None | None | None | None |
| PALB2 | None | None | None | None |
| ALDOB | None | None | None | None |
| ALDOB | None | None | None | None |
| ALDOB | None | None | None | None |
| ALDOB | None | None | None | None |
| ALDOB | None | None | None | None |
| ALDOB | None | None | None | None |
| ALDOB | None | None | None | None |
| GNE | None | None | None | None |
| GNE | None | None | None | None |
| GNE | None | None | None | None |
| GNE | None | None | None | None |
| GNE | None | None | None | None |
| GNE | None | None | None | None |
| GNE | None | None | None | None |
| GNE | None | None | None | None |
| GNE | None | None | None | None |
| GNE | None | None | None | None |
| GNE | None | None | None | None |
| GNE | None | None | None | None |
| CHRNB2 | None | None | None | None |
| CHRNB2 | None | None | None | None |
| ERCC3 | None | None | None | None |
| ERCC3 | None | None | None | None |
| ERCC3 | None | None | None | None |
| ERCC3 | None | None | None | None |
| ERCC3 | None | None | None | None |
| UNC119 | None | None | None | None |
| ATL1 | None | None | None | None |
| ATL1 | None | None | None | None |
| ATL1 | None | None | None | None |
| ATL1 | 0.8 | 1 | 3q5d | A |
| ATL1 | None | None | None | None |
| ATL1 | None | None | None | None |
| ATL1 | None | None | None | None |
| ATL1 | None | None | None | None |
| ATL1 | None | None | None | None |
| ATL1 | None | None | None | None |
| NR3C2 | None | None | None | None |
| TBK1 | None | None | None | None |
| HYAL1 | None | None | None | None |
| HYAL1 | None | None | None | None |
| HYAL1 | None | None | None | None |
| HYAL1 | None | None | None | None |
| CTSD | None | None | None | None |

| | | | | |
|---|---|---|---|---|
| PRKCG | None | None | None | None |
| TGFBR1 | None | None | None | None |
| XDH | None | None | None | None |
| XDH | None | None | None | None |
| XDH | None | None | None | None |
| XDH | None | None | None | None |
| XDH | None | None | None | None |
| XDH | None | None | None | None |
| NMNAT1 | 0.833 | 1 | 1kqn | E |
| NMNAT1 | None | None | None | None |
| NMNAT1 | None | None | None | None |
| NMNAT1 | None | None | None | None |
| NMNAT1 | None | None | None | None |
| NMNAT1 | None | None | None | None |
| SERPING1 | None | None | None | None |
| SERPING1 | None | None | None | None |
| SERPING1 | None | None | None | None |
| SERPING1 | None | None | None | None |
| SERPING1 | None | None | None | None |
| PDZRN3 | None | None | None | None |
| FABP2 | None | None | None | None |
| IFNGR1 | None | None | None | None |
| IFNGR1 | None | None | None | None |
| TDP1 | None | None | None | None |
| TDP1 | None | None | None | None |
| INS | None | None | None | None |
| INS | None | None | None | None |
| INS | None | None | None | None |
| PPOX | 0.857 | 1 | 4ivm | B |
| PPOX | None | None | None | None |
| PPOX | None | None | None | None |
| PPOX | None | None | None | None |
| PPOX | None | None | None | None |
| PPOX | None | None | None | None |
| PPOX | None | None | None | None |
| PPOX | None | None | None | None |
| PPOX | None | None | None | None |
| GFER | None | None | None | None |
| GFER | None | None | None | None |
| GFER | None | None | None | None |
| GFER | None | None | None | None |
| LIFR | None | None | None | None |
| EGLN1 | None | None | None | None |
| EGLN1 | 0.889 | 6 | 5v18 | A |
| EGLN1 | 0.944 | 2 | 5a3u | C |
| GP1BA | None | None | None | None |
| GP1BA | None | None | None | None |
| GP1BA | None | None | None | None |
| GP1BA | None | None | None | None |
| GP1BA | None | None | None | None |
| MMADHC | None | None | None | None |
| MMADHC | None | None | None | None |
| MMADHC | None | None | None | None |
| MMADHC | None | None | None | None |

| | | | | |
|---|---|---|---|---|
| MMADHC | None | None | None | None |
| MMADHC | None | None | None | None |
| MMADHC | None | None | None | None |
| ACVRL1 | None | None | None | None |
| ACVRL1 | None | None | None | None |
| RFXANK | None | None | None | None |
| CHIT1 | None | None | None | None |
| UBR1 | None | None | None | None |
| BLM | None | None | None | None |
| BLM | None | None | None | None |
| BLM | None | None | None | None |
| BLM | None | None | None | None |
| BLM | None | None | None | None |
| BLM | None | None | None | None |
| BLM | None | None | None | None |
| BLM | None | None | None | None |
| BLM | None | None | None | None |
| BLM | None | None | None | None |
| BLM | None | None | None | None |
| BLM | None | None | None | None |
| BLM | None | None | None | None |
| VDR | None | None | None | None |
| VDR | None | None | None | None |
| VDR | None | None | None | None |
| VDR | None | None | None | None |
| VDR | None | None | None | None |
| VDR | None | None | None | None |
| EPHA2 | None | None | None | None |
| EPHA2 | None | None | None | None |
| EPHA2 | None | None | None | None |
| EPHA2 | None | None | None | None |
| EPHA2 | None | None | None | None |
| EPHA2 | None | None | None | None |
| EPHA2 | None | None | None | None |
| EPHA2 | None | None | None | None |
| EPHA2 | None | None | None | None |
| EPHA2 | None | None | None | None |
| UBA1 | None | None | None | None |
| STAT1 | None | None | None | None |
| STAT1 | None | None | None | None |
| STAT1 | None | None | None | None |
| STAT1 | None | None | None | None |
| STAT1 | None | None | None | None |
| STAT1 | None | None | None | None |
| STAT1 | None | None | None | None |
| STAT1 | None | None | None | None |
| DNM1L | None | None | None | None |
| DNM1L | None | None | None | None |
| OTC | None | None | None | None |
| OTC | None | None | None | None |
| OTC | None | None | None | None |
| OTC | None | None | None | None |
| OTC | None | None | None | None |
| OTC | None | None | None | None |

| | | | | |
|---|---|---|---|---|
| OTC | None | None | None | None |
| OTC | None | None | None | None |
| OTC | None | None | None | None |
| OTC | None | None | None | None |
| OTC | None | None | None | None |
| OTC | None | None | None | None |
| OTC | None | None | None | None |
| OTC | None | None | None | None |
| OTC | None | None | None | None |
| OTC | None | None | None | None |
| OTC | None | None | None | None |
| RARS | None | None | None | None |
| RARS | None | None | None | None |
| RARS | None | None | None | None |
| SEPSECS | None | None | None | None |
| SEPSECS | None | None | None | None |
| SEPSECS | None | None | None | None |
| SEPSECS | None | None | None | None |
| SEPSECS | None | None | None | None |
| BRCA1 | None | None | None | None |
| BRCA1 | None | None | None | None |
| BRCA1 | None | None | None | None |
| BRCA1 | None | None | None | None |
| BRCA1 | None | None | None | None |
| BRCA1 | None | None | None | None |
| BRCA1 | None | None | None | None |
| BRCA1 | None | None | None | None |
| BRCA1 | None | None | None | None |
| BRCA1 | None | None | None | None |
| BRCA1 | None | None | None | None |
| BRCA1 | None | None | None | None |
| BRCA1 | None | None | None | None |
| BRCA1 | None | None | None | None |
| BRCA1 | None | None | None | None |
| BRCA1 | None | None | None | None |
| BRCA1 | None | None | None | None |
| BRCA1 | None | None | None | None |
| BRCA1 | None | None | None | None |
| BRCA1 | None | None | None | None |
| BRCA1 | None | None | None | None |
| BRCA1 | None | None | None | None |
| CUBN | None | None | None | None |
| ITGB3 | None | None | None | None |
| MTRR | None | None | None | None |
| MTRR | None | None | None | None |
| MTRR | None | None | None | None |
| MTRR | None | None | None | None |
| MTRR | None | None | None | None |
| MTRR | None | None | None | None |
| MTRR | None | None | None | None |
| MTRR | None | None | None | None |
| IL7R | None | None | None | None |
| IL7R | None | None | None | None |
| RPS24 | None | None | None | None |

| | | | | |
|---|---|---|---|---|
| RPS24 | None | None | None | None |
| HOGA1 | None | None | None | None |
| HOGA1 | None | None | None | None |
| HOGA1 | None | None | None | None |
| HOGA1 | None | None | None | None |
| HOGA1 | None | None | None | None |
| HOGA1 | None | None | None | None |
| HOGA1 | None | None | None | None |
| HOGA1 | None | None | None | None |
| HOGA1 | None | None | None | None |
| HOGA1 | None | None | None | None |
| HOGA1 | None | None | None | None |
| HOGA1 | None | None | None | None |
| HOGA1 | None | None | None | None |
| GJB2 | None | None | None | None |
| GJB2 | None | None | None | None |
| GJB2 | None | None | None | None |
| GJB2 | None | None | None | None |
| GJB2 | None | None | None | None |
| GJB2 | None | None | None | None |
| GJB2 | None | None | None | None |
| GJB2 | None | None | None | None |
| GJB2 | None | None | None | None |
| GJB2 | None | None | None | None |
| GJB2 | None | None | None | None |
| GJB2 | None | None | None | None |
| GJB2 | None | None | None | None |
| GJB2 | None | None | None | None |
| GJB2 | None | None | None | None |
| GJB2 | None | None | None | None |
| GJB2 | None | None | None | None |
| GJB2 | None | None | None | None |
| GJB2 | None | None | None | None |
| GJB2 | None | None | None | None |
| GJB2 | None | None | None | None |
| GJB2 | None | None | None | None |
| GJB2 | None | None | None | None |
| GJB2 | None | None | None | None |
| GJB2 | None | None | None | None |
| GJB2 | None | None | None | None |
| GJB2 | None | None | None | None |
| GJB2 | None | None | None | None |
| GJB2 | None | None | None | None |
| GJB2 | None | None | None | None |
| GJB2 | None | None | None | None |
| GJB2 | None | None | None | None |
| GJB2 | None | None | None | None |
| GJB2 | None | None | None | None |
| GJB2 | None | None | None | None |
| GJB2 | None | None | None | None |
| GJB2 | None | None | None | None |
| GJB2 | None | None | None | None |
| GJB2 | None | None | None | None |
| GJB2 | None | None | None | None |

| | | | | |
|---|---|---|---|---|
| GJB2 | None | None | None | None |
| GJB2 | None | None | None | None |
| GJB2 | None | None | None | None |
| GJB2 | None | None | None | None |
| GJB2 | None | None | None | None |
| GJB2 | None | None | None | None |
| GJB2 | None | None | None | None |
| GJB2 | None | None | None | None |
| GJB2 | None | None | None | None |
| GJB2 | None | None | None | None |
| GJB2 | None | None | None | None |
| GJB2 | None | None | None | None |
| GJB2 | None | None | None | None |
| GJB2 | None | None | None | None |
| CPS1 | None | None | None | None |
| CPS1 | None | None | None | None |
| CPS1 | None | None | None | None |
| CPS1 | None | None | None | None |
| CPS1 | None | None | None | None |
| CPS1 | None | None | None | None |
| CPS1 | None | None | None | None |
| CPS1 | None | None | None | None |
| CPS1 | None | None | None | None |
| CPS1 | None | None | None | None |
| PSPH | None | None | None | None |
| PSPH | None | None | None | None |
| PSPH | None | None | None | None |
| PSPH | None | None | None | None |
| PSPH | None | None | None | None |
| PSPH | None | None | None | None |
| PEPD | None | None | None | None |
| PEPD | None | None | None | None |
| PEPD | None | None | None | None |
| PEPD | None | None | None | None |
| PEPD | None | None | None | None |
| PEPD | None | None | None | None |
| PEPD | None | None | None | None |
| PEPD | None | None | None | None |
| PEPD | None | None | None | None |
| KDM5C | None | None | None | None |
| KDM5C | None | None | None | None |
| KDM5C | None | None | None | None |
| ITGB6 | None | None | None | None |
| ITGB6 | None | None | None | None |
| ITGB6 | None | None | None | None |
| DARS | None | None | None | None |
| DARS | None | None | None | None |
| DARS | None | None | None | None |
| DARS | None | None | None | None |
| BTK | None | None | None | None |
| BTK | None | None | None | None |
| BTK | None | None | None | None |
| HDAC4 | None | None | None | None |
| ITGA2B | None | None | None | None |

| | | | | |
|---|---|---|---|---|
| ITGA2B | None | None | None | None |
| ITGA2B | None | None | None | None |
| ITGA2B | None | None | None | None |
| PCSK9 | None | None | None | None |
| PCSK9 | None | None | None | None |
| PCSK9 | None | None | None | None |
| PCSK9 | None | None | None | None |
| MUTYH | None | None | None | None |
| MUTYH | None | None | None | None |
| MUTYH | None | None | None | None |
| MUTYH | None | None | None | None |
| MUTYH | None | None | None | None |
| MUTYH | None | None | None | None |
| MUTYH | None | None | None | None |
| MUTYH | None | None | None | None |
| MUTYH | None | None | None | None |
| MUTYH | None | None | None | None |
| MUTYH | None | None | None | None |
| MUTYH | None | None | None | None |
| MUTYH | None | None | None | None |
| MUTYH | None | None | None | None |
| MUTYH | None | None | None | None |
| MUTYH | None | None | None | None |
| MUTYH | None | None | None | None |
| MUTYH | None | None | None | None |
| MUTYH | None | None | None | None |
| MUTYH | None | None | None | None |
| MUTYH | None | None | None | None |
| MUTYH | None | None | None | None |
| MUTYH | None | None | None | None |
| MUTYH | None | None | None | None |
| SLC25A12 | None | None | None | None |
| GSN | None | None | None | None |
| IHH | 0.909 | 1 | 3n1p | B |
| IHH | None | None | None | None |
| IHH | None | None | None | None |
| IHH | 0.818 | 2 | 3n1p | B |
| IHH | None | None | None | None |
| IHH | None | None | None | None |
| IHH | 0.818 | 7 | 3n1p | B |
| CD247 | None | None | None | None |
| EP300 | None | None | None | None |
| TTN | None | None | None | None |
| MRI1 | None | None | None | None |
| SGSH | None | None | None | None |
| SGSH | None | None | None | None |
| SGSH | None | None | None | None |
| SGSH | None | None | None | None |
| SGSH | None | None | None | None |
| SGSH | None | None | None | None |
| SGSH | None | None | None | None |
| SGSH | None | None | None | None |
| SGSH | None | None | None | None |
| SGSH | None | None | None | None |

| | | | | |
|---|---|---|---|---|
| SGSH | None | None | None | None |
| SGSH | None | None | None | None |
| SGSH | None | None | None | None |
| SGSH | None | None | None | None |
| SGSH | None | None | None | None |
| SGSH | None | None | None | None |
| SGSH | None | None | None | None |
| SGSH | None | None | None | None |
| SGSH | None | None | None | None |
| SGSH | None | None | None | None |
| SGSH | None | None | None | None |
| SGSH | None | None | None | None |
| CBL | None | None | None | None |
| CANT1 | None | None | None | None |
| CANT1 | None | None | None | None |
| CANT1 | None | None | None | None |
| CANT1 | None | None | None | None |
| CANT1 | None | None | None | None |
| CANT1 | None | None | None | None |
| CANT1 | None | None | None | None |
| SLC1A3 | None | None | None | None |
| SLC1A3 | None | None | None | None |
| SLC1A5 | None | None | None | None |
| SLC1A3 | None | None | None | None |
| SLC1A5 | None | None | None | None |
| SLC1A3 | None | None | None | None |
| SLC1A3 | None | None | None | None |
| CTSD | None | None | None | None |
| CTSD | None | None | None | None |
| CTSD | None | None | None | None |
| CTSD | None | None | None | None |
| CTSD | None | None | None | None |
| MTOR | None | None | None | None |
| ILK | None | None | None | None |
| PCDH15 | None | None | None | None |
| PCDH15 | None | None | None | None |
| PCDH15 | None | None | None | None |
| PCDH15 | None | None | None | None |
| PCDH15 | None | None | None | None |
| PCDH15 | None | None | None | None |
| KLHL40 | None | None | None | None |
| KLHL40 | None | None | None | None |
| KLHL40 | None | None | None | None |
| KLHL40 | None | None | None | None |
| CSTA | None | None | None | None |
| CSTA | None | None | None | None |
| HNMT | None | None | None | None |
| HNMT | None | None | None | None |
| IFNGR2 | None | None | None | None |
| IFNGR2 | None | None | None | None |
| PEX5 | None | None | None | None |
| KIF7 | None | None | None | None |
| KIF7 | None | None | None | None |
| MYL3 | None | None | None | None |

| | | | | |
|---|---|---|---|---|
| MYL3 | None | None | None | None |
| MYL3 | None | None | None | None |
| MYL3 | None | None | None | None |
| MYL3 | None | None | None | None |
| MYL3 | None | None | None | None |
| MYL3 | None | None | None | None |
| NAGLU | None | None | None | None |
| NAGLU | None | None | None | None |
| NAGLU | None | None | None | None |
| NAGLU | None | None | None | None |
| NAGLU | None | None | None | None |
| NAGLU | None | None | None | None |
| NAGLU | None | None | None | None |
| NAGLU | None | None | None | None |
| NAGLU | None | None | None | None |
| NAGLU | None | None | None | None |
| NAGLU | None | None | None | None |
| NAGLU | None | None | None | None |
| NAGLU | None | None | None | None |
| TNFRSF13C | None | None | None | None |
| KIF21A | None | None | None | None |
| KIF21A | None | None | None | None |
| KIF21A | None | None | None | None |
| KIF21A | None | None | None | None |
| KIF21A | None | None | None | None |
| DOCK2 | None | None | None | None |
| DSP | None | None | None | None |
| CDKN2A | None | None | None | None |
| CDKN2A | None | None | None | None |
| CDKN2A | None | None | None | None |
| CDKN2A | None | None | None | None |
| CDKN2A | None | None | None | None |
| CDKN2A | None | None | None | None |
| CDKN2A | None | None | None | None |
| CDKN2A | None | None | None | None |
| CDKN2A | None | None | None | None |
| CDKN2A | None | None | None | None |
| CDKN2A | None | None | None | None |
| CDKN2A | None | None | None | None |
| CDKN2A | None | None | None | None |
| CDKN2A | None | None | None | None |
| VCL | None | None | None | None |
| TPP1 | None | None | None | None |
| TPP1 | None | None | None | None |
| TPP1 | None | None | None | None |
| TPP1 | None | None | None | None |
| TPP1 | None | None | None | None |
| TPP1 | None | None | None | None |
| TPP1 | None | None | None | None |
| TPP1 | None | None | None | None |
| TPP1 | None | None | None | None |
| TPP1 | None | None | None | None |
| TPP1 | None | None | None | None |
| TPP1 | None | None | None | None |

| | | | | |
|---|---|---|---|---|
| TPP1 | None | None | None | None |
| TPP1 | None | None | None | None |
| TPP1 | None | None | None | None |
| TPP1 | None | None | None | None |
| TPP1 | None | None | None | None |
| TPP1 | None | None | None | None |
| TPP1 | None | None | None | None |
| TPP1 | None | None | None | None |
| TPP1 | None | None | None | None |
| TPP1 | None | None | None | None |
| IL10RB | None | None | None | None |
| IL10RB | None | None | None | None |
| CCM2 | None | None | None | None |
| CCM2 | None | None | None | None |
| CCM2 | None | None | None | None |
| DCX | None | None | None | None |
| DCX | None | None | None | None |
| DCX | None | None | None | None |
| DCX | None | None | None | None |
| DCX | None | None | None | None |
| DCX | None | None | None | None |
| DCX | None | None | None | None |
| DCX | None | None | None | None |
| BRCA2 | None | None | None | None |
| NNT | None | None | None | None |
| C2 | None | None | None | None |
| C2 | None | None | None | None |
| C2 | None | None | None | None |
| C2 | None | None | None | None |
| WRN | None | None | None | None |
| WRN | None | None | None | None |
| F13A1 | None | None | None | None |
| F13A1 | None | None | None | None |
| F13A1 | None | None | None | None |
| F13A1 | None | None | None | None |
| TF | None | None | None | None |
| TF | None | None | None | None |
| TF | None | None | None | None |
| TF | None | None | None | None |
| TF | None | None | None | None |
| ITPA | None | None | None | None |
| ITPA | None | None | None | None |
| SNCA | None | None | None | None |
| ITGB4 | None | None | None | None |
| ITGB4 | None | None | None | None |
| ITGB4 | None | None | None | None |
| LRP6 | None | None | None | None |
| LRP6 | None | None | None | None |
| LRP6 | None | None | None | None |
| PLG | None | None | None | None |
| PLG | None | None | None | None |
| TYROBP | None | None | None | None |
| UQCRC2 | None | None | None | None |
| GPSM2 | None | None | None | None |

| | | | | |
|---|---|---|---|---|
| GPSM2 | None | None | None | None |
| GPSM2 | None | None | None | None |
| SCN5A | None | None | None | None |
| SCN5A | None | None | None | None |
| SCN5A | None | None | None | None |
| SCN5A | None | None | None | None |
| SCN5A | None | None | None | None |
| ATIC | None | None | None | None |
| EZH2 | None | None | None | None |
| CPA1 | None | None | None | None |
| SCN4B | None | None | None | None |
| SCN4B | None | None | None | None |
| MAN1B1 | None | None | None | None |
| MAN1B1 | None | None | None | None |
| MAN1B1 | None | None | None | None |
| MAN1B1 | None | None | None | None |
| CYCS | None | None | None | None |
| IDUA | None | None | None | None |
| IDUA | None | None | None | None |
| IDUA | None | None | None | None |
| IDUA | None | None | None | None |
| IDUA | None | None | None | None |
| IDUA | None | None | None | None |
| IDUA | None | None | None | None |
| IDUA | None | None | None | None |
| IDUA | None | None | None | None |
| IDUA | None | None | None | None |
| IDUA | None | None | None | None |
| IDUA | None | None | None | None |
| IDUA | None | None | None | None |
| IDUA | None | None | None | None |
| IDUA | None | None | None | None |
| IDUA | None | None | None | None |
| IDUA | None | None | None | None |
| IDUA | None | None | None | None |
| IDUA | None | None | None | None |
| IDUA | None | None | None | None |
| IDUA | None | None | None | None |
| IDUA | None | None | None | None |
| IDUA | None | None | None | None |
| IDUA | None | None | None | None |
| IDUA | None | None | None | None |
| HIKESHI | None | None | None | None |
| APC | None | None | None | None |
| APC | None | None | None | None |
| APC | None | None | None | None |
| APP | None | None | None | None |
| CEP152 | None | None | None | None |
| GAA | None | None | None | None |
| GAA | None | None | None | None |
| GAA | None | None | None | None |
| GAA | None | None | None | None |
| GAA | None | None | None | None |
| GAA | None | None | None | None |

| | | | | |
|---|---|---|---|---|
| GAA | None | None | None | None |
| GAA | None | None | None | None |
| GAA | None | None | None | None |
| GAA | None | None | None | None |
| GAA | None | None | None | None |
| GAA | None | None | None | None |
| GAA | None | None | None | None |
| GAA | None | None | None | None |
| GAA | None | None | None | None |
| GAA | None | None | None | None |
| GAA | None | None | None | None |
| GAA | None | None | None | None |
| GAA | None | None | None | None |
| GAA | None | None | None | None |
| GAA | None | None | None | None |
| GAA | None | None | None | None |
| GAA | None | None | None | None |
| GAA | None | None | None | None |
| GAA | None | None | None | None |
| GAA | None | None | None | None |
| GAA | None | None | None | None |
| GAA | None | None | None | None |
| GAA | None | None | None | None |
| GAA | None | None | None | None |
| GAA | None | None | None | None |
| GAA | None | None | None | None |
| GAA | None | None | None | None |
| GAA | None | None | None | None |
| GAA | None | None | None | None |
| GAA | None | None | None | None |
| GAA | None | None | None | None |
| GAA | None | None | None | None |
| GAA | None | None | None | None |
| GAA | None | None | None | None |
| GAA | None | None | None | None |
| GAA | None | None | None | None |
| GAA | None | None | None | None |
| GAA | None | None | None | None |
| GAA | None | None | None | None |
| GAA | None | None | None | None |
| GAA | None | None | None | None |
| GAA | None | None | None | None |
| GAA | None | None | None | None |
| FOLR1 | None | None | None | None |
| DSP | None | None | None | None |
| DSP | None | None | None | None |
| DSP | None | None | None | None |
| DSP | None | None | None | None |
| DSP | None | None | None | None |
| CCM2 | None | None | None | None |
| FXN | None | None | None | None |
| FXN | None | None | None | None |
| COMP | None | None | None | None |
| COMP | None | None | None | None |

| | | | | |
|---|---|---|---|---|
| COMP | None | None | None | None |
| COMP | None | None | None | None |
| COMP | None | None | None | None |
| COMP | None | None | None | None |
| COMP | None | None | None | None |
| COMP | None | None | None | None |
| COMP | None | None | None | None |
| COMP | None | None | None | None |
| COMP | None | None | None | None |
| COMP | None | None | None | None |
| AIFM1 | None | None | None | None |
| AIFM1 | None | None | None | None |
| AIFM1 | None | None | None | None |
| AIFM1 | None | None | None | None |
| AIFM1 | None | None | None | None |
| AIFM1 | None | None | None | None |
| AIFM1 | None | None | None | None |
| MYH9 | None | None | None | None |
| FHL2 | None | None | None | None |
| FHL2 | None | None | None | None |
| TNFRSF4 | None | None | None | None |
| CFB | None | None | None | None |
| CFB | None | None | None | None |
| CFB | None | None | None | None |
| CFB | None | None | None | None |
| CFB | None | None | None | None |
| CFB | None | None | None | None |
| CFB | None | None | None | None |
| CFB | None | None | None | None |
| CFB | None | None | None | None |
| CFB | None | None | None | None |
| MVD | None | None | None | None |
| MVD | None | None | None | None |
| HSD17B4 | None | None | None | None |
| HSD17B4 | None | None | None | None |
| HSD17B4 | None | None | None | None |
| HSD17B4 | None | None | None | None |
| HSD17B4 | None | None | None | None |
| HSD17B4 | None | None | None | None |
| F11 | None | None | None | None |
| F11 | None | None | None | None |
| F11 | None | None | None | None |
| F11 | None | None | None | None |
| F11 | None | None | None | None |
| F11 | None | None | None | None |
| F11 | None | None | None | None |
| F11 | None | None | None | None |
| F11 | None | None | None | None |
| SPTA1 | None | None | None | None |
| SPTA1 | None | None | None | None |
| SPTA1 | None | None | None | None |
| SPTA1 | None | None | None | None |
| SPTA1 | None | None | None | None |
| SPTA1 | None | None | None | None |

| | | | | |
|---|---|---|---|---|
| GMPR | None | None | None | None |
| MITF | None | None | None | None |
| NPC1 | None | None | None | None |
| NPC1 | None | None | None | None |
| NPC1 | None | None | None | None |
| NPC1 | None | None | None | None |
| NPC1 | None | None | None | None |
| NPC1 | None | None | None | None |
| NPC1 | None | None | None | None |
| NPC1 | None | None | None | None |
| NPC1 | None | None | None | None |
| NPC1 | None | None | None | None |
| NPC1 | None | None | None | None |
| NPC1 | None | None | None | None |
| NPC1 | None | None | None | None |
| NPC1 | None | None | None | None |
| NPC1 | None | None | None | None |
| NPC1 | None | None | None | None |
| NPC1 | None | None | None | None |
| NPC1 | None | None | None | None |
| NPC1 | None | None | None | None |
| NPC1 | None | None | None | None |
| NPC1 | None | None | None | None |
| NPC1 | None | None | None | None |
| NPC1 | None | None | None | None |
| NPC1 | None | None | None | None |
| NPC1 | None | None | None | None |
| NPC1 | None | None | None | None |
| NPC1 | None | None | None | None |
| NPC1 | None | None | None | None |
| NPC1 | None | None | None | None |
| NPC1 | None | None | None | None |
| NPC1 | None | None | None | None |
| NPC1 | None | None | None | None |
| NPC1 | None | None | None | None |
| NPC1 | None | None | None | None |
| NPC1 | None | None | None | None |
| NPC1 | None | None | None | None |
| NPC1 | None | None | None | None |
| NPC1 | None | None | None | None |
| NPC1 | None | None | None | None |
| NPC1 | None | None | None | None |
| NPC1 | None | None | None | None |
| NPC1 | None | None | None | None |
| NPC1 | None | None | None | None |
| C3 | None | None | None | None |
| NLRX1 | None | None | None | None |
| IGLL1 | None | None | None | None |
| APRT | None | None | None | None |
| APRT | None | None | None | None |
| APRT | None | None | None | None |
| APRT | 0.923 | 1 | 1zn8 | A |
| APRT | None | None | None | None |
| ZIC3 | None | None | None | None |

| | | | | |
|---|---|---|---|---|
| ZIC3 | None | None | None | None |
| KLHL3 | None | None | None | None |
| RPS6KA3 | None | None | None | None |
| RPS6KA3 | None | None | None | None |
| RPS6KA3 | None | None | None | None |
| F8 | None | None | None | None |
| F8 | None | None | None | None |
| F8 | None | None | None | None |
| F8 | None | None | None | None |
| F8 | None | None | None | None |
| F8 | None | None | None | None |
| F8 | None | None | None | None |
| F8 | None | None | None | None |
| F8 | None | None | None | None |
| F8 | None | None | None | None |
| F8 | None | None | None | None |
| F8 | None | None | None | None |
| F8 | None | None | None | None |
| F8 | None | None | None | None |
| F8 | None | None | None | None |
| F8 | None | None | None | None |
| F8 | None | None | None | None |
| ISG15 | None | None | None | None |
| EXOSC3 | None | None | None | None |
| EXOSC3 | None | None | None | None |
| EXOSC3 | None | None | None | None |
| NKX2-5 | None | None | None | None |
| NKX2-5 | None | None | None | None |
| NKX2-5 | None | None | None | None |
| NKX2-5 | None | None | None | None |
| FH | None | None | None | None |
| FH | None | None | None | None |
| FH | None | None | None | None |
| FH | None | None | None | None |
| FH | None | None | None | None |
| FH | None | None | None | None |
| FH | None | None | None | None |
| FH | None | None | None | None |
| FH | None | None | None | None |
| FH | None | None | None | None |
| FH | None | None | None | None |
| FH | None | None | None | None |
| SNRNP200 | None | None | None | None |
| SNRNP200 | None | None | None | None |
| SNRNP200 | None | None | None | None |
| SNRNP200 | None | None | None | None |
| CHRNA1 | None | None | None | None |
| CHRNA1 | None | None | None | None |
| MYOM1 | None | None | None | None |
| MYOM1 | None | None | None | None |
| MYOM1 | None | None | None | None |
| MYOM1 | None | None | None | None |
| MYOM1 | None | None | None | None |
| MYOM1 | None | None | None | None |

| | | | | |
|---|---|---|---|---|
| LDLR | None | None | None | None |
| LDLR | None | None | None | None |
| LDLR | None | None | None | None |
| LDLR | None | None | None | None |
| LDLR | None | None | None | None |
| LDLR | None | None | None | None |
| EHMT1 | None | None | None | None |
| EHMT1 | None | None | None | None |
| EHMT1 | None | None | None | None |
| COL9A2 | None | None | None | None |
| HFE2 | None | None | None | None |
| HFE2 | None | None | None | None |
| CD3G | None | None | None | None |
| MMAB | None | None | None | None |
| MMAB | None | None | None | None |
| MMAB | None | None | None | None |
| MMAB | None | None | None | None |
| MMAB | None | None | None | None |
| MMAB | None | None | None | None |
| MMAB | None | None | None | None |
| MMAB | None | None | None | None |
| MMAB | None | None | None | None |
| MMAB | None | None | None | None |
| TAF1 | None | None | None | None |
| TAF1 | None | None | None | None |
| CD4 | None | None | None | None |
| QARS | None | None | None | None |
| QARS | None | None | None | None |
| QARS | None | None | None | None |
| QARS | None | None | None | None |
| QARS | None | None | None | None |
| FGFR3 | None | None | None | None |
| FGFR3 | None | None | None | None |
| FGFR3 | None | None | None | None |
| FGFR3 | None | None | None | None |
| FGFR3 | None | None | None | None |
| FGFR3 | None | None | None | None |
| FLNB | None | None | None | None |
| FLNB | None | None | None | None |
| FLNB | None | None | None | None |
| FLNB | None | None | None | None |
| FLNB | None | None | None | None |
| FLNB | None | None | None | None |
| FLNB | None | None | None | None |
| FLNB | None | None | None | None |
| IL21R | None | None | None | None |
| RB1 | None | None | None | None |
| RB1 | None | None | None | None |
| RB1 | None | None | None | None |
| NTRK1 | None | None | None | None |
| NOTCH1 | None | None | None | None |
| MYOM1 | None | None | None | None |
| MYOM1 | None | None | None | None |
| MYOM1 | None | None | None | None |

| | | | | |
|---|---|---|---|---|
| MYOM1 | None | None | None | None |
| MYOM1 | None | None | None | None |
| MYOM1 | None | None | None | None |
| COQ9 | None | None | None | None |
| COQ9 | None | None | None | None |
| CRBN | None | None | None | None |
| DCTN1 | None | None | None | None |
| DCTN1 | None | None | None | None |
| DCTN1 | None | None | None | None |
| DCTN1 | None | None | None | None |
| DCTN1 | None | None | None | None |
| DCTN1 | None | None | None | None |
| THAP1 | None | None | None | None |
| THAP1 | None | None | None | None |
| GNAS | None | None | None | None |
| GNAS | None | None | None | None |
| HNF4A | None | None | None | None |
| HNF4A | None | None | None | None |
| ALDH3A2 | None | None | None | None |
| ALDH3A2 | None | None | None | None |
| ALDH3A2 | None | None | None | None |
| ALDH3A2 | None | None | None | None |
| ALDH3A2 | None | None | None | None |
| ALDH3A2 | None | None | None | None |
| ALDH3A2 | None | None | None | None |
| ALDH3A2 | None | None | None | None |
| ALDH3A2 | None | None | None | None |
| HCN1 | None | None | None | None |
| PLEC | None | None | None | None |
| SCN5A | None | None | None | None |
| SCN5A | None | None | None | None |
| STAMBP | None | None | None | None |
| STAMBP | None | None | None | None |
| RNF125 | None | None | None | None |
| RNF125 | None | None | None | None |
| USH1C | None | None | None | None |
| OBSL1 | None | None | None | None |
| GRIN2B | None | None | None | None |
| GRIN2B | None | None | None | None |
| HCN4 | None | None | None | None |
| HCN4 | None | None | None | None |
| TALDO1 | None | None | None | None |
| ITGB3 | None | None | None | None |
| CHKB | None | None | None | None |
| CHKB | None | None | None | None |
| CHKB | None | None | None | None |
| CHKB | None | None | None | None |
| ACTN4 | None | None | None | None |
| ACTN4 | None | None | None | None |
| ACTN4 | None | None | None | None |
| ACTN4 | None | None | None | None |
| CDT1 | None | None | None | None |
| CDT1 | None | None | None | None |
| CDT1 | None | None | None | None |

| | | | | |
|---|---|---|---|---|
| CDT1 | None | None | None | None |
| CDT1 | None | None | None | None |
| CDT1 | None | None | None | None |
| MAPT | None | None | None | None |
| F5 | None | None | None | None |
| F5 | None | None | None | None |
| F5 | None | None | None | None |
| ESRRB | None | None | None | None |
| DGUOK | None | None | None | None |
| DGUOK | None | None | None | None |
| DGUOK | None | None | None | None |
| DGUOK | None | None | None | None |
| DGUOK | None | None | None | None |
| DGUOK | None | None | None | None |
| DGUOK | None | None | None | None |
| DGUOK | None | None | None | None |
| DGUOK | None | None | None | None |
| CENPJ | None | None | None | None |
| TTN | None | None | None | None |
| TTN | None | None | None | None |
| TTN | None | None | None | None |
| TTN | None | None | None | None |
| TNFRSF1A | None | None | None | None |
| TNFRSF1A | None | None | None | None |
| TNFRSF1A | None | None | None | None |
| TNFRSF1A | None | None | None | None |
| TNFRSF1A | None | None | None | None |
| TNFRSF1A | None | None | None | None |
| TNFRSF1A | None | None | None | None |
| TNFRSF1A | None | None | None | None |
| TNFRSF1A | None | None | None | None |
| TNFRSF1A | None | None | None | None |
| TNFRSF1A | None | None | None | None |
| TNFRSF1A | None | None | None | None |
| ASL | None | None | None | None |
| ASL | None | None | None | None |
| ASL | None | None | None | None |
| ASL | None | None | None | None |
| ASL | None | None | None | None |
| ASL | None | None | None | None |
| ASL | None | None | None | None |
| ASL | None | None | None | None |
| ASL | None | None | None | None |
| ASL | None | None | None | None |
| MYO7A | None | None | None | None |
| MYO7A | None | None | None | None |
| MYO7A | None | None | None | None |
| MYO7A | None | None | None | None |
| MYO7A | None | None | None | None |
| MYO7A | None | None | None | None |
| MYO7A | None | None | None | None |
| MYO7A | None | None | None | None |
| MYO7A | None | None | None | None |
| MYO7A | None | None | None | None |

| | | | | |
|---|---|---|---|---|
| MYO7A | None | None | None | None |
| MYO7A | None | None | None | None |
| MYO7A | None | None | None | None |
| MYO7A | None | None | None | None |
| MYO7A | None | None | None | None |
| MYO7A | None | None | None | None |
| MYO7A | None | None | None | None |
| MYO7A | None | None | None | None |
| MYO7A | None | None | None | None |
| MYO7A | None | None | None | None |
| APC | None | None | None | None |
| ALAD | None | None | None | None |
| ALAD | None | None | None | None |
| ALAD | None | None | None | None |
| ALAD | None | None | None | None |
| ALAD | None | None | None | None |
| ALAD | None | None | None | None |
| POR | None | None | None | None |
| CSF1R | None | None | None | None |
| CSF1R | None | None | None | None |
| CSF1R | None | None | None | None |
| CSF1R | None | None | None | None |
| CSF1R | None | None | None | None |
| CSF1R | None | None | None | None |
| CSF1R | None | None | None | None |
| CSF1R | None | None | None | None |
| CSF1R | None | None | None | None |
| CSF1R | None | None | None | None |
| CSF1R | None | None | None | None |
| VWF | None | None | None | None |
| VWF | None | None | None | None |
| PHF8 | None | None | None | None |
| ACY1 | None | None | None | None |
| SMN1 | None | None | None | None |
| SMN1 | None | None | None | None |
| DLAT | None | None | None | None |
| DLAT | None | None | None | None |
| B2M | None | None | None | None |
| SOX9 | None | None | None | None |
| SOX9 | None | None | None | None |
| SOX9 | None | None | None | None |
| SOX9 | None | None | None | None |
| SOX9 | None | None | None | None |
| CFH | None | None | None | None |
| FLNC | None | None | None | None |
| FLNC | None | None | None | None |
| FLNC | None | None | None | None |
| ITGB2 | None | None | None | None |
| ITGB2 | None | None | None | None |
| ITGB2 | None | None | None | None |
| ITGB2 | None | None | None | None |
| ITGB2 | None | None | None | None |
| ITGB2 | None | None | None | None |
| ITGB2 | None | None | None | None |

| | | | | |
|---|---|---|---|---|
| ITGB2 | None | None | None | None |
| ITGB2 | None | None | None | None |
| PSEN1 | None | None | None | None |
| PSEN1 | None | None | None | None |
| PSEN1 | None | None | None | None |
| PSEN1 | None | None | None | None |
| PSEN1 | None | None | None | None |
| PSEN1 | None | None | None | None |
| OXCT1 | None | None | None | None |
| OXCT1 | None | None | None | None |
| OXCT1 | None | None | None | None |
| OXCT1 | None | None | None | None |
| OXCT1 | None | None | None | None |
| OXCT1 | None | None | None | None |
| OXCT1 | None | None | None | None |
| DHH | None | None | None | None |
| PRKN | None | None | None | None |
| PRKN | None | None | None | None |
| PRKN | None | None | None | None |
| PRKN | None | None | None | None |
| PRKN | None | None | None | None |
| PRKN | None | None | None | None |
| PRKN | None | None | None | None |
| PRKN | None | None | None | None |
| PRKN | None | None | None | None |
| PRKN | None | None | None | None |
| PRKN | None | None | None | None |
| LCAT | None | None | None | None |
| LCAT | None | None | None | None |
| LCAT | None | None | None | None |
| LCAT | None | None | None | None |
| LCAT | None | None | None | None |
| LCAT | None | None | None | None |
| LCAT | None | None | None | None |
| LCAT | None | None | None | None |
| LCAT | None | None | None | None |
| LCAT | None | None | None | None |
| LCAT | None | None | None | None |
| LCAT | None | None | None | None |
| FOXP3 | None | None | None | None |
| FOXP3 | None | None | None | None |
| FOXP3 | None | None | None | None |
| FOXP3 | None | None | None | None |
| SUFU | None | None | None | None |
| HSPA9 | None | None | None | None |
| HSPA9 | None | None | None | None |
| SQSTM1 | None | None | None | None |
| DARS2 | None | None | None | None |
| DARS2 | None | None | None | None |
| DARS2 | None | None | None | None |
| DARS2 | None | None | None | None |
| DARS2 | None | None | None | None |
| DARS2 | None | None | None | None |
| DARS2 | None | None | None | None |

| | | | | |
|---|---|---|---|---|
| DARS2 | None | None | None | None |
| DARS2 | None | None | None | None |
| DARS2 | None | None | None | None |
| DARS2 | None | None | None | None |
| DARS2 | None | None | None | None |
| NT5C3A | None | None | None | None |
| NT5C3A | None | None | None | None |
| NT5C3A | None | None | None | None |
| NT5C3A | None | None | None | None |
| ETFA | None | None | None | None |
| ETFA | None | None | None | None |
| ETFA | None | None | None | None |
| ETFA | None | None | None | None |
| ETFA | None | None | None | None |
| ETFA | None | None | None | None |
| ETFA | None | None | None | None |
| TTN | None | None | None | None |
| TTN | None | None | None | None |
| FLT4 | None | None | None | None |
| FGB | None | None | None | None |
| HPCA | None | None | None | None |
| TAP2 | None | None | None | None |
| TAP2 | None | None | None | None |
| PTPRC | None | None | None | None |
| SLC4A1 | None | None | None | None |
| SLC4A1 | None | None | None | None |
| SLC4A1 | None | None | None | None |
| SLC4A1 | None | None | None | None |
| SLC4A1 | None | None | None | None |
| SLC4A1 | None | None | None | None |
| MYH7 | None | None | None | None |
| MYH7 | None | None | None | None |
| MYH7 | None | None | None | None |
| MYH7 | None | None | None | None |
| MYH7 | None | None | None | None |
| MYH7 | None | None | None | None |
| MYH7 | None | None | None | None |
| MYH7 | None | None | None | None |
| MYH7 | None | None | None | None |
| PARN | None | None | None | None |
| PARN | None | None | None | None |
| NDUFS6 | None | None | None | None |
| VRK1 | None | None | None | None |
| VRK1 | None | None | None | None |
| VRK1 | None | None | None | None |
| VRK1 | None | None | None | None |
| VRK1 | None | None | None | None |
| VRK1 | None | None | None | None |
| COLEC11 | None | None | None | None |
| VHL | None | None | None | None |
| TTN | None | None | None | None |
| NCF2 | None | None | None | None |
| NCF2 | None | None | None | None |
| NCF2 | None | None | None | None |

| | | | | |
|---|---|---|---|---|
| NCF2 | None | None | None | None |
| NCF2 | None | None | None | None |
| RPS26 | None | None | None | None |
| CETP | None | None | None | None |
| CETP | None | None | None | None |
| CETP | None | None | None | None |
| CETP | None | None | None | None |
| CETP | None | None | None | None |
| CETP | None | None | None | None |
| CETP | None | None | None | None |
| CETP | None | None | None | None |
| CETP | None | None | None | None |
| CETP | None | None | None | None |
| CETP | None | None | None | None |
| CETP | None | None | None | None |
| CETP | None | None | None | None |
| FLNA | None | None | None | None |
| FLNA | None | None | None | None |
| FLNA | None | None | None | None |
| FLNA | None | None | None | None |
| FLNA | None | None | None | None |
| FLNA | None | None | None | None |
| FLNA | None | None | None | None |
| MYBPC3 | None | None | None | None |
| MYBPC3 | None | None | None | None |
| MYBPC3 | None | None | None | None |
| MYBPC3 | None | None | None | None |
| MYBPC3 | None | None | None | None |
| MYBPC3 | None | None | None | None |
| MYBPC3 | None | None | None | None |
| MYBPC3 | None | None | None | None |
| MYBPC3 | None | None | None | None |
| MYBPC3 | None | None | None | None |
| LSS | None | None | None | None |
| C6 | None | None | None | None |
| C6 | None | None | None | None |
| RLBP1 | None | None | None | None |
| RLBP1 | None | None | None | None |
| RLBP1 | None | None | None | None |
| RLBP1 | None | None | None | None |
| RLBP1 | None | None | None | None |
| RLBP1 | None | None | None | None |
| RLBP1 | None | None | None | None |
| RLBP1 | None | None | None | None |
| MMAA | None | None | None | None |
| MMAA | None | None | None | None |
| MMAA | None | None | None | None |
| MMAA | None | None | None | None |
| MMAA | None | None | None | None |
| MMAA | None | None | None | None |
| IL1RN | None | None | None | None |
| MASTL | None | None | None | None |
| FHL1 | None | None | None | None |
| FHL1 | None | None | None | None |

| | | | | |
|---|---|---|---|---|
| FHL1 | None | None | None | None |
| FHL1 | None | None | None | None |
| FHL1 | None | None | None | None |
| FHL1 | None | None | None | None |
| FHL1 | None | None | None | None |
| FHL1 | None | None | None | None |
| FHL1 | None | None | None | None |
| EIF2B1 | None | None | None | None |
| EIF2B1 | None | None | None | None |
| COQ8A | None | None | None | None |
| COQ8A | None | None | None | None |
| COQ8A | None | None | None | None |
| COQ8A | None | None | None | None |
| COQ8A | None | None | None | None |
| COQ8A | None | None | None | None |
| COQ8A | None | None | None | None |
| COQ8A | None | None | None | None |
| COQ8A | None | None | None | None |
| COQ8A | None | None | None | None |
| COQ8A | None | None | None | None |
| COQ8A | None | None | None | None |
| COQ8A | None | None | None | None |
| COQ8A | None | None | None | None |
| COQ8A | None | None | None | None |
| COQ8A | None | None | None | None |
| UNC119 | None | None | None | None |
| IGHMBP2 | None | None | None | None |
| JAG1 | None | None | None | None |
| JAG1 | None | None | None | None |
| JAG1 | None | None | None | None |
| GNB1 | None | None | None | None |
| GNB1 | None | None | None | None |
| FLNA | None | None | None | None |
| FLNA | None | None | None | None |
| FLNA | None | None | None | None |
| FLNA | None | None | None | None |
| FLNA | None | None | None | None |
| CREBBP | None | None | None | None |
| CSF3R | None | None | None | None |
| CSF3R | None | None | None | None |
| HIBCH | None | None | None | None |
| HIBCH | None | None | None | None |
| HIBCH | None | None | None | None |
| MMACHC | None | None | None | None |
| MMACHC | None | None | None | None |
| MMACHC | None | None | None | None |
| MMACHC | None | None | None | None |
| MMACHC | None | None | None | None |
| MMACHC | None | None | None | None |
| MMACHC | None | None | None | None |
| MMACHC | None | None | None | None |
| MMACHC | None | None | None | None |
| MMACHC | None | None | None | None |
| MMACHC | None | None | None | None |

| | | | | |
|---|---|---|---|---|
| KLF1 | None | None | None | None |
| NAGS | None | None | None | None |
| NAGS | None | None | None | None |
| NAGS | None | None | None | None |
| PIK3R2 | None | None | None | None |
| EDNRB | None | None | None | None |
| EDNRB | None | None | None | None |
| EDNRB | None | None | None | None |
| EDNRB | None | None | None | None |
| E | None | None | None | None |
| RUNX1 | None | None | None | None |
| RUNX1 | None | None | None | None |
| CTNNB1 | None | None | None | None |
| CTNNB1 | None | None | None | None |
| CTNNB1 | None | None | None | None |
| CTNNB1 | None | None | None | None |
| CTNNB1 | None | None | None | None |
| CTNNB1 | None | None | None | None |
| TTN | None | None | None | None |
| TTN | None | None | None | None |
| TYK2 | None | None | None | None |
| TYK2 | None | None | None | None |
| TYK2 | None | None | None | None |
| ERCC4 | None | None | None | None |
| ERCC4 | None | None | None | None |
| ERCC4 | None | None | None | None |
| ERCC4 | None | None | None | None |
| CFI | None | None | None | None |
| CFI | None | None | None | None |
| CFI | None | None | None | None |
| CFI | None | None | None | None |
| CFI | None | None | None | None |
| CFI | None | None | None | None |
| CFI | None | None | None | None |
| CFI | None | None | None | None |
| CFI | None | None | None | None |
| CFI | None | None | None | None |
| CFI | None | None | None | None |
| CFI | None | None | None | None |
| CFI | None | None | None | None |
| CFI | None | None | None | None |
| NOTCH3 | None | None | None | None |
| NOTCH3 | None | None | None | None |
| NOTCH3 | None | None | None | None |
| NOTCH3 | None | None | None | None |
| TBX1 | None | None | None | None |
| TBX1 | None | None | None | None |
| DDB2 | None | None | None | None |
| DDB2 | None | None | None | None |
| TP53 | None | None | None | None |
| POMGNT1 | None | None | None | None |
| POMGNT1 | None | None | None | None |
| POMGNT1 | None | None | None | None |
| POMGNT1 | None | None | None | None |

| | | | | |
|---|---|---|---|---|
| POMGNT1 | None | None | None | None |
| POMGNT1 | None | None | None | None |
| POMGNT1 | None | None | None | None |
| POMGNT1 | None | None | None | None |
| POMGNT1 | None | None | None | None |
| POMGNT1 | None | None | None | None |
| POMGNT1 | None | None | None | None |
| POMGNT1 | None | None | None | None |
| POMGNT1 | None | None | None | None |
| FMR1 | None | None | None | None |
| FMR1 | None | None | None | None |
| SLC9A3R1 | None | None | None | None |
| HSPB1 | None | None | None | None |
| HSPB1 | None | None | None | None |
| HSPB1 | None | None | None | None |
| HSPB1 | None | None | None | None |
| HSPB1 | None | None | None | None |
| HSPB1 | None | None | None | None |
| HSPB1 | None | None | None | None |
| SOD3 | None | None | None | None |
| AGA | 1 | 2 | 1apz | D |
| AGA | None | None | None | None |
| AGA | None | None | None | None |
| NDUFS7 | None | None | None | None |
| NDUFS7 | None | None | None | None |
| SOX2 | None | None | None | None |
| CSRP3 | None | None | None | None |
| CSRP3 | None | None | None | None |
| PAH | None | None | None | None |
| PAH | None | None | None | None |
| PAH | None | None | None | None |
| PAH | None | None | None | None |
| CACNA1S | None | None | None | None |
| TRPV4 | None | None | None | None |
| TRPV4 | None | None | None | None |
| TRPV4 | None | None | None | None |
| TRPV4 | None | None | None | None |
| TRPV4 | None | None | None | None |
| TRPV4 | None | None | None | None |
| TRPV4 | None | None | None | None |
| TRPV4 | None | None | None | None |
| TRPV4 | None | None | None | None |
| TRPV4 | None | None | None | None |
| TRPV4 | None | None | None | None |
| TRPV4 | None | None | None | None |
| TRPV4 | None | None | None | None |
| TNNI3 | None | None | None | None |
| MLYCD | None | None | None | None |
| MLYCD | None | None | None | None |
| MLYCD | None | None | None | None |
| MLYCD | None | None | None | None |
| AIP | None | None | None | None |
| AIP | None | None | None | None |
| ERCC2 | None | None | None | None |

| | | | | |
|---|---|---|---|---|
| ERCC2 | None | None | None | None |
| ERCC2 | None | None | None | None |
| ERCC2 | None | None | None | None |
| ERCC2 | None | None | None | None |
| ERCC2 | None | None | None | None |
| ERCC2 | None | None | None | None |
| ERCC2 | None | None | None | None |
| ERCC2 | None | None | None | None |
| ERCC2 | None | None | None | None |
| ERCC2 | None | None | None | None |
| ERCC2 | None | None | None | None |
| ERCC2 | None | None | None | None |
| ERCC2 | None | None | None | None |
| PUS1 | None | None | None | None |
| PUS1 | None | None | None | None |
| PUS1 | None | None | None | None |
| CASK | None | None | None | None |
| CASK | None | None | None | None |
| BLVRA | None | None | None | None |
| DICER1 | None | None | None | None |
| DICER1 | None | None | None | None |
| DICER1 | None | None | None | None |
| EIF2AK3 | None | None | None | None |
| EIF2AK3 | None | None | None | None |
| EIF2AK3 | None | None | None | None |
| KIT | None | None | None | None |
| KIT | None | None | None | None |
| KIT | None | None | None | None |
| KIT | None | None | None | None |
| KIT | None | None | None | None |
| KIT | None | None | None | None |
| DNMT3B | None | None | None | None |
| TGM1 | None | None | None | None |
| NDUFA10 | None | None | None | None |
| NDUFA10 | None | None | None | None |
| PRPF31 | None | None | None | None |
| PRPF31 | None | None | None | None |
| ECE1 | None | None | None | None |
| ECE1 | None | None | None | None |
| ECE1 | None | None | None | None |
| NDUFV2 | None | None | None | None |
| NDUFV2 | None | None | None | None |
| NDUFV2 | None | None | None | None |
| GHRHR | None | None | None | None |
| GHRHR | None | None | None | None |
| GHRHR | None | None | None | None |
| SLC4A1 | None | None | None | None |
| FGF9 | None | None | None | None |
| BARD1 | None | None | None | None |
| BARD1 | None | None | None | None |
| BARD1 | None | None | None | None |
| PTPN12 | None | None | None | None |
| MASP2 | None | None | None | None |
| MASP2 | None | None | None | None |

| | | | | |
|---|---|---|---|---|
| MASP2 | None | None | None | None |
| MASP2 | None | None | None | None |
| HOXB1 | None | None | None | None |
| SLC4A1 | None | None | None | None |
| SLC4A1 | None | None | None | None |
| SLC4A1 | None | None | None | None |
| SLC4A1 | None | None | None | None |
| SLC4A1 | None | None | None | None |
| SLC4A1 | None | None | None | None |
| SLC4A1 | None | None | None | None |
| SLC4A1 | None | None | None | None |
| SLC4A1 | None | None | None | None |
| SLC4A1 | None | None | None | None |
| SLC4A1 | None | None | None | None |
| SLC4A1 | None | None | None | None |
| SLC4A1 | None | None | None | None |
| SLC4A1 | None | None | None | None |
| SLC4A1 | None | None | None | None |
| SLC4A1 | None | None | None | None |
| SLC4A1 | None | None | None | None |
| CBL | None | None | None | None |
| FAS | None | None | None | None |
| MTMR2 | None | None | None | None |
| MTMR2 | None | None | None | None |
| MTMR2 | None | None | None | None |
| MTMR2 | None | None | None | None |
| MTMR2 | None | None | None | None |
| MTMR2 | None | None | None | None |
| MTMR2 | None | None | None | None |
| MTMR2 | None | None | None | None |
| KLHL3 | None | None | None | None |
| KLHL3 | None | None | None | None |
| KLHL3 | None | None | None | None |
| KLHL3 | None | None | None | None |
| KLHL3 | None | None | None | None |
| KLHL3 | None | None | None | None |
| RNASEH1 | None | None | None | None |
| ACTN1 | None | None | None | None |
| PHKG2 | None | None | None | None |
| PHKG2 | None | None | None | None |
| PHKG2 | None | None | None | None |
| NTRK1 | None | None | None | None |
| APOH | None | None | None | None |
| PMS2 | None | None | None | None |
| PMS2 | None | None | None | None |
| PMS2 | None | None | None | None |
| PMS2 | None | None | None | None |
| PMS2 | None | None | None | None |
| PMS2 | None | None | None | None |
| PMS2 | None | None | None | None |
| PMS2 | None | None | None | None |
| PMS2 | None | None | None | None |
| PMS2 | None | None | None | None |
| PMS2 | None | None | None | None |

| | | | | |
|---|---|---|---|---|
| PMS2 | None | None | None | None |
| PMS2 | None | None | None | None |
| PMS2 | None | None | None | None |
| PMS2 | None | None | None | None |
| PMS2 | None | None | None | None |
| PMS2 | None | None | None | None |
| PMS2 | None | None | None | None |
| PMS2 | None | None | None | None |
| PMS2 | None | None | None | None |
| PMS2 | None | None | None | None |
| PMS2 | None | None | None | None |
| MOCS2 | None | None | None | None |
| MOCS2 | None | None | None | None |
| RPGRIP1 | None | None | None | None |
| RPGRIP1 | None | None | None | None |
| LIG4 | None | None | None | None |
| PROC | None | None | None | None |
| PROC | None | None | None | None |
| NAT1 | None | None | None | None |
| NAT1 | None | None | None | None |
| PNPO | None | None | None | None |
| PNPO | None | None | None | None |
| PNPO | None | None | None | None |
| PNPO | None | None | None | None |
| PNPO | None | None | None | None |
| MRE11 | None | None | None | None |
| MRE11 | None | None | None | None |
| MRE11 | None | None | None | None |
| MRE11 | None | None | None | None |
| DDC | None | None | None | None |
| DDC | None | None | None | None |
| DDC | None | None | None | None |
| DDC | None | None | None | None |
| DDC | None | None | None | None |
| DDC | 0.909 | 7 | 3rch | B |
| DDC | 0.909 | 7 | 3rch | B |
| DDC | None | None | None | None |
| DDC | None | None | None | None |
| DDC | None | None | None | None |
| DDC | None | None | None | None |
| ANK2 | None | None | None | None |
| ANK2 | None | None | None | None |
| ANK2 | None | None | None | None |
| AK2 | None | None | None | None |
| AK2 | None | None | None | None |
| AK2 | None | None | None | None |
| AK2 | None | None | None | None |
| AK2 | None | None | None | None |
| AK2 | None | None | None | None |
| GPHN | None | None | None | None |
| HSD17B4 | None | None | None | None |
| HSD17B4 | None | None | None | None |
| HSD17B4 | None | None | None | None |
| HSD17B4 | None | None | None | None |

| | | | | |
|---|---|---|---|---|
| HSD17B4 | None | None | None | None |
| HSD17B4 | None | None | None | None |
| APTX | None | None | None | None |
| APTX | None | None | None | None |
| APTX | None | None | None | None |
| APTX | None | None | None | None |
| APTX | None | None | None | None |
| APTX | None | None | None | None |
| APTX | None | None | None | None |
| FANCL | None | None | None | None |
| FANCL | None | None | None | None |
| EMG1 | None | None | None | None |
| EMG1 | None | None | None | None |
| EZH2 | None | None | None | None |
| ABCB6 | None | None | None | None |
| NCF2 | None | None | None | None |
| NCF2 | None | None | None | None |
| COL4A2 | None | None | None | None |
| COL4A2 | 1 | 1 | 1li1 | F |
| COL4A2 | None | None | None | None |
| COL4A2 | None | None | None | None |
| COL4A2 | None | None | None | None |
| SLX4 | None | None | None | None |
| SERPIND1 | None | None | None | None |
| FGFR3 | None | None | None | None |
| FGFR3 | None | None | None | None |
| FGFR3 | None | None | None | None |
| FGFR3 | None | None | None | None |
| FGFR3 | None | None | None | None |
| FGFR3 | None | None | None | None |
| TREM2 | None | None | None | None |
| TREM2 | None | None | None | None |
| TREM2 | None | None | None | None |
| TREM2 | None | None | None | None |
| TREM2 | None | None | None | None |
| TREM2 | None | None | None | None |
| TREM2 | None | None | None | None |
| LIG4 | None | None | None | None |
| LIG4 | None | None | None | None |
| LIG4 | None | None | None | None |
| LIG4 | None | None | None | None |
| LIG4 | None | None | None | None |
| LIG4 | None | None | None | None |
| ECHS1 | None | None | None | None |
| ECHS1 | None | None | None | None |
| ECHS1 | None | None | None | None |
| MTR | None | None | None | None |
| MTR | None | None | None | None |
| MTR | None | None | None | None |
| MTR | None | None | None | None |
| MTR | None | None | None | None |
| PMVK | None | None | None | None |
| PRNP | None | None | None | None |
| CHN1 | None | None | None | None |

| | | | | |
|---|---|---|---|---|
| CHN1 | None | None | None | None |
| CHN1 | None | None | None | None |
| CHN1 | None | None | None | None |
| CHN1 | None | None | None | None |
| CHN1 | None | None | None | None |
| CHN1 | None | None | None | None |
| CHN1 | None | None | None | None |
| SLC2A1 | None | None | None | None |
| SLC2A1 | None | None | None | None |
| SLC2A1 | None | None | None | None |
| SLC2A1 | None | None | None | None |
| SLC2A1 | None | None | None | None |
| SLC2A1 | None | None | None | None |
| SLC2A1 | None | None | None | None |
| SLC2A1 | None | None | None | None |
| SLC2A1 | None | None | None | None |
| SLC2A1 | None | None | None | None |
| SLC2A1 | None | None | None | None |
| SLC2A1 | None | None | None | None |
| SLC2A1 | None | None | None | None |
| SLC2A1 | None | None | None | None |
| SLC2A1 | None | None | None | None |
| SLC2A1 | None | None | None | None |
| SLC2A1 | None | None | None | None |
| PMM2 | None | None | None | None |
| PMM2 | None | None | None | None |
| PMM2 | None | None | None | None |
| PMM2 | None | None | None | None |
| PMM2 | None | None | None | None |
| PMM2 | None | None | None | None |
| PMM2 | None | None | None | None |
| PMM2 | None | None | None | None |
| PMM2 | None | None | None | None |
| PMM2 | None | None | None | None |
| PMM2 | None | None | None | None |
| PMM2 | None | None | None | None |
| PMM2 | None | None | None | None |
| PMM2 | None | None | None | None |
| PMM2 | None | None | None | None |
| PMM2 | None | None | None | None |
| PMM2 | None | None | None | None |
| PMM2 | None | None | None | None |
| PMM2 | None | None | None | None |
| PMM2 | None | None | None | None |
| PMM2 | None | None | None | None |
| PMM2 | None | None | None | None |
| PMM2 | None | None | None | None |
| PMM2 | None | None | None | None |
| PMM2 | None | None | None | None |
| PMM2 | None | None | None | None |
| PMM2 | None | None | None | None |
| PMM2 | None | None | None | None |
| CCKAR | None | None | None | None |
| MAN1B1 | None | None | None | None |

| | | | | |
|---|---|---|---|---|
| MAN1B1 | None | None | None | None |
| LAMA2 | None | None | None | None |
| LAMA2 | None | None | None | None |
| TBC1D4 | None | None | None | None |
| TBC1D4 | None | None | None | None |
| LDLR | None | None | None | None |
| LDLR | None | None | None | None |
| LDLR | None | None | None | None |
| LDLR | None | None | None | None |
| LDLR | None | None | None | None |
| LDLR | None | None | None | None |
| LDLR | None | None | None | None |
| LDLR | None | None | None | None |
| LDLR | None | None | None | None |
| LDLR | None | None | None | None |
| LDLR | None | None | None | None |
| LDLR | None | None | None | None |
| LDLR | None | None | None | None |
| LDLR | None | None | None | None |
| LDLR | None | None | None | None |
| LDLR | None | None | None | None |
| LDLR | None | None | None | None |
| LDLR | None | None | None | None |
| LDLR | None | None | None | None |
| LDLR | None | None | None | None |
| LDLR | None | None | None | None |
| LDLR | None | None | None | None |
| LDLR | None | None | None | None |
| LDLR | None | None | None | None |
| LDLR | None | None | None | None |
| LDLR | None | None | None | None |
| LDLR | None | None | None | None |
| LDLR | None | None | None | None |
| LDLR | None | None | None | None |
| LDLR | None | None | None | None |
| LDLR | None | None | None | None |
| LDLR | None | None | None | None |
| LDLR | None | None | None | None |
| LDLR | None | None | None | None |
| LDLR | None | None | None | None |
| LDLR | None | None | None | None |
| LDLR | None | None | None | None |
| LDLR | None | None | None | None |
| LDLR | None | None | None | None |
| LDLR | None | None | None | None |
| LDLR | None | None | None | None |
| LDLR | None | None | None | None |
| LDLR | None | None | None | None |
| FANCM | None | None | None | None |
| HAMP | None | None | None | None |
| GUSB | None | None | None | None |
| GUSB | None | None | None | None |
| GUSB | None | None | None | None |
| GUSB | None | None | None | None |

| | | | | |
|---|---|---|---|---|
| GUSB | None | None | None | None |
| GUSB | None | None | None | None |
| GUSB | None | None | None | None |
| GUSB | None | None | None | None |
| GUSB | None | None | None | None |
| GUSB | None | None | None | None |
| GUSB | None | None | None | None |
| GUSB | None | None | None | None |
| GUSB | None | None | None | None |
| LIG1 | None | None | None | None |
| LIG1 | None | None | None | None |
| MYH7 | None | None | None | None |
| MYH7 | None | None | None | None |
| MYH7 | None | None | None | None |
| MYH7 | None | None | None | None |
| MYH7 | None | None | None | None |
| MYH7 | None | None | None | None |
| MYH7 | None | None | None | None |
| MYH7 | None | None | None | None |
| MYH7 | None | None | None | None |
| MYH7 | None | None | None | None |
| MYH7 | None | None | None | None |
| MYH7 | None | None | None | None |
| MYH7 | None | None | None | None |
| MYH7 | None | None | None | None |
| MYH7 | None | None | None | None |
| MYH7 | None | None | None | None |
| MYH7 | None | None | None | None |
| MYH7 | None | None | None | None |
| MYH7 | None | None | None | None |
| MYH7 | None | None | None | None |
| MYH7 | None | None | None | None |
| MYH7 | None | None | None | None |
| MYH7 | None | None | None | None |
| MYH7 | None | None | None | None |
| MYH7 | None | None | None | None |
| MYH7 | None | None | None | None |
| MYH7 | None | None | None | None |
| MYH7 | None | None | None | None |
| MYH7 | None | None | None | None |
| MYH7 | None | None | None | None |
| MYH7 | None | None | None | None |
| MYH7 | None | None | None | None |
| MYH7 | None | None | None | None |
| MYH7 | None | None | None | None |
| HCN4 | None | None | None | None |
| RECQL4 | None | None | None | None |
| RECQL4 | None | None | None | None |
| RECQL4 | None | None | None | None |
| RECQL4 | None | None | None | None |
| RECQL4 | None | None | None | None |
| RECQL4 | None | None | None | None |
| RECQL4 | None | None | None | None |
| RECQL4 | None | None | None | None |

| | | | | |
|---|---|---|---|---|
| RECQL4 | None | None | None | None |
| RECQL4 | None | None | None | None |
| RECQL4 | None | None | None | None |
| RECQL4 | None | None | None | None |
| RECQL4 | None | None | None | None |
| CHEK2 | None | None | None | None |
| PPARG | None | None | None | None |
| ACADVL | None | None | None | None |
| ACADVL | None | None | None | None |
| ACADVL | None | None | None | None |
| ACADVL | None | None | None | None |
| ACADVL | None | None | None | None |
| ACADVL | None | None | None | None |
| ACADVL | None | None | None | None |
| ACADVL | None | None | None | None |
| ACADVL | None | None | None | None |
| ACADVL | None | None | None | None |
| ACADVL | None | None | None | None |
| ACADVL | None | None | None | None |
| ACADVL | None | None | None | None |
| ACADVL | None | None | None | None |
| ACADVL | None | None | None | None |
| ACADVL | None | None | None | None |
| ACADVL | None | None | None | None |
| ACADVL | None | None | None | None |
| ACADVL | None | None | None | None |
| ACADVL | None | None | None | None |
| ACADVL | None | None | None | None |
| ACADVL | None | None | None | None |
| APOA1 | None | None | None | None |
| APOA1 | None | None | None | None |
| APOA1 | None | None | None | None |
| APOA1 | None | None | None | None |
| APOA1 | None | None | None | None |
| DNMT3A | None | None | None | None |
| DNMT3A | None | None | None | None |
| APC | None | None | None | None |
| TMOD1 | None | None | None | None |
| USH1C | None | None | None | None |
| USH1C | None | None | None | None |
| USH1C | None | None | None | None |
| USH1C | None | None | None | None |
| ERCC6 | None | None | None | None |
| ERCC6 | None | None | None | None |
| ERCC6 | None | None | None | None |
| ERCC8 | None | None | None | None |
| ERCC8 | None | None | None | None |
| ERCC8 | None | None | None | None |
| DPAGT1 | None | None | None | None |
| DPAGT1 | None | None | None | None |
| DPAGT1 | None | None | None | None |
| DPAGT1 | None | None | None | None |
| DPAGT1 | None | None | None | None |
| DPAGT1 | None | None | None | None |

| | | | | |
|---|---|---|---|---|
| DPAGT1 | None | None | None | None |
| IL2RG | None | None | None | None |
| HNF1A | None | None | None | None |
| HNF4A | None | None | None | None |
| FLT4 | None | None | None | None |
| FLT4 | None | None | None | None |
| FLT4 | None | None | None | None |
| CD3D | None | None | None | None |
| CD3D | None | None | None | None |
| MCOLN1 | None | None | None | None |
| MCOLN1 | None | None | None | None |
| MOCS2 | None | None | None | None |
| MOCS2 | None | None | None | None |
| ABCG8 | None | None | None | None |
| ABCG8 | None | None | None | None |
| ABCG8 | None | None | None | None |
| ABCG8 | None | None | None | None |
| ABCG8 | None | None | None | None |
| ABCG8 | None | None | None | None |
| ABCG8 | None | None | None | None |
| ABCG8 | None | None | None | None |
| ABCG8 | None | None | None | None |
| ABCG8 | None | None | None | None |
| ABCG8 | None | None | None | None |
| ABCG8 | None | None | None | None |
| ABCG8 | None | None | None | None |
| ABCG8 | None | None | None | None |
| CTNNA1 | None | None | None | None |
| TYMP | None | None | None | None |
| TYMP | None | None | None | None |
| TYMP | None | None | None | None |
| TYMP | None | None | None | None |
| TYMP | None | None | None | None |
| TYMP | None | None | None | None |
| TYMP | None | None | None | None |
| TYMP | None | None | None | None |
| TYMP | None | None | None | None |
| TYMP | None | None | None | None |
| TYMP | None | None | None | None |
| TYMP | None | None | None | None |
| TYMP | None | None | None | None |
| GAMT | None | None | None | None |
| GAMT | None | None | None | None |
| GAMT | None | None | None | None |
| GAMT | None | None | None | None |
| GAMT | None | None | None | None |
| GAMT | None | None | None | None |
| GAMT | None | None | None | None |
| GAMT | None | None | None | None |
| GAMT | None | None | None | None |
| GAMT | None | None | None | None |
| GAMT | None | None | None | None |
| GLRA1 | None | None | None | None |
| GLRA1 | None | None | None | None |

| | | | | |
|---|---|---|---|---|
| GLRA1 | None | None | None | None |
| GLRA1 | None | None | None | None |
| GLRA1 | None | None | None | None |
| GLRA1 | None | None | None | None |
| GLRA1 | None | None | None | None |
| GLRA1 | None | None | None | None |
| GLRA1 | None | None | None | None |
| ADK | None | None | None | None |
| ACTN2 | None | None | None | None |
| UBE2T | None | None | None | None |
| TTN | None | None | None | None |
| ATP7B | None | None | None | None |
| ATP7B | None | None | None | None |
| ATP7B | None | None | None | None |
| ATP7B | None | None | None | None |
| ATP7B | None | None | None | None |
| ATP7B | None | None | None | None |
| ATP7B | None | None | None | None |
| ATP7B | None | None | None | None |
| ATP7B | None | None | None | None |
| ATP7B | None | None | None | None |
| ATP7B | None | None | None | None |
| ATP7B | None | None | None | None |
| ATP7B | None | None | None | None |
| CDKN1B | None | None | None | None |
| CDKN1B | None | None | None | None |
| CDKN1B | None | None | None | None |
| CDKN1B | None | None | None | None |
| CDKN1B | None | None | None | None |
| CDKN1B | None | None | None | None |
| MYOC | None | None | None | None |
| MYOC | None | None | None | None |
| MYOC | None | None | None | None |
| MYOC | None | None | None | None |
| MYOC | None | None | None | None |
| MYOC | None | None | None | None |
| MYOC | None | None | None | None |
| MYOC | None | None | None | None |
| MYOC | None | None | None | None |
| MYOC | None | None | None | None |
| MYOC | None | None | None | None |
| MYOC | None | None | None | None |
| MYOC | None | None | None | None |
| MYOC | None | None | None | None |
| MYOC | None | None | None | None |
| SPTAN1 | None | None | None | None |
| GC | None | None | None | None |
| GC | None | None | None | None |
| MAD1L1 | None | None | None | None |
| NDST1 | None | None | None | None |
| ARL2BP | None | None | None | None |
| TOR1A | None | None | None | None |
| C1S | None | None | None | None |
| ARL6 | None | None | None | None |

| | | | | |
|---|---|---|---|---|
| ARL6 | None | None | None | None |
| ARL6 | None | None | None | None |
| ARL6 | None | None | None | None |
| ARL6 | None | None | None | None |
| ARL6 | None | None | None | None |
| FGFR1 | None | None | None | None |
| KLHL7 | None | None | None | None |
| FLNC | None | None | None | None |
| PTPRC | None | None | None | None |
| GCH1 | None | None | None | None |
| GCH1 | None | None | None | None |
| GCH1 | None | None | None | None |
| GCH1 | None | None | None | None |
| GCH1 | None | None | None | None |
| GCH1 | None | None | None | None |
| GCH1 | None | None | None | None |
| GCH1 | None | None | None | None |
| GCH1 | None | None | None | None |
| GCH1 | None | None | None | None |
| GCH1 | None | None | None | None |
| GCH1 | None | None | None | None |
| GCH1 | None | None | None | None |
| GCH1 | None | None | None | None |
| CFH | None | None | None | None |
| COL2A1 | None | None | None | None |
| COL2A1 | None | None | None | None |
| OCRL | None | None | None | None |
| WRN | None | None | None | None |
| WRN | None | None | None | None |
| WRN | None | None | None | None |
| WRN | None | None | None | None |
| MASP1 | None | None | None | None |
| MASP1 | None | None | None | None |
| NSD1 | None | None | None | None |
| PITX2 | None | None | None | None |
| PITX2 | None | None | None | None |
| PITX2 | None | None | None | None |
| PITX2 | None | None | None | None |
| PITX2 | None | None | None | None |
| PITX2 | None | None | None | None |
| B4GALT7 | 0.917 | 1 | 4irq | A |
| B4GALT7 | None | None | None | None |
| B4GALT7 | None | None | None | None |
| B4GALT7 | None | None | None | None |
| B4GALT7 | None | None | None | None |
| FSHR | None | None | None | None |
| IGHMBP2 | None | None | None | None |
| IGHMBP2 | None | None | None | None |
| IGHMBP2 | None | None | None | None |
| IGHMBP2 | None | None | None | None |
| IGHMBP2 | None | None | None | None |
| IGHMBP2 | None | None | None | None |
| IGHMBP2 | None | None | None | None |
| IGHMBP2 | None | None | None | None |

| | | | | |
|---|---|---|---|---|
| IGHMBP2 | None | None | None | None |
| IGHMBP2 | None | None | None | None |
| IGHMBP2 | None | None | None | None |
| IGHMBP2 | None | None | None | None |
| IGHMBP2 | None | None | None | None |
| IGHMBP2 | None | None | None | None |
| IGHMBP2 | None | None | None | None |
| IGHMBP2 | None | None | None | None |
| TTN | None | None | None | None |
| TTN | None | None | None | None |
| IRAK4 | None | None | None | None |
| PROC | None | None | None | None |
| PROC | None | None | None | None |
| RAD50 | None | None | None | None |
| RAD50 | None | None | None | None |
| DYNC1H1 | None | None | None | None |
| DYNC1H1 | None | None | None | None |
| DYNC1H1 | None | None | None | None |
| DYNC1H1 | None | None | None | None |
| DYNC1H1 | None | None | None | None |
| DYNC1H1 | None | None | None | None |
| DYNC1H1 | None | None | None | None |
| DYNC1H1 | None | None | None | None |
| DYNC1H1 | None | None | None | None |
| DYNC1H1 | None | None | None | None |
| DYNC1H1 | None | None | None | None |
| CYB5R3 | None | None | None | None |
| CYB5R3 | None | None | None | None |
| CYB5R3 | None | None | None | None |
| CYB5R3 | None | None | None | None |
| CYB5R3 | None | None | None | None |
| CYB5R3 | None | None | None | None |
| CYB5R3 | None | None | None | None |
| CYB5R3 | None | None | None | None |
| CYB5R3 | None | None | None | None |
| CYB5R3 | None | None | None | None |
| CYB5R3 | None | None | None | None |
| SPTB | None | None | None | None |
| FANCL | None | None | None | None |
| FAM126A | None | None | None | None |
| SEPT9 | None | None | None | None |
| TTN | None | None | None | None |
| TTN | None | None | None | None |
| TTN | None | None | None | None |
| CNTN1 | None | None | None | None |
| C1QTNF5 | None | None | None | None |
| VCL | None | None | None | None |
| VCL | None | None | None | None |
| VCL | None | None | None | None |
| VCL | None | None | None | None |
| VCL | None | None | None | None |
| VCL | None | None | None | None |
| PPT1 | None | None | None | None |
| PPT1 | None | None | None | None |

| | | | | |
|---|---|---|---|---|
| PPT1 | None | None | None | None |
| PPT1 | None | None | None | None |
| PPT1 | None | None | None | None |
| PPT1 | None | None | None | None |
| PPT1 | None | None | None | None |
| PPT1 | None | None | None | None |
| PPT1 | None | None | None | None |
| PPT1 | None | None | None | None |
| PPT1 | None | None | None | None |
| PPT1 | None | None | None | None |
| PPT1 | None | None | None | None |
| PPT1 | None | None | None | None |
| PPT1 | None | None | None | None |
| PPT1 | None | None | None | None |
| PPT1 | None | None | None | None |
| PPT1 | None | None | None | None |
| PPT1 | None | None | None | None |
| NOTCH2 | None | None | None | None |
| NOTCH2 | None | None | None | None |
| IFIH1 | None | None | None | None |
| IFIH1 | None | None | None | None |
| IKBKG | None | None | None | None |
| DBH | None | None | None | None |
| DBH | None | None | None | None |
| DBH | None | None | None | None |
| DBH | None | None | None | None |
| DBH | None | None | None | None |
| DBH | None | None | None | None |
| DBH | None | None | None | None |
| DBH | None | None | None | None |
| DBH | None | None | None | None |
| DBH | None | None | None | None |
| IL10RA | None | None | None | None |
| IL10RA | None | None | None | None |
| IL10RA | None | None | None | None |
| IL10RA | None | None | None | None |
| IL10RA | None | None | None | None |
| IL10RA | None | None | None | None |
| IL10RA | None | None | None | None |
| ILK | None | None | None | None |
| HPD | None | None | None | None |
| HPD | None | None | None | None |
| HPD | None | None | None | None |
| MECP2 | None | None | None | None |
| MECP2 | None | None | None | None |
| MECP2 | None | None | None | None |
| MECP2 | None | None | None | None |
| MECP2 | None | None | None | None |
| MECP2 | None | None | None | None |
| MECP2 | None | None | None | None |
| MECP2 | None | None | None | None |
| KYNU | None | None | None | None |
| FAS | None | None | None | None |
| FAS | None | None | None | None |

| | | | | |
|---|---|---|---|---|
| ELP1 | None | None | None | None |
| ELP1 | None | None | None | None |
| ELP1 | None | None | None | None |
| ELP1 | None | None | None | None |
| ELP1 | None | None | None | None |
| ELP1 | None | None | None | None |
| ELP1 | None | None | None | None |
| ELP1 | None | None | None | None |
| HESX1 | None | None | None | None |
| HESX1 | None | None | None | None |
| HESX1 | None | None | None | None |
| HESX1 | None | None | None | None |
| GAD1 | None | None | None | None |
| GAD1 | None | None | None | None |
| NDUFA1 | None | None | None | None |
| NDUFA1 | None | None | None | None |
| NDUFA1 | None | None | None | None |
| PNPT1 | None | None | None | None |
| PNPT1 | None | None | None | None |
| PNPT1 | None | None | None | None |
| PNPT1 | None | None | None | None |
| PNPT1 | None | None | None | None |
| PNPT1 | None | None | None | None |
| PNPT1 | None | None | None | None |
| SLC25A13 | None | None | None | None |
| SLC25A13 | None | None | None | None |
| SLC25A13 | None | None | None | None |
| AP4M1 | None | None | None | None |
| AP4M1 | None | None | None | None |
| MEFV | None | None | None | None |
| MEFV | None | None | None | None |
| MEFV | None | None | None | None |
| MEFV | None | None | None | None |
| MEFV | None | None | None | None |
| MEFV | None | None | None | None |
| MEFV | None | None | None | None |
| MEFV | None | None | None | None |
| MEFV | None | None | None | None |
| CAPN3 | None | None | None | None |
| CAPN3 | None | None | None | None |
| CAPN3 | None | None | None | None |
| CAPN3 | None | None | None | None |
| CAPN3 | None | None | None | None |
| CAPN3 | None | None | None | None |
| CAPN3 | None | None | None | None |
| GP6 | None | None | None | None |
| FAS | None | None | None | None |
| FBN1 | None | None | None | None |
| FBN1 | None | None | None | None |
| BMPR2 | None | None | None | None |
| BMPR2 | None | None | None | None |
| BMPR2 | None | None | None | None |
| BMPR2 | None | None | None | None |
| BMPR2 | None | None | None | None |

| | | | | |
|---|---|---|---|---|
| RET | None | None | None | None |
| RET | None | None | None | None |
| RET | None | None | None | None |
| RET | None | None | None | None |
| RET | None | None | None | None |
| RET | None | None | None | None |
| RET | None | None | None | None |
| RET | None | None | None | None |
| RET | None | None | None | None |
| RET | None | None | None | None |
| RET | None | None | None | None |
| RET | None | None | None | None |
| RET | None | None | None | None |
| RET | None | None | None | None |
| RET | None | None | None | None |
| RET | None | None | None | None |
| RET | None | None | None | None |
| RET | None | None | None | None |
| RET | None | None | None | None |
| RET | None | None | None | None |
| RET | None | None | None | None |
| RET | None | None | None | None |
| RET | None | None | None | None |
| RET | None | None | None | None |
| RET | None | None | None | None |
| RET | None | None | None | None |
| RET | None | None | None | None |
| RET | None | None | None | None |
| CSF2RA | None | None | None | None |
| CSF2RA | None | None | None | None |
| CSF2RA | None | None | None | None |
| BRCA1 | None | None | None | None |
| ZAP70 | None | None | None | None |
| ZAP70 | None | None | None | None |
| ZAP70 | None | None | None | None |
| PIK3R2 | None | None | None | None |
| TCN2 | None | None | None | None |
| TCN2 | None | None | None | None |
| TCN2 | None | None | None | None |
| TCN2 | None | None | None | None |
| TCN2 | None | None | None | None |
| TCN2 | None | None | None | None |
| TCN2 | None | None | None | None |
| TCN2 | None | None | None | None |
| EPM2A | None | None | None | None |
| EPM2A | None | None | None | None |
| EPM2A | None | None | None | None |
| EPM2A | None | None | None | None |
| AARS | None | None | None | None |
| AARS | None | None | None | None |
| AARS | None | None | None | None |
| PCDH15 | None | None | None | None |
| PCDH15 | None | None | None | None |
| PCDH15 | None | None | None | None |

| | | | | |
|---|---|---|---|---|
| PCDH15 | None | None | None | None |
| PCDH15 | None | None | None | None |
| PCDH15 | None | None | None | None |
| PCDH15 | None | None | None | None |
| PCDH15 | None | None | None | None |
| PCDH15 | None | None | None | None |
| PCDH15 | None | None | None | None |
| NDUFS4 | None | None | None | None |
| NDUFS4 | None | None | None | None |
| NDUFS4 | None | None | None | None |
| FGF10 | None | None | None | None |
| FGF10 | None | None | None | None |
| FGF10 | None | None | None | None |
| FGF10 | None | None | None | None |
| CDON | None | None | None | None |
| STAMBP | 0.909 | 2 | 3rzv | A |
| BRCA1 | None | None | None | None |
| BRCA1 | None | None | None | None |
| BRCA1 | None | None | None | None |
| BRCA1 | None | None | None | None |
| BRCA1 | None | None | None | None |
| BRCA1 | None | None | None | None |
| BRCA1 | None | None | None | None |
| BRCA1 | None | None | None | None |
| BRCA1 | None | None | None | None |
| BRCA1 | None | None | None | None |
| PLXNC1 | None | None | None | None |
| RNASEH2C | None | None | None | None |
| RNASEH2C | None | None | None | None |
| APC | None | None | None | None |
| PLA2G4A | None | None | None | None |
| PLA2G4A | None | None | None | None |
| PSAT1 | None | None | None | None |
| PSAT1 | None | None | None | None |
| TRAPPC3 | None | None | None | None |
| QDPR | None | None | None | None |
| QDPR | None | None | None | None |
| QDPR | None | None | None | None |
| QDPR | None | None | None | None |
| KRT5 | None | None | None | None |
| KRT5 | None | None | None | None |
| KRT5 | None | None | None | None |
| MLH1 | None | None | None | None |
| MLH1 | None | None | None | None |
| MLH1 | None | None | None | None |
| MLH1 | None | None | None | None |
| MLH1 | None | None | None | None |
| MLH1 | None | None | None | None |
| MLH1 | None | None | None | None |
| MLH1 | None | None | None | None |
| MLH1 | None | None | None | None |
| MLH1 | None | None | None | None |
| MLH1 | None | None | None | None |
| MLH1 | None | None | None | None |

| | | | | |
|---|---|---|---|---|
| MLH1 | None | None | None | None |
| MLH1 | None | None | None | None |
| MLH1 | None | None | None | None |
| MLH1 | None | None | None | None |
| MLH1 | None | None | None | None |
| KRIT1 | None | None | None | None |
| KRIT1 | None | None | None | None |
| KRIT1 | None | None | None | None |
| ALX4 | None | None | None | None |
| ALX4 | None | None | None | None |
| ALX4 | None | None | None | None |
| TJP2 | None | None | None | None |
| TJP2 | None | None | None | None |
| PANK2 | None | None | None | None |
| PANK2 | None | None | None | None |
| PANK2 | None | None | None | None |
| PANK2 | None | None | None | None |
| PANK2 | None | None | None | None |
| PANK2 | None | None | None | None |
| PANK2 | None | None | None | None |
| NDUFA6 | None | None | None | None |
| TSHR | None | None | None | None |
| TSHR | None | None | None | None |
| TSHR | None | None | None | None |
| CFTR | None | None | None | None |
| CFTR | None | None | None | None |
| MYOM1 | None | None | None | None |
| MYOM1 | None | None | None | None |
| SPTB | None | None | None | None |
| GPT2 | None | None | None | None |
| NDUFS2 | None | None | None | None |
| NDUFS2 | None | None | None | None |
| NDUFS2 | None | None | None | None |
| NDUFS2 | None | None | None | None |
| NDUFS2 | None | None | None | None |
| CASQ2 | None | None | None | None |
| CASQ2 | None | None | None | None |
| CASQ2 | None | None | None | None |
| CASQ2 | None | None | None | None |
| CASQ2 | None | None | None | None |
| CASQ2 | None | None | None | None |
| CASQ2 | None | None | None | None |
| CASQ2 | None | None | None | None |
| CASQ2 | None | None | None | None |
| CPOX | None | None | None | None |
| CPOX | None | None | None | None |
| CPOX | 0.857 | 2 | 2aex | A |
| CPOX | None | None | None | None |
| CPOX | None | None | None | None |
| CPOX | None | None | None | None |
| CPOX | None | None | None | None |
| CPOX | None | None | None | None |
| CPOX | None | None | None | None |
| MVK | None | None | None | None |

| | | | | |
|---|---|---|---|---|
| MVK | None | None | None | None |
| MVK | None | None | None | None |
| MVK | None | None | None | None |
| MVK | None | None | None | None |
| MVK | None | None | None | None |
| MVK | None | None | None | None |
| MVK | None | None | None | None |
| MVK | None | None | None | None |
| MVK | None | None | None | None |
| MVK | None | None | None | None |
| MVK | None | None | None | None |
| MVK | None | None | None | None |
| MVK | None | None | None | None |
| MVK | None | None | None | None |
| GSS | None | None | None | None |
| GSS | None | None | None | None |
| GSS | None | None | None | None |
| GSS | None | None | None | None |
| GSS | None | None | None | None |
| DSG2 | None | None | None | None |
| DSG2 | None | None | None | None |
| DSG2 | None | None | None | None |
| DSG2 | None | None | None | None |
| DSG2 | None | None | None | None |
| DSG2 | None | None | None | None |
| DSG2 | None | None | None | None |
| DSG2 | None | None | None | None |
| DSG2 | None | None | None | None |
| DSG2 | None | None | None | None |
| DSG2 | None | None | None | None |
| DSG2 | None | None | None | None |
| DSG2 | None | None | None | None |
| DSG2 | None | None | None | None |
| PAPSS2 | None | None | None | None |
| PLCB2 | None | None | None | None |
| HMGCS2 | None | None | None | None |
| HMGCS2 | None | None | None | None |
| HMGCS2 | 0.938 | 1 | 2wya | C |
| HMGCS2 | None | None | None | None |
| HMGCS2 | None | None | None | None |
| HMGCS2 | None | None | None | None |
| HMGCS2 | None | None | None | None |
| HMGCS2 | None | None | None | None |
| MPDZ | None | None | None | None |
| GATA4 | None | None | None | None |
| GATA4 | None | None | None | None |
| WARS2 | None | None | None | None |
| WARS2 | None | None | None | None |
| HGD | 1 | 47 | 1eyb | A |
| HGD | 1 | 47 | 1eyb | A |
| HGD | 1 | 70 | 1eyb | A |
| HGD | None | None | None | None |
| HGD | None | None | None | None |
| HGD | None | None | None | None |

| | | | | |
|---|---|---|---|---|
| HGD | None | None | None | None |
| HGD | None | None | None | None |
| HGD | None | None | None | None |
| HGD | 1 | 1 | 1ey2 | A |
| HGD | 1 | 9 | 1ey2 | A |
| HGD | None | None | None | None |
| HGD | 1 | 10 | 1eyb | A |
| HGD | 1 | 7 | 1ey2 | A |
| PAX8 | None | None | None | None |
| PAX8 | None | None | None | None |
| PAX8 | None | None | None | None |
| PAX8 | None | None | None | None |
| SPTB | None | None | None | None |
| NCF4 | None | None | None | None |
| CFH | None | None | None | None |
| CFH | None | None | None | None |
| CSF1R | None | None | None | None |
| CSF1R | None | None | None | None |
| CA8 | None | None | None | None |
| CA8 | None | None | None | None |
| CA8 | None | None | None | None |
| APC | None | None | None | None |
| APC | None | None | None | None |
| APC | None | None | None | None |
| APC | None | None | None | None |
| APC | None | None | None | None |
| NDUFA2 | None | None | None | None |
| GIF | None | None | None | None |
| GIF | None | None | None | None |
| GIF | None | None | None | None |
| GIF | None | None | None | None |
| C2 | None | None | None | None |
| ACP5 | None | None | None | None |
| ACP5 | None | None | None | None |
| ACP5 | None | None | None | None |
| ACP5 | None | None | None | None |
| ACP5 | None | None | None | None |
| RNASEH2A | None | None | None | None |
| RNASEH2A | None | None | None | None |
| RNASEH2A | None | None | None | None |
| RNASEH2A | None | None | None | None |
| RNASEH2A | None | None | None | None |
| RNASEH2A | None | None | None | None |
| RNASEH2A | None | None | None | None |
| DSG2 | None | None | None | None |
| USH1G | None | None | None | None |
| FUS | None | None | None | None |
| FUS | None | None | None | None |
| FUS | None | None | None | None |
| FUS | None | None | None | None |
| FUS | None | None | None | None |
| MRPL44 | None | None | None | None |
| MRPL44 | None | None | None | None |
| MRPL44 | None | None | None | None |

| | | | | |
|---|---|---|---|---|
| MRPL44 | None | None | None | None |
| DMGDH | None | None | None | None |
| AARS | None | None | None | None |
| STAR | None | None | None | None |
| STAR | None | None | None | None |
| STAR | None | None | None | None |
| STAR | None | None | None | None |
| STAR | None | None | None | None |
| STAR | None | None | None | None |
| STAR | None | None | None | None |
| STAR | None | None | None | None |
| LARP7 | None | None | None | None |
| PRPF8 | None | None | None | None |
| PRPF8 | None | None | None | None |
| PRPF8 | None | None | None | None |
| PRPF8 | None | None | None | None |
| PSAP | None | None | None | None |
| PSAP | None | None | None | None |
| PSAP | None | None | None | None |
| DMPK | None | None | None | None |
| EPCAM | None | None | None | None |
| EPCAM | None | None | None | None |
| EPCAM | None | None | None | None |
| EPCAM | None | None | None | None |
| EPCAM | None | None | None | None |
| EPCAM | None | None | None | None |
| EPCAM | None | None | None | None |
| EPCAM | None | None | None | None |
| ASS1 | None | None | None | None |
| ASS1 | None | None | None | None |
| ASS1 | None | None | None | None |
| ASS1 | None | None | None | None |
| ASS1 | None | None | None | None |
| ASS1 | 0.857 | 20 | 2nz2 | A |
| ASS1 | 0.857 | 9 | 2nz2 | A |
| ASS1 | None | None | None | None |
| ASS1 | None | None | None | None |
| ASS1 | None | None | None | None |
| ASS1 | 0.857 | 9 | 2nz2 | A |
| ASS1 | 0.857 | 9 | 2nz2 | A |
| ASS1 | 0.857 | 9 | 2nz2 | A |
| ASS1 | None | None | None | None |
| ASS1 | None | None | None | None |
| ASS1 | None | None | None | None |
| ASS1 | None | None | None | None |
| ASS1 | None | None | None | None |
| ASS1 | None | None | None | None |
| ASS1 | None | None | None | None |
| POLE | None | None | None | None |
| POLE | None | None | None | None |
| POLE | None | None | None | None |
| RNASET2 | None | None | None | None |
| RNASET2 | None | None | None | None |
| RNASET2 | None | None | None | None |

| | | | | |
|---|---|---|---|---|
| FLG | None | None | None | None |
| F10 | None | None | None | None |
| AGT | None | None | None | None |
| AGT | None | None | None | None |
| PLN | None | None | None | None |
| PLN | None | None | None | None |
| PLN | None | None | None | None |
| PTHLH | None | None | None | None |
| IGF1R | None | None | None | None |
| IGF1R | None | None | None | None |
| IGF1R | None | None | None | None |
| JAG1 | None | None | None | None |
| JAG1 | None | None | None | None |
| JAG1 | None | None | None | None |
| JAG1 | None | None | None | None |
| JAG1 | None | None | None | None |
| JAG1 | None | None | None | None |
| AP1S3 | None | None | None | None |
| SPTA1 | None | None | None | None |
| SPTA1 | None | None | None | None |
| SPTA1 | None | None | None | None |
| SPTA1 | None | None | None | None |
| SPG7 | None | None | None | None |
| SPG7 | None | None | None | None |
| SPG7 | None | None | None | None |
| SPG7 | None | None | None | None |
| SPG7 | None | None | None | None |
| SPG7 | None | None | None | None |
| FN1 | None | None | None | None |
| FN1 | None | None | None | None |
| NSMCE3 | None | None | None | None |
| PCCA | None | None | None | None |
| PCCA | None | None | None | None |
| SPTB | None | None | None | None |
| RYR2 | None | None | None | None |
| RYR2 | None | None | None | None |
| MMP9 | None | None | None | None |
| MMP9 | None | None | None | None |
| GLMN | None | None | None | None |
| GLMN | None | None | None | None |
| GLMN | None | None | None | None |
| GLMN | None | None | None | None |
| GLMN | None | None | None | None |
| GLMN | None | None | None | None |
| GLMN | None | None | None | None |
| GLMN | None | None | None | None |
| GLMN | None | None | None | None |
| GLMN | None | None | None | None |
| MARS | None | None | None | None |
| MARS | None | None | None | None |
| MARS | None | None | None | None |
| MARS | None | None | None | None |
| MARS | None | None | None | None |
| MARS | None | None | None | None |

| | | | | |
|---|---|---|---|---|
| SNAP29 | None | None | None | None |
| SNAP29 | None | None | None | None |
| ETHE1 | None | None | None | None |
| ETHE1 | None | None | None | None |
| ETHE1 | None | None | None | None |
| ETHE1 | None | None | None | None |
| ETHE1 | None | None | None | None |
| ETHE1 | None | None | None | None |
| ETHE1 | None | None | None | None |
| SAMHD1 | None | None | None | None |
| KCNQ4 | None | None | None | None |
| FXYD2 | None | None | None | None |
| FXYD2 | None | None | None | None |
| FXYD2 | None | None | None | None |
| LRRK2 | None | None | None | None |
| LRRK2 | None | None | None | None |
| NDUFS1 | None | None | None | None |
| NDUFS1 | None | None | None | None |
| NDUFS1 | None | None | None | None |
| NDUFS1 | None | None | None | None |
| NDUFS1 | None | None | None | None |
| NDUFS1 | None | None | None | None |
| NDUFS1 | None | None | None | None |
| NDUFS1 | None | None | None | None |
| NDUFS1 | None | None | None | None |
| NDUFS1 | None | None | None | None |
| NDUFS1 | None | None | None | None |
| EZH2 | None | None | None | None |
| EZH2 | None | None | None | None |
| LARS | None | None | None | None |
| TLL1 | None | None | None | None |
| CRYBB2 | None | None | None | None |
| CRYBB2 | None | None | None | None |
| EXOSC8 | None | None | None | None |
| SRP72 | None | None | None | None |
| FOXC2 | None | None | None | None |
| HMBS | None | None | None | None |
| HMBS | None | None | None | None |
| HMBS | None | None | None | None |
| HMBS | None | None | None | None |
| HMBS | None | None | None | None |
| HMBS | None | None | None | None |
| HMBS | None | None | None | None |
| HMBS | None | None | None | None |
| HMBS | None | None | None | None |
| HMBS | None | None | None | None |
| HMBS | None | None | None | None |
| HMBS | None | None | None | None |
| HMBS | None | None | None | None |
| HMBS | None | None | None | None |
| HMBS | None | None | None | None |
| HMBS | None | None | None | None |
| HMBS | None | None | None | None |
| HMBS | None | None | None | None |

| | | | | |
|---|---|---|---|---|
| HMBS | None | None | None | None |
| HMBS | None | None | None | None |
| HMBS | None | None | None | None |
| HMBS | None | None | None | None |
| HMBS | None | None | None | None |
| HMBS | None | None | None | None |
| HMBS | None | None | None | None |
| OGT | None | None | None | None |
| NOG | None | None | None | None |
| NOG | None | None | None | None |
| NOG | None | None | None | None |
| NOG | None | None | None | None |
| NOG | None | None | None | None |
| NOG | None | None | None | None |
| NOG | None | None | None | None |
| NOG | None | None | None | None |
| NOG | None | None | None | None |
| NOG | None | None | None | None |
| NOG | None | None | None | None |
| DLG3 | None | None | None | None |
| SCN2A | None | None | None | None |
| STAC3 | None | None | None | None |
| PFKM | None | None | None | None |
| PFKM | None | None | None | None |
| PFKM | None | None | None | None |
| PFKM | None | None | None | None |
| PFKM | None | None | None | None |
| DNMT1 | None | None | None | None |
| DNMT1 | None | None | None | None |
| TNNI3 | None | None | None | None |
| TNNI3 | None | None | None | None |
| TNNI3 | None | None | None | None |
| STXBP2 | None | None | None | None |
| STXBP2 | None | None | None | None |
| STXBP2 | None | None | None | None |
| STXBP2 | None | None | None | None |
| STXBP2 | None | None | None | None |
| STXBP2 | None | None | None | None |
| STXBP2 | None | None | None | None |
| STXBP2 | None | None | None | None |
| REN | None | None | None | None |
| DSP | None | None | None | None |
| DSP | None | None | None | None |
| HNF1B | None | None | None | None |
| ADA | None | None | None | None |
| ADA | None | None | None | None |
| ADA | None | None | None | None |
| ADA | None | None | None | None |
| ADA | None | None | None | None |
| ADA | None | None | None | None |
| ADA | None | None | None | None |
| ADA | None | None | None | None |
| ADA | None | None | None | None |
| ADA | None | None | None | None |

| | | | | |
|---|---|---|---|---|
| ADA | None | None | None | None |
| ADA | None | None | None | None |
| ADA | None | None | None | None |
| ADA | None | None | None | None |
| ADA | None | None | None | None |
| DCX | None | None | None | None |
| DCX | None | None | None | None |
| DCX | None | None | None | None |
| DCX | None | None | None | None |
| DCX | None | None | None | None |
| ARSB | None | None | None | None |
| ARSB | None | None | None | None |
| ARSB | None | None | None | None |
| ARSB | None | None | None | None |
| ARSB | None | None | None | None |
| ARSB | None | None | None | None |
| ARSB | None | None | None | None |
| ARSB | None | None | None | None |
| ARSB | None | None | None | None |
| ARSB | None | None | None | None |
| ARSB | None | None | None | None |
| ARSB | None | None | None | None |
| ARSB | None | None | None | None |
| WWOX | None | None | None | None |
| TNNI3 | None | None | None | None |
| TNNI3 | None | None | None | None |
| TNNI3 | None | None | None | None |
| TNNI3 | None | None | None | None |
| CYBA | None | None | None | None |
| SPAST | None | None | None | None |
| SPAST | None | None | None | None |
| SPAST | None | None | None | None |
| SPAST | None | None | None | None |
| SPAST | None | None | None | None |
| SPAST | None | None | None | None |
| SPAST | None | None | None | None |
| SPAST | None | None | None | None |
| KRIT1 | None | None | None | None |
| KRIT1 | None | None | None | None |
| TP53 | None | None | None | None |
| ISPD | None | None | None | None |
| ISPD | None | None | None | None |
| ISPD | None | None | None | None |
| ISPD | None | None | None | None |
| KCNMA1 | None | None | None | None |
| STK11 | None | None | None | None |
| STK11 | None | None | None | None |
| STK11 | None | None | None | None |
| STK11 | None | None | None | None |
| STK11 | None | None | None | None |
| STK11 | None | None | None | None |
| NDUFA13 | None | None | None | None |
| PHF6 | None | None | None | None |
| PHF6 | None | None | None | None |

| | | | | |
|---|---|---|---|---|
| PHF6 | None | None | None | None |
| PHYH | None | None | None | None |
| PHYH | None | None | None | None |
| PHYH | None | None | None | None |
| PHYH | None | None | None | None |
| PHYH | None | None | None | None |
| NDUFS3 | None | None | None | None |
| NDUFS3 | None | None | None | None |
| ALDH18A1 | None | None | None | None |
| ALDH18A1 | None | None | None | None |
| ALDH18A1 | None | None | None | None |
| ALDH18A1 | None | None | None | None |
| ALDH18A1 | None | None | None | None |
| PAX6 | None | None | None | None |
| PAX6 | None | None | None | None |
| INPP5E | None | None | None | None |
| INPP5E | None | None | None | None |
| INPP5E | None | None | None | None |
| INPP5E | None | None | None | None |
| INPP5E | None | None | None | None |
| INPP5E | None | None | None | None |
| INPP5E | None | None | None | None |
| INPP5E | None | None | None | None |
| INPP5E | None | None | None | None |
| TNNT2 | None | None | None | None |
| TNNT2 | None | None | None | None |
| TNNT2 | None | None | None | None |
| TNNT2 | None | None | None | None |
| TNNT2 | None | None | None | None |
| TNNT2 | None | None | None | None |
| TNNT2 | None | None | None | None |
| TNNT2 | None | None | None | None |
| TYK2 | None | None | None | None |
| TYK2 | None | None | None | None |
| TYK2 | None | None | None | None |
| CFTR | None | None | None | None |
| CFTR | None | None | None | None |
| CFTR | None | None | None | None |
| CFTR | None | None | None | None |
| CFTR | None | None | None | None |
| CFTR | None | None | None | None |
| CFTR | None | None | None | None |
| CFTR | None | None | None | None |
| CFTR | None | None | None | None |
| CFTR | None | None | None | None |
| CFTR | None | None | None | None |
| CFTR | None | None | None | None |
| CFTR | None | None | None | None |
| CFTR | None | None | None | None |
| CFTR | None | None | None | None |
| CFTR | None | None | None | None |
| CFTR | None | None | None | None |
| CFTR | None | None | None | None |
| CFTR | None | None | None | None |

| | | | | |
|---|---|---|---|---|
| CFTR | None | None | None | None |
| CFTR | None | None | None | None |
| CFTR | None | None | None | None |
| CFTR | None | None | None | None |
| CFTR | None | None | None | None |
| CFTR | None | None | None | None |
| CFTR | None | None | None | None |
| CFTR | None | None | None | None |
| CFTR | None | None | None | None |
| CFTR | None | None | None | None |
| CFTR | None | None | None | None |
| CFTR | None | None | None | None |
| CFTR | None | None | None | None |
| CFTR | None | None | None | None |
| CFTR | None | None | None | None |
| CFTR | None | None | None | None |
| CFTR | None | None | None | None |
| CFTR | None | None | None | None |
| CFTR | None | None | None | None |
| CFTR | None | None | None | None |
| CFTR | None | None | None | None |
| CFTR | None | None | None | None |
| CFTR | None | None | None | None |
| CFTR | None | None | None | None |
| CFTR | None | None | None | None |
| CFTR | None | None | None | None |
| CFTR | None | None | None | None |
| CFTR | None | None | None | None |
| CFTR | None | None | None | None |
| CFTR | None | None | None | None |
| CFTR | None | None | None | None |
| CFTR | None | None | None | None |
| CFTR | None | None | None | None |
| CFTR | None | None | None | None |
| SERPINF1 | None | None | None | None |
| SERPINF1 | None | None | None | None |
| IDS | None | None | None | None |
| IDS | None | None | None | None |
| IDS | None | None | None | None |
| IDS | None | None | None | None |
| IDS | None | None | None | None |
| IDS | None | None | None | None |
| IDS | None | None | None | None |
| IDS | None | None | None | None |
| IDS | None | None | None | None |
| NFKBIA | None | None | None | None |
| NFKBIA | None | None | None | None |
| SPAST | None | None | None | None |
| FGF23 | None | None | None | None |
| FGF23 | None | None | None | None |
| FGF23 | None | None | None | None |
| FGF23 | None | None | None | None |
| FGF23 | None | None | None | None |
| FGF23 | None | None | None | None |

| | | | | |
|---|---|---|---|---|
| FGF23 | None | None | None | None |
| DLAT | None | None | None | None |
| STUB1 | None | None | None | None |
| HSD17B4 | None | None | None | None |
| HSD17B4 | None | None | None | None |
| DYSF | None | None | None | None |
| CEP135 | None | None | None | None |
| CEP135 | None | None | None | None |
| PRKCG | None | None | None | None |
| CSF1R | None | None | None | None |
| PEX5 | None | None | None | None |
| PAX6 | None | None | None | None |
| PAX6 | None | None | None | None |
| PAX6 | None | None | None | None |
| PAX6 | None | None | None | None |
| PAX6 | None | None | None | None |
| PAX6 | None | None | None | None |
| NF2 | None | None | None | None |
| NF2 | None | None | None | None |
| NF2 | None | None | None | None |
| REM1 | None | None | None | None |

| source_id | classification | binding_site | binding_site_single_water | protein_binding_site |
|---|---|---|---|---|
| rs121918343 | pathogenic | 1 | 1 | 0 |
| rs267606649 | pathogenic | 1 | 1 | 1 |
| rs121918342 | pathogenic | 0 | 1 | 0 |
| rs267606650 | pathogenic | 1 | 1 | 1 |
| rs1130335 | benign | 1 | 1 | 1 |
| rs199476087 | pathogenic | 1 | 1 | 1 |
| rs772163112 | benign | 0 | 0 | 0 |
| rs755793 | benign | 0 | 0 | 0 |
| rs121918155 | pathogenic | 1 | 1 | 0 |
| rs121918156 | pathogenic | 1 | 1 | 0 |
| rs121918154 | pathogenic | 1 | 1 | 1 |
| rs121918151 | pathogenic | 1 | 1 | 1 |
| rs121918144 | pathogenic | 0 | 1 | 0 |
| rs121918146 | pathogenic | 1 | 1 | 1 |
| rs121918160 | pathogenic | 1 | 1 | 1 |
| rs121918150 | pathogenic | 1 | 1 | 1 |
| rs121918141 | pathogenic | 1 | 1 | 1 |
| rs142742242 | pathogenic | 1 | 1 | 1 |
| rs121918142 | pathogenic | 1 | 1 | 1 |
| rs121918157 | pathogenic | 1 | 1 | 1 |
| rs121918101 | pathogenic | 1 | 1 | 1 |
| rs121908260 | pathogenic | 1 | 1 | 1 |
| rs80356667 | pathogenic | 1 | 1 | 1 |
| rs80356668 | pathogenic | 1 | 1 | 1 |
| rs397514540 | pathogenic | 1 | 1 | 1 |
| rs104894685 | pathogenic | 0 | 1 | 0 |
| rs199869995 | pathogenic | 1 | 1 | 1 |
| rs121909072 | pathogenic | 0 | 0 | 0 |
| rs143669310 | benign | 1 | 1 | 1 |
| rs747347498 | benign | 0 | 0 | 0 |
| rs138283222 | benign | 0 | 0 | 0 |
| rs118203931 | pathogenic | 1 | 1 | 1 |
| rs118203933 | pathogenic | 1 | 1 | 0 |
| rs118203932 | pathogenic | 1 | 1 | 0 |
| rs2228063 | benign | 1 | 1 | 1 |
| rs377767360 | pathogenic | 0 | 0 | 0 |
| rs397518413 | pathogenic | 1 | 1 | 1 |
| rs1800458 | benign | 1 | 1 | 0 |
| rs121918083 | pathogenic | 1 | 1 | 1 |
| rs121918094 | pathogenic | 1 | 1 | 1 |
| rs121918098 | pathogenic | 1 | 1 | 1 |
| rs121918093 | pathogenic | 1 | 1 | 1 |
| rs79977247 | pathogenic | 1 | 1 | 1 |
| rs28933979 | pathogenic | 1 | 1 | 1 |
| rs121918068 | pathogenic | 1 | 1 | 1 |
| rs121918077 | pathogenic | 1 | 1 | 1 |
| rs11541796 | pathogenic | 1 | 1 | 1 |
| rs104894665 | pathogenic | 1 | 1 | 1 |
| rs121918090 | pathogenic | 1 | 1 | 1 |
| rs121918081 | pathogenic | 1 | 1 | 1 |
| rs121918080 | pathogenic | 1 | 1 | 1 |
| rs121918076 | pathogenic | 1 | 1 | 1 |
| rs121918097 | pathogenic | 1 | 1 | 1 |

| | | | | |
|---|---|---|---|---|
| rs121918079 | pathogenic | 1 | 1 | 1 |
| rs121918069 | pathogenic | 1 | 1 | 1 |
| rs121918086 | pathogenic | 1 | 1 | 1 |
| rs121918091 | pathogenic | 1 | 1 | 1 |
| rs121918085 | pathogenic | 1 | 1 | 1 |
| rs121918100 | pathogenic | 1 | 1 | 1 |
| rs121918084 | pathogenic | 1 | 1 | 1 |
| rs121918071 | pathogenic | 1 | 1 | 1 |
| rs121918072 | pathogenic | 1 | 1 | 1 |
| rs121918082 | pathogenic | 1 | 1 | 1 |
| rs121918087 | pathogenic | 1 | 1 | 1 |
| rs76410435 | benign | 1 | 1 | 1 |
| rs121918089 | pathogenic | 1 | 1 | 1 |
| rs121918073 | pathogenic | 1 | 1 | 1 |
| rs121918075 | pathogenic | 1 | 1 | 1 |
| rs121918088 | pathogenic | 1 | 1 | 1 |
| rs76992529 | pathogenic | 1 | 1 | 1 |
| rs536294863 | benign | 1 | 1 | 1 |
| rs730881162 | benign | 1 | 1 | 1 |
| rs113994087 | pathogenic | 1 | 1 | 1 |
| rs74716434 | benign | 1 | 1 | 1 |
| rs121918694 | pathogenic | 1 | 1 | 1 |
| rs121918707 | pathogenic | 1 | 1 | 1 |
| rs121918695 | pathogenic | 1 | 1 | 0 |
| rs121918690 | pathogenic | 1 | 1 | 0 |
| rs121918696 | pathogenic | 1 | 1 | 0 |
| rs121918693 | pathogenic | 1 | 1 | 0 |
| rs28999969 | pathogenic | 1 | 1 | 1 |
| rs121918697 | pathogenic | 1 | 1 | 1 |
| rs121918688 | pathogenic | 1 | 1 | 0 |
| rs28999970 | pathogenic | 1 | 1 | 0 |
| rs121918686 | pathogenic | 1 | 1 | 0 |
| rs28999971 | pathogenic | 1 | 1 | 1 |
| rs121918698 | pathogenic | 1 | 1 | 1 |
| rs121918691 | pathogenic | 1 | 1 | 0 |
| rs121918692 | pathogenic | 1 | 1 | 1 |
| rs121918703 | pathogenic | 1 | 1 | 1 |
| rs121918687 | pathogenic | 1 | 1 | 1 |
| rs28933408 | pathogenic | 1 | 1 | 1 |
| rs121918702 | pathogenic | 1 | 1 | 1 |
| rs17875899 | benign | 1 | 1 | 1 |
| rs111033620 | pathogenic | 1 | 1 | 1 |
| rs111033622 | pathogenic | 1 | 1 | 1 |
| rs111033621 | pathogenic | 1 | 1 | 1 |
| rs111033618 | pathogenic | 1 | 1 | 1 |
| rs5918 | benign | 0 | 0 | 0 |
| rs121918452 | pathogenic | 0 | 0 | 0 |
| rs121918445 | pathogenic | 1 | 1 | 1 |
| rs5917 | benign | 1 | 1 | 1 |
| rs121918444 | pathogenic | 1 | 1 | 1 |
| rs121918446 | pathogenic | 1 | 1 | 1 |
| rs121918449 | pathogenic | 0 | 0 | 0 |
| rs121918448 | benign | 0 | 0 | 0 |
| rs13306487 | benign | 1 | 1 | 1 |

| | | | | |
|---|---|---|---|---|
| rs1805044 | benign | 0 | 0 | 0 |
| rs117078377 | benign | 0 | 0 | 0 |
| rs138132026 | benign | 1 | 1 | 1 |
| rs75762896 | benign | 1 | 1 | 0 |
| rs121918043 | pathogenic | 1 | 1 | 0 |
| rs143581690 | benign | 1 | 1 | 0 |
| rs147654263 | benign | 0 | 0 | 0 |
| rs587782529 | pathogenic | 1 | 1 | 1 |
| rs121912664 | pathogenic | 1 | 1 | 1 |
| rs150293825 | benign | 1 | 1 | 1 |
| rs17882252 | benign | 1 | 1 | 1 |
| rs121912662 | pathogenic | 1 | 1 | 1 |
| rs35993958 | benign | 1 | 1 | 1 |
| rs606231253 | pathogenic | 1 | 1 | 1 |
| rs2227983 | benign | 1 | 1 | 1 |
| rs587779351 | pathogenic | 0 | 0 | 0 |
| rs779773463 | pathogenic | 1 | 1 | 1 |
| rs713040 | benign | 1 | 1 | 1 |
| rs334 | pathogenic | 1 | 1 | 1 |
| rs33930165 | pathogenic | 1 | 1 | 1 |
| rs33918131 | benign | 1 | 1 | 1 |
| rs63750783 | pathogenic | 1 | 1 | 1 |
| rs33986703 | pathogenic | 1 | 1 | 1 |
| rs33972047 | pathogenic | 1 | 1 | 1 |
| rs33936254 | pathogenic | 1 | 1 | 1 |
| rs33959855 | benign | 1 | 1 | 1 |
| rs33950507 | pathogenic | 1 | 1 | 1 |
| rs33916412 | pathogenic | 1 | 1 | 0 |
| rs1135071 | pathogenic | 1 | 1 | 1 |
| rs33960103 | pathogenic | 1 | 1 | 1 |
| rs11549407 | pathogenic | 1 | 1 | 1 |
| rs33922842 | pathogenic | 1 | 1 | 1 |
| rs35278874 | benign | 1 | 1 | 1 |
| rs34621955 | benign | 1 | 1 | 1 |
| rs33995148 | pathogenic | 1 | 1 | 1 |
| rs33945705 | benign | 1 | 1 | 1 |
| rs33993568 | benign | 1 | 1 | 1 |
| rs35553496 | benign | 1 | 1 | 1 |
| rs33913712 | pathogenic | 1 | 1 | 1 |
| rs33917785 | pathogenic | 1 | 1 | 1 |
| rs34515413 | benign | 1 | 1 | 1 |
| rs35256489 | pathogenic | 1 | 1 | 0 |
| rs34484056 | pathogenic | 1 | 1 | 1 |
| rs33947020 | benign | 1 | 1 | 1 |
| rs33946267 | pathogenic | 1 | 1 | 1 |
| rs33966761 | pathogenic | 1 | 1 | 1 |
| rs36020563 | benign | 1 | 1 | 1 |
| rs122454124 | pathogenic | 1 | 1 | 1 |
| rs122454126 | pathogenic | 1 | 1 | 0 |
| rs122454127 | pathogenic | 1 | 1 | 1 |
| rs122454130 | pathogenic | 1 | 1 | 1 |
| rs122454125 | pathogenic | 1 | 1 | 1 |
| rs122454131 | pathogenic | 1 | 1 | 0 |
| rs121908302 | pathogenic | 0 | 1 | 0 |

| | | | | |
|---|---|---|---|---|
| rs77829017 | pathogenic | 0 | 1 | 0 |
| rs1141814 | pathogenic | 0 | 1 | 0 |
| rs121908312 | pathogenic | 0 | 1 | 0 |
| rs79653797 | pathogenic | 1 | 1 | 0 |
| rs439898 | pathogenic | 1 | 1 | 0 |
| rs121908299 | pathogenic | 0 | 1 | 0 |
| rs80356763 | pathogenic | 1 | 1 | 1 |
| rs781152868 | pathogenic | 0 | 1 | 0 |
| rs121908297 | pathogenic | 0 | 1 | 0 |
| rs398123531 | benign | 0 | 1 | 0 |
| rs398123532 | pathogenic | 0 | 1 | 0 |
| rs364897 | pathogenic | 1 | 1 | 1 |
| rs1064644 | pathogenic | 0 | 1 | 0 |
| rs409652 | pathogenic | 1 | 1 | 1 |
| rs121908300 | pathogenic | 0 | 1 | 0 |
| rs381737 | pathogenic | 0 | 1 | 0 |
| rs74500255 | pathogenic | 0 | 1 | 0 |
| rs121908313 | pathogenic | 0 | 1 | 0 |
| rs367968666 | pathogenic | 0 | 1 | 0 |
| rs78973108 | pathogenic | 1 | 1 | 1 |
| rs121908298 | pathogenic | 1 | 1 | 1 |
| rs78396650 | pathogenic | 0 | 0 | 0 |
| rs78198234 | pathogenic | 1 | 1 | 0 |
| rs121908304 | pathogenic | 1 | 1 | 1 |
| rs76539814 | pathogenic | 1 | 1 | 1 |
| rs121908305 | pathogenic | 0 | 1 | 0 |
| rs121908306 | pathogenic | 1 | 1 | 0 |
| rs121908308 | pathogenic | 1 | 1 | 1 |
| rs121908307 | pathogenic | 0 | 1 | 0 |
| rs75548401 | benign | 0 | 1 | 0 |
| rs76763715 | pathogenic | 0 | 0 | 0 |
| rs121908314 | pathogenic | 0 | 1 | 0 |
| rs398123528 | pathogenic | 0 | 1 | 0 |
| rs121908311 | pathogenic | 0 | 0 | 0 |
| rs121908310 | pathogenic | 1 | 1 | 0 |
| rs74598136 | pathogenic | 0 | 1 | 0 |
| rs1064651 | pathogenic | 1 | 1 | 1 |
| rs77369218 | pathogenic | 1 | 1 | 1 |
| rs121908295 | pathogenic | 0 | 1 | 0 |
| rs421016 | pathogenic | 0 | 1 | 0 |
| rs368060 | pathogenic | 0 | 1 | 0 |
| rs80356771 | pathogenic | 1 | 1 | 1 |
| rs80356772 | pathogenic | 1 | 1 | 1 |
| rs121908301 | pathogenic | 0 | 0 | 0 |
| rs747506979 | pathogenic | 0 | 1 | 0 |
| rs75822236 | pathogenic | 0 | 1 | 0 |
| rs1990760 | benign | 1 | 1 | 0 |
| rs28934568 | pathogenic | 1 | 1 | 0 |
| rs34833812 | benign | 1 | 1 | 0 |
| rs193922666 | benign | 1 | 1 | 0 |
| rs104893812 | pathogenic | 1 | 1 | 0 |
| rs144701411 | benign | 0 | 0 | 0 |
| rs104893813 | pathogenic | 0 | 1 | 0 |
| rs199660234 | pathogenic | 1 | 1 | 0 |

| | | | | |
|---|---|---|---|---|
| rs104893814 | pathogenic | 1 | 1 | 0 |
| rs143095746 | benign | 1 | 1 | 1 |
| rs104893807 | pathogenic | 1 | 1 | 0 |
| rs104893811 | pathogenic | 0 | 0 | 0 |
| rs104893816 | pathogenic | 0 | 0 | 0 |
| rs121918714 | pathogenic | 0 | 1 | 0 |
| rs104893810 | pathogenic | 1 | 1 | 0 |
| rs104893815 | pathogenic | 1 | 1 | 0 |
| rs4303 | benign | 0 | 1 | 0 |
| rs121912704 | pathogenic | 0 | 1 | 0 |
| rs56394458 | benign | 1 | 1 | 0 |
| rs397514688 | pathogenic | 0 | 0 | 0 |
| rs367797185 | pathogenic | 1 | 1 | 0 |
| rs12709426 | benign | 0 | 1 | 0 |
| rs2228547 | benign | 0 | 0 | 0 |
| rs111033546 | pathogenic | 0 | 0 | 0 |
| rs111033553 | pathogenic | 0 | 0 | 0 |
| rs111033554 | pathogenic | 1 | 1 | 1 |
| rs111033544 | pathogenic | 1 | 1 | 1 |
| rs111033555 | pathogenic | 1 | 1 | 1 |
| rs111033545 | pathogenic | 1 | 1 | 1 |
| rs111033549 | pathogenic | 0 | 1 | 0 |
| rs111033552 | pathogenic | 1 | 1 | 1 |
| rs752204035 | benign | 0 | 0 | 0 |
| rs11569534 | benign | 0 | 0 | 0 |
| rs28937571 | pathogenic | 1 | 1 | 1 |
| rs104894215 | pathogenic | 1 | 1 | 1 |
| rs3888798 | benign | 1 | 1 | 1 |
| rs764263034 | benign | 1 | 1 | 1 |
| rs1800973 | benign | 1 | 1 | 1 |
| rs367623154 | benign | 1 | 1 | 1 |
| rs201230446 | pathogenic | 1 | 1 | 1 |
| rs267606766 | pathogenic | 0 | 1 | 0 |
| rs267606769 | pathogenic | 0 | 1 | 0 |
| rs267606767 | pathogenic | 1 | 1 | 0 |
| rs267606768 | pathogenic | 0 | 1 | 0 |
| rs61733129 | benign | 0 | 1 | 0 |
| rs201947120 | pathogenic | 1 | 1 | 1 |
| rs283413 | benign | 0 | 1 | 0 |
| rs36119840 | benign | 0 | 0 | 0 |
| rs281865419 | pathogenic | 1 | 1 | 1 |
| rs137852937 | pathogenic | 0 | 1 | 0 |
| rs2230709 | benign | 0 | 1 | 0 |
| rs6695033 | benign | 0 | 1 | 0 |
| rs180177032 | pathogenic | 1 | 1 | 1 |
| rs180177033 | pathogenic | 1 | 1 | 1 |
| rs121913351 | pathogenic | 1 | 1 | 1 |
| rs397507473 | pathogenic | 1 | 1 | 1 |
| rs121913355 | pathogenic | 1 | 1 | 1 |
| rs121913357 | pathogenic | 1 | 1 | 1 |
| rs180177036 | pathogenic | 1 | 1 | 1 |
| rs180177039 | pathogenic | 1 | 1 | 1 |
| rs180177038 | pathogenic | 1 | 1 | 1 |
| rs180177040 | pathogenic | 1 | 1 | 1 |

| | | | | |
|---|---|---|---|---|
| rs121913338 | pathogenic | 1 | 1 | 1 |
| rs121913366 | pathogenic | 1 | 1 | 1 |
| rs121913369 | pathogenic | 1 | 1 | 1 |
| rs113488022 | pathogenic | 1 | 1 | 1 |
| rs121913364 | pathogenic | 1 | 1 | 1 |
| rs180177042 | pathogenic | 1 | 1 | 1 |
| rs200461617 | benign | 0 | 0 | 0 |
| rs141609745 | benign | 0 | 0 | 0 |
| rs104894383 | pathogenic | 1 | 1 | 1 |
| rs104894384 | pathogenic | 1 | 1 | 1 |
| rs104894381 | pathogenic | 1 | 1 | 0 |
| rs147710408 | benign | 1 | 1 | 1 |
| rs77357563 | benign | 1 | 1 | 1 |
| rs104894378 | pathogenic | 0 | 0 | 0 |
| rs104894382 | pathogenic | 0 | 0 | 0 |
| rs137853292 | pathogenic | 1 | 1 | 1 |
| rs587778834 | pathogenic | 0 | 0 | 0 |
| rs137853294 | pathogenic | 1 | 1 | 1 |
| rs551747882 | benign | 0 | 0 | 0 |
| rs3092903 | benign | 1 | 1 | 1 |
| rs137853296 | pathogenic | 0 | 0 | 0 |
| rs398122917 | pathogenic | 0 | 1 | 0 |
| rs41304992 | benign | 0 | 0 | 0 |
| rs121912708 | pathogenic | 1 | 1 | 1 |
| rs140102105 | pathogenic | 1 | 1 | 1 |
| rs376917645 | pathogenic | 1 | 1 | 0 |
| rs372660425 | pathogenic | 1 | 1 | 0 |
| rs555896752 | pathogenic | 1 | 1 | 1 |
| rs777829351 | pathogenic | 1 | 1 | 0 |
| rs121912711 | pathogenic | 1 | 1 | 0 |
| rs754449549 | pathogenic | 0 | 1 | 0 |
| rs141701364 | benign | 0 | 1 | 0 |
| rs761011399 | benign | 1 | 1 | 1 |
| rs121912710 | benign | 1 | 1 | 0 |
| rs121912707 | pathogenic | 1 | 1 | 0 |
| rs747597620 | pathogenic | 0 | 1 | 0 |
| rs186558364 | pathogenic | 1 | 1 | 1 |
| rs556400964 | pathogenic | 1 | 1 | 1 |
| rs61757684 | benign | 1 | 1 | 1 |
| rs121917766 | pathogenic | 1 | 1 | 1 |
| rs121918248 | pathogenic | 0 | 0 | 0 |
| rs398123278 | pathogenic | 0 | 0 | 0 |
| rs121918251 | pathogenic | 1 | 1 | 1 |
| rs727504022 | pathogenic | 0 | 0 | 0 |
| rs190834116 | pathogenic | 1 | 1 | 1 |
| rs121918249 | pathogenic | 1 | 1 | 0 |
| rs121918257 | pathogenic | 1 | 1 | 0 |
| rs142068898 | benign | 0 | 0 | 0 |
| rs757000253 | pathogenic | 0 | 0 | 0 |
| rs200908035 | pathogenic | 1 | 1 | 0 |
| rs760782399 | pathogenic | 0 | 0 | 0 |
| rs778702777 | pathogenic | 0 | 0 | 0 |
| rs121918258 | pathogenic | 1 | 1 | 0 |
| rs121918256 | pathogenic | 0 | 0 | 0 |

| | | | | |
|---|---|---|---|---|
| rs200596762 | pathogenic | 1 | 1 | 0 |
| rs547709692 | pathogenic | 0 | 0 | 0 |
| rs758577372 | pathogenic | 0 | 1 | 0 |
| rs564069299 | pathogenic | 1 | 1 | 1 |
| rs772552898 | pathogenic | 1 | 1 | 1 |
| rs368790885 | pathogenic | 1 | 1 | 1 |
| rs121918250 | pathogenic | 0 | 0 | 0 |
| rs533755473 | pathogenic | 1 | 1 | 1 |
| rs753288303 | pathogenic | 1 | 1 | 1 |
| rs2229385 | benign | 0 | 0 | 0 |
| rs1141321 | benign | 0 | 0 | 0 |
| rs753564352 | pathogenic | 0 | 0 | 0 |
| rs775593146 | pathogenic | 0 | 0 | 0 |
| rs121918254 | pathogenic | 0 | 0 | 0 |
| rs8589 | benign | 1 | 1 | 0 |
| rs777758903 | pathogenic | 0 | 0 | 0 |
| rs140600746 | pathogenic | 1 | 1 | 0 |
| rs121918255 | pathogenic | 1 | 1 | 0 |
| rs121918252 | pathogenic | 0 | 0 | 0 |
| rs779990936 | pathogenic | 1 | 1 | 0 |
| rs121912455 | pathogenic | 1 | 1 | 1 |
| rs121912459 | pathogenic | 1 | 1 | 1 |
| rs74315452 | pathogenic | 1 | 1 | 1 |
| rs121912451 | pathogenic | 1 | 1 | 1 |
| rs79006549 | pathogenic | 0 | 0 | 0 |
| rs28934889 | benign | 0 | 1 | 0 |
| rs111033557 | benign | 1 | 1 | 1 |
| rs28934597 | pathogenic | 1 | 1 | 1 |
| rs28934596 | pathogenic | 1 | 1 | 1 |
| rs28934595 | pathogenic | 1 | 1 | 1 |
| rs146519482 | pathogenic | 1 | 1 | 1 |
| rs111033563 | pathogenic | 1 | 1 | 1 |
| rs75385605 | benign | 1 | 1 | 1 |
| rs121908569 | pathogenic | 1 | 1 | 1 |
| rs35719940 | benign | 0 | 0 | 0 |
| rs121918664 | pathogenic | 0 | 0 | 0 |
| rs104893665 | pathogenic | 1 | 1 | 1 |
| rs104893666 | pathogenic | 1 | 1 | 1 |
| rs200845457 | benign | 1 | 1 | 1 |
| rs779204655 | pathogenic | 1 | 1 | 1 |
| rs121917747 | pathogenic | 1 | 1 | 1 |
| rs6042 | benign | 1 | 1 | 1 |
| rs104894380 | pathogenic | 0 | 0 | 0 |
| rs369083541 | pathogenic | 1 | 1 | 1 |
| rs145243532 | benign | 1 | 1 | 1 |
| rs121909500 | pathogenic | 1 | 1 | 1 |
| rs140993310 | benign | 1 | 1 | 1 |
| rs200837270 | pathogenic | 1 | 1 | 1 |
| rs121918484 | pathogenic | 1 | 1 | 1 |
| rs121918306 | pathogenic | 0 | 0 | 0 |
| rs118204089 | pathogenic | 1 | 1 | 0 |
| rs118204083 | pathogenic | 1 | 1 | 0 |
| rs201953584 | pathogenic | 1 | 1 | 1 |
| rs74315362 | pathogenic | 1 | 1 | 1 |

| | | | | |
|---|---|---|---|---|
| rs118204084 | pathogenic | 1 | 1 | 0 |
| rs118204085 | pathogenic | 1 | 1 | 1 |
| rs113403872 | pathogenic | 1 | 1 | 0 |
| rs1052176 | benign | 1 | 1 | 1 |
| rs2031640 | benign | 0 | 0 | 0 |
| rs104894230 | pathogenic | 1 | 1 | 1 |
| rs104894229 | pathogenic | 1 | 1 | 1 |
| rs104894228 | pathogenic | 1 | 1 | 1 |
| rs104894226 | pathogenic | 1 | 1 | 1 |
| rs121917757 | pathogenic | 1 | 1 | 1 |
| rs121917758 | pathogenic | 1 | 1 | 1 |
| rs28933406 | pathogenic | 1 | 1 | 1 |
| rs121917756 | pathogenic | 1 | 1 | 1 |
| rs104894227 | pathogenic | 1 | 1 | 1 |
| rs369106578 | pathogenic | 1 | 1 | 1 |
| rs397517142 | benign | 1 | 1 | 1 |
| rs104894231 | pathogenic | 1 | 1 | 1 |
| rs121917759 | pathogenic | 1 | 1 | 1 |
| rs104893827 | pathogenic | 0 | 0 | 0 |
| rs104893826 | pathogenic | 1 | 1 | 1 |
| rs5030802 | pathogenic | 1 | 1 | 1 |
| rs5030827 | pathogenic | 1 | 1 | 0 |
| rs119103277 | pathogenic | 1 | 1 | 1 |
| rs5030808 | pathogenic | 0 | 1 | 0 |
| rs5030809 | pathogenic | 1 | 1 | 1 |
| rs765978945 | pathogenic | 1 | 1 | 1 |
| rs104893824 | pathogenic | 1 | 1 | 1 |
| rs104893831 | pathogenic | 0 | 1 | 0 |
| rs104893830 | pathogenic | 0 | 1 | 0 |
| rs119103278 | pathogenic | 0 | 1 | 0 |
| rs5030818 | pathogenic | 1 | 1 | 1 |
| rs397516444 | pathogenic | 1 | 1 | 1 |
| rs28940297 | pathogenic | 1 | 1 | 1 |
| rs267607170 | pathogenic | 1 | 1 | 1 |
| rs104893825 | pathogenic | 1 | 1 | 1 |
| rs5030820 | pathogenic | 0 | 1 | 0 |
| rs5030821 | pathogenic | 0 | 1 | 0 |
| rs5030823 | pathogenic | 1 | 1 | 1 |
| rs5030824 | pathogenic | 1 | 1 | 1 |
| rs28940301 | pathogenic | 0 | 1 | 0 |
| rs28940300 | pathogenic | 0 | 1 | 0 |
| rs28940298 | pathogenic | 0 | 1 | 0 |
| rs200019083 | benign | 0 | 0 | 0 |
| rs759748655 | pathogenic | 0 | 0 | 0 |
| rs121917739 | pathogenic | 1 | 1 | 1 |
| rs75353611 | pathogenic | 1 | 1 | 1 |
| rs79744198 | benign | 1 | 1 | 1 |
| rs79228041 | pathogenic | 1 | 1 | 1 |
| rs80106970 | benign | 1 | 1 | 1 |
| rs375353223 | benign | 0 | 0 | 0 |
| rs56311081 | benign | 0 | 0 | 0 |
| rs55985569 | benign | 1 | 1 | 1 |
| rs45483396 | benign | 0 | 0 | 0 |
| rs35284565 | benign | 1 | 1 | 1 |

| | | | | |
|---|---|---|---|---|
| rs34349517 | benign | 0 | 0 | 0 |
| rs35776110 | benign | 0 | 0 | 0 |
| rs33917957 | benign | 1 | 1 | 1 |
| rs587778445 | benign | 0 | 0 | 0 |
| rs34624984 | benign | 0 | 0 | 0 |
| rs701753 | benign | 0 | 0 | 0 |
| rs855314 | benign | 0 | 0 | 0 |
| rs121918371 | pathogenic | 1 | 1 | 0 |
| rs398122912 | pathogenic | 0 | 1 | 0 |
| rs1126728 | benign | 0 | 0 | 0 |
| rs587777404 | pathogenic | 1 | 1 | 0 |
| rs11208257 | benign | 0 | 0 | 0 |
| rs6676290 | benign | 0 | 0 | 0 |
| rs104894118 | pathogenic | 0 | 0 | 0 |
| rs121918655 | pathogenic | 1 | 1 | 1 |
| rs104894120 | pathogenic | 1 | 1 | 1 |
| rs137852238 | pathogenic | 0 | 0 | 0 |
| rs137852237 | pathogenic | 0 | 0 | 0 |
| rs137852499 | pathogenic | 1 | 1 | 1 |
| rs137852480 | pathogenic | 1 | 1 | 1 |
| rs137852494 | pathogenic | 1 | 1 | 1 |
| rs137852502 | pathogenic | 1 | 1 | 1 |
| rs137852500 | pathogenic | 1 | 1 | 1 |
| rs137852487 | pathogenic | 1 | 1 | 1 |
| rs137852481 | pathogenic | 1 | 1 | 1 |
| rs137852501 | pathogenic | 0 | 0 | 0 |
| rs137852478 | pathogenic | 1 | 1 | 1 |
| rs137852485 | pathogenic | 1 | 1 | 1 |
| rs137852482 | pathogenic | 1 | 1 | 0 |
| rs137852483 | pathogenic | 0 | 1 | 0 |
| rs137852477 | pathogenic | 1 | 1 | 0 |
| rs137852496 | pathogenic | 1 | 1 | 0 |
| rs137852484 | pathogenic | 0 | 1 | 0 |
| rs137852498 | pathogenic | 1 | 1 | 0 |
| rs137852497 | pathogenic | 1 | 1 | 0 |
| rs137852493 | pathogenic | 1 | 1 | 0 |
| rs137852504 | pathogenic | 1 | 1 | 1 |
| rs267606863 | pathogenic | 1 | 1 | 1 |
| rs137852486 | pathogenic | 1 | 1 | 1 |
| rs137852479 | pathogenic | 1 | 1 | 1 |
| rs112122950 | benign | 0 | 0 | 0 |
| rs72546648 | benign | 1 | 1 | 1 |
| rs72552291 | pathogenic | 0 | 0 | 0 |
| rs79184941 | pathogenic | 1 | 1 | 1 |
| rs77543610 | pathogenic | 1 | 1 | 1 |
| rs121918505 | pathogenic | 1 | 1 | 1 |
| rs776587763 | pathogenic | 1 | 1 | 1 |
| rs121918497 | pathogenic | 1 | 1 | 1 |
| rs121918499 | pathogenic | 1 | 1 | 1 |
| rs121918501 | pathogenic | 1 | 1 | 1 |
| rs121918500 | pathogenic | 0 | 0 | 0 |
| rs121918504 | pathogenic | 1 | 1 | 1 |
| rs121918510 | pathogenic | 1 | 1 | 1 |
| rs121918493 | pathogenic | 1 | 1 | 1 |

| | | | | |
|---|---|---|---|---|
| rs121918489 | pathogenic | 0 | 0 | 0 |
| rs121918495 | pathogenic | 1 | 1 | 1 |
| rs121918488 | pathogenic | 0 | 0 | 0 |
| rs121918496 | pathogenic | 0 | 0 | 0 |
| rs121918492 | pathogenic | 1 | 1 | 1 |
| rs121918494 | pathogenic | 1 | 1 | 1 |
| rs121918502 | pathogenic | 1 | 1 | 1 |
| rs121918490 | pathogenic | 1 | 1 | 1 |
| rs72466451 | pathogenic | 1 | 1 | 1 |
| rs66468541 | pathogenic | 0 | 0 | 0 |
| rs137852662 | pathogenic | 0 | 0 | 0 |
| rs137852663 | pathogenic | 0 | 0 | 0 |
| rs606231407 | pathogenic | 0 | 0 | 0 |
| rs137852233 | pathogenic | 1 | 1 | 1 |
| rs112306225 | benign | 0 | 0 | 0 |
| rs387906808 | pathogenic | 1 | 1 | 0 |
| rs730882094 | pathogenic | 1 | 1 | 1 |
| rs373869746 | pathogenic | 1 | 1 | 1 |
| rs777524402 | pathogenic | 1 | 1 | 1 |
| rs769737896 | pathogenic | 0 | 1 | 0 |
| rs13306515 | benign | 1 | 1 | 0 |
| rs746982741 | pathogenic | 0 | 0 | 0 |
| rs2230200 | benign | 1 | 1 | 1 |
| rs1047286 | benign | 0 | 0 | 0 |
| rs141737564 | benign | 0 | 0 | 0 |
| rs774826179 | benign | 0 | 0 | 0 |
| rs11569422 | benign | 1 | 1 | 1 |
| rs146613648 | benign | 0 | 0 | 0 |
| rs144432231 | benign | 0 | 0 | 0 |
| rs140655115 | benign | 0 | 0 | 0 |
| rs387906837 | pathogenic | 1 | 1 | 0 |
| rs121909076 | pathogenic | 1 | 1 | 0 |
| rs121909077 | pathogenic | 0 | 0 | 0 |
| rs121909074 | pathogenic | 0 | 0 | 0 |
| rs137853330 | pathogenic | 1 | 1 | 1 |
| rs137853331 | pathogenic | 1 | 1 | 1 |
| rs137853332 | pathogenic | 1 | 1 | 1 |
| rs28931586 | pathogenic | 1 | 1 | 0 |
| rs11569321 | benign | 1 | 1 | 1 |
| rs119486097 | benign | 0 | 0 | 0 |
| rs1799807 | pathogenic | 1 | 1 | 1 |
| rs201820739 | pathogenic | 1 | 1 | 0 |
| rs114706984 | pathogenic | 1 | 1 | 0 |
| rs140080572 | pathogenic | 1 | 1 | 0 |
| rs28933389 | pathogenic | 0 | 1 | 0 |
| rs16849700 | benign | 1 | 1 | 0 |
| rs115624085 | pathogenic | 1 | 1 | 1 |
| rs747196387 | pathogenic | 1 | 1 | 1 |
| rs104893684 | pathogenic | 1 | 1 | 0 |
| rs121918557 | pathogenic | 1 | 1 | 0 |
| rs121918556 | pathogenic | 1 | 1 | 1 |
| rs398124571 | pathogenic | 1 | 1 | 1 |
| rs776631396 | pathogenic | 0 | 1 | 0 |
| rs398124576 | pathogenic | 1 | 1 | 1 |

| | | | | |
|---|---|---|---|---|
| rs398124581 | pathogenic | 1 | 1 | 1 |
| rs398124582 | pathogenic | 1 | 1 | 1 |
| rs149766077 | pathogenic | 0 | 1 | 0 |
| rs79761867 | pathogenic | 0 | 1 | 0 |
| rs148905512 | pathogenic | 0 | 1 | 0 |
| rs121965004 | pathogenic | 1 | 1 | 1 |
| rs386834233 | pathogenic | 0 | 1 | 0 |
| rs398124598 | pathogenic | 0 | 1 | 0 |
| rs398124603 | pathogenic | 1 | 1 | 1 |
| rs546716073 | benign | 1 | 1 | 1 |
| rs200310662 | benign | 1 | 1 | 1 |
| rs115494727 | benign | 1 | 1 | 1 |
| rs781513008 | benign | 0 | 0 | 0 |
| rs587776398 | benign | 1 | 1 | 0 |
| rs35288908 | benign | 1 | 1 | 1 |
| rs150634562 | pathogenic | 1 | 1 | 0 |
| rs142948132 | pathogenic | 1 | 1 | 1 |
| rs132630321 | pathogenic | 1 | 1 | 1 |
| rs132630317 | pathogenic | 1 | 1 | 1 |
| rs132630320 | pathogenic | 0 | 0 | 0 |
| rs781394318 | pathogenic | 1 | 1 | 1 |
| rs111033624 | pathogenic | 1 | 1 | 1 |
| rs111033630 | pathogenic | 1 | 1 | 1 |
| rs111033627 | pathogenic | 1 | 1 | 1 |
| rs193922287 | pathogenic | 1 | 1 | 1 |
| rs759072800 | pathogenic | 0 | 1 | 0 |
| rs104894010 | pathogenic | 0 | 0 | 0 |
| rs193922296 | pathogenic | 0 | 0 | 0 |
| rs104894006 | pathogenic | 1 | 1 | 1 |
| rs193922311 | pathogenic | 1 | 1 | 0 |
| rs80356654 | pathogenic | 1 | 1 | 0 |
| rs144723656 | benign | 1 | 1 | 1 |
| rs148311934 | pathogenic | 1 | 1 | 0 |
| rs80356655 | pathogenic | 1 | 1 | 1 |
| rs769268803 | pathogenic | 1 | 1 | 0 |
| rs104894008 | pathogenic | 0 | 1 | 0 |
| rs104894005 | pathogenic | 1 | 1 | 1 |
| rs104894009 | pathogenic | 1 | 1 | 0 |
| rs193922340 | pathogenic | 0 | 0 | 0 |
| rs104894016 | pathogenic | 0 | 1 | 0 |
| rs193921338 | pathogenic | 0 | 0 | 0 |
| rs193922272 | pathogenic | 1 | 1 | 1 |
| rs104894012 | pathogenic | 1 | 1 | 0 |
| rs193922285 | pathogenic | 1 | 1 | 1 |
| rs104894336 | pathogenic | 1 | 1 | 1 |
| rs28931580 | pathogenic | 1 | 1 | 1 |
| rs104894326 | pathogenic | 1 | 1 | 1 |
| rs104894331 | pathogenic | 0 | 0 | 0 |
| rs104894338 | pathogenic | 0 | 0 | 0 |
| rs104894333 | pathogenic | 1 | 1 | 1 |
| rs104894330 | pathogenic | 1 | 1 | 1 |
| rs143886391 | benign | 1 | 1 | 1 |
| rs104894335 | pathogenic | 1 | 1 | 1 |
| rs104894337 | pathogenic | 0 | 0 | 0 |

| | | | | |
|---|---|---|---|---|
| rs104894328 | pathogenic | 0 | 0 | 0 |
| rs104894341 | pathogenic | 1 | 1 | 0 |
| rs104894329 | pathogenic | 0 | 0 | 0 |
| rs137852850 | pathogenic | 1 | 1 | 1 |
| rs137852854 | pathogenic | 0 | 1 | 0 |
| rs137852849 | pathogenic | 0 | 1 | 0 |
| rs137852848 | pathogenic | 1 | 1 | 1 |
| rs137852852 | pathogenic | 0 | 1 | 0 |
| rs137852853 | pathogenic | 0 | 1 | 0 |
| rs137852847 | pathogenic | 0 | 1 | 0 |
| rs137852846 | pathogenic | 0 | 1 | 0 |
| rs137852844 | pathogenic | 0 | 1 | 0 |
| rs144411158 | pathogenic | 0 | 0 | 0 |
| rs2302234 | benign | 0 | 0 | 0 |
| rs139620139 | pathogenic | 0 | 0 | 0 |
| rs146689929 | benign | 0 | 0 | 0 |
| rs373328681 | pathogenic | 1 | 1 | 0 |
| rs387906620 | pathogenic | 1 | 1 | 1 |
| rs1799983 | benign | 0 | 1 | 0 |
| rs28999111 | pathogenic | 1 | 1 | 1 |
| rs104894544 | pathogenic | 1 | 1 | 1 |
| rs104894546 | pathogenic | 1 | 1 | 1 |
| rs28999112 | pathogenic | 0 | 1 | 0 |
| rs104894545 | pathogenic | 1 | 1 | 1 |
| rs104894547 | pathogenic | 1 | 1 | 1 |
| rs74315310 | pathogenic | 0 | 0 | 0 |
| rs267606852 | pathogenic | 0 | 1 | 0 |
| rs137853586 | pathogenic | 0 | 1 | 0 |
| rs137853582 | pathogenic | 1 | 1 | 1 |
| rs61754634 | pathogenic | 1 | 1 | 1 |
| rs137853587 | pathogenic | 0 | 1 | 0 |
| rs267606851 | pathogenic | 1 | 1 | 1 |
| rs267606853 | pathogenic | 1 | 1 | 1 |
| rs137853584 | pathogenic | 1 | 1 | 1 |
| rs137853585 | pathogenic | 1 | 1 | 1 |
| rs460184 | pathogenic | 1 | 1 | 1 |
| rs104893751 | pathogenic | 0 | 1 | 0 |
| rs77358162 | benign | 0 | 0 | 0 |
| rs387906936 | pathogenic | 0 | 0 | 0 |
| rs769525399 | pathogenic | 0 | 0 | 0 |
| rs371271054 | pathogenic | 1 | 1 | 0 |
| rs118204000 | pathogenic | 0 | 0 | 0 |
| rs138568622 | pathogenic | 1 | 1 | 1 |
| rs34509359 | benign | 0 | 0 | 0 |
| rs104894130 | pathogenic | 1 | 1 | 0 |
| rs281865424 | pathogenic | 0 | 0 | 0 |
| rs121912778 | pathogenic | 1 | 1 | 0 |
| rs771055145 | pathogenic | 1 | 1 | 0 |
| rs776174514 | pathogenic | 1 | 1 | 0 |
| rs545986367 | pathogenic | 1 | 1 | 1 |
| rs570768038 | benign | 1 | 1 | 1 |
| rs74315442 | pathogenic | 1 | 1 | 1 |
| rs79716074 | benign | 1 | 1 | 1 |
| rs748486078 | pathogenic | 1 | 1 | 1 |

| | | | | |
|---|---|---|---|---|
| rs2397084 | benign | 1 | 1 | 1 |
| rs11465553 | benign | 1 | 1 | 1 |
| rs144576902 | benign | 1 | 1 | 1 |
| rs763780 | benign | 1 | 1 | 1 |
| rs267606870 | pathogenic | 1 | 1 | 0 |
| rs121913502 | pathogenic | 1 | 1 | 0 |
| rs118101777 | benign | 1 | 1 | 1 |
| rs118053940 | benign | 0 | 0 | 0 |
| rs368771578 | benign | 1 | 1 | 1 |
| rs11540654 | pathogenic | 1 | 1 | 1 |
| rs121912658 | pathogenic | 1 | 1 | 1 |
| rs28934873 | pathogenic | 0 | 1 | 0 |
| rs780442292 | pathogenic | 1 | 1 | 1 |
| rs758781593 | benign | 1 | 1 | 1 |
| rs28934875 | pathogenic | 1 | 1 | 1 |
| rs28934874 | pathogenic | 1 | 1 | 0 |
| rs587782705 | pathogenic | 1 | 1 | 1 |
| rs587780068 | pathogenic | 1 | 1 | 1 |
| rs587782144 | pathogenic | 1 | 1 | 1 |
| rs138729528 | pathogenic | 1 | 1 | 1 |
| rs28934578 | pathogenic | 1 | 1 | 1 |
| rs397514495 | pathogenic | 1 | 1 | 1 |
| rs121912665 | pathogenic | 1 | 1 | 1 |
| rs397516435 | pathogenic | 1 | 1 | 1 |
| rs587778720 | pathogenic | 1 | 1 | 1 |
| rs397516436 | pathogenic | 1 | 1 | 1 |
| rs587781386 | benign | 1 | 1 | 1 |
| rs121912666 | pathogenic | 1 | 1 | 0 |
| rs587780073 | pathogenic | 1 | 1 | 1 |
| rs587782664 | pathogenic | 1 | 1 | 1 |
| rs730882005 | pathogenic | 1 | 1 | 1 |
| rs28934573 | pathogenic | 1 | 1 | 1 |
| rs121912655 | pathogenic | 1 | 1 | 1 |
| rs28934575 | pathogenic | 1 | 1 | 1 |
| rs121912656 | pathogenic | 1 | 1 | 1 |
| rs11540652 | pathogenic | 1 | 1 | 1 |
| rs121912651 | pathogenic | 1 | 1 | 1 |
| rs730882027 | pathogenic | 1 | 1 | 1 |
| rs121912653 | pathogenic | 1 | 1 | 1 |
| rs121912652 | pathogenic | 1 | 1 | 1 |
| rs121912657 | pathogenic | 0 | 1 | 0 |
| rs121913343 | pathogenic | 1 | 1 | 1 |
| rs28934576 | pathogenic | 1 | 1 | 1 |
| rs763098116 | pathogenic | 1 | 1 | 1 |
| rs121912660 | pathogenic | 1 | 1 | 1 |
| rs28934574 | pathogenic | 1 | 1 | 1 |
| rs371409680 | pathogenic | 1 | 1 | 1 |
| rs121912667 | pathogenic | 1 | 1 | 1 |
| rs121912663 | pathogenic | 1 | 1 | 1 |
| rs201744589 | pathogenic | 0 | 0 | 0 |
| rs33926449 | benign | 0 | 0 | 0 |
| rs74315302 | pathogenic | 0 | 1 | 0 |
| rs202040269 | pathogenic | 1 | 1 | 1 |
| rs375958814 | pathogenic | 1 | 1 | 1 |

| | | | | |
|---|---|---|---|---|
| rs121909624 | pathogenic | 1 | 1 | 1 |
| rs121909621 | pathogenic | 1 | 1 | 1 |
| rs121909622 | pathogenic | 1 | 1 | 0 |
| rs4220 | benign | 1 | 1 | 1 |
| rs147559466 | pathogenic | 1 | 1 | 1 |
| rs757434857 | pathogenic | 1 | 1 | 1 |
| rs398123072 | pathogenic | 1 | 1 | 1 |
| rs121434280 | pathogenic | 0 | 1 | 0 |
| rs398123074 | pathogenic | 0 | 1 | 0 |
| rs762114560 | pathogenic | 1 | 1 | 1 |
| rs121434283 | pathogenic | 1 | 1 | 0 |
| rs121434277 | pathogenic | 0 | 1 | 0 |
| rs745844469 | pathogenic | 1 | 1 | 1 |
| rs121434279 | pathogenic | 1 | 1 | 0 |
| rs121434278 | pathogenic | 1 | 1 | 1 |
| rs373715782 | pathogenic | 1 | 1 | 1 |
| rs200724875 | pathogenic | 1 | 1 | 1 |
| rs149678400 | pathogenic | 0 | 1 | 0 |
| rs770273135 | pathogenic | 1 | 1 | 1 |
| rs121434276 | pathogenic | 0 | 1 | 0 |
| rs121434281 | pathogenic | 1 | 1 | 0 |
| rs768884003 | pathogenic | 1 | 1 | 1 |
| rs201375579 | pathogenic | 0 | 0 | 0 |
| rs121434274 | pathogenic | 1 | 1 | 0 |
| rs121434282 | pathogenic | 1 | 1 | 0 |
| rs779759347 | pathogenic | 0 | 1 | 0 |
| rs77931234 | pathogenic | 1 | 1 | 1 |
| rs148207467 | pathogenic | 1 | 1 | 1 |
| rs766140986 | pathogenic | 1 | 1 | 1 |
| rs121434275 | pathogenic | 1 | 1 | 1 |
| rs748110745 | pathogenic | 1 | 1 | 1 |
| rs104894274 | pathogenic | 1 | 1 | 1 |
| rs104894273 | pathogenic | 0 | 1 | 0 |
| rs104894278 | pathogenic | 1 | 1 | 1 |
| rs104894275 | pathogenic | 1 | 1 | 1 |
| rs104894277 | pathogenic | 1 | 1 | 1 |
| rs104894276 | pathogenic | 0 | 0 | 0 |
| rs104894280 | pathogenic | 1 | 1 | 1 |
| rs145882709 | pathogenic | 1 | 1 | 1 |
| rs104894279 | pathogenic | 1 | 1 | 1 |
| rs80356670 | pathogenic | 1 | 1 | 1 |
| rs121918102 | pathogenic | 1 | 1 | 1 |
| rs80356671 | pathogenic | 1 | 1 | 1 |
| rs80356672 | pathogenic | 1 | 1 | 1 |
| rs758609113 | pathogenic | 1 | 1 | 0 |
| rs121918188 | pathogenic | 0 | 1 | 0 |
| rs121918189 | pathogenic | 1 | 1 | 0 |
| rs121918191 | pathogenic | 1 | 1 | 1 |
| rs1769259 | benign | 1 | 1 | 1 |
| rs140222720 | benign | 1 | 1 | 1 |
| rs2297085 | benign | 1 | 1 | 1 |
| rs200679026 | pathogenic | 1 | 1 | 1 |
| rs121918192 | pathogenic | 0 | 1 | 0 |
| rs145771944 | benign | 0 | 1 | 0 |

| | | | | |
|---|---|---|---|---|
| rs201508880 | pathogenic | 0 | 1 | 0 |
| rs17221854 | pathogenic | 0 | 0 | 0 |
| rs120074189 | pathogenic | 1 | 1 | 1 |
| rs120074190 | pathogenic | 1 | 1 | 1 |
| rs758600283 | benign | 1 | 1 | 1 |
| rs200305064 | pathogenic | 1 | 1 | 1 |
| rs121917891 | pathogenic | 1 | 1 | 1 |
| rs3772809 | benign | 0 | 1 | 0 |
| rs387906619 | pathogenic | 1 | 1 | 0 |
| rs2304681 | benign | 0 | 1 | 0 |
| rs121917849 | pathogenic | 0 | 0 | 0 |
| rs143010236 | pathogenic | 0 | 0 | 0 |
| rs121917850 | pathogenic | 1 | 1 | 0 |
| rs35916840 | pathogenic | 1 | 1 | 0 |
| rs28939688 | pathogenic | 1 | 1 | 1 |
| rs11258194 | benign | 1 | 1 | 1 |
| rs199840952 | pathogenic | 1 | 1 | 1 |
| rs121918159 | pathogenic | 1 | 1 | 1 |
| rs200396166 | pathogenic | 0 | 0 | 0 |
| rs757966746 | pathogenic | 0 | 1 | 0 |
| rs121918608 | pathogenic | 0 | 1 | 0 |
| rs74315401 | pathogenic | 1 | 1 | 1 |
| rs11538758 | pathogenic | 0 | 0 | 0 |
| rs74315402 | pathogenic | 1 | 1 | 1 |
| rs1799990 | pathogenic | 1 | 1 | 1 |
| rs74315410 | pathogenic | 1 | 1 | 1 |
| rs150351644 | benign | 1 | 1 | 1 |
| rs16990018 | benign | 1 | 1 | 1 |
| rs74315403 | pathogenic | 1 | 1 | 1 |
| rs74315408 | pathogenic | 1 | 1 | 1 |
| rs74315411 | pathogenic | 1 | 1 | 1 |
| rs74315413 | pathogenic | 1 | 1 | 1 |
| rs74315405 | pathogenic | 1 | 1 | 1 |
| rs28933385 | pathogenic | 1 | 1 | 1 |
| rs74315412 | pathogenic | 1 | 1 | 1 |
| rs74315407 | pathogenic | 1 | 1 | 1 |
| rs74315406 | pathogenic | 1 | 1 | 1 |
| rs11571117 | benign | 1 | 1 | 1 |
| rs121917742 | pathogenic | 1 | 1 | 1 |
| rs149782619 | pathogenic | 1 | 1 | 1 |
| rs397514492 | pathogenic | 1 | 1 | 1 |
| rs397514489 | pathogenic | 0 | 1 | 0 |
| rs397514493 | pathogenic | 1 | 1 | 1 |
| rs373849532 | pathogenic | 1 | 1 | 1 |
| rs121965048 | pathogenic | 1 | 1 | 1 |
| rs121965037 | pathogenic | 1 | 1 | 1 |
| rs121965060 | pathogenic | 0 | 1 | 0 |
| rs386833603 | pathogenic | 1 | 1 | 1 |
| rs121965038 | pathogenic | 1 | 1 | 1 |
| rs121965039 | pathogenic | 1 | 1 | 1 |
| rs121965040 | pathogenic | 1 | 1 | 1 |
| rs267606925 | pathogenic | 1 | 1 | 1 |
| rs121965056 | pathogenic | 0 | 1 | 0 |
| rs121965059 | pathogenic | 0 | 0 | 0 |

| | | | | |
|---|---|---|---|---|
| rs386833615 | pathogenic | 1 | 1 | 0 |
| rs121965051 | pathogenic | 1 | 1 | 0 |
| rs121965046 | pathogenic | 0 | 1 | 0 |
| rs121965052 | pathogenic | 0 | 1 | 0 |
| rs386833617 | pathogenic | 0 | 1 | 0 |
| rs386833618 | pathogenic | 1 | 1 | 0 |
| rs121965041 | pathogenic | 0 | 1 | 0 |
| rs121965042 | pathogenic | 0 | 1 | 0 |
| rs121965057 | pathogenic | 1 | 1 | 1 |
| rs386833621 | pathogenic | 1 | 1 | 1 |
| rs121965049 | pathogenic | 1 | 1 | 1 |
| rs386833623 | pathogenic | 1 | 1 | 1 |
| rs121965047 | pathogenic | 1 | 1 | 1 |
| rs121965053 | pathogenic | 0 | 1 | 0 |
| rs121965045 | pathogenic | 0 | 1 | 0 |
| rs121965054 | pathogenic | 0 | 1 | 0 |
| rs121965043 | pathogenic | 0 | 1 | 0 |
| rs386833598 | pathogenic | 0 | 0 | 0 |
| rs137854545 | pathogenic | 1 | 1 | 1 |
| rs137854544 | pathogenic | 1 | 1 | 1 |
| rs104894455 | pathogenic | 0 | 1 | 0 |
| rs1049564 | benign | 0 | 1 | 0 |
| rs104894460 | pathogenic | 1 | 1 | 1 |
| rs104894453 | pathogenic | 1 | 1 | 1 |
| rs104894450 | pathogenic | 0 | 1 | 0 |
| rs104894454 | pathogenic | 0 | 1 | 0 |
| rs104894452 | pathogenic | 1 | 1 | 1 |
| rs104894451 | pathogenic | 0 | 1 | 0 |
| rs104894368 | pathogenic | 1 | 1 | 1 |
| rs730880950 | pathogenic | 1 | 1 | 1 |
| rs121909535 | pathogenic | 1 | 1 | 1 |
| rs121909537 | pathogenic | 1 | 1 | 1 |
| rs121909536 | pathogenic | 1 | 1 | 1 |
| rs121909542 | pathogenic | 1 | 1 | 1 |
| rs121909538 | pathogenic | 1 | 1 | 1 |
| rs121909539 | pathogenic | 1 | 1 | 1 |
| rs121909540 | pathogenic | 1 | 1 | 1 |
| rs17560 | benign | 1 | 1 | 1 |
| rs121909543 | pathogenic | 0 | 1 | 0 |
| rs121909544 | pathogenic | 1 | 1 | 1 |
| rs2307456 | benign | 0 | 1 | 0 |
| rs745778317 | pathogenic | 0 | 1 | 0 |
| rs759607901 | pathogenic | 0 | 1 | 0 |
| rs201188114 | benign | 0 | 0 | 0 |
| rs1800472 | benign | 0 | 0 | 0 |
| rs587780728 | benign | 1 | 1 | 1 |
| rs144386518 | benign | 1 | 1 | 1 |
| rs281865439 | pathogenic | 1 | 1 | 1 |
| rs140175796 | pathogenic | 1 | 1 | 1 |
| rs5030843 | pathogenic | 1 | 1 | 0 |
| rs75166491 | pathogenic | 1 | 1 | 0 |
| rs199475679 | pathogenic | 1 | 1 | 1 |
| rs281865440 | pathogenic | 1 | 1 | 1 |
| rs74486803 | pathogenic | 0 | 1 | 0 |

| | | | | |
|---|---|---|---|---|
| rs199475602 | pathogenic | 0 | 1 | 0 |
| rs5030844 | pathogenic | 1 | 1 | 0 |
| rs62514927 | pathogenic | 1 | 1 | 1 |
| rs62514928 | pathogenic | 1 | 1 | 1 |
| rs62516152 | pathogenic | 0 | 1 | 0 |
| rs62507348 | pathogenic | 1 | 1 | 1 |
| rs76687508 | pathogenic | 1 | 1 | 1 |
| rs62508730 | pathogenic | 1 | 1 | 1 |
| rs62508588 | pathogenic | 0 | 1 | 0 |
| rs5030846 | pathogenic | 0 | 1 | 0 |
| rs74503222 | pathogenic | 1 | 1 | 0 |
| rs5030847 | pathogenic | 1 | 1 | 0 |
| rs5030850 | pathogenic | 1 | 1 | 1 |
| rs778154939 | pathogenic | 0 | 1 | 0 |
| rs62514952 | pathogenic | 1 | 1 | 1 |
| rs78655458 | pathogenic | 1 | 1 | 1 |
| rs62508698 | pathogenic | 1 | 1 | 0 |
| rs5030851 | pathogenic | 1 | 1 | 0 |
| rs62642933 | pathogenic | 1 | 1 | 1 |
| rs5030853 | pathogenic | 1 | 1 | 1 |
| rs62642935 | pathogenic | 1 | 1 | 1 |
| rs199475615 | pathogenic | 0 | 0 | 0 |
| rs62516062 | pathogenic | 1 | 1 | 0 |
| rs62516092 | pathogenic | 0 | 0 | 0 |
| rs62516095 | pathogenic | 1 | 1 | 1 |
| rs62642937 | pathogenic | 1 | 1 | 1 |
| rs62516141 | pathogenic | 0 | 1 | 0 |
| rs5030856 | pathogenic | 1 | 1 | 1 |
| rs5030857 | pathogenic | 1 | 1 | 1 |
| rs5030858 | pathogenic | 1 | 1 | 1 |
| rs79931499 | pathogenic | 1 | 1 | 1 |
| rs5030860 | pathogenic | 1 | 1 | 1 |
| rs281865437 | pathogenic | 1 | 1 | 1 |
| rs62644499 | pathogenic | 1 | 1 | 1 |
| rs147689487 | benign | 0 | 0 | 0 |
| rs397515417 | pathogenic | 1 | 1 | 1 |
| rs121909332 | pathogenic | 1 | 1 | 1 |
| rs121909330 | pathogenic | 1 | 1 | 0 |
| rs121909329 | pathogenic | 1 | 1 | 0 |
| rs121909335 | pathogenic | 1 | 1 | 1 |
| rs387906789 | pathogenic | 1 | 1 | 1 |
| rs121909334 | pathogenic | 1 | 1 | 1 |
| rs121909331 | pathogenic | 1 | 1 | 1 |
| rs372487178 | pathogenic | 1 | 1 | 1 |
| rs387906937 | pathogenic | 1 | 1 | 0 |
| rs16845 | benign | 1 | 1 | 1 |
| rs672601313 | pathogenic | 1 | 1 | 1 |
| rs117805308 | benign | 0 | 0 | 0 |
| rs121434261 | pathogenic | 1 | 1 | 1 |
| rs141629512 | benign | 0 | 0 | 0 |
| rs199957176 | benign | 1 | 1 | 1 |
| rs2230461 | benign | 1 | 1 | 1 |
| rs764618040 | pathogenic | 1 | 1 | 1 |
| rs267606949 | pathogenic | 1 | 1 | 1 |

| | | | | |
|---|---|---|---|---|
| rs587777483 | pathogenic | 1 | 1 | 1 |
| rs267606947 | pathogenic | 1 | 1 | 1 |
| rs6941583 | benign | 0 | 0 | 0 |
| rs749597959 | benign | 1 | 1 | 1 |
| rs182340753 | benign | 1 | 1 | 1 |
| rs146678221 | benign | 0 | 0 | 0 |
| rs11539202 | benign | 0 | 0 | 0 |
| rs148723565 | benign | 0 | 0 | 0 |
| rs61752925 | benign | 1 | 1 | 1 |
| rs113309941 | pathogenic | 0 | 0 | 0 |
| rs121434520 | pathogenic | 1 | 1 | 1 |
| rs121434517 | pathogenic | 1 | 1 | 1 |
| rs267607027 | pathogenic | 1 | 1 | 1 |
| rs138603088 | pathogenic | 0 | 0 | 0 |
| rs121434516 | pathogenic | 1 | 1 | 0 |
| rs121434521 | pathogenic | 0 | 0 | 0 |
| rs145735112 | benign | 1 | 1 | 1 |
| rs121434519 | pathogenic | 1 | 1 | 1 |
| rs75326924 | pathogenic | 1 | 1 | 0 |
| rs142186404 | pathogenic | 0 | 0 | 0 |
| rs121918035 | pathogenic | 0 | 0 | 0 |
| rs374385878 | pathogenic | 1 | 1 | 0 |
| rs11557652 | benign | 0 | 0 | 0 |
| rs118203986 | pathogenic | 1 | 1 | 1 |
| rs141675330 | benign | 1 | 1 | 1 |
| rs104893927 | pathogenic | 0 | 0 | 0 |
| rs104893935 | pathogenic | 0 | 0 | 0 |
| rs104893933 | pathogenic | 1 | 1 | 0 |
| rs104893934 | pathogenic | 0 | 0 | 0 |
| rs145183203 | benign | 1 | 1 | 0 |
| rs2230806 | benign | 0 | 0 | 0 |
| rs9282541 | benign | 0 | 0 | 0 |
| rs559110055 | benign | 0 | 0 | 0 |
| rs200030513 | benign | 0 | 0 | 0 |
| rs9282543 | benign | 0 | 0 | 0 |
| rs147675550 | benign | 0 | 0 | 0 |
| rs137854496 | pathogenic | 0 | 0 | 0 |
| rs374190304 | benign | 0 | 0 | 0 |
| rs114620717 | benign | 0 | 0 | 0 |
| rs2066718 | benign | 0 | 0 | 0 |
| rs35819696 | benign | 0 | 0 | 0 |
| rs138880920 | benign | 0 | 0 | 0 |
| rs2066715 | benign | 0 | 0 | 0 |
| rs35207495 | benign | 0 | 0 | 0 |
| rs2066714 | benign | 0 | 0 | 0 |
| rs28937313 | pathogenic | 0 | 0 | 0 |
| rs28937314 | pathogenic | 0 | 0 | 0 |
| rs137854495 | pathogenic | 0 | 0 | 0 |
| rs140365800 | benign | 0 | 0 | 0 |
| rs13306073 | benign | 0 | 0 | 0 |
| rs28933692 | pathogenic | 0 | 0 | 0 |
| rs33918808 | benign | 0 | 0 | 0 |
| rs76881554 | benign | 0 | 0 | 0 |
| rs143180998 | benign | 0 | 0 | 0 |

| | | | | |
|---|---|---|---|---|
| rs137854500 | pathogenic | 0 | 0 | 0 |
| rs538614702 | benign | 0 | 0 | 0 |
| rs201879964 | benign | 0 | 0 | 0 |
| rs2230808 | benign | 0 | 0 | 0 |
| rs138422574 | benign | 0 | 0 | 0 |
| rs137854498 | pathogenic | 0 | 0 | 0 |
| rs150125857 | benign | 0 | 0 | 0 |
| rs146292819 | pathogenic | 0 | 0 | 0 |
| rs142688906 | benign | 0 | 0 | 0 |
| rs137854499 | pathogenic | 0 | 0 | 0 |
| rs34879708 | benign | 0 | 0 | 0 |
| rs121913028 | pathogenic | 1 | 1 | 1 |
| rs1041951 | benign | 1 | 1 | 0 |
| rs138714094 | benign | 1 | 1 | 0 |
| rs200650502 | benign | 0 | 1 | 0 |
| rs370708663 | pathogenic | 1 | 1 | 1 |
| rs118204040 | pathogenic | 0 | 1 | 0 |
| rs146899669 | pathogenic | 1 | 1 | 0 |
| rs118204039 | pathogenic | 0 | 1 | 0 |
| rs121909233 | pathogenic | 0 | 0 | 0 |
| rs121909225 | pathogenic | 1 | 1 | 0 |
| rs121909236 | pathogenic | 0 | 0 | 0 |
| rs398123317 | pathogenic | 0 | 0 | 0 |
| rs121909226 | pathogenic | 1 | 1 | 0 |
| rs121909238 | pathogenic | 1 | 1 | 0 |
| rs121909230 | pathogenic | 0 | 0 | 0 |
| rs121909237 | pathogenic | 0 | 0 | 0 |
| rs121909222 | pathogenic | 1 | 1 | 0 |
| rs121909223 | pathogenic | 1 | 1 | 0 |
| rs121909218 | pathogenic | 1 | 1 | 0 |
| rs121909229 | pathogenic | 1 | 1 | 0 |
| rs121909224 | pathogenic | 1 | 1 | 0 |
| rs121909241 | pathogenic | 0 | 0 | 0 |
| rs121909221 | pathogenic | 0 | 0 | 0 |
| rs121913294 | pathogenic | 0 | 0 | 0 |
| rs121909234 | pathogenic | 0 | 0 | 0 |
| rs121909240 | pathogenic | 0 | 0 | 0 |
| rs767623493 | benign | 0 | 0 | 0 |
| rs121909239 | pathogenic | 0 | 0 | 0 |
| rs120074145 | pathogenic | 1 | 1 | 1 |
| rs120074148 | pathogenic | 1 | 1 | 1 |
| rs762991875 | pathogenic | 0 | 0 | 0 |
| rs148639841 | pathogenic | 1 | 1 | 1 |
| rs199524907 | pathogenic | 1 | 1 | 1 |
| rs120074141 | pathogenic | 1 | 1 | 1 |
| rs120074144 | pathogenic | 0 | 1 | 0 |
| rs120074146 | pathogenic | 1 | 1 | 1 |
| rs120074147 | pathogenic | 0 | 1 | 0 |
| rs120074143 | pathogenic | 0 | 1 | 0 |
| rs120074140 | pathogenic | 1 | 1 | 0 |
| rs28937315 | pathogenic | 1 | 1 | 1 |
| rs76066357 | benign | 1 | 1 | 1 |
| rs137852911 | pathogenic | 0 | 1 | 0 |
| rs137852907 | pathogenic | 1 | 1 | 0 |

| | | | | |
|---|---|---|---|---|
| rs137852910 | pathogenic | 1 | 1 | 1 |
| rs537585211 | benign | 0 | 0 | 0 |
| rs202134424 | pathogenic | 0 | 0 | 0 |
| rs200930463 | pathogenic | 0 | 0 | 0 |
| rs587777241 | pathogenic | 1 | 1 | 0 |
| rs77563738 | pathogenic | 0 | 0 | 0 |
| rs750868279 | pathogenic | 1 | 1 | 0 |
| rs148936893 | pathogenic | 0 | 0 | 0 |
| rs146597836 | pathogenic | 0 | 0 | 0 |
| rs775440641 | pathogenic | 1 | 1 | 1 |
| rs770689762 | pathogenic | 0 | 1 | 0 |
| rs376785840 | pathogenic | 0 | 0 | 0 |
| rs137852531 | pathogenic | 0 | 0 | 0 |
| rs137852538 | pathogenic | 1 | 1 | 0 |
| rs137852529 | pathogenic | 0 | 1 | 0 |
| rs137852528 | pathogenic | 0 | 1 | 0 |
| rs137852535 | pathogenic | 1 | 1 | 0 |
| rs137853045 | pathogenic | 0 | 1 | 0 |
| rs137853046 | pathogenic | 1 | 1 | 1 |
| rs137853043 | pathogenic | 1 | 1 | 1 |
| rs137853044 | pathogenic | 1 | 1 | 1 |
| rs137853047 | pathogenic | 1 | 1 | 1 |
| rs267606990 | pathogenic | 1 | 1 | 1 |
| rs751437780 | pathogenic | 1 | 1 | 0 |
| rs121918461 | pathogenic | 1 | 1 | 1 |
| rs121918460 | pathogenic | 1 | 1 | 1 |
| rs121918459 | pathogenic | 1 | 1 | 0 |
| rs397507511 | pathogenic | 1 | 1 | 1 |
| rs121918454 | pathogenic | 1 | 1 | 1 |
| rs121918453 | pathogenic | 1 | 1 | 1 |
| rs121918462 | pathogenic | 1 | 1 | 0 |
| rs121918465 | pathogenic | 1 | 1 | 0 |
| rs121918464 | pathogenic | 1 | 1 | 0 |
| rs121918466 | pathogenic | 1 | 1 | 1 |
| rs397507521 | pathogenic | 1 | 1 | 1 |
| rs397507527 | pathogenic | 1 | 1 | 1 |
| rs777603059 | pathogenic | 0 | 1 | 0 |
| rs121918456 | pathogenic | 1 | 1 | 1 |
| rs121918463 | pathogenic | 1 | 1 | 1 |
| rs121918455 | pathogenic | 0 | 1 | 0 |
| rs28933386 | pathogenic | 0 | 1 | 0 |
| rs774356443 | benign | 1 | 1 | 1 |
| rs121918467 | pathogenic | 1 | 1 | 1 |
| rs121918468 | pathogenic | 1 | 1 | 1 |
| rs121918469 | pathogenic | 1 | 1 | 1 |
| rs397507539 | pathogenic | 1 | 1 | 0 |
| rs121918458 | pathogenic | 1 | 1 | 1 |
| rs121918470 | pathogenic | 1 | 1 | 1 |
| rs35289681 | benign | 0 | 1 | 0 |
| rs4302331 | benign | 0 | 1 | 0 |
| rs11544803 | pathogenic | 1 | 1 | 1 |
| rs116254156 | benign | 0 | 0 | 0 |
| rs104893909 | pathogenic | 1 | 1 | 1 |
| rs104893911 | pathogenic | 1 | 1 | 1 |

| | | | | |
|---|---|---|---|---|
| rs104893908 | pathogenic | 1 | 1 | 0 |
| rs258751 | benign | 1 | 1 | 1 |
| rs104893914 | pathogenic | 1 | 1 | 0 |
| rs121909727 | pathogenic | 1 | 1 | 1 |
| rs104893910 | pathogenic | 1 | 1 | 1 |
| rs121909726 | pathogenic | 1 | 1 | 1 |
| rs104893912 | pathogenic | 1 | 1 | 1 |
| rs138407155 | benign | 1 | 1 | 1 |
| rs574061789 | benign | 1 | 1 | 1 |
| rs121912718 | pathogenic | 1 | 1 | 1 |
| rs121912719 | pathogenic | 1 | 1 | 1 |
| rs140770089 | benign | 1 | 1 | 1 |
| rs28931573 | pathogenic | 1 | 1 | 1 |
| rs121912717 | pathogenic | 1 | 1 | 1 |
| rs376456598 | pathogenic | 0 | 0 | 0 |
| rs398123499 | pathogenic | 1 | 1 | 1 |
| rs773048903 | pathogenic | 1 | 1 | 0 |
| rs398123503 | pathogenic | 1 | 1 | 1 |
| rs375785084 | pathogenic | 0 | 1 | 0 |
| rs137852874 | pathogenic | 1 | 1 | 1 |
| rs373713279 | pathogenic | 1 | 1 | 1 |
| rs137852873 | pathogenic | 1 | 1 | 0 |
| rs367823977 | pathogenic | 1 | 1 | 1 |
| rs398123508 | pathogenic | 1 | 1 | 1 |
| rs137852871 | pathogenic | 1 | 1 | 1 |
| rs753698250 | pathogenic | 0 | 1 | 0 |
| rs182923857 | pathogenic | 1 | 1 | 1 |
| rs148090804 | benign | 1 | 1 | 1 |
| rs137852870 | pathogenic | 1 | 1 | 1 |
| rs267607131 | pathogenic | 1 | 1 | 0 |
| rs137852902 | pathogenic | 0 | 0 | 0 |
| rs61741646 | benign | 1 | 1 | 1 |
| rs137852905 | pathogenic | 0 | 0 | 0 |
| rs121918681 | pathogenic | 0 | 1 | 0 |
| rs8177238 | benign | 1 | 1 | 0 |
| rs5030776 | pathogenic | 1 | 1 | 1 |
| rs104894572 | pathogenic | 0 | 0 | 0 |
| rs104894576 | pathogenic | 0 | 0 | 0 |
| rs80084721 | pathogenic | 0 | 0 | 0 |
| rs104894146 | pathogenic | 0 | 1 | 0 |
| rs104894138 | pathogenic | 1 | 1 | 0 |
| rs104894153 | pathogenic | 1 | 1 | 0 |
| rs104894135 | pathogenic | 1 | 1 | 0 |
| rs104894147 | pathogenic | 1 | 1 | 0 |
| rs104894148 | pathogenic | 1 | 1 | 0 |
| rs104894154 | pathogenic | 1 | 1 | 0 |
| rs104894150 | pathogenic | 1 | 1 | 0 |
| rs104894136 | pathogenic | 1 | 1 | 0 |
| rs104894144 | pathogenic | 0 | 0 | 0 |
| rs104894137 | pathogenic | 0 | 0 | 0 |
| rs61754278 | pathogenic | 1 | 1 | 0 |
| rs104894149 | pathogenic | 1 | 1 | 0 |
| rs104894139 | pathogenic | 1 | 1 | 0 |
| rs104894142 | pathogenic | 0 | 0 | 0 |

| | | | | |
|---|---|---|---|---|
| rs104894143 | pathogenic | 0 | 0 | 0 |
| rs104894145 | pathogenic | 0 | 0 | 0 |
| rs121909545 | benign | 0 | 1 | 0 |
| rs782308462 | pathogenic | 1 | 1 | 1 |
| rs375002796 | pathogenic | 0 | 1 | 0 |
| rs201727685 | pathogenic | 0 | 0 | 0 |
| rs768050261 | pathogenic | 1 | 1 | 1 |
| rs780851340 | benign | 0 | 0 | 0 |
| rs139824241 | benign | 1 | 1 | 0 |
| rs757486575 | pathogenic | 1 | 1 | 0 |
| rs750831299 | pathogenic | 1 | 1 | 1 |
| rs767602961 | pathogenic | 1 | 1 | 1 |
| rs121913661 | pathogenic | 1 | 1 | 1 |
| rs777692271 | pathogenic | 1 | 1 | 1 |
| rs121913659 | pathogenic | 1 | 1 | 1 |
| rs587777533 | pathogenic | 1 | 1 | 1 |
| rs104894210 | pathogenic | 0 | 0 | 0 |
| rs104894216 | pathogenic | 1 | 1 | 1 |
| rs104894207 | pathogenic | 1 | 1 | 1 |
| rs104894208 | pathogenic | 0 | 1 | 0 |
| rs104894214 | pathogenic | 1 | 1 | 1 |
| rs104894211 | pathogenic | 1 | 1 | 1 |
| rs61749384 | pathogenic | 0 | 0 | 0 |
| rs61750071 | pathogenic | 1 | 1 | 1 |
| rs61750074 | pathogenic | 0 | 0 | 0 |
| rs11063988 | benign | 0 | 0 | 0 |
| rs216311 | benign | 0 | 0 | 0 |
| rs1800382 | benign | 1 | 1 | 1 |
| rs11063987 | benign | 1 | 1 | 1 |
| rs387906847 | pathogenic | 1 | 1 | 1 |
| rs73003466 | benign | 0 | 0 | 0 |
| rs149787233 | benign | 1 | 1 | 1 |
| rs144451841 | pathogenic | 1 | 1 | 1 |
| rs141638421 | pathogenic | 1 | 1 | 1 |
| rs121434497 | pathogenic | 0 | 0 | 0 |
| rs121434498 | pathogenic | 0 | 0 | 0 |
| rs267607230 | pathogenic | 1 | 1 | 0 |
| rs121434499 | pathogenic | 1 | 1 | 0 |
| rs546749336 | benign | 1 | 1 | 1 |
| rs138873805 | benign | 0 | 0 | 0 |
| rs730880522 | pathogenic | 0 | 0 | 0 |
| rs200371894 | benign | 0 | 0 | 0 |
| rs730880509 | benign | 0 | 0 | 0 |
| rs267606704 | pathogenic | 1 | 1 | 1 |
| rs267606706 | pathogenic | 0 | 0 | 0 |
| rs267606705 | pathogenic | 1 | 1 | 1 |
| rs267606707 | pathogenic | 0 | 0 | 0 |
| rs387906665 | pathogenic | 1 | 1 | 1 |
| rs267606708 | pathogenic | 1 | 1 | 1 |
| rs148368481 | benign | 0 | 0 | 0 |
| rs13204 | benign | 0 | 0 | 0 |
| rs140547520 | pathogenic | 1 | 1 | 1 |
| rs755234990 | benign | 1 | 1 | 0 |
| rs151184052 | benign | 0 | 0 | 0 |

| | | | | |
|---|---|---|---|---|
| rs3736358 | benign | 0 | 0 | 0 |
| rs3736360 | benign | 0 | 0 | 0 |
| rs1138469 | benign | 0 | 0 | 0 |
| rs74315351 | pathogenic | 0 | 1 | 0 |
| rs137853051 | pathogenic | 1 | 1 | 0 |
| rs74315353 | pathogenic | 0 | 1 | 0 |
| rs74315352 | pathogenic | 0 | 1 | 0 |
| rs74315354 | pathogenic | 1 | 1 | 1 |
| rs28938172 | pathogenic | 1 | 1 | 1 |
| rs52836744 | pathogenic | 0 | 0 | 0 |
| rs121913159 | pathogenic | 1 | 1 | 1 |
| rs121913145 | pathogenic | 1 | 1 | 1 |
| rs891087 | benign | 1 | 1 | 1 |
| rs9282757 | benign | 1 | 1 | 1 |
| rs397517404 | pathogenic | 1 | 1 | 1 |
| rs199918720 | benign | 1 | 1 | 0 |
| rs368299411 | benign | 1 | 1 | 0 |
| rs139399951 | benign | 1 | 1 | 0 |
| rs763981974 | benign | 1 | 1 | 0 |
| rs144242114 | benign | 0 | 0 | 0 |
| rs750965801 | benign | 0 | 0 | 0 |
| rs561310777 | benign | 0 | 0 | 0 |
| rs148185335 | benign | 0 | 0 | 0 |
| rs147742157 | benign | 0 | 0 | 0 |
| rs375664373 | pathogenic | 1 | 1 | 1 |
| rs777327250 | pathogenic | 0 | 0 | 0 |
| rs121908471 | pathogenic | 1 | 1 | 0 |
| rs2301612 | pathogenic | 1 | 1 | 0 |
| rs121908476 | pathogenic | 0 | 0 | 0 |
| rs36220239 | benign | 0 | 0 | 0 |
| rs121908473 | pathogenic | 0 | 0 | 0 |
| rs28647808 | benign | 0 | 0 | 0 |
| rs36090624 | benign | 0 | 0 | 0 |
| rs121908002 | pathogenic | 1 | 1 | 1 |
| rs4251545 | benign | 1 | 1 | 1 |
| rs78705382 | pathogenic | 0 | 0 | 0 |
| rs75184679 | pathogenic | 1 | 1 | 1 |
| rs74555752 | pathogenic | 0 | 0 | 0 |
| rs56320207 | benign | 1 | 1 | 1 |
| rs28730837 | pathogenic | 1 | 1 | 1 |
| rs119469014 | pathogenic | 1 | 1 | 1 |
| rs119468010 | pathogenic | 1 | 1 | 1 |
| rs35220041 | benign | 0 | 0 | 0 |
| rs762321137 | pathogenic | 1 | 1 | 1 |
| rs28931570 | pathogenic | 1 | 1 | 1 |
| rs709932 | benign | 1 | 1 | 1 |
| rs746531546 | benign | 1 | 1 | 1 |
| rs6647 | benign | 1 | 1 | 1 |
| rs141095970 | benign | 1 | 1 | 1 |
| rs143370956 | benign | 1 | 1 | 1 |
| rs28929474 | pathogenic | 1 | 1 | 1 |
| rs61761869 | pathogenic | 1 | 1 | 1 |
| rs1303 | benign | 1 | 1 | 1 |
| rs149447502 | benign | 0 | 0 | 0 |

| | | | | |
|---|---|---|---|---|
| rs267607180 | pathogenic | 1 | 1 | 1 |
| rs150304121 | benign | 0 | 0 | 0 |
| rs147630375 | benign | 1 | 1 | 1 |
| rs11539445 | benign | 1 | 1 | 1 |
| rs372098364 | pathogenic | 1 | 1 | 0 |
| rs760072286 | benign | 0 | 0 | 0 |
| rs745595833 | pathogenic | 1 | 1 | 1 |
| rs587777214 | pathogenic | 0 | 0 | 0 |
| rs150318528 | benign | 0 | 0 | 0 |
| rs137853254 | pathogenic | 1 | 1 | 1 |
| rs137853255 | pathogenic | 0 | 0 | 0 |
| rs137853253 | pathogenic | 0 | 0 | 0 |
| rs202166915 | benign | 0 | 0 | 0 |
| rs137853259 | pathogenic | 0 | 0 | 0 |
| rs2229137 | benign | 0 | 0 | 0 |
| rs137853258 | pathogenic | 1 | 1 | 0 |
| rs137853252 | pathogenic | 1 | 1 | 1 |
| rs137853256 | pathogenic | 0 | 0 | 0 |
| rs137853250 | pathogenic | 1 | 1 | 1 |
| rs121434615 | pathogenic | 0 | 0 | 0 |
| rs115014558 | benign | 1 | 1 | 1 |
| rs34116584 | benign | 1 | 1 | 1 |
| rs34885252 | benign | 1 | 1 | 1 |
| rs180177262 | pathogenic | 1 | 1 | 1 |
| rs180177162 | pathogenic | 1 | 1 | 1 |
| rs121908523 | pathogenic | 1 | 1 | 1 |
| rs180177170 | pathogenic | 1 | 1 | 1 |
| rs180177180 | pathogenic | 1 | 1 | 1 |
| rs767586362 | pathogenic | 1 | 1 | 1 |
| rs121908521 | pathogenic | 1 | 1 | 1 |
| rs180177203 | pathogenic | 1 | 1 | 1 |
| rs138025751 | pathogenic | 1 | 1 | 1 |
| rs180177210 | pathogenic | 0 | 1 | 0 |
| rs180177211 | pathogenic | 0 | 1 | 0 |
| rs121908524 | pathogenic | 1 | 1 | 0 |
| rs121908530 | pathogenic | 1 | 1 | 0 |
| rs180177227 | pathogenic | 0 | 1 | 0 |
| rs180177230 | pathogenic | 1 | 1 | 1 |
| rs121908529 | pathogenic | 0 | 1 | 0 |
| rs117195882 | benign | 1 | 1 | 0 |
| rs180177238 | pathogenic | 1 | 1 | 0 |
| rs34664134 | benign | 1 | 1 | 1 |
| rs180177246 | pathogenic | 0 | 1 | 0 |
| rs536352238 | pathogenic | 0 | 0 | 0 |
| rs121908520 | pathogenic | 1 | 1 | 0 |
| rs180177248 | pathogenic | 1 | 1 | 0 |
| rs180177252 | pathogenic | 1 | 1 | 1 |
| rs121908526 | pathogenic | 1 | 1 | 1 |
| rs121908525 | pathogenic | 1 | 1 | 1 |
| rs147106773 | benign | 1 | 1 | 1 |
| rs180177264 | pathogenic | 0 | 1 | 0 |
| rs146525143 | pathogenic | 1 | 1 | 1 |
| rs140992177 | benign | 1 | 1 | 1 |
| rs13408961 | benign | 0 | 1 | 0 |

| | | | | |
|---|---|---|---|---|
| rs4426527 | benign | 1 | 1 | 1 |
| rs3088308 | benign | 0 | 1 | 0 |
| rs4469829 | benign | 1 | 1 | 0 |
| rs104886003 | pathogenic | 1 | 1 | 1 |
| rs121913286 | pathogenic | 1 | 1 | 1 |
| rs121913279 | pathogenic | 1 | 1 | 1 |
| rs121434636 | pathogenic | 1 | 1 | 1 |
| rs755299132 | pathogenic | 1 | 1 | 1 |
| rs748318386 | pathogenic | 1 | 1 | 1 |
| rs387906757 | pathogenic | 1 | 1 | 1 |
| rs121965064 | pathogenic | 1 | 1 | 1 |
| rs28934608 | pathogenic | 0 | 0 | 0 |
| rs5967 | benign | 1 | 1 | 1 |
| rs773905328 | pathogenic | 0 | 0 | 0 |
| rs104894256 | pathogenic | 0 | 1 | 0 |
| rs28931612 | pathogenic | 0 | 1 | 0 |
| rs104894263 | pathogenic | 1 | 1 | 1 |
| rs607969 | benign | 1 | 1 | 0 |
| rs104894262 | pathogenic | 0 | 1 | 0 |
| rs104894268 | pathogenic | 1 | 1 | 1 |
| rs104894264 | pathogenic | 0 | 1 | 0 |
| rs104894259 | pathogenic | 1 | 1 | 0 |
| rs2959656 | benign | 0 | 0 | 0 |
| rs121913035 | pathogenic | 1 | 1 | 0 |
| rs139605474 | benign | 0 | 0 | 0 |
| rs3817552 | benign | 0 | 0 | 0 |
| rs150541115 | benign | 0 | 0 | 0 |
| rs562763010 | benign | 1 | 1 | 1 |
| rs200053119 | pathogenic | 0 | 0 | 0 |
| rs73826386 | benign | 0 | 0 | 0 |
| rs143655258 | benign | 0 | 0 | 0 |
| rs758857853 | pathogenic | 1 | 1 | 0 |
| rs2228380 | benign | 0 | 0 | 0 |
| rs141078733 | benign | 1 | 1 | 1 |
| rs55824172 | pathogenic | 1 | 1 | 1 |
| rs756751089 | pathogenic | 1 | 1 | 1 |
| rs121964990 | pathogenic | 0 | 0 | 0 |
| rs143555186 | benign | 0 | 0 | 0 |
| rs121964993 | pathogenic | 1 | 1 | 0 |
| rs121964992 | pathogenic | 1 | 1 | 1 |
| rs121964991 | pathogenic | 1 | 1 | 0 |
| rs121964988 | pathogenic | 1 | 1 | 1 |
| rs4687 | benign | 1 | 1 | 1 |
| rs267607125 | pathogenic | 1 | 1 | 1 |
| rs751682878 | benign | 1 | 1 | 1 |
| rs267607126 | pathogenic | 1 | 1 | 1 |
| rs768713596 | pathogenic | 0 | 0 | 0 |
| rs147288996 | benign | 0 | 0 | 0 |
| rs387906639 | pathogenic | 1 | 1 | 1 |
| rs730881069 | pathogenic | 1 | 1 | 1 |
| rs730881071 | pathogenic | 1 | 1 | 1 |
| rs397516348 | pathogenic | 0 | 0 | 0 |
| rs121917760 | pathogenic | 0 | 0 | 0 |
| rs104894724 | pathogenic | 1 | 1 | 1 |

| | | | | |
|---|---|---|---|---|
| rs397516349 | pathogenic | 1 | 1 | 1 |
| rs12075 | benign | 0 | 0 | 0 |
| rs147549987 | benign | 0 | 0 | 0 |
| rs56079734 | benign | 0 | 0 | 0 |
| rs1017842 | benign | 1 | 1 | 1 |
| rs137852593 | pathogenic | 1 | 1 | 1 |
| rs137852571 | pathogenic | 1 | 1 | 1 |
| rs137852591 | pathogenic | 1 | 1 | 1 |
| rs9332971 | pathogenic | 1 | 1 | 1 |
| rs755226547 | pathogenic | 1 | 1 | 1 |
| rs79206939 | benign | 0 | 0 | 0 |
| rs121918214 | pathogenic | 1 | 1 | 0 |
| rs757075712 | pathogenic | 1 | 1 | 0 |
| rs150483902 | benign | 1 | 1 | 1 |
| rs113993973 | pathogenic | 0 | 1 | 0 |
| rs946616 | benign | 0 | 0 | 0 |
| rs113993976 | pathogenic | 1 | 1 | 0 |
| rs151022264 | benign | 0 | 1 | 0 |
| rs113993977 | pathogenic | 1 | 1 | 0 |
| rs113993978 | pathogenic | 1 | 1 | 0 |
| rs2228499 | benign | 0 | 1 | 0 |
| rs113993979 | pathogenic | 1 | 1 | 0 |
| rs113993980 | pathogenic | 1 | 1 | 0 |
| rs113993983 | pathogenic | 0 | 1 | 0 |
| rs150547274 | pathogenic | 1 | 1 | 0 |
| rs113993984 | pathogenic | 1 | 1 | 0 |
| rs113993986 | pathogenic | 1 | 1 | 0 |
| rs34313873 | benign | 0 | 1 | 0 |
| rs113993988 | pathogenic | 1 | 1 | 0 |
| rs199638097 | pathogenic | 1 | 1 | 1 |
| rs118204447 | pathogenic | 1 | 1 | 1 |
| rs11862754 | benign | 0 | 0 | 0 |
| rs118204445 | pathogenic | 0 | 0 | 0 |
| rs118204441 | pathogenic | 0 | 0 | 0 |
| rs118204438 | pathogenic | 0 | 0 | 0 |
| rs118204436 | pathogenic | 0 | 0 | 0 |
| rs146093755 | pathogenic | 0 | 0 | 0 |
| rs781439830 | pathogenic | 0 | 0 | 0 |
| rs398123438 | pathogenic | 0 | 0 | 0 |
| rs398123439 | pathogenic | 0 | 0 | 0 |
| rs118204444 | pathogenic | 0 | 0 | 0 |
| rs7187889 | benign | 0 | 0 | 0 |
| rs118204435 | pathogenic | 1 | 1 | 0 |
| rs34745339 | benign | 0 | 0 | 0 |
| rs398123440 | pathogenic | 0 | 0 | 0 |
| rs118204442 | pathogenic | 0 | 0 | 0 |
| rs770053354 | pathogenic | 0 | 0 | 0 |
| rs118204448 | pathogenic | 0 | 0 | 0 |
| rs118204443 | pathogenic | 0 | 0 | 0 |
| rs118204446 | pathogenic | 0 | 0 | 0 |
| rs746756997 | pathogenic | 0 | 0 | 0 |
| rs267606838 | pathogenic | 0 | 0 | 0 |
| rs118204437 | pathogenic | 0 | 0 | 0 |
| rs398123429 | pathogenic | 0 | 0 | 0 |

| | | | | |
|---|---|---|---|---|
| rs398123430 | pathogenic | 0 | 0 | 0 |
| rs2303269 | benign | 0 | 0 | 0 |
| rs114703967 | benign | 0 | 0 | 0 |
| rs118204439 | pathogenic | 0 | 0 | 0 |
| rs151296605 | benign | 0 | 0 | 0 |
| rs118204440 | pathogenic | 0 | 0 | 0 |
| rs78127134 | benign | 0 | 0 | 0 |
| rs372893383 | pathogenic | 0 | 0 | 0 |
| rs61753266 | pathogenic | 1 | 1 | 1 |
| rs139579928 | benign | 1 | 1 | 1 |
| rs56275071 | pathogenic | 1 | 1 | 1 |
| rs121909736 | pathogenic | 1 | 1 | 0 |
| rs121909737 | pathogenic | 1 | 1 | 0 |
| rs121909735 | pathogenic | 1 | 1 | 0 |
| rs121909731 | pathogenic | 1 | 1 | 1 |
| rs121909734 | pathogenic | 1 | 1 | 1 |
| rs121909733 | pathogenic | 1 | 1 | 1 |
| rs121909732 | pathogenic | 1 | 1 | 1 |
| rs121909730 | pathogenic | 1 | 1 | 1 |
| rs202198360 | benign | 1 | 1 | 1 |
| rs587777294 | pathogenic | 1 | 1 | 0 |
| rs587777297 | pathogenic | 1 | 1 | 0 |
| rs104893913 | pathogenic | 1 | 1 | 0 |
| rs17860424 | pathogenic | 1 | 1 | 1 |
| rs1045485 | benign | 1 | 1 | 1 |
| rs13926 | benign | 0 | 0 | 0 |
| rs79343837 | benign | 0 | 0 | 0 |
| rs45516306 | benign | 1 | 1 | 1 |
| rs139849355 | benign | 1 | 1 | 1 |
| rs104894874 | pathogenic | 1 | 1 | 1 |
| rs104894870 | pathogenic | 1 | 1 | 1 |
| rs137852221 | pathogenic | 1 | 1 | 1 |
| rs104894869 | pathogenic | 1 | 1 | 1 |
| rs104894880 | pathogenic | 1 | 1 | 1 |
| rs104894872 | pathogenic | 1 | 1 | 1 |
| rs104894868 | pathogenic | 1 | 1 | 1 |
| rs104894867 | pathogenic | 1 | 1 | 1 |
| rs104894871 | pathogenic | 1 | 1 | 1 |
| rs104894877 | pathogenic | 1 | 1 | 1 |
| rs104894883 | pathogenic | 1 | 1 | 1 |
| rs104894875 | pathogenic | 1 | 1 | 1 |
| rs104894876 | pathogenic | 1 | 1 | 1 |
| rs104894878 | pathogenic | 1 | 1 | 1 |
| rs137852220 | pathogenic | 1 | 1 | 1 |
| rs28933684 | pathogenic | 1 | 1 | 1 |
| rs104894873 | pathogenic | 1 | 1 | 1 |
| rs138409264 | benign | 0 | 0 | 0 |
| rs2236722 | benign | 0 | 0 | 0 |
| rs28757184 | benign | 0 | 0 | 0 |
| rs121434538 | pathogenic | 0 | 0 | 0 |
| rs700519 | benign | 0 | 0 | 0 |
| rs80051519 | pathogenic | 0 | 0 | 0 |
| rs121434536 | pathogenic | 1 | 1 | 0 |
| rs121434534 | pathogenic | 1 | 1 | 0 |

| | | | | |
|---|---|---|---|---|
| rs78310315 | pathogenic | 1 | 1 | 0 |
| rs540839115 | pathogenic | 1 | 1 | 1 |
| rs80338731 | pathogenic | 1 | 1 | 1 |
| rs180177151 | pathogenic | 1 | 1 | 1 |
| rs180177152 | pathogenic | 0 | 0 | 0 |
| rs80338732 | pathogenic | 1 | 1 | 1 |
| rs80338675 | pathogenic | 1 | 1 | 1 |
| rs80338676 | pathogenic | 1 | 1 | 1 |
| rs180177153 | pathogenic | 0 | 0 | 0 |
| rs180177154 | pathogenic | 1 | 1 | 1 |
| rs5896 | benign | 1 | 1 | 1 |
| rs121918477 | pathogenic | 0 | 0 | 0 |
| rs121918428 | pathogenic | 1 | 1 | 1 |
| rs121918427 | pathogenic | 1 | 1 | 1 |
| rs544300820 | benign | 1 | 1 | 1 |
| rs121918110 | pathogenic | 1 | 1 | 1 |
| rs1057141 | benign | 0 | 0 | 0 |
| rs1135216 | benign | 1 | 1 | 1 |
| rs121917702 | pathogenic | 1 | 1 | 1 |
| rs28941474 | pathogenic | 0 | 0 | 0 |
| rs104893944 | pathogenic | 1 | 1 | 1 |
| rs104893947 | pathogenic | 1 | 1 | 0 |
| rs104893943 | pathogenic | 1 | 1 | 1 |
| rs104893942 | pathogenic | 1 | 1 | 1 |
| rs104893940 | pathogenic | 1 | 1 | 1 |
| rs34045013 | benign | 0 | 0 | 0 |
| rs143083947 | pathogenic | 0 | 0 | 0 |
| rs145786986 | benign | 0 | 0 | 0 |
| rs119450942 | pathogenic | 0 | 0 | 0 |
| rs374259530 | pathogenic | 0 | 0 | 0 |
| rs756210458 | pathogenic | 0 | 0 | 0 |
| rs11089991 | benign | 0 | 0 | 0 |
| rs28941471 | pathogenic | 0 | 1 | 0 |
| rs750614500 | pathogenic | 0 | 1 | 0 |
| rs199761158 | benign | 1 | 1 | 1 |
| rs119450944 | pathogenic | 1 | 1 | 1 |
| rs373458753 | pathogenic | 1 | 1 | 1 |
| rs202064195 | pathogenic | 1 | 1 | 1 |
| rs776496275 | pathogenic | 1 | 1 | 1 |
| rs761493155 | pathogenic | 1 | 1 | 1 |
| rs763542069 | pathogenic | 0 | 0 | 0 |
| rs119450941 | pathogenic | 0 | 0 | 0 |
| rs119450940 | pathogenic | 0 | 0 | 0 |
| rs777821034 | pathogenic | 1 | 1 | 1 |
| rs771121666 | pathogenic | 1 | 1 | 1 |
| rs372895468 | pathogenic | 1 | 1 | 1 |
| rs61750603 | pathogenic | 0 | 0 | 0 |
| rs76863441 | pathogenic | 0 | 0 | 0 |
| rs142601143 | benign | 1 | 1 | 1 |
| rs185832993 | benign | 1 | 1 | 0 |
| rs17860568 | benign | 0 | 0 | 0 |
| rs142064825 | benign | 0 | 0 | 0 |
| rs564076183 | benign | 0 | 0 | 0 |
| rs386834034 | pathogenic | 0 | 0 | 0 |

| | | | | |
|---|---|---|---|---|
| rs147663769 | benign | 0 | 0 | 0 |
| rs776478974 | benign | 0 | 0 | 0 |
| rs111840783 | benign | 0 | 0 | 0 |
| rs1516446 | benign | 1 | 1 | 1 |
| rs143749154 | benign | 1 | 1 | 1 |
| rs80257412 | benign | 0 | 0 | 0 |
| rs142943120 | benign | 0 | 0 | 0 |
| rs144553482 | benign | 0 | 0 | 0 |
| rs35997569 | benign | 0 | 0 | 0 |
| rs772909106 | pathogenic | 0 | 0 | 0 |
| rs149433837 | benign | 1 | 1 | 1 |
| rs532178791 | pathogenic | 0 | 0 | 0 |
| rs145588689 | pathogenic | 1 | 1 | 0 |
| rs17843776 | benign | 1 | 1 | 0 |
| rs121917890 | pathogenic | 1 | 1 | 1 |
| rs121917892 | pathogenic | 0 | 0 | 0 |
| rs606231289 | pathogenic | 0 | 0 | 0 |
| rs370011798 | pathogenic | 0 | 0 | 0 |
| rs34689457 | benign | 0 | 0 | 0 |
| rs118204451 | pathogenic | 0 | 0 | 0 |
| rs118204452 | pathogenic | 1 | 1 | 1 |
| rs652785 | benign | 0 | 0 | 0 |
| rs35378126 | benign | 0 | 0 | 0 |
| rs80358214 | pathogenic | 1 | 1 | 1 |
| rs80358215 | pathogenic | 0 | 1 | 0 |
| rs267606858 | pathogenic | 1 | 1 | 1 |
| rs143137713 | pathogenic | 1 | 1 | 1 |
| rs782168634 | benign | 0 | 0 | 0 |
| rs143873938 | benign | 0 | 0 | 0 |
| rs782776511 | benign | 0 | 0 | 0 |
| rs201093148 | benign | 0 | 0 | 0 |
| rs267606815 | pathogenic | 0 | 0 | 0 |
| rs137853311 | pathogenic | 0 | 0 | 0 |
| rs267606817 | pathogenic | 0 | 0 | 0 |
| rs72650006 | benign | 0 | 0 | 0 |
| rs727503638 | benign | 1 | 1 | 0 |
| rs189286381 | benign | 0 | 0 | 0 |
| rs376644553 | benign | 0 | 0 | 0 |
| rs121434610 | pathogenic | 1 | 1 | 1 |
| rs267607076 | pathogenic | 1 | 1 | 1 |
| rs201275730 | benign | 0 | 0 | 0 |
| rs121909587 | pathogenic | 0 | 0 | 0 |
| rs121909558 | pathogenic | 1 | 1 | 0 |
| rs28929469 | pathogenic | 1 | 1 | 0 |
| rs121909547 | pathogenic | 1 | 1 | 0 |
| rs121909552 | pathogenic | 1 | 1 | 0 |
| rs121909573 | pathogenic | 1 | 1 | 1 |
| rs121909567 | pathogenic | 1 | 1 | 0 |
| rs121909569 | pathogenic | 0 | 0 | 0 |
| rs121909563 | pathogenic | 1 | 1 | 0 |
| rs121909570 | pathogenic | 1 | 1 | 1 |
| rs121909571 | pathogenic | 1 | 1 | 0 |
| rs121909572 | pathogenic | 1 | 1 | 1 |
| rs121909565 | pathogenic | 1 | 1 | 1 |

| | | | | |
|---|---|---|---|---|
| rs121909557 | pathogenic | 1 | 1 | 1 |
| rs121909568 | pathogenic | 1 | 1 | 1 |
| rs121909566 | pathogenic | 1 | 1 | 1 |
| rs121909549 | pathogenic | 1 | 1 | 1 |
| rs121909554 | pathogenic | 1 | 1 | 1 |
| rs121909550 | pathogenic | 1 | 1 | 1 |
| rs121909546 | pathogenic | 1 | 1 | 0 |
| rs121909555 | pathogenic | 1 | 1 | 1 |
| rs121909564 | pathogenic | 1 | 1 | 1 |
| rs548046212 | pathogenic | 0 | 0 | 0 |
| rs147353781 | benign | 0 | 0 | 0 |
| rs104894678 | pathogenic | 1 | 1 | 1 |
| rs1130426 | benign | 0 | 0 | 0 |
| rs104894677 | pathogenic | 1 | 1 | 1 |
| rs141917423 | benign | 1 | 1 | 1 |
| rs574777883 | benign | 0 | 0 | 0 |
| rs150199226 | benign | 1 | 1 | 1 |
| rs199618701 | benign | 0 | 1 | 0 |
| rs121964923 | pathogenic | 0 | 0 | 0 |
| rs121964924 | pathogenic | 0 | 0 | 0 |
| rs201258823 | pathogenic | 0 | 1 | 0 |
| rs267606773 | pathogenic | 0 | 0 | 0 |
| rs35638832 | benign | 0 | 0 | 0 |
| rs121908632 | pathogenic | 1 | 1 | 1 |
| rs121908625 | pathogenic | 1 | 1 | 1 |
| rs748949603 | pathogenic | 0 | 0 | 0 |
| rs757523840 | pathogenic | 0 | 0 | 0 |
| rs62541771 | pathogenic | 0 | 0 | 0 |
| rs776384541 | pathogenic | 1 | 1 | 0 |
| rs121908627 | pathogenic | 1 | 1 | 0 |
| rs28937594 | pathogenic | 0 | 0 | 0 |
| rs121907888 | pathogenic | 0 | 0 | 0 |
| rs121907889 | pathogenic | 0 | 1 | 0 |
| rs28941785 | pathogenic | 1 | 1 | 1 |
| rs28941786 | pathogenic | 1 | 1 | 1 |
| rs1021737 | benign | 0 | 0 | 0 |
| rs121908370 | pathogenic | 1 | 1 | 1 |
| rs63750083 | pathogenic | 0 | 0 | 0 |
| rs17883862 | benign | 0 | 0 | 0 |
| rs587782070 | pathogenic | 1 | 1 | 1 |
| rs587781269 | pathogenic | 1 | 1 | 1 |
| rs28909982 | pathogenic | 1 | 1 | 1 |
| rs373648967 | benign | 1 | 1 | 1 |
| rs368570187 | benign | 1 | 1 | 1 |
| rs730881701 | pathogenic | 1 | 1 | 1 |
| rs137853007 | pathogenic | 0 | 0 | 0 |
| rs575910805 | benign | 0 | 0 | 0 |
| rs104893875 | pathogenic | 1 | 1 | 1 |
| rs104893877 | pathogenic | 1 | 1 | 1 |
| rs119485093 | pathogenic | 1 | 1 | 1 |
| rs119485092 | pathogenic | 0 | 0 | 0 |
| rs119485090 | pathogenic | 0 | 0 | 0 |
| rs28942099 | pathogenic | 1 | 1 | 1 |
| rs373730800 | pathogenic | 1 | 1 | 1 |

| | | | | |
|---|---|---|---|---|
| rs113993995 | pathogenic | 0 | 0 | 0 |
| rs201070132 | pathogenic | 0 | 0 | 0 |
| rs113993996 | pathogenic | 0 | 0 | 0 |
| rs113993998 | pathogenic | 0 | 0 | 0 |
| rs78950939 | pathogenic | 1 | 1 | 1 |
| rs56378716 | pathogenic | 1 | 1 | 1 |
| rs754921704 | benign | 0 | 1 | 0 |
| rs766379103 | pathogenic | 1 | 1 | 0 |
| rs121912515 | pathogenic | 0 | 1 | 0 |
| rs764831888 | benign | 1 | 1 | 0 |
| rs572527340 | benign | 0 | 0 | 0 |
| rs35969327 | benign | 0 | 0 | 0 |
| rs35814893 | benign | 0 | 0 | 0 |
| rs34032300 | benign | 0 | 0 | 0 |
| rs80358260 | pathogenic | 0 | 0 | 0 |
| rs151220873 | benign | 0 | 0 | 0 |
| rs80358261 | pathogenic | 1 | 1 | 0 |
| rs11694 | pathogenic | 0 | 0 | 0 |
| rs80358264 | pathogenic | 0 | 0 | 0 |
| rs80358266 | pathogenic | 1 | 1 | 0 |
| rs104894458 | pathogenic | 1 | 1 | 1 |
| rs121918028 | pathogenic | 0 | 0 | 0 |
| rs11583680 | pathogenic | 0 | 0 | 0 |
| rs4715 | benign | 1 | 1 | 1 |
| rs34957318 | benign | 0 | 0 | 0 |
| rs1124 | benign | 1 | 1 | 1 |
| rs104894778 | pathogenic | 1 | 1 | 1 |
| rs104894777 | pathogenic | 1 | 1 | 1 |
| rs104894769 | pathogenic | 1 | 1 | 1 |
| rs104894768 | pathogenic | 1 | 1 | 1 |
| rs104894771 | pathogenic | 1 | 1 | 1 |
| rs575372249 | benign | 1 | 1 | 1 |
| rs118203914 | pathogenic | 0 | 0 | 0 |
| rs118203916 | pathogenic | 1 | 1 | 0 |
| rs140154690 | benign | 0 | 0 | 0 |
| rs34382405 | benign | 0 | 0 | 0 |
| rs73718082 | benign | 0 | 0 | 0 |
| rs764932724 | benign | 0 | 0 | 0 |
| rs554931092 | pathogenic | 1 | 1 | 0 |
| rs202060864 | pathogenic | 0 | 0 | 0 |
| rs116567033 | benign | 0 | 0 | 0 |
| rs11243011 | benign | 0 | 0 | 0 |
| rs142073519 | pathogenic | 0 | 0 | 0 |
| rs397514611 | pathogenic | 1 | 1 | 0 |
| rs397514612 | pathogenic | 0 | 0 | 0 |
| rs372054960 | pathogenic | 0 | 0 | 0 |
| rs775690041 | pathogenic | 0 | 0 | 0 |
| rs148921184 | pathogenic | 0 | 0 | 0 |
| rs12711521 | benign | 0 | 0 | 0 |
| rs2273346 | benign | 0 | 0 | 0 |
| rs12085877 | benign | 1 | 1 | 1 |
| rs144471433 | benign | 1 | 1 | 1 |
| rs397514671 | pathogenic | 1 | 1 | 1 |
| rs76394680 | benign | 0 | 0 | 0 |

| | | | | |
|---|---|---|---|---|
| rs78289603 | pathogenic | 1 | 1 | 1 |
| rs137852913 | pathogenic | 1 | 1 | 1 |
| rs137852914 | pathogenic | 1 | 1 | 1 |
| rs121918653 | pathogenic | 1 | 1 | 1 |
| rs35515200 | benign | 1 | 1 | 0 |
| rs17876030 | benign | 1 | 1 | 1 |
| rs556623443 | benign | 1 | 1 | 1 |
| rs121434365 | pathogenic | 1 | 1 | 1 |
| rs121434364 | pathogenic | 0 | 0 | 0 |
| rs41400249 | benign | 0 | 0 | 0 |
| rs149440376 | benign | 1 | 1 | 1 |
| rs181590179 | benign | 0 | 0 | 0 |
| rs113993968 | pathogenic | 0 | 0 | 0 |
| rs587776445 | pathogenic | 1 | 1 | 1 |
| rs113993969 | pathogenic | 1 | 1 | 1 |
| rs113993970 | pathogenic | 1 | 1 | 1 |
| rs587776449 | pathogenic | 1 | 1 | 1 |
| rs113993971 | pathogenic | 1 | 1 | 1 |
| rs267607194 | pathogenic | 1 | 1 | 1 |
| rs397514745 | pathogenic | 0 | 0 | 0 |
| rs201650281 | pathogenic | 0 | 0 | 0 |
| rs150529876 | benign | 1 | 1 | 1 |
| rs77573084 | benign | 0 | 0 | 0 |
| rs398122379 | pathogenic | 1 | 1 | 0 |
| rs765478968 | benign | 0 | 0 | 0 |
| rs201399681 | benign | 0 | 0 | 0 |
| rs17159287 | benign | 0 | 0 | 0 |
| rs538571144 | benign | 0 | 0 | 0 |
| rs137852647 | pathogenic | 0 | 0 | 0 |
| rs201358272 | pathogenic | 1 | 1 | 1 |
| rs181251337 | benign | 0 | 0 | 0 |
| rs3213119 | benign | 0 | 0 | 0 |
| rs137853102 | pathogenic | 1 | 1 | 0 |
| rs4956145 | benign | 0 | 0 | 0 |
| rs146036912 | pathogenic | 1 | 1 | 1 |
| rs375717077 | pathogenic | 1 | 1 | 1 |
| rs137853103 | pathogenic | 1 | 1 | 1 |
| rs776184830 | pathogenic | 1 | 1 | 1 |
| rs200273673 | pathogenic | 1 | 1 | 1 |
| rs267607025 | pathogenic | 1 | 1 | 1 |
| rs267607024 | pathogenic | 1 | 1 | 1 |
| rs515726188 | pathogenic | 1 | 1 | 1 |
| rs147315735 | benign | 0 | 0 | 0 |
| rs121918308 | pathogenic | 1 | 1 | 0 |
| rs515726195 | pathogenic | 0 | 0 | 0 |
| rs515726196 | pathogenic | 1 | 1 | 0 |
| rs121918311 | pathogenic | 0 | 1 | 0 |
| rs121918309 | pathogenic | 0 | 0 | 0 |
| rs387906891 | pathogenic | 0 | 0 | 0 |
| rs182614164 | pathogenic | 1 | 1 | 0 |
| rs121918307 | pathogenic | 0 | 0 | 0 |
| rs201440849 | benign | 0 | 0 | 0 |
| rs118203957 | pathogenic | 0 | 0 | 0 |
| rs4986790 | benign | 0 | 0 | 0 |

| | | | | |
|---|---|---|---|---|
| rs281864928 | pathogenic | 1 | 1 | 0 |
| rs267607156 | pathogenic | 0 | 0 | 0 |
| rs55916807 | benign | 0 | 0 | 0 |
| rs749750052 | benign | 0 | 0 | 0 |
| rs74382477 | benign | 0 | 0 | 0 |
| rs201016638 | benign | 0 | 0 | 0 |
| rs376683989 | benign | 0 | 0 | 0 |
| rs144439703 | pathogenic | 0 | 0 | 0 |
| rs779122022 | benign | 0 | 0 | 0 |
| rs121918056 | pathogenic | 0 | 0 | 0 |
| rs113994093 | pathogenic | 1 | 1 | 1 |
| rs141367015 | pathogenic | 0 | 0 | 0 |
| rs775538075 | pathogenic | 0 | 0 | 0 |
| rs121918055 | pathogenic | 0 | 0 | 0 |
| rs754246294 | benign | 1 | 1 | 1 |
| rs2307447 | benign | 1 | 1 | 1 |
| rs113994096 | pathogenic | 1 | 1 | 1 |
| rs367610201 | pathogenic | 1 | 1 | 1 |
| rs758438414 | pathogenic | 1 | 1 | 0 |
| rs121918046 | pathogenic | 0 | 0 | 0 |
| rs201871736 | benign | 0 | 0 | 0 |
| rs773994204 | benign | 0 | 0 | 0 |
| rs2307450 | benign | 1 | 1 | 1 |
| rs200257554 | benign | 0 | 0 | 0 |
| rs185645212 | benign | 0 | 0 | 0 |
| rs121918054 | pathogenic | 0 | 0 | 0 |
| rs113994097 | pathogenic | 0 | 0 | 0 |
| rs202037973 | pathogenic | 0 | 0 | 0 |
| rs769827124 | pathogenic | 1 | 1 | 0 |
| rs41549716 | benign | 1 | 1 | 1 |
| rs113994098 | pathogenic | 1 | 1 | 0 |
| rs144500145 | pathogenic | 1 | 1 | 0 |
| rs121918053 | pathogenic | 1 | 1 | 0 |
| rs121918050 | pathogenic | 1 | 1 | 0 |
| rs368587966 | pathogenic | 0 | 0 | 0 |
| rs139590686 | pathogenic | 0 | 0 | 0 |
| rs121918048 | pathogenic | 1 | 1 | 0 |
| rs768653086 | pathogenic | 1 | 1 | 0 |
| rs113994099 | pathogenic | 1 | 1 | 0 |
| rs121918051 | pathogenic | 1 | 1 | 0 |
| rs753160398 | pathogenic | 1 | 1 | 0 |
| rs201477273 | pathogenic | 1 | 1 | 0 |
| rs142347031 | pathogenic | 0 | 0 | 0 |
| rs121918049 | pathogenic | 0 | 0 | 0 |
| rs267606959 | pathogenic | 1 | 1 | 0 |
| rs201732356 | pathogenic | 0 | 0 | 0 |
| rs368435864 | pathogenic | 0 | 0 | 0 |
| rs764036283 | pathogenic | 0 | 0 | 0 |
| rs2307442 | benign | 0 | 0 | 0 |
| rs2307440 | benign | 0 | 0 | 0 |
| rs548076633 | pathogenic | 0 | 1 | 0 |
| rs750792237 | benign | 0 | 0 | 0 |
| rs3087374 | benign | 0 | 0 | 0 |
| rs121918060 | pathogenic | 0 | 1 | 0 |

| | | | | |
|---|---|---|---|---|
| rs397514765 | pathogenic | 1 | 1 | 1 |
| rs121918063 | pathogenic | 0 | 0 | 0 |
| rs121918058 | pathogenic | 1 | 1 | 1 |
| rs121918064 | pathogenic | 0 | 0 | 0 |
| rs36033115 | benign | 0 | 1 | 0 |
| rs121918057 | pathogenic | 0 | 1 | 0 |
| rs121918065 | pathogenic | 1 | 1 | 1 |
| rs121918061 | pathogenic | 1 | 1 | 1 |
| rs121918066 | pathogenic | 0 | 1 | 0 |
| rs707555 | benign | 0 | 1 | 0 |
| rs8192708 | benign | 0 | 1 | 0 |
| rs11552145 | benign | 0 | 0 | 0 |
| rs1804160 | benign | 0 | 1 | 0 |
| rs34103538 | benign | 0 | 1 | 0 |
| rs751393950 | pathogenic | 1 | 1 | 1 |
| rs121907979 | pathogenic | 1 | 1 | 1 |
| rs121907975 | pathogenic | 1 | 1 | 1 |
| rs121907962 | pathogenic | 1 | 1 | 1 |
| rs121907957 | pathogenic | 0 | 0 | 0 |
| rs121907953 | pathogenic | 1 | 1 | 1 |
| rs28941770 | pathogenic | 1 | 1 | 1 |
| rs28941771 | pathogenic | 1 | 1 | 1 |
| rs121907969 | pathogenic | 1 | 1 | 1 |
| rs121907973 | pathogenic | 1 | 1 | 1 |
| rs1800429 | pathogenic | 0 | 0 | 0 |
| rs121907976 | pathogenic | 1 | 1 | 0 |
| rs121907961 | pathogenic | 0 | 0 | 0 |
| rs121907974 | pathogenic | 0 | 0 | 0 |
| rs121907959 | pathogenic | 0 | 0 | 0 |
| rs121907971 | pathogenic | 1 | 1 | 0 |
| rs121907954 | pathogenic | 0 | 0 | 0 |
| rs121907977 | pathogenic | 0 | 0 | 0 |
| rs121907963 | pathogenic | 1 | 1 | 0 |
| rs370266293 | pathogenic | 1 | 1 | 0 |
| rs1800430 | benign | 0 | 0 | 0 |
| rs121907958 | pathogenic | 0 | 0 | 0 |
| rs202173526 | pathogenic | 0 | 0 | 0 |
| rs121907978 | pathogenic | 0 | 0 | 0 |
| rs121907981 | pathogenic | 0 | 1 | 0 |
| rs145012038 | benign | 1 | 1 | 1 |
| rs121907968 | pathogenic | 1 | 1 | 1 |
| rs121907966 | pathogenic | 1 | 1 | 1 |
| rs121907956 | pathogenic | 1 | 1 | 1 |
| rs28942071 | pathogenic | 1 | 1 | 1 |
| rs121907955 | pathogenic | 1 | 1 | 1 |
| rs4727853 | benign | 0 | 0 | 0 |
| rs201880593 | benign | 0 | 0 | 0 |
| rs201434579 | benign | 1 | 1 | 1 |
| rs74551128 | pathogenic | 1 | 1 | 0 |
| rs193922500 | pathogenic | 0 | 0 | 0 |
| rs121909009 | pathogenic | 1 | 1 | 0 |
| rs121908805 | pathogenic | 1 | 1 | 0 |
| rs139573311 | pathogenic | 0 | 0 | 0 |
| rs213950 | benign | 0 | 0 | 0 |

| | | | | |
|---|---|---|---|---|
| rs79282516 | pathogenic | 0 | 0 | 0 |
| rs121909017 | pathogenic | 1 | 1 | 1 |
| rs77101217 | pathogenic | 1 | 1 | 1 |
| rs1800092 | pathogenic | 0 | 1 | 0 |
| rs77646904 | pathogenic | 0 | 0 | 0 |
| rs1800095 | benign | 1 | 1 | 1 |
| rs387906368 | pathogenic | 1 | 1 | 1 |
| rs113993959 | pathogenic | 1 | 1 | 1 |
| rs121908755 | pathogenic | 1 | 1 | 1 |
| rs121909005 | pathogenic | 1 | 1 | 1 |
| rs121908757 | pathogenic | 1 | 1 | 1 |
| rs75527207 | pathogenic | 1 | 1 | 0 |
| rs121909013 | pathogenic | 1 | 1 | 0 |
| rs121909044 | pathogenic | 1 | 1 | 0 |
| rs74597325 | pathogenic | 1 | 1 | 0 |
| rs75549581 | pathogenic | 0 | 0 | 0 |
| rs80055610 | pathogenic | 1 | 1 | 1 |
| rs1800097 | pathogenic | 0 | 0 | 0 |
| rs121909006 | pathogenic | 0 | 1 | 0 |
| rs397508276 | pathogenic | 0 | 0 | 0 |
| rs397508296 | pathogenic | 1 | 1 | 1 |
| rs139468767 | pathogenic | 0 | 0 | 0 |
| rs121908759 | pathogenic | 1 | 1 | 0 |
| rs121909033 | pathogenic | 1 | 1 | 0 |
| rs397508331 | pathogenic | 0 | 0 | 0 |
| rs387907279 | pathogenic | 1 | 1 | 1 |
| rs546530618 | benign | 0 | 0 | 0 |
| rs558259739 | benign | 0 | 0 | 0 |
| rs201827340 | benign | 0 | 0 | 0 |
| rs148865119 | pathogenic | 1 | 1 | 0 |
| rs2229413 | benign | 1 | 1 | 1 |
| rs34040148 | benign | 1 | 1 | 1 |
| rs121964964 | pathogenic | 1 | 1 | 1 |
| rs760214620 | pathogenic | 1 | 1 | 1 |
| rs775992753 | pathogenic | 1 | 1 | 1 |
| rs770095972 | pathogenic | 1 | 1 | 1 |
| rs781444670 | pathogenic | 1 | 1 | 1 |
| rs147474549 | pathogenic | 0 | 0 | 0 |
| rs121964965 | pathogenic | 0 | 1 | 0 |
| rs121964963 | pathogenic | 1 | 1 | 1 |
| rs755952006 | pathogenic | 1 | 1 | 1 |
| rs121964970 | pathogenic | 1 | 1 | 1 |
| rs121964973 | pathogenic | 1 | 1 | 1 |
| rs773734233 | pathogenic | 1 | 1 | 1 |
| rs758236584 | pathogenic | 1 | 1 | 1 |
| rs149119723 | pathogenic | 1 | 1 | 0 |
| rs5742905 | pathogenic | 0 | 0 | 0 |
| rs121964962 | pathogenic | 1 | 1 | 1 |
| rs781567152 | pathogenic | 1 | 1 | 0 |
| rs398123151 | pathogenic | 1 | 1 | 1 |
| rs760417941 | pathogenic | 1 | 1 | 1 |
| rs774926464 | pathogenic | 0 | 0 | 0 |
| rs121964972 | pathogenic | 1 | 1 | 0 |
| rs372010465 | pathogenic | 1 | 1 | 1 |

| | | | | |
|---|---|---|---|---|
| rs763036586 | pathogenic | 1 | 1 | 1 |
| rs769080151 | pathogenic | 1 | 1 | 1 |
| rs121964967 | pathogenic | 1 | 1 | 1 |
| rs28934892 | pathogenic | 1 | 1 | 0 |
| rs28934891 | pathogenic | 1 | 1 | 1 |
| rs121964971 | pathogenic | 1 | 1 | 1 |
| rs121964968 | pathogenic | 1 | 1 | 1 |
| rs150828989 | benign | 1 | 1 | 1 |
| rs3775291 | benign | 1 | 1 | 0 |
| rs104894419 | pathogenic | 1 | 1 | 1 |
| rs2232642 | benign | 0 | 0 | 0 |
| rs104894551 | pathogenic | 1 | 1 | 0 |
| rs766328537 | pathogenic | 0 | 1 | 0 |
| rs104894553 | pathogenic | 1 | 1 | 0 |
| rs80099330 | benign | 0 | 0 | 0 |
| rs199565861 | pathogenic | 0 | 0 | 0 |
| rs104894548 | pathogenic | 0 | 0 | 0 |
| rs104894549 | pathogenic | 0 | 0 | 0 |
| rs104894550 | pathogenic | 1 | 1 | 1 |
| rs12948217 | pathogenic | 1 | 1 | 1 |
| rs761064915 | pathogenic | 0 | 0 | 0 |
| rs28940279 | pathogenic | 1 | 1 | 1 |
| rs774323189 | pathogenic | 1 | 1 | 1 |
| rs28940574 | pathogenic | 0 | 0 | 0 |
| rs515299 | benign | 1 | 1 | 1 |
| rs10127939 | benign | 1 | 1 | 1 |
| rs396991 | benign | 1 | 1 | 1 |
| rs145956175 | benign | 0 | 1 | 0 |
| rs747838255 | pathogenic | 1 | 1 | 1 |
| rs762659685 | pathogenic | 1 | 1 | 0 |
| rs104894831 | pathogenic | 0 | 1 | 0 |
| rs104894848 | pathogenic | 1 | 1 | 0 |
| rs104894833 | pathogenic | 0 | 1 | 0 |
| rs781838005 | pathogenic | 0 | 1 | 0 |
| rs104894834 | pathogenic | 1 | 1 | 0 |
| rs372966991 | pathogenic | 1 | 1 | 0 |
| rs138886989 | benign | 1 | 1 | 1 |
| rs28935195 | pathogenic | 0 | 1 | 0 |
| rs28935196 | pathogenic | 0 | 1 | 0 |
| rs372416832 | pathogenic | 1 | 1 | 1 |
| rs28935197 | pathogenic | 1 | 1 | 0 |
| rs28935486 | pathogenic | 1 | 1 | 0 |
| rs28935487 | pathogenic | 1 | 1 | 0 |
| rs28935488 | pathogenic | 1 | 1 | 1 |
| rs28935495 | pathogenic | 1 | 1 | 1 |
| rs28935485 | pathogenic | 1 | 1 | 1 |
| rs140329381 | pathogenic | 0 | 1 | 0 |
| rs104894830 | pathogenic | 0 | 1 | 0 |
| rs104894846 | pathogenic | 0 | 1 | 0 |
| rs28935489 | pathogenic | 0 | 1 | 0 |
| rs104894828 | pathogenic | 1 | 1 | 0 |
| rs28935491 | pathogenic | 1 | 1 | 0 |
| rs104894832 | pathogenic | 1 | 1 | 0 |
| rs28935492 | pathogenic | 1 | 1 | 0 |

| | | | | |
|---|---|---|---|---|
| rs730880437 | benign | 1 | 1 | 0 |
| rs28935493 | pathogenic | 0 | 1 | 0 |
| rs28935494 | pathogenic | 1 | 1 | 1 |
| rs111422676 | pathogenic | 1 | 1 | 1 |
| rs397515869 | benign | 1 | 1 | 0 |
| rs782449839 | pathogenic | 0 | 1 | 0 |
| rs104894852 | pathogenic | 1 | 1 | 1 |
| rs116929575 | pathogenic | 1 | 1 | 1 |
| rs7023954 | benign | 0 | 0 | 0 |
| rs184520335 | benign | 1 | 1 | 1 |
| rs143080527 | benign | 0 | 0 | 0 |
| rs1050565 | benign | 0 | 0 | 0 |
| rs11545035 | benign | 0 | 0 | 0 |
| rs267607190 | pathogenic | 1 | 1 | 1 |
| rs200283987 | benign | 0 | 0 | 0 |
| rs372556807 | benign | 1 | 1 | 0 |
| rs563073392 | benign | 0 | 0 | 0 |
| rs4539 | benign | 0 | 0 | 0 |
| rs28931609 | pathogenic | 1 | 1 | 1 |
| rs121912978 | pathogenic | 1 | 1 | 1 |
| rs104894072 | pathogenic | 1 | 1 | 1 |
| rs551933154 | benign | 1 | 1 | 1 |
| rs752962897 | benign | 1 | 1 | 0 |
| rs121912977 | pathogenic | 1 | 1 | 0 |
| rs4537 | benign | 1 | 1 | 1 |
| rs201830462 | benign | 0 | 0 | 0 |
| rs4544 | benign | 0 | 0 | 0 |
| rs562670189 | benign | 0 | 0 | 0 |
| rs4538 | benign | 0 | 1 | 0 |
| rs61757294 | pathogenic | 0 | 0 | 0 |
| rs5313 | benign | 0 | 0 | 0 |
| rs4545 | benign | 0 | 0 | 0 |
| rs72554627 | pathogenic | 1 | 1 | 1 |
| rs72554626 | pathogenic | 1 | 1 | 1 |
| rs387906861 | pathogenic | 1 | 1 | 1 |
| rs749655461 | pathogenic | 1 | 1 | 1 |
| rs121909606 | pathogenic | 1 | 1 | 1 |
| rs121909607 | pathogenic | 1 | 1 | 1 |
| rs200203992 | benign | 0 | 0 | 0 |
| rs6050 | benign | 0 | 0 | 0 |
| rs184635235 | benign | 0 | 0 | 0 |
| rs139146037 | benign | 0 | 0 | 0 |
| rs121909612 | pathogenic | 0 | 0 | 0 |
| rs78506343 | pathogenic | 0 | 0 | 0 |
| rs121918138 | pathogenic | 1 | 1 | 1 |
| rs121918139 | pathogenic | 1 | 1 | 1 |
| rs121918140 | pathogenic | 0 | 0 | 0 |
| rs36063028 | benign | 1 | 1 | 1 |
| rs45495192 | benign | 1 | 1 | 1 |
| rs45458802 | benign | 1 | 1 | 1 |
| rs552865793 | pathogenic | 1 | 1 | 1 |
| rs753744791 | pathogenic | 1 | 1 | 1 |
| rs777518512 | benign | 1 | 1 | 1 |
| rs142708991 | benign | 1 | 1 | 0 |

| | | | | |
|---|---|---|---|---|
| rs746069957 | benign | 0 | 0 | 0 |
| rs1334811 | benign | 0 | 0 | 0 |
| rs138894008 | benign | 1 | 1 | 0 |
| rs35030851 | benign | 1 | 1 | 0 |
| rs35378598 | benign | 0 | 0 | 0 |
| rs35818764 | benign | 1 | 1 | 1 |
| rs4631561 | benign | 1 | 1 | 1 |
| rs121918660 | pathogenic | 0 | 0 | 0 |
| rs373175587 | benign | 0 | 0 | 0 |
| rs61731956 | pathogenic | 1 | 1 | 1 |
| rs180177305 | pathogenic | 0 | 0 | 0 |
| rs119490108 | pathogenic | 0 | 0 | 0 |
| rs180177312 | pathogenic | 1 | 1 | 0 |
| rs180177314 | pathogenic | 1 | 1 | 0 |
| rs180177322 | pathogenic | 1 | 1 | 1 |
| rs180177323 | pathogenic | 1 | 1 | 1 |
| rs180177325 | pathogenic | 0 | 0 | 0 |
| rs35196441 | benign | 0 | 0 | 0 |
| rs28763904 | benign | 0 | 0 | 0 |
| rs2229519 | benign | 0 | 0 | 0 |
| rs137852886 | pathogenic | 0 | 0 | 0 |
| rs137852892 | pathogenic | 0 | 0 | 0 |
| rs770427750 | pathogenic | 0 | 0 | 0 |
| rs137852887 | pathogenic | 0 | 0 | 0 |
| rs137852893 | pathogenic | 0 | 0 | 0 |
| rs80338671 | pathogenic | 1 | 1 | 0 |
| rs2228389 | benign | 0 | 0 | 0 |
| rs80338672 | pathogenic | 0 | 0 | 0 |
| rs201958741 | pathogenic | 0 | 0 | 0 |
| rs80338673 | pathogenic | 0 | 0 | 0 |
| rs137852888 | pathogenic | 0 | 0 | 0 |
| rs137852889 | pathogenic | 0 | 0 | 0 |
| rs137852894 | pathogenic | 0 | 0 | 0 |
| rs137852890 | pathogenic | 0 | 0 | 0 |
| rs137852891 | pathogenic | 0 | 0 | 0 |
| rs766935302 | pathogenic | 0 | 0 | 0 |
| rs193074572 | benign | 0 | 0 | 0 |
| rs555210692 | benign | 1 | 1 | 0 |
| rs119103242 | pathogenic | 0 | 1 | 0 |
| rs143725101 | benign | 0 | 1 | 0 |
| rs28940589 | pathogenic | 1 | 1 | 1 |
| rs113994145 | pathogenic | 0 | 0 | 0 |
| rs773872994 | benign | 0 | 0 | 0 |
| rs28940590 | pathogenic | 1 | 1 | 0 |
| rs145400751 | benign | 1 | 1 | 1 |
| rs587782331 | benign | 0 | 0 | 0 |
| rs730881787 | benign | 0 | 0 | 0 |
| rs587778531 | benign | 0 | 0 | 0 |
| rs781652274 | benign | 0 | 0 | 0 |
| rs55882234 | benign | 0 | 0 | 0 |
| rs2020908 | benign | 0 | 0 | 0 |
| rs63750741 | pathogenic | 1 | 1 | 1 |
| rs41295268 | benign | 1 | 1 | 1 |
| rs63750897 | benign | 1 | 1 | 0 |

| | | | | |
|---|---|---|---|---|
| rs63751005 | benign | 1 | 1 | 0 |
| rs587779223 | benign | 0 | 0 | 0 |
| rs202127474 | benign | 0 | 0 | 0 |
| rs374230313 | benign | 0 | 0 | 0 |
| rs2020912 | benign | 0 | 0 | 0 |
| rs63751017 | pathogenic | 0 | 0 | 0 |
| rs587779246 | pathogenic | 0 | 0 | 0 |
| rs63749919 | pathogenic | 0 | 0 | 0 |
| rs63750563 | pathogenic | 1 | 1 | 1 |
| rs63749999 | pathogenic | 0 | 0 | 0 |
| rs200492211 | pathogenic | 0 | 0 | 0 |
| rs576269342 | benign | 0 | 0 | 0 |
| rs267608075 | benign | 0 | 0 | 0 |
| rs149605979 | benign | 0 | 0 | 0 |
| rs63750252 | benign | 0 | 0 | 0 |
| rs754289472 | benign | 0 | 0 | 0 |
| rs63750119 | pathogenic | 0 | 0 | 0 |
| rs587782109 | benign | 0 | 0 | 0 |
| rs34625968 | benign | 1 | 1 | 1 |
| rs764786814 | benign | 1 | 1 | 1 |
| rs199594809 | benign | 1 | 1 | 1 |
| rs74315364 | pathogenic | 1 | 1 | 1 |
| rs201253919 | pathogenic | 0 | 0 | 0 |
| rs387906828 | pathogenic | 1 | 1 | 1 |
| rs34521609 | benign | 1 | 1 | 1 |
| rs199472894 | pathogenic | 0 | 0 | 0 |
| rs199472902 | pathogenic | 0 | 0 | 0 |
| rs121912505 | pathogenic | 0 | 0 | 0 |
| rs28928905 | pathogenic | 0 | 0 | 0 |
| rs374376640 | benign | 0 | 0 | 0 |
| rs202102276 | benign | 0 | 0 | 0 |
| rs199472918 | pathogenic | 0 | 0 | 0 |
| rs121912516 | pathogenic | 1 | 1 | 1 |
| rs121912504 | pathogenic | 0 | 0 | 0 |
| rs9333649 | pathogenic | 0 | 0 | 0 |
| rs121912508 | pathogenic | 0 | 0 | 0 |
| rs199473428 | pathogenic | 1 | 1 | 1 |
| rs104894021 | pathogenic | 0 | 0 | 0 |
| rs28928904 | pathogenic | 1 | 1 | 1 |
| rs199473039 | benign | 1 | 1 | 1 |
| rs199472958 | pathogenic | 1 | 1 | 1 |
| rs1137617 | pathogenic | 1 | 1 | 1 |
| rs199473532 | pathogenic | 1 | 1 | 1 |
| rs199472988 | pathogenic | 0 | 0 | 0 |
| rs189014161 | pathogenic | 0 | 0 | 0 |
| rs12720441 | pathogenic | 0 | 0 | 0 |
| rs199473001 | pathogenic | 1 | 1 | 0 |
| rs121912506 | pathogenic | 1 | 1 | 0 |
| rs773724817 | pathogenic | 1 | 1 | 1 |
| rs118204002 | pathogenic | 1 | 1 | 1 |
| rs138556525 | pathogenic | 1 | 1 | 1 |
| rs72558181 | pathogenic | 1 | 1 | 1 |
| rs118204005 | pathogenic | 1 | 1 | 1 |
| rs118204004 | pathogenic | 0 | 0 | 0 |

| | | | | |
|---|---|---|---|---|
| rs118204001 | pathogenic | 1 | 1 | 1 |
| rs118204006 | pathogenic | 1 | 1 | 1 |
| rs118204003 | pathogenic | 0 | 0 | 0 |
| rs121913147 | pathogenic | 1 | 1 | 0 |
| rs2229429 | benign | 1 | 1 | 0 |
| rs182552223 | benign | 0 | 0 | 0 |
| rs146588336 | benign | 0 | 0 | 0 |
| rs121964983 | pathogenic | 0 | 1 | 0 |
| rs121964982 | pathogenic | 0 | 0 | 0 |
| rs386833679 | pathogenic | 0 | 0 | 0 |
| rs386833680 | pathogenic | 0 | 0 | 0 |
| rs121964981 | pathogenic | 1 | 1 | 0 |
| rs121964984 | pathogenic | 0 | 0 | 0 |
| rs144971200 | benign | 0 | 0 | 0 |
| rs121964985 | pathogenic | 1 | 1 | 0 |
| rs141246107 | benign | 0 | 0 | 0 |
| rs104894561 | pathogenic | 1 | 1 | 0 |
| rs377023736 | pathogenic | 0 | 0 | 0 |
| rs121964907 | pathogenic | 1 | 1 | 1 |
| rs121964909 | pathogenic | 1 | 1 | 1 |
| rs765070743 | pathogenic | 0 | 0 | 0 |
| rs2228119 | benign | 0 | 0 | 0 |
| rs745976989 | pathogenic | 1 | 1 | 1 |
| rs121964904 | pathogenic | 0 | 0 | 0 |
| rs386833430 | pathogenic | 1 | 1 | 1 |
| rs267606992 | pathogenic | 0 | 0 | 0 |
| rs61752063 | pathogenic | 0 | 0 | 0 |
| rs61752067 | pathogenic | 1 | 1 | 1 |
| rs104894934 | pathogenic | 0 | 0 | 0 |
| rs104894929 | pathogenic | 1 | 1 | 1 |
| rs80189640 | benign | 0 | 0 | 0 |
| rs137854547 | pathogenic | 1 | 1 | 1 |
| rs137854540 | pathogenic | 1 | 1 | 1 |
| rs782092363 | pathogenic | 1 | 1 | 1 |
| rs147783915 | benign | 0 | 0 | 0 |
| rs28933687 | pathogenic | 1 | 1 | 1 |
| rs149503319 | benign | 0 | 0 | 0 |
| rs1805147 | benign | 0 | 0 | 0 |
| rs587777218 | pathogenic | 1 | 1 | 1 |
| rs200786705 | benign | 0 | 0 | 0 |
| rs56219475 | benign | 0 | 0 | 0 |
| rs587781172 | benign | 0 | 0 | 0 |
| rs757094189 | pathogenic | 0 | 0 | 0 |
| rs140811086 | benign | 1 | 1 | 1 |
| rs574088829 | pathogenic | 1 | 1 | 1 |
| rs121918525 | pathogenic | 0 | 0 | 0 |
| rs121918527 | pathogenic | 0 | 0 | 0 |
| rs121964848 | pathogenic | 0 | 1 | 0 |
| rs121964845 | pathogenic | 1 | 1 | 1 |
| rs121964846 | pathogenic | 0 | 1 | 0 |
| rs17217716 | benign | 1 | 1 | 1 |
| rs141711342 | benign | 0 | 0 | 0 |
| rs372619120 | benign | 0 | 0 | 0 |
| rs63750002 | benign | 1 | 1 | 1 |

| | | | | |
|---|---|---|---|---|
| rs193922373 | benign | 0 | 0 | 0 |
| rs41295286 | benign | 0 | 0 | 0 |
| rs17217772 | benign | 0 | 0 | 0 |
| rs145649774 | benign | 0 | 0 | 0 |
| rs63750214 | pathogenic | 0 | 0 | 0 |
| rs63750716 | benign | 0 | 0 | 0 |
| rs202026056 | benign | 0 | 0 | 0 |
| rs4987188 | benign | 1 | 1 | 0 |
| rs267607938 | benign | 0 | 0 | 0 |
| rs17224367 | benign | 0 | 0 | 0 |
| rs63750006 | benign | 0 | 0 | 0 |
| rs587781373 | benign | 0 | 0 | 0 |
| rs63751207 | pathogenic | 1 | 1 | 0 |
| rs730881759 | benign | 0 | 0 | 0 |
| rs372350768 | benign | 1 | 1 | 1 |
| rs55778204 | benign | 1 | 1 | 0 |
| rs63749910 | benign | 0 | 0 | 0 |
| rs63750844 | benign | 0 | 0 | 0 |
| rs28929483 | pathogenic | 0 | 0 | 0 |
| rs61756468 | benign | 0 | 0 | 0 |
| rs63750875 | pathogenic | 1 | 1 | 0 |
| rs776528054 | benign | 1 | 1 | 0 |
| rs751939698 | benign | 0 | 0 | 0 |
| rs587780685 | benign | 0 | 0 | 0 |
| rs730881763 | benign | 1 | 1 | 0 |
| rs373226409 | benign | 0 | 0 | 0 |
| rs202145681 | benign | 1 | 1 | 1 |
| rs63749841 | benign | 1 | 1 | 1 |
| rs63750757 | benign | 1 | 1 | 1 |
| rs587778527 | benign | 1 | 1 | 1 |
| rs63750849 | benign | 0 | 0 | 0 |
| rs587779969 | benign | 0 | 0 | 0 |
| rs121908046 | pathogenic | 1 | 1 | 1 |
| rs28940882 | pathogenic | 1 | 1 | 0 |
| rs28940883 | pathogenic | 1 | 1 | 1 |
| rs727503943 | pathogenic | 1 | 1 | 1 |
| rs137853859 | pathogenic | 1 | 1 | 1 |
| rs121908045 | pathogenic | 1 | 1 | 0 |
| rs137853861 | pathogenic | 1 | 1 | 0 |
| rs3180383 | pathogenic | 0 | 1 | 0 |
| rs144431814 | benign | 0 | 0 | 0 |
| rs139502866 | benign | 0 | 0 | 0 |
| rs17524437 | benign | 0 | 0 | 0 |
| rs760960779 | benign | 0 | 0 | 0 |
| rs148960463 | pathogenic | 0 | 0 | 0 |
| rs111033647 | pathogenic | 1 | 1 | 1 |
| rs111033648 | pathogenic | 0 | 0 | 0 |
| rs1800461 | benign | 0 | 0 | 0 |
| rs111033656 | pathogenic | 0 | 0 | 0 |
| rs111033663 | pathogenic | 1 | 1 | 0 |
| rs111033669 | pathogenic | 1 | 1 | 1 |
| rs111033690 | pathogenic | 0 | 0 | 0 |
| rs111033686 | pathogenic | 0 | 0 | 0 |
| rs111033687 | pathogenic | 0 | 0 | 0 |

| | | | | |
|---|---|---|---|---|
| rs111033695 | pathogenic | 0 | 0 | 0 |
| rs111033697 | pathogenic | 0 | 0 | 0 |
| rs111033693 | pathogenic | 0 | 0 | 0 |
| rs111033694 | pathogenic | 0 | 0 | 0 |
| rs111033721 | pathogenic | 1 | 1 | 1 |
| rs111033722 | pathogenic | 0 | 0 | 0 |
| rs111033725 | pathogenic | 1 | 1 | 0 |
| rs75391579 | pathogenic | 1 | 1 | 1 |
| rs111033728 | pathogenic | 1 | 1 | 1 |
| rs111033735 | pathogenic | 1 | 1 | 1 |
| rs111033739 | pathogenic | 1 | 1 | 1 |
| rs111033736 | pathogenic | 0 | 0 | 0 |
| rs111033744 | pathogenic | 1 | 1 | 1 |
| rs111033750 | pathogenic | 1 | 1 | 1 |
| rs111033752 | pathogenic | 0 | 0 | 0 |
| rs111033751 | pathogenic | 0 | 0 | 0 |
| rs111033846 | pathogenic | 0 | 0 | 0 |
| rs368166217 | pathogenic | 0 | 0 | 0 |
| rs111033764 | pathogenic | 0 | 0 | 0 |
| rs111033847 | benign | 0 | 0 | 0 |
| rs111033778 | pathogenic | 0 | 0 | 0 |
| rs111033773 | pathogenic | 0 | 0 | 0 |
| rs111033792 | pathogenic | 0 | 0 | 0 |
| rs367543266 | pathogenic | 1 | 1 | 1 |
| rs111033794 | pathogenic | 0 | 0 | 0 |
| rs111033802 | pathogenic | 0 | 0 | 0 |
| rs111033803 | pathogenic | 0 | 0 | 0 |
| rs111033809 | pathogenic | 1 | 1 | 0 |
| rs121918376 | pathogenic | 0 | 1 | 0 |
| rs121918377 | pathogenic | 0 | 1 | 0 |
| rs121918378 | pathogenic | 1 | 1 | 1 |
| rs121918374 | pathogenic | 1 | 1 | 1 |
| rs137852379 | pathogenic | 0 | 0 | 0 |
| rs137852388 | pathogenic | 1 | 1 | 1 |
| rs137852368 | pathogenic | 0 | 0 | 0 |
| rs28935499 | pathogenic | 0 | 0 | 0 |
| rs28933668 | pathogenic | 0 | 0 | 0 |
| rs28933669 | pathogenic | 0 | 0 | 0 |
| rs28933670 | pathogenic | 0 | 0 | 0 |
| rs28933671 | pathogenic | 0 | 0 | 0 |
| rs28933672 | pathogenic | 0 | 0 | 0 |
| rs28937272 | pathogenic | 0 | 0 | 0 |
| rs35383156 | benign | 0 | 0 | 0 |
| rs137852418 | pathogenic | 0 | 0 | 0 |
| rs137852419 | pathogenic | 0 | 0 | 0 |
| rs137852428 | pathogenic | 0 | 0 | 0 |
| rs137852436 | pathogenic | 0 | 0 | 0 |
| rs28937285 | pathogenic | 0 | 0 | 0 |
| rs35213384 | benign | 1 | 1 | 1 |
| rs3810950 | benign | 0 | 0 | 0 |
| rs201479289 | pathogenic | 0 | 0 | 0 |
| rs121912820 | pathogenic | 0 | 1 | 0 |
| rs121912815 | pathogenic | 0 | 1 | 0 |
| rs115510708 | benign | 0 | 0 | 0 |

| | | | | |
|---|---|---|---|---|
| rs868749 | benign | 0 | 0 | 0 |
| rs121912823 | pathogenic | 0 | 0 | 0 |
| rs115212829 | benign | 0 | 0 | 0 |
| rs121912822 | pathogenic | 0 | 0 | 0 |
| rs121912816 | pathogenic | 0 | 1 | 0 |
| rs4838544 | benign | 0 | 0 | 0 |
| rs121912818 | pathogenic | 0 | 0 | 0 |
| rs121912821 | pathogenic | 0 | 0 | 0 |
| rs121912817 | pathogenic | 0 | 0 | 0 |
| rs8178992 | benign | 0 | 0 | 0 |
| rs121912819 | pathogenic | 1 | 1 | 0 |
| rs80097077 | benign | 1 | 1 | 0 |
| rs121909359 | pathogenic | 1 | 1 | 0 |
| rs121909358 | pathogenic | 1 | 1 | 1 |
| rs121909361 | pathogenic | 1 | 1 | 1 |
| rs121909357 | pathogenic | 0 | 1 | 0 |
| rs121909365 | pathogenic | 1 | 1 | 1 |
| rs6413484 | pathogenic | 1 | 1 | 1 |
| rs121909373 | pathogenic | 1 | 1 | 1 |
| rs121909366 | pathogenic | 1 | 1 | 1 |
| rs121909367 | pathogenic | 1 | 1 | 1 |
| rs121909368 | pathogenic | 1 | 1 | 1 |
| rs121909369 | pathogenic | 0 | 0 | 0 |
| rs121909363 | pathogenic | 1 | 1 | 1 |
| rs45588036 | pathogenic | 1 | 1 | 1 |
| rs121909364 | pathogenic | 1 | 1 | 1 |
| rs80358282 | benign | 0 | 0 | 0 |
| rs80358284 | pathogenic | 1 | 1 | 1 |
| rs61735303 | benign | 0 | 0 | 0 |
| rs2229274 | benign | 0 | 0 | 0 |
| rs145006491 | benign | 0 | 0 | 0 |
| rs121913580 | pathogenic | 1 | 1 | 0 |
| rs730882196 | pathogenic | 0 | 0 | 0 |
| rs153477 | benign | 0 | 0 | 0 |
| rs153478 | benign | 1 | 1 | 0 |
| rs61740602 | benign | 1 | 1 | 0 |
| rs104893892 | pathogenic | 1 | 1 | 0 |
| rs121909595 | pathogenic | 1 | 1 | 1 |
| rs28931605 | pathogenic | 1 | 1 | 1 |
| rs121909597 | pathogenic | 0 | 1 | 0 |
| rs202233735 | pathogenic | 0 | 1 | 0 |
| rs121909596 | pathogenic | 1 | 1 | 1 |
| rs75156162 | benign | 0 | 1 | 0 |
| rs2305430 | benign | 1 | 1 | 1 |
| rs150318966 | benign | 0 | 0 | 0 |
| rs121909598 | pathogenic | 0 | 1 | 0 |
| rs5744175 | benign | 0 | 0 | 0 |
| rs28935476 | pathogenic | 1 | 1 | 1 |
| rs28935475 | pathogenic | 1 | 1 | 1 |
| rs122461163 | pathogenic | 1 | 1 | 1 |
| rs62626305 | pathogenic | 1 | 1 | 1 |
| rs587777157 | pathogenic | 0 | 0 | 0 |
| rs137854478 | pathogenic | 1 | 1 | 0 |
| rs137854465 | pathogenic | 0 | 0 | 0 |

| | | | | |
|---|---|---|---|---|
| rs137854470 | pathogenic | 0 | 0 | 0 |
| rs137854482 | pathogenic | 0 | 0 | 0 |
| rs137854473 | pathogenic | 0 | 0 | 0 |
| rs137854456 | pathogenic | 0 | 0 | 0 |
| rs140598 | benign | 0 | 0 | 0 |
| rs104894105 | pathogenic | 1 | 1 | 1 |
| rs104894107 | pathogenic | 1 | 1 | 1 |
| rs104894106 | pathogenic | 1 | 1 | 1 |
| rs74621026 | benign | 1 | 1 | 1 |
| rs2290332 | benign | 0 | 0 | 0 |
| rs780202604 | pathogenic | 1 | 1 | 1 |
| rs74315305 | pathogenic | 1 | 1 | 0 |
| rs766751878 | pathogenic | 1 | 1 | 1 |
| rs137854603 | pathogenic | 1 | 1 | 1 |
| rs139592595 | benign | 1 | 1 | 0 |
| rs150565592 | benign | 1 | 1 | 1 |
| rs763520126 | benign | 0 | 0 | 0 |
| rs55980502 | benign | 0 | 0 | 0 |
| rs137852812 | pathogenic | 0 | 0 | 0 |
| rs137852813 | pathogenic | 0 | 0 | 0 |
| rs190222208 | benign | 0 | 0 | 0 |
| rs138459502 | pathogenic | 0 | 0 | 0 |
| rs730881041 | pathogenic | 0 | 0 | 0 |
| rs267607080 | pathogenic | 0 | 0 | 0 |
| rs137852814 | pathogenic | 0 | 0 | 0 |
| rs267607079 | pathogenic | 0 | 0 | 0 |
| rs192378417 | benign | 0 | 0 | 0 |
| rs1288775 | benign | 0 | 0 | 0 |
| rs80338737 | pathogenic | 1 | 1 | 1 |
| rs397514708 | pathogenic | 0 | 0 | 0 |
| rs775933965 | pathogenic | 0 | 0 | 0 |
| rs747608698 | benign | 0 | 1 | 0 |
| rs1145086 | benign | 0 | 1 | 0 |
| rs121912953 | pathogenic | 1 | 1 | 0 |
| rs243849 | benign | 0 | 0 | 0 |
| rs121912955 | pathogenic | 1 | 1 | 1 |
| rs17859943 | benign | 0 | 1 | 0 |
| rs755557988 | benign | 1 | 1 | 1 |
| rs10195 | benign | 0 | 0 | 0 |
| rs61735357 | benign | 1 | 1 | 1 |
| rs137882374 | benign | 0 | 0 | 0 |
| rs5388 | benign | 0 | 0 | 0 |
| rs6352 | benign | 0 | 0 | 0 |
| rs76645461 | pathogenic | 1 | 1 | 0 |
| rs1050828 | pathogenic | 0 | 0 | 0 |
| rs1050829 | pathogenic | 0 | 0 | 0 |
| rs137852314 | pathogenic | 0 | 0 | 0 |
| rs137852331 | pathogenic | 0 | 0 | 0 |
| rs5030872 | pathogenic | 0 | 0 | 0 |
| rs267606836 | pathogenic | 0 | 0 | 0 |
| rs5030868 | pathogenic | 0 | 0 | 0 |
| rs137852330 | pathogenic | 0 | 1 | 0 |
| rs137852326 | pathogenic | 1 | 1 | 1 |
| rs137852318 | pathogenic | 1 | 1 | 1 |

| | | | | |
|---|---|---|---|---|
| rs74575103 | benign | 0 | 0 | 0 |
| rs137852327 | pathogenic | 1 | 1 | 1 |
| rs76723693 | pathogenic | 0 | 0 | 0 |
| rs5030869 | pathogenic | 0 | 0 | 0 |
| rs34193178 | pathogenic | 1 | 1 | 1 |
| rs137852316 | pathogenic | 1 | 1 | 0 |
| rs398123546 | pathogenic | 1 | 1 | 1 |
| rs72554665 | pathogenic | 0 | 0 | 0 |
| rs72554664 | pathogenic | 0 | 0 | 0 |
| rs28939068 | pathogenic | 1 | 1 | 1 |
| rs59301204 | pathogenic | 1 | 1 | 1 |
| rs143715750 | benign | 1 | 1 | 1 |
| rs35263902 | pathogenic | 1 | 1 | 1 |
| rs202214518 | benign | 1 | 1 | 1 |
| rs34076756 | benign | 0 | 0 | 0 |
| rs199592341 | pathogenic | 1 | 1 | 0 |
| rs121908833 | pathogenic | 1 | 1 | 1 |
| rs121908834 | pathogenic | 1 | 1 | 0 |
| rs138454151 | benign | 0 | 0 | 0 |
| rs141323599 | benign | 0 | 0 | 0 |
| rs137853200 | pathogenic | 0 | 0 | 0 |
| rs137853199 | pathogenic | 0 | 0 | 0 |
| rs139787163 | benign | 0 | 0 | 0 |
| rs61731097 | benign | 1 | 1 | 1 |
| rs61729795 | benign | 0 | 0 | 0 |
| rs149242330 | benign | 0 | 0 | 0 |
| rs587777490 | pathogenic | 1 | 1 | 1 |
| rs398123011 | pathogenic | 1 | 1 | 0 |
| rs775186117 | pathogenic | 0 | 0 | 0 |
| rs72677237 | benign | 0 | 0 | 0 |
| rs104894163 | pathogenic | 1 | 1 | 1 |
| rs121909588 | pathogenic | 0 | 0 | 0 |
| rs1042579 | benign | 0 | 0 | 0 |
| rs121909273 | pathogenic | 0 | 0 | 0 |
| rs121909274 | pathogenic | 0 | 0 | 0 |
| rs119473034 | pathogenic | 1 | 1 | 1 |
| rs398123065 | pathogenic | 0 | 0 | 0 |
| rs587777042 | pathogenic | 1 | 1 | 0 |
| rs761413864 | pathogenic | 1 | 1 | 1 |
| rs118203965 | pathogenic | 0 | 0 | 0 |
| rs118203966 | pathogenic | 0 | 0 | 0 |
| rs1801201 | benign | 1 | 1 | 0 |
| rs1136201 | benign | 1 | 1 | 1 |
| rs144148008 | benign | 0 | 0 | 0 |
| rs200118378 | benign | 1 | 1 | 1 |
| rs112984118 | benign | 1 | 1 | 1 |
| rs149446102 | benign | 1 | 1 | 1 |
| rs1427463 | benign | 0 | 0 | 0 |
| rs201936720 | benign | 1 | 1 | 1 |
| rs61751983 | benign | 1 | 1 | 1 |
| rs104894632 | pathogenic | 1 | 1 | 1 |
| rs149202905 | benign | 0 | 0 | 0 |
| rs2230210 | benign | 1 | 1 | 1 |
| rs111631988 | benign | 0 | 0 | 0 |

| | | | | |
|---|---|---|---|---|
| rs137852550 | pathogenic | 0 | 0 | 0 |
| rs5963403 | benign | 0 | 0 | 0 |
| rs138018739 | benign | 0 | 0 | 0 |
| rs62640590 | benign | 0 | 0 | 0 |
| rs121909581 | pathogenic | 1 | 1 | 1 |
| rs202017913 | benign | 1 | 1 | 1 |
| rs143163513 | benign | 0 | 0 | 0 |
| rs238238 | benign | 0 | 1 | 0 |
| rs143945974 | benign | 0 | 0 | 0 |
| rs238239 | benign | 1 | 1 | 1 |
| rs121918403 | pathogenic | 0 | 1 | 0 |
| rs61735456 | benign | 0 | 1 | 0 |
| rs121918404 | pathogenic | 1 | 1 | 0 |
| rs12674488 | benign | 0 | 0 | 0 |
| rs201599657 | pathogenic | 0 | 0 | 0 |
| rs1057090 | benign | 0 | 0 | 0 |
| rs1057091 | benign | 0 | 0 | 0 |
| rs587783740 | benign | 0 | 0 | 0 |
| rs75956769 | benign | 0 | 0 | 0 |
| rs201691299 | benign | 0 | 0 | 0 |
| rs62621087 | benign | 0 | 0 | 0 |
| rs201308454 | benign | 0 | 0 | 0 |
| rs145657360 | benign | 0 | 0 | 0 |
| rs121434592 | pathogenic | 1 | 1 | 0 |
| rs375650413 | benign | 0 | 0 | 0 |
| rs121434437 | pathogenic | 1 | 1 | 0 |
| rs763494292 | pathogenic | 0 | 0 | 0 |
| rs121964997 | pathogenic | 1 | 1 | 0 |
| rs121964996 | pathogenic | 0 | 0 | 0 |
| rs41283425 | benign | 1 | 1 | 1 |
| rs376289528 | benign | 1 | 1 | 1 |
| rs144171604 | benign | 1 | 1 | 1 |
| rs782440692 | pathogenic | 1 | 1 | 1 |
| rs782058451 | pathogenic | 1 | 1 | 1 |
| rs78437817 | benign | 0 | 0 | 0 |
| rs200327969 | benign | 1 | 1 | 1 |
| rs143043662 | benign | 0 | 0 | 0 |
| rs111966833 | benign | 1 | 1 | 1 |
| rs35523678 | benign | 0 | 0 | 0 |
| rs515726208 | pathogenic | 0 | 0 | 0 |
| rs397517239 | pathogenic | 1 | 1 | 1 |
| rs148966602 | benign | 1 | 1 | 1 |
| rs76371972 | benign | 1 | 1 | 1 |
| rs2229968 | benign | 1 | 1 | 1 |
| rs61751538 | benign | 1 | 1 | 1 |
| rs200800978 | pathogenic | 1 | 1 | 1 |
| rs148695964 | benign | 1 | 1 | 1 |
| rs140042418 | benign | 1 | 1 | 1 |
| rs116185403 | benign | 0 | 0 | 0 |
| rs142828669 | benign | 1 | 1 | 1 |
| rs139832048 | benign | 0 | 1 | 0 |
| rs58750568 | benign | 0 | 1 | 0 |
| rs77681311 | benign | 0 | 0 | 0 |
| rs151208452 | benign | 0 | 0 | 0 |

| | | | | |
|---|---|---|---|---|
| rs76463933 | benign | 0 | 0 | 0 |
| rs144434647 | benign | 0 | 0 | 0 |
| rs117361485 | benign | 0 | 0 | 0 |
| rs587777588 | pathogenic | 0 | 0 | 0 |
| rs80338796 | pathogenic | 0 | 0 | 0 |
| rs121434594 | pathogenic | 0 | 0 | 0 |
| rs146766138 | pathogenic | 1 | 1 | 1 |
| rs28939679 | pathogenic | 1 | 1 | 1 |
| rs748754134 | pathogenic | 1 | 1 | 0 |
| rs764276946 | pathogenic | 1 | 1 | 1 |
| rs111033588 | pathogenic | 1 | 1 | 1 |
| rs201633414 | pathogenic | 1 | 1 | 1 |
| rs121909262 | pathogenic | 1 | 1 | 1 |
| rs104893702 | pathogenic | 1 | 1 | 1 |
| rs121909265 | pathogenic | 1 | 1 | 1 |
| rs121909266 | pathogenic | 1 | 1 | 1 |
| rs193922439 | pathogenic | 1 | 1 | 0 |
| rs104893691 | pathogenic | 1 | 1 | 1 |
| rs104893695 | pathogenic | 0 | 0 | 0 |
| rs104893708 | pathogenic | 1 | 1 | 1 |
| rs121909260 | pathogenic | 0 | 0 | 0 |
| rs104893696 | pathogenic | 0 | 0 | 0 |
| rs121909267 | pathogenic | 0 | 0 | 0 |
| rs121909263 | pathogenic | 0 | 0 | 0 |
| rs121909264 | pathogenic | 0 | 0 | 0 |
| rs104893694 | pathogenic | 1 | 1 | 1 |
| rs121909268 | pathogenic | 1 | 1 | 1 |
| rs104893689 | pathogenic | 1 | 1 | 0 |
| rs104893707 | pathogenic | 1 | 1 | 0 |
| rs104893697 | pathogenic | 1 | 1 | 1 |
| rs28936684 | pathogenic | 1 | 1 | 1 |
| rs62269092 | benign | 1 | 1 | 0 |
| rs121909259 | pathogenic | 1 | 1 | 0 |
| rs548403340 | benign | 1 | 1 | 0 |
| rs142818334 | benign | 0 | 0 | 0 |
| rs12493789 | benign | 1 | 1 | 1 |
| rs193922423 | pathogenic | 1 | 1 | 0 |
| rs104893719 | pathogenic | 1 | 1 | 1 |
| rs117375173 | benign | 1 | 1 | 0 |
| rs104893712 | pathogenic | 0 | 0 | 0 |
| rs121917776 | pathogenic | 1 | 1 | 1 |
| rs767325003 | benign | 1 | 1 | 1 |
| rs35824453 | benign | 0 | 0 | 0 |
| rs141641266 | benign | 0 | 0 | 0 |
| rs1050239 | benign | 0 | 0 | 0 |
| rs141303384 | benign | 0 | 0 | 0 |
| rs12720458 | pathogenic | 0 | 0 | 0 |
| rs368507376 | pathogenic | 1 | 1 | 1 |
| rs17215500 | pathogenic | 1 | 1 | 1 |
| rs120074188 | pathogenic | 1 | 1 | 1 |
| rs397508097 | pathogenic | 1 | 1 | 1 |
| rs138551008 | pathogenic | 0 | 0 | 0 |
| rs387907139 | pathogenic | 1 | 1 | 1 |
| rs765487627 | pathogenic | 0 | 0 | 0 |

| | | | | |
|---|---|---|---|---|
| rs587778587 | benign | 0 | 0 | 0 |
| rs45568339 | benign | 0 | 0 | 0 |
| rs78179744 | benign | 0 | 0 | 0 |
| rs45624036 | benign | 0 | 0 | 0 |
| rs61755166 | benign | 0 | 0 | 0 |
| rs180177132 | pathogenic | 1 | 1 | 1 |
| rs587776527 | pathogenic | 0 | 0 | 0 |
| rs373783514 | benign | 0 | 0 | 0 |
| rs730881896 | benign | 0 | 0 | 0 |
| rs118203998 | pathogenic | 0 | 0 | 0 |
| rs41281039 | pathogenic | 1 | 1 | 1 |
| rs118204430 | pathogenic | 1 | 1 | 1 |
| rs1800546 | pathogenic | 1 | 1 | 0 |
| rs76917243 | pathogenic | 0 | 1 | 0 |
| rs78340951 | pathogenic | 0 | 1 | 0 |
| rs77718928 | pathogenic | 0 | 0 | 0 |
| rs369586696 | pathogenic | 0 | 1 | 0 |
| rs745517517 | pathogenic | 1 | 1 | 0 |
| rs372872777 | pathogenic | 0 | 0 | 0 |
| rs748704459 | pathogenic | 0 | 0 | 0 |
| rs769215411 | pathogenic | 0 | 0 | 0 |
| rs139425890 | pathogenic | 0 | 0 | 0 |
| rs779694939 | pathogenic | 1 | 1 | 0 |
| rs773729410 | pathogenic | 0 | 0 | 0 |
| rs121908623 | pathogenic | 1 | 1 | 1 |
| rs121908622 | pathogenic | 1 | 1 | 1 |
| rs121908621 | pathogenic | 1 | 1 | 1 |
| rs762106720 | pathogenic | 1 | 1 | 0 |
| rs199877522 | pathogenic | 0 | 0 | 0 |
| rs74315291 | pathogenic | 1 | 1 | 1 |
| rs79137415 | benign | 0 | 0 | 0 |
| rs121913045 | pathogenic | 0 | 0 | 0 |
| rs34295337 | pathogenic | 0 | 0 | 0 |
| rs121913046 | pathogenic | 0 | 0 | 0 |
| rs121913047 | pathogenic | 0 | 0 | 0 |
| rs121913048 | pathogenic | 0 | 0 | 0 |
| rs267607166 | pathogenic | 0 | 0 | 0 |
| rs17850684 | benign | 1 | 1 | 1 |
| rs119476051 | pathogenic | 1 | 1 | 1 |
| rs17850683 | benign | 1 | 1 | 1 |
| rs119476049 | pathogenic | 1 | 1 | 1 |
| rs119476046 | pathogenic | 1 | 1 | 1 |
| rs119476048 | pathogenic | 0 | 0 | 0 |
| rs119476047 | pathogenic | 1 | 1 | 1 |
| rs28939094 | pathogenic | 1 | 1 | 1 |
| rs119476050 | pathogenic | 0 | 0 | 0 |
| rs387906941 | pathogenic | 1 | 1 | 1 |
| rs41511344 | pathogenic | 1 | 1 | 0 |
| rs748112833 | pathogenic | 1 | 1 | 1 |
| rs74342080 | benign | 1 | 1 | 0 |
| rs116482870 | benign | 0 | 0 | 0 |
| rs104893743 | pathogenic | 0 | 0 | 0 |
| rs117179004 | benign | 0 | 0 | 0 |
| rs587779409 | pathogenic | 1 | 1 | 1 |

| | | | | |
|---|---|---|---|---|
| rs3745406 | benign | 1 | 1 | 0 |
| rs387906696 | pathogenic | 0 | 0 | 0 |
| rs72549369 | pathogenic | 1 | 1 | 0 |
| rs45523133 | benign | 0 | 0 | 0 |
| rs119460972 | pathogenic | 0 | 1 | 0 |
| rs34929837 | benign | 0 | 1 | 0 |
| rs17323225 | benign | 0 | 1 | 0 |
| rs17011368 | benign | 0 | 0 | 0 |
| rs748902766 | pathogenic | 1 | 1 | 1 |
| rs387907292 | pathogenic | 0 | 0 | 0 |
| rs387907293 | pathogenic | 1 | 1 | 0 |
| rs371526758 | pathogenic | 1 | 1 | 1 |
| rs142968179 | pathogenic | 1 | 1 | 1 |
| rs387907291 | pathogenic | 1 | 1 | 0 |
| rs121907949 | pathogenic | 0 | 0 | 0 |
| rs121907947 | pathogenic | 0 | 0 | 0 |
| rs121907950 | pathogenic | 1 | 1 | 1 |
| rs121907948 | pathogenic | 1 | 1 | 1 |
| rs4926 | benign | 0 | 0 | 0 |
| rs748809996 | pathogenic | 0 | 0 | 0 |
| rs1799883 | benign | 1 | 1 | 0 |
| rs104893974 | pathogenic | 0 | 1 | 0 |
| rs104893973 | pathogenic | 0 | 1 | 0 |
| rs34452707 | benign | 0 | 0 | 0 |
| rs119467003 | pathogenic | 1 | 1 | 1 |
| rs121908261 | pathogenic | 0 | 1 | 0 |
| rs28933985 | pathogenic | 1 | 1 | 1 |
| rs80356669 | pathogenic | 1 | 1 | 1 |
| rs28936677 | pathogenic | 1 | 1 | 0 |
| rs121918326 | pathogenic | 0 | 0 | 0 |
| rs121918324 | pathogenic | 1 | 1 | 0 |
| rs121918325 | pathogenic | 1 | 1 | 0 |
| rs41270025 | pathogenic | 1 | 1 | 0 |
| rs121918323 | pathogenic | 0 | 0 | 0 |
| rs12735723 | pathogenic | 1 | 1 | 0 |
| rs36013429 | benign | 0 | 0 | 0 |
| rs28936676 | pathogenic | 1 | 1 | 0 |
| rs771809901 | pathogenic | 1 | 1 | 1 |
| rs758350837 | benign | 1 | 1 | 0 |
| rs121908192 | pathogenic | 1 | 1 | 1 |
| rs147568500 | pathogenic | 1 | 1 | 1 |
| rs3729734 | benign | 0 | 0 | 0 |
| rs371536347 | benign | 1 | 1 | 1 |
| rs80358193 | pathogenic | 1 | 1 | 1 |
| rs119476044 | pathogenic | 1 | 1 | 1 |
| rs111292798 | benign | 1 | 1 | 1 |
| rs121908063 | pathogenic | 1 | 1 | 0 |
| rs121908065 | pathogenic | 1 | 1 | 1 |
| rs121908062 | pathogenic | 1 | 1 | 1 |
| rs121908064 | pathogenic | 1 | 1 | 1 |
| rs61746421 | benign | 1 | 1 | 1 |
| rs34886916 | benign | 1 | 1 | 1 |
| rs118204045 | pathogenic | 0 | 0 | 0 |
| rs138607412 | benign | 1 | 1 | 1 |

| | | | | |
|---|---|---|---|---|
| rs118204046 | pathogenic | 0 | 0 | 0 |
| rs118204048 | pathogenic | 0 | 0 | 0 |
| rs118204044 | pathogenic | 0 | 0 | 0 |
| rs149664056 | benign | 0 | 0 | 0 |
| rs121909285 | pathogenic | 0 | 0 | 0 |
| rs1802498 | benign | 0 | 0 | 0 |
| rs1065761 | benign | 0 | 0 | 0 |
| rs119477054 | pathogenic | 0 | 0 | 0 |
| rs12720097 | benign | 0 | 0 | 0 |
| rs747281324 | pathogenic | 1 | 1 | 0 |
| rs367543028 | pathogenic | 1 | 1 | 0 |
| rs146077918 | benign | 0 | 0 | 0 |
| rs142551229 | benign | 0 | 0 | 0 |
| rs367543039 | pathogenic | 0 | 0 | 0 |
| rs587779884 | pathogenic | 1 | 1 | 0 |
| rs137853153 | pathogenic | 0 | 0 | 0 |
| rs367543029 | pathogenic | 0 | 0 | 0 |
| rs587783037 | pathogenic | 1 | 1 | 0 |
| rs142928725 | benign | 0 | 0 | 0 |
| rs28385141 | benign | 0 | 0 | 0 |
| rs587779888 | pathogenic | 0 | 0 | 0 |
| rs121909801 | pathogenic | 1 | 1 | 1 |
| rs121909790 | pathogenic | 1 | 1 | 0 |
| rs121909797 | pathogenic | 1 | 1 | 0 |
| rs121909794 | pathogenic | 1 | 1 | 1 |
| rs121909791 | pathogenic | 1 | 1 | 1 |
| rs121909793 | pathogenic | 1 | 1 | 1 |
| rs147977279 | benign | 1 | 1 | 1 |
| rs147352564 | benign | 0 | 0 | 0 |
| rs368758045 | benign | 1 | 1 | 1 |
| rs114498261 | benign | 1 | 1 | 0 |
| rs35484156 | benign | 0 | 0 | 0 |
| rs139176878 | benign | 0 | 0 | 0 |
| rs11543934 | benign | 0 | 0 | 0 |
| rs34192549 | benign | 0 | 0 | 0 |
| rs568736295 | benign | 0 | 0 | 0 |
| rs55747232 | benign | 0 | 0 | 0 |
| rs781887272 | benign | 0 | 0 | 0 |
| rs146273341 | benign | 0 | 0 | 0 |
| rs41473544 | benign | 0 | 0 | 0 |
| rs137852680 | pathogenic | 0 | 0 | 0 |
| rs55891000 | benign | 0 | 0 | 0 |
| rs140351189 | benign | 0 | 0 | 0 |
| rs137852679 | pathogenic | 1 | 1 | 0 |
| rs137852678 | pathogenic | 0 | 0 | 0 |
| rs137852677 | pathogenic | 1 | 1 | 1 |
| rs138620818 | benign | 0 | 0 | 0 |
| rs121908531 | pathogenic | 0 | 0 | 0 |
| rs72554307 | pathogenic | 0 | 0 | 0 |
| rs72554308 | pathogenic | 0 | 0 | 0 |
| rs72554312 | pathogenic | 1 | 1 | 0 |
| rs67939655 | pathogenic | 0 | 0 | 0 |
| rs67486158 | pathogenic | 0 | 0 | 0 |
| rs72554338 | pathogenic | 1 | 1 | 1 |

| | | | | |
|---|---|---|---|---|
| rs1800324 | pathogenic | 1 | 1 | 1 |
| rs72554356 | pathogenic | 1 | 1 | 1 |
| rs66656800 | pathogenic | 1 | 1 | 1 |
| rs68026851 | pathogenic | 1 | 1 | 1 |
| rs72558412 | pathogenic | 0 | 0 | 0 |
| rs72558415 | pathogenic | 0 | 0 | 0 |
| rs72558423 | pathogenic | 0 | 0 | 0 |
| rs67120076 | pathogenic | 1 | 1 | 1 |
| rs1800328 | pathogenic | 1 | 1 | 1 |
| rs72558451 | pathogenic | 1 | 1 | 1 |
| rs72558454 | pathogenic | 1 | 1 | 1 |
| rs672601372 | pathogenic | 0 | 0 | 0 |
| rs139644798 | pathogenic | 0 | 0 | 0 |
| rs369398935 | pathogenic | 0 | 0 | 0 |
| rs267607035 | pathogenic | 0 | 0 | 0 |
| rs267607036 | pathogenic | 1 | 1 | 1 |
| rs2302566 | benign | 1 | 1 | 0 |
| rs115959591 | benign | 0 | 0 | 0 |
| rs773876739 | pathogenic | 0 | 0 | 0 |
| rs1799967 | benign | 0 | 0 | 0 |
| rs80356968 | benign | 0 | 0 | 0 |
| rs28897695 | benign | 1 | 1 | 1 |
| rs80357314 | benign | 0 | 0 | 0 |
| rs80357169 | benign | 0 | 0 | 0 |
| rs397507239 | benign | 1 | 1 | 1 |
| rs80187739 | pathogenic | 1 | 1 | 1 |
| rs41293459 | pathogenic | 1 | 1 | 1 |
| rs55770810 | pathogenic | 1 | 1 | 1 |
| rs80356860 | pathogenic | 0 | 0 | 0 |
| rs56195342 | benign | 0 | 0 | 0 |
| rs80357442 | benign | 0 | 0 | 0 |
| rs41293463 | pathogenic | 1 | 1 | 1 |
| rs80357069 | pathogenic | 0 | 0 | 0 |
| rs397509275 | benign | 0 | 0 | 0 |
| rs80356920 | benign | 0 | 0 | 0 |
| rs397509284 | pathogenic | 0 | 0 | 0 |
| rs80357286 | benign | 0 | 0 | 0 |
| rs80357274 | benign | 0 | 0 | 0 |
| rs765656957 | benign | 0 | 0 | 0 |
| rs755427809 | benign | 0 | 0 | 0 |
| rs80357322 | benign | 0 | 0 | 0 |
| rs121434430 | pathogenic | 0 | 0 | 0 |
| rs121918450 | pathogenic | 1 | 1 | 1 |
| rs1532268 | benign | 0 | 0 | 0 |
| rs2303080 | benign | 0 | 0 | 0 |
| rs10064631 | benign | 0 | 0 | 0 |
| rs162036 | benign | 0 | 0 | 0 |
| rs2287780 | benign | 0 | 0 | 0 |
| rs16879334 | benign | 0 | 0 | 0 |
| rs137853061 | pathogenic | 1 | 1 | 0 |
| rs10380 | benign | 0 | 0 | 0 |
| rs1494558 | benign | 0 | 0 | 0 |
| rs1494555 | benign | 0 | 0 | 0 |
| rs192994060 | benign | 0 | 0 | 0 |

| | | | | |
|---|---|---|---|---|
| rs147128920 | benign | 1 | 1 | 0 |
| rs201803986 | pathogenic | 0 | 0 | 0 |
| rs746419489 | pathogenic | 0 | 1 | 0 |
| rs764396564 | pathogenic | 0 | 0 | 0 |
| rs758304537 | pathogenic | 0 | 0 | 0 |
| rs267606763 | pathogenic | 0 | 0 | 0 |
| rs267606762 | pathogenic | 0 | 0 | 0 |
| rs150702945 | pathogenic | 1 | 1 | 1 |
| rs202047589 | pathogenic | 0 | 0 | 0 |
| rs755562733 | pathogenic | 0 | 0 | 0 |
| rs267606764 | pathogenic | 1 | 1 | 1 |
| rs138207257 | pathogenic | 1 | 1 | 0 |
| rs149150736 | pathogenic | 1 | 1 | 1 |
| rs777046879 | pathogenic | 0 | 0 | 0 |
| rs104894408 | pathogenic | 1 | 1 | 1 |
| rs1801002 | pathogenic | 1 | 1 | 1 |
| rs28929485 | pathogenic | 1 | 1 | 1 |
| rs104894396 | pathogenic | 1 | 1 | 1 |
| rs371024165 | pathogenic | 0 | 0 | 0 |
| rs35887622 | pathogenic | 1 | 1 | 1 |
| rs587783644 | pathogenic | 1 | 1 | 1 |
| rs72474224 | pathogenic | 1 | 1 | 1 |
| rs111033296 | pathogenic | 1 | 1 | 1 |
| rs104894407 | pathogenic | 1 | 1 | 1 |
| rs72561723 | pathogenic | 1 | 1 | 1 |
| rs104894398 | pathogenic | 1 | 1 | 1 |
| rs28931594 | pathogenic | 1 | 1 | 1 |
| rs104894412 | pathogenic | 1 | 1 | 1 |
| rs111033297 | pathogenic | 1 | 1 | 1 |
| rs104894404 | pathogenic | 1 | 1 | 1 |
| rs104894410 | pathogenic | 1 | 1 | 1 |
| rs104894403 | pathogenic | 1 | 1 | 1 |
| rs397516870 | benign | 0 | 0 | 0 |
| rs200023879 | pathogenic | 1 | 1 | 1 |
| rs121912968 | pathogenic | 0 | 0 | 0 |
| rs104894402 | pathogenic | 1 | 1 | 1 |
| rs28931593 | pathogenic | 1 | 1 | 1 |
| rs104894397 | pathogenic | 0 | 0 | 0 |
| rs80338944 | pathogenic | 0 | 0 | 0 |
| rs199883710 | pathogenic | 0 | 0 | 0 |
| rs727504302 | pathogenic | 0 | 0 | 0 |
| rs145216882 | benign | 1 | 1 | 1 |
| rs781534323 | pathogenic | 1 | 1 | 1 |
| rs111033218 | benign | 1 | 1 | 1 |
| rs104894409 | pathogenic | 0 | 0 | 0 |
| rs80338945 | pathogenic | 1 | 1 | 1 |
| rs111033299 | pathogenic | 0 | 0 | 0 |
| rs143343083 | pathogenic | 1 | 1 | 1 |
| rs80338946 | pathogenic | 0 | 0 | 0 |
| rs150529554 | pathogenic | 0 | 0 | 0 |
| rs111033295 | pathogenic | 0 | 0 | 0 |
| rs111033188 | benign | 0 | 0 | 0 |
| rs397516874 | pathogenic | 0 | 0 | 0 |
| rs76434661 | pathogenic | 0 | 0 | 0 |

| | | | | |
|---|---|---|---|---|
| rs80338948 | pathogenic | 0 | 0 | 0 |
| rs104894401 | pathogenic | 0 | 0 | 0 |
| rs767178508 | pathogenic | 0 | 0 | 0 |
| rs772264564 | pathogenic | 0 | 0 | 0 |
| rs28931592 | pathogenic | 0 | 0 | 0 |
| rs80338949 | pathogenic | 1 | 1 | 1 |
| rs376898963 | benign | 1 | 1 | 1 |
| rs111033360 | benign | 0 | 0 | 0 |
| rs774518779 | pathogenic | 1 | 1 | 1 |
| rs28931595 | pathogenic | 1 | 1 | 1 |
| rs771748289 | pathogenic | 0 | 0 | 0 |
| rs104894406 | pathogenic | 0 | 0 | 0 |
| rs76838169 | benign | 0 | 0 | 0 |
| rs111033294 | pathogenic | 0 | 0 | 0 |
| rs200214298 | pathogenic | 0 | 0 | 0 |
| rs28940283 | pathogenic | 0 | 1 | 0 |
| rs1047883 | benign | 0 | 0 | 0 |
| rs138254425 | benign | 0 | 0 | 0 |
| rs121912592 | pathogenic | 1 | 1 | 0 |
| rs369061090 | pathogenic | 1 | 1 | 0 |
| rs758724746 | pathogenic | 0 | 1 | 0 |
| rs121912595 | pathogenic | 0 | 0 | 0 |
| rs76340296 | benign | 0 | 0 | 0 |
| rs140578009 | benign | 1 | 1 | 1 |
| rs74445297 | benign | 0 | 0 | 0 |
| rs78599516 | benign | 1 | 1 | 0 |
| rs104894035 | pathogenic | 1 | 1 | 0 |
| rs79451216 | benign | 0 | 0 | 0 |
| rs104894036 | pathogenic | 0 | 0 | 0 |
| rs73343757 | benign | 0 | 0 | 0 |
| rs121917722 | pathogenic | 1 | 1 | 1 |
| rs747700126 | pathogenic | 0 | 1 | 0 |
| rs121917725 | pathogenic | 1 | 1 | 1 |
| rs121917721 | pathogenic | 1 | 1 | 1 |
| rs121917723 | pathogenic | 1 | 1 | 0 |
| rs17569 | benign | 1 | 1 | 1 |
| rs2230062 | benign | 1 | 1 | 1 |
| rs17570 | benign | 0 | 0 | 0 |
| rs121917724 | pathogenic | 0 | 1 | 0 |
| rs199422235 | pathogenic | 0 | 0 | 0 |
| rs199422237 | pathogenic | 1 | 1 | 1 |
| rs199422234 | pathogenic | 1 | 1 | 0 |
| rs140015315 | pathogenic | 1 | 1 | 1 |
| rs730880298 | pathogenic | 1 | 1 | 1 |
| rs779692470 | pathogenic | 0 | 0 | 0 |
| rs369152939 | pathogenic | 0 | 0 | 0 |
| rs527236040 | pathogenic | 0 | 0 | 0 |
| rs587776985 | pathogenic | 1 | 1 | 1 |
| rs147077598 | pathogenic | 1 | 1 | 1 |
| rs128620185 | pathogenic | 1 | 1 | 1 |
| rs128620189 | pathogenic | 0 | 0 | 0 |
| rs128621190 | pathogenic | 0 | 0 | 0 |
| rs61754648 | benign | 0 | 0 | 0 |
| rs137852906 | pathogenic | 0 | 0 | 0 |

| | | | | |
|---|---|---|---|---|
| rs76811038 | pathogenic | 0 | 0 | 0 |
| rs149468422 | benign | 0 | 0 | 0 |
| rs5911 | benign | 0 | 0 | 0 |
| rs28942112 | pathogenic | 1 | 1 | 1 |
| rs143275858 | pathogenic | 1 | 1 | 0 |
| rs28362261 | benign | 0 | 0 | 0 |
| rs28362263 | benign | 1 | 1 | 1 |
| rs558707786 | pathogenic | 0 | 0 | 0 |
| rs138775799 | pathogenic | 0 | 0 | 0 |
| rs748170941 | pathogenic | 0 | 0 | 0 |
| rs121908380 | pathogenic | 0 | 0 | 0 |
| rs765123255 | pathogenic | 1 | 1 | 1 |
| rs730881832 | pathogenic | 0 | 1 | 0 |
| rs587782165 | benign | 1 | 1 | 1 |
| rs762307622 | pathogenic | 0 | 0 | 0 |
| rs34612342 | pathogenic | 1 | 1 | 0 |
| rs143353451 | pathogenic | 0 | 1 | 0 |
| rs747993448 | pathogenic | 0 | 1 | 0 |
| rs376561094 | pathogenic | 0 | 0 | 0 |
| rs776487884 | pathogenic | 1 | 1 | 0 |
| rs34126013 | pathogenic | 1 | 1 | 0 |
| rs200495564 | pathogenic | 1 | 1 | 0 |
| rs140342925 | pathogenic | 1 | 1 | 0 |
| rs587780749 | pathogenic | 1 | 1 | 0 |
| rs773087549 | pathogenic | 1 | 1 | 0 |
| rs730881833 | pathogenic | 0 | 0 | 0 |
| rs374950566 | pathogenic | 1 | 1 | 0 |
| rs587780752 | benign | 0 | 0 | 0 |
| rs3219489 | benign | 0 | 0 | 0 |
| rs199742231 | benign | 0 | 0 | 0 |
| rs587780082 | pathogenic | 0 | 0 | 0 |
| rs35881803 | benign | 0 | 0 | 0 |
| rs41305623 | benign | 1 | 1 | 1 |
| rs121917856 | pathogenic | 1 | 1 | 1 |
| rs576284122 | benign | 0 | 0 | 0 |
| rs142036701 | benign | 0 | 0 | 0 |
| rs121917852 | pathogenic | 1 | 1 | 1 |
| rs121917854 | pathogenic | 1 | 1 | 1 |
| rs121917853 | pathogenic | 1 | 1 | 1 |
| rs121917857 | pathogenic | 0 | 1 | 0 |
| rs193922740 | pathogenic | 1 | 1 | 1 |
| rs565779970 | pathogenic | 1 | 1 | 1 |
| rs6723526 | benign | 0 | 0 | 0 |
| rs141094096 | pathogenic | 1 | 1 | 1 |
| rs104894637 | pathogenic | 0 | 0 | 0 |
| rs104894636 | pathogenic | 0 | 0 | 0 |
| rs104894642 | pathogenic | 0 | 0 | 0 |
| rs104894638 | pathogenic | 0 | 0 | 0 |
| rs753666460 | pathogenic | 0 | 0 | 0 |
| rs104894643 | pathogenic | 1 | 1 | 0 |
| rs34520362 | pathogenic | 0 | 0 | 0 |
| rs374621913 | pathogenic | 0 | 0 | 0 |
| rs113641837 | benign | 0 | 0 | 0 |
| rs104894635 | pathogenic | 0 | 0 | 0 |

| | | | | |
|---|---|---|---|---|
| rs143947056 | pathogenic | 0 | 0 | 0 |
| rs138504221 | pathogenic | 0 | 0 | 0 |
| rs9894254 | benign | 0 | 0 | 0 |
| rs104894640 | pathogenic | 1 | 1 | 1 |
| rs58786455 | benign | 0 | 0 | 0 |
| rs144143780 | pathogenic | 0 | 0 | 0 |
| rs62620232 | benign | 0 | 0 | 0 |
| rs34297805 | benign | 1 | 1 | 1 |
| rs104894641 | pathogenic | 0 | 0 | 0 |
| rs148876719 | benign | 0 | 0 | 0 |
| rs104894639 | pathogenic | 0 | 0 | 0 |
| rs7503034 | benign | 0 | 0 | 0 |
| rs201631570 | benign | 0 | 0 | 0 |
| rs150181226 | pathogenic | 0 | 0 | 0 |
| rs377546036 | pathogenic | 0 | 0 | 0 |
| rs267606700 | pathogenic | 0 | 0 | 0 |
| rs267606701 | pathogenic | 1 | 1 | 0 |
| rs267606699 | pathogenic | 1 | 1 | 0 |
| rs9903215 | benign | 0 | 0 | 0 |
| rs139486406 | benign | 0 | 0 | 0 |
| rs143791614 | benign | 0 | 0 | 0 |
| rs573483474 | benign | 0 | 0 | 0 |
| rs2032892 | benign | 0 | 0 | 0 |
| rs137852619 | pathogenic | 0 | 0 | 0 |
| rs200243548 | benign | 0 | 0 | 0 |
| rs115702388 | benign | 0 | 0 | 0 |
| rs138085358 | pathogenic | 0 | 0 | 0 |
| rs121912789 | pathogenic | 1 | 1 | 1 |
| rs763407972 | pathogenic | 1 | 1 | 1 |
| rs147278302 | benign | 0 | 0 | 0 |
| rs374540411 | benign | 0 | 0 | 0 |
| rs121912790 | pathogenic | 1 | 1 | 0 |
| rs752049917 | benign | 0 | 0 | 0 |
| rs139640491 | benign | 1 | 1 | 1 |
| rs138010738 | benign | 1 | 1 | 0 |
| rs201328768 | pathogenic | 1 | 1 | 0 |
| rs2135720 | benign | 1 | 1 | 0 |
| rs201816080 | benign | 1 | 1 | 0 |
| rs754391973 | pathogenic | 0 | 0 | 0 |
| rs139087859 | benign | 0 | 0 | 0 |
| rs6805421 | benign | 0 | 0 | 0 |
| rs146266900 | benign | 0 | 0 | 0 |
| rs397509419 | pathogenic | 0 | 0 | 0 |
| rs149980411 | benign | 0 | 0 | 0 |
| rs747711488 | pathogenic | 1 | 1 | 1 |
| rs149474339 | pathogenic | 1 | 1 | 1 |
| rs758252808 | pathogenic | 1 | 1 | 0 |
| rs745756308 | pathogenic | 0 | 0 | 0 |
| rs4986958 | benign | 0 | 0 | 0 |
| rs74315444 | pathogenic | 0 | 0 | 0 |
| rs61752137 | pathogenic | 0 | 0 | 0 |
| rs8179065 | benign | 1 | 1 | 1 |
| rs758361736 | pathogenic | 1 | 1 | 0 |
| rs139794067 | pathogenic | 1 | 1 | 1 |

| | | | | |
|---|---|---|---|---|
| rs730880954 | pathogenic | 1 | 1 | 1 |
| rs730880955 | pathogenic | 1 | 1 | 1 |
| rs150634297 | pathogenic | 0 | 0 | 0 |
| rs104893750 | pathogenic | 1 | 1 | 1 |
| rs104893748 | pathogenic | 1 | 1 | 1 |
| rs104893749 | pathogenic | 1 | 1 | 1 |
| rs753520553 | pathogenic | 1 | 1 | 0 |
| rs61737294 | benign | 0 | 0 | 0 |
| rs104894601 | pathogenic | 0 | 0 | 0 |
| rs376090795 | pathogenic | 0 | 0 | 0 |
| rs104894596 | pathogenic | 0 | 0 | 0 |
| rs104894595 | pathogenic | 0 | 0 | 0 |
| rs104894598 | pathogenic | 0 | 0 | 0 |
| rs104894597 | pathogenic | 0 | 0 | 0 |
| rs104894591 | pathogenic | 0 | 0 | 0 |
| rs104894593 | pathogenic | 0 | 0 | 0 |
| rs104894594 | pathogenic | 0 | 0 | 0 |
| rs104894590 | pathogenic | 0 | 0 | 0 |
| rs86312 | benign | 0 | 0 | 0 |
| rs77874543 | benign | 0 | 0 | 0 |
| rs121912590 | pathogenic | 1 | 1 | 1 |
| rs121912589 | pathogenic | 1 | 1 | 1 |
| rs121912586 | pathogenic | 1 | 1 | 1 |
| rs121912585 | pathogenic | 1 | 1 | 1 |
| rs121912587 | pathogenic | 1 | 1 | 1 |
| rs762909359 | pathogenic | 0 | 0 | 0 |
| rs28931610 | pathogenic | 0 | 0 | 0 |
| rs104894097 | pathogenic | 1 | 1 | 1 |
| rs775176191 | benign | 0 | 0 | 0 |
| rs199907548 | pathogenic | 0 | 0 | 0 |
| rs104894095 | pathogenic | 1 | 1 | 1 |
| rs104894109 | pathogenic | 1 | 1 | 1 |
| rs372266620 | benign | 1 | 1 | 1 |
| rs104894099 | pathogenic | 0 | 0 | 0 |
| rs137854599 | pathogenic | 1 | 1 | 1 |
| rs104894094 | pathogenic | 0 | 0 | 0 |
| rs104894104 | pathogenic | 0 | 0 | 0 |
| rs6413463 | benign | 1 | 1 | 1 |
| rs104894098 | pathogenic | 0 | 0 | 0 |
| rs6413464 | benign | 0 | 0 | 0 |
| rs3731249 | benign | 0 | 0 | 0 |
| rs16931179 | benign | 0 | 0 | 0 |
| rs2734715 | benign | 0 | 0 | 0 |
| rs759080581 | pathogenic | 0 | 0 | 0 |
| rs1800746 | benign | 0 | 0 | 0 |
| rs202189057 | pathogenic | 0 | 0 | 0 |
| rs121908204 | pathogenic | 0 | 0 | 0 |
| rs756564767 | pathogenic | 0 | 0 | 0 |
| rs34758634 | benign | 1 | 1 | 1 |
| rs28940573 | pathogenic | 0 | 0 | 0 |
| rs119455955 | pathogenic | 1 | 1 | 0 |
| rs752164603 | pathogenic | 1 | 1 | 0 |
| rs763162812 | pathogenic | 0 | 0 | 0 |
| rs119455957 | pathogenic | 0 | 0 | 0 |

| | | | | |
|---|---|---|---|---|
| rs119455958 | pathogenic | 1 | 1 | 0 |
| rs750428882 | pathogenic | 0 | 0 | 0 |
| rs141482368 | benign | 0 | 0 | 0 |
| rs119455953 | pathogenic | 0 | 0 | 0 |
| rs119455954 | pathogenic | 0 | 0 | 0 |
| rs112812685 | benign | 1 | 1 | 0 |
| rs138744051 | benign | 1 | 1 | 0 |
| rs121908200 | pathogenic | 0 | 0 | 0 |
| rs119455956 | pathogenic | 0 | 0 | 0 |
| rs78970155 | benign | 0 | 0 | 0 |
| rs2834167 | benign | 0 | 0 | 0 |
| rs387907326 | pathogenic | 1 | 1 | 1 |
| rs2107732 | benign | 0 | 0 | 0 |
| rs11552377 | benign | 0 | 0 | 0 |
| rs137852843 | pathogenic | 0 | 0 | 0 |
| rs104894783 | pathogenic | 0 | 0 | 0 |
| rs122457137 | pathogenic | 1 | 1 | 1 |
| rs104894779 | pathogenic | 1 | 1 | 1 |
| rs104894786 | pathogenic | 0 | 0 | 0 |
| rs104894784 | pathogenic | 0 | 0 | 0 |
| rs104894785 | pathogenic | 0 | 0 | 0 |
| rs201870761 | pathogenic | 0 | 0 | 0 |
| rs104894781 | pathogenic | 0 | 0 | 0 |
| rs28897728 | benign | 0 | 0 | 0 |
| rs370273690 | pathogenic | 1 | 1 | 0 |
| rs9332739 | benign | 0 | 0 | 0 |
| rs150878060 | benign | 0 | 0 | 0 |
| rs573509224 | benign | 0 | 0 | 0 |
| rs36221133 | benign | 0 | 0 | 0 |
| rs2230009 | benign | 0 | 0 | 0 |
| rs267607008 | pathogenic | 0 | 0 | 0 |
| rs5985 | benign | 1 | 1 | 1 |
| rs5982 | benign | 1 | 1 | 1 |
| rs5987 | benign | 0 | 0 | 0 |
| rs5988 | benign | 0 | 0 | 0 |
| rs121918680 | pathogenic | 1 | 1 | 0 |
| rs121918679 | pathogenic | 1 | 1 | 0 |
| rs7845 | benign | 1 | 1 | 0 |
| rs1049296 | benign | 1 | 1 | 1 |
| rs121918677 | pathogenic | 1 | 1 | 0 |
| rs200086262 | pathogenic | 1 | 1 | 0 |
| rs746930990 | pathogenic | 1 | 1 | 0 |
| rs104893878 | pathogenic | 1 | 1 | 1 |
| rs121912468 | pathogenic | 1 | 1 | 1 |
| rs121912467 | pathogenic | 1 | 1 | 1 |
| rs145351926 | benign | 1 | 1 | 1 |
| rs141212743 | pathogenic | 1 | 1 | 1 |
| rs397515473 | pathogenic | 1 | 1 | 0 |
| rs121918313 | pathogenic | 0 | 1 | 0 |
| rs73015965 | pathogenic | 0 | 0 | 0 |
| rs121918030 | pathogenic | 0 | 1 | 0 |
| rs77782321 | benign | 0 | 0 | 0 |
| rs374661051 | pathogenic | 1 | 1 | 1 |
| rs267606854 | pathogenic | 0 | 0 | 0 |

| | | | | |
|---|---|---|---|---|
| rs41279678 | benign | 0 | 0 | 0 |
| rs61754640 | benign | 0 | 0 | 0 |
| rs199473314 | pathogenic | 0 | 0 | 0 |
| rs137854601 | pathogenic | 0 | 0 | 0 |
| rs137854614 | pathogenic | 0 | 0 | 0 |
| rs137854615 | pathogenic | 0 | 0 | 0 |
| rs199473635 | pathogenic | 0 | 0 | 0 |
| rs121434478 | pathogenic | 1 | 1 | 1 |
| rs775407864 | pathogenic | 1 | 1 | 1 |
| rs12706927 | benign | 0 | 1 | 0 |
| rs775614261 | benign | 0 | 0 | 0 |
| rs768926261 | benign | 0 | 0 | 0 |
| rs114057640 | benign | 0 | 0 | 0 |
| rs387906886 | pathogenic | 1 | 1 | 0 |
| rs138658585 | benign | 0 | 0 | 0 |
| rs4880091 | benign | 0 | 0 | 0 |
| rs121918552 | pathogenic | 1 | 1 | 1 |
| rs10794537 | benign | 0 | 0 | 0 |
| rs121965022 | pathogenic | 1 | 1 | 0 |
| rs121965020 | pathogenic | 0 | 0 | 0 |
| rs758452450 | pathogenic | 0 | 0 | 0 |
| rs747981483 | benign | 0 | 0 | 0 |
| rs58037052 | benign | 0 | 0 | 0 |
| rs121965029 | pathogenic | 1 | 1 | 0 |
| rs147490060 | benign | 0 | 0 | 0 |
| rs3755955 | benign | 0 | 0 | 0 |
| rs148789453 | pathogenic | 0 | 0 | 0 |
| rs121965030 | pathogenic | 0 | 0 | 0 |
| rs76722191 | benign | 0 | 0 | 0 |
| rs199801029 | pathogenic | 0 | 0 | 0 |
| rs121965033 | pathogenic | 0 | 0 | 0 |
| rs368454909 | pathogenic | 1 | 1 | 0 |
| rs6831280 | benign | 0 | 0 | 0 |
| rs121965024 | pathogenic | 0 | 0 | 0 |
| rs121965019 | pathogenic | 0 | 0 | 0 |
| rs11934801 | benign | 0 | 0 | 0 |
| rs73066479 | benign | 0 | 0 | 0 |
| rs121965027 | pathogenic | 0 | 0 | 0 |
| rs121965026 | pathogenic | 1 | 1 | 0 |
| rs121965021 | pathogenic | 0 | 0 | 0 |
| rs398123257 | benign | 0 | 0 | 0 |
| rs121965031 | pathogenic | 0 | 0 | 0 |
| rs202003795 | pathogenic | 0 | 0 | 0 |
| rs559510809 | pathogenic | 0 | 0 | 0 |
| rs762936223 | benign | 0 | 0 | 0 |
| rs137854580 | pathogenic | 1 | 1 | 1 |
| rs112263157 | benign | 0 | 0 | 0 |
| rs2289181 | benign | 0 | 0 | 0 |
| rs138034915 | benign | 0 | 1 | 0 |
| rs528367092 | pathogenic | 0 | 1 | 0 |
| rs1042393 | benign | 0 | 0 | 0 |
| rs370950728 | pathogenic | 0 | 1 | 0 |
| rs530478036 | benign | 0 | 1 | 0 |
| rs374569672 | benign | 0 | 1 | 0 |

| | | | | |
|---|---|---|---|---|
| rs121907944 | pathogenic | 0 | 1 | 0 |
| rs200856561 | pathogenic | 0 | 1 | 0 |
| rs577915581 | pathogenic | 0 | 1 | 0 |
| rs201896815 | pathogenic | 0 | 1 | 0 |
| rs121907945 | pathogenic | 0 | 1 | 0 |
| rs121907940 | pathogenic | 0 | 1 | 0 |
| rs543300039 | pathogenic | 0 | 1 | 0 |
| rs121907936 | pathogenic | 0 | 1 | 0 |
| rs766074609 | pathogenic | 0 | 1 | 0 |
| rs755253527 | pathogenic | 0 | 1 | 0 |
| rs560575383 | pathogenic | 0 | 1 | 0 |
| rs528369909 | benign | 0 | 1 | 0 |
| rs770610356 | pathogenic | 0 | 1 | 0 |
| rs747610090 | pathogenic | 0 | 1 | 0 |
| rs145712232 | benign | 0 | 1 | 0 |
| rs7215458 | benign | 1 | 1 | 0 |
| rs535644999 | pathogenic | 0 | 1 | 0 |
| rs772883420 | pathogenic | 1 | 1 | 0 |
| rs398123169 | pathogenic | 0 | 1 | 0 |
| rs121907937 | pathogenic | 0 | 1 | 0 |
| rs121907941 | pathogenic | 1 | 1 | 0 |
| rs121907942 | pathogenic | 0 | 1 | 0 |
| rs779556619 | pathogenic | 0 | 1 | 0 |
| rs1800307 | benign | 0 | 1 | 0 |
| rs377544304 | pathogenic | 1 | 1 | 0 |
| rs549029029 | pathogenic | 0 | 1 | 0 |
| rs753269119 | pathogenic | 1 | 1 | 1 |
| rs757617999 | pathogenic | 0 | 1 | 0 |
| rs28937909 | pathogenic | 0 | 1 | 0 |
| rs28940868 | pathogenic | 0 | 1 | 0 |
| rs368438393 | pathogenic | 0 | 1 | 0 |
| rs536906561 | pathogenic | 0 | 1 | 0 |
| rs374143224 | pathogenic | 0 | 1 | 0 |
| rs778418246 | pathogenic | 0 | 1 | 0 |
| rs398123172 | pathogenic | 0 | 1 | 0 |
| rs121907938 | pathogenic | 0 | 1 | 0 |
| rs1800312 | pathogenic | 0 | 1 | 0 |
| rs1126690 | benign | 0 | 1 | 0 |
| rs1800314 | benign | 0 | 1 | 0 |
| rs369532274 | pathogenic | 0 | 1 | 0 |
| rs121907943 | pathogenic | 0 | 1 | 0 |
| rs780321415 | pathogenic | 0 | 1 | 0 |
| rs377286472 | benign | 0 | 1 | 0 |
| rs121918406 | pathogenic | 1 | 1 | 1 |
| rs147484870 | benign | 0 | 0 | 0 |
| rs193922671 | pathogenic | 0 | 0 | 0 |
| rs201397978 | benign | 1 | 1 | 1 |
| rs34884895 | benign | 1 | 1 | 1 |
| rs138329459 | benign | 0 | 0 | 0 |
| rs2289366 | benign | 1 | 1 | 1 |
| rs141858334 | benign | 1 | 1 | 1 |
| rs59907886 | benign | 1 | 1 | 1 |
| rs137852653 | pathogenic | 0 | 0 | 0 |
| rs137852652 | pathogenic | 0 | 0 | 0 |

| | | | | |
|---|---|---|---|---|
| rs137852656 | pathogenic | 0 | 0 | 0 |
| rs61739916 | benign | 0 | 0 | 0 |
| rs28936668 | pathogenic | 0 | 0 | 0 |
| rs137852651 | pathogenic | 0 | 0 | 0 |
| rs137852650 | pathogenic | 0 | 0 | 0 |
| rs28936669 | pathogenic | 0 | 0 | 0 |
| rs137852654 | pathogenic | 0 | 0 | 0 |
| rs28936368 | pathogenic | 1 | 1 | 0 |
| rs137852655 | pathogenic | 0 | 0 | 0 |
| rs61752496 | benign | 0 | 0 | 0 |
| rs750098055 | benign | 0 | 0 | 0 |
| rs764149793 | benign | 0 | 0 | 0 |
| rs724160017 | pathogenic | 0 | 1 | 0 |
| rs724160019 | pathogenic | 0 | 0 | 0 |
| rs202219398 | pathogenic | 1 | 1 | 0 |
| rs724160023 | pathogenic | 1 | 1 | 0 |
| rs761953453 | pathogenic | 0 | 0 | 0 |
| rs80338835 | pathogenic | 1 | 1 | 1 |
| rs56284940 | benign | 0 | 0 | 0 |
| rs74575749 | benign | 0 | 0 | 0 |
| rs587777075 | pathogenic | 1 | 1 | 1 |
| rs641153 | benign | 0 | 0 | 0 |
| rs12614 | benign | 0 | 0 | 0 |
| rs117314762 | benign | 0 | 0 | 0 |
| rs537478097 | benign | 1 | 1 | 1 |
| rs144812066 | benign | 0 | 0 | 0 |
| rs4151651 | benign | 0 | 0 | 0 |
| rs138207668 | benign | 1 | 1 | 1 |
| rs149101394 | benign | 1 | 1 | 1 |
| rs4151659 | benign | 1 | 1 | 1 |
| rs45484591 | benign | 1 | 1 | 1 |
| rs761991070 | pathogenic | 0 | 0 | 0 |
| rs755948940 | pathogenic | 0 | 0 | 0 |
| rs137853097 | pathogenic | 0 | 0 | 0 |
| rs775297352 | benign | 0 | 0 | 0 |
| rs28943591 | benign | 0 | 0 | 0 |
| rs191468413 | pathogenic | 1 | 1 | 0 |
| rs11539471 | benign | 1 | 1 | 1 |
| rs11205 | benign | 0 | 0 | 0 |
| rs768409400 | pathogenic | 0 | 0 | 0 |
| rs762013077 | pathogenic | 0 | 0 | 0 |
| rs768474112 | pathogenic | 0 | 0 | 0 |
| rs121965063 | pathogenic | 1 | 1 | 1 |
| rs143648758 | pathogenic | 0 | 0 | 0 |
| rs121965066 | pathogenic | 0 | 0 | 0 |
| rs752907087 | pathogenic | 0 | 0 | 0 |
| rs747702749 | pathogenic | 0 | 0 | 0 |
| rs754087775 | pathogenic | 0 | 0 | 0 |
| rs121918641 | pathogenic | 1 | 1 | 1 |
| rs121918642 | pathogenic | 1 | 1 | 1 |
| rs121918640 | pathogenic | 1 | 1 | 1 |
| rs121918637 | pathogenic | 1 | 1 | 1 |
| rs121918644 | pathogenic | 1 | 1 | 1 |
| rs16840544 | benign | 0 | 0 | 0 |

| | | | | |
|---|---|---|---|---|
| rs1042391 | benign | 1 | 1 | 1 |
| rs104893744 | pathogenic | 1 | 1 | 1 |
| rs120074136 | pathogenic | 0 | 0 | 0 |
| rs1805081 | benign | 1 | 1 | 0 |
| rs80358251 | benign | 0 | 0 | 0 |
| rs774588125 | benign | 0 | 0 | 0 |
| rs120074134 | pathogenic | 0 | 0 | 0 |
| rs139751448 | pathogenic | 0 | 0 | 0 |
| rs77080672 | benign | 0 | 0 | 0 |
| rs372445155 | pathogenic | 1 | 1 | 0 |
| rs13381670 | benign | 0 | 0 | 0 |
| rs1788799 | benign | 0 | 0 | 0 |
| rs376213990 | pathogenic | 0 | 0 | 0 |
| rs757475924 | pathogenic | 0 | 0 | 0 |
| rs80358253 | pathogenic | 0 | 0 | 0 |
| rs1805082 | benign | 0 | 0 | 0 |
| rs120074130 | pathogenic | 0 | 0 | 0 |
| rs768999208 | pathogenic | 0 | 0 | 0 |
| rs34302553 | benign | 0 | 0 | 0 |
| rs28940897 | pathogenic | 0 | 0 | 0 |
| rs1140458 | benign | 0 | 0 | 0 |
| rs143124972 | pathogenic | 0 | 0 | 0 |
| rs120074135 | pathogenic | 0 | 0 | 0 |
| rs543206298 | pathogenic | 0 | 0 | 0 |
| rs120074132 | pathogenic | 0 | 0 | 0 |
| rs759826138 | pathogenic | 0 | 0 | 0 |
| rs28942108 | pathogenic | 0 | 0 | 0 |
| rs80358254 | pathogenic | 0 | 0 | 0 |
| rs80358257 | pathogenic | 0 | 0 | 0 |
| rs781261962 | pathogenic | 0 | 0 | 0 |
| rs28942107 | pathogenic | 0 | 0 | 0 |
| rs80358258 | pathogenic | 0 | 0 | 0 |
| rs80358259 | pathogenic | 1 | 1 | 0 |
| rs141440861 | benign | 0 | 0 | 0 |
| rs28942106 | pathogenic | 0 | 0 | 0 |
| rs778878523 | pathogenic | 0 | 0 | 0 |
| rs28942105 | pathogenic | 0 | 0 | 0 |
| rs61731969 | benign | 0 | 0 | 0 |
| rs200444084 | pathogenic | 0 | 0 | 0 |
| rs113371321 | pathogenic | 0 | 0 | 0 |
| rs369098773 | pathogenic | 0 | 0 | 0 |
| rs35248744 | benign | 0 | 0 | 0 |
| rs758902805 | pathogenic | 0 | 0 | 0 |
| rs120074131 | pathogenic | 0 | 0 | 0 |
| rs1805084 | benign | 0 | 0 | 0 |
| rs117793540 | benign | 0 | 0 | 0 |
| rs530058063 | benign | 0 | 0 | 0 |
| rs1064422 | pathogenic | 1 | 1 | 1 |
| rs104894506 | pathogenic | 1 | 1 | 1 |
| rs104894507 | pathogenic | 1 | 1 | 1 |
| rs104894508 | pathogenic | 1 | 1 | 0 |
| rs28999113 | pathogenic | 1 | 1 | 0 |
| rs281860266 | pathogenic | 0 | 0 | 0 |
| rs104894961 | pathogenic | 1 | 1 | 0 |

| | | | | |
|---|---|---|---|---|
| rs122463168 | pathogenic | 0 | 0 | 0 |
| rs199469624 | pathogenic | 1 | 1 | 1 |
| rs150107747 | benign | 0 | 0 | 0 |
| rs144984628 | pathogenic | 0 | 0 | 0 |
| rs28935171 | pathogenic | 0 | 0 | 0 |
| rs137852443 | pathogenic | 1 | 1 | 0 |
| rs28933674 | pathogenic | 1 | 1 | 1 |
| rs28933675 | pathogenic | 1 | 1 | 1 |
| rs28933676 | pathogenic | 1 | 1 | 1 |
| rs28933677 | pathogenic | 1 | 1 | 1 |
| rs28933678 | pathogenic | 0 | 0 | 0 |
| rs28933679 | pathogenic | 1 | 1 | 1 |
| rs28933680 | pathogenic | 0 | 0 | 0 |
| rs28933681 | pathogenic | 0 | 0 | 0 |
| rs28933682 | pathogenic | 0 | 0 | 0 |
| rs28937294 | pathogenic | 1 | 1 | 0 |
| rs137852459 | pathogenic | 0 | 0 | 0 |
| rs137852461 | pathogenic | 0 | 0 | 0 |
| rs137852464 | pathogenic | 1 | 1 | 1 |
| rs137852465 | pathogenic | 1 | 1 | 1 |
| rs1800297 | benign | 1 | 1 | 0 |
| rs137852374 | pathogenic | 1 | 1 | 1 |
| rs672601312 | pathogenic | 1 | 1 | 1 |
| rs387907196 | pathogenic | 1 | 1 | 1 |
| rs148348866 | benign | 0 | 0 | 0 |
| rs141138948 | pathogenic | 1 | 1 | 1 |
| rs137852685 | pathogenic | 1 | 1 | 0 |
| rs104893900 | pathogenic | 1 | 1 | 0 |
| rs137852686 | pathogenic | 1 | 1 | 1 |
| rs104893906 | pathogenic | 1 | 1 | 0 |
| rs587782207 | pathogenic | 1 | 1 | 1 |
| rs75086406 | pathogenic | 0 | 0 | 0 |
| rs199822819 | pathogenic | 0 | 0 | 0 |
| rs398123166 | pathogenic | 1 | 1 | 1 |
| rs121913123 | pathogenic | 1 | 1 | 1 |
| rs772190176 | pathogenic | 0 | 1 | 0 |
| rs587782216 | pathogenic | 1 | 1 | 1 |
| rs398123159 | pathogenic | 1 | 1 | 1 |
| rs121913122 | pathogenic | 1 | 1 | 1 |
| rs755436052 | pathogenic | 1 | 1 | 1 |
| rs200796606 | pathogenic | 1 | 1 | 1 |
| rs2070080 | pathogenic | 0 | 1 | 0 |
| rs149616320 | benign | 0 | 0 | 0 |
| rs143529458 | benign | 0 | 0 | 0 |
| rs267607077 | pathogenic | 1 | 1 | 1 |
| rs143898031 | benign | 1 | 1 | 0 |
| rs148304857 | benign | 1 | 1 | 1 |
| rs137852809 | pathogenic | 0 | 0 | 0 |
| rs3765623 | benign | 0 | 0 | 0 |
| rs16944397 | benign | 0 | 0 | 0 |
| rs730880167 | benign | 0 | 0 | 0 |
| rs187108957 | benign | 0 | 0 | 0 |
| rs117342470 | benign | 0 | 0 | 0 |
| rs751200138 | pathogenic | 1 | 1 | 1 |

| | | | | |
|---|---|---|---|---|
| rs755799528 | pathogenic | 0 | 0 | 0 |
| rs537484504 | benign | 0 | 0 | 0 |
| rs748944640 | pathogenic | 0 | 0 | 0 |
| rs199774121 | pathogenic | 0 | 0 | 0 |
| rs766094434 | pathogenic | 1 | 1 | 0 |
| rs777321035 | pathogenic | 1 | 1 | 1 |
| rs146269774 | benign | 0 | 1 | 0 |
| rs144085805 | benign | 0 | 1 | 0 |
| rs367643028 | benign | 0 | 0 | 0 |
| rs42524 | benign | 0 | 0 | 0 |
| rs28940586 | pathogenic | 0 | 0 | 0 |
| rs74315327 | pathogenic | 1 | 1 | 1 |
| rs199676861 | pathogenic | 0 | 0 | 0 |
| rs115802744 | benign | 1 | 1 | 0 |
| rs752866643 | pathogenic | 1 | 1 | 1 |
| rs28941784 | pathogenic | 1 | 1 | 1 |
| rs398124434 | pathogenic | 1 | 1 | 1 |
| rs756414548 | pathogenic | 1 | 1 | 1 |
| rs376128990 | pathogenic | 1 | 1 | 1 |
| rs746219370 | pathogenic | 1 | 1 | 1 |
| rs756195708 | pathogenic | 1 | 1 | 1 |
| rs369296618 | pathogenic | 1 | 1 | 1 |
| rs9593 | benign | 1 | 1 | 1 |
| rs139510844 | benign | 0 | 0 | 0 |
| rs28382158 | benign | 0 | 0 | 0 |
| rs28919570 | pathogenic | 0 | 0 | 0 |
| rs587777331 | pathogenic | 0 | 0 | 0 |
| rs587777333 | pathogenic | 0 | 0 | 0 |
| rs587777332 | pathogenic | 0 | 0 | 0 |
| rs774980346 | pathogenic | 0 | 0 | 0 |
| rs370934093 | pathogenic | 0 | 0 | 0 |
| rs121913479 | pathogenic | 1 | 1 | 1 |
| rs121913484 | pathogenic | 0 | 0 | 0 |
| rs121913485 | pathogenic | 1 | 1 | 1 |
| rs28931614 | pathogenic | 1 | 1 | 1 |
| rs17881656 | benign | 1 | 1 | 1 |
| rs28931615 | pathogenic | 1 | 1 | 1 |
| rs147846832 | benign | 0 | 0 | 0 |
| rs80356506 | pathogenic | 0 | 0 | 0 |
| rs80356494 | pathogenic | 0 | 0 | 0 |
| rs121908894 | pathogenic | 0 | 0 | 0 |
| rs121908895 | pathogenic | 0 | 0 | 0 |
| rs80356508 | pathogenic | 0 | 0 | 0 |
| rs80356507 | pathogenic | 0 | 0 | 0 |
| rs121908896 | pathogenic | 0 | 0 | 0 |
| rs745372589 | pathogenic | 1 | 1 | 0 |
| rs183898408 | benign | 0 | 0 | 0 |
| rs200844292 | benign | 0 | 0 | 0 |
| rs146897002 | benign | 0 | 0 | 0 |
| rs121964869 | pathogenic | 1 | 1 | 1 |
| rs201768800 | benign | 1 | 1 | 1 |
| rs9807556 | benign | 1 | 1 | 1 |
| rs184721031 | benign | 0 | 0 | 0 |
| rs201104206 | benign | 0 | 0 | 0 |

| | | | | |
|---|---|---|---|---|
| rs188677538 | benign | 0 | 0 | 0 |
| rs149528866 | benign | 0 | 0 | 0 |
| rs115382168 | benign | 0 | 0 | 0 |
| rs78846023 | benign | 1 | 1 | 1 |
| rs61730662 | benign | 0 | 0 | 0 |
| rs121918368 | pathogenic | 0 | 0 | 0 |
| rs566433112 | pathogenic | 1 | 1 | 1 |
| rs121909342 | pathogenic | 0 | 0 | 0 |
| rs67586389 | pathogenic | 0 | 0 | 0 |
| rs72466485 | pathogenic | 0 | 0 | 0 |
| rs72466486 | pathogenic | 1 | 1 | 1 |
| rs72466487 | pathogenic | 1 | 1 | 1 |
| rs267607112 | pathogenic | 0 | 0 | 0 |
| rs118204013 | pathogenic | 0 | 0 | 0 |
| rs121913495 | pathogenic | 0 | 0 | 0 |
| rs11554273 | pathogenic | 0 | 0 | 0 |
| rs193922477 | pathogenic | 1 | 1 | 1 |
| rs193922479 | pathogenic | 1 | 1 | 1 |
| rs72547554 | pathogenic | 0 | 0 | 0 |
| rs72547561 | pathogenic | 1 | 1 | 1 |
| rs72547562 | pathogenic | 1 | 1 | 0 |
| rs115977487 | benign | 1 | 1 | 0 |
| rs72547564 | pathogenic | 0 | 0 | 0 |
| rs72547568 | pathogenic | 1 | 1 | 0 |
| rs72547569 | pathogenic | 1 | 1 | 1 |
| rs72547571 | pathogenic | 0 | 0 | 0 |
| rs72547575 | pathogenic | 1 | 1 | 1 |
| rs544994462 | benign | 0 | 0 | 0 |
| rs190470017 | benign | 0 | 0 | 0 |
| rs199473266 | pathogenic | 1 | 1 | 1 |
| rs137854602 | pathogenic | 0 | 0 | 0 |
| rs143739249 | pathogenic | 0 | 0 | 0 |
| rs397514697 | pathogenic | 1 | 1 | 1 |
| rs373764886 | pathogenic | 0 | 0 | 0 |
| rs370242930 | pathogenic | 0 | 0 | 0 |
| rs377145777 | pathogenic | 1 | 1 | 1 |
| rs191256186 | benign | 0 | 0 | 0 |
| rs150070901 | benign | 1 | 1 | 1 |
| rs138098032 | benign | 0 | 0 | 0 |
| rs117731813 | benign | 1 | 1 | 1 |
| rs104894488 | pathogenic | 1 | 1 | 1 |
| rs11302 | benign | 0 | 1 | 0 |
| rs121918447 | pathogenic | 1 | 1 | 1 |
| rs138205828 | benign | 1 | 1 | 0 |
| rs149858290 | benign | 0 | 0 | 0 |
| rs750764003 | pathogenic | 0 | 1 | 0 |
| rs141381896 | benign | 0 | 0 | 0 |
| rs149027682 | benign | 0 | 0 | 0 |
| rs121908415 | pathogenic | 0 | 0 | 0 |
| rs121908416 | pathogenic | 0 | 0 | 0 |
| rs121908417 | pathogenic | 0 | 0 | 0 |
| rs587780306 | benign | 0 | 0 | 0 |
| rs3218725 | benign | 0 | 0 | 0 |
| rs3218729 | benign | 0 | 0 | 0 |

| | | | | |
|---|---|---|---|---|
| rs387906917 | pathogenic | 0 | 0 | 0 |
| rs200652608 | pathogenic | 0 | 0 | 0 |
| rs3218721 | benign | 0 | 0 | 0 |
| rs63750869 | pathogenic | 1 | 1 | 1 |
| rs118203910 | pathogenic | 0 | 1 | 0 |
| rs9332701 | benign | 1 | 1 | 0 |
| rs6027 | benign | 0 | 0 | 0 |
| rs121909110 | pathogenic | 0 | 1 | 0 |
| rs762550967 | pathogenic | 1 | 1 | 0 |
| rs763615602 | pathogenic | 1 | 1 | 0 |
| rs144907403 | benign | 0 | 0 | 0 |
| rs104893631 | pathogenic | 1 | 1 | 0 |
| rs144181978 | pathogenic | 1 | 1 | 0 |
| rs74874677 | benign | 1 | 1 | 1 |
| rs104893632 | pathogenic | 0 | 1 | 0 |
| rs371342464 | benign | 1 | 1 | 1 |
| rs104893633 | pathogenic | 1 | 1 | 0 |
| rs201088712 | pathogenic | 0 | 0 | 0 |
| rs201857541 | benign | 1 | 1 | 1 |
| rs139517732 | pathogenic | 1 | 1 | 0 |
| rs191057824 | benign | 1 | 1 | 0 |
| rs35683768 | benign | 1 | 1 | 0 |
| rs104895271 | pathogenic | 0 | 0 | 0 |
| rs104895217 | pathogenic | 1 | 1 | 1 |
| rs104895223 | pathogenic | 1 | 1 | 1 |
| rs104895218 | pathogenic | 1 | 1 | 1 |
| rs104895225 | pathogenic | 1 | 1 | 1 |
| rs4149637 | benign | 1 | 1 | 1 |
| rs104895219 | pathogenic | 1 | 1 | 1 |
| rs104895220 | pathogenic | 1 | 1 | 1 |
| rs104895228 | pathogenic | 1 | 1 | 1 |
| rs104895221 | pathogenic | 1 | 1 | 1 |
| rs104895222 | pathogenic | 1 | 1 | 1 |
| rs104895247 | pathogenic | 0 | 0 | 0 |
| rs28940585 | pathogenic | 0 | 0 | 0 |
| rs561367199 | pathogenic | 1 | 1 | 1 |
| rs143793815 | benign | 0 | 0 | 0 |
| rs28941473 | pathogenic | 1 | 1 | 1 |
| rs751590073 | pathogenic | 1 | 1 | 1 |
| rs111407265 | benign | 1 | 1 | 1 |
| rs373697663 | pathogenic | 1 | 1 | 1 |
| rs369879957 | pathogenic | 0 | 0 | 0 |
| rs28941472 | pathogenic | 1 | 1 | 1 |
| rs28940287 | pathogenic | 1 | 1 | 1 |
| rs77625410 | benign | 0 | 0 | 0 |
| rs111033477 | benign | 0 | 0 | 0 |
| rs115123584 | benign | 0 | 0 | 0 |
| rs374576916 | benign | 0 | 0 | 0 |
| rs397516317 | pathogenic | 0 | 0 | 0 |
| rs368657015 | pathogenic | 0 | 0 | 0 |
| rs397516321 | pathogenic | 0 | 0 | 0 |
| rs397516322 | pathogenic | 0 | 0 | 0 |
| rs111033215 | pathogenic | 0 | 0 | 0 |
| rs199606180 | pathogenic | 0 | 0 | 0 |

| | | | | |
|---|---|---|---|---|
| rs111033192 | pathogenic | 0 | 0 | 0 |
| rs948962 | benign | 1 | 1 | 1 |
| rs142293185 | benign | 0 | 0 | 0 |
| rs376764423 | pathogenic | 0 | 0 | 0 |
| rs773844428 | pathogenic | 0 | 0 | 0 |
| rs186644871 | benign | 0 | 0 | 0 |
| rs200313391 | benign | 0 | 0 | 0 |
| rs377670513 | pathogenic | 0 | 0 | 0 |
| rs1132036 | benign | 0 | 0 | 0 |
| rs111033231 | benign | 0 | 0 | 0 |
| rs145945630 | pathogenic | 0 | 0 | 0 |
| rs121912984 | pathogenic | 1 | 1 | 1 |
| rs1800435 | benign | 0 | 1 | 0 |
| rs121912980 | pathogenic | 0 | 0 | 0 |
| rs121912982 | pathogenic | 1 | 1 | 1 |
| rs121912983 | pathogenic | 0 | 0 | 0 |
| rs121912981 | pathogenic | 0 | 0 | 0 |
| rs1057868 | benign | 0 | 0 | 0 |
| rs56048668 | benign | 0 | 0 | 0 |
| rs55865465 | benign | 0 | 0 | 0 |
| rs748096324 | benign | 1 | 1 | 0 |
| rs150475750 | benign | 0 | 0 | 0 |
| rs143025739 | benign | 1 | 1 | 0 |
| rs79702016 | benign | 1 | 1 | 0 |
| rs41338945 | benign | 0 | 0 | 0 |
| rs146406037 | benign | 1 | 1 | 1 |
| rs143274491 | benign | 0 | 0 | 0 |
| rs3829986 | benign | 0 | 0 | 0 |
| rs149168939 | benign | 0 | 0 | 0 |
| rs1800385 | benign | 0 | 0 | 0 |
| rs1800386 | pathogenic | 0 | 0 | 0 |
| rs121918524 | pathogenic | 1 | 1 | 1 |
| rs121912700 | pathogenic | 0 | 0 | 0 |
| rs104893930 | pathogenic | 1 | 1 | 1 |
| rs104893931 | pathogenic | 0 | 0 | 0 |
| rs116125936 | benign | 0 | 0 | 0 |
| rs627441 | benign | 0 | 0 | 0 |
| rs104894481 | pathogenic | 0 | 0 | 0 |
| rs137853129 | pathogenic | 0 | 0 | 0 |
| rs137853130 | pathogenic | 1 | 1 | 0 |
| rs28940282 | pathogenic | 0 | 0 | 0 |
| rs2229989 | benign | 0 | 0 | 0 |
| rs104894647 | pathogenic | 0 | 0 | 0 |
| rs1061170 | benign | 1 | 1 | 1 |
| rs576402053 | benign | 0 | 0 | 0 |
| rs34972246 | benign | 0 | 0 | 0 |
| rs192725607 | benign | 0 | 0 | 0 |
| rs137852615 | pathogenic | 1 | 1 | 0 |
| rs137852617 | pathogenic | 1 | 1 | 1 |
| rs137852611 | pathogenic | 0 | 0 | 0 |
| rs137852612 | pathogenic | 0 | 0 | 0 |
| rs137852614 | pathogenic | 1 | 1 | 1 |
| rs137852610 | pathogenic | 1 | 1 | 1 |
| rs137852618 | pathogenic | 1 | 1 | 1 |

| | | | | |
|---|---|---|---|---|
| rs137852613 | pathogenic | 1 | 1 | 0 |
| rs137852609 | pathogenic | 0 | 0 | 0 |
| rs63750306 | pathogenic | 1 | 1 | 1 |
| rs63750265 | pathogenic | 1 | 1 | 1 |
| rs63751287 | pathogenic | 1 | 1 | 1 |
| rs199723282 | benign | 0 | 0 | 0 |
| rs121917807 | pathogenic | 0 | 0 | 0 |
| rs121917809 | pathogenic | 0 | 0 | 0 |
| rs75134564 | pathogenic | 0 | 0 | 0 |
| rs267606930 | pathogenic | 1 | 1 | 0 |
| rs121909302 | pathogenic | 0 | 0 | 0 |
| rs121909303 | pathogenic | 0 | 0 | 0 |
| rs121909299 | pathogenic | 0 | 0 | 0 |
| rs121909301 | pathogenic | 0 | 0 | 0 |
| rs121909300 | pathogenic | 1 | 1 | 0 |
| rs111033589 | pathogenic | 0 | 0 | 0 |
| rs137853059 | pathogenic | 0 | 0 | 0 |
| rs137853057 | pathogenic | 1 | 1 | 1 |
| rs1801474 | benign | 0 | 0 | 0 |
| rs137853060 | pathogenic | 1 | 1 | 1 |
| rs137853058 | pathogenic | 1 | 1 | 1 |
| rs34424986 | pathogenic | 0 | 0 | 0 |
| rs199657839 | benign | 1 | 1 | 1 |
| rs56092260 | benign | 0 | 0 | 0 |
| rs1801582 | benign | 1 | 1 | 1 |
| rs1801334 | benign | 1 | 1 | 1 |
| rs191486604 | pathogenic | 0 | 0 | 0 |
| rs121908051 | pathogenic | 0 | 0 | 0 |
| rs121908055 | pathogenic | 1 | 1 | 0 |
| rs28940886 | pathogenic | 0 | 0 | 0 |
| rs121908050 | pathogenic | 0 | 0 | 0 |
| rs28940887 | pathogenic | 0 | 0 | 0 |
| rs387906300 | pathogenic | 0 | 0 | 0 |
| rs28942087 | pathogenic | 0 | 0 | 0 |
| rs121908049 | pathogenic | 1 | 1 | 0 |
| rs121908054 | pathogenic | 1 | 1 | 0 |
| rs121908048 | pathogenic | 0 | 0 | 0 |
| rs28940888 | pathogenic | 1 | 1 | 1 |
| rs121908053 | pathogenic | 1 | 1 | 0 |
| rs122467169 | pathogenic | 1 | 1 | 1 |
| rs122467172 | pathogenic | 1 | 1 | 1 |
| rs122467170 | pathogenic | 1 | 1 | 1 |
| rs28935477 | pathogenic | 1 | 1 | 0 |
| rs28942088 | pathogenic | 0 | 0 | 0 |
| rs751478142 | pathogenic | 1 | 1 | 0 |
| rs765368797 | pathogenic | 0 | 0 | 0 |
| rs147810437 | benign | 0 | 0 | 0 |
| rs121918209 | pathogenic | 1 | 1 | 1 |
| rs121918208 | pathogenic | 1 | 1 | 1 |
| rs121918210 | pathogenic | 1 | 1 | 1 |
| rs35515638 | benign | 1 | 1 | 1 |
| rs121918207 | pathogenic | 1 | 1 | 1 |
| rs141298312 | benign | 0 | 0 | 0 |
| rs121918211 | pathogenic | 0 | 0 | 0 |

| | | | | |
|---|---|---|---|---|
| rs149660059 | benign | 0 | 0 | 0 |
| rs200670286 | pathogenic | 1 | 1 | 1 |
| rs121918212 | pathogenic | 1 | 1 | 1 |
| rs121918213 | pathogenic | 1 | 1 | 1 |
| rs121918205 | pathogenic | 1 | 1 | 1 |
| rs104894025 | pathogenic | 1 | 1 | 1 |
| rs104894026 | pathogenic | 1 | 1 | 1 |
| rs104894028 | pathogenic | 0 | 1 | 0 |
| rs104894029 | pathogenic | 0 | 1 | 0 |
| rs119458971 | pathogenic | 0 | 0 | 0 |
| rs119458969 | pathogenic | 1 | 1 | 1 |
| rs1801591 | benign | 1 | 1 | 1 |
| rs140169311 | benign | 0 | 0 | 0 |
| rs753363278 | benign | 1 | 1 | 1 |
| rs199763682 | pathogenic | 0 | 0 | 0 |
| rs119458970 | pathogenic | 1 | 1 | 1 |
| rs763722868 | pathogenic | 0 | 0 | 0 |
| rs56404770 | benign | 0 | 0 | 0 |
| rs34221241 | benign | 0 | 0 | 0 |
| rs121909625 | pathogenic | 0 | 0 | 0 |
| rs550921485 | pathogenic | 0 | 0 | 0 |
| rs1800454 | benign | 0 | 0 | 0 |
| rs241447 | benign | 0 | 0 | 0 |
| rs2230606 | benign | 1 | 1 | 0 |
| rs5036 | pathogenic | 1 | 1 | 1 |
| rs13306788 | benign | 0 | 0 | 0 |
| rs28929480 | pathogenic | 0 | 0 | 0 |
| rs121912749 | pathogenic | 0 | 0 | 0 |
| rs56361140 | pathogenic | 0 | 0 | 0 |
| rs28931583 | pathogenic | 1 | 1 | 1 |
| rs36211715 | pathogenic | 1 | 1 | 1 |
| rs138049878 | pathogenic | 1 | 1 | 1 |
| rs727503253 | pathogenic | 1 | 1 | 1 |
| rs267606908 | pathogenic | 0 | 0 | 0 |
| rs121913631 | pathogenic | 1 | 1 | 1 |
| rs36211716 | benign | 1 | 1 | 1 |
| rs121913639 | pathogenic | 1 | 1 | 1 |
| rs730880761 | pathogenic | 1 | 1 | 1 |
| rs121913629 | pathogenic | 1 | 1 | 1 |
| rs754368658 | pathogenic | 0 | 0 | 0 |
| rs777090017 | pathogenic | 0 | 0 | 0 |
| rs267606913 | pathogenic | 1 | 1 | 1 |
| rs773138218 | pathogenic | 0 | 0 | 0 |
| rs371295780 | pathogenic | 0 | 0 | 0 |
| rs387906830 | pathogenic | 1 | 1 | 0 |
| rs771364038 | pathogenic | 0 | 1 | 0 |
| rs772731615 | pathogenic | 0 | 0 | 0 |
| rs137853063 | pathogenic | 0 | 0 | 0 |
| rs387907076 | pathogenic | 0 | 1 | 0 |
| rs35460768 | benign | 0 | 0 | 0 |
| rs2278196 | benign | 1 | 1 | 1 |
| rs119103275 | pathogenic | 0 | 0 | 0 |
| rs119103276 | pathogenic | 1 | 1 | 1 |
| rs374402066 | pathogenic | 1 | 1 | 1 |

| | | | | |
|---|---|---|---|---|
| rs119103274 | pathogenic | 0 | 0 | 0 |
| rs2274064 | benign | 0 | 0 | 0 |
| rs267607023 | pathogenic | 1 | 1 | 0 |
| rs142117489 | benign | 1 | 1 | 0 |
| rs371258270 | benign | 0 | 0 | 0 |
| rs34716057 | benign | 0 | 0 | 0 |
| rs201778606 | benign | 0 | 0 | 0 |
| rs201438792 | benign | 0 | 0 | 0 |
| rs140547417 | benign | 0 | 0 | 0 |
| rs5881 | benign | 0 | 0 | 0 |
| rs752298084 | benign | 0 | 0 | 0 |
| rs34855278 | benign | 0 | 0 | 0 |
| rs5880 | benign | 1 | 1 | 0 |
| rs5882 | benign | 0 | 0 | 0 |
| rs1800777 | benign | 0 | 0 | 0 |
| rs577301331 | benign | 0 | 0 | 0 |
| rs137853313 | pathogenic | 0 | 0 | 0 |
| rs28935169 | pathogenic | 0 | 0 | 0 |
| rs137853315 | pathogenic | 0 | 0 | 0 |
| rs137853314 | pathogenic | 0 | 0 | 0 |
| rs28935469 | pathogenic | 0 | 0 | 0 |
| rs137853318 | pathogenic | 0 | 0 | 0 |
| rs28935470 | pathogenic | 0 | 0 | 0 |
| rs3729986 | benign | 0 | 0 | 0 |
| rs730880619 | pathogenic | 0 | 0 | 0 |
| rs201012766 | benign | 0 | 0 | 0 |
| rs727503218 | pathogenic | 0 | 0 | 0 |
| rs11570052 | benign | 0 | 0 | 0 |
| rs730880623 | pathogenic | 0 | 0 | 0 |
| rs138753870 | benign | 0 | 0 | 0 |
| rs397516068 | pathogenic | 0 | 0 | 0 |
| rs3729989 | benign | 1 | 1 | 1 |
| rs397516074 | pathogenic | 0 | 0 | 0 |
| rs561449819 | pathogenic | 0 | 0 | 0 |
| rs121917779 | benign | 0 | 0 | 0 |
| rs142881576 | pathogenic | 0 | 0 | 0 |
| rs77384282 | benign | 0 | 0 | 0 |
| rs201865787 | benign | 0 | 0 | 0 |
| rs142244640 | benign | 0 | 0 | 0 |
| rs200488706 | benign | 0 | 0 | 0 |
| rs137853291 | pathogenic | 1 | 1 | 0 |
| rs28933990 | pathogenic | 1 | 1 | 0 |
| rs140570189 | benign | 0 | 0 | 0 |
| rs201588178 | benign | 0 | 0 | 0 |
| rs104893846 | pathogenic | 0 | 0 | 0 |
| rs754545360 | pathogenic | 0 | 1 | 0 |
| rs104893851 | pathogenic | 0 | 1 | 0 |
| rs104893849 | pathogenic | 0 | 0 | 0 |
| rs571038432 | pathogenic | 1 | 1 | 0 |
| rs2270655 | benign | 1 | 1 | 1 |
| rs315952 | benign | 0 | 0 | 0 |
| rs151176548 | benign | 0 | 0 | 0 |
| rs122459147 | pathogenic | 1 | 1 | 0 |
| rs122458140 | pathogenic | 0 | 0 | 0 |

| | | | | |
|---|---|---|---|---|
| rs122458142 | pathogenic | 1 | 1 | 0 |
| rs267606812 | pathogenic | 1 | 1 | 0 |
| rs267606813 | pathogenic | 1 | 1 | 0 |
| rs122458143 | pathogenic | 0 | 0 | 0 |
| rs122459146 | pathogenic | 0 | 0 | 0 |
| rs122458144 | pathogenic | 0 | 0 | 0 |
| rs122458145 | pathogenic | 0 | 0 | 0 |
| rs113994007 | pathogenic | 1 | 1 | 0 |
| rs758746181 | pathogenic | 1 | 1 | 1 |
| rs145034527 | pathogenic | 1 | 1 | 0 |
| rs119468006 | pathogenic | 1 | 1 | 0 |
| rs201908721 | pathogenic | 0 | 0 | 0 |
| rs755933881 | pathogenic | 1 | 1 | 0 |
| rs771578775 | pathogenic | 0 | 0 | 0 |
| rs17849927 | benign | 0 | 0 | 0 |
| rs139133094 | benign | 0 | 0 | 0 |
| rs12593 | benign | 0 | 0 | 0 |
| rs578189699 | pathogenic | 1 | 1 | 0 |
| rs119468008 | pathogenic | 0 | 0 | 0 |
| rs768524626 | benign | 0 | 0 | 0 |
| rs119468009 | pathogenic | 0 | 0 | 0 |
| rs119468004 | pathogenic | 0 | 0 | 0 |
| rs199874519 | pathogenic | 0 | 0 | 0 |
| rs376478331 | pathogenic | 0 | 0 | 0 |
| rs61995958 | pathogenic | 0 | 0 | 0 |
| rs199714731 | benign | 0 | 0 | 0 |
| rs149185954 | benign | 0 | 0 | 0 |
| rs121918353 | pathogenic | 0 | 0 | 0 |
| rs1131695 | benign | 0 | 0 | 0 |
| rs28939668 | pathogenic | 0 | 0 | 0 |
| rs758432471 | pathogenic | 1 | 1 | 1 |
| rs752746786 | pathogenic | 1 | 1 | 1 |
| rs28935471 | pathogenic | 0 | 0 | 0 |
| rs80338837 | pathogenic | 0 | 0 | 0 |
| rs137853312 | pathogenic | 0 | 0 | 0 |
| rs28935472 | pathogenic | 0 | 0 | 0 |
| rs28935473 | pathogenic | 0 | 0 | 0 |
| rs267606752 | pathogenic | 0 | 0 | 0 |
| rs3917973 | benign | 0 | 0 | 0 |
| rs3918018 | benign | 0 | 0 | 0 |
| rs757976755 | pathogenic | 0 | 0 | 0 |
| rs121918329 | pathogenic | 1 | 1 | 0 |
| rs770114459 | pathogenic | 0 | 0 | 0 |
| rs6662272 | benign | 0 | 0 | 0 |
| rs121918242 | pathogenic | 1 | 1 | 0 |
| rs121918240 | pathogenic | 1 | 1 | 0 |
| rs200094982 | pathogenic | 0 | 0 | 0 |
| rs121918241 | pathogenic | 0 | 0 | 0 |
| rs140522266 | pathogenic | 1 | 1 | 0 |
| rs121918243 | pathogenic | 1 | 1 | 0 |
| rs370596113 | pathogenic | 1 | 1 | 0 |
| rs398124295 | pathogenic | 1 | 1 | 0 |
| rs587776889 | pathogenic | 1 | 1 | 0 |
| rs747527726 | pathogenic | 1 | 1 | 0 |

| | | | | |
|---|---|---|---|---|
| rs112631212 | benign | 0 | 0 | 0 |
| rs104894605 | pathogenic | 0 | 0 | 0 |
| rs104894607 | pathogenic | 0 | 0 | 0 |
| rs104894606 | pathogenic | 0 | 0 | 0 |
| rs143873873 | benign | 0 | 0 | 0 |
| rs104894388 | pathogenic | 1 | 1 | 0 |
| rs104894391 | pathogenic | 1 | 1 | 0 |
| rs5350 | benign | 1 | 1 | 1 |
| rs104894390 | pathogenic | 1 | 1 | 1 |
| rs5346 | benign | 1 | 1 | 1 |
| rs121912500 | pathogenic | 0 | 0 | 0 |
| rs74315450 | pathogenic | 1 | 1 | 0 |
| rs121913396 | pathogenic | 1 | 1 | 1 |
| rs28931588 | pathogenic | 1 | 1 | 1 |
| rs121913400 | pathogenic | 0 | 0 | 0 |
| rs28931589 | pathogenic | 1 | 1 | 1 |
| rs121913403 | pathogenic | 0 | 0 | 0 |
| rs121913413 | pathogenic | 0 | 0 | 0 |
| rs760107623 | pathogenic | 1 | 1 | 0 |
| rs16866378 | benign | 1 | 1 | 0 |
| rs201025290 | pathogenic | 1 | 1 | 0 |
| rs12720356 | benign | 1 | 1 | 0 |
| rs55882956 | benign | 0 | 1 | 0 |
| rs4986933 | benign | 1 | 1 | 0 |
| rs2020957 | benign | 1 | 1 | 1 |
| rs1800124 | benign | 1 | 1 | 1 |
| rs150077735 | benign | 1 | 1 | 1 |
| rs201419000 | benign | 0 | 0 | 0 |
| rs772044176 | pathogenic | 0 | 0 | 0 |
| rs138346388 | benign | 0 | 0 | 0 |
| rs121964916 | pathogenic | 0 | 0 | 0 |
| rs112534524 | pathogenic | 0 | 0 | 0 |
| rs182078921 | benign | 1 | 1 | 1 |
| rs113273712 | benign | 0 | 0 | 0 |
| rs74817407 | benign | 1 | 1 | 1 |
| rs61733901 | benign | 1 | 1 | 1 |
| rs121964912 | pathogenic | 1 | 1 | 1 |
| rs41278047 | benign | 1 | 1 | 1 |
| rs7437875 | benign | 1 | 1 | 1 |
| rs113460688 | benign | 1 | 1 | 1 |
| rs754572081 | benign | 1 | 1 | 1 |
| rs201082692 | benign | 1 | 1 | 1 |
| rs202085803 | benign | 1 | 1 | 1 |
| rs141320511 | benign | 0 | 0 | 0 |
| rs78501403 | benign | 0 | 0 | 0 |
| rs28939675 | pathogenic | 0 | 0 | 0 |
| rs74315522 | pathogenic | 0 | 0 | 0 |
| rs121434639 | pathogenic | 1 | 1 | 0 |
| rs121434640 | pathogenic | 1 | 1 | 1 |
| rs757282628 | benign | 1 | 1 | 1 |
| rs386834010 | pathogenic | 0 | 0 | 0 |
| rs200863680 | pathogenic | 0 | 0 | 0 |
| rs193919336 | pathogenic | 1 | 1 | 0 |
| rs386834011 | pathogenic | 0 | 0 | 0 |

| | | | | |
|---|---|---|---|---|
| rs28940869 | pathogenic | 0 | 0 | 0 |
| rs386834014 | pathogenic | 1 | 1 | 0 |
| rs267606960 | pathogenic | 0 | 0 | 0 |
| rs28942068 | pathogenic | 0 | 0 | 0 |
| rs193919335 | pathogenic | 0 | 0 | 0 |
| rs74374973 | pathogenic | 0 | 0 | 0 |
| rs386834018 | pathogenic | 0 | 0 | 0 |
| rs267606962 | pathogenic | 0 | 0 | 0 |
| rs6659553 | benign | 0 | 0 | 0 |
| rs29281 | benign | 0 | 0 | 0 |
| rs782265090 | benign | 0 | 0 | 0 |
| rs35910969 | pathogenic | 0 | 0 | 0 |
| rs121909113 | pathogenic | 1 | 1 | 1 |
| rs29001571 | pathogenic | 1 | 1 | 1 |
| rs587781250 | pathogenic | 1 | 1 | 1 |
| rs28939680 | pathogenic | 1 | 1 | 1 |
| rs28939681 | pathogenic | 1 | 1 | 1 |
| rs121909112 | pathogenic | 1 | 1 | 1 |
| rs28937568 | pathogenic | 1 | 1 | 1 |
| rs1799895 | pathogenic | 0 | 0 | 0 |
| rs386833431 | pathogenic | 1 | 1 | 1 |
| rs121964905 | pathogenic | 1 | 1 | 1 |
| rs121964906 | pathogenic | 1 | 1 | 1 |
| rs11551664 | pathogenic | 1 | 1 | 1 |
| rs104894705 | pathogenic | 1 | 1 | 1 |
| rs104893805 | pathogenic | 0 | 0 | 0 |
| rs104894205 | pathogenic | 0 | 0 | 0 |
| rs104894204 | pathogenic | 1 | 1 | 0 |
| rs62642926 | pathogenic | 0 | 0 | 0 |
| rs5030841 | pathogenic | 1 | 1 | 1 |
| rs75193786 | pathogenic | 1 | 1 | 1 |
| rs62508682 | pathogenic | 0 | 0 | 0 |
| rs3850625 | benign | 1 | 1 | 1 |
| rs141908793 | benign | 0 | 0 | 0 |
| rs187864727 | benign | 0 | 0 | 0 |
| rs548909101 | benign | 0 | 0 | 0 |
| rs3825394 | benign | 0 | 1 | 0 |
| rs143548402 | benign | 0 | 0 | 0 |
| rs267607146 | pathogenic | 0 | 0 | 0 |
| rs267607144 | pathogenic | 0 | 0 | 0 |
| rs267607148 | pathogenic | 1 | 1 | 1 |
| rs267607143 | pathogenic | 0 | 0 | 0 |
| rs267607145 | pathogenic | 0 | 0 | 0 |
| rs121912636 | pathogenic | 1 | 1 | 1 |
| rs121912634 | pathogenic | 1 | 1 | 1 |
| rs570675468 | benign | 0 | 0 | 0 |
| rs368861241 | pathogenic | 1 | 1 | 1 |
| rs104894528 | pathogenic | 1 | 1 | 1 |
| rs77493891 | benign | 0 | 0 | 0 |
| rs62640903 | benign | 0 | 0 | 0 |
| rs149926091 | benign | 1 | 1 | 1 |
| rs641081 | benign | 0 | 0 | 0 |
| rs4930199 | benign | 0 | 0 | 0 |
| rs121913020 | pathogenic | 0 | 0 | 0 |

| | | | | |
|---|---|---|---|---|
| rs1799793 | benign | 0 | 0 | 0 |
| rs121913016 | pathogenic | 0 | 0 | 0 |
| rs121913025 | pathogenic | 0 | 0 | 0 |
| rs121913019 | pathogenic | 0 | 0 | 0 |
| rs376556895 | pathogenic | 1 | 1 | 1 |
| rs121913024 | pathogenic | 1 | 1 | 1 |
| rs121913021 | pathogenic | 0 | 0 | 0 |
| rs121913023 | pathogenic | 0 | 0 | 0 |
| rs41556519 | pathogenic | 0 | 0 | 0 |
| rs121913022 | pathogenic | 0 | 0 | 0 |
| rs144564120 | pathogenic | 0 | 0 | 0 |
| rs121913026 | pathogenic | 1 | 1 | 1 |
| rs121913018 | pathogenic | 1 | 1 | 1 |
| rs142954643 | benign | 0 | 0 | 0 |
| rs76655496 | benign | 0 | 0 | 0 |
| rs104894371 | pathogenic | 1 | 1 | 0 |
| rs137964936 | benign | 0 | 0 | 0 |
| rs759736132 | pathogenic | 0 | 0 | 0 |
| rs387906595 | pathogenic | 1 | 1 | 0 |
| rs768248216 | pathogenic | 0 | 0 | 0 |
| rs548231008 | benign | 0 | 0 | 0 |
| rs757094384 | benign | 0 | 0 | 0 |
| rs867529 | benign | 0 | 0 | 0 |
| rs13045 | benign | 1 | 1 | 1 |
| rs121908570 | pathogenic | 0 | 0 | 0 |
| rs201872586 | benign | 1 | 1 | 0 |
| rs149092990 | benign | 0 | 0 | 0 |
| rs115585711 | benign | 0 | 0 | 0 |
| rs137909416 | benign | 1 | 1 | 0 |
| rs143388949 | benign | 1 | 1 | 1 |
| rs73137716 | benign | 1 | 1 | 1 |
| rs121908947 | pathogenic | 1 | 1 | 1 |
| rs398122904 | pathogenic | 1 | 1 | 1 |
| rs35715497 | benign | 1 | 1 | 0 |
| rs35462421 | benign | 0 | 0 | 0 |
| rs119475043 | pathogenic | 0 | 0 | 0 |
| rs119475042 | pathogenic | 0 | 0 | 0 |
| rs1076669 | benign | 1 | 1 | 0 |
| rs141146885 | benign | 0 | 0 | 0 |
| rs3026906 | pathogenic | 0 | 0 | 0 |
| rs75362221 | benign | 1 | 1 | 1 |
| rs138472116 | benign | 1 | 1 | 1 |
| rs371040282 | pathogenic | 1 | 1 | 1 |
| rs4988496 | benign | 0 | 0 | 0 |
| rs121918117 | pathogenic | 1 | 1 | 1 |
| rs4988498 | benign | 0 | 0 | 0 |
| rs5035 | benign | 0 | 0 | 0 |
| rs121918322 | pathogenic | 1 | 1 | 1 |
| rs370359540 | benign | 0 | 0 | 0 |
| rs758972589 | pathogenic | 0 | 0 | 0 |
| rs144856889 | benign | 0 | 0 | 0 |
| rs121434623 | pathogenic | 1 | 1 | 1 |
| rs61735600 | benign | 0 | 0 | 0 |
| rs72550870 | benign | 1 | 1 | 1 |

| | | | | |
|---|---|---|---|---|
| rs56392418 | benign | 0 | 0 | 0 |
| rs41307788 | benign | 1 | 1 | 1 |
| rs387907239 | pathogenic | 1 | 1 | 0 |
| rs28931584 | pathogenic | 0 | 0 | 0 |
| rs121912745 | pathogenic | 0 | 0 | 0 |
| rs121912744 | pathogenic | 0 | 0 | 0 |
| rs121912754 | pathogenic | 1 | 1 | 1 |
| rs121912746 | pathogenic | 0 | 0 | 0 |
| rs121912757 | pathogenic | 0 | 0 | 0 |
| rs121912758 | pathogenic | 0 | 0 | 0 |
| rs121912748 | pathogenic | 0 | 0 | 0 |
| rs121912755 | pathogenic | 0 | 0 | 0 |
| rs121912741 | pathogenic | 0 | 0 | 0 |
| rs121912753 | pathogenic | 0 | 0 | 0 |
| rs121912750 | pathogenic | 0 | 0 | 0 |
| rs2285644 | pathogenic | 0 | 0 | 0 |
| rs121912751 | pathogenic | 0 | 0 | 0 |
| rs5026 | benign | 0 | 0 | 0 |
| rs121912759 | pathogenic | 0 | 0 | 0 |
| rs28931585 | pathogenic | 0 | 0 | 0 |
| rs17122769 | benign | 0 | 0 | 0 |
| rs28362322 | benign | 1 | 1 | 1 |
| rs3824874 | benign | 0 | 0 | 0 |
| rs146004831 | benign | 0 | 0 | 0 |
| rs142155860 | benign | 0 | 0 | 0 |
| rs121434404 | pathogenic | 0 | 0 | 0 |
| rs61735578 | benign | 1 | 1 | 1 |
| rs558018 | benign | 0 | 0 | 0 |
| rs61735577 | benign | 1 | 1 | 1 |
| rs76784113 | benign | 0 | 0 | 0 |
| rs199469640 | pathogenic | 0 | 0 | 0 |
| rs199469628 | pathogenic | 1 | 1 | 0 |
| rs199469629 | pathogenic | 0 | 1 | 0 |
| rs387907156 | pathogenic | 0 | 1 | 0 |
| rs199469643 | pathogenic | 0 | 0 | 0 |
| rs199469646 | pathogenic | 0 | 0 | 0 |
| rs766294940 | pathogenic | 0 | 0 | 0 |
| rs387907347 | pathogenic | 0 | 0 | 0 |
| rs137853590 | pathogenic | 1 | 1 | 1 |
| rs137853589 | pathogenic | 1 | 1 | 1 |
| rs137853588 | pathogenic | 1 | 1 | 1 |
| rs55909005 | benign | 0 | 0 | 0 |
| rs4581 | benign | 0 | 0 | 0 |
| rs63750123 | benign | 0 | 0 | 0 |
| rs10254120 | benign | 0 | 0 | 0 |
| rs121434629 | pathogenic | 1 | 1 | 0 |
| rs200313585 | benign | 0 | 0 | 0 |
| rs143162541 | benign | 1 | 1 | 0 |
| rs760615315 | benign | 0 | 0 | 0 |
| rs768488890 | benign | 1 | 1 | 1 |
| rs116373169 | benign | 1 | 1 | 0 |
| rs63750871 | pathogenic | 1 | 1 | 0 |
| rs116349687 | benign | 0 | 0 | 0 |
| rs375289386 | benign | 0 | 0 | 0 |

| | | | | |
|---|---|---|---|---|
| rs760228510 | pathogenic | 1 | 1 | 1 |
| rs587779343 | pathogenic | 1 | 1 | 1 |
| rs573125799 | benign | 0 | 0 | 0 |
| rs1805322 | benign | 0 | 0 | 0 |
| rs267608153 | pathogenic | 0 | 0 | 0 |
| rs139194813 | benign | 0 | 0 | 0 |
| rs530021751 | benign | 0 | 0 | 0 |
| rs200640585 | pathogenic | 0 | 0 | 0 |
| rs139438201 | benign | 0 | 0 | 0 |
| rs200513014 | benign | 0 | 0 | 0 |
| rs567102013 | benign | 0 | 0 | 0 |
| rs121908608 | pathogenic | 0 | 0 | 0 |
| rs121908606 | pathogenic | 0 | 0 | 0 |
| rs17103671 | benign | 0 | 0 | 0 |
| rs34725281 | benign | 0 | 0 | 0 |
| rs104894421 | pathogenic | 1 | 1 | 0 |
| rs121918145 | pathogenic | 1 | 1 | 1 |
| rs121918152 | pathogenic | 1 | 1 | 1 |
| rs56379106 | benign | 0 | 0 | 0 |
| rs4987076 | benign | 0 | 0 | 0 |
| rs370243877 | pathogenic | 0 | 0 | 0 |
| rs17679445 | benign | 1 | 1 | 1 |
| rs550423482 | pathogenic | 1 | 1 | 1 |
| rs104894629 | pathogenic | 1 | 1 | 1 |
| rs773450573 | pathogenic | 1 | 1 | 1 |
| rs116679717 | benign | 0 | 0 | 0 |
| rs372000848 | pathogenic | 0 | 0 | 0 |
| rs138595973 | benign | 0 | 0 | 0 |
| rs115244417 | benign | 0 | 0 | 0 |
| rs11575292 | benign | 0 | 0 | 0 |
| rs137853211 | pathogenic | 1 | 1 | 1 |
| rs137853207 | pathogenic | 0 | 0 | 0 |
| rs137853210 | pathogenic | 1 | 1 | 1 |
| rs6262 | benign | 0 | 0 | 0 |
| rs6263 | benign | 0 | 1 | 0 |
| rs137853208 | pathogenic | 0 | 1 | 0 |
| rs137853212 | pathogenic | 1 | 1 | 0 |
| rs137853209 | pathogenic | 1 | 1 | 1 |
| rs201951824 | pathogenic | 0 | 0 | 0 |
| rs11575542 | benign | 0 | 0 | 0 |
| rs199681788 | benign | 0 | 1 | 0 |
| rs748835902 | benign | 1 | 1 | 1 |
| rs753719949 | benign | 0 | 0 | 0 |
| rs267606647 | pathogenic | 0 | 0 | 0 |
| rs267606648 | pathogenic | 1 | 1 | 1 |
| rs267606643 | pathogenic | 0 | 1 | 0 |
| rs559947967 | pathogenic | 1 | 1 | 0 |
| rs267606645 | pathogenic | 1 | 1 | 1 |
| rs12116440 | benign | 0 | 0 | 0 |
| rs121908539 | pathogenic | 0 | 0 | 0 |
| rs137853096 | pathogenic | 1 | 1 | 0 |
| rs148363262 | benign | 1 | 1 | 0 |
| rs587777442 | pathogenic | 0 | 0 | 0 |
| rs25640 | benign | 1 | 1 | 1 |

| | | | | |
|---|---|---|---|---|
| rs28943589 | benign | 0 | 0 | 0 |
| rs1143650 | benign | 1 | 1 | 1 |
| rs150886026 | benign | 0 | 1 | 0 |
| rs121908133 | pathogenic | 1 | 1 | 0 |
| rs121908131 | pathogenic | 0 | 0 | 0 |
| rs754477154 | pathogenic | 0 | 0 | 0 |
| rs747334007 | benign | 0 | 0 | 0 |
| rs121908132 | pathogenic | 0 | 0 | 0 |
| rs104894103 | pathogenic | 1 | 1 | 0 |
| rs140088149 | benign | 0 | 0 | 0 |
| rs149803148 | benign | 0 | 0 | 0 |
| rs11064480 | benign | 0 | 0 | 0 |
| rs74435397 | pathogenic | 1 | 1 | 1 |
| rs587783626 | pathogenic | 1 | 1 | 1 |
| rs148211042 | pathogenic | 1 | 1 | 1 |
| rs17849502 | benign | 0 | 0 | 0 |
| rs13306575 | pathogenic | 1 | 1 | 1 |
| rs201426733 | benign | 0 | 0 | 0 |
| rs182278784 | benign | 1 | 1 | 1 |
| rs77786415 | benign | 1 | 1 | 1 |
| rs200192119 | benign | 1 | 1 | 1 |
| rs12877501 | benign | 1 | 1 | 1 |
| rs77985244 | benign | 1 | 1 | 1 |
| rs5907 | pathogenic | 1 | 1 | 0 |
| rs2305178 | benign | 0 | 0 | 0 |
| rs3135867 | benign | 0 | 0 | 0 |
| rs121913482 | pathogenic | 1 | 1 | 1 |
| rs121913483 | pathogenic | 1 | 1 | 1 |
| rs4647924 | pathogenic | 1 | 1 | 1 |
| rs121913111 | pathogenic | 1 | 1 | 1 |
| rs104894002 | pathogenic | 0 | 0 | 0 |
| rs104894001 | pathogenic | 0 | 0 | 0 |
| rs75932628 | benign | 0 | 0 | 0 |
| rs201258663 | pathogenic | 0 | 0 | 0 |
| rs104893998 | pathogenic | 0 | 0 | 0 |
| rs2234253 | benign | 1 | 1 | 0 |
| rs121908402 | pathogenic | 1 | 1 | 0 |
| rs1805389 | benign | 0 | 0 | 0 |
| rs1805388 | benign | 0 | 0 | 0 |
| rs748385144 | pathogenic | 0 | 0 | 0 |
| rs104894420 | pathogenic | 0 | 0 | 0 |
| rs104894418 | pathogenic | 0 | 0 | 0 |
| rs61731910 | benign | 0 | 0 | 0 |
| rs758723288 | pathogenic | 1 | 1 | 0 |
| rs201865375 | pathogenic | 1 | 1 | 0 |
| rs565090080 | pathogenic | 1 | 1 | 1 |
| rs113042166 | benign | 0 | 0 | 0 |
| rs61736326 | benign | 1 | 1 | 1 |
| rs12030699 | benign | 1 | 1 | 1 |
| rs121913578 | pathogenic | 1 | 1 | 1 |
| rs61739582 | benign | 0 | 0 | 0 |
| rs745983207 | pathogenic | 0 | 0 | 0 |
| rs763524380 | benign | 0 | 0 | 0 |
| rs121912792 | pathogenic | 0 | 0 | 0 |

| | | | | |
|---|---|---|---|---|
| rs121912793 | pathogenic | 0 | 0 | 0 |
| rs387906599 | pathogenic | 0 | 0 | 0 |
| rs121912794 | pathogenic | 0 | 1 | 0 |
| rs121912795 | pathogenic | 0 | 0 | 0 |
| rs121912796 | pathogenic | 1 | 1 | 0 |
| rs121912797 | pathogenic | 0 | 1 | 0 |
| rs121912798 | pathogenic | 1 | 1 | 1 |
| rs80359812 | pathogenic | 1 | 1 | 0 |
| rs776583130 | benign | 1 | 1 | 0 |
| rs80359814 | pathogenic | 1 | 1 | 0 |
| rs202060209 | pathogenic | 1 | 1 | 0 |
| rs267607061 | pathogenic | 1 | 1 | 0 |
| rs267607060 | pathogenic | 0 | 0 | 0 |
| rs80359816 | pathogenic | 1 | 1 | 0 |
| rs80359818 | pathogenic | 1 | 1 | 0 |
| rs387907313 | pathogenic | 0 | 0 | 0 |
| rs587784395 | benign | 0 | 0 | 0 |
| rs5811 | benign | 0 | 0 | 0 |
| rs121909740 | pathogenic | 1 | 1 | 0 |
| rs148518827 | benign | 1 | 1 | 0 |
| rs121909739 | pathogenic | 1 | 1 | 0 |
| rs780528770 | benign | 1 | 1 | 0 |
| rs13306758 | pathogenic | 0 | 0 | 0 |
| rs267607059 | pathogenic | 0 | 0 | 0 |
| rs61730638 | benign | 0 | 0 | 0 |
| rs104894533 | pathogenic | 0 | 0 | 0 |
| rs104894527 | pathogenic | 0 | 0 | 0 |
| rs200503569 | pathogenic | 1 | 1 | 1 |
| rs80338700 | pathogenic | 1 | 1 | 1 |
| rs765433263 | pathogenic | 1 | 1 | 1 |
| rs104894530 | pathogenic | 1 | 1 | 1 |
| rs80338701 | pathogenic | 1 | 1 | 1 |
| rs141498002 | pathogenic | 1 | 1 | 1 |
| rs104894525 | pathogenic | 0 | 0 | 0 |
| rs80338703 | pathogenic | 0 | 0 | 0 |
| rs28936415 | pathogenic | 1 | 1 | 0 |
| rs150719105 | pathogenic | 0 | 0 | 0 |
| rs148032587 | pathogenic | 0 | 0 | 0 |
| rs150577656 | pathogenic | 0 | 0 | 0 |
| rs190521996 | pathogenic | 0 | 0 | 0 |
| rs104894526 | pathogenic | 1 | 1 | 1 |
| rs80338704 | pathogenic | 0 | 0 | 0 |
| rs199562225 | pathogenic | 1 | 1 | 1 |
| rs532870929 | pathogenic | 0 | 0 | 0 |
| rs398123309 | pathogenic | 0 | 0 | 0 |
| rs78290141 | pathogenic | 0 | 0 | 0 |
| rs80338705 | pathogenic | 0 | 0 | 0 |
| rs104894531 | pathogenic | 1 | 1 | 1 |
| rs80338706 | pathogenic | 0 | 0 | 0 |
| rs80338707 | pathogenic | 0 | 0 | 0 |
| rs80338708 | pathogenic | 0 | 0 | 0 |
| rs80338709 | pathogenic | 0 | 0 | 0 |
| rs104893833 | benign | 0 | 0 | 0 |
| rs61744585 | benign | 0 | 0 | 0 |

| | | | | |
|---|---|---|---|---|
| rs73569515 | benign | 0 | 0 | 0 |
| rs398123373 | pathogenic | 0 | 0 | 0 |
| rs756854513 | pathogenic | 0 | 0 | 0 |
| rs58232698 | benign | 0 | 0 | 0 |
| rs9600455 | benign | 0 | 0 | 0 |
| rs201016593 | pathogenic | 0 | 0 | 0 |
| rs756039188 | pathogenic | 0 | 0 | 0 |
| rs121908024 | pathogenic | 0 | 0 | 0 |
| rs373144619 | benign | 0 | 0 | 0 |
| rs138078086 | benign | 0 | 0 | 0 |
| rs376207800 | pathogenic | 0 | 0 | 0 |
| rs749038326 | pathogenic | 1 | 1 | 0 |
| rs372845091 | benign | 0 | 0 | 0 |
| rs377271627 | pathogenic | 0 | 0 | 0 |
| rs563390335 | pathogenic | 0 | 0 | 0 |
| rs13306510 | benign | 0 | 0 | 0 |
| rs764042910 | pathogenic | 0 | 0 | 0 |
| rs201384282 | benign | 1 | 1 | 1 |
| rs730882086 | pathogenic | 1 | 1 | 1 |
| rs756613387 | pathogenic | 1 | 1 | 1 |
| rs121908028 | benign | 1 | 1 | 0 |
| rs773328511 | benign | 0 | 0 | 0 |
| rs749220643 | pathogenic | 1 | 1 | 0 |
| rs730882089 | pathogenic | 0 | 0 | 0 |
| rs146651743 | benign | 0 | 0 | 0 |
| rs148698650 | pathogenic | 0 | 0 | 0 |
| rs140241383 | benign | 0 | 0 | 0 |
| rs757252110 | pathogenic | 0 | 0 | 0 |
| rs13306512 | pathogenic | 1 | 1 | 1 |
| rs121908043 | pathogenic | 1 | 1 | 1 |
| rs773658037 | pathogenic | 0 | 0 | 0 |
| rs774439908 | pathogenic | 0 | 0 | 0 |
| rs779732323 | pathogenic | 1 | 1 | 0 |
| rs749780672 | pathogenic | 0 | 0 | 0 |
| rs375651668 | benign | 1 | 1 | 1 |
| rs193922568 | pathogenic | 1 | 1 | 0 |
| rs750363970 | benign | 1 | 1 | 0 |
| rs5932 | benign | 0 | 0 | 0 |
| rs755154048 | pathogenic | 1 | 1 | 0 |
| rs759876319 | pathogenic | 0 | 0 | 0 |
| rs764550980 | pathogenic | 1 | 1 | 0 |
| rs779940524 | pathogenic | 0 | 0 | 0 |
| rs563382937 | pathogenic | 1 | 1 | 1 |
| rs28942083 | pathogenic | 0 | 1 | 0 |
| rs770744861 | pathogenic | 0 | 1 | 0 |
| rs121908031 | pathogenic | 0 | 1 | 0 |
| rs776217028 | pathogenic | 0 | 1 | 0 |
| rs763625913 | pathogenic | 0 | 0 | 0 |
| rs3736772 | benign | 0 | 0 | 0 |
| rs104894696 | benign | 1 | 1 | 1 |
| rs121918177 | pathogenic | 0 | 1 | 0 |
| rs121918181 | pathogenic | 1 | 1 | 0 |
| rs121918174 | pathogenic | 0 | 1 | 0 |
| rs121918182 | pathogenic | 0 | 0 | 0 |

| | | | | |
|---|---|---|---|---|
| rs121918175 | pathogenic | 1 | 1 | 1 |
| rs121918185 | pathogenic | 1 | 1 | 1 |
| rs121918173 | pathogenic | 1 | 1 | 0 |
| rs121918178 | pathogenic | 1 | 1 | 1 |
| rs121918179 | pathogenic | 1 | 1 | 1 |
| rs121918183 | pathogenic | 0 | 0 | 0 |
| rs121918176 | pathogenic | 1 | 1 | 1 |
| rs121918172 | pathogenic | 0 | 1 | 0 |
| rs121918184 | pathogenic | 0 | 0 | 0 |
| rs121434560 | pathogenic | 1 | 1 | 0 |
| rs121434561 | pathogenic | 1 | 1 | 0 |
| rs3218716 | pathogenic | 1 | 1 | 1 |
| rs145532615 | benign | 0 | 0 | 0 |
| rs149840927 | pathogenic | 0 | 0 | 0 |
| rs730880762 | pathogenic | 0 | 0 | 0 |
| rs730880764 | pathogenic | 0 | 0 | 0 |
| rs730880765 | pathogenic | 0 | 0 | 0 |
| rs587782962 | pathogenic | 0 | 0 | 0 |
| rs141735183 | pathogenic | 0 | 0 | 0 |
| rs397516187 | pathogenic | 0 | 0 | 0 |
| rs727505026 | pathogenic | 0 | 0 | 0 |
| rs529700838 | pathogenic | 0 | 0 | 0 |
| rs540263945 | pathogenic | 0 | 0 | 0 |
| rs149103761 | benign | 0 | 0 | 0 |
| rs730880906 | pathogenic | 0 | 0 | 0 |
| rs730880785 | pathogenic | 0 | 0 | 0 |
| rs397516199 | pathogenic | 0 | 0 | 0 |
| rs730880725 | benign | 0 | 0 | 0 |
| rs730880796 | pathogenic | 0 | 0 | 0 |
| rs201307101 | benign | 0 | 0 | 0 |
| rs397516214 | pathogenic | 0 | 0 | 0 |
| rs3729823 | benign | 0 | 0 | 0 |
| rs397516218 | pathogenic | 0 | 0 | 0 |
| rs121913647 | pathogenic | 0 | 0 | 0 |
| rs373514686 | pathogenic | 0 | 0 | 0 |
| rs763538103 | pathogenic | 0 | 0 | 0 |
| rs121913650 | pathogenic | 0 | 0 | 0 |
| rs139222507 | pathogenic | 0 | 0 | 0 |
| rs369940645 | pathogenic | 0 | 0 | 0 |
| rs763073072 | pathogenic | 0 | 0 | 0 |
| rs145734640 | pathogenic | 0 | 0 | 0 |
| rs730880821 | pathogenic | 0 | 0 | 0 |
| rs28933098 | pathogenic | 0 | 0 | 0 |
| rs372381770 | pathogenic | 0 | 0 | 0 |
| rs146796870 | benign | 0 | 0 | 0 |
| rs104894485 | pathogenic | 1 | 1 | 1 |
| rs35407712 | benign | 0 | 0 | 0 |
| rs35842750 | benign | 0 | 0 | 0 |
| rs2721191 | benign | 0 | 0 | 0 |
| rs35098923 | benign | 0 | 0 | 0 |
| rs34293591 | benign | 1 | 1 | 0 |
| rs36023964 | benign | 0 | 0 | 0 |
| rs4251691 | benign | 0 | 0 | 0 |
| rs34666647 | benign | 0 | 0 | 0 |

| | | | | |
|---|---|---|---|---|
| rs4925828 | benign | 0 | 0 | 0 |
| rs35348691 | benign | 0 | 0 | 0 |
| rs36078464 | benign | 0 | 0 | 0 |
| rs34915097 | benign | 0 | 0 | 0 |
| rs35101495 | benign | 0 | 0 | 0 |
| rs141568342 | pathogenic | 0 | 0 | 0 |
| rs1800571 | pathogenic | 0 | 0 | 0 |
| rs140566084 | benign | 1 | 1 | 1 |
| rs727503791 | pathogenic | 1 | 1 | 0 |
| rs545215807 | pathogenic | 0 | 0 | 0 |
| rs113994168 | pathogenic | 0 | 0 | 0 |
| rs113994167 | pathogenic | 0 | 0 | 0 |
| rs200788251 | pathogenic | 0 | 0 | 0 |
| rs200573371 | pathogenic | 0 | 0 | 0 |
| rs771874163 | pathogenic | 1 | 1 | 1 |
| rs112406105 | pathogenic | 1 | 1 | 1 |
| rs118204015 | pathogenic | 1 | 1 | 1 |
| rs113994169 | pathogenic | 0 | 0 | 0 |
| rs118204018 | pathogenic | 0 | 1 | 0 |
| rs138834083 | pathogenic | 0 | 0 | 0 |
| rs533055438 | pathogenic | 1 | 1 | 1 |
| rs2309689 | pathogenic | 1 | 1 | 1 |
| rs138058572 | pathogenic | 1 | 1 | 1 |
| rs118204017 | pathogenic | 1 | 1 | 0 |
| rs766742117 | pathogenic | 1 | 1 | 1 |
| rs751995154 | pathogenic | 1 | 1 | 1 |
| rs200771970 | pathogenic | 0 | 0 | 0 |
| rs118204014 | pathogenic | 0 | 0 | 0 |
| rs148584617 | pathogenic | 1 | 1 | 1 |
| rs121912720 | pathogenic | 0 | 0 | 0 |
| rs121912721 | pathogenic | 0 | 0 | 0 |
| rs28929476 | pathogenic | 0 | 0 | 0 |
| rs28931574 | pathogenic | 1 | 1 | 1 |
| rs12718465 | benign | 0 | 0 | 0 |
| rs144689354 | pathogenic | 0 | 0 | 0 |
| rs149095705 | pathogenic | 0 | 0 | 0 |
| rs572839648 | benign | 1 | 1 | 1 |
| rs121909715 | pathogenic | 1 | 1 | 0 |
| rs34077456 | benign | 1 | 1 | 1 |
| rs397517876 | benign | 0 | 0 | 0 |
| rs1064074 | benign | 0 | 0 | 0 |
| rs142751309 | benign | 0 | 0 | 0 |
| rs116275562 | benign | 0 | 0 | 0 |
| rs148095899 | benign | 0 | 0 | 0 |
| rs751838040 | pathogenic | 0 | 0 | 0 |
| rs121434324 | pathogenic | 1 | 1 | 1 |
| rs143367518 | pathogenic | 0 | 0 | 0 |
| rs121434325 | pathogenic | 0 | 0 | 0 |
| rs397515328 | pathogenic | 0 | 0 | 0 |
| rs376039938 | pathogenic | 0 | 0 | 0 |
| rs387907243 | pathogenic | 0 | 0 | 0 |
| rs28934876 | pathogenic | 0 | 0 | 0 |
| rs387907245 | pathogenic | 0 | 0 | 0 |
| rs138544311 | benign | 0 | 0 | 0 |

| | | | | |
|---|---|---|---|---|
| rs643788 | benign | 0 | 0 | 0 |
| rs749977756 | benign | 0 | 0 | 0 |
| rs1800574 | benign | 0 | 0 | 0 |
| rs137853336 | pathogenic | 1 | 1 | 1 |
| rs111916034 | benign | 0 | 0 | 0 |
| rs307826 | benign | 0 | 0 | 0 |
| rs35874891 | benign | 0 | 0 | 0 |
| rs111033580 | pathogenic | 1 | 1 | 1 |
| rs111033581 | pathogenic | 1 | 1 | 1 |
| rs121908373 | pathogenic | 0 | 0 | 0 |
| rs767122713 | pathogenic | 0 | 0 | 0 |
| rs2233213 | benign | 0 | 0 | 0 |
| rs121908605 | pathogenic | 1 | 1 | 0 |
| rs11887534 | benign | 0 | 0 | 0 |
| rs4148211 | benign | 0 | 0 | 0 |
| rs80025980 | benign | 0 | 0 | 0 |
| rs137854891 | pathogenic | 1 | 1 | 0 |
| rs74358901 | benign | 0 | 0 | 0 |
| rs9282574 | benign | 0 | 0 | 0 |
| rs137852993 | pathogenic | 0 | 0 | 0 |
| rs137852990 | pathogenic | 0 | 0 | 0 |
| rs137852987 | pathogenic | 0 | 0 | 0 |
| rs4148217 | benign | 0 | 0 | 0 |
| rs140690030 | benign | 0 | 0 | 0 |
| rs137852992 | pathogenic | 0 | 0 | 0 |
| rs6544718 | benign | 0 | 0 | 0 |
| rs137852989 | pathogenic | 0 | 0 | 0 |
| rs150893072 | benign | 1 | 1 | 1 |
| rs28931613 | pathogenic | 1 | 1 | 1 |
| rs143789597 | benign | 1 | 1 | 1 |
| rs121913037 | pathogenic | 1 | 1 | 0 |
| rs188802138 | pathogenic | 1 | 1 | 0 |
| rs121913038 | pathogenic | 1 | 1 | 0 |
| rs146835206 | benign | 0 | 0 | 0 |
| rs121913041 | pathogenic | 1 | 1 | 1 |
| rs121913039 | pathogenic | 1 | 1 | 1 |
| rs149977726 | pathogenic | 1 | 1 | 0 |
| rs121913036 | pathogenic | 0 | 0 | 0 |
| rs121913040 | pathogenic | 0 | 0 | 0 |
| rs765007158 | benign | 0 | 0 | 0 |
| rs112723255 | benign | 0 | 0 | 0 |
| rs574164748 | benign | 0 | 0 | 0 |
| rs80338734 | pathogenic | 1 | 1 | 0 |
| rs104894694 | pathogenic | 1 | 1 | 0 |
| rs753198836 | pathogenic | 0 | 0 | 0 |
| rs760101382 | pathogenic | 0 | 0 | 0 |
| rs121909272 | pathogenic | 1 | 1 | 0 |
| rs370421531 | pathogenic | 0 | 0 | 0 |
| rs565109128 | benign | 0 | 0 | 0 |
| rs17851582 | benign | 0 | 0 | 0 |
| rs753228876 | benign | 0 | 0 | 0 |
| rs141471799 | benign | 0 | 0 | 0 |
| rs281864916 | pathogenic | 1 | 1 | 0 |
| rs121918409 | pathogenic | 1 | 1 | 1 |

| | | | | |
|---|---|---|---|---|
| rs121918413 | pathogenic | 1 | 1 | 1 |
| rs121918411 | pathogenic | 1 | 1 | 1 |
| rs121918408 | pathogenic | 1 | 1 | 1 |
| rs121918412 | pathogenic | 1 | 1 | 1 |
| rs121918410 | pathogenic | 1 | 1 | 1 |
| rs116474260 | benign | 1 | 1 | 1 |
| rs281864919 | pathogenic | 1 | 1 | 1 |
| rs397514452 | pathogenic | 1 | 1 | 0 |
| rs747843638 | benign | 0 | 0 | 0 |
| rs774357609 | pathogenic | 1 | 1 | 1 |
| rs201804005 | benign | 0 | 0 | 0 |
| rs746485916 | pathogenic | 0 | 0 | 0 |
| rs764131178 | pathogenic | 0 | 0 | 0 |
| rs374094065 | pathogenic | 0 | 0 | 0 |
| rs76151636 | pathogenic | 1 | 1 | 0 |
| rs753250853 | pathogenic | 0 | 0 | 0 |
| rs761632029 | pathogenic | 0 | 0 | 0 |
| rs775541743 | pathogenic | 0 | 0 | 0 |
| rs778749563 | pathogenic | 0 | 0 | 0 |
| rs60431989 | pathogenic | 1 | 1 | 0 |
| rs755554442 | pathogenic | 1 | 1 | 0 |
| rs749085322 | pathogenic | 0 | 0 | 0 |
| rs756029120 | pathogenic | 1 | 1 | 0 |
| rs137853279 | pathogenic | 1 | 1 | 0 |
| rs777354267 | pathogenic | 1 | 1 | 1 |
| rs142940486 | benign | 1 | 1 | 1 |
| rs140167393 | benign | 1 | 1 | 1 |
| rs2066827 | benign | 0 | 0 | 0 |
| rs765681672 | benign | 0 | 0 | 0 |
| rs137985549 | benign | 0 | 0 | 0 |
| rs74315340 | pathogenic | 1 | 1 | 1 |
| rs74315341 | pathogenic | 0 | 0 | 0 |
| rs74315335 | pathogenic | 0 | 0 | 0 |
| rs61730974 | benign | 0 | 0 | 0 |
| rs121909193 | pathogenic | 0 | 0 | 0 |
| rs74315334 | pathogenic | 0 | 0 | 0 |
| rs74315329 | pathogenic | 0 | 0 | 0 |
| rs74315330 | pathogenic | 0 | 0 | 0 |
| rs121909194 | pathogenic | 0 | 0 | 0 |
| rs28936694 | pathogenic | 0 | 0 | 0 |
| rs74315336 | pathogenic | 0 | 0 | 0 |
| rs74315338 | pathogenic | 1 | 1 | 1 |
| rs74315328 | pathogenic | 0 | 0 | 0 |
| rs74315331 | pathogenic | 0 | 0 | 0 |
| rs74315332 | pathogenic | 0 | 0 | 0 |
| rs141696651 | benign | 0 | 0 | 0 |
| rs7041 | benign | 0 | 0 | 0 |
| rs4588 | benign | 0 | 0 | 0 |
| rs121908981 | pathogenic | 1 | 1 | 1 |
| rs606231459 | pathogenic | 0 | 0 | 0 |
| rs398123053 | pathogenic | 0 | 0 | 0 |
| rs774259802 | benign | 0 | 0 | 0 |
| rs121909582 | pathogenic | 1 | 1 | 1 |
| rs104893680 | pathogenic | 1 | 1 | 1 |

| | | | | |
|---|---|---|---|---|
| rs137854907 | pathogenic | 0 | 0 | 0 |
| rs765715798 | pathogenic | 0 | 0 | 0 |
| rs104893678 | pathogenic | 0 | 0 | 0 |
| rs104893679 | pathogenic | 0 | 1 | 0 |
| rs104893681 | pathogenic | 1 | 1 | 0 |
| rs121909640 | pathogenic | 1 | 1 | 0 |
| rs780705654 | pathogenic | 0 | 0 | 0 |
| rs201672146 | benign | 1 | 1 | 0 |
| rs12136658 | benign | 0 | 0 | 0 |
| rs56127440 | benign | 0 | 0 | 0 |
| rs104894433 | pathogenic | 1 | 1 | 1 |
| rs104894435 | pathogenic | 0 | 1 | 0 |
| rs104894437 | pathogenic | 1 | 1 | 1 |
| rs104894441 | pathogenic | 1 | 1 | 1 |
| rs104894440 | pathogenic | 1 | 1 | 1 |
| rs104894445 | pathogenic | 1 | 1 | 1 |
| rs104894436 | pathogenic | 0 | 0 | 0 |
| rs104894438 | pathogenic | 0 | 0 | 0 |
| rs200891969 | pathogenic | 1 | 1 | 0 |
| rs104894443 | pathogenic | 1 | 1 | 1 |
| rs104894434 | pathogenic | 1 | 1 | 1 |
| rs41298442 | pathogenic | 1 | 1 | 1 |
| rs104894442 | pathogenic | 1 | 1 | 1 |
| rs35453854 | benign | 0 | 0 | 0 |
| rs121912898 | pathogenic | 0 | 0 | 0 |
| rs121912897 | pathogenic | 1 | 1 | 0 |
| rs138260625 | benign | 0 | 0 | 0 |
| rs147802438 | benign | 1 | 1 | 0 |
| rs146443310 | benign | 1 | 1 | 0 |
| rs747319628 | pathogenic | 1 | 1 | 0 |
| rs1801195 | benign | 0 | 0 | 0 |
| rs145045341 | benign | 1 | 1 | 0 |
| rs116001173 | benign | 1 | 1 | 0 |
| rs587784175 | pathogenic | 0 | 0 | 0 |
| rs104893857 | pathogenic | 0 | 0 | 0 |
| rs104893862 | pathogenic | 1 | 1 | 0 |
| rs104893858 | pathogenic | 1 | 1 | 1 |
| rs121909249 | pathogenic | 0 | 1 | 0 |
| rs121909248 | pathogenic | 1 | 1 | 1 |
| rs104893859 | pathogenic | 1 | 1 | 0 |
| rs187063864 | pathogenic | 1 | 1 | 0 |
| rs121917817 | pathogenic | 1 | 1 | 0 |
| rs121917818 | pathogenic | 0 | 0 | 0 |
| rs753594601 | pathogenic | 0 | 0 | 0 |
| rs28937869 | pathogenic | 1 | 1 | 0 |
| rs6165 | benign | 0 | 0 | 0 |
| rs200089714 | pathogenic | 1 | 1 | 0 |
| rs372000714 | pathogenic | 0 | 0 | 0 |
| rs2228206 | benign | 0 | 0 | 0 |
| rs560096 | benign | 0 | 0 | 0 |
| rs137852666 | pathogenic | 0 | 0 | 0 |
| rs10896380 | benign | 0 | 0 | 0 |
| rs773690764 | pathogenic | 0 | 0 | 0 |
| rs201060167 | pathogenic | 0 | 0 | 0 |

| | | | | |
|---|---|---|---|---|
| rs137852670 | pathogenic | 0 | 0 | 0 |
| rs780594709 | pathogenic | 0 | 0 | 0 |
| rs145226920 | pathogenic | 0 | 0 | 0 |
| rs137852665 | pathogenic | 0 | 0 | 0 |
| rs724159960 | pathogenic | 0 | 0 | 0 |
| rs756985703 | pathogenic | 0 | 0 | 0 |
| rs137852667 | pathogenic | 0 | 0 | 0 |
| rs151079750 | pathogenic | 1 | 1 | 0 |
| rs2306636 | benign | 0 | 0 | 0 |
| rs72647885 | benign | 1 | 1 | 0 |
| rs377584435 | pathogenic | 1 | 1 | 1 |
| rs757583846 | pathogenic | 1 | 1 | 0 |
| rs121918148 | pathogenic | 0 | 0 | 0 |
| rs149201802 | benign | 0 | 0 | 0 |
| rs587781904 | pathogenic | 1 | 1 | 1 |
| rs143421325 | benign | 0 | 0 | 0 |
| rs587781253 | pathogenic | 0 | 0 | 0 |
| rs587780331 | benign | 0 | 0 | 0 |
| rs151001016 | benign | 0 | 0 | 0 |
| rs150888094 | benign | 0 | 0 | 0 |
| rs149581331 | benign | 0 | 0 | 0 |
| rs774840535 | benign | 0 | 0 | 0 |
| rs138428684 | benign | 0 | 0 | 0 |
| rs149300055 | benign | 0 | 0 | 0 |
| rs571005578 | benign | 0 | 0 | 0 |
| rs200731839 | benign | 0 | 0 | 0 |
| rs121965007 | pathogenic | 0 | 0 | 0 |
| rs121965013 | pathogenic | 0 | 1 | 0 |
| rs121965009 | pathogenic | 0 | 0 | 0 |
| rs1800457 | benign | 1 | 1 | 0 |
| rs121965006 | pathogenic | 1 | 1 | 0 |
| rs121965008 | pathogenic | 0 | 0 | 0 |
| rs61732609 | pathogenic | 0 | 0 | 0 |
| rs121965011 | pathogenic | 0 | 0 | 0 |
| rs121965015 | pathogenic | 0 | 0 | 0 |
| rs121965018 | pathogenic | 1 | 1 | 0 |
| rs121965016 | pathogenic | 0 | 0 | 0 |
| rs77806 | benign | 0 | 0 | 0 |
| rs149731356 | benign | 0 | 0 | 0 |
| rs72549407 | pathogenic | 0 | 0 | 0 |
| rs200363929 | benign | 0 | 0 | 0 |
| rs376403708 | benign | 1 | 1 | 0 |
| rs56376197 | benign | 1 | 1 | 1 |
| rs200237973 | benign | 1 | 1 | 1 |
| rs548330075 | benign | 0 | 0 | 0 |
| rs148291350 | benign | 0 | 0 | 0 |
| rs71579353 | pathogenic | 0 | 0 | 0 |
| rs146278697 | benign | 0 | 0 | 0 |
| rs141033098 | benign | 0 | 0 | 0 |
| rs397517234 | pathogenic | 0 | 0 | 0 |
| rs71579374 | benign | 0 | 0 | 0 |
| rs140381835 | benign | 0 | 0 | 0 |
| rs137852702 | pathogenic | 0 | 0 | 0 |
| rs137852696 | pathogenic | 1 | 1 | 1 |

| | | | | |
|---|---|---|---|---|
| rs137852697 | pathogenic | 0 | 0 | 0 |
| rs386833639 | pathogenic | 0 | 0 | 0 |
| rs386833640 | pathogenic | 0 | 0 | 0 |
| rs386833641 | benign | 0 | 0 | 0 |
| rs137852701 | pathogenic | 0 | 0 | 0 |
| rs386833642 | pathogenic | 0 | 0 | 0 |
| rs137852695 | pathogenic | 0 | 0 | 0 |
| rs386833646 | pathogenic | 0 | 0 | 0 |
| rs137852700 | pathogenic | 1 | 1 | 0 |
| rs386833649 | pathogenic | 1 | 1 | 0 |
| rs386833650 | pathogenic | 0 | 0 | 0 |
| rs148412181 | pathogenic | 0 | 0 | 0 |
| rs386833655 | pathogenic | 0 | 0 | 0 |
| rs386833656 | pathogenic | 1 | 1 | 0 |
| rs137852698 | pathogenic | 0 | 0 | 0 |
| rs386833661 | pathogenic | 0 | 0 | 0 |
| rs386833670 | pathogenic | 0 | 0 | 0 |
| rs111033632 | pathogenic | 1 | 1 | 1 |
| rs34308573 | benign | 1 | 1 | 1 |
| rs376048533 | pathogenic | 1 | 1 | 0 |
| rs3747517 | benign | 1 | 1 | 0 |
| rs137853328 | pathogenic | 0 | 0 | 0 |
| rs77273740 | benign | 0 | 0 | 0 |
| rs267606760 | pathogenic | 0 | 0 | 0 |
| rs77576840 | pathogenic | 0 | 0 | 0 |
| rs45465204 | benign | 1 | 1 | 0 |
| rs5320 | benign | 0 | 0 | 0 |
| rs5324 | benign | 0 | 0 | 0 |
| rs4531 | benign | 0 | 0 | 0 |
| rs267606761 | pathogenic | 0 | 0 | 0 |
| rs75215331 | pathogenic | 0 | 0 | 0 |
| rs6271 | benign | 0 | 0 | 0 |
| rs201643277 | pathogenic | 0 | 0 | 0 |
| rs4252250 | benign | 0 | 0 | 0 |
| rs137853580 | pathogenic | 0 | 0 | 0 |
| rs368287711 | pathogenic | 0 | 0 | 0 |
| rs137853579 | pathogenic | 0 | 0 | 0 |
| rs3135932 | benign | 0 | 0 | 0 |
| rs2228055 | benign | 0 | 0 | 0 |
| rs140322345 | benign | 1 | 1 | 1 |
| rs1154510 | pathogenic | 0 | 0 | 0 |
| rs11833399 | benign | 0 | 0 | 0 |
| rs36023382 | benign | 1 | 1 | 1 |
| rs61754444 | benign | 0 | 0 | 0 |
| rs28935168 | pathogenic | 0 | 0 | 0 |
| rs28934907 | pathogenic | 0 | 0 | 0 |
| rs28934904 | pathogenic | 1 | 1 | 0 |
| rs61748392 | pathogenic | 1 | 1 | 0 |
| rs28934908 | pathogenic | 0 | 0 | 0 |
| rs28934905 | pathogenic | 0 | 0 | 0 |
| rs28934906 | pathogenic | 1 | 1 | 0 |
| rs606231307 | pathogenic | 0 | 0 | 0 |
| rs9333296 | benign | 0 | 0 | 0 |
| rs3218614 | benign | 0 | 0 | 0 |

| | | | | |
|---|---|---|---|---|
| rs2230792 | benign | 0 | 0 | 0 |
| rs2230793 | benign | 0 | 0 | 0 |
| rs2230794 | benign | 0 | 0 | 0 |
| rs749052963 | pathogenic | 0 | 0 | 0 |
| rs2230798 | benign | 0 | 0 | 0 |
| rs2230795 | benign | 0 | 0 | 0 |
| rs3204145 | benign | 0 | 0 | 0 |
| rs1538660 | benign | 0 | 0 | 0 |
| rs9878928 | benign | 0 | 0 | 0 |
| rs104893742 | pathogenic | 0 | 0 | 0 |
| rs28936702 | pathogenic | 0 | 0 | 0 |
| rs28936703 | pathogenic | 0 | 0 | 0 |
| rs769392 | benign | 0 | 0 | 0 |
| rs769402 | benign | 1 | 1 | 1 |
| rs104894884 | pathogenic | 1 | 1 | 1 |
| rs1801316 | benign | 1 | 1 | 1 |
| rs104894885 | pathogenic | 1 | 1 | 1 |
| rs782572 | benign | 0 | 0 | 0 |
| rs151166046 | pathogenic | 0 | 0 | 0 |
| rs772153760 | pathogenic | 1 | 1 | 0 |
| rs35405862 | benign | 0 | 0 | 0 |
| rs146571352 | pathogenic | 0 | 1 | 0 |
| rs374698153 | pathogenic | 0 | 0 | 0 |
| rs7594497 | benign | 0 | 0 | 0 |
| rs80338716 | pathogenic | 0 | 0 | 0 |
| rs80338719 | pathogenic | 1 | 1 | 1 |
| rs80338721 | pathogenic | 0 | 0 | 0 |
| rs730882249 | pathogenic | 0 | 0 | 0 |
| rs146262009 | pathogenic | 0 | 0 | 0 |
| rs104895083 | pathogenic | 1 | 1 | 1 |
| rs104895100 | benign | 1 | 1 | 1 |
| rs104895085 | pathogenic | 0 | 0 | 0 |
| rs28940580 | pathogenic | 0 | 0 | 0 |
| rs104895098 | pathogenic | 0 | 0 | 0 |
| rs28940578 | pathogenic | 0 | 0 | 0 |
| rs61752717 | pathogenic | 0 | 0 | 0 |
| rs28940579 | pathogenic | 0 | 0 | 0 |
| rs104895097 | pathogenic | 0 | 0 | 0 |
| rs140425651 | benign | 1 | 1 | 0 |
| rs768090444 | pathogenic | 0 | 0 | 0 |
| rs764459544 | pathogenic | 0 | 0 | 0 |
| rs80338802 | pathogenic | 1 | 1 | 1 |
| rs149969786 | pathogenic | 1 | 1 | 0 |
| rs115311625 | pathogenic | 1 | 1 | 1 |
| rs778768583 | pathogenic | 1 | 1 | 1 |
| rs892090 | benign | 0 | 0 | 0 |
| rs28929498 | pathogenic | 1 | 1 | 1 |
| rs137854467 | pathogenic | 0 | 0 | 0 |
| rs137854480 | pathogenic | 0 | 0 | 0 |
| rs148682262 | benign | 0 | 0 | 0 |
| rs137852744 | pathogenic | 0 | 0 | 0 |
| rs137852745 | pathogenic | 0 | 1 | 0 |
| rs137852749 | pathogenic | 0 | 1 | 0 |
| rs137852746 | pathogenic | 0 | 1 | 0 |

| | | | | |
|---|---|---|---|---|
| rs145633958 | benign | 0 | 0 | 0 |
| rs141679950 | benign | 0 | 0 | 0 |
| rs76397662 | benign | 1 | 1 | 1 |
| rs370736139 | benign | 0 | 0 | 0 |
| rs781750106 | benign | 0 | 0 | 0 |
| rs546866208 | benign | 0 | 0 | 0 |
| rs142338976 | benign | 0 | 0 | 0 |
| rs115272158 | benign | 0 | 0 | 0 |
| rs115423919 | benign | 0 | 0 | 0 |
| rs537874538 | benign | 0 | 0 | 0 |
| rs9282834 | benign | 0 | 0 | 0 |
| rs201745826 | benign | 0 | 0 | 0 |
| rs75873440 | pathogenic | 0 | 0 | 0 |
| rs147219360 | benign | 0 | 0 | 0 |
| rs77939446 | pathogenic | 0 | 0 | 0 |
| rs77558292 | pathogenic | 0 | 0 | 0 |
| rs80069458 | pathogenic | 0 | 0 | 0 |
| rs76262710 | pathogenic | 0 | 0 | 0 |
| rs79781594 | pathogenic | 0 | 0 | 0 |
| rs77316810 | pathogenic | 0 | 0 | 0 |
| rs77503355 | pathogenic | 0 | 0 | 0 |
| rs79890926 | pathogenic | 0 | 0 | 0 |
| rs377767402 | benign | 0 | 0 | 0 |
| rs121913308 | pathogenic | 0 | 0 | 0 |
| rs55846256 | benign | 0 | 0 | 0 |
| rs75076352 | pathogenic | 0 | 0 | 0 |
| rs75996173 | pathogenic | 0 | 0 | 0 |
| rs77709286 | pathogenic | 0 | 0 | 0 |
| rs151058706 | benign | 0 | 0 | 0 |
| rs138964358 | benign | 0 | 0 | 0 |
| rs137852353 | pathogenic | 1 | 1 | 0 |
| rs201196020 | benign | 0 | 0 | 0 |
| rs137853201 | pathogenic | 1 | 1 | 1 |
| rs113994174 | pathogenic | 1 | 1 | 1 |
| rs104893674 | pathogenic | 1 | 1 | 1 |
| rs779519249 | benign | 1 | 1 | 1 |
| rs9606756 | benign | 0 | 0 | 0 |
| rs75680863 | benign | 1 | 1 | 1 |
| rs35838082 | benign | 0 | 0 | 0 |
| rs1801198 | benign | 0 | 0 | 0 |
| rs117458738 | benign | 1 | 1 | 0 |
| rs9621049 | benign | 1 | 1 | 0 |
| rs1131603 | benign | 1 | 1 | 0 |
| rs4820889 | benign | 0 | 0 | 0 |
| rs777767978 | pathogenic | 1 | 1 | 0 |
| rs147399860 | benign | 1 | 1 | 1 |
| rs104893950 | pathogenic | 0 | 0 | 0 |
| rs777308612 | benign | 0 | 0 | 0 |
| rs151091410 | benign | 0 | 0 | 0 |
| rs11537667 | benign | 0 | 0 | 0 |
| rs267606621 | pathogenic | 1 | 1 | 0 |
| rs143058902 | benign | 0 | 0 | 0 |
| rs370261904 | pathogenic | 0 | 0 | 0 |
| rs141169746 | benign | 0 | 0 | 0 |

| | | | | |
|---|---|---|---|---|
| rs111033436 | benign | 0 | 0 | 0 |
| rs148533341 | benign | 0 | 0 | 0 |
| rs143827620 | benign | 0 | 0 | 0 |
| rs4935502 | benign | 0 | 0 | 0 |
| rs61735473 | benign | 0 | 0 | 0 |
| rs267606932 | pathogenic | 1 | 1 | 0 |
| rs61730754 | benign | 0 | 0 | 0 |
| rs374491359 | pathogenic | 1 | 1 | 1 |
| rs761800077 | benign | 1 | 1 | 1 |
| rs760391381 | benign | 1 | 1 | 1 |
| rs104893885 | pathogenic | 0 | 0 | 0 |
| rs104893886 | pathogenic | 1 | 1 | 1 |
| rs104893884 | pathogenic | 0 | 0 | 0 |
| rs147715509 | benign | 0 | 0 | 0 |
| rs115533243 | benign | 0 | 0 | 0 |
| rs397509390 | pathogenic | 0 | 1 | 0 |
| rs80357134 | pathogenic | 0 | 0 | 0 |
| rs766004110 | benign | 1 | 1 | 1 |
| rs769650474 | benign | 0 | 0 | 0 |
| rs398122635 | benign | 1 | 1 | 0 |
| rs199522616 | pathogenic | 0 | 0 | 0 |
| rs28897672 | pathogenic | 1 | 1 | 0 |
| rs80357086 | pathogenic | 1 | 1 | 1 |
| rs80357064 | pathogenic | 1 | 1 | 0 |
| rs80356936 | benign | 0 | 0 | 0 |
| rs28897673 | benign | 0 | 0 | 0 |
| rs146747324 | benign | 0 | 0 | 0 |
| rs78635798 | pathogenic | 0 | 0 | 0 |
| rs75146158 | pathogenic | 1 | 1 | 1 |
| rs587782886 | benign | 0 | 0 | 0 |
| rs121434634 | pathogenic | 0 | 0 | 0 |
| rs121434635 | pathogenic | 0 | 0 | 0 |
| rs587777778 | pathogenic | 0 | 0 | 0 |
| rs118203967 | pathogenic | 0 | 0 | 0 |
| rs751375244 | pathogenic | 1 | 1 | 1 |
| rs104893863 | pathogenic | 1 | 1 | 0 |
| rs104893865 | pathogenic | 0 | 0 | 0 |
| rs104893864 | pathogenic | 1 | 1 | 1 |
| rs104893866 | pathogenic | 1 | 1 | 0 |
| rs561738497 | benign | 0 | 0 | 0 |
| rs57599352 | pathogenic | 1 | 1 | 1 |
| rs267607448 | pathogenic | 1 | 1 | 1 |
| rs63751596 | pathogenic | 0 | 0 | 0 |
| rs63751612 | benign | 0 | 0 | 0 |
| rs63751684 | benign | 0 | 0 | 0 |
| rs267607861 | benign | 0 | 0 | 0 |
| rs35831931 | benign | 0 | 0 | 0 |
| rs63750876 | benign | 0 | 0 | 0 |
| rs41295284 | benign | 0 | 0 | 0 |
| rs35502531 | benign | 0 | 0 | 0 |
| rs63750449 | benign | 0 | 0 | 0 |
| rs63750899 | pathogenic | 0 | 0 | 0 |
| rs55907433 | benign | 0 | 0 | 0 |
| rs63750217 | pathogenic | 0 | 0 | 0 |

| | | | | |
|---|---|---|---|---|
| rs63751275 | pathogenic | 0 | 0 | 0 |
| rs63750702 | benign | 1 | 1 | 1 |
| rs63750114 | benign | 0 | 0 | 0 |
| rs2020873 | benign | 0 | 0 | 0 |
| rs140195825 | benign | 1 | 1 | 1 |
| rs182763805 | benign | 0 | 0 | 0 |
| rs533696377 | benign | 0 | 0 | 0 |
| rs41278788 | benign | 0 | 0 | 0 |
| rs104894193 | pathogenic | 1 | 1 | 0 |
| rs3824915 | benign | 1 | 1 | 1 |
| rs104894196 | pathogenic | 0 | 0 | 0 |
| rs78681604 | benign | 1 | 1 | 1 |
| rs77321498 | benign | 1 | 1 | 1 |
| rs137852965 | pathogenic | 0 | 0 | 0 |
| rs137852961 | pathogenic | 0 | 0 | 0 |
| rs137852962 | pathogenic | 1 | 1 | 0 |
| rs753376100 | pathogenic | 0 | 0 | 0 |
| rs137852968 | pathogenic | 1 | 1 | 1 |
| rs137852959 | pathogenic | 1 | 1 | 0 |
| rs137852967 | pathogenic | 0 | 0 | 0 |
| rs113437301 | benign | 1 | 1 | 1 |
| rs61747482 | benign | 1 | 1 | 1 |
| rs2234919 | benign | 1 | 1 | 0 |
| rs121908863 | pathogenic | 0 | 0 | 0 |
| rs140455771 | pathogenic | 0 | 0 | 0 |
| rs121908810 | pathogenic | 1 | 1 | 1 |
| rs143879853 | benign | 0 | 0 | 0 |
| rs188319622 | benign | 0 | 0 | 0 |
| rs121918645 | pathogenic | 0 | 0 | 0 |
| rs115352435 | pathogenic | 0 | 0 | 0 |
| rs121434427 | pathogenic | 1 | 1 | 1 |
| rs121434428 | pathogenic | 0 | 0 | 0 |
| rs35086265 | benign | 1 | 1 | 1 |
| rs11576415 | benign | 0 | 0 | 0 |
| rs121434429 | pathogenic | 1 | 1 | 1 |
| rs4074536 | benign | 1 | 1 | 1 |
| rs10801999 | benign | 0 | 0 | 0 |
| rs121434550 | pathogenic | 0 | 0 | 0 |
| rs148057999 | pathogenic | 0 | 0 | 0 |
| rs28730716 | benign | 0 | 0 | 0 |
| rs142036299 | benign | 0 | 0 | 0 |
| rs762381137 | pathogenic | 0 | 0 | 0 |
| rs121434549 | pathogenic | 0 | 0 | 0 |
| rs139228801 | pathogenic | 0 | 0 | 0 |
| rs147219463 | benign | 0 | 0 | 0 |
| rs1131857 | benign | 0 | 0 | 0 |
| rs121917874 | pathogenic | 0 | 1 | 0 |
| rs2228056 | benign | 0 | 0 | 0 |
| rs121917870 | pathogenic | 0 | 0 | 0 |
| rs121917866 | pathogenic | 0 | 0 | 0 |
| rs11921054 | benign | 0 | 0 | 0 |
| rs121917868 | pathogenic | 0 | 0 | 0 |
| rs28931603 | pathogenic | 0 | 0 | 0 |
| rs104895295 | pathogenic | 1 | 1 | 0 |

| | | | | |
|---|---|---|---|---|
| rs104895382 | pathogenic | 0 | 0 | 0 |
| rs104895298 | pathogenic | 0 | 0 | 0 |
| rs104895300 | pathogenic | 0 | 0 | 0 |
| rs104895332 | pathogenic | 0 | 0 | 0 |
| rs758026399 | pathogenic | 1 | 1 | 1 |
| rs104895366 | pathogenic | 0 | 0 | 0 |
| rs104895304 | pathogenic | 0 | 0 | 0 |
| rs759997079 | pathogenic | 0 | 0 | 0 |
| rs121917789 | pathogenic | 0 | 0 | 0 |
| rs104895319 | pathogenic | 0 | 0 | 0 |
| rs104895317 | pathogenic | 1 | 1 | 0 |
| rs28934897 | pathogenic | 0 | 0 | 0 |
| rs104895380 | benign | 0 | 0 | 0 |
| rs104895360 | pathogenic | 0 | 0 | 0 |
| rs28936396 | pathogenic | 1 | 1 | 0 |
| rs121909307 | pathogenic | 1 | 1 | 1 |
| rs28938472 | pathogenic | 1 | 1 | 0 |
| rs121909308 | pathogenic | 1 | 1 | 0 |
| rs121909309 | pathogenic | 0 | 0 | 0 |
| rs121913013 | benign | 1 | 1 | 1 |
| rs201855245 | benign | 1 | 1 | 1 |
| rs2230232 | benign | 1 | 1 | 0 |
| rs113451409 | benign | 0 | 0 | 0 |
| rs121913011 | pathogenic | 1 | 1 | 0 |
| rs185821167 | pathogenic | 1 | 1 | 0 |
| rs2230234 | benign | 0 | 0 | 0 |
| rs193922639 | benign | 1 | 1 | 0 |
| rs774863785 | pathogenic | 1 | 1 | 0 |
| rs576404380 | pathogenic | 1 | 1 | 0 |
| rs2230235 | benign | 1 | 1 | 0 |
| rs200484060 | pathogenic | 1 | 1 | 0 |
| rs199761749 | benign | 0 | 0 | 0 |
| rs199681901 | benign | 0 | 0 | 0 |
| rs121908951 | pathogenic | 0 | 0 | 0 |
| rs769251460 | benign | 0 | 0 | 0 |
| rs28937320 | pathogenic | 0 | 0 | 0 |
| rs200607527 | pathogenic | 0 | 0 | 0 |
| rs137852640 | pathogenic | 1 | 1 | 1 |
| rs137852636 | pathogenic | 1 | 1 | 1 |
| rs137852638 | pathogenic | 0 | 0 | 0 |
| rs142637231 | pathogenic | 0 | 0 | 0 |
| rs151328418 | benign | 0 | 1 | 0 |
| rs137852639 | pathogenic | 1 | 1 | 1 |
| rs34605667 | benign | 0 | 0 | 0 |
| rs104894073 | pathogenic | 1 | 1 | 0 |
| rs56298569 | pathogenic | 0 | 0 | 0 |
| rs757600616 | pathogenic | 0 | 0 | 0 |
| rs139194636 | pathogenic | 0 | 0 | 0 |
| rs369517993 | pathogenic | 1 | 1 | 1 |
| rs138846036 | benign | 1 | 1 | 1 |
| rs200808744 | pathogenic | 1 | 1 | 1 |
| rs2255543 | benign | 0 | 0 | 0 |
| rs149165166 | pathogenic | 0 | 0 | 0 |
| rs544956641 | pathogenic | 1 | 1 | 1 |

| | | | | |
|---|---|---|---|---|
| rs28941783 | pathogenic | 0 | 0 | 0 |
| rs562853291 | pathogenic | 1 | 1 | 1 |
| rs28942100 | pathogenic | 0 | 0 | 0 |
| rs120074174 | pathogenic | 1 | 1 | 1 |
| rs120074171 | pathogenic | 1 | 1 | 1 |
| rs120074173 | pathogenic | 0 | 0 | 0 |
| rs120074172 | pathogenic | 1 | 1 | 1 |
| rs767159114 | pathogenic | 1 | 1 | 1 |
| rs104893657 | pathogenic | 0 | 0 | 0 |
| rs104893656 | pathogenic | 0 | 0 | 0 |
| rs104893659 | pathogenic | 0 | 0 | 0 |
| rs104893658 | pathogenic | 0 | 0 | 0 |
| rs200386310 | pathogenic | 0 | 0 | 0 |
| rs2072712 | benign | 0 | 0 | 0 |
| rs35274867 | benign | 0 | 0 | 0 |
| rs35343172 | benign | 0 | 0 | 0 |
| rs10079250 | benign | 0 | 0 | 0 |
| rs34951517 | benign | 0 | 0 | 0 |
| rs267606695 | pathogenic | 0 | 0 | 0 |
| rs149391728 | benign | 0 | 0 | 0 |
| rs387906598 | pathogenic | 1 | 1 | 0 |
| rs137854578 | pathogenic | 0 | 0 | 0 |
| rs141519952 | benign | 0 | 0 | 0 |
| rs146572883 | benign | 0 | 0 | 0 |
| rs111866410 | benign | 0 | 0 | 0 |
| rs139387758 | benign | 1 | 1 | 1 |
| rs35672074 | benign | 1 | 1 | 1 |
| rs121434322 | pathogenic | 0 | 0 | 0 |
| rs150884181 | benign | 0 | 0 | 0 |
| rs35867471 | benign | 0 | 0 | 0 |
| rs2867802 | benign | 0 | 0 | 0 |
| rs137902889 | benign | 0 | 0 | 0 |
| rs369804864 | pathogenic | 0 | 0 | 0 |
| rs781050795 | pathogenic | 0 | 0 | 0 |
| rs2305799 | benign | 0 | 0 | 0 |
| rs2229531 | benign | 0 | 0 | 0 |
| rs147025508 | benign | 0 | 0 | 0 |
| rs76857106 | pathogenic | 0 | 0 | 0 |
| rs7247284 | benign | 0 | 0 | 0 |
| rs62619782 | benign | 0 | 0 | 0 |
| rs377244188 | pathogenic | 0 | 0 | 0 |
| rs75718910 | pathogenic | 0 | 0 | 0 |
| rs79843600 | pathogenic | 0 | 0 | 0 |
| rs75037667 | pathogenic | 1 | 1 | 1 |
| rs121913006 | pathogenic | 0 | 0 | 0 |
| rs397517925 | pathogenic | 1 | 1 | 1 |
| rs267606831 | pathogenic | 1 | 1 | 1 |
| rs121909667 | pathogenic | 1 | 1 | 1 |
| rs121909669 | pathogenic | 1 | 1 | 1 |
| rs121909668 | pathogenic | 1 | 1 | 1 |
| rs121909671 | pathogenic | 1 | 1 | 1 |
| rs534596036 | benign | 0 | 0 | 0 |
| rs144037629 | pathogenic | 0 | 0 | 0 |
| rs11546406 | benign | 0 | 0 | 0 |

| | | | | |
|---|---|---|---|---|
| rs143697995 | pathogenic | 0 | 0 | 0 |
| rs121908331 | pathogenic | 0 | 0 | 0 |
| rs149377346 | benign | 0 | 0 | 0 |
| rs104894086 | pathogenic | 1 | 1 | 0 |
| rs104894089 | pathogenic | 0 | 0 | 0 |
| rs104894090 | pathogenic | 1 | 1 | 0 |
| rs387907235 | pathogenic | 0 | 0 | 0 |
| rs137852689 | pathogenic | 0 | 0 | 0 |
| rs137852690 | pathogenic | 1 | 1 | 0 |
| rs104894087 | pathogenic | 0 | 0 | 0 |
| rs104894085 | pathogenic | 0 | 0 | 0 |
| rs79383654 | benign | 0 | 0 | 0 |
| rs121434239 | pathogenic | 0 | 0 | 0 |
| rs121434240 | pathogenic | 0 | 0 | 0 |
| rs121434236 | pathogenic | 0 | 0 | 0 |
| rs121434238 | pathogenic | 1 | 1 | 1 |
| rs121918107 | pathogenic | 1 | 1 | 1 |
| rs121918103 | pathogenic | 1 | 1 | 1 |
| rs121918104 | pathogenic | 1 | 1 | 1 |
| rs61747614 | benign | 1 | 1 | 1 |
| rs555329870 | benign | 0 | 0 | 0 |
| rs186094931 | benign | 0 | 0 | 0 |
| rs267606785 | pathogenic | 0 | 0 | 0 |
| rs34474955 | benign | 0 | 0 | 0 |
| rs1126497 | benign | 1 | 1 | 1 |
| rs397514661 | pathogenic | 0 | 0 | 0 |
| rs189732445 | benign | 0 | 0 | 0 |
| rs146685071 | benign | 0 | 0 | 0 |
| rs121908636 | pathogenic | 1 | 1 | 0 |
| rs121908643 | pathogenic | 0 | 0 | 0 |
| rs121908644 | pathogenic | 1 | 1 | 1 |
| rs575001023 | pathogenic | 1 | 1 | 1 |
| rs121908646 | pathogenic | 1 | 1 | 0 |
| rs121908638 | pathogenic | 1 | 1 | 1 |
| rs777828000 | pathogenic | 1 | 1 | 1 |
| rs74923032 | benign | 0 | 0 | 0 |
| rs192838388 | pathogenic | 1 | 1 | 1 |
| rs148918985 | pathogenic | 1 | 1 | 1 |
| rs773540764 | benign | 1 | 1 | 1 |
| rs370595480 | pathogenic | 1 | 1 | 1 |
| rs762387914 | pathogenic | 1 | 1 | 1 |
| rs121908642 | pathogenic | 1 | 1 | 1 |
| rs183276875 | pathogenic | 1 | 1 | 1 |
| rs121908648 | pathogenic | 1 | 1 | 1 |
| rs121908647 | pathogenic | 1 | 1 | 1 |
| rs121908640 | pathogenic | 1 | 1 | 1 |
| rs771937610 | pathogenic | 1 | 1 | 1 |
| rs121908641 | pathogenic | 1 | 1 | 1 |
| rs5745068 | benign | 0 | 0 | 0 |
| rs73155056 | benign | 1 | 1 | 1 |
| rs148788180 | benign | 0 | 0 | 0 |
| rs121918137 | pathogenic | 0 | 0 | 0 |
| rs41269593 | benign | 0 | 0 | 0 |
| rs11159 | benign | 1 | 1 | 0 |

| | | | | |
|---|---|---|---|---|
| rs114733570 | pathogenic | 1 | 1 | 0 |
| rs3211783 | benign | 0 | 0 | 0 |
| rs4762 | benign | 0 | 0 | 0 |
| rs74315283 | pathogenic | 0 | 0 | 0 |
| rs111033559 | pathogenic | 0 | 0 | 0 |
| rs111033560 | pathogenic | 1 | 1 | 1 |
| rs749962743 | benign | 0 | 0 | 0 |
| rs267606986 | pathogenic | 0 | 0 | 0 |
| rs34516635 | benign | 0 | 0 | 0 |
| rs117440569 | benign | 0 | 0 | 0 |
| rs61740877 | benign | 0 | 0 | 0 |
| rs121918352 | pathogenic | 0 | 0 | 0 |
| rs183974372 | benign | 0 | 0 | 0 |
| rs140511204 | benign | 1 | 1 | 0 |
| rs6040067 | benign | 0 | 0 | 0 |
| rs199674138 | benign | 0 | 0 | 0 |
| rs121918350 | pathogenic | 0 | 0 | 0 |
| rs138292988 | benign | 1 | 1 | 1 |
| rs145288947 | benign | 0 | 0 | 0 |
| rs369714529 | benign | 0 | 0 | 0 |
| rs200945419 | benign | 0 | 0 | 0 |
| rs116959874 | benign | 0 | 0 | 0 |
| rs141659620 | pathogenic | 1 | 1 | 0 |
| rs111475461 | benign | 0 | 0 | 0 |
| rs61755320 | pathogenic | 0 | 0 | 0 |
| rs748600162 | pathogenic | 1 | 1 | 1 |
| rs369227537 | pathogenic | 1 | 1 | 1 |
| rs372981030 | pathogenic | 1 | 1 | 1 |
| rs137854486 | pathogenic | 0 | 0 | 0 |
| rs137854487 | pathogenic | 0 | 0 | 0 |
| rs199905054 | pathogenic | 0 | 0 | 0 |
| rs771438170 | pathogenic | 0 | 0 | 0 |
| rs774949844 | pathogenic | 0 | 0 | 0 |
| rs4899145 | benign | 0 | 0 | 0 |
| rs200642525 | benign | 0 | 0 | 0 |
| rs190140598 | pathogenic | 0 | 0 | 0 |
| rs2250889 | benign | 0 | 0 | 0 |
| rs17577 | benign | 0 | 0 | 0 |
| rs142032681 | benign | 0 | 0 | 0 |
| rs144577963 | benign | 0 | 0 | 0 |
| rs61754623 | benign | 0 | 0 | 0 |
| rs139873200 | benign | 0 | 0 | 0 |
| rs532530906 | benign | 0 | 0 | 0 |
| rs145762716 | benign | 0 | 0 | 0 |
| rs34062020 | benign | 0 | 0 | 0 |
| rs35258161 | benign | 0 | 0 | 0 |
| rs149792649 | benign | 0 | 0 | 0 |
| rs78101774 | benign | 0 | 0 | 0 |
| rs766466297 | pathogenic | 0 | 1 | 0 |
| rs140467171 | pathogenic | 0 | 0 | 0 |
| rs141340466 | pathogenic | 0 | 0 | 0 |
| rs201555303 | pathogenic | 0 | 0 | 0 |
| rs143592405 | pathogenic | 0 | 0 | 0 |
| rs781249411 | pathogenic | 0 | 0 | 0 |

| | | | | |
|---|---|---|---|---|
| rs116644127 | benign | 0 | 0 | 0 |
| rs151146863 | benign | 1 | 1 | 1 |
| rs116440799 | benign | 0 | 0 | 0 |
| rs138958351 | benign | 0 | 0 | 0 |
| rs143319354 | benign | 0 | 0 | 0 |
| rs768669208 | pathogenic | 1 | 1 | 1 |
| rs28940289 | pathogenic | 1 | 1 | 0 |
| rs745656120 | pathogenic | 1 | 1 | 0 |
| rs387906987 | pathogenic | 0 | 0 | 0 |
| rs147240777 | benign | 0 | 0 | 0 |
| rs140945833 | benign | 0 | 0 | 0 |
| rs146614981 | benign | 0 | 0 | 0 |
| rs149878562 | benign | 0 | 0 | 0 |
| rs28938168 | pathogenic | 0 | 0 | 0 |
| rs33939927 | pathogenic | 1 | 1 | 1 |
| rs34995376 | pathogenic | 1 | 1 | 1 |
| rs201034481 | pathogenic | 1 | 1 | 1 |
| rs774069983 | benign | 0 | 0 | 0 |
| rs199422226 | pathogenic | 1 | 1 | 0 |
| rs199422225 | pathogenic | 1 | 1 | 1 |
| rs199422224 | pathogenic | 1 | 1 | 0 |
| rs149271416 | pathogenic | 1 | 1 | 0 |
| rs1801318 | benign | 1 | 1 | 1 |
| rs372691318 | pathogenic | 1 | 1 | 1 |
| rs147685849 | benign | 0 | 0 | 0 |
| rs387907199 | pathogenic | 0 | 0 | 0 |
| rs767122069 | pathogenic | 0 | 0 | 0 |
| rs2302427 | benign | 0 | 0 | 0 |
| rs201135441 | benign | 0 | 0 | 0 |
| rs201861847 | pathogenic | 0 | 0 | 0 |
| rs137852952 | pathogenic | 0 | 0 | 0 |
| rs16986560 | benign | 0 | 0 | 0 |
| rs147344332 | pathogenic | 1 | 1 | 1 |
| rs36027220 | pathogenic | 0 | 0 | 0 |
| rs192401229 | benign | 0 | 0 | 0 |
| rs121909106 | pathogenic | 1 | 1 | 0 |
| rs118204103 | pathogenic | 1 | 1 | 0 |
| rs118204104 | pathogenic | 1 | 1 | 0 |
| rs118204105 | pathogenic | 1 | 1 | 0 |
| rs118204106 | pathogenic | 0 | 0 | 0 |
| rs118204119 | pathogenic | 0 | 0 | 0 |
| rs118204107 | pathogenic | 0 | 0 | 0 |
| rs118204094 | pathogenic | 0 | 0 | 0 |
| rs118204098 | pathogenic | 1 | 1 | 0 |
| rs118204101 | pathogenic | 1 | 1 | 0 |
| rs118204096 | pathogenic | 1 | 1 | 0 |
| rs118204108 | pathogenic | 0 | 0 | 0 |
| rs536814318 | pathogenic | 0 | 0 | 0 |
| rs34413634 | pathogenic | 1 | 1 | 0 |
| rs118204100 | pathogenic | 0 | 0 | 0 |
| rs118204109 | pathogenic | 0 | 0 | 0 |
| rs118204116 | pathogenic | 1 | 1 | 0 |
| rs118204110 | pathogenic | 0 | 0 | 0 |
| rs118204099 | pathogenic | 0 | 0 | 0 |

| | | | | |
|---|---|---|---|---|
| rs118204111 | pathogenic | 0 | 0 | 0 |
| rs118204112 | pathogenic | 0 | 0 | 0 |
| rs118204113 | pathogenic | 0 | 0 | 0 |
| rs118204114 | pathogenic | 0 | 0 | 0 |
| rs118204115 | pathogenic | 0 | 0 | 0 |
| rs150428209 | benign | 0 | 0 | 0 |
| rs144949995 | benign | 0 | 0 | 0 |
| rs190023981 | benign | 0 | 0 | 0 |
| rs104894611 | pathogenic | 1 | 1 | 1 |
| rs28937580 | pathogenic | 1 | 1 | 1 |
| rs121908949 | pathogenic | 0 | 0 | 0 |
| rs104894612 | pathogenic | 1 | 1 | 1 |
| rs104894609 | pathogenic | 1 | 1 | 1 |
| rs104894610 | pathogenic | 1 | 1 | 1 |
| rs104894615 | pathogenic | 0 | 0 | 0 |
| rs104894603 | pathogenic | 0 | 0 | 0 |
| rs104894602 | pathogenic | 1 | 1 | 1 |
| rs121908948 | pathogenic | 1 | 1 | 1 |
| rs104894608 | pathogenic | 1 | 1 | 1 |
| rs371545673 | benign | 0 | 0 | 0 |
| rs199925238 | benign | 0 | 0 | 0 |
| rs146313451 | benign | 0 | 0 | 0 |
| rs121918193 | pathogenic | 1 | 1 | 0 |
| rs2228500 | benign | 0 | 0 | 0 |
| rs121918194 | pathogenic | 1 | 1 | 0 |
| rs121918196 | pathogenic | 0 | 0 | 0 |
| rs41291971 | benign | 0 | 0 | 0 |
| rs199473690 | pathogenic | 0 | 0 | 0 |
| rs2228613 | benign | 0 | 0 | 0 |
| rs267607130 | pathogenic | 1 | 1 | 1 |
| rs77615401 | benign | 1 | 1 | 1 |
| rs374618872 | benign | 1 | 1 | 1 |
| rs146165014 | benign | 0 | 0 | 0 |
| rs144586070 | benign | 0 | 0 | 0 |
| rs121918541 | pathogenic | 0 | 0 | 0 |
| rs142105943 | benign | 0 | 0 | 0 |
| rs121918540 | pathogenic | 0 | 0 | 0 |
| rs6791 | benign | 0 | 0 | 0 |
| rs61736587 | pathogenic | 0 | 0 | 0 |
| rs61736586 | benign | 0 | 0 | 0 |
| rs121917741 | pathogenic | 1 | 1 | 1 |
| rs41302885 | benign | 0 | 0 | 0 |
| rs141026028 | benign | 0 | 0 | 0 |
| rs193922493 | pathogenic | 0 | 0 | 0 |
| rs73598374 | benign | 0 | 0 | 0 |
| rs121908725 | pathogenic | 1 | 1 | 0 |
| rs199422328 | pathogenic | 0 | 1 | 0 |
| rs11555566 | benign | 1 | 1 | 1 |
| rs267606634 | pathogenic | 0 | 0 | 0 |
| rs121908714 | pathogenic | 1 | 1 | 0 |
| rs121908717 | pathogenic | 1 | 1 | 0 |
| rs267606635 | pathogenic | 1 | 1 | 0 |
| rs121908739 | pathogenic | 0 | 0 | 0 |
| rs121908735 | pathogenic | 0 | 1 | 0 |

| | | | | |
|---|---|---|---|---|
| rs121908716 | pathogenic | 0 | 1 | 0 |
| rs121908723 | pathogenic | 0 | 1 | 0 |
| rs121908721 | pathogenic | 0 | 0 | 0 |
| rs199422327 | pathogenic | 0 | 0 | 0 |
| rs121908715 | pathogenic | 0 | 0 | 0 |
| rs587783563 | pathogenic | 0 | 0 | 0 |
| rs104894780 | pathogenic | 0 | 0 | 0 |
| rs56030372 | pathogenic | 1 | 1 | 1 |
| rs104894782 | pathogenic | 1 | 1 | 1 |
| rs149495971 | benign | 0 | 0 | 0 |
| rs118203942 | pathogenic | 1 | 1 | 0 |
| rs118203939 | pathogenic | 0 | 0 | 0 |
| rs118203938 | pathogenic | 0 | 0 | 0 |
| rs118203943 | pathogenic | 0 | 0 | 0 |
| rs118203940 | pathogenic | 0 | 0 | 0 |
| rs727503809 | pathogenic | 0 | 0 | 0 |
| rs1065757 | benign | 0 | 0 | 0 |
| rs1071598 | benign | 0 | 0 | 0 |
| rs25414 | benign | 0 | 0 | 0 |
| rs118203944 | pathogenic | 0 | 0 | 0 |
| rs118203941 | pathogenic | 0 | 0 | 0 |
| rs144222167 | benign | 0 | 0 | 0 |
| rs201101343 | pathogenic | 0 | 0 | 0 |
| rs144601717 | benign | 0 | 0 | 0 |
| rs267607129 | pathogenic | 0 | 0 | 0 |
| rs104894729 | pathogenic | 0 | 0 | 0 |
| rs104894727 | pathogenic | 0 | 0 | 0 |
| rs104894725 | pathogenic | 0 | 0 | 0 |
| rs104894515 | pathogenic | 1 | 1 | 1 |
| rs74688377 | benign | 0 | 0 | 0 |
| rs121908513 | pathogenic | 1 | 1 | 0 |
| rs121908509 | pathogenic | 0 | 0 | 0 |
| rs121908514 | pathogenic | 1 | 1 | 0 |
| rs121908512 | pathogenic | 1 | 1 | 0 |
| rs121908510 | pathogenic | 0 | 0 | 0 |
| rs121908511 | pathogenic | 1 | 1 | 1 |
| rs121908518 | pathogenic | 1 | 1 | 1 |
| rs34358665 | benign | 0 | 0 | 0 |
| rs147568834 | benign | 0 | 0 | 0 |
| rs201717599 | benign | 0 | 0 | 0 |
| rs61744487 | benign | 0 | 0 | 0 |
| rs370627877 | pathogenic | 1 | 1 | 1 |
| rs368593151 | pathogenic | 1 | 1 | 1 |
| rs114363936 | benign | 0 | 0 | 0 |
| rs137853333 | pathogenic | 0 | 0 | 0 |
| rs137853080 | pathogenic | 1 | 1 | 1 |
| rs137853077 | pathogenic | 1 | 1 | 1 |
| rs780749732 | benign | 1 | 1 | 1 |
| rs137853081 | pathogenic | 1 | 1 | 0 |
| rs137853078 | pathogenic | 0 | 0 | 0 |
| rs121913315 | pathogenic | 1 | 1 | 1 |
| rs752513525 | pathogenic | 1 | 1 | 1 |
| rs104894918 | pathogenic | 0 | 0 | 0 |
| rs104894917 | pathogenic | 0 | 0 | 0 |

| | | | | |
|---|---|---|---|---|
| rs104894919 | pathogenic | 0 | 0 | 0 |
| rs28939672 | pathogenic | 0 | 0 | 0 |
| rs104894173 | pathogenic | 0 | 0 | 0 |
| rs104894179 | pathogenic | 0 | 0 | 0 |
| rs104894174 | pathogenic | 1 | 1 | 0 |
| rs104894178 | pathogenic | 1 | 1 | 0 |
| rs28939714 | pathogenic | 1 | 1 | 1 |
| rs104894270 | pathogenic | 1 | 1 | 1 |
| rs3765571 | benign | 0 | 0 | 0 |
| rs752669339 | pathogenic | 1 | 1 | 1 |
| rs537043237 | pathogenic | 1 | 1 | 1 |
| rs774047299 | pathogenic | 1 | 1 | 1 |
| rs121434583 | pathogenic | 1 | 1 | 1 |
| rs121907927 | pathogenic | 0 | 0 | 0 |
| rs121907925 | pathogenic | 0 | 1 | 0 |
| rs746212325 | pathogenic | 0 | 0 | 0 |
| rs754637179 | pathogenic | 0 | 0 | 0 |
| rs780882740 | pathogenic | 0 | 0 | 0 |
| rs121918130 | pathogenic | 0 | 0 | 0 |
| rs121918129 | pathogenic | 0 | 0 | 0 |
| rs13297509 | pathogenic | 0 | 0 | 0 |
| rs746867724 | pathogenic | 0 | 0 | 0 |
| rs121918128 | pathogenic | 0 | 0 | 0 |
| rs752300607 | pathogenic | 0 | 0 | 0 |
| rs121964860 | pathogenic | 0 | 0 | 0 |
| rs45466197 | pathogenic | 1 | 1 | 1 |
| rs3730238 | benign | 1 | 1 | 1 |
| rs730881125 | pathogenic | 1 | 1 | 1 |
| rs121964861 | pathogenic | 1 | 1 | 1 |
| rs397516484 | pathogenic | 0 | 0 | 0 |
| rs367785431 | pathogenic | 0 | 0 | 0 |
| rs141121678 | pathogenic | 0 | 0 | 0 |
| rs12720263 | benign | 0 | 0 | 0 |
| rs2304256 | benign | 0 | 0 | 0 |
| rs2304255 | benign | 0 | 0 | 0 |
| rs121909045 | pathogenic | 0 | 0 | 0 |
| rs397508796 | pathogenic | 0 | 0 | 0 |
| rs151020603 | pathogenic | 0 | 0 | 0 |
| rs77284892 | pathogenic | 0 | 0 | 0 |
| rs368505753 | pathogenic | 0 | 0 | 0 |
| rs115545701 | pathogenic | 0 | 0 | 0 |
| rs121908749 | pathogenic | 0 | 0 | 0 |
| rs75961395 | pathogenic | 0 | 0 | 0 |
| rs121908750 | pathogenic | 0 | 0 | 0 |
| rs121908751 | pathogenic | 0 | 0 | 0 |
| rs397508464 | pathogenic | 0 | 0 | 0 |
| rs397508461 | pathogenic | 0 | 0 | 0 |
| rs121909031 | pathogenic | 0 | 0 | 0 |
| rs113993958 | pathogenic | 0 | 0 | 0 |
| rs77834169 | pathogenic | 0 | 0 | 0 |
| rs78655421 | pathogenic | 0 | 0 | 0 |
| rs141723617 | benign | 0 | 0 | 0 |
| rs35516286 | benign | 0 | 0 | 0 |
| rs1800079 | pathogenic | 1 | 1 | 0 |

| | | | | |
|---|---|---|---|---|
| rs80282562 | pathogenic | 0 | 0 | 0 |
| rs121908752 | pathogenic | 0 | 0 | 0 |
| rs556880586 | pathogenic | 0 | 0 | 0 |
| rs121908760 | pathogenic | 0 | 0 | 0 |
| rs397508350 | pathogenic | 0 | 0 | 0 |
| rs374946172 | pathogenic | 0 | 0 | 0 |
| rs397508387 | pathogenic | 0 | 0 | 0 |
| rs201386642 | pathogenic | 0 | 0 | 0 |
| rs121909012 | pathogenic | 0 | 0 | 0 |
| rs121909008 | pathogenic | 0 | 0 | 0 |
| rs149790377 | pathogenic | 0 | 0 | 0 |
| rs397508442 | pathogenic | 0 | 0 | 0 |
| rs121909035 | pathogenic | 0 | 0 | 0 |
| rs386134230 | pathogenic | 0 | 0 | 0 |
| rs1800112 | benign | 0 | 0 | 0 |
| rs144055758 | pathogenic | 0 | 0 | 0 |
| rs142394380 | pathogenic | 0 | 0 | 0 |
| rs121909036 | pathogenic | 0 | 0 | 0 |
| rs78194216 | pathogenic | 0 | 0 | 0 |
| rs121909019 | pathogenic | 0 | 0 | 0 |
| rs121909020 | pathogenic | 0 | 0 | 0 |
| rs200321110 | pathogenic | 0 | 0 | 0 |
| rs202179988 | pathogenic | 0 | 0 | 0 |
| rs121909037 | pathogenic | 0 | 0 | 0 |
| rs186045772 | pathogenic | 0 | 0 | 0 |
| rs139304906 | pathogenic | 0 | 0 | 0 |
| rs78802634 | pathogenic | 0 | 0 | 0 |
| rs121908761 | pathogenic | 0 | 0 | 0 |
| rs36210737 | pathogenic | 0 | 0 | 0 |
| rs139729994 | pathogenic | 0 | 0 | 0 |
| rs79850223 | pathogenic | 0 | 0 | 0 |
| rs1800120 | benign | 0 | 0 | 0 |
| rs74767530 | pathogenic | 0 | 0 | 0 |
| rs374705585 | pathogenic | 0 | 0 | 0 |
| rs1136287 | benign | 0 | 0 | 0 |
| rs6828 | benign | 0 | 0 | 0 |
| rs104894856 | pathogenic | 0 | 0 | 0 |
| rs104894860 | pathogenic | 0 | 0 | 0 |
| rs61736892 | pathogenic | 0 | 0 | 0 |
| rs145807417 | benign | 0 | 0 | 0 |
| rs104894853 | pathogenic | 0 | 0 | 0 |
| rs199422227 | pathogenic | 0 | 0 | 0 |
| rs113993946 | pathogenic | 0 | 0 | 0 |
| rs199422231 | pathogenic | 0 | 0 | 0 |
| rs199422228 | pathogenic | 0 | 0 | 0 |
| rs142195196 | benign | 0 | 1 | 0 |
| rs148656104 | benign | 1 | 1 | 1 |
| rs141944844 | benign | 0 | 0 | 0 |
| rs104894342 | pathogenic | 0 | 0 | 0 |
| rs104894343 | pathogenic | 0 | 0 | 0 |
| rs104894344 | pathogenic | 0 | 0 | 0 |
| rs573322878 | benign | 0 | 0 | 0 |
| rs104894347 | pathogenic | 0 | 0 | 0 |
| rs28937882 | pathogenic | 0 | 0 | 0 |

| | | | | |
|---|---|---|---|---|
| rs193922702 | pathogenic | 0 | 0 | 0 |
| rs200786104 | benign | 0 | 0 | 0 |
| rs690016544 | pathogenic | 1 | 1 | 1 |
| rs28943592 | benign | 0 | 0 | 0 |
| rs28943594 | benign | 0 | 0 | 0 |
| rs121908957 | pathogenic | 0 | 0 | 0 |
| rs115408010 | benign | 1 | 1 | 1 |
| rs77591659 | benign | 0 | 0 | 0 |
| rs121918511 | pathogenic | 0 | 0 | 0 |
| rs538359948 | benign | 0 | 0 | 0 |
| rs76708142 | benign | 0 | 0 | 0 |
| rs121907913 | pathogenic | 1 | 1 | 0 |
| rs121907920 | pathogenic | 0 | 0 | 0 |
| rs121907923 | pathogenic | 1 | 1 | 0 |
| rs121907928 | pathogenic | 1 | 1 | 0 |
| rs121907919 | pathogenic | 1 | 1 | 0 |
| rs121907918 | pathogenic | 1 | 1 | 0 |
| rs74315493 | pathogenic | 1 | 1 | 1 |
| rs74315494 | pathogenic | 1 | 1 | 1 |
| rs145446060 | benign | 1 | 1 | 1 |
| rs550409143 | benign | 1 | 1 | 0 |

| compound_binding_site | cofactor_binding_site | nucleic_binding_site | ion_binding_site |
|---|---|---|---|
| 1 | 0 | 0 | 0 |
| 1 | 0 | 0 | 0 |
| 0 | 0 | 0 | 0 |
| 1 | 0 | 0 | 0 |
| 0 | 0 | 0 | 0 |
| 0 | 0 | 0 | 0 |
| 0 | 0 | 0 | 0 |
| 0 | 0 | 0 | 0 |
| 1 | 0 | 0 | 1 |
| 1 | 0 | 0 | 0 |
| 0 | 0 | 0 | 1 |
| 1 | 0 | 1 | 0 |
| 0 | 0 | 0 | 0 |
| 0 | 0 | 0 | 0 |
| 0 | 0 | 0 | 0 |
| 0 | 0 | 0 | 0 |
| 0 | 0 | 0 | 0 |
| 1 | 0 | 0 | 1 |
| 1 | 0 | 1 | 0 |
| 0 | 0 | 0 | 0 |
| 1 | 0 | 0 | 1 |
| 1 | 0 | 0 | 0 |
| 1 | 0 | 0 | 0 |
| 1 | 0 | 0 | 0 |
| 0 | 0 | 0 | 0 |
| 0 | 0 | 0 | 0 |
| 1 | 0 | 0 | 1 |
| 0 | 0 | 0 | 0 |
| 0 | 0 | 0 | 0 |
| 0 | 0 | 0 | 0 |
| 0 | 0 | 0 | 0 |
| 1 | 0 | 0 | 0 |
| 1 | 0 | 0 | 1 |
| 1 | 0 | 0 | 0 |
| 0 | 0 | 0 | 0 |
| 0 | 0 | 0 | 0 |
| 0 | 0 | 0 | 0 |
| 1 | 0 | 0 | 1 |
| 1 | 0 | 0 | 1 |
| 1 | 0 | 0 | 1 |
| 1 | 0 | 0 | 0 |
| 1 | 0 | 0 | 0 |
| 1 | 0 | 0 | 0 |
| 1 | 0 | 0 | 0 |
| 1 | 0 | 0 | 0 |
| 1 | 0 | 0 | 0 |
| 1 | 0 | 0 | 0 |
| 1 | 0 | 0 | 0 |
| 1 | 0 | 0 | 0 |
| 1 | 0 | 0 | 0 |
| 1 | 0 | 0 | 0 |
| 1 | 0 | 0 | 0 |
| 1 | 0 | 0 | 0 |

| | | | |
|---|---|---|---|
| 1 | 0 | 0 | 1 |
| 1 | 0 | 0 | 0 |
| 1 | 0 | 0 | 0 |
| 1 | 0 | 0 | 0 |
| 0 | 0 | 0 | 0 |
| 1 | 0 | 0 | 0 |
| 0 | 0 | 0 | 0 |
| 1 | 0 | 0 | 0 |
| 1 | 0 | 0 | 0 |
| 1 | 0 | 0 | 1 |
| 1 | 0 | 0 | 0 |
| 1 | 0 | 0 | 0 |
| 1 | 0 | 0 | 0 |
| 1 | 0 | 0 | 0 |
| 1 | 0 | 0 | 0 |
| 1 | 0 | 0 | 0 |
| 1 | 0 | 0 | 0 |
| 1 | 0 | 0 | 1 |
| 1 | 0 | 0 | 1 |
| 1 | 1 | 0 | 1 |
| 1 | 0 | 0 | 0 |
| 1 | 0 | 0 | 0 |
| 1 | 0 | 0 | 0 |
| 1 | 0 | 0 | 0 |
| 1 | 1 | 0 | 0 |
| 1 | 0 | 0 | 0 |
| 1 | 0 | 0 | 0 |
| 1 | 0 | 0 | 0 |
| 1 | 0 | 0 | 0 |
| 1 | 0 | 0 | 0 |
| 1 | 1 | 0 | 0 |
| 1 | 1 | 0 | 0 |
| 1 | 1 | 0 | 0 |
| 1 | 1 | 0 | 0 |
| 1 | 1 | 0 | 0 |
| 1 | 0 | 0 | 0 |
| 1 | 1 | 0 | 0 |
| 1 | 0 | 0 | 0 |
| 1 | 0 | 0 | 0 |
| 1 | 1 | 0 | 0 |
| 0 | 0 | 0 | 0 |
| 0 | 0 | 0 | 0 |
| 0 | 0 | 0 | 0 |
| 0 | 0 | 0 | 0 |
| 0 | 0 | 0 | 0 |
| 0 | 0 | 0 | 0 |
| 0 | 0 | 0 | 0 |
| 1 | 0 | 0 | 1 |
| 0 | 0 | 0 | 0 |
| 1 | 0 | 0 | 0 |
| 1 | 0 | 0 | 0 |
| 0 | 0 | 0 | 0 |
| 0 | 0 | 0 | 0 |
| 0 | 0 | 0 | 0 |

| | | | |
|---|---|---|---|
| 0 | 0 | 0 | 0 |
| 0 | 0 | 0 | 0 |
| 0 | 0 | 0 | 0 |
| 1 | 0 | 0 | 0 |
| 1 | 0 | 0 | 0 |
| 1 | 0 | 0 | 0 |
| 0 | 0 | 0 | 0 |
| 0 | 0 | 1 | 0 |
| 0 | 0 | 1 | 0 |
| 0 | 0 | 1 | 0 |
| 0 | 0 | 1 | 0 |
| 0 | 0 | 1 | 0 |
| 0 | 0 | 0 | 0 |
| 0 | 0 | 0 | 0 |
| 1 | 0 | 0 | 0 |
| 0 | 0 | 0 | 0 |
| 0 | 0 | 0 | 0 |
| 1 | 0 | 0 | 0 |
| 1 | 0 | 0 | 0 |
| 1 | 0 | 0 | 0 |
| 1 | 0 | 0 | 0 |
| 1 | 1 | 0 | 0 |
| 1 | 0 | 0 | 0 |
| 1 | 0 | 0 | 0 |
| 0 | 0 | 0 | 0 |
| 0 | 0 | 0 | 0 |
| 1 | 0 | 0 | 0 |
| 1 | 1 | 0 | 0 |
| 1 | 0 | 0 | 0 |
| 1 | 0 | 0 | 0 |
| 1 | 1 | 0 | 0 |
| 1 | 1 | 0 | 0 |
| 1 | 1 | 0 | 0 |
| 1 | 1 | 0 | 0 |
| 1 | 1 | 0 | 0 |
| 1 | 1 | 0 | 0 |
| 1 | 1 | 0 | 0 |
| 1 | 1 | 0 | 0 |
| 1 | 1 | 0 | 0 |
| 1 | 1 | 0 | 0 |
| 1 | 1 | 0 | 0 |
| 1 | 1 | 0 | 0 |
| 1 | 0 | 0 | 0 |
| 1 | 0 | 0 | 0 |
| 1 | 0 | 0 | 0 |
| 1 | 1 | 0 | 0 |
| 1 | 0 | 0 | 0 |
| 1 | 1 | 1 | 0 |
| 1 | 1 | 1 | 0 |
| 1 | 1 | 0 | 0 |
| 1 | 1 | 0 | 0 |
| 1 | 0 | 0 | 0 |
| 1 | 0 | 0 | 0 |
| 0 | 0 | 0 | 0 |

| | | | |
|---|---|---|---|
| 0 | 0 | 0 | 0 |
| 0 | 0 | 0 | 0 |
| 0 | 0 | 0 | 0 |
| 1 | 0 | 0 | 0 |
| 1 | 0 | 0 | 0 |
| 0 | 0 | 0 | 0 |
| 0 | 0 | 0 | 0 |
| 0 | 0 | 0 | 0 |
| 0 | 0 | 0 | 0 |
| 0 | 0 | 0 | 0 |
| 0 | 0 | 0 | 0 |
| 0 | 0 | 0 | 0 |
| 0 | 0 | 0 | 0 |
| 0 | 0 | 0 | 0 |
| 0 | 0 | 0 | 0 |
| 0 | 0 | 0 | 0 |
| 0 | 0 | 0 | 0 |
| 0 | 0 | 0 | 0 |
| 0 | 0 | 0 | 0 |
| 0 | 0 | 0 | 0 |
| 0 | 0 | 0 | 0 |
| 0 | 0 | 0 | 0 |
| 1 | 0 | 0 | 0 |
| 1 | 0 | 0 | 0 |
| 0 | 0 | 0 | 0 |
| 0 | 0 | 0 | 0 |
| 1 | 0 | 0 | 0 |
| 0 | 0 | 0 | 0 |
| 0 | 0 | 0 | 0 |
| 0 | 0 | 0 | 0 |
| 0 | 0 | 0 | 0 |
| 0 | 0 | 0 | 0 |
| 0 | 0 | 0 | 0 |
| 0 | 0 | 0 | 0 |
| 1 | 0 | 0 | 0 |
| 0 | 0 | 0 | 0 |
| 0 | 0 | 0 | 0 |
| 0 | 0 | 0 | 0 |
| 0 | 0 | 0 | 0 |
| 0 | 0 | 0 | 0 |
| 0 | 0 | 0 | 0 |
| 0 | 0 | 0 | 0 |
| 0 | 0 | 0 | 0 |
| 0 | 0 | 0 | 0 |
| 0 | 0 | 0 | 0 |
| 0 | 0 | 0 | 0 |
| 0 | 0 | 1 | 0 |
| 1 | 0 | 0 | 0 |
| 1 | 0 | 0 | 0 |
| 1 | 1 | 0 | 0 |
| 1 | 0 | 0 | 0 |
| 0 | 0 | 0 | 0 |
| 0 | 0 | 0 | 0 |
| 1 | 0 | 0 | 0 |

| | | | |
|---|---|---|---|
| 1 | 0 | 0 | 0 |
| 1 | 0 | 0 | 0 |
| 1 | 0 | 0 | 0 |
| 0 | 0 | 0 | 0 |
| 0 | 0 | 0 | 0 |
| 0 | 0 | 0 | 0 |
| 1 | 0 | 0 | 0 |
| 1 | 0 | 0 | 0 |
| 0 | 0 | 0 | 0 |
| 0 | 0 | 0 | 0 |
| 1 | 0 | 0 | 0 |
| 0 | 0 | 0 | 0 |
| 1 | 0 | 0 | 0 |
| 0 | 0 | 0 | 0 |
| 0 | 0 | 0 | 0 |
| 0 | 0 | 0 | 0 |
| 0 | 0 | 0 | 0 |
| 0 | 0 | 0 | 0 |
| 0 | 0 | 0 | 0 |
| 0 | 0 | 0 | 0 |
| 0 | 0 | 0 | 0 |
| 0 | 0 | 0 | 0 |
| 0 | 0 | 0 | 0 |
| 0 | 0 | 0 | 0 |
| 0 | 0 | 0 | 0 |
| 0 | 0 | 0 | 0 |
| 1 | 0 | 0 | 0 |
| 1 | 0 | 0 | 0 |
| 1 | 0 | 0 | 0 |
| 1 | 0 | 0 | 0 |
| 1 | 0 | 0 | 0 |
| 1 | 0 | 0 | 0 |
| 0 | 0 | 0 | 0 |
| 0 | 0 | 0 | 0 |
| 1 | 0 | 0 | 0 |
| 0 | 0 | 0 | 0 |
| 0 | 0 | 0 | 0 |
| 0 | 0 | 0 | 0 |
| 0 | 0 | 0 | 0 |
| 0 | 0 | 0 | 0 |
| 0 | 0 | 0 | 0 |
| 0 | 0 | 0 | 0 |
| 0 | 0 | 0 | 0 |
| 0 | 0 | 0 | 0 |
| 1 | 1 | 0 | 0 |
| 1 | 1 | 0 | 0 |
| 1 | 1 | 0 | 1 |
| 1 | 1 | 0 | 1 |
| 1 | 1 | 0 | 1 |
| 1 | 1 | 0 | 1 |
| 1 | 1 | 0 | 0 |
| 1 | 1 | 0 | 1 |
| 1 | 1 | 0 | 1 |
| 1 | 1 | 0 | 1 |

| | | | |
|---|---|---|---|
| 1 | 1 | 0 | 1 |
| 1 | 1 | 0 | 1 |
| 1 | 1 | 0 | 1 |
| 1 | 1 | 0 | 1 |
| 1 | 1 | 0 | 1 |
| 1 | 0 | 0 | 0 |
| 0 | 0 | 0 | 0 |
| 0 | 0 | 0 | 0 |
| 0 | 0 | 0 | 0 |
| 1 | 0 | 0 | 0 |
| 0 | 0 | 1 | 0 |
| 0 | 0 | 0 | 0 |
| 0 | 0 | 0 | 0 |
| 0 | 0 | 0 | 0 |
| 0 | 0 | 0 | 0 |
| 0 | 0 | 0 | 0 |
| 0 | 0 | 0 | 0 |
| 0 | 0 | 0 | 0 |
| 0 | 0 | 0 | 0 |
| 0 | 0 | 0 | 0 |
| 0 | 0 | 0 | 0 |
| 0 | 0 | 0 | 0 |
| 0 | 0 | 0 | 0 |
| 0 | 0 | 0 | 0 |
| 0 | 0 | 0 | 0 |
| 1 | 1 | 0 | 0 |
| 1 | 1 | 0 | 0 |
| 1 | 1 | 0 | 0 |
| 0 | 1 | 0 | 0 |
| 1 | 0 | 0 | 0 |
| 0 | 0 | 0 | 0 |
| 0 | 0 | 0 | 0 |
| 0 | 0 | 0 | 0 |
| 1 | 0 | 0 | 0 |
| 1 | 1 | 0 | 0 |
| 0 | 0 | 0 | 0 |
| 0 | 0 | 0 | 0 |
| 0 | 0 | 0 | 0 |
| 0 | 0 | 0 | 0 |
| 1 | 0 | 0 | 0 |
| 0 | 0 | 0 | 0 |
| 0 | 0 | 0 | 0 |
| 0 | 0 | 0 | 0 |
| 0 | 0 | 0 | 0 |
| 0 | 0 | 0 | 0 |
| 0 | 1 | 0 | 0 |
| 0 | 1 | 0 | 0 |
| 0 | 0 | 0 | 0 |
| 0 | 0 | 0 | 0 |
| 0 | 1 | 0 | 0 |
| 0 | 0 | 0 | 0 |
| 0 | 0 | 0 | 0 |
| 0 | 1 | 0 | 0 |
| 0 | 0 | 0 | 0 |

| | | | |
|---|---|---|---|
| 1 | 1 | 0 | 0 |
| 0 | 0 | 0 | 0 |
| 0 | 0 | 0 | 0 |
| 0 | 0 | 0 | 0 |
| 0 | 0 | 0 | 0 |
| 0 | 0 | 0 | 0 |
| 0 | 0 | 0 | 0 |
| 0 | 0 | 0 | 0 |
| 0 | 0 | 0 | 0 |
| 0 | 0 | 0 | 0 |
| 0 | 0 | 0 | 0 |
| 0 | 0 | 0 | 0 |
| 0 | 0 | 0 | 0 |
| 0 | 0 | 0 | 0 |
| 0 | 1 | 0 | 0 |
| 0 | 0 | 0 | 0 |
| 0 | 1 | 0 | 0 |
| 0 | 1 | 0 | 0 |
| 0 | 0 | 0 | 0 |
| 0 | 1 | 0 | 0 |
| 1 | 0 | 0 | 1 |
| 1 | 1 | 0 | 0 |
| 1 | 0 | 0 | 0 |
| 1 | 0 | 0 | 1 |
| 0 | 0 | 0 | 0 |
| 0 | 0 | 0 | 0 |
| 0 | 0 | 0 | 0 |
| 0 | 0 | 0 | 0 |
| 1 | 0 | 0 | 0 |
| 0 | 0 | 0 | 0 |
| 1 | 0 | 0 | 0 |
| 0 | 0 | 0 | 0 |
| 0 | 0 | 0 | 0 |
| 0 | 0 | 0 | 0 |
| 0 | 0 | 0 | 0 |
| 0 | 0 | 0 | 0 |
| 0 | 0 | 0 | 0 |
| 1 | 1 | 0 | 0 |
| 1 | 1 | 0 | 0 |
| 1 | 1 | 0 | 0 |
| 0 | 0 | 0 | 0 |
| 0 | 0 | 0 | 0 |
| 0 | 0 | 0 | 0 |
| 1 | 0 | 0 | 0 |
| 1 | 0 | 0 | 1 |
| 1 | 0 | 0 | 1 |
| 1 | 0 | 0 | 0 |
| 1 | 0 | 0 | 1 |
| 1 | 0 | 0 | 0 |
| 0 | 0 | 0 | 0 |
| 1 | 1 | 0 | 0 |
| 1 | 1 | 0 | 0 |
| 0 | 0 | 0 | 0 |
| 0 | 0 | 0 | 0 |

| | | | |
|---|---|---|---|
| 1 | 0 | 0 | 0 |
| 1 | 0 | 0 | 0 |
| 1 | 0 | 0 | 0 |
| 0 | 0 | 0 | 0 |
| 0 | 0 | 0 | 0 |
| 1 | 1 | 0 | 1 |
| 1 | 1 | 0 | 1 |
| 1 | 1 | 0 | 1 |
| 1 | 1 | 0 | 1 |
| 1 | 1 | 0 | 1 |
| 1 | 1 | 0 | 1 |
| 1 | 1 | 0 | 1 |
| 1 | 1 | 0 | 0 |
| 1 | 1 | 0 | 0 |
| 1 | 1 | 1 | 1 |
| 0 | 0 | 0 | 0 |
| 1 | 1 | 0 | 0 |
| 1 | 1 | 0 | 0 |
| 0 | 0 | 0 | 0 |
| 0 | 0 | 0 | 0 |
| 0 | 0 | 0 | 0 |
| 1 | 0 | 0 | 0 |
| 1 | 0 | 0 | 0 |
| 0 | 0 | 0 | 0 |
| 1 | 0 | 0 | 0 |
| 1 | 0 | 0 | 0 |
| 1 | 0 | 0 | 0 |
| 0 | 0 | 0 | 0 |
| 0 | 0 | 0 | 0 |
| 0 | 0 | 0 | 0 |
| 0 | 0 | 0 | 0 |
| 0 | 0 | 0 | 0 |
| 0 | 0 | 0 | 0 |
| 0 | 0 | 0 | 0 |
| 0 | 0 | 0 | 0 |
| 0 | 0 | 0 | 0 |
| 0 | 0 | 0 | 0 |
| 0 | 0 | 0 | 0 |
| 0 | 0 | 0 | 0 |
| 0 | 0 | 0 | 0 |
| 0 | 0 | 0 | 0 |
| 0 | 0 | 0 | 0 |
| 0 | 0 | 0 | 0 |
| 0 | 0 | 0 | 0 |
| 0 | 0 | 0 | 0 |
| 1 | 0 | 0 | 0 |
| 1 | 0 | 0 | 0 |
| 1 | 0 | 0 | 0 |
| 1 | 0 | 0 | 0 |
| 0 | 0 | 0 | 0 |
| 0 | 0 | 0 | 0 |
| 0 | 0 | 0 | 0 |
| 0 | 0 | 0 | 0 |
| 0 | 0 | 0 | 0 |

| | | | |
|---|---|---|---|
| 0 | 0 | 0 | 0 |
| 0 | 0 | 0 | 0 |
| 0 | 0 | 0 | 0 |
| 0 | 0 | 0 | 0 |
| 0 | 0 | 0 | 0 |
| 0 | 0 | 0 | 0 |
| 0 | 0 | 0 | 0 |
| 1 | 0 | 0 | 1 |
| 0 | 0 | 0 | 0 |
| 0 | 0 | 0 | 0 |
| 1 | 0 | 0 | 0 |
| 0 | 0 | 0 | 0 |
| 0 | 0 | 0 | 0 |
| 0 | 0 | 0 | 0 |
| 0 | 0 | 0 | 0 |
| 1 | 0 | 0 | 0 |
| 0 | 0 | 0 | 0 |
| 0 | 0 | 0 | 0 |
| 0 | 0 | 0 | 0 |
| 0 | 0 | 0 | 0 |
| 0 | 0 | 0 | 0 |
| 0 | 0 | 0 | 0 |
| 0 | 0 | 0 | 0 |
| 1 | 1 | 0 | 1 |
| 0 | 0 | 0 | 0 |
| 0 | 0 | 0 | 0 |
| 0 | 0 | 0 | 0 |
| 1 | 1 | 0 | 1 |
| 1 | 1 | 0 | 0 |
| 0 | 0 | 0 | 0 |
| 1 | 0 | 0 | 0 |
| 1 | 1 | 0 | 0 |
| 0 | 0 | 0 | 0 |
| 1 | 0 | 0 | 0 |
| 1 | 1 | 0 | 0 |
| 1 | 0 | 0 | 0 |
| 1 | 1 | 0 | 1 |
| 1 | 1 | 0 | 1 |
| 1 | 0 | 0 | 0 |
| 0 | 0 | 0 | 0 |
| 0 | 0 | 0 | 0 |
| 0 | 0 | 0 | 0 |
| 0 | 0 | 0 | 0 |
| 0 | 0 | 0 | 0 |
| 0 | 0 | 0 | 0 |
| 0 | 0 | 0 | 0 |
| 0 | 0 | 0 | 0 |
| 0 | 0 | 0 | 0 |
| 0 | 0 | 0 | 0 |
| 0 | 0 | 0 | 0 |
| 0 | 0 | 0 | 0 |
| 0 | 0 | 0 | 0 |
| 0 | 0 | 0 | 0 |
| 0 | 0 | 0 | 0 |

| | | | |
|---|---|---|---|
| 0 | 0 | 0 | 0 |
| 0 | 0 | 0 | 0 |
| 0 | 0 | 0 | 0 |
| 0 | 0 | 0 | 0 |
| 0 | 0 | 0 | 0 |
| 0 | 0 | 0 | 0 |
| 0 | 0 | 0 | 0 |
| 0 | 0 | 0 | 0 |
| 0 | 0 | 0 | 0 |
| 0 | 0 | 0 | 0 |
| 0 | 0 | 0 | 0 |
| 0 | 0 | 0 | 0 |
| 0 | 0 | 0 | 0 |
| 1 | 1 | 0 | 0 |
| 0 | 0 | 0 | 0 |
| 1 | 0 | 0 | 0 |
| 0 | 0 | 0 | 1 |
| 0 | 0 | 0 | 0 |
| 0 | 0 | 0 | 0 |
| 0 | 0 | 0 | 0 |
| 0 | 0 | 0 | 0 |
| 0 | 0 | 0 | 0 |
| 0 | 0 | 0 | 0 |
| 0 | 0 | 0 | 0 |
| 0 | 0 | 0 | 0 |
| 0 | 0 | 0 | 0 |
| 0 | 0 | 0 | 0 |
| 0 | 0 | 0 | 0 |
| 0 | 0 | 0 | 0 |
| 0 | 0 | 0 | 0 |
| 1 | 0 | 0 | 0 |
| 1 | 0 | 0 | 0 |
| 0 | 0 | 0 | 0 |
| 0 | 0 | 0 | 0 |
| 0 | 0 | 0 | 0 |
| 0 | 0 | 0 | 0 |
| 0 | 0 | 0 | 0 |
| 0 | 0 | 0 | 0 |
| 0 | 0 | 0 | 0 |
| 0 | 0 | 0 | 0 |
| 1 | 0 | 0 | 0 |
| 1 | 0 | 0 | 0 |
| 1 | 0 | 0 | 0 |
| 1 | 0 | 0 | 0 |
| 0 | 0 | 0 | 0 |
| 1 | 0 | 0 | 0 |
| 0 | 0 | 0 | 0 |
| 0 | 0 | 0 | 0 |
| 1 | 0 | 0 | 0 |
| 1 | 0 | 0 | 0 |
| 0 | 0 | 0 | 0 |
| 0 | 0 | 0 | 0 |
| 0 | 0 | 0 | 0 |
| 0 | 0 | 0 | 0 |

| | | | |
|---|---|---|---|
| 0 | 0 | 0 | 0 |
| 0 | 0 | 0 | 0 |
| 0 | 0 | 0 | 0 |
| 0 | 0 | 0 | 0 |
| 0 | 0 | 0 | 0 |
| 0 | 0 | 0 | 0 |
| 0 | 0 | 0 | 0 |
| 0 | 0 | 0 | 0 |
| 0 | 0 | 0 | 0 |
| 0 | 0 | 0 | 0 |
| 0 | 0 | 0 | 0 |
| 0 | 0 | 0 | 0 |
| 0 | 0 | 0 | 0 |
| 0 | 0 | 0 | 0 |
| 0 | 0 | 0 | 0 |
| 1 | 0 | 0 | 1 |
| 0 | 0 | 0 | 0 |
| 0 | 0 | 0 | 0 |
| 0 | 0 | 0 | 0 |
| 0 | 0 | 0 | 0 |
| 0 | 0 | 0 | 0 |
| 1 | 1 | 0 | 0 |
| 1 | 1 | 0 | 0 |
| 1 | 0 | 0 | 0 |
| 1 | 0 | 0 | 0 |
| 0 | 0 | 0 | 0 |
| 0 | 0 | 0 | 0 |
| 0 | 0 | 0 | 0 |
| 0 | 0 | 0 | 0 |
| 1 | 0 | 0 | 0 |
| 1 | 0 | 0 | 0 |
| 1 | 0 | 0 | 0 |
| 1 | 1 | 0 | 0 |
| 1 | 1 | 0 | 0 |
| 1 | 0 | 0 | 0 |
| 0 | 0 | 0 | 0 |
| 0 | 0 | 0 | 0 |
| 0 | 1 | 0 | 0 |
| 0 | 0 | 0 | 0 |
| 0 | 0 | 0 | 0 |
| 0 | 0 | 0 | 0 |
| 1 | 1 | 0 | 0 |
| 1 | 0 | 0 | 0 |
| 1 | 0 | 0 | 0 |
| 0 | 0 | 0 | 0 |
| 0 | 0 | 0 | 0 |
| 0 | 0 | 0 | 0 |
| 0 | 0 | 0 | 0 |
| 0 | 0 | 0 | 0 |
| 0 | 0 | 0 | 0 |
| 0 | 0 | 0 | 0 |
| 0 | 0 | 0 | 0 |
| 0 | 0 | 0 | 0 |
| 0 | 0 | 0 | 0 |

| | | | |
|---|---|---|---|
| 0 | 0 | 0 | 0 |
| 1 | 0 | 0 | 0 |
| 0 | 0 | 0 | 0 |
| 0 | 0 | 0 | 0 |
| 0 | 0 | 0 | 0 |
| 0 | 0 | 0 | 0 |
| 1 | 0 | 0 | 0 |
| 0 | 0 | 0 | 0 |
| 0 | 0 | 0 | 0 |
| 0 | 0 | 0 | 0 |
| 0 | 0 | 0 | 0 |
| 0 | 0 | 0 | 0 |
| 0 | 0 | 0 | 0 |
| 0 | 0 | 0 | 0 |
| 0 | 0 | 0 | 0 |
| 0 | 0 | 0 | 0 |
| 0 | 1 | 0 | 0 |
| 0 | 0 | 0 | 0 |
| 0 | 0 | 0 | 0 |
| 0 | 0 | 0 | 0 |
| 1 | 0 | 0 | 0 |
| 0 | 0 | 0 | 0 |
| 0 | 0 | 0 | 0 |
| 1 | 1 | 0 | 0 |
| 0 | 0 | 0 | 0 |
| 0 | 0 | 0 | 0 |
| 0 | 0 | 0 | 0 |
| 0 | 0 | 0 | 0 |
| 1 | 0 | 0 | 0 |
| 0 | 0 | 0 | 0 |
| 0 | 0 | 0 | 0 |
| 0 | 0 | 0 | 0 |
| 0 | 0 | 0 | 0 |
| 0 | 0 | 0 | 0 |
| 0 | 0 | 0 | 0 |
| 1 | 0 | 0 | 0 |
| 0 | 0 | 0 | 0 |
| 0 | 0 | 0 | 0 |
| 0 | 0 | 0 | 0 |
| 0 | 0 | 0 | 0 |
| 1 | 1 | 0 | 0 |
| 0 | 0 | 0 | 0 |
| 0 | 0 | 0 | 0 |
| 0 | 0 | 0 | 0 |
| 0 | 0 | 0 | 0 |
| 0 | 0 | 0 | 0 |
| 1 | 0 | 0 | 0 |
| 1 | 0 | 0 | 1 |
| 1 | 0 | 0 | 0 |
| 0 | 0 | 0 | 0 |
| 0 | 0 | 0 | 0 |
| 0 | 0 | 0 | 0 |
| 1 | 0 | 0 | 0 |
| 1 | 0 | 0 | 0 |

| | | | |
|---|---|---|---|
| 1 | 0 | 0 | 0 |
| 0 | 0 | 0 | 0 |
| 0 | 0 | 0 | 0 |
| 0 | 0 | 0 | 0 |
| 1 | 1 | 0 | 0 |
| 1 | 1 | 0 | 0 |
| 0 | 0 | 0 | 0 |
| 0 | 0 | 0 | 0 |
| 1 | 0 | 0 | 0 |
| 1 | 0 | 0 | 0 |
| 1 | 0 | 1 | 0 |
| 0 | 0 | 0 | 0 |
| 0 | 0 | 1 | 0 |
| 0 | 0 | 1 | 0 |
| 0 | 0 | 1 | 0 |
| 1 | 0 | 0 | 0 |
| 1 | 0 | 0 | 0 |
| 0 | 0 | 0 | 0 |
| 0 | 0 | 0 | 0 |
| 1 | 0 | 0 | 1 |
| 1 | 0 | 0 | 1 |
| 1 | 0 | 0 | 0 |
| 0 | 0 | 0 | 0 |
| 0 | 0 | 0 | 0 |
| 0 | 0 | 0 | 0 |
| 0 | 0 | 0 | 0 |
| 0 | 0 | 0 | 0 |
| 1 | 0 | 0 | 0 |
| 0 | 0 | 0 | 0 |
| 0 | 0 | 0 | 1 |
| 0 | 0 | 0 | 1 |
| 0 | 0 | 1 | 1 |
| 0 | 0 | 1 | 1 |
| 0 | 0 | 1 | 1 |
| 0 | 0 | 1 | 1 |
| 0 | 0 | 1 | 0 |
| 0 | 0 | 1 | 0 |
| 0 | 0 | 0 | 0 |
| 0 | 0 | 0 | 0 |
| 0 | 0 | 0 | 0 |
| 0 | 0 | 0 | 0 |
| 0 | 0 | 1 | 0 |
| 0 | 0 | 1 | 0 |
| 0 | 0 | 1 | 0 |
| 0 | 0 | 1 | 0 |
| 0 | 0 | 1 | 0 |
| 0 | 0 | 1 | 0 |
| 0 | 0 | 1 | 0 |
| 0 | 0 | 1 | 0 |
| 0 | 0 | 0 | 0 |
| 0 | 0 | 0 | 0 |
| 0 | 0 | 0 | 0 |
| 0 | 0 | 0 | 0 |
| 1 | 0 | 0 | 0 |

| | | | |
|---|---|---|---|
| 0 | 0 | 0 | 0 |
| 0 | 0 | 0 | 0 |
| 0 | 0 | 0 | 0 |
| 1 | 0 | 0 | 0 |
| 0 | 0 | 0 | 0 |
| 0 | 0 | 0 | 0 |
| 0 | 0 | 0 | 0 |
| 0 | 0 | 0 | 0 |
| 0 | 0 | 0 | 0 |
| 0 | 0 | 0 | 0 |
| 1 | 1 | 0 | 0 |
| 0 | 0 | 0 | 0 |
| 1 | 1 | 0 | 0 |
| 1 | 1 | 0 | 0 |
| 0 | 1 | 0 | 0 |
| 0 | 0 | 0 | 0 |
| 0 | 0 | 0 | 0 |
| 0 | 0 | 0 | 0 |
| 0 | 0 | 0 | 0 |
| 0 | 0 | 0 | 0 |
| 1 | 1 | 0 | 0 |
| 0 | 1 | 0 | 0 |
| 0 | 0 | 0 | 0 |
| 0 | 1 | 0 | 0 |
| 1 | 1 | 0 | 0 |
| 0 | 0 | 0 | 0 |
| 0 | 0 | 0 | 0 |
| 1 | 1 | 0 | 0 |
| 1 | 1 | 0 | 0 |
| 0 | 1 | 0 | 0 |
| 0 | 0 | 0 | 0 |
| 0 | 0 | 0 | 0 |
| 0 | 0 | 0 | 0 |
| 0 | 0 | 0 | 0 |
| 0 | 0 | 0 | 0 |
| 0 | 0 | 0 | 0 |
| 0 | 0 | 0 | 0 |
| 0 | 0 | 0 | 0 |
| 0 | 0 | 0 | 0 |
| 0 | 0 | 0 | 0 |
| 1 | 0 | 0 | 0 |
| 1 | 0 | 0 | 0 |
| 1 | 0 | 0 | 0 |
| 1 | 0 | 0 | 0 |
| 1 | 1 | 0 | 1 |
| 0 | 0 | 0 | 0 |
| 1 | 0 | 0 | 0 |
| 0 | 0 | 0 | 0 |
| 1 | 0 | 0 | 0 |
| 0 | 0 | 0 | 0 |
| 1 | 0 | 0 | 0 |
| 1 | 0 | 0 | 0 |
| 0 | 0 | 0 | 0 |
| 0 | 0 | 0 | 0 |

| | | | |
|---|---|---|---|
| 0 | 0 | 0 | 0 |
| 0 | 0 | 0 | 0 |
| 0 | 0 | 0 | 0 |
| 0 | 0 | 0 | 0 |
| 1 | 0 | 0 | 0 |
| 1 | 0 | 0 | 0 |
| 1 | 0 | 0 | 0 |
| 0 | 0 | 0 | 0 |
| 1 | 1 | 0 | 0 |
| 0 | 0 | 0 | 0 |
| 0 | 0 | 0 | 0 |
| 0 | 0 | 0 | 0 |
| 1 | 1 | 0 | 0 |
| 1 | 0 | 0 | 0 |
| 0 | 0 | 0 | 0 |
| 0 | 0 | 0 | 0 |
| 1 | 1 | 0 | 0 |
| 0 | 0 | 0 | 0 |
| 0 | 0 | 0 | 0 |
| 0 | 0 | 0 | 0 |
| 0 | 0 | 0 | 0 |
| 0 | 0 | 0 | 0 |
| 0 | 0 | 0 | 0 |
| 0 | 0 | 0 | 0 |
| 1 | 0 | 0 | 0 |
| 0 | 0 | 0 | 0 |
| 1 | 0 | 0 | 0 |
| 0 | 0 | 0 | 0 |
| 0 | 0 | 0 | 0 |
| 0 | 0 | 0 | 0 |
| 1 | 0 | 0 | 0 |
| 1 | 0 | 0 | 0 |
| 0 | 0 | 0 | 0 |
| 0 | 0 | 0 | 0 |
| 1 | 0 | 0 | 0 |
| 0 | 0 | 0 | 0 |
| 0 | 0 | 0 | 0 |
| 1 | 0 | 0 | 0 |
| 0 | 0 | 0 | 0 |
| 1 | 0 | 0 | 0 |
| 1 | 0 | 0 | 0 |
| 0 | 0 | 0 | 0 |
| 0 | 0 | 0 | 0 |
| 1 | 1 | 0 | 0 |
| 1 | 0 | 0 | 0 |
| 1 | 1 | 0 | 0 |
| 0 | 0 | 0 | 0 |
| 0 | 0 | 0 | 0 |
| 0 | 0 | 0 | 0 |
| 0 | 0 | 0 | 0 |
| 1 | 1 | 0 | 0 |
| 0 | 0 | 0 | 0 |
| 0 | 0 | 0 | 0 |
| 0 | 0 | 0 | 0 |

| | | | |
|---|---|---|---|
| 0 | 1 | 0 | 0 |
| 1 | 0 | 0 | 0 |
| 0 | 0 | 0 | 0 |
| 0 | 0 | 0 | 0 |
| 0 | 0 | 0 | 0 |
| 1 | 1 | 0 | 0 |
| 0 | 0 | 0 | 0 |
| 0 | 0 | 0 | 0 |
| 0 | 0 | 0 | 0 |
| 1 | 0 | 0 | 0 |
| 1 | 0 | 0 | 0 |
| 0 | 0 | 0 | 0 |
| 0 | 0 | 0 | 0 |
| 0 | 0 | 0 | 0 |
| 0 | 0 | 0 | 0 |
| 0 | 0 | 0 | 0 |
| 0 | 0 | 0 | 0 |
| 0 | 0 | 0 | 0 |
| 0 | 0 | 0 | 0 |
| 0 | 0 | 0 | 0 |
| 0 | 0 | 0 | 0 |
| 0 | 0 | 0 | 0 |
| 0 | 0 | 0 | 0 |
| 1 | 1 | 0 | 0 |
| 0 | 0 | 0 | 0 |
| 0 | 0 | 0 | 0 |
| 1 | 1 | 1 | 0 |
| 0 | 0 | 0 | 0 |
| 0 | 0 | 0 | 0 |
| 0 | 0 | 0 | 0 |
| 1 | 1 | 1 | 0 |
| 1 | 0 | 0 | 0 |
| 1 | 0 | 0 | 0 |
| 1 | 0 | 0 | 0 |
| 1 | 0 | 0 | 0 |
| 1 | 1 | 1 | 0 |
| 1 | 1 | 1 | 0 |
| 0 | 0 | 0 | 0 |
| 0 | 0 | 0 | 0 |
| 1 | 1 | 1 | 0 |
| 0 | 0 | 0 | 0 |
| 0 | 0 | 0 | 0 |
| 0 | 0 | 0 | 0 |
| 0 | 0 | 0 | 0 |
| 0 | 0 | 0 | 0 |
| 0 | 0 | 0 | 0 |
| 0 | 0 | 0 | 0 |
| 0 | 0 | 0 | 0 |
| 1 | 0 | 0 | 0 |
| 1 | 0 | 0 | 0 |
| 1 | 0 | 0 | 0 |
| 0 | 0 | 0 | 0 |
| 0 | 0 | 0 | 0 |
| 0 | 0 | 0 | 0 |

| | | | |
|---|---|---|---|
| 0 | 0 | 0 | 0 |
| 1 | 0 | 0 | 0 |
| 0 | 0 | 0 | 0 |
| 0 | 0 | 0 | 0 |
| 0 | 0 | 0 | 0 |
| 0 | 0 | 0 | 0 |
| 0 | 0 | 0 | 0 |
| 0 | 0 | 0 | 0 |
| 0 | 0 | 0 | 0 |
| 0 | 0 | 0 | 0 |
| 1 | 1 | 0 | 0 |
| 1 | 1 | 0 | 0 |
| 0 | 0 | 0 | 0 |
| 0 | 0 | 0 | 0 |
| 0 | 0 | 0 | 0 |
| 1 | 0 | 0 | 0 |
| 1 | 0 | 0 | 0 |
| 1 | 1 | 0 | 1 |
| 0 | 0 | 0 | 0 |
| 0 | 0 | 0 | 0 |
| 0 | 0 | 0 | 0 |
| 0 | 0 | 0 | 0 |
| 1 | 0 | 0 | 1 |
| 0 | 0 | 0 | 0 |
| 0 | 0 | 0 | 0 |
| 1 | 0 | 0 | 0 |
| 0 | 0 | 0 | 0 |
| 0 | 0 | 0 | 0 |
| 0 | 0 | 0 | 0 |
| 0 | 0 | 0 | 0 |
| 0 | 0 | 0 | 0 |
| 0 | 0 | 0 | 0 |
| 0 | 0 | 0 | 0 |
| 0 | 0 | 0 | 0 |
| 0 | 0 | 0 | 0 |
| 1 | 0 | 0 | 0 |
| 1 | 0 | 0 | 0 |
| 1 | 1 | 0 | 0 |
| 1 | 1 | 0 | 0 |
| 0 | 0 | 0 | 0 |
| 0 | 0 | 0 | 0 |
| 1 | 0 | 0 | 0 |
| 0 | 0 | 0 | 0 |
| 1 | 0 | 0 | 0 |
| 1 | 1 | 0 | 0 |
| 0 | 0 | 0 | 0 |
| 0 | 0 | 0 | 0 |
| 0 | 0 | 0 | 0 |
| 0 | 0 | 0 | 0 |
| 0 | 0 | 0 | 0 |
| 0 | 0 | 0 | 0 |
| 1 | 0 | 0 | 0 |
| 0 | 0 | 0 | 0 |
| 0 | 0 | 0 | 0 |

| | | | |
|---|---|---|---|
| 0 | 0 | 0 | 0 |
| 1 | 1 | 0 | 0 |
| 0 | 0 | 0 | 0 |
| 1 | 0 | 0 | 0 |
| 1 | 0 | 0 | 0 |
| 0 | 0 | 0 | 0 |
| 0 | 0 | 0 | 0 |
| 0 | 0 | 0 | 0 |
| 0 | 0 | 0 | 0 |
| 0 | 0 | 0 | 0 |
| 0 | 0 | 0 | 0 |
| 1 | 1 | 0 | 0 |
| 1 | 1 | 0 | 1 |
| 0 | 0 | 0 | 0 |
| 0 | 1 | 0 | 0 |
| 0 | 0 | 0 | 0 |
| 0 | 0 | 0 | 0 |
| 0 | 0 | 0 | 0 |
| 0 | 0 | 0 | 0 |
| 0 | 0 | 0 | 0 |
| 0 | 0 | 0 | 0 |
| 1 | 0 | 0 | 0 |
| 0 | 0 | 0 | 0 |
| 0 | 0 | 0 | 0 |
| 0 | 0 | 0 | 0 |
| 0 | 0 | 0 | 0 |
| 0 | 0 | 0 | 0 |
| 1 | 0 | 0 | 0 |
| 0 | 0 | 0 | 0 |
| 0 | 0 | 0 | 0 |
| 0 | 0 | 0 | 0 |
| 0 | 0 | 0 | 0 |
| 0 | 0 | 0 | 0 |
| 0 | 0 | 0 | 0 |
| 0 | 0 | 0 | 0 |
| 0 | 0 | 0 | 0 |
| 0 | 0 | 0 | 0 |
| 0 | 0 | 0 | 0 |
| 0 | 0 | 0 | 0 |
| 0 | 0 | 0 | 0 |
| 0 | 0 | 0 | 0 |
| 0 | 0 | 0 | 0 |
| 0 | 0 | 0 | 0 |
| 0 | 0 | 0 | 0 |
| 0 | 0 | 0 | 0 |
| 0 | 0 | 0 | 0 |
| 0 | 0 | 0 | 0 |
| 0 | 0 | 0 | 0 |
| 0 | 0 | 0 | 0 |
| 0 | 0 | 0 | 0 |
| 0 | 0 | 0 | 0 |
| 0 | 0 | 0 | 0 |
| 0 | 0 | 0 | 0 |
| 0 | 0 | 0 | 0 |

| | | | |
|---|---|---|---|
| 0 | 0 | 0 | 0 |
| 0 | 0 | 0 | 0 |
| 0 | 0 | 0 | 0 |
| 0 | 0 | 0 | 0 |
| 0 | 0 | 0 | 0 |
| 0 | 0 | 0 | 0 |
| 0 | 0 | 0 | 0 |
| 0 | 0 | 0 | 0 |
| 0 | 0 | 0 | 0 |
| 0 | 0 | 0 | 0 |
| 0 | 0 | 0 | 0 |
| 0 | 0 | 0 | 0 |
| 1 | 0 | 0 | 0 |
| 1 | 1 | 0 | 0 |
| 0 | 0 | 0 | 0 |
| 1 | 0 | 0 | 0 |
| 0 | 0 | 0 | 0 |
| 1 | 0 | 0 | 0 |
| 0 | 0 | 0 | 0 |
| 0 | 0 | 0 | 0 |
| 1 | 0 | 0 | 0 |
| 0 | 0 | 0 | 0 |
| 0 | 0 | 0 | 0 |
| 1 | 0 | 0 | 0 |
| 1 | 0 | 0 | 0 |
| 0 | 0 | 0 | 0 |
| 0 | 0 | 0 | 0 |
| 1 | 0 | 0 | 0 |
| 1 | 0 | 0 | 0 |
| 1 | 0 | 0 | 0 |
| 1 | 0 | 0 | 0 |
| 1 | 0 | 0 | 0 |
| 0 | 0 | 0 | 0 |
| 0 | 0 | 0 | 0 |
| 0 | 0 | 0 | 0 |
| 0 | 0 | 0 | 0 |
| 0 | 0 | 0 | 0 |
| 0 | 0 | 0 | 0 |
| 0 | 0 | 0 | 0 |
| 0 | 0 | 0 | 0 |
| 0 | 0 | 0 | 0 |
| 0 | 0 | 0 | 0 |
| 0 | 0 | 0 | 0 |
| 0 | 0 | 0 | 0 |
| 1 | 1 | 0 | 0 |
| 0 | 0 | 0 | 0 |
| 0 | 0 | 0 | 0 |
| 0 | 0 | 0 | 0 |
| 0 | 0 | 0 | 0 |
| 1 | 1 | 0 | 0 |
| 1 | 0 | 0 | 0 |
| 0 | 0 | 0 | 0 |
| 0 | 0 | 0 | 0 |
| 0 | 0 | 0 | 1 |

| | | | |
|---|---|---|---|
| 0 | 0 | 0 | 0 |
| 0 | 0 | 0 | 0 |
| 0 | 0 | 0 | 0 |
| 0 | 0 | 0 | 0 |
| 1 | 0 | 0 | 1 |
| 0 | 0 | 0 | 0 |
| 1 | 0 | 0 | 0 |
| 0 | 0 | 0 | 0 |
| 0 | 0 | 0 | 0 |
| 0 | 0 | 0 | 0 |
| 0 | 0 | 0 | 0 |
| 0 | 0 | 0 | 0 |
| 0 | 0 | 0 | 0 |
| 1 | 0 | 0 | 0 |
| 0 | 0 | 0 | 0 |
| 0 | 0 | 0 | 0 |
| 1 | 1 | 0 | 0 |
| 0 | 0 | 0 | 0 |
| 1 | 0 | 0 | 0 |
| 1 | 0 | 0 | 0 |
| 0 | 0 | 0 | 0 |
| 0 | 0 | 0 | 0 |
| 0 | 0 | 0 | 0 |
| 1 | 0 | 0 | 0 |
| 1 | 0 | 0 | 0 |
| 1 | 0 | 0 | 0 |
| 1 | 0 | 0 | 0 |
| 1 | 0 | 0 | 0 |
| 1 | 0 | 0 | 0 |
| 1 | 0 | 0 | 0 |
| 1 | 0 | 0 | 0 |
| 1 | 0 | 0 | 0 |
| 1 | 0 | 0 | 0 |
| 1 | 0 | 0 | 0 |
| 0 | 0 | 0 | 0 |
| 0 | 0 | 0 | 0 |
| 0 | 0 | 0 | 0 |
| 1 | 0 | 0 | 0 |
| 1 | 0 | 0 | 0 |
| 0 | 0 | 0 | 0 |
| 0 | 0 | 0 | 0 |
| 1 | 0 | 0 | 0 |
| 0 | 0 | 0 | 0 |
| 1 | 0 | 0 | 0 |
| 1 | 0 | 0 | 0 |
| 1 | 0 | 0 | 0 |
| 1 | 0 | 0 | 0 |
| 1 | 0 | 0 | 0 |
| 0 | 0 | 0 | 0 |
| 0 | 0 | 0 | 0 |
| 1 | 0 | 0 | 0 |
| 0 | 0 | 0 | 0 |
| 1 | 0 | 0 | 0 |
| 1 | 0 | 0 | 0 |

| | | | |
|---|---|---|---|
| 1 | 0 | 0 | 0 |
| 1 | 0 | 0 | 0 |
| 1 | 0 | 0 | 0 |
| 1 | 0 | 0 | 0 |
| 1 | 0 | 0 | 0 |
| 1 | 0 | 0 | 0 |
| 0 | 0 | 0 | 0 |
| 1 | 0 | 0 | 0 |
| 1 | 0 | 0 | 0 |
| 1 | 0 | 0 | 0 |
| 1 | 0 | 0 | 0 |
| 1 | 0 | 0 | 0 |
| 1 | 0 | 0 | 0 |
| 1 | 0 | 0 | 0 |
| 0 | 0 | 0 | 0 |
| 0 | 0 | 0 | 0 |
| 1 | 1 | 0 | 0 |
| 1 | 1 | 0 | 0 |
| 0 | 0 | 0 | 0 |
| 0 | 0 | 0 | 0 |
| 0 | 0 | 0 | 0 |
| 1 | 1 | 0 | 1 |
| 0 | 0 | 0 | 0 |
| 0 | 0 | 0 | 0 |
| 0 | 0 | 0 | 0 |
| 0 | 0 | 0 | 0 |
| 0 | 0 | 0 | 0 |
| 0 | 0 | 0 | 0 |
| 0 | 0 | 0 | 0 |
| 1 | 0 | 1 | 0 |
| 0 | 0 | 0 | 0 |
| 0 | 0 | 0 | 0 |
| 0 | 0 | 0 | 0 |
| 0 | 0 | 0 | 0 |
| 0 | 0 | 0 | 0 |
| 0 | 0 | 0 | 0 |
| 0 | 0 | 0 | 0 |
| 0 | 0 | 0 | 0 |
| 0 | 0 | 0 | 0 |
| 0 | 0 | 0 | 0 |
| 1 | 1 | 0 | 0 |
| 1 | 1 | 0 | 0 |
| 1 | 0 | 0 | 0 |
| 1 | 1 | 0 | 0 |
| 1 | 0 | 0 | 0 |
| 1 | 1 | 0 | 0 |
| 1 | 0 | 0 | 0 |
| 1 | 0 | 0 | 0 |
| 0 | 0 | 0 | 0 |
| 0 | 0 | 0 | 0 |
| 1 | 0 | 0 | 0 |
| 1 | 0 | 0 | 0 |
| 1 | 0 | 0 | 0 |
| 0 | 0 | 0 | 0 |

| | | | |
|---|---|---|---|
| 0 | 0 | 0 | 0 |
| 0 | 0 | 0 | 0 |
| 0 | 0 | 0 | 0 |
| 1 | 1 | 1 | 0 |
| 0 | 0 | 0 | 0 |
| 0 | 0 | 0 | 0 |
| 0 | 0 | 0 | 0 |
| 0 | 0 | 0 | 0 |
| 0 | 1 | 0 | 0 |
| 0 | 1 | 0 | 0 |
| 0 | 1 | 0 | 0 |
| 0 | 0 | 0 | 0 |
| 0 | 0 | 0 | 0 |
| 0 | 0 | 0 | 0 |
| 1 | 0 | 0 | 0 |
| 1 | 0 | 0 | 0 |
| 0 | 0 | 0 | 0 |
| 0 | 0 | 0 | 0 |
| 1 | 0 | 0 | 0 |
| 0 | 0 | 0 | 0 |
| 1 | 0 | 0 | 0 |
| 1 | 0 | 0 | 0 |
| 0 | 0 | 0 | 0 |
| 0 | 0 | 0 | 0 |
| 0 | 0 | 0 | 0 |
| 0 | 0 | 0 | 0 |
| 0 | 0 | 0 | 0 |
| 0 | 0 | 1 | 0 |
| 0 | 0 | 0 | 0 |
| 0 | 1 | 0 | 0 |
| 0 | 0 | 0 | 0 |
| 0 | 0 | 0 | 0 |
| 0 | 0 | 0 | 0 |
| 0 | 0 | 0 | 0 |
| 0 | 0 | 0 | 0 |
| 0 | 0 | 0 | 0 |
| 1 | 0 | 0 | 0 |
| 1 | 0 | 0 | 0 |
| 1 | 0 | 0 | 0 |
| 0 | 0 | 0 | 0 |
| 0 | 0 | 0 | 0 |
| 0 | 0 | 0 | 0 |
| 0 | 0 | 0 | 0 |
| 0 | 0 | 0 | 0 |
| 0 | 0 | 0 | 0 |
| 0 | 0 | 0 | 0 |
| 0 | 0 | 0 | 0 |
| 0 | 0 | 0 | 1 |
| 0 | 0 | 0 | 0 |
| 0 | 0 | 0 | 0 |
| 0 | 0 | 0 | 0 |
| 0 | 0 | 0 | 0 |
| 1 | 1 | 0 | 0 |
| 0 | 0 | 0 | 0 |

| | | | |
|---|---|---|---|
| 0 | 0 | 0 | 0 |
| 0 | 0 | 0 | 0 |
| 0 | 0 | 0 | 0 |
| 0 | 0 | 0 | 0 |
| 1 | 0 | 0 | 0 |
| 0 | 0 | 0 | 0 |
| 0 | 0 | 0 | 0 |
| 0 | 0 | 0 | 0 |
| 0 | 0 | 0 | 0 |
| 0 | 0 | 0 | 0 |
| 0 | 0 | 0 | 0 |
| 0 | 0 | 0 | 0 |
| 0 | 0 | 0 | 0 |
| 0 | 0 | 0 | 0 |
| 0 | 0 | 0 | 0 |
| 0 | 0 | 0 | 0 |
| 0 | 0 | 0 | 0 |
| 0 | 0 | 0 | 0 |
| 0 | 0 | 0 | 0 |
| 0 | 0 | 0 | 0 |
| 0 | 0 | 0 | 0 |
| 0 | 0 | 0 | 0 |
| 0 | 0 | 0 | 0 |
| 0 | 0 | 0 | 0 |
| 0 | 0 | 0 | 0 |
| 0 | 0 | 0 | 0 |
| 0 | 0 | 0 | 0 |
| 0 | 0 | 0 | 0 |
| 0 | 0 | 0 | 0 |
| 0 | 0 | 0 | 0 |
| 0 | 0 | 0 | 0 |
| 0 | 0 | 0 | 0 |
| 0 | 0 | 0 | 0 |
| 0 | 0 | 0 | 0 |
| 0 | 0 | 0 | 0 |
| 0 | 0 | 0 | 0 |
| 0 | 0 | 0 | 0 |
| 0 | 0 | 0 | 0 |
| 0 | 0 | 0 | 0 |
| 0 | 0 | 0 | 0 |
| 1 | 1 | 0 | 0 |
| 0 | 0 | 0 | 0 |
| 0 | 0 | 0 | 0 |
| 0 | 0 | 0 | 0 |
| 0 | 0 | 0 | 0 |
| 1 | 0 | 0 | 0 |
| 1 | 0 | 0 | 0 |
| 0 | 0 | 0 | 0 |
| 1 | 0 | 0 | 0 |
| 0 | 0 | 0 | 0 |
| 0 | 0 | 0 | 0 |
| 1 | 0 | 0 | 0 |
| 1 | 0 | 0 | 0 |
| 0 | 0 | 0 | 0 |

| | | | |
|---|---|---|---|
| 0 | 0 | 0 | 0 |
| 0 | 0 | 0 | 0 |
| 0 | 0 | 0 | 0 |
| 0 | 0 | 0 | 0 |
| 1 | 0 | 0 | 0 |
| 0 | 0 | 0 | 0 |
| 0 | 0 | 0 | 0 |
| 0 | 0 | 0 | 0 |
| 0 | 0 | 0 | 0 |
| 0 | 0 | 0 | 0 |
| 0 | 0 | 0 | 0 |
| 0 | 0 | 0 | 0 |
| 0 | 0 | 0 | 0 |
| 0 | 0 | 0 | 0 |
| 0 | 0 | 0 | 0 |
| 1 | 1 | 0 | 1 |
| 0 | 0 | 0 | 0 |
| 0 | 0 | 0 | 0 |
| 0 | 0 | 0 | 0 |
| 0 | 0 | 0 | 0 |
| 0 | 0 | 0 | 0 |
| 0 | 0 | 0 | 0 |
| 0 | 0 | 0 | 0 |
| 0 | 0 | 0 | 0 |
| 0 | 0 | 0 | 0 |
| 0 | 0 | 0 | 0 |
| 0 | 0 | 0 | 0 |
| 1 | 0 | 0 | 0 |
| 1 | 0 | 0 | 0 |
| 0 | 0 | 0 | 0 |
| 1 | 1 | 0 | 0 |
| 0 | 0 | 0 | 0 |
| 0 | 0 | 0 | 0 |
| 0 | 0 | 0 | 0 |
| 1 | 1 | 0 | 0 |
| 1 | 1 | 0 | 0 |
| 0 | 0 | 0 | 0 |
| 1 | 0 | 0 | 0 |
| 0 | 0 | 0 | 0 |
| 1 | 1 | 0 | 0 |
| 0 | 1 | 0 | 0 |
| 0 | 0 | 0 | 0 |
| 0 | 0 | 0 | 0 |
| 0 | 0 | 0 | 0 |
| 1 | 1 | 0 | 0 |
| 1 | 1 | 0 | 0 |
| 1 | 1 | 0 | 0 |
| 0 | 0 | 0 | 0 |
| 0 | 0 | 0 | 0 |
| 0 | 0 | 0 | 0 |
| 0 | 0 | 0 | 0 |
| 0 | 0 | 0 | 0 |
| 0 | 0 | 0 | 0 |
| 0 | 0 | 0 | 0 |

| | | | |
|---|---|---|---|
| 0 | 0 | 0 | 0 |
| 0 | 0 | 0 | 0 |
| 1 | 0 | 0 | 0 |
| 1 | 0 | 0 | 0 |
| 0 | 0 | 0 | 0 |
| 1 | 0 | 0 | 0 |
| 0 | 0 | 0 | 0 |
| 0 | 0 | 0 | 0 |
| 0 | 0 | 0 | 0 |
| 0 | 1 | 0 | 0 |
| 0 | 0 | 0 | 0 |
| 0 | 0 | 0 | 0 |
| 0 | 0 | 0 | 0 |
| 0 | 0 | 0 | 0 |
| 0 | 0 | 0 | 0 |
| 0 | 0 | 0 | 0 |
| 0 | 0 | 0 | 0 |
| 1 | 0 | 0 | 0 |
| 0 | 0 | 0 | 0 |
| 1 | 0 | 0 | 0 |
| 0 | 0 | 0 | 0 |
| 1 | 0 | 0 | 0 |
| 0 | 0 | 0 | 0 |
| 1 | 0 | 0 | 0 |
| 0 | 0 | 0 | 0 |
| 0 | 0 | 0 | 0 |
| 0 | 0 | 0 | 0 |
| 0 | 0 | 0 | 0 |
| 0 | 0 | 0 | 0 |
| 0 | 0 | 0 | 0 |
| 0 | 0 | 0 | 0 |
| 1 | 0 | 0 | 0 |
| 0 | 0 | 0 | 0 |
| 0 | 0 | 0 | 0 |
| 0 | 0 | 0 | 0 |
| 0 | 0 | 0 | 0 |
| 0 | 0 | 0 | 0 |
| 0 | 0 | 0 | 0 |
| 1 | 1 | 0 | 0 |
| 1 | 1 | 0 | 0 |
| 1 | 1 | 0 | 0 |
| 1 | 1 | 0 | 0 |
| 1 | 1 | 0 | 0 |
| 1 | 0 | 0 | 0 |
| 1 | 0 | 0 | 0 |
| 1 | 0 | 0 | 0 |
| 0 | 0 | 0 | 0 |
| 0 | 0 | 0 | 0 |
| 0 | 0 | 0 | 0 |
| 0 | 0 | 0 | 0 |
| 0 | 0 | 0 | 0 |
| 0 | 0 | 0 | 0 |
| 0 | 0 | 0 | 0 |
| 0 | 0 | 0 | 0 |

| | | | |
|---|---|---|---|
| 0 | 0 | 0 | 0 |
| 0 | 0 | 0 | 0 |
| 0 | 0 | 0 | 0 |
| 0 | 0 | 0 | 0 |
| 0 | 0 | 0 | 0 |
| 1 | 0 | 0 | 0 |
| 1 | 0 | 0 | 0 |
| 0 | 0 | 0 | 0 |
| 0 | 0 | 0 | 0 |
| 0 | 0 | 0 | 0 |
| 0 | 0 | 0 | 0 |
| 1 | 0 | 0 | 0 |
| 0 | 0 | 1 | 0 |
| 0 | 0 | 0 | 0 |
| 0 | 0 | 0 | 0 |
| 0 | 0 | 0 | 0 |
| 1 | 0 | 0 | 0 |
| 0 | 0 | 0 | 0 |
| 1 | 0 | 0 | 0 |
| 1 | 0 | 0 | 0 |
| 0 | 0 | 0 | 0 |
| 1 | 0 | 0 | 0 |
| 1 | 1 | 0 | 0 |
| 0 | 0 | 0 | 0 |
| 1 | 1 | 0 | 0 |
| 1 | 1 | 0 | 0 |
| 1 | 1 | 0 | 0 |
| 0 | 0 | 0 | 0 |
| 1 | 0 | 0 | 0 |
| 0 | 0 | 0 | 0 |
| 0 | 0 | 0 | 0 |
| 0 | 0 | 0 | 0 |
| 0 | 0 | 0 | 0 |
| 0 | 0 | 0 | 0 |
| 0 | 0 | 0 | 0 |
| 0 | 0 | 0 | 0 |
| 0 | 0 | 0 | 0 |
| 0 | 0 | 0 | 0 |
| 0 | 0 | 0 | 0 |
| 0 | 0 | 0 | 0 |
| 0 | 0 | 0 | 0 |
| 0 | 0 | 0 | 0 |
| 0 | 0 | 0 | 0 |
| 0 | 0 | 0 | 0 |
| 0 | 0 | 0 | 0 |
| 0 | 0 | 0 | 0 |
| 0 | 0 | 0 | 0 |
| 0 | 0 | 0 | 0 |
| 0 | 0 | 0 | 0 |
| 0 | 0 | 0 | 0 |
| 0 | 0 | 0 | 0 |
| 0 | 0 | 0 | 0 |
| 0 | 0 | 0 | 0 |
| 0 | 0 | 0 | 0 |

| | | | |
|---|---|---|---|
| 0 | 0 | 0 | 0 |
| 0 | 0 | 0 | 0 |
| 0 | 0 | 0 | 0 |
| 0 | 0 | 0 | 0 |
| 0 | 0 | 0 | 0 |
| 0 | 0 | 0 | 0 |
| 0 | 0 | 0 | 0 |
| 0 | 0 | 0 | 0 |
| 0 | 0 | 0 | 0 |
| 0 | 0 | 0 | 0 |
| 0 | 1 | 0 | 0 |
| 0 | 1 | 0 | 0 |
| 0 | 1 | 0 | 0 |
| 0 | 1 | 0 | 0 |
| 0 | 0 | 0 | 0 |
| 0 | 1 | 0 | 0 |
| 0 | 1 | 0 | 0 |
| 1 | 1 | 0 | 0 |
| 0 | 1 | 0 | 0 |
| 1 | 0 | 0 | 0 |
| 1 | 1 | 0 | 0 |
| 1 | 1 | 0 | 0 |
| 0 | 0 | 1 | 1 |
| 0 | 0 | 0 | 0 |
| 1 | 0 | 0 | 0 |
| 0 | 0 | 0 | 0 |
| 0 | 0 | 0 | 0 |
| 0 | 0 | 0 | 0 |
| 0 | 0 | 0 | 0 |
| 0 | 0 | 0 | 0 |
| 0 | 0 | 0 | 0 |
| 0 | 0 | 0 | 0 |
| 0 | 0 | 0 | 0 |
| 0 | 0 | 0 | 0 |
| 0 | 0 | 0 | 0 |
| 0 | 0 | 0 | 0 |
| 0 | 0 | 0 | 0 |
| 0 | 0 | 0 | 0 |
| 0 | 0 | 0 | 0 |
| 0 | 0 | 0 | 0 |
| 1 | 0 | 0 | 0 |
| 0 | 0 | 0 | 0 |
| 1 | 0 | 0 | 0 |
| 1 | 0 | 0 | 0 |
| 0 | 0 | 0 | 0 |
| 1 | 0 | 0 | 0 |
| 0 | 0 | 0 | 0 |
| 0 | 0 | 0 | 0 |
| 0 | 0 | 0 | 0 |
| 0 | 0 | 0 | 0 |
| 0 | 0 | 0 | 0 |
| 0 | 0 | 0 | 0 |
| 1 | 1 | 0 | 0 |
| 1 | 1 | 0 | 0 |

| | | | |
|---|---|---|---|
| 1 | 1 | 0 | 0 |
| 0 | 0 | 0 | 0 |
| 0 | 0 | 0 | 0 |
| 0 | 0 | 0 | 0 |
| 0 | 0 | 0 | 0 |
| 0 | 0 | 0 | 0 |
| 0 | 1 | 0 | 0 |
| 0 | 0 | 0 | 0 |
| 0 | 0 | 0 | 0 |
| 0 | 1 | 0 | 0 |
| 1 | 0 | 0 | 0 |
| 0 | 0 | 0 | 0 |
| 0 | 0 | 1 | 0 |
| 0 | 0 | 1 | 0 |
| 0 | 0 | 0 | 0 |
| 0 | 0 | 0 | 0 |
| 0 | 0 | 0 | 0 |
| 0 | 1 | 0 | 0 |
| 0 | 0 | 0 | 0 |
| 0 | 0 | 0 | 0 |
| 1 | 0 | 0 | 0 |
| 0 | 0 | 0 | 1 |
| 0 | 0 | 0 | 0 |
| 0 | 0 | 0 | 0 |
| 0 | 0 | 0 | 0 |
| 0 | 0 | 0 | 0 |
| 0 | 0 | 0 | 0 |
| 0 | 0 | 0 | 0 |
| 0 | 0 | 0 | 0 |
| 0 | 0 | 0 | 0 |
| 0 | 0 | 0 | 0 |
| 0 | 0 | 0 | 0 |
| 0 | 0 | 0 | 0 |
| 0 | 0 | 0 | 0 |
| 0 | 0 | 0 | 0 |
| 0 | 0 | 0 | 0 |
| 1 | 0 | 0 | 0 |
| 0 | 0 | 0 | 0 |
| 1 | 0 | 0 | 0 |
| 0 | 0 | 0 | 0 |
| 0 | 0 | 0 | 0 |
| 0 | 0 | 0 | 0 |
| 0 | 0 | 0 | 0 |
| 0 | 0 | 0 | 0 |
| 0 | 0 | 0 | 0 |
| 0 | 0 | 0 | 0 |
| 0 | 0 | 0 | 0 |
| 0 | 0 | 0 | 0 |
| 0 | 0 | 0 | 0 |
| 1 | 0 | 0 | 0 |
| 0 | 0 | 0 | 0 |
| 0 | 0 | 0 | 0 |
| 0 | 0 | 0 | 0 |
| 0 | 0 | 0 | 0 |

| | | | |
|---|---|---|---|
| 0 | 0 | 0 | 0 |
| 0 | 0 | 0 | 0 |
| 0 | 0 | 0 | 0 |
| 0 | 0 | 0 | 0 |
| 0 | 0 | 0 | 0 |
| 0 | 0 | 0 | 0 |
| 0 | 0 | 0 | 0 |
| 0 | 0 | 0 | 0 |
| 0 | 0 | 0 | 0 |
| 0 | 0 | 0 | 0 |
| 0 | 0 | 0 | 0 |
| 0 | 0 | 0 | 0 |
| 0 | 0 | 1 | 0 |
| 1 | 0 | 0 | 0 |
| 1 | 0 | 0 | 0 |
| 0 | 0 | 0 | 0 |
| 0 | 0 | 0 | 0 |
| 0 | 0 | 0 | 0 |
| 0 | 0 | 0 | 0 |
| 0 | 0 | 0 | 0 |
| 0 | 0 | 0 | 0 |
| 0 | 0 | 0 | 0 |
| 0 | 0 | 0 | 0 |
| 1 | 1 | 0 | 1 |
| 0 | 0 | 0 | 0 |
| 1 | 1 | 0 | 0 |
| 1 | 1 | 0 | 1 |
| 0 | 0 | 0 | 0 |
| 0 | 0 | 0 | 0 |
| 0 | 0 | 0 | 0 |
| 0 | 0 | 0 | 0 |
| 0 | 0 | 0 | 0 |
| 0 | 0 | 0 | 0 |
| 0 | 0 | 0 | 0 |
| 0 | 0 | 0 | 0 |
| 1 | 0 | 0 | 0 |
| 0 | 0 | 0 | 0 |
| 0 | 0 | 0 | 0 |
| 0 | 0 | 0 | 0 |
| 0 | 0 | 0 | 0 |
| 0 | 0 | 0 | 0 |
| 0 | 0 | 0 | 0 |
| 1 | 0 | 0 | 0 |
| 1 | 0 | 0 | 0 |
| 1 | 0 | 0 | 0 |
| 1 | 0 | 0 | 0 |
| 1 | 0 | 0 | 0 |
| 0 | 0 | 0 | 0 |
| 0 | 0 | 0 | 0 |
| 1 | 0 | 0 | 0 |
| 1 | 0 | 0 | 0 |
| 0 | 0 | 0 | 0 |
| 1 | 0 | 0 | 0 |
| 1 | 0 | 0 | 0 |

| | | | |
|---|---|---|---|
| 0 | 0 | 0 | 0 |
| 1 | 0 | 0 | 0 |
| 0 | 0 | 0 | 0 |
| 0 | 0 | 0 | 0 |
| 0 | 0 | 0 | 0 |
| 0 | 0 | 0 | 0 |
| 1 | 0 | 0 | 0 |
| 0 | 0 | 0 | 0 |
| 0 | 0 | 0 | 0 |
| 0 | 0 | 0 | 0 |
| 0 | 0 | 0 | 0 |
| 1 | 1 | 0 | 0 |
| 0 | 0 | 0 | 0 |
| 0 | 0 | 0 | 0 |
| 0 | 0 | 0 | 0 |
| 0 | 0 | 0 | 0 |
| 0 | 0 | 0 | 0 |
| 0 | 0 | 0 | 0 |
| 0 | 0 | 0 | 0 |
| 0 | 0 | 0 | 0 |
| 0 | 0 | 0 | 0 |
| 0 | 0 | 0 | 0 |
| 0 | 0 | 0 | 0 |
| 0 | 0 | 0 | 0 |
| 0 | 0 | 0 | 0 |
| 0 | 0 | 0 | 0 |
| 0 | 0 | 0 | 0 |
| 0 | 0 | 0 | 0 |
| 1 | 1 | 0 | 0 |
| 1 | 0 | 0 | 0 |
| 0 | 0 | 0 | 0 |
| 0 | 0 | 0 | 0 |
| 0 | 0 | 0 | 0 |
| 0 | 0 | 0 | 0 |
| 0 | 0 | 0 | 0 |
| 0 | 0 | 0 | 0 |
| 0 | 0 | 0 | 0 |
| 0 | 0 | 0 | 0 |
| 0 | 0 | 0 | 0 |
| 0 | 0 | 0 | 0 |
| 0 | 0 | 0 | 0 |
| 0 | 0 | 0 | 0 |
| 0 | 0 | 0 | 0 |
| 0 | 0 | 0 | 0 |
| 0 | 0 | 0 | 0 |
| 0 | 0 | 0 | 0 |
| 0 | 0 | 0 | 0 |
| 0 | 0 | 0 | 0 |
| 0 | 0 | 0 | 0 |
| 0 | 0 | 0 | 0 |
| 0 | 0 | 0 | 0 |
| 0 | 0 | 0 | 0 |
| 0 | 0 | 1 | 0 |
| 0 | 0 | 0 | 0 |

| | | | |
|---|---|---|---|
| 0 | 0 | 0 | 0 |
| 0 | 0 | 0 | 0 |
| 0 | 0 | 0 | 0 |
| 0 | 0 | 0 | 0 |
| 1 | 0 | 0 | 0 |
| 0 | 0 | 0 | 0 |
| 0 | 0 | 0 | 0 |
| 1 | 1 | 0 | 0 |
| 0 | 0 | 0 | 0 |
| 1 | 1 | 0 | 0 |
| 0 | 0 | 0 | 0 |
| 0 | 0 | 0 | 0 |
| 0 | 0 | 0 | 0 |
| 0 | 0 | 0 | 0 |
| 0 | 0 | 0 | 0 |
| 0 | 0 | 0 | 0 |
| 1 | 0 | 0 | 0 |
| 0 | 0 | 0 | 0 |
| 0 | 0 | 0 | 0 |
| 1 | 0 | 0 | 0 |
| 1 | 0 | 0 | 0 |
| 0 | 0 | 0 | 0 |
| 0 | 0 | 0 | 0 |
| 0 | 0 | 0 | 0 |
| 0 | 0 | 0 | 0 |
| 0 | 0 | 0 | 0 |
| 1 | 0 | 0 | 0 |
| 0 | 0 | 0 | 0 |
| 0 | 0 | 0 | 0 |
| 1 | 0 | 0 | 0 |
| 0 | 0 | 0 | 0 |
| 0 | 0 | 0 | 0 |
| 0 | 0 | 0 | 0 |
| 1 | 1 | 0 | 0 |
| 0 | 0 | 0 | 0 |
| 0 | 0 | 0 | 0 |
| 0 | 0 | 0 | 0 |
| 0 | 0 | 0 | 0 |
| 0 | 1 | 0 | 0 |
| 0 | 0 | 0 | 0 |
| 0 | 0 | 0 | 0 |
| 0 | 0 | 0 | 0 |
| 0 | 0 | 0 | 0 |
| 1 | 1 | 0 | 0 |
| 0 | 0 | 0 | 0 |
| 0 | 0 | 0 | 0 |
| 0 | 0 | 0 | 0 |
| 0 | 0 | 0 | 0 |
| 0 | 0 | 0 | 0 |
| 0 | 0 | 0 | 0 |
| 0 | 0 | 0 | 0 |
| 1 | 0 | 0 | 0 |
| 0 | 0 | 0 | 0 |
| 0 | 0 | 0 | 0 |

| | | | |
|---|---|---|---|
| 1 | 0 | 0 | 1 |
| 1 | 0 | 0 | 1 |
| 1 | 0 | 0 | 1 |
| 0 | 0 | 0 | 0 |
| 0 | 0 | 0 | 0 |
| 0 | 0 | 0 | 0 |
| 1 | 0 | 0 | 0 |
| 0 | 0 | 0 | 0 |
| 0 | 0 | 0 | 0 |
| 0 | 0 | 0 | 0 |
| 0 | 0 | 1 | 0 |
| 0 | 0 | 0 | 0 |
| 0 | 0 | 0 | 0 |
| 0 | 0 | 0 | 0 |
| 0 | 0 | 0 | 0 |
| 0 | 0 | 0 | 0 |
| 1 | 0 | 0 | 0 |
| 0 | 0 | 0 | 0 |
| 0 | 0 | 0 | 0 |
| 0 | 0 | 0 | 0 |
| 0 | 0 | 0 | 0 |
| 0 | 0 | 0 | 0 |
| 0 | 0 | 0 | 0 |
| 1 | 1 | 0 | 0 |
| 0 | 0 | 0 | 0 |
| 0 | 0 | 0 | 0 |
| 0 | 0 | 0 | 0 |
| 0 | 0 | 0 | 0 |
| 0 | 0 | 0 | 0 |
| 0 | 0 | 0 | 0 |
| 0 | 0 | 0 | 0 |
| 0 | 0 | 0 | 0 |
| 0 | 1 | 0 | 0 |
| 0 | 0 | 0 | 0 |
| 1 | 1 | 0 | 0 |
| 0 | 0 | 0 | 0 |
| 0 | 0 | 0 | 0 |
| 0 | 0 | 0 | 0 |
| 0 | 0 | 0 | 0 |
| 0 | 0 | 0 | 0 |
| 0 | 0 | 0 | 0 |
| 1 | 0 | 0 | 1 |
| 0 | 0 | 0 | 0 |
| 1 | 0 | 0 | 1 |
| 0 | 0 | 0 | 0 |
| 1 | 0 | 0 | 1 |
| 0 | 0 | 0 | 0 |
| 0 | 0 | 0 | 0 |
| 0 | 0 | 0 | 0 |
| 1 | 0 | 0 | 0 |
| 0 | 0 | 0 | 0 |
| 0 | 0 | 0 | 0 |
| 0 | 0 | 0 | 0 |
| 0 | 0 | 0 | 0 |

| | | | |
|---|---|---|---|
| 0 | 0 | 0 | 0 |
| 0 | 0 | 0 | 0 |
| 0 | 0 | 0 | 0 |
| 0 | 0 | 0 | 0 |
| 0 | 0 | 0 | 0 |
| 0 | 0 | 0 | 0 |
| 0 | 0 | 0 | 0 |
| 0 | 0 | 0 | 0 |
| 0 | 0 | 0 | 0 |
| 0 | 0 | 0 | 0 |
| 0 | 0 | 0 | 0 |
| 0 | 0 | 0 | 0 |
| 0 | 0 | 0 | 0 |
| 0 | 0 | 0 | 0 |
| 0 | 0 | 0 | 0 |
| 0 | 0 | 0 | 0 |
| 0 | 0 | 0 | 0 |
| 0 | 0 | 0 | 0 |
| 0 | 0 | 1 | 0 |
| 0 | 0 | 0 | 0 |
| 0 | 0 | 0 | 0 |
| 0 | 0 | 0 | 0 |
| 0 | 0 | 0 | 0 |
| 0 | 0 | 0 | 0 |
| 0 | 0 | 0 | 0 |
| 0 | 0 | 0 | 0 |
| 0 | 0 | 0 | 0 |
| 0 | 0 | 0 | 0 |
| 0 | 0 | 1 | 0 |
| 0 | 0 | 0 | 0 |
| 0 | 0 | 1 | 0 |
| 1 | 0 | 0 | 0 |
| 1 | 1 | 1 | 0 |
| 1 | 1 | 1 | 0 |
| 0 | 0 | 0 | 0 |
| 0 | 0 | 0 | 0 |
| 0 | 1 | 0 | 0 |
| 1 | 1 | 1 | 0 |
| 1 | 1 | 1 | 0 |
| 0 | 0 | 1 | 0 |
| 0 | 0 | 1 | 0 |
| 1 | 0 | 0 | 0 |
| 0 | 0 | 0 | 0 |
| 0 | 0 | 0 | 0 |
| 1 | 1 | 0 | 0 |
| 0 | 0 | 0 | 0 |
| 0 | 0 | 0 | 0 |
| 0 | 0 | 0 | 0 |
| 0 | 0 | 0 | 0 |
| 0 | 0 | 0 | 0 |
| 0 | 0 | 0 | 0 |
| 0 | 0 | 0 | 0 |
| 0 | 0 | 0 | 0 |
| 0 | 0 | 0 | 0 |

| | | | |
|---|---|---|---|
| 0 | 0 | 0 | 0 |
| 0 | 0 | 0 | 0 |
| 0 | 0 | 0 | 0 |
| 0 | 0 | 0 | 0 |
| 0 | 0 | 0 | 0 |
| 0 | 0 | 0 | 0 |
| 0 | 0 | 0 | 0 |
| 0 | 0 | 0 | 0 |
| 0 | 0 | 0 | 0 |
| 0 | 0 | 0 | 0 |
| 0 | 0 | 0 | 0 |
| 0 | 0 | 0 | 0 |
| 0 | 0 | 0 | 0 |
| 0 | 0 | 0 | 0 |
| 0 | 0 | 0 | 0 |
| 0 | 0 | 0 | 0 |
| 0 | 0 | 0 | 0 |
| 0 | 0 | 0 | 0 |
| 0 | 0 | 0 | 0 |
| 1 | 0 | 0 | 0 |
| 1 | 0 | 0 | 0 |
| 0 | 0 | 0 | 0 |
| 0 | 0 | 0 | 0 |
| 0 | 0 | 0 | 0 |
| 0 | 0 | 0 | 0 |
| 1 | 0 | 0 | 0 |
| 0 | 0 | 0 | 0 |
| 0 | 0 | 0 | 0 |
| 0 | 0 | 0 | 0 |
| 1 | 0 | 0 | 0 |
| 0 | 0 | 0 | 0 |
| 0 | 0 | 0 | 0 |
| 1 | 0 | 0 | 0 |
| 1 | 0 | 0 | 0 |
| 0 | 0 | 0 | 0 |
| 0 | 0 | 0 | 0 |
| 0 | 0 | 0 | 0 |
| 0 | 0 | 0 | 0 |
| 0 | 0 | 0 | 0 |
| 0 | 0 | 0 | 0 |
| 0 | 0 | 0 | 0 |
| 0 | 0 | 0 | 0 |
| 0 | 0 | 0 | 0 |
| 0 | 0 | 0 | 0 |
| 0 | 0 | 0 | 0 |
| 0 | 0 | 0 | 0 |
| 0 | 0 | 0 | 0 |
| 0 | 1 | 0 | 0 |
| 0 | 1 | 0 | 0 |
| 0 | 0 | 0 | 0 |
| 0 | 1 | 0 | 0 |
| 0 | 1 | 0 | 0 |
| 0 | 0 | 0 | 0 |
| 0 | 0 | 0 | 0 |

| | | | |
|---|---|---|---|
| 0 | 0 | 0 | 0 |
| 0 | 0 | 0 | 0 |
| 1 | 1 | 0 | 1 |
| 0 | 0 | 0 | 0 |
| 0 | 0 | 0 | 0 |
| 0 | 0 | 0 | 0 |
| 0 | 0 | 0 | 0 |
| 0 | 1 | 0 | 0 |
| 1 | 1 | 0 | 0 |
| 1 | 1 | 0 | 0 |
| 1 | 1 | 0 | 0 |
| 1 | 1 | 0 | 0 |
| 1 | 1 | 0 | 0 |
| 0 | 1 | 0 | 0 |
| 0 | 1 | 0 | 0 |
| 0 | 0 | 0 | 0 |
| 0 | 0 | 0 | 0 |
| 0 | 0 | 0 | 0 |
| 0 | 0 | 0 | 0 |
| 0 | 0 | 0 | 0 |
| 0 | 0 | 0 | 0 |
| 0 | 0 | 0 | 0 |
| 0 | 1 | 0 | 0 |
| 0 | 1 | 0 | 0 |
| 0 | 0 | 0 | 0 |
| 0 | 0 | 1 | 0 |
| 0 | 0 | 0 | 0 |
| 0 | 0 | 0 | 0 |
| 0 | 0 | 0 | 0 |
| 0 | 1 | 0 | 0 |
| 0 | 0 | 0 | 0 |
| 0 | 0 | 0 | 0 |
| 0 | 0 | 0 | 0 |
| 0 | 0 | 0 | 0 |
| 0 | 0 | 0 | 0 |
| 0 | 0 | 0 | 0 |
| 0 | 0 | 0 | 0 |
| 0 | 0 | 0 | 0 |
| 0 | 0 | 0 | 0 |
| 1 | 0 | 0 | 0 |
| 1 | 1 | 0 | 0 |
| 1 | 0 | 0 | 0 |
| 0 | 0 | 0 | 0 |
| 0 | 1 | 0 | 0 |
| 1 | 1 | 0 | 0 |
| 0 | 1 | 0 | 0 |
| 0 | 0 | 0 | 0 |
| 1 | 0 | 0 | 0 |
| 0 | 1 | 0 | 0 |
| 0 | 0 | 0 | 0 |
| 0 | 0 | 0 | 0 |
| 0 | 0 | 0 | 0 |
| 1 | 1 | 0 | 0 |
| 0 | 0 | 0 | 0 |

| | | | |
|---|---|---|---|
| 0 | 0 | 0 | 0 |
| 0 | 0 | 0 | 0 |
| 1 | 0 | 0 | 0 |
| 1 | 1 | 0 | 0 |
| 1 | 1 | 0 | 0 |
| 0 | 0 | 0 | 0 |
| 0 | 0 | 0 | 0 |
| 0 | 0 | 0 | 0 |
| 1 | 0 | 0 | 0 |
| 0 | 0 | 0 | 0 |
| 0 | 0 | 0 | 0 |
| 1 | 0 | 0 | 1 |
| 0 | 0 | 0 | 0 |
| 1 | 0 | 0 | 0 |
| 0 | 0 | 0 | 0 |
| 0 | 0 | 0 | 0 |
| 0 | 0 | 0 | 0 |
| 0 | 0 | 0 | 0 |
| 0 | 0 | 0 | 0 |
| 0 | 0 | 0 | 0 |
| 0 | 0 | 0 | 0 |
| 1 | 0 | 0 | 1 |
| 1 | 0 | 0 | 0 |
| 0 | 0 | 0 | 0 |
| 0 | 0 | 0 | 0 |
| 0 | 0 | 0 | 0 |
| 0 | 0 | 0 | 0 |
| 0 | 0 | 0 | 0 |
| 1 | 0 | 0 | 0 |
| 1 | 0 | 0 | 0 |
| 0 | 0 | 0 | 0 |
| 0 | 0 | 0 | 0 |
| 0 | 0 | 0 | 0 |
| 0 | 0 | 0 | 0 |
| 0 | 0 | 0 | 0 |
| 0 | 0 | 0 | 0 |
| 1 | 0 | 0 | 0 |
| 0 | 0 | 0 | 0 |
| 0 | 0 | 0 | 0 |
| 1 | 0 | 0 | 0 |
| 1 | 0 | 0 | 0 |
| 1 | 0 | 0 | 0 |
| 1 | 0 | 0 | 0 |
| 0 | 0 | 0 | 0 |
| 0 | 0 | 0 | 0 |
| 1 | 0 | 0 | 0 |
| 0 | 0 | 0 | 0 |
| 0 | 0 | 0 | 0 |
| 0 | 0 | 0 | 0 |
| 0 | 0 | 0 | 0 |
| 0 | 0 | 0 | 0 |
| 1 | 0 | 0 | 0 |
| 1 | 0 | 0 | 0 |
| 1 | 0 | 0 | 0 |

| | | | |
|---|---|---|---|
| 1 | 0 | 0 | 0 |
| 0 | 0 | 0 | 0 |
| 0 | 0 | 0 | 0 |
| 0 | 0 | 0 | 0 |
| 1 | 0 | 0 | 0 |
| 0 | 0 | 0 | 0 |
| 1 | 0 | 0 | 0 |
| 0 | 0 | 0 | 0 |
| 0 | 0 | 0 | 0 |
| 0 | 0 | 0 | 0 |
| 0 | 0 | 0 | 0 |
| 0 | 0 | 0 | 0 |
| 0 | 0 | 0 | 0 |
| 0 | 0 | 0 | 0 |
| 0 | 0 | 0 | 0 |
| 1 | 0 | 0 | 0 |
| 0 | 0 | 0 | 0 |
| 0 | 0 | 0 | 0 |
| 0 | 0 | 0 | 0 |
| 0 | 0 | 0 | 0 |
| 0 | 0 | 0 | 0 |
| 0 | 0 | 0 | 0 |
| 1 | 0 | 0 | 0 |
| 1 | 0 | 0 | 0 |
| 0 | 0 | 0 | 0 |
| 0 | 0 | 0 | 0 |
| 0 | 0 | 0 | 0 |
| 0 | 0 | 0 | 0 |
| 0 | 0 | 0 | 0 |
| 0 | 0 | 0 | 0 |
| 0 | 0 | 0 | 0 |
| 0 | 0 | 0 | 0 |
| 1 | 0 | 0 | 0 |
| 0 | 0 | 0 | 0 |
| 0 | 0 | 0 | 0 |
| 0 | 0 | 0 | 0 |
| 0 | 0 | 0 | 0 |
| 0 | 0 | 0 | 0 |
| 0 | 0 | 0 | 0 |
| 0 | 0 | 0 | 0 |
| 0 | 0 | 0 | 0 |
| 0 | 0 | 0 | 0 |
| 0 | 0 | 0 | 0 |
| 0 | 0 | 0 | 0 |
| 0 | 1 | 0 | 0 |
| 0 | 0 | 0 | 0 |
| 0 | 0 | 0 | 0 |
| 0 | 0 | 0 | 0 |
| 0 | 0 | 0 | 0 |
| 0 | 0 | 0 | 0 |
| 0 | 0 | 0 | 0 |
| 0 | 0 | 0 | 0 |
| 0 | 0 | 0 | 0 |
| 1 | 0 | 0 | 0 |

| | | | |
|---|---|---|---|
| 0 | 0 | 0 | 0 |
| 0 | 0 | 0 | 0 |
| 0 | 0 | 0 | 0 |
| 0 | 0 | 0 | 0 |
| 0 | 0 | 0 | 0 |
| 0 | 0 | 0 | 0 |
| 0 | 0 | 0 | 0 |
| 0 | 0 | 0 | 0 |
| 0 | 0 | 0 | 0 |
| 1 | 0 | 0 | 0 |
| 0 | 0 | 0 | 0 |
| 0 | 0 | 0 | 0 |
| 1 | 1 | 0 | 0 |
| 1 | 1 | 0 | 0 |
| 1 | 1 | 0 | 0 |
| 1 | 1 | 0 | 0 |
| 0 | 0 | 0 | 0 |
| 0 | 0 | 0 | 0 |
| 0 | 0 | 0 | 0 |
| 0 | 0 | 0 | 0 |
| 0 | 0 | 0 | 0 |
| 0 | 0 | 0 | 0 |
| 0 | 0 | 0 | 0 |
| 0 | 0 | 0 | 0 |
| 0 | 0 | 0 | 0 |
| 0 | 0 | 0 | 1 |
| 0 | 0 | 0 | 0 |
| 0 | 0 | 0 | 0 |
| 0 | 0 | 0 | 0 |
| 0 | 0 | 0 | 0 |
| 0 | 0 | 0 | 0 |
| 0 | 0 | 0 | 0 |
| 0 | 0 | 0 | 0 |
| 0 | 0 | 0 | 0 |
| 0 | 0 | 0 | 0 |
| 0 | 0 | 0 | 0 |
| 0 | 0 | 0 | 0 |
| 1 | 1 | 0 | 0 |
| 0 | 0 | 0 | 0 |
| 0 | 0 | 0 | 0 |
| 1 | 1 | 0 | 0 |
| 0 | 0 | 0 | 0 |
| 0 | 0 | 0 | 0 |
| 1 | 0 | 0 | 1 |
| 0 | 0 | 0 | 0 |
| 0 | 0 | 0 | 0 |
| 0 | 0 | 0 | 0 |
| 0 | 0 | 0 | 0 |
| 0 | 0 | 0 | 0 |
| 0 | 0 | 0 | 0 |
| 0 | 0 | 0 | 0 |
| 0 | 0 | 1 | 0 |
| 0 | 0 | 1 | 0 |
| 0 | 0 | 1 | 0 |

| | | | |
|---|---|---|---|
| 0 | 0 | 1 | 0 |
| 0 | 0 | 0 | 0 |
| 0 | 0 | 0 | 0 |
| 0 | 0 | 0 | 0 |
| 0 | 0 | 0 | 0 |
| 0 | 0 | 0 | 0 |
| 0 | 0 | 0 | 0 |
| 0 | 0 | 0 | 0 |
| 0 | 0 | 0 | 0 |
| 0 | 0 | 0 | 0 |
| 0 | 0 | 0 | 0 |
| 0 | 0 | 0 | 0 |
| 0 | 0 | 0 | 0 |
| 0 | 0 | 0 | 0 |
| 0 | 0 | 0 | 0 |
| 0 | 0 | 0 | 0 |
| 0 | 0 | 0 | 0 |
| 0 | 0 | 0 | 0 |
| 0 | 0 | 0 | 0 |
| 0 | 0 | 0 | 0 |
| 0 | 0 | 0 | 0 |
| 0 | 0 | 0 | 0 |
| 0 | 0 | 0 | 0 |
| 0 | 1 | 0 | 1 |
| 0 | 0 | 0 | 0 |
| 0 | 0 | 0 | 0 |
| 0 | 0 | 0 | 0 |
| 0 | 0 | 0 | 0 |
| 0 | 0 | 0 | 0 |
| 0 | 0 | 0 | 0 |
| 0 | 0 | 0 | 0 |
| 0 | 0 | 0 | 0 |
| 0 | 0 | 0 | 0 |
| 0 | 0 | 0 | 0 |
| 0 | 0 | 0 | 0 |
| 0 | 0 | 0 | 0 |
| 0 | 0 | 0 | 0 |
| 0 | 0 | 0 | 0 |
| 0 | 0 | 0 | 0 |
| 0 | 0 | 0 | 0 |
| 0 | 0 | 0 | 0 |
| 0 | 0 | 0 | 0 |
| 0 | 0 | 0 | 0 |
| 0 | 0 | 0 | 0 |
| 0 | 0 | 0 | 0 |
| 0 | 0 | 0 | 0 |
| 1 | 0 | 0 | 0 |
| 1 | 0 | 0 | 0 |
| 0 | 0 | 0 | 0 |
| 0 | 1 | 0 | 0 |
| 1 | 1 | 0 | 1 |
| 1 | 1 | 0 | 1 |
| 1 | 1 | 0 | 1 |
| 0 | 0 | 0 | 0 |

| | | | |
|---|---|---|---|
| 1 | 1 | 0 | 0 |
| 0 | 0 | 0 | 0 |
| 0 | 0 | 0 | 0 |
| 0 | 0 | 0 | 0 |
| 0 | 0 | 0 | 0 |
| 0 | 0 | 0 | 0 |
| 0 | 0 | 0 | 0 |
| 0 | 0 | 0 | 0 |
| 0 | 0 | 0 | 0 |
| 0 | 0 | 0 | 0 |
| 0 | 0 | 0 | 0 |
| 0 | 1 | 0 | 0 |
| 0 | 0 | 0 | 0 |
| 0 | 0 | 0 | 0 |
| 0 | 1 | 0 | 0 |
| 0 | 0 | 0 | 0 |
| 1 | 1 | 0 | 0 |
| 0 | 0 | 0 | 0 |
| 0 | 0 | 0 | 0 |
| 1 | 0 | 0 | 0 |
| 0 | 0 | 0 | 0 |
| 0 | 0 | 0 | 0 |
| 0 | 0 | 0 | 0 |
| 0 | 0 | 0 | 0 |
| 0 | 0 | 0 | 0 |
| 0 | 0 | 0 | 0 |
| 0 | 0 | 0 | 0 |
| 0 | 0 | 0 | 0 |
| 0 | 0 | 0 | 0 |
| 0 | 0 | 0 | 0 |
| 0 | 0 | 0 | 0 |
| 0 | 0 | 0 | 0 |
| 0 | 0 | 0 | 0 |
| 1 | 0 | 0 | 0 |
| 0 | 0 | 0 | 0 |
| 0 | 1 | 0 | 0 |
| 0 | 0 | 0 | 0 |
| 0 | 0 | 0 | 0 |
| 0 | 0 | 0 | 0 |
| 0 | 0 | 0 | 0 |
| 0 | 0 | 0 | 0 |
| 0 | 0 | 0 | 0 |
| 0 | 0 | 0 | 0 |
| 0 | 0 | 0 | 0 |
| 0 | 0 | 0 | 0 |
| 0 | 0 | 0 | 0 |
| 0 | 0 | 0 | 0 |
| 0 | 0 | 0 | 0 |
| 1 | 1 | 0 | 0 |
| 0 | 0 | 0 | 0 |
| 0 | 0 | 0 | 0 |
| 0 | 0 | 0 | 0 |
| 0 | 0 | 0 | 0 |
| 0 | 0 | 0 | 0 |

| | | | |
|---|---|---|---|
| 0 | 0 | 0 | 0 |
| 0 | 0 | 0 | 0 |
| 0 | 0 | 0 | 0 |
| 0 | 0 | 0 | 0 |
| 0 | 0 | 0 | 0 |
| 0 | 0 | 0 | 0 |
| 0 | 0 | 0 | 0 |
| 0 | 1 | 0 | 0 |
| 0 | 0 | 0 | 0 |
| 0 | 0 | 0 | 0 |
| 0 | 0 | 0 | 0 |
| 0 | 0 | 0 | 0 |
| 0 | 0 | 1 | 0 |
| 0 | 0 | 0 | 0 |
| 0 | 0 | 1 | 0 |
| 0 | 0 | 1 | 0 |
| 0 | 0 | 0 | 0 |
| 0 | 0 | 0 | 0 |
| 0 | 0 | 0 | 0 |
| 0 | 0 | 0 | 0 |
| 0 | 1 | 0 | 0 |
| 0 | 1 | 0 | 0 |
| 0 | 0 | 0 | 0 |
| 0 | 0 | 0 | 0 |
| 0 | 1 | 0 | 0 |
| 0 | 0 | 0 | 0 |
| 0 | 0 | 0 | 0 |
| 0 | 1 | 0 | 0 |
| 0 | 0 | 0 | 0 |
| 0 | 0 | 0 | 0 |
| 0 | 0 | 0 | 0 |
| 0 | 0 | 0 | 0 |
| 1 | 1 | 0 | 0 |
| 1 | 1 | 0 | 0 |
| 1 | 1 | 0 | 0 |
| 0 | 1 | 0 | 0 |
| 1 | 0 | 0 | 0 |
| 0 | 1 | 0 | 0 |
| 1 | 1 | 0 | 0 |
| 0 | 0 | 0 | 0 |
| 0 | 0 | 0 | 0 |
| 0 | 0 | 0 | 0 |
| 0 | 0 | 0 | 0 |
| 0 | 0 | 0 | 0 |
| 0 | 0 | 0 | 0 |
| 0 | 0 | 0 | 0 |
| 0 | 0 | 0 | 0 |
| 0 | 0 | 0 | 0 |
| 0 | 0 | 0 | 0 |
| 1 | 1 | 0 | 0 |
| 1 | 1 | 0 | 0 |
| 0 | 0 | 0 | 0 |
| 0 | 0 | 0 | 0 |
| 0 | 0 | 0 | 0 |

| | | | |
|---|---|---|---|
| 0 | 0 | 0 | 0 |
| 0 | 0 | 0 | 0 |
| 0 | 0 | 0 | 0 |
| 0 | 0 | 0 | 0 |
| 0 | 0 | 0 | 0 |
| 0 | 0 | 0 | 0 |
| 1 | 1 | 0 | 0 |
| 1 | 1 | 0 | 0 |
| 0 | 0 | 0 | 0 |
| 0 | 0 | 0 | 0 |
| 0 | 0 | 0 | 0 |
| 0 | 0 | 0 | 0 |
| 0 | 0 | 0 | 0 |
| 0 | 0 | 0 | 0 |
| 0 | 0 | 0 | 0 |
| 0 | 0 | 0 | 0 |
| 0 | 0 | 0 | 0 |
| 0 | 0 | 0 | 0 |
| 0 | 0 | 0 | 0 |
| 0 | 0 | 0 | 0 |
| 0 | 0 | 0 | 0 |
| 0 | 0 | 0 | 0 |
| 0 | 0 | 0 | 0 |
| 0 | 0 | 0 | 0 |
| 0 | 0 | 0 | 0 |
| 0 | 0 | 0 | 0 |
| 0 | 0 | 0 | 0 |
| 1 | 1 | 0 | 0 |
| 0 | 0 | 0 | 0 |
| 0 | 0 | 0 | 0 |
| 0 | 0 | 0 | 0 |
| 0 | 0 | 0 | 0 |
| 0 | 0 | 0 | 0 |
| 0 | 0 | 0 | 0 |
| 0 | 0 | 0 | 0 |
| 0 | 0 | 0 | 0 |
| 0 | 0 | 0 | 0 |
| 0 | 0 | 0 | 0 |
| 0 | 0 | 0 | 0 |
| 0 | 0 | 0 | 0 |
| 0 | 0 | 0 | 0 |
| 0 | 0 | 0 | 0 |
| 0 | 0 | 0 | 0 |
| 0 | 0 | 0 | 0 |
| 0 | 0 | 0 | 0 |
| 0 | 0 | 0 | 0 |
| 0 | 0 | 0 | 0 |
| 0 | 0 | 0 | 0 |
| 0 | 0 | 0 | 0 |
| 0 | 0 | 0 | 0 |
| 0 | 0 | 0 | 0 |
| 0 | 0 | 0 | 0 |
| 0 | 0 | 0 | 0 |
| 0 | 0 | 0 | 0 |

| | | | |
|---|---|---|---|
| 0 | 0 | 0 | 0 |
| 0 | 0 | 0 | 0 |
| 0 | 0 | 0 | 0 |
| 0 | 0 | 0 | 0 |
| 0 | 0 | 0 | 0 |
| 0 | 0 | 0 | 0 |
| 0 | 0 | 0 | 0 |
| 0 | 0 | 0 | 0 |
| 0 | 0 | 0 | 0 |
| 0 | 0 | 0 | 0 |
| 1 | 0 | 0 | 0 |
| 1 | 1 | 0 | 0 |
| 1 | 0 | 0 | 0 |
| 0 | 0 | 0 | 0 |
| 0 | 0 | 0 | 0 |
| 0 | 0 | 0 | 0 |
| 0 | 0 | 0 | 0 |
| 0 | 0 | 0 | 0 |
| 0 | 0 | 0 | 0 |
| 0 | 0 | 0 | 0 |
| 0 | 0 | 0 | 0 |
| 0 | 0 | 0 | 0 |
| 0 | 0 | 0 | 0 |
| 0 | 0 | 0 | 0 |
| 0 | 0 | 0 | 0 |
| 0 | 0 | 0 | 0 |
| 0 | 0 | 0 | 0 |
| 0 | 0 | 0 | 0 |
| 0 | 0 | 0 | 0 |
| 0 | 0 | 0 | 0 |
| 0 | 0 | 0 | 0 |
| 0 | 1 | 0 | 0 |
| 0 | 0 | 0 | 0 |
| 0 | 0 | 0 | 0 |
| 1 | 0 | 0 | 0 |
| 1 | 0 | 0 | 0 |
| 1 | 0 | 0 | 0 |
| 0 | 0 | 0 | 0 |
| 0 | 0 | 0 | 0 |
| 0 | 0 | 0 | 0 |
| 0 | 0 | 0 | 0 |
| 0 | 0 | 0 | 0 |
| 0 | 0 | 0 | 0 |
| 0 | 0 | 0 | 0 |
| 0 | 0 | 0 | 0 |
| 0 | 0 | 0 | 0 |
| 0 | 0 | 0 | 0 |
| 0 | 0 | 0 | 0 |
| 0 | 0 | 0 | 0 |
| 0 | 0 | 0 | 0 |
| 0 | 0 | 0 | 0 |
| 0 | 0 | 0 | 0 |
| 0 | 0 | 0 | 1 |
| 0 | 0 | 0 | 0 |

| | | | |
|---|---|---|---|
| 0 | 0 | 0 | 0 |
| 0 | 0 | 0 | 0 |
| 0 | 0 | 0 | 0 |
| 0 | 0 | 0 | 0 |
| 0 | 0 | 0 | 0 |
| 0 | 0 | 0 | 0 |
| 0 | 0 | 0 | 0 |
| 0 | 0 | 0 | 0 |
| 0 | 0 | 0 | 0 |
| 0 | 0 | 0 | 0 |
| 0 | 0 | 0 | 0 |
| 1 | 0 | 0 | 0 |
| 0 | 0 | 0 | 0 |
| 0 | 0 | 0 | 0 |
| 0 | 0 | 1 | 0 |
| 0 | 0 | 0 | 0 |
| 0 | 0 | 0 | 0 |
| 0 | 0 | 0 | 0 |
| 0 | 0 | 0 | 0 |
| 0 | 0 | 0 | 0 |
| 0 | 0 | 0 | 0 |
| 0 | 0 | 0 | 0 |
| 0 | 0 | 0 | 0 |
| 0 | 0 | 0 | 0 |
| 0 | 0 | 0 | 0 |
| 0 | 0 | 0 | 0 |
| 0 | 0 | 0 | 0 |
| 0 | 0 | 0 | 0 |
| 0 | 0 | 0 | 0 |
| 0 | 0 | 0 | 0 |
| 0 | 0 | 0 | 0 |
| 0 | 0 | 0 | 0 |
| 0 | 0 | 0 | 0 |
| 1 | 0 | 0 | 0 |
| 0 | 0 | 0 | 0 |
| 1 | 0 | 0 | 1 |
| 0 | 0 | 0 | 0 |
| 0 | 0 | 0 | 0 |
| 0 | 0 | 0 | 0 |
| 0 | 0 | 0 | 0 |
| 0 | 0 | 0 | 0 |
| 0 | 0 | 0 | 0 |
| 0 | 0 | 0 | 0 |
| 0 | 1 | 0 | 0 |
| 0 | 0 | 0 | 0 |
| 0 | 0 | 0 | 0 |
| 0 | 0 | 0 | 0 |
| 0 | 0 | 0 | 0 |
| 0 | 0 | 0 | 0 |
| 0 | 0 | 0 | 0 |
| 0 | 0 | 0 | 0 |
| 0 | 0 | 0 | 0 |
| 0 | 0 | 0 | 0 |
| 0 | 0 | 0 | 0 |

| | | | |
|---|---|---|---|
| 0 | 0 | 0 | 0 |
| 0 | 0 | 0 | 0 |
| 0 | 0 | 0 | 0 |
| 0 | 0 | 0 | 0 |
| 0 | 0 | 0 | 0 |
| 0 | 1 | 0 | 0 |
| 0 | 0 | 0 | 0 |
| 0 | 0 | 0 | 0 |
| 0 | 0 | 0 | 0 |
| 0 | 0 | 0 | 0 |
| 0 | 0 | 0 | 0 |
| 0 | 0 | 0 | 0 |
| 0 | 0 | 0 | 0 |
| 0 | 0 | 0 | 0 |
| 0 | 0 | 0 | 0 |
| 0 | 1 | 0 | 0 |
| 1 | 1 | 0 | 0 |
| 0 | 0 | 1 | 0 |
| 0 | 0 | 0 | 0 |
| 0 | 0 | 0 | 0 |
| 0 | 0 | 0 | 0 |
| 0 | 0 | 0 | 0 |
| 0 | 0 | 0 | 0 |
| 0 | 0 | 0 | 0 |
| 0 | 0 | 0 | 0 |
| 0 | 0 | 0 | 0 |
| 0 | 0 | 0 | 0 |
| 1 | 0 | 0 | 0 |
| 0 | 0 | 0 | 0 |
| 0 | 0 | 0 | 0 |
| 0 | 0 | 1 | 0 |
| 0 | 0 | 0 | 0 |
| 0 | 0 | 0 | 0 |
| 0 | 0 | 0 | 0 |
| 0 | 0 | 0 | 0 |
| 0 | 0 | 0 | 0 |
| 0 | 0 | 0 | 0 |
| 1 | 1 | 0 | 0 |
| 0 | 0 | 0 | 0 |
| 0 | 0 | 0 | 0 |
| 0 | 0 | 0 | 0 |
| 1 | 0 | 0 | 0 |
| 1 | 0 | 0 | 0 |
| 0 | 0 | 0 | 0 |
| 0 | 0 | 0 | 0 |
| 0 | 0 | 0 | 0 |
| 0 | 0 | 0 | 0 |
| 0 | 0 | 0 | 0 |
| 0 | 0 | 0 | 0 |
| 0 | 0 | 0 | 0 |
| 0 | 0 | 0 | 0 |
| 0 | 0 | 0 | 0 |
| 0 | 0 | 0 | 0 |
| 0 | 0 | 0 | 0 |

| | | | |
|---|---|---|---|
| 0 | 0 | 0 | 0 |
| 0 | 0 | 0 | 0 |
| 0 | 0 | 0 | 0 |
| 0 | 0 | 0 | 0 |
| 0 | 0 | 0 | 0 |
| 0 | 0 | 0 | 0 |
| 0 | 0 | 0 | 0 |
| 0 | 0 | 0 | 0 |
| 0 | 0 | 0 | 0 |
| 0 | 0 | 0 | 0 |
| 0 | 0 | 0 | 0 |
| 0 | 0 | 0 | 0 |
| 1 | 0 | 0 | 0 |
| 0 | 0 | 0 | 0 |
| 0 | 0 | 0 | 0 |
| 0 | 0 | 0 | 0 |
| 0 | 0 | 0 | 0 |
| 0 | 0 | 0 | 0 |
| 0 | 0 | 0 | 0 |
| 0 | 0 | 0 | 0 |
| 0 | 0 | 0 | 0 |
| 0 | 0 | 0 | 0 |
| 0 | 0 | 0 | 0 |
| 1 | 0 | 0 | 0 |
| 0 | 0 | 0 | 0 |
| 0 | 0 | 0 | 0 |
| 0 | 0 | 0 | 0 |
| 1 | 1 | 0 | 1 |
| 0 | 0 | 0 | 0 |
| 0 | 0 | 0 | 0 |
| 0 | 0 | 0 | 0 |
| 0 | 0 | 0 | 0 |
| 0 | 0 | 0 | 0 |
| 0 | 0 | 0 | 0 |
| 0 | 0 | 0 | 0 |
| 0 | 0 | 0 | 0 |
| 0 | 0 | 0 | 0 |
| 0 | 0 | 0 | 0 |
| 0 | 0 | 0 | 0 |
| 0 | 0 | 0 | 0 |
| 0 | 0 | 0 | 0 |
| 0 | 0 | 0 | 0 |
| 0 | 0 | 0 | 0 |
| 0 | 0 | 0 | 0 |
| 0 | 0 | 0 | 0 |
| 0 | 0 | 0 | 0 |
| 0 | 0 | 0 | 0 |
| 0 | 0 | 0 | 0 |
| 0 | 0 | 0 | 0 |
| 0 | 0 | 0 | 0 |
| 0 | 0 | 0 | 0 |
| 0 | 0 | 0 | 0 |
| 0 | 0 | 0 | 0 |
| 0 | 0 | 0 | 0 |

| | | | |
|---|---|---|---|
| 0 | 0 | 0 | 0 |
| 0 | 0 | 0 | 0 |
| 0 | 0 | 0 | 0 |
| 0 | 0 | 0 | 0 |
| 0 | 0 | 0 | 0 |
| 0 | 0 | 0 | 0 |
| 0 | 0 | 0 | 0 |
| 0 | 0 | 0 | 0 |
| 1 | 0 | 0 | 0 |
| 1 | 0 | 0 | 0 |
| 1 | 0 | 0 | 0 |
| 0 | 0 | 0 | 0 |
| 1 | 0 | 0 | 0 |
| 0 | 0 | 0 | 0 |
| 0 | 0 | 0 | 0 |
| 1 | 0 | 0 | 0 |
| 0 | 0 | 0 | 0 |
| 0 | 0 | 0 | 0 |
| 0 | 0 | 0 | 0 |
| 0 | 0 | 0 | 0 |
| 0 | 0 | 0 | 0 |
| 0 | 0 | 0 | 0 |
| 0 | 0 | 0 | 0 |
| 0 | 0 | 0 | 0 |
| 0 | 0 | 0 | 0 |
| 0 | 0 | 0 | 0 |
| 0 | 0 | 0 | 0 |
| 1 | 0 | 0 | 0 |
| 1 | 0 | 0 | 0 |
| 0 | 0 | 0 | 0 |
| 0 | 0 | 0 | 0 |
| 0 | 0 | 0 | 0 |
| 1 | 0 | 0 | 0 |
| 0 | 0 | 0 | 0 |
| 0 | 0 | 0 | 0 |
| 0 | 0 | 0 | 0 |
| 0 | 0 | 0 | 0 |
| 0 | 0 | 0 | 0 |
| 0 | 0 | 0 | 0 |
| 0 | 0 | 0 | 0 |
| 1 | 0 | 0 | 0 |
| 1 | 0 | 0 | 0 |
| 0 | 0 | 0 | 0 |
| 0 | 0 | 0 | 0 |
| 0 | 0 | 0 | 0 |
| 0 | 0 | 0 | 0 |
| 0 | 0 | 0 | 0 |
| 0 | 0 | 0 | 0 |
| 0 | 0 | 0 | 0 |
| 0 | 0 | 0 | 0 |
| 0 | 0 | 0 | 0 |
| 0 | 0 | 0 | 0 |
| 0 | 0 | 0 | 0 |
| 0 | 0 | 0 | 0 |

| | | | |
|---|---|---|---|
| 0 | 0 | 0 | 0 |
| 0 | 0 | 0 | 0 |
| 0 | 0 | 0 | 0 |
| 0 | 0 | 0 | 0 |
| 0 | 0 | 0 | 0 |
| 0 | 0 | 0 | 0 |
| 0 | 0 | 0 | 0 |
| 0 | 0 | 0 | 0 |
| 0 | 0 | 0 | 0 |
| 0 | 0 | 0 | 0 |
| 1 | 0 | 0 | 0 |
| 0 | 0 | 0 | 0 |
| 1 | 0 | 0 | 0 |
| 0 | 0 | 0 | 0 |
| 0 | 0 | 0 | 0 |
| 0 | 0 | 0 | 0 |
| 0 | 0 | 0 | 0 |
| 0 | 1 | 0 | 0 |
| 0 | 0 | 0 | 0 |
| 0 | 0 | 0 | 0 |
| 0 | 0 | 0 | 0 |
| 0 | 0 | 0 | 0 |
| 1 | 0 | 0 | 0 |
| 0 | 0 | 0 | 0 |
| 0 | 0 | 0 | 0 |
| 1 | 0 | 0 | 0 |
| 1 | 0 | 0 | 0 |
| 1 | 0 | 0 | 0 |
| 0 | 0 | 0 | 0 |
| 0 | 0 | 0 | 0 |
| 0 | 0 | 0 | 0 |
| 0 | 0 | 0 | 0 |
| 0 | 0 | 0 | 0 |
| 0 | 0 | 0 | 0 |
| 0 | 0 | 0 | 0 |
| 0 | 0 | 0 | 0 |
| 0 | 0 | 0 | 0 |
| 0 | 0 | 0 | 0 |
| 0 | 0 | 0 | 0 |
| 0 | 0 | 0 | 0 |
| 1 | 1 | 0 | 0 |
| 0 | 0 | 0 | 0 |
| 0 | 0 | 0 | 0 |
| 0 | 0 | 0 | 0 |
| 0 | 0 | 0 | 0 |
| 0 | 0 | 0 | 0 |
| 0 | 0 | 0 | 0 |
| 1 | 0 | 0 | 0 |
| 0 | 0 | 0 | 0 |
| 0 | 0 | 0 | 0 |
| 0 | 0 | 0 | 0 |
| 0 | 0 | 0 | 0 |
| 0 | 0 | 0 | 0 |
| 0 | 0 | 0 | 0 |

| | | | |
|---|---|---|---|
| 1 | 0 | 0 | 0 |
| 0 | 0 | 0 | 0 |
| 1 | 0 | 0 | 0 |
| 0 | 0 | 0 | 0 |
| 0 | 0 | 0 | 0 |
| 0 | 0 | 0 | 0 |
| 0 | 0 | 0 | 0 |
| 0 | 0 | 0 | 0 |
| 1 | 1 | 0 | 0 |
| 0 | 0 | 0 | 0 |
| 1 | 1 | 0 | 0 |
| 1 | 1 | 0 | 0 |
| 0 | 0 | 0 | 0 |
| 1 | 1 | 0 | 0 |
| 0 | 0 | 0 | 0 |
| 0 | 0 | 0 | 0 |
| 0 | 0 | 0 | 0 |
| 0 | 0 | 0 | 0 |
| 0 | 0 | 0 | 0 |
| 0 | 0 | 0 | 0 |
| 1 | 0 | 0 | 0 |
| 0 | 0 | 0 | 0 |
| 0 | 0 | 0 | 0 |
| 0 | 0 | 0 | 0 |
| 1 | 0 | 0 | 0 |
| 0 | 0 | 0 | 0 |
| 1 | 0 | 0 | 0 |
| 1 | 0 | 0 | 0 |
| 1 | 1 | 0 | 0 |
| 0 | 0 | 0 | 0 |
| 1 | 1 | 0 | 0 |
| 1 | 1 | 0 | 0 |
| 1 | 1 | 0 | 0 |
| 0 | 0 | 0 | 0 |
| 1 | 1 | 0 | 0 |
| 0 | 0 | 0 | 0 |
| 1 | 0 | 0 | 0 |
| 0 | 0 | 0 | 0 |
| 1 | 1 | 0 | 0 |
| 1 | 1 | 0 | 0 |
| 1 | 0 | 0 | 0 |
| 0 | 0 | 0 | 0 |
| 0 | 0 | 0 | 0 |
| 1 | 0 | 0 | 0 |
| 0 | 0 | 0 | 0 |
| 0 | 0 | 0 | 0 |
| 1 | 0 | 0 | 0 |
| 0 | 0 | 0 | 0 |
| 1 | 0 | 0 | 0 |
| 0 | 0 | 0 | 0 |
| 0 | 0 | 0 | 0 |
| 0 | 0 | 0 | 0 |
| 0 | 0 | 0 | 0 |
| 0 | 0 | 0 | 0 |

| | | | |
|---|---|---|---|
| 0 | 0 | 0 | 0 |
| 0 | 0 | 0 | 0 |
| 0 | 0 | 0 | 0 |
| 0 | 0 | 0 | 0 |
| 0 | 0 | 0 | 0 |
| 0 | 0 | 0 | 0 |
| 0 | 0 | 0 | 0 |
| 0 | 0 | 0 | 0 |
| 0 | 0 | 0 | 0 |
| 1 | 1 | 0 | 0 |
| 1 | 1 | 0 | 0 |
| 0 | 0 | 0 | 0 |
| 0 | 0 | 0 | 0 |
| 0 | 0 | 0 | 0 |
| 1 | 0 | 1 | 0 |
| 0 | 0 | 0 | 0 |
| 0 | 0 | 0 | 0 |
| 1 | 0 | 1 | 0 |
| 0 | 0 | 0 | 0 |
| 0 | 0 | 0 | 0 |
| 0 | 0 | 0 | 0 |
| 0 | 0 | 1 | 0 |
| 1 | 0 | 1 | 0 |
| 0 | 0 | 1 | 0 |
| 0 | 0 | 1 | 0 |
| 0 | 0 | 1 | 1 |
| 0 | 0 | 1 | 1 |
| 0 | 0 | 0 | 0 |
| 0 | 0 | 0 | 0 |
| 0 | 0 | 0 | 0 |
| 0 | 0 | 0 | 0 |
| 0 | 0 | 0 | 0 |
| 0 | 0 | 0 | 0 |
| 0 | 0 | 0 | 0 |
| 0 | 0 | 0 | 0 |
| 0 | 0 | 0 | 0 |
| 0 | 0 | 0 | 0 |
| 0 | 0 | 0 | 0 |
| 0 | 0 | 0 | 0 |
| 0 | 0 | 0 | 0 |
| 0 | 0 | 0 | 0 |
| 0 | 0 | 0 | 0 |
| 0 | 0 | 0 | 0 |
| 0 | 0 | 1 | 0 |
| 0 | 0 | 0 | 0 |
| 0 | 0 | 0 | 0 |
| 0 | 0 | 0 | 0 |
| 0 | 0 | 0 | 0 |
| 0 | 0 | 0 | 0 |
| 0 | 0 | 0 | 0 |
| 1 | 0 | 0 | 0 |
| 0 | 0 | 0 | 0 |
| 0 | 0 | 0 | 0 |
| 0 | 0 | 0 | 0 |

| | | | |
|---|---|---|---|
| 0 | 0 | 0 | 0 |
| 0 | 0 | 0 | 0 |
| 0 | 0 | 0 | 0 |
| 1 | 0 | 0 | 0 |
| 0 | 0 | 0 | 0 |
| 0 | 0 | 0 | 0 |
| 0 | 0 | 0 | 0 |
| 0 | 0 | 0 | 0 |
| 1 | 0 | 0 | 0 |
| 1 | 0 | 0 | 0 |
| 0 | 0 | 0 | 0 |
| 0 | 0 | 0 | 0 |
| 0 | 0 | 0 | 0 |
| 0 | 0 | 0 | 0 |
| 0 | 0 | 0 | 0 |
| 0 | 0 | 0 | 0 |
| 0 | 0 | 1 | 0 |
| 0 | 0 | 0 | 0 |
| 0 | 0 | 0 | 0 |
| 0 | 0 | 0 | 0 |
| 0 | 0 | 0 | 0 |
| 0 | 0 | 0 | 0 |
| 0 | 0 | 0 | 0 |
| 0 | 0 | 0 | 0 |
| 0 | 0 | 0 | 0 |
| 0 | 0 | 0 | 0 |
| 0 | 0 | 0 | 0 |
| 0 | 0 | 0 | 0 |
| 0 | 0 | 0 | 0 |
| 0 | 0 | 0 | 0 |
| 0 | 0 | 0 | 0 |
| 0 | 0 | 0 | 0 |
| 0 | 0 | 0 | 0 |
| 0 | 0 | 0 | 0 |
| 0 | 0 | 0 | 0 |
| 0 | 0 | 0 | 0 |
| 0 | 0 | 0 | 0 |
| 0 | 0 | 0 | 0 |
| 0 | 0 | 0 | 0 |
| 0 | 0 | 0 | 0 |
| 0 | 0 | 0 | 0 |
| 0 | 0 | 0 | 0 |
| 0 | 0 | 0 | 0 |
| 0 | 0 | 0 | 0 |
| 0 | 0 | 0 | 0 |
| 0 | 0 | 0 | 0 |
| 0 | 0 | 0 | 0 |
| 0 | 0 | 0 | 0 |
| 0 | 0 | 0 | 0 |
| 0 | 1 | 0 | 0 |
| 0 | 0 | 0 | 0 |
| 0 | 0 | 0 | 0 |
| 0 | 0 | 0 | 0 |
| 0 | 0 | 0 | 0 |

| | | | |
|---|---|---|---|
| 0 | 0 | 1 | 0 |
| 0 | 0 | 0 | 0 |
| 0 | 0 | 0 | 0 |
| 0 | 0 | 0 | 0 |
| 0 | 0 | 0 | 0 |
| 0 | 0 | 0 | 0 |
| 0 | 0 | 0 | 0 |
| 1 | 0 | 0 | 0 |
| 0 | 0 | 0 | 0 |
| 0 | 0 | 0 | 0 |
| 0 | 0 | 0 | 0 |
| 1 | 0 | 0 | 0 |
| 0 | 0 | 0 | 0 |
| 0 | 0 | 0 | 0 |
| 0 | 0 | 0 | 0 |
| 0 | 0 | 0 | 0 |
| 0 | 0 | 0 | 0 |
| 0 | 0 | 0 | 0 |
| 0 | 0 | 0 | 0 |
| 0 | 0 | 0 | 0 |
| 0 | 0 | 0 | 0 |
| 0 | 0 | 0 | 0 |
| 0 | 0 | 0 | 0 |
| 0 | 0 | 0 | 0 |
| 0 | 0 | 0 | 0 |
| 0 | 0 | 0 | 0 |
| 0 | 0 | 0 | 0 |
| 0 | 0 | 0 | 0 |
| 0 | 0 | 0 | 0 |
| 0 | 0 | 0 | 0 |
| 0 | 0 | 0 | 0 |
| 0 | 0 | 0 | 0 |
| 0 | 0 | 0 | 0 |
| 0 | 0 | 0 | 0 |
| 0 | 0 | 0 | 0 |
| 0 | 0 | 0 | 0 |
| 0 | 0 | 0 | 0 |
| 0 | 0 | 0 | 0 |
| 0 | 0 | 0 | 0 |
| 0 | 0 | 0 | 0 |
| 0 | 0 | 0 | 0 |
| 0 | 0 | 0 | 0 |
| 0 | 0 | 0 | 0 |
| 0 | 0 | 0 | 0 |
| 0 | 0 | 0 | 0 |
| 0 | 0 | 0 | 0 |
| 0 | 0 | 0 | 0 |
| 0 | 0 | 0 | 0 |
| 0 | 0 | 0 | 0 |
| 0 | 0 | 0 | 0 |
| 0 | 0 | 0 | 0 |
| 0 | 0 | 0 | 0 |
| 0 | 0 | 0 | 0 |
| 0 | 0 | 0 | 0 |

| | | | |
|---|---|---|---|
| 0 | 0 | 0 | 0 |
| 0 | 0 | 0 | 0 |
| 0 | 0 | 0 | 0 |
| 0 | 0 | 0 | 0 |
| 0 | 0 | 0 | 0 |
| 0 | 0 | 0 | 0 |
| 0 | 0 | 0 | 0 |
| 0 | 0 | 0 | 0 |
| 0 | 0 | 0 | 0 |
| 0 | 0 | 0 | 0 |
| 0 | 0 | 0 | 0 |
| 0 | 0 | 0 | 0 |
| 0 | 0 | 0 | 0 |
| 0 | 0 | 0 | 0 |
| 0 | 0 | 0 | 0 |
| 0 | 0 | 0 | 0 |
| 0 | 0 | 0 | 0 |
| 0 | 0 | 0 | 0 |
| 0 | 0 | 0 | 1 |
| 0 | 1 | 0 | 1 |
| 0 | 0 | 0 | 0 |
| 0 | 0 | 0 | 0 |
| 0 | 0 | 0 | 0 |
| 0 | 0 | 0 | 0 |
| 0 | 0 | 0 | 0 |
| 1 | 0 | 0 | 0 |
| 1 | 0 | 0 | 0 |
| 0 | 0 | 0 | 0 |
| 0 | 0 | 0 | 0 |
| 0 | 0 | 0 | 0 |
| 0 | 0 | 0 | 0 |
| 0 | 0 | 0 | 0 |
| 0 | 0 | 0 | 0 |
| 1 | 0 | 0 | 1 |
| 1 | 0 | 0 | 0 |
| 1 | 0 | 0 | 1 |
| 1 | 0 | 0 | 0 |
| 0 | 0 | 0 | 0 |
| 0 | 0 | 0 | 0 |
| 0 | 0 | 0 | 0 |
| 0 | 0 | 0 | 0 |
| 1 | 0 | 0 | 0 |
| 1 | 0 | 0 | 1 |
| 1 | 0 | 0 | 0 |
| 0 | 0 | 0 | 0 |
| 0 | 0 | 0 | 0 |
| 0 | 0 | 0 | 0 |
| 0 | 0 | 0 | 0 |
| 0 | 0 | 0 | 0 |
| 1 | 0 | 0 | 0 |
| 0 | 0 | 0 | 0 |
| 0 | 0 | 0 | 0 |
| 0 | 0 | 0 | 0 |
| 0 | 0 | 0 | 0 |

| | | | |
|---|---|---|---|
| 0 | 0 | 0 | 0 |
| 0 | 0 | 0 | 0 |
| 0 | 0 | 0 | 0 |
| 1 | 0 | 0 | 0 |
| 1 | 0 | 0 | 0 |
| 0 | 0 | 0 | 0 |
| 1 | 0 | 0 | 0 |
| 0 | 0 | 0 | 0 |
| 0 | 0 | 0 | 0 |
| 0 | 0 | 0 | 0 |
| 0 | 0 | 0 | 0 |
| 1 | 0 | 1 | 0 |
| 0 | 0 | 0 | 0 |
| 0 | 0 | 0 | 0 |
| 0 | 0 | 0 | 0 |
| 0 | 0 | 1 | 0 |
| 0 | 0 | 0 | 0 |
| 0 | 0 | 0 | 0 |
| 0 | 0 | 0 | 0 |
| 0 | 0 | 1 | 0 |
| 1 | 0 | 1 | 0 |
| 1 | 0 | 0 | 0 |
| 1 | 0 | 0 | 0 |
| 1 | 0 | 0 | 0 |
| 0 | 0 | 1 | 0 |
| 0 | 0 | 0 | 0 |
| 1 | 0 | 0 | 0 |
| 0 | 0 | 0 | 0 |
| 0 | 0 | 0 | 0 |
| 0 | 0 | 0 | 0 |
| 0 | 0 | 0 | 0 |
| 0 | 0 | 0 | 0 |
| 0 | 0 | 0 | 0 |
| 0 | 0 | 0 | 0 |
| 0 | 0 | 0 | 0 |
| 0 | 0 | 0 | 0 |
| 0 | 0 | 0 | 1 |
| 0 | 0 | 0 | 1 |
| 0 | 0 | 0 | 1 |
| 0 | 0 | 0 | 0 |
| 0 | 0 | 0 | 0 |
| 0 | 0 | 0 | 0 |
| 0 | 0 | 0 | 0 |
| 0 | 0 | 0 | 0 |
| 0 | 0 | 0 | 0 |
| 0 | 0 | 0 | 0 |
| 0 | 0 | 0 | 0 |
| 0 | 0 | 0 | 0 |
| 0 | 0 | 0 | 0 |
| 0 | 0 | 0 | 0 |
| 0 | 0 | 0 | 0 |
| 0 | 0 | 0 | 0 |
| 0 | 0 | 0 | 0 |
| 0 | 0 | 0 | 0 |

| | | | |
|---|---|---|---|
| 0 | 0 | 0 | 0 |
| 0 | 0 | 0 | 0 |
| 0 | 0 | 0 | 0 |
| 0 | 0 | 0 | 0 |
| 0 | 0 | 0 | 0 |
| 0 | 0 | 0 | 0 |
| 0 | 0 | 0 | 0 |
| 0 | 0 | 0 | 0 |
| 0 | 0 | 0 | 0 |
| 0 | 0 | 0 | 0 |
| 0 | 0 | 0 | 0 |
| 0 | 0 | 0 | 0 |
| 0 | 0 | 0 | 0 |
| 0 | 0 | 0 | 0 |
| 0 | 0 | 0 | 0 |
| 0 | 0 | 0 | 0 |
| 0 | 1 | 0 | 0 |
| 0 | 1 | 0 | 0 |
| 0 | 0 | 0 | 0 |
| 0 | 0 | 0 | 0 |
| 0 | 0 | 0 | 0 |
| 0 | 0 | 0 | 0 |
| 0 | 0 | 0 | 0 |
| 0 | 0 | 0 | 0 |
| 0 | 0 | 0 | 0 |
| 0 | 0 | 0 | 0 |
| 0 | 0 | 0 | 0 |
| 0 | 0 | 0 | 0 |
| 0 | 0 | 0 | 0 |
| 0 | 0 | 0 | 0 |
| 0 | 0 | 0 | 0 |
| 1 | 0 | 0 | 0 |
| 0 | 0 | 0 | 0 |
| 0 | 0 | 0 | 0 |
| 0 | 0 | 0 | 0 |
| 0 | 0 | 0 | 0 |
| 0 | 0 | 0 | 0 |
| 0 | 0 | 0 | 0 |
| 0 | 0 | 0 | 0 |
| 0 | 0 | 0 | 0 |
| 0 | 0 | 0 | 0 |
| 0 | 0 | 0 | 0 |
| 0 | 0 | 0 | 0 |
| 0 | 0 | 0 | 0 |
| 0 | 0 | 0 | 0 |
| 0 | 0 | 0 | 0 |
| 1 | 1 | 0 | 0 |
| 0 | 0 | 0 | 0 |
| 0 | 0 | 0 | 0 |
| 0 | 0 | 0 | 0 |
| 0 | 0 | 0 | 0 |
| 0 | 0 | 0 | 0 |
| 1 | 0 | 0 | 0 |
| 0 | 0 | 0 | 0 |

| | | | |
|---|---|---|---|
| 0 | 0 | 0 | 0 |
| 0 | 0 | 0 | 0 |
| 0 | 0 | 0 | 0 |
| 0 | 0 | 0 | 0 |
| 0 | 0 | 0 | 0 |
| 0 | 0 | 0 | 0 |
| 1 | 0 | 0 | 0 |
| 0 | 0 | 0 | 0 |
| 0 | 0 | 0 | 0 |
| 0 | 0 | 0 | 0 |
| 0 | 0 | 0 | 0 |
| 0 | 0 | 0 | 0 |
| 0 | 0 | 0 | 0 |
| 0 | 0 | 0 | 0 |
| 0 | 0 | 0 | 0 |
| 0 | 0 | 0 | 0 |
| 0 | 0 | 0 | 0 |
| 0 | 0 | 0 | 0 |
| 0 | 0 | 0 | 0 |
| 0 | 0 | 0 | 0 |
| 0 | 0 | 0 | 0 |
| 0 | 0 | 0 | 0 |
| 0 | 0 | 0 | 0 |
| 0 | 0 | 0 | 0 |
| 0 | 0 | 0 | 0 |
| 0 | 0 | 0 | 0 |
| 0 | 0 | 0 | 0 |
| 0 | 0 | 0 | 0 |
| 0 | 0 | 0 | 0 |
| 0 | 0 | 0 | 0 |
| 0 | 0 | 0 | 0 |
| 0 | 0 | 0 | 0 |
| 0 | 0 | 0 | 0 |
| 0 | 0 | 0 | 0 |
| 0 | 0 | 0 | 0 |
| 0 | 0 | 0 | 0 |
| 0 | 0 | 0 | 0 |
| 0 | 0 | 0 | 0 |
| 0 | 0 | 0 | 0 |
| 0 | 0 | 0 | 0 |
| 0 | 0 | 0 | 0 |
| 0 | 0 | 0 | 0 |
| 0 | 0 | 0 | 0 |
| 0 | 0 | 0 | 0 |
| 0 | 0 | 0 | 0 |
| 0 | 0 | 0 | 0 |
| 0 | 0 | 0 | 0 |
| 0 | 0 | 0 | 0 |
| 0 | 0 | 0 | 0 |
| 0 | 0 | 0 | 0 |
| 0 | 0 | 0 | 0 |
| 0 | 0 | 0 | 0 |
| 0 | 0 | 0 | 0 |
| 0 | 0 | 0 | 0 |

| | | | |
|---|---|---|---|
| 0 | 0 | 0 | 0 |
| 0 | 0 | 0 | 0 |
| 0 | 0 | 0 | 0 |
| 0 | 0 | 0 | 0 |
| 0 | 0 | 0 | 0 |
| 0 | 0 | 0 | 0 |
| 0 | 0 | 0 | 0 |
| 0 | 0 | 0 | 0 |
| 0 | 0 | 0 | 0 |
| 0 | 0 | 0 | 0 |
| 0 | 0 | 0 | 0 |
| 0 | 0 | 0 | 0 |
| 0 | 0 | 0 | 0 |
| 0 | 0 | 0 | 0 |
| 0 | 0 | 0 | 0 |
| 0 | 0 | 0 | 0 |
| 0 | 0 | 0 | 0 |
| 0 | 0 | 0 | 0 |
| 0 | 0 | 0 | 0 |
| 0 | 0 | 0 | 0 |
| 0 | 0 | 0 | 0 |
| 0 | 0 | 0 | 0 |
| 0 | 0 | 0 | 0 |
| 0 | 0 | 0 | 0 |
| 1 | 1 | 0 | 0 |
| 0 | 0 | 0 | 0 |
| 0 | 0 | 0 | 0 |
| 0 | 0 | 0 | 0 |
| 0 | 0 | 0 | 0 |
| 0 | 0 | 0 | 0 |
| 0 | 0 | 0 | 0 |
| 0 | 0 | 0 | 0 |
| 0 | 0 | 0 | 0 |
| 0 | 0 | 0 | 0 |
| 0 | 0 | 0 | 0 |
| 1 | 0 | 0 | 0 |
| 1 | 0 | 0 | 1 |
| 1 | 0 | 0 | 0 |
| 0 | 0 | 0 | 0 |
| 1 | 0 | 0 | 0 |
| 1 | 1 | 0 | 0 |
| 1 | 1 | 0 | 0 |
| 0 | 0 | 0 | 0 |
| 0 | 0 | 0 | 0 |
| 0 | 0 | 0 | 0 |
| 0 | 0 | 0 | 0 |
| 0 | 0 | 0 | 0 |
| 0 | 0 | 0 | 0 |
| 0 | 0 | 0 | 0 |
| 0 | 0 | 0 | 0 |
| 0 | 0 | 0 | 0 |
| 0 | 0 | 0 | 0 |
| 0 | 0 | 0 | 0 |
| 0 | 0 | 0 | 0 |

| | | | |
|---|---|---|---|
| 0 | 0 | 0 | 0 |
| 0 | 0 | 0 | 0 |
| 0 | 0 | 0 | 0 |
| 0 | 0 | 0 | 0 |
| 0 | 0 | 0 | 0 |
| 0 | 0 | 0 | 0 |
| 0 | 0 | 0 | 0 |
| 0 | 0 | 0 | 0 |
| 0 | 0 | 0 | 0 |
| 0 | 0 | 0 | 0 |
| 0 | 0 | 0 | 0 |
| 0 | 0 | 0 | 0 |
| 0 | 0 | 0 | 0 |
| 1 | 0 | 0 | 0 |
| 0 | 0 | 0 | 0 |
| 0 | 0 | 0 | 0 |
| 1 | 1 | 0 | 0 |
| 0 | 0 | 0 | 0 |
| 0 | 0 | 0 | 0 |
| 0 | 0 | 0 | 0 |
| 0 | 0 | 0 | 0 |
| 0 | 0 | 0 | 0 |
| 0 | 0 | 0 | 0 |
| 1 | 0 | 0 | 0 |
| 0 | 0 | 0 | 0 |
| 0 | 0 | 0 | 0 |
| 0 | 0 | 0 | 0 |
| 0 | 0 | 0 | 0 |
| 0 | 0 | 0 | 0 |
| 0 | 0 | 0 | 0 |
| 0 | 0 | 0 | 0 |
| 1 | 0 | 0 | 0 |
| 0 | 0 | 0 | 0 |
| 0 | 0 | 0 | 0 |
| 0 | 0 | 0 | 0 |
| 0 | 0 | 0 | 0 |
| 0 | 0 | 0 | 0 |
| 0 | 0 | 0 | 0 |
| 0 | 0 | 0 | 0 |
| 0 | 0 | 0 | 0 |
| 0 | 0 | 0 | 0 |
| 0 | 0 | 0 | 0 |
| 0 | 0 | 0 | 0 |
| 0 | 0 | 0 | 0 |
| 0 | 0 | 0 | 0 |
| 0 | 0 | 0 | 0 |
| 0 | 0 | 0 | 0 |
| 0 | 0 | 0 | 0 |
| 0 | 0 | 0 | 0 |
| 0 | 0 | 0 | 0 |
| 0 | 0 | 0 | 0 |
| 0 | 0 | 0 | 0 |
| 0 | 0 | 0 | 0 |
| 0 | 0 | 0 | 0 |

| | | | |
|---|---|---|---|
| 0 | 0 | 0 | 0 |
| 0 | 0 | 0 | 0 |
| 0 | 0 | 0 | 0 |
| 0 | 0 | 0 | 0 |
| 0 | 0 | 0 | 0 |
| 0 | 0 | 0 | 0 |
| 0 | 0 | 0 | 0 |
| 0 | 0 | 0 | 0 |
| 0 | 0 | 0 | 0 |
| 0 | 0 | 0 | 0 |
| 0 | 0 | 0 | 0 |
| 0 | 0 | 0 | 0 |
| 0 | 0 | 0 | 0 |
| 0 | 0 | 0 | 0 |
| 0 | 0 | 0 | 0 |
| 0 | 0 | 0 | 0 |
| 0 | 0 | 0 | 0 |
| 1 | 0 | 0 | 0 |
| 0 | 0 | 0 | 0 |
| 0 | 0 | 0 | 0 |
| 0 | 0 | 0 | 0 |
| 0 | 0 | 0 | 0 |
| 0 | 0 | 0 | 0 |
| 0 | 0 | 0 | 0 |
| 1 | 0 | 0 | 0 |
| 0 | 0 | 0 | 0 |
| 0 | 0 | 0 | 0 |
| 0 | 0 | 0 | 0 |
| 0 | 0 | 0 | 0 |
| 0 | 0 | 0 | 0 |
| 0 | 0 | 0 | 0 |
| 0 | 0 | 0 | 0 |
| 0 | 0 | 0 | 0 |
| 0 | 0 | 0 | 0 |
| 0 | 0 | 0 | 0 |
| 0 | 0 | 0 | 0 |
| 0 | 0 | 0 | 0 |
| 0 | 0 | 0 | 0 |
| 0 | 0 | 0 | 0 |
| 0 | 0 | 0 | 0 |
| 0 | 0 | 0 | 0 |
| 0 | 0 | 0 | 0 |
| 0 | 0 | 0 | 0 |
| 0 | 0 | 0 | 0 |
| 0 | 0 | 0 | 0 |
| 0 | 0 | 0 | 0 |
| 0 | 0 | 0 | 0 |
| 0 | 0 | 0 | 0 |
| 0 | 0 | 0 | 0 |
| 0 | 0 | 0 | 0 |
| 0 | 0 | 0 | 0 |
| 0 | 0 | 0 | 0 |
| 0 | 0 | 0 | 0 |
| 0 | 0 | 0 | 0 |

| | | | |
|---|---|---|---|
| 0 | 0 | 0 | 0 |
| 0 | 0 | 0 | 0 |
| 0 | 0 | 0 | 0 |
| 0 | 0 | 0 | 0 |
| 0 | 0 | 0 | 0 |
| 0 | 0 | 0 | 0 |
| 0 | 0 | 0 | 0 |
| 0 | 0 | 0 | 0 |
| 0 | 0 | 0 | 0 |
| 0 | 0 | 0 | 0 |
| 0 | 0 | 0 | 0 |
| 0 | 0 | 0 | 0 |
| 0 | 0 | 0 | 0 |
| 0 | 0 | 0 | 0 |
| 1 | 1 | 0 | 0 |
| 1 | 1 | 0 | 0 |
| 0 | 0 | 0 | 0 |
| 0 | 0 | 0 | 0 |
| 0 | 0 | 0 | 0 |
| 0 | 0 | 0 | 0 |
| 0 | 0 | 0 | 0 |
| 0 | 0 | 0 | 0 |
| 0 | 0 | 0 | 0 |
| 0 | 0 | 0 | 0 |
| 0 | 0 | 0 | 0 |
| 0 | 0 | 0 | 0 |
| 0 | 0 | 0 | 0 |
| 1 | 0 | 0 | 0 |
| 0 | 0 | 0 | 0 |
| 1 | 0 | 0 | 0 |
| 0 | 0 | 0 | 0 |
| 0 | 0 | 0 | 0 |
| 0 | 0 | 0 | 0 |
| 0 | 0 | 0 | 0 |
| 0 | 0 | 0 | 0 |
| 0 | 0 | 0 | 0 |
| 1 | 0 | 0 | 0 |
| 0 | 0 | 0 | 0 |
| 0 | 0 | 0 | 0 |
| 0 | 0 | 0 | 0 |
| 0 | 0 | 0 | 0 |
| 0 | 0 | 0 | 0 |
| 0 | 0 | 0 | 0 |
| 0 | 0 | 0 | 0 |
| 0 | 0 | 0 | 0 |
| 0 | 0 | 0 | 0 |
| 0 | 0 | 0 | 0 |
| 0 | 0 | 0 | 0 |
| 0 | 0 | 0 | 0 |
| 0 | 0 | 0 | 0 |
| 0 | 0 | 0 | 0 |
| 0 | 0 | 0 | 0 |
| 0 | 0 | 0 | 0 |
| 0 | 0 | 0 | 0 |

| | | | |
|---|---|---|---|
| 0 | 0 | 0 | 0 |
| 0 | 0 | 0 | 0 |
| 0 | 0 | 0 | 0 |
| 0 | 0 | 0 | 0 |
| 0 | 0 | 0 | 0 |
| 0 | 0 | 0 | 0 |
| 0 | 0 | 0 | 0 |
| 0 | 0 | 0 | 0 |
| 0 | 0 | 0 | 0 |
| 0 | 0 | 0 | 0 |
| 0 | 0 | 0 | 0 |
| 0 | 0 | 0 | 0 |
| 0 | 0 | 0 | 0 |
| 0 | 0 | 0 | 0 |
| 0 | 0 | 0 | 0 |
| 0 | 0 | 0 | 0 |
| 0 | 0 | 0 | 0 |
| 0 | 0 | 0 | 0 |
| 0 | 0 | 0 | 0 |
| 0 | 0 | 0 | 0 |
| 0 | 0 | 0 | 0 |
| 0 | 0 | 0 | 0 |
| 0 | 0 | 0 | 0 |
| 0 | 0 | 0 | 0 |
| 0 | 0 | 0 | 0 |
| 0 | 0 | 0 | 0 |
| 0 | 0 | 0 | 0 |
| 0 | 0 | 0 | 0 |
| 0 | 0 | 0 | 0 |
| 0 | 0 | 0 | 0 |
| 0 | 0 | 0 | 0 |
| 0 | 0 | 0 | 0 |
| 0 | 0 | 0 | 0 |
| 0 | 0 | 0 | 0 |
| 0 | 0 | 0 | 0 |
| 0 | 0 | 0 | 0 |
| 0 | 0 | 0 | 0 |
| 0 | 0 | 0 | 0 |
| 0 | 0 | 0 | 0 |
| 0 | 0 | 0 | 0 |
| 0 | 0 | 0 | 0 |
| 0 | 0 | 0 | 0 |
| 0 | 0 | 0 | 0 |
| 0 | 0 | 0 | 0 |
| 0 | 0 | 0 | 0 |
| 0 | 0 | 0 | 0 |
| 0 | 0 | 0 | 0 |
| 0 | 0 | 0 | 0 |
| 1 | 0 | 0 | 1 |
| 0 | 0 | 0 | 0 |
| 1 | 0 | 0 | 0 |
| 1 | 0 | 0 | 0 |
| 0 | 0 | 0 | 0 |
| 0 | 0 | 0 | 1 |

| | | | |
|---|---|---|---|
| 0 | 0 | 0 | 0 |
| 0 | 0 | 0 | 0 |
| 0 | 0 | 0 | 0 |
| 0 | 0 | 0 | 0 |
| 0 | 0 | 0 | 0 |
| 0 | 0 | 0 | 0 |
| 0 | 0 | 0 | 0 |
| 0 | 0 | 0 | 0 |
| 0 | 0 | 0 | 0 |
| 0 | 0 | 0 | 0 |
| 0 | 0 | 0 | 0 |
| 0 | 0 | 0 | 0 |
| 0 | 0 | 0 | 0 |
| 0 | 0 | 0 | 0 |
| 0 | 0 | 0 | 0 |
| 0 | 0 | 0 | 0 |
| 0 | 0 | 0 | 0 |
| 0 | 0 | 0 | 0 |
| 0 | 0 | 0 | 0 |
| 0 | 0 | 0 | 0 |
| 0 | 0 | 0 | 0 |
| 0 | 0 | 0 | 0 |
| 0 | 0 | 0 | 0 |
| 0 | 0 | 0 | 0 |
| 0 | 0 | 0 | 0 |
| 0 | 0 | 0 | 0 |
| 0 | 0 | 1 | 0 |
| 0 | 0 | 1 | 0 |
| 0 | 0 | 1 | 0 |
| 0 | 0 | 1 | 0 |
| 0 | 0 | 0 | 0 |
| 0 | 0 | 0 | 0 |
| 0 | 0 | 0 | 0 |
| 1 | 0 | 0 | 0 |
| 1 | 0 | 0 | 0 |
| 0 | 0 | 0 | 0 |
| 0 | 0 | 0 | 0 |
| 0 | 0 | 0 | 0 |
| 0 | 0 | 0 | 0 |
| 1 | 0 | 0 | 0 |
| 0 | 0 | 0 | 0 |
| 0 | 0 | 0 | 0 |
| 0 | 0 | 0 | 0 |
| 0 | 0 | 0 | 0 |
| 0 | 0 | 0 | 0 |
| 0 | 1 | 0 | 0 |
| 0 | 0 | 0 | 0 |
| 0 | 0 | 0 | 0 |
| 0 | 0 | 0 | 0 |
| 0 | 0 | 0 | 0 |
| 0 | 0 | 0 | 0 |
| 0 | 0 | 0 | 0 |
| 0 | 0 | 0 | 0 |
| 0 | 0 | 0 | 0 |

| | | | |
|---|---|---|---|
| 0 | 0 | 0 | 0 |
| 0 | 0 | 0 | 0 |
| 0 | 0 | 0 | 0 |
| 0 | 0 | 0 | 0 |
| 0 | 0 | 0 | 0 |
| 0 | 0 | 0 | 1 |
| 0 | 0 | 0 | 0 |
| 0 | 0 | 0 | 0 |
| 0 | 0 | 0 | 0 |
| 0 | 0 | 0 | 0 |
| 0 | 0 | 0 | 0 |
| 0 | 0 | 0 | 0 |
| 0 | 0 | 0 | 0 |
| 0 | 1 | 0 | 0 |
| 0 | 1 | 0 | 0 |
| 0 | 1 | 0 | 0 |
| 0 | 1 | 0 | 0 |
| 0 | 1 | 0 | 0 |
| 0 | 0 | 0 | 0 |
| 0 | 0 | 0 | 0 |
| 0 | 0 | 0 | 0 |
| 0 | 0 | 0 | 0 |
| 0 | 0 | 0 | 0 |
| 0 | 0 | 0 | 0 |
| 0 | 0 | 0 | 0 |
| 0 | 0 | 0 | 0 |
| 0 | 0 | 0 | 0 |
| 0 | 0 | 0 | 0 |
| 0 | 0 | 0 | 0 |
| 0 | 0 | 0 | 0 |
| 0 | 0 | 0 | 0 |
| 0 | 0 | 0 | 0 |
| 0 | 0 | 0 | 0 |
| 0 | 0 | 0 | 0 |
| 0 | 0 | 0 | 0 |
| 0 | 0 | 0 | 0 |
| 0 | 0 | 0 | 0 |
| 0 | 0 | 0 | 0 |
| 0 | 0 | 0 | 0 |
| 0 | 0 | 0 | 0 |
| 0 | 0 | 0 | 0 |
| 0 | 0 | 0 | 0 |
| 0 | 0 | 0 | 0 |
| 0 | 0 | 0 | 0 |
| 0 | 0 | 0 | 0 |
| 0 | 0 | 0 | 0 |
| 0 | 0 | 0 | 0 |
| 0 | 0 | 0 | 0 |
| 0 | 0 | 0 | 0 |
| 0 | 0 | 0 | 0 |
| 0 | 0 | 0 | 0 |
| 0 | 0 | 0 | 0 |
| 0 | 0 | 0 | 0 |
| 0 | 0 | 0 | 0 |

| | | | |
|---|---|---|---|
| 0 | 0 | 0 | 0 |
| 0 | 0 | 0 | 0 |
| 0 | 0 | 0 | 0 |
| 1 | 0 | 0 | 0 |
| 0 | 0 | 0 | 0 |
| 0 | 0 | 0 | 0 |
| 0 | 0 | 0 | 0 |
| 0 | 0 | 0 | 0 |
| 0 | 0 | 0 | 0 |
| 0 | 0 | 0 | 0 |
| 0 | 0 | 0 | 0 |
| 0 | 0 | 0 | 0 |
| 0 | 0 | 0 | 0 |
| 0 | 0 | 0 | 0 |
| 0 | 0 | 0 | 0 |
| 0 | 0 | 0 | 0 |
| 0 | 0 | 0 | 0 |
| 0 | 0 | 0 | 0 |
| 0 | 0 | 0 | 0 |
| 0 | 0 | 0 | 0 |
| 1 | 1 | 0 | 0 |
| 1 | 1 | 0 | 0 |
| 0 | 0 | 0 | 0 |
| 0 | 1 | 0 | 0 |
| 0 | 0 | 0 | 0 |
| 0 | 0 | 0 | 0 |
| 0 | 0 | 0 | 0 |
| 0 | 0 | 0 | 0 |
| 0 | 0 | 0 | 0 |
| 0 | 0 | 0 | 0 |
| 0 | 0 | 0 | 0 |
| 0 | 0 | 0 | 0 |
| 0 | 0 | 0 | 0 |
| 0 | 0 | 0 | 0 |
| 0 | 0 | 0 | 0 |
| 0 | 0 | 0 | 0 |
| 0 | 0 | 0 | 0 |
| 0 | 0 | 0 | 0 |
| 0 | 0 | 0 | 0 |
| 1 | 0 | 0 | 0 |
| 1 | 0 | 0 | 0 |
| 0 | 0 | 0 | 0 |
| 0 | 0 | 0 | 0 |
| 1 | 0 | 0 | 0 |
| 0 | 0 | 0 | 0 |
| 0 | 0 | 0 | 0 |
| 0 | 0 | 0 | 0 |
| 0 | 0 | 0 | 0 |
| 0 | 0 | 0 | 0 |
| 0 | 0 | 0 | 0 |
| 0 | 0 | 0 | 0 |
| 0 | 0 | 0 | 0 |
| 0 | 0 | 0 | 0 |
| 0 | 0 | 0 | 0 |

| | | | |
|---|---|---|---|
| 0 | 0 | 0 | 0 |
| 0 | 0 | 0 | 0 |
| 0 | 0 | 0 | 0 |
| 0 | 0 | 0 | 0 |
| 0 | 0 | 0 | 0 |
| 1 | 0 | 0 | 0 |
| 0 | 0 | 0 | 0 |
| 0 | 0 | 0 | 0 |
| 0 | 1 | 0 | 0 |
| 1 | 1 | 0 | 0 |
| 0 | 0 | 0 | 0 |
| 1 | 1 | 0 | 0 |
| 1 | 0 | 0 | 0 |
| 0 | 0 | 0 | 0 |
| 0 | 0 | 0 | 0 |
| 0 | 0 | 0 | 0 |
| 1 | 1 | 0 | 0 |
| 0 | 0 | 0 | 0 |
| 0 | 0 | 0 | 0 |
| 1 | 0 | 0 | 0 |
| 0 | 0 | 0 | 0 |
| 0 | 0 | 0 | 0 |
| 0 | 0 | 0 | 0 |
| 0 | 0 | 0 | 0 |
| 0 | 0 | 0 | 0 |
| 0 | 0 | 0 | 0 |
| 0 | 0 | 0 | 0 |
| 0 | 0 | 0 | 0 |
| 0 | 0 | 0 | 0 |
| 0 | 0 | 0 | 0 |
| 1 | 0 | 0 | 0 |
| 0 | 0 | 0 | 0 |
| 0 | 0 | 0 | 0 |
| 0 | 0 | 0 | 0 |
| 0 | 0 | 0 | 0 |
| 1 | 0 | 0 | 0 |
| 0 | 0 | 0 | 0 |
| 0 | 0 | 0 | 0 |
| 0 | 0 | 0 | 0 |
| 0 | 0 | 0 | 0 |
| 0 | 0 | 0 | 0 |
| 0 | 0 | 0 | 0 |
| 0 | 0 | 0 | 0 |
| 0 | 0 | 0 | 0 |
| 0 | 0 | 0 | 0 |
| 0 | 0 | 0 | 0 |
| 0 | 0 | 0 | 0 |
| 0 | 0 | 0 | 0 |
| 0 | 0 | 0 | 0 |
| 0 | 0 | 0 | 0 |
| 0 | 0 | 0 | 0 |
| 0 | 0 | 0 | 0 |
| 0 | 0 | 0 | 0 |

| | | | |
|---|---|---|---|
| 0 | 0 | 0 | 0 |
| 0 | 0 | 0 | 0 |
| 0 | 0 | 0 | 0 |
| 0 | 0 | 0 | 0 |
| 0 | 0 | 0 | 0 |
| 0 | 0 | 0 | 0 |
| 0 | 0 | 0 | 0 |
| 0 | 0 | 0 | 0 |
| 0 | 0 | 0 | 0 |
| 0 | 0 | 0 | 0 |
| 0 | 0 | 0 | 0 |
| 0 | 0 | 0 | 0 |
| 0 | 0 | 0 | 0 |
| 0 | 0 | 0 | 0 |
| 0 | 0 | 0 | 1 |
| 0 | 0 | 0 | 0 |
| 0 | 0 | 0 | 0 |
| 0 | 0 | 0 | 0 |
| 0 | 0 | 0 | 0 |
| 0 | 0 | 0 | 0 |
| 0 | 0 | 0 | 0 |
| 0 | 0 | 0 | 0 |
| 0 | 0 | 0 | 0 |
| 0 | 0 | 0 | 0 |
| 0 | 0 | 0 | 0 |
| 0 | 0 | 0 | 0 |
| 0 | 0 | 0 | 0 |
| 0 | 0 | 0 | 0 |
| 0 | 0 | 0 | 0 |
| 0 | 0 | 0 | 0 |
| 0 | 0 | 0 | 0 |
| 0 | 0 | 0 | 0 |
| 0 | 0 | 0 | 0 |
| 0 | 0 | 0 | 0 |
| 0 | 0 | 0 | 0 |
| 0 | 0 | 0 | 0 |
| 0 | 0 | 0 | 0 |
| 0 | 0 | 0 | 0 |
| 0 | 0 | 0 | 0 |
| 0 | 0 | 1 | 0 |
| 0 | 0 | 0 | 0 |
| 0 | 0 | 0 | 0 |
| 0 | 0 | 0 | 0 |
| 1 | 0 | 0 | 0 |
| 0 | 0 | 0 | 0 |
| 0 | 0 | 0 | 0 |
| 0 | 0 | 0 | 0 |
| 0 | 0 | 0 | 0 |
| 1 | 0 | 0 | 1 |
| 0 | 0 | 0 | 0 |
| 0 | 0 | 0 | 0 |
| 0 | 0 | 0 | 0 |
| 0 | 0 | 0 | 0 |
| 0 | 0 | 0 | 0 |

| | | | |
|---|---|---|---|
| 0 | 0 | 0 | 0 |
| 0 | 0 | 0 | 0 |
| 0 | 0 | 0 | 0 |
| 0 | 0 | 0 | 0 |
| 0 | 0 | 0 | 0 |
| 0 | 0 | 0 | 0 |
| 0 | 0 | 0 | 0 |
| 0 | 0 | 0 | 0 |
| 0 | 0 | 0 | 0 |
| 1 | 0 | 0 | 0 |
| 0 | 0 | 0 | 0 |
| 0 | 0 | 0 | 0 |
| 0 | 0 | 0 | 0 |
| 0 | 0 | 0 | 0 |
| 1 | 0 | 0 | 0 |
| 0 | 0 | 0 | 0 |
| 0 | 0 | 0 | 0 |
| 0 | 0 | 0 | 0 |
| 0 | 0 | 0 | 0 |
| 0 | 0 | 0 | 0 |
| 0 | 0 | 0 | 0 |
| 0 | 0 | 0 | 0 |
| 0 | 0 | 0 | 0 |
| 0 | 0 | 0 | 0 |
| 1 | 0 | 0 | 0 |
| 0 | 0 | 0 | 0 |
| 0 | 0 | 0 | 0 |
| 0 | 0 | 0 | 0 |
| 0 | 0 | 0 | 0 |
| 0 | 0 | 0 | 0 |
| 0 | 0 | 0 | 0 |
| 0 | 0 | 0 | 0 |
| 0 | 0 | 0 | 0 |
| 0 | 0 | 0 | 0 |
| 1 | 0 | 0 | 0 |
| 1 | 0 | 0 | 0 |
| 0 | 0 | 0 | 0 |
| 0 | 0 | 0 | 0 |
| 1 | 0 | 0 | 0 |
| 0 | 0 | 1 | 0 |
| 0 | 0 | 1 | 0 |
| 0 | 0 | 1 | 0 |
| 0 | 0 | 1 | 0 |
| 0 | 0 | 0 | 0 |
| 1 | 0 | 0 | 0 |
| 0 | 0 | 0 | 0 |
| 0 | 0 | 0 | 0 |
| 0 | 0 | 0 | 0 |
| 0 | 0 | 1 | 0 |
| 0 | 0 | 0 | 0 |
| 0 | 0 | 0 | 0 |
| 0 | 0 | 0 | 0 |
| 0 | 0 | 0 | 0 |
| 0 | 0 | 0 | 0 |

| | | | |
|---|---|---|---|
| 0 | 0 | 0 | 0 |
| 0 | 0 | 0 | 0 |
| 0 | 0 | 0 | 0 |
| 0 | 0 | 0 | 0 |
| 0 | 0 | 0 | 0 |
| 0 | 0 | 0 | 0 |
| 0 | 0 | 0 | 0 |
| 0 | 0 | 0 | 0 |
| 0 | 0 | 0 | 0 |
| 0 | 0 | 0 | 0 |
| 0 | 1 | 0 | 0 |
| 0 | 0 | 0 | 0 |
| 0 | 0 | 0 | 0 |
| 0 | 0 | 0 | 0 |
| 0 | 0 | 0 | 0 |
| 0 | 1 | 0 | 0 |
| 0 | 0 | 0 | 0 |
| 0 | 0 | 0 | 0 |
| 0 | 0 | 0 | 0 |
| 0 | 0 | 0 | 0 |
| 0 | 0 | 0 | 0 |
| 0 | 0 | 0 | 0 |
| 0 | 0 | 0 | 0 |
| 0 | 0 | 0 | 0 |
| 0 | 0 | 0 | 0 |
| 0 | 0 | 0 | 0 |
| 0 | 0 | 0 | 0 |
| 0 | 0 | 0 | 0 |
| 0 | 0 | 0 | 0 |
| 0 | 0 | 0 | 0 |
| 0 | 0 | 0 | 0 |
| 0 | 0 | 0 | 0 |
| 0 | 0 | 0 | 0 |
| 0 | 0 | 0 | 0 |
| 0 | 0 | 0 | 0 |
| 0 | 0 | 0 | 0 |
| 0 | 0 | 0 | 0 |
| 0 | 0 | 0 | 0 |
| 0 | 0 | 0 | 0 |
| 0 | 0 | 0 | 0 |
| 0 | 0 | 0 | 0 |
| 0 | 0 | 0 | 0 |
| 0 | 0 | 0 | 0 |
| 0 | 0 | 0 | 0 |
| 1 | 1 | 0 | 0 |
| 0 | 0 | 0 | 0 |
| 0 | 0 | 0 | 0 |
| 0 | 0 | 0 | 0 |
| 0 | 0 | 0 | 0 |
| 0 | 0 | 0 | 0 |
| 0 | 0 | 0 | 0 |
| 0 | 0 | 0 | 0 |
| 0 | 0 | 0 | 0 |
| 0 | 0 | 0 | 0 |

| | | | |
|---|---|---|---|
| 0 | 0 | 0 | 0 |
| 0 | 0 | 0 | 0 |
| 0 | 0 | 1 | 0 |
| 1 | 0 | 0 | 0 |
| 0 | 0 | 0 | 0 |
| 0 | 0 | 0 | 0 |
| 0 | 0 | 0 | 0 |
| 0 | 0 | 0 | 0 |
| 0 | 0 | 0 | 0 |
| 0 | 0 | 0 | 0 |
| 0 | 0 | 0 | 0 |
| 0 | 0 | 0 | 0 |
| 1 | 0 | 0 | 0 |
| 0 | 0 | 0 | 0 |
| 0 | 0 | 0 | 0 |
| 0 | 0 | 0 | 0 |
| 0 | 0 | 0 | 0 |
| 0 | 0 | 0 | 0 |
| 0 | 0 | 0 | 0 |
| 0 | 0 | 0 | 0 |
| 0 | 0 | 0 | 0 |
| 0 | 0 | 0 | 0 |
| 0 | 0 | 0 | 0 |
| 0 | 0 | 0 | 0 |
| 0 | 0 | 0 | 0 |
| 0 | 0 | 0 | 0 |
| 0 | 0 | 0 | 0 |
| 0 | 0 | 0 | 0 |
| 0 | 0 | 0 | 0 |
| 0 | 0 | 0 | 0 |
| 0 | 0 | 0 | 0 |
| 0 | 0 | 0 | 0 |
| 0 | 0 | 0 | 0 |
| 0 | 0 | 0 | 0 |
| 0 | 0 | 0 | 0 |
| 0 | 0 | 0 | 0 |
| 0 | 0 | 0 | 0 |
| 0 | 0 | 0 | 0 |
| 0 | 0 | 0 | 0 |
| 0 | 0 | 0 | 0 |
| 0 | 1 | 0 | 0 |
| 1 | 1 | 0 | 0 |
| 0 | 0 | 0 | 0 |
| 0 | 0 | 0 | 0 |
| 0 | 0 | 0 | 0 |
| 0 | 0 | 0 | 0 |
| 0 | 0 | 0 | 0 |
| 0 | 0 | 0 | 0 |
| 0 | 1 | 0 | 0 |
| 0 | 0 | 0 | 0 |
| 0 | 0 | 0 | 0 |
| 0 | 0 | 0 | 0 |
| 0 | 0 | 0 | 1 |
| 0 | 0 | 0 | 0 |

| | | | |
|---|---|---|---|
| 0 | 0 | 0 | 1 |
| 0 | 0 | 0 | 1 |
| 0 | 0 | 0 | 1 |
| 0 | 0 | 0 | 0 |
| 0 | 0 | 0 | 0 |
| 0 | 0 | 0 | 0 |
| 0 | 0 | 0 | 0 |
| 1 | 0 | 0 | 0 |
| 0 | 0 | 0 | 0 |
| 1 | 1 | 0 | 0 |
| 0 | 1 | 0 | 0 |
| 0 | 0 | 0 | 0 |
| 1 | 1 | 0 | 0 |
| 0 | 0 | 0 | 0 |
| 0 | 0 | 0 | 0 |
| 0 | 0 | 0 | 0 |
| 0 | 0 | 0 | 0 |
| 1 | 0 | 0 | 0 |
| 0 | 0 | 0 | 0 |
| 0 | 0 | 0 | 0 |
| 0 | 0 | 0 | 0 |
| 0 | 0 | 0 | 0 |
| 0 | 0 | 0 | 0 |
| 0 | 0 | 0 | 0 |
| 0 | 0 | 0 | 0 |
| 0 | 0 | 0 | 0 |
| 0 | 0 | 0 | 0 |
| 0 | 0 | 0 | 0 |
| 0 | 0 | 0 | 0 |
| 0 | 0 | 0 | 0 |
| 0 | 0 | 0 | 0 |
| 1 | 0 | 0 | 0 |
| 0 | 0 | 0 | 0 |
| 0 | 0 | 0 | 0 |
| 0 | 0 | 0 | 0 |
| 0 | 0 | 0 | 0 |
| 0 | 0 | 0 | 0 |
| 0 | 0 | 0 | 0 |
| 0 | 0 | 0 | 0 |
| 0 | 0 | 0 | 0 |
| 0 | 0 | 0 | 0 |
| 1 | 1 | 0 | 0 |
| 0 | 0 | 0 | 0 |
| 0 | 0 | 0 | 0 |
| 1 | 0 | 0 | 0 |
| 0 | 1 | 0 | 0 |
| 0 | 0 | 0 | 0 |
| 0 | 0 | 0 | 0 |
| 0 | 1 | 0 | 0 |
| 1 | 1 | 0 | 0 |
| 1 | 1 | 0 | 0 |
| 1 | 1 | 0 | 0 |
| 1 | 1 | 0 | 0 |
| 1 | 1 | 0 | 0 |

| | | | |
|---|---|---|---|
| 0 | 0 | 0 | 0 |
| 0 | 0 | 0 | 0 |
| 0 | 0 | 0 | 0 |
| 0 | 0 | 0 | 0 |
| 0 | 0 | 0 | 0 |
| 1 | 0 | 0 | 0 |
| 1 | 0 | 0 | 0 |
| 0 | 0 | 0 | 0 |
| 0 | 0 | 0 | 0 |
| 1 | 0 | 0 | 0 |
| 0 | 0 | 0 | 0 |
| 0 | 0 | 1 | 0 |
| 0 | 0 | 0 | 0 |
| 0 | 0 | 0 | 0 |
| 0 | 0 | 0 | 0 |
| 0 | 0 | 0 | 0 |
| 0 | 0 | 0 | 0 |
| 0 | 0 | 0 | 0 |
| 0 | 0 | 0 | 0 |
| 0 | 0 | 0 | 0 |
| 1 | 1 | 0 | 0 |
| 1 | 0 | 0 | 0 |
| 0 | 0 | 0 | 0 |
| 0 | 0 | 1 | 0 |
| 0 | 0 | 0 | 0 |
| 0 | 0 | 1 | 0 |
| 0 | 0 | 0 | 0 |
| 0 | 0 | 0 | 0 |
| 0 | 0 | 0 | 0 |
| 0 | 0 | 0 | 0 |
| 0 | 0 | 0 | 0 |
| 0 | 0 | 0 | 0 |
| 0 | 0 | 0 | 0 |
| 0 | 0 | 0 | 0 |
| 0 | 0 | 0 | 0 |
| 0 | 0 | 0 | 0 |
| 1 | 0 | 0 | 0 |
| 0 | 0 | 0 | 0 |
| 1 | 0 | 0 | 0 |
| 1 | 0 | 0 | 0 |
| 1 | 0 | 0 | 0 |
| 0 | 0 | 0 | 0 |
| 0 | 0 | 0 | 0 |
| 0 | 0 | 0 | 0 |
| 0 | 0 | 0 | 0 |
| 0 | 0 | 0 | 0 |
| 0 | 0 | 0 | 0 |
| 1 | 0 | 1 | 0 |
| 0 | 0 | 0 | 0 |
| 0 | 0 | 0 | 0 |
| 0 | 0 | 0 | 0 |
| 0 | 0 | 0 | 0 |
| 1 | 1 | 0 | 0 |
| 0 | 0 | 0 | 0 |

| | | | |
|---|---|---|---|
| 0 | 0 | 0 | 0 |
| 0 | 1 | 0 | 0 |
| 0 | 0 | 0 | 0 |
| 0 | 0 | 0 | 0 |
| 0 | 0 | 0 | 0 |
| 0 | 0 | 0 | 0 |
| 0 | 0 | 0 | 0 |
| 0 | 0 | 0 | 0 |
| 0 | 0 | 0 | 0 |
| 0 | 0 | 0 | 0 |
| 0 | 0 | 0 | 0 |
| 0 | 0 | 0 | 0 |
| 0 | 0 | 0 | 0 |
| 0 | 0 | 0 | 0 |
| 0 | 0 | 0 | 0 |
| 0 | 0 | 0 | 0 |
| 0 | 0 | 0 | 0 |
| 0 | 0 | 0 | 0 |
| 0 | 0 | 0 | 0 |
| 0 | 0 | 0 | 0 |
| 1 | 0 | 0 | 0 |
| 0 | 0 | 0 | 0 |
| 0 | 0 | 0 | 0 |
| 0 | 0 | 0 | 0 |
| 1 | 0 | 0 | 0 |
| 0 | 0 | 0 | 0 |
| 0 | 0 | 0 | 0 |
| 0 | 0 | 0 | 1 |
| 0 | 0 | 0 | 0 |
| 0 | 0 | 0 | 0 |
| 0 | 0 | 0 | 0 |
| 0 | 0 | 0 | 0 |
| 0 | 0 | 0 | 0 |
| 0 | 0 | 0 | 0 |
| 0 | 0 | 0 | 0 |
| 0 | 0 | 0 | 0 |
| 0 | 0 | 0 | 0 |
| 0 | 0 | 0 | 0 |
| 0 | 0 | 0 | 0 |
| 0 | 0 | 0 | 0 |
| 0 | 0 | 0 | 0 |
| 0 | 0 | 0 | 0 |
| 0 | 0 | 0 | 0 |
| 0 | 0 | 0 | 0 |
| 0 | 0 | 0 | 0 |
| 0 | 0 | 0 | 0 |
| 0 | 0 | 0 | 0 |
| 0 | 0 | 0 | 0 |
| 0 | 0 | 0 | 0 |
| 0 | 0 | 0 | 0 |
| 0 | 0 | 0 | 0 |
| 0 | 0 | 0 | 0 |
| 0 | 0 | 0 | 0 |
| 0 | 0 | 0 | 0 |

| | | | |
|---|---|---|---|
| 0 | 0 | 0 | 0 |
| 0 | 0 | 0 | 0 |
| 0 | 0 | 0 | 0 |
| 0 | 0 | 0 | 0 |
| 0 | 0 | 0 | 0 |
| 0 | 0 | 0 | 0 |
| 0 | 0 | 0 | 0 |
| 0 | 0 | 0 | 0 |
| 0 | 0 | 0 | 0 |
| 0 | 0 | 0 | 0 |
| 0 | 0 | 0 | 0 |
| 0 | 0 | 0 | 0 |
| 0 | 0 | 0 | 0 |
| 0 | 0 | 0 | 0 |
| 0 | 0 | 0 | 0 |
| 1 | 0 | 0 | 0 |
| 0 | 0 | 0 | 0 |
| 0 | 0 | 0 | 0 |
| 0 | 1 | 0 | 0 |
| 0 | 0 | 0 | 0 |
| 0 | 0 | 0 | 0 |
| 0 | 0 | 0 | 0 |
| 0 | 0 | 0 | 0 |
| 0 | 0 | 0 | 0 |
| 0 | 0 | 0 | 0 |
| 1 | 0 | 0 | 0 |
| 0 | 0 | 0 | 0 |
| 0 | 0 | 0 | 0 |
| 0 | 0 | 0 | 0 |
| 0 | 0 | 0 | 0 |
| 0 | 0 | 0 | 0 |
| 0 | 0 | 0 | 0 |
| 0 | 0 | 0 | 0 |
| 1 | 1 | 0 | 0 |
| 0 | 0 | 0 | 0 |
| 0 | 0 | 0 | 0 |
| 0 | 0 | 0 | 0 |
| 0 | 0 | 0 | 0 |
| 0 | 0 | 0 | 0 |
| 0 | 0 | 0 | 0 |
| 0 | 0 | 0 | 0 |
| 0 | 0 | 0 | 0 |
| 0 | 0 | 0 | 0 |
| 0 | 0 | 0 | 0 |
| 0 | 0 | 0 | 0 |
| 0 | 0 | 0 | 0 |
| 0 | 0 | 0 | 0 |
| 1 | 0 | 0 | 0 |
| 0 | 0 | 0 | 0 |
| 0 | 0 | 0 | 0 |
| 0 | 0 | 0 | 0 |
| 0 | 0 | 0 | 0 |
| 0 | 0 | 0 | 0 |
| 1 | 0 | 0 | 1 |

| | | | |
|---|---|---|---|
| 0 | 0 | 0 | 0 |
| 0 | 0 | 0 | 0 |
| 0 | 0 | 1 | 0 |
| 0 | 0 | 0 | 0 |
| 0 | 0 | 0 | 0 |
| 0 | 0 | 0 | 0 |
| 0 | 0 | 0 | 0 |
| 0 | 0 | 0 | 0 |
| 0 | 0 | 0 | 0 |
| 0 | 0 | 0 | 0 |
| 0 | 0 | 0 | 0 |
| 0 | 0 | 0 | 0 |
| 0 | 0 | 0 | 0 |
| 0 | 0 | 0 | 0 |
| 0 | 0 | 0 | 0 |
| 0 | 0 | 0 | 0 |
| 0 | 0 | 0 | 0 |
| 0 | 0 | 0 | 0 |
| 0 | 0 | 0 | 0 |
| 0 | 0 | 0 | 0 |
| 0 | 0 | 0 | 0 |
| 0 | 0 | 0 | 0 |
| 0 | 0 | 0 | 0 |
| 0 | 0 | 0 | 0 |
| 0 | 0 | 0 | 0 |
| 0 | 0 | 0 | 0 |
| 0 | 0 | 0 | 0 |
| 0 | 0 | 0 | 0 |
| 0 | 0 | 0 | 0 |
| 0 | 0 | 0 | 0 |
| 0 | 0 | 0 | 0 |
| 1 | 0 | 0 | 0 |
| 0 | 0 | 0 | 0 |
| 0 | 0 | 0 | 0 |
| 0 | 0 | 0 | 0 |
| 0 | 0 | 0 | 0 |
| 0 | 0 | 0 | 0 |
| 0 | 0 | 0 | 0 |
| 0 | 0 | 0 | 0 |
| 0 | 0 | 0 | 0 |
| 1 | 1 | 0 | 0 |
| 0 | 0 | 0 | 0 |
| 0 | 0 | 0 | 0 |
| 0 | 0 | 0 | 0 |
| 0 | 0 | 0 | 0 |
| 1 | 1 | 0 | 0 |
| 0 | 0 | 0 | 0 |
| 1 | 1 | 0 | 0 |
| 0 | 0 | 0 | 0 |
| 1 | 0 | 0 | 0 |
| 1 | 0 | 0 | 0 |
| 1 | 0 | 0 | 0 |
| 0 | 0 | 0 | 0 |
| 0 | 0 | 0 | 0 |

| | | | |
|---|---|---|---|
| 1 | 0 | 0 | 0 |
| 0 | 0 | 0 | 0 |
| 0 | 0 | 0 | 0 |
| 0 | 0 | 0 | 0 |
| 0 | 0 | 0 | 0 |
| 0 | 0 | 0 | 0 |
| 0 | 0 | 0 | 0 |
| 0 | 0 | 0 | 0 |
| 0 | 0 | 0 | 0 |
| 0 | 0 | 0 | 0 |
| 0 | 0 | 0 | 0 |
| 0 | 0 | 0 | 0 |
| 0 | 0 | 0 | 0 |
| 0 | 0 | 0 | 0 |
| 0 | 0 | 0 | 0 |
| 1 | 1 | 1 | 0 |
| 0 | 0 | 0 | 0 |
| 1 | 0 | 0 | 0 |
| 0 | 0 | 0 | 0 |
| 0 | 0 | 0 | 0 |
| 0 | 0 | 0 | 0 |
| 0 | 1 | 0 | 0 |
| 0 | 0 | 0 | 0 |
| 0 | 1 | 0 | 0 |
| 0 | 1 | 0 | 0 |
| 0 | 0 | 0 | 0 |
| 0 | 0 | 0 | 0 |
| 0 | 0 | 0 | 0 |
| 0 | 0 | 0 | 0 |
| 0 | 0 | 0 | 0 |
| 0 | 0 | 0 | 0 |
| 0 | 0 | 0 | 0 |
| 1 | 1 | 0 | 0 |
| 0 | 0 | 0 | 0 |
| 0 | 0 | 0 | 0 |
| 0 | 0 | 0 | 0 |
| 0 | 1 | 0 | 0 |
| 1 | 1 | 0 | 0 |
| 0 | 0 | 0 | 0 |
| 0 | 0 | 0 | 0 |
| 0 | 0 | 0 | 0 |
| 0 | 0 | 0 | 0 |
| 0 | 0 | 0 | 0 |
| 0 | 0 | 0 | 0 |
| 1 | 1 | 0 | 0 |
| 0 | 0 | 0 | 0 |
| 1 | 0 | 0 | 0 |
| 1 | 1 | 0 | 0 |
| 0 | 0 | 0 | 0 |
| 0 | 0 | 0 | 0 |
| 1 | 1 | 0 | 0 |
| 1 | 1 | 0 | 0 |
| 0 | 0 | 0 | 0 |
| 1 | 1 | 0 | 0 |

| | | | |
|---|---|---|---|
| 0 | 0 | 0 | 0 |
| 0 | 0 | 0 | 0 |
| 0 | 0 | 0 | 0 |
| 1 | 1 | 0 | 0 |
| 0 | 0 | 0 | 0 |
| 0 | 0 | 0 | 0 |
| 0 | 0 | 0 | 0 |
| 0 | 0 | 0 | 0 |
| 1 | 0 | 0 | 0 |
| 0 | 0 | 0 | 0 |
| 0 | 0 | 0 | 0 |
| 0 | 0 | 0 | 0 |
| 1 | 0 | 1 | 0 |
| 1 | 0 | 0 | 0 |
| 0 | 0 | 0 | 0 |
| 0 | 0 | 0 | 0 |
| 0 | 0 | 0 | 0 |
| 0 | 0 | 0 | 0 |
| 0 | 0 | 0 | 0 |
| 0 | 0 | 0 | 0 |
| 0 | 0 | 0 | 0 |
| 0 | 0 | 0 | 0 |
| 0 | 0 | 0 | 0 |
| 1 | 0 | 0 | 0 |
| 0 | 0 | 0 | 0 |
| 0 | 0 | 0 | 0 |
| 0 | 0 | 0 | 0 |
| 0 | 0 | 0 | 0 |
| 0 | 0 | 0 | 0 |
| 0 | 0 | 0 | 0 |
| 0 | 0 | 0 | 0 |
| 0 | 0 | 0 | 0 |
| 0 | 0 | 0 | 0 |
| 0 | 0 | 0 | 0 |
| 0 | 0 | 0 | 0 |
| 0 | 0 | 0 | 0 |
| 0 | 0 | 0 | 0 |
| 0 | 0 | 0 | 0 |
| 0 | 0 | 0 | 0 |
| 0 | 0 | 0 | 0 |
| 0 | 0 | 0 | 0 |
| 0 | 0 | 0 | 0 |
| 0 | 0 | 0 | 0 |
| 1 | 1 | 0 | 0 |
| 1 | 1 | 0 | 0 |
| 0 | 0 | 0 | 0 |
| 0 | 0 | 0 | 0 |
| 0 | 1 | 0 | 0 |
| 0 | 1 | 0 | 0 |
| 0 | 1 | 0 | 0 |
| 0 | 0 | 0 | 0 |
| 0 | 0 | 0 | 0 |
| 0 | 0 | 0 | 0 |
| 0 | 0 | 0 | 0 |

| | | | |
|---|---|---|---|
| 0 | 0 | 0 | 0 |
| 0 | 0 | 0 | 0 |
| 0 | 0 | 0 | 0 |
| 0 | 0 | 0 | 0 |
| 1 | 0 | 0 | 0 |
| 0 | 0 | 0 | 0 |
| 0 | 0 | 0 | 0 |
| 1 | 0 | 0 | 0 |
| 1 | 0 | 0 | 0 |
| 1 | 0 | 0 | 0 |
| 1 | 0 | 0 | 0 |
| 1 | 0 | 0 | 0 |
| 0 | 0 | 0 | 0 |
| 1 | 0 | 0 | 0 |
| 1 | 0 | 0 | 0 |
| 0 | 0 | 0 | 0 |
| 0 | 0 | 0 | 0 |
| 0 | 0 | 0 | 0 |
| 1 | 0 | 0 | 0 |
| 1 | 0 | 0 | 0 |
| 1 | 0 | 0 | 0 |
| 1 | 0 | 0 | 0 |
| 0 | 0 | 0 | 0 |
| 0 | 0 | 0 | 0 |
| 0 | 0 | 0 | 0 |
| 0 | 0 | 0 | 0 |
| 0 | 0 | 0 | 0 |
| 0 | 0 | 0 | 0 |
| 0 | 0 | 0 | 0 |
| 0 | 0 | 0 | 0 |
| 0 | 0 | 0 | 0 |
| 0 | 0 | 0 | 0 |
| 1 | 0 | 0 | 0 |
| 0 | 0 | 0 | 0 |
| 0 | 0 | 0 | 0 |
| 1 | 0 | 0 | 0 |
| 0 | 0 | 0 | 0 |
| 0 | 0 | 0 | 0 |
| 0 | 0 | 0 | 0 |
| 0 | 0 | 0 | 0 |
| 0 | 0 | 0 | 0 |
| 0 | 0 | 0 | 0 |
| 0 | 0 | 0 | 0 |
| 0 | 0 | 0 | 0 |
| 0 | 0 | 0 | 0 |
| 0 | 0 | 0 | 0 |
| 0 | 0 | 0 | 0 |
| 0 | 0 | 0 | 0 |
| 0 | 0 | 0 | 0 |
| 0 | 0 | 0 | 0 |
| 0 | 0 | 0 | 0 |
| 0 | 0 | 0 | 0 |
| 0 | 0 | 0 | 0 |
| 0 | 0 | 0 | 0 |

| | | | |
|---|---|---|---|
| 0 | 0 | 0 | 0 |
| 0 | 0 | 0 | 0 |
| 0 | 0 | 0 | 0 |
| 0 | 0 | 0 | 0 |
| 0 | 0 | 0 | 0 |
| 0 | 0 | 0 | 0 |
| 0 | 0 | 0 | 0 |
| 0 | 0 | 0 | 0 |
| 0 | 0 | 0 | 0 |
| 0 | 0 | 0 | 0 |
| 0 | 0 | 0 | 0 |
| 0 | 0 | 0 | 1 |
| 0 | 0 | 0 | 0 |
| 0 | 0 | 0 | 0 |
| 0 | 0 | 0 | 0 |
| 0 | 0 | 0 | 0 |
| 0 | 0 | 0 | 0 |
| 0 | 0 | 0 | 1 |
| 0 | 0 | 0 | 0 |
| 0 | 0 | 0 | 0 |
| 0 | 0 | 0 | 1 |
| 0 | 0 | 0 | 0 |
| 0 | 0 | 0 | 0 |
| 0 | 0 | 0 | 0 |
| 0 | 0 | 0 | 0 |
| 0 | 0 | 0 | 0 |
| 0 | 0 | 0 | 0 |
| 0 | 0 | 0 | 0 |
| 0 | 0 | 0 | 1 |
| 0 | 0 | 0 | 0 |
| 0 | 0 | 0 | 0 |
| 0 | 0 | 0 | 0 |
| 0 | 0 | 0 | 0 |
| 0 | 0 | 0 | 0 |
| 0 | 0 | 0 | 0 |
| 0 | 0 | 0 | 0 |
| 0 | 0 | 0 | 0 |
| 0 | 0 | 0 | 0 |
| 0 | 0 | 0 | 0 |
| 0 | 0 | 0 | 0 |
| 0 | 0 | 0 | 0 |
| 0 | 0 | 0 | 0 |
| 0 | 0 | 0 | 0 |
| 0 | 0 | 0 | 0 |
| 0 | 0 | 0 | 0 |
| 0 | 0 | 0 | 0 |
| 0 | 0 | 0 | 0 |
| 0 | 0 | 0 | 0 |
| 0 | 0 | 0 | 0 |
| 0 | 0 | 0 | 0 |
| 0 | 0 | 0 | 0 |
| 0 | 0 | 0 | 0 |
| 0 | 0 | 0 | 0 |
| 0 | 0 | 0 | 0 |

| | | | |
|---|---|---|---|
| 0 | 0 | 0 | 0 |
| 0 | 0 | 0 | 0 |
| 1 | 0 | 0 | 0 |
| 0 | 0 | 0 | 0 |
| 1 | 0 | 0 | 0 |
| 0 | 0 | 0 | 0 |
| 0 | 0 | 0 | 0 |
| 0 | 0 | 0 | 0 |
| 0 | 0 | 0 | 0 |
| 1 | 1 | 0 | 0 |
| 0 | 0 | 1 | 0 |
| 0 | 0 | 0 | 0 |
| 0 | 0 | 0 | 0 |
| 0 | 0 | 0 | 0 |
| 0 | 0 | 0 | 0 |
| 0 | 0 | 0 | 0 |
| 0 | 0 | 0 | 0 |
| 0 | 0 | 0 | 0 |
| 0 | 0 | 0 | 0 |
| 0 | 0 | 0 | 0 |
| 0 | 0 | 0 | 0 |
| 0 | 0 | 0 | 0 |
| 0 | 0 | 0 | 0 |
| 0 | 0 | 0 | 0 |
| 0 | 0 | 0 | 0 |
| 0 | 0 | 0 | 0 |
| 0 | 0 | 0 | 0 |
| 0 | 0 | 0 | 0 |
| 0 | 0 | 0 | 0 |
| 0 | 0 | 0 | 0 |
| 0 | 0 | 0 | 0 |
| 0 | 0 | 0 | 0 |
| 0 | 0 | 0 | 0 |
| 0 | 0 | 0 | 0 |
| 0 | 0 | 0 | 0 |
| 0 | 0 | 0 | 0 |
| 0 | 0 | 0 | 0 |
| 0 | 0 | 0 | 0 |
| 0 | 0 | 0 | 0 |
| 0 | 0 | 0 | 0 |
| 0 | 0 | 0 | 0 |
| 0 | 0 | 0 | 0 |
| 0 | 0 | 0 | 0 |
| 0 | 0 | 0 | 0 |
| 0 | 0 | 0 | 0 |
| 1 | 0 | 0 | 0 |
| 0 | 0 | 0 | 0 |
| 0 | 0 | 0 | 0 |
| 0 | 0 | 0 | 0 |
| 0 | 0 | 0 | 0 |
| 1 | 0 | 0 | 0 |
| 0 | 0 | 0 | 0 |
| 0 | 0 | 0 | 0 |
| 0 | 0 | 0 | 0 |

| | | | |
|---|---|---|---|
| 0 | 0 | 0 | 0 |
| 0 | 0 | 0 | 0 |
| 0 | 0 | 0 | 0 |
| 0 | 0 | 0 | 0 |
| 0 | 0 | 0 | 0 |
| 0 | 0 | 0 | 0 |
| 0 | 0 | 0 | 0 |
| 0 | 0 | 0 | 0 |
| 1 | 1 | 0 | 0 |
| 0 | 0 | 0 | 0 |
| 0 | 0 | 0 | 0 |
| 0 | 0 | 0 | 0 |
| 0 | 0 | 0 | 0 |
| 0 | 0 | 0 | 0 |
| 0 | 1 | 0 | 0 |
| 0 | 1 | 0 | 0 |
| 0 | 1 | 0 | 0 |
| 0 | 0 | 0 | 0 |
| 0 | 0 | 0 | 0 |
| 0 | 0 | 0 | 0 |
| 0 | 1 | 0 | 0 |
| 0 | 0 | 0 | 0 |
| 0 | 0 | 0 | 0 |
| 1 | 1 | 0 | 0 |
| 0 | 1 | 0 | 0 |
| 0 | 1 | 0 | 0 |
| 0 | 0 | 0 | 0 |
| 0 | 0 | 0 | 0 |
| 0 | 0 | 0 | 0 |
| 0 | 0 | 0 | 0 |
| 0 | 0 | 0 | 0 |
| 0 | 0 | 0 | 0 |
| 0 | 0 | 0 | 0 |
| 0 | 0 | 0 | 0 |
| 0 | 0 | 0 | 0 |
| 0 | 0 | 0 | 0 |
| 0 | 0 | 0 | 0 |
| 1 | 0 | 0 | 0 |
| 0 | 0 | 0 | 0 |
| 0 | 0 | 0 | 0 |
| 0 | 0 | 0 | 0 |
| 0 | 0 | 0 | 0 |
| 0 | 0 | 0 | 0 |
| 0 | 0 | 0 | 0 |
| 0 | 0 | 0 | 0 |
| 0 | 0 | 0 | 0 |
| 0 | 0 | 0 | 0 |
| 0 | 0 | 0 | 0 |
| 0 | 0 | 0 | 0 |
| 0 | 0 | 0 | 0 |
| 0 | 0 | 0 | 0 |
| 0 | 0 | 0 | 0 |
| 0 | 0 | 0 | 0 |
| 0 | 0 | 0 | 0 |

| | | | |
|---|---|---|---|
| 0 | 0 | 0 | 0 |
| 0 | 0 | 0 | 0 |
| 0 | 0 | 0 | 0 |
| 0 | 0 | 1 | 0 |
| 0 | 0 | 0 | 0 |
| 0 | 0 | 0 | 0 |
| 0 | 0 | 0 | 0 |
| 0 | 0 | 0 | 0 |
| 0 | 0 | 0 | 0 |
| 0 | 0 | 0 | 0 |
| 0 | 0 | 0 | 0 |
| 0 | 0 | 0 | 0 |
| 1 | 0 | 0 | 0 |
| 0 | 0 | 0 | 0 |
| 0 | 0 | 0 | 0 |
| 0 | 0 | 0 | 0 |
| 0 | 1 | 0 | 0 |
| 0 | 0 | 0 | 0 |
| 0 | 0 | 0 | 0 |
| 0 | 0 | 0 | 0 |
| 0 | 0 | 0 | 0 |
| 0 | 0 | 0 | 0 |
| 0 | 0 | 0 | 0 |
| 0 | 0 | 0 | 0 |
| 0 | 0 | 0 | 0 |
| 0 | 0 | 0 | 0 |
| 0 | 0 | 0 | 0 |
| 0 | 0 | 0 | 0 |
| 0 | 0 | 0 | 0 |
| 0 | 0 | 0 | 0 |
| 1 | 0 | 0 | 0 |
| 1 | 0 | 0 | 0 |
| 1 | 0 | 0 | 0 |
| 0 | 0 | 0 | 0 |
| 1 | 0 | 0 | 0 |
| 1 | 0 | 0 | 0 |
| 1 | 0 | 0 | 0 |
| 0 | 0 | 0 | 0 |
| 0 | 0 | 0 | 0 |
| 0 | 0 | 0 | 0 |
| 0 | 0 | 0 | 0 |
| 0 | 0 | 0 | 0 |
| 0 | 1 | 0 | 0 |
| 0 | 1 | 0 | 0 |
| 0 | 0 | 0 | 0 |
| 0 | 0 | 0 | 0 |
| 1 | 0 | 0 | 0 |
| 0 | 0 | 0 | 0 |
| 0 | 0 | 0 | 0 |
| 0 | 0 | 0 | 0 |
| 0 | 0 | 0 | 0 |
| 0 | 0 | 0 | 0 |
| 1 | 0 | 0 | 0 |
| 0 | 0 | 0 | 0 |

| | | | |
|---|---|---|---|
| 0 | 0 | 0 | 0 |
| 1 | 0 | 0 | 0 |
| 1 | 0 | 0 | 0 |
| 0 | 0 | 0 | 0 |
| 0 | 0 | 0 | 0 |
| 0 | 0 | 0 | 0 |
| 0 | 0 | 0 | 0 |
| 0 | 1 | 0 | 0 |
| 0 | 0 | 0 | 0 |
| 0 | 0 | 0 | 0 |
| 0 | 0 | 0 | 0 |
| 0 | 0 | 0 | 0 |
| 0 | 0 | 0 | 0 |
| 0 | 0 | 0 | 0 |
| 1 | 0 | 0 | 0 |
| 0 | 0 | 0 | 0 |
| 0 | 0 | 0 | 0 |
| 0 | 0 | 0 | 0 |
| 0 | 0 | 0 | 0 |
| 1 | 0 | 0 | 0 |
| 1 | 0 | 0 | 0 |
| 0 | 0 | 0 | 0 |
| 1 | 0 | 0 | 0 |
| 1 | 0 | 0 | 0 |
| 0 | 0 | 0 | 0 |
| 0 | 0 | 0 | 0 |
| 0 | 0 | 0 | 0 |
| 0 | 0 | 0 | 0 |
| 0 | 0 | 0 | 0 |
| 0 | 0 | 0 | 0 |
| 0 | 0 | 0 | 0 |
| 0 | 0 | 0 | 0 |
| 0 | 0 | 0 | 0 |
| 0 | 0 | 0 | 0 |
| 0 | 0 | 0 | 0 |
| 0 | 0 | 0 | 0 |
| 0 | 0 | 0 | 0 |
| 0 | 0 | 0 | 0 |
| 0 | 0 | 0 | 0 |
| 0 | 0 | 0 | 0 |
| 0 | 0 | 0 | 0 |
| 0 | 0 | 0 | 0 |
| 0 | 0 | 0 | 0 |
| 0 | 0 | 0 | 0 |
| 0 | 0 | 0 | 0 |
| 0 | 0 | 0 | 0 |
| 0 | 0 | 0 | 0 |
| 0 | 0 | 0 | 0 |
| 0 | 0 | 0 | 0 |
| 0 | 0 | 0 | 0 |
| 0 | 0 | 0 | 0 |
| 0 | 0 | 0 | 0 |
| 0 | 0 | 0 | 0 |
| 1 | 1 | 0 | 1 |

| | | | |
|---|---|---|---|
| 0 | 0 | 0 | 0 |
| 0 | 0 | 0 | 0 |
| 0 | 0 | 0 | 0 |
| 0 | 0 | 0 | 0 |
| 0 | 1 | 0 | 0 |
| 0 | 0 | 0 | 0 |
| 0 | 0 | 0 | 0 |
| 0 | 1 | 0 | 0 |
| 0 | 0 | 0 | 0 |
| 0 | 0 | 0 | 0 |
| 0 | 0 | 0 | 0 |
| 0 | 0 | 0 | 0 |
| 0 | 0 | 0 | 0 |
| 0 | 0 | 0 | 0 |
| 0 | 1 | 0 | 1 |
| 0 | 1 | 0 | 0 |
| 0 | 0 | 0 | 0 |
| 0 | 0 | 0 | 0 |
| 0 | 1 | 0 | 0 |
| 0 | 1 | 0 | 1 |
| 0 | 0 | 0 | 0 |
| 0 | 0 | 0 | 0 |
| 0 | 0 | 0 | 0 |
| 0 | 0 | 0 | 0 |
| 0 | 0 | 0 | 0 |
| 1 | 0 | 0 | 0 |
| 0 | 0 | 0 | 0 |
| 0 | 0 | 1 | 0 |
| 0 | 0 | 1 | 0 |
| 0 | 0 | 1 | 0 |
| 0 | 0 | 0 | 0 |
| 0 | 0 | 0 | 0 |
| 0 | 0 | 0 | 0 |
| 0 | 0 | 0 | 0 |
| 0 | 0 | 0 | 0 |
| 0 | 0 | 1 | 0 |
| 0 | 0 | 0 | 0 |
| 0 | 0 | 0 | 0 |
| 1 | 0 | 1 | 0 |
| 0 | 0 | 1 | 0 |
| 1 | 1 | 0 | 0 |
| 0 | 1 | 0 | 0 |
| 0 | 0 | 0 | 0 |
| 0 | 0 | 0 | 0 |
| 1 | 0 | 0 | 0 |
| 0 | 0 | 0 | 0 |
| 0 | 0 | 1 | 0 |
| 0 | 0 | 0 | 0 |
| 0 | 0 | 0 | 0 |
| 0 | 0 | 0 | 0 |
| 0 | 0 | 0 | 0 |
| 0 | 0 | 0 | 0 |
| 0 | 0 | 0 | 0 |
| 0 | 0 | 0 | 0 |

| | | | |
|---|---|---|---|
| 0 | 0 | 0 | 0 |
| 0 | 0 | 0 | 0 |
| 0 | 0 | 0 | 0 |
| 0 | 0 | 0 | 0 |
| 0 | 0 | 0 | 0 |
| 0 | 0 | 0 | 0 |
| 0 | 0 | 0 | 0 |
| 1 | 1 | 0 | 0 |
| 0 | 0 | 0 | 0 |
| 0 | 0 | 0 | 0 |
| 0 | 0 | 0 | 0 |
| 0 | 0 | 0 | 1 |
| 0 | 0 | 0 | 0 |
| 0 | 0 | 0 | 0 |
| 0 | 0 | 0 | 0 |
| 0 | 0 | 0 | 0 |
| 0 | 0 | 0 | 0 |
| 0 | 0 | 0 | 0 |
| 0 | 0 | 0 | 0 |
| 0 | 0 | 0 | 0 |
| 0 | 0 | 0 | 0 |
| 0 | 0 | 0 | 0 |
| 0 | 0 | 0 | 0 |
| 0 | 0 | 0 | 0 |
| 0 | 0 | 0 | 0 |
| 0 | 0 | 0 | 0 |
| 0 | 0 | 0 | 0 |
| 0 | 0 | 0 | 0 |
| 0 | 0 | 0 | 0 |
| 0 | 1 | 0 | 0 |
| 0 | 1 | 0 | 0 |
| 0 | 0 | 0 | 0 |
| 0 | 0 | 0 | 0 |
| 0 | 0 | 0 | 0 |
| 0 | 0 | 0 | 0 |
| 0 | 1 | 0 | 0 |
| 0 | 0 | 0 | 0 |
| 0 | 0 | 0 | 0 |
| 0 | 0 | 0 | 0 |
| 0 | 0 | 0 | 0 |
| 0 | 0 | 0 | 0 |
| 1 | 1 | 0 | 0 |
| 1 | 0 | 0 | 0 |
| 0 | 0 | 0 | 0 |
| 0 | 0 | 0 | 0 |
| 0 | 0 | 0 | 0 |
| 0 | 0 | 0 | 0 |
| 0 | 0 | 0 | 0 |
| 0 | 0 | 0 | 0 |
| 0 | 0 | 0 | 0 |
| 0 | 0 | 0 | 0 |
| 0 | 0 | 0 | 0 |
| 0 | 0 | 0 | 0 |
| 0 | 0 | 0 | 0 |

| | | | |
|---|---|---|---|
| 0 | 0 | 0 | 0 |
| 0 | 0 | 0 | 0 |
| 0 | 0 | 0 | 0 |
| 0 | 0 | 0 | 0 |
| 0 | 0 | 0 | 0 |
| 0 | 0 | 0 | 0 |
| 0 | 0 | 0 | 0 |
| 0 | 0 | 0 | 0 |
| 1 | 0 | 0 | 0 |
| 1 | 0 | 0 | 0 |
| 0 | 0 | 0 | 0 |
| 0 | 0 | 0 | 0 |
| 0 | 0 | 0 | 0 |
| 1 | 0 | 0 | 0 |
| 0 | 0 | 0 | 0 |
| 0 | 0 | 0 | 0 |
| 0 | 0 | 0 | 0 |
| 0 | 0 | 0 | 0 |
| 0 | 0 | 0 | 0 |
| 1 | 1 | 0 | 0 |
| 0 | 0 | 1 | 0 |
| 0 | 0 | 0 | 0 |
| 0 | 0 | 0 | 0 |
| 0 | 0 | 0 | 0 |
| 0 | 0 | 0 | 0 |
| 0 | 0 | 0 | 0 |
| 0 | 0 | 0 | 0 |
| 0 | 0 | 0 | 0 |
| 0 | 0 | 0 | 0 |
| 0 | 0 | 0 | 0 |
| 0 | 0 | 0 | 0 |
| 0 | 0 | 0 | 0 |
| 0 | 0 | 0 | 0 |
| 0 | 0 | 0 | 0 |
| 0 | 0 | 0 | 0 |
| 0 | 0 | 0 | 0 |
| 0 | 0 | 0 | 0 |
| 0 | 0 | 0 | 0 |
| 0 | 0 | 0 | 0 |
| 0 | 0 | 0 | 0 |
| 0 | 0 | 0 | 0 |
| 0 | 0 | 0 | 0 |
| 0 | 0 | 0 | 0 |
| 0 | 0 | 0 | 0 |
| 0 | 0 | 0 | 0 |
| 0 | 0 | 0 | 0 |
| 1 | 0 | 1 | 0 |
| 1 | 0 | 0 | 0 |
| 0 | 0 | 0 | 0 |
| 0 | 0 | 0 | 0 |
| 0 | 0 | 1 | 0 |
| 0 | 0 | 0 | 0 |
| 0 | 0 | 0 | 0 |
| 0 | 0 | 0 | 0 |

| | | | |
|---|---|---|---|
| 0 | 0 | 0 | 0 |
| 0 | 0 | 0 | 0 |
| 0 | 0 | 0 | 0 |
| 0 | 0 | 0 | 0 |
| 0 | 0 | 0 | 0 |
| 0 | 0 | 0 | 0 |
| 0 | 0 | 0 | 0 |
| 0 | 0 | 0 | 0 |
| 0 | 0 | 0 | 0 |
| 0 | 0 | 0 | 0 |
| 0 | 0 | 0 | 0 |
| 0 | 0 | 0 | 0 |
| 0 | 0 | 0 | 0 |
| 0 | 0 | 0 | 0 |
| 0 | 0 | 0 | 0 |
| 0 | 0 | 0 | 0 |
| 0 | 0 | 0 | 0 |
| 0 | 0 | 0 | 0 |
| 0 | 0 | 0 | 0 |
| 1 | 0 | 0 | 0 |
| 0 | 0 | 0 | 0 |
| 0 | 0 | 0 | 0 |
| 0 | 0 | 0 | 0 |
| 0 | 0 | 0 | 0 |
| 0 | 0 | 0 | 0 |
| 0 | 0 | 0 | 0 |
| 0 | 0 | 0 | 0 |
| 0 | 0 | 0 | 0 |
| 0 | 0 | 0 | 0 |
| 0 | 0 | 0 | 0 |
| 0 | 0 | 0 | 0 |
| 0 | 0 | 0 | 0 |
| 0 | 0 | 0 | 0 |
| 0 | 0 | 0 | 0 |
| 0 | 0 | 0 | 0 |
| 0 | 0 | 0 | 0 |
| 0 | 0 | 0 | 0 |
| 0 | 0 | 0 | 0 |
| 1 | 0 | 0 | 1 |
| 0 | 0 | 0 | 0 |
| 0 | 0 | 0 | 0 |
| 0 | 0 | 0 | 0 |
| 0 | 0 | 0 | 1 |
| 0 | 0 | 0 | 1 |
| 0 | 0 | 0 | 1 |
| 0 | 0 | 0 | 0 |
| 0 | 0 | 0 | 0 |
| 0 | 0 | 0 | 0 |
| 0 | 0 | 0 | 0 |
| 0 | 0 | 0 | 0 |
| 0 | 0 | 0 | 0 |
| 0 | 0 | 0 | 0 |
| 0 | 0 | 0 | 0 |
| 0 | 0 | 0 | 0 |

| | | | |
|---|---|---|---|
| 0 | 0 | 0 | 0 |
| 0 | 0 | 0 | 0 |
| 0 | 0 | 0 | 0 |
| 0 | 0 | 0 | 0 |
| 0 | 0 | 0 | 0 |
| 0 | 0 | 0 | 0 |
| 0 | 0 | 0 | 0 |
| 0 | 0 | 0 | 0 |
| 0 | 0 | 0 | 0 |
| 0 | 0 | 0 | 0 |
| 0 | 0 | 0 | 0 |
| 0 | 0 | 0 | 0 |
| 0 | 0 | 0 | 0 |
| 0 | 0 | 0 | 0 |
| 0 | 0 | 0 | 0 |
| 0 | 0 | 0 | 0 |
| 0 | 0 | 0 | 0 |
| 0 | 0 | 0 | 0 |
| 0 | 0 | 0 | 0 |
| 0 | 0 | 0 | 0 |
| 0 | 0 | 0 | 0 |
| 0 | 0 | 0 | 0 |
| 0 | 0 | 0 | 0 |
| 0 | 0 | 0 | 0 |
| 0 | 0 | 0 | 0 |
| 0 | 0 | 0 | 0 |
| 0 | 0 | 0 | 0 |
| 0 | 0 | 0 | 0 |
| 0 | 0 | 0 | 0 |
| 0 | 0 | 0 | 0 |
| 0 | 0 | 0 | 0 |
| 0 | 0 | 0 | 0 |
| 1 | 1 | 0 | 1 |
| 1 | 1 | 0 | 1 |
| 1 | 0 | 0 | 0 |
| 0 | 0 | 0 | 0 |
| 0 | 0 | 0 | 0 |
| 0 | 0 | 0 | 0 |
| 0 | 0 | 0 | 0 |
| 0 | 0 | 0 | 0 |
| 0 | 0 | 0 | 0 |
| 0 | 0 | 0 | 0 |
| 1 | 1 | 0 | 0 |
| 0 | 0 | 0 | 0 |
| 1 | 0 | 0 | 0 |
| 0 | 0 | 0 | 0 |
| 0 | 0 | 0 | 0 |
| 0 | 0 | 0 | 0 |
| 0 | 0 | 0 | 0 |
| 0 | 0 | 0 | 0 |
| 1 | 0 | 0 | 0 |
| 0 | 0 | 0 | 0 |
| 0 | 0 | 0 | 0 |
| 0 | 0 | 0 | 0 |

| | | | |
|---|---|---|---|
| 0 | 0 | 0 | 0 |
| 0 | 0 | 0 | 0 |
| 0 | 0 | 0 | 0 |
| 0 | 0 | 0 | 0 |
| 0 | 0 | 0 | 0 |
| 0 | 0 | 0 | 0 |
| 0 | 0 | 0 | 0 |
| 0 | 0 | 0 | 0 |
| 0 | 0 | 0 | 0 |
| 0 | 0 | 0 | 0 |
| 0 | 0 | 0 | 0 |
| 0 | 0 | 0 | 0 |
| 0 | 0 | 0 | 0 |
| 0 | 0 | 0 | 0 |
| 0 | 0 | 0 | 0 |
| 0 | 0 | 0 | 0 |
| 0 | 0 | 0 | 0 |
| 0 | 0 | 0 | 0 |
| 0 | 0 | 0 | 0 |
| 0 | 0 | 0 | 1 |
| 0 | 0 | 0 | 0 |
| 0 | 0 | 0 | 1 |
| 0 | 0 | 0 | 1 |
| 0 | 0 | 0 | 1 |
| 0 | 0 | 0 | 0 |
| 0 | 0 | 0 | 0 |
| 0 | 0 | 0 | 0 |
| 0 | 0 | 0 | 0 |
| 0 | 0 | 0 | 0 |
| 0 | 0 | 0 | 0 |
| 0 | 0 | 0 | 0 |
| 0 | 0 | 0 | 0 |
| 0 | 0 | 0 | 0 |
| 0 | 0 | 0 | 0 |
| 0 | 0 | 0 | 0 |
| 0 | 1 | 0 | 0 |
| 0 | 0 | 0 | 0 |
| 0 | 0 | 0 | 0 |
| 0 | 1 | 0 | 0 |
| 0 | 0 | 0 | 0 |
| 0 | 0 | 0 | 0 |
| 0 | 0 | 0 | 0 |
| 0 | 0 | 0 | 0 |
| 0 | 0 | 0 | 0 |
| 0 | 0 | 0 | 0 |
| 0 | 0 | 0 | 0 |
| 0 | 0 | 0 | 0 |
| 0 | 0 | 0 | 0 |
| 0 | 0 | 0 | 0 |
| 0 | 0 | 0 | 0 |
| 0 | 0 | 0 | 0 |
| 0 | 0 | 0 | 0 |
| 0 | 0 | 0 | 0 |
| 0 | 0 | 0 | 0 |

| | | | |
|---|---|---|---|
| 0 | 0 | 0 | 0 |
| 0 | 0 | 0 | 0 |
| 0 | 0 | 0 | 0 |
| 0 | 0 | 0 | 0 |
| 0 | 0 | 0 | 0 |
| 0 | 0 | 0 | 0 |
| 0 | 0 | 0 | 0 |
| 0 | 0 | 0 | 0 |
| 0 | 0 | 1 | 0 |
| 0 | 0 | 1 | 0 |
| 0 | 0 | 0 | 0 |
| 0 | 0 | 0 | 0 |
| 0 | 0 | 0 | 0 |
| 0 | 0 | 0 | 0 |
| 0 | 0 | 0 | 0 |
| 0 | 1 | 0 | 0 |
| 0 | 0 | 0 | 0 |
| 0 | 0 | 0 | 0 |
| 1 | 1 | 0 | 0 |
| 0 | 0 | 0 | 0 |
| 0 | 0 | 0 | 0 |
| 0 | 0 | 0 | 0 |
| 0 | 0 | 0 | 0 |
| 0 | 0 | 0 | 0 |
| 0 | 0 | 0 | 0 |
| 0 | 0 | 0 | 0 |
| 0 | 0 | 0 | 0 |
| 0 | 0 | 0 | 0 |
| 0 | 0 | 0 | 0 |
| 0 | 0 | 0 | 0 |
| 0 | 0 | 0 | 0 |
| 0 | 0 | 0 | 0 |
| 0 | 0 | 0 | 0 |
| 0 | 0 | 0 | 0 |
| 0 | 0 | 0 | 0 |
| 0 | 0 | 0 | 0 |
| 0 | 0 | 0 | 0 |
| 0 | 0 | 0 | 0 |
| 0 | 0 | 0 | 0 |
| 0 | 0 | 0 | 0 |
| 0 | 0 | 0 | 0 |
| 0 | 0 | 0 | 0 |
| 0 | 0 | 0 | 0 |
| 0 | 0 | 0 | 0 |
| 0 | 0 | 0 | 0 |
| 0 | 0 | 0 | 0 |
| 0 | 0 | 0 | 0 |
| 0 | 0 | 0 | 0 |
| 0 | 0 | 0 | 0 |
| 0 | 0 | 0 | 0 |
| 0 | 0 | 0 | 0 |
| 0 | 0 | 0 | 0 |
| 0 | 0 | 0 | 0 |
| 1 | 0 | 0 | 0 |

| | | | |
|---|---|---|---|
| 0 | 0 | 0 | 0 |
| 0 | 0 | 0 | 0 |
| 0 | 0 | 0 | 0 |
| 0 | 0 | 0 | 0 |
| 0 | 0 | 0 | 0 |
| 0 | 0 | 0 | 0 |
| 0 | 0 | 0 | 0 |
| 0 | 0 | 0 | 0 |
| 0 | 0 | 0 | 0 |
| 0 | 0 | 0 | 0 |
| 1 | 0 | 0 | 0 |
| 0 | 0 | 0 | 0 |
| 0 | 0 | 0 | 0 |
| 0 | 0 | 0 | 0 |
| 1 | 1 | 0 | 1 |
| 0 | 0 | 0 | 0 |
| 1 | 0 | 0 | 0 |
| 1 | 1 | 0 | 0 |
| 0 | 0 | 0 | 0 |
| 0 | 0 | 0 | 0 |
| 0 | 0 | 0 | 0 |
| 0 | 0 | 0 | 0 |
| 0 | 0 | 0 | 0 |
| 0 | 0 | 0 | 1 |
| 0 | 0 | 0 | 0 |
| 0 | 0 | 0 | 0 |
| 0 | 0 | 0 | 0 |
| 0 | 0 | 0 | 1 |
| 0 | 0 | 0 | 0 |
| 0 | 0 | 0 | 0 |
| 0 | 0 | 0 | 0 |
| 0 | 0 | 0 | 0 |
| 0 | 0 | 0 | 0 |
| 0 | 0 | 0 | 0 |
| 0 | 0 | 0 | 0 |
| 0 | 0 | 0 | 0 |
| 0 | 0 | 0 | 0 |
| 0 | 0 | 0 | 0 |
| 0 | 0 | 0 | 0 |
| 0 | 0 | 0 | 0 |
| 0 | 0 | 0 | 0 |
| 0 | 0 | 0 | 0 |
| 0 | 0 | 0 | 0 |
| 0 | 0 | 0 | 0 |
| 0 | 0 | 1 | 0 |
| 0 | 0 | 0 | 0 |
| 0 | 0 | 0 | 0 |
| 0 | 0 | 0 | 0 |
| 0 | 0 | 0 | 0 |
| 0 | 0 | 0 | 0 |
| 0 | 0 | 0 | 0 |
| 0 | 0 | 0 | 0 |
| 0 | 0 | 0 | 0 |
| 0 | 0 | 0 | 0 |

| | | | |
|---|---|---|---|
| 0 | 0 | 0 | 0 |
| 0 | 0 | 0 | 0 |
| 0 | 0 | 0 | 0 |
| 0 | 0 | 0 | 0 |
| 0 | 0 | 0 | 0 |
| 0 | 0 | 0 | 0 |
| 1 | 0 | 0 | 1 |
| 1 | 0 | 0 | 0 |
| 0 | 0 | 0 | 0 |
| 0 | 0 | 0 | 0 |
| 0 | 0 | 0 | 0 |
| 0 | 0 | 0 | 0 |
| 0 | 0 | 0 | 0 |
| 0 | 0 | 0 | 0 |
| 0 | 0 | 0 | 0 |
| 0 | 0 | 0 | 0 |
| 0 | 0 | 0 | 0 |
| 0 | 0 | 0 | 0 |
| 0 | 0 | 0 | 0 |
| 0 | 0 | 0 | 0 |
| 1 | 0 | 0 | 0 |
| 0 | 0 | 0 | 0 |
| 0 | 0 | 0 | 0 |
| 0 | 0 | 0 | 0 |
| 0 | 0 | 0 | 0 |
| 0 | 0 | 0 | 0 |
| 0 | 0 | 0 | 0 |
| 0 | 0 | 0 | 0 |
| 0 | 0 | 0 | 0 |
| 0 | 0 | 0 | 0 |
| 0 | 0 | 0 | 0 |
| 0 | 0 | 0 | 0 |
| 0 | 0 | 0 | 0 |
| 0 | 0 | 0 | 0 |
| 0 | 0 | 0 | 0 |
| 0 | 0 | 0 | 0 |
| 0 | 0 | 0 | 0 |
| 0 | 0 | 0 | 0 |
| 0 | 0 | 0 | 0 |
| 0 | 0 | 0 | 0 |
| 0 | 0 | 0 | 0 |
| 0 | 0 | 0 | 0 |
| 0 | 0 | 0 | 0 |
| 0 | 0 | 0 | 0 |
| 0 | 0 | 0 | 0 |
| 0 | 0 | 0 | 0 |
| 0 | 0 | 0 | 0 |
| 0 | 0 | 0 | 0 |
| 0 | 0 | 0 | 0 |
| 0 | 0 | 0 | 0 |
| 0 | 0 | 0 | 0 |
| 0 | 0 | 0 | 0 |
| 0 | 0 | 0 | 0 |
| 0 | 0 | 0 | 0 |

| | | | |
|---|---|---|---|
| 0 | 0 | 0 | 0 |
| 0 | 0 | 0 | 0 |
| 0 | 0 | 0 | 0 |
| 1 | 0 | 0 | 0 |
| 0 | 0 | 0 | 0 |
| 1 | 0 | 0 | 0 |
| 0 | 0 | 0 | 0 |
| 0 | 0 | 0 | 0 |
| 1 | 0 | 0 | 0 |
| 0 | 0 | 0 | 0 |
| 0 | 0 | 0 | 0 |
| 0 | 0 | 0 | 0 |
| 0 | 0 | 0 | 0 |
| 0 | 0 | 0 | 0 |
| 0 | 0 | 0 | 0 |
| 0 | 0 | 0 | 0 |
| 0 | 0 | 0 | 0 |
| 0 | 0 | 0 | 0 |
| 1 | 0 | 0 | 0 |
| 0 | 0 | 0 | 0 |
| 0 | 0 | 0 | 0 |
| 0 | 0 | 0 | 0 |
| 0 | 0 | 0 | 0 |
| 0 | 0 | 0 | 0 |
| 0 | 0 | 0 | 0 |
| 0 | 0 | 0 | 0 |
| 0 | 0 | 0 | 0 |
| 0 | 0 | 0 | 0 |
| 0 | 1 | 0 | 0 |
| 0 | 0 | 0 | 0 |
| 0 | 0 | 0 | 0 |
| 0 | 0 | 0 | 0 |
| 0 | 1 | 0 | 0 |
| 1 | 1 | 0 | 0 |
| 1 | 0 | 0 | 0 |
| 0 | 0 | 0 | 0 |
| 0 | 0 | 0 | 0 |
| 0 | 0 | 0 | 0 |
| 0 | 0 | 0 | 0 |
| 0 | 0 | 0 | 0 |
| 1 | 0 | 0 | 0 |
| 0 | 0 | 0 | 0 |
| 0 | 0 | 0 | 0 |
| 0 | 0 | 0 | 0 |
| 0 | 0 | 0 | 0 |
| 0 | 0 | 0 | 0 |
| 0 | 0 | 0 | 0 |
| 0 | 0 | 0 | 0 |
| 0 | 0 | 0 | 0 |
| 0 | 0 | 0 | 0 |
| 0 | 0 | 0 | 0 |
| 0 | 0 | 0 | 0 |
| 0 | 0 | 0 | 0 |
| 1 | 0 | 0 | 0 |

| | | | |
|---|---|---|---|
| 1 | 0 | 0 | 1 |
| 0 | 0 | 0 | 0 |
| 0 | 0 | 0 | 0 |
| 0 | 0 | 0 | 0 |
| 0 | 0 | 0 | 0 |
| 0 | 0 | 0 | 0 |
| 0 | 0 | 0 | 0 |
| 0 | 0 | 0 | 0 |
| 0 | 0 | 0 | 0 |
| 0 | 0 | 0 | 0 |
| 0 | 0 | 0 | 0 |
| 0 | 0 | 0 | 0 |
| 0 | 0 | 0 | 0 |
| 0 | 0 | 0 | 0 |
| 0 | 0 | 0 | 0 |
| 0 | 0 | 0 | 0 |
| 0 | 0 | 0 | 0 |
| 0 | 0 | 0 | 0 |
| 0 | 0 | 0 | 0 |
| 0 | 0 | 0 | 0 |
| 0 | 0 | 0 | 0 |
| 0 | 0 | 0 | 0 |
| 0 | 1 | 0 | 0 |
| 0 | 0 | 0 | 0 |
| 0 | 0 | 0 | 0 |
| 0 | 0 | 0 | 0 |
| 0 | 0 | 0 | 0 |
| 0 | 0 | 0 | 0 |
| 0 | 0 | 0 | 0 |
| 0 | 0 | 0 | 0 |
| 0 | 0 | 0 | 0 |
| 0 | 0 | 0 | 0 |
| 0 | 0 | 0 | 0 |
| 0 | 0 | 0 | 0 |
| 0 | 0 | 0 | 0 |
| 0 | 0 | 0 | 0 |
| 0 | 0 | 0 | 0 |
| 0 | 0 | 0 | 0 |
| 0 | 0 | 0 | 0 |
| 0 | 0 | 0 | 0 |
| 0 | 0 | 0 | 0 |
| 0 | 0 | 0 | 0 |
| 0 | 0 | 0 | 0 |
| 0 | 0 | 0 | 0 |
| 0 | 0 | 0 | 0 |
| 0 | 0 | 0 | 0 |
| 0 | 0 | 0 | 0 |
| 0 | 0 | 0 | 0 |
| 0 | 0 | 0 | 0 |
| 0 | 0 | 0 | 0 |
| 0 | 0 | 0 | 0 |
| 0 | 0 | 0 | 0 |
| 0 | 0 | 0 | 0 |
| 0 | 0 | 0 | 0 |

| | | | |
|---|---|---|---|
| 0 | 0 | 0 | 0 |
| 0 | 0 | 0 | 0 |
| 0 | 0 | 0 | 0 |
| 0 | 0 | 0 | 0 |
| 0 | 0 | 0 | 0 |
| 0 | 0 | 0 | 0 |
| 0 | 1 | 0 | 0 |
| 0 | 1 | 0 | 0 |
| 0 | 0 | 0 | 0 |
| 0 | 0 | 0 | 0 |
| 0 | 0 | 0 | 0 |
| 0 | 0 | 0 | 0 |
| 0 | 0 | 0 | 0 |
| 0 | 0 | 0 | 0 |
| 0 | 0 | 0 | 0 |
| 0 | 0 | 0 | 0 |
| 1 | 0 | 0 | 0 |
| 0 | 0 | 0 | 0 |
| 1 | 0 | 0 | 0 |
| 1 | 0 | 0 | 0 |
| 1 | 0 | 0 | 0 |
| 1 | 0 | 0 | 0 |
| 0 | 0 | 0 | 0 |
| 0 | 0 | 0 | 0 |
| 0 | 0 | 0 | 0 |
| 0 | 0 | 0 | 0 |
| 0 | 0 | 0 | 0 |
| 0 | 0 | 0 | 0 |
| 0 | 0 | 0 | 0 |
| 0 | 0 | 0 | 0 |
| 0 | 0 | 0 | 0 |
| 0 | 0 | 0 | 0 |
| 0 | 0 | 0 | 0 |
| 0 | 0 | 0 | 0 |
| 0 | 0 | 0 | 0 |
| 0 | 0 | 1 | 0 |
| 1 | 0 | 0 | 0 |
| 1 | 0 | 0 | 0 |
| 1 | 0 | 0 | 0 |
| 0 | 0 | 0 | 0 |
| 0 | 0 | 0 | 0 |
| 0 | 0 | 0 | 0 |
| 0 | 0 | 0 | 0 |
| 1 | 0 | 0 | 0 |
| 1 | 0 | 0 | 0 |
| 1 | 0 | 0 | 0 |
| 0 | 0 | 0 | 0 |
| 0 | 0 | 0 | 0 |
| 1 | 0 | 0 | 0 |
| 0 | 0 | 0 | 0 |
| 0 | 0 | 0 | 0 |
| 1 | 0 | 0 | 0 |
| 0 | 0 | 0 | 0 |
| 0 | 0 | 0 | 0 |

| | | | |
|---|---|---|---|
| 0 | 0 | 0 | 0 |
| 0 | 0 | 0 | 0 |
| 0 | 0 | 0 | 0 |
| 0 | 0 | 0 | 0 |
| 0 | 0 | 0 | 0 |
| 0 | 0 | 0 | 0 |
| 0 | 0 | 0 | 0 |
| 0 | 0 | 0 | 0 |
| 0 | 0 | 0 | 0 |
| 0 | 0 | 0 | 0 |
| 0 | 0 | 0 | 0 |
| 0 | 0 | 0 | 0 |
| 0 | 0 | 0 | 0 |
| 0 | 0 | 0 | 0 |
| 0 | 0 | 0 | 0 |
| 0 | 0 | 0 | 0 |
| 0 | 0 | 0 | 0 |
| 0 | 0 | 0 | 0 |
| 0 | 0 | 0 | 0 |
| 0 | 0 | 0 | 0 |
| 0 | 0 | 0 | 0 |
| 0 | 0 | 0 | 0 |
| 1 | 1 | 0 | 0 |
| 0 | 0 | 0 | 0 |
| 0 | 1 | 0 | 0 |
| 0 | 0 | 0 | 0 |
| 0 | 0 | 0 | 0 |
| 0 | 0 | 0 | 0 |
| 0 | 0 | 0 | 0 |
| 0 | 0 | 0 | 0 |
| 0 | 0 | 0 | 0 |
| 0 | 0 | 0 | 0 |
| 0 | 0 | 0 | 0 |
| 0 | 0 | 0 | 0 |
| 0 | 0 | 0 | 0 |
| 0 | 0 | 0 | 0 |
| 0 | 0 | 0 | 0 |
| 0 | 0 | 0 | 0 |
| 0 | 0 | 0 | 0 |
| 0 | 0 | 0 | 0 |
| 0 | 0 | 0 | 0 |
| 0 | 0 | 0 | 0 |
| 0 | 0 | 0 | 0 |
| 0 | 0 | 0 | 0 |
| 0 | 0 | 0 | 0 |
| 1 | 0 | 0 | 1 |
| 0 | 0 | 0 | 0 |
| 0 | 0 | 0 | 0 |
| 0 | 0 | 0 | 0 |
| 1 | 0 | 0 | 0 |
| 1 | 0 | 0 | 0 |
| 1 | 0 | 0 | 0 |
| 0 | 0 | 0 | 0 |
| 0 | 0 | 0 | 0 |

| | | | |
|---|---|---|---|
| 0 | 0 | 0 | 0 |
| 0 | 0 | 0 | 0 |
| 0 | 0 | 0 | 0 |
| 0 | 0 | 0 | 0 |
| 0 | 0 | 0 | 0 |
| 0 | 0 | 0 | 0 |
| 0 | 0 | 0 | 0 |
| 0 | 0 | 0 | 0 |
| 0 | 0 | 0 | 0 |
| 0 | 0 | 0 | 0 |
| 0 | 0 | 0 | 1 |
| 0 | 0 | 0 | 0 |
| 0 | 0 | 0 | 0 |
| 0 | 0 | 0 | 0 |
| 0 | 0 | 0 | 0 |
| 0 | 0 | 0 | 0 |
| 0 | 0 | 0 | 0 |
| 0 | 0 | 0 | 0 |
| 0 | 0 | 0 | 0 |
| 0 | 0 | 0 | 0 |
| 0 | 0 | 0 | 0 |
| 0 | 0 | 0 | 0 |
| 0 | 0 | 0 | 0 |
| 0 | 0 | 0 | 0 |
| 0 | 0 | 0 | 0 |
| 0 | 0 | 0 | 0 |
| 0 | 0 | 0 | 0 |
| 0 | 0 | 0 | 0 |
| 0 | 0 | 0 | 0 |
| 0 | 0 | 0 | 0 |
| 0 | 1 | 0 | 0 |
| 0 | 0 | 0 | 0 |
| 1 | 1 | 0 | 0 |
| 1 | 1 | 0 | 0 |
| 0 | 0 | 0 | 0 |
| 0 | 0 | 0 | 0 |
| 0 | 0 | 0 | 0 |
| 0 | 0 | 0 | 0 |
| 0 | 0 | 0 | 0 |
| 0 | 0 | 0 | 0 |
| 0 | 0 | 0 | 0 |
| 0 | 1 | 0 | 0 |
| 0 | 0 | 0 | 0 |
| 0 | 0 | 0 | 0 |
| 0 | 0 | 0 | 0 |
| 1 | 0 | 0 | 0 |
| 1 | 0 | 0 | 0 |
| 1 | 0 | 0 | 0 |
| 1 | 1 | 1 | 0 |
| 0 | 0 | 0 | 0 |
| 1 | 1 | 1 | 1 |
| 0 | 0 | 0 | 0 |
| 0 | 0 | 0 | 0 |
| 0 | 0 | 0 | 0 |

| | | | |
|---|---|---|---|
| 0 | 0 | 0 | 0 |
| 0 | 0 | 0 | 0 |
| 0 | 0 | 0 | 0 |
| 0 | 0 | 0 | 0 |
| 1 | 0 | 0 | 0 |
| 1 | 0 | 0 | 0 |
| 0 | 0 | 0 | 0 |
| 0 | 0 | 0 | 0 |
| 0 | 0 | 0 | 0 |
| 0 | 0 | 0 | 0 |
| 0 | 0 | 0 | 0 |
| 0 | 0 | 0 | 0 |
| 0 | 0 | 0 | 0 |
| 0 | 0 | 0 | 0 |
| 0 | 0 | 0 | 0 |
| 0 | 0 | 0 | 0 |
| 0 | 0 | 0 | 0 |
| 0 | 0 | 0 | 0 |
| 0 | 0 | 0 | 0 |
| 0 | 0 | 0 | 0 |
| 0 | 0 | 0 | 0 |
| 0 | 0 | 0 | 0 |
| 0 | 0 | 0 | 0 |
| 0 | 0 | 0 | 0 |
| 0 | 0 | 0 | 0 |
| 0 | 0 | 0 | 0 |
| 0 | 0 | 0 | 0 |
| 0 | 0 | 0 | 0 |
| 0 | 0 | 0 | 0 |
| 0 | 0 | 0 | 0 |
| 0 | 0 | 0 | 0 |
| 0 | 0 | 0 | 0 |
| 0 | 0 | 0 | 0 |
| 0 | 0 | 0 | 0 |
| 0 | 0 | 0 | 0 |
| 0 | 0 | 0 | 0 |
| 0 | 0 | 0 | 0 |
| 0 | 0 | 0 | 0 |
| 0 | 0 | 0 | 0 |
| 0 | 0 | 0 | 0 |
| 0 | 0 | 0 | 0 |
| 0 | 0 | 0 | 0 |
| 0 | 0 | 0 | 0 |
| 0 | 0 | 0 | 0 |
| 0 | 0 | 0 | 0 |
| 0 | 0 | 0 | 0 |
| 0 | 0 | 0 | 0 |
| 0 | 0 | 0 | 0 |
| 0 | 0 | 0 | 0 |
| 0 | 0 | 0 | 0 |
| 0 | 0 | 0 | 0 |
| 0 | 0 | 0 | 0 |
| 0 | 0 | 0 | 0 |
| 0 | 1 | 0 | 0 |

| | | | |
|---|---|---|---|
| 0 | 0 | 0 | 0 |
| 0 | 0 | 0 | 0 |
| 0 | 0 | 0 | 0 |
| 0 | 0 | 0 | 0 |
| 0 | 0 | 0 | 0 |
| 0 | 0 | 0 | 0 |
| 0 | 0 | 0 | 0 |
| 0 | 0 | 0 | 0 |
| 0 | 0 | 0 | 0 |
| 0 | 0 | 0 | 0 |
| 0 | 0 | 0 | 0 |
| 0 | 0 | 0 | 0 |
| 0 | 0 | 0 | 0 |
| 0 | 0 | 0 | 0 |
| 0 | 0 | 0 | 0 |
| 0 | 0 | 0 | 0 |
| 0 | 0 | 0 | 0 |
| 0 | 0 | 0 | 0 |
| 0 | 0 | 0 | 0 |
| 0 | 0 | 0 | 0 |
| 0 | 0 | 0 | 0 |
| 0 | 0 | 0 | 0 |
| 0 | 0 | 0 | 0 |
| 0 | 0 | 0 | 0 |
| 0 | 0 | 0 | 0 |
| 0 | 0 | 0 | 0 |
| 0 | 0 | 0 | 0 |
| 0 | 0 | 0 | 0 |
| 0 | 0 | 0 | 0 |
| 0 | 0 | 0 | 0 |
| 0 | 0 | 0 | 0 |
| 0 | 0 | 0 | 0 |
| 0 | 0 | 0 | 0 |
| 0 | 0 | 0 | 0 |
| 0 | 0 | 0 | 0 |
| 0 | 0 | 0 | 0 |
| 0 | 0 | 0 | 0 |
| 0 | 0 | 0 | 0 |
| 0 | 0 | 0 | 0 |
| 0 | 0 | 0 | 0 |
| 0 | 0 | 0 | 0 |
| 0 | 0 | 0 | 0 |
| 0 | 0 | 0 | 0 |
| 0 | 0 | 0 | 0 |
| 0 | 0 | 0 | 0 |
| 0 | 0 | 0 | 0 |
| 0 | 0 | 0 | 0 |
| 0 | 0 | 0 | 0 |
| 0 | 0 | 0 | 0 |
| 0 | 0 | 0 | 0 |
| 0 | 0 | 0 | 0 |
| 0 | 0 | 0 | 0 |
| 0 | 0 | 0 | 0 |
| 0 | 0 | 0 | 0 |

| | | | |
|---|---|---|---|
| 0 | 0 | 0 | 0 |
| 0 | 0 | 0 | 0 |
| 0 | 0 | 0 | 0 |
| 0 | 0 | 0 | 0 |
| 0 | 0 | 0 | 0 |
| 0 | 0 | 0 | 0 |
| 0 | 0 | 0 | 0 |
| 0 | 0 | 0 | 0 |
| 0 | 0 | 0 | 0 |
| 0 | 0 | 0 | 0 |
| 0 | 0 | 0 | 0 |
| 0 | 0 | 1 | 0 |
| 0 | 0 | 0 | 0 |
| 0 | 0 | 1 | 0 |
| 0 | 0 | 1 | 0 |
| 0 | 0 | 1 | 0 |
| 0 | 0 | 1 | 0 |
| 0 | 0 | 0 | 0 |
| 0 | 0 | 0 | 0 |
| 0 | 0 | 0 | 0 |
| 1 | 0 | 0 | 0 |

| glycan_binding_site | water_binding_site | water_only_binding_site | non_binding_site |
|---|---|---|---|
| 0 | 0 | 0 | 0 |
| 0 | 0 | 0 | 0 |
| 0 | 0 | 0 | 1 |
| 0 | 0 | 0 | 0 |
| 0 | 0 | 0 | 0 |
| 0 | 0 | 0 | 0 |
| 0 | 0 | 0 | 1 |
| 0 | 0 | 0 | 1 |
| 0 | 0 | 0 | 0 |
| 0 | 0 | 0 | 0 |
| 0 | 0 | 0 | 0 |
| 0 | 0 | 0 | 0 |
| 0 | 0 | 0 | 1 |
| 1 | 0 | 0 | 0 |
| 0 | 0 | 0 | 0 |
| 0 | 0 | 0 | 0 |
| 0 | 0 | 0 | 0 |
| 0 | 0 | 0 | 0 |
| 0 | 0 | 0 | 0 |
| 0 | 0 | 0 | 0 |
| 0 | 1 | 0 | 0 |
| 1 | 0 | 0 | 0 |
| 1 | 1 | 0 | 0 |
| 0 | 1 | 0 | 0 |
| 0 | 0 | 0 | 0 |
| 0 | 0 | 0 | 1 |
| 0 | 0 | 0 | 0 |
| 0 | 0 | 0 | 1 |
| 0 | 0 | 0 | 0 |
| 0 | 0 | 0 | 1 |
| 0 | 0 | 0 | 1 |
| 0 | 1 | 0 | 0 |
| 0 | 1 | 0 | 0 |
| 0 | 1 | 0 | 0 |
| 0 | 1 | 0 | 0 |
| 0 | 0 | 0 | 1 |
| 0 | 0 | 0 | 0 |
| 0 | 0 | 0 | 0 |
| 0 | 1 | 0 | 0 |
| 0 | 0 | 0 | 0 |
| 0 | 1 | 0 | 0 |
| 0 | 1 | 0 | 0 |
| 0 | 0 | 0 | 0 |
| 0 | 0 | 0 | 0 |
| 0 | 0 | 0 | 0 |
| 0 | 1 | 0 | 0 |
| 0 | 0 | 0 | 0 |
| 0 | 0 | 0 | 0 |
| 0 | 0 | 0 | 0 |
| 0 | 1 | 0 | 0 |
| 0 | 1 | 0 | 0 |
| 0 | 1 | 0 | 0 |
| 0 | 0 | 0 | 0 |

| | | | |
|---|---|---|---|
| 0 | 1 | 0 | 0 |
| 0 | 0 | 0 | 0 |
| 0 | 0 | 0 | 0 |
| 0 | 0 | 0 | 0 |
| 0 | 1 | 0 | 0 |
| 0 | 1 | 0 | 0 |
| 0 | 0 | 0 | 0 |
| 0 | 1 | 0 | 0 |
| 0 | 1 | 0 | 0 |
| 0 | 1 | 0 | 0 |
| 0 | 0 | 0 | 0 |
| 0 | 0 | 0 | 0 |
| 0 | 0 | 0 | 0 |
| 0 | 1 | 0 | 0 |
| 0 | 1 | 0 | 0 |
| 0 | 1 | 0 | 0 |
| 0 | 1 | 0 | 0 |
| 0 | 0 | 0 | 0 |
| 0 | 0 | 0 | 0 |
| 0 | 0 | 0 | 0 |
| 0 | 0 | 0 | 0 |
| 0 | 0 | 0 | 0 |
| 0 | 0 | 0 | 0 |
| 0 | 0 | 0 | 0 |
| 0 | 0 | 0 | 0 |
| 0 | 0 | 0 | 0 |
| 0 | 0 | 0 | 0 |
| 0 | 0 | 0 | 0 |
| 0 | 0 | 0 | 0 |
| 0 | 0 | 0 | 0 |
| 0 | 0 | 0 | 0 |
| 0 | 0 | 0 | 0 |
| 0 | 0 | 0 | 0 |
| 0 | 0 | 0 | 0 |
| 0 | 0 | 0 | 0 |
| 0 | 0 | 0 | 0 |
| 0 | 0 | 0 | 0 |
| 0 | 0 | 0 | 0 |
| 0 | 0 | 0 | 0 |
| 0 | 0 | 0 | 0 |
| 1 | 0 | 0 | 0 |
| 0 | 0 | 0 | 0 |
| 1 | 0 | 0 | 0 |
| 1 | 0 | 0 | 0 |
| 0 | 0 | 0 | 0 |
| 0 | 0 | 0 | 1 |
| 0 | 0 | 0 | 1 |
| 0 | 0 | 0 | 0 |
| 1 | 0 | 0 | 0 |
| 0 | 0 | 0 | 0 |
| 0 | 0 | 0 | 0 |
| 0 | 0 | 0 | 1 |
| 0 | 0 | 0 | 1 |
| 0 | 0 | 0 | 0 |

| | | | |
|---|---|---|---|
| 0 | 0 | 0 | 1 |
| 0 | 0 | 0 | 1 |
| 0 | 0 | 0 | 0 |
| 0 | 0 | 0 | 0 |
| 0 | 0 | 0 | 0 |
| 0 | 0 | 0 | 0 |
| 0 | 0 | 0 | 1 |
| 0 | 0 | 0 | 0 |
| 0 | 0 | 0 | 0 |
| 0 | 0 | 0 | 0 |
| 0 | 0 | 0 | 0 |
| 0 | 0 | 0 | 0 |
| 0 | 0 | 0 | 0 |
| 1 | 0 | 0 | 0 |
| 1 | 0 | 0 | 0 |
| 0 | 0 | 0 | 1 |
| 0 | 0 | 0 | 0 |
| 0 | 0 | 0 | 0 |
| 0 | 0 | 0 | 0 |
| 0 | 0 | 0 | 0 |
| 0 | 0 | 0 | 0 |
| 0 | 0 | 0 | 0 |
| 0 | 0 | 0 | 0 |
| 0 | 0 | 0 | 0 |
| 0 | 0 | 0 | 0 |
| 0 | 0 | 0 | 0 |
| 0 | 0 | 0 | 0 |
| 0 | 0 | 0 | 0 |
| 0 | 0 | 0 | 0 |
| 0 | 0 | 0 | 0 |
| 0 | 0 | 0 | 0 |
| 0 | 0 | 0 | 0 |
| 0 | 0 | 0 | 0 |
| 0 | 0 | 0 | 0 |
| 0 | 0 | 0 | 0 |
| 0 | 0 | 0 | 0 |
| 0 | 0 | 0 | 0 |
| 0 | 0 | 0 | 0 |
| 0 | 0 | 0 | 0 |
| 0 | 0 | 0 | 0 |
| 0 | 0 | 0 | 0 |
| 0 | 0 | 0 | 0 |
| 0 | 0 | 0 | 0 |
| 0 | 0 | 0 | 0 |
| 0 | 0 | 0 | 0 |
| 0 | 0 | 0 | 0 |
| 1 | 0 | 0 | 0 |
| 0 | 0 | 0 | 0 |
| 0 | 0 | 0 | 0 |
| 0 | 0 | 0 | 0 |
| 0 | 0 | 0 | 0 |
| 0 | 0 | 0 | 0 |
| 0 | 0 | 0 | 0 |
| 0 | 1 | 1 | 1 |

| | | | |
|---|---|---|---|
| 0 | 1 | 1 | 1 |
| 0 | 1 | 1 | 1 |
| 0 | 0 | 0 | 1 |
| 0 | 0 | 0 | 0 |
| 0 | 0 | 0 | 0 |
| 0 | 1 | 1 | 1 |
| 0 | 0 | 0 | 0 |
| 0 | 1 | 1 | 1 |
| 0 | 1 | 1 | 1 |
| 0 | 0 | 0 | 1 |
| 0 | 0 | 0 | 1 |
| 0 | 1 | 0 | 0 |
| 0 | 1 | 1 | 1 |
| 0 | 0 | 0 | 0 |
| 0 | 1 | 1 | 1 |
| 0 | 0 | 0 | 1 |
| 0 | 0 | 0 | 1 |
| 0 | 1 | 1 | 1 |
| 0 | 1 | 1 | 1 |
| 0 | 0 | 0 | 0 |
| 0 | 1 | 0 | 0 |
| 0 | 0 | 0 | 1 |
| 0 | 0 | 0 | 0 |
| 0 | 0 | 0 | 0 |
| 0 | 0 | 0 | 0 |
| 0 | 0 | 0 | 1 |
| 0 | 0 | 0 | 0 |
| 0 | 0 | 0 | 0 |
| 0 | 0 | 0 | 1 |
| 0 | 0 | 0 | 1 |
| 0 | 0 | 0 | 1 |
| 0 | 0 | 0 | 1 |
| 0 | 0 | 0 | 1 |
| 0 | 0 | 0 | 1 |
| 0 | 0 | 0 | 0 |
| 0 | 0 | 0 | 1 |
| 1 | 0 | 0 | 0 |
| 1 | 0 | 0 | 0 |
| 0 | 1 | 1 | 1 |
| 0 | 1 | 1 | 1 |
| 0 | 0 | 0 | 1 |
| 0 | 1 | 0 | 0 |
| 0 | 1 | 0 | 0 |
| 0 | 0 | 0 | 1 |
| 0 | 0 | 0 | 1 |
| 0 | 0 | 0 | 1 |
| 0 | 0 | 0 | 0 |
| 0 | 0 | 0 | 0 |
| 0 | 0 | 0 | 0 |
| 0 | 0 | 0 | 0 |
| 0 | 0 | 0 | 0 |
| 0 | 0 | 0 | 1 |
| 0 | 0 | 0 | 1 |
| 0 | 0 | 0 | 0 |

| | | | |
|---|---|---|---|
| 0 | 0 | 0 | 0 |
| 0 | 0 | 0 | 0 |
| 0 | 0 | 0 | 0 |
| 0 | 0 | 0 | 1 |
| 0 | 0 | 0 | 1 |
| 0 | 0 | 0 | 1 |
| 0 | 0 | 0 | 0 |
| 0 | 0 | 0 | 0 |
| 0 | 0 | 0 | 1 |
| 0 | 0 | 0 | 1 |
| 1 | 0 | 0 | 0 |
| 0 | 0 | 0 | 1 |
| 1 | 0 | 0 | 0 |
| 0 | 0 | 0 | 1 |
| 0 | 0 | 0 | 1 |
| 0 | 0 | 0 | 1 |
| 0 | 0 | 0 | 1 |
| 0 | 0 | 0 | 0 |
| 0 | 0 | 0 | 0 |
| 0 | 0 | 0 | 0 |
| 0 | 0 | 0 | 0 |
| 0 | 0 | 0 | 1 |
| 0 | 0 | 0 | 0 |
| 0 | 0 | 0 | 1 |
| 0 | 0 | 0 | 1 |
| 0 | 0 | 0 | 0 |
| 0 | 1 | 0 | 0 |
| 0 | 0 | 0 | 0 |
| 0 | 0 | 0 | 0 |
| 0 | 1 | 0 | 0 |
| 0 | 0 | 0 | 0 |
| 0 | 0 | 0 | 0 |
| 0 | 0 | 0 | 1 |
| 0 | 0 | 0 | 1 |
| 0 | 0 | 0 | 0 |
| 0 | 0 | 0 | 1 |
| 0 | 0 | 0 | 1 |
| 0 | 0 | 0 | 0 |
| 0 | 0 | 0 | 1 |
| 0 | 0 | 0 | 1 |
| 0 | 0 | 0 | 0 |
| 0 | 0 | 0 | 1 |
| 0 | 0 | 0 | 1 |
| 0 | 0 | 0 | 1 |
| 0 | 0 | 0 | 0 |
| 0 | 0 | 0 | 0 |
| 0 | 0 | 0 | 0 |
| 0 | 0 | 0 | 0 |
| 0 | 0 | 0 | 0 |
| 0 | 0 | 0 | 0 |
| 0 | 0 | 0 | 0 |
| 0 | 0 | 0 | 0 |
| 0 | 0 | 0 | 0 |
| 0 | 0 | 0 | 0 |

| | | | |
|---|---|---|---|
| 0 | 0 | 0 | 0 |
| 0 | 0 | 0 | 0 |
| 0 | 0 | 0 | 0 |
| 0 | 0 | 0 | 0 |
| 0 | 0 | 0 | 0 |
| 0 | 0 | 0 | 0 |
| 0 | 0 | 0 | 1 |
| 0 | 0 | 0 | 1 |
| 0 | 0 | 0 | 0 |
| 0 | 0 | 0 | 0 |
| 0 | 0 | 0 | 0 |
| 0 | 0 | 0 | 0 |
| 0 | 0 | 0 | 0 |
| 0 | 0 | 0 | 1 |
| 0 | 0 | 0 | 1 |
| 0 | 0 | 0 | 0 |
| 0 | 0 | 0 | 1 |
| 0 | 0 | 0 | 0 |
| 0 | 0 | 0 | 1 |
| 0 | 0 | 0 | 0 |
| 0 | 0 | 0 | 1 |
| 0 | 0 | 0 | 1 |
| 0 | 0 | 0 | 1 |
| 0 | 0 | 0 | 0 |
| 0 | 0 | 0 | 0 |
| 0 | 0 | 0 | 0 |
| 0 | 0 | 0 | 0 |
| 0 | 0 | 0 | 0 |
| 0 | 0 | 0 | 0 |
| 0 | 0 | 0 | 0 |
| 0 | 0 | 0 | 1 |
| 0 | 0 | 0 | 1 |
| 0 | 0 | 0 | 0 |
| 0 | 0 | 0 | 0 |
| 0 | 0 | 0 | 0 |
| 0 | 0 | 0 | 1 |
| 0 | 0 | 0 | 0 |
| 0 | 0 | 0 | 0 |
| 0 | 0 | 0 | 0 |
| 0 | 0 | 0 | 0 |
| 0 | 0 | 0 | 1 |
| 0 | 0 | 0 | 1 |
| 0 | 0 | 0 | 0 |
| 0 | 0 | 0 | 1 |
| 0 | 0 | 0 | 0 |
| 0 | 0 | 0 | 0 |
| 0 | 0 | 0 | 0 |
| 0 | 0 | 0 | 1 |
| 0 | 0 | 0 | 1 |
| 0 | 0 | 0 | 0 |
| 0 | 0 | 0 | 1 |
| 0 | 0 | 0 | 1 |
| 0 | 0 | 0 | 0 |
| 0 | 0 | 0 | 1 |

| | | | |
|---|---|---|---|
| 0 | 0 | 0 | 0 |
| 0 | 0 | 0 | 1 |
| 0 | 0 | 0 | 1 |
| 0 | 0 | 0 | 0 |
| 0 | 0 | 0 | 0 |
| 0 | 0 | 0 | 0 |
| 0 | 0 | 0 | 1 |
| 0 | 0 | 0 | 0 |
| 0 | 0 | 0 | 0 |
| 0 | 0 | 0 | 1 |
| 0 | 0 | 0 | 1 |
| 0 | 0 | 0 | 1 |
| 0 | 0 | 0 | 1 |
| 0 | 0 | 0 | 1 |
| 0 | 0 | 0 | 0 |
| 0 | 0 | 0 | 1 |
| 0 | 0 | 0 | 0 |
| 0 | 0 | 0 | 0 |
| 0 | 0 | 0 | 1 |
| 0 | 0 | 0 | 0 |
| 0 | 0 | 0 | 0 |
| 0 | 0 | 0 | 0 |
| 0 | 0 | 0 | 0 |
| 0 | 0 | 0 | 0 |
| 0 | 0 | 0 | 1 |
| 0 | 0 | 0 | 1 |
| 0 | 0 | 0 | 0 |
| 0 | 0 | 0 | 0 |
| 0 | 0 | 0 | 0 |
| 0 | 0 | 0 | 0 |
| 0 | 0 | 0 | 0 |
| 0 | 0 | 0 | 0 |
| 0 | 0 | 0 | 0 |
| 0 | 0 | 0 | 0 |
| 0 | 0 | 0 | 1 |
| 0 | 0 | 0 | 1 |
| 0 | 0 | 0 | 0 |
| 0 | 0 | 0 | 0 |
| 0 | 0 | 0 | 0 |
| 0 | 0 | 0 | 0 |
| 0 | 0 | 0 | 0 |
| 0 | 0 | 0 | 0 |
| 0 | 0 | 0 | 1 |
| 0 | 0 | 0 | 0 |
| 0 | 0 | 0 | 0 |
| 0 | 0 | 0 | 0 |
| 0 | 0 | 0 | 0 |
| 0 | 0 | 0 | 0 |
| 1 | 1 | 0 | 0 |
| 0 | 0 | 0 | 1 |
| 0 | 0 | 0 | 0 |
| 0 | 0 | 0 | 0 |
| 0 | 0 | 0 | 0 |
| 0 | 0 | 0 | 0 |

| | | | |
|---|---|---|---|
| 0 | 0 | 0 | 0 |
| 0 | 0 | 0 | 0 |
| 0 | 0 | 0 | 0 |
| 0 | 0 | 0 | 0 |
| 0 | 0 | 0 | 1 |
| 0 | 0 | 0 | 0 |
| 0 | 0 | 0 | 0 |
| 0 | 0 | 0 | 0 |
| 0 | 0 | 0 | 0 |
| 0 | 0 | 0 | 0 |
| 0 | 0 | 0 | 0 |
| 0 | 0 | 0 | 0 |
| 0 | 0 | 0 | 0 |
| 0 | 0 | 0 | 0 |
| 0 | 0 | 0 | 0 |
| 0 | 0 | 0 | 0 |
| 0 | 0 | 0 | 0 |
| 0 | 0 | 0 | 0 |
| 0 | 0 | 0 | 1 |
| 0 | 0 | 0 | 0 |
| 0 | 0 | 0 | 0 |
| 0 | 0 | 0 | 0 |
| 0 | 0 | 0 | 0 |
| 0 | 0 | 0 | 1 |
| 0 | 0 | 0 | 0 |
| 0 | 0 | 0 | 0 |
| 0 | 0 | 0 | 0 |
| 0 | 0 | 0 | 1 |
| 0 | 0 | 0 | 1 |
| 0 | 0 | 0 | 1 |
| 0 | 0 | 0 | 0 |
| 0 | 0 | 0 | 0 |
| 0 | 0 | 0 | 0 |
| 0 | 0 | 0 | 0 |
| 0 | 0 | 0 | 0 |
| 0 | 0 | 0 | 1 |
| 0 | 0 | 0 | 1 |
| 0 | 0 | 0 | 0 |
| 0 | 0 | 0 | 0 |
| 0 | 0 | 0 | 1 |
| 0 | 0 | 0 | 1 |
| 0 | 0 | 0 | 1 |
| 0 | 0 | 0 | 1 |
| 0 | 0 | 0 | 1 |
| 0 | 0 | 0 | 0 |
| 0 | 0 | 0 | 0 |
| 0 | 0 | 0 | 0 |
| 0 | 0 | 0 | 0 |
| 0 | 0 | 0 | 0 |
| 0 | 0 | 0 | 1 |
| 0 | 0 | 0 | 1 |
| 0 | 0 | 0 | 0 |
| 0 | 0 | 0 | 1 |
| 0 | 0 | 0 | 0 |

| | | | |
|---|---|---|---|
| 0 | 0 | 0 | 1 |
| 0 | 0 | 0 | 1 |
| 0 | 0 | 0 | 0 |
| 0 | 0 | 0 | 1 |
| 0 | 0 | 0 | 1 |
| 0 | 0 | 0 | 1 |
| 0 | 0 | 0 | 1 |
| 0 | 0 | 0 | 0 |
| 0 | 0 | 0 | 1 |
| 0 | 0 | 0 | 1 |
| 0 | 0 | 0 | 0 |
| 0 | 0 | 0 | 1 |
| 0 | 0 | 0 | 1 |
| 0 | 0 | 0 | 1 |
| 0 | 0 | 0 | 0 |
| 0 | 0 | 0 | 0 |
| 0 | 0 | 0 | 1 |
| 0 | 0 | 0 | 1 |
| 0 | 0 | 0 | 0 |
| 0 | 0 | 0 | 0 |
| 0 | 0 | 0 | 0 |
| 0 | 0 | 0 | 0 |
| 0 | 0 | 0 | 0 |
| 0 | 0 | 0 | 0 |
| 0 | 0 | 0 | 0 |
| 0 | 0 | 0 | 1 |
| 0 | 0 | 0 | 0 |
| 0 | 0 | 0 | 0 |
| 0 | 0 | 0 | 0 |
| 0 | 0 | 0 | 1 |
| 0 | 0 | 0 | 0 |
| 0 | 0 | 0 | 0 |
| 0 | 0 | 0 | 1 |
| 0 | 0 | 0 | 0 |
| 0 | 0 | 0 | 0 |
| 0 | 0 | 0 | 0 |
| 0 | 0 | 0 | 0 |
| 0 | 0 | 0 | 0 |
| 0 | 0 | 0 | 0 |
| 0 | 0 | 0 | 0 |
| 0 | 0 | 0 | 1 |
| 0 | 0 | 0 | 0 |
| 0 | 0 | 0 | 1 |
| 0 | 0 | 0 | 0 |
| 0 | 0 | 0 | 0 |
| 1 | 0 | 0 | 0 |
| 0 | 0 | 0 | 0 |
| 1 | 0 | 0 | 0 |
| 0 | 0 | 0 | 0 |
| 0 | 0 | 0 | 0 |
| 0 | 0 | 0 | 1 |
| 0 | 0 | 0 | 0 |
| 0 | 0 | 0 | 0 |
| 0 | 0 | 0 | 0 |

| | | | |
|---|---|---|---|
| 0 | 0 | 0 | 1 |
| 0 | 0 | 0 | 0 |
| 0 | 0 | 0 | 1 |
| 0 | 0 | 0 | 1 |
| 1 | 0 | 0 | 0 |
| 0 | 0 | 0 | 0 |
| 1 | 0 | 0 | 0 |
| 1 | 0 | 0 | 0 |
| 0 | 0 | 0 | 0 |
| 0 | 0 | 0 | 1 |
| 0 | 0 | 0 | 1 |
| 0 | 0 | 0 | 1 |
| 0 | 0 | 0 | 1 |
| 1 | 0 | 0 | 0 |
| 0 | 0 | 0 | 1 |
| 0 | 0 | 0 | 0 |
| 0 | 0 | 0 | 0 |
| 0 | 0 | 0 | 0 |
| 1 | 0 | 0 | 0 |
| 0 | 0 | 0 | 1 |
| 1 | 0 | 0 | 0 |
| 0 | 0 | 0 | 1 |
| 0 | 0 | 0 | 0 |
| 0 | 0 | 0 | 1 |
| 0 | 0 | 0 | 1 |
| 0 | 0 | 0 | 1 |
| 0 | 0 | 0 | 0 |
| 0 | 0 | 0 | 1 |
| 0 | 0 | 0 | 1 |
| 0 | 0 | 0 | 1 |
| 0 | 0 | 0 | 0 |
| 0 | 0 | 0 | 0 |
| 0 | 0 | 0 | 1 |
| 0 | 0 | 0 | 1 |
| 0 | 0 | 0 | 0 |
| 0 | 0 | 0 | 0 |
| 0 | 0 | 0 | 0 |
| 1 | 0 | 0 | 0 |
| 0 | 0 | 0 | 0 |
| 0 | 0 | 0 | 1 |
| 0 | 0 | 0 | 0 |
| 0 | 0 | 0 | 0 |
| 0 | 0 | 0 | 0 |
| 0 | 0 | 0 | 0 |
| 0 | 0 | 0 | 1 |
| 1 | 0 | 0 | 0 |
| 0 | 0 | 0 | 0 |
| 0 | 0 | 0 | 0 |
| 0 | 0 | 0 | 0 |
| 1 | 0 | 0 | 0 |
| 0 | 0 | 0 | 0 |
| 0 | 0 | 0 | 0 |
| 0 | 0 | 0 | 1 |
| 0 | 0 | 0 | 0 |

| | | | |
|---|---|---|---|
| 0 | 0 | 0 | 0 |
| 0 | 0 | 0 | 0 |
| 0 | 0 | 0 | 1 |
| 0 | 0 | 0 | 1 |
| 0 | 0 | 0 | 1 |
| 0 | 0 | 0 | 0 |
| 0 | 0 | 0 | 1 |
| 0 | 0 | 0 | 1 |
| 0 | 0 | 0 | 0 |
| 0 | 0 | 0 | 0 |
| 1 | 0 | 0 | 0 |
| 1 | 0 | 0 | 0 |
| 0 | 0 | 0 | 1 |
| 1 | 0 | 0 | 0 |
| 0 | 0 | 0 | 0 |
| 0 | 0 | 0 | 0 |
| 0 | 1 | 0 | 0 |
| 0 | 0 | 0 | 0 |
| 0 | 0 | 0 | 0 |
| 0 | 0 | 0 | 1 |
| 0 | 0 | 0 | 0 |
| 0 | 0 | 0 | 0 |
| 0 | 0 | 0 | 0 |
| 0 | 0 | 0 | 0 |
| 0 | 0 | 0 | 0 |
| 0 | 0 | 0 | 1 |
| 0 | 0 | 0 | 1 |
| 0 | 0 | 0 | 1 |
| 0 | 0 | 0 | 0 |
| 0 | 0 | 0 | 0 |
| 0 | 0 | 0 | 0 |
| 0 | 0 | 0 | 0 |
| 0 | 0 | 0 | 0 |
| 0 | 0 | 0 | 0 |
| 0 | 0 | 0 | 0 |
| 0 | 0 | 0 | 1 |
| 0 | 0 | 0 | 0 |
| 0 | 0 | 0 | 0 |
| 0 | 0 | 0 | 1 |
| 0 | 0 | 0 | 1 |
| 0 | 0 | 0 | 1 |
| 0 | 0 | 0 | 0 |
| 0 | 0 | 0 | 0 |
| 0 | 0 | 0 | 0 |
| 0 | 0 | 0 | 0 |
| 0 | 0 | 0 | 0 |
| 0 | 0 | 0 | 0 |
| 0 | 0 | 0 | 1 |
| 0 | 0 | 0 | 1 |
| 0 | 0 | 0 | 0 |
| 0 | 0 | 0 | 0 |
| 0 | 0 | 0 | 0 |
| 0 | 0 | 0 | 0 |
| 0 | 0 | 0 | 1 |

| | | | |
|---|---|---|---|
| 0 | 0 | 0 | 1 |
| 0 | 0 | 0 | 0 |
| 0 | 0 | 0 | 1 |
| 0 | 0 | 0 | 0 |
| 0 | 0 | 0 | 1 |
| 0 | 0 | 0 | 1 |
| 0 | 0 | 0 | 0 |
| 0 | 0 | 0 | 1 |
| 0 | 0 | 0 | 1 |
| 0 | 0 | 0 | 1 |
| 0 | 0 | 0 | 1 |
| 0 | 0 | 0 | 1 |
| 0 | 0 | 0 | 1 |
| 0 | 0 | 0 | 1 |
| 0 | 0 | 0 | 1 |
| 0 | 0 | 0 | 1 |
| 0 | 0 | 0 | 0 |
| 0 | 0 | 0 | 0 |
| 0 | 0 | 0 | 1 |
| 0 | 0 | 0 | 0 |
| 0 | 0 | 0 | 0 |
| 0 | 0 | 0 | 0 |
| 0 | 0 | 0 | 1 |
| 0 | 0 | 0 | 0 |
| 0 | 0 | 0 | 0 |
| 0 | 0 | 0 | 1 |
| 0 | 0 | 0 | 1 |
| 0 | 0 | 0 | 1 |
| 0 | 1 | 0 | 0 |
| 0 | 0 | 0 | 0 |
| 0 | 0 | 0 | 1 |
| 0 | 0 | 0 | 0 |
| 0 | 0 | 0 | 0 |
| 0 | 0 | 0 | 0 |
| 0 | 0 | 0 | 0 |
| 0 | 0 | 0 | 0 |
| 0 | 0 | 0 | 1 |
| 0 | 0 | 0 | 1 |
| 0 | 0 | 0 | 1 |
| 0 | 0 | 0 | 1 |
| 0 | 0 | 0 | 0 |
| 0 | 0 | 0 | 1 |
| 0 | 0 | 0 | 0 |
| 0 | 0 | 0 | 1 |
| 1 | 0 | 0 | 0 |
| 0 | 0 | 0 | 1 |
| 0 | 0 | 0 | 0 |
| 0 | 0 | 0 | 0 |
| 1 | 0 | 0 | 0 |
| 0 | 0 | 0 | 0 |
| 0 | 0 | 0 | 0 |
| 0 | 0 | 0 | 0 |
| 0 | 0 | 0 | 0 |
| 0 | 0 | 0 | 0 |

| | | | |
|---|---|---|---|
| 0 | 0 | 0 | 0 |
| 0 | 0 | 0 | 0 |
| 0 | 0 | 0 | 0 |
| 0 | 0 | 0 | 0 |
| 0 | 0 | 0 | 0 |
| 0 | 0 | 0 | 0 |
| 0 | 0 | 0 | 0 |
| 0 | 0 | 0 | 1 |
| 0 | 1 | 0 | 0 |
| 0 | 0 | 0 | 0 |
| 0 | 0 | 0 | 0 |
| 0 | 0 | 0 | 1 |
| 0 | 0 | 0 | 0 |
| 0 | 1 | 0 | 0 |
| 0 | 1 | 0 | 0 |
| 0 | 1 | 0 | 0 |
| 0 | 0 | 0 | 0 |
| 0 | 0 | 0 | 0 |
| 0 | 0 | 0 | 0 |
| 0 | 1 | 0 | 0 |
| 0 | 1 | 0 | 0 |
| 0 | 0 | 0 | 0 |
| 0 | 1 | 0 | 0 |
| 0 | 1 | 0 | 0 |
| 0 | 1 | 0 | 0 |
| 0 | 1 | 0 | 0 |
| 0 | 1 | 0 | 0 |
| 0 | 1 | 0 | 0 |
| 0 | 1 | 0 | 0 |
| 0 | 0 | 0 | 0 |
| 0 | 1 | 0 | 0 |
| 0 | 0 | 0 | 0 |
| 0 | 0 | 0 | 0 |
| 0 | 0 | 0 | 0 |
| 0 | 0 | 0 | 0 |
| 0 | 0 | 0 | 0 |
| 0 | 0 | 0 | 0 |
| 0 | 0 | 0 | 0 |
| 0 | 0 | 0 | 0 |
| 0 | 0 | 0 | 0 |
| 0 | 0 | 0 | 1 |
| 0 | 0 | 0 | 0 |
| 0 | 0 | 0 | 0 |
| 0 | 0 | 0 | 0 |
| 0 | 0 | 0 | 0 |
| 0 | 0 | 0 | 0 |
| 0 | 0 | 0 | 0 |
| 0 | 0 | 0 | 0 |
| 0 | 0 | 0 | 0 |
| 0 | 0 | 0 | 1 |
| 0 | 0 | 0 | 1 |
| 0 | 0 | 0 | 1 |
| 1 | 0 | 0 | 0 |
| 0 | 0 | 0 | 0 |

| | | | |
|---|---|---|---|
| 0 | 0 | 0 | 0 |
| 0 | 0 | 0 | 0 |
| 1 | 0 | 0 | 0 |
| 1 | 0 | 0 | 0 |
| 0 | 0 | 0 | 0 |
| 0 | 0 | 0 | 0 |
| 0 | 0 | 0 | 0 |
| 0 | 0 | 0 | 1 |
| 0 | 0 | 0 | 1 |
| 0 | 0 | 0 | 0 |
| 0 | 0 | 0 | 0 |
| 0 | 0 | 0 | 1 |
| 0 | 0 | 0 | 0 |
| 0 | 0 | 0 | 0 |
| 0 | 0 | 0 | 0 |
| 0 | 0 | 0 | 0 |
| 0 | 0 | 0 | 0 |
| 0 | 0 | 0 | 1 |
| 0 | 0 | 0 | 0 |
| 0 | 0 | 0 | 1 |
| 0 | 0 | 0 | 0 |
| 0 | 0 | 0 | 0 |
| 0 | 0 | 0 | 1 |
| 0 | 0 | 0 | 0 |
| 0 | 0 | 0 | 0 |
| 0 | 0 | 0 | 1 |
| 0 | 0 | 0 | 0 |
| 0 | 0 | 0 | 0 |
| 0 | 0 | 0 | 0 |
| 0 | 0 | 0 | 0 |
| 0 | 0 | 0 | 0 |
| 0 | 0 | 0 | 0 |
| 0 | 0 | 0 | 1 |
| 0 | 0 | 0 | 0 |
| 0 | 0 | 0 | 0 |
| 0 | 0 | 0 | 0 |
| 0 | 0 | 0 | 1 |
| 0 | 0 | 0 | 0 |
| 0 | 0 | 0 | 0 |
| 0 | 0 | 0 | 0 |
| 0 | 0 | 0 | 0 |
| 0 | 1 | 0 | 0 |
| 0 | 1 | 0 | 0 |
| 0 | 1 | 0 | 0 |
| 0 | 0 | 0 | 0 |
| 0 | 0 | 0 | 1 |
| 0 | 0 | 0 | 0 |
| 0 | 0 | 0 | 0 |
| 0 | 0 | 0 | 0 |
| 0 | 0 | 0 | 0 |
| 0 | 0 | 0 | 0 |
| 0 | 0 | 0 | 0 |
| 0 | 0 | 0 | 1 |
| 0 | 0 | 0 | 1 |

| | | | |
|---|---|---|---|
| 0 | 0 | 0 | 1 |
| 0 | 0 | 0 | 1 |
| 0 | 0 | 0 | 0 |
| 0 | 0 | 0 | 0 |
| 0 | 0 | 0 | 0 |
| 0 | 1 | 0 | 0 |
| 0 | 0 | 0 | 0 |
| 0 | 0 | 0 | 1 |
| 0 | 0 | 0 | 0 |
| 0 | 0 | 0 | 1 |
| 0 | 0 | 0 | 1 |
| 0 | 0 | 0 | 1 |
| 0 | 0 | 0 | 0 |
| 0 | 0 | 0 | 0 |
| 0 | 0 | 0 | 0 |
| 0 | 0 | 0 | 0 |
| 0 | 0 | 0 | 0 |
| 1 | 0 | 0 | 0 |
| 0 | 0 | 0 | 1 |
| 0 | 0 | 0 | 1 |
| 0 | 0 | 0 | 1 |
| 0 | 0 | 0 | 0 |
| 0 | 0 | 0 | 1 |
| 0 | 0 | 0 | 0 |
| 0 | 0 | 0 | 0 |
| 0 | 0 | 0 | 0 |
| 0 | 0 | 0 | 0 |
| 0 | 0 | 0 | 0 |
| 0 | 0 | 0 | 0 |
| 0 | 0 | 0 | 0 |
| 0 | 0 | 0 | 0 |
| 0 | 0 | 0 | 0 |
| 0 | 0 | 0 | 0 |
| 0 | 0 | 0 | 0 |
| 0 | 0 | 0 | 0 |
| 0 | 0 | 0 | 0 |
| 0 | 0 | 0 | 0 |
| 0 | 0 | 0 | 0 |
| 1 | 0 | 0 | 0 |
| 0 | 0 | 0 | 0 |
| 0 | 0 | 0 | 0 |
| 0 | 0 | 0 | 1 |
| 0 | 0 | 0 | 0 |
| 0 | 1 | 0 | 0 |
| 0 | 0 | 0 | 0 |
| 0 | 0 | 0 | 0 |
| 0 | 0 | 0 | 1 |
| 0 | 0 | 0 | 0 |
| 0 | 0 | 0 | 0 |
| 0 | 0 | 0 | 0 |
| 0 | 0 | 0 | 0 |
| 0 | 0 | 0 | 0 |
| 0 | 0 | 0 | 1 |
| 0 | 0 | 0 | 1 |

| | | | |
|---|---|---|---|
| 0 | 0 | 0 | 0 |
| 0 | 0 | 0 | 0 |
| 0 | 0 | 0 | 1 |
| 0 | 0 | 0 | 1 |
| 0 | 0 | 0 | 1 |
| 0 | 0 | 0 | 0 |
| 0 | 0 | 0 | 1 |
| 0 | 0 | 0 | 1 |
| 0 | 0 | 0 | 0 |
| 0 | 0 | 0 | 0 |
| 0 | 0 | 0 | 0 |
| 0 | 0 | 0 | 0 |
| 0 | 0 | 0 | 0 |
| 0 | 0 | 0 | 1 |
| 0 | 0 | 0 | 1 |
| 0 | 0 | 0 | 1 |
| 0 | 0 | 0 | 1 |
| 0 | 0 | 0 | 1 |
| 0 | 0 | 0 | 0 |
| 0 | 0 | 0 | 0 |
| 0 | 0 | 0 | 1 |
| 0 | 0 | 0 | 1 |
| 0 | 0 | 0 | 0 |
| 0 | 0 | 0 | 0 |
| 0 | 0 | 0 | 1 |
| 0 | 0 | 0 | 1 |
| 0 | 1 | 0 | 0 |
| 0 | 0 | 0 | 1 |
| 0 | 0 | 0 | 0 |
| 0 | 0 | 0 | 0 |
| 0 | 0 | 0 | 0 |
| 0 | 0 | 0 | 0 |
| 0 | 0 | 0 | 0 |
| 1 | 0 | 0 | 0 |
| 1 | 0 | 0 | 0 |
| 0 | 0 | 0 | 0 |
| 0 | 0 | 0 | 0 |
| 0 | 0 | 0 | 0 |
| 0 | 0 | 0 | 1 |
| 0 | 0 | 0 | 0 |
| 0 | 0 | 0 | 1 |
| 0 | 0 | 0 | 1 |
| 0 | 0 | 0 | 1 |
| 0 | 0 | 0 | 1 |
| 0 | 0 | 0 | 1 |
| 0 | 0 | 0 | 0 |
| 0 | 0 | 0 | 0 |
| 0 | 0 | 0 | 0 |
| 0 | 0 | 0 | 0 |
| 0 | 0 | 0 | 0 |
| 0 | 0 | 0 | 0 |
| 0 | 0 | 0 | 0 |
| 0 | 0 | 0 | 0 |
| 0 | 0 | 0 | 1 |

| | | | |
|---|---|---|---|
| 0 | 0 | 0 | 1 |
| 0 | 0 | 0 | 0 |
| 0 | 0 | 0 | 0 |
| 0 | 0 | 0 | 0 |
| 0 | 0 | 0 | 1 |
| 0 | 0 | 0 | 0 |
| 0 | 0 | 0 | 0 |
| 0 | 0 | 0 | 0 |
| 0 | 0 | 0 | 1 |
| 0 | 0 | 0 | 1 |
| 0 | 0 | 0 | 0 |
| 0 | 0 | 0 | 0 |
| 0 | 0 | 0 | 0 |
| 0 | 0 | 0 | 1 |
| 0 | 0 | 0 | 0 |
| 0 | 0 | 0 | 0 |
| 0 | 0 | 0 | 0 |
| 0 | 0 | 0 | 0 |
| 0 | 0 | 0 | 0 |
| 0 | 0 | 0 | 0 |
| 0 | 0 | 0 | 0 |
| 0 | 0 | 0 | 1 |
| 0 | 0 | 0 | 0 |
| 0 | 0 | 0 | 1 |
| 0 | 0 | 0 | 0 |
| 0 | 0 | 0 | 0 |
| 0 | 0 | 0 | 1 |
| 0 | 0 | 0 | 0 |
| 0 | 0 | 0 | 0 |
| 0 | 0 | 0 | 0 |
| 0 | 0 | 0 | 0 |
| 0 | 0 | 0 | 0 |
| 0 | 0 | 0 | 0 |
| 0 | 0 | 0 | 0 |
| 0 | 0 | 0 | 1 |
| 0 | 0 | 0 | 0 |
| 0 | 0 | 0 | 0 |
| 0 | 0 | 0 | 0 |
| 0 | 0 | 0 | 0 |
| 0 | 0 | 0 | 0 |
| 0 | 0 | 0 | 0 |
| 0 | 0 | 0 | 0 |
| 0 | 0 | 0 | 0 |
| 0 | 0 | 0 | 0 |
| 0 | 0 | 0 | 0 |
| 0 | 0 | 0 | 0 |
| 0 | 0 | 0 | 0 |
| 0 | 0 | 0 | 1 |
| 0 | 0 | 0 | 0 |
| 0 | 0 | 0 | 1 |
| 0 | 0 | 0 | 0 |
| 0 | 0 | 0 | 0 |
| 0 | 0 | 0 | 0 |
| 0 | 0 | 0 | 0 |

| | | | |
|---|---|---|---|
| 0 | 0 | 0 | 0 |
| 0 | 0 | 0 | 0 |
| 0 | 0 | 0 | 1 |
| 0 | 0 | 0 | 0 |
| 0 | 0 | 0 | 0 |
| 0 | 0 | 0 | 1 |
| 0 | 0 | 0 | 1 |
| 0 | 0 | 0 | 1 |
| 0 | 0 | 0 | 0 |
| 0 | 0 | 0 | 1 |
| 0 | 0 | 0 | 0 |
| 0 | 0 | 0 | 0 |
| 0 | 0 | 0 | 0 |
| 0 | 0 | 0 | 1 |
| 0 | 0 | 0 | 0 |
| 0 | 0 | 0 | 1 |
| 0 | 0 | 0 | 0 |
| 0 | 0 | 0 | 0 |
| 1 | 0 | 0 | 0 |
| 0 | 0 | 0 | 1 |
| 0 | 0 | 0 | 1 |
| 0 | 0 | 0 | 0 |
| 0 | 0 | 0 | 1 |
| 0 | 0 | 0 | 0 |
| 0 | 0 | 0 | 0 |
| 0 | 0 | 0 | 1 |
| 0 | 0 | 0 | 1 |
| 0 | 0 | 0 | 0 |
| 0 | 0 | 0 | 1 |
| 1 | 0 | 0 | 0 |
| 0 | 0 | 0 | 1 |
| 0 | 0 | 0 | 1 |
| 0 | 0 | 0 | 1 |
| 0 | 0 | 0 | 1 |
| 0 | 0 | 0 | 1 |
| 0 | 0 | 0 | 1 |
| 0 | 0 | 0 | 1 |
| 0 | 0 | 0 | 1 |
| 0 | 0 | 0 | 1 |
| 0 | 0 | 0 | 1 |
| 0 | 0 | 0 | 1 |
| 0 | 0 | 0 | 1 |
| 0 | 0 | 0 | 1 |
| 0 | 0 | 0 | 1 |
| 0 | 0 | 0 | 1 |
| 0 | 0 | 0 | 1 |
| 0 | 0 | 0 | 1 |
| 0 | 0 | 0 | 1 |
| 0 | 0 | 0 | 1 |
| 0 | 0 | 0 | 1 |
| 0 | 0 | 0 | 1 |
| 0 | 0 | 0 | 1 |
| 0 | 0 | 0 | 1 |

| | | | |
|---|---|---|---|
| 0 | 0 | 0 | 1 |
| 0 | 0 | 0 | 1 |
| 0 | 0 | 0 | 1 |
| 0 | 0 | 0 | 1 |
| 0 | 0 | 0 | 1 |
| 0 | 0 | 0 | 1 |
| 0 | 0 | 0 | 1 |
| 0 | 0 | 0 | 1 |
| 0 | 0 | 0 | 1 |
| 0 | 0 | 0 | 1 |
| 0 | 0 | 0 | 1 |
| 0 | 0 | 0 | 0 |
| 0 | 0 | 0 | 0 |
| 0 | 0 | 0 | 0 |
| 0 | 0 | 0 | 1 |
| 0 | 0 | 0 | 0 |
| 0 | 0 | 0 | 1 |
| 0 | 0 | 0 | 0 |
| 0 | 0 | 0 | 1 |
| 0 | 0 | 0 | 1 |
| 0 | 0 | 0 | 0 |
| 0 | 0 | 0 | 1 |
| 0 | 0 | 0 | 1 |
| 0 | 0 | 0 | 0 |
| 0 | 0 | 0 | 0 |
| 0 | 0 | 0 | 1 |
| 0 | 0 | 0 | 1 |
| 0 | 0 | 0 | 0 |
| 0 | 0 | 0 | 0 |
| 0 | 0 | 0 | 0 |
| 0 | 0 | 0 | 0 |
| 0 | 0 | 0 | 0 |
| 0 | 0 | 0 | 1 |
| 0 | 0 | 0 | 1 |
| 0 | 0 | 0 | 1 |
| 0 | 0 | 0 | 1 |
| 0 | 0 | 0 | 1 |
| 0 | 0 | 0 | 1 |
| 0 | 0 | 0 | 1 |
| 0 | 0 | 0 | 0 |
| 0 | 0 | 0 | 0 |
| 0 | 0 | 0 | 1 |
| 0 | 0 | 0 | 0 |
| 0 | 0 | 0 | 0 |
| 0 | 0 | 0 | 0 |
| 0 | 0 | 0 | 1 |
| 0 | 0 | 0 | 0 |
| 0 | 0 | 0 | 1 |
| 0 | 0 | 0 | 1 |
| 0 | 0 | 0 | 0 |
| 0 | 0 | 0 | 0 |
| 0 | 0 | 0 | 0 |
| 0 | 0 | 0 | 1 |
| 0 | 0 | 0 | 0 |

| | | | |
|---|---|---|---|
| 0 | 0 | 0 | 0 |
| 0 | 0 | 0 | 1 |
| 0 | 0 | 0 | 1 |
| 0 | 0 | 0 | 1 |
| 0 | 0 | 0 | 0 |
| 0 | 0 | 0 | 1 |
| 0 | 0 | 0 | 0 |
| 0 | 0 | 0 | 1 |
| 0 | 0 | 0 | 1 |
| 0 | 0 | 0 | 0 |
| 0 | 1 | 1 | 1 |
| 0 | 0 | 0 | 1 |
| 0 | 0 | 0 | 1 |
| 0 | 0 | 0 | 0 |
| 0 | 0 | 0 | 1 |
| 0 | 0 | 0 | 1 |
| 0 | 0 | 0 | 0 |
| 0 | 0 | 0 | 1 |
| 0 | 0 | 0 | 0 |
| 0 | 0 | 0 | 0 |
| 0 | 0 | 0 | 0 |
| 0 | 0 | 0 | 0 |
| 0 | 0 | 0 | 0 |
| 0 | 0 | 0 | 0 |
| 0 | 0 | 0 | 0 |
| 0 | 0 | 0 | 0 |
| 0 | 0 | 0 | 0 |
| 0 | 0 | 0 | 0 |
| 0 | 0 | 0 | 0 |
| 0 | 0 | 0 | 0 |
| 0 | 0 | 0 | 0 |
| 0 | 0 | 0 | 0 |
| 0 | 0 | 0 | 0 |
| 0 | 0 | 0 | 0 |
| 0 | 0 | 0 | 0 |
| 0 | 0 | 0 | 0 |
| 0 | 0 | 0 | 1 |
| 0 | 0 | 0 | 0 |
| 0 | 0 | 0 | 0 |
| 0 | 0 | 0 | 1 |
| 0 | 0 | 0 | 1 |
| 0 | 0 | 0 | 0 |
| 0 | 0 | 0 | 0 |
| 0 | 0 | 0 | 0 |
| 0 | 0 | 0 | 0 |
| 0 | 0 | 0 | 0 |
| 0 | 0 | 0 | 0 |
| 0 | 0 | 0 | 0 |
| 0 | 0 | 0 | 1 |
| 0 | 0 | 0 | 1 |
| 0 | 0 | 0 | 0 |
| 0 | 0 | 0 | 1 |
| 0 | 0 | 0 | 0 |
| 0 | 0 | 0 | 0 |

| | | | |
|---|---|---|---|
| 0 | 0 | 0 | 0 |
| 0 | 0 | 0 | 0 |
| 0 | 0 | 0 | 0 |
| 0 | 0 | 0 | 0 |
| 0 | 0 | 0 | 0 |
| 0 | 0 | 0 | 0 |
| 0 | 0 | 0 | 0 |
| 0 | 0 | 0 | 0 |
| 0 | 0 | 0 | 0 |
| 0 | 0 | 0 | 0 |
| 0 | 0 | 0 | 0 |
| 0 | 0 | 0 | 0 |
| 0 | 0 | 0 | 0 |
| 0 | 0 | 0 | 0 |
| 0 | 0 | 0 | 1 |
| 0 | 0 | 0 | 0 |
| 0 | 1 | 0 | 0 |
| 0 | 0 | 0 | 0 |
| 0 | 0 | 0 | 1 |
| 0 | 0 | 0 | 0 |
| 0 | 0 | 0 | 0 |
| 0 | 1 | 0 | 0 |
| 0 | 0 | 0 | 0 |
| 0 | 0 | 0 | 0 |
| 0 | 0 | 0 | 0 |
| 0 | 0 | 0 | 1 |
| 0 | 0 | 0 | 0 |
| 0 | 0 | 0 | 0 |
| 0 | 0 | 0 | 0 |
| 0 | 0 | 0 | 0 |
| 0 | 0 | 0 | 1 |
| 0 | 0 | 0 | 0 |
| 0 | 0 | 0 | 1 |
| 0 | 0 | 0 | 1 |
| 1 | 0 | 0 | 0 |
| 0 | 0 | 0 | 0 |
| 0 | 0 | 0 | 1 |
| 0 | 0 | 0 | 1 |
| 0 | 0 | 0 | 1 |
| 0 | 0 | 0 | 1 |
| 0 | 0 | 0 | 0 |
| 0 | 0 | 0 | 0 |
| 0 | 0 | 0 | 0 |
| 0 | 0 | 0 | 0 |
| 0 | 0 | 0 | 0 |
| 0 | 0 | 0 | 0 |
| 0 | 0 | 0 | 0 |
| 0 | 0 | 0 | 0 |
| 0 | 0 | 0 | 1 |
| 0 | 0 | 0 | 1 |
| 0 | 0 | 0 | 0 |
| 0 | 0 | 0 | 0 |
| 0 | 0 | 0 | 0 |
| 0 | 0 | 0 | 1 |

| | | | |
|---|---|---|---|
| 0 | 0 | 0 | 1 |
| 0 | 0 | 0 | 1 |
| 0 | 0 | 0 | 1 |
| 0 | 0 | 0 | 0 |
| 0 | 0 | 0 | 1 |
| 0 | 0 | 0 | 1 |
| 0 | 0 | 0 | 0 |
| 0 | 0 | 0 | 1 |
| 0 | 0 | 0 | 0 |
| 0 | 0 | 0 | 0 |
| 0 | 0 | 0 | 0 |
| 0 | 0 | 0 | 0 |
| 0 | 0 | 0 | 0 |
| 0 | 0 | 0 | 0 |
| 0 | 0 | 0 | 0 |
| 0 | 0 | 0 | 0 |
| 0 | 0 | 0 | 1 |
| 0 | 0 | 0 | 0 |
| 0 | 0 | 0 | 0 |
| 0 | 0 | 0 | 1 |
| 0 | 0 | 0 | 0 |
| 0 | 0 | 0 | 0 |
| 0 | 0 | 0 | 1 |
| 0 | 0 | 0 | 0 |
| 0 | 0 | 0 | 1 |
| 0 | 0 | 0 | 1 |
| 0 | 0 | 0 | 1 |
| 0 | 0 | 0 | 0 |
| 0 | 0 | 0 | 0 |
| 0 | 0 | 0 | 0 |
| 0 | 0 | 0 | 1 |
| 0 | 0 | 0 | 0 |
| 0 | 0 | 0 | 0 |
| 0 | 0 | 0 | 0 |
| 0 | 0 | 0 | 1 |
| 0 | 0 | 0 | 1 |
| 0 | 0 | 0 | 0 |
| 0 | 0 | 0 | 0 |
| 0 | 0 | 0 | 0 |
| 0 | 0 | 0 | 1 |
| 0 | 0 | 0 | 1 |
| 0 | 0 | 0 | 1 |
| 0 | 0 | 0 | 1 |
| 0 | 0 | 0 | 0 |
| 0 | 0 | 0 | 1 |
| 0 | 0 | 0 | 0 |
| 0 | 0 | 0 | 1 |
| 0 | 0 | 0 | 0 |
| 0 | 0 | 0 | 0 |
| 0 | 0 | 0 | 1 |
| 0 | 0 | 0 | 1 |
| 0 | 0 | 0 | 0 |
| 0 | 0 | 0 | 0 |
| 0 | 0 | 0 | 1 |

| | | | |
|---|---|---|---|
| 0 | 0 | 0 | 1 |
| 0 | 0 | 0 | 1 |
| 0 | 0 | 0 | 1 |
| 0 | 0 | 0 | 1 |
| 0 | 0 | 0 | 0 |
| 0 | 0 | 0 | 1 |
| 0 | 0 | 0 | 1 |
| 0 | 0 | 0 | 0 |
| 0 | 0 | 0 | 0 |
| 0 | 0 | 0 | 1 |
| 0 | 0 | 0 | 0 |
| 0 | 0 | 0 | 0 |
| 1 | 0 | 0 | 0 |
| 1 | 0 | 0 | 0 |
| 0 | 0 | 0 | 0 |
| 1 | 0 | 0 | 0 |
| 1 | 0 | 0 | 0 |
| 1 | 0 | 0 | 0 |
| 1 | 0 | 0 | 0 |
| 0 | 0 | 0 | 1 |
| 0 | 0 | 0 | 1 |
| 0 | 0 | 0 | 1 |
| 0 | 0 | 0 | 1 |
| 0 | 0 | 0 | 1 |
| 0 | 0 | 0 | 0 |
| 0 | 0 | 0 | 1 |
| 1 | 0 | 0 | 0 |
| 1 | 0 | 0 | 0 |
| 0 | 0 | 0 | 1 |
| 0 | 0 | 0 | 1 |
| 0 | 0 | 0 | 1 |
| 0 | 0 | 0 | 1 |
| 0 | 0 | 0 | 1 |
| 0 | 0 | 0 | 0 |
| 0 | 0 | 0 | 0 |
| 0 | 0 | 0 | 1 |
| 0 | 0 | 0 | 0 |
| 0 | 0 | 0 | 1 |
| 0 | 0 | 0 | 0 |
| 0 | 0 | 0 | 0 |
| 0 | 0 | 0 | 0 |
| 0 | 0 | 0 | 0 |
| 0 | 0 | 0 | 1 |
| 0 | 0 | 0 | 0 |
| 0 | 0 | 0 | 0 |
| 1 | 0 | 0 | 0 |
| 1 | 0 | 0 | 0 |
| 1 | 0 | 0 | 0 |
| 0 | 0 | 0 | 0 |
| 0 | 0 | 0 | 0 |
| 0 | 0 | 0 | 0 |
| 1 | 0 | 0 | 0 |
| 1 | 0 | 0 | 0 |
| 0 | 0 | 0 | 1 |

| | | | |
|---|---|---|---|
| 0 | 0 | 0 | 0 |
| 0 | 0 | 0 | 1 |
| 0 | 0 | 0 | 0 |
| 0 | 0 | 0 | 0 |
| 0 | 0 | 0 | 0 |
| 0 | 0 | 0 | 1 |
| 0 | 0 | 0 | 0 |
| 0 | 0 | 0 | 1 |
| 0 | 0 | 0 | 1 |
| 0 | 0 | 0 | 0 |
| 0 | 0 | 0 | 1 |
| 0 | 0 | 0 | 1 |
| 0 | 0 | 0 | 1 |
| 0 | 0 | 0 | 1 |
| 0 | 0 | 0 | 1 |
| 0 | 1 | 0 | 0 |
| 0 | 0 | 0 | 0 |
| 0 | 0 | 0 | 1 |
| 0 | 0 | 0 | 0 |
| 0 | 0 | 0 | 1 |
| 0 | 0 | 0 | 0 |
| 0 | 0 | 0 | 0 |
| 0 | 0 | 0 | 0 |
| 0 | 0 | 0 | 0 |
| 0 | 0 | 0 | 0 |
| 0 | 0 | 0 | 0 |
| 0 | 0 | 0 | 0 |
| 0 | 0 | 0 | 0 |
| 0 | 0 | 0 | 0 |
| 0 | 0 | 0 | 0 |
| 0 | 0 | 0 | 0 |
| 0 | 0 | 0 | 0 |
| 0 | 0 | 0 | 1 |
| 0 | 0 | 0 | 1 |
| 0 | 0 | 0 | 0 |
| 0 | 0 | 0 | 0 |
| 0 | 0 | 0 | 1 |
| 0 | 0 | 0 | 0 |
| 0 | 0 | 0 | 1 |
| 0 | 0 | 0 | 0 |
| 0 | 0 | 0 | 0 |
| 0 | 0 | 0 | 0 |
| 0 | 0 | 0 | 1 |
| 0 | 0 | 0 | 1 |
| 0 | 0 | 0 | 0 |
| 0 | 0 | 0 | 0 |
| 0 | 0 | 0 | 0 |
| 0 | 0 | 0 | 0 |
| 0 | 0 | 0 | 0 |
| 0 | 0 | 0 | 0 |
| 0 | 0 | 0 | 1 |
| 0 | 0 | 0 | 0 |
| 0 | 0 | 0 | 0 |
| 0 | 0 | 0 | 1 |

| | | | |
|---|---|---|---|
| 0 | 0 | 0 | 0 |
| 0 | 0 | 0 | 1 |
| 0 | 0 | 0 | 0 |
| 0 | 0 | 0 | 0 |
| 0 | 0 | 0 | 0 |
| 0 | 0 | 0 | 0 |
| 0 | 0 | 0 | 0 |
| 0 | 0 | 0 | 0 |
| 0 | 0 | 0 | 0 |
| 0 | 0 | 0 | 0 |
| 0 | 0 | 0 | 0 |
| 0 | 0 | 0 | 1 |
| 0 | 0 | 0 | 0 |
| 0 | 0 | 0 | 1 |
| 0 | 0 | 0 | 1 |
| 0 | 0 | 0 | 1 |
| 0 | 0 | 0 | 0 |
| 0 | 0 | 0 | 0 |
| 0 | 1 | 1 | 1 |
| 0 | 0 | 0 | 0 |
| 0 | 0 | 0 | 1 |
| 0 | 0 | 0 | 0 |
| 0 | 0 | 0 | 1 |
| 0 | 0 | 0 | 0 |
| 0 | 0 | 0 | 1 |
| 0 | 0 | 0 | 1 |
| 0 | 0 | 0 | 1 |
| 0 | 0 | 0 | 0 |
| 0 | 0 | 0 | 1 |
| 0 | 0 | 0 | 1 |
| 0 | 0 | 0 | 1 |
| 1 | 0 | 0 | 0 |
| 0 | 0 | 0 | 1 |
| 0 | 0 | 0 | 0 |
| 0 | 0 | 0 | 0 |
| 0 | 0 | 0 | 0 |
| 0 | 0 | 0 | 1 |
| 0 | 0 | 0 | 1 |
| 0 | 0 | 0 | 0 |
| 0 | 0 | 0 | 0 |
| 0 | 0 | 0 | 0 |
| 0 | 0 | 0 | 0 |
| 0 | 0 | 0 | 0 |
| 0 | 0 | 0 | 0 |
| 0 | 0 | 0 | 0 |
| 0 | 0 | 0 | 0 |
| 0 | 0 | 0 | 1 |
| 0 | 0 | 0 | 1 |
| 0 | 0 | 0 | 0 |
| 0 | 0 | 0 | 0 |
| 0 | 0 | 0 | 0 |
| 0 | 0 | 0 | 1 |
| 0 | 0 | 0 | 1 |
| 0 | 0 | 0 | 0 |

| | | | |
|---|---|---|---|
| 0 | 0 | 0 | 0 |
| 0 | 0 | 0 | 1 |
| 0 | 0 | 0 | 1 |
| 0 | 0 | 0 | 1 |
| 0 | 0 | 0 | 0 |
| 0 | 0 | 0 | 0 |
| 0 | 0 | 0 | 0 |
| 0 | 0 | 0 | 0 |
| 0 | 0 | 0 | 0 |
| 0 | 0 | 0 | 0 |
| 0 | 0 | 0 | 1 |
| 0 | 0 | 0 | 0 |
| 0 | 0 | 0 | 0 |
| 0 | 0 | 0 | 0 |
| 0 | 0 | 0 | 1 |
| 0 | 0 | 0 | 1 |
| 0 | 0 | 0 | 0 |
| 0 | 0 | 0 | 1 |
| 0 | 0 | 0 | 0 |
| 0 | 0 | 0 | 0 |
| 0 | 0 | 0 | 1 |
| 0 | 0 | 0 | 0 |
| 0 | 0 | 0 | 0 |
| 0 | 0 | 0 | 1 |
| 0 | 0 | 0 | 0 |
| 0 | 0 | 0 | 0 |
| 0 | 0 | 0 | 0 |
| 0 | 0 | 0 | 1 |
| 0 | 0 | 0 | 0 |
| 0 | 0 | 0 | 0 |
| 0 | 0 | 0 | 0 |
| 0 | 0 | 0 | 1 |
| 0 | 0 | 0 | 1 |
| 0 | 0 | 0 | 1 |
| 0 | 0 | 0 | 1 |
| 0 | 0 | 0 | 1 |
| 0 | 0 | 0 | 1 |
| 0 | 0 | 0 | 1 |
| 0 | 0 | 0 | 1 |
| 0 | 0 | 0 | 1 |
| 0 | 0 | 0 | 1 |
| 0 | 0 | 0 | 1 |
| 1 | 0 | 0 | 0 |
| 0 | 0 | 0 | 1 |
| 0 | 0 | 0 | 1 |
| 0 | 0 | 0 | 1 |
| 0 | 0 | 0 | 1 |
| 0 | 0 | 0 | 1 |
| 0 | 0 | 0 | 1 |
| 0 | 0 | 0 | 1 |
| 0 | 0 | 0 | 1 |
| 0 | 0 | 0 | 1 |
| 0 | 0 | 0 | 1 |
| 0 | 0 | 0 | 1 |

| | | | |
|---|---|---|---|
| 0 | 0 | 0 | 1 |
| 0 | 0 | 0 | 1 |
| 0 | 0 | 0 | 1 |
| 0 | 0 | 0 | 1 |
| 0 | 0 | 0 | 1 |
| 0 | 0 | 0 | 1 |
| 0 | 0 | 0 | 1 |
| 0 | 0 | 0 | 1 |
| 0 | 0 | 0 | 0 |
| 0 | 0 | 0 | 0 |
| 0 | 0 | 0 | 0 |
| 0 | 0 | 0 | 0 |
| 0 | 0 | 0 | 0 |
| 0 | 0 | 0 | 0 |
| 0 | 0 | 0 | 0 |
| 0 | 0 | 0 | 0 |
| 0 | 0 | 0 | 0 |
| 0 | 0 | 0 | 0 |
| 0 | 0 | 0 | 0 |
| 0 | 0 | 0 | 0 |
| 1 | 0 | 0 | 0 |
| 1 | 0 | 0 | 0 |
| 0 | 0 | 0 | 0 |
| 0 | 0 | 0 | 0 |
| 0 | 0 | 0 | 0 |
| 0 | 0 | 0 | 1 |
| 0 | 0 | 0 | 1 |
| 0 | 0 | 0 | 0 |
| 0 | 0 | 0 | 0 |
| 1 | 0 | 0 | 0 |
| 1 | 0 | 0 | 0 |
| 0 | 0 | 0 | 0 |
| 1 | 0 | 0 | 0 |
| 1 | 0 | 0 | 0 |
| 0 | 0 | 0 | 0 |
| 0 | 0 | 0 | 0 |
| 0 | 0 | 0 | 0 |
| 0 | 0 | 0 | 0 |
| 0 | 0 | 0 | 0 |
| 0 | 0 | 0 | 0 |
| 0 | 0 | 0 | 0 |
| 0 | 0 | 0 | 0 |
| 1 | 0 | 0 | 0 |
| 1 | 0 | 0 | 0 |
| 1 | 0 | 0 | 0 |
| 0 | 0 | 0 | 0 |
| 0 | 0 | 0 | 1 |
| 0 | 0 | 0 | 1 |
| 0 | 0 | 0 | 1 |
| 0 | 0 | 0 | 1 |
| 0 | 0 | 0 | 1 |
| 0 | 0 | 0 | 1 |
| 0 | 0 | 0 | 0 |
| 0 | 0 | 0 | 0 |

| | | | |
|---|---|---|---|
| 0 | 0 | 0 | 0 |
| 0 | 0 | 0 | 0 |
| 0 | 0 | 0 | 0 |
| 0 | 0 | 0 | 0 |
| 0 | 0 | 0 | 1 |
| 0 | 0 | 0 | 0 |
| 0 | 0 | 0 | 0 |
| 0 | 0 | 0 | 0 |
| 0 | 0 | 0 | 1 |
| 0 | 0 | 0 | 0 |
| 1 | 0 | 0 | 0 |
| 0 | 0 | 0 | 1 |
| 0 | 0 | 0 | 0 |
| 0 | 0 | 0 | 0 |
| 0 | 0 | 0 | 0 |
| 0 | 0 | 0 | 0 |
| 0 | 0 | 0 | 1 |
| 0 | 0 | 0 | 0 |
| 0 | 0 | 0 | 0 |
| 0 | 0 | 0 | 1 |
| 0 | 0 | 0 | 0 |
| 0 | 0 | 0 | 0 |
| 0 | 0 | 0 | 0 |
| 0 | 0 | 0 | 0 |
| 0 | 0 | 0 | 0 |
| 0 | 0 | 0 | 1 |
| 0 | 0 | 0 | 1 |
| 0 | 0 | 0 | 1 |
| 0 | 0 | 0 | 1 |
| 0 | 0 | 0 | 1 |
| 0 | 0 | 0 | 1 |
| 0 | 0 | 0 | 1 |
| 0 | 0 | 0 | 1 |
| 0 | 0 | 0 | 1 |
| 0 | 0 | 0 | 0 |
| 0 | 0 | 0 | 0 |
| 0 | 0 | 0 | 0 |
| 0 | 0 | 0 | 0 |
| 0 | 0 | 0 | 0 |
| 0 | 0 | 0 | 0 |
| 0 | 0 | 0 | 1 |
| 0 | 0 | 0 | 1 |
| 0 | 0 | 0 | 1 |
| 0 | 0 | 0 | 0 |
| 0 | 0 | 0 | 0 |
| 0 | 0 | 0 | 0 |
| 0 | 0 | 0 | 1 |
| 0 | 0 | 0 | 1 |
| 0 | 0 | 0 | 0 |
| 0 | 0 | 0 | 0 |
| 0 | 0 | 0 | 1 |
| 0 | 0 | 0 | 1 |
| 0 | 0 | 0 | 1 |
| 0 | 0 | 0 | 1 |

| | | | |
|---|---|---|---|
| 0 | 0 | 0 | 1 |
| 0 | 0 | 0 | 1 |
| 0 | 0 | 0 | 1 |
| 0 | 0 | 0 | 0 |
| 0 | 0 | 0 | 0 |
| 0 | 0 | 0 | 1 |
| 0 | 0 | 0 | 1 |
| 0 | 0 | 0 | 1 |
| 0 | 0 | 0 | 1 |
| 0 | 0 | 0 | 1 |
| 0 | 0 | 0 | 0 |
| 0 | 0 | 0 | 1 |
| 0 | 0 | 0 | 0 |
| 0 | 0 | 0 | 0 |
| 0 | 0 | 0 | 0 |
| 0 | 0 | 0 | 1 |
| 0 | 0 | 0 | 1 |
| 0 | 0 | 0 | 1 |
| 0 | 0 | 0 | 1 |
| 0 | 0 | 0 | 1 |
| 0 | 0 | 0 | 0 |
| 0 | 0 | 0 | 1 |
| 0 | 0 | 0 | 1 |
| 0 | 0 | 0 | 0 |
| 0 | 0 | 0 | 1 |
| 0 | 0 | 0 | 0 |
| 0 | 1 | 0 | 0 |
| 0 | 0 | 0 | 1 |
| 0 | 0 | 0 | 1 |
| 0 | 0 | 0 | 1 |
| 0 | 0 | 0 | 1 |
| 0 | 0 | 0 | 1 |
| 0 | 0 | 0 | 1 |
| 0 | 0 | 0 | 1 |
| 0 | 0 | 0 | 1 |
| 0 | 0 | 0 | 0 |
| 0 | 0 | 0 | 1 |
| 0 | 0 | 0 | 1 |
| 0 | 0 | 0 | 0 |
| 0 | 0 | 0 | 0 |
| 0 | 0 | 0 | 1 |
| 0 | 0 | 0 | 1 |
| 1 | 0 | 0 | 0 |
| 1 | 0 | 0 | 0 |
| 0 | 0 | 0 | 0 |
| 0 | 0 | 0 | 0 |
| 1 | 0 | 0 | 0 |
| 1 | 0 | 0 | 0 |
| 0 | 0 | 0 | 1 |
| 1 | 0 | 0 | 0 |
| 1 | 0 | 0 | 0 |
| 1 | 0 | 0 | 0 |
| 1 | 0 | 0 | 0 |
| 1 | 0 | 0 | 0 |

| | | | |
|---|---|---|---|
| 0 | 0 | 0 | 0 |
| 0 | 0 | 0 | 0 |
| 1 | 0 | 0 | 0 |
| 1 | 0 | 0 | 0 |
| 1 | 0 | 0 | 0 |
| 1 | 0 | 0 | 0 |
| 0 | 0 | 0 | 0 |
| 0 | 0 | 0 | 0 |
| 0 | 0 | 0 | 0 |
| 0 | 0 | 0 | 1 |
| 0 | 0 | 0 | 1 |
| 0 | 0 | 0 | 0 |
| 0 | 0 | 0 | 1 |
| 0 | 0 | 0 | 0 |
| 0 | 0 | 0 | 0 |
| 0 | 0 | 0 | 1 |
| 0 | 0 | 0 | 0 |
| 0 | 0 | 0 | 1 |
| 0 | 0 | 0 | 1 |
| 0 | 0 | 0 | 1 |
| 0 | 0 | 0 | 1 |
| 0 | 0 | 0 | 1 |
| 0 | 0 | 0 | 1 |
| 0 | 0 | 0 | 0 |
| 0 | 0 | 0 | 0 |
| 0 | 0 | 0 | 1 |
| 0 | 0 | 0 | 1 |
| 0 | 0 | 0 | 1 |
| 0 | 0 | 0 | 0 |
| 0 | 0 | 0 | 0 |
| 0 | 0 | 0 | 1 |
| 0 | 0 | 0 | 1 |
| 0 | 0 | 0 | 1 |
| 0 | 0 | 0 | 0 |
| 0 | 0 | 0 | 0 |
| 0 | 0 | 0 | 1 |
| 0 | 0 | 0 | 0 |
| 0 | 0 | 0 | 1 |
| 0 | 0 | 0 | 1 |
| 0 | 0 | 0 | 0 |
| 0 | 0 | 0 | 0 |
| 0 | 0 | 0 | 0 |
| 0 | 0 | 0 | 0 |
| 0 | 0 | 0 | 0 |
| 0 | 0 | 0 | 0 |
| 0 | 0 | 0 | 1 |
| 0 | 0 | 0 | 1 |
| 0 | 0 | 0 | 0 |
| 0 | 0 | 0 | 0 |
| 0 | 0 | 0 | 0 |
| 0 | 0 | 0 | 1 |
| 0 | 0 | 0 | 1 |
| 0 | 0 | 0 | 0 |
| 0 | 0 | 0 | 0 |

| | | | |
|---|---|---|---|
| 0 | 0 | 0 | 1 |
| 0 | 0 | 0 | 1 |
| 0 | 0 | 0 | 1 |
| 0 | 0 | 0 | 1 |
| 0 | 1 | 0 | 0 |
| 0 | 0 | 0 | 0 |
| 0 | 0 | 0 | 1 |
| 0 | 0 | 0 | 0 |
| 0 | 0 | 0 | 1 |
| 0 | 0 | 0 | 0 |
| 0 | 0 | 0 | 1 |
| 0 | 0 | 0 | 1 |
| 0 | 0 | 0 | 1 |
| 0 | 0 | 0 | 1 |
| 0 | 0 | 0 | 1 |
| 0 | 0 | 0 | 1 |
| 0 | 0 | 0 | 0 |
| 0 | 0 | 0 | 1 |
| 0 | 0 | 0 | 1 |
| 0 | 0 | 0 | 0 |
| 0 | 0 | 0 | 0 |
| 0 | 0 | 0 | 1 |
| 0 | 0 | 0 | 1 |
| 0 | 0 | 0 | 0 |
| 0 | 0 | 0 | 1 |
| 0 | 0 | 0 | 0 |
| 0 | 0 | 0 | 0 |
| 0 | 0 | 0 | 0 |
| 0 | 0 | 0 | 0 |
| 0 | 0 | 0 | 0 |
| 0 | 0 | 0 | 0 |
| 0 | 0 | 0 | 0 |
| 0 | 0 | 0 | 1 |
| 0 | 0 | 0 | 0 |
| 0 | 0 | 0 | 1 |
| 0 | 0 | 0 | 1 |
| 0 | 0 | 0 | 1 |
| 0 | 0 | 0 | 1 |
| 0 | 0 | 0 | 0 |
| 0 | 0 | 0 | 1 |
| 0 | 0 | 0 | 1 |
| 0 | 0 | 0 | 1 |
| 0 | 0 | 0 | 1 |
| 0 | 0 | 0 | 0 |
| 0 | 0 | 0 | 1 |
| 0 | 0 | 0 | 1 |
| 0 | 0 | 0 | 1 |
| 0 | 0 | 0 | 1 |
| 0 | 0 | 0 | 1 |
| 0 | 0 | 0 | 1 |
| 0 | 0 | 0 | 0 |
| 0 | 0 | 0 | 0 |
| 0 | 0 | 0 | 0 |
| 0 | 0 | 0 | 1 |

| | | | |
|---|---|---|---|
| 0 | 0 | 0 | 0 |
| 0 | 0 | 0 | 0 |
| 0 | 0 | 0 | 0 |
| 0 | 0 | 0 | 0 |
| 1 | 0 | 0 | 0 |
| 1 | 0 | 0 | 0 |
| 0 | 0 | 0 | 0 |
| 0 | 0 | 0 | 0 |
| 0 | 0 | 0 | 1 |
| 0 | 0 | 0 | 1 |
| 0 | 0 | 0 | 0 |
| 0 | 0 | 0 | 1 |
| 0 | 0 | 0 | 1 |
| 0 | 0 | 0 | 0 |
| 0 | 0 | 0 | 0 |
| 0 | 0 | 0 | 0 |
| 0 | 0 | 0 | 0 |
| 0 | 0 | 0 | 0 |
| 0 | 0 | 0 | 0 |
| 0 | 0 | 0 | 1 |
| 0 | 0 | 0 | 1 |
| 0 | 0 | 0 | 0 |
| 0 | 0 | 0 | 1 |
| 0 | 0 | 0 | 0 |
| 0 | 0 | 0 | 1 |
| 0 | 0 | 0 | 1 |
| 0 | 0 | 0 | 1 |
| 0 | 0 | 0 | 1 |
| 0 | 0 | 0 | 1 |
| 0 | 0 | 0 | 0 |
| 0 | 0 | 0 | 1 |
| 0 | 0 | 0 | 1 |
| 0 | 0 | 0 | 0 |
| 0 | 0 | 0 | 1 |
| 0 | 0 | 0 | 0 |
| 0 | 0 | 0 | 0 |
| 0 | 0 | 0 | 0 |
| 0 | 0 | 0 | 0 |
| 0 | 0 | 0 | 0 |
| 0 | 0 | 0 | 0 |
| 0 | 0 | 0 | 0 |
| 0 | 0 | 0 | 0 |
| 0 | 0 | 0 | 1 |
| 0 | 0 | 0 | 0 |
| 0 | 0 | 0 | 1 |
| 0 | 0 | 0 | 0 |
| 0 | 0 | 0 | 1 |
| 0 | 0 | 0 | 1 |
| 0 | 0 | 0 | 1 |
| 0 | 0 | 0 | 0 |
| 0 | 0 | 0 | 1 |
| 0 | 0 | 0 | 1 |
| 0 | 0 | 0 | 1 |
| 0 | 0 | 0 | 1 |

| | | | |
|---|---|---|---|
| 1 | 0 | 0 | 0 |
| 0 | 0 | 0 | 1 |
| 0 | 0 | 0 | 1 |
| 0 | 0 | 0 | 1 |
| 0 | 0 | 0 | 1 |
| 0 | 0 | 0 | 1 |
| 0 | 0 | 0 | 1 |
| 0 | 0 | 0 | 1 |
| 0 | 0 | 0 | 1 |
| 0 | 0 | 0 | 1 |
| 0 | 0 | 0 | 0 |
| 0 | 0 | 0 | 1 |
| 0 | 0 | 0 | 1 |
| 0 | 0 | 0 | 1 |
| 0 | 0 | 0 | 0 |
| 0 | 0 | 0 | 0 |
| 0 | 0 | 0 | 0 |
| 0 | 0 | 0 | 0 |
| 0 | 0 | 0 | 0 |
| 0 | 0 | 0 | 1 |
| 0 | 0 | 0 | 1 |
| 0 | 0 | 0 | 1 |
| 0 | 0 | 0 | 0 |
| 0 | 0 | 0 | 1 |
| 0 | 0 | 0 | 1 |
| 0 | 0 | 0 | 1 |
| 0 | 0 | 0 | 1 |
| 0 | 0 | 0 | 1 |
| 0 | 0 | 0 | 0 |
| 0 | 0 | 0 | 0 |
| 0 | 0 | 0 | 0 |
| 0 | 0 | 0 | 0 |
| 0 | 0 | 0 | 0 |
| 0 | 0 | 0 | 0 |
| 0 | 0 | 0 | 1 |
| 0 | 0 | 0 | 1 |
| 0 | 0 | 0 | 0 |
| 0 | 0 | 0 | 0 |
| 0 | 0 | 0 | 0 |
| 0 | 0 | 0 | 0 |
| 0 | 0 | 0 | 0 |
| 0 | 0 | 0 | 0 |
| 0 | 0 | 0 | 1 |
| 0 | 0 | 0 | 1 |
| 0 | 0 | 0 | 0 |
| 0 | 0 | 0 | 1 |
| 0 | 0 | 0 | 1 |
| 0 | 0 | 0 | 1 |
| 0 | 0 | 0 | 1 |
| 0 | 0 | 0 | 1 |
| 0 | 0 | 0 | 1 |
| 0 | 0 | 0 | 1 |
| 0 | 0 | 0 | 1 |
| 0 | 0 | 0 | 1 |

| | | | |
|---|---|---|---|
| 0 | 0 | 0 | 0 |
| 0 | 0 | 0 | 1 |
| 0 | 0 | 0 | 0 |
| 0 | 0 | 0 | 1 |
| 0 | 0 | 0 | 1 |
| 0 | 0 | 0 | 1 |
| 0 | 0 | 0 | 0 |
| 0 | 0 | 0 | 0 |
| 0 | 0 | 0 | 1 |
| 0 | 0 | 0 | 1 |
| 0 | 0 | 0 | 1 |
| 0 | 0 | 0 | 1 |
| 0 | 0 | 0 | 1 |
| 0 | 0 | 0 | 1 |
| 0 | 0 | 0 | 0 |
| 0 | 0 | 0 | 0 |
| 0 | 0 | 0 | 0 |
| 0 | 0 | 0 | 0 |
| 0 | 0 | 0 | 1 |
| 0 | 0 | 0 | 0 |
| 0 | 0 | 0 | 0 |
| 0 | 0 | 0 | 0 |
| 0 | 0 | 0 | 0 |
| 0 | 0 | 0 | 0 |
| 0 | 0 | 0 | 1 |
| 0 | 0 | 0 | 0 |
| 0 | 0 | 0 | 1 |
| 0 | 0 | 0 | 1 |
| 0 | 0 | 0 | 1 |
| 0 | 0 | 0 | 0 |
| 0 | 0 | 0 | 1 |
| 0 | 0 | 0 | 1 |
| 0 | 0 | 0 | 0 |
| 0 | 0 | 0 | 0 |
| 0 | 0 | 0 | 1 |
| 0 | 0 | 0 | 1 |
| 0 | 0 | 0 | 1 |
| 0 | 0 | 0 | 1 |
| 0 | 0 | 0 | 1 |
| 0 | 0 | 0 | 0 |
| 0 | 0 | 0 | 0 |
| 0 | 0 | 0 | 0 |
| 0 | 0 | 0 | 0 |
| 0 | 0 | 0 | 0 |
| 0 | 0 | 0 | 0 |
| 0 | 0 | 0 | 1 |
| 0 | 0 | 0 | 1 |
| 0 | 0 | 0 | 0 |
| 0 | 0 | 0 | 0 |
| 0 | 0 | 0 | 1 |
| 0 | 0 | 0 | 0 |
| 0 | 0 | 0 | 0 |
| 0 | 0 | 0 | 1 |
| 0 | 0 | 0 | 1 |

| | | | |
|---|---|---|---|
| 0 | 0 | 0 | 1 |
| 0 | 0 | 0 | 0 |
| 0 | 0 | 0 | 0 |
| 0 | 0 | 0 | 1 |
| 0 | 0 | 0 | 1 |
| 0 | 0 | 0 | 0 |
| 0 | 0 | 0 | 0 |
| 0 | 0 | 0 | 0 |
| 0 | 0 | 0 | 0 |
| 0 | 0 | 0 | 0 |
| 0 | 0 | 0 | 0 |
| 0 | 0 | 0 | 0 |
| 0 | 0 | 0 | 0 |
| 0 | 0 | 0 | 0 |
| 0 | 0 | 0 | 0 |
| 0 | 0 | 0 | 1 |
| 0 | 0 | 0 | 0 |
| 0 | 0 | 0 | 1 |
| 0 | 0 | 0 | 1 |
| 0 | 0 | 0 | 1 |
| 0 | 0 | 0 | 0 |
| 0 | 0 | 0 | 1 |
| 0 | 0 | 0 | 0 |
| 0 | 0 | 0 | 0 |
| 0 | 0 | 0 | 1 |
| 0 | 0 | 0 | 0 |
| 0 | 0 | 0 | 1 |
| 0 | 0 | 0 | 1 |
| 0 | 0 | 0 | 1 |
| 0 | 0 | 0 | 0 |
| 0 | 0 | 0 | 0 |
| 0 | 0 | 0 | 0 |
| 0 | 0 | 0 | 0 |
| 0 | 0 | 0 | 0 |
| 0 | 0 | 0 | 0 |
| 0 | 0 | 0 | 0 |
| 0 | 0 | 0 | 0 |
| 0 | 0 | 0 | 1 |
| 0 | 0 | 0 | 1 |
| 0 | 0 | 0 | 0 |
| 0 | 0 | 0 | 0 |
| 0 | 0 | 0 | 0 |
| 0 | 0 | 0 | 0 |
| 0 | 0 | 0 | 0 |
| 0 | 0 | 0 | 0 |
| 0 | 0 | 0 | 0 |
| 0 | 0 | 0 | 1 |
| 0 | 0 | 0 | 0 |
| 0 | 0 | 0 | 0 |
| 0 | 0 | 0 | 0 |
| 0 | 0 | 0 | 0 |
| 0 | 0 | 0 | 1 |
| 0 | 0 | 0 | 0 |
| 0 | 0 | 0 | 0 |

| | | | |
|---|---|---|---|
| 0 | 0 | 0 | 0 |
| 0 | 0 | 0 | 0 |
| 0 | 0 | 0 | 0 |
| 0 | 0 | 0 | 0 |
| 0 | 0 | 0 | 0 |
| 0 | 0 | 0 | 0 |
| 0 | 0 | 0 | 0 |
| 0 | 0 | 0 | 0 |
| 1 | 0 | 0 | 0 |
| 0 | 0 | 0 | 0 |
| 0 | 0 | 0 | 1 |
| 0 | 0 | 0 | 0 |
| 0 | 0 | 0 | 1 |
| 0 | 0 | 0 | 0 |
| 0 | 0 | 0 | 1 |
| 0 | 0 | 0 | 1 |
| 0 | 0 | 0 | 1 |
| 0 | 0 | 0 | 1 |
| 0 | 0 | 0 | 0 |
| 0 | 0 | 0 | 0 |
| 0 | 0 | 0 | 1 |
| 0 | 0 | 0 | 0 |
| 0 | 0 | 0 | 0 |
| 0 | 0 | 0 | 1 |
| 0 | 0 | 0 | 0 |
| 0 | 0 | 0 | 0 |
| 0 | 0 | 0 | 0 |
| 0 | 0 | 0 | 1 |
| 0 | 0 | 0 | 0 |
| 0 | 0 | 0 | 0 |
| 0 | 0 | 0 | 1 |
| 1 | 0 | 0 | 0 |
| 0 | 0 | 0 | 1 |
| 0 | 0 | 0 | 1 |
| 1 | 0 | 0 | 0 |
| 1 | 0 | 0 | 0 |
| 1 | 0 | 0 | 0 |
| 0 | 0 | 0 | 1 |
| 0 | 0 | 0 | 1 |
| 0 | 0 | 0 | 0 |
| 1 | 0 | 0 | 0 |
| 0 | 0 | 0 | 0 |
| 0 | 0 | 0 | 0 |
| 1 | 0 | 0 | 0 |
| 1 | 0 | 0 | 0 |
| 0 | 0 | 0 | 0 |
| 0 | 0 | 0 | 1 |
| 0 | 0 | 0 | 1 |
| 0 | 0 | 0 | 1 |
| 0 | 0 | 0 | 1 |
| 1 | 0 | 0 | 0 |
| 0 | 0 | 0 | 0 |
| 0 | 0 | 0 | 0 |
| 0 | 0 | 0 | 0 |

| | | | |
|---|---|---|---|
| 0 | 0 | 0 | 0 |
| 0 | 0 | 0 | 1 |
| 0 | 0 | 0 | 0 |
| 0 | 0 | 0 | 0 |
| 0 | 0 | 0 | 0 |
| 0 | 0 | 0 | 1 |
| 0 | 0 | 0 | 0 |
| 0 | 0 | 0 | 0 |
| 0 | 0 | 0 | 1 |
| 0 | 0 | 0 | 0 |
| 0 | 0 | 0 | 1 |
| 0 | 0 | 0 | 1 |
| 0 | 0 | 0 | 1 |
| 0 | 0 | 0 | 0 |
| 0 | 0 | 0 | 1 |
| 0 | 0 | 0 | 0 |
| 0 | 0 | 0 | 1 |
| 0 | 0 | 0 | 1 |
| 0 | 0 | 0 | 0 |
| 0 | 0 | 0 | 0 |
| 0 | 0 | 0 | 0 |
| 0 | 0 | 0 | 0 |
| 0 | 0 | 0 | 0 |
| 0 | 0 | 0 | 0 |
| 0 | 0 | 0 | 0 |
| 0 | 0 | 0 | 1 |
| 0 | 0 | 0 | 1 |
| 0 | 0 | 0 | 1 |
| 0 | 0 | 0 | 1 |
| 0 | 0 | 0 | 1 |
| 0 | 0 | 0 | 1 |
| 0 | 0 | 0 | 1 |
| 0 | 0 | 0 | 0 |
| 0 | 0 | 0 | 0 |
| 0 | 0 | 0 | 0 |
| 0 | 0 | 0 | 0 |
| 0 | 0 | 0 | 0 |
| 0 | 0 | 0 | 0 |
| 0 | 0 | 0 | 1 |
| 0 | 0 | 0 | 1 |
| 0 | 0 | 0 | 1 |
| 0 | 0 | 0 | 1 |
| 0 | 0 | 0 | 1 |
| 0 | 0 | 0 | 1 |
| 0 | 1 | 0 | 0 |
| 0 | 0 | 0 | 0 |
| 0 | 0 | 0 | 1 |
| 0 | 0 | 0 | 0 |
| 0 | 0 | 0 | 0 |
| 0 | 0 | 0 | 0 |
| 0 | 0 | 0 | 0 |
| 0 | 0 | 0 | 0 |
| 0 | 0 | 0 | 0 |
| 0 | 0 | 0 | 0 |

| | | | |
|---|---|---|---|
| 0 | 0 | 0 | 1 |
| 0 | 0 | 0 | 1 |
| 1 | 0 | 0 | 0 |
| 1 | 0 | 0 | 0 |
| 0 | 0 | 0 | 1 |
| 0 | 0 | 0 | 0 |
| 0 | 0 | 0 | 0 |
| 0 | 0 | 0 | 1 |
| 0 | 0 | 0 | 1 |
| 0 | 0 | 0 | 0 |
| 0 | 0 | 0 | 1 |
| 0 | 0 | 0 | 1 |
| 0 | 0 | 0 | 0 |
| 0 | 0 | 0 | 0 |
| 0 | 0 | 0 | 0 |
| 0 | 0 | 0 | 0 |
| 0 | 0 | 0 | 1 |
| 0 | 0 | 0 | 1 |
| 0 | 0 | 0 | 1 |
| 0 | 0 | 0 | 1 |
| 0 | 0 | 0 | 1 |
| 0 | 0 | 0 | 1 |
| 0 | 0 | 0 | 1 |
| 0 | 0 | 0 | 1 |
| 0 | 0 | 0 | 1 |
| 0 | 0 | 0 | 0 |
| 0 | 0 | 0 | 1 |
| 0 | 0 | 0 | 1 |
| 0 | 0 | 0 | 1 |
| 0 | 0 | 0 | 1 |
| 0 | 0 | 0 | 1 |
| 0 | 0 | 0 | 1 |
| 0 | 0 | 0 | 1 |
| 0 | 0 | 0 | 1 |
| 0 | 0 | 0 | 1 |
| 0 | 0 | 0 | 1 |
| 0 | 0 | 0 | 1 |
| 0 | 0 | 0 | 0 |
| 0 | 0 | 0 | 1 |
| 0 | 0 | 0 | 1 |
| 0 | 0 | 0 | 0 |
| 0 | 0 | 0 | 1 |
| 0 | 0 | 0 | 1 |
| 0 | 0 | 0 | 0 |
| 0 | 0 | 0 | 0 |
| 0 | 0 | 0 | 1 |
| 0 | 0 | 0 | 1 |
| 0 | 0 | 0 | 1 |
| 0 | 0 | 0 | 1 |
| 0 | 0 | 0 | 1 |
| 0 | 0 | 0 | 1 |
| 0 | 0 | 0 | 0 |
| 0 | 0 | 0 | 0 |
| 0 | 0 | 0 | 0 |

| | | | |
|---|---|---|---|
| 0 | 0 | 0 | 0 |
| 0 | 0 | 0 | 1 |
| 0 | 0 | 0 | 1 |
| 0 | 0 | 0 | 1 |
| 0 | 0 | 0 | 1 |
| 0 | 0 | 0 | 1 |
| 0 | 0 | 0 | 1 |
| 0 | 0 | 0 | 1 |
| 0 | 0 | 0 | 0 |
| 0 | 0 | 0 | 1 |
| 0 | 0 | 0 | 1 |
| 0 | 0 | 0 | 1 |
| 0 | 0 | 0 | 1 |
| 0 | 0 | 0 | 1 |
| 0 | 0 | 0 | 1 |
| 0 | 0 | 0 | 1 |
| 0 | 0 | 0 | 1 |
| 0 | 0 | 0 | 1 |
| 0 | 0 | 0 | 0 |
| 0 | 0 | 0 | 0 |
| 0 | 0 | 0 | 0 |
| 0 | 0 | 0 | 0 |
| 0 | 0 | 0 | 1 |
| 0 | 0 | 0 | 0 |
| 0 | 0 | 0 | 0 |
| 0 | 0 | 0 | 1 |
| 0 | 0 | 0 | 1 |
| 0 | 0 | 0 | 1 |
| 0 | 0 | 0 | 1 |
| 0 | 0 | 0 | 1 |
| 0 | 0 | 0 | 1 |
| 0 | 0 | 0 | 1 |
| 0 | 0 | 0 | 0 |
| 0 | 0 | 0 | 1 |
| 0 | 0 | 0 | 1 |
| 0 | 0 | 0 | 1 |
| 0 | 0 | 0 | 0 |
| 0 | 0 | 0 | 1 |
| 0 | 0 | 0 | 0 |
| 0 | 0 | 0 | 0 |
| 0 | 0 | 0 | 0 |
| 0 | 0 | 0 | 0 |
| 0 | 0 | 0 | 0 |
| 0 | 0 | 0 | 1 |
| 0 | 0 | 0 | 1 |
| 0 | 0 | 0 | 1 |
| 0 | 0 | 0 | 0 |
| 0 | 0 | 0 | 0 |
| 0 | 0 | 0 | 0 |
| 0 | 0 | 0 | 0 |
| 0 | 0 | 0 | 0 |
| 0 | 0 | 0 | 0 |
| 0 | 0 | 0 | 0 |
| 0 | 0 | 0 | 1 |

| | | | |
|---|---|---|---|
| 0 | 0 | 0 | 0 |
| 0 | 0 | 0 | 0 |
| 0 | 0 | 0 | 1 |
| 1 | 0 | 0 | 0 |
| 1 | 0 | 0 | 0 |
| 0 | 0 | 0 | 1 |
| 0 | 0 | 0 | 1 |
| 0 | 0 | 0 | 1 |
| 0 | 0 | 0 | 1 |
| 0 | 0 | 0 | 1 |
| 0 | 0 | 0 | 1 |
| 0 | 0 | 0 | 0 |
| 0 | 0 | 0 | 1 |
| 0 | 0 | 0 | 1 |
| 0 | 0 | 0 | 0 |
| 0 | 0 | 0 | 1 |
| 0 | 0 | 0 | 0 |
| 0 | 0 | 0 | 1 |
| 0 | 0 | 0 | 0 |
| 1 | 0 | 0 | 0 |
| 0 | 0 | 0 | 1 |
| 0 | 0 | 0 | 1 |
| 0 | 0 | 0 | 0 |
| 0 | 0 | 0 | 1 |
| 0 | 0 | 0 | 0 |
| 0 | 0 | 0 | 1 |
| 0 | 0 | 0 | 1 |
| 0 | 0 | 0 | 0 |
| 0 | 0 | 0 | 1 |
| 0 | 0 | 0 | 0 |
| 0 | 0 | 0 | 1 |
| 0 | 0 | 0 | 0 |
| 0 | 0 | 0 | 0 |
| 0 | 0 | 0 | 0 |
| 0 | 0 | 0 | 1 |
| 0 | 0 | 0 | 0 |
| 0 | 0 | 0 | 1 |
| 0 | 0 | 0 | 1 |
| 0 | 0 | 0 | 0 |
| 0 | 0 | 0 | 1 |
| 0 | 0 | 0 | 1 |
| 0 | 0 | 0 | 1 |
| 0 | 0 | 0 | 1 |
| 0 | 0 | 0 | 0 |
| 0 | 0 | 0 | 0 |
| 0 | 0 | 0 | 1 |
| 0 | 0 | 0 | 1 |
| 0 | 0 | 0 | 1 |
| 0 | 0 | 0 | 0 |
| 0 | 0 | 0 | 1 |
| 0 | 0 | 0 | 0 |
| 0 | 0 | 0 | 1 |
| 0 | 0 | 0 | 1 |
| 0 | 0 | 0 | 0 |

| | | | |
|---|---|---|---|
| 0 | 0 | 0 | 1 |
| 0 | 0 | 0 | 1 |
| 0 | 0 | 0 | 1 |
| 0 | 0 | 0 | 1 |
| 0 | 0 | 0 | 1 |
| 0 | 0 | 0 | 1 |
| 0 | 0 | 0 | 1 |
| 0 | 0 | 0 | 0 |
| 0 | 0 | 0 | 1 |
| 0 | 0 | 0 | 1 |
| 0 | 0 | 0 | 1 |
| 0 | 0 | 0 | 1 |
| 0 | 0 | 0 | 0 |
| 0 | 0 | 0 | 1 |
| 0 | 0 | 0 | 0 |
| 0 | 0 | 0 | 0 |
| 0 | 0 | 0 | 1 |
| 0 | 0 | 0 | 1 |
| 0 | 0 | 0 | 1 |
| 0 | 0 | 0 | 1 |
| 0 | 0 | 0 | 0 |
| 0 | 0 | 0 | 0 |
| 0 | 0 | 0 | 1 |
| 0 | 0 | 0 | 1 |
| 0 | 0 | 0 | 0 |
| 0 | 0 | 0 | 1 |
| 0 | 0 | 0 | 0 |
| 0 | 0 | 0 | 0 |
| 0 | 0 | 0 | 0 |
| 0 | 0 | 0 | 0 |
| 0 | 0 | 0 | 1 |
| 0 | 0 | 0 | 1 |
| 0 | 0 | 0 | 0 |
| 0 | 0 | 0 | 0 |
| 0 | 0 | 0 | 0 |
| 0 | 0 | 0 | 0 |
| 0 | 0 | 0 | 0 |
| 0 | 0 | 0 | 0 |
| 0 | 0 | 0 | 0 |
| 0 | 0 | 0 | 1 |
| 0 | 0 | 0 | 1 |
| 0 | 0 | 0 | 1 |
| 0 | 0 | 0 | 1 |
| 0 | 0 | 0 | 1 |
| 0 | 0 | 0 | 1 |
| 0 | 0 | 0 | 0 |
| 0 | 0 | 0 | 1 |
| 0 | 0 | 0 | 1 |
| 0 | 0 | 0 | 1 |
| 0 | 0 | 0 | 0 |
| 0 | 0 | 0 | 0 |
| 0 | 0 | 0 | 1 |
| 0 | 0 | 0 | 1 |
| 0 | 0 | 0 | 1 |

| | | | |
|---|---|---|---|
| 0 | 0 | 0 | 1 |
| 0 | 0 | 0 | 1 |
| 0 | 0 | 0 | 1 |
| 0 | 0 | 0 | 1 |
| 0 | 0 | 0 | 0 |
| 0 | 0 | 0 | 1 |
| 0 | 0 | 0 | 0 |
| 0 | 0 | 0 | 0 |
| 0 | 0 | 0 | 0 |
| 0 | 0 | 0 | 0 |
| 0 | 0 | 0 | 0 |
| 0 | 0 | 0 | 1 |
| 0 | 0 | 0 | 0 |
| 0 | 0 | 0 | 0 |
| 0 | 0 | 0 | 1 |
| 0 | 0 | 0 | 1 |
| 0 | 0 | 0 | 1 |
| 0 | 0 | 0 | 1 |
| 0 | 0 | 0 | 1 |
| 0 | 0 | 0 | 1 |
| 0 | 0 | 0 | 1 |
| 0 | 0 | 0 | 1 |
| 0 | 0 | 0 | 1 |
| 0 | 0 | 0 | 0 |
| 0 | 0 | 0 | 1 |
| 0 | 0 | 0 | 1 |
| 0 | 0 | 0 | 1 |
| 0 | 0 | 0 | 0 |
| 0 | 0 | 0 | 1 |
| 0 | 0 | 0 | 1 |
| 0 | 0 | 0 | 0 |
| 0 | 0 | 0 | 0 |
| 0 | 0 | 0 | 1 |
| 0 | 0 | 0 | 0 |
| 0 | 0 | 0 | 1 |
| 0 | 0 | 0 | 1 |
| 0 | 0 | 0 | 1 |
| 0 | 0 | 0 | 1 |
| 0 | 0 | 0 | 1 |
| 0 | 0 | 0 | 1 |
| 0 | 0 | 0 | 1 |
| 0 | 0 | 0 | 1 |
| 0 | 0 | 0 | 1 |
| 0 | 0 | 0 | 1 |
| 0 | 0 | 0 | 1 |
| 0 | 0 | 0 | 1 |
| 0 | 0 | 0 | 1 |
| 0 | 0 | 0 | 1 |
| 0 | 0 | 0 | 0 |
| 0 | 0 | 0 | 1 |
| 0 | 0 | 0 | 1 |
| 0 | 0 | 0 | 1 |
| 0 | 0 | 0 | 1 |
| 0 | 0 | 0 | 1 |

| | | | |
|---|---|---|---|
| 0 | 0 | 0 | 1 |
| 0 | 0 | 0 | 1 |
| 0 | 0 | 0 | 1 |
| 0 | 0 | 0 | 1 |
| 0 | 0 | 0 | 1 |
| 0 | 0 | 0 | 1 |
| 0 | 0 | 0 | 1 |
| 0 | 0 | 0 | 1 |
| 0 | 0 | 0 | 1 |
| 0 | 0 | 0 | 1 |
| 0 | 0 | 0 | 0 |
| 0 | 0 | 0 | 0 |
| 0 | 0 | 0 | 0 |
| 1 | 0 | 0 | 0 |
| 1 | 0 | 0 | 0 |
| 0 | 0 | 0 | 1 |
| 1 | 0 | 0 | 0 |
| 0 | 0 | 0 | 0 |
| 0 | 0 | 0 | 0 |
| 0 | 0 | 0 | 0 |
| 0 | 0 | 0 | 0 |
| 1 | 0 | 0 | 0 |
| 0 | 0 | 0 | 1 |
| 0 | 0 | 0 | 0 |
| 1 | 0 | 0 | 0 |
| 1 | 0 | 0 | 0 |
| 0 | 0 | 0 | 1 |
| 0 | 0 | 0 | 0 |
| 0 | 0 | 0 | 1 |
| 0 | 0 | 0 | 1 |
| 0 | 0 | 0 | 1 |
| 0 | 0 | 0 | 0 |
| 0 | 0 | 0 | 1 |
| 0 | 0 | 0 | 1 |
| 0 | 0 | 0 | 0 |
| 0 | 0 | 0 | 0 |
| 0 | 0 | 0 | 0 |
| 0 | 0 | 0 | 0 |
| 0 | 0 | 0 | 0 |
| 0 | 0 | 0 | 1 |
| 0 | 0 | 0 | 1 |
| 0 | 0 | 0 | 0 |
| 0 | 0 | 0 | 1 |
| 0 | 0 | 0 | 0 |
| 0 | 0 | 0 | 1 |
| 0 | 0 | 0 | 1 |
| 0 | 0 | 0 | 1 |
| 0 | 0 | 0 | 0 |
| 0 | 0 | 0 | 0 |
| 0 | 0 | 0 | 0 |
| 0 | 0 | 0 | 0 |
| 0 | 0 | 0 | 1 |
| 0 | 0 | 0 | 0 |
| 0 | 0 | 0 | 1 |

| | | | |
|---|---|---|---|
| 0 | 0 | 0 | 1 |
| 0 | 0 | 0 | 1 |
| 0 | 0 | 0 | 1 |
| 0 | 0 | 0 | 1 |
| 0 | 0 | 0 | 1 |
| 0 | 0 | 0 | 0 |
| 0 | 0 | 0 | 0 |
| 0 | 0 | 0 | 0 |
| 0 | 0 | 0 | 0 |
| 0 | 0 | 0 | 1 |
| 1 | 0 | 0 | 0 |
| 1 | 0 | 0 | 0 |
| 0 | 0 | 0 | 0 |
| 0 | 0 | 0 | 0 |
| 0 | 0 | 0 | 0 |
| 0 | 0 | 0 | 0 |
| 0 | 0 | 0 | 1 |
| 0 | 0 | 0 | 1 |
| 0 | 0 | 0 | 1 |
| 0 | 0 | 0 | 1 |
| 0 | 0 | 0 | 1 |
| 0 | 0 | 0 | 1 |
| 0 | 0 | 0 | 1 |
| 0 | 0 | 0 | 1 |
| 0 | 0 | 0 | 1 |
| 0 | 0 | 0 | 1 |
| 0 | 0 | 0 | 1 |
| 0 | 0 | 0 | 1 |
| 0 | 0 | 0 | 0 |
| 0 | 0 | 0 | 1 |
| 0 | 0 | 0 | 1 |
| 0 | 0 | 0 | 1 |
| 0 | 0 | 0 | 1 |
| 0 | 0 | 0 | 0 |
| 0 | 0 | 0 | 1 |
| 0 | 0 | 0 | 0 |
| 0 | 0 | 0 | 1 |
| 0 | 0 | 0 | 0 |
| 0 | 0 | 0 | 1 |
| 0 | 0 | 0 | 0 |
| 0 | 0 | 0 | 1 |
| 0 | 0 | 0 | 1 |
| 0 | 0 | 0 | 1 |
| 0 | 0 | 0 | 0 |
| 0 | 0 | 0 | 1 |
| 0 | 0 | 0 | 1 |
| 0 | 0 | 0 | 1 |
| 0 | 0 | 0 | 1 |
| 0 | 0 | 0 | 1 |
| 0 | 0 | 0 | 1 |
| 0 | 0 | 0 | 1 |
| 0 | 0 | 0 | 1 |
| 0 | 0 | 0 | 0 |
| 0 | 0 | 0 | 0 |

| | | | |
|---|---|---|---|
| 0 | 0 | 0 | 1 |
| 0 | 0 | 0 | 0 |
| 0 | 0 | 0 | 1 |
| 0 | 0 | 0 | 1 |
| 0 | 0 | 0 | 0 |
| 0 | 0 | 0 | 0 |
| 0 | 0 | 0 | 0 |
| 0 | 0 | 0 | 1 |
| 0 | 0 | 0 | 1 |
| 0 | 0 | 0 | 0 |
| 0 | 0 | 0 | 0 |
| 0 | 0 | 0 | 0 |
| 0 | 0 | 0 | 0 |
| 0 | 0 | 0 | 0 |
| 0 | 0 | 0 | 1 |
| 0 | 0 | 0 | 0 |
| 0 | 0 | 0 | 0 |
| 0 | 0 | 0 | 0 |
| 0 | 0 | 0 | 1 |
| 0 | 0 | 0 | 1 |
| 0 | 0 | 0 | 1 |
| 0 | 0 | 0 | 1 |
| 0 | 0 | 0 | 1 |
| 0 | 0 | 0 | 0 |
| 0 | 0 | 0 | 1 |
| 0 | 0 | 0 | 1 |
| 0 | 0 | 0 | 0 |
| 0 | 0 | 0 | 0 |
| 0 | 0 | 0 | 1 |
| 0 | 0 | 0 | 1 |
| 0 | 0 | 0 | 0 |
| 0 | 0 | 0 | 1 |
| 0 | 0 | 0 | 1 |
| 0 | 0 | 0 | 1 |
| 0 | 0 | 0 | 1 |
| 0 | 0 | 0 | 0 |
| 0 | 0 | 0 | 1 |
| 0 | 0 | 0 | 0 |
| 0 | 0 | 0 | 0 |
| 0 | 0 | 0 | 1 |
| 0 | 0 | 0 | 1 |
| 0 | 0 | 0 | 0 |
| 0 | 0 | 0 | 0 |
| 0 | 0 | 0 | 1 |
| 0 | 0 | 0 | 0 |
| 0 | 0 | 0 | 0 |
| 0 | 0 | 0 | 0 |
| 0 | 0 | 0 | 1 |
| 0 | 0 | 0 | 0 |
| 0 | 0 | 0 | 0 |
| 0 | 0 | 0 | 0 |
| 0 | 0 | 0 | 1 |
| 0 | 0 | 0 | 0 |
| 0 | 0 | 0 | 1 |

| | | | |
|---|---|---|---|
| 0 | 0 | 0 | 1 |
| 0 | 0 | 0 | 1 |
| 0 | 0 | 0 | 1 |
| 0 | 0 | 0 | 1 |
| 0 | 0 | 0 | 0 |
| 0 | 0 | 0 | 0 |
| 0 | 0 | 0 | 1 |
| 0 | 0 | 0 | 1 |
| 0 | 0 | 0 | 1 |
| 0 | 0 | 0 | 0 |
| 0 | 0 | 0 | 1 |
| 0 | 0 | 0 | 1 |
| 0 | 0 | 0 | 0 |
| 0 | 0 | 0 | 1 |
| 0 | 0 | 0 | 1 |
| 0 | 0 | 0 | 1 |
| 0 | 0 | 0 | 1 |
| 0 | 0 | 0 | 1 |
| 0 | 0 | 0 | 1 |
| 0 | 0 | 0 | 1 |
| 0 | 0 | 0 | 1 |
| 0 | 0 | 0 | 1 |
| 0 | 0 | 0 | 1 |
| 0 | 0 | 0 | 0 |
| 0 | 0 | 0 | 1 |
| 1 | 0 | 0 | 0 |
| 0 | 0 | 0 | 1 |
| 0 | 0 | 0 | 0 |
| 0 | 0 | 0 | 1 |
| 0 | 0 | 0 | 0 |
| 0 | 0 | 0 | 0 |
| 0 | 0 | 0 | 0 |
| 0 | 0 | 0 | 0 |
| 0 | 0 | 0 | 0 |
| 0 | 0 | 0 | 1 |
| 0 | 0 | 0 | 0 |
| 0 | 0 | 0 | 1 |
| 0 | 0 | 0 | 0 |
| 0 | 0 | 0 | 1 |
| 0 | 0 | 0 | 1 |
| 0 | 0 | 0 | 0 |
| 0 | 0 | 0 | 0 |
| 0 | 0 | 0 | 0 |
| 0 | 0 | 0 | 0 |
| 0 | 0 | 0 | 0 |
| 0 | 0 | 0 | 0 |
| 0 | 0 | 0 | 0 |
| 0 | 0 | 0 | 0 |
| 0 | 0 | 0 | 1 |
| 0 | 0 | 0 | 0 |
| 0 | 0 | 0 | 1 |
| 0 | 0 | 0 | 1 |
| 0 | 0 | 0 | 1 |
| 0 | 0 | 0 | 1 |

| | | | |
|---|---|---|---|
| 0 | 0 | 0 | 1 |
| 0 | 0 | 0 | 1 |
| 0 | 0 | 0 | 1 |
| 0 | 0 | 0 | 1 |
| 0 | 0 | 0 | 1 |
| 0 | 0 | 0 | 1 |
| 0 | 0 | 0 | 0 |
| 0 | 0 | 0 | 0 |
| 0 | 0 | 0 | 0 |
| 0 | 0 | 0 | 0 |
| 0 | 0 | 0 | 0 |
| 0 | 0 | 0 | 0 |
| 0 | 0 | 0 | 0 |
| 0 | 0 | 0 | 0 |
| 0 | 0 | 0 | 0 |
| 0 | 0 | 0 | 0 |
| 1 | 0 | 0 | 0 |
| 0 | 0 | 0 | 0 |
| 0 | 0 | 0 | 1 |
| 0 | 0 | 0 | 0 |
| 0 | 0 | 0 | 1 |
| 0 | 0 | 0 | 1 |
| 0 | 0 | 0 | 1 |
| 0 | 0 | 0 | 1 |
| 0 | 0 | 0 | 1 |
| 0 | 0 | 0 | 0 |
| 0 | 0 | 0 | 0 |
| 0 | 0 | 0 | 0 |
| 0 | 0 | 0 | 0 |
| 0 | 0 | 0 | 0 |
| 0 | 0 | 0 | 0 |
| 1 | 0 | 0 | 0 |
| 0 | 0 | 0 | 0 |
| 1 | 0 | 0 | 0 |
| 0 | 0 | 0 | 1 |
| 1 | 0 | 0 | 0 |
| 1 | 0 | 0 | 0 |
| 0 | 0 | 0 | 0 |
| 1 | 0 | 0 | 0 |
| 0 | 0 | 0 | 1 |
| 0 | 0 | 0 | 0 |
| 0 | 0 | 0 | 0 |
| 0 | 0 | 0 | 1 |
| 0 | 0 | 0 | 1 |
| 0 | 0 | 0 | 1 |
| 0 | 0 | 0 | 1 |
| 0 | 0 | 0 | 1 |
| 0 | 0 | 0 | 0 |
| 0 | 0 | 0 | 0 |
| 0 | 0 | 0 | 0 |
| 0 | 0 | 0 | 0 |
| 0 | 0 | 0 | 1 |
| 0 | 0 | 0 | 0 |
| 0 | 0 | 0 | 1 |

| | | | |
|---|---|---|---|
| 0 | 0 | 0 | 1 |
| 0 | 0 | 0 | 1 |
| 0 | 0 | 0 | 1 |
| 0 | 0 | 0 | 1 |
| 0 | 0 | 0 | 1 |
| 0 | 0 | 0 | 0 |
| 0 | 0 | 0 | 1 |
| 0 | 0 | 0 | 1 |
| 0 | 0 | 0 | 1 |
| 0 | 0 | 0 | 1 |
| 0 | 0 | 0 | 0 |
| 0 | 0 | 0 | 0 |
| 0 | 0 | 0 | 0 |
| 0 | 0 | 0 | 1 |
| 0 | 0 | 0 | 1 |
| 0 | 0 | 0 | 1 |
| 0 | 0 | 0 | 1 |
| 0 | 0 | 0 | 0 |
| 0 | 0 | 0 | 1 |
| 0 | 0 | 0 | 1 |
| 0 | 0 | 0 | 1 |
| 0 | 0 | 0 | 1 |
| 0 | 0 | 0 | 0 |
| 0 | 0 | 0 | 1 |
| 0 | 0 | 0 | 0 |
| 0 | 0 | 0 | 0 |
| 0 | 0 | 0 | 0 |
| 0 | 0 | 0 | 0 |
| 0 | 0 | 0 | 1 |
| 0 | 0 | 0 | 0 |
| 0 | 0 | 0 | 1 |
| 0 | 0 | 0 | 1 |
| 0 | 0 | 0 | 1 |
| 0 | 0 | 0 | 1 |
| 0 | 0 | 0 | 1 |
| 0 | 0 | 0 | 1 |
| 0 | 0 | 0 | 1 |
| 0 | 0 | 0 | 0 |
| 0 | 0 | 0 | 0 |
| 0 | 0 | 0 | 0 |
| 0 | 0 | 0 | 0 |
| 0 | 0 | 0 | 0 |
| 0 | 0 | 0 | 1 |
| 0 | 0 | 0 | 0 |
| 0 | 0 | 0 | 0 |
| 0 | 0 | 0 | 1 |
| 0 | 0 | 0 | 0 |
| 0 | 0 | 0 | 0 |
| 0 | 0 | 0 | 0 |
| 1 | 0 | 0 | 0 |
| 0 | 0 | 0 | 1 |
| 0 | 0 | 0 | 1 |
| 0 | 0 | 0 | 1 |
| 0 | 0 | 0 | 0 |

| | | | |
|---|---|---|---|
| 0 | 0 | 0 | 0 |
| 0 | 0 | 0 | 1 |
| 0 | 0 | 0 | 0 |
| 0 | 0 | 0 | 1 |
| 0 | 0 | 0 | 1 |
| 0 | 0 | 0 | 1 |
| 0 | 0 | 0 | 1 |
| 0 | 0 | 0 | 1 |
| 0 | 0 | 0 | 0 |
| 0 | 0 | 0 | 1 |
| 0 | 0 | 0 | 0 |
| 0 | 0 | 0 | 0 |
| 0 | 0 | 0 | 0 |
| 0 | 0 | 0 | 0 |
| 0 | 0 | 0 | 1 |
| 0 | 0 | 0 | 1 |
| 0 | 0 | 0 | 0 |
| 0 | 0 | 0 | 0 |
| 0 | 0 | 0 | 1 |
| 0 | 0 | 0 | 1 |
| 0 | 0 | 0 | 0 |
| 0 | 0 | 0 | 1 |
| 0 | 0 | 0 | 1 |
| 0 | 0 | 0 | 1 |
| 0 | 0 | 0 | 0 |
| 0 | 0 | 0 | 1 |
| 0 | 0 | 0 | 0 |
| 0 | 0 | 0 | 0 |
| 0 | 0 | 0 | 0 |
| 0 | 0 | 0 | 1 |
| 0 | 0 | 0 | 0 |
| 0 | 0 | 0 | 0 |
| 0 | 0 | 0 | 0 |
| 0 | 0 | 0 | 1 |
| 0 | 0 | 0 | 0 |
| 0 | 0 | 0 | 1 |
| 0 | 0 | 0 | 0 |
| 0 | 0 | 0 | 0 |
| 0 | 0 | 0 | 0 |
| 0 | 0 | 0 | 0 |
| 0 | 0 | 0 | 0 |
| 0 | 0 | 0 | 1 |
| 0 | 0 | 0 | 0 |
| 0 | 0 | 0 | 0 |
| 0 | 1 | 0 | 0 |
| 1 | 0 | 0 | 0 |
| 0 | 0 | 0 | 0 |
| 1 | 0 | 0 | 0 |
| 1 | 0 | 0 | 0 |
| 1 | 0 | 0 | 0 |
| 0 | 0 | 0 | 0 |
| 0 | 0 | 0 | 0 |
| 0 | 0 | 0 | 1 |
| 0 | 0 | 0 | 0 |

| | | | |
|---|---|---|---|
| 0 | 0 | 0 | 1 |
| 0 | 0 | 0 | 1 |
| 0 | 0 | 0 | 1 |
| 0 | 0 | 0 | 1 |
| 0 | 0 | 0 | 1 |
| 0 | 0 | 0 | 1 |
| 0 | 0 | 0 | 1 |
| 0 | 0 | 0 | 1 |
| 0 | 0 | 0 | 1 |
| 0 | 0 | 0 | 0 |
| 0 | 0 | 0 | 0 |
| 0 | 0 | 0 | 1 |
| 0 | 0 | 0 | 1 |
| 0 | 0 | 0 | 1 |
| 0 | 0 | 0 | 0 |
| 0 | 0 | 0 | 1 |
| 0 | 0 | 0 | 1 |
| 0 | 0 | 0 | 0 |
| 0 | 0 | 0 | 1 |
| 0 | 0 | 0 | 1 |
| 0 | 0 | 0 | 1 |
| 0 | 0 | 0 | 0 |
| 0 | 0 | 0 | 0 |
| 0 | 0 | 0 | 0 |
| 0 | 0 | 0 | 0 |
| 0 | 0 | 0 | 0 |
| 0 | 0 | 0 | 0 |
| 0 | 0 | 0 | 0 |
| 0 | 0 | 0 | 1 |
| 0 | 0 | 0 | 0 |
| 1 | 0 | 0 | 0 |
| 0 | 0 | 0 | 1 |
| 0 | 0 | 0 | 1 |
| 0 | 0 | 0 | 1 |
| 0 | 0 | 0 | 1 |
| 0 | 0 | 0 | 1 |
| 0 | 0 | 0 | 1 |
| 0 | 0 | 0 | 1 |
| 0 | 0 | 0 | 1 |
| 0 | 0 | 0 | 1 |
| 0 | 0 | 0 | 1 |
| 0 | 0 | 0 | 1 |
| 0 | 0 | 0 | 1 |
| 0 | 0 | 0 | 0 |
| 0 | 0 | 0 | 1 |
| 0 | 0 | 0 | 0 |
| 0 | 0 | 0 | 1 |
| 0 | 0 | 0 | 1 |
| 0 | 0 | 0 | 1 |
| 0 | 0 | 0 | 1 |
| 0 | 0 | 0 | 0 |
| 0 | 0 | 0 | 1 |
| 0 | 0 | 0 | 1 |
| 0 | 0 | 0 | 0 |

| | | | |
|---|---|---|---|
| 0 | 0 | 0 | 0 |
| 0 | 0 | 0 | 0 |
| 0 | 0 | 0 | 0 |
| 0 | 0 | 0 | 0 |
| 0 | 0 | 0 | 1 |
| 0 | 0 | 0 | 1 |
| 0 | 0 | 0 | 1 |
| 0 | 0 | 0 | 0 |
| 0 | 0 | 0 | 0 |
| 0 | 0 | 0 | 0 |
| 0 | 0 | 0 | 0 |
| 0 | 0 | 0 | 1 |
| 0 | 0 | 0 | 1 |
| 0 | 0 | 0 | 1 |
| 0 | 0 | 0 | 1 |
| 0 | 0 | 0 | 0 |
| 0 | 0 | 0 | 0 |
| 0 | 0 | 0 | 1 |
| 0 | 0 | 0 | 1 |
| 0 | 0 | 0 | 1 |
| 0 | 0 | 0 | 1 |
| 0 | 0 | 0 | 0 |
| 0 | 0 | 0 | 1 |
| 0 | 0 | 0 | 1 |
| 0 | 0 | 0 | 0 |
| 0 | 0 | 0 | 0 |
| 0 | 0 | 0 | 0 |
| 0 | 0 | 0 | 0 |
| 0 | 0 | 0 | 1 |
| 0 | 0 | 0 | 1 |
| 0 | 0 | 0 | 1 |
| 0 | 0 | 0 | 0 |
| 0 | 0 | 0 | 1 |
| 0 | 0 | 0 | 1 |
| 0 | 0 | 0 | 1 |
| 0 | 0 | 0 | 1 |
| 0 | 0 | 0 | 1 |
| 0 | 0 | 0 | 1 |
| 0 | 0 | 0 | 1 |
| 0 | 0 | 0 | 1 |
| 0 | 0 | 0 | 1 |
| 0 | 0 | 0 | 1 |
| 0 | 0 | 0 | 0 |
| 0 | 0 | 0 | 1 |
| 0 | 0 | 0 | 1 |
| 0 | 0 | 0 | 1 |
| 0 | 0 | 0 | 1 |
| 0 | 0 | 0 | 1 |
| 0 | 0 | 0 | 1 |
| 0 | 0 | 0 | 0 |
| 0 | 0 | 0 | 1 |
| 0 | 0 | 0 | 1 |
| 0 | 0 | 0 | 1 |
| 0 | 0 | 0 | 1 |

| | | | |
|---|---|---|---|
| 0 | 0 | 0 | 0 |
| 0 | 0 | 0 | 1 |
| 0 | 0 | 0 | 1 |
| 0 | 0 | 0 | 1 |
| 0 | 0 | 0 | 1 |
| 0 | 0 | 0 | 1 |
| 0 | 0 | 0 | 1 |
| 0 | 0 | 0 | 0 |
| 0 | 0 | 0 | 1 |
| 0 | 0 | 0 | 1 |
| 0 | 0 | 0 | 0 |
| 0 | 0 | 0 | 0 |
| 0 | 0 | 0 | 0 |
| 0 | 0 | 0 | 1 |
| 0 | 0 | 0 | 0 |
| 0 | 0 | 0 | 0 |
| 0 | 0 | 0 | 0 |
| 0 | 0 | 0 | 0 |
| 0 | 0 | 0 | 1 |
| 0 | 0 | 0 | 0 |
| 0 | 0 | 0 | 0 |
| 0 | 0 | 0 | 0 |
| 0 | 0 | 0 | 0 |
| 0 | 0 | 0 | 0 |
| 0 | 0 | 0 | 0 |
| 0 | 0 | 0 | 0 |
| 0 | 0 | 0 | 0 |
| 0 | 0 | 0 | 0 |
| 0 | 0 | 0 | 0 |
| 0 | 0 | 0 | 0 |
| 0 | 0 | 0 | 0 |
| 0 | 0 | 0 | 0 |
| 0 | 0 | 0 | 1 |
| 0 | 0 | 0 | 0 |
| 0 | 0 | 0 | 1 |
| 0 | 0 | 0 | 0 |
| 0 | 0 | 0 | 0 |
| 0 | 0 | 0 | 1 |
| 0 | 0 | 0 | 1 |
| 0 | 0 | 0 | 1 |
| 0 | 0 | 0 | 1 |
| 0 | 0 | 0 | 0 |
| 0 | 0 | 0 | 0 |
| 0 | 0 | 0 | 0 |
| 0 | 0 | 0 | 1 |
| 0 | 0 | 0 | 0 |
| 0 | 0 | 0 | 1 |
| 0 | 0 | 0 | 0 |
| 0 | 0 | 0 | 1 |
| 0 | 0 | 0 | 1 |
| 0 | 0 | 0 | 1 |
| 0 | 0 | 0 | 1 |
| 0 | 0 | 0 | 1 |
| 0 | 0 | 0 | 1 |

| | | | |
|---|---|---|---|
| 0 | 0 | 0 | 1 |
| 0 | 0 | 0 | 1 |
| 0 | 0 | 0 | 1 |
| 0 | 0 | 0 | 1 |
| 0 | 0 | 0 | 1 |
| 0 | 0 | 0 | 0 |
| 0 | 0 | 0 | 0 |
| 0 | 0 | 0 | 1 |
| 0 | 0 | 0 | 0 |
| 0 | 0 | 0 | 0 |
| 0 | 0 | 0 | 1 |
| 0 | 0 | 0 | 1 |
| 0 | 0 | 0 | 1 |
| 0 | 0 | 0 | 1 |
| 0 | 0 | 0 | 1 |
| 0 | 0 | 0 | 1 |
| 0 | 0 | 0 | 1 |
| 0 | 0 | 0 | 1 |
| 0 | 0 | 0 | 0 |
| 0 | 0 | 0 | 0 |
| 0 | 0 | 0 | 1 |
| 0 | 0 | 0 | 1 |
| 0 | 0 | 0 | 1 |
| 0 | 0 | 0 | 0 |
| 0 | 0 | 0 | 1 |
| 0 | 0 | 0 | 0 |
| 0 | 0 | 0 | 0 |
| 0 | 0 | 0 | 1 |
| 0 | 0 | 0 | 1 |
| 0 | 0 | 0 | 1 |
| 0 | 0 | 0 | 0 |
| 0 | 0 | 0 | 1 |
| 0 | 0 | 0 | 0 |
| 0 | 0 | 0 | 0 |
| 0 | 0 | 0 | 0 |
| 0 | 0 | 0 | 0 |
| 0 | 0 | 0 | 0 |
| 0 | 0 | 0 | 1 |
| 0 | 0 | 0 | 1 |
| 0 | 0 | 0 | 1 |
| 0 | 0 | 0 | 0 |
| 0 | 0 | 0 | 0 |
| 0 | 0 | 0 | 0 |
| 0 | 0 | 0 | 0 |
| 0 | 0 | 0 | 1 |
| 0 | 0 | 0 | 1 |
| 0 | 0 | 0 | 1 |
| 0 | 0 | 0 | 0 |
| 0 | 0 | 0 | 0 |
| 0 | 0 | 0 | 0 |
| 0 | 0 | 0 | 1 |
| 0 | 0 | 0 | 1 |
| 0 | 0 | 0 | 1 |
| 0 | 0 | 0 | 1 |

| | | | |
|---|---|---|---|
| 0 | 0 | 0 | 1 |
| 0 | 0 | 0 | 1 |
| 0 | 0 | 0 | 1 |
| 0 | 0 | 0 | 0 |
| 0 | 0 | 0 | 0 |
| 0 | 0 | 0 | 1 |
| 0 | 0 | 0 | 0 |
| 0 | 0 | 0 | 1 |
| 0 | 0 | 0 | 1 |
| 0 | 0 | 0 | 1 |
| 0 | 0 | 0 | 1 |
| 0 | 0 | 0 | 0 |
| 0 | 0 | 0 | 1 |
| 0 | 0 | 0 | 0 |
| 0 | 0 | 0 | 1 |
| 0 | 0 | 0 | 0 |
| 0 | 0 | 0 | 1 |
| 0 | 0 | 0 | 1 |
| 0 | 0 | 0 | 1 |
| 0 | 0 | 0 | 0 |
| 0 | 0 | 0 | 0 |
| 0 | 0 | 0 | 0 |
| 0 | 0 | 0 | 0 |
| 0 | 0 | 0 | 0 |
| 0 | 0 | 0 | 0 |
| 0 | 0 | 0 | 1 |
| 0 | 0 | 0 | 0 |
| 0 | 0 | 0 | 1 |
| 0 | 0 | 0 | 1 |
| 0 | 0 | 0 | 1 |
| 0 | 0 | 0 | 1 |
| 0 | 0 | 0 | 1 |
| 0 | 0 | 0 | 0 |
| 0 | 1 | 0 | 0 |
| 0 | 0 | 0 | 1 |
| 0 | 0 | 0 | 1 |
| 0 | 0 | 0 | 0 |
| 0 | 0 | 0 | 0 |
| 0 | 0 | 0 | 0 |
| 0 | 0 | 0 | 1 |
| 0 | 0 | 0 | 0 |
| 0 | 0 | 0 | 0 |
| 0 | 0 | 0 | 1 |
| 0 | 0 | 0 | 0 |
| 0 | 0 | 0 | 1 |
| 0 | 0 | 0 | 1 |
| 0 | 0 | 0 | 1 |
| 0 | 0 | 0 | 1 |
| 0 | 0 | 0 | 1 |
| 1 | 0 | 0 | 0 |
| 0 | 0 | 0 | 1 |
| 0 | 0 | 0 | 1 |
| 0 | 0 | 0 | 1 |
| 0 | 0 | 0 | 1 |

| | | | |
|---|---|---|---|
| 0 | 0 | 0 | 1 |
| 0 | 0 | 0 | 1 |
| 0 | 0 | 0 | 1 |
| 0 | 0 | 0 | 0 |
| 0 | 0 | 0 | 1 |
| 0 | 0 | 0 | 1 |
| 0 | 0 | 0 | 1 |
| 0 | 0 | 0 | 0 |
| 0 | 0 | 0 | 1 |
| 0 | 0 | 0 | 1 |
| 0 | 0 | 0 | 1 |
| 0 | 0 | 0 | 1 |
| 0 | 0 | 0 | 1 |
| 0 | 0 | 0 | 1 |
| 0 | 0 | 0 | 1 |
| 0 | 0 | 0 | 1 |
| 0 | 0 | 0 | 0 |
| 0 | 0 | 0 | 0 |
| 0 | 0 | 0 | 1 |
| 0 | 0 | 0 | 1 |
| 0 | 0 | 0 | 1 |
| 0 | 0 | 0 | 1 |
| 0 | 0 | 0 | 1 |
| 0 | 0 | 0 | 1 |
| 0 | 0 | 0 | 1 |
| 0 | 0 | 0 | 1 |
| 0 | 0 | 0 | 1 |
| 0 | 0 | 0 | 0 |
| 0 | 0 | 0 | 0 |
| 0 | 0 | 0 | 1 |
| 0 | 0 | 0 | 1 |
| 0 | 0 | 0 | 0 |
| 0 | 0 | 0 | 1 |
| 0 | 0 | 0 | 0 |
| 1 | 0 | 0 | 0 |
| 1 | 0 | 0 | 0 |
| 1 | 0 | 0 | 0 |
| 1 | 0 | 0 | 0 |
| 0 | 0 | 0 | 1 |
| 0 | 0 | 0 | 1 |
| 0 | 0 | 0 | 1 |
| 0 | 0 | 0 | 1 |
| 0 | 0 | 0 | 1 |
| 0 | 0 | 0 | 1 |
| 0 | 0 | 0 | 0 |
| 0 | 0 | 0 | 0 |
| 0 | 0 | 0 | 0 |
| 0 | 0 | 0 | 1 |
| 0 | 0 | 0 | 1 |
| 0 | 0 | 0 | 1 |
| 0 | 0 | 0 | 1 |
| 0 | 0 | 0 | 0 |
| 0 | 0 | 0 | 0 |
| 0 | 0 | 0 | 0 |

| | | | |
|---|---|---|---|
| 0 | 0 | 0 | 0 |
| 0 | 0 | 0 | 0 |
| 0 | 0 | 0 | 1 |
| 0 | 0 | 0 | 0 |
| 0 | 0 | 0 | 0 |
| 0 | 0 | 0 | 0 |
| 0 | 0 | 0 | 0 |
| 0 | 0 | 0 | 1 |
| 0 | 0 | 0 | 1 |
| 0 | 0 | 0 | 1 |
| 0 | 0 | 0 | 1 |
| 0 | 0 | 0 | 1 |
| 0 | 0 | 0 | 1 |
| 0 | 0 | 0 | 1 |
| 0 | 0 | 0 | 1 |
| 0 | 0 | 0 | 1 |
| 0 | 0 | 0 | 1 |
| 0 | 0 | 0 | 1 |
| 0 | 0 | 0 | 1 |
| 0 | 0 | 0 | 1 |
| 0 | 0 | 0 | 0 |
| 0 | 0 | 0 | 0 |
| 0 | 0 | 0 | 0 |
| 0 | 0 | 0 | 0 |
| 0 | 0 | 0 | 0 |
| 0 | 0 | 0 | 1 |
| 0 | 0 | 0 | 1 |
| 0 | 0 | 0 | 0 |
| 0 | 0 | 0 | 1 |
| 0 | 0 | 0 | 1 |
| 0 | 0 | 0 | 0 |
| 0 | 0 | 0 | 0 |
| 0 | 0 | 0 | 0 |
| 0 | 0 | 0 | 1 |
| 0 | 0 | 0 | 0 |
| 0 | 0 | 0 | 1 |
| 0 | 0 | 0 | 1 |
| 0 | 0 | 0 | 0 |
| 0 | 0 | 0 | 1 |
| 0 | 0 | 0 | 1 |
| 0 | 0 | 0 | 1 |
| 0 | 0 | 0 | 1 |
| 0 | 0 | 0 | 1 |
| 0 | 0 | 0 | 1 |
| 0 | 0 | 0 | 1 |
| 0 | 0 | 0 | 1 |
| 0 | 0 | 0 | 1 |
| 0 | 0 | 0 | 1 |
| 0 | 0 | 0 | 0 |
| 0 | 0 | 0 | 1 |
| 1 | 0 | 0 | 0 |
| 1 | 0 | 0 | 0 |
| 0 | 0 | 0 | 1 |
| 0 | 0 | 0 | 1 |

| | | | |
|---|---|---|---|
| 1 | 0 | 0 | 0 |
| 0 | 0 | 0 | 1 |
| 0 | 0 | 0 | 1 |
| 0 | 0 | 0 | 1 |
| 0 | 0 | 0 | 1 |
| 1 | 0 | 0 | 0 |
| 1 | 0 | 0 | 0 |
| 0 | 0 | 0 | 1 |
| 0 | 0 | 0 | 1 |
| 0 | 0 | 0 | 1 |
| 0 | 0 | 0 | 1 |
| 0 | 0 | 0 | 0 |
| 0 | 0 | 0 | 1 |
| 0 | 0 | 0 | 1 |
| 0 | 0 | 0 | 1 |
| 0 | 0 | 0 | 1 |
| 0 | 0 | 0 | 0 |
| 0 | 0 | 0 | 0 |
| 0 | 0 | 0 | 1 |
| 0 | 0 | 0 | 1 |
| 0 | 0 | 0 | 1 |
| 0 | 0 | 0 | 1 |
| 0 | 0 | 0 | 1 |
| 0 | 0 | 0 | 1 |
| 0 | 0 | 0 | 0 |
| 0 | 0 | 0 | 1 |
| 0 | 0 | 0 | 1 |
| 0 | 0 | 0 | 1 |
| 0 | 0 | 0 | 1 |
| 0 | 0 | 0 | 1 |
| 0 | 0 | 0 | 1 |
| 0 | 0 | 0 | 0 |
| 0 | 0 | 0 | 0 |
| 0 | 0 | 0 | 1 |
| 0 | 0 | 0 | 1 |
| 0 | 0 | 0 | 0 |
| 1 | 0 | 0 | 0 |
| 0 | 0 | 0 | 0 |
| 1 | 0 | 0 | 0 |
| 0 | 0 | 0 | 0 |
| 0 | 0 | 0 | 0 |
| 0 | 0 | 0 | 0 |
| 0 | 0 | 0 | 0 |
| 0 | 0 | 0 | 0 |
| 0 | 0 | 0 | 0 |
| 0 | 0 | 0 | 0 |
| 0 | 0 | 0 | 0 |
| 1 | 0 | 0 | 0 |
| 0 | 0 | 0 | 1 |
| 0 | 0 | 0 | 1 |
| 0 | 0 | 0 | 1 |
| 0 | 0 | 0 | 1 |
| 0 | 0 | 0 | 0 |
| 0 | 0 | 0 | 1 |

| | | | |
|---|---|---|---|
| 0 | 0 | 0 | 1 |
| 0 | 0 | 0 | 1 |
| 0 | 0 | 0 | 1 |
| 0 | 0 | 0 | 1 |
| 0 | 0 | 0 | 1 |
| 0 | 0 | 0 | 1 |
| 0 | 0 | 0 | 1 |
| 0 | 0 | 0 | 0 |
| 0 | 0 | 0 | 0 |
| 0 | 0 | 0 | 1 |
| 0 | 0 | 0 | 1 |
| 0 | 0 | 0 | 1 |
| 0 | 0 | 0 | 1 |
| 0 | 0 | 0 | 0 |
| 0 | 0 | 0 | 1 |
| 0 | 0 | 0 | 1 |
| 0 | 0 | 0 | 0 |
| 0 | 0 | 0 | 1 |
| 1 | 0 | 0 | 0 |
| 0 | 0 | 0 | 1 |
| 0 | 0 | 0 | 1 |
| 0 | 0 | 0 | 1 |
| 0 | 0 | 0 | 1 |
| 0 | 0 | 0 | 0 |
| 0 | 0 | 0 | 1 |
| 0 | 0 | 0 | 1 |
| 0 | 0 | 0 | 1 |
| 0 | 0 | 0 | 1 |
| 0 | 0 | 0 | 1 |
| 0 | 0 | 0 | 1 |
| 0 | 0 | 0 | 1 |
| 1 | 0 | 0 | 0 |
| 0 | 0 | 0 | 1 |
| 0 | 0 | 0 | 1 |
| 0 | 0 | 0 | 1 |
| 0 | 0 | 0 | 1 |
| 0 | 0 | 0 | 1 |
| 0 | 0 | 0 | 1 |
| 1 | 0 | 0 | 0 |
| 0 | 0 | 0 | 1 |
| 0 | 0 | 0 | 1 |
| 0 | 0 | 0 | 1 |
| 0 | 0 | 0 | 1 |
| 0 | 0 | 0 | 1 |
| 0 | 0 | 0 | 1 |
| 0 | 0 | 0 | 0 |
| 0 | 0 | 0 | 1 |
| 0 | 0 | 0 | 1 |
| 0 | 0 | 0 | 1 |
| 0 | 0 | 0 | 1 |
| 0 | 0 | 0 | 1 |
| 0 | 0 | 0 | 1 |
| 0 | 0 | 0 | 1 |
| 0 | 0 | 0 | 1 |

| | | | |
|---|---|---|---|
| 0 | 0 | 0 | 1 |
| 0 | 0 | 0 | 1 |
| 0 | 0 | 0 | 1 |
| 0 | 0 | 0 | 1 |
| 0 | 0 | 0 | 1 |
| 0 | 0 | 0 | 1 |
| 0 | 0 | 0 | 1 |
| 0 | 0 | 0 | 1 |
| 0 | 0 | 0 | 1 |
| 0 | 0 | 0 | 1 |
| 0 | 0 | 0 | 1 |
| 0 | 0 | 0 | 1 |
| 0 | 0 | 0 | 1 |
| 0 | 0 | 0 | 1 |
| 0 | 0 | 0 | 1 |
| 1 | 0 | 0 | 0 |
| 0 | 0 | 0 | 1 |
| 0 | 0 | 0 | 0 |
| 0 | 0 | 0 | 1 |
| 0 | 0 | 0 | 1 |
| 1 | 0 | 0 | 0 |
| 0 | 0 | 0 | 1 |
| 0 | 0 | 0 | 1 |
| 0 | 0 | 0 | 1 |
| 0 | 0 | 0 | 0 |
| 0 | 0 | 0 | 1 |
| 0 | 0 | 0 | 0 |
| 0 | 0 | 0 | 1 |
| 0 | 0 | 0 | 1 |
| 0 | 0 | 0 | 1 |
| 0 | 0 | 0 | 1 |
| 0 | 0 | 0 | 1 |
| 0 | 0 | 0 | 1 |
| 0 | 0 | 0 | 1 |
| 0 | 0 | 0 | 1 |
| 0 | 0 | 0 | 1 |
| 0 | 0 | 0 | 1 |
| 0 | 0 | 0 | 1 |
| 0 | 0 | 0 | 1 |
| 0 | 0 | 0 | 1 |
| 0 | 0 | 0 | 1 |
| 0 | 0 | 0 | 1 |
| 0 | 0 | 0 | 1 |
| 0 | 0 | 0 | 0 |
| 0 | 0 | 0 | 1 |
| 0 | 0 | 0 | 1 |
| 0 | 0 | 0 | 0 |
| 0 | 0 | 0 | 0 |
| 0 | 0 | 0 | 1 |
| 0 | 0 | 0 | 0 |
| 0 | 0 | 0 | 0 |
| 0 | 0 | 0 | 0 |
| 0 | 0 | 0 | 1 |
| 0 | 0 | 0 | 1 |

| | | | |
|---|---|---|---|
| 0 | 0 | 0 | 1 |
| 0 | 0 | 0 | 1 |
| 0 | 0 | 0 | 1 |
| 0 | 0 | 0 | 1 |
| 0 | 0 | 0 | 1 |
| 0 | 0 | 0 | 1 |
| 0 | 0 | 0 | 1 |
| 1 | 0 | 0 | 0 |
| 0 | 0 | 0 | 1 |
| 0 | 0 | 0 | 1 |
| 0 | 0 | 0 | 1 |
| 0 | 0 | 0 | 1 |
| 0 | 0 | 0 | 1 |
| 0 | 0 | 0 | 1 |
| 0 | 0 | 0 | 0 |
| 0 | 0 | 0 | 0 |
| 0 | 0 | 0 | 1 |
| 0 | 0 | 0 | 0 |
| 0 | 0 | 0 | 1 |
| 0 | 0 | 0 | 1 |
| 0 | 0 | 0 | 0 |
| 0 | 0 | 0 | 1 |
| 0 | 0 | 0 | 1 |
| 0 | 0 | 0 | 1 |
| 0 | 0 | 0 | 0 |
| 0 | 0 | 0 | 1 |
| 0 | 0 | 0 | 1 |
| 1 | 0 | 0 | 0 |
| 0 | 0 | 0 | 0 |
| 1 | 0 | 0 | 0 |
| 1 | 0 | 0 | 0 |
| 0 | 0 | 0 | 1 |
| 0 | 0 | 0 | 1 |
| 0 | 0 | 0 | 1 |
| 0 | 0 | 0 | 1 |
| 0 | 0 | 0 | 1 |
| 0 | 0 | 0 | 0 |
| 0 | 0 | 0 | 0 |
| 0 | 0 | 0 | 1 |
| 0 | 0 | 0 | 1 |
| 0 | 0 | 0 | 1 |
| 0 | 0 | 0 | 1 |
| 0 | 0 | 0 | 0 |
| 0 | 0 | 0 | 1 |
| 0 | 0 | 0 | 1 |
| 0 | 0 | 0 | 1 |
| 0 | 0 | 0 | 1 |
| 0 | 0 | 0 | 1 |
| 0 | 0 | 0 | 0 |
| 0 | 0 | 0 | 0 |
| 0 | 0 | 0 | 0 |
| 0 | 0 | 0 | 0 |
| 0 | 0 | 0 | 0 |
| 0 | 0 | 0 | 1 |

| | | | |
|---|---|---|---|
| 0 | 0 | 0 | 0 |
| 0 | 0 | 0 | 0 |
| 0 | 0 | 0 | 1 |
| 1 | 0 | 0 | 0 |
| 0 | 0 | 0 | 1 |
| 0 | 0 | 0 | 1 |
| 0 | 0 | 0 | 1 |
| 0 | 0 | 0 | 1 |
| 0 | 0 | 0 | 1 |
| 1 | 0 | 0 | 0 |
| 0 | 0 | 0 | 1 |
| 0 | 0 | 0 | 1 |
| 0 | 0 | 0 | 1 |
| 0 | 0 | 0 | 1 |
| 0 | 0 | 0 | 1 |
| 0 | 0 | 0 | 1 |
| 0 | 0 | 0 | 1 |
| 0 | 0 | 0 | 1 |
| 0 | 0 | 0 | 1 |
| 0 | 0 | 0 | 1 |
| 0 | 0 | 0 | 1 |
| 0 | 0 | 0 | 1 |
| 0 | 0 | 0 | 1 |
| 0 | 0 | 0 | 1 |
| 0 | 0 | 0 | 1 |
| 0 | 0 | 0 | 1 |
| 0 | 0 | 0 | 1 |
| 0 | 0 | 0 | 1 |
| 0 | 0 | 0 | 1 |
| 0 | 0 | 0 | 1 |
| 0 | 0 | 0 | 1 |
| 0 | 0 | 0 | 1 |
| 1 | 0 | 0 | 0 |
| 0 | 0 | 0 | 1 |
| 0 | 0 | 0 | 1 |
| 0 | 0 | 0 | 1 |
| 0 | 0 | 0 | 1 |
| 0 | 0 | 0 | 1 |
| 0 | 0 | 0 | 1 |
| 0 | 0 | 0 | 1 |
| 0 | 0 | 0 | 1 |
| 0 | 0 | 0 | 1 |
| 0 | 0 | 0 | 1 |
| 0 | 0 | 0 | 1 |
| 0 | 0 | 0 | 1 |
| 0 | 0 | 0 | 1 |
| 0 | 0 | 0 | 1 |
| 0 | 0 | 0 | 0 |
| 0 | 0 | 0 | 0 |
| 0 | 0 | 0 | 0 |
| 0 | 0 | 0 | 0 |
| 0 | 1 | 0 | 0 |
| 0 | 0 | 0 | 1 |
| 0 | 0 | 0 | 0 |

| | | | |
|---|---|---|---|
| 0 | 0 | 0 | 1 |
| 0 | 0 | 0 | 0 |
| 0 | 0 | 0 | 1 |
| 0 | 0 | 0 | 1 |
| 0 | 0 | 0 | 1 |
| 1 | 0 | 0 | 0 |
| 0 | 0 | 0 | 0 |
| 0 | 0 | 0 | 0 |
| 1 | 0 | 0 | 0 |
| 0 | 0 | 0 | 0 |
| 0 | 0 | 0 | 1 |
| 0 | 0 | 0 | 0 |
| 0 | 0 | 0 | 1 |
| 0 | 0 | 0 | 1 |
| 0 | 0 | 0 | 1 |
| 1 | 0 | 0 | 0 |
| 0 | 0 | 0 | 1 |
| 0 | 0 | 0 | 1 |
| 0 | 0 | 0 | 0 |
| 0 | 0 | 0 | 0 |
| 1 | 0 | 0 | 0 |
| 1 | 0 | 0 | 0 |
| 0 | 0 | 0 | 0 |
| 0 | 0 | 0 | 0 |
| 0 | 0 | 0 | 1 |
| 0 | 0 | 0 | 0 |
| 0 | 0 | 0 | 0 |
| 0 | 0 | 0 | 0 |
| 0 | 0 | 0 | 0 |
| 0 | 0 | 0 | 0 |
| 0 | 0 | 0 | 0 |
| 0 | 0 | 0 | 1 |
| 0 | 0 | 0 | 1 |
| 0 | 0 | 0 | 0 |
| 0 | 0 | 0 | 0 |
| 0 | 0 | 0 | 1 |
| 0 | 0 | 0 | 0 |
| 0 | 0 | 0 | 0 |
| 0 | 0 | 0 | 0 |
| 0 | 0 | 0 | 0 |
| 0 | 0 | 0 | 0 |
| 0 | 0 | 0 | 1 |
| 0 | 0 | 0 | 1 |
| 0 | 0 | 0 | 1 |
| 0 | 0 | 0 | 0 |
| 0 | 0 | 0 | 0 |
| 0 | 0 | 0 | 0 |
| 0 | 0 | 0 | 1 |
| 0 | 0 | 0 | 1 |
| 0 | 0 | 0 | 1 |
| 0 | 0 | 0 | 1 |
| 0 | 0 | 0 | 1 |
| 0 | 0 | 0 | 1 |
| 0 | 0 | 0 | 0 |

| | | | |
|---|---|---|---|
| 0 | 0 | 0 | 1 |
| 0 | 0 | 0 | 1 |
| 0 | 0 | 0 | 1 |
| 0 | 0 | 0 | 1 |
| 1 | 0 | 0 | 0 |
| 0 | 0 | 0 | 0 |
| 0 | 0 | 0 | 1 |
| 0 | 0 | 0 | 1 |
| 0 | 0 | 0 | 1 |
| 0 | 0 | 0 | 1 |
| 0 | 0 | 0 | 1 |
| 0 | 0 | 0 | 0 |
| 0 | 0 | 0 | 1 |
| 0 | 0 | 0 | 0 |
| 0 | 0 | 0 | 0 |
| 0 | 0 | 0 | 0 |
| 0 | 0 | 0 | 0 |
| 0 | 0 | 0 | 0 |
| 0 | 0 | 0 | 0 |
| 0 | 0 | 0 | 0 |
| 0 | 0 | 0 | 0 |
| 0 | 0 | 0 | 0 |
| 0 | 0 | 0 | 0 |
| 0 | 0 | 0 | 1 |
| 0 | 0 | 0 | 1 |
| 0 | 0 | 0 | 1 |
| 0 | 0 | 0 | 1 |
| 0 | 0 | 0 | 1 |
| 0 | 0 | 0 | 1 |
| 0 | 0 | 0 | 1 |
| 0 | 0 | 0 | 1 |
| 0 | 0 | 0 | 0 |
| 0 | 0 | 0 | 1 |
| 0 | 0 | 0 | 0 |
| 0 | 0 | 0 | 0 |
| 0 | 0 | 0 | 0 |
| 0 | 0 | 0 | 0 |
| 0 | 0 | 0 | 1 |
| 0 | 0 | 0 | 1 |
| 0 | 0 | 0 | 1 |
| 0 | 0 | 0 | 1 |
| 0 | 0 | 0 | 1 |
| 0 | 0 | 0 | 1 |
| 0 | 0 | 0 | 1 |
| 0 | 0 | 0 | 1 |
| 1 | 0 | 0 | 0 |
| 0 | 0 | 0 | 1 |
| 0 | 0 | 0 | 1 |
| 0 | 0 | 0 | 1 |
| 1 | 0 | 0 | 0 |
| 1 | 0 | 0 | 0 |
| 0 | 0 | 0 | 0 |
| 0 | 0 | 0 | 1 |
| 0 | 0 | 0 | 1 |

| | | | |
|---|---|---|---|
| 0 | 0 | 0 | 1 |
| 0 | 0 | 0 | 1 |
| 0 | 0 | 0 | 1 |
| 0 | 0 | 0 | 0 |
| 0 | 0 | 0 | 1 |
| 0 | 0 | 0 | 1 |
| 0 | 0 | 0 | 0 |
| 0 | 0 | 0 | 1 |
| 0 | 0 | 0 | 1 |
| 0 | 0 | 0 | 1 |
| 0 | 0 | 0 | 0 |
| 0 | 0 | 0 | 0 |
| 0 | 0 | 0 | 1 |
| 0 | 0 | 0 | 1 |
| 0 | 0 | 0 | 1 |
| 0 | 0 | 0 | 1 |
| 0 | 0 | 0 | 0 |
| 0 | 0 | 0 | 0 |
| 0 | 0 | 0 | 1 |
| 0 | 0 | 0 | 0 |
| 0 | 0 | 0 | 0 |
| 0 | 0 | 0 | 0 |
| 0 | 0 | 0 | 1 |
| 0 | 0 | 0 | 0 |
| 0 | 0 | 0 | 0 |
| 0 | 0 | 0 | 1 |
| 0 | 0 | 0 | 0 |
| 0 | 0 | 0 | 1 |
| 0 | 0 | 0 | 1 |
| 0 | 0 | 0 | 0 |
| 0 | 0 | 0 | 1 |
| 0 | 0 | 0 | 1 |
| 0 | 0 | 0 | 0 |
| 0 | 0 | 0 | 1 |
| 0 | 0 | 0 | 1 |
| 0 | 0 | 0 | 0 |
| 0 | 0 | 0 | 1 |
| 0 | 0 | 0 | 0 |
| 0 | 0 | 0 | 1 |
| 0 | 0 | 0 | 0 |
| 0 | 0 | 0 | 0 |
| 0 | 0 | 0 | 1 |
| 0 | 0 | 0 | 0 |
| 0 | 0 | 0 | 0 |
| 0 | 0 | 0 | 1 |
| 0 | 0 | 0 | 1 |
| 0 | 0 | 0 | 1 |
| 0 | 0 | 0 | 1 |
| 0 | 0 | 0 | 1 |
| 0 | 0 | 0 | 1 |
| 0 | 0 | 0 | 1 |
| 0 | 0 | 0 | 1 |
| 0 | 0 | 0 | 1 |
| 0 | 0 | 0 | 1 |

| | | | |
|---|---|---|---|
| 0 | 0 | 0 | 1 |
| 0 | 0 | 0 | 1 |
| 0 | 0 | 0 | 1 |
| 0 | 0 | 0 | 0 |
| 0 | 0 | 0 | 1 |
| 0 | 0 | 0 | 0 |
| 0 | 0 | 0 | 1 |
| 0 | 0 | 0 | 1 |
| 0 | 0 | 0 | 0 |
| 0 | 0 | 0 | 0 |
| 0 | 0 | 0 | 1 |
| 0 | 0 | 0 | 0 |
| 0 | 0 | 0 | 0 |
| 0 | 0 | 0 | 0 |
| 0 | 0 | 0 | 1 |
| 0 | 0 | 0 | 0 |
| 0 | 0 | 0 | 0 |
| 0 | 0 | 0 | 1 |
| 1 | 0 | 0 | 0 |
| 1 | 0 | 0 | 0 |
| 1 | 0 | 0 | 0 |
| 1 | 0 | 0 | 0 |
| 0 | 0 | 0 | 1 |
| 0 | 0 | 0 | 0 |
| 0 | 0 | 0 | 0 |
| 0 | 0 | 0 | 0 |
| 0 | 0 | 0 | 0 |
| 0 | 0 | 0 | 0 |
| 0 | 0 | 0 | 0 |
| 0 | 0 | 0 | 0 |
| 0 | 0 | 0 | 0 |
| 0 | 0 | 0 | 0 |
| 0 | 0 | 0 | 0 |
| 0 | 0 | 0 | 1 |
| 0 | 0 | 0 | 1 |
| 0 | 0 | 0 | 0 |
| 0 | 0 | 0 | 1 |
| 0 | 0 | 0 | 0 |
| 0 | 0 | 0 | 0 |
| 0 | 0 | 0 | 0 |
| 0 | 0 | 0 | 0 |
| 0 | 0 | 0 | 1 |
| 0 | 0 | 0 | 0 |
| 0 | 0 | 0 | 0 |
| 0 | 0 | 0 | 1 |
| 0 | 0 | 0 | 1 |
| 0 | 0 | 0 | 1 |
| 0 | 0 | 0 | 1 |
| 0 | 0 | 0 | 1 |
| 0 | 0 | 0 | 1 |
| 0 | 0 | 0 | 1 |
| 0 | 0 | 0 | 1 |
| 0 | 0 | 0 | 1 |
| 0 | 0 | 0 | 1 |

| | | | |
|---|---|---|---|
| 0 | 0 | 0 | 1 |
| 0 | 0 | 0 | 0 |
| 0 | 0 | 0 | 1 |
| 0 | 0 | 0 | 1 |
| 0 | 0 | 0 | 1 |
| 0 | 0 | 0 | 1 |
| 0 | 0 | 0 | 1 |
| 0 | 0 | 0 | 1 |
| 0 | 0 | 0 | 1 |
| 0 | 0 | 0 | 1 |
| 0 | 0 | 0 | 1 |
| 0 | 0 | 0 | 0 |
| 0 | 0 | 0 | 1 |
| 0 | 0 | 0 | 1 |
| 0 | 0 | 0 | 0 |
| 0 | 0 | 0 | 1 |
| 0 | 0 | 0 | 1 |
| 0 | 0 | 0 | 1 |
| 0 | 0 | 0 | 1 |
| 0 | 0 | 0 | 1 |
| 1 | 0 | 0 | 0 |
| 0 | 0 | 0 | 1 |
| 1 | 0 | 0 | 0 |
| 1 | 0 | 0 | 0 |
| 0 | 0 | 0 | 1 |
| 0 | 0 | 0 | 0 |
| 0 | 0 | 0 | 1 |
| 0 | 0 | 0 | 1 |
| 0 | 0 | 0 | 1 |
| 0 | 0 | 0 | 1 |
| 0 | 0 | 0 | 1 |
| 0 | 0 | 0 | 0 |
| 0 | 0 | 0 | 1 |
| 0 | 0 | 0 | 0 |
| 0 | 0 | 0 | 1 |
| 0 | 0 | 0 | 1 |
| 0 | 0 | 0 | 1 |
| 0 | 0 | 0 | 1 |
| 0 | 0 | 0 | 1 |
| 0 | 0 | 0 | 0 |
| 0 | 0 | 0 | 1 |
| 0 | 0 | 0 | 1 |
| 0 | 0 | 0 | 1 |
| 0 | 0 | 0 | 0 |
| 0 | 0 | 0 | 1 |
| 0 | 0 | 0 | 1 |
| 0 | 0 | 0 | 1 |
| 1 | 0 | 0 | 0 |
| 0 | 0 | 0 | 0 |
| 0 | 0 | 0 | 1 |
| 0 | 0 | 0 | 1 |
| 0 | 0 | 0 | 0 |
| 1 | 0 | 0 | 0 |
| 0 | 0 | 0 | 0 |

| | | | |
|---|---|---|---|
| 1 | 0 | 0 | 0 |
| 0 | 0 | 0 | 1 |
| 0 | 0 | 0 | 0 |
| 0 | 0 | 0 | 0 |
| 0 | 0 | 0 | 0 |
| 0 | 0 | 0 | 1 |
| 0 | 0 | 0 | 1 |
| 0 | 0 | 0 | 1 |
| 0 | 0 | 0 | 1 |
| 0 | 0 | 0 | 0 |
| 0 | 0 | 0 | 1 |
| 0 | 0 | 0 | 1 |
| 0 | 0 | 0 | 1 |
| 0 | 0 | 0 | 1 |
| 0 | 0 | 0 | 0 |
| 0 | 0 | 0 | 1 |
| 0 | 0 | 0 | 1 |
| 0 | 0 | 0 | 0 |
| 0 | 0 | 0 | 1 |
| 0 | 0 | 0 | 0 |
| 0 | 0 | 0 | 0 |
| 0 | 0 | 0 | 1 |
| 0 | 0 | 0 | 0 |
| 0 | 0 | 0 | 1 |
| 0 | 0 | 0 | 0 |
| 0 | 0 | 0 | 0 |
| 0 | 0 | 0 | 1 |
| 0 | 0 | 0 | 1 |
| 1 | 0 | 0 | 0 |
| 0 | 0 | 0 | 1 |
| 0 | 0 | 0 | 1 |
| 0 | 0 | 0 | 1 |
| 0 | 0 | 0 | 1 |
| 0 | 0 | 0 | 1 |
| 0 | 0 | 0 | 0 |
| 0 | 0 | 0 | 0 |
| 0 | 0 | 0 | 1 |
| 0 | 0 | 0 | 0 |
| 0 | 0 | 0 | 0 |
| 0 | 0 | 0 | 0 |
| 0 | 0 | 0 | 0 |
| 0 | 0 | 0 | 0 |
| 0 | 0 | 0 | 0 |
| 0 | 0 | 0 | 1 |
| 0 | 0 | 0 | 0 |
| 0 | 0 | 0 | 1 |
| 0 | 0 | 0 | 1 |
| 0 | 0 | 0 | 0 |
| 0 | 0 | 0 | 0 |
| 0 | 0 | 0 | 0 |
| 0 | 0 | 0 | 0 |
| 0 | 0 | 0 | 0 |
| 0 | 0 | 0 | 1 |
| 0 | 0 | 0 | 1 |

| | | | |
|---|---|---|---|
| 0 | 0 | 0 | 1 |
| 0 | 0 | 0 | 0 |
| 0 | 0 | 0 | 0 |
| 0 | 0 | 0 | 0 |
| 0 | 0 | 0 | 0 |
| 0 | 0 | 0 | 0 |
| 0 | 0 | 0 | 0 |
| 0 | 0 | 0 | 1 |
| 0 | 0 | 0 | 1 |
| 0 | 0 | 0 | 1 |
| 0 | 0 | 0 | 0 |
| 0 | 0 | 0 | 0 |
| 0 | 0 | 0 | 1 |
| 0 | 0 | 0 | 0 |
| 0 | 0 | 0 | 1 |
| 0 | 0 | 0 | 0 |
| 0 | 0 | 0 | 1 |
| 0 | 0 | 0 | 1 |
| 0 | 0 | 0 | 1 |
| 0 | 0 | 0 | 1 |
| 0 | 0 | 0 | 1 |
| 0 | 0 | 0 | 1 |
| 0 | 0 | 0 | 1 |
| 1 | 0 | 0 | 0 |
| 0 | 0 | 0 | 0 |
| 0 | 0 | 0 | 1 |
| 0 | 0 | 0 | 1 |
| 0 | 0 | 0 | 1 |
| 0 | 0 | 0 | 1 |
| 0 | 0 | 0 | 0 |
| 0 | 0 | 0 | 0 |
| 0 | 0 | 0 | 0 |
| 0 | 0 | 0 | 0 |
| 0 | 0 | 0 | 1 |
| 0 | 0 | 0 | 0 |
| 0 | 0 | 0 | 0 |
| 0 | 0 | 0 | 0 |
| 0 | 0 | 0 | 0 |
| 0 | 0 | 0 | 0 |
| 0 | 0 | 0 | 1 |
| 0 | 0 | 0 | 1 |
| 0 | 0 | 0 | 0 |
| 0 | 0 | 0 | 1 |
| 0 | 0 | 0 | 1 |
| 0 | 0 | 0 | 0 |
| 0 | 0 | 0 | 1 |
| 0 | 0 | 0 | 1 |
| 0 | 0 | 0 | 1 |
| 0 | 0 | 0 | 1 |
| 0 | 0 | 0 | 1 |
| 0 | 0 | 0 | 0 |
| 0 | 0 | 0 | 1 |
| 0 | 0 | 0 | 0 |
| 0 | 0 | 0 | 0 |

| | | | |
|---|---|---|---|
| 0 | 0 | 0 | 1 |
| 0 | 0 | 0 | 1 |
| 0 | 0 | 0 | 0 |
| 0 | 0 | 0 | 0 |
| 0 | 0 | 0 | 1 |
| 0 | 0 | 0 | 1 |
| 0 | 0 | 0 | 1 |
| 0 | 0 | 0 | 1 |
| 0 | 0 | 0 | 1 |
| 0 | 0 | 0 | 1 |
| 0 | 0 | 0 | 1 |
| 0 | 0 | 0 | 1 |
| 0 | 0 | 0 | 0 |
| 0 | 0 | 0 | 1 |
| 0 | 0 | 0 | 1 |
| 0 | 0 | 0 | 1 |
| 0 | 0 | 0 | 1 |
| 0 | 0 | 0 | 1 |
| 0 | 0 | 0 | 1 |
| 0 | 0 | 0 | 1 |
| 0 | 0 | 0 | 1 |
| 0 | 0 | 0 | 1 |
| 0 | 0 | 0 | 1 |
| 0 | 0 | 0 | 1 |
| 0 | 0 | 0 | 1 |
| 0 | 0 | 0 | 1 |
| 0 | 0 | 0 | 1 |
| 0 | 0 | 0 | 1 |
| 0 | 0 | 0 | 1 |
| 0 | 0 | 0 | 1 |
| 0 | 0 | 0 | 1 |
| 0 | 0 | 0 | 0 |
| 0 | 0 | 0 | 1 |
| 0 | 0 | 0 | 1 |
| 0 | 0 | 0 | 1 |
| 0 | 0 | 0 | 1 |
| 0 | 0 | 0 | 1 |
| 0 | 0 | 0 | 1 |
| 0 | 0 | 0 | 1 |
| 0 | 0 | 0 | 1 |
| 0 | 0 | 0 | 0 |
| 0 | 0 | 0 | 0 |
| 0 | 0 | 0 | 1 |
| 0 | 0 | 0 | 1 |
| 0 | 0 | 0 | 1 |
| 0 | 0 | 0 | 1 |
| 0 | 0 | 0 | 1 |
| 0 | 0 | 0 | 1 |
| 0 | 0 | 0 | 0 |
| 0 | 0 | 0 | 0 |
| 0 | 0 | 0 | 1 |
| 0 | 0 | 0 | 1 |
| 0 | 0 | 0 | 0 |
| 0 | 0 | 0 | 1 |

| | | | |
|---|---|---|---|
| 0 | 0 | 0 | 0 |
| 0 | 0 | 0 | 0 |
| 0 | 0 | 0 | 0 |
| 0 | 0 | 0 | 1 |
| 0 | 0 | 0 | 1 |
| 0 | 0 | 0 | 1 |
| 0 | 0 | 0 | 1 |
| 0 | 0 | 0 | 0 |
| 0 | 0 | 0 | 0 |
| 0 | 0 | 0 | 0 |
| 0 | 0 | 0 | 0 |
| 0 | 0 | 0 | 1 |
| 0 | 0 | 0 | 0 |
| 0 | 0 | 0 | 1 |
| 0 | 0 | 0 | 1 |
| 0 | 0 | 0 | 1 |
| 0 | 0 | 0 | 1 |
| 0 | 0 | 0 | 0 |
| 0 | 0 | 0 | 1 |
| 0 | 0 | 0 | 1 |
| 0 | 0 | 0 | 1 |
| 0 | 0 | 0 | 1 |
| 0 | 0 | 0 | 1 |
| 0 | 0 | 0 | 1 |
| 0 | 0 | 0 | 1 |
| 0 | 0 | 0 | 1 |
| 0 | 0 | 0 | 1 |
| 0 | 0 | 0 | 1 |
| 0 | 0 | 0 | 1 |
| 0 | 0 | 0 | 1 |
| 0 | 0 | 0 | 0 |
| 0 | 0 | 0 | 0 |
| 0 | 0 | 0 | 1 |
| 0 | 0 | 0 | 1 |
| 0 | 0 | 0 | 1 |
| 0 | 0 | 0 | 1 |
| 0 | 0 | 0 | 1 |
| 0 | 0 | 0 | 1 |
| 0 | 0 | 0 | 1 |
| 0 | 0 | 0 | 1 |
| 0 | 0 | 0 | 1 |
| 0 | 0 | 0 | 0 |
| 0 | 0 | 0 | 1 |
| 0 | 0 | 0 | 1 |
| 0 | 0 | 0 | 0 |
| 0 | 0 | 0 | 0 |
| 0 | 0 | 0 | 1 |
| 0 | 0 | 0 | 1 |
| 0 | 0 | 0 | 0 |
| 0 | 0 | 0 | 0 |
| 0 | 0 | 0 | 0 |
| 0 | 0 | 0 | 0 |
| 0 | 0 | 0 | 0 |
| 0 | 0 | 0 | 0 |

| | | | |
|---|---|---|---|
| 0 | 0 | 0 | 1 |
| 0 | 0 | 0 | 1 |
| 0 | 0 | 0 | 1 |
| 0 | 0 | 0 | 1 |
| 0 | 0 | 0 | 1 |
| 0 | 0 | 0 | 0 |
| 0 | 0 | 0 | 0 |
| 0 | 0 | 0 | 0 |
| 0 | 0 | 0 | 0 |
| 0 | 0 | 0 | 0 |
| 0 | 0 | 0 | 1 |
| 0 | 0 | 0 | 0 |
| 0 | 0 | 0 | 0 |
| 0 | 0 | 0 | 0 |
| 0 | 0 | 0 | 1 |
| 0 | 0 | 0 | 0 |
| 0 | 0 | 0 | 1 |
| 0 | 0 | 0 | 1 |
| 1 | 0 | 0 | 0 |
| 1 | 0 | 0 | 0 |
| 0 | 0 | 0 | 0 |
| 0 | 0 | 0 | 0 |
| 0 | 0 | 0 | 1 |
| 0 | 0 | 0 | 0 |
| 0 | 0 | 0 | 0 |
| 0 | 0 | 0 | 0 |
| 0 | 0 | 0 | 0 |
| 0 | 0 | 0 | 1 |
| 0 | 0 | 0 | 1 |
| 0 | 0 | 0 | 1 |
| 0 | 0 | 0 | 1 |
| 0 | 0 | 0 | 1 |
| 0 | 0 | 0 | 0 |
| 0 | 0 | 0 | 1 |
| 0 | 0 | 0 | 0 |
| 0 | 0 | 0 | 0 |
| 0 | 0 | 0 | 0 |
| 0 | 0 | 0 | 0 |
| 0 | 0 | 0 | 0 |
| 0 | 0 | 0 | 0 |
| 0 | 0 | 0 | 0 |
| 1 | 0 | 0 | 0 |
| 0 | 0 | 0 | 0 |
| 0 | 0 | 0 | 1 |
| 0 | 0 | 0 | 1 |
| 0 | 0 | 0 | 1 |
| 0 | 0 | 0 | 1 |
| 0 | 0 | 0 | 0 |
| 0 | 0 | 0 | 0 |
| 0 | 0 | 0 | 0 |
| 0 | 0 | 0 | 1 |
| 0 | 0 | 0 | 1 |
| 0 | 0 | 0 | 0 |
| 0 | 0 | 0 | 1 |

| | | | |
|---|---|---|---|
| 0 | 0 | 0 | 1 |
| 0 | 0 | 0 | 0 |
| 0 | 0 | 0 | 1 |
| 0 | 0 | 0 | 1 |
| 0 | 0 | 0 | 1 |
| 0 | 0 | 0 | 1 |
| 0 | 0 | 0 | 1 |
| 0 | 0 | 0 | 1 |
| 0 | 0 | 0 | 1 |
| 0 | 0 | 0 | 1 |
| 0 | 0 | 0 | 1 |
| 0 | 0 | 0 | 1 |
| 0 | 0 | 0 | 0 |
| 0 | 0 | 0 | 0 |
| 0 | 0 | 0 | 0 |
| 0 | 0 | 0 | 0 |
| 0 | 0 | 0 | 0 |
| 0 | 0 | 0 | 0 |
| 0 | 0 | 0 | 0 |
| 0 | 0 | 0 | 1 |
| 1 | 1 | 0 | 0 |
| 0 | 0 | 0 | 0 |
| 0 | 0 | 0 | 0 |
| 0 | 0 | 0 | 0 |
| 0 | 0 | 0 | 0 |
| 0 | 0 | 0 | 1 |
| 0 | 0 | 0 | 1 |
| 0 | 0 | 0 | 0 |
| 0 | 0 | 0 | 1 |
| 0 | 0 | 0 | 0 |
| 0 | 0 | 0 | 0 |
| 0 | 0 | 0 | 1 |
| 0 | 0 | 0 | 0 |
| 0 | 0 | 0 | 1 |
| 0 | 0 | 0 | 1 |
| 0 | 0 | 0 | 1 |
| 0 | 0 | 0 | 1 |
| 0 | 0 | 0 | 1 |
| 0 | 0 | 0 | 1 |
| 0 | 0 | 0 | 1 |
| 0 | 0 | 0 | 0 |
| 0 | 0 | 0 | 1 |
| 0 | 0 | 0 | 1 |
| 0 | 0 | 0 | 0 |
| 0 | 0 | 0 | 0 |
| 0 | 0 | 0 | 1 |
| 0 | 0 | 0 | 0 |
| 0 | 0 | 0 | 0 |
| 0 | 0 | 0 | 1 |
| 0 | 0 | 0 | 1 |
| 0 | 0 | 0 | 0 |
| 0 | 0 | 0 | 1 |
| 0 | 0 | 0 | 1 |
| 0 | 0 | 0 | 1 |

| | | | |
|---|---|---|---|
| 0 | 0 | 0 | 1 |
| 0 | 0 | 0 | 1 |
| 0 | 0 | 0 | 1 |
| 0 | 0 | 0 | 1 |
| 0 | 0 | 0 | 0 |
| 0 | 0 | 0 | 0 |
| 0 | 0 | 0 | 1 |
| 0 | 0 | 0 | 1 |
| 0 | 0 | 0 | 1 |
| 0 | 0 | 0 | 1 |
| 0 | 0 | 0 | 1 |
| 0 | 0 | 0 | 0 |
| 0 | 0 | 0 | 0 |
| 0 | 0 | 0 | 1 |
| 0 | 0 | 0 | 1 |
| 0 | 0 | 0 | 0 |
| 0 | 0 | 0 | 1 |
| 0 | 0 | 0 | 1 |
| 0 | 0 | 0 | 0 |
| 0 | 0 | 0 | 1 |
| 0 | 0 | 0 | 1 |
| 0 | 0 | 0 | 1 |
| 0 | 0 | 0 | 1 |
| 0 | 0 | 0 | 0 |
| 0 | 0 | 0 | 1 |
| 0 | 0 | 0 | 0 |
| 0 | 0 | 0 | 1 |
| 0 | 0 | 0 | 1 |
| 1 | 0 | 0 | 0 |
| 1 | 0 | 0 | 0 |
| 0 | 0 | 0 | 0 |
| 0 | 0 | 0 | 0 |
| 0 | 0 | 0 | 0 |
| 0 | 0 | 0 | 0 |
| 0 | 0 | 0 | 1 |
| 0 | 0 | 0 | 1 |
| 0 | 0 | 0 | 1 |
| 1 | 0 | 0 | 0 |
| 0 | 0 | 0 | 1 |
| 0 | 0 | 0 | 1 |
| 0 | 0 | 0 | 0 |
| 0 | 0 | 0 | 0 |
| 0 | 0 | 0 | 0 |
| 0 | 0 | 0 | 1 |
| 0 | 0 | 0 | 0 |
| 0 | 0 | 0 | 1 |
| 0 | 0 | 0 | 1 |
| 0 | 0 | 0 | 0 |
| 0 | 0 | 0 | 1 |
| 0 | 0 | 0 | 1 |
| 0 | 0 | 0 | 1 |
| 0 | 0 | 0 | 0 |
| 0 | 0 | 0 | 1 |
| 0 | 0 | 0 | 0 |

| | | | |
|---|---|---|---|
| 0 | 0 | 0 | 1 |
| 0 | 0 | 0 | 0 |
| 0 | 0 | 0 | 0 |
| 0 | 0 | 0 | 1 |
| 0 | 0 | 0 | 1 |
| 0 | 0 | 0 | 1 |
| 0 | 0 | 0 | 0 |
| 0 | 0 | 0 | 1 |
| 0 | 0 | 0 | 1 |
| 0 | 0 | 0 | 1 |
| 0 | 0 | 0 | 1 |
| 0 | 0 | 0 | 1 |
| 0 | 0 | 0 | 1 |
| 0 | 0 | 0 | 1 |
| 0 | 0 | 0 | 1 |
| 0 | 0 | 0 | 1 |
| 0 | 0 | 0 | 1 |
| 0 | 0 | 0 | 1 |
| 0 | 0 | 0 | 1 |
| 0 | 0 | 0 | 1 |
| 0 | 0 | 0 | 1 |
| 0 | 0 | 0 | 0 |
| 0 | 0 | 0 | 1 |
| 0 | 0 | 0 | 1 |
| 0 | 0 | 0 | 1 |
| 0 | 0 | 0 | 1 |
| 0 | 0 | 0 | 0 |
| 0 | 0 | 0 | 1 |
| 0 | 0 | 0 | 0 |
| 0 | 0 | 0 | 1 |
| 0 | 0 | 0 | 1 |
| 0 | 0 | 0 | 0 |
| 0 | 0 | 0 | 1 |
| 0 | 0 | 0 | 1 |
| 0 | 0 | 0 | 1 |
| 0 | 0 | 0 | 1 |
| 0 | 0 | 0 | 1 |
| 0 | 0 | 0 | 1 |
| 0 | 0 | 0 | 0 |
| 0 | 0 | 0 | 0 |
| 0 | 0 | 0 | 0 |
| 0 | 0 | 0 | 1 |
| 0 | 0 | 0 | 1 |
| 0 | 0 | 0 | 1 |
| 0 | 0 | 0 | 1 |
| 0 | 0 | 0 | 0 |
| 0 | 0 | 0 | 1 |
| 0 | 0 | 0 | 0 |
| 0 | 0 | 0 | 1 |
| 0 | 0 | 0 | 0 |
| 0 | 0 | 0 | 0 |
| 0 | 0 | 0 | 0 |
| 0 | 0 | 0 | 1 |
| 0 | 0 | 0 | 1 |

| | | | |
|---|---|---|---|
| 0 | 0 | 0 | 0 |
| 0 | 0 | 0 | 0 |
| 0 | 0 | 0 | 1 |
| 0 | 0 | 0 | 1 |
| 0 | 0 | 0 | 1 |
| 0 | 0 | 0 | 1 |
| 0 | 0 | 0 | 1 |
| 0 | 0 | 0 | 1 |
| 0 | 0 | 0 | 1 |
| 0 | 0 | 0 | 1 |
| 0 | 0 | 0 | 1 |
| 0 | 0 | 0 | 1 |
| 0 | 0 | 0 | 1 |
| 0 | 0 | 0 | 1 |
| 0 | 0 | 0 | 1 |
| 0 | 0 | 0 | 0 |
| 0 | 0 | 0 | 0 |
| 0 | 0 | 0 | 0 |
| 0 | 0 | 0 | 1 |
| 0 | 0 | 0 | 1 |
| 0 | 0 | 0 | 1 |
| 0 | 0 | 0 | 0 |
| 0 | 0 | 0 | 0 |
| 0 | 0 | 0 | 0 |
| 0 | 0 | 0 | 0 |
| 0 | 0 | 0 | 1 |
| 0 | 0 | 0 | 1 |
| 0 | 0 | 0 | 1 |
| 0 | 0 | 0 | 1 |
| 0 | 0 | 0 | 1 |
| 0 | 0 | 0 | 0 |
| 0 | 0 | 0 | 1 |
| 0 | 0 | 0 | 0 |
| 0 | 0 | 0 | 1 |
| 0 | 1 | 1 | 1 |
| 0 | 1 | 1 | 1 |
| 0 | 0 | 0 | 0 |
| 0 | 0 | 0 | 0 |
| 0 | 0 | 0 | 1 |
| 0 | 0 | 0 | 1 |
| 0 | 0 | 0 | 1 |
| 0 | 0 | 0 | 0 |
| 0 | 0 | 0 | 1 |
| 0 | 0 | 0 | 1 |
| 0 | 0 | 0 | 0 |
| 0 | 0 | 0 | 1 |
| 0 | 0 | 0 | 0 |
| 0 | 0 | 0 | 0 |
| 0 | 0 | 0 | 1 |
| 0 | 0 | 0 | 1 |
| 0 | 0 | 0 | 0 |
| 0 | 0 | 0 | 0 |
| 0 | 0 | 0 | 1 |
| 0 | 0 | 0 | 0 |

| | | | |
|---|---|---|---|
| 0 | 0 | 0 | 1 |
| 0 | 0 | 0 | 0 |
| 0 | 0 | 0 | 1 |
| 0 | 0 | 0 | 0 |
| 0 | 0 | 0 | 1 |
| 0 | 0 | 0 | 1 |
| 0 | 0 | 0 | 1 |
| 0 | 0 | 0 | 1 |
| 0 | 0 | 0 | 0 |
| 0 | 0 | 0 | 1 |
| 0 | 0 | 0 | 1 |
| 0 | 0 | 0 | 1 |
| 0 | 0 | 0 | 0 |
| 0 | 0 | 0 | 0 |
| 0 | 0 | 0 | 0 |
| 0 | 0 | 0 | 1 |
| 0 | 0 | 0 | 0 |
| 0 | 0 | 0 | 1 |
| 0 | 1 | 0 | 0 |
| 0 | 0 | 0 | 0 |
| 0 | 0 | 0 | 0 |
| 0 | 0 | 0 | 0 |
| 0 | 0 | 0 | 0 |
| 1 | 0 | 0 | 0 |
| 0 | 0 | 0 | 1 |
| 0 | 0 | 0 | 1 |
| 0 | 0 | 0 | 0 |
| 0 | 0 | 0 | 0 |
| 0 | 0 | 0 | 0 |
| 0 | 0 | 0 | 0 |
| 0 | 0 | 0 | 1 |
| 0 | 0 | 0 | 1 |
| 0 | 0 | 0 | 1 |
| 0 | 0 | 0 | 1 |
| 0 | 0 | 0 | 1 |
| 1 | 0 | 0 | 0 |
| 1 | 0 | 0 | 0 |
| 0 | 0 | 0 | 1 |
| 0 | 0 | 0 | 1 |
| 0 | 0 | 0 | 1 |
| 0 | 0 | 0 | 1 |
| 0 | 0 | 0 | 1 |
| 0 | 0 | 0 | 1 |
| 0 | 0 | 0 | 0 |
| 0 | 0 | 0 | 0 |
| 0 | 0 | 0 | 0 |
| 0 | 0 | 0 | 1 |
| 0 | 0 | 0 | 0 |
| 0 | 0 | 0 | 0 |
| 0 | 0 | 0 | 0 |
| 0 | 0 | 0 | 1 |
| 0 | 0 | 0 | 1 |
| 0 | 0 | 0 | 1 |
| 0 | 0 | 0 | 1 |

| | | | |
|---|---|---|---|
| 0 | 0 | 0 | 1 |
| 0 | 0 | 0 | 1 |
| 0 | 0 | 0 | 1 |
| 0 | 0 | 0 | 1 |
| 0 | 0 | 0 | 0 |
| 0 | 0 | 0 | 1 |
| 0 | 0 | 0 | 0 |
| 0 | 0 | 0 | 0 |
| 0 | 0 | 0 | 0 |
| 0 | 0 | 0 | 0 |
| 0 | 0 | 0 | 0 |
| 0 | 0 | 0 | 0 |
| 0 | 0 | 0 | 1 |
| 0 | 0 | 0 | 0 |
| 0 | 0 | 0 | 0 |
| 0 | 0 | 0 | 1 |
| 0 | 0 | 0 | 1 |
| 0 | 0 | 0 | 1 |
| 0 | 0 | 0 | 0 |
| 0 | 0 | 0 | 0 |
| 0 | 0 | 0 | 0 |
| 0 | 0 | 0 | 0 |
| 0 | 0 | 0 | 1 |
| 0 | 0 | 0 | 1 |
| 0 | 0 | 0 | 1 |
| 0 | 0 | 0 | 1 |
| 0 | 0 | 0 | 1 |
| 0 | 0 | 0 | 0 |
| 0 | 0 | 0 | 0 |
| 0 | 0 | 0 | 0 |
| 0 | 0 | 0 | 0 |
| 0 | 0 | 0 | 0 |
| 0 | 0 | 0 | 0 |
| 0 | 0 | 0 | 1 |
| 0 | 0 | 0 | 1 |
| 0 | 0 | 0 | 0 |
| 0 | 0 | 0 | 1 |
| 0 | 0 | 0 | 1 |
| 0 | 0 | 0 | 1 |
| 0 | 0 | 0 | 1 |
| 0 | 0 | 0 | 0 |
| 0 | 0 | 0 | 1 |
| 0 | 0 | 0 | 0 |
| 0 | 0 | 0 | 1 |
| 0 | 0 | 0 | 1 |
| 0 | 0 | 0 | 1 |
| 0 | 0 | 0 | 1 |
| 0 | 0 | 0 | 0 |
| 0 | 0 | 0 | 1 |
| 0 | 0 | 0 | 1 |
| 0 | 0 | 0 | 1 |
| 0 | 0 | 0 | 1 |
| 0 | 0 | 0 | 1 |
| 0 | 0 | 0 | 1 |

| | | | |
|---|---|---|---|
| 0 | 0 | 0 | 1 |
| 0 | 0 | 0 | 1 |
| 0 | 0 | 0 | 1 |
| 0 | 0 | 0 | 1 |
| 0 | 0 | 0 | 1 |
| 0 | 0 | 0 | 1 |
| 0 | 0 | 0 | 1 |
| 0 | 0 | 0 | 1 |
| 0 | 0 | 0 | 1 |
| 0 | 0 | 0 | 1 |
| 0 | 0 | 0 | 1 |
| 0 | 0 | 0 | 0 |
| 0 | 0 | 0 | 1 |
| 0 | 0 | 0 | 1 |
| 0 | 0 | 0 | 1 |
| 0 | 0 | 0 | 1 |
| 0 | 0 | 0 | 1 |
| 0 | 0 | 0 | 0 |
| 0 | 0 | 0 | 0 |
| 0 | 0 | 0 | 0 |
| 0 | 0 | 0 | 0 |
| 0 | 0 | 0 | 1 |
| 1 | 0 | 0 | 0 |
| 0 | 0 | 0 | 1 |
| 0 | 0 | 0 | 1 |
| 0 | 0 | 0 | 1 |
| 0 | 0 | 0 | 1 |
| 0 | 0 | 0 | 1 |
| 0 | 0 | 0 | 0 |
| 0 | 0 | 0 | 0 |
| 0 | 0 | 0 | 1 |
| 0 | 0 | 0 | 1 |
| 1 | 0 | 0 | 0 |
| 0 | 0 | 0 | 1 |
| 0 | 0 | 0 | 0 |
| 1 | 0 | 0 | 0 |
| 1 | 0 | 0 | 0 |
| 0 | 0 | 0 | 1 |
| 1 | 0 | 0 | 0 |
| 0 | 0 | 0 | 1 |
| 1 | 0 | 0 | 0 |
| 0 | 0 | 0 | 1 |
| 0 | 0 | 0 | 0 |
| 0 | 0 | 0 | 1 |
| 0 | 0 | 0 | 1 |
| 0 | 0 | 0 | 1 |
| 0 | 0 | 0 | 1 |
| 0 | 0 | 0 | 1 |
| 0 | 0 | 0 | 1 |
| 0 | 0 | 0 | 0 |
| 0 | 0 | 0 | 1 |
| 1 | 0 | 0 | 0 |
| 0 | 0 | 0 | 1 |
| 0 | 0 | 0 | 1 |

| | | | |
|---|---|---|---|
| 0 | 0 | 0 | 0 |
| 0 | 0 | 0 | 0 |
| 0 | 0 | 0 | 0 |
| 0 | 0 | 0 | 0 |
| 0 | 0 | 0 | 0 |
| 0 | 0 | 0 | 1 |
| 0 | 0 | 0 | 0 |
| 0 | 0 | 0 | 1 |
| 0 | 0 | 0 | 1 |
| 0 | 0 | 0 | 0 |
| 0 | 0 | 0 | 0 |
| 0 | 0 | 0 | 0 |
| 0 | 0 | 0 | 1 |
| 0 | 0 | 0 | 1 |
| 0 | 0 | 0 | 1 |
| 0 | 0 | 0 | 1 |
| 0 | 0 | 0 | 1 |
| 0 | 0 | 0 | 1 |
| 0 | 0 | 0 | 1 |
| 0 | 0 | 0 | 1 |
| 0 | 0 | 0 | 1 |
| 0 | 0 | 0 | 1 |
| 0 | 0 | 0 | 1 |
| 0 | 0 | 0 | 1 |
| 0 | 0 | 0 | 1 |
| 0 | 0 | 0 | 1 |
| 0 | 0 | 0 | 1 |
| 0 | 0 | 0 | 1 |
| 0 | 0 | 0 | 1 |
| 0 | 0 | 0 | 1 |
| 0 | 0 | 0 | 1 |
| 0 | 0 | 0 | 1 |
| 0 | 0 | 0 | 1 |
| 0 | 0 | 0 | 1 |
| 0 | 0 | 0 | 1 |
| 0 | 0 | 0 | 1 |
| 0 | 0 | 0 | 1 |
| 0 | 0 | 0 | 1 |
| 0 | 0 | 0 | 1 |
| 0 | 0 | 0 | 1 |
| 0 | 0 | 0 | 1 |
| 0 | 0 | 0 | 1 |
| 0 | 0 | 0 | 1 |
| 0 | 0 | 0 | 1 |
| 0 | 0 | 0 | 1 |
| 0 | 0 | 0 | 0 |
| 0 | 0 | 0 | 1 |
| 0 | 0 | 0 | 1 |
| 0 | 0 | 0 | 1 |
| 0 | 0 | 0 | 1 |
| 0 | 0 | 0 | 0 |
| 0 | 0 | 0 | 1 |
| 0 | 0 | 0 | 1 |
| 0 | 0 | 0 | 1 |

| | | | |
|---|---|---|---|
| 0 | 0 | 0 | 1 |
| 0 | 0 | 0 | 1 |
| 0 | 0 | 0 | 1 |
| 0 | 0 | 0 | 1 |
| 0 | 0 | 0 | 1 |
| 0 | 0 | 0 | 1 |
| 0 | 0 | 0 | 1 |
| 0 | 0 | 0 | 0 |
| 0 | 0 | 0 | 0 |
| 0 | 0 | 0 | 1 |
| 0 | 0 | 0 | 1 |
| 0 | 0 | 0 | 1 |
| 0 | 0 | 0 | 1 |
| 0 | 0 | 0 | 1 |
| 0 | 0 | 0 | 0 |
| 0 | 0 | 0 | 0 |
| 0 | 0 | 0 | 0 |
| 0 | 0 | 0 | 1 |
| 0 | 0 | 0 | 1 |
| 0 | 0 | 0 | 1 |
| 0 | 0 | 0 | 0 |
| 0 | 0 | 0 | 0 |
| 0 | 0 | 0 | 0 |
| 0 | 0 | 0 | 0 |
| 0 | 0 | 0 | 0 |
| 0 | 0 | 0 | 0 |
| 0 | 0 | 0 | 1 |
| 0 | 0 | 0 | 1 |
| 0 | 0 | 0 | 0 |
| 0 | 0 | 0 | 1 |
| 0 | 0 | 0 | 1 |
| 0 | 0 | 0 | 1 |
| 0 | 0 | 0 | 0 |
| 0 | 0 | 0 | 1 |
| 0 | 0 | 0 | 1 |
| 0 | 0 | 0 | 1 |
| 0 | 0 | 0 | 0 |
| 0 | 0 | 0 | 0 |
| 0 | 0 | 0 | 0 |
| 0 | 0 | 0 | 1 |
| 0 | 0 | 0 | 1 |
| 0 | 0 | 0 | 1 |
| 0 | 0 | 0 | 1 |
| 0 | 0 | 0 | 1 |
| 0 | 0 | 0 | 1 |
| 0 | 0 | 0 | 0 |
| 0 | 0 | 0 | 1 |
| 0 | 0 | 0 | 1 |
| 0 | 0 | 0 | 1 |
| 0 | 0 | 0 | 1 |
| 0 | 0 | 0 | 1 |
| 0 | 0 | 0 | 1 |
| 0 | 0 | 0 | 1 |
| 0 | 0 | 0 | 1 |

| | | | |
|---|---|---|---|
| 0 | 0 | 0 | 1 |
| 0 | 0 | 0 | 1 |
| 0 | 0 | 0 | 1 |
| 0 | 0 | 0 | 0 |
| 0 | 0 | 0 | 1 |
| 0 | 0 | 0 | 1 |
| 0 | 0 | 0 | 1 |
| 0 | 0 | 0 | 0 |
| 0 | 0 | 0 | 0 |
| 0 | 0 | 0 | 1 |
| 0 | 0 | 0 | 1 |
| 0 | 0 | 0 | 1 |
| 0 | 0 | 0 | 0 |
| 0 | 0 | 0 | 1 |
| 0 | 0 | 0 | 1 |
| 0 | 0 | 0 | 1 |
| 0 | 0 | 0 | 0 |
| 0 | 0 | 0 | 1 |
| 0 | 0 | 0 | 1 |
| 0 | 0 | 0 | 1 |
| 0 | 0 | 0 | 1 |
| 0 | 0 | 0 | 1 |
| 0 | 0 | 0 | 1 |
| 0 | 0 | 0 | 1 |
| 0 | 0 | 0 | 1 |
| 0 | 0 | 0 | 1 |
| 0 | 0 | 0 | 1 |
| 0 | 0 | 0 | 0 |
| 0 | 0 | 0 | 0 |
| 0 | 0 | 0 | 0 |
| 0 | 0 | 0 | 0 |
| 0 | 0 | 0 | 0 |
| 0 | 0 | 0 | 0 |
| 0 | 0 | 0 | 1 |
| 0 | 0 | 0 | 0 |
| 0 | 0 | 0 | 0 |
| 0 | 0 | 0 | 0 |
| 0 | 0 | 0 | 1 |
| 0 | 0 | 0 | 1 |
| 0 | 0 | 0 | 1 |
| 0 | 0 | 0 | 1 |
| 0 | 0 | 0 | 1 |
| 0 | 0 | 0 | 0 |
| 0 | 0 | 0 | 0 |
| 0 | 0 | 0 | 1 |
| 0 | 0 | 0 | 1 |
| 0 | 0 | 0 | 0 |
| 0 | 0 | 0 | 1 |
| 0 | 0 | 0 | 1 |
| 0 | 0 | 0 | 1 |
| 0 | 0 | 0 | 1 |
| 0 | 0 | 0 | 1 |
| 0 | 0 | 0 | 0 |
| 0 | 0 | 0 | 0 |

| | | | |
|---|---|---|---|
| 0 | 0 | 0 | 0 |
| 0 | 0 | 0 | 0 |
| 0 | 0 | 0 | 0 |
| 0 | 0 | 0 | 0 |
| 0 | 0 | 0 | 0 |
| 0 | 0 | 0 | 0 |
| 0 | 0 | 0 | 0 |
| 0 | 0 | 0 | 0 |
| 0 | 0 | 0 | 1 |
| 0 | 0 | 0 | 0 |
| 0 | 0 | 0 | 1 |
| 0 | 0 | 0 | 1 |
| 0 | 0 | 0 | 1 |
| 0 | 0 | 0 | 1 |
| 0 | 0 | 0 | 0 |
| 0 | 0 | 0 | 1 |
| 0 | 0 | 0 | 1 |
| 0 | 0 | 0 | 1 |
| 0 | 0 | 0 | 1 |
| 0 | 0 | 0 | 0 |
| 0 | 0 | 0 | 0 |
| 0 | 0 | 0 | 1 |
| 0 | 0 | 0 | 0 |
| 0 | 0 | 0 | 0 |
| 0 | 0 | 0 | 0 |
| 0 | 0 | 0 | 0 |
| 0 | 0 | 0 | 0 |
| 0 | 0 | 0 | 1 |
| 0 | 0 | 0 | 1 |
| 0 | 0 | 0 | 1 |
| 0 | 0 | 0 | 0 |
| 0 | 0 | 0 | 1 |
| 0 | 0 | 0 | 1 |
| 0 | 0 | 0 | 1 |
| 0 | 0 | 0 | 1 |
| 0 | 0 | 0 | 1 |
| 0 | 0 | 0 | 1 |
| 0 | 0 | 0 | 1 |
| 0 | 0 | 0 | 1 |
| 0 | 0 | 0 | 1 |
| 0 | 0 | 0 | 1 |
| 0 | 0 | 0 | 0 |
| 0 | 0 | 0 | 1 |
| 0 | 0 | 0 | 1 |
| 0 | 0 | 0 | 1 |
| 0 | 0 | 0 | 1 |
| 0 | 0 | 0 | 1 |
| 0 | 0 | 0 | 1 |
| 0 | 0 | 0 | 0 |
| 0 | 0 | 0 | 1 |
| 0 | 0 | 0 | 1 |
| 0 | 0 | 0 | 1 |
| 0 | 0 | 0 | 0 |
| 0 | 0 | 0 | 0 |

| | | | |
|---|---|---|---|
| 0 | 0 | 0 | 1 |
| 0 | 0 | 0 | 1 |
| 0 | 0 | 0 | 1 |
| 0 | 0 | 0 | 1 |
| 0 | 0 | 0 | 0 |
| 1 | 0 | 0 | 0 |
| 0 | 0 | 0 | 1 |
| 0 | 0 | 0 | 0 |
| 0 | 0 | 0 | 1 |
| 0 | 0 | 0 | 1 |
| 0 | 0 | 0 | 0 |
| 0 | 0 | 0 | 1 |
| 0 | 0 | 0 | 0 |
| 0 | 0 | 0 | 0 |
| 0 | 0 | 0 | 0 |
| 0 | 0 | 0 | 0 |
| 0 | 0 | 0 | 1 |
| 0 | 0 | 0 | 1 |
| 0 | 0 | 0 | 0 |
| 0 | 0 | 0 | 0 |
| 0 | 0 | 0 | 0 |
| 0 | 0 | 0 | 0 |
| 0 | 0 | 0 | 0 |
| 0 | 0 | 0 | 1 |
| 0 | 0 | 0 | 1 |
| 0 | 0 | 0 | 0 |
| 0 | 0 | 0 | 1 |
| 0 | 0 | 0 | 0 |
| 0 | 0 | 0 | 0 |
| 0 | 0 | 0 | 0 |
| 0 | 0 | 0 | 1 |
| 1 | 0 | 0 | 0 |
| 1 | 0 | 0 | 0 |
| 0 | 0 | 0 | 1 |
| 0 | 0 | 0 | 1 |
| 0 | 0 | 0 | 0 |
| 0 | 0 | 0 | 0 |
| 0 | 0 | 0 | 1 |
| 0 | 0 | 0 | 0 |
| 0 | 0 | 0 | 0 |
| 0 | 1 | 0 | 0 |
| 0 | 0 | 0 | 0 |
| 0 | 0 | 0 | 1 |
| 0 | 0 | 0 | 1 |
| 0 | 0 | 0 | 0 |
| 0 | 0 | 0 | 1 |
| 0 | 0 | 0 | 0 |
| 0 | 0 | 0 | 1 |
| 0 | 0 | 0 | 1 |
| 0 | 0 | 0 | 1 |
| 0 | 0 | 0 | 1 |
| 0 | 0 | 0 | 1 |
| 0 | 0 | 0 | 1 |
| 0 | 0 | 0 | 1 |

| | | | |
|---|---|---|---|
| 0 | 0 | 0 | 1 |
| 0 | 0 | 0 | 1 |
| 0 | 0 | 0 | 1 |
| 0 | 0 | 0 | 1 |
| 0 | 0 | 0 | 1 |
| 0 | 0 | 0 | 1 |
| 0 | 0 | 0 | 1 |
| 0 | 0 | 0 | 0 |
| 0 | 0 | 0 | 1 |
| 1 | 0 | 0 | 0 |
| 0 | 0 | 0 | 0 |
| 0 | 0 | 0 | 0 |
| 0 | 0 | 0 | 1 |
| 0 | 0 | 0 | 1 |
| 0 | 0 | 0 | 0 |
| 0 | 0 | 0 | 1 |
| 0 | 0 | 0 | 1 |
| 0 | 0 | 0 | 1 |
| 0 | 0 | 0 | 1 |
| 0 | 0 | 0 | 1 |
| 0 | 0 | 0 | 1 |
| 0 | 0 | 0 | 1 |
| 0 | 0 | 0 | 1 |
| 0 | 0 | 0 | 1 |
| 0 | 0 | 0 | 1 |
| 0 | 0 | 0 | 1 |
| 0 | 0 | 0 | 1 |
| 0 | 0 | 0 | 1 |
| 0 | 0 | 0 | 1 |
| 0 | 0 | 0 | 0 |
| 0 | 0 | 0 | 0 |
| 0 | 0 | 0 | 1 |
| 0 | 0 | 0 | 1 |
| 0 | 0 | 0 | 1 |
| 0 | 0 | 0 | 1 |
| 0 | 0 | 0 | 0 |
| 0 | 0 | 0 | 1 |
| 0 | 0 | 0 | 1 |
| 0 | 0 | 0 | 1 |
| 0 | 0 | 0 | 1 |
| 0 | 0 | 0 | 1 |
| 0 | 0 | 0 | 0 |
| 0 | 0 | 0 | 0 |
| 0 | 0 | 0 | 0 |
| 0 | 0 | 0 | 1 |
| 0 | 0 | 0 | 1 |
| 0 | 0 | 0 | 1 |
| 0 | 0 | 0 | 1 |
| 0 | 0 | 0 | 1 |
| 0 | 0 | 0 | 1 |
| 0 | 0 | 0 | 1 |
| 0 | 0 | 0 | 1 |
| 0 | 0 | 0 | 1 |
| 0 | 0 | 0 | 0 |

| | | | |
|---|---|---|---|
| 0 | 0 | 0 | 1 |
| 0 | 0 | 0 | 1 |
| 0 | 0 | 0 | 1 |
| 0 | 0 | 0 | 1 |
| 0 | 0 | 0 | 1 |
| 0 | 0 | 0 | 1 |
| 0 | 0 | 0 | 1 |
| 0 | 0 | 0 | 1 |
| 0 | 0 | 0 | 0 |
| 0 | 0 | 0 | 0 |
| 0 | 0 | 0 | 1 |
| 0 | 0 | 0 | 1 |
| 0 | 0 | 0 | 1 |
| 0 | 0 | 0 | 0 |
| 0 | 0 | 0 | 1 |
| 0 | 0 | 0 | 1 |
| 0 | 0 | 0 | 1 |
| 1 | 0 | 0 | 0 |
| 1 | 0 | 0 | 0 |
| 0 | 0 | 0 | 0 |
| 0 | 0 | 0 | 0 |
| 0 | 0 | 0 | 1 |
| 0 | 0 | 0 | 1 |
| 0 | 0 | 0 | 1 |
| 0 | 0 | 0 | 1 |
| 1 | 0 | 0 | 0 |
| 0 | 0 | 0 | 1 |
| 0 | 0 | 0 | 1 |
| 0 | 0 | 0 | 1 |
| 0 | 0 | 0 | 1 |
| 0 | 0 | 0 | 1 |
| 0 | 0 | 0 | 1 |
| 0 | 0 | 0 | 1 |
| 0 | 0 | 0 | 1 |
| 0 | 0 | 0 | 1 |
| 0 | 0 | 0 | 1 |
| 0 | 0 | 0 | 1 |
| 0 | 0 | 0 | 1 |
| 0 | 0 | 0 | 1 |
| 0 | 0 | 0 | 0 |
| 0 | 0 | 0 | 1 |
| 0 | 0 | 0 | 1 |
| 0 | 0 | 0 | 0 |
| 0 | 0 | 0 | 1 |
| 0 | 0 | 0 | 1 |
| 0 | 0 | 0 | 1 |
| 0 | 0 | 0 | 0 |
| 0 | 0 | 0 | 0 |
| 0 | 0 | 0 | 1 |
| 0 | 0 | 0 | 1 |
| 0 | 0 | 0 | 0 |
| 0 | 0 | 0 | 1 |
| 0 | 0 | 0 | 1 |
| 0 | 0 | 0 | 1 |

| | | | |
|---|---|---|---|
| 0 | 0 | 0 | 1 |
| 0 | 0 | 0 | 1 |
| 0 | 0 | 0 | 1 |
| 0 | 0 | 0 | 1 |
| 0 | 0 | 0 | 1 |
| 0 | 0 | 0 | 1 |
| 0 | 0 | 0 | 1 |
| 0 | 0 | 0 | 1 |
| 0 | 0 | 0 | 1 |
| 0 | 0 | 0 | 1 |
| 0 | 0 | 0 | 1 |
| 0 | 0 | 0 | 1 |
| 0 | 0 | 0 | 1 |
| 0 | 0 | 0 | 0 |
| 0 | 0 | 0 | 0 |
| 0 | 0 | 0 | 0 |
| 0 | 0 | 0 | 0 |
| 0 | 0 | 0 | 1 |
| 0 | 0 | 0 | 1 |
| 0 | 0 | 0 | 0 |
| 0 | 0 | 0 | 1 |
| 0 | 0 | 0 | 1 |
| 0 | 0 | 0 | 1 |
| 0 | 0 | 0 | 1 |
| 0 | 0 | 0 | 1 |
| 0 | 0 | 0 | 0 |
| 0 | 0 | 0 | 1 |
| 0 | 0 | 0 | 1 |
| 0 | 0 | 0 | 1 |
| 0 | 0 | 0 | 0 |
| 0 | 0 | 0 | 0 |
| 0 | 0 | 0 | 1 |
| 0 | 0 | 0 | 1 |
| 0 | 0 | 0 | 1 |
| 0 | 0 | 0 | 1 |
| 0 | 0 | 0 | 1 |
| 0 | 0 | 0 | 1 |
| 0 | 0 | 0 | 1 |
| 0 | 0 | 0 | 0 |
| 0 | 0 | 0 | 1 |
| 0 | 0 | 0 | 1 |
| 0 | 0 | 0 | 0 |
| 0 | 0 | 0 | 0 |
| 0 | 0 | 0 | 0 |
| 0 | 0 | 0 | 0 |
| 0 | 0 | 0 | 1 |
| 0 | 0 | 0 | 0 |
| 0 | 0 | 0 | 1 |
| 0 | 0 | 0 | 1 |
| 0 | 0 | 0 | 1 |
| 0 | 0 | 0 | 1 |
| 0 | 0 | 0 | 1 |
| 0 | 0 | 0 | 1 |
| 0 | 0 | 0 | 1 |

| | | | |
|---|---|---|---|
| 0 | 0 | 0 | 1 |
| 0 | 0 | 0 | 1 |
| 0 | 0 | 0 | 0 |
| 0 | 0 | 0 | 1 |
| 0 | 0 | 0 | 1 |
| 0 | 0 | 0 | 1 |
| 0 | 0 | 0 | 1 |
| 0 | 0 | 0 | 1 |
| 0 | 0 | 0 | 1 |
| 0 | 0 | 0 | 1 |
| 0 | 0 | 0 | 1 |
| 0 | 0 | 0 | 1 |
| 0 | 0 | 0 | 1 |
| 0 | 0 | 0 | 1 |
| 0 | 0 | 0 | 1 |
| 0 | 0 | 0 | 1 |
| 0 | 0 | 0 | 1 |
| 0 | 0 | 0 | 1 |
| 0 | 0 | 0 | 1 |
| 0 | 0 | 0 | 1 |
| 0 | 0 | 0 | 1 |
| 0 | 0 | 0 | 1 |
| 0 | 0 | 0 | 1 |
| 0 | 0 | 0 | 1 |
| 0 | 0 | 0 | 1 |
| 0 | 0 | 0 | 1 |
| 0 | 0 | 0 | 1 |
| 0 | 0 | 0 | 1 |
| 0 | 0 | 0 | 1 |
| 0 | 0 | 0 | 1 |
| 1 | 0 | 0 | 0 |
| 0 | 0 | 0 | 1 |
| 0 | 0 | 0 | 0 |
| 0 | 0 | 0 | 0 |
| 0 | 0 | 0 | 0 |
| 0 | 0 | 0 | 0 |
| 0 | 0 | 0 | 1 |
| 0 | 0 | 0 | 0 |
| 0 | 0 | 0 | 1 |
| 0 | 0 | 0 | 1 |
| 1 | 0 | 0 | 0 |
| 1 | 0 | 0 | 0 |
| 0 | 0 | 0 | 0 |
| 0 | 0 | 0 | 1 |
| 0 | 0 | 0 | 0 |
| 0 | 0 | 0 | 0 |
| 0 | 0 | 0 | 1 |
| 0 | 0 | 0 | 1 |
| 0 | 0 | 0 | 1 |
| 0 | 0 | 0 | 1 |
| 0 | 0 | 0 | 0 |
| 0 | 0 | 0 | 1 |
| 0 | 0 | 0 | 1 |
| 0 | 0 | 0 | 1 |

| | | | |
|---|---|---|---|
| 0 | 0 | 0 | 1 |
| 0 | 0 | 0 | 1 |
| 0 | 0 | 0 | 1 |
| 0 | 0 | 0 | 1 |
| 0 | 0 | 0 | 1 |
| 1 | 0 | 0 | 0 |
| 0 | 0 | 0 | 1 |
| 0 | 0 | 0 | 0 |
| 0 | 0 | 0 | 0 |
| 0 | 0 | 0 | 0 |
| 0 | 0 | 0 | 1 |
| 0 | 0 | 0 | 0 |
| 0 | 0 | 0 | 1 |
| 0 | 0 | 0 | 1 |
| 0 | 0 | 0 | 1 |
| 0 | 1 | 1 | 1 |
| 0 | 0 | 0 | 1 |
| 0 | 0 | 0 | 0 |
| 0 | 0 | 0 | 1 |
| 0 | 0 | 0 | 0 |
| 0 | 0 | 0 | 1 |
| 0 | 0 | 0 | 0 |
| 0 | 0 | 0 | 0 |
| 0 | 0 | 0 | 0 |
| 0 | 0 | 0 | 1 |
| 0 | 0 | 0 | 1 |
| 0 | 0 | 0 | 1 |
| 0 | 0 | 0 | 1 |
| 0 | 0 | 0 | 0 |
| 0 | 0 | 0 | 1 |
| 0 | 0 | 0 | 1 |
| 0 | 0 | 0 | 1 |
| 0 | 0 | 0 | 1 |
| 0 | 0 | 0 | 1 |
| 0 | 0 | 0 | 0 |
| 0 | 0 | 0 | 0 |
| 0 | 0 | 0 | 1 |
| 0 | 0 | 0 | 0 |
| 0 | 0 | 0 | 0 |
| 0 | 0 | 0 | 1 |
| 0 | 0 | 0 | 0 |
| 0 | 0 | 0 | 0 |
| 0 | 0 | 0 | 1 |
| 0 | 0 | 0 | 1 |
| 0 | 0 | 0 | 1 |
| 0 | 0 | 0 | 1 |
| 0 | 0 | 0 | 1 |
| 0 | 0 | 0 | 1 |
| 0 | 0 | 0 | 1 |
| 0 | 0 | 0 | 1 |
| 0 | 0 | 0 | 1 |
| 0 | 0 | 0 | 1 |
| 0 | 0 | 0 | 1 |
| 0 | 0 | 0 | 1 |

| | | | |
|---|---|---|---|
| 0 | 0 | 0 | 1 |
| 0 | 0 | 0 | 0 |
| 0 | 0 | 0 | 1 |
| 0 | 0 | 0 | 1 |
| 0 | 0 | 0 | 0 |
| 0 | 0 | 0 | 1 |
| 0 | 0 | 0 | 1 |
| 0 | 0 | 0 | 1 |
| 0 | 0 | 0 | 0 |
| 0 | 0 | 0 | 0 |
| 0 | 0 | 0 | 1 |
| 0 | 0 | 0 | 0 |
| 0 | 0 | 0 | 0 |
| 0 | 0 | 0 | 1 |
| 0 | 0 | 0 | 1 |
| 0 | 0 | 0 | 0 |
| 0 | 0 | 0 | 1 |
| 0 | 0 | 0 | 0 |
| 0 | 0 | 0 | 0 |
| 0 | 0 | 0 | 1 |
| 0 | 0 | 0 | 0 |
| 0 | 0 | 0 | 0 |
| 1 | 0 | 0 | 0 |
| 0 | 0 | 0 | 1 |
| 0 | 0 | 0 | 1 |
| 0 | 0 | 0 | 0 |
| 0 | 0 | 0 | 1 |
| 0 | 0 | 0 | 1 |
| 0 | 0 | 0 | 1 |
| 0 | 0 | 0 | 1 |
| 0 | 0 | 0 | 0 |
| 0 | 0 | 0 | 1 |
| 0 | 0 | 0 | 0 |
| 0 | 0 | 0 | 1 |
| 0 | 0 | 0 | 0 |
| 0 | 0 | 0 | 0 |
| 0 | 0 | 0 | 1 |
| 0 | 0 | 0 | 1 |
| 0 | 0 | 0 | 1 |
| 0 | 0 | 0 | 1 |
| 0 | 0 | 0 | 1 |
| 0 | 0 | 0 | 1 |
| 0 | 0 | 0 | 1 |
| 0 | 0 | 0 | 1 |
| 0 | 0 | 0 | 1 |
| 0 | 0 | 0 | 1 |
| 0 | 0 | 0 | 1 |
| 0 | 0 | 0 | 1 |
| 0 | 0 | 0 | 1 |
| 0 | 0 | 0 | 1 |
| 0 | 0 | 0 | 1 |
| 0 | 0 | 0 | 1 |
| 0 | 0 | 0 | 1 |
| 0 | 0 | 0 | 0 |

| | | | |
|---|---|---|---|
| 0 | 0 | 0 | 1 |
| 0 | 0 | 0 | 1 |
| 0 | 0 | 0 | 1 |
| 0 | 0 | 0 | 1 |
| 0 | 0 | 0 | 0 |
| 0 | 0 | 0 | 1 |
| 0 | 0 | 0 | 1 |
| 0 | 0 | 0 | 1 |
| 0 | 0 | 0 | 1 |
| 0 | 0 | 0 | 1 |
| 0 | 0 | 0 | 0 |
| 0 | 0 | 0 | 1 |
| 0 | 0 | 0 | 1 |
| 0 | 0 | 0 | 1 |
| 0 | 0 | 0 | 0 |
| 0 | 0 | 0 | 0 |
| 0 | 0 | 0 | 0 |
| 0 | 0 | 0 | 0 |
| 0 | 0 | 0 | 1 |
| 0 | 0 | 0 | 0 |
| 0 | 0 | 0 | 0 |
| 1 | 0 | 0 | 0 |
| 0 | 0 | 0 | 1 |
| 0 | 0 | 0 | 0 |
| 1 | 0 | 0 | 0 |
| 0 | 0 | 0 | 1 |
| 1 | 0 | 0 | 0 |
| 0 | 0 | 0 | 0 |
| 1 | 0 | 0 | 0 |
| 1 | 0 | 0 | 0 |
| 1 | 0 | 0 | 0 |
| 0 | 0 | 0 | 1 |
| 0 | 0 | 0 | 1 |
| 0 | 0 | 0 | 1 |
| 0 | 0 | 0 | 1 |
| 0 | 0 | 0 | 1 |
| 0 | 0 | 0 | 1 |
| 0 | 1 | 0 | 0 |
| 0 | 0 | 0 | 0 |
| 0 | 0 | 0 | 1 |
| 0 | 0 | 0 | 1 |
| 0 | 0 | 0 | 1 |
| 0 | 0 | 0 | 0 |
| 0 | 0 | 0 | 1 |
| 0 | 0 | 0 | 0 |
| 0 | 0 | 0 | 1 |
| 0 | 0 | 0 | 1 |
| 0 | 0 | 0 | 1 |
| 0 | 1 | 0 | 0 |
| 0 | 1 | 0 | 0 |
| 0 | 1 | 0 | 0 |
| 0 | 0 | 0 | 1 |
| 0 | 0 | 0 | 1 |
| 0 | 0 | 0 | 0 |

| | | | |
|---|---|---|---|
| 0 | 0 | 0 | 1 |
| 0 | 0 | 0 | 0 |
| 0 | 0 | 0 | 1 |
| 0 | 1 | 0 | 0 |
| 0 | 1 | 0 | 0 |
| 0 | 0 | 0 | 1 |
| 0 | 1 | 0 | 0 |
| 0 | 1 | 0 | 0 |
| 0 | 0 | 0 | 1 |
| 0 | 0 | 0 | 1 |
| 0 | 0 | 0 | 1 |
| 0 | 0 | 0 | 1 |
| 0 | 0 | 0 | 1 |
| 0 | 0 | 0 | 1 |
| 0 | 0 | 0 | 1 |
| 0 | 0 | 0 | 1 |
| 0 | 0 | 0 | 1 |
| 0 | 0 | 0 | 1 |
| 0 | 0 | 0 | 1 |
| 0 | 0 | 0 | 1 |
| 1 | 0 | 0 | 0 |
| 0 | 0 | 0 | 1 |
| 0 | 0 | 0 | 1 |
| 0 | 0 | 0 | 1 |
| 0 | 0 | 0 | 1 |
| 0 | 0 | 0 | 0 |
| 0 | 0 | 0 | 0 |
| 0 | 0 | 0 | 1 |
| 0 | 0 | 0 | 1 |
| 0 | 0 | 0 | 1 |
| 0 | 0 | 0 | 1 |
| 0 | 0 | 0 | 1 |
| 0 | 0 | 0 | 1 |
| 0 | 0 | 0 | 1 |
| 0 | 0 | 0 | 1 |
| 0 | 0 | 0 | 1 |
| 0 | 0 | 0 | 1 |
| 0 | 0 | 0 | 1 |
| 0 | 0 | 0 | 1 |
| 0 | 0 | 0 | 1 |
| 0 | 0 | 0 | 1 |
| 0 | 0 | 0 | 1 |
| 0 | 0 | 0 | 1 |
| 0 | 0 | 0 | 0 |
| 0 | 0 | 0 | 1 |
| 0 | 0 | 0 | 0 |
| 0 | 0 | 0 | 0 |
| 0 | 0 | 0 | 0 |
| 0 | 0 | 0 | 0 |
| 0 | 0 | 0 | 0 |
| 0 | 0 | 0 | 0 |
| 0 | 0 | 0 | 1 |
| 0 | 0 | 0 | 1 |
| 0 | 0 | 0 | 1 |

| | | | |
|---|---|---|---|
| 0 | 0 | 0 | 1 |
| 0 | 0 | 0 | 1 |
| 0 | 0 | 0 | 1 |
| 0 | 0 | 0 | 0 |
| 0 | 0 | 0 | 1 |
| 0 | 0 | 0 | 0 |
| 0 | 0 | 0 | 1 |
| 0 | 0 | 0 | 1 |
| 0 | 0 | 0 | 0 |
| 0 | 0 | 0 | 1 |
| 0 | 0 | 0 | 1 |
| 0 | 0 | 0 | 1 |
| 0 | 0 | 0 | 1 |
| 0 | 0 | 0 | 1 |
| 0 | 0 | 0 | 1 |
| 0 | 0 | 0 | 0 |
| 0 | 0 | 0 | 0 |
| 0 | 0 | 0 | 0 |
| 0 | 0 | 0 | 0 |
| 0 | 0 | 0 | 0 |
| 0 | 0 | 0 | 1 |
| 0 | 0 | 0 | 1 |
| 0 | 0 | 0 | 1 |
| 0 | 0 | 0 | 1 |
| 0 | 0 | 0 | 0 |
| 0 | 0 | 0 | 1 |
| 0 | 0 | 0 | 1 |
| 0 | 0 | 0 | 1 |
| 0 | 0 | 0 | 0 |
| 0 | 0 | 0 | 1 |
| 0 | 0 | 0 | 0 |
| 0 | 0 | 0 | 0 |
| 0 | 0 | 0 | 0 |
| 0 | 0 | 0 | 0 |
| 0 | 0 | 0 | 0 |
| 0 | 0 | 0 | 1 |
| 0 | 0 | 0 | 0 |
| 0 | 0 | 0 | 0 |
| 0 | 0 | 0 | 0 |
| 0 | 0 | 0 | 0 |
| 0 | 0 | 0 | 0 |
| 0 | 0 | 0 | 0 |
| 0 | 0 | 0 | 0 |
| 0 | 0 | 0 | 0 |
| 0 | 0 | 0 | 0 |
| 0 | 0 | 0 | 0 |
| 0 | 0 | 0 | 0 |
| 0 | 0 | 0 | 0 |
| 0 | 0 | 0 | 1 |
| 0 | 0 | 0 | 0 |
| 0 | 0 | 0 | 1 |
| 0 | 0 | 0 | 1 |
| 0 | 0 | 0 | 1 |
| 0 | 0 | 0 | 0 |

| | | | |
|---|---|---|---|
| 0 | 0 | 0 | 0 |
| 0 | 0 | 0 | 1 |
| 0 | 0 | 0 | 1 |
| 0 | 0 | 0 | 1 |
| 0 | 0 | 0 | 1 |
| 0 | 0 | 0 | 0 |
| 0 | 0 | 0 | 1 |
| 0 | 0 | 0 | 1 |
| 0 | 0 | 0 | 1 |
| 0 | 0 | 0 | 1 |
| 0 | 0 | 0 | 1 |
| 0 | 0 | 0 | 1 |
| 0 | 0 | 0 | 1 |
| 1 | 0 | 0 | 0 |
| 0 | 0 | 0 | 1 |
| 0 | 0 | 0 | 1 |
| 0 | 0 | 0 | 1 |
| 0 | 0 | 0 | 0 |
| 0 | 0 | 0 | 1 |
| 0 | 0 | 0 | 1 |
| 0 | 0 | 0 | 1 |
| 0 | 0 | 0 | 1 |
| 0 | 0 | 0 | 0 |
| 0 | 0 | 0 | 1 |
| 0 | 0 | 0 | 1 |
| 0 | 0 | 0 | 0 |
| 0 | 0 | 0 | 0 |
| 0 | 0 | 0 | 0 |
| 0 | 0 | 0 | 1 |
| 0 | 0 | 0 | 1 |
| 0 | 0 | 0 | 1 |
| 0 | 0 | 0 | 1 |
| 0 | 0 | 0 | 1 |
| 0 | 0 | 0 | 1 |
| 0 | 0 | 0 | 1 |
| 0 | 0 | 0 | 1 |
| 0 | 0 | 0 | 1 |
| 0 | 0 | 0 | 1 |
| 0 | 0 | 0 | 1 |
| 0 | 0 | 0 | 1 |
| 0 | 0 | 0 | 1 |
| 0 | 0 | 0 | 1 |
| 0 | 0 | 0 | 1 |
| 0 | 0 | 0 | 1 |
| 0 | 0 | 0 | 1 |
| 0 | 0 | 0 | 1 |
| 0 | 0 | 0 | 1 |
| 0 | 0 | 0 | 1 |
| 0 | 0 | 0 | 1 |
| 0 | 0 | 0 | 1 |
| 0 | 0 | 0 | 1 |
| 0 | 0 | 0 | 1 |
| 0 | 0 | 0 | 1 |
| 0 | 0 | 0 | 1 |

| | | | |
|---|---|---|---|
| 0 | 0 | 0 | 1 |
| 0 | 0 | 0 | 0 |
| 0 | 0 | 0 | 1 |
| 0 | 0 | 0 | 1 |
| 0 | 0 | 0 | 1 |
| 0 | 0 | 0 | 0 |
| 0 | 0 | 0 | 0 |
| 0 | 0 | 0 | 0 |
| 0 | 0 | 0 | 1 |
| 0 | 0 | 0 | 1 |
| 0 | 0 | 0 | 1 |
| 0 | 0 | 0 | 1 |
| 0 | 0 | 0 | 1 |
| 0 | 0 | 0 | 1 |
| 0 | 0 | 0 | 0 |
| 0 | 0 | 0 | 0 |
| 0 | 0 | 0 | 0 |
| 0 | 0 | 0 | 1 |
| 0 | 0 | 0 | 0 |
| 0 | 0 | 0 | 0 |
| 0 | 0 | 0 | 0 |
| 0 | 0 | 0 | 0 |
| 0 | 0 | 0 | 0 |
| 0 | 0 | 0 | 0 |
| 0 | 0 | 0 | 1 |
| 0 | 0 | 0 | 1 |
| 0 | 0 | 0 | 1 |
| 0 | 0 | 0 | 1 |
| 0 | 0 | 0 | 1 |
| 0 | 0 | 0 | 1 |
| 0 | 0 | 0 | 1 |
| 0 | 0 | 0 | 1 |
| 0 | 0 | 0 | 0 |
| 0 | 0 | 0 | 1 |
| 0 | 0 | 0 | 1 |
| 0 | 0 | 0 | 0 |
| 0 | 0 | 0 | 0 |
| 0 | 0 | 0 | 0 |
| 0 | 0 | 0 | 0 |
| 0 | 0 | 0 | 1 |
| 0 | 0 | 0 | 1 |
| 0 | 0 | 0 | 1 |
| 0 | 0 | 0 | 1 |
| 0 | 0 | 0 | 0 |
| 0 | 0 | 0 | 0 |
| 0 | 0 | 0 | 0 |
| 0 | 0 | 0 | 1 |
| 0 | 0 | 0 | 1 |
| 0 | 0 | 0 | 0 |
| 0 | 0 | 0 | 1 |
| 0 | 0 | 0 | 1 |
| 0 | 0 | 0 | 0 |
| 0 | 0 | 0 | 1 |
| 0 | 0 | 0 | 1 |

| | | | |
|---|---|---|---|
| 0 | 0 | 0 | 1 |
| 0 | 0 | 0 | 1 |
| 0 | 0 | 0 | 1 |
| 0 | 0 | 0 | 1 |
| 0 | 0 | 0 | 1 |
| 0 | 0 | 0 | 1 |
| 0 | 0 | 0 | 1 |
| 0 | 0 | 0 | 1 |
| 0 | 0 | 0 | 0 |
| 0 | 0 | 0 | 0 |
| 0 | 0 | 0 | 1 |
| 0 | 0 | 0 | 0 |
| 0 | 0 | 0 | 0 |
| 0 | 0 | 0 | 0 |
| 0 | 0 | 0 | 1 |
| 0 | 0 | 0 | 1 |
| 0 | 0 | 0 | 0 |
| 0 | 0 | 0 | 0 |
| 0 | 0 | 0 | 0 |
| 0 | 0 | 0 | 1 |
| 0 | 0 | 0 | 1 |
| 0 | 0 | 0 | 1 |
| 0 | 0 | 0 | 0 |
| 0 | 0 | 0 | 1 |
| 0 | 0 | 0 | 0 |
| 0 | 0 | 0 | 1 |
| 0 | 0 | 0 | 1 |
| 0 | 0 | 0 | 1 |
| 0 | 0 | 0 | 1 |
| 0 | 0 | 0 | 0 |
| 0 | 0 | 0 | 0 |
| 0 | 0 | 0 | 0 |
| 0 | 0 | 0 | 1 |
| 0 | 0 | 0 | 1 |
| 0 | 0 | 0 | 1 |
| 0 | 0 | 0 | 1 |
| 0 | 0 | 0 | 1 |
| 0 | 0 | 0 | 1 |
| 0 | 0 | 0 | 1 |
| 0 | 0 | 0 | 1 |
| 0 | 0 | 0 | 0 |
| 0 | 0 | 0 | 1 |
| 0 | 0 | 0 | 1 |
| 0 | 0 | 0 | 1 |
| 0 | 0 | 0 | 1 |
| 0 | 0 | 0 | 0 |
| 0 | 0 | 0 | 1 |
| 0 | 0 | 0 | 0 |
| 0 | 0 | 0 | 1 |
| 0 | 0 | 0 | 0 |
| 0 | 0 | 0 | 0 |
| 0 | 0 | 0 | 0 |
| 0 | 0 | 0 | 1 |
| 0 | 0 | 0 | 1 |

| | | | |
|---|---|---|---|
| 0 | 0 | 0 | 1 |
| 0 | 0 | 0 | 1 |
| 0 | 0 | 0 | 1 |
| 0 | 0 | 0 | 1 |
| 0 | 0 | 0 | 1 |
| 0 | 0 | 0 | 1 |
| 0 | 0 | 0 | 1 |
| 0 | 0 | 0 | 0 |
| 0 | 0 | 0 | 0 |
| 0 | 0 | 0 | 1 |
| 0 | 0 | 0 | 0 |
| 0 | 0 | 0 | 1 |
| 0 | 0 | 0 | 1 |
| 0 | 0 | 0 | 1 |
| 0 | 0 | 0 | 1 |
| 0 | 0 | 0 | 1 |
| 0 | 0 | 0 | 1 |
| 0 | 0 | 0 | 1 |
| 0 | 0 | 0 | 1 |
| 0 | 0 | 0 | 1 |
| 0 | 0 | 0 | 1 |
| 0 | 0 | 0 | 1 |
| 0 | 0 | 0 | 1 |
| 0 | 0 | 0 | 1 |
| 0 | 0 | 0 | 1 |
| 0 | 0 | 0 | 1 |
| 0 | 0 | 0 | 1 |
| 0 | 0 | 0 | 1 |
| 0 | 0 | 0 | 0 |
| 0 | 0 | 0 | 1 |
| 0 | 0 | 0 | 0 |
| 0 | 0 | 0 | 1 |
| 0 | 0 | 0 | 0 |
| 0 | 0 | 0 | 0 |
| 0 | 0 | 0 | 1 |
| 0 | 0 | 0 | 0 |
| 0 | 0 | 0 | 0 |
| 0 | 0 | 0 | 1 |
| 0 | 0 | 0 | 1 |
| 0 | 0 | 0 | 1 |
| 0 | 0 | 0 | 1 |
| 0 | 0 | 0 | 0 |
| 0 | 0 | 0 | 0 |
| 0 | 0 | 0 | 1 |
| 0 | 0 | 0 | 1 |
| 0 | 0 | 0 | 0 |
| 0 | 0 | 0 | 0 |
| 0 | 0 | 0 | 0 |
| 0 | 0 | 0 | 0 |
| 0 | 0 | 0 | 1 |
| 0 | 0 | 0 | 0 |
| 0 | 0 | 0 | 0 |
| 0 | 0 | 0 | 1 |
| 0 | 0 | 0 | 1 |

| | | | |
|---|---|---|---|
| 0 | 0 | 0 | 1 |
| 0 | 0 | 0 | 1 |
| 0 | 0 | 0 | 1 |
| 0 | 0 | 0 | 1 |
| 0 | 0 | 0 | 0 |
| 0 | 0 | 0 | 0 |
| 0 | 0 | 0 | 0 |
| 0 | 0 | 0 | 0 |
| 0 | 0 | 0 | 1 |
| 0 | 0 | 0 | 0 |
| 0 | 0 | 0 | 0 |
| 0 | 0 | 0 | 0 |
| 0 | 0 | 0 | 0 |
| 0 | 0 | 0 | 1 |
| 0 | 0 | 0 | 1 |
| 0 | 0 | 0 | 1 |
| 0 | 0 | 0 | 1 |
| 0 | 0 | 0 | 1 |
| 0 | 0 | 0 | 1 |
| 0 | 0 | 0 | 1 |
| 0 | 0 | 0 | 1 |
| 0 | 0 | 0 | 1 |
| 0 | 0 | 0 | 1 |
| 0 | 0 | 0 | 1 |
| 0 | 0 | 0 | 1 |
| 0 | 0 | 0 | 0 |
| 0 | 0 | 0 | 0 |
| 0 | 0 | 0 | 0 |
| 0 | 0 | 0 | 0 |
| 0 | 0 | 0 | 1 |
| 0 | 0 | 0 | 1 |
| 0 | 0 | 0 | 1 |
| 0 | 0 | 0 | 1 |
| 0 | 0 | 0 | 1 |
| 0 | 0 | 0 | 1 |
| 0 | 0 | 0 | 1 |
| 0 | 0 | 0 | 1 |
| 0 | 0 | 0 | 1 |
| 0 | 0 | 0 | 1 |
| 0 | 0 | 0 | 1 |
| 0 | 0 | 0 | 1 |
| 0 | 0 | 0 | 1 |
| 0 | 0 | 0 | 1 |
| 0 | 0 | 0 | 1 |
| 0 | 0 | 0 | 1 |
| 0 | 0 | 0 | 1 |
| 0 | 0 | 0 | 1 |
| 0 | 0 | 0 | 1 |
| 0 | 0 | 0 | 1 |
| 0 | 0 | 0 | 1 |
| 0 | 0 | 0 | 1 |
| 0 | 0 | 0 | 1 |
| 0 | 0 | 0 | 1 |
| 0 | 0 | 0 | 0 |

| | | | |
|---|---|---|---|
| 0 | 0 | 0 | 1 |
| 0 | 0 | 0 | 1 |
| 0 | 0 | 0 | 1 |
| 0 | 0 | 0 | 1 |
| 0 | 0 | 0 | 1 |
| 0 | 0 | 0 | 1 |
| 0 | 0 | 0 | 1 |
| 0 | 0 | 0 | 1 |
| 0 | 0 | 0 | 1 |
| 0 | 0 | 0 | 1 |
| 0 | 0 | 0 | 1 |
| 0 | 0 | 0 | 1 |
| 0 | 0 | 0 | 1 |
| 0 | 0 | 0 | 1 |
| 0 | 0 | 0 | 1 |
| 0 | 0 | 0 | 1 |
| 0 | 0 | 0 | 1 |
| 0 | 0 | 0 | 1 |
| 0 | 0 | 0 | 1 |
| 0 | 0 | 0 | 1 |
| 0 | 0 | 0 | 1 |
| 0 | 0 | 0 | 1 |
| 0 | 0 | 0 | 1 |
| 0 | 0 | 0 | 1 |
| 0 | 0 | 0 | 1 |
| 0 | 0 | 0 | 1 |
| 0 | 0 | 0 | 1 |
| 0 | 0 | 0 | 1 |
| 0 | 0 | 0 | 1 |
| 0 | 0 | 0 | 1 |
| 0 | 0 | 0 | 1 |
| 0 | 0 | 0 | 1 |
| 0 | 0 | 0 | 1 |
| 0 | 0 | 0 | 1 |
| 0 | 0 | 0 | 1 |
| 0 | 0 | 0 | 1 |
| 0 | 0 | 0 | 1 |
| 0 | 0 | 0 | 1 |
| 0 | 0 | 0 | 1 |
| 0 | 0 | 0 | 1 |
| 0 | 0 | 0 | 1 |
| 0 | 0 | 0 | 1 |
| 0 | 0 | 0 | 1 |
| 0 | 0 | 0 | 1 |
| 0 | 0 | 0 | 1 |
| 0 | 0 | 0 | 1 |
| 0 | 0 | 0 | 0 |
| 0 | 0 | 0 | 1 |
| 0 | 0 | 0 | 1 |
| 0 | 0 | 0 | 1 |
| 0 | 0 | 0 | 1 |
| 0 | 0 | 0 | 1 |
| 0 | 0 | 0 | 1 |
| 0 | 0 | 0 | 1 |

| | | | |
|---|---|---|---|
| 0 | 0 | 0 | 1 |
| 0 | 0 | 0 | 1 |
| 0 | 0 | 0 | 0 |
| 0 | 0 | 0 | 1 |
| 0 | 0 | 0 | 1 |
| 0 | 0 | 0 | 1 |
| 0 | 0 | 0 | 0 |
| 0 | 0 | 0 | 1 |
| 0 | 0 | 0 | 1 |
| 0 | 0 | 0 | 1 |
| 0 | 0 | 0 | 1 |
| 0 | 0 | 0 | 0 |
| 0 | 0 | 0 | 1 |
| 0 | 0 | 0 | 0 |
| 0 | 0 | 0 | 0 |
| 0 | 0 | 0 | 0 |
| 0 | 0 | 0 | 0 |
| 0 | 0 | 0 | 0 |
| 0 | 0 | 0 | 0 |
| 0 | 0 | 0 | 0 |
| 0 | 0 | 0 | 0 |

| surface_non_binding_site | non_binding_site_single_water | surface_non_binding_site_single_water |
|---|---|---|
| 0 | 0 | 0 |
| 0 | 0 | 0 |
| 0 | 0 | 0 |
| 0 | 0 | 0 |
| 0 | 0 | 0 |
| 0 | 0 | 0 |
| 1 | 1 | 1 |
| 1 | 1 | 1 |
| 0 | 0 | 0 |
| 0 | 0 | 0 |
| 0 | 0 | 0 |
| 0 | 0 | 0 |
| 0 | 0 | 0 |
| 0 | 0 | 0 |
| 0 | 0 | 0 |
| 0 | 0 | 0 |
| 0 | 0 | 0 |
| 0 | 0 | 0 |
| 0 | 0 | 0 |
| 0 | 0 | 0 |
| 0 | 0 | 0 |
| 0 | 0 | 0 |
| 0 | 0 | 0 |
| 0 | 0 | 0 |
| 0 | 0 | 0 |
| 1 | 0 | 0 |
| 0 | 0 | 0 |
| 1 | 1 | 1 |
| 0 | 0 | 0 |
| 0 | 1 | 0 |
| 0 | 1 | 0 |
| 0 | 0 | 0 |
| 0 | 0 | 0 |
| 0 | 0 | 0 |
| 0 | 0 | 0 |
| 1 | 1 | 1 |
| 0 | 0 | 0 |
| 0 | 0 | 0 |
| 0 | 0 | 0 |
| 0 | 0 | 0 |
| 0 | 0 | 0 |
| 0 | 0 | 0 |
| 0 | 0 | 0 |
| 0 | 0 | 0 |
| 0 | 0 | 0 |
| 0 | 0 | 0 |
| 0 | 0 | 0 |
| 0 | 0 | 0 |
| 0 | 0 | 0 |
| 0 | 0 | 0 |
| 0 | 0 | 0 |
| 0 | 0 | 0 |
| 0 | 0 | 0 |

| | | |
|---|---|---|
| 0 | 0 | 0 |
| 0 | 0 | 0 |
| 0 | 0 | 0 |
| 0 | 0 | 0 |
| 0 | 0 | 0 |
| 0 | 0 | 0 |
| 0 | 0 | 0 |
| 0 | 0 | 0 |
| 0 | 0 | 0 |
| 0 | 0 | 0 |
| 0 | 0 | 0 |
| 0 | 0 | 0 |
| 0 | 0 | 0 |
| 0 | 0 | 0 |
| 0 | 0 | 0 |
| 0 | 0 | 0 |
| 0 | 0 | 0 |
| 0 | 0 | 0 |
| 0 | 0 | 0 |
| 0 | 0 | 0 |
| 0 | 0 | 0 |
| 0 | 0 | 0 |
| 0 | 0 | 0 |
| 0 | 0 | 0 |
| 0 | 0 | 0 |
| 0 | 0 | 0 |
| 0 | 0 | 0 |
| 0 | 0 | 0 |
| 0 | 0 | 0 |
| 0 | 0 | 0 |
| 0 | 0 | 0 |
| 0 | 0 | 0 |
| 0 | 0 | 0 |
| 0 | 0 | 0 |
| 0 | 0 | 0 |
| 0 | 0 | 0 |
| 0 | 0 | 0 |
| 0 | 0 | 0 |
| 0 | 0 | 0 |
| 0 | 0 | 0 |
| 0 | 0 | 0 |
| 0 | 0 | 0 |
| 0 | 0 | 0 |
| 0 | 0 | 0 |
| 0 | 0 | 0 |
| 1 | 1 | 1 |
| 0 | 1 | 0 |
| 0 | 0 | 0 |
| 0 | 0 | 0 |
| 0 | 0 | 0 |
| 0 | 0 | 0 |
| 1 | 1 | 1 |
| 1 | 1 | 1 |
| 0 | 0 | 0 |

| | | |
|---|---|---|
| 0 | 1 | 0 |
| 0 | 1 | 0 |
| 0 | 0 | 0 |
| 0 | 0 | 0 |
| 0 | 0 | 0 |
| 0 | 0 | 0 |
| 0 | 1 | 0 |
| 0 | 0 | 0 |
| 0 | 0 | 0 |
| 0 | 0 | 0 |
| 0 | 0 | 0 |
| 0 | 0 | 0 |
| 0 | 0 | 0 |
| 0 | 0 | 0 |
| 0 | 0 | 0 |
| 1 | 1 | 1 |
| 0 | 0 | 0 |
| 0 | 0 | 0 |
| 0 | 0 | 0 |
| 0 | 0 | 0 |
| 0 | 0 | 0 |
| 0 | 0 | 0 |
| 0 | 0 | 0 |
| 0 | 0 | 0 |
| 0 | 0 | 0 |
| 0 | 0 | 0 |
| 0 | 0 | 0 |
| 0 | 0 | 0 |
| 0 | 0 | 0 |
| 0 | 0 | 0 |
| 0 | 0 | 0 |
| 0 | 0 | 0 |
| 0 | 0 | 0 |
| 0 | 0 | 0 |
| 0 | 0 | 0 |
| 0 | 0 | 0 |
| 0 | 0 | 0 |
| 0 | 0 | 0 |
| 0 | 0 | 0 |
| 0 | 0 | 0 |
| 0 | 0 | 0 |
| 0 | 0 | 0 |
| 0 | 0 | 0 |
| 0 | 0 | 0 |
| 0 | 0 | 0 |
| 0 | 0 | 0 |
| 0 | 0 | 0 |
| 0 | 0 | 0 |
| 0 | 0 | 0 |
| 0 | 0 | 0 |
| 0 | 0 | 0 |
| 0 | 0 | 0 |
| 0 | 0 | 0 |
| 0 | 0 | 0 |

| | | |
|---|---|---|
| 1 | 0 | 0 |
| 1 | 0 | 0 |
| 1 | 0 | 0 |
| 0 | 0 | 0 |
| 0 | 0 | 0 |
| 0 | 0 | 0 |
| 0 | 0 | 0 |
| 1 | 0 | 0 |
| 0 | 0 | 0 |
| 1 | 0 | 0 |
| 1 | 0 | 0 |
| 0 | 0 | 0 |
| 1 | 0 | 0 |
| 0 | 0 | 0 |
| 0 | 0 | 0 |
| 1 | 0 | 0 |
| 1 | 0 | 0 |
| 1 | 0 | 0 |
| 1 | 0 | 0 |
| 0 | 0 | 0 |
| 0 | 0 | 0 |
| 0 | 1 | 0 |
| 0 | 0 | 0 |
| 0 | 0 | 0 |
| 0 | 0 | 0 |
| 1 | 0 | 0 |
| 0 | 0 | 0 |
| 0 | 0 | 0 |
| 1 | 0 | 0 |
| 1 | 0 | 0 |
| 1 | 1 | 1 |
| 0 | 0 | 0 |
| 0 | 0 | 0 |
| 0 | 1 | 0 |
| 0 | 0 | 0 |
| 0 | 0 | 0 |
| 0 | 0 | 0 |
| 0 | 0 | 0 |
| 0 | 0 | 0 |
| 1 | 0 | 0 |
| 1 | 0 | 0 |
| 0 | 0 | 0 |
| 0 | 0 | 0 |
| 1 | 1 | 1 |
| 1 | 0 | 0 |
| 1 | 0 | 0 |
| 0 | 0 | 0 |
| 0 | 0 | 0 |
| 0 | 0 | 0 |
| 0 | 0 | 0 |
| 0 | 0 | 0 |
| 0 | 1 | 0 |
| 1 | 0 | 0 |
| 0 | 0 | 0 |

| | | |
|---|---|---|
| 0 | 0 | 0 |
| 0 | 0 | 0 |
| 0 | 0 | 0 |
| 1 | 1 | 1 |
| 1 | 1 | 1 |
| 1 | 0 | 0 |
| 0 | 0 | 0 |
| 0 | 0 | 0 |
| 1 | 0 | 0 |
| 1 | 0 | 0 |
| 0 | 0 | 0 |
| 0 | 1 | 0 |
| 0 | 0 | 0 |
| 1 | 0 | 0 |
| 0 | 1 | 0 |
| 1 | 1 | 1 |
| 1 | 1 | 1 |
| 0 | 0 | 0 |
| 0 | 0 | 0 |
| 0 | 0 | 0 |
| 0 | 0 | 0 |
| 1 | 0 | 0 |
| 0 | 0 | 0 |
| 0 | 1 | 0 |
| 1 | 1 | 1 |
| 0 | 0 | 0 |
| 0 | 0 | 0 |
| 0 | 0 | 0 |
| 0 | 0 | 0 |
| 0 | 0 | 0 |
| 0 | 0 | 0 |
| 0 | 0 | 0 |
| 1 | 0 | 0 |
| 1 | 0 | 0 |
| 0 | 0 | 0 |
| 1 | 0 | 0 |
| 0 | 0 | 0 |
| 0 | 0 | 0 |
| 1 | 0 | 0 |
| 0 | 1 | 0 |
| 0 | 0 | 0 |
| 1 | 0 | 0 |
| 1 | 0 | 0 |
| 1 | 0 | 0 |
| 0 | 0 | 0 |
| 0 | 0 | 0 |
| 0 | 0 | 0 |
| 0 | 0 | 0 |
| 0 | 0 | 0 |
| 0 | 0 | 0 |
| 0 | 0 | 0 |
| 0 | 0 | 0 |
| 0 | 0 | 0 |
| 0 | 0 | 0 |

| | | |
|---|---|---|
| 0 | 0 | 0 |
| 0 | 0 | 0 |
| 0 | 0 | 0 |
| 0 | 0 | 0 |
| 0 | 0 | 0 |
| 0 | 0 | 0 |
| 0 | 1 | 0 |
| 0 | 1 | 0 |
| 0 | 0 | 0 |
| 0 | 0 | 0 |
| 0 | 0 | 0 |
| 0 | 0 | 0 |
| 0 | 0 | 0 |
| 0 | 1 | 0 |
| 0 | 1 | 0 |
| 0 | 0 | 0 |
| 0 | 1 | 0 |
| 0 | 0 | 0 |
| 1 | 1 | 1 |
| 0 | 0 | 0 |
| 1 | 1 | 1 |
| 1 | 0 | 0 |
| 1 | 1 | 1 |
| 0 | 0 | 0 |
| 0 | 0 | 0 |
| 0 | 0 | 0 |
| 0 | 0 | 0 |
| 0 | 0 | 0 |
| 0 | 0 | 0 |
| 0 | 0 | 0 |
| 0 | 0 | 0 |
| 1 | 0 | 0 |
| 0 | 0 | 0 |
| 0 | 0 | 0 |
| 0 | 0 | 0 |
| 0 | 0 | 0 |
| 0 | 0 | 0 |
| 0 | 0 | 0 |
| 0 | 0 | 0 |
| 0 | 0 | 0 |
| 0 | 1 | 0 |
| 0 | 1 | 0 |
| 0 | 0 | 0 |
| 0 | 1 | 0 |
| 0 | 0 | 0 |
| 0 | 0 | 0 |
| 0 | 0 | 0 |
| 1 | 1 | 1 |
| 1 | 1 | 1 |
| 0 | 0 | 0 |
| 0 | 1 | 0 |
| 1 | 1 | 1 |
| 0 | 0 | 0 |
| 0 | 1 | 0 |

| | | |
|---|---|---|
| 0 | 0 | 0 |
| 1 | 1 | 1 |
| 0 | 0 | 0 |
| 0 | 0 | 0 |
| 0 | 0 | 0 |
| 0 | 0 | 0 |
| 0 | 1 | 0 |
| 0 | 0 | 0 |
| 0 | 0 | 0 |
| 1 | 1 | 1 |
| 1 | 1 | 1 |
| 1 | 1 | 1 |
| 1 | 1 | 1 |
| 1 | 1 | 1 |
| 0 | 0 | 0 |
| 1 | 1 | 1 |
| 0 | 0 | 0 |
| 0 | 0 | 0 |
| 1 | 1 | 1 |
| 0 | 0 | 0 |
| 0 | 0 | 0 |
| 0 | 0 | 0 |
| 0 | 0 | 0 |
| 0 | 0 | 0 |
| 1 | 1 | 1 |
| 1 | 0 | 0 |
| 0 | 0 | 0 |
| 0 | 0 | 0 |
| 0 | 0 | 0 |
| 0 | 0 | 0 |
| 0 | 0 | 0 |
| 0 | 0 | 0 |
| 0 | 0 | 0 |
| 0 | 0 | 0 |
| 1 | 1 | 1 |
| 1 | 1 | 1 |
| 0 | 0 | 0 |
| 0 | 0 | 0 |
| 0 | 0 | 0 |
| 0 | 0 | 0 |
| 0 | 0 | 0 |
| 0 | 0 | 0 |
| 0 | 1 | 0 |
| 0 | 0 | 0 |
| 0 | 0 | 0 |
| 0 | 0 | 0 |
| 0 | 0 | 0 |
| 0 | 0 | 0 |
| 0 | 0 | 0 |
| 1 | 1 | 1 |
| 0 | 0 | 0 |
| 0 | 0 | 0 |
| 0 | 0 | 0 |
| 0 | 0 | 0 |

| | | |
|---|---|---|
| 0 | 0 | 0 |
| 0 | 0 | 0 |
| 0 | 0 | 0 |
| 0 | 0 | 0 |
| 1 | 1 | 1 |
| 0 | 0 | 0 |
| 0 | 0 | 0 |
| 0 | 0 | 0 |
| 0 | 0 | 0 |
| 0 | 0 | 0 |
| 0 | 0 | 0 |
| 0 | 0 | 0 |
| 0 | 0 | 0 |
| 0 | 0 | 0 |
| 0 | 0 | 0 |
| 0 | 0 | 0 |
| 0 | 0 | 0 |
| 0 | 0 | 0 |
| 1 | 1 | 1 |
| 0 | 0 | 0 |
| 0 | 0 | 0 |
| 0 | 0 | 0 |
| 0 | 0 | 0 |
| 1 | 0 | 0 |
| 0 | 0 | 0 |
| 0 | 0 | 0 |
| 0 | 0 | 0 |
| 1 | 0 | 0 |
| 1 | 0 | 0 |
| 1 | 0 | 0 |
| 0 | 0 | 0 |
| 0 | 0 | 0 |
| 0 | 0 | 0 |
| 0 | 0 | 0 |
| 0 | 0 | 0 |
| 1 | 0 | 0 |
| 1 | 0 | 0 |
| 0 | 0 | 0 |
| 0 | 0 | 0 |
| 1 | 0 | 0 |
| 1 | 0 | 0 |
| 1 | 0 | 0 |
| 0 | 1 | 0 |
| 1 | 1 | 1 |
| 0 | 0 | 0 |
| 0 | 0 | 0 |
| 0 | 0 | 0 |
| 0 | 0 | 0 |
| 0 | 0 | 0 |
| 0 | 1 | 0 |
| 1 | 1 | 1 |
| 0 | 0 | 0 |
| 1 | 1 | 1 |
| 0 | 0 | 0 |

| | | |
|---|---|---|
| 1 | 1 | 1 |
| 1 | 1 | 1 |
| 0 | 0 | 0 |
| 1 | 1 | 1 |
| 1 | 1 | 1 |
| 1 | 1 | 1 |
| 1 | 1 | 1 |
| 0 | 0 | 0 |
| 1 | 0 | 0 |
| 1 | 1 | 1 |
| 0 | 0 | 0 |
| 1 | 1 | 1 |
| 1 | 1 | 1 |
| 0 | 1 | 0 |
| 0 | 0 | 0 |
| 0 | 0 | 0 |
| 1 | 1 | 1 |
| 1 | 1 | 1 |
| 0 | 0 | 0 |
| 0 | 0 | 0 |
| 0 | 0 | 0 |
| 0 | 0 | 0 |
| 0 | 0 | 0 |
| 0 | 0 | 0 |
| 0 | 0 | 0 |
| 1 | 1 | 1 |
| 0 | 0 | 0 |
| 0 | 0 | 0 |
| 0 | 0 | 0 |
| 1 | 0 | 0 |
| 0 | 0 | 0 |
| 0 | 0 | 0 |
| 0 | 0 | 0 |
| 0 | 0 | 0 |
| 0 | 0 | 0 |
| 0 | 0 | 0 |
| 0 | 0 | 0 |
| 0 | 0 | 0 |
| 0 | 0 | 0 |
| 0 | 0 | 0 |
| 0 | 1 | 0 |
| 0 | 0 | 0 |
| 1 | 1 | 1 |
| 0 | 0 | 0 |
| 0 | 0 | 0 |
| 0 | 0 | 0 |
| 0 | 0 | 0 |
| 0 | 0 | 0 |
| 0 | 0 | 0 |
| 0 | 0 | 0 |
| 0 | 1 | 0 |
| 0 | 0 | 0 |
| 0 | 0 | 0 |
| 0 | 0 | 0 |

| | | |
|---|---|---|
| 1 | 1 | 1 |
| 0 | 0 | 0 |
| 0 | 1 | 0 |
| 0 | 1 | 0 |
| 0 | 0 | 0 |
| 0 | 0 | 0 |
| 0 | 0 | 0 |
| 0 | 0 | 0 |
| 0 | 0 | 0 |
| 0 | 1 | 0 |
| 1 | 1 | 1 |
| 1 | 1 | 1 |
| 1 | 1 | 1 |
| 0 | 0 | 0 |
| 1 | 1 | 1 |
| 0 | 0 | 0 |
| 0 | 0 | 0 |
| 0 | 0 | 0 |
| 0 | 0 | 0 |
| 1 | 0 | 0 |
| 0 | 0 | 0 |
| 1 | 1 | 1 |
| 0 | 0 | 0 |
| 0 | 1 | 0 |
| 1 | 1 | 1 |
| 1 | 1 | 1 |
| 0 | 0 | 0 |
| 1 | 1 | 1 |
| 1 | 1 | 1 |
| 1 | 1 | 1 |
| 0 | 0 | 0 |
| 0 | 0 | 0 |
| 1 | 1 | 1 |
| 1 | 1 | 1 |
| 0 | 0 | 0 |
| 0 | 0 | 0 |
| 0 | 0 | 0 |
| 0 | 0 | 0 |
| 0 | 0 | 0 |
| 1 | 1 | 1 |
| 0 | 0 | 0 |
| 0 | 0 | 0 |
| 0 | 0 | 0 |
| 0 | 0 | 0 |
| 1 | 0 | 0 |
| 0 | 0 | 0 |
| 0 | 0 | 0 |
| 0 | 0 | 0 |
| 0 | 0 | 0 |
| 0 | 0 | 0 |
| 0 | 0 | 0 |
| 0 | 0 | 0 |
| 1 | 0 | 0 |
| 0 | 0 | 0 |

| | | |
|---|---|---|
| 0 | 0 | 0 |
| 0 | 0 | 0 |
| 0 | 0 | 0 |
| 0 | 0 | 0 |
| 0 | 0 | 0 |
| 0 | 0 | 0 |
| 0 | 0 | 0 |
| 1 | 0 | 0 |
| 0 | 0 | 0 |
| 0 | 0 | 0 |
| 0 | 0 | 0 |
| 0 | 0 | 0 |
| 1 | 1 | 1 |
| 0 | 0 | 0 |
| 0 | 0 | 0 |
| 0 | 0 | 0 |
| 0 | 0 | 0 |
| 0 | 0 | 0 |
| 0 | 0 | 0 |
| 1 | 1 | 1 |
| 0 | 0 | 0 |
| 0 | 0 | 0 |
| 0 | 0 | 0 |
| 0 | 0 | 0 |
| 0 | 0 | 0 |
| 1 | 0 | 0 |
| 1 | 1 | 1 |
| 0 | 1 | 0 |
| 0 | 0 | 0 |
| 0 | 0 | 0 |
| 0 | 0 | 0 |
| 0 | 0 | 0 |
| 0 | 0 | 0 |
| 0 | 0 | 0 |
| 0 | 0 | 0 |
| 1 | 0 | 0 |
| 0 | 0 | 0 |
| 0 | 0 | 0 |
| 1 | 1 | 1 |
| 1 | 0 | 0 |
| 0 | 1 | 0 |
| 0 | 0 | 0 |
| 0 | 0 | 0 |
| 0 | 0 | 0 |
| 0 | 0 | 0 |
| 0 | 0 | 0 |
| 0 | 0 | 0 |
| 1 | 1 | 1 |
| 1 | 1 | 1 |
| 0 | 0 | 0 |
| 0 | 0 | 0 |
| 0 | 0 | 0 |
| 0 | 0 | 0 |
| 1 | 1 | 1 |

| | | |
|---|---|---|
| 1 | 1 | 1 |
| 0 | 0 | 0 |
| 0 | 1 | 0 |
| 0 | 0 | 0 |
| 1 | 0 | 0 |
| 1 | 0 | 0 |
| 0 | 0 | 0 |
| 1 | 0 | 0 |
| 1 | 0 | 0 |
| 1 | 0 | 0 |
| 1 | 0 | 0 |
| 1 | 0 | 0 |
| 1 | 1 | 1 |
| 1 | 1 | 1 |
| 1 | 1 | 1 |
| 1 | 1 | 1 |
| 0 | 0 | 0 |
| 0 | 0 | 0 |
| 1 | 0 | 0 |
| 0 | 0 | 0 |
| 0 | 0 | 0 |
| 0 | 0 | 0 |
| 1 | 0 | 0 |
| 0 | 0 | 0 |
| 0 | 0 | 0 |
| 1 | 1 | 1 |
| 1 | 0 | 0 |
| 1 | 0 | 0 |
| 0 | 0 | 0 |
| 0 | 0 | 0 |
| 0 | 0 | 0 |
| 0 | 0 | 0 |
| 0 | 0 | 0 |
| 0 | 0 | 0 |
| 0 | 0 | 0 |
| 0 | 0 | 0 |
| 1 | 0 | 0 |
| 1 | 1 | 1 |
| 1 | 1 | 1 |
| 1 | 1 | 1 |
| 0 | 0 | 0 |
| 1 | 1 | 1 |
| 0 | 0 | 0 |
| 1 | 1 | 1 |
| 0 | 0 | 0 |
| 0 | 1 | 0 |
| 0 | 0 | 0 |
| 0 | 0 | 0 |
| 0 | 0 | 0 |
| 0 | 0 | 0 |
| 0 | 0 | 0 |
| 0 | 0 | 0 |
| 0 | 0 | 0 |
| 0 | 0 | 0 |

| | | |
|---|---|---|
| 0 | 0 | 0 |
| 0 | 0 | 0 |
| 0 | 0 | 0 |
| 0 | 0 | 0 |
| 0 | 0 | 0 |
| 0 | 0 | 0 |
| 0 | 0 | 0 |
| 1 | 1 | 1 |
| 0 | 0 | 0 |
| 0 | 0 | 0 |
| 0 | 0 | 0 |
| 1 | 0 | 0 |
| 0 | 0 | 0 |
| 0 | 0 | 0 |
| 0 | 0 | 0 |
| 0 | 0 | 0 |
| 0 | 0 | 0 |
| 0 | 0 | 0 |
| 0 | 0 | 0 |
| 0 | 0 | 0 |
| 0 | 0 | 0 |
| 0 | 0 | 0 |
| 0 | 0 | 0 |
| 0 | 0 | 0 |
| 0 | 0 | 0 |
| 0 | 0 | 0 |
| 0 | 0 | 0 |
| 0 | 0 | 0 |
| 0 | 0 | 0 |
| 0 | 0 | 0 |
| 0 | 0 | 0 |
| 0 | 0 | 0 |
| 0 | 0 | 0 |
| 0 | 0 | 0 |
| 0 | 0 | 0 |
| 0 | 0 | 0 |
| 0 | 0 | 0 |
| 0 | 0 | 0 |
| 0 | 0 | 0 |
| 0 | 0 | 0 |
| 0 | 0 | 0 |
| 1 | 0 | 0 |
| 0 | 0 | 0 |
| 0 | 0 | 0 |
| 0 | 0 | 0 |
| 0 | 0 | 0 |
| 0 | 0 | 0 |
| 0 | 0 | 0 |
| 0 | 0 | 0 |
| 0 | 0 | 0 |
| 0 | 1 | 0 |
| 0 | 1 | 0 |
| 1 | 0 | 0 |
| 0 | 0 | 0 |
| 0 | 0 | 0 |

| | | |
|---|---|---|
| 0 | 0 | 0 |
| 0 | 0 | 0 |
| 0 | 0 | 0 |
| 0 | 0 | 0 |
| 0 | 0 | 0 |
| 0 | 0 | 0 |
| 0 | 0 | 0 |
| 1 | 0 | 0 |
| 1 | 0 | 0 |
| 0 | 0 | 0 |
| 0 | 0 | 0 |
| 1 | 0 | 0 |
| 0 | 0 | 0 |
| 0 | 0 | 0 |
| 0 | 0 | 0 |
| 0 | 0 | 0 |
| 0 | 0 | 0 |
| 1 | 0 | 0 |
| 0 | 0 | 0 |
| 1 | 0 | 0 |
| 0 | 0 | 0 |
| 0 | 0 | 0 |
| 1 | 1 | 1 |
| 0 | 0 | 0 |
| 0 | 0 | 0 |
| 1 | 0 | 0 |
| 0 | 0 | 0 |
| 0 | 0 | 0 |
| 0 | 0 | 0 |
| 0 | 0 | 0 |
| 0 | 0 | 0 |
| 0 | 0 | 0 |
| 1 | 0 | 0 |
| 0 | 0 | 0 |
| 0 | 0 | 0 |
| 0 | 0 | 0 |
| 1 | 1 | 1 |
| 0 | 0 | 0 |
| 0 | 0 | 0 |
| 0 | 0 | 0 |
| 0 | 0 | 0 |
| 0 | 0 | 0 |
| 0 | 0 | 0 |
| 0 | 0 | 0 |
| 0 | 0 | 0 |
| 0 | 0 | 0 |
| 0 | 0 | 0 |
| 0 | 0 | 0 |
| 0 | 0 | 0 |
| 0 | 0 | 0 |
| 0 | 0 | 0 |
| 0 | 0 | 0 |
| 0 | 0 | 0 |
| 0 | 0 | 0 |

| | | |
|---|---|---|
| 1 | 0 | 0 |
| 0 | 1 | 0 |
| 0 | 0 | 0 |
| 0 | 0 | 0 |
| 0 | 0 | 0 |
| 0 | 0 | 0 |
| 0 | 0 | 0 |
| 0 | 0 | 0 |
| 0 | 0 | 0 |
| 1 | 0 | 0 |
| 0 | 1 | 0 |
| 1 | 1 | 1 |
| 0 | 0 | 0 |
| 0 | 0 | 0 |
| 0 | 0 | 0 |
| 0 | 0 | 0 |
| 0 | 0 | 0 |
| 0 | 0 | 0 |
| 1 | 1 | 1 |
| 1 | 0 | 0 |
| 1 | 0 | 0 |
| 0 | 0 | 0 |
| 0 | 1 | 0 |
| 0 | 0 | 0 |
| 0 | 0 | 0 |
| 0 | 0 | 0 |
| 0 | 0 | 0 |
| 0 | 0 | 0 |
| 0 | 0 | 0 |
| 0 | 0 | 0 |
| 0 | 0 | 0 |
| 0 | 0 | 0 |
| 0 | 0 | 0 |
| 0 | 0 | 0 |
| 0 | 0 | 0 |
| 0 | 0 | 0 |
| 0 | 0 | 0 |
| 0 | 0 | 0 |
| 0 | 0 | 0 |
| 0 | 0 | 0 |
| 0 | 0 | 0 |
| 1 | 0 | 0 |
| 0 | 0 | 0 |
| 0 | 0 | 0 |
| 0 | 0 | 0 |
| 0 | 0 | 0 |
| 1 | 0 | 0 |
| 0 | 0 | 0 |
| 0 | 0 | 0 |
| 0 | 0 | 0 |
| 0 | 0 | 0 |
| 0 | 0 | 0 |
| 1 | 0 | 0 |
| 0 | 1 | 0 |

| | | |
|---|---|---|
| 0 | 0 | 0 |
| 0 | 0 | 0 |
| 1 | 0 | 0 |
| 1 | 0 | 0 |
| 1 | 0 | 0 |
| 0 | 0 | 0 |
| 1 | 0 | 0 |
| 0 | 0 | 0 |
| 0 | 0 | 0 |
| 0 | 0 | 0 |
| 0 | 0 | 0 |
| 0 | 0 | 0 |
| 0 | 0 | 0 |
| 1 | 0 | 0 |
| 1 | 0 | 0 |
| 1 | 0 | 0 |
| 0 | 0 | 0 |
| 1 | 1 | 1 |
| 0 | 0 | 0 |
| 0 | 0 | 0 |
| 1 | 0 | 0 |
| 1 | 0 | 0 |
| 0 | 0 | 0 |
| 0 | 0 | 0 |
| 1 | 0 | 0 |
| 0 | 0 | 0 |
| 0 | 0 | 0 |
| 1 | 0 | 0 |
| 0 | 0 | 0 |
| 0 | 0 | 0 |
| 0 | 0 | 0 |
| 0 | 0 | 0 |
| 0 | 0 | 0 |
| 0 | 0 | 0 |
| 0 | 0 | 0 |
| 0 | 0 | 0 |
| 0 | 0 | 0 |
| 0 | 0 | 0 |
| 0 | 0 | 0 |
| 0 | 0 | 0 |
| 1 | 0 | 0 |
| 1 | 0 | 0 |
| 1 | 0 | 0 |
| 1 | 1 | 1 |
| 1 | 1 | 1 |
| 0 | 0 | 0 |
| 0 | 0 | 0 |
| 0 | 0 | 0 |
| 0 | 0 | 0 |
| 0 | 0 | 0 |
| 0 | 0 | 0 |
| 0 | 0 | 0 |
| 0 | 0 | 0 |
| 1 | 0 | 0 |

| | | |
|---|---|---|
| 1 | 0 | 0 |
| 0 | 0 | 0 |
| 0 | 0 | 0 |
| 0 | 0 | 0 |
| 1 | 0 | 0 |
| 0 | 0 | 0 |
| 0 | 0 | 0 |
| 0 | 0 | 0 |
| 1 | 0 | 0 |
| 1 | 0 | 0 |
| 0 | 0 | 0 |
| 0 | 0 | 0 |
| 0 | 0 | 0 |
| 1 | 0 | 0 |
| 0 | 0 | 0 |
| 0 | 0 | 0 |
| 0 | 0 | 0 |
| 0 | 0 | 0 |
| 0 | 0 | 0 |
| 0 | 0 | 0 |
| 0 | 0 | 0 |
| 1 | 1 | 1 |
| 0 | 0 | 0 |
| 0 | 1 | 0 |
| 0 | 0 | 0 |
| 0 | 0 | 0 |
| 0 | 0 | 0 |
| 0 | 0 | 0 |
| 0 | 0 | 0 |
| 0 | 0 | 0 |
| 0 | 0 | 0 |
| 0 | 0 | 0 |
| 0 | 0 | 0 |
| 0 | 0 | 0 |
| 0 | 1 | 0 |
| 0 | 0 | 0 |
| 0 | 0 | 0 |
| 0 | 0 | 0 |
| 0 | 0 | 0 |
| 0 | 0 | 0 |
| 0 | 0 | 0 |
| 0 | 0 | 0 |
| 0 | 0 | 0 |
| 0 | 0 | 0 |
| 0 | 0 | 0 |
| 0 | 0 | 0 |
| 0 | 0 | 0 |
| 1 | 1 | 1 |
| 0 | 0 | 0 |
| 1 | 1 | 1 |
| 0 | 0 | 0 |
| 0 | 0 | 0 |
| 0 | 0 | 0 |
| 0 | 0 | 0 |

| | | |
|---|---|---|
| 0 | 0 | 0 |
| 0 | 0 | 0 |
| 1 | 1 | 1 |
| 0 | 0 | 0 |
| 0 | 0 | 0 |
| 0 | 1 | 0 |
| 0 | 1 | 0 |
| 0 | 1 | 0 |
| 0 | 0 | 0 |
| 0 | 1 | 0 |
| 0 | 0 | 0 |
| 0 | 0 | 0 |
| 0 | 0 | 0 |
| 1 | 1 | 1 |
| 0 | 0 | 0 |
| 0 | 1 | 0 |
| 0 | 0 | 0 |
| 0 | 0 | 0 |
| 0 | 0 | 0 |
| 1 | 1 | 1 |
| 1 | 1 | 1 |
| 0 | 0 | 0 |
| 1 | 1 | 1 |
| 0 | 0 | 0 |
| 0 | 0 | 0 |
| 1 | 1 | 1 |
| 1 | 1 | 1 |
| 0 | 0 | 0 |
| 1 | 1 | 1 |
| 0 | 0 | 0 |
| 0 | 1 | 0 |
| 0 | 1 | 0 |
| 1 | 1 | 1 |
| 0 | 1 | 0 |
| 1 | 1 | 1 |
| 1 | 1 | 1 |
| 1 | 1 | 1 |
| 1 | 1 | 1 |
| 1 | 1 | 1 |
| 1 | 1 | 1 |
| 1 | 1 | 1 |
| 1 | 1 | 1 |
| 0 | 1 | 0 |
| 0 | 1 | 0 |
| 0 | 1 | 0 |
| 0 | 1 | 0 |
| 0 | 1 | 0 |
| 0 | 1 | 0 |
| 0 | 1 | 0 |
| 0 | 1 | 0 |
| 0 | 1 | 0 |
| 0 | 1 | 0 |
| 0 | 1 | 0 |
| 0 | 1 | 0 |

| | | |
|---|---|---|
| 0 | 1 | 0 |
| 1 | 1 | 1 |
| 1 | 1 | 1 |
| 1 | 1 | 1 |
| 1 | 1 | 1 |
| 1 | 1 | 1 |
| 1 | 1 | 1 |
| 0 | 1 | 0 |
| 0 | 1 | 0 |
| 0 | 1 | 0 |
| 0 | 1 | 0 |
| 0 | 0 | 0 |
| 0 | 0 | 0 |
| 0 | 0 | 0 |
| 1 | 0 | 0 |
| 0 | 0 | 0 |
| 1 | 0 | 0 |
| 0 | 0 | 0 |
| 1 | 0 | 0 |
| 1 | 1 | 1 |
| 0 | 0 | 0 |
| 1 | 1 | 1 |
| 1 | 1 | 1 |
| 0 | 0 | 0 |
| 0 | 0 | 0 |
| 1 | 1 | 1 |
| 1 | 1 | 1 |
| 0 | 0 | 0 |
| 0 | 0 | 0 |
| 0 | 0 | 0 |
| 0 | 0 | 0 |
| 0 | 0 | 0 |
| 0 | 1 | 0 |
| 0 | 1 | 0 |
| 1 | 1 | 1 |
| 1 | 1 | 1 |
| 1 | 1 | 1 |
| 1 | 1 | 1 |
| 1 | 1 | 1 |
| 0 | 0 | 0 |
| 0 | 0 | 0 |
| 0 | 1 | 0 |
| 0 | 0 | 0 |
| 0 | 0 | 0 |
| 0 | 0 | 0 |
| 1 | 0 | 0 |
| 0 | 0 | 0 |
| 1 | 0 | 0 |
| 1 | 0 | 0 |
| 0 | 0 | 0 |
| 0 | 0 | 0 |
| 0 | 0 | 0 |
| 0 | 0 | 0 |
| 0 | 0 | 0 |

| | | |
|---|---|---|
| 0 | 0 | 0 |
| 0 | 1 | 0 |
| 1 | 1 | 1 |
| 1 | 1 | 1 |
| 0 | 0 | 0 |
| 1 | 1 | 1 |
| 0 | 0 | 0 |
| 1 | 1 | 1 |
| 1 | 1 | 1 |
| 0 | 0 | 0 |
| 1 | 0 | 0 |
| 1 | 1 | 1 |
| 1 | 1 | 1 |
| 0 | 0 | 0 |
| 1 | 0 | 0 |
| 1 | 0 | 0 |
| 0 | 0 | 0 |
| 1 | 0 | 0 |
| 0 | 0 | 0 |
| 0 | 0 | 0 |
| 0 | 0 | 0 |
| 0 | 0 | 0 |
| 0 | 0 | 0 |
| 0 | 0 | 0 |
| 0 | 0 | 0 |
| 0 | 0 | 0 |
| 0 | 0 | 0 |
| 0 | 0 | 0 |
| 0 | 0 | 0 |
| 0 | 0 | 0 |
| 0 | 0 | 0 |
| 0 | 0 | 0 |
| 0 | 0 | 0 |
| 0 | 0 | 0 |
| 0 | 0 | 0 |
| 0 | 0 | 0 |
| 1 | 0 | 0 |
| 0 | 0 | 0 |
| 0 | 0 | 0 |
| 0 | 0 | 0 |
| 0 | 0 | 0 |
| 0 | 0 | 0 |
| 0 | 0 | 0 |
| 0 | 0 | 0 |
| 0 | 0 | 0 |
| 0 | 0 | 0 |
| 0 | 0 | 0 |
| 0 | 0 | 0 |
| 1 | 0 | 0 |
| 1 | 0 | 0 |
| 0 | 0 | 0 |
| 1 | 1 | 1 |
| 0 | 0 | 0 |
| 0 | 0 | 0 |

| | | |
|---|---|---|
| 0 | 0 | 0 |
| 0 | 0 | 0 |
| 0 | 0 | 0 |
| 0 | 0 | 0 |
| 0 | 0 | 0 |
| 0 | 0 | 0 |
| 0 | 0 | 0 |
| 0 | 0 | 0 |
| 0 | 0 | 0 |
| 0 | 0 | 0 |
| 0 | 0 | 0 |
| 0 | 0 | 0 |
| 0 | 0 | 0 |
| 0 | 0 | 0 |
| 0 | 1 | 0 |
| 0 | 0 | 0 |
| 0 | 0 | 0 |
| 0 | 0 | 0 |
| 1 | 0 | 0 |
| 0 | 0 | 0 |
| 0 | 0 | 0 |
| 0 | 0 | 0 |
| 0 | 0 | 0 |
| 0 | 0 | 0 |
| 0 | 0 | 0 |
| 1 | 0 | 0 |
| 0 | 0 | 0 |
| 0 | 0 | 0 |
| 0 | 0 | 0 |
| 0 | 0 | 0 |
| 1 | 1 | 1 |
| 0 | 0 | 0 |
| 1 | 1 | 1 |
| 1 | 0 | 0 |
| 0 | 0 | 0 |
| 0 | 0 | 0 |
| 1 | 1 | 1 |
| 1 | 1 | 1 |
| 1 | 1 | 1 |
| 1 | 0 | 0 |
| 0 | 0 | 0 |
| 0 | 0 | 0 |
| 0 | 0 | 0 |
| 0 | 0 | 0 |
| 0 | 0 | 0 |
| 0 | 0 | 0 |
| 0 | 0 | 0 |
| 0 | 0 | 0 |
| 1 | 1 | 1 |
| 1 | 1 | 1 |
| 0 | 0 | 0 |
| 0 | 0 | 0 |
| 0 | 0 | 0 |
| 0 | 1 | 0 |

| | | |
|---|---|---|
| 1 | 1 | 1 |
| 1 | 1 | 1 |
| 1 | 0 | 0 |
| 0 | 0 | 0 |
| 1 | 0 | 0 |
| 1 | 1 | 1 |
| 0 | 0 | 0 |
| 1 | 1 | 1 |
| 0 | 0 | 0 |
| 0 | 0 | 0 |
| 0 | 0 | 0 |
| 0 | 0 | 0 |
| 0 | 0 | 0 |
| 0 | 0 | 0 |
| 0 | 0 | 0 |
| 0 | 0 | 0 |
| 0 | 1 | 0 |
| 0 | 0 | 0 |
| 0 | 0 | 0 |
| 1 | 0 | 0 |
| 0 | 0 | 0 |
| 0 | 0 | 0 |
| 1 | 1 | 1 |
| 0 | 0 | 0 |
| 1 | 1 | 1 |
| 1 | 1 | 1 |
| 0 | 1 | 0 |
| 0 | 0 | 0 |
| 0 | 0 | 0 |
| 0 | 0 | 0 |
| 1 | 1 | 1 |
| 0 | 0 | 0 |
| 0 | 0 | 0 |
| 0 | 0 | 0 |
| 0 | 1 | 0 |
| 0 | 1 | 0 |
| 0 | 0 | 0 |
| 0 | 0 | 0 |
| 0 | 0 | 0 |
| 1 | 1 | 1 |
| 1 | 1 | 1 |
| 0 | 1 | 0 |
| 0 | 1 | 0 |
| 0 | 0 | 0 |
| 1 | 1 | 1 |
| 0 | 0 | 0 |
| 1 | 1 | 1 |
| 0 | 0 | 0 |
| 0 | 0 | 0 |
| 1 | 1 | 1 |
| 0 | 1 | 0 |
| 0 | 0 | 0 |
| 0 | 0 | 0 |
| 1 | 1 | 1 |

| | | |
|---|---|---|
| 1 | 1 | 1 |
| 1 | 1 | 1 |
| 0 | 1 | 0 |
| 0 | 0 | 0 |
| 0 | 0 | 0 |
| 1 | 0 | 0 |
| 1 | 0 | 0 |
| 0 | 0 | 0 |
| 0 | 0 | 0 |
| 1 | 1 | 1 |
| 0 | 0 | 0 |
| 0 | 0 | 0 |
| 0 | 0 | 0 |
| 0 | 0 | 0 |
| 0 | 0 | 0 |
| 0 | 0 | 0 |
| 0 | 0 | 0 |
| 0 | 0 | 0 |
| 0 | 0 | 0 |
| 0 | 1 | 0 |
| 1 | 1 | 1 |
| 1 | 1 | 1 |
| 1 | 1 | 1 |
| 1 | 1 | 1 |
| 0 | 0 | 0 |
| 0 | 1 | 0 |
| 0 | 0 | 0 |
| 0 | 0 | 0 |
| 1 | 1 | 1 |
| 1 | 1 | 1 |
| 1 | 1 | 1 |
| 1 | 1 | 1 |
| 1 | 1 | 1 |
| 0 | 0 | 0 |
| 0 | 0 | 0 |
| 1 | 1 | 1 |
| 0 | 0 | 0 |
| 0 | 1 | 0 |
| 0 | 0 | 0 |
| 0 | 0 | 0 |
| 0 | 0 | 0 |
| 0 | 0 | 0 |
| 0 | 1 | 0 |
| 0 | 0 | 0 |
| 0 | 0 | 0 |
| 0 | 0 | 0 |
| 0 | 0 | 0 |
| 0 | 0 | 0 |
| 0 | 0 | 0 |
| 0 | 0 | 0 |
| 0 | 0 | 0 |
| 0 | 0 | 0 |
| 0 | 0 | 0 |
| 0 | 1 | 0 |

| | | |
|---|---|---|
| 0 | 0 | 0 |
| 0 | 1 | 0 |
| 0 | 0 | 0 |
| 0 | 0 | 0 |
| 0 | 0 | 0 |
| 1 | 1 | 1 |
| 0 | 0 | 0 |
| 1 | 1 | 1 |
| 0 | 1 | 0 |
| 0 | 0 | 0 |
| 0 | 1 | 0 |
| 0 | 1 | 0 |
| 0 | 1 | 0 |
| 0 | 1 | 0 |
| 0 | 1 | 0 |
| 0 | 0 | 0 |
| 0 | 0 | 0 |
| 0 | 1 | 0 |
| 0 | 0 | 0 |
| 1 | 1 | 1 |
| 0 | 0 | 0 |
| 0 | 0 | 0 |
| 0 | 0 | 0 |
| 0 | 0 | 0 |
| 0 | 0 | 0 |
| 0 | 0 | 0 |
| 0 | 0 | 0 |
| 0 | 0 | 0 |
| 0 | 0 | 0 |
| 0 | 0 | 0 |
| 0 | 0 | 0 |
| 0 | 0 | 0 |
| 1 | 0 | 0 |
| 1 | 0 | 0 |
| 0 | 0 | 0 |
| 0 | 0 | 0 |
| 0 | 0 | 0 |
| 0 | 0 | 0 |
| 1 | 0 | 0 |
| 0 | 0 | 0 |
| 0 | 0 | 0 |
| 0 | 0 | 0 |
| 1 | 0 | 0 |
| 0 | 1 | 0 |
| 0 | 0 | 0 |
| 0 | 0 | 0 |
| 0 | 0 | 0 |
| 0 | 0 | 0 |
| 0 | 0 | 0 |
| 0 | 0 | 0 |
| 1 | 0 | 0 |
| 0 | 0 | 0 |
| 0 | 0 | 0 |
| 1 | 0 | 0 |

| | | |
|---|---|---|
| 0 | 0 | 0 |
| 1 | 0 | 0 |
| 0 | 0 | 0 |
| 0 | 0 | 0 |
| 0 | 0 | 0 |
| 0 | 0 | 0 |
| 0 | 0 | 0 |
| 0 | 0 | 0 |
| 0 | 0 | 0 |
| 0 | 0 | 0 |
| 0 | 0 | 0 |
| 1 | 1 | 1 |
| 0 | 0 | 0 |
| 1 | 1 | 1 |
| 1 | 0 | 0 |
| 1 | 0 | 0 |
| 0 | 0 | 0 |
| 0 | 0 | 0 |
| 0 | 0 | 0 |
| 0 | 0 | 0 |
| 1 | 0 | 0 |
| 0 | 0 | 0 |
| 0 | 1 | 0 |
| 0 | 0 | 0 |
| 1 | 1 | 1 |
| 1 | 1 | 1 |
| 1 | 1 | 1 |
| 0 | 0 | 0 |
| 1 | 1 | 1 |
| 1 | 1 | 1 |
| 1 | 1 | 1 |
| 0 | 0 | 0 |
| 1 | 1 | 1 |
| 0 | 0 | 0 |
| 0 | 0 | 0 |
| 0 | 0 | 0 |
| 0 | 1 | 0 |
| 1 | 1 | 1 |
| 0 | 0 | 0 |
| 0 | 0 | 0 |
| 0 | 0 | 0 |
| 0 | 0 | 0 |
| 0 | 0 | 0 |
| 0 | 0 | 0 |
| 0 | 0 | 0 |
| 0 | 0 | 0 |
| 1 | 1 | 1 |
| 1 | 1 | 1 |
| 0 | 0 | 0 |
| 0 | 0 | 0 |
| 0 | 0 | 0 |
| 1 | 1 | 1 |
| 1 | 1 | 1 |
| 0 | 0 | 0 |

| | | |
|---|---|---|
| 0 | 0 | 0 |
| 0 | 1 | 0 |
| 0 | 1 | 0 |
| 1 | 1 | 1 |
| 0 | 0 | 0 |
| 0 | 0 | 0 |
| 0 | 0 | 0 |
| 0 | 0 | 0 |
| 0 | 0 | 0 |
| 0 | 0 | 0 |
| 1 | 1 | 1 |
| 0 | 0 | 0 |
| 0 | 0 | 0 |
| 0 | 0 | 0 |
| 1 | 0 | 0 |
| 1 | 1 | 1 |
| 0 | 0 | 0 |
| 1 | 0 | 0 |
| 0 | 0 | 0 |
| 0 | 0 | 0 |
| 1 | 0 | 0 |
| 0 | 0 | 0 |
| 0 | 0 | 0 |
| 1 | 0 | 0 |
| 0 | 0 | 0 |
| 0 | 0 | 0 |
| 0 | 0 | 0 |
| 1 | 0 | 0 |
| 0 | 0 | 0 |
| 0 | 0 | 0 |
| 0 | 0 | 0 |
| 1 | 1 | 1 |
| 1 | 1 | 1 |
| 1 | 1 | 1 |
| 0 | 1 | 0 |
| 0 | 1 | 0 |
| 0 | 1 | 0 |
| 1 | 1 | 1 |
| 1 | 1 | 1 |
| 1 | 1 | 1 |
| 0 | 1 | 0 |
| 1 | 1 | 1 |
| 0 | 0 | 0 |
| 0 | 1 | 0 |
| 1 | 1 | 1 |
| 1 | 1 | 1 |
| 0 | 1 | 0 |
| 1 | 1 | 1 |
| 1 | 1 | 1 |
| 1 | 1 | 1 |
| 1 | 1 | 1 |
| 1 | 1 | 1 |
| 1 | 1 | 1 |
| 1 | 1 | 1 |

| | | |
|---|---|---|
| 0 | 1 | 0 |
| 0 | 1 | 0 |
| 1 | 1 | 1 |
| 1 | 1 | 1 |
| 1 | 1 | 1 |
| 1 | 1 | 1 |
| 1 | 1 | 1 |
| 1 | 1 | 1 |
| 0 | 0 | 0 |
| 0 | 0 | 0 |
| 0 | 0 | 0 |
| 0 | 0 | 0 |
| 0 | 0 | 0 |
| 0 | 0 | 0 |
| 0 | 0 | 0 |
| 0 | 0 | 0 |
| 0 | 0 | 0 |
| 0 | 0 | 0 |
| 0 | 0 | 0 |
| 0 | 0 | 0 |
| 0 | 0 | 0 |
| 0 | 0 | 0 |
| 0 | 0 | 0 |
| 0 | 0 | 0 |
| 0 | 0 | 0 |
| 1 | 1 | 1 |
| 1 | 1 | 1 |
| 0 | 0 | 0 |
| 0 | 0 | 0 |
| 0 | 0 | 0 |
| 0 | 0 | 0 |
| 0 | 0 | 0 |
| 0 | 0 | 0 |
| 0 | 0 | 0 |
| 0 | 0 | 0 |
| 0 | 0 | 0 |
| 0 | 0 | 0 |
| 0 | 0 | 0 |
| 0 | 0 | 0 |
| 0 | 0 | 0 |
| 0 | 0 | 0 |
| 0 | 0 | 0 |
| 0 | 0 | 0 |
| 0 | 0 | 0 |
| 0 | 0 | 0 |
| 0 | 0 | 0 |
| 1 | 1 | 1 |
| 0 | 1 | 0 |
| 1 | 1 | 1 |
| 1 | 1 | 1 |
| 1 | 1 | 1 |
| 1 | 1 | 1 |
| 0 | 0 | 0 |
| 0 | 0 | 0 |

| | | |
|---|---|---|
| 0 | 0 | 0 |
| 0 | 0 | 0 |
| 0 | 0 | 0 |
| 0 | 0 | 0 |
| 0 | 1 | 0 |
| 0 | 0 | 0 |
| 0 | 0 | 0 |
| 0 | 0 | 0 |
| 1 | 1 | 1 |
| 0 | 0 | 0 |
| 0 | 0 | 0 |
| 0 | 1 | 0 |
| 0 | 0 | 0 |
| 0 | 0 | 0 |
| 0 | 0 | 0 |
| 0 | 0 | 0 |
| 1 | 1 | 1 |
| 0 | 0 | 0 |
| 0 | 0 | 0 |
| 1 | 1 | 1 |
| 0 | 0 | 0 |
| 0 | 0 | 0 |
| 0 | 0 | 0 |
| 0 | 0 | 0 |
| 0 | 0 | 0 |
| 0 | 1 | 0 |
| 0 | 1 | 0 |
| 1 | 1 | 1 |
| 1 | 1 | 1 |
| 0 | 1 | 0 |
| 1 | 1 | 1 |
| 1 | 1 | 1 |
| 1 | 0 | 0 |
| 1 | 0 | 0 |
| 0 | 0 | 0 |
| 0 | 0 | 0 |
| 0 | 0 | 0 |
| 0 | 0 | 0 |
| 0 | 0 | 0 |
| 0 | 0 | 0 |
| 1 | 1 | 1 |
| 1 | 1 | 1 |
| 1 | 1 | 1 |
| 0 | 0 | 0 |
| 0 | 0 | 0 |
| 0 | 0 | 0 |
| 1 | 1 | 1 |
| 0 | 1 | 0 |
| 0 | 0 | 0 |
| 0 | 0 | 0 |
| 1 | 1 | 1 |
| 1 | 1 | 1 |
| 1 | 1 | 1 |
| 1 | 1 | 1 |

| | | |
|---|---|---|
| 0 | 1 | 0 |
| 1 | 1 | 1 |
| 0 | 1 | 0 |
| 0 | 0 | 0 |
| 0 | 0 | 0 |
| 1 | 1 | 1 |
| 1 | 1 | 1 |
| 1 | 1 | 1 |
| 1 | 1 | 1 |
| 1 | 1 | 1 |
| 0 | 0 | 0 |
| 0 | 1 | 0 |
| 0 | 0 | 0 |
| 0 | 0 | 0 |
| 0 | 0 | 0 |
| 1 | 1 | 1 |
| 1 | 1 | 1 |
| 0 | 1 | 0 |
| 1 | 1 | 1 |
| 1 | 1 | 1 |
| 0 | 0 | 0 |
| 1 | 1 | 1 |
| 1 | 1 | 1 |
| 0 | 0 | 0 |
| 1 | 0 | 0 |
| 0 | 0 | 0 |
| 0 | 0 | 0 |
| 1 | 1 | 1 |
| 1 | 1 | 1 |
| 1 | 1 | 1 |
| 1 | 1 | 1 |
| 1 | 1 | 1 |
| 1 | 1 | 1 |
| 1 | 1 | 1 |
| 1 | 1 | 1 |
| 0 | 0 | 0 |
| 1 | 1 | 1 |
| 1 | 1 | 1 |
| 0 | 0 | 0 |
| 0 | 0 | 0 |
| 1 | 1 | 1 |
| 0 | 1 | 0 |
| 0 | 0 | 0 |
| 0 | 0 | 0 |
| 0 | 0 | 0 |
| 0 | 0 | 0 |
| 0 | 0 | 0 |
| 0 | 0 | 0 |
| 1 | 1 | 1 |
| 0 | 0 | 0 |
| 0 | 0 | 0 |
| 0 | 0 | 0 |
| 0 | 0 | 0 |
| 0 | 0 | 0 |

| | | |
|---|---|---|
| 0 | 0 | 0 |
| 0 | 0 | 0 |
| 0 | 0 | 0 |
| 0 | 0 | 0 |
| 0 | 0 | 0 |
| 0 | 0 | 0 |
| 0 | 0 | 0 |
| 0 | 0 | 0 |
| 0 | 0 | 0 |
| 0 | 1 | 0 |
| 0 | 1 | 0 |
| 0 | 0 | 0 |
| 1 | 1 | 1 |
| 0 | 0 | 0 |
| 0 | 0 | 0 |
| 0 | 1 | 0 |
| 0 | 0 | 0 |
| 1 | 0 | 0 |
| 1 | 1 | 1 |
| 1 | 1 | 1 |
| 0 | 0 | 0 |
| 0 | 1 | 0 |
| 0 | 1 | 0 |
| 0 | 0 | 0 |
| 0 | 0 | 0 |
| 0 | 1 | 0 |
| 0 | 1 | 0 |
| 1 | 1 | 1 |
| 0 | 0 | 0 |
| 0 | 0 | 0 |
| 1 | 1 | 1 |
| 1 | 1 | 1 |
| 0 | 0 | 0 |
| 0 | 0 | 0 |
| 0 | 0 | 0 |
| 0 | 1 | 0 |
| 0 | 0 | 0 |
| 0 | 1 | 0 |
| 0 | 1 | 0 |
| 0 | 0 | 0 |
| 0 | 0 | 0 |
| 0 | 0 | 0 |
| 0 | 0 | 0 |
| 0 | 0 | 0 |
| 0 | 0 | 0 |
| 1 | 1 | 1 |
| 0 | 1 | 0 |
| 0 | 0 | 0 |
| 0 | 0 | 0 |
| 0 | 0 | 0 |
| 1 | 1 | 1 |
| 1 | 1 | 1 |
| 0 | 0 | 0 |
| 0 | 0 | 0 |

| | | |
|---|---|---|
| 1 | 1 | 1 |
| 1 | 1 | 1 |
| 1 | 1 | 1 |
| 1 | 1 | 1 |
| 0 | 0 | 0 |
| 0 | 0 | 0 |
| 1 | 0 | 0 |
| 0 | 0 | 0 |
| 1 | 0 | 0 |
| 0 | 0 | 0 |
| 1 | 1 | 1 |
| 1 | 1 | 1 |
| 1 | 1 | 1 |
| 1 | 1 | 1 |
| 1 | 1 | 1 |
| 1 | 1 | 1 |
| 0 | 0 | 0 |
| 1 | 1 | 1 |
| 1 | 1 | 1 |
| 0 | 0 | 0 |
| 0 | 0 | 0 |
| 0 | 1 | 0 |
| 0 | 1 | 0 |
| 0 | 0 | 0 |
| 0 | 1 | 0 |
| 0 | 0 | 0 |
| 0 | 0 | 0 |
| 0 | 0 | 0 |
| 0 | 0 | 0 |
| 0 | 0 | 0 |
| 0 | 0 | 0 |
| 0 | 0 | 0 |
| 0 | 1 | 0 |
| 0 | 0 | 0 |
| 1 | 1 | 1 |
| 1 | 1 | 1 |
| 1 | 1 | 1 |
| 1 | 1 | 1 |
| 0 | 0 | 0 |
| 1 | 1 | 1 |
| 1 | 1 | 1 |
| 1 | 1 | 1 |
| 1 | 1 | 1 |
| 0 | 0 | 0 |
| 1 | 1 | 1 |
| 1 | 1 | 1 |
| 1 | 1 | 1 |
| 1 | 1 | 1 |
| 1 | 1 | 1 |
| 1 | 1 | 1 |
| 0 | 0 | 0 |
| 0 | 0 | 0 |
| 0 | 0 | 0 |
| 0 | 1 | 0 |

| | | |
|---|---|---|
| 0 | 0 | 0 |
| 0 | 0 | 0 |
| 0 | 0 | 0 |
| 0 | 0 | 0 |
| 0 | 0 | 0 |
| 0 | 0 | 0 |
| 0 | 0 | 0 |
| 0 | 0 | 0 |
| 1 | 1 | 1 |
| 1 | 1 | 1 |
| 0 | 0 | 0 |
| 1 | 1 | 1 |
| 1 | 1 | 1 |
| 0 | 0 | 0 |
| 0 | 0 | 0 |
| 0 | 0 | 0 |
| 0 | 0 | 0 |
| 0 | 0 | 0 |
| 0 | 0 | 0 |
| 1 | 1 | 1 |
| 1 | 1 | 1 |
| 0 | 0 | 0 |
| 1 | 1 | 1 |
| 0 | 0 | 0 |
| 1 | 1 | 1 |
| 1 | 1 | 1 |
| 1 | 1 | 1 |
| 0 | 1 | 0 |
| 0 | 1 | 0 |
| 0 | 0 | 0 |
| 0 | 1 | 0 |
| 1 | 1 | 1 |
| 0 | 0 | 0 |
| 1 | 1 | 1 |
| 0 | 0 | 0 |
| 0 | 0 | 0 |
| 0 | 0 | 0 |
| 0 | 0 | 0 |
| 0 | 0 | 0 |
| 0 | 0 | 0 |
| 0 | 0 | 0 |
| 0 | 0 | 0 |
| 1 | 1 | 1 |
| 0 | 0 | 0 |
| 1 | 1 | 1 |
| 0 | 0 | 0 |
| 1 | 0 | 0 |
| 1 | 1 | 1 |
| 1 | 1 | 1 |
| 0 | 0 | 0 |
| 1 | 1 | 1 |
| 1 | 1 | 1 |
| 0 | 1 | 0 |
| 1 | 1 | 1 |

| | | |
|---|---|---|
| 0 | 0 | 0 |
| 0 | 1 | 0 |
| 1 | 1 | 1 |
| 0 | 1 | 0 |
| 0 | 1 | 0 |
| 0 | 1 | 0 |
| 0 | 1 | 0 |
| 1 | 1 | 1 |
| 1 | 1 | 1 |
| 1 | 1 | 1 |
| 0 | 0 | 0 |
| 1 | 1 | 1 |
| 1 | 1 | 1 |
| 0 | 1 | 0 |
| 0 | 0 | 0 |
| 0 | 0 | 0 |
| 0 | 0 | 0 |
| 0 | 0 | 0 |
| 0 | 0 | 0 |
| 0 | 1 | 0 |
| 0 | 1 | 0 |
| 1 | 1 | 1 |
| 0 | 0 | 0 |
| 0 | 1 | 0 |
| 0 | 1 | 0 |
| 0 | 1 | 0 |
| 1 | 1 | 1 |
| 1 | 1 | 1 |
| 0 | 0 | 0 |
| 0 | 0 | 0 |
| 0 | 0 | 0 |
| 0 | 0 | 0 |
| 0 | 0 | 0 |
| 0 | 0 | 0 |
| 1 | 1 | 1 |
| 1 | 1 | 1 |
| 0 | 0 | 0 |
| 0 | 0 | 0 |
| 0 | 0 | 0 |
| 0 | 0 | 0 |
| 0 | 0 | 0 |
| 0 | 0 | 0 |
| 1 | 1 | 1 |
| 1 | 1 | 1 |
| 0 | 0 | 0 |
| 1 | 1 | 1 |
| 1 | 1 | 1 |
| 1 | 1 | 1 |
| 1 | 1 | 1 |
| 1 | 1 | 1 |
| 1 | 0 | 0 |
| 0 | 1 | 0 |
| 0 | 1 | 0 |
| 1 | 0 | 0 |

| | | |
|---|---|---|
| 0 | 0 | 0 |
| 1 | 1 | 1 |
| 0 | 0 | 0 |
| 0 | 1 | 0 |
| 1 | 0 | 0 |
| 0 | 0 | 0 |
| 0 | 0 | 0 |
| 0 | 0 | 0 |
| 1 | 0 | 0 |
| 1 | 0 | 0 |
| 0 | 0 | 0 |
| 1 | 1 | 1 |
| 1 | 0 | 0 |
| 1 | 0 | 0 |
| 0 | 0 | 0 |
| 0 | 0 | 0 |
| 0 | 0 | 0 |
| 0 | 0 | 0 |
| 1 | 1 | 1 |
| 0 | 0 | 0 |
| 0 | 0 | 0 |
| 0 | 0 | 0 |
| 0 | 0 | 0 |
| 0 | 0 | 0 |
| 1 | 1 | 1 |
| 0 | 0 | 0 |
| 1 | 1 | 1 |
| 0 | 1 | 0 |
| 1 | 1 | 1 |
| 0 | 0 | 0 |
| 1 | 1 | 1 |
| 1 | 1 | 1 |
| 0 | 0 | 0 |
| 0 | 0 | 0 |
| 1 | 1 | 1 |
| 1 | 1 | 1 |
| 1 | 1 | 1 |
| 0 | 1 | 0 |
| 1 | 0 | 0 |
| 0 | 0 | 0 |
| 0 | 0 | 0 |
| 0 | 0 | 0 |
| 0 | 0 | 0 |
| 0 | 0 | 0 |
| 0 | 0 | 0 |
| 0 | 1 | 0 |
| 0 | 1 | 0 |
| 0 | 0 | 0 |
| 0 | 0 | 0 |
| 1 | 1 | 1 |
| 0 | 0 | 0 |
| 0 | 0 | 0 |
| 0 | 1 | 0 |
| 1 | 1 | 1 |

| | | |
|---|---|---|
| 1 | 1 | 1 |
| 0 | 0 | 0 |
| 0 | 0 | 0 |
| 1 | 0 | 0 |
| 1 | 1 | 1 |
| 0 | 0 | 0 |
| 0 | 0 | 0 |
| 0 | 0 | 0 |
| 0 | 0 | 0 |
| 0 | 0 | 0 |
| 0 | 0 | 0 |
| 0 | 0 | 0 |
| 0 | 0 | 0 |
| 0 | 0 | 0 |
| 0 | 0 | 0 |
| 0 | 1 | 0 |
| 0 | 0 | 0 |
| 1 | 1 | 1 |
| 1 | 0 | 0 |
| 1 | 1 | 1 |
| 0 | 0 | 0 |
| 1 | 1 | 1 |
| 0 | 0 | 0 |
| 0 | 0 | 0 |
| 1 | 1 | 1 |
| 0 | 0 | 0 |
| 0 | 1 | 0 |
| 0 | 1 | 0 |
| 0 | 1 | 0 |
| 0 | 0 | 0 |
| 0 | 0 | 0 |
| 0 | 0 | 0 |
| 0 | 0 | 0 |
| 0 | 0 | 0 |
| 0 | 0 | 0 |
| 0 | 0 | 0 |
| 0 | 0 | 0 |
| 1 | 1 | 1 |
| 1 | 0 | 0 |
| 0 | 0 | 0 |
| 0 | 0 | 0 |
| 0 | 0 | 0 |
| 0 | 0 | 0 |
| 0 | 0 | 0 |
| 0 | 0 | 0 |
| 0 | 0 | 0 |
| 1 | 1 | 1 |
| 0 | 0 | 0 |
| 0 | 0 | 0 |
| 0 | 0 | 0 |
| 0 | 0 | 0 |
| 0 | 1 | 0 |
| 0 | 0 | 0 |
| 0 | 0 | 0 |

| | | |
|---|---|---|
| 0 | 0 | 0 |
| 0 | 0 | 0 |
| 0 | 0 | 0 |
| 0 | 0 | 0 |
| 0 | 0 | 0 |
| 0 | 0 | 0 |
| 0 | 0 | 0 |
| 0 | 0 | 0 |
| 0 | 0 | 0 |
| 0 | 0 | 0 |
| 1 | 1 | 1 |
| 0 | 0 | 0 |
| 1 | 0 | 0 |
| 0 | 0 | 0 |
| 0 | 1 | 0 |
| 1 | 1 | 1 |
| 1 | 1 | 1 |
| 1 | 1 | 1 |
| 0 | 0 | 0 |
| 0 | 0 | 0 |
| 1 | 1 | 1 |
| 0 | 0 | 0 |
| 0 | 0 | 0 |
| 1 | 1 | 1 |
| 0 | 0 | 0 |
| 0 | 0 | 0 |
| 0 | 0 | 0 |
| 1 | 0 | 0 |
| 0 | 0 | 0 |
| 0 | 0 | 0 |
| 1 | 0 | 0 |
| 0 | 0 | 0 |
| 1 | 0 | 0 |
| 1 | 0 | 0 |
| 0 | 0 | 0 |
| 0 | 0 | 0 |
| 0 | 0 | 0 |
| 1 | 0 | 0 |
| 1 | 0 | 0 |
| 0 | 0 | 0 |
| 0 | 0 | 0 |
| 0 | 0 | 0 |
| 0 | 0 | 0 |
| 0 | 0 | 0 |
| 0 | 0 | 0 |
| 0 | 0 | 0 |
| 0 | 0 | 0 |
| 0 | 0 | 0 |
| 0 | 0 | 0 |
| 0 | 0 | 0 |
| 0 | 0 | 0 |
| 0 | 0 | 0 |
| 0 | 0 | 0 |
| 0 | 0 | 0 |

| | | |
|---|---|---|
| 0 | 0 | 0 |
| 1 | 0 | 0 |
| 0 | 0 | 0 |
| 0 | 0 | 0 |
| 0 | 0 | 0 |
| 1 | 0 | 0 |
| 0 | 0 | 0 |
| 0 | 0 | 0 |
| 1 | 1 | 1 |
| 0 | 0 | 0 |
| 0 | 1 | 0 |
| 1 | 1 | 1 |
| 1 | 1 | 1 |
| 0 | 0 | 0 |
| 1 | 1 | 1 |
| 0 | 0 | 0 |
| 1 | 1 | 1 |
| 1 | 1 | 1 |
| 0 | 0 | 0 |
| 0 | 0 | 0 |
| 0 | 0 | 0 |
| 0 | 0 | 0 |
| 0 | 0 | 0 |
| 0 | 0 | 0 |
| 0 | 0 | 0 |
| 0 | 1 | 0 |
| 1 | 1 | 1 |
| 1 | 1 | 1 |
| 1 | 0 | 0 |
| 1 | 1 | 1 |
| 1 | 1 | 1 |
| 0 | 1 | 0 |
| 0 | 0 | 0 |
| 0 | 0 | 0 |
| 0 | 0 | 0 |
| 0 | 0 | 0 |
| 0 | 0 | 0 |
| 0 | 0 | 0 |
| 0 | 1 | 0 |
| 0 | 1 | 0 |
| 0 | 1 | 0 |
| 0 | 1 | 0 |
| 0 | 1 | 0 |
| 0 | 1 | 0 |
| 0 | 0 | 0 |
| 0 | 0 | 0 |
| 0 | 1 | 0 |
| 0 | 0 | 0 |
| 0 | 0 | 0 |
| 0 | 0 | 0 |
| 0 | 0 | 0 |
| 0 | 0 | 0 |
| 0 | 0 | 0 |
| 0 | 0 | 0 |

| | | |
|---|---|---|
| 1 | 1 | 1 |
| 1 | 1 | 1 |
| 0 | 0 | 0 |
| 0 | 0 | 0 |
| 1 | 1 | 1 |
| 0 | 0 | 0 |
| 0 | 0 | 0 |
| 1 | 1 | 1 |
| 1 | 1 | 1 |
| 0 | 0 | 0 |
| 1 | 1 | 1 |
| 1 | 1 | 1 |
| 0 | 0 | 0 |
| 0 | 0 | 0 |
| 0 | 0 | 0 |
| 0 | 0 | 0 |
| 0 | 1 | 0 |
| 0 | 1 | 0 |
| 1 | 1 | 1 |
| 1 | 1 | 1 |
| 1 | 1 | 1 |
| 0 | 1 | 0 |
| 1 | 1 | 1 |
| 1 | 1 | 1 |
| 1 | 1 | 1 |
| 0 | 0 | 0 |
| 1 | 1 | 1 |
| 1 | 1 | 1 |
| 1 | 1 | 1 |
| 0 | 1 | 0 |
| 0 | 1 | 0 |
| 1 | 1 | 1 |
| 1 | 1 | 1 |
| 1 | 1 | 1 |
| 1 | 1 | 1 |
| 1 | 1 | 1 |
| 1 | 1 | 1 |
| 0 | 0 | 0 |
| 0 | 0 | 0 |
| 1 | 0 | 0 |
| 0 | 0 | 0 |
| 1 | 1 | 1 |
| 1 | 1 | 1 |
| 0 | 0 | 0 |
| 0 | 0 | 0 |
| 0 | 1 | 0 |
| 0 | 1 | 0 |
| 0 | 1 | 0 |
| 1 | 1 | 1 |
| 1 | 1 | 1 |
| 1 | 1 | 1 |
| 0 | 0 | 0 |
| 0 | 0 | 0 |
| 0 | 0 | 0 |

| | | |
|---|---|---|
| 0 | 0 | 0 |
| 1 | 1 | 1 |
| 1 | 1 | 1 |
| 1 | 1 | 1 |
| 1 | 1 | 1 |
| 1 | 1 | 1 |
| 1 | 1 | 1 |
| 1 | 1 | 1 |
| 0 | 0 | 0 |
| 1 | 1 | 1 |
| 1 | 1 | 1 |
| 1 | 1 | 1 |
| 0 | 1 | 0 |
| 0 | 1 | 0 |
| 1 | 1 | 1 |
| 1 | 1 | 1 |
| 0 | 1 | 0 |
| 0 | 1 | 0 |
| 0 | 0 | 0 |
| 0 | 0 | 0 |
| 0 | 0 | 0 |
| 0 | 0 | 0 |
| 0 | 1 | 0 |
| 0 | 0 | 0 |
| 0 | 0 | 0 |
| 1 | 1 | 1 |
| 1 | 1 | 1 |
| 1 | 1 | 1 |
| 1 | 1 | 1 |
| 1 | 1 | 1 |
| 1 | 1 | 1 |
| 1 | 1 | 1 |
| 0 | 0 | 0 |
| 1 | 1 | 1 |
| 1 | 1 | 1 |
| 0 | 1 | 0 |
| 0 | 0 | 0 |
| 1 | 1 | 1 |
| 0 | 0 | 0 |
| 0 | 0 | 0 |
| 0 | 0 | 0 |
| 0 | 0 | 0 |
| 0 | 0 | 0 |
| 1 | 1 | 1 |
| 1 | 1 | 1 |
| 1 | 1 | 1 |
| 0 | 0 | 0 |
| 0 | 0 | 0 |
| 0 | 0 | 0 |
| 0 | 0 | 0 |
| 0 | 0 | 0 |
| 0 | 0 | 0 |
| 0 | 0 | 0 |
| 0 | 1 | 0 |

| | | |
|---|---|---|
| 0 | 0 | 0 |
| 0 | 0 | 0 |
| 0 | 1 | 0 |
| 0 | 0 | 0 |
| 0 | 0 | 0 |
| 1 | 1 | 1 |
| 1 | 1 | 1 |
| 1 | 0 | 0 |
| 1 | 1 | 1 |
| 1 | 1 | 1 |
| 1 | 1 | 1 |
| 0 | 0 | 0 |
| 1 | 1 | 1 |
| 1 | 1 | 1 |
| 0 | 0 | 0 |
| 1 | 1 | 1 |
| 0 | 0 | 0 |
| 1 | 1 | 1 |
| 0 | 0 | 0 |
| 0 | 0 | 0 |
| 1 | 1 | 1 |
| 1 | 1 | 1 |
| 0 | 0 | 0 |
| 1 | 1 | 1 |
| 0 | 0 | 0 |
| 1 | 1 | 1 |
| 1 | 1 | 1 |
| 0 | 0 | 0 |
| 1 | 1 | 1 |
| 0 | 0 | 0 |
| 0 | 1 | 0 |
| 0 | 0 | 0 |
| 0 | 0 | 0 |
| 0 | 0 | 0 |
| 1 | 1 | 1 |
| 0 | 0 | 0 |
| 1 | 1 | 1 |
| 1 | 1 | 1 |
| 0 | 0 | 0 |
| 1 | 1 | 1 |
| 1 | 1 | 1 |
| 1 | 1 | 1 |
| 1 | 1 | 1 |
| 0 | 0 | 0 |
| 0 | 0 | 0 |
| 1 | 1 | 1 |
| 1 | 1 | 1 |
| 1 | 0 | 0 |
| 0 | 0 | 0 |
| 1 | 0 | 0 |
| 0 | 0 | 0 |
| 1 | 1 | 1 |
| 1 | 1 | 1 |
| 0 | 0 | 0 |

| | | |
|---|---|---|
| 1 | 1 | 1 |
| 1 | 1 | 1 |
| 1 | 1 | 1 |
| 1 | 1 | 1 |
| 0 | 1 | 0 |
| 1 | 1 | 1 |
| 1 | 1 | 1 |
| 0 | 0 | 0 |
| 1 | 1 | 1 |
| 1 | 1 | 1 |
| 1 | 1 | 1 |
| 1 | 1 | 1 |
| 0 | 0 | 0 |
| 1 | 1 | 1 |
| 0 | 0 | 0 |
| 0 | 0 | 0 |
| 1 | 1 | 1 |
| 1 | 1 | 1 |
| 1 | 1 | 1 |
| 1 | 1 | 1 |
| 0 | 0 | 0 |
| 0 | 0 | 0 |
| 1 | 1 | 1 |
| 1 | 1 | 1 |
| 0 | 0 | 0 |
| 1 | 1 | 1 |
| 0 | 0 | 0 |
| 0 | 0 | 0 |
| 0 | 0 | 0 |
| 0 | 0 | 0 |
| 0 | 1 | 0 |
| 0 | 1 | 0 |
| 0 | 0 | 0 |
| 0 | 0 | 0 |
| 0 | 0 | 0 |
| 0 | 0 | 0 |
| 0 | 0 | 0 |
| 0 | 0 | 0 |
| 0 | 0 | 0 |
| 1 | 0 | 0 |
| 1 | 1 | 1 |
| 0 | 1 | 0 |
| 0 | 1 | 0 |
| 0 | 1 | 0 |
| 0 | 1 | 0 |
| 0 | 0 | 0 |
| 0 | 1 | 0 |
| 0 | 1 | 0 |
| 1 | 1 | 1 |
| 0 | 0 | 0 |
| 0 | 0 | 0 |
| 1 | 1 | 1 |
| 1 | 1 | 1 |
| 1 | 1 | 1 |

| | | |
|---|---|---|
| 1 | 1 | 1 |
| 1 | 1 | 1 |
| 1 | 1 | 1 |
| 1 | 1 | 1 |
| 0 | 0 | 0 |
| 1 | 1 | 1 |
| 0 | 0 | 0 |
| 0 | 0 | 0 |
| 0 | 0 | 0 |
| 0 | 0 | 0 |
| 0 | 0 | 0 |
| 1 | 1 | 1 |
| 0 | 0 | 0 |
| 0 | 0 | 0 |
| 1 | 1 | 1 |
| 1 | 1 | 1 |
| 1 | 1 | 1 |
| 1 | 1 | 1 |
| 1 | 1 | 1 |
| 1 | 1 | 1 |
| 1 | 1 | 1 |
| 1 | 1 | 1 |
| 1 | 1 | 1 |
| 0 | 0 | 0 |
| 1 | 1 | 1 |
| 1 | 1 | 1 |
| 1 | 1 | 1 |
| 0 | 0 | 0 |
| 1 | 0 | 0 |
| 1 | 0 | 0 |
| 0 | 0 | 0 |
| 0 | 0 | 0 |
| 1 | 1 | 1 |
| 0 | 0 | 0 |
| 0 | 1 | 0 |
| 1 | 1 | 1 |
| 1 | 1 | 1 |
| 1 | 1 | 1 |
| 1 | 1 | 1 |
| 1 | 1 | 1 |
| 1 | 1 | 1 |
| 1 | 1 | 1 |
| 1 | 1 | 1 |
| 1 | 1 | 1 |
| 1 | 1 | 1 |
| 1 | 1 | 1 |
| 1 | 1 | 1 |
| 1 | 1 | 1 |
| 0 | 0 | 0 |
| 0 | 1 | 0 |
| 1 | 1 | 1 |
| 1 | 0 | 0 |
| 1 | 0 | 0 |
| 1 | 1 | 1 |

| | | |
|---|---|---|
| 1 | 1 | 1 |
| 0 | 1 | 0 |
| 1 | 1 | 1 |
| 0 | 1 | 0 |
| 1 | 0 | 0 |
| 1 | 1 | 1 |
| 1 | 1 | 1 |
| 1 | 1 | 1 |
| 1 | 1 | 1 |
| 1 | 1 | 1 |
| 0 | 0 | 0 |
| 0 | 0 | 0 |
| 0 | 0 | 0 |
| 0 | 0 | 0 |
| 0 | 0 | 0 |
| 1 | 0 | 0 |
| 0 | 0 | 0 |
| 0 | 0 | 0 |
| 0 | 0 | 0 |
| 0 | 0 | 0 |
| 0 | 0 | 0 |
| 0 | 0 | 0 |
| 0 | 1 | 0 |
| 0 | 0 | 0 |
| 0 | 0 | 0 |
| 0 | 0 | 0 |
| 1 | 1 | 1 |
| 0 | 0 | 0 |
| 0 | 1 | 0 |
| 0 | 1 | 0 |
| 0 | 1 | 0 |
| 0 | 0 | 0 |
| 1 | 1 | 1 |
| 1 | 1 | 1 |
| 0 | 0 | 0 |
| 0 | 0 | 0 |
| 0 | 0 | 0 |
| 0 | 0 | 0 |
| 0 | 0 | 0 |
| 1 | 0 | 0 |
| 1 | 0 | 0 |
| 0 | 0 | 0 |
| 1 | 0 | 0 |
| 0 | 0 | 0 |
| 1 | 1 | 1 |
| 1 | 0 | 0 |
| 1 | 1 | 1 |
| 0 | 0 | 0 |
| 0 | 0 | 0 |
| 0 | 0 | 0 |
| 0 | 0 | 0 |
| 1 | 1 | 1 |
| 0 | 0 | 0 |
| 1 | 1 | 1 |

| | | |
|---|---|---|
| 1 | 1 | 1 |
| 1 | 1 | 1 |
| 1 | 1 | 1 |
| 1 | 1 | 1 |
| 1 | 1 | 1 |
| 0 | 0 | 0 |
| 0 | 0 | 0 |
| 0 | 0 | 0 |
| 0 | 0 | 0 |
| 0 | 1 | 0 |
| 0 | 0 | 0 |
| 0 | 0 | 0 |
| 0 | 0 | 0 |
| 0 | 0 | 0 |
| 0 | 0 | 0 |
| 0 | 0 | 0 |
| 1 | 1 | 1 |
| 0 | 1 | 0 |
| 1 | 1 | 1 |
| 1 | 1 | 1 |
| 1 | 1 | 1 |
| 1 | 1 | 1 |
| 1 | 1 | 1 |
| 1 | 1 | 1 |
| 1 | 1 | 1 |
| 1 | 1 | 1 |
| 1 | 1 | 1 |
| 1 | 1 | 1 |
| 0 | 0 | 0 |
| 0 | 1 | 0 |
| 0 | 1 | 0 |
| 1 | 0 | 0 |
| 1 | 0 | 0 |
| 0 | 0 | 0 |
| 1 | 1 | 1 |
| 0 | 0 | 0 |
| 1 | 0 | 0 |
| 0 | 0 | 0 |
| 1 | 1 | 1 |
| 0 | 0 | 0 |
| 1 | 1 | 1 |
| 1 | 1 | 1 |
| 0 | 1 | 0 |
| 0 | 0 | 0 |
| 1 | 1 | 1 |
| 1 | 1 | 1 |
| 1 | 1 | 1 |
| 1 | 1 | 1 |
| 1 | 1 | 1 |
| 1 | 1 | 1 |
| 1 | 1 | 1 |
| 1 | 0 | 0 |
| 0 | 0 | 0 |
| 0 | 0 | 0 |

| | | |
|---|---|---|
| 1 | 1 | 1 |
| 0 | 0 | 0 |
| 1 | 1 | 1 |
| 1 | 1 | 1 |
| 0 | 0 | 0 |
| 0 | 0 | 0 |
| 0 | 0 | 0 |
| 1 | 1 | 1 |
| 1 | 1 | 1 |
| 0 | 0 | 0 |
| 0 | 0 | 0 |
| 0 | 0 | 0 |
| 0 | 0 | 0 |
| 0 | 0 | 0 |
| 1 | 1 | 1 |
| 0 | 0 | 0 |
| 0 | 0 | 0 |
| 0 | 0 | 0 |
| 1 | 1 | 1 |
| 1 | 1 | 1 |
| 1 | 1 | 1 |
| 1 | 1 | 1 |
| 1 | 1 | 1 |
| 0 | 0 | 0 |
| 0 | 1 | 0 |
| 0 | 1 | 0 |
| 0 | 0 | 0 |
| 0 | 0 | 0 |
| 1 | 1 | 1 |
| 1 | 1 | 1 |
| 0 | 0 | 0 |
| 1 | 1 | 1 |
| 1 | 1 | 1 |
| 0 | 1 | 0 |
| 1 | 1 | 1 |
| 0 | 0 | 0 |
| 0 | 1 | 0 |
| 0 | 0 | 0 |
| 0 | 0 | 0 |
| 0 | 1 | 0 |
| 0 | 1 | 0 |
| 0 | 0 | 0 |
| 0 | 0 | 0 |
| 0 | 1 | 0 |
| 0 | 0 | 0 |
| 0 | 0 | 0 |
| 0 | 0 | 0 |
| 0 | 1 | 0 |
| 0 | 0 | 0 |
| 0 | 0 | 0 |
| 0 | 0 | 0 |
| 1 | 1 | 1 |
| 0 | 0 | 0 |
| 1 | 1 | 1 |

| | | |
|---|---|---|
| 1 | 1 | 1 |
| 1 | 1 | 1 |
| 1 | 1 | 1 |
| 1 | 1 | 1 |
| 0 | 0 | 0 |
| 0 | 0 | 0 |
| 1 | 1 | 1 |
| 1 | 0 | 0 |
| 1 | 1 | 1 |
| 0 | 0 | 0 |
| 1 | 0 | 0 |
| 1 | 0 | 0 |
| 0 | 0 | 0 |
| 1 | 1 | 1 |
| 1 | 1 | 1 |
| 1 | 1 | 1 |
| 1 | 1 | 1 |
| 1 | 1 | 1 |
| 1 | 1 | 1 |
| 1 | 1 | 1 |
| 1 | 1 | 1 |
| 0 | 1 | 0 |
| 1 | 1 | 1 |
| 0 | 0 | 0 |
| 1 | 1 | 1 |
| 0 | 0 | 0 |
| 1 | 1 | 1 |
| 0 | 0 | 0 |
| 1 | 1 | 1 |
| 0 | 0 | 0 |
| 0 | 0 | 0 |
| 0 | 0 | 0 |
| 0 | 0 | 0 |
| 0 | 0 | 0 |
| 0 | 1 | 0 |
| 0 | 0 | 0 |
| 1 | 1 | 1 |
| 0 | 0 | 0 |
| 1 | 1 | 1 |
| 1 | 1 | 1 |
| 0 | 0 | 0 |
| 0 | 0 | 0 |
| 0 | 0 | 0 |
| 0 | 0 | 0 |
| 0 | 0 | 0 |
| 0 | 0 | 0 |
| 0 | 0 | 0 |
| 0 | 0 | 0 |
| 1 | 1 | 1 |
| 0 | 0 | 0 |
| 1 | 0 | 0 |
| 1 | 0 | 0 |
| 1 | 1 | 1 |
| 1 | 1 | 1 |

| | | |
|---|---|---|
| 1 | 1 | 1 |
| 1 | 1 | 1 |
| 1 | 1 | 1 |
| 0 | 1 | 0 |
| 0 | 1 | 0 |
| 0 | 1 | 0 |
| 0 | 0 | 0 |
| 0 | 0 | 0 |
| 0 | 0 | 0 |
| 0 | 0 | 0 |
| 0 | 0 | 0 |
| 0 | 0 | 0 |
| 0 | 0 | 0 |
| 0 | 0 | 0 |
| 0 | 0 | 0 |
| 0 | 0 | 0 |
| 0 | 0 | 0 |
| 0 | 0 | 0 |
| 1 | 1 | 1 |
| 0 | 0 | 0 |
| 0 | 1 | 0 |
| 0 | 1 | 0 |
| 0 | 1 | 0 |
| 1 | 1 | 1 |
| 0 | 1 | 0 |
| 0 | 0 | 0 |
| 0 | 0 | 0 |
| 0 | 0 | 0 |
| 0 | 0 | 0 |
| 0 | 0 | 0 |
| 0 | 0 | 0 |
| 0 | 0 | 0 |
| 0 | 0 | 0 |
| 0 | 0 | 0 |
| 1 | 1 | 1 |
| 0 | 0 | 0 |
| 0 | 0 | 0 |
| 0 | 0 | 0 |
| 0 | 0 | 0 |
| 0 | 1 | 0 |
| 0 | 0 | 0 |
| 0 | 0 | 0 |
| 1 | 1 | 1 |
| 0 | 1 | 0 |
| 1 | 1 | 1 |
| 0 | 1 | 0 |
| 0 | 1 | 0 |
| 0 | 0 | 0 |
| 0 | 0 | 0 |
| 0 | 0 | 0 |
| 0 | 0 | 0 |
| 0 | 1 | 0 |
| 0 | 0 | 0 |
| 1 | 1 | 1 |

| | | |
|---|---|---|
| 0 | 1 | 0 |
| 1 | 1 | 1 |
| 1 | 1 | 1 |
| 1 | 1 | 1 |
| 1 | 1 | 1 |
| 0 | 0 | 0 |
| 0 | 1 | 0 |
| 1 | 1 | 1 |
| 1 | 1 | 1 |
| 1 | 1 | 1 |
| 0 | 0 | 0 |
| 0 | 0 | 0 |
| 0 | 0 | 0 |
| 0 | 0 | 0 |
| 0 | 0 | 0 |
| 0 | 1 | 0 |
| 1 | 0 | 0 |
| 0 | 0 | 0 |
| 1 | 1 | 1 |
| 1 | 1 | 1 |
| 1 | 1 | 1 |
| 1 | 1 | 1 |
| 0 | 0 | 0 |
| 1 | 1 | 1 |
| 0 | 0 | 0 |
| 0 | 0 | 0 |
| 0 | 0 | 0 |
| 0 | 0 | 0 |
| 1 | 1 | 1 |
| 0 | 0 | 0 |
| 1 | 1 | 1 |
| 0 | 1 | 0 |
| 0 | 1 | 0 |
| 0 | 1 | 0 |
| 1 | 1 | 1 |
| 1 | 1 | 1 |
| 0 | 1 | 0 |
| 0 | 0 | 0 |
| 0 | 0 | 0 |
| 0 | 0 | 0 |
| 0 | 0 | 0 |
| 0 | 0 | 0 |
| 1 | 1 | 1 |
| 0 | 0 | 0 |
| 0 | 0 | 0 |
| 1 | 1 | 1 |
| 0 | 0 | 0 |
| 0 | 0 | 0 |
| 0 | 0 | 0 |
| 0 | 0 | 0 |
| 1 | 1 | 1 |
| 1 | 1 | 1 |
| 1 | 1 | 1 |
| 0 | 0 | 0 |

| | | |
|---|---|---|
| 0 | 0 | 0 |
| 1 | 1 | 1 |
| 0 | 0 | 0 |
| 0 | 1 | 0 |
| 1 | 0 | 0 |
| 1 | 0 | 0 |
| 1 | 0 | 0 |
| 1 | 1 | 1 |
| 0 | 0 | 0 |
| 1 | 1 | 1 |
| 0 | 0 | 0 |
| 0 | 0 | 0 |
| 0 | 0 | 0 |
| 0 | 0 | 0 |
| 1 | 1 | 1 |
| 1 | 1 | 1 |
| 0 | 0 | 0 |
| 0 | 0 | 0 |
| 0 | 1 | 0 |
| 1 | 1 | 1 |
| 0 | 0 | 0 |
| 1 | 0 | 0 |
| 1 | 0 | 0 |
| 1 | 1 | 1 |
| 0 | 0 | 0 |
| 1 | 0 | 0 |
| 0 | 0 | 0 |
| 0 | 0 | 0 |
| 0 | 0 | 0 |
| 1 | 1 | 1 |
| 0 | 0 | 0 |
| 0 | 0 | 0 |
| 0 | 0 | 0 |
| 1 | 1 | 1 |
| 0 | 0 | 0 |
| 1 | 1 | 1 |
| 0 | 0 | 0 |
| 0 | 0 | 0 |
| 0 | 0 | 0 |
| 0 | 0 | 0 |
| 0 | 0 | 0 |
| 1 | 1 | 1 |
| 0 | 0 | 0 |
| 0 | 0 | 0 |
| 0 | 0 | 0 |
| 0 | 0 | 0 |
| 0 | 0 | 0 |
| 0 | 0 | 0 |
| 0 | 0 | 0 |
| 0 | 0 | 0 |
| 0 | 0 | 0 |
| 0 | 0 | 0 |
| 1 | 1 | 1 |
| 0 | 0 | 0 |

| | | |
|---|---|---|
| 0 | 1 | 0 |
| 1 | 1 | 1 |
| 1 | 1 | 1 |
| 1 | 1 | 1 |
| 1 | 1 | 1 |
| 1 | 1 | 1 |
| 1 | 1 | 1 |
| 1 | 1 | 1 |
| 1 | 1 | 1 |
| 0 | 0 | 0 |
| 0 | 0 | 0 |
| 1 | 1 | 1 |
| 1 | 1 | 1 |
| 1 | 1 | 1 |
| 0 | 0 | 0 |
| 1 | 1 | 1 |
| 1 | 1 | 1 |
| 0 | 0 | 0 |
| 0 | 1 | 0 |
| 0 | 1 | 0 |
| 0 | 1 | 0 |
| 0 | 0 | 0 |
| 0 | 0 | 0 |
| 0 | 0 | 0 |
| 0 | 0 | 0 |
| 0 | 0 | 0 |
| 0 | 0 | 0 |
| 0 | 0 | 0 |
| 1 | 1 | 1 |
| 0 | 0 | 0 |
| 0 | 0 | 0 |
| 1 | 1 | 1 |
| 1 | 1 | 1 |
| 1 | 1 | 1 |
| 1 | 1 | 1 |
| 1 | 1 | 1 |
| 1 | 1 | 1 |
| 1 | 1 | 1 |
| 1 | 1 | 1 |
| 1 | 1 | 1 |
| 1 | 1 | 1 |
| 1 | 1 | 1 |
| 1 | 1 | 1 |
| 0 | 0 | 0 |
| 1 | 1 | 1 |
| 0 | 0 | 0 |
| 1 | 1 | 1 |
| 1 | 1 | 1 |
| 1 | 1 | 1 |
| 1 | 1 | 1 |
| 0 | 0 | 0 |
| 1 | 1 | 1 |
| 1 | 1 | 1 |
| 0 | 0 | 0 |

| | | |
|---|---|---|
| 0 | 0 | 0 |
| 0 | 0 | 0 |
| 0 | 0 | 0 |
| 0 | 0 | 0 |
| 1 | 1 | 1 |
| 0 | 1 | 0 |
| 1 | 1 | 1 |
| 0 | 0 | 0 |
| 0 | 0 | 0 |
| 0 | 0 | 0 |
| 0 | 0 | 0 |
| 1 | 1 | 1 |
| 1 | 1 | 1 |
| 1 | 1 | 1 |
| 1 | 1 | 1 |
| 0 | 0 | 0 |
| 0 | 0 | 0 |
| 0 | 1 | 0 |
| 0 | 1 | 0 |
| 1 | 1 | 1 |
| 1 | 1 | 1 |
| 0 | 0 | 0 |
| 1 | 1 | 1 |
| 0 | 1 | 0 |
| 0 | 0 | 0 |
| 0 | 0 | 0 |
| 0 | 0 | 0 |
| 0 | 0 | 0 |
| 1 | 1 | 1 |
| 1 | 1 | 1 |
| 1 | 1 | 1 |
| 0 | 0 | 0 |
| 1 | 1 | 1 |
| 1 | 1 | 1 |
| 1 | 1 | 1 |
| 1 | 1 | 1 |
| 1 | 1 | 1 |
| 1 | 1 | 1 |
| 1 | 1 | 1 |
| 1 | 1 | 1 |
| 1 | 1 | 1 |
| 1 | 1 | 1 |
| 0 | 0 | 0 |
| 0 | 1 | 0 |
| 0 | 1 | 0 |
| 1 | 1 | 1 |
| 0 | 1 | 0 |
| 1 | 1 | 1 |
| 1 | 1 | 1 |
| 0 | 0 | 0 |
| 1 | 1 | 1 |
| 1 | 1 | 1 |
| 1 | 1 | 1 |
| 1 | 1 | 1 |

| | | |
|---|---|---|
| 0 | 0 | 0 |
| 1 | 1 | 1 |
| 0 | 0 | 0 |
| 0 | 1 | 0 |
| 1 | 1 | 1 |
| 1 | 1 | 1 |
| 1 | 1 | 1 |
| 0 | 0 | 0 |
| 1 | 1 | 1 |
| 1 | 1 | 1 |
| 0 | 0 | 0 |
| 0 | 0 | 0 |
| 0 | 0 | 0 |
| 1 | 1 | 1 |
| 0 | 0 | 0 |
| 0 | 0 | 0 |
| 0 | 0 | 0 |
| 0 | 0 | 0 |
| 0 | 1 | 0 |
| 0 | 0 | 0 |
| 0 | 0 | 0 |
| 0 | 0 | 0 |
| 0 | 0 | 0 |
| 0 | 0 | 0 |
| 0 | 0 | 0 |
| 0 | 0 | 0 |
| 0 | 0 | 0 |
| 0 | 0 | 0 |
| 0 | 0 | 0 |
| 0 | 0 | 0 |
| 0 | 0 | 0 |
| 0 | 0 | 0 |
| 1 | 1 | 1 |
| 0 | 0 | 0 |
| 1 | 1 | 1 |
| 0 | 0 | 0 |
| 0 | 0 | 0 |
| 1 | 1 | 1 |
| 1 | 1 | 1 |
| 0 | 1 | 0 |
| 0 | 1 | 0 |
| 0 | 0 | 0 |
| 0 | 0 | 0 |
| 0 | 0 | 0 |
| 1 | 1 | 1 |
| 0 | 0 | 0 |
| 1 | 1 | 1 |
| 0 | 0 | 0 |
| 0 | 1 | 0 |
| 0 | 1 | 0 |
| 0 | 1 | 0 |
| 0 | 1 | 0 |
| 0 | 1 | 0 |
| 1 | 1 | 1 |

| | | |
|---|---|---|
| 1 | 1 | 1 |
| 1 | 1 | 1 |
| 0 | 1 | 0 |
| 1 | 1 | 1 |
| 1 | 1 | 1 |
| 0 | 0 | 0 |
| 0 | 0 | 0 |
| 1 | 1 | 1 |
| 0 | 0 | 0 |
| 0 | 0 | 0 |
| 1 | 1 | 1 |
| 0 | 1 | 0 |
| 1 | 1 | 1 |
| 1 | 1 | 1 |
| 1 | 1 | 1 |
| 0 | 0 | 0 |
| 1 | 1 | 1 |
| 1 | 1 | 1 |
| 0 | 0 | 0 |
| 0 | 0 | 0 |
| 0 | 0 | 0 |
| 1 | 1 | 1 |
| 1 | 1 | 1 |
| 0 | 0 | 0 |
| 1 | 1 | 1 |
| 0 | 0 | 0 |
| 0 | 0 | 0 |
| 1 | 1 | 1 |
| 1 | 1 | 1 |
| 1 | 1 | 1 |
| 0 | 0 | 0 |
| 1 | 0 | 0 |
| 0 | 0 | 0 |
| 0 | 0 | 0 |
| 0 | 0 | 0 |
| 0 | 0 | 0 |
| 0 | 0 | 0 |
| 1 | 1 | 1 |
| 0 | 0 | 0 |
| 0 | 1 | 0 |
| 0 | 0 | 0 |
| 0 | 0 | 0 |
| 0 | 0 | 0 |
| 0 | 0 | 0 |
| 1 | 1 | 1 |
| 0 | 1 | 0 |
| 0 | 1 | 0 |
| 0 | 0 | 0 |
| 0 | 0 | 0 |
| 0 | 0 | 0 |
| 1 | 1 | 1 |
| 0 | 1 | 0 |
| 1 | 1 | 1 |
| 1 | 1 | 1 |

| | | |
|---|---|---|
| 1 | 1 | 1 |
| 0 | 1 | 0 |
| 0 | 1 | 0 |
| 0 | 0 | 0 |
| 0 | 0 | 0 |
| 1 | 1 | 1 |
| 0 | 0 | 0 |
| 1 | 1 | 1 |
| 1 | 1 | 1 |
| 1 | 1 | 1 |
| 1 | 1 | 1 |
| 0 | 0 | 0 |
| 1 | 0 | 0 |
| 0 | 0 | 0 |
| 1 | 1 | 1 |
| 0 | 0 | 0 |
| 0 | 0 | 0 |
| 0 | 0 | 0 |
| 1 | 1 | 1 |
| 0 | 0 | 0 |
| 0 | 0 | 0 |
| 0 | 0 | 0 |
| 0 | 0 | 0 |
| 0 | 0 | 0 |
| 0 | 0 | 0 |
| 1 | 1 | 1 |
| 0 | 0 | 0 |
| 0 | 1 | 0 |
| 0 | 1 | 0 |
| 0 | 1 | 0 |
| 0 | 1 | 0 |
| 0 | 1 | 0 |
| 0 | 0 | 0 |
| 0 | 0 | 0 |
| 1 | 1 | 1 |
| 1 | 1 | 1 |
| 0 | 0 | 0 |
| 0 | 0 | 0 |
| 0 | 0 | 0 |
| 1 | 0 | 0 |
| 0 | 0 | 0 |
| 0 | 0 | 0 |
| 1 | 1 | 1 |
| 0 | 0 | 0 |
| 1 | 1 | 1 |
| 1 | 1 | 1 |
| 1 | 1 | 1 |
| 1 | 1 | 1 |
| 1 | 1 | 1 |
| 0 | 0 | 0 |
| 1 | 1 | 1 |
| 1 | 1 | 1 |
| 1 | 1 | 1 |
| 0 | 1 | 0 |

| | | |
|---|---|---|
| 1 | 1 | 1 |
| 1 | 1 | 1 |
| 1 | 1 | 1 |
| 0 | 0 | 0 |
| 1 | 1 | 1 |
| 1 | 1 | 1 |
| 0 | 1 | 0 |
| 0 | 0 | 0 |
| 1 | 1 | 1 |
| 1 | 1 | 1 |
| 1 | 1 | 1 |
| 1 | 1 | 1 |
| 1 | 1 | 1 |
| 0 | 1 | 0 |
| 0 | 1 | 0 |
| 0 | 1 | 0 |
| 0 | 0 | 0 |
| 0 | 0 | 0 |
| 0 | 1 | 0 |
| 1 | 1 | 1 |
| 0 | 1 | 0 |
| 1 | 1 | 1 |
| 0 | 1 | 0 |
| 1 | 1 | 1 |
| 1 | 1 | 1 |
| 1 | 1 | 1 |
| 1 | 1 | 1 |
| 0 | 0 | 0 |
| 0 | 0 | 0 |
| 1 | 1 | 1 |
| 1 | 1 | 1 |
| 0 | 0 | 0 |
| 0 | 1 | 0 |
| 0 | 0 | 0 |
| 0 | 0 | 0 |
| 0 | 0 | 0 |
| 0 | 0 | 0 |
| 0 | 0 | 0 |
| 1 | 1 | 1 |
| 1 | 1 | 1 |
| 1 | 1 | 1 |
| 0 | 1 | 0 |
| 1 | 1 | 1 |
| 1 | 1 | 1 |
| 0 | 0 | 0 |
| 0 | 0 | 0 |
| 0 | 0 | 0 |
| 1 | 1 | 1 |
| 1 | 1 | 1 |
| 1 | 1 | 1 |
| 1 | 1 | 1 |
| 0 | 0 | 0 |
| 0 | 0 | 0 |
| 0 | 0 | 0 |

| | | |
|---|---|---|
| 0 | 0 | 0 |
| 0 | 0 | 0 |
| 1 | 1 | 1 |
| 0 | 0 | 0 |
| 0 | 0 | 0 |
| 0 | 0 | 0 |
| 0 | 0 | 0 |
| 1 | 1 | 1 |
| 1 | 1 | 1 |
| 1 | 1 | 1 |
| 1 | 1 | 1 |
| 1 | 1 | 1 |
| 0 | 1 | 0 |
| 0 | 1 | 0 |
| 1 | 1 | 1 |
| 1 | 1 | 1 |
| 1 | 1 | 1 |
| 1 | 1 | 1 |
| 1 | 1 | 1 |
| 1 | 1 | 1 |
| 0 | 0 | 0 |
| 0 | 0 | 0 |
| 0 | 0 | 0 |
| 0 | 0 | 0 |
| 0 | 0 | 0 |
| 1 | 1 | 1 |
| 1 | 1 | 1 |
| 0 | 0 | 0 |
| 0 | 1 | 0 |
| 0 | 1 | 0 |
| 0 | 0 | 0 |
| 0 | 0 | 0 |
| 0 | 0 | 0 |
| 1 | 1 | 1 |
| 0 | 0 | 0 |
| 1 | 1 | 1 |
| 0 | 1 | 0 |
| 0 | 0 | 0 |
| 1 | 1 | 1 |
| 1 | 1 | 1 |
| 0 | 1 | 0 |
| 0 | 1 | 0 |
| 1 | 1 | 1 |
| 1 | 1 | 1 |
| 1 | 1 | 1 |
| 1 | 1 | 1 |
| 1 | 1 | 1 |
| 1 | 1 | 1 |
| 0 | 0 | 0 |
| 1 | 1 | 1 |
| 0 | 0 | 0 |
| 0 | 0 | 0 |
| 1 | 1 | 1 |
| 0 | 1 | 0 |

| | | |
|---|---|---|
| 0 | 0 | 0 |
| 1 | 1 | 1 |
| 1 | 1 | 1 |
| 1 | 1 | 1 |
| 1 | 1 | 1 |
| 0 | 0 | 0 |
| 0 | 0 | 0 |
| 1 | 1 | 1 |
| 0 | 1 | 0 |
| 1 | 1 | 1 |
| 1 | 1 | 1 |
| 0 | 0 | 0 |
| 0 | 1 | 0 |
| 1 | 1 | 1 |
| 0 | 1 | 0 |
| 1 | 1 | 1 |
| 0 | 0 | 0 |
| 0 | 0 | 0 |
| 1 | 1 | 1 |
| 1 | 1 | 1 |
| 1 | 1 | 1 |
| 1 | 1 | 1 |
| 1 | 1 | 1 |
| 1 | 1 | 1 |
| 0 | 0 | 0 |
| 1 | 1 | 1 |
| 1 | 1 | 1 |
| 1 | 1 | 1 |
| 1 | 1 | 1 |
| 1 | 1 | 1 |
| 0 | 1 | 0 |
| 0 | 0 | 0 |
| 0 | 0 | 0 |
| 0 | 1 | 0 |
| 1 | 1 | 1 |
| 0 | 0 | 0 |
| 0 | 0 | 0 |
| 0 | 0 | 0 |
| 0 | 0 | 0 |
| 0 | 0 | 0 |
| 0 | 0 | 0 |
| 0 | 0 | 0 |
| 0 | 0 | 0 |
| 0 | 0 | 0 |
| 0 | 0 | 0 |
| 0 | 0 | 0 |
| 0 | 0 | 0 |
| 0 | 0 | 0 |
| 1 | 0 | 0 |
| 1 | 1 | 1 |
| 1 | 0 | 0 |
| 1 | 1 | 1 |
| 0 | 0 | 0 |
| 1 | 1 | 1 |

| | | |
|---|---|---|
| 1 | 1 | 1 |
| 1 | 1 | 1 |
| 1 | 1 | 1 |
| 1 | 1 | 1 |
| 1 | 1 | 1 |
| 1 | 1 | 1 |
| 1 | 1 | 1 |
| 0 | 0 | 0 |
| 0 | 0 | 0 |
| 1 | 0 | 0 |
| 1 | 1 | 1 |
| 1 | 1 | 1 |
| 1 | 1 | 1 |
| 0 | 0 | 0 |
| 1 | 1 | 1 |
| 1 | 1 | 1 |
| 0 | 0 | 0 |
| 1 | 1 | 1 |
| 0 | 0 | 0 |
| 0 | 1 | 0 |
| 1 | 1 | 1 |
| 1 | 1 | 1 |
| 1 | 1 | 1 |
| 0 | 0 | 0 |
| 1 | 1 | 1 |
| 1 | 1 | 1 |
| 0 | 1 | 0 |
| 0 | 1 | 0 |
| 0 | 1 | 0 |
| 0 | 1 | 0 |
| 1 | 1 | 1 |
| 0 | 0 | 0 |
| 1 | 1 | 1 |
| 0 | 1 | 0 |
| 1 | 1 | 1 |
| 1 | 1 | 1 |
| 1 | 1 | 1 |
| 1 | 1 | 1 |
| 0 | 0 | 0 |
| 1 | 1 | 1 |
| 1 | 1 | 1 |
| 1 | 1 | 1 |
| 0 | 1 | 0 |
| 1 | 1 | 1 |
| 1 | 1 | 1 |
| 0 | 0 | 0 |
| 0 | 1 | 0 |
| 0 | 1 | 0 |
| 1 | 0 | 0 |
| 1 | 0 | 0 |
| 0 | 1 | 0 |
| 1 | 0 | 0 |
| 0 | 0 | 0 |
| 1 | 0 | 0 |

| | | |
|---|---|---|
| 1 | 0 | 0 |
| 1 | 0 | 0 |
| 1 | 0 | 0 |
| 1 | 0 | 0 |
| 0 | 0 | 0 |
| 0 | 0 | 0 |
| 0 | 0 | 0 |
| 0 | 0 | 0 |
| 1 | 0 | 0 |
| 1 | 0 | 0 |
| 1 | 0 | 0 |
| 1 | 0 | 0 |
| 1 | 0 | 0 |
| 0 | 0 | 0 |
| 1 | 0 | 0 |
| 0 | 0 | 0 |
| 1 | 0 | 0 |
| 0 | 0 | 0 |
| 1 | 0 | 0 |
| 0 | 0 | 0 |
| 0 | 0 | 0 |
| 1 | 0 | 0 |
| 1 | 0 | 0 |
| 0 | 0 | 0 |
| 0 | 0 | 0 |
| 1 | 0 | 0 |
| 0 | 0 | 0 |
| 0 | 0 | 0 |
| 0 | 0 | 0 |
| 1 | 0 | 0 |
| 1 | 0 | 0 |
| 0 | 0 | 0 |
| 1 | 0 | 0 |
| 1 | 0 | 0 |
| 1 | 0 | 0 |
| 1 | 0 | 0 |
| 1 | 0 | 0 |
| 1 | 0 | 0 |
| 1 | 0 | 0 |
| 1 | 0 | 0 |
| 1 | 0 | 0 |
| 1 | 0 | 0 |
| 1 | 0 | 0 |
| 0 | 0 | 0 |
| 1 | 1 | 1 |
| 1 | 1 | 1 |
| 0 | 0 | 0 |
| 0 | 0 | 0 |
| 0 | 1 | 0 |
| 0 | 0 | 0 |
| 0 | 0 | 0 |
| 0 | 0 | 0 |
| 1 | 1 | 1 |
| 1 | 1 | 1 |

| | | |
|---|---|---|
| 1 | 1 | 1 |
| 1 | 1 | 1 |
| 1 | 1 | 1 |
| 1 | 1 | 1 |
| 1 | 1 | 1 |
| 1 | 1 | 1 |
| 1 | 1 | 1 |
| 0 | 0 | 0 |
| 1 | 1 | 1 |
| 1 | 1 | 1 |
| 0 | 1 | 0 |
| 0 | 1 | 0 |
| 1 | 0 | 0 |
| 1 | 1 | 1 |
| 0 | 0 | 0 |
| 0 | 0 | 0 |
| 0 | 1 | 0 |
| 0 | 0 | 0 |
| 1 | 1 | 1 |
| 1 | 1 | 1 |
| 0 | 0 | 0 |
| 0 | 1 | 0 |
| 0 | 1 | 0 |
| 1 | 1 | 1 |
| 0 | 0 | 0 |
| 1 | 1 | 1 |
| 0 | 1 | 0 |
| 0 | 0 | 0 |
| 0 | 0 | 0 |
| 0 | 0 | 0 |
| 0 | 0 | 0 |
| 1 | 1 | 1 |
| 1 | 1 | 1 |
| 0 | 1 | 0 |
| 1 | 1 | 1 |
| 1 | 1 | 1 |
| 0 | 0 | 0 |
| 0 | 0 | 0 |
| 1 | 1 | 1 |
| 1 | 1 | 1 |
| 1 | 1 | 1 |
| 1 | 1 | 1 |
| 0 | 0 | 0 |
| 0 | 1 | 0 |
| 1 | 1 | 1 |
| 1 | 1 | 1 |
| 1 | 1 | 1 |
| 1 | 1 | 1 |
| 0 | 0 | 0 |
| 0 | 0 | 0 |
| 0 | 0 | 0 |
| 0 | 0 | 0 |
| 0 | 0 | 0 |
| 1 | 1 | 1 |

| | | |
|---|---|---|
| 0 | 0 | 0 |
| 0 | 0 | 0 |
| 0 | 1 | 0 |
| 0 | 0 | 0 |
| 1 | 1 | 1 |
| 0 | 1 | 0 |
| 0 | 1 | 0 |
| 1 | 1 | 1 |
| 1 | 1 | 1 |
| 0 | 0 | 0 |
| 1 | 1 | 1 |
| 0 | 1 | 0 |
| 1 | 1 | 1 |
| 0 | 1 | 0 |
| 0 | 1 | 0 |
| 0 | 1 | 0 |
| 0 | 1 | 0 |
| 1 | 1 | 1 |
| 1 | 1 | 1 |
| 0 | 1 | 0 |
| 0 | 1 | 0 |
| 0 | 1 | 0 |
| 0 | 1 | 0 |
| 0 | 1 | 0 |
| 0 | 1 | 0 |
| 0 | 1 | 0 |
| 0 | 1 | 0 |
| 0 | 1 | 0 |
| 0 | 1 | 0 |
| 0 | 1 | 0 |
| 1 | 1 | 1 |
| 1 | 1 | 1 |
| 0 | 0 | 0 |
| 1 | 1 | 1 |
| 1 | 1 | 1 |
| 1 | 1 | 1 |
| 1 | 1 | 1 |
| 0 | 1 | 0 |
| 1 | 1 | 1 |
| 1 | 1 | 1 |
| 1 | 1 | 1 |
| 1 | 1 | 1 |
| 1 | 1 | 1 |
| 0 | 1 | 0 |
| 0 | 1 | 0 |
| 1 | 1 | 1 |
| 1 | 1 | 1 |
| 0 | 0 | 0 |
| 0 | 0 | 0 |
| 0 | 0 | 0 |
| 0 | 0 | 0 |
| 0 | 0 | 0 |
| 1 | 1 | 1 |
| 0 | 0 | 0 |

| | | |
|---|---|---|
| 1 | 1 | 1 |
| 0 | 0 | 0 |
| 1 | 1 | 1 |
| 0 | 1 | 0 |
| 0 | 1 | 0 |
| 0 | 0 | 0 |
| 0 | 0 | 0 |
| 0 | 0 | 0 |
| 0 | 0 | 0 |
| 0 | 0 | 0 |
| 0 | 1 | 0 |
| 0 | 0 | 0 |
| 0 | 1 | 0 |
| 1 | 1 | 1 |
| 0 | 1 | 0 |
| 0 | 0 | 0 |
| 1 | 1 | 1 |
| 1 | 1 | 1 |
| 0 | 0 | 0 |
| 0 | 0 | 0 |
| 0 | 0 | 0 |
| 0 | 0 | 0 |
| 0 | 0 | 0 |
| 0 | 0 | 0 |
| 0 | 1 | 0 |
| 0 | 0 | 0 |
| 0 | 0 | 0 |
| 0 | 0 | 0 |
| 0 | 0 | 0 |
| 0 | 0 | 0 |
| 0 | 0 | 0 |
| 1 | 1 | 1 |
| 1 | 1 | 1 |
| 0 | 0 | 0 |
| 0 | 0 | 0 |
| 1 | 0 | 0 |
| 0 | 0 | 0 |
| 0 | 0 | 0 |
| 0 | 0 | 0 |
| 0 | 0 | 0 |
| 0 | 0 | 0 |
| 1 | 0 | 0 |
| 1 | 1 | 1 |
| 1 | 1 | 1 |
| 0 | 0 | 0 |
| 0 | 0 | 0 |
| 0 | 0 | 0 |
| 1 | 1 | 1 |
| 1 | 1 | 1 |
| 1 | 1 | 1 |
| 1 | 1 | 1 |
| 1 | 1 | 1 |
| 1 | 1 | 1 |
| 0 | 0 | 0 |

| | | |
|---|---|---|
| 1 | 1 | 1 |
| 1 | 1 | 1 |
| 1 | 1 | 1 |
| 1 | 1 | 1 |
| 0 | 0 | 0 |
| 0 | 0 | 0 |
| 1 | 0 | 0 |
| 1 | 0 | 0 |
| 0 | 1 | 0 |
| 1 | 1 | 1 |
| 1 | 1 | 1 |
| 0 | 0 | 0 |
| 1 | 1 | 1 |
| 0 | 0 | 0 |
| 0 | 0 | 0 |
| 0 | 0 | 0 |
| 0 | 0 | 0 |
| 0 | 0 | 0 |
| 0 | 0 | 0 |
| 0 | 0 | 0 |
| 0 | 0 | 0 |
| 0 | 0 | 0 |
| 0 | 0 | 0 |
| 0 | 1 | 0 |
| 0 | 1 | 0 |
| 0 | 1 | 0 |
| 1 | 1 | 1 |
| 1 | 1 | 1 |
| 1 | 1 | 1 |
| 1 | 1 | 1 |
| 1 | 1 | 1 |
| 0 | 0 | 0 |
| 1 | 1 | 1 |
| 0 | 0 | 0 |
| 0 | 0 | 0 |
| 0 | 0 | 0 |
| 0 | 0 | 0 |
| 1 | 1 | 1 |
| 1 | 1 | 1 |
| 0 | 1 | 0 |
| 1 | 1 | 1 |
| 1 | 1 | 1 |
| 1 | 1 | 1 |
| 0 | 1 | 0 |
| 1 | 1 | 1 |
| 0 | 0 | 0 |
| 1 | 1 | 1 |
| 1 | 1 | 1 |
| 1 | 1 | 1 |
| 0 | 0 | 0 |
| 0 | 0 | 0 |
| 0 | 0 | 0 |
| 0 | 1 | 0 |
| 0 | 1 | 0 |

| | | |
|---|---|---|
| 0 | 1 | 0 |
| 0 | 1 | 0 |
| 0 | 1 | 0 |
| 0 | 0 | 0 |
| 0 | 1 | 0 |
| 1 | 1 | 1 |
| 0 | 0 | 0 |
| 0 | 1 | 0 |
| 1 | 1 | 1 |
| 1 | 1 | 1 |
| 0 | 0 | 0 |
| 0 | 0 | 0 |
| 1 | 1 | 1 |
| 1 | 1 | 1 |
| 0 | 1 | 0 |
| 0 | 1 | 0 |
| 0 | 0 | 0 |
| 0 | 0 | 0 |
| 1 | 1 | 1 |
| 0 | 0 | 0 |
| 0 | 0 | 0 |
| 0 | 0 | 0 |
| 0 | 1 | 0 |
| 0 | 0 | 0 |
| 0 | 0 | 0 |
| 1 | 1 | 1 |
| 0 | 0 | 0 |
| 0 | 1 | 0 |
| 1 | 1 | 1 |
| 0 | 0 | 0 |
| 0 | 1 | 0 |
| 1 | 1 | 1 |
| 0 | 0 | 0 |
| 0 | 1 | 0 |
| 0 | 1 | 0 |
| 0 | 0 | 0 |
| 1 | 1 | 1 |
| 0 | 0 | 0 |
| 1 | 1 | 1 |
| 0 | 0 | 0 |
| 0 | 0 | 0 |
| 1 | 0 | 0 |
| 0 | 0 | 0 |
| 0 | 0 | 0 |
| 1 | 1 | 1 |
| 1 | 0 | 0 |
| 0 | 1 | 0 |
| 1 | 1 | 1 |
| 1 | 1 | 1 |
| 1 | 1 | 1 |
| 1 | 1 | 1 |
| 0 | 1 | 0 |
| 0 | 1 | 0 |
| 0 | 1 | 0 |

| | | |
|---|---|---|
| 0 | 1 | 0 |
| 0 | 1 | 0 |
| 0 | 1 | 0 |
| 0 | 0 | 0 |
| 1 | 0 | 0 |
| 0 | 0 | 0 |
| 1 | 1 | 1 |
| 1 | 0 | 0 |
| 0 | 0 | 0 |
| 0 | 0 | 0 |
| 1 | 1 | 1 |
| 0 | 0 | 0 |
| 0 | 0 | 0 |
| 0 | 0 | 0 |
| 1 | 0 | 0 |
| 0 | 0 | 0 |
| 0 | 0 | 0 |
| 0 | 1 | 0 |
| 0 | 0 | 0 |
| 0 | 0 | 0 |
| 0 | 0 | 0 |
| 0 | 0 | 0 |
| 1 | 1 | 1 |
| 0 | 0 | 0 |
| 0 | 0 | 0 |
| 0 | 0 | 0 |
| 0 | 0 | 0 |
| 0 | 0 | 0 |
| 0 | 0 | 0 |
| 0 | 0 | 0 |
| 0 | 0 | 0 |
| 0 | 0 | 0 |
| 0 | 0 | 0 |
| 0 | 1 | 0 |
| 0 | 1 | 0 |
| 0 | 0 | 0 |
| 1 | 1 | 1 |
| 0 | 0 | 0 |
| 0 | 0 | 0 |
| 0 | 0 | 0 |
| 0 | 0 | 0 |
| 1 | 1 | 1 |
| 0 | 0 | 0 |
| 0 | 0 | 0 |
| 1 | 1 | 1 |
| 0 | 1 | 0 |
| 1 | 1 | 1 |
| 1 | 1 | 1 |
| 1 | 1 | 1 |
| 0 | 1 | 0 |
| 1 | 1 | 1 |
| 1 | 1 | 1 |
| 1 | 1 | 1 |
| 1 | 1 | 1 |

| | | |
|---|---|---|
| 1 | 1 | 1 |
| 0 | 0 | 0 |
| 1 | 1 | 1 |
| 1 | 1 | 1 |
| 1 | 1 | 1 |
| 1 | 1 | 1 |
| 1 | 1 | 1 |
| 1 | 1 | 1 |
| 1 | 1 | 1 |
| 0 | 1 | 0 |
| 1 | 1 | 1 |
| 0 | 0 | 0 |
| 1 | 0 | 0 |
| 0 | 1 | 0 |
| 0 | 0 | 0 |
| 0 | 1 | 0 |
| 0 | 1 | 0 |
| 1 | 1 | 1 |
| 1 | 1 | 1 |
| 1 | 1 | 1 |
| 0 | 0 | 0 |
| 0 | 1 | 0 |
| 0 | 0 | 0 |
| 0 | 0 | 0 |
| 1 | 1 | 1 |
| 0 | 0 | 0 |
| 1 | 1 | 1 |
| 1 | 1 | 1 |
| 1 | 1 | 1 |
| 1 | 1 | 1 |
| 1 | 1 | 1 |
| 0 | 0 | 0 |
| 1 | 1 | 1 |
| 0 | 0 | 0 |
| 1 | 1 | 1 |
| 0 | 1 | 0 |
| 0 | 1 | 0 |
| 0 | 1 | 0 |
| 1 | 1 | 1 |
| 0 | 0 | 0 |
| 1 | 1 | 1 |
| 1 | 1 | 1 |
| 0 | 1 | 0 |
| 0 | 0 | 0 |
| 1 | 1 | 1 |
| 1 | 1 | 1 |
| 1 | 1 | 1 |
| 0 | 0 | 0 |
| 0 | 0 | 0 |
| 1 | 1 | 1 |
| 1 | 1 | 1 |
| 0 | 0 | 0 |
| 0 | 0 | 0 |
| 0 | 0 | 0 |

| | | |
|---|---|---|
| 0 | 0 | 0 |
| 1 | 1 | 1 |
| 0 | 0 | 0 |
| 0 | 0 | 0 |
| 0 | 0 | 0 |
| 1 | 1 | 1 |
| 1 | 1 | 1 |
| 0 | 1 | 0 |
| 1 | 1 | 1 |
| 0 | 0 | 0 |
| 1 | 1 | 1 |
| 0 | 1 | 0 |
| 1 | 1 | 1 |
| 1 | 1 | 1 |
| 0 | 0 | 0 |
| 0 | 1 | 0 |
| 1 | 1 | 1 |
| 0 | 0 | 0 |
| 1 | 1 | 1 |
| 0 | 0 | 0 |
| 0 | 0 | 0 |
| 1 | 1 | 1 |
| 0 | 0 | 0 |
| 1 | 1 | 1 |
| 0 | 0 | 0 |
| 0 | 0 | 0 |
| 1 | 1 | 1 |
| 0 | 1 | 0 |
| 0 | 0 | 0 |
| 1 | 1 | 1 |
| 1 | 1 | 1 |
| 1 | 1 | 1 |
| 1 | 1 | 1 |
| 0 | 1 | 0 |
| 0 | 0 | 0 |
| 0 | 0 | 0 |
| 1 | 1 | 1 |
| 0 | 0 | 0 |
| 0 | 0 | 0 |
| 0 | 0 | 0 |
| 0 | 0 | 0 |
| 0 | 0 | 0 |
| 0 | 0 | 0 |
| 0 | 1 | 0 |
| 0 | 0 | 0 |
| 1 | 1 | 1 |
| 0 | 1 | 0 |
| 0 | 0 | 0 |
| 0 | 0 | 0 |
| 0 | 0 | 0 |
| 0 | 0 | 0 |
| 0 | 0 | 0 |
| 1 | 1 | 1 |
| 1 | 1 | 1 |

| | | |
|---|---|---|
| 1 | 1 | 1 |
| 0 | 0 | 0 |
| 0 | 0 | 0 |
| 0 | 0 | 0 |
| 0 | 0 | 0 |
| 0 | 0 | 0 |
| 0 | 0 | 0 |
| 1 | 0 | 0 |
| 1 | 0 | 0 |
| 1 | 1 | 1 |
| 0 | 0 | 0 |
| 0 | 0 | 0 |
| 1 | 1 | 1 |
| 0 | 0 | 0 |
| 1 | 1 | 1 |
| 0 | 0 | 0 |
| 1 | 1 | 1 |
| 1 | 1 | 1 |
| 0 | 1 | 0 |
| 0 | 1 | 0 |
| 0 | 1 | 0 |
| 1 | 1 | 1 |
| 0 | 1 | 0 |
| 0 | 0 | 0 |
| 0 | 0 | 0 |
| 1 | 1 | 1 |
| 1 | 1 | 1 |
| 1 | 1 | 1 |
| 1 | 1 | 1 |
| 0 | 0 | 0 |
| 0 | 0 | 0 |
| 0 | 0 | 0 |
| 0 | 0 | 0 |
| 1 | 1 | 1 |
| 0 | 0 | 0 |
| 0 | 0 | 0 |
| 0 | 0 | 0 |
| 0 | 0 | 0 |
| 0 | 0 | 0 |
| 1 | 1 | 1 |
| 1 | 1 | 1 |
| 0 | 0 | 0 |
| 1 | 1 | 1 |
| 1 | 1 | 1 |
| 0 | 0 | 0 |
| 1 | 0 | 0 |
| 1 | 1 | 1 |
| 0 | 1 | 0 |
| 1 | 0 | 0 |
| 0 | 1 | 0 |
| 0 | 0 | 0 |
| 1 | 1 | 1 |
| 0 | 0 | 0 |
| 0 | 0 | 0 |

| | | |
|---|---|---|
| 0 | 1 | 0 |
| 1 | 1 | 1 |
| 0 | 0 | 0 |
| 0 | 0 | 0 |
| 1 | 1 | 1 |
| 1 | 1 | 1 |
| 1 | 1 | 1 |
| 1 | 1 | 1 |
| 1 | 1 | 1 |
| 1 | 1 | 1 |
| 1 | 1 | 1 |
| 1 | 1 | 1 |
| 0 | 0 | 0 |
| 1 | 1 | 1 |
| 1 | 1 | 1 |
| 1 | 1 | 1 |
| 1 | 1 | 1 |
| 1 | 1 | 1 |
| 1 | 1 | 1 |
| 1 | 1 | 1 |
| 1 | 1 | 1 |
| 1 | 1 | 1 |
| 1 | 1 | 1 |
| 1 | 1 | 1 |
| 1 | 1 | 1 |
| 1 | 1 | 1 |
| 0 | 1 | 0 |
| 1 | 1 | 1 |
| 1 | 1 | 1 |
| 1 | 1 | 1 |
| 1 | 1 | 1 |
| 0 | 0 | 0 |
| 1 | 1 | 1 |
| 0 | 1 | 0 |
| 1 | 1 | 1 |
| 1 | 1 | 1 |
| 0 | 1 | 0 |
| 1 | 1 | 1 |
| 1 | 1 | 1 |
| 1 | 1 | 1 |
| 0 | 0 | 0 |
| 0 | 0 | 0 |
| 1 | 1 | 1 |
| 1 | 1 | 1 |
| 1 | 1 | 1 |
| 1 | 0 | 0 |
| 1 | 0 | 0 |
| 1 | 1 | 1 |
| 0 | 0 | 0 |
| 0 | 0 | 0 |
| 1 | 1 | 1 |
| 0 | 1 | 0 |
| 0 | 0 | 0 |
| 1 | 1 | 1 |

| | | |
|---|---|---|
| 0 | 0 | 0 |
| 0 | 0 | 0 |
| 0 | 0 | 0 |
| 1 | 1 | 1 |
| 1 | 1 | 1 |
| 1 | 1 | 1 |
| 1 | 1 | 1 |
| 0 | 0 | 0 |
| 0 | 0 | 0 |
| 0 | 0 | 0 |
| 0 | 0 | 0 |
| 1 | 1 | 1 |
| 0 | 0 | 0 |
| 1 | 1 | 1 |
| 1 | 1 | 1 |
| 1 | 1 | 1 |
| 1 | 1 | 1 |
| 0 | 0 | 0 |
| 1 | 1 | 1 |
| 1 | 1 | 1 |
| 1 | 1 | 1 |
| 1 | 1 | 1 |
| 0 | 1 | 0 |
| 1 | 1 | 1 |
| 1 | 1 | 1 |
| 0 | 1 | 0 |
| 1 | 1 | 1 |
| 1 | 1 | 1 |
| 1 | 1 | 1 |
| 1 | 1 | 1 |
| 0 | 0 | 0 |
| 0 | 0 | 0 |
| 1 | 1 | 1 |
| 1 | 1 | 1 |
| 1 | 1 | 1 |
| 1 | 1 | 1 |
| 1 | 1 | 1 |
| 1 | 1 | 1 |
| 1 | 1 | 1 |
| 1 | 1 | 1 |
| 1 | 1 | 1 |
| 0 | 0 | 0 |
| 1 | 1 | 1 |
| 1 | 1 | 1 |
| 0 | 0 | 0 |
| 0 | 0 | 0 |
| 1 | 1 | 1 |
| 1 | 1 | 1 |
| 0 | 0 | 0 |
| 0 | 0 | 0 |
| 0 | 0 | 0 |
| 0 | 0 | 0 |
| 0 | 0 | 0 |
| 0 | 0 | 0 |

| | | |
|---|---|---|
| 1 | 1 | 1 |
| 1 | 1 | 1 |
| 1 | 1 | 1 |
| 1 | 1 | 1 |
| 1 | 1 | 1 |
| 0 | 0 | 0 |
| 0 | 0 | 0 |
| 0 | 0 | 0 |
| 0 | 0 | 0 |
| 0 | 0 | 0 |
| 0 | 1 | 0 |
| 0 | 0 | 0 |
| 0 | 0 | 0 |
| 0 | 0 | 0 |
| 0 | 1 | 0 |
| 0 | 0 | 0 |
| 0 | 1 | 0 |
| 0 | 1 | 0 |
| 0 | 0 | 0 |
| 0 | 0 | 0 |
| 0 | 0 | 0 |
| 0 | 0 | 0 |
| 1 | 0 | 0 |
| 0 | 0 | 0 |
| 0 | 0 | 0 |
| 0 | 0 | 0 |
| 0 | 0 | 0 |
| 1 | 1 | 1 |
| 1 | 1 | 1 |
| 1 | 1 | 1 |
| 1 | 1 | 1 |
| 1 | 1 | 1 |
| 0 | 0 | 0 |
| 0 | 1 | 0 |
| 0 | 0 | 0 |
| 0 | 0 | 0 |
| 0 | 0 | 0 |
| 0 | 0 | 0 |
| 0 | 0 | 0 |
| 0 | 0 | 0 |
| 0 | 0 | 0 |
| 0 | 0 | 0 |
| 0 | 0 | 0 |
| 1 | 1 | 1 |
| 1 | 1 | 1 |
| 1 | 1 | 1 |
| 1 | 1 | 1 |
| 0 | 0 | 0 |
| 0 | 0 | 0 |
| 0 | 0 | 0 |
| 1 | 1 | 1 |
| 1 | 1 | 1 |
| 0 | 0 | 0 |
| 1 | 1 | 1 |

| | | |
|---|---|---|
| 1 | 1 | 1 |
| 0 | 0 | 0 |
| 0 | 1 | 0 |
| 0 | 1 | 0 |
| 1 | 1 | 1 |
| 1 | 1 | 1 |
| 0 | 1 | 0 |
| 1 | 1 | 1 |
| 0 | 1 | 0 |
| 1 | 1 | 1 |
| 1 | 1 | 1 |
| 1 | 1 | 1 |
| 0 | 0 | 0 |
| 0 | 0 | 0 |
| 0 | 0 | 0 |
| 0 | 0 | 0 |
| 0 | 0 | 0 |
| 0 | 0 | 0 |
| 0 | 0 | 0 |
| 0 | 1 | 0 |
| 0 | 0 | 0 |
| 0 | 0 | 0 |
| 0 | 0 | 0 |
| 0 | 0 | 0 |
| 0 | 0 | 0 |
| 1 | 1 | 1 |
| 1 | 1 | 1 |
| 0 | 0 | 0 |
| 0 | 1 | 0 |
| 0 | 0 | 0 |
| 0 | 0 | 0 |
| 1 | 1 | 1 |
| 0 | 0 | 0 |
| 1 | 1 | 1 |
| 1 | 1 | 1 |
| 1 | 1 | 1 |
| 1 | 0 | 0 |
| 1 | 1 | 1 |
| 1 | 1 | 1 |
| 1 | 1 | 1 |
| 0 | 0 | 0 |
| 1 | 1 | 1 |
| 1 | 1 | 1 |
| 0 | 0 | 0 |
| 0 | 0 | 0 |
| 1 | 1 | 1 |
| 0 | 0 | 0 |
| 0 | 0 | 0 |
| 1 | 1 | 1 |
| 1 | 1 | 1 |
| 0 | 0 | 0 |
| 1 | 1 | 1 |
| 1 | 1 | 1 |
| 1 | 1 | 1 |

| | | |
|---|---|---|
| 1 | 1 | 1 |
| 1 | 1 | 1 |
| 1 | 1 | 1 |
| 1 | 1 | 1 |
| 0 | 0 | 0 |
| 0 | 0 | 0 |
| 1 | 1 | 1 |
| 1 | 1 | 1 |
| 1 | 1 | 1 |
| 0 | 1 | 0 |
| 1 | 1 | 1 |
| 0 | 0 | 0 |
| 0 | 0 | 0 |
| 1 | 1 | 1 |
| 1 | 1 | 1 |
| 0 | 0 | 0 |
| 0 | 1 | 0 |
| 1 | 1 | 1 |
| 0 | 0 | 0 |
| 1 | 1 | 1 |
| 1 | 1 | 1 |
| 1 | 1 | 1 |
| 1 | 1 | 1 |
| 0 | 0 | 0 |
| 1 | 1 | 1 |
| 0 | 0 | 0 |
| 1 | 1 | 1 |
| 1 | 1 | 1 |
| 0 | 0 | 0 |
| 0 | 0 | 0 |
| 0 | 0 | 0 |
| 0 | 0 | 0 |
| 0 | 0 | 0 |
| 0 | 0 | 0 |
| 1 | 1 | 1 |
| 1 | 1 | 1 |
| 1 | 1 | 1 |
| 0 | 0 | 0 |
| 1 | 1 | 1 |
| 1 | 1 | 1 |
| 0 | 0 | 0 |
| 0 | 0 | 0 |
| 0 | 0 | 0 |
| 1 | 1 | 1 |
| 0 | 0 | 0 |
| 0 | 1 | 0 |
| 0 | 1 | 0 |
| 0 | 0 | 0 |
| 1 | 1 | 1 |
| 1 | 1 | 1 |
| 1 | 1 | 1 |
| 0 | 0 | 0 |
| 1 | 1 | 1 |
| 0 | 0 | 0 |

| | | |
|---|---|---|
| 1 | 1 | 1 |
| 0 | 0 | 0 |
| 0 | 0 | 0 |
| 0 | 1 | 0 |
| 1 | 1 | 1 |
| 1 | 1 | 1 |
| 0 | 0 | 0 |
| 1 | 1 | 1 |
| 0 | 1 | 0 |
| 0 | 1 | 0 |
| 1 | 1 | 1 |
| 1 | 1 | 1 |
| 1 | 1 | 1 |
| 1 | 1 | 1 |
| 1 | 1 | 1 |
| 1 | 1 | 1 |
| 1 | 1 | 1 |
| 1 | 1 | 1 |
| 1 | 1 | 1 |
| 1 | 1 | 1 |
| 1 | 1 | 1 |
| 0 | 0 | 0 |
| 0 | 1 | 0 |
| 0 | 1 | 0 |
| 1 | 1 | 1 |
| 1 | 1 | 1 |
| 0 | 0 | 0 |
| 1 | 1 | 1 |
| 0 | 0 | 0 |
| 0 | 1 | 0 |
| 1 | 1 | 1 |
| 0 | 0 | 0 |
| 0 | 0 | 0 |
| 1 | 0 | 0 |
| 1 | 1 | 1 |
| 1 | 1 | 1 |
| 1 | 1 | 1 |
| 1 | 1 | 1 |
| 0 | 0 | 0 |
| 0 | 0 | 0 |
| 0 | 0 | 0 |
| 1 | 1 | 1 |
| 1 | 1 | 1 |
| 0 | 1 | 0 |
| 0 | 1 | 0 |
| 0 | 0 | 0 |
| 1 | 1 | 1 |
| 0 | 0 | 0 |
| 1 | 1 | 1 |
| 0 | 0 | 0 |
| 0 | 0 | 0 |
| 0 | 0 | 0 |
| 1 | 1 | 1 |
| 1 | 1 | 1 |

| | | |
|---|---|---|
| 0 | 0 | 0 |
| 0 | 0 | 0 |
| 1 | 1 | 1 |
| 1 | 1 | 1 |
| 1 | 1 | 1 |
| 1 | 1 | 1 |
| 1 | 1 | 1 |
| 1 | 1 | 1 |
| 1 | 1 | 1 |
| 1 | 1 | 1 |
| 1 | 1 | 1 |
| 1 | 1 | 1 |
| 1 | 1 | 1 |
| 1 | 1 | 1 |
| 1 | 1 | 1 |
| 0 | 0 | 0 |
| 0 | 0 | 0 |
| 0 | 0 | 0 |
| 1 | 1 | 1 |
| 1 | 1 | 1 |
| 0 | 1 | 0 |
| 0 | 0 | 0 |
| 0 | 0 | 0 |
| 0 | 0 | 0 |
| 0 | 0 | 0 |
| 1 | 1 | 1 |
| 1 | 1 | 1 |
| 1 | 1 | 1 |
| 1 | 1 | 1 |
| 1 | 1 | 1 |
| 0 | 0 | 0 |
| 0 | 1 | 0 |
| 0 | 0 | 0 |
| 1 | 1 | 1 |
| 1 | 0 | 0 |
| 1 | 0 | 0 |
| 0 | 0 | 0 |
| 0 | 0 | 0 |
| 0 | 1 | 0 |
| 1 | 1 | 1 |
| 1 | 0 | 0 |
| 0 | 0 | 0 |
| 1 | 1 | 1 |
| 0 | 1 | 0 |
| 0 | 0 | 0 |
| 1 | 0 | 0 |
| 0 | 0 | 0 |
| 0 | 0 | 0 |
| 1 | 1 | 1 |
| 0 | 1 | 0 |
| 0 | 0 | 0 |
| 0 | 0 | 0 |
| 1 | 1 | 1 |
| 0 | 0 | 0 |

| | | |
|---|---|---|
| 0 | 1 | 0 |
| 0 | 0 | 0 |
| 1 | 0 | 0 |
| 0 | 0 | 0 |
| 1 | 1 | 1 |
| 1 | 1 | 1 |
| 1 | 1 | 1 |
| 0 | 1 | 0 |
| 0 | 0 | 0 |
| 1 | 1 | 1 |
| 0 | 1 | 0 |
| 0 | 1 | 0 |
| 0 | 0 | 0 |
| 0 | 0 | 0 |
| 0 | 0 | 0 |
| 1 | 1 | 1 |
| 0 | 0 | 0 |
| 0 | 1 | 0 |
| 0 | 0 | 0 |
| 0 | 0 | 0 |
| 0 | 0 | 0 |
| 0 | 0 | 0 |
| 0 | 0 | 0 |
| 0 | 0 | 0 |
| 0 | 1 | 0 |
| 0 | 1 | 0 |
| 0 | 0 | 0 |
| 0 | 0 | 0 |
| 0 | 0 | 0 |
| 0 | 0 | 0 |
| 1 | 1 | 1 |
| 1 | 1 | 1 |
| 1 | 1 | 1 |
| 1 | 1 | 1 |
| 1 | 1 | 1 |
| 0 | 0 | 0 |
| 0 | 0 | 0 |
| 0 | 1 | 0 |
| 1 | 1 | 1 |
| 1 | 1 | 1 |
| 0 | 1 | 0 |
| 1 | 1 | 1 |
| 1 | 1 | 1 |
| 0 | 0 | 0 |
| 0 | 0 | 0 |
| 0 | 0 | 0 |
| 1 | 1 | 1 |
| 0 | 0 | 0 |
| 0 | 0 | 0 |
| 0 | 0 | 0 |
| 1 | 1 | 1 |
| 1 | 1 | 1 |
| 0 | 1 | 0 |
| 0 | 1 | 0 |

| | | |
|---|---|---|
| 0 | 1 | 0 |
| 1 | 1 | 1 |
| 1 | 0 | 0 |
| 1 | 1 | 1 |
| 0 | 0 | 0 |
| 1 | 0 | 0 |
| 0 | 0 | 0 |
| 0 | 0 | 0 |
| 0 | 0 | 0 |
| 0 | 0 | 0 |
| 0 | 0 | 0 |
| 0 | 0 | 0 |
| 0 | 1 | 0 |
| 0 | 0 | 0 |
| 0 | 0 | 0 |
| 1 | 1 | 1 |
| 1 | 1 | 1 |
| 1 | 1 | 1 |
| 0 | 0 | 0 |
| 0 | 0 | 0 |
| 0 | 0 | 0 |
| 0 | 0 | 0 |
| 0 | 1 | 0 |
| 0 | 1 | 0 |
| 1 | 1 | 1 |
| 1 | 1 | 1 |
| 1 | 1 | 1 |
| 0 | 0 | 0 |
| 0 | 0 | 0 |
| 0 | 0 | 0 |
| 0 | 0 | 0 |
| 0 | 0 | 0 |
| 0 | 0 | 0 |
| 0 | 1 | 0 |
| 1 | 1 | 1 |
| 0 | 0 | 0 |
| 1 | 1 | 1 |
| 1 | 1 | 1 |
| 1 | 1 | 1 |
| 1 | 1 | 1 |
| 0 | 0 | 0 |
| 1 | 1 | 1 |
| 0 | 0 | 0 |
| 0 | 1 | 0 |
| 1 | 1 | 1 |
| 1 | 1 | 1 |
| 1 | 1 | 1 |
| 0 | 0 | 0 |
| 1 | 1 | 1 |
| 1 | 1 | 1 |
| 1 | 1 | 1 |
| 1 | 1 | 1 |
| 1 | 1 | 1 |
| 0 | 1 | 0 |

| | | |
|---|---|---|
| 0 | 1 | 0 |
| 1 | 1 | 1 |
| 1 | 1 | 1 |
| 0 | 1 | 0 |
| 1 | 1 | 1 |
| 0 | 1 | 0 |
| 0 | 1 | 0 |
| 0 | 1 | 0 |
| 0 | 1 | 0 |
| 0 | 1 | 0 |
| 0 | 1 | 0 |
| 0 | 0 | 0 |
| 0 | 1 | 0 |
| 0 | 1 | 0 |
| 0 | 1 | 0 |
| 0 | 1 | 0 |
| 0 | 1 | 0 |
| 0 | 0 | 0 |
| 0 | 0 | 0 |
| 0 | 0 | 0 |
| 0 | 0 | 0 |
| 0 | 1 | 0 |
| 0 | 0 | 0 |
| 1 | 1 | 1 |
| 1 | 1 | 1 |
| 1 | 1 | 1 |
| 1 | 1 | 1 |
| 0 | 1 | 0 |
| 0 | 0 | 0 |
| 0 | 0 | 0 |
| 1 | 1 | 1 |
| 1 | 1 | 1 |
| 0 | 0 | 0 |
| 1 | 1 | 1 |
| 0 | 0 | 0 |
| 0 | 0 | 0 |
| 0 | 0 | 0 |
| 1 | 1 | 1 |
| 0 | 0 | 0 |
| 1 | 1 | 1 |
| 0 | 0 | 0 |
| 1 | 1 | 1 |
| 0 | 0 | 0 |
| 1 | 0 | 0 |
| 1 | 0 | 0 |
| 1 | 0 | 0 |
| 1 | 0 | 0 |
| 0 | 1 | 0 |
| 0 | 1 | 0 |
| 0 | 0 | 0 |
| 1 | 0 | 0 |
| 0 | 0 | 0 |
| 1 | 0 | 0 |
| 1 | 1 | 1 |

| | | |
|---|---|---|
| 0 | 0 | 0 |
| 0 | 0 | 0 |
| 0 | 0 | 0 |
| 0 | 0 | 0 |
| 0 | 0 | 0 |
| 1 | 1 | 1 |
| 0 | 0 | 0 |
| 0 | 0 | 0 |
| 1 | 1 | 1 |
| 0 | 0 | 0 |
| 0 | 0 | 0 |
| 0 | 0 | 0 |
| 0 | 1 | 0 |
| 0 | 1 | 0 |
| 0 | 1 | 0 |
| 0 | 1 | 0 |
| 0 | 1 | 0 |
| 0 | 1 | 0 |
| 0 | 1 | 0 |
| 0 | 1 | 0 |
| 0 | 1 | 0 |
| 0 | 1 | 0 |
| 0 | 1 | 0 |
| 0 | 1 | 0 |
| 0 | 1 | 0 |
| 0 | 1 | 0 |
| 0 | 1 | 0 |
| 0 | 1 | 0 |
| 0 | 1 | 0 |
| 0 | 1 | 0 |
| 0 | 1 | 0 |
| 0 | 1 | 0 |
| 0 | 1 | 0 |
| 0 | 1 | 0 |
| 0 | 1 | 0 |
| 0 | 1 | 0 |
| 0 | 1 | 0 |
| 0 | 1 | 0 |
| 0 | 1 | 0 |
| 0 | 1 | 0 |
| 0 | 1 | 0 |
| 0 | 1 | 0 |
| 0 | 1 | 0 |
| 0 | 1 | 0 |
| 0 | 1 | 0 |
| 0 | 0 | 0 |
| 1 | 1 | 1 |
| 1 | 1 | 1 |
| 1 | 1 | 1 |
| 1 | 1 | 1 |
| 0 | 0 | 0 |
| 1 | 1 | 1 |
| 1 | 1 | 1 |
| 0 | 1 | 0 |

| | | |
|---|---|---|
| 1 | 1 | 1 |
| 1 | 1 | 1 |
| 1 | 1 | 1 |
| 1 | 1 | 1 |
| 0 | 1 | 0 |
| 0 | 1 | 0 |
| 0 | 1 | 0 |
| 0 | 0 | 0 |
| 0 | 0 | 0 |
| 1 | 1 | 1 |
| 0 | 1 | 0 |
| 1 | 1 | 1 |
| 1 | 1 | 1 |
| 1 | 1 | 1 |
| 0 | 0 | 0 |
| 0 | 0 | 0 |
| 0 | 0 | 0 |
| 1 | 1 | 1 |
| 1 | 0 | 0 |
| 1 | 1 | 1 |
| 0 | 0 | 0 |
| 0 | 0 | 0 |
| 0 | 0 | 0 |
| 0 | 0 | 0 |
| 0 | 0 | 0 |
| 0 | 0 | 0 |
| 0 | 1 | 0 |
| 1 | 1 | 1 |
| 0 | 0 | 0 |
| 1 | 1 | 1 |
| 1 | 1 | 1 |
| 1 | 1 | 1 |
| 0 | 0 | 0 |
| 1 | 1 | 1 |
| 1 | 1 | 1 |
| 1 | 1 | 1 |
| 0 | 0 | 0 |
| 0 | 0 | 0 |
| 0 | 0 | 0 |
| 1 | 1 | 1 |
| 1 | 1 | 1 |
| 1 | 1 | 1 |
| 1 | 1 | 1 |
| 1 | 1 | 1 |
| 0 | 1 | 0 |
| 0 | 0 | 0 |
| 1 | 1 | 1 |
| 0 | 1 | 0 |
| 1 | 1 | 1 |
| 0 | 1 | 0 |
| 1 | 1 | 1 |
| 1 | 1 | 1 |
| 0 | 1 | 0 |
| 1 | 1 | 1 |

| | | |
|---|---|---|
| 1 | 1 | 1 |
| 0 | 1 | 0 |
| 1 | 1 | 1 |
| 0 | 0 | 0 |
| 1 | 1 | 1 |
| 1 | 1 | 1 |
| 1 | 1 | 1 |
| 0 | 0 | 0 |
| 0 | 0 | 0 |
| 1 | 1 | 1 |
| 1 | 1 | 1 |
| 1 | 1 | 1 |
| 0 | 0 | 0 |
| 0 | 1 | 0 |
| 0 | 1 | 0 |
| 0 | 1 | 0 |
| 0 | 0 | 0 |
| 1 | 1 | 1 |
| 1 | 1 | 1 |
| 1 | 1 | 1 |
| 1 | 1 | 1 |
| 1 | 1 | 1 |
| 1 | 1 | 1 |
| 1 | 1 | 1 |
| 0 | 1 | 0 |
| 1 | 1 | 1 |
| 0 | 1 | 0 |
| 0 | 0 | 0 |
| 0 | 0 | 0 |
| 0 | 0 | 0 |
| 0 | 0 | 0 |
| 0 | 0 | 0 |
| 0 | 0 | 0 |
| 0 | 1 | 0 |
| 0 | 0 | 0 |
| 0 | 0 | 0 |
| 0 | 0 | 0 |
| 1 | 1 | 1 |
| 1 | 1 | 1 |
| 1 | 1 | 1 |
| 1 | 1 | 1 |
| 0 | 1 | 0 |
| 0 | 0 | 0 |
| 0 | 0 | 0 |
| 1 | 1 | 1 |
| 1 | 1 | 1 |
| 0 | 0 | 0 |
| 1 | 1 | 1 |
| 1 | 1 | 1 |
| 1 | 1 | 1 |
| 1 | 1 | 1 |
| 1 | 1 | 1 |
| 0 | 0 | 0 |
| 0 | 0 | 0 |

| | | |
|---|---|---|
| 0 | 0 | 0 |
| 0 | 0 | 0 |
| 0 | 0 | 0 |
| 0 | 0 | 0 |
| 0 | 0 | 0 |
| 0 | 0 | 0 |
| 0 | 0 | 0 |
| 0 | 0 | 0 |
| 1 | 1 | 1 |
| 0 | 0 | 0 |
| 1 | 1 | 1 |
| 1 | 1 | 1 |
| 1 | 1 | 1 |
| 1 | 1 | 1 |
| 0 | 0 | 0 |
| 1 | 1 | 1 |
| 1 | 1 | 1 |
| 1 | 1 | 1 |
| 1 | 1 | 1 |
| 0 | 0 | 0 |
| 0 | 0 | 0 |
| 1 | 1 | 1 |
| 0 | 0 | 0 |
| 0 | 0 | 0 |
| 0 | 0 | 0 |
| 0 | 0 | 0 |
| 0 | 0 | 0 |
| 0 | 1 | 0 |
| 0 | 1 | 0 |
| 0 | 1 | 0 |
| 0 | 0 | 0 |
| 1 | 1 | 1 |
| 1 | 1 | 1 |
| 1 | 1 | 1 |
| 1 | 1 | 1 |
| 1 | 1 | 1 |
| 1 | 1 | 1 |
| 1 | 1 | 1 |
| 1 | 1 | 1 |
| 1 | 1 | 1 |
| 0 | 1 | 0 |
| 0 | 0 | 0 |
| 1 | 1 | 1 |
| 1 | 1 | 1 |
| 1 | 1 | 1 |
| 0 | 1 | 0 |
| 1 | 1 | 1 |
| 1 | 1 | 1 |
| 0 | 0 | 0 |
| 0 | 1 | 0 |
| 1 | 1 | 1 |
| 1 | 1 | 1 |
| 0 | 0 | 0 |
| 0 | 0 | 0 |

| | | |
|---|---|---|
| 1 | 1 | 1 |
| 1 | 1 | 1 |
| 0 | 1 | 0 |
| 1 | 0 | 0 |
| 0 | 0 | 0 |
| 0 | 0 | 0 |
| 0 | 1 | 0 |
| 0 | 0 | 0 |
| 1 | 1 | 1 |
| 1 | 1 | 1 |
| 0 | 0 | 0 |
| 1 | 0 | 0 |
| 0 | 0 | 0 |
| 0 | 0 | 0 |
| 0 | 0 | 0 |
| 0 | 0 | 0 |
| 1 | 1 | 1 |
| 1 | 1 | 1 |
| 0 | 0 | 0 |
| 0 | 0 | 0 |
| 0 | 0 | 0 |
| 0 | 0 | 0 |
| 0 | 0 | 0 |
| 1 | 1 | 1 |
| 1 | 1 | 1 |
| 0 | 0 | 0 |
| 1 | 1 | 1 |
| 0 | 0 | 0 |
| 0 | 0 | 0 |
| 0 | 0 | 0 |
| 0 | 1 | 0 |
| 0 | 0 | 0 |
| 0 | 0 | 0 |
| 1 | 1 | 1 |
| 1 | 1 | 1 |
| 0 | 0 | 0 |
| 0 | 0 | 0 |
| 1 | 0 | 0 |
| 0 | 0 | 0 |
| 0 | 0 | 0 |
| 0 | 0 | 0 |
| 0 | 0 | 0 |
| 0 | 1 | 0 |
| 1 | 1 | 1 |
| 0 | 0 | 0 |
| 0 | 1 | 0 |
| 0 | 0 | 0 |
| 1 | 1 | 1 |
| 1 | 1 | 1 |
| 1 | 1 | 1 |
| 1 | 1 | 1 |
| 1 | 1 | 1 |
| 0 | 1 | 0 |
| 1 | 1 | 1 |

| | | |
|---|---|---|
| 1 | 1 | 1 |
| 1 | 1 | 1 |
| 1 | 1 | 1 |
| 0 | 1 | 0 |
| 1 | 1 | 1 |
| 1 | 1 | 1 |
| 0 | 1 | 0 |
| 0 | 0 | 0 |
| 1 | 1 | 1 |
| 0 | 0 | 0 |
| 0 | 0 | 0 |
| 0 | 0 | 0 |
| 0 | 1 | 0 |
| 1 | 1 | 1 |
| 0 | 0 | 0 |
| 0 | 1 | 0 |
| 0 | 1 | 0 |
| 0 | 1 | 0 |
| 0 | 1 | 0 |
| 0 | 1 | 0 |
| 1 | 1 | 1 |
| 1 | 1 | 1 |
| 1 | 1 | 1 |
| 0 | 1 | 0 |
| 1 | 1 | 1 |
| 1 | 1 | 1 |
| 1 | 1 | 1 |
| 1 | 0 | 0 |
| 1 | 1 | 1 |
| 0 | 0 | 0 |
| 0 | 0 | 0 |
| 1 | 1 | 1 |
| 1 | 1 | 1 |
| 0 | 1 | 0 |
| 0 | 1 | 0 |
| 0 | 0 | 0 |
| 1 | 1 | 1 |
| 1 | 1 | 1 |
| 1 | 1 | 1 |
| 1 | 1 | 1 |
| 1 | 1 | 1 |
| 0 | 0 | 0 |
| 0 | 0 | 0 |
| 0 | 0 | 0 |
| 1 | 1 | 1 |
| 1 | 1 | 1 |
| 1 | 1 | 1 |
| 1 | 1 | 1 |
| 1 | 1 | 1 |
| 1 | 1 | 1 |
| 1 | 1 | 1 |
| 1 | 1 | 1 |
| 1 | 1 | 1 |
| 0 | 0 | 0 |

| | | |
|---|---|---|
| 1 | 1 | 1 |
| 1 | 1 | 1 |
| 1 | 1 | 1 |
| 1 | 1 | 1 |
| 1 | 1 | 1 |
| 1 | 1 | 1 |
| 1 | 1 | 1 |
| 0 | 1 | 0 |
| 0 | 0 | 0 |
| 0 | 0 | 0 |
| 1 | 1 | 1 |
| 0 | 1 | 0 |
| 0 | 1 | 0 |
| 0 | 0 | 0 |
| 1 | 1 | 1 |
| 1 | 1 | 1 |
| 1 | 1 | 1 |
| 0 | 0 | 0 |
| 0 | 0 | 0 |
| 0 | 0 | 0 |
| 0 | 0 | 0 |
| 0 | 1 | 0 |
| 1 | 1 | 1 |
| 1 | 1 | 1 |
| 1 | 1 | 1 |
| 0 | 0 | 0 |
| 1 | 1 | 1 |
| 0 | 1 | 0 |
| 1 | 1 | 1 |
| 1 | 1 | 1 |
| 1 | 1 | 1 |
| 1 | 1 | 1 |
| 1 | 1 | 1 |
| 1 | 1 | 1 |
| 1 | 1 | 1 |
| 1 | 1 | 1 |
| 1 | 1 | 1 |
| 1 | 1 | 1 |
| 1 | 1 | 1 |
| 0 | 0 | 0 |
| 1 | 1 | 1 |
| 1 | 1 | 1 |
| 0 | 0 | 0 |
| 1 | 1 | 1 |
| 1 | 1 | 1 |
| 1 | 1 | 1 |
| 0 | 0 | 0 |
| 0 | 0 | 0 |
| 1 | 1 | 1 |
| 1 | 1 | 1 |
| 0 | 0 | 0 |
| 1 | 1 | 1 |
| 0 | 1 | 0 |
| 1 | 1 | 1 |

| | | |
|---|---|---|
| 1 | 1 | 1 |
| 1 | 1 | 1 |
| 1 | 1 | 1 |
| 1 | 1 | 1 |
| 1 | 1 | 1 |
| 1 | 1 | 1 |
| 1 | 1 | 1 |
| 0 | 1 | 0 |
| 1 | 1 | 1 |
| 1 | 1 | 1 |
| 1 | 1 | 1 |
| 1 | 1 | 1 |
| 1 | 1 | 1 |
| 0 | 0 | 0 |
| 0 | 0 | 0 |
| 0 | 0 | 0 |
| 0 | 0 | 0 |
| 0 | 1 | 0 |
| 1 | 1 | 1 |
| 0 | 0 | 0 |
| 1 | 1 | 1 |
| 1 | 0 | 0 |
| 1 | 1 | 1 |
| 1 | 1 | 1 |
| 1 | 1 | 1 |
| 0 | 0 | 0 |
| 0 | 1 | 0 |
| 1 | 1 | 1 |
| 1 | 1 | 1 |
| 0 | 0 | 0 |
| 0 | 0 | 0 |
| 1 | 1 | 1 |
| 1 | 1 | 1 |
| 1 | 1 | 1 |
| 1 | 1 | 1 |
| 1 | 1 | 1 |
| 1 | 1 | 1 |
| 1 | 1 | 1 |
| 0 | 0 | 0 |
| 1 | 1 | 1 |
| 1 | 1 | 1 |
| 0 | 0 | 0 |
| 0 | 0 | 0 |
| 0 | 0 | 0 |
| 0 | 0 | 0 |
| 1 | 1 | 1 |
| 0 | 0 | 0 |
| 1 | 1 | 1 |
| 1 | 1 | 1 |
| 1 | 1 | 1 |
| 0 | 1 | 0 |
| 1 | 0 | 0 |
| 1 | 0 | 0 |
| 1 | 0 | 0 |

| | | |
|---|---|---|
| 1 | 1 | 1 |
| 1 | 1 | 1 |
| 0 | 0 | 0 |
| 1 | 1 | 1 |
| 1 | 1 | 1 |
| 1 | 1 | 1 |
| 1 | 1 | 1 |
| 0 | 1 | 0 |
| 1 | 1 | 1 |
| 1 | 1 | 1 |
| 1 | 1 | 1 |
| 0 | 1 | 0 |
| 0 | 1 | 0 |
| 0 | 1 | 0 |
| 0 | 1 | 0 |
| 0 | 1 | 0 |
| 0 | 1 | 0 |
| 0 | 1 | 0 |
| 0 | 1 | 0 |
| 0 | 1 | 0 |
| 0 | 1 | 0 |
| 0 | 1 | 0 |
| 0 | 1 | 0 |
| 0 | 1 | 0 |
| 0 | 1 | 0 |
| 0 | 1 | 0 |
| 0 | 1 | 0 |
| 0 | 1 | 0 |
| 1 | 1 | 1 |
| 1 | 1 | 1 |
| 0 | 0 | 0 |
| 1 | 1 | 1 |
| 0 | 0 | 0 |
| 0 | 0 | 0 |
| 0 | 0 | 0 |
| 0 | 0 | 0 |
| 1 | 1 | 1 |
| 0 | 0 | 0 |
| 1 | 1 | 1 |
| 1 | 1 | 1 |
| 0 | 0 | 0 |
| 0 | 0 | 0 |
| 0 | 0 | 0 |
| 1 | 1 | 1 |
| 0 | 0 | 0 |
| 0 | 0 | 0 |
| 1 | 1 | 1 |
| 1 | 1 | 1 |
| 1 | 1 | 1 |
| 1 | 1 | 1 |
| 0 | 0 | 0 |
| 1 | 1 | 1 |
| 1 | 1 | 1 |
| 1 | 1 | 1 |

| | | |
|---|---|---|
| 1 | 1 | 1 |
| 1 | 1 | 1 |
| 1 | 1 | 1 |
| 1 | 1 | 1 |
| 1 | 1 | 1 |
| 0 | 0 | 0 |
| 1 | 1 | 1 |
| 0 | 0 | 0 |
| 0 | 0 | 0 |
| 0 | 0 | 0 |
| 1 | 1 | 1 |
| 0 | 0 | 0 |
| 1 | 1 | 1 |
| 1 | 1 | 1 |
| 1 | 1 | 1 |
| 1 | 0 | 0 |
| 1 | 1 | 1 |
| 0 | 0 | 0 |
| 1 | 1 | 1 |
| 0 | 0 | 0 |
| 1 | 1 | 1 |
| 0 | 0 | 0 |
| 0 | 0 | 0 |
| 0 | 0 | 0 |
| 1 | 1 | 1 |
| 0 | 1 | 0 |
| 1 | 1 | 1 |
| 1 | 1 | 1 |
| 0 | 0 | 0 |
| 1 | 1 | 1 |
| 1 | 1 | 1 |
| 1 | 1 | 1 |
| 1 | 1 | 1 |
| 1 | 1 | 1 |
| 0 | 0 | 0 |
| 0 | 0 | 0 |
| 1 | 1 | 1 |
| 0 | 0 | 0 |
| 0 | 0 | 0 |
| 0 | 1 | 0 |
| 0 | 0 | 0 |
| 0 | 0 | 0 |
| 0 | 1 | 0 |
| 0 | 1 | 0 |
| 1 | 1 | 1 |
| 0 | 1 | 0 |
| 1 | 1 | 1 |
| 1 | 1 | 1 |
| 1 | 1 | 1 |
| 1 | 1 | 1 |
| 1 | 1 | 1 |
| 1 | 1 | 1 |
| 1 | 1 | 1 |
| 1 | 1 | 1 |

| | | |
|---|---|---|
| 1 | 1 | 1 |
| 0 | 0 | 0 |
| 0 | 1 | 0 |
| 1 | 1 | 1 |
| 0 | 0 | 0 |
| 1 | 1 | 1 |
| 1 | 1 | 1 |
| 1 | 1 | 1 |
| 0 | 0 | 0 |
| 0 | 0 | 0 |
| 1 | 1 | 1 |
| 0 | 0 | 0 |
| 0 | 0 | 0 |
| 1 | 1 | 1 |
| 1 | 1 | 1 |
| 0 | 0 | 0 |
| 0 | 1 | 0 |
| 0 | 0 | 0 |
| 0 | 0 | 0 |
| 1 | 1 | 1 |
| 0 | 0 | 0 |
| 0 | 0 | 0 |
| 0 | 0 | 0 |
| 1 | 1 | 1 |
| 0 | 1 | 0 |
| 0 | 0 | 0 |
| 1 | 1 | 1 |
| 1 | 1 | 1 |
| 1 | 1 | 1 |
| 1 | 1 | 1 |
| 0 | 0 | 0 |
| 1 | 1 | 1 |
| 0 | 0 | 0 |
| 1 | 1 | 1 |
| 0 | 0 | 0 |
| 0 | 0 | 0 |
| 1 | 1 | 1 |
| 1 | 1 | 1 |
| 1 | 1 | 1 |
| 1 | 1 | 1 |
| 1 | 1 | 1 |
| 1 | 1 | 1 |
| 1 | 1 | 1 |
| 1 | 1 | 1 |
| 1 | 1 | 1 |
| 1 | 1 | 1 |
| 1 | 0 | 0 |
| 1 | 1 | 1 |
| 1 | 1 | 1 |
| 1 | 1 | 1 |
| 1 | 1 | 1 |
| 1 | 1 | 1 |
| 1 | 1 | 1 |
| 0 | 0 | 0 |

| | | |
|---|---|---|
| 1 | 1 | 1 |
| 1 | 1 | 1 |
| 1 | 1 | 1 |
| 1 | 1 | 1 |
| 0 | 0 | 0 |
| 1 | 1 | 1 |
| 0 | 1 | 0 |
| 0 | 1 | 0 |
| 0 | 1 | 0 |
| 1 | 1 | 1 |
| 0 | 0 | 0 |
| 1 | 1 | 1 |
| 1 | 1 | 1 |
| 1 | 1 | 1 |
| 0 | 0 | 0 |
| 0 | 0 | 0 |
| 0 | 0 | 0 |
| 0 | 0 | 0 |
| 0 | 1 | 0 |
| 0 | 0 | 0 |
| 0 | 0 | 0 |
| 0 | 0 | 0 |
| 1 | 1 | 1 |
| 0 | 0 | 0 |
| 0 | 0 | 0 |
| 1 | 1 | 1 |
| 0 | 0 | 0 |
| 0 | 0 | 0 |
| 0 | 0 | 0 |
| 0 | 0 | 0 |
| 0 | 0 | 0 |
| 1 | 1 | 1 |
| 1 | 1 | 1 |
| 0 | 1 | 0 |
| 1 | 1 | 1 |
| 1 | 1 | 1 |
| 1 | 1 | 1 |
| 0 | 0 | 0 |
| 0 | 0 | 0 |
| 0 | 1 | 0 |
| 1 | 1 | 1 |
| 0 | 0 | 0 |
| 0 | 0 | 0 |
| 0 | 1 | 0 |
| 0 | 0 | 0 |
| 1 | 1 | 1 |
| 1 | 1 | 1 |
| 1 | 1 | 1 |
| 0 | 0 | 0 |
| 0 | 0 | 0 |
| 0 | 0 | 0 |
| 1 | 1 | 1 |
| 0 | 1 | 0 |
| 0 | 0 | 0 |

| | | |
|---|---|---|
| 1 | 1 | 1 |
| 0 | 0 | 0 |
| 1 | 1 | 1 |
| 0 | 0 | 0 |
| 0 | 0 | 0 |
| 0 | 1 | 0 |
| 0 | 0 | 0 |
| 0 | 0 | 0 |
| 1 | 1 | 1 |
| 1 | 1 | 1 |
| 1 | 1 | 1 |
| 1 | 1 | 1 |
| 1 | 1 | 1 |
| 1 | 1 | 1 |
| 1 | 1 | 1 |
| 1 | 1 | 1 |
| 1 | 1 | 1 |
| 0 | 1 | 0 |
| 1 | 1 | 1 |
| 1 | 1 | 1 |
| 0 | 0 | 0 |
| 0 | 1 | 0 |
| 0 | 1 | 0 |
| 0 | 1 | 0 |
| 0 | 1 | 0 |
| 0 | 0 | 0 |
| 0 | 0 | 0 |
| 1 | 1 | 1 |
| 1 | 1 | 1 |
| 1 | 1 | 1 |
| 1 | 1 | 1 |
| 1 | 1 | 1 |
| 1 | 1 | 1 |
| 0 | 1 | 0 |
| 1 | 1 | 1 |
| 1 | 1 | 1 |
| 1 | 1 | 1 |
| 1 | 1 | 1 |
| 0 | 1 | 0 |
| 1 | 1 | 1 |
| 1 | 1 | 1 |
| 0 | 1 | 0 |
| 1 | 1 | 1 |
| 0 | 0 | 0 |
| 1 | 1 | 1 |
| 0 | 0 | 0 |
| 0 | 0 | 0 |
| 0 | 0 | 0 |
| 0 | 0 | 0 |
| 0 | 0 | 0 |
| 0 | 0 | 0 |
| 0 | 1 | 0 |
| 1 | 1 | 1 |
| 1 | 1 | 1 |

| | | |
|---|---|---|
| 1 | 1 | 1 |
| 1 | 1 | 1 |
| 1 | 1 | 1 |
| 0 | 0 | 0 |
| 1 | 1 | 1 |
| 0 | 0 | 0 |
| 1 | 1 | 1 |
| 1 | 1 | 1 |
| 0 | 0 | 0 |
| 1 | 1 | 1 |
| 1 | 1 | 1 |
| 0 | 1 | 0 |
| 1 | 1 | 1 |
| 1 | 1 | 1 |
| 1 | 1 | 1 |
| 0 | 0 | 0 |
| 0 | 0 | 0 |
| 0 | 0 | 0 |
| 0 | 0 | 0 |
| 0 | 0 | 0 |
| 1 | 1 | 1 |
| 1 | 1 | 1 |
| 1 | 1 | 1 |
| 1 | 1 | 1 |
| 0 | 0 | 0 |
| 1 | 1 | 1 |
| 1 | 1 | 1 |
| 1 | 1 | 1 |
| 0 | 0 | 0 |
| 1 | 1 | 1 |
| 0 | 0 | 0 |
| 0 | 0 | 0 |
| 0 | 0 | 0 |
| 0 | 0 | 0 |
| 0 | 0 | 0 |
| 1 | 1 | 1 |
| 0 | 0 | 0 |
| 0 | 0 | 0 |
| 0 | 0 | 0 |
| 0 | 0 | 0 |
| 0 | 0 | 0 |
| 0 | 0 | 0 |
| 0 | 0 | 0 |
| 0 | 0 | 0 |
| 0 | 0 | 0 |
| 0 | 0 | 0 |
| 0 | 0 | 0 |
| 0 | 0 | 0 |
| 1 | 1 | 1 |
| 0 | 0 | 0 |
| 1 | 1 | 1 |
| 1 | 1 | 1 |
| 0 | 1 | 0 |
| 0 | 0 | 0 |

| | | |
|---|---|---|
| 0 | 0 | 0 |
| 0 | 1 | 0 |
| 1 | 1 | 1 |
| 1 | 1 | 1 |
| 1 | 1 | 1 |
| 0 | 0 | 0 |
| 1 | 1 | 1 |
| 0 | 1 | 0 |
| 1 | 1 | 1 |
| 0 | 1 | 0 |
| 0 | 1 | 0 |
| 0 | 1 | 0 |
| 1 | 1 | 1 |
| 0 | 0 | 0 |
| 0 | 1 | 0 |
| 1 | 1 | 1 |
| 1 | 1 | 1 |
| 0 | 0 | 0 |
| 1 | 1 | 1 |
| 1 | 1 | 1 |
| 1 | 1 | 1 |
| 1 | 1 | 1 |
| 0 | 0 | 0 |
| 1 | 1 | 1 |
| 1 | 1 | 1 |
| 0 | 0 | 0 |
| 0 | 0 | 0 |
| 0 | 0 | 0 |
| 1 | 1 | 1 |
| 0 | 1 | 0 |
| 1 | 1 | 1 |
| 1 | 1 | 1 |
| 1 | 1 | 1 |
| 1 | 1 | 1 |
| 1 | 1 | 1 |
| 1 | 1 | 1 |
| 1 | 1 | 1 |
| 1 | 1 | 1 |
| 1 | 1 | 1 |
| 1 | 1 | 1 |
| 1 | 1 | 1 |
| 1 | 1 | 1 |
| 1 | 1 | 1 |
| 1 | 1 | 1 |
| 1 | 1 | 1 |
| 1 | 1 | 1 |
| 1 | 1 | 1 |
| 1 | 1 | 1 |
| 1 | 0 | 0 |
| 1 | 1 | 1 |
| 1 | 1 | 1 |
| 1 | 1 | 1 |
| 1 | 1 | 1 |
| 1 | 1 | 1 |

| | | |
|---|---|---|
| 1 | 1 | 1 |
| 0 | 0 | 0 |
| 1 | 1 | 1 |
| 1 | 1 | 1 |
| 1 | 1 | 1 |
| 0 | 0 | 0 |
| 0 | 0 | 0 |
| 0 | 0 | 0 |
| 1 | 1 | 1 |
| 1 | 1 | 1 |
| 0 | 1 | 0 |
| 1 | 1 | 1 |
| 1 | 1 | 1 |
| 1 | 1 | 1 |
| 0 | 0 | 0 |
| 0 | 0 | 0 |
| 0 | 0 | 0 |
| 1 | 1 | 1 |
| 0 | 0 | 0 |
| 0 | 0 | 0 |
| 0 | 0 | 0 |
| 0 | 0 | 0 |
| 0 | 0 | 0 |
| 0 | 0 | 0 |
| 0 | 1 | 0 |
| 1 | 1 | 1 |
| 1 | 1 | 1 |
| 0 | 1 | 0 |
| 0 | 1 | 0 |
| 1 | 1 | 1 |
| 1 | 1 | 1 |
| 1 | 1 | 1 |
| 0 | 0 | 0 |
| 1 | 1 | 1 |
| 0 | 1 | 0 |
| 0 | 0 | 0 |
| 0 | 0 | 0 |
| 0 | 0 | 0 |
| 0 | 0 | 0 |
| 1 | 1 | 1 |
| 0 | 1 | 0 |
| 1 | 1 | 1 |
| 1 | 1 | 1 |
| 0 | 0 | 0 |
| 0 | 0 | 0 |
| 0 | 0 | 0 |
| 0 | 1 | 0 |
| 1 | 1 | 1 |
| 0 | 0 | 0 |
| 1 | 1 | 1 |
| 1 | 1 | 1 |
| 0 | 0 | 0 |
| 1 | 1 | 1 |
| 0 | 1 | 0 |

| | | |
|---|---|---|
| 1 | 1 | 1 |
| 0 | 1 | 0 |
| 1 | 1 | 1 |
| 1 | 1 | 1 |
| 1 | 1 | 1 |
| 1 | 1 | 1 |
| 0 | 1 | 0 |
| 1 | 1 | 1 |
| 0 | 0 | 0 |
| 0 | 0 | 0 |
| 1 | 1 | 1 |
| 0 | 0 | 0 |
| 0 | 0 | 0 |
| 0 | 0 | 0 |
| 1 | 1 | 1 |
| 1 | 1 | 1 |
| 0 | 0 | 0 |
| 0 | 0 | 0 |
| 0 | 0 | 0 |
| 1 | 1 | 1 |
| 1 | 1 | 1 |
| 1 | 1 | 1 |
| 0 | 0 | 0 |
| 1 | 1 | 1 |
| 0 | 0 | 0 |
| 1 | 1 | 1 |
| 1 | 1 | 1 |
| 1 | 1 | 1 |
| 1 | 1 | 1 |
| 0 | 0 | 0 |
| 0 | 0 | 0 |
| 0 | 0 | 0 |
| 1 | 1 | 1 |
| 0 | 1 | 0 |
| 1 | 1 | 1 |
| 1 | 1 | 1 |
| 1 | 1 | 1 |
| 0 | 1 | 0 |
| 0 | 1 | 0 |
| 0 | 1 | 0 |
| 0 | 0 | 0 |
| 1 | 1 | 1 |
| 1 | 1 | 1 |
| 1 | 1 | 1 |
| 1 | 1 | 1 |
| 0 | 0 | 0 |
| 1 | 0 | 0 |
| 0 | 0 | 0 |
| 1 | 1 | 1 |
| 0 | 0 | 0 |
| 0 | 0 | 0 |
| 0 | 0 | 0 |
| 0 | 1 | 0 |
| 1 | 0 | 0 |

| | | |
|---|---|---|
| 1 | 0 | 0 |
| 0 | 0 | 0 |
| 0 | 1 | 0 |
| 1 | 1 | 1 |
| 0 | 1 | 0 |
| 0 | 1 | 0 |
| 0 | 1 | 0 |
| 0 | 0 | 0 |
| 0 | 0 | 0 |
| 0 | 1 | 0 |
| 0 | 0 | 0 |
| 1 | 1 | 1 |
| 1 | 1 | 1 |
| 1 | 1 | 1 |
| 0 | 1 | 0 |
| 1 | 1 | 1 |
| 1 | 1 | 1 |
| 1 | 1 | 1 |
| 1 | 1 | 1 |
| 0 | 1 | 0 |
| 1 | 1 | 1 |
| 1 | 1 | 1 |
| 1 | 1 | 1 |
| 1 | 1 | 1 |
| 0 | 1 | 0 |
| 0 | 1 | 0 |
| 0 | 1 | 0 |
| 0 | 1 | 0 |
| 0 | 0 | 0 |
| 0 | 1 | 0 |
| 0 | 0 | 0 |
| 1 | 1 | 1 |
| 0 | 0 | 0 |
| 0 | 0 | 0 |
| 0 | 1 | 0 |
| 0 | 0 | 0 |
| 0 | 0 | 0 |
| 1 | 1 | 1 |
| 0 | 1 | 0 |
| 0 | 1 | 0 |
| 1 | 1 | 1 |
| 0 | 0 | 0 |
| 0 | 0 | 0 |
| 1 | 1 | 1 |
| 1 | 1 | 1 |
| 0 | 0 | 0 |
| 0 | 0 | 0 |
| 0 | 0 | 0 |
| 0 | 0 | 0 |
| 0 | 1 | 0 |
| 0 | 0 | 0 |
| 0 | 0 | 0 |
| 1 | 1 | 1 |
| 1 | 1 | 1 |

| | | |
|---|---|---|
| 1 | 1 | 1 |
| 1 | 1 | 1 |
| 1 | 1 | 1 |
| 1 | 1 | 1 |
| 0 | 0 | 0 |
| 0 | 0 | 0 |
| 0 | 0 | 0 |
| 0 | 0 | 0 |
| 1 | 1 | 1 |
| 0 | 0 | 0 |
| 0 | 0 | 0 |
| 0 | 0 | 0 |
| 0 | 0 | 0 |
| 1 | 1 | 1 |
| 1 | 0 | 0 |
| 0 | 1 | 0 |
| 1 | 1 | 1 |
| 1 | 1 | 1 |
| 1 | 1 | 1 |
| 0 | 1 | 0 |
| 1 | 1 | 1 |
| 1 | 1 | 1 |
| 1 | 1 | 1 |
| 1 | 1 | 1 |
| 1 | 1 | 1 |
| 0 | 0 | 0 |
| 0 | 0 | 0 |
| 0 | 0 | 0 |
| 0 | 0 | 0 |
| 0 | 1 | 0 |
| 0 | 1 | 0 |
| 0 | 1 | 0 |
| 1 | 1 | 1 |
| 0 | 1 | 0 |
| 0 | 1 | 0 |
| 1 | 1 | 1 |
| 1 | 1 | 1 |
| 1 | 1 | 1 |
| 1 | 1 | 1 |
| 1 | 1 | 1 |
| 1 | 1 | 1 |
| 1 | 1 | 1 |
| 1 | 1 | 1 |
| 1 | 1 | 1 |
| 1 | 1 | 1 |
| 1 | 1 | 1 |
| 1 | 1 | 1 |
| 1 | 1 | 1 |
| 1 | 1 | 1 |
| 1 | 1 | 1 |
| 1 | 1 | 1 |
| 1 | 1 | 1 |
| 1 | 1 | 1 |
| 0 | 0 | 0 |

| | | |
|---|---|---|
| 1 | 1 | 1 |
| 1 | 1 | 1 |
| 1 | 1 | 1 |
| 0 | 1 | 0 |
| 0 | 1 | 0 |
| 0 | 1 | 0 |
| 0 | 1 | 0 |
| 0 | 1 | 0 |
| 1 | 1 | 1 |
| 1 | 1 | 1 |
| 1 | 1 | 1 |
| 1 | 1 | 1 |
| 1 | 1 | 1 |
| 1 | 1 | 1 |
| 1 | 1 | 1 |
| 1 | 1 | 1 |
| 1 | 1 | 1 |
| 1 | 1 | 1 |
| 1 | 1 | 1 |
| 1 | 1 | 1 |
| 1 | 1 | 1 |
| 1 | 1 | 1 |
| 1 | 1 | 1 |
| 1 | 1 | 1 |
| 1 | 1 | 1 |
| 1 | 1 | 1 |
| 1 | 1 | 1 |
| 1 | 1 | 1 |
| 1 | 1 | 1 |
| 1 | 1 | 1 |
| 1 | 1 | 1 |
| 1 | 1 | 1 |
| 1 | 1 | 1 |
| 0 | 1 | 0 |
| 1 | 1 | 1 |
| 1 | 1 | 1 |
| 1 | 1 | 1 |
| 1 | 1 | 1 |
| 1 | 1 | 1 |
| 1 | 1 | 1 |
| 0 | 1 | 0 |
| 1 | 1 | 1 |
| 1 | 1 | 1 |
| 1 | 1 | 1 |
| 0 | 1 | 0 |
| 1 | 0 | 0 |
| 0 | 0 | 0 |
| 1 | 1 | 1 |
| 0 | 1 | 0 |
| 1 | 1 | 1 |
| 1 | 1 | 1 |
| 0 | 1 | 0 |
| 0 | 1 | 0 |
| 0 | 1 | 0 |

| | | |
|---|---|---|
| 0 | 1 | 0 |
| 1 | 1 | 1 |
| 0 | 0 | 0 |
| 1 | 1 | 1 |
| 1 | 1 | 1 |
| 1 | 1 | 1 |
| 0 | 0 | 0 |
| 1 | 1 | 1 |
| 1 | 1 | 1 |
| 0 | 1 | 0 |
| 0 | 1 | 0 |
| 0 | 0 | 0 |
| 1 | 1 | 1 |
| 0 | 0 | 0 |
| 0 | 0 | 0 |
| 0 | 0 | 0 |
| 0 | 0 | 0 |
| 0 | 0 | 0 |
| 0 | 0 | 0 |
| 0 | 0 | 0 |
| 0 | 0 | 0 |

| organism_scientific | evo_cons | cluster_id | pdb_id | chain_id | resi | resn | change | uniprot_id |
|---|---|---|---|---|---|---|---|---|
| HOMO SAPIENS | 5 | 87 | 3uzw | A | 106 | LEU | p.Leu106Phe | P51857 |
| HOMO SAPIENS | 3 | 87 | 3uzw | A | 133 | PRO | p.Pro133Arg | P51857 |
| HOMO SAPIENS | 0 | 87 | 3uzw | A | 198 | PRO | p.Pro198Leu | P51857 |
| HOMO SAPIENS | 7 | 87 | 3uzw | A | 261 | ARG | p.Arg261Cys | P51857 |
| HOMO SAPIENS | 0 | 4693 | 3mk1 | A | 3 | PRO | p.Pro25Leu | P05187 |
| HOMO SAPIENS | 8 | 4830 | 2h64 | B | 101 | CYS | p.Cys124Arg | P36894 |
| HOMO SAPIENS | 2 | 4830 | 2h64 | B | 116 | PRO | p.Pro139Ser | P36894 |
| HOMO SAPIENS | 0 | 6937 | 3dar | A | 186 | MET | p.Met186Thr | P21802 |
| HOMO SAPIENS | 3 | 5 | 1aut | C | 30 | GLN | p.Gln226His | P04070 |
| HOMO SAPIENS | 1 | 5 | 1aut | C | 68 | LEU | p.Leu265Phe | P04070 |
| HOMO SAPIENS | 0 | 5 | 1aut | C | 75 | ARG | p.Arg272Cys | P04070 |
| HOMO SAPIENS | 6 | 5 | 1aut | C | 92 | PRO | p.Pro289Leu | P04070 |
| HOMO SAPIENS | 0 | 5 | 1aut | C | 104 | ALA | p.Ala301Val | P04070 |
| HOMO SAPIENS | 1 | 5 | 1aut | C | 112 | ALA | p.Ala309Thr | P04070 |
| HOMO SAPIENS | 2 | 5 | 1aut | C | 115 | SER | p.Ser312Leu | P04070 |
| HOMO SAPIENS | 1 | 5 | 1aut | C | 133 | GLY | p.Gly334Ser | P04070 |
| HOMO SAPIENS | 0 | 5 | 1aut | C | 147 | ARG | p.Arg348Ter | P04070 |
| HOMO SAPIENS | 8 | 5 | 1aut | C | 194 | ASP | p.Asp401Asn | P04070 |
| HOMO SAPIENS | 7 | 5 | 1aut | C | 237 | TRP | p.Trp444Cys | P04070 |
| HOMO SAPIENS | 0 | 5 | 1aut | C | 238 | ILE | p.Ile445Met | P04070 |
| HOMO SAPIENS | 8 | 43 | 1aiy | F | 10 | HIS | p.His34Asp | P01308 |
| HOMO SAPIENS | 2 | 43 | 1aiy | F | 22 | ARG | p.Arg46Gln | P01308 |
| HOMO SAPIENS | 4 | 43 | 1aiy | F | 23 | GLY | p.Gly47Val | P01308 |
| HOMO SAPIENS | 7 | 43 | 1aiy | F | 24 | PHE | p.Phe48Ser | P01308 |
| HOMO SAPIENS | 5 | 24 | 2fg8 | A | 33 | THR | p.Thr30Ile | P02792 |
| HOMO SAPIENS | 4 | 24 | 2fg8 | A | 99 | ALA | p.Ala96Thr | P02792 |
| HOMO SAPIENS | 5 | 24 | 2fg8 | A | 107 | GLU | p.Glu104Ter | P02792 |
| HOMO SAPIENS | 3 | 5409 | 5bnq | A | 199 | MET | p.Met199Lys | O14788 |
| HOMO SAPIENS | -1 | 839 | 3swc | A | 959 | GLN | p.Gln990His | Q9H9B1 |
| HOMO SAPIENS | -1 | 839 | 3swc | A | 968 | ALA | p.Ala999Val | Q9H9B1 |
| HOMO SAPIENS | -1 | 839 | 3swc | A | 973 | SER | p.Ser1004Ile | Q9H9B1 |
| HOMO SAPIENS | 8 | 17 | 1eou | A | 18 | LYS | p.Lys18Glu | P00918 |
| HOMO SAPIENS | 2 | 17 | 1eou | A | 107 | HIS | p.His107Tyr | P00918 |
| HOMO SAPIENS | 1 | 17 | 1eou | A | 236 | PRO | p.Pro236His | P00918 |
| HOMO SAPIENS | 6 | 17 | 1eou | A | 252 | ASN | p.Asn252Asp | P00918 |
| HOMO SAPIENS | 2 | 9140 | 5c4v | A | 445 | ARG | p.Arg445Ter | Q13485 |
| HOMO SAPIENS | 7 | 9140 | 5c4v | A | 496 | ARG | p.Arg496Cys | Q13485 |
| HOMO SAPIENS | -1 | 42 | 4pwi | A | 6 | GLY | p.Gly26Ser | P02766 |
| HOMO SAPIENS | 6 | 42 | 4pwi | A | 10 | CYS | p.Cys30Arg | P02766 |
| HOMO SAPIENS | 8 | 42 | 4pwi | A | 12 | LEU | p.Leu32Pro | P02766 |
| HOMO SAPIENS | 9 | 42 | 4pwi | A | 18 | ASP | p.Asp38Gly | P02766 |
| HOMO SAPIENS | 8 | 42 | 4pwi | A | 20 | VAL | p.Val40Ile | P02766 |
| HOMO SAPIENS | 4 | 42 | 4pwi | A | 30 | MET | p.Val50Ala | P02766 |
| HOMO SAPIENS | 4 | 42 | 4pwi | A | 30 | MET | p.Val50Leu | P02766 |
| HOMO SAPIENS | 6 | 42 | 4pwi | A | 33 | PHE | p.Phe53Ile | P02766 |
| HOMO SAPIENS | 7 | 42 | 4pwi | A | 36 | ALA | p.Ala56Pro | P02766 |
| HOMO SAPIENS | 7 | 42 | 4pwi | A | 42 | GLU | p.Glu62Gly | P02766 |
| HOMO SAPIENS | 8 | 42 | 4pwi | A | 44 | PHE | p.Phe64Ser | P02766 |
| HOMO SAPIENS | 4 | 42 | 4pwi | A | 47 | GLY | p.Gly67Ala | P02766 |
| HOMO SAPIENS | 6 | 42 | 4pwi | A | 49 | THR | p.Thr69Ala | P02766 |
| HOMO SAPIENS | 6 | 42 | 4pwi | A | 50 | SER | p.Ser70Ile | P02766 |
| HOMO SAPIENS | 6 | 42 | 4pwi | A | 50 | SER | p.Ser70Arg | P02766 |
| HOMO SAPIENS | 0 | 42 | 4pwi | A | 53 | GLY | p.Gly73Glu | P02766 |

| | | | | | | | | |
|---|---|---|---|---|---|---|---|---|
| HOMO SAPIENS | 7 | 42 | 4pwi | A | 55 | LEU | p.Leu75Pro | P02766 |
| HOMO SAPIENS | 6 | 42 | 4pwi | A | 58 | LEU | p.Leu78His | P02766 |
| HOMO SAPIENS | 7 | 42 | 4pwi | A | 61 | GLU | p.Glu81Lys | P02766 |
| HOMO SAPIENS | 8 | 42 | 4pwi | A | 64 | PHE | p.Phe84Leu | P02766 |
| HOMO SAPIENS | 8 | 42 | 4pwi | A | 68 | ILE | p.Ile88Leu | P02766 |
| HOMO SAPIENS | 8 | 42 | 4pwi | A | 69 | TYR | p.Tyr89His | P02766 |
| HOMO SAPIENS | 4 | 42 | 4pwi | A | 71 | VAL | p.Val91Ala | P02766 |
| HOMO SAPIENS | 6 | 42 | 4pwi | A | 77 | SER | p.Ser97Tyr | P02766 |
| HOMO SAPIENS | 4 | 42 | 4pwi | A | 84 | ILE | p.Ile104Ser | P02766 |
| HOMO SAPIENS | 8 | 42 | 4pwi | A | 89 | GLU | p.Glu109Gln | P02766 |
| HOMO SAPIENS | 8 | 42 | 4pwi | A | 97 | ALA | p.Ala117Gly | P02766 |
| HOMO SAPIENS | 7 | 42 | 4pwi | A | 99 | ASP | p.Asp119Asn | P02766 |
| HOMO SAPIENS | 9 | 42 | 4pwi | A | 107 | ILE | p.Ile127Val | P02766 |
| HOMO SAPIENS | 9 | 42 | 4pwi | A | 111 | LEU | p.Leu131Met | P02766 |
| HOMO SAPIENS | 7 | 42 | 4pwi | A | 114 | TYR | p.Tyr134Cys | P02766 |
| HOMO SAPIENS | 7 | 42 | 4pwi | A | 114 | TYR | p.Tyr134His | P02766 |
| HOMO SAPIENS | 7 | 42 | 4pwi | A | 122 | VAL | p.Val142Ile | P02766 |
| HOMO SAPIENS | -1 | 42 | 4pwi | A | 126 | LYS | p.Lys146Arg | P02766 |
| HOMO SAPIENS | -1 | 42 | 4pwi | A | 127 | GLU | p.Glu147Ter | P02766 |
| HOMO SAPIENS | -1 | 1312 | 4dce | A | 1275 | ARG | p.Arg1275Gln | Q9UM73 |
| HOMO SAPIENS | -1 | 1312 | 4dce | A | 1280 | ALA | p.Ala1280Val | Q9UM73 |
| HOMO SAPIENS | 1 | 20 | 1y0x | X | 234 | ALA | p.Ala234Thr | P10828 |
| HOMO SAPIENS | 0 | 20 | 1y0x | X | 243 | ARG | p.Arg243Trp | P10828 |
| HOMO SAPIENS | 3 | 20 | 1y0x | X | 316 | ARG | p.Arg316His | P10828 |
| HOMO SAPIENS | 1 | 20 | 1y0x | X | 317 | ALA | p.Ala317Thr | P10828 |
| HOMO SAPIENS | 2 | 20 | 1y0x | X | 320 | ARG | p.Arg320Cys | P10828 |
| HOMO SAPIENS | 2 | 20 | 1y0x | X | 320 | ARG | p.Arg320His | P10828 |
| HOMO SAPIENS | 0 | 20 | 1y0x | X | 332 | GLY | p.Gly332Arg | P10828 |
| HOMO SAPIENS | 1 | 20 | 1y0x | X | 338 | ARG | p.Arg338Trp | P10828 |
| HOMO SAPIENS | 0 | 20 | 1y0x | X | 340 | GLN | p.Gln340His | P10828 |
| HOMO SAPIENS | 0 | 20 | 1y0x | X | 345 | GLY | p.Gly345Val | P10828 |
| HOMO SAPIENS | 0 | 20 | 1y0x | X | 345 | GLY | p.Gly345Ser | P10828 |
| HOMO SAPIENS | 0 | 20 | 1y0x | X | 347 | GLY | p.Gly347Glu | P10828 |
| HOMO SAPIENS | 1 | 20 | 1y0x | X | 438 | ARG | p.Arg438His | P10828 |
| HOMO SAPIENS | 0 | 20 | 1y0x | X | 442 | MET | p.Met442Val | P10828 |
| HOMO SAPIENS | 0 | 20 | 1y0x | X | 443 | LYS | p.Lys443Glu | P10828 |
| HOMO SAPIENS | 0 | 20 | 1y0x | X | 446 | CYS | p.Cys446Arg | P10828 |
| HOMO SAPIENS | 2 | 20 | 1y0x | X | 453 | PRO | p.Pro453His | P10828 |
| HOMO SAPIENS | 2 | 20 | 1y0x | X | 453 | PRO | p.Pro453Thr | P10828 |
| HOMO SAPIENS | 3 | 20 | 1y0x | X | 459 | PHE | p.Phe459Cys | P10828 |
| HOMO SAPIENS | 0 | 4409 | 3qb7 | D | 87 | GLU | p.Glu109Lys | P31785 |
| HOMO SAPIENS | 1 | 4409 | 3qb7 | D | 92 | GLY | p.Gly114Asp | P31785 |
| HOMO SAPIENS | 1 | 4409 | 3qb7 | D | 93 | CYS | p.Cys115Arg | P31785 |
| HOMO SAPIENS | 0 | 4409 | 3qb7 | D | 131 | ILE | p.Ile153Asn | P31785 |
| HOMO SAPIENS | 9 | 4409 | 3qb7 | D | 200 | ARG | p.Arg222Cys | P31785 |
| HOMO SAPIENS | 0 | 3891 | 3ije | B | 33 | LEU | p.Leu59Pro | P05106 |
| HOMO SAPIENS | 0 | 3891 | 3ije | B | 117 | LEU | p.Leu143Trp | P05106 |
| HOMO SAPIENS | 3 | 3891 | 3ije | B | 119 | ASP | p.Asp145Tyr | P05106 |
| HOMO SAPIENS | 2 | 3891 | 3ije | B | 143 | ARG | p.Arg169Gln | P05106 |
| HOMO SAPIENS | 2 | 3891 | 3ije | B | 214 | ARG | p.Arg240Gln | P05106 |
| HOMO SAPIENS | 2 | 3891 | 3ije | B | 214 | ARG | p.Arg240Trp | P05106 |
| HOMO SAPIENS | 2 | 3891 | 3ije | B | 374 | CYS | p.Cys400Tyr | P05106 |
| HOMO SAPIENS | 1 | 3891 | 3ije | B | 407 | PRO | p.Pro433Ala | P05106 |
| HOMO SAPIENS | 0 | 3891 | 3ije | B | 489 | ARG | p.Arg515Gln | P05106 |

| | | | | | | | | |
|---|---|---|---|---|---|---|---|---|
| HOMO SAPIENS | -1 | 9613 | 5mke | B | 28 | ARG | p.Arg28Pro | Q13563 |
| HOMO SAPIENS | -1 | 9613 | 5mke | B | 190 | ALA | p.Ala190Thr | Q13563 |
| HOMO SAPIENS | 4 | 9613 | 5mke | B | 262 | VAL | p.Val262Met | Q13563 |
| HOMO SAPIENS | 0 | 9613 | 5mke | B | 482 | PHE | p.Phe482Cys | Q13563 |
| HOMO SAPIENS | 4 | 9613 | 5mke | B | 511 | ASP | p.Asp511Val | Q13563 |
| HOMO SAPIENS | 5 | 9613 | 5mke | B | 516 | VAL | p.Val516Leu | Q13563 |
| HOMO SAPIENS | -1 | 9613 | 5mke | B | 807 | ARG | p.Arg807Gln | Q13563 |
| HOMO SAPIENS | 4 | 2428 | 1olh | C | 337 | ARG | p.Arg337Cys | P04637 |
| HOMO SAPIENS | 4 | 2428 | 1olh | C | 337 | ARG | p.Arg337His | P04637 |
| HOMO SAPIENS | 8 | 2428 | 1olh | C | 338 | PHE | p.Phe338Leu | P04637 |
| HOMO SAPIENS | 8 | 2428 | 1olh | C | 339 | GLU | p.Glu339Lys | P04637 |
| HOMO SAPIENS | 7 | 2428 | 1olh | C | 344 | LEU | p.Leu344Pro | P04637 |
| HOMO SAPIENS | 0 | 2428 | 1olh | C | 360 | GLY | p.Gly360Ala | P04637 |
| HOMO SAPIENS | 7 | 4867 | 1mox | B | 404 | GLY | p.Gly428Asp | P00533 |
| HOMO SAPIENS | 5 | 4867 | 1mox | B | 497 | ARG | p.Arg521Lys | P00533 |
| HOMO SAPIENS | 1 | 4630 | 1fu1 | A | 43 | TRP | p.Trp43Arg | Q13426 |
| HOMO SAPIENS | 2 | 4630 | 1fu1 | A | 161 | ARG | p.Arg161Ter | Q13426 |
| HOMO SAPIENS | 2 | 15 | 1lfy | B | 2 | HIS | p.His3Gln | P68871 |
| HOMO SAPIENS | 2 | 15 | 1lfy | B | 6 | GLU | p.Glu7Ala | P68871 |
| HOMO SAPIENS | 2 | 15 | 1lfy | B | 6 | GLU | p.Glu7Lys | P68871 |
| HOMO SAPIENS | 4 | 15 | 1lfy | B | 9 | SER | p.Ser10Cys | P68871 |
| HOMO SAPIENS | 6 | 15 | 1lfy | B | 15 | TRP | p.Trp16Ter | P68871 |
| HOMO SAPIENS | 3 | 15 | 1lfy | B | 17 | LYS | p.Lys18Glu | P68871 |
| HOMO SAPIENS | 6 | 15 | 1lfy | B | 19 | ASN | p.Asn20Ser | P68871 |
| HOMO SAPIENS | 6 | 15 | 1lfy | B | 22 | GLU | p.Glu23Ala | P68871 |
| HOMO SAPIENS | 6 | 15 | 1lfy | B | 22 | GLU | p.Glu23Lys | P68871 |
| HOMO SAPIENS | 8 | 15 | 1lfy | B | 26 | GLU | p.Glu27Lys | P68871 |
| HOMO SAPIENS | 8 | 15 | 1lfy | B | 28 | LEU | p.Leu29Pro | P68871 |
| HOMO SAPIENS | 8 | 15 | 1lfy | B | 30 | ARG | p.Arg31Ser | P68871 |
| HOMO SAPIENS | 8 | 15 | 1lfy | B | 30 | ARG | p.Arg31Lys | P68871 |
| HOMO SAPIENS | 6 | 15 | 1lfy | B | 39 | GLN | p.Gln40Ter | P68871 |
| HOMO SAPIENS | 2 | 15 | 1lfy | B | 43 | GLU | p.Glu44Gln | P68871 |
| HOMO SAPIENS | 6 | 15 | 1lfy | B | 57 | ASN | p.Asn58Lys | P68871 |
| HOMO SAPIENS | 6 | 15 | 1lfy | B | 59 | LYS | p.Lys60Asn | P68871 |
| HOMO SAPIENS | 8 | 15 | 1lfy | B | 61 | LYS | p.Lys62Glu | P68871 |
| HOMO SAPIENS | 4 | 15 | 1lfy | B | 73 | ASP | p.Asp74Tyr | P68871 |
| HOMO SAPIENS | 4 | 15 | 1lfy | B | 87 | THR | p.Thr88Ile | P68871 |
| HOMO SAPIENS | 4 | 15 | 1lfy | B | 87 | THR | p.Thr88Pro | P68871 |
| HOMO SAPIENS | 5 | 15 | 1lfy | B | 90 | GLU | p.Glu91Lys | P68871 |
| HOMO SAPIENS | 6 | 15 | 1lfy | B | 91 | LEU | p.Leu92Pro | P68871 |
| HOMO SAPIENS | 5 | 15 | 1lfy | B | 97 | HIS | p.His98Gln | P68871 |
| HOMO SAPIENS | 6 | 15 | 1lfy | B | 110 | LEU | p.Leu111Pro | P68871 |
| HOMO SAPIENS | 6 | 15 | 1lfy | B | 113 | VAL | p.Val114Glu | P68871 |
| HOMO SAPIENS | 2 | 15 | 1lfy | B | 119 | GLY | p.Gly120Ala | P68871 |
| HOMO SAPIENS | 4 | 15 | 1lfy | B | 121 | GLU | p.Glu122Gln | P68871 |
| HOMO SAPIENS | 5 | 15 | 1lfy | B | 134 | VAL | p.Val135Glu | P68871 |
| HOMO SAPIENS | 3 | 15 | 1lfy | B | 143 | HIS | p.His144Gln | P68871 |
| HOMO SAPIENS | 4 | 91 | 5d9k | A | 75 | GLY | p.Gly75Val | P51812 |
| HOMO SAPIENS | 5 | 91 | 5d9k | A | 82 | VAL | p.Val82Phe | P51812 |
| HOMO SAPIENS | 0 | 91 | 5d9k | A | 114 | ARG | p.Arg114Trp | P51812 |
| HOMO SAPIENS | 4 | 91 | 5d9k | A | 189 | ILE | p.Ile189Lys | P51812 |
| HOMO SAPIENS | -1 | 91 | 5d9k | A | 227 | SER | p.Ser227Ala | P51812 |
| HOMO SAPIENS | 1 | 91 | 5d9k | A | 268 | PHE | p.Phe268Ser | P51812 |
| HOMO SAPIENS | 0 | 954 | 2vt0 | A | 15 | VAL | p.Val54Leu | P04062 |

| | | | | | | | | |
|---|---|---|---|---|---|---|---|---|
| HOMO SAPIENS | 5 | 954 | 2vt0 | A | 46 | GLY | p.Gly85Glu | P04062 |
| HOMO SAPIENS | 5 | 954 | 2vt0 | A | 48 | ARG | p.Arg87Trp | P04062 |
| HOMO SAPIENS | 6 | 954 | 2vt0 | A | 79 | LYS | p.Lys118Asn | P04062 |
| HOMO SAPIENS | 0 | 954 | 2vt0 | A | 120 | ARG | p.Arg159Gln | P04062 |
| HOMO SAPIENS | 0 | 954 | 2vt0 | A | 120 | ARG | p.Arg159Trp | P04062 |
| HOMO SAPIENS | 0 | 954 | 2vt0 | A | 122 | PRO | p.Pro161Ser | P04062 |
| HOMO SAPIENS | 5 | 954 | 2vt0 | A | 131 | ARG | p.Arg170Leu | P04062 |
| HOMO SAPIENS | 5 | 954 | 2vt0 | A | 135 | TYR | p.Tyr174Cys | P04062 |
| HOMO SAPIENS | 0 | 954 | 2vt0 | A | 157 | LYS | p.Lys196Gln | P04062 |
| HOMO SAPIENS | 5 | 954 | 2vt0 | A | 163 | ARG | p.Arg202Gln | P04062 |
| HOMO SAPIENS | 5 | 954 | 2vt0 | A | 170 | ARG | p.Arg209Cys | P04062 |
| HOMO SAPIENS | 5 | 954 | 2vt0 | A | 188 | ASN | p.Asn227Ser | P04062 |
| HOMO SAPIENS | 3 | 954 | 2vt0 | A | 196 | SER | p.Ser235Pro | P04062 |
| HOMO SAPIENS | 5 | 954 | 2vt0 | A | 202 | GLY | p.Gly241Arg | P04062 |
| HOMO SAPIENS | 0 | 954 | 2vt0 | A | 212 | TYR | p.Tyr251His | P04062 |
| HOMO SAPIENS | 0 | 954 | 2vt0 | A | 213 | PHE | p.Phe252Ile | P04062 |
| HOMO SAPIENS | 5 | 954 | 2vt0 | A | 216 | PHE | p.Phe255Tyr | P04062 |
| HOMO SAPIENS | 6 | 954 | 2vt0 | A | 251 | PHE | p.Phe290Leu | P04062 |
| HOMO SAPIENS | 5 | 954 | 2vt0 | A | 255 | HIS | p.His294Gln | P04062 |
| HOMO SAPIENS | 6 | 954 | 2vt0 | A | 257 | ARG | p.Arg296Gln | P04062 |
| HOMO SAPIENS | 5 | 954 | 2vt0 | A | 289 | PRO | p.Pro328Leu | P04062 |
| HOMO SAPIENS | 0 | 954 | 2vt0 | A | 309 | ALA | p.Ala348Val | P04062 |
| HOMO SAPIENS | 0 | 954 | 2vt0 | A | 311 | HIS | p.His350Arg | P04062 |
| HOMO SAPIENS | 6 | 954 | 2vt0 | A | 312 | TRP | p.Trp351Cys | P04062 |
| HOMO SAPIENS | 5 | 954 | 2vt0 | A | 323 | THR | p.Thr362Ile | P04062 |
| HOMO SAPIENS | 5 | 954 | 2vt0 | A | 325 | GLY | p.Gly364Arg | P04062 |
| HOMO SAPIENS | 6 | 954 | 2vt0 | A | 342 | CYS | p.Cys381Gly | P04062 |
| HOMO SAPIENS | 5 | 954 | 2vt0 | A | 353 | ARG | p.Arg392Gly | P04062 |
| HOMO SAPIENS | 3 | 954 | 2vt0 | A | 364 | SER | p.Ser403Thr | P04062 |
| HOMO SAPIENS | 5 | 954 | 2vt0 | A | 369 | THR | p.Thr408Met | P04062 |
| HOMO SAPIENS | 3 | 954 | 2vt0 | A | 370 | ASN | p.Asn409Ser | P04062 |
| HOMO SAPIENS | 0 | 954 | 2vt0 | A | 371 | LEU | p.Leu410Val | P04062 |
| HOMO SAPIENS | 0 | 954 | 2vt0 | A | 375 | VAL | p.Val414Leu | P04062 |
| HOMO SAPIENS | 0 | 954 | 2vt0 | A | 377 | GLY | p.Gly416Ser | P04062 |
| HOMO SAPIENS | 8 | 954 | 2vt0 | A | 398 | VAL | p.Val437Phe | P04062 |
| HOMO SAPIENS | 0 | 954 | 2vt0 | A | 401 | PRO | p.Pro440Leu | P04062 |
| HOMO SAPIENS | 3 | 954 | 2vt0 | A | 409 | ASP | p.Asp448His | P04062 |
| HOMO SAPIENS | 3 | 954 | 2vt0 | A | 409 | ASP | p.Asp448Val | P04062 |
| HOMO SAPIENS | 0 | 954 | 2vt0 | A | 415 | PRO | p.Pro454Arg | P04062 |
| HOMO SAPIENS | 5 | 954 | 2vt0 | A | 444 | LEU | p.Leu483Arg | P04062 |
| HOMO SAPIENS | 5 | 954 | 2vt0 | A | 456 | ALA | p.Ala495Pro | P04062 |
| HOMO SAPIENS | 5 | 954 | 2vt0 | A | 463 | ARG | p.Arg502Cys | P04062 |
| HOMO SAPIENS | 5 | 954 | 2vt0 | A | 463 | ARG | p.Arg502His | P04062 |
| HOMO SAPIENS | 5 | 954 | 2vt0 | A | 478 | GLY | p.Gly517Ser | P04062 |
| HOMO SAPIENS | 5 | 954 | 2vt0 | A | 496 | ARG | p.Arg535Cys | P04062 |
| HOMO SAPIENS | 5 | 954 | 2vt0 | A | 496 | ARG | p.Arg535His | P04062 |
| HOMO SAPIENS | 1 | 9037 | 2rqb | A | 946 | ALA | p.Ala946Thr | Q9BYX4 |
| HOMO SAPIENS | 0 | 253 | 5E+092 | A | 308 | LEU | p.Leu308Pro | P37173 |
| HOMO SAPIENS | 0 | 253 | 5E+092 | A | 315 | THR | p.Thr315Met | P37173 |
| HOMO SAPIENS | 0 | 253 | 5E+092 | A | 328 | HIS | p.His328Gln | P37173 |
| HOMO SAPIENS | 0 | 253 | 5E+092 | A | 336 | TYR | p.Tyr336Asn | P37173 |
| HOMO SAPIENS | 0 | 253 | 5E+092 | A | 348 | ARG | p.Arg348Cys | P37173 |
| HOMO SAPIENS | 0 | 253 | 5E+092 | A | 355 | ALA | p.Ala355Pro | P37173 |
| HOMO SAPIENS | 1 | 253 | 5E+092 | A | 356 | ARG | p.Arg356Gly | P37173 |

| | | | | | | | | |
|---|---|---|---|---|---|---|---|---|
| HOMO SAPIENS | 0 | 253 | 5E+092 | A | 357 | GLY | p.Gly357Trp | P37173 |
| HOMO SAPIENS | 1 | 253 | 5E+092 | A | 403 | ARG | p.Arg403His | P37173 |
| HOMO SAPIENS | 5 | 253 | 5E+092 | A | 449 | SER | p.Ser449Phe | P37173 |
| HOMO SAPIENS | 0 | 253 | 5E+092 | A | 460 | ARG | p.Arg460Cys | P37173 |
| HOMO SAPIENS | 0 | 253 | 5E+092 | A | 460 | ARG | p.Arg460His | P37173 |
| HOMO SAPIENS | 2 | 253 | 5E+092 | A | 526 | GLU | p.Glu526Gln | P37173 |
| HOMO SAPIENS | 5 | 253 | 5E+092 | A | 528 | ARG | p.Arg528Cys | P37173 |
| HOMO SAPIENS | 5 | 253 | 5E+092 | A | 528 | ARG | p.Arg528His | P37173 |
| HOMO SAPIENS | 6 | 158 | 5amc | A | 231 | ARG | p.Arg260Leu | P12821 |
| HOMO SAPIENS | 7 | 158 | 5amc | A | 237 | TYR | p.Tyr266Ter | P12821 |
| HOMO SAPIENS | 6 | 158 | 5amc | A | 325 | GLY | p.Gly354Arg | P12821 |
| HOMO SAPIENS | 0 | 158 | 5amc | A | 467 | ARG | p.Arg496Ter | P12821 |
| HOMO SAPIENS | 4 | 158 | 5amc | A | 479 | ARG | p.Arg508Ter | P12821 |
| HOMO SAPIENS | 0 | 158 | 5amc | A | 563 | ASP | p.Asp592Gly | P12821 |
| HOMO SAPIENS | -1 | 33062 | 1gr3 | A | 545 | GLY | p.Gly545Arg | Q03692 |
| HOMO SAPIENS | 5 | 33062 | 1gr3 | A | 591 | CYS | p.Cys591Arg | Q03692 |
| HOMO SAPIENS | 2 | 33062 | 1gr3 | A | 595 | GLY | p.Gly595Glu | Q03692 |
| HOMO SAPIENS | 0 | 33062 | 1gr3 | A | 597 | TYR | p.Tyr597Cys | Q03692 |
| HOMO SAPIENS | 5 | 33062 | 1gr3 | A | 598 | TYR | p.Tyr598Asp | Q03692 |
| HOMO SAPIENS | 7 | 33062 | 1gr3 | A | 600 | SER | p.Ser600Pro | Q03692 |
| HOMO SAPIENS | 1 | 33062 | 1gr3 | A | 614 | LEU | p.Leu614Pro | Q03692 |
| HOMO SAPIENS | 6 | 33062 | 1gr3 | A | 651 | TRP | p.Trp651Arg | Q03692 |
| HOMO SAPIENS | 7 | 33062 | 1gr3 | A | 671 | SER | p.Ser671Pro | Q03692 |
| HOMO SAPIENS | -1 | 38257 | 2l2i | B | 87 | PRO | p.Pro87Ala | Q13351 |
| HOMO SAPIENS | 2 | 2036 | 1ghq | A | 231 | GLY | p.Gly1224Asp | P01024 |
| HOMO SAPIENS | 5 | 3594 | 4cde | F | 388 | TYR | p.Tyr412Cys | P53634 |
| HOMO SAPIENS | 9 | 3594 | 4cde | F | 405 | TRP | p.Trp429Cys | P53634 |
| HOMO SAPIENS | 0 | 3594 | 4cde | F | 429 | ILE | p.Ile453Val | P53634 |
| HOMO SAPIENS | 7 | 18 | 1jkb | A | 34 | TRP | p.Trp52Ter | P61626 |
| HOMO SAPIENS | 9 | 18 | 1jkb | A | 70 | THR | p.Thr88Asn | P61626 |
| HOMO SAPIENS | 8 | 18 | 1jkb | A | 121 | VAL | p.Val139Ile | P61626 |
| HOMO SAPIENS | 2 | 414 | 3fj6 | A | 136 | ARG | p.Arg135Cys | Q02127 |
| HOMO SAPIENS | 1 | 414 | 3fj6 | A | 153 | GLY | p.Gly152Arg | Q02127 |
| HOMO SAPIENS | 0 | 414 | 3fj6 | A | 200 | ARG | p.Arg199Cys | Q02127 |
| HOMO SAPIENS | 0 | 414 | 3fj6 | A | 203 | GLY | p.Gly202Ala | Q02127 |
| HOMO SAPIENS | 0 | 414 | 3fj6 | A | 245 | ARG | p.Arg244Trp | Q02127 |
| HOMO SAPIENS | 0 | 414 | 3fj6 | A | 342 | ALA | p.Ala341Val | Q02127 |
| HOMO SAPIENS | 1 | 414 | 3fj6 | A | 347 | ARG | p.Arg346Trp | Q02127 |
| HOMO SAPIENS | 6 | 103 | 1u3w | A | 77 | GLY | p.Gly78Ter | P00326 |
| HOMO SAPIENS | -1 | 8369 | 4ux8 | F | 16 | ARG | p.Arg93Trp | P39905 |
| HOMO SAPIENS | 0 | 366 | 2zjc | A | 32 | GLY | p.Arg108Trp | P01375 |
| HOMO SAPIENS | 0 | 1457 | 3v9i | A | 352 | LEU | p.Ser352Leu | P30038 |
| HOMO SAPIENS | 0 | 1457 | 3v9i | A | 470 | VAL | p.Val470Ile | P30038 |
| HOMO SAPIENS | 0 | 1457 | 3v9i | A | 473 | THR | p.Thr473Ala | P30038 |
| HOMO SAPIENS | 1 | 2 | 4wo5 | A | 462 | ARG | p.Arg462Ile | P15056 |
| HOMO SAPIENS | 3 | 2 | 4wo5 | A | 463 | ILE | p.Ile463Ser | P15056 |
| HOMO SAPIENS | 1 | 2 | 4wo5 | A | 466 | GLY | p.Gly466Val | P15056 |
| HOMO SAPIENS | 1 | 2 | 4wo5 | A | 468 | PHE | p.Phe468Ser | P15056 |
| HOMO SAPIENS | 2 | 2 | 4wo5 | A | 469 | GLY | p.Gly469Ala | P15056 |
| HOMO SAPIENS | 2 | 2 | 4wo5 | A | 469 | GLY | p.Gly469Arg | P15056 |
| HOMO SAPIENS | 2 | 2 | 4wo5 | A | 485 | LEU | p.Leu485Phe | P15056 |
| HOMO SAPIENS | 0 | 2 | 4wo5 | A | 501 | GLU | p.Glu501Gly | P15056 |
| HOMO SAPIENS | 0 | 2 | 4wo5 | A | 501 | GLU | p.Glu501Lys | P15056 |
| HOMO SAPIENS | 7 | 2 | 4wo5 | A | 581 | ASN | p.Asn581Asp | P15056 |

| | | | | | | | | |
|---|---|---|---|---|---|---|---|---|
| HOMO SAPIENS | 8 | 2 | 4wo5 | A | 594 | ASP | p.Asp594Gly | P15056 |
| HOMO SAPIENS | 0 | 2 | 4wo5 | A | 597 | LEU | p.Leu597Arg | P15056 |
| HOMO SAPIENS | 0 | 2 | 4wo5 | A | 597 | LEU | p.Leu597Val | P15056 |
| HOMO SAPIENS | 0 | 2 | 4wo5 | A | 600 | VAL | p.Val600Glu | P15056 |
| HOMO SAPIENS | 2 | 2 | 4wo5 | A | 601 | LYS | p.Lys601Glu | P15056 |
| HOMO SAPIENS | 8 | 2 | 4wo5 | A | 638 | ASP | p.Asp638Glu | P15056 |
| HOMO SAPIENS | -1 | 30366 | 5bqd | B | 19 | ALA | p.Ala19Ser | Q99593 |
| HOMO SAPIENS | -1 | 30366 | 5bqd | B | 23 | PRO | p.Pro23His | Q99593 |
| HOMO SAPIENS | 1 | 30366 | 5bqd | B | 49 | GLN | p.Gln49Lys | Q99593 |
| HOMO SAPIENS | 2 | 30366 | 5bqd | B | 54 | ILE | p.Ile54Thr | Q99593 |
| HOMO SAPIENS | 8 | 30366 | 5bqd | B | 80 | GLY | p.Gly80Arg | Q99593 |
| HOMO SAPIENS | 6 | 30366 | 5bqd | B | 106 | ILE | p.Ile106Val | Q99593 |
| HOMO SAPIENS | 7 | 30366 | 5bqd | B | 111 | ASP | p.Asp111Tyr | Q99593 |
| HOMO SAPIENS | -1 | 30366 | 5bqd | B | 237 | ARG | p.Arg237Gln | Q99593 |
| HOMO SAPIENS | -1 | 30366 | 5bqd | B | 237 | ARG | p.Arg237Trp | Q99593 |
| HOMO SAPIENS | 5 | 4789 | 1n4m | A | 567 | SER | p.Ser567Leu | P06400 |
| HOMO SAPIENS | 0 | 4789 | 1n4m | A | 658 | ALA | p.Ala658Asp | P06400 |
| HOMO SAPIENS | 7 | 4789 | 1n4m | A | 661 | ARG | p.Arg661Trp | P06400 |
| HOMO SAPIENS | 6 | 4789 | 1n4m | A | 668 | ARG | p.Arg668His | P06400 |
| HOMO SAPIENS | 8 | 4789 | 1n4m | A | 697 | ASP | p.Asp697Glu | P06400 |
| HOMO SAPIENS | 6 | 4789 | 1n4m | A | 712 | CYS | p.Cys712Arg | P06400 |
| HOMO SAPIENS | 1 | 1798 | 2r5g | A | 101 | HIS | p.His101Gln | O15247 |
| HOMO SAPIENS | 0 | 1798 | 2r5g | A | 160 | PRO | p.Pro160Ala | O15247 |
| HOMO SAPIENS | 2 | 22 | 4x0t | A | 82 | ARG | p.Arg110Ter | P49419 |
| HOMO SAPIENS | 0 | 22 | 4x0t | A | 149 | ALA | p.Ala177Pro | P49419 |
| HOMO SAPIENS | 4 | 22 | 4x0t | A | 164 | THR | p.Thr192Met | P49419 |
| HOMO SAPIENS | 9 | 22 | 4x0t | A | 167 | ASN | p.Asn195Ser | P49419 |
| HOMO SAPIENS | 0 | 22 | 4x0t | A | 175 | TRP | p.Trp203Gly | P49419 |
| HOMO SAPIENS | 0 | 22 | 4x0t | A | 194 | THR | p.Thr222Ala | P49419 |
| HOMO SAPIENS | 0 | 22 | 4x0t | A | 273 | ASN | p.Asn301Ile | P49419 |
| HOMO SAPIENS | 0 | 22 | 4x0t | A | 307 | ARG | p.Arg335Gln | P49419 |
| HOMO SAPIENS | 3 | 22 | 4x0t | A | 313 | SER | p.Ser341Asn | P49419 |
| HOMO SAPIENS | 0 | 22 | 4x0t | A | 329 | ILE | p.Ile357Val | P49419 |
| HOMO SAPIENS | 1 | 22 | 4x0t | A | 380 | TYR | p.Tyr408Ter | P49419 |
| HOMO SAPIENS | 9 | 22 | 4x0t | A | 399 | GLU | p.Glu427Gln | P49419 |
| HOMO SAPIENS | 0 | 22 | 4x0t | A | 406 | TYR | p.Tyr434Cys | P49419 |
| HOMO SAPIENS | 0 | 22 | 4x0t | A | 431 | ILE | p.Ile459Phe | P49419 |
| HOMO SAPIENS | 4 | 22 | 4x0t | A | 477 | GLY | p.Gly505Arg | P49419 |
| HOMO SAPIENS | 0 | 22 | 4x0t | A | 495 | THR | p.Thr523Ala | P49419 |
| HOMO SAPIENS | 8 | 11042 | 1y6b | A | 1145 | PRO | p.Pro1147Ser | P35968 |
| HOMO SAPIENS | -1 | 3867 | 2xij | A | 18 | GLN | p.Gln18Ter | P22033 |
| HOMO SAPIENS | -1 | 3867 | 2xij | A | 31 | ARG | p.Arg31Ter | P22033 |
| HOMO SAPIENS | 2 | 3867 | 2xij | A | 93 | ARG | p.Arg93His | P22033 |
| HOMO SAPIENS | 0 | 3867 | 2xij | A | 94 | GLY | p.Gly94Arg | P22033 |
| HOMO SAPIENS | 2 | 3867 | 2xij | A | 95 | PRO | p.Pro95Leu | P22033 |
| HOMO SAPIENS | 2 | 3867 | 2xij | A | 105 | TRP | p.Trp105Arg | P22033 |
| HOMO SAPIENS | 2 | 3867 | 2xij | A | 108 | ARG | p.Arg108Cys | P22033 |
| HOMO SAPIENS | 3 | 3867 | 2xij | A | 154 | ARG | p.Arg154His | P22033 |
| HOMO SAPIENS | 3 | 3867 | 2xij | A | 156 | ASP | p.Asp156Val | P22033 |
| HOMO SAPIENS | 0 | 3867 | 2xij | A | 189 | ASN | p.Asn189Ile | P22033 |
| HOMO SAPIENS | 0 | 3867 | 2xij | A | 191 | ALA | p.Ala191Glu | P22033 |
| HOMO SAPIENS | 0 | 3867 | 2xij | A | 203 | GLY | p.Gly203Arg | P22033 |
| HOMO SAPIENS | 2 | 3867 | 2xij | A | 215 | GLY | p.Gly215Ser | P22033 |
| HOMO SAPIENS | 0 | 3867 | 2xij | A | 219 | ASN | p.Asn219Tyr | P22033 |

| | | | | | | | | |
|---|---|---|---|---|---|---|---|---|
| HOMO SAPIENS | 3 | 3867 | 2xij | A | 228 | ARG | p.Arg228Ter | P22033 |
| HOMO SAPIENS | 1 | 3867 | 2xij | A | 297 | THR | p.Thr297Ile | P22033 |
| HOMO SAPIENS | 0 | 3867 | 2xij | A | 326 | ARG | p.Arg326Lys | P22033 |
| HOMO SAPIENS | 3 | 3867 | 2xij | A | 369 | ARG | p.Arg369His | P22033 |
| HOMO SAPIENS | 3 | 3867 | 2xij | A | 369 | ARG | p.Arg369Cys | P22033 |
| HOMO SAPIENS | 0 | 3867 | 2xij | A | 370 | THR | p.Thr370Pro | P22033 |
| HOMO SAPIENS | 0 | 3867 | 2xij | A | 377 | ALA | p.Ala377Glu | P22033 |
| HOMO SAPIENS | 2 | 3867 | 2xij | A | 426 | GLY | p.Gly426Ala | P22033 |
| HOMO SAPIENS | 2 | 3867 | 2xij | A | 427 | GLY | p.Gly427Asp | P22033 |
| HOMO SAPIENS | 1 | 3867 | 2xij | A | 499 | THR | p.Ala499Thr | P22033 |
| HOMO SAPIENS | 2 | 3867 | 2xij | A | 532 | ARG | p.Arg532His | P22033 |
| HOMO SAPIENS | 3 | 3867 | 2xij | A | 555 | ALA | p.Ala555Thr | P22033 |
| HOMO SAPIENS | 0 | 3867 | 2xij | A | 573 | PHE | p.Phe573Ser | P22033 |
| HOMO SAPIENS | 3 | 3867 | 2xij | A | 623 | GLY | p.Gly623Arg | P22033 |
| HOMO SAPIENS | 5 | 3867 | 2xij | A | 671 | VAL | p.Ile671Val | P22033 |
| HOMO SAPIENS | 4 | 3867 | 2xij | A | 694 | ARG | p.Arg694Trp | P22033 |
| HOMO SAPIENS | 1 | 3867 | 2xij | A | 700 | MET | p.Met700Lys | P22033 |
| HOMO SAPIENS | 8 | 3867 | 2xij | A | 703 | GLY | p.Gly703Arg | P22033 |
| HOMO SAPIENS | 0 | 3867 | 2xij | A | 717 | GLY | p.Gly717Val | P22033 |
| HOMO SAPIENS | 1 | 3867 | 2xij | A | 727 | ARG | p.Arg727Ter | P22033 |
| HOMO SAPIENS | -1 | 52 | 3k91 | B | 72 | GLY | p.Gly73Ser | P00441 |
| HOMO SAPIENS | 6 | 52 | 3k91 | B | 96 | ASP | p.Asp97Asn | P00441 |
| HOMO SAPIENS | 5 | 52 | 3k91 | B | 112 | ILE | p.Ile113Thr | P00441 |
| HOMO SAPIENS | -1 | 52 | 3k91 | B | 134 | SER | p.Ser135Asn | P00441 |
| HOMO SAPIENS | 0 | 3950 | 4fom | A | 296 | ASN | p.Asn296His | Q9NQS3 |
| HOMO SAPIENS | 1 | 10 | 1de4 | A | 31 | VAL | p.Val53Met | Q30201 |
| HOMO SAPIENS | 0 | 10 | 1de4 | A | 37 | VAL | p.Val59Met | Q30201 |
| HOMO SAPIENS | 0 | 10 | 1de4 | A | 71 | GLY | p.Gly93Arg | Q30201 |
| HOMO SAPIENS | 0 | 10 | 1de4 | A | 83 | ILE | p.Ile105Thr | Q30201 |
| HOMO SAPIENS | 0 | 10 | 1de4 | A | 105 | GLN | p.Gln127His | Q30201 |
| HOMO SAPIENS | 1 | 10 | 1de4 | A | 146 | GLU | p.Glu168Ter | Q30201 |
| HOMO SAPIENS | 2 | 10 | 1de4 | A | 261 | GLN | p.Gln283Pro | Q30201 |
| HOMO SAPIENS | 0 | 8038 | 4x7n | A | 586 | ILE | p.Ile586Leu | Q9NZJ5 |
| HOMO SAPIENS | 0 | 8038 | 4x7n | A | 588 | ARG | p.Arg588Gln | Q9NZJ5 |
| HOMO SAPIENS | 0 | 35175 | 5ugw | A | 1062 | ALA | p.Ala1062Thr | O14746 |
| HOMO SAPIENS | 2 | 35175 | 5ugw | A | 1090 | VAL | p.Val1090Met | O14746 |
| HOMO SAPIENS | 0 | 1722 | 4z3k | C | 147 | ARG | p.Arg150Gly | P35270 |
| HOMO SAPIENS | 0 | 1722 | 4z3k | C | 160 | PRO | p.Pro163Leu | P35270 |
| HOMO SAPIENS | 0 | 1722 | 4z3k | C | 207 | ARG | p.Arg210Trp | P35270 |
| HOMO SAPIENS | 0 | 1722 | 4z3k | C | 216 | ARG | p.Arg219Ter | P35270 |
| HOMO SAPIENS | 0 | 1722 | 4z3k | C | 248 | LYS | p.Lys251Ter | P35270 |
| HOMO SAPIENS | 1 | 2832 | 4x8v | L | 115 | HIS | p.His175Gln | P08709 |
| HOMO SAPIENS | 0 | 825 | 1emh | A | 242 | PHE | p.Phe251Ser | P13051 |
| HOMO SAPIENS | 0 | 122 | 5uwk | A | 109 | ARG | p.Arg109Ter | P45452 |
| HOMO SAPIENS | 1 | 122 | 5uwk | A | 170 | LYS | p.Lys170Thr | P45452 |
| HOMO SAPIENS | 9 | 122 | 5uwk | A | 232 | HIS | p.His232Asn | P45452 |
| HOMO SAPIENS | 5 | 122 | 5uwk | A | 257 | ASP | p.Asp257Val | P45452 |
| HOMO SAPIENS | 9 | 1074 | 3p5i | A | 264 | TRP | p.Trp264Arg | Q9UJ71 |
| HOMO SAPIENS | 0 | 146 | 3r3g | A | 14C | GLU | p.Glu352Lys | P00734 |
| HOMO SAPIENS | 3 | 9522 | 1wyq | A | 83 | LEU | p.Leu253Pro | O15020 |
| HOMO SAPIENS | 8 | 67 | 2vgb | A | 130 | SER | p.Ser130Tyr | P30613 |
| HOMO SAPIENS | 8 | 67 | 2vgb | A | 163 | ARG | p.Arg163Cys | P30613 |
| HOMO SAPIENS | 4 | 67 | 2vgb | A | 241 | GLU | p.Glu241Ter | P30613 |
| HOMO SAPIENS | 9 | 67 | 2vgb | A | 384 | THR | p.Thr384Met | P30613 |

| | | | | | | | | |
|---|---|---|---|---|---|---|---|---|
| HOMO SAPIENS | 0 | 67 | 2vgb | A | 421 | GLN | p.Gln421Lys | P30613 |
| HOMO SAPIENS | 6 | 67 | 2vgb | A | 479 | ARG | p.Arg479His | P30613 |
| HOMO SAPIENS | 8 | 67 | 2vgb | A | 510 | ARG | p.Arg510Gln | P30613 |
| HOMO SAPIENS | 4 | 67 | 2vgb | A | 569 | ARG | p.Arg569Trp | P30613 |
| HOMO SAPIENS | 0 | 9088 | 5v44 | A | 232 | ASN | p.Asn232Lys | Q9NZJ4 |
| HOMO SAPIENS | 4 | 16 | 3lbn | A | 12 | GLY | p.Gly12Val | P01112 |
| HOMO SAPIENS | 4 | 16 | 3lbn | A | 12 | GLY | p.Gly12Ser | P01112 |
| HOMO SAPIENS | 4 | 16 | 3lbn | A | 13 | GLY | p.Gly13Cys | P01112 |
| HOMO SAPIENS | 4 | 16 | 3lbn | A | 13 | GLY | p.Gly13Asp | P01112 |
| HOMO SAPIENS | 5 | 16 | 3lbn | A | 22 | GLN | p.Gln22Lys | P01112 |
| HOMO SAPIENS | 5 | 16 | 3lbn | A | 58 | THR | p.Thr58Ile | P01112 |
| HOMO SAPIENS | 3 | 16 | 3lbn | A | 61 | GLN | p.Gln61Lys | P01112 |
| HOMO SAPIENS | 1 | 16 | 3lbn | A | 63 | GLU | p.Glu63Lys | P01112 |
| HOMO SAPIENS | 9 | 16 | 3lbn | A | 117 | LYS | p.Lys117Arg | P01112 |
| HOMO SAPIENS | 6 | 16 | 3lbn | A | 123 | ARG | p.Arg123Cys | P01112 |
| HOMO SAPIENS | 1 | 16 | 3lbn | A | 138 | GLY | p.Gly138Ser | P01112 |
| HOMO SAPIENS | 9 | 16 | 3lbn | A | 146 | ALA | p.Ala146Thr | P01112 |
| HOMO SAPIENS | 9 | 16 | 3lbn | A | 146 | ALA | p.Ala146Val | P01112 |
| HOMO SAPIENS | 1 | 359 | 5nvy | C | 63 | LEU | p.Leu63Pro | P40337 |
| HOMO SAPIENS | 1 | 359 | 5nvy | C | 64 | ARG | p.Arg64Pro | P40337 |
| HOMO SAPIENS | 1 | 359 | 5nvy | C | 70 | GLU | p.Glu70Lys | P40337 |
| HOMO SAPIENS | 0 | 359 | 5nvy | C | 84 | VAL | p.Val84Leu | P40337 |
| HOMO SAPIENS | 1 | 359 | 5nvy | C | 88 | TRP | p.Trp88Ser | P40337 |
| HOMO SAPIENS | 1 | 359 | 5nvy | C | 93 | GLY | p.Gly93Cys | P40337 |
| HOMO SAPIENS | 1 | 359 | 5nvy | C | 98 | TYR | p.Tyr98His | P40337 |
| HOMO SAPIENS | 1 | 359 | 5nvy | C | 111 | SER | p.Ser111Arg | P40337 |
| HOMO SAPIENS | 1 | 359 | 5nvy | C | 112 | TYR | p.Tyr112His | P40337 |
| HOMO SAPIENS | 1 | 359 | 5nvy | C | 126 | ASP | p.Asp126Tyr | P40337 |
| HOMO SAPIENS | 1 | 359 | 5nvy | C | 130 | VAL | p.Val130Leu | P40337 |
| HOMO SAPIENS | 1 | 359 | 5nvy | C | 135 | LEU | p.Leu135Phe | P40337 |
| HOMO SAPIENS | 9 | 359 | 5nvy | C | 161 | ARG | p.Arg161Gly | P40337 |
| HOMO SAPIENS | 7 | 359 | 5nvy | C | 162 | CYS | p.Cys162Phe | P40337 |
| HOMO SAPIENS | 4 | 359 | 5nvy | C | 163 | LEU | p.Leu163Pro | P40337 |
| HOMO SAPIENS | 3 | 359 | 5nvy | C | 164 | GLN | p.Gln164Arg | P40337 |
| HOMO SAPIENS | 9 | 359 | 5nvy | C | 166 | VAL | p.Val166Phe | P40337 |
| HOMO SAPIENS | 7 | 359 | 5nvy | C | 167 | ARG | p.Arg167Gly | P40337 |
| HOMO SAPIENS | 7 | 359 | 5nvy | C | 167 | ARG | p.Arg167Gln | P40337 |
| HOMO SAPIENS | 1 | 359 | 5nvy | C | 183 | SER | p.Ser183Ter | P40337 |
| HOMO SAPIENS | 2 | 359 | 5nvy | C | 188 | LEU | p.Leu188Val | P40337 |
| HOMO SAPIENS | 1 | 359 | 5nvy | C | 191 | HIS | p.His191Asp | P40337 |
| HOMO SAPIENS | 1 | 359 | 5nvy | C | 192 | PRO | p.Pro192Ser | P40337 |
| HOMO SAPIENS | 1 | 359 | 5nvy | C | 200 | ARG | p.Arg200Trp | P40337 |
| HOMO SAPIENS | -1 | 359 | 5nvy | C | 211 | MET | p.Met211Leu | P40337 |
| HOMO SAPIENS | 0 | 14099 | 4btm | A | 282 | ARG | p.Arg258Gln | Q5TCY1 |
| HOMO SAPIENS | 0 | 696 | 5h1b | A | 150 | ARG | p.Arg150Gln | Q06609 |
| HOMO SAPIENS | -1 | 208 | 2bxl | A | 1 | ASP | p.Asp25Val | P02768 |
| HOMO SAPIENS | 0 | 208 | 2bxl | A | 269 | ASP | p.Asp293Gly | P02768 |
| HOMO SAPIENS | 0 | 208 | 2bxl | A | 570 | GLU | p.Glu594Lys | P02768 |
| HOMO SAPIENS | 0 | 208 | 2bxl | A | 573 | LYS | p.Lys597Glu | P02768 |
| HOMO SAPIENS | -1 | 12813 | 2uzx | B | 35 | MET | p.Met35Leu | P08581 |
| HOMO SAPIENS | 1 | 12813 | 2uzx | B | 156 | SER | p.Ser156Leu | P08581 |
| HOMO SAPIENS | 0 | 12813 | 2uzx | B | 168 | GLU | p.Glu168Asp | P08581 |
| HOMO SAPIENS | 2 | 12813 | 2uzx | B | 211 | LEU | p.Leu211Trp | P08581 |
| HOMO SAPIENS | 2 | 12813 | 2uzx | B | 218 | ARG | p.Arg218Ser | P08581 |

| | | | | | | | | |
|---|---|---|---|---|---|---|---|---|
| HOMO SAPIENS | 1 | 12813 | 2uzx | B | 238 | LEU | p.Leu238Ser | P08581 |
| HOMO SAPIENS | 4 | 12813 | 2uzx | B | 320 | ALA | p.Ala320Val | P08581 |
| HOMO SAPIENS | 1 | 12813 | 2uzx | B | 375 | ASN | p.Asn375Ser | P08581 |
| HOMO SAPIENS | 0 | 12813 | 2uzx | B | 575 | LEU | p.Leu575Ile | P08581 |
| HOMO SAPIENS | 1 | 10780 | 4ejx | A | 348 | ARG | p.Arg367Cys | P00450 |
| HOMO SAPIENS | 2 | 10780 | 4ejx | A | 525 | ASP | p.Asp544Glu | P00450 |
| HOMO SAPIENS | 1 | 1407 | 5jn5 | A | 88 | ILE | p.Ile88Val | P36871 |
| HOMO SAPIENS | 7 | 1407 | 5jn5 | A | 115 | THR | p.Thr115Ala | P36871 |
| HOMO SAPIENS | 3 | 1407 | 5jn5 | A | 121 | GLY | p.Gly121Arg | P36871 |
| HOMO SAPIENS | 2 | 1407 | 5jn5 | A | 221 | ARG | p.Arg221Cys | P36871 |
| HOMO SAPIENS | 6 | 1407 | 5jn5 | A | 263 | TYR | p.Asp263Tyr | P36871 |
| HOMO SAPIENS | 3 | 1407 | 5jn5 | A | 420 | TYR | p.Tyr420His | P36871 |
| HOMO SAPIENS | 0 | 1407 | 5jn5 | A | 501 | VAL | p.Val501Ile | P36871 |
| HOMO SAPIENS | -1 | 1878 | 1yow | A | 255 | ARG | p.Arg255Leu | Q13285 |
| HOMO SAPIENS | 6 | 1878 | 1yow | A | 293 | ASP | p.Asp293Asn | Q13285 |
| HOMO SAPIENS | 1 | 1878 | 1yow | A | 437 | LEU | p.Leu437Gln | Q13285 |
| HOMO SAPIENS | 0 | 3071 | 2wpm | E | 145 | ARG | p.Arg191His | P00740 |
| HOMO SAPIENS | 0 | 3071 | 2wpm | E | 145 | ARG | p.Arg191Cys | P00740 |
| HOMO SAPIENS | 3 | 589 | 3ggc | B | 15 | GLY | p.Gly16Ser | P00492 |
| HOMO SAPIENS | 4 | 589 | 3ggc | B | 40 | LEU | p.Leu41Pro | P00492 |
| HOMO SAPIENS | 2 | 589 | 3ggc | B | 50 | ARG | p.Arg51Gly | P00492 |
| HOMO SAPIENS | 5 | 589 | 3ggc | B | 51 | ASP | p.Asp52Gly | P00492 |
| HOMO SAPIENS | 0 | 589 | 3ggc | B | 57 | GLY | p.Gly58Arg | P00492 |
| HOMO SAPIENS | 6 | 589 | 3ggc | B | 69 | GLY | p.Gly70Glu | P00492 |
| HOMO SAPIENS | 1 | 589 | 3ggc | B | 73 | PHE | p.Phe74Leu | P00492 |
| HOMO SAPIENS | 0 | 589 | 3ggc | B | 77 | LEU | p.Leu78Val | P00492 |
| HOMO SAPIENS | 3 | 589 | 3ggc | B | 79 | ASP | p.Asp80Val | P00492 |
| HOMO SAPIENS | 0 | 589 | 3ggc | B | 103 | SER | p.Ser104Arg | P00492 |
| HOMO SAPIENS | 0 | 589 | 3ggc | B | 109 | SER | p.Ser110Leu | P00492 |
| HOMO SAPIENS | 1 | 589 | 3ggc | B | 129 | VAL | p.Val130Asp | P00492 |
| HOMO SAPIENS | 4 | 589 | 3ggc | B | 131 | ILE | p.Ile132Met | P00492 |
| HOMO SAPIENS | 7 | 589 | 3ggc | B | 142 | MET | p.Met143Lys | P00492 |
| HOMO SAPIENS | 0 | 589 | 3ggc | B | 160 | ALA | p.Ala161Ser | P00492 |
| HOMO SAPIENS | 6 | 589 | 3ggc | B | 167 | THR | p.Thr168Ile | P00492 |
| HOMO SAPIENS | 0 | 589 | 3ggc | B | 169 | ARG | p.Arg170Ter | P00492 |
| HOMO SAPIENS | 5 | 589 | 3ggc | B | 175 | PRO | p.Pro176Leu | P00492 |
| HOMO SAPIENS | 8 | 589 | 3ggc | B | 193 | ASP | p.Asp194Glu | P00492 |
| HOMO SAPIENS | 8 | 589 | 3ggc | B | 193 | ASP | p.Asp194Asn | P00492 |
| HOMO SAPIENS | 8 | 589 | 3ggc | B | 198 | PHE | p.Phe199Val | P00492 |
| HOMO SAPIENS | 8 | 589 | 3ggc | B | 200 | ASP | p.Asp201Gly | P00492 |
| HOMO SAPIENS | 0 | 4269 | 2pla | B | 174 | GLU | p.Glu174Lys | Q8N335 |
| HOMO SAPIENS | 1 | 4269 | 2pla | B | 220 | ARG | p.Arg220His | Q8N335 |
| HOMO SAPIENS | 4 | 4269 | 2pla | B | 280 | ALA | p.Ala280Val | Q8N335 |
| HOMO SAPIENS | 0 | 2712 | 1ii4 | E | 252 | TRP | p.Ser252Leu | P21802 |
| HOMO SAPIENS | 0 | 2712 | 1ii4 | E | 253 | PRO | p.Pro253Arg | P21802 |
| HOMO SAPIENS | 5 | 2712 | 1ii4 | E | 267 | SER | p.Ser267Pro | P21802 |
| HOMO SAPIENS | 0 | 2712 | 1ii4 | E | 278 | CYS | p.Cys278Phe | P21802 |
| HOMO SAPIENS | 0 | 2712 | 1ii4 | E | 289 | GLN | p.Gln289Pro | P21802 |
| HOMO SAPIENS | 0 | 2712 | 1ii4 | E | 290 | TRP | p.Trp290Cys | P21802 |
| HOMO SAPIENS | 0 | 2712 | 1ii4 | E | 290 | TRP | p.Trp290Arg | P21802 |
| HOMO SAPIENS | 0 | 2712 | 1ii4 | E | 292 | LYS | p.Lys292Glu | P21802 |
| HOMO SAPIENS | 0 | 2712 | 1ii4 | E | 315 | ALA | p.Ala315Ser | P21802 |
| HOMO SAPIENS | 0 | 2712 | 1ii4 | E | 321 | ASP | p.Asp321Ala | P21802 |
| HOMO SAPIENS | 1 | 2712 | 1ii4 | E | 328 | TYR | p.Tyr328Cys | P21802 |

| | | | | | | | | |
|---|---|---|---|---|---|---|---|---|
| HOMO SAPIENS | 9 | 2712 | 1ii4 | E | 340 | TYR | p.Tyr340His | P21802 |
| HOMO SAPIENS | 1 | 2712 | 1ii4 | E | 341 | THR | p.Thr341Pro | P21802 |
| HOMO SAPIENS | 0 | 2712 | 1ii4 | E | 342 | CYS | p.Cys342Arg | P21802 |
| HOMO SAPIENS | 0 | 2712 | 1ii4 | E | 342 | CYS | p.Cys342Trp | P21802 |
| HOMO SAPIENS | 0 | 2712 | 1ii4 | E | 344 | ALA | p.Ala344Gly | P21802 |
| HOMO SAPIENS | 0 | 2712 | 1ii4 | E | 347 | SER | p.Ser347Cys | P21802 |
| HOMO SAPIENS | 0 | 2712 | 1ii4 | E | 351 | SER | p.Ser351Cys | P21802 |
| HOMO SAPIENS | 1 | 2712 | 1ii4 | E | 354 | SER | p.Ser354Cys | P21802 |
| HOMO SAPIENS | 2 | 53 | 4pj1 | H | 5 | ASP | p.Asp29Gly | P10809 |
| HOMO SAPIENS | 0 | 53 | 4pj1 | H | 74 | VAL | p.Val98Ile | P10809 |
| HOMO SAPIENS | 0 | 29968 | 2fdb | M | 71 | LYS | p.Lys89Glu | P55075 |
| HOMO SAPIENS | 2 | 29968 | 2fdb | M | 98 | ARG | p.Arg116Gly | P55075 |
| HOMO SAPIENS | 2 | 29968 | 2fdb | M | 122 | GLY | p.Gly140Ser | P55075 |
| HOMO SAPIENS | 8 | 5061 | 5f85 | B | 60 | GLY | p.Gly106Ser | P00740 |
| HOMO SAPIENS | 0 | 35944 | 1h6h | A | 85 | THR | p.Thr85Asn | Q15080 |
| HOMO SAPIENS | 8 | 35944 | 1h6h | A | 105 | ARG | p.Arg105Gln | Q15080 |
| HOMO SAPIENS | 7 | 12016 | 1hj7 | A | 295 | ASN | p.Asn316Ser | P01130 |
| HOMO SAPIENS | 5 | 12016 | 1hj7 | A | 302 | GLY | p.Gly323Ser | P01130 |
| HOMO SAPIENS | 6 | 12016 | 1hj7 | A | 319 | CYS | p.Cys340Trp | P01130 |
| HOMO SAPIENS | 7 | 12016 | 1hj7 | A | 329 | ARG | p.Arg350Ter | P01130 |
| HOMO SAPIENS | 5 | 12016 | 1hj7 | A | 331 | CYS | p.Cys352Ter | P01130 |
| HOMO SAPIENS | 2 | 12016 | 1hj7 | A | 345 | GLN | p.Gln366Arg | P01130 |
| HOMO SAPIENS | 2 | 1481 | 2i07 | A | 239 | TYR | p.Tyr261Ter | P01024 |
| HOMO SAPIENS | -1 | 1481 | 2i07 | A | 292 | LEU | p.Pro314Leu | P01024 |
| HOMO SAPIENS | 2 | 1481 | 2i07 | A | 326 | ILE | p.Ile348Val | P01024 |
| HOMO SAPIENS | 2 | 1481 | 2i07 | A | 413 | GLU | p.Glu435Lys | P01024 |
| HOMO SAPIENS | 4 | 1481 | 2i07 | A | 447 | GLU | p.Glu469Asp | P01024 |
| HOMO SAPIENS | 2 | 1481 | 2i07 | A | 597 | VAL | p.Val619Met | P01024 |
| HOMO SAPIENS | 5 | 1481 | 2i07 | A | 603 | ILE | p.Ile625Phe | P01024 |
| HOMO SAPIENS | 0 | 1481 | 2i07 | A | 611 | LYS | p.Lys633Arg | P01024 |
| HOMO SAPIENS | 0 | 11123 | 3c5n | A | 315 | GLY | p.Gly368Trp | O00294 |
| HOMO SAPIENS | 8 | 11123 | 3c5n | A | 329 | PHE | p.Phe382Ser | O00294 |
| HOMO SAPIENS | 5 | 11123 | 3c5n | A | 429 | ARG | p.Arg482Trp | O00294 |
| HOMO SAPIENS | 8 | 11123 | 3c5n | A | 438 | PHE | p.Phe491Leu | O00294 |
| HOMO SAPIENS | 3 | 4398 | 3fx0 | B | 1288 | ALA | p.Ala288Gly | Q9Y6K9 |
| HOMO SAPIENS | 7 | 4398 | 3fx0 | B | 1315 | GLU | p.Glu315Ala | Q9Y6K9 |
| HOMO SAPIENS | 8 | 4398 | 3fx0 | B | 1319 | ARG | p.Arg319Gln | Q9Y6K9 |
| HOMO SAPIENS | 6 | 3989 | 3qd6 | S | 83 | CYS | p.Cys83Arg | P25942 |
| HOMO SAPIENS | 1 | 3989 | 3qd6 | S | 124 | SER | p.Ser124Leu | P25942 |
| HOMO SAPIENS | 2 | 24003 | 2m0v | A | 225 | GLU | p.Glu225Lys | O14745 |
| HOMO SAPIENS | 2 | 46 | 4aqd | A | 70 | ASP | p.Asp98Gly | P06276 |
| HOMO SAPIENS | 4 | 46 | 4aqd | A | 115 | GLY | p.Gly143Asp | P06276 |
| HOMO SAPIENS | 3 | 46 | 4aqd | A | 184 | ALA | p.Ala212Val | P06276 |
| HOMO SAPIENS | 0 | 46 | 4aqd | A | 225 | GLY | p.Gly253Ter | P06276 |
| HOMO SAPIENS | 2 | 46 | 4aqd | A | 243 | THR | p.Thr271Met | P06276 |
| HOMO SAPIENS | 2 | 46 | 4aqd | A | 255 | GLU | p.Glu283Asp | P06276 |
| HOMO SAPIENS | 5 | 46 | 4aqd | A | 267 | LYS | p.Lys295Arg | P06276 |
| HOMO SAPIENS | 3 | 46 | 4aqd | A | 271 | GLU | p.Glu299Ter | P06276 |
| HOMO SAPIENS | 2 | 46 | 4aqd | A | 307 | LEU | p.Leu335Pro | P06276 |
| HOMO SAPIENS | 1 | 46 | 4aqd | A | 330 | LEU | p.Leu358Ile | P06276 |
| HOMO SAPIENS | 0 | 46 | 4aqd | A | 497 | GLU | p.Glu525Val | P06276 |
| HOMO SAPIENS | 3 | 786 | 1ols | B | 51 | GLY | p.Gly101Asp | P21953 |
| HOMO SAPIENS | 2 | 786 | 1ols | B | 87 | ALA | p.Ala137Val | P21953 |
| HOMO SAPIENS | 0 | 786 | 1ols | B | 110 | ILE | p.Ile160Ser | P21953 |

| | | | | | | | | |
|---|---|---|---|---|---|---|---|---|
| HOMO SAPIENS | 4 | 786 | 1ols | B | 120 | ARG | p.Arg170Cys | P21953 |
| HOMO SAPIENS | 2 | 786 | 1ols | B | 126 | ASN | p.Asn176Tyr | P21953 |
| HOMO SAPIENS | 0 | 786 | 1ols | B | 133 | ARG | p.Arg183Gly | P21953 |
| HOMO SAPIENS | 0 | 786 | 1ols | B | 133 | ARG | p.Arg183Pro | P21953 |
| HOMO SAPIENS | 0 | 786 | 1ols | B | 135 | PRO | p.Pro185Leu | P21953 |
| HOMO SAPIENS | 3 | 786 | 1ols | B | 156 | HIS | p.His206Tyr | P21953 |
| HOMO SAPIENS | 0 | 786 | 1ols | B | 228 | GLY | p.Gly278Ser | P21953 |
| HOMO SAPIENS | 1 | 786 | 1ols | B | 235 | ARG | p.Arg285Ter | P21953 |
| HOMO SAPIENS | 6 | 786 | 1ols | B | 274 | ARG | p.Arg324Ter | P21953 |
| HOMO SAPIENS | 1 | 831 | 4zt1 | B | 48 | VAL | p.Val202Ile | P12830 |
| HOMO SAPIENS | 6 | 831 | 4zt1 | B | 70 | ARG | p.Arg224Cys | P12830 |
| HOMO SAPIENS | 5 | 831 | 4zt1 | B | 79 | HIS | p.His233Gln | P12830 |
| HOMO SAPIENS | 4 | 831 | 4zt1 | B | 120 | GLY | p.Gly274Ser | P12830 |
| HOMO SAPIENS | 7 | 831 | 4zt1 | B | 187 | TYR | p.Tyr341Ter | P12830 |
| HOMO SAPIENS | 0 | 8497 | 3sla | A | 287 | ASN | p.Asn287Ser | P35222 |
| HOMO SAPIENS | 7 | 23986 | 2ke1 | A | 301 | VAL | p.Val301Met | O43918 |
| HOMO SAPIENS | 0 | 4967 | 1rj7 | B | 334 | ARG | p.Arg334His | Q92838 |
| HOMO SAPIENS | 0 | 4967 | 1rj7 | B | 338 | THR | p.Thr338Met | Q92838 |
| HOMO SAPIENS | 3 | 4967 | 1rj7 | B | 349 | ALA | p.Ala349Thr | Q92838 |
| HOMO SAPIENS | 2 | 4967 | 1rj7 | B | 358 | GLN | p.Gln358Glu | Q92838 |
| HOMO SAPIENS | 1 | 4967 | 1rj7 | B | 375 | LYS | p.Lys375Glu | Q92838 |
| HOMO SAPIENS | 9 | 7623 | 1d4t | A | 32 | ARG | p.Arg32Thr | O60880 |
| HOMO SAPIENS | 6 | 7623 | 1d4t | A | 55 | ARG | p.Arg55Leu | O60880 |
| HOMO SAPIENS | 0 | 7623 | 1d4t | A | 68 | THR | p.Thr68Ile | O60880 |
| HOMO SAPIENS | 6 | 705 | 3idh | A | 59 | PRO | p.Pro59Ser | P35557 |
| HOMO SAPIENS | 5 | 705 | 3idh | A | 124 | ASP | p.Asp124Asn | P35557 |
| HOMO SAPIENS | 4 | 705 | 3idh | A | 131 | SER | p.Ser131Pro | P35557 |
| HOMO SAPIENS | 0 | 705 | 3idh | A | 147 | GLY | p.Gly147Asp | P35557 |
| HOMO SAPIENS | 5 | 705 | 3idh | A | 186 | ARG | p.Arg186Ter | P35557 |
| HOMO SAPIENS | 0 | 705 | 3idh | A | 202 | MET | p.Met202Thr | P35557 |
| HOMO SAPIENS | 2 | 705 | 3idh | A | 210 | MET | p.Met210Lys | P35557 |
| HOMO SAPIENS | 1 | 705 | 3idh | A | 215 | TYR | p.Tyr215Ter | P35557 |
| HOMO SAPIENS | 0 | 705 | 3idh | A | 226 | VAL | p.Val226Met | P35557 |
| HOMO SAPIENS | 9 | 705 | 3idh | A | 228 | THR | p.Thr228Met | P35557 |
| HOMO SAPIENS | 9 | 705 | 3idh | A | 256 | GLU | p.Glu256Lys | P35557 |
| HOMO SAPIENS | 2 | 705 | 3idh | A | 261 | GLY | p.Gly261Arg | P35557 |
| HOMO SAPIENS | 6 | 705 | 3idh | A | 279 | GLU | p.Glu279Gln | P35557 |
| HOMO SAPIENS | 7 | 705 | 3idh | A | 299 | GLY | p.Gly299Arg | P35557 |
| HOMO SAPIENS | 2 | 705 | 3idh | A | 318 | GLY | p.Gly318Trp | P35557 |
| HOMO SAPIENS | 4 | 705 | 3idh | A | 378 | ALA | p.Ala378Thr | P35557 |
| HOMO SAPIENS | 0 | 705 | 3idh | A | 387 | ALA | p.Ala387Glu | P35557 |
| HOMO SAPIENS | 5 | 705 | 3idh | A | 414 | LYS | p.Lys414Glu | P35557 |
| HOMO SAPIENS | 0 | 705 | 3idh | A | 455 | VAL | p.Val455Met | P35557 |
| HOMO SAPIENS | -1 | 705 | 3idh | A | 462 | MET | p.Met462Ile | P35557 |
| HOMO SAPIENS | 1 | 509 | 4nef | A | 22 | LEU | p.Leu22Val | P41181 |
| HOMO SAPIENS | 4 | 509 | 4nef | A | 57 | GLN | p.Gln57Pro | P41181 |
| HOMO SAPIENS | 8 | 509 | 4nef | A | 64 | GLY | p.Gly64Arg | P41181 |
| HOMO SAPIENS | 7 | 509 | 4nef | A | 68 | ASN | p.Asn68Ser | P41181 |
| HOMO SAPIENS | 1 | 509 | 4nef | A | 100 | GLY | p.Gly100Val | P41181 |
| HOMO SAPIENS | 0 | 509 | 4nef | A | 125 | THR | p.Thr125Met | P41181 |
| HOMO SAPIENS | 0 | 509 | 4nef | A | 126 | THR | p.Thr126Met | P41181 |
| HOMO SAPIENS | 4 | 509 | 4nef | A | 146 | PHE | p.Phe146Leu | P41181 |
| HOMO SAPIENS | 0 | 509 | 4nef | A | 175 | GLY | p.Gly175Arg | P41181 |
| HOMO SAPIENS | 5 | 509 | 4nef | A | 181 | CYS | p.Cys181Trp | P41181 |

| | | | | | | | | | |
|---|---|---|---|---|---|---|---|---|---|
| HOMO SAPIENS | 9 | 509 | 4nef | A | | 187 | ARG | p.Arg187Cys | P41181 |
| HOMO SAPIENS | 0 | 509 | 4nef | A | | 190 | ALA | p.Ala190Thr | P41181 |
| HOMO SAPIENS | 0 | 509 | 4nef | A | | 216 | SER | p.Ser216Pro | P41181 |
| HOMO SAPIENS | 4 | 2242 | 1z70 | X | | 1155 | SER | p.Ser155Pro | Q8NBK3 |
| HOMO SAPIENS | 3 | 2242 | 1z70 | X | | 1218 | CYS | p.Cys218Tyr | Q8NBK3 |
| HOMO SAPIENS | 0 | 2242 | 1z70 | X | | 1279 | ALA | p.Ala279Val | Q8NBK3 |
| HOMO SAPIENS | -1 | 2242 | 1z70 | X | | 1336 | OCS | p.Cys336Arg | Q8NBK3 |
| HOMO SAPIENS | 4 | 2242 | 1z70 | X | | 1345 | ARG | p.Arg345Cys | Q8NBK3 |
| HOMO SAPIENS | 2 | 2242 | 1z70 | X | | 1348 | ALA | p.Ala348Pro | Q8NBK3 |
| HOMO SAPIENS | 5 | 2242 | 1z70 | X | | 1349 | ARG | p.Arg349Gln | Q8NBK3 |
| HOMO SAPIENS | 5 | 2242 | 1z70 | X | | 1349 | ARG | p.Arg349Trp | Q8NBK3 |
| HOMO SAPIENS | 3 | 2242 | 1z70 | X | | 1359 | SER | p.Ser359Ter | Q8NBK3 |
| HOMO SAPIENS | 5 | 8269 | 5wrr | B | | 136 | ARG | p.Arg136Ter | Q96MK3 |
| HOMO SAPIENS | 1 | 8269 | 5wrr | B | | 332 | LYS | p.Asn332Lys | Q96MK3 |
| HOMO SAPIENS | 3 | 8269 | 5wrr | B | | 478 | ARG | p.Arg478Ter | Q96MK3 |
| HOMO SAPIENS | 5 | 8269 | 5wrr | B | | 504 | THR | p.Thr504Ile | Q96MK3 |
| HOMO SAPIENS | 5 | 1411 | 4h1s | B | | 221 | SER | p.Ser221Ter | P21589 |
| HOMO SAPIENS | 1 | 1411 | 4h1s | B | | 358 | CYS | p.Cys358Tyr | P21589 |
| HOMO SAPIENS | 2 | 23 | 5vvd | B | | 298 | GLU | p.Glu298Asp | P29474 |
| HOMO SAPIENS | 0 | 19 | 5jlh | E | | 88 | THR | p.Thr89Ile | P63261 |
| HOMO SAPIENS | 6 | 19 | 5jlh | E | | 117 | LYS | p.Lys118Met | P63261 |
| HOMO SAPIENS | 5 | 19 | 5jlh | E | | 263 | PRO | p.Pro264Leu | P63261 |
| HOMO SAPIENS | 1 | 19 | 5jlh | E | | 277 | THR | p.Thr278Ile | P63261 |
| HOMO SAPIENS | 6 | 19 | 5jlh | E | | 331 | PRO | p.Pro332Ala | P63261 |
| HOMO SAPIENS | 5 | 19 | 5jlh | E | | 369 | VAL | p.Val370Ala | P63261 |
| HOMO SAPIENS | 0 | 14306 | 4w4o | C | | 92 | ARG | p.Arg92Ter | P12314 |
| HOMO SAPIENS | 3 | 603 | 1nuh | A | | 4 | THR | p.Thr5Ile | P06744 |
| HOMO SAPIENS | 2 | 603 | 1nuh | A | | 19 | HIS | p.His20Pro | P06744 |
| HOMO SAPIENS | 8 | 603 | 1nuh | A | | 158 | GLY | p.Gly159Ser | P06744 |
| HOMO SAPIENS | 6 | 603 | 1nuh | A | | 223 | THR | p.Thr224Met | P06744 |
| HOMO SAPIENS | 0 | 603 | 1nuh | A | | 338 | LEU | p.Leu339Pro | P06744 |
| HOMO SAPIENS | 6 | 603 | 1nuh | A | | 342 | GLN | p.Gln343Arg | P06744 |
| HOMO SAPIENS | 6 | 603 | 1nuh | A | | 374 | THR | p.Thr375Arg | P06744 |
| HOMO SAPIENS | 4 | 603 | 1nuh | A | | 524 | ILE | p.Ile525Thr | P06744 |
| HOMO SAPIENS | 4 | 603 | 1nuh | A | | 538 | ASP | p.Asp539Asn | P06744 |
| HOMO SAPIENS | 1 | 3107 | 3kzj | A | | 1197 | VAL | p.Val1197Ala | P08603 |
| HOMO SAPIENS | 9 | 1748 | 2noh | A | | 46 | ARG | p.Arg46Gln | O15527 |
| HOMO SAPIENS | 2 | 11750 | 3s4y | A | | 33 | ARG | p.Arg33His | Q9H3S4 |
| HOMO SAPIENS | 2 | 11750 | 3s4y | A | | 40 | LEU | p.Leu40Pro | Q9H3S4 |
| HOMO SAPIENS | 4 | 11750 | 3s4y | A | | 142 | LEU | p.Leu142Phe | Q9H3S4 |
| HOMO SAPIENS | 5 | 11750 | 3s4y | A | | 219 | ASN | p.Asn219Ser | Q9H3S4 |
| HOMO SAPIENS | 1 | 2591 | 5vnj | A | | 382 | PHE | p.Phe382Leu | Q15436 |
| HOMO SAPIENS | 2 | 2591 | 5vnj | A | | 702 | MET | p.Met702Val | Q15436 |
| HOMO SAPIENS | 1 | 2426 | 5m8t | A | | 87 | ARG | p.Arg87Trp | P17643 |
| HOMO SAPIENS | 1 | 2426 | 5m8t | A | | 166 | SER | p.Ser166Ter | P17643 |
| HOMO SAPIENS | 0 | 2426 | 5m8t | A | | 356 | ARG | p.Arg356Gln | P17643 |
| HOMO SAPIENS | 2 | 2426 | 5m8t | A | | 374 | SER | p.Arg374Ter | P17643 |
| HOMO SAPIENS | 6 | 2426 | 5m8t | A | | 378 | ASN | p.Asn378Ser | P17643 |
| HOMO SAPIENS | 6 | 2426 | 5m8t | A | | 382 | LEU | p.Leu382Pro | P17643 |
| HOMO SAPIENS | 7 | 10097 | 4n6v | | 1 | 46 | GLN | p.Gln46Ter | P04080 |
| HOMO SAPIENS | 3 | 10097 | 4n6v | | 1 | 65 | VAL | p.Val65Ile | P04080 |
| HOMO SAPIENS | 8 | 10097 | 4n6v | | 1 | 68 | ARG | p.Arg68Ter | P04080 |
| HOMO SAPIENS | 2 | 859 | 4z9a | A | | 105 | GLN | p.Gln106Arg | P24666 |
| HOMO SAPIENS | 8 | 1420 | 5nan | F | | 95 | SER | p.Ser95Leu | Q96PD4 |

| | | | | | | | | |
|---|---|---|---|---|---|---|---|---|
| HOMO SAPIENS | 6 | 1420 | 5nan | F | 126 | GLU | p.Glu126Gly | Q96PD4 |
| HOMO SAPIENS | 8 | 1420 | 5nan | F | 155 | VAL | p.Val155Ile | Q96PD4 |
| HOMO SAPIENS | 6 | 1420 | 5nan | F | 157 | PRO | p.Pro157Ser | Q96PD4 |
| HOMO SAPIENS | -1 | 1420 | 5nan | F | 161 | HIS | p.His161Arg | Q96PD4 |
| HOMO SAPIENS | 3 | 288 | 5i96 | B | 140 | GLN | p.Arg140Gly | P48735 |
| HOMO SAPIENS | 3 | 288 | 5i96 | B | 140 | GLN | p.Arg140Gln | P48735 |
| HOMO SAPIENS | 2 | 288 | 5i96 | B | 261 | ARG | p.Arg261His | P48735 |
| HOMO SAPIENS | 2 | 288 | 5i96 | B | 435 | THR | p.Thr435Met | P48735 |
| HOMO SAPIENS | 9 | 120 | 1tup | B | 107 | TYR | p.Tyr107His | P04637 |
| HOMO SAPIENS | 8 | 120 | 1tup | B | 110 | ARG | p.Arg110Leu | P04637 |
| HOMO SAPIENS | 5 | 120 | 1tup | B | 120 | LYS | p.Lys120Glu | P04637 |
| HOMO SAPIENS | 4 | 120 | 1tup | B | 133 | MET | p.Met133Thr | P04637 |
| HOMO SAPIENS | 3 | 120 | 1tup | B | 134 | PHE | p.Phe134Cys | P04637 |
| HOMO SAPIENS | 8 | 120 | 1tup | B | 136 | GLN | p.Gln136His | P04637 |
| HOMO SAPIENS | 9 | 120 | 1tup | B | 138 | ALA | p.Ala138Pro | P04637 |
| HOMO SAPIENS | 9 | 120 | 1tup | B | 151 | PRO | p.Pro151Ser | P04637 |
| HOMO SAPIENS | 9 | 120 | 1tup | B | 152 | PRO | p.Pro152Leu | P04637 |
| HOMO SAPIENS | 9 | 120 | 1tup | B | 158 | ARG | p.Arg158Cys | P04637 |
| HOMO SAPIENS | 9 | 120 | 1tup | B | 158 | ARG | p.Arg158His | P04637 |
| HOMO SAPIENS | 9 | 120 | 1tup | B | 175 | ARG | p.Arg175Cys | P04637 |
| HOMO SAPIENS | 9 | 120 | 1tup | B | 175 | ARG | p.Arg175Leu | P04637 |
| HOMO SAPIENS | 8 | 120 | 1tup | B | 181 | ARG | p.Arg181Leu | P04637 |
| HOMO SAPIENS | 9 | 120 | 1tup | B | 189 | ALA | p.Ala189Val | P04637 |
| HOMO SAPIENS | 9 | 120 | 1tup | B | 196 | ARG | p.Arg196Gly | P04637 |
| HOMO SAPIENS | 9 | 120 | 1tup | B | 213 | ARG | p.Arg213Gln | P04637 |
| HOMO SAPIENS | 9 | 120 | 1tup | B | 213 | ARG | p.Arg213Ter | P04637 |
| HOMO SAPIENS | 9 | 120 | 1tup | B | 214 | HIS | p.His214Gln | P04637 |
| HOMO SAPIENS | 9 | 120 | 1tup | B | 220 | TYR | p.Tyr220Cys | P04637 |
| HOMO SAPIENS | 0 | 120 | 1tup | B | 234 | TYR | p.Tyr234Cys | P04637 |
| HOMO SAPIENS | 9 | 120 | 1tup | B | 237 | MET | p.Met237Ile | P04637 |
| HOMO SAPIENS | 0 | 120 | 1tup | B | 238 | CYS | p.Cys238Tyr | P04637 |
| HOMO SAPIENS | 9 | 120 | 1tup | B | 241 | SER | p.Ser241Phe | P04637 |
| HOMO SAPIENS | 9 | 120 | 1tup | B | 242 | CYS | p.Cys242Ser | P04637 |
| HOMO SAPIENS | 9 | 120 | 1tup | B | 245 | GLY | p.Gly245Cys | P04637 |
| HOMO SAPIENS | 9 | 120 | 1tup | B | 245 | GLY | p.Gly245Asp | P04637 |
| HOMO SAPIENS | 9 | 120 | 1tup | B | 248 | ARG | p.Arg248Leu | P04637 |
| HOMO SAPIENS | 9 | 120 | 1tup | B | 248 | ARG | p.Arg248Trp | P04637 |
| HOMO SAPIENS | 9 | 120 | 1tup | B | 251 | ILE | p.Ile251Ser | P04637 |
| HOMO SAPIENS | 9 | 120 | 1tup | B | 252 | LEU | p.Leu252Pro | P04637 |
| HOMO SAPIENS | 9 | 120 | 1tup | B | 258 | GLU | p.Glu258Lys | P04637 |
| HOMO SAPIENS | 8 | 120 | 1tup | B | 272 | VAL | p.Val272Leu | P04637 |
| HOMO SAPIENS | 9 | 120 | 1tup | B | 273 | ARG | p.Arg273Cys | P04637 |
| HOMO SAPIENS | 9 | 120 | 1tup | B | 273 | ARG | p.Arg273His | P04637 |
| HOMO SAPIENS | 9 | 120 | 1tup | B | 277 | CYS | p.Cys277Phe | P04637 |
| HOMO SAPIENS | 9 | 120 | 1tup | B | 280 | ARG | p.Arg280Ile | P04637 |
| HOMO SAPIENS | 9 | 120 | 1tup | B | 282 | ARG | p.Arg282Trp | P04637 |
| HOMO SAPIENS | 9 | 120 | 1tup | B | 283 | ARG | p.Arg283His | P04637 |
| HOMO SAPIENS | 9 | 120 | 1tup | B | 285 | GLU | p.Glu285Val | P04637 |
| HOMO SAPIENS | -1 | 120 | 1tup | B | 292 | LYS | p.Lys292Ile | P04637 |
| HOMO SAPIENS | -1 | 120 | 1tup | B | 298 | GLU | p.Glu298Lys | P04637 |
| HOMO SAPIENS | -1 | 2791 | 4ygb | A | 39 | VAL | p.Val39Ala | P49257 |
| HOMO SAPIENS | 2 | 106 | 4x6j | A | 32 | GLY | p.Gly146Arg | P43235 |
| HOMO SAPIENS | 0 | 106 | 4x6j | A | 76 | GLN | p.Gln190Ter | P43235 |
| HOMO SAPIENS | 7 | 106 | 4x6j | A | 198 | ARG | p.Arg312Ter | P43235 |

| | | | | | | | | |
|---|---|---|---|---|---|---|---|---|
| HOMO SAPIENS | 5 | 490 | 2hlo | B | 172 | LEU | p.Leu202Gln | P02675 |
| HOMO SAPIENS | 0 | 490 | 2hlo | B | 353 | LEU | p.Leu383Arg | P02675 |
| HOMO SAPIENS | 4 | 490 | 2hlo | B | 400 | GLY | p.Gly430Asp | P02675 |
| HOMO SAPIENS | 7 | 490 | 2hlo | B | 448 | ARG | p.Arg478Lys | P02675 |
| HOMO SAPIENS | 1 | 289 | 4p13 | B | 18 | GLU | p.Glu43Lys | P11310 |
| HOMO SAPIENS | 2 | 289 | 4p13 | B | 20 | GLN | p.Gln45Arg | P11310 |
| HOMO SAPIENS | 5 | 289 | 4p13 | B | 28 | ARG | p.Arg53Cys | P11310 |
| HOMO SAPIENS | 1 | 289 | 4p13 | B | 42 | TYR | p.Tyr67His | P11310 |
| HOMO SAPIENS | 1 | 289 | 4p13 | B | 53 | ILE | p.Ile78Thr | P11310 |
| HOMO SAPIENS | 1 | 289 | 4p13 | B | 59 | LEU | p.Leu84Phe | P11310 |
| HOMO SAPIENS | 2 | 289 | 4p13 | B | 96 | THR | p.Thr121Ile | P11310 |
| HOMO SAPIENS | 0 | 289 | 4p13 | B | 124 | MET | p.Met149Ile | P11310 |
| HOMO SAPIENS | 5 | 289 | 4p13 | B | 143 | ASP | p.Asp168Ala | P11310 |
| HOMO SAPIENS | 7 | 289 | 4p13 | B | 168 | THR | p.Thr193Ala | P11310 |
| HOMO SAPIENS | 1 | 289 | 4p13 | B | 170 | GLY | p.Gly195Arg | P11310 |
| HOMO SAPIENS | 2 | 289 | 4p13 | B | 181 | ARG | p.Arg206Cys | P11310 |
| HOMO SAPIENS | 2 | 289 | 4p13 | B | 181 | ARG | p.Arg206His | P11310 |
| HOMO SAPIENS | 8 | 289 | 4p13 | B | 203 | THR | p.Thr228Ile | P11310 |
| HOMO SAPIENS | 2 | 289 | 4p13 | B | 208 | ILE | p.Ile233Asn | P11310 |
| HOMO SAPIENS | 1 | 289 | 4p13 | B | 219 | CYS | p.Cys244Arg | P11310 |
| HOMO SAPIENS | 6 | 289 | 4p13 | B | 220 | SER | p.Ser245Leu | P11310 |
| HOMO SAPIENS | 5 | 289 | 4p13 | B | 228 | GLU | p.Glu253Lys | P11310 |
| HOMO SAPIENS | 2 | 289 | 4p13 | B | 241 | ASP | p.Asp266Gly | P11310 |
| HOMO SAPIENS | 1 | 289 | 4p13 | B | 242 | GLY | p.Gly267Arg | P11310 |
| HOMO SAPIENS | 7 | 289 | 4p13 | B | 256 | ARG | p.Arg281Thr | P11310 |
| HOMO SAPIENS | 4 | 289 | 4p13 | B | 269 | ARG | p.Arg294Thr | P11310 |
| HOMO SAPIENS | 4 | 289 | 4p13 | B | 304 | GLU | p.Lys329Glu | P11310 |
| HOMO SAPIENS | 1 | 289 | 4p13 | B | 324 | ARG | p.Arg349Ter | P11310 |
| HOMO SAPIENS | 3 | 289 | 4p13 | B | 326 | THR | p.Thr351Ile | P11310 |
| HOMO SAPIENS | 7 | 289 | 4p13 | B | 350 | ILE | p.Ile375Thr | P11310 |
| HOMO SAPIENS | 3 | 289 | 4p13 | B | 359 | GLU | p.Glu384Ter | P11310 |
| HOMO SAPIENS | 1 | 4563 | 3i2b | A | 16 | ARG | p.Arg16Cys | Q03393 |
| HOMO SAPIENS | 5 | 4563 | 3i2b | A | 25 | ARG | p.Arg25Gln | Q03393 |
| HOMO SAPIENS | 1 | 4563 | 3i2b | A | 47 | ASN | p.Asn47Asp | Q03393 |
| HOMO SAPIENS | 8 | 4563 | 3i2b | A | 52 | ASN | p.Asn52Ser | Q03393 |
| HOMO SAPIENS | 1 | 4563 | 3i2b | A | 56 | VAL | p.Val56Met | Q03393 |
| HOMO SAPIENS | 1 | 4563 | 3i2b | A | 87 | PRO | p.Pro87Ser | Q03393 |
| HOMO SAPIENS | 1 | 4563 | 3i2b | A | 96 | ASP | p.Asp96Asn | Q03393 |
| HOMO SAPIENS | 0 | 4563 | 3i2b | A | 99 | TYR | p.Tyr99Ter | Q03393 |
| HOMO SAPIENS | 2 | 4563 | 3i2b | A | 116 | ASP | p.Asp116Gly | Q03393 |
| HOMO SAPIENS | 2 | 41 | 3exx | A | 1 | GLY | p.Gly90Cys | P01308 |
| HOMO SAPIENS | 5 | 41 | 3exx | A | 3 | VAL | p.Val92Leu | P01308 |
| HOMO SAPIENS | 4 | 41 | 3exx | A | 7 | CYS | p.Cys96Ser | P01308 |
| HOMO SAPIENS | 9 | 41 | 3exx | A | 19 | TYR | p.Tyr108Cys | P01308 |
| HOMO SAPIENS | 9 | 131 | 4mjo | D | 118 | ASP | p.Asp119Asn | P09467 |
| HOMO SAPIENS | 0 | 131 | 4mjo | D | 163 | GLY | p.Gly164Ser | P09467 |
| HOMO SAPIENS | 0 | 131 | 4mjo | D | 176 | ALA | p.Ala177Asp | P09467 |
| HOMO SAPIENS | 5 | 131 | 4mjo | D | 193 | PHE | p.Phe194Ser | P09467 |
| HOMO SAPIENS | 4 | 131 | 4mjo | D | 217 | ARG | p.Arg218Lys | P09467 |
| HOMO SAPIENS | 4 | 131 | 4mjo | D | 223 | VAL | p.Val224Ile | P09467 |
| HOMO SAPIENS | 4 | 131 | 4mjo | D | 232 | PHE | p.Phe233Ile | P09467 |
| HOMO SAPIENS | 5 | 131 | 4mjo | D | 243 | ARG | p.Arg244Trp | P09467 |
| HOMO SAPIENS | 0 | 131 | 4mjo | D | 283 | PRO | p.Pro284Arg | P09467 |
| HOMO SAPIENS | 0 | 131 | 4mjo | D | 288 | MET | p.Met289Leu | P09467 |

| | | | | | | | | |
|---|---|---|---|---|---|---|---|---|
| HOMO SAPIENS | 0 | 131 | 4mjo | D | 323 | ASP | p.Asp324Asn | P09467 |
| HOMO SAPIENS | -1 | 13122 | 3hfc | A | 583 | ARG | p.Arg583Cys | P51787 |
| HOMO SAPIENS | 1 | 13122 | 3hfc | A | 587 | THR | p.Thr587Met | P51787 |
| HOMO SAPIENS | 0 | 13122 | 3hfc | A | 589 | GLY | p.Gly589Asp | P51787 |
| HOMO SAPIENS | 4 | 3002 | 3nbq | B | 94 | ARG | p.Arg94Cys | Q16831 |
| HOMO SAPIENS | 1 | 3616 | 3dbp | A | 97 | ILE | p.Ile286Asn | P11172 |
| HOMO SAPIENS | 0 | 3616 | 3dbp | A | 240 | GLY | p.Gly429Arg | P11172 |
| HOMO SAPIENS | 0 | 3616 | 3dbp | A | 257 | ILE | p.Ile446Val | P11172 |
| HOMO SAPIENS | 1 | 584 | 1drf | A | 79 | LEU | p.Leu80Phe | P00374 |
| HOMO SAPIENS | 3 | 1609 | 3qa2 | B | 49 | VAL | p.Val49Ile | P50053 |
| HOMO SAPIENS | 0 | 20560 | 5mug | A | 101 | HIS | p.His101Gln | P49638 |
| HOMO SAPIENS | 2 | 20560 | 5mug | A | 120 | ALA | p.Ala120Thr | P49638 |
| HOMO SAPIENS | 3 | 20560 | 5mug | A | 192 | ARG | p.Arg192His | P49638 |
| HOMO SAPIENS | 3 | 20560 | 5mug | A | 221 | ARG | p.Arg221Trp | P49638 |
| HOMO SAPIENS | 4 | 256 | 3vtv | A | 65 | THR | p.Glu50Lys | Q9GZQ8 |
| HOMO SAPIENS | 4 | 256 | 3vtv | A | 113 | VAL | p.Met98Lys | Q9GZQ8 |
| HOMO SAPIENS | 4 | 5397 | 1m61 | A | 192 | ARG | p.Arg192Trp | P43403 |
| HOMO SAPIENS | 5 | 39881 | 1aut | L | 107 | HIS | p.His149Pro | P04070 |
| HOMO SAPIENS | 5 | 4010 | 4uf8 | I | 33 | ALA | p.Ala33Val | Q13501 |
| HOMO SAPIENS | 3 | 313 | 5w49 | A | 98 | PRO | p.Pro98Leu | P23526 |
| HOMO SAPIENS | 2 | 313 | 5w49 | A | 143 | TYR | p.Tyr143Cys | P23526 |
| HOMO SAPIENS | -1 | 1643 | 2m8t | A | 102 | PRO | p.Pro102Leu | P04156 |
| HOMO SAPIENS | -1 | 1643 | 2m8t | A | 105 | PRO | p.Pro105Leu | P04156 |
| HOMO SAPIENS | -1 | 1643 | 2m8t | A | 117 | ALA | p.Ala117Val | P04156 |
| HOMO SAPIENS | 5 | 1643 | 2m8t | A | 129 | MET | p.Met129Val | P04156 |
| HOMO SAPIENS | 5 | 1643 | 2m8t | A | 131 | GLY | p.Gly131Val | P04156 |
| HOMO SAPIENS | 2 | 1643 | 2m8t | A | 142 | GLY | p.Gly142Ser | P04156 |
| HOMO SAPIENS | 3 | 1643 | 2m8t | A | 171 | ASN | p.Asn171Ser | P04156 |
| HOMO SAPIENS | 9 | 1643 | 2m8t | A | 178 | ASP | p.Asp178Asn | P04156 |
| HOMO SAPIENS | 9 | 1643 | 2m8t | A | 180 | VAL | p.Val180Ile | P04156 |
| HOMO SAPIENS | 9 | 1643 | 2m8t | A | 183 | THR | p.Thr183Ala | P04156 |
| HOMO SAPIENS | 9 | 1643 | 2m8t | A | 187 | HIS | p.His187Arg | P04156 |
| HOMO SAPIENS | 8 | 1643 | 2m8t | A | 198 | PHE | p.Phe198Ser | P04156 |
| HOMO SAPIENS | 9 | 1643 | 2m8t | A | 200 | GLU | p.Glu200Lys | P04156 |
| HOMO SAPIENS | 9 | 1643 | 2m8t | A | 208 | ARG | p.Arg208His | P04156 |
| HOMO SAPIENS | 3 | 1643 | 2m8t | A | 210 | VAL | p.Val210Ile | P04156 |
| HOMO SAPIENS | 6 | 1643 | 2m8t | A | 217 | GLN | p.Gln217Arg | P04156 |
| HOMO SAPIENS | 1 | 29 | 5kos | A | 151 | GLY | p.Gly217Arg | P00797 |
| HOMO SAPIENS | 1 | 29 | 5kos | A | 164 | ARG | p.Arg230Lys | P00797 |
| HOMO SAPIENS | 3 | 504 | 4zkv | B | 37 | ARG | p.Arg37Pro | P49773 |
| HOMO SAPIENS | 2 | 504 | 4zkv | B | 62 | GLN | p.Gln62Ter | P49773 |
| HOMO SAPIENS | 2 | 504 | 4zkv | B | 84 | CYS | p.Cys84Arg | P49773 |
| HOMO SAPIENS | 4 | 504 | 4zkv | B | 93 | GLY | p.Gly93Asp | P49773 |
| HOMO SAPIENS | 9 | 504 | 4zkv | B | 112 | HIS | p.His112Asn | P49773 |
| HOMO SAPIENS | 0 | 74 | 5vwo | C | 54 | ASN | p.Asn54Lys | P04181 |
| HOMO SAPIENS | 0 | 74 | 5vwo | C | 55 | TYR | p.Tyr55His | P04181 |
| HOMO SAPIENS | 2 | 74 | 5vwo | C | 90 | GLN | p.Gln90Glu | P04181 |
| HOMO SAPIENS | 6 | 74 | 5vwo | C | 91 | GLY | p.Gly91Glu | P04181 |
| HOMO SAPIENS | 2 | 74 | 5vwo | C | 93 | CYS | p.Cys93Phe | P04181 |
| HOMO SAPIENS | 1 | 74 | 5vwo | C | 154 | ARG | p.Arg154Leu | P04181 |
| HOMO SAPIENS | 0 | 74 | 5vwo | C | 180 | ARG | p.Arg180Thr | P04181 |
| HOMO SAPIENS | 0 | 74 | 5vwo | C | 199 | PRO | p.Pro199Gln | P04181 |
| HOMO SAPIENS | 4 | 74 | 5vwo | C | 209 | TYR | p.Tyr209Ter | P04181 |
| HOMO SAPIENS | 0 | 74 | 5vwo | C | 226 | ALA | p.Ala226Val | P04181 |

| | | | | | | | | |
|---|---|---|---|---|---|---|---|---|
| HOMO SAPIENS | 5 | 74 | 5vwo | C | 233 | GLN | p.Gln233Arg | P04181 |
| HOMO SAPIENS | 5 | 74 | 5vwo | C | 241 | PRO | p.Pro241Leu | P04181 |
| HOMO SAPIENS | 2 | 74 | 5vwo | C | 245 | TYR | p.Tyr245Cys | P04181 |
| HOMO SAPIENS | 6 | 74 | 5vwo | C | 250 | ARG | p.Arg250Pro | P04181 |
| HOMO SAPIENS | 6 | 74 | 5vwo | C | 250 | ARG | p.Arg250Ter | P04181 |
| HOMO SAPIENS | 8 | 74 | 5vwo | C | 267 | THR | p.Thr267Ile | P04181 |
| HOMO SAPIENS | 0 | 74 | 5vwo | C | 270 | ALA | p.Ala270Pro | P04181 |
| HOMO SAPIENS | 0 | 74 | 5vwo | C | 271 | ARG | p.Arg271Lys | P04181 |
| HOMO SAPIENS | 1 | 74 | 5vwo | C | 299 | TYR | p.Tyr299Ter | P04181 |
| HOMO SAPIENS | 0 | 74 | 5vwo | C | 318 | GLU | p.Glu318Lys | P04181 |
| HOMO SAPIENS | 1 | 74 | 5vwo | C | 319 | HIS | p.His319Tyr | P04181 |
| HOMO SAPIENS | 1 | 74 | 5vwo | C | 331 | ARG | p.Arg331Ter | P04181 |
| HOMO SAPIENS | 0 | 74 | 5vwo | C | 332 | VAL | p.Val332Met | P04181 |
| HOMO SAPIENS | 1 | 74 | 5vwo | C | 353 | GLY | p.Gly353Asp | P04181 |
| HOMO SAPIENS | 2 | 74 | 5vwo | C | 375 | GLY | p.Gly375Ala | P04181 |
| HOMO SAPIENS | 0 | 74 | 5vwo | C | 394 | CYS | p.Cys394Arg | P04181 |
| HOMO SAPIENS | 0 | 74 | 5vwo | C | 402 | LEU | p.Leu402Pro | P04181 |
| HOMO SAPIENS | 0 | 74 | 5vwo | C | 436 | ILE | p.Ile436Asn | P04181 |
| HOMO SAPIENS | 7 | 24356 | 4az0 | A | 104 | VAL | p.Val132Met | P10619 |
| HOMO SAPIENS | 4 | 24356 | 4az0 | A | 221 | TYR | p.Tyr249Asn | P10619 |
| HOMO SAPIENS | 3 | 245 | 1rsz | A | 24 | ARG | p.Arg24Ter | P00491 |
| HOMO SAPIENS | 3 | 245 | 1rsz | A | 51 | SER | p.Gly51Ser | P00491 |
| HOMO SAPIENS | 1 | 245 | 1rsz | A | 58 | ARG | p.Arg58Ter | P00491 |
| HOMO SAPIENS | 4 | 245 | 1rsz | A | 89 | GLU | p.Glu89Lys | P00491 |
| HOMO SAPIENS | 6 | 245 | 1rsz | A | 128 | ASP | p.Asp128Gly | P00491 |
| HOMO SAPIENS | 0 | 245 | 1rsz | A | 174 | ALA | p.Ala174Pro | P00491 |
| HOMO SAPIENS | 8 | 245 | 1rsz | A | 192 | TYR | p.Tyr192Cys | P00491 |
| HOMO SAPIENS | 8 | 245 | 1rsz | A | 234 | ARG | p.Arg234Pro | P00491 |
| HOMO SAPIENS | 0 | 37 | 5tby | E | 22 | GLU | p.Glu22Lys | P10916 |
| HOMO SAPIENS | 1 | 37 | 5tby | E | 86 | PHE | p.Phe86Ser | P10916 |
| HOMO SAPIENS | 9 | 71 | 5m9j | A | 12 | GLN | p.Gln36Leu | P03950 |
| HOMO SAPIENS | 3 | 71 | 5m9j | A | 17 | LYS | p.Lys41Glu | P03950 |
| HOMO SAPIENS | 3 | 71 | 5m9j | A | 17 | LYS | p.Lys41Ile | P03950 |
| HOMO SAPIENS | 4 | 71 | 5m9j | A | 28 | SER | p.Ser52Asn | P03950 |
| HOMO SAPIENS | 7 | 71 | 5m9j | A | 31 | ARG | p.Arg55Lys | P03950 |
| HOMO SAPIENS | 9 | 71 | 5m9j | A | 39 | CYS | p.Cys63Trp | P03950 |
| HOMO SAPIENS | 9 | 71 | 5m9j | A | 40 | LYS | p.Lys64Ile | P03950 |
| HOMO SAPIENS | 2 | 71 | 5m9j | A | 60 | GLU | p.Lys84Glu | P03950 |
| HOMO SAPIENS | 0 | 71 | 5m9j | A | 112 | PRO | p.Pro136Leu | P03950 |
| HOMO SAPIENS | 9 | 71 | 5m9j | A | 113 | VAL | p.Val137Ile | P03950 |
| HOMO SAPIENS | 3 | 326 | 5dqi | A | 209 | GLY | p.Gly209Val | Q9Y253 |
| HOMO SAPIENS | 1 | 326 | 5dqi | A | 242 | SER | p.Ser242Ter | Q9Y253 |
| HOMO SAPIENS | 1 | 326 | 5dqi | A | 303 | ARG | p.Arg303Ter | Q9Y253 |
| HOMO SAPIENS | 1 | 8203 | 3iu3 | K | 132 | VAL | p.Val153Ile | P01589 |
| HOMO SAPIENS | 0 | 4995 | 5ffo | H | 234 | THR | p.Thr263Ile | P01137 |
| HOMO SAPIENS | 5 | 38609 | 2ruk | A | 49 | ASP | p.Asp49His | P04637 |
| HOMO SAPIENS | 5 | 38609 | 2ruk | A | 58 | PRO | p.Pro58Arg | P04637 |
| HOMO SAPIENS | 0 | 1607 | 6pah | A | 117 | THR | p.Thr117Ile | P00439 |
| HOMO SAPIENS | 3 | 1607 | 6pah | A | 145 | ASP | p.Asp145Val | P00439 |
| HOMO SAPIENS | 5 | 1607 | 6pah | A | 158 | ARG | p.Arg158Gln | P00439 |
| HOMO SAPIENS | 5 | 1607 | 6pah | A | 158 | ARG | p.Arg158Trp | P00439 |
| HOMO SAPIENS | 5 | 1607 | 6pah | A | 169 | ARG | p.Arg169His | P00439 |
| HOMO SAPIENS | 5 | 1607 | 6pah | A | 169 | ARG | p.Arg169Cys | P00439 |
| HOMO SAPIENS | 3 | 1607 | 6pah | A | 176 | ARG | p.Arg176Leu | P00439 |

| | | | | | | | | |
|---|---|---|---|---|---|---|---|---|
| HOMO SAPIENS | 3 | 1607 | 6pah | A | 177 | VAL | p.Val177Leu | P00439 |
| HOMO SAPIENS | 1 | 1607 | 6pah | A | 194 | LEU | p.Leu194Pro | P00439 |
| HOMO SAPIENS | 1 | 1607 | 6pah | A | 204 | TYR | p.Tyr204Cys | P00439 |
| HOMO SAPIENS | 1 | 1607 | 6pah | A | 204 | TYR | p.Tyr204Ter | P00439 |
| HOMO SAPIENS | 2 | 1607 | 6pah | A | 230 | VAL | p.Val230Ile | P00439 |
| HOMO SAPIENS | 4 | 1607 | 6pah | A | 232 | GLN | p.Gln232Ter | P00439 |
| HOMO SAPIENS | 4 | 1607 | 6pah | A | 241 | ARG | p.Arg241Cys | P00439 |
| HOMO SAPIENS | 4 | 1607 | 6pah | A | 241 | ARG | p.Arg241His | P00439 |
| HOMO SAPIENS | 5 | 1607 | 6pah | A | 243 | ARG | p.Arg243Gln | P00439 |
| HOMO SAPIENS | 5 | 1607 | 6pah | A | 243 | ARG | p.Arg243Ter | P00439 |
| HOMO SAPIENS | 7 | 1607 | 6pah | A | 249 | LEU | p.Leu249Phe | P00439 |
| HOMO SAPIENS | 5 | 1607 | 6pah | A | 252 | ARG | p.Arg252Gly | P00439 |
| HOMO SAPIENS | 5 | 1607 | 6pah | A | 261 | ARG | p.Arg261Ter | P00439 |
| HOMO SAPIENS | 4 | 1607 | 6pah | A | 267 | GLN | p.Gln267Leu | P00439 |
| HOMO SAPIENS | 1 | 1607 | 6pah | A | 272 | GLY | p.Gly272Ter | P00439 |
| HOMO SAPIENS | 2 | 1607 | 6pah | A | 277 | TYR | p.Tyr277Asp | P00439 |
| HOMO SAPIENS | 7 | 1607 | 6pah | A | 280 | GLU | p.Glu280Lys | P00439 |
| HOMO SAPIENS | 8 | 1607 | 6pah | A | 281 | PRO | p.Pro281Leu | P00439 |
| HOMO SAPIENS | 0 | 1607 | 6pah | A | 299 | PHE | p.Phe299Cys | P00439 |
| HOMO SAPIENS | 5 | 1607 | 6pah | A | 300 | ALA | p.Ala300Ser | P00439 |
| HOMO SAPIENS | 5 | 1607 | 6pah | A | 309 | ALA | p.Ala309Asp | P00439 |
| HOMO SAPIENS | 4 | 1607 | 6pah | A | 320 | LYS | p.Lys320Asn | P00439 |
| HOMO SAPIENS | 0 | 1607 | 6pah | A | 345 | ALA | p.Ala345Ser | P00439 |
| HOMO SAPIENS | 0 | 1607 | 6pah | A | 348 | LEU | p.Leu348Val | P00439 |
| HOMO SAPIENS | 6 | 1607 | 6pah | A | 356 | TYR | p.Tyr356Ter | P00439 |
| HOMO SAPIENS | 2 | 1607 | 6pah | A | 380 | THR | p.Thr380Met | P00439 |
| HOMO SAPIENS | 0 | 1607 | 6pah | A | 386 | TYR | p.Tyr386Cys | P00439 |
| HOMO SAPIENS | 5 | 1607 | 6pah | A | 390 | GLU | p.Glu390Gly | P00439 |
| HOMO SAPIENS | 5 | 1607 | 6pah | A | 403 | ALA | p.Ala403Val | P00439 |
| HOMO SAPIENS | 4 | 1607 | 6pah | A | 408 | ARG | p.Arg408Trp | P00439 |
| HOMO SAPIENS | 3 | 1607 | 6pah | A | 413 | ARG | p.Arg413Pro | P00439 |
| HOMO SAPIENS | 4 | 1607 | 6pah | A | 414 | TYR | p.Tyr414Cys | P00439 |
| HOMO SAPIENS | 4 | 1607 | 6pah | A | 414 | TYR | p.Tyr414His | P00439 |
| HOMO SAPIENS | 4 | 1607 | 6pah | A | 415 | ASP | p.Asp415Asn | P00439 |
| HOMO SAPIENS | -1 | 676 | 5dc5 | A | 7 | PRO | p.Pro7Leu | Q9BY41 |
| HOMO SAPIENS | 0 | 676 | 5dc5 | A | 311 | THR | p.Thr311Met | Q9BY41 |
| HOMO SAPIENS | 1 | 124 | 5ftk | E | 95 | ARG | p.Arg95Gly | P55072 |
| HOMO SAPIENS | 1 | 124 | 5ftk | E | 155 | ARG | p.Arg155Cys | P55072 |
| HOMO SAPIENS | 1 | 124 | 5ftk | E | 155 | ARG | p.Arg155His | P55072 |
| HOMO SAPIENS | 1 | 124 | 5ftk | E | 159 | ARG | p.Arg159His | P55072 |
| HOMO SAPIENS | 1 | 124 | 5ftk | E | 159 | ARG | p.Arg159Gly | P55072 |
| HOMO SAPIENS | 0 | 124 | 5ftk | E | 191 | ARG | p.Arg191Gln | P55072 |
| HOMO SAPIENS | 0 | 124 | 5ftk | E | 232 | ALA | p.Ala232Glu | P55072 |
| HOMO SAPIENS | 5 | 5442 | 1fgg | B | 223 | GLY | p.Gly223Ser | O94766 |
| HOMO SAPIENS | 0 | 5442 | 1fgg | B | 277 | ARG | p.Arg277Gln | O94766 |
| HOMO SAPIENS | 0 | 9072 | 2gys | B | 225 | GLU | p.Glu249Gln | P32927 |
| HOMO SAPIENS | 0 | 9072 | 2gys | B | 247 | SER | p.Ser271Ter | P32927 |
| HOMO SAPIENS | 0 | 9072 | 2gys | B | 294 | ASP | p.Asp312Asn | P32927 |
| HOMO SAPIENS | 0 | 1484 | 4zrj | A | 62 | PHE | p.Phe62Ser | P35240 |
| HOMO SAPIENS | 3 | 1484 | 4zrj | A | 205 | MET | p.Met205Val | P35240 |
| HOMO SAPIENS | 3 | 1484 | 4zrj | A | 301 | ILE | p.Ile301Met | P35240 |
| HOMO SAPIENS | 4 | 40923 | 2enq | A | 68 | ILE | p.Ile391Met | P42336 |
| HOMO SAPIENS | 1 | 6179 | 5ofv | A | 125 | THR | p.Thr125Met | O43175 |
| HOMO SAPIENS | 0 | 6179 | 5ofv | A | 135 | ARG | p.Arg135Trp | O43175 |

| | | | | | | | | |
|---|---|---|---|---|---|---|---|---|
| HOMO SAPIENS | 1 | 6179 | 5ofv | A | 163 | ARG | p.Arg163Gln | O43175 |
| HOMO SAPIENS | 9 | 6179 | 5ofv | A | 261 | VAL | p.Val261Met | O43175 |
| HOMO SAPIENS | 7 | 34118 | 2i5o | A | 24 | MET | p.Met647Leu | Q9Y253 |
| HOMO SAPIENS | 8 | 3234 | 5c84 | A | 311 | LYS | p.Lys311Asn | P98170 |
| HOMO SAPIENS | 3 | 3234 | 5c84 | A | 321 | ALA | p.Ala321Gly | P98170 |
| HOMO SAPIENS | -1 | 15129 | 1zy8 | K | 1 | GLY | p.Gly54Ser | O00330 |
| HOMO SAPIENS | -1 | 15129 | 1zy8 | K | 48 | THR | p.Thr101Ala | O00330 |
| HOMO SAPIENS | -1 | 15129 | 1zy8 | K | 105 | GLU | p.Glu158Asp | O00330 |
| HOMO SAPIENS | 8 | 15129 | 1zy8 | K | 136 | ARG | p.Arg189His | O00330 |
| HOMO SAPIENS | -1 | 15129 | 1zy8 | K | 195 | GLN | p.Gln248Ter | O00330 |
| HOMO SAPIENS | 8 | 202 | 4bzb | A | 123 | HIS | p.His123Pro | Q9Y3Z3 |
| HOMO SAPIENS | 4 | 202 | 4bzb | A | 145 | ARG | p.Arg145Ter | Q9Y3Z3 |
| HOMO SAPIENS | 6 | 202 | 4bzb | A | 164 | ARG | p.Arg164Ter | Q9Y3Z3 |
| HOMO SAPIENS | 3 | 202 | 4bzb | A | 201 | ILE | p.Ile201Asn | Q9Y3Z3 |
| HOMO SAPIENS | 0 | 202 | 4bzb | A | 209 | GLY | p.Gly209Ser | Q9Y3Z3 |
| HOMO SAPIENS | 0 | 202 | 4bzb | A | 254 | MET | p.Met254Val | Q9Y3Z3 |
| HOMO SAPIENS | 5 | 202 | 4bzb | A | 531 | ARG | p.Arg531Ser | Q9Y3Z3 |
| HOMO SAPIENS | 6 | 202 | 4bzb | A | 548 | GLN | p.Gln548Ter | Q9Y3Z3 |
| HOMO SAPIENS | 3 | 30947 | 5lgd | A | 90 | PRO | p.Pro90Ser | P16671 |
| HOMO SAPIENS | 6 | 30947 | 5lgd | A | 254 | PHE | p.Phe254Leu | P16671 |
| HOMO SAPIENS | 0 | 30947 | 5lgd | A | 413 | ILE | p.Ile413Leu | P16671 |
| HOMO SAPIENS | 2 | 3906 | 5jtc | A | 735 | ARG | p.Arg735Trp | Q12797 |
| HOMO SAPIENS | 1 | 45 | 5uyx | C | 146 | GLU | p.Glu146Val | P48643 |
| HOMO SAPIENS | 1 | 45 | 5uyx | C | 147 | HIS | p.His147Arg | P48643 |
| HOMO SAPIENS | 0 | 45 | 5uyx | C | 362 | ILE | p.Ile362Met | P48643 |
| HOMO SAPIENS | 8 | 39596 | 1mhn | A | 95 | GLY | p.Gly95Arg | Q16637 |
| HOMO SAPIENS | 4 | 39596 | 1mhn | A | 111 | ALA | p.Ala111Gly | Q16637 |
| HOMO SAPIENS | 9 | 39596 | 1mhn | A | 116 | ILE | p.Ile116Phe | Q16637 |
| HOMO SAPIENS | 1 | 39596 | 1mhn | A | 136 | GLN | p.Gln136Glu | Q16637 |
| HOMO SAPIENS | 1 | 6408 | 5xjy | A | 85 | PRO | p.Pro85Leu | O95477 |
| HOMO SAPIENS | 0 | 6408 | 5xjy | A | 219 | ARG | p.Arg219Lys | O95477 |
| HOMO SAPIENS | 0 | 6408 | 5xjy | A | 230 | ARG | p.Arg230Cys | O95477 |
| HOMO SAPIENS | 1 | 6408 | 5xjy | A | 281 | MET | p.Met281Thr | O95477 |
| HOMO SAPIENS | -1 | 6408 | 5xjy | A | 343 | ALA | p.Ala343Gly | O95477 |
| HOMO SAPIENS | 1 | 6408 | 5xjy | A | 399 | VAL | p.Val399Ala | O95477 |
| HOMO SAPIENS | 1 | 6408 | 5xjy | A | 496 | ARG | p.Arg496Trp | O95477 |
| HOMO SAPIENS | 1 | 6408 | 5xjy | A | 590 | TRP | p.Trp590Ser | O95477 |
| HOMO SAPIENS | 1 | 6408 | 5xjy | A | 638 | ARG | p.Arg638Gln | O95477 |
| HOMO SAPIENS | 4 | 6408 | 5xjy | A | 697 | ALA | p.Ala697Thr | O95477 |
| HOMO SAPIENS | 1 | 6408 | 5xjy | A | 771 | VAL | p.Val771Met | O95477 |
| HOMO SAPIENS | 1 | 6408 | 5xjy | A | 774 | THR | p.Thr774Pro | O95477 |
| HOMO SAPIENS | 1 | 6408 | 5xjy | A | 776 | LYS | p.Lys776Asn | O95477 |
| HOMO SAPIENS | 0 | 6408 | 5xjy | A | 825 | VAL | p.Val825Ile | O95477 |
| HOMO SAPIENS | -1 | 6408 | 5xjy | A | 868 | GLU | p.Glu868Lys | O95477 |
| HOMO SAPIENS | -1 | 6408 | 5xjy | A | 883 | ILE | p.Ile883Met | O95477 |
| HOMO SAPIENS | 0 | 6408 | 5xjy | A | 935 | ASN | p.Asn935Ser | O95477 |
| HOMO SAPIENS | 0 | 6408 | 5xjy | A | 935 | ASN | p.Asn935His | O95477 |
| HOMO SAPIENS | 0 | 6408 | 5xjy | A | 937 | ALA | p.Ala937Val | O95477 |
| HOMO SAPIENS | 0 | 6408 | 5xjy | A | 1018 | ASP | p.Asp1018Gly | O95477 |
| HOMO SAPIENS | 0 | 6408 | 5xjy | A | 1096 | VAL | p.Val1096Ile | O95477 |
| HOMO SAPIENS | 0 | 6408 | 5xjy | A | 1099 | ASP | p.Asp1099Tyr | O95477 |
| HOMO SAPIENS | 0 | 6408 | 5xjy | A | 1172 | GLU | p.Glu1172Asp | O95477 |
| HOMO SAPIENS | 0 | 6408 | 5xjy | A | 1181 | SER | p.Ser1181Phe | O95477 |
| HOMO SAPIENS | 0 | 6408 | 5xjy | A | 1182 | ALA | p.Ala1182Thr | O95477 |

| | | | | | | | | |
|---|---|---|---|---|---|---|---|---|
| HOMO SAPIENS | -1 | 6408 | 5xjy | A | 1289 | ASP | p.Asp1289Asn | O95477 |
| HOMO SAPIENS | 0 | 6408 | 5xjy | A | 1399 | THR | p.Thr1399Ser | O95477 |
| HOMO SAPIENS | 1 | 6408 | 5xjy | A | 1408 | LEU | p.Leu1408Phe | O95477 |
| HOMO SAPIENS | 0 | 6408 | 5xjy | A | 1587 | LYS | p.Lys1587Arg | O95477 |
| HOMO SAPIENS | 1 | 6408 | 5xjy | A | 1674 | VAL | p.Val1674Ile | O95477 |
| HOMO SAPIENS | 1 | 6408 | 5xjy | A | 1680 | ARG | p.Arg1680Trp | O95477 |
| HOMO SAPIENS | 1 | 6408 | 5xjy | A | 1680 | ARG | p.Arg1680Gln | O95477 |
| HOMO SAPIENS | 0 | 6408 | 5xjy | A | 1800 | ASN | p.Asn1800His | O95477 |
| HOMO SAPIENS | -1 | 6408 | 5xjy | A | 1925 | ARG | p.Arg1925Gln | O95477 |
| HOMO SAPIENS | 0 | 6408 | 5xjy | A | 2009 | PHE | p.Phe2009Ser | O95477 |
| HOMO SAPIENS | -1 | 6408 | 5xjy | A | 2243 | ASP | p.Asp2243Glu | O95477 |
| HOMO SAPIENS | 4 | 23760 | 2mut | A | 231 | LEU | p.Phe231Leu | P07992 |
| HOMO SAPIENS | 6 | 1460 | 2hre | B | 96 | ARG | p.Arg96Gln | P22830 |
| HOMO SAPIENS | 8 | 1460 | 2hre | B | 129 | GLY | p.Gly129Arg | P22830 |
| HOMO SAPIENS | 9 | 1460 | 2hre | B | 172 | GLU | p.Glu172Lys | P22830 |
| HOMO SAPIENS | 4 | 1460 | 2hre | B | 285 | GLN | p.Gln285Arg | P22830 |
| HOMO SAPIENS | 5 | 1460 | 2hre | B | 362 | VAL | p.Val362Gly | P22830 |
| HOMO SAPIENS | 6 | 1460 | 2hre | B | 411 | CYS | p.Cys411Gly | P22830 |
| HOMO SAPIENS | 6 | 1460 | 2hre | B | 417 | PHE | p.Phe417Ser | P22830 |
| HOMO SAPIENS | 0 | 2210 | 5bug | A | 19 | ASP | p.Asp19Asn | P60484 |
| HOMO SAPIENS | 0 | 2210 | 5bug | A | 35 | MET | p.Met35Arg | P60484 |
| HOMO SAPIENS | 1 | 2210 | 5bug | A | 61 | HIS | p.His61Asp | P60484 |
| HOMO SAPIENS | 0 | 2210 | 5bug | A | 68 | TYR | p.Tyr68His | P60484 |
| HOMO SAPIENS | 8 | 2210 | 5bug | A | 70 | LEU | p.Leu70Pro | P60484 |
| HOMO SAPIENS | 6 | 2210 | 5bug | A | 93 | HIS | p.His93Arg | P60484 |
| HOMO SAPIENS | 1 | 2210 | 5bug | A | 112 | LEU | p.Leu112Pro | P60484 |
| HOMO SAPIENS | 0 | 2210 | 5bug | A | 121 | ALA | p.Ala121Gly | P60484 |
| HOMO SAPIENS | 3 | 2210 | 5bug | A | 123 | HIS | p.His123Arg | P60484 |
| HOMO SAPIENS | 7 | 2210 | 5bug | A | 124 | CYS | p.Cys124Arg | P60484 |
| HOMO SAPIENS | 9 | 2210 | 5bug | A | 129 | GLY | p.Gly129Glu | P60484 |
| HOMO SAPIENS | 9 | 2210 | 5bug | A | 130 | ARG | p.Arg130Gln | P60484 |
| HOMO SAPIENS | 9 | 2210 | 5bug | A | 130 | ARG | p.Arg130Gly | P60484 |
| HOMO SAPIENS | 0 | 2210 | 5bug | A | 132 | GLY | p.Gly132Val | P60484 |
| HOMO SAPIENS | 0 | 2210 | 5bug | A | 170 | SER | p.Ser170Arg | P60484 |
| HOMO SAPIENS | 1 | 2210 | 5bug | A | 173 | ARG | p.Arg173His | P60484 |
| HOMO SAPIENS | 0 | 2210 | 5bug | A | 217 | VAL | p.Val217Ile | P60484 |
| HOMO SAPIENS | 0 | 2210 | 5bug | A | 241 | PHE | p.Phe241Ser | P60484 |
| HOMO SAPIENS | 0 | 2210 | 5bug | A | 250 | CYS | p.Cys250Ter | P60484 |
| HOMO SAPIENS | 1 | 2210 | 5bug | A | 252 | ASP | p.Asp252Gly | P60484 |
| HOMO SAPIENS | 3 | 123 | 2ibu | A | 93 | ASN | p.Asn93Ser | P24752 |
| HOMO SAPIENS | 0 | 123 | 2ibu | A | 145 | GLN | p.Gln145Glu | P24752 |
| HOMO SAPIENS | 0 | 123 | 2ibu | A | 152 | GLY | p.Gly152Ala | P24752 |
| HOMO SAPIENS | 2 | 123 | 2ibu | A | 158 | ASN | p.Asn158Asp | P24752 |
| HOMO SAPIENS | 2 | 123 | 2ibu | A | 158 | ASN | p.Asn158Ser | P24752 |
| HOMO SAPIENS | 5 | 123 | 2ibu | A | 183 | GLY | p.Gly183Arg | P24752 |
| HOMO SAPIENS | 3 | 123 | 2ibu | A | 272 | GLN | p.Gln272Ter | P24752 |
| HOMO SAPIENS | 3 | 123 | 2ibu | A | 312 | ILE | p.Ile312Thr | P24752 |
| HOMO SAPIENS | 1 | 123 | 2ibu | A | 333 | ALA | p.Ala333Pro | P24752 |
| HOMO SAPIENS | 5 | 123 | 2ibu | A | 379 | GLY | p.Gly379Val | P24752 |
| HOMO SAPIENS | 0 | 123 | 2ibu | A | 380 | ALA | p.Ala380Thr | P24752 |
| HOMO SAPIENS | 2 | 369 | 5nrw | A | 1175 | TYR | p.Tyr1175Cys | Q92793 |
| HOMO SAPIENS | 2 | 830 | 2vc2 | A | 116 | LEU | p.Leu147Val | P08514 |
| HOMO SAPIENS | 0 | 830 | 2vc2 | A | 183 | LEU | p.Leu214Pro | P08514 |
| HOMO SAPIENS | 4 | 830 | 2vc2 | A | 242 | GLY | p.Gly273Asp | P08514 |

| | | | | | | | | |
|---|---|---|---|---|---|---|---|---|
| HOMO SAPIENS | 6 | 830 | 2vc2 | A | 324 | GLU | p.Glu355Lys | P08514 |
| HOMO SAPIENS | -1 | 10353 | 5ejc | E | 989 | ALA | p.Ala989Ser | Q92574 |
| HOMO SAPIENS | 0 | 15448 | 3lgg | A | 21 | GLY | p.Gly47Arg | Q9NZK5 |
| HOMO SAPIENS | 0 | 15448 | 3lgg | A | 21 | GLY | p.Gly47Val | Q9NZK5 |
| HOMO SAPIENS | 9 | 15448 | 3lgg | A | 86 | HIS | p.His112Gln | Q9NZK5 |
| HOMO SAPIENS | 0 | 15448 | 3lgg | A | 143 | ARG | p.Arg169Gln | Q9NZK5 |
| HOMO SAPIENS | 3 | 15448 | 3lgg | A | 221 | ALA | p.Ala247Val | Q9NZK5 |
| HOMO SAPIENS | 0 | 15448 | 3lgg | A | 225 | PRO | p.Pro251Leu | Q9NZK5 |
| HOMO SAPIENS | 0 | 15448 | 3lgg | A | 283 | MET | p.Met309Ile | Q9NZK5 |
| HOMO SAPIENS | 2 | 15448 | 3lgg | A | 334 | THR | p.Thr360Ala | Q9NZK5 |
| HOMO SAPIENS | 7 | 15448 | 3lgg | A | 357 | GLY | p.Gly383Ser | Q9NZK5 |
| HOMO SAPIENS | 0 | 15448 | 3lgg | A | 427 | TYR | p.Tyr453Cys | Q9NZK5 |
| HOMO SAPIENS | 0 | 872 | 2y3i | A | 87 | LEU | p.Leu88Pro | P00558 |
| HOMO SAPIENS | 1 | 872 | 2y3i | A | 163 | ASP | p.Asp164Val | P00558 |
| HOMO SAPIENS | 4 | 872 | 2y3i | A | 205 | ARG | p.Arg206Pro | P00558 |
| HOMO SAPIENS | 4 | 872 | 2y3i | A | 267 | ASP | p.Asp268Asn | P00558 |
| HOMO SAPIENS | 8 | 872 | 2y3i | A | 284 | ASP | p.Asp285Val | P00558 |
| HOMO SAPIENS | 4 | 8 | 5jco | B | 188 | ILE | p.Ile188Leu | Q71U36 |
| HOMO SAPIENS | 7 | 8 | 5jco | B | 263 | PRO | p.Pro263Thr | Q71U36 |
| HOMO SAPIENS | 6 | 8 | 5jco | B | 264 | ARG | p.Arg264Cys | Q71U36 |
| HOMO SAPIENS | 7 | 8 | 5jco | B | 402 | ARG | p.Arg402His | Q71U36 |
| HOMO SAPIENS | 7 | 8 | 5jco | B | 419 | SER | p.Ser419Leu | Q71U36 |
| HOMO SAPIENS | -1 | 880 | 4nwf | A | 2 | THR | p.Thr2Ile | Q06124 |
| HOMO SAPIENS | 2 | 880 | 4nwf | A | 58 | ASN | p.Asn58Ser | Q06124 |
| HOMO SAPIENS | 4 | 880 | 4nwf | A | 61 | ASP | p.Asp61Gly | Q06124 |
| HOMO SAPIENS | 2 | 880 | 4nwf | A | 62 | TYR | p.Tyr62Asp | Q06124 |
| HOMO SAPIENS | 3 | 880 | 4nwf | A | 63 | TYR | p.Tyr63Cys | Q06124 |
| HOMO SAPIENS | 2 | 880 | 4nwf | A | 69 | GLU | p.Glu69Gln | Q06124 |
| HOMO SAPIENS | 2 | 880 | 4nwf | A | 72 | ALA | p.Ala72Gly | Q06124 |
| HOMO SAPIENS | 2 | 880 | 4nwf | A | 72 | ALA | p.Ala72Ser | Q06124 |
| HOMO SAPIENS | 3 | 880 | 4nwf | A | 73 | THR | p.Thr73Ile | Q06124 |
| HOMO SAPIENS | 5 | 880 | 4nwf | A | 76 | GLU | p.Glu76Ala | Q06124 |
| HOMO SAPIENS | 5 | 880 | 4nwf | A | 76 | GLU | p.Glu76Lys | Q06124 |
| HOMO SAPIENS | 3 | 880 | 4nwf | A | 79 | GLN | p.Gln79Arg | Q06124 |
| HOMO SAPIENS | 5 | 880 | 4nwf | A | 152 | ARG | p.Arg152His | Q06124 |
| HOMO SAPIENS | 2 | 880 | 4nwf | A | 268 | GLY | p.Gly268Cys | Q06124 |
| HOMO SAPIENS | 9 | 880 | 4nwf | A | 276 | LYS | p.Lys276Arg | Q06124 |
| HOMO SAPIENS | 9 | 880 | 4nwf | A | 279 | TYR | p.Tyr279Cys | Q06124 |
| HOMO SAPIENS | 5 | 880 | 4nwf | A | 285 | PHE | p.Phe285Ser | Q06124 |
| HOMO SAPIENS | 0 | 880 | 4nwf | A | 308 | ASP | p.Asn308Ser | Q06124 |
| HOMO SAPIENS | 0 | 880 | 4nwf | A | 308 | ASP | p.Asn308Asp | Q06124 |
| HOMO SAPIENS | 5 | 880 | 4nwf | A | 392 | ALA | p.Ala392Thr | Q06124 |
| HOMO SAPIENS | 1 | 880 | 4nwf | A | 411 | THR | p.Thr415Met | Q06124 |
| HOMO SAPIENS | 0 | 880 | 4nwf | A | 461 | ALA | p.Ala465Thr | Q06124 |
| HOMO SAPIENS | 0 | 880 | 4nwf | A | 464 | GLY | p.Gly468Ala | Q06124 |
| HOMO SAPIENS | 1 | 880 | 4nwf | A | 491 | PRO | p.Pro495Ser | Q06124 |
| HOMO SAPIENS | 1 | 880 | 4nwf | A | 502 | SER | p.Ser506Thr | Q06124 |
| HOMO SAPIENS | 8 | 880 | 4nwf | A | 510 | GLN | p.Gln514Pro | Q06124 |
| HOMO SAPIENS | 4 | 1872 | 3wf1 | D | 109 | ARG | p.Arg109Trp | P16278 |
| HOMO SAPIENS | 0 | 1872 | 3wf1 | D | 521 | ARG | p.Arg521Cys | P16278 |
| HOMO SAPIENS | 5 | 910 | 5xte | A | 45 | SER | p.Ser45Phe | O14949 |
| HOMO SAPIENS | 2 | 9017 | 5f0k | A | 382 | LYS | p.Lys382Arg | Q96QK1 |
| HOMO SAPIENS | 1 | 710 | 4lsj | A | 559 | ILE | p.Ile559Asn | P04150 |
| HOMO SAPIENS | 5 | 710 | 4lsj | A | 571 | VAL | p.Val571Ala | P04150 |

| | | | | | | | | |
|---|---|---|---|---|---|---|---|---|
| HOMO SAPIENS | 0 | 710 | 4lsj | A | 641 | ASP | p.Asp641Val | P04150 |
| HOMO SAPIENS | 1 | 710 | 4lsj | A | 678 | ASP | p.Asp678Glu | P04150 |
| HOMO SAPIENS | 2 | 710 | 4lsj | A | 679 | GLY | p.Gly679Ser | P04150 |
| HOMO SAPIENS | 1 | 710 | 4lsj | A | 737 | PHE | p.Phe737Leu | P04150 |
| HOMO SAPIENS | 1 | 710 | 4lsj | A | 747 | ILE | p.Ile747Met | P04150 |
| HOMO SAPIENS | 1 | 710 | 4lsj | A | 753 | LEU | p.Leu753Phe | P04150 |
| HOMO SAPIENS | 1 | 710 | 4lsj | A | 773 | LEU | p.Leu773Pro | P04150 |
| HOMO SAPIENS | 1 | 2145 | 1av1 | B | 71 | PHE | p.Phe95Tyr | P02647 |
| HOMO SAPIENS | 1 | 2145 | 1av1 | B | 128 | GLU | p.Glu152Lys | P02647 |
| HOMO SAPIENS | 1 | 2145 | 1av1 | B | 136 | GLU | p.Glu160Lys | P02647 |
| HOMO SAPIENS | 2 | 2145 | 1av1 | B | 143 | PRO | p.Pro167Arg | P02647 |
| HOMO SAPIENS | 6 | 2145 | 1av1 | B | 164 | ALA | p.Ala188Ser | P02647 |
| HOMO SAPIENS | 7 | 2145 | 1av1 | B | 173 | ARG | p.Arg197Cys | P02647 |
| HOMO SAPIENS | 2 | 2145 | 1av1 | B | 198 | GLU | p.Glu222Lys | P02647 |
| HOMO SAPIENS | -1 | 3432 | 2beu | A | 1 | SER | p.Ser46Ter | P12694 |
| HOMO SAPIENS | 5 | 3432 | 2beu | A | 79 | ARG | p.Arg124Trp | P12694 |
| HOMO SAPIENS | 7 | 3432 | 2beu | A | 114 | ARG | p.Arg159Gln | P12694 |
| HOMO SAPIENS | 6 | 3432 | 2beu | A | 166 | THR | p.Thr211Met | P12694 |
| HOMO SAPIENS | 4 | 3432 | 2beu | A | 175 | ALA | p.Ala220Val | P12694 |
| HOMO SAPIENS | 5 | 3432 | 2beu | A | 204 | GLY | p.Gly249Ser | P12694 |
| HOMO SAPIENS | 4 | 3432 | 2beu | A | 209 | ALA | p.Ala254Asp | P12694 |
| HOMO SAPIENS | 6 | 3432 | 2beu | A | 220 | ARG | p.Arg265Trp | P12694 |
| HOMO SAPIENS | 6 | 3432 | 2beu | A | 234 | TYR | p.Tyr279Cys | P12694 |
| HOMO SAPIENS | 5 | 3432 | 2beu | A | 240 | ALA | p.Ala285Pro | P12694 |
| HOMO SAPIENS | 4 | 3432 | 2beu | A | 245 | GLY | p.Gly290Arg | P12694 |
| HOMO SAPIENS | 5 | 3432 | 2beu | A | 269 | ARG | p.Arg314Ter | P12694 |
| HOMO SAPIENS | 3 | 3432 | 2beu | A | 301 | ARG | p.Arg346Cys | P12694 |
| HOMO SAPIENS | 5 | 3432 | 2beu | A | 359 | ASN | p.Asn404Ser | P12694 |
| HOMO SAPIENS | 4 | 3432 | 2beu | A | 393 | TYR | p.Tyr438Asn | P12694 |
| HOMO SAPIENS | 8 | 4541 | 1rr8 | C | 533 | ASP | p.Asp533Gly | P11387 |
| HOMO SAPIENS | 2 | 3247 | 1tzn | d | 1105 | GLY | p.Gly105Asp | P58335 |
| HOMO SAPIENS | 3 | 3247 | 1tzn | d | 1111 | ARG | p.Arg111Cys | P58335 |
| HOMO SAPIENS | 2 | 3247 | 1tzn | d | 1189 | ILE | p.Ile189Thr | P58335 |
| HOMO SAPIENS | 9 | 467 | 1suv | C | 58 | ASP | p.Asp77Asn | P02787 |
| HOMO SAPIENS | 2 | 467 | 1suv | C | 277 | ASP | p.Asp296Gly | P02787 |
| HOMO SAPIENS | 9 | 6485 | 1xwd | B | 51 | CYS | p.Cys69Gly | P01225 |
| HOMO SAPIENS | 1 | 694 | 1wuu | A | 28 | PRO | p.Pro28Thr | P51570 |
| HOMO SAPIENS | 0 | 694 | 1wuu | A | 32 | VAL | p.Val32Met | P51570 |
| HOMO SAPIENS | 0 | 694 | 1wuu | A | 198 | ALA | p.Ala198Val | P51570 |
| HOMO SAPIENS | 0 | 143 | 4nkx | A | 93 | PHE | p.Phe93Cys | P05093 |
| HOMO SAPIENS | 3 | 143 | 4nkx | A | 96 | ARG | p.Arg96Trp | P05093 |
| HOMO SAPIENS | 3 | 143 | 4nkx | A | 96 | ARG | p.Arg96Gln | P05093 |
| HOMO SAPIENS | 0 | 143 | 4nkx | A | 106 | SER | p.Ser106Pro | P05093 |
| HOMO SAPIENS | 1 | 143 | 4nkx | A | 114 | PHE | p.Phe114Val | P05093 |
| HOMO SAPIENS | 1 | 143 | 4nkx | A | 116 | ASP | p.Asp116Val | P05093 |
| HOMO SAPIENS | 3 | 143 | 4nkx | A | 125 | ARG | p.Arg125Gln | P05093 |
| HOMO SAPIENS | 0 | 143 | 4nkx | A | 201 | TYR | p.Tyr201Asp | P05093 |
| HOMO SAPIENS | 0 | 143 | 4nkx | A | 239 | ARG | p.Arg239Ter | P05093 |
| HOMO SAPIENS | 0 | 143 | 4nkx | A | 329 | TYR | p.Tyr329Asp | P05093 |
| HOMO SAPIENS | 1 | 143 | 4nkx | A | 342 | PRO | p.Pro342Thr | P05093 |
| HOMO SAPIENS | 2 | 143 | 4nkx | A | 347 | ARG | p.Arg347His | P05093 |
| HOMO SAPIENS | 2 | 143 | 4nkx | A | 347 | ARG | p.Arg347Cys | P05093 |
| HOMO SAPIENS | 1 | 143 | 4nkx | A | 358 | ARG | p.Arg358Gln | P05093 |
| HOMO SAPIENS | 0 | 143 | 4nkx | A | 362 | ARG | p.Arg362Cys | P05093 |

| | | | | | | | | |
|---|---|---|---|---|---|---|---|---|
| HOMO SAPIENS | 1 | 143 | 4nkx | A | 406 | TRP | p.Trp406Arg | P05093 |
| HOMO SAPIENS | 0 | 143 | 4nkx | A | 428 | PRO | p.Pro428Leu | P05093 |
| HOMO SAPIENS | 6 | 1441 | 1uuh | A | 46 | ARG | p.Arg46Gly | P16070 |
| HOMO SAPIENS | 3 | 82 | 4wlv | D | 37 | GLY | p.Gly37Arg | P40926 |
| HOMO SAPIENS | 0 | 82 | 4wlv | D | 133 | PRO | p.Pro133Leu | P40926 |
| HOMO SAPIENS | 2 | 392 | 5xtd | A | 56 | SER | p.Ser56Pro | P49821 |
| HOMO SAPIENS | 2 | 392 | 5xtd | A | 59 | ARG | p.Arg59Ter | P49821 |
| HOMO SAPIENS | 2 | 392 | 5xtd | A | 74 | ASP | p.Asp74Asn | P49821 |
| HOMO SAPIENS | 4 | 392 | 5xtd | A | 116 | ASN | p.Asn116Lys | P49821 |
| HOMO SAPIENS | 4 | 392 | 5xtd | A | 117 | ALA | p.Ala117Thr | P49821 |
| HOMO SAPIENS | 5 | 392 | 5xtd | A | 122 | PRO | p.Pro122Leu | P49821 |
| HOMO SAPIENS | 3 | 392 | 5xtd | A | 160 | GLY | p.Gly160Glu | P49821 |
| HOMO SAPIENS | 4 | 392 | 5xtd | A | 214 | GLU | p.Glu214Lys | P49821 |
| HOMO SAPIENS | 2 | 392 | 5xtd | A | 357 | MET | p.Met357Thr | P49821 |
| HOMO SAPIENS | 4 | 392 | 5xtd | A | 423 | THR | p.Thr423Met | P49821 |
| HOMO SAPIENS | 0 | 22593 | 3pdf | A | 8 | TYR | p.Tyr32Ter | P53634 |
| HOMO SAPIENS | 0 | 22593 | 3pdf | A | 15 | TRP | p.Trp39Ser | P53634 |
| HOMO SAPIENS | 0 | 22593 | 3pdf | A | 103 | HIS | p.His127Pro | P53634 |
| HOMO SAPIENS | 9 | 22593 | 3pdf | A | 228 | GLN | p.Gln252Leu | P53634 |
| HOMO SAPIENS | 3 | 22593 | 3pdf | A | 262 | GLN | p.Gln286Arg | P53634 |
| HOMO SAPIENS | 4 | 22593 | 3pdf | A | 277 | GLY | p.Gly301Ser | P53634 |
| HOMO SAPIENS | 0 | 22593 | 3pdf | A | 323 | TYR | p.Tyr347Cys | P53634 |
| HOMO SAPIENS | 5 | 3461 | 4c2a | A | 1306 | GLN | p.Arg1306Trp | P04275 |
| HOMO SAPIENS | 5 | 3461 | 4c2a | A | 1374 | ARG | p.Arg1374Cys | P04275 |
| HOMO SAPIENS | 5 | 3461 | 4c2a | A | 1379 | ARG | p.Arg1379Cys | P04275 |
| HOMO SAPIENS | 4 | 3461 | 4c2a | A | 1380 | ILE | p.Ile1380Val | P04275 |
| HOMO SAPIENS | 0 | 3461 | 4c2a | A | 1381 | ALA | p.Thr1381Ala | P04275 |
| HOMO SAPIENS | 5 | 3461 | 4c2a | A | 1399 | ARG | p.Arg1399His | P04275 |
| HOMO SAPIENS | 5 | 3461 | 4c2a | A | 1435 | ASN | p.Asn1435Ser | P04275 |
| HOMO SAPIENS | 0 | 4533 | 5uj7 | C | 174 | TYR | p.Tyr174Cys | O43929 |
| HOMO SAPIENS | 0 | 4533 | 5uj7 | C | 202 | LEU | p.Leu202Val | O43929 |
| HOMO SAPIENS | 0 | 1374 | 3j07 | O | 39 | PRO | p.Pro39Leu | P02511 |
| HOMO SAPIENS | 0 | 1374 | 3j07 | O | 107 | ARG | p.Arg107His | P02511 |
| HOMO SAPIENS | 0 | 1374 | 3j07 | O | 157 | ARG | p.Arg157His | P02511 |
| HOMO SAPIENS | -1 | 942 | 1s9i | B | 57 | PHE | p.Phe57Cys | P36507 |
| HOMO SAPIENS | -1 | 942 | 1s9i | B | 57 | PHE | p.Phe57Val | P36507 |
| HOMO SAPIENS | 2 | 942 | 1s9i | B | 128 | PRO | p.Pro128Gln | P36507 |
| HOMO SAPIENS | 1 | 942 | 1s9i | B | 134 | TYR | p.Tyr134His | P36507 |
| HOMO SAPIENS | 1 | 942 | 1s9i | B | 188 | HIS | p.His188Gln | P36507 |
| HOMO SAPIENS | 0 | 942 | 1s9i | B | 262 | VAL | p.Val262Phe | P36507 |
| HOMO SAPIENS | 0 | 942 | 1s9i | B | 263 | GLY | p.Gly263Arg | P36507 |
| HOMO SAPIENS | -1 | 942 | 1s9i | B | 298 | PRO | p.Pro298Leu | P36507 |
| HOMO SAPIENS | -1 | 942 | 1s9i | B | 305 | VAL | p.Val305Ile | P36507 |
| HOMO SAPIENS | 0 | 3245 | 1fbv | A | 367 | GLN | p.Gln367Pro | P22681 |
| HOMO SAPIENS | 0 | 3245 | 1fbv | A | 371 | TYR | p.Tyr371His | P22681 |
| HOMO SAPIENS | 1 | 3245 | 1fbv | A | 382 | LYS | p.Lys382Glu | P22681 |
| HOMO SAPIENS | 2 | 3245 | 1fbv | A | 390 | ASP | p.Asp390Tyr | P22681 |
| HOMO SAPIENS | 9 | 3245 | 1fbv | A | 396 | CYS | p.Cys396Arg | P22681 |
| HOMO SAPIENS | 7 | 3245 | 1fbv | A | 420 | ARG | p.Arg420Gln | P22681 |
| HOMO SAPIENS | 1 | 3245 | 1fbv | A | 429 | ILE | p.Ile429Met | P22681 |
| HOMO SAPIENS | 0 | 2604 | 2pbd | P | 111 | LEU | p.Leu112Met | P07737 |
| HOMO SAPIENS | 3 | 2604 | 2pbd | P | 116 | GLU | p.Glu117Gly | P07737 |
| HOMO SAPIENS | 4 | 2013 | 3b7w | A | 472 | ARG | p.Arg472Gly | Q08AH3 |
| HOMO SAPIENS | 0 | 24346 | 3sh5 | A | 85 | ARG | p.Arg4281His | P98160 |

| | | | | | | | | |
|---|---|---|---|---|---|---|---|---|
| HOMO SAPIENS | 0 | 24346 | 3sh5 | A | 122 | ARG | p.Arg4318Trp | P98160 |
| HOMO SAPIENS | 0 | 24346 | 3sh5 | A | 135 | SER | p.Ser4331Asn | P98160 |
| HOMO SAPIENS | 0 | 24346 | 3sh5 | A | 136 | VAL | p.Val4332Ile | P98160 |
| HOMO SAPIENS | 0 | 707 | 4p2g | A | 26 | MET | p.Met26Ile | Q99497 |
| HOMO SAPIENS | 5 | 707 | 4p2g | A | 39 | ALA | p.Ala39Ser | Q99497 |
| HOMO SAPIENS | 5 | 707 | 4p2g | A | 64 | GLU | p.Glu64Asp | Q99497 |
| HOMO SAPIENS | 4 | 707 | 4p2g | A | 149 | ASP | p.Asp149Ala | Q99497 |
| HOMO SAPIENS | 6 | 707 | 4p2g | A | 163 | GLU | p.Glu163Lys | Q99497 |
| HOMO SAPIENS | 6 | 707 | 4p2g | A | 166 | LEU | p.Leu166Pro | Q99497 |
| HOMO SAPIENS | 3 | 5121 | 4xss | E | 31 | GLY | p.Gly58Arg | P06213 |
| HOMO SAPIENS | 2 | 5121 | 4xss | E | 119 | ILE | p.Ile146Met | P06213 |
| HOMO SAPIENS | 8 | 5121 | 4xss | E | 209 | HIS | p.His236Arg | P06213 |
| HOMO SAPIENS | 2 | 5121 | 4xss | E | 234 | ASP | p.Asp261Glu | P06213 |
| HOMO SAPIENS | 1 | 5121 | 4xss | E | 276 | GLN | p.Gln303His | P06213 |
| HOMO SAPIENS | 1 | 27790 | 5erp | B | 140 | THR | p.Thr275Met | Q02487 |
| HOMO SAPIENS | 0 | 27790 | 5erp | B | 150 | ILE | p.Ile285Thr | Q02487 |
| HOMO SAPIENS | 1 | 27790 | 5erp | B | 205 | THR | p.Thr340Ala | Q02487 |
| HOMO SAPIENS | 1 | 27790 | 5erp | B | 223 | THR | p.Thr358Ile | Q02487 |
| HOMO SAPIENS | 0 | 27790 | 5erp | B | 301 | GLY | p.Gly436Ala | Q02487 |
| HOMO SAPIENS | -1 | 27790 | 5erp | B | 315 | ARG | p.Arg450Ser | Q02487 |
| HOMO SAPIENS | 0 | 27790 | 5erp | B | 369 | SER | p.Ser504Gly | Q02487 |
| HOMO SAPIENS | 0 | 27790 | 5erp | B | 385 | ILE | p.Ile520Thr | Q02487 |
| HOMO SAPIENS | 0 | 27790 | 5erp | B | 461 | ALA | p.Ala596Val | Q02487 |
| HOMO SAPIENS | 0 | 27790 | 5erp | B | 503 | GLN | p.Gln638His | Q02487 |
| HOMO SAPIENS | 5 | 6599 | 4nsc | A | 129 | ARG | p.Arg129Leu | Q9BPX6 |
| HOMO SAPIENS | -1 | 6599 | 4nsc | A | 183 | GLN | p.Gln183Ter | Q9BPX6 |
| HOMO SAPIENS | 0 | 20420 | 3ghm | A | 398 | ARG | p.Arg398His | Q76LX8 |
| HOMO SAPIENS | 0 | 20420 | 3ghm | A | 448 | GLN | p.Gln448Glu | Q76LX8 |
| HOMO SAPIENS | 1 | 20420 | 3ghm | A | 449 | GLN | p.Gln449Ter | Q76LX8 |
| HOMO SAPIENS | 0 | 20420 | 3ghm | A | 456 | GLN | p.Gln456His | Q76LX8 |
| HOMO SAPIENS | 1 | 20420 | 3ghm | A | 528 | ARG | p.Arg528Gly | Q76LX8 |
| HOMO SAPIENS | 0 | 20420 | 3ghm | A | 618 | PRO | p.Pro618Ala | Q76LX8 |
| HOMO SAPIENS | 0 | 20420 | 3ghm | A | 625 | ARG | p.Arg625His | Q76LX8 |
| HOMO SAPIENS | 0 | 275 | 4rmz | A | 293 | GLN | p.Gln293Ter | Q9NWZ3 |
| HOMO SAPIENS | 0 | 275 | 4rmz | A | 428 | ALA | p.Ala428Thr | Q9NWZ3 |
| HOMO SAPIENS | 2 | 10458 | 3puf | B | 138 | LEU | p.Leu138Phe | Q5TBB1 |
| HOMO SAPIENS | 6 | 10458 | 3puf | B | 177 | ALA | p.Ala177Thr | Q5TBB1 |
| HOMO SAPIENS | -1 | 10458 | 3puf | B | 185 | VAL | p.Val185Gly | Q5TBB1 |
| HOMO SAPIENS | 0 | 26648 | 2n90 | A | 36 | ARG | p.Arg444Gln | P04629 |
| HOMO SAPIENS | 2 | 977 | 1dnw | D | 166 | ALA | p.Ala332Val | P05164 |
| HOMO SAPIENS | 4 | 977 | 1dnw | D | 333 | ARG | p.Arg499Cys | P05164 |
| HOMO SAPIENS | 1 | 977 | 1dnw | D | 403 | ARG | p.Arg569Trp | P05164 |
| HOMO SAPIENS | 0 | 25704 | 3ajb | A | 82 | GLN | p.Gln82Arg | P56589 |
| HOMO SAPIENS | 3 | 255 | 3t1p | A | 38 | TYR | p.Tyr62Ter | P01009 |
| HOMO SAPIENS | 3 | 255 | 3t1p | A | 39 | ARG | p.Arg63Cys | P01009 |
| HOMO SAPIENS | 0 | 255 | 3t1p | A | 101 | ARG | p.Arg125His | P01009 |
| HOMO SAPIENS | 0 | 255 | 3t1p | A | 212 | GLN | p.Gln236Arg | P01009 |
| HOMO SAPIENS | 0 | 255 | 3t1p | A | 213 | VAL | p.Val237Ala | P01009 |
| HOMO SAPIENS | 1 | 255 | 3t1p | A | 269 | HIS | p.His293Gln | P01009 |
| HOMO SAPIENS | 8 | 255 | 3t1p | A | 341 | ASP | p.Asp365Asn | P01009 |
| HOMO SAPIENS | 9 | 255 | 3t1p | A | 342 | GLU | p.Glu366Lys | P01009 |
| HOMO SAPIENS | 0 | 255 | 3t1p | A | 369 | PRO | p.Pro393Ser | P01009 |
| HOMO SAPIENS | 0 | 255 | 3t1p | A | 376 | GLU | p.Glu400Asp | P01009 |
| HOMO SAPIENS | -1 | 5175 | 3zxi | A | 35 | ALA | p.Ala35Asp | Q9Y2Z4 |

| | | | | | | | | | |
|---|---|---|---|---|---|---|---|---|---|
| HOMO SAPIENS | 0 | 5175 | 3zxi | A | | 52 | PHE | p.Phe52Leu | Q9Y2Z4 |
| HOMO SAPIENS | -1 | 5175 | 3zxi | A | | 68 | GLY | p.Gly68Ser | Q9Y2Z4 |
| HOMO SAPIENS | 0 | 5175 | 3zxi | A | | 179 | LYS | p.Lys179Gln | Q9Y2Z4 |
| HOMO SAPIENS | 0 | 5175 | 3zxi | A | | 191 | GLY | p.Gly191Val | Q9Y2Z4 |
| HOMO SAPIENS | 0 | 5175 | 3zxi | A | | 251 | ILE | p.Ile251Val | Q9Y2Z4 |
| HOMO SAPIENS | 2 | 5175 | 3zxi | A | | 295 | ASP | p.Asp295Asn | Q9Y2Z4 |
| HOMO SAPIENS | 0 | 5175 | 3zxi | A | | 311 | ASP | p.Asp311Glu | Q9Y2Z4 |
| HOMO SAPIENS | 0 | 5175 | 3zxi | A | | 360 | ARG | p.Arg360Ter | Q9Y2Z4 |
| HOMO SAPIENS | 1 | 3016 | 3exg | | 3 | 4 | ASP | p.Asp33His | P08559 |
| HOMO SAPIENS | 1 | 3016 | 3exg | | 3 | 176 | PHE | p.Phe205Leu | P08559 |
| HOMO SAPIENS | 1 | 3016 | 3exg | | 3 | 214 | TYR | p.Tyr243Asn | P08559 |
| HOMO SAPIENS | 1 | 3016 | 3exg | | 3 | 229 | ASP | p.Asp258Ala | P08559 |
| HOMO SAPIENS | 0 | 3016 | 3exg | | 3 | 233 | VAL | p.Val262Ile | P08559 |
| HOMO SAPIENS | 1 | 3016 | 3exg | | 3 | 234 | ARG | p.Arg263Gly | P08559 |
| HOMO SAPIENS | 0 | 3016 | 3exg | | 3 | 253 | MET | p.Met282Leu | P08559 |
| HOMO SAPIENS | 7 | 3016 | 3exg | | 3 | 259 | ARG | p.Arg288His | P08559 |
| HOMO SAPIENS | -1 | 3016 | 3exg | | 3 | 273 | ARG | p.Arg302Cys | P08559 |
| HOMO SAPIENS | 2 | 3016 | 3exg | | 3 | 286 | ASP | p.Asp315Asn | P08559 |
| HOMO SAPIENS | 1 | 3016 | 3exg | | 3 | 349 | ARG | p.Arg378His | P08559 |
| HOMO SAPIENS | 5 | 6729 | 4a0l | E | | 572 | ARG | p.Arg572Cys | Q13620 |
| HOMO SAPIENS | 0 | 1067 | 4kyo | A | | 9 | THR | p.Thr9Asn | P21549 |
| HOMO SAPIENS | 0 | 1067 | 4kyo | A | | 11 | PRO | p.Pro11Leu | P21549 |
| HOMO SAPIENS | 0 | 1067 | 4kyo | A | | 22 | ASN | p.Asn22Ser | P21549 |
| HOMO SAPIENS | 4 | 1067 | 4kyo | A | | 25 | LEU | p.Leu25Arg | P21549 |
| HOMO SAPIENS | 7 | 1067 | 4kyo | A | | 36 | ARG | p.Arg36His | P21549 |
| HOMO SAPIENS | 3 | 1067 | 4kyo | A | | 41 | GLY | p.Gly41Arg | P21549 |
| HOMO SAPIENS | 0 | 1067 | 4kyo | A | | 42 | GLY | p.Gly42Glu | P21549 |
| HOMO SAPIENS | 3 | 1067 | 4kyo | A | | 56 | ILE | p.Ile56Asn | P21549 |
| HOMO SAPIENS | 3 | 1067 | 4kyo | A | | 59 | GLU | p.Glu59Lys | P21549 |
| HOMO SAPIENS | 3 | 1067 | 4kyo | A | | 66 | TYR | p.Tyr66Ter | P21549 |
| HOMO SAPIENS | 3 | 1067 | 4kyo | A | | 111 | ARG | p.Arg111Leu | P21549 |
| HOMO SAPIENS | 7 | 1067 | 4kyo | A | | 118 | ARG | p.Arg118His | P21549 |
| HOMO SAPIENS | 3 | 1067 | 4kyo | A | | 122 | ARG | p.Arg122Ter | P21549 |
| HOMO SAPIENS | 3 | 1067 | 4kyo | A | | 124 | HIS | p.His124Pro | P21549 |
| HOMO SAPIENS | 0 | 1067 | 4kyo | A | | 152 | PHE | p.Phe152Ile | P21549 |
| HOMO SAPIENS | 1 | 1067 | 4kyo | A | | 156 | GLY | p.Gly156Arg | P21549 |
| HOMO SAPIENS | 0 | 1067 | 4kyo | A | | 161 | GLY | p.Gly161Arg | P21549 |
| HOMO SAPIENS | 3 | 1067 | 4kyo | A | | 166 | LEU | p.Leu166Pro | P21549 |
| HOMO SAPIENS | 5 | 1067 | 4kyo | A | | 170 | GLY | p.Gly170Arg | P21549 |
| HOMO SAPIENS | 5 | 1067 | 4kyo | A | | 186 | ALA | p.Ala186Val | P21549 |
| HOMO SAPIENS | 3 | 1067 | 4kyo | A | | 187 | SER | p.Ser187Phe | P21549 |
| HOMO SAPIENS | 3 | 1067 | 4kyo | A | | 197 | ARG | p.Arg197Pro | P21549 |
| HOMO SAPIENS | 5 | 1067 | 4kyo | A | | 201 | ASP | p.Asp201Glu | P21549 |
| HOMO SAPIENS | 0 | 1067 | 4kyo | A | | 202 | ILE | p.Ile202Asn | P21549 |
| HOMO SAPIENS | 1 | 1067 | 4kyo | A | | 205 | SER | p.Ser205Pro | P21549 |
| HOMO SAPIENS | 1 | 1067 | 4kyo | A | | 205 | SER | p.Ser205Ter | P21549 |
| HOMO SAPIENS | 2 | 1067 | 4kyo | A | | 216 | GLY | p.Gly216Arg | P21549 |
| HOMO SAPIENS | 5 | 1067 | 4kyo | A | | 233 | ARG | p.Arg233Cys | P21549 |
| HOMO SAPIENS | 5 | 1067 | 4kyo | A | | 244 | ILE | p.Ile244Thr | P21549 |
| HOMO SAPIENS | 5 | 1067 | 4kyo | A | | 244 | ILE | p.Ile244Met | P21549 |
| HOMO SAPIENS | 4 | 1067 | 4kyo | A | | 253 | CYS | p.Cys253Arg | P21549 |
| HOMO SAPIENS | 7 | 1067 | 4kyo | A | | 274 | GLU | p.Glu274Asp | P21549 |
| HOMO SAPIENS | 7 | 1067 | 4kyo | A | | 279 | ILE | p.Ile279Thr | P21549 |
| HOMO SAPIENS | 6 | 1067 | 4kyo | A | | 295 | ALA | p.Ala295Thr | P21549 |

| | | | | | | | | |
|---|---|---|---|---|---|---|---|---|
| HOMO SAPIENS | 5 | 1067 | 4kyo | A | 340 | ILE | p.Ile340Met | P21549 |
| HOMO SAPIENS | 1 | 790 | 2ric | C | 270 | SER | p.Ser290Thr | P35247 |
| HOMO SAPIENS | 0 | 790 | 2ric | C | 289 | GLU | p.Glu309Lys | P35247 |
| HOMO SAPIENS | 1 | 212 | 5sxe | A | 545 | GLU | p.Glu545Lys | P42336 |
| HOMO SAPIENS | 3 | 212 | 5sxe | A | 546 | GLN | p.Gln546Glu | P42336 |
| HOMO SAPIENS | 2 | 212 | 5sxe | A | 1047 | HIS | p.His1047Leu | P42336 |
| HOMO SAPIENS | 7 | 152 | 2zqq | F | 197 | ARG | p.Arg197Ter | Q13825 |
| HOMO SAPIENS | 0 | 152 | 2zqq | F | 248 | ALA | p.Ala248Thr | Q13825 |
| HOMO SAPIENS | 0 | 152 | 2zqq | F | 275 | ALA | p.Ala275Val | Q13825 |
| HOMO SAPIENS | 5 | 152 | 2zqq | F | 331 | LYS | p.Lys331Ter | Q13825 |
| HOMO SAPIENS | 6 | 17801 | 2j8l | A | 283 | PHE | p.Phe301Leu | P03951 |
| HOMO SAPIENS | 3 | 17801 | 2j8l | A | 308 | ARG | p.Arg326Cys | P03951 |
| HOMO SAPIENS | 0 | 17801 | 2j8l | A | 321 | CYS | p.Cys339Phe | P03951 |
| HOMO SAPIENS | 3 | 17801 | 2j8l | A | 351 | TYR | p.Tyr369Ter | P03951 |
| HOMO SAPIENS | 7 | 1802 | 5db0 | A | 22 | LEU | p.Leu22Arg | O00255 |
| HOMO SAPIENS | 7 | 1802 | 5db0 | A | 26 | GLU | p.Glu26Lys | O00255 |
| HOMO SAPIENS | 7 | 1802 | 5db0 | A | 139 | HIS | p.His139Asp | O00255 |
| HOMO SAPIENS | 0 | 1802 | 5db0 | A | 171 | ARG | p.Arg176Gln | O00255 |
| HOMO SAPIENS | 0 | 1802 | 5db0 | A | 184 | VAL | p.Val189Glu | O00255 |
| HOMO SAPIENS | 7 | 1802 | 5db0 | A | 255 | GLU | p.Glu260Lys | O00255 |
| HOMO SAPIENS | 4 | 1802 | 5db0 | A | 418 | ASP | p.Asp423Asn | O00255 |
| HOMO SAPIENS | 0 | 1802 | 5db0 | A | 436 | TRP | p.Trp441Arg | O00255 |
| HOMO SAPIENS | -1 | 1802 | 5db0 | A | 541 | ALA | p.Thr546Ala | O00255 |
| HOMO SAPIENS | 4 | 1802 | 5db0 | A | 552 | THR | p.Thr557Ser | O00255 |
| HOMO SAPIENS | 0 | 3450 | 2yuz | A | 36 | PRO | p.Pro461Gln | Q00872 |
| HOMO SAPIENS | 1 | 3450 | 2yuz | A | 56 | GLN | p.His481Gln | Q00872 |
| HOMO SAPIENS | 2 | 3450 | 2yuz | A | 68 | ALA | p.Ala493Val | Q00872 |
| HOMO SAPIENS | 0 | 3211 | 3rqg | A | 72 | VAL | p.Val72Leu | Q9BUL8 |
| HOMO SAPIENS | 2 | 4168 | 4q4f | A | 121 | ARG | p.Arg121Ter | Q14108 |
| HOMO SAPIENS | 2 | 4168 | 4q4f | A | 121 | ARG | p.Arg121Gln | Q14108 |
| HOMO SAPIENS | 2 | 4168 | 4q4f | A | 159 | MET | p.Met159Val | Q14108 |
| HOMO SAPIENS | 4 | 4168 | 4q4f | A | 363 | HIS | p.His363Asn | Q14108 |
| HOMO SAPIENS | 2 | 4168 | 4q4f | A | 396 | VAL | p.Val396Ile | Q14108 |
| HOMO SAPIENS | 0 | 1448 | 1y97 | A | 87 | ARG | p.Arg130Ter | Q9BQ50 |
| HOMO SAPIENS | 0 | 3444 | 4iwp | A | 151 | SER | p.Ser151Phe | Q9UHD2 |
| HOMO SAPIENS | 0 | 3444 | 4iwp | A | 401 | LYS | p.Lys401Glu | Q9UHD2 |
| HOMO SAPIENS | 0 | 116 | 1zy8 | B | 194 | GLY | p.Gly229Cys | P09622 |
| HOMO SAPIENS | 0 | 116 | 1zy8 | B | 255 | ASP | p.Asp290Gly | P09622 |
| HOMO SAPIENS | 2 | 116 | 1zy8 | B | 326 | MET | p.Met361Val | P09622 |
| HOMO SAPIENS | 0 | 116 | 1zy8 | B | 340 | GLU | p.Glu375Lys | P09622 |
| HOMO SAPIENS | 2 | 116 | 1zy8 | B | 358 | ILE | p.Ile393Thr | P09622 |
| HOMO SAPIENS | 1 | 116 | 1zy8 | B | 453 | PRO | p.Pro488Leu | P09622 |
| HOMO SAPIENS | 7 | 127 | 4qo7 | B | 160 | SER | p.Ser161Arg | P00338 |
| HOMO SAPIENS | 0 | 2025 | 1wrl | A | 8 | ALA | p.Ala8Val | P63316 |
| HOMO SAPIENS | 8 | 2025 | 1wrl | A | 24 | PHE | p.Phe24Leu | P63316 |
| HOMO SAPIENS | 0 | 2025 | 1wrl | A | 84 | SER | p.Cys84Tyr | P63316 |
| HOMO SAPIENS | 8 | 3791 | 1mhd | B | 93 | ARG | p.Arg93Ter | P84022 |
| HOMO SAPIENS | 0 | 1575 | 4phc | A | 205 | GLY | p.Gly205Asp | P12081 |
| HOMO SAPIENS | 0 | 1575 | 4phc | A | 454 | TYR | p.Tyr454Ser | P12081 |
| HOMO SAPIENS | 0 | 39620 | 1ozs | B | 135 | ARG | p.Arg136Gln | P19429 |
| HOMO SAPIENS | 5 | 39620 | 1ozs | B | 140 | ARG | p.Arg141Trp | P19429 |
| HOMO SAPIENS | 5 | 39620 | 1ozs | B | 142 | THR | p.Thr143Asn | P19429 |
| HOMO SAPIENS | 5 | 39620 | 1ozs | B | 143 | LEU | p.Leu144Gln | P19429 |
| HOMO SAPIENS | 5 | 39620 | 1ozs | B | 144 | ARG | p.Arg145Gly | P19429 |

| | | | | | | | | |
|---|---|---|---|---|---|---|---|---|
| HOMO SAPIENS | 5 | 39620 | 1ozs | B | 144 | ARG | p.Arg145Gln | P19429 |
| HOMO SAPIENS | -1 | 20934 | 4nuv | D | 42 | GLY | p.Gly42Asp | Q16570 |
| HOMO SAPIENS | -1 | 40565 | 5lcw | S | 4 | VAL | p.Val4Met | O60566 |
| HOMO SAPIENS | 0 | 40565 | 5lcw | S | 40 | THR | p.Thr40Met | O60566 |
| HOMO SAPIENS | 4 | 40565 | 5lcw | S | 198 | ALA | p.Glu390Asp | O60566 |
| HOMO SAPIENS | 4 | 758 | 4oiu | A | 726 | ARG | p.Arg727Leu | P10275 |
| HOMO SAPIENS | 4 | 758 | 4oiu | A | 730 | VAL | p.Val731Met | P10275 |
| HOMO SAPIENS | 2 | 758 | 4oiu | A | 798 | GLN | p.Gln799Glu | P10275 |
| HOMO SAPIENS | 3 | 758 | 4oiu | A | 855 | ARG | p.Arg856His | P10275 |
| HOMO SAPIENS | 1 | 758 | 4oiu | A | 886 | MET | p.Met887Val | P10275 |
| HOMO SAPIENS | 0 | 3303 | 4ie0 | A | 134 | ALA | p.Ala134Thr | Q9C0B1 |
| HOMO SAPIENS | 9 | 3303 | 4ie0 | A | 316 | ARG | p.Arg316Gln | Q9C0B1 |
| HOMO SAPIENS | 1 | 41502 | 3fgh | A | 136 | PRO | p.Pro178Leu | Q00059 |
| HOMO SAPIENS | 5 | 175 | 3ceh | B | 58 | THR | p.Thr59Lys | P06737 |
| HOMO SAPIENS | 8 | 175 | 3ceh | B | 93 | ARG | p.Arg94Ter | P06737 |
| HOMO SAPIENS | 5 | 175 | 3ceh | B | 221 | VAL | p.Val222Ile | P06737 |
| HOMO SAPIENS | 0 | 175 | 3ceh | B | 338 | ASN | p.Asn339Ser | P06737 |
| HOMO SAPIENS | 1 | 175 | 3ceh | B | 372 | PHE | p.Phe373Leu | P06737 |
| HOMO SAPIENS | 9 | 175 | 3ceh | B | 376 | ASN | p.Asn377Lys | P06737 |
| HOMO SAPIENS | 8 | 175 | 3ceh | B | 398 | ARG | p.Arg399Ter | P06737 |
| HOMO SAPIENS | 5 | 175 | 3ceh | B | 424 | ARG | p.Arg425Pro | P06737 |
| HOMO SAPIENS | 9 | 175 | 3ceh | B | 455 | VAL | p.Val456Met | P06737 |
| HOMO SAPIENS | 8 | 175 | 3ceh | B | 490 | ARG | p.Arg491Cys | P06737 |
| HOMO SAPIENS | 8 | 175 | 3ceh | B | 631 | ASN | p.Asn632Ile | P06737 |
| HOMO SAPIENS | 9 | 175 | 3ceh | B | 648 | TYR | p.Tyr649Ter | P06737 |
| HOMO SAPIENS | 9 | 175 | 3ceh | B | 672 | GLU | p.Glu673Lys | P06737 |
| HOMO SAPIENS | 9 | 175 | 3ceh | B | 674 | SER | p.Ser675Leu | P06737 |
| HOMO SAPIENS | 0 | 175 | 3ceh | B | 805 | ILE | p.Ile806Leu | P06737 |
| HOMO SAPIENS | 0 | 175 | 3ceh | B | 820 | TYR | p.Tyr821His | P06737 |
| HOMO SAPIENS | 1 | 6750 | 4fdj | A | 47 | GLY | p.Gly47Arg | P34059 |
| HOMO SAPIENS | 9 | 6750 | 4fdj | A | 60 | ASP | p.Asp60Asn | P34059 |
| HOMO SAPIENS | 5 | 6750 | 4fdj | A | 67 | LEU | p.Leu67Met | P34059 |
| HOMO SAPIENS | 0 | 6750 | 4fdj | A | 69 | PHE | p.Phe69Val | P34059 |
| HOMO SAPIENS | 3 | 6750 | 4fdj | A | 94 | ARG | p.Arg94Gly | P34059 |
| HOMO SAPIENS | 0 | 6750 | 4fdj | A | 113 | ILE | p.Ile113Phe | P34059 |
| HOMO SAPIENS | 0 | 6750 | 4fdj | A | 138 | VAL | p.Val138Ala | P34059 |
| HOMO SAPIENS | 0 | 6750 | 4fdj | A | 139 | GLY | p.Gly139Ser | P34059 |
| HOMO SAPIENS | 0 | 6750 | 4fdj | A | 151 | PRO | p.Pro151Thr | P34059 |
| HOMO SAPIENS | 0 | 6750 | 4fdj | A | 155 | GLY | p.Gly155Arg | P34059 |
| HOMO SAPIENS | 0 | 6750 | 4fdj | A | 159 | TRP | p.Trp159Ter | P34059 |
| HOMO SAPIENS | 0 | 6750 | 4fdj | A | 162 | SER | p.Ser162Phe | P34059 |
| HOMO SAPIENS | 0 | 6750 | 4fdj | A | 200 | THR | p.Thr200Met | P34059 |
| HOMO SAPIENS | 0 | 6750 | 4fdj | A | 204 | ASN | p.Asn204Lys | P34059 |
| HOMO SAPIENS | 0 | 6750 | 4fdj | A | 231 | ALA | p.Ala231Gly | P34059 |
| HOMO SAPIENS | 2 | 6750 | 4fdj | A | 235 | THR | p.Thr235Lys | P34059 |
| HOMO SAPIENS | 1 | 6750 | 4fdj | A | 259 | ARG | p.Arg259Gln | P34059 |
| HOMO SAPIENS | 0 | 6750 | 4fdj | A | 287 | SER | p.Ser287Leu | P34059 |
| HOMO SAPIENS | 0 | 6750 | 4fdj | A | 291 | ALA | p.Ala291Thr | P34059 |
| HOMO SAPIENS | 3 | 6750 | 4fdj | A | 301 | GLY | p.Gly301Cys | P34059 |
| HOMO SAPIENS | 0 | 6750 | 4fdj | A | 312 | THR | p.Thr312Ser | P34059 |
| HOMO SAPIENS | 0 | 6750 | 4fdj | A | 318 | MET | p.Met318Arg | P34059 |
| HOMO SAPIENS | 0 | 6750 | 4fdj | A | 340 | GLY | p.Gly340Asp | P34059 |
| HOMO SAPIENS | 0 | 6750 | 4fdj | A | 386 | ARG | p.Arg386Cys | P34059 |
| HOMO SAPIENS | 0 | 6750 | 4fdj | A | 391 | MET | p.Met391Val | P34059 |

| | | | | | | | | |
|---|---|---|---|---|---|---|---|---|
| HOMO SAPIENS | 0 | 6750 | 4fdj | A | 392 | ALA | p.Ala392Val | P34059 |
| HOMO SAPIENS | 0 | 6750 | 4fdj | A | 393 | ALA | p.Ala393Ser | P34059 |
| HOMO SAPIENS | 0 | 6750 | 4fdj | A | 459 | ALA | p.Ala459Val | P34059 |
| HOMO SAPIENS | 0 | 6750 | 4fdj | A | 473 | GLN | p.Gln473Ter | P34059 |
| HOMO SAPIENS | 2 | 6750 | 4fdj | A | 480 | VAL | p.Val480Phe | P34059 |
| HOMO SAPIENS | 0 | 6750 | 4fdj | A | 487 | ASN | p.Asn487Ser | P34059 |
| HOMO SAPIENS | 0 | 6750 | 4fdj | A | 488 | VAL | p.Val488Met | P34059 |
| HOMO SAPIENS | 0 | 6750 | 4fdj | A | 520 | TRP | p.Trp520Ter | P34059 |
| HOMO SAPIENS | 4 | 1088 | 2p3u | A | 102 | GLU | p.Glu142Lys | P00742 |
| HOMO SAPIENS | 6 | 370 | 1nr1 | D | 73 | ASP | p.Asp126His | P00367 |
| HOMO SAPIENS | 3 | 370 | 1nr1 | D | 221 | ARG | p.Arg274Cys | P00367 |
| HOMO SAPIENS | 2 | 370 | 1nr1 | D | 265 | ARG | p.Arg318Lys | P00367 |
| HOMO SAPIENS | 1 | 370 | 1nr1 | D | 269 | ARG | p.Arg322His | P00367 |
| HOMO SAPIENS | 1 | 370 | 1nr1 | D | 296 | GLU | p.Glu349Ala | P00367 |
| HOMO SAPIENS | 1 | 370 | 1nr1 | D | 445 | SER | p.Ser498Leu | P00367 |
| HOMO SAPIENS | 1 | 370 | 1nr1 | D | 446 | GLY | p.Gly499Asp | P00367 |
| HOMO SAPIENS | 1 | 370 | 1nr1 | D | 446 | GLY | p.Gly499Ser | P00367 |
| HOMO SAPIENS | 1 | 370 | 1nr1 | D | 448 | SER | p.Ser501Pro | P00367 |
| HOMO SAPIENS | 2 | 370 | 1nr1 | D | 454 | HIS | p.His507Tyr | P00367 |
| HOMO SAPIENS | 1 | 7720 | 2he0 | A | 147 | HIS | p.His2018Gln | P46531 |
| HOMO SAPIENS | 2 | 11844 | 5l0v | A | 218 | ARG | p.Arg218Ter | Q8NBL1 |
| HOMO SAPIENS | 2 | 11844 | 5l0v | A | 279 | ARG | p.Arg279Trp | Q8NBL1 |
| HOMO SAPIENS | 9 | 1485 | 5va0 | A | 477 | ARG | p.Arg477His | P04150 |
| HOMO SAPIENS | 0 | 235 | 2fun | B | 2248 | ARG | p.Arg248Trp | Q14790 |
| HOMO SAPIENS | 4 | 235 | 2fun | B | 2285 | HIS | p.Asp285His | Q14790 |
| HOMO SAPIENS | 0 | 4822 | 5y3n | A | 307 | ARG | p.Arg307Gly | Q12931 |
| HOMO SAPIENS | 8 | 32928 | 4m6w | A | 1886 | HIS | p.His1886Tyr | Q8IYD8 |
| HOMO SAPIENS | 0 | 2711 | 4y5v | F | 22 | GLY | p.Gly46Glu | P19235 |
| HOMO SAPIENS | -1 | 2711 | 4y5v | F | 166 | ALA | p.Ala190Ser | P19235 |
| HOMO SAPIENS | 4 | 5161 | 5bpu | A | 42 | HIS | p.His42Arg | Q00604 |
| HOMO SAPIENS | 0 | 5161 | 5bpu | A | 44 | TYR | p.Tyr44Cys | Q00604 |
| HOMO SAPIENS | 0 | 5161 | 5bpu | A | 45 | VAL | p.Val45Glu | Q00604 |
| HOMO SAPIENS | 0 | 5161 | 5bpu | A | 60 | VAL | p.Val60Glu | Q00604 |
| HOMO SAPIENS | 0 | 5161 | 5bpu | A | 61 | LEU | p.Leu61Phe | Q00604 |
| HOMO SAPIENS | 7 | 5161 | 5bpu | A | 69 | CYS | p.Cys69Ser | Q00604 |
| HOMO SAPIENS | 0 | 5161 | 5bpu | A | 75 | SER | p.Ser75Cys | Q00604 |
| HOMO SAPIENS | 0 | 5161 | 5bpu | A | 90 | ARG | p.Arg90Pro | Q00604 |
| HOMO SAPIENS | 9 | 5161 | 5bpu | A | 96 | CYS | p.Cys96Tyr | Q00604 |
| HOMO SAPIENS | 9 | 5161 | 5bpu | A | 96 | CYS | p.Cys96Trp | Q00604 |
| HOMO SAPIENS | 0 | 5161 | 5bpu | A | 101 | SER | p.Ser101Phe | Q00604 |
| HOMO SAPIENS | 0 | 5161 | 5bpu | A | 105 | ALA | p.Ala105Thr | Q00604 |
| HOMO SAPIENS | 0 | 5161 | 5bpu | A | 110 | CYS | p.Cys110Gly | Q00604 |
| HOMO SAPIENS | 0 | 5161 | 5bpu | A | 121 | ARG | p.Arg121Trp | Q00604 |
| HOMO SAPIENS | 0 | 5161 | 5bpu | A | 121 | ARG | p.Arg121Leu | Q00604 |
| HOMO SAPIENS | 0 | 5161 | 5bpu | A | 124 | LEU | p.Leu124Phe | Q00604 |
| HOMO SAPIENS | 8 | 5161 | 5bpu | A | 128 | CYS | p.Cys128Ter | Q00604 |
| HOMO SAPIENS | 1 | 13944 | 5xqh | A | 238 | ARG | p.Arg238His | Q9GZN7 |
| HOMO SAPIENS | -1 | 7508 | 5jl6 | A | 39 | TRP | p.Trp39Arg | P11511 |
| HOMO SAPIENS | 0 | 7508 | 5jl6 | A | 201 | THR | p.Thr201Met | P11511 |
| HOMO SAPIENS | 0 | 7508 | 5jl6 | A | 210 | GLU | p.Glu210Lys | P11511 |
| HOMO SAPIENS | 1 | 7508 | 5jl6 | A | 264 | ARG | p.Arg264Cys | P11511 |
| HOMO SAPIENS | 2 | 7508 | 5jl6 | A | 365 | ARG | p.Arg365Gln | P11511 |
| HOMO SAPIENS | 7 | 7508 | 5jl6 | A | 375 | ARG | p.Arg375Cys | P11511 |
| HOMO SAPIENS | 8 | 7508 | 5jl6 | A | 435 | ARG | p.Arg435Cys | P11511 |

| | | | | | | | | |
|---|---|---|---|---|---|---|---|---|
| HOMO SAPIENS | 9 | 7508 | 5jl6 | A | 437 | CYS | p.Cys437Tyr | P11511 |
| HOMO SAPIENS | -1 | 30206 | 5l95 | B | 246 | ARG | p.Arg246Ter | Q9GZZ9 |
| HOMO SAPIENS | 4 | 875 | 3efh | B | 43 | GLU | p.Glu43Asp | P60891 |
| HOMO SAPIENS | 5 | 875 | 3efh | B | 65 | ASP | p.Asp65Asn | P60891 |
| HOMO SAPIENS | 0 | 875 | 3efh | B | 87 | ALA | p.Ala87Thr | P60891 |
| HOMO SAPIENS | 2 | 875 | 3efh | B | 115 | MET | p.Met115Thr | P60891 |
| HOMO SAPIENS | 8 | 875 | 3efh | B | 133 | GLN | p.Gln133Pro | P60891 |
| HOMO SAPIENS | 8 | 875 | 3efh | B | 152 | LEU | p.Leu152Pro | P60891 |
| HOMO SAPIENS | 7 | 875 | 3efh | B | 290 | ILE | p.Ile290Thr | P60891 |
| HOMO SAPIENS | 6 | 875 | 3efh | B | 306 | GLY | p.Gly306Arg | P60891 |
| HOMO SAPIENS | 0 | 7978 | 5edm | A | 122 | MET | p.Thr165Met | P00734 |
| HOMO SAPIENS | -1 | 7978 | 5edm | A | 257 | ARG | p.Arg314Cys | P00734 |
| HOMO SAPIENS | 0 | 1716 | 1b98 | M | 126 | ARG | p.Arg206Gln | P34130 |
| HOMO SAPIENS | 0 | 1716 | 1b98 | M | 126 | ARG | p.Arg206Trp | P34130 |
| HOMO SAPIENS | 9 | 3960 | 2qyp | B | 28 | ILE | p.Ile338Val | P07602 |
| HOMO SAPIENS | 1 | 3960 | 2qyp | B | 39 | LEU | p.Leu349Pro | P07602 |
| HOMO SAPIENS | 0 | 31414 | 5u1d | A | 333 | ILE | p.Ile393Val | Q03518 |
| HOMO SAPIENS | 0 | 31414 | 5u1d | A | 637 | ASP | p.Asp697Gly | Q03518 |
| HOMO SAPIENS | 0 | 31414 | 5u1d | A | 659 | ARG | p.Arg719Gln | Q03518 |
| HOMO SAPIENS | 3 | 119 | 2pho | A | 11 | ILE | p.Ile11Thr | P05089 |
| HOMO SAPIENS | 8 | 119 | 2pho | A | 21 | ARG | p.Arg21Ter | P05089 |
| HOMO SAPIENS | 0 | 119 | 2pho | A | 122 | TRP | p.Trp122Ter | P05089 |
| HOMO SAPIENS | 6 | 119 | 2pho | A | 138 | GLY | p.Gly138Val | P05089 |
| HOMO SAPIENS | 5 | 119 | 2pho | A | 290 | THR | p.Thr290Ser | P05089 |
| HOMO SAPIENS | 5 | 119 | 2pho | A | 291 | ARG | p.Arg291Ter | P05089 |
| HOMO SAPIENS | 0 | 3732 | 2i99 | A | 247 | TYR | p.Tyr247Ter | Q14894 |
| HOMO SAPIENS | -1 | 1382 | 4ffx | A | 2 | ALA | p.Ala2Val | P30566 |
| HOMO SAPIENS | 1 | 1382 | 4ffx | A | 72 | ILE | p.Ile72Met | P30566 |
| HOMO SAPIENS | 1 | 1382 | 4ffx | A | 100 | PRO | p.Pro100Ala | P30566 |
| HOMO SAPIENS | 0 | 1382 | 4ffx | A | 114 | TYR | p.Tyr114His | P30566 |
| HOMO SAPIENS | 1 | 1382 | 4ffx | A | 141 | ARG | p.Arg141Trp | P30566 |
| HOMO SAPIENS | 0 | 1382 | 4ffx | A | 147 | LYS | p.Lys147Met | P30566 |
| HOMO SAPIENS | 3 | 1382 | 4ffx | A | 190 | ARG | p.Arg190Gln | P30566 |
| HOMO SAPIENS | 3 | 1382 | 4ffx | A | 190 | ARG | p.Arg190Ter | P30566 |
| HOMO SAPIENS | 0 | 1382 | 4ffx | A | 217 | HIS | p.His217Asp | P30566 |
| HOMO SAPIENS | 2 | 1382 | 4ffx | A | 246 | LYS | p.Lys246Glu | P30566 |
| HOMO SAPIENS | 5 | 1382 | 4ffx | A | 303 | ARG | p.Arg303Cys | P30566 |
| HOMO SAPIENS | 1 | 1382 | 4ffx | A | 318 | PRO | p.Pro318Leu | P30566 |
| HOMO SAPIENS | 6 | 1382 | 4ffx | A | 332 | ASP | p.Asp332His | P30566 |
| HOMO SAPIENS | 6 | 1382 | 4ffx | A | 337 | ARG | p.Arg337Ter | P30566 |
| HOMO SAPIENS | 1 | 1382 | 4ffx | A | 396 | ARG | p.Arg396His | P30566 |
| HOMO SAPIENS | 1 | 1382 | 4ffx | A | 426 | ARG | p.Arg426His | P30566 |
| HOMO SAPIENS | 0 | 1382 | 4ffx | A | 438 | SER | p.Ser438Pro | P30566 |
| HOMO SAPIENS | 1 | 1382 | 4ffx | A | 447 | SER | p.Ser447Pro | P30566 |
| HOMO SAPIENS | 2 | 1382 | 4ffx | A | 448 | SER | p.Ser448Pro | P30566 |
| HOMO SAPIENS | 2 | 1382 | 4ffx | A | 450 | THR | p.Thr450Ser | P30566 |
| HOMO SAPIENS | 0 | 6212 | 1ao3 | A | 46 | SER | p.Ser1731Thr | P04275 |
| HOMO SAPIENS | 0 | 1711 | 3f9c | A | 279 | VAL | p.Val279Phe | Q13093 |
| HOMO SAPIENS | 0 | 6112 | 4fvl | A | 312 | HIS | p.His312Arg | P45452 |
| HOMO SAPIENS | 0 | 6112 | 4fvl | A | 323 | THR | p.Thr323Met | P45452 |
| HOMO SAPIENS | 0 | 6112 | 4fvl | A | 390 | ASP | p.Asp390Gly | P45452 |
| HOMO SAPIENS | 0 | 6112 | 4fvl | A | 458 | ARG | p.Arg458Cys | P45452 |
| HOMO SAPIENS | 2 | 6180 | 5ggn | A | 97 | ARG | p.Arg97Trp | Q8WZA1 |
| HOMO SAPIENS | 2 | 6180 | 5ggn | A | 215 | ARG | p.Arg215Ter | Q8WZA1 |

| | | | | | | | | |
|---|---|---|---|---|---|---|---|---|
| HOMO SAPIENS | -1 | 5526 | 4ak3 | A | 11 | ASP | p.Asp1232Asn | P02461 |
| HOMO SAPIENS | 6 | 5526 | 4ak3 | A | 38 | ALA | p.Ala1259Thr | P02461 |
| HOMO SAPIENS | 0 | 5526 | 4ak3 | A | 92 | LYS | p.Lys1313Arg | P02461 |
| HOMO SAPIENS | 0 | 5526 | 4ak3 | A | 132 | GLN | p.His1353Gln | P02461 |
| HOMO SAPIENS | 4 | 16616 | 1hci | B | 433 | SER | p.Ser433Leu | P35609 |
| HOMO SAPIENS | 1 | 16616 | 1hci | B | 475 | ASP | p.Asp475Asn | P35609 |
| HOMO SAPIENS | 1 | 16616 | 1hci | B | 476 | ALA | p.Ala476Ser | P35609 |
| HOMO SAPIENS | 2 | 16616 | 1hci | B | 507 | MET | p.Met507Val | P35609 |
| HOMO SAPIENS | 1 | 16616 | 1hci | B | 604 | MET | p.Met604Val | P35609 |
| HOMO SAPIENS | 2 | 16616 | 1hci | B | 640 | ARG | p.Arg640His | P35609 |
| HOMO SAPIENS | 1 | 16616 | 1hci | B | 721 | ARG | p.Arg721Cys | P35609 |
| HOMO SAPIENS | -1 | 3402 | 5iaa | B | 57 | MET | p.Met57Val | Q9GZZ9 |
| HOMO SAPIENS | 7 | 3250 | 3irq | B | 193 | PRO | p.Pro193Ala | P55265 |
| HOMO SAPIENS | 2 | 15200 | 2wns | A | 30 | SER | p.Ser30Gly | P11172 |
| HOMO SAPIENS | 0 | 15200 | 2wns | A | 96 | ARG | p.Arg96Gly | P11172 |
| HOMO SAPIENS | 0 | 15200 | 2wns | A | 109 | VAL | p.Val109Gly | P11172 |
| HOMO SAPIENS | 2 | 15759 | 4x4w | B | 140 | LEU | p.Leu166Ser | Q96Q11 |
| HOMO SAPIENS | 0 | 15759 | 4x4w | B | 197 | ILE | p.Ile223Thr | Q96Q11 |
| HOMO SAPIENS | 0 | 3156 | 3w03 | B | 14 | ALA | p.Ala14Thr | Q9H9Q4 |
| HOMO SAPIENS | 0 | 3156 | 3w03 | B | 57 | ARG | p.Arg57Gly | Q9H9Q4 |
| HOMO SAPIENS | 0 | 3156 | 3w03 | B | 123 | CYS | p.Cys123Arg | Q9H9Q4 |
| HOMO SAPIENS | 0 | 11693 | 3ojy | A | 63 | GLN | p.Gln93Lys | P07357 |
| HOMO SAPIENS | 0 | 1421 | 2qc8 | E | 275 | GLU | p.Glu275Asp | P15104 |
| HOMO SAPIENS | 9 | 1421 | 2qc8 | E | 324 | ARG | p.Arg324Cys | P15104 |
| HOMO SAPIENS | 6 | 1421 | 2qc8 | E | 341 | ARG | p.Arg341Cys | P15104 |
| HOMO SAPIENS | 6 | 2850 | 3qvb | A | 83 | THR | p.Thr83Met | P46976 |
| HOMO SAPIENS | 8 | 2850 | 3qvb | A | 102 | ASP | p.Asp102His | P46976 |
| HOMO SAPIENS | 1 | 24314 | 4m9p | A | 480 | PRO | p.Pro480Leu | P21333 |
| HOMO SAPIENS | 0 | 24314 | 4m9p | A | 528 | VAL | p.Val528Met | P21333 |
| HOMO SAPIENS | 0 | 24314 | 4m9p | A | 590 | VAL | p.Val590Ile | P21333 |
| HOMO SAPIENS | 0 | 24314 | 4m9p | A | 605 | ASP | p.Asp605Asn | P21333 |
| HOMO SAPIENS | 1 | 24314 | 4m9p | A | 637 | PRO | p.Pro637Gln | P21333 |
| HOMO SAPIENS | 0 | 24314 | 4m9p | A | 656 | LEU | p.Leu656Phe | P21333 |
| HOMO SAPIENS | 2 | 24314 | 4m9p | A | 711 | VAL | p.Val711Asp | P21333 |
| HOMO SAPIENS | 0 | 4951 | 5joe | A | 40 | THR | p.Thr9621Ser | Q8WZ42 |
| HOMO SAPIENS | 1 | 4951 | 5joe | A | 41 | GLU | p.Glu9622Gln | Q8WZ42 |
| HOMO SAPIENS | 0 | 4951 | 5joe | A | 82 | ALA | p.Ala9663Ser | Q8WZ42 |
| HOMO SAPIENS | 4 | 4951 | 5joe | A | 89 | ILE | p.Ile9670Asn | Q8WZ42 |
| HOMO SAPIENS | 0 | 9664 | 3c6m | A | 56 | GLY | p.Gly56Ser | P52788 |
| HOMO SAPIENS | 0 | 9664 | 3c6m | A | 132 | VAL | p.Val132Gly | P52788 |
| HOMO SAPIENS | 1 | 9664 | 3c6m | A | 190 | THR | p.Thr190Ile | P52788 |
| HOMO SAPIENS | -1 | 12118 | 5hcd | B | 19 | GLN | p.Gln19Ter | P01031 |
| HOMO SAPIENS | 0 | 1583 | 2b4x | L | 7 | ILE | p.Ile39Asn | P01008 |
| HOMO SAPIENS | 0 | 1583 | 2b4x | L | 24 | ARG | p.Arg56Cys | P01008 |
| HOMO SAPIENS | 0 | 1583 | 2b4x | L | 47 | ARG | p.Arg79Cys | P01008 |
| HOMO SAPIENS | 0 | 1583 | 2b4x | L | 47 | ARG | p.Arg79His | P01008 |
| HOMO SAPIENS | 1 | 1583 | 2b4x | L | 95 | CYS | p.Cys127Arg | P01008 |
| HOMO SAPIENS | 1 | 1583 | 2b4x | L | 99 | LEU | p.Leu131Phe | P01008 |
| HOMO SAPIENS | 0 | 1583 | 2b4x | L | 116 | SER | p.Ser148Pro | P01008 |
| HOMO SAPIENS | 0 | 1583 | 2b4x | L | 129 | ARG | p.Arg161Gln | P01008 |
| HOMO SAPIENS | 0 | 1583 | 2b4x | L | 135 | ASN | p.Asn167Thr | P01008 |
| HOMO SAPIENS | 2 | 1583 | 2b4x | L | 187 | ASN | p.Asn219Asp | P01008 |
| HOMO SAPIENS | 0 | 1583 | 2b4x | L | 191 | SER | p.Ser223Pro | P01008 |
| HOMO SAPIENS | 2 | 1583 | 2b4x | L | 349 | SER | p.Ser381Pro | P01008 |

| | | | | | | | | |
|---|---|---|---|---|---|---|---|---|
| HOMO SAPIENS | 0 | 1583 | 2b4x | L | 382 | ALA | p.Ala414Thr | P01008 |
| HOMO SAPIENS | 0 | 1583 | 2b4x | L | 387 | ALA | p.Ala419Val | P01008 |
| HOMO SAPIENS | 0 | 1583 | 2b4x | L | 392 | GLY | p.Gly424Asp | P01008 |
| HOMO SAPIENS | 0 | 1583 | 2b4x | L | 393 | ARG | p.Arg425His | P01008 |
| HOMO SAPIENS | 0 | 1583 | 2b4x | L | 393 | ARG | p.Arg425Cys | P01008 |
| HOMO SAPIENS | 0 | 1583 | 2b4x | L | 394 | SER | p.Ser426Leu | P01008 |
| HOMO SAPIENS | 0 | 1583 | 2b4x | L | 404 | ALA | p.Ala436Thr | P01008 |
| HOMO SAPIENS | 2 | 1583 | 2b4x | L | 407 | PRO | p.Pro439Leu | P01008 |
| HOMO SAPIENS | 2 | 1583 | 2b4x | L | 429 | PRO | p.Pro461Leu | P01008 |
| HOMO SAPIENS | 0 | 11647 | 2a1t | S | 78 | ALA | p.Ala78Ser | P38117 |
| HOMO SAPIENS | 0 | 11647 | 2a1t | S | 98 | ARG | p.Arg98Cys | P38117 |
| HOMO SAPIENS | 8 | 11647 | 2a1t | S | 128 | ASP | p.Asp128Asn | P38117 |
| HOMO SAPIENS | 5 | 11647 | 2a1t | S | 154 | THR | p.Thr154Met | P38117 |
| HOMO SAPIENS | 6 | 11647 | 2a1t | S | 164 | ARG | p.Arg164Gln | P38117 |
| HOMO SAPIENS | 5 | 11647 | 2a1t | S | 174 | ARG | p.Arg174His | P38117 |
| HOMO SAPIENS | -1 | 11200 | 4tx5 | A | 4 | ILE | p.Ile59Val | Q9NR28 |
| HOMO SAPIENS | 0 | 11200 | 4tx5 | A | 169 | GLY | p.Gly224Arg | Q9NR28 |
| HOMO SAPIENS | 3 | 417 | 2vr2 | A | 6 | ARG | p.Arg6Gln | Q14117 |
| HOMO SAPIENS | 1 | 417 | 2vr2 | A | 334 | GLN | p.Gln334Arg | Q14117 |
| HOMO SAPIENS | 2 | 417 | 2vr2 | A | 360 | TRP | p.Trp360Arg | Q14117 |
| HOMO SAPIENS | 0 | 417 | 2vr2 | A | 379 | SER | p.Ser379Arg | Q14117 |
| HOMO SAPIENS | 0 | 417 | 2vr2 | A | 435 | GLY | p.Gly435Arg | Q14117 |
| HOMO SAPIENS | 0 | 12498 | 3eo3 | B | 423 | ILE | p.Ile423Val | Q9Y223 |
| HOMO SAPIENS | 5 | 12498 | 3eo3 | B | 572 | VAL | p.Val572Leu | Q9Y223 |
| HOMO SAPIENS | 0 | 12498 | 3eo3 | B | 576 | GLY | p.Gly576Glu | Q9Y223 |
| HOMO SAPIENS | 0 | 12498 | 3eo3 | B | 587 | ILE | p.Ile587Thr | Q9Y223 |
| HOMO SAPIENS | 0 | 12498 | 3eo3 | B | 615 | SER | p.Ser615Ter | Q9Y223 |
| HOMO SAPIENS | 1 | 12498 | 3eo3 | B | 631 | ALA | p.Ala631Val | Q9Y223 |
| HOMO SAPIENS | 1 | 12498 | 3eo3 | B | 669 | GLY | p.Gly669Arg | Q9Y223 |
| HOMO SAPIENS | 1 | 12498 | 3eo3 | B | 696 | VAL | p.Val696Met | Q9Y223 |
| HOMO SAPIENS | 2 | 12498 | 3eo3 | B | 712 | MET | p.Met712Thr | Q9Y223 |
| HOMO SAPIENS | 0 | 1568 | 1r74 | A | 49 | LEU | p.Leu50Pro | Q14749 |
| HOMO SAPIENS | 0 | 1568 | 1r74 | A | 176 | HIS | p.His177Asn | Q14749 |
| HOMO SAPIENS | 3 | 147 | 5eig | A | 67 | THR | p.Thr67Ile | P32929 |
| HOMO SAPIENS | 3 | 147 | 5eig | A | 240 | GLN | p.Gln240Glu | P32929 |
| HOMO SAPIENS | -1 | 147 | 5eig | A | 403 | SER | p.Ser403Ile | P32929 |
| HOMO SAPIENS | 8 | 20036 | 2lsr | A | 31 | ARG | p.Arg31Ter | Q9Y6N9 |
| HOMO SAPIENS | -1 | 41382 | 4uis | B | 431 | UNK | p.Ala431Glu | P49768 |
| HOMO SAPIENS | -1 | 8536 | 3i6w | B | 85 | PRO | p.Pro85Leu | O96017 |
| HOMO SAPIENS | 0 | 8536 | 3i6w | B | 93 | TRP | p.Trp93Ter | O96017 |
| HOMO SAPIENS | 0 | 8536 | 3i6w | B | 95 | ARG | p.Arg95Ter | O96017 |
| HOMO SAPIENS | 0 | 8536 | 3i6w | B | 117 | ARG | p.Arg117Gly | O96017 |
| HOMO SAPIENS | 0 | 8536 | 3i6w | B | 119 | LYS | p.Lys119Arg | O96017 |
| HOMO SAPIENS | 0 | 8536 | 3i6w | B | 137 | ARG | p.Arg137Gln | O96017 |
| HOMO SAPIENS | 0 | 8536 | 3i6w | B | 137 | ARG | p.Arg137Ter | O96017 |
| HOMO SAPIENS | 0 | 8536 | 3i6w | B | 145 | ARG | p.Arg145Trp | O96017 |
| HOMO SAPIENS | 0 | 8536 | 3i6w | B | 160 | ILE | p.Ile160Met | O96017 |
| HOMO SAPIENS | 0 | 41322 | 4bxl | C | 46 | GLU | p.Glu46Lys | P37840 |
| HOMO SAPIENS | 5 | 41322 | 4bxl | C | 53 | ALA | p.Ala53Thr | P37840 |
| HOMO SAPIENS | 2 | 24966 | 3hve | A | 15 | ARG | p.Arg15Ser | Q9H2C0 |
| HOMO SAPIENS | 1 | 24966 | 3hve | A | 138 | ARG | p.Arg138His | Q9H2C0 |
| HOMO SAPIENS | 0 | 24966 | 3hve | A | 201 | ARG | p.Arg201Ter | Q9H2C0 |
| HOMO SAPIENS | 0 | 11680 | 5an9 | J | 8 | ASN | p.Asn8Lys | Q9Y3A5 |
| HOMO SAPIENS | 1 | 11680 | 5an9 | J | 33 | LYS | p.Lys33Arg | Q9Y3A5 |

| | | | | | | | | |
|---|---|---|---|---|---|---|---|---|
| HOMO SAPIENS | 1 | 11680 | 5an9 | J | 126 | ARG | p.Arg126Thr | Q9Y3A5 |
| HOMO SAPIENS | 1 | 11680 | 5an9 | J | 130 | VAL | p.Val130Met | Q9Y3A5 |
| HOMO SAPIENS | 1 | 11680 | 5an9 | J | 169 | ARG | p.Arg169Cys | Q9Y3A5 |
| HOMO SAPIENS | 1 | 11680 | 5an9 | J | 218 | ARG | p.Arg218Ter | Q9Y3A5 |
| HOMO SAPIENS | 4 | 978 | 3zs0 | B | 7 | TYR | p.Tyr173Cys | P05164 |
| HOMO SAPIENS | 1 | 978 | 3zs0 | B | 85 | MET | p.Met251Thr | P05164 |
| HOMO SAPIENS | 4 | 5140 | 2l4r | A | 52 | CYS | p.Cys52Ter | Q12809 |
| HOMO SAPIENS | 4 | 5140 | 2l4r | A | 63 | PRO | p.Pro63His | Q12809 |
| HOMO SAPIENS | 0 | 5140 | 2l4r | A | 100 | ARG | p.Arg100Gly | Q12809 |
| HOMO SAPIENS | 4 | 5140 | 2l4r | A | 129 | PHE | p.Phe129Leu | Q12809 |
| HOMO SAPIENS | 0 | 25836 | 2gy7 | B | 103 | GLU | p.Glu103Asp | Q02763 |
| HOMO SAPIENS | 0 | 25836 | 2gy7 | B | 148 | ILE | p.Ile148Thr | Q02763 |
| HOMO SAPIENS | 0 | 25836 | 2gy7 | B | 226 | ALA | p.Ala226Val | Q02763 |
| HOMO SAPIENS | 0 | 25836 | 2gy7 | B | 391 | THR | p.Thr391Ile | Q02763 |
| HOMO SAPIENS | 4 | 12563 | 5kwy | D | 20 | GLU | p.Glu20Ter | P61916 |
| HOMO SAPIENS | 2 | 12563 | 5kwy | D | 30 | VAL | p.Val30Met | P61916 |
| HOMO SAPIENS | 4 | 12563 | 5kwy | D | 39 | VAL | p.Val39Met | P61916 |
| HOMO SAPIENS | 8 | 12563 | 5kwy | D | 67 | SER | p.Ser67Pro | P61916 |
| HOMO SAPIENS | 8 | 12563 | 5kwy | D | 99 | CYS | p.Cys99Arg | P61916 |
| HOMO SAPIENS | 2 | 12563 | 5kwy | D | 118 | GLU | p.Glu118Ter | P61916 |
| HOMO SAPIENS | 2 | 12563 | 5kwy | D | 120 | PRO | p.Pro120Ser | P61916 |
| HOMO SAPIENS | -1 | 1679 | 1pmk | A | -2 | VAL | p.Val374Phe | P00747 |
| HOMO SAPIENS | -1 | 5315 | 2w2n | P | 53 | ALA | p.Ala53Val | Q8NBP7 |
| HOMO SAPIENS | 8 | 13496 | 2yad | A | 138 | THR | p.Asn138Thr | P11686 |
| HOMO SAPIENS | -1 | 13496 | 2yad | A | 173 | ARG | p.Arg167Gln | P11686 |
| HOMO SAPIENS | 2 | 13496 | 2yad | A | 186 | SER | p.Asn186Ser | P11686 |
| HOMO SAPIENS | 0 | 4426 | 3qd6 | C | 123 | ALA | p.Ala123Glu | P29965 |
| HOMO SAPIENS | 1 | 4426 | 3qd6 | C | 140 | TRP | p.Trp140Gly | P29965 |
| HOMO SAPIENS | 0 | 4426 | 3qd6 | C | 155 | LEU | p.Leu155Pro | P29965 |
| HOMO SAPIENS | 6 | 4426 | 3qd6 | C | 227 | GLY | p.Gly227Val | P29965 |
| HOMO SAPIENS | 0 | 4426 | 3qd6 | C | 235 | ALA | p.Ala235Pro | P29965 |
| HOMO SAPIENS | 0 | 10734 | 3dyd | B | 45 | PRO | p.Pro45Ser | P17735 |
| HOMO SAPIENS | -1 | 10734 | 3dyd | B | 57 | ARG | p.Arg57Ter | P17735 |
| HOMO SAPIENS | 8 | 10734 | 3dyd | B | 417 | ARG | p.Arg417Ter | P17735 |
| HOMO SAPIENS | 0 | 6956 | 4pzs | A | 1600 | LEU | p.Leu1600Phe | Q09472 |
| HOMO SAPIENS | 0 | 9370 | 3hfv | A | 21 | SER | p.Ser57Cys | O95363 |
| HOMO SAPIENS | 1 | 9370 | 3hfv | A | 25 | ASP | p.Asp61Glu | O95363 |
| HOMO SAPIENS | 1 | 9370 | 3hfv | A | 130 | ALA | p.Ala166Thr | O95363 |
| HOMO SAPIENS | 5 | 9370 | 3hfv | A | 148 | ASP | p.Asp184Asn | O95363 |
| HOMO SAPIENS | 0 | 9370 | 3hfv | A | 187 | ARG | p.Arg223Cys | O95363 |
| HOMO SAPIENS | 0 | 9370 | 3hfv | A | 210 | THR | p.Thr246Met | O95363 |
| HOMO SAPIENS | 1 | 9370 | 3hfv | A | 244 | ASN | p.Asn280Ser | O95363 |
| HOMO SAPIENS | 2 | 9370 | 3hfv | A | 288 | TYR | p.Tyr324Cys | O95363 |
| HOMO SAPIENS | 5 | 9370 | 3hfv | A | 293 | ILE | p.Ile329Thr | O95363 |
| HOMO SAPIENS | 1 | 9370 | 3hfv | A | 355 | ASP | p.Asp391Val | O95363 |
| HOMO SAPIENS | 1 | 9370 | 3hfv | A | 371 | THR | p.Thr407Met | O95363 |
| HOMO SAPIENS | 1 | 9370 | 3hfv | A | 383 | ARG | p.Arg419Cys | O95363 |
| HOMO SAPIENS | 1 | 9370 | 3hfv | A | 387 | ARG | p.Arg423Gln | O95363 |
| HOMO SAPIENS | 0 | 14401 | 1q3x | A | 371 | ASP | p.Asp371Tyr | O00187 |
| HOMO SAPIENS | 0 | 14401 | 1q3x | A | 377 | VAL | p.Val377Ala | O00187 |
| HOMO SAPIENS | 0 | 14401 | 1q3x | A | 439 | ARG | p.Arg439His | O00187 |
| HOMO SAPIENS | 0 | 14401 | 1q3x | A | 577 | GLN | p.Gln577His | O00187 |
| HOMO SAPIENS | 2 | 18270 | 3teq | A | 429 | ARG | p.Arg429Cys | Q13586 |
| HOMO SAPIENS | -1 | 16081 | 4mqt | A | 231 | VAL | p.Val231Ile | P08172 |

| | | | | | | | | |
|---|---|---|---|---|---|---|---|---|
| HOMO SAPIENS | 9 | 6607 | 3whu | B | 81 | ASP | p.Asp81His | Q8NI22 |
| HOMO SAPIENS | 9 | 6607 | 3whu | B | 129 | ASP | p.Asp129Glu | Q8NI22 |
| HOMO SAPIENS | 9 | 6607 | 3whu | B | 136 | ILE | p.Ile136Thr | Q8NI22 |
| HOMO SAPIENS | 0 | 7208 | 2e9w | D | 11 | ASN | p.Asn36Ser | P21583 |
| HOMO SAPIENS | 0 | 24566 | 4bdx | A | 10 | LEU | p.Leu140Val | P00748 |
| HOMO SAPIENS | 5 | 24566 | 4bdx | A | 77 | PRO | p.Ala207Pro | P00748 |
| HOMO SAPIENS | 7 | 606 | 3qnf | A | 319 | MET | p.Met319Val | Q9NZ08 |
| HOMO SAPIENS | 8 | 4097 | 1ydl | A | 21 | LEU | p.Leu21Pro | Q6ZYL4 |
| HOMO SAPIENS | 6 | 4097 | 1ydl | A | 56 | ARG | p.Arg56Ter | Q6ZYL4 |
| HOMO SAPIENS | 8 | 36691 | 1dqb | A | 42 | ARG | p.Arg403Lys | P07204 |
| HOMO SAPIENS | 0 | 5642 | 3j7y | X | 59 | ARG | p.Arg59Pro | Q13084 |
| HOMO SAPIENS | 1 | 5642 | 3j7y | X | 204 | VAL | p.Val204Met | Q13084 |
| HOMO SAPIENS | 1 | 1119 | 3nwu | B | 252 | ALA | p.Ala252Thr | Q92743 |
| HOMO SAPIENS | 3 | 1119 | 3nwu | B | 274 | ARG | p.Arg274Gln | Q92743 |
| HOMO SAPIENS | 9 | 1119 | 3nwu | B | 297 | VAL | p.Val297Met | Q92743 |
| HOMO SAPIENS | -1 | 1119 | 3nwu | B | 302 | ARG | p.Arg302Ter | Q92743 |
| HOMO SAPIENS | 9 | 1119 | 3nwu | B | 321 | ALA | p.Ala321Thr | Q92743 |
| HOMO SAPIENS | -1 | 1119 | 3nwu | B | 370 | ARG | p.Arg370Ter | Q92743 |
| HOMO SAPIENS | 0 | 416 | 4ycu | B | 105 | LEU | p.Leu105His | Q15046 |
| HOMO SAPIENS | 0 | 416 | 4ycu | B | 145 | TYR | p.Tyr145His | Q15046 |
| HOMO SAPIENS | 0 | 416 | 4ycu | B | 200 | PRO | p.Pro200Leu | Q15046 |
| HOMO SAPIENS | 0 | 416 | 4ycu | B | 201 | TYR | p.Tyr201His | Q15046 |
| HOMO SAPIENS | 0 | 416 | 4ycu | B | 420 | ARG | p.Arg420Cys | Q15046 |
| HOMO SAPIENS | 9 | 3297 | 4fpb | A | 680 | THR | p.Thr680Ser | P19367 |
| HOMO SAPIENS | 0 | 6771 | 2zt6 | A | 201 | ALA | p.Ala255Val | P41250 |
| HOMO SAPIENS | 0 | 6771 | 2zt6 | A | 202 | ASP | p.Asp256His | P41250 |
| HOMO SAPIENS | 0 | 6771 | 2zt6 | A | 334 | ARG | p.Arg388Gln | P41250 |
| HOMO SAPIENS | -1 | 6771 | 2zt6 | A | 439 | ASN | p.Asn493Ser | P41250 |
| HOMO SAPIENS | -1 | 6771 | 2zt6 | A | 500 | ASP | p.Asp554Asn | P41250 |
| HOMO SAPIENS | 1 | 6771 | 2zt6 | A | 581 | SER | p.Ser635Leu | P41250 |
| HOMO SAPIENS | -1 | 6771 | 2zt6 | A | 684 | GLU | p.Glu738Lys | P41250 |
| HOMO SAPIENS | 0 | 2687 | 4grw | B | 276 | VAL | p.Val298Phe | P29460 |
| HOMO SAPIENS | 6 | 913 | 1m75 | B | 45 | ASP | p.Asp57Glu | Q16836 |
| HOMO SAPIENS | 2 | 913 | 1m75 | B | 74 | PRO | p.Leu86Pro | Q16836 |
| HOMO SAPIENS | 3 | 913 | 1m75 | B | 214 | TYR | p.Tyr226His | Q16836 |
| HOMO SAPIENS | 3 | 913 | 1m75 | B | 224 | ARG | p.Arg236Ter | Q16836 |
| HOMO SAPIENS | 4 | 913 | 1m75 | B | 246 | PRO | p.Pro258Leu | Q16836 |
| HOMO SAPIENS | 2 | 4713 | 4djn | B | 40 | ARG | p.Arg40Cys | Q7LG56 |
| HOMO SAPIENS | 1 | 4713 | 4djn | B | 41 | ARG | p.Arg41Pro | Q7LG56 |
| HOMO SAPIENS | 2 | 4713 | 4djn | B | 110 | ARG | p.Arg110His | Q7LG56 |
| HOMO SAPIENS | 2 | 4713 | 4djn | B | 121 | ARG | p.Arg121His | Q7LG56 |
| HOMO SAPIENS | 9 | 4713 | 4djn | B | 131 | GLU | p.Glu131Lys | Q7LG56 |
| HOMO SAPIENS | 2 | 4713 | 4djn | B | 182 | THR | p.Thr182Ala | Q7LG56 |
| HOMO SAPIENS | 6 | 4713 | 4djn | B | 194 | GLU | p.Glu194Lys | Q7LG56 |
| HOMO SAPIENS | 2 | 4713 | 4djn | B | 211 | ARG | p.Arg211Lys | Q7LG56 |
| HOMO SAPIENS | 0 | 4713 | 4djn | B | 224 | ILE | p.Ile224Ser | Q7LG56 |
| HOMO SAPIENS | 3 | 4713 | 4djn | B | 229 | GLY | p.Gly229Val | Q7LG56 |
| HOMO SAPIENS | 2 | 4713 | 4djn | B | 236 | CYS | p.Cys236Phe | Q7LG56 |
| HOMO SAPIENS | 0 | 4713 | 4djn | B | 273 | GLY | p.Gly273Ser | Q7LG56 |
| HOMO SAPIENS | -1 | 4713 | 4djn | B | 282 | MSE | p.Met282Ile | Q7LG56 |
| HOMO SAPIENS | 3 | 4713 | 4djn | B | 284 | GLN | p.Gln284Ter | Q7LG56 |
| HOMO SAPIENS | 3 | 4713 | 4djn | B | 305 | ALA | p.Ala305Ser | Q7LG56 |
| HOMO SAPIENS | -1 | 8974 | 5n7i | A | 500 | ARG | p.Arg500Ter | Q8N4S9 |
| HOMO SAPIENS | 0 | 17072 | 3fxi | B | 299 | ASP | p.Asp299Gly | O00206 |

| | | | | | | | | |
|---|---|---|---|---|---|---|---|---|
| HOMO SAPIENS | 1 | 14355 | 2wwk | T | 55 | ILE | p.Ile34306Asn | Q8WZ42 |
| HOMO SAPIENS | 0 | 14355 | 2wwk | T | 64 | LEU | p.Leu34315Pro | Q8WZ42 |
| HOMO SAPIENS | 2 | 17797 | 1xu1 | S | 72 | ARG | p.Arg72His | O14836 |
| HOMO SAPIENS | -1 | 10836 | 4ztu | A | 39 | GLN | p.Gln39Arg | P54098 |
| HOMO SAPIENS | -1 | 10836 | 4ztu | A | 42 | ARG | p.Arg42Gln | P54098 |
| HOMO SAPIENS | -1 | 10836 | 4ztu | A | 45 | GLN | p.Gln45Arg | P54098 |
| HOMO SAPIENS | -1 | 10836 | 4ztu | A | 52 | GLN | p.Gln52Glu | P54098 |
| HOMO SAPIENS | 0 | 10836 | 4ztu | A | 88 | PHE | p.Phe88Leu | P54098 |
| HOMO SAPIENS | 1 | 10836 | 4ztu | A | 194 | ALA | p.Ala194Val | P54098 |
| HOMO SAPIENS | 1 | 10836 | 4ztu | A | 227 | ARG | p.Arg227Trp | P54098 |
| HOMO SAPIENS | 0 | 10836 | 4ztu | A | 232 | ARG | p.Arg232His | P54098 |
| HOMO SAPIENS | 1 | 10836 | 4ztu | A | 285 | GLN | p.Gln285His | P54098 |
| HOMO SAPIENS | 1 | 10836 | 4ztu | A | 434 | TYR | p.Tyr434His | P54098 |
| HOMO SAPIENS | -1 | 10836 | 4ztu | A | 511 | SER | p.Ser511Asn | P54098 |
| HOMO SAPIENS | 0 | 10836 | 4ztu | A | 533 | CYS | p.Cys533Trp | P54098 |
| HOMO SAPIENS | 1 | 10836 | 4ztu | A | 546 | ARG | p.Arg546Cys | P54098 |
| HOMO SAPIENS | 1 | 10836 | 4ztu | A | 587 | PRO | p.Pro587Leu | P54098 |
| HOMO SAPIENS | 0 | 10836 | 4ztu | A | 603 | MET | p.Met603Thr | P54098 |
| HOMO SAPIENS | 0 | 10836 | 4ztu | A | 623 | LEU | p.Leu623Trp | P54098 |
| HOMO SAPIENS | -1 | 10836 | 4ztu | A | 627 | ARG | p.Arg627Trp | P54098 |
| HOMO SAPIENS | -1 | 10836 | 4ztu | A | 628 | ARG | p.Arg628Pro | P54098 |
| HOMO SAPIENS | 0 | 10836 | 4ztu | A | 635 | PRO | p.Pro635Leu | P54098 |
| HOMO SAPIENS | 0 | 10836 | 4ztu | A | 662 | GLU | p.Glu662Lys | P54098 |
| HOMO SAPIENS | -1 | 10836 | 4ztu | A | 674 | GLY | p.Gly674Asp | P54098 |
| HOMO SAPIENS | -1 | 10836 | 4ztu | A | 722 | ARG | p.Arg722His | P54098 |
| HOMO SAPIENS | -1 | 10836 | 4ztu | A | 737 | GLY | p.Gly737Arg | P54098 |
| HOMO SAPIENS | 0 | 10836 | 4ztu | A | 748 | TRP | p.Trp748Ser | P54098 |
| HOMO SAPIENS | 0 | 10836 | 4ztu | A | 749 | PHE | p.Phe749Ser | P54098 |
| HOMO SAPIENS | 1 | 10836 | 4ztu | A | 807 | ARG | p.Arg807Cys | P54098 |
| HOMO SAPIENS | 1 | 10836 | 4ztu | A | 831 | TYR | p.Tyr831Cys | P54098 |
| HOMO SAPIENS | 8 | 10836 | 4ztu | A | 848 | GLY | p.Gly848Ser | P54098 |
| HOMO SAPIENS | 1 | 10836 | 4ztu | A | 852 | ARG | p.Arg852Cys | P54098 |
| HOMO SAPIENS | 9 | 10836 | 4ztu | A | 853 | ARG | p.Arg853Trp | P54098 |
| HOMO SAPIENS | 1 | 10836 | 4ztu | A | 864 | ASN | p.Asn864Ser | P54098 |
| HOMO SAPIENS | 1 | 10836 | 4ztu | A | 879 | GLN | p.Gln879Arg | P54098 |
| HOMO SAPIENS | 1 | 10836 | 4ztu | A | 914 | THR | p.Thr914Pro | P54098 |
| HOMO SAPIENS | 6 | 10836 | 4ztu | A | 932 | HIS | p.His932Tyr | P54098 |
| HOMO SAPIENS | 2 | 10836 | 4ztu | A | 944 | GLU | p.Glu944Lys | P54098 |
| HOMO SAPIENS | 6 | 10836 | 4ztu | A | 955 | TYR | p.Tyr955Cys | P54098 |
| HOMO SAPIENS | 1 | 10836 | 4ztu | A | 957 | ALA | p.Ala957Ser | P54098 |
| HOMO SAPIENS | 1 | 10836 | 4ztu | A | 957 | ALA | p.Ala957Val | P54098 |
| HOMO SAPIENS | 1 | 10836 | 4ztu | A | 964 | ARG | p.Arg964Cys | P54098 |
| HOMO SAPIENS | 1 | 10836 | 4ztu | A | 966 | LEU | p.Leu966Arg | P54098 |
| HOMO SAPIENS | 0 | 10836 | 4ztu | A | 1051 | GLY | p.Gly1051Arg | P54098 |
| HOMO SAPIENS | 1 | 10836 | 4ztu | A | 1073 | PRO | p.Pro1073Leu | P54098 |
| HOMO SAPIENS | 1 | 10836 | 4ztu | A | 1096 | ARG | p.Arg1096Cys | P54098 |
| HOMO SAPIENS | 1 | 10836 | 4ztu | A | 1096 | ARG | p.Arg1096His | P54098 |
| HOMO SAPIENS | 0 | 10836 | 4ztu | A | 1116 | MET | p.Met1116Thr | P54098 |
| HOMO SAPIENS | 1 | 10836 | 4ztu | A | 1142 | ARG | p.Arg1142Trp | P54098 |
| HOMO SAPIENS | 1 | 10836 | 4ztu | A | 1146 | ARG | p.Arg1146Cys | P54098 |
| HOMO SAPIENS | 1 | 10836 | 4ztu | A | 1157 | ASN | p.Asn1157Ser | P54098 |
| HOMO SAPIENS | -1 | 10836 | 4ztu | A | 1234 | ARG | p.Arg1234Gln | P54098 |
| HOMO SAPIENS | -1 | 10836 | 4ztu | A | 1236 | GLN | p.Gln1236His | P54098 |
| HOMO SAPIENS | 5 | 2666 | 1jph | A | 62 | PRO | p.Pro62Leu | P06132 |

| | | | | | | | | |
|---|---|---|---|---|---|---|---|---|
| HOMO SAPIENS | 5 | 2666 | 1jph | A | 116 | GLN | p.Gln116Ter | P06132 |
| HOMO SAPIENS | 0 | 2666 | 1jph | A | 165 | MET | p.Met165Arg | P06132 |
| HOMO SAPIENS | 6 | 2666 | 1jph | A | 167 | GLU | p.Glu167Lys | P06132 |
| HOMO SAPIENS | 0 | 2666 | 1jph | A | 195 | LEU | p.Leu195Phe | P06132 |
| HOMO SAPIENS | 7 | 2666 | 1jph | A | 253 | LEU | p.Leu253Gln | P06132 |
| HOMO SAPIENS | 0 | 2666 | 1jph | A | 281 | GLY | p.Gly281Glu | P06132 |
| HOMO SAPIENS | 9 | 2666 | 1jph | A | 304 | ASN | p.Asn304Lys | P06132 |
| HOMO SAPIENS | 8 | 2666 | 1jph | A | 311 | TYR | p.Tyr311Cys | P06132 |
| HOMO SAPIENS | 5 | 2666 | 1jph | A | 332 | ARG | p.Arg332His | P06132 |
| HOMO SAPIENS | 1 | 889 | 1m51 | A | 184 | LEU | p.Val184Leu | P35558 |
| HOMO SAPIENS | 0 | 889 | 1m51 | A | 267 | VAL | p.Ile267Val | P35558 |
| HOMO SAPIENS | 2 | 889 | 1m51 | A | 276 | GLU | p.Glu276Lys | P35558 |
| HOMO SAPIENS | 6 | 889 | 1m51 | A | 368 | VAL | p.Val368Ile | P35558 |
| HOMO SAPIENS | 0 | 889 | 1m51 | A | 579 | SER | p.Ser579Arg | P35558 |
| HOMO SAPIENS | 1 | 4084 | 2gjx | A | 28 | GLN | p.Gln28Ter | P06865 |
| HOMO SAPIENS | 0 | 4084 | 2gjx | A | 39 | LEU | p.Leu39Arg | P06865 |
| HOMO SAPIENS | 0 | 4084 | 2gjx | A | 127 | LEU | p.Leu127Arg | P06865 |
| HOMO SAPIENS | 6 | 4084 | 2gjx | A | 137 | ARG | p.Arg137Ter | P06865 |
| HOMO SAPIENS | 6 | 4084 | 2gjx | A | 170 | ARG | p.Arg170Gln | P06865 |
| HOMO SAPIENS | 9 | 4084 | 2gjx | A | 178 | ARG | p.Arg178Cys | P06865 |
| HOMO SAPIENS | 9 | 4084 | 2gjx | A | 178 | ARG | p.Arg178His | P06865 |
| HOMO SAPIENS | 5 | 4084 | 2gjx | A | 180 | TYR | p.Tyr180His | P06865 |
| HOMO SAPIENS | 5 | 4084 | 2gjx | A | 180 | TYR | p.Tyr180Ter | P06865 |
| HOMO SAPIENS | 1 | 4084 | 2gjx | A | 197 | LYS | p.Lys197Thr | P06865 |
| HOMO SAPIENS | 0 | 4084 | 2gjx | A | 200 | VAL | p.Val200Met | P06865 |
| HOMO SAPIENS | 9 | 4084 | 2gjx | A | 204 | HIS | p.His204Arg | P06865 |
| HOMO SAPIENS | 5 | 4084 | 2gjx | A | 210 | SER | p.Ser210Phe | P06865 |
| HOMO SAPIENS | 0 | 4084 | 2gjx | A | 211 | PHE | p.Phe211Ser | P06865 |
| HOMO SAPIENS | 2 | 4084 | 2gjx | A | 250 | GLY | p.Gly250Asp | P06865 |
| HOMO SAPIENS | 9 | 4084 | 2gjx | A | 258 | ASP | p.Asp258His | P06865 |
| HOMO SAPIENS | 0 | 4084 | 2gjx | A | 269 | GLY | p.Gly269Ser | P06865 |
| HOMO SAPIENS | 0 | 4084 | 2gjx | A | 301 | MET | p.Met301Arg | P06865 |
| HOMO SAPIENS | 3 | 4084 | 2gjx | A | 393 | ARG | p.Arg393Ter | P06865 |
| HOMO SAPIENS | 3 | 4084 | 2gjx | A | 393 | ARG | p.Arg393Pro | P06865 |
| HOMO SAPIENS | 0 | 4084 | 2gjx | A | 399 | ASN | p.Asn399Asp | P06865 |
| HOMO SAPIENS | 2 | 4084 | 2gjx | A | 420 | TRP | p.Trp420Cys | P06865 |
| HOMO SAPIENS | 1 | 4084 | 2gjx | A | 434 | PHE | p.Phe434Leu | P06865 |
| HOMO SAPIENS | 0 | 4084 | 2gjx | A | 454 | GLY | p.Gly454Ser | P06865 |
| HOMO SAPIENS | 1 | 4084 | 2gjx | A | 474 | TRP | p.Trp474Cys | P06865 |
| HOMO SAPIENS | 0 | 4084 | 2gjx | A | 479 | ALA | p.Ala479Thr | P06865 |
| HOMO SAPIENS | 5 | 4084 | 2gjx | A | 485 | TRP | p.Trp485Arg | P06865 |
| HOMO SAPIENS | 4 | 4084 | 2gjx | A | 499 | ARG | p.Arg499Cys | P06865 |
| HOMO SAPIENS | 4 | 4084 | 2gjx | A | 499 | ARG | p.Arg499His | P06865 |
| HOMO SAPIENS | 2 | 4084 | 2gjx | A | 504 | ARG | p.Arg504Cys | P06865 |
| HOMO SAPIENS | 2 | 4084 | 2gjx | A | 504 | ARG | p.Arg504His | P06865 |
| HOMO SAPIENS | -1 | 1943 | 1xmj | A | 417 | ASN | p.Asn417Lys | P13569 |
| HOMO SAPIENS | -1 | 1943 | 1xmj | A | 422 | SER | p.Ser422Phe | P13569 |
| HOMO SAPIENS | 1 | 1943 | 1xmj | A | 438 | THR | p.Thr438Ala | P13569 |
| HOMO SAPIENS | 0 | 1943 | 1xmj | A | 455 | ALA | p.Ala455Glu | P13569 |
| HOMO SAPIENS | 4 | 1943 | 1xmj | A | 456 | VAL | p.Val456Ala | P13569 |
| HOMO SAPIENS | 9 | 1943 | 1xmj | A | 458 | GLY | p.Gly458Val | P13569 |
| HOMO SAPIENS | 8 | 1943 | 1xmj | A | 466 | SER | p.Ser466Ter | P13569 |
| HOMO SAPIENS | 0 | 1943 | 1xmj | A | 467 | LEU | p.Leu467Pro | P13569 |
| HOMO SAPIENS | 1 | 1943 | 1xmj | A | 470 | MET | p.Val470Met | P13569 |

| | | | | | | | | |
|---|---|---|---|---|---|---|---|---|
| HOMO SAPIENS | 5 | 1943 | 1xmj | A | 480 | GLY | p.Gly480Cys | P13569 |
| HOMO SAPIENS | 3 | 1943 | 1xmj | A | 492 | SER | p.Ser492Phe | P13569 |
| HOMO SAPIENS | 8 | 1943 | 1xmj | A | 493 | GLN | p.Gln493Ter | P13569 |
| HOMO SAPIENS | 5 | 1943 | 1xmj | A | 506 | ILE | p.Ile506Met | P13569 |
| HOMO SAPIENS | 3 | 1943 | 1xmj | A | 520 | VAL | p.Val520Phe | P13569 |
| HOMO SAPIENS | 1 | 1943 | 1xmj | A | 528 | GLU | p.Glu528Asp | P13569 |
| HOMO SAPIENS | 1 | 1943 | 1xmj | A | 534 | ALA | p.Ala534Glu | P13569 |
| HOMO SAPIENS | 4 | 1943 | 1xmj | A | 542 | GLY | p.Gly542Ter | P13569 |
| HOMO SAPIENS | 6 | 1943 | 1xmj | A | 549 | SER | p.Ser549Asn | P13569 |
| HOMO SAPIENS | 6 | 1943 | 1xmj | A | 549 | SER | p.Ser549Arg | P13569 |
| HOMO SAPIENS | 6 | 1943 | 1xmj | A | 549 | SER | p.Ser549Arg | P13569 |
| HOMO SAPIENS | 6 | 1943 | 1xmj | A | 551 | GLY | p.Gly551Asp | P13569 |
| HOMO SAPIENS | 6 | 1943 | 1xmj | A | 551 | GLY | p.Gly551Ser | P13569 |
| HOMO SAPIENS | 6 | 1943 | 1xmj | A | 553 | GLN | p.Arg553Gln | P13569 |
| HOMO SAPIENS | 6 | 1943 | 1xmj | A | 553 | GLN | p.Arg553Ter | P13569 |
| HOMO SAPIENS | 0 | 1943 | 1xmj | A | 559 | ALA | p.Ala559Thr | P13569 |
| HOMO SAPIENS | 7 | 1943 | 1xmj | A | 560 | ARG | p.Arg560Lys | P13569 |
| HOMO SAPIENS | 4 | 1943 | 1xmj | A | 562 | VAL | p.Val562Ile | P13569 |
| HOMO SAPIENS | 3 | 1943 | 1xmj | A | 563 | TYR | p.Tyr563Asn | P13569 |
| HOMO SAPIENS | 3 | 1943 | 1xmj | A | 569 | TYR | p.Tyr569Asp | P13569 |
| HOMO SAPIENS | 3 | 1943 | 1xmj | A | 585 | GLU | p.Glu585Ter | P13569 |
| HOMO SAPIENS | 1 | 1943 | 1xmj | A | 618 | ILE | p.Ile618Thr | P13569 |
| HOMO SAPIENS | 9 | 1943 | 1xmj | A | 622 | GLY | p.Gly622Asp | P13569 |
| HOMO SAPIENS | -1 | 1943 | 1xmj | A | 648 | ASP | p.Asp648Val | P13569 |
| HOMO SAPIENS | 0 | 1943 | 1xmj | A | 673 | GLY | p.Gly673Ter | P13569 |
| HOMO SAPIENS | 5 | 5111 | 4rec | A | 749 | ARG | p.Arg749Ter | Q9Y2M0 |
| HOMO SAPIENS | -1 | 4580 | 4l0d | B | 2 | GLY | p.Pro2Leu | P35520 |
| HOMO SAPIENS | -1 | 4580 | 4l0d | B | 12 | PRO | p.Pro12Ser | P35520 |
| HOMO SAPIENS | -1 | 4580 | 4l0d | B | 18 | ARG | p.Arg18Cys | P35520 |
| HOMO SAPIENS | 1 | 4580 | 4l0d | B | 49 | PRO | p.Pro49Leu | P35520 |
| HOMO SAPIENS | 2 | 4580 | 4l0d | B | 70 | PRO | p.Pro70Leu | P35520 |
| HOMO SAPIENS | 1 | 4580 | 4l0d | B | 102 | LYS | p.Lys102Gln | P35520 |
| HOMO SAPIENS | 7 | 4580 | 4l0d | B | 114 | ALA | p.Ala114Val | P35520 |
| HOMO SAPIENS | 5 | 4580 | 4l0d | B | 116 | GLY | p.Gly116Arg | P35520 |
| HOMO SAPIENS | 8 | 4580 | 4l0d | B | 121 | ARG | p.Arg121Cys | P35520 |
| HOMO SAPIENS | 8 | 4580 | 4l0d | B | 121 | ARG | p.Arg121His | P35520 |
| HOMO SAPIENS | 5 | 4580 | 4l0d | B | 125 | ARG | p.Arg125Gln | P35520 |
| HOMO SAPIENS | 4 | 4580 | 4l0d | B | 134 | GLY | p.Gly134Arg | P35520 |
| HOMO SAPIENS | 4 | 4580 | 4l0d | B | 139 | GLY | p.Gly139Arg | P35520 |
| HOMO SAPIENS | 0 | 4580 | 4l0d | B | 145 | PRO | p.Pro145Leu | P35520 |
| HOMO SAPIENS | 0 | 4580 | 4l0d | B | 148 | GLY | p.Gly148Arg | P35520 |
| HOMO SAPIENS | 0 | 4580 | 4l0d | B | 168 | VAL | p.Val168Met | P35520 |
| HOMO SAPIENS | 2 | 4580 | 4l0d | B | 191 | THR | p.Thr191Met | P35520 |
| HOMO SAPIENS | 5 | 4580 | 4l0d | B | 234 | ASP | p.Asp234Asn | P35520 |
| HOMO SAPIENS | 0 | 4580 | 4l0d | B | 257 | THR | p.Thr257Met | P35520 |
| HOMO SAPIENS | 4 | 4580 | 4l0d | B | 262 | THR | p.Thr262Met | P35520 |
| HOMO SAPIENS | 0 | 4580 | 4l0d | B | 278 | ILE | p.Ile278Thr | P35520 |
| HOMO SAPIENS | 2 | 4580 | 4l0d | B | 307 | GLY | p.Gly307Ser | P35520 |
| HOMO SAPIENS | 4 | 4580 | 4l0d | B | 320 | VAL | p.Val320Ala | P35520 |
| HOMO SAPIENS | 2 | 4580 | 4l0d | B | 336 | ARG | p.Arg336Cys | P35520 |
| HOMO SAPIENS | 2 | 4580 | 4l0d | B | 336 | ARG | p.Arg336His | P35520 |
| HOMO SAPIENS | 0 | 4580 | 4l0d | B | 351 | GLY | p.Gly351Arg | P35520 |
| HOMO SAPIENS | 0 | 4580 | 4l0d | B | 353 | THR | p.Thr353Met | P35520 |
| HOMO SAPIENS | 0 | 4580 | 4l0d | B | 371 | VAL | p.Val371Met | P35520 |

| | | | | | | | | |
|---|---|---|---|---|---|---|---|---|
| HOMO SAPIENS | 2 | 4580 | 4l0d | B | 379 | ARG | p.Arg379Gln | P35520 |
| HOMO SAPIENS | 2 | 4580 | 4l0d | B | 379 | ARG | p.Arg379Trp | P35520 |
| HOMO SAPIENS | 2 | 4580 | 4l0d | B | 384 | LYS | p.Lys384Glu | P35520 |
| HOMO SAPIENS | 0 | 4580 | 4l0d | B | 422 | PRO | p.Pro422Leu | P35520 |
| HOMO SAPIENS | 0 | 4580 | 4l0d | B | 444 | ASP | p.Asp444Asn | P35520 |
| HOMO SAPIENS | 0 | 4580 | 4l0d | B | 466 | SER | p.Ser466Leu | P35520 |
| HOMO SAPIENS | 0 | 4580 | 4l0d | B | 539 | LEU | p.Leu539Ser | P35520 |
| HOMO SAPIENS | 0 | 4580 | 4l0d | B | 548 | ARG | p.Arg548Gln | P35520 |
| HOMO SAPIENS | 1 | 6536 | 3ulu | A | 412 | LEU | p.Leu412Phe | O15455 |
| HOMO SAPIENS | 1 | 28493 | 3ii6 | X | 814 | ARG | p.Arg814Ter | P49917 |
| HOMO SAPIENS | 1 | 28493 | 3ii6 | X | 857 | ALA | p.Ala857Thr | P49917 |
| HOMO SAPIENS | 9 | 3543 | 2q51 | A | 24 | GLU | p.Glu24Gly | P45381 |
| HOMO SAPIENS | 1 | 3543 | 2q51 | A | 27 | GLY | p.Gly27Arg | P45381 |
| HOMO SAPIENS | 9 | 3543 | 2q51 | A | 71 | ARG | p.Arg71His | P45381 |
| HOMO SAPIENS | -1 | 3543 | 2q51 | A | 82 | MSE | p.Met82Thr | P45381 |
| HOMO SAPIENS | 0 | 3543 | 2q51 | A | 143 | ILE | p.Ile143Phe | P45381 |
| HOMO SAPIENS | 1 | 3543 | 2q51 | A | 152 | CYS | p.Cys152Arg | P45381 |
| HOMO SAPIENS | 2 | 3543 | 2q51 | A | 218 | CYS | p.Cys218Ter | P45381 |
| HOMO SAPIENS | 1 | 3543 | 2q51 | A | 231 | TYR | p.Tyr231Cys | P45381 |
| HOMO SAPIENS | 1 | 3543 | 2q51 | A | 231 | TYR | p.Tyr231Ter | P45381 |
| HOMO SAPIENS | 2 | 3543 | 2q51 | A | 274 | GLY | p.Gly274Arg | P45381 |
| HOMO SAPIENS | 6 | 3543 | 2q51 | A | 285 | GLU | p.Glu285Ala | P45381 |
| HOMO SAPIENS | 5 | 3543 | 2q51 | A | 287 | ALA | p.Ala287Thr | P45381 |
| HOMO SAPIENS | 2 | 3543 | 2q51 | A | 305 | ALA | p.Ala305Glu | P45381 |
| HOMO SAPIENS | 2 | 24687 | 1hfi | A | 25 | SER | p.Ser890Ile | P08603 |
| HOMO SAPIENS | 0 | 1508 | 3wn5 | C | 45 | LEU | p.Leu66Arg | P08637 |
| HOMO SAPIENS | 0 | 1508 | 3wn5 | C | 155 | VAL | p.Phe176Val | P08637 |
| HOMO SAPIENS | 0 | 750 | 5tpa | A | 54 | VAL | p.Val452Met | Q12879 |
| HOMO SAPIENS | 9 | 750 | 5tpa | A | 120 | ARG | p.Arg518Cys | Q12879 |
| HOMO SAPIENS | 0 | 750 | 5tpa | A | 199 | ALA | p.Ala716Thr | Q12879 |
| HOMO SAPIENS | 4 | 1363 | 3hg2 | B | 40 | PRO | p.Pro40Ser | P06280 |
| HOMO SAPIENS | 3 | 1363 | 3hg2 | B | 65 | SER | p.Ser65Thr | P06280 |
| HOMO SAPIENS | 4 | 1363 | 3hg2 | B | 66 | GLU | p.Glu66Gln | P06280 |
| HOMO SAPIENS | 2 | 1363 | 3hg2 | B | 80 | GLY | p.Gly80Asp | P06280 |
| HOMO SAPIENS | 3 | 1363 | 3hg2 | B | 112 | ARG | p.Arg112Cys | P06280 |
| HOMO SAPIENS | 3 | 1363 | 3hg2 | B | 112 | ARG | p.Arg112His | P06280 |
| HOMO SAPIENS | 0 | 1363 | 3hg2 | B | 139 | ASN | p.Asn139Ser | P06280 |
| HOMO SAPIENS | 7 | 1363 | 3hg2 | B | 156 | ALA | p.Ala156Thr | P06280 |
| HOMO SAPIENS | 5 | 1363 | 3hg2 | B | 162 | TRP | p.Trp162Arg | P06280 |
| HOMO SAPIENS | 5 | 1363 | 3hg2 | B | 207 | TYR | p.Tyr207His | P06280 |
| HOMO SAPIENS | 0 | 1363 | 3hg2 | B | 215 | ASN | p.Asn215Ser | P06280 |
| HOMO SAPIENS | 0 | 1363 | 3hg2 | B | 264 | ASP | p.Asp264Val | P06280 |
| HOMO SAPIENS | 5 | 1363 | 3hg2 | B | 266 | ASP | p.Asp266Val | P06280 |
| HOMO SAPIENS | 2 | 1363 | 3hg2 | B | 269 | VAL | p.Val269Ala | P06280 |
| HOMO SAPIENS | 3 | 1363 | 3hg2 | B | 272 | ASN | p.Asn272Ser | P06280 |
| HOMO SAPIENS | 3 | 1363 | 3hg2 | B | 279 | GLN | p.Gln279Glu | P06280 |
| HOMO SAPIENS | 0 | 1363 | 3hg2 | B | 289 | ILE | p.Ile289Val | P06280 |
| HOMO SAPIENS | 0 | 1363 | 3hg2 | B | 296 | MET | p.Met296Val | P06280 |
| HOMO SAPIENS | 0 | 1363 | 3hg2 | B | 296 | MET | p.Met296Ile | P06280 |
| HOMO SAPIENS | 0 | 1363 | 3hg2 | B | 297 | SER | p.Ser297Phe | P06280 |
| HOMO SAPIENS | 2 | 1363 | 3hg2 | B | 301 | ARG | p.Arg301Gln | P06280 |
| HOMO SAPIENS | 3 | 1363 | 3hg2 | B | 327 | GLN | p.Gln327Lys | P06280 |
| HOMO SAPIENS | 0 | 1363 | 3hg2 | B | 328 | GLY | p.Gly328Arg | P06280 |
| HOMO SAPIENS | 0 | 1363 | 3hg2 | B | 328 | GLY | p.Gly328Ala | P06280 |

| | | | | | | | | |
|---|---|---|---|---|---|---|---|---|
| HOMO SAPIENS | 2 | 1363 | 3hg2 | B | 331 | LEU | p.Leu331Phe | P06280 |
| HOMO SAPIENS | 0 | 1363 | 3hg2 | B | 342 | ARG | p.Arg342Gln | P06280 |
| HOMO SAPIENS | 1 | 1363 | 3hg2 | B | 361 | GLY | p.Gly361Arg | P06280 |
| HOMO SAPIENS | 1 | 1363 | 3hg2 | B | 363 | ARG | p.Arg363His | P06280 |
| HOMO SAPIENS | 1 | 1363 | 3hg2 | B | 385 | THR | p.Thr385Ala | P06280 |
| HOMO SAPIENS | 1 | 1363 | 3hg2 | B | 399 | TRP | p.Trp399Ser | P06280 |
| HOMO SAPIENS | 1 | 1363 | 3hg2 | B | 410 | THR | p.Thr410Ala | P06280 |
| HOMO SAPIENS | 2 | 3904 | 3caa | B | 385 | MET | p.Met414Val | P01011 |
| HOMO SAPIENS | 0 | 558 | 5eub | A | 56 | VAL | p.Val56Ile | Q13126 |
| HOMO SAPIENS | 0 | 558 | 5eub | A | 189 | TRP | p.Trp189Leu | Q13126 |
| HOMO SAPIENS | 0 | 558 | 5eub | A | 218 | ALA | p.Ala218Gly | Q13126 |
| HOMO SAPIENS | 1 | 3865 | 1cb5 | B | 443 | ILE | p.Ile443Val | Q13867 |
| HOMO SAPIENS | 2 | 14329 | 4m92 | A | 24 | ILE | p.Ile65Val | Q13454 |
| HOMO SAPIENS | 1 | 7381 | 5kz5 | k | 50 | GLY | p.Gly50Glu | Q9H1K1 |
| HOMO SAPIENS | 0 | 1102 | 4fdh | E | 86 | PRO | p.Pro86Ala | P19099 |
| HOMO SAPIENS | 0 | 1102 | 4fdh | E | 118 | ALA | p.Ala118Thr | P19099 |
| HOMO SAPIENS | 0 | 1102 | 4fdh | E | 159 | PRO | p.Pro159Leu | P19099 |
| HOMO SAPIENS | 0 | 1102 | 4fdh | E | 173 | LYS | p.Lys173Arg | P19099 |
| HOMO SAPIENS | 0 | 1102 | 4fdh | E | 181 | ARG | p.Arg181Trp | P19099 |
| HOMO SAPIENS | 0 | 1102 | 4fdh | E | 185 | THR | p.Thr185Ile | P19099 |
| HOMO SAPIENS | 0 | 1102 | 4fdh | E | 198 | GLU | p.Glu198Asp | P19099 |
| HOMO SAPIENS | 0 | 1102 | 4fdh | E | 214 | HIS | p.His214Asp | P19099 |
| HOMO SAPIENS | 0 | 1102 | 4fdh | E | 251 | LYS | p.Lys251Arg | P19099 |
| HOMO SAPIENS | 0 | 1102 | 4fdh | E | 255 | GLU | p.Glu255Ter | P19099 |
| HOMO SAPIENS | 0 | 1102 | 4fdh | E | 281 | ASN | p.Asn281Ser | P19099 |
| HOMO SAPIENS | 0 | 1102 | 4fdh | E | 320 | ALA | p.Ala320Val | P19099 |
| HOMO SAPIENS | 0 | 1102 | 4fdh | E | 339 | ILE | p.Ile339Thr | P19099 |
| HOMO SAPIENS | 0 | 1102 | 4fdh | E | 341 | ARG | p.Arg341Cys | P19099 |
| HOMO SAPIENS | 2 | 1102 | 4fdh | E | 374 | ARG | p.Arg374Trp | P19099 |
| HOMO SAPIENS | 0 | 1102 | 4fdh | E | 386 | VAL | p.Val386Ala | P19099 |
| HOMO SAPIENS | 0 | 1102 | 4fdh | E | 390 | LEU | p.Leu390Phe | P19099 |
| HOMO SAPIENS | -1 | 1102 | 4fdh | E | 435 | GLY | p.Gly435Ser | P19099 |
| HOMO SAPIENS | 0 | 1102 | 4fdh | E | 461 | LEU | p.Leu461Pro | P19099 |
| HOMO SAPIENS | 0 | 1102 | 4fdh | E | 498 | THR | p.Thr498Ala | P19099 |
| HOMO SAPIENS | 6 | 10292 | 2of5 | F | 128 | GLY | p.Gly128Arg | P78560 |
| HOMO SAPIENS | 6 | 10292 | 2of5 | F | 170 | ARG | p.Arg170Cys | P78560 |
| HOMO SAPIENS | -1 | 18965 | 3ghg | A | 16 | ARG | p.Arg35Cys | P02671 |
| HOMO SAPIENS | -1 | 18965 | 3ghg | A | 16 | ARG | p.Arg35His | P02671 |
| HOMO SAPIENS | -1 | 18965 | 3ghg | A | 283 | PRO | p.Pro302Ala | P02671 |
| HOMO SAPIENS | -1 | 18965 | 3ghg | A | 312 | THR | p.Thr331Ala | P02671 |
| HOMO SAPIENS | -1 | 18965 | 3ghg | A | 381 | SER | p.Ser400Phe | P02671 |
| HOMO SAPIENS | -1 | 18965 | 3ghg | A | 463 | VAL | p.Val482Met | P02671 |
| HOMO SAPIENS | -1 | 18965 | 3ghg | A | 526 | GLU | p.Glu545Val | P02671 |
| HOMO SAPIENS | -1 | 18965 | 3ghg | A | 554 | ARG | p.Arg573Leu | P02671 |
| HOMO SAPIENS | 8 | 3023 | 4ttb | A | 101 | ARG | p.Arg101Trp | Q6PHW0 |
| HOMO SAPIENS | 2 | 3023 | 4ttb | A | 116 | ILE | p.Ile116Thr | Q6PHW0 |
| HOMO SAPIENS | 0 | 3023 | 4ttb | A | 220 | ALA | p.Ala220Thr | Q6PHW0 |
| HOMO SAPIENS | 2 | 3023 | 4ttb | A | 271 | GLU | p.Glu271Lys | Q6PHW0 |
| HOMO SAPIENS | 3 | 33226 | 1ya5 | T | 11 | SER | p.Ser11Ter | O15273 |
| HOMO SAPIENS | 6 | 33226 | 1ya5 | T | 64 | SER | p.Ser64Leu | O15273 |
| HOMO SAPIENS | 6 | 33226 | 1ya5 | T | 70 | ARG | p.Arg70Gln | O15273 |
| HOMO SAPIENS | 3 | 33226 | 1ya5 | T | 75 | GLY | p.Gly75Ser | O15273 |
| HOMO SAPIENS | 0 | 33226 | 1ya5 | T | 87 | ARG | p.Arg87Trp | O15273 |
| HOMO SAPIENS | 2 | 5667 | 3dax | A | 398 | PRO | p.Pro398Ala | P22680 |

| | | | | | | | | |
|---|---|---|---|---|---|---|---|---|
| HOMO SAPIENS | 0 | 12861 | 5mya | A | 464 | ASN | p.Asn464His | Q02763 |
| HOMO SAPIENS | 2 | 12861 | 5mya | A | 486 | VAL | p.Val486Ile | Q02763 |
| HOMO SAPIENS | 0 | 12861 | 5mya | A | 522 | ARG | p.Arg522His | Q02763 |
| HOMO SAPIENS | 2 | 12861 | 5mya | A | 600 | VAL | p.Val600Leu | Q02763 |
| HOMO SAPIENS | 0 | 12861 | 5mya | A | 634 | LEU | p.Leu634Phe | Q02763 |
| HOMO SAPIENS | 0 | 12861 | 5mya | A | 684 | HIS | p.His684Gln | Q02763 |
| HOMO SAPIENS | 0 | 12861 | 5mya | A | 724 | ALA | p.Ala724Thr | Q02763 |
| HOMO SAPIENS | 0 | 7743 | 3qzw | G | 90 | GLY | p.Gly111Ser | P01732 |
| HOMO SAPIENS | 2 | 9498 | 3kci | A | 4267 | ASP | p.Asp4267Glu | O95714 |
| HOMO SAPIENS | 5 | 201 | 3ips | B | 415 | ARG | p.Arg415Gln | Q13133 |
| HOMO SAPIENS | 3 | 229 | 2q50 | A | 68 | LEU | p.Leu68Pro | Q9UBQ7 |
| HOMO SAPIENS | 2 | 229 | 2q50 | A | 99 | ARG | p.Arg99Ter | Q9UBQ7 |
| HOMO SAPIENS | 3 | 229 | 2q50 | A | 160 | GLY | p.Gly160Arg | Q9UBQ7 |
| HOMO SAPIENS | 5 | 229 | 2q50 | A | 165 | GLY | p.Gly165Asp | Q9UBQ7 |
| HOMO SAPIENS | 4 | 229 | 2q50 | A | 302 | ARG | p.Arg302Cys | Q9UBQ7 |
| HOMO SAPIENS | 4 | 229 | 2q50 | A | 302 | ARG | p.Arg302His | Q9UBQ7 |
| HOMO SAPIENS | -1 | 229 | 2q50 | A | 322 | MSE | p.Met322Arg | Q9UBQ7 |
| HOMO SAPIENS | -1 | 4519 | 5clw | A | 40 | PRO | p.Pro40Thr | Q04446 |
| HOMO SAPIENS | 1 | 4519 | 5clw | A | 59 | ILE | p.Ile59Thr | Q04446 |
| HOMO SAPIENS | 0 | 4519 | 5clw | A | 190 | ARG | p.Arg190Gly | Q04446 |
| HOMO SAPIENS | 1 | 4519 | 5clw | A | 224 | LEU | p.Leu224Pro | Q04446 |
| HOMO SAPIENS | 0 | 4519 | 5clw | A | 236 | GLN | p.Gln236His | Q04446 |
| HOMO SAPIENS | 1 | 4519 | 5clw | A | 254 | THR | p.Thr254Ala | Q04446 |
| HOMO SAPIENS | 0 | 4519 | 5clw | A | 257 | PHE | p.Phe257Leu | Q04446 |
| HOMO SAPIENS | 1 | 4519 | 5clw | A | 262 | ARG | p.Arg262Cys | Q04446 |
| HOMO SAPIENS | 3 | 4519 | 5clw | A | 329 | TYR | p.Tyr329Ser | Q04446 |
| HOMO SAPIENS | 0 | 4519 | 5clw | A | 507 | THR | p.Thr507Ala | Q04446 |
| HOMO SAPIENS | 0 | 4519 | 5clw | A | 515 | ARG | p.Arg515Cys | Q04446 |
| HOMO SAPIENS | 0 | 4519 | 5clw | A | 515 | ARG | p.Arg515His | Q04446 |
| HOMO SAPIENS | 0 | 4519 | 5clw | A | 524 | ARG | p.Arg524Gln | Q04446 |
| HOMO SAPIENS | 0 | 4519 | 5clw | A | 524 | ARG | p.Arg524Ter | Q04446 |
| HOMO SAPIENS | 0 | 4519 | 5clw | A | 545 | HIS | p.His545Arg | Q04446 |
| HOMO SAPIENS | 2 | 4519 | 5clw | A | 548 | TRP | p.Trp548Ter | Q04446 |
| HOMO SAPIENS | 0 | 4519 | 5clw | A | 592 | GLU | p.Glu592Ter | Q04446 |
| HOMO SAPIENS | 0 | 4519 | 5clw | A | 628 | HIS | p.His628Arg | Q04446 |
| HOMO SAPIENS | 0 | 4519 | 5clw | A | 637 | ARG | p.Arg637Ter | Q04446 |
| HOMO SAPIENS | 0 | 4519 | 5clw | A | 673 | ALA | p.Ala673Thr | Q04446 |
| HOMO SAPIENS | 1 | 10405 | 3bg3 | D | 507 | VAL | p.Val507Ile | P11498 |
| HOMO SAPIENS | 0 | 10405 | 3bg3 | D | 583 | ARG | p.Arg583Leu | P11498 |
| HOMO SAPIENS | 1 | 10405 | 3bg3 | D | 589 | LYS | p.Lys589Asn | P11498 |
| HOMO SAPIENS | 6 | 10405 | 3bg3 | D | 610 | ALA | p.Ala610Thr | P11498 |
| HOMO SAPIENS | 2 | 10405 | 3bg3 | D | 631 | ARG | p.Arg631Gln | P11498 |
| HOMO SAPIENS | 1 | 10405 | 3bg3 | D | 732 | ARG | p.Arg732Gln | P11498 |
| HOMO SAPIENS | 6 | 10405 | 3bg3 | D | 743 | MET | p.Met743Ile | P11498 |
| HOMO SAPIENS | 0 | 10405 | 3bg3 | D | 750 | THR | p.Thr750Met | P11498 |
| HOMO SAPIENS | -1 | 13751 | 2o8d | B | 350 | ALA | p.Ala350Val | P52701 |
| HOMO SAPIENS | -1 | 13751 | 2o8d | B | 352 | VAL | p.Val352Ile | P52701 |
| HOMO SAPIENS | -1 | 13751 | 2o8d | B | 355 | GLY | p.Gly355Ser | P52701 |
| HOMO SAPIENS | 0 | 13751 | 2o8d | B | 385 | ARG | p.Arg385Trp | P52701 |
| HOMO SAPIENS | 0 | 13751 | 2o8d | B | 390 | ASP | p.Asp390Glu | P52701 |
| HOMO SAPIENS | 0 | 13751 | 2o8d | B | 396 | LEU | p.Leu396Val | P52701 |
| HOMO SAPIENS | 0 | 13751 | 2o8d | B | 449 | LEU | p.Leu449Pro | P52701 |
| HOMO SAPIENS | 0 | 13751 | 2o8d | B | 468 | ARG | p.Arg468His | P52701 |
| HOMO SAPIENS | 0 | 13751 | 2o8d | B | 503 | SER | p.Ser503Cys | P52701 |

| | | | | | | | | |
|---|---|---|---|---|---|---|---|---|
| HOMO SAPIENS | 1 | 13751 | 2o8d | B | 509 | VAL | p.Val509Ala | P52701 |
| HOMO SAPIENS | 0 | 13751 | 2o8d | B | 675 | GLU | p.Glu675Asp | P52701 |
| HOMO SAPIENS | 0 | 13751 | 2o8d | B | 795 | ILE | p.Ile795Thr | P52701 |
| HOMO SAPIENS | 0 | 13751 | 2o8d | B | 850 | TYR | p.Tyr850Ter | P52701 |
| HOMO SAPIENS | 0 | 13751 | 2o8d | B | 878 | VAL | p.Val878Ala | P52701 |
| HOMO SAPIENS | 0 | 13751 | 2o8d | B | 911 | ARG | p.Arg911Ter | P52701 |
| HOMO SAPIENS | 0 | 13751 | 2o8d | B | 922 | ARG | p.Arg922Ter | P52701 |
| HOMO SAPIENS | 0 | 13751 | 2o8d | B | 969 | TYR | p.Tyr969Cys | P52701 |
| HOMO SAPIENS | 0 | 13751 | 2o8d | B | 1005 | ARG | p.Arg1005Ter | P52701 |
| HOMO SAPIENS | 0 | 13751 | 2o8d | B | 1035 | ARG | p.Arg1035Ter | P52701 |
| HOMO SAPIENS | 1 | 13751 | 2o8d | B | 1048 | GLN | p.Gln1048Ter | P52701 |
| HOMO SAPIENS | 1 | 13751 | 2o8d | B | 1051 | VAL | p.Val1051Ile | P52701 |
| HOMO SAPIENS | 0 | 13751 | 2o8d | B | 1054 | ILE | p.Ile1054Phe | P52701 |
| HOMO SAPIENS | 0 | 13751 | 2o8d | B | 1054 | ILE | p.Ile1054Met | P52701 |
| HOMO SAPIENS | 0 | 13751 | 2o8d | B | 1163 | GLU | p.Glu1163Val | P52701 |
| HOMO SAPIENS | 0 | 13751 | 2o8d | B | 1237 | GLU | p.Glu1237Asp | P52701 |
| HOMO SAPIENS | 0 | 13751 | 2o8d | B | 1242 | ARG | p.Arg1242His | P52701 |
| HOMO SAPIENS | -1 | 13751 | 2o8d | B | 1278 | PRO | p.Pro1278Thr | P52701 |
| HOMO SAPIENS | 0 | 13751 | 2o8d | B | 1304 | ARG | p.Arg1304Lys | P52701 |
| HOMO SAPIENS | 0 | 13751 | 2o8d | B | 1317 | HIS | p.His1317Gln | P52701 |
| HOMO SAPIENS | 0 | 13751 | 2o8d | B | 1329 | SER | p.Ser1329Leu | P52701 |
| HOMO SAPIENS | 3 | 13273 | 4g8k | A | 265 | GLU | p.Glu265Ter | Q05823 |
| HOMO SAPIENS | -1 | 11088 | 5uj7 | A | 666 | ARG | p.Arg666Trp | Q13415 |
| HOMO SAPIENS | 4 | 11088 | 5uj7 | A | 720 | ARG | p.Arg720Gln | Q13415 |
| HOMO SAPIENS | 0 | 11088 | 5uj7 | A | 816 | MET | p.Met816Thr | Q13415 |
| HOMO SAPIENS | 1 | 4352 | 5va1 | A | 421 | THR | p.Thr421Met | Q12809 |
| HOMO SAPIENS | 0 | 4352 | 5va1 | A | 451 | PRO | p.Pro451Leu | Q12809 |
| HOMO SAPIENS | 0 | 4352 | 5va1 | A | 470 | ASN | p.Asn470Asp | Q12809 |
| HOMO SAPIENS | 0 | 4352 | 5va1 | A | 490 | ALA | p.Ala490Thr | Q12809 |
| HOMO SAPIENS | 0 | 4352 | 5va1 | A | 491 | VAL | p.Val491Ile | Q12809 |
| HOMO SAPIENS | 1 | 4352 | 5va1 | A | 533 | VAL | p.Val533Gly | Q12809 |
| HOMO SAPIENS | 0 | 4352 | 5va1 | A | 552 | LEU | p.Leu552Ser | Q12809 |
| HOMO SAPIENS | 0 | 4352 | 5va1 | A | 558 | ALA | p.Ala558Pro | Q12809 |
| HOMO SAPIENS | 1 | 4352 | 5va1 | A | 561 | ALA | p.Ala561Val | Q12809 |
| HOMO SAPIENS | 0 | 4352 | 5va1 | A | 572 | GLY | p.Gly572Arg | Q12809 |
| HOMO SAPIENS | -1 | 4352 | 5va1 | A | 582 | ARG | p.Arg582Cys | Q12809 |
| HOMO SAPIENS | 0 | 4352 | 5va1 | A | 584 | GLY | p.Gly584Ser | Q12809 |
| HOMO SAPIENS | 0 | 4352 | 5va1 | A | 588 | ASN | p.Asn588Lys | Q12809 |
| HOMO SAPIENS | 0 | 4352 | 5va1 | A | 593 | ILE | p.Ile593Arg | Q12809 |
| HOMO SAPIENS | 1 | 4352 | 5va1 | A | 627 | PHE | p.Phe627Leu | Q12809 |
| HOMO SAPIENS | 0 | 4352 | 5va1 | A | 630 | VAL | p.Val630Leu | Q12809 |
| HOMO SAPIENS | 0 | 4352 | 5va1 | A | 652 | TYR | p.Tyr652Ter | Q12809 |
| HOMO SAPIENS | 0 | 4352 | 5va1 | A | 711 | ILE | p.Ile711Val | Q12809 |
| HOMO SAPIENS | 0 | 4352 | 5va1 | A | 735 | SER | p.Ser735Leu | Q12809 |
| HOMO SAPIENS | 0 | 4352 | 5va1 | A | 744 | ARG | p.Arg744Ter | Q12809 |
| HOMO SAPIENS | 0 | 4352 | 5va1 | A | 784 | ARG | p.Arg784Trp | Q12809 |
| HOMO SAPIENS | 1 | 4352 | 5va1 | A | 820 | GLY | p.Gly820Arg | Q12809 |
| HOMO SAPIENS | 5 | 4352 | 5va1 | A | 822 | VAL | p.Val822Met | Q12809 |
| HOMO SAPIENS | 1 | 4352 | 5va1 | A | 863 | ARG | p.Arg863Ter | Q12809 |
| HOMO SAPIENS | 5 | 303 | 2obv | A | 55 | ALA | p.Ala55Asp | Q00266 |
| HOMO SAPIENS | 7 | 303 | 2obv | A | 259 | ALA | p.Ala259Gly | Q00266 |
| HOMO SAPIENS | 9 | 303 | 2obv | A | 264 | ARG | p.Arg264His | Q00266 |
| HOMO SAPIENS | 9 | 303 | 2obv | A | 264 | ARG | p.Arg264Cys | Q00266 |
| HOMO SAPIENS | 0 | 303 | 2obv | A | 305 | LEU | p.Leu305Pro | Q00266 |

| | | | | | | | | |
|---|---|---|---|---|---|---|---|---|
| HOMO SAPIENS | 7 | 303 | 2obv | A | 322 | ILE | p.Ile322Met | Q00266 |
| HOMO SAPIENS | 0 | 303 | 2obv | A | 336 | GLY | p.Gly336Arg | Q00266 |
| HOMO SAPIENS | 0 | 303 | 2obv | A | 357 | PRO | p.Pro357Leu | Q00266 |
| HOMO SAPIENS | 0 | 2077 | 4zxb | E | 462 | ASN | p.Asn489Ser | P06213 |
| HOMO SAPIENS | -1 | 2077 | 4zxb | E | 519 | ASP | p.Asp546Glu | P06213 |
| HOMO SAPIENS | 2 | 2077 | 4zxb | E | 819 | THR | p.Thr858Ala | P06213 |
| HOMO SAPIENS | 0 | 2077 | 4zxb | E | 907 | ASP | p.Asp946Glu | P06213 |
| HOMO SAPIENS | 7 | 3274 | 1wsv | B | 14 | HIS | p.His42Arg | P48728 |
| HOMO SAPIENS | 0 | 3274 | 1wsv | B | 19 | GLY | p.Gly47Arg | P48728 |
| HOMO SAPIENS | 4 | 3274 | 1wsv | B | 45 | ARG | p.Arg73Cys | P48728 |
| HOMO SAPIENS | 1 | 3274 | 1wsv | B | 49 | SER | p.Ser77Leu | P48728 |
| HOMO SAPIENS | 3 | 3274 | 1wsv | B | 241 | GLY | p.Gly269Asp | P48728 |
| HOMO SAPIENS | 7 | 3274 | 1wsv | B | 248 | ASP | p.Asp276His | P48728 |
| HOMO SAPIENS | 0 | 3274 | 1wsv | B | 272 | MET | p.Met300Val | P48728 |
| HOMO SAPIENS | 4 | 3274 | 1wsv | B | 292 | ARG | p.Arg320His | P48728 |
| HOMO SAPIENS | 0 | 3274 | 1wsv | B | 354 | ARG | p.Arg382Gln | P48728 |
| HOMO SAPIENS | -1 | 435 | 3uyt | A | 44 | THR | p.Thr44Ala | P48730 |
| HOMO SAPIENS | 1 | 4698 | 5ife | C | 666 | LYS | p.Lys666Asn | O75533 |
| HOMO SAPIENS | 1 | 16917 | 1apy | C | 37 | GLY | p.Gly60Asp | P20933 |
| HOMO SAPIENS | 0 | 16917 | 1apy | C | 49 | SER | p.Ser72Pro | P20933 |
| HOMO SAPIENS | 0 | 16917 | 1apy | C | 84 | ARG | p.Arg107Ter | P20933 |
| HOMO SAPIENS | 0 | 16917 | 1apy | C | 126 | SER | p.Thr149Ser | P20933 |
| HOMO SAPIENS | 0 | 16917 | 1apy | C | 135 | TRP | p.Trp158Ter | P20933 |
| HOMO SAPIENS | 0 | 16917 | 1apy | C | 140 | CYS | p.Cys163Ser | P20933 |
| HOMO SAPIENS | 0 | 16917 | 1apy | C | 145 | TRP | p.Trp168Ter | P20933 |
| HOMO SAPIENS | 0 | 10413 | 4frw | A | 185 | THR | p.Thr185Met | Q96NY8 |
| HOMO SAPIENS | 0 | 5217 | 5n6w | G | 96 | TRP | p.Trp96Arg | O15537 |
| HOMO SAPIENS | 1 | 5217 | 5n6w | G | 102 | ARG | p.Arg102Trp | O15537 |
| HOMO SAPIENS | 0 | 5217 | 5n6w | G | 109 | GLY | p.Gly109Arg | O15537 |
| HOMO SAPIENS | -1 | 5217 | 5n6w | G | 223 | CYS | p.Cys223Arg | O15537 |
| HOMO SAPIENS | 0 | 9485 | 4z6y | E | 138 | ALA | p.Ala138Asp | Q9P0N9 |
| HOMO SAPIENS | 3 | 4504 | 4mwt | B | 411 | GLY | p.Gly439Ser | P10619 |
| HOMO SAPIENS | 3 | 4504 | 4mwt | B | 412 | PHE | p.Phe440Val | P10619 |
| HOMO SAPIENS | 6 | 6391 | 4kxy | A | 318 | ARG | p.Arg318Cys | P29401 |
| HOMO SAPIENS | 6 | 19546 | 2bx6 | A | 87 | THR | p.Thr87Ile | O75695 |
| HOMO SAPIENS | 1 | 19546 | 2bx6 | A | 118 | ARG | p.Arg118His | O75695 |
| HOMO SAPIENS | 1 | 19546 | 2bx6 | A | 201 | ILE | p.Ile201Val | O75695 |
| HOMO SAPIENS | 1 | 19546 | 2bx6 | A | 282 | ARG | p.Arg282Trp | O75695 |
| HOMO SAPIENS | 1 | 5265 | 5usr | H | 68 | ARG | p.Arg68Leu | Q9HD34 |
| HOMO SAPIENS | 2 | 1012 | 4nyi | S | 569 | LEU | p.Leu569Val | Q07889 |
| HOMO SAPIENS | 2 | 1012 | 4nyi | S | 655 | PRO | p.Pro655Leu | Q07889 |
| HOMO SAPIENS | 4 | 1012 | 4nyi | S | 663 | ILE | p.Ile663Met | Q07889 |
| HOMO SAPIENS | 2 | 1012 | 4nyi | S | 702 | TYR | p.Tyr702Cys | Q07889 |
| HOMO SAPIENS | 2 | 1012 | 4nyi | S | 708 | ALA | p.Ala708Thr | Q07889 |
| HOMO SAPIENS | 4 | 1012 | 4nyi | S | 733 | ILE | p.Ile733Phe | Q07889 |
| HOMO SAPIENS | 1 | 1848 | 3g1n | A | 4013 | ARG | p.Arg4013Trp | Q7Z6Z7 |
| HOMO SAPIENS | 1 | 1848 | 3g1n | A | 4187 | ARG | p.Arg4187Cys | Q7Z6Z7 |
| HOMO SAPIENS | 1 | 77 | 4pod | A | 41 | CYS | p.Cys79Tyr | P60174 |
| HOMO SAPIENS | 9 | 77 | 4pod | A | 104 | GLU | p.Glu142Asp | P60174 |
| HOMO SAPIENS | 1 | 77 | 4pod | A | 122 | GLY | p.Gly160Arg | P60174 |
| HOMO SAPIENS | -1 | 8672 | 3thz | A | 8 | THR | p.Thr8Met | P43246 |
| HOMO SAPIENS | 0 | 8672 | 3thz | A | 19 | PHE | p.Phe19Leu | P43246 |
| HOMO SAPIENS | 0 | 8672 | 3thz | A | 23 | PHE | p.Phe23Leu | P43246 |
| HOMO SAPIENS | 0 | 8672 | 3thz | A | 96 | ARG | p.Arg96His | P43246 |

| | | | | | | | | |
|---|---|---|---|---|---|---|---|---|
| HOMO SAPIENS | 0 | 8672 | 3thz | A | 102 | VAL | p.Val102Ile | P43246 |
| HOMO SAPIENS | 0 | 8672 | 3thz | A | 106 | ARG | p.Arg106Lys | P43246 |
| HOMO SAPIENS | 0 | 8672 | 3thz | A | 127 | ASN | p.Asn127Ser | P43246 |
| HOMO SAPIENS | 0 | 8672 | 3thz | A | 128 | LEU | p.Leu128Val | P43246 |
| HOMO SAPIENS | 0 | 8672 | 3thz | A | 163 | VAL | p.Val163Gly | P43246 |
| HOMO SAPIENS | 0 | 8672 | 3thz | A | 169 | ILE | p.Ile169Val | P43246 |
| HOMO SAPIENS | 0 | 8672 | 3thz | A | 315 | GLY | p.Gly315Asp | P43246 |
| HOMO SAPIENS | -1 | 8672 | 3thz | A | 322 | GLY | p.Gly322Asp | P43246 |
| HOMO SAPIENS | 0 | 8672 | 3thz | A | 331 | ASN | p.Asn331Asp | P43246 |
| HOMO SAPIENS | 1 | 8672 | 3thz | A | 390 | LEU | p.Leu390Phe | P43246 |
| HOMO SAPIENS | 0 | 8672 | 3thz | A | 419 | GLN | p.Gln419Lys | P43246 |
| HOMO SAPIENS | 0 | 8672 | 3thz | A | 447 | PHE | p.Phe447Leu | P43246 |
| HOMO SAPIENS | 0 | 8672 | 3thz | A | 524 | ARG | p.Arg524Pro | P43246 |
| HOMO SAPIENS | 0 | 8672 | 3thz | A | 539 | PHE | p.Phe539Leu | P43246 |
| HOMO SAPIENS | -1 | 8672 | 3thz | A | 546 | LYS | p.Lys546Asn | P43246 |
| HOMO SAPIENS | 0 | 8672 | 3thz | A | 564 | THR | p.Thr564Ala | P43246 |
| HOMO SAPIENS | 0 | 8672 | 3thz | A | 577 | ILE | p.Ile577Thr | P43246 |
| HOMO SAPIENS | 0 | 8672 | 3thz | A | 588 | TYR | p.Tyr588Ter | P43246 |
| HOMO SAPIENS | 0 | 8672 | 3thz | A | 622 | PRO | p.Pro622Leu | P43246 |
| HOMO SAPIENS | 0 | 8672 | 3thz | A | 629 | GLN | p.Gln629Arg | P43246 |
| HOMO SAPIENS | 0 | 8672 | 3thz | A | 636 | ALA | p.Ala636Pro | P43246 |
| HOMO SAPIENS | 2 | 8672 | 3thz | A | 642 | VAL | p.Val642Ile | P43246 |
| HOMO SAPIENS | 1 | 8672 | 3thz | A | 654 | ASP | p.Asp654Glu | P43246 |
| HOMO SAPIENS | 0 | 8672 | 3thz | A | 662 | GLN | p.Gln662His | P43246 |
| HOMO SAPIENS | 2 | 8672 | 3thz | A | 681 | GLN | p.Gln681His | P43246 |
| HOMO SAPIENS | 0 | 8672 | 3thz | A | 707 | CYS | p.Cys707Ser | P43246 |
| HOMO SAPIENS | 0 | 8672 | 3thz | A | 809 | GLU | p.Glu809Lys | P43246 |
| HOMO SAPIENS | 0 | 8672 | 3thz | A | 813 | MET | p.Met813Val | P43246 |
| HOMO SAPIENS | 1 | 8672 | 3thz | A | 834 | ALA | p.Ala834Thr | P43246 |
| HOMO SAPIENS | 0 | 8672 | 3thz | A | 849 | LEU | p.Leu849Val | P43246 |
| HOMO SAPIENS | -1 | 8672 | 3thz | A | 860 | SER | p.Ser860Leu | P43246 |
| HOMO SAPIENS | -1 | 8672 | 3thz | A | 934 | THR | p.Thr934Lys | P43246 |
| HOMO SAPIENS | 2 | 292 | 1i3k | A | 34 | ASN | p.Asn34Ser | Q14376 |
| HOMO SAPIENS | 7 | 292 | 1i3k | A | 90 | GLY | p.Gly90Glu | Q14376 |
| HOMO SAPIENS | 2 | 292 | 1i3k | A | 103 | ASP | p.Asp103Gly | Q14376 |
| HOMO SAPIENS | 2 | 292 | 1i3k | A | 136 | TYR | p.Tyr136Ter | Q14376 |
| HOMO SAPIENS | 0 | 292 | 1i3k | A | 169 | ARG | p.Arg169Trp | Q14376 |
| HOMO SAPIENS | 0 | 292 | 1i3k | A | 183 | LEU | p.Leu183Pro | Q14376 |
| HOMO SAPIENS | 0 | 292 | 1i3k | A | 302 | GLY | p.Gly302Asp | Q14376 |
| HOMO SAPIENS | 0 | 292 | 1i3k | A | 313 | LEU | p.Leu313Met | Q14376 |
| HOMO SAPIENS | 1 | 2084 | 5vc8 | A | 295 | GLN | p.Gln295Leu | O96028 |
| HOMO SAPIENS | -1 | 10157 | 5wrw | A | 7 | GLY | p.Gly7Ala | O76094 |
| HOMO SAPIENS | -1 | 10157 | 5wrw | A | 7 | GLY | p.Gly7Arg | O76094 |
| HOMO SAPIENS | -1 | 10157 | 5wrw | A | 8 | GLY | p.Gly8Ala | O76094 |
| HOMO SAPIENS | 0 | 3113 | 3in5 | A | 29 | GLU | p.Glu29Lys | Q9UBT6 |
| HOMO SAPIENS | 5 | 4546 | 5in3 | B | 44 | VAL | p.Val44Leu | P07902 |
| HOMO SAPIENS | -1 | 4546 | 5in3 | B | 51 | ARG | p.Arg51Leu | P07902 |
| HOMO SAPIENS | -1 | 4546 | 5in3 | B | 62 | LEU | p.Leu62Met | P07902 |
| HOMO SAPIENS | 4 | 4546 | 5in3 | B | 66 | PRO | p.Pro66His | P07902 |
| HOMO SAPIENS | 2 | 4546 | 5in3 | B | 74 | LEU | p.Leu74Pro | P07902 |
| HOMO SAPIENS | 5 | 4546 | 5in3 | B | 97 | ASN | p.Asn97Ser | P07902 |
| HOMO SAPIENS | 1 | 4546 | 5in3 | B | 135 | SER | p.Ser135Leu | P07902 |
| HOMO SAPIENS | 5 | 4546 | 5in3 | B | 138 | THR | p.Thr138Met | P07902 |
| HOMO SAPIENS | 7 | 4546 | 5in3 | B | 139 | LEU | p.Leu139Pro | P07902 |

| | | | | | | | | |
|---|---|---|---|---|---|---|---|---|
| HOMO SAPIENS | 2 | 4546 | 5in3 | B | 142 | MET | p.Met142Lys | P07902 |
| HOMO SAPIENS | 1 | 4546 | 5in3 | B | 143 | SER | p.Ser143Leu | P07902 |
| HOMO SAPIENS | 2 | 4546 | 5in3 | B | 148 | ARG | p.Arg148Gly | P07902 |
| HOMO SAPIENS | 2 | 4546 | 5in3 | B | 148 | ARG | p.Arg148Gln | P07902 |
| HOMO SAPIENS | 5 | 4546 | 5in3 | B | 183 | PRO | p.Pro183Thr | P07902 |
| HOMO SAPIENS | 5 | 4546 | 5in3 | B | 185 | PRO | p.Pro185His | P07902 |
| HOMO SAPIENS | 5 | 4546 | 5in3 | B | 186 | HIS | p.His186Tyr | P07902 |
| HOMO SAPIENS | 7 | 4546 | 5in3 | B | 188 | GLN | p.Gln188Arg | P07902 |
| HOMO SAPIENS | 5 | 4546 | 5in3 | B | 195 | LEU | p.Leu195Pro | P07902 |
| HOMO SAPIENS | 4 | 4546 | 5in3 | B | 201 | ARG | p.Arg201His | P07902 |
| HOMO SAPIENS | 4 | 4546 | 5in3 | B | 201 | ARG | p.Arg201Cys | P07902 |
| HOMO SAPIENS | 4 | 4546 | 5in3 | B | 203 | GLU | p.Glu203Lys | P07902 |
| HOMO SAPIENS | 2 | 4546 | 5in3 | B | 209 | TYR | p.Tyr209Cys | P07902 |
| HOMO SAPIENS | 5 | 4546 | 5in3 | B | 223 | ARG | p.Arg223Ser | P07902 |
| HOMO SAPIENS | 5 | 4546 | 5in3 | B | 226 | LEU | p.Leu226Pro | P07902 |
| HOMO SAPIENS | 5 | 4546 | 5in3 | B | 226 | LEU | p.Leu226Val | P07902 |
| HOMO SAPIENS | 2 | 4546 | 5in3 | B | 227 | LEU | p.Leu227Pro | P07902 |
| HOMO SAPIENS | 8 | 4546 | 5in3 | B | 258 | ARG | p.Arg258Cys | P07902 |
| HOMO SAPIENS | 1 | 4546 | 5in3 | B | 265 | PRO | p.Pro265Ala | P07902 |
| HOMO SAPIENS | 8 | 4546 | 5in3 | B | 268 | THR | p.Thr268Asn | P07902 |
| HOMO SAPIENS | 7 | 4546 | 5in3 | B | 278 | ILE | p.Ile278Asn | P07902 |
| HOMO SAPIENS | 5 | 4546 | 5in3 | B | 285 | LYS | p.Lys285Asn | P07902 |
| HOMO SAPIENS | 1 | 4546 | 5in3 | B | 319 | HIS | p.His319Gln | P07902 |
| HOMO SAPIENS | 4 | 4546 | 5in3 | B | 321 | HIS | p.His321Tyr | P07902 |
| HOMO SAPIENS | 4 | 4546 | 5in3 | B | 325 | PRO | p.Pro325Leu | P07902 |
| HOMO SAPIENS | 2 | 4546 | 5in3 | B | 328 | ARG | p.Arg328His | P07902 |
| HOMO SAPIENS | 2 | 4546 | 5in3 | B | 329 | SER | p.Ser329Phe | P07902 |
| HOMO SAPIENS | 5 | 4546 | 5in3 | B | 334 | LYS | p.Lys334Arg | P07902 |
| HOMO SAPIENS | 6 | 1290 | 5uav | C | 119 | ARG | p.Arg119Gly | P32322 |
| HOMO SAPIENS | 6 | 1290 | 5uav | C | 119 | ARG | p.Arg119His | P32322 |
| HOMO SAPIENS | 3 | 1290 | 5uav | C | 251 | ARG | p.Arg251His | P32322 |
| HOMO SAPIENS | 7 | 1290 | 5uav | C | 266 | ARG | p.Arg266Gln | P32322 |
| HOMO SAPIENS | 0 | 9618 | 2r7e | A | 22 | GLY | p.Gly41Cys | P00451 |
| HOMO SAPIENS | 2 | 9618 | 2r7e | A | 113 | GLU | p.Glu132Asp | P00451 |
| HOMO SAPIENS | -1 | 9618 | 2r7e | A | 336 | ARG | p.Arg355Ter | P00451 |
| HOMO SAPIENS | 0 | 9618 | 2r7e | A | 372 | ARG | p.Arg391His | P00451 |
| HOMO SAPIENS | 0 | 9618 | 2r7e | A | 373 | SER | p.Ser392Leu | P00451 |
| HOMO SAPIENS | 0 | 9618 | 2r7e | A | 373 | SER | p.Ser392Pro | P00451 |
| HOMO SAPIENS | 3 | 9618 | 2r7e | A | 386 | ILE | p.Ile405Ser | P00451 |
| HOMO SAPIENS | 6 | 9618 | 2r7e | A | 390 | GLU | p.Glu409Gly | P00451 |
| HOMO SAPIENS | 2 | 9618 | 2r7e | A | 412 | LEU | p.Leu431Phe | P00451 |
| HOMO SAPIENS | 5 | 9618 | 2r7e | A | 425 | LYS | p.Lys444Arg | P00451 |
| HOMO SAPIENS | 0 | 9618 | 2r7e | A | 484 | ARG | p.Arg503His | P00451 |
| HOMO SAPIENS | 2 | 9618 | 2r7e | A | 531 | ARG | p.Arg550His | P00451 |
| HOMO SAPIENS | 3 | 9618 | 2r7e | A | 535 | SER | p.Ser554Gly | P00451 |
| HOMO SAPIENS | 2 | 9618 | 2r7e | A | 593 | ARG | p.Arg612Cys | P00451 |
| HOMO SAPIENS | 0 | 9618 | 2r7e | A | 704 | ALA | p.Ala723Thr | P00451 |
| HOMO SAPIENS | 0 | 9618 | 2r7e | A | 720 | GLU | p.Glu739Lys | P00451 |
| HOMO SAPIENS | 1 | 13085 | 3ud1 | A | 1075 | ILE | p.Thr1075Ile | P16157 |
| HOMO SAPIENS | -1 | 3289 | 2fy4 | A | 2 | ALA | p.Ala120Thr | P28329 |
| HOMO SAPIENS | 7 | 3289 | 2fy4 | A | 18 | VAL | p.Val136Met | P28329 |
| HOMO SAPIENS | 5 | 3289 | 2fy4 | A | 92 | LEU | p.Leu210Pro | P28329 |
| HOMO SAPIENS | 2 | 3289 | 2fy4 | A | 93 | PRO | p.Pro211Ala | P28329 |
| HOMO SAPIENS | 4 | 3289 | 2fy4 | A | 131 | LEU | p.Leu249Val | P28329 |

| | | | | | | | | |
|---|---|---|---|---|---|---|---|---|
| HOMO SAPIENS | 2 | 3289 | 2fy4 | A | 181 | PRO | p.Pro299Leu | P28329 |
| HOMO SAPIENS | 0 | 3289 | 2fy4 | A | 218 | ILE | p.Ile336Thr | P28329 |
| HOMO SAPIENS | 2 | 3289 | 2fy4 | A | 261 | ASP | p.Asp379His | P28329 |
| HOMO SAPIENS | 0 | 3289 | 2fy4 | A | 302 | ARG | p.Arg420Cys | P28329 |
| HOMO SAPIENS | 5 | 3289 | 2fy4 | A | 323 | GLU | p.Glu441Lys | P28329 |
| HOMO SAPIENS | 0 | 3289 | 2fy4 | A | 343 | MET | p.Val461Met | P28329 |
| HOMO SAPIENS | 2 | 3289 | 2fy4 | A | 364 | ARG | p.Arg482Gly | P28329 |
| HOMO SAPIENS | 3 | 3289 | 2fy4 | A | 380 | SER | p.Ser498Leu | P28329 |
| HOMO SAPIENS | 6 | 3289 | 2fy4 | A | 388 | VAL | p.Val506Leu | P28329 |
| HOMO SAPIENS | 7 | 3289 | 2fy4 | A | 429 | HIS | p.His547Gln | P28329 |
| HOMO SAPIENS | 8 | 3289 | 2fy4 | A | 442 | ARG | p.Arg560His | P28329 |
| HOMO SAPIENS | 3 | 3289 | 2fy4 | A | 443 | ARG | p.Arg561Gln | P28329 |
| HOMO SAPIENS | 0 | 4137 | 3hhr | B | 38 | CYS | p.Cys56Ter | P10912 |
| HOMO SAPIENS | 0 | 4137 | 3hhr | B | 43 | ARG | p.Arg61Ter | P10912 |
| HOMO SAPIENS | 0 | 4137 | 3hhr | B | 44 | GLU | p.Glu62Lys | P10912 |
| HOMO SAPIENS | 0 | 4137 | 3hhr | B | 96 | PHE | p.Phe114Ser | P10912 |
| HOMO SAPIENS | 5 | 4137 | 3hhr | B | 131 | PRO | p.Pro149Gln | P10912 |
| HOMO SAPIENS | 0 | 4137 | 3hhr | B | 144 | VAL | p.Val162Ile | P10912 |
| HOMO SAPIENS | 0 | 4137 | 3hhr | B | 150 | HIS | p.His168Gln | P10912 |
| HOMO SAPIENS | 0 | 4137 | 3hhr | B | 152 | ASP | p.Asp170His | P10912 |
| HOMO SAPIENS | 0 | 4137 | 3hhr | B | 153 | ILE | p.Ile171Thr | P10912 |
| HOMO SAPIENS | 0 | 4137 | 3hhr | B | 154 | GLN | p.Gln172Pro | P10912 |
| HOMO SAPIENS | 0 | 4137 | 3hhr | B | 155 | VAL | p.Val173Gly | P10912 |
| HOMO SAPIENS | 0 | 4137 | 3hhr | B | 217 | ARG | p.Arg235Ter | P10912 |
| HOMO SAPIENS | 5 | 4137 | 3hhr | B | 224 | GLU | p.Glu242Asp | P10912 |
| HOMO SAPIENS | 5 | 4137 | 3hhr | B | 224 | GLU | p.Glu242Ter | P10912 |
| HOMO SAPIENS | 6 | 2493 | 5bpb | A | 69 | HIS | p.His69Tyr | Q9ULV1 |
| HOMO SAPIENS | 6 | 2493 | 5bpb | A | 105 | MET | p.Met105Val | Q9ULV1 |
| HOMO SAPIENS | -1 | 2493 | 5bpb | A | 168 | PRO | p.Pro168Ser | Q9ULV1 |
| HOMO SAPIENS | 0 | 32123 | 4ccz | A | 314 | ASP | p.Asp314Asn | Q99707 |
| HOMO SAPIENS | 0 | 32123 | 4ccz | A | 345 | VAL | p.Val345Phe | Q99707 |
| HOMO SAPIENS | 1 | 32123 | 4ccz | A | 585 | ARG | p.Arg585Ter | Q99707 |
| HOMO SAPIENS | 4 | 4618 | 2ag4 | A | 26 | PRO | p.Pro55Leu | P17900 |
| HOMO SAPIENS | 4 | 4618 | 2ag4 | A | 30 | VAL | p.Ile59Val | P17900 |
| HOMO SAPIENS | 2 | 4618 | 2ag4 | A | 40 | VAL | p.Met69Val | P17900 |
| HOMO SAPIENS | 2 | 4618 | 2ag4 | A | 124 | VAL | p.Val153Ala | P17900 |
| HOMO SAPIENS | 8 | 4618 | 2ag4 | A | 140 | ARG | p.Arg169Pro | P17900 |
| HOMO SAPIENS | 5 | 1082 | 1hk0 | X | 14 | ARG | p.Arg15Cys | P07320 |
| HOMO SAPIENS | 7 | 1082 | 1hk0 | X | 23 | PRO | p.Pro24Thr | P07320 |
| HOMO SAPIENS | 6 | 1082 | 1hk0 | X | 36 | ARG | p.Arg37Ser | P07320 |
| HOMO SAPIENS | 5 | 1082 | 1hk0 | X | 55 | TYR | p.Tyr56Ter | P07320 |
| HOMO SAPIENS | 3 | 1082 | 1hk0 | X | 58 | ARG | p.Arg59His | P07320 |
| HOMO SAPIENS | 7 | 1082 | 1hk0 | X | 88 | HIS | p.His88Gln | P07320 |
| HOMO SAPIENS | 6 | 1082 | 1hk0 | X | 95 | ARG | p.Arg95Ser | P07320 |
| HOMO SAPIENS | 5 | 1082 | 1hk0 | X | 126 | VAL | p.Val126Met | P07320 |
| HOMO SAPIENS | 8 | 1082 | 1hk0 | X | 157 | TRP | p.Trp157Ter | P07320 |
| HOMO SAPIENS | 0 | 22879 | 3j0a | B | 644 | ILE | p.Ile644Phe | O60602 |
| HOMO SAPIENS | 8 | 6 | 2o23 | B | 122 | LEU | p.Leu122Val | Q99714 |
| HOMO SAPIENS | 1 | 6 | 2o23 | B | 130 | ARG | p.Arg130Cys | Q99714 |
| HOMO SAPIENS | 1 | 6 | 2o23 | B | 247 | ASN | p.Asn247Ser | Q99714 |
| HOMO SAPIENS | 0 | 6 | 2o23 | B | 249 | GLU | p.Glu249Gln | Q99714 |
| HOMO SAPIENS | 1 | 12910 | 4hl4 | A | 67 | ARG | p.Arg67Ter | Q96BZ9 |
| HOMO SAPIENS | 8 | 16105 | 1lmj | A | 7 | GLU | p.Glu1073Lys | P35555 |
| HOMO SAPIENS | 8 | 16105 | 1lmj | A | 8 | CYS | p.Cys1074Arg | P35555 |

| | | | | | | | | |
|---|---|---|---|---|---|---|---|---|
| HOMO SAPIENS | 9 | 16105 | 1lmj | A | 51 | CYS | p.Cys1117Tyr | P35555 |
| HOMO SAPIENS | 9 | 16105 | 1lmj | A | 63 | CYS | p.Cys1129Tyr | P35555 |
| HOMO SAPIENS | 9 | 16105 | 1lmj | A | 65 | ASN | p.Asn1131Tyr | P35555 |
| HOMO SAPIENS | 4 | 16105 | 1lmj | A | 71 | ARG | p.Arg1137Pro | P35555 |
| HOMO SAPIENS | 1 | 16105 | 1lmj | A | 82 | PRO | p.Pro1148Ala | P35555 |
| HOMO SAPIENS | 0 | 4795 | 3s5f | A | 106 | LEU | p.Leu106Ser | Q16595 |
| HOMO SAPIENS | 0 | 4795 | 3s5f | A | 130 | GLY | p.Gly130Val | Q16595 |
| HOMO SAPIENS | 0 | 4795 | 3s5f | A | 154 | ILE | p.Ile154Phe | Q16595 |
| HOMO SAPIENS | -1 | 4795 | 3s5f | A | 209 | ASP | p.Asp209Gly | Q16595 |
| HOMO SAPIENS | -1 | 2884 | 4lli | A | 1655 | ASP | p.Asp1655Glu | Q9Y4I1 |
| HOMO SAPIENS | 0 | 1786 | 7pck | C | 56P | TYR | p.Tyr71Ter | P43235 |
| HOMO SAPIENS | 0 | 1786 | 7pck | C | 64P | GLY | p.Gly79Glu | P43235 |
| HOMO SAPIENS | 7 | 38286 | 2l53 | B | 1914 | ARG | p.Arg1914Gly | Q14524 |
| HOMO SAPIENS | 1 | 38286 | 2l53 | B | 1924 | ALA | p.Ala1924Thr | Q14524 |
| HOMO SAPIENS | 0 | 31747 | 3ksy | A | 25 | PRO | p.Pro25Ser | Q07889 |
| HOMO SAPIENS | 3 | 31747 | 3ksy | A | 37 | THR | p.Thr37Ala | Q07889 |
| HOMO SAPIENS | 0 | 31747 | 3ksy | A | 54 | GLN | p.Gln54His | Q07889 |
| HOMO SAPIENS | -1 | 31747 | 3ksy | A | 190 | ASP | p.Asp190Glu | Q07889 |
| HOMO SAPIENS | 2 | 31747 | 3ksy | A | 266 | THR | p.Thr266Lys | Q07889 |
| HOMO SAPIENS | 1 | 31747 | 3ksy | A | 269 | MET | p.Met269Arg | Q07889 |
| HOMO SAPIENS | 1 | 31747 | 3ksy | A | 340 | PRO | p.Pro340Ser | Q07889 |
| HOMO SAPIENS | 1 | 31747 | 3ksy | A | 423 | ASN | p.Asn423Lys | Q07889 |
| HOMO SAPIENS | 1 | 31747 | 3ksy | A | 424 | GLU | p.Glu424Lys | Q07889 |
| HOMO SAPIENS | 1 | 31747 | 3ksy | A | 432 | TRP | p.Trp432Arg | Q07889 |
| HOMO SAPIENS | 1 | 31747 | 3ksy | A | 552 | ARG | p.Arg552Gly | Q07889 |
| HOMO SAPIENS | 1 | 31747 | 3ksy | A | 552 | ARG | p.Arg552Ser | Q07889 |
| HOMO SAPIENS | 3 | 7529 | 6jdw | A | 93 | ILE | p.Ile93Met | P50440 |
| HOMO SAPIENS | 2 | 7529 | 6jdw | A | 110 | GLN | p.Gln110His | P50440 |
| HOMO SAPIENS | 5 | 7529 | 6jdw | A | 149 | TRP | p.Trp149Ter | P50440 |
| HOMO SAPIENS | 0 | 7529 | 6jdw | A | 169 | ARG | p.Arg169Ter | P50440 |
| HOMO SAPIENS | 0 | 7529 | 6jdw | A | 260 | ARG | p.Arg260Ter | P50440 |
| HOMO SAPIENS | 3 | 7529 | 6jdw | A | 344 | ILE | p.Ile344Val | P50440 |
| HOMO SAPIENS | 5 | 7529 | 6jdw | A | 418 | LEU | p.Leu418Ile | P50440 |
| HOMO SAPIENS | 1 | 14788 | 1eak | A | 101 | ARG | p.Arg101His | P08253 |
| HOMO SAPIENS | 0 | 14788 | 1eak | A | 383 | ASP | p.Asp383Glu | P08253 |
| HOMO SAPIENS | 7 | 14788 | 1eak | A | 404 | GLN | p.Glu404Lys | P08253 |
| HOMO SAPIENS | 2 | 14788 | 1eak | A | 447 | ALA | p.Ala447Val | P08253 |
| HOMO SAPIENS | 0 | 1255 | 5xtc | p | 8 | PRO | p.Pro8His | Q9Y6M9 |
| HOMO SAPIENS | 0 | 1255 | 5xtc | p | 146 | PRO | p.Pro146Ser | Q9Y6M9 |
| HOMO SAPIENS | 3 | 6721 | 1a22 | A | 25 | PHE | p.Phe51Tyr | P01241 |
| HOMO SAPIENS | 2 | 6721 | 1a22 | A | 95 | SER | p.Ser121Arg | P01241 |
| HOMO SAPIENS | 2 | 6721 | 1a22 | A | 110 | VAL | p.Val136Ile | P01241 |
| HOMO SAPIENS | 0 | 1380 | 5i6z | A | 605 | LYS | p.Lys605Asn | P31645 |
| HOMO SAPIENS | 0 | 941 | 1qki | D | 48 | ILE | p.Ile48Thr | P11413 |
| HOMO SAPIENS | 3 | 941 | 1qki | D | 68 | VAL | p.Val68Met | P11413 |
| HOMO SAPIENS | 2 | 941 | 1qki | D | 126 | ASN | p.Asn126Asp | P11413 |
| HOMO SAPIENS | 8 | 941 | 1qki | D | 163 | GLY | p.Gly163Ser | P11413 |
| HOMO SAPIENS | 6 | 941 | 1qki | D | 165 | ASN | p.Asn165Asp | P11413 |
| HOMO SAPIENS | 4 | 941 | 1qki | D | 181 | ASP | p.Asp181Val | P11413 |
| HOMO SAPIENS | 4 | 941 | 1qki | D | 182 | ARG | p.Arg182Trp | P11413 |
| HOMO SAPIENS | 5 | 941 | 1qki | D | 188 | SER | p.Ser188Phe | P11413 |
| HOMO SAPIENS | 9 | 941 | 1qki | D | 198 | ARG | p.Arg198Cys | P11413 |
| HOMO SAPIENS | 8 | 941 | 1qki | D | 213 | VAL | p.Val213Leu | P11413 |
| HOMO SAPIENS | 5 | 941 | 1qki | D | 282 | ASP | p.Asp282His | P11413 |

| | | | | | | | | |
|---|---|---|---|---|---|---|---|---|
| HOMO SAPIENS | 7 | 941 | 1qki | D | 285 | ARG | p.Arg285His | P11413 |
| HOMO SAPIENS | 5 | 941 | 1qki | D | 291 | VAL | p.Val291Met | P11413 |
| HOMO SAPIENS | 8 | 941 | 1qki | D | 323 | LEU | p.Leu323Pro | P11413 |
| HOMO SAPIENS | 3 | 941 | 1qki | D | 335 | ALA | p.Ala335Thr | P11413 |
| HOMO SAPIENS | 6 | 941 | 1qki | D | 350 | ASP | p.Asp350His | P11413 |
| HOMO SAPIENS | 5 | 941 | 1qki | D | 393 | ARG | p.Arg393His | P11413 |
| HOMO SAPIENS | 5 | 941 | 1qki | D | 454 | ARG | p.Arg454Cys | P11413 |
| HOMO SAPIENS | 2 | 941 | 1qki | D | 459 | LEU | p.Arg459Leu | P11413 |
| HOMO SAPIENS | 4 | 941 | 1qki | D | 463 | ARG | p.Arg463His | P11413 |
| HOMO SAPIENS | 4 | 2483 | 1tij | A | 68 | LEU | p.Leu94Gln | P01034 |
| HOMO SAPIENS | 2 | 20632 | 3v5b | A | 331 | ARG | p.Arg331Pro | P02545 |
| HOMO SAPIENS | 2 | 20632 | 3v5b | A | 374 | HIS | p.His374Gln | P02545 |
| HOMO SAPIENS | 4 | 13929 | 5xtd | J | 75 | ARG | p.Arg75His | Q16795 |
| HOMO SAPIENS | 3 | 13929 | 5xtd | J | 243 | VAL | p.Val243Ala | Q16795 |
| HOMO SAPIENS | 1 | 13929 | 5xtd | J | 294 | PRO | p.Pro294Leu | Q16795 |
| HOMO SAPIENS | 2 | 13929 | 5xtd | J | 321 | ARG | p.Arg321Pro | Q16795 |
| HOMO SAPIENS | 0 | 9411 | 5thl | A | 41 | ARG | p.Gly41Arg | P54577 |
| HOMO SAPIENS | 0 | 9411 | 5thl | A | 196 | GLU | p.Glu196Lys | P54577 |
| HOMO SAPIENS | 0 | 9411 | 5thl | A | 316 | ALA | p.Ala316Ser | P54577 |
| HOMO SAPIENS | 0 | 9411 | 5thl | A | 339 | ALA | p.Ala339Thr | P54577 |
| HOMO SAPIENS | 4 | 16273 | 2e8n | A | 46 | THR | p.Thr940Ile | P29317 |
| HOMO SAPIENS | 7 | 16273 | 2e8n | A | 54 | GLY | p.Gly948Trp | P29317 |
| HOMO SAPIENS | 6 | 16273 | 2e8n | A | 65 | ALA | p.Ala959Thr | P29317 |
| HOMO SAPIENS | 7 | 30387 | 2k9y | A | 541 | VAL | p.Val541Met | P29317 |
| HOMO SAPIENS | 0 | 6961 | 4nha | A | 843 | ILE | p.Ile853Val | Q9UPY3 |
| HOMO SAPIENS | -1 | 6961 | 4nha | A | 862 | ALA | p.Ala872Thr | Q9UPY3 |
| HOMO SAPIENS | 3 | 15708 | 4pzg | A | 16 | ARG | p.Arg16Ter | Q9Y5X0 |
| HOMO SAPIENS | 4 | 15708 | 4pzg | A | 51 | ARG | p.Arg51Gln | Q9Y5X0 |
| HOMO SAPIENS | 3 | 30166 | 4o00 | A | 44 | ARG | p.Arg14257Ter | Q8WZ42 |
| HOMO SAPIENS | 2 | 30166 | 4o00 | A | 54 | ARG | p.Arg14267His | Q8WZ42 |
| HOMO SAPIENS | 8 | 11158 | 4hca | A | 275 | TRP | p.Trp274Arg | P23771 |
| HOMO SAPIENS | 2 | 4328 | 4e0s | A | 1476 | ARG | p.Arg1476Ter | P01031 |
| HOMO SAPIENS | 0 | 11462 | 1dx5 | I | 455 | ALA | p.Ala473Val | P07204 |
| HOMO SAPIENS | -1 | 11150 | 2fic | A | 45 | LYS | p.Lys35Asn | O00499 |
| HOMO SAPIENS | 6 | 11150 | 2fic | A | 161 | ASP | p.Asp151Asn | O00499 |
| HOMO SAPIENS | 8 | 27382 | 4mqv | B | 17 | ARG | p.Arg17Ter | Q9NZC9 |
| HOMO SAPIENS | -1 | 37991 | 1q0p | A | 231 | GLN | p.Gln256Ter | P00751 |
| HOMO SAPIENS | 1 | 443 | 5xte | U | 215 | LEU | p.Leu215Phe | P08574 |
| HOMO SAPIENS | 8 | 21471 | 5y4e | A | 1461 | ASN | p.Thr692Ala | Q01484 |
| HOMO SAPIENS | -1 | 35443 | 3um3 | B | 129 | ASP | p.Asp129Val | Q9H444 |
| HOMO SAPIENS | -1 | 35443 | 3um3 | B | 161 | GLU | p.Glu161Lys | Q9H444 |
| HOMO SAPIENS | 6 | 28781 | 2n2a | B | 111 | ILE | p.Ile654Val | P04626 |
| HOMO SAPIENS | 7 | 28781 | 2n2a | B | 112 | ILE | p.Ile655Val | P04626 |
| HOMO SAPIENS | -1 | 3346 | 4ztz | C | 61 | GLY | p.Gly61Arg | Q9UHN1 |
| HOMO SAPIENS | 4 | 3346 | 4ztz | C | 103 | GLY | p.Gly103Ser | Q9UHN1 |
| HOMO SAPIENS | 4 | 3346 | 4ztz | C | 133 | HIS | p.His133Tyr | Q9UHN1 |
| HOMO SAPIENS | -1 | 3346 | 4ztz | C | 153 | LEU | p.Leu153Val | Q9UHN1 |
| HOMO SAPIENS | -1 | 3346 | 4ztz | C | 169 | ALA | p.Ala169Thr | Q9UHN1 |
| HOMO SAPIENS | 7 | 3346 | 4ztz | C | 369 | ARG | p.Arg369Gly | Q9UHN1 |
| HOMO SAPIENS | 4 | 3346 | 4ztz | C | 386 | ASP | p.Asp386Glu | Q9UHN1 |
| HOMO SAPIENS | 4 | 3346 | 4ztz | C | 451 | GLY | p.Gly451Glu | Q9UHN1 |
| HOMO SAPIENS | 4 | 2131 | 5fo9 | E | 1549 | LEU | p.Leu1549Met | P01024 |
| HOMO SAPIENS | 2 | 2131 | 5fo9 | E | 1619 | SER | p.Ser1619Arg | P01024 |
| HOMO SAPIENS | 3 | 12662 | 4jhp | C | 75 | ILE | p.Ile75Val | Q92834 |

| | | | | | | | | |
|---|---|---|---|---|---|---|---|---|
| HOMO SAPIENS | 1 | 12662 | 4jhp | C | 173 | GLY | p.Gly173Arg | Q92834 |
| HOMO SAPIENS | 0 | 12662 | 4jhp | C | 184 | GLN | p.Gln184His | Q92834 |
| HOMO SAPIENS | 3 | 12662 | 4jhp | C | 262 | ALA | p.Ala262Gly | Q92834 |
| HOMO SAPIENS | 3 | 12662 | 4jhp | C | 302 | CYS | p.Cys302Tyr | Q92834 |
| HOMO SAPIENS | 5 | 1093 | 2jg9 | A | 186 | GLN | p.Gln208Ter | P02745 |
| HOMO SAPIENS | 0 | 13181 | 4r4l | A | 12 | MET | p.Met13Ile | Q13976 |
| HOMO SAPIENS | 8 | 25939 | 4iyp | A | 106 | ARG | p.Arg106Gln | P78318 |
| HOMO SAPIENS | 1 | 217 | 2xsx | A | 71 | SER | p.Asn71Ser | P13929 |
| HOMO SAPIENS | 0 | 217 | 2xsx | A | 76 | ALA | p.Ala76Pro | P13929 |
| HOMO SAPIENS | 1 | 217 | 2xsx | A | 85 | ALA | p.Val85Ala | P13929 |
| HOMO SAPIENS | 8 | 217 | 2xsx | A | 156 | GLY | p.Gly156Asp | P13929 |
| HOMO SAPIENS | 5 | 217 | 2xsx | A | 332 | VAL | p.Val332Ile | P13929 |
| HOMO SAPIENS | 7 | 217 | 2xsx | A | 374 | GLY | p.Gly374Glu | P13929 |
| HOMO SAPIENS | 1 | 5508 | 3szm | A | 682 | THR | p.Thr682Asn | Q8NEM0 |
| HOMO SAPIENS | 0 | 5508 | 3szm | A | 715 | TRP | p.Trp715Ter | Q8NEM0 |
| HOMO SAPIENS | 1 | 5508 | 3szm | A | 761 | VAL | p.Ala761Val | Q8NEM0 |
| HOMO SAPIENS | 1 | 5508 | 3szm | A | 828 | PRO | p.Pro828Ser | Q8NEM0 |
| HOMO SAPIENS | 4 | 5508 | 3szm | A | 829 | GLU | p.Glu829Asp | Q8NEM0 |
| HOMO SAPIENS | 2 | 16353 | 2q0z | X | 171 | GLY | p.Gly1978Ala | O75643 |
| HOMO SAPIENS | 2 | 16353 | 2q0z | X | 188 | ALA | p.Ala1995Thr | O75643 |
| HOMO SAPIENS | 0 | 4127 | 3swr | A | 860 | GLY | p.Gly860Arg | P26358 |
| HOMO SAPIENS | -1 | 4127 | 3swr | A | 1122 | SER | p.Ser1122Cys | P26358 |
| HOMO SAPIENS | 0 | 4127 | 3swr | A | 1216 | ASP | p.Asp1216Glu | P26358 |
| HOMO SAPIENS | 0 | 6382 | 3o96 | A | 17 | GLU | p.Glu17Lys | P31749 |
| HOMO SAPIENS | 6 | 14314 | 5xtc | Q | 71 | PRO | p.Pro71Leu | O75306 |
| HOMO SAPIENS | 2 | 5886 | 4fao | E | 22 | ARG | p.Arg40His | Q13705 |
| HOMO SAPIENS | 1 | 2534 | 3mp4 | A | 37 | GLU | p.Glu37Ter | P35914 |
| HOMO SAPIENS | 3 | 2534 | 3mp4 | A | 41 | MET | p.Arg41Gln | P35914 |
| HOMO SAPIENS | 1 | 2534 | 3mp4 | A | 70 | VAL | p.Val70Leu | P35914 |
| HOMO SAPIENS | 2 | 2196 | 3ifq | A | 142 | ARG | p.Arg142His | P14923 |
| HOMO SAPIENS | 2 | 2196 | 3ifq | A | 153 | ASP | p.Asp153Glu | P14923 |
| HOMO SAPIENS | 3 | 2196 | 3ifq | A | 176 | ARG | p.Arg176Gln | P14923 |
| HOMO SAPIENS | 3 | 2196 | 3ifq | A | 265 | ARG | p.Arg265His | P14923 |
| HOMO SAPIENS | 2 | 2196 | 3ifq | A | 301 | GLU | p.Glu301Gly | P14923 |
| HOMO SAPIENS | 0 | 2196 | 3ifq | A | 456 | VAL | p.Val456Ile | P14923 |
| HOMO SAPIENS | 3 | 2196 | 3ifq | A | 603 | VAL | p.Val603Leu | P14923 |
| HOMO SAPIENS | 0 | 2196 | 3ifq | A | 648 | VAL | p.Val648Ile | P14923 |
| HOMO SAPIENS | 1 | 8746 | 1cgj | I | 32 | PRO | p.Pro55Ser | P00995 |
| HOMO SAPIENS | 1 | 8746 | 1cgj | I | 44 | ARG | p.Arg67His | P00995 |
| HOMO SAPIENS | 1 | 8746 | 1cgj | I | 44 | ARG | p.Arg67Cys | P00995 |
| HOMO SAPIENS | 3 | 3212 | 1syq | A | 105 | ARG | p.Arg105Ter | P18206 |
| HOMO SAPIENS | 3 | 3212 | 1syq | A | 126 | PHE | p.Phe126Leu | P18206 |
| HOMO SAPIENS | 4 | 10028 | 3eto | A | 1609 | ARG | p.Arg1608His | P46531 |
| HOMO SAPIENS | 1 | 10028 | 3eto | A | 1672 | VAL | p.Val1671Ile | P46531 |
| HOMO SAPIENS | 5 | 10028 | 3eto | A | 1692 | GLN | p.Gln1691His | P46531 |
| HOMO SAPIENS | 3 | 3749 | 5xtc | Z | 70 | GLY | p.Gly70Ter | O43676 |
| HOMO SAPIENS | 2 | 17039 | 3brv | B | 57 | GLU | p.Glu57Lys | Q9Y6K9 |
| HOMO SAPIENS | 8 | 3589 | 2fh3 | A | 433 | VAL | p.Val460Leu | P06396 |
| HOMO SAPIENS | 1 | 3589 | 2fh3 | A | 454 | ARG | p.Arg481Cys | P06396 |
| HOMO SAPIENS | 4 | 3589 | 2fh3 | A | 498 | ARG | p.Arg525His | P06396 |
| HOMO SAPIENS | 2 | 3589 | 2fh3 | A | 520 | TYR | p.Tyr547Cys | P06396 |
| HOMO SAPIENS | 1 | 3589 | 2fh3 | A | 529 | GLY | p.Gly556Arg | P06396 |
| HOMO SAPIENS | 2 | 3589 | 2fh3 | A | 536 | THR | p.Thr563Ser | P06396 |
| HOMO SAPIENS | 2 | 3589 | 2fh3 | A | 579 | VAL | p.Val606Met | P06396 |

| | | | | | | | | |
|---|---|---|---|---|---|---|---|---|
| HOMO SAPIENS | 1 | 3589 | 2fh3 | A | 589 | THR | p.Thr616Met | P06396 |
| HOMO SAPIENS | 5 | 3589 | 2fh3 | A | 668 | THR | p.Thr695Met | P06396 |
| HOMO SAPIENS | 0 | 148 | 1lcl | A | 130 | ILE | p.Ile130Ser | Q05315 |
| HOMO SAPIENS | -1 | 24727 | 4fj3 | P | 237 | ALA | p.Ala237Thr | P04049 |
| HOMO SAPIENS | 0 | 24727 | 4fj3 | P | 257 | SER | p.Ser257Leu | P04049 |
| HOMO SAPIENS | 0 | 24727 | 4fj3 | P | 261 | PRO | p.Pro261Ala | P04049 |
| HOMO SAPIENS | 0 | 2949 | 5xtb | B | 77 | ARG | p.Arg77Gly | O00217 |
| HOMO SAPIENS | 0 | 2949 | 5xtb | B | 79 | PRO | p.Pro79Leu | O00217 |
| HOMO SAPIENS | 0 | 2949 | 5xtb | B | 100 | ALA | p.Ala100Val | O00217 |
| HOMO SAPIENS | 1 | 2949 | 5xtb | B | 115 | LYS | p.Lys115Glu | O00217 |
| HOMO SAPIENS | 0 | 2949 | 5xtb | B | 138 | ARG | p.Arg138His | O00217 |
| HOMO SAPIENS | 1 | 3959 | 5k5s | B | 25 | ARG | p.Arg25Ter | P41180 |
| HOMO SAPIENS | 4 | 3959 | 5k5s | B | 39 | PRO | p.Pro39Ala | P41180 |
| HOMO SAPIENS | 0 | 3959 | 5k5s | B | 47 | LYS | p.Lys47Asn | P41180 |
| HOMO SAPIENS | 2 | 3959 | 5k5s | B | 62 | ARG | p.Arg62Met | P41180 |
| HOMO SAPIENS | 2 | 3959 | 5k5s | B | 66 | ARG | p.Arg66Cys | P41180 |
| HOMO SAPIENS | 4 | 3959 | 5k5s | B | 90 | ASN | p.Asn90Thr | P41180 |
| HOMO SAPIENS | 1 | 3959 | 5k5s | B | 116 | ALA | p.Ala116Thr | P41180 |
| HOMO SAPIENS | 1 | 3959 | 5k5s | B | 118 | ASN | p.Asn118Lys | P41180 |
| HOMO SAPIENS | -1 | 3959 | 5k5s | B | 125 | LEU | p.Leu125Pro | P41180 |
| HOMO SAPIENS | -1 | 3959 | 5k5s | B | 127 | GLU | p.Glu127Ala | P41180 |
| HOMO SAPIENS | -1 | 3959 | 5k5s | B | 128 | PHE | p.Phe128Leu | P41180 |
| HOMO SAPIENS | -1 | 3959 | 5k5s | B | 131 | CYS | p.Cys131Trp | P41180 |
| HOMO SAPIENS | 1 | 3959 | 5k5s | B | 138 | THR | p.Thr138Met | P41180 |
| HOMO SAPIENS | 0 | 3959 | 5k5s | B | 143 | GLY | p.Gly143Glu | P41180 |
| HOMO SAPIENS | 5 | 3959 | 5k5s | B | 151 | THR | p.Thr151Met | P41180 |
| HOMO SAPIENS | 4 | 3959 | 5k5s | B | 180 | PHE | p.Phe180Cys | P41180 |
| HOMO SAPIENS | 0 | 3959 | 5k5s | B | 185 | ARG | p.Arg185Gln | P41180 |
| HOMO SAPIENS | 0 | 3959 | 5k5s | B | 185 | ARG | p.Arg185Ter | P41180 |
| HOMO SAPIENS | 2 | 3959 | 5k5s | B | 191 | GLU | p.Glu191Lys | P41180 |
| HOMO SAPIENS | 2 | 3959 | 5k5s | B | 227 | ARG | p.Arg227Leu | P41180 |
| HOMO SAPIENS | 2 | 3959 | 5k5s | B | 250 | GLU | p.Glu250Lys | P41180 |
| HOMO SAPIENS | 6 | 3959 | 5k5s | B | 297 | GLU | p.Glu297Lys | P41180 |
| HOMO SAPIENS | 1 | 3959 | 5k5s | B | 336 | LYS | p.Lys336Asn | P41180 |
| HOMO SAPIENS | 6 | 3959 | 5k5s | B | 429 | HIS | p.His429Tyr | P41180 |
| HOMO SAPIENS | 0 | 3959 | 5k5s | B | 445 | THR | p.Thr445Ala | P41180 |
| HOMO SAPIENS | 3 | 3959 | 5k5s | B | 509 | GLY | p.Gly509Arg | P41180 |
| HOMO SAPIENS | 1 | 3959 | 5k5s | B | 553 | GLY | p.Gly553Arg | P41180 |
| HOMO SAPIENS | 3 | 3959 | 5k5s | B | 592 | ASN | p.Asn592Ser | P41180 |
| HOMO SAPIENS | -1 | 3959 | 5k5s | B | 604 | GLU | p.Glu604Lys | P41180 |
| HOMO SAPIENS | 4 | 2564 | 5l0c | C | 975 | ARG | p.Arg975Trp | P18206 |
| HOMO SAPIENS | 4 | 2564 | 5l0c | C | 975 | ARG | p.Arg975Leu | P18206 |
| HOMO SAPIENS | 1 | 5695 | 5i8r | A | 294 | ARG | p.Arg294Gln | P17405 |
| HOMO SAPIENS | 0 | 5695 | 5i8r | A | 485 | ALA | p.Ala485Val | P17405 |
| HOMO SAPIENS | 1 | 5695 | 5i8r | A | 506 | GLY | p.Gly506Arg | P17405 |
| HOMO SAPIENS | -1 | 1978 | 3avs | A | 1060 | HIS | p.His1060Leu | O15550 |
| HOMO SAPIENS | 0 | 11173 | 4v0c | B | 362 | LYS | p.Lys362Arg | P51787 |
| HOMO SAPIENS | 3 | 11173 | 4v0c | B | 381 | CYS | p.Cys381Tyr | P51787 |
| HOMO SAPIENS | 7 | 11173 | 4v0c | B | 518 | ARG | p.Arg518Gly | P51787 |
| HOMO SAPIENS | 2 | 11173 | 4v0c | B | 525 | ALA | p.Ala525Thr | P51787 |
| HOMO SAPIENS | 0 | 11173 | 4v0c | B | 530 | GLN | p.Gln530Ter | P51787 |
| HOMO SAPIENS | -1 | 11173 | 4v0c | B | 536 | TYR | p.Tyr536Ter | P51787 |
| HOMO SAPIENS | 2 | 1113 | 5xtd | N | 60 | ARG | p.Arg60Ter | Q9UI09 |
| HOMO SAPIENS | 3 | 6061 | 2mmz | A | 148 | LEU | p.Leu148Ser | Q86SX6 |

| | | | | | | | | |
|---|---|---|---|---|---|---|---|---|
| HOMO SAPIENS | -1 | 26081 | 3eu7 | A | 837 | GLU | p.Glu837Ter | Q86YC2 |
| HOMO SAPIENS | 0 | 26081 | 3eu7 | A | 864 | PRO | p.Pro864Ser | Q86YC2 |
| HOMO SAPIENS | 5 | 26081 | 3eu7 | A | 891 | CYS | p.Cys891Trp | Q86YC2 |
| HOMO SAPIENS | 0 | 26081 | 3eu7 | A | 932 | VAL | p.Val932Met | Q86YC2 |
| HOMO SAPIENS | 5 | 26081 | 3eu7 | A | 961 | LEU | p.Leu961Met | Q86YC2 |
| HOMO SAPIENS | 5 | 26081 | 3eu7 | A | 1038 | TRP | p.Trp1038Ter | Q86YC2 |
| HOMO SAPIENS | -1 | 26081 | 3eu7 | A | 1086 | ARG | p.Arg1086Ter | Q86YC2 |
| HOMO SAPIENS | 0 | 26081 | 3eu7 | A | 1122 | ASP | p.Asp1122Glu | Q86YC2 |
| HOMO SAPIENS | 5 | 26081 | 3eu7 | A | 1158 | HIS | p.His1158Gln | Q86YC2 |
| HOMO SAPIENS | 5 | 26081 | 3eu7 | A | 1183 | TYR | p.Tyr1183Ter | Q86YC2 |
| HOMO SAPIENS | 4 | 199 | 1qo5 | C | 45 | ARG | p.Arg46Trp | P05062 |
| HOMO SAPIENS | 5 | 199 | 1qo5 | C | 147 | TRP | p.Trp148Arg | P05062 |
| HOMO SAPIENS | 0 | 199 | 1qo5 | C | 149 | ALA | p.Ala150Pro | P05062 |
| HOMO SAPIENS | 0 | 199 | 1qo5 | C | 174 | ALA | p.Ala175Asp | P05062 |
| HOMO SAPIENS | 0 | 199 | 1qo5 | C | 334 | ASN | p.Asn335Lys | P05062 |
| HOMO SAPIENS | 0 | 199 | 1qo5 | C | 337 | ALA | p.Ala338Val | P05062 |
| HOMO SAPIENS | 5 | 199 | 1qo5 | C | 342 | TYR | p.Tyr343His | P05062 |
| HOMO SAPIENS | 0 | 6910 | 4zht | B | 59 | ARG | p.Arg59Ter | Q9Y223 |
| HOMO SAPIENS | 4 | 6910 | 4zht | B | 129 | ARG | p.Arg129Ter | Q9Y223 |
| HOMO SAPIENS | 4 | 6910 | 4zht | B | 129 | ARG | p.Arg129Gln | Q9Y223 |
| HOMO SAPIENS | 1 | 6910 | 4zht | B | 162 | ARG | p.Arg162Cys | Q9Y223 |
| HOMO SAPIENS | 0 | 6910 | 4zht | B | 176 | ASP | p.Asp176Val | Q9Y223 |
| HOMO SAPIENS | 0 | 6910 | 4zht | B | 216 | VAL | p.Val216Ala | Q9Y223 |
| HOMO SAPIENS | 0 | 6910 | 4zht | B | 246 | ARG | p.Arg246Trp | Q9Y223 |
| HOMO SAPIENS | 0 | 6910 | 4zht | B | 263 | ARG | p.Arg263Leu | Q9Y223 |
| HOMO SAPIENS | 0 | 6910 | 4zht | B | 266 | ARG | p.Arg266Gln | Q9Y223 |
| HOMO SAPIENS | 0 | 6910 | 4zht | B | 266 | ARG | p.Arg266Trp | Q9Y223 |
| HOMO SAPIENS | 0 | 6910 | 4zht | B | 277 | ARG | p.Arg277Cys | Q9Y223 |
| HOMO SAPIENS | 0 | 6910 | 4zht | B | 378 | ASP | p.Asp378Tyr | Q9Y223 |
| HOMO SAPIENS | 3 | 211 | 5kxi | C | 262 | VAL | p.Val287Leu | P17787 |
| HOMO SAPIENS | 0 | 211 | 5kxi | C | 369 | PHE | p.Phe478Leu | P17787 |
| HOMO SAPIENS | 0 | 2454 | 5iy9 | V | null | PHE | p.Phe99Ser | P19447 |
| HOMO SAPIENS | 0 | 2454 | 5iy9 | V | null | ARG | p.Arg109Ter | P19447 |
| HOMO SAPIENS | 0 | 2454 | 5iy9 | V | null | THR | p.Thr119Pro | P19447 |
| HOMO SAPIENS | 0 | 2454 | 5iy9 | V | 425 | ARG | p.Arg425Ter | P19447 |
| HOMO SAPIENS | 1 | 2454 | 5iy9 | V | 545 | GLN | p.Gln545Ter | P19447 |
| HOMO SAPIENS | -1 | 3805 | 3rbq | A | 57 | LYS | p.Lys57Ter | Q13432 |
| HOMO SAPIENS | 2 | 2649 | 4ido | A | 43 | ASP | p.Asp43Glu | Q8WXF7 |
| HOMO SAPIENS | 2 | 2649 | 4ido | A | 157 | LEU | p.Leu157Trp | Q8WXF7 |
| HOMO SAPIENS | 2 | 2649 | 4ido | A | 193 | PHE | p.Phe193Cys | Q8WXF7 |
| HOMO SAPIENS | 7 | 2649 | 4ido | A | 217 | ARG | p.Arg217Gln | Q8WXF7 |
| HOMO SAPIENS | 4 | 2649 | 4ido | A | 239 | ARG | p.Arg239Cys | Q8WXF7 |
| HOMO SAPIENS | 3 | 2649 | 4ido | A | 258 | HIS | p.His258Arg | Q8WXF7 |
| HOMO SAPIENS | 4 | 2649 | 4ido | A | 259 | SER | p.Ser259Tyr | Q8WXF7 |
| HOMO SAPIENS | 4 | 2649 | 4ido | A | 408 | MET | p.Met408Val | Q8WXF7 |
| HOMO SAPIENS | 2 | 2649 | 4ido | A | 415 | ARG | p.Arg415Trp | Q8WXF7 |
| HOMO SAPIENS | 3 | 2649 | 4ido | A | 416 | ARG | p.Arg416Cys | Q8WXF7 |
| HOMO SAPIENS | 5 | 2203 | 3wff | A | 810 | LEU | p.Ser810Leu | P08235 |
| HOMO SAPIENS | 4 | 12648 | 5ep6 | B | 696 | GLU | p.Glu696Lys | Q9UHD2 |
| HOMO SAPIENS | 0 | 30622 | 2pe4 | A | 90 | GLU | p.Glu90Asp | Q12794 |
| HOMO SAPIENS | 0 | 30622 | 2pe4 | A | 256 | GLY | p.Gly256Arg | Q12794 |
| HOMO SAPIENS | 1 | 30622 | 2pe4 | A | 268 | GLU | p.Glu268Lys | Q12794 |
| HOMO SAPIENS | 0 | 30622 | 2pe4 | A | 392 | THR | p.Thr392Met | Q12794 |
| HOMO SAPIENS | 5 | 5411 | 1lya | A | 93 | SER | p.Ser157Leu | P07339 |

| | | | | | | | | |
|---|---|---|---|---|---|---|---|---|
| HOMO SAPIENS | 3 | 5743 | 2uzp | A | 189 | ASN | p.Asn189Lys | P05129 |
| HOMO SAPIENS | 1 | 12529 | 3kfd | J | 21 | ASN | p.Asn45Ser | P36897 |
| HOMO SAPIENS | 1 | 1063 | 2e1q | B | 149 | ARG | p.Arg149Cys | P47989 |
| HOMO SAPIENS | -1 | 1063 | 2e1q | B | 172 | GLY | p.Gly172Arg | P47989 |
| HOMO SAPIENS | 4 | 1063 | 2e1q | B | 228 | ARG | p.Arg228Ter | P47989 |
| HOMO SAPIENS | 5 | 1063 | 2e1q | B | 395 | LYS | p.Lys395Met | P47989 |
| HOMO SAPIENS | 2 | 1063 | 2e1q | B | 646 | ILE | p.Ile646Val | P47989 |
| HOMO SAPIENS | 4 | 1063 | 2e1q | B | 703 | ILE | p.Ile703Val | P47989 |
| HOMO SAPIENS | 8 | 2525 | 1kqn | F | 18 | ASN | p.Asn18Ser | Q9HAN9 |
| HOMO SAPIENS | 0 | 2525 | 1kqn | F | 151 | VAL | p.Val151Phe | Q9HAN9 |
| HOMO SAPIENS | 0 | 2525 | 1kqn | F | 153 | LEU | p.Leu153Val | Q9HAN9 |
| HOMO SAPIENS | 0 | 2525 | 1kqn | F | 169 | TRP | p.Trp169Ter | Q9HAN9 |
| HOMO SAPIENS | 0 | 2525 | 1kqn | F | 207 | ARG | p.Arg207Trp | Q9HAN9 |
| HOMO SAPIENS | 0 | 2525 | 1kqn | F | 273 | ASN | p.Asn273Asp | Q9HAN9 |
| HOMO SAPIENS | 0 | 12299 | 5duq | A | 432 | VAL | p.Val454Glu | P05155 |
| HOMO SAPIENS | 0 | 12299 | 5duq | A | 436 | ALA | p.Ala458Thr | P05155 |
| HOMO SAPIENS | 0 | 12299 | 5duq | A | 443 | ALA | p.Ala465Val | P05155 |
| HOMO SAPIENS | 0 | 12299 | 5duq | A | 444 | ARG | p.Arg466His | P05155 |
| HOMO SAPIENS | 0 | 12299 | 5duq | A | 458 | MET | p.Val480Met | P05155 |
| HOMO SAPIENS | 0 | 327 | 1uhp | A | 36 | ASP | p.Asp274Asn | Q9UPQ7 |
| HOMO SAPIENS | 0 | 64 | 2mo5 | A | 54 | ALA | p.Ala55Thr | P12104 |
| HOMO SAPIENS | 2 | 35969 | 1jrh | I | 60 | CYS | p.Cys77Tyr | P15260 |
| HOMO SAPIENS | 4 | 35969 | 1jrh | I | 70 | ILE | p.Ile87Thr | P15260 |
| HOMO SAPIENS | 2 | 1068 | 1mu7 | A | 304 | ARG | p.Arg304Gln | Q9NUW8 |
| HOMO SAPIENS | 8 | 1068 | 1mu7 | A | 493 | HIS | p.His493Arg | Q9NUW8 |
| HOMO SAPIENS | 0 | 32360 | 2kqp | A | 31 | ARG | p.Arg55Cys | P01308 |
| HOMO SAPIENS | 0 | 32360 | 2kqp | A | 65 | ARG | p.Arg89His | P01308 |
| HOMO SAPIENS | 0 | 32360 | 2kqp | A | 65 | ARG | p.Arg89Cys | P01308 |
| HOMO SAPIENS | 6 | 10805 | 3nks | A | 12 | ILE | p.Ile12Thr | P50336 |
| HOMO SAPIENS | 0 | 10805 | 3nks | A | 20 | HIS | p.His20Pro | P50336 |
| HOMO SAPIENS | 0 | 10805 | 3nks | A | 59 | ARG | p.Arg59Trp | P50336 |
| HOMO SAPIENS | 0 | 10805 | 3nks | A | 168 | ARG | p.Arg168Cys | P50336 |
| HOMO SAPIENS | 0 | 10805 | 3nks | A | 168 | ARG | p.Arg168His | P50336 |
| HOMO SAPIENS | 2 | 10805 | 3nks | A | 232 | GLY | p.Gly232Arg | P50336 |
| HOMO SAPIENS | 3 | 10805 | 3nks | A | 256 | PRO | p.Pro256Arg | P50336 |
| HOMO SAPIENS | 0 | 10805 | 3nks | A | 304 | ARG | p.Arg304His | P50336 |
| HOMO SAPIENS | 0 | 10805 | 3nks | A | 349 | ASP | p.Asp349Ala | P50336 |
| HOMO SAPIENS | 9 | 7008 | 3u2l | A | 125 | GLN | p.Gln125Ter | P55789 |
| HOMO SAPIENS | 4 | 7008 | 3u2l | A | 172 | HIS | p.His172Asn | P55789 |
| HOMO SAPIENS | 4 | 7008 | 3u2l | A | 194 | ARG | p.Arg194His | P55789 |
| HOMO SAPIENS | 3 | 7008 | 3u2l | A | 198 | GLY | p.Gly198Ser | P55789 |
| HOMO SAPIENS | 0 | 20821 | 3e0g | A | 65 | HIS | p.His116Tyr | P42702 |
| HOMO SAPIENS | 3 | 1151 | 5l9v | A | 278 | ASP | p.Asp278Asn | Q9GZT9 |
| HOMO SAPIENS | 4 | 1151 | 5l9v | A | 317 | CYS | p.Pro317Arg | Q9GZT9 |
| HOMO SAPIENS | 5 | 1151 | 5l9v | A | 371 | ARG | p.Arg371His | Q9GZT9 |
| HOMO SAPIENS | 2 | 3935 | 1gwb | A | 36 | ARG | p.Arg36Gly | P07359 |
| HOMO SAPIENS | 1 | 3935 | 1gwb | A | 73 | LEU | p.Leu73Phe | P07359 |
| HOMO SAPIENS | 1 | 3935 | 1gwb | A | 172 | ALA | p.Ala172Val | P07359 |
| HOMO SAPIENS | 0 | 3935 | 1gwb | A | 249 | GLY | p.Gly249Val | P07359 |
| HOMO SAPIENS | 0 | 3935 | 1gwb | A | 255 | MET | p.Met255Val | P07359 |
| HOMO SAPIENS | 2 | 14541 | 5cuz | A | 138 | GLU | p.Glu138Lys | Q9H3L0 |
| HOMO SAPIENS | 2 | 14541 | 5cuz | A | 143 | SER | p.Ser143Ile | Q9H3L0 |
| HOMO SAPIENS | 4 | 14541 | 5cuz | A | 182 | THR | p.Thr182Asn | Q9H3L0 |
| HOMO SAPIENS | 6 | 14541 | 5cuz | A | 206 | ASN | p.Asn206Ile | Q9H3L0 |

| | | | | | | | | |
|---|---|---|---|---|---|---|---|---|
| HOMO SAPIENS | 0 | 14541 | 5cuz | A | 249 | TYR | p.Tyr249Cys | Q9H3L0 |
| HOMO SAPIENS | 4 | 14541 | 5cuz | A | 250 | ARG | p.Arg250Ter | Q9H3L0 |
| HOMO SAPIENS | 2 | 14541 | 5cuz | A | 259 | LEU | p.Leu259Pro | Q9H3L0 |
| HOMO SAPIENS | 0 | 4206 | 4fao | D | 30 | PRO | p.Pro30Ser | P37023 |
| HOMO SAPIENS | 1 | 4206 | 4fao | D | 50 | TRP | p.Trp50Cys | P37023 |
| HOMO SAPIENS | 0 | 16553 | 3v30 | A | 251 | GLN | p.Gln251Glu | O14593 |
| HOMO SAPIENS | 0 | 5012 | 5hbf | A | 442 | ALA | p.Ala442Gly | Q13231 |
| HOMO SAPIENS | 5 | 4844 | 5tdc | A | 136 | HIS | p.His136Arg | Q8IWV7 |
| HOMO SAPIENS | 0 | 15435 | 4cgz | A | 643 | ARG | p.Arg643His | P54132 |
| HOMO SAPIENS | 1 | 15435 | 4cgz | A | 672 | GLN | p.Gln672Arg | P54132 |
| HOMO SAPIENS | 1 | 15435 | 4cgz | A | 700 | GLN | p.Gln700Ter | P54132 |
| HOMO SAPIENS | 0 | 15435 | 4cgz | A | 707 | PRO | p.Pro707Ser | P54132 |
| HOMO SAPIENS | 0 | 15435 | 4cgz | A | 755 | LYS | p.Lys755Glu | P54132 |
| HOMO SAPIENS | 0 | 15435 | 4cgz | A | 881 | TRP | p.Trp881Ter | P54132 |
| HOMO SAPIENS | 2 | 15435 | 4cgz | A | 899 | ARG | p.Arg899Ter | P54132 |
| HOMO SAPIENS | 0 | 15435 | 4cgz | A | 1036 | CYS | p.Cys1036Phe | P54132 |
| HOMO SAPIENS | 0 | 15435 | 4cgz | A | 1055 | CYS | p.Cys1055Ser | P54132 |
| HOMO SAPIENS | 0 | 15435 | 4cgz | A | 1139 | ARG | p.Arg1139Ter | P54132 |
| HOMO SAPIENS | -1 | 15435 | 4cgz | A | 1198 | VAL | p.Val1198Met | P54132 |
| HOMO SAPIENS | -1 | 15435 | 4cgz | A | 1205 | VAL | p.Val1205Ile | P54132 |
| HOMO SAPIENS | 0 | 15435 | 4cgz | A | 1251 | GLU | p.Glu1251Gln | P54132 |
| HOMO SAPIENS | 0 | 11185 | 1kb4 | A | 30 | ARG | p.Arg30Ter | P11473 |
| HOMO SAPIENS | 5 | 11185 | 1kb4 | A | 33 | GLY | p.Gly33Asp | P11473 |
| HOMO SAPIENS | 5 | 11185 | 1kb4 | A | 46 | GLY | p.Gly46Asp | P11473 |
| HOMO SAPIENS | 6 | 11185 | 1kb4 | A | 50 | ARG | p.Arg50Gln | P11473 |
| HOMO SAPIENS | 9 | 11185 | 1kb4 | A | 73 | ARG | p.Arg73Gln | P11473 |
| HOMO SAPIENS | 9 | 11185 | 1kb4 | A | 80 | ARG | p.Arg80Gln | P11473 |
| HOMO SAPIENS | 0 | 5382 | 3fl7 | A | 41 | LEU | p.Leu41Val | P29317 |
| HOMO SAPIENS | 2 | 5382 | 3fl7 | A | 175 | ARG | p.Arg175Cys | P29317 |
| HOMO SAPIENS | 1 | 5382 | 3fl7 | A | 195 | ARG | p.Arg195Cys | P29317 |
| HOMO SAPIENS | 0 | 5382 | 3fl7 | A | 232 | ASP | p.Asp232Gly | P29317 |
| HOMO SAPIENS | 0 | 5382 | 3fl7 | A | 277 | SER | p.Ser277Leu | P29317 |
| HOMO SAPIENS | 0 | 5382 | 3fl7 | A | 315 | ARG | p.Arg315Gln | P29317 |
| HOMO SAPIENS | 7 | 5382 | 3fl7 | A | 350 | PRO | p.Pro350Thr | P29317 |
| HOMO SAPIENS | 0 | 5382 | 3fl7 | A | 391 | GLY | p.Gly391Arg | P29317 |
| HOMO SAPIENS | 0 | 5382 | 3fl7 | A | 418 | VAL | p.Val418Ile | P29317 |
| HOMO SAPIENS | 0 | 5382 | 3fl7 | A | 511 | THR | p.Thr511Met | P29317 |
| HOMO SAPIENS | 2 | 29168 | 4p22 | B | 144 | VAL | p.Val144Ile | P22314 |
| HOMO SAPIENS | 5 | 4348 | 1bf5 | A | 241 | ARG | p.Arg241Gln | P42224 |
| HOMO SAPIENS | 2 | 4348 | 1bf5 | A | 266 | VAL | p.Val266Ile | P42224 |
| HOMO SAPIENS | 4 | 4348 | 1bf5 | A | 320 | GLU | p.Glu320Gln | P42224 |
| HOMO SAPIENS | 1 | 4348 | 1bf5 | A | 372 | ASN | p.Asn372Lys | P42224 |
| HOMO SAPIENS | 1 | 4348 | 1bf5 | A | 447 | ASP | p.Asp447Glu | P42224 |
| HOMO SAPIENS | 2 | 4348 | 1bf5 | A | 463 | GLN | p.Gln463His | P42224 |
| HOMO SAPIENS | 4 | 4348 | 1bf5 | A | 600 | LEU | p.Leu600Pro | P42224 |
| HOMO SAPIENS | 0 | 4348 | 1bf5 | A | 706 | LEU | p.Leu706Ser | P42224 |
| HOMO SAPIENS | 4 | 4991 | 4bej | B | 141 | ASN | p.Asn141Ser | O00429 |
| HOMO SAPIENS | 0 | 4991 | 4bej | B | 395 | ALA | p.Ala395Asp | O00429 |
| HOMO SAPIENS | 2 | 11792 | 1fvo | A | 40 | ARG | p.Arg40Cys | P00480 |
| HOMO SAPIENS | 2 | 11792 | 1fvo | A | 40 | ARG | p.Arg40His | P00480 |
| HOMO SAPIENS | 0 | 11792 | 1fvo | A | 45 | LEU | p.Leu45Pro | P00480 |
| HOMO SAPIENS | 1 | 11792 | 1fvo | A | 47 | ASN | p.Asn47Ile | P00480 |
| HOMO SAPIENS | 0 | 11792 | 1fvo | A | 50 | GLY | p.Gly50Arg | P00480 |
| HOMO SAPIENS | 5 | 11792 | 1fvo | A | 87 | GLU | p.Glu87Lys | P00480 |

| | | | | | | | | |
|---|---|---|---|---|---|---|---|---|
| HOMO SAPIENS | 3 | 11792 | 1fvo | A | 111 | LEU | p.Leu111Pro | P00480 |
| HOMO SAPIENS | 1 | 11792 | 1fvo | A | 125 | THR | p.Thr125Met | P00480 |
| HOMO SAPIENS | 3 | 11792 | 1fvo | A | 129 | ARG | p.Arg129His | P00480 |
| HOMO SAPIENS | 9 | 11792 | 1fvo | A | 141 | ARG | p.Arg141Pro | P00480 |
| HOMO SAPIENS | 0 | 11792 | 1fvo | A | 206 | MET | p.Met206Arg | P00480 |
| HOMO SAPIENS | 0 | 11792 | 1fvo | A | 207 | SER | p.Ser207Arg | P00480 |
| HOMO SAPIENS | 0 | 11792 | 1fvo | A | 216 | GLN | p.Gln216Glu | P00480 |
| HOMO SAPIENS | 1 | 11792 | 1fvo | A | 225 | PRO | p.Pro225Leu | P00480 |
| HOMO SAPIENS | 0 | 11792 | 1fvo | A | 270 | ARG | p.Gln270Arg | P00480 |
| HOMO SAPIENS | 0 | 11792 | 1fvo | A | 270 | ARG | p.Gln270Ter | P00480 |
| HOMO SAPIENS | 2 | 11792 | 1fvo | A | 277 | ARG | p.Arg277Trp | P00480 |
| HOMO SAPIENS | 0 | 3721 | 4r3z | B | 2 | ASP | p.Asp2Gly | P54136 |
| HOMO SAPIENS | 0 | 3721 | 4r3z | B | 456 | SER | p.Ser456Leu | P54136 |
| HOMO SAPIENS | 5 | 3721 | 4r3z | B | 512 | ARG | p.Arg512Gln | P54136 |
| HOMO SAPIENS | 3 | 5749 | 3hl2 | A | 239 | ALA | p.Ala239Thr | Q9HD40 |
| HOMO SAPIENS | 1 | 5749 | 3hl2 | A | 334 | TYR | p.Tyr334Cys | Q9HD40 |
| HOMO SAPIENS | 1 | 5749 | 3hl2 | A | 452 | LYS | p.Lys452Asn | Q9HD40 |
| HOMO SAPIENS | -1 | 5749 | 3hl2 | A | 465 | ARG | p.Arg465Leu | Q9HD40 |
| HOMO SAPIENS | -1 | 5749 | 3hl2 | A | 489 | ASP | p.Asp489Gly | Q9HD40 |
| HOMO SAPIENS | 5 | 1721 | 3k15 | A | 1652 | MET | p.Met1652Ile | P38398 |
| HOMO SAPIENS | 5 | 1721 | 3k15 | A | 1652 | MET | p.Met1652Lys | P38398 |
| HOMO SAPIENS | 5 | 1721 | 3k15 | A | 1662 | PHE | p.Phe1662Cys | P38398 |
| HOMO SAPIENS | 5 | 1721 | 3k15 | A | 1664 | LEU | p.Leu1664Pro | P38398 |
| HOMO SAPIENS | 0 | 1721 | 3k15 | A | 1665 | VAL | p.Val1665Met | P38398 |
| HOMO SAPIENS | 5 | 1721 | 3k15 | A | 1690 | LYS | p.Lys1690Gln | P38398 |
| HOMO SAPIENS | 5 | 1721 | 3k15 | A | 1692 | ASP | p.Asp1692Asn | P38398 |
| HOMO SAPIENS | 5 | 1721 | 3k15 | A | 1699 | ARG | p.Arg1699Gln | P38398 |
| HOMO SAPIENS | 5 | 1721 | 3k15 | A | 1699 | ARG | p.Arg1699Trp | P38398 |
| HOMO SAPIENS | 3 | 1721 | 3k15 | A | 1706 | GLY | p.Gly1706Glu | P38398 |
| HOMO SAPIENS | 5 | 1721 | 3k15 | A | 1720 | THR | p.Thr1720Ala | P38398 |
| HOMO SAPIENS | 5 | 1721 | 3k15 | A | 1751 | ARG | p.Arg1751Pro | P38398 |
| HOMO SAPIENS | 4 | 1721 | 3k15 | A | 1775 | MET | p.Met1775Arg | P38398 |
| HOMO SAPIENS | 2 | 1721 | 3k15 | A | 1788 | GLY | p.Gly1788Val | P38398 |
| HOMO SAPIENS | 5 | 1721 | 3k15 | A | 1794 | GLU | p.Glu1794Asp | P38398 |
| HOMO SAPIENS | 5 | 1721 | 3k15 | A | 1804 | VAL | p.Val1804Asp | P38398 |
| HOMO SAPIENS | 3 | 1721 | 3k15 | A | 1815 | TRP | p.Trp1815Ter | P38398 |
| HOMO SAPIENS | 1 | 1721 | 3k15 | A | 1819 | ASN | p.Asn1819Ser | P38398 |
| HOMO SAPIENS | 5 | 1721 | 3k15 | A | 1856 | PRO | p.Pro1856Ser | P38398 |
| HOMO SAPIENS | 5 | 1721 | 3k15 | A | 1858 | ILE | p.Ile1858Leu | P38398 |
| HOMO SAPIENS | 5 | 1721 | 3k15 | A | 1858 | ILE | p.Ile1858Thr | P38398 |
| HOMO SAPIENS | 5 | 1721 | 3k15 | A | 1859 | PRO | p.Pro1859Arg | P38398 |
| HOMO SAPIENS | 2 | 22086 | 3kq4 | B | 1297 | PRO | p.Pro1297Leu | O60494 |
| HOMO SAPIENS | 1 | 12135 | 2k9j | B | 724 | ARG | p.Arg750Ter | P05106 |
| HOMO SAPIENS | -1 | 24027 | 2qtl | A | 175 | LEU | p.Ser202Leu | Q9UBK8 |
| HOMO SAPIENS | 0 | 24027 | 2qtl | A | 257 | SER | p.Ser284Thr | Q9UBK8 |
| HOMO SAPIENS | 0 | 24027 | 2qtl | A | 333 | LEU | p.Leu360Val | Q9UBK8 |
| HOMO SAPIENS | 0 | 24027 | 2qtl | A | 350 | LYS | p.Lys377Arg | Q9UBK8 |
| HOMO SAPIENS | 0 | 24027 | 2qtl | A | 415 | ARG | p.Arg442Cys | Q9UBK8 |
| HOMO SAPIENS | 2 | 24027 | 2qtl | A | 450 | PRO | p.Pro477Arg | Q9UBK8 |
| HOMO SAPIENS | 4 | 24027 | 2qtl | A | 487 | GLY | p.Gly514Arg | Q9UBK8 |
| HOMO SAPIENS | 1 | 24027 | 2qtl | A | 595 | HIS | p.His622Tyr | Q9UBK8 |
| HOMO SAPIENS | 0 | 5758 | 3di2 | B | 46 | THR | p.Thr66Ile | P16871 |
| HOMO SAPIENS | 1 | 5758 | 3di2 | B | 118 | VAL | p.Ile138Val | P16871 |
| HOMO SAPIENS | 4 | 848 | 5oa3 | Y | 79 | LEU | p.Leu79Val | P62847 |

| | | | | | | | | |
|---|---|---|---|---|---|---|---|---|
| HOMO SAPIENS | 2 | 848 | 5oa3 | Y | 124 | ASN | p.Asn124Ser | P62847 |
| HOMO SAPIENS | 0 | 81 | 3s5o | A | 36 | ALA | p.Ala36Glu | Q86XE5 |
| HOMO SAPIENS | 0 | 81 | 3s5o | A | 39 | TYR | p.Tyr39Ter | Q86XE5 |
| HOMO SAPIENS | 0 | 81 | 3s5o | A | 45 | PRO | p.Pro45Leu | Q86XE5 |
| HOMO SAPIENS | 0 | 81 | 3s5o | A | 70 | ARG | p.Arg70Ter | Q86XE5 |
| HOMO SAPIENS | 0 | 81 | 3s5o | A | 70 | ARG | p.Arg70Pro | Q86XE5 |
| HOMO SAPIENS | 1 | 81 | 3s5o | A | 97 | ARG | p.Arg97Cys | Q86XE5 |
| HOMO SAPIENS | 4 | 81 | 3s5o | A | 113 | GLU | p.Glu113Lys | Q86XE5 |
| HOMO SAPIENS | 1 | 81 | 3s5o | A | 190 | PRO | p.Pro190Leu | Q86XE5 |
| HOMO SAPIENS | 0 | 81 | 3s5o | A | 245 | VAL | p.Val245Phe | Q86XE5 |
| HOMO SAPIENS | 0 | 81 | 3s5o | A | 257 | CYS | p.Cys257Gly | Q86XE5 |
| HOMO SAPIENS | 0 | 81 | 3s5o | A | 287 | GLY | p.Gly287Val | Q86XE5 |
| HOMO SAPIENS | 7 | 81 | 3s5o | A | 303 | ARG | p.Arg303Cys | Q86XE5 |
| HOMO SAPIENS | 0 | 81 | 3s5o | A | 325 | GLY | p.Gly325Ser | Q86XE5 |
| HOMO SAPIENS | -1 | 622 | 5era | A | 12 | GLY | p.Gly12Arg | P29033 |
| HOMO SAPIENS | -1 | 622 | 5era | A | 12 | GLY | p.Gly12Asp | P29033 |
| HOMO SAPIENS | -1 | 622 | 5era | A | 17 | SER | p.Ser17Phe | P29033 |
| HOMO SAPIENS | 5 | 622 | 5era | A | 24 | TRP | p.Trp24Ter | P29033 |
| HOMO SAPIENS | 0 | 622 | 5era | A | 32 | ARG | p.Arg32Cys | P29033 |
| HOMO SAPIENS | 5 | 622 | 5era | A | 34 | MET | p.Met34Thr | P29033 |
| HOMO SAPIENS | 5 | 622 | 5era | A | 36 | LEU | p.Leu36Pro | P29033 |
| HOMO SAPIENS | 5 | 622 | 5era | A | 37 | VAL | p.Val37Ile | P29033 |
| HOMO SAPIENS | 5 | 622 | 5era | A | 40 | ALA | p.Ala40Glu | P29033 |
| HOMO SAPIENS | 5 | 622 | 5era | A | 44 | TRP | p.Trp44Cys | P29033 |
| HOMO SAPIENS | 5 | 622 | 5era | A | 45 | GLY | p.Gly45Glu | P29033 |
| HOMO SAPIENS | 5 | 622 | 5era | A | 47 | GLU | p.Glu47Ter | P29033 |
| HOMO SAPIENS | 5 | 622 | 5era | A | 50 | ASP | p.Asp50Asn | P29033 |
| HOMO SAPIENS | 5 | 622 | 5era | A | 54 | ASN | p.Asn54Lys | P29033 |
| HOMO SAPIENS | 5 | 622 | 5era | A | 57 | GLN | p.Gln57Ter | P29033 |
| HOMO SAPIENS | 5 | 622 | 5era | A | 59 | GLY | p.Gly59Ala | P29033 |
| HOMO SAPIENS | 5 | 622 | 5era | A | 59 | GLY | p.Gly59Ser | P29033 |
| HOMO SAPIENS | 5 | 622 | 5era | A | 66 | ASP | p.Asp66His | P29033 |
| HOMO SAPIENS | 5 | 622 | 5era | A | 68 | TYR | p.Tyr68Cys | P29033 |
| HOMO SAPIENS | 5 | 622 | 5era | A | 70 | PRO | p.Pro70Ala | P29033 |
| HOMO SAPIENS | 5 | 622 | 5era | A | 73 | HIS | p.His73Arg | P29033 |
| HOMO SAPIENS | 5 | 622 | 5era | A | 75 | ARG | p.Arg75Trp | P29033 |
| HOMO SAPIENS | 5 | 622 | 5era | A | 75 | ARG | p.Arg75Gln | P29033 |
| HOMO SAPIENS | 2 | 622 | 5era | A | 77 | TRP | p.Trp77Arg | P29033 |
| HOMO SAPIENS | 2 | 622 | 5era | A | 77 | TRP | p.Trp77Ter | P29033 |
| HOMO SAPIENS | 0 | 622 | 5era | A | 80 | GLN | p.Gln80Ter | P29033 |
| HOMO SAPIENS | 0 | 622 | 5era | A | 80 | GLN | p.Gln80Pro | P29033 |
| HOMO SAPIENS | 5 | 622 | 5era | A | 81 | LEU | p.Leu81Val | P29033 |
| HOMO SAPIENS | 5 | 622 | 5era | A | 82 | ILE | p.Ile82Met | P29033 |
| HOMO SAPIENS | 5 | 622 | 5era | A | 83 | PHE | p.Phe83Leu | P29033 |
| HOMO SAPIENS | 5 | 622 | 5era | A | 84 | VAL | p.Val84Leu | P29033 |
| HOMO SAPIENS | 5 | 622 | 5era | A | 90 | LEU | p.Leu90Pro | P29033 |
| HOMO SAPIENS | 5 | 622 | 5era | A | 95 | VAL | p.Val95Met | P29033 |
| HOMO SAPIENS | -1 | 622 | 5era | A | 100 | HIS | p.His100Tyr | P29033 |
| HOMO SAPIENS | -1 | 622 | 5era | A | 113 | SER | p.Ser113Arg | P29033 |
| HOMO SAPIENS | -1 | 622 | 5era | A | 119 | GLU | p.Glu119Lys | P29033 |
| HOMO SAPIENS | -1 | 622 | 5era | A | 122 | LYS | p.Lys122Ile | P29033 |
| HOMO SAPIENS | -1 | 622 | 5era | A | 123 | THR | p.Thr123Asn | P29033 |
| HOMO SAPIENS | -1 | 622 | 5era | A | 124 | GLN | p.Gln124Ter | P29033 |
| HOMO SAPIENS | 5 | 622 | 5era | A | 139 | SER | p.Ser139Asn | P29033 |

| | | | | | | | | |
|---|---|---|---|---|---|---|---|---|
| HOMO SAPIENS | 5 | 622 | 5era | A | 143 | ARG | p.Arg143Trp | P29033 |
| HOMO SAPIENS | 5 | 622 | 5era | A | 143 | ARG | p.Arg143Gln | P29033 |
| HOMO SAPIENS | 0 | 622 | 5era | A | 147 | GLU | p.Glu147Lys | P29033 |
| HOMO SAPIENS | 7 | 622 | 5era | A | 155 | TYR | p.Tyr155Ter | P29033 |
| HOMO SAPIENS | 2 | 622 | 5era | A | 159 | ASP | p.Asp159Val | P29033 |
| HOMO SAPIENS | 5 | 622 | 5era | A | 163 | MET | p.Met163Val | P29033 |
| HOMO SAPIENS | 5 | 622 | 5era | A | 165 | ARG | p.Arg165Trp | P29033 |
| HOMO SAPIENS | 5 | 622 | 5era | A | 167 | VAL | p.Val167Met | P29033 |
| HOMO SAPIENS | 5 | 622 | 5era | A | 169 | CYS | p.Cys169Tyr | P29033 |
| HOMO SAPIENS | 5 | 622 | 5era | A | 179 | ASP | p.Asp179Asn | P29033 |
| HOMO SAPIENS | 0 | 622 | 5era | A | 199 | SER | p.Ser199Phe | P29033 |
| HOMO SAPIENS | 0 | 622 | 5era | A | 202 | CYS | p.Cys202Phe | P29033 |
| HOMO SAPIENS | 5 | 622 | 5era | A | 203 | ILE | p.Ile203Thr | P29033 |
| HOMO SAPIENS | 5 | 622 | 5era | A | 206 | ASN | p.Asn206Ser | P29033 |
| HOMO SAPIENS | 0 | 8103 | 5dot | A | 163 | GLY | p.Gly163Ter | P31327 |
| HOMO SAPIENS | 0 | 8103 | 5dot | A | 337 | HIS | p.His337Arg | P31327 |
| HOMO SAPIENS | 3 | 8103 | 5dot | A | 344 | THR | p.Thr344Ala | P31327 |
| HOMO SAPIENS | 1 | 8103 | 5dot | A | 405 | THR | p.Thr405Asn | P31327 |
| HOMO SAPIENS | 5 | 8103 | 5dot | A | 544 | THR | p.Thr544Met | P31327 |
| HOMO SAPIENS | 5 | 8103 | 5dot | A | 716 | ASN | p.Asn716Lys | P31327 |
| HOMO SAPIENS | -1 | 8103 | 5dot | A | 780 | ARG | p.Arg780His | P31327 |
| HOMO SAPIENS | 0 | 8103 | 5dot | A | 982 | GLY | p.Gly982Asp | P31327 |
| HOMO SAPIENS | 3 | 8103 | 5dot | A | 1119 | ALA | p.Ala1119Ser | P31327 |
| HOMO SAPIENS | 1 | 8103 | 5dot | A | 1376 | GLY | p.Gly1376Ser | P31327 |
| HOMO SAPIENS | 4 | 6833 | 1l8l | B | 27 | ARG | p.Arg27Ser | P78330 |
| HOMO SAPIENS | 3 | 6833 | 1l8l | B | 32 | ASP | p.Asp32Gly | P78330 |
| HOMO SAPIENS | 3 | 6833 | 1l8l | B | 32 | ASP | p.Asp32Asn | P78330 |
| HOMO SAPIENS | 1 | 6833 | 1l8l | B | 49 | ARG | p.Arg49Trp | P78330 |
| HOMO SAPIENS | 0 | 6833 | 1l8l | B | 52 | MET | p.Met52Thr | P78330 |
| HOMO SAPIENS | 0 | 6833 | 1l8l | B | 83 | GLN | p.Gln83His | P78330 |
| HOMO SAPIENS | 6 | 810 | 2iw2 | A | 185 | ARG | p.Arg184Gln | P12955 |
| HOMO SAPIENS | 0 | 810 | 2iw2 | A | 213 | ALA | p.Ala212Pro | P12955 |
| HOMO SAPIENS | 1 | 810 | 2iw2 | A | 266 | ARG | p.Arg265Ter | P12955 |
| HOMO SAPIENS | 9 | 810 | 2iw2 | A | 277 | ASP | p.Asp276Asn | P12955 |
| HOMO SAPIENS | 0 | 810 | 2iw2 | A | 279 | GLY | p.Gly278Asp | P12955 |
| HOMO SAPIENS | 9 | 810 | 2iw2 | A | 378 | HIS | p.His377Gln | P12955 |
| HOMO SAPIENS | 0 | 810 | 2iw2 | A | 389 | ARG | p.Arg388His | P12955 |
| HOMO SAPIENS | 0 | 810 | 2iw2 | A | 436 | PHE | p.Leu435Phe | P12955 |
| HOMO SAPIENS | 0 | 810 | 2iw2 | A | 449 | GLY | p.Gly448Arg | P12955 |
| HOMO SAPIENS | -1 | 1530 | 5fwj | A | 388 | ALA | p.Ala388Pro | P41229 |
| HOMO SAPIENS | 4 | 1530 | 5fwj | A | 451 | SER | p.Ser451Arg | P41229 |
| HOMO SAPIENS | 3 | 1530 | 5fwj | A | 731 | LEU | p.Leu731Phe | P41229 |
| HOMO SAPIENS | 3 | 578 | 5net | B | 123 | ALA | p.Ala143Thr | P18564 |
| HOMO SAPIENS | 6 | 578 | 5net | B | 176 | PRO | p.Pro196Thr | P18564 |
| HOMO SAPIENS | 3 | 578 | 5net | B | 280 | GLU | p.Glu300Lys | P18564 |
| HOMO SAPIENS | -1 | 5247 | 4j15 | A | 274 | ALA | p.Ala274Val | P14868 |
| HOMO SAPIENS | -1 | 5247 | 4j15 | A | 280 | HIS | p.His280Leu | P14868 |
| HOMO SAPIENS | 2 | 5247 | 4j15 | A | 460 | ARG | p.Arg460His | P14868 |
| HOMO SAPIENS | 5 | 5247 | 4j15 | A | 494 | ARG | p.Arg494Cys | P14868 |
| HOMO SAPIENS | 7 | 2566 | 1b55 | A | 28 | ARG | p.Arg28His | Q06187 |
| HOMO SAPIENS | 5 | 2566 | 1b55 | A | 33 | THR | p.Thr33Pro | Q06187 |
| HOMO SAPIENS | 0 | 2566 | 1b55 | A | 113 | VAL | p.Val113Asp | Q06187 |
| HOMO SAPIENS | 0 | 356 | 5a2s | A | 786 | ALA | p.Ala786Thr | P56524 |
| HOMO SAPIENS | 5 | 2960 | 4cak | A | 553 | ARG | p.Arg584Ter | P08514 |

| | | | | | | | | |
|---|---|---|---|---|---|---|---|---|
| HOMO SAPIENS | 5 | 2960 | 4cak | A | 565 | ILE | p.Ile596Thr | P08514 |
| HOMO SAPIENS | -1 | 2960 | 4cak | A | 841 | LEU | p.Leu872Met | P08514 |
| HOMO SAPIENS | -1 | 2960 | 4cak | A | 843 | ILE | p.Ile874Ser | P08514 |
| HOMO SAPIENS | 0 | 5054 | 4k8r | B | 216 | PHE | p.Phe216Leu | Q8NBP7 |
| HOMO SAPIENS | 1 | 5054 | 4k8r | B | 417 | HIS | p.His417Gln | Q8NBP7 |
| HOMO SAPIENS | 1 | 5054 | 4k8r | B | 425 | ASN | p.Asn425Ser | Q8NBP7 |
| HOMO SAPIENS | 1 | 5054 | 4k8r | B | 443 | ALA | p.Ala443Thr | Q8NBP7 |
| HOMO SAPIENS | 0 | 27283 | 3n5n | X | 71 | HIS | p.His82Arg | Q9UIF7 |
| HOMO SAPIENS | 1 | 27283 | 3n5n | X | 83 | ARG | p.Arg94Ter | Q9UIF7 |
| HOMO SAPIENS | 8 | 27283 | 3n5n | X | 89 | TRP | p.Trp100Ter | Q9UIF7 |
| HOMO SAPIENS | 8 | 27283 | 3n5n | X | 90 | TYR | p.Tyr101Ter | Q9UIF7 |
| HOMO SAPIENS | 7 | 27283 | 3n5n | X | 95 | ARG | p.Arg106Trp | Q9UIF7 |
| HOMO SAPIENS | 7 | 27283 | 3n5n | X | 117 | TRP | p.Trp128Arg | Q9UIF7 |
| HOMO SAPIENS | 0 | 27283 | 3n5n | X | 137 | GLY | p.Gly148Arg | Q9UIF7 |
| HOMO SAPIENS | 5 | 27283 | 3n5n | X | 142 | TRP | p.Trp153Ter | Q9UIF7 |
| HOMO SAPIENS | 0 | 27283 | 3n5n | X | 165 | TYR | p.Tyr176Cys | Q9UIF7 |
| HOMO SAPIENS | 0 | 27283 | 3n5n | X | 168 | ARG | p.Arg179His | Q9UIF7 |
| HOMO SAPIENS | 0 | 27283 | 3n5n | X | 168 | ARG | p.Arg179Cys | Q9UIF7 |
| HOMO SAPIENS | 0 | 27283 | 3n5n | X | 196 | GLN | p.Gln207Ter | Q9UIF7 |
| HOMO SAPIENS | 8 | 27283 | 3n5n | X | 201 | VAL | p.Val212Met | Q9UIF7 |
| HOMO SAPIENS | 9 | 27283 | 3n5n | X | 227 | ARG | p.Arg238Trp | Q9UIF7 |
| HOMO SAPIENS | 9 | 27283 | 3n5n | X | 231 | ARG | p.Arg242Cys | Q9UIF7 |
| HOMO SAPIENS | 9 | 27283 | 3n5n | X | 231 | ARG | p.Arg242His | Q9UIF7 |
| HOMO SAPIENS | 3 | 27283 | 3n5n | X | 232 | VAL | p.Val243Phe | Q9UIF7 |
| HOMO SAPIENS | 3 | 27283 | 3n5n | X | 246 | GLN | p.Gln257Ter | Q9UIF7 |
| HOMO SAPIENS | 9 | 27283 | 3n5n | X | 272 | GLY | p.Gly283Glu | Q9UIF7 |
| HOMO SAPIENS | 9 | 27283 | 3n5n | X | 281 | PRO | p.Pro292Leu | Q9UIF7 |
| HOMO SAPIENS | -1 | 27283 | 3n5n | X | 314 | ASP | p.Asp325Glu | Q9UIF7 |
| HOMO SAPIENS | 0 | 27283 | 3n5n | X | 324 | GLN | p.Gln335His | Q9UIF7 |
| HOMO SAPIENS | 0 | 27283 | 3n5n | X | 324 | GLN | p.Gln335Arg | Q9UIF7 |
| HOMO SAPIENS | 0 | 27283 | 3n5n | X | 324 | GLN | p.Gln335Ter | Q9UIF7 |
| HOMO SAPIENS | -1 | 20852 | 4p60 | B | 243 | ARG | p.Arg243Lys | O75746 |
| HOMO SAPIENS | -1 | 1875 | 5dd2 | G | 152 | VAL | p.Val179Leu | P06396 |
| HOMO SAPIENS | 9 | 2126 | 3n1p | B | 46 | PRO | p.Pro46Leu | Q14623 |
| HOMO SAPIENS | 8 | 2126 | 3n1p | B | 76 | GLU | p.Glu76Ala | Q14623 |
| HOMO SAPIENS | 9 | 2126 | 3n1p | B | 77 | ARG | p.Arg77Ser | Q14623 |
| HOMO SAPIENS | 6 | 2126 | 3n1p | B | 95 | GLU | p.Glu95Lys | Q14623 |
| HOMO SAPIENS | 9 | 2126 | 3n1p | B | 100 | ASP | p.Asp100Glu | Q14623 |
| HOMO SAPIENS | 9 | 2126 | 3n1p | B | 131 | GLU | p.Glu131Lys | Q14623 |
| HOMO SAPIENS | 0 | 2126 | 3n1p | B | 190 | VAL | p.Val190Ala | Q14623 |
| HOMO SAPIENS | 5 | 40410 | 3ioz | B | 70 | GLN | p.Gln70Leu | P20963 |
| HOMO SAPIENS | 1 | 5754 | 5lkz | A | 1108 | THR | p.Tyr1191Ter | Q09472 |
| HOMO SAPIENS | 3 | 8159 | 1tiu | A | 3 | GLU | p.Lys12679Glu | Q8WZ42 |
| HOMO SAPIENS | 4 | 3275 | 4ldq | A | 210 | ARG | p.Arg210Gln | Q9BV20 |
| HOMO SAPIENS | 0 | 8143 | 4miv | B | 66 | SER | p.Ser66Trp | P51688 |
| HOMO SAPIENS | 1 | 8143 | 4miv | B | 74 | ARG | p.Arg74Cys | P51688 |
| HOMO SAPIENS | 0 | 8143 | 4miv | B | 128 | PRO | p.Pro128Leu | P51688 |
| HOMO SAPIENS | 1 | 8143 | 4miv | B | 150 | ARG | p.Arg150Gln | P51688 |
| HOMO SAPIENS | 0 | 8143 | 4miv | B | 191 | GLY | p.Gly191Arg | P51688 |
| HOMO SAPIENS | 0 | 8143 | 4miv | B | 206 | ARG | p.Arg206Pro | P51688 |
| HOMO SAPIENS | 0 | 8143 | 4miv | B | 225 | PHE | p.Phe225Leu | P51688 |
| HOMO SAPIENS | 0 | 8143 | 4miv | B | 233 | ARG | p.Arg233Ter | P51688 |
| HOMO SAPIENS | 0 | 8143 | 4miv | B | 234 | ALA | p.Ala234Gly | P51688 |
| HOMO SAPIENS | 0 | 8143 | 4miv | B | 245 | ARG | p.Arg245His | P51688 |

| | | | | | | | | |
|---|---|---|---|---|---|---|---|---|
| HOMO SAPIENS | 0 | 8143 | 4miv | B | 293 | PRO | p.Pro293Ser | P51688 |
| HOMO SAPIENS | 1 | 8143 | 4miv | B | 298 | SER | p.Ser298Pro | P51688 |
| HOMO SAPIENS | 0 | 8143 | 4miv | B | 361 | VAL | p.Val361Ile | P51688 |
| HOMO SAPIENS | 0 | 8143 | 4miv | B | 369 | GLU | p.Glu369Lys | P51688 |
| HOMO SAPIENS | 0 | 8143 | 4miv | B | 372 | MET | p.Met372Ile | P51688 |
| HOMO SAPIENS | 1 | 8143 | 4miv | B | 380 | GLN | p.Gln380Arg | P51688 |
| HOMO SAPIENS | 0 | 8143 | 4miv | B | 387 | VAL | p.Val387Met | P51688 |
| HOMO SAPIENS | 0 | 8143 | 4miv | B | 394 | MET | p.Met394Ile | P51688 |
| HOMO SAPIENS | 0 | 8143 | 4miv | B | 433 | ARG | p.Arg433Gln | P51688 |
| HOMO SAPIENS | 1 | 8143 | 4miv | B | 446 | HIS | p.His446Arg | P51688 |
| HOMO SAPIENS | 5 | 8143 | 4miv | B | 447 | GLU | p.Glu447Lys | P51688 |
| HOMO SAPIENS | 2 | 8143 | 4miv | B | 456 | ARG | p.Arg456His | P51688 |
| HOMO SAPIENS | 1 | 21880 | 2oo9 | B | 857 | LEU | p.Leu857Phe | P22681 |
| HOMO SAPIENS | 0 | 9467 | 1s1d | A | 154 | LEU | p.Leu224Pro | Q8WVQ1 |
| HOMO SAPIENS | 0 | 9467 | 1s1d | A | 156 | VAL | p.Val226Met | Q8WVQ1 |
| HOMO SAPIENS | 0 | 9467 | 1s1d | A | 229 | PRO | p.Pro299Leu | Q8WVQ1 |
| HOMO SAPIENS | 4 | 9467 | 1s1d | A | 230 | ARG | p.Arg300Cys | Q8WVQ1 |
| HOMO SAPIENS | 4 | 9467 | 1s1d | A | 230 | ARG | p.Arg300His | Q8WVQ1 |
| HOMO SAPIENS | 0 | 9467 | 1s1d | A | 253 | ALA | p.Ala323Thr | Q8WVQ1 |
| HOMO SAPIENS | 2 | 9467 | 1s1d | A | 305 | MET | p.Met375Leu | Q8WVQ1 |
| HOMO SAPIENS | -1 | 2361 | 5llm | A | 5 | ASN | p.Asn5Lys | P43003 |
| HOMO SAPIENS | 0 | 2361 | 5llm | A | 71 | TYR | p.Tyr71Cys | P43003 |
| HOMO SAPIENS | -1 | 2361 | 5llm | A | 211 | LYS | p.Glu219Asp | Q15758 |
| HOMO SAPIENS | 8 | 2361 | 5llm | A | 270 | PRO | p.Pro290Arg | P43003 |
| HOMO SAPIENS | 1 | 2361 | 5llm | A | 321 | THR | p.Ala329Thr | Q15758 |
| HOMO SAPIENS | 1 | 2361 | 5llm | A | 365 | ARG | p.Arg385His | P43003 |
| HOMO SAPIENS | 1 | 2361 | 5llm | A | 479 | ARG | p.Arg499Gln | P43003 |
| HOMO SAPIENS | 3 | 5412 | 4obz | D | 165 | PHE | p.Phe229Ile | P07339 |
| HOMO SAPIENS | 2 | 5412 | 4obz | D | 187 | ASP | p.Asp251Asn | P07339 |
| HOMO SAPIENS | 0 | 5412 | 4obz | D | 218 | GLY | p.Gly282Arg | P07339 |
| HOMO SAPIENS | 0 | 5412 | 4obz | D | 245 | ARG | p.Arg309His | P07339 |
| HOMO SAPIENS | 2 | 5412 | 4obz | D | 319 | TRP | p.Trp383Cys | P07339 |
| HOMO SAPIENS | 0 | 3544 | 5h64 | A | 277 | MET | p.Met277Ile | P42345 |
| HOMO SAPIENS | 0 | 14230 | 2kbx | A | 145 | ARG | p.Arg145Ser | Q13418 |
| HOMO SAPIENS | 2 | 34970 | 4xhz | A | 834 | ARG | p.Arg855Trp | Q96QU1 |
| HOMO SAPIENS | 4 | 34970 | 4xhz | A | 854 | SER | p.Ser875Ter | Q96QU1 |
| HOMO SAPIENS | 1 | 34970 | 4xhz | A | 908 | ARG | p.Arg929Gln | Q96QU1 |
| HOMO SAPIENS | 1 | 34970 | 4xhz | A | 941 | ARG | p.Arg962Cys | Q96QU1 |
| HOMO SAPIENS | 0 | 34970 | 4xhz | A | 970 | ARG | p.Arg991Ter | Q96QU1 |
| HOMO SAPIENS | 0 | 34970 | 4xhz | A | 976 | GLU | p.Glu997Gly | Q96QU1 |
| HOMO SAPIENS | 0 | 25734 | 4asc | A | 345 | SER | p.Asn345Ser | Q2TBA0 |
| HOMO SAPIENS | 0 | 25734 | 4asc | A | 514 | VAL | p.Val514Met | Q2TBA0 |
| HOMO SAPIENS | 6 | 25734 | 4asc | A | 528 | GLU | p.Glu528Lys | Q2TBA0 |
| HOMO SAPIENS | 0 | 25734 | 4asc | A | 580 | LEU | p.Leu580Ile | Q2TBA0 |
| HOMO SAPIENS | 8 | 2356 | 3kse | D | 22 | LYS | p.Lys22Ter | P01040 |
| HOMO SAPIENS | 1 | 2356 | 3kse | D | 58 | ARG | p.Arg58Ter | P01040 |
| HOMO SAPIENS | 2 | 6000 | 1jqd | A | 60 | GLY | p.Gly60Asp | P50135 |
| HOMO SAPIENS | 9 | 6000 | 1jqd | A | 208 | LEU | p.Leu208Pro | P50135 |
| HOMO SAPIENS | 0 | 14741 | 5eh1 | A | 58 | THR | p.Thr58Arg | P38484 |
| HOMO SAPIENS | 0 | 14741 | 5eh1 | A | 168 | THR | p.Thr168Asn | P38484 |
| HOMO SAPIENS | 1 | 4814 | 4kxk | D | 427 | ARG | p.Arg427Ter | P50542 |
| HOMO SAPIENS | 0 | 188 | 2xt3 | A | 49 | ASP | p.Asp52Asn | Q2M1P5 |
| HOMO SAPIENS | 3 | 188 | 2xt3 | A | 142 | TYR | p.Tyr145Ser | Q2M1P5 |
| HOMO SAPIENS | 9 | 8681 | 5tby | D | 57 | ALA | p.Ala57Asp | P08590 |

| | | | | | | | | |
|---|---|---|---|---|---|---|---|---|
| HOMO SAPIENS | 3 | 8681 | 5tby | D | 62 | ASP | p.Asp62Asn | P08590 |
| HOMO SAPIENS | 0 | 8681 | 5tby | D | 65 | PRO | p.Pro65Arg | P08590 |
| HOMO SAPIENS | 7 | 8681 | 5tby | D | 79 | VAL | p.Val79Ile | P08590 |
| HOMO SAPIENS | 0 | 8681 | 5tby | D | 143 | GLU | p.Glu143Lys | P08590 |
| HOMO SAPIENS | 0 | 8681 | 5tby | D | 149 | MET | p.Met149Val | P08590 |
| HOMO SAPIENS | 3 | 8681 | 5tby | D | 154 | ARG | p.Arg154His | P08590 |
| HOMO SAPIENS | 6 | 30984 | 4xwh | A | 140 | TYR | p.Tyr140Cys | P54802 |
| HOMO SAPIENS | 4 | 30984 | 4xwh | A | 141 | SER | p.Ser141Thr | P54802 |
| HOMO SAPIENS | 2 | 30984 | 4xwh | A | 234 | ARG | p.Arg234Cys | P54802 |
| HOMO SAPIENS | 2 | 30984 | 4xwh | A | 336 | GLU | p.Glu336Ter | P54802 |
| HOMO SAPIENS | 6 | 30984 | 4xwh | A | 482 | ARG | p.Arg482Trp | P54802 |
| HOMO SAPIENS | 2 | 30984 | 4xwh | A | 521 | PRO | p.Pro521Leu | P54802 |
| HOMO SAPIENS | 0 | 30984 | 4xwh | A | 565 | ARG | p.Arg565Gln | P54802 |
| HOMO SAPIENS | 0 | 30984 | 4xwh | A | 565 | ARG | p.Arg565Trp | P54802 |
| HOMO SAPIENS | 6 | 30984 | 4xwh | A | 626 | ARG | p.Arg626Ter | P54802 |
| HOMO SAPIENS | 6 | 30984 | 4xwh | A | 643 | ARG | p.Arg643His | P54802 |
| HOMO SAPIENS | 6 | 30984 | 4xwh | A | 643 | ARG | p.Arg643Cys | P54802 |
| HOMO SAPIENS | 2 | 30984 | 4xwh | A | 674 | ARG | p.Arg674His | P54802 |
| HOMO SAPIENS | 2 | 30984 | 4xwh | A | 737 | ARG | p.Arg737Gly | P54802 |
| HOMO SAPIENS | 5 | 10545 | 1oqe | N | 6 | PRO | p.Pro21Arg | Q96RJ3 |
| HOMO SAPIENS | 1 | 16995 | 5d3a | B | 12 | MET | p.Met947Arg | Q7Z4S6 |
| HOMO SAPIENS | 1 | 16995 | 5d3a | B | 12 | MET | p.Met947Val | Q7Z4S6 |
| HOMO SAPIENS | 2 | 16995 | 5d3a | B | 19 | ARG | p.Arg954Gln | Q7Z4S6 |
| HOMO SAPIENS | 2 | 16995 | 5d3a | B | 19 | ARG | p.Arg954Trp | Q7Z4S6 |
| HOMO SAPIENS | 0 | 16995 | 5d3a | B | 75 | ILE | p.Ile1010Thr | Q7Z4S6 |
| HOMO SAPIENS | 0 | 17091 | 2yin | B | 1324 | GLN | p.Gln1324Ter | Q92608 |
| HOMO SAPIENS | 5 | 27606 | 1lm7 | B | 2366 | ARG | p.Arg2366Cys | P15924 |
| HOMO SAPIENS | 2 | 15731 | 1bi7 | B | 24 | ARG | p.Arg24Pro | P42771 |
| HOMO SAPIENS | 0 | 15731 | 1bi7 | B | 28 | VAL | p.Val28Gly | P42771 |
| HOMO SAPIENS | 0 | 15731 | 1bi7 | B | 49 | ILE | p.Ile49Thr | P42771 |
| HOMO SAPIENS | 0 | 15731 | 1bi7 | B | 53 | MET | p.Met53Ile | P42771 |
| HOMO SAPIENS | 2 | 15731 | 1bi7 | B | 56 | SER | p.Ser56Ile | P42771 |
| HOMO SAPIENS | 1 | 15731 | 1bi7 | B | 57 | ALA | p.Ala57Val | P42771 |
| HOMO SAPIENS | 4 | 15731 | 1bi7 | B | 59 | VAL | p.Val59Gly | P42771 |
| HOMO SAPIENS | 9 | 15731 | 1bi7 | B | 89 | GLY | p.Gly89Asp | P42771 |
| HOMO SAPIENS | 0 | 15731 | 1bi7 | B | 101 | GLY | p.Gly101Trp | P42771 |
| HOMO SAPIENS | 0 | 15731 | 1bi7 | B | 114 | PRO | p.Pro114Ser | P42771 |
| HOMO SAPIENS | 9 | 15731 | 1bi7 | B | 123 | HIS | p.His123Gln | P42771 |
| HOMO SAPIENS | 8 | 15731 | 1bi7 | B | 126 | VAL | p.Val126Asp | P42771 |
| HOMO SAPIENS | 7 | 15731 | 1bi7 | B | 127 | ALA | p.Ala127Ser | P42771 |
| HOMO SAPIENS | -1 | 15731 | 1bi7 | B | 148 | ALA | p.Ala148Thr | P42771 |
| HOMO SAPIENS | -1 | 19858 | 3jbk | M | 866 | ALA | p.Ala934Val | P18206 |
| HOMO SAPIENS | 0 | 19484 | 3ee6 | B | 62 | SER | p.Ser62Leu | O14773 |
| HOMO SAPIENS | 1 | 19484 | 3ee6 | B | 66 | GLN | p.Gln66Ter | O14773 |
| HOMO SAPIENS | 0 | 19484 | 3ee6 | B | 100 | GLN | p.Gln100Arg | O14773 |
| HOMO SAPIENS | 0 | 19484 | 3ee6 | B | 104 | LEU | p.Leu104Ter | O14773 |
| HOMO SAPIENS | 0 | 19484 | 3ee6 | B | 127 | ARG | p.Arg127Gln | O14773 |
| HOMO SAPIENS | 0 | 19484 | 3ee6 | B | 127 | ARG | p.Arg127Ter | O14773 |
| HOMO SAPIENS | -1 | 19484 | 3ee6 | B | 185 | ARG | p.Arg185Cys | O14773 |
| HOMO SAPIENS | 0 | 19484 | 3ee6 | B | 206 | ARG | p.Arg206Cys | O14773 |
| HOMO SAPIENS | 0 | 19484 | 3ee6 | B | 208 | ARG | p.Arg208Ter | O14773 |
| HOMO SAPIENS | 0 | 19484 | 3ee6 | B | 214 | GLN | p.Gln214Ter | O14773 |
| HOMO SAPIENS | 1 | 19484 | 3ee6 | B | 276 | ASP | p.Asp276Val | O14773 |
| HOMO SAPIENS | 0 | 19484 | 3ee6 | B | 284 | GLY | p.Gly284Val | O14773 |

| | | | | | | | | |
|---|---|---|---|---|---|---|---|---|
| HOMO SAPIENS | 0 | 19484 | 3ee6 | B | 286 | ASN | p.Asn286Ser | O14773 |
| HOMO SAPIENS | 0 | 19484 | 3ee6 | B | 339 | ARG | p.Arg339Gly | O14773 |
| HOMO SAPIENS | 2 | 19484 | 3ee6 | B | 345 | MET | p.Met345Leu | O14773 |
| HOMO SAPIENS | 0 | 19484 | 3ee6 | B | 365 | CYS | p.Cys365Arg | O14773 |
| HOMO SAPIENS | 0 | 19484 | 3ee6 | B | 365 | CYS | p.Cys365Tyr | O14773 |
| HOMO SAPIENS | 0 | 19484 | 3ee6 | B | 373 | GLN | p.Gln373Glu | O14773 |
| HOMO SAPIENS | 6 | 19484 | 3ee6 | B | 418 | ARG | p.Arg418Gln | O14773 |
| HOMO SAPIENS | 7 | 19484 | 3ee6 | B | 422 | GLN | p.Gln422His | O14773 |
| HOMO SAPIENS | 0 | 19484 | 3ee6 | B | 447 | ARG | p.Arg447His | O14773 |
| HOMO SAPIENS | 0 | 19484 | 3ee6 | B | 466 | VAL | p.Val466Met | O14773 |
| HOMO SAPIENS | 0 | 4654 | 5t5w | A | 47 | LYS | p.Lys47Glu | Q08334 |
| HOMO SAPIENS | 0 | 4654 | 5t5w | A | 141 | GLU | p.Glu141Ter | Q08334 |
| HOMO SAPIENS | -1 | 18815 | 4wj7 | A | 53 | VAL | p.Val53Ile | Q9BSQ5 |
| HOMO SAPIENS | 0 | 18815 | 4wj7 | A | 120 | VAL | p.Val120Ile | Q9BSQ5 |
| HOMO SAPIENS | 0 | 18815 | 4wj7 | A | 198 | LEU | p.Leu198Arg | Q9BSQ5 |
| HOMO SAPIENS | 0 | 4602 | 1mjd | A | 47 | SER | p.Ser47Arg | O43602 |
| HOMO SAPIENS | 6 | 4602 | 1mjd | A | 59 | ARG | p.Arg59Leu | O43602 |
| HOMO SAPIENS | 7 | 4602 | 1mjd | A | 62 | ASP | p.Asp62Asn | O43602 |
| HOMO SAPIENS | 7 | 4602 | 1mjd | A | 71 | ALA | p.Ala71Ser | O43602 |
| HOMO SAPIENS | 7 | 4602 | 1mjd | A | 78 | ARG | p.Arg78Leu | O43602 |
| HOMO SAPIENS | 7 | 4602 | 1mjd | A | 89 | ARG | p.Arg89Gly | O43602 |
| HOMO SAPIENS | 8 | 4602 | 1mjd | A | 104 | ILE | p.Ile104Phe | O43602 |
| HOMO SAPIENS | 7 | 4602 | 1mjd | A | 125 | TYR | p.Tyr125His | O43602 |
| HOMO SAPIENS | 5 | 34467 | 1n0w | B | 1529 | GLY | p.Gly1529Arg | P51587 |
| HOMO SAPIENS | 8 | 5738 | 1u31 | A | 130 | ASN | p.Asn1009Lys | Q13423 |
| HOMO SAPIENS | 0 | 5760 | 2i6s | A | 298 | GLU | p.Glu318Asp | P06681 |
| HOMO SAPIENS | 0 | 5760 | 2i6s | A | 573 | ARG | p.Arg593Gln | P06681 |
| HOMO SAPIENS | 0 | 5760 | 2i6s | A | 592 | SER | p.Ser612Asn | P06681 |
| HOMO SAPIENS | 0 | 5760 | 2i6s | A | 621 | VAL | p.Val641Ala | P06681 |
| HOMO SAPIENS | 1 | 10977 | 2fby | A | 114 | VAL | p.Val114Ile | Q14191 |
| HOMO SAPIENS | 0 | 10977 | 2fby | A | 135 | LYS | p.Lys135Glu | Q14191 |
| HOMO SAPIENS | -1 | 1175 | 1qrk | A | 34 | VAL | p.Val35Leu | P00488 |
| HOMO SAPIENS | 4 | 1175 | 1qrk | A | 564 | PRO | p.Pro565Leu | P00488 |
| HOMO SAPIENS | 0 | 1175 | 1qrk | A | 650 | VAL | p.Val651Ile | P00488 |
| HOMO SAPIENS | 0 | 1175 | 1qrk | A | 651 | GLU | p.Gln652Glu | P00488 |
| HOMO SAPIENS | 3 | 909 | 4h0w | A | 375 | GLU | p.Glu394Lys | P02787 |
| HOMO SAPIENS | 0 | 909 | 4h0w | A | 458 | ALA | p.Ala477Pro | P02787 |
| HOMO SAPIENS | 6 | 909 | 4h0w | A | 515 | TYR | p.Tyr534Ter | P02787 |
| HOMO SAPIENS | 7 | 909 | 4h0w | A | 570 | PRO | p.Pro589Ser | P02787 |
| HOMO SAPIENS | 3 | 909 | 4h0w | A | 652 | GLY | p.Gly671Glu | P02787 |
| HOMO SAPIENS | 5 | 1964 | 2car | A | 151 | TRP | p.Trp151Ter | Q9BY32 |
| HOMO SAPIENS | 7 | 1964 | 2car | A | 178 | ARG | p.Arg178Cys | Q9BY32 |
| HOMO SAPIENS | 0 | 144 | 2n0a | J | 30 | ALA | p.Ala30Pro | P37840 |
| HOMO SAPIENS | 1 | 12288 | 4q58 | D | 1225 | ARG | p.Arg1225His | P16144 |
| HOMO SAPIENS | 2 | 12288 | 4q58 | D | 1281 | ARG | p.Arg1281Trp | P16144 |
| HOMO SAPIENS | 2 | 12288 | 4q58 | D | 1287 | ASN | p.Asn1287Lys | P16144 |
| HOMO SAPIENS | 0 | 5085 | 3s94 | A | 360 | ARG | p.Arg360His | O75581 |
| HOMO SAPIENS | 0 | 5085 | 3s94 | A | 433 | ASN | p.Asn433Ser | O75581 |
| HOMO SAPIENS | 2 | 5085 | 3s94 | A | 611 | ARG | p.Arg611Cys | O75581 |
| HOMO SAPIENS | 0 | 15452 | 4dur | A | 19 | LYS | p.Lys38Glu | P00747 |
| HOMO SAPIENS | 2 | 15452 | 4dur | A | 216 | ARG | p.Arg235His | P00747 |
| HOMO SAPIENS | 7 | 8406 | 4wo1 | D | 28 | VAL | p.Val55Leu | O43914 |
| HOMO SAPIENS | 2 | 895 | 5xte | K | 183 | ARG | p.Arg183Trp | P22695 |
| HOMO SAPIENS | 2 | 4191 | 4wnd | A | 120 | ARG | p.Arg127Ter | P81274 |

| | | | | | | | | |
|---|---|---|---|---|---|---|---|---|
| HOMO SAPIENS | 2 | 4191 | 4wnd | A | 120 | ARG | p.Arg127Gln | P81274 |
| HOMO SAPIENS | 1 | 4191 | 4wnd | A | 349 | GLY | p.Gly356Arg | P81274 |
| HOMO SAPIENS | 0 | 41734 | 2kbi | A | 1777 | VAL | p.Val1777Met | Q14524 |
| HOMO SAPIENS | 1 | 41734 | 2kbi | A | 1784 | GLU | p.Glu1784Lys | Q14524 |
| HOMO SAPIENS | 5 | 41734 | 2kbi | A | 1795 | TYR | p.Tyr1795Cys | Q14524 |
| HOMO SAPIENS | 5 | 41734 | 2kbi | A | 1795 | TYR | p.Tyr1795His | Q14524 |
| HOMO SAPIENS | 7 | 41734 | 2kbi | A | 1826 | ARG | p.Arg1826Cys | Q14524 |
| HOMO SAPIENS | 3 | 2119 | 1pl0 | A | 426 | LYS | p.Lys426Arg | P31939 |
| HOMO SAPIENS | 5 | 3475 | 5u62 | C | 50 | LEU | p.Leu50Ser | Q15910 |
| HOMO SAPIENS | 3 | 379 | 4uez | A | 200 | TYR | p.Tyr200Ter | P15085 |
| HOMO SAPIENS | 0 | 8195 | 5xaw | B | 67 | ARG | p.Arg67Trp | Q8IWT1 |
| HOMO SAPIENS | 0 | 8195 | 5xaw | B | 94 | THR | p.Thr94Met | Q8IWT1 |
| HOMO SAPIENS | 1 | 5711 | 1fo3 | A | 265 | ARG | p.Arg265His | Q9UKM7 |
| HOMO SAPIENS | 4 | 5711 | 1fo3 | A | 334 | ARG | p.Arg334Cys | Q9UKM7 |
| HOMO SAPIENS | 1 | 5711 | 1fo3 | A | 478 | LYS | p.Lys478Arg | Q9UKM7 |
| HOMO SAPIENS | 1 | 5711 | 1fo3 | A | 682 | ASP | p.Asp682Glu | Q9UKM7 |
| HOMO SAPIENS | 0 | 170 | 2n3y | A | 41 | GLY | p.Gly42Ser | P99999 |
| HOMO SAPIENS | 0 | 4122 | 3w81 | A | 33 | HIS | p.His33Gln | P35475 |
| HOMO SAPIENS | 0 | 4122 | 3w81 | A | 64 | TYR | p.Tyr64Ter | P35475 |
| HOMO SAPIENS | 0 | 4122 | 3w81 | A | 70 | GLN | p.Gln70Ter | P35475 |
| HOMO SAPIENS | 0 | 4122 | 3w81 | A | 75 | ALA | p.Ala75Thr | P35475 |
| HOMO SAPIENS | 0 | 4122 | 3w81 | A | 79 | ALA | p.Ala79Val | P35475 |
| HOMO SAPIENS | 0 | 4122 | 3w81 | A | 79 | ALA | p.Ala79Thr | P35475 |
| HOMO SAPIENS | 0 | 4122 | 3w81 | A | 89 | ARG | p.Arg89Gln | P35475 |
| HOMO SAPIENS | 0 | 4122 | 3w81 | A | 99 | THR | p.Thr99Ile | P35475 |
| HOMO SAPIENS | 0 | 4122 | 3w81 | A | 105 | ARG | p.Arg105Gln | P35475 |
| HOMO SAPIENS | 0 | 4122 | 3w81 | A | 238 | LEU | p.Leu238Gln | P35475 |
| HOMO SAPIENS | 0 | 4122 | 3w81 | A | 300 | ALA | p.Ala300Thr | P35475 |
| HOMO SAPIENS | 0 | 4122 | 3w81 | A | 322 | VAL | p.Val322Glu | P35475 |
| HOMO SAPIENS | 0 | 4122 | 3w81 | A | 327 | ALA | p.Ala327Pro | P35475 |
| HOMO SAPIENS | 0 | 4122 | 3w81 | A | 346 | LEU | p.Leu346Arg | P35475 |
| HOMO SAPIENS | 0 | 4122 | 3w81 | A | 349 | ASP | p.Asp349Asn | P35475 |
| HOMO SAPIENS | 0 | 4122 | 3w81 | A | 361 | ALA | p.Ala361Thr | P35475 |
| HOMO SAPIENS | 0 | 4122 | 3w81 | A | 366 | THR | p.Thr366Pro | P35475 |
| HOMO SAPIENS | 0 | 4122 | 3w81 | A | 402 | TRP | p.Trp402Ter | P35475 |
| HOMO SAPIENS | 0 | 4122 | 3w81 | A | 409 | GLY | p.Gly409Arg | P35475 |
| HOMO SAPIENS | 0 | 4122 | 3w81 | A | 454 | VAL | p.Val454Ile | P35475 |
| HOMO SAPIENS | 1 | 4122 | 3w81 | A | 490 | LEU | p.Leu490Pro | P35475 |
| HOMO SAPIENS | 1 | 4122 | 3w81 | A | 492 | ARG | p.Arg492Pro | P35475 |
| HOMO SAPIENS | 0 | 4122 | 3w81 | A | 533 | PRO | p.Pro533Arg | P35475 |
| HOMO SAPIENS | 0 | 4122 | 3w81 | A | 591 | ALA | p.Ala591Thr | P35475 |
| HOMO SAPIENS | 7 | 4122 | 3w81 | A | 619 | ARG | p.Arg619Gly | P35475 |
| HOMO SAPIENS | 0 | 15325 | 3wvz | B | 54 | VAL | p.Val54Leu | Q53FT3 |
| HOMO SAPIENS | 3 | 3228 | 5iz8 | A | 421 | TRP | p.Trp421Ter | P25054 |
| HOMO SAPIENS | 2 | 3228 | 5iz8 | A | 437 | PRO | p.Pro437Arg | P25054 |
| HOMO SAPIENS | 3 | 3228 | 5iz8 | A | 499 | ARG | p.Arg499Ter | P25054 |
| HOMO SAPIENS | -1 | 29229 | 5buo | A | 539 | SER | p.Ser614Gly | P05067 |
| HOMO SAPIENS | -1 | 34931 | 4n7v | C | 54 | SER | p.Ser54Leu | O94986 |
| HOMO SAPIENS | 0 | 2450 | 5kzx | A | 123 | GLY | p.Gly123Glu | P10253 |
| HOMO SAPIENS | 1 | 2450 | 5kzx | A | 190 | ARG | p.Arg190His | P10253 |
| HOMO SAPIENS | -1 | 2450 | 5kzx | A | 199 | HIS | p.His199Arg | P10253 |
| HOMO SAPIENS | 0 | 2450 | 5kzx | A | 219 | GLY | p.Gly219Arg | P10253 |
| HOMO SAPIENS | 0 | 2450 | 5kzx | A | 220 | VAL | p.Val220Leu | P10253 |
| HOMO SAPIENS | 0 | 2450 | 5kzx | A | 222 | VAL | p.Val222Met | P10253 |

| | | | | | | | | |
|---|---|---|---|---|---|---|---|---|
| HOMO SAPIENS | 0 | 2450 | 5kzx | A | 237 | ALA | p.Ala237Val | P10253 |
| HOMO SAPIENS | 0 | 2450 | 5kzx | A | 251 | SER | p.Ser251Leu | P10253 |
| HOMO SAPIENS | 0 | 2450 | 5kzx | A | 254 | SER | p.Ser254Leu | P10253 |
| HOMO SAPIENS | 0 | 2450 | 5kzx | A | 262 | GLU | p.Glu262Lys | P10253 |
| HOMO SAPIENS | 0 | 2450 | 5kzx | A | 293 | GLY | p.Gly293Arg | P10253 |
| HOMO SAPIENS | 0 | 2450 | 5kzx | A | 299 | LEU | p.Leu299Arg | P10253 |
| HOMO SAPIENS | 0 | 2450 | 5kzx | A | 309 | GLY | p.Gly309Arg | P10253 |
| HOMO SAPIENS | 0 | 2450 | 5kzx | A | 318 | MET | p.Met318Thr | P10253 |
| HOMO SAPIENS | 0 | 2450 | 5kzx | A | 355 | LEU | p.Leu355Pro | P10253 |
| HOMO SAPIENS | 1 | 2450 | 5kzx | A | 361 | PRO | p.Pro361Leu | P10253 |
| HOMO SAPIENS | 4 | 2450 | 5kzx | A | 408 | MET | p.Met408Val | P10253 |
| HOMO SAPIENS | 1 | 2450 | 5kzx | A | 429 | GLN | p.Gln429Glu | P10253 |
| HOMO SAPIENS | 4 | 2450 | 5kzx | A | 437 | ARG | p.Arg437Cys | P10253 |
| HOMO SAPIENS | 0 | 2450 | 5kzx | A | 439 | MET | p.Met439Lys | P10253 |
| HOMO SAPIENS | 1 | 2450 | 5kzx | A | 448 | SER | p.Ser448Thr | P10253 |
| HOMO SAPIENS | 0 | 2450 | 5kzx | A | 451 | PRO | p.Pro451Arg | P10253 |
| HOMO SAPIENS | 1 | 2450 | 5kzx | A | 459 | ASP | p.Asp459His | P10253 |
| HOMO SAPIENS | 8 | 2450 | 5kzx | A | 481 | TRP | p.Trp481Arg | P10253 |
| HOMO SAPIENS | 1 | 2450 | 5kzx | A | 489 | ASP | p.Asp489Asn | P10253 |
| HOMO SAPIENS | 0 | 2450 | 5kzx | A | 521 | GLU | p.Glu521Lys | P10253 |
| HOMO SAPIENS | 0 | 2450 | 5kzx | A | 529 | SER | p.Ser529Val | P10253 |
| HOMO SAPIENS | 0 | 2450 | 5kzx | A | 545 | PRO | p.Pro545Leu | P10253 |
| HOMO SAPIENS | 0 | 2450 | 5kzx | A | 552 | LEU | p.Leu552Pro | P10253 |
| HOMO SAPIENS | 0 | 2450 | 5kzx | A | 576 | GLY | p.Gly576Ser | P10253 |
| HOMO SAPIENS | 9 | 2450 | 5kzx | A | 600 | ARG | p.Arg600His | P10253 |
| HOMO SAPIENS | 8 | 2450 | 5kzx | A | 615 | GLY | p.Gly615Arg | P10253 |
| HOMO SAPIENS | 3 | 2450 | 5kzx | A | 619 | SER | p.Ser619Asn | P10253 |
| HOMO SAPIENS | 0 | 2450 | 5kzx | A | 638 | GLY | p.Gly638Trp | P10253 |
| HOMO SAPIENS | 0 | 2450 | 5kzx | A | 643 | GLY | p.Gly643Arg | P10253 |
| HOMO SAPIENS | 9 | 2450 | 5kzx | A | 645 | ASP | p.Asp645Glu | P10253 |
| HOMO SAPIENS | 9 | 2450 | 5kzx | A | 645 | ASP | p.Asp645His | P10253 |
| HOMO SAPIENS | 0 | 2450 | 5kzx | A | 648 | GLY | p.Gly648Ser | P10253 |
| HOMO SAPIENS | 5 | 2450 | 5kzx | A | 660 | ARG | p.Arg660His | P10253 |
| HOMO SAPIENS | 8 | 2450 | 5kzx | A | 672 | ARG | p.Arg672Gln | P10253 |
| HOMO SAPIENS | 0 | 2450 | 5kzx | A | 702 | ARG | p.Arg702Leu | P10253 |
| HOMO SAPIENS | 3 | 2450 | 5kzx | A | 725 | ARG | p.Arg725Trp | P10253 |
| HOMO SAPIENS | 0 | 2450 | 5kzx | A | 746 | TRP | p.Trp746Cys | P10253 |
| HOMO SAPIENS | 0 | 2450 | 5kzx | A | 780 | ILE | p.Val780Ile | P10253 |
| HOMO SAPIENS | 1 | 2450 | 5kzx | A | 816 | VAL | p.Val816Ile | P10253 |
| HOMO SAPIENS | 0 | 2450 | 5kzx | A | 838 | GLN | p.Gln838Ter | P10253 |
| HOMO SAPIENS | 2 | 2450 | 5kzx | A | 854 | ARG | p.Arg854Ter | P10253 |
| HOMO SAPIENS | 3 | 2450 | 5kzx | A | 870 | ARG | p.Arg870Ter | P10253 |
| HOMO SAPIENS | 0 | 2450 | 5kzx | A | 890 | VAL | p.Val890Leu | P10253 |
| HOMO SAPIENS | 6 | 1882 | 4km6 | A | 175 | CYS | p.Cys175Ter | P15328 |
| HOMO SAPIENS | 3 | 14767 | 1lm5 | A | 2628 | LEU | p.Leu2628Pro | P15924 |
| HOMO SAPIENS | 5 | 14767 | 1lm5 | A | 2655 | ALA | p.Ala2655Gly | P15924 |
| HOMO SAPIENS | 3 | 14767 | 1lm5 | A | 2731 | TYR | p.Tyr2731His | P15924 |
| HOMO SAPIENS | 1 | 14767 | 1lm5 | A | 2767 | THR | p.Thr2767Asn | P15924 |
| HOMO SAPIENS | -1 | 14767 | 1lm5 | A | 2819 | MET | p.Met2819Leu | P15924 |
| HOMO SAPIENS | -1 | 11246 | 4y5o | A | 289 | SER | p.Ser289Asn | Q9BSQ5 |
| HOMO SAPIENS | 0 | 7380 | 5kz5 | I | 60 | ARG | p.Arg60His | Q16595 |
| HOMO SAPIENS | 0 | 7380 | 5kz5 | I | 76 | MET | p.Met76Val | Q16595 |
| HOMO SAPIENS | 2 | 22187 | 3fby | A | 328 | CYS | p.Cys328Arg | P49747 |
| HOMO SAPIENS | 2 | 22187 | 3fby | A | 342 | ASP | p.Asp342Tyr | P49747 |

| | | | | | | | | |
|---|---|---|---|---|---|---|---|---|
| HOMO SAPIENS | 2 | 22187 | 3fby | A | 348 | CYS | p.Cys348Arg | P49747 |
| HOMO SAPIENS | 2 | 22187 | 3fby | A | 386 | ASN | p.Asn386Asp | P49747 |
| HOMO SAPIENS | 2 | 22187 | 3fby | A | 453 | ASN | p.Asn453Ser | P49747 |
| HOMO SAPIENS | 2 | 22187 | 3fby | A | 468 | CYS | p.Cys468Tyr | P49747 |
| HOMO SAPIENS | 3 | 22187 | 3fby | A | 472 | ASP | p.Asp472Tyr | P49747 |
| HOMO SAPIENS | 3 | 22187 | 3fby | A | 473 | ASP | p.Asp473Gly | P49747 |
| HOMO SAPIENS | 2 | 22187 | 3fby | A | 523 | ASN | p.Asn523Lys | P49747 |
| HOMO SAPIENS | 1 | 22187 | 3fby | A | 718 | ARG | p.Arg718Trp | P49747 |
| HOMO SAPIENS | 1 | 22187 | 3fby | A | 719 | GLY | p.Gly719Asp | P49747 |
| HOMO SAPIENS | 0 | 22187 | 3fby | A | 756 | GLN | p.Gln756Arg | P49747 |
| HOMO SAPIENS | -1 | 2784 | 5kvh | A | 88 | MET | p.Met88Val | O95831 |
| HOMO SAPIENS | -1 | 2784 | 5kvh | A | 96 | ASN | p.Asn96Ser | O95831 |
| HOMO SAPIENS | 0 | 2784 | 5kvh | A | 282 | THR | p.Thr282Met | O95831 |
| HOMO SAPIENS | 1 | 2784 | 5kvh | A | 366 | ASN | p.Asn366Ser | O95831 |
| HOMO SAPIENS | 3 | 2784 | 5kvh | A | 463 | ARG | p.Arg463Ile | O95831 |
| HOMO SAPIENS | 0 | 2784 | 5kvh | A | 498 | VAL | p.Val498Met | O95831 |
| HOMO SAPIENS | -1 | 2784 | 5kvh | A | 549 | ALA | p.Ala549Val | O95831 |
| HOMO SAPIENS | 0 | 38316 | 2lnk | C | 1933 | ARG | p.Arg1933Ter | P35579 |
| HOMO SAPIENS | 0 | 33808 | 2miu | A | 29 | VAL | p.Val29Met | Q14192 |
| HOMO SAPIENS | 1 | 33808 | 2miu | A | 37 | ALA | p.Ala37Ser | Q14192 |
| HOMO SAPIENS | 2 | 20994 | 2hey | R | 65 | ARG | p.Arg65Cys | P43489 |
| HOMO SAPIENS | -1 | 5687 | 2xwj | I | 7 | ARG | p.Arg32Gln | P00751 |
| HOMO SAPIENS | -1 | 5687 | 2xwj | I | 7 | ARG | p.Arg32Trp | P00751 |
| HOMO SAPIENS | 0 | 5687 | 2xwj | I | 49 | ARG | p.Arg74His | P00751 |
| HOMO SAPIENS | 0 | 5687 | 2xwj | I | 177 | ARG | p.Arg202Gly | P00751 |
| HOMO SAPIENS | 0 | 5687 | 2xwj | I | 217 | ILE | p.Ile242Leu | P00751 |
| HOMO SAPIENS | -1 | 5687 | 2xwj | I | 227 | GLY | p.Gly252Ser | P00751 |
| HOMO SAPIENS | 0 | 5687 | 2xwj | I | 483 | HIS | p.His508Gln | P00751 |
| HOMO SAPIENS | 0 | 5687 | 2xwj | I | 508 | LYS | p.Lys533Arg | P00751 |
| HOMO SAPIENS | 0 | 5687 | 2xwj | I | 540 | LYS | p.Lys565Glu | P00751 |
| HOMO SAPIENS | 0 | 5687 | 2xwj | I | 541 | GLU | p.Glu566Ala | P00751 |
| HOMO SAPIENS | 2 | 953 | 3d4j | A | 249 | PHE | p.Phe249Ser | P53602 |
| HOMO SAPIENS | 1 | 953 | 3d4j | A | 292 | ASN | p.Asn292Ser | P53602 |
| HOMO SAPIENS | 0 | 3300 | 1s9c | A | 140 | ASN | p.Asn457Tyr | P51659 |
| HOMO SAPIENS | 2 | 3300 | 1s9c | A | 167 | ILE | p.Ile484Val | P51659 |
| HOMO SAPIENS | 0 | 3300 | 1s9c | A | 174 | ALA | p.Ala491Thr | P51659 |
| HOMO SAPIENS | 8 | 3300 | 1s9c | A | 193 | ASP | p.Asp510Asn | P51659 |
| HOMO SAPIENS | 3 | 3300 | 1s9c | A | 194 | TRP | p.Trp511Arg | P51659 |
| HOMO SAPIENS | 0 | 3300 | 1s9c | A | 242 | VAL | p.Ile559Val | P51659 |
| HOMO SAPIENS | 0 | 10810 | 5i25 | A | 5 | GLN | p.Gln23Ter | P03951 |
| HOMO SAPIENS | 0 | 10810 | 5i25 | A | 55 | TRP | p.Trp73Ter | P03951 |
| HOMO SAPIENS | 0 | 10810 | 5i25 | A | 91 | ALA | p.Ala109Thr | P03951 |
| HOMO SAPIENS | 0 | 10810 | 5i25 | A | 117 | GLU | p.Glu135Ter | P03951 |
| HOMO SAPIENS | 0 | 10810 | 5i25 | A | 118 | CYS | p.Cys136Ter | P03951 |
| HOMO SAPIENS | 0 | 10810 | 5i25 | A | 128 | CYS | p.Cys146Ter | P03951 |
| HOMO SAPIENS | 0 | 10810 | 5i25 | A | 210 | ARG | p.Arg228Ter | P03951 |
| HOMO SAPIENS | 0 | 10810 | 5i25 | A | 226 | GLN | p.Gln244Ter | P03951 |
| HOMO SAPIENS | 0 | 10810 | 5i25 | A | 233 | GLN | p.Gln251Ter | P03951 |
| HOMO SAPIENS | 0 | 16668 | 3lbx | A | 28 | ARG | p.Arg28Leu | P02549 |
| HOMO SAPIENS | 0 | 16668 | 3lbx | A | 28 | ARG | p.Arg28Cys | P02549 |
| HOMO SAPIENS | 1 | 16668 | 3lbx | A | 41 | ARG | p.Arg41Trp | P02549 |
| HOMO SAPIENS | 1 | 16668 | 3lbx | A | 45 | ARG | p.Arg45Ser | P02549 |
| HOMO SAPIENS | 2 | 16668 | 3lbx | A | 48 | LYS | p.Lys48Arg | P02549 |
| HOMO SAPIENS | 1 | 16668 | 3lbx | A | 152 | ASP | p.Asp152Asn | P02549 |

| | | | | | | | | |
|---|---|---|---|---|---|---|---|---|
| HOMO SAPIENS | 0 | 226 | 2ble | A | 256 | ILE | p.Phe256Ile | P36959 |
| HOMO SAPIENS | 2 | 36808 | 4c7n | A | 3 | SER | p.Ser357Pro | O75030 |
| HOMO SAPIENS | 0 | 4092 | 5jnx | A | 113 | CYS | p.Cys113Arg | O15118 |
| HOMO SAPIENS | 4 | 4092 | 5jnx | A | 215 | HIS | p.His215Arg | O15118 |
| HOMO SAPIENS | 6 | 4092 | 5jnx | A | 237 | PRO | p.Pro237Ser | O15118 |
| HOMO SAPIENS | 0 | 4092 | 5jnx | A | 363 | ALA | p.Ala363Val | O15118 |
| HOMO SAPIENS | 0 | 4092 | 5jnx | A | 378 | VAL | p.Val378Ala | O15118 |
| HOMO SAPIENS | 8 | 4092 | 5jnx | A | 404 | ARG | p.Arg404Gln | O15118 |
| HOMO SAPIENS | 8 | 4092 | 5jnx | A | 411 | ARG | p.Arg411Gln | O15118 |
| HOMO SAPIENS | 2 | 4092 | 5jnx | A | 474 | PRO | p.Pro474Leu | O15118 |
| HOMO SAPIENS | 8 | 4092 | 5jnx | A | 511 | THR | p.Thr511Met | O15118 |
| HOMO SAPIENS | 0 | 4092 | 5jnx | A | 642 | MET | p.Met642Ile | O15118 |
| HOMO SAPIENS | 2 | 4092 | 5jnx | A | 664 | VAL | p.Val664Met | O15118 |
| HOMO SAPIENS | 0 | 4092 | 5jnx | A | 734 | SER | p.Ser734Ile | O15118 |
| HOMO SAPIENS | -1 | 4092 | 5jnx | A | 775 | GLN | p.Gln775Pro | O15118 |
| HOMO SAPIENS | 0 | 4092 | 5jnx | A | 858 | ILE | p.Ile858Val | O15118 |
| HOMO SAPIENS | 0 | 4092 | 5jnx | A | 889 | VAL | p.Val889Met | O15118 |
| HOMO SAPIENS | 0 | 4092 | 5jnx | A | 910 | GLY | p.Gly910Ser | O15118 |
| HOMO SAPIENS | 0 | 4092 | 5jnx | A | 911 | GLY | p.Gly911Ser | O15118 |
| HOMO SAPIENS | 0 | 4092 | 5jnx | A | 928 | GLN | p.Gln928Pro | O15118 |
| HOMO SAPIENS | 0 | 4092 | 5jnx | A | 931 | ASN | p.Asn931Lys | O15118 |
| HOMO SAPIENS | 0 | 4092 | 5jnx | A | 940 | SER | p.Ser940Leu | O15118 |
| HOMO SAPIENS | 0 | 4092 | 5jnx | A | 950 | VAL | p.Val950Met | O15118 |
| HOMO SAPIENS | 0 | 4092 | 5jnx | A | 954 | SER | p.Ser954Leu | O15118 |
| HOMO SAPIENS | 0 | 4092 | 5jnx | A | 958 | ARG | p.Arg958Gln | O15118 |
| HOMO SAPIENS | 0 | 4092 | 5jnx | A | 958 | ARG | p.Arg958Ter | O15118 |
| HOMO SAPIENS | -1 | 4092 | 5jnx | A | 978 | ARG | p.Arg978Cys | O15118 |
| HOMO SAPIENS | 0 | 4092 | 5jnx | A | 992 | GLY | p.Gly992Trp | O15118 |
| HOMO SAPIENS | 0 | 4092 | 5jnx | A | 1007 | PRO | p.Pro1007Ala | O15118 |
| HOMO SAPIENS | 0 | 4092 | 5jnx | A | 1019 | TYR | p.Tyr1019Cys | O15118 |
| HOMO SAPIENS | 2 | 4092 | 5jnx | A | 1035 | ALA | p.Ala1035Val | O15118 |
| HOMO SAPIENS | 2 | 4092 | 5jnx | A | 1054 | ALA | p.Ala1054Thr | O15118 |
| HOMO SAPIENS | 2 | 4092 | 5jnx | A | 1061 | ILE | p.Ile1061Thr | O15118 |
| HOMO SAPIENS | 0 | 4092 | 5jnx | A | 1073 | GLY | p.Gly1073Ser | O15118 |
| HOMO SAPIENS | 2 | 4092 | 5jnx | A | 1088 | TYR | p.Tyr1088Cys | O15118 |
| HOMO SAPIENS | 2 | 4092 | 5jnx | A | 1142 | MET | p.Met1142Thr | O15118 |
| HOMO SAPIENS | 2 | 4092 | 5jnx | A | 1156 | ASN | p.Asn1156Ser | O15118 |
| HOMO SAPIENS | 0 | 4092 | 5jnx | A | 1179 | MET | p.Met1179Val | O15118 |
| HOMO SAPIENS | 0 | 4092 | 5jnx | A | 1186 | ARG | p.Arg1186His | O15118 |
| HOMO SAPIENS | 0 | 4092 | 5jnx | A | 1187 | ALA | p.Ala1187Gly | O15118 |
| HOMO SAPIENS | 2 | 4092 | 5jnx | A | 1189 | GLU | p.Glu1189Gly | O15118 |
| HOMO SAPIENS | 2 | 4092 | 5jnx | A | 1200 | SER | p.Ser1200Gly | O15118 |
| HOMO SAPIENS | 2 | 4092 | 5jnx | A | 1205 | THR | p.Thr1205Lys | O15118 |
| HOMO SAPIENS | 0 | 4092 | 5jnx | A | 1213 | LEU | p.Leu1213Phe | O15118 |
| HOMO SAPIENS | -1 | 4092 | 5jnx | A | 1266 | ARG | p.Arg1266Gln | O15118 |
| HOMO SAPIENS | 9 | 16498 | 4i6o | A | 64 | ARG | p.Arg735Trp | P01024 |
| HOMO SAPIENS | 1 | 21902 | 3un9 | C | 751 | ARG | p.Arg751His | Q86UT6 |
| HOMO SAPIENS | 7 | 20391 | 2h32 | B | 105 | PRO | p.Pro142Leu | P15814 |
| HOMO SAPIENS | 6 | 1635 | 4x45 | A | 65 | ASP | p.Asp65Val | P07741 |
| HOMO SAPIENS | 1 | 1635 | 4x45 | A | 98 | TRP | p.Trp98Ter | P07741 |
| HOMO SAPIENS | 0 | 1635 | 4x45 | A | 110 | LEU | p.Leu110Pro | P07741 |
| HOMO SAPIENS | 7 | 1635 | 4x45 | A | 136 | MET | p.Met136Thr | P07741 |
| HOMO SAPIENS | 1 | 1635 | 4x45 | A | 150 | VAL | p.Val150Phe | P07741 |
| HOMO SAPIENS | 4 | 34810 | 2rpc | A | 16 | CYS | p.Cys253Ser | O60481 |

| | | | | | | | | |
|---|---|---|---|---|---|---|---|---|
| HOMO SAPIENS | 1 | 34810 | 2rpc | A | 18 | TRP | p.Trp255Gly | O60481 |
| HOMO SAPIENS | 1 | 14359 | 4hxi | A | 78 | MET | p.Met78Val | Q9UH77 |
| HOMO SAPIENS | 0 | 5342 | 4d9t | A | 628 | GLU | p.Glu628Asp | P51812 |
| HOMO SAPIENS | -1 | 5342 | 4d9t | A | 723 | ARG | p.Arg723His | P51812 |
| HOMO SAPIENS | -1 | 5342 | 4d9t | A | 729 | ARG | p.Arg729Gln | P51812 |
| HOMO SAPIENS | 1 | 9619 | 3j2s | B | 1781 | ARG | p.Arg1800Cys | P00451 |
| HOMO SAPIENS | 0 | 9619 | 3j2s | B | 1823 | MET | p.Met1842Ile | P00451 |
| HOMO SAPIENS | 1 | 9619 | 3j2s | B | 1825 | PRO | p.Pro1844Ser | P00451 |
| HOMO SAPIENS | 0 | 9619 | 3j2s | B | 1826 | THR | p.Thr1845Pro | P00451 |
| HOMO SAPIENS | 0 | 9619 | 3j2s | B | 1834 | ALA | p.Ala1853Val | P00451 |
| HOMO SAPIENS | 0 | 9619 | 3j2s | B | 1846 | ASP | p.Asp1865Asn | P00451 |
| HOMO SAPIENS | 1 | 9619 | 3j2s | B | 1848 | HIS | p.His1867Arg | P00451 |
| HOMO SAPIENS | 0 | 9619 | 3j2s | B | 1854 | PRO | p.Pro1873Arg | P00451 |
| HOMO SAPIENS | 1 | 9619 | 3j2s | B | 1885 | GLU | p.Glu1904Lys | P00451 |
| HOMO SAPIENS | 0 | 9619 | 3j2s | B | 1922 | ASN | p.Asn1941Ser | P00451 |
| HOMO SAPIENS | 1 | 9619 | 3j2s | B | 1941 | ARG | p.Arg1960Gln | P00451 |
| HOMO SAPIENS | 4 | 9619 | 3j2s | B | 2105 | TYR | p.Tyr2124Cys | P00451 |
| HOMO SAPIENS | 6 | 9619 | 3j2s | B | 2150 | ARG | p.Arg2169His | P00451 |
| HOMO SAPIENS | 0 | 9619 | 3j2s | B | 2159 | ARG | p.Arg2178Cys | P00451 |
| HOMO SAPIENS | 0 | 9619 | 3j2s | B | 2159 | ARG | p.Arg2178His | P00451 |
| HOMO SAPIENS | 2 | 9619 | 3j2s | B | 2238 | MET | p.Met2257Val | P00451 |
| HOMO SAPIENS | 2 | 9619 | 3j2s | B | 2300 | PRO | p.Pro2319Ser | P00451 |
| HOMO SAPIENS | 1 | 667 | 1z2m | A | 127 | GLU | p.Glu127Ter | P05161 |
| HOMO SAPIENS | 0 | 15717 | 2nn6 | G | 30 | GLY | p.Gly31Ala | Q9NQT5 |
| HOMO SAPIENS | -1 | 15717 | 2nn6 | G | 55 | ASN | p.Asn56His | Q9NQT5 |
| HOMO SAPIENS | 7 | 15717 | 2nn6 | G | 131 | ASP | p.Asp132Ala | Q9NQT5 |
| HOMO SAPIENS | 7 | 784 | 3rkq | A | 161 | ARG | p.Arg161Pro | P52952 |
| HOMO SAPIENS | 6 | 784 | 3rkq | A | 178 | THR | p.Thr178Met | P52952 |
| HOMO SAPIENS | 7 | 784 | 3rkq | A | 183 | LYS | p.Lys183Glu | P52952 |
| HOMO SAPIENS | 8 | 784 | 3rkq | A | 190 | ARG | p.Arg190Cys | P52952 |
| HOMO SAPIENS | 6 | 385 | 30000 | C | 70 | ALA | p.Ala70Thr | P07954 |
| HOMO SAPIENS | 2 | 385 | 30000 | C | 101 | ARG | p.Arg101Pro | P07954 |
| HOMO SAPIENS | 2 | 385 | 30000 | C | 174 | PRO | p.Pro174Arg | P07954 |
| HOMO SAPIENS | 6 | 385 | 30000 | C | 187 | SER | p.Ser187Ter | P07954 |
| HOMO SAPIENS | 9 | 385 | 30000 | C | 233 | ARG | p.Arg233His | P07954 |
| HOMO SAPIENS | 3 | 385 | 30000 | C | 273 | ALA | p.Ala273Thr | P07954 |
| HOMO SAPIENS | 4 | 385 | 30000 | C | 334 | SER | p.Ser334Arg | P07954 |
| HOMO SAPIENS | 6 | 385 | 30000 | C | 340 | ASN | p.Asn340Lys | P07954 |
| HOMO SAPIENS | 8 | 385 | 30000 | C | 343 | ARG | p.Arg343Ter | P07954 |
| HOMO SAPIENS | 3 | 385 | 30000 | C | 350 | ARG | p.Arg350Trp | P07954 |
| HOMO SAPIENS | 4 | 385 | 30000 | C | 376 | GLN | p.Gln376Pro | P07954 |
| HOMO SAPIENS | 6 | 385 | 30000 | C | 434 | CYS | p.Cys434Ter | P07954 |
| HOMO SAPIENS | 0 | 3316 | 4f91 | B | 934 | THR | p.Thr934Ala | O75643 |
| HOMO SAPIENS | 0 | 3316 | 4f91 | B | 1002 | ASN | p.Asn1002Ser | O75643 |
| HOMO SAPIENS | 0 | 3316 | 4f91 | B | 1087 | SER | p.Ser1087Leu | O75643 |
| HOMO SAPIENS | 4 | 3316 | 4f91 | B | 1389 | VAL | p.Val1389Ile | O75643 |
| HOMO SAPIENS | 0 | 619 | 5hbt | B | 195 | ASP | p.Asp240His | P02708 |
| HOMO SAPIENS | 1 | 619 | 5hbt | B | 209 | ARG | p.Arg254Leu | P02708 |
| HOMO SAPIENS | 0 | 18171 | 2y25 | B | 1408 | ASP | p.Asp1408Asn | P52179 |
| HOMO SAPIENS | 0 | 18171 | 2y25 | B | 1453 | MET | p.Met1453Thr | P52179 |
| HOMO SAPIENS | 0 | 18171 | 2y25 | B | 1459 | ILE | p.Ile1459Lys | P52179 |
| HOMO SAPIENS | 0 | 18171 | 2y25 | B | 1558 | PHE | p.Phe1558Leu | P52179 |
| HOMO SAPIENS | 0 | 18171 | 2y25 | B | 1573 | ARG | p.Arg1573Gln | P52179 |
| HOMO SAPIENS | 7 | 18171 | 2y25 | B | 1663 | VAL | p.Val1663Met | P52179 |

| | | | | | | | | |
|---|---|---|---|---|---|---|---|---|
| HOMO SAPIENS | 8 | 14262 | 2fcw | B | 117 | SER | p.Ser138Ter | P01130 |
| HOMO SAPIENS | 8 | 14262 | 2fcw | B | 118 | ASP | p.Asp139Glu | P01130 |
| HOMO SAPIENS | 9 | 14262 | 2fcw | B | 119 | GLU | p.Glu140Lys | P01130 |
| HOMO SAPIENS | 7 | 14262 | 2fcw | B | 122 | CYS | p.Cys143Ter | P01130 |
| HOMO SAPIENS | 6 | 14262 | 2fcw | B | 134 | CYS | p.Cys155Ter | P01130 |
| HOMO SAPIENS | 7 | 14262 | 2fcw | B | 147 | ASP | p.Asp168Glu | P01130 |
| HOMO SAPIENS | 4 | 2624 | 3b7b | A | 855 | VAL | p.Val886Met | Q9H9B1 |
| HOMO SAPIENS | 7 | 2624 | 3b7b | A | 868 | ILE | p.Ile899Val | Q9H9B1 |
| HOMO SAPIENS | 0 | 2624 | 3b7b | A | 892 | GLU | p.Glu923Lys | Q9H9B1 |
| HOMO SAPIENS | 0 | 4791 | 5ctd | B | 68 | LEU | p.Pro549Ala | Q14055 |
| HOMO SAPIENS | 5 | 460 | 4ui1 | D | 80 | CYS | p.Cys80Arg | Q6ZVN8 |
| HOMO SAPIENS | 3 | 460 | 4ui1 | D | 101 | LEU | p.Leu101Pro | Q6ZVN8 |
| HOMO SAPIENS | 0 | 40782 | 1sy6 | A | 47 | LYS | p.Lys69Ter | P09693 |
| HOMO SAPIENS | -1 | 1092 | 2idx | A | 62 | THR | p.Thr62Met | Q96EY8 |
| HOMO SAPIENS | 6 | 1092 | 2idx | A | 183 | HIS | p.His183Leu | Q96EY8 |
| HOMO SAPIENS | 7 | 1092 | 2idx | A | 186 | ARG | p.Arg186Trp | Q96EY8 |
| HOMO SAPIENS | 6 | 1092 | 2idx | A | 190 | ARG | p.Arg190Cys | Q96EY8 |
| HOMO SAPIENS | 6 | 1092 | 2idx | A | 190 | ARG | p.Arg190His | Q96EY8 |
| HOMO SAPIENS | 7 | 1092 | 2idx | A | 191 | ARG | p.Arg191Trp | Q96EY8 |
| HOMO SAPIENS | 7 | 1092 | 2idx | A | 191 | ARG | p.Arg191Gln | Q96EY8 |
| HOMO SAPIENS | 6 | 1092 | 2idx | A | 195 | ARG | p.Arg195His | Q96EY8 |
| HOMO SAPIENS | 1 | 1092 | 2idx | A | 234 | GLN | p.Gln234Ter | Q96EY8 |
| HOMO SAPIENS | 0 | 1092 | 2idx | A | 239 | MET | p.Met239Lys | Q96EY8 |
| HOMO SAPIENS | -1 | 6418 | 5fur | G | 11 | THR | p.Thr11Ile | P21675 |
| HOMO SAPIENS | -1 | 6418 | 5fur | G | 290 | LEU | p.Leu269Val | P21675 |
| HOMO SAPIENS | 0 | 6100 | 1wiq | A | 240 | ARG | p.Arg265Trp | P01730 |
| HOMO SAPIENS | 0 | 13757 | 4ye6 | A | 45 | GLY | p.Gly45Val | P47897 |
| HOMO SAPIENS | 1 | 13757 | 4ye6 | A | 57 | TYR | p.Tyr57His | P47897 |
| HOMO SAPIENS | 8 | 13757 | 4ye6 | A | 403 | ARG | p.Arg403Trp | P47897 |
| HOMO SAPIENS | 7 | 13757 | 4ye6 | A | 524 | ARG | p.Arg524Trp | P47897 |
| HOMO SAPIENS | 6 | 13757 | 4ye6 | A | 690 | ARG | p.Arg690Cys | P47897 |
| HOMO SAPIENS | 0 | 26958 | 2lzl | A | 370 | GLY | p.Gly370Cys | P22607 |
| HOMO SAPIENS | 0 | 26958 | 2lzl | A | 371 | SER | p.Ser371Cys | P22607 |
| HOMO SAPIENS | 0 | 26958 | 2lzl | A | 373 | TYR | p.Tyr373Cys | P22607 |
| HOMO SAPIENS | 0 | 26958 | 2lzl | A | 380 | GLY | p.Gly380Arg | P22607 |
| HOMO SAPIENS | 0 | 26958 | 2lzl | A | 384 | PHE | p.Phe384Leu | P22607 |
| HOMO SAPIENS | 5 | 26958 | 2lzl | A | 391 | ALA | p.Ala391Glu | P22607 |
| HOMO SAPIENS | 6 | 11325 | 2wa6 | A | 121 | TYR | p.Tyr121Phe | O75369 |
| HOMO SAPIENS | 2 | 11325 | 2wa6 | A | 161 | PHE | p.Phe161Cys | O75369 |
| HOMO SAPIENS | 0 | 11325 | 2wa6 | A | 171 | LEU | p.Leu171Arg | O75369 |
| HOMO SAPIENS | 2 | 11325 | 2wa6 | A | 173 | ALA | p.Ala173Val | O75369 |
| HOMO SAPIENS | 0 | 11325 | 2wa6 | A | 202 | MET | p.Met202Val | O75369 |
| HOMO SAPIENS | 3 | 11325 | 2wa6 | A | 227 | GLU | p.Glu227Lys | O75369 |
| HOMO SAPIENS | 0 | 11325 | 2wa6 | A | 234 | LEU | p.Leu234Val | O75369 |
| HOMO SAPIENS | 2 | 11325 | 2wa6 | A | 235 | SER | p.Ser235Pro | O75369 |
| HOMO SAPIENS | 9 | 7240 | 3tgx | A | 137 | TYR | p.Tyr156Cys | Q9HBE5 |
| HOMO SAPIENS | 0 | 35479 | 4elj | A | 307 | THR | p.Thr307Ile | P06400 |
| HOMO SAPIENS | 0 | 35479 | 4elj | A | 310 | GLY | p.Gly310Glu | P06400 |
| HOMO SAPIENS | 0 | 35479 | 4elj | A | 377 | THR | p.Thr377Ser | P06400 |
| HOMO SAPIENS | 9 | 36706 | 1he7 | A | 359 | TYR | p.Tyr359Cys | P04629 |
| HOMO SAPIENS | 4 | 6538 | 4d0e | A | 504 | ARG | p.Arg504His | P46531 |
| HOMO SAPIENS | 1 | 26518 | 5fm5 | M | 600 | GLU | p.Glu600Val | P52179 |
| HOMO SAPIENS | -1 | 26518 | 5fm5 | M | 651 | ARG | p.Arg651Gln | P52179 |
| HOMO SAPIENS | -1 | 26518 | 5fm5 | M | 660 | PRO | p.Pro660Ala | P52179 |

| | | | | | | | | |
|---|---|---|---|---|---|---|---|---|
| HOMO SAPIENS | -1 | 26518 | 5fm5 | M | 688 | THR | p.Thr688Ser | P52179 |
| HOMO SAPIENS | -1 | 26518 | 5fm5 | M | 704 | GLU | p.Glu704Lys | P52179 |
| HOMO SAPIENS | -1 | 26518 | 5fm5 | M | 727 | THR | p.Thr727Ala | P52179 |
| HOMO SAPIENS | 2 | 27054 | 4rhp | B | 209 | LEU | p.Leu209Val | O75208 |
| HOMO SAPIENS | -1 | 27054 | 4rhp | B | 288 | LYS | p.Lys288Asn | O75208 |
| HOMO SAPIENS | 5 | 2384 | 5fqd | B | 419 | ARG | p.Arg419Ter | Q96SW2 |
| HOMO SAPIENS | 4 | 5154 | 2hl3 | A | 56 | LYS | p.Lys56Arg | Q14203 |
| HOMO SAPIENS | 0 | 5154 | 2hl3 | A | 59 | GLY | p.Gly59Ser | Q14203 |
| HOMO SAPIENS | 0 | 5154 | 2hl3 | A | 71 | GLY | p.Gly71Ala | Q14203 |
| HOMO SAPIENS | 0 | 5154 | 2hl3 | A | 71 | GLY | p.Gly71Arg | Q14203 |
| HOMO SAPIENS | 7 | 5154 | 2hl3 | A | 72 | THR | p.Thr72Pro | Q14203 |
| HOMO SAPIENS | 2 | 5154 | 2hl3 | A | 74 | GLN | p.Gln74Pro | Q14203 |
| HOMO SAPIENS | 2 | 20039 | 2l1g | A | 9 | GLY | p.Gly9Cys | Q9NVV9 |
| HOMO SAPIENS | 5 | 20039 | 2l1g | A | 81 | PHE | p.Phe81Leu | Q9NVV9 |
| HOMO SAPIENS | -1 | 97 | 5uz7 | A | 201 | ARG | p.Arg201His | P63092 |
| HOMO SAPIENS | -1 | 97 | 5uz7 | A | 201 | ARG | p.Arg201Cys | P63092 |
| HOMO SAPIENS | 0 | 3257 | 4b7w | B | 278 | GLU | p.Glu278Asp | P41235 |
| HOMO SAPIENS | 1 | 3257 | 4b7w | B | 331 | ARG | p.Arg331Cys | P41235 |
| HOMO SAPIENS | 0 | 2814 | 4qgk | B | 10 | GLN | p.Gln10Ter | P51648 |
| HOMO SAPIENS | 0 | 2814 | 4qgk | B | 177 | ARG | p.Arg177Ter | P51648 |
| HOMO SAPIENS | 9 | 2814 | 4qgk | B | 184 | THR | p.Thr184Met | P51648 |
| HOMO SAPIENS | 1 | 2814 | 4qgk | B | 188 | ALA | p.Ala188Val | P51648 |
| HOMO SAPIENS | 0 | 2814 | 4qgk | B | 214 | CYS | p.Cys214Tyr | P51648 |
| HOMO SAPIENS | 0 | 2814 | 4qgk | B | 245 | ASP | p.Asp245Asn | P51648 |
| HOMO SAPIENS | 0 | 2814 | 4qgk | B | 266 | LYS | p.Lys266Asn | P51648 |
| HOMO SAPIENS | 0 | 2814 | 4qgk | B | 315 | PRO | p.Pro315Ser | P51648 |
| HOMO SAPIENS | 0 | 2814 | 4qgk | B | 386 | ASN | p.Asn386Ser | P51648 |
| HOMO SAPIENS | -1 | 5486 | 5u6o | B | 47 | GLY | p.Gly47Val | O60741 |
| HOMO SAPIENS | 1 | 21475 | 3pdy | A | 679 | THR | p.Thr789Ile | Q15149 |
| HOMO SAPIENS | 0 | 32890 | 4djc | B | 1501 | LEU | p.Leu1501Val | Q14524 |
| HOMO SAPIENS | -1 | 32890 | 4djc | B | 1512 | ARG | p.Arg1512Trp | Q14524 |
| HOMO SAPIENS | 0 | 26683 | 2xze | B | 38 | ARG | p.Arg38Cys | O95630 |
| HOMO SAPIENS | 0 | 26683 | 2xze | B | 100 | PHE | p.Phe100Tyr | O95630 |
| HOMO SAPIENS | -1 | 23175 | 5dka | A | 163 | SER | p.Ser163Leu | Q96EQ8 |
| HOMO SAPIENS | -1 | 23175 | 5dka | A | 174 | ARG | p.Arg174Cys | Q96EQ8 |
| HOMO SAPIENS | 6 | 21177 | 3k1r | A | 155 | ARG | p.Arg155Ter | Q9Y6N9 |
| HOMO SAPIENS | 0 | 1237 | 2wwm | O | 40 | PRO | p.Pro40Arg | O75147 |
| HOMO SAPIENS | 1 | 2543 | 5ewl | B | 64 | VAL | p.Val64Leu | Q13224 |
| HOMO SAPIENS | 2 | 2543 | 5ewl | B | 271 | ALA | p.Ala271Glu | Q13224 |
| HOMO SAPIENS | 3 | 34262 | 2mng | A | 613 | PHE | p.Phe613Leu | Q9Y3Q4 |
| HOMO SAPIENS | 7 | 34262 | 2mng | A | 672 | SER | p.Ser672Arg | Q9Y3Q4 |
| HOMO SAPIENS | 4 | 1667 | 1f05 | A | 321 | LYS | p.Lys321Met | P37837 |
| HOMO SAPIENS | 0 | 25179 | 2knc | B | 752 | SER | p.Ser778Pro | P05106 |
| HOMO SAPIENS | 0 | 2185 | 2ig7 | A | 50 | TYR | p.Tyr50Cys | Q9Y259 |
| HOMO SAPIENS | 0 | 2185 | 2ig7 | A | 224 | ASN | p.Asn224His | Q9Y259 |
| HOMO SAPIENS | 0 | 2185 | 2ig7 | A | 270 | TYR | p.Tyr270Ter | Q9Y259 |
| HOMO SAPIENS | 0 | 2185 | 2ig7 | A | 328 | GLN | p.Gln328Arg | Q9Y259 |
| HOMO SAPIENS | 3 | 2201 | 2r0o | A | 179 | PRO | p.Pro179Leu | O43707 |
| HOMO SAPIENS | 3 | 2201 | 2r0o | A | 255 | GLU | p.Lys255Glu | O43707 |
| HOMO SAPIENS | 3 | 2201 | 2r0o | A | 259 | THR | p.Thr259Ile | O43707 |
| HOMO SAPIENS | 4 | 2201 | 2r0o | A | 262 | SER | p.Ser262Pro | O43707 |
| HOMO SAPIENS | -1 | 31221 | 2wvr | C | 44 | THR | p.Thr44Pro | Q9H211 |
| HOMO SAPIENS | -1 | 31221 | 2wvr | C | 135 | ALA | p.Ala135Val | Q9H211 |
| HOMO SAPIENS | -1 | 31221 | 2wvr | C | 456 | GLU | p.Glu456Ala | Q9H211 |

| | | | | | | | | |
|---|---|---|---|---|---|---|---|---|
| HOMO SAPIENS | -1 | 31221 | 2wvr | C | 462 | ARG | p.Arg462Gln | Q9H211 |
| HOMO SAPIENS | -1 | 31221 | 2wvr | C | 468 | GLU | p.Glu468Lys | Q9H211 |
| HOMO SAPIENS | -1 | 31221 | 2wvr | C | 537 | ALA | p.Ala537Val | Q9H211 |
| HOMO SAPIENS | 5 | 2576 | 5o3l | C | 363 | VAL | p.Val680Ile | P10636 |
| HOMO SAPIENS | 5 | 1345 | 1czt | A | 37 | ARG | p.Arg2102Cys | P12259 |
| HOMO SAPIENS | 1 | 1345 | 1czt | A | 83 | MET | p.Met2148Thr | P12259 |
| HOMO SAPIENS | 1 | 1345 | 1czt | A | 157 | ASP | p.Asp2222Gly | P12259 |
| HOMO SAPIENS | 8 | 35985 | 1lo1 | A | 110 | ALA | p.Ala110Val | O95718 |
| HOMO SAPIENS | 1 | 617 | 2ocp | B | 44 | GLU | p.Glu44Lys | Q16854 |
| HOMO SAPIENS | 5 | 617 | 2ocp | B | 46 | ASN | p.Asn46Ser | Q16854 |
| HOMO SAPIENS | 0 | 617 | 2ocp | B | 59 | THR | p.Thr59Met | Q16854 |
| HOMO SAPIENS | 7 | 617 | 2ocp | B | 142 | ARG | p.Arg142Lys | Q16854 |
| HOMO SAPIENS | 3 | 617 | 2ocp | B | 154 | ASN | p.Asn154Lys | Q16854 |
| HOMO SAPIENS | 3 | 617 | 2ocp | B | 170 | GLN | p.Gln170Arg | Q16854 |
| HOMO SAPIENS | 3 | 617 | 2ocp | B | 227 | GLU | p.Glu227Lys | Q16854 |
| HOMO SAPIENS | 3 | 617 | 2ocp | B | 233 | LYS | p.Lys233Glu | Q16854 |
| HOMO SAPIENS | 3 | 617 | 2ocp | B | 255 | ASP | p.Asp255Tyr | Q16854 |
| HOMO SAPIENS | -1 | 38975 | 5itz | D | 421 | GLN | p.Gln421His | Q9HC77 |
| HOMO SAPIENS | 2 | 6235 | 2f8v | A | 15 | VAL | p.Val15Ile | Q8WZ42 |
| HOMO SAPIENS | 0 | 6235 | 2f8v | A | 54 | VAL | p.Val54Met | Q8WZ42 |
| HOMO SAPIENS | 0 | 6235 | 2f8v | A | 59 | SER | p.Ser59Arg | Q8WZ42 |
| HOMO SAPIENS | 0 | 6235 | 2f8v | A | 60 | ASP | p.Asp60Tyr | Q8WZ42 |
| HOMO SAPIENS | 0 | 12630 | 1ext | B | 12 | ASP | p.Asp41Glu | P19438 |
| HOMO SAPIENS | 3 | 12630 | 1ext | B | 30 | CYS | p.Cys59Arg | P19438 |
| HOMO SAPIENS | 3 | 12630 | 1ext | B | 30 | CYS | p.Cys59Ser | P19438 |
| HOMO SAPIENS | 5 | 12630 | 1ext | B | 33 | CYS | p.Cys62Tyr | P19438 |
| HOMO SAPIENS | 5 | 12630 | 1ext | B | 33 | CYS | p.Cys62Gly | P19438 |
| HOMO SAPIENS | 1 | 12630 | 1ext | B | 46 | PRO | p.Pro75Leu | P19438 |
| HOMO SAPIENS | 3 | 12630 | 1ext | B | 50 | THR | p.Thr79Met | P19438 |
| HOMO SAPIENS | 6 | 12630 | 1ext | B | 52 | CYS | p.Cys81Phe | P19438 |
| HOMO SAPIENS | 6 | 12630 | 1ext | B | 70 | CYS | p.Cys99Ser | P19438 |
| HOMO SAPIENS | 5 | 12630 | 1ext | B | 88 | CYS | p.Cys117Arg | P19438 |
| HOMO SAPIENS | 5 | 12630 | 1ext | B | 88 | CYS | p.Cys117Tyr | P19438 |
| HOMO SAPIENS | -1 | 12630 | 1ext | B | 170 | ILE | p.Ile199Asn | P19438 |
| HOMO SAPIENS | 0 | 994 | 1aos | B | 95 | ARG | p.Arg95Cys | P04424 |
| HOMO SAPIENS | 0 | 994 | 1aos | B | 111 | ARG | p.Arg111Gln | P04424 |
| HOMO SAPIENS | 0 | 994 | 1aos | B | 131 | THR | p.Thr131Met | P04424 |
| HOMO SAPIENS | 6 | 994 | 1aos | B | 178 | VAL | p.Val178Met | P04424 |
| HOMO SAPIENS | 9 | 994 | 1aos | B | 182 | ARG | p.Arg182Gln | P04424 |
| HOMO SAPIENS | 9 | 994 | 1aos | B | 186 | ARG | p.Arg186Trp | P04424 |
| HOMO SAPIENS | 8 | 994 | 1aos | B | 193 | ARG | p.Arg193Gln | P04424 |
| HOMO SAPIENS | 5 | 994 | 1aos | B | 217 | ARG | p.Arg217Ter | P04424 |
| HOMO SAPIENS | 0 | 994 | 1aos | B | 286 | GLN | p.Gln286Arg | P04424 |
| HOMO SAPIENS | 5 | 994 | 1aos | B | 379 | ARG | p.Arg379Cys | P04424 |
| HOMO SAPIENS | 4 | 16066 | 5mv9 | A | 1719 | TYR | p.Tyr1719Cys | Q13402 |
| HOMO SAPIENS | -1 | 16066 | 5mv9 | A | 1739 | ARG | p.Arg1739Ter | Q13402 |
| HOMO SAPIENS | 4 | 16066 | 5mv9 | A | 1775 | ILE | p.Ile1775Thr | Q13402 |
| HOMO SAPIENS | 2 | 16066 | 5mv9 | A | 1786 | SER | p.Ser1786Thr | Q13402 |
| HOMO SAPIENS | 4 | 16066 | 5mv9 | A | 1798 | GLN | p.Gln1798Ter | Q13402 |
| HOMO SAPIENS | 0 | 16066 | 5mv9 | A | 1858 | LEU | p.Leu1858Pro | Q13402 |
| HOMO SAPIENS | 6 | 16066 | 5mv9 | A | 1873 | ARG | p.Arg1873Trp | Q13402 |
| HOMO SAPIENS | 6 | 16066 | 5mv9 | A | 1873 | ARG | p.Arg1873Leu | Q13402 |
| HOMO SAPIENS | 6 | 16066 | 5mv9 | A | 1883 | ARG | p.Arg1883Gln | Q13402 |
| HOMO SAPIENS | 4 | 16066 | 5mv9 | A | 1887 | PRO | p.Pro1887Leu | Q13402 |

| | | | | | | | | |
|---|---|---|---|---|---|---|---|---|
| HOMO SAPIENS | 5 | 16066 | 5mv9 | A | 1942 | GLY | p.Gly1942Ter | Q13402 |
| HOMO SAPIENS | 5 | 16066 | 5mv9 | A | 1954 | LEU | p.Leu1954Ile | Q13402 |
| HOMO SAPIENS | 5 | 16066 | 5mv9 | A | 1956 | VAL | p.Val1956Ile | Q13402 |
| HOMO SAPIENS | 5 | 16066 | 5mv9 | A | 1967 | ARG | p.Arg1967Ter | Q13402 |
| HOMO SAPIENS | 5 | 16066 | 5mv9 | A | 1990 | GLN | p.Gln1990Ter | Q13402 |
| HOMO SAPIENS | 2 | 16066 | 5mv9 | A | 2005 | LYS | p.Lys2005Glu | Q13402 |
| HOMO SAPIENS | 1 | 16066 | 5mv9 | A | 2072 | VAL | p.Val2072Ile | Q13402 |
| HOMO SAPIENS | 4 | 16066 | 5mv9 | A | 2109 | THR | p.Thr2109Ile | Q13402 |
| HOMO SAPIENS | 4 | 16066 | 5mv9 | A | 2142 | ASP | p.Asp2142Asn | Q13402 |
| HOMO SAPIENS | -1 | 16066 | 5mv9 | A | 2214 | GLY | p.Gly2214Ser | Q13402 |
| HOMO SAPIENS | 7 | 27942 | 1deb | A | 24 | ARG | p.Arg24Ter | P25054 |
| HOMO SAPIENS | 0 | 876 | 5hnr | B | 12 | PHE | p.Phe12Leu | P13716 |
| HOMO SAPIENS | 0 | 876 | 5hnr | B | 59 | LYS | p.Lys59Asn | P13716 |
| HOMO SAPIENS | -1 | 876 | 5hnr | B | 133 | GLY | p.Gly133Arg | P13716 |
| HOMO SAPIENS | 2 | 876 | 5hnr | B | 240 | ARG | p.Arg240Trp | P13716 |
| HOMO SAPIENS | 0 | 876 | 5hnr | B | 274 | ALA | p.Ala274Thr | P13716 |
| HOMO SAPIENS | 0 | 876 | 5hnr | B | 275 | VAL | p.Val275Met | P13716 |
| HOMO SAPIENS | 0 | 1142 | 3qe2 | A | 503 | VAL | p.Ala500Val | P16435 |
| HOMO SAPIENS | 8 | 14358 | 4dkd | C | 32 | VAL | p.Val32Gly | P07333 |
| HOMO SAPIENS | 0 | 14358 | 4dkd | C | 60 | LEU | p.Leu60Val | P07333 |
| HOMO SAPIENS | 7 | 14358 | 4dkd | C | 75 | THR | p.Thr75Ile | P07333 |
| HOMO SAPIENS | -1 | 14358 | 4dkd | C | 90 | PRO | p.Pro90Thr | P07333 |
| HOMO SAPIENS | 0 | 14358 | 4dkd | C | 106 | ARG | p.Arg106Trp | P07333 |
| HOMO SAPIENS | 0 | 14358 | 4dkd | C | 241 | ASN | p.Asn241Asp | P07333 |
| HOMO SAPIENS | 0 | 14358 | 4dkd | C | 245 | ALA | p.Ala245Ser | P07333 |
| HOMO SAPIENS | 0 | 14358 | 4dkd | C | 255 | ASN | p.Asn255Ile | P07333 |
| HOMO SAPIENS | 0 | 14358 | 4dkd | C | 268 | VAL | p.Val268Ile | P07333 |
| HOMO SAPIENS | 0 | 14358 | 4dkd | C | 279 | VAL | p.Val279Met | P07333 |
| HOMO SAPIENS | 0 | 14358 | 4dkd | C | 294 | ARG | p.Arg294Gln | P07333 |
| HOMO SAPIENS | 5 | 8968 | 3ppy | A | 1565 | VAL | p.Val1565Leu | P04275 |
| HOMO SAPIENS | 4 | 8968 | 3ppy | A | 1584 | TYR | p.Tyr1584Cys | P04275 |
| HOMO SAPIENS | 5 | 5341 | 3k3n | A | 279 | PHE | p.Phe315Ser | Q9UPP1 |
| HOMO SAPIENS | 0 | 29463 | 1q7l | A | 197 | ARG | p.Arg197Trp | Q03154 |
| HOMO SAPIENS | 0 | 33761 | 2leh | B | 30 | ASP | p.Asp30Asn | Q16637 |
| HOMO SAPIENS | 6 | 33761 | 2leh | B | 44 | ASP | p.Asp44Val | Q16637 |
| HOMO SAPIENS | -1 | 12249 | 2pnr | C | 224 | GLU | p.Glu310Lys | P10515 |
| HOMO SAPIENS | -1 | 12249 | 2pnr | C | 232 | VAL | p.Val318Ala | P10515 |
| HOMO SAPIENS | -1 | 7 | 5whk | B | -9 | ALA | p.Ala11Pro | P61769 |
| HOMO SAPIENS | 8 | 20298 | 4euw | A | 154 | PHE | p.Phe154Leu | P48436 |
| HOMO SAPIENS | 9 | 20298 | 4euw | A | 158 | ALA | p.Ala158Thr | P48436 |
| HOMO SAPIENS | 5 | 20298 | 4euw | A | 165 | HIS | p.His165Tyr | P48436 |
| HOMO SAPIENS | 3 | 20298 | 4euw | A | 169 | HIS | p.His169Gln | P48436 |
| HOMO SAPIENS | 0 | 20298 | 4euw | A | 173 | LYS | p.Lys173Glu | P48436 |
| HOMO SAPIENS | 1 | 16465 | 2jgx | A | 402 | TYR | p.Tyr402His | P08603 |
| HOMO SAPIENS | 0 | 8993 | 3v8o | A | 659 | LEU | p.Leu659Arg | Q14315 |
| HOMO SAPIENS | 6 | 8993 | 3v8o | A | 693 | ASP | p.Asp693Ala | Q14315 |
| HOMO SAPIENS | 0 | 8993 | 3v8o | A | 709 | ALA | p.Ala709Thr | Q14315 |
| HOMO SAPIENS | 2 | 3852 | 5es4 | B | 106 | ASP | p.Asp128Asn | P05107 |
| HOMO SAPIENS | 2 | 3852 | 5es4 | B | 116 | SER | p.Ser138Pro | P05107 |
| HOMO SAPIENS | 1 | 3852 | 5es4 | B | 127 | LEU | p.Leu149Pro | P05107 |
| HOMO SAPIENS | 0 | 3852 | 5es4 | B | 147 | GLY | p.Gly169Arg | P05107 |
| HOMO SAPIENS | 3 | 3852 | 5es4 | B | 156 | PRO | p.Pro178Leu | P05107 |
| HOMO SAPIENS | 0 | 3852 | 5es4 | B | 174 | LYS | p.Lys196Thr | P05107 |
| HOMO SAPIENS | 2 | 3852 | 5es4 | B | 251 | GLY | p.Gly273Arg | P05107 |

| | | | | | | | | |
|---|---|---|---|---|---|---|---|---|
| HOMO SAPIENS | 2 | 3852 | 5es4 | B | 329 | ASN | p.Asn351Ser | P05107 |
| HOMO SAPIENS | 2 | 3852 | 5es4 | B | 571 | ARG | p.Arg593Cys | P05107 |
| HOMO SAPIENS | 7 | 5962 | 5fn2 | B | 146 | MET | p.Met146Leu | P49768 |
| HOMO SAPIENS | 3 | 5962 | 5fn2 | B | 166 | LEU | p.Leu166Pro | P49768 |
| HOMO SAPIENS | 1 | 5962 | 5fn2 | B | 233 | MET | p.Met233Leu | P49768 |
| HOMO SAPIENS | 1 | 5962 | 5fn2 | B | 261 | VAL | p.Val261Gly | P49768 |
| HOMO SAPIENS | 1 | 5962 | 5fn2 | B | 266 | GLY | p.Gly266Ser | P49768 |
| HOMO SAPIENS | -1 | 5962 | 5fn2 | B | 333 | ASP | p.Asp333Gly | P49768 |
| HOMO SAPIENS | 6 | 1918 | 3dlx | A | 58 | THR | p.Thr58Met | P55809 |
| HOMO SAPIENS | 5 | 1918 | 3dlx | A | 133 | VAL | p.Val133Glu | P55809 |
| HOMO SAPIENS | 5 | 1918 | 3dlx | A | 219 | GLY | p.Gly219Glu | P55809 |
| HOMO SAPIENS | 0 | 1918 | 3dlx | A | 221 | VAL | p.Val221Met | P55809 |
| HOMO SAPIENS | 5 | 1918 | 3dlx | A | 283 | SER | p.Ser283Ter | P55809 |
| HOMO SAPIENS | 4 | 1918 | 3dlx | A | 324 | GLY | p.Gly324Glu | P55809 |
| HOMO SAPIENS | 0 | 1918 | 3dlx | A | 456 | CYS | p.Cys456Phe | P55809 |
| HOMO SAPIENS | 0 | 9010 | 2wfr | A | 162 | LEU | p.Leu162Pro | O43323 |
| HOMO SAPIENS | 5 | 3770 | 5n38 | A | 56 | VAL | p.Val56Glu | O60260 |
| HOMO SAPIENS | 1 | 3770 | 5n38 | A | 161 | LYS | p.Lys161Asn | O60260 |
| HOMO SAPIENS | 1 | 3770 | 5n38 | A | 167 | SER | p.Ser167Asn | O60260 |
| HOMO SAPIENS | 1 | 3770 | 5n38 | A | 211 | LYS | p.Lys211Asn | O60260 |
| HOMO SAPIENS | 1 | 3770 | 5n38 | A | 212 | CYS | p.Cys212Tyr | O60260 |
| HOMO SAPIENS | 1 | 3770 | 5n38 | A | 275 | ARG | p.Arg275Trp | O60260 |
| HOMO SAPIENS | 2 | 3770 | 5n38 | A | 334 | ARG | p.Arg334Cys | O60260 |
| HOMO SAPIENS | 1 | 3770 | 5n38 | A | 366 | ARG | p.Arg366Trp | O60260 |
| HOMO SAPIENS | -1 | 3770 | 5n38 | A | 380 | VAL | p.Val380Leu | O60260 |
| HOMO SAPIENS | 1 | 3770 | 5n38 | A | 394 | ASP | p.Asp394Asn | O60260 |
| HOMO SAPIENS | 1 | 3770 | 5n38 | A | 430 | GLY | p.Gly430Asp | O60260 |
| HOMO SAPIENS | -1 | 2467 | 4xx1 | A | 10 | PRO | p.Pro34Leu | P04180 |
| HOMO SAPIENS | 4 | 2467 | 4xx1 | A | 83 | TYR | p.Tyr107Ter | P04180 |
| HOMO SAPIENS | 4 | 2467 | 4xx1 | A | 93 | ALA | p.Ala117Thr | P04180 |
| HOMO SAPIENS | 4 | 2467 | 4xx1 | A | 123 | THR | p.Thr147Ile | P04180 |
| HOMO SAPIENS | 4 | 2467 | 4xx1 | A | 135 | ARG | p.Arg159Trp | P04180 |
| HOMO SAPIENS | 3 | 2467 | 4xx1 | A | 158 | ARG | p.Arg182Cys | P04180 |
| HOMO SAPIENS | 0 | 2467 | 4xx1 | A | 209 | LEU | p.Leu233Pro | P04180 |
| HOMO SAPIENS | 1 | 2467 | 4xx1 | A | 228 | ASN | p.Asn252Lys | P04180 |
| HOMO SAPIENS | 4 | 2467 | 4xx1 | A | 252 | MET | p.Met276Lys | P04180 |
| HOMO SAPIENS | 3 | 2467 | 4xx1 | A | 293 | MET | p.Met317Ile | P04180 |
| HOMO SAPIENS | 5 | 2467 | 4xx1 | A | 321 | THR | p.Thr345Met | P04180 |
| HOMO SAPIENS | 9 | 2467 | 4xx1 | A | 347 | THR | p.Thr371Met | P04180 |
| HOMO SAPIENS | 1 | 5834 | 3qrf | G | 371 | PHE | p.Phe371Cys | Q9BZS1 |
| HOMO SAPIENS | 1 | 5834 | 3qrf | G | 373 | PHE | p.Phe373Ala | Q9BZS1 |
| HOMO SAPIENS | 9 | 5834 | 3qrf | G | 384 | ALA | p.Ala384Thr | Q9BZS1 |
| HOMO SAPIENS | 6 | 5834 | 3qrf | G | 397 | ARG | p.Arg397Trp | Q9BZS1 |
| HOMO SAPIENS | -1 | 11869 | 4km8 | A | 15 | PRO | p.Pro15Leu | Q9UMX1 |
| HOMO SAPIENS | 5 | 262 | 4kbo | A | 126 | ARG | p.Arg126Trp | P38646 |
| HOMO SAPIENS | 3 | 262 | 4kbo | A | 128 | TYR | p.Tyr128Cys | P38646 |
| HOMO SAPIENS | -1 | 11347 | 4uf9 | D | 117 | ALA | p.Ala117Val | Q13501 |
| HOMO SAPIENS | 0 | 2173 | 4ah6 | A | 45 | SER | p.Ser45Gly | Q6PI48 |
| HOMO SAPIENS | 2 | 2173 | 4ah6 | A | 152 | CYS | p.Cys152Phe | Q6PI48 |
| HOMO SAPIENS | 4 | 2173 | 4ah6 | A | 179 | ARG | p.Arg179His | Q6PI48 |
| HOMO SAPIENS | 0 | 2173 | 4ah6 | A | 196 | LYS | p.Lys196Arg | Q6PI48 |
| HOMO SAPIENS | 8 | 2173 | 4ah6 | A | 263 | ARG | p.Arg263Gln | Q6PI48 |
| HOMO SAPIENS | 1 | 2173 | 4ah6 | A | 338 | GLY | p.Gly338Glu | Q6PI48 |
| HOMO SAPIENS | 0 | 2173 | 4ah6 | A | 425 | GLU | p.Glu425Ter | Q6PI48 |

| | | | | | | | | |
|---|---|---|---|---|---|---|---|---|
| HOMO SAPIENS | 1 | 2173 | 4ah6 | A | 548 | LEU | p.Leu548Met | Q6PI48 |
| HOMO SAPIENS | 0 | 2173 | 4ah6 | A | 609 | ARG | p.Arg609Trp | Q6PI48 |
| HOMO SAPIENS | 2 | 2173 | 4ah6 | A | 613 | LEU | p.Leu613Phe | Q6PI48 |
| HOMO SAPIENS | 0 | 2173 | 4ah6 | A | 626 | LEU | p.Leu626Gln | Q6PI48 |
| HOMO SAPIENS | 0 | 2173 | 4ah6 | A | 626 | LEU | p.Leu626Val | Q6PI48 |
| HOMO SAPIENS | 7 | 3203 | 2cn1 | A | 87 | ASP | p.Asp137Val | Q9H0P0 |
| HOMO SAPIENS | 8 | 3203 | 2cn1 | A | 166 | GLN | p.Gln216Ter | Q9H0P0 |
| HOMO SAPIENS | 8 | 3203 | 2cn1 | A | 179 | ASN | p.Asn229Ser | Q9H0P0 |
| HOMO SAPIENS | 7 | 3203 | 2cn1 | A | 230 | GLY | p.Gly280Arg | Q9H0P0 |
| HOMO SAPIENS | 2 | 3424 | 2a1t | R | 116 | GLY | p.Gly116Arg | P13804 |
| HOMO SAPIENS | 2 | 3424 | 2a1t | R | 157 | VAL | p.Val157Gly | P13804 |
| HOMO SAPIENS | 2 | 3424 | 2a1t | R | 171 | THR | p.Thr171Ile | P13804 |
| HOMO SAPIENS | 0 | 3424 | 2a1t | R | 178 | THR | p.Thr178Arg | P13804 |
| HOMO SAPIENS | 0 | 3424 | 2a1t | R | 194 | VAL | p.Val194Met | P13804 |
| HOMO SAPIENS | 1 | 3424 | 2a1t | R | 209 | ARG | p.Arg209Ter | P13804 |
| HOMO SAPIENS | 6 | 3424 | 2a1t | R | 266 | THR | p.Thr266Met | P13804 |
| HOMO SAPIENS | 0 | 27598 | 1g1c | A | 26 | ARG | p.Arg2098Gly | Q8WZ42 |
| HOMO SAPIENS | 1 | 27598 | 1g1c | A | 46 | ILE | p.Ile2118Thr | Q8WZ42 |
| HOMO SAPIENS | 0 | 32885 | 4bsk | A | 149 | ASN | p.Asn149Asp | P35916 |
| HOMO SAPIENS | -1 | 8074 | 3ghg | B | 17 | ARG | p.Arg47Ter | P02675 |
| HOMO SAPIENS | -1 | 447 | 5g4p | A | 190 | ALA | p.Ala190Thr | P84074 |
| HOMO SAPIENS | 0 | 10761 | 5u1d | B | 379 | VAL | p.Val379Ile | Q03519 |
| HOMO SAPIENS | 0 | 10761 | 5u1d | B | 665 | THR | p.Thr665Ala | Q03519 |
| HOMO SAPIENS | 5 | 29251 | 5fn7 | A | 79 | ILE | p.Ile294Leu | P08575 |
| HOMO SAPIENS | -1 | 27727 | 4ky9 | A | 56 | LYS | p.Lys56Glu | P02730 |
| HOMO SAPIENS | 2 | 27727 | 4ky9 | A | 72 | GLU | p.Glu72Asp | P02730 |
| HOMO SAPIENS | 2 | 27727 | 4ky9 | A | 90 | GLU | p.Glu90Lys | P02730 |
| HOMO SAPIENS | 1 | 27727 | 4ky9 | A | 130 | GLY | p.Gly130Arg | P02730 |
| HOMO SAPIENS | 2 | 27727 | 4ky9 | A | 150 | ARG | p.Arg150Ter | P02730 |
| HOMO SAPIENS | 0 | 27727 | 4ky9 | A | 327 | PRO | p.Pro327Arg | P02730 |
| HOMO SAPIENS | 2 | 13202 | 2fxo | B | 870 | ARG | p.Arg870His | P12883 |
| HOMO SAPIENS | 2 | 13202 | 2fxo | B | 870 | ARG | p.Arg870Cys | P12883 |
| HOMO SAPIENS | 4 | 13202 | 2fxo | B | 904 | ARG | p.Arg904Cys | P12883 |
| HOMO SAPIENS | 4 | 13202 | 2fxo | B | 906 | ASP | p.Asp906Gly | P12883 |
| HOMO SAPIENS | 2 | 13202 | 2fxo | B | 908 | LEU | p.Leu908Val | P12883 |
| HOMO SAPIENS | 2 | 13202 | 2fxo | B | 923 | ASN | p.Asn923Lys | P12883 |
| HOMO SAPIENS | 7 | 13202 | 2fxo | B | 935 | GLU | p.Glu935Lys | P12883 |
| HOMO SAPIENS | 7 | 13202 | 2fxo | B | 935 | GLU | p.Glu935Val | P12883 |
| HOMO SAPIENS | 8 | 13202 | 2fxo | B | 949 | GLU | p.Glu949Lys | P12883 |
| HOMO SAPIENS | 0 | 13200 | 2a1s | A | 349 | ARG | p.Arg349Trp | O95453 |
| HOMO SAPIENS | 0 | 13200 | 2a1s | A | 421 | LYS | p.Lys421Arg | O95453 |
| HOMO SAPIENS | 6 | 3196 | 5xtb | T | 115 | CYS | p.Cys115Tyr | O75380 |
| HOMO SAPIENS | 0 | 1108 | 5ukf | A | 89 | ARG | p.Arg89Gln | Q99986 |
| HOMO SAPIENS | 0 | 1108 | 5ukf | A | 119 | HIS | p.His119Arg | Q99986 |
| HOMO SAPIENS | 0 | 1108 | 5ukf | A | 133 | ARG | p.Arg133Cys | Q99986 |
| HOMO SAPIENS | 0 | 1108 | 5ukf | A | 236 | VAL | p.Val236Leu | Q99986 |
| HOMO SAPIENS | 0 | 1108 | 5ukf | A | 321 | ARG | p.Arg321Cys | Q99986 |
| HOMO SAPIENS | -1 | 1108 | 5ukf | A | 358 | ARG | p.Arg358Ter | Q99986 |
| HOMO SAPIENS | 0 | 5842 | 4yli | C | 204 | GLY | p.Gly204Ser | Q9BWP8 |
| HOMO SAPIENS | -1 | 31362 | 4wqo | A | 25 | PRO | p.Pro25Leu | P40337 |
| HOMO SAPIENS | 2 | 10107 | 2j8h | A | 34 | VAL | p.Val31886Ile | Q8WZ42 |
| HOMO SAPIENS | 8 | 20190 | 1wm5 | A | 77 | ARG | p.Arg77Gln | P19878 |
| HOMO SAPIENS | 1 | 20190 | 1wm5 | A | 100 | GLN | p.Gln100Ter | P19878 |
| HOMO SAPIENS | 0 | 20190 | 1wm5 | A | 102 | ARG | p.Arg102Ter | P19878 |

| | | | | | | | | | |
|---|---|---|---|---|---|---|---|---|---|
| HOMO SAPIENS | 0 | 20190 | 1wm5 | A | | 128 | ALA | p.Ala128Val | P19878 |
| HOMO SAPIENS | 0 | 20190 | 1wm5 | A | | 181 | LYS | p.Lys181Arg | P19878 |
| HOMO SAPIENS | 7 | 1278 | 5a2q | a | | 33 | ASP | p.Asp33Asn | P62854 |
| HOMO SAPIENS | 3 | 19515 | 2obd | A | | 11 | ILE | p.Ile28Met | P11597 |
| HOMO SAPIENS | 3 | 19515 | 2obd | A | | 89 | VAL | p.Val106Met | P11597 |
| HOMO SAPIENS | 3 | 19515 | 2obd | A | | 137 | ARG | p.Arg154Trp | P11597 |
| HOMO SAPIENS | 3 | 19515 | 2obd | A | | 139 | ASP | p.Asp156Asn | P11597 |
| HOMO SAPIENS | 3 | 19515 | 2obd | A | | 204 | SER | p.Ser221Arg | P11597 |
| HOMO SAPIENS | 0 | 19515 | 2obd | A | | 297 | GLU | p.Glu314Lys | P11597 |
| HOMO SAPIENS | 0 | 19515 | 2obd | A | | 314 | GLY | p.Gly331Ser | P11597 |
| HOMO SAPIENS | 0 | 19515 | 2obd | A | | 332 | SER | p.Ser349Tyr | P11597 |
| HOMO SAPIENS | 0 | 19515 | 2obd | A | | 368 | VAL | p.Val385Met | P11597 |
| HOMO SAPIENS | 0 | 19515 | 2obd | A | | 373 | ALA | p.Ala390Pro | P11597 |
| HOMO SAPIENS | 3 | 19515 | 2obd | A | | 405 | ILE | p.Val422Ile | P11597 |
| HOMO SAPIENS | 0 | 19515 | 2obd | A | | 451 | ARG | p.Arg468Gln | P11597 |
| HOMO SAPIENS | 0 | 19515 | 2obd | A | | 451 | ARG | p.Arg468Ter | P11597 |
| HOMO SAPIENS | 0 | 6626 | 3hop | A | | 39 | ALA | p.Ala39Gly | P21333 |
| HOMO SAPIENS | 1 | 6626 | 3hop | A | | 82 | GLU | p.Glu82Val | P21333 |
| HOMO SAPIENS | 1 | 6626 | 3hop | A | | 128 | ALA | p.Ala128Val | P21333 |
| HOMO SAPIENS | 2 | 6626 | 3hop | A | | 203 | ASP | p.Asp203Tyr | P21333 |
| HOMO SAPIENS | 1 | 6626 | 3hop | A | | 207 | PRO | p.Pro207Leu | P21333 |
| HOMO SAPIENS | 2 | 6626 | 3hop | A | | 210 | CYS | p.Cys210Phe | P21333 |
| HOMO SAPIENS | 7 | 6626 | 3hop | A | | 254 | GLU | p.Glu254Lys | P21333 |
| HOMO SAPIENS | 4 | 4210 | 2avg | A | | 8 | VAL | p.Val158Met | Q14896 |
| HOMO SAPIENS | 6 | 4210 | 2avg | A | | 15 | GLU | p.Glu165Asp | Q14896 |
| HOMO SAPIENS | 4 | 4210 | 2avg | A | | 27 | ARG | p.Arg177His | Q14896 |
| HOMO SAPIENS | 0 | 4210 | 2avg | A | | 28 | VAL | p.Val178Met | Q14896 |
| HOMO SAPIENS | 6 | 4210 | 2avg | A | | 39 | VAL | p.Val189Ile | Q14896 |
| HOMO SAPIENS | 2 | 4210 | 2avg | A | | 52 | LYS | p.Lys202Gln | Q14896 |
| HOMO SAPIENS | 4 | 4210 | 2avg | A | | 67 | SER | p.Ser217Gly | Q14896 |
| HOMO SAPIENS | 6 | 4210 | 2avg | A | | 69 | VAL | p.Val219Leu | Q14896 |
| HOMO SAPIENS | 2 | 4210 | 2avg | A | | 86 | GLY | p.Ser236Gly | Q14896 |
| HOMO SAPIENS | 4 | 4210 | 2avg | A | | 108 | GLU | p.Glu258Lys | Q14896 |
| HOMO SAPIENS | 0 | 22237 | 1w6k | A | | 588 | GLY | p.Gly588Ser | P48449 |
| HOMO SAPIENS | 0 | 10765 | 4a5w | B | | 98 | GLU | p.Glu98Ala | P13671 |
| HOMO SAPIENS | 0 | 10765 | 4a5w | B | | 596 | ARG | p.Arg596Ter | P13671 |
| HOMO SAPIENS | -1 | 15515 | 3hx3 | A | | 11 | MSE | p.Met10Lys | P12271 |
| HOMO SAPIENS | 8 | 15515 | 3hx3 | A | | 64 | ARG | p.Arg64Gln | P12271 |
| HOMO SAPIENS | 0 | 15515 | 3hx3 | A | | 182 | PHE | p.Phe182Cys | P12271 |
| HOMO SAPIENS | 0 | 15515 | 3hx3 | A | | 216 | ARG | p.Arg216Gln | P12271 |
| HOMO SAPIENS | -1 | 15515 | 3hx3 | A | | 226 | MSE | p.Met226Lys | P12271 |
| HOMO SAPIENS | 0 | 15515 | 3hx3 | A | | 234 | TRP | p.Arg234Trp | P12271 |
| HOMO SAPIENS | 0 | 15515 | 3hx3 | A | | 273 | LEU | p.Leu273Phe | P12271 |
| HOMO SAPIENS | 0 | 15515 | 3hx3 | A | | 292 | THR | p.Thr292Met | P12271 |
| HOMO SAPIENS | 0 | 5152 | 2www | D | | 95 | GLN | p.Gln95Ter | Q8IVH4 |
| HOMO SAPIENS | 0 | 5152 | 2www | D | | 133 | GLN | p.Gln133Ter | Q8IVH4 |
| HOMO SAPIENS | 0 | 5152 | 2www | D | | 145 | ARG | p.Arg145Ter | Q8IVH4 |
| HOMO SAPIENS | 0 | 5152 | 2www | D | | 207 | TYR | p.Tyr207Cys | Q8IVH4 |
| HOMO SAPIENS | 8 | 5152 | 2www | D | | 330 | ARG | p.Arg330Ter | Q8IVH4 |
| HOMO SAPIENS | 0 | 5152 | 2www | D | | 363 | GLN | p.Gln363His | Q8IVH4 |
| HOMO SAPIENS | 1 | 1626 | 1ilr | | 1 | 105 | SER | p.Ser130Arg | P18510 |
| HOMO SAPIENS | -1 | 929 | 5loh | A | | 12 | GLY | p.Gly12Arg | Q96GX5 |
| HOMO SAPIENS | 9 | 3206 | 1x63 | A | | 21 | CYS | p.Cys104Arg | Q13642 |
| HOMO SAPIENS | 8 | 3206 | 1x63 | A | | 39 | TRP | p.Trp122Ser | Q13642 |

| | | | | | | | | |
|---|---|---|---|---|---|---|---|---|
| HOMO SAPIENS | 9 | 3206 | 1x63 | A | 40 | HIS | p.His123Tyr | Q13642 |
| HOMO SAPIENS | 9 | 3206 | 1x63 | A | 40 | HIS | p.His123Leu | Q13642 |
| HOMO SAPIENS | 9 | 3206 | 1x63 | A | 40 | HIS | p.His123Gln | Q13642 |
| HOMO SAPIENS | 3 | 3206 | 1x63 | A | 49 | CYS | p.Cys132Phe | Q13642 |
| HOMO SAPIENS | 3 | 3206 | 1x63 | A | 67 | CYS | p.Cys150Tyr | Q13642 |
| HOMO SAPIENS | 3 | 3206 | 1x63 | A | 70 | CYS | p.Cys153Arg | Q13642 |
| HOMO SAPIENS | 3 | 3206 | 1x63 | A | 70 | CYS | p.Cys153Tyr | Q13642 |
| HOMO SAPIENS | 5 | 1052 | 3ecs | E | 208 | ASN | p.Asn208Tyr | Q14232 |
| HOMO SAPIENS | 2 | 1052 | 3ecs | E | 275 | TYR | p.Tyr275Cys | Q14232 |
| HOMO SAPIENS | 0 | 24253 | 5i35 | A | 271 | ARG | p.Arg271Cys | Q8NI60 |
| HOMO SAPIENS | 0 | 24253 | 5i35 | A | 272 | GLY | p.Gly272Asp | Q8NI60 |
| HOMO SAPIENS | 0 | 24253 | 5i35 | A | 299 | ARG | p.Arg299Trp | Q8NI60 |
| HOMO SAPIENS | 1 | 24253 | 5i35 | A | 339 | ALA | p.Ala339Thr | Q8NI60 |
| HOMO SAPIENS | 0 | 24253 | 5i35 | A | 348 | ARG | p.Arg348Ter | Q8NI60 |
| HOMO SAPIENS | 0 | 24253 | 5i35 | A | 395 | ILE | p.Ile395Met | Q8NI60 |
| HOMO SAPIENS | 0 | 24253 | 5i35 | A | 396 | ASP | p.Asp396Glu | Q8NI60 |
| HOMO SAPIENS | 0 | 24253 | 5i35 | A | 480 | PHE | p.Phe480Leu | Q8NI60 |
| HOMO SAPIENS | 2 | 24253 | 5i35 | A | 511 | THR | p.Thr511Met | Q8NI60 |
| HOMO SAPIENS | 0 | 24253 | 5i35 | A | 514 | TYR | p.Tyr514Cys | Q8NI60 |
| HOMO SAPIENS | 0 | 24253 | 5i35 | A | 538 | ARG | p.Arg538Gln | Q8NI60 |
| HOMO SAPIENS | 0 | 24253 | 5i35 | A | 549 | GLY | p.Gly549Ser | Q8NI60 |
| HOMO SAPIENS | 0 | 24253 | 5i35 | A | 551 | GLU | p.Glu551Lys | Q8NI60 |
| HOMO SAPIENS | -1 | 24253 | 5i35 | A | 555 | MSE | p.Met555Ile | Q8NI60 |
| HOMO SAPIENS | 0 | 24253 | 5i35 | A | 568 | GLU | p.Glu568Ter | Q8NI60 |
| HOMO SAPIENS | 0 | 24253 | 5i35 | A | 602 | PRO | p.Pro602Arg | Q8NI60 |
| HOMO SAPIENS | -1 | 15505 | 4goj | C | 22 | GLY | p.Gly22Val | Q13432 |
| HOMO SAPIENS | 5 | 23902 | 2lrr | A | 765 | HIS | p.His765Gln | P38935 |
| HOMO SAPIENS | 1 | 29650 | 2vj2 | A | 234 | CYS | p.Cys234Tyr | P78504 |
| HOMO SAPIENS | 0 | 29650 | 2vj2 | A | 255 | TYR | p.Tyr255Ter | P78504 |
| HOMO SAPIENS | 3 | 29650 | 2vj2 | A | 274 | GLY | p.Gly274Asp | P78504 |
| HOMO SAPIENS | 1 | 105 | 5he0 | B | 77 | GLY | p.Gly77Ser | P62873 |
| HOMO SAPIENS | 2 | 105 | 5he0 | B | 80 | ILE | p.Ile80Thr | P62873 |
| HOMO SAPIENS | 3 | 4587 | 3rgh | A | 159 | ASP | p.Asp1159Ala | P21333 |
| HOMO SAPIENS | 3 | 4587 | 3rgh | A | 184 | ASP | p.Asp1184Glu | P21333 |
| HOMO SAPIENS | 2 | 4587 | 3rgh | A | 186 | SER | p.Ser1186Leu | P21333 |
| HOMO SAPIENS | 4 | 4587 | 3rgh | A | 188 | ALA | p.Ala1188Thr | P21333 |
| HOMO SAPIENS | 2 | 4587 | 3rgh | A | 199 | SER | p.Ser1199Leu | P21333 |
| HOMO SAPIENS | 0 | 20283 | 4n3w | A | 1278 | GLU | p.Glu1278Lys | Q92793 |
| HOMO SAPIENS | 1 | 14294 | 2d9q | B | 208 | MET | p.Met231Thr | Q99062 |
| HOMO SAPIENS | 6 | 14294 | 2d9q | B | 297 | ASP | p.Asp320Asn | Q99062 |
| HOMO SAPIENS | 0 | 11642 | 3bpt | A | 66 | ARG | p.Arg66Trp | Q6NVY1 |
| HOMO SAPIENS | 0 | 11642 | 3bpt | A | 122 | TYR | p.Tyr122Cys | Q6NVY1 |
| HOMO SAPIENS | 0 | 11642 | 3bpt | A | 345 | GLY | p.Gly345Ser | Q6NVY1 |
| HOMO SAPIENS | 1 | 3942 | 3sby | A | 60 | ASP | p.Asp60His | Q9Y4U1 |
| HOMO SAPIENS | 0 | 3942 | 3sby | A | 111 | ARG | p.Arg111Ter | Q9Y4U1 |
| HOMO SAPIENS | 1 | 3942 | 3sby | A | 116 | LEU | p.Leu116Pro | Q9Y4U1 |
| HOMO SAPIENS | 1 | 3942 | 3sby | A | 130 | TYR | p.Tyr130Cys | Q9Y4U1 |
| HOMO SAPIENS | 1 | 3942 | 3sby | A | 132 | ARG | p.Arg132Ter | Q9Y4U1 |
| HOMO SAPIENS | 1 | 3942 | 3sby | A | 147 | GLY | p.Gly147Ala | Q9Y4U1 |
| HOMO SAPIENS | 1 | 3942 | 3sby | A | 161 | ARG | p.Arg161Gln | Q9Y4U1 |
| HOMO SAPIENS | 1 | 3942 | 3sby | A | 161 | ARG | p.Arg161Ter | Q9Y4U1 |
| HOMO SAPIENS | 1 | 3942 | 3sby | A | 203 | TRP | p.Trp203Ter | Q9Y4U1 |
| HOMO SAPIENS | 1 | 3942 | 3sby | A | 203 | TRP | p.Trp203Ter | Q9Y4U1 |
| HOMO SAPIENS | 1 | 3942 | 3sby | A | 205 | TYR | p.Tyr205Ter | Q9Y4U1 |

| | | | | | | | | | |
|---|---|---|---|---|---|---|---|---|---|
| HOMO SAPIENS | 0 | 38340 | 2mbh | B | | 39 | MET | p.Met39Leu | Q13351 |
| HOMO SAPIENS | 2 | 7360 | 4k30 | A | | 430 | LEU | p.Leu430Pro | Q8N159 |
| HOMO SAPIENS | 2 | 7360 | 4k30 | A | | 433 | GLU | p.Glu433Asp | Q8N159 |
| HOMO SAPIENS | 7 | 7360 | 4k30 | A | | 484 | TRP | p.Trp484Arg | Q8N159 |
| HOMO SAPIENS | 1 | 41903 | 2xs6 | A | | 151 | PRO | p.Pro151Ser | O00459 |
| HOMO SAPIENS | 0 | 172 | 5glh | A | | 183 | ALA | p.Ala183Gly | P24530 |
| HOMO SAPIENS | 0 | 172 | 5glh | A | | 201 | ARG | p.Arg201Ter | P24530 |
| HOMO SAPIENS | 0 | 172 | 5glh | A | | 244 | THR | p.Thr244Met | P24530 |
| HOMO SAPIENS | 0 | 172 | 5glh | A | | 253 | ARG | p.Arg253Ter | P24530 |
| HOMO SAPIENS | 9 | 172 | 5glh | A | | 1017 | ILE | p.Leu17Phe | P00720 |
| HOMO SAPIENS | -1 | 3342 | 1ljm | A | | 1058 | HIS | p.His58Asn | Q01196 |
| HOMO SAPIENS | 4 | 3342 | 1ljm | A | | 1174 | ARG | p.Arg174Gln | Q01196 |
| HOMO SAPIENS | 5 | 25607 | 1p22 | C | | 32 | ASP | p.Asp32Ala | P35222 |
| HOMO SAPIENS | 5 | 25607 | 1p22 | C | | 32 | ASP | p.Asp32Tyr | P35222 |
| HOMO SAPIENS | -1 | 25607 | 1p22 | C | | 33 | SEP | p.Ser33Phe | P35222 |
| HOMO SAPIENS | 5 | 25607 | 1p22 | C | | 34 | GLY | p.Gly34Glu | P35222 |
| HOMO SAPIENS | -1 | 25607 | 1p22 | C | | 37 | SEP | p.Ser37Cys | P35222 |
| HOMO SAPIENS | -1 | 25607 | 1p22 | C | | 41 | THR | p.Thr41Ile | P35222 |
| HOMO SAPIENS | 1 | 48 | 4uow | | 9 | 60 | GLN | p.Arg34111His | Q8WZ42 |
| HOMO SAPIENS | 0 | 48 | 4uow | | 9 | 64 | LEU | p.Val34115Ala | Q8WZ42 |
| HOMO SAPIENS | 0 | 1127 | 5c01 | A | | 638 | ARG | p.Arg638Ter | P29597 |
| HOMO SAPIENS | 0 | 1127 | 5c01 | A | | 684 | ILE | p.Ile684Ser | P29597 |
| HOMO SAPIENS | 0 | 1127 | 5c01 | A | | 703 | ARG | p.Arg703Trp | P29597 |
| HOMO SAPIENS | 5 | 16389 | 2aq0 | B | | 128 | ALA | p.Ala860Asp | Q92889 |
| HOMO SAPIENS | 2 | 16389 | 2aq0 | B | | 141 | ILE | p.Ile873Val | Q92889 |
| HOMO SAPIENS | 5 | 16389 | 2aq0 | B | | 143 | GLU | p.Glu875Gly | Q92889 |
| HOMO SAPIENS | 0 | 16389 | 2aq0 | B | | 180 | GLY | p.Gly912Arg | Q92889 |
| HOMO SAPIENS | 0 | 15816 | 2xrc | A | | 89 | THR | p.Thr107Ala | P05156 |
| HOMO SAPIENS | 0 | 15816 | 2xrc | A | | 133 | ASN | p.Asn151Ser | P05156 |
| HOMO SAPIENS | 0 | 15816 | 2xrc | A | | 185 | THR | p.Thr203Ile | P05156 |
| HOMO SAPIENS | 0 | 15816 | 2xrc | A | | 225 | GLY | p.Gly243Asp | P05156 |
| HOMO SAPIENS | 0 | 15816 | 2xrc | A | | 243 | GLY | p.Gly261Ala | P05156 |
| HOMO SAPIENS | 0 | 15816 | 2xrc | A | | 269 | GLY | p.Gly287Arg | P05156 |
| HOMO SAPIENS | -1 | 15816 | 2xrc | A | | 288 | ILE | p.Ile306Val | P05156 |
| HOMO SAPIENS | -1 | 15816 | 2xrc | A | | 388 | ARG | p.Arg406His | P05156 |
| HOMO SAPIENS | 7 | 15816 | 2xrc | A | | 398 | ILE | p.Ile416Leu | P05156 |
| HOMO SAPIENS | 9 | 15816 | 2xrc | A | | 400 | HIS | p.His418Leu | P05156 |
| HOMO SAPIENS | 0 | 15816 | 2xrc | A | | 423 | LYS | p.Lys441Arg | P05156 |
| HOMO SAPIENS | -1 | 15816 | 2xrc | A | | 530 | GLU | p.Glu548Gln | P05156 |
| HOMO SAPIENS | 0 | 15816 | 2xrc | A | | 535 | PRO | p.Pro553Ala | P05156 |
| HOMO SAPIENS | 0 | 15816 | 2xrc | A | | 536 | GLU | p.Glu554Gly | P05156 |
| HOMO SAPIENS | 1 | 12385 | 4zlp | B | | 1450 | ALA | p.Ala1450Pro | Q9UM47 |
| HOMO SAPIENS | 7 | 12385 | 4zlp | B | | 1482 | GLY | p.Gly1482Arg | Q9UM47 |
| HOMO SAPIENS | 2 | 12385 | 4zlp | B | | 1518 | LEU | p.Leu1518Met | Q9UM47 |
| HOMO SAPIENS | 0 | 12385 | 4zlp | B | | 1560 | ARG | p.Arg1560Pro | Q9UM47 |
| HOMO SAPIENS | 0 | 1227 | 4a04 | A | | 148 | PHE | p.Phe148Tyr | O43435 |
| HOMO SAPIENS | 4 | 1227 | 4a04 | A | | 194 | HIS | p.His194Gln | O43435 |
| HOMO SAPIENS | 2 | 8017 | 3ei4 | B | | 244 | LYS | p.Lys244Glu | Q92466 |
| HOMO SAPIENS | 1 | 8017 | 3ei4 | B | | 273 | ARG | p.Arg273His | Q92466 |
| HOMO SAPIENS | 3 | 36112 | 2k8f | B | | 9 | SER | p.Ser9Arg | P04637 |
| HOMO SAPIENS | 2 | 12880 | 5ggf | A | | 265 | ARG | p.Arg265His | Q8WZA1 |
| HOMO SAPIENS | 2 | 12880 | 5ggf | A | | 287 | ILE | p.Ile287Ser | Q8WZA1 |
| HOMO SAPIENS | 2 | 12880 | 5ggf | A | | 311 | ARG | p.Arg311Gln | Q8WZA1 |
| HOMO SAPIENS | 2 | 12880 | 5ggf | A | | 425 | TRP | p.Trp425Ser | Q8WZA1 |

| | | | | | | | | |
|---|---|---|---|---|---|---|---|---|
| HOMO SAPIENS | 2 | 12880 | 5ggf | A | 442 | ARG | p.Arg442Cys | Q8WZA1 |
| HOMO SAPIENS | 2 | 12880 | 5ggf | A | 448 | GLY | p.Gly448Arg | Q8WZA1 |
| HOMO SAPIENS | 0 | 12880 | 5ggf | A | 490 | CYS | p.Cys490Tyr | Q8WZA1 |
| HOMO SAPIENS | 0 | 12880 | 5ggf | A | 493 | PRO | p.Pro493Arg | Q8WZA1 |
| HOMO SAPIENS | 3 | 12880 | 5ggf | A | 550 | SER | p.Ser550Asn | Q8WZA1 |
| HOMO SAPIENS | 1 | 12880 | 5ggf | A | 556 | ASP | p.Asp556Asn | Q8WZA1 |
| HOMO SAPIENS | 0 | 12880 | 5ggf | A | 580 | ARG | p.Arg580Ter | Q8WZA1 |
| HOMO SAPIENS | 2 | 12880 | 5ggf | A | 605 | ARG | p.Arg605Pro | Q8WZA1 |
| HOMO SAPIENS | 0 | 12880 | 5ggf | A | 623 | VAL | p.Met623Val | Q8WZA1 |
| HOMO SAPIENS | 1 | 10187 | 4qw2 | A | 145 | ALA | p.Ala145Ser | Q06787 |
| HOMO SAPIENS | 1 | 10187 | 4qw2 | A | 174 | ASN | p.Asn174Lys | Q06787 |
| HOMO SAPIENS | 0 | 24002 | 2m0t | A | 110 | LEU | p.Leu110Val | O14745 |
| HOMO SAPIENS | 9 | 3126 | 4mjh | A | 99 | LEU | p.Leu99Met | P04792 |
| HOMO SAPIENS | 1 | 3126 | 4mjh | A | 127 | ARG | p.Arg127Trp | P04792 |
| HOMO SAPIENS | 1 | 3126 | 4mjh | A | 127 | ARG | p.Arg127Leu | P04792 |
| HOMO SAPIENS | 1 | 3126 | 4mjh | A | 135 | SER | p.Ser135Phe | P04792 |
| HOMO SAPIENS | 2 | 3126 | 4mjh | A | 136 | ARG | p.Arg136Trp | P04792 |
| HOMO SAPIENS | 1 | 3126 | 4mjh | A | 140 | ARG | p.Arg140Gly | P04792 |
| HOMO SAPIENS | 0 | 3126 | 4mjh | A | 151 | THR | p.Thr151Ile | P04792 |
| HOMO SAPIENS | -1 | 15780 | 2jlp | A | 213 | ARG | p.Arg231Gly | P08294 |
| HOMO SAPIENS | 8 | 3578 | 1apz | D | 203 | GLY | p.Gly226Asp | P20933 |
| HOMO SAPIENS | 1 | 3578 | 1apz | D | 279 | GLY | p.Gly302Arg | P20933 |
| HOMO SAPIENS | 0 | 3578 | 1apz | D | 283 | CYS | p.Cys306Arg | P20933 |
| HOMO SAPIENS | 5 | 1335 | 5xtd | C | 115 | ARG | p.Arg115Cys | O75251 |
| HOMO SAPIENS | 0 | 1335 | 5xtd | C | 122 | VAL | p.Val122Met | O75251 |
| HOMO SAPIENS | 2 | 5063 | 2le4 | A | 37 | ARG | p.Arg74Pro | P48431 |
| HOMO SAPIENS | 1 | 25870 | 2o10 | A | 44 | LEU | p.Leu44Pro | P50461 |
| HOMO SAPIENS | 4 | 25870 | 2o10 | A | 58 | CYS | p.Cys58Gly | P50461 |
| HOMO SAPIENS | 0 | 18748 | 5fii | C | 39 | PHE | p.Phe39Leu | P00439 |
| HOMO SAPIENS | 6 | 18748 | 5fii | C | 48 | LEU | p.Leu48Ser | P00439 |
| HOMO SAPIENS | 2 | 18748 | 5fii | C | 65 | ILE | p.Ile65Asn | P00439 |
| HOMO SAPIENS | 3 | 18748 | 5fii | C | 95 | ILE | p.Ile95Phe | P00439 |
| HOMO SAPIENS | 1 | 14261 | 2vay | B | 1539 | ARG | p.Arg1539Cys | Q13698 |
| HOMO SAPIENS | 0 | 2494 | 4dx1 | A | 183 | GLU | p.Glu183Asp | Q9HBA0 |
| HOMO SAPIENS | 0 | 2494 | 4dx1 | A | 217 | ALA | p.Ala217Ser | Q9HBA0 |
| HOMO SAPIENS | 0 | 2494 | 4dx1 | A | 217 | ALA | p.Ala217Val | Q9HBA0 |
| HOMO SAPIENS | 4 | 2494 | 4dx1 | A | 224 | ARG | p.Arg224Trp | Q9HBA0 |
| HOMO SAPIENS | 4 | 2494 | 4dx1 | A | 254 | VAL | p.Val254Leu | Q9HBA0 |
| HOMO SAPIENS | 0 | 2494 | 4dx1 | A | 269 | ARG | p.Arg269Cys | Q9HBA0 |
| HOMO SAPIENS | 0 | 2494 | 4dx1 | A | 269 | ARG | p.Arg269His | Q9HBA0 |
| HOMO SAPIENS | 0 | 2494 | 4dx1 | A | 278 | GLU | p.Glu278Lys | Q9HBA0 |
| HOMO SAPIENS | 0 | 2494 | 4dx1 | A | 315 | ARG | p.Arg315Trp | Q9HBA0 |
| HOMO SAPIENS | 0 | 2494 | 4dx1 | A | 316 | ARG | p.Arg316Cys | Q9HBA0 |
| HOMO SAPIENS | 0 | 2494 | 4dx1 | A | 331 | ILE | p.Ile331Phe | Q9HBA0 |
| HOMO SAPIENS | 0 | 2494 | 4dx1 | A | 333 | ASP | p.Asp333Gly | Q9HBA0 |
| HOMO SAPIENS | 1 | 2494 | 4dx1 | A | 365 | VAL | p.Val365Met | Q9HBA0 |
| HOMO SAPIENS | 0 | 23998 | 2l1r | B | 161 | ARG | p.Arg162Trp | P19429 |
| HOMO SAPIENS | 0 | 8354 | 4f0x | D | 187 | SER | p.Ser187Ter | O95822 |
| HOMO SAPIENS | 2 | 8354 | 4f0x | D | 259 | GLY | p.Gly259Ala | O95822 |
| HOMO SAPIENS | 2 | 8354 | 4f0x | D | 284 | THR | p.Thr284Ala | O95822 |
| HOMO SAPIENS | 3 | 8354 | 4f0x | D | 309 | LYS | p.Lys309Gln | O95822 |
| HOMO SAPIENS | 0 | 21182 | 4apo | A | 228 | GLN | p.Gln228Lys | O00170 |
| HOMO SAPIENS | 1 | 21182 | 4apo | A | 307 | GLN | p.Gln307Arg | O00170 |
| HOMO SAPIENS | 0 | 2287 | 5of4 | B | 112 | ARG | p.Arg112His | P18074 |

| | | | | | | | | |
|---|---|---|---|---|---|---|---|---|
| HOMO SAPIENS | 0 | 2287 | 5of4 | B | 312 | ASP | p.Asp312Asn | P18074 |
| HOMO SAPIENS | 3 | 2287 | 5of4 | B | 461 | LEU | p.Leu461Val | P18074 |
| HOMO SAPIENS | 0 | 2287 | 5of4 | B | 485 | LEU | p.Leu485Pro | P18074 |
| HOMO SAPIENS | 1 | 2287 | 5of4 | B | 541 | SER | p.Ser541Arg | P18074 |
| HOMO SAPIENS | 0 | 2287 | 5of4 | B | 616 | ARG | p.Arg616Pro | P18074 |
| HOMO SAPIENS | 0 | 2287 | 5of4 | B | 616 | ARG | p.Arg616Trp | P18074 |
| HOMO SAPIENS | 1 | 2287 | 5of4 | B | 658 | ARG | p.Arg658Cys | P18074 |
| HOMO SAPIENS | 1 | 2287 | 5of4 | B | 681 | ASP | p.Asp681Asn | P18074 |
| HOMO SAPIENS | 1 | 2287 | 5of4 | B | 683 | ARG | p.Arg683Trp | P18074 |
| HOMO SAPIENS | 0 | 2287 | 5of4 | B | 713 | GLY | p.Gly713Arg | P18074 |
| HOMO SAPIENS | 0 | 2287 | 5of4 | B | 717 | ALA | p.Ala717Gly | P18074 |
| HOMO SAPIENS | 0 | 2287 | 5of4 | B | 722 | ARG | p.Arg722Trp | P18074 |
| HOMO SAPIENS | 0 | 2287 | 5of4 | B | 725 | ALA | p.Ala725Pro | P18074 |
| HOMO SAPIENS | 4 | 12188 | 4its | A | 122 | ARG | p.Arg122Gly | Q9Y606 |
| HOMO SAPIENS | 4 | 12188 | 4its | A | 133 | ASP | p.Asp133Asn | Q9Y606 |
| HOMO SAPIENS | 4 | 12188 | 4its | A | 144 | ARG | p.Arg144Trp | Q9Y606 |
| HOMO SAPIENS | 0 | 31529 | 1kgd | A | 761 | ARG | p.Arg766Gln | O14936 |
| HOMO SAPIENS | 1 | 31529 | 1kgd | A | 824 | GLN | p.Gln829Arg | O14936 |
| HOMO SAPIENS | 0 | 10906 | 2h63 | C | 18 | ARG | p.Arg18Ter | P53004 |
| HOMO SAPIENS | 9 | 27767 | 4wyq | A | 280 | LEU | p.Leu278Phe | Q9UPY3 |
| HOMO SAPIENS | 0 | 27767 | 4wyq | A | 297 | SER | p.Ser295Cys | Q9UPY3 |
| HOMO SAPIENS | 1 | 27767 | 4wyq | A | 334 | TYR | p.Tyr332Ter | Q9UPY3 |
| HOMO SAPIENS | 7 | 7775 | 5sv7 | C | 136 | SER | p.Ser136Cys | Q9NZJ5 |
| HOMO SAPIENS | 2 | 7775 | 5sv7 | C | 166 | GLN | p.Gln166Arg | Q9NZJ5 |
| HOMO SAPIENS | 2 | 7775 | 5sv7 | C | 332 | GLU | p.Glu332Ter | Q9NZJ5 |
| HOMO SAPIENS | 0 | 21449 | 2e9w | B | 84 | THR | p.Thr84Met | P10721 |
| HOMO SAPIENS | 0 | 21449 | 2e9w | B | 168 | ALA | p.Ala168Glu | P10721 |
| HOMO SAPIENS | 0 | 21449 | 2e9w | B | 178 | ALA | p.Ala178Thr | P10721 |
| HOMO SAPIENS | 0 | 21449 | 2e9w | B | 293 | ASN | p.Asn293Ser | P10721 |
| HOMO SAPIENS | -1 | 21449 | 2e9w | B | 318 | MET | p.Met318Leu | P10721 |
| HOMO SAPIENS | 0 | 21449 | 2e9w | B | 374 | VAL | p.Val374Ile | P10721 |
| HOMO SAPIENS | 6 | 3010 | 3flg | A | 66 | SER | p.Ser270Pro | Q9UBC3 |
| HOMO SAPIENS | 1 | 33918 | 2xzz | A | 760 | ARG | p.Arg760Ter | P22735 |
| HOMO SAPIENS | 2 | 1254 | 5xtc | w | 65 | ASN | p.Asn65Ser | O95299 |
| HOMO SAPIENS | 1 | 1254 | 5xtc | w | 238 | GLU | p.Glu238Lys | O95299 |
| HOMO SAPIENS | 3 | 10245 | 3siv | B | 194 | ALA | p.Ala194Glu | Q8WWY3 |
| HOMO SAPIENS | 4 | 10245 | 3siv | B | 216 | ALA | p.Ala216Pro | Q8WWY3 |
| HOMO SAPIENS | 0 | 2670 | 3dwb | A | 341 | THR | p.Thr341Ile | P42892 |
| HOMO SAPIENS | 1 | 2670 | 3dwb | A | 462 | SER | p.Ser462Asn | P42892 |
| HOMO SAPIENS | 1 | 2670 | 3dwb | A | 754 | ARG | p.Arg754Cys | P42892 |
| HOMO SAPIENS | 1 | 2961 | 5xtb | O | 40 | VAL | p.Val40Met | P19404 |
| HOMO SAPIENS | 5 | 2961 | 5xtb | O | 172 | ILE | p.Ile172Thr | P19404 |
| HOMO SAPIENS | 5 | 2961 | 5xtb | O | 190 | ASP | p.Asp190Gly | P19404 |
| HOMO SAPIENS | 5 | 29655 | 2xdg | A | 57 | ALA | p.Ala57Thr | Q02643 |
| HOMO SAPIENS | 4 | 29655 | 2xdg | A | 72 | GLU | p.Glu72Ter | Q02643 |
| HOMO SAPIENS | 0 | 29655 | 2xdg | A | 121 | GLU | p.Glu121Asp | Q02643 |
| HOMO SAPIENS | -1 | 17935 | 1hyn | P | 38 | ASP | p.Asp38Ala | P02730 |
| HOMO SAPIENS | 7 | 12825 | 1g82 | D | 99 | SER | p.Ser99Asn | P31371 |
| HOMO SAPIENS | 6 | 24927 | 1jm7 | B | 85 | VAL | p.Val85Leu | Q99728 |
| HOMO SAPIENS | 1 | 24927 | 1jm7 | B | 112 | ARG | p.Arg112Ter | Q99728 |
| HOMO SAPIENS | 0 | 24927 | 1jm7 | B | 116 | HIS | p.His116Tyr | Q99728 |
| HOMO SAPIENS | 7 | 92 | 5hde | A | 61 | LYS | p.Lys61Arg | Q05209 |
| HOMO SAPIENS | 0 | 17246 | 1szb | A | 84 | ARG | p.Arg99Gln | O00187 |
| HOMO SAPIENS | 1 | 17246 | 1szb | A | 105 | ASP | p.Asp120Gly | O00187 |

| | | | | | | | | |
|---|---|---|---|---|---|---|---|---|
| HOMO SAPIENS | 0 | 17246 | 1szb | A | 111 | PRO | p.Pro126Leu | O00187 |
| HOMO SAPIENS | 9 | 17246 | 1szb | A | 141 | CYS | p.Cys156Tyr | O00187 |
| HOMO SAPIENS | 0 | 32473 | 1b72 | A | 207 | ARG | p.Arg207Cys | P14653 |
| HOMO SAPIENS | 0 | 1920 | 4yzf | B | 488 | VAL | p.Val488Met | P02730 |
| HOMO SAPIENS | 0 | 1920 | 4yzf | B | 589 | ARG | p.Arg589Cys | P02730 |
| HOMO SAPIENS | 0 | 1920 | 4yzf | B | 589 | ARG | p.Arg589His | P02730 |
| HOMO SAPIENS | 0 | 1920 | 4yzf | B | 602 | ARG | p.Arg602Pro | P02730 |
| HOMO SAPIENS | 0 | 1920 | 4yzf | B | 613 | SER | p.Ser613Phe | P02730 |
| HOMO SAPIENS | -1 | 1920 | 4yzf | B | 646 | ARG | p.Arg646Gln | P02730 |
| HOMO SAPIENS | -1 | 1920 | 4yzf | B | 646 | ARG | p.Arg646Trp | P02730 |
| HOMO SAPIENS | 0 | 1920 | 4yzf | B | 701 | GLY | p.Gly701Asp | P02730 |
| HOMO SAPIENS | 0 | 1920 | 4yzf | B | 760 | ARG | p.Arg760Gln | P02730 |
| HOMO SAPIENS | 0 | 1920 | 4yzf | B | 771 | GLY | p.Gly771Asp | P02730 |
| HOMO SAPIENS | 0 | 1920 | 4yzf | B | 773 | SER | p.Ser773Pro | P02730 |
| HOMO SAPIENS | 0 | 1920 | 4yzf | B | 837 | THR | p.Thr837Ala | P02730 |
| HOMO SAPIENS | 0 | 1920 | 4yzf | B | 854 | PRO | p.Pro854Leu | P02730 |
| HOMO SAPIENS | 0 | 1920 | 4yzf | B | 858 | ALA | p.Ala858Asp | P02730 |
| HOMO SAPIENS | 0 | 1920 | 4yzf | B | 862 | VAL | p.Val862Ile | P02730 |
| HOMO SAPIENS | 0 | 1920 | 4yzf | B | 868 | PRO | p.Pro868Leu | P02730 |
| HOMO SAPIENS | 0 | 1920 | 4yzf | B | 870 | ARG | p.Arg870Trp | P02730 |
| HOMO SAPIENS | 0 | 9863 | 2juj | A | 54 | VAL | p.Val904Ile | P22681 |
| HOMO SAPIENS | 0 | 21934 | 2na7 | A | 184 | ILE | p.Ile184Val | P25445 |
| HOMO SAPIENS | -1 | 6746 | 1lw3 | A | 3 | THR | p.Lys3Thr | Q13614 |
| HOMO SAPIENS | -1 | 6746 | 1lw3 | A | 44 | VAL | p.Val44Ile | Q13614 |
| HOMO SAPIENS | 3 | 6746 | 1lw3 | A | 183 | ALA | p.Ala183Ser | Q13614 |
| HOMO SAPIENS | 4 | 6746 | 1lw3 | A | 482 | GLN | p.Gln482Ter | Q13614 |
| HOMO SAPIENS | 4 | 6746 | 1lw3 | A | 502 | GLU | p.Glu502Gln | Q13614 |
| HOMO SAPIENS | 3 | 6746 | 1lw3 | A | 545 | ASN | p.Asn545Ser | Q13614 |
| HOMO SAPIENS | 0 | 6746 | 1lw3 | A | 586 | ARG | p.Arg586Trp | Q13614 |
| HOMO SAPIENS | -1 | 6746 | 1lw3 | A | 602 | ALA | p.Ala602Gly | Q13614 |
| HOMO SAPIENS | 1 | 319 | 4ch9 | B | 336 | ARG | p.Arg336Ile | Q9UH77 |
| HOMO SAPIENS | 3 | 319 | 4ch9 | B | 340 | ALA | p.Ala340Val | Q9UH77 |
| HOMO SAPIENS | 0 | 319 | 4ch9 | B | 384 | ARG | p.Arg384Gln | Q9UH77 |
| HOMO SAPIENS | 5 | 319 | 4ch9 | B | 426 | PRO | p.Pro426Leu | Q9UH77 |
| HOMO SAPIENS | 3 | 319 | 4ch9 | B | 431 | ARG | p.Arg431Gln | Q9UH77 |
| HOMO SAPIENS | 2 | 319 | 4ch9 | B | 575 | ARG | p.Arg575Trp | Q9UH77 |
| HOMO SAPIENS | 0 | 574 | 2qkk | A | 142 | VAL | p.Val142Ile | O60930 |
| HOMO SAPIENS | 0 | 24021 | 2n8y | A | 752 | ARG | p.Arg752Gln | P12814 |
| HOMO SAPIENS | 0 | 18693 | 2y7j | B | 44 | ARG | p.Arg44Ter | P15735 |
| HOMO SAPIENS | 0 | 18693 | 2y7j | B | 106 | VAL | p.Val106Glu | P15735 |
| HOMO SAPIENS | 3 | 18693 | 2y7j | B | 189 | GLY | p.Gly189Glu | P15735 |
| HOMO SAPIENS | 0 | 17087 | 2ifg | A | 237 | THR | p.Thr237Met | P04629 |
| HOMO SAPIENS | 0 | 7980 | 1qub | A | 247 | VAL | p.Val266Leu | P02749 |
| HOMO SAPIENS | -1 | 2205 | 1h7u | A | 18 | ILE | p.Ile18Val | P54278 |
| HOMO SAPIENS | -1 | 2205 | 1h7u | A | 20 | ARG | p.Arg20Gln | P54278 |
| HOMO SAPIENS | 9 | 2205 | 1h7u | A | 46 | SER | p.Ser46Ile | P54278 |
| HOMO SAPIENS | 0 | 2205 | 1h7u | A | 60 | ASP | p.Asp60Glu | P54278 |
| HOMO SAPIENS | 0 | 2205 | 1h7u | A | 80 | PHE | p.Phe80Leu | P54278 |
| HOMO SAPIENS | 0 | 2205 | 1h7u | A | 118 | SER | p.Ser118Arg | P54278 |
| HOMO SAPIENS | 0 | 2205 | 1h7u | A | 126 | HIS | p.His126Gln | P54278 |
| HOMO SAPIENS | 0 | 2205 | 1h7u | A | 128 | SER | p.Ser128Leu | P54278 |
| HOMO SAPIENS | 1 | 2205 | 1h7u | A | 134 | ARG | p.Arg134Ter | P54278 |
| HOMO SAPIENS | 0 | 2205 | 1h7u | A | 166 | LEU | p.Leu166Pro | P54278 |
| HOMO SAPIENS | 1 | 2205 | 1h7u | A | 191 | TYR | p.Tyr191Cys | P54278 |

| | | | | | | | | |
|---|---|---|---|---|---|---|---|---|
| HOMO SAPIENS | 0 | 2205 | 1h7u | A | 211 | ARG | p.Arg211Ter | P54278 |
| HOMO SAPIENS | 0 | 2205 | 1h7u | A | 233 | GLN | p.Gln233Ter | P54278 |
| HOMO SAPIENS | 0 | 2205 | 1h7u | A | 255 | TYR | p.Tyr255Ter | P54278 |
| HOMO SAPIENS | 0 | 2205 | 1h7u | A | 277 | THR | p.Thr277Lys | P54278 |
| HOMO SAPIENS | 0 | 2205 | 1h7u | A | 301 | LYS | p.Lys301Asn | P54278 |
| HOMO SAPIENS | 0 | 2205 | 1h7u | A | 312 | MET | p.Met312Ile | P54278 |
| HOMO SAPIENS | 0 | 2205 | 1h7u | A | 312 | MET | p.Met312Thr | P54278 |
| HOMO SAPIENS | 0 | 2205 | 1h7u | A | 315 | ARG | p.Arg315Ter | P54278 |
| HOMO SAPIENS | 0 | 2205 | 1h7u | A | 318 | TYR | p.Tyr318Cys | P54278 |
| HOMO SAPIENS | 0 | 2205 | 1h7u | A | 335 | ASN | p.Asn335Ile | P54278 |
| HOMO SAPIENS | 0 | 2205 | 1h7u | A | 360 | ILE | p.Ile360Met | P54278 |
| HOMO SAPIENS | 6 | 20754 | 5mpo | B | 7 | VAL | p.Val7Phe | O96033 |
| HOMO SAPIENS | 4 | 20754 | 5mpo | B | 64 | VAL | p.Val64Ile | O96033 |
| HOMO SAPIENS | 5 | 34944 | 4qam | B | 1114 | ASP | p.Asp1114Gly | Q96KN7 |
| HOMO SAPIENS | 5 | 34944 | 4qam | B | 1240 | GLY | p.Gly1240Glu | Q96KN7 |
| HOMO SAPIENS | 9 | 40496 | 3vnn | A | 278 | ARG | p.Arg278His | P49917 |
| HOMO SAPIENS | 6 | 30131 | 4dt7 | E | 40 | PRO | p.Pro210Leu | P04070 |
| HOMO SAPIENS | -1 | 30131 | 4dt7 | E | 50 | ARG | p.Arg220Trp | P04070 |
| HOMO SAPIENS | 7 | 15311 | 2pqt | A | 64 | ARG | p.Arg64Trp | P18440 |
| HOMO SAPIENS | 2 | 15311 | 2pqt | A | 149 | VAL | p.Val149Ile | P18440 |
| HOMO SAPIENS | -1 | 19581 | 3hy8 | A | 33 | ASP | p.Asp33Val | Q9NVS9 |
| HOMO SAPIENS | 5 | 19581 | 3hy8 | A | 116 | ARG | p.Arg116Gln | Q9NVS9 |
| HOMO SAPIENS | 4 | 19581 | 3hy8 | A | 225 | ARG | p.Arg225His | Q9NVS9 |
| HOMO SAPIENS | 2 | 19581 | 3hy8 | A | 229 | TRP | p.Arg229Trp | Q9NVS9 |
| HOMO SAPIENS | 2 | 19581 | 3hy8 | A | 229 | TRP | p.Arg229Gln | Q9NVS9 |
| HOMO SAPIENS | 0 | 10266 | 3t1i | A | 41 | ASP | p.Asp41His | P49959 |
| HOMO SAPIENS | 1 | 10266 | 3t1i | A | 305 | ARG | p.Arg305Trp | P49959 |
| HOMO SAPIENS | 0 | 10266 | 3t1i | A | 308 | PHE | p.Phe308Leu | P49959 |
| HOMO SAPIENS | 0 | 10266 | 3t1i | A | 334 | SER | p.Ser334Arg | P49959 |
| HOMO SAPIENS | 1 | 1600 | 3rbf | A | 61 | GLU | p.Glu61Asp | P20711 |
| HOMO SAPIENS | 0 | 1600 | 3rbf | A | 91 | ALA | p.Ala91Val | P20711 |
| HOMO SAPIENS | -1 | 1600 | 3rbf | A | 102 | GLY | p.Gly102Ser | P20711 |
| HOMO SAPIENS | 3 | 1600 | 3rbf | A | 147 | SER | p.Ser147Arg | P20711 |
| HOMO SAPIENS | 1 | 1600 | 3rbf | A | 210 | PRO | p.Pro210Leu | P20711 |
| HOMO SAPIENS | 0 | 1600 | 3rbf | A | 217 | MET | p.Met217Val | P20711 |
| HOMO SAPIENS | 4 | 1600 | 3rbf | A | 250 | SER | p.Ser250Phe | P20711 |
| HOMO SAPIENS | 2 | 1600 | 3rbf | A | 275 | ALA | p.Ala275Thr | P20711 |
| HOMO SAPIENS | 1 | 1600 | 3rbf | A | 309 | PHE | p.Phe309Leu | P20711 |
| HOMO SAPIENS | -1 | 1600 | 3rbf | A | 347 | ARG | p.Arg347Gln | P20711 |
| HOMO SAPIENS | 0 | 1600 | 3rbf | A | 462 | ARG | p.Arg462Gln | P20711 |
| HOMO SAPIENS | 6 | 20830 | 5y4f | A | 714 | THR | p.Thr715Ile | Q01484 |
| HOMO SAPIENS | 7 | 20830 | 5y4f | A | 719 | ASP | p.Asp720Glu | Q01484 |
| HOMO SAPIENS | 0 | 20830 | 5y4f | A | 814 | VAL | p.Val815Ile | Q01484 |
| HOMO SAPIENS | -1 | 302 | 2c9y | A | 9 | GLU | p.Glu9Ter | P54819 |
| HOMO SAPIENS | 0 | 302 | 2c9y | A | 103 | ARG | p.Arg103Trp | P54819 |
| HOMO SAPIENS | 3 | 302 | 2c9y | A | 165 | ASP | p.Asp165Gly | P54819 |
| HOMO SAPIENS | 0 | 302 | 2c9y | A | 182 | ALA | p.Ala182Asp | P54819 |
| HOMO SAPIENS | 2 | 302 | 2c9y | A | 186 | ARG | p.Arg186Cys | P54819 |
| HOMO SAPIENS | 0 | 302 | 2c9y | A | 209 | ALA | p.Ala209Thr | P54819 |
| HOMO SAPIENS | -1 | 16920 | 1jlj | B | 10 | ASN | p.Asn10Tyr | Q9NQX3 |
| HOMO SAPIENS | 9 | 8462 | 1zbq | B | 16 | GLY | p.Gly16Ser | P51659 |
| HOMO SAPIENS | 5 | 8462 | 1zbq | B | 19 | ALA | p.Ala19Gly | P51659 |
| HOMO SAPIENS | 1 | 8462 | 1zbq | B | 34 | ALA | p.Ala34Val | P51659 |
| HOMO SAPIENS | 1 | 8462 | 1zbq | B | 106 | ARG | p.Arg106His | P51659 |

| | | | | | | | | |
|---|---|---|---|---|---|---|---|---|
| HOMO SAPIENS | 0 | 8462 | 1zbq | B | 140 | LYS | p.Lys140Asn | P51659 |
| HOMO SAPIENS | 0 | 8462 | 1zbq | B | 292 | THR | p.Thr292Ser | P51659 |
| HOMO SAPIENS | 0 | 4375 | 4ndi | A | 199 | ARG | p.Arg213His | Q7Z2E3 |
| HOMO SAPIENS | 3 | 4375 | 4ndi | A | 201 | HIS | p.His215Arg | Q7Z2E3 |
| HOMO SAPIENS | 0 | 4375 | 4ndi | A | 206 | PRO | p.Pro220Leu | Q7Z2E3 |
| HOMO SAPIENS | 2 | 4375 | 4ndi | A | 223 | LEU | p.Leu237Phe | Q7Z2E3 |
| HOMO SAPIENS | 0 | 4375 | 4ndi | A | 245 | ARG | p.Arg259His | Q7Z2E3 |
| HOMO SAPIENS | 0 | 4375 | 4ndi | A | 263 | VAL | p.Val277Gly | Q7Z2E3 |
| HOMO SAPIENS | 0 | 4375 | 4ndi | A | 279 | TRP | p.Trp293Ter | Q7Z2E3 |
| HOMO SAPIENS | 0 | 27004 | 4ccg | Y | 321 | ASP | p.Asp321Glu | Q9NW38 |
| HOMO SAPIENS | -1 | 27004 | 4ccg | Y | 372 | GLY | p.Gly372Ala | Q9NW38 |
| HOMO SAPIENS | -1 | 6383 | 5fai | A | 34 | ALA | p.Ala34Gly | Q92979 |
| HOMO SAPIENS | 8 | 6383 | 5fai | A | 86 | ASP | p.Asp86Gly | Q92979 |
| HOMO SAPIENS | 2 | 24585 | 4mi5 | A | 684 | ARG | p.Arg679Cys | Q15910 |
| HOMO SAPIENS | 0 | 15967 | 3nhb | A | 723 | ARG | p.Arg723Gln | Q9NP58 |
| HOMO SAPIENS | 5 | 18152 | 1oey | A | 389 | HIS | p.His389Gln | P19878 |
| HOMO SAPIENS | 5 | 18152 | 1oey | A | 395 | ARG | p.Arg395Trp | P19878 |
| HOMO SAPIENS | -1 | 1044 | 1li1 | F | 2 | VAL | p.Val1486Ile | P08572 |
| HOMO SAPIENS | 6 | 1044 | 1li1 | F | 37 | GLU | p.Glu1521Lys | P08572 |
| HOMO SAPIENS | 5 | 1044 | 1li1 | F | 157 | SER | p.Ser1639Leu | P08572 |
| HOMO SAPIENS | 5 | 1044 | 1li1 | F | 179 | ARG | p.Arg1662His | P08572 |
| HOMO SAPIENS | 5 | 1044 | 1li1 | F | 180 | GLY | p.Gly1663Arg | P08572 |
| HOMO SAPIENS | 6 | 20076 | 4uyi | A | 671 | LEU | p.Leu671Ser | Q8IY92 |
| HOMO SAPIENS | 0 | 19588 | 1jmj | A | 189 | ARG | p.Arg208His | P05546 |
| HOMO SAPIENS | -1 | 38023 | 1ry7 | B | 65 | GLY | p.Gly65Arg | P22607 |
| HOMO SAPIENS | -1 | 38023 | 1ry7 | B | 139 | ASP | p.Asp139Glu | P22607 |
| HOMO SAPIENS | 0 | 38023 | 1ry7 | B | 248 | ARG | p.Arg248Cys | P22607 |
| HOMO SAPIENS | 0 | 38023 | 1ry7 | B | 249 | SER | p.Ser249Cys | P22607 |
| HOMO SAPIENS | 0 | 38023 | 1ry7 | B | 250 | PRO | p.Pro250Arg | P22607 |
| HOMO SAPIENS | 0 | 38023 | 1ry7 | B | 322 | GLU | p.Glu322Lys | P22607 |
| HOMO SAPIENS | 5 | 16884 | 5eli | A | 33 | GLN | p.Gln33Ter | Q9NZC2 |
| HOMO SAPIENS | 0 | 16884 | 5eli | A | 44 | TRP | p.Trp44Ter | Q9NZC2 |
| HOMO SAPIENS | 0 | 16884 | 5eli | A | 47 | ARG | p.Arg47His | Q9NZC2 |
| HOMO SAPIENS | 0 | 16884 | 5eli | A | 66 | THR | p.Thr66Met | Q9NZC2 |
| HOMO SAPIENS | 0 | 16884 | 5eli | A | 78 | TRP | p.Trp78Ter | Q9NZC2 |
| HOMO SAPIENS | 1 | 16884 | 5eli | A | 96 | THR | p.Thr96Lys | Q9NZC2 |
| HOMO SAPIENS | 9 | 16884 | 5eli | A | 126 | VAL | p.Val126Gly | Q9NZC2 |
| HOMO SAPIENS | -1 | 12267 | 3w5o | A | 3 | ALA | p.Ala3Val | P49917 |
| HOMO SAPIENS | 1 | 12267 | 3w5o | A | 9 | THR | p.Thr9Ile | P49917 |
| HOMO SAPIENS | 2 | 12267 | 3w5o | A | 188 | ARG | p.Arg188Gln | P49917 |
| HOMO SAPIENS | 0 | 12267 | 3w5o | A | 469 | GLY | p.Gly469Glu | P49917 |
| HOMO SAPIENS | 1 | 12267 | 3w5o | A | 580 | ARG | p.Arg580Ter | P49917 |
| HOMO SAPIENS | 0 | 12267 | 3w5o | A | 600 | GLU | p.Glu600Lys | P49917 |
| HOMO SAPIENS | 9 | 2839 | 2hw5 | B | 54 | ARG | p.Arg54Cys | P30084 |
| HOMO SAPIENS | 9 | 2839 | 2hw5 | B | 59 | ASN | p.Asn59Ser | P30084 |
| HOMO SAPIENS | 0 | 2839 | 2hw5 | B | 273 | LYS | p.Lys273Glu | P30084 |
| HOMO SAPIENS | 1 | 12266 | 2o2k | B | 939 | GLY | p.Gly939Cys | Q99707 |
| HOMO SAPIENS | 5 | 12266 | 2o2k | B | 1164 | ARG | p.Arg1164His | Q99707 |
| HOMO SAPIENS | 1 | 12266 | 2o2k | B | 1166 | LEU | p.Leu1166Met | Q99707 |
| HOMO SAPIENS | 5 | 12266 | 2o2k | B | 1173 | PRO | p.Pro1173Leu | Q99707 |
| HOMO SAPIENS | 5 | 12266 | 2o2k | B | 1222 | ASN | p.Asn1222Ser | Q99707 |
| HOMO SAPIENS | 0 | 30891 | 3ch4 | B | 138 | ARG | p.Arg138Ter | Q15126 |
| HOMO SAPIENS | -1 | 4433 | 1qlz | A | 54 | GLY | p.Gly54Ser | P04156 |
| HOMO SAPIENS | 0 | 22568 | 3cxl | A | 20 | LEU | p.Leu20Phe | P15882 |

| | | | | | | | | |
|---|---|---|---|---|---|---|---|---|
| HOMO SAPIENS | 0 | 22568 | 3cxl | A | 126 | ILE | p.Ile126Met | P15882 |
| HOMO SAPIENS | 0 | 22568 | 3cxl | A | 141 | PRO | p.Pro141Leu | P15882 |
| HOMO SAPIENS | 0 | 22568 | 3cxl | A | 143 | TYR | p.Tyr143His | P15882 |
| HOMO SAPIENS | 0 | 22568 | 3cxl | A | 223 | ALA | p.Ala223Val | P15882 |
| HOMO SAPIENS | 0 | 22568 | 3cxl | A | 228 | GLY | p.Gly228Ser | P15882 |
| HOMO SAPIENS | 0 | 22568 | 3cxl | A | 252 | PRO | p.Pro252Gln | P15882 |
| HOMO SAPIENS | 0 | 22568 | 3cxl | A | 313 | GLU | p.Glu313Lys | P15882 |
| HOMO SAPIENS | 3 | 2668 | 5eqg | A | 34 | ASN | p.Asn34Ile | P11166 |
| HOMO SAPIENS | 3 | 2668 | 5eqg | A | 77 | MET | p.Met77Val | P11166 |
| HOMO SAPIENS | 6 | 2668 | 5eqg | A | 91 | GLY | p.Gly91Asp | P11166 |
| HOMO SAPIENS | 6 | 2668 | 5eqg | A | 92 | ARG | p.Arg92Trp | P11166 |
| HOMO SAPIENS | 6 | 2668 | 5eqg | A | 93 | ARG | p.Arg93Trp | P11166 |
| HOMO SAPIENS | 0 | 2668 | 5eqg | A | 95 | SER | p.Ser95Ile | P11166 |
| HOMO SAPIENS | 3 | 2668 | 5eqg | A | 126 | ARG | p.Arg126His | P11166 |
| HOMO SAPIENS | 3 | 2668 | 5eqg | A | 126 | ARG | p.Arg126Cys | P11166 |
| HOMO SAPIENS | 4 | 2668 | 5eqg | A | 232 | ARG | p.Arg232Cys | P11166 |
| HOMO SAPIENS | 3 | 2668 | 5eqg | A | 249 | ARG | p.Arg249Gln | P11166 |
| HOMO SAPIENS | 3 | 2668 | 5eqg | A | 255 | LYS | p.Lys255Arg | P11166 |
| HOMO SAPIENS | 0 | 2668 | 5eqg | A | 275 | ALA | p.Ala275Thr | P11166 |
| HOMO SAPIENS | 4 | 2668 | 5eqg | A | 299 | GLU | p.Glu299Lys | P11166 |
| HOMO SAPIENS | 6 | 2668 | 5eqg | A | 314 | GLY | p.Gly314Ser | P11166 |
| HOMO SAPIENS | 7 | 2668 | 5eqg | A | 315 | ILE | p.Ile315Met | P11166 |
| HOMO SAPIENS | -1 | 2668 | 5eqg | A | 458 | ARG | p.Arg458Trp | P11166 |
| HOMO SAPIENS | -1 | 2668 | 5eqg | A | 468 | ARG | p.Arg468Trp | P11166 |
| HOMO SAPIENS | 3 | 2291 | 2amy | A | 31 | PHE | p.Phe31Leu | O15305 |
| HOMO SAPIENS | 7 | 2291 | 2amy | A | 32 | LEU | p.Leu32Arg | O15305 |
| HOMO SAPIENS | 7 | 2291 | 2amy | A | 65 | ASP | p.Asp65Tyr | O15305 |
| HOMO SAPIENS | 1 | 2291 | 2amy | A | 108 | ALA | p.Ala108Val | O15305 |
| HOMO SAPIENS | 3 | 2291 | 2amy | A | 113 | PRO | p.Pro113Leu | O15305 |
| HOMO SAPIENS | 3 | 2291 | 2amy | A | 113 | PRO | p.Pro113Ser | O15305 |
| HOMO SAPIENS | 6 | 2291 | 2amy | A | 117 | GLY | p.Gly117Arg | O15305 |
| HOMO SAPIENS | 6 | 2291 | 2amy | A | 119 | PHE | p.Phe119Leu | O15305 |
| HOMO SAPIENS | 6 | 2291 | 2amy | A | 123 | ARG | p.Arg123Gln | O15305 |
| HOMO SAPIENS | 0 | 2291 | 2amy | A | 129 | VAL | p.Val129Met | O15305 |
| HOMO SAPIENS | 7 | 2291 | 2amy | A | 139 | GLU | p.Glu139Lys | O15305 |
| HOMO SAPIENS | 5 | 2291 | 2amy | A | 141 | ARG | p.Arg141His | O15305 |
| HOMO SAPIENS | 7 | 2291 | 2amy | A | 144 | PHE | p.Phe144Leu | O15305 |
| HOMO SAPIENS | 6 | 2291 | 2amy | A | 148 | ASP | p.Asp148Asn | O15305 |
| HOMO SAPIENS | 6 | 2291 | 2amy | A | 153 | ILE | p.Ile153Thr | O15305 |
| HOMO SAPIENS | 2 | 2291 | 2amy | A | 157 | PHE | p.Phe157Ser | O15305 |
| HOMO SAPIENS | 1 | 2291 | 2amy | A | 162 | ARG | p.Arg162Trp | O15305 |
| HOMO SAPIENS | 6 | 2291 | 2amy | A | 188 | ASP | p.Asp188Gly | O15305 |
| HOMO SAPIENS | 5 | 2291 | 2amy | A | 194 | ARG | p.Arg194Ter | O15305 |
| HOMO SAPIENS | 0 | 2291 | 2amy | A | 207 | PHE | p.Phe207Ser | O15305 |
| HOMO SAPIENS | 5 | 2291 | 2amy | A | 208 | GLY | p.Gly208Ala | O15305 |
| HOMO SAPIENS | 1 | 2291 | 2amy | A | 216 | ASN | p.Asn216Ile | O15305 |
| HOMO SAPIENS | 5 | 2291 | 2amy | A | 218 | HIS | p.His218Leu | O15305 |
| HOMO SAPIENS | 6 | 2291 | 2amy | A | 223 | ASP | p.Asp223Glu | O15305 |
| HOMO SAPIENS | 6 | 2291 | 2amy | A | 226 | THR | p.Thr226Ser | O15305 |
| HOMO SAPIENS | 5 | 2291 | 2amy | A | 231 | VAL | p.Val231Met | O15305 |
| HOMO SAPIENS | 1 | 2291 | 2amy | A | 237 | THR | p.Thr237Met | O15305 |
| HOMO SAPIENS | 1 | 2291 | 2amy | A | 241 | CYS | p.Cys241Ser | O15305 |
| HOMO SAPIENS | 5 | 35907 | 1d6g | A | 21 | GLY | p.Gly21Arg | P32238 |
| HOMO SAPIENS | -1 | 33209 | 1x9d | A | 197 | PRO | p.Pro197Leu | Q9UKM7 |

| | | | | | | | | |
|---|---|---|---|---|---|---|---|---|
| HOMO SAPIENS | -1 | 33209 | 1x9d | A | 201 | PRO | p.Pro201Leu | Q9UKM7 |
| HOMO SAPIENS | 6 | 21305 | 4yep | B | 1326 | ARG | p.Arg1326Ter | P24043 |
| HOMO SAPIENS | 6 | 21305 | 4yep | B | 1350 | ARG | p.Arg1350Ter | P24043 |
| HOMO SAPIENS | 0 | 21553 | 3qyb | A | 1120 | VAL | p.Val1119Ala | O60343 |
| HOMO SAPIENS | 2 | 21553 | 3qyb | A | 1148 | THR | p.Thr1147Met | O60343 |
| HOMO SAPIENS | -1 | 30645 | 3m0c | C | -17 | TRP | p.Trp4Ter | P01130 |
| HOMO SAPIENS | -1 | 30645 | 3m0c | C | -17 | TRP | p.Trp4Ter | P01130 |
| HOMO SAPIENS | -1 | 30645 | 3m0c | C | 12 | GLN | p.Gln33Ter | P01130 |
| HOMO SAPIENS | -1 | 30645 | 3m0c | C | 15 | ASP | p.Asp36Glu | P01130 |
| HOMO SAPIENS | -1 | 30645 | 3m0c | C | 28 | SER | p.Ser49Arg | P01130 |
| HOMO SAPIENS | -1 | 30645 | 3m0c | C | 41 | THR | p.Thr62Met | P01130 |
| HOMO SAPIENS | -1 | 30645 | 3m0c | C | 69 | ASP | p.Asp90Asn | P01130 |
| HOMO SAPIENS | -1 | 30645 | 3m0c | C | 76 | ASN | p.Asn97Lys | P01130 |
| HOMO SAPIENS | -1 | 30645 | 3m0c | C | 78 | SER | p.Ser99Ter | P01130 |
| HOMO SAPIENS | -1 | 30645 | 3m0c | C | 81 | GLN | p.Gln102Ter | P01130 |
| HOMO SAPIENS | -1 | 30645 | 3m0c | C | 84 | PRO | p.Pro105Ser | P01130 |
| HOMO SAPIENS | -1 | 30645 | 3m0c | C | 194 | ARG | p.Arg215Cys | P01130 |
| HOMO SAPIENS | -1 | 30645 | 3m0c | C | 198 | GLY | p.Gly219Asp | P01130 |
| HOMO SAPIENS | -1 | 30645 | 3m0c | C | 201 | CYS | p.Cys222Tyr | P01130 |
| HOMO SAPIENS | -1 | 30645 | 3m0c | C | 201 | CYS | p.Cys222Ter | P01130 |
| HOMO SAPIENS | -1 | 30645 | 3m0c | C | 206 | ASP | p.Asp227Glu | P01130 |
| HOMO SAPIENS | -1 | 30645 | 3m0c | C | 249 | CYS | p.Cys270Ter | P01130 |
| HOMO SAPIENS | -1 | 30645 | 3m0c | C | 250 | VAL | p.Val271Ile | P01130 |
| HOMO SAPIENS | 0 | 30645 | 3m0c | C | 255 | CYS | p.Cys276Tyr | P01130 |
| HOMO SAPIENS | 0 | 30645 | 3m0c | C | 255 | CYS | p.Cys276Ter | P01130 |
| HOMO SAPIENS | 0 | 30645 | 3m0c | C | 256 | GLU | p.Glu277Lys | P01130 |
| HOMO SAPIENS | 0 | 30645 | 3m0c | C | 265 | SER | p.Ser286Arg | P01130 |
| HOMO SAPIENS | 0 | 30645 | 3m0c | C | 284 | TRP | p.Trp305Ter | P01130 |
| HOMO SAPIENS | 0 | 30645 | 3m0c | C | 292 | CYS | p.Cys313Ter | P01130 |
| HOMO SAPIENS | 1 | 30645 | 3m0c | C | 385 | ARG | p.Arg406Trp | P01130 |
| HOMO SAPIENS | 0 | 30645 | 3m0c | C | 395 | ARG | p.Arg416Gln | P01130 |
| HOMO SAPIENS | 0 | 30645 | 3m0c | C | 398 | TYR | p.Tyr419Ter | P01130 |
| HOMO SAPIENS | 1 | 30645 | 3m0c | C | 415 | VAL | p.Val436Ala | P01130 |
| HOMO SAPIENS | 1 | 30645 | 3m0c | C | 424 | ASP | p.Asp445Glu | P01130 |
| HOMO SAPIENS | 1 | 30645 | 3m0c | C | 425 | LEU | p.Leu446Val | P01130 |
| HOMO SAPIENS | -1 | 30645 | 3m0c | C | 440 | GLY | p.Gly461Cys | P01130 |
| HOMO SAPIENS | -1 | 30645 | 3m0c | C | 441 | VAL | p.Val462Ile | P01130 |
| HOMO SAPIENS | 1 | 30645 | 3m0c | C | 447 | VAL | p.Val468Ile | P01130 |
| HOMO SAPIENS | 1 | 30645 | 3m0c | C | 489 | THR | p.Thr510Met | P01130 |
| HOMO SAPIENS | 0 | 30645 | 3m0c | C | 524 | GLY | p.Gly545Glu | P01130 |
| HOMO SAPIENS | 1 | 30645 | 3m0c | C | 615 | GLY | p.Gly636Ser | P01130 |
| HOMO SAPIENS | 1 | 30645 | 3m0c | C | 625 | LEU | p.Leu646Ile | P01130 |
| HOMO SAPIENS | 1 | 30645 | 3m0c | C | 630 | ASP | p.Asp651Gly | P01130 |
| HOMO SAPIENS | 3 | 30645 | 3m0c | C | 646 | CYS | p.Cys667Tyr | P01130 |
| HOMO SAPIENS | 4 | 30645 | 3m0c | C | 654 | GLY | p.Gly675Ser | P01130 |
| HOMO SAPIENS | 6 | 30645 | 3m0c | C | 660 | CYS | p.Cys681Ter | P01130 |
| HOMO SAPIENS | 5 | 30645 | 3m0c | C | 676 | ALA | p.Ala697Pro | P01130 |
| HOMO SAPIENS | -1 | 30645 | 3m0c | C | 749 | GLN | p.Gln770Ter | P01130 |
| HOMO SAPIENS | -1 | 41323 | 4bxo | A | 1812 | PRO | p.Pro1812Ala | Q8IYD8 |
| HOMO SAPIENS | 5 | 8335 | 3h0t | C | 12 | GLY | p.Gly71Asp | P81172 |
| HOMO SAPIENS | 6 | 609 | 1bhg | B | 148 | PRO | p.Pro148Ser | P08236 |
| HOMO SAPIENS | 1 | 609 | 1bhg | B | 176 | LEU | p.Leu176Phe | P08236 |
| HOMO SAPIENS | 6 | 609 | 1bhg | B | 216 | ARG | p.Arg216Trp | P08236 |
| HOMO SAPIENS | 0 | 609 | 1bhg | B | 350 | LYS | p.Lys350Asn | P08236 |

| | | | | | | | | |
|---|---|---|---|---|---|---|---|---|
| HOMO SAPIENS | 1 | 609 | 1bhg | B | 354 | ALA | p.Ala354Val | P08236 |
| HOMO SAPIENS | 1 | 609 | 1bhg | B | 357 | ARG | p.Arg357Ter | P08236 |
| HOMO SAPIENS | 0 | 609 | 1bhg | B | 382 | ARG | p.Arg382Cys | P08236 |
| HOMO SAPIENS | 0 | 609 | 1bhg | B | 495 | TYR | p.Tyr495Cys | P08236 |
| HOMO SAPIENS | 2 | 609 | 1bhg | B | 507 | TRP | p.Trp507Ter | P08236 |
| HOMO SAPIENS | 0 | 609 | 1bhg | B | 577 | ARG | p.Arg577Leu | P08236 |
| HOMO SAPIENS | 1 | 609 | 1bhg | B | 611 | ARG | p.Arg611Trp | P08236 |
| HOMO SAPIENS | 0 | 609 | 1bhg | B | 619 | ALA | p.Ala619Val | P08236 |
| HOMO SAPIENS | 0 | 609 | 1bhg | B | 627 | TRP | p.Trp627Cys | P08236 |
| HOMO SAPIENS | 5 | 3422 | 1x9n | A | 566 | GLU | p.Glu566Lys | P18858 |
| HOMO SAPIENS | 2 | 3422 | 1x9n | A | 771 | ARG | p.Arg771Trp | P18858 |
| HOMO SAPIENS | 0 | 3279 | 5tby | B | 797 | ALA | p.Ala797Thr | P12883 |
| HOMO SAPIENS | -1 | 3279 | 5tby | B | 982 | MET | p.Met982Thr | P12883 |
| HOMO SAPIENS | -1 | 3279 | 5tby | B | 992 | LEU | p.Leu992Met | P12883 |
| HOMO SAPIENS | -1 | 3279 | 5tby | B | 1004 | GLN | p.Gln1004Glu | P12883 |
| HOMO SAPIENS | -1 | 3279 | 5tby | B | 1013 | GLU | p.Glu1013Lys | P12883 |
| HOMO SAPIENS | -1 | 3279 | 5tby | B | 1032 | ASP | p.Asp1032Asn | P12883 |
| HOMO SAPIENS | -1 | 3279 | 5tby | B | 1053 | ARG | p.Arg1053Gln | P12883 |
| HOMO SAPIENS | -1 | 3279 | 5tby | B | 1117 | GLU | p.Glu1117Ter | P12883 |
| HOMO SAPIENS | -1 | 3279 | 5tby | B | 1193 | ARG | p.Arg1193His | P12883 |
| HOMO SAPIENS | -1 | 3279 | 5tby | B | 1205 | GLU | p.Glu1205Lys | P12883 |
| HOMO SAPIENS | -1 | 3279 | 5tby | B | 1207 | ILE | p.Ile1207Met | P12883 |
| HOMO SAPIENS | -1 | 3279 | 5tby | B | 1250 | ARG | p.Arg1250Pro | P12883 |
| HOMO SAPIENS | -1 | 3279 | 5tby | B | 1259 | HIS | p.His1259Gln | P12883 |
| HOMO SAPIENS | -1 | 3279 | 5tby | B | 1272 | ASP | p.Asp1272Asn | P12883 |
| HOMO SAPIENS | -1 | 3279 | 5tby | B | 1295 | GLU | p.Glu1295Lys | P12883 |
| HOMO SAPIENS | -1 | 3279 | 5tby | B | 1335 | SER | p.Ser1335Leu | P12883 |
| HOMO SAPIENS | -1 | 3279 | 5tby | B | 1399 | GLU | p.Glu1399Asp | P12883 |
| HOMO SAPIENS | -1 | 3279 | 5tby | B | 1413 | SER | p.Ser1413Ter | P12883 |
| HOMO SAPIENS | -1 | 3279 | 5tby | B | 1459 | LYS | p.Lys1459Asn | P12883 |
| HOMO SAPIENS | -1 | 3279 | 5tby | B | 1467 | LEU | p.Leu1467Val | P12883 |
| HOMO SAPIENS | -1 | 3279 | 5tby | B | 1491 | SER | p.Ser1491Cys | P12883 |
| HOMO SAPIENS | -1 | 3279 | 5tby | B | 1496 | GLU | p.Glu1496Ala | P12883 |
| HOMO SAPIENS | -1 | 3279 | 5tby | B | 1500 | ARG | p.Arg1500Pro | P12883 |
| HOMO SAPIENS | -1 | 3279 | 5tby | B | 1606 | ARG | p.Arg1606His | P12883 |
| HOMO SAPIENS | -1 | 3279 | 5tby | B | 1664 | ASN | p.Asn1664Lys | P12883 |
| HOMO SAPIENS | -1 | 3279 | 5tby | B | 1712 | ARG | p.Arg1712Trp | P12883 |
| HOMO SAPIENS | -1 | 3279 | 5tby | B | 1769 | LEU | p.Leu1769Met | P12883 |
| HOMO SAPIENS | -1 | 3279 | 5tby | B | 1808 | GLY | p.Gly1808Ser | P12883 |
| HOMO SAPIENS | -1 | 3279 | 5tby | B | 1818 | ARG | p.Arg1818Trp | P12883 |
| HOMO SAPIENS | -1 | 3279 | 5tby | B | 1820 | ARG | p.Arg1820Trp | P12883 |
| HOMO SAPIENS | -1 | 3279 | 5tby | B | 1844 | GLU | p.Glu1844Lys | P12883 |
| HOMO SAPIENS | -1 | 3279 | 5tby | B | 1845 | ARG | p.Arg1845Trp | P12883 |
| HOMO SAPIENS | -1 | 3279 | 5tby | B | 1854 | THR | p.Thr1854Met | P12883 |
| HOMO SAPIENS | -1 | 3279 | 5tby | B | 1909 | ARG | p.Arg1909Trp | P12883 |
| HOMO SAPIENS | 2 | 2260 | 4kl1 | A | 553 | ASP | p.Asp553Asn | Q9Y3Q4 |
| HOMO SAPIENS | 0 | 40570 | 5lst | A | 522 | ARG | p.Arg522Cys | O94761 |
| HOMO SAPIENS | 0 | 40570 | 5lst | A | 522 | ARG | p.Arg522His | O94761 |
| HOMO SAPIENS | 0 | 40570 | 5lst | A | 591 | PRO | p.Pro591Leu | O94761 |
| HOMO SAPIENS | 0 | 40570 | 5lst | A | 793 | PRO | p.Pro793Ser | O94761 |
| HOMO SAPIENS | 4 | 40570 | 5lst | A | 799 | VAL | p.Val799Met | O94761 |
| HOMO SAPIENS | 0 | 40570 | 5lst | A | 1004 | ARG | p.Arg1004Trp | O94761 |
| HOMO SAPIENS | 0 | 40570 | 5lst | A | 1005 | ARG | p.Arg1005Gln | O94761 |
| HOMO SAPIENS | 0 | 40570 | 5lst | A | 1021 | ARG | p.Arg1021Gln | O94761 |

| | | | | | | | | |
|---|---|---|---|---|---|---|---|---|
| HOMO SAPIENS | 0 | 40570 | 5lst | A | 1043 | LEU | p.Leu1043Pro | O94761 |
| HOMO SAPIENS | 0 | 40570 | 5lst | A | 1045 | ALA | p.Ala1045Thr | O94761 |
| HOMO SAPIENS | 0 | 40570 | 5lst | A | 1105 | GLY | p.Gly1105Asp | O94761 |
| HOMO SAPIENS | 0 | 40570 | 5lst | A | 1105 | GLY | p.Gly1105Ser | O94761 |
| HOMO SAPIENS | -1 | 40570 | 5lst | A | 1113 | GLY | p.Gly1113Arg | O94761 |
| HOMO SAPIENS | -1 | 18865 | 1gxc | A | 64 | GLU | p.Glu64Lys | O96017 |
| HOMO SAPIENS | -1 | 11673 | 3dzu | D | 85 | PRO | p.Pro113Gln | P37231 |
| HOMO SAPIENS | 0 | 19383 | 3b96 | A | 63 | LYS | p.Lys103Arg | P49748 |
| HOMO SAPIENS | 4 | 19383 | 3b96 | A | 140 | ALA | p.Ala180Thr | P49748 |
| HOMO SAPIENS | 1 | 19383 | 3b96 | A | 145 | GLY | p.Gly185Ser | P49748 |
| HOMO SAPIENS | 0 | 19383 | 3b96 | A | 220 | THR | p.Thr260Met | P49748 |
| HOMO SAPIENS | 5 | 19383 | 3b96 | A | 243 | VAL | p.Val283Ala | P49748 |
| HOMO SAPIENS | 0 | 19383 | 3b96 | A | 249 | GLY | p.Gly289Arg | P49748 |
| HOMO SAPIENS | 0 | 19383 | 3b96 | A | 254 | GLY | p.Gly294Glu | P49748 |
| HOMO SAPIENS | 7 | 19383 | 3b96 | A | 326 | ARG | p.Arg366Cys | P49748 |
| HOMO SAPIENS | 7 | 19383 | 3b96 | A | 326 | ARG | p.Arg366His | P49748 |
| HOMO SAPIENS | 5 | 19383 | 3b96 | A | 342 | LYS | p.Lys382Gln | P49748 |
| HOMO SAPIENS | 0 | 19383 | 3b96 | A | 369 | THR | p.Thr409Met | P49748 |
| HOMO SAPIENS | 5 | 19383 | 3b96 | A | 376 | ALA | p.Ala416Ser | P49748 |
| HOMO SAPIENS | 0 | 19383 | 3b96 | A | 385 | ALA | p.Ala425Thr | P49748 |
| HOMO SAPIENS | 5 | 19383 | 3b96 | A | 399 | GLY | p.Gly439Asp | P49748 |
| HOMO SAPIENS | 7 | 19383 | 3b96 | A | 401 | GLY | p.Gly441Asp | P49748 |
| HOMO SAPIENS | 9 | 19383 | 3b96 | A | 413 | ARG | p.Arg453Gln | P49748 |
| HOMO SAPIENS | 2 | 19383 | 3b96 | A | 418 | PHE | p.Phe458Leu | P49748 |
| HOMO SAPIENS | 6 | 19383 | 3b96 | A | 419 | ARG | p.Arg459Trp | P49748 |
| HOMO SAPIENS | 6 | 19383 | 3b96 | A | 419 | ARG | p.Arg459Gln | P49748 |
| HOMO SAPIENS | -1 | 19383 | 3b96 | A | 471 | ARG | p.Arg511Gln | P49748 |
| HOMO SAPIENS | 0 | 19383 | 3b96 | A | 573 | ARG | p.Arg613Trp | P49748 |
| HOMO SAPIENS | 0 | 19383 | 3b96 | A | 575 | ARG | p.Arg615Gln | P49748 |
| HOMO SAPIENS | 0 | 22286 | 3k2s | A | 3 | PRO | p.Pro27His | P02647 |
| HOMO SAPIENS | 0 | 22286 | 3k2s | A | 4 | PRO | p.Pro28Arg | P02647 |
| HOMO SAPIENS | 0 | 22286 | 3k2s | A | 10 | ARG | p.Arg34Leu | P02647 |
| HOMO SAPIENS | 0 | 22286 | 3k2s | A | 26 | GLY | p.Gly50Arg | P02647 |
| HOMO SAPIENS | 0 | 22286 | 3k2s | A | 37 | ALA | p.Ala61Thr | P02647 |
| HOMO SAPIENS | 1 | 1622 | 2qrv | D | 631 | ARG | p.Arg635Trp | Q9Y6K1 |
| HOMO SAPIENS | 1 | 1622 | 2qrv | D | 900 | PRO | p.Pro904Leu | Q9Y6K1 |
| HOMO SAPIENS | 6 | 40796 | 1v18 | B | 1509 | LEU | p.Leu1509Val | P25054 |
| HOMO SAPIENS | 1 | 22375 | 4pki | G | 1214 | LYS | p.Asp214Asn | P28289 |
| HOMO SAPIENS | 1 | 16307 | 5mv8 | B | 483 | ALA | p.Ala483Ser | Q9Y6N9 |
| HOMO SAPIENS | 6 | 16307 | 5mv8 | B | 506 | ASN | p.Asn506Lys | Q9Y6N9 |
| HOMO SAPIENS | 4 | 16307 | 5mv8 | B | 519 | ASP | p.Glu519Asp | Q9Y6N9 |
| HOMO SAPIENS | 0 | 16307 | 5mv8 | B | 530 | GLY | p.Gly530Arg | Q9Y6N9 |
| HOMO SAPIENS | 4 | 36221 | 4cvo | A | 127 | VAL | p.Val127Ile | Q03468 |
| HOMO SAPIENS | 7 | 36221 | 4cvo | A | 134 | ARG | p.Arg134Trp | Q03468 |
| HOMO SAPIENS | 0 | 36221 | 4cvo | A | 156 | GLN | p.Gln156Ter | Q03468 |
| HOMO SAPIENS | 0 | 10829 | 4a11 | B | 13 | GLU | p.Glu13Ter | Q13216 |
| HOMO SAPIENS | 1 | 10829 | 4a11 | B | 100 | TYR | p.Tyr100Ter | Q13216 |
| HOMO SAPIENS | 0 | 10829 | 4a11 | B | 160 | ALA | p.Ala160Val | Q13216 |
| HOMO SAPIENS | 1 | 6759 | 5lev | A | 29 | ILE | p.Ile29Phe | Q9H3H5 |
| HOMO SAPIENS | 0 | 6759 | 5lev | A | 108 | MET | p.Met108Ile | Q9H3H5 |
| HOMO SAPIENS | 2 | 6759 | 5lev | A | 117 | VAL | p.Val117Ile | Q9H3H5 |
| HOMO SAPIENS | 1 | 6759 | 5lev | A | 170 | TYR | p.Tyr170Cys | Q9H3H5 |
| HOMO SAPIENS | 0 | 6759 | 5lev | A | 264 | GLY | p.Val264Gly | Q9H3H5 |
| HOMO SAPIENS | 0 | 6759 | 5lev | A | 332 | PHE | p.Phe332Val | Q9H3H5 |

| | | | | | | | | |
|---|---|---|---|---|---|---|---|---|
| HOMO SAPIENS | 7 | 6759 | 5lev | A | 393 | ILE | p.Ile393Val | Q9H3H5 |
| HOMO SAPIENS | -1 | 33473 | 5m5e | C | 9 | ASN | p.Asn31Ser | P31785 |
| HOMO SAPIENS | 0 | 26155 | 1ic8 | B | 98 | ALA | p.Ala98Val | P20823 |
| HOMO SAPIENS | 0 | 15329 | 4iqr | B | 127 | ARG | p.Arg136Trp | P41235 |
| HOMO SAPIENS | 0 | 20725 | 4bsj | A | 350 | GLU | p.Glu350Val | P35916 |
| HOMO SAPIENS | 0 | 20725 | 4bsj | A | 494 | THR | p.Thr494Ala | P35916 |
| HOMO SAPIENS | 0 | 20725 | 4bsj | A | 527 | ASN | p.Asn527Ser | P35916 |
| HOMO SAPIENS | 6 | 29959 | 1xiw | B | 47 | ARG | p.Arg68Ter | P04234 |
| HOMO SAPIENS | 2 | 29959 | 1xiw | B | 72 | CYS | p.Cys93Ter | P04234 |
| HOMO SAPIENS | 3 | 20242 | 5tjc | A | 102 | ARG | p.Arg102Ter | Q9GZU1 |
| HOMO SAPIENS | 3 | 20242 | 5tjc | A | 232 | THR | p.Thr232Pro | Q9GZU1 |
| HOMO SAPIENS | 7 | 9590 | 4ap8 | A | 50 | THR | p.Thr50Ala | O96007 |
| HOMO SAPIENS | 4 | 9590 | 4ap8 | A | 168 | GLU | p.Glu168Lys | O96007 |
| HOMO SAPIENS | -1 | 4297 | 5do7 | B | 19 | ASP | p.Asp19His | Q9H221 |
| HOMO SAPIENS | -1 | 4297 | 5do7 | B | 54 | TYR | p.Tyr54Cys | Q9H221 |
| HOMO SAPIENS | -1 | 4297 | 5do7 | B | 80 | CYS | p.Cys80Tyr | Q9H221 |
| HOMO SAPIENS | 5 | 4297 | 5do7 | B | 107 | SER | p.Ser107Ter | Q9H221 |
| HOMO SAPIENS | 1 | 4297 | 5do7 | B | 164 | ARG | p.Arg164Pro | Q9H221 |
| HOMO SAPIENS | 0 | 4297 | 5do7 | B | 210 | VAL | p.Val210Met | Q9H221 |
| HOMO SAPIENS | 0 | 4297 | 5do7 | B | 231 | PRO | p.Pro231Thr | Q9H221 |
| HOMO SAPIENS | 0 | 4297 | 5do7 | B | 263 | ARG | p.Arg263Gln | Q9H221 |
| HOMO SAPIENS | 0 | 4297 | 5do7 | B | 361 | TRP | p.Trp361Ter | Q9H221 |
| HOMO SAPIENS | 0 | 4297 | 5do7 | B | 400 | THR | p.Thr400Lys | Q9H221 |
| HOMO SAPIENS | 0 | 4297 | 5do7 | B | 549 | ALA | p.Ala549Pro | Q9H221 |
| HOMO SAPIENS | 0 | 4297 | 5do7 | B | 596 | LEU | p.Leu596Arg | Q9H221 |
| HOMO SAPIENS | 0 | 4297 | 5do7 | B | 632 | VAL | p.Val632Ala | Q9H221 |
| HOMO SAPIENS | 0 | 4297 | 5do7 | B | 658 | TYR | p.Tyr658Ter | Q9H221 |
| HOMO SAPIENS | 2 | 9356 | 4igg | B | 206 | GLN | p.Gln206His | P35221 |
| HOMO SAPIENS | 6 | 2179 | 2wk6 | A | 44 | ARG | p.Arg44Gln | P19971 |
| HOMO SAPIENS | 6 | 2179 | 2wk6 | A | 81 | ARG | p.Arg81Gln | P19971 |
| HOMO SAPIENS | 6 | 2179 | 2wk6 | A | 145 | GLY | p.Gly145Arg | P19971 |
| HOMO SAPIENS | 7 | 2179 | 2wk6 | A | 146 | ARG | p.Arg146His | P19971 |
| HOMO SAPIENS | 7 | 2179 | 2wk6 | A | 153 | GLY | p.Gly153Ser | P19971 |
| HOMO SAPIENS | 0 | 2179 | 2wk6 | A | 173 | MET | p.Met173Ile | P19971 |
| HOMO SAPIENS | 6 | 2179 | 2wk6 | A | 202 | ARG | p.Arg202Lys | P19971 |
| HOMO SAPIENS | 6 | 2179 | 2wk6 | A | 208 | VAL | p.Val208Met | P19971 |
| HOMO SAPIENS | 5 | 2179 | 2wk6 | A | 222 | LYS | p.Lys222Arg | P19971 |
| HOMO SAPIENS | 5 | 2179 | 2wk6 | A | 289 | GLU | p.Glu289Ala | P19971 |
| HOMO SAPIENS | 2 | 2179 | 2wk6 | A | 311 | GLY | p.Gly311Arg | P19971 |
| HOMO SAPIENS | 2 | 2179 | 2wk6 | A | 376 | ARG | p.Arg376Pro | P19971 |
| HOMO SAPIENS | 2 | 2179 | 2wk6 | A | 465 | ALA | p.Ala465Thr | P19971 |
| HOMO SAPIENS | -1 | 11856 | 3orh | A | 3 | ALA | p.Ala3Ser | Q14353 |
| HOMO SAPIENS | 1 | 11856 | 3orh | A | 20 | TRP | p.Trp20Ser | Q14353 |
| HOMO SAPIENS | 1 | 11856 | 3orh | A | 50 | MET | p.Met50Leu | Q14353 |
| HOMO SAPIENS | 1 | 11856 | 3orh | A | 110 | VAL | p.Val110Phe | Q14353 |
| HOMO SAPIENS | 8 | 11856 | 3orh | A | 164 | GLY | p.Gly164Asp | Q14353 |
| HOMO SAPIENS | 1 | 11856 | 3orh | A | 169 | CYS | p.Cys169Tyr | Q14353 |
| HOMO SAPIENS | 1 | 11856 | 3orh | A | 174 | TRP | p.Trp174Ter | Q14353 |
| HOMO SAPIENS | 2 | 11856 | 3orh | A | 196 | ALA | p.Ala196Val | Q14353 |
| HOMO SAPIENS | 1 | 11856 | 3orh | A | 209 | THR | p.Thr209Met | Q14353 |
| HOMO SAPIENS | 1 | 11856 | 3orh | A | 219 | ASP | p.Asp219Asn | Q14353 |
| HOMO SAPIENS | 1 | 11856 | 3orh | A | 224 | ALA | p.Ala224Thr | Q14353 |
| HOMO SAPIENS | 1 | 13244 | 2m6i | D | 218 | ARG | p.Arg246Leu | P23415 |
| HOMO SAPIENS | 1 | 13244 | 2m6i | D | 244 | ILE | p.Ile272Asn | P23415 |

| | | | | | | | | |
|---|---|---|---|---|---|---|---|---|
| HOMO SAPIENS | 5 | 13244 | 2m6i | D | 250 | PRO | p.Pro278Thr | P23415 |
| HOMO SAPIENS | 6 | 13244 | 2m6i | D | 266 | GLN | p.Gln294His | P23415 |
| HOMO SAPIENS | 1 | 13244 | 2m6i | D | 271 | ARG | p.Arg299Leu | P23415 |
| HOMO SAPIENS | 1 | 13244 | 2m6i | D | 276 | LYS | p.Lys304Glu | P23415 |
| HOMO SAPIENS | 1 | 13244 | 2m6i | D | 279 | TYR | p.Tyr307Cys | P23415 |
| HOMO SAPIENS | 0 | 13244 | 2m6i | D | 316 | GLY | p.Gly378Ser | P23415 |
| HOMO SAPIENS | 1 | 13244 | 2m6i | D | 400 | ARG | p.Arg428His | P23415 |
| HOMO SAPIENS | 3 | 1967 | 2i6b | A | 301 | ALA | p.Ala318Glu | P55263 |
| HOMO SAPIENS | 4 | 11096 | 1h8b | A | 61 | ALA | p.Ala882Thr | P35609 |
| HOMO SAPIENS | 0 | 15423 | 5ngz | A | 2 | GLN | p.Gln2Glu | Q9NPD8 |
| HOMO SAPIENS | 3 | 30475 | 3lpw | A | 21 | SER | p.Ser22894Tyr | Q8WZ42 |
| HOMO SAPIENS | 6 | 32687 | 2arf | A | 1041 | ARG | p.Arg1041Trp | P35670 |
| HOMO SAPIENS | 8 | 32687 | 2arf | A | 1061 | GLY | p.Gly1061Glu | P35670 |
| HOMO SAPIENS | 6 | 32687 | 2arf | A | 1064 | GLU | p.Glu1064Ala | P35670 |
| HOMO SAPIENS | 2 | 32687 | 2arf | A | 1069 | HIS | p.His1069Gln | P35670 |
| HOMO SAPIENS | 2 | 32687 | 2arf | A | 1088 | LEU | p.Leu1088Ter | P35670 |
| HOMO SAPIENS | 0 | 32687 | 2arf | A | 1099 | GLY | p.Gly1099Ser | P35670 |
| HOMO SAPIENS | 4 | 32687 | 2arf | A | 1106 | VAL | p.Val1106Asp | P35670 |
| HOMO SAPIENS | 2 | 32687 | 2arf | A | 1142 | GLN | p.Gln1142His | P35670 |
| HOMO SAPIENS | 8 | 32687 | 2arf | A | 1148 | ILE | p.Ile1148Thr | P35670 |
| HOMO SAPIENS | 6 | 32687 | 2arf | A | 1151 | ARG | p.Arg1151Cys | P35670 |
| HOMO SAPIENS | 4 | 32687 | 2arf | A | 1169 | MET | p.Met1169Val | P35670 |
| HOMO SAPIENS | 4 | 32687 | 2arf | A | 1173 | GLU | p.Glu1173Lys | P35670 |
| HOMO SAPIENS | 2 | 32687 | 2arf | A | 1176 | GLY | p.Gly1176Arg | P35670 |
| HOMO SAPIENS | 3 | 15544 | 5uq3 | C | 575 | PRO | p.Pro69Leu | P46527 |
| HOMO SAPIENS | 6 | 15544 | 5uq3 | C | 592 | GLU | p.Glu86Asp | P46527 |
| HOMO SAPIENS | 6 | 15544 | 5uq3 | C | 599 | ARG | p.Arg93Trp | P46527 |
| HOMO SAPIENS | -1 | 15544 | 5uq3 | C | 615 | VAL | p.Val109Gly | P46527 |
| HOMO SAPIENS | -1 | 15544 | 5uq3 | C | 631 | SER | p.Ser125Pro | P46527 |
| HOMO SAPIENS | -1 | 15544 | 5uq3 | C | 639 | PRO | p.Pro133Ser | P46527 |
| HOMO SAPIENS | 2 | 2703 | 4wxq | A | 245 | CYS | p.Cys245Tyr | Q99972 |
| HOMO SAPIENS | 0 | 2703 | 4wxq | A | 252 | GLY | p.Gly252Arg | Q99972 |
| HOMO SAPIENS | 1 | 2703 | 4wxq | A | 337 | GLN | p.Gln337Arg | Q99972 |
| HOMO SAPIENS | 5 | 2703 | 4wxq | A | 347 | TYR | p.Tyr347Ter | Q99972 |
| HOMO SAPIENS | 3 | 2703 | 4wxq | A | 364 | GLY | p.Gly364Val | Q99972 |
| HOMO SAPIENS | 2 | 2703 | 4wxq | A | 367 | GLY | p.Gly367Arg | Q99972 |
| HOMO SAPIENS | 2 | 2703 | 4wxq | A | 368 | GLN | p.Gln368Ter | Q99972 |
| HOMO SAPIENS | 6 | 2703 | 4wxq | A | 370 | PRO | p.Pro370Leu | Q99972 |
| HOMO SAPIENS | 1 | 2703 | 4wxq | A | 380 | ASP | p.Asp380His | Q99972 |
| HOMO SAPIENS | 8 | 2703 | 4wxq | A | 399 | GLY | p.Gly399Val | Q99972 |
| HOMO SAPIENS | 0 | 2703 | 4wxq | A | 423 | LYS | p.Lys423Glu | Q99972 |
| HOMO SAPIENS | 3 | 2703 | 4wxq | A | 433 | CYS | p.Cys433Arg | Q99972 |
| HOMO SAPIENS | 3 | 2703 | 4wxq | A | 437 | TYR | p.Tyr437His | Q99972 |
| HOMO SAPIENS | 2 | 2703 | 4wxq | A | 477 | ILE | p.Ile477Ser | Q99972 |
| HOMO SAPIENS | 3 | 2703 | 4wxq | A | 480 | ASN | p.Asn480Lys | Q99972 |
| HOMO SAPIENS | -1 | 34100 | 2fot | C | 1171 | GLN | p.Gln1173Pro | Q13813 |
| HOMO SAPIENS | 0 | 8773 | 1kw2 | A | 432 | ASP | p.Asp432Glu | P02774 |
| HOMO SAPIENS | 0 | 8773 | 1kw2 | A | 436 | LYS | p.Lys436Thr | P02774 |
| HOMO SAPIENS | 0 | 8402 | 4dzo | B | 649 | TYR | p.Tyr649Ter | Q9Y6D9 |
| HOMO SAPIENS | 0 | 39962 | 1nst | A | 611 | GLY | p.Gly611Ser | P52848 |
| HOMO SAPIENS | 2 | 22578 | 3dof | B | 45 | MET | p.Met45Arg | Q9Y2Y0 |
| HOMO SAPIENS | 1 | 23290 | 5j1s | A | 292 | ILE | p.Ile292Thr | O14656 |
| HOMO SAPIENS | 4 | 9505 | 4j1y | A | 534 | ARG | p.Arg534Trp | P09871 |
| HOMO SAPIENS | 9 | 260 | 2h57 | A | 31 | THR | p.Thr31Met | Q9H0F7 |

| | | | | | | | | |
|---|---|---|---|---|---|---|---|---|
| HOMO SAPIENS | 0 | 260 | 2h57 | A | 91 | ILE | p.Ile91Thr | Q9H0F7 |
| HOMO SAPIENS | 0 | 260 | 2h57 | A | 121 | ARG | p.Arg121His | Q9H0F7 |
| HOMO SAPIENS | 0 | 260 | 2h57 | A | 122 | ARG | p.Arg122Ter | Q9H0F7 |
| HOMO SAPIENS | 2 | 260 | 2h57 | A | 169 | GLY | p.Gly169Ala | Q9H0F7 |
| HOMO SAPIENS | 4 | 260 | 2h57 | A | 170 | LEU | p.Leu170Trp | Q9H0F7 |
| HOMO SAPIENS | 4 | 34023 | 2cr3 | A | 18 | GLY | p.Gly48Ser | P11362 |
| HOMO SAPIENS | 0 | 37667 | 3ii7 | A | 420 | ARG | p.Arg420Cys | Q8IXQ5 |
| HOMO SAPIENS | 0 | 41720 | 2k9u | A | 68 | ARG | p.Arg2364His | Q14315 |
| HOMO SAPIENS | 1 | 29250 | 5fmv | A | 568 | HIS | p.His568Gln | P08575 |
| HOMO SAPIENS | 1 | 1280 | 1fb1 | A | 69 | PRO | p.Pro69Leu | P30793 |
| HOMO SAPIENS | 3 | 1280 | 1fb1 | A | 88 | ARG | p.Arg88Trp | P30793 |
| HOMO SAPIENS | 2 | 1280 | 1fb1 | A | 108 | GLY | p.Gly108Asp | P30793 |
| HOMO SAPIENS | 4 | 1280 | 1fb1 | A | 134 | ASP | p.Asp134Val | P30793 |
| HOMO SAPIENS | 6 | 1280 | 1fb1 | A | 135 | ILE | p.Ile135Lys | P30793 |
| HOMO SAPIENS | 9 | 1280 | 1fb1 | A | 144 | HIS | p.His144Pro | P30793 |
| HOMO SAPIENS | 9 | 1280 | 1fb1 | A | 184 | ARG | p.Arg184His | P30793 |
| HOMO SAPIENS | 1 | 1280 | 1fb1 | A | 196 | ALA | p.Ala196Ser | P30793 |
| HOMO SAPIENS | 3 | 1280 | 1fb1 | A | 201 | GLY | p.Gly201Glu | P30793 |
| HOMO SAPIENS | 2 | 1280 | 1fb1 | A | 204 | VAL | p.Val204Ile | P30793 |
| HOMO SAPIENS | 2 | 1280 | 1fb1 | A | 211 | MET | p.Met211Ile | P30793 |
| HOMO SAPIENS | 0 | 1280 | 1fb1 | A | 221 | MET | p.Met221Thr | P30793 |
| HOMO SAPIENS | 3 | 1280 | 1fb1 | A | 224 | LYS | p.Lys224Arg | P30793 |
| HOMO SAPIENS | 2 | 1280 | 1fb1 | A | 249 | ARG | p.Arg249Ser | P30793 |
| HOMO SAPIENS | 8 | 41352 | 4k12 | A | 47 | ILE | p.Ile551Thr | P08603 |
| HOMO SAPIENS | 9 | 20703 | 5nir | A | 57 | CYS | p.Cys57Tyr | P02458 |
| HOMO SAPIENS | 9 | 20703 | 5nir | A | 64 | CYS | p.Cys64Ter | P02458 |
| HOMO SAPIENS | 0 | 20087 | 4cmn | A | 299 | MET | p.Met299Ile | Q01968 |
| HOMO SAPIENS | 1 | 20819 | 3aaf | A | 979 | ILE | p.Ile979Met | Q14191 |
| HOMO SAPIENS | 1 | 20819 | 3aaf | A | 984 | LEU | p.Leu984Ile | Q14191 |
| HOMO SAPIENS | 3 | 20819 | 3aaf | A | 987 | ARG | p.Arg987Ter | Q14191 |
| HOMO SAPIENS | -1 | 20819 | 3aaf | A | 1074 | PHE | p.Leu1074Phe | Q14191 |
| HOMO SAPIENS | 0 | 3619 | 3dem | A | 206 | ASN | p.Asn225Ser | P48740 |
| HOMO SAPIENS | 1 | 3619 | 3dem | A | 269 | ARG | p.Arg288Leu | P48740 |
| HOMO SAPIENS | 0 | 3105 | 3ooi | A | 164 | TYR | p.Tyr2015His | Q96L73 |
| HOMO SAPIENS | 6 | 15828 | 2lkx | A | 16 | LEU | p.Leu100Gln | Q99697 |
| HOMO SAPIENS | 8 | 15828 | 2lkx | A | 24 | ARG | p.Arg108His | Q99697 |
| HOMO SAPIENS | 2 | 15828 | 2lkx | A | 30 | THR | p.Thr114Pro | Q99697 |
| HOMO SAPIENS | 8 | 15828 | 2lkx | A | 45 | VAL | p.Val129Leu | Q99697 |
| HOMO SAPIENS | 8 | 15828 | 2lkx | A | 46 | ARG | p.Arg130Trp | Q99697 |
| HOMO SAPIENS | 9 | 15828 | 2lkx | A | 53 | ARG | p.Arg137Pro | Q99697 |
| HOMO SAPIENS | 9 | 13017 | 4irq | A | 141 | ARG | p.Arg141Trp | Q9UBV7 |
| HOMO SAPIENS | 0 | 13017 | 4irq | A | 186 | ALA | p.Ala186Asp | Q9UBV7 |
| HOMO SAPIENS | 0 | 13017 | 4irq | A | 206 | LEU | p.Leu206Pro | Q9UBV7 |
| HOMO SAPIENS | 1 | 13017 | 4irq | A | 214 | CYS | p.Cys214Tyr | Q9UBV7 |
| HOMO SAPIENS | 0 | 13017 | 4irq | A | 270 | ARG | p.Arg270Cys | Q9UBV7 |
| HOMO SAPIENS | -1 | 13053 | 4mqw | X | 307 | ALA | p.Ala307Thr | P23945 |
| HOMO SAPIENS | 0 | 19406 | 4b3g | A | 43 | ARG | p.Arg43Ter | P38935 |
| HOMO SAPIENS | 0 | 19406 | 4b3g | A | 46 | CYS | p.Cys46Ter | P38935 |
| HOMO SAPIENS | 0 | 19406 | 4b3g | A | 75 | ALA | p.Ala75Thr | P38935 |
| HOMO SAPIENS | 0 | 19406 | 4b3g | A | 201 | LEU | p.Leu201Ser | P38935 |
| HOMO SAPIENS | 3 | 19406 | 4b3g | A | 213 | HIS | p.His213Arg | P38935 |
| HOMO SAPIENS | 0 | 19406 | 4b3g | A | 275 | ILE | p.Ile275Val | P38935 |
| HOMO SAPIENS | -1 | 19406 | 4b3g | A | 320 | ARG | p.Arg320Ter | P38935 |
| HOMO SAPIENS | 0 | 19406 | 4b3g | A | 361 | LEU | p.Leu361Pro | P38935 |

| | | | | | | | | |
|---|---|---|---|---|---|---|---|---|
| HOMO SAPIENS | 0 | 19406 | 4b3g | A | 369 | PHE | p.Phe369Leu | P38935 |
| HOMO SAPIENS | 0 | 19406 | 4b3g | A | 493 | THR | p.Thr493Ile | P38935 |
| HOMO SAPIENS | 0 | 19406 | 4b3g | A | 496 | CYS | p.Cys496Ter | P38935 |
| HOMO SAPIENS | 0 | 19406 | 4b3g | A | 514 | GLU | p.Glu514Lys | P38935 |
| HOMO SAPIENS | 0 | 19406 | 4b3g | A | 528 | ALA | p.Ala528Thr | P38935 |
| HOMO SAPIENS | 0 | 19406 | 4b3g | A | 531 | PRO | p.Pro531Thr | P38935 |
| HOMO SAPIENS | 0 | 19406 | 4b3g | A | 580 | VAL | p.Val580Ile | P38935 |
| HOMO SAPIENS | 6 | 19406 | 4b3g | A | 603 | ARG | p.Arg603His | P38935 |
| HOMO SAPIENS | 0 | 21501 | 5jde | A | 40 | SER | p.Ser2831Asn | Q8WZ42 |
| HOMO SAPIENS | 5 | 21501 | 5jde | A | 189 | ALA | p.Ala2980Thr | Q8WZ42 |
| HOMO SAPIENS | 6 | 15008 | 3mop | H | 12 | ARG | p.Arg12Cys | Q9NWZ3 |
| HOMO SAPIENS | 8 | 17455 | 3jtc | C | 15 | ARG | p.Arg57Trp | P04070 |
| HOMO SAPIENS | -1 | 17455 | 3jtc | C | 20 | CGU | p.Glu62Ala | P04070 |
| HOMO SAPIENS | 0 | 27795 | 5gox | B | 625 | TYR | p.Tyr625Ter | Q92878 |
| HOMO SAPIENS | 0 | 27795 | 5gox | B | 653 | SER | p.Ser653Ter | Q92878 |
| HOMO SAPIENS | -1 | 6558 | 5nug | A | 137 | VAL | p.Val137Ile | Q14204 |
| HOMO SAPIENS | -1 | 6558 | 5nug | A | 567 | ARG | p.Arg567His | Q14204 |
| HOMO SAPIENS | -1 | 6558 | 5nug | A | 1140 | THR | p.Thr1140Met | Q14204 |
| HOMO SAPIENS | -1 | 6558 | 5nug | A | 1991 | ASP | p.Asp1991Asn | Q14204 |
| HOMO SAPIENS | -1 | 6558 | 5nug | A | 2401 | LYS | p.Lys2401Asn | Q14204 |
| HOMO SAPIENS | 0 | 6558 | 5nug | A | 2488 | ARG | p.Arg2488His | Q14204 |
| HOMO SAPIENS | 0 | 6558 | 5nug | A | 3907 | HIS | p.His3907Gln | Q14204 |
| HOMO SAPIENS | 0 | 6558 | 5nug | A | 3981 | THR | p.Thr3981Arg | Q14204 |
| HOMO SAPIENS | 0 | 6558 | 5nug | A | 4394 | THR | p.Thr4394Met | Q14204 |
| HOMO SAPIENS | 0 | 6558 | 5nug | A | 4573 | ASN | p.Asn4573Ser | Q14204 |
| HOMO SAPIENS | 0 | 6558 | 5nug | A | 4603 | SER | p.Ser4603Ile | Q14204 |
| HOMO SAPIENS | 1 | 4244 | 1umk | A | 57 | ARG | p.Arg58Gln | P00387 |
| HOMO SAPIENS | 1 | 4244 | 1umk | A | 72 | LEU | p.Leu73Pro | P00387 |
| HOMO SAPIENS | 1 | 4244 | 1umk | A | 105 | VAL | p.Val106Met | P00387 |
| HOMO SAPIENS | 0 | 4244 | 1umk | A | 116 | THR | p.Thr117Ser | P00387 |
| HOMO SAPIENS | 8 | 4244 | 1umk | A | 127 | SER | p.Ser128Pro | P00387 |
| HOMO SAPIENS | 0 | 4244 | 1umk | A | 148 | LEU | p.Leu149Pro | P00387 |
| HOMO SAPIENS | 0 | 4244 | 1umk | A | 159 | ARG | p.Arg160Ter | P00387 |
| HOMO SAPIENS | 0 | 4244 | 1umk | A | 203 | CYS | p.Cys204Arg | P00387 |
| HOMO SAPIENS | 0 | 4244 | 1umk | A | 203 | CYS | p.Cys204Tyr | P00387 |
| HOMO SAPIENS | 2 | 4244 | 1umk | A | 239 | ASP | p.Asp240Gly | P00387 |
| HOMO SAPIENS | 0 | 4244 | 1umk | A | 291 | GLY | p.Gly292Asp | P00387 |
| HOMO SAPIENS | 4 | 34495 | 1s35 | A | 1151 | ASP | p.Asn1151Asp | P11277 |
| HOMO SAPIENS | 3 | 30110 | 3zqs | A | 224 | THR | p.Thr224Ala | Q9NW38 |
| HOMO SAPIENS | 3 | 17588 | 5dse | B | 53 | LEU | p.Leu53Pro | Q9BYI3 |
| HOMO SAPIENS | 0 | 1564 | 4yqf | B | 72 | HIS | p.His347Gln | Q9UHD8 |
| HOMO SAPIENS | 0 | 17673 | 4jnw | A | 32 | ARG | p.Arg32186His | Q8WZ42 |
| HOMO SAPIENS | 3 | 17673 | 4jnw | A | 140 | ILE | p.Ile32294Val | Q8WZ42 |
| HOMO SAPIENS | 5 | 17673 | 4jnw | A | 269 | ALA | p.Ala32423Thr | Q8WZ42 |
| HOMO SAPIENS | 3 | 12858 | 3s97 | C | 302 | LEU | p.Leu302Pro | Q12860 |
| HOMO SAPIENS | 0 | 6147 | 4nn0 | A | 114 | ARG | p.Arg114Gln | Q9BXJ0 |
| HOMO SAPIENS | 0 | 7549 | 1tr2 | A | 277 | LEU | p.Leu277Met | P18206 |
| HOMO SAPIENS | 1 | 7549 | 1tr2 | A | 413 | ALA | p.Ala413Thr | P18206 |
| HOMO SAPIENS | 1 | 7549 | 1tr2 | A | 519 | ILE | p.Ile519Leu | P18206 |
| HOMO SAPIENS | 5 | 7549 | 1tr2 | A | 547 | ARG | p.Arg547Ter | P18206 |
| HOMO SAPIENS | 2 | 7549 | 1tr2 | A | 636 | HIS | p.His636Arg | P18206 |
| HOMO SAPIENS | 2 | 7549 | 1tr2 | A | 824 | ILE | p.Ile824Met | P18206 |
| HOMO SAPIENS | 2 | 13900 | 3gro | A | 45 | CYS | p.Cys45Tyr | P50897 |
| HOMO SAPIENS | 2 | 13900 | 3gro | A | 75 | THR | p.Thr75Pro | P50897 |

| | | | | | | | | |
|---|---|---|---|---|---|---|---|---|
| HOMO SAPIENS | 2 | 13900 | 3gro | A | 79 | ASP | p.Asp79Gly | P50897 |
| HOMO SAPIENS | 5 | 13900 | 3gro | A | 91 | GLN | p.Gln91Pro | P50897 |
| HOMO SAPIENS | 5 | 13900 | 3gro | A | 96 | CYS | p.Cys96Tyr | P50897 |
| HOMO SAPIENS | 0 | 13900 | 3gro | A | 104 | LYS | p.Lys104Ter | P50897 |
| HOMO SAPIENS | 2 | 13900 | 3gro | A | 108 | GLY | p.Gly108Arg | P50897 |
| HOMO SAPIENS | 2 | 13900 | 3gro | A | 109 | TYR | p.Tyr109Asp | P50897 |
| HOMO SAPIENS | 0 | 13900 | 3gro | A | 122 | ARG | p.Arg122Trp | P50897 |
| HOMO SAPIENS | 0 | 13900 | 3gro | A | 138 | SER | p.Ser138Leu | P50897 |
| HOMO SAPIENS | 5 | 13900 | 3gro | A | 151 | ARG | p.Arg151Ter | P50897 |
| HOMO SAPIENS | 7 | 13900 | 3gro | A | 164 | ARG | p.Arg164Ter | P50897 |
| HOMO SAPIENS | 0 | 13900 | 3gro | A | 177 | GLN | p.Gln177Glu | P50897 |
| HOMO SAPIENS | 0 | 13900 | 3gro | A | 181 | VAL | p.Val181Leu | P50897 |
| HOMO SAPIENS | 0 | 13900 | 3gro | A | 184 | GLU | p.Glu184Lys | P50897 |
| HOMO SAPIENS | 5 | 13900 | 3gro | A | 186 | TRP | p.Trp186Ter | P50897 |
| HOMO SAPIENS | 0 | 13900 | 3gro | A | 219 | LEU | p.Leu219Gln | P50897 |
| HOMO SAPIENS | 0 | 13900 | 3gro | A | 222 | LEU | p.Leu222Pro | P50897 |
| HOMO SAPIENS | 5 | 13900 | 3gro | A | 296 | TRP | p.Trp296Ter | P50897 |
| HOMO SAPIENS | 6 | 35028 | 5mwb | A | 444 | CYS | p.Cys444Tyr | Q04721 |
| HOMO SAPIENS | 0 | 35028 | 5mwb | A | 514 | ASN | p.Asn514Ser | Q04721 |
| HOMO SAPIENS | 5 | 1483 | 4gl2 | A | 822 | ARG | p.Arg822Gln | Q9BYX4 |
| HOMO SAPIENS | 0 | 1483 | 4gl2 | A | 843 | ARG | p.His843Arg | Q9BYX4 |
| HOMO SAPIENS | -1 | 28838 | 3cl3 | D | 153 | LEU | p.Leu153Arg | Q9Y6K9 |
| HOMO SAPIENS | 2 | 22563 | 4zel | B | 79 | ARG | p.Arg79Trp | P09172 |
| HOMO SAPIENS | 2 | 22563 | 4zel | B | 101 | VAL | p.Val101Met | P09172 |
| HOMO SAPIENS | 2 | 22563 | 4zel | B | 114 | ASP | p.Asp114Glu | P09172 |
| HOMO SAPIENS | 0 | 22563 | 4zel | B | 201 | ASN | p.Asn201Ser | P09172 |
| HOMO SAPIENS | 2 | 22563 | 4zel | B | 211 | ALA | p.Ala211Thr | P09172 |
| HOMO SAPIENS | -1 | 22563 | 4zel | B | 290 | ASP | p.Asp290Asn | P09172 |
| HOMO SAPIENS | 2 | 22563 | 4zel | B | 318 | ALA | p.Ala318Ser | P09172 |
| HOMO SAPIENS | 2 | 22563 | 4zel | B | 345 | ASP | p.Asp345Asn | P09172 |
| HOMO SAPIENS | 2 | 22563 | 4zel | B | 362 | ALA | p.Ala362Glu | P09172 |
| HOMO SAPIENS | 2 | 22563 | 4zel | B | 549 | ARG | p.Arg549Cys | P09172 |
| HOMO SAPIENS | 9 | 9780 | 1y6k | R | 36 | TYR | p.Tyr57Cys | Q13651 |
| HOMO SAPIENS | 1 | 9780 | 1y6k | R | 40 | LEU | p.Leu61Val | Q13651 |
| HOMO SAPIENS | 1 | 9780 | 1y6k | R | 63 | THR | p.Thr84Ile | Q13651 |
| HOMO SAPIENS | 9 | 9780 | 1y6k | R | 80 | ARG | p.Arg101Trp | Q13651 |
| HOMO SAPIENS | 0 | 9780 | 1y6k | R | 120 | GLY | p.Gly141Arg | Q13651 |
| HOMO SAPIENS | 0 | 9780 | 1y6k | R | 138 | SER | p.Ser159Gly | Q13651 |
| HOMO SAPIENS | 0 | 9780 | 1y6k | R | 203 | ILE | p.Ile224Val | Q13651 |
| HOMO SAPIENS | 0 | 16019 | 3rep | A | 211 | ARG | p.Arg211Cys | Q13418 |
| HOMO SAPIENS | 3 | 7879 | 5ec3 | A | 33 | ALA | p.Ala33Thr | P32754 |
| HOMO SAPIENS | 4 | 7879 | 5ec3 | A | 113 | ARG | p.Arg113Gln | P32754 |
| HOMO SAPIENS | 3 | 7879 | 5ec3 | A | 340 | VAL | p.Val340Leu | P32754 |
| HOMO SAPIENS | 0 | 8904 | 1qk9 | A | 6 | LYS | p.Lys82Arg | P51608 |
| HOMO SAPIENS | 4 | 8904 | 1qk9 | A | 24 | LEU | p.Leu100Val | P51608 |
| HOMO SAPIENS | 7 | 8904 | 1qk9 | A | 30 | ARG | p.Arg106Trp | P51608 |
| HOMO SAPIENS | 6 | 8904 | 1qk9 | A | 57 | ARG | p.Arg133Cys | P51608 |
| HOMO SAPIENS | 3 | 8904 | 1qk9 | A | 61 | GLU | p.Glu137Gly | P51608 |
| HOMO SAPIENS | 2 | 8904 | 1qk9 | A | 64 | ALA | p.Ala140Val | P51608 |
| HOMO SAPIENS | 5 | 8904 | 1qk9 | A | 79 | PHE | p.Phe155Ser | P51608 |
| HOMO SAPIENS | 3 | 8904 | 1qk9 | A | 82 | THR | p.Thr158Met | P51608 |
| HOMO SAPIENS | 1 | 23121 | 3e9k | A | 198 | THR | p.Thr198Ala | Q16719 |
| HOMO SAPIENS | -1 | 26090 | 3tje | F | 19 | LEU | p.Leu35Met | P25445 |
| HOMO SAPIENS | 8 | 26090 | 3tje | F | 106 | THR | p.Thr122Ile | P25445 |

| | | | | | | | | |
|---|---|---|---|---|---|---|---|---|
| HOMO SAPIENS | 2 | 36404 | 5cqr | A | 765 | GLY | p.Gly765Glu | O95163 |
| HOMO SAPIENS | 2 | 36404 | 5cqr | A | 816 | ILE | p.Ile816Leu | O95163 |
| HOMO SAPIENS | 2 | 36404 | 5cqr | A | 830 | ILE | p.Ile830Met | O95163 |
| HOMO SAPIENS | 5 | 36404 | 5cqr | A | 939 | TYR | p.Tyr939Ter | O95163 |
| HOMO SAPIENS | 2 | 36404 | 5cqr | A | 952 | LYS | p.Lys952Ile | O95163 |
| HOMO SAPIENS | 5 | 36404 | 5cqr | A | 1013 | GLY | p.Gly1013Ser | O95163 |
| HOMO SAPIENS | 5 | 36404 | 5cqr | A | 1072 | CYS | p.Cys1072Ser | O95163 |
| HOMO SAPIENS | -1 | 36404 | 5cqr | A | 1158 | PRO | p.Pro1158Leu | O95163 |
| HOMO SAPIENS | 1 | 34149 | 2k40 | A | 18 | ASN | p.Asn125Ser | Q9UBX0 |
| HOMO SAPIENS | 0 | 34149 | 2k40 | A | 42 | LEU | p.Glu149Lys | Q9UBX0 |
| HOMO SAPIENS | 9 | 34149 | 2k40 | A | 53 | ARG | p.Arg160Cys | Q9UBX0 |
| HOMO SAPIENS | 0 | 34149 | 2k40 | A | 63 | SER | p.Ser170Leu | Q9UBX0 |
| HOMO SAPIENS | 2 | 2468 | 3vp6 | A | 484 | LEU | p.Leu484Met | Q99259 |
| HOMO SAPIENS | 0 | 2468 | 3vp6 | A | 532 | ARG | p.Arg532Gln | Q99259 |
| HOMO SAPIENS | 5 | 1253 | 5xtd | S | 8 | GLY | p.Gly8Arg | O15239 |
| HOMO SAPIENS | 7 | 1253 | 5xtd | S | 32 | GLY | p.Gly32Arg | O15239 |
| HOMO SAPIENS | 7 | 1253 | 5xtd | S | 37 | ARG | p.Arg37Ser | O15239 |
| HOMO SAPIENS | -1 | 13292 | 3u1k | A | 121 | VAL | p.Ile121Val | Q8TCS8 |
| HOMO SAPIENS | 2 | 13292 | 3u1k | A | 165 | PRO | p.Pro165Ser | Q8TCS8 |
| HOMO SAPIENS | 2 | 13292 | 3u1k | A | 192 | ARG | p.Arg192Gln | Q8TCS8 |
| HOMO SAPIENS | 1 | 13292 | 3u1k | A | 315 | LYS | p.Lys315Arg | Q8TCS8 |
| HOMO SAPIENS | 4 | 13292 | 3u1k | A | 509 | VAL | p.Val509Ile | Q8TCS8 |
| HOMO SAPIENS | 7 | 13292 | 3u1k | A | 531 | THR | p.Thr531Arg | Q8TCS8 |
| HOMO SAPIENS | 0 | 13292 | 3u1k | A | 590 | ASN | p.Asn590Asp | Q8TCS8 |
| HOMO SAPIENS | 0 | 27048 | 4p5w | B | 184 | ARG | p.Arg184Ter | Q9UJS0 |
| HOMO SAPIENS | 0 | 27048 | 4p5w | B | 225 | SER | p.Ser225Ter | Q9UJS0 |
| HOMO SAPIENS | -1 | 27048 | 4p5w | B | 631 | ALA | p.Arg360Ter | Q9UJS0 |
| HOMO SAPIENS | 0 | 24240 | 4mdr | A | 318 | ARG | p.Arg318Ter | O00189 |
| HOMO SAPIENS | 0 | 24240 | 4mdr | A | 338 | ARG | p.Arg338Ter | O00189 |
| HOMO SAPIENS | 0 | 13429 | 4cg4 | A | 479 | PHE | p.Phe479Leu | O15553 |
| HOMO SAPIENS | 0 | 13429 | 4cg4 | A | 487 | VAL | p.Val487Leu | O15553 |
| HOMO SAPIENS | 3 | 13429 | 4cg4 | A | 653 | ARG | p.Arg653His | O15553 |
| HOMO SAPIENS | 1 | 13429 | 4cg4 | A | 680 | MET | p.Met680Ile | O15553 |
| HOMO SAPIENS | 1 | 13429 | 4cg4 | A | 688 | TYR | p.Tyr688Ter | O15553 |
| HOMO SAPIENS | 0 | 13429 | 4cg4 | A | 694 | MET | p.Met694Ile | O15553 |
| HOMO SAPIENS | 0 | 13429 | 4cg4 | A | 694 | MET | p.Met694Val | O15553 |
| HOMO SAPIENS | 1 | 13429 | 4cg4 | A | 726 | VAL | p.Val726Ala | O15553 |
| HOMO SAPIENS | 0 | 13429 | 4cg4 | A | 761 | ARG | p.Arg761His | O15553 |
| HOMO SAPIENS | 0 | 2079 | 4okh | A | 691 | GLY | p.Gly691Arg | P20807 |
| HOMO SAPIENS | 2 | 2079 | 4okh | A | 748 | ARG | p.Arg748Ter | P20807 |
| HOMO SAPIENS | 0 | 2079 | 4okh | A | 763 | TYR | p.Tyr763Cys | P20807 |
| HOMO SAPIENS | 1 | 2079 | 4okh | A | 769 | ARG | p.Arg769Gln | P20807 |
| HOMO SAPIENS | 9 | 2079 | 4okh | A | 777 | ILE | p.Ile777Val | P20807 |
| HOMO SAPIENS | 8 | 2079 | 4okh | A | 778 | ASP | p.Asp778His | P20807 |
| HOMO SAPIENS | 5 | 2079 | 4okh | A | 780 | ASP | p.Asp780His | P20807 |
| HOMO SAPIENS | 1 | 1200 | 2gi7 | A | 142 | ARG | p.Arg162Trp | Q9HCN6 |
| HOMO SAPIENS | 4 | 9199 | 3ezq | C | 260 | ASP | p.Asp260Val | P25445 |
| HOMO SAPIENS | 0 | 36522 | 5ms9 | A | 122 | ARG | p.Arg122Cys | P35555 |
| HOMO SAPIENS | 0 | 36522 | 5ms9 | A | 240 | ARG | p.Arg240Cys | P35555 |
| HOMO SAPIENS | 0 | 17448 | 3g2f | A | 225 | ARG | p.Arg225His | Q13873 |
| HOMO SAPIENS | 0 | 17448 | 3g2f | A | 347 | CYS | p.Cys347Tyr | Q13873 |
| HOMO SAPIENS | 3 | 17448 | 3g2f | A | 485 | ASP | p.Asp485Gly | Q13873 |
| HOMO SAPIENS | 7 | 17448 | 3g2f | A | 491 | ARG | p.Arg491Gln | Q13873 |
| HOMO SAPIENS | 7 | 17448 | 3g2f | A | 491 | ARG | p.Arg491Trp | Q13873 |

| | | | | | | | | |
|---|---|---|---|---|---|---|---|---|
| HOMO SAPIENS | 0 | 4861 | 4ux8 | A | 56 | LEU | p.Leu56Met | P07949 |
| HOMO SAPIENS | 1 | 4861 | 4ux8 | A | 88 | ILE | p.Ile88Leu | P07949 |
| HOMO SAPIENS | 0 | 4861 | 4ux8 | A | 114 | ARG | p.Arg114His | P07949 |
| HOMO SAPIENS | 5 | 4861 | 4ux8 | A | 180 | ARG | p.Arg180Gln | P07949 |
| HOMO SAPIENS | 0 | 4861 | 4ux8 | A | 208 | GLU | p.Glu208Asp | P07949 |
| HOMO SAPIENS | 0 | 4861 | 4ux8 | A | 373 | ALA | p.Ala373Val | P07949 |
| HOMO SAPIENS | 0 | 4861 | 4ux8 | A | 375 | LEU | p.Leu375Gln | P07949 |
| HOMO SAPIENS | -1 | 4861 | 4ux8 | A | 386 | ALA | p.Ala386Val | P07949 |
| HOMO SAPIENS | 0 | 4861 | 4ux8 | A | 446 | GLY | p.Gly446Arg | P07949 |
| HOMO SAPIENS | 0 | 4861 | 4ux8 | A | 480 | GLU | p.Glu480Lys | P07949 |
| HOMO SAPIENS | 0 | 4861 | 4ux8 | A | 489 | ASP | p.Asp489Asn | P07949 |
| HOMO SAPIENS | -1 | 4861 | 4ux8 | A | 510 | ALA | p.Ala510Val | P07949 |
| HOMO SAPIENS | -1 | 4861 | 4ux8 | A | 533 | GLY | p.Gly533Cys | P07949 |
| HOMO SAPIENS | -1 | 4861 | 4ux8 | A | 567 | ASP | p.Asp567Asn | P07949 |
| HOMO SAPIENS | -1 | 4861 | 4ux8 | A | 609 | CYS | p.Cys609Tyr | P07949 |
| HOMO SAPIENS | -1 | 4861 | 4ux8 | A | 609 | CYS | p.Cys609Gly | P07949 |
| HOMO SAPIENS | -1 | 4861 | 4ux8 | A | 611 | CYS | p.Cys611Trp | P07949 |
| HOMO SAPIENS | -1 | 4861 | 4ux8 | A | 618 | CYS | p.Cys618Gly | P07949 |
| HOMO SAPIENS | -1 | 4861 | 4ux8 | A | 618 | CYS | p.Cys618Phe | P07949 |
| HOMO SAPIENS | -1 | 4861 | 4ux8 | A | 620 | CYS | p.Cys620Gly | P07949 |
| HOMO SAPIENS | -1 | 4861 | 4ux8 | A | 620 | CYS | p.Cys620Ser | P07949 |
| HOMO SAPIENS | -1 | 4861 | 4ux8 | A | 620 | CYS | p.Cys620Trp | P07949 |
| HOMO SAPIENS | -1 | 4861 | 4ux8 | A | 623 | GLU | p.Glu623Lys | P07949 |
| HOMO SAPIENS | -1 | 4861 | 4ux8 | A | 631 | ASP | p.Asp631Gly | P07949 |
| HOMO SAPIENS | -1 | 4861 | 4ux8 | A | 631 | ASP | p.Asp631Glu | P07949 |
| HOMO SAPIENS | -1 | 4861 | 4ux8 | A | 634 | CYS | p.Cys634Gly | P07949 |
| HOMO SAPIENS | -1 | 4861 | 4ux8 | A | 634 | CYS | p.Cys634Phe | P07949 |
| HOMO SAPIENS | -1 | 4861 | 4ux8 | A | 634 | CYS | p.Cys634Trp | P07949 |
| HOMO SAPIENS | 0 | 36308 | 4nkq | B | 123 | PRO | p.Pro142Leu | P15509 |
| HOMO SAPIENS | 0 | 36308 | 4nkq | B | 145 | ARG | p.Arg164Gln | P15509 |
| HOMO SAPIENS | -1 | 36308 | 4nkq | B | 177 | GLY | p.Gly196Arg | P15509 |
| HOMO SAPIENS | 0 | 36006 | 1oqa | A | 107 | HIS | p.His1860Pro | P38398 |
| HOMO SAPIENS | 7 | 15394 | 2ozo | A | 465 | ARG | p.Arg465His | P43403 |
| HOMO SAPIENS | 7 | 15394 | 2ozo | A | 465 | ARG | p.Arg465Cys | P43403 |
| HOMO SAPIENS | 4 | 15394 | 2ozo | A | 518 | SER | p.Ser518Arg | P43403 |
| HOMO SAPIENS | 1 | 20813 | 3o5z | A | 19 | ARG | p.Arg19Trp | O00459 |
| HOMO SAPIENS | 2 | 4329 | 2bb5 | A | 5 | ILE | p.Ile23Val | P20062 |
| HOMO SAPIENS | 4 | 4329 | 2bb5 | A | 59 | LYS | p.Lys77Met | P20062 |
| HOMO SAPIENS | 4 | 4329 | 2bb5 | A | 197 | ARG | p.Arg215Trp | P20062 |
| HOMO SAPIENS | 1 | 4329 | 2bb5 | A | 241 | PRO | p.Arg259Pro | P20062 |
| HOMO SAPIENS | 2 | 4329 | 2bb5 | A | 315 | THR | p.Thr333Lys | P20062 |
| HOMO SAPIENS | 5 | 4329 | 2bb5 | A | 330 | SER | p.Ser348Phe | P20062 |
| HOMO SAPIENS | 4 | 4329 | 2bb5 | A | 358 | LEU | p.Leu376Ser | P20062 |
| HOMO SAPIENS | 5 | 4329 | 2bb5 | A | 381 | ARG | p.Arg399Gln | P20062 |
| HOMO SAPIENS | 4 | 18294 | 4r30 | A | 163 | ASN | p.Asn163Asp | O95278 |
| HOMO SAPIENS | 6 | 18294 | 4r30 | A | 227 | ALA | p.Ala227Val | O95278 |
| HOMO SAPIENS | 4 | 18294 | 4r30 | A | 241 | ARG | p.Arg241Ter | O95278 |
| HOMO SAPIENS | 4 | 18294 | 4r30 | A | 325 | ARG | p.Arg325His | O95278 |
| HOMO SAPIENS | 0 | 2632 | 4xem | A | 22 | GLU | p.Glu22Lys | P49588 |
| HOMO SAPIENS | 0 | 2632 | 4xem | A | 275 | GLY | p.Gly275Asp | P49588 |
| HOMO SAPIENS | 1 | 2632 | 4xem | A | 329 | ARG | p.Arg329His | P49588 |
| HOMO SAPIENS | 0 | 11431 | 5t4n | A | 284 | GLN | p.Gln305Glu | Q96QU1 |
| HOMO SAPIENS | 0 | 11431 | 5t4n | A | 315 | ARG | p.Arg336Ter | Q96QU1 |
| HOMO SAPIENS | 0 | 11431 | 5t4n | A | 322 | ARG | p.Arg343Lys | Q96QU1 |

| | | | | | | | | |
|---|---|---|---|---|---|---|---|---|
| HOMO SAPIENS | 1 | 11431 | 5t4n | A | 326 | LEU | p.Leu347Phe | Q96QU1 |
| HOMO SAPIENS | 0 | 11431 | 5t4n | A | 373 | TYR | p.Tyr394Cys | Q96QU1 |
| HOMO SAPIENS | 0 | 11431 | 5t4n | A | 398 | ASN | p.Asn419Ser | Q96QU1 |
| HOMO SAPIENS | 0 | 11431 | 5t4n | A | 414 | ALA | p.Asp435Ala | Q96QU1 |
| HOMO SAPIENS | 0 | 11431 | 5t4n | A | 433 | VAL | p.Val454Phe | Q96QU1 |
| HOMO SAPIENS | 1 | 11431 | 5t4n | A | 507 | VAL | p.Val528Asp | Q96QU1 |
| HOMO SAPIENS | 0 | 11431 | 5t4n | A | 547 | ALA | p.Ala568Ser | Q96QU1 |
| HOMO SAPIENS | 3 | 3194 | 5xtd | L | 84 | ARG | p.Arg84His | O43181 |
| HOMO SAPIENS | 1 | 3194 | 5xtd | L | 130 | THR | p.Thr130Ser | O43181 |
| HOMO SAPIENS | 8 | 3194 | 5xtd | L | 136 | SER | p.Ser136Ala | O43181 |
| HOMO SAPIENS | 0 | 40749 | 1nun | A | 106 | CYS | p.Cys106Phe | O15520 |
| HOMO SAPIENS | 9 | 40749 | 1nun | A | 156 | ILE | p.Ile156Arg | O15520 |
| HOMO SAPIENS | 1 | 40749 | 1nun | A | 193 | ARG | p.Arg193Ter | O15520 |
| HOMO SAPIENS | 0 | 40749 | 1nun | A | 207 | HIS | p.His207Pro | O15520 |
| HOMO SAPIENS | 0 | 7026 | 3n1f | D | 875 | SER | p.Ser875Gly | Q4KMG0 |
| HOMO SAPIENS | 6 | 23802 | 3rzv | A | 424 | ARG | p.Arg424Ter | O95630 |
| HOMO SAPIENS | 0 | 36720 | 1jm7 | A | 12 | GLN | p.Gln12Ter | P38398 |
| HOMO SAPIENS | 3 | 36720 | 1jm7 | A | 23 | GLU | p.Glu23Asp | P38398 |
| HOMO SAPIENS | 1 | 36720 | 1jm7 | A | 45 | LYS | p.Lys45Gln | P38398 |
| HOMO SAPIENS | 8 | 36720 | 1jm7 | A | 47 | CYS | p.Cys47Ter | P38398 |
| HOMO SAPIENS | 0 | 36720 | 1jm7 | A | 59 | SER | p.Ser59Ter | P38398 |
| HOMO SAPIENS | 8 | 36720 | 1jm7 | A | 61 | CYS | p.Cys61Gly | P38398 |
| HOMO SAPIENS | 8 | 36720 | 1jm7 | A | 63 | LEU | p.Leu63Ter | P38398 |
| HOMO SAPIENS | 7 | 36720 | 1jm7 | A | 64 | CYS | p.Cys64Gly | P38398 |
| HOMO SAPIENS | 0 | 36720 | 1jm7 | A | 101 | TYR | p.Tyr101Ter | P38398 |
| HOMO SAPIENS | 0 | 36720 | 1jm7 | A | null | TYR | p.Tyr105Cys | P38398 |
| HOMO SAPIENS | 0 | 28500 | 3kuz | A | 1217 | CYS | p.Cys1217Arg | O60486 |
| HOMO SAPIENS | 5 | 8097 | 3p56 | C | 69 | ARG | p.Arg69Trp | Q8TDP1 |
| HOMO SAPIENS | 5 | 8097 | 3p56 | C | 143 | LYS | p.Lys143Ile | Q8TDP1 |
| HOMO SAPIENS | 2 | 30073 | 3nmz | A | 380 | ARG | p.Arg380Gln | P25054 |
| HOMO SAPIENS | 1 | 22573 | 1cjy | A | 111 | SER | p.Ser111Pro | P47712 |
| HOMO SAPIENS | 0 | 22573 | 1cjy | A | 485 | ARG | p.Arg485His | P47712 |
| HOMO SAPIENS | 2 | 965 | 3E+077 | A | 99 | ALA | p.Ala99Val | Q9Y617 |
| HOMO SAPIENS | 2 | 965 | 3E+077 | A | 100 | ASP | p.Asp100Ala | Q9Y617 |
| HOMO SAPIENS | 4 | 1503 | 3kxc | A | 62 | ARG | p.Arg62Trp | O43617 |
| HOMO SAPIENS | 6 | 9418 | 1hdr | A | 22 | GLY | p.Gly23Asp | P09417 |
| HOMO SAPIENS | 2 | 9418 | 1hdr | A | 35 | TRP | p.Trp36Arg | P09417 |
| HOMO SAPIENS | 2 | 9418 | 1hdr | A | 107 | TRP | p.Trp108Gly | P09417 |
| HOMO SAPIENS | 6 | 9418 | 1hdr | A | 149 | TYR | p.Tyr150Cys | P09417 |
| HOMO SAPIENS | -1 | 12079 | 3tnu | B | 373 | GLY | p.Gly373Ala | P13647 |
| HOMO SAPIENS | 1 | 12079 | 3tnu | B | 463 | LEU | p.Leu463Pro | P13647 |
| HOMO SAPIENS | 1 | 12079 | 3tnu | B | 471 | ARG | p.Arg471Cys | P13647 |
| HOMO SAPIENS | 0 | 8135 | 3rbn | B | 89 | SER | p.Ser556Ile | P40692 |
| HOMO SAPIENS | 0 | 8135 | 3rbn | B | 111 | GLU | p.Glu578Gly | P40692 |
| HOMO SAPIENS | 2 | 8135 | 3rbn | B | 114 | PRO | p.Pro581Leu | P40692 |
| HOMO SAPIENS | 0 | 8135 | 3rbn | B | 133 | GLU | p.Glu600Gly | P40692 |
| HOMO SAPIENS | 2 | 8135 | 3rbn | B | 136 | PRO | p.Pro603Arg | P40692 |
| HOMO SAPIENS | 2 | 8135 | 3rbn | B | 136 | PRO | p.Pro603Leu | P40692 |
| HOMO SAPIENS | 7 | 8135 | 3rbn | B | 140 | LEU | p.Leu607His | P40692 |
| HOMO SAPIENS | 2 | 8135 | 3rbn | B | 151 | LYS | p.Lys618Ala | P40692 |
| HOMO SAPIENS | 2 | 8135 | 3rbn | B | 151 | LYS | p.Lys618Thr | P40692 |
| HOMO SAPIENS | 5 | 8135 | 3rbn | B | 181 | PRO | p.Pro648Ser | P40692 |
| HOMO SAPIENS | 2 | 8135 | 3rbn | B | 188 | ILE | p.Ile655Val | P40692 |
| HOMO SAPIENS | 2 | 8135 | 3rbn | B | 214 | ALA | p.Ala681Thr | P40692 |

| | | | | | | | | |
|---|---|---|---|---|---|---|---|---|
| HOMO SAPIENS | 2 | 8135 | 3rbn | B | 220 | ARG | p.Arg687Trp | P40692 |
| HOMO SAPIENS | 2 | 8135 | 3rbn | B | 222 | GLN | p.Gln689Arg | P40692 |
| HOMO SAPIENS | -1 | 8135 | 3rbn | B | 236 | GLN | p.Gln701Ter | P40692 |
| HOMO SAPIENS | 7 | 8135 | 3rbn | B | 251 | HIS | p.His718Tyr | P40692 |
| HOMO SAPIENS | 0 | 8135 | 3rbn | B | 284 | LYS | p.Lys751Arg | P40692 |
| HOMO SAPIENS | 1 | 21998 | 5d68 | A | 355 | LEU | p.Leu355Phe | O00522 |
| HOMO SAPIENS | 3 | 21998 | 5d68 | A | 380 | THR | p.Thr380Met | O00522 |
| HOMO SAPIENS | 0 | 21998 | 5d68 | A | 415 | ILE | p.Ile415Met | O00522 |
| HOMO SAPIENS | 0 | 33789 | 2m0c | A | 13 | ARG | p.Arg218Gln | Q9H161 |
| HOMO SAPIENS | 7 | 33789 | 2m0c | A | 52 | ARG | p.Arg257Thr | Q9H161 |
| HOMO SAPIENS | 0 | 33789 | 2m0c | A | 67 | ARG | p.Arg272Pro | Q9H161 |
| HOMO SAPIENS | 6 | 8263 | 2osg | A | 51 | GLY | p.Gly355Arg | Q9UDY2 |
| HOMO SAPIENS | 5 | 8263 | 2osg | A | 60 | THR | p.Thr364Met | Q9UDY2 |
| HOMO SAPIENS | 1 | 2475 | 5E+026 | B | 234 | THR | p.Thr234Ala | Q9BZ23 |
| HOMO SAPIENS | 2 | 2475 | 5E+026 | B | 264 | ARG | p.Arg264Trp | Q9BZ23 |
| HOMO SAPIENS | 4 | 2475 | 5E+026 | B | 286 | ARG | p.Arg286Cys | Q9BZ23 |
| HOMO SAPIENS | 0 | 2475 | 5E+026 | B | 357 | ARG | p.Arg357Trp | Q9BZ23 |
| HOMO SAPIENS | 2 | 2475 | 5E+026 | B | 481 | ARG | p.Arg481Ter | Q9BZ23 |
| HOMO SAPIENS | 8 | 2475 | 5E+026 | B | 521 | GLY | p.Gly521Arg | Q9BZ23 |
| HOMO SAPIENS | 2 | 2475 | 5E+026 | B | 528 | THR | p.Thr528Met | Q9BZ23 |
| HOMO SAPIENS | 1 | 3742 | 5xtd | E | 108 | HIS | p.His134Tyr | P56556 |
| HOMO SAPIENS | 1 | 16725 | 3g04 | C | 36 | ASP | p.Asp36His | P16473 |
| HOMO SAPIENS | 3 | 16725 | 3g04 | C | 52 | PRO | p.Pro52Thr | P16473 |
| HOMO SAPIENS | 1 | 16725 | 3g04 | C | 162 | PRO | p.Pro162Ala | P16473 |
| HOMO SAPIENS | -1 | 12366 | 5d2d | C | 750 | PRO | p.Pro750Leu | P13569 |
| HOMO SAPIENS | 0 | 12366 | 5d2d | C | 764 | ARG | p.Arg764Ter | P13569 |
| HOMO SAPIENS | 0 | 34845 | 2y23 | A | 1151 | ASP | p.Asp1151Glu | P52179 |
| HOMO SAPIENS | 0 | 34845 | 2y23 | A | 1180 | ASP | p.Asp1180Ala | P52179 |
| HOMO SAPIENS | 6 | 36740 | 3lbx | B | 2053 | ALA | p.Ala2053Pro | P11277 |
| HOMO SAPIENS | 0 | 19289 | 3ihj | A | 404 | ARG | p.Arg404Ter | Q8TD30 |
| HOMO SAPIENS | 5 | 1034 | 5xtd | Q | 228 | ARG | p.Arg228Gln | O75306 |
| HOMO SAPIENS | 5 | 1034 | 5xtd | Q | 229 | PRO | p.Pro229Gln | O75306 |
| HOMO SAPIENS | 4 | 1034 | 5xtd | Q | 323 | ARG | p.Arg323Gln | O75306 |
| HOMO SAPIENS | 4 | 1034 | 5xtd | Q | 352 | PRO | p.Pro352Ala | O75306 |
| HOMO SAPIENS | 5 | 1034 | 5xtd | Q | 413 | SER | p.Ser413Pro | O75306 |
| HOMO SAPIENS | 1 | 2772 | 2vaf | A | 1047 | THR | p.Thr66Ala | O14958 |
| HOMO SAPIENS | 5 | 2772 | 2vaf | A | 1057 | VAL | p.Val76Met | O14958 |
| HOMO SAPIENS | 0 | 2772 | 2vaf | A | 1148 | LEU | p.Leu167His | O14958 |
| HOMO SAPIENS | 1 | 2772 | 2vaf | A | 1149 | ILE | p.Ile168Thr | O14958 |
| HOMO SAPIENS | 8 | 2772 | 2vaf | A | 1225 | HIS | p.His244Arg | O14958 |
| HOMO SAPIENS | 8 | 2772 | 2vaf | A | 1225 | HIS | p.His244Tyr | O14958 |
| HOMO SAPIENS | 7 | 2772 | 2vaf | A | 1251 | ILE | p.Ile270Thr | O14958 |
| HOMO SAPIENS | 3 | 2772 | 2vaf | A | 1288 | ASP | p.Asp307His | O14958 |
| HOMO SAPIENS | 5 | 2772 | 2vaf | A | 1289 | PRO | p.Pro308Leu | O14958 |
| HOMO SAPIENS | 1 | 31885 | 2aex | A | 132 | ALA | p.Ala132Val | P36551 |
| HOMO SAPIENS | 1 | 31885 | 2aex | A | 272 | ASN | p.Asn272His | P36551 |
| HOMO SAPIENS | 8 | 31885 | 2aex | A | 279 | GLY | p.Gly279Arg | P36551 |
| HOMO SAPIENS | 5 | 31885 | 2aex | A | 294 | VAL | p.Val294Ile | P36551 |
| HOMO SAPIENS | 5 | 31885 | 2aex | A | 295 | HIS | p.His295Asp | P36551 |
| HOMO SAPIENS | 1 | 31885 | 2aex | A | 331 | ARG | p.Arg331Trp | P36551 |
| HOMO SAPIENS | 3 | 31885 | 2aex | A | 352 | ARG | p.Arg352Cys | P36551 |
| HOMO SAPIENS | 1 | 31885 | 2aex | A | 404 | LYS | p.Lys404Glu | P36551 |
| HOMO SAPIENS | 2 | 31885 | 2aex | A | 447 | ARG | p.Arg447Cys | P36551 |
| HOMO SAPIENS | 5 | 11721 | 2r3v | A | 20 | HIS | p.His20Pro | Q03426 |

| | | | | | | | | |
|---|---|---|---|---|---|---|---|---|
| HOMO SAPIENS | 1 | 11721 | 2r3v | A | 116 | TYR | p.Tyr116His | Q03426 |
| HOMO SAPIENS | 1 | 11721 | 2r3v | A | 148 | ALA | p.Ala148Thr | Q03426 |
| HOMO SAPIENS | 1 | 11721 | 2r3v | A | 167 | PRO | p.Pro167Leu | Q03426 |
| HOMO SAPIENS | 6 | 11721 | 2r3v | A | 203 | VAL | p.Val203Ala | Q03426 |
| HOMO SAPIENS | 2 | 11721 | 2r3v | A | 215 | ARG | p.Arg215Ter | Q03426 |
| HOMO SAPIENS | 2 | 11721 | 2r3v | A | 237 | THR | p.Thr237Ser | Q03426 |
| HOMO SAPIENS | 0 | 11721 | 2r3v | A | 268 | ILE | p.Ile268Thr | Q03426 |
| HOMO SAPIENS | 0 | 11721 | 2r3v | A | 268 | ILE | p.Ile268Val | Q03426 |
| HOMO SAPIENS | 0 | 11721 | 2r3v | A | 301 | ASN | p.Asn301Ser | Q03426 |
| HOMO SAPIENS | 6 | 11721 | 2r3v | A | 310 | VAL | p.Val310Met | Q03426 |
| HOMO SAPIENS | 5 | 11721 | 2r3v | A | 334 | ALA | p.Ala334Thr | Q03426 |
| HOMO SAPIENS | 2 | 11721 | 2r3v | A | 377 | VAL | p.Val377Ile | Q03426 |
| HOMO SAPIENS | 1 | 11721 | 2r3v | A | 386 | ASP | p.Asp386Asn | Q03426 |
| HOMO SAPIENS | 1 | 11721 | 2r3v | A | 388 | ARG | p.Arg388Ter | Q03426 |
| HOMO SAPIENS | 8 | 5010 | 2hgs | A | 125 | ARG | p.Arg125Cys | P48637 |
| HOMO SAPIENS | 1 | 5010 | 2hgs | A | 164 | ARG | p.Arg164Gln | P48637 |
| HOMO SAPIENS | 0 | 5010 | 2hgs | A | 219 | ASP | p.Asp219Gly | P48637 |
| HOMO SAPIENS | 7 | 5010 | 2hgs | A | 267 | ARG | p.Arg267Trp | P48637 |
| HOMO SAPIENS | 0 | 5010 | 2hgs | A | 283 | ARG | p.Arg283Cys | P48637 |
| HOMO SAPIENS | 0 | 6892 | 5erd | B | 7 | VAL | p.Val56Met | Q14126 |
| HOMO SAPIENS | 1 | 6892 | 5erd | B | 25 | HIS | p.His74Arg | Q14126 |
| HOMO SAPIENS | 0 | 6892 | 5erd | B | 40 | TYR | p.Tyr89Cys | Q14126 |
| HOMO SAPIENS | 0 | 6892 | 5erd | B | 97 | ARG | p.Arg146Leu | Q14126 |
| HOMO SAPIENS | 7 | 6892 | 5erd | B | 217 | ASN | p.Asn266Ser | Q14126 |
| HOMO SAPIENS | 0 | 6892 | 5erd | B | 243 | ARG | p.Arg292His | Q14126 |
| HOMO SAPIENS | 0 | 6892 | 5erd | B | 244 | ILE | p.Ile293Val | Q14126 |
| HOMO SAPIENS | 0 | 6892 | 5erd | B | 343 | VAL | p.Val392Ile | Q14126 |
| HOMO SAPIENS | 4 | 6892 | 5erd | B | 350 | GLU | p.Glu399Lys | Q14126 |
| HOMO SAPIENS | 0 | 6892 | 5erd | B | 410 | TYR | p.Tyr459Cys | Q14126 |
| HOMO SAPIENS | 0 | 6892 | 5erd | B | 466 | VAL | p.Val515Ile | Q14126 |
| HOMO SAPIENS | 0 | 6892 | 5erd | B | 482 | PHE | p.Phe531Cys | Q14126 |
| HOMO SAPIENS | 0 | 6892 | 5erd | B | 484 | VAL | p.Val533Ile | Q14126 |
| HOMO SAPIENS | 0 | 6892 | 5erd | B | 545 | LEU | p.Leu594Pro | Q14126 |
| HOMO SAPIENS | 0 | 1338 | 2ax4 | A | 58 | THR | p.Thr48Arg | O95340 |
| HOMO SAPIENS | 7 | 8675 | 2zkm | X | 385 | LYS | p.Lys385Arg | Q00722 |
| HOMO SAPIENS | 1 | 11769 | 2wya | B | 54 | VAL | p.Val54Met | P54868 |
| HOMO SAPIENS | 1 | 11769 | 2wya | B | 116 | ARG | p.Arg116Cys | P54868 |
| HOMO SAPIENS | 5 | 11769 | 2wya | B | 167 | TYR | p.Tyr167Cys | P54868 |
| HOMO SAPIENS | 1 | 11769 | 2wya | B | 174 | PHE | p.Phe174Leu | P54868 |
| HOMO SAPIENS | 0 | 11769 | 2wya | B | 212 | GLY | p.Gly212Arg | P54868 |
| HOMO SAPIENS | 1 | 11769 | 2wya | B | 288 | ARG | p.Arg288Ter | P54868 |
| HOMO SAPIENS | 0 | 11769 | 2wya | B | 375 | SER | p.Ser375Cys | P54868 |
| HOMO SAPIENS | 4 | 11769 | 2wya | B | 500 | ARG | p.Arg500His | P54868 |
| HOMO SAPIENS | 0 | 3102 | 2iwo | A | 1880 | ARG | p.Arg1908Lys | O75970 |
| HOMO SAPIENS | 6 | 14296 | 2m9w | A | 38 | GLY | p.Gly296Ser | P43694 |
| HOMO SAPIENS | 0 | 14296 | 2m9w | A | 58 | GLN | p.Gln316Glu | P43694 |
| HOMO SAPIENS | 2 | 704 | 5ekd | B | 239 | ARG | p.Arg239Ter | Q9UGM6 |
| HOMO SAPIENS | 2 | 704 | 5ekd | B | 264 | TYR | p.Tyr264Cys | Q9UGM6 |
| HOMO SAPIENS | 6 | 2228 | 1ey2 | A | 47 | SER | p.Ser47Ter | Q93099 |
| HOMO SAPIENS | 6 | 2228 | 1ey2 | A | 48 | ALA | p.Ala48Ser | Q93099 |
| HOMO SAPIENS | 6 | 2228 | 1ey2 | A | 53 | ARG | p.Arg53Gln | Q93099 |
| HOMO SAPIENS | 3 | 2228 | 1ey2 | A | 80 | HIS | p.Gln80His | Q93099 |
| HOMO SAPIENS | 0 | 2228 | 1ey2 | A | 120 | CYS | p.Cys120Trp | Q93099 |
| HOMO SAPIENS | 3 | 2228 | 1ey2 | A | 122 | ALA | p.Ala122Val | Q93099 |

| | | | | | | | | |
|---|---|---|---|---|---|---|---|---|
| HOMO SAPIENS | 3 | 2228 | 1ey2 | A | 161 | GLY | p.Gly161Arg | Q93099 |
| HOMO SAPIENS | 6 | 2228 | 1ey2 | A | 225 | ARG | p.Arg225His | Q93099 |
| HOMO SAPIENS | 0 | 2228 | 1ey2 | A | 230 | PRO | p.Pro230Ser | Q93099 |
| HOMO SAPIENS | 6 | 2228 | 1ey2 | A | 270 | GLY | p.Gly270Arg | Q93099 |
| HOMO SAPIENS | 6 | 2228 | 1ey2 | A | 330 | ARG | p.Arg330Ser | Q93099 |
| HOMO SAPIENS | -1 | 2228 | 1ey2 | A | 368 | MSE | p.Met368Val | Q93099 |
| HOMO SAPIENS | 6 | 2228 | 1ey2 | A | 371 | HIS | p.His371Arg | Q93099 |
| HOMO SAPIENS | 6 | 2228 | 1ey2 | A | 401 | GLU | p.Glu401Gln | Q93099 |
| HOMO SAPIENS | 5 | 34648 | 2k27 | A | 31 | ARG | p.Arg31His | Q06710 |
| HOMO SAPIENS | 7 | 34648 | 2k27 | A | 40 | GLN | p.Gln40Pro | Q06710 |
| HOMO SAPIENS | 5 | 34648 | 2k27 | A | 57 | CYS | p.Cys57Tyr | Q06710 |
| HOMO SAPIENS | 5 | 34648 | 2k27 | A | 62 | LEU | p.Leu62Arg | Q06710 |
| HOMO SAPIENS | 1 | 21399 | 3edu | A | 1819 | GLU | p.Glu1819Ter | P11277 |
| HOMO SAPIENS | 1 | 18153 | 1oey | J | 245 | TYR | p.Tyr245Ter | Q15080 |
| HOMO SAPIENS | 2 | 41567 | 3sw0 | X | 1050 | ASN | p.Asn1050Tyr | P08603 |
| HOMO SAPIENS | 5 | 41567 | 3sw0 | X | 1059 | ILE | p.Ile1059Thr | P08603 |
| HOMO SAPIENS | 0 | 36362 | 4wrm | A | 362 | HIS | p.His362Arg | P07333 |
| HOMO SAPIENS | -1 | 36362 | 4wrm | A | 413 | GLY | p.Gly413Ser | P07333 |
| HOMO SAPIENS | 0 | 31255 | 2w2j | A | 100 | SER | p.Ser100Pro | P35219 |
| HOMO SAPIENS | 0 | 31255 | 2w2j | A | 100 | SER | p.Ser100Leu | P35219 |
| HOMO SAPIENS | 5 | 31255 | 2w2j | A | 237 | ARG | p.Arg237Gln | P35219 |
| HOMO SAPIENS | -1 | 26450 | 1th1 | C | 1395 | SER | p.Ser1395Cys | P25054 |
| HOMO SAPIENS | -1 | 26450 | 1th1 | C | 1413 | MET | p.Met1413Val | P25054 |
| HOMO SAPIENS | -1 | 26450 | 1th1 | C | 1446 | ALA | p.Ala1446Thr | P25054 |
| HOMO SAPIENS | -1 | 26450 | 1th1 | C | 1454 | LYS | p.Lys1454Glu | P25054 |
| HOMO SAPIENS | 0 | 26450 | 1th1 | C | 1474 | ALA | p.Ala1474Thr | P25054 |
| HOMO SAPIENS | 1 | 981 | 5xtb | F | 14 | LEU | p.Leu14Val | O43678 |
| HOMO SAPIENS | 1 | 11069 | 3kq4 | A | 46 | SER | p.Ser46Leu | P27352 |
| HOMO SAPIENS | 1 | 11069 | 3kq4 | A | 97 | MET | p.Met97Thr | P27352 |
| HOMO SAPIENS | 1 | 11069 | 3kq4 | A | 255 | ASN | p.Asn255Ser | P27352 |
| HOMO SAPIENS | 1 | 11069 | 3kq4 | A | 330 | ASN | p.Asn330Lys | P27352 |
| HOMO SAPIENS | 0 | 41489 | 3erb | A | 53 | PRO | p.Pro73Leu | P06681 |
| HOMO SAPIENS | 6 | 8715 | 2bq8 | X | 23 | THR | p.Thr44Met | P13686 |
| HOMO SAPIENS | 0 | 8715 | 2bq8 | X | 88 | GLY | p.Gly109Arg | P13686 |
| HOMO SAPIENS | 6 | 8715 | 2bq8 | X | 127 | VAL | p.Val148Met | P13686 |
| HOMO SAPIENS | 2 | 8715 | 2bq8 | X | 179 | VAL | p.Val200Met | P13686 |
| HOMO SAPIENS | 6 | 8715 | 2bq8 | X | 251 | ARG | p.Arg272Cys | P13686 |
| HOMO SAPIENS | 7 | 8112 | 3p56 | A | 37 | GLY | p.Gly37Ser | O75792 |
| HOMO SAPIENS | -1 | 8112 | 3p56 | A | 202 | LEU | p.Leu202Ser | O75792 |
| HOMO SAPIENS | 1 | 8112 | 3p56 | A | 205 | ASP | p.Asp205Glu | O75792 |
| HOMO SAPIENS | 4 | 8112 | 3p56 | A | 212 | ASN | p.Asn212Ile | O75792 |
| HOMO SAPIENS | 0 | 8112 | 3p56 | A | 235 | ARG | p.Arg235Gln | O75792 |
| HOMO SAPIENS | 4 | 8112 | 3p56 | A | 240 | THR | p.Thr240Met | O75792 |
| HOMO SAPIENS | 0 | 8112 | 3p56 | A | 291 | ARG | p.Arg291His | O75792 |
| HOMO SAPIENS | 0 | 37603 | 2yqg | A | 10 | ARG | p.Arg49His | Q14126 |
| HOMO SAPIENS | 0 | 21178 | 3k1r | B | 458 | ASP | p.Asp458Val | Q495M9 |
| HOMO SAPIENS | -1 | 36828 | 4fq3 | B | 507 | GLY | p.Gly507Asp | P35637 |
| HOMO SAPIENS | 5 | 36828 | 4fq3 | B | 517 | HIS | p.His517Gln | P35637 |
| HOMO SAPIENS | 8 | 36828 | 4fq3 | B | 518 | ARG | p.Arg518Lys | P35637 |
| HOMO SAPIENS | 6 | 36828 | 4fq3 | B | 521 | ARG | p.Arg521Cys | P35637 |
| HOMO SAPIENS | 6 | 36828 | 4fq3 | B | 521 | ARG | p.Arg521His | P35637 |
| HOMO SAPIENS | 0 | 6363 | 3j7y | c | null | HIS | p.His13Arg | Q9H9J2 |
| HOMO SAPIENS | 2 | 6363 | 3j7y | c | 105 | ARG | p.Arg105Cys | Q9H9J2 |
| HOMO SAPIENS | 2 | 6363 | 3j7y | c | 138 | THR | p.Thr138Ile | Q9H9J2 |

| | | | | | | | | |
|---|---|---|---|---|---|---|---|---|
| HOMO SAPIENS | 1 | 6363 | 3j7y | c | 156 | LEU | p.Leu156Arg | Q9H9J2 |
| HOMO SAPIENS | 1 | 6782 | 5l46 | A | 109 | HIS | p.His109Arg | Q9UI17 |
| HOMO SAPIENS | 0 | 23951 | 5t76 | A | 175 | GLY | p.Gly931Ser | P49588 |
| HOMO SAPIENS | 8 | 12604 | 3p0l | A | 182 | ARG | p.Arg182Leu | P49675 |
| HOMO SAPIENS | 2 | 12604 | 3p0l | A | 187 | VAL | p.Val187Met | P49675 |
| HOMO SAPIENS | 2 | 12604 | 3p0l | A | 188 | ARG | p.Arg188Cys | P49675 |
| HOMO SAPIENS | 0 | 12604 | 3p0l | A | 193 | ARG | p.Arg193Ter | P49675 |
| HOMO SAPIENS | 9 | 12604 | 3p0l | A | 217 | ARG | p.Arg217Thr | P49675 |
| HOMO SAPIENS | 4 | 12604 | 3p0l | A | 218 | ALA | p.Ala218Val | P49675 |
| HOMO SAPIENS | 0 | 12604 | 3p0l | A | 250 | TRP | p.Trp250Ter | P49675 |
| HOMO SAPIENS | 1 | 12604 | 3p0l | A | 258 | GLN | p.Gln258Ter | P49675 |
| HOMO SAPIENS | -1 | 27765 | 4wkr | A | 4 | GLU | p.Glu4Lys | Q4G0J3 |
| HOMO SAPIENS | 2 | 3587 | 4kit | C | 2301 | PRO | p.Pro2301Thr | Q6P2Q9 |
| HOMO SAPIENS | 3 | 3587 | 4kit | C | 2304 | PHE | p.Phe2304Leu | Q6P2Q9 |
| HOMO SAPIENS | 3 | 3587 | 4kit | C | 2309 | HIS | p.His2309Pro | Q6P2Q9 |
| HOMO SAPIENS | 3 | 3587 | 4kit | C | 2310 | ARG | p.Arg2310Lys | Q6P2Q9 |
| HOMO SAPIENS | 0 | 9073 | 1n69 | B | 21 | ASN | p.Asn215His | P07602 |
| HOMO SAPIENS | 0 | 9073 | 1n69 | B | 23 | THR | p.Thr217Ile | P07602 |
| HOMO SAPIENS | 0 | 9073 | 1n69 | B | 47 | CYS | p.Cys241Ser | P07602 |
| HOMO SAPIENS | 0 | 475 | 2vd5 | A | 225 | THR | p.Thr225Met | Q09013 |
| HOMO SAPIENS | 1 | 9066 | 4mzv | A | 31 | ASN | p.Asn31Lys | P16422 |
| HOMO SAPIENS | 1 | 9066 | 4mzv | A | 47 | GLN | p.Gln47His | P16422 |
| HOMO SAPIENS | 3 | 9066 | 4mzv | A | 66 | CYS | p.Cys66Tyr | P16422 |
| HOMO SAPIENS | 1 | 9066 | 4mzv | A | 102 | SER | p.Ser102Gly | P16422 |
| HOMO SAPIENS | 5 | 9066 | 4mzv | A | 115 | THR | p.Met115Thr | P16422 |
| HOMO SAPIENS | 1 | 9066 | 4mzv | A | 138 | ARG | p.Arg138Ter | P16422 |
| HOMO SAPIENS | 1 | 9066 | 4mzv | A | 153 | ARG | p.Arg153Thr | P16422 |
| HOMO SAPIENS | 1 | 9066 | 4mzv | A | 165 | ALA | p.Ala165Thr | P16422 |
| HOMO SAPIENS | 8 | 1827 | 2nz2 | A | 14 | GLY | p.Gly14Ser | P00966 |
| HOMO SAPIENS | 5 | 1827 | 2nz2 | A | 18 | SER | p.Ser18Leu | P00966 |
| HOMO SAPIENS | 3 | 1827 | 2nz2 | A | 86 | ARG | p.Arg86Cys | P00966 |
| HOMO SAPIENS | 3 | 1827 | 2nz2 | A | 86 | ARG | p.Arg86His | P00966 |
| HOMO SAPIENS | 1 | 1827 | 2nz2 | A | 179 | TRP | p.Trp179Arg | P00966 |
| HOMO SAPIENS | 7 | 1827 | 2nz2 | A | 180 | SER | p.Ser180Asn | P00966 |
| HOMO SAPIENS | 7 | 1827 | 2nz2 | A | 191 | GLU | p.Glu191Lys | P00966 |
| HOMO SAPIENS | 4 | 1827 | 2nz2 | A | 256 | GLU | p.Glu256Lys | P00966 |
| HOMO SAPIENS | 4 | 1827 | 2nz2 | A | 263 | VAL | p.Val263Met | P00966 |
| HOMO SAPIENS | 5 | 1827 | 2nz2 | A | 265 | ARG | p.Arg265Cys | P00966 |
| HOMO SAPIENS | 5 | 1827 | 2nz2 | A | 268 | ILE | p.Ile268Met | P00966 |
| HOMO SAPIENS | 6 | 1827 | 2nz2 | A | 269 | VAL | p.Val269Met | P00966 |
| HOMO SAPIENS | 7 | 1827 | 2nz2 | A | 272 | ARG | p.Arg272Cys | P00966 |
| HOMO SAPIENS | 3 | 1827 | 2nz2 | A | 304 | ARG | p.Arg304Trp | P00966 |
| HOMO SAPIENS | 3 | 1827 | 2nz2 | A | 307 | ARG | p.Arg307Cys | P00966 |
| HOMO SAPIENS | 3 | 1827 | 2nz2 | A | 310 | LYS | p.Lys310Gln | P00966 |
| HOMO SAPIENS | 2 | 1827 | 2nz2 | A | 362 | GLY | p.Gly362Val | P00966 |
| HOMO SAPIENS | 2 | 1827 | 2nz2 | A | 363 | ARG | p.Arg363Trp | P00966 |
| HOMO SAPIENS | 2 | 1827 | 2nz2 | A | 363 | ARG | p.Arg363Gln | P00966 |
| HOMO SAPIENS | 1 | 1827 | 2nz2 | A | 390 | GLY | p.Gly390Arg | P00966 |
| HOMO SAPIENS | 0 | 29524 | 5vbn | B | 2165 | ARG | p.Arg2165His | Q07864 |
| HOMO SAPIENS | 2 | 29524 | 5vbn | B | 2255 | ILE | p.Ile2255Phe | Q07864 |
| HOMO SAPIENS | 2 | 29524 | 5vbn | B | 2274 | LEU | p.Leu2274Val | Q07864 |
| HOMO SAPIENS | 0 | 31897 | 3t0o | A | 184 | CYS | p.Cys184Arg | O00584 |
| HOMO SAPIENS | -1 | 31897 | 3t0o | A | 215 | GLU | p.Glu215Lys | O00584 |
| HOMO SAPIENS | 0 | 31897 | 3t0o | A | 236 | ARG | p.Arg236Trp | O00584 |

| | | | | | | | | |
|---|---|---|---|---|---|---|---|---|
| HOMO SAPIENS | 7 | 18490 | 4pcw | B | 31 | GLU | p.Glu32Ter | P20930 |
| HOMO SAPIENS | -1 | 39865 | 2h9e | L | 152 | GLY | p.Gly192Arg | P00742 |
| HOMO SAPIENS | 2 | 4309 | 2x0b | F | 174 | THR | p.Thr207Met | P01019 |
| HOMO SAPIENS | 3 | 4309 | 2x0b | F | 342 | ARG | p.Arg375Gln | P01019 |
| HOMO SAPIENS | 2 | 3094 | 1zll | B | 9 | ARG | p.Arg9Cys | P26678 |
| HOMO SAPIENS | 9 | 3094 | 1zll | B | 39 | LEU | p.Leu39Ter | P26678 |
| HOMO SAPIENS | 5 | 3094 | 1zll | B | 49 | VAL | p.Val49Met | P26678 |
| HOMO SAPIENS | -1 | 41498 | 3ffd | P | 8 | LEU | p.Leu44Pro | P12272 |
| HOMO SAPIENS | 1 | 20697 | 1igr | A | 407 | ARG | p.Arg437His | P08069 |
| HOMO SAPIENS | 0 | 20697 | 1igr | A | 416 | MET | p.Met446Val | P08069 |
| HOMO SAPIENS | -1 | 20697 | 1igr | A | 462 | VAL | p.Val492Ile | P08069 |
| HOMO SAPIENS | 0 | 3973 | 4cbz | A | 37 | LEU | p.Leu37Ser | P78504 |
| HOMO SAPIENS | 0 | 3973 | 4cbz | A | 45 | VAL | p.Val45Leu | P78504 |
| HOMO SAPIENS | 0 | 3973 | 4cbz | A | 134 | LEU | p.Leu134Met | P78504 |
| HOMO SAPIENS | -1 | 3973 | 4cbz | A | 146 | VAL | p.Val146Ile | P78504 |
| HOMO SAPIENS | 0 | 3973 | 4cbz | A | 176 | VAL | p.Val176Ile | P78504 |
| HOMO SAPIENS | 0 | 3973 | 4cbz | A | 184 | ARG | p.Arg184Cys | P78504 |
| HOMO SAPIENS | 6 | 1697 | 4hmy | S | 33 | ARG | p.Arg33Trp | Q96PC3 |
| HOMO SAPIENS | 2 | 23752 | 5j4o | A | 13 | ARG | p.Arg1611Ser | P02549 |
| HOMO SAPIENS | 0 | 23752 | 5j4o | A | 42 | ALA | p.Ala1640Gly | P02549 |
| HOMO SAPIENS | 0 | 23752 | 5j4o | A | 108 | HIS | p.His1706Gln | P02549 |
| HOMO SAPIENS | 6 | 23752 | 5j4o | A | 206 | LEU | p.Leu1804Phe | P02549 |
| HOMO SAPIENS | 8 | 531 | 2qz4 | A | 349 | GLY | p.Gly349Ser | Q9UQ90 |
| HOMO SAPIENS | 0 | 531 | 2qz4 | A | 486 | ARG | p.Arg486Gln | Q9UQ90 |
| HOMO SAPIENS | 1 | 531 | 2qz4 | A | 510 | ALA | p.Ala510Val | Q9UQ90 |
| HOMO SAPIENS | 0 | 531 | 2qz4 | A | 529 | ALA | p.Ala529Val | Q9UQ90 |
| HOMO SAPIENS | 0 | 531 | 2qz4 | A | 558 | LYS | p.Lys558Ter | Q9UQ90 |
| HOMO SAPIENS | 1 | 531 | 2qz4 | A | 559 | LYS | p.Lys559Ter | Q9UQ90 |
| HOMO SAPIENS | 2 | 7736 | 1fnh | A | 114 | TRP | p.Trp1834Arg | P02751 |
| HOMO SAPIENS | 0 | 7736 | 1fnh | A | 163 | LEU | p.Leu1883Arg | P02751 |
| HOMO SAPIENS | 3 | 20337 | 5wy5 | B | 264 | LEU | p.Leu264Phe | Q96MG7 |
| HOMO SAPIENS | 8 | 36096 | 2jku | A | 643 | GLY | p.Gly668Arg | P05165 |
| HOMO SAPIENS | 6 | 36096 | 2jku | A | 663 | CYS | p.Cys688Arg | P05165 |
| HOMO SAPIENS | 0 | 9850 | 3kbt | A | 1606 | TYR | p.Tyr1606Ter | P11277 |
| HOMO SAPIENS | 4 | 6876 | 4jkq | A | 217 | ILE | p.Ile217Val | Q92736 |
| HOMO SAPIENS | 5 | 6876 | 4jkq | A | 420 | ARG | p.Arg420Trp | Q92736 |
| HOMO SAPIENS | 2 | 27603 | 1itv | B | 62 | PRO | p.Arg574Pro | P14780 |
| HOMO SAPIENS | 4 | 27603 | 1itv | B | 156 | ARG | p.Arg668Gln | P14780 |
| HOMO SAPIENS | 2 | 22954 | 4f52 | E | 48 | ILE | p.Ile48Met | Q92990 |
| HOMO SAPIENS | 0 | 22954 | 4f52 | E | 91 | ASP | p.Asp91Asn | Q92990 |
| HOMO SAPIENS | 0 | 22954 | 4f52 | E | 146 | ALA | p.Ala146Thr | Q92990 |
| HOMO SAPIENS | 0 | 22954 | 4f52 | E | 181 | ILE | p.Ile181Lys | Q92990 |
| HOMO SAPIENS | 0 | 22954 | 4f52 | E | 194 | ASN | p.Asn194Ser | Q92990 |
| HOMO SAPIENS | 2 | 22954 | 4f52 | E | 254 | PRO | p.Pro254Arg | Q92990 |
| HOMO SAPIENS | 2 | 22954 | 4f52 | E | 300 | ILE | p.Ile300Thr | Q92990 |
| HOMO SAPIENS | 2 | 22954 | 4f52 | E | 336 | LEU | p.Leu336Ser | Q92990 |
| HOMO SAPIENS | 0 | 22954 | 4f52 | E | 353 | TYR | p.Tyr353His | Q92990 |
| HOMO SAPIENS | 0 | 22954 | 4f52 | E | 387 | MET | p.Met387Val | Q92990 |
| HOMO SAPIENS | 4 | 2946 | 5gl7 | A | 344 | TYR | p.Tyr344Cys | P56192 |
| HOMO SAPIENS | 0 | 2946 | 5gl7 | A | 370 | PHE | p.Phe370Leu | P56192 |
| HOMO SAPIENS | 0 | 2946 | 5gl7 | A | 393 | ALA | p.Ala393Thr | P56192 |
| HOMO SAPIENS | 0 | 2946 | 5gl7 | A | 523 | ILE | p.Ile523Thr | P56192 |
| HOMO SAPIENS | 0 | 2946 | 5gl7 | A | 567 | SER | p.Ser567Leu | P56192 |
| HOMO SAPIENS | 0 | 2946 | 5gl7 | A | 800 | PRO | p.Pro800Thr | P56192 |

| | | | | | | | | |
|---|---|---|---|---|---|---|---|---|
| HOMO SAPIENS | 3 | 34961 | 4wy4 | C | 44 | TYR | p.Tyr44His | O95721 |
| HOMO SAPIENS | 8 | 34961 | 4wy4 | C | 89 | GLU | p.Glu89Lys | O95721 |
| HOMO SAPIENS | 0 | 9427 | 4chl | B | 21 | ALA | p.Ala21Ser | O95571 |
| HOMO SAPIENS | 2 | 9427 | 4chl | B | 62 | ALA | p.Ala62Thr | O95571 |
| HOMO SAPIENS | 0 | 9427 | 4chl | B | 114 | ILE | p.Ile114Phe | O95571 |
| HOMO SAPIENS | 2 | 9427 | 4chl | B | 159 | ARG | p.Arg159His | O95571 |
| HOMO SAPIENS | 8 | 9427 | 4chl | B | 163 | ARG | p.Arg163Trp | O95571 |
| HOMO SAPIENS | 8 | 9427 | 4chl | B | 163 | ARG | p.Arg163Gln | O95571 |
| HOMO SAPIENS | 2 | 9427 | 4chl | B | 185 | LEU | p.Leu185Arg | O95571 |
| HOMO SAPIENS | 0 | 40917 | 2e8o | A | 11 | PRO | p.Pro26Leu | Q9Y3Z3 |
| HOMO SAPIENS | -1 | 39190 | 4gow | A | 556 | TYR | p.Tyr556Ter | P56696 |
| HOMO SAPIENS | 0 | 2399 | 2mkv | A | 8 | GLY | p.Gly10Ser | P54710 |
| HOMO SAPIENS | 1 | 2399 | 2mkv | A | 24 | VAL | p.Val26Ile | P54710 |
| HOMO SAPIENS | 4 | 2399 | 2mkv | A | 39 | GLY | p.Gly41Arg | P54710 |
| HOMO SAPIENS | 0 | 20801 | 2zej | B | 1441 | ARG | p.Arg1441Cys | Q5S007 |
| HOMO SAPIENS | 0 | 20801 | 2zej | B | 1441 | ARG | p.Arg1441His | Q5S007 |
| HOMO SAPIENS | 5 | 1019 | 5xtb | M | 62 | ARG | p.Arg62Gln | P28331 |
| HOMO SAPIENS | 1 | 1019 | 5xtb | M | 85 | ALA | p.Ala85Thr | P28331 |
| HOMO SAPIENS | 5 | 1019 | 5xtb | M | 231 | LEU | p.Leu231Val | P28331 |
| HOMO SAPIENS | 2 | 1019 | 5xtb | M | 241 | ARG | p.Arg241Trp | P28331 |
| HOMO SAPIENS | 1 | 1019 | 5xtb | M | 252 | ASP | p.Asp252Gly | P28331 |
| HOMO SAPIENS | 1 | 1019 | 5xtb | M | 408 | ARG | p.Arg408Cys | P28331 |
| HOMO SAPIENS | 2 | 1019 | 5xtb | M | 417 | ARG | p.Arg417Ser | P28331 |
| HOMO SAPIENS | 1 | 1019 | 5xtb | M | 557 | ARG | p.Arg557Ter | P28331 |
| HOMO SAPIENS | 0 | 1019 | 5xtb | M | 567 | ILE | p.Ile567Val | P28331 |
| HOMO SAPIENS | 1 | 1019 | 5xtb | M | 595 | THR | p.Thr595Ala | P28331 |
| HOMO SAPIENS | 1 | 1019 | 5xtb | M | 703 | ALA | p.Ala703Thr | P28331 |
| HOMO SAPIENS | -1 | 3437 | 5hyn | A | 185 | ASP | p.Asp185His | Q15910 |
| HOMO SAPIENS | -1 | 3437 | 5hyn | A | 482 | ALA | p.Ala482Thr | Q15910 |
| HOMO SAPIENS | 0 | 8201 | 2wfd | A | 373 | TYR | p.Tyr373Cys | Q9P2J5 |
| HOMO SAPIENS | 1 | 3590 | 3edi | A | 90 | VAL | p.Val238Ala | O43897 |
| HOMO SAPIENS | 6 | 4555 | 1ytq | A | 64 | ALA | p.Ala65Ser | P43320 |
| HOMO SAPIENS | 0 | 4555 | 1ytq | A | 194 | TRP | p.Trp195Gly | P43320 |
| HOMO SAPIENS | 0 | 1013 | 2nn6 | C | 272 | SER | p.Ser272Thr | Q96B26 |
| HOMO SAPIENS | -1 | 29026 | 5m73 | D | 530 | ILE | p.Ile530Asn | O76094 |
| HOMO SAPIENS | 8 | 2252 | 1d5v | A | 57 | SER | p.Ser125Leu | Q99958 |
| HOMO SAPIENS | 1 | 4337 | 3ecr | A | 29 | ARG | p.Arg26His | P08397 |
| HOMO SAPIENS | 2 | 4337 | 3ecr | A | 34 | ALA | p.Ala31Thr | P08397 |
| HOMO SAPIENS | 2 | 4337 | 3ecr | A | 37 | GLN | p.Gln34Lys | P08397 |
| HOMO SAPIENS | 1 | 4337 | 3ecr | A | 58 | ALA | p.Ala55Ser | P08397 |
| HOMO SAPIENS | 0 | 4337 | 3ecr | A | 84 | LEU | p.Leu81Pro | P08397 |
| HOMO SAPIENS | 0 | 4337 | 3ecr | A | 114 | GLY | p.Gly111Arg | P08397 |
| HOMO SAPIENS | 1 | 4337 | 3ecr | A | 119 | ARG | p.Arg116Trp | P08397 |
| HOMO SAPIENS | 1 | 4337 | 3ecr | A | 152 | ARG | p.Arg149Gln | P08397 |
| HOMO SAPIENS | 1 | 4337 | 3ecr | A | 170 | ARG | p.Arg167Trp | P08397 |
| HOMO SAPIENS | 1 | 4337 | 3ecr | A | 176 | ARG | p.Arg173Gln | P08397 |
| HOMO SAPIENS | 0 | 4337 | 3ecr | A | 180 | LEU | p.Leu177Arg | P08397 |
| HOMO SAPIENS | 0 | 4337 | 3ecr | A | 181 | ASP | p.Asp178Asn | P08397 |
| HOMO SAPIENS | 1 | 4337 | 3ecr | A | 198 | ARG | p.Arg195Cys | P08397 |
| HOMO SAPIENS | 0 | 4337 | 3ecr | A | 201 | TRP | p.Trp198Ter | P08397 |
| HOMO SAPIENS | 1 | 4337 | 3ecr | A | 204 | ARG | p.Arg201Trp | P08397 |
| HOMO SAPIENS | 1 | 4337 | 3ecr | A | 219 | GLY | p.Gly216Asp | P08397 |
| HOMO SAPIENS | 1 | 4337 | 3ecr | A | 226 | GLU | p.Glu223Lys | P08397 |
| HOMO SAPIENS | 0 | 4337 | 3ecr | A | 248 | LEU | p.Leu245Arg | P08397 |

| | | | | | | | | |
|---|---|---|---|---|---|---|---|---|
| HOMO SAPIENS | 1 | 4337 | 3ecr | A | 250 | CYS | p.Cys247Arg | P08397 |
| HOMO SAPIENS | 0 | 4337 | 3ecr | A | 253 | GLU | p.Glu250Lys | P08397 |
| HOMO SAPIENS | 0 | 4337 | 3ecr | A | 255 | ALA | p.Ala252Thr | P08397 |
| HOMO SAPIENS | 0 | 4337 | 3ecr | A | 255 | ALA | p.Ala252Val | P08397 |
| HOMO SAPIENS | 0 | 4337 | 3ecr | A | 259 | HIS | p.His256Asn | P08397 |
| HOMO SAPIENS | 0 | 4337 | 3ecr | A | 324 | ARG | p.Arg321His | P08397 |
| HOMO SAPIENS | -1 | 4337 | 3ecr | A | 362 | ASP | p.Asp359Asn | P08397 |
| HOMO SAPIENS | 1 | 1204 | 5lwv | A | 854 | ILE | p.Ala864Thr | O15294 |
| HOMO SAPIENS | 1 | 25016 | 1m4u | A | 35 | PRO | p.Pro35Arg | Q13253 |
| HOMO SAPIENS | 1 | 25016 | 1m4u | A | 35 | PRO | p.Pro35Ser | Q13253 |
| HOMO SAPIENS | 1 | 25016 | 1m4u | A | 167 | ARG | p.Arg167Gly | Q13253 |
| HOMO SAPIENS | 1 | 25016 | 1m4u | A | 184 | CYS | p.Cys184Tyr | Q13253 |
| HOMO SAPIENS | 0 | 25016 | 1m4u | A | 189 | GLY | p.Gly189Cys | Q13253 |
| HOMO SAPIENS | 1 | 25016 | 1m4u | A | 204 | ARG | p.Arg204Leu | Q13253 |
| HOMO SAPIENS | 1 | 25016 | 1m4u | A | 205 | TRP | p.Trp205Cys | Q13253 |
| HOMO SAPIENS | 1 | 25016 | 1m4u | A | 217 | TRP | p.Trp217Gly | Q13253 |
| HOMO SAPIENS | 0 | 25016 | 1m4u | A | 222 | TYR | p.Tyr222Cys | Q13253 |
| HOMO SAPIENS | 0 | 25016 | 1m4u | A | 222 | TYR | p.Tyr222Asp | Q13253 |
| HOMO SAPIENS | 1 | 25016 | 1m4u | A | 223 | PRO | p.Pro223Leu | Q13253 |
| HOMO SAPIENS | 0 | 1394 | 1um7 | A | 95 | ALA | p.Ala463Val | Q92796 |
| HOMO SAPIENS | 0 | 21294 | 4jpz | B | 1788 | GLU | p.Glu1788Asp | Q99250 |
| HOMO SAPIENS | 9 | 24068 | 2db6 | A | 47 | ARG | p.Arg119Cys | Q96MF2 |
| HOMO SAPIENS | 5 | 1869 | 4omt | A | 39 | ARG | p.Arg39Leu | P08237 |
| HOMO SAPIENS | 2 | 1869 | 4omt | A | 100 | ARG | p.Arg100Gln | P08237 |
| HOMO SAPIENS | 7 | 1869 | 4omt | A | 543 | ASP | p.Asp543Ala | P08237 |
| HOMO SAPIENS | 1 | 1869 | 4omt | A | 686 | TRP | p.Trp686Cys | P08237 |
| HOMO SAPIENS | 1 | 1869 | 4omt | A | 696 | ARG | p.Arg696His | P08237 |
| HOMO SAPIENS | 0 | 12163 | 4wxx | A | 495 | TYR | p.Tyr495Cys | P26358 |
| HOMO SAPIENS | 0 | 12163 | 4wxx | A | 528 | ILE | p.Ile528Met | P26358 |
| HOMO SAPIENS | 0 | 12391 | 1j1d | C | 36 | LYS | p.Lys36Gln | P19429 |
| HOMO SAPIENS | 0 | 12391 | 1j1d | C | 82 | PRO | p.Pro82Ser | P19429 |
| HOMO SAPIENS | 7 | 12391 | 1j1d | C | 102 | ALA | p.Ala102Thr | P19429 |
| HOMO SAPIENS | 1 | 5000 | 4cca | A | 17 | GLY | p.Gly17Arg | Q15833 |
| HOMO SAPIENS | 0 | 5000 | 4cca | A | 205 | VAL | p.Val205Ile | Q15833 |
| HOMO SAPIENS | 0 | 5000 | 4cca | A | 209 | LEU | p.Leu209Pro | Q15833 |
| HOMO SAPIENS | 1 | 5000 | 4cca | A | 459 | ARG | p.Arg459Trp | Q15833 |
| HOMO SAPIENS | 7 | 5000 | 4cca | A | 477 | PRO | p.Pro477Leu | Q15833 |
| HOMO SAPIENS | -1 | 5000 | 4cca | A | 526 | ILE | p.Ile526Val | Q15833 |
| HOMO SAPIENS | 0 | 5000 | 4cca | A | 541 | GLY | p.Gly541Ser | Q15833 |
| HOMO SAPIENS | 0 | 5000 | 4cca | A | 555 | ARG | p.Arg555Gly | Q15833 |
| HOMO SAPIENS | 5 | 28551 | 3vcm | P | 26 | ARG | p.Arg49Ter | P00797 |
| HOMO SAPIENS | 6 | 36453 | 5dzz | A | 2070 | ASP | p.Asp2070His | P15924 |
| HOMO SAPIENS | 0 | 36453 | 5dzz | A | 2166 | ARG | p.Arg2166Ter | P15924 |
| HOMO SAPIENS | 0 | 20268 | 2da6 | A | 96 | ASN | p.Asn321Ser | P35680 |
| HOMO SAPIENS | 1 | 1405 | 3iar | A | 8 | ASP | p.Asp8Asn | P00813 |
| HOMO SAPIENS | 8 | 1405 | 3iar | A | 15 | HIS | p.His15Asp | P00813 |
| HOMO SAPIENS | 6 | 1405 | 3iar | A | 74 | GLY | p.Gly74Val | P00813 |
| HOMO SAPIENS | 6 | 1405 | 3iar | A | 80 | LYS | p.Lys80Arg | P00813 |
| HOMO SAPIENS | 5 | 1405 | 3iar | A | 97 | TYR | p.Tyr97Cys | P00813 |
| HOMO SAPIENS | 5 | 1405 | 3iar | A | 101 | ARG | p.Arg101Leu | P00813 |
| HOMO SAPIENS | 5 | 1405 | 3iar | A | 101 | ARG | p.Arg101Trp | P00813 |
| HOMO SAPIENS | 5 | 1405 | 3iar | A | 106 | LEU | p.Leu106Val | P00813 |
| HOMO SAPIENS | 0 | 1405 | 3iar | A | 107 | LEU | p.Leu107Pro | P00813 |
| HOMO SAPIENS | 9 | 1405 | 3iar | A | 156 | ARG | p.Arg156Cys | P00813 |

| | | | | | | | | |
|---|---|---|---|---|---|---|---|---|
| HOMO SAPIENS | 6 | 1405 | 3iar | A | 211 | ARG | p.Arg211His | P00813 |
| HOMO SAPIENS | 0 | 1405 | 3iar | A | 216 | GLY | p.Gly216Arg | P00813 |
| HOMO SAPIENS | 0 | 1405 | 3iar | A | 291 | SER | p.Ser291Leu | P00813 |
| HOMO SAPIENS | 6 | 1405 | 3iar | A | 304 | LEU | p.Leu304Arg | P00813 |
| HOMO SAPIENS | 0 | 1405 | 3iar | A | 329 | ALA | p.Ala329Val | P00813 |
| HOMO SAPIENS | -1 | 31340 | 4atu | I | 186 | ARG | p.Arg186His | O43602 |
| HOMO SAPIENS | -1 | 31340 | 4atu | I | 192 | ARG | p.Arg192Trp | O43602 |
| HOMO SAPIENS | -1 | 31340 | 4atu | I | 196 | ARG | p.Arg196His | O43602 |
| HOMO SAPIENS | -1 | 31340 | 4atu | I | 203 | THR | p.Thr203Arg | O43602 |
| HOMO SAPIENS | -1 | 31340 | 4atu | I | 286 | ALA | p.Ala286Ser | O43602 |
| HOMO SAPIENS | 4 | 30734 | 1fsu | A | 95 | ARG | p.Arg95Gln | P15848 |
| HOMO SAPIENS | 0 | 30734 | 1fsu | A | 117 | CYS | p.Cys117Arg | P15848 |
| HOMO SAPIENS | 0 | 30734 | 1fsu | A | 137 | GLY | p.Gly137Val | P15848 |
| HOMO SAPIENS | 0 | 30734 | 1fsu | A | 210 | TYR | p.Tyr210Cys | P15848 |
| HOMO SAPIENS | 0 | 30734 | 1fsu | A | 236 | LEU | p.Leu236Pro | P15848 |
| HOMO SAPIENS | 1 | 30734 | 1fsu | A | 315 | ARG | p.Arg315Gln | P15848 |
| HOMO SAPIENS | 0 | 30734 | 1fsu | A | 358 | VAL | p.Val358Met | P15848 |
| HOMO SAPIENS | 2 | 30734 | 1fsu | A | 376 | VAL | p.Val376Met | P15848 |
| HOMO SAPIENS | 0 | 30734 | 1fsu | A | 384 | SER | p.Ser384Asn | P15848 |
| HOMO SAPIENS | 0 | 30734 | 1fsu | A | 393 | HIS | p.His393Pro | P15848 |
| HOMO SAPIENS | 0 | 30734 | 1fsu | A | 405 | CYS | p.Cys405Tyr | P15848 |
| HOMO SAPIENS | 0 | 30734 | 1fsu | A | 458 | ASN | p.Asn458Ser | P15848 |
| HOMO SAPIENS | 1 | 30734 | 1fsu | A | 484 | ARG | p.Arg484Gly | P15848 |
| HOMO SAPIENS | 0 | 38072 | 1wmv | A | 51 | PRO | p.Pro98Leu | Q9NZC7 |
| HOMO SAPIENS | -1 | 5674 | 1j1e | C | 185 | ASN | p.Asn185Lys | P19429 |
| HOMO SAPIENS | -1 | 5674 | 1j1e | C | 192 | ARG | p.Arg192His | P19429 |
| HOMO SAPIENS | -1 | 5674 | 1j1e | C | 196 | ASP | p.Asp196Asn | P19429 |
| HOMO SAPIENS | -1 | 5674 | 1j1e | C | 206 | LYS | p.Lys206Gln | P19429 |
| HOMO SAPIENS | 0 | 38067 | 1wlp | A | 13 | PRO | p.Pro156Gln | P13498 |
| HOMO SAPIENS | -1 | 20441 | 3vfd | A | 289 | HIS | p.His289Tyr | Q9UBP0 |
| HOMO SAPIENS | 2 | 20441 | 3vfd | A | 344 | ILE | p.Ile344Lys | Q9UBP0 |
| HOMO SAPIENS | 0 | 20441 | 3vfd | A | 362 | SER | p.Ser362Cys | Q9UBP0 |
| HOMO SAPIENS | 1 | 20441 | 3vfd | A | 386 | ASN | p.Asn386Ser | Q9UBP0 |
| HOMO SAPIENS | 9 | 20441 | 3vfd | A | 441 | ASP | p.Asp441Gly | Q9UBP0 |
| HOMO SAPIENS | -1 | 20441 | 3vfd | A | 448 | CYS | p.Cys448Tyr | Q9UBP0 |
| HOMO SAPIENS | 2 | 20441 | 3vfd | A | 500 | ARG | p.Arg499Cys | Q9UBP0 |
| HOMO SAPIENS | 1 | 20441 | 3vfd | A | 563 | ARG | p.Arg562Gly | Q9UBP0 |
| HOMO SAPIENS | 3 | 18471 | 4dx8 | J | 26 | ARG | p.Arg26Gln | O00522 |
| HOMO SAPIENS | -1 | 18471 | 4dx8 | J | 101 | GLY | p.Gly101Val | O00522 |
| HOMO SAPIENS | -1 | 34735 | 2ly4 | B | 83 | ALA | p.Ala83Glu | P04637 |
| HOMO SAPIENS | 0 | 31128 | 4cvh | A | 116 | ARG | p.Arg116Cys | A4D126 |
| HOMO SAPIENS | 8 | 31128 | 4cvh | A | 215 | GLN | p.Gln215Ter | A4D126 |
| HOMO SAPIENS | 0 | 31128 | 4cvh | A | 268 | ARG | p.Arg268Ter | A4D126 |
| HOMO SAPIENS | 0 | 31128 | 4cvh | A | 316 | THR | p.Thr316Lys | A4D126 |
| HOMO SAPIENS | 1 | 7164 | 3mt5 | A | 369 | ASP | p.Asp434Gly | Q12791 |
| HOMO SAPIENS | 2 | 26812 | 2wtk | F | 49 | TYR | p.Tyr49Asp | Q15831 |
| HOMO SAPIENS | 0 | 26812 | 2wtk | F | 67 | LEU | p.Leu67Pro | Q15831 |
| HOMO SAPIENS | 4 | 26812 | 2wtk | F | 104 | ARG | p.Arg104Ser | Q15831 |
| HOMO SAPIENS | 0 | 26812 | 2wtk | F | 135 | GLY | p.Gly135Arg | Q15831 |
| HOMO SAPIENS | 0 | 26812 | 2wtk | F | 163 | GLY | p.Gly163Asp | Q15831 |
| HOMO SAPIENS | 2 | 26812 | 2wtk | F | 194 | ALA | p.Asp194Asn | Q15831 |
| HOMO SAPIENS | 2 | 1357 | 5xtd | W | 57 | ARG | p.Arg57His | Q9P0J0 |
| HOMO SAPIENS | 0 | 28086 | 4nn2 | A | 229 | HIS | p.His229Arg | Q8IWS0 |
| HOMO SAPIENS | 0 | 28086 | 4nn2 | A | 234 | LYS | p.Lys234Glu | Q8IWS0 |

| | | | | | | | | |
|---|---|---|---|---|---|---|---|---|
| HOMO SAPIENS | 0 | 28086 | 4nn2 | A | 257 | ARG | p.Arg257Gly | Q8IWS0 |
| HOMO SAPIENS | 5 | 31182 | 2a1x | A | 176 | GLN | p.Gln176Lys | O14832 |
| HOMO SAPIENS | 1 | 31182 | 2a1x | A | 204 | GLY | p.Gly204Ser | O14832 |
| HOMO SAPIENS | 0 | 31182 | 2a1x | A | 269 | ASN | p.Asn269His | O14832 |
| HOMO SAPIENS | 3 | 31182 | 2a1x | A | 275 | ARG | p.Arg275Gln | O14832 |
| HOMO SAPIENS | 3 | 31182 | 2a1x | A | 275 | ARG | p.Arg275Trp | O14832 |
| HOMO SAPIENS | 4 | 2823 | 5xtb | P | 145 | THR | p.Thr145Ile | O75489 |
| HOMO SAPIENS | 5 | 2823 | 5xtb | P | 199 | ARG | p.Arg199Trp | O75489 |
| HOMO SAPIENS | 0 | 8645 | 2h5g | A | 372 | SER | p.Ser372Tyr | P54886 |
| HOMO SAPIENS | 3 | 8645 | 2h5g | A | 715 | ASP | p.Asp715His | P54886 |
| HOMO SAPIENS | 0 | 8645 | 2h5g | A | 765 | ARG | p.Arg765Gln | P54886 |
| HOMO SAPIENS | 0 | 8645 | 2h5g | A | 782 | TYR | p.Tyr782Cys | P54886 |
| HOMO SAPIENS | 0 | 8645 | 2h5g | A | 784 | HIS | p.His784Tyr | P54886 |
| HOMO SAPIENS | 1 | 13992 | 2cue | A | 39 | ARG | p.Arg242Thr | P26367 |
| HOMO SAPIENS | 9 | 13992 | 2cue | A | 55 | PHE | p.Phe258Ser | P26367 |
| HOMO SAPIENS | 0 | 30418 | 2xsw | A | 303 | VAL | p.Val303Met | Q9NRR6 |
| HOMO SAPIENS | 0 | 30418 | 2xsw | A | 315 | PRO | p.Pro315Leu | Q9NRR6 |
| HOMO SAPIENS | 0 | 30418 | 2xsw | A | 341 | GLY | p.Gly341Ser | Q9NRR6 |
| HOMO SAPIENS | 0 | 30418 | 2xsw | A | 378 | ARG | p.Arg378Cys | Q9NRR6 |
| HOMO SAPIENS | 0 | 30418 | 2xsw | A | 435 | ARG | p.Arg435Gln | Q9NRR6 |
| HOMO SAPIENS | 0 | 30418 | 2xsw | A | 515 | ARG | p.Arg515Trp | Q9NRR6 |
| HOMO SAPIENS | 1 | 30418 | 2xsw | A | 526 | PRO | p.Pro526Leu | Q9NRR6 |
| HOMO SAPIENS | 0 | 30418 | 2xsw | A | 563 | ARG | p.Arg563His | Q9NRR6 |
| HOMO SAPIENS | 2 | 30418 | 2xsw | A | 585 | ARG | p.Arg585His | Q9NRR6 |
| HOMO SAPIENS | 2 | 11167 | 4y99 | B | 205 | ARG | p.Arg215Leu | P45379 |
| HOMO SAPIENS | 5 | 11167 | 4y99 | B | 244 | GLU | p.Glu254Asp | P45379 |
| HOMO SAPIENS | 1 | 11167 | 4y99 | B | 253 | LYS | p.Lys263Arg | P45379 |
| HOMO SAPIENS | 8 | 11167 | 4y99 | B | 265 | ARG | p.Arg275Gln | P45379 |
| HOMO SAPIENS | 3 | 11167 | 4y99 | B | 270 | ASP | p.Asp280Asn | P45379 |
| HOMO SAPIENS | -1 | 11167 | 4y99 | B | 278 | ARG | p.Arg288Pro | P45379 |
| HOMO SAPIENS | -1 | 11167 | 4y99 | B | 286 | ARG | p.Arg296Cys | P45379 |
| HOMO SAPIENS | -1 | 11167 | 4y99 | B | 286 | ARG | p.Arg296His | P45379 |
| HOMO SAPIENS | 0 | 15993 | 4po6 | A | 197 | ARG | p.Arg197His | P29597 |
| HOMO SAPIENS | -1 | 15993 | 4po6 | A | 362 | VAL | p.Val362Phe | P29597 |
| HOMO SAPIENS | -1 | 15993 | 4po6 | A | 363 | GLY | p.Gly363Ser | P29597 |
| HOMO SAPIENS | 0 | 5646 | 5uak | A | 7 | GLU | p.Glu7Ter | P13569 |
| HOMO SAPIENS | 0 | 5646 | 5uak | A | 27 | GLY | p.Gly27Ter | P13569 |
| HOMO SAPIENS | 0 | 5646 | 5uak | A | 46 | ALA | p.Ala46Asp | P13569 |
| HOMO SAPIENS | 0 | 5646 | 5uak | A | 60 | GLU | p.Glu60Ter | P13569 |
| HOMO SAPIENS | 0 | 5646 | 5uak | A | 67 | PRO | p.Pro67Leu | P13569 |
| HOMO SAPIENS | 0 | 5646 | 5uak | A | 74 | ARG | p.Arg74Trp | P13569 |
| HOMO SAPIENS | 0 | 5646 | 5uak | A | 75 | ARG | p.Arg75Ter | P13569 |
| HOMO SAPIENS | 0 | 5646 | 5uak | A | 85 | GLY | p.Gly85Glu | P13569 |
| HOMO SAPIENS | 0 | 5646 | 5uak | A | 91 | GLY | p.Gly91Arg | P13569 |
| HOMO SAPIENS | 0 | 5646 | 5uak | A | 92 | GLU | p.Glu92Lys | P13569 |
| HOMO SAPIENS | 0 | 5646 | 5uak | A | 98 | GLN | p.Gln98Arg | P13569 |
| HOMO SAPIENS | 0 | 5646 | 5uak | A | 98 | GLN | p.Gln98Ter | P13569 |
| HOMO SAPIENS | 0 | 5646 | 5uak | A | 109 | TYR | p.Tyr109Cys | P13569 |
| HOMO SAPIENS | 0 | 5646 | 5uak | A | 110 | ASP | p.Asp110His | P13569 |
| HOMO SAPIENS | 0 | 5646 | 5uak | A | 117 | ARG | p.Arg117Cys | P13569 |
| HOMO SAPIENS | 0 | 5646 | 5uak | A | 117 | ARG | p.Arg117His | P13569 |
| HOMO SAPIENS | 0 | 5646 | 5uak | A | 125 | ILE | p.Ile125Thr | P13569 |
| HOMO SAPIENS | 0 | 5646 | 5uak | A | 148 | ILE | p.Ile148Thr | P13569 |
| HOMO SAPIENS | 0 | 5646 | 5uak | A | 170 | ARG | p.Arg170His | P13569 |

| | | | | | | | | |
|---|---|---|---|---|---|---|---|---|
| HOMO SAPIENS | 0 | 5646 | 5uak | A | 178 | GLY | p.Gly178Arg | P13569 |
| HOMO SAPIENS | 0 | 5646 | 5uak | A | 206 | LEU | p.Leu206Trp | P13569 |
| HOMO SAPIENS | 0 | 5646 | 5uak | A | 373 | ASP | p.Asp373Asn | P13569 |
| HOMO SAPIENS | -1 | 5646 | 5uak | A | 709 | ARG | p.Arg709Ter | P13569 |
| HOMO SAPIENS | -1 | 5646 | 5uak | A | 732 | LEU | p.Leu732Ter | P13569 |
| HOMO SAPIENS | -1 | 5646 | 5uak | A | 785 | ARG | p.Arg785Ter | P13569 |
| HOMO SAPIENS | -1 | 5646 | 5uak | A | 831 | GLU | p.Glu831Ter | P13569 |
| HOMO SAPIENS | -1 | 5646 | 5uak | A | 836 | ASP | p.Asp836Tyr | P13569 |
| HOMO SAPIENS | 0 | 5646 | 5uak | A | 851 | ARG | p.Arg851Ter | P13569 |
| HOMO SAPIENS | 0 | 5646 | 5uak | A | 913 | TYR | p.Tyr913Cys | P13569 |
| HOMO SAPIENS | 0 | 5646 | 5uak | A | 913 | TYR | p.Tyr913Ter | P13569 |
| HOMO SAPIENS | 0 | 5646 | 5uak | A | 945 | SER | p.Ser945Leu | P13569 |
| HOMO SAPIENS | 0 | 5646 | 5uak | A | 949 | HIS | p.His949Tyr | P13569 |
| HOMO SAPIENS | 0 | 5646 | 5uak | A | 970 | GLY | p.Gly970Asp | P13569 |
| HOMO SAPIENS | 0 | 5646 | 5uak | A | 1027 | ILE | p.Ile1027Thr | P13569 |
| HOMO SAPIENS | 0 | 5646 | 5uak | A | 1032 | TYR | p.Tyr1032Cys | P13569 |
| HOMO SAPIENS | 0 | 5646 | 5uak | A | 1061 | GLY | p.Gly1061Arg | P13569 |
| HOMO SAPIENS | 1 | 5646 | 5uak | A | 1065 | LEU | p.Leu1065Pro | P13569 |
| HOMO SAPIENS | 1 | 5646 | 5uak | A | 1066 | ARG | p.Arg1066Cys | P13569 |
| HOMO SAPIENS | 1 | 5646 | 5uak | A | 1066 | ARG | p.Arg1066His | P13569 |
| HOMO SAPIENS | 1 | 5646 | 5uak | A | 1067 | ALA | p.Ala1067Thr | P13569 |
| HOMO SAPIENS | 0 | 5646 | 5uak | A | 1069 | GLY | p.Gly1069Arg | P13569 |
| HOMO SAPIENS | 0 | 5646 | 5uak | A | 1070 | ARG | p.Arg1070Trp | P13569 |
| HOMO SAPIENS | 1 | 5646 | 5uak | A | 1071 | GLN | p.Gln1071Pro | P13569 |
| HOMO SAPIENS | 0 | 5646 | 5uak | A | 1074 | PHE | p.Phe1074Leu | P13569 |
| HOMO SAPIENS | 0 | 5646 | 5uak | A | 1077 | LEU | p.Leu1077Pro | P13569 |
| HOMO SAPIENS | 0 | 5646 | 5uak | A | 1089 | TRP | p.Trp1089Ter | P13569 |
| HOMO SAPIENS | 0 | 5646 | 5uak | A | 1092 | TYR | p.Tyr1092Ter | P13569 |
| HOMO SAPIENS | 0 | 5646 | 5uak | A | 1101 | MET | p.Met1101Lys | P13569 |
| HOMO SAPIENS | 0 | 5646 | 5uak | A | 1156 | LEU | p.Leu1156Phe | P13569 |
| HOMO SAPIENS | 0 | 5646 | 5uak | A | 1158 | ARG | p.Arg1158Ter | P13569 |
| HOMO SAPIENS | 0 | 5646 | 5uak | A | 1162 | ARG | p.Arg1162Leu | P13569 |
| HOMO SAPIENS | 0 | 5646 | 5uak | A | 1162 | ARG | p.Arg1162Ter | P13569 |
| HOMO SAPIENS | -1 | 5646 | 5uak | A | 1476 | GLN | p.Gln1476Ter | P13569 |
| HOMO SAPIENS | 0 | 22763 | 1imv | A | 52 | MET | p.Thr72Met | P36955 |
| HOMO SAPIENS | 0 | 22763 | 1imv | A | 301 | TYR | p.Tyr321Ter | P36955 |
| HOMO SAPIENS | 0 | 23179 | 5fql | A | 160 | PRO | p.Pro160Arg | P22304 |
| HOMO SAPIENS | 0 | 23179 | 5fql | A | 172 | ARG | p.Arg172Ter | P22304 |
| HOMO SAPIENS | 0 | 23179 | 5fql | A | 214 | THR | p.Thr214Met | P22304 |
| HOMO SAPIENS | 0 | 23179 | 5fql | A | 309 | THR | p.Thr309Ala | P22304 |
| HOMO SAPIENS | 0 | 23179 | 5fql | A | 333 | SER | p.Ser333Leu | P22304 |
| HOMO SAPIENS | 0 | 23179 | 5fql | A | 443 | ARG | p.Arg443Ter | P22304 |
| HOMO SAPIENS | 0 | 23179 | 5fql | A | 468 | ARG | p.Arg468Leu | P22304 |
| HOMO SAPIENS | 0 | 23179 | 5fql | A | 468 | ARG | p.Arg468Trp | P22304 |
| HOMO SAPIENS | 0 | 23179 | 5fql | A | 502 | TRP | p.Trp502Ser | P22304 |
| HOMO SAPIENS | 9 | 20989 | 1nfi | F | 185 | THR | p.Thr185Met | P25963 |
| HOMO SAPIENS | 4 | 20989 | 1nfi | F | 194 | GLY | p.Gly194Ala | P25963 |
| HOMO SAPIENS | 0 | 11538 | 3eab | A | 162 | VAL | p.Val162Ile | Q9UBP0 |
| HOMO SAPIENS | 0 | 33399 | 2p39 | A | 71 | SER | p.Ser71Gly | Q9GZV9 |
| HOMO SAPIENS | 3 | 33399 | 2p39 | A | 96 | MET | p.Met96Thr | Q9GZV9 |
| HOMO SAPIENS | 0 | 33399 | 2p39 | A | 129 | SER | p.Ser129Phe | Q9GZV9 |
| HOMO SAPIENS | -1 | 33399 | 2p39 | A | 172 | PRO | p.Pro172Leu | Q9GZV9 |
| HOMO SAPIENS | -1 | 33399 | 2p39 | A | 176 | ARG | p.Arg176Gln | Q9GZV9 |
| HOMO SAPIENS | -1 | 33399 | 2p39 | A | 179 | ARG | p.Arg179Trp | Q9GZV9 |

| | | | | | | | | |
|---|---|---|---|---|---|---|---|---|
| HOMO SAPIENS | -1 | 33399 | 2p39 | A | 179 | ARG | p.Arg179Gln | Q9GZV9 |
| HOMO SAPIENS | 0 | 16217 | 2dne | A | 79 | ILE | p.Ile163Leu | P10515 |
| HOMO SAPIENS | 8 | 7666 | 4kbq | A | 65 | ASN | p.Asn65Ser | Q9UNE7 |
| HOMO SAPIENS | 1 | 6407 | 1ikt | A | 71 | THR | p.Thr687Ile | P51659 |
| HOMO SAPIENS | 0 | 6407 | 1ikt | A | 112 | MET | p.Met728Val | P51659 |
| HOMO SAPIENS | 0 | 5849 | 4ihb | A | 67 | VAL | p.Val67Asp | O75923 |
| HOMO SAPIENS | 6 | 17703 | 5fcn | A | 20 | HIS | p.His99Gln | Q66GS9 |
| HOMO SAPIENS | 8 | 17703 | 5fcn | A | 33 | ARG | p.Arg112His | Q66GS9 |
| HOMO SAPIENS | 0 | 34095 | 2E+073 | A | 101 | HIS | p.His101Tyr | P05129 |
| HOMO SAPIENS | -1 | 41358 | 4liq | E | 507 | THR | p.Thr507Met | P07333 |
| HOMO SAPIENS | -1 | 21290 | 1fch | A | 235 | MET | p.Met272Arg | P50542 |
| HOMO SAPIENS | 5 | 23431 | 6pax | A | 23 | ARG | p.Arg26Gly | P26367 |
| HOMO SAPIENS | 1 | 23431 | 6pax | A | 61 | GLY | p.Gly64Val | P26367 |
| HOMO SAPIENS | 1 | 23431 | 6pax | A | 65 | PRO | p.Pro68Ser | P26367 |
| HOMO SAPIENS | 1 | 23431 | 6pax | A | 116 | SER | p.Ser119Arg | P26367 |
| HOMO SAPIENS | 9 | 23431 | 6pax | A | 123 | VAL | p.Val126Asp | P26367 |
| HOMO SAPIENS | 9 | 23431 | 6pax | A | 125 | ARG | p.Arg128Cys | P26367 |
| HOMO SAPIENS | 9 | 33003 | 4zrj | B | 535 | LEU | p.Leu535Pro | P35240 |
| HOMO SAPIENS | 7 | 33003 | 4zrj | B | 538 | GLN | p.Gln538Pro | P35240 |
| HOMO SAPIENS | 0 | 33003 | 4zrj | B | 585 | TRP | p.Ala585Thr | P35240 |
| HOMO SAPIENS | 0 | 3800 | 2nzj | A | 193 | VAL | p.Val193Met | O75628 |

| gene_name | max_cons | max_bs_id | max_pdb_id | max_chain_id |
|---|---|---|---|---|
| AKR1D1 | None | None | None | None |
| AKR1D1 | None | None | None | None |
| AKR1D1 | None | None | None | None |
| AKR1D1 | None | None | None | None |
| ALPP | None | None | None | None |
| BMPR1A | None | None | None | None |
| BMPR1A | None | None | None | None |
| FGFR2 | None | None | None | None |
| PROC | None | None | None | None |
| PROC | None | None | None | None |
| PROC | None | None | None | None |
| PROC | None | None | None | None |
| PROC | None | None | None | None |
| PROC | None | None | None | None |
| PROC | None | None | None | None |
| PROC | None | None | None | None |
| PROC | None | None | None | None |
| PROC | None | None | None | None |
| PROC | None | None | None | None |
| PROC | None | None | None | None |
| INS | 1 | 2 | 3zs2 | J |
| INS | None | None | None | None |
| INS | 1 | 1 | 3zs2 | J |
| INS | 1 | 1 | 3zs2 | J |
| FTL | None | None | None | None |
| FTL | None | None | None | None |
| FTL | None | None | None | None |
| TNFSF11 | None | None | None | None |
| EHMT1 | None | None | None | None |
| EHMT1 | None | None | None | None |
| EHMT1 | None | None | None | None |
| CA2 | 0.939 | 1 | 4idr | X |
| CA2 | 0.939 | 1 | 4idr | X |
| CA2 | 0.902 | 5 | 4idr | X |
| CA2 | 0.902 | 5 | 4idr | X |
| SMAD4 | None | None | None | None |
| SMAD4 | None | None | None | None |
| TTR | None | None | None | None |
| TTR | 0.961 | 7 | 3grb | A |
| TTR | None | None | None | None |
| TTR | 0.947 | 3 | 1tta | A |
| TTR | 0.923 | 1 | 3grb | B |
| TTR | None | None | None | None |
| TTR | None | None | None | None |
| TTR | None | None | None | None |
| TTR | 0.909 | 5 | 4tl4 | A |
| TTR | None | None | None | None |
| TTR | None | None | None | None |
| TTR | None | None | None | None |
| TTR | 0.902 | 9 | 3m1o | B |
| TTR | 0.902 | 9 | 3m1o | B |
| TTR | 0.902 | 9 | 3m1o | B |
| TTR | None | None | None | None |

| | | | | |
|---|---|---|---|---|
| TTR | 0.902 | 9 | 3m1o | B |
| TTR | None | None | None | None |
| TTR | None | None | None | None |
| TTR | None | None | None | None |
| TTR | 0.909 | 5 | 4tl4 | A |
| TTR | 0.909 | 5 | 4tl4 | A |
| TTR | None | None | None | None |
| TTR | 0.945 | 4 | 4deu | A |
| TTR | 0.941 | 3 | 3did | B |
| TTR | 0.941 | 3 | 3did | B |
| TTR | None | None | None | None |
| TTR | None | None | None | None |
| TTR | None | None | None | None |
| TTR | 0.941 | 3 | 3did | B |
| TTR | 0.923 | 1 | 4hiq | A |
| TTR | 0.923 | 1 | 4hiq | A |
| TTR | 0.961 | 7 | 3grb | A |
| TTR | None | None | None | None |
| TTR | None | None | None | None |
| ALK | None | None | None | None |
| ALK | None | None | None | None |
| THRB | None | None | None | None |
| THRB | None | None | None | None |
| THRB | None | None | None | None |
| THRB | None | None | None | None |
| THRB | None | None | None | None |
| THRB | None | None | None | None |
| THRB | None | None | None | None |
| THRB | None | None | None | None |
| THRB | None | None | None | None |
| THRB | None | None | None | None |
| THRB | None | None | None | None |
| THRB | None | None | None | None |
| THRB | None | None | None | None |
| THRB | None | None | None | None |
| THRB | None | None | None | None |
| THRB | None | None | None | None |
| THRB | None | None | None | None |
| THRB | None | None | None | None |
| IL2RG | None | None | None | None |
| IL2RG | None | None | None | None |
| IL2RG | None | None | None | None |
| IL2RG | None | None | None | None |
| IL2RG | None | None | None | None |
| ITGB3 | None | None | None | None |
| ITGB3 | None | None | None | None |
| ITGB3 | None | None | None | None |
| ITGB3 | None | None | None | None |
| ITGB3 | None | None | None | None |
| ITGB3 | None | None | None | None |
| ITGB3 | None | None | None | None |
| ITGB3 | None | None | None | None |
| ITGB3 | None | None | None | None |

| | | | | |
|---|---|---|---|---|
| PKD2 | None | None | None | None |
| PKD2 | None | None | None | None |
| PKD2 | None | None | None | None |
| PKD2 | None | None | None | None |
| PKD2 | None | None | None | None |
| PKD2 | None | None | None | None |
| PKD2 | None | None | None | None |
| TP53 | None | None | None | None |
| TP53 | None | None | None | None |
| TP53 | None | None | None | None |
| TP53 | None | None | None | None |
| TP53 | None | None | None | None |
| TP53 | None | None | None | None |
| EGFR | None | None | None | None |
| EGFR | None | None | None | None |
| XRCC4 | None | None | None | None |
| XRCC4 | None | None | None | None |
| HBB | None | None | None | None |
| HBB | None | None | None | None |
| HBB | None | None | None | None |
| HBB | None | None | None | None |
| HBB | None | None | None | None |
| HBB | None | None | None | None |
| HBB | None | None | None | None |
| HBB | None | None | None | None |
| HBB | None | None | None | None |
| HBB | None | None | None | None |
| HBB | None | None | None | None |
| HBB | None | None | None | None |
| HBB | None | None | None | None |
| HBB | None | None | None | None |
| HBB | None | None | None | None |
| HBB | None | None | None | None |
| HBB | None | None | None | None |
| HBB | None | None | None | None |
| HBB | None | None | None | None |
| HBB | None | None | None | None |
| HBB | None | None | None | None |
| HBB | None | None | None | None |
| HBB | None | None | None | None |
| HBB | None | None | None | None |
| HBB | None | None | None | None |
| HBB | None | None | None | None |
| HBB | None | None | None | None |
| HBB | None | None | None | None |
| HBB | None | None | None | None |
| HBB | None | None | None | None |
| RPS6KA3 | None | None | None | None |
| RPS6KA3 | None | None | None | None |
| RPS6KA3 | None | None | None | None |
| RPS6KA3 | None | None | None | None |
| RPS6KA3 | None | None | None | None |
| RPS6KA3 | None | None | None | None |
| GBA | 1 | 15 | 3gxd | B |

| | | | | | |
|---|---|---|---|---|---|
| GBA | | 1 | 55 | 3gxd | B |
| GBA | | 1 | 15 | 3gxd | B |
| GBA | None | None | | None | None |
| GBA | None | None | | None | None |
| GBA | None | None | | None | None |
| GBA | | 1 | 12 | 1ogs | A |
| GBA | None | None | | None | None |
| GBA | | 1 | 2 | 1ogs | A |
| GBA | | 1 | 11 | 1ogs | A |
| GBA | None | None | | None | None |
| GBA | None | None | | None | None |
| GBA | | 1 | 18 | 1ogs | A |
| GBA | | 1 | 3 | 1ogs | A |
| GBA | None | None | | None | None |
| GBA | | 1 | 2 | 1ogs | A |
| GBA | None | None | | None | None |
| GBA | None | None | | None | None |
| GBA | | 1 | 3 | 1ogs | A |
| GBA | | 1 | 3 | 1ogs | A |
| GBA | None | None | | None | None |
| GBA | | 1 | 28 | 1ogs | A |
| GBA | None | None | | None | None |
| GBA | None | None | | None | None |
| GBA | None | None | | None | None |
| GBA | None | None | | None | None |
| GBA | None | None | | None | None |
| GBA | None | None | | None | None |
| GBA | None | None | | None | None |
| GBA | None | None | | None | None |
| GBA | None | None | | None | None |
| GBA | None | None | | None | None |
| GBA | None | None | | None | None |
| GBA | None | None | | None | None |
| GBA | None | None | | None | None |
| GBA | None | None | | None | None |
| GBA | None | None | | None | None |
| GBA | None | None | | None | None |
| GBA | None | None | | None | None |
| GBA | | 1 | 15 | 3gxd | B |
| GBA | | 1 | 44 | 3gxd | B |
| GBA | None | None | | None | None |
| GBA | | 1 | 44 | 3gxd | B |
| GBA | | 1 | 44 | 3gxd | B |
| GBA | None | None | | None | None |
| GBA | None | None | | None | None |
| GBA | None | None | | None | None |
| IFIH1 | None | None | | None | None |
| TGFBR2 | None | None | | None | None |
| TGFBR2 | None | None | | None | None |
| TGFBR2 | None | None | | None | None |
| TGFBR2 | None | None | | None | None |
| TGFBR2 | None | None | | None | None |
| TGFBR2 | None | None | | None | None |
| TGFBR2 | None | None | | None | None |

| | | | | |
|---|---|---|---|---|
| TGFBR2 | None | None | None | None |
| TGFBR2 | None | None | None | None |
| TGFBR2 | None | None | None | None |
| TGFBR2 | None | None | None | None |
| TGFBR2 | None | None | None | None |
| TGFBR2 | None | None | None | None |
| TGFBR2 | None | None | None | None |
| TGFBR2 | None | None | None | None |
| ACE | None | None | None | None |
| ACE | None | None | None | None |
| ACE | None | None | None | None |
| ACE | None | None | None | None |
| ACE | None | None | None | None |
| ACE | None | None | None | None |
| COL10A1 | None | None | None | None |
| COL10A1 | None | None | None | None |
| COL10A1 | None | None | None | None |
| COL10A1 | None | None | None | None |
| COL10A1 | None | None | None | None |
| COL10A1 | None | None | None | None |
| COL10A1 | None | None | None | None |
| COL10A1 | None | None | None | None |
| COL10A1 | None | None | None | None |
| KLF1 | None | None | None | None |
| C3 | None | None | None | None |
| CTSC | None | None | None | None |
| CTSC | 0.934 | 1 | 2djg | C |
| CTSC | None | None | None | None |
| LYZ | None | None | None | None |
| LYZ | 0.923 | 1 | 1lz4 | A |
| LYZ | None | None | None | None |
| DHODH | None | None | None | None |
| DHODH | None | None | None | None |
| DHODH | None | None | None | None |
| DHODH | None | None | None | None |
| DHODH | None | None | None | None |
| DHODH | None | None | None | None |
| DHODH | None | None | None | None |
| ADH1C | None | None | None | None |
| GDNF | None | None | None | None |
| TNF | None | None | None | None |
| ALDH4A1 | None | None | None | None |
| ALDH4A1 | None | None | None | None |
| ALDH4A1 | None | None | None | None |
| BRAF | None | None | None | None |
| BRAF | None | None | None | None |
| BRAF | None | None | None | None |
| BRAF | None | None | None | None |
| BRAF | None | None | None | None |
| BRAF | None | None | None | None |
| BRAF | None | None | None | None |
| BRAF | None | None | None | None |
| BRAF | None | None | None | None |
| BRAF | None | None | None | None |

| | | | | |
|---|---|---|---|---|
| BRAF | None | None | None | None |
| BRAF | None | None | None | None |
| BRAF | None | None | None | None |
| BRAF | None | None | None | None |
| BRAF | None | None | None | None |
| BRAF | None | None | None | None |
| TBX5 | None | None | None | None |
| TBX5 | None | None | None | None |
| TBX5 | None | None | None | None |
| TBX5 | None | None | None | None |
| TBX5 | None | None | None | None |
| TBX5 | None | None | None | None |
| TBX5 | None | None | None | None |
| TBX5 | None | None | None | None |
| TBX5 | None | None | None | None |
| RB1 | None | None | None | None |
| RB1 | None | None | None | None |
| RB1 | None | None | None | None |
| RB1 | None | None | None | None |
| RB1 | None | None | None | None |
| RB1 | None | None | None | None |
| CLIC2 | None | None | None | None |
| CLIC2 | None | None | None | None |
| ALDH7A1 | None | None | None | None |
| ALDH7A1 | None | None | None | None |
| ALDH7A1 | None | None | None | None |
| ALDH7A1 | None | None | None | None |
| ALDH7A1 | None | None | None | None |
| ALDH7A1 | None | None | None | None |
| ALDH7A1 | None | None | None | None |
| ALDH7A1 | None | None | None | None |
| ALDH7A1 | None | None | None | None |
| ALDH7A1 | None | None | None | None |
| ALDH7A1 | None | None | None | None |
| ALDH7A1 | None | None | None | None |
| ALDH7A1 | None | None | None | None |
| ALDH7A1 | None | None | None | None |
| ALDH7A1 | None | None | None | None |
| ALDH7A1 | None | None | None | None |
| KDR | None | None | None | None |
| MUT | None | None | None | None |
| MUT | None | None | None | None |
| MUT | None | None | None | None |
| MUT | None | None | None | None |
| MUT | None | None | None | None |
| MUT | None | None | None | None |
| MUT | None | None | None | None |
| MUT | None | None | None | None |
| MUT | None | None | None | None |
| MUT | None | None | None | None |
| MUT | None | None | None | None |
| MUT | None | None | None | None |
| MUT | None | None | None | None |
| MUT | None | None | None | None |

| | | | | |
|---|---|---|---|---|
| MUT | None | None | None | None |
| MUT | None | None | None | None |
| MUT | None | None | None | None |
| MUT | None | None | None | None |
| MUT | None | None | None | None |
| MUT | None | None | None | None |
| MUT | None | None | None | None |
| MUT | None | None | None | None |
| MUT | None | None | None | None |
| MUT | None | None | None | None |
| MUT | None | None | None | None |
| MUT | None | None | None | None |
| MUT | None | None | None | None |
| MUT | None | None | None | None |
| MUT | None | None | None | None |
| MUT | None | None | None | None |
| MUT | None | None | None | None |
| MUT | None | None | None | None |
| MUT | None | None | None | None |
| MUT | None | None | None | None |
| SOD1 | None | None | None | None |
| SOD1 | None | None | None | None |
| SOD1 | None | None | None | None |
| SOD1 | None | None | None | None |
| NECTIN3 | None | None | None | None |
| HFE | None | None | None | None |
| HFE | None | None | None | None |
| HFE | None | None | None | None |
| HFE | None | None | None | None |
| HFE | None | None | None | None |
| HFE | None | None | None | None |
| HFE | None | None | None | None |
| EIF2AK3 | None | None | None | None |
| EIF2AK3 | None | None | None | None |
| TERT | None | None | None | None |
| TERT | None | None | None | None |
| SPR | None | None | None | None |
| SPR | None | None | None | None |
| SPR | None | None | None | None |
| SPR | None | None | None | None |
| SPR | None | None | None | None |
| F7 | None | None | None | None |
| UNG | None | None | None | None |
| MMP13 | None | None | None | None |
| MMP13 | None | None | None | None |
| MMP13 | None | None | None | None |
| MMP13 | None | None | None | None |
| CD207 | None | None | None | None |
| F2 | 1 | 2 | 1nrq | L |
| SPTBN2 | None | None | None | None |
| PKLR | None | None | None | None |
| PKLR | None | None | None | None |
| PKLR | None | None | None | None |
| PKLR | None | None | None | None |

| | | | | |
|---|---|---|---|---|
| PKLR | None | None | None | None |
| PKLR | None | None | None | None |
| PKLR | None | None | None | None |
| PKLR | None | None | None | None |
| SACS | None | None | None | None |
| HRAS | None | None | None | None |
| HRAS | None | None | None | None |
| HRAS | None | None | None | None |
| HRAS | None | None | None | None |
| HRAS | None | None | None | None |
| HRAS | None | None | None | None |
| HRAS | None | None | None | None |
| HRAS | None | None | None | None |
| HRAS | None | None | None | None |
| HRAS | None | None | None | None |
| HRAS | None | None | None | None |
| HRAS | None | None | None | None |
| HRAS | None | None | None | None |
| VHL | None | None | None | None |
| VHL | None | None | None | None |
| VHL | None | None | None | None |
| VHL | None | None | None | None |
| VHL | None | None | None | None |
| VHL | None | None | None | None |
| VHL | None | None | None | None |
| VHL | None | None | None | None |
| VHL | None | None | None | None |
| VHL | None | None | None | None |
| VHL | None | None | None | None |
| VHL | None | None | None | None |
| VHL | None | None | None | None |
| VHL | None | None | None | None |
| VHL | None | None | None | None |
| VHL | None | None | None | None |
| VHL | None | None | None | None |
| VHL | None | None | None | None |
| VHL | None | None | None | None |
| VHL | None | None | None | None |
| VHL | None | None | None | None |
| VHL | None | None | None | None |
| VHL | None | None | None | None |
| VHL | None | None | None | None |
| VHL | None | None | None | None |
| TTBK1 | None | None | None | None |
| RAD51 | None | None | None | None |
| ALB | None | None | None | None |
| ALB | None | None | None | None |
| ALB | None | None | None | None |
| ALB | None | None | None | None |
| MET | None | None | None | None |
| MET | None | None | None | None |
| MET | None | None | None | None |
| MET | None | None | None | None |
| MET | None | None | None | None |

| | | | | |
|---|---|---|---|---|
| MET | None | None | None | None |
| MET | None | None | None | None |
| MET | None | None | None | None |
| MET | None | None | None | None |
| CP | None | None | None | None |
| CP | None | None | None | None |
| PGM1 | None | None | None | None |
| PGM1 | None | None | None | None |
| PGM1 | None | None | None | None |
| PGM1 | None | None | None | None |
| PGM1 | None | None | None | None |
| PGM1 | None | None | None | None |
| PGM1 | None | None | None | None |
| NR5A1 | None | None | None | None |
| NR5A1 | None | None | None | None |
| NR5A1 | None | None | None | None |
| F9 | None | None | None | None |
| F9 | None | None | None | None |
| HPRT1 | None | None | None | None |
| HPRT1 | None | None | None | None |
| HPRT1 | None | None | None | None |
| HPRT1 | None | None | None | None |
| HPRT1 | None | None | None | None |
| HPRT1 | None | None | None | None |
| HPRT1 | None | None | None | None |
| HPRT1 | None | None | None | None |
| HPRT1 | None | None | None | None |
| HPRT1 | None | None | None | None |
| HPRT1 | None | None | None | None |
| HPRT1 | None | None | None | None |
| HPRT1 | None | None | None | None |
| HPRT1 | None | None | None | None |
| HPRT1 | None | None | None | None |
| HPRT1 | None | None | None | None |
| HPRT1 | None | None | None | None |
| HPRT1 | None | None | None | None |
| HPRT1 | None | None | None | None |
| HPRT1 | None | None | None | None |
| HPRT1 | None | None | None | None |
| HPRT1 | None | None | None | None |
| GPD1L | None | None | None | None |
| GPD1L | None | None | None | None |
| GPD1L | None | None | None | None |
| FGFR2 | None | None | None | None |
| FGFR2 | None | None | None | None |
| FGFR2 | None | None | None | None |
| FGFR2 | None | None | None | None |
| FGFR2 | None | None | None | None |
| FGFR2 | None | None | None | None |
| FGFR2 | None | None | None | None |
| FGFR2 | None | None | None | None |
| FGFR2 | None | None | None | None |
| FGFR2 | None | None | None | None |
| FGFR2 | None | None | None | None |

| | | | | |
|---|---|---|---|---|
| FGFR2 | None | None | None | None |
| FGFR2 | None | None | None | None |
| FGFR2 | None | None | None | None |
| FGFR2 | None | None | None | None |
| FGFR2 | None | None | None | None |
| FGFR2 | None | None | None | None |
| FGFR2 | None | None | None | None |
| FGFR2 | None | None | None | None |
| HSPD1 | None | None | None | None |
| HSPD1 | None | None | None | None |
| FGF8 | None | None | None | None |
| FGF8 | None | None | None | None |
| FGF8 | None | None | None | None |
| F9 | None | None | None | None |
| NCF4 | None | None | None | None |
| NCF4 | None | None | None | None |
| LDLR | None | None | None | None |
| LDLR | None | None | None | None |
| LDLR | None | None | None | None |
| LDLR | None | None | None | None |
| LDLR | None | None | None | None |
| LDLR | None | None | None | None |
| C3 | None | None | None | None |
| C3 | None | None | None | None |
| C3 | None | None | None | None |
| C3 | None | None | None | None |
| C3 | None | None | None | None |
| C3 | None | None | None | None |
| C3 | None | None | None | None |
| C3 | None | None | None | None |
| TULP1 | None | None | None | None |
| TULP1 | None | None | None | None |
| TULP1 | None | None | None | None |
| TULP1 | None | None | None | None |
| IKBKG | None | None | None | None |
| IKBKG | None | None | None | None |
| IKBKG | None | None | None | None |
| CD40 | None | None | None | None |
| CD40 | None | None | None | None |
| SLC9A3R1 | None | None | None | None |
| BCHE | None | None | None | None |
| BCHE | None | None | None | None |
| BCHE | None | None | None | None |
| BCHE | None | None | None | None |
| BCHE | None | None | None | None |
| BCHE | None | None | None | None |
| BCHE | None | None | None | None |
| BCHE | None | None | None | None |
| BCHE | None | None | None | None |
| BCHE | None | None | None | None |
| BCHE | None | None | None | None |
| BCKDHB | None | None | None | None |
| BCKDHB | None | None | None | None |
| BCKDHB | None | None | None | None |

| | | | | |
|---|---|---|---|---|
| BCKDHB | None | None | None | None |
| BCKDHB | None | None | None | None |
| BCKDHB | None | None | None | None |
| BCKDHB | None | None | None | None |
| BCKDHB | None | None | None | None |
| BCKDHB | None | None | None | None |
| BCKDHB | None | None | None | None |
| BCKDHB | None | None | None | None |
| BCKDHB | None | None | None | None |
| CDH1 | None | None | None | None |
| CDH1 | None | None | None | None |
| CDH1 | None | None | None | None |
| CDH1 | None | None | None | None |
| CDH1 | None | None | None | None |
| CTNNB1 | None | None | None | None |
| AIRE | None | None | None | None |
| EDA | 1 | 1 | 1rj8 | A |
| EDA | None | None | None | None |
| EDA | None | None | None | None |
| EDA | None | None | None | None |
| EDA | None | None | None | None |
| SH2D1A | None | None | None | None |
| SH2D1A | None | None | None | None |
| SH2D1A | None | None | None | None |
| GCK | None | None | None | None |
| GCK | None | None | None | None |
| GCK | None | None | None | None |
| GCK | None | None | None | None |
| GCK | None | None | None | None |
| GCK | None | None | None | None |
| GCK | None | None | None | None |
| GCK | None | None | None | None |
| GCK | None | None | None | None |
| GCK | None | None | None | None |
| GCK | None | None | None | None |
| GCK | None | None | None | None |
| GCK | None | None | None | None |
| GCK | None | None | None | None |
| GCK | None | None | None | None |
| GCK | None | None | None | None |
| GCK | None | None | None | None |
| GCK | None | None | None | None |
| GCK | None | None | None | None |
| GCK | None | None | None | None |
| AQP2 | None | None | None | None |
| AQP2 | None | None | None | None |
| AQP2 | None | None | None | None |
| AQP2 | None | None | None | None |
| AQP2 | None | None | None | None |
| AQP2 | None | None | None | None |
| AQP2 | None | None | None | None |
| AQP2 | None | None | None | None |
| AQP2 | None | None | None | None |
| AQP2 | None | None | None | None |

| | | | | |
|---|---|---|---|---|
| AQP2 | None | None | None | None |
| AQP2 | None | None | None | None |
| AQP2 | None | None | None | None |
| SUMF1 | None | None | None | None |
| SUMF1 | None | None | None | None |
| SUMF1 | None | None | None | None |
| SUMF1 | None | None | None | None |
| SUMF1 | None | None | None | None |
| SUMF1 | None | None | None | None |
| SUMF1 | None | None | None | None |
| SUMF1 | None | None | None | None |
| SUMF1 | None | None | None | None |
| FAM20A | None | None | None | None |
| FAM20A | None | None | None | None |
| FAM20A | None | None | None | None |
| FAM20A | None | None | None | None |
| NT5E | None | None | None | None |
| NT5E | None | None | None | None |
| NOS3 | None | None | None | None |
| ACTG1 | None | None | None | None |
| ACTG1 | None | None | None | None |
| ACTG1 | None | None | None | None |
| ACTG1 | None | None | None | None |
| ACTG1 | None | None | None | None |
| ACTG1 | None | None | None | None |
| FCGR1A | None | None | None | None |
| GPI | None | None | None | None |
| GPI | None | None | None | None |
| GPI | 1 | 3 | 1jiq | A |
| GPI | None | None | None | None |
| GPI | None | None | None | None |
| GPI | None | None | None | None |
| GPI | None | None | None | None |
| GPI | None | None | None | None |
| GPI | None | None | None | None |
| CFH | None | None | None | None |
| OGG1 | None | None | None | None |
| TPK1 | None | None | None | None |
| TPK1 | None | None | None | None |
| TPK1 | None | None | None | None |
| TPK1 | None | None | None | None |
| SEC23A | None | None | None | None |
| SEC23A | None | None | None | None |
| TYRP1 | None | None | None | None |
| TYRP1 | None | None | None | None |
| TYRP1 | None | None | None | None |
| TYRP1 | None | None | None | None |
| TYRP1 | None | None | None | None |
| TYRP1 | None | None | None | None |
| CSTB | None | None | None | None |
| CSTB | None | None | None | None |
| CSTB | None | None | None | None |
| ACP1 | None | None | None | None |
| IL17F | None | None | None | None |

| | | | | |
|---|---|---|---|---|
| IL17F | None | None | None | None |
| IL17F | None | None | None | None |
| IL17F | None | None | None | None |
| IL17F | None | None | None | None |
| IDH2 | None | None | None | None |
| IDH2 | None | None | None | None |
| IDH2 | None | None | None | None |
| IDH2 | None | None | None | None |
| TP53 | 0.925 | 11 | 2ocj | A |
| TP53 | None | None | None | None |
| TP53 | None | None | None | None |
| TP53 | None | None | None | None |
| TP53 | None | None | None | None |
| TP53 | 0.9 | 1 | 4agp | A |
| TP53 | 0.9 | 1 | 4agp | A |
| TP53 | 0.925 | 10 | 4hje | A |
| TP53 | None | None | None | None |
| TP53 | None | None | None | None |
| TP53 | None | None | None | None |
| TP53 | 0.925 | 6 | 3kz8 | B |
| TP53 | 0.925 | 6 | 3kz8 | B |
| TP53 | None | None | None | None |
| TP53 | 0.925 | 5 | 3kmd | B |
| TP53 | 0.925 | 5 | 3kmd | B |
| TP53 | 0.925 | 8 | 3kz8 | B |
| TP53 | 0.925 | 8 | 3kz8 | B |
| TP53 | 0.925 | 1 | 3kmd | B |
| TP53 | 0.9 | 11 | 2pcx | A |
| TP53 | 0.9 | 1 | 5aol | A |
| TP53 | None | None | None | None |
| TP53 | 0.9 | 1 | 3q06 | B |
| TP53 | None | None | None | None |
| TP53 | None | None | None | None |
| TP53 | None | None | None | None |
| TP53 | None | None | None | None |
| TP53 | None | None | None | None |
| TP53 | None | None | None | None |
| TP53 | None | None | None | None |
| TP53 | None | None | None | None |
| TP53 | None | None | None | None |
| TP53 | None | None | None | None |
| TP53 | None | None | None | None |
| TP53 | None | None | None | None |
| TP53 | None | None | None | None |
| TP53 | None | None | None | None |
| TP53 | None | None | None | None |
| TP53 | None | None | None | None |
| TP53 | None | None | None | None |
| TP53 | None | None | None | None |
| TP53 | None | None | None | None |
| LMAN1 | None | None | None | None |
| CTSK | None | None | None | None |
| CTSK | None | None | None | None |
| CTSK | None | None | None | None |

| | | | | |
|---|---|---|---|---|
| FGB | None | None | None | None |
| FGB | None | None | None | None |
| FGB | None | None | None | None |
| FGB | None | None | None | None |
| ACADM | None | None | None | None |
| ACADM | None | None | None | None |
| ACADM | None | None | None | None |
| ACADM | None | None | None | None |
| ACADM | None | None | None | None |
| ACADM | None | None | None | None |
| ACADM | None | None | None | None |
| ACADM | None | None | None | None |
| ACADM | None | None | None | None |
| ACADM | None | None | None | None |
| ACADM | None | None | None | None |
| ACADM | None | None | None | None |
| ACADM | None | None | None | None |
| ACADM | None | None | None | None |
| ACADM | None | None | None | None |
| ACADM | None | None | None | None |
| ACADM | None | None | None | None |
| ACADM | None | None | None | None |
| ACADM | None | None | None | None |
| ACADM | None | None | None | None |
| ACADM | None | None | None | None |
| ACADM | None | None | None | None |
| ACADM | None | None | None | None |
| ACADM | None | None | None | None |
| ACADM | None | None | None | None |
| ACADM | None | None | None | None |
| ACADM | None | None | None | None |
| PTS | None | None | None | None |
| PTS | None | None | None | None |
| PTS | None | None | None | None |
| PTS | None | None | None | None |
| PTS | None | None | None | None |
| PTS | None | None | None | None |
| PTS | None | None | None | None |
| PTS | None | None | None | None |
| PTS | None | None | None | None |
| INS | None | None | None | None |
| INS | 1 | 5 | 2l1z | A |
| INS | 1 | 5 | 2l1z | A |
| INS | 1 | 1 | 1his | A |
| FBP1 | None | None | None | None |
| FBP1 | None | None | None | None |
| FBP1 | None | None | None | None |
| FBP1 | None | None | None | None |
| FBP1 | None | None | None | None |
| FBP1 | None | None | None | None |
| FBP1 | None | None | None | None |
| FBP1 | None | None | None | None |
| FBP1 | None | None | None | None |
| FBP1 | None | None | None | None |

| | | | | |
|---|---|---|---|---|
| FBP1 | None | None | None | None |
| KCNQ1 | None | None | None | None |
| KCNQ1 | None | None | None | None |
| KCNQ1 | None | None | None | None |
| UPP1 | None | None | None | None |
| UMPS | 1 | 3 | 2qcl | A |
| UMPS | None | None | None | None |
| UMPS | None | None | None | None |
| DHFR | None | None | None | None |
| KHK | None | None | None | None |
| TTPA | None | None | None | None |
| TTPA | None | None | None | None |
| TTPA | None | None | None | None |
| TTPA | None | None | None | None |
| MAP1LC3B | None | None | None | None |
| MAP1LC3B | None | None | None | None |
| ZAP70 | None | None | None | None |
| PROC | None | None | None | None |
| SQSTM1 | None | None | None | None |
| AHCY | None | None | None | None |
| AHCY | None | None | None | None |
| PRNP | None | None | None | None |
| PRNP | None | None | None | None |
| PRNP | None | None | None | None |
| PRNP | None | None | None | None |
| PRNP | None | None | None | None |
| PRNP | None | None | None | None |
| PRNP | None | None | None | None |
| PRNP | None | None | None | None |
| PRNP | None | None | None | None |
| PRNP | None | None | None | None |
| PRNP | None | None | None | None |
| PRNP | None | None | None | None |
| PRNP | None | None | None | None |
| PRNP | None | None | None | None |
| PRNP | None | None | None | None |
| PRNP | None | None | None | None |
| REN | None | None | None | None |
| REN | None | None | None | None |
| HINT1 | None | None | None | None |
| HINT1 | None | None | None | None |
| HINT1 | None | None | None | None |
| HINT1 | None | None | None | None |
| HINT1 | 0.923 | 1 | 5ipc | A |
| OAT | None | None | None | None |
| OAT | None | None | None | None |
| OAT | None | None | None | None |
| OAT | None | None | None | None |
| OAT | None | None | None | None |
| OAT | None | None | None | None |
| OAT | None | None | None | None |
| OAT | None | None | None | None |
| OAT | None | None | None | None |
| OAT | None | None | None | None |

| | | | | |
|---|---|---|---|---|
| OAT | None | None | None | None |
| OAT | None | None | None | None |
| OAT | None | None | None | None |
| OAT | None | None | None | None |
| OAT | None | None | None | None |
| OAT | None | None | None | None |
| OAT | None | None | None | None |
| OAT | None | None | None | None |
| OAT | None | None | None | None |
| OAT | None | None | None | None |
| OAT | None | None | None | None |
| OAT | None | None | None | None |
| OAT | None | None | None | None |
| OAT | None | None | None | None |
| OAT | None | None | None | None |
| OAT | None | None | None | None |
| OAT | None | None | None | None |
| OAT | None | None | None | None |
| CTSA | None | None | None | None |
| CTSA | None | None | None | None |
| PNP | None | None | None | None |
| PNP | None | None | None | None |
| PNP | None | None | None | None |
| PNP | None | None | None | None |
| PNP | None | None | None | None |
| PNP | None | None | None | None |
| PNP | 0.933 | 1 | 3ggs | C |
| PNP | None | None | None | None |
| MYL2 | None | None | None | None |
| MYL2 | None | None | None | None |
| ANG | None | None | None | None |
| ANG | None | None | None | None |
| ANG | None | None | None | None |
| ANG | None | None | None | None |
| ANG | None | None | None | None |
| ANG | None | None | None | None |
| ANG | None | None | None | None |
| ANG | None | None | None | None |
| ANG | None | None | None | None |
| ANG | None | None | None | None |
| POLH | None | None | None | None |
| POLH | None | None | None | None |
| POLH | None | None | None | None |
| IL2RA | None | None | None | None |
| TGFB1 | None | None | None | None |
| TP53 | None | None | None | None |
| TP53 | None | None | None | None |
| PAH | None | None | None | None |
| PAH | None | None | None | None |
| PAH | None | None | None | None |
| PAH | None | None | None | None |
| PAH | None | None | None | None |
| PAH | None | None | None | None |
| PAH | None | None | None | None |

| | | | | |
|---|---|---|---|---|
| PAH | None | None | None | None |
| PAH | None | None | None | None |
| PAH | None | None | None | None |
| PAH | None | None | None | None |
| PAH | None | None | None | None |
| PAH | None | None | None | None |
| PAH | None | None | None | None |
| PAH | None | None | None | None |
| PAH | None | None | None | None |
| PAH | None | None | None | None |
| PAH | None | None | None | None |
| PAH | None | None | None | None |
| PAH | None | None | None | None |
| PAH | None | None | None | None |
| PAH | None | None | None | None |
| PAH | None | None | None | None |
| PAH | None | None | None | None |
| PAH | None | None | None | None |
| PAH | None | None | None | None |
| PAH | None | None | None | None |
| PAH | None | None | None | None |
| PAH | None | None | None | None |
| PAH | None | None | None | None |
| PAH | None | None | None | None |
| PAH | None | None | None | None |
| PAH | None | None | None | None |
| PAH | None | None | None | None |
| PAH | None | None | None | None |
| PAH | None | None | None | None |
| PAH | None | None | None | None |
| PAH | None | None | None | None |
| PAH | None | None | None | None |
| PAH | None | None | None | None |
| PAH | None | None | None | None |
| HDAC8 | None | None | None | None |
| HDAC8 | None | None | None | None |
| VCP | None | None | None | None |
| VCP | None | None | None | None |
| VCP | None | None | None | None |
| VCP | None | None | None | None |
| VCP | None | None | None | None |
| VCP | None | None | None | None |
| VCP | None | None | None | None |
| B3GAT3 | None | None | None | None |
| B3GAT3 | None | None | None | None |
| CSF2RB | None | None | None | None |
| CSF2RB | None | None | None | None |
| CSF2RB | None | None | None | None |
| NF2 | None | None | None | None |
| NF2 | None | None | None | None |
| NF2 | None | None | None | None |
| PIK3CA | None | None | None | None |
| PHGDH | None | None | None | None |
| PHGDH | None | None | None | None |

| | | | | |
|---|---|---|---|---|
| PHGDH | None | None | None | None |
| PHGDH | None | None | None | None |
| POLH | None | None | None | None |
| XIAP | None | None | None | None |
| XIAP | None | None | None | None |
| PDHX | None | None | None | None |
| PDHX | None | None | None | None |
| PDHX | None | None | None | None |
| PDHX | None | None | None | None |
| PDHX | None | None | None | None |
| SAMHD1 | None | None | None | None |
| SAMHD1 | None | None | None | None |
| SAMHD1 | None | None | None | None |
| SAMHD1 | None | None | None | None |
| SAMHD1 | None | None | None | None |
| SAMHD1 | None | None | None | None |
| SAMHD1 | None | None | None | None |
| SAMHD1 | None | None | None | None |
| CD36 | None | None | None | None |
| CD36 | None | None | None | None |
| CD36 | None | None | None | None |
| ASPH | None | None | None | None |
| CCT5 | None | None | None | None |
| CCT5 | None | None | None | None |
| CCT5 | None | None | None | None |
| SMN1 | None | None | None | None |
| SMN1 | None | None | None | None |
| SMN1 | None | None | None | None |
| SMN1 | None | None | None | None |
| ABCA1 | None | None | None | None |
| ABCA1 | None | None | None | None |
| ABCA1 | None | None | None | None |
| ABCA1 | None | None | None | None |
| ABCA1 | None | None | None | None |
| ABCA1 | None | None | None | None |
| ABCA1 | None | None | None | None |
| ABCA1 | None | None | None | None |
| ABCA1 | None | None | None | None |
| ABCA1 | None | None | None | None |
| ABCA1 | None | None | None | None |
| ABCA1 | None | None | None | None |
| ABCA1 | None | None | None | None |
| ABCA1 | None | None | None | None |
| ABCA1 | None | None | None | None |
| ABCA1 | None | None | None | None |
| ABCA1 | None | None | None | None |
| ABCA1 | None | None | None | None |
| ABCA1 | None | None | None | None |
| ABCA1 | None | None | None | None |
| ABCA1 | None | None | None | None |
| ABCA1 | None | None | None | None |
| ABCA1 | None | None | None | None |
| ABCA1 | None | None | None | None |
| ABCA1 | None | None | None | None |
| ABCA1 | None | None | None | None |

| | | | | |
|---|---|---|---|---|
| ABCA1 | None | None | None | None |
| ABCA1 | None | None | None | None |
| ABCA1 | None | None | None | None |
| ABCA1 | None | None | None | None |
| ABCA1 | None | None | None | None |
| ABCA1 | None | None | None | None |
| ABCA1 | None | None | None | None |
| ABCA1 | None | None | None | None |
| ABCA1 | None | None | None | None |
| ABCA1 | None | None | None | None |
| ABCA1 | None | None | None | None |
| ERCC1 | None | None | None | None |
| FECH | None | None | None | None |
| FECH | None | None | None | None |
| FECH | None | None | None | None |
| FECH | None | None | None | None |
| FECH | None | None | None | None |
| FECH | None | None | None | None |
| FECH | None | None | None | None |
| PTEN | None | None | None | None |
| PTEN | None | None | None | None |
| PTEN | None | None | None | None |
| PTEN | None | None | None | None |
| PTEN | None | None | None | None |
| PTEN | None | None | None | None |
| PTEN | None | None | None | None |
| PTEN | None | None | None | None |
| PTEN | None | None | None | None |
| PTEN | None | None | None | None |
| PTEN | None | None | None | None |
| PTEN | None | None | None | None |
| PTEN | None | None | None | None |
| PTEN | None | None | None | None |
| PTEN | None | None | None | None |
| PTEN | None | None | None | None |
| PTEN | None | None | None | None |
| PTEN | None | None | None | None |
| PTEN | None | None | None | None |
| PTEN | None | None | None | None |
| ACAT1 | None | None | None | None |
| ACAT1 | None | None | None | None |
| ACAT1 | None | None | None | None |
| ACAT1 | None | None | None | None |
| ACAT1 | None | None | None | None |
| ACAT1 | None | None | None | None |
| ACAT1 | None | None | None | None |
| ACAT1 | None | None | None | None |
| ACAT1 | None | None | None | None |
| ACAT1 | None | None | None | None |
| ACAT1 | None | None | None | None |
| CREBBP | None | None | None | None |
| ITGA2B | None | None | None | None |
| ITGA2B | None | None | None | None |
| ITGA2B | None | None | None | None |

| | | | | |
|---|---|---|---|---|
| ITGA2B | None | None | None | None |
| TSC1 | None | None | None | None |
| ADA2 | None | None | None | None |
| ADA2 | None | None | None | None |
| ADA2 | None | None | None | None |
| ADA2 | None | None | None | None |
| ADA2 | None | None | None | None |
| ADA2 | None | None | None | None |
| ADA2 | None | None | None | None |
| ADA2 | None | None | None | None |
| ADA2 | 1 | 1 | 3lgd | B |
| ADA2 | None | None | None | None |
| PGK1 | None | None | None | None |
| PGK1 | None | None | None | None |
| PGK1 | None | None | None | None |
| PGK1 | None | None | None | None |
| PGK1 | None | None | None | None |
| TUBA1A | None | None | None | None |
| TUBA1A | None | None | None | None |
| TUBA1A | None | None | None | None |
| TUBA1A | None | None | None | None |
| TUBA1A | None | None | None | None |
| PTPN11 | None | None | None | None |
| PTPN11 | None | None | None | None |
| PTPN11 | None | None | None | None |
| PTPN11 | None | None | None | None |
| PTPN11 | None | None | None | None |
| PTPN11 | None | None | None | None |
| PTPN11 | None | None | None | None |
| PTPN11 | None | None | None | None |
| PTPN11 | None | None | None | None |
| PTPN11 | None | None | None | None |
| PTPN11 | None | None | None | None |
| PTPN11 | None | None | None | None |
| PTPN11 | None | None | None | None |
| PTPN11 | None | None | None | None |
| PTPN11 | None | None | None | None |
| PTPN11 | None | None | None | None |
| PTPN11 | None | None | None | None |
| PTPN11 | None | None | None | None |
| PTPN11 | None | None | None | None |
| PTPN11 | None | None | None | None |
| PTPN11 | None | None | None | None |
| PTPN11 | None | None | None | None |
| PTPN11 | None | None | None | None |
| PTPN11 | None | None | None | None |
| PTPN11 | None | None | None | None |
| PTPN11 | None | None | None | None |
| GLB1 | None | None | None | None |
| GLB1 | None | None | None | None |
| UQCRQ | None | None | None | None |
| VPS35 | None | None | None | None |
| NR3C1 | None | None | None | None |
| NR3C1 | None | None | None | None |

| | | | | |
|---|---|---|---|---|
| NR3C1 | None | None | None | None |
| NR3C1 | None | None | None | None |
| NR3C1 | None | None | None | None |
| NR3C1 | None | None | None | None |
| NR3C1 | None | None | None | None |
| NR3C1 | None | None | None | None |
| NR3C1 | None | None | None | None |
| APOA1 | None | None | None | None |
| APOA1 | None | None | None | None |
| APOA1 | None | None | None | None |
| APOA1 | None | None | None | None |
| APOA1 | None | None | None | None |
| APOA1 | None | None | None | None |
| APOA1 | None | None | None | None |
| BCKDHA | None | None | None | None |
| BCKDHA | None | None | None | None |
| BCKDHA | 0.963 | 1 | 1ols | A |
| BCKDHA | None | None | None | None |
| BCKDHA | None | None | None | None |
| BCKDHA | None | None | None | None |
| BCKDHA | None | None | None | None |
| BCKDHA | 1 | 1 | 2beu | A |
| BCKDHA | None | None | None | None |
| BCKDHA | None | None | None | None |
| BCKDHA | None | None | None | None |
| BCKDHA | None | None | None | None |
| BCKDHA | None | None | None | None |
| BCKDHA | None | None | None | None |
| BCKDHA | None | None | None | None |
| TOP1 | None | None | None | None |
| ANTXR2 | None | None | None | None |
| ANTXR2 | None | None | None | None |
| ANTXR2 | None | None | None | None |
| TF | None | None | None | None |
| TF | None | None | None | None |
| FSHB | None | None | None | None |
| GALK1 | None | None | None | None |
| GALK1 | None | None | None | None |
| GALK1 | None | None | None | None |
| CYP17A1 | None | None | None | None |
| CYP17A1 | None | None | None | None |
| CYP17A1 | None | None | None | None |
| CYP17A1 | None | None | None | None |
| CYP17A1 | None | None | None | None |
| CYP17A1 | None | None | None | None |
| CYP17A1 | None | None | None | None |
| CYP17A1 | None | None | None | None |
| CYP17A1 | None | None | None | None |
| CYP17A1 | None | None | None | None |
| CYP17A1 | None | None | None | None |
| CYP17A1 | None | None | None | None |
| CYP17A1 | None | None | None | None |
| CYP17A1 | None | None | None | None |
| CYP17A1 | None | None | None | None |

| | | | | |
|---|---|---|---|---|
| CYP17A1 | None | None | None | None |
| CYP17A1 | None | None | None | None |
| CD44 | None | None | None | None |
| MDH2 | None | None | None | None |
| MDH2 | None | None | None | None |
| NDUFV1 | None | None | None | None |
| NDUFV1 | None | None | None | None |
| NDUFV1 | None | None | None | None |
| NDUFV1 | None | None | None | None |
| NDUFV1 | None | None | None | None |
| NDUFV1 | None | None | None | None |
| NDUFV1 | None | None | None | None |
| NDUFV1 | None | None | None | None |
| NDUFV1 | None | None | None | None |
| NDUFV1 | None | None | None | None |
| CTSC | None | None | None | None |
| CTSC | None | None | None | None |
| CTSC | None | None | None | None |
| CTSC | None | None | None | None |
| CTSC | None | None | None | None |
| CTSC | None | None | None | None |
| CTSC | None | None | None | None |
| VWF | None | None | None | None |
| VWF | None | None | None | None |
| VWF | None | None | None | None |
| VWF | None | None | None | None |
| VWF | None | None | None | None |
| VWF | None | None | None | None |
| VWF | None | None | None | None |
| ORC4 | None | None | None | None |
| ORC4 | None | None | None | None |
| CRYAB | None | None | None | None |
| CRYAB | None | None | None | None |
| CRYAB | None | None | None | None |
| MAP2K2 | None | None | None | None |
| MAP2K2 | None | None | None | None |
| MAP2K2 | None | None | None | None |
| MAP2K2 | None | None | None | None |
| MAP2K2 | None | None | None | None |
| MAP2K2 | None | None | None | None |
| MAP2K2 | None | None | None | None |
| MAP2K2 | None | None | None | None |
| MAP2K2 | None | None | None | None |
| CBL | None | None | None | None |
| CBL | None | None | None | None |
| CBL | None | None | None | None |
| CBL | None | None | None | None |
| CBL | None | None | None | None |
| CBL | None | None | None | None |
| CBL | None | None | None | None |
| PFN1 | None | None | None | None |
| PFN1 | None | None | None | None |
| ACSM2A | None | None | None | None |
| HSPG2 | None | None | None | None |

| | | | | |
|---|---|---|---|---|
| HSPG2 | None | None | None | None |
| HSPG2 | None | None | None | None |
| HSPG2 | None | None | None | None |
| PARK7 | None | None | None | None |
| PARK7 | None | None | None | None |
| PARK7 | None | None | None | None |
| PARK7 | None | None | None | None |
| PARK7 | None | None | None | None |
| PARK7 | None | None | None | None |
| INSR | None | None | None | None |
| INSR | None | None | None | None |
| INSR | None | None | None | None |
| INSR | None | None | None | None |
| INSR | None | None | None | None |
| DSC2 | None | None | None | None |
| DSC2 | None | None | None | None |
| DSC2 | None | None | None | None |
| DSC2 | None | None | None | None |
| DSC2 | None | None | None | None |
| DSC2 | None | None | None | None |
| DSC2 | None | None | None | None |
| DSC2 | None | None | None | None |
| DSC2 | None | None | None | None |
| DSC2 | None | None | None | None |
| MICU1 | None | None | None | None |
| MICU1 | None | None | None | None |
| ADAMTS13 | None | None | None | None |
| ADAMTS13 | None | None | None | None |
| ADAMTS13 | None | None | None | None |
| ADAMTS13 | None | None | None | None |
| ADAMTS13 | None | None | None | None |
| ADAMTS13 | None | None | None | None |
| ADAMTS13 | None | None | None | None |
| IRAK4 | None | None | None | None |
| IRAK4 | None | None | None | None |
| RNASEH2B | None | None | None | None |
| RNASEH2B | None | None | None | None |
| RNASEH2B | None | None | None | None |
| NTRK1 | None | None | None | None |
| MPO | None | None | None | None |
| MPO | None | None | None | None |
| MPO | None | None | None | None |
| PEX3 | None | None | None | None |
| SERPINA1 | None | None | None | None |
| SERPINA1 | None | None | None | None |
| SERPINA1 | None | None | None | None |
| SERPINA1 | None | None | None | None |
| SERPINA1 | None | None | None | None |
| SERPINA1 | None | None | None | None |
| SERPINA1 | None | None | None | None |
| SERPINA1 | None | None | None | None |
| SERPINA1 | None | None | None | None |
| SERPINA1 | None | None | None | None |
| YARS2 | None | None | None | None |

| | | | | |
|---|---|---|---|---|
| YARS2 | None | None | None | None |
| YARS2 | None | None | None | None |
| YARS2 | None | None | None | None |
| YARS2 | None | None | None | None |
| YARS2 | None | None | None | None |
| YARS2 | None | None | None | None |
| YARS2 | None | None | None | None |
| YARS2 | None | None | None | None |
| PDHA1 | None | None | None | None |
| PDHA1 | None | None | None | None |
| PDHA1 | None | None | None | None |
| PDHA1 | None | None | None | None |
| PDHA1 | None | None | None | None |
| PDHA1 | None | None | None | None |
| PDHA1 | None | None | None | None |
| PDHA1 | 0.96 | 1 | 3exh | C |
| PDHA1 | None | None | None | None |
| PDHA1 | None | None | None | None |
| PDHA1 | None | None | None | None |
| CUL4B | None | None | None | None |
| AGXT | None | None | None | None |
| AGXT | None | None | None | None |
| AGXT | None | None | None | None |
| AGXT | None | None | None | None |
| AGXT | None | None | None | None |
| AGXT | None | None | None | None |
| AGXT | None | None | None | None |
| AGXT | None | None | None | None |
| AGXT | None | None | None | None |
| AGXT | None | None | None | None |
| AGXT | None | None | None | None |
| AGXT | None | None | None | None |
| AGXT | None | None | None | None |
| AGXT | None | None | None | None |
| AGXT | None | None | None | None |
| AGXT | None | None | None | None |
| AGXT | None | None | None | None |
| AGXT | None | None | None | None |
| AGXT | None | None | None | None |
| AGXT | None | None | None | None |
| AGXT | None | None | None | None |
| AGXT | None | None | None | None |
| AGXT | None | None | None | None |
| AGXT | None | None | None | None |
| AGXT | None | None | None | None |
| AGXT | None | None | None | None |
| AGXT | None | None | None | None |
| AGXT | None | None | None | None |
| AGXT | None | None | None | None |
| AGXT | None | None | None | None |
| AGXT | None | None | None | None |
| AGXT | None | None | None | None |
| AGXT | None | None | None | None |
| AGXT | None | None | None | None |

| | | | | |
|---|---|---|---|---|
| AGXT | None | None | None | None |
| SFTPD | None | None | None | None |
| SFTPD | None | None | None | None |
| PIK3CA | None | None | None | None |
| PIK3CA | None | None | None | None |
| PIK3CA | None | None | None | None |
| AUH | None | None | None | None |
| AUH | None | None | None | None |
| AUH | None | None | None | None |
| AUH | None | None | None | None |
| F11 | None | None | None | None |
| F11 | None | None | None | None |
| F11 | None | None | None | None |
| F11 | None | None | None | None |
| MEN1 | None | None | None | None |
| MEN1 | None | None | None | None |
| MEN1 | None | None | None | None |
| MEN1 | None | None | None | None |
| MEN1 | 1 | 2 | 4i80 | A |
| MEN1 | None | None | None | None |
| MEN1 | None | None | None | None |
| MEN1 | None | None | None | None |
| MEN1 | None | None | None | None |
| MEN1 | None | None | None | None |
| MYBPC1 | None | None | None | None |
| MYBPC1 | None | None | None | None |
| MYBPC1 | None | None | None | None |
| PDCD10 | None | None | None | None |
| SCARB2 | None | None | None | None |
| SCARB2 | None | None | None | None |
| SCARB2 | None | None | None | None |
| SCARB2 | None | None | None | None |
| SCARB2 | None | None | None | None |
| TREX2 | None | None | None | None |
| TBK1 | None | None | None | None |
| TBK1 | None | None | None | None |
| DLD | None | None | None | None |
| DLD | None | None | None | None |
| DLD | None | None | None | None |
| DLD | None | None | None | None |
| DLD | None | None | None | None |
| DLD | None | None | None | None |
| LDHA | None | None | None | None |
| TNNC1 | None | None | None | None |
| TNNC1 | None | None | None | None |
| TNNC1 | None | None | None | None |
| SMAD3 | None | None | None | None |
| HARS | None | None | None | None |
| HARS | None | None | None | None |
| TNNI3 | None | None | None | None |
| TNNI3 | None | None | None | None |
| TNNI3 | None | None | None | None |
| TNNI3 | None | None | None | None |
| TNNI3 | None | None | None | None |

| | | | | |
|---|---|---|---|---|
| TNNI3 | None | None | None | None |
| ACKR1 | None | None | None | None |
| BUB1B | None | None | None | None |
| BUB1B | None | None | None | None |
| BUB1B | None | None | None | None |
| AR | None | None | None | None |
| AR | None | None | None | None |
| AR | None | None | None | None |
| AR | None | None | None | None |
| AR | None | None | None | None |
| FTO | None | None | None | None |
| FTO | None | None | None | None |
| TFAM | None | None | None | None |
| PYGL | None | None | None | None |
| PYGL | None | None | None | None |
| PYGL | None | None | None | None |
| PYGL | None | None | None | None |
| PYGL | None | None | None | None |
| PYGL | None | None | None | None |
| PYGL | None | None | None | None |
| PYGL | None | None | None | None |
| PYGL | None | None | None | None |
| PYGL | None | None | None | None |
| PYGL | None | None | None | None |
| PYGL | None | None | None | None |
| PYGL | None | None | None | None |
| PYGL | None | None | None | None |
| PYGL | None | None | None | None |
| PYGL | None | None | None | None |
| GALNS | None | None | None | None |
| GALNS | None | None | None | None |
| GALNS | None | None | None | None |
| GALNS | None | None | None | None |
| GALNS | None | None | None | None |
| GALNS | None | None | None | None |
| GALNS | None | None | None | None |
| GALNS | None | None | None | None |
| GALNS | None | None | None | None |
| GALNS | None | None | None | None |
| GALNS | None | None | None | None |
| GALNS | None | None | None | None |
| GALNS | None | None | None | None |
| GALNS | None | None | None | None |
| GALNS | None | None | None | None |
| GALNS | None | None | None | None |
| GALNS | None | None | None | None |
| GALNS | None | None | None | None |
| GALNS | None | None | None | None |
| GALNS | None | None | None | None |
| GALNS | None | None | None | None |
| GALNS | None | None | None | None |
| GALNS | None | None | None | None |
| GALNS | None | None | None | None |

| | | | | |
|---|---|---|---|---|
| GALNS | None | None | None | None |
| GALNS | None | None | None | None |
| GALNS | None | None | None | None |
| GALNS | None | None | None | None |
| GALNS | None | None | None | None |
| GALNS | None | None | None | None |
| GALNS | None | None | None | None |
| GALNS | None | None | None | None |
| F10 | None | None | None | None |
| GLUD1 | None | None | None | None |
| GLUD1 | None | None | None | None |
| GLUD1 | None | None | None | None |
| GLUD1 | None | None | None | None |
| GLUD1 | None | None | None | None |
| GLUD1 | None | None | None | None |
| GLUD1 | None | None | None | None |
| GLUD1 | None | None | None | None |
| GLUD1 | None | None | None | None |
| GLUD1 | None | None | None | None |
| NOTCH1 | None | None | None | None |
| POGLUT1 | None | None | None | None |
| POGLUT1 | None | None | None | None |
| NR3C1 | None | None | None | None |
| CASP8 | None | None | None | None |
| CASP8 | None | None | None | None |
| TRAP1 | None | None | None | None |
| FANCM | None | None | None | None |
| EPOR | None | None | None | None |
| EPOR | None | None | None | None |
| NDP | None | None | None | None |
| NDP | None | None | None | None |
| NDP | None | None | None | None |
| NDP | None | None | None | None |
| NDP | None | None | None | None |
| NDP | None | None | None | None |
| NDP | None | None | None | None |
| NDP | None | None | None | None |
| NDP | None | None | None | None |
| NDP | None | None | None | None |
| NDP | None | None | None | None |
| NDP | None | None | None | None |
| NDP | None | None | None | None |
| NDP | None | None | None | None |
| NDP | None | None | None | None |
| NDP | None | None | None | None |
| NDP | None | None | None | None |
| ROGDI | None | None | None | None |
| CYP19A1 | None | None | None | None |
| CYP19A1 | None | None | None | None |
| CYP19A1 | None | None | None | None |
| CYP19A1 | None | None | None | None |
| CYP19A1 | None | None | None | None |
| CYP19A1 | None | None | None | None |
| CYP19A1 | None | None | None | None |

| | | | | |
|---|---|---|---|---|
| CYP19A1 | None | None | None | None |
| UBA5 | None | None | None | None |
| PRPS1 | None | None | None | None |
| PRPS1 | None | None | None | None |
| PRPS1 | None | None | None | None |
| PRPS1 | None | None | None | None |
| PRPS1 | None | None | None | None |
| PRPS1 | None | None | None | None |
| PRPS1 | None | None | None | None |
| PRPS1 | None | None | None | None |
| F2 | None | None | None | None |
| F2 | None | None | None | None |
| NTF4 | None | None | None | None |
| NTF4 | None | None | None | None |
| PSAP | None | None | None | None |
| PSAP | None | None | None | None |
| TAP1 | None | None | None | None |
| TAP1 | None | None | None | None |
| TAP1 | None | None | None | None |
| ARG1 | None | None | None | None |
| ARG1 | None | None | None | None |
| ARG1 | None | None | None | None |
| ARG1 | None | None | None | None |
| ARG1 | None | None | None | None |
| ARG1 | None | None | None | None |
| CRYM | None | None | None | None |
| ADSL | None | None | None | None |
| ADSL | None | None | None | None |
| ADSL | None | None | None | None |
| ADSL | None | None | None | None |
| ADSL | None | None | None | None |
| ADSL | None | None | None | None |
| ADSL | None | None | None | None |
| ADSL | None | None | None | None |
| ADSL | None | None | None | None |
| ADSL | None | None | None | None |
| ADSL | None | None | None | None |
| ADSL | None | None | None | None |
| ADSL | None | None | None | None |
| ADSL | None | None | None | None |
| ADSL | None | None | None | None |
| ADSL | None | None | None | None |
| ADSL | None | None | None | None |
| ADSL | None | None | None | None |
| ADSL | None | None | None | None |
| ADSL | None | None | None | None |
| VWF | None | None | None | None |
| PLA2G7 | None | None | None | None |
| MMP13 | None | None | None | None |
| MMP13 | None | None | None | None |
| MMP13 | None | None | None | None |
| MMP13 | None | None | None | None |
| POMGNT1 | None | None | None | None |
| POMGNT1 | None | None | None | None |

| | | | | |
|---|---|---|---|---|
| COL3A1 | None | None | None | None |
| COL3A1 | None | None | None | None |
| COL3A1 | None | None | None | None |
| COL3A1 | None | None | None | None |
| ACTN2 | None | None | None | None |
| ACTN2 | None | None | None | None |
| ACTN2 | None | None | None | None |
| ACTN2 | None | None | None | None |
| ACTN2 | None | None | None | None |
| ACTN2 | None | None | None | None |
| ACTN2 | None | None | None | None |
| UBA5 | None | None | None | None |
| ADAR | None | None | None | None |
| UMPS | None | None | None | None |
| UMPS | None | None | None | None |
| UMPS | None | None | None | None |
| TRNT1 | None | None | None | None |
| TRNT1 | None | None | None | None |
| NHEJ1 | None | None | None | None |
| NHEJ1 | None | None | None | None |
| NHEJ1 | None | None | None | None |
| C8A | None | None | None | None |
| GLUL | None | None | None | None |
| GLUL | None | None | None | None |
| GLUL | None | None | None | None |
| GYG1 | None | None | None | None |
| GYG1 | 0.923 | 1 | 3u2u | B |
| FLNA | None | None | None | None |
| FLNA | None | None | None | None |
| FLNA | None | None | None | None |
| FLNA | None | None | None | None |
| FLNA | None | None | None | None |
| FLNA | None | None | None | None |
| FLNA | None | None | None | None |
| TTN | None | None | None | None |
| TTN | None | None | None | None |
| TTN | None | None | None | None |
| TTN | None | None | None | None |
| SMS | None | None | None | None |
| SMS | None | None | None | None |
| SMS | None | None | None | None |
| C5 | None | None | None | None |
| SERPINC1 | None | None | None | None |
| SERPINC1 | None | None | None | None |
| SERPINC1 | None | None | None | None |
| SERPINC1 | None | None | None | None |
| SERPINC1 | None | None | None | None |
| SERPINC1 | None | None | None | None |
| SERPINC1 | None | None | None | None |
| SERPINC1 | None | None | None | None |
| SERPINC1 | None | None | None | None |
| SERPINC1 | None | None | None | None |
| SERPINC1 | None | None | None | None |
| SERPINC1 | None | None | None | None |

| | | | | |
|---|---|---|---|---|
| SERPINC1 | None | None | None | None |
| SERPINC1 | None | None | None | None |
| SERPINC1 | None | None | None | None |
| SERPINC1 | None | None | None | None |
| SERPINC1 | None | None | None | None |
| SERPINC1 | None | None | None | None |
| SERPINC1 | None | None | None | None |
| SERPINC1 | None | None | None | None |
| SERPINC1 | None | None | None | None |
| ETFB | None | None | None | None |
| ETFB | None | None | None | None |
| ETFB | None | None | None | None |
| ETFB | None | None | None | None |
| ETFB | None | None | None | None |
| ETFB | None | None | None | None |
| DIABLO | None | None | None | None |
| DIABLO | None | None | None | None |
| DPYS | None | None | None | None |
| DPYS | None | None | None | None |
| DPYS | None | None | None | None |
| DPYS | None | None | None | None |
| DPYS | None | None | None | None |
| GNE | None | None | None | None |
| GNE | None | None | None | None |
| GNE | None | None | None | None |
| GNE | None | None | None | None |
| GNE | None | None | None | None |
| GNE | None | None | None | None |
| GNE | None | None | None | None |
| GNE | None | None | None | None |
| GNE | None | None | None | None |
| GNMT | None | None | None | None |
| GNMT | None | None | None | None |
| CTH | None | None | None | None |
| CTH | None | None | None | None |
| CTH | None | None | None | None |
| USH1C | None | None | None | None |
| PSEN1 | None | None | None | None |
| CHEK2 | None | None | None | None |
| CHEK2 | None | None | None | None |
| CHEK2 | None | None | None | None |
| CHEK2 | None | None | None | None |
| CHEK2 | None | None | None | None |
| CHEK2 | None | None | None | None |
| CHEK2 | None | None | None | None |
| CHEK2 | None | None | None | None |
| CHEK2 | None | None | None | None |
| SNCA | None | None | None | None |
| SNCA | None | None | None | None |
| GAN | None | None | None | None |
| GAN | None | None | None | None |
| GAN | None | None | None | None |
| SBDS | None | None | None | None |
| SBDS | None | None | None | None |

| | | | | |
|---|---|---|---|---|
| SBDS | None | None | None | None |
| SBDS | None | None | None | None |
| SBDS | None | None | None | None |
| SBDS | None | None | None | None |
| MPO | 1 | 3 | 3f9p | A |
| MPO | None | None | None | None |
| KCNH2 | None | None | None | None |
| KCNH2 | None | None | None | None |
| KCNH2 | None | None | None | None |
| KCNH2 | None | None | None | None |
| TEK | None | None | None | None |
| TEK | None | None | None | None |
| TEK | None | None | None | None |
| TEK | None | None | None | None |
| NPC2 | None | None | None | None |
| NPC2 | None | None | None | None |
| NPC2 | None | None | None | None |
| NPC2 | None | None | None | None |
| NPC2 | None | None | None | None |
| NPC2 | None | None | None | None |
| NPC2 | None | None | None | None |
| PLG | None | None | None | None |
| PCSK9 | None | None | None | None |
| SFTPC | None | None | None | None |
| SFTPC | None | None | None | None |
| SFTPC | None | None | None | None |
| CD40LG | None | None | None | None |
| CD40LG | None | None | None | None |
| CD40LG | None | None | None | None |
| CD40LG | None | None | None | None |
| CD40LG | None | None | None | None |
| TAT | None | None | None | None |
| TAT | None | None | None | None |
| TAT | None | None | None | None |
| EP300 | None | None | None | None |
| FARS2 | None | None | None | None |
| FARS2 | None | None | None | None |
| FARS2 | None | None | None | None |
| FARS2 | None | None | None | None |
| FARS2 | None | None | None | None |
| FARS2 | None | None | None | None |
| FARS2 | None | None | None | None |
| FARS2 | None | None | None | None |
| FARS2 | None | None | None | None |
| FARS2 | None | None | None | None |
| FARS2 | None | None | None | None |
| FARS2 | None | None | None | None |
| FARS2 | None | None | None | None |
| MASP2 | None | None | None | None |
| MASP2 | None | None | None | None |
| MASP2 | None | None | None | None |
| MASP2 | None | None | None | None |
| STIM1 | None | None | None | None |
| CHRM2 | None | None | None | None |

| | | | | |
|---|---|---|---|---|
| MCFD2 | None | None | None | None |
| MCFD2 | None | None | None | None |
| MCFD2 | None | None | None | None |
| KITLG | None | None | None | None |
| F12 | None | None | None | None |
| F12 | None | None | None | None |
| ERAP1 | None | None | None | None |
| GTF2H5 | None | None | None | None |
| GTF2H5 | None | None | None | None |
| THBD | None | None | None | None |
| MRPL28 | None | None | None | None |
| MRPL28 | None | None | None | None |
| HTRA1 | None | None | None | None |
| HTRA1 | None | None | None | None |
| HTRA1 | None | None | None | None |
| HTRA1 | None | None | None | None |
| HTRA1 | None | None | None | None |
| HTRA1 | None | None | None | None |
| KARS | None | None | None | None |
| KARS | None | None | None | None |
| KARS | None | None | None | None |
| KARS | None | None | None | None |
| KARS | None | None | None | None |
| HK1 | None | None | None | None |
| GARS | None | None | None | None |
| GARS | None | None | None | None |
| GARS | None | None | None | None |
| GARS | None | None | None | None |
| GARS | None | None | None | None |
| GARS | None | None | None | None |
| GARS | None | None | None | None |
| IL12B | None | None | None | None |
| HADH | None | None | None | None |
| HADH | None | None | None | None |
| HADH | None | None | None | None |
| HADH | None | None | None | None |
| HADH | None | None | None | None |
| RRM2B | None | None | None | None |
| RRM2B | None | None | None | None |
| RRM2B | None | None | None | None |
| RRM2B | None | None | None | None |
| RRM2B | None | None | None | None |
| RRM2B | None | None | None | None |
| RRM2B | None | None | None | None |
| RRM2B | None | None | None | None |
| RRM2B | None | None | None | None |
| RRM2B | None | None | None | None |
| RRM2B | None | None | None | None |
| RRM2B | None | None | None | None |
| RRM2B | None | None | None | None |
| RRM2B | None | None | None | None |
| RRM2B | None | None | None | None |
| MARVELD2 | None | None | None | None |
| TLR4 | None | None | None | None |

| | | | | |
|---|---|---|---|---|
| TTN | None | None | None | None |
| TTN | None | None | None | None |
| TNFRSF13B | None | None | None | None |
| POLG | None | None | None | None |
| POLG | None | None | None | None |
| POLG | None | None | None | None |
| POLG | None | None | None | None |
| POLG | None | None | None | None |
| POLG | None | None | None | None |
| POLG | None | None | None | None |
| POLG | None | None | None | None |
| POLG | None | None | None | None |
| POLG | None | None | None | None |
| POLG | None | None | None | None |
| POLG | None | None | None | None |
| POLG | None | None | None | None |
| POLG | None | None | None | None |
| POLG | None | None | None | None |
| POLG | None | None | None | None |
| POLG | None | None | None | None |
| POLG | None | None | None | None |
| POLG | None | None | None | None |
| POLG | None | None | None | None |
| POLG | None | None | None | None |
| POLG | None | None | None | None |
| POLG | None | None | None | None |
| POLG | None | None | None | None |
| POLG | None | None | None | None |
| POLG | None | None | None | None |
| POLG | None | None | None | None |
| POLG | None | None | None | None |
| POLG | None | None | None | None |
| POLG | None | None | None | None |
| POLG | None | None | None | None |
| POLG | None | None | None | None |
| POLG | None | None | None | None |
| POLG | None | None | None | None |
| POLG | None | None | None | None |
| POLG | None | None | None | None |
| POLG | None | None | None | None |
| POLG | None | None | None | None |
| POLG | None | None | None | None |
| POLG | None | None | None | None |
| POLG | None | None | None | None |
| POLG | None | None | None | None |
| POLG | None | None | None | None |
| POLG | None | None | None | None |
| POLG | None | None | None | None |
| POLG | None | None | None | None |
| POLG | None | None | None | None |
| POLG | None | None | None | None |
| POLG | None | None | None | None |
| POLG | None | None | None | None |
| POLG | None | None | None | None |
| UROD | None | None | None | None |

| | | | | |
|---|---|---|---|---|
| UROD | None | None | None | None |
| UROD | None | None | None | None |
| UROD | None | None | None | None |
| UROD | None | None | None | None |
| UROD | None | None | None | None |
| UROD | None | None | None | None |
| UROD | None | None | None | None |
| UROD | None | None | None | None |
| UROD | None | None | None | None |
| PCK1 | None | None | None | None |
| PCK1 | None | None | None | None |
| PCK1 | None | None | None | None |
| PCK1 | None | None | None | None |
| PCK1 | None | None | None | None |
| HEXA | None | None | None | None |
| HEXA | None | None | None | None |
| HEXA | None | None | None | None |
| HEXA | None | None | None | None |
| HEXA | None | None | None | None |
| HEXA | None | None | None | None |
| HEXA | None | None | None | None |
| HEXA | None | None | None | None |
| HEXA | None | None | None | None |
| HEXA | None | None | None | None |
| HEXA | None | None | None | None |
| HEXA | None | None | None | None |
| HEXA | None | None | None | None |
| HEXA | None | None | None | None |
| HEXA | None | None | None | None |
| HEXA | None | None | None | None |
| HEXA | None | None | None | None |
| HEXA | None | None | None | None |
| HEXA | None | None | None | None |
| HEXA | None | None | None | None |
| HEXA | None | None | None | None |
| HEXA | None | None | None | None |
| HEXA | None | None | None | None |
| HEXA | None | None | None | None |
| HEXA | None | None | None | None |
| HEXA | None | None | None | None |
| HEXA | None | None | None | None |
| HEXA | None | None | None | None |
| HEXA | None | None | None | None |
| HEXA | None | None | None | None |
| HEXA | None | None | None | None |
| CFTR | None | None | None | None |
| CFTR | None | None | None | None |
| CFTR | None | None | None | None |
| CFTR | None | None | None | None |
| CFTR | None | None | None | None |
| CFTR | None | None | None | None |
| CFTR | None | None | None | None |
| CFTR | None | None | None | None |
| CFTR | None | None | None | None |

| | | | | |
|---|---|---|---|---|
| CFTR | None | None | None | None |
| CFTR | None | None | None | None |
| CFTR | None | None | None | None |
| CFTR | None | None | None | None |
| CFTR | None | None | None | None |
| CFTR | None | None | None | None |
| CFTR | None | None | None | None |
| CFTR | None | None | None | None |
| CFTR | None | None | None | None |
| CFTR | None | None | None | None |
| CFTR | None | None | None | None |
| CFTR | None | None | None | None |
| CFTR | None | None | None | None |
| CFTR | None | None | None | None |
| CFTR | None | None | None | None |
| CFTR | None | None | None | None |
| CFTR | None | None | None | None |
| CFTR | None | None | None | None |
| CFTR | None | None | None | None |
| CFTR | None | None | None | None |
| CFTR | None | None | None | None |
| CFTR | None | None | None | None |
| CFTR | None | None | None | None |
| CFTR | None | None | None | None |
| CFTR | None | None | None | None |
| FAN1 | None | None | None | None |
| CBS | None | None | None | None |
| CBS | None | None | None | None |
| CBS | None | None | None | None |
| CBS | None | None | None | None |
| CBS | None | None | None | None |
| CBS | None | None | None | None |
| CBS | None | None | None | None |
| CBS | None | None | None | None |
| CBS | None | None | None | None |
| CBS | None | None | None | None |
| CBS | None | None | None | None |
| CBS | None | None | None | None |
| CBS | None | None | None | None |
| CBS | None | None | None | None |
| CBS | None | None | None | None |
| CBS | None | None | None | None |
| CBS | None | None | None | None |
| CBS | None | None | None | None |
| CBS | None | None | None | None |
| CBS | None | None | None | None |
| CBS | None | None | None | None |
| CBS | None | None | None | None |
| CBS | None | None | None | None |
| CBS | None | None | None | None |
| CBS | None | None | None | None |
| CBS | None | None | None | None |
| CBS | None | None | None | None |
| CBS | None | None | None | None |

| | | | | |
|---|---|---|---|---|
| CBS | None | None | None | None |
| CBS | None | None | None | None |
| CBS | None | None | None | None |
| CBS | None | None | None | None |
| CBS | None | None | None | None |
| CBS | None | None | None | None |
| CBS | None | None | None | None |
| CBS | None | None | None | None |
| TLR3 | None | None | None | None |
| LIG4 | None | None | None | None |
| LIG4 | None | None | None | None |
| ASPA | None | None | None | None |
| ASPA | None | None | None | None |
| ASPA | None | None | None | None |
| ASPA | None | None | None | None |
| ASPA | None | None | None | None |
| ASPA | None | None | None | None |
| ASPA | None | None | None | None |
| ASPA | None | None | None | None |
| ASPA | None | None | None | None |
| ASPA | None | None | None | None |
| ASPA | None | None | None | None |
| ASPA | None | None | None | None |
| ASPA | None | None | None | None |
| CFH | None | None | None | None |
| FCGR3A | None | None | None | None |
| FCGR3A | None | None | None | None |
| GRIN2A | None | None | None | None |
| GRIN2A | None | None | None | None |
| GRIN2A | None | None | None | None |
| GLA | None | None | None | None |
| GLA | None | None | None | None |
| GLA | None | None | None | None |
| GLA | None | None | None | None |
| GLA | None | None | None | None |
| GLA | None | None | None | None |
| GLA | None | None | None | None |
| GLA | None | None | None | None |
| GLA | None | None | None | None |
| GLA | None | None | None | None |
| GLA | None | None | None | None |
| GLA | None | None | None | None |
| GLA | None | None | None | None |
| GLA | None | None | None | None |
| GLA | None | None | None | None |
| GLA | None | None | None | None |
| GLA | None | None | None | None |
| GLA | None | None | None | None |
| GLA | None | None | None | None |
| GLA | None | None | None | None |
| GLA | None | None | None | None |
| GLA | None | None | None | None |
| GLA | None | None | None | None |
| GLA | None | None | None | None |

| | | | | |
|---|---|---|---|---|
| GLA | None | None | None | None |
| GLA | None | None | None | None |
| GLA | None | None | None | None |
| GLA | None | None | None | None |
| GLA | None | None | None | None |
| GLA | None | None | None | None |
| GLA | None | None | None | None |
| SERPINA3 | None | None | None | None |
| MTAP | None | None | None | None |
| MTAP | None | None | None | None |
| MTAP | None | None | None | None |
| BLMH | None | None | None | None |
| TUSC3 | None | None | None | None |
| ISCU | None | None | None | None |
| CYP11B2 | None | None | None | None |
| CYP11B2 | None | None | None | None |
| CYP11B2 | None | None | None | None |
| CYP11B2 | None | None | None | None |
| CYP11B2 | None | None | None | None |
| CYP11B2 | None | None | None | None |
| CYP11B2 | None | None | None | None |
| CYP11B2 | None | None | None | None |
| CYP11B2 | None | None | None | None |
| CYP11B2 | None | None | None | None |
| CYP11B2 | None | None | None | None |
| CYP11B2 | None | None | None | None |
| CYP11B2 | None | None | None | None |
| CYP11B2 | None | None | None | None |
| CYP11B2 | None | None | None | None |
| CYP11B2 | None | None | None | None |
| CYP11B2 | None | None | None | None |
| CYP11B2 | None | None | None | None |
| CYP11B2 | None | None | None | None |
| CRADD | None | None | None | None |
| CRADD | None | None | None | None |
| FGA | None | None | None | None |
| FGA | None | None | None | None |
| FGA | None | None | None | None |
| FGA | None | None | None | None |
| FGA | None | None | None | None |
| FGA | None | None | None | None |
| FGA | None | None | None | None |
| FGA | None | None | None | None |
| IYD | 1 | 1 | 4ttb | A |
| IYD | None | None | None | None |
| IYD | None | None | None | None |
| IYD | None | None | None | None |
| TCAP | None | None | None | None |
| TCAP | None | None | None | None |
| TCAP | None | None | None | None |
| TCAP | None | None | None | None |
| TCAP | None | None | None | None |
| CYP7A1 | None | None | None | None |

| | | | | |
|---|---|---|---|---|
| TEK | None | None | None | None |
| TEK | None | None | None | None |
| TEK | None | None | None | None |
| TEK | None | None | None | None |
| TEK | None | None | None | None |
| TEK | None | None | None | None |
| TEK | None | None | None | None |
| CD8A | None | None | None | None |
| HERC2 | None | None | None | None |
| NR1H3 | None | None | None | None |
| GRHPR | None | None | None | None |
| GRHPR | None | None | None | None |
| GRHPR | None | None | None | None |
| GRHPR | None | None | None | None |
| GRHPR | None | None | None | None |
| GRHPR | None | None | None | None |
| GRHPR | None | None | None | None |
| GBE1 | None | None | None | None |
| GBE1 | None | None | None | None |
| GBE1 | None | None | None | None |
| GBE1 | None | None | None | None |
| GBE1 | None | None | None | None |
| GBE1 | None | None | None | None |
| GBE1 | None | None | None | None |
| GBE1 | None | None | None | None |
| GBE1 | None | None | None | None |
| GBE1 | None | None | None | None |
| GBE1 | None | None | None | None |
| GBE1 | None | None | None | None |
| GBE1 | None | None | None | None |
| GBE1 | None | None | None | None |
| GBE1 | None | None | None | None |
| GBE1 | None | None | None | None |
| GBE1 | None | None | None | None |
| GBE1 | None | None | None | None |
| GBE1 | None | None | None | None |
| GBE1 | None | None | None | None |
| PC | None | None | None | None |
| PC | None | None | None | None |
| PC | None | None | None | None |
| PC | None | None | None | None |
| PC | None | None | None | None |
| PC | None | None | None | None |
| PC | None | None | None | None |
| PC | None | None | None | None |
| MSH6 | None | None | None | None |
| MSH6 | None | None | None | None |
| MSH6 | None | None | None | None |
| MSH6 | None | None | None | None |
| MSH6 | None | None | None | None |
| MSH6 | None | None | None | None |
| MSH6 | None | None | None | None |
| MSH6 | None | None | None | None |
| MSH6 | None | None | None | None |

| | | | | |
|---|---|---|---|---|
| MSH6 | None | None | None | None |
| MSH6 | None | None | None | None |
| MSH6 | None | None | None | None |
| MSH6 | None | None | None | None |
| MSH6 | None | None | None | None |
| MSH6 | None | None | None | None |
| MSH6 | None | None | None | None |
| MSH6 | None | None | None | None |
| MSH6 | None | None | None | None |
| MSH6 | None | None | None | None |
| MSH6 | None | None | None | None |
| MSH6 | None | None | None | None |
| MSH6 | None | None | None | None |
| MSH6 | None | None | None | None |
| MSH6 | None | None | None | None |
| MSH6 | None | None | None | None |
| MSH6 | None | None | None | None |
| MSH6 | None | None | None | None |
| MSH6 | None | None | None | None |
| MSH6 | None | None | None | None |
| MSH6 | None | None | None | None |
| RNASEL | None | None | None | None |
| ORC1 | None | None | None | None |
| ORC1 | None | None | None | None |
| ORC1 | None | None | None | None |
| KCNH2 | None | None | None | None |
| KCNH2 | None | None | None | None |
| KCNH2 | None | None | None | None |
| KCNH2 | None | None | None | None |
| KCNH2 | None | None | None | None |
| KCNH2 | None | None | None | None |
| KCNH2 | None | None | None | None |
| KCNH2 | None | None | None | None |
| KCNH2 | None | None | None | None |
| KCNH2 | None | None | None | None |
| KCNH2 | None | None | None | None |
| KCNH2 | None | None | None | None |
| KCNH2 | None | None | None | None |
| KCNH2 | None | None | None | None |
| KCNH2 | None | None | None | None |
| KCNH2 | None | None | None | None |
| KCNH2 | None | None | None | None |
| KCNH2 | None | None | None | None |
| KCNH2 | None | None | None | None |
| KCNH2 | None | None | None | None |
| KCNH2 | None | None | None | None |
| KCNH2 | None | None | None | None |
| KCNH2 | None | None | None | None |
| KCNH2 | None | None | None | None |
| MAT1A | None | None | None | None |
| MAT1A | None | None | None | None |
| MAT1A | None | None | None | None |
| MAT1A | None | None | None | None |
| MAT1A | None | None | None | None |

| | | | | |
|---|---|---|---|---|
| MAT1A | None | None | None | None |
| MAT1A | None | None | None | None |
| MAT1A | None | None | None | None |
| INSR | None | None | None | None |
| INSR | None | None | None | None |
| INSR | None | None | None | None |
| INSR | None | None | None | None |
| AMT | None | None | None | None |
| AMT | None | None | None | None |
| AMT | None | None | None | None |
| AMT | None | None | None | None |
| AMT | None | None | None | None |
| AMT | None | None | None | None |
| AMT | None | None | None | None |
| AMT | None | None | None | None |
| AMT | None | None | None | None |
| CSNK1D | None | None | None | None |
| SF3B1 | None | None | None | None |
| AGA | None | None | None | None |
| AGA | None | None | None | None |
| AGA | None | None | None | None |
| AGA | None | None | None | None |
| AGA | None | None | None | None |
| AGA | None | None | None | None |
| AGA | None | None | None | None |
| NECTIN4 | None | None | None | None |
| RS1 | None | None | None | None |
| RS1 | None | None | None | None |
| RS1 | None | None | None | None |
| RS1 | None | None | None | None |
| TBC1D7 | None | None | None | None |
| CTSA | None | None | None | None |
| CTSA | None | None | None | None |
| TKT | None | None | None | None |
| RP2 | None | None | None | None |
| RP2 | None | None | None | None |
| RP2 | None | None | None | None |
| RP2 | None | None | None | None |
| LYRM4 | None | None | None | None |
| SOS1 | None | None | None | None |
| SOS1 | None | None | None | None |
| SOS1 | None | None | None | None |
| SOS1 | None | None | None | None |
| SOS1 | None | None | None | None |
| SOS1 | None | None | None | None |
| HUWE1 | None | None | None | None |
| HUWE1 | None | None | None | None |
| TPI1 | None | None | None | None |
| TPI1 | None | None | None | None |
| TPI1 | None | None | None | None |
| MSH2 | None | None | None | None |
| MSH2 | None | None | None | None |
| MSH2 | None | None | None | None |
| MSH2 | None | None | None | None |

| | | | | |
|---|---|---|---|---|
| MSH2 | None | None | None | None |
| MSH2 | None | None | None | None |
| MSH2 | None | None | None | None |
| MSH2 | None | None | None | None |
| MSH2 | None | None | None | None |
| MSH2 | None | None | None | None |
| MSH2 | None | None | None | None |
| MSH2 | None | None | None | None |
| MSH2 | None | None | None | None |
| MSH2 | None | None | None | None |
| MSH2 | None | None | None | None |
| MSH2 | None | None | None | None |
| MSH2 | None | None | None | None |
| MSH2 | None | None | None | None |
| MSH2 | None | None | None | None |
| MSH2 | None | None | None | None |
| MSH2 | None | None | None | None |
| MSH2 | None | None | None | None |
| MSH2 | None | None | None | None |
| MSH2 | None | None | None | None |
| MSH2 | None | None | None | None |
| MSH2 | None | None | None | None |
| MSH2 | None | None | None | None |
| MSH2 | None | None | None | None |
| MSH2 | None | None | None | None |
| MSH2 | None | None | None | None |
| MSH2 | None | None | None | None |
| MSH2 | None | None | None | None |
| MSH2 | None | None | None | None |
| MSH2 | None | None | None | None |
| MSH2 | None | None | None | None |
| MSH2 | None | None | None | None |
| GALE | None | None | None | None |
| GALE | None | None | None | None |
| GALE | None | None | None | None |
| GALE | None | None | None | None |
| GALE | None | None | None | None |
| GALE | None | None | None | None |
| GALE | None | None | None | None |
| GALE | None | None | None | None |
| NSD2 | None | None | None | None |
| SRP72 | None | None | None | None |
| SRP72 | None | None | None | None |
| SRP72 | None | None | None | None |
| POLK | None | None | None | None |
| GALT | None | None | None | None |
| GALT | None | None | None | None |
| GALT | None | None | None | None |
| GALT | None | None | None | None |
| GALT | None | None | None | None |
| GALT | None | None | None | None |
| GALT | None | None | None | None |
| GALT | None | None | None | None |
| GALT | None | None | None | None |

| | | | | |
|---|---|---|---|---|
| GALT | None | None | None | None |
| GALT | None | None | None | None |
| GALT | None | None | None | None |
| GALT | None | None | None | None |
| GALT | None | None | None | None |
| GALT | None | None | None | None |
| GALT | None | None | None | None |
| GALT | None | None | None | None |
| GALT | None | None | None | None |
| GALT | None | None | None | None |
| GALT | None | None | None | None |
| GALT | None | None | None | None |
| GALT | None | None | None | None |
| GALT | None | None | None | None |
| GALT | None | None | None | None |
| GALT | None | None | None | None |
| GALT | None | None | None | None |
| GALT | None | None | None | None |
| GALT | None | None | None | None |
| GALT | None | None | None | None |
| GALT | None | None | None | None |
| GALT | None | None | None | None |
| GALT | None | None | None | None |
| GALT | None | None | None | None |
| GALT | None | None | None | None |
| GALT | None | None | None | None |
| GALT | None | None | None | None |
| GALT | None | None | None | None |
| PYCR1 | None | None | None | None |
| PYCR1 | None | None | None | None |
| PYCR1 | None | None | None | None |
| PYCR1 | None | None | None | None |
| F8 | None | None | None | None |
| F8 | None | None | None | None |
| F8 | None | None | None | None |
| F8 | None | None | None | None |
| F8 | None | None | None | None |
| F8 | None | None | None | None |
| F8 | None | None | None | None |
| F8 | None | None | None | None |
| F8 | None | None | None | None |
| F8 | None | None | None | None |
| F8 | None | None | None | None |
| F8 | None | None | None | None |
| F8 | None | None | None | None |
| F8 | None | None | None | None |
| F8 | None | None | None | None |
| F8 | None | None | None | None |
| ANK1 | None | None | None | None |
| CHAT | None | None | None | None |
| CHAT | None | None | None | None |
| CHAT | None | None | None | None |
| CHAT | None | None | None | None |
| CHAT | None | None | None | None |

| | | | | |
|---|---|---|---|---|
| CHAT | None | None | None | None |
| CHAT | None | None | None | None |
| CHAT | None | None | None | None |
| CHAT | None | None | None | None |
| CHAT | None | None | None | None |
| CHAT | None | None | None | None |
| CHAT | None | None | None | None |
| CHAT | None | None | None | None |
| CHAT | None | None | None | None |
| CHAT | None | None | None | None |
| CHAT | None | None | None | None |
| CHAT | None | None | None | None |
| GHR | None | None | None | None |
| GHR | None | None | None | None |
| GHR | None | None | None | None |
| GHR | None | None | None | None |
| GHR | None | None | None | None |
| GHR | None | None | None | None |
| GHR | None | None | None | None |
| GHR | None | None | None | None |
| GHR | None | None | None | None |
| GHR | None | None | None | None |
| GHR | None | None | None | None |
| GHR | None | None | None | None |
| GHR | None | None | None | None |
| GHR | None | None | None | None |
| FZD4 | None | None | None | None |
| FZD4 | None | None | None | None |
| FZD4 | None | None | None | None |
| MTR | None | None | None | None |
| MTR | None | None | None | None |
| MTR | None | None | None | None |
| GM2A | None | None | None | None |
| GM2A | None | None | None | None |
| GM2A | None | None | None | None |
| GM2A | None | None | None | None |
| GM2A | None | None | None | None |
| CRYGD | None | None | None | None |
| CRYGD | None | None | None | None |
| CRYGD | None | None | None | None |
| CRYGD | None | None | None | None |
| CRYGD | None | None | None | None |
| CRYGD | None | None | None | None |
| CRYGD | None | None | None | None |
| CRYGD | None | None | None | None |
| CRYGD | None | None | None | None |
| TLR5 | None | None | None | None |
| HSD17B10 | None | None | None | None |
| HSD17B10 | None | None | None | None |
| HSD17B10 | None | None | None | None |
| HSD17B10 | None | None | None | None |
| TBC1D20 | None | None | None | None |
| FBN1 | None | None | None | None |
| FBN1 | None | None | None | None |

| | | | | |
|---|---|---|---|---|
| FBN1 | None | None | None | None |
| FBN1 | None | None | None | None |
| FBN1 | None | None | None | None |
| FBN1 | None | None | None | None |
| FBN1 | None | None | None | None |
| FXN | None | None | None | None |
| FXN | None | None | None | None |
| FXN | None | None | None | None |
| FXN | None | None | None | None |
| MYO5A | None | None | None | None |
| CTSK | None | None | None | None |
| CTSK | None | None | None | None |
| SCN5A | None | None | None | None |
| SCN5A | None | None | None | None |
| SOS1 | None | None | None | None |
| SOS1 | None | None | None | None |
| SOS1 | None | None | None | None |
| SOS1 | None | None | None | None |
| SOS1 | None | None | None | None |
| SOS1 | None | None | None | None |
| SOS1 | None | None | None | None |
| SOS1 | None | None | None | None |
| SOS1 | None | None | None | None |
| SOS1 | None | None | None | None |
| SOS1 | None | None | None | None |
| SOS1 | None | None | None | None |
| GATM | None | None | None | None |
| GATM | None | None | None | None |
| GATM | None | None | None | None |
| GATM | None | None | None | None |
| GATM | None | None | None | None |
| GATM | None | None | None | None |
| GATM | None | None | None | None |
| MMP2 | None | None | None | None |
| MMP2 | None | None | None | None |
| MMP2 | None | None | None | None |
| MMP2 | None | None | None | None |
| NDUFB9 | None | None | None | None |
| NDUFB9 | None | None | None | None |
| GH1 | None | None | None | None |
| GH1 | None | None | None | None |
| GH1 | None | None | None | None |
| SLC6A4 | None | None | None | None |
| G6PD | None | None | None | None |
| G6PD | None | None | None | None |
| G6PD | None | None | None | None |
| G6PD | None | None | None | None |
| G6PD | None | None | None | None |
| G6PD | None | None | None | None |
| G6PD | None | None | None | None |
| G6PD | None | None | None | None |
| G6PD | None | None | None | None |
| G6PD | None | None | None | None |
| G6PD | None | None | None | None |

| | | | | |
|---|---|---|---|---|
| G6PD | None | None | None | None |
| G6PD | None | None | None | None |
| G6PD | None | None | None | None |
| G6PD | None | None | None | None |
| G6PD | None | None | None | None |
| G6PD | None | None | None | None |
| G6PD | None | None | None | None |
| G6PD | None | None | None | None |
| G6PD | None | None | None | None |
| CST3 | None | None | None | None |
| LMNA | None | None | None | None |
| LMNA | None | None | None | None |
| NDUFA9 | None | None | None | None |
| NDUFA9 | None | None | None | None |
| NDUFA9 | None | None | None | None |
| NDUFA9 | None | None | None | None |
| YARS | None | None | None | None |
| YARS | None | None | None | None |
| YARS | None | None | None | None |
| YARS | None | None | None | None |
| EPHA2 | None | None | None | None |
| EPHA2 | None | None | None | None |
| EPHA2 | None | None | None | None |
| EPHA2 | None | None | None | None |
| DICER1 | None | None | None | None |
| DICER1 | None | None | None | None |
| SNX10 | None | None | None | None |
| SNX10 | None | None | None | None |
| TTN | None | None | None | None |
| TTN | None | None | None | None |
| GATA3 | None | None | None | None |
| C5 | None | None | None | None |
| THBD | None | None | None | None |
| BIN1 | None | None | None | None |
| BIN1 | None | None | None | None |
| SMARCAL1 | None | None | None | None |
| CFB | None | None | None | None |
| CYC1 | None | None | None | None |
| ANK2 | None | None | None | None |
| CHMP4B | None | None | None | None |
| CHMP4B | None | None | None | None |
| ERBB2 | None | None | None | None |
| ERBB2 | None | None | None | None |
| POLG2 | None | None | None | None |
| POLG2 | None | None | None | None |
| POLG2 | None | None | None | None |
| POLG2 | None | None | None | None |
| POLG2 | None | None | None | None |
| POLG2 | None | None | None | None |
| POLG2 | None | None | None | None |
| POLG2 | None | None | None | None |
| C3 | None | None | None | None |
| C3 | None | None | None | None |
| RPGR | None | None | None | None |

| | | | | |
|---|---|---|---|---|
| RPGR | None | None | None | None |
| RPGR | None | None | None | None |
| RPGR | None | None | None | None |
| RPGR | None | None | None | None |
| C1QA | None | None | None | None |
| PRKG1 | None | None | None | None |
| IGBP1 | None | None | None | None |
| ENO3 | None | None | None | None |
| ENO3 | None | None | None | None |
| ENO3 | None | None | None | None |
| ENO3 | None | None | None | None |
| ENO3 | None | None | None | None |
| ENO3 | None | None | None | None |
| MCPH1 | None | None | None | None |
| MCPH1 | None | None | None | None |
| MCPH1 | None | None | None | None |
| MCPH1 | None | None | None | None |
| MCPH1 | None | None | None | None |
| SNRNP200 | None | None | None | None |
| SNRNP200 | None | None | None | None |
| DNMT1 | None | None | None | None |
| DNMT1 | None | None | None | None |
| DNMT1 | None | None | None | None |
| AKT1 | None | None | None | None |
| NDUFS2 | None | None | None | None |
| ACVR2B | None | None | None | None |
| HMGCL | None | None | None | None |
| HMGCL | None | None | None | None |
| HMGCL | None | None | None | None |
| JUP | None | None | None | None |
| JUP | None | None | None | None |
| JUP | None | None | None | None |
| JUP | None | None | None | None |
| JUP | None | None | None | None |
| JUP | None | None | None | None |
| JUP | None | None | None | None |
| JUP | None | None | None | None |
| SPINK1 | None | None | None | None |
| SPINK1 | None | None | None | None |
| SPINK1 | None | None | None | None |
| VCL | None | None | None | None |
| VCL | None | None | None | None |
| NOTCH1 | None | None | None | None |
| NOTCH1 | None | None | None | None |
| NOTCH1 | None | None | None | None |
| NDUFB3 | None | None | None | None |
| IKBKG | None | None | None | None |
| GSN | None | None | None | None |
| GSN | None | None | None | None |
| GSN | None | None | None | None |
| GSN | None | None | None | None |
| GSN | None | None | None | None |
| GSN | None | None | None | None |
| GSN | None | None | None | None |

| | | | | |
|---|---|---|---|---|
| GSN | None | None | None | None |
| GSN | None | None | None | None |
| CLC | None | None | None | None |
| RAF1 | None | None | None | None |
| RAF1 | None | None | None | None |
| RAF1 | None | None | None | None |
| NDUFS8 | None | None | None | None |
| NDUFS8 | None | None | None | None |
| NDUFS8 | None | None | None | None |
| NDUFS8 | None | None | None | None |
| NDUFS8 | None | None | None | None |
| CASR | None | None | None | None |
| CASR | None | None | None | None |
| CASR | None | None | None | None |
| CASR | None | None | None | None |
| CASR | None | None | None | None |
| CASR | None | None | None | None |
| CASR | None | None | None | None |
| CASR | None | None | None | None |
| CASR | None | None | None | None |
| CASR | None | None | None | None |
| CASR | None | None | None | None |
| CASR | None | None | None | None |
| CASR | None | None | None | None |
| CASR | None | None | None | None |
| CASR | None | None | None | None |
| CASR | None | None | None | None |
| CASR | None | None | None | None |
| CASR | None | None | None | None |
| CASR | None | None | None | None |
| CASR | None | None | None | None |
| CASR | None | None | None | None |
| CASR | None | None | None | None |
| CASR | None | None | None | None |
| CASR | None | None | None | None |
| CASR | None | None | None | None |
| CASR | None | None | None | None |
| CASR | None | None | None | None |
| CASR | None | None | None | None |
| CASR | None | None | None | None |
| VCL | None | None | None | None |
| VCL | None | None | None | None |
| SMPD1 | None | None | None | None |
| SMPD1 | None | None | None | None |
| SMPD1 | None | None | None | None |
| KDM6A | None | None | None | None |
| KCNQ1 | None | None | None | None |
| KCNQ1 | None | None | None | None |
| KCNQ1 | None | None | None | None |
| KCNQ1 | None | None | None | None |
| KCNQ1 | None | None | None | None |
| KCNQ1 | None | None | None | None |
| NDUFA12 | None | None | None | None |
| GLRX5 | None | None | None | None |

| | | | | |
|---|---|---|---|---|
| PALB2 | None | None | None | None |
| PALB2 | None | None | None | None |
| PALB2 | None | None | None | None |
| PALB2 | None | None | None | None |
| PALB2 | None | None | None | None |
| PALB2 | None | None | None | None |
| PALB2 | None | None | None | None |
| PALB2 | None | None | None | None |
| PALB2 | None | None | None | None |
| PALB2 | None | None | None | None |
| ALDOB | None | None | None | None |
| ALDOB | None | None | None | None |
| ALDOB | None | None | None | None |
| ALDOB | None | None | None | None |
| ALDOB | None | None | None | None |
| ALDOB | None | None | None | None |
| ALDOB | None | None | None | None |
| GNE | None | None | None | None |
| GNE | None | None | None | None |
| GNE | None | None | None | None |
| GNE | None | None | None | None |
| GNE | None | None | None | None |
| GNE | None | None | None | None |
| GNE | None | None | None | None |
| GNE | None | None | None | None |
| GNE | None | None | None | None |
| GNE | None | None | None | None |
| GNE | None | None | None | None |
| GNE | None | None | None | None |
| CHRNB2 | None | None | None | None |
| CHRNB2 | None | None | None | None |
| ERCC3 | None | None | None | None |
| ERCC3 | None | None | None | None |
| ERCC3 | None | None | None | None |
| ERCC3 | None | None | None | None |
| ERCC3 | None | None | None | None |
| UNC119 | None | None | None | None |
| ATL1 | None | None | None | None |
| ATL1 | None | None | None | None |
| ATL1 | None | None | None | None |
| ATL1 | None | None | None | None |
| ATL1 | None | None | None | None |
| ATL1 | None | None | None | None |
| ATL1 | None | None | None | None |
| ATL1 | None | None | None | None |
| ATL1 | None | None | None | None |
| ATL1 | None | None | None | None |
| NR3C2 | None | None | None | None |
| TBK1 | None | None | None | None |
| HYAL1 | None | None | None | None |
| HYAL1 | None | None | None | None |
| HYAL1 | None | None | None | None |
| HYAL1 | None | None | None | None |
| CTSD | None | None | None | None |

| | | | | |
|---|---|---|---|---|
| PRKCG | None | None | None | None |
| TGFBR1 | None | None | None | None |
| XDH | None | None | None | None |
| XDH | None | None | None | None |
| XDH | None | None | None | None |
| XDH | None | None | None | None |
| XDH | None | None | None | None |
| XDH | None | None | None | None |
| NMNAT1 | None | None | None | None |
| NMNAT1 | None | None | None | None |
| NMNAT1 | None | None | None | None |
| NMNAT1 | None | None | None | None |
| NMNAT1 | None | None | None | None |
| NMNAT1 | None | None | None | None |
| SERPING1 | None | None | None | None |
| SERPING1 | None | None | None | None |
| SERPING1 | None | None | None | None |
| SERPING1 | None | None | None | None |
| SERPING1 | None | None | None | None |
| PDZRN3 | None | None | None | None |
| FABP2 | None | None | None | None |
| IFNGR1 | None | None | None | None |
| IFNGR1 | None | None | None | None |
| TDP1 | None | None | None | None |
| TDP1 | None | None | None | None |
| INS | None | None | None | None |
| INS | None | None | None | None |
| INS | None | None | None | None |
| PPOX | None | None | None | None |
| PPOX | None | None | None | None |
| PPOX | None | None | None | None |
| PPOX | None | None | None | None |
| PPOX | None | None | None | None |
| PPOX | None | None | None | None |
| PPOX | None | None | None | None |
| PPOX | None | None | None | None |
| PPOX | None | None | None | None |
| GFER | None | None | None | None |
| GFER | None | None | None | None |
| GFER | None | None | None | None |
| GFER | None | None | None | None |
| LIFR | None | None | None | None |
| EGLN1 | None | None | None | None |
| EGLN1 | None | None | None | None |
| EGLN1 | 0.944 | 2 | 5a3u | C |
| GP1BA | None | None | None | None |
| GP1BA | None | None | None | None |
| GP1BA | None | None | None | None |
| GP1BA | None | None | None | None |
| GP1BA | None | None | None | None |
| MMADHC | None | None | None | None |
| MMADHC | None | None | None | None |
| MMADHC | None | None | None | None |
| MMADHC | None | None | None | None |

| | | | | |
|---|---|---|---|---|
| MMADHC | None | None | None | None |
| MMADHC | None | None | None | None |
| MMADHC | None | None | None | None |
| ACVRL1 | None | None | None | None |
| ACVRL1 | None | None | None | None |
| RFXANK | None | None | None | None |
| CHIT1 | None | None | None | None |
| UBR1 | None | None | None | None |
| BLM | None | None | None | None |
| BLM | None | None | None | None |
| BLM | None | None | None | None |
| BLM | None | None | None | None |
| BLM | None | None | None | None |
| BLM | None | None | None | None |
| BLM | None | None | None | None |
| BLM | None | None | None | None |
| BLM | None | None | None | None |
| BLM | None | None | None | None |
| BLM | None | None | None | None |
| BLM | None | None | None | None |
| BLM | None | None | None | None |
| VDR | None | None | None | None |
| VDR | None | None | None | None |
| VDR | None | None | None | None |
| VDR | None | None | None | None |
| VDR | None | None | None | None |
| VDR | None | None | None | None |
| EPHA2 | None | None | None | None |
| EPHA2 | None | None | None | None |
| EPHA2 | None | None | None | None |
| EPHA2 | None | None | None | None |
| EPHA2 | None | None | None | None |
| EPHA2 | None | None | None | None |
| EPHA2 | None | None | None | None |
| EPHA2 | None | None | None | None |
| EPHA2 | None | None | None | None |
| EPHA2 | None | None | None | None |
| UBA1 | None | None | None | None |
| STAT1 | None | None | None | None |
| STAT1 | None | None | None | None |
| STAT1 | None | None | None | None |
| STAT1 | None | None | None | None |
| STAT1 | None | None | None | None |
| STAT1 | None | None | None | None |
| STAT1 | None | None | None | None |
| STAT1 | None | None | None | None |
| DNM1L | None | None | None | None |
| DNM1L | None | None | None | None |
| OTC | None | None | None | None |
| OTC | None | None | None | None |
| OTC | None | None | None | None |
| OTC | None | None | None | None |
| OTC | None | None | None | None |
| OTC | None | None | None | None |

| | | | | |
|---|---|---|---|---|
| OTC | None | None | None | None |
| OTC | None | None | None | None |
| OTC | None | None | None | None |
| OTC | None | None | None | None |
| OTC | None | None | None | None |
| OTC | None | None | None | None |
| OTC | None | None | None | None |
| OTC | None | None | None | None |
| OTC | None | None | None | None |
| OTC | None | None | None | None |
| OTC | None | None | None | None |
| RARS | None | None | None | None |
| RARS | None | None | None | None |
| RARS | None | None | None | None |
| SEPSECS | None | None | None | None |
| SEPSECS | None | None | None | None |
| SEPSECS | None | None | None | None |
| SEPSECS | None | None | None | None |
| SEPSECS | None | None | None | None |
| BRCA1 | None | None | None | None |
| BRCA1 | None | None | None | None |
| BRCA1 | None | None | None | None |
| BRCA1 | None | None | None | None |
| BRCA1 | None | None | None | None |
| BRCA1 | None | None | None | None |
| BRCA1 | None | None | None | None |
| BRCA1 | None | None | None | None |
| BRCA1 | None | None | None | None |
| BRCA1 | None | None | None | None |
| BRCA1 | None | None | None | None |
| BRCA1 | None | None | None | None |
| BRCA1 | None | None | None | None |
| BRCA1 | None | None | None | None |
| BRCA1 | None | None | None | None |
| BRCA1 | None | None | None | None |
| BRCA1 | None | None | None | None |
| BRCA1 | None | None | None | None |
| BRCA1 | None | None | None | None |
| BRCA1 | None | None | None | None |
| BRCA1 | None | None | None | None |
| BRCA1 | None | None | None | None |
| CUBN | None | None | None | None |
| ITGB3 | None | None | None | None |
| MTRR | None | None | None | None |
| MTRR | None | None | None | None |
| MTRR | None | None | None | None |
| MTRR | None | None | None | None |
| MTRR | None | None | None | None |
| MTRR | None | None | None | None |
| MTRR | None | None | None | None |
| MTRR | None | None | None | None |
| IL7R | None | None | None | None |
| IL7R | None | None | None | None |
| RPS24 | None | None | None | None |

| | | | | |
|---|---|---|---|---|
| RPS24 | None | None | None | None |
| HOGA1 | None | None | None | None |
| HOGA1 | None | None | None | None |
| HOGA1 | None | None | None | None |
| HOGA1 | None | None | None | None |
| HOGA1 | None | None | None | None |
| HOGA1 | None | None | None | None |
| HOGA1 | None | None | None | None |
| HOGA1 | None | None | None | None |
| HOGA1 | None | None | None | None |
| HOGA1 | None | None | None | None |
| HOGA1 | None | None | None | None |
| HOGA1 | None | None | None | None |
| HOGA1 | None | None | None | None |
| GJB2 | None | None | None | None |
| GJB2 | None | None | None | None |
| GJB2 | None | None | None | None |
| GJB2 | None | None | None | None |
| GJB2 | None | None | None | None |
| GJB2 | None | None | None | None |
| GJB2 | None | None | None | None |
| GJB2 | None | None | None | None |
| GJB2 | None | None | None | None |
| GJB2 | None | None | None | None |
| GJB2 | None | None | None | None |
| GJB2 | None | None | None | None |
| GJB2 | None | None | None | None |
| GJB2 | None | None | None | None |
| GJB2 | None | None | None | None |
| GJB2 | None | None | None | None |
| GJB2 | None | None | None | None |
| GJB2 | None | None | None | None |
| GJB2 | None | None | None | None |
| GJB2 | None | None | None | None |
| GJB2 | None | None | None | None |
| GJB2 | None | None | None | None |
| GJB2 | None | None | None | None |
| GJB2 | None | None | None | None |
| GJB2 | None | None | None | None |
| GJB2 | None | None | None | None |
| GJB2 | None | None | None | None |
| GJB2 | None | None | None | None |
| GJB2 | None | None | None | None |
| GJB2 | None | None | None | None |
| GJB2 | None | None | None | None |
| GJB2 | None | None | None | None |
| GJB2 | None | None | None | None |
| GJB2 | None | None | None | None |
| GJB2 | None | None | None | None |
| GJB2 | None | None | None | None |
| GJB2 | None | None | None | None |
| GJB2 | None | None | None | None |
| GJB2 | None | None | None | None |
| GJB2 | None | None | None | None |

| | | | | |
|---|---|---|---|---|
| GJB2 | None | None | None | None |
| GJB2 | None | None | None | None |
| GJB2 | None | None | None | None |
| GJB2 | None | None | None | None |
| GJB2 | None | None | None | None |
| GJB2 | None | None | None | None |
| GJB2 | None | None | None | None |
| GJB2 | None | None | None | None |
| GJB2 | None | None | None | None |
| GJB2 | None | None | None | None |
| GJB2 | None | None | None | None |
| GJB2 | None | None | None | None |
| GJB2 | None | None | None | None |
| GJB2 | None | None | None | None |
| CPS1 | None | None | None | None |
| CPS1 | None | None | None | None |
| CPS1 | None | None | None | None |
| CPS1 | None | None | None | None |
| CPS1 | None | None | None | None |
| CPS1 | None | None | None | None |
| CPS1 | None | None | None | None |
| CPS1 | None | None | None | None |
| CPS1 | None | None | None | None |
| CPS1 | None | None | None | None |
| PSPH | None | None | None | None |
| PSPH | None | None | None | None |
| PSPH | None | None | None | None |
| PSPH | None | None | None | None |
| PSPH | None | None | None | None |
| PSPH | None | None | None | None |
| PEPD | None | None | None | None |
| PEPD | None | None | None | None |
| PEPD | None | None | None | None |
| PEPD | None | None | None | None |
| PEPD | None | None | None | None |
| PEPD | None | None | None | None |
| PEPD | None | None | None | None |
| PEPD | None | None | None | None |
| PEPD | None | None | None | None |
| KDM5C | None | None | None | None |
| KDM5C | None | None | None | None |
| KDM5C | None | None | None | None |
| ITGB6 | None | None | None | None |
| ITGB6 | None | None | None | None |
| ITGB6 | None | None | None | None |
| DARS | None | None | None | None |
| DARS | None | None | None | None |
| DARS | None | None | None | None |
| DARS | None | None | None | None |
| BTK | None | None | None | None |
| BTK | None | None | None | None |
| BTK | None | None | None | None |
| HDAC4 | None | None | None | None |
| ITGA2B | None | None | None | None |

| | | | | |
|---|---|---|---|---|
| ITGA2B | None | None | None | None |
| ITGA2B | None | None | None | None |
| ITGA2B | None | None | None | None |
| PCSK9 | None | None | None | None |
| PCSK9 | None | None | None | None |
| PCSK9 | None | None | None | None |
| PCSK9 | None | None | None | None |
| MUTYH | None | None | None | None |
| MUTYH | None | None | None | None |
| MUTYH | None | None | None | None |
| MUTYH | None | None | None | None |
| MUTYH | None | None | None | None |
| MUTYH | None | None | None | None |
| MUTYH | None | None | None | None |
| MUTYH | None | None | None | None |
| MUTYH | None | None | None | None |
| MUTYH | None | None | None | None |
| MUTYH | None | None | None | None |
| MUTYH | None | None | None | None |
| MUTYH | None | None | None | None |
| MUTYH | None | None | None | None |
| MUTYH | None | None | None | None |
| MUTYH | None | None | None | None |
| MUTYH | None | None | None | None |
| MUTYH | None | None | None | None |
| MUTYH | None | None | None | None |
| MUTYH | None | None | None | None |
| MUTYH | None | None | None | None |
| MUTYH | None | None | None | None |
| MUTYH | None | None | None | None |
| MUTYH | None | None | None | None |
| SLC25A12 | None | None | None | None |
| GSN | None | None | None | None |
| IHH | 0.909 | 1 | 3n1p | B |
| IHH | None | None | None | None |
| IHH | None | None | None | None |
| IHH | None | None | None | None |
| IHH | None | None | None | None |
| IHH | None | None | None | None |
| IHH | None | None | None | None |
| CD247 | None | None | None | None |
| EP300 | None | None | None | None |
| TTN | None | None | None | None |
| MRI1 | None | None | None | None |
| SGSH | None | None | None | None |
| SGSH | None | None | None | None |
| SGSH | None | None | None | None |
| SGSH | None | None | None | None |
| SGSH | None | None | None | None |
| SGSH | None | None | None | None |
| SGSH | None | None | None | None |
| SGSH | None | None | None | None |
| SGSH | None | None | None | None |
| SGSH | None | None | None | None |

| | | | | |
|---|---|---|---|---|
| SGSH | None | None | None | None |
| SGSH | None | None | None | None |
| SGSH | None | None | None | None |
| SGSH | None | None | None | None |
| SGSH | None | None | None | None |
| SGSH | None | None | None | None |
| SGSH | None | None | None | None |
| SGSH | None | None | None | None |
| SGSH | None | None | None | None |
| SGSH | None | None | None | None |
| SGSH | None | None | None | None |
| SGSH | None | None | None | None |
| CBL | None | None | None | None |
| CANT1 | None | None | None | None |
| CANT1 | None | None | None | None |
| CANT1 | None | None | None | None |
| CANT1 | None | None | None | None |
| CANT1 | None | None | None | None |
| CANT1 | None | None | None | None |
| CANT1 | None | None | None | None |
| SLC1A3 | None | None | None | None |
| SLC1A3 | None | None | None | None |
| SLC1A5 | None | None | None | None |
| SLC1A3 | None | None | None | None |
| SLC1A5 | None | None | None | None |
| SLC1A3 | None | None | None | None |
| SLC1A3 | None | None | None | None |
| CTSD | None | None | None | None |
| CTSD | None | None | None | None |
| CTSD | None | None | None | None |
| CTSD | None | None | None | None |
| CTSD | None | None | None | None |
| MTOR | None | None | None | None |
| ILK | None | None | None | None |
| PCDH15 | None | None | None | None |
| PCDH15 | None | None | None | None |
| PCDH15 | None | None | None | None |
| PCDH15 | None | None | None | None |
| PCDH15 | None | None | None | None |
| PCDH15 | None | None | None | None |
| KLHL40 | None | None | None | None |
| KLHL40 | None | None | None | None |
| KLHL40 | None | None | None | None |
| KLHL40 | None | None | None | None |
| CSTA | None | None | None | None |
| CSTA | None | None | None | None |
| HNMT | None | None | None | None |
| HNMT | None | None | None | None |
| IFNGR2 | None | None | None | None |
| IFNGR2 | None | None | None | None |
| PEX5 | None | None | None | None |
| KIF7 | None | None | None | None |
| KIF7 | None | None | None | None |
| MYL3 | None | None | None | None |

| | | | | |
|---|---|---|---|---|
| MYL3 | None | None | None | None |
| MYL3 | None | None | None | None |
| MYL3 | None | None | None | None |
| MYL3 | None | None | None | None |
| MYL3 | None | None | None | None |
| MYL3 | None | None | None | None |
| NAGLU | None | None | None | None |
| NAGLU | None | None | None | None |
| NAGLU | None | None | None | None |
| NAGLU | None | None | None | None |
| NAGLU | None | None | None | None |
| NAGLU | None | None | None | None |
| NAGLU | None | None | None | None |
| NAGLU | None | None | None | None |
| NAGLU | None | None | None | None |
| NAGLU | None | None | None | None |
| NAGLU | None | None | None | None |
| NAGLU | None | None | None | None |
| NAGLU | None | None | None | None |
| TNFRSF13C | None | None | None | None |
| KIF21A | None | None | None | None |
| KIF21A | None | None | None | None |
| KIF21A | None | None | None | None |
| KIF21A | None | None | None | None |
| KIF21A | None | None | None | None |
| DOCK2 | None | None | None | None |
| DSP | None | None | None | None |
| CDKN2A | None | None | None | None |
| CDKN2A | None | None | None | None |
| CDKN2A | None | None | None | None |
| CDKN2A | None | None | None | None |
| CDKN2A | None | None | None | None |
| CDKN2A | None | None | None | None |
| CDKN2A | None | None | None | None |
| CDKN2A | None | None | None | None |
| CDKN2A | None | None | None | None |
| CDKN2A | None | None | None | None |
| CDKN2A | None | None | None | None |
| CDKN2A | None | None | None | None |
| CDKN2A | None | None | None | None |
| CDKN2A | None | None | None | None |
| VCL | None | None | None | None |
| TPP1 | None | None | None | None |
| TPP1 | None | None | None | None |
| TPP1 | None | None | None | None |
| TPP1 | None | None | None | None |
| TPP1 | None | None | None | None |
| TPP1 | None | None | None | None |
| TPP1 | None | None | None | None |
| TPP1 | None | None | None | None |
| TPP1 | None | None | None | None |
| TPP1 | None | None | None | None |
| TPP1 | None | None | None | None |
| TPP1 | None | None | None | None |

| | | | | |
|---|---|---|---|---|
| TPP1 | None | None | None | None |
| TPP1 | None | None | None | None |
| TPP1 | None | None | None | None |
| TPP1 | None | None | None | None |
| TPP1 | None | None | None | None |
| TPP1 | None | None | None | None |
| TPP1 | None | None | None | None |
| TPP1 | None | None | None | None |
| TPP1 | None | None | None | None |
| TPP1 | None | None | None | None |
| IL10RB | None | None | None | None |
| IL10RB | None | None | None | None |
| CCM2 | None | None | None | None |
| CCM2 | None | None | None | None |
| CCM2 | None | None | None | None |
| DCX | None | None | None | None |
| DCX | None | None | None | None |
| DCX | None | None | None | None |
| DCX | None | None | None | None |
| DCX | None | None | None | None |
| DCX | None | None | None | None |
| DCX | None | None | None | None |
| DCX | None | None | None | None |
| BRCA2 | None | None | None | None |
| NNT | None | None | None | None |
| C2 | None | None | None | None |
| C2 | None | None | None | None |
| C2 | None | None | None | None |
| C2 | None | None | None | None |
| WRN | None | None | None | None |
| WRN | None | None | None | None |
| F13A1 | None | None | None | None |
| F13A1 | None | None | None | None |
| F13A1 | None | None | None | None |
| F13A1 | None | None | None | None |
| TF | None | None | None | None |
| TF | None | None | None | None |
| TF | None | None | None | None |
| TF | None | None | None | None |
| TF | None | None | None | None |
| ITPA | None | None | None | None |
| ITPA | None | None | None | None |
| SNCA | None | None | None | None |
| ITGB4 | None | None | None | None |
| ITGB4 | None | None | None | None |
| ITGB4 | None | None | None | None |
| LRP6 | None | None | None | None |
| LRP6 | None | None | None | None |
| LRP6 | None | None | None | None |
| PLG | None | None | None | None |
| PLG | None | None | None | None |
| TYROBP | None | None | None | None |
| UQCRC2 | None | None | None | None |
| GPSM2 | None | None | None | None |

| | | | | |
|---|---|---|---|---|
| GPSM2 | None | None | None | None |
| GPSM2 | None | None | None | None |
| SCN5A | None | None | None | None |
| SCN5A | None | None | None | None |
| SCN5A | None | None | None | None |
| SCN5A | None | None | None | None |
| SCN5A | None | None | None | None |
| ATIC | None | None | None | None |
| EZH2 | None | None | None | None |
| CPA1 | None | None | None | None |
| SCN4B | None | None | None | None |
| SCN4B | None | None | None | None |
| MAN1B1 | None | None | None | None |
| MAN1B1 | None | None | None | None |
| MAN1B1 | None | None | None | None |
| MAN1B1 | None | None | None | None |
| CYCS | None | None | None | None |
| IDUA | None | None | None | None |
| IDUA | None | None | None | None |
| IDUA | None | None | None | None |
| IDUA | None | None | None | None |
| IDUA | None | None | None | None |
| IDUA | None | None | None | None |
| IDUA | None | None | None | None |
| IDUA | None | None | None | None |
| IDUA | None | None | None | None |
| IDUA | None | None | None | None |
| IDUA | None | None | None | None |
| IDUA | None | None | None | None |
| IDUA | None | None | None | None |
| IDUA | None | None | None | None |
| IDUA | None | None | None | None |
| IDUA | None | None | None | None |
| IDUA | None | None | None | None |
| IDUA | None | None | None | None |
| IDUA | None | None | None | None |
| IDUA | None | None | None | None |
| IDUA | None | None | None | None |
| IDUA | None | None | None | None |
| IDUA | None | None | None | None |
| IDUA | None | None | None | None |
| IDUA | None | None | None | None |
| HIKESHI | None | None | None | None |
| APC | None | None | None | None |
| APC | None | None | None | None |
| APC | None | None | None | None |
| APP | None | None | None | None |
| CEP152 | None | None | None | None |
| GAA | None | None | None | None |
| GAA | None | None | None | None |
| GAA | None | None | None | None |
| GAA | None | None | None | None |
| GAA | None | None | None | None |
| GAA | None | None | None | None |

| | | | | |
|---|---|---|---|---|
| GAA | None | None | None | None |
| GAA | None | None | None | None |
| GAA | None | None | None | None |
| GAA | None | None | None | None |
| GAA | None | None | None | None |
| GAA | None | None | None | None |
| GAA | None | None | None | None |
| GAA | None | None | None | None |
| GAA | None | None | None | None |
| GAA | None | None | None | None |
| GAA | None | None | None | None |
| GAA | None | None | None | None |
| GAA | None | None | None | None |
| GAA | None | None | None | None |
| GAA | None | None | None | None |
| GAA | None | None | None | None |
| GAA | None | None | None | None |
| GAA | None | None | None | None |
| GAA | None | None | None | None |
| GAA | None | None | None | None |
| GAA | None | None | None | None |
| GAA | None | None | None | None |
| GAA | None | None | None | None |
| GAA | None | None | None | None |
| GAA | None | None | None | None |
| GAA | None | None | None | None |
| GAA | None | None | None | None |
| GAA | None | None | None | None |
| GAA | None | None | None | None |
| GAA | None | None | None | None |
| GAA | None | None | None | None |
| GAA | None | None | None | None |
| GAA | None | None | None | None |
| GAA | None | None | None | None |
| GAA | None | None | None | None |
| GAA | None | None | None | None |
| GAA | None | None | None | None |
| GAA | None | None | None | None |
| GAA | None | None | None | None |
| GAA | None | None | None | None |
| GAA | None | None | None | None |
| GAA | None | None | None | None |
| GAA | None | None | None | None |
| GAA | None | None | None | None |
| FOLR1 | None | None | None | None |
| DSP | None | None | None | None |
| DSP | None | None | None | None |
| DSP | None | None | None | None |
| DSP | None | None | None | None |
| DSP | None | None | None | None |
| CCM2 | None | None | None | None |
| FXN | None | None | None | None |
| FXN | None | None | None | None |
| COMP | None | None | None | None |
| COMP | None | None | None | None |

| | | | | |
|---|---|---|---|---|
| COMP | None | None | None | None |
| COMP | None | None | None | None |
| COMP | None | None | None | None |
| COMP | None | None | None | None |
| COMP | None | None | None | None |
| COMP | None | None | None | None |
| COMP | None | None | None | None |
| COMP | None | None | None | None |
| COMP | None | None | None | None |
| COMP | None | None | None | None |
| AIFM1 | None | None | None | None |
| AIFM1 | None | None | None | None |
| AIFM1 | None | None | None | None |
| AIFM1 | None | None | None | None |
| AIFM1 | None | None | None | None |
| AIFM1 | None | None | None | None |
| AIFM1 | None | None | None | None |
| MYH9 | None | None | None | None |
| FHL2 | None | None | None | None |
| FHL2 | None | None | None | None |
| TNFRSF4 | None | None | None | None |
| CFB | None | None | None | None |
| CFB | None | None | None | None |
| CFB | None | None | None | None |
| CFB | None | None | None | None |
| CFB | None | None | None | None |
| CFB | None | None | None | None |
| CFB | None | None | None | None |
| CFB | None | None | None | None |
| CFB | None | None | None | None |
| CFB | None | None | None | None |
| MVD | None | None | None | None |
| MVD | None | None | None | None |
| HSD17B4 | None | None | None | None |
| HSD17B4 | None | None | None | None |
| HSD17B4 | None | None | None | None |
| HSD17B4 | None | None | None | None |
| HSD17B4 | None | None | None | None |
| HSD17B4 | None | None | None | None |
| F11 | None | None | None | None |
| F11 | None | None | None | None |
| F11 | None | None | None | None |
| F11 | None | None | None | None |
| F11 | None | None | None | None |
| F11 | None | None | None | None |
| F11 | None | None | None | None |
| F11 | None | None | None | None |
| F11 | None | None | None | None |
| SPTA1 | None | None | None | None |
| SPTA1 | None | None | None | None |
| SPTA1 | None | None | None | None |
| SPTA1 | None | None | None | None |
| SPTA1 | None | None | None | None |
| SPTA1 | None | None | None | None |

| | | | | |
|---|---|---|---|---|
| GMPR | None | None | None | None |
| MITF | None | None | None | None |
| NPC1 | None | None | None | None |
| NPC1 | None | None | None | None |
| NPC1 | None | None | None | None |
| NPC1 | None | None | None | None |
| NPC1 | None | None | None | None |
| NPC1 | None | None | None | None |
| NPC1 | None | None | None | None |
| NPC1 | None | None | None | None |
| NPC1 | None | None | None | None |
| NPC1 | None | None | None | None |
| NPC1 | None | None | None | None |
| NPC1 | None | None | None | None |
| NPC1 | None | None | None | None |
| NPC1 | None | None | None | None |
| NPC1 | None | None | None | None |
| NPC1 | None | None | None | None |
| NPC1 | None | None | None | None |
| NPC1 | None | None | None | None |
| NPC1 | None | None | None | None |
| NPC1 | None | None | None | None |
| NPC1 | None | None | None | None |
| NPC1 | None | None | None | None |
| NPC1 | None | None | None | None |
| NPC1 | None | None | None | None |
| NPC1 | None | None | None | None |
| NPC1 | None | None | None | None |
| NPC1 | None | None | None | None |
| NPC1 | None | None | None | None |
| NPC1 | None | None | None | None |
| NPC1 | None | None | None | None |
| NPC1 | None | None | None | None |
| NPC1 | None | None | None | None |
| NPC1 | None | None | None | None |
| NPC1 | None | None | None | None |
| NPC1 | None | None | None | None |
| NPC1 | None | None | None | None |
| NPC1 | None | None | None | None |
| NPC1 | None | None | None | None |
| NPC1 | None | None | None | None |
| NPC1 | None | None | None | None |
| NPC1 | None | None | None | None |
| NPC1 | None | None | None | None |
| NPC1 | None | None | None | None |
| C3 | None | None | None | None |
| NLRX1 | None | None | None | None |
| IGLL1 | None | None | None | None |
| APRT | None | None | None | None |
| APRT | None | None | None | None |
| APRT | None | None | None | None |
| APRT | 0.923 | 1 | 1zn8 | A |
| APRT | None | None | None | None |
| ZIC3 | None | None | None | None |

| | | | | |
|---|---|---|---|---|
| ZIC3 | None | None | None | None |
| KLHL3 | None | None | None | None |
| RPS6KA3 | None | None | None | None |
| RPS6KA3 | None | None | None | None |
| RPS6KA3 | None | None | None | None |
| F8 | None | None | None | None |
| F8 | None | None | None | None |
| F8 | None | None | None | None |
| F8 | None | None | None | None |
| F8 | None | None | None | None |
| F8 | None | None | None | None |
| F8 | None | None | None | None |
| F8 | None | None | None | None |
| F8 | None | None | None | None |
| F8 | None | None | None | None |
| F8 | None | None | None | None |
| F8 | None | None | None | None |
| F8 | None | None | None | None |
| F8 | None | None | None | None |
| F8 | None | None | None | None |
| F8 | None | None | None | None |
| F8 | None | None | None | None |
| ISG15 | None | None | None | None |
| EXOSC3 | None | None | None | None |
| EXOSC3 | None | None | None | None |
| EXOSC3 | None | None | None | None |
| NKX2-5 | None | None | None | None |
| NKX2-5 | None | None | None | None |
| NKX2-5 | None | None | None | None |
| NKX2-5 | None | None | None | None |
| FH | None | None | None | None |
| FH | None | None | None | None |
| FH | None | None | None | None |
| FH | None | None | None | None |
| FH | None | None | None | None |
| FH | None | None | None | None |
| FH | None | None | None | None |
| FH | None | None | None | None |
| FH | None | None | None | None |
| FH | None | None | None | None |
| FH | None | None | None | None |
| FH | None | None | None | None |
| SNRNP200 | None | None | None | None |
| SNRNP200 | None | None | None | None |
| SNRNP200 | None | None | None | None |
| SNRNP200 | None | None | None | None |
| CHRNA1 | None | None | None | None |
| CHRNA1 | None | None | None | None |
| MYOM1 | None | None | None | None |
| MYOM1 | None | None | None | None |
| MYOM1 | None | None | None | None |
| MYOM1 | None | None | None | None |
| MYOM1 | None | None | None | None |
| MYOM1 | None | None | None | None |

| | | | | |
|---|---|---|---|---|
| LDLR | None | None | None | None |
| LDLR | None | None | None | None |
| LDLR | None | None | None | None |
| LDLR | None | None | None | None |
| LDLR | None | None | None | None |
| LDLR | None | None | None | None |
| EHMT1 | None | None | None | None |
| EHMT1 | None | None | None | None |
| EHMT1 | None | None | None | None |
| COL9A2 | None | None | None | None |
| HFE2 | None | None | None | None |
| HFE2 | None | None | None | None |
| CD3G | None | None | None | None |
| MMAB | None | None | None | None |
| MMAB | None | None | None | None |
| MMAB | None | None | None | None |
| MMAB | None | None | None | None |
| MMAB | None | None | None | None |
| MMAB | None | None | None | None |
| MMAB | None | None | None | None |
| MMAB | None | None | None | None |
| MMAB | None | None | None | None |
| MMAB | None | None | None | None |
| TAF1 | None | None | None | None |
| TAF1 | None | None | None | None |
| CD4 | None | None | None | None |
| QARS | None | None | None | None |
| QARS | None | None | None | None |
| QARS | None | None | None | None |
| QARS | None | None | None | None |
| QARS | None | None | None | None |
| FGFR3 | None | None | None | None |
| FGFR3 | None | None | None | None |
| FGFR3 | None | None | None | None |
| FGFR3 | None | None | None | None |
| FGFR3 | None | None | None | None |
| FGFR3 | None | None | None | None |
| FLNB | None | None | None | None |
| FLNB | None | None | None | None |
| FLNB | None | None | None | None |
| FLNB | None | None | None | None |
| FLNB | None | None | None | None |
| FLNB | None | None | None | None |
| FLNB | None | None | None | None |
| FLNB | None | None | None | None |
| IL21R | None | None | None | None |
| RB1 | None | None | None | None |
| RB1 | None | None | None | None |
| RB1 | None | None | None | None |
| NTRK1 | None | None | None | None |
| NOTCH1 | None | None | None | None |
| MYOM1 | None | None | None | None |
| MYOM1 | None | None | None | None |
| MYOM1 | None | None | None | None |

| | | | | |
|---|---|---|---|---|
| MYOM1 | None | None | None | None |
| MYOM1 | None | None | None | None |
| MYOM1 | None | None | None | None |
| COQ9 | None | None | None | None |
| COQ9 | None | None | None | None |
| CRBN | None | None | None | None |
| DCTN1 | None | None | None | None |
| DCTN1 | None | None | None | None |
| DCTN1 | None | None | None | None |
| DCTN1 | None | None | None | None |
| DCTN1 | None | None | None | None |
| DCTN1 | None | None | None | None |
| THAP1 | None | None | None | None |
| THAP1 | None | None | None | None |
| GNAS | None | None | None | None |
| GNAS | None | None | None | None |
| HNF4A | None | None | None | None |
| HNF4A | None | None | None | None |
| ALDH3A2 | None | None | None | None |
| ALDH3A2 | None | None | None | None |
| ALDH3A2 | None | None | None | None |
| ALDH3A2 | None | None | None | None |
| ALDH3A2 | None | None | None | None |
| ALDH3A2 | None | None | None | None |
| ALDH3A2 | None | None | None | None |
| ALDH3A2 | None | None | None | None |
| ALDH3A2 | None | None | None | None |
| HCN1 | None | None | None | None |
| PLEC | None | None | None | None |
| SCN5A | None | None | None | None |
| SCN5A | None | None | None | None |
| STAMBP | None | None | None | None |
| STAMBP | None | None | None | None |
| RNF125 | None | None | None | None |
| RNF125 | None | None | None | None |
| USH1C | None | None | None | None |
| OBSL1 | None | None | None | None |
| GRIN2B | None | None | None | None |
| GRIN2B | None | None | None | None |
| HCN4 | None | None | None | None |
| HCN4 | None | None | None | None |
| TALDO1 | None | None | None | None |
| ITGB3 | None | None | None | None |
| CHKB | None | None | None | None |
| CHKB | None | None | None | None |
| CHKB | None | None | None | None |
| CHKB | None | None | None | None |
| ACTN4 | None | None | None | None |
| ACTN4 | None | None | None | None |
| ACTN4 | None | None | None | None |
| ACTN4 | None | None | None | None |
| CDT1 | None | None | None | None |
| CDT1 | None | None | None | None |
| CDT1 | None | None | None | None |

| | | | | |
|---|---|---|---|---|
| CDT1 | None | None | None | None |
| CDT1 | None | None | None | None |
| CDT1 | None | None | None | None |
| MAPT | None | None | None | None |
| F5 | None | None | None | None |
| F5 | None | None | None | None |
| F5 | None | None | None | None |
| ESRRB | None | None | None | None |
| DGUOK | None | None | None | None |
| DGUOK | None | None | None | None |
| DGUOK | None | None | None | None |
| DGUOK | None | None | None | None |
| DGUOK | None | None | None | None |
| DGUOK | None | None | None | None |
| DGUOK | None | None | None | None |
| DGUOK | None | None | None | None |
| DGUOK | None | None | None | None |
| CENPJ | None | None | None | None |
| TTN | None | None | None | None |
| TTN | None | None | None | None |
| TTN | None | None | None | None |
| TTN | None | None | None | None |
| TNFRSF1A | None | None | None | None |
| TNFRSF1A | None | None | None | None |
| TNFRSF1A | None | None | None | None |
| TNFRSF1A | None | None | None | None |
| TNFRSF1A | None | None | None | None |
| TNFRSF1A | None | None | None | None |
| TNFRSF1A | None | None | None | None |
| TNFRSF1A | None | None | None | None |
| TNFRSF1A | None | None | None | None |
| TNFRSF1A | None | None | None | None |
| TNFRSF1A | None | None | None | None |
| TNFRSF1A | None | None | None | None |
| ASL | None | None | None | None |
| ASL | None | None | None | None |
| ASL | None | None | None | None |
| ASL | None | None | None | None |
| ASL | None | None | None | None |
| ASL | None | None | None | None |
| ASL | None | None | None | None |
| ASL | None | None | None | None |
| ASL | None | None | None | None |
| ASL | None | None | None | None |
| MYO7A | None | None | None | None |
| MYO7A | None | None | None | None |
| MYO7A | None | None | None | None |
| MYO7A | None | None | None | None |
| MYO7A | None | None | None | None |
| MYO7A | None | None | None | None |
| MYO7A | None | None | None | None |
| MYO7A | None | None | None | None |
| MYO7A | None | None | None | None |
| MYO7A | None | None | None | None |

| | | | | |
|---|---|---|---|---|
| MYO7A | None | None | None | None |
| MYO7A | None | None | None | None |
| MYO7A | None | None | None | None |
| MYO7A | None | None | None | None |
| MYO7A | None | None | None | None |
| MYO7A | None | None | None | None |
| MYO7A | None | None | None | None |
| MYO7A | None | None | None | None |
| MYO7A | None | None | None | None |
| MYO7A | None | None | None | None |
| APC | None | None | None | None |
| ALAD | None | None | None | None |
| ALAD | None | None | None | None |
| ALAD | None | None | None | None |
| ALAD | None | None | None | None |
| ALAD | None | None | None | None |
| ALAD | None | None | None | None |
| POR | None | None | None | None |
| CSF1R | None | None | None | None |
| CSF1R | None | None | None | None |
| CSF1R | None | None | None | None |
| CSF1R | None | None | None | None |
| CSF1R | None | None | None | None |
| CSF1R | None | None | None | None |
| CSF1R | None | None | None | None |
| CSF1R | None | None | None | None |
| CSF1R | None | None | None | None |
| CSF1R | None | None | None | None |
| CSF1R | None | None | None | None |
| VWF | None | None | None | None |
| VWF | None | None | None | None |
| PHF8 | None | None | None | None |
| ACY1 | None | None | None | None |
| SMN1 | None | None | None | None |
| SMN1 | None | None | None | None |
| DLAT | None | None | None | None |
| DLAT | None | None | None | None |
| B2M | None | None | None | None |
| SOX9 | None | None | None | None |
| SOX9 | None | None | None | None |
| SOX9 | None | None | None | None |
| SOX9 | None | None | None | None |
| SOX9 | None | None | None | None |
| CFH | None | None | None | None |
| FLNC | None | None | None | None |
| FLNC | None | None | None | None |
| FLNC | None | None | None | None |
| ITGB2 | None | None | None | None |
| ITGB2 | None | None | None | None |
| ITGB2 | None | None | None | None |
| ITGB2 | None | None | None | None |
| ITGB2 | None | None | None | None |
| ITGB2 | None | None | None | None |
| ITGB2 | None | None | None | None |

| | | | | |
|---|---|---|---|---|
| ITGB2 | None | None | None | None |
| ITGB2 | None | None | None | None |
| PSEN1 | None | None | None | None |
| PSEN1 | None | None | None | None |
| PSEN1 | None | None | None | None |
| PSEN1 | None | None | None | None |
| PSEN1 | None | None | None | None |
| PSEN1 | None | None | None | None |
| OXCT1 | None | None | None | None |
| OXCT1 | None | None | None | None |
| OXCT1 | None | None | None | None |
| OXCT1 | None | None | None | None |
| OXCT1 | None | None | None | None |
| OXCT1 | None | None | None | None |
| OXCT1 | None | None | None | None |
| DHH | None | None | None | None |
| PRKN | None | None | None | None |
| PRKN | None | None | None | None |
| PRKN | None | None | None | None |
| PRKN | None | None | None | None |
| PRKN | None | None | None | None |
| PRKN | None | None | None | None |
| PRKN | None | None | None | None |
| PRKN | None | None | None | None |
| PRKN | None | None | None | None |
| PRKN | None | None | None | None |
| PRKN | None | None | None | None |
| LCAT | None | None | None | None |
| LCAT | None | None | None | None |
| LCAT | None | None | None | None |
| LCAT | None | None | None | None |
| LCAT | None | None | None | None |
| LCAT | None | None | None | None |
| LCAT | None | None | None | None |
| LCAT | None | None | None | None |
| LCAT | None | None | None | None |
| LCAT | None | None | None | None |
| LCAT | None | None | None | None |
| LCAT | None | None | None | None |
| FOXP3 | None | None | None | None |
| FOXP3 | None | None | None | None |
| FOXP3 | None | None | None | None |
| FOXP3 | None | None | None | None |
| SUFU | None | None | None | None |
| HSPA9 | None | None | None | None |
| HSPA9 | None | None | None | None |
| SQSTM1 | None | None | None | None |
| DARS2 | None | None | None | None |
| DARS2 | None | None | None | None |
| DARS2 | None | None | None | None |
| DARS2 | None | None | None | None |
| DARS2 | None | None | None | None |
| DARS2 | None | None | None | None |
| DARS2 | None | None | None | None |

| | | | | |
|---|---|---|---|---|
| DARS2 | None | None | None | None |
| DARS2 | None | None | None | None |
| DARS2 | None | None | None | None |
| DARS2 | None | None | None | None |
| DARS2 | None | None | None | None |
| NT5C3A | None | None | None | None |
| NT5C3A | None | None | None | None |
| NT5C3A | None | None | None | None |
| NT5C3A | None | None | None | None |
| ETFA | None | None | None | None |
| ETFA | None | None | None | None |
| ETFA | None | None | None | None |
| ETFA | None | None | None | None |
| ETFA | None | None | None | None |
| ETFA | None | None | None | None |
| ETFA | None | None | None | None |
| TTN | None | None | None | None |
| TTN | None | None | None | None |
| FLT4 | None | None | None | None |
| FGB | None | None | None | None |
| HPCA | None | None | None | None |
| TAP2 | None | None | None | None |
| TAP2 | None | None | None | None |
| PTPRC | None | None | None | None |
| SLC4A1 | None | None | None | None |
| SLC4A1 | None | None | None | None |
| SLC4A1 | None | None | None | None |
| SLC4A1 | None | None | None | None |
| SLC4A1 | None | None | None | None |
| SLC4A1 | None | None | None | None |
| MYH7 | None | None | None | None |
| MYH7 | None | None | None | None |
| MYH7 | None | None | None | None |
| MYH7 | None | None | None | None |
| MYH7 | None | None | None | None |
| MYH7 | None | None | None | None |
| MYH7 | None | None | None | None |
| MYH7 | None | None | None | None |
| MYH7 | None | None | None | None |
| PARN | None | None | None | None |
| PARN | None | None | None | None |
| NDUFS6 | None | None | None | None |
| VRK1 | None | None | None | None |
| VRK1 | None | None | None | None |
| VRK1 | None | None | None | None |
| VRK1 | None | None | None | None |
| VRK1 | None | None | None | None |
| VRK1 | None | None | None | None |
| COLEC11 | None | None | None | None |
| VHL | None | None | None | None |
| TTN | None | None | None | None |
| NCF2 | None | None | None | None |
| NCF2 | None | None | None | None |
| NCF2 | None | None | None | None |

| | | | | |
|---|---|---|---|---|
| NCF2 | None | None | None | None |
| NCF2 | None | None | None | None |
| RPS26 | None | None | None | None |
| CETP | None | None | None | None |
| CETP | None | None | None | None |
| CETP | None | None | None | None |
| CETP | None | None | None | None |
| CETP | None | None | None | None |
| CETP | None | None | None | None |
| CETP | None | None | None | None |
| CETP | None | None | None | None |
| CETP | None | None | None | None |
| CETP | None | None | None | None |
| CETP | None | None | None | None |
| CETP | None | None | None | None |
| CETP | None | None | None | None |
| FLNA | None | None | None | None |
| FLNA | None | None | None | None |
| FLNA | None | None | None | None |
| FLNA | None | None | None | None |
| FLNA | None | None | None | None |
| FLNA | None | None | None | None |
| FLNA | None | None | None | None |
| MYBPC3 | None | None | None | None |
| MYBPC3 | None | None | None | None |
| MYBPC3 | None | None | None | None |
| MYBPC3 | None | None | None | None |
| MYBPC3 | None | None | None | None |
| MYBPC3 | None | None | None | None |
| MYBPC3 | None | None | None | None |
| MYBPC3 | None | None | None | None |
| MYBPC3 | None | None | None | None |
| MYBPC3 | None | None | None | None |
| LSS | None | None | None | None |
| C6 | None | None | None | None |
| C6 | None | None | None | None |
| RLBP1 | None | None | None | None |
| RLBP1 | None | None | None | None |
| RLBP1 | None | None | None | None |
| RLBP1 | None | None | None | None |
| RLBP1 | None | None | None | None |
| RLBP1 | None | None | None | None |
| RLBP1 | None | None | None | None |
| RLBP1 | None | None | None | None |
| MMAA | None | None | None | None |
| MMAA | None | None | None | None |
| MMAA | None | None | None | None |
| MMAA | None | None | None | None |
| MMAA | None | None | None | None |
| MMAA | None | None | None | None |
| IL1RN | None | None | None | None |
| MASTL | None | None | None | None |
| FHL1 | None | None | None | None |
| FHL1 | None | None | None | None |

| | | | | |
|---|---|---|---|---|
| FHL1 | None | None | None | None |
| FHL1 | None | None | None | None |
| FHL1 | None | None | None | None |
| FHL1 | None | None | None | None |
| FHL1 | None | None | None | None |
| FHL1 | None | None | None | None |
| FHL1 | None | None | None | None |
| EIF2B1 | None | None | None | None |
| EIF2B1 | None | None | None | None |
| COQ8A | None | None | None | None |
| COQ8A | None | None | None | None |
| COQ8A | None | None | None | None |
| COQ8A | None | None | None | None |
| COQ8A | None | None | None | None |
| COQ8A | None | None | None | None |
| COQ8A | None | None | None | None |
| COQ8A | None | None | None | None |
| COQ8A | None | None | None | None |
| COQ8A | None | None | None | None |
| COQ8A | None | None | None | None |
| COQ8A | None | None | None | None |
| COQ8A | None | None | None | None |
| COQ8A | None | None | None | None |
| COQ8A | None | None | None | None |
| COQ8A | None | None | None | None |
| UNC119 | None | None | None | None |
| IGHMBP2 | None | None | None | None |
| JAG1 | None | None | None | None |
| JAG1 | None | None | None | None |
| JAG1 | None | None | None | None |
| GNB1 | None | None | None | None |
| GNB1 | None | None | None | None |
| FLNA | None | None | None | None |
| FLNA | None | None | None | None |
| FLNA | None | None | None | None |
| FLNA | None | None | None | None |
| FLNA | None | None | None | None |
| CREBBP | None | None | None | None |
| CSF3R | None | None | None | None |
| CSF3R | None | None | None | None |
| HIBCH | None | None | None | None |
| HIBCH | None | None | None | None |
| HIBCH | None | None | None | None |
| MMACHC | None | None | None | None |
| MMACHC | None | None | None | None |
| MMACHC | None | None | None | None |
| MMACHC | None | None | None | None |
| MMACHC | None | None | None | None |
| MMACHC | None | None | None | None |
| MMACHC | None | None | None | None |
| MMACHC | None | None | None | None |
| MMACHC | None | None | None | None |
| MMACHC | None | None | None | None |
| MMACHC | None | None | None | None |

| | | | | |
|---|---|---|---|---|
| KLF1 | None | None | None | None |
| NAGS | None | None | None | None |
| NAGS | None | None | None | None |
| NAGS | None | None | None | None |
| PIK3R2 | None | None | None | None |
| EDNRB | None | None | None | None |
| EDNRB | None | None | None | None |
| EDNRB | None | None | None | None |
| EDNRB | None | None | None | None |
| E | None | None | None | None |
| RUNX1 | None | None | None | None |
| RUNX1 | None | None | None | None |
| CTNNB1 | None | None | None | None |
| CTNNB1 | None | None | None | None |
| CTNNB1 | None | None | None | None |
| CTNNB1 | None | None | None | None |
| CTNNB1 | None | None | None | None |
| CTNNB1 | None | None | None | None |
| TTN | None | None | None | None |
| TTN | None | None | None | None |
| TYK2 | None | None | None | None |
| TYK2 | None | None | None | None |
| TYK2 | None | None | None | None |
| ERCC4 | None | None | None | None |
| ERCC4 | None | None | None | None |
| ERCC4 | None | None | None | None |
| ERCC4 | None | None | None | None |
| CFI | None | None | None | None |
| CFI | None | None | None | None |
| CFI | None | None | None | None |
| CFI | None | None | None | None |
| CFI | None | None | None | None |
| CFI | None | None | None | None |
| CFI | None | None | None | None |
| CFI | None | None | None | None |
| CFI | None | None | None | None |
| CFI | None | None | None | None |
| CFI | None | None | None | None |
| CFI | None | None | None | None |
| CFI | None | None | None | None |
| CFI | None | None | None | None |
| NOTCH3 | None | None | None | None |
| NOTCH3 | None | None | None | None |
| NOTCH3 | None | None | None | None |
| NOTCH3 | None | None | None | None |
| TBX1 | None | None | None | None |
| TBX1 | None | None | None | None |
| DDB2 | None | None | None | None |
| DDB2 | None | None | None | None |
| TP53 | None | None | None | None |
| POMGNT1 | None | None | None | None |
| POMGNT1 | None | None | None | None |
| POMGNT1 | None | None | None | None |
| POMGNT1 | None | None | None | None |

| | | | | |
|---|---|---|---|---|
| POMGNT1 | None | None | None | None |
| POMGNT1 | None | None | None | None |
| POMGNT1 | None | None | None | None |
| POMGNT1 | None | None | None | None |
| POMGNT1 | None | None | None | None |
| POMGNT1 | None | None | None | None |
| POMGNT1 | None | None | None | None |
| POMGNT1 | None | None | None | None |
| POMGNT1 | None | None | None | None |
| FMR1 | None | None | None | None |
| FMR1 | None | None | None | None |
| SLC9A3R1 | None | None | None | None |
| HSPB1 | None | None | None | None |
| HSPB1 | None | None | None | None |
| HSPB1 | None | None | None | None |
| HSPB1 | None | None | None | None |
| HSPB1 | None | None | None | None |
| HSPB1 | None | None | None | None |
| HSPB1 | None | None | None | None |
| SOD3 | None | None | None | None |
| AGA | 1 | 2 | 1apz | D |
| AGA | None | None | None | None |
| AGA | None | None | None | None |
| NDUFS7 | None | None | None | None |
| NDUFS7 | None | None | None | None |
| SOX2 | None | None | None | None |
| CSRP3 | None | None | None | None |
| CSRP3 | None | None | None | None |
| PAH | None | None | None | None |
| PAH | None | None | None | None |
| PAH | None | None | None | None |
| PAH | None | None | None | None |
| CACNA1S | None | None | None | None |
| TRPV4 | None | None | None | None |
| TRPV4 | None | None | None | None |
| TRPV4 | None | None | None | None |
| TRPV4 | None | None | None | None |
| TRPV4 | None | None | None | None |
| TRPV4 | None | None | None | None |
| TRPV4 | None | None | None | None |
| TRPV4 | None | None | None | None |
| TRPV4 | None | None | None | None |
| TRPV4 | None | None | None | None |
| TRPV4 | None | None | None | None |
| TRPV4 | None | None | None | None |
| TRPV4 | None | None | None | None |
| TNNI3 | None | None | None | None |
| MLYCD | None | None | None | None |
| MLYCD | None | None | None | None |
| MLYCD | None | None | None | None |
| MLYCD | None | None | None | None |
| AIP | None | None | None | None |
| AIP | None | None | None | None |
| ERCC2 | None | None | None | None |

| | | | | |
|---|---|---|---|---|
| ERCC2 | None | None | None | None |
| ERCC2 | None | None | None | None |
| ERCC2 | None | None | None | None |
| ERCC2 | None | None | None | None |
| ERCC2 | None | None | None | None |
| ERCC2 | None | None | None | None |
| ERCC2 | None | None | None | None |
| ERCC2 | None | None | None | None |
| ERCC2 | None | None | None | None |
| ERCC2 | None | None | None | None |
| ERCC2 | None | None | None | None |
| ERCC2 | None | None | None | None |
| ERCC2 | None | None | None | None |
| PUS1 | None | None | None | None |
| PUS1 | None | None | None | None |
| PUS1 | None | None | None | None |
| CASK | None | None | None | None |
| CASK | None | None | None | None |
| BLVRA | None | None | None | None |
| DICER1 | None | None | None | None |
| DICER1 | None | None | None | None |
| DICER1 | None | None | None | None |
| EIF2AK3 | None | None | None | None |
| EIF2AK3 | None | None | None | None |
| EIF2AK3 | None | None | None | None |
| KIT | None | None | None | None |
| KIT | None | None | None | None |
| KIT | None | None | None | None |
| KIT | None | None | None | None |
| KIT | None | None | None | None |
| KIT | None | None | None | None |
| DNMT3B | None | None | None | None |
| TGM1 | None | None | None | None |
| NDUFA10 | None | None | None | None |
| NDUFA10 | None | None | None | None |
| PRPF31 | None | None | None | None |
| PRPF31 | None | None | None | None |
| ECE1 | None | None | None | None |
| ECE1 | None | None | None | None |
| ECE1 | None | None | None | None |
| NDUFV2 | None | None | None | None |
| NDUFV2 | None | None | None | None |
| NDUFV2 | None | None | None | None |
| GHRHR | None | None | None | None |
| GHRHR | None | None | None | None |
| GHRHR | None | None | None | None |
| SLC4A1 | None | None | None | None |
| FGF9 | None | None | None | None |
| BARD1 | None | None | None | None |
| BARD1 | None | None | None | None |
| BARD1 | None | None | None | None |
| PTPN12 | None | None | None | None |
| MASP2 | None | None | None | None |
| MASP2 | None | None | None | None |

| | | | | |
|---|---|---|---|---|
| MASP2 | None | None | None | None |
| MASP2 | None | None | None | None |
| HOXB1 | None | None | None | None |
| SLC4A1 | None | None | None | None |
| SLC4A1 | None | None | None | None |
| SLC4A1 | None | None | None | None |
| SLC4A1 | None | None | None | None |
| SLC4A1 | None | None | None | None |
| SLC4A1 | None | None | None | None |
| SLC4A1 | None | None | None | None |
| SLC4A1 | None | None | None | None |
| SLC4A1 | None | None | None | None |
| SLC4A1 | None | None | None | None |
| SLC4A1 | None | None | None | None |
| SLC4A1 | None | None | None | None |
| SLC4A1 | None | None | None | None |
| SLC4A1 | None | None | None | None |
| SLC4A1 | None | None | None | None |
| SLC4A1 | None | None | None | None |
| SLC4A1 | None | None | None | None |
| CBL | None | None | None | None |
| FAS | None | None | None | None |
| MTMR2 | None | None | None | None |
| MTMR2 | None | None | None | None |
| MTMR2 | None | None | None | None |
| MTMR2 | None | None | None | None |
| MTMR2 | None | None | None | None |
| MTMR2 | None | None | None | None |
| MTMR2 | None | None | None | None |
| MTMR2 | None | None | None | None |
| KLHL3 | None | None | None | None |
| KLHL3 | None | None | None | None |
| KLHL3 | None | None | None | None |
| KLHL3 | None | None | None | None |
| KLHL3 | None | None | None | None |
| KLHL3 | None | None | None | None |
| RNASEH1 | None | None | None | None |
| ACTN1 | None | None | None | None |
| PHKG2 | None | None | None | None |
| PHKG2 | None | None | None | None |
| PHKG2 | None | None | None | None |
| NTRK1 | None | None | None | None |
| APOH | None | None | None | None |
| PMS2 | None | None | None | None |
| PMS2 | None | None | None | None |
| PMS2 | None | None | None | None |
| PMS2 | None | None | None | None |
| PMS2 | None | None | None | None |
| PMS2 | None | None | None | None |
| PMS2 | None | None | None | None |
| PMS2 | None | None | None | None |
| PMS2 | None | None | None | None |
| PMS2 | None | None | None | None |
| PMS2 | None | None | None | None |

| | | | | |
|---|---|---|---|---|
| PMS2 | None | None | None | None |
| PMS2 | None | None | None | None |
| PMS2 | None | None | None | None |
| PMS2 | None | None | None | None |
| PMS2 | None | None | None | None |
| PMS2 | None | None | None | None |
| PMS2 | None | None | None | None |
| PMS2 | None | None | None | None |
| PMS2 | None | None | None | None |
| PMS2 | None | None | None | None |
| PMS2 | None | None | None | None |
| MOCS2 | None | None | None | None |
| MOCS2 | None | None | None | None |
| RPGRIP1 | None | None | None | None |
| RPGRIP1 | None | None | None | None |
| LIG4 | None | None | None | None |
| PROC | None | None | None | None |
| PROC | None | None | None | None |
| NAT1 | None | None | None | None |
| NAT1 | None | None | None | None |
| PNPO | None | None | None | None |
| PNPO | None | None | None | None |
| PNPO | None | None | None | None |
| PNPO | None | None | None | None |
| PNPO | None | None | None | None |
| MRE11 | None | None | None | None |
| MRE11 | None | None | None | None |
| MRE11 | None | None | None | None |
| MRE11 | None | None | None | None |
| DDC | None | None | None | None |
| DDC | None | None | None | None |
| DDC | None | None | None | None |
| DDC | None | None | None | None |
| DDC | None | None | None | None |
| DDC | 0.909 | 7 | 3rch | B |
| DDC | 0.909 | 7 | 3rch | B |
| DDC | None | None | None | None |
| DDC | None | None | None | None |
| DDC | None | None | None | None |
| DDC | None | None | None | None |
| ANK2 | None | None | None | None |
| ANK2 | None | None | None | None |
| ANK2 | None | None | None | None |
| AK2 | None | None | None | None |
| AK2 | None | None | None | None |
| AK2 | None | None | None | None |
| AK2 | None | None | None | None |
| AK2 | None | None | None | None |
| AK2 | None | None | None | None |
| GPHN | None | None | None | None |
| HSD17B4 | None | None | None | None |
| HSD17B4 | None | None | None | None |
| HSD17B4 | None | None | None | None |
| HSD17B4 | None | None | None | None |

| | | | | |
|---|---|---|---|---|
| HSD17B4 | None | None | None | None |
| HSD17B4 | None | None | None | None |
| APTX | None | None | None | None |
| APTX | None | None | None | None |
| APTX | None | None | None | None |
| APTX | None | None | None | None |
| APTX | None | None | None | None |
| APTX | None | None | None | None |
| APTX | None | None | None | None |
| FANCL | None | None | None | None |
| FANCL | None | None | None | None |
| EMG1 | None | None | None | None |
| EMG1 | None | None | None | None |
| EZH2 | None | None | None | None |
| ABCB6 | None | None | None | None |
| NCF2 | None | None | None | None |
| NCF2 | None | None | None | None |
| COL4A2 | None | None | None | None |
| COL4A2 | 1 | 1 | 1li1 | F |
| COL4A2 | None | None | None | None |
| COL4A2 | None | None | None | None |
| COL4A2 | None | None | None | None |
| SLX4 | None | None | None | None |
| SERPIND1 | None | None | None | None |
| FGFR3 | None | None | None | None |
| FGFR3 | None | None | None | None |
| FGFR3 | None | None | None | None |
| FGFR3 | None | None | None | None |
| FGFR3 | None | None | None | None |
| FGFR3 | None | None | None | None |
| TREM2 | None | None | None | None |
| TREM2 | None | None | None | None |
| TREM2 | None | None | None | None |
| TREM2 | None | None | None | None |
| TREM2 | None | None | None | None |
| TREM2 | None | None | None | None |
| TREM2 | None | None | None | None |
| LIG4 | None | None | None | None |
| LIG4 | None | None | None | None |
| LIG4 | None | None | None | None |
| LIG4 | None | None | None | None |
| LIG4 | None | None | None | None |
| LIG4 | None | None | None | None |
| ECHS1 | None | None | None | None |
| ECHS1 | None | None | None | None |
| ECHS1 | None | None | None | None |
| MTR | None | None | None | None |
| MTR | None | None | None | None |
| MTR | None | None | None | None |
| MTR | None | None | None | None |
| MTR | None | None | None | None |
| PMVK | None | None | None | None |
| PRNP | None | None | None | None |
| CHN1 | None | None | None | None |

| | | | | |
|---|---|---|---|---|
| CHN1 | None | None | None | None |
| CHN1 | None | None | None | None |
| CHN1 | None | None | None | None |
| CHN1 | None | None | None | None |
| CHN1 | None | None | None | None |
| CHN1 | None | None | None | None |
| CHN1 | None | None | None | None |
| SLC2A1 | None | None | None | None |
| SLC2A1 | None | None | None | None |
| SLC2A1 | None | None | None | None |
| SLC2A1 | None | None | None | None |
| SLC2A1 | None | None | None | None |
| SLC2A1 | None | None | None | None |
| SLC2A1 | None | None | None | None |
| SLC2A1 | None | None | None | None |
| SLC2A1 | None | None | None | None |
| SLC2A1 | None | None | None | None |
| SLC2A1 | None | None | None | None |
| SLC2A1 | None | None | None | None |
| SLC2A1 | None | None | None | None |
| SLC2A1 | None | None | None | None |
| SLC2A1 | None | None | None | None |
| SLC2A1 | None | None | None | None |
| SLC2A1 | None | None | None | None |
| PMM2 | None | None | None | None |
| PMM2 | None | None | None | None |
| PMM2 | None | None | None | None |
| PMM2 | None | None | None | None |
| PMM2 | None | None | None | None |
| PMM2 | None | None | None | None |
| PMM2 | None | None | None | None |
| PMM2 | None | None | None | None |
| PMM2 | None | None | None | None |
| PMM2 | None | None | None | None |
| PMM2 | None | None | None | None |
| PMM2 | None | None | None | None |
| PMM2 | None | None | None | None |
| PMM2 | None | None | None | None |
| PMM2 | None | None | None | None |
| PMM2 | None | None | None | None |
| PMM2 | None | None | None | None |
| PMM2 | None | None | None | None |
| PMM2 | None | None | None | None |
| PMM2 | None | None | None | None |
| PMM2 | None | None | None | None |
| PMM2 | None | None | None | None |
| PMM2 | None | None | None | None |
| PMM2 | None | None | None | None |
| PMM2 | None | None | None | None |
| PMM2 | None | None | None | None |
| PMM2 | None | None | None | None |
| PMM2 | None | None | None | None |
| CCKAR | None | None | None | None |
| MAN1B1 | None | None | None | None |

| | | | | |
|---|---|---|---|---|
| MAN1B1 | None | None | None | None |
| LAMA2 | None | None | None | None |
| LAMA2 | None | None | None | None |
| TBC1D4 | None | None | None | None |
| TBC1D4 | None | None | None | None |
| LDLR | None | None | None | None |
| LDLR | None | None | None | None |
| LDLR | None | None | None | None |
| LDLR | None | None | None | None |
| LDLR | None | None | None | None |
| LDLR | None | None | None | None |
| LDLR | None | None | None | None |
| LDLR | None | None | None | None |
| LDLR | None | None | None | None |
| LDLR | None | None | None | None |
| LDLR | None | None | None | None |
| LDLR | None | None | None | None |
| LDLR | None | None | None | None |
| LDLR | None | None | None | None |
| LDLR | None | None | None | None |
| LDLR | None | None | None | None |
| LDLR | None | None | None | None |
| LDLR | None | None | None | None |
| LDLR | None | None | None | None |
| LDLR | None | None | None | None |
| LDLR | None | None | None | None |
| LDLR | None | None | None | None |
| LDLR | None | None | None | None |
| LDLR | None | None | None | None |
| LDLR | None | None | None | None |
| LDLR | None | None | None | None |
| LDLR | None | None | None | None |
| LDLR | None | None | None | None |
| LDLR | None | None | None | None |
| LDLR | None | None | None | None |
| LDLR | None | None | None | None |
| LDLR | None | None | None | None |
| LDLR | None | None | None | None |
| LDLR | None | None | None | None |
| LDLR | None | None | None | None |
| LDLR | None | None | None | None |
| LDLR | None | None | None | None |
| LDLR | None | None | None | None |
| LDLR | None | None | None | None |
| LDLR | None | None | None | None |
| LDLR | None | None | None | None |
| LDLR | None | None | None | None |
| LDLR | None | None | None | None |
| FANCM | None | None | None | None |
| HAMP | None | None | None | None |
| GUSB | None | None | None | None |
| GUSB | None | None | None | None |
| GUSB | None | None | None | None |
| GUSB | None | None | None | None |

| | | | | |
|---|---|---|---|---|
| GUSB | None | None | None | None |
| GUSB | None | None | None | None |
| GUSB | None | None | None | None |
| GUSB | None | None | None | None |
| GUSB | None | None | None | None |
| GUSB | None | None | None | None |
| GUSB | None | None | None | None |
| GUSB | None | None | None | None |
| GUSB | None | None | None | None |
| LIG1 | None | None | None | None |
| LIG1 | None | None | None | None |
| MYH7 | None | None | None | None |
| MYH7 | None | None | None | None |
| MYH7 | None | None | None | None |
| MYH7 | None | None | None | None |
| MYH7 | None | None | None | None |
| MYH7 | None | None | None | None |
| MYH7 | None | None | None | None |
| MYH7 | None | None | None | None |
| MYH7 | None | None | None | None |
| MYH7 | None | None | None | None |
| MYH7 | None | None | None | None |
| MYH7 | None | None | None | None |
| MYH7 | None | None | None | None |
| MYH7 | None | None | None | None |
| MYH7 | None | None | None | None |
| MYH7 | None | None | None | None |
| MYH7 | None | None | None | None |
| MYH7 | None | None | None | None |
| MYH7 | None | None | None | None |
| MYH7 | None | None | None | None |
| MYH7 | None | None | None | None |
| MYH7 | None | None | None | None |
| MYH7 | None | None | None | None |
| MYH7 | None | None | None | None |
| MYH7 | None | None | None | None |
| MYH7 | None | None | None | None |
| MYH7 | None | None | None | None |
| MYH7 | None | None | None | None |
| MYH7 | None | None | None | None |
| MYH7 | None | None | None | None |
| MYH7 | None | None | None | None |
| MYH7 | None | None | None | None |
| MYH7 | None | None | None | None |
| HCN4 | None | None | None | None |
| RECQL4 | None | None | None | None |
| RECQL4 | None | None | None | None |
| RECQL4 | None | None | None | None |
| RECQL4 | None | None | None | None |
| RECQL4 | None | None | None | None |
| RECQL4 | None | None | None | None |
| RECQL4 | None | None | None | None |
| RECQL4 | None | None | None | None |

| | | | | |
|---|---|---|---|---|
| RECQL4 | None | None | None | None |
| RECQL4 | None | None | None | None |
| RECQL4 | None | None | None | None |
| RECQL4 | None | None | None | None |
| RECQL4 | None | None | None | None |
| CHEK2 | None | None | None | None |
| PPARG | None | None | None | None |
| ACADVL | None | None | None | None |
| ACADVL | None | None | None | None |
| ACADVL | None | None | None | None |
| ACADVL | None | None | None | None |
| ACADVL | None | None | None | None |
| ACADVL | None | None | None | None |
| ACADVL | None | None | None | None |
| ACADVL | None | None | None | None |
| ACADVL | None | None | None | None |
| ACADVL | None | None | None | None |
| ACADVL | None | None | None | None |
| ACADVL | None | None | None | None |
| ACADVL | None | None | None | None |
| ACADVL | None | None | None | None |
| ACADVL | None | None | None | None |
| ACADVL | None | None | None | None |
| ACADVL | None | None | None | None |
| ACADVL | None | None | None | None |
| ACADVL | None | None | None | None |
| ACADVL | None | None | None | None |
| ACADVL | None | None | None | None |
| ACADVL | None | None | None | None |
| APOA1 | None | None | None | None |
| APOA1 | None | None | None | None |
| APOA1 | None | None | None | None |
| APOA1 | None | None | None | None |
| APOA1 | None | None | None | None |
| DNMT3A | None | None | None | None |
| DNMT3A | None | None | None | None |
| APC | None | None | None | None |
| TMOD1 | None | None | None | None |
| USH1C | None | None | None | None |
| USH1C | None | None | None | None |
| USH1C | None | None | None | None |
| USH1C | None | None | None | None |
| ERCC6 | None | None | None | None |
| ERCC6 | None | None | None | None |
| ERCC6 | None | None | None | None |
| ERCC8 | None | None | None | None |
| ERCC8 | None | None | None | None |
| ERCC8 | None | None | None | None |
| DPAGT1 | None | None | None | None |
| DPAGT1 | None | None | None | None |
| DPAGT1 | None | None | None | None |
| DPAGT1 | None | None | None | None |
| DPAGT1 | None | None | None | None |
| DPAGT1 | None | None | None | None |

| | | | | |
|---|---|---|---|---|
| DPAGT1 | None | None | None | None |
| IL2RG | None | None | None | None |
| HNF1A | None | None | None | None |
| HNF4A | None | None | None | None |
| FLT4 | None | None | None | None |
| FLT4 | None | None | None | None |
| FLT4 | None | None | None | None |
| CD3D | None | None | None | None |
| CD3D | None | None | None | None |
| MCOLN1 | None | None | None | None |
| MCOLN1 | None | None | None | None |
| MOCS2 | None | None | None | None |
| MOCS2 | None | None | None | None |
| ABCG8 | None | None | None | None |
| ABCG8 | None | None | None | None |
| ABCG8 | None | None | None | None |
| ABCG8 | None | None | None | None |
| ABCG8 | None | None | None | None |
| ABCG8 | None | None | None | None |
| ABCG8 | None | None | None | None |
| ABCG8 | None | None | None | None |
| ABCG8 | None | None | None | None |
| ABCG8 | None | None | None | None |
| ABCG8 | None | None | None | None |
| ABCG8 | None | None | None | None |
| ABCG8 | None | None | None | None |
| ABCG8 | None | None | None | None |
| CTNNA1 | None | None | None | None |
| TYMP | None | None | None | None |
| TYMP | None | None | None | None |
| TYMP | None | None | None | None |
| TYMP | None | None | None | None |
| TYMP | None | None | None | None |
| TYMP | None | None | None | None |
| TYMP | None | None | None | None |
| TYMP | None | None | None | None |
| TYMP | None | None | None | None |
| TYMP | None | None | None | None |
| TYMP | None | None | None | None |
| TYMP | None | None | None | None |
| TYMP | None | None | None | None |
| GAMT | None | None | None | None |
| GAMT | None | None | None | None |
| GAMT | None | None | None | None |
| GAMT | None | None | None | None |
| GAMT | None | None | None | None |
| GAMT | None | None | None | None |
| GAMT | None | None | None | None |
| GAMT | None | None | None | None |
| GAMT | None | None | None | None |
| GAMT | None | None | None | None |
| GAMT | None | None | None | None |
| GLRA1 | None | None | None | None |
| GLRA1 | None | None | None | None |

| | | | | |
|---|---|---|---|---|
| GLRA1 | None | None | None | None |
| GLRA1 | None | None | None | None |
| GLRA1 | None | None | None | None |
| GLRA1 | None | None | None | None |
| GLRA1 | None | None | None | None |
| GLRA1 | None | None | None | None |
| GLRA1 | None | None | None | None |
| ADK | None | None | None | None |
| ACTN2 | None | None | None | None |
| UBE2T | None | None | None | None |
| TTN | None | None | None | None |
| ATP7B | None | None | None | None |
| ATP7B | None | None | None | None |
| ATP7B | None | None | None | None |
| ATP7B | None | None | None | None |
| ATP7B | None | None | None | None |
| ATP7B | None | None | None | None |
| ATP7B | None | None | None | None |
| ATP7B | None | None | None | None |
| ATP7B | None | None | None | None |
| ATP7B | None | None | None | None |
| ATP7B | None | None | None | None |
| ATP7B | None | None | None | None |
| ATP7B | None | None | None | None |
| CDKN1B | None | None | None | None |
| CDKN1B | None | None | None | None |
| CDKN1B | None | None | None | None |
| CDKN1B | None | None | None | None |
| CDKN1B | None | None | None | None |
| CDKN1B | None | None | None | None |
| MYOC | None | None | None | None |
| MYOC | None | None | None | None |
| MYOC | None | None | None | None |
| MYOC | None | None | None | None |
| MYOC | None | None | None | None |
| MYOC | None | None | None | None |
| MYOC | None | None | None | None |
| MYOC | None | None | None | None |
| MYOC | None | None | None | None |
| MYOC | None | None | None | None |
| MYOC | None | None | None | None |
| MYOC | None | None | None | None |
| MYOC | None | None | None | None |
| MYOC | None | None | None | None |
| MYOC | None | None | None | None |
| SPTAN1 | None | None | None | None |
| GC | None | None | None | None |
| GC | None | None | None | None |
| MAD1L1 | None | None | None | None |
| NDST1 | None | None | None | None |
| ARL2BP | None | None | None | None |
| TOR1A | None | None | None | None |
| C1S | None | None | None | None |
| ARL6 | None | None | None | None |

| | | | | |
|---|---|---|---|---|
| ARL6 | None | None | None | None |
| ARL6 | None | None | None | None |
| ARL6 | None | None | None | None |
| ARL6 | None | None | None | None |
| ARL6 | None | None | None | None |
| FGFR1 | None | None | None | None |
| KLHL7 | None | None | None | None |
| FLNC | None | None | None | None |
| PTPRC | None | None | None | None |
| GCH1 | None | None | None | None |
| GCH1 | None | None | None | None |
| GCH1 | None | None | None | None |
| GCH1 | None | None | None | None |
| GCH1 | None | None | None | None |
| GCH1 | None | None | None | None |
| GCH1 | None | None | None | None |
| GCH1 | None | None | None | None |
| GCH1 | None | None | None | None |
| GCH1 | None | None | None | None |
| GCH1 | None | None | None | None |
| GCH1 | None | None | None | None |
| GCH1 | None | None | None | None |
| GCH1 | None | None | None | None |
| CFH | None | None | None | None |
| COL2A1 | None | None | None | None |
| COL2A1 | None | None | None | None |
| OCRL | None | None | None | None |
| WRN | None | None | None | None |
| WRN | None | None | None | None |
| WRN | None | None | None | None |
| WRN | None | None | None | None |
| MASP1 | None | None | None | None |
| MASP1 | None | None | None | None |
| NSD1 | None | None | None | None |
| PITX2 | None | None | None | None |
| PITX2 | None | None | None | None |
| PITX2 | None | None | None | None |
| PITX2 | None | None | None | None |
| PITX2 | None | None | None | None |
| PITX2 | None | None | None | None |
| B4GALT7 | 0.917 | 1 | 4irq | A |
| B4GALT7 | None | None | None | None |
| B4GALT7 | None | None | None | None |
| B4GALT7 | None | None | None | None |
| B4GALT7 | None | None | None | None |
| FSHR | None | None | None | None |
| IGHMBP2 | None | None | None | None |
| IGHMBP2 | None | None | None | None |
| IGHMBP2 | None | None | None | None |
| IGHMBP2 | None | None | None | None |
| IGHMBP2 | None | None | None | None |
| IGHMBP2 | None | None | None | None |
| IGHMBP2 | None | None | None | None |
| IGHMBP2 | None | None | None | None |

| | | | | |
|---|---|---|---|---|
| IGHMBP2 | None | None | None | None |
| IGHMBP2 | None | None | None | None |
| IGHMBP2 | None | None | None | None |
| IGHMBP2 | None | None | None | None |
| IGHMBP2 | None | None | None | None |
| IGHMBP2 | None | None | None | None |
| IGHMBP2 | None | None | None | None |
| IGHMBP2 | None | None | None | None |
| TTN | None | None | None | None |
| TTN | None | None | None | None |
| IRAK4 | None | None | None | None |
| PROC | None | None | None | None |
| PROC | None | None | None | None |
| RAD50 | None | None | None | None |
| RAD50 | None | None | None | None |
| DYNC1H1 | None | None | None | None |
| DYNC1H1 | None | None | None | None |
| DYNC1H1 | None | None | None | None |
| DYNC1H1 | None | None | None | None |
| DYNC1H1 | None | None | None | None |
| DYNC1H1 | None | None | None | None |
| DYNC1H1 | None | None | None | None |
| DYNC1H1 | None | None | None | None |
| DYNC1H1 | None | None | None | None |
| DYNC1H1 | None | None | None | None |
| DYNC1H1 | None | None | None | None |
| CYB5R3 | None | None | None | None |
| CYB5R3 | None | None | None | None |
| CYB5R3 | None | None | None | None |
| CYB5R3 | None | None | None | None |
| CYB5R3 | None | None | None | None |
| CYB5R3 | None | None | None | None |
| CYB5R3 | None | None | None | None |
| CYB5R3 | None | None | None | None |
| CYB5R3 | None | None | None | None |
| CYB5R3 | None | None | None | None |
| CYB5R3 | None | None | None | None |
| SPTB | None | None | None | None |
| FANCL | None | None | None | None |
| FAM126A | None | None | None | None |
| SEPT9 | None | None | None | None |
| TTN | None | None | None | None |
| TTN | None | None | None | None |
| TTN | None | None | None | None |
| CNTN1 | None | None | None | None |
| C1QTNF5 | None | None | None | None |
| VCL | None | None | None | None |
| VCL | None | None | None | None |
| VCL | None | None | None | None |
| VCL | None | None | None | None |
| VCL | None | None | None | None |
| VCL | None | None | None | None |
| PPT1 | None | None | None | None |
| PPT1 | None | None | None | None |

| | | | | |
|---|---|---|---|---|
| PPT1 | None | None | None | None |
| PPT1 | None | None | None | None |
| PPT1 | None | None | None | None |
| PPT1 | None | None | None | None |
| PPT1 | None | None | None | None |
| PPT1 | None | None | None | None |
| PPT1 | None | None | None | None |
| PPT1 | None | None | None | None |
| PPT1 | None | None | None | None |
| PPT1 | None | None | None | None |
| PPT1 | None | None | None | None |
| PPT1 | None | None | None | None |
| PPT1 | None | None | None | None |
| PPT1 | None | None | None | None |
| PPT1 | None | None | None | None |
| PPT1 | None | None | None | None |
| PPT1 | None | None | None | None |
| NOTCH2 | None | None | None | None |
| NOTCH2 | None | None | None | None |
| IFIH1 | None | None | None | None |
| IFIH1 | None | None | None | None |
| IKBKG | None | None | None | None |
| DBH | None | None | None | None |
| DBH | None | None | None | None |
| DBH | None | None | None | None |
| DBH | None | None | None | None |
| DBH | None | None | None | None |
| DBH | None | None | None | None |
| DBH | None | None | None | None |
| DBH | None | None | None | None |
| DBH | None | None | None | None |
| DBH | None | None | None | None |
| IL10RA | None | None | None | None |
| IL10RA | None | None | None | None |
| IL10RA | None | None | None | None |
| IL10RA | None | None | None | None |
| IL10RA | None | None | None | None |
| IL10RA | None | None | None | None |
| IL10RA | None | None | None | None |
| ILK | None | None | None | None |
| HPD | None | None | None | None |
| HPD | None | None | None | None |
| HPD | None | None | None | None |
| MECP2 | None | None | None | None |
| MECP2 | None | None | None | None |
| MECP2 | None | None | None | None |
| MECP2 | None | None | None | None |
| MECP2 | None | None | None | None |
| MECP2 | None | None | None | None |
| MECP2 | None | None | None | None |
| MECP2 | None | None | None | None |
| KYNU | None | None | None | None |
| FAS | None | None | None | None |
| FAS | None | None | None | None |

| | | | | |
|---|---|---|---|---|
| ELP1 | None | None | None | None |
| ELP1 | None | None | None | None |
| ELP1 | None | None | None | None |
| ELP1 | None | None | None | None |
| ELP1 | None | None | None | None |
| ELP1 | None | None | None | None |
| ELP1 | None | None | None | None |
| ELP1 | None | None | None | None |
| HESX1 | None | None | None | None |
| HESX1 | None | None | None | None |
| HESX1 | None | None | None | None |
| HESX1 | None | None | None | None |
| GAD1 | None | None | None | None |
| GAD1 | None | None | None | None |
| NDUFA1 | None | None | None | None |
| NDUFA1 | None | None | None | None |
| NDUFA1 | None | None | None | None |
| PNPT1 | None | None | None | None |
| PNPT1 | None | None | None | None |
| PNPT1 | None | None | None | None |
| PNPT1 | None | None | None | None |
| PNPT1 | None | None | None | None |
| PNPT1 | None | None | None | None |
| PNPT1 | None | None | None | None |
| SLC25A13 | None | None | None | None |
| SLC25A13 | None | None | None | None |
| SLC25A13 | None | None | None | None |
| AP4M1 | None | None | None | None |
| AP4M1 | None | None | None | None |
| MEFV | None | None | None | None |
| MEFV | None | None | None | None |
| MEFV | None | None | None | None |
| MEFV | None | None | None | None |
| MEFV | None | None | None | None |
| MEFV | None | None | None | None |
| MEFV | None | None | None | None |
| MEFV | None | None | None | None |
| MEFV | None | None | None | None |
| CAPN3 | None | None | None | None |
| CAPN3 | None | None | None | None |
| CAPN3 | None | None | None | None |
| CAPN3 | None | None | None | None |
| CAPN3 | None | None | None | None |
| CAPN3 | None | None | None | None |
| CAPN3 | None | None | None | None |
| GP6 | None | None | None | None |
| FAS | None | None | None | None |
| FBN1 | None | None | None | None |
| FBN1 | None | None | None | None |
| BMPR2 | None | None | None | None |
| BMPR2 | None | None | None | None |
| BMPR2 | None | None | None | None |
| BMPR2 | None | None | None | None |
| BMPR2 | None | None | None | None |

| | | | | |
|---|---|---|---|---|
| RET | None | None | None | None |
| RET | None | None | None | None |
| RET | None | None | None | None |
| RET | None | None | None | None |
| RET | None | None | None | None |
| RET | None | None | None | None |
| RET | None | None | None | None |
| RET | None | None | None | None |
| RET | None | None | None | None |
| RET | None | None | None | None |
| RET | None | None | None | None |
| RET | None | None | None | None |
| RET | None | None | None | None |
| RET | None | None | None | None |
| RET | None | None | None | None |
| RET | None | None | None | None |
| RET | None | None | None | None |
| RET | None | None | None | None |
| RET | None | None | None | None |
| RET | None | None | None | None |
| RET | None | None | None | None |
| RET | None | None | None | None |
| RET | None | None | None | None |
| RET | None | None | None | None |
| RET | None | None | None | None |
| RET | None | None | None | None |
| RET | None | None | None | None |
| RET | None | None | None | None |
| CSF2RA | None | None | None | None |
| CSF2RA | None | None | None | None |
| CSF2RA | None | None | None | None |
| BRCA1 | None | None | None | None |
| ZAP70 | None | None | None | None |
| ZAP70 | None | None | None | None |
| ZAP70 | None | None | None | None |
| PIK3R2 | None | None | None | None |
| TCN2 | None | None | None | None |
| TCN2 | None | None | None | None |
| TCN2 | None | None | None | None |
| TCN2 | None | None | None | None |
| TCN2 | None | None | None | None |
| TCN2 | None | None | None | None |
| TCN2 | None | None | None | None |
| TCN2 | None | None | None | None |
| EPM2A | None | None | None | None |
| EPM2A | None | None | None | None |
| EPM2A | None | None | None | None |
| EPM2A | None | None | None | None |
| AARS | None | None | None | None |
| AARS | None | None | None | None |
| AARS | None | None | None | None |
| PCDH15 | None | None | None | None |
| PCDH15 | None | None | None | None |
| PCDH15 | None | None | None | None |

| | | | | |
|---|---|---|---|---|
| PCDH15 | None | None | None | None |
| PCDH15 | None | None | None | None |
| PCDH15 | None | None | None | None |
| PCDH15 | None | None | None | None |
| PCDH15 | None | None | None | None |
| PCDH15 | None | None | None | None |
| PCDH15 | None | None | None | None |
| NDUFS4 | None | None | None | None |
| NDUFS4 | None | None | None | None |
| NDUFS4 | None | None | None | None |
| FGF10 | None | None | None | None |
| FGF10 | None | None | None | None |
| FGF10 | None | None | None | None |
| FGF10 | None | None | None | None |
| CDON | None | None | None | None |
| STAMBP | 0.909 | 2 | 3rzv | A |
| BRCA1 | None | None | None | None |
| BRCA1 | None | None | None | None |
| BRCA1 | None | None | None | None |
| BRCA1 | None | None | None | None |
| BRCA1 | None | None | None | None |
| BRCA1 | None | None | None | None |
| BRCA1 | None | None | None | None |
| BRCA1 | None | None | None | None |
| BRCA1 | None | None | None | None |
| BRCA1 | None | None | None | None |
| PLXNC1 | None | None | None | None |
| RNASEH2C | None | None | None | None |
| RNASEH2C | None | None | None | None |
| APC | None | None | None | None |
| PLA2G4A | None | None | None | None |
| PLA2G4A | None | None | None | None |
| PSAT1 | None | None | None | None |
| PSAT1 | None | None | None | None |
| TRAPPC3 | None | None | None | None |
| QDPR | None | None | None | None |
| QDPR | None | None | None | None |
| QDPR | None | None | None | None |
| QDPR | None | None | None | None |
| KRT5 | None | None | None | None |
| KRT5 | None | None | None | None |
| KRT5 | None | None | None | None |
| MLH1 | None | None | None | None |
| MLH1 | None | None | None | None |
| MLH1 | None | None | None | None |
| MLH1 | None | None | None | None |
| MLH1 | None | None | None | None |
| MLH1 | None | None | None | None |
| MLH1 | None | None | None | None |
| MLH1 | None | None | None | None |
| MLH1 | None | None | None | None |
| MLH1 | None | None | None | None |
| MLH1 | None | None | None | None |
| MLH1 | None | None | None | None |

| | | | | |
|---|---|---|---|---|
| MLH1 | None | None | None | None |
| MLH1 | None | None | None | None |
| MLH1 | None | None | None | None |
| MLH1 | None | None | None | None |
| MLH1 | None | None | None | None |
| KRIT1 | None | None | None | None |
| KRIT1 | None | None | None | None |
| KRIT1 | None | None | None | None |
| ALX4 | None | None | None | None |
| ALX4 | None | None | None | None |
| ALX4 | None | None | None | None |
| TJP2 | None | None | None | None |
| TJP2 | None | None | None | None |
| PANK2 | None | None | None | None |
| PANK2 | None | None | None | None |
| PANK2 | None | None | None | None |
| PANK2 | None | None | None | None |
| PANK2 | None | None | None | None |
| PANK2 | None | None | None | None |
| PANK2 | None | None | None | None |
| NDUFA6 | None | None | None | None |
| TSHR | None | None | None | None |
| TSHR | None | None | None | None |
| TSHR | None | None | None | None |
| CFTR | None | None | None | None |
| CFTR | None | None | None | None |
| MYOM1 | None | None | None | None |
| MYOM1 | None | None | None | None |
| SPTB | None | None | None | None |
| GPT2 | None | None | None | None |
| NDUFS2 | None | None | None | None |
| NDUFS2 | None | None | None | None |
| NDUFS2 | None | None | None | None |
| NDUFS2 | None | None | None | None |
| NDUFS2 | None | None | None | None |
| CASQ2 | None | None | None | None |
| CASQ2 | None | None | None | None |
| CASQ2 | None | None | None | None |
| CASQ2 | None | None | None | None |
| CASQ2 | None | None | None | None |
| CASQ2 | None | None | None | None |
| CASQ2 | None | None | None | None |
| CASQ2 | None | None | None | None |
| CASQ2 | None | None | None | None |
| CPOX | None | None | None | None |
| CPOX | None | None | None | None |
| CPOX | None | None | None | None |
| CPOX | None | None | None | None |
| CPOX | None | None | None | None |
| CPOX | None | None | None | None |
| CPOX | None | None | None | None |
| CPOX | None | None | None | None |
| CPOX | None | None | None | None |
| MVK | None | None | None | None |

| | | | | |
|---|---|---|---|---|
| MVK | None | None | None | None |
| MVK | None | None | None | None |
| MVK | None | None | None | None |
| MVK | None | None | None | None |
| MVK | None | None | None | None |
| MVK | None | None | None | None |
| MVK | None | None | None | None |
| MVK | None | None | None | None |
| MVK | None | None | None | None |
| MVK | None | None | None | None |
| MVK | None | None | None | None |
| MVK | None | None | None | None |
| MVK | None | None | None | None |
| MVK | None | None | None | None |
| GSS | None | None | None | None |
| GSS | None | None | None | None |
| GSS | None | None | None | None |
| GSS | None | None | None | None |
| GSS | None | None | None | None |
| DSG2 | None | None | None | None |
| DSG2 | None | None | None | None |
| DSG2 | None | None | None | None |
| DSG2 | None | None | None | None |
| DSG2 | None | None | None | None |
| DSG2 | None | None | None | None |
| DSG2 | None | None | None | None |
| DSG2 | None | None | None | None |
| DSG2 | None | None | None | None |
| DSG2 | None | None | None | None |
| DSG2 | None | None | None | None |
| DSG2 | None | None | None | None |
| DSG2 | None | None | None | None |
| DSG2 | None | None | None | None |
| PAPSS2 | None | None | None | None |
| PLCB2 | None | None | None | None |
| HMGCS2 | None | None | None | None |
| HMGCS2 | None | None | None | None |
| HMGCS2 | 0.938 | 1 | 2wya | C |
| HMGCS2 | None | None | None | None |
| HMGCS2 | None | None | None | None |
| HMGCS2 | None | None | None | None |
| HMGCS2 | None | None | None | None |
| HMGCS2 | None | None | None | None |
| MPDZ | None | None | None | None |
| GATA4 | None | None | None | None |
| GATA4 | None | None | None | None |
| WARS2 | None | None | None | None |
| WARS2 | None | None | None | None |
| HGD | 1 | 47 | 1eyb | A |
| HGD | 1 | 47 | 1eyb | A |
| HGD | 1 | 70 | 1eyb | A |
| HGD | None | None | None | None |
| HGD | None | None | None | None |
| HGD | None | None | None | None |

| | | | | |
|---|---|---|---|---|
| HGD | None | None | None | None |
| HGD | None | None | None | None |
| HGD | None | None | None | None |
| HGD | | 1 | 1 | 1ey2 | A |
| HGD | | 1 | 9 | 1ey2 | A |
| HGD | None | None | None | None |
| HGD | | 1 | 10 | 1eyb | A |
| HGD | | 1 | 7 | 1ey2 | A |
| PAX8 | None | None | None | None |
| PAX8 | None | None | None | None |
| PAX8 | None | None | None | None |
| PAX8 | None | None | None | None |
| SPTB | None | None | None | None |
| NCF4 | None | None | None | None |
| CFH | None | None | None | None |
| CFH | None | None | None | None |
| CSF1R | None | None | None | None |
| CSF1R | None | None | None | None |
| CA8 | None | None | None | None |
| CA8 | None | None | None | None |
| CA8 | None | None | None | None |
| APC | None | None | None | None |
| APC | None | None | None | None |
| APC | None | None | None | None |
| APC | None | None | None | None |
| APC | None | None | None | None |
| NDUFA2 | None | None | None | None |
| GIF | None | None | None | None |
| GIF | None | None | None | None |
| GIF | None | None | None | None |
| GIF | None | None | None | None |
| C2 | None | None | None | None |
| ACP5 | None | None | None | None |
| ACP5 | None | None | None | None |
| ACP5 | None | None | None | None |
| ACP5 | None | None | None | None |
| ACP5 | None | None | None | None |
| RNASEH2A | None | None | None | None |
| RNASEH2A | None | None | None | None |
| RNASEH2A | None | None | None | None |
| RNASEH2A | None | None | None | None |
| RNASEH2A | None | None | None | None |
| RNASEH2A | None | None | None | None |
| RNASEH2A | None | None | None | None |
| DSG2 | None | None | None | None |
| USH1G | None | None | None | None |
| FUS | None | None | None | None |
| FUS | None | None | None | None |
| FUS | None | None | None | None |
| FUS | None | None | None | None |
| FUS | None | None | None | None |
| MRPL44 | None | None | None | None |
| MRPL44 | None | None | None | None |
| MRPL44 | None | None | None | None |

| | | | | |
|---|---|---|---|---|
| MRPL44 | None | None | None | None |
| DMGDH | None | None | None | None |
| AARS | None | None | None | None |
| STAR | None | None | None | None |
| STAR | None | None | None | None |
| STAR | None | None | None | None |
| STAR | None | None | None | None |
| STAR | None | None | None | None |
| STAR | None | None | None | None |
| STAR | None | None | None | None |
| STAR | None | None | None | None |
| LARP7 | None | None | None | None |
| PRPF8 | None | None | None | None |
| PRPF8 | None | None | None | None |
| PRPF8 | None | None | None | None |
| PRPF8 | None | None | None | None |
| PSAP | None | None | None | None |
| PSAP | None | None | None | None |
| PSAP | None | None | None | None |
| DMPK | None | None | None | None |
| EPCAM | None | None | None | None |
| EPCAM | None | None | None | None |
| EPCAM | None | None | None | None |
| EPCAM | None | None | None | None |
| EPCAM | None | None | None | None |
| EPCAM | None | None | None | None |
| EPCAM | None | None | None | None |
| EPCAM | None | None | None | None |
| ASS1 | None | None | None | None |
| ASS1 | None | None | None | None |
| ASS1 | None | None | None | None |
| ASS1 | None | None | None | None |
| ASS1 | None | None | None | None |
| ASS1 | None | None | None | None |
| ASS1 | None | None | None | None |
| ASS1 | None | None | None | None |
| ASS1 | None | None | None | None |
| ASS1 | None | None | None | None |
| ASS1 | None | None | None | None |
| ASS1 | None | None | None | None |
| ASS1 | None | None | None | None |
| ASS1 | None | None | None | None |
| ASS1 | None | None | None | None |
| ASS1 | None | None | None | None |
| ASS1 | None | None | None | None |
| ASS1 | None | None | None | None |
| ASS1 | None | None | None | None |
| ASS1 | None | None | None | None |
| POLE | None | None | None | None |
| POLE | None | None | None | None |
| POLE | None | None | None | None |
| RNASET2 | None | None | None | None |
| RNASET2 | None | None | None | None |
| RNASET2 | None | None | None | None |

| | | | | |
|---|---|---|---|---|
| FLG | None | None | None | None |
| F10 | None | None | None | None |
| AGT | None | None | None | None |
| AGT | None | None | None | None |
| PLN | None | None | None | None |
| PLN | None | None | None | None |
| PLN | None | None | None | None |
| PTHLH | None | None | None | None |
| IGF1R | None | None | None | None |
| IGF1R | None | None | None | None |
| IGF1R | None | None | None | None |
| JAG1 | None | None | None | None |
| JAG1 | None | None | None | None |
| JAG1 | None | None | None | None |
| JAG1 | None | None | None | None |
| JAG1 | None | None | None | None |
| JAG1 | None | None | None | None |
| AP1S3 | None | None | None | None |
| SPTA1 | None | None | None | None |
| SPTA1 | None | None | None | None |
| SPTA1 | None | None | None | None |
| SPTA1 | None | None | None | None |
| SPG7 | None | None | None | None |
| SPG7 | None | None | None | None |
| SPG7 | None | None | None | None |
| SPG7 | None | None | None | None |
| SPG7 | None | None | None | None |
| SPG7 | None | None | None | None |
| FN1 | None | None | None | None |
| FN1 | None | None | None | None |
| NSMCE3 | None | None | None | None |
| PCCA | None | None | None | None |
| PCCA | None | None | None | None |
| SPTB | None | None | None | None |
| RYR2 | None | None | None | None |
| RYR2 | None | None | None | None |
| MMP9 | None | None | None | None |
| MMP9 | None | None | None | None |
| GLMN | None | None | None | None |
| GLMN | None | None | None | None |
| GLMN | None | None | None | None |
| GLMN | None | None | None | None |
| GLMN | None | None | None | None |
| GLMN | None | None | None | None |
| GLMN | None | None | None | None |
| GLMN | None | None | None | None |
| GLMN | None | None | None | None |
| GLMN | None | None | None | None |
| MARS | None | None | None | None |
| MARS | None | None | None | None |
| MARS | None | None | None | None |
| MARS | None | None | None | None |
| MARS | None | None | None | None |
| MARS | None | None | None | None |

| | | | | |
|---|---|---|---|---|
| SNAP29 | None | None | None | None |
| SNAP29 | None | None | None | None |
| ETHE1 | None | None | None | None |
| ETHE1 | None | None | None | None |
| ETHE1 | None | None | None | None |
| ETHE1 | None | None | None | None |
| ETHE1 | None | None | None | None |
| ETHE1 | None | None | None | None |
| ETHE1 | None | None | None | None |
| SAMHD1 | None | None | None | None |
| KCNQ4 | None | None | None | None |
| FXYD2 | None | None | None | None |
| FXYD2 | None | None | None | None |
| FXYD2 | None | None | None | None |
| LRRK2 | None | None | None | None |
| LRRK2 | None | None | None | None |
| NDUFS1 | None | None | None | None |
| NDUFS1 | None | None | None | None |
| NDUFS1 | None | None | None | None |
| NDUFS1 | None | None | None | None |
| NDUFS1 | None | None | None | None |
| NDUFS1 | None | None | None | None |
| NDUFS1 | None | None | None | None |
| NDUFS1 | None | None | None | None |
| NDUFS1 | None | None | None | None |
| NDUFS1 | None | None | None | None |
| NDUFS1 | None | None | None | None |
| EZH2 | None | None | None | None |
| EZH2 | None | None | None | None |
| LARS | None | None | None | None |
| TLL1 | None | None | None | None |
| CRYBB2 | None | None | None | None |
| CRYBB2 | None | None | None | None |
| EXOSC8 | None | None | None | None |
| SRP72 | None | None | None | None |
| FOXC2 | None | None | None | None |
| HMBS | None | None | None | None |
| HMBS | None | None | None | None |
| HMBS | None | None | None | None |
| HMBS | None | None | None | None |
| HMBS | None | None | None | None |
| HMBS | None | None | None | None |
| HMBS | None | None | None | None |
| HMBS | None | None | None | None |
| HMBS | None | None | None | None |
| HMBS | None | None | None | None |
| HMBS | None | None | None | None |
| HMBS | None | None | None | None |
| HMBS | None | None | None | None |
| HMBS | None | None | None | None |
| HMBS | None | None | None | None |
| HMBS | None | None | None | None |
| HMBS | None | None | None | None |
| HMBS | None | None | None | None |

| | | | | |
|---|---|---|---|---|
| HMBS | None | None | None | None |
| HMBS | None | None | None | None |
| HMBS | None | None | None | None |
| HMBS | None | None | None | None |
| HMBS | None | None | None | None |
| HMBS | None | None | None | None |
| HMBS | None | None | None | None |
| OGT | None | None | None | None |
| NOG | None | None | None | None |
| NOG | None | None | None | None |
| NOG | None | None | None | None |
| NOG | None | None | None | None |
| NOG | None | None | None | None |
| NOG | None | None | None | None |
| NOG | None | None | None | None |
| NOG | None | None | None | None |
| NOG | None | None | None | None |
| NOG | None | None | None | None |
| NOG | None | None | None | None |
| DLG3 | None | None | None | None |
| SCN2A | None | None | None | None |
| STAC3 | None | None | None | None |
| PFKM | None | None | None | None |
| PFKM | None | None | None | None |
| PFKM | None | None | None | None |
| PFKM | None | None | None | None |
| PFKM | None | None | None | None |
| DNMT1 | None | None | None | None |
| DNMT1 | None | None | None | None |
| TNNI3 | None | None | None | None |
| TNNI3 | None | None | None | None |
| TNNI3 | None | None | None | None |
| STXBP2 | None | None | None | None |
| STXBP2 | None | None | None | None |
| STXBP2 | None | None | None | None |
| STXBP2 | None | None | None | None |
| STXBP2 | None | None | None | None |
| STXBP2 | None | None | None | None |
| STXBP2 | None | None | None | None |
| STXBP2 | None | None | None | None |
| REN | None | None | None | None |
| DSP | None | None | None | None |
| DSP | None | None | None | None |
| HNF1B | None | None | None | None |
| ADA | None | None | None | None |
| ADA | None | None | None | None |
| ADA | None | None | None | None |
| ADA | None | None | None | None |
| ADA | None | None | None | None |
| ADA | None | None | None | None |
| ADA | None | None | None | None |
| ADA | None | None | None | None |
| ADA | None | None | None | None |
| ADA | None | None | None | None |

| | | | | |
|---|---|---|---|---|
| ADA | None | None | None | None |
| ADA | None | None | None | None |
| ADA | None | None | None | None |
| ADA | None | None | None | None |
| ADA | None | None | None | None |
| DCX | None | None | None | None |
| DCX | None | None | None | None |
| DCX | None | None | None | None |
| DCX | None | None | None | None |
| DCX | None | None | None | None |
| ARSB | None | None | None | None |
| ARSB | None | None | None | None |
| ARSB | None | None | None | None |
| ARSB | None | None | None | None |
| ARSB | None | None | None | None |
| ARSB | None | None | None | None |
| ARSB | None | None | None | None |
| ARSB | None | None | None | None |
| ARSB | None | None | None | None |
| ARSB | None | None | None | None |
| ARSB | None | None | None | None |
| ARSB | None | None | None | None |
| ARSB | None | None | None | None |
| WWOX | None | None | None | None |
| TNNI3 | None | None | None | None |
| TNNI3 | None | None | None | None |
| TNNI3 | None | None | None | None |
| TNNI3 | None | None | None | None |
| CYBA | None | None | None | None |
| SPAST | None | None | None | None |
| SPAST | None | None | None | None |
| SPAST | None | None | None | None |
| SPAST | None | None | None | None |
| SPAST | None | None | None | None |
| SPAST | None | None | None | None |
| SPAST | None | None | None | None |
| SPAST | None | None | None | None |
| KRIT1 | None | None | None | None |
| KRIT1 | None | None | None | None |
| TP53 | None | None | None | None |
| ISPD | None | None | None | None |
| ISPD | None | None | None | None |
| ISPD | None | None | None | None |
| ISPD | None | None | None | None |
| KCNMA1 | None | None | None | None |
| STK11 | None | None | None | None |
| STK11 | None | None | None | None |
| STK11 | None | None | None | None |
| STK11 | None | None | None | None |
| STK11 | None | None | None | None |
| STK11 | None | None | None | None |
| NDUFA13 | None | None | None | None |
| PHF6 | None | None | None | None |
| PHF6 | None | None | None | None |

| | | | | |
|---|---|---|---|---|
| PHF6 | None | None | None | None |
| PHYH | None | None | None | None |
| PHYH | None | None | None | None |
| PHYH | None | None | None | None |
| PHYH | None | None | None | None |
| PHYH | None | None | None | None |
| NDUFS3 | None | None | None | None |
| NDUFS3 | None | None | None | None |
| ALDH18A1 | None | None | None | None |
| ALDH18A1 | None | None | None | None |
| ALDH18A1 | None | None | None | None |
| ALDH18A1 | None | None | None | None |
| ALDH18A1 | None | None | None | None |
| PAX6 | None | None | None | None |
| PAX6 | None | None | None | None |
| INPP5E | None | None | None | None |
| INPP5E | None | None | None | None |
| INPP5E | None | None | None | None |
| INPP5E | None | None | None | None |
| INPP5E | None | None | None | None |
| INPP5E | None | None | None | None |
| INPP5E | None | None | None | None |
| INPP5E | None | None | None | None |
| INPP5E | None | None | None | None |
| TNNT2 | None | None | None | None |
| TNNT2 | None | None | None | None |
| TNNT2 | None | None | None | None |
| TNNT2 | None | None | None | None |
| TNNT2 | None | None | None | None |
| TNNT2 | None | None | None | None |
| TNNT2 | None | None | None | None |
| TNNT2 | None | None | None | None |
| TYK2 | None | None | None | None |
| TYK2 | None | None | None | None |
| TYK2 | None | None | None | None |
| CFTR | None | None | None | None |
| CFTR | None | None | None | None |
| CFTR | None | None | None | None |
| CFTR | None | None | None | None |
| CFTR | None | None | None | None |
| CFTR | None | None | None | None |
| CFTR | None | None | None | None |
| CFTR | None | None | None | None |
| CFTR | None | None | None | None |
| CFTR | None | None | None | None |
| CFTR | None | None | None | None |
| CFTR | None | None | None | None |
| CFTR | None | None | None | None |
| CFTR | None | None | None | None |
| CFTR | None | None | None | None |
| CFTR | None | None | None | None |
| CFTR | None | None | None | None |
| CFTR | None | None | None | None |
| CFTR | None | None | None | None |

| | | | | |
|---|---|---|---|---|
| CFTR | None | None | None | None |
| CFTR | None | None | None | None |
| CFTR | None | None | None | None |
| CFTR | None | None | None | None |
| CFTR | None | None | None | None |
| CFTR | None | None | None | None |
| CFTR | None | None | None | None |
| CFTR | None | None | None | None |
| CFTR | None | None | None | None |
| CFTR | None | None | None | None |
| CFTR | None | None | None | None |
| CFTR | None | None | None | None |
| CFTR | None | None | None | None |
| CFTR | None | None | None | None |
| CFTR | None | None | None | None |
| CFTR | None | None | None | None |
| CFTR | None | None | None | None |
| CFTR | None | None | None | None |
| CFTR | None | None | None | None |
| CFTR | None | None | None | None |
| CFTR | None | None | None | None |
| CFTR | None | None | None | None |
| CFTR | None | None | None | None |
| CFTR | None | None | None | None |
| CFTR | None | None | None | None |
| CFTR | None | None | None | None |
| CFTR | None | None | None | None |
| CFTR | None | None | None | None |
| CFTR | None | None | None | None |
| CFTR | None | None | None | None |
| CFTR | None | None | None | None |
| CFTR | None | None | None | None |
| CFTR | None | None | None | None |
| CFTR | None | None | None | None |
| SERPINF1 | None | None | None | None |
| SERPINF1 | None | None | None | None |
| IDS | None | None | None | None |
| IDS | None | None | None | None |
| IDS | None | None | None | None |
| IDS | None | None | None | None |
| IDS | None | None | None | None |
| IDS | None | None | None | None |
| IDS | None | None | None | None |
| IDS | None | None | None | None |
| IDS | None | None | None | None |
| NFKBIA | None | None | None | None |
| NFKBIA | None | None | None | None |
| SPAST | None | None | None | None |
| FGF23 | None | None | None | None |
| FGF23 | None | None | None | None |
| FGF23 | None | None | None | None |
| FGF23 | None | None | None | None |
| FGF23 | None | None | None | None |
| FGF23 | None | None | None | None |

| | | | | |
|---|---|---|---|---|
| FGF23 | None | None | None | None |
| DLAT | None | None | None | None |
| STUB1 | None | None | None | None |
| HSD17B4 | None | None | None | None |
| HSD17B4 | None | None | None | None |
| DYSF | None | None | None | None |
| CEP135 | None | None | None | None |
| CEP135 | None | None | None | None |
| PRKCG | None | None | None | None |
| CSF1R | None | None | None | None |
| PEX5 | None | None | None | None |
| PAX6 | None | None | None | None |
| PAX6 | None | None | None | None |
| PAX6 | None | None | None | None |
| PAX6 | None | None | None | None |
| PAX6 | None | None | None | None |
| PAX6 | None | None | None | None |
| NF2 | None | None | None | None |
| NF2 | None | None | None | None |
| NF2 | None | None | None | None |
| REM1 | None | None | None | None |

| source_id | classification | binding_site | binding_site_single_water | protein_binding_site |
|---|---|---|---|---|
| rs121918343 | pathogenic | 1 | 1 | 0 |
| rs267606649 | pathogenic | 1 | 1 | 1 |
| rs121918342 | pathogenic | 0 | 1 | 0 |
| rs267606650 | pathogenic | 1 | 1 | 1 |
| rs1130335 | benign | 1 | 1 | 1 |
| rs199476087 | pathogenic | 1 | 1 | 1 |
| rs772163112 | benign | 0 | 0 | 0 |
| rs755793 | benign | 0 | 0 | 0 |
| rs121918155 | pathogenic | 1 | 1 | 0 |
| rs121918156 | pathogenic | 1 | 1 | 0 |
| rs121918154 | pathogenic | 1 | 1 | 1 |
| rs121918151 | pathogenic | 1 | 1 | 1 |
| rs121918144 | pathogenic | 0 | 1 | 0 |
| rs121918146 | pathogenic | 1 | 1 | 1 |
| rs121918160 | pathogenic | 1 | 1 | 1 |
| rs121918150 | pathogenic | 1 | 1 | 1 |
| rs121918141 | pathogenic | 1 | 1 | 1 |
| rs142742242 | pathogenic | 1 | 1 | 1 |
| rs121918142 | pathogenic | 1 | 1 | 1 |
| rs121918157 | pathogenic | 1 | 1 | 1 |
| rs121918101 | pathogenic | 1 | 1 | 1 |
| rs121908260 | pathogenic | 1 | 1 | 1 |
| rs80356667 | pathogenic | 1 | 1 | 1 |
| rs80356668 | pathogenic | 1 | 1 | 1 |
| rs397514540 | pathogenic | 1 | 1 | 1 |
| rs104894685 | pathogenic | 0 | 1 | 0 |
| rs199869995 | pathogenic | 1 | 1 | 1 |
| rs121909072 | pathogenic | 0 | 0 | 0 |
| rs143669310 | benign | 1 | 1 | 1 |
| rs747347498 | benign | 0 | 0 | 0 |
| rs138283222 | benign | 0 | 0 | 0 |
| rs118203931 | pathogenic | 1 | 1 | 1 |
| rs118203933 | pathogenic | 1 | 1 | 0 |
| rs118203932 | pathogenic | 1 | 1 | 0 |
| rs2228063 | benign | 1 | 1 | 1 |
| rs377767360 | pathogenic | 0 | 0 | 0 |
| rs397518413 | pathogenic | 1 | 1 | 1 |
| rs1800458 | benign | 1 | 1 | 0 |
| rs121918083 | pathogenic | 1 | 1 | 1 |
| rs121918094 | pathogenic | 1 | 1 | 1 |
| rs121918098 | pathogenic | 1 | 1 | 1 |
| rs121918093 | pathogenic | 1 | 1 | 1 |
| rs79977247 | pathogenic | 1 | 1 | 1 |
| rs28933979 | pathogenic | 1 | 1 | 1 |
| rs121918068 | pathogenic | 1 | 1 | 1 |
| rs121918077 | pathogenic | 1 | 1 | 1 |
| rs11541796 | pathogenic | 1 | 1 | 1 |
| rs104894665 | pathogenic | 1 | 1 | 1 |
| rs121918090 | pathogenic | 1 | 1 | 1 |
| rs121918081 | pathogenic | 1 | 1 | 1 |
| rs121918080 | pathogenic | 1 | 1 | 1 |
| rs121918076 | pathogenic | 1 | 1 | 1 |
| rs121918097 | pathogenic | 1 | 1 | 1 |

| | | | | |
|---|---|---|---|---|
| rs121918079 | pathogenic | 1 | 1 | 1 |
| rs121918069 | pathogenic | 1 | 1 | 1 |
| rs121918086 | pathogenic | 1 | 1 | 1 |
| rs121918091 | pathogenic | 1 | 1 | 1 |
| rs121918085 | pathogenic | 1 | 1 | 1 |
| rs121918100 | pathogenic | 1 | 1 | 1 |
| rs121918084 | pathogenic | 1 | 1 | 1 |
| rs121918071 | pathogenic | 1 | 1 | 1 |
| rs121918072 | pathogenic | 1 | 1 | 1 |
| rs121918082 | pathogenic | 1 | 1 | 1 |
| rs121918087 | pathogenic | 1 | 1 | 1 |
| rs76410435 | benign | 1 | 1 | 1 |
| rs121918089 | pathogenic | 1 | 1 | 1 |
| rs121918073 | pathogenic | 1 | 1 | 1 |
| rs121918075 | pathogenic | 1 | 1 | 1 |
| rs121918088 | pathogenic | 1 | 1 | 1 |
| rs76992529 | pathogenic | 1 | 1 | 1 |
| rs536294863 | benign | 1 | 1 | 1 |
| rs730881162 | benign | 1 | 1 | 1 |
| rs113994087 | pathogenic | 1 | 1 | 1 |
| rs74716434 | benign | 1 | 1 | 1 |
| rs121918694 | pathogenic | 1 | 1 | 1 |
| rs121918707 | pathogenic | 1 | 1 | 1 |
| rs121918695 | pathogenic | 1 | 1 | 0 |
| rs121918690 | pathogenic | 1 | 1 | 0 |
| rs121918696 | pathogenic | 1 | 1 | 0 |
| rs121918693 | pathogenic | 1 | 1 | 0 |
| rs28999969 | pathogenic | 1 | 1 | 1 |
| rs121918697 | pathogenic | 1 | 1 | 1 |
| rs121918688 | pathogenic | 1 | 1 | 0 |
| rs28999970 | pathogenic | 1 | 1 | 0 |
| rs121918686 | pathogenic | 1 | 1 | 0 |
| rs28999971 | pathogenic | 1 | 1 | 1 |
| rs121918698 | pathogenic | 1 | 1 | 1 |
| rs121918691 | pathogenic | 1 | 1 | 0 |
| rs121918692 | pathogenic | 1 | 1 | 1 |
| rs121918703 | pathogenic | 1 | 1 | 1 |
| rs121918687 | pathogenic | 1 | 1 | 1 |
| rs28933408 | pathogenic | 1 | 1 | 1 |
| rs121918702 | pathogenic | 1 | 1 | 1 |
| rs17875899 | benign | 1 | 1 | 1 |
| rs111033620 | pathogenic | 1 | 1 | 1 |
| rs111033622 | pathogenic | 1 | 1 | 1 |
| rs111033621 | pathogenic | 1 | 1 | 1 |
| rs111033618 | pathogenic | 1 | 1 | 1 |
| rs5918 | benign | 0 | 0 | 0 |
| rs121918452 | pathogenic | 0 | 0 | 0 |
| rs121918445 | pathogenic | 1 | 1 | 1 |
| rs5917 | benign | 1 | 1 | 1 |
| rs121918444 | pathogenic | 1 | 1 | 1 |
| rs121918446 | pathogenic | 1 | 1 | 1 |
| rs121918449 | pathogenic | 0 | 0 | 0 |
| rs121918448 | benign | 0 | 0 | 0 |
| rs13306487 | benign | 1 | 1 | 1 |

| | | | | |
|---|---|---|---|---|
| rs1805044 | benign | 0 | 0 | 0 |
| rs117078377 | benign | 0 | 0 | 0 |
| rs138132026 | benign | 1 | 1 | 1 |
| rs75762896 | benign | 1 | 1 | 0 |
| rs121918043 | pathogenic | 1 | 1 | 0 |
| rs143581690 | benign | 1 | 1 | 0 |
| rs147654263 | benign | 0 | 0 | 0 |
| rs587782529 | pathogenic | 1 | 1 | 1 |
| rs121912664 | pathogenic | 1 | 1 | 1 |
| rs150293825 | benign | 1 | 1 | 1 |
| rs17882252 | benign | 1 | 1 | 1 |
| rs121912662 | pathogenic | 1 | 1 | 1 |
| rs35993958 | benign | 1 | 1 | 1 |
| rs606231253 | pathogenic | 1 | 1 | 1 |
| rs2227983 | benign | 1 | 1 | 1 |
| rs587779351 | pathogenic | 0 | 0 | 0 |
| rs779773463 | pathogenic | 1 | 1 | 1 |
| rs713040 | benign | 1 | 1 | 1 |
| rs334 | pathogenic | 1 | 1 | 1 |
| rs33930165 | pathogenic | 1 | 1 | 1 |
| rs33918131 | benign | 1 | 1 | 1 |
| rs63750783 | pathogenic | 1 | 1 | 1 |
| rs33986703 | pathogenic | 1 | 1 | 1 |
| rs33972047 | pathogenic | 1 | 1 | 1 |
| rs33936254 | pathogenic | 1 | 1 | 1 |
| rs33959855 | benign | 1 | 1 | 1 |
| rs33950507 | pathogenic | 1 | 1 | 1 |
| rs33916412 | pathogenic | 1 | 1 | 0 |
| rs1135071 | pathogenic | 1 | 1 | 1 |
| rs33960103 | pathogenic | 1 | 1 | 1 |
| rs11549407 | pathogenic | 1 | 1 | 1 |
| rs33922842 | pathogenic | 1 | 1 | 1 |
| rs35278874 | benign | 1 | 1 | 1 |
| rs34621955 | benign | 1 | 1 | 1 |
| rs33995148 | pathogenic | 1 | 1 | 1 |
| rs33945705 | benign | 1 | 1 | 1 |
| rs33993568 | benign | 1 | 1 | 1 |
| rs35553496 | benign | 1 | 1 | 1 |
| rs33913712 | pathogenic | 1 | 1 | 1 |
| rs33917785 | pathogenic | 1 | 1 | 1 |
| rs34515413 | benign | 1 | 1 | 1 |
| rs35256489 | pathogenic | 1 | 1 | 0 |
| rs34484056 | pathogenic | 1 | 1 | 1 |
| rs33947020 | benign | 1 | 1 | 1 |
| rs33946267 | pathogenic | 1 | 1 | 1 |
| rs33966761 | pathogenic | 1 | 1 | 1 |
| rs36020563 | benign | 1 | 1 | 1 |
| rs122454124 | pathogenic | 1 | 1 | 1 |
| rs122454126 | pathogenic | 1 | 1 | 0 |
| rs122454127 | pathogenic | 1 | 1 | 1 |
| rs122454130 | pathogenic | 1 | 1 | 1 |
| rs122454125 | pathogenic | 1 | 1 | 1 |
| rs122454131 | pathogenic | 1 | 1 | 0 |
| rs121908302 | pathogenic | 0 | 1 | 0 |

| | | | | |
|---|---|---|---|---|
| rs77829017 | pathogenic | 0 | 1 | 0 |
| rs1141814 | pathogenic | 0 | 1 | 0 |
| rs121908312 | pathogenic | 0 | 1 | 0 |
| rs79653797 | pathogenic | 1 | 1 | 0 |
| rs439898 | pathogenic | 1 | 1 | 0 |
| rs121908299 | pathogenic | 0 | 1 | 0 |
| rs80356763 | pathogenic | 1 | 1 | 1 |
| rs781152868 | pathogenic | 0 | 1 | 0 |
| rs121908297 | pathogenic | 0 | 1 | 0 |
| rs398123531 | benign | 0 | 1 | 0 |
| rs398123532 | pathogenic | 0 | 1 | 0 |
| rs364897 | pathogenic | 1 | 1 | 1 |
| rs1064644 | pathogenic | 0 | 1 | 0 |
| rs409652 | pathogenic | 1 | 1 | 1 |
| rs121908300 | pathogenic | 0 | 1 | 0 |
| rs381737 | pathogenic | 0 | 1 | 0 |
| rs74500255 | pathogenic | 0 | 1 | 0 |
| rs121908313 | pathogenic | 0 | 1 | 0 |
| rs367968666 | pathogenic | 0 | 1 | 0 |
| rs78973108 | pathogenic | 1 | 1 | 1 |
| rs121908298 | pathogenic | 1 | 1 | 1 |
| rs78396650 | pathogenic | 0 | 0 | 0 |
| rs78198234 | pathogenic | 1 | 1 | 0 |
| rs121908304 | pathogenic | 1 | 1 | 1 |
| rs76539814 | pathogenic | 1 | 1 | 1 |
| rs121908305 | pathogenic | 0 | 1 | 0 |
| rs121908306 | pathogenic | 1 | 1 | 0 |
| rs121908308 | pathogenic | 1 | 1 | 1 |
| rs121908307 | pathogenic | 0 | 1 | 0 |
| rs75548401 | benign | 0 | 1 | 0 |
| rs76763715 | pathogenic | 0 | 0 | 0 |
| rs121908314 | pathogenic | 0 | 1 | 0 |
| rs398123528 | pathogenic | 0 | 1 | 0 |
| rs121908311 | pathogenic | 0 | 0 | 0 |
| rs121908310 | pathogenic | 1 | 1 | 0 |
| rs74598136 | pathogenic | 0 | 1 | 0 |
| rs1064651 | pathogenic | 1 | 1 | 1 |
| rs77369218 | pathogenic | 1 | 1 | 1 |
| rs121908295 | pathogenic | 0 | 1 | 0 |
| rs421016 | pathogenic | 0 | 1 | 0 |
| rs368060 | pathogenic | 0 | 1 | 0 |
| rs80356771 | pathogenic | 1 | 1 | 1 |
| rs80356772 | pathogenic | 1 | 1 | 1 |
| rs121908301 | pathogenic | 0 | 0 | 0 |
| rs747506979 | pathogenic | 0 | 1 | 0 |
| rs75822236 | pathogenic | 0 | 1 | 0 |
| rs1990760 | benign | 1 | 1 | 0 |
| rs28934568 | pathogenic | 1 | 1 | 0 |
| rs34833812 | benign | 1 | 1 | 0 |
| rs193922666 | benign | 1 | 1 | 0 |
| rs104893812 | pathogenic | 1 | 1 | 0 |
| rs144701411 | benign | 0 | 0 | 0 |
| rs104893813 | pathogenic | 0 | 1 | 0 |
| rs199660234 | pathogenic | 1 | 1 | 0 |

| | | | | |
|---|---|---|---|---|
| rs104893814 | pathogenic | 1 | 1 | 0 |
| rs143095746 | benign | 1 | 1 | 1 |
| rs104893807 | pathogenic | 1 | 1 | 0 |
| rs104893811 | pathogenic | 0 | 0 | 0 |
| rs104893816 | pathogenic | 0 | 0 | 0 |
| rs121918714 | pathogenic | 0 | 1 | 0 |
| rs104893810 | pathogenic | 1 | 1 | 0 |
| rs104893815 | pathogenic | 1 | 1 | 0 |
| rs4303 | benign | 0 | 1 | 0 |
| rs121912704 | pathogenic | 0 | 1 | 0 |
| rs56394458 | benign | 1 | 1 | 0 |
| rs397514688 | pathogenic | 0 | 0 | 0 |
| rs367797185 | pathogenic | 1 | 1 | 0 |
| rs12709426 | benign | 0 | 1 | 0 |
| rs2228547 | benign | 0 | 0 | 0 |
| rs111033546 | pathogenic | 0 | 0 | 0 |
| rs111033553 | pathogenic | 0 | 0 | 0 |
| rs111033554 | pathogenic | 1 | 1 | 1 |
| rs111033544 | pathogenic | 1 | 1 | 1 |
| rs111033555 | pathogenic | 1 | 1 | 1 |
| rs111033545 | pathogenic | 1 | 1 | 1 |
| rs111033549 | pathogenic | 0 | 1 | 0 |
| rs111033552 | pathogenic | 1 | 1 | 1 |
| rs752204035 | benign | 0 | 0 | 0 |
| rs11569534 | benign | 0 | 0 | 0 |
| rs28937571 | pathogenic | 1 | 1 | 1 |
| rs104894215 | pathogenic | 1 | 1 | 1 |
| rs3888798 | benign | 1 | 1 | 1 |
| rs764263034 | benign | 1 | 1 | 1 |
| rs1800973 | benign | 1 | 1 | 1 |
| rs367623154 | benign | 1 | 1 | 1 |
| rs201230446 | pathogenic | 1 | 1 | 1 |
| rs267606766 | pathogenic | 0 | 1 | 0 |
| rs267606769 | pathogenic | 0 | 1 | 0 |
| rs267606767 | pathogenic | 1 | 1 | 0 |
| rs267606768 | pathogenic | 0 | 1 | 0 |
| rs61733129 | benign | 0 | 1 | 0 |
| rs201947120 | pathogenic | 1 | 1 | 1 |
| rs283413 | benign | 0 | 1 | 0 |
| rs36119840 | benign | 0 | 0 | 0 |
| rs281865419 | pathogenic | 1 | 1 | 1 |
| rs137852937 | pathogenic | 0 | 1 | 0 |
| rs2230709 | benign | 0 | 1 | 0 |
| rs6695033 | benign | 0 | 1 | 0 |
| rs180177032 | pathogenic | 1 | 1 | 1 |
| rs180177033 | pathogenic | 1 | 1 | 1 |
| rs121913351 | pathogenic | 1 | 1 | 1 |
| rs397507473 | pathogenic | 1 | 1 | 1 |
| rs121913355 | pathogenic | 1 | 1 | 1 |
| rs121913357 | pathogenic | 1 | 1 | 1 |
| rs180177036 | pathogenic | 1 | 1 | 1 |
| rs180177039 | pathogenic | 1 | 1 | 1 |
| rs180177038 | pathogenic | 1 | 1 | 1 |
| rs180177040 | pathogenic | 1 | 1 | 1 |

| | | | | |
|---|---|---|---|---|
| rs121913338 | pathogenic | 1 | 1 | 1 |
| rs121913366 | pathogenic | 1 | 1 | 1 |
| rs121913369 | pathogenic | 1 | 1 | 1 |
| rs113488022 | pathogenic | 1 | 1 | 1 |
| rs121913364 | pathogenic | 1 | 1 | 1 |
| rs180177042 | pathogenic | 1 | 1 | 1 |
| rs200461617 | benign | 0 | 0 | 0 |
| rs141609745 | benign | 0 | 0 | 0 |
| rs104894383 | pathogenic | 1 | 1 | 1 |
| rs104894384 | pathogenic | 1 | 1 | 1 |
| rs104894381 | pathogenic | 1 | 1 | 0 |
| rs147710408 | benign | 1 | 1 | 1 |
| rs77357563 | benign | 1 | 1 | 1 |
| rs104894378 | pathogenic | 0 | 0 | 0 |
| rs104894382 | pathogenic | 0 | 0 | 0 |
| rs137853292 | pathogenic | 1 | 1 | 1 |
| rs587778834 | pathogenic | 0 | 0 | 0 |
| rs137853294 | pathogenic | 1 | 1 | 1 |
| rs551747882 | benign | 0 | 0 | 0 |
| rs3092903 | benign | 1 | 1 | 1 |
| rs137853296 | pathogenic | 0 | 0 | 0 |
| rs398122917 | pathogenic | 0 | 1 | 0 |
| rs41304992 | benign | 0 | 0 | 0 |
| rs121912708 | pathogenic | 1 | 1 | 1 |
| rs140102105 | pathogenic | 1 | 1 | 1 |
| rs376917645 | pathogenic | 1 | 1 | 0 |
| rs372660425 | pathogenic | 1 | 1 | 0 |
| rs555896752 | pathogenic | 1 | 1 | 1 |
| rs777829351 | pathogenic | 1 | 1 | 0 |
| rs121912711 | pathogenic | 1 | 1 | 0 |
| rs754449549 | pathogenic | 0 | 1 | 0 |
| rs141701364 | benign | 0 | 1 | 0 |
| rs761011399 | benign | 1 | 1 | 1 |
| rs121912710 | benign | 1 | 1 | 0 |
| rs121912707 | pathogenic | 1 | 1 | 0 |
| rs747597620 | pathogenic | 0 | 1 | 0 |
| rs186558364 | pathogenic | 1 | 1 | 1 |
| rs556400964 | pathogenic | 1 | 1 | 1 |
| rs61757684 | benign | 1 | 1 | 1 |
| rs121917766 | pathogenic | 1 | 1 | 1 |
| rs121918248 | pathogenic | 0 | 0 | 0 |
| rs398123278 | pathogenic | 0 | 0 | 0 |
| rs121918251 | pathogenic | 1 | 1 | 1 |
| rs727504022 | pathogenic | 0 | 0 | 0 |
| rs190834116 | pathogenic | 1 | 1 | 1 |
| rs121918249 | pathogenic | 1 | 1 | 0 |
| rs121918257 | pathogenic | 1 | 1 | 0 |
| rs142068898 | benign | 0 | 0 | 0 |
| rs757000253 | pathogenic | 0 | 0 | 0 |
| rs200908035 | pathogenic | 1 | 1 | 0 |
| rs760782399 | pathogenic | 0 | 0 | 0 |
| rs778702777 | pathogenic | 0 | 0 | 0 |
| rs121918258 | pathogenic | 1 | 1 | 0 |
| rs121918256 | pathogenic | 0 | 0 | 0 |

| | | | | |
|---|---|---|---|---|
| rs200596762 | pathogenic | 1 | 1 | 0 |
| rs547709692 | pathogenic | 0 | 0 | 0 |
| rs758577372 | pathogenic | 0 | 1 | 0 |
| rs564069299 | pathogenic | 1 | 1 | 1 |
| rs772552898 | pathogenic | 1 | 1 | 1 |
| rs368790885 | pathogenic | 1 | 1 | 1 |
| rs121918250 | pathogenic | 0 | 0 | 0 |
| rs533755473 | pathogenic | 1 | 1 | 1 |
| rs753288303 | pathogenic | 1 | 1 | 1 |
| rs2229385 | benign | 0 | 0 | 0 |
| rs1141321 | benign | 0 | 0 | 0 |
| rs753564352 | pathogenic | 0 | 0 | 0 |
| rs775593146 | pathogenic | 0 | 0 | 0 |
| rs121918254 | pathogenic | 0 | 0 | 0 |
| rs8589 | benign | 1 | 1 | 0 |
| rs777758903 | pathogenic | 0 | 0 | 0 |
| rs140600746 | pathogenic | 1 | 1 | 0 |
| rs121918255 | pathogenic | 1 | 1 | 0 |
| rs121918252 | pathogenic | 0 | 0 | 0 |
| rs779990936 | pathogenic | 1 | 1 | 0 |
| rs121912455 | pathogenic | 1 | 1 | 1 |
| rs121912459 | pathogenic | 1 | 1 | 1 |
| rs74315452 | pathogenic | 1 | 1 | 1 |
| rs121912451 | pathogenic | 1 | 1 | 1 |
| rs79006549 | pathogenic | 0 | 0 | 0 |
| rs28934889 | benign | 0 | 1 | 0 |
| rs111033557 | benign | 1 | 1 | 1 |
| rs28934597 | pathogenic | 1 | 1 | 1 |
| rs28934596 | pathogenic | 1 | 1 | 1 |
| rs28934595 | pathogenic | 1 | 1 | 1 |
| rs146519482 | pathogenic | 1 | 1 | 1 |
| rs111033563 | pathogenic | 1 | 1 | 1 |
| rs75385605 | benign | 1 | 1 | 1 |
| rs121908569 | pathogenic | 1 | 1 | 1 |
| rs35719940 | benign | 0 | 0 | 0 |
| rs121918664 | pathogenic | 0 | 0 | 0 |
| rs104893665 | pathogenic | 1 | 1 | 1 |
| rs104893666 | pathogenic | 1 | 1 | 1 |
| rs200845457 | benign | 1 | 1 | 1 |
| rs779204655 | pathogenic | 1 | 1 | 1 |
| rs121917747 | pathogenic | 1 | 1 | 1 |
| rs6042 | benign | 1 | 1 | 1 |
| rs104894380 | pathogenic | 0 | 0 | 0 |
| rs369083541 | pathogenic | 1 | 1 | 1 |
| rs145243532 | benign | 1 | 1 | 1 |
| rs121909500 | pathogenic | 1 | 1 | 1 |
| rs140993310 | benign | 1 | 1 | 1 |
| rs200837270 | pathogenic | 1 | 1 | 1 |
| rs121918484 | pathogenic | 1 | 1 | 1 |
| rs121918306 | pathogenic | 0 | 0 | 0 |
| rs118204089 | pathogenic | 1 | 1 | 0 |
| rs118204083 | pathogenic | 1 | 1 | 0 |
| rs201953584 | pathogenic | 1 | 1 | 1 |
| rs74315362 | pathogenic | 1 | 1 | 1 |

| | | | | |
|---|---|---|---|---|
| rs118204084 | pathogenic | 1 | 1 | 0 |
| rs118204085 | pathogenic | 1 | 1 | 1 |
| rs113403872 | pathogenic | 1 | 1 | 0 |
| rs1052176 | benign | 1 | 1 | 1 |
| rs2031640 | benign | 0 | 0 | 0 |
| rs104894230 | pathogenic | 1 | 1 | 1 |
| rs104894229 | pathogenic | 1 | 1 | 1 |
| rs104894228 | pathogenic | 1 | 1 | 1 |
| rs104894226 | pathogenic | 1 | 1 | 1 |
| rs121917757 | pathogenic | 1 | 1 | 1 |
| rs121917758 | pathogenic | 1 | 1 | 1 |
| rs28933406 | pathogenic | 1 | 1 | 1 |
| rs121917756 | pathogenic | 1 | 1 | 1 |
| rs104894227 | pathogenic | 1 | 1 | 1 |
| rs369106578 | pathogenic | 1 | 1 | 1 |
| rs397517142 | benign | 1 | 1 | 1 |
| rs104894231 | pathogenic | 1 | 1 | 1 |
| rs121917759 | pathogenic | 1 | 1 | 1 |
| rs104893827 | pathogenic | 0 | 0 | 0 |
| rs104893826 | pathogenic | 1 | 1 | 1 |
| rs5030802 | pathogenic | 1 | 1 | 1 |
| rs5030827 | pathogenic | 1 | 1 | 0 |
| rs119103277 | pathogenic | 1 | 1 | 1 |
| rs5030808 | pathogenic | 0 | 1 | 0 |
| rs5030809 | pathogenic | 1 | 1 | 1 |
| rs765978945 | pathogenic | 1 | 1 | 1 |
| rs104893824 | pathogenic | 1 | 1 | 1 |
| rs104893831 | pathogenic | 0 | 1 | 0 |
| rs104893830 | pathogenic | 0 | 1 | 0 |
| rs119103278 | pathogenic | 0 | 1 | 0 |
| rs5030818 | pathogenic | 1 | 1 | 1 |
| rs397516444 | pathogenic | 1 | 1 | 1 |
| rs28940297 | pathogenic | 1 | 1 | 1 |
| rs267607170 | pathogenic | 1 | 1 | 1 |
| rs104893825 | pathogenic | 1 | 1 | 1 |
| rs5030820 | pathogenic | 0 | 1 | 0 |
| rs5030821 | pathogenic | 0 | 1 | 0 |
| rs5030823 | pathogenic | 1 | 1 | 1 |
| rs5030824 | pathogenic | 1 | 1 | 1 |
| rs28940301 | pathogenic | 0 | 1 | 0 |
| rs28940300 | pathogenic | 0 | 1 | 0 |
| rs28940298 | pathogenic | 0 | 1 | 0 |
| rs200019083 | benign | 0 | 0 | 0 |
| rs759748655 | pathogenic | 0 | 0 | 0 |
| rs121917739 | pathogenic | 1 | 1 | 1 |
| rs75353611 | pathogenic | 1 | 1 | 1 |
| rs79744198 | benign | 1 | 1 | 1 |
| rs79228041 | pathogenic | 1 | 1 | 1 |
| rs80106970 | benign | 1 | 1 | 1 |
| rs375353223 | benign | 0 | 0 | 0 |
| rs56311081 | benign | 0 | 0 | 0 |
| rs55985569 | benign | 1 | 1 | 1 |
| rs45483396 | benign | 0 | 0 | 0 |
| rs35284565 | benign | 1 | 1 | 1 |

| | | | | |
|---|---|---|---|---|
| rs34349517 | benign | 0 | 0 | 0 |
| rs35776110 | benign | 0 | 0 | 0 |
| rs33917957 | benign | 1 | 1 | 1 |
| rs587778445 | benign | 0 | 0 | 0 |
| rs34624984 | benign | 0 | 0 | 0 |
| rs701753 | benign | 0 | 0 | 0 |
| rs855314 | benign | 0 | 0 | 0 |
| rs121918371 | pathogenic | 1 | 1 | 0 |
| rs398122912 | pathogenic | 0 | 1 | 0 |
| rs1126728 | benign | 0 | 0 | 0 |
| rs587777404 | pathogenic | 1 | 1 | 0 |
| rs11208257 | benign | 0 | 0 | 0 |
| rs6676290 | benign | 0 | 0 | 0 |
| rs104894118 | pathogenic | 0 | 0 | 0 |
| rs121918655 | pathogenic | 1 | 1 | 1 |
| rs104894120 | pathogenic | 1 | 1 | 1 |
| rs137852238 | pathogenic | 0 | 0 | 0 |
| rs137852237 | pathogenic | 0 | 0 | 0 |
| rs137852499 | pathogenic | 1 | 1 | 1 |
| rs137852480 | pathogenic | 1 | 1 | 1 |
| rs137852494 | pathogenic | 1 | 1 | 1 |
| rs137852502 | pathogenic | 1 | 1 | 1 |
| rs137852500 | pathogenic | 1 | 1 | 1 |
| rs137852487 | pathogenic | 1 | 1 | 1 |
| rs137852481 | pathogenic | 1 | 1 | 1 |
| rs137852501 | pathogenic | 0 | 0 | 0 |
| rs137852478 | pathogenic | 1 | 1 | 1 |
| rs137852485 | pathogenic | 1 | 1 | 1 |
| rs137852482 | pathogenic | 1 | 1 | 0 |
| rs137852483 | pathogenic | 0 | 1 | 0 |
| rs137852477 | pathogenic | 1 | 1 | 0 |
| rs137852496 | pathogenic | 1 | 1 | 0 |
| rs137852484 | pathogenic | 0 | 1 | 0 |
| rs137852498 | pathogenic | 1 | 1 | 0 |
| rs137852497 | pathogenic | 1 | 1 | 0 |
| rs137852493 | pathogenic | 1 | 1 | 0 |
| rs137852504 | pathogenic | 1 | 1 | 1 |
| rs267606863 | pathogenic | 1 | 1 | 1 |
| rs137852486 | pathogenic | 1 | 1 | 1 |
| rs137852479 | pathogenic | 1 | 1 | 1 |
| rs112122950 | benign | 0 | 0 | 0 |
| rs72546648 | benign | 1 | 1 | 1 |
| rs72552291 | pathogenic | 0 | 0 | 0 |
| rs79184941 | pathogenic | 1 | 1 | 1 |
| rs77543610 | pathogenic | 1 | 1 | 1 |
| rs121918505 | pathogenic | 1 | 1 | 1 |
| rs776587763 | pathogenic | 1 | 1 | 1 |
| rs121918497 | pathogenic | 1 | 1 | 1 |
| rs121918499 | pathogenic | 1 | 1 | 1 |
| rs121918501 | pathogenic | 1 | 1 | 1 |
| rs121918500 | pathogenic | 0 | 0 | 0 |
| rs121918504 | pathogenic | 1 | 1 | 1 |
| rs121918510 | pathogenic | 1 | 1 | 1 |
| rs121918493 | pathogenic | 1 | 1 | 1 |

| | | | | |
|---|---|---|---|---|
| rs121918489 | pathogenic | 0 | 0 | 0 |
| rs121918495 | pathogenic | 1 | 1 | 1 |
| rs121918488 | pathogenic | 0 | 0 | 0 |
| rs121918496 | pathogenic | 0 | 0 | 0 |
| rs121918492 | pathogenic | 1 | 1 | 1 |
| rs121918494 | pathogenic | 1 | 1 | 1 |
| rs121918502 | pathogenic | 1 | 1 | 1 |
| rs121918490 | pathogenic | 1 | 1 | 1 |
| rs72466451 | pathogenic | 1 | 1 | 1 |
| rs66468541 | pathogenic | 0 | 0 | 0 |
| rs137852662 | pathogenic | 0 | 0 | 0 |
| rs137852663 | pathogenic | 0 | 0 | 0 |
| rs606231407 | pathogenic | 0 | 0 | 0 |
| rs137852233 | pathogenic | 1 | 1 | 1 |
| rs112306225 | benign | 0 | 0 | 0 |
| rs387906808 | pathogenic | 1 | 1 | 0 |
| rs730882094 | pathogenic | 1 | 1 | 1 |
| rs373869746 | pathogenic | 1 | 1 | 1 |
| rs777524402 | pathogenic | 1 | 1 | 1 |
| rs769737896 | pathogenic | 0 | 1 | 0 |
| rs13306515 | benign | 1 | 1 | 0 |
| rs746982741 | pathogenic | 0 | 0 | 0 |
| rs2230200 | benign | 1 | 1 | 1 |
| rs1047286 | benign | 0 | 0 | 0 |
| rs141737564 | benign | 0 | 0 | 0 |
| rs774826179 | benign | 0 | 0 | 0 |
| rs11569422 | benign | 1 | 1 | 1 |
| rs146613648 | benign | 0 | 0 | 0 |
| rs144432231 | benign | 0 | 0 | 0 |
| rs140655115 | benign | 0 | 0 | 0 |
| rs387906837 | pathogenic | 1 | 1 | 0 |
| rs121909076 | pathogenic | 1 | 1 | 0 |
| rs121909077 | pathogenic | 0 | 0 | 0 |
| rs121909074 | pathogenic | 0 | 0 | 0 |
| rs137853330 | pathogenic | 1 | 1 | 1 |
| rs137853331 | pathogenic | 1 | 1 | 1 |
| rs137853332 | pathogenic | 1 | 1 | 1 |
| rs28931586 | pathogenic | 1 | 1 | 0 |
| rs11569321 | benign | 1 | 1 | 1 |
| rs119486097 | benign | 0 | 0 | 0 |
| rs1799807 | pathogenic | 1 | 1 | 1 |
| rs201820739 | pathogenic | 1 | 1 | 0 |
| rs114706984 | pathogenic | 1 | 1 | 0 |
| rs140080572 | pathogenic | 1 | 1 | 0 |
| rs28933389 | pathogenic | 0 | 1 | 0 |
| rs16849700 | benign | 1 | 1 | 0 |
| rs115624085 | pathogenic | 1 | 1 | 1 |
| rs747196387 | pathogenic | 1 | 1 | 1 |
| rs104893684 | pathogenic | 1 | 1 | 0 |
| rs121918557 | pathogenic | 1 | 1 | 0 |
| rs121918556 | pathogenic | 1 | 1 | 1 |
| rs398124571 | pathogenic | 1 | 1 | 1 |
| rs776631396 | pathogenic | 0 | 1 | 0 |
| rs398124576 | pathogenic | 1 | 1 | 1 |

| | | | | |
|---|---|---|---|---|
| rs398124581 | pathogenic | 1 | 1 | 1 |
| rs398124582 | pathogenic | 1 | 1 | 1 |
| rs149766077 | pathogenic | 0 | 1 | 0 |
| rs79761867 | pathogenic | 0 | 1 | 0 |
| rs148905512 | pathogenic | 0 | 1 | 0 |
| rs121965004 | pathogenic | 1 | 1 | 1 |
| rs386834233 | pathogenic | 0 | 1 | 0 |
| rs398124598 | pathogenic | 0 | 1 | 0 |
| rs398124603 | pathogenic | 1 | 1 | 1 |
| rs546716073 | benign | 1 | 1 | 1 |
| rs200310662 | benign | 1 | 1 | 1 |
| rs115494727 | benign | 1 | 1 | 1 |
| rs781513008 | benign | 0 | 0 | 0 |
| rs587776398 | benign | 1 | 1 | 0 |
| rs35288908 | benign | 1 | 1 | 1 |
| rs150634562 | pathogenic | 1 | 1 | 0 |
| rs142948132 | pathogenic | 1 | 1 | 1 |
| rs132630321 | pathogenic | 1 | 1 | 1 |
| rs132630317 | pathogenic | 1 | 1 | 1 |
| rs132630320 | pathogenic | 0 | 0 | 0 |
| rs781394318 | pathogenic | 1 | 1 | 1 |
| rs111033624 | pathogenic | 1 | 1 | 1 |
| rs111033630 | pathogenic | 1 | 1 | 1 |
| rs111033627 | pathogenic | 1 | 1 | 1 |
| rs193922287 | pathogenic | 1 | 1 | 1 |
| rs759072800 | pathogenic | 0 | 1 | 0 |
| rs104894010 | pathogenic | 0 | 0 | 0 |
| rs193922296 | pathogenic | 0 | 0 | 0 |
| rs104894006 | pathogenic | 1 | 1 | 1 |
| rs193922311 | pathogenic | 1 | 1 | 0 |
| rs80356654 | pathogenic | 1 | 1 | 0 |
| rs144723656 | benign | 1 | 1 | 1 |
| rs148311934 | pathogenic | 1 | 1 | 0 |
| rs80356655 | pathogenic | 1 | 1 | 1 |
| rs769268803 | pathogenic | 1 | 1 | 0 |
| rs104894008 | pathogenic | 0 | 1 | 0 |
| rs104894005 | pathogenic | 1 | 1 | 1 |
| rs104894009 | pathogenic | 1 | 1 | 0 |
| rs193922340 | pathogenic | 0 | 0 | 0 |
| rs104894016 | pathogenic | 0 | 1 | 0 |
| rs193921338 | pathogenic | 0 | 0 | 0 |
| rs193922272 | pathogenic | 1 | 1 | 1 |
| rs104894012 | pathogenic | 1 | 1 | 0 |
| rs193922285 | pathogenic | 1 | 1 | 1 |
| rs104894336 | pathogenic | 1 | 1 | 1 |
| rs28931580 | pathogenic | 1 | 1 | 1 |
| rs104894326 | pathogenic | 1 | 1 | 1 |
| rs104894331 | pathogenic | 0 | 0 | 0 |
| rs104894338 | pathogenic | 0 | 0 | 0 |
| rs104894333 | pathogenic | 1 | 1 | 1 |
| rs104894330 | pathogenic | 1 | 1 | 1 |
| rs143886391 | benign | 1 | 1 | 1 |
| rs104894335 | pathogenic | 1 | 1 | 1 |
| rs104894337 | pathogenic | 0 | 0 | 0 |

| | | | | |
|---|---|---|---|---|
| rs104894328 | pathogenic | 0 | 0 | 0 |
| rs104894341 | pathogenic | 1 | 1 | 0 |
| rs104894329 | pathogenic | 0 | 0 | 0 |
| rs137852850 | pathogenic | 1 | 1 | 1 |
| rs137852854 | pathogenic | 0 | 1 | 0 |
| rs137852849 | pathogenic | 0 | 1 | 0 |
| rs137852848 | pathogenic | 1 | 1 | 1 |
| rs137852852 | pathogenic | 0 | 1 | 0 |
| rs137852853 | pathogenic | 0 | 1 | 0 |
| rs137852847 | pathogenic | 0 | 1 | 0 |
| rs137852846 | pathogenic | 0 | 1 | 0 |
| rs137852844 | pathogenic | 0 | 1 | 0 |
| rs144411158 | pathogenic | 0 | 0 | 0 |
| rs2302234 | benign | 0 | 0 | 0 |
| rs139620139 | pathogenic | 0 | 0 | 0 |
| rs146689929 | benign | 0 | 0 | 0 |
| rs373328681 | pathogenic | 1 | 1 | 0 |
| rs387906620 | pathogenic | 1 | 1 | 1 |
| rs1799983 | benign | 0 | 1 | 0 |
| rs28999111 | pathogenic | 1 | 1 | 1 |
| rs104894544 | pathogenic | 1 | 1 | 1 |
| rs104894546 | pathogenic | 1 | 1 | 1 |
| rs28999112 | pathogenic | 0 | 1 | 0 |
| rs104894545 | pathogenic | 1 | 1 | 1 |
| rs104894547 | pathogenic | 1 | 1 | 1 |
| rs74315310 | pathogenic | 0 | 0 | 0 |
| rs267606852 | pathogenic | 0 | 1 | 0 |
| rs137853586 | pathogenic | 0 | 1 | 0 |
| rs137853582 | pathogenic | 1 | 1 | 1 |
| rs61754634 | pathogenic | 1 | 1 | 1 |
| rs137853587 | pathogenic | 0 | 1 | 0 |
| rs267606851 | pathogenic | 1 | 1 | 1 |
| rs267606853 | pathogenic | 1 | 1 | 1 |
| rs137853584 | pathogenic | 1 | 1 | 1 |
| rs137853585 | pathogenic | 1 | 1 | 1 |
| rs460184 | pathogenic | 1 | 1 | 1 |
| rs104893751 | pathogenic | 0 | 1 | 0 |
| rs77358162 | benign | 0 | 0 | 0 |
| rs387906936 | pathogenic | 0 | 0 | 0 |
| rs769525399 | pathogenic | 0 | 0 | 0 |
| rs371271054 | pathogenic | 1 | 1 | 0 |
| rs118204000 | pathogenic | 0 | 0 | 0 |
| rs138568622 | pathogenic | 1 | 1 | 1 |
| rs34509359 | benign | 0 | 0 | 0 |
| rs104894130 | pathogenic | 1 | 1 | 0 |
| rs281865424 | pathogenic | 0 | 0 | 0 |
| rs121912778 | pathogenic | 1 | 1 | 0 |
| rs771055145 | pathogenic | 1 | 1 | 0 |
| rs776174514 | pathogenic | 1 | 1 | 0 |
| rs545986367 | pathogenic | 1 | 1 | 1 |
| rs570768038 | benign | 1 | 1 | 1 |
| rs74315442 | pathogenic | 1 | 1 | 1 |
| rs79716074 | benign | 1 | 1 | 1 |
| rs748486078 | pathogenic | 1 | 1 | 1 |

| | | | | |
|---|---|---|---|---|
| rs2397084 | benign | 1 | 1 | 1 |
| rs11465553 | benign | 1 | 1 | 1 |
| rs144576902 | benign | 1 | 1 | 1 |
| rs763780 | benign | 1 | 1 | 1 |
| rs267606870 | pathogenic | 1 | 1 | 0 |
| rs121913502 | pathogenic | 1 | 1 | 0 |
| rs118101777 | benign | 1 | 1 | 1 |
| rs118053940 | benign | 0 | 0 | 0 |
| rs368771578 | benign | 1 | 1 | 1 |
| rs11540654 | pathogenic | 1 | 1 | 1 |
| rs121912658 | pathogenic | 1 | 1 | 1 |
| rs28934873 | pathogenic | 0 | 1 | 0 |
| rs780442292 | pathogenic | 1 | 1 | 1 |
| rs758781593 | benign | 1 | 1 | 1 |
| rs28934875 | pathogenic | 1 | 1 | 1 |
| rs28934874 | pathogenic | 1 | 1 | 0 |
| rs587782705 | pathogenic | 1 | 1 | 1 |
| rs587780068 | pathogenic | 1 | 1 | 1 |
| rs587782144 | pathogenic | 1 | 1 | 1 |
| rs138729528 | pathogenic | 1 | 1 | 1 |
| rs28934578 | pathogenic | 1 | 1 | 1 |
| rs397514495 | pathogenic | 1 | 1 | 1 |
| rs121912665 | pathogenic | 1 | 1 | 1 |
| rs397516435 | pathogenic | 1 | 1 | 1 |
| rs587778720 | pathogenic | 1 | 1 | 1 |
| rs397516436 | pathogenic | 1 | 1 | 1 |
| rs587781386 | benign | 1 | 1 | 1 |
| rs121912666 | pathogenic | 1 | 1 | 0 |
| rs587780073 | pathogenic | 1 | 1 | 1 |
| rs587782664 | pathogenic | 1 | 1 | 1 |
| rs730882005 | pathogenic | 1 | 1 | 1 |
| rs28934573 | pathogenic | 1 | 1 | 1 |
| rs121912655 | pathogenic | 1 | 1 | 1 |
| rs28934575 | pathogenic | 1 | 1 | 1 |
| rs121912656 | pathogenic | 1 | 1 | 1 |
| rs11540652 | pathogenic | 1 | 1 | 1 |
| rs121912651 | pathogenic | 1 | 1 | 1 |
| rs730882027 | pathogenic | 1 | 1 | 1 |
| rs121912653 | pathogenic | 1 | 1 | 1 |
| rs121912652 | pathogenic | 1 | 1 | 1 |
| rs121912657 | pathogenic | 0 | 1 | 0 |
| rs121913343 | pathogenic | 1 | 1 | 1 |
| rs28934576 | pathogenic | 1 | 1 | 1 |
| rs763098116 | pathogenic | 1 | 1 | 1 |
| rs121912660 | pathogenic | 1 | 1 | 1 |
| rs28934574 | pathogenic | 1 | 1 | 1 |
| rs371409680 | pathogenic | 1 | 1 | 1 |
| rs121912667 | pathogenic | 1 | 1 | 1 |
| rs121912663 | pathogenic | 1 | 1 | 1 |
| rs201744589 | pathogenic | 0 | 0 | 0 |
| rs33926449 | benign | 0 | 0 | 0 |
| rs74315302 | pathogenic | 0 | 1 | 0 |
| rs202040269 | pathogenic | 1 | 1 | 1 |
| rs375958814 | pathogenic | 1 | 1 | 1 |

| | | | | |
|---|---|---|---|---|
| rs121909624 | pathogenic | 1 | 1 | 1 |
| rs121909621 | pathogenic | 1 | 1 | 1 |
| rs121909622 | pathogenic | 1 | 1 | 0 |
| rs4220 | benign | 1 | 1 | 1 |
| rs147559466 | pathogenic | 1 | 1 | 1 |
| rs757434857 | pathogenic | 1 | 1 | 1 |
| rs398123072 | pathogenic | 1 | 1 | 1 |
| rs121434280 | pathogenic | 0 | 1 | 0 |
| rs398123074 | pathogenic | 0 | 1 | 0 |
| rs762114560 | pathogenic | 1 | 1 | 1 |
| rs121434283 | pathogenic | 1 | 1 | 0 |
| rs121434277 | pathogenic | 0 | 1 | 0 |
| rs745844469 | pathogenic | 1 | 1 | 1 |
| rs121434279 | pathogenic | 1 | 1 | 0 |
| rs121434278 | pathogenic | 1 | 1 | 1 |
| rs373715782 | pathogenic | 1 | 1 | 1 |
| rs200724875 | pathogenic | 1 | 1 | 1 |
| rs149678400 | pathogenic | 0 | 1 | 0 |
| rs770273135 | pathogenic | 1 | 1 | 1 |
| rs121434276 | pathogenic | 0 | 1 | 0 |
| rs121434281 | pathogenic | 1 | 1 | 0 |
| rs768884003 | pathogenic | 1 | 1 | 1 |
| rs201375579 | pathogenic | 0 | 0 | 0 |
| rs121434274 | pathogenic | 1 | 1 | 0 |
| rs121434282 | pathogenic | 1 | 1 | 0 |
| rs779759347 | pathogenic | 0 | 1 | 0 |
| rs77931234 | pathogenic | 1 | 1 | 1 |
| rs148207467 | pathogenic | 1 | 1 | 1 |
| rs766140986 | pathogenic | 1 | 1 | 1 |
| rs121434275 | pathogenic | 1 | 1 | 1 |
| rs748110745 | pathogenic | 1 | 1 | 1 |
| rs104894274 | pathogenic | 1 | 1 | 1 |
| rs104894273 | pathogenic | 0 | 1 | 0 |
| rs104894278 | pathogenic | 1 | 1 | 1 |
| rs104894275 | pathogenic | 1 | 1 | 1 |
| rs104894277 | pathogenic | 1 | 1 | 1 |
| rs104894276 | pathogenic | 0 | 0 | 0 |
| rs104894280 | pathogenic | 1 | 1 | 1 |
| rs145882709 | pathogenic | 1 | 1 | 1 |
| rs104894279 | pathogenic | 1 | 1 | 1 |
| rs80356670 | pathogenic | 1 | 1 | 1 |
| rs121918102 | pathogenic | 1 | 1 | 1 |
| rs80356671 | pathogenic | 1 | 1 | 1 |
| rs80356672 | pathogenic | 1 | 1 | 1 |
| rs758609113 | pathogenic | 1 | 1 | 0 |
| rs121918188 | pathogenic | 0 | 1 | 0 |
| rs121918189 | pathogenic | 1 | 1 | 0 |
| rs121918191 | pathogenic | 1 | 1 | 1 |
| rs1769259 | benign | 1 | 1 | 1 |
| rs140222720 | benign | 1 | 1 | 1 |
| rs2297085 | benign | 1 | 1 | 1 |
| rs200679026 | pathogenic | 1 | 1 | 1 |
| rs121918192 | pathogenic | 0 | 1 | 0 |
| rs145771944 | benign | 0 | 1 | 0 |

| | | | | |
|---|---|---|---|---|
| rs201508880 | pathogenic | 0 | 1 | 0 |
| rs17221854 | pathogenic | 0 | 0 | 0 |
| rs120074189 | pathogenic | 1 | 1 | 1 |
| rs120074190 | pathogenic | 1 | 1 | 1 |
| rs758600283 | benign | 1 | 1 | 1 |
| rs200305064 | pathogenic | 1 | 1 | 1 |
| rs121917891 | pathogenic | 1 | 1 | 1 |
| rs3772809 | benign | 0 | 1 | 0 |
| rs387906619 | pathogenic | 1 | 1 | 0 |
| rs2304681 | benign | 0 | 1 | 0 |
| rs121917849 | pathogenic | 0 | 0 | 0 |
| rs143010236 | pathogenic | 0 | 0 | 0 |
| rs121917850 | pathogenic | 1 | 1 | 0 |
| rs35916840 | pathogenic | 1 | 1 | 0 |
| rs28939688 | pathogenic | 1 | 1 | 1 |
| rs11258194 | benign | 1 | 1 | 1 |
| rs199840952 | pathogenic | 1 | 1 | 1 |
| rs121918159 | pathogenic | 1 | 1 | 1 |
| rs200396166 | pathogenic | 0 | 0 | 0 |
| rs757966746 | pathogenic | 0 | 1 | 0 |
| rs121918608 | pathogenic | 0 | 1 | 0 |
| rs74315401 | pathogenic | 1 | 1 | 1 |
| rs11538758 | pathogenic | 0 | 0 | 0 |
| rs74315402 | pathogenic | 1 | 1 | 1 |
| rs1799990 | pathogenic | 1 | 1 | 1 |
| rs74315410 | pathogenic | 1 | 1 | 1 |
| rs150351644 | benign | 1 | 1 | 1 |
| rs16990018 | benign | 1 | 1 | 1 |
| rs74315403 | pathogenic | 1 | 1 | 1 |
| rs74315408 | pathogenic | 1 | 1 | 1 |
| rs74315411 | pathogenic | 1 | 1 | 1 |
| rs74315413 | pathogenic | 1 | 1 | 1 |
| rs74315405 | pathogenic | 1 | 1 | 1 |
| rs28933385 | pathogenic | 1 | 1 | 1 |
| rs74315412 | pathogenic | 1 | 1 | 1 |
| rs74315407 | pathogenic | 1 | 1 | 1 |
| rs74315406 | pathogenic | 1 | 1 | 1 |
| rs11571117 | benign | 1 | 1 | 1 |
| rs121917742 | pathogenic | 1 | 1 | 1 |
| rs149782619 | pathogenic | 1 | 1 | 1 |
| rs397514492 | pathogenic | 1 | 1 | 1 |
| rs397514489 | pathogenic | 0 | 1 | 0 |
| rs397514493 | pathogenic | 1 | 1 | 1 |
| rs373849532 | pathogenic | 1 | 1 | 1 |
| rs121965048 | pathogenic | 1 | 1 | 1 |
| rs121965037 | pathogenic | 1 | 1 | 1 |
| rs121965060 | pathogenic | 0 | 1 | 0 |
| rs386833603 | pathogenic | 1 | 1 | 1 |
| rs121965038 | pathogenic | 1 | 1 | 1 |
| rs121965039 | pathogenic | 1 | 1 | 1 |
| rs121965040 | pathogenic | 1 | 1 | 1 |
| rs267606925 | pathogenic | 1 | 1 | 1 |
| rs121965056 | pathogenic | 0 | 1 | 0 |
| rs121965059 | pathogenic | 0 | 0 | 0 |

| | | | | |
|---|---|---|---|---|
| rs386833615 | pathogenic | 1 | 1 | 0 |
| rs121965051 | pathogenic | 1 | 1 | 0 |
| rs121965046 | pathogenic | 0 | 1 | 0 |
| rs121965052 | pathogenic | 0 | 1 | 0 |
| rs386833617 | pathogenic | 0 | 1 | 0 |
| rs386833618 | pathogenic | 1 | 1 | 0 |
| rs121965041 | pathogenic | 0 | 1 | 0 |
| rs121965042 | pathogenic | 0 | 1 | 0 |
| rs121965057 | pathogenic | 1 | 1 | 1 |
| rs386833621 | pathogenic | 1 | 1 | 1 |
| rs121965049 | pathogenic | 1 | 1 | 1 |
| rs386833623 | pathogenic | 1 | 1 | 1 |
| rs121965047 | pathogenic | 1 | 1 | 1 |
| rs121965053 | pathogenic | 0 | 1 | 0 |
| rs121965045 | pathogenic | 0 | 1 | 0 |
| rs121965054 | pathogenic | 0 | 1 | 0 |
| rs121965043 | pathogenic | 0 | 1 | 0 |
| rs386833598 | pathogenic | 0 | 0 | 0 |
| rs137854545 | pathogenic | 1 | 1 | 1 |
| rs137854544 | pathogenic | 1 | 1 | 1 |
| rs104894455 | pathogenic | 0 | 1 | 0 |
| rs1049564 | benign | 0 | 1 | 0 |
| rs104894460 | pathogenic | 1 | 1 | 1 |
| rs104894453 | pathogenic | 1 | 1 | 1 |
| rs104894450 | pathogenic | 0 | 1 | 0 |
| rs104894454 | pathogenic | 0 | 1 | 0 |
| rs104894452 | pathogenic | 1 | 1 | 1 |
| rs104894451 | pathogenic | 0 | 1 | 0 |
| rs104894368 | pathogenic | 1 | 1 | 1 |
| rs730880950 | pathogenic | 1 | 1 | 1 |
| rs121909535 | pathogenic | 1 | 1 | 1 |
| rs121909537 | pathogenic | 1 | 1 | 1 |
| rs121909536 | pathogenic | 1 | 1 | 1 |
| rs121909542 | pathogenic | 1 | 1 | 1 |
| rs121909538 | pathogenic | 1 | 1 | 1 |
| rs121909539 | pathogenic | 1 | 1 | 1 |
| rs121909540 | pathogenic | 1 | 1 | 1 |
| rs17560 | benign | 1 | 1 | 1 |
| rs121909543 | pathogenic | 0 | 1 | 0 |
| rs121909544 | pathogenic | 1 | 1 | 1 |
| rs2307456 | benign | 0 | 1 | 0 |
| rs745778317 | pathogenic | 0 | 1 | 0 |
| rs759607901 | pathogenic | 0 | 1 | 0 |
| rs201188114 | benign | 0 | 0 | 0 |
| rs1800472 | benign | 0 | 0 | 0 |
| rs587780728 | benign | 1 | 1 | 1 |
| rs144386518 | benign | 1 | 1 | 1 |
| rs281865439 | pathogenic | 1 | 1 | 1 |
| rs140175796 | pathogenic | 1 | 1 | 1 |
| rs5030843 | pathogenic | 1 | 1 | 0 |
| rs75166491 | pathogenic | 1 | 1 | 0 |
| rs199475679 | pathogenic | 1 | 1 | 1 |
| rs281865440 | pathogenic | 1 | 1 | 1 |
| rs74486803 | pathogenic | 0 | 1 | 0 |

| | | | | |
|---|---|---|---|---|
| rs199475602 | pathogenic | 0 | 1 | 0 |
| rs5030844 | pathogenic | 1 | 1 | 0 |
| rs62514927 | pathogenic | 1 | 1 | 1 |
| rs62514928 | pathogenic | 1 | 1 | 1 |
| rs62516152 | pathogenic | 0 | 1 | 0 |
| rs62507348 | pathogenic | 1 | 1 | 1 |
| rs76687508 | pathogenic | 1 | 1 | 1 |
| rs62508730 | pathogenic | 1 | 1 | 1 |
| rs62508588 | pathogenic | 0 | 1 | 0 |
| rs5030846 | pathogenic | 0 | 1 | 0 |
| rs74503222 | pathogenic | 1 | 1 | 0 |
| rs5030847 | pathogenic | 1 | 1 | 0 |
| rs5030850 | pathogenic | 1 | 1 | 1 |
| rs778154939 | pathogenic | 0 | 1 | 0 |
| rs62514952 | pathogenic | 1 | 1 | 1 |
| rs78655458 | pathogenic | 1 | 1 | 1 |
| rs62508698 | pathogenic | 1 | 1 | 0 |
| rs5030851 | pathogenic | 1 | 1 | 0 |
| rs62642933 | pathogenic | 1 | 1 | 1 |
| rs5030853 | pathogenic | 1 | 1 | 1 |
| rs62642935 | pathogenic | 1 | 1 | 1 |
| rs199475615 | pathogenic | 0 | 0 | 0 |
| rs62516062 | pathogenic | 1 | 1 | 0 |
| rs62516092 | pathogenic | 0 | 0 | 0 |
| rs62516095 | pathogenic | 1 | 1 | 1 |
| rs62642937 | pathogenic | 1 | 1 | 1 |
| rs62516141 | pathogenic | 0 | 1 | 0 |
| rs5030856 | pathogenic | 1 | 1 | 1 |
| rs5030857 | pathogenic | 1 | 1 | 1 |
| rs5030858 | pathogenic | 1 | 1 | 1 |
| rs79931499 | pathogenic | 1 | 1 | 1 |
| rs5030860 | pathogenic | 1 | 1 | 1 |
| rs281865437 | pathogenic | 1 | 1 | 1 |
| rs62644499 | pathogenic | 1 | 1 | 1 |
| rs147689487 | benign | 0 | 0 | 0 |
| rs397515417 | pathogenic | 1 | 1 | 1 |
| rs121909332 | pathogenic | 1 | 1 | 1 |
| rs121909330 | pathogenic | 1 | 1 | 0 |
| rs121909329 | pathogenic | 1 | 1 | 0 |
| rs121909335 | pathogenic | 1 | 1 | 1 |
| rs387906789 | pathogenic | 1 | 1 | 1 |
| rs121909334 | pathogenic | 1 | 1 | 1 |
| rs121909331 | pathogenic | 1 | 1 | 1 |
| rs372487178 | pathogenic | 1 | 1 | 1 |
| rs387906937 | pathogenic | 1 | 1 | 0 |
| rs16845 | benign | 1 | 1 | 1 |
| rs672601313 | pathogenic | 1 | 1 | 1 |
| rs117805308 | benign | 0 | 0 | 0 |
| rs121434261 | pathogenic | 1 | 1 | 1 |
| rs141629512 | benign | 0 | 0 | 0 |
| rs199957176 | benign | 1 | 1 | 1 |
| rs2230461 | benign | 1 | 1 | 1 |
| rs764618040 | pathogenic | 1 | 1 | 1 |
| rs267606949 | pathogenic | 1 | 1 | 1 |

| | | | | |
|---|---|---|---|---|
| rs587777483 | pathogenic | 1 | 1 | 1 |
| rs267606947 | pathogenic | 1 | 1 | 1 |
| rs6941583 | benign | 0 | 0 | 0 |
| rs749597959 | benign | 1 | 1 | 1 |
| rs182340753 | benign | 1 | 1 | 1 |
| rs146678221 | benign | 0 | 0 | 0 |
| rs11539202 | benign | 0 | 0 | 0 |
| rs148723565 | benign | 0 | 0 | 0 |
| rs61752925 | benign | 1 | 1 | 1 |
| rs113309941 | pathogenic | 0 | 0 | 0 |
| rs121434520 | pathogenic | 1 | 1 | 1 |
| rs121434517 | pathogenic | 1 | 1 | 1 |
| rs267607027 | pathogenic | 1 | 1 | 1 |
| rs138603088 | pathogenic | 0 | 0 | 0 |
| rs121434516 | pathogenic | 1 | 1 | 0 |
| rs121434521 | pathogenic | 0 | 0 | 0 |
| rs145735112 | benign | 1 | 1 | 1 |
| rs121434519 | pathogenic | 1 | 1 | 1 |
| rs75326924 | pathogenic | 1 | 1 | 0 |
| rs142186404 | pathogenic | 0 | 0 | 0 |
| rs121918035 | pathogenic | 0 | 0 | 0 |
| rs374385878 | pathogenic | 1 | 1 | 0 |
| rs11557652 | benign | 0 | 0 | 0 |
| rs118203986 | pathogenic | 1 | 1 | 1 |
| rs141675330 | benign | 1 | 1 | 1 |
| rs104893927 | pathogenic | 0 | 0 | 0 |
| rs104893935 | pathogenic | 0 | 0 | 0 |
| rs104893933 | pathogenic | 1 | 1 | 0 |
| rs104893934 | pathogenic | 0 | 0 | 0 |
| rs145183203 | benign | 1 | 1 | 0 |
| rs2230806 | benign | 0 | 0 | 0 |
| rs9282541 | benign | 0 | 0 | 0 |
| rs559110055 | benign | 0 | 0 | 0 |
| rs200030513 | benign | 0 | 0 | 0 |
| rs9282543 | benign | 0 | 0 | 0 |
| rs147675550 | benign | 0 | 0 | 0 |
| rs137854496 | pathogenic | 0 | 0 | 0 |
| rs374190304 | benign | 0 | 0 | 0 |
| rs114620717 | benign | 0 | 0 | 0 |
| rs2066718 | benign | 0 | 0 | 0 |
| rs35819696 | benign | 0 | 0 | 0 |
| rs138880920 | benign | 0 | 0 | 0 |
| rs2066715 | benign | 0 | 0 | 0 |
| rs35207495 | benign | 0 | 0 | 0 |
| rs2066714 | benign | 0 | 0 | 0 |
| rs28937313 | pathogenic | 0 | 0 | 0 |
| rs28937314 | pathogenic | 0 | 0 | 0 |
| rs137854495 | pathogenic | 0 | 0 | 0 |
| rs140365800 | benign | 0 | 0 | 0 |
| rs13306073 | benign | 0 | 0 | 0 |
| rs28933692 | pathogenic | 0 | 0 | 0 |
| rs33918808 | benign | 0 | 0 | 0 |
| rs76881554 | benign | 0 | 0 | 0 |
| rs143180998 | benign | 0 | 0 | 0 |

| | | | | |
|---|---|---|---|---|
| rs137854500 | pathogenic | 0 | 0 | 0 |
| rs538614702 | benign | 0 | 0 | 0 |
| rs201879964 | benign | 0 | 0 | 0 |
| rs2230808 | benign | 0 | 0 | 0 |
| rs138422574 | benign | 0 | 0 | 0 |
| rs137854498 | pathogenic | 0 | 0 | 0 |
| rs150125857 | benign | 0 | 0 | 0 |
| rs146292819 | pathogenic | 0 | 0 | 0 |
| rs142688906 | benign | 0 | 0 | 0 |
| rs137854499 | pathogenic | 0 | 0 | 0 |
| rs34879708 | benign | 0 | 0 | 0 |
| rs121913028 | pathogenic | 1 | 1 | 1 |
| rs1041951 | benign | 1 | 1 | 0 |
| rs138714094 | benign | 1 | 1 | 0 |
| rs200650502 | benign | 0 | 1 | 0 |
| rs370708663 | pathogenic | 1 | 1 | 1 |
| rs118204040 | pathogenic | 0 | 1 | 0 |
| rs146899669 | pathogenic | 1 | 1 | 0 |
| rs118204039 | pathogenic | 0 | 1 | 0 |
| rs121909233 | pathogenic | 0 | 0 | 0 |
| rs121909225 | pathogenic | 1 | 1 | 0 |
| rs121909236 | pathogenic | 0 | 0 | 0 |
| rs398123317 | pathogenic | 0 | 0 | 0 |
| rs121909226 | pathogenic | 1 | 1 | 0 |
| rs121909238 | pathogenic | 1 | 1 | 0 |
| rs121909230 | pathogenic | 0 | 0 | 0 |
| rs121909237 | pathogenic | 0 | 0 | 0 |
| rs121909222 | pathogenic | 1 | 1 | 0 |
| rs121909223 | pathogenic | 1 | 1 | 0 |
| rs121909218 | pathogenic | 1 | 1 | 0 |
| rs121909229 | pathogenic | 1 | 1 | 0 |
| rs121909224 | pathogenic | 1 | 1 | 0 |
| rs121909241 | pathogenic | 0 | 0 | 0 |
| rs121909221 | pathogenic | 0 | 0 | 0 |
| rs121913294 | pathogenic | 0 | 0 | 0 |
| rs121909234 | pathogenic | 0 | 0 | 0 |
| rs121909240 | pathogenic | 0 | 0 | 0 |
| rs767623493 | benign | 0 | 0 | 0 |
| rs121909239 | pathogenic | 0 | 0 | 0 |
| rs120074145 | pathogenic | 1 | 1 | 1 |
| rs120074148 | pathogenic | 1 | 1 | 1 |
| rs762991875 | pathogenic | 0 | 0 | 0 |
| rs148639841 | pathogenic | 1 | 1 | 1 |
| rs199524907 | pathogenic | 1 | 1 | 1 |
| rs120074141 | pathogenic | 1 | 1 | 1 |
| rs120074144 | pathogenic | 0 | 1 | 0 |
| rs120074146 | pathogenic | 1 | 1 | 1 |
| rs120074147 | pathogenic | 0 | 1 | 0 |
| rs120074143 | pathogenic | 0 | 1 | 0 |
| rs120074140 | pathogenic | 1 | 1 | 0 |
| rs28937315 | pathogenic | 1 | 1 | 1 |
| rs76066357 | benign | 1 | 1 | 1 |
| rs137852911 | pathogenic | 0 | 1 | 0 |
| rs137852907 | pathogenic | 1 | 1 | 0 |

| | | | | |
|---|---|---|---|---|
| rs137852910 | pathogenic | 1 | 1 | 1 |
| rs537585211 | benign | 0 | 0 | 0 |
| rs202134424 | pathogenic | 0 | 0 | 0 |
| rs200930463 | pathogenic | 0 | 0 | 0 |
| rs587777241 | pathogenic | 1 | 1 | 0 |
| rs77563738 | pathogenic | 0 | 0 | 0 |
| rs750868279 | pathogenic | 1 | 1 | 0 |
| rs148936893 | pathogenic | 0 | 0 | 0 |
| rs146597836 | pathogenic | 0 | 0 | 0 |
| rs775440641 | pathogenic | 1 | 1 | 1 |
| rs770689762 | pathogenic | 0 | 1 | 0 |
| rs376785840 | pathogenic | 0 | 0 | 0 |
| rs137852531 | pathogenic | 0 | 0 | 0 |
| rs137852538 | pathogenic | 1 | 1 | 0 |
| rs137852529 | pathogenic | 0 | 1 | 0 |
| rs137852528 | pathogenic | 0 | 1 | 0 |
| rs137852535 | pathogenic | 1 | 1 | 0 |
| rs137853045 | pathogenic | 0 | 1 | 0 |
| rs137853046 | pathogenic | 1 | 1 | 1 |
| rs137853043 | pathogenic | 1 | 1 | 1 |
| rs137853044 | pathogenic | 1 | 1 | 1 |
| rs137853047 | pathogenic | 1 | 1 | 1 |
| rs267606990 | pathogenic | 1 | 1 | 1 |
| rs751437780 | pathogenic | 1 | 1 | 0 |
| rs121918461 | pathogenic | 1 | 1 | 1 |
| rs121918460 | pathogenic | 1 | 1 | 1 |
| rs121918459 | pathogenic | 1 | 1 | 0 |
| rs397507511 | pathogenic | 1 | 1 | 1 |
| rs121918454 | pathogenic | 1 | 1 | 1 |
| rs121918453 | pathogenic | 1 | 1 | 1 |
| rs121918462 | pathogenic | 1 | 1 | 0 |
| rs121918465 | pathogenic | 1 | 1 | 0 |
| rs121918464 | pathogenic | 1 | 1 | 0 |
| rs121918466 | pathogenic | 1 | 1 | 1 |
| rs397507521 | pathogenic | 1 | 1 | 1 |
| rs397507527 | pathogenic | 1 | 1 | 1 |
| rs777603059 | pathogenic | 0 | 1 | 0 |
| rs121918456 | pathogenic | 1 | 1 | 1 |
| rs121918463 | pathogenic | 1 | 1 | 1 |
| rs121918455 | pathogenic | 0 | 1 | 0 |
| rs28933386 | pathogenic | 0 | 1 | 0 |
| rs774356443 | benign | 1 | 1 | 1 |
| rs121918467 | pathogenic | 1 | 1 | 1 |
| rs121918468 | pathogenic | 1 | 1 | 1 |
| rs121918469 | pathogenic | 1 | 1 | 1 |
| rs397507539 | pathogenic | 1 | 1 | 0 |
| rs121918458 | pathogenic | 1 | 1 | 1 |
| rs121918470 | pathogenic | 1 | 1 | 1 |
| rs35289681 | benign | 0 | 1 | 0 |
| rs4302331 | benign | 0 | 1 | 0 |
| rs11544803 | pathogenic | 1 | 1 | 1 |
| rs116254156 | benign | 0 | 0 | 0 |
| rs104893909 | pathogenic | 1 | 1 | 1 |
| rs104893911 | pathogenic | 1 | 1 | 1 |

| | | | | |
|---|---|---|---|---|
| rs104893908 | pathogenic | 1 | 1 | 0 |
| rs258751 | benign | 1 | 1 | 1 |
| rs104893914 | pathogenic | 1 | 1 | 0 |
| rs121909727 | pathogenic | 1 | 1 | 1 |
| rs104893910 | pathogenic | 1 | 1 | 1 |
| rs121909726 | pathogenic | 1 | 1 | 1 |
| rs104893912 | pathogenic | 1 | 1 | 1 |
| rs138407155 | benign | 1 | 1 | 1 |
| rs574061789 | benign | 1 | 1 | 1 |
| rs121912718 | pathogenic | 1 | 1 | 1 |
| rs121912719 | pathogenic | 1 | 1 | 1 |
| rs140770089 | benign | 1 | 1 | 1 |
| rs28931573 | pathogenic | 1 | 1 | 1 |
| rs121912717 | pathogenic | 1 | 1 | 1 |
| rs376456598 | pathogenic | 0 | 0 | 0 |
| rs398123499 | pathogenic | 1 | 1 | 1 |
| rs773048903 | pathogenic | 1 | 1 | 0 |
| rs398123503 | pathogenic | 1 | 1 | 1 |
| rs375785084 | pathogenic | 0 | 1 | 0 |
| rs137852874 | pathogenic | 1 | 1 | 1 |
| rs373713279 | pathogenic | 1 | 1 | 1 |
| rs137852873 | pathogenic | 1 | 1 | 0 |
| rs367823977 | pathogenic | 1 | 1 | 1 |
| rs398123508 | pathogenic | 1 | 1 | 1 |
| rs137852871 | pathogenic | 1 | 1 | 1 |
| rs753698250 | pathogenic | 0 | 1 | 0 |
| rs182923857 | pathogenic | 1 | 1 | 1 |
| rs148090804 | benign | 1 | 1 | 1 |
| rs137852870 | pathogenic | 1 | 1 | 1 |
| rs267607131 | pathogenic | 1 | 1 | 0 |
| rs137852902 | pathogenic | 0 | 0 | 0 |
| rs61741646 | benign | 1 | 1 | 1 |
| rs137852905 | pathogenic | 0 | 0 | 0 |
| rs121918681 | pathogenic | 0 | 1 | 0 |
| rs8177238 | benign | 1 | 1 | 0 |
| rs5030776 | pathogenic | 1 | 1 | 1 |
| rs104894572 | pathogenic | 0 | 0 | 0 |
| rs104894576 | pathogenic | 0 | 0 | 0 |
| rs80084721 | pathogenic | 0 | 0 | 0 |
| rs104894146 | pathogenic | 0 | 1 | 0 |
| rs104894138 | pathogenic | 1 | 1 | 0 |
| rs104894153 | pathogenic | 1 | 1 | 0 |
| rs104894135 | pathogenic | 1 | 1 | 0 |
| rs104894147 | pathogenic | 1 | 1 | 0 |
| rs104894148 | pathogenic | 1 | 1 | 0 |
| rs104894154 | pathogenic | 1 | 1 | 0 |
| rs104894150 | pathogenic | 1 | 1 | 0 |
| rs104894136 | pathogenic | 1 | 1 | 0 |
| rs104894144 | pathogenic | 0 | 0 | 0 |
| rs104894137 | pathogenic | 0 | 0 | 0 |
| rs61754278 | pathogenic | 1 | 1 | 0 |
| rs104894149 | pathogenic | 1 | 1 | 0 |
| rs104894139 | pathogenic | 1 | 1 | 0 |
| rs104894142 | pathogenic | 0 | 0 | 0 |

| | | | | |
|---|---|---|---|---|
| rs104894143 | pathogenic | 0 | 0 | 0 |
| rs104894145 | pathogenic | 0 | 0 | 0 |
| rs121909545 | benign | 0 | 1 | 0 |
| rs782308462 | pathogenic | 1 | 1 | 1 |
| rs375002796 | pathogenic | 0 | 1 | 0 |
| rs201727685 | pathogenic | 0 | 0 | 0 |
| rs768050261 | pathogenic | 1 | 1 | 1 |
| rs780851340 | benign | 0 | 0 | 0 |
| rs139824241 | benign | 1 | 1 | 0 |
| rs757486575 | pathogenic | 1 | 1 | 0 |
| rs750831299 | pathogenic | 1 | 1 | 1 |
| rs767602961 | pathogenic | 1 | 1 | 1 |
| rs121913661 | pathogenic | 1 | 1 | 1 |
| rs777692271 | pathogenic | 1 | 1 | 1 |
| rs121913659 | pathogenic | 1 | 1 | 1 |
| rs587777533 | pathogenic | 1 | 1 | 1 |
| rs104894210 | pathogenic | 0 | 0 | 0 |
| rs104894216 | pathogenic | 1 | 1 | 1 |
| rs104894207 | pathogenic | 1 | 1 | 1 |
| rs104894208 | pathogenic | 0 | 1 | 0 |
| rs104894214 | pathogenic | 1 | 1 | 1 |
| rs104894211 | pathogenic | 1 | 1 | 1 |
| rs61749384 | pathogenic | 0 | 0 | 0 |
| rs61750071 | pathogenic | 1 | 1 | 1 |
| rs61750074 | pathogenic | 0 | 0 | 0 |
| rs11063988 | benign | 0 | 0 | 0 |
| rs216311 | benign | 0 | 0 | 0 |
| rs1800382 | benign | 1 | 1 | 1 |
| rs11063987 | benign | 1 | 1 | 1 |
| rs387906847 | pathogenic | 1 | 1 | 1 |
| rs73003466 | benign | 0 | 0 | 0 |
| rs149787233 | benign | 1 | 1 | 1 |
| rs144451841 | pathogenic | 1 | 1 | 1 |
| rs141638421 | pathogenic | 1 | 1 | 1 |
| rs121434497 | pathogenic | 0 | 0 | 0 |
| rs121434498 | pathogenic | 0 | 0 | 0 |
| rs267607230 | pathogenic | 1 | 1 | 0 |
| rs121434499 | pathogenic | 1 | 1 | 0 |
| rs546749336 | benign | 1 | 1 | 1 |
| rs138873805 | benign | 0 | 0 | 0 |
| rs730880522 | pathogenic | 0 | 0 | 0 |
| rs200371894 | benign | 0 | 0 | 0 |
| rs730880509 | benign | 0 | 0 | 0 |
| rs267606704 | pathogenic | 1 | 1 | 1 |
| rs267606706 | pathogenic | 0 | 0 | 0 |
| rs267606705 | pathogenic | 1 | 1 | 1 |
| rs267606707 | pathogenic | 0 | 0 | 0 |
| rs387906665 | pathogenic | 1 | 1 | 1 |
| rs267606708 | pathogenic | 1 | 1 | 1 |
| rs148368481 | benign | 0 | 0 | 0 |
| rs13204 | benign | 0 | 0 | 0 |
| rs140547520 | pathogenic | 1 | 1 | 1 |
| rs755234990 | benign | 1 | 1 | 0 |
| rs151184052 | benign | 0 | 0 | 0 |

| | | | | |
|---|---|---|---|---|
| rs3736358 | benign | 0 | 0 | 0 |
| rs3736360 | benign | 0 | 0 | 0 |
| rs1138469 | benign | 0 | 0 | 0 |
| rs74315351 | pathogenic | 0 | 1 | 0 |
| rs137853051 | pathogenic | 1 | 1 | 0 |
| rs74315353 | pathogenic | 0 | 1 | 0 |
| rs74315352 | pathogenic | 0 | 1 | 0 |
| rs74315354 | pathogenic | 1 | 1 | 1 |
| rs28938172 | pathogenic | 1 | 1 | 1 |
| rs52836744 | pathogenic | 0 | 0 | 0 |
| rs121913159 | pathogenic | 1 | 1 | 1 |
| rs121913145 | pathogenic | 1 | 1 | 1 |
| rs891087 | benign | 1 | 1 | 1 |
| rs9282757 | benign | 1 | 1 | 1 |
| rs397517404 | pathogenic | 1 | 1 | 1 |
| rs199918720 | benign | 1 | 1 | 0 |
| rs368299411 | benign | 1 | 1 | 0 |
| rs139399951 | benign | 1 | 1 | 0 |
| rs763981974 | benign | 1 | 1 | 0 |
| rs144242114 | benign | 0 | 0 | 0 |
| rs750965801 | benign | 0 | 0 | 0 |
| rs561310777 | benign | 0 | 0 | 0 |
| rs148185335 | benign | 0 | 0 | 0 |
| rs147742157 | benign | 0 | 0 | 0 |
| rs375664373 | pathogenic | 1 | 1 | 1 |
| rs777327250 | pathogenic | 0 | 0 | 0 |
| rs121908471 | pathogenic | 1 | 1 | 0 |
| rs2301612 | pathogenic | 1 | 1 | 0 |
| rs121908476 | pathogenic | 0 | 0 | 0 |
| rs36220239 | benign | 0 | 0 | 0 |
| rs121908473 | pathogenic | 0 | 0 | 0 |
| rs28647808 | benign | 0 | 0 | 0 |
| rs36090624 | benign | 0 | 0 | 0 |
| rs121908002 | pathogenic | 1 | 1 | 1 |
| rs4251545 | benign | 1 | 1 | 1 |
| rs78705382 | pathogenic | 0 | 0 | 0 |
| rs75184679 | pathogenic | 1 | 1 | 1 |
| rs74555752 | pathogenic | 0 | 0 | 0 |
| rs56320207 | benign | 1 | 1 | 1 |
| rs28730837 | pathogenic | 1 | 1 | 1 |
| rs119469014 | pathogenic | 1 | 1 | 1 |
| rs119468010 | pathogenic | 1 | 1 | 1 |
| rs35220041 | benign | 0 | 0 | 0 |
| rs762321137 | pathogenic | 1 | 1 | 1 |
| rs28931570 | pathogenic | 1 | 1 | 1 |
| rs709932 | benign | 1 | 1 | 1 |
| rs746531546 | benign | 1 | 1 | 1 |
| rs6647 | benign | 1 | 1 | 1 |
| rs141095970 | benign | 1 | 1 | 1 |
| rs143370956 | benign | 1 | 1 | 1 |
| rs28929474 | pathogenic | 1 | 1 | 1 |
| rs61761869 | pathogenic | 1 | 1 | 1 |
| rs1303 | benign | 1 | 1 | 1 |
| rs149447502 | benign | 0 | 0 | 0 |

| | | | | |
|---|---|---|---|---|
| rs267607180 | pathogenic | 1 | 1 | 1 |
| rs150304121 | benign | 0 | 0 | 0 |
| rs147630375 | benign | 1 | 1 | 1 |
| rs11539445 | benign | 1 | 1 | 1 |
| rs372098364 | pathogenic | 1 | 1 | 0 |
| rs760072286 | benign | 0 | 0 | 0 |
| rs745595833 | pathogenic | 1 | 1 | 1 |
| rs587777214 | pathogenic | 0 | 0 | 0 |
| rs150318528 | benign | 0 | 0 | 0 |
| rs137853254 | pathogenic | 1 | 1 | 1 |
| rs137853255 | pathogenic | 0 | 0 | 0 |
| rs137853253 | pathogenic | 0 | 0 | 0 |
| rs202166915 | benign | 0 | 0 | 0 |
| rs137853259 | pathogenic | 0 | 0 | 0 |
| rs2229137 | benign | 0 | 0 | 0 |
| rs137853258 | pathogenic | 1 | 1 | 0 |
| rs137853252 | pathogenic | 1 | 1 | 1 |
| rs137853256 | pathogenic | 0 | 0 | 0 |
| rs137853250 | pathogenic | 1 | 1 | 1 |
| rs121434615 | pathogenic | 0 | 0 | 0 |
| rs115014558 | benign | 1 | 1 | 1 |
| rs34116584 | benign | 1 | 1 | 1 |
| rs34885252 | benign | 1 | 1 | 1 |
| rs180177262 | pathogenic | 1 | 1 | 1 |
| rs180177162 | pathogenic | 1 | 1 | 1 |
| rs121908523 | pathogenic | 1 | 1 | 1 |
| rs180177170 | pathogenic | 1 | 1 | 1 |
| rs180177180 | pathogenic | 1 | 1 | 1 |
| rs767586362 | pathogenic | 1 | 1 | 1 |
| rs121908521 | pathogenic | 1 | 1 | 1 |
| rs180177203 | pathogenic | 1 | 1 | 1 |
| rs138025751 | pathogenic | 1 | 1 | 1 |
| rs180177210 | pathogenic | 0 | 1 | 0 |
| rs180177211 | pathogenic | 0 | 1 | 0 |
| rs121908524 | pathogenic | 1 | 1 | 0 |
| rs121908530 | pathogenic | 1 | 1 | 0 |
| rs180177227 | pathogenic | 0 | 1 | 0 |
| rs180177230 | pathogenic | 1 | 1 | 1 |
| rs121908529 | pathogenic | 0 | 1 | 0 |
| rs117195882 | benign | 1 | 1 | 0 |
| rs180177238 | pathogenic | 1 | 1 | 0 |
| rs34664134 | benign | 1 | 1 | 1 |
| rs180177246 | pathogenic | 0 | 1 | 0 |
| rs536352238 | pathogenic | 0 | 0 | 0 |
| rs121908520 | pathogenic | 1 | 1 | 0 |
| rs180177248 | pathogenic | 1 | 1 | 0 |
| rs180177252 | pathogenic | 1 | 1 | 1 |
| rs121908526 | pathogenic | 1 | 1 | 1 |
| rs121908525 | pathogenic | 1 | 1 | 1 |
| rs147106773 | benign | 1 | 1 | 1 |
| rs180177264 | pathogenic | 0 | 1 | 0 |
| rs146525143 | pathogenic | 1 | 1 | 1 |
| rs140992177 | benign | 1 | 1 | 1 |
| rs13408961 | benign | 0 | 1 | 0 |

| | | | | |
|---|---|---|---|---|
| rs4426527 | benign | 1 | 1 | 1 |
| rs3088308 | benign | 0 | 1 | 0 |
| rs4469829 | benign | 1 | 1 | 0 |
| rs104886003 | pathogenic | 1 | 1 | 1 |
| rs121913286 | pathogenic | 1 | 1 | 1 |
| rs121913279 | pathogenic | 1 | 1 | 1 |
| rs121434636 | pathogenic | 1 | 1 | 1 |
| rs755299132 | pathogenic | 1 | 1 | 1 |
| rs748318386 | pathogenic | 1 | 1 | 1 |
| rs387906757 | pathogenic | 1 | 1 | 1 |
| rs121965064 | pathogenic | 1 | 1 | 1 |
| rs28934608 | pathogenic | 0 | 0 | 0 |
| rs5967 | benign | 1 | 1 | 1 |
| rs773905328 | pathogenic | 0 | 0 | 0 |
| rs104894256 | pathogenic | 0 | 1 | 0 |
| rs28931612 | pathogenic | 0 | 1 | 0 |
| rs104894263 | pathogenic | 1 | 1 | 1 |
| rs607969 | benign | 1 | 1 | 0 |
| rs104894262 | pathogenic | 0 | 1 | 0 |
| rs104894268 | pathogenic | 1 | 1 | 1 |
| rs104894264 | pathogenic | 0 | 1 | 0 |
| rs104894259 | pathogenic | 1 | 1 | 0 |
| rs2959656 | benign | 0 | 0 | 0 |
| rs121913035 | pathogenic | 1 | 1 | 0 |
| rs139605474 | benign | 0 | 0 | 0 |
| rs3817552 | benign | 0 | 0 | 0 |
| rs150541115 | benign | 0 | 0 | 0 |
| rs562763010 | benign | 1 | 1 | 1 |
| rs200053119 | pathogenic | 0 | 0 | 0 |
| rs73826386 | benign | 0 | 0 | 0 |
| rs143655258 | benign | 0 | 0 | 0 |
| rs758857853 | pathogenic | 1 | 1 | 0 |
| rs2228380 | benign | 0 | 0 | 0 |
| rs141078733 | benign | 1 | 1 | 1 |
| rs55824172 | pathogenic | 1 | 1 | 1 |
| rs756751089 | pathogenic | 1 | 1 | 1 |
| rs121964990 | pathogenic | 0 | 0 | 0 |
| rs143555186 | benign | 0 | 0 | 0 |
| rs121964993 | pathogenic | 1 | 1 | 0 |
| rs121964992 | pathogenic | 1 | 1 | 1 |
| rs121964991 | pathogenic | 1 | 1 | 0 |
| rs121964988 | pathogenic | 1 | 1 | 1 |
| rs4687 | benign | 1 | 1 | 1 |
| rs267607125 | pathogenic | 1 | 1 | 1 |
| rs751682878 | benign | 1 | 1 | 1 |
| rs267607126 | pathogenic | 1 | 1 | 1 |
| rs768713596 | pathogenic | 0 | 0 | 0 |
| rs147288996 | benign | 0 | 0 | 0 |
| rs387906639 | pathogenic | 1 | 1 | 1 |
| rs730881069 | pathogenic | 1 | 1 | 1 |
| rs730881071 | pathogenic | 1 | 1 | 1 |
| rs397516348 | pathogenic | 0 | 0 | 0 |
| rs121917760 | pathogenic | 0 | 0 | 0 |
| rs104894724 | pathogenic | 1 | 1 | 1 |

| | | | | |
|---|---|---|---|---|
| rs397516349 | pathogenic | 1 | 1 | 1 |
| rs12075 | benign | 0 | 0 | 0 |
| rs147549987 | benign | 0 | 0 | 0 |
| rs56079734 | benign | 0 | 0 | 0 |
| rs1017842 | benign | 1 | 1 | 1 |
| rs137852593 | pathogenic | 1 | 1 | 1 |
| rs137852571 | pathogenic | 1 | 1 | 1 |
| rs137852591 | pathogenic | 1 | 1 | 1 |
| rs9332971 | pathogenic | 1 | 1 | 1 |
| rs755226547 | pathogenic | 1 | 1 | 1 |
| rs79206939 | benign | 0 | 0 | 0 |
| rs121918214 | pathogenic | 1 | 1 | 0 |
| rs757075712 | pathogenic | 1 | 1 | 0 |
| rs150483902 | benign | 1 | 1 | 1 |
| rs113993973 | pathogenic | 0 | 1 | 0 |
| rs946616 | benign | 0 | 0 | 0 |
| rs113993976 | pathogenic | 1 | 1 | 0 |
| rs151022264 | benign | 0 | 1 | 0 |
| rs113993977 | pathogenic | 1 | 1 | 0 |
| rs113993978 | pathogenic | 1 | 1 | 0 |
| rs2228499 | benign | 0 | 1 | 0 |
| rs113993979 | pathogenic | 1 | 1 | 0 |
| rs113993980 | pathogenic | 1 | 1 | 0 |
| rs113993983 | pathogenic | 0 | 1 | 0 |
| rs150547274 | pathogenic | 1 | 1 | 0 |
| rs113993984 | pathogenic | 1 | 1 | 0 |
| rs113993986 | pathogenic | 1 | 1 | 0 |
| rs34313873 | benign | 0 | 1 | 0 |
| rs113993988 | pathogenic | 1 | 1 | 0 |
| rs199638097 | pathogenic | 1 | 1 | 1 |
| rs118204447 | pathogenic | 1 | 1 | 1 |
| rs11862754 | benign | 0 | 0 | 0 |
| rs118204445 | pathogenic | 0 | 0 | 0 |
| rs118204441 | pathogenic | 0 | 0 | 0 |
| rs118204438 | pathogenic | 0 | 0 | 0 |
| rs118204436 | pathogenic | 0 | 0 | 0 |
| rs146093755 | pathogenic | 0 | 0 | 0 |
| rs781439830 | pathogenic | 0 | 0 | 0 |
| rs398123438 | pathogenic | 0 | 0 | 0 |
| rs398123439 | pathogenic | 0 | 0 | 0 |
| rs118204444 | pathogenic | 0 | 0 | 0 |
| rs7187889 | benign | 0 | 0 | 0 |
| rs118204435 | pathogenic | 1 | 1 | 0 |
| rs34745339 | benign | 0 | 0 | 0 |
| rs398123440 | pathogenic | 0 | 0 | 0 |
| rs118204442 | pathogenic | 0 | 0 | 0 |
| rs770053354 | pathogenic | 0 | 0 | 0 |
| rs118204448 | pathogenic | 0 | 0 | 0 |
| rs118204443 | pathogenic | 0 | 0 | 0 |
| rs118204446 | pathogenic | 0 | 0 | 0 |
| rs746756997 | pathogenic | 0 | 0 | 0 |
| rs267606838 | pathogenic | 0 | 0 | 0 |
| rs118204437 | pathogenic | 0 | 0 | 0 |
| rs398123429 | pathogenic | 0 | 0 | 0 |

| | | | | |
|---|---|---|---|---|
| rs398123430 | pathogenic | 0 | 0 | 0 |
| rs2303269 | benign | 0 | 0 | 0 |
| rs114703967 | benign | 0 | 0 | 0 |
| rs118204439 | pathogenic | 0 | 0 | 0 |
| rs151296605 | benign | 0 | 0 | 0 |
| rs118204440 | pathogenic | 0 | 0 | 0 |
| rs78127134 | benign | 0 | 0 | 0 |
| rs372893383 | pathogenic | 0 | 0 | 0 |
| rs61753266 | pathogenic | 1 | 1 | 1 |
| rs139579928 | benign | 1 | 1 | 1 |
| rs56275071 | pathogenic | 1 | 1 | 1 |
| rs121909736 | pathogenic | 1 | 1 | 0 |
| rs121909737 | pathogenic | 1 | 1 | 0 |
| rs121909735 | pathogenic | 1 | 1 | 0 |
| rs121909731 | pathogenic | 1 | 1 | 1 |
| rs121909734 | pathogenic | 1 | 1 | 1 |
| rs121909733 | pathogenic | 1 | 1 | 1 |
| rs121909732 | pathogenic | 1 | 1 | 1 |
| rs121909730 | pathogenic | 1 | 1 | 1 |
| rs202198360 | benign | 1 | 1 | 1 |
| rs587777294 | pathogenic | 1 | 1 | 0 |
| rs587777297 | pathogenic | 1 | 1 | 0 |
| rs104893913 | pathogenic | 1 | 1 | 0 |
| rs17860424 | pathogenic | 1 | 1 | 1 |
| rs1045485 | benign | 1 | 1 | 1 |
| rs13926 | benign | 0 | 0 | 0 |
| rs79343837 | benign | 0 | 0 | 0 |
| rs45516306 | benign | 1 | 1 | 1 |
| rs139849355 | benign | 1 | 1 | 1 |
| rs104894874 | pathogenic | 1 | 1 | 1 |
| rs104894870 | pathogenic | 1 | 1 | 1 |
| rs137852221 | pathogenic | 1 | 1 | 1 |
| rs104894869 | pathogenic | 1 | 1 | 1 |
| rs104894880 | pathogenic | 1 | 1 | 1 |
| rs104894872 | pathogenic | 1 | 1 | 1 |
| rs104894868 | pathogenic | 1 | 1 | 1 |
| rs104894867 | pathogenic | 1 | 1 | 1 |
| rs104894871 | pathogenic | 1 | 1 | 1 |
| rs104894877 | pathogenic | 1 | 1 | 1 |
| rs104894883 | pathogenic | 1 | 1 | 1 |
| rs104894875 | pathogenic | 1 | 1 | 1 |
| rs104894876 | pathogenic | 1 | 1 | 1 |
| rs104894878 | pathogenic | 1 | 1 | 1 |
| rs137852220 | pathogenic | 1 | 1 | 1 |
| rs28933684 | pathogenic | 1 | 1 | 1 |
| rs104894873 | pathogenic | 1 | 1 | 1 |
| rs138409264 | benign | 0 | 0 | 0 |
| rs2236722 | benign | 0 | 0 | 0 |
| rs28757184 | benign | 0 | 0 | 0 |
| rs121434538 | pathogenic | 0 | 0 | 0 |
| rs700519 | benign | 0 | 0 | 0 |
| rs80051519 | pathogenic | 0 | 0 | 0 |
| rs121434536 | pathogenic | 1 | 1 | 0 |
| rs121434534 | pathogenic | 1 | 1 | 0 |

| | | | | |
|---|---|---|---|---|
| rs78310315 | pathogenic | 1 | 1 | 0 |
| rs540839115 | pathogenic | 1 | 1 | 1 |
| rs80338731 | pathogenic | 1 | 1 | 1 |
| rs180177151 | pathogenic | 1 | 1 | 1 |
| rs180177152 | pathogenic | 0 | 0 | 0 |
| rs80338732 | pathogenic | 1 | 1 | 1 |
| rs80338675 | pathogenic | 1 | 1 | 1 |
| rs80338676 | pathogenic | 1 | 1 | 1 |
| rs180177153 | pathogenic | 0 | 0 | 0 |
| rs180177154 | pathogenic | 1 | 1 | 1 |
| rs5896 | benign | 1 | 1 | 1 |
| rs121918477 | pathogenic | 0 | 0 | 0 |
| rs121918428 | pathogenic | 1 | 1 | 1 |
| rs121918427 | pathogenic | 1 | 1 | 1 |
| rs544300820 | benign | 1 | 1 | 1 |
| rs121918110 | pathogenic | 1 | 1 | 1 |
| rs1057141 | benign | 0 | 0 | 0 |
| rs1135216 | benign | 1 | 1 | 1 |
| rs121917702 | pathogenic | 1 | 1 | 1 |
| rs28941474 | pathogenic | 0 | 0 | 0 |
| rs104893944 | pathogenic | 1 | 1 | 1 |
| rs104893947 | pathogenic | 1 | 1 | 0 |
| rs104893943 | pathogenic | 1 | 1 | 1 |
| rs104893942 | pathogenic | 1 | 1 | 1 |
| rs104893940 | pathogenic | 1 | 1 | 1 |
| rs34045013 | benign | 0 | 0 | 0 |
| rs143083947 | pathogenic | 0 | 0 | 0 |
| rs145786986 | benign | 0 | 0 | 0 |
| rs119450942 | pathogenic | 0 | 0 | 0 |
| rs374259530 | pathogenic | 0 | 0 | 0 |
| rs756210458 | pathogenic | 0 | 0 | 0 |
| rs11089991 | benign | 0 | 0 | 0 |
| rs28941471 | pathogenic | 0 | 1 | 0 |
| rs750614500 | pathogenic | 0 | 1 | 0 |
| rs199761158 | benign | 1 | 1 | 1 |
| rs119450944 | pathogenic | 1 | 1 | 1 |
| rs373458753 | pathogenic | 1 | 1 | 1 |
| rs202064195 | pathogenic | 1 | 1 | 1 |
| rs776496275 | pathogenic | 1 | 1 | 1 |
| rs761493155 | pathogenic | 1 | 1 | 1 |
| rs763542069 | pathogenic | 0 | 0 | 0 |
| rs119450941 | pathogenic | 0 | 0 | 0 |
| rs119450940 | pathogenic | 0 | 0 | 0 |
| rs777821034 | pathogenic | 1 | 1 | 1 |
| rs771121666 | pathogenic | 1 | 1 | 1 |
| rs372895468 | pathogenic | 1 | 1 | 1 |
| rs61750603 | pathogenic | 0 | 0 | 0 |
| rs76863441 | pathogenic | 0 | 0 | 0 |
| rs142601143 | benign | 1 | 1 | 1 |
| rs185832993 | benign | 1 | 1 | 0 |
| rs17860568 | benign | 0 | 0 | 0 |
| rs142064825 | benign | 0 | 0 | 0 |
| rs564076183 | benign | 0 | 0 | 0 |
| rs386834034 | pathogenic | 0 | 0 | 0 |

| | | | | |
|---|---|---|---|---|
| rs147663769 | benign | 0 | 0 | 0 |
| rs776478974 | benign | 0 | 0 | 0 |
| rs111840783 | benign | 0 | 0 | 0 |
| rs1516446 | benign | 1 | 1 | 1 |
| rs143749154 | benign | 1 | 1 | 1 |
| rs80257412 | benign | 0 | 0 | 0 |
| rs142943120 | benign | 0 | 0 | 0 |
| rs144553482 | benign | 0 | 0 | 0 |
| rs35997569 | benign | 0 | 0 | 0 |
| rs772909106 | pathogenic | 0 | 0 | 0 |
| rs149433837 | benign | 1 | 1 | 1 |
| rs532178791 | pathogenic | 0 | 0 | 0 |
| rs145588689 | pathogenic | 1 | 1 | 0 |
| rs17843776 | benign | 1 | 1 | 0 |
| rs121917890 | pathogenic | 1 | 1 | 1 |
| rs121917892 | pathogenic | 0 | 0 | 0 |
| rs606231289 | pathogenic | 0 | 0 | 0 |
| rs370011798 | pathogenic | 0 | 0 | 0 |
| rs34689457 | benign | 0 | 0 | 0 |
| rs118204451 | pathogenic | 0 | 0 | 0 |
| rs118204452 | pathogenic | 1 | 1 | 1 |
| rs652785 | benign | 0 | 0 | 0 |
| rs35378126 | benign | 0 | 0 | 0 |
| rs80358214 | pathogenic | 1 | 1 | 1 |
| rs80358215 | pathogenic | 0 | 1 | 0 |
| rs267606858 | pathogenic | 1 | 1 | 1 |
| rs143137713 | pathogenic | 1 | 1 | 1 |
| rs782168634 | benign | 0 | 0 | 0 |
| rs143873938 | benign | 0 | 0 | 0 |
| rs782776511 | benign | 0 | 0 | 0 |
| rs201093148 | benign | 0 | 0 | 0 |
| rs267606815 | pathogenic | 0 | 0 | 0 |
| rs137853311 | pathogenic | 0 | 0 | 0 |
| rs267606817 | pathogenic | 0 | 0 | 0 |
| rs72650006 | benign | 0 | 0 | 0 |
| rs727503638 | benign | 1 | 1 | 0 |
| rs189286381 | benign | 0 | 0 | 0 |
| rs376644553 | benign | 0 | 0 | 0 |
| rs121434610 | pathogenic | 1 | 1 | 1 |
| rs267607076 | pathogenic | 1 | 1 | 1 |
| rs201275730 | benign | 0 | 0 | 0 |
| rs121909587 | pathogenic | 0 | 0 | 0 |
| rs121909558 | pathogenic | 1 | 1 | 0 |
| rs28929469 | pathogenic | 1 | 1 | 0 |
| rs121909547 | pathogenic | 1 | 1 | 0 |
| rs121909552 | pathogenic | 1 | 1 | 0 |
| rs121909573 | pathogenic | 1 | 1 | 1 |
| rs121909567 | pathogenic | 1 | 1 | 0 |
| rs121909569 | pathogenic | 0 | 0 | 0 |
| rs121909563 | pathogenic | 1 | 1 | 0 |
| rs121909570 | pathogenic | 1 | 1 | 1 |
| rs121909571 | pathogenic | 1 | 1 | 0 |
| rs121909572 | pathogenic | 1 | 1 | 1 |
| rs121909565 | pathogenic | 1 | 1 | 1 |

| | | | | |
|---|---|---|---|---|
| rs121909557 | pathogenic | 1 | 1 | 1 |
| rs121909568 | pathogenic | 1 | 1 | 1 |
| rs121909566 | pathogenic | 1 | 1 | 1 |
| rs121909549 | pathogenic | 1 | 1 | 1 |
| rs121909554 | pathogenic | 1 | 1 | 1 |
| rs121909550 | pathogenic | 1 | 1 | 1 |
| rs121909546 | pathogenic | 1 | 1 | 0 |
| rs121909555 | pathogenic | 1 | 1 | 1 |
| rs121909564 | pathogenic | 1 | 1 | 1 |
| rs548046212 | pathogenic | 0 | 0 | 0 |
| rs147353781 | benign | 0 | 0 | 0 |
| rs104894678 | pathogenic | 1 | 1 | 1 |
| rs1130426 | benign | 0 | 0 | 0 |
| rs104894677 | pathogenic | 1 | 1 | 1 |
| rs141917423 | benign | 1 | 1 | 1 |
| rs574777883 | benign | 0 | 0 | 0 |
| rs150199226 | benign | 1 | 1 | 1 |
| rs199618701 | benign | 0 | 1 | 0 |
| rs121964923 | pathogenic | 0 | 0 | 0 |
| rs121964924 | pathogenic | 0 | 0 | 0 |
| rs201258823 | pathogenic | 0 | 1 | 0 |
| rs267606773 | pathogenic | 0 | 0 | 0 |
| rs35638832 | benign | 0 | 0 | 0 |
| rs121908632 | pathogenic | 1 | 1 | 1 |
| rs121908625 | pathogenic | 1 | 1 | 1 |
| rs748949603 | pathogenic | 0 | 0 | 0 |
| rs757523840 | pathogenic | 0 | 0 | 0 |
| rs62541771 | pathogenic | 0 | 0 | 0 |
| rs776384541 | pathogenic | 1 | 1 | 0 |
| rs121908627 | pathogenic | 1 | 1 | 0 |
| rs28937594 | pathogenic | 0 | 0 | 0 |
| rs121907888 | pathogenic | 0 | 0 | 0 |
| rs121907889 | pathogenic | 0 | 1 | 0 |
| rs28941785 | pathogenic | 1 | 1 | 1 |
| rs28941786 | pathogenic | 1 | 1 | 1 |
| rs1021737 | benign | 0 | 0 | 0 |
| rs121908370 | pathogenic | 1 | 1 | 1 |
| rs63750083 | pathogenic | 0 | 0 | 0 |
| rs17883862 | benign | 0 | 0 | 0 |
| rs587782070 | pathogenic | 1 | 1 | 1 |
| rs587781269 | pathogenic | 1 | 1 | 1 |
| rs28909982 | pathogenic | 1 | 1 | 1 |
| rs373648967 | benign | 1 | 1 | 1 |
| rs368570187 | benign | 1 | 1 | 1 |
| rs730881701 | pathogenic | 1 | 1 | 1 |
| rs137853007 | pathogenic | 0 | 0 | 0 |
| rs575910805 | benign | 0 | 0 | 0 |
| rs104893875 | pathogenic | 1 | 1 | 1 |
| rs104893877 | pathogenic | 1 | 1 | 1 |
| rs119485093 | pathogenic | 1 | 1 | 1 |
| rs119485092 | pathogenic | 0 | 0 | 0 |
| rs119485090 | pathogenic | 0 | 0 | 0 |
| rs28942099 | pathogenic | 1 | 1 | 1 |
| rs373730800 | pathogenic | 1 | 1 | 1 |

| | | | | |
|---|---|---|---|---|
| rs113993995 | pathogenic | 0 | 0 | 0 |
| rs201070132 | pathogenic | 0 | 0 | 0 |
| rs113993996 | pathogenic | 0 | 0 | 0 |
| rs113993998 | pathogenic | 0 | 0 | 0 |
| rs78950939 | pathogenic | 1 | 1 | 1 |
| rs56378716 | pathogenic | 1 | 1 | 1 |
| rs754921704 | benign | 0 | 1 | 0 |
| rs766379103 | pathogenic | 1 | 1 | 0 |
| rs121912515 | pathogenic | 0 | 1 | 0 |
| rs764831888 | benign | 1 | 1 | 0 |
| rs572527340 | benign | 0 | 0 | 0 |
| rs35969327 | benign | 0 | 0 | 0 |
| rs35814893 | benign | 0 | 0 | 0 |
| rs34032300 | benign | 0 | 0 | 0 |
| rs80358260 | pathogenic | 0 | 0 | 0 |
| rs151220873 | benign | 0 | 0 | 0 |
| rs80358261 | pathogenic | 1 | 1 | 0 |
| rs11694 | pathogenic | 0 | 0 | 0 |
| rs80358264 | pathogenic | 0 | 0 | 0 |
| rs80358266 | pathogenic | 1 | 1 | 0 |
| rs104894458 | pathogenic | 1 | 1 | 1 |
| rs121918028 | pathogenic | 0 | 0 | 0 |
| rs11583680 | pathogenic | 0 | 0 | 0 |
| rs4715 | benign | 1 | 1 | 1 |
| rs34957318 | benign | 0 | 0 | 0 |
| rs1124 | benign | 1 | 1 | 1 |
| rs104894778 | pathogenic | 1 | 1 | 1 |
| rs104894777 | pathogenic | 1 | 1 | 1 |
| rs104894769 | pathogenic | 1 | 1 | 1 |
| rs104894768 | pathogenic | 1 | 1 | 1 |
| rs104894771 | pathogenic | 1 | 1 | 1 |
| rs575372249 | benign | 1 | 1 | 1 |
| rs118203914 | pathogenic | 0 | 0 | 0 |
| rs118203916 | pathogenic | 1 | 1 | 0 |
| rs140154690 | benign | 0 | 0 | 0 |
| rs34382405 | benign | 0 | 0 | 0 |
| rs73718082 | benign | 0 | 0 | 0 |
| rs764932724 | benign | 0 | 0 | 0 |
| rs554931092 | pathogenic | 1 | 1 | 0 |
| rs202060864 | pathogenic | 0 | 0 | 0 |
| rs116567033 | benign | 0 | 0 | 0 |
| rs11243011 | benign | 0 | 0 | 0 |
| rs142073519 | pathogenic | 0 | 0 | 0 |
| rs397514611 | pathogenic | 1 | 1 | 0 |
| rs397514612 | pathogenic | 0 | 0 | 0 |
| rs372054960 | pathogenic | 0 | 0 | 0 |
| rs775690041 | pathogenic | 0 | 0 | 0 |
| rs148921184 | pathogenic | 0 | 0 | 0 |
| rs12711521 | benign | 0 | 0 | 0 |
| rs2273346 | benign | 0 | 0 | 0 |
| rs12085877 | benign | 1 | 1 | 1 |
| rs144471433 | benign | 1 | 1 | 1 |
| rs397514671 | pathogenic | 1 | 1 | 1 |
| rs76394680 | benign | 0 | 0 | 0 |

| | | | | |
|---|---|---|---|---|
| rs78289603 | pathogenic | 1 | 1 | 1 |
| rs137852913 | pathogenic | 1 | 1 | 1 |
| rs137852914 | pathogenic | 1 | 1 | 1 |
| rs121918653 | pathogenic | 1 | 1 | 1 |
| rs35515200 | benign | 1 | 1 | 0 |
| rs17876030 | benign | 1 | 1 | 1 |
| rs556623443 | benign | 1 | 1 | 1 |
| rs121434365 | pathogenic | 1 | 1 | 1 |
| rs121434364 | pathogenic | 0 | 0 | 0 |
| rs41400249 | benign | 0 | 0 | 0 |
| rs149440376 | benign | 1 | 1 | 1 |
| rs181590179 | benign | 0 | 0 | 0 |
| rs113993968 | pathogenic | 0 | 0 | 0 |
| rs587776445 | pathogenic | 1 | 1 | 1 |
| rs113993969 | pathogenic | 1 | 1 | 1 |
| rs113993970 | pathogenic | 1 | 1 | 1 |
| rs587776449 | pathogenic | 1 | 1 | 1 |
| rs113993971 | pathogenic | 1 | 1 | 1 |
| rs267607194 | pathogenic | 1 | 1 | 1 |
| rs397514745 | pathogenic | 0 | 0 | 0 |
| rs201650281 | pathogenic | 0 | 0 | 0 |
| rs150529876 | benign | 1 | 1 | 1 |
| rs77573084 | benign | 0 | 0 | 0 |
| rs398122379 | pathogenic | 1 | 1 | 0 |
| rs765478968 | benign | 0 | 0 | 0 |
| rs201399681 | benign | 0 | 0 | 0 |
| rs17159287 | benign | 0 | 0 | 0 |
| rs538571144 | benign | 0 | 0 | 0 |
| rs137852647 | pathogenic | 0 | 0 | 0 |
| rs201358272 | pathogenic | 1 | 1 | 1 |
| rs181251337 | benign | 0 | 0 | 0 |
| rs3213119 | benign | 0 | 0 | 0 |
| rs137853102 | pathogenic | 1 | 1 | 0 |
| rs4956145 | benign | 0 | 0 | 0 |
| rs146036912 | pathogenic | 1 | 1 | 1 |
| rs375717077 | pathogenic | 1 | 1 | 1 |
| rs137853103 | pathogenic | 1 | 1 | 1 |
| rs776184830 | pathogenic | 1 | 1 | 1 |
| rs200273673 | pathogenic | 1 | 1 | 1 |
| rs267607025 | pathogenic | 1 | 1 | 1 |
| rs267607024 | pathogenic | 1 | 1 | 1 |
| rs515726188 | pathogenic | 1 | 1 | 1 |
| rs147315735 | benign | 0 | 0 | 0 |
| rs121918308 | pathogenic | 1 | 1 | 0 |
| rs515726195 | pathogenic | 0 | 0 | 0 |
| rs515726196 | pathogenic | 1 | 1 | 0 |
| rs121918311 | pathogenic | 0 | 1 | 0 |
| rs121918309 | pathogenic | 0 | 0 | 0 |
| rs387906891 | pathogenic | 0 | 0 | 0 |
| rs182614164 | pathogenic | 1 | 1 | 0 |
| rs121918307 | pathogenic | 0 | 0 | 0 |
| rs201440849 | benign | 0 | 0 | 0 |
| rs118203957 | pathogenic | 0 | 0 | 0 |
| rs4986790 | benign | 0 | 0 | 0 |

| | | | | |
|---|---|---|---|---|
| rs281864928 | pathogenic | 1 | 1 | 0 |
| rs267607156 | pathogenic | 0 | 0 | 0 |
| rs55916807 | benign | 0 | 0 | 0 |
| rs749750052 | benign | 0 | 0 | 0 |
| rs74382477 | benign | 0 | 0 | 0 |
| rs201016638 | benign | 0 | 0 | 0 |
| rs376683989 | benign | 0 | 0 | 0 |
| rs144439703 | pathogenic | 0 | 0 | 0 |
| rs779122022 | benign | 0 | 0 | 0 |
| rs121918056 | pathogenic | 0 | 0 | 0 |
| rs113994093 | pathogenic | 1 | 1 | 1 |
| rs141367015 | pathogenic | 0 | 0 | 0 |
| rs775538075 | pathogenic | 0 | 0 | 0 |
| rs121918055 | pathogenic | 0 | 0 | 0 |
| rs754246294 | benign | 1 | 1 | 1 |
| rs2307447 | benign | 1 | 1 | 1 |
| rs113994096 | pathogenic | 1 | 1 | 1 |
| rs367610201 | pathogenic | 1 | 1 | 1 |
| rs758438414 | pathogenic | 1 | 1 | 0 |
| rs121918046 | pathogenic | 0 | 0 | 0 |
| rs201871736 | benign | 0 | 0 | 0 |
| rs773994204 | benign | 0 | 0 | 0 |
| rs2307450 | benign | 1 | 1 | 1 |
| rs200257554 | benign | 0 | 0 | 0 |
| rs185645212 | benign | 0 | 0 | 0 |
| rs121918054 | pathogenic | 0 | 0 | 0 |
| rs113994097 | pathogenic | 0 | 0 | 0 |
| rs202037973 | pathogenic | 0 | 0 | 0 |
| rs769827124 | pathogenic | 1 | 1 | 0 |
| rs41549716 | benign | 1 | 1 | 1 |
| rs113994098 | pathogenic | 1 | 1 | 0 |
| rs144500145 | pathogenic | 1 | 1 | 0 |
| rs121918053 | pathogenic | 1 | 1 | 0 |
| rs121918050 | pathogenic | 1 | 1 | 0 |
| rs368587966 | pathogenic | 0 | 0 | 0 |
| rs139590686 | pathogenic | 0 | 0 | 0 |
| rs121918048 | pathogenic | 1 | 1 | 0 |
| rs768653086 | pathogenic | 1 | 1 | 0 |
| rs113994099 | pathogenic | 1 | 1 | 0 |
| rs121918051 | pathogenic | 1 | 1 | 0 |
| rs753160398 | pathogenic | 1 | 1 | 0 |
| rs201477273 | pathogenic | 1 | 1 | 0 |
| rs142347031 | pathogenic | 0 | 0 | 0 |
| rs121918049 | pathogenic | 0 | 0 | 0 |
| rs267606959 | pathogenic | 1 | 1 | 0 |
| rs201732356 | pathogenic | 0 | 0 | 0 |
| rs368435864 | pathogenic | 0 | 0 | 0 |
| rs764036283 | pathogenic | 0 | 0 | 0 |
| rs2307442 | benign | 0 | 0 | 0 |
| rs2307440 | benign | 0 | 0 | 0 |
| rs548076633 | pathogenic | 0 | 1 | 0 |
| rs750792237 | benign | 0 | 0 | 0 |
| rs3087374 | benign | 0 | 0 | 0 |
| rs121918060 | pathogenic | 0 | 1 | 0 |

| | | | | |
|---|---|---|---|---|
| rs397514765 | pathogenic | 1 | 1 | 1 |
| rs121918063 | pathogenic | 0 | 0 | 0 |
| rs121918058 | pathogenic | 1 | 1 | 1 |
| rs121918064 | pathogenic | 0 | 0 | 0 |
| rs36033115 | benign | 0 | 1 | 0 |
| rs121918057 | pathogenic | 0 | 1 | 0 |
| rs121918065 | pathogenic | 1 | 1 | 1 |
| rs121918061 | pathogenic | 1 | 1 | 1 |
| rs121918066 | pathogenic | 0 | 1 | 0 |
| rs707555 | benign | 0 | 1 | 0 |
| rs8192708 | benign | 0 | 1 | 0 |
| rs11552145 | benign | 0 | 0 | 0 |
| rs1804160 | benign | 0 | 1 | 0 |
| rs34103538 | benign | 0 | 1 | 0 |
| rs751393950 | pathogenic | 1 | 1 | 1 |
| rs121907979 | pathogenic | 1 | 1 | 1 |
| rs121907975 | pathogenic | 1 | 1 | 1 |
| rs121907962 | pathogenic | 1 | 1 | 1 |
| rs121907957 | pathogenic | 0 | 0 | 0 |
| rs121907953 | pathogenic | 1 | 1 | 1 |
| rs28941770 | pathogenic | 1 | 1 | 1 |
| rs28941771 | pathogenic | 1 | 1 | 1 |
| rs121907969 | pathogenic | 1 | 1 | 1 |
| rs121907973 | pathogenic | 1 | 1 | 1 |
| rs1800429 | pathogenic | 0 | 0 | 0 |
| rs121907976 | pathogenic | 1 | 1 | 0 |
| rs121907961 | pathogenic | 0 | 0 | 0 |
| rs121907974 | pathogenic | 0 | 0 | 0 |
| rs121907959 | pathogenic | 0 | 0 | 0 |
| rs121907971 | pathogenic | 1 | 1 | 0 |
| rs121907954 | pathogenic | 0 | 0 | 0 |
| rs121907977 | pathogenic | 0 | 0 | 0 |
| rs121907963 | pathogenic | 1 | 1 | 0 |
| rs370266293 | pathogenic | 1 | 1 | 0 |
| rs1800430 | benign | 0 | 0 | 0 |
| rs121907958 | pathogenic | 0 | 0 | 0 |
| rs202173526 | pathogenic | 0 | 0 | 0 |
| rs121907978 | pathogenic | 0 | 0 | 0 |
| rs121907981 | pathogenic | 0 | 1 | 0 |
| rs145012038 | benign | 1 | 1 | 1 |
| rs121907968 | pathogenic | 1 | 1 | 1 |
| rs121907966 | pathogenic | 1 | 1 | 1 |
| rs121907956 | pathogenic | 1 | 1 | 1 |
| rs28942071 | pathogenic | 1 | 1 | 1 |
| rs121907955 | pathogenic | 1 | 1 | 1 |
| rs4727853 | benign | 0 | 0 | 0 |
| rs201880593 | benign | 0 | 0 | 0 |
| rs201434579 | benign | 1 | 1 | 1 |
| rs74551128 | pathogenic | 1 | 1 | 0 |
| rs193922500 | pathogenic | 0 | 0 | 0 |
| rs121909009 | pathogenic | 1 | 1 | 0 |
| rs121908805 | pathogenic | 1 | 1 | 0 |
| rs139573311 | pathogenic | 0 | 0 | 0 |
| rs213950 | benign | 0 | 0 | 0 |

| | | | | |
|---|---|---|---|---|
| rs79282516 | pathogenic | 0 | 0 | 0 |
| rs121909017 | pathogenic | 1 | 1 | 1 |
| rs77101217 | pathogenic | 1 | 1 | 1 |
| rs1800092 | pathogenic | 0 | 1 | 0 |
| rs77646904 | pathogenic | 0 | 0 | 0 |
| rs1800095 | benign | 1 | 1 | 1 |
| rs387906368 | pathogenic | 1 | 1 | 1 |
| rs113993959 | pathogenic | 1 | 1 | 1 |
| rs121908755 | pathogenic | 1 | 1 | 1 |
| rs121909005 | pathogenic | 1 | 1 | 1 |
| rs121908757 | pathogenic | 1 | 1 | 1 |
| rs75527207 | pathogenic | 1 | 1 | 0 |
| rs121909013 | pathogenic | 1 | 1 | 0 |
| rs121909044 | pathogenic | 1 | 1 | 0 |
| rs74597325 | pathogenic | 1 | 1 | 0 |
| rs75549581 | pathogenic | 0 | 0 | 0 |
| rs80055610 | pathogenic | 1 | 1 | 1 |
| rs1800097 | pathogenic | 0 | 0 | 0 |
| rs121909006 | pathogenic | 0 | 1 | 0 |
| rs397508276 | pathogenic | 0 | 0 | 0 |
| rs397508296 | pathogenic | 1 | 1 | 1 |
| rs139468767 | pathogenic | 0 | 0 | 0 |
| rs121908759 | pathogenic | 1 | 1 | 0 |
| rs121909033 | pathogenic | 1 | 1 | 0 |
| rs397508331 | pathogenic | 0 | 0 | 0 |
| rs387907279 | pathogenic | 1 | 1 | 1 |
| rs546530618 | benign | 0 | 0 | 0 |
| rs558259739 | benign | 0 | 0 | 0 |
| rs201827340 | benign | 0 | 0 | 0 |
| rs148865119 | pathogenic | 1 | 1 | 0 |
| rs2229413 | benign | 1 | 1 | 1 |
| rs34040148 | benign | 1 | 1 | 1 |
| rs121964964 | pathogenic | 1 | 1 | 1 |
| rs760214620 | pathogenic | 1 | 1 | 1 |
| rs775992753 | pathogenic | 1 | 1 | 1 |
| rs770095972 | pathogenic | 1 | 1 | 1 |
| rs781444670 | pathogenic | 1 | 1 | 1 |
| rs147474549 | pathogenic | 0 | 0 | 0 |
| rs121964965 | pathogenic | 0 | 1 | 0 |
| rs121964963 | pathogenic | 1 | 1 | 1 |
| rs755952006 | pathogenic | 1 | 1 | 1 |
| rs121964970 | pathogenic | 1 | 1 | 1 |
| rs121964973 | pathogenic | 1 | 1 | 1 |
| rs773734233 | pathogenic | 1 | 1 | 1 |
| rs758236584 | pathogenic | 1 | 1 | 1 |
| rs149119723 | pathogenic | 1 | 1 | 0 |
| rs5742905 | pathogenic | 0 | 0 | 0 |
| rs121964962 | pathogenic | 1 | 1 | 1 |
| rs781567152 | pathogenic | 1 | 1 | 0 |
| rs398123151 | pathogenic | 1 | 1 | 1 |
| rs760417941 | pathogenic | 1 | 1 | 1 |
| rs774926464 | pathogenic | 0 | 0 | 0 |
| rs121964972 | pathogenic | 1 | 1 | 0 |
| rs372010465 | pathogenic | 1 | 1 | 1 |

| | | | | |
|---|---|---|---|---|
| rs763036586 | pathogenic | 1 | 1 | 1 |
| rs769080151 | pathogenic | 1 | 1 | 1 |
| rs121964967 | pathogenic | 1 | 1 | 1 |
| rs28934892 | pathogenic | 1 | 1 | 0 |
| rs28934891 | pathogenic | 1 | 1 | 1 |
| rs121964971 | pathogenic | 1 | 1 | 1 |
| rs121964968 | pathogenic | 1 | 1 | 1 |
| rs150828989 | benign | 1 | 1 | 1 |
| rs3775291 | benign | 1 | 1 | 0 |
| rs104894419 | pathogenic | 1 | 1 | 1 |
| rs2232642 | benign | 0 | 0 | 0 |
| rs104894551 | pathogenic | 1 | 1 | 0 |
| rs766328537 | pathogenic | 0 | 1 | 0 |
| rs104894553 | pathogenic | 1 | 1 | 0 |
| rs80099330 | benign | 0 | 0 | 0 |
| rs199565861 | pathogenic | 0 | 0 | 0 |
| rs104894548 | pathogenic | 0 | 0 | 0 |
| rs104894549 | pathogenic | 0 | 0 | 0 |
| rs104894550 | pathogenic | 1 | 1 | 1 |
| rs12948217 | pathogenic | 1 | 1 | 1 |
| rs761064915 | pathogenic | 0 | 0 | 0 |
| rs28940279 | pathogenic | 1 | 1 | 1 |
| rs774323189 | pathogenic | 1 | 1 | 1 |
| rs28940574 | pathogenic | 0 | 0 | 0 |
| rs515299 | benign | 1 | 1 | 1 |
| rs10127939 | benign | 1 | 1 | 1 |
| rs396991 | benign | 1 | 1 | 1 |
| rs145956175 | benign | 0 | 1 | 0 |
| rs747838255 | pathogenic | 1 | 1 | 1 |
| rs762659685 | pathogenic | 1 | 1 | 0 |
| rs104894831 | pathogenic | 0 | 1 | 0 |
| rs104894848 | pathogenic | 1 | 1 | 0 |
| rs104894833 | pathogenic | 0 | 1 | 0 |
| rs781838005 | pathogenic | 0 | 1 | 0 |
| rs104894834 | pathogenic | 1 | 1 | 0 |
| rs372966991 | pathogenic | 1 | 1 | 0 |
| rs138886989 | benign | 1 | 1 | 1 |
| rs28935195 | pathogenic | 0 | 1 | 0 |
| rs28935196 | pathogenic | 0 | 1 | 0 |
| rs372416832 | pathogenic | 1 | 1 | 1 |
| rs28935197 | pathogenic | 1 | 1 | 0 |
| rs28935486 | pathogenic | 1 | 1 | 0 |
| rs28935487 | pathogenic | 1 | 1 | 0 |
| rs28935488 | pathogenic | 1 | 1 | 1 |
| rs28935495 | pathogenic | 1 | 1 | 1 |
| rs28935485 | pathogenic | 1 | 1 | 1 |
| rs140329381 | pathogenic | 0 | 1 | 0 |
| rs104894830 | pathogenic | 0 | 1 | 0 |
| rs104894846 | pathogenic | 0 | 1 | 0 |
| rs28935489 | pathogenic | 0 | 1 | 0 |
| rs104894828 | pathogenic | 1 | 1 | 0 |
| rs28935491 | pathogenic | 1 | 1 | 0 |
| rs104894832 | pathogenic | 1 | 1 | 0 |
| rs28935492 | pathogenic | 1 | 1 | 0 |

| | | | | |
|---|---|---|---|---|
| rs730880437 | benign | 1 | 1 | 0 |
| rs28935493 | pathogenic | 0 | 1 | 0 |
| rs28935494 | pathogenic | 1 | 1 | 1 |
| rs111422676 | pathogenic | 1 | 1 | 1 |
| rs397515869 | benign | 1 | 1 | 0 |
| rs782449839 | pathogenic | 0 | 1 | 0 |
| rs104894852 | pathogenic | 1 | 1 | 1 |
| rs116929575 | pathogenic | 1 | 1 | 1 |
| rs7023954 | benign | 0 | 0 | 0 |
| rs184520335 | benign | 1 | 1 | 1 |
| rs143080527 | benign | 0 | 0 | 0 |
| rs1050565 | benign | 0 | 0 | 0 |
| rs11545035 | benign | 0 | 0 | 0 |
| rs267607190 | pathogenic | 1 | 1 | 1 |
| rs200283987 | benign | 0 | 0 | 0 |
| rs372556807 | benign | 1 | 1 | 0 |
| rs563073392 | benign | 0 | 0 | 0 |
| rs4539 | benign | 0 | 0 | 0 |
| rs28931609 | pathogenic | 1 | 1 | 1 |
| rs121912978 | pathogenic | 1 | 1 | 1 |
| rs104894072 | pathogenic | 1 | 1 | 1 |
| rs551933154 | benign | 1 | 1 | 1 |
| rs752962897 | benign | 1 | 1 | 0 |
| rs121912977 | pathogenic | 1 | 1 | 0 |
| rs4537 | benign | 1 | 1 | 1 |
| rs201830462 | benign | 0 | 0 | 0 |
| rs4544 | benign | 0 | 0 | 0 |
| rs562670189 | benign | 0 | 0 | 0 |
| rs4538 | benign | 0 | 1 | 0 |
| rs61757294 | pathogenic | 0 | 0 | 0 |
| rs5313 | benign | 0 | 0 | 0 |
| rs4545 | benign | 0 | 0 | 0 |
| rs72554627 | pathogenic | 1 | 1 | 1 |
| rs72554626 | pathogenic | 1 | 1 | 1 |
| rs387906861 | pathogenic | 1 | 1 | 1 |
| rs749655461 | pathogenic | 1 | 1 | 1 |
| rs121909606 | pathogenic | 1 | 1 | 1 |
| rs121909607 | pathogenic | 1 | 1 | 1 |
| rs200203992 | benign | 0 | 0 | 0 |
| rs6050 | benign | 0 | 0 | 0 |
| rs184635235 | benign | 0 | 0 | 0 |
| rs139146037 | benign | 0 | 0 | 0 |
| rs121909612 | pathogenic | 0 | 0 | 0 |
| rs78506343 | pathogenic | 0 | 0 | 0 |
| rs121918138 | pathogenic | 1 | 1 | 1 |
| rs121918139 | pathogenic | 1 | 1 | 1 |
| rs121918140 | pathogenic | 0 | 0 | 0 |
| rs36063028 | benign | 1 | 1 | 1 |
| rs45495192 | benign | 1 | 1 | 1 |
| rs45458802 | benign | 1 | 1 | 1 |
| rs552865793 | pathogenic | 1 | 1 | 1 |
| rs753744791 | pathogenic | 1 | 1 | 1 |
| rs777518512 | benign | 1 | 1 | 1 |
| rs142708991 | benign | 1 | 1 | 0 |

| | | | | |
|---|---|---|---|---|
| rs746069957 | benign | 0 | 0 | 0 |
| rs1334811 | benign | 0 | 0 | 0 |
| rs138894008 | benign | 1 | 1 | 0 |
| rs35030851 | benign | 1 | 1 | 0 |
| rs35378598 | benign | 0 | 0 | 0 |
| rs35818764 | benign | 1 | 1 | 1 |
| rs4631561 | benign | 1 | 1 | 1 |
| rs121918660 | pathogenic | 0 | 0 | 0 |
| rs373175587 | benign | 0 | 0 | 0 |
| rs61731956 | pathogenic | 1 | 1 | 1 |
| rs180177305 | pathogenic | 0 | 0 | 0 |
| rs119490108 | pathogenic | 0 | 0 | 0 |
| rs180177312 | pathogenic | 1 | 1 | 0 |
| rs180177314 | pathogenic | 1 | 1 | 0 |
| rs180177322 | pathogenic | 1 | 1 | 1 |
| rs180177323 | pathogenic | 1 | 1 | 1 |
| rs180177325 | pathogenic | 0 | 0 | 0 |
| rs35196441 | benign | 0 | 0 | 0 |
| rs28763904 | benign | 0 | 0 | 0 |
| rs2229519 | benign | 0 | 0 | 0 |
| rs137852886 | pathogenic | 0 | 0 | 0 |
| rs137852892 | pathogenic | 0 | 0 | 0 |
| rs770427750 | pathogenic | 0 | 0 | 0 |
| rs137852887 | pathogenic | 0 | 0 | 0 |
| rs137852893 | pathogenic | 0 | 0 | 0 |
| rs80338671 | pathogenic | 1 | 1 | 0 |
| rs2228389 | benign | 0 | 0 | 0 |
| rs80338672 | pathogenic | 0 | 0 | 0 |
| rs201958741 | pathogenic | 0 | 0 | 0 |
| rs80338673 | pathogenic | 0 | 0 | 0 |
| rs137852888 | pathogenic | 0 | 0 | 0 |
| rs137852889 | pathogenic | 0 | 0 | 0 |
| rs137852894 | pathogenic | 0 | 0 | 0 |
| rs137852890 | pathogenic | 0 | 0 | 0 |
| rs137852891 | pathogenic | 0 | 0 | 0 |
| rs766935302 | pathogenic | 0 | 0 | 0 |
| rs193074572 | benign | 0 | 0 | 0 |
| rs555210692 | benign | 1 | 1 | 0 |
| rs119103242 | pathogenic | 0 | 1 | 0 |
| rs143725101 | benign | 0 | 1 | 0 |
| rs28940589 | pathogenic | 1 | 1 | 1 |
| rs113994145 | pathogenic | 0 | 0 | 0 |
| rs773872994 | benign | 0 | 0 | 0 |
| rs28940590 | pathogenic | 1 | 1 | 0 |
| rs145400751 | benign | 1 | 1 | 1 |
| rs587782331 | benign | 0 | 0 | 0 |
| rs730881787 | benign | 0 | 0 | 0 |
| rs587778531 | benign | 0 | 0 | 0 |
| rs781652274 | benign | 0 | 0 | 0 |
| rs55882234 | benign | 0 | 0 | 0 |
| rs2020908 | benign | 0 | 0 | 0 |
| rs63750741 | pathogenic | 1 | 1 | 1 |
| rs41295268 | benign | 1 | 1 | 1 |
| rs63750897 | benign | 1 | 1 | 0 |

| | | | | |
|---|---|---|---|---|
| rs63751005 | benign | 1 | 1 | 0 |
| rs587779223 | benign | 0 | 0 | 0 |
| rs202127474 | benign | 0 | 0 | 0 |
| rs374230313 | benign | 0 | 0 | 0 |
| rs2020912 | benign | 0 | 0 | 0 |
| rs63751017 | pathogenic | 0 | 0 | 0 |
| rs587779246 | pathogenic | 0 | 0 | 0 |
| rs63749919 | pathogenic | 0 | 0 | 0 |
| rs63750563 | pathogenic | 1 | 1 | 1 |
| rs63749999 | pathogenic | 0 | 0 | 0 |
| rs200492211 | pathogenic | 0 | 0 | 0 |
| rs576269342 | benign | 0 | 0 | 0 |
| rs267608075 | benign | 0 | 0 | 0 |
| rs149605979 | benign | 0 | 0 | 0 |
| rs63750252 | benign | 0 | 0 | 0 |
| rs754289472 | benign | 0 | 0 | 0 |
| rs63750119 | pathogenic | 0 | 0 | 0 |
| rs587782109 | benign | 0 | 0 | 0 |
| rs34625968 | benign | 1 | 1 | 1 |
| rs764786814 | benign | 1 | 1 | 1 |
| rs199594809 | benign | 1 | 1 | 1 |
| rs74315364 | pathogenic | 1 | 1 | 1 |
| rs201253919 | pathogenic | 0 | 0 | 0 |
| rs387906828 | pathogenic | 1 | 1 | 1 |
| rs34521609 | benign | 1 | 1 | 1 |
| rs199472894 | pathogenic | 0 | 0 | 0 |
| rs199472902 | pathogenic | 0 | 0 | 0 |
| rs121912505 | pathogenic | 0 | 0 | 0 |
| rs28928905 | pathogenic | 0 | 0 | 0 |
| rs374376640 | benign | 0 | 0 | 0 |
| rs202102276 | benign | 0 | 0 | 0 |
| rs199472918 | pathogenic | 0 | 0 | 0 |
| rs121912516 | pathogenic | 1 | 1 | 1 |
| rs121912504 | pathogenic | 0 | 0 | 0 |
| rs9333649 | pathogenic | 0 | 0 | 0 |
| rs121912508 | pathogenic | 0 | 0 | 0 |
| rs199473428 | pathogenic | 1 | 1 | 1 |
| rs104894021 | pathogenic | 0 | 0 | 0 |
| rs28928904 | pathogenic | 1 | 1 | 1 |
| rs199473039 | benign | 1 | 1 | 1 |
| rs199472958 | pathogenic | 1 | 1 | 1 |
| rs1137617 | pathogenic | 1 | 1 | 1 |
| rs199473532 | pathogenic | 1 | 1 | 1 |
| rs199472988 | pathogenic | 0 | 0 | 0 |
| rs189014161 | pathogenic | 0 | 0 | 0 |
| rs12720441 | pathogenic | 0 | 0 | 0 |
| rs199473001 | pathogenic | 1 | 1 | 0 |
| rs121912506 | pathogenic | 1 | 1 | 0 |
| rs773724817 | pathogenic | 1 | 1 | 1 |
| rs118204002 | pathogenic | 1 | 1 | 1 |
| rs138556525 | pathogenic | 1 | 1 | 1 |
| rs72558181 | pathogenic | 1 | 1 | 1 |
| rs118204005 | pathogenic | 1 | 1 | 1 |
| rs118204004 | pathogenic | 0 | 0 | 0 |

| | | | | |
|---|---|---|---|---|
| rs118204001 | pathogenic | 1 | 1 | 1 |
| rs118204006 | pathogenic | 1 | 1 | 1 |
| rs118204003 | pathogenic | 0 | 0 | 0 |
| rs121913147 | pathogenic | 1 | 1 | 0 |
| rs2229429 | benign | 1 | 1 | 0 |
| rs182552223 | benign | 0 | 0 | 0 |
| rs146588336 | benign | 0 | 0 | 0 |
| rs121964983 | pathogenic | 0 | 1 | 0 |
| rs121964982 | pathogenic | 0 | 0 | 0 |
| rs386833679 | pathogenic | 0 | 0 | 0 |
| rs386833680 | pathogenic | 0 | 0 | 0 |
| rs121964981 | pathogenic | 1 | 1 | 0 |
| rs121964984 | pathogenic | 0 | 0 | 0 |
| rs144971200 | benign | 0 | 0 | 0 |
| rs121964985 | pathogenic | 1 | 1 | 0 |
| rs141246107 | benign | 0 | 0 | 0 |
| rs104894561 | pathogenic | 1 | 1 | 0 |
| rs377023736 | pathogenic | 0 | 0 | 0 |
| rs121964907 | pathogenic | 1 | 1 | 1 |
| rs121964909 | pathogenic | 1 | 1 | 1 |
| rs765070743 | pathogenic | 0 | 0 | 0 |
| rs2228119 | benign | 0 | 0 | 0 |
| rs745976989 | pathogenic | 1 | 1 | 1 |
| rs121964904 | pathogenic | 0 | 0 | 0 |
| rs386833430 | pathogenic | 1 | 1 | 1 |
| rs267606992 | pathogenic | 0 | 0 | 0 |
| rs61752063 | pathogenic | 0 | 0 | 0 |
| rs61752067 | pathogenic | 1 | 1 | 1 |
| rs104894934 | pathogenic | 0 | 0 | 0 |
| rs104894929 | pathogenic | 1 | 1 | 1 |
| rs80189640 | benign | 0 | 0 | 0 |
| rs137854547 | pathogenic | 1 | 1 | 1 |
| rs137854540 | pathogenic | 1 | 1 | 1 |
| rs782092363 | pathogenic | 1 | 1 | 1 |
| rs147783915 | benign | 0 | 0 | 0 |
| rs28933687 | pathogenic | 1 | 1 | 1 |
| rs149503319 | benign | 0 | 0 | 0 |
| rs1805147 | benign | 0 | 0 | 0 |
| rs587777218 | pathogenic | 1 | 1 | 1 |
| rs200786705 | benign | 0 | 0 | 0 |
| rs56219475 | benign | 0 | 0 | 0 |
| rs587781172 | benign | 0 | 0 | 0 |
| rs757094189 | pathogenic | 0 | 0 | 0 |
| rs140811086 | benign | 1 | 1 | 1 |
| rs574088829 | pathogenic | 1 | 1 | 1 |
| rs121918525 | pathogenic | 0 | 0 | 0 |
| rs121918527 | pathogenic | 0 | 0 | 0 |
| rs121964848 | pathogenic | 0 | 1 | 0 |
| rs121964845 | pathogenic | 1 | 1 | 1 |
| rs121964846 | pathogenic | 0 | 1 | 0 |
| rs17217716 | benign | 1 | 1 | 1 |
| rs141711342 | benign | 0 | 0 | 0 |
| rs372619120 | benign | 0 | 0 | 0 |
| rs63750002 | benign | 1 | 1 | 1 |

| | | | | |
|---|---|---|---|---|
| rs193922373 | benign | 0 | 0 | 0 |
| rs41295286 | benign | 0 | 0 | 0 |
| rs17217772 | benign | 0 | 0 | 0 |
| rs145649774 | benign | 0 | 0 | 0 |
| rs63750214 | pathogenic | 0 | 0 | 0 |
| rs63750716 | benign | 0 | 0 | 0 |
| rs202026056 | benign | 0 | 0 | 0 |
| rs4987188 | benign | 1 | 1 | 0 |
| rs267607938 | benign | 0 | 0 | 0 |
| rs17224367 | benign | 0 | 0 | 0 |
| rs63750006 | benign | 0 | 0 | 0 |
| rs587781373 | benign | 0 | 0 | 0 |
| rs63751207 | pathogenic | 1 | 1 | 0 |
| rs730881759 | benign | 0 | 0 | 0 |
| rs372350768 | benign | 1 | 1 | 1 |
| rs55778204 | benign | 1 | 1 | 0 |
| rs63749910 | benign | 0 | 0 | 0 |
| rs63750844 | benign | 0 | 0 | 0 |
| rs28929483 | pathogenic | 0 | 0 | 0 |
| rs61756468 | benign | 0 | 0 | 0 |
| rs63750875 | pathogenic | 1 | 1 | 0 |
| rs776528054 | benign | 1 | 1 | 0 |
| rs751939698 | benign | 0 | 0 | 0 |
| rs587780685 | benign | 0 | 0 | 0 |
| rs730881763 | benign | 1 | 1 | 0 |
| rs373226409 | benign | 0 | 0 | 0 |
| rs202145681 | benign | 1 | 1 | 1 |
| rs63749841 | benign | 1 | 1 | 1 |
| rs63750757 | benign | 1 | 1 | 1 |
| rs587778527 | benign | 1 | 1 | 1 |
| rs63750849 | benign | 0 | 0 | 0 |
| rs587779969 | benign | 0 | 0 | 0 |
| rs121908046 | pathogenic | 1 | 1 | 1 |
| rs28940882 | pathogenic | 1 | 1 | 0 |
| rs28940883 | pathogenic | 1 | 1 | 1 |
| rs727503943 | pathogenic | 1 | 1 | 1 |
| rs137853859 | pathogenic | 1 | 1 | 1 |
| rs121908045 | pathogenic | 1 | 1 | 0 |
| rs137853861 | pathogenic | 1 | 1 | 0 |
| rs3180383 | pathogenic | 0 | 1 | 0 |
| rs144431814 | benign | 0 | 0 | 0 |
| rs139502866 | benign | 0 | 0 | 0 |
| rs17524437 | benign | 0 | 0 | 0 |
| rs760960779 | benign | 0 | 0 | 0 |
| rs148960463 | pathogenic | 0 | 0 | 0 |
| rs111033647 | pathogenic | 1 | 1 | 1 |
| rs111033648 | pathogenic | 0 | 0 | 0 |
| rs1800461 | benign | 0 | 0 | 0 |
| rs111033656 | pathogenic | 0 | 0 | 0 |
| rs111033663 | pathogenic | 1 | 1 | 0 |
| rs111033669 | pathogenic | 1 | 1 | 1 |
| rs111033690 | pathogenic | 0 | 0 | 0 |
| rs111033686 | pathogenic | 0 | 0 | 0 |
| rs111033687 | pathogenic | 0 | 0 | 0 |

| | | | | |
|---|---|---|---|---|
| rs111033695 | pathogenic | 0 | 0 | 0 |
| rs111033697 | pathogenic | 0 | 0 | 0 |
| rs111033693 | pathogenic | 0 | 0 | 0 |
| rs111033694 | pathogenic | 0 | 0 | 0 |
| rs111033721 | pathogenic | 1 | 1 | 1 |
| rs111033722 | pathogenic | 0 | 0 | 0 |
| rs111033725 | pathogenic | 1 | 1 | 0 |
| rs75391579 | pathogenic | 1 | 1 | 1 |
| rs111033728 | pathogenic | 1 | 1 | 1 |
| rs111033735 | pathogenic | 1 | 1 | 1 |
| rs111033739 | pathogenic | 1 | 1 | 1 |
| rs111033736 | pathogenic | 0 | 0 | 0 |
| rs111033744 | pathogenic | 1 | 1 | 1 |
| rs111033750 | pathogenic | 1 | 1 | 1 |
| rs111033752 | pathogenic | 0 | 0 | 0 |
| rs111033751 | pathogenic | 0 | 0 | 0 |
| rs111033846 | pathogenic | 0 | 0 | 0 |
| rs368166217 | pathogenic | 0 | 0 | 0 |
| rs111033764 | pathogenic | 0 | 0 | 0 |
| rs111033847 | benign | 0 | 0 | 0 |
| rs111033778 | pathogenic | 0 | 0 | 0 |
| rs111033773 | pathogenic | 0 | 0 | 0 |
| rs111033792 | pathogenic | 0 | 0 | 0 |
| rs367543266 | pathogenic | 1 | 1 | 1 |
| rs111033794 | pathogenic | 0 | 0 | 0 |
| rs111033802 | pathogenic | 0 | 0 | 0 |
| rs111033803 | pathogenic | 0 | 0 | 0 |
| rs111033809 | pathogenic | 1 | 1 | 0 |
| rs121918376 | pathogenic | 0 | 1 | 0 |
| rs121918377 | pathogenic | 0 | 1 | 0 |
| rs121918378 | pathogenic | 1 | 1 | 1 |
| rs121918374 | pathogenic | 1 | 1 | 1 |
| rs137852379 | pathogenic | 0 | 0 | 0 |
| rs137852388 | pathogenic | 1 | 1 | 1 |
| rs137852368 | pathogenic | 0 | 0 | 0 |
| rs28935499 | pathogenic | 0 | 0 | 0 |
| rs28933668 | pathogenic | 0 | 0 | 0 |
| rs28933669 | pathogenic | 0 | 0 | 0 |
| rs28933670 | pathogenic | 0 | 0 | 0 |
| rs28933671 | pathogenic | 0 | 0 | 0 |
| rs28933672 | pathogenic | 0 | 0 | 0 |
| rs28937272 | pathogenic | 0 | 0 | 0 |
| rs35383156 | benign | 0 | 0 | 0 |
| rs137852418 | pathogenic | 0 | 0 | 0 |
| rs137852419 | pathogenic | 0 | 0 | 0 |
| rs137852428 | pathogenic | 0 | 0 | 0 |
| rs137852436 | pathogenic | 0 | 0 | 0 |
| rs28937285 | pathogenic | 0 | 0 | 0 |
| rs35213384 | benign | 1 | 1 | 1 |
| rs3810950 | benign | 0 | 0 | 0 |
| rs201479289 | pathogenic | 0 | 0 | 0 |
| rs121912820 | pathogenic | 0 | 1 | 0 |
| rs121912815 | pathogenic | 0 | 1 | 0 |
| rs115510708 | benign | 0 | 0 | 0 |

| | | | | |
|---|---|---|---|---|
| rs868749 | benign | 0 | 0 | 0 |
| rs121912823 | pathogenic | 0 | 0 | 0 |
| rs115212829 | benign | 0 | 0 | 0 |
| rs121912822 | pathogenic | 0 | 0 | 0 |
| rs121912816 | pathogenic | 0 | 1 | 0 |
| rs4838544 | benign | 0 | 0 | 0 |
| rs121912818 | pathogenic | 0 | 0 | 0 |
| rs121912821 | pathogenic | 0 | 0 | 0 |
| rs121912817 | pathogenic | 0 | 0 | 0 |
| rs8178992 | benign | 0 | 0 | 0 |
| rs121912819 | pathogenic | 1 | 1 | 0 |
| rs80097077 | benign | 1 | 1 | 0 |
| rs121909359 | pathogenic | 1 | 1 | 0 |
| rs121909358 | pathogenic | 1 | 1 | 1 |
| rs121909361 | pathogenic | 1 | 1 | 1 |
| rs121909357 | pathogenic | 0 | 1 | 0 |
| rs121909365 | pathogenic | 1 | 1 | 1 |
| rs6413484 | pathogenic | 1 | 1 | 1 |
| rs121909373 | pathogenic | 1 | 1 | 1 |
| rs121909366 | pathogenic | 1 | 1 | 1 |
| rs121909367 | pathogenic | 1 | 1 | 1 |
| rs121909368 | pathogenic | 1 | 1 | 1 |
| rs121909369 | pathogenic | 0 | 0 | 0 |
| rs121909363 | pathogenic | 1 | 1 | 1 |
| rs45588036 | pathogenic | 1 | 1 | 1 |
| rs121909364 | pathogenic | 1 | 1 | 1 |
| rs80358282 | benign | 0 | 0 | 0 |
| rs80358284 | pathogenic | 1 | 1 | 1 |
| rs61735303 | benign | 0 | 0 | 0 |
| rs2229274 | benign | 0 | 0 | 0 |
| rs145006491 | benign | 0 | 0 | 0 |
| rs121913580 | pathogenic | 1 | 1 | 0 |
| rs730882196 | pathogenic | 0 | 0 | 0 |
| rs153477 | benign | 0 | 0 | 0 |
| rs153478 | benign | 1 | 1 | 0 |
| rs61740602 | benign | 1 | 1 | 0 |
| rs104893892 | pathogenic | 1 | 1 | 0 |
| rs121909595 | pathogenic | 1 | 1 | 1 |
| rs28931605 | pathogenic | 1 | 1 | 1 |
| rs121909597 | pathogenic | 0 | 1 | 0 |
| rs202233735 | pathogenic | 0 | 1 | 0 |
| rs121909596 | pathogenic | 1 | 1 | 1 |
| rs75156162 | benign | 0 | 1 | 0 |
| rs2305430 | benign | 1 | 1 | 1 |
| rs150318966 | benign | 0 | 0 | 0 |
| rs121909598 | pathogenic | 0 | 1 | 0 |
| rs5744175 | benign | 0 | 0 | 0 |
| rs28935476 | pathogenic | 1 | 1 | 1 |
| rs28935475 | pathogenic | 1 | 1 | 1 |
| rs122461163 | pathogenic | 1 | 1 | 1 |
| rs62626305 | pathogenic | 1 | 1 | 1 |
| rs587777157 | pathogenic | 0 | 0 | 0 |
| rs137854478 | pathogenic | 1 | 1 | 0 |
| rs137854465 | pathogenic | 0 | 0 | 0 |

| | | | | |
|---|---|---|---|---|
| rs137854470 | pathogenic | 0 | 0 | 0 |
| rs137854482 | pathogenic | 0 | 0 | 0 |
| rs137854473 | pathogenic | 0 | 0 | 0 |
| rs137854456 | pathogenic | 0 | 0 | 0 |
| rs140598 | benign | 0 | 0 | 0 |
| rs104894105 | pathogenic | 1 | 1 | 1 |
| rs104894107 | pathogenic | 1 | 1 | 1 |
| rs104894106 | pathogenic | 1 | 1 | 1 |
| rs74621026 | benign | 1 | 1 | 1 |
| rs2290332 | benign | 0 | 0 | 0 |
| rs780202604 | pathogenic | 1 | 1 | 1 |
| rs74315305 | pathogenic | 1 | 1 | 0 |
| rs766751878 | pathogenic | 1 | 1 | 1 |
| rs137854603 | pathogenic | 1 | 1 | 1 |
| rs139592595 | benign | 1 | 1 | 0 |
| rs150565592 | benign | 1 | 1 | 1 |
| rs763520126 | benign | 0 | 0 | 0 |
| rs55980502 | benign | 0 | 0 | 0 |
| rs137852812 | pathogenic | 0 | 0 | 0 |
| rs137852813 | pathogenic | 0 | 0 | 0 |
| rs190222208 | benign | 0 | 0 | 0 |
| rs138459502 | pathogenic | 0 | 0 | 0 |
| rs730881041 | pathogenic | 0 | 0 | 0 |
| rs267607080 | pathogenic | 0 | 0 | 0 |
| rs137852814 | pathogenic | 0 | 0 | 0 |
| rs267607079 | pathogenic | 0 | 0 | 0 |
| rs192378417 | benign | 0 | 0 | 0 |
| rs1288775 | benign | 0 | 0 | 0 |
| rs80338737 | pathogenic | 1 | 1 | 1 |
| rs397514708 | pathogenic | 0 | 0 | 0 |
| rs775933965 | pathogenic | 0 | 0 | 0 |
| rs747608698 | benign | 0 | 1 | 0 |
| rs1145086 | benign | 0 | 1 | 0 |
| rs121912953 | pathogenic | 1 | 1 | 0 |
| rs243849 | benign | 0 | 0 | 0 |
| rs121912955 | pathogenic | 1 | 1 | 1 |
| rs17859943 | benign | 0 | 1 | 0 |
| rs755557988 | benign | 1 | 1 | 1 |
| rs10195 | benign | 0 | 0 | 0 |
| rs61735357 | benign | 1 | 1 | 1 |
| rs137882374 | benign | 0 | 0 | 0 |
| rs5388 | benign | 0 | 0 | 0 |
| rs6352 | benign | 0 | 0 | 0 |
| rs76645461 | pathogenic | 1 | 1 | 0 |
| rs1050828 | pathogenic | 0 | 0 | 0 |
| rs1050829 | pathogenic | 0 | 0 | 0 |
| rs137852314 | pathogenic | 0 | 0 | 0 |
| rs137852331 | pathogenic | 0 | 0 | 0 |
| rs5030872 | pathogenic | 0 | 0 | 0 |
| rs267606836 | pathogenic | 0 | 0 | 0 |
| rs5030868 | pathogenic | 0 | 0 | 0 |
| rs137852330 | pathogenic | 0 | 1 | 0 |
| rs137852326 | pathogenic | 1 | 1 | 1 |
| rs137852318 | pathogenic | 1 | 1 | 1 |

| | | | | |
|---|---|---|---|---|
| rs74575103 | benign | 0 | 0 | 0 |
| rs137852327 | pathogenic | 1 | 1 | 1 |
| rs76723693 | pathogenic | 0 | 0 | 0 |
| rs5030869 | pathogenic | 0 | 0 | 0 |
| rs34193178 | pathogenic | 1 | 1 | 1 |
| rs137852316 | pathogenic | 1 | 1 | 0 |
| rs398123546 | pathogenic | 1 | 1 | 1 |
| rs72554665 | pathogenic | 0 | 0 | 0 |
| rs72554664 | pathogenic | 0 | 0 | 0 |
| rs28939068 | pathogenic | 1 | 1 | 1 |
| rs59301204 | pathogenic | 1 | 1 | 1 |
| rs143715750 | benign | 1 | 1 | 1 |
| rs35263902 | pathogenic | 1 | 1 | 1 |
| rs202214518 | benign | 1 | 1 | 1 |
| rs34076756 | benign | 0 | 0 | 0 |
| rs199592341 | pathogenic | 1 | 1 | 0 |
| rs121908833 | pathogenic | 1 | 1 | 1 |
| rs121908834 | pathogenic | 1 | 1 | 0 |
| rs138454151 | benign | 0 | 0 | 0 |
| rs141323599 | benign | 0 | 0 | 0 |
| rs137853200 | pathogenic | 0 | 0 | 0 |
| rs137853199 | pathogenic | 0 | 0 | 0 |
| rs139787163 | benign | 0 | 0 | 0 |
| rs61731097 | benign | 1 | 1 | 1 |
| rs61729795 | benign | 0 | 0 | 0 |
| rs149242330 | benign | 0 | 0 | 0 |
| rs587777490 | pathogenic | 1 | 1 | 1 |
| rs398123011 | pathogenic | 1 | 1 | 0 |
| rs775186117 | pathogenic | 0 | 0 | 0 |
| rs72677237 | benign | 0 | 0 | 0 |
| rs104894163 | pathogenic | 1 | 1 | 1 |
| rs121909588 | pathogenic | 0 | 0 | 0 |
| rs1042579 | benign | 0 | 0 | 0 |
| rs121909273 | pathogenic | 0 | 0 | 0 |
| rs121909274 | pathogenic | 0 | 0 | 0 |
| rs119473034 | pathogenic | 1 | 1 | 1 |
| rs398123065 | pathogenic | 0 | 0 | 0 |
| rs587777042 | pathogenic | 1 | 1 | 0 |
| rs761413864 | pathogenic | 1 | 1 | 1 |
| rs118203965 | pathogenic | 0 | 0 | 0 |
| rs118203966 | pathogenic | 0 | 0 | 0 |
| rs1801201 | benign | 1 | 1 | 0 |
| rs1136201 | benign | 1 | 1 | 1 |
| rs144148008 | benign | 0 | 0 | 0 |
| rs200118378 | benign | 1 | 1 | 1 |
| rs112984118 | benign | 1 | 1 | 1 |
| rs149446102 | benign | 1 | 1 | 1 |
| rs1427463 | benign | 0 | 0 | 0 |
| rs201936720 | benign | 1 | 1 | 1 |
| rs61751983 | benign | 1 | 1 | 1 |
| rs104894632 | pathogenic | 1 | 1 | 1 |
| rs149202905 | benign | 0 | 0 | 0 |
| rs2230210 | benign | 1 | 1 | 1 |
| rs111631988 | benign | 0 | 0 | 0 |

| | | | | |
|---|---|---|---|---|
| rs137852550 | pathogenic | 0 | 0 | 0 |
| rs5963403 | benign | 0 | 0 | 0 |
| rs138018739 | benign | 0 | 0 | 0 |
| rs62640590 | benign | 0 | 0 | 0 |
| rs121909581 | pathogenic | 1 | 1 | 1 |
| rs202017913 | benign | 1 | 1 | 1 |
| rs143163513 | benign | 0 | 0 | 0 |
| rs238238 | benign | 0 | 1 | 0 |
| rs143945974 | benign | 0 | 0 | 0 |
| rs238239 | benign | 1 | 1 | 1 |
| rs121918403 | pathogenic | 0 | 1 | 0 |
| rs61735456 | benign | 0 | 1 | 0 |
| rs121918404 | pathogenic | 1 | 1 | 0 |
| rs12674488 | benign | 0 | 0 | 0 |
| rs201599657 | pathogenic | 0 | 0 | 0 |
| rs1057090 | benign | 0 | 0 | 0 |
| rs1057091 | benign | 0 | 0 | 0 |
| rs587783740 | benign | 0 | 0 | 0 |
| rs75956769 | benign | 0 | 0 | 0 |
| rs201691299 | benign | 0 | 0 | 0 |
| rs62621087 | benign | 0 | 0 | 0 |
| rs201308454 | benign | 0 | 0 | 0 |
| rs145657360 | benign | 0 | 0 | 0 |
| rs121434592 | pathogenic | 1 | 1 | 0 |
| rs375650413 | benign | 0 | 0 | 0 |
| rs121434437 | pathogenic | 1 | 1 | 0 |
| rs763494292 | pathogenic | 0 | 0 | 0 |
| rs121964997 | pathogenic | 1 | 1 | 0 |
| rs121964996 | pathogenic | 0 | 0 | 0 |
| rs41283425 | benign | 1 | 1 | 1 |
| rs376289528 | benign | 1 | 1 | 1 |
| rs144171604 | benign | 1 | 1 | 1 |
| rs782440692 | pathogenic | 1 | 1 | 1 |
| rs782058451 | pathogenic | 1 | 1 | 1 |
| rs78437817 | benign | 0 | 0 | 0 |
| rs200327969 | benign | 1 | 1 | 1 |
| rs143043662 | benign | 0 | 0 | 0 |
| rs111966833 | benign | 1 | 1 | 1 |
| rs35523678 | benign | 0 | 0 | 0 |
| rs515726208 | pathogenic | 0 | 0 | 0 |
| rs397517239 | pathogenic | 1 | 1 | 1 |
| rs148966602 | benign | 1 | 1 | 1 |
| rs76371972 | benign | 1 | 1 | 1 |
| rs2229968 | benign | 1 | 1 | 1 |
| rs61751538 | benign | 1 | 1 | 1 |
| rs200800978 | pathogenic | 1 | 1 | 1 |
| rs148695964 | benign | 1 | 1 | 1 |
| rs140042418 | benign | 1 | 1 | 1 |
| rs116185403 | benign | 0 | 0 | 0 |
| rs142828669 | benign | 1 | 1 | 1 |
| rs139832048 | benign | 0 | 1 | 0 |
| rs58750568 | benign | 0 | 1 | 0 |
| rs77681311 | benign | 0 | 0 | 0 |
| rs151208452 | benign | 0 | 0 | 0 |

| | | | | |
|---|---|---|---|---|
| rs76463933 | benign | 0 | 0 | 0 |
| rs144434647 | benign | 0 | 0 | 0 |
| rs117361485 | benign | 0 | 0 | 0 |
| rs587777588 | pathogenic | 0 | 0 | 0 |
| rs80338796 | pathogenic | 0 | 0 | 0 |
| rs121434594 | pathogenic | 0 | 0 | 0 |
| rs146766138 | pathogenic | 1 | 1 | 1 |
| rs28939679 | pathogenic | 1 | 1 | 1 |
| rs748754134 | pathogenic | 1 | 1 | 0 |
| rs764276946 | pathogenic | 1 | 1 | 1 |
| rs111033588 | pathogenic | 1 | 1 | 1 |
| rs201633414 | pathogenic | 1 | 1 | 1 |
| rs121909262 | pathogenic | 1 | 1 | 1 |
| rs104893702 | pathogenic | 1 | 1 | 1 |
| rs121909265 | pathogenic | 1 | 1 | 1 |
| rs121909266 | pathogenic | 1 | 1 | 1 |
| rs193922439 | pathogenic | 1 | 1 | 0 |
| rs104893691 | pathogenic | 1 | 1 | 1 |
| rs104893695 | pathogenic | 0 | 0 | 0 |
| rs104893708 | pathogenic | 1 | 1 | 1 |
| rs121909260 | pathogenic | 0 | 0 | 0 |
| rs104893696 | pathogenic | 0 | 0 | 0 |
| rs121909267 | pathogenic | 0 | 0 | 0 |
| rs121909263 | pathogenic | 0 | 0 | 0 |
| rs121909264 | pathogenic | 0 | 0 | 0 |
| rs104893694 | pathogenic | 1 | 1 | 1 |
| rs121909268 | pathogenic | 1 | 1 | 1 |
| rs104893689 | pathogenic | 1 | 1 | 0 |
| rs104893707 | pathogenic | 1 | 1 | 0 |
| rs104893697 | pathogenic | 1 | 1 | 1 |
| rs28936684 | pathogenic | 1 | 1 | 1 |
| rs62269092 | benign | 1 | 1 | 0 |
| rs121909259 | pathogenic | 1 | 1 | 0 |
| rs548403340 | benign | 1 | 1 | 0 |
| rs142818334 | benign | 0 | 0 | 0 |
| rs12493789 | benign | 1 | 1 | 1 |
| rs193922423 | pathogenic | 1 | 1 | 0 |
| rs104893719 | pathogenic | 1 | 1 | 1 |
| rs117375173 | benign | 1 | 1 | 0 |
| rs104893712 | pathogenic | 0 | 0 | 0 |
| rs121917776 | pathogenic | 1 | 1 | 1 |
| rs767325003 | benign | 1 | 1 | 1 |
| rs35824453 | benign | 0 | 0 | 0 |
| rs141641266 | benign | 0 | 0 | 0 |
| rs1050239 | benign | 0 | 0 | 0 |
| rs141303384 | benign | 0 | 0 | 0 |
| rs12720458 | pathogenic | 0 | 0 | 0 |
| rs368507376 | pathogenic | 1 | 1 | 1 |
| rs17215500 | pathogenic | 1 | 1 | 1 |
| rs120074188 | pathogenic | 1 | 1 | 1 |
| rs397508097 | pathogenic | 1 | 1 | 1 |
| rs138551008 | pathogenic | 0 | 0 | 0 |
| rs387907139 | pathogenic | 1 | 1 | 1 |
| rs765487627 | pathogenic | 0 | 0 | 0 |

| | | | | |
|---|---|---|---|---|
| rs587778587 | benign | 0 | 0 | 0 |
| rs45568339 | benign | 0 | 0 | 0 |
| rs78179744 | benign | 0 | 0 | 0 |
| rs45624036 | benign | 0 | 0 | 0 |
| rs61755166 | benign | 0 | 0 | 0 |
| rs180177132 | pathogenic | 1 | 1 | 1 |
| rs587776527 | pathogenic | 0 | 0 | 0 |
| rs373783514 | benign | 0 | 0 | 0 |
| rs730881896 | benign | 0 | 0 | 0 |
| rs118203998 | pathogenic | 0 | 0 | 0 |
| rs41281039 | pathogenic | 1 | 1 | 1 |
| rs118204430 | pathogenic | 1 | 1 | 1 |
| rs1800546 | pathogenic | 1 | 1 | 0 |
| rs76917243 | pathogenic | 0 | 1 | 0 |
| rs78340951 | pathogenic | 0 | 1 | 0 |
| rs77718928 | pathogenic | 0 | 0 | 0 |
| rs369586696 | pathogenic | 0 | 1 | 0 |
| rs745517517 | pathogenic | 1 | 1 | 0 |
| rs372872777 | pathogenic | 0 | 0 | 0 |
| rs748704459 | pathogenic | 0 | 0 | 0 |
| rs769215411 | pathogenic | 0 | 0 | 0 |
| rs139425890 | pathogenic | 0 | 0 | 0 |
| rs779694939 | pathogenic | 1 | 1 | 0 |
| rs773729410 | pathogenic | 0 | 0 | 0 |
| rs121908623 | pathogenic | 1 | 1 | 1 |
| rs121908622 | pathogenic | 1 | 1 | 1 |
| rs121908621 | pathogenic | 1 | 1 | 1 |
| rs762106720 | pathogenic | 1 | 1 | 0 |
| rs199877522 | pathogenic | 0 | 0 | 0 |
| rs74315291 | pathogenic | 1 | 1 | 1 |
| rs79137415 | benign | 0 | 0 | 0 |
| rs121913045 | pathogenic | 0 | 0 | 0 |
| rs34295337 | pathogenic | 0 | 0 | 0 |
| rs121913046 | pathogenic | 0 | 0 | 0 |
| rs121913047 | pathogenic | 0 | 0 | 0 |
| rs121913048 | pathogenic | 0 | 0 | 0 |
| rs267607166 | pathogenic | 0 | 0 | 0 |
| rs17850684 | benign | 1 | 1 | 1 |
| rs119476051 | pathogenic | 1 | 1 | 1 |
| rs17850683 | benign | 1 | 1 | 1 |
| rs119476049 | pathogenic | 1 | 1 | 1 |
| rs119476046 | pathogenic | 1 | 1 | 1 |
| rs119476048 | pathogenic | 0 | 0 | 0 |
| rs119476047 | pathogenic | 1 | 1 | 1 |
| rs28939094 | pathogenic | 1 | 1 | 1 |
| rs119476050 | pathogenic | 0 | 0 | 0 |
| rs387906941 | pathogenic | 1 | 1 | 1 |
| rs41511344 | pathogenic | 1 | 1 | 0 |
| rs748112833 | pathogenic | 1 | 1 | 1 |
| rs74342080 | benign | 1 | 1 | 0 |
| rs116482870 | benign | 0 | 0 | 0 |
| rs104893743 | pathogenic | 0 | 0 | 0 |
| rs117179004 | benign | 0 | 0 | 0 |
| rs587779409 | pathogenic | 1 | 1 | 1 |

| | | | | |
|---|---|---|---|---|
| rs3745406 | benign | 1 | 1 | 0 |
| rs387906696 | pathogenic | 0 | 0 | 0 |
| rs72549369 | pathogenic | 1 | 1 | 0 |
| rs45523133 | benign | 0 | 0 | 0 |
| rs119460972 | pathogenic | 0 | 1 | 0 |
| rs34929837 | benign | 0 | 1 | 0 |
| rs17323225 | benign | 0 | 1 | 0 |
| rs17011368 | benign | 0 | 0 | 0 |
| rs748902766 | pathogenic | 1 | 1 | 1 |
| rs387907292 | pathogenic | 0 | 0 | 0 |
| rs387907293 | pathogenic | 1 | 1 | 0 |
| rs371526758 | pathogenic | 1 | 1 | 1 |
| rs142968179 | pathogenic | 1 | 1 | 1 |
| rs387907291 | pathogenic | 1 | 1 | 0 |
| rs121907949 | pathogenic | 0 | 0 | 0 |
| rs121907947 | pathogenic | 0 | 0 | 0 |
| rs121907950 | pathogenic | 1 | 1 | 1 |
| rs121907948 | pathogenic | 1 | 1 | 1 |
| rs4926 | benign | 0 | 0 | 0 |
| rs748809996 | pathogenic | 0 | 0 | 0 |
| rs1799883 | benign | 1 | 1 | 0 |
| rs104893974 | pathogenic | 0 | 1 | 0 |
| rs104893973 | pathogenic | 0 | 1 | 0 |
| rs34452707 | benign | 0 | 0 | 0 |
| rs119467003 | pathogenic | 1 | 1 | 1 |
| rs121908261 | pathogenic | 0 | 1 | 0 |
| rs28933985 | pathogenic | 1 | 1 | 1 |
| rs80356669 | pathogenic | 1 | 1 | 1 |
| rs28936677 | pathogenic | 1 | 1 | 0 |
| rs121918326 | pathogenic | 0 | 0 | 0 |
| rs121918324 | pathogenic | 1 | 1 | 0 |
| rs121918325 | pathogenic | 1 | 1 | 0 |
| rs41270025 | pathogenic | 1 | 1 | 0 |
| rs121918323 | pathogenic | 0 | 0 | 0 |
| rs12735723 | pathogenic | 1 | 1 | 0 |
| rs36013429 | benign | 0 | 0 | 0 |
| rs28936676 | pathogenic | 1 | 1 | 0 |
| rs771809901 | pathogenic | 1 | 1 | 1 |
| rs758350837 | benign | 1 | 1 | 0 |
| rs121908192 | pathogenic | 1 | 1 | 1 |
| rs147568500 | pathogenic | 1 | 1 | 1 |
| rs3729734 | benign | 0 | 0 | 0 |
| rs371536347 | benign | 1 | 1 | 1 |
| rs80358193 | pathogenic | 1 | 1 | 1 |
| rs119476044 | pathogenic | 1 | 1 | 1 |
| rs111292798 | benign | 1 | 1 | 1 |
| rs121908063 | pathogenic | 1 | 1 | 0 |
| rs121908065 | pathogenic | 1 | 1 | 1 |
| rs121908062 | pathogenic | 1 | 1 | 1 |
| rs121908064 | pathogenic | 1 | 1 | 1 |
| rs61746421 | benign | 1 | 1 | 1 |
| rs34886916 | benign | 1 | 1 | 1 |
| rs118204045 | pathogenic | 0 | 0 | 0 |
| rs138607412 | benign | 1 | 1 | 1 |

| | | | | |
|---|---|---|---|---|
| rs118204046 | pathogenic | 0 | 0 | 0 |
| rs118204048 | pathogenic | 0 | 0 | 0 |
| rs118204044 | pathogenic | 0 | 0 | 0 |
| rs149664056 | benign | 0 | 0 | 0 |
| rs121909285 | pathogenic | 0 | 0 | 0 |
| rs1802498 | benign | 0 | 0 | 0 |
| rs1065761 | benign | 0 | 0 | 0 |
| rs119477054 | pathogenic | 0 | 0 | 0 |
| rs12720097 | benign | 0 | 0 | 0 |
| rs747281324 | pathogenic | 1 | 1 | 0 |
| rs367543028 | pathogenic | 1 | 1 | 0 |
| rs146077918 | benign | 0 | 0 | 0 |
| rs142551229 | benign | 0 | 0 | 0 |
| rs367543039 | pathogenic | 0 | 0 | 0 |
| rs587779884 | pathogenic | 1 | 1 | 0 |
| rs137853153 | pathogenic | 0 | 0 | 0 |
| rs367543029 | pathogenic | 0 | 0 | 0 |
| rs587783037 | pathogenic | 1 | 1 | 0 |
| rs142928725 | benign | 0 | 0 | 0 |
| rs28385141 | benign | 0 | 0 | 0 |
| rs587779888 | pathogenic | 0 | 0 | 0 |
| rs121909801 | pathogenic | 1 | 1 | 1 |
| rs121909790 | pathogenic | 1 | 1 | 0 |
| rs121909797 | pathogenic | 1 | 1 | 0 |
| rs121909794 | pathogenic | 1 | 1 | 1 |
| rs121909791 | pathogenic | 1 | 1 | 1 |
| rs121909793 | pathogenic | 1 | 1 | 1 |
| rs147977279 | benign | 1 | 1 | 1 |
| rs147352564 | benign | 0 | 0 | 0 |
| rs368758045 | benign | 1 | 1 | 1 |
| rs114498261 | benign | 1 | 1 | 0 |
| rs35484156 | benign | 0 | 0 | 0 |
| rs139176878 | benign | 0 | 0 | 0 |
| rs11543934 | benign | 0 | 0 | 0 |
| rs34192549 | benign | 0 | 0 | 0 |
| rs568736295 | benign | 0 | 0 | 0 |
| rs55747232 | benign | 0 | 0 | 0 |
| rs781887272 | benign | 0 | 0 | 0 |
| rs146273341 | benign | 0 | 0 | 0 |
| rs41473544 | benign | 0 | 0 | 0 |
| rs137852680 | pathogenic | 0 | 0 | 0 |
| rs55891000 | benign | 0 | 0 | 0 |
| rs140351189 | benign | 0 | 0 | 0 |
| rs137852679 | pathogenic | 1 | 1 | 0 |
| rs137852678 | pathogenic | 0 | 0 | 0 |
| rs137852677 | pathogenic | 1 | 1 | 1 |
| rs138620818 | benign | 0 | 0 | 0 |
| rs121908531 | pathogenic | 0 | 0 | 0 |
| rs72554307 | pathogenic | 0 | 0 | 0 |
| rs72554308 | pathogenic | 0 | 0 | 0 |
| rs72554312 | pathogenic | 1 | 1 | 0 |
| rs67939655 | pathogenic | 0 | 0 | 0 |
| rs67486158 | pathogenic | 0 | 0 | 0 |
| rs72554338 | pathogenic | 1 | 1 | 1 |

| | | | | |
|---|---|---|---|---|
| rs1800324 | pathogenic | 1 | 1 | 1 |
| rs72554356 | pathogenic | 1 | 1 | 1 |
| rs66656800 | pathogenic | 1 | 1 | 1 |
| rs68026851 | pathogenic | 1 | 1 | 1 |
| rs72558412 | pathogenic | 0 | 0 | 0 |
| rs72558415 | pathogenic | 0 | 0 | 0 |
| rs72558423 | pathogenic | 0 | 0 | 0 |
| rs67120076 | pathogenic | 1 | 1 | 1 |
| rs1800328 | pathogenic | 1 | 1 | 1 |
| rs72558451 | pathogenic | 1 | 1 | 1 |
| rs72558454 | pathogenic | 1 | 1 | 1 |
| rs672601372 | pathogenic | 0 | 0 | 0 |
| rs139644798 | pathogenic | 0 | 0 | 0 |
| rs369398935 | pathogenic | 0 | 0 | 0 |
| rs267607035 | pathogenic | 0 | 0 | 0 |
| rs267607036 | pathogenic | 1 | 1 | 1 |
| rs2302566 | benign | 1 | 1 | 0 |
| rs115959591 | benign | 0 | 0 | 0 |
| rs773876739 | pathogenic | 0 | 0 | 0 |
| rs1799967 | benign | 0 | 0 | 0 |
| rs80356968 | benign | 0 | 0 | 0 |
| rs28897695 | benign | 1 | 1 | 1 |
| rs80357314 | benign | 0 | 0 | 0 |
| rs80357169 | benign | 0 | 0 | 0 |
| rs397507239 | benign | 1 | 1 | 1 |
| rs80187739 | pathogenic | 1 | 1 | 1 |
| rs41293459 | pathogenic | 1 | 1 | 1 |
| rs55770810 | pathogenic | 1 | 1 | 1 |
| rs80356860 | pathogenic | 0 | 0 | 0 |
| rs56195342 | benign | 0 | 0 | 0 |
| rs80357442 | benign | 0 | 0 | 0 |
| rs41293463 | pathogenic | 1 | 1 | 1 |
| rs80357069 | pathogenic | 0 | 0 | 0 |
| rs397509275 | benign | 0 | 0 | 0 |
| rs80356920 | benign | 0 | 0 | 0 |
| rs397509284 | pathogenic | 0 | 0 | 0 |
| rs80357286 | benign | 0 | 0 | 0 |
| rs80357274 | benign | 0 | 0 | 0 |
| rs765656957 | benign | 0 | 0 | 0 |
| rs755427809 | benign | 0 | 0 | 0 |
| rs80357322 | benign | 0 | 0 | 0 |
| rs121434430 | pathogenic | 0 | 0 | 0 |
| rs121918450 | pathogenic | 1 | 1 | 1 |
| rs1532268 | benign | 0 | 0 | 0 |
| rs2303080 | benign | 0 | 0 | 0 |
| rs10064631 | benign | 0 | 0 | 0 |
| rs162036 | benign | 0 | 0 | 0 |
| rs2287780 | benign | 0 | 0 | 0 |
| rs16879334 | benign | 0 | 0 | 0 |
| rs137853061 | pathogenic | 1 | 1 | 0 |
| rs10380 | benign | 0 | 0 | 0 |
| rs1494558 | benign | 0 | 0 | 0 |
| rs1494555 | benign | 0 | 0 | 0 |
| rs192994060 | benign | 0 | 0 | 0 |

| | | | | |
|---|---|---|---|---|
| rs147128920 | benign | 1 | 1 | 0 |
| rs201803986 | pathogenic | 0 | 0 | 0 |
| rs746419489 | pathogenic | 0 | 1 | 0 |
| rs764396564 | pathogenic | 0 | 0 | 0 |
| rs758304537 | pathogenic | 0 | 0 | 0 |
| rs267606763 | pathogenic | 0 | 0 | 0 |
| rs267606762 | pathogenic | 0 | 0 | 0 |
| rs150702945 | pathogenic | 1 | 1 | 1 |
| rs202047589 | pathogenic | 0 | 0 | 0 |
| rs755562733 | pathogenic | 0 | 0 | 0 |
| rs267606764 | pathogenic | 1 | 1 | 1 |
| rs138207257 | pathogenic | 1 | 1 | 0 |
| rs149150736 | pathogenic | 1 | 1 | 1 |
| rs777046879 | pathogenic | 0 | 0 | 0 |
| rs104894408 | pathogenic | 1 | 1 | 1 |
| rs1801002 | pathogenic | 1 | 1 | 1 |
| rs28929485 | pathogenic | 1 | 1 | 1 |
| rs104894396 | pathogenic | 1 | 1 | 1 |
| rs371024165 | pathogenic | 0 | 0 | 0 |
| rs35887622 | pathogenic | 1 | 1 | 1 |
| rs587783644 | pathogenic | 1 | 1 | 1 |
| rs72474224 | pathogenic | 1 | 1 | 1 |
| rs111033296 | pathogenic | 1 | 1 | 1 |
| rs104894407 | pathogenic | 1 | 1 | 1 |
| rs72561723 | pathogenic | 1 | 1 | 1 |
| rs104894398 | pathogenic | 1 | 1 | 1 |
| rs28931594 | pathogenic | 1 | 1 | 1 |
| rs104894412 | pathogenic | 1 | 1 | 1 |
| rs111033297 | pathogenic | 1 | 1 | 1 |
| rs104894404 | pathogenic | 1 | 1 | 1 |
| rs104894410 | pathogenic | 1 | 1 | 1 |
| rs104894403 | pathogenic | 1 | 1 | 1 |
| rs397516870 | benign | 0 | 0 | 0 |
| rs200023879 | pathogenic | 1 | 1 | 1 |
| rs121912968 | pathogenic | 0 | 0 | 0 |
| rs104894402 | pathogenic | 1 | 1 | 1 |
| rs28931593 | pathogenic | 1 | 1 | 1 |
| rs104894397 | pathogenic | 0 | 0 | 0 |
| rs80338944 | pathogenic | 0 | 0 | 0 |
| rs199883710 | pathogenic | 0 | 0 | 0 |
| rs727504302 | pathogenic | 0 | 0 | 0 |
| rs145216882 | benign | 1 | 1 | 1 |
| rs781534323 | pathogenic | 1 | 1 | 1 |
| rs111033218 | benign | 1 | 1 | 1 |
| rs104894409 | pathogenic | 0 | 0 | 0 |
| rs80338945 | pathogenic | 1 | 1 | 1 |
| rs111033299 | pathogenic | 0 | 0 | 0 |
| rs143343083 | pathogenic | 1 | 1 | 1 |
| rs80338946 | pathogenic | 0 | 0 | 0 |
| rs150529554 | pathogenic | 0 | 0 | 0 |
| rs111033295 | pathogenic | 0 | 0 | 0 |
| rs111033188 | benign | 0 | 0 | 0 |
| rs397516874 | pathogenic | 0 | 0 | 0 |
| rs76434661 | pathogenic | 0 | 0 | 0 |

| | | | | |
|---|---|---|---|---|
| rs80338948 | pathogenic | 0 | 0 | 0 |
| rs104894401 | pathogenic | 0 | 0 | 0 |
| rs767178508 | pathogenic | 0 | 0 | 0 |
| rs772264564 | pathogenic | 0 | 0 | 0 |
| rs28931592 | pathogenic | 0 | 0 | 0 |
| rs80338949 | pathogenic | 1 | 1 | 1 |
| rs376898963 | benign | 1 | 1 | 1 |
| rs111033360 | benign | 0 | 0 | 0 |
| rs774518779 | pathogenic | 1 | 1 | 1 |
| rs28931595 | pathogenic | 1 | 1 | 1 |
| rs771748289 | pathogenic | 0 | 0 | 0 |
| rs104894406 | pathogenic | 0 | 0 | 0 |
| rs76838169 | benign | 0 | 0 | 0 |
| rs111033294 | pathogenic | 0 | 0 | 0 |
| rs200214298 | pathogenic | 0 | 0 | 0 |
| rs28940283 | pathogenic | 0 | 1 | 0 |
| rs1047883 | benign | 0 | 0 | 0 |
| rs138254425 | benign | 0 | 0 | 0 |
| rs121912592 | pathogenic | 1 | 1 | 0 |
| rs369061090 | pathogenic | 1 | 1 | 0 |
| rs758724746 | pathogenic | 0 | 1 | 0 |
| rs121912595 | pathogenic | 0 | 0 | 0 |
| rs76340296 | benign | 0 | 0 | 0 |
| rs140578009 | benign | 1 | 1 | 1 |
| rs74445297 | benign | 0 | 0 | 0 |
| rs78599516 | benign | 1 | 1 | 0 |
| rs104894035 | pathogenic | 1 | 1 | 0 |
| rs79451216 | benign | 0 | 0 | 0 |
| rs104894036 | pathogenic | 0 | 0 | 0 |
| rs73343757 | benign | 0 | 0 | 0 |
| rs121917722 | pathogenic | 1 | 1 | 1 |
| rs747700126 | pathogenic | 0 | 1 | 0 |
| rs121917725 | pathogenic | 1 | 1 | 1 |
| rs121917721 | pathogenic | 1 | 1 | 1 |
| rs121917723 | pathogenic | 1 | 1 | 0 |
| rs17569 | benign | 1 | 1 | 1 |
| rs2230062 | benign | 1 | 1 | 1 |
| rs17570 | benign | 0 | 0 | 0 |
| rs121917724 | pathogenic | 0 | 1 | 0 |
| rs199422235 | pathogenic | 0 | 0 | 0 |
| rs199422237 | pathogenic | 1 | 1 | 1 |
| rs199422234 | pathogenic | 1 | 1 | 0 |
| rs140015315 | pathogenic | 1 | 1 | 1 |
| rs730880298 | pathogenic | 1 | 1 | 1 |
| rs779692470 | pathogenic | 0 | 0 | 0 |
| rs369152939 | pathogenic | 0 | 0 | 0 |
| rs527236040 | pathogenic | 0 | 0 | 0 |
| rs587776985 | pathogenic | 1 | 1 | 1 |
| rs147077598 | pathogenic | 1 | 1 | 1 |
| rs128620185 | pathogenic | 1 | 1 | 1 |
| rs128620189 | pathogenic | 0 | 0 | 0 |
| rs128621190 | pathogenic | 0 | 0 | 0 |
| rs61754648 | benign | 0 | 0 | 0 |
| rs137852906 | pathogenic | 0 | 0 | 0 |

| | | | | |
|---|---|---|---|---|
| rs76811038 | pathogenic | 0 | 0 | 0 |
| rs149468422 | benign | 0 | 0 | 0 |
| rs5911 | benign | 0 | 0 | 0 |
| rs28942112 | pathogenic | 1 | 1 | 1 |
| rs143275858 | pathogenic | 1 | 1 | 0 |
| rs28362261 | benign | 0 | 0 | 0 |
| rs28362263 | benign | 1 | 1 | 1 |
| rs558707786 | pathogenic | 0 | 0 | 0 |
| rs138775799 | pathogenic | 0 | 0 | 0 |
| rs748170941 | pathogenic | 0 | 0 | 0 |
| rs121908380 | pathogenic | 0 | 0 | 0 |
| rs765123255 | pathogenic | 1 | 1 | 1 |
| rs730881832 | pathogenic | 0 | 1 | 0 |
| rs587782165 | benign | 1 | 1 | 1 |
| rs762307622 | pathogenic | 0 | 0 | 0 |
| rs34612342 | pathogenic | 1 | 1 | 0 |
| rs143353451 | pathogenic | 0 | 1 | 0 |
| rs747993448 | pathogenic | 0 | 1 | 0 |
| rs376561094 | pathogenic | 0 | 0 | 0 |
| rs776487884 | pathogenic | 1 | 1 | 0 |
| rs34126013 | pathogenic | 1 | 1 | 0 |
| rs200495564 | pathogenic | 1 | 1 | 0 |
| rs140342925 | pathogenic | 1 | 1 | 0 |
| rs587780749 | pathogenic | 1 | 1 | 0 |
| rs773087549 | pathogenic | 1 | 1 | 0 |
| rs730881833 | pathogenic | 0 | 0 | 0 |
| rs374950566 | pathogenic | 1 | 1 | 0 |
| rs587780752 | benign | 0 | 0 | 0 |
| rs3219489 | benign | 0 | 0 | 0 |
| rs199742231 | benign | 0 | 0 | 0 |
| rs587780082 | pathogenic | 0 | 0 | 0 |
| rs35881803 | benign | 0 | 0 | 0 |
| rs41305623 | benign | 1 | 1 | 1 |
| rs121917856 | pathogenic | 1 | 1 | 1 |
| rs576284122 | benign | 0 | 0 | 0 |
| rs142036701 | benign | 0 | 0 | 0 |
| rs121917852 | pathogenic | 1 | 1 | 1 |
| rs121917854 | pathogenic | 1 | 1 | 1 |
| rs121917853 | pathogenic | 1 | 1 | 1 |
| rs121917857 | pathogenic | 0 | 1 | 0 |
| rs193922740 | pathogenic | 1 | 1 | 1 |
| rs565779970 | pathogenic | 1 | 1 | 1 |
| rs6723526 | benign | 0 | 0 | 0 |
| rs141094096 | pathogenic | 1 | 1 | 1 |
| rs104894637 | pathogenic | 0 | 0 | 0 |
| rs104894636 | pathogenic | 0 | 0 | 0 |
| rs104894642 | pathogenic | 0 | 0 | 0 |
| rs104894638 | pathogenic | 0 | 0 | 0 |
| rs753666460 | pathogenic | 0 | 0 | 0 |
| rs104894643 | pathogenic | 1 | 1 | 0 |
| rs34520362 | pathogenic | 0 | 0 | 0 |
| rs374621913 | pathogenic | 0 | 0 | 0 |
| rs113641837 | benign | 0 | 0 | 0 |
| rs104894635 | pathogenic | 0 | 0 | 0 |

| | | | | |
|---|---|---|---|---|
| rs143947056 | pathogenic | 0 | 0 | 0 |
| rs138504221 | pathogenic | 0 | 0 | 0 |
| rs9894254 | benign | 0 | 0 | 0 |
| rs104894640 | pathogenic | 1 | 1 | 1 |
| rs58786455 | benign | 0 | 0 | 0 |
| rs144143780 | pathogenic | 0 | 0 | 0 |
| rs62620232 | benign | 0 | 0 | 0 |
| rs34297805 | benign | 1 | 1 | 1 |
| rs104894641 | pathogenic | 0 | 0 | 0 |
| rs148876719 | benign | 0 | 0 | 0 |
| rs104894639 | pathogenic | 0 | 0 | 0 |
| rs7503034 | benign | 0 | 0 | 0 |
| rs201631570 | benign | 0 | 0 | 0 |
| rs150181226 | pathogenic | 0 | 0 | 0 |
| rs377546036 | pathogenic | 0 | 0 | 0 |
| rs267606700 | pathogenic | 0 | 0 | 0 |
| rs267606701 | pathogenic | 1 | 1 | 0 |
| rs267606699 | pathogenic | 1 | 1 | 0 |
| rs9903215 | benign | 0 | 0 | 0 |
| rs139486406 | benign | 0 | 0 | 0 |
| rs143791614 | benign | 0 | 0 | 0 |
| rs573483474 | benign | 0 | 0 | 0 |
| rs2032892 | benign | 0 | 0 | 0 |
| rs137852619 | pathogenic | 0 | 0 | 0 |
| rs200243548 | benign | 0 | 0 | 0 |
| rs115702388 | benign | 0 | 0 | 0 |
| rs138085358 | pathogenic | 0 | 0 | 0 |
| rs121912789 | pathogenic | 1 | 1 | 1 |
| rs763407972 | pathogenic | 1 | 1 | 1 |
| rs147278302 | benign | 0 | 0 | 0 |
| rs374540411 | benign | 0 | 0 | 0 |
| rs121912790 | pathogenic | 1 | 1 | 0 |
| rs752049917 | benign | 0 | 0 | 0 |
| rs139640491 | benign | 1 | 1 | 1 |
| rs138010738 | benign | 1 | 1 | 0 |
| rs201328768 | pathogenic | 1 | 1 | 0 |
| rs2135720 | benign | 1 | 1 | 0 |
| rs201816080 | benign | 1 | 1 | 0 |
| rs754391973 | pathogenic | 0 | 0 | 0 |
| rs139087859 | benign | 0 | 0 | 0 |
| rs6805421 | benign | 0 | 0 | 0 |
| rs146266900 | benign | 0 | 0 | 0 |
| rs397509419 | pathogenic | 0 | 0 | 0 |
| rs149980411 | benign | 0 | 0 | 0 |
| rs747711488 | pathogenic | 1 | 1 | 1 |
| rs149474339 | pathogenic | 1 | 1 | 1 |
| rs758252808 | pathogenic | 1 | 1 | 0 |
| rs745756308 | pathogenic | 0 | 0 | 0 |
| rs4986958 | benign | 0 | 0 | 0 |
| rs74315444 | pathogenic | 0 | 0 | 0 |
| rs61752137 | pathogenic | 0 | 0 | 0 |
| rs8179065 | benign | 1 | 1 | 1 |
| rs758361736 | pathogenic | 1 | 1 | 0 |
| rs139794067 | pathogenic | 1 | 1 | 1 |

| | | | | |
|---|---|---|---|---|
| rs730880954 | pathogenic | 1 | 1 | 1 |
| rs730880955 | pathogenic | 1 | 1 | 1 |
| rs150634297 | pathogenic | 0 | 0 | 0 |
| rs104893750 | pathogenic | 1 | 1 | 1 |
| rs104893748 | pathogenic | 1 | 1 | 1 |
| rs104893749 | pathogenic | 1 | 1 | 1 |
| rs753520553 | pathogenic | 1 | 1 | 0 |
| rs61737294 | benign | 0 | 0 | 0 |
| rs104894601 | pathogenic | 0 | 0 | 0 |
| rs376090795 | pathogenic | 0 | 0 | 0 |
| rs104894596 | pathogenic | 0 | 0 | 0 |
| rs104894595 | pathogenic | 0 | 0 | 0 |
| rs104894598 | pathogenic | 0 | 0 | 0 |
| rs104894597 | pathogenic | 0 | 0 | 0 |
| rs104894591 | pathogenic | 0 | 0 | 0 |
| rs104894593 | pathogenic | 0 | 0 | 0 |
| rs104894594 | pathogenic | 0 | 0 | 0 |
| rs104894590 | pathogenic | 0 | 0 | 0 |
| rs86312 | benign | 0 | 0 | 0 |
| rs77874543 | benign | 0 | 0 | 0 |
| rs121912590 | pathogenic | 1 | 1 | 1 |
| rs121912589 | pathogenic | 1 | 1 | 1 |
| rs121912586 | pathogenic | 1 | 1 | 1 |
| rs121912585 | pathogenic | 1 | 1 | 1 |
| rs121912587 | pathogenic | 1 | 1 | 1 |
| rs762909359 | pathogenic | 0 | 0 | 0 |
| rs28931610 | pathogenic | 0 | 0 | 0 |
| rs104894097 | pathogenic | 1 | 1 | 1 |
| rs775176191 | benign | 0 | 0 | 0 |
| rs199907548 | pathogenic | 0 | 0 | 0 |
| rs104894095 | pathogenic | 1 | 1 | 1 |
| rs104894109 | pathogenic | 1 | 1 | 1 |
| rs372266620 | benign | 1 | 1 | 1 |
| rs104894099 | pathogenic | 0 | 0 | 0 |
| rs137854599 | pathogenic | 1 | 1 | 1 |
| rs104894094 | pathogenic | 0 | 0 | 0 |
| rs104894104 | pathogenic | 0 | 0 | 0 |
| rs6413463 | benign | 1 | 1 | 1 |
| rs104894098 | pathogenic | 0 | 0 | 0 |
| rs6413464 | benign | 0 | 0 | 0 |
| rs3731249 | benign | 0 | 0 | 0 |
| rs16931179 | benign | 0 | 0 | 0 |
| rs2734715 | benign | 0 | 0 | 0 |
| rs759080581 | pathogenic | 0 | 0 | 0 |
| rs1800746 | benign | 0 | 0 | 0 |
| rs202189057 | pathogenic | 0 | 0 | 0 |
| rs121908204 | pathogenic | 0 | 0 | 0 |
| rs756564767 | pathogenic | 0 | 0 | 0 |
| rs34758634 | benign | 1 | 1 | 1 |
| rs28940573 | pathogenic | 0 | 0 | 0 |
| rs119455955 | pathogenic | 1 | 1 | 0 |
| rs752164603 | pathogenic | 1 | 1 | 0 |
| rs763162812 | pathogenic | 0 | 0 | 0 |
| rs119455957 | pathogenic | 0 | 0 | 0 |

| | | | | |
|---|---|---|---|---|
| rs119455958 | pathogenic | 1 | 1 | 0 |
| rs750428882 | pathogenic | 0 | 0 | 0 |
| rs141482368 | benign | 0 | 0 | 0 |
| rs119455953 | pathogenic | 0 | 0 | 0 |
| rs119455954 | pathogenic | 0 | 0 | 0 |
| rs112812685 | benign | 1 | 1 | 0 |
| rs138744051 | benign | 1 | 1 | 0 |
| rs121908200 | pathogenic | 0 | 0 | 0 |
| rs119455956 | pathogenic | 0 | 0 | 0 |
| rs78970155 | benign | 0 | 0 | 0 |
| rs2834167 | benign | 0 | 0 | 0 |
| rs387907326 | pathogenic | 1 | 1 | 1 |
| rs2107732 | benign | 0 | 0 | 0 |
| rs11552377 | benign | 0 | 0 | 0 |
| rs137852843 | pathogenic | 0 | 0 | 0 |
| rs104894783 | pathogenic | 0 | 0 | 0 |
| rs122457137 | pathogenic | 1 | 1 | 1 |
| rs104894779 | pathogenic | 1 | 1 | 1 |
| rs104894786 | pathogenic | 0 | 0 | 0 |
| rs104894784 | pathogenic | 0 | 0 | 0 |
| rs104894785 | pathogenic | 0 | 0 | 0 |
| rs201870761 | pathogenic | 0 | 0 | 0 |
| rs104894781 | pathogenic | 0 | 0 | 0 |
| rs28897728 | benign | 0 | 0 | 0 |
| rs370273690 | pathogenic | 1 | 1 | 0 |
| rs9332739 | benign | 0 | 0 | 0 |
| rs150878060 | benign | 0 | 0 | 0 |
| rs573509224 | benign | 0 | 0 | 0 |
| rs36221133 | benign | 0 | 0 | 0 |
| rs2230009 | benign | 0 | 0 | 0 |
| rs267607008 | pathogenic | 0 | 0 | 0 |
| rs5985 | benign | 1 | 1 | 1 |
| rs5982 | benign | 1 | 1 | 1 |
| rs5987 | benign | 0 | 0 | 0 |
| rs5988 | benign | 0 | 0 | 0 |
| rs121918680 | pathogenic | 1 | 1 | 0 |
| rs121918679 | pathogenic | 1 | 1 | 0 |
| rs7845 | benign | 1 | 1 | 0 |
| rs1049296 | benign | 1 | 1 | 1 |
| rs121918677 | pathogenic | 1 | 1 | 0 |
| rs200086262 | pathogenic | 1 | 1 | 0 |
| rs746930990 | pathogenic | 1 | 1 | 0 |
| rs104893878 | pathogenic | 1 | 1 | 1 |
| rs121912468 | pathogenic | 1 | 1 | 1 |
| rs121912467 | pathogenic | 1 | 1 | 1 |
| rs145351926 | benign | 1 | 1 | 1 |
| rs141212743 | pathogenic | 1 | 1 | 1 |
| rs397515473 | pathogenic | 1 | 1 | 0 |
| rs121918313 | pathogenic | 0 | 1 | 0 |
| rs73015965 | pathogenic | 0 | 0 | 0 |
| rs121918030 | pathogenic | 0 | 1 | 0 |
| rs77782321 | benign | 0 | 0 | 0 |
| rs374661051 | pathogenic | 1 | 1 | 1 |
| rs267606854 | pathogenic | 0 | 0 | 0 |

| | | | | |
|---|---|---|---|---|
| rs41279678 | benign | 0 | 0 | 0 |
| rs61754640 | benign | 0 | 0 | 0 |
| rs199473314 | pathogenic | 0 | 0 | 0 |
| rs137854601 | pathogenic | 0 | 0 | 0 |
| rs137854614 | pathogenic | 0 | 0 | 0 |
| rs137854615 | pathogenic | 0 | 0 | 0 |
| rs199473635 | pathogenic | 0 | 0 | 0 |
| rs121434478 | pathogenic | 1 | 1 | 1 |
| rs775407864 | pathogenic | 1 | 1 | 1 |
| rs12706927 | benign | 0 | 1 | 0 |
| rs775614261 | benign | 0 | 0 | 0 |
| rs768926261 | benign | 0 | 0 | 0 |
| rs114057640 | benign | 0 | 0 | 0 |
| rs387906886 | pathogenic | 1 | 1 | 0 |
| rs138658585 | benign | 0 | 0 | 0 |
| rs4880091 | benign | 0 | 0 | 0 |
| rs121918552 | pathogenic | 1 | 1 | 1 |
| rs10794537 | benign | 0 | 0 | 0 |
| rs121965022 | pathogenic | 1 | 1 | 0 |
| rs121965020 | pathogenic | 0 | 0 | 0 |
| rs758452450 | pathogenic | 0 | 0 | 0 |
| rs747981483 | benign | 0 | 0 | 0 |
| rs58037052 | benign | 0 | 0 | 0 |
| rs121965029 | pathogenic | 1 | 1 | 0 |
| rs147490060 | benign | 0 | 0 | 0 |
| rs3755955 | benign | 0 | 0 | 0 |
| rs148789453 | pathogenic | 0 | 0 | 0 |
| rs121965030 | pathogenic | 0 | 0 | 0 |
| rs76722191 | benign | 0 | 0 | 0 |
| rs199801029 | pathogenic | 0 | 0 | 0 |
| rs121965033 | pathogenic | 0 | 0 | 0 |
| rs368454909 | pathogenic | 1 | 1 | 0 |
| rs6831280 | benign | 0 | 0 | 0 |
| rs121965024 | pathogenic | 0 | 0 | 0 |
| rs121965019 | pathogenic | 0 | 0 | 0 |
| rs11934801 | benign | 0 | 0 | 0 |
| rs73066479 | benign | 0 | 0 | 0 |
| rs121965027 | pathogenic | 0 | 0 | 0 |
| rs121965026 | pathogenic | 1 | 1 | 0 |
| rs121965021 | pathogenic | 0 | 0 | 0 |
| rs398123257 | benign | 0 | 0 | 0 |
| rs121965031 | pathogenic | 0 | 0 | 0 |
| rs202003795 | pathogenic | 0 | 0 | 0 |
| rs559510809 | pathogenic | 0 | 0 | 0 |
| rs762936223 | benign | 0 | 0 | 0 |
| rs137854580 | pathogenic | 1 | 1 | 1 |
| rs112263157 | benign | 0 | 0 | 0 |
| rs2289181 | benign | 0 | 0 | 0 |
| rs138034915 | benign | 0 | 1 | 0 |
| rs528367092 | pathogenic | 0 | 1 | 0 |
| rs1042393 | benign | 0 | 0 | 0 |
| rs370950728 | pathogenic | 0 | 1 | 0 |
| rs530478036 | benign | 0 | 1 | 0 |
| rs374569672 | benign | 0 | 1 | 0 |

| | | | | |
|---|---|---|---|---|
| rs121907944 | pathogenic | 0 | 1 | 0 |
| rs200856561 | pathogenic | 0 | 1 | 0 |
| rs577915581 | pathogenic | 0 | 1 | 0 |
| rs201896815 | pathogenic | 0 | 1 | 0 |
| rs121907945 | pathogenic | 0 | 1 | 0 |
| rs121907940 | pathogenic | 0 | 1 | 0 |
| rs543300039 | pathogenic | 0 | 1 | 0 |
| rs121907936 | pathogenic | 0 | 1 | 0 |
| rs766074609 | pathogenic | 0 | 1 | 0 |
| rs755253527 | pathogenic | 0 | 1 | 0 |
| rs560575383 | pathogenic | 0 | 1 | 0 |
| rs528369909 | benign | 0 | 1 | 0 |
| rs770610356 | pathogenic | 0 | 1 | 0 |
| rs747610090 | pathogenic | 0 | 1 | 0 |
| rs145712232 | benign | 0 | 1 | 0 |
| rs7215458 | benign | 1 | 1 | 0 |
| rs535644999 | pathogenic | 0 | 1 | 0 |
| rs772883420 | pathogenic | 1 | 1 | 0 |
| rs398123169 | pathogenic | 0 | 1 | 0 |
| rs121907937 | pathogenic | 0 | 1 | 0 |
| rs121907941 | pathogenic | 1 | 1 | 0 |
| rs121907942 | pathogenic | 0 | 1 | 0 |
| rs779556619 | pathogenic | 0 | 1 | 0 |
| rs1800307 | benign | 0 | 1 | 0 |
| rs377544304 | pathogenic | 1 | 1 | 0 |
| rs549029029 | pathogenic | 0 | 1 | 0 |
| rs753269119 | pathogenic | 1 | 1 | 1 |
| rs757617999 | pathogenic | 0 | 1 | 0 |
| rs28937909 | pathogenic | 0 | 1 | 0 |
| rs28940868 | pathogenic | 0 | 1 | 0 |
| rs368438393 | pathogenic | 0 | 1 | 0 |
| rs536906561 | pathogenic | 0 | 1 | 0 |
| rs374143224 | pathogenic | 0 | 1 | 0 |
| rs778418246 | pathogenic | 0 | 1 | 0 |
| rs398123172 | pathogenic | 0 | 1 | 0 |
| rs121907938 | pathogenic | 0 | 1 | 0 |
| rs1800312 | pathogenic | 0 | 1 | 0 |
| rs1126690 | benign | 0 | 1 | 0 |
| rs1800314 | benign | 0 | 1 | 0 |
| rs369532274 | pathogenic | 0 | 1 | 0 |
| rs121907943 | pathogenic | 0 | 1 | 0 |
| rs780321415 | pathogenic | 0 | 1 | 0 |
| rs377286472 | benign | 0 | 1 | 0 |
| rs121918406 | pathogenic | 1 | 1 | 1 |
| rs147484870 | benign | 0 | 0 | 0 |
| rs193922671 | pathogenic | 0 | 0 | 0 |
| rs201397978 | benign | 1 | 1 | 1 |
| rs34884895 | benign | 1 | 1 | 1 |
| rs138329459 | benign | 0 | 0 | 0 |
| rs2289366 | benign | 1 | 1 | 1 |
| rs141858334 | benign | 1 | 1 | 1 |
| rs59907886 | benign | 1 | 1 | 1 |
| rs137852653 | pathogenic | 0 | 0 | 0 |
| rs137852652 | pathogenic | 0 | 0 | 0 |

| | | | | |
|---|---|---|---|---|
| rs137852656 | pathogenic | 0 | 0 | 0 |
| rs61739916 | benign | 0 | 0 | 0 |
| rs28936668 | pathogenic | 0 | 0 | 0 |
| rs137852651 | pathogenic | 0 | 0 | 0 |
| rs137852650 | pathogenic | 0 | 0 | 0 |
| rs28936669 | pathogenic | 0 | 0 | 0 |
| rs137852654 | pathogenic | 0 | 0 | 0 |
| rs28936368 | pathogenic | 1 | 1 | 0 |
| rs137852655 | pathogenic | 0 | 0 | 0 |
| rs61752496 | benign | 0 | 0 | 0 |
| rs750098055 | benign | 0 | 0 | 0 |
| rs764149793 | benign | 0 | 0 | 0 |
| rs724160017 | pathogenic | 0 | 1 | 0 |
| rs724160019 | pathogenic | 0 | 0 | 0 |
| rs202219398 | pathogenic | 1 | 1 | 0 |
| rs724160023 | pathogenic | 1 | 1 | 0 |
| rs761953453 | pathogenic | 0 | 0 | 0 |
| rs80338835 | pathogenic | 1 | 1 | 1 |
| rs56284940 | benign | 0 | 0 | 0 |
| rs74575749 | benign | 0 | 0 | 0 |
| rs587777075 | pathogenic | 1 | 1 | 1 |
| rs641153 | benign | 0 | 0 | 0 |
| rs12614 | benign | 0 | 0 | 0 |
| rs117314762 | benign | 0 | 0 | 0 |
| rs537478097 | benign | 1 | 1 | 1 |
| rs144812066 | benign | 0 | 0 | 0 |
| rs4151651 | benign | 0 | 0 | 0 |
| rs138207668 | benign | 1 | 1 | 1 |
| rs149101394 | benign | 1 | 1 | 1 |
| rs4151659 | benign | 1 | 1 | 1 |
| rs45484591 | benign | 1 | 1 | 1 |
| rs761991070 | pathogenic | 0 | 0 | 0 |
| rs755948940 | pathogenic | 0 | 0 | 0 |
| rs137853097 | pathogenic | 0 | 0 | 0 |
| rs775297352 | benign | 0 | 0 | 0 |
| rs28943591 | benign | 0 | 0 | 0 |
| rs191468413 | pathogenic | 1 | 1 | 0 |
| rs11539471 | benign | 1 | 1 | 1 |
| rs11205 | benign | 0 | 0 | 0 |
| rs768409400 | pathogenic | 0 | 0 | 0 |
| rs762013077 | pathogenic | 0 | 0 | 0 |
| rs768474112 | pathogenic | 0 | 0 | 0 |
| rs121965063 | pathogenic | 1 | 1 | 1 |
| rs143648758 | pathogenic | 0 | 0 | 0 |
| rs121965066 | pathogenic | 0 | 0 | 0 |
| rs752907087 | pathogenic | 0 | 0 | 0 |
| rs747702749 | pathogenic | 0 | 0 | 0 |
| rs754087775 | pathogenic | 0 | 0 | 0 |
| rs121918641 | pathogenic | 1 | 1 | 1 |
| rs121918642 | pathogenic | 1 | 1 | 1 |
| rs121918640 | pathogenic | 1 | 1 | 1 |
| rs121918637 | pathogenic | 1 | 1 | 1 |
| rs121918644 | pathogenic | 1 | 1 | 1 |
| rs16840544 | benign | 0 | 0 | 0 |

| | | | | |
|---|---|---|---|---|
| rs1042391 | benign | 1 | 1 | 1 |
| rs104893744 | pathogenic | 1 | 1 | 1 |
| rs120074136 | pathogenic | 0 | 0 | 0 |
| rs1805081 | benign | 1 | 1 | 0 |
| rs80358251 | benign | 0 | 0 | 0 |
| rs774588125 | benign | 0 | 0 | 0 |
| rs120074134 | pathogenic | 0 | 0 | 0 |
| rs139751448 | pathogenic | 0 | 0 | 0 |
| rs77080672 | benign | 0 | 0 | 0 |
| rs372445155 | pathogenic | 1 | 1 | 0 |
| rs13381670 | benign | 0 | 0 | 0 |
| rs1788799 | benign | 0 | 0 | 0 |
| rs376213990 | pathogenic | 0 | 0 | 0 |
| rs757475924 | pathogenic | 0 | 0 | 0 |
| rs80358253 | pathogenic | 0 | 0 | 0 |
| rs1805082 | benign | 0 | 0 | 0 |
| rs120074130 | pathogenic | 0 | 0 | 0 |
| rs768999208 | pathogenic | 0 | 0 | 0 |
| rs34302553 | benign | 0 | 0 | 0 |
| rs28940897 | pathogenic | 0 | 0 | 0 |
| rs1140458 | benign | 0 | 0 | 0 |
| rs143124972 | pathogenic | 0 | 0 | 0 |
| rs120074135 | pathogenic | 0 | 0 | 0 |
| rs543206298 | pathogenic | 0 | 0 | 0 |
| rs120074132 | pathogenic | 0 | 0 | 0 |
| rs759826138 | pathogenic | 0 | 0 | 0 |
| rs28942108 | pathogenic | 0 | 0 | 0 |
| rs80358254 | pathogenic | 0 | 0 | 0 |
| rs80358257 | pathogenic | 0 | 0 | 0 |
| rs781261962 | pathogenic | 0 | 0 | 0 |
| rs28942107 | pathogenic | 0 | 0 | 0 |
| rs80358258 | pathogenic | 0 | 0 | 0 |
| rs80358259 | pathogenic | 1 | 1 | 0 |
| rs141440861 | benign | 0 | 0 | 0 |
| rs28942106 | pathogenic | 0 | 0 | 0 |
| rs778878523 | pathogenic | 0 | 0 | 0 |
| rs28942105 | pathogenic | 0 | 0 | 0 |
| rs61731969 | benign | 0 | 0 | 0 |
| rs200444084 | pathogenic | 0 | 0 | 0 |
| rs113371321 | pathogenic | 0 | 0 | 0 |
| rs369098773 | pathogenic | 0 | 0 | 0 |
| rs35248744 | benign | 0 | 0 | 0 |
| rs758902805 | pathogenic | 0 | 0 | 0 |
| rs120074131 | pathogenic | 0 | 0 | 0 |
| rs1805084 | benign | 0 | 0 | 0 |
| rs117793540 | benign | 0 | 0 | 0 |
| rs530058063 | benign | 0 | 0 | 0 |
| rs1064422 | pathogenic | 1 | 1 | 1 |
| rs104894506 | pathogenic | 1 | 1 | 1 |
| rs104894507 | pathogenic | 1 | 1 | 1 |
| rs104894508 | pathogenic | 1 | 1 | 0 |
| rs28999113 | pathogenic | 1 | 1 | 0 |
| rs281860266 | pathogenic | 0 | 0 | 0 |
| rs104894961 | pathogenic | 1 | 1 | 0 |

| | | | | |
|---|---|---|---|---|
| rs122463168 | pathogenic | 0 | 0 | 0 |
| rs199469624 | pathogenic | 1 | 1 | 1 |
| rs150107747 | benign | 0 | 0 | 0 |
| rs144984628 | pathogenic | 0 | 0 | 0 |
| rs28935171 | pathogenic | 0 | 0 | 0 |
| rs137852443 | pathogenic | 1 | 1 | 0 |
| rs28933674 | pathogenic | 1 | 1 | 1 |
| rs28933675 | pathogenic | 1 | 1 | 1 |
| rs28933676 | pathogenic | 1 | 1 | 1 |
| rs28933677 | pathogenic | 1 | 1 | 1 |
| rs28933678 | pathogenic | 0 | 0 | 0 |
| rs28933679 | pathogenic | 1 | 1 | 1 |
| rs28933680 | pathogenic | 0 | 0 | 0 |
| rs28933681 | pathogenic | 0 | 0 | 0 |
| rs28933682 | pathogenic | 0 | 0 | 0 |
| rs28937294 | pathogenic | 1 | 1 | 0 |
| rs137852459 | pathogenic | 0 | 0 | 0 |
| rs137852461 | pathogenic | 0 | 0 | 0 |
| rs137852464 | pathogenic | 1 | 1 | 1 |
| rs137852465 | pathogenic | 1 | 1 | 1 |
| rs1800297 | benign | 1 | 1 | 0 |
| rs137852374 | pathogenic | 1 | 1 | 1 |
| rs672601312 | pathogenic | 1 | 1 | 1 |
| rs387907196 | pathogenic | 1 | 1 | 1 |
| rs148348866 | benign | 0 | 0 | 0 |
| rs141138948 | pathogenic | 1 | 1 | 1 |
| rs137852685 | pathogenic | 1 | 1 | 0 |
| rs104893900 | pathogenic | 1 | 1 | 0 |
| rs137852686 | pathogenic | 1 | 1 | 1 |
| rs104893906 | pathogenic | 1 | 1 | 0 |
| rs587782207 | pathogenic | 1 | 1 | 1 |
| rs75086406 | pathogenic | 0 | 0 | 0 |
| rs199822819 | pathogenic | 0 | 0 | 0 |
| rs398123166 | pathogenic | 1 | 1 | 1 |
| rs121913123 | pathogenic | 1 | 1 | 1 |
| rs772190176 | pathogenic | 0 | 1 | 0 |
| rs587782216 | pathogenic | 1 | 1 | 1 |
| rs398123159 | pathogenic | 1 | 1 | 1 |
| rs121913122 | pathogenic | 1 | 1 | 1 |
| rs755436052 | pathogenic | 1 | 1 | 1 |
| rs200796606 | pathogenic | 1 | 1 | 1 |
| rs2070080 | pathogenic | 0 | 1 | 0 |
| rs149616320 | benign | 0 | 0 | 0 |
| rs143529458 | benign | 0 | 0 | 0 |
| rs267607077 | pathogenic | 1 | 1 | 1 |
| rs143898031 | benign | 1 | 1 | 0 |
| rs148304857 | benign | 1 | 1 | 1 |
| rs137852809 | pathogenic | 0 | 0 | 0 |
| rs3765623 | benign | 0 | 0 | 0 |
| rs16944397 | benign | 0 | 0 | 0 |
| rs730880167 | benign | 0 | 0 | 0 |
| rs187108957 | benign | 0 | 0 | 0 |
| rs117342470 | benign | 0 | 0 | 0 |
| rs751200138 | pathogenic | 1 | 1 | 1 |

| | | | | |
|---|---|---|---|---|
| rs755799528 | pathogenic | 0 | 0 | 0 |
| rs537484504 | benign | 0 | 0 | 0 |
| rs748944640 | pathogenic | 0 | 0 | 0 |
| rs199774121 | pathogenic | 0 | 0 | 0 |
| rs766094434 | pathogenic | 1 | 1 | 0 |
| rs777321035 | pathogenic | 1 | 1 | 1 |
| rs146269774 | benign | 0 | 1 | 0 |
| rs144085805 | benign | 0 | 1 | 0 |
| rs367643028 | benign | 0 | 0 | 0 |
| rs42524 | benign | 0 | 0 | 0 |
| rs28940586 | pathogenic | 0 | 0 | 0 |
| rs74315327 | pathogenic | 1 | 1 | 1 |
| rs199676861 | pathogenic | 0 | 0 | 0 |
| rs115802744 | benign | 1 | 1 | 0 |
| rs752866643 | pathogenic | 1 | 1 | 1 |
| rs28941784 | pathogenic | 1 | 1 | 1 |
| rs398124434 | pathogenic | 1 | 1 | 1 |
| rs756414548 | pathogenic | 1 | 1 | 1 |
| rs376128990 | pathogenic | 1 | 1 | 1 |
| rs746219370 | pathogenic | 1 | 1 | 1 |
| rs756195708 | pathogenic | 1 | 1 | 1 |
| rs369296618 | pathogenic | 1 | 1 | 1 |
| rs9593 | benign | 1 | 1 | 1 |
| rs139510844 | benign | 0 | 0 | 0 |
| rs28382158 | benign | 0 | 0 | 0 |
| rs28919570 | pathogenic | 0 | 0 | 0 |
| rs587777331 | pathogenic | 0 | 0 | 0 |
| rs587777333 | pathogenic | 0 | 0 | 0 |
| rs587777332 | pathogenic | 0 | 0 | 0 |
| rs774980346 | pathogenic | 0 | 0 | 0 |
| rs370934093 | pathogenic | 0 | 0 | 0 |
| rs121913479 | pathogenic | 1 | 1 | 1 |
| rs121913484 | pathogenic | 0 | 0 | 0 |
| rs121913485 | pathogenic | 1 | 1 | 1 |
| rs28931614 | pathogenic | 1 | 1 | 1 |
| rs17881656 | benign | 1 | 1 | 1 |
| rs28931615 | pathogenic | 1 | 1 | 1 |
| rs147846832 | benign | 0 | 0 | 0 |
| rs80356506 | pathogenic | 0 | 0 | 0 |
| rs80356494 | pathogenic | 0 | 0 | 0 |
| rs121908894 | pathogenic | 0 | 0 | 0 |
| rs121908895 | pathogenic | 0 | 0 | 0 |
| rs80356508 | pathogenic | 0 | 0 | 0 |
| rs80356507 | pathogenic | 0 | 0 | 0 |
| rs121908896 | pathogenic | 0 | 0 | 0 |
| rs745372589 | pathogenic | 1 | 1 | 0 |
| rs183898408 | benign | 0 | 0 | 0 |
| rs200844292 | benign | 0 | 0 | 0 |
| rs146897002 | benign | 0 | 0 | 0 |
| rs121964869 | pathogenic | 1 | 1 | 1 |
| rs201768800 | benign | 1 | 1 | 1 |
| rs9807556 | benign | 1 | 1 | 1 |
| rs184721031 | benign | 0 | 0 | 0 |
| rs201104206 | benign | 0 | 0 | 0 |

| | | | | |
|---|---|---|---|---|
| rs188677538 | benign | 0 | 0 | 0 |
| rs149528866 | benign | 0 | 0 | 0 |
| rs115382168 | benign | 0 | 0 | 0 |
| rs78846023 | benign | 1 | 1 | 1 |
| rs61730662 | benign | 0 | 0 | 0 |
| rs121918368 | pathogenic | 0 | 0 | 0 |
| rs566433112 | pathogenic | 1 | 1 | 1 |
| rs121909342 | pathogenic | 0 | 0 | 0 |
| rs67586389 | pathogenic | 0 | 0 | 0 |
| rs72466485 | pathogenic | 0 | 0 | 0 |
| rs72466486 | pathogenic | 1 | 1 | 1 |
| rs72466487 | pathogenic | 1 | 1 | 1 |
| rs267607112 | pathogenic | 0 | 0 | 0 |
| rs118204013 | pathogenic | 0 | 0 | 0 |
| rs121913495 | pathogenic | 0 | 0 | 0 |
| rs11554273 | pathogenic | 0 | 0 | 0 |
| rs193922477 | pathogenic | 1 | 1 | 1 |
| rs193922479 | pathogenic | 1 | 1 | 1 |
| rs72547554 | pathogenic | 0 | 0 | 0 |
| rs72547561 | pathogenic | 1 | 1 | 1 |
| rs72547562 | pathogenic | 1 | 1 | 0 |
| rs115977487 | benign | 1 | 1 | 0 |
| rs72547564 | pathogenic | 0 | 0 | 0 |
| rs72547568 | pathogenic | 1 | 1 | 0 |
| rs72547569 | pathogenic | 1 | 1 | 1 |
| rs72547571 | pathogenic | 0 | 0 | 0 |
| rs72547575 | pathogenic | 1 | 1 | 1 |
| rs544994462 | benign | 0 | 0 | 0 |
| rs190470017 | benign | 0 | 0 | 0 |
| rs199473266 | pathogenic | 1 | 1 | 1 |
| rs137854602 | pathogenic | 0 | 0 | 0 |
| rs143739249 | pathogenic | 0 | 0 | 0 |
| rs397514697 | pathogenic | 1 | 1 | 1 |
| rs373764886 | pathogenic | 0 | 0 | 0 |
| rs370242930 | pathogenic | 0 | 0 | 0 |
| rs377145777 | pathogenic | 1 | 1 | 1 |
| rs191256186 | benign | 0 | 0 | 0 |
| rs150070901 | benign | 1 | 1 | 1 |
| rs138098032 | benign | 0 | 0 | 0 |
| rs117731813 | benign | 1 | 1 | 1 |
| rs104894488 | pathogenic | 1 | 1 | 1 |
| rs11302 | benign | 0 | 1 | 0 |
| rs121918447 | pathogenic | 1 | 1 | 1 |
| rs138205828 | benign | 1 | 1 | 0 |
| rs149858290 | benign | 0 | 0 | 0 |
| rs750764003 | pathogenic | 0 | 1 | 0 |
| rs141381896 | benign | 0 | 0 | 0 |
| rs149027682 | benign | 0 | 0 | 0 |
| rs121908415 | pathogenic | 0 | 0 | 0 |
| rs121908416 | pathogenic | 0 | 0 | 0 |
| rs121908417 | pathogenic | 0 | 0 | 0 |
| rs587780306 | benign | 0 | 0 | 0 |
| rs3218725 | benign | 0 | 0 | 0 |
| rs3218729 | benign | 0 | 0 | 0 |

| | | | | |
|---|---|---|---|---|
| rs387906917 | pathogenic | 0 | 0 | 0 |
| rs200652608 | pathogenic | 0 | 0 | 0 |
| rs3218721 | benign | 0 | 0 | 0 |
| rs63750869 | pathogenic | 1 | 1 | 1 |
| rs118203910 | pathogenic | 0 | 1 | 0 |
| rs9332701 | benign | 1 | 1 | 0 |
| rs6027 | benign | 0 | 0 | 0 |
| rs121909110 | pathogenic | 0 | 1 | 0 |
| rs762550967 | pathogenic | 1 | 1 | 0 |
| rs763615602 | pathogenic | 1 | 1 | 0 |
| rs144907403 | benign | 0 | 0 | 0 |
| rs104893631 | pathogenic | 1 | 1 | 0 |
| rs144181978 | pathogenic | 1 | 1 | 0 |
| rs74874677 | benign | 1 | 1 | 1 |
| rs104893632 | pathogenic | 0 | 1 | 0 |
| rs371342464 | benign | 1 | 1 | 1 |
| rs104893633 | pathogenic | 1 | 1 | 0 |
| rs201088712 | pathogenic | 0 | 0 | 0 |
| rs201857541 | benign | 1 | 1 | 1 |
| rs139517732 | pathogenic | 1 | 1 | 0 |
| rs191057824 | benign | 1 | 1 | 0 |
| rs35683768 | benign | 1 | 1 | 0 |
| rs104895271 | pathogenic | 0 | 0 | 0 |
| rs104895217 | pathogenic | 1 | 1 | 1 |
| rs104895223 | pathogenic | 1 | 1 | 1 |
| rs104895218 | pathogenic | 1 | 1 | 1 |
| rs104895225 | pathogenic | 1 | 1 | 1 |
| rs4149637 | benign | 1 | 1 | 1 |
| rs104895219 | pathogenic | 1 | 1 | 1 |
| rs104895220 | pathogenic | 1 | 1 | 1 |
| rs104895228 | pathogenic | 1 | 1 | 1 |
| rs104895221 | pathogenic | 1 | 1 | 1 |
| rs104895222 | pathogenic | 1 | 1 | 1 |
| rs104895247 | pathogenic | 0 | 0 | 0 |
| rs28940585 | pathogenic | 0 | 0 | 0 |
| rs561367199 | pathogenic | 1 | 1 | 1 |
| rs143793815 | benign | 0 | 0 | 0 |
| rs28941473 | pathogenic | 1 | 1 | 1 |
| rs751590073 | pathogenic | 1 | 1 | 1 |
| rs111407265 | benign | 1 | 1 | 1 |
| rs373697663 | pathogenic | 1 | 1 | 1 |
| rs369879957 | pathogenic | 0 | 0 | 0 |
| rs28941472 | pathogenic | 1 | 1 | 1 |
| rs28940287 | pathogenic | 1 | 1 | 1 |
| rs77625410 | benign | 0 | 0 | 0 |
| rs111033477 | benign | 0 | 0 | 0 |
| rs115123584 | benign | 0 | 0 | 0 |
| rs374576916 | benign | 0 | 0 | 0 |
| rs397516317 | pathogenic | 0 | 0 | 0 |
| rs368657015 | pathogenic | 0 | 0 | 0 |
| rs397516321 | pathogenic | 0 | 0 | 0 |
| rs397516322 | pathogenic | 0 | 0 | 0 |
| rs111033215 | pathogenic | 0 | 0 | 0 |
| rs199606180 | pathogenic | 0 | 0 | 0 |

| | | | | |
|---|---|---|---|---|
| rs111033192 | pathogenic | 0 | 0 | 0 |
| rs948962 | benign | 1 | 1 | 1 |
| rs142293185 | benign | 0 | 0 | 0 |
| rs376764423 | pathogenic | 0 | 0 | 0 |
| rs773844428 | pathogenic | 0 | 0 | 0 |
| rs186644871 | benign | 0 | 0 | 0 |
| rs200313391 | benign | 0 | 0 | 0 |
| rs377670513 | pathogenic | 0 | 0 | 0 |
| rs1132036 | benign | 0 | 0 | 0 |
| rs111033231 | benign | 0 | 0 | 0 |
| rs145945630 | pathogenic | 0 | 0 | 0 |
| rs121912984 | pathogenic | 1 | 1 | 1 |
| rs1800435 | benign | 0 | 1 | 0 |
| rs121912980 | pathogenic | 0 | 0 | 0 |
| rs121912982 | pathogenic | 1 | 1 | 1 |
| rs121912983 | pathogenic | 0 | 0 | 0 |
| rs121912981 | pathogenic | 0 | 0 | 0 |
| rs1057868 | benign | 0 | 0 | 0 |
| rs56048668 | benign | 0 | 0 | 0 |
| rs55865465 | benign | 0 | 0 | 0 |
| rs748096324 | benign | 1 | 1 | 0 |
| rs150475750 | benign | 0 | 0 | 0 |
| rs143025739 | benign | 1 | 1 | 0 |
| rs79702016 | benign | 1 | 1 | 0 |
| rs41338945 | benign | 0 | 0 | 0 |
| rs146406037 | benign | 1 | 1 | 1 |
| rs143274491 | benign | 0 | 0 | 0 |
| rs3829986 | benign | 0 | 0 | 0 |
| rs149168939 | benign | 0 | 0 | 0 |
| rs1800385 | benign | 0 | 0 | 0 |
| rs1800386 | pathogenic | 0 | 0 | 0 |
| rs121918524 | pathogenic | 1 | 1 | 1 |
| rs121912700 | pathogenic | 0 | 0 | 0 |
| rs104893930 | pathogenic | 1 | 1 | 1 |
| rs104893931 | pathogenic | 0 | 0 | 0 |
| rs116125936 | benign | 0 | 0 | 0 |
| rs627441 | benign | 0 | 0 | 0 |
| rs104894481 | pathogenic | 0 | 0 | 0 |
| rs137853129 | pathogenic | 0 | 0 | 0 |
| rs137853130 | pathogenic | 1 | 1 | 0 |
| rs28940282 | pathogenic | 0 | 0 | 0 |
| rs2229989 | benign | 0 | 0 | 0 |
| rs104894647 | pathogenic | 0 | 0 | 0 |
| rs1061170 | benign | 1 | 1 | 1 |
| rs576402053 | benign | 0 | 0 | 0 |
| rs34972246 | benign | 0 | 0 | 0 |
| rs192725607 | benign | 0 | 0 | 0 |
| rs137852615 | pathogenic | 1 | 1 | 0 |
| rs137852617 | pathogenic | 1 | 1 | 1 |
| rs137852611 | pathogenic | 0 | 0 | 0 |
| rs137852612 | pathogenic | 0 | 0 | 0 |
| rs137852614 | pathogenic | 1 | 1 | 1 |
| rs137852610 | pathogenic | 1 | 1 | 1 |
| rs137852618 | pathogenic | 1 | 1 | 1 |

| | | | | |
|---|---|---|---|---|
| rs137852613 | pathogenic | 1 | 1 | 0 |
| rs137852609 | pathogenic | 0 | 0 | 0 |
| rs63750306 | pathogenic | 1 | 1 | 1 |
| rs63750265 | pathogenic | 1 | 1 | 1 |
| rs63751287 | pathogenic | 1 | 1 | 1 |
| rs199723282 | benign | 0 | 0 | 0 |
| rs121917807 | pathogenic | 0 | 0 | 0 |
| rs121917809 | pathogenic | 0 | 0 | 0 |
| rs75134564 | pathogenic | 0 | 0 | 0 |
| rs267606930 | pathogenic | 1 | 1 | 0 |
| rs121909302 | pathogenic | 0 | 0 | 0 |
| rs121909303 | pathogenic | 0 | 0 | 0 |
| rs121909299 | pathogenic | 0 | 0 | 0 |
| rs121909301 | pathogenic | 0 | 0 | 0 |
| rs121909300 | pathogenic | 1 | 1 | 0 |
| rs111033589 | pathogenic | 0 | 0 | 0 |
| rs137853059 | pathogenic | 0 | 0 | 0 |
| rs137853057 | pathogenic | 1 | 1 | 1 |
| rs1801474 | benign | 0 | 0 | 0 |
| rs137853060 | pathogenic | 1 | 1 | 1 |
| rs137853058 | pathogenic | 1 | 1 | 1 |
| rs34424986 | pathogenic | 0 | 0 | 0 |
| rs199657839 | benign | 1 | 1 | 1 |
| rs56092260 | benign | 0 | 0 | 0 |
| rs1801582 | benign | 1 | 1 | 1 |
| rs1801334 | benign | 1 | 1 | 1 |
| rs191486604 | pathogenic | 0 | 0 | 0 |
| rs121908051 | pathogenic | 0 | 0 | 0 |
| rs121908055 | pathogenic | 1 | 1 | 0 |
| rs28940886 | pathogenic | 0 | 0 | 0 |
| rs121908050 | pathogenic | 0 | 0 | 0 |
| rs28940887 | pathogenic | 0 | 0 | 0 |
| rs387906300 | pathogenic | 0 | 0 | 0 |
| rs28942087 | pathogenic | 0 | 0 | 0 |
| rs121908049 | pathogenic | 1 | 1 | 0 |
| rs121908054 | pathogenic | 1 | 1 | 0 |
| rs121908048 | pathogenic | 0 | 0 | 0 |
| rs28940888 | pathogenic | 1 | 1 | 1 |
| rs121908053 | pathogenic | 1 | 1 | 0 |
| rs122467169 | pathogenic | 1 | 1 | 1 |
| rs122467172 | pathogenic | 1 | 1 | 1 |
| rs122467170 | pathogenic | 1 | 1 | 1 |
| rs28935477 | pathogenic | 1 | 1 | 0 |
| rs28942088 | pathogenic | 0 | 0 | 0 |
| rs751478142 | pathogenic | 1 | 1 | 0 |
| rs765368797 | pathogenic | 0 | 0 | 0 |
| rs147810437 | benign | 0 | 0 | 0 |
| rs121918209 | pathogenic | 1 | 1 | 1 |
| rs121918208 | pathogenic | 1 | 1 | 1 |
| rs121918210 | pathogenic | 1 | 1 | 1 |
| rs35515638 | benign | 1 | 1 | 1 |
| rs121918207 | pathogenic | 1 | 1 | 1 |
| rs141298312 | benign | 0 | 0 | 0 |
| rs121918211 | pathogenic | 0 | 0 | 0 |

| | | | | |
|---|---|---|---|---|
| rs149660059 | benign | 0 | 0 | 0 |
| rs200670286 | pathogenic | 1 | 1 | 1 |
| rs121918212 | pathogenic | 1 | 1 | 1 |
| rs121918213 | pathogenic | 1 | 1 | 1 |
| rs121918205 | pathogenic | 1 | 1 | 1 |
| rs104894025 | pathogenic | 1 | 1 | 1 |
| rs104894026 | pathogenic | 1 | 1 | 1 |
| rs104894028 | pathogenic | 0 | 1 | 0 |
| rs104894029 | pathogenic | 0 | 1 | 0 |
| rs119458971 | pathogenic | 0 | 0 | 0 |
| rs119458969 | pathogenic | 1 | 1 | 1 |
| rs1801591 | benign | 1 | 1 | 1 |
| rs140169311 | benign | 0 | 0 | 0 |
| rs753363278 | benign | 1 | 1 | 1 |
| rs199763682 | pathogenic | 0 | 0 | 0 |
| rs119458970 | pathogenic | 1 | 1 | 1 |
| rs763722868 | pathogenic | 0 | 0 | 0 |
| rs56404770 | benign | 0 | 0 | 0 |
| rs34221241 | benign | 0 | 0 | 0 |
| rs121909625 | pathogenic | 0 | 0 | 0 |
| rs550921485 | pathogenic | 0 | 0 | 0 |
| rs1800454 | benign | 0 | 0 | 0 |
| rs241447 | benign | 0 | 0 | 0 |
| rs2230606 | benign | 1 | 1 | 0 |
| rs5036 | pathogenic | 1 | 1 | 1 |
| rs13306788 | benign | 0 | 0 | 0 |
| rs28929480 | pathogenic | 0 | 0 | 0 |
| rs121912749 | pathogenic | 0 | 0 | 0 |
| rs56361140 | pathogenic | 0 | 0 | 0 |
| rs28931583 | pathogenic | 1 | 1 | 1 |
| rs36211715 | pathogenic | 1 | 1 | 1 |
| rs138049878 | pathogenic | 1 | 1 | 1 |
| rs727503253 | pathogenic | 1 | 1 | 1 |
| rs267606908 | pathogenic | 0 | 0 | 0 |
| rs121913631 | pathogenic | 1 | 1 | 1 |
| rs36211716 | benign | 1 | 1 | 1 |
| rs121913639 | pathogenic | 1 | 1 | 1 |
| rs730880761 | pathogenic | 1 | 1 | 1 |
| rs121913629 | pathogenic | 1 | 1 | 1 |
| rs754368658 | pathogenic | 0 | 0 | 0 |
| rs777090017 | pathogenic | 0 | 0 | 0 |
| rs267606913 | pathogenic | 1 | 1 | 1 |
| rs773138218 | pathogenic | 0 | 0 | 0 |
| rs371295780 | pathogenic | 0 | 0 | 0 |
| rs387906830 | pathogenic | 1 | 1 | 0 |
| rs771364038 | pathogenic | 0 | 1 | 0 |
| rs772731615 | pathogenic | 0 | 0 | 0 |
| rs137853063 | pathogenic | 0 | 0 | 0 |
| rs387907076 | pathogenic | 0 | 1 | 0 |
| rs35460768 | benign | 0 | 0 | 0 |
| rs2278196 | benign | 1 | 1 | 1 |
| rs119103275 | pathogenic | 0 | 0 | 0 |
| rs119103276 | pathogenic | 1 | 1 | 1 |
| rs374402066 | pathogenic | 1 | 1 | 1 |

| | | | | |
|---|---|---|---|---|
| rs119103274 | pathogenic | 0 | 0 | 0 |
| rs2274064 | benign | 0 | 0 | 0 |
| rs267607023 | pathogenic | 1 | 1 | 0 |
| rs142117489 | benign | 1 | 1 | 0 |
| rs371258270 | benign | 0 | 0 | 0 |
| rs34716057 | benign | 0 | 0 | 0 |
| rs201778606 | benign | 0 | 0 | 0 |
| rs201438792 | benign | 0 | 0 | 0 |
| rs140547417 | benign | 0 | 0 | 0 |
| rs5881 | benign | 0 | 0 | 0 |
| rs752298084 | benign | 0 | 0 | 0 |
| rs34855278 | benign | 0 | 0 | 0 |
| rs5880 | benign | 1 | 1 | 0 |
| rs5882 | benign | 0 | 0 | 0 |
| rs1800777 | benign | 0 | 0 | 0 |
| rs577301331 | benign | 0 | 0 | 0 |
| rs137853313 | pathogenic | 0 | 0 | 0 |
| rs28935169 | pathogenic | 0 | 0 | 0 |
| rs137853315 | pathogenic | 0 | 0 | 0 |
| rs137853314 | pathogenic | 0 | 0 | 0 |
| rs28935469 | pathogenic | 0 | 0 | 0 |
| rs137853318 | pathogenic | 0 | 0 | 0 |
| rs28935470 | pathogenic | 0 | 0 | 0 |
| rs3729986 | benign | 0 | 0 | 0 |
| rs730880619 | pathogenic | 0 | 0 | 0 |
| rs201012766 | benign | 0 | 0 | 0 |
| rs727503218 | pathogenic | 0 | 0 | 0 |
| rs11570052 | benign | 0 | 0 | 0 |
| rs730880623 | pathogenic | 0 | 0 | 0 |
| rs138753870 | benign | 0 | 0 | 0 |
| rs397516068 | pathogenic | 0 | 0 | 0 |
| rs3729989 | benign | 1 | 1 | 1 |
| rs397516074 | pathogenic | 0 | 0 | 0 |
| rs561449819 | pathogenic | 0 | 0 | 0 |
| rs121917779 | benign | 0 | 0 | 0 |
| rs142881576 | pathogenic | 0 | 0 | 0 |
| rs77384282 | benign | 0 | 0 | 0 |
| rs201865787 | benign | 0 | 0 | 0 |
| rs142244640 | benign | 0 | 0 | 0 |
| rs200488706 | benign | 0 | 0 | 0 |
| rs137853291 | pathogenic | 1 | 1 | 0 |
| rs28933990 | pathogenic | 1 | 1 | 0 |
| rs140570189 | benign | 0 | 0 | 0 |
| rs201588178 | benign | 0 | 0 | 0 |
| rs104893846 | pathogenic | 0 | 0 | 0 |
| rs754545360 | pathogenic | 0 | 1 | 0 |
| rs104893851 | pathogenic | 0 | 1 | 0 |
| rs104893849 | pathogenic | 0 | 0 | 0 |
| rs571038432 | pathogenic | 1 | 1 | 0 |
| rs2270655 | benign | 1 | 1 | 1 |
| rs315952 | benign | 0 | 0 | 0 |
| rs151176548 | benign | 0 | 0 | 0 |
| rs122459147 | pathogenic | 1 | 1 | 0 |
| rs122458140 | pathogenic | 0 | 0 | 0 |

| | | | | |
|---|---|---|---|---|
| rs122458142 | pathogenic | 1 | 1 | 0 |
| rs267606812 | pathogenic | 1 | 1 | 0 |
| rs267606813 | pathogenic | 1 | 1 | 0 |
| rs122458143 | pathogenic | 0 | 0 | 0 |
| rs122459146 | pathogenic | 0 | 0 | 0 |
| rs122458144 | pathogenic | 0 | 0 | 0 |
| rs122458145 | pathogenic | 0 | 0 | 0 |
| rs113994007 | pathogenic | 1 | 1 | 0 |
| rs758746181 | pathogenic | 1 | 1 | 1 |
| rs145034527 | pathogenic | 1 | 1 | 0 |
| rs119468006 | pathogenic | 1 | 1 | 0 |
| rs201908721 | pathogenic | 0 | 0 | 0 |
| rs755933881 | pathogenic | 1 | 1 | 0 |
| rs771578775 | pathogenic | 0 | 0 | 0 |
| rs17849927 | benign | 0 | 0 | 0 |
| rs139133094 | benign | 0 | 0 | 0 |
| rs12593 | benign | 0 | 0 | 0 |
| rs578189699 | pathogenic | 1 | 1 | 0 |
| rs119468008 | pathogenic | 0 | 0 | 0 |
| rs768524626 | benign | 0 | 0 | 0 |
| rs119468009 | pathogenic | 0 | 0 | 0 |
| rs119468004 | pathogenic | 0 | 0 | 0 |
| rs199874519 | pathogenic | 0 | 0 | 0 |
| rs376478331 | pathogenic | 0 | 0 | 0 |
| rs61995958 | pathogenic | 0 | 0 | 0 |
| rs199714731 | benign | 0 | 0 | 0 |
| rs149185954 | benign | 0 | 0 | 0 |
| rs121918353 | pathogenic | 0 | 0 | 0 |
| rs1131695 | benign | 0 | 0 | 0 |
| rs28939668 | pathogenic | 0 | 0 | 0 |
| rs758432471 | pathogenic | 1 | 1 | 1 |
| rs752746786 | pathogenic | 1 | 1 | 1 |
| rs28935471 | pathogenic | 0 | 0 | 0 |
| rs80338837 | pathogenic | 0 | 0 | 0 |
| rs137853312 | pathogenic | 0 | 0 | 0 |
| rs28935472 | pathogenic | 0 | 0 | 0 |
| rs28935473 | pathogenic | 0 | 0 | 0 |
| rs267606752 | pathogenic | 0 | 0 | 0 |
| rs3917973 | benign | 0 | 0 | 0 |
| rs3918018 | benign | 0 | 0 | 0 |
| rs757976755 | pathogenic | 0 | 0 | 0 |
| rs121918329 | pathogenic | 1 | 1 | 0 |
| rs770114459 | pathogenic | 0 | 0 | 0 |
| rs6662272 | benign | 0 | 0 | 0 |
| rs121918242 | pathogenic | 1 | 1 | 0 |
| rs121918240 | pathogenic | 1 | 1 | 0 |
| rs200094982 | pathogenic | 0 | 0 | 0 |
| rs121918241 | pathogenic | 0 | 0 | 0 |
| rs140522266 | pathogenic | 1 | 1 | 0 |
| rs121918243 | pathogenic | 1 | 1 | 0 |
| rs370596113 | pathogenic | 1 | 1 | 0 |
| rs398124295 | pathogenic | 1 | 1 | 0 |
| rs587776889 | pathogenic | 1 | 1 | 0 |
| rs747527726 | pathogenic | 1 | 1 | 0 |

| | | | | |
|---|---|---|---|---|
| rs112631212 | benign | 0 | 0 | 0 |
| rs104894605 | pathogenic | 0 | 0 | 0 |
| rs104894607 | pathogenic | 0 | 0 | 0 |
| rs104894606 | pathogenic | 0 | 0 | 0 |
| rs143873873 | benign | 0 | 0 | 0 |
| rs104894388 | pathogenic | 1 | 1 | 0 |
| rs104894391 | pathogenic | 1 | 1 | 0 |
| rs5350 | benign | 1 | 1 | 1 |
| rs104894390 | pathogenic | 1 | 1 | 1 |
| rs5346 | benign | 1 | 1 | 1 |
| rs121912500 | pathogenic | 0 | 0 | 0 |
| rs74315450 | pathogenic | 1 | 1 | 0 |
| rs121913396 | pathogenic | 1 | 1 | 1 |
| rs28931588 | pathogenic | 1 | 1 | 1 |
| rs121913400 | pathogenic | 0 | 0 | 0 |
| rs28931589 | pathogenic | 1 | 1 | 1 |
| rs121913403 | pathogenic | 0 | 0 | 0 |
| rs121913413 | pathogenic | 0 | 0 | 0 |
| rs760107623 | pathogenic | 1 | 1 | 0 |
| rs16866378 | benign | 1 | 1 | 0 |
| rs201025290 | pathogenic | 1 | 1 | 0 |
| rs12720356 | benign | 1 | 1 | 0 |
| rs55882956 | benign | 0 | 1 | 0 |
| rs4986933 | benign | 1 | 1 | 0 |
| rs2020957 | benign | 1 | 1 | 1 |
| rs1800124 | benign | 1 | 1 | 1 |
| rs150077735 | benign | 1 | 1 | 1 |
| rs201419000 | benign | 0 | 0 | 0 |
| rs772044176 | pathogenic | 0 | 0 | 0 |
| rs138346388 | benign | 0 | 0 | 0 |
| rs121964916 | pathogenic | 0 | 0 | 0 |
| rs112534524 | pathogenic | 0 | 0 | 0 |
| rs182078921 | benign | 1 | 1 | 1 |
| rs113273712 | benign | 0 | 0 | 0 |
| rs74817407 | benign | 1 | 1 | 1 |
| rs61733901 | benign | 1 | 1 | 1 |
| rs121964912 | pathogenic | 1 | 1 | 1 |
| rs41278047 | benign | 1 | 1 | 1 |
| rs7437875 | benign | 1 | 1 | 1 |
| rs113460688 | benign | 1 | 1 | 1 |
| rs754572081 | benign | 1 | 1 | 1 |
| rs201082692 | benign | 1 | 1 | 1 |
| rs202085803 | benign | 1 | 1 | 1 |
| rs141320511 | benign | 0 | 0 | 0 |
| rs78501403 | benign | 0 | 0 | 0 |
| rs28939675 | pathogenic | 0 | 0 | 0 |
| rs74315522 | pathogenic | 0 | 0 | 0 |
| rs121434639 | pathogenic | 1 | 1 | 0 |
| rs121434640 | pathogenic | 1 | 1 | 1 |
| rs757282628 | benign | 1 | 1 | 1 |
| rs386834010 | pathogenic | 0 | 0 | 0 |
| rs200863680 | pathogenic | 0 | 0 | 0 |
| rs193919336 | pathogenic | 1 | 1 | 0 |
| rs386834011 | pathogenic | 0 | 0 | 0 |

| | | | | |
|---|---|---|---|---|
| rs28940869 | pathogenic | 0 | 0 | 0 |
| rs386834014 | pathogenic | 1 | 1 | 0 |
| rs267606960 | pathogenic | 0 | 0 | 0 |
| rs28942068 | pathogenic | 0 | 0 | 0 |
| rs193919335 | pathogenic | 0 | 0 | 0 |
| rs74374973 | pathogenic | 0 | 0 | 0 |
| rs386834018 | pathogenic | 0 | 0 | 0 |
| rs267606962 | pathogenic | 0 | 0 | 0 |
| rs6659553 | benign | 0 | 0 | 0 |
| rs29281 | benign | 0 | 0 | 0 |
| rs782265090 | benign | 0 | 0 | 0 |
| rs35910969 | pathogenic | 0 | 0 | 0 |
| rs121909113 | pathogenic | 1 | 1 | 1 |
| rs29001571 | pathogenic | 1 | 1 | 1 |
| rs587781250 | pathogenic | 1 | 1 | 1 |
| rs28939680 | pathogenic | 1 | 1 | 1 |
| rs28939681 | pathogenic | 1 | 1 | 1 |
| rs121909112 | pathogenic | 1 | 1 | 1 |
| rs28937568 | pathogenic | 1 | 1 | 1 |
| rs1799895 | pathogenic | 0 | 0 | 0 |
| rs386833431 | pathogenic | 1 | 1 | 1 |
| rs121964905 | pathogenic | 1 | 1 | 1 |
| rs121964906 | pathogenic | 1 | 1 | 1 |
| rs11551664 | pathogenic | 1 | 1 | 1 |
| rs104894705 | pathogenic | 1 | 1 | 1 |
| rs104893805 | pathogenic | 0 | 0 | 0 |
| rs104894205 | pathogenic | 0 | 0 | 0 |
| rs104894204 | pathogenic | 1 | 1 | 0 |
| rs62642926 | pathogenic | 0 | 0 | 0 |
| rs5030841 | pathogenic | 1 | 1 | 1 |
| rs75193786 | pathogenic | 1 | 1 | 1 |
| rs62508682 | pathogenic | 0 | 0 | 0 |
| rs3850625 | benign | 1 | 1 | 1 |
| rs141908793 | benign | 0 | 0 | 0 |
| rs187864727 | benign | 0 | 0 | 0 |
| rs548909101 | benign | 0 | 0 | 0 |
| rs3825394 | benign | 0 | 1 | 0 |
| rs143548402 | benign | 0 | 0 | 0 |
| rs267607146 | pathogenic | 0 | 0 | 0 |
| rs267607144 | pathogenic | 0 | 0 | 0 |
| rs267607148 | pathogenic | 1 | 1 | 1 |
| rs267607143 | pathogenic | 0 | 0 | 0 |
| rs267607145 | pathogenic | 0 | 0 | 0 |
| rs121912636 | pathogenic | 1 | 1 | 1 |
| rs121912634 | pathogenic | 1 | 1 | 1 |
| rs570675468 | benign | 0 | 0 | 0 |
| rs368861241 | pathogenic | 1 | 1 | 1 |
| rs104894528 | pathogenic | 1 | 1 | 1 |
| rs77493891 | benign | 0 | 0 | 0 |
| rs62640903 | benign | 0 | 0 | 0 |
| rs149926091 | benign | 1 | 1 | 1 |
| rs641081 | benign | 0 | 0 | 0 |
| rs4930199 | benign | 0 | 0 | 0 |
| rs121913020 | pathogenic | 0 | 0 | 0 |

| | | | | |
|---|---|---|---|---|
| rs1799793 | benign | 0 | 0 | 0 |
| rs121913016 | pathogenic | 0 | 0 | 0 |
| rs121913025 | pathogenic | 0 | 0 | 0 |
| rs121913019 | pathogenic | 0 | 0 | 0 |
| rs376556895 | pathogenic | 1 | 1 | 1 |
| rs121913024 | pathogenic | 1 | 1 | 1 |
| rs121913021 | pathogenic | 0 | 0 | 0 |
| rs121913023 | pathogenic | 0 | 0 | 0 |
| rs41556519 | pathogenic | 0 | 0 | 0 |
| rs121913022 | pathogenic | 0 | 0 | 0 |
| rs144564120 | pathogenic | 0 | 0 | 0 |
| rs121913026 | pathogenic | 1 | 1 | 1 |
| rs121913018 | pathogenic | 1 | 1 | 1 |
| rs142954643 | benign | 0 | 0 | 0 |
| rs76655496 | benign | 0 | 0 | 0 |
| rs104894371 | pathogenic | 1 | 1 | 0 |
| rs137964936 | benign | 0 | 0 | 0 |
| rs759736132 | pathogenic | 0 | 0 | 0 |
| rs387906595 | pathogenic | 1 | 1 | 0 |
| rs768248216 | pathogenic | 0 | 0 | 0 |
| rs548231008 | benign | 0 | 0 | 0 |
| rs757094384 | benign | 0 | 0 | 0 |
| rs867529 | benign | 0 | 0 | 0 |
| rs13045 | benign | 1 | 1 | 1 |
| rs121908570 | pathogenic | 0 | 0 | 0 |
| rs201872586 | benign | 1 | 1 | 0 |
| rs149092990 | benign | 0 | 0 | 0 |
| rs115585711 | benign | 0 | 0 | 0 |
| rs137909416 | benign | 1 | 1 | 0 |
| rs143388949 | benign | 1 | 1 | 1 |
| rs73137716 | benign | 1 | 1 | 1 |
| rs121908947 | pathogenic | 1 | 1 | 1 |
| rs398122904 | pathogenic | 1 | 1 | 1 |
| rs35715497 | benign | 1 | 1 | 0 |
| rs35462421 | benign | 0 | 0 | 0 |
| rs119475043 | pathogenic | 0 | 0 | 0 |
| rs119475042 | pathogenic | 0 | 0 | 0 |
| rs1076669 | benign | 1 | 1 | 0 |
| rs141146885 | benign | 0 | 0 | 0 |
| rs3026906 | pathogenic | 0 | 0 | 0 |
| rs75362221 | benign | 1 | 1 | 1 |
| rs138472116 | benign | 1 | 1 | 1 |
| rs371040282 | pathogenic | 1 | 1 | 1 |
| rs4988496 | benign | 0 | 0 | 0 |
| rs121918117 | pathogenic | 1 | 1 | 1 |
| rs4988498 | benign | 0 | 0 | 0 |
| rs5035 | benign | 0 | 0 | 0 |
| rs121918322 | pathogenic | 1 | 1 | 1 |
| rs370359540 | benign | 0 | 0 | 0 |
| rs758972589 | pathogenic | 0 | 0 | 0 |
| rs144856889 | benign | 0 | 0 | 0 |
| rs121434623 | pathogenic | 1 | 1 | 1 |
| rs61735600 | benign | 0 | 0 | 0 |
| rs72550870 | benign | 1 | 1 | 1 |

| | | | | |
|---|---|---|---|---|
| rs56392418 | benign | 0 | 0 | 0 |
| rs41307788 | benign | 1 | 1 | 1 |
| rs387907239 | pathogenic | 1 | 1 | 0 |
| rs28931584 | pathogenic | 0 | 0 | 0 |
| rs121912745 | pathogenic | 0 | 0 | 0 |
| rs121912744 | pathogenic | 0 | 0 | 0 |
| rs121912754 | pathogenic | 1 | 1 | 1 |
| rs121912746 | pathogenic | 0 | 0 | 0 |
| rs121912757 | pathogenic | 0 | 0 | 0 |
| rs121912758 | pathogenic | 0 | 0 | 0 |
| rs121912748 | pathogenic | 0 | 0 | 0 |
| rs121912755 | pathogenic | 0 | 0 | 0 |
| rs121912741 | pathogenic | 0 | 0 | 0 |
| rs121912753 | pathogenic | 0 | 0 | 0 |
| rs121912750 | pathogenic | 0 | 0 | 0 |
| rs2285644 | pathogenic | 0 | 0 | 0 |
| rs121912751 | pathogenic | 0 | 0 | 0 |
| rs5026 | benign | 0 | 0 | 0 |
| rs121912759 | pathogenic | 0 | 0 | 0 |
| rs28931585 | pathogenic | 0 | 0 | 0 |
| rs17122769 | benign | 0 | 0 | 0 |
| rs28362322 | benign | 1 | 1 | 1 |
| rs3824874 | benign | 0 | 0 | 0 |
| rs146004831 | benign | 0 | 0 | 0 |
| rs142155860 | benign | 0 | 0 | 0 |
| rs121434404 | pathogenic | 0 | 0 | 0 |
| rs61735578 | benign | 1 | 1 | 1 |
| rs558018 | benign | 0 | 0 | 0 |
| rs61735577 | benign | 1 | 1 | 1 |
| rs76784113 | benign | 0 | 0 | 0 |
| rs199469640 | pathogenic | 0 | 0 | 0 |
| rs199469628 | pathogenic | 1 | 1 | 0 |
| rs199469629 | pathogenic | 0 | 1 | 0 |
| rs387907156 | pathogenic | 0 | 1 | 0 |
| rs199469643 | pathogenic | 0 | 0 | 0 |
| rs199469646 | pathogenic | 0 | 0 | 0 |
| rs766294940 | pathogenic | 0 | 0 | 0 |
| rs387907347 | pathogenic | 0 | 0 | 0 |
| rs137853590 | pathogenic | 1 | 1 | 1 |
| rs137853589 | pathogenic | 1 | 1 | 1 |
| rs137853588 | pathogenic | 1 | 1 | 1 |
| rs55909005 | benign | 0 | 0 | 0 |
| rs4581 | benign | 0 | 0 | 0 |
| rs63750123 | benign | 0 | 0 | 0 |
| rs10254120 | benign | 0 | 0 | 0 |
| rs121434629 | pathogenic | 1 | 1 | 0 |
| rs200313585 | benign | 0 | 0 | 0 |
| rs143162541 | benign | 1 | 1 | 0 |
| rs760615315 | benign | 0 | 0 | 0 |
| rs768488890 | benign | 1 | 1 | 1 |
| rs116373169 | benign | 1 | 1 | 0 |
| rs63750871 | pathogenic | 1 | 1 | 0 |
| rs116349687 | benign | 0 | 0 | 0 |
| rs375289386 | benign | 0 | 0 | 0 |

| | | | | |
|---|---|---|---|---|
| rs760228510 | pathogenic | 1 | 1 | 1 |
| rs587779343 | pathogenic | 1 | 1 | 1 |
| rs573125799 | benign | 0 | 0 | 0 |
| rs1805322 | benign | 0 | 0 | 0 |
| rs267608153 | pathogenic | 0 | 0 | 0 |
| rs139194813 | benign | 0 | 0 | 0 |
| rs530021751 | benign | 0 | 0 | 0 |
| rs200640585 | pathogenic | 0 | 0 | 0 |
| rs139438201 | benign | 0 | 0 | 0 |
| rs200513014 | benign | 0 | 0 | 0 |
| rs567102013 | benign | 0 | 0 | 0 |
| rs121908608 | pathogenic | 0 | 0 | 0 |
| rs121908606 | pathogenic | 0 | 0 | 0 |
| rs17103671 | benign | 0 | 0 | 0 |
| rs34725281 | benign | 0 | 0 | 0 |
| rs104894421 | pathogenic | 1 | 1 | 0 |
| rs121918145 | pathogenic | 1 | 1 | 1 |
| rs121918152 | pathogenic | 1 | 1 | 1 |
| rs56379106 | benign | 0 | 0 | 0 |
| rs4987076 | benign | 0 | 0 | 0 |
| rs370243877 | pathogenic | 0 | 0 | 0 |
| rs17679445 | benign | 1 | 1 | 1 |
| rs550423482 | pathogenic | 1 | 1 | 1 |
| rs104894629 | pathogenic | 1 | 1 | 1 |
| rs773450573 | pathogenic | 1 | 1 | 1 |
| rs116679717 | benign | 0 | 0 | 0 |
| rs372000848 | pathogenic | 0 | 0 | 0 |
| rs138595973 | benign | 0 | 0 | 0 |
| rs115244417 | benign | 0 | 0 | 0 |
| rs11575292 | benign | 0 | 0 | 0 |
| rs137853211 | pathogenic | 1 | 1 | 1 |
| rs137853207 | pathogenic | 0 | 0 | 0 |
| rs137853210 | pathogenic | 1 | 1 | 1 |
| rs6262 | benign | 0 | 0 | 0 |
| rs6263 | benign | 0 | 1 | 0 |
| rs137853208 | pathogenic | 0 | 1 | 0 |
| rs137853212 | pathogenic | 1 | 1 | 0 |
| rs137853209 | pathogenic | 1 | 1 | 1 |
| rs201951824 | pathogenic | 0 | 0 | 0 |
| rs11575542 | benign | 0 | 0 | 0 |
| rs199681788 | benign | 0 | 1 | 0 |
| rs748835902 | benign | 1 | 1 | 1 |
| rs753719949 | benign | 0 | 0 | 0 |
| rs267606647 | pathogenic | 0 | 0 | 0 |
| rs267606648 | pathogenic | 1 | 1 | 1 |
| rs267606643 | pathogenic | 0 | 1 | 0 |
| rs559947967 | pathogenic | 1 | 1 | 0 |
| rs267606645 | pathogenic | 1 | 1 | 1 |
| rs12116440 | benign | 0 | 0 | 0 |
| rs121908539 | pathogenic | 0 | 0 | 0 |
| rs137853096 | pathogenic | 1 | 1 | 0 |
| rs148363262 | benign | 1 | 1 | 0 |
| rs587777442 | pathogenic | 0 | 0 | 0 |
| rs25640 | benign | 1 | 1 | 1 |

| | | | | |
|---|---|---|---|---|
| rs28943589 | benign | 0 | 0 | 0 |
| rs1143650 | benign | 1 | 1 | 1 |
| rs150886026 | benign | 0 | 1 | 0 |
| rs121908133 | pathogenic | 1 | 1 | 0 |
| rs121908131 | pathogenic | 0 | 0 | 0 |
| rs754477154 | pathogenic | 0 | 0 | 0 |
| rs747334007 | benign | 0 | 0 | 0 |
| rs121908132 | pathogenic | 0 | 0 | 0 |
| rs104894103 | pathogenic | 1 | 1 | 0 |
| rs140088149 | benign | 0 | 0 | 0 |
| rs149803148 | benign | 0 | 0 | 0 |
| rs11064480 | benign | 0 | 0 | 0 |
| rs74435397 | pathogenic | 1 | 1 | 1 |
| rs587783626 | pathogenic | 1 | 1 | 1 |
| rs148211042 | pathogenic | 1 | 1 | 1 |
| rs17849502 | benign | 0 | 0 | 0 |
| rs13306575 | pathogenic | 1 | 1 | 1 |
| rs201426733 | benign | 0 | 0 | 0 |
| rs182278784 | benign | 1 | 1 | 1 |
| rs77786415 | benign | 1 | 1 | 1 |
| rs200192119 | benign | 1 | 1 | 1 |
| rs12877501 | benign | 1 | 1 | 1 |
| rs77985244 | benign | 1 | 1 | 1 |
| rs5907 | pathogenic | 1 | 1 | 0 |
| rs2305178 | benign | 0 | 0 | 0 |
| rs3135867 | benign | 0 | 0 | 0 |
| rs121913482 | pathogenic | 1 | 1 | 1 |
| rs121913483 | pathogenic | 1 | 1 | 1 |
| rs4647924 | pathogenic | 1 | 1 | 1 |
| rs121913111 | pathogenic | 1 | 1 | 1 |
| rs104894002 | pathogenic | 0 | 0 | 0 |
| rs104894001 | pathogenic | 0 | 0 | 0 |
| rs75932628 | benign | 0 | 0 | 0 |
| rs201258663 | pathogenic | 0 | 0 | 0 |
| rs104893998 | pathogenic | 0 | 0 | 0 |
| rs2234253 | benign | 1 | 1 | 0 |
| rs121908402 | pathogenic | 1 | 1 | 0 |
| rs1805389 | benign | 0 | 0 | 0 |
| rs1805388 | benign | 0 | 0 | 0 |
| rs748385144 | pathogenic | 0 | 0 | 0 |
| rs104894420 | pathogenic | 0 | 0 | 0 |
| rs104894418 | pathogenic | 0 | 0 | 0 |
| rs61731910 | benign | 0 | 0 | 0 |
| rs758723288 | pathogenic | 1 | 1 | 0 |
| rs201865375 | pathogenic | 1 | 1 | 0 |
| rs565090080 | pathogenic | 1 | 1 | 1 |
| rs113042166 | benign | 0 | 0 | 0 |
| rs61736326 | benign | 1 | 1 | 1 |
| rs12030699 | benign | 1 | 1 | 1 |
| rs121913578 | pathogenic | 1 | 1 | 1 |
| rs61739582 | benign | 0 | 0 | 0 |
| rs745983207 | pathogenic | 0 | 0 | 0 |
| rs763524380 | benign | 0 | 0 | 0 |
| rs121912792 | pathogenic | 0 | 0 | 0 |

| | | | | |
|---|---|---|---|---|
| rs121912793 | pathogenic | 0 | 0 | 0 |
| rs387906599 | pathogenic | 0 | 0 | 0 |
| rs121912794 | pathogenic | 0 | 1 | 0 |
| rs121912795 | pathogenic | 0 | 0 | 0 |
| rs121912796 | pathogenic | 1 | 1 | 0 |
| rs121912797 | pathogenic | 0 | 1 | 0 |
| rs121912798 | pathogenic | 1 | 1 | 1 |
| rs80359812 | pathogenic | 1 | 1 | 0 |
| rs776583130 | benign | 1 | 1 | 0 |
| rs80359814 | pathogenic | 1 | 1 | 0 |
| rs202060209 | pathogenic | 1 | 1 | 0 |
| rs267607061 | pathogenic | 1 | 1 | 0 |
| rs267607060 | pathogenic | 0 | 0 | 0 |
| rs80359816 | pathogenic | 1 | 1 | 0 |
| rs80359818 | pathogenic | 1 | 1 | 0 |
| rs387907313 | pathogenic | 0 | 0 | 0 |
| rs587784395 | benign | 0 | 0 | 0 |
| rs5811 | benign | 0 | 0 | 0 |
| rs121909740 | pathogenic | 1 | 1 | 0 |
| rs148518827 | benign | 1 | 1 | 0 |
| rs121909739 | pathogenic | 1 | 1 | 0 |
| rs780528770 | benign | 1 | 1 | 0 |
| rs13306758 | pathogenic | 0 | 0 | 0 |
| rs267607059 | pathogenic | 0 | 0 | 0 |
| rs61730638 | benign | 0 | 0 | 0 |
| rs104894533 | pathogenic | 0 | 0 | 0 |
| rs104894527 | pathogenic | 0 | 0 | 0 |
| rs200503569 | pathogenic | 1 | 1 | 1 |
| rs80338700 | pathogenic | 1 | 1 | 1 |
| rs765433263 | pathogenic | 1 | 1 | 1 |
| rs104894530 | pathogenic | 1 | 1 | 1 |
| rs80338701 | pathogenic | 1 | 1 | 1 |
| rs141498002 | pathogenic | 1 | 1 | 1 |
| rs104894525 | pathogenic | 0 | 0 | 0 |
| rs80338703 | pathogenic | 0 | 0 | 0 |
| rs28936415 | pathogenic | 1 | 1 | 0 |
| rs150719105 | pathogenic | 0 | 0 | 0 |
| rs148032587 | pathogenic | 0 | 0 | 0 |
| rs150577656 | pathogenic | 0 | 0 | 0 |
| rs190521996 | pathogenic | 0 | 0 | 0 |
| rs104894526 | pathogenic | 1 | 1 | 1 |
| rs80338704 | pathogenic | 0 | 0 | 0 |
| rs199562225 | pathogenic | 1 | 1 | 1 |
| rs532870929 | pathogenic | 0 | 0 | 0 |
| rs398123309 | pathogenic | 0 | 0 | 0 |
| rs78290141 | pathogenic | 0 | 0 | 0 |
| rs80338705 | pathogenic | 0 | 0 | 0 |
| rs104894531 | pathogenic | 1 | 1 | 1 |
| rs80338706 | pathogenic | 0 | 0 | 0 |
| rs80338707 | pathogenic | 0 | 0 | 0 |
| rs80338708 | pathogenic | 0 | 0 | 0 |
| rs80338709 | pathogenic | 0 | 0 | 0 |
| rs104893833 | benign | 0 | 0 | 0 |
| rs61744585 | benign | 0 | 0 | 0 |

| | | | | |
|---|---|---|---|---|
| rs73569515 | benign | 0 | 0 | 0 |
| rs398123373 | pathogenic | 0 | 0 | 0 |
| rs756854513 | pathogenic | 0 | 0 | 0 |
| rs58232698 | benign | 0 | 0 | 0 |
| rs9600455 | benign | 0 | 0 | 0 |
| rs201016593 | pathogenic | 0 | 0 | 0 |
| rs756039188 | pathogenic | 0 | 0 | 0 |
| rs121908024 | pathogenic | 0 | 0 | 0 |
| rs373144619 | benign | 0 | 0 | 0 |
| rs138078086 | benign | 0 | 0 | 0 |
| rs376207800 | pathogenic | 0 | 0 | 0 |
| rs749038326 | pathogenic | 1 | 1 | 0 |
| rs372845091 | benign | 0 | 0 | 0 |
| rs377271627 | pathogenic | 0 | 0 | 0 |
| rs563390335 | pathogenic | 0 | 0 | 0 |
| rs13306510 | benign | 0 | 0 | 0 |
| rs764042910 | pathogenic | 0 | 0 | 0 |
| rs201384282 | benign | 1 | 1 | 1 |
| rs730882086 | pathogenic | 1 | 1 | 1 |
| rs756613387 | pathogenic | 1 | 1 | 1 |
| rs121908028 | benign | 1 | 1 | 0 |
| rs773328511 | benign | 0 | 0 | 0 |
| rs749220643 | pathogenic | 1 | 1 | 0 |
| rs730882089 | pathogenic | 0 | 0 | 0 |
| rs146651743 | benign | 0 | 0 | 0 |
| rs148698650 | pathogenic | 0 | 0 | 0 |
| rs140241383 | benign | 0 | 0 | 0 |
| rs757252110 | pathogenic | 0 | 0 | 0 |
| rs13306512 | pathogenic | 1 | 1 | 1 |
| rs121908043 | pathogenic | 1 | 1 | 1 |
| rs773658037 | pathogenic | 0 | 0 | 0 |
| rs774439908 | pathogenic | 0 | 0 | 0 |
| rs779732323 | pathogenic | 1 | 1 | 0 |
| rs749780672 | pathogenic | 0 | 0 | 0 |
| rs375651668 | benign | 1 | 1 | 1 |
| rs193922568 | pathogenic | 1 | 1 | 0 |
| rs750363970 | benign | 1 | 1 | 0 |
| rs5932 | benign | 0 | 0 | 0 |
| rs755154048 | pathogenic | 1 | 1 | 0 |
| rs759876319 | pathogenic | 0 | 0 | 0 |
| rs764550980 | pathogenic | 1 | 1 | 0 |
| rs779940524 | pathogenic | 0 | 0 | 0 |
| rs563382937 | pathogenic | 1 | 1 | 1 |
| rs28942083 | pathogenic | 0 | 1 | 0 |
| rs770744861 | pathogenic | 0 | 1 | 0 |
| rs121908031 | pathogenic | 0 | 1 | 0 |
| rs776217028 | pathogenic | 0 | 1 | 0 |
| rs763625913 | pathogenic | 0 | 0 | 0 |
| rs3736772 | benign | 0 | 0 | 0 |
| rs104894696 | benign | 1 | 1 | 1 |
| rs121918177 | pathogenic | 0 | 1 | 0 |
| rs121918181 | pathogenic | 1 | 1 | 0 |
| rs121918174 | pathogenic | 0 | 1 | 0 |
| rs121918182 | pathogenic | 0 | 0 | 0 |

| | | | | |
|---|---|---|---|---|
| rs121918175 | pathogenic | 1 | 1 | 1 |
| rs121918185 | pathogenic | 1 | 1 | 1 |
| rs121918173 | pathogenic | 1 | 1 | 0 |
| rs121918178 | pathogenic | 1 | 1 | 1 |
| rs121918179 | pathogenic | 1 | 1 | 1 |
| rs121918183 | pathogenic | 0 | 0 | 0 |
| rs121918176 | pathogenic | 1 | 1 | 1 |
| rs121918172 | pathogenic | 0 | 1 | 0 |
| rs121918184 | pathogenic | 0 | 0 | 0 |
| rs121434560 | pathogenic | 1 | 1 | 0 |
| rs121434561 | pathogenic | 1 | 1 | 0 |
| rs3218716 | pathogenic | 1 | 1 | 1 |
| rs145532615 | benign | 0 | 0 | 0 |
| rs149840927 | pathogenic | 0 | 0 | 0 |
| rs730880762 | pathogenic | 0 | 0 | 0 |
| rs730880764 | pathogenic | 0 | 0 | 0 |
| rs730880765 | pathogenic | 0 | 0 | 0 |
| rs587782962 | pathogenic | 0 | 0 | 0 |
| rs141735183 | pathogenic | 0 | 0 | 0 |
| rs397516187 | pathogenic | 0 | 0 | 0 |
| rs727505026 | pathogenic | 0 | 0 | 0 |
| rs529700838 | pathogenic | 0 | 0 | 0 |
| rs540263945 | pathogenic | 0 | 0 | 0 |
| rs149103761 | benign | 0 | 0 | 0 |
| rs730880906 | pathogenic | 0 | 0 | 0 |
| rs730880785 | pathogenic | 0 | 0 | 0 |
| rs397516199 | pathogenic | 0 | 0 | 0 |
| rs730880725 | benign | 0 | 0 | 0 |
| rs730880796 | pathogenic | 0 | 0 | 0 |
| rs201307101 | benign | 0 | 0 | 0 |
| rs397516214 | pathogenic | 0 | 0 | 0 |
| rs3729823 | benign | 0 | 0 | 0 |
| rs397516218 | pathogenic | 0 | 0 | 0 |
| rs121913647 | pathogenic | 0 | 0 | 0 |
| rs373514686 | pathogenic | 0 | 0 | 0 |
| rs763538103 | pathogenic | 0 | 0 | 0 |
| rs121913650 | pathogenic | 0 | 0 | 0 |
| rs139222507 | pathogenic | 0 | 0 | 0 |
| rs369940645 | pathogenic | 0 | 0 | 0 |
| rs763073072 | pathogenic | 0 | 0 | 0 |
| rs145734640 | pathogenic | 0 | 0 | 0 |
| rs730880821 | pathogenic | 0 | 0 | 0 |
| rs28933098 | pathogenic | 0 | 0 | 0 |
| rs372381770 | pathogenic | 0 | 0 | 0 |
| rs146796870 | benign | 0 | 0 | 0 |
| rs104894485 | pathogenic | 1 | 1 | 1 |
| rs35407712 | benign | 0 | 0 | 0 |
| rs35842750 | benign | 0 | 0 | 0 |
| rs2721191 | benign | 0 | 0 | 0 |
| rs35098923 | benign | 0 | 0 | 0 |
| rs34293591 | benign | 1 | 1 | 0 |
| rs36023964 | benign | 0 | 0 | 0 |
| rs4251691 | benign | 0 | 0 | 0 |
| rs34666647 | benign | 0 | 0 | 0 |

| | | | | |
|---|---|---|---|---|
| rs4925828 | benign | 0 | 0 | 0 |
| rs35348691 | benign | 0 | 0 | 0 |
| rs36078464 | benign | 0 | 0 | 0 |
| rs34915097 | benign | 0 | 0 | 0 |
| rs35101495 | benign | 0 | 0 | 0 |
| rs141568342 | pathogenic | 0 | 0 | 0 |
| rs1800571 | pathogenic | 0 | 0 | 0 |
| rs140566084 | benign | 1 | 1 | 1 |
| rs727503791 | pathogenic | 1 | 1 | 0 |
| rs545215807 | pathogenic | 0 | 0 | 0 |
| rs113994168 | pathogenic | 0 | 0 | 0 |
| rs113994167 | pathogenic | 0 | 0 | 0 |
| rs200788251 | pathogenic | 0 | 0 | 0 |
| rs200573371 | pathogenic | 0 | 0 | 0 |
| rs771874163 | pathogenic | 1 | 1 | 1 |
| rs112406105 | pathogenic | 1 | 1 | 1 |
| rs118204015 | pathogenic | 1 | 1 | 1 |
| rs113994169 | pathogenic | 0 | 0 | 0 |
| rs118204018 | pathogenic | 0 | 1 | 0 |
| rs138834083 | pathogenic | 0 | 0 | 0 |
| rs533055438 | pathogenic | 1 | 1 | 1 |
| rs2309689 | pathogenic | 1 | 1 | 1 |
| rs138058572 | pathogenic | 1 | 1 | 1 |
| rs118204017 | pathogenic | 1 | 1 | 0 |
| rs766742117 | pathogenic | 1 | 1 | 1 |
| rs751995154 | pathogenic | 1 | 1 | 1 |
| rs200771970 | pathogenic | 0 | 0 | 0 |
| rs118204014 | pathogenic | 0 | 0 | 0 |
| rs148584617 | pathogenic | 1 | 1 | 1 |
| rs121912720 | pathogenic | 0 | 0 | 0 |
| rs121912721 | pathogenic | 0 | 0 | 0 |
| rs28929476 | pathogenic | 0 | 0 | 0 |
| rs28931574 | pathogenic | 1 | 1 | 1 |
| rs12718465 | benign | 0 | 0 | 0 |
| rs144689354 | pathogenic | 0 | 0 | 0 |
| rs149095705 | pathogenic | 0 | 0 | 0 |
| rs572839648 | benign | 1 | 1 | 1 |
| rs121909715 | pathogenic | 1 | 1 | 0 |
| rs34077456 | benign | 1 | 1 | 1 |
| rs397517876 | benign | 0 | 0 | 0 |
| rs1064074 | benign | 0 | 0 | 0 |
| rs142751309 | benign | 0 | 0 | 0 |
| rs116275562 | benign | 0 | 0 | 0 |
| rs148095899 | benign | 0 | 0 | 0 |
| rs751838040 | pathogenic | 0 | 0 | 0 |
| rs121434324 | pathogenic | 1 | 1 | 1 |
| rs143367518 | pathogenic | 0 | 0 | 0 |
| rs121434325 | pathogenic | 0 | 0 | 0 |
| rs397515328 | pathogenic | 0 | 0 | 0 |
| rs376039938 | pathogenic | 0 | 0 | 0 |
| rs387907243 | pathogenic | 0 | 0 | 0 |
| rs28934876 | pathogenic | 0 | 0 | 0 |
| rs387907245 | pathogenic | 0 | 0 | 0 |
| rs138544311 | benign | 0 | 0 | 0 |

| | | | | |
|---|---|---|---|---|
| rs643788 | benign | 0 | 0 | 0 |
| rs749977756 | benign | 0 | 0 | 0 |
| rs1800574 | benign | 0 | 0 | 0 |
| rs137853336 | pathogenic | 1 | 1 | 1 |
| rs111916034 | benign | 0 | 0 | 0 |
| rs307826 | benign | 0 | 0 | 0 |
| rs35874891 | benign | 0 | 0 | 0 |
| rs111033580 | pathogenic | 1 | 1 | 1 |
| rs111033581 | pathogenic | 1 | 1 | 1 |
| rs121908373 | pathogenic | 0 | 0 | 0 |
| rs767122713 | pathogenic | 0 | 0 | 0 |
| rs2233213 | benign | 0 | 0 | 0 |
| rs121908605 | pathogenic | 1 | 1 | 0 |
| rs11887534 | benign | 0 | 0 | 0 |
| rs4148211 | benign | 0 | 0 | 0 |
| rs80025980 | benign | 0 | 0 | 0 |
| rs137854891 | pathogenic | 1 | 1 | 0 |
| rs74358901 | benign | 0 | 0 | 0 |
| rs9282574 | benign | 0 | 0 | 0 |
| rs137852993 | pathogenic | 0 | 0 | 0 |
| rs137852990 | pathogenic | 0 | 0 | 0 |
| rs137852987 | pathogenic | 0 | 0 | 0 |
| rs4148217 | benign | 0 | 0 | 0 |
| rs140690030 | benign | 0 | 0 | 0 |
| rs137852992 | pathogenic | 0 | 0 | 0 |
| rs6544718 | benign | 0 | 0 | 0 |
| rs137852989 | pathogenic | 0 | 0 | 0 |
| rs150893072 | benign | 1 | 1 | 1 |
| rs28931613 | pathogenic | 1 | 1 | 1 |
| rs143789597 | benign | 1 | 1 | 1 |
| rs121913037 | pathogenic | 1 | 1 | 0 |
| rs188802138 | pathogenic | 1 | 1 | 0 |
| rs121913038 | pathogenic | 1 | 1 | 0 |
| rs146835206 | benign | 0 | 0 | 0 |
| rs121913041 | pathogenic | 1 | 1 | 1 |
| rs121913039 | pathogenic | 1 | 1 | 1 |
| rs149977726 | pathogenic | 1 | 1 | 0 |
| rs121913036 | pathogenic | 0 | 0 | 0 |
| rs121913040 | pathogenic | 0 | 0 | 0 |
| rs765007158 | benign | 0 | 0 | 0 |
| rs112723255 | benign | 0 | 0 | 0 |
| rs574164748 | benign | 0 | 0 | 0 |
| rs80338734 | pathogenic | 1 | 1 | 0 |
| rs104894694 | pathogenic | 1 | 1 | 0 |
| rs753198836 | pathogenic | 0 | 0 | 0 |
| rs760101382 | pathogenic | 0 | 0 | 0 |
| rs121909272 | pathogenic | 1 | 1 | 0 |
| rs370421531 | pathogenic | 0 | 0 | 0 |
| rs565109128 | benign | 0 | 0 | 0 |
| rs17851582 | benign | 0 | 0 | 0 |
| rs753228876 | benign | 0 | 0 | 0 |
| rs141471799 | benign | 0 | 0 | 0 |
| rs281864916 | pathogenic | 1 | 1 | 0 |
| rs121918409 | pathogenic | 1 | 1 | 1 |

| | | | | |
|---|---|---|---|---|
| rs121918413 | pathogenic | 1 | 1 | 1 |
| rs121918411 | pathogenic | 1 | 1 | 1 |
| rs121918408 | pathogenic | 1 | 1 | 1 |
| rs121918412 | pathogenic | 1 | 1 | 1 |
| rs121918410 | pathogenic | 1 | 1 | 1 |
| rs116474260 | benign | 1 | 1 | 1 |
| rs281864919 | pathogenic | 1 | 1 | 1 |
| rs397514452 | pathogenic | 1 | 1 | 0 |
| rs747843638 | benign | 0 | 0 | 0 |
| rs774357609 | pathogenic | 1 | 1 | 1 |
| rs201804005 | benign | 0 | 0 | 0 |
| rs746485916 | pathogenic | 0 | 0 | 0 |
| rs764131178 | pathogenic | 0 | 0 | 0 |
| rs374094065 | pathogenic | 0 | 0 | 0 |
| rs76151636 | pathogenic | 1 | 1 | 0 |
| rs753250853 | pathogenic | 0 | 0 | 0 |
| rs761632029 | pathogenic | 0 | 0 | 0 |
| rs775541743 | pathogenic | 0 | 0 | 0 |
| rs778749563 | pathogenic | 0 | 0 | 0 |
| rs60431989 | pathogenic | 1 | 1 | 0 |
| rs755554442 | pathogenic | 1 | 1 | 0 |
| rs749085322 | pathogenic | 0 | 0 | 0 |
| rs756029120 | pathogenic | 1 | 1 | 0 |
| rs137853279 | pathogenic | 1 | 1 | 0 |
| rs777354267 | pathogenic | 1 | 1 | 1 |
| rs142940486 | benign | 1 | 1 | 1 |
| rs140167393 | benign | 1 | 1 | 1 |
| rs2066827 | benign | 0 | 0 | 0 |
| rs765681672 | benign | 0 | 0 | 0 |
| rs137985549 | benign | 0 | 0 | 0 |
| rs74315340 | pathogenic | 1 | 1 | 1 |
| rs74315341 | pathogenic | 0 | 0 | 0 |
| rs74315335 | pathogenic | 0 | 0 | 0 |
| rs61730974 | benign | 0 | 0 | 0 |
| rs121909193 | pathogenic | 0 | 0 | 0 |
| rs74315334 | pathogenic | 0 | 0 | 0 |
| rs74315329 | pathogenic | 0 | 0 | 0 |
| rs74315330 | pathogenic | 0 | 0 | 0 |
| rs121909194 | pathogenic | 0 | 0 | 0 |
| rs28936694 | pathogenic | 0 | 0 | 0 |
| rs74315336 | pathogenic | 0 | 0 | 0 |
| rs74315338 | pathogenic | 1 | 1 | 1 |
| rs74315328 | pathogenic | 0 | 0 | 0 |
| rs74315331 | pathogenic | 0 | 0 | 0 |
| rs74315332 | pathogenic | 0 | 0 | 0 |
| rs141696651 | benign | 0 | 0 | 0 |
| rs7041 | benign | 0 | 0 | 0 |
| rs4588 | benign | 0 | 0 | 0 |
| rs121908981 | pathogenic | 1 | 1 | 1 |
| rs606231459 | pathogenic | 0 | 0 | 0 |
| rs398123053 | pathogenic | 0 | 0 | 0 |
| rs774259802 | benign | 0 | 0 | 0 |
| rs121909582 | pathogenic | 1 | 1 | 1 |
| rs104893680 | pathogenic | 1 | 1 | 1 |

| | | | | |
|---|---|---|---|---|
| rs137854907 | pathogenic | 0 | 0 | 0 |
| rs765715798 | pathogenic | 0 | 0 | 0 |
| rs104893678 | pathogenic | 0 | 0 | 0 |
| rs104893679 | pathogenic | 0 | 1 | 0 |
| rs104893681 | pathogenic | 1 | 1 | 0 |
| rs121909640 | pathogenic | 1 | 1 | 0 |
| rs780705654 | pathogenic | 0 | 0 | 0 |
| rs201672146 | benign | 1 | 1 | 0 |
| rs12136658 | benign | 0 | 0 | 0 |
| rs56127440 | benign | 0 | 0 | 0 |
| rs104894433 | pathogenic | 1 | 1 | 1 |
| rs104894435 | pathogenic | 0 | 1 | 0 |
| rs104894437 | pathogenic | 1 | 1 | 1 |
| rs104894441 | pathogenic | 1 | 1 | 1 |
| rs104894440 | pathogenic | 1 | 1 | 1 |
| rs104894445 | pathogenic | 1 | 1 | 1 |
| rs104894436 | pathogenic | 0 | 0 | 0 |
| rs104894438 | pathogenic | 0 | 0 | 0 |
| rs200891969 | pathogenic | 1 | 1 | 0 |
| rs104894443 | pathogenic | 1 | 1 | 1 |
| rs104894434 | pathogenic | 1 | 1 | 1 |
| rs41298442 | pathogenic | 1 | 1 | 1 |
| rs104894442 | pathogenic | 1 | 1 | 1 |
| rs35453854 | benign | 0 | 0 | 0 |
| rs121912898 | pathogenic | 0 | 0 | 0 |
| rs121912897 | pathogenic | 1 | 1 | 0 |
| rs138260625 | benign | 0 | 0 | 0 |
| rs147802438 | benign | 1 | 1 | 0 |
| rs146443310 | benign | 1 | 1 | 0 |
| rs747319628 | pathogenic | 1 | 1 | 0 |
| rs1801195 | benign | 0 | 0 | 0 |
| rs145045341 | benign | 1 | 1 | 0 |
| rs116001173 | benign | 1 | 1 | 0 |
| rs587784175 | pathogenic | 0 | 0 | 0 |
| rs104893857 | pathogenic | 0 | 0 | 0 |
| rs104893862 | pathogenic | 1 | 1 | 0 |
| rs104893858 | pathogenic | 1 | 1 | 1 |
| rs121909249 | pathogenic | 0 | 1 | 0 |
| rs121909248 | pathogenic | 1 | 1 | 1 |
| rs104893859 | pathogenic | 1 | 1 | 0 |
| rs187063864 | pathogenic | 1 | 1 | 0 |
| rs121917817 | pathogenic | 1 | 1 | 0 |
| rs121917818 | pathogenic | 0 | 0 | 0 |
| rs753594601 | pathogenic | 0 | 0 | 0 |
| rs28937869 | pathogenic | 1 | 1 | 0 |
| rs6165 | benign | 0 | 0 | 0 |
| rs200089714 | pathogenic | 1 | 1 | 0 |
| rs372000714 | pathogenic | 0 | 0 | 0 |
| rs2228206 | benign | 0 | 0 | 0 |
| rs560096 | benign | 0 | 0 | 0 |
| rs137852666 | pathogenic | 0 | 0 | 0 |
| rs10896380 | benign | 0 | 0 | 0 |
| rs773690764 | pathogenic | 0 | 0 | 0 |
| rs201060167 | pathogenic | 0 | 0 | 0 |

| | | | | |
|---|---|---|---|---|
| rs137852670 | pathogenic | 0 | 0 | 0 |
| rs780594709 | pathogenic | 0 | 0 | 0 |
| rs145226920 | pathogenic | 0 | 0 | 0 |
| rs137852665 | pathogenic | 0 | 0 | 0 |
| rs724159960 | pathogenic | 0 | 0 | 0 |
| rs756985703 | pathogenic | 0 | 0 | 0 |
| rs137852667 | pathogenic | 0 | 0 | 0 |
| rs151079750 | pathogenic | 1 | 1 | 0 |
| rs2306636 | benign | 0 | 0 | 0 |
| rs72647885 | benign | 1 | 1 | 0 |
| rs377584435 | pathogenic | 1 | 1 | 1 |
| rs757583846 | pathogenic | 1 | 1 | 0 |
| rs121918148 | pathogenic | 0 | 0 | 0 |
| rs149201802 | benign | 0 | 0 | 0 |
| rs587781904 | pathogenic | 1 | 1 | 1 |
| rs143421325 | benign | 0 | 0 | 0 |
| rs587781253 | pathogenic | 0 | 0 | 0 |
| rs587780331 | benign | 0 | 0 | 0 |
| rs151001016 | benign | 0 | 0 | 0 |
| rs150888094 | benign | 0 | 0 | 0 |
| rs149581331 | benign | 0 | 0 | 0 |
| rs774840535 | benign | 0 | 0 | 0 |
| rs138428684 | benign | 0 | 0 | 0 |
| rs149300055 | benign | 0 | 0 | 0 |
| rs571005578 | benign | 0 | 0 | 0 |
| rs200731839 | benign | 0 | 0 | 0 |
| rs121965007 | pathogenic | 0 | 0 | 0 |
| rs121965013 | pathogenic | 0 | 1 | 0 |
| rs121965009 | pathogenic | 0 | 0 | 0 |
| rs1800457 | benign | 1 | 1 | 0 |
| rs121965006 | pathogenic | 1 | 1 | 0 |
| rs121965008 | pathogenic | 0 | 0 | 0 |
| rs61732609 | pathogenic | 0 | 0 | 0 |
| rs121965011 | pathogenic | 0 | 0 | 0 |
| rs121965015 | pathogenic | 0 | 0 | 0 |
| rs121965018 | pathogenic | 1 | 1 | 0 |
| rs121965016 | pathogenic | 0 | 0 | 0 |
| rs77806 | benign | 0 | 0 | 0 |
| rs149731356 | benign | 0 | 0 | 0 |
| rs72549407 | pathogenic | 0 | 0 | 0 |
| rs200363929 | benign | 0 | 0 | 0 |
| rs376403708 | benign | 1 | 1 | 0 |
| rs56376197 | benign | 1 | 1 | 1 |
| rs200237973 | benign | 1 | 1 | 1 |
| rs548330075 | benign | 0 | 0 | 0 |
| rs148291350 | benign | 0 | 0 | 0 |
| rs71579353 | pathogenic | 0 | 0 | 0 |
| rs146278697 | benign | 0 | 0 | 0 |
| rs141033098 | benign | 0 | 0 | 0 |
| rs397517234 | pathogenic | 0 | 0 | 0 |
| rs71579374 | benign | 0 | 0 | 0 |
| rs140381835 | benign | 0 | 0 | 0 |
| rs137852702 | pathogenic | 0 | 0 | 0 |
| rs137852696 | pathogenic | 1 | 1 | 1 |

| | | | | |
|---|---|---|---|---|
| rs137852697 | pathogenic | 0 | 0 | 0 |
| rs386833639 | pathogenic | 0 | 0 | 0 |
| rs386833640 | pathogenic | 0 | 0 | 0 |
| rs386833641 | benign | 0 | 0 | 0 |
| rs137852701 | pathogenic | 0 | 0 | 0 |
| rs386833642 | pathogenic | 0 | 0 | 0 |
| rs137852695 | pathogenic | 0 | 0 | 0 |
| rs386833646 | pathogenic | 0 | 0 | 0 |
| rs137852700 | pathogenic | 1 | 1 | 0 |
| rs386833649 | pathogenic | 1 | 1 | 0 |
| rs386833650 | pathogenic | 0 | 0 | 0 |
| rs148412181 | pathogenic | 0 | 0 | 0 |
| rs386833655 | pathogenic | 0 | 0 | 0 |
| rs386833656 | pathogenic | 1 | 1 | 0 |
| rs137852698 | pathogenic | 0 | 0 | 0 |
| rs386833661 | pathogenic | 0 | 0 | 0 |
| rs386833670 | pathogenic | 0 | 0 | 0 |
| rs111033632 | pathogenic | 1 | 1 | 1 |
| rs34308573 | benign | 1 | 1 | 1 |
| rs376048533 | pathogenic | 1 | 1 | 0 |
| rs3747517 | benign | 1 | 1 | 0 |
| rs137853328 | pathogenic | 0 | 0 | 0 |
| rs77273740 | benign | 0 | 0 | 0 |
| rs267606760 | pathogenic | 0 | 0 | 0 |
| rs77576840 | pathogenic | 0 | 0 | 0 |
| rs45465204 | benign | 1 | 1 | 0 |
| rs5320 | benign | 0 | 0 | 0 |
| rs5324 | benign | 0 | 0 | 0 |
| rs4531 | benign | 0 | 0 | 0 |
| rs267606761 | pathogenic | 0 | 0 | 0 |
| rs75215331 | pathogenic | 0 | 0 | 0 |
| rs6271 | benign | 0 | 0 | 0 |
| rs201643277 | pathogenic | 0 | 0 | 0 |
| rs4252250 | benign | 0 | 0 | 0 |
| rs137853580 | pathogenic | 0 | 0 | 0 |
| rs368287711 | pathogenic | 0 | 0 | 0 |
| rs137853579 | pathogenic | 0 | 0 | 0 |
| rs3135932 | benign | 0 | 0 | 0 |
| rs2228055 | benign | 0 | 0 | 0 |
| rs140322345 | benign | 1 | 1 | 1 |
| rs1154510 | pathogenic | 0 | 0 | 0 |
| rs11833399 | benign | 0 | 0 | 0 |
| rs36023382 | benign | 1 | 1 | 1 |
| rs61754444 | benign | 0 | 0 | 0 |
| rs28935168 | pathogenic | 0 | 0 | 0 |
| rs28934907 | pathogenic | 0 | 0 | 0 |
| rs28934904 | pathogenic | 1 | 1 | 0 |
| rs61748392 | pathogenic | 1 | 1 | 0 |
| rs28934908 | pathogenic | 0 | 0 | 0 |
| rs28934905 | pathogenic | 0 | 0 | 0 |
| rs28934906 | pathogenic | 1 | 1 | 0 |
| rs606231307 | pathogenic | 0 | 0 | 0 |
| rs9333296 | benign | 0 | 0 | 0 |
| rs3218614 | benign | 0 | 0 | 0 |

| | | | | |
|---|---|---|---|---|
| rs2230792 | benign | 0 | 0 | 0 |
| rs2230793 | benign | 0 | 0 | 0 |
| rs2230794 | benign | 0 | 0 | 0 |
| rs749052963 | pathogenic | 0 | 0 | 0 |
| rs2230798 | benign | 0 | 0 | 0 |
| rs2230795 | benign | 0 | 0 | 0 |
| rs3204145 | benign | 0 | 0 | 0 |
| rs1538660 | benign | 0 | 0 | 0 |
| rs9878928 | benign | 0 | 0 | 0 |
| rs104893742 | pathogenic | 0 | 0 | 0 |
| rs28936702 | pathogenic | 0 | 0 | 0 |
| rs28936703 | pathogenic | 0 | 0 | 0 |
| rs769392 | benign | 0 | 0 | 0 |
| rs769402 | benign | 1 | 1 | 1 |
| rs104894884 | pathogenic | 1 | 1 | 1 |
| rs1801316 | benign | 1 | 1 | 1 |
| rs104894885 | pathogenic | 1 | 1 | 1 |
| rs782572 | benign | 0 | 0 | 0 |
| rs151166046 | pathogenic | 0 | 0 | 0 |
| rs772153760 | pathogenic | 1 | 1 | 0 |
| rs35405862 | benign | 0 | 0 | 0 |
| rs146571352 | pathogenic | 0 | 1 | 0 |
| rs374698153 | pathogenic | 0 | 0 | 0 |
| rs7594497 | benign | 0 | 0 | 0 |
| rs80338716 | pathogenic | 0 | 0 | 0 |
| rs80338719 | pathogenic | 1 | 1 | 1 |
| rs80338721 | pathogenic | 0 | 0 | 0 |
| rs730882249 | pathogenic | 0 | 0 | 0 |
| rs146262009 | pathogenic | 0 | 0 | 0 |
| rs104895083 | pathogenic | 1 | 1 | 1 |
| rs104895100 | benign | 1 | 1 | 1 |
| rs104895085 | pathogenic | 0 | 0 | 0 |
| rs28940580 | pathogenic | 0 | 0 | 0 |
| rs104895098 | pathogenic | 0 | 0 | 0 |
| rs28940578 | pathogenic | 0 | 0 | 0 |
| rs61752717 | pathogenic | 0 | 0 | 0 |
| rs28940579 | pathogenic | 0 | 0 | 0 |
| rs104895097 | pathogenic | 0 | 0 | 0 |
| rs140425651 | benign | 1 | 1 | 0 |
| rs768090444 | pathogenic | 0 | 0 | 0 |
| rs764459544 | pathogenic | 0 | 0 | 0 |
| rs80338802 | pathogenic | 1 | 1 | 1 |
| rs149969786 | pathogenic | 1 | 1 | 0 |
| rs115311625 | pathogenic | 1 | 1 | 1 |
| rs778768583 | pathogenic | 1 | 1 | 1 |
| rs892090 | benign | 0 | 0 | 0 |
| rs28929498 | pathogenic | 1 | 1 | 1 |
| rs137854467 | pathogenic | 0 | 0 | 0 |
| rs137854480 | pathogenic | 0 | 0 | 0 |
| rs148682262 | benign | 0 | 0 | 0 |
| rs137852744 | pathogenic | 0 | 0 | 0 |
| rs137852745 | pathogenic | 0 | 1 | 0 |
| rs137852749 | pathogenic | 0 | 1 | 0 |
| rs137852746 | pathogenic | 0 | 1 | 0 |

| | | | | |
|---|---|---|---|---|
| rs145633958 | benign | 0 | 0 | 0 |
| rs141679950 | benign | 0 | 0 | 0 |
| rs76397662 | benign | 1 | 1 | 1 |
| rs370736139 | benign | 0 | 0 | 0 |
| rs781750106 | benign | 0 | 0 | 0 |
| rs546866208 | benign | 0 | 0 | 0 |
| rs142338976 | benign | 0 | 0 | 0 |
| rs115272158 | benign | 0 | 0 | 0 |
| rs115423919 | benign | 0 | 0 | 0 |
| rs537874538 | benign | 0 | 0 | 0 |
| rs9282834 | benign | 0 | 0 | 0 |
| rs201745826 | benign | 0 | 0 | 0 |
| rs75873440 | pathogenic | 0 | 0 | 0 |
| rs147219360 | benign | 0 | 0 | 0 |
| rs77939446 | pathogenic | 0 | 0 | 0 |
| rs77558292 | pathogenic | 0 | 0 | 0 |
| rs80069458 | pathogenic | 0 | 0 | 0 |
| rs76262710 | pathogenic | 0 | 0 | 0 |
| rs79781594 | pathogenic | 0 | 0 | 0 |
| rs77316810 | pathogenic | 0 | 0 | 0 |
| rs77503355 | pathogenic | 0 | 0 | 0 |
| rs79890926 | pathogenic | 0 | 0 | 0 |
| rs377767402 | benign | 0 | 0 | 0 |
| rs121913308 | pathogenic | 0 | 0 | 0 |
| rs55846256 | benign | 0 | 0 | 0 |
| rs75076352 | pathogenic | 0 | 0 | 0 |
| rs75996173 | pathogenic | 0 | 0 | 0 |
| rs77709286 | pathogenic | 0 | 0 | 0 |
| rs151058706 | benign | 0 | 0 | 0 |
| rs138964358 | benign | 0 | 0 | 0 |
| rs137852353 | pathogenic | 1 | 1 | 0 |
| rs201196020 | benign | 0 | 0 | 0 |
| rs137853201 | pathogenic | 1 | 1 | 1 |
| rs113994174 | pathogenic | 1 | 1 | 1 |
| rs104893674 | pathogenic | 1 | 1 | 1 |
| rs779519249 | benign | 1 | 1 | 1 |
| rs9606756 | benign | 0 | 0 | 0 |
| rs75680863 | benign | 1 | 1 | 1 |
| rs35838082 | benign | 0 | 0 | 0 |
| rs1801198 | benign | 0 | 0 | 0 |
| rs117458738 | benign | 1 | 1 | 0 |
| rs9621049 | benign | 1 | 1 | 0 |
| rs1131603 | benign | 1 | 1 | 0 |
| rs4820889 | benign | 0 | 0 | 0 |
| rs777767978 | pathogenic | 1 | 1 | 0 |
| rs147399860 | benign | 1 | 1 | 1 |
| rs104893950 | pathogenic | 0 | 0 | 0 |
| rs777308612 | benign | 0 | 0 | 0 |
| rs151091410 | benign | 0 | 0 | 0 |
| rs11537667 | benign | 0 | 0 | 0 |
| rs267606621 | pathogenic | 1 | 1 | 0 |
| rs143058902 | benign | 0 | 0 | 0 |
| rs370261904 | pathogenic | 0 | 0 | 0 |
| rs141169746 | benign | 0 | 0 | 0 |

| | | | | |
|---|---|---|---|---|
| rs111033436 | benign | 0 | 0 | 0 |
| rs148533341 | benign | 0 | 0 | 0 |
| rs143827620 | benign | 0 | 0 | 0 |
| rs4935502 | benign | 0 | 0 | 0 |
| rs61735473 | benign | 0 | 0 | 0 |
| rs267606932 | pathogenic | 1 | 1 | 0 |
| rs61730754 | benign | 0 | 0 | 0 |
| rs374491359 | pathogenic | 1 | 1 | 1 |
| rs761800077 | benign | 1 | 1 | 1 |
| rs760391381 | benign | 1 | 1 | 1 |
| rs104893885 | pathogenic | 0 | 0 | 0 |
| rs104893886 | pathogenic | 1 | 1 | 1 |
| rs104893884 | pathogenic | 0 | 0 | 0 |
| rs147715509 | benign | 0 | 0 | 0 |
| rs115533243 | benign | 0 | 0 | 0 |
| rs397509390 | pathogenic | 0 | 1 | 0 |
| rs80357134 | pathogenic | 0 | 0 | 0 |
| rs766004110 | benign | 1 | 1 | 1 |
| rs769650474 | benign | 0 | 0 | 0 |
| rs398122635 | benign | 1 | 1 | 0 |
| rs199522616 | pathogenic | 0 | 0 | 0 |
| rs28897672 | pathogenic | 1 | 1 | 0 |
| rs80357086 | pathogenic | 1 | 1 | 1 |
| rs80357064 | pathogenic | 1 | 1 | 0 |
| rs80356936 | benign | 0 | 0 | 0 |
| rs28897673 | benign | 0 | 0 | 0 |
| rs146747324 | benign | 0 | 0 | 0 |
| rs78635798 | pathogenic | 0 | 0 | 0 |
| rs75146158 | pathogenic | 1 | 1 | 1 |
| rs587782886 | benign | 0 | 0 | 0 |
| rs121434634 | pathogenic | 0 | 0 | 0 |
| rs121434635 | pathogenic | 0 | 0 | 0 |
| rs587777778 | pathogenic | 0 | 0 | 0 |
| rs118203967 | pathogenic | 0 | 0 | 0 |
| rs751375244 | pathogenic | 1 | 1 | 1 |
| rs104893863 | pathogenic | 1 | 1 | 0 |
| rs104893865 | pathogenic | 0 | 0 | 0 |
| rs104893864 | pathogenic | 1 | 1 | 1 |
| rs104893866 | pathogenic | 1 | 1 | 0 |
| rs561738497 | benign | 0 | 0 | 0 |
| rs57599352 | pathogenic | 1 | 1 | 1 |
| rs267607448 | pathogenic | 1 | 1 | 1 |
| rs63751596 | pathogenic | 0 | 0 | 0 |
| rs63751612 | benign | 0 | 0 | 0 |
| rs63751684 | benign | 0 | 0 | 0 |
| rs267607861 | benign | 0 | 0 | 0 |
| rs35831931 | benign | 0 | 0 | 0 |
| rs63750876 | benign | 0 | 0 | 0 |
| rs41295284 | benign | 0 | 0 | 0 |
| rs35502531 | benign | 0 | 0 | 0 |
| rs63750449 | benign | 0 | 0 | 0 |
| rs63750899 | pathogenic | 0 | 0 | 0 |
| rs55907433 | benign | 0 | 0 | 0 |
| rs63750217 | pathogenic | 0 | 0 | 0 |

| | | | | |
|---|---|---|---|---|
| rs63751275 | pathogenic | 0 | 0 | 0 |
| rs63750702 | benign | 1 | 1 | 1 |
| rs63750114 | benign | 0 | 0 | 0 |
| rs2020873 | benign | 0 | 0 | 0 |
| rs140195825 | benign | 1 | 1 | 1 |
| rs182763805 | benign | 0 | 0 | 0 |
| rs533696377 | benign | 0 | 0 | 0 |
| rs41278788 | benign | 0 | 0 | 0 |
| rs104894193 | pathogenic | 1 | 1 | 0 |
| rs3824915 | benign | 1 | 1 | 1 |
| rs104894196 | pathogenic | 0 | 0 | 0 |
| rs78681604 | benign | 1 | 1 | 1 |
| rs77321498 | benign | 1 | 1 | 1 |
| rs137852965 | pathogenic | 0 | 0 | 0 |
| rs137852961 | pathogenic | 0 | 0 | 0 |
| rs137852962 | pathogenic | 1 | 1 | 0 |
| rs753376100 | pathogenic | 0 | 0 | 0 |
| rs137852968 | pathogenic | 1 | 1 | 1 |
| rs137852959 | pathogenic | 1 | 1 | 0 |
| rs137852967 | pathogenic | 0 | 0 | 0 |
| rs113437301 | benign | 1 | 1 | 1 |
| rs61747482 | benign | 1 | 1 | 1 |
| rs2234919 | benign | 1 | 1 | 0 |
| rs121908863 | pathogenic | 0 | 0 | 0 |
| rs140455771 | pathogenic | 0 | 0 | 0 |
| rs121908810 | pathogenic | 1 | 1 | 1 |
| rs143879853 | benign | 0 | 0 | 0 |
| rs188319622 | benign | 0 | 0 | 0 |
| rs121918645 | pathogenic | 0 | 0 | 0 |
| rs115352435 | pathogenic | 0 | 0 | 0 |
| rs121434427 | pathogenic | 1 | 1 | 1 |
| rs121434428 | pathogenic | 0 | 0 | 0 |
| rs35086265 | benign | 1 | 1 | 1 |
| rs11576415 | benign | 0 | 0 | 0 |
| rs121434429 | pathogenic | 1 | 1 | 1 |
| rs4074536 | benign | 1 | 1 | 1 |
| rs10801999 | benign | 0 | 0 | 0 |
| rs121434550 | pathogenic | 0 | 0 | 0 |
| rs148057999 | pathogenic | 0 | 0 | 0 |
| rs28730716 | benign | 0 | 0 | 0 |
| rs142036299 | benign | 0 | 0 | 0 |
| rs762381137 | pathogenic | 0 | 0 | 0 |
| rs121434549 | pathogenic | 0 | 0 | 0 |
| rs139228801 | pathogenic | 0 | 0 | 0 |
| rs147219463 | benign | 0 | 0 | 0 |
| rs1131857 | benign | 0 | 0 | 0 |
| rs121917874 | pathogenic | 0 | 1 | 0 |
| rs2228056 | benign | 0 | 0 | 0 |
| rs121917870 | pathogenic | 0 | 0 | 0 |
| rs121917866 | pathogenic | 0 | 0 | 0 |
| rs11921054 | benign | 0 | 0 | 0 |
| rs121917868 | pathogenic | 0 | 0 | 0 |
| rs28931603 | pathogenic | 0 | 0 | 0 |
| rs104895295 | pathogenic | 1 | 1 | 0 |

| | | | | |
|---|---|---|---|---|
| rs104895382 | pathogenic | 0 | 0 | 0 |
| rs104895298 | pathogenic | 0 | 0 | 0 |
| rs104895300 | pathogenic | 0 | 0 | 0 |
| rs104895332 | pathogenic | 0 | 0 | 0 |
| rs758026399 | pathogenic | 1 | 1 | 1 |
| rs104895366 | pathogenic | 0 | 0 | 0 |
| rs104895304 | pathogenic | 0 | 0 | 0 |
| rs759997079 | pathogenic | 0 | 0 | 0 |
| rs121917789 | pathogenic | 0 | 0 | 0 |
| rs104895319 | pathogenic | 0 | 0 | 0 |
| rs104895317 | pathogenic | 1 | 1 | 0 |
| rs28934897 | pathogenic | 0 | 0 | 0 |
| rs104895380 | benign | 0 | 0 | 0 |
| rs104895360 | pathogenic | 0 | 0 | 0 |
| rs28936396 | pathogenic | 1 | 1 | 0 |
| rs121909307 | pathogenic | 1 | 1 | 1 |
| rs28938472 | pathogenic | 1 | 1 | 0 |
| rs121909308 | pathogenic | 1 | 1 | 0 |
| rs121909309 | pathogenic | 0 | 0 | 0 |
| rs121913013 | benign | 1 | 1 | 1 |
| rs201855245 | benign | 1 | 1 | 1 |
| rs2230232 | benign | 1 | 1 | 0 |
| rs113451409 | benign | 0 | 0 | 0 |
| rs121913011 | pathogenic | 1 | 1 | 0 |
| rs185821167 | pathogenic | 1 | 1 | 0 |
| rs2230234 | benign | 0 | 0 | 0 |
| rs193922639 | benign | 1 | 1 | 0 |
| rs774863785 | pathogenic | 1 | 1 | 0 |
| rs576404380 | pathogenic | 1 | 1 | 0 |
| rs2230235 | benign | 1 | 1 | 0 |
| rs200484060 | pathogenic | 1 | 1 | 0 |
| rs199761749 | benign | 0 | 0 | 0 |
| rs199681901 | benign | 0 | 0 | 0 |
| rs121908951 | pathogenic | 0 | 0 | 0 |
| rs769251460 | benign | 0 | 0 | 0 |
| rs28937320 | pathogenic | 0 | 0 | 0 |
| rs200607527 | pathogenic | 0 | 0 | 0 |
| rs137852640 | pathogenic | 1 | 1 | 1 |
| rs137852636 | pathogenic | 1 | 1 | 1 |
| rs137852638 | pathogenic | 0 | 0 | 0 |
| rs142637231 | pathogenic | 0 | 0 | 0 |
| rs151328418 | benign | 0 | 1 | 0 |
| rs137852639 | pathogenic | 1 | 1 | 1 |
| rs34605667 | benign | 0 | 0 | 0 |
| rs104894073 | pathogenic | 1 | 1 | 0 |
| rs56298569 | pathogenic | 0 | 0 | 0 |
| rs757600616 | pathogenic | 0 | 0 | 0 |
| rs139194636 | pathogenic | 0 | 0 | 0 |
| rs369517993 | pathogenic | 1 | 1 | 1 |
| rs138846036 | benign | 1 | 1 | 1 |
| rs200808744 | pathogenic | 1 | 1 | 1 |
| rs2255543 | benign | 0 | 0 | 0 |
| rs149165166 | pathogenic | 0 | 0 | 0 |
| rs544956641 | pathogenic | 1 | 1 | 1 |

| | | | | |
|---|---|---|---|---|
| rs28941783 | pathogenic | 0 | 0 | 0 |
| rs562853291 | pathogenic | 1 | 1 | 1 |
| rs28942100 | pathogenic | 0 | 0 | 0 |
| rs120074174 | pathogenic | 1 | 1 | 1 |
| rs120074171 | pathogenic | 1 | 1 | 1 |
| rs120074173 | pathogenic | 0 | 0 | 0 |
| rs120074172 | pathogenic | 1 | 1 | 1 |
| rs767159114 | pathogenic | 1 | 1 | 1 |
| rs104893657 | pathogenic | 0 | 0 | 0 |
| rs104893656 | pathogenic | 0 | 0 | 0 |
| rs104893659 | pathogenic | 0 | 0 | 0 |
| rs104893658 | pathogenic | 0 | 0 | 0 |
| rs200386310 | pathogenic | 0 | 0 | 0 |
| rs2072712 | benign | 0 | 0 | 0 |
| rs35274867 | benign | 0 | 0 | 0 |
| rs35343172 | benign | 0 | 0 | 0 |
| rs10079250 | benign | 0 | 0 | 0 |
| rs34951517 | benign | 0 | 0 | 0 |
| rs267606695 | pathogenic | 0 | 0 | 0 |
| rs149391728 | benign | 0 | 0 | 0 |
| rs387906598 | pathogenic | 1 | 1 | 0 |
| rs137854578 | pathogenic | 0 | 0 | 0 |
| rs141519952 | benign | 0 | 0 | 0 |
| rs146572883 | benign | 0 | 0 | 0 |
| rs111866410 | benign | 0 | 0 | 0 |
| rs139387758 | benign | 1 | 1 | 1 |
| rs35672074 | benign | 1 | 1 | 1 |
| rs121434322 | pathogenic | 0 | 0 | 0 |
| rs150884181 | benign | 0 | 0 | 0 |
| rs35867471 | benign | 0 | 0 | 0 |
| rs2867802 | benign | 0 | 0 | 0 |
| rs137902889 | benign | 0 | 0 | 0 |
| rs369804864 | pathogenic | 0 | 0 | 0 |
| rs781050795 | pathogenic | 0 | 0 | 0 |
| rs2305799 | benign | 0 | 0 | 0 |
| rs2229531 | benign | 0 | 0 | 0 |
| rs147025508 | benign | 0 | 0 | 0 |
| rs76857106 | pathogenic | 0 | 0 | 0 |
| rs7247284 | benign | 0 | 0 | 0 |
| rs62619782 | benign | 0 | 0 | 0 |
| rs377244188 | pathogenic | 0 | 0 | 0 |
| rs75718910 | pathogenic | 0 | 0 | 0 |
| rs79843600 | pathogenic | 0 | 0 | 0 |
| rs75037667 | pathogenic | 1 | 1 | 1 |
| rs121913006 | pathogenic | 0 | 0 | 0 |
| rs397517925 | pathogenic | 1 | 1 | 1 |
| rs267606831 | pathogenic | 1 | 1 | 1 |
| rs121909667 | pathogenic | 1 | 1 | 1 |
| rs121909669 | pathogenic | 1 | 1 | 1 |
| rs121909668 | pathogenic | 1 | 1 | 1 |
| rs121909671 | pathogenic | 1 | 1 | 1 |
| rs534596036 | benign | 0 | 0 | 0 |
| rs144037629 | pathogenic | 0 | 0 | 0 |
| rs11546406 | benign | 0 | 0 | 0 |

| | | | | |
|---|---|---|---|---|
| rs143697995 | pathogenic | 0 | 0 | 0 |
| rs121908331 | pathogenic | 0 | 0 | 0 |
| rs149377346 | benign | 0 | 0 | 0 |
| rs104894086 | pathogenic | 1 | 1 | 0 |
| rs104894089 | pathogenic | 0 | 0 | 0 |
| rs104894090 | pathogenic | 1 | 1 | 0 |
| rs387907235 | pathogenic | 0 | 0 | 0 |
| rs137852689 | pathogenic | 0 | 0 | 0 |
| rs137852690 | pathogenic | 1 | 1 | 0 |
| rs104894087 | pathogenic | 0 | 0 | 0 |
| rs104894085 | pathogenic | 0 | 0 | 0 |
| rs79383654 | benign | 0 | 0 | 0 |
| rs121434239 | pathogenic | 0 | 0 | 0 |
| rs121434240 | pathogenic | 0 | 0 | 0 |
| rs121434236 | pathogenic | 0 | 0 | 0 |
| rs121434238 | pathogenic | 1 | 1 | 1 |
| rs121918107 | pathogenic | 1 | 1 | 1 |
| rs121918103 | pathogenic | 1 | 1 | 1 |
| rs121918104 | pathogenic | 1 | 1 | 1 |
| rs61747614 | benign | 1 | 1 | 1 |
| rs555329870 | benign | 0 | 0 | 0 |
| rs186094931 | benign | 0 | 0 | 0 |
| rs267606785 | pathogenic | 0 | 0 | 0 |
| rs34474955 | benign | 0 | 0 | 0 |
| rs1126497 | benign | 1 | 1 | 1 |
| rs397514661 | pathogenic | 0 | 0 | 0 |
| rs189732445 | benign | 0 | 0 | 0 |
| rs146685071 | benign | 0 | 0 | 0 |
| rs121908636 | pathogenic | 1 | 1 | 0 |
| rs121908643 | pathogenic | 0 | 0 | 0 |
| rs121908644 | pathogenic | 1 | 1 | 1 |
| rs575001023 | pathogenic | 1 | 1 | 1 |
| rs121908646 | pathogenic | 1 | 1 | 0 |
| rs121908638 | pathogenic | 1 | 1 | 1 |
| rs777828000 | pathogenic | 1 | 1 | 1 |
| rs74923032 | benign | 0 | 0 | 0 |
| rs192838388 | pathogenic | 1 | 1 | 1 |
| rs148918985 | pathogenic | 1 | 1 | 1 |
| rs773540764 | benign | 1 | 1 | 1 |
| rs370595480 | pathogenic | 1 | 1 | 1 |
| rs762387914 | pathogenic | 1 | 1 | 1 |
| rs121908642 | pathogenic | 1 | 1 | 1 |
| rs183276875 | pathogenic | 1 | 1 | 1 |
| rs121908648 | pathogenic | 1 | 1 | 1 |
| rs121908647 | pathogenic | 1 | 1 | 1 |
| rs121908640 | pathogenic | 1 | 1 | 1 |
| rs771937610 | pathogenic | 1 | 1 | 1 |
| rs121908641 | pathogenic | 1 | 1 | 1 |
| rs5745068 | benign | 0 | 0 | 0 |
| rs73155056 | benign | 1 | 1 | 1 |
| rs148788180 | benign | 0 | 0 | 0 |
| rs121918137 | pathogenic | 0 | 0 | 0 |
| rs41269593 | benign | 0 | 0 | 0 |
| rs11159 | benign | 1 | 1 | 0 |

| | | | | |
|---|---|---|---|---|
| rs114733570 | pathogenic | 1 | 1 | 0 |
| rs3211783 | benign | 0 | 0 | 0 |
| rs4762 | benign | 0 | 0 | 0 |
| rs74315283 | pathogenic | 0 | 0 | 0 |
| rs111033559 | pathogenic | 0 | 0 | 0 |
| rs111033560 | pathogenic | 1 | 1 | 1 |
| rs749962743 | benign | 0 | 0 | 0 |
| rs267606986 | pathogenic | 0 | 0 | 0 |
| rs34516635 | benign | 0 | 0 | 0 |
| rs117440569 | benign | 0 | 0 | 0 |
| rs61740877 | benign | 0 | 0 | 0 |
| rs121918352 | pathogenic | 0 | 0 | 0 |
| rs183974372 | benign | 0 | 0 | 0 |
| rs140511204 | benign | 1 | 1 | 0 |
| rs6040067 | benign | 0 | 0 | 0 |
| rs199674138 | benign | 0 | 0 | 0 |
| rs121918350 | pathogenic | 0 | 0 | 0 |
| rs138292988 | benign | 1 | 1 | 1 |
| rs145288947 | benign | 0 | 0 | 0 |
| rs369714529 | benign | 0 | 0 | 0 |
| rs200945419 | benign | 0 | 0 | 0 |
| rs116959874 | benign | 0 | 0 | 0 |
| rs141659620 | pathogenic | 1 | 1 | 0 |
| rs111475461 | benign | 0 | 0 | 0 |
| rs61755320 | pathogenic | 0 | 0 | 0 |
| rs748600162 | pathogenic | 1 | 1 | 1 |
| rs369227537 | pathogenic | 1 | 1 | 1 |
| rs372981030 | pathogenic | 1 | 1 | 1 |
| rs137854486 | pathogenic | 0 | 0 | 0 |
| rs137854487 | pathogenic | 0 | 0 | 0 |
| rs199905054 | pathogenic | 0 | 0 | 0 |
| rs771438170 | pathogenic | 0 | 0 | 0 |
| rs774949844 | pathogenic | 0 | 0 | 0 |
| rs4899145 | benign | 0 | 0 | 0 |
| rs200642525 | benign | 0 | 0 | 0 |
| rs190140598 | pathogenic | 0 | 0 | 0 |
| rs2250889 | benign | 0 | 0 | 0 |
| rs17577 | benign | 0 | 0 | 0 |
| rs142032681 | benign | 0 | 0 | 0 |
| rs144577963 | benign | 0 | 0 | 0 |
| rs61754623 | benign | 0 | 0 | 0 |
| rs139873200 | benign | 0 | 0 | 0 |
| rs532530906 | benign | 0 | 0 | 0 |
| rs145762716 | benign | 0 | 0 | 0 |
| rs34062020 | benign | 0 | 0 | 0 |
| rs35258161 | benign | 0 | 0 | 0 |
| rs149792649 | benign | 0 | 0 | 0 |
| rs78101774 | benign | 0 | 0 | 0 |
| rs766466297 | pathogenic | 0 | 1 | 0 |
| rs140467171 | pathogenic | 0 | 0 | 0 |
| rs141340466 | pathogenic | 0 | 0 | 0 |
| rs201555303 | pathogenic | 0 | 0 | 0 |
| rs143592405 | pathogenic | 0 | 0 | 0 |
| rs781249411 | pathogenic | 0 | 0 | 0 |

| | | | | |
|---|---|---|---|---|
| rs116644127 | benign | 0 | 0 | 0 |
| rs151146863 | benign | 1 | 1 | 1 |
| rs116440799 | benign | 0 | 0 | 0 |
| rs138958351 | benign | 0 | 0 | 0 |
| rs143319354 | benign | 0 | 0 | 0 |
| rs768669208 | pathogenic | 1 | 1 | 1 |
| rs28940289 | pathogenic | 1 | 1 | 0 |
| rs745656120 | pathogenic | 1 | 1 | 0 |
| rs387906987 | pathogenic | 0 | 0 | 0 |
| rs147240777 | benign | 0 | 0 | 0 |
| rs140945833 | benign | 0 | 0 | 0 |
| rs146614981 | benign | 0 | 0 | 0 |
| rs149878562 | benign | 0 | 0 | 0 |
| rs28938168 | pathogenic | 0 | 0 | 0 |
| rs33939927 | pathogenic | 1 | 1 | 1 |
| rs34995376 | pathogenic | 1 | 1 | 1 |
| rs201034481 | pathogenic | 1 | 1 | 1 |
| rs774069983 | benign | 0 | 0 | 0 |
| rs199422226 | pathogenic | 1 | 1 | 0 |
| rs199422225 | pathogenic | 1 | 1 | 1 |
| rs199422224 | pathogenic | 1 | 1 | 0 |
| rs149271416 | pathogenic | 1 | 1 | 0 |
| rs1801318 | benign | 1 | 1 | 1 |
| rs372691318 | pathogenic | 1 | 1 | 1 |
| rs147685849 | benign | 0 | 0 | 0 |
| rs387907199 | pathogenic | 0 | 0 | 0 |
| rs767122069 | pathogenic | 0 | 0 | 0 |
| rs2302427 | benign | 0 | 0 | 0 |
| rs201135441 | benign | 0 | 0 | 0 |
| rs201861847 | pathogenic | 0 | 0 | 0 |
| rs137852952 | pathogenic | 0 | 0 | 0 |
| rs16986560 | benign | 0 | 0 | 0 |
| rs147344332 | pathogenic | 1 | 1 | 1 |
| rs36027220 | pathogenic | 0 | 0 | 0 |
| rs192401229 | benign | 0 | 0 | 0 |
| rs121909106 | pathogenic | 1 | 1 | 0 |
| rs118204103 | pathogenic | 1 | 1 | 0 |
| rs118204104 | pathogenic | 1 | 1 | 0 |
| rs118204105 | pathogenic | 1 | 1 | 0 |
| rs118204106 | pathogenic | 0 | 0 | 0 |
| rs118204119 | pathogenic | 0 | 0 | 0 |
| rs118204107 | pathogenic | 0 | 0 | 0 |
| rs118204094 | pathogenic | 0 | 0 | 0 |
| rs118204098 | pathogenic | 1 | 1 | 0 |
| rs118204101 | pathogenic | 1 | 1 | 0 |
| rs118204096 | pathogenic | 1 | 1 | 0 |
| rs118204108 | pathogenic | 0 | 0 | 0 |
| rs536814318 | pathogenic | 0 | 0 | 0 |
| rs34413634 | pathogenic | 1 | 1 | 0 |
| rs118204100 | pathogenic | 0 | 0 | 0 |
| rs118204109 | pathogenic | 0 | 0 | 0 |
| rs118204116 | pathogenic | 1 | 1 | 0 |
| rs118204110 | pathogenic | 0 | 0 | 0 |
| rs118204099 | pathogenic | 0 | 0 | 0 |

| | | | | |
|---|---|---|---|---|
| rs118204111 | pathogenic | 0 | 0 | 0 |
| rs118204112 | pathogenic | 0 | 0 | 0 |
| rs118204113 | pathogenic | 0 | 0 | 0 |
| rs118204114 | pathogenic | 0 | 0 | 0 |
| rs118204115 | pathogenic | 0 | 0 | 0 |
| rs150428209 | benign | 0 | 0 | 0 |
| rs144949995 | benign | 0 | 0 | 0 |
| rs190023981 | benign | 0 | 0 | 0 |
| rs104894611 | pathogenic | 1 | 1 | 1 |
| rs28937580 | pathogenic | 1 | 1 | 1 |
| rs121908949 | pathogenic | 0 | 0 | 0 |
| rs104894612 | pathogenic | 1 | 1 | 1 |
| rs104894609 | pathogenic | 1 | 1 | 1 |
| rs104894610 | pathogenic | 1 | 1 | 1 |
| rs104894615 | pathogenic | 0 | 0 | 0 |
| rs104894603 | pathogenic | 0 | 0 | 0 |
| rs104894602 | pathogenic | 1 | 1 | 1 |
| rs121908948 | pathogenic | 1 | 1 | 1 |
| rs104894608 | pathogenic | 1 | 1 | 1 |
| rs371545673 | benign | 0 | 0 | 0 |
| rs199925238 | benign | 0 | 0 | 0 |
| rs146313451 | benign | 0 | 0 | 0 |
| rs121918193 | pathogenic | 1 | 1 | 0 |
| rs2228500 | benign | 0 | 0 | 0 |
| rs121918194 | pathogenic | 1 | 1 | 0 |
| rs121918196 | pathogenic | 0 | 0 | 0 |
| rs41291971 | benign | 0 | 0 | 0 |
| rs199473690 | pathogenic | 0 | 0 | 0 |
| rs2228613 | benign | 0 | 0 | 0 |
| rs267607130 | pathogenic | 1 | 1 | 1 |
| rs77615401 | benign | 1 | 1 | 1 |
| rs374618872 | benign | 1 | 1 | 1 |
| rs146165014 | benign | 0 | 0 | 0 |
| rs144586070 | benign | 0 | 0 | 0 |
| rs121918541 | pathogenic | 0 | 0 | 0 |
| rs142105943 | benign | 0 | 0 | 0 |
| rs121918540 | pathogenic | 0 | 0 | 0 |
| rs6791 | benign | 0 | 0 | 0 |
| rs61736587 | pathogenic | 0 | 0 | 0 |
| rs61736586 | benign | 0 | 0 | 0 |
| rs121917741 | pathogenic | 1 | 1 | 1 |
| rs41302885 | benign | 0 | 0 | 0 |
| rs141026028 | benign | 0 | 0 | 0 |
| rs193922493 | pathogenic | 0 | 0 | 0 |
| rs73598374 | benign | 0 | 0 | 0 |
| rs121908725 | pathogenic | 1 | 1 | 0 |
| rs199422328 | pathogenic | 0 | 1 | 0 |
| rs11555566 | benign | 1 | 1 | 1 |
| rs267606634 | pathogenic | 0 | 0 | 0 |
| rs121908714 | pathogenic | 1 | 1 | 0 |
| rs121908717 | pathogenic | 1 | 1 | 0 |
| rs267606635 | pathogenic | 1 | 1 | 0 |
| rs121908739 | pathogenic | 0 | 0 | 0 |
| rs121908735 | pathogenic | 0 | 1 | 0 |

| | | | | |
|---|---|---|---|---|
| rs121908716 | pathogenic | 0 | 1 | 0 |
| rs121908723 | pathogenic | 0 | 1 | 0 |
| rs121908721 | pathogenic | 0 | 0 | 0 |
| rs199422327 | pathogenic | 0 | 0 | 0 |
| rs121908715 | pathogenic | 0 | 0 | 0 |
| rs587783563 | pathogenic | 0 | 0 | 0 |
| rs104894780 | pathogenic | 0 | 0 | 0 |
| rs56030372 | pathogenic | 1 | 1 | 1 |
| rs104894782 | pathogenic | 1 | 1 | 1 |
| rs149495971 | benign | 0 | 0 | 0 |
| rs118203942 | pathogenic | 1 | 1 | 0 |
| rs118203939 | pathogenic | 0 | 0 | 0 |
| rs118203938 | pathogenic | 0 | 0 | 0 |
| rs118203943 | pathogenic | 0 | 0 | 0 |
| rs118203940 | pathogenic | 0 | 0 | 0 |
| rs727503809 | pathogenic | 0 | 0 | 0 |
| rs1065757 | benign | 0 | 0 | 0 |
| rs1071598 | benign | 0 | 0 | 0 |
| rs25414 | benign | 0 | 0 | 0 |
| rs118203944 | pathogenic | 0 | 0 | 0 |
| rs118203941 | pathogenic | 0 | 0 | 0 |
| rs144222167 | benign | 0 | 0 | 0 |
| rs201101343 | pathogenic | 0 | 0 | 0 |
| rs144601717 | benign | 0 | 0 | 0 |
| rs267607129 | pathogenic | 0 | 0 | 0 |
| rs104894729 | pathogenic | 0 | 0 | 0 |
| rs104894727 | pathogenic | 0 | 0 | 0 |
| rs104894725 | pathogenic | 0 | 0 | 0 |
| rs104894515 | pathogenic | 1 | 1 | 1 |
| rs74688377 | benign | 0 | 0 | 0 |
| rs121908513 | pathogenic | 1 | 1 | 0 |
| rs121908509 | pathogenic | 0 | 0 | 0 |
| rs121908514 | pathogenic | 1 | 1 | 0 |
| rs121908512 | pathogenic | 1 | 1 | 0 |
| rs121908510 | pathogenic | 0 | 0 | 0 |
| rs121908511 | pathogenic | 1 | 1 | 1 |
| rs121908518 | pathogenic | 1 | 1 | 1 |
| rs34358665 | benign | 0 | 0 | 0 |
| rs147568834 | benign | 0 | 0 | 0 |
| rs201717599 | benign | 0 | 0 | 0 |
| rs61744487 | benign | 0 | 0 | 0 |
| rs370627877 | pathogenic | 1 | 1 | 1 |
| rs368593151 | pathogenic | 1 | 1 | 1 |
| rs114363936 | benign | 0 | 0 | 0 |
| rs137853333 | pathogenic | 0 | 0 | 0 |
| rs137853080 | pathogenic | 1 | 1 | 1 |
| rs137853077 | pathogenic | 1 | 1 | 1 |
| rs780749732 | benign | 1 | 1 | 1 |
| rs137853081 | pathogenic | 1 | 1 | 0 |
| rs137853078 | pathogenic | 0 | 0 | 0 |
| rs121913315 | pathogenic | 1 | 1 | 1 |
| rs752513525 | pathogenic | 1 | 1 | 1 |
| rs104894918 | pathogenic | 0 | 0 | 0 |
| rs104894917 | pathogenic | 0 | 0 | 0 |

| | | | | |
|---|---|---|---|---|
| rs104894919 | pathogenic | 0 | 0 | 0 |
| rs28939672 | pathogenic | 0 | 0 | 0 |
| rs104894173 | pathogenic | 0 | 0 | 0 |
| rs104894179 | pathogenic | 0 | 0 | 0 |
| rs104894174 | pathogenic | 1 | 1 | 0 |
| rs104894178 | pathogenic | 1 | 1 | 0 |
| rs28939714 | pathogenic | 1 | 1 | 1 |
| rs104894270 | pathogenic | 1 | 1 | 1 |
| rs3765571 | benign | 0 | 0 | 0 |
| rs752669339 | pathogenic | 1 | 1 | 1 |
| rs537043237 | pathogenic | 1 | 1 | 1 |
| rs774047299 | pathogenic | 1 | 1 | 1 |
| rs121434583 | pathogenic | 1 | 1 | 1 |
| rs121907927 | pathogenic | 0 | 0 | 0 |
| rs121907925 | pathogenic | 0 | 1 | 0 |
| rs746212325 | pathogenic | 0 | 0 | 0 |
| rs754637179 | pathogenic | 0 | 0 | 0 |
| rs780882740 | pathogenic | 0 | 0 | 0 |
| rs121918130 | pathogenic | 0 | 0 | 0 |
| rs121918129 | pathogenic | 0 | 0 | 0 |
| rs13297509 | pathogenic | 0 | 0 | 0 |
| rs746867724 | pathogenic | 0 | 0 | 0 |
| rs121918128 | pathogenic | 0 | 0 | 0 |
| rs752300607 | pathogenic | 0 | 0 | 0 |
| rs121964860 | pathogenic | 0 | 0 | 0 |
| rs45466197 | pathogenic | 1 | 1 | 1 |
| rs3730238 | benign | 1 | 1 | 1 |
| rs730881125 | pathogenic | 1 | 1 | 1 |
| rs121964861 | pathogenic | 1 | 1 | 1 |
| rs397516484 | pathogenic | 0 | 0 | 0 |
| rs367785431 | pathogenic | 0 | 0 | 0 |
| rs141121678 | pathogenic | 0 | 0 | 0 |
| rs12720263 | benign | 0 | 0 | 0 |
| rs2304256 | benign | 0 | 0 | 0 |
| rs2304255 | benign | 0 | 0 | 0 |
| rs121909045 | pathogenic | 0 | 0 | 0 |
| rs397508796 | pathogenic | 0 | 0 | 0 |
| rs151020603 | pathogenic | 0 | 0 | 0 |
| rs77284892 | pathogenic | 0 | 0 | 0 |
| rs368505753 | pathogenic | 0 | 0 | 0 |
| rs115545701 | pathogenic | 0 | 0 | 0 |
| rs121908749 | pathogenic | 0 | 0 | 0 |
| rs75961395 | pathogenic | 0 | 0 | 0 |
| rs121908750 | pathogenic | 0 | 0 | 0 |
| rs121908751 | pathogenic | 0 | 0 | 0 |
| rs397508464 | pathogenic | 0 | 0 | 0 |
| rs397508461 | pathogenic | 0 | 0 | 0 |
| rs121909031 | pathogenic | 0 | 0 | 0 |
| rs113993958 | pathogenic | 0 | 0 | 0 |
| rs77834169 | pathogenic | 0 | 0 | 0 |
| rs78655421 | pathogenic | 0 | 0 | 0 |
| rs141723617 | benign | 0 | 0 | 0 |
| rs35516286 | benign | 0 | 0 | 0 |
| rs1800079 | pathogenic | 1 | 1 | 0 |

| | | | | |
|---|---|---|---|---|
| rs80282562 | pathogenic | 0 | 0 | 0 |
| rs121908752 | pathogenic | 0 | 0 | 0 |
| rs556880586 | pathogenic | 0 | 0 | 0 |
| rs121908760 | pathogenic | 0 | 0 | 0 |
| rs397508350 | pathogenic | 0 | 0 | 0 |
| rs374946172 | pathogenic | 0 | 0 | 0 |
| rs397508387 | pathogenic | 0 | 0 | 0 |
| rs201386642 | pathogenic | 0 | 0 | 0 |
| rs121909012 | pathogenic | 0 | 0 | 0 |
| rs121909008 | pathogenic | 0 | 0 | 0 |
| rs149790377 | pathogenic | 0 | 0 | 0 |
| rs397508442 | pathogenic | 0 | 0 | 0 |
| rs121909035 | pathogenic | 0 | 0 | 0 |
| rs386134230 | pathogenic | 0 | 0 | 0 |
| rs1800112 | benign | 0 | 0 | 0 |
| rs144055758 | pathogenic | 0 | 0 | 0 |
| rs142394380 | pathogenic | 0 | 0 | 0 |
| rs121909036 | pathogenic | 0 | 0 | 0 |
| rs78194216 | pathogenic | 0 | 0 | 0 |
| rs121909019 | pathogenic | 0 | 0 | 0 |
| rs121909020 | pathogenic | 0 | 0 | 0 |
| rs200321110 | pathogenic | 0 | 0 | 0 |
| rs202179988 | pathogenic | 0 | 0 | 0 |
| rs121909037 | pathogenic | 0 | 0 | 0 |
| rs186045772 | pathogenic | 0 | 0 | 0 |
| rs139304906 | pathogenic | 0 | 0 | 0 |
| rs78802634 | pathogenic | 0 | 0 | 0 |
| rs121908761 | pathogenic | 0 | 0 | 0 |
| rs36210737 | pathogenic | 0 | 0 | 0 |
| rs139729994 | pathogenic | 0 | 0 | 0 |
| rs79850223 | pathogenic | 0 | 0 | 0 |
| rs1800120 | benign | 0 | 0 | 0 |
| rs74767530 | pathogenic | 0 | 0 | 0 |
| rs374705585 | pathogenic | 0 | 0 | 0 |
| rs1136287 | benign | 0 | 0 | 0 |
| rs6828 | benign | 0 | 0 | 0 |
| rs104894856 | pathogenic | 0 | 0 | 0 |
| rs104894860 | pathogenic | 0 | 0 | 0 |
| rs61736892 | pathogenic | 0 | 0 | 0 |
| rs145807417 | benign | 0 | 0 | 0 |
| rs104894853 | pathogenic | 0 | 0 | 0 |
| rs199422227 | pathogenic | 0 | 0 | 0 |
| rs113993946 | pathogenic | 0 | 0 | 0 |
| rs199422231 | pathogenic | 0 | 0 | 0 |
| rs199422228 | pathogenic | 0 | 0 | 0 |
| rs142195196 | benign | 0 | 1 | 0 |
| rs148656104 | benign | 1 | 1 | 1 |
| rs141944844 | benign | 0 | 0 | 0 |
| rs104894342 | pathogenic | 0 | 0 | 0 |
| rs104894343 | pathogenic | 0 | 0 | 0 |
| rs104894344 | pathogenic | 0 | 0 | 0 |
| rs573322878 | benign | 0 | 0 | 0 |
| rs104894347 | pathogenic | 0 | 0 | 0 |
| rs28937882 | pathogenic | 0 | 0 | 0 |

| | | | | |
|---|---|---|---|---|
| rs193922702 | pathogenic | 0 | 0 | 0 |
| rs200786104 | benign | 0 | 0 | 0 |
| rs690016544 | pathogenic | 1 | 1 | 1 |
| rs28943592 | benign | 0 | 0 | 0 |
| rs28943594 | benign | 0 | 0 | 0 |
| rs121908957 | pathogenic | 0 | 0 | 0 |
| rs115408010 | benign | 1 | 1 | 1 |
| rs77591659 | benign | 0 | 0 | 0 |
| rs121918511 | pathogenic | 0 | 0 | 0 |
| rs538359948 | benign | 0 | 0 | 0 |
| rs76708142 | benign | 0 | 0 | 0 |
| rs121907913 | pathogenic | 1 | 1 | 0 |
| rs121907920 | pathogenic | 0 | 0 | 0 |
| rs121907923 | pathogenic | 1 | 1 | 0 |
| rs121907928 | pathogenic | 1 | 1 | 0 |
| rs121907919 | pathogenic | 1 | 1 | 0 |
| rs121907918 | pathogenic | 1 | 1 | 0 |
| rs74315493 | pathogenic | 1 | 1 | 1 |
| rs74315494 | pathogenic | 1 | 1 | 1 |
| rs145446060 | benign | 1 | 1 | 1 |
| rs550409143 | benign | 1 | 1 | 0 |

| compound_binding_site | cofactor_binding_site | nucleic_binding_site | ion_binding_site |
|---|---|---|---|
| 1 | 0 | 0 | 0 |
| 1 | 0 | 0 | 0 |
| 0 | 0 | 0 | 0 |
| 1 | 0 | 0 | 0 |
| 0 | 0 | 0 | 0 |
| 0 | 0 | 0 | 0 |
| 0 | 0 | 0 | 0 |
| 0 | 0 | 0 | 0 |
| 1 | 0 | 0 | 1 |
| 1 | 0 | 0 | 0 |
| 0 | 0 | 0 | 1 |
| 1 | 0 | 1 | 0 |
| 0 | 0 | 0 | 0 |
| 0 | 0 | 0 | 0 |
| 0 | 0 | 0 | 0 |
| 0 | 0 | 0 | 0 |
| 0 | 0 | 0 | 0 |
| 1 | 0 | 0 | 1 |
| 1 | 0 | 1 | 0 |
| 0 | 0 | 0 | 0 |
| 1 | 0 | 0 | 1 |
| 1 | 0 | 0 | 0 |
| 1 | 0 | 0 | 0 |
| 1 | 0 | 0 | 0 |
| 0 | 0 | 0 | 0 |
| 0 | 0 | 0 | 0 |
| 1 | 0 | 0 | 1 |
| 0 | 0 | 0 | 0 |
| 0 | 0 | 0 | 0 |
| 0 | 0 | 0 | 0 |
| 0 | 0 | 0 | 0 |
| 1 | 0 | 0 | 0 |
| 1 | 0 | 0 | 1 |
| 1 | 0 | 0 | 0 |
| 0 | 0 | 0 | 0 |
| 0 | 0 | 0 | 0 |
| 0 | 0 | 0 | 0 |
| 1 | 0 | 0 | 1 |
| 1 | 0 | 0 | 1 |
| 1 | 0 | 0 | 1 |
| 1 | 0 | 0 | 0 |
| 1 | 0 | 0 | 0 |
| 1 | 0 | 0 | 0 |
| 1 | 0 | 0 | 0 |
| 1 | 0 | 0 | 0 |
| 1 | 0 | 0 | 0 |
| 1 | 0 | 0 | 0 |
| 1 | 0 | 0 | 0 |
| 1 | 0 | 0 | 0 |
| 1 | 0 | 0 | 0 |
| 1 | 0 | 0 | 0 |
| 1 | 0 | 0 | 0 |
| 1 | 0 | 0 | 0 |

| | | | |
|---|---|---|---|
| 1 | 0 | 0 | 1 |
| 1 | 0 | 0 | 0 |
| 1 | 0 | 0 | 0 |
| 1 | 0 | 0 | 0 |
| 0 | 0 | 0 | 0 |
| 1 | 0 | 0 | 0 |
| 0 | 0 | 0 | 0 |
| 1 | 0 | 0 | 0 |
| 1 | 0 | 0 | 0 |
| 1 | 0 | 0 | 1 |
| 1 | 0 | 0 | 0 |
| 1 | 0 | 0 | 0 |
| 1 | 0 | 0 | 0 |
| 1 | 0 | 0 | 0 |
| 1 | 0 | 0 | 0 |
| 1 | 0 | 0 | 0 |
| 1 | 0 | 0 | 0 |
| 1 | 0 | 0 | 1 |
| 1 | 0 | 0 | 1 |
| 1 | 1 | 0 | 1 |
| 1 | 0 | 0 | 0 |
| 1 | 0 | 0 | 0 |
| 1 | 0 | 0 | 0 |
| 1 | 0 | 0 | 0 |
| 1 | 1 | 0 | 0 |
| 1 | 0 | 0 | 0 |
| 1 | 0 | 0 | 0 |
| 1 | 0 | 0 | 0 |
| 1 | 0 | 0 | 0 |
| 1 | 0 | 0 | 0 |
| 1 | 1 | 0 | 0 |
| 1 | 1 | 0 | 0 |
| 1 | 1 | 0 | 0 |
| 1 | 1 | 0 | 0 |
| 1 | 1 | 0 | 0 |
| 1 | 0 | 0 | 0 |
| 1 | 1 | 0 | 0 |
| 1 | 0 | 0 | 0 |
| 1 | 0 | 0 | 0 |
| 1 | 1 | 0 | 0 |
| 0 | 0 | 0 | 0 |
| 0 | 0 | 0 | 0 |
| 0 | 0 | 0 | 0 |
| 0 | 0 | 0 | 0 |
| 0 | 0 | 0 | 0 |
| 0 | 0 | 0 | 0 |
| 0 | 0 | 0 | 0 |
| 1 | 0 | 0 | 1 |
| 0 | 0 | 0 | 0 |
| 1 | 0 | 0 | 0 |
| 1 | 0 | 0 | 0 |
| 0 | 0 | 0 | 0 |
| 0 | 0 | 0 | 0 |
| 0 | 0 | 0 | 0 |

| | | | |
|---|---|---|---|
| 0 | 0 | 0 | 0 |
| 0 | 0 | 0 | 0 |
| 0 | 0 | 0 | 0 |
| 1 | 0 | 0 | 0 |
| 1 | 0 | 0 | 0 |
| 1 | 0 | 0 | 0 |
| 0 | 0 | 0 | 0 |
| 0 | 0 | 1 | 0 |
| 0 | 0 | 1 | 0 |
| 0 | 0 | 1 | 0 |
| 0 | 0 | 1 | 0 |
| 0 | 0 | 1 | 0 |
| 0 | 0 | 0 | 0 |
| 0 | 0 | 0 | 0 |
| 1 | 0 | 0 | 0 |
| 0 | 0 | 0 | 0 |
| 0 | 0 | 0 | 0 |
| 1 | 0 | 0 | 0 |
| 1 | 0 | 0 | 0 |
| 1 | 0 | 0 | 0 |
| 1 | 0 | 0 | 0 |
| 1 | 1 | 0 | 0 |
| 1 | 0 | 0 | 0 |
| 1 | 0 | 0 | 0 |
| 0 | 0 | 0 | 0 |
| 0 | 0 | 0 | 0 |
| 1 | 0 | 0 | 0 |
| 1 | 1 | 0 | 0 |
| 1 | 0 | 0 | 0 |
| 1 | 0 | 0 | 0 |
| 1 | 1 | 0 | 0 |
| 1 | 1 | 0 | 0 |
| 1 | 1 | 0 | 0 |
| 1 | 1 | 0 | 0 |
| 1 | 1 | 0 | 0 |
| 1 | 1 | 0 | 0 |
| 1 | 1 | 0 | 0 |
| 1 | 1 | 0 | 0 |
| 1 | 1 | 0 | 0 |
| 1 | 1 | 0 | 0 |
| 1 | 1 | 0 | 0 |
| 1 | 1 | 0 | 0 |
| 1 | 0 | 0 | 0 |
| 1 | 0 | 0 | 0 |
| 1 | 0 | 0 | 0 |
| 1 | 1 | 0 | 0 |
| 1 | 0 | 0 | 0 |
| 1 | 1 | 1 | 0 |
| 1 | 1 | 1 | 0 |
| 1 | 1 | 0 | 0 |
| 1 | 1 | 0 | 0 |
| 1 | 0 | 0 | 0 |
| 1 | 0 | 0 | 0 |
| 0 | 0 | 0 | 0 |

| | | | |
|---|---|---|---|
| 0 | 0 | 0 | 0 |
| 0 | 0 | 0 | 0 |
| 0 | 0 | 0 | 0 |
| 1 | 0 | 0 | 0 |
| 1 | 0 | 0 | 0 |
| 0 | 0 | 0 | 0 |
| 0 | 0 | 0 | 0 |
| 0 | 0 | 0 | 0 |
| 0 | 0 | 0 | 0 |
| 0 | 0 | 0 | 0 |
| 0 | 0 | 0 | 0 |
| 0 | 0 | 0 | 0 |
| 0 | 0 | 0 | 0 |
| 0 | 0 | 0 | 0 |
| 0 | 0 | 0 | 0 |
| 0 | 0 | 0 | 0 |
| 0 | 0 | 0 | 0 |
| 0 | 0 | 0 | 0 |
| 0 | 0 | 0 | 0 |
| 0 | 0 | 0 | 0 |
| 0 | 0 | 0 | 0 |
| 0 | 0 | 0 | 0 |
| 1 | 0 | 0 | 0 |
| 1 | 0 | 0 | 0 |
| 0 | 0 | 0 | 0 |
| 0 | 0 | 0 | 0 |
| 1 | 0 | 0 | 0 |
| 0 | 0 | 0 | 0 |
| 0 | 0 | 0 | 0 |
| 0 | 0 | 0 | 0 |
| 0 | 0 | 0 | 0 |
| 0 | 0 | 0 | 0 |
| 0 | 0 | 0 | 0 |
| 0 | 0 | 0 | 0 |
| 1 | 0 | 0 | 0 |
| 0 | 0 | 0 | 0 |
| 0 | 0 | 0 | 0 |
| 0 | 0 | 0 | 0 |
| 0 | 0 | 0 | 0 |
| 0 | 0 | 0 | 0 |
| 0 | 0 | 0 | 0 |
| 0 | 0 | 0 | 0 |
| 0 | 0 | 0 | 0 |
| 0 | 0 | 0 | 0 |
| 0 | 0 | 0 | 0 |
| 0 | 0 | 0 | 0 |
| 0 | 0 | 1 | 0 |
| 1 | 0 | 0 | 0 |
| 1 | 0 | 0 | 0 |
| 1 | 1 | 0 | 0 |
| 1 | 0 | 0 | 0 |
| 0 | 0 | 0 | 0 |
| 0 | 0 | 0 | 0 |
| 1 | 0 | 0 | 0 |

| | | | |
|---|---|---|---|
| 1 | 0 | 0 | 0 |
| 1 | 0 | 0 | 0 |
| 1 | 0 | 0 | 0 |
| 0 | 0 | 0 | 0 |
| 0 | 0 | 0 | 0 |
| 0 | 0 | 0 | 0 |
| 1 | 0 | 0 | 0 |
| 1 | 0 | 0 | 0 |
| 0 | 0 | 0 | 0 |
| 0 | 0 | 0 | 0 |
| 1 | 0 | 0 | 0 |
| 0 | 0 | 0 | 0 |
| 1 | 0 | 0 | 0 |
| 0 | 0 | 0 | 0 |
| 0 | 0 | 0 | 0 |
| 0 | 0 | 0 | 0 |
| 0 | 0 | 0 | 0 |
| 0 | 0 | 0 | 0 |
| 0 | 0 | 0 | 0 |
| 0 | 0 | 0 | 0 |
| 0 | 0 | 0 | 0 |
| 0 | 0 | 0 | 0 |
| 0 | 0 | 0 | 0 |
| 0 | 0 | 0 | 0 |
| 0 | 0 | 0 | 0 |
| 0 | 0 | 0 | 0 |
| 1 | 0 | 0 | 0 |
| 1 | 0 | 0 | 0 |
| 1 | 0 | 0 | 0 |
| 1 | 0 | 0 | 0 |
| 1 | 0 | 0 | 0 |
| 1 | 0 | 0 | 0 |
| 0 | 0 | 0 | 0 |
| 0 | 0 | 0 | 0 |
| 1 | 0 | 0 | 0 |
| 0 | 0 | 0 | 0 |
| 0 | 0 | 0 | 0 |
| 0 | 0 | 0 | 0 |
| 0 | 0 | 0 | 0 |
| 0 | 0 | 0 | 0 |
| 0 | 0 | 0 | 0 |
| 0 | 0 | 0 | 0 |
| 0 | 0 | 0 | 0 |
| 0 | 0 | 0 | 0 |
| 1 | 1 | 0 | 0 |
| 1 | 1 | 0 | 0 |
| 1 | 1 | 0 | 1 |
| 1 | 1 | 0 | 1 |
| 1 | 1 | 0 | 1 |
| 1 | 1 | 0 | 1 |
| 1 | 1 | 0 | 0 |
| 1 | 1 | 0 | 1 |
| 1 | 1 | 0 | 1 |
| 1 | 1 | 0 | 1 |

| | | | |
|---|---|---|---|
| 1 | 1 | 0 | 1 |
| 1 | 1 | 0 | 1 |
| 1 | 1 | 0 | 1 |
| 1 | 1 | 0 | 1 |
| 1 | 1 | 0 | 1 |
| 1 | 0 | 0 | 0 |
| 0 | 0 | 0 | 0 |
| 0 | 0 | 0 | 0 |
| 0 | 0 | 0 | 0 |
| 1 | 0 | 0 | 0 |
| 0 | 0 | 1 | 0 |
| 0 | 0 | 0 | 0 |
| 0 | 0 | 0 | 0 |
| 0 | 0 | 0 | 0 |
| 0 | 0 | 0 | 0 |
| 0 | 0 | 0 | 0 |
| 0 | 0 | 0 | 0 |
| 0 | 0 | 0 | 0 |
| 0 | 0 | 0 | 0 |
| 0 | 0 | 0 | 0 |
| 0 | 0 | 0 | 0 |
| 0 | 0 | 0 | 0 |
| 0 | 0 | 0 | 0 |
| 0 | 0 | 0 | 0 |
| 0 | 0 | 0 | 0 |
| 1 | 1 | 0 | 0 |
| 1 | 1 | 0 | 0 |
| 1 | 1 | 0 | 0 |
| 0 | 1 | 0 | 0 |
| 1 | 0 | 0 | 0 |
| 0 | 0 | 0 | 0 |
| 0 | 0 | 0 | 0 |
| 0 | 0 | 0 | 0 |
| 1 | 0 | 0 | 0 |
| 1 | 1 | 0 | 0 |
| 0 | 0 | 0 | 0 |
| 0 | 0 | 0 | 0 |
| 0 | 0 | 0 | 0 |
| 0 | 0 | 0 | 0 |
| 1 | 0 | 0 | 0 |
| 0 | 0 | 0 | 0 |
| 0 | 0 | 0 | 0 |
| 0 | 0 | 0 | 0 |
| 0 | 0 | 0 | 0 |
| 0 | 0 | 0 | 0 |
| 0 | 1 | 0 | 0 |
| 0 | 1 | 0 | 0 |
| 0 | 0 | 0 | 0 |
| 0 | 0 | 0 | 0 |
| 0 | 1 | 0 | 0 |
| 0 | 0 | 0 | 0 |
| 0 | 0 | 0 | 0 |
| 0 | 1 | 0 | 0 |
| 0 | 0 | 0 | 0 |

| | | | |
|---|---|---|---|
| 1 | 1 | 0 | 0 |
| 0 | 0 | 0 | 0 |
| 0 | 0 | 0 | 0 |
| 0 | 0 | 0 | 0 |
| 0 | 0 | 0 | 0 |
| 0 | 0 | 0 | 0 |
| 0 | 0 | 0 | 0 |
| 0 | 0 | 0 | 0 |
| 0 | 0 | 0 | 0 |
| 0 | 0 | 0 | 0 |
| 0 | 0 | 0 | 0 |
| 0 | 0 | 0 | 0 |
| 0 | 0 | 0 | 0 |
| 0 | 0 | 0 | 0 |
| 0 | 1 | 0 | 0 |
| 0 | 0 | 0 | 0 |
| 0 | 1 | 0 | 0 |
| 0 | 1 | 0 | 0 |
| 0 | 0 | 0 | 0 |
| 0 | 1 | 0 | 0 |
| 1 | 0 | 0 | 1 |
| 1 | 1 | 0 | 0 |
| 1 | 0 | 0 | 0 |
| 1 | 0 | 0 | 1 |
| 0 | 0 | 0 | 0 |
| 0 | 0 | 0 | 0 |
| 0 | 0 | 0 | 0 |
| 0 | 0 | 0 | 0 |
| 1 | 0 | 0 | 0 |
| 0 | 0 | 0 | 0 |
| 1 | 0 | 0 | 0 |
| 0 | 0 | 0 | 0 |
| 0 | 0 | 0 | 0 |
| 0 | 0 | 0 | 0 |
| 0 | 0 | 0 | 0 |
| 0 | 0 | 0 | 0 |
| 0 | 0 | 0 | 0 |
| 1 | 1 | 0 | 0 |
| 1 | 1 | 0 | 0 |
| 1 | 1 | 0 | 0 |
| 0 | 0 | 0 | 0 |
| 0 | 0 | 0 | 0 |
| 0 | 0 | 0 | 0 |
| 1 | 0 | 0 | 0 |
| 1 | 0 | 0 | 1 |
| 1 | 0 | 0 | 1 |
| 1 | 0 | 0 | 0 |
| 1 | 0 | 0 | 1 |
| 1 | 0 | 0 | 0 |
| 0 | 0 | 0 | 0 |
| 1 | 1 | 0 | 0 |
| 1 | 1 | 0 | 0 |
| 0 | 0 | 0 | 0 |
| 0 | 0 | 0 | 0 |

| | | | |
|---|---|---|---|
| 1 | 0 | 0 | 0 |
| 1 | 0 | 0 | 0 |
| 1 | 0 | 0 | 0 |
| 0 | 0 | 0 | 0 |
| 0 | 0 | 0 | 0 |
| 1 | 1 | 0 | 1 |
| 1 | 1 | 0 | 1 |
| 1 | 1 | 0 | 1 |
| 1 | 1 | 0 | 1 |
| 1 | 1 | 0 | 1 |
| 1 | 1 | 0 | 1 |
| 1 | 1 | 0 | 1 |
| 1 | 1 | 0 | 0 |
| 1 | 1 | 0 | 0 |
| 1 | 1 | 1 | 1 |
| 0 | 0 | 0 | 0 |
| 1 | 1 | 0 | 0 |
| 1 | 1 | 0 | 0 |
| 0 | 0 | 0 | 0 |
| 0 | 0 | 0 | 0 |
| 0 | 0 | 0 | 0 |
| 1 | 0 | 0 | 0 |
| 1 | 0 | 0 | 0 |
| 0 | 0 | 0 | 0 |
| 1 | 0 | 0 | 0 |
| 1 | 0 | 0 | 0 |
| 1 | 0 | 0 | 0 |
| 0 | 0 | 0 | 0 |
| 0 | 0 | 0 | 0 |
| 0 | 0 | 0 | 0 |
| 0 | 0 | 0 | 0 |
| 0 | 0 | 0 | 0 |
| 0 | 0 | 0 | 0 |
| 0 | 0 | 0 | 0 |
| 0 | 0 | 0 | 0 |
| 0 | 0 | 0 | 0 |
| 0 | 0 | 0 | 0 |
| 0 | 0 | 0 | 0 |
| 0 | 0 | 0 | 0 |
| 0 | 0 | 0 | 0 |
| 0 | 0 | 0 | 0 |
| 0 | 0 | 0 | 0 |
| 0 | 0 | 0 | 0 |
| 0 | 0 | 0 | 0 |
| 0 | 0 | 0 | 0 |
| 1 | 0 | 0 | 0 |
| 1 | 0 | 0 | 0 |
| 1 | 0 | 0 | 0 |
| 1 | 0 | 0 | 0 |
| 0 | 0 | 0 | 0 |
| 0 | 0 | 0 | 0 |
| 0 | 0 | 0 | 0 |
| 0 | 0 | 0 | 0 |
| 0 | 0 | 0 | 0 |

| | | | |
|---|---|---|---|
| 0 | 0 | 0 | 0 |
| 0 | 0 | 0 | 0 |
| 0 | 0 | 0 | 0 |
| 0 | 0 | 0 | 0 |
| 0 | 0 | 0 | 0 |
| 0 | 0 | 0 | 0 |
| 0 | 0 | 0 | 0 |
| 1 | 0 | 0 | 1 |
| 0 | 0 | 0 | 0 |
| 0 | 0 | 0 | 0 |
| 1 | 0 | 0 | 0 |
| 0 | 0 | 0 | 0 |
| 0 | 0 | 0 | 0 |
| 0 | 0 | 0 | 0 |
| 0 | 0 | 0 | 0 |
| 1 | 0 | 0 | 0 |
| 0 | 0 | 0 | 0 |
| 0 | 0 | 0 | 0 |
| 0 | 0 | 0 | 0 |
| 0 | 0 | 0 | 0 |
| 0 | 0 | 0 | 0 |
| 0 | 0 | 0 | 0 |
| 0 | 0 | 0 | 0 |
| 1 | 1 | 0 | 1 |
| 0 | 0 | 0 | 0 |
| 0 | 0 | 0 | 0 |
| 0 | 0 | 0 | 0 |
| 1 | 1 | 0 | 1 |
| 1 | 1 | 0 | 0 |
| 0 | 0 | 0 | 0 |
| 1 | 0 | 0 | 0 |
| 1 | 1 | 0 | 0 |
| 0 | 0 | 0 | 0 |
| 1 | 0 | 0 | 0 |
| 1 | 1 | 0 | 0 |
| 1 | 0 | 0 | 0 |
| 1 | 1 | 0 | 1 |
| 1 | 1 | 0 | 1 |
| 1 | 0 | 0 | 0 |
| 0 | 0 | 0 | 0 |
| 0 | 0 | 0 | 0 |
| 0 | 0 | 0 | 0 |
| 0 | 0 | 0 | 0 |
| 0 | 0 | 0 | 0 |
| 0 | 0 | 0 | 0 |
| 0 | 0 | 0 | 0 |
| 0 | 0 | 0 | 0 |
| 0 | 0 | 0 | 0 |
| 0 | 0 | 0 | 0 |
| 0 | 0 | 0 | 0 |
| 0 | 0 | 0 | 0 |
| 0 | 0 | 0 | 0 |
| 0 | 0 | 0 | 0 |
| 0 | 0 | 0 | 0 |

| | | | |
|---|---|---|---|
| 0 | 0 | 0 | 0 |
| 0 | 0 | 0 | 0 |
| 0 | 0 | 0 | 0 |
| 0 | 0 | 0 | 0 |
| 0 | 0 | 0 | 0 |
| 0 | 0 | 0 | 0 |
| 0 | 0 | 0 | 0 |
| 0 | 0 | 0 | 0 |
| 0 | 0 | 0 | 0 |
| 0 | 0 | 0 | 0 |
| 0 | 0 | 0 | 0 |
| 0 | 0 | 0 | 0 |
| 0 | 0 | 0 | 0 |
| 1 | 1 | 0 | 0 |
| 0 | 0 | 0 | 0 |
| 1 | 0 | 0 | 0 |
| 0 | 0 | 0 | 1 |
| 0 | 0 | 0 | 0 |
| 0 | 0 | 0 | 0 |
| 0 | 0 | 0 | 0 |
| 0 | 0 | 0 | 0 |
| 0 | 0 | 0 | 0 |
| 0 | 0 | 0 | 0 |
| 0 | 0 | 0 | 0 |
| 0 | 0 | 0 | 0 |
| 0 | 0 | 0 | 0 |
| 0 | 0 | 0 | 0 |
| 0 | 0 | 0 | 0 |
| 0 | 0 | 0 | 0 |
| 0 | 0 | 0 | 0 |
| 1 | 0 | 0 | 0 |
| 1 | 0 | 0 | 0 |
| 0 | 0 | 0 | 0 |
| 0 | 0 | 0 | 0 |
| 0 | 0 | 0 | 0 |
| 0 | 0 | 0 | 0 |
| 0 | 0 | 0 | 0 |
| 0 | 0 | 0 | 0 |
| 0 | 0 | 0 | 0 |
| 0 | 0 | 0 | 0 |
| 1 | 0 | 0 | 0 |
| 1 | 0 | 0 | 0 |
| 1 | 0 | 0 | 0 |
| 1 | 0 | 0 | 0 |
| 0 | 0 | 0 | 0 |
| 1 | 0 | 0 | 0 |
| 0 | 0 | 0 | 0 |
| 0 | 0 | 0 | 0 |
| 1 | 0 | 0 | 0 |
| 1 | 0 | 0 | 0 |
| 0 | 0 | 0 | 0 |
| 0 | 0 | 0 | 0 |
| 0 | 0 | 0 | 0 |
| 0 | 0 | 0 | 0 |

| | | | |
|---|---|---|---|
| 0 | 0 | 0 | 0 |
| 0 | 0 | 0 | 0 |
| 0 | 0 | 0 | 0 |
| 0 | 0 | 0 | 0 |
| 0 | 0 | 0 | 0 |
| 0 | 0 | 0 | 0 |
| 0 | 0 | 0 | 0 |
| 0 | 0 | 0 | 0 |
| 0 | 0 | 0 | 0 |
| 0 | 0 | 0 | 0 |
| 0 | 0 | 0 | 0 |
| 0 | 0 | 0 | 0 |
| 0 | 0 | 0 | 0 |
| 0 | 0 | 0 | 0 |
| 0 | 0 | 0 | 0 |
| 1 | 0 | 0 | 1 |
| 0 | 0 | 0 | 0 |
| 0 | 0 | 0 | 0 |
| 0 | 0 | 0 | 0 |
| 0 | 0 | 0 | 0 |
| 0 | 0 | 0 | 0 |
| 1 | 1 | 0 | 0 |
| 1 | 1 | 0 | 0 |
| 1 | 0 | 0 | 0 |
| 1 | 0 | 0 | 0 |
| 0 | 0 | 0 | 0 |
| 0 | 0 | 0 | 0 |
| 0 | 0 | 0 | 0 |
| 0 | 0 | 0 | 0 |
| 1 | 0 | 0 | 0 |
| 1 | 0 | 0 | 0 |
| 1 | 0 | 0 | 0 |
| 1 | 1 | 0 | 0 |
| 1 | 1 | 0 | 0 |
| 1 | 0 | 0 | 0 |
| 0 | 0 | 0 | 0 |
| 0 | 0 | 0 | 0 |
| 0 | 1 | 0 | 0 |
| 0 | 0 | 0 | 0 |
| 0 | 0 | 0 | 0 |
| 0 | 0 | 0 | 0 |
| 1 | 1 | 0 | 0 |
| 1 | 0 | 0 | 0 |
| 1 | 0 | 0 | 0 |
| 0 | 0 | 0 | 0 |
| 0 | 0 | 0 | 0 |
| 0 | 0 | 0 | 0 |
| 0 | 0 | 0 | 0 |
| 0 | 0 | 0 | 0 |
| 0 | 0 | 0 | 0 |
| 0 | 0 | 0 | 0 |
| 0 | 0 | 0 | 0 |
| 0 | 0 | 0 | 0 |
| 0 | 0 | 0 | 0 |

| | | | |
|---|---|---|---|
| 0 | 0 | 0 | 0 |
| 1 | 0 | 0 | 0 |
| 0 | 0 | 0 | 0 |
| 0 | 0 | 0 | 0 |
| 0 | 0 | 0 | 0 |
| 0 | 0 | 0 | 0 |
| 1 | 0 | 0 | 0 |
| 0 | 0 | 0 | 0 |
| 0 | 0 | 0 | 0 |
| 0 | 0 | 0 | 0 |
| 0 | 0 | 0 | 0 |
| 0 | 0 | 0 | 0 |
| 0 | 0 | 0 | 0 |
| 0 | 0 | 0 | 0 |
| 0 | 0 | 0 | 0 |
| 0 | 0 | 0 | 0 |
| 0 | 1 | 0 | 0 |
| 0 | 0 | 0 | 0 |
| 0 | 0 | 0 | 0 |
| 0 | 0 | 0 | 0 |
| 1 | 0 | 0 | 0 |
| 0 | 0 | 0 | 0 |
| 0 | 0 | 0 | 0 |
| 1 | 1 | 0 | 0 |
| 0 | 0 | 0 | 0 |
| 0 | 0 | 0 | 0 |
| 0 | 0 | 0 | 0 |
| 0 | 0 | 0 | 0 |
| 1 | 0 | 0 | 0 |
| 0 | 0 | 0 | 0 |
| 0 | 0 | 0 | 0 |
| 0 | 0 | 0 | 0 |
| 0 | 0 | 0 | 0 |
| 0 | 0 | 0 | 0 |
| 0 | 0 | 0 | 0 |
| 1 | 0 | 0 | 0 |
| 0 | 0 | 0 | 0 |
| 0 | 0 | 0 | 0 |
| 0 | 0 | 0 | 0 |
| 0 | 0 | 0 | 0 |
| 1 | 1 | 0 | 0 |
| 0 | 0 | 0 | 0 |
| 0 | 0 | 0 | 0 |
| 0 | 0 | 0 | 0 |
| 0 | 0 | 0 | 0 |
| 0 | 0 | 0 | 0 |
| 1 | 0 | 0 | 0 |
| 1 | 0 | 0 | 1 |
| 1 | 0 | 0 | 0 |
| 0 | 0 | 0 | 0 |
| 0 | 0 | 0 | 0 |
| 0 | 0 | 0 | 0 |
| 1 | 0 | 0 | 0 |
| 1 | 0 | 0 | 0 |

| | | | |
|---|---|---|---|
| 1 | 0 | 0 | 0 |
| 0 | 0 | 0 | 0 |
| 0 | 0 | 0 | 0 |
| 0 | 0 | 0 | 0 |
| 1 | 1 | 0 | 0 |
| 1 | 1 | 0 | 0 |
| 0 | 0 | 0 | 0 |
| 0 | 0 | 0 | 0 |
| 1 | 0 | 0 | 0 |
| 1 | 0 | 0 | 0 |
| 1 | 0 | 1 | 0 |
| 0 | 0 | 0 | 0 |
| 0 | 0 | 1 | 0 |
| 0 | 0 | 1 | 0 |
| 0 | 0 | 1 | 0 |
| 1 | 0 | 0 | 0 |
| 1 | 0 | 0 | 0 |
| 0 | 0 | 0 | 0 |
| 0 | 0 | 0 | 0 |
| 1 | 0 | 0 | 1 |
| 1 | 0 | 0 | 1 |
| 1 | 0 | 0 | 0 |
| 0 | 0 | 0 | 0 |
| 0 | 0 | 0 | 0 |
| 0 | 0 | 0 | 0 |
| 0 | 0 | 0 | 0 |
| 0 | 0 | 0 | 0 |
| 1 | 0 | 0 | 0 |
| 0 | 0 | 0 | 0 |
| 0 | 0 | 0 | 1 |
| 0 | 0 | 0 | 1 |
| 0 | 0 | 1 | 1 |
| 0 | 0 | 1 | 1 |
| 0 | 0 | 1 | 1 |
| 0 | 0 | 1 | 1 |
| 0 | 0 | 1 | 0 |
| 0 | 0 | 1 | 0 |
| 0 | 0 | 0 | 0 |
| 0 | 0 | 0 | 0 |
| 0 | 0 | 0 | 0 |
| 0 | 0 | 0 | 0 |
| 0 | 0 | 1 | 0 |
| 0 | 0 | 1 | 0 |
| 0 | 0 | 1 | 0 |
| 0 | 0 | 1 | 0 |
| 0 | 0 | 1 | 0 |
| 0 | 0 | 1 | 0 |
| 0 | 0 | 1 | 0 |
| 0 | 0 | 1 | 0 |
| 0 | 0 | 0 | 0 |
| 0 | 0 | 0 | 0 |
| 0 | 0 | 0 | 0 |
| 0 | 0 | 0 | 0 |
| 1 | 0 | 0 | 0 |

| | | | |
|---|---|---|---|
| 0 | 0 | 0 | 0 |
| 0 | 0 | 0 | 0 |
| 0 | 0 | 0 | 0 |
| 1 | 0 | 0 | 0 |
| 0 | 0 | 0 | 0 |
| 0 | 0 | 0 | 0 |
| 0 | 0 | 0 | 0 |
| 0 | 0 | 0 | 0 |
| 0 | 0 | 0 | 0 |
| 0 | 0 | 0 | 0 |
| 1 | 1 | 0 | 0 |
| 0 | 0 | 0 | 0 |
| 1 | 1 | 0 | 0 |
| 1 | 1 | 0 | 0 |
| 0 | 1 | 0 | 0 |
| 0 | 0 | 0 | 0 |
| 0 | 0 | 0 | 0 |
| 0 | 0 | 0 | 0 |
| 0 | 0 | 0 | 0 |
| 0 | 0 | 0 | 0 |
| 1 | 1 | 0 | 0 |
| 0 | 1 | 0 | 0 |
| 0 | 0 | 0 | 0 |
| 0 | 1 | 0 | 0 |
| 1 | 1 | 0 | 0 |
| 0 | 0 | 0 | 0 |
| 0 | 0 | 0 | 0 |
| 1 | 1 | 0 | 0 |
| 1 | 1 | 0 | 0 |
| 0 | 1 | 0 | 0 |
| 0 | 0 | 0 | 0 |
| 0 | 0 | 0 | 0 |
| 0 | 0 | 0 | 0 |
| 0 | 0 | 0 | 0 |
| 0 | 0 | 0 | 0 |
| 0 | 0 | 0 | 0 |
| 0 | 0 | 0 | 0 |
| 0 | 0 | 0 | 0 |
| 0 | 0 | 0 | 0 |
| 0 | 0 | 0 | 0 |
| 1 | 0 | 0 | 0 |
| 1 | 0 | 0 | 0 |
| 1 | 0 | 0 | 0 |
| 1 | 0 | 0 | 0 |
| 1 | 1 | 0 | 1 |
| 0 | 0 | 0 | 0 |
| 1 | 0 | 0 | 0 |
| 0 | 0 | 0 | 0 |
| 1 | 0 | 0 | 0 |
| 0 | 0 | 0 | 0 |
| 1 | 0 | 0 | 0 |
| 1 | 0 | 0 | 0 |
| 0 | 0 | 0 | 0 |
| 0 | 0 | 0 | 0 |

| | | | |
|---|---|---|---|
| 0 | 0 | 0 | 0 |
| 0 | 0 | 0 | 0 |
| 0 | 0 | 0 | 0 |
| 0 | 0 | 0 | 0 |
| 1 | 0 | 0 | 0 |
| 1 | 0 | 0 | 0 |
| 1 | 0 | 0 | 0 |
| 0 | 0 | 0 | 0 |
| 1 | 1 | 0 | 0 |
| 0 | 0 | 0 | 0 |
| 0 | 0 | 0 | 0 |
| 0 | 0 | 0 | 0 |
| 1 | 1 | 0 | 0 |
| 1 | 0 | 0 | 0 |
| 0 | 0 | 0 | 0 |
| 0 | 0 | 0 | 0 |
| 1 | 1 | 0 | 0 |
| 0 | 0 | 0 | 0 |
| 0 | 0 | 0 | 0 |
| 0 | 0 | 0 | 0 |
| 0 | 0 | 0 | 0 |
| 0 | 0 | 0 | 0 |
| 0 | 0 | 0 | 0 |
| 0 | 0 | 0 | 0 |
| 1 | 0 | 0 | 0 |
| 0 | 0 | 0 | 0 |
| 1 | 0 | 0 | 0 |
| 0 | 0 | 0 | 0 |
| 0 | 0 | 0 | 0 |
| 0 | 0 | 0 | 0 |
| 1 | 0 | 0 | 0 |
| 1 | 0 | 0 | 0 |
| 0 | 0 | 0 | 0 |
| 0 | 0 | 0 | 0 |
| 1 | 0 | 0 | 0 |
| 0 | 0 | 0 | 0 |
| 0 | 0 | 0 | 0 |
| 1 | 0 | 0 | 0 |
| 0 | 0 | 0 | 0 |
| 1 | 0 | 0 | 0 |
| 1 | 0 | 0 | 0 |
| 0 | 0 | 0 | 0 |
| 0 | 0 | 0 | 0 |
| 1 | 1 | 0 | 0 |
| 1 | 0 | 0 | 0 |
| 1 | 1 | 0 | 0 |
| 0 | 0 | 0 | 0 |
| 0 | 0 | 0 | 0 |
| 0 | 0 | 0 | 0 |
| 0 | 0 | 0 | 0 |
| 1 | 1 | 0 | 0 |
| 0 | 0 | 0 | 0 |
| 0 | 0 | 0 | 0 |
| 0 | 0 | 0 | 0 |

| | | | |
|---|---|---|---|
| 0 | 1 | 0 | 0 |
| 1 | 0 | 0 | 0 |
| 0 | 0 | 0 | 0 |
| 0 | 0 | 0 | 0 |
| 0 | 0 | 0 | 0 |
| 1 | 1 | 0 | 0 |
| 0 | 0 | 0 | 0 |
| 0 | 0 | 0 | 0 |
| 0 | 0 | 0 | 0 |
| 1 | 0 | 0 | 0 |
| 1 | 0 | 0 | 0 |
| 0 | 0 | 0 | 0 |
| 0 | 0 | 0 | 0 |
| 0 | 0 | 0 | 0 |
| 0 | 0 | 0 | 0 |
| 0 | 0 | 0 | 0 |
| 0 | 0 | 0 | 0 |
| 0 | 0 | 0 | 0 |
| 0 | 0 | 0 | 0 |
| 0 | 0 | 0 | 0 |
| 0 | 0 | 0 | 0 |
| 0 | 0 | 0 | 0 |
| 0 | 0 | 0 | 0 |
| 1 | 1 | 0 | 0 |
| 0 | 0 | 0 | 0 |
| 0 | 0 | 0 | 0 |
| 1 | 1 | 1 | 0 |
| 0 | 0 | 0 | 0 |
| 0 | 0 | 0 | 0 |
| 0 | 0 | 0 | 0 |
| 1 | 1 | 1 | 0 |
| 1 | 0 | 0 | 0 |
| 1 | 0 | 0 | 0 |
| 1 | 0 | 0 | 0 |
| 1 | 0 | 0 | 0 |
| 1 | 1 | 1 | 0 |
| 1 | 1 | 1 | 0 |
| 0 | 0 | 0 | 0 |
| 0 | 0 | 0 | 0 |
| 1 | 1 | 1 | 0 |
| 0 | 0 | 0 | 0 |
| 0 | 0 | 0 | 0 |
| 0 | 0 | 0 | 0 |
| 0 | 0 | 0 | 0 |
| 0 | 0 | 0 | 0 |
| 0 | 0 | 0 | 0 |
| 0 | 0 | 0 | 0 |
| 0 | 0 | 0 | 0 |
| 1 | 0 | 0 | 0 |
| 1 | 0 | 0 | 0 |
| 1 | 0 | 0 | 0 |
| 0 | 0 | 0 | 0 |
| 0 | 0 | 0 | 0 |
| 0 | 0 | 0 | 0 |

| | | | |
|---|---|---|---|
| 0 | 0 | 0 | 0 |
| 1 | 0 | 0 | 0 |
| 0 | 0 | 0 | 0 |
| 0 | 0 | 0 | 0 |
| 0 | 0 | 0 | 0 |
| 0 | 0 | 0 | 0 |
| 0 | 0 | 0 | 0 |
| 0 | 0 | 0 | 0 |
| 0 | 0 | 0 | 0 |
| 0 | 0 | 0 | 0 |
| 1 | 1 | 0 | 0 |
| 1 | 1 | 0 | 0 |
| 0 | 0 | 0 | 0 |
| 0 | 0 | 0 | 0 |
| 0 | 0 | 0 | 0 |
| 1 | 0 | 0 | 0 |
| 1 | 0 | 0 | 0 |
| 1 | 1 | 0 | 1 |
| 0 | 0 | 0 | 0 |
| 0 | 0 | 0 | 0 |
| 0 | 0 | 0 | 0 |
| 0 | 0 | 0 | 0 |
| 1 | 0 | 0 | 1 |
| 0 | 0 | 0 | 0 |
| 0 | 0 | 0 | 0 |
| 1 | 0 | 0 | 0 |
| 0 | 0 | 0 | 0 |
| 0 | 0 | 0 | 0 |
| 0 | 0 | 0 | 0 |
| 0 | 0 | 0 | 0 |
| 0 | 0 | 0 | 0 |
| 0 | 0 | 0 | 0 |
| 0 | 0 | 0 | 0 |
| 0 | 0 | 0 | 0 |
| 0 | 0 | 0 | 0 |
| 1 | 0 | 0 | 0 |
| 1 | 0 | 0 | 0 |
| 1 | 1 | 0 | 0 |
| 1 | 1 | 0 | 0 |
| 0 | 0 | 0 | 0 |
| 0 | 0 | 0 | 0 |
| 1 | 0 | 0 | 0 |
| 0 | 0 | 0 | 0 |
| 1 | 0 | 0 | 0 |
| 1 | 1 | 0 | 0 |
| 0 | 0 | 0 | 0 |
| 0 | 0 | 0 | 0 |
| 0 | 0 | 0 | 0 |
| 0 | 0 | 0 | 0 |
| 0 | 0 | 0 | 0 |
| 0 | 0 | 0 | 0 |
| 1 | 0 | 0 | 0 |
| 0 | 0 | 0 | 0 |
| 0 | 0 | 0 | 0 |

| | | | |
|---|---|---|---|
| 0 | 0 | 0 | 0 |
| 1 | 1 | 0 | 0 |
| 0 | 0 | 0 | 0 |
| 1 | 0 | 0 | 0 |
| 1 | 0 | 0 | 0 |
| 0 | 0 | 0 | 0 |
| 0 | 0 | 0 | 0 |
| 0 | 0 | 0 | 0 |
| 0 | 0 | 0 | 0 |
| 0 | 0 | 0 | 0 |
| 0 | 0 | 0 | 0 |
| 1 | 1 | 0 | 0 |
| 1 | 1 | 0 | 1 |
| 0 | 0 | 0 | 0 |
| 0 | 1 | 0 | 0 |
| 0 | 0 | 0 | 0 |
| 0 | 0 | 0 | 0 |
| 0 | 0 | 0 | 0 |
| 0 | 0 | 0 | 0 |
| 0 | 0 | 0 | 0 |
| 0 | 0 | 0 | 0 |
| 1 | 0 | 0 | 0 |
| 0 | 0 | 0 | 0 |
| 0 | 0 | 0 | 0 |
| 0 | 0 | 0 | 0 |
| 0 | 0 | 0 | 0 |
| 0 | 0 | 0 | 0 |
| 1 | 0 | 0 | 0 |
| 0 | 0 | 0 | 0 |
| 0 | 0 | 0 | 0 |
| 0 | 0 | 0 | 0 |
| 0 | 0 | 0 | 0 |
| 0 | 0 | 0 | 0 |
| 0 | 0 | 0 | 0 |
| 0 | 0 | 0 | 0 |
| 0 | 0 | 0 | 0 |
| 0 | 0 | 0 | 0 |
| 0 | 0 | 0 | 0 |
| 0 | 0 | 0 | 0 |
| 0 | 0 | 0 | 0 |
| 0 | 0 | 0 | 0 |
| 0 | 0 | 0 | 0 |
| 0 | 0 | 0 | 0 |
| 0 | 0 | 0 | 0 |
| 0 | 0 | 0 | 0 |
| 0 | 0 | 0 | 0 |
| 0 | 0 | 0 | 0 |
| 0 | 0 | 0 | 0 |
| 0 | 0 | 0 | 0 |
| 0 | 0 | 0 | 0 |
| 0 | 0 | 0 | 0 |
| 0 | 0 | 0 | 0 |
| 0 | 0 | 0 | 0 |
| 0 | 0 | 0 | 0 |

| | | | |
|---|---|---|---|
| 0 | 0 | 0 | 0 |
| 0 | 0 | 0 | 0 |
| 0 | 0 | 0 | 0 |
| 0 | 0 | 0 | 0 |
| 0 | 0 | 0 | 0 |
| 0 | 0 | 0 | 0 |
| 0 | 0 | 0 | 0 |
| 0 | 0 | 0 | 0 |
| 0 | 0 | 0 | 0 |
| 0 | 0 | 0 | 0 |
| 0 | 0 | 0 | 0 |
| 0 | 0 | 0 | 0 |
| 1 | 0 | 0 | 0 |
| 1 | 1 | 0 | 0 |
| 0 | 0 | 0 | 0 |
| 1 | 0 | 0 | 0 |
| 0 | 0 | 0 | 0 |
| 1 | 0 | 0 | 0 |
| 0 | 0 | 0 | 0 |
| 0 | 0 | 0 | 0 |
| 1 | 0 | 0 | 0 |
| 0 | 0 | 0 | 0 |
| 0 | 0 | 0 | 0 |
| 1 | 0 | 0 | 0 |
| 1 | 0 | 0 | 0 |
| 0 | 0 | 0 | 0 |
| 0 | 0 | 0 | 0 |
| 1 | 0 | 0 | 0 |
| 1 | 0 | 0 | 0 |
| 1 | 0 | 0 | 0 |
| 1 | 0 | 0 | 0 |
| 1 | 0 | 0 | 0 |
| 0 | 0 | 0 | 0 |
| 0 | 0 | 0 | 0 |
| 0 | 0 | 0 | 0 |
| 0 | 0 | 0 | 0 |
| 0 | 0 | 0 | 0 |
| 0 | 0 | 0 | 0 |
| 0 | 0 | 0 | 0 |
| 0 | 0 | 0 | 0 |
| 0 | 0 | 0 | 0 |
| 0 | 0 | 0 | 0 |
| 0 | 0 | 0 | 0 |
| 0 | 0 | 0 | 0 |
| 1 | 1 | 0 | 0 |
| 0 | 0 | 0 | 0 |
| 0 | 0 | 0 | 0 |
| 0 | 0 | 0 | 0 |
| 0 | 0 | 0 | 0 |
| 1 | 1 | 0 | 0 |
| 1 | 0 | 0 | 0 |
| 0 | 0 | 0 | 0 |
| 0 | 0 | 0 | 0 |
| 0 | 0 | 0 | 1 |

| | | | |
|---|---|---|---|
| 0 | 0 | 0 | 0 |
| 0 | 0 | 0 | 0 |
| 0 | 0 | 0 | 0 |
| 0 | 0 | 0 | 0 |
| 1 | 0 | 0 | 1 |
| 0 | 0 | 0 | 0 |
| 1 | 0 | 0 | 0 |
| 0 | 0 | 0 | 0 |
| 0 | 0 | 0 | 0 |
| 0 | 0 | 0 | 0 |
| 0 | 0 | 0 | 0 |
| 0 | 0 | 0 | 0 |
| 0 | 0 | 0 | 0 |
| 1 | 0 | 0 | 0 |
| 0 | 0 | 0 | 0 |
| 0 | 0 | 0 | 0 |
| 1 | 1 | 0 | 0 |
| 0 | 0 | 0 | 0 |
| 1 | 0 | 0 | 0 |
| 1 | 0 | 0 | 0 |
| 0 | 0 | 0 | 0 |
| 0 | 0 | 0 | 0 |
| 0 | 0 | 0 | 0 |
| 1 | 0 | 0 | 0 |
| 1 | 0 | 0 | 0 |
| 1 | 0 | 0 | 0 |
| 1 | 0 | 0 | 0 |
| 1 | 0 | 0 | 0 |
| 1 | 0 | 0 | 0 |
| 1 | 0 | 0 | 0 |
| 1 | 0 | 0 | 0 |
| 1 | 0 | 0 | 0 |
| 1 | 0 | 0 | 0 |
| 1 | 0 | 0 | 0 |
| 0 | 0 | 0 | 0 |
| 0 | 0 | 0 | 0 |
| 0 | 0 | 0 | 0 |
| 1 | 0 | 0 | 0 |
| 1 | 0 | 0 | 0 |
| 0 | 0 | 0 | 0 |
| 0 | 0 | 0 | 0 |
| 1 | 0 | 0 | 0 |
| 0 | 0 | 0 | 0 |
| 1 | 0 | 0 | 0 |
| 1 | 0 | 0 | 0 |
| 1 | 0 | 0 | 0 |
| 1 | 0 | 0 | 0 |
| 1 | 0 | 0 | 0 |
| 0 | 0 | 0 | 0 |
| 0 | 0 | 0 | 0 |
| 1 | 0 | 0 | 0 |
| 0 | 0 | 0 | 0 |
| 1 | 0 | 0 | 0 |
| 1 | 0 | 0 | 0 |

| | | | |
|---|---|---|---|
| 1 | 0 | 0 | 0 |
| 1 | 0 | 0 | 0 |
| 1 | 0 | 0 | 0 |
| 1 | 0 | 0 | 0 |
| 1 | 0 | 0 | 0 |
| 1 | 0 | 0 | 0 |
| 0 | 0 | 0 | 0 |
| 1 | 0 | 0 | 0 |
| 1 | 0 | 0 | 0 |
| 1 | 0 | 0 | 0 |
| 1 | 0 | 0 | 0 |
| 1 | 0 | 0 | 0 |
| 1 | 0 | 0 | 0 |
| 1 | 0 | 0 | 0 |
| 0 | 0 | 0 | 0 |
| 0 | 0 | 0 | 0 |
| 1 | 1 | 0 | 0 |
| 1 | 1 | 0 | 0 |
| 0 | 0 | 0 | 0 |
| 0 | 0 | 0 | 0 |
| 0 | 0 | 0 | 0 |
| 1 | 1 | 0 | 1 |
| 0 | 0 | 0 | 0 |
| 0 | 0 | 0 | 0 |
| 0 | 0 | 0 | 0 |
| 0 | 0 | 0 | 0 |
| 0 | 0 | 0 | 0 |
| 0 | 0 | 0 | 0 |
| 0 | 0 | 0 | 0 |
| 1 | 0 | 1 | 0 |
| 0 | 0 | 0 | 0 |
| 0 | 0 | 0 | 0 |
| 0 | 0 | 0 | 0 |
| 0 | 0 | 0 | 0 |
| 0 | 0 | 0 | 0 |
| 0 | 0 | 0 | 0 |
| 0 | 0 | 0 | 0 |
| 0 | 0 | 0 | 0 |
| 0 | 0 | 0 | 0 |
| 0 | 0 | 0 | 0 |
| 1 | 1 | 0 | 0 |
| 1 | 1 | 0 | 0 |
| 1 | 0 | 0 | 0 |
| 1 | 1 | 0 | 0 |
| 1 | 0 | 0 | 0 |
| 1 | 1 | 0 | 0 |
| 1 | 0 | 0 | 0 |
| 1 | 0 | 0 | 0 |
| 0 | 0 | 0 | 0 |
| 0 | 0 | 0 | 0 |
| 1 | 0 | 0 | 0 |
| 1 | 0 | 0 | 0 |
| 1 | 0 | 0 | 0 |
| 0 | 0 | 0 | 0 |

| | | | |
|---|---|---|---|
| 0 | 0 | 0 | 0 |
| 0 | 0 | 0 | 0 |
| 0 | 0 | 0 | 0 |
| 1 | 1 | 1 | 0 |
| 0 | 0 | 0 | 0 |
| 0 | 0 | 0 | 0 |
| 0 | 0 | 0 | 0 |
| 0 | 0 | 0 | 0 |
| 0 | 1 | 0 | 0 |
| 0 | 1 | 0 | 0 |
| 0 | 1 | 0 | 0 |
| 0 | 0 | 0 | 0 |
| 0 | 0 | 0 | 0 |
| 0 | 0 | 0 | 0 |
| 1 | 0 | 0 | 0 |
| 1 | 0 | 0 | 0 |
| 0 | 0 | 0 | 0 |
| 0 | 0 | 0 | 0 |
| 1 | 0 | 0 | 0 |
| 0 | 0 | 0 | 0 |
| 1 | 0 | 0 | 0 |
| 1 | 0 | 0 | 0 |
| 0 | 0 | 0 | 0 |
| 0 | 0 | 0 | 0 |
| 0 | 0 | 0 | 0 |
| 0 | 0 | 0 | 0 |
| 0 | 0 | 0 | 0 |
| 0 | 0 | 1 | 0 |
| 0 | 0 | 0 | 0 |
| 0 | 1 | 0 | 0 |
| 0 | 0 | 0 | 0 |
| 0 | 0 | 0 | 0 |
| 0 | 0 | 0 | 0 |
| 0 | 0 | 0 | 0 |
| 0 | 0 | 0 | 0 |
| 0 | 0 | 0 | 0 |
| 1 | 0 | 0 | 0 |
| 1 | 0 | 0 | 0 |
| 1 | 0 | 0 | 0 |
| 0 | 0 | 0 | 0 |
| 0 | 0 | 0 | 0 |
| 0 | 0 | 0 | 0 |
| 0 | 0 | 0 | 0 |
| 0 | 0 | 0 | 0 |
| 0 | 0 | 0 | 0 |
| 0 | 0 | 0 | 0 |
| 0 | 0 | 0 | 0 |
| 0 | 0 | 0 | 1 |
| 0 | 0 | 0 | 0 |
| 0 | 0 | 0 | 0 |
| 0 | 0 | 0 | 0 |
| 0 | 0 | 0 | 0 |
| 1 | 1 | 0 | 0 |
| 0 | 0 | 0 | 0 |

| | | | |
|---|---|---|---|
| 0 | 0 | 0 | 0 |
| 0 | 0 | 0 | 0 |
| 0 | 0 | 0 | 0 |
| 0 | 0 | 0 | 0 |
| 1 | 0 | 0 | 0 |
| 0 | 0 | 0 | 0 |
| 0 | 0 | 0 | 0 |
| 0 | 0 | 0 | 0 |
| 0 | 0 | 0 | 0 |
| 0 | 0 | 0 | 0 |
| 0 | 0 | 0 | 0 |
| 0 | 0 | 0 | 0 |
| 0 | 0 | 0 | 0 |
| 0 | 0 | 0 | 0 |
| 0 | 0 | 0 | 0 |
| 0 | 0 | 0 | 0 |
| 0 | 0 | 0 | 0 |
| 0 | 0 | 0 | 0 |
| 0 | 0 | 0 | 0 |
| 0 | 0 | 0 | 0 |
| 0 | 0 | 0 | 0 |
| 0 | 0 | 0 | 0 |
| 0 | 0 | 0 | 0 |
| 0 | 0 | 0 | 0 |
| 0 | 0 | 0 | 0 |
| 0 | 0 | 0 | 0 |
| 0 | 0 | 0 | 0 |
| 0 | 0 | 0 | 0 |
| 0 | 0 | 0 | 0 |
| 0 | 0 | 0 | 0 |
| 0 | 0 | 0 | 0 |
| 0 | 0 | 0 | 0 |
| 0 | 0 | 0 | 0 |
| 0 | 0 | 0 | 0 |
| 0 | 0 | 0 | 0 |
| 0 | 0 | 0 | 0 |
| 0 | 0 | 0 | 0 |
| 0 | 0 | 0 | 0 |
| 0 | 0 | 0 | 0 |
| 0 | 0 | 0 | 0 |
| 1 | 1 | 0 | 0 |
| 0 | 0 | 0 | 0 |
| 0 | 0 | 0 | 0 |
| 0 | 0 | 0 | 0 |
| 0 | 0 | 0 | 0 |
| 1 | 0 | 0 | 0 |
| 1 | 0 | 0 | 0 |
| 0 | 0 | 0 | 0 |
| 1 | 0 | 0 | 0 |
| 0 | 0 | 0 | 0 |
| 0 | 0 | 0 | 0 |
| 1 | 0 | 0 | 0 |
| 1 | 0 | 0 | 0 |
| 0 | 0 | 0 | 0 |

| | | | |
|---|---|---|---|
| 0 | 0 | 0 | 0 |
| 0 | 0 | 0 | 0 |
| 0 | 0 | 0 | 0 |
| 0 | 0 | 0 | 0 |
| 1 | 0 | 0 | 0 |
| 0 | 0 | 0 | 0 |
| 0 | 0 | 0 | 0 |
| 0 | 0 | 0 | 0 |
| 0 | 0 | 0 | 0 |
| 0 | 0 | 0 | 0 |
| 0 | 0 | 0 | 0 |
| 0 | 0 | 0 | 0 |
| 0 | 0 | 0 | 0 |
| 0 | 0 | 0 | 0 |
| 0 | 0 | 0 | 0 |
| 1 | 1 | 0 | 1 |
| 0 | 0 | 0 | 0 |
| 0 | 0 | 0 | 0 |
| 0 | 0 | 0 | 0 |
| 0 | 0 | 0 | 0 |
| 0 | 0 | 0 | 0 |
| 0 | 0 | 0 | 0 |
| 0 | 0 | 0 | 0 |
| 0 | 0 | 0 | 0 |
| 0 | 0 | 0 | 0 |
| 0 | 0 | 0 | 0 |
| 0 | 0 | 0 | 0 |
| 1 | 0 | 0 | 0 |
| 1 | 0 | 0 | 0 |
| 0 | 0 | 0 | 0 |
| 1 | 1 | 0 | 0 |
| 0 | 0 | 0 | 0 |
| 0 | 0 | 0 | 0 |
| 0 | 0 | 0 | 0 |
| 1 | 1 | 0 | 0 |
| 1 | 1 | 0 | 0 |
| 0 | 0 | 0 | 0 |
| 1 | 0 | 0 | 0 |
| 0 | 0 | 0 | 0 |
| 1 | 1 | 0 | 0 |
| 0 | 1 | 0 | 0 |
| 0 | 0 | 0 | 0 |
| 0 | 0 | 0 | 0 |
| 0 | 0 | 0 | 0 |
| 1 | 1 | 0 | 0 |
| 1 | 1 | 0 | 0 |
| 1 | 1 | 0 | 0 |
| 0 | 0 | 0 | 0 |
| 0 | 0 | 0 | 0 |
| 0 | 0 | 0 | 0 |
| 0 | 0 | 0 | 0 |
| 0 | 0 | 0 | 0 |
| 0 | 0 | 0 | 0 |
| 0 | 0 | 0 | 0 |

| | | | |
|---|---|---|---|
| 0 | 0 | 0 | 0 |
| 0 | 0 | 0 | 0 |
| 1 | 0 | 0 | 0 |
| 1 | 0 | 0 | 0 |
| 0 | 0 | 0 | 0 |
| 1 | 0 | 0 | 0 |
| 0 | 0 | 0 | 0 |
| 0 | 0 | 0 | 0 |
| 0 | 0 | 0 | 0 |
| 0 | 1 | 0 | 0 |
| 0 | 0 | 0 | 0 |
| 0 | 0 | 0 | 0 |
| 0 | 0 | 0 | 0 |
| 0 | 0 | 0 | 0 |
| 0 | 0 | 0 | 0 |
| 0 | 0 | 0 | 0 |
| 0 | 0 | 0 | 0 |
| 1 | 0 | 0 | 0 |
| 0 | 0 | 0 | 0 |
| 1 | 0 | 0 | 0 |
| 0 | 0 | 0 | 0 |
| 1 | 0 | 0 | 0 |
| 0 | 0 | 0 | 0 |
| 1 | 0 | 0 | 0 |
| 0 | 0 | 0 | 0 |
| 0 | 0 | 0 | 0 |
| 0 | 0 | 0 | 0 |
| 0 | 0 | 0 | 0 |
| 0 | 0 | 0 | 0 |
| 0 | 0 | 0 | 0 |
| 0 | 0 | 0 | 0 |
| 1 | 0 | 0 | 0 |
| 0 | 0 | 0 | 0 |
| 0 | 0 | 0 | 0 |
| 0 | 0 | 0 | 0 |
| 0 | 0 | 0 | 0 |
| 0 | 0 | 0 | 0 |
| 0 | 0 | 0 | 0 |
| 1 | 1 | 0 | 0 |
| 1 | 1 | 0 | 0 |
| 1 | 1 | 0 | 0 |
| 1 | 1 | 0 | 0 |
| 1 | 1 | 0 | 0 |
| 1 | 0 | 0 | 0 |
| 1 | 0 | 0 | 0 |
| 1 | 0 | 0 | 0 |
| 0 | 0 | 0 | 0 |
| 0 | 0 | 0 | 0 |
| 0 | 0 | 0 | 0 |
| 0 | 0 | 0 | 0 |
| 0 | 0 | 0 | 0 |
| 0 | 0 | 0 | 0 |
| 0 | 0 | 0 | 0 |
| 0 | 0 | 0 | 0 |

| | | | |
|---|---|---|---|
| 0 | 0 | 0 | 0 |
| 0 | 0 | 0 | 0 |
| 0 | 0 | 0 | 0 |
| 0 | 0 | 0 | 0 |
| 0 | 0 | 0 | 0 |
| 1 | 0 | 0 | 0 |
| 1 | 0 | 0 | 0 |
| 0 | 0 | 0 | 0 |
| 0 | 0 | 0 | 0 |
| 0 | 0 | 0 | 0 |
| 0 | 0 | 0 | 0 |
| 1 | 0 | 0 | 0 |
| 0 | 0 | 1 | 0 |
| 0 | 0 | 0 | 0 |
| 0 | 0 | 0 | 0 |
| 0 | 0 | 0 | 0 |
| 1 | 0 | 0 | 0 |
| 0 | 0 | 0 | 0 |
| 1 | 0 | 0 | 0 |
| 1 | 0 | 0 | 0 |
| 0 | 0 | 0 | 0 |
| 1 | 0 | 0 | 0 |
| 1 | 1 | 0 | 0 |
| 0 | 0 | 0 | 0 |
| 1 | 1 | 0 | 0 |
| 1 | 1 | 0 | 0 |
| 1 | 1 | 0 | 0 |
| 0 | 0 | 0 | 0 |
| 1 | 0 | 0 | 0 |
| 0 | 0 | 0 | 0 |
| 0 | 0 | 0 | 0 |
| 0 | 0 | 0 | 0 |
| 0 | 0 | 0 | 0 |
| 0 | 0 | 0 | 0 |
| 0 | 0 | 0 | 0 |
| 0 | 0 | 0 | 0 |
| 0 | 0 | 0 | 0 |
| 0 | 0 | 0 | 0 |
| 0 | 0 | 0 | 0 |
| 0 | 0 | 0 | 0 |
| 0 | 0 | 0 | 0 |
| 0 | 0 | 0 | 0 |
| 0 | 0 | 0 | 0 |
| 0 | 0 | 0 | 0 |
| 0 | 0 | 0 | 0 |
| 0 | 0 | 0 | 0 |
| 0 | 0 | 0 | 0 |
| 0 | 0 | 0 | 0 |
| 0 | 0 | 0 | 0 |
| 0 | 0 | 0 | 0 |
| 0 | 0 | 0 | 0 |
| 0 | 0 | 0 | 0 |
| 0 | 0 | 0 | 0 |
| 0 | 0 | 0 | 0 |

| | | | |
|---|---|---|---|
| 0 | 0 | 0 | 0 |
| 0 | 0 | 0 | 0 |
| 0 | 0 | 0 | 0 |
| 0 | 0 | 0 | 0 |
| 0 | 0 | 0 | 0 |
| 0 | 0 | 0 | 0 |
| 0 | 0 | 0 | 0 |
| 0 | 0 | 0 | 0 |
| 0 | 0 | 0 | 0 |
| 0 | 0 | 0 | 0 |
| 0 | 1 | 0 | 0 |
| 0 | 1 | 0 | 0 |
| 0 | 1 | 0 | 0 |
| 0 | 1 | 0 | 0 |
| 0 | 0 | 0 | 0 |
| 0 | 1 | 0 | 0 |
| 0 | 1 | 0 | 0 |
| 1 | 1 | 0 | 0 |
| 0 | 1 | 0 | 0 |
| 1 | 0 | 0 | 0 |
| 1 | 1 | 0 | 0 |
| 1 | 1 | 0 | 0 |
| 0 | 0 | 1 | 1 |
| 0 | 0 | 0 | 0 |
| 1 | 0 | 0 | 0 |
| 0 | 0 | 0 | 0 |
| 0 | 0 | 0 | 0 |
| 0 | 0 | 0 | 0 |
| 0 | 0 | 0 | 0 |
| 0 | 0 | 0 | 0 |
| 0 | 0 | 0 | 0 |
| 0 | 0 | 0 | 0 |
| 0 | 0 | 0 | 0 |
| 0 | 0 | 0 | 0 |
| 0 | 0 | 0 | 0 |
| 0 | 0 | 0 | 0 |
| 0 | 0 | 0 | 0 |
| 0 | 0 | 0 | 0 |
| 0 | 0 | 0 | 0 |
| 0 | 0 | 0 | 0 |
| 1 | 0 | 0 | 0 |
| 0 | 0 | 0 | 0 |
| 1 | 0 | 0 | 0 |
| 1 | 0 | 0 | 0 |
| 0 | 0 | 0 | 0 |
| 1 | 0 | 0 | 0 |
| 0 | 0 | 0 | 0 |
| 0 | 0 | 0 | 0 |
| 0 | 0 | 0 | 0 |
| 0 | 0 | 0 | 0 |
| 0 | 0 | 0 | 0 |
| 0 | 0 | 0 | 0 |
| 1 | 1 | 0 | 0 |
| 1 | 1 | 0 | 0 |

| | | | |
|---|---|---|---|
| 1 | 1 | 0 | 0 |
| 0 | 0 | 0 | 0 |
| 0 | 0 | 0 | 0 |
| 0 | 0 | 0 | 0 |
| 0 | 0 | 0 | 0 |
| 0 | 0 | 0 | 0 |
| 0 | 1 | 0 | 0 |
| 0 | 0 | 0 | 0 |
| 0 | 0 | 0 | 0 |
| 0 | 1 | 0 | 0 |
| 1 | 0 | 0 | 0 |
| 0 | 0 | 0 | 0 |
| 0 | 0 | 1 | 0 |
| 0 | 0 | 1 | 0 |
| 0 | 0 | 0 | 0 |
| 0 | 0 | 0 | 0 |
| 0 | 0 | 0 | 0 |
| 0 | 1 | 0 | 0 |
| 0 | 0 | 0 | 0 |
| 0 | 0 | 0 | 0 |
| 1 | 0 | 0 | 0 |
| 0 | 0 | 0 | 1 |
| 0 | 0 | 0 | 0 |
| 0 | 0 | 0 | 0 |
| 0 | 0 | 0 | 0 |
| 0 | 0 | 0 | 0 |
| 0 | 0 | 0 | 0 |
| 0 | 0 | 0 | 0 |
| 0 | 0 | 0 | 0 |
| 0 | 0 | 0 | 0 |
| 0 | 0 | 0 | 0 |
| 0 | 0 | 0 | 0 |
| 0 | 0 | 0 | 0 |
| 0 | 0 | 0 | 0 |
| 0 | 0 | 0 | 0 |
| 0 | 0 | 0 | 0 |
| 1 | 0 | 0 | 0 |
| 0 | 0 | 0 | 0 |
| 1 | 0 | 0 | 0 |
| 0 | 0 | 0 | 0 |
| 0 | 0 | 0 | 0 |
| 0 | 0 | 0 | 0 |
| 0 | 0 | 0 | 0 |
| 0 | 0 | 0 | 0 |
| 0 | 0 | 0 | 0 |
| 0 | 0 | 0 | 0 |
| 0 | 0 | 0 | 0 |
| 0 | 0 | 0 | 0 |
| 0 | 0 | 0 | 0 |
| 1 | 0 | 0 | 0 |
| 0 | 0 | 0 | 0 |
| 0 | 0 | 0 | 0 |
| 0 | 0 | 0 | 0 |
| 0 | 0 | 0 | 0 |

| | | | |
|---|---|---|---|
| 0 | 0 | 0 | 0 |
| 0 | 0 | 0 | 0 |
| 0 | 0 | 0 | 0 |
| 0 | 0 | 0 | 0 |
| 0 | 0 | 0 | 0 |
| 0 | 0 | 0 | 0 |
| 0 | 0 | 0 | 0 |
| 0 | 0 | 0 | 0 |
| 0 | 0 | 0 | 0 |
| 0 | 0 | 0 | 0 |
| 0 | 0 | 0 | 0 |
| 0 | 0 | 0 | 0 |
| 0 | 0 | 1 | 0 |
| 1 | 0 | 0 | 0 |
| 1 | 0 | 0 | 0 |
| 0 | 0 | 0 | 0 |
| 0 | 0 | 0 | 0 |
| 0 | 0 | 0 | 0 |
| 0 | 0 | 0 | 0 |
| 0 | 0 | 0 | 0 |
| 0 | 0 | 0 | 0 |
| 0 | 0 | 0 | 0 |
| 0 | 0 | 0 | 0 |
| 1 | 1 | 0 | 1 |
| 0 | 0 | 0 | 0 |
| 1 | 1 | 0 | 0 |
| 1 | 1 | 0 | 1 |
| 0 | 0 | 0 | 0 |
| 0 | 0 | 0 | 0 |
| 0 | 0 | 0 | 0 |
| 0 | 0 | 0 | 0 |
| 0 | 0 | 0 | 0 |
| 0 | 0 | 0 | 0 |
| 0 | 0 | 0 | 0 |
| 0 | 0 | 0 | 0 |
| 1 | 0 | 0 | 0 |
| 0 | 0 | 0 | 0 |
| 0 | 0 | 0 | 0 |
| 0 | 0 | 0 | 0 |
| 0 | 0 | 0 | 0 |
| 0 | 0 | 0 | 0 |
| 0 | 0 | 0 | 0 |
| 1 | 0 | 0 | 0 |
| 1 | 0 | 0 | 0 |
| 1 | 0 | 0 | 0 |
| 1 | 0 | 0 | 0 |
| 1 | 0 | 0 | 0 |
| 0 | 0 | 0 | 0 |
| 0 | 0 | 0 | 0 |
| 1 | 0 | 0 | 0 |
| 1 | 0 | 0 | 0 |
| 0 | 0 | 0 | 0 |
| 1 | 0 | 0 | 0 |
| 1 | 0 | 0 | 0 |

| | | | |
|---|---|---|---|
| 0 | 0 | 0 | 0 |
| 1 | 0 | 0 | 0 |
| 0 | 0 | 0 | 0 |
| 0 | 0 | 0 | 0 |
| 0 | 0 | 0 | 0 |
| 0 | 0 | 0 | 0 |
| 1 | 0 | 0 | 0 |
| 0 | 0 | 0 | 0 |
| 0 | 0 | 0 | 0 |
| 0 | 0 | 0 | 0 |
| 0 | 0 | 0 | 0 |
| 1 | 1 | 0 | 0 |
| 0 | 0 | 0 | 0 |
| 0 | 0 | 0 | 0 |
| 0 | 0 | 0 | 0 |
| 0 | 0 | 0 | 0 |
| 0 | 0 | 0 | 0 |
| 0 | 0 | 0 | 0 |
| 0 | 0 | 0 | 0 |
| 0 | 0 | 0 | 0 |
| 0 | 0 | 0 | 0 |
| 0 | 0 | 0 | 0 |
| 0 | 0 | 0 | 0 |
| 0 | 0 | 0 | 0 |
| 0 | 0 | 0 | 0 |
| 0 | 0 | 0 | 0 |
| 0 | 0 | 0 | 0 |
| 0 | 0 | 0 | 0 |
| 1 | 1 | 0 | 0 |
| 1 | 0 | 0 | 0 |
| 0 | 0 | 0 | 0 |
| 0 | 0 | 0 | 0 |
| 0 | 0 | 0 | 0 |
| 0 | 0 | 0 | 0 |
| 0 | 0 | 0 | 0 |
| 0 | 0 | 0 | 0 |
| 0 | 0 | 0 | 0 |
| 0 | 0 | 0 | 0 |
| 0 | 0 | 0 | 0 |
| 0 | 0 | 0 | 0 |
| 0 | 0 | 0 | 0 |
| 0 | 0 | 0 | 0 |
| 0 | 0 | 0 | 0 |
| 0 | 0 | 0 | 0 |
| 0 | 0 | 0 | 0 |
| 0 | 0 | 0 | 0 |
| 0 | 0 | 0 | 0 |
| 0 | 0 | 0 | 0 |
| 0 | 0 | 0 | 0 |
| 0 | 0 | 0 | 0 |
| 0 | 0 | 0 | 0 |
| 0 | 0 | 0 | 0 |
| 0 | 0 | 1 | 0 |
| 0 | 0 | 0 | 0 |

| | | | |
|---|---|---|---|
| 0 | 0 | 0 | 0 |
| 0 | 0 | 0 | 0 |
| 0 | 0 | 0 | 0 |
| 0 | 0 | 0 | 0 |
| 1 | 0 | 0 | 0 |
| 0 | 0 | 0 | 0 |
| 0 | 0 | 0 | 0 |
| 1 | 1 | 0 | 0 |
| 0 | 0 | 0 | 0 |
| 1 | 1 | 0 | 0 |
| 0 | 0 | 0 | 0 |
| 0 | 0 | 0 | 0 |
| 0 | 0 | 0 | 0 |
| 0 | 0 | 0 | 0 |
| 0 | 0 | 0 | 0 |
| 0 | 0 | 0 | 0 |
| 1 | 0 | 0 | 0 |
| 0 | 0 | 0 | 0 |
| 0 | 0 | 0 | 0 |
| 1 | 0 | 0 | 0 |
| 1 | 0 | 0 | 0 |
| 0 | 0 | 0 | 0 |
| 0 | 0 | 0 | 0 |
| 0 | 0 | 0 | 0 |
| 0 | 0 | 0 | 0 |
| 0 | 0 | 0 | 0 |
| 1 | 0 | 0 | 0 |
| 0 | 0 | 0 | 0 |
| 0 | 0 | 0 | 0 |
| 1 | 0 | 0 | 0 |
| 0 | 0 | 0 | 0 |
| 0 | 0 | 0 | 0 |
| 0 | 0 | 0 | 0 |
| 1 | 1 | 0 | 0 |
| 0 | 0 | 0 | 0 |
| 0 | 0 | 0 | 0 |
| 0 | 0 | 0 | 0 |
| 0 | 0 | 0 | 0 |
| 0 | 1 | 0 | 0 |
| 0 | 0 | 0 | 0 |
| 0 | 0 | 0 | 0 |
| 0 | 0 | 0 | 0 |
| 0 | 0 | 0 | 0 |
| 1 | 1 | 0 | 0 |
| 0 | 0 | 0 | 0 |
| 0 | 0 | 0 | 0 |
| 0 | 0 | 0 | 0 |
| 0 | 0 | 0 | 0 |
| 0 | 0 | 0 | 0 |
| 0 | 0 | 0 | 0 |
| 0 | 0 | 0 | 0 |
| 1 | 0 | 0 | 0 |
| 0 | 0 | 0 | 0 |
| 0 | 0 | 0 | 0 |

| | | | |
|---|---|---|---|
| 1 | 0 | 0 | 1 |
| 1 | 0 | 0 | 1 |
| 1 | 0 | 0 | 1 |
| 0 | 0 | 0 | 0 |
| 0 | 0 | 0 | 0 |
| 0 | 0 | 0 | 0 |
| 1 | 0 | 0 | 0 |
| 0 | 0 | 0 | 0 |
| 0 | 0 | 0 | 0 |
| 0 | 0 | 0 | 0 |
| 0 | 0 | 1 | 0 |
| 0 | 0 | 0 | 0 |
| 0 | 0 | 0 | 0 |
| 0 | 0 | 0 | 0 |
| 0 | 0 | 0 | 0 |
| 0 | 0 | 0 | 0 |
| 1 | 0 | 0 | 0 |
| 0 | 0 | 0 | 0 |
| 0 | 0 | 0 | 0 |
| 0 | 0 | 0 | 0 |
| 0 | 0 | 0 | 0 |
| 0 | 0 | 0 | 0 |
| 0 | 0 | 0 | 0 |
| 1 | 1 | 0 | 0 |
| 0 | 0 | 0 | 0 |
| 0 | 0 | 0 | 0 |
| 0 | 0 | 0 | 0 |
| 0 | 0 | 0 | 0 |
| 0 | 0 | 0 | 0 |
| 0 | 0 | 0 | 0 |
| 0 | 0 | 0 | 0 |
| 0 | 0 | 0 | 0 |
| 0 | 1 | 0 | 0 |
| 0 | 0 | 0 | 0 |
| 1 | 1 | 0 | 0 |
| 0 | 0 | 0 | 0 |
| 0 | 0 | 0 | 0 |
| 0 | 0 | 0 | 0 |
| 0 | 0 | 0 | 0 |
| 0 | 0 | 0 | 0 |
| 0 | 0 | 0 | 0 |
| 1 | 0 | 0 | 1 |
| 0 | 0 | 0 | 0 |
| 1 | 0 | 0 | 1 |
| 0 | 0 | 0 | 0 |
| 1 | 0 | 0 | 1 |
| 0 | 0 | 0 | 0 |
| 0 | 0 | 0 | 0 |
| 0 | 0 | 0 | 0 |
| 1 | 0 | 0 | 0 |
| 0 | 0 | 0 | 0 |
| 0 | 0 | 0 | 0 |
| 0 | 0 | 0 | 0 |
| 0 | 0 | 0 | 0 |

| | | | |
|---|---|---|---|
| 0 | 0 | 0 | 0 |
| 0 | 0 | 0 | 0 |
| 0 | 0 | 0 | 0 |
| 0 | 0 | 0 | 0 |
| 0 | 0 | 0 | 0 |
| 0 | 0 | 0 | 0 |
| 0 | 0 | 0 | 0 |
| 0 | 0 | 0 | 0 |
| 0 | 0 | 0 | 0 |
| 0 | 0 | 0 | 0 |
| 0 | 0 | 0 | 0 |
| 0 | 0 | 0 | 0 |
| 0 | 0 | 0 | 0 |
| 0 | 0 | 0 | 0 |
| 0 | 0 | 0 | 0 |
| 0 | 0 | 0 | 0 |
| 0 | 0 | 0 | 0 |
| 0 | 0 | 0 | 0 |
| 0 | 0 | 1 | 0 |
| 0 | 0 | 0 | 0 |
| 0 | 0 | 0 | 0 |
| 0 | 0 | 0 | 0 |
| 0 | 0 | 0 | 0 |
| 0 | 0 | 0 | 0 |
| 0 | 0 | 0 | 0 |
| 0 | 0 | 0 | 0 |
| 0 | 0 | 0 | 0 |
| 0 | 0 | 0 | 0 |
| 0 | 0 | 1 | 0 |
| 0 | 0 | 0 | 0 |
| 0 | 0 | 1 | 0 |
| 1 | 0 | 0 | 0 |
| 1 | 1 | 1 | 0 |
| 1 | 1 | 1 | 0 |
| 0 | 0 | 0 | 0 |
| 0 | 0 | 0 | 0 |
| 0 | 1 | 0 | 0 |
| 1 | 1 | 1 | 0 |
| 1 | 1 | 1 | 0 |
| 0 | 0 | 1 | 0 |
| 0 | 0 | 1 | 0 |
| 1 | 0 | 0 | 0 |
| 0 | 0 | 0 | 0 |
| 0 | 0 | 0 | 0 |
| 1 | 1 | 0 | 0 |
| 0 | 0 | 0 | 0 |
| 0 | 0 | 0 | 0 |
| 0 | 0 | 0 | 0 |
| 0 | 0 | 0 | 0 |
| 0 | 0 | 0 | 0 |
| 0 | 0 | 0 | 0 |
| 0 | 0 | 0 | 0 |
| 0 | 0 | 0 | 0 |
| 0 | 0 | 0 | 0 |

| | | | |
|---|---|---|---|
| 0 | 0 | 0 | 0 |
| 0 | 0 | 0 | 0 |
| 0 | 0 | 0 | 0 |
| 0 | 0 | 0 | 0 |
| 0 | 0 | 0 | 0 |
| 0 | 0 | 0 | 0 |
| 0 | 0 | 0 | 0 |
| 0 | 0 | 0 | 0 |
| 0 | 0 | 0 | 0 |
| 0 | 0 | 0 | 0 |
| 0 | 0 | 0 | 0 |
| 0 | 0 | 0 | 0 |
| 0 | 0 | 0 | 0 |
| 0 | 0 | 0 | 0 |
| 0 | 0 | 0 | 0 |
| 0 | 0 | 0 | 0 |
| 0 | 0 | 0 | 0 |
| 0 | 0 | 0 | 0 |
| 0 | 0 | 0 | 0 |
| 1 | 0 | 0 | 0 |
| 1 | 0 | 0 | 0 |
| 0 | 0 | 0 | 0 |
| 0 | 0 | 0 | 0 |
| 0 | 0 | 0 | 0 |
| 0 | 0 | 0 | 0 |
| 1 | 0 | 0 | 0 |
| 0 | 0 | 0 | 0 |
| 0 | 0 | 0 | 0 |
| 0 | 0 | 0 | 0 |
| 1 | 0 | 0 | 0 |
| 0 | 0 | 0 | 0 |
| 0 | 0 | 0 | 0 |
| 1 | 0 | 0 | 0 |
| 1 | 0 | 0 | 0 |
| 0 | 0 | 0 | 0 |
| 0 | 0 | 0 | 0 |
| 0 | 0 | 0 | 0 |
| 0 | 0 | 0 | 0 |
| 0 | 0 | 0 | 0 |
| 0 | 0 | 0 | 0 |
| 0 | 0 | 0 | 0 |
| 0 | 0 | 0 | 0 |
| 0 | 0 | 0 | 0 |
| 0 | 0 | 0 | 0 |
| 0 | 0 | 0 | 0 |
| 0 | 0 | 0 | 0 |
| 0 | 0 | 0 | 0 |
| 0 | 1 | 0 | 0 |
| 0 | 1 | 0 | 0 |
| 0 | 0 | 0 | 0 |
| 0 | 1 | 0 | 0 |
| 0 | 1 | 0 | 0 |
| 0 | 0 | 0 | 0 |
| 0 | 0 | 0 | 0 |

| | | | |
|---|---|---|---|
| 0 | 0 | 0 | 0 |
| 0 | 0 | 0 | 0 |
| 1 | 1 | 0 | 1 |
| 0 | 0 | 0 | 0 |
| 0 | 0 | 0 | 0 |
| 0 | 0 | 0 | 0 |
| 0 | 0 | 0 | 0 |
| 0 | 1 | 0 | 0 |
| 1 | 1 | 0 | 0 |
| 1 | 1 | 0 | 0 |
| 1 | 1 | 0 | 0 |
| 1 | 1 | 0 | 0 |
| 1 | 1 | 0 | 0 |
| 0 | 1 | 0 | 0 |
| 0 | 1 | 0 | 0 |
| 0 | 0 | 0 | 0 |
| 0 | 0 | 0 | 0 |
| 0 | 0 | 0 | 0 |
| 0 | 0 | 0 | 0 |
| 0 | 0 | 0 | 0 |
| 0 | 0 | 0 | 0 |
| 0 | 0 | 0 | 0 |
| 0 | 1 | 0 | 0 |
| 0 | 1 | 0 | 0 |
| 0 | 0 | 0 | 0 |
| 0 | 0 | 1 | 0 |
| 0 | 0 | 0 | 0 |
| 0 | 0 | 0 | 0 |
| 0 | 0 | 0 | 0 |
| 0 | 1 | 0 | 0 |
| 0 | 0 | 0 | 0 |
| 0 | 0 | 0 | 0 |
| 0 | 0 | 0 | 0 |
| 0 | 0 | 0 | 0 |
| 0 | 0 | 0 | 0 |
| 0 | 0 | 0 | 0 |
| 0 | 0 | 0 | 0 |
| 0 | 0 | 0 | 0 |
| 0 | 0 | 0 | 0 |
| 1 | 0 | 0 | 0 |
| 1 | 1 | 0 | 0 |
| 1 | 0 | 0 | 0 |
| 0 | 0 | 0 | 0 |
| 0 | 1 | 0 | 0 |
| 1 | 1 | 0 | 0 |
| 0 | 1 | 0 | 0 |
| 0 | 0 | 0 | 0 |
| 1 | 0 | 0 | 0 |
| 0 | 1 | 0 | 0 |
| 0 | 0 | 0 | 0 |
| 0 | 0 | 0 | 0 |
| 0 | 0 | 0 | 0 |
| 1 | 1 | 0 | 0 |
| 0 | 0 | 0 | 0 |

| | | | |
|---|---|---|---|
| 0 | 0 | 0 | 0 |
| 0 | 0 | 0 | 0 |
| 1 | 0 | 0 | 0 |
| 1 | 1 | 0 | 0 |
| 1 | 1 | 0 | 0 |
| 0 | 0 | 0 | 0 |
| 0 | 0 | 0 | 0 |
| 0 | 0 | 0 | 0 |
| 1 | 0 | 0 | 0 |
| 0 | 0 | 0 | 0 |
| 0 | 0 | 0 | 0 |
| 1 | 0 | 0 | 1 |
| 0 | 0 | 0 | 0 |
| 1 | 0 | 0 | 0 |
| 0 | 0 | 0 | 0 |
| 0 | 0 | 0 | 0 |
| 0 | 0 | 0 | 0 |
| 0 | 0 | 0 | 0 |
| 0 | 0 | 0 | 0 |
| 0 | 0 | 0 | 0 |
| 0 | 0 | 0 | 0 |
| 1 | 0 | 0 | 1 |
| 1 | 0 | 0 | 0 |
| 0 | 0 | 0 | 0 |
| 0 | 0 | 0 | 0 |
| 0 | 0 | 0 | 0 |
| 0 | 0 | 0 | 0 |
| 0 | 0 | 0 | 0 |
| 1 | 0 | 0 | 0 |
| 1 | 0 | 0 | 0 |
| 0 | 0 | 0 | 0 |
| 0 | 0 | 0 | 0 |
| 0 | 0 | 0 | 0 |
| 0 | 0 | 0 | 0 |
| 0 | 0 | 0 | 0 |
| 0 | 0 | 0 | 0 |
| 1 | 0 | 0 | 0 |
| 0 | 0 | 0 | 0 |
| 0 | 0 | 0 | 0 |
| 1 | 0 | 0 | 0 |
| 1 | 0 | 0 | 0 |
| 1 | 0 | 0 | 0 |
| 1 | 0 | 0 | 0 |
| 0 | 0 | 0 | 0 |
| 0 | 0 | 0 | 0 |
| 1 | 0 | 0 | 0 |
| 0 | 0 | 0 | 0 |
| 0 | 0 | 0 | 0 |
| 0 | 0 | 0 | 0 |
| 0 | 0 | 0 | 0 |
| 0 | 0 | 0 | 0 |
| 1 | 0 | 0 | 0 |
| 1 | 0 | 0 | 0 |
| 1 | 0 | 0 | 0 |

| | | | |
|---|---|---|---|
| 1 | 0 | 0 | 0 |
| 0 | 0 | 0 | 0 |
| 0 | 0 | 0 | 0 |
| 0 | 0 | 0 | 0 |
| 1 | 0 | 0 | 0 |
| 0 | 0 | 0 | 0 |
| 1 | 0 | 0 | 0 |
| 0 | 0 | 0 | 0 |
| 0 | 0 | 0 | 0 |
| 0 | 0 | 0 | 0 |
| 0 | 0 | 0 | 0 |
| 0 | 0 | 0 | 0 |
| 0 | 0 | 0 | 0 |
| 0 | 0 | 0 | 0 |
| 0 | 0 | 0 | 0 |
| 1 | 0 | 0 | 0 |
| 0 | 0 | 0 | 0 |
| 0 | 0 | 0 | 0 |
| 0 | 0 | 0 | 0 |
| 0 | 0 | 0 | 0 |
| 0 | 0 | 0 | 0 |
| 0 | 0 | 0 | 0 |
| 1 | 0 | 0 | 0 |
| 1 | 0 | 0 | 0 |
| 0 | 0 | 0 | 0 |
| 0 | 0 | 0 | 0 |
| 0 | 0 | 0 | 0 |
| 0 | 0 | 0 | 0 |
| 0 | 0 | 0 | 0 |
| 0 | 0 | 0 | 0 |
| 0 | 0 | 0 | 0 |
| 0 | 0 | 0 | 0 |
| 1 | 0 | 0 | 0 |
| 0 | 0 | 0 | 0 |
| 0 | 0 | 0 | 0 |
| 0 | 0 | 0 | 0 |
| 0 | 0 | 0 | 0 |
| 0 | 0 | 0 | 0 |
| 0 | 0 | 0 | 0 |
| 0 | 0 | 0 | 0 |
| 0 | 0 | 0 | 0 |
| 0 | 0 | 0 | 0 |
| 0 | 0 | 0 | 0 |
| 0 | 0 | 0 | 0 |
| 0 | 1 | 0 | 0 |
| 0 | 0 | 0 | 0 |
| 0 | 0 | 0 | 0 |
| 0 | 0 | 0 | 0 |
| 0 | 0 | 0 | 0 |
| 0 | 0 | 0 | 0 |
| 0 | 0 | 0 | 0 |
| 0 | 0 | 0 | 0 |
| 0 | 0 | 0 | 0 |
| 1 | 0 | 0 | 0 |

| | | | |
|---|---|---|---|
| 0 | 0 | 0 | 0 |
| 0 | 0 | 0 | 0 |
| 0 | 0 | 0 | 0 |
| 0 | 0 | 0 | 0 |
| 0 | 0 | 0 | 0 |
| 0 | 0 | 0 | 0 |
| 0 | 0 | 0 | 0 |
| 0 | 0 | 0 | 0 |
| 0 | 0 | 0 | 0 |
| 1 | 0 | 0 | 0 |
| 0 | 0 | 0 | 0 |
| 0 | 0 | 0 | 0 |
| 1 | 1 | 0 | 0 |
| 1 | 1 | 0 | 0 |
| 1 | 1 | 0 | 0 |
| 1 | 1 | 0 | 0 |
| 0 | 0 | 0 | 0 |
| 0 | 0 | 0 | 0 |
| 0 | 0 | 0 | 0 |
| 0 | 0 | 0 | 0 |
| 0 | 0 | 0 | 0 |
| 0 | 0 | 0 | 0 |
| 0 | 0 | 0 | 0 |
| 0 | 0 | 0 | 0 |
| 0 | 0 | 0 | 0 |
| 0 | 0 | 0 | 1 |
| 0 | 0 | 0 | 0 |
| 0 | 0 | 0 | 0 |
| 0 | 0 | 0 | 0 |
| 0 | 0 | 0 | 0 |
| 0 | 0 | 0 | 0 |
| 0 | 0 | 0 | 0 |
| 0 | 0 | 0 | 0 |
| 0 | 0 | 0 | 0 |
| 0 | 0 | 0 | 0 |
| 0 | 0 | 0 | 0 |
| 0 | 0 | 0 | 0 |
| 1 | 1 | 0 | 0 |
| 0 | 0 | 0 | 0 |
| 0 | 0 | 0 | 0 |
| 1 | 1 | 0 | 0 |
| 0 | 0 | 0 | 0 |
| 0 | 0 | 0 | 0 |
| 1 | 0 | 0 | 1 |
| 0 | 0 | 0 | 0 |
| 0 | 0 | 0 | 0 |
| 0 | 0 | 0 | 0 |
| 0 | 0 | 0 | 0 |
| 0 | 0 | 0 | 0 |
| 0 | 0 | 0 | 0 |
| 0 | 0 | 0 | 0 |
| 0 | 0 | 1 | 0 |
| 0 | 0 | 1 | 0 |
| 0 | 0 | 1 | 0 |

| | | | |
|---|---|---|---|
| 0 | 0 | 1 | 0 |
| 0 | 0 | 0 | 0 |
| 0 | 0 | 0 | 0 |
| 0 | 0 | 0 | 0 |
| 0 | 0 | 0 | 0 |
| 0 | 0 | 0 | 0 |
| 0 | 0 | 0 | 0 |
| 0 | 0 | 0 | 0 |
| 0 | 0 | 0 | 0 |
| 0 | 0 | 0 | 0 |
| 0 | 0 | 0 | 0 |
| 0 | 0 | 0 | 0 |
| 0 | 0 | 0 | 0 |
| 0 | 0 | 0 | 0 |
| 0 | 0 | 0 | 0 |
| 0 | 0 | 0 | 0 |
| 0 | 0 | 0 | 0 |
| 0 | 0 | 0 | 0 |
| 0 | 0 | 0 | 0 |
| 0 | 0 | 0 | 0 |
| 0 | 0 | 0 | 0 |
| 0 | 0 | 0 | 0 |
| 0 | 0 | 0 | 0 |
| 0 | 1 | 0 | 1 |
| 0 | 0 | 0 | 0 |
| 0 | 0 | 0 | 0 |
| 0 | 0 | 0 | 0 |
| 0 | 0 | 0 | 0 |
| 0 | 0 | 0 | 0 |
| 0 | 0 | 0 | 0 |
| 0 | 0 | 0 | 0 |
| 0 | 0 | 0 | 0 |
| 0 | 0 | 0 | 0 |
| 0 | 0 | 0 | 0 |
| 0 | 0 | 0 | 0 |
| 0 | 0 | 0 | 0 |
| 0 | 0 | 0 | 0 |
| 0 | 0 | 0 | 0 |
| 0 | 0 | 0 | 0 |
| 0 | 0 | 0 | 0 |
| 0 | 0 | 0 | 0 |
| 0 | 0 | 0 | 0 |
| 0 | 0 | 0 | 0 |
| 0 | 0 | 0 | 0 |
| 0 | 0 | 0 | 0 |
| 0 | 0 | 0 | 0 |
| 1 | 0 | 0 | 0 |
| 1 | 0 | 0 | 0 |
| 0 | 0 | 0 | 0 |
| 0 | 1 | 0 | 0 |
| 1 | 1 | 0 | 1 |
| 1 | 1 | 0 | 1 |
| 1 | 1 | 0 | 1 |
| 0 | 0 | 0 | 0 |

| | | | |
|---|---|---|---|
| 1 | 1 | 0 | 0 |
| 0 | 0 | 0 | 0 |
| 0 | 0 | 0 | 0 |
| 0 | 0 | 0 | 0 |
| 0 | 0 | 0 | 0 |
| 0 | 0 | 0 | 0 |
| 0 | 0 | 0 | 0 |
| 0 | 0 | 0 | 0 |
| 0 | 0 | 0 | 0 |
| 0 | 0 | 0 | 0 |
| 0 | 0 | 0 | 0 |
| 0 | 1 | 0 | 0 |
| 0 | 0 | 0 | 0 |
| 0 | 0 | 0 | 0 |
| 0 | 1 | 0 | 0 |
| 0 | 0 | 0 | 0 |
| 1 | 1 | 0 | 0 |
| 0 | 0 | 0 | 0 |
| 0 | 0 | 0 | 0 |
| 1 | 0 | 0 | 0 |
| 0 | 0 | 0 | 0 |
| 0 | 0 | 0 | 0 |
| 0 | 0 | 0 | 0 |
| 0 | 0 | 0 | 0 |
| 0 | 0 | 0 | 0 |
| 0 | 0 | 0 | 0 |
| 0 | 0 | 0 | 0 |
| 0 | 0 | 0 | 0 |
| 0 | 0 | 0 | 0 |
| 0 | 0 | 0 | 0 |
| 0 | 0 | 0 | 0 |
| 0 | 0 | 0 | 0 |
| 0 | 0 | 0 | 0 |
| 1 | 0 | 0 | 0 |
| 0 | 0 | 0 | 0 |
| 0 | 1 | 0 | 0 |
| 0 | 0 | 0 | 0 |
| 0 | 0 | 0 | 0 |
| 0 | 0 | 0 | 0 |
| 0 | 0 | 0 | 0 |
| 0 | 0 | 0 | 0 |
| 0 | 0 | 0 | 0 |
| 0 | 0 | 0 | 0 |
| 0 | 0 | 0 | 0 |
| 0 | 0 | 0 | 0 |
| 0 | 0 | 0 | 0 |
| 0 | 0 | 0 | 0 |
| 0 | 0 | 0 | 0 |
| 1 | 1 | 0 | 0 |
| 0 | 0 | 0 | 0 |
| 0 | 0 | 0 | 0 |
| 0 | 0 | 0 | 0 |
| 0 | 0 | 0 | 0 |
| 0 | 0 | 0 | 0 |

| | | | |
|---|---|---|---|
| 0 | 0 | 0 | 0 |
| 0 | 0 | 0 | 0 |
| 0 | 0 | 0 | 0 |
| 0 | 0 | 0 | 0 |
| 0 | 0 | 0 | 0 |
| 0 | 0 | 0 | 0 |
| 0 | 0 | 0 | 0 |
| 0 | 1 | 0 | 0 |
| 0 | 0 | 0 | 0 |
| 0 | 0 | 0 | 0 |
| 0 | 0 | 0 | 0 |
| 0 | 0 | 0 | 0 |
| 0 | 0 | 1 | 0 |
| 0 | 0 | 0 | 0 |
| 0 | 0 | 1 | 0 |
| 0 | 0 | 1 | 0 |
| 0 | 0 | 0 | 0 |
| 0 | 0 | 0 | 0 |
| 0 | 0 | 0 | 0 |
| 0 | 0 | 0 | 0 |
| 0 | 1 | 0 | 0 |
| 0 | 1 | 0 | 0 |
| 0 | 0 | 0 | 0 |
| 0 | 0 | 0 | 0 |
| 0 | 1 | 0 | 0 |
| 0 | 0 | 0 | 0 |
| 0 | 0 | 0 | 0 |
| 0 | 1 | 0 | 0 |
| 0 | 0 | 0 | 0 |
| 0 | 0 | 0 | 0 |
| 0 | 0 | 0 | 0 |
| 0 | 0 | 0 | 0 |
| 1 | 1 | 0 | 0 |
| 1 | 1 | 0 | 0 |
| 1 | 1 | 0 | 0 |
| 0 | 1 | 0 | 0 |
| 1 | 0 | 0 | 0 |
| 0 | 1 | 0 | 0 |
| 1 | 1 | 0 | 0 |
| 0 | 0 | 0 | 0 |
| 0 | 0 | 0 | 0 |
| 0 | 0 | 0 | 0 |
| 0 | 0 | 0 | 0 |
| 0 | 0 | 0 | 0 |
| 0 | 0 | 0 | 0 |
| 0 | 0 | 0 | 0 |
| 0 | 0 | 0 | 0 |
| 0 | 0 | 0 | 0 |
| 0 | 0 | 0 | 0 |
| 1 | 1 | 0 | 0 |
| 1 | 1 | 0 | 0 |
| 0 | 0 | 0 | 0 |
| 0 | 0 | 0 | 0 |
| 0 | 0 | 0 | 0 |

| | | | |
|---|---|---|---|
| 0 | 0 | 0 | 0 |
| 0 | 0 | 0 | 0 |
| 0 | 0 | 0 | 0 |
| 0 | 0 | 0 | 0 |
| 0 | 0 | 0 | 0 |
| 0 | 0 | 0 | 0 |
| 1 | 1 | 0 | 0 |
| 1 | 1 | 0 | 0 |
| 0 | 0 | 0 | 0 |
| 0 | 0 | 0 | 0 |
| 0 | 0 | 0 | 0 |
| 0 | 0 | 0 | 0 |
| 0 | 0 | 0 | 0 |
| 0 | 0 | 0 | 0 |
| 0 | 0 | 0 | 0 |
| 0 | 0 | 0 | 0 |
| 0 | 0 | 0 | 0 |
| 0 | 0 | 0 | 0 |
| 0 | 0 | 0 | 0 |
| 0 | 0 | 0 | 0 |
| 0 | 0 | 0 | 0 |
| 0 | 0 | 0 | 0 |
| 0 | 0 | 0 | 0 |
| 0 | 0 | 0 | 0 |
| 0 | 0 | 0 | 0 |
| 0 | 0 | 0 | 0 |
| 0 | 0 | 0 | 0 |
| 1 | 1 | 0 | 0 |
| 0 | 0 | 0 | 0 |
| 0 | 0 | 0 | 0 |
| 0 | 0 | 0 | 0 |
| 0 | 0 | 0 | 0 |
| 0 | 0 | 0 | 0 |
| 0 | 0 | 0 | 0 |
| 0 | 0 | 0 | 0 |
| 0 | 0 | 0 | 0 |
| 0 | 0 | 0 | 0 |
| 0 | 0 | 0 | 0 |
| 0 | 0 | 0 | 0 |
| 0 | 0 | 0 | 0 |
| 0 | 0 | 0 | 0 |
| 0 | 0 | 0 | 0 |
| 0 | 0 | 0 | 0 |
| 0 | 0 | 0 | 0 |
| 0 | 0 | 0 | 0 |
| 0 | 0 | 0 | 0 |
| 0 | 0 | 0 | 0 |
| 0 | 0 | 0 | 0 |
| 0 | 0 | 0 | 0 |
| 0 | 0 | 0 | 0 |
| 0 | 0 | 0 | 0 |
| 0 | 0 | 0 | 0 |
| 0 | 0 | 0 | 0 |
| 0 | 0 | 0 | 0 |

| | | | |
|---|---|---|---|
| 0 | 0 | 0 | 0 |
| 0 | 0 | 0 | 0 |
| 0 | 0 | 0 | 0 |
| 0 | 0 | 0 | 0 |
| 0 | 0 | 0 | 0 |
| 0 | 0 | 0 | 0 |
| 0 | 0 | 0 | 0 |
| 0 | 0 | 0 | 0 |
| 0 | 0 | 0 | 0 |
| 0 | 0 | 0 | 0 |
| 1 | 0 | 0 | 0 |
| 1 | 1 | 0 | 0 |
| 1 | 0 | 0 | 0 |
| 0 | 0 | 0 | 0 |
| 0 | 0 | 0 | 0 |
| 0 | 0 | 0 | 0 |
| 0 | 0 | 0 | 0 |
| 0 | 0 | 0 | 0 |
| 0 | 0 | 0 | 0 |
| 0 | 0 | 0 | 0 |
| 0 | 0 | 0 | 0 |
| 0 | 0 | 0 | 0 |
| 0 | 0 | 0 | 0 |
| 0 | 0 | 0 | 0 |
| 0 | 0 | 0 | 0 |
| 0 | 0 | 0 | 0 |
| 0 | 0 | 0 | 0 |
| 0 | 0 | 0 | 0 |
| 0 | 0 | 0 | 0 |
| 0 | 0 | 0 | 0 |
| 0 | 0 | 0 | 0 |
| 0 | 1 | 0 | 0 |
| 0 | 0 | 0 | 0 |
| 0 | 0 | 0 | 0 |
| 1 | 0 | 0 | 0 |
| 1 | 0 | 0 | 0 |
| 1 | 0 | 0 | 0 |
| 0 | 0 | 0 | 0 |
| 0 | 0 | 0 | 0 |
| 0 | 0 | 0 | 0 |
| 0 | 0 | 0 | 0 |
| 0 | 0 | 0 | 0 |
| 0 | 0 | 0 | 0 |
| 0 | 0 | 0 | 0 |
| 0 | 0 | 0 | 0 |
| 0 | 0 | 0 | 0 |
| 0 | 0 | 0 | 0 |
| 0 | 0 | 0 | 0 |
| 0 | 0 | 0 | 0 |
| 0 | 0 | 0 | 0 |
| 0 | 0 | 0 | 0 |
| 0 | 0 | 0 | 0 |
| 0 | 0 | 0 | 1 |
| 0 | 0 | 0 | 0 |

| | | | |
|---|---|---|---|
| 0 | 0 | 0 | 0 |
| 0 | 0 | 0 | 0 |
| 0 | 0 | 0 | 0 |
| 0 | 0 | 0 | 0 |
| 0 | 0 | 0 | 0 |
| 0 | 0 | 0 | 0 |
| 0 | 0 | 0 | 0 |
| 0 | 0 | 0 | 0 |
| 0 | 0 | 0 | 0 |
| 0 | 0 | 0 | 0 |
| 0 | 0 | 0 | 0 |
| 1 | 0 | 0 | 0 |
| 0 | 0 | 0 | 0 |
| 0 | 0 | 0 | 0 |
| 0 | 0 | 1 | 0 |
| 0 | 0 | 0 | 0 |
| 0 | 0 | 0 | 0 |
| 0 | 0 | 0 | 0 |
| 0 | 0 | 0 | 0 |
| 0 | 0 | 0 | 0 |
| 0 | 0 | 0 | 0 |
| 0 | 0 | 0 | 0 |
| 0 | 0 | 0 | 0 |
| 0 | 0 | 0 | 0 |
| 0 | 0 | 0 | 0 |
| 0 | 0 | 0 | 0 |
| 0 | 0 | 0 | 0 |
| 0 | 0 | 0 | 0 |
| 0 | 0 | 0 | 0 |
| 0 | 0 | 0 | 0 |
| 0 | 0 | 0 | 0 |
| 0 | 0 | 0 | 0 |
| 0 | 0 | 0 | 0 |
| 1 | 0 | 0 | 0 |
| 0 | 0 | 0 | 0 |
| 1 | 0 | 0 | 1 |
| 0 | 0 | 0 | 0 |
| 0 | 0 | 0 | 0 |
| 0 | 0 | 0 | 0 |
| 0 | 0 | 0 | 0 |
| 0 | 0 | 0 | 0 |
| 0 | 0 | 0 | 0 |
| 0 | 0 | 0 | 0 |
| 0 | 1 | 0 | 0 |
| 0 | 0 | 0 | 0 |
| 0 | 0 | 0 | 0 |
| 0 | 0 | 0 | 0 |
| 0 | 0 | 0 | 0 |
| 0 | 0 | 0 | 0 |
| 0 | 0 | 0 | 0 |
| 0 | 0 | 0 | 0 |
| 0 | 0 | 0 | 0 |
| 0 | 0 | 0 | 0 |
| 0 | 0 | 0 | 0 |

| | | | |
|---|---|---|---|
| 0 | 0 | 0 | 0 |
| 0 | 0 | 0 | 0 |
| 0 | 0 | 0 | 0 |
| 0 | 0 | 0 | 0 |
| 0 | 0 | 0 | 0 |
| 0 | 1 | 0 | 0 |
| 0 | 0 | 0 | 0 |
| 0 | 0 | 0 | 0 |
| 0 | 0 | 0 | 0 |
| 0 | 0 | 0 | 0 |
| 0 | 0 | 0 | 0 |
| 0 | 0 | 0 | 0 |
| 0 | 0 | 0 | 0 |
| 0 | 0 | 0 | 0 |
| 0 | 0 | 0 | 0 |
| 0 | 1 | 0 | 0 |
| 1 | 1 | 0 | 0 |
| 0 | 0 | 1 | 0 |
| 0 | 0 | 0 | 0 |
| 0 | 0 | 0 | 0 |
| 0 | 0 | 0 | 0 |
| 0 | 0 | 0 | 0 |
| 0 | 0 | 0 | 0 |
| 0 | 0 | 0 | 0 |
| 0 | 0 | 0 | 0 |
| 0 | 0 | 0 | 0 |
| 0 | 0 | 0 | 0 |
| 1 | 0 | 0 | 0 |
| 0 | 0 | 0 | 0 |
| 0 | 0 | 0 | 0 |
| 0 | 0 | 1 | 0 |
| 0 | 0 | 0 | 0 |
| 0 | 0 | 0 | 0 |
| 0 | 0 | 0 | 0 |
| 0 | 0 | 0 | 0 |
| 0 | 0 | 0 | 0 |
| 0 | 0 | 0 | 0 |
| 1 | 1 | 0 | 0 |
| 0 | 0 | 0 | 0 |
| 0 | 0 | 0 | 0 |
| 0 | 0 | 0 | 0 |
| 1 | 0 | 0 | 0 |
| 1 | 0 | 0 | 0 |
| 0 | 0 | 0 | 0 |
| 0 | 0 | 0 | 0 |
| 0 | 0 | 0 | 0 |
| 0 | 0 | 0 | 0 |
| 0 | 0 | 0 | 0 |
| 0 | 0 | 0 | 0 |
| 0 | 0 | 0 | 0 |
| 0 | 0 | 0 | 0 |
| 0 | 0 | 0 | 0 |
| 0 | 0 | 0 | 0 |
| 0 | 0 | 0 | 0 |

| | | | |
|---|---|---|---|
| 0 | 0 | 0 | 0 |
| 0 | 0 | 0 | 0 |
| 0 | 0 | 0 | 0 |
| 0 | 0 | 0 | 0 |
| 0 | 0 | 0 | 0 |
| 0 | 0 | 0 | 0 |
| 0 | 0 | 0 | 0 |
| 0 | 0 | 0 | 0 |
| 0 | 0 | 0 | 0 |
| 0 | 0 | 0 | 0 |
| 0 | 0 | 0 | 0 |
| 0 | 0 | 0 | 0 |
| 1 | 0 | 0 | 0 |
| 0 | 0 | 0 | 0 |
| 0 | 0 | 0 | 0 |
| 0 | 0 | 0 | 0 |
| 0 | 0 | 0 | 0 |
| 0 | 0 | 0 | 0 |
| 0 | 0 | 0 | 0 |
| 0 | 0 | 0 | 0 |
| 0 | 0 | 0 | 0 |
| 0 | 0 | 0 | 0 |
| 0 | 0 | 0 | 0 |
| 1 | 0 | 0 | 0 |
| 0 | 0 | 0 | 0 |
| 0 | 0 | 0 | 0 |
| 0 | 0 | 0 | 0 |
| 1 | 1 | 0 | 1 |
| 0 | 0 | 0 | 0 |
| 0 | 0 | 0 | 0 |
| 0 | 0 | 0 | 0 |
| 0 | 0 | 0 | 0 |
| 0 | 0 | 0 | 0 |
| 0 | 0 | 0 | 0 |
| 0 | 0 | 0 | 0 |
| 0 | 0 | 0 | 0 |
| 0 | 0 | 0 | 0 |
| 0 | 0 | 0 | 0 |
| 0 | 0 | 0 | 0 |
| 0 | 0 | 0 | 0 |
| 0 | 0 | 0 | 0 |
| 0 | 0 | 0 | 0 |
| 0 | 0 | 0 | 0 |
| 0 | 0 | 0 | 0 |
| 0 | 0 | 0 | 0 |
| 0 | 0 | 0 | 0 |
| 0 | 0 | 0 | 0 |
| 0 | 0 | 0 | 0 |
| 0 | 0 | 0 | 0 |
| 0 | 0 | 0 | 0 |
| 0 | 0 | 0 | 0 |
| 0 | 0 | 0 | 0 |
| 0 | 0 | 0 | 0 |
| 0 | 0 | 0 | 0 |

| | | | |
|---|---|---|---|
| 0 | 0 | 0 | 0 |
| 0 | 0 | 0 | 0 |
| 0 | 0 | 0 | 0 |
| 0 | 0 | 0 | 0 |
| 0 | 0 | 0 | 0 |
| 0 | 0 | 0 | 0 |
| 0 | 0 | 0 | 0 |
| 0 | 0 | 0 | 0 |
| 1 | 0 | 0 | 0 |
| 1 | 0 | 0 | 0 |
| 1 | 0 | 0 | 0 |
| 0 | 0 | 0 | 0 |
| 1 | 0 | 0 | 0 |
| 0 | 0 | 0 | 0 |
| 0 | 0 | 0 | 0 |
| 1 | 0 | 0 | 0 |
| 0 | 0 | 0 | 0 |
| 0 | 0 | 0 | 0 |
| 0 | 0 | 0 | 0 |
| 0 | 0 | 0 | 0 |
| 0 | 0 | 0 | 0 |
| 0 | 0 | 0 | 0 |
| 0 | 0 | 0 | 0 |
| 0 | 0 | 0 | 0 |
| 0 | 0 | 0 | 0 |
| 0 | 0 | 0 | 0 |
| 0 | 0 | 0 | 0 |
| 1 | 0 | 0 | 0 |
| 1 | 0 | 0 | 0 |
| 0 | 0 | 0 | 0 |
| 0 | 0 | 0 | 0 |
| 0 | 0 | 0 | 0 |
| 1 | 0 | 0 | 0 |
| 0 | 0 | 0 | 0 |
| 0 | 0 | 0 | 0 |
| 0 | 0 | 0 | 0 |
| 0 | 0 | 0 | 0 |
| 0 | 0 | 0 | 0 |
| 0 | 0 | 0 | 0 |
| 0 | 0 | 0 | 0 |
| 1 | 0 | 0 | 0 |
| 1 | 0 | 0 | 0 |
| 0 | 0 | 0 | 0 |
| 0 | 0 | 0 | 0 |
| 0 | 0 | 0 | 0 |
| 0 | 0 | 0 | 0 |
| 0 | 0 | 0 | 0 |
| 0 | 0 | 0 | 0 |
| 0 | 0 | 0 | 0 |
| 0 | 0 | 0 | 0 |
| 0 | 0 | 0 | 0 |
| 0 | 0 | 0 | 0 |
| 0 | 0 | 0 | 0 |
| 0 | 0 | 0 | 0 |

| | | | |
|---|---|---|---|
| 0 | 0 | 0 | 0 |
| 0 | 0 | 0 | 0 |
| 0 | 0 | 0 | 0 |
| 0 | 0 | 0 | 0 |
| 0 | 0 | 0 | 0 |
| 0 | 0 | 0 | 0 |
| 0 | 0 | 0 | 0 |
| 0 | 0 | 0 | 0 |
| 0 | 0 | 0 | 0 |
| 0 | 0 | 0 | 0 |
| 1 | 0 | 0 | 0 |
| 0 | 0 | 0 | 0 |
| 1 | 0 | 0 | 0 |
| 0 | 0 | 0 | 0 |
| 0 | 0 | 0 | 0 |
| 0 | 0 | 0 | 0 |
| 0 | 0 | 0 | 0 |
| 0 | 1 | 0 | 0 |
| 0 | 0 | 0 | 0 |
| 0 | 0 | 0 | 0 |
| 0 | 0 | 0 | 0 |
| 0 | 0 | 0 | 0 |
| 1 | 0 | 0 | 0 |
| 0 | 0 | 0 | 0 |
| 0 | 0 | 0 | 0 |
| 1 | 0 | 0 | 0 |
| 1 | 0 | 0 | 0 |
| 1 | 0 | 0 | 0 |
| 0 | 0 | 0 | 0 |
| 0 | 0 | 0 | 0 |
| 0 | 0 | 0 | 0 |
| 0 | 0 | 0 | 0 |
| 0 | 0 | 0 | 0 |
| 0 | 0 | 0 | 0 |
| 0 | 0 | 0 | 0 |
| 0 | 0 | 0 | 0 |
| 0 | 0 | 0 | 0 |
| 0 | 0 | 0 | 0 |
| 0 | 0 | 0 | 0 |
| 0 | 0 | 0 | 0 |
| 1 | 1 | 0 | 0 |
| 0 | 0 | 0 | 0 |
| 0 | 0 | 0 | 0 |
| 0 | 0 | 0 | 0 |
| 0 | 0 | 0 | 0 |
| 0 | 0 | 0 | 0 |
| 0 | 0 | 0 | 0 |
| 1 | 0 | 0 | 0 |
| 0 | 0 | 0 | 0 |
| 0 | 0 | 0 | 0 |
| 0 | 0 | 0 | 0 |
| 0 | 0 | 0 | 0 |
| 0 | 0 | 0 | 0 |
| 0 | 0 | 0 | 0 |

| | | | |
|---|---|---|---|
| 1 | 0 | 0 | 0 |
| 0 | 0 | 0 | 0 |
| 1 | 0 | 0 | 0 |
| 0 | 0 | 0 | 0 |
| 0 | 0 | 0 | 0 |
| 0 | 0 | 0 | 0 |
| 0 | 0 | 0 | 0 |
| 0 | 0 | 0 | 0 |
| 1 | 1 | 0 | 0 |
| 0 | 0 | 0 | 0 |
| 1 | 1 | 0 | 0 |
| 1 | 1 | 0 | 0 |
| 0 | 0 | 0 | 0 |
| 1 | 1 | 0 | 0 |
| 0 | 0 | 0 | 0 |
| 0 | 0 | 0 | 0 |
| 0 | 0 | 0 | 0 |
| 0 | 0 | 0 | 0 |
| 0 | 0 | 0 | 0 |
| 0 | 0 | 0 | 0 |
| 1 | 0 | 0 | 0 |
| 0 | 0 | 0 | 0 |
| 0 | 0 | 0 | 0 |
| 0 | 0 | 0 | 0 |
| 1 | 0 | 0 | 0 |
| 0 | 0 | 0 | 0 |
| 1 | 0 | 0 | 0 |
| 1 | 0 | 0 | 0 |
| 1 | 1 | 0 | 0 |
| 0 | 0 | 0 | 0 |
| 1 | 1 | 0 | 0 |
| 1 | 1 | 0 | 0 |
| 1 | 1 | 0 | 0 |
| 0 | 0 | 0 | 0 |
| 1 | 1 | 0 | 0 |
| 0 | 0 | 0 | 0 |
| 1 | 0 | 0 | 0 |
| 0 | 0 | 0 | 0 |
| 1 | 1 | 0 | 0 |
| 1 | 1 | 0 | 0 |
| 1 | 0 | 0 | 0 |
| 0 | 0 | 0 | 0 |
| 0 | 0 | 0 | 0 |
| 1 | 0 | 0 | 0 |
| 0 | 0 | 0 | 0 |
| 0 | 0 | 0 | 0 |
| 1 | 0 | 0 | 0 |
| 0 | 0 | 0 | 0 |
| 1 | 0 | 0 | 0 |
| 0 | 0 | 0 | 0 |
| 0 | 0 | 0 | 0 |
| 0 | 0 | 0 | 0 |
| 0 | 0 | 0 | 0 |
| 0 | 0 | 0 | 0 |

| | | | |
|---|---|---|---|
| 0 | 0 | 0 | 0 |
| 0 | 0 | 0 | 0 |
| 0 | 0 | 0 | 0 |
| 0 | 0 | 0 | 0 |
| 0 | 0 | 0 | 0 |
| 0 | 0 | 0 | 0 |
| 0 | 0 | 0 | 0 |
| 0 | 0 | 0 | 0 |
| 0 | 0 | 0 | 0 |
| 1 | 1 | 0 | 0 |
| 1 | 1 | 0 | 0 |
| 0 | 0 | 0 | 0 |
| 0 | 0 | 0 | 0 |
| 0 | 0 | 0 | 0 |
| 1 | 0 | 1 | 0 |
| 0 | 0 | 0 | 0 |
| 0 | 0 | 0 | 0 |
| 1 | 0 | 1 | 0 |
| 0 | 0 | 0 | 0 |
| 0 | 0 | 0 | 0 |
| 0 | 0 | 0 | 0 |
| 0 | 0 | 1 | 0 |
| 1 | 0 | 1 | 0 |
| 0 | 0 | 1 | 0 |
| 0 | 0 | 1 | 0 |
| 0 | 0 | 1 | 1 |
| 0 | 0 | 1 | 1 |
| 0 | 0 | 0 | 0 |
| 0 | 0 | 0 | 0 |
| 0 | 0 | 0 | 0 |
| 0 | 0 | 0 | 0 |
| 0 | 0 | 0 | 0 |
| 0 | 0 | 0 | 0 |
| 0 | 0 | 0 | 0 |
| 0 | 0 | 0 | 0 |
| 0 | 0 | 0 | 0 |
| 0 | 0 | 0 | 0 |
| 0 | 0 | 0 | 0 |
| 0 | 0 | 0 | 0 |
| 0 | 0 | 0 | 0 |
| 0 | 0 | 0 | 0 |
| 0 | 0 | 0 | 0 |
| 0 | 0 | 0 | 0 |
| 0 | 0 | 1 | 0 |
| 0 | 0 | 0 | 0 |
| 0 | 0 | 0 | 0 |
| 0 | 0 | 0 | 0 |
| 0 | 0 | 0 | 0 |
| 0 | 0 | 0 | 0 |
| 0 | 0 | 0 | 0 |
| 1 | 0 | 0 | 0 |
| 0 | 0 | 0 | 0 |
| 0 | 0 | 0 | 0 |
| 0 | 0 | 0 | 0 |

| | | | |
|---|---|---|---|
| 0 | 0 | 0 | 0 |
| 0 | 0 | 0 | 0 |
| 0 | 0 | 0 | 0 |
| 1 | 0 | 0 | 0 |
| 0 | 0 | 0 | 0 |
| 0 | 0 | 0 | 0 |
| 0 | 0 | 0 | 0 |
| 0 | 0 | 0 | 0 |
| 1 | 0 | 0 | 0 |
| 1 | 0 | 0 | 0 |
| 0 | 0 | 0 | 0 |
| 0 | 0 | 0 | 0 |
| 0 | 0 | 0 | 0 |
| 0 | 0 | 0 | 0 |
| 0 | 0 | 0 | 0 |
| 0 | 0 | 0 | 0 |
| 0 | 0 | 1 | 0 |
| 0 | 0 | 0 | 0 |
| 0 | 0 | 0 | 0 |
| 0 | 0 | 0 | 0 |
| 0 | 0 | 0 | 0 |
| 0 | 0 | 0 | 0 |
| 0 | 0 | 0 | 0 |
| 0 | 0 | 0 | 0 |
| 0 | 0 | 0 | 0 |
| 0 | 0 | 0 | 0 |
| 0 | 0 | 0 | 0 |
| 0 | 0 | 0 | 0 |
| 0 | 0 | 0 | 0 |
| 0 | 0 | 0 | 0 |
| 0 | 0 | 0 | 0 |
| 0 | 0 | 0 | 0 |
| 0 | 0 | 0 | 0 |
| 0 | 0 | 0 | 0 |
| 0 | 0 | 0 | 0 |
| 0 | 0 | 0 | 0 |
| 0 | 0 | 0 | 0 |
| 0 | 0 | 0 | 0 |
| 0 | 0 | 0 | 0 |
| 0 | 0 | 0 | 0 |
| 0 | 0 | 0 | 0 |
| 0 | 0 | 0 | 0 |
| 0 | 0 | 0 | 0 |
| 0 | 0 | 0 | 0 |
| 0 | 0 | 0 | 0 |
| 0 | 0 | 0 | 0 |
| 0 | 0 | 0 | 0 |
| 0 | 0 | 0 | 0 |
| 0 | 0 | 0 | 0 |
| 0 | 1 | 0 | 0 |
| 0 | 0 | 0 | 0 |
| 0 | 0 | 0 | 0 |
| 0 | 0 | 0 | 0 |
| 0 | 0 | 0 | 0 |

| | | | |
|---|---|---|---|
| 0 | 0 | 1 | 0 |
| 0 | 0 | 0 | 0 |
| 0 | 0 | 0 | 0 |
| 0 | 0 | 0 | 0 |
| 0 | 0 | 0 | 0 |
| 0 | 0 | 0 | 0 |
| 0 | 0 | 0 | 0 |
| 1 | 0 | 0 | 0 |
| 0 | 0 | 0 | 0 |
| 0 | 0 | 0 | 0 |
| 0 | 0 | 0 | 0 |
| 1 | 0 | 0 | 0 |
| 0 | 0 | 0 | 0 |
| 0 | 0 | 0 | 0 |
| 0 | 0 | 0 | 0 |
| 0 | 0 | 0 | 0 |
| 0 | 0 | 0 | 0 |
| 0 | 0 | 0 | 0 |
| 0 | 0 | 0 | 0 |
| 0 | 0 | 0 | 0 |
| 0 | 0 | 0 | 0 |
| 0 | 0 | 0 | 0 |
| 0 | 0 | 0 | 0 |
| 0 | 0 | 0 | 0 |
| 0 | 0 | 0 | 0 |
| 0 | 0 | 0 | 0 |
| 0 | 0 | 0 | 0 |
| 0 | 0 | 0 | 0 |
| 0 | 0 | 0 | 0 |
| 0 | 0 | 0 | 0 |
| 0 | 0 | 0 | 0 |
| 0 | 0 | 0 | 0 |
| 0 | 0 | 0 | 0 |
| 0 | 0 | 0 | 0 |
| 0 | 0 | 0 | 0 |
| 0 | 0 | 0 | 0 |
| 0 | 0 | 0 | 0 |
| 0 | 0 | 0 | 0 |
| 0 | 0 | 0 | 0 |
| 0 | 0 | 0 | 0 |
| 0 | 0 | 0 | 0 |
| 0 | 0 | 0 | 0 |
| 0 | 0 | 0 | 0 |
| 0 | 0 | 0 | 0 |
| 0 | 0 | 0 | 0 |
| 0 | 0 | 0 | 0 |
| 0 | 0 | 0 | 0 |
| 0 | 0 | 0 | 0 |
| 0 | 0 | 0 | 0 |
| 0 | 0 | 0 | 0 |
| 0 | 0 | 0 | 0 |
| 0 | 0 | 0 | 0 |
| 0 | 0 | 0 | 0 |
| 0 | 0 | 0 | 0 |

| | | | |
|---|---|---|---|
| 0 | 0 | 0 | 0 |
| 0 | 0 | 0 | 0 |
| 0 | 0 | 0 | 0 |
| 0 | 0 | 0 | 0 |
| 0 | 0 | 0 | 0 |
| 0 | 0 | 0 | 0 |
| 0 | 0 | 0 | 0 |
| 0 | 0 | 0 | 0 |
| 0 | 0 | 0 | 0 |
| 0 | 0 | 0 | 0 |
| 0 | 0 | 0 | 0 |
| 0 | 0 | 0 | 0 |
| 0 | 0 | 0 | 0 |
| 0 | 0 | 0 | 0 |
| 0 | 0 | 0 | 0 |
| 0 | 0 | 0 | 0 |
| 0 | 0 | 0 | 0 |
| 0 | 0 | 0 | 0 |
| 0 | 0 | 0 | 1 |
| 0 | 1 | 0 | 1 |
| 0 | 0 | 0 | 0 |
| 0 | 0 | 0 | 0 |
| 0 | 0 | 0 | 0 |
| 0 | 0 | 0 | 0 |
| 0 | 0 | 0 | 0 |
| 1 | 0 | 0 | 0 |
| 1 | 0 | 0 | 0 |
| 0 | 0 | 0 | 0 |
| 0 | 0 | 0 | 0 |
| 0 | 0 | 0 | 0 |
| 0 | 0 | 0 | 0 |
| 0 | 0 | 0 | 0 |
| 0 | 0 | 0 | 0 |
| 1 | 0 | 0 | 1 |
| 1 | 0 | 0 | 0 |
| 1 | 0 | 0 | 1 |
| 1 | 0 | 0 | 0 |
| 0 | 0 | 0 | 0 |
| 0 | 0 | 0 | 0 |
| 0 | 0 | 0 | 0 |
| 0 | 0 | 0 | 0 |
| 1 | 0 | 0 | 0 |
| 1 | 0 | 0 | 1 |
| 1 | 0 | 0 | 0 |
| 0 | 0 | 0 | 0 |
| 0 | 0 | 0 | 0 |
| 0 | 0 | 0 | 0 |
| 0 | 0 | 0 | 0 |
| 0 | 0 | 0 | 0 |
| 1 | 0 | 0 | 0 |
| 0 | 0 | 0 | 0 |
| 0 | 0 | 0 | 0 |
| 0 | 0 | 0 | 0 |
| 0 | 0 | 0 | 0 |

| | | | |
|---|---|---|---|
| 0 | 0 | 0 | 0 |
| 0 | 0 | 0 | 0 |
| 0 | 0 | 0 | 0 |
| 1 | 0 | 0 | 0 |
| 1 | 0 | 0 | 0 |
| 0 | 0 | 0 | 0 |
| 1 | 0 | 0 | 0 |
| 0 | 0 | 0 | 0 |
| 0 | 0 | 0 | 0 |
| 0 | 0 | 0 | 0 |
| 0 | 0 | 0 | 0 |
| 1 | 0 | 1 | 0 |
| 0 | 0 | 0 | 0 |
| 0 | 0 | 0 | 0 |
| 0 | 0 | 0 | 0 |
| 0 | 0 | 1 | 0 |
| 0 | 0 | 0 | 0 |
| 0 | 0 | 0 | 0 |
| 0 | 0 | 0 | 0 |
| 0 | 0 | 1 | 0 |
| 1 | 0 | 1 | 0 |
| 1 | 0 | 0 | 0 |
| 1 | 0 | 0 | 0 |
| 1 | 0 | 0 | 0 |
| 0 | 0 | 1 | 0 |
| 0 | 0 | 0 | 0 |
| 1 | 0 | 0 | 0 |
| 0 | 0 | 0 | 0 |
| 0 | 0 | 0 | 0 |
| 0 | 0 | 0 | 0 |
| 0 | 0 | 0 | 0 |
| 0 | 0 | 0 | 0 |
| 0 | 0 | 0 | 0 |
| 0 | 0 | 0 | 0 |
| 0 | 0 | 0 | 0 |
| 0 | 0 | 0 | 0 |
| 0 | 0 | 0 | 1 |
| 0 | 0 | 0 | 1 |
| 0 | 0 | 0 | 1 |
| 0 | 0 | 0 | 0 |
| 0 | 0 | 0 | 0 |
| 0 | 0 | 0 | 0 |
| 0 | 0 | 0 | 0 |
| 0 | 0 | 0 | 0 |
| 0 | 0 | 0 | 0 |
| 0 | 0 | 0 | 0 |
| 0 | 0 | 0 | 0 |
| 0 | 0 | 0 | 0 |
| 0 | 0 | 0 | 0 |
| 0 | 0 | 0 | 0 |
| 0 | 0 | 0 | 0 |
| 0 | 0 | 0 | 0 |
| 0 | 0 | 0 | 0 |
| 0 | 0 | 0 | 0 |

| | | | |
|---|---|---|---|
| 0 | 0 | 0 | 0 |
| 0 | 0 | 0 | 0 |
| 0 | 0 | 0 | 0 |
| 0 | 0 | 0 | 0 |
| 0 | 0 | 0 | 0 |
| 0 | 0 | 0 | 0 |
| 0 | 0 | 0 | 0 |
| 0 | 0 | 0 | 0 |
| 0 | 0 | 0 | 0 |
| 0 | 0 | 0 | 0 |
| 0 | 0 | 0 | 0 |
| 0 | 0 | 0 | 0 |
| 0 | 0 | 0 | 0 |
| 0 | 0 | 0 | 0 |
| 0 | 0 | 0 | 0 |
| 0 | 0 | 0 | 0 |
| 0 | 1 | 0 | 0 |
| 0 | 1 | 0 | 0 |
| 0 | 0 | 0 | 0 |
| 0 | 0 | 0 | 0 |
| 0 | 0 | 0 | 0 |
| 0 | 0 | 0 | 0 |
| 0 | 0 | 0 | 0 |
| 0 | 0 | 0 | 0 |
| 0 | 0 | 0 | 0 |
| 0 | 0 | 0 | 0 |
| 0 | 0 | 0 | 0 |
| 0 | 0 | 0 | 0 |
| 0 | 0 | 0 | 0 |
| 0 | 0 | 0 | 0 |
| 0 | 0 | 0 | 0 |
| 1 | 0 | 0 | 0 |
| 0 | 0 | 0 | 0 |
| 0 | 0 | 0 | 0 |
| 0 | 0 | 0 | 0 |
| 0 | 0 | 0 | 0 |
| 0 | 0 | 0 | 0 |
| 0 | 0 | 0 | 0 |
| 0 | 0 | 0 | 0 |
| 0 | 0 | 0 | 0 |
| 0 | 0 | 0 | 0 |
| 0 | 0 | 0 | 0 |
| 0 | 0 | 0 | 0 |
| 0 | 0 | 0 | 0 |
| 0 | 0 | 0 | 0 |
| 0 | 0 | 0 | 0 |
| 1 | 1 | 0 | 0 |
| 0 | 0 | 0 | 0 |
| 0 | 0 | 0 | 0 |
| 0 | 0 | 0 | 0 |
| 0 | 0 | 0 | 0 |
| 0 | 0 | 0 | 0 |
| 1 | 0 | 0 | 0 |
| 0 | 0 | 0 | 0 |

| | | | |
|---|---|---|---|
| 0 | 0 | 0 | 0 |
| 0 | 0 | 0 | 0 |
| 0 | 0 | 0 | 0 |
| 0 | 0 | 0 | 0 |
| 0 | 0 | 0 | 0 |
| 0 | 0 | 0 | 0 |
| 1 | 0 | 0 | 0 |
| 0 | 0 | 0 | 0 |
| 0 | 0 | 0 | 0 |
| 0 | 0 | 0 | 0 |
| 0 | 0 | 0 | 0 |
| 0 | 0 | 0 | 0 |
| 0 | 0 | 0 | 0 |
| 0 | 0 | 0 | 0 |
| 0 | 0 | 0 | 0 |
| 0 | 0 | 0 | 0 |
| 0 | 0 | 0 | 0 |
| 0 | 0 | 0 | 0 |
| 0 | 0 | 0 | 0 |
| 0 | 0 | 0 | 0 |
| 0 | 0 | 0 | 0 |
| 0 | 0 | 0 | 0 |
| 0 | 0 | 0 | 0 |
| 0 | 0 | 0 | 0 |
| 0 | 0 | 0 | 0 |
| 0 | 0 | 0 | 0 |
| 0 | 0 | 0 | 0 |
| 0 | 0 | 0 | 0 |
| 0 | 0 | 0 | 0 |
| 0 | 0 | 0 | 0 |
| 0 | 0 | 0 | 0 |
| 0 | 0 | 0 | 0 |
| 0 | 0 | 0 | 0 |
| 0 | 0 | 0 | 0 |
| 0 | 0 | 0 | 0 |
| 0 | 0 | 0 | 0 |
| 0 | 0 | 0 | 0 |
| 0 | 0 | 0 | 0 |
| 0 | 0 | 0 | 0 |
| 0 | 0 | 0 | 0 |
| 0 | 0 | 0 | 0 |
| 0 | 0 | 0 | 0 |
| 0 | 0 | 0 | 0 |
| 0 | 0 | 0 | 0 |
| 0 | 0 | 0 | 0 |
| 0 | 0 | 0 | 0 |
| 0 | 0 | 0 | 0 |
| 0 | 0 | 0 | 0 |
| 0 | 0 | 0 | 0 |
| 0 | 0 | 0 | 0 |
| 0 | 0 | 0 | 0 |
| 0 | 0 | 0 | 0 |
| 0 | 0 | 0 | 0 |
| 0 | 0 | 0 | 0 |

| | | | |
|---|---|---|---|
| 0 | 0 | 0 | 0 |
| 0 | 0 | 0 | 0 |
| 0 | 0 | 0 | 0 |
| 0 | 0 | 0 | 0 |
| 0 | 0 | 0 | 0 |
| 0 | 0 | 0 | 0 |
| 0 | 0 | 0 | 0 |
| 0 | 0 | 0 | 0 |
| 0 | 0 | 0 | 0 |
| 0 | 0 | 0 | 0 |
| 0 | 0 | 0 | 0 |
| 0 | 0 | 0 | 0 |
| 0 | 0 | 0 | 0 |
| 0 | 0 | 0 | 0 |
| 0 | 0 | 0 | 0 |
| 0 | 0 | 0 | 0 |
| 0 | 0 | 0 | 0 |
| 0 | 0 | 0 | 0 |
| 0 | 0 | 0 | 0 |
| 0 | 0 | 0 | 0 |
| 0 | 0 | 0 | 0 |
| 0 | 0 | 0 | 0 |
| 0 | 0 | 0 | 0 |
| 0 | 0 | 0 | 0 |
| 1 | 1 | 0 | 0 |
| 0 | 0 | 0 | 0 |
| 0 | 0 | 0 | 0 |
| 0 | 0 | 0 | 0 |
| 0 | 0 | 0 | 0 |
| 0 | 0 | 0 | 0 |
| 0 | 0 | 0 | 0 |
| 0 | 0 | 0 | 0 |
| 0 | 0 | 0 | 0 |
| 0 | 0 | 0 | 0 |
| 0 | 0 | 0 | 0 |
| 1 | 0 | 0 | 0 |
| 1 | 0 | 0 | 1 |
| 1 | 0 | 0 | 0 |
| 0 | 0 | 0 | 0 |
| 1 | 0 | 0 | 0 |
| 1 | 1 | 0 | 0 |
| 1 | 1 | 0 | 0 |
| 0 | 0 | 0 | 0 |
| 0 | 0 | 0 | 0 |
| 0 | 0 | 0 | 0 |
| 0 | 0 | 0 | 0 |
| 0 | 0 | 0 | 0 |
| 0 | 0 | 0 | 0 |
| 0 | 0 | 0 | 0 |
| 0 | 0 | 0 | 0 |
| 0 | 0 | 0 | 0 |
| 0 | 0 | 0 | 0 |
| 0 | 0 | 0 | 0 |
| 0 | 0 | 0 | 0 |

| | | | |
|---|---|---|---|
| 0 | 0 | 0 | 0 |
| 0 | 0 | 0 | 0 |
| 0 | 0 | 0 | 0 |
| 0 | 0 | 0 | 0 |
| 0 | 0 | 0 | 0 |
| 0 | 0 | 0 | 0 |
| 0 | 0 | 0 | 0 |
| 0 | 0 | 0 | 0 |
| 0 | 0 | 0 | 0 |
| 0 | 0 | 0 | 0 |
| 0 | 0 | 0 | 0 |
| 0 | 0 | 0 | 0 |
| 0 | 0 | 0 | 0 |
| 1 | 0 | 0 | 0 |
| 0 | 0 | 0 | 0 |
| 0 | 0 | 0 | 0 |
| 1 | 1 | 0 | 0 |
| 0 | 0 | 0 | 0 |
| 0 | 0 | 0 | 0 |
| 0 | 0 | 0 | 0 |
| 0 | 0 | 0 | 0 |
| 0 | 0 | 0 | 0 |
| 0 | 0 | 0 | 0 |
| 1 | 0 | 0 | 0 |
| 0 | 0 | 0 | 0 |
| 0 | 0 | 0 | 0 |
| 0 | 0 | 0 | 0 |
| 0 | 0 | 0 | 0 |
| 0 | 0 | 0 | 0 |
| 0 | 0 | 0 | 0 |
| 0 | 0 | 0 | 0 |
| 1 | 0 | 0 | 0 |
| 0 | 0 | 0 | 0 |
| 0 | 0 | 0 | 0 |
| 0 | 0 | 0 | 0 |
| 0 | 0 | 0 | 0 |
| 0 | 0 | 0 | 0 |
| 0 | 0 | 0 | 0 |
| 0 | 0 | 0 | 0 |
| 0 | 0 | 0 | 0 |
| 0 | 0 | 0 | 0 |
| 0 | 0 | 0 | 0 |
| 0 | 0 | 0 | 0 |
| 0 | 0 | 0 | 0 |
| 0 | 0 | 0 | 0 |
| 0 | 0 | 0 | 0 |
| 0 | 0 | 0 | 0 |
| 0 | 0 | 0 | 0 |
| 0 | 0 | 0 | 0 |
| 0 | 0 | 0 | 0 |
| 0 | 0 | 0 | 0 |
| 0 | 0 | 0 | 0 |
| 0 | 0 | 0 | 0 |
| 0 | 0 | 0 | 0 |

| | | | |
|---|---|---|---|
| 0 | 0 | 0 | 0 |
| 0 | 0 | 0 | 0 |
| 0 | 0 | 0 | 0 |
| 0 | 0 | 0 | 0 |
| 0 | 0 | 0 | 0 |
| 0 | 0 | 0 | 0 |
| 0 | 0 | 0 | 0 |
| 0 | 0 | 0 | 0 |
| 0 | 0 | 0 | 0 |
| 0 | 0 | 0 | 0 |
| 0 | 0 | 0 | 0 |
| 0 | 0 | 0 | 0 |
| 0 | 0 | 0 | 0 |
| 0 | 0 | 0 | 0 |
| 0 | 0 | 0 | 0 |
| 0 | 0 | 0 | 0 |
| 0 | 0 | 0 | 0 |
| 1 | 0 | 0 | 0 |
| 0 | 0 | 0 | 0 |
| 0 | 0 | 0 | 0 |
| 0 | 0 | 0 | 0 |
| 0 | 0 | 0 | 0 |
| 0 | 0 | 0 | 0 |
| 0 | 0 | 0 | 0 |
| 1 | 0 | 0 | 0 |
| 0 | 0 | 0 | 0 |
| 0 | 0 | 0 | 0 |
| 0 | 0 | 0 | 0 |
| 0 | 0 | 0 | 0 |
| 0 | 0 | 0 | 0 |
| 0 | 0 | 0 | 0 |
| 0 | 0 | 0 | 0 |
| 0 | 0 | 0 | 0 |
| 0 | 0 | 0 | 0 |
| 0 | 0 | 0 | 0 |
| 0 | 0 | 0 | 0 |
| 0 | 0 | 0 | 0 |
| 0 | 0 | 0 | 0 |
| 0 | 0 | 0 | 0 |
| 0 | 0 | 0 | 0 |
| 0 | 0 | 0 | 0 |
| 0 | 0 | 0 | 0 |
| 0 | 0 | 0 | 0 |
| 0 | 0 | 0 | 0 |
| 0 | 0 | 0 | 0 |
| 0 | 0 | 0 | 0 |
| 0 | 0 | 0 | 0 |
| 0 | 0 | 0 | 0 |
| 0 | 0 | 0 | 0 |
| 0 | 0 | 0 | 0 |
| 0 | 0 | 0 | 0 |
| 0 | 0 | 0 | 0 |
| 0 | 0 | 0 | 0 |
| 0 | 0 | 0 | 0 |

| | | | |
|---|---|---|---|
| 0 | 0 | 0 | 0 |
| 0 | 0 | 0 | 0 |
| 0 | 0 | 0 | 0 |
| 0 | 0 | 0 | 0 |
| 0 | 0 | 0 | 0 |
| 0 | 0 | 0 | 0 |
| 0 | 0 | 0 | 0 |
| 0 | 0 | 0 | 0 |
| 0 | 0 | 0 | 0 |
| 0 | 0 | 0 | 0 |
| 0 | 0 | 0 | 0 |
| 0 | 0 | 0 | 0 |
| 0 | 0 | 0 | 0 |
| 0 | 0 | 0 | 0 |
| 1 | 1 | 0 | 0 |
| 1 | 1 | 0 | 0 |
| 0 | 0 | 0 | 0 |
| 0 | 0 | 0 | 0 |
| 0 | 0 | 0 | 0 |
| 0 | 0 | 0 | 0 |
| 0 | 0 | 0 | 0 |
| 0 | 0 | 0 | 0 |
| 0 | 0 | 0 | 0 |
| 0 | 0 | 0 | 0 |
| 0 | 0 | 0 | 0 |
| 0 | 0 | 0 | 0 |
| 0 | 0 | 0 | 0 |
| 1 | 0 | 0 | 0 |
| 0 | 0 | 0 | 0 |
| 1 | 0 | 0 | 0 |
| 0 | 0 | 0 | 0 |
| 0 | 0 | 0 | 0 |
| 0 | 0 | 0 | 0 |
| 0 | 0 | 0 | 0 |
| 0 | 0 | 0 | 0 |
| 0 | 0 | 0 | 0 |
| 1 | 0 | 0 | 0 |
| 0 | 0 | 0 | 0 |
| 0 | 0 | 0 | 0 |
| 0 | 0 | 0 | 0 |
| 0 | 0 | 0 | 0 |
| 0 | 0 | 0 | 0 |
| 0 | 0 | 0 | 0 |
| 0 | 0 | 0 | 0 |
| 0 | 0 | 0 | 0 |
| 0 | 0 | 0 | 0 |
| 0 | 0 | 0 | 0 |
| 0 | 0 | 0 | 0 |
| 0 | 0 | 0 | 0 |
| 0 | 0 | 0 | 0 |
| 0 | 0 | 0 | 0 |
| 0 | 0 | 0 | 0 |
| 0 | 0 | 0 | 0 |
| 0 | 0 | 0 | 0 |

| | | | |
|---|---|---|---|
| 0 | 0 | 0 | 0 |
| 0 | 0 | 0 | 0 |
| 0 | 0 | 0 | 0 |
| 0 | 0 | 0 | 0 |
| 0 | 0 | 0 | 0 |
| 0 | 0 | 0 | 0 |
| 0 | 0 | 0 | 0 |
| 0 | 0 | 0 | 0 |
| 0 | 0 | 0 | 0 |
| 0 | 0 | 0 | 0 |
| 0 | 0 | 0 | 0 |
| 0 | 0 | 0 | 0 |
| 0 | 0 | 0 | 0 |
| 0 | 0 | 0 | 0 |
| 0 | 0 | 0 | 0 |
| 0 | 0 | 0 | 0 |
| 0 | 0 | 0 | 0 |
| 0 | 0 | 0 | 0 |
| 0 | 0 | 0 | 0 |
| 0 | 0 | 0 | 0 |
| 0 | 0 | 0 | 0 |
| 0 | 0 | 0 | 0 |
| 0 | 0 | 0 | 0 |
| 0 | 0 | 0 | 0 |
| 0 | 0 | 0 | 0 |
| 0 | 0 | 0 | 0 |
| 0 | 0 | 0 | 0 |
| 0 | 0 | 0 | 0 |
| 0 | 0 | 0 | 0 |
| 0 | 0 | 0 | 0 |
| 0 | 0 | 0 | 0 |
| 0 | 0 | 0 | 0 |
| 0 | 0 | 0 | 0 |
| 0 | 0 | 0 | 0 |
| 0 | 0 | 0 | 0 |
| 0 | 0 | 0 | 0 |
| 0 | 0 | 0 | 0 |
| 0 | 0 | 0 | 0 |
| 0 | 0 | 0 | 0 |
| 0 | 0 | 0 | 0 |
| 0 | 0 | 0 | 0 |
| 0 | 0 | 0 | 0 |
| 0 | 0 | 0 | 0 |
| 0 | 0 | 0 | 0 |
| 0 | 0 | 0 | 0 |
| 0 | 0 | 0 | 0 |
| 0 | 0 | 0 | 0 |
| 0 | 0 | 0 | 0 |
| 1 | 0 | 0 | 1 |
| 0 | 0 | 0 | 0 |
| 1 | 0 | 0 | 0 |
| 1 | 0 | 0 | 0 |
| 0 | 0 | 0 | 0 |
| 0 | 0 | 0 | 1 |

| | | | |
|---|---|---|---|
| 0 | 0 | 0 | 0 |
| 0 | 0 | 0 | 0 |
| 0 | 0 | 0 | 0 |
| 0 | 0 | 0 | 0 |
| 0 | 0 | 0 | 0 |
| 0 | 0 | 0 | 0 |
| 0 | 0 | 0 | 0 |
| 0 | 0 | 0 | 0 |
| 0 | 0 | 0 | 0 |
| 0 | 0 | 0 | 0 |
| 0 | 0 | 0 | 0 |
| 0 | 0 | 0 | 0 |
| 0 | 0 | 0 | 0 |
| 0 | 0 | 0 | 0 |
| 0 | 0 | 0 | 0 |
| 0 | 0 | 0 | 0 |
| 0 | 0 | 0 | 0 |
| 0 | 0 | 0 | 0 |
| 0 | 0 | 0 | 0 |
| 0 | 0 | 0 | 0 |
| 0 | 0 | 0 | 0 |
| 0 | 0 | 0 | 0 |
| 0 | 0 | 0 | 0 |
| 0 | 0 | 0 | 0 |
| 0 | 0 | 0 | 0 |
| 0 | 0 | 0 | 0 |
| 0 | 0 | 1 | 0 |
| 0 | 0 | 1 | 0 |
| 0 | 0 | 1 | 0 |
| 0 | 0 | 1 | 0 |
| 0 | 0 | 0 | 0 |
| 0 | 0 | 0 | 0 |
| 0 | 0 | 0 | 0 |
| 1 | 0 | 0 | 0 |
| 1 | 0 | 0 | 0 |
| 0 | 0 | 0 | 0 |
| 0 | 0 | 0 | 0 |
| 0 | 0 | 0 | 0 |
| 0 | 0 | 0 | 0 |
| 1 | 0 | 0 | 0 |
| 0 | 0 | 0 | 0 |
| 0 | 0 | 0 | 0 |
| 0 | 0 | 0 | 0 |
| 0 | 0 | 0 | 0 |
| 0 | 0 | 0 | 0 |
| 0 | 1 | 0 | 0 |
| 0 | 0 | 0 | 0 |
| 0 | 0 | 0 | 0 |
| 0 | 0 | 0 | 0 |
| 0 | 0 | 0 | 0 |
| 0 | 0 | 0 | 0 |
| 0 | 0 | 0 | 0 |
| 0 | 0 | 0 | 0 |
| 0 | 0 | 0 | 0 |

| | | | |
|---|---|---|---|
| 0 | 0 | 0 | 0 |
| 0 | 0 | 0 | 0 |
| 0 | 0 | 0 | 0 |
| 0 | 0 | 0 | 0 |
| 0 | 0 | 0 | 0 |
| 0 | 0 | 0 | 1 |
| 0 | 0 | 0 | 0 |
| 0 | 0 | 0 | 0 |
| 0 | 0 | 0 | 0 |
| 0 | 0 | 0 | 0 |
| 0 | 0 | 0 | 0 |
| 0 | 0 | 0 | 0 |
| 0 | 0 | 0 | 0 |
| 0 | 1 | 0 | 0 |
| 0 | 1 | 0 | 0 |
| 0 | 1 | 0 | 0 |
| 0 | 1 | 0 | 0 |
| 0 | 1 | 0 | 0 |
| 0 | 0 | 0 | 0 |
| 0 | 0 | 0 | 0 |
| 0 | 0 | 0 | 0 |
| 0 | 0 | 0 | 0 |
| 0 | 0 | 0 | 0 |
| 0 | 0 | 0 | 0 |
| 0 | 0 | 0 | 0 |
| 0 | 0 | 0 | 0 |
| 0 | 0 | 0 | 0 |
| 0 | 0 | 0 | 0 |
| 0 | 0 | 0 | 0 |
| 0 | 0 | 0 | 0 |
| 0 | 0 | 0 | 0 |
| 0 | 0 | 0 | 0 |
| 0 | 0 | 0 | 0 |
| 0 | 0 | 0 | 0 |
| 0 | 0 | 0 | 0 |
| 0 | 0 | 0 | 0 |
| 0 | 0 | 0 | 0 |
| 0 | 0 | 0 | 0 |
| 0 | 0 | 0 | 0 |
| 0 | 0 | 0 | 0 |
| 0 | 0 | 0 | 0 |
| 0 | 0 | 0 | 0 |
| 0 | 0 | 0 | 0 |
| 0 | 0 | 0 | 0 |
| 0 | 0 | 0 | 0 |
| 0 | 0 | 0 | 0 |
| 0 | 0 | 0 | 0 |
| 0 | 0 | 0 | 0 |
| 0 | 0 | 0 | 0 |
| 0 | 0 | 0 | 0 |
| 0 | 0 | 0 | 0 |
| 0 | 0 | 0 | 0 |
| 0 | 0 | 0 | 0 |
| 0 | 0 | 0 | 0 |

| | | | |
|---|---|---|---|
| 0 | 0 | 0 | 0 |
| 0 | 0 | 0 | 0 |
| 0 | 0 | 0 | 0 |
| 1 | 0 | 0 | 0 |
| 0 | 0 | 0 | 0 |
| 0 | 0 | 0 | 0 |
| 0 | 0 | 0 | 0 |
| 0 | 0 | 0 | 0 |
| 0 | 0 | 0 | 0 |
| 0 | 0 | 0 | 0 |
| 0 | 0 | 0 | 0 |
| 0 | 0 | 0 | 0 |
| 0 | 0 | 0 | 0 |
| 0 | 0 | 0 | 0 |
| 0 | 0 | 0 | 0 |
| 0 | 0 | 0 | 0 |
| 0 | 0 | 0 | 0 |
| 0 | 0 | 0 | 0 |
| 0 | 0 | 0 | 0 |
| 0 | 0 | 0 | 0 |
| 1 | 1 | 0 | 0 |
| 1 | 1 | 0 | 0 |
| 0 | 0 | 0 | 0 |
| 0 | 1 | 0 | 0 |
| 0 | 0 | 0 | 0 |
| 0 | 0 | 0 | 0 |
| 0 | 0 | 0 | 0 |
| 0 | 0 | 0 | 0 |
| 0 | 0 | 0 | 0 |
| 0 | 0 | 0 | 0 |
| 0 | 0 | 0 | 0 |
| 0 | 0 | 0 | 0 |
| 0 | 0 | 0 | 0 |
| 0 | 0 | 0 | 0 |
| 0 | 0 | 0 | 0 |
| 0 | 0 | 0 | 0 |
| 0 | 0 | 0 | 0 |
| 0 | 0 | 0 | 0 |
| 0 | 0 | 0 | 0 |
| 1 | 0 | 0 | 0 |
| 1 | 0 | 0 | 0 |
| 0 | 0 | 0 | 0 |
| 0 | 0 | 0 | 0 |
| 1 | 0 | 0 | 0 |
| 0 | 0 | 0 | 0 |
| 0 | 0 | 0 | 0 |
| 0 | 0 | 0 | 0 |
| 0 | 0 | 0 | 0 |
| 0 | 0 | 0 | 0 |
| 0 | 0 | 0 | 0 |
| 0 | 0 | 0 | 0 |
| 0 | 0 | 0 | 0 |
| 0 | 0 | 0 | 0 |
| 0 | 0 | 0 | 0 |

| | | | |
|---|---|---|---|
| 0 | 0 | 0 | 0 |
| 0 | 0 | 0 | 0 |
| 0 | 0 | 0 | 0 |
| 0 | 0 | 0 | 0 |
| 0 | 0 | 0 | 0 |
| 1 | 0 | 0 | 0 |
| 0 | 0 | 0 | 0 |
| 0 | 0 | 0 | 0 |
| 0 | 1 | 0 | 0 |
| 1 | 1 | 0 | 0 |
| 0 | 0 | 0 | 0 |
| 1 | 1 | 0 | 0 |
| 1 | 0 | 0 | 0 |
| 0 | 0 | 0 | 0 |
| 0 | 0 | 0 | 0 |
| 0 | 0 | 0 | 0 |
| 1 | 1 | 0 | 0 |
| 0 | 0 | 0 | 0 |
| 0 | 0 | 0 | 0 |
| 1 | 0 | 0 | 0 |
| 0 | 0 | 0 | 0 |
| 0 | 0 | 0 | 0 |
| 0 | 0 | 0 | 0 |
| 0 | 0 | 0 | 0 |
| 0 | 0 | 0 | 0 |
| 0 | 0 | 0 | 0 |
| 0 | 0 | 0 | 0 |
| 0 | 0 | 0 | 0 |
| 0 | 0 | 0 | 0 |
| 0 | 0 | 0 | 0 |
| 1 | 0 | 0 | 0 |
| 0 | 0 | 0 | 0 |
| 0 | 0 | 0 | 0 |
| 0 | 0 | 0 | 0 |
| 0 | 0 | 0 | 0 |
| 1 | 0 | 0 | 0 |
| 0 | 0 | 0 | 0 |
| 0 | 0 | 0 | 0 |
| 0 | 0 | 0 | 0 |
| 0 | 0 | 0 | 0 |
| 0 | 0 | 0 | 0 |
| 0 | 0 | 0 | 0 |
| 0 | 0 | 0 | 0 |
| 0 | 0 | 0 | 0 |
| 0 | 0 | 0 | 0 |
| 0 | 0 | 0 | 0 |
| 0 | 0 | 0 | 0 |
| 0 | 0 | 0 | 0 |
| 0 | 0 | 0 | 0 |
| 0 | 0 | 0 | 0 |
| 0 | 0 | 0 | 0 |
| 0 | 0 | 0 | 0 |
| 0 | 0 | 0 | 0 |
| 0 | 0 | 0 | 0 |

| | | | |
|---|---|---|---|
| 0 | 0 | 0 | 0 |
| 0 | 0 | 0 | 0 |
| 0 | 0 | 0 | 0 |
| 0 | 0 | 0 | 0 |
| 0 | 0 | 0 | 0 |
| 0 | 0 | 0 | 0 |
| 0 | 0 | 0 | 0 |
| 0 | 0 | 0 | 0 |
| 0 | 0 | 0 | 0 |
| 0 | 0 | 0 | 0 |
| 0 | 0 | 0 | 0 |
| 0 | 0 | 0 | 0 |
| 0 | 0 | 0 | 0 |
| 0 | 0 | 0 | 0 |
| 0 | 0 | 0 | 1 |
| 0 | 0 | 0 | 0 |
| 0 | 0 | 0 | 0 |
| 0 | 0 | 0 | 0 |
| 0 | 0 | 0 | 0 |
| 0 | 0 | 0 | 0 |
| 0 | 0 | 0 | 0 |
| 0 | 0 | 0 | 0 |
| 0 | 0 | 0 | 0 |
| 0 | 0 | 0 | 0 |
| 0 | 0 | 0 | 0 |
| 0 | 0 | 0 | 0 |
| 0 | 0 | 0 | 0 |
| 0 | 0 | 0 | 0 |
| 0 | 0 | 0 | 0 |
| 0 | 0 | 0 | 0 |
| 0 | 0 | 0 | 0 |
| 0 | 0 | 0 | 0 |
| 0 | 0 | 0 | 0 |
| 0 | 0 | 0 | 0 |
| 0 | 0 | 0 | 0 |
| 0 | 0 | 0 | 0 |
| 0 | 0 | 0 | 0 |
| 0 | 0 | 0 | 0 |
| 0 | 0 | 0 | 0 |
| 0 | 0 | 1 | 0 |
| 0 | 0 | 0 | 0 |
| 0 | 0 | 0 | 0 |
| 0 | 0 | 0 | 0 |
| 1 | 0 | 0 | 0 |
| 0 | 0 | 0 | 0 |
| 0 | 0 | 0 | 0 |
| 0 | 0 | 0 | 0 |
| 0 | 0 | 0 | 0 |
| 1 | 0 | 0 | 1 |
| 0 | 0 | 0 | 0 |
| 0 | 0 | 0 | 0 |
| 0 | 0 | 0 | 0 |
| 0 | 0 | 0 | 0 |
| 0 | 0 | 0 | 0 |

| | | | |
|---|---|---|---|
| 0 | 0 | 0 | 0 |
| 0 | 0 | 0 | 0 |
| 0 | 0 | 0 | 0 |
| 0 | 0 | 0 | 0 |
| 0 | 0 | 0 | 0 |
| 0 | 0 | 0 | 0 |
| 0 | 0 | 0 | 0 |
| 0 | 0 | 0 | 0 |
| 0 | 0 | 0 | 0 |
| 1 | 0 | 0 | 0 |
| 0 | 0 | 0 | 0 |
| 0 | 0 | 0 | 0 |
| 0 | 0 | 0 | 0 |
| 0 | 0 | 0 | 0 |
| 1 | 0 | 0 | 0 |
| 0 | 0 | 0 | 0 |
| 0 | 0 | 0 | 0 |
| 0 | 0 | 0 | 0 |
| 0 | 0 | 0 | 0 |
| 0 | 0 | 0 | 0 |
| 0 | 0 | 0 | 0 |
| 0 | 0 | 0 | 0 |
| 0 | 0 | 0 | 0 |
| 0 | 0 | 0 | 0 |
| 1 | 0 | 0 | 0 |
| 0 | 0 | 0 | 0 |
| 0 | 0 | 0 | 0 |
| 0 | 0 | 0 | 0 |
| 0 | 0 | 0 | 0 |
| 0 | 0 | 0 | 0 |
| 0 | 0 | 0 | 0 |
| 0 | 0 | 0 | 0 |
| 0 | 0 | 0 | 0 |
| 0 | 0 | 0 | 0 |
| 1 | 0 | 0 | 0 |
| 1 | 0 | 0 | 0 |
| 0 | 0 | 0 | 0 |
| 0 | 0 | 0 | 0 |
| 1 | 0 | 0 | 0 |
| 0 | 0 | 1 | 0 |
| 0 | 0 | 1 | 0 |
| 0 | 0 | 1 | 0 |
| 0 | 0 | 1 | 0 |
| 0 | 0 | 0 | 0 |
| 1 | 0 | 0 | 0 |
| 0 | 0 | 0 | 0 |
| 0 | 0 | 0 | 0 |
| 0 | 0 | 0 | 0 |
| 0 | 0 | 1 | 0 |
| 0 | 0 | 0 | 0 |
| 0 | 0 | 0 | 0 |
| 0 | 0 | 0 | 0 |
| 0 | 0 | 0 | 0 |
| 0 | 0 | 0 | 0 |

| | | | |
|---|---|---|---|
| 0 | 0 | 0 | 0 |
| 0 | 0 | 0 | 0 |
| 0 | 0 | 0 | 0 |
| 0 | 0 | 0 | 0 |
| 0 | 0 | 0 | 0 |
| 0 | 0 | 0 | 0 |
| 0 | 0 | 0 | 0 |
| 0 | 0 | 0 | 0 |
| 0 | 0 | 0 | 0 |
| 0 | 0 | 0 | 0 |
| 0 | 1 | 0 | 0 |
| 0 | 0 | 0 | 0 |
| 0 | 0 | 0 | 0 |
| 0 | 0 | 0 | 0 |
| 0 | 0 | 0 | 0 |
| 0 | 1 | 0 | 0 |
| 0 | 0 | 0 | 0 |
| 0 | 0 | 0 | 0 |
| 0 | 0 | 0 | 0 |
| 0 | 0 | 0 | 0 |
| 0 | 0 | 0 | 0 |
| 0 | 0 | 0 | 0 |
| 0 | 0 | 0 | 0 |
| 0 | 0 | 0 | 0 |
| 0 | 0 | 0 | 0 |
| 0 | 0 | 0 | 0 |
| 0 | 0 | 0 | 0 |
| 0 | 0 | 0 | 0 |
| 0 | 0 | 0 | 0 |
| 0 | 0 | 0 | 0 |
| 0 | 0 | 0 | 0 |
| 0 | 0 | 0 | 0 |
| 0 | 0 | 0 | 0 |
| 0 | 0 | 0 | 0 |
| 0 | 0 | 0 | 0 |
| 0 | 0 | 0 | 0 |
| 0 | 0 | 0 | 0 |
| 0 | 0 | 0 | 0 |
| 0 | 0 | 0 | 0 |
| 0 | 0 | 0 | 0 |
| 0 | 0 | 0 | 0 |
| 0 | 0 | 0 | 0 |
| 0 | 0 | 0 | 0 |
| 0 | 0 | 0 | 0 |
| 1 | 1 | 0 | 0 |
| 0 | 0 | 0 | 0 |
| 0 | 0 | 0 | 0 |
| 0 | 0 | 0 | 0 |
| 0 | 0 | 0 | 0 |
| 0 | 0 | 0 | 0 |
| 0 | 0 | 0 | 0 |
| 0 | 0 | 0 | 0 |
| 0 | 0 | 0 | 0 |
| 0 | 0 | 0 | 0 |

| | | | |
|---|---|---|---|
| 0 | 0 | 0 | 0 |
| 0 | 0 | 0 | 0 |
| 0 | 0 | 1 | 0 |
| 1 | 0 | 0 | 0 |
| 0 | 0 | 0 | 0 |
| 0 | 0 | 0 | 0 |
| 0 | 0 | 0 | 0 |
| 0 | 0 | 0 | 0 |
| 0 | 0 | 0 | 0 |
| 0 | 0 | 0 | 0 |
| 0 | 0 | 0 | 0 |
| 0 | 0 | 0 | 0 |
| 1 | 0 | 0 | 0 |
| 0 | 0 | 0 | 0 |
| 0 | 0 | 0 | 0 |
| 0 | 0 | 0 | 0 |
| 0 | 0 | 0 | 0 |
| 0 | 0 | 0 | 0 |
| 0 | 0 | 0 | 0 |
| 0 | 0 | 0 | 0 |
| 0 | 0 | 0 | 0 |
| 0 | 0 | 0 | 0 |
| 0 | 0 | 0 | 0 |
| 0 | 0 | 0 | 0 |
| 0 | 0 | 0 | 0 |
| 0 | 0 | 0 | 0 |
| 0 | 0 | 0 | 0 |
| 0 | 0 | 0 | 0 |
| 0 | 0 | 0 | 0 |
| 0 | 0 | 0 | 0 |
| 0 | 0 | 0 | 0 |
| 0 | 0 | 0 | 0 |
| 0 | 0 | 0 | 0 |
| 0 | 0 | 0 | 0 |
| 0 | 0 | 0 | 0 |
| 0 | 0 | 0 | 0 |
| 0 | 0 | 0 | 0 |
| 0 | 0 | 0 | 0 |
| 0 | 0 | 0 | 0 |
| 0 | 0 | 0 | 0 |
| 0 | 1 | 0 | 0 |
| 1 | 1 | 0 | 0 |
| 0 | 0 | 0 | 0 |
| 0 | 0 | 0 | 0 |
| 0 | 0 | 0 | 0 |
| 0 | 0 | 0 | 0 |
| 0 | 0 | 0 | 0 |
| 0 | 0 | 0 | 0 |
| 0 | 1 | 0 | 0 |
| 0 | 0 | 0 | 0 |
| 0 | 0 | 0 | 0 |
| 0 | 0 | 0 | 0 |
| 0 | 0 | 0 | 1 |
| 0 | 0 | 0 | 0 |

| | | | |
|---|---|---|---|
| 0 | 0 | 0 | 1 |
| 0 | 0 | 0 | 1 |
| 0 | 0 | 0 | 1 |
| 0 | 0 | 0 | 0 |
| 0 | 0 | 0 | 0 |
| 0 | 0 | 0 | 0 |
| 0 | 0 | 0 | 0 |
| 1 | 0 | 0 | 0 |
| 0 | 0 | 0 | 0 |
| 1 | 1 | 0 | 0 |
| 0 | 1 | 0 | 0 |
| 0 | 0 | 0 | 0 |
| 1 | 1 | 0 | 0 |
| 0 | 0 | 0 | 0 |
| 0 | 0 | 0 | 0 |
| 0 | 0 | 0 | 0 |
| 0 | 0 | 0 | 0 |
| 1 | 0 | 0 | 0 |
| 0 | 0 | 0 | 0 |
| 0 | 0 | 0 | 0 |
| 0 | 0 | 0 | 0 |
| 0 | 0 | 0 | 0 |
| 0 | 0 | 0 | 0 |
| 0 | 0 | 0 | 0 |
| 0 | 0 | 0 | 0 |
| 0 | 0 | 0 | 0 |
| 0 | 0 | 0 | 0 |
| 0 | 0 | 0 | 0 |
| 0 | 0 | 0 | 0 |
| 0 | 0 | 0 | 0 |
| 0 | 0 | 0 | 0 |
| 1 | 0 | 0 | 0 |
| 0 | 0 | 0 | 0 |
| 0 | 0 | 0 | 0 |
| 0 | 0 | 0 | 0 |
| 0 | 0 | 0 | 0 |
| 0 | 0 | 0 | 0 |
| 0 | 0 | 0 | 0 |
| 0 | 0 | 0 | 0 |
| 0 | 0 | 0 | 0 |
| 0 | 0 | 0 | 0 |
| 1 | 1 | 0 | 0 |
| 0 | 0 | 0 | 0 |
| 0 | 0 | 0 | 0 |
| 1 | 0 | 0 | 0 |
| 0 | 1 | 0 | 0 |
| 0 | 0 | 0 | 0 |
| 0 | 0 | 0 | 0 |
| 0 | 1 | 0 | 0 |
| 1 | 1 | 0 | 0 |
| 1 | 1 | 0 | 0 |
| 1 | 1 | 0 | 0 |
| 1 | 1 | 0 | 0 |
| 1 | 1 | 0 | 0 |

| | | | |
|---|---|---|---|
| 0 | 0 | 0 | 0 |
| 0 | 0 | 0 | 0 |
| 0 | 0 | 0 | 0 |
| 0 | 0 | 0 | 0 |
| 0 | 0 | 0 | 0 |
| 1 | 0 | 0 | 0 |
| 1 | 0 | 0 | 0 |
| 0 | 0 | 0 | 0 |
| 0 | 0 | 0 | 0 |
| 1 | 0 | 0 | 0 |
| 0 | 0 | 0 | 0 |
| 0 | 0 | 1 | 0 |
| 0 | 0 | 0 | 0 |
| 0 | 0 | 0 | 0 |
| 0 | 0 | 0 | 0 |
| 0 | 0 | 0 | 0 |
| 0 | 0 | 0 | 0 |
| 0 | 0 | 0 | 0 |
| 0 | 0 | 0 | 0 |
| 0 | 0 | 0 | 0 |
| 1 | 1 | 0 | 0 |
| 1 | 0 | 0 | 0 |
| 0 | 0 | 0 | 0 |
| 0 | 0 | 1 | 0 |
| 0 | 0 | 0 | 0 |
| 0 | 0 | 1 | 0 |
| 0 | 0 | 0 | 0 |
| 0 | 0 | 0 | 0 |
| 0 | 0 | 0 | 0 |
| 0 | 0 | 0 | 0 |
| 0 | 0 | 0 | 0 |
| 0 | 0 | 0 | 0 |
| 0 | 0 | 0 | 0 |
| 0 | 0 | 0 | 0 |
| 0 | 0 | 0 | 0 |
| 0 | 0 | 0 | 0 |
| 1 | 0 | 0 | 0 |
| 0 | 0 | 0 | 0 |
| 1 | 0 | 0 | 0 |
| 1 | 0 | 0 | 0 |
| 1 | 0 | 0 | 0 |
| 0 | 0 | 0 | 0 |
| 0 | 0 | 0 | 0 |
| 0 | 0 | 0 | 0 |
| 0 | 0 | 0 | 0 |
| 0 | 0 | 0 | 0 |
| 0 | 0 | 0 | 0 |
| 1 | 0 | 1 | 0 |
| 0 | 0 | 0 | 0 |
| 0 | 0 | 0 | 0 |
| 0 | 0 | 0 | 0 |
| 0 | 0 | 0 | 0 |
| 1 | 1 | 0 | 0 |
| 0 | 0 | 0 | 0 |

| | | | |
|---|---|---|---|
| 0 | 0 | 0 | 0 |
| 0 | 1 | 0 | 0 |
| 0 | 0 | 0 | 0 |
| 0 | 0 | 0 | 0 |
| 0 | 0 | 0 | 0 |
| 0 | 0 | 0 | 0 |
| 0 | 0 | 0 | 0 |
| 0 | 0 | 0 | 0 |
| 0 | 0 | 0 | 0 |
| 0 | 0 | 0 | 0 |
| 0 | 0 | 0 | 0 |
| 0 | 0 | 0 | 0 |
| 0 | 0 | 0 | 0 |
| 0 | 0 | 0 | 0 |
| 0 | 0 | 0 | 0 |
| 0 | 0 | 0 | 0 |
| 0 | 0 | 0 | 0 |
| 0 | 0 | 0 | 0 |
| 0 | 0 | 0 | 0 |
| 0 | 0 | 0 | 0 |
| 1 | 0 | 0 | 0 |
| 0 | 0 | 0 | 0 |
| 0 | 0 | 0 | 0 |
| 0 | 0 | 0 | 0 |
| 1 | 0 | 0 | 0 |
| 0 | 0 | 0 | 0 |
| 0 | 0 | 0 | 0 |
| 0 | 0 | 0 | 1 |
| 0 | 0 | 0 | 0 |
| 0 | 0 | 0 | 0 |
| 0 | 0 | 0 | 0 |
| 0 | 0 | 0 | 0 |
| 0 | 0 | 0 | 0 |
| 0 | 0 | 0 | 0 |
| 0 | 0 | 0 | 0 |
| 0 | 0 | 0 | 0 |
| 0 | 0 | 0 | 0 |
| 0 | 0 | 0 | 0 |
| 0 | 0 | 0 | 0 |
| 0 | 0 | 0 | 0 |
| 0 | 0 | 0 | 0 |
| 0 | 0 | 0 | 0 |
| 0 | 0 | 0 | 0 |
| 0 | 0 | 0 | 0 |
| 0 | 0 | 0 | 0 |
| 0 | 0 | 0 | 0 |
| 0 | 0 | 0 | 0 |
| 0 | 0 | 0 | 0 |
| 0 | 0 | 0 | 0 |
| 0 | 0 | 0 | 0 |
| 0 | 0 | 0 | 0 |
| 0 | 0 | 0 | 0 |
| 0 | 0 | 0 | 0 |
| 0 | 0 | 0 | 0 |

| | | | |
|---|---|---|---|
| 0 | 0 | 0 | 0 |
| 0 | 0 | 0 | 0 |
| 0 | 0 | 0 | 0 |
| 0 | 0 | 0 | 0 |
| 0 | 0 | 0 | 0 |
| 0 | 0 | 0 | 0 |
| 0 | 0 | 0 | 0 |
| 0 | 0 | 0 | 0 |
| 0 | 0 | 0 | 0 |
| 0 | 0 | 0 | 0 |
| 0 | 0 | 0 | 0 |
| 0 | 0 | 0 | 0 |
| 0 | 0 | 0 | 0 |
| 0 | 0 | 0 | 0 |
| 0 | 0 | 0 | 0 |
| 1 | 0 | 0 | 0 |
| 0 | 0 | 0 | 0 |
| 0 | 0 | 0 | 0 |
| 0 | 1 | 0 | 0 |
| 0 | 0 | 0 | 0 |
| 0 | 0 | 0 | 0 |
| 0 | 0 | 0 | 0 |
| 0 | 0 | 0 | 0 |
| 0 | 0 | 0 | 0 |
| 0 | 0 | 0 | 0 |
| 1 | 0 | 0 | 0 |
| 0 | 0 | 0 | 0 |
| 0 | 0 | 0 | 0 |
| 0 | 0 | 0 | 0 |
| 0 | 0 | 0 | 0 |
| 0 | 0 | 0 | 0 |
| 0 | 0 | 0 | 0 |
| 0 | 0 | 0 | 0 |
| 1 | 1 | 0 | 0 |
| 0 | 0 | 0 | 0 |
| 0 | 0 | 0 | 0 |
| 0 | 0 | 0 | 0 |
| 0 | 0 | 0 | 0 |
| 0 | 0 | 0 | 0 |
| 0 | 0 | 0 | 0 |
| 0 | 0 | 0 | 0 |
| 0 | 0 | 0 | 0 |
| 0 | 0 | 0 | 0 |
| 0 | 0 | 0 | 0 |
| 0 | 0 | 0 | 0 |
| 0 | 0 | 0 | 0 |
| 0 | 0 | 0 | 0 |
| 1 | 0 | 0 | 0 |
| 0 | 0 | 0 | 0 |
| 0 | 0 | 0 | 0 |
| 0 | 0 | 0 | 0 |
| 0 | 0 | 0 | 0 |
| 0 | 0 | 0 | 0 |
| 1 | 0 | 0 | 1 |

| | | | |
|---|---|---|---|
| 0 | 0 | 0 | 0 |
| 0 | 0 | 0 | 0 |
| 0 | 0 | 1 | 0 |
| 0 | 0 | 0 | 0 |
| 0 | 0 | 0 | 0 |
| 0 | 0 | 0 | 0 |
| 0 | 0 | 0 | 0 |
| 0 | 0 | 0 | 0 |
| 0 | 0 | 0 | 0 |
| 0 | 0 | 0 | 0 |
| 0 | 0 | 0 | 0 |
| 0 | 0 | 0 | 0 |
| 0 | 0 | 0 | 0 |
| 0 | 0 | 0 | 0 |
| 0 | 0 | 0 | 0 |
| 0 | 0 | 0 | 0 |
| 0 | 0 | 0 | 0 |
| 0 | 0 | 0 | 0 |
| 0 | 0 | 0 | 0 |
| 0 | 0 | 0 | 0 |
| 0 | 0 | 0 | 0 |
| 0 | 0 | 0 | 0 |
| 0 | 0 | 0 | 0 |
| 0 | 0 | 0 | 0 |
| 0 | 0 | 0 | 0 |
| 0 | 0 | 0 | 0 |
| 0 | 0 | 0 | 0 |
| 0 | 0 | 0 | 0 |
| 0 | 0 | 0 | 0 |
| 0 | 0 | 0 | 0 |
| 0 | 0 | 0 | 0 |
| 1 | 0 | 0 | 0 |
| 0 | 0 | 0 | 0 |
| 0 | 0 | 0 | 0 |
| 0 | 0 | 0 | 0 |
| 0 | 0 | 0 | 0 |
| 0 | 0 | 0 | 0 |
| 0 | 0 | 0 | 0 |
| 0 | 0 | 0 | 0 |
| 0 | 0 | 0 | 0 |
| 1 | 1 | 0 | 0 |
| 0 | 0 | 0 | 0 |
| 0 | 0 | 0 | 0 |
| 0 | 0 | 0 | 0 |
| 0 | 0 | 0 | 0 |
| 1 | 1 | 0 | 0 |
| 0 | 0 | 0 | 0 |
| 1 | 1 | 0 | 0 |
| 0 | 0 | 0 | 0 |
| 1 | 0 | 0 | 0 |
| 1 | 0 | 0 | 0 |
| 1 | 0 | 0 | 0 |
| 0 | 0 | 0 | 0 |
| 0 | 0 | 0 | 0 |

| | | | |
|---|---|---|---|
| 1 | 0 | 0 | 0 |
| 0 | 0 | 0 | 0 |
| 0 | 0 | 0 | 0 |
| 0 | 0 | 0 | 0 |
| 0 | 0 | 0 | 0 |
| 0 | 0 | 0 | 0 |
| 0 | 0 | 0 | 0 |
| 0 | 0 | 0 | 0 |
| 0 | 0 | 0 | 0 |
| 0 | 0 | 0 | 0 |
| 0 | 0 | 0 | 0 |
| 0 | 0 | 0 | 0 |
| 0 | 0 | 0 | 0 |
| 0 | 0 | 0 | 0 |
| 0 | 0 | 0 | 0 |
| 1 | 1 | 1 | 0 |
| 0 | 0 | 0 | 0 |
| 1 | 0 | 0 | 0 |
| 0 | 0 | 0 | 0 |
| 0 | 0 | 0 | 0 |
| 0 | 0 | 0 | 0 |
| 0 | 1 | 0 | 0 |
| 0 | 0 | 0 | 0 |
| 0 | 1 | 0 | 0 |
| 0 | 1 | 0 | 0 |
| 0 | 0 | 0 | 0 |
| 0 | 0 | 0 | 0 |
| 0 | 0 | 0 | 0 |
| 0 | 0 | 0 | 0 |
| 0 | 0 | 0 | 0 |
| 0 | 0 | 0 | 0 |
| 0 | 0 | 0 | 0 |
| 1 | 1 | 0 | 0 |
| 0 | 0 | 0 | 0 |
| 0 | 0 | 0 | 0 |
| 0 | 0 | 0 | 0 |
| 0 | 1 | 0 | 0 |
| 1 | 1 | 0 | 0 |
| 0 | 0 | 0 | 0 |
| 0 | 0 | 0 | 0 |
| 0 | 0 | 0 | 0 |
| 0 | 0 | 0 | 0 |
| 0 | 0 | 0 | 0 |
| 0 | 0 | 0 | 0 |
| 1 | 1 | 0 | 0 |
| 0 | 0 | 0 | 0 |
| 1 | 0 | 0 | 0 |
| 1 | 1 | 0 | 0 |
| 0 | 0 | 0 | 0 |
| 0 | 0 | 0 | 0 |
| 1 | 1 | 0 | 0 |
| 1 | 1 | 0 | 0 |
| 0 | 0 | 0 | 0 |
| 1 | 1 | 0 | 0 |

| | | | |
|---|---|---|---|
| 0 | 0 | 0 | 0 |
| 0 | 0 | 0 | 0 |
| 0 | 0 | 0 | 0 |
| 1 | 1 | 0 | 0 |
| 0 | 0 | 0 | 0 |
| 0 | 0 | 0 | 0 |
| 0 | 0 | 0 | 0 |
| 0 | 0 | 0 | 0 |
| 1 | 0 | 0 | 0 |
| 0 | 0 | 0 | 0 |
| 0 | 0 | 0 | 0 |
| 0 | 0 | 0 | 0 |
| 1 | 0 | 1 | 0 |
| 1 | 0 | 0 | 0 |
| 0 | 0 | 0 | 0 |
| 0 | 0 | 0 | 0 |
| 0 | 0 | 0 | 0 |
| 0 | 0 | 0 | 0 |
| 0 | 0 | 0 | 0 |
| 0 | 0 | 0 | 0 |
| 0 | 0 | 0 | 0 |
| 0 | 0 | 0 | 0 |
| 0 | 0 | 0 | 0 |
| 1 | 0 | 0 | 0 |
| 0 | 0 | 0 | 0 |
| 0 | 0 | 0 | 0 |
| 0 | 0 | 0 | 0 |
| 0 | 0 | 0 | 0 |
| 0 | 0 | 0 | 0 |
| 0 | 0 | 0 | 0 |
| 0 | 0 | 0 | 0 |
| 0 | 0 | 0 | 0 |
| 0 | 0 | 0 | 0 |
| 0 | 0 | 0 | 0 |
| 0 | 0 | 0 | 0 |
| 0 | 0 | 0 | 0 |
| 0 | 0 | 0 | 0 |
| 0 | 0 | 0 | 0 |
| 0 | 0 | 0 | 0 |
| 0 | 0 | 0 | 0 |
| 0 | 0 | 0 | 0 |
| 0 | 0 | 0 | 0 |
| 0 | 0 | 0 | 0 |
| 1 | 1 | 0 | 0 |
| 1 | 1 | 0 | 0 |
| 0 | 0 | 0 | 0 |
| 0 | 0 | 0 | 0 |
| 0 | 1 | 0 | 0 |
| 0 | 1 | 0 | 0 |
| 0 | 1 | 0 | 0 |
| 0 | 0 | 0 | 0 |
| 0 | 0 | 0 | 0 |
| 0 | 0 | 0 | 0 |
| 0 | 0 | 0 | 0 |

| | | | |
|---|---|---|---|
| 0 | 0 | 0 | 0 |
| 0 | 0 | 0 | 0 |
| 0 | 0 | 0 | 0 |
| 0 | 0 | 0 | 0 |
| 1 | 0 | 0 | 0 |
| 0 | 0 | 0 | 0 |
| 0 | 0 | 0 | 0 |
| 1 | 0 | 0 | 0 |
| 1 | 0 | 0 | 0 |
| 1 | 0 | 0 | 0 |
| 1 | 0 | 0 | 0 |
| 1 | 0 | 0 | 0 |
| 0 | 0 | 0 | 0 |
| 1 | 0 | 0 | 0 |
| 1 | 0 | 0 | 0 |
| 0 | 0 | 0 | 0 |
| 0 | 0 | 0 | 0 |
| 0 | 0 | 0 | 0 |
| 1 | 0 | 0 | 0 |
| 1 | 0 | 0 | 0 |
| 1 | 0 | 0 | 0 |
| 1 | 0 | 0 | 0 |
| 0 | 0 | 0 | 0 |
| 0 | 0 | 0 | 0 |
| 0 | 0 | 0 | 0 |
| 0 | 0 | 0 | 0 |
| 0 | 0 | 0 | 0 |
| 0 | 0 | 0 | 0 |
| 0 | 0 | 0 | 0 |
| 0 | 0 | 0 | 0 |
| 0 | 0 | 0 | 0 |
| 0 | 0 | 0 | 0 |
| 1 | 0 | 0 | 0 |
| 0 | 0 | 0 | 0 |
| 0 | 0 | 0 | 0 |
| 1 | 0 | 0 | 0 |
| 0 | 0 | 0 | 0 |
| 0 | 0 | 0 | 0 |
| 0 | 0 | 0 | 0 |
| 0 | 0 | 0 | 0 |
| 0 | 0 | 0 | 0 |
| 0 | 0 | 0 | 0 |
| 0 | 0 | 0 | 0 |
| 0 | 0 | 0 | 0 |
| 0 | 0 | 0 | 0 |
| 0 | 0 | 0 | 0 |
| 0 | 0 | 0 | 0 |
| 0 | 0 | 0 | 0 |
| 0 | 0 | 0 | 0 |
| 0 | 0 | 0 | 0 |
| 0 | 0 | 0 | 0 |
| 0 | 0 | 0 | 0 |
| 0 | 0 | 0 | 0 |
| 0 | 0 | 0 | 0 |

| | | | |
|---|---|---|---|
| 0 | 0 | 0 | 0 |
| 0 | 0 | 0 | 0 |
| 0 | 0 | 0 | 0 |
| 0 | 0 | 0 | 0 |
| 0 | 0 | 0 | 0 |
| 0 | 0 | 0 | 0 |
| 0 | 0 | 0 | 0 |
| 0 | 0 | 0 | 0 |
| 0 | 0 | 0 | 0 |
| 0 | 0 | 0 | 0 |
| 0 | 0 | 0 | 0 |
| 0 | 0 | 0 | 1 |
| 0 | 0 | 0 | 0 |
| 0 | 0 | 0 | 0 |
| 0 | 0 | 0 | 0 |
| 0 | 0 | 0 | 0 |
| 0 | 0 | 0 | 0 |
| 0 | 0 | 0 | 1 |
| 0 | 0 | 0 | 0 |
| 0 | 0 | 0 | 0 |
| 0 | 0 | 0 | 1 |
| 0 | 0 | 0 | 0 |
| 0 | 0 | 0 | 0 |
| 0 | 0 | 0 | 0 |
| 0 | 0 | 0 | 0 |
| 0 | 0 | 0 | 0 |
| 0 | 0 | 0 | 0 |
| 0 | 0 | 0 | 0 |
| 0 | 0 | 0 | 1 |
| 0 | 0 | 0 | 0 |
| 0 | 0 | 0 | 0 |
| 0 | 0 | 0 | 0 |
| 0 | 0 | 0 | 0 |
| 0 | 0 | 0 | 0 |
| 0 | 0 | 0 | 0 |
| 0 | 0 | 0 | 0 |
| 0 | 0 | 0 | 0 |
| 0 | 0 | 0 | 0 |
| 0 | 0 | 0 | 0 |
| 0 | 0 | 0 | 0 |
| 0 | 0 | 0 | 0 |
| 0 | 0 | 0 | 0 |
| 0 | 0 | 0 | 0 |
| 0 | 0 | 0 | 0 |
| 0 | 0 | 0 | 0 |
| 0 | 0 | 0 | 0 |
| 0 | 0 | 0 | 0 |
| 0 | 0 | 0 | 0 |
| 0 | 0 | 0 | 0 |
| 0 | 0 | 0 | 0 |
| 0 | 0 | 0 | 0 |
| 0 | 0 | 0 | 0 |
| 0 | 0 | 0 | 0 |
| 0 | 0 | 0 | 0 |

| | | | |
|---|---|---|---|
| 0 | 0 | 0 | 0 |
| 0 | 0 | 0 | 0 |
| 1 | 0 | 0 | 0 |
| 0 | 0 | 0 | 0 |
| 1 | 0 | 0 | 0 |
| 0 | 0 | 0 | 0 |
| 0 | 0 | 0 | 0 |
| 0 | 0 | 0 | 0 |
| 0 | 0 | 0 | 0 |
| 1 | 1 | 0 | 0 |
| 0 | 0 | 1 | 0 |
| 0 | 0 | 0 | 0 |
| 0 | 0 | 0 | 0 |
| 0 | 0 | 0 | 0 |
| 0 | 0 | 0 | 0 |
| 0 | 0 | 0 | 0 |
| 0 | 0 | 0 | 0 |
| 0 | 0 | 0 | 0 |
| 0 | 0 | 0 | 0 |
| 0 | 0 | 0 | 0 |
| 0 | 0 | 0 | 0 |
| 0 | 0 | 0 | 0 |
| 0 | 0 | 0 | 0 |
| 0 | 0 | 0 | 0 |
| 0 | 0 | 0 | 0 |
| 0 | 0 | 0 | 0 |
| 0 | 0 | 0 | 0 |
| 0 | 0 | 0 | 0 |
| 0 | 0 | 0 | 0 |
| 0 | 0 | 0 | 0 |
| 0 | 0 | 0 | 0 |
| 0 | 0 | 0 | 0 |
| 0 | 0 | 0 | 0 |
| 0 | 0 | 0 | 0 |
| 0 | 0 | 0 | 0 |
| 0 | 0 | 0 | 0 |
| 0 | 0 | 0 | 0 |
| 0 | 0 | 0 | 0 |
| 0 | 0 | 0 | 0 |
| 0 | 0 | 0 | 0 |
| 0 | 0 | 0 | 0 |
| 0 | 0 | 0 | 0 |
| 0 | 0 | 0 | 0 |
| 0 | 0 | 0 | 0 |
| 0 | 0 | 0 | 0 |
| 1 | 0 | 0 | 0 |
| 0 | 0 | 0 | 0 |
| 0 | 0 | 0 | 0 |
| 0 | 0 | 0 | 0 |
| 0 | 0 | 0 | 0 |
| 1 | 0 | 0 | 0 |
| 0 | 0 | 0 | 0 |
| 0 | 0 | 0 | 0 |
| 0 | 0 | 0 | 0 |

| | | | |
|---|---|---|---|
| 0 | 0 | 0 | 0 |
| 0 | 0 | 0 | 0 |
| 0 | 0 | 0 | 0 |
| 0 | 0 | 0 | 0 |
| 0 | 0 | 0 | 0 |
| 0 | 0 | 0 | 0 |
| 0 | 0 | 0 | 0 |
| 0 | 0 | 0 | 0 |
| 1 | 1 | 0 | 0 |
| 0 | 0 | 0 | 0 |
| 0 | 0 | 0 | 0 |
| 0 | 0 | 0 | 0 |
| 0 | 0 | 0 | 0 |
| 0 | 0 | 0 | 0 |
| 0 | 1 | 0 | 0 |
| 0 | 1 | 0 | 0 |
| 0 | 1 | 0 | 0 |
| 0 | 0 | 0 | 0 |
| 0 | 0 | 0 | 0 |
| 0 | 0 | 0 | 0 |
| 0 | 1 | 0 | 0 |
| 0 | 0 | 0 | 0 |
| 0 | 0 | 0 | 0 |
| 1 | 1 | 0 | 0 |
| 0 | 1 | 0 | 0 |
| 0 | 1 | 0 | 0 |
| 0 | 0 | 0 | 0 |
| 0 | 0 | 0 | 0 |
| 0 | 0 | 0 | 0 |
| 0 | 0 | 0 | 0 |
| 0 | 0 | 0 | 0 |
| 0 | 0 | 0 | 0 |
| 0 | 0 | 0 | 0 |
| 0 | 0 | 0 | 0 |
| 0 | 0 | 0 | 0 |
| 0 | 0 | 0 | 0 |
| 0 | 0 | 0 | 0 |
| 1 | 0 | 0 | 0 |
| 0 | 0 | 0 | 0 |
| 0 | 0 | 0 | 0 |
| 0 | 0 | 0 | 0 |
| 0 | 0 | 0 | 0 |
| 0 | 0 | 0 | 0 |
| 0 | 0 | 0 | 0 |
| 0 | 0 | 0 | 0 |
| 0 | 0 | 0 | 0 |
| 0 | 0 | 0 | 0 |
| 0 | 0 | 0 | 0 |
| 0 | 0 | 0 | 0 |
| 0 | 0 | 0 | 0 |
| 0 | 0 | 0 | 0 |
| 0 | 0 | 0 | 0 |
| 0 | 0 | 0 | 0 |
| 0 | 0 | 0 | 0 |

| | | | |
|---|---|---|---|
| 0 | 0 | 0 | 0 |
| 0 | 0 | 0 | 0 |
| 0 | 0 | 0 | 0 |
| 0 | 0 | 1 | 0 |
| 0 | 0 | 0 | 0 |
| 0 | 0 | 0 | 0 |
| 0 | 0 | 0 | 0 |
| 0 | 0 | 0 | 0 |
| 0 | 0 | 0 | 0 |
| 0 | 0 | 0 | 0 |
| 0 | 0 | 0 | 0 |
| 0 | 0 | 0 | 0 |
| 1 | 0 | 0 | 0 |
| 0 | 0 | 0 | 0 |
| 0 | 0 | 0 | 0 |
| 0 | 0 | 0 | 0 |
| 0 | 1 | 0 | 0 |
| 0 | 0 | 0 | 0 |
| 0 | 0 | 0 | 0 |
| 0 | 0 | 0 | 0 |
| 0 | 0 | 0 | 0 |
| 0 | 0 | 0 | 0 |
| 0 | 0 | 0 | 0 |
| 0 | 0 | 0 | 0 |
| 0 | 0 | 0 | 0 |
| 0 | 0 | 0 | 0 |
| 0 | 0 | 0 | 0 |
| 0 | 0 | 0 | 0 |
| 0 | 0 | 0 | 0 |
| 0 | 0 | 0 | 0 |
| 1 | 0 | 0 | 0 |
| 1 | 0 | 0 | 0 |
| 1 | 0 | 0 | 0 |
| 0 | 0 | 0 | 0 |
| 1 | 0 | 0 | 0 |
| 1 | 0 | 0 | 0 |
| 1 | 0 | 0 | 0 |
| 0 | 0 | 0 | 0 |
| 0 | 0 | 0 | 0 |
| 0 | 0 | 0 | 0 |
| 0 | 0 | 0 | 0 |
| 0 | 0 | 0 | 0 |
| 0 | 1 | 0 | 0 |
| 0 | 1 | 0 | 0 |
| 0 | 0 | 0 | 0 |
| 0 | 0 | 0 | 0 |
| 1 | 0 | 0 | 0 |
| 0 | 0 | 0 | 0 |
| 0 | 0 | 0 | 0 |
| 0 | 0 | 0 | 0 |
| 0 | 0 | 0 | 0 |
| 0 | 0 | 0 | 0 |
| 1 | 0 | 0 | 0 |
| 0 | 0 | 0 | 0 |

| | | | |
|---|---|---|---|
| 0 | 0 | 0 | 0 |
| 1 | 0 | 0 | 0 |
| 1 | 0 | 0 | 0 |
| 0 | 0 | 0 | 0 |
| 0 | 0 | 0 | 0 |
| 0 | 0 | 0 | 0 |
| 0 | 0 | 0 | 0 |
| 0 | 1 | 0 | 0 |
| 0 | 0 | 0 | 0 |
| 0 | 0 | 0 | 0 |
| 0 | 0 | 0 | 0 |
| 0 | 0 | 0 | 0 |
| 0 | 0 | 0 | 0 |
| 0 | 0 | 0 | 0 |
| 1 | 0 | 0 | 0 |
| 0 | 0 | 0 | 0 |
| 0 | 0 | 0 | 0 |
| 0 | 0 | 0 | 0 |
| 0 | 0 | 0 | 0 |
| 1 | 0 | 0 | 0 |
| 1 | 0 | 0 | 0 |
| 0 | 0 | 0 | 0 |
| 1 | 0 | 0 | 0 |
| 1 | 0 | 0 | 0 |
| 0 | 0 | 0 | 0 |
| 0 | 0 | 0 | 0 |
| 0 | 0 | 0 | 0 |
| 0 | 0 | 0 | 0 |
| 0 | 0 | 0 | 0 |
| 0 | 0 | 0 | 0 |
| 0 | 0 | 0 | 0 |
| 0 | 0 | 0 | 0 |
| 0 | 0 | 0 | 0 |
| 0 | 0 | 0 | 0 |
| 0 | 0 | 0 | 0 |
| 0 | 0 | 0 | 0 |
| 0 | 0 | 0 | 0 |
| 0 | 0 | 0 | 0 |
| 0 | 0 | 0 | 0 |
| 0 | 0 | 0 | 0 |
| 0 | 0 | 0 | 0 |
| 0 | 0 | 0 | 0 |
| 0 | 0 | 0 | 0 |
| 0 | 0 | 0 | 0 |
| 0 | 0 | 0 | 0 |
| 0 | 0 | 0 | 0 |
| 0 | 0 | 0 | 0 |
| 0 | 0 | 0 | 0 |
| 0 | 0 | 0 | 0 |
| 0 | 0 | 0 | 0 |
| 0 | 0 | 0 | 0 |
| 0 | 0 | 0 | 0 |
| 0 | 0 | 0 | 0 |
| 1 | 1 | 0 | 1 |

| | | | |
|---|---|---|---|
| 0 | 0 | 0 | 0 |
| 0 | 0 | 0 | 0 |
| 0 | 0 | 0 | 0 |
| 0 | 0 | 0 | 0 |
| 0 | 1 | 0 | 0 |
| 0 | 0 | 0 | 0 |
| 0 | 0 | 0 | 0 |
| 0 | 1 | 0 | 0 |
| 0 | 0 | 0 | 0 |
| 0 | 0 | 0 | 0 |
| 0 | 0 | 0 | 0 |
| 0 | 0 | 0 | 0 |
| 0 | 0 | 0 | 0 |
| 0 | 0 | 0 | 0 |
| 0 | 1 | 0 | 1 |
| 0 | 1 | 0 | 0 |
| 0 | 0 | 0 | 0 |
| 0 | 0 | 0 | 0 |
| 0 | 1 | 0 | 0 |
| 0 | 1 | 0 | 1 |
| 0 | 0 | 0 | 0 |
| 0 | 0 | 0 | 0 |
| 0 | 0 | 0 | 0 |
| 0 | 0 | 0 | 0 |
| 0 | 0 | 0 | 0 |
| 1 | 0 | 0 | 0 |
| 0 | 0 | 0 | 0 |
| 0 | 0 | 1 | 0 |
| 0 | 0 | 1 | 0 |
| 0 | 0 | 1 | 0 |
| 0 | 0 | 0 | 0 |
| 0 | 0 | 0 | 0 |
| 0 | 0 | 0 | 0 |
| 0 | 0 | 0 | 0 |
| 0 | 0 | 0 | 0 |
| 0 | 0 | 1 | 0 |
| 0 | 0 | 0 | 0 |
| 0 | 0 | 0 | 0 |
| 1 | 0 | 1 | 0 |
| 0 | 0 | 1 | 0 |
| 1 | 1 | 0 | 0 |
| 0 | 1 | 0 | 0 |
| 0 | 0 | 0 | 0 |
| 0 | 0 | 0 | 0 |
| 1 | 0 | 0 | 0 |
| 0 | 0 | 0 | 0 |
| 0 | 0 | 1 | 0 |
| 0 | 0 | 0 | 0 |
| 0 | 0 | 0 | 0 |
| 0 | 0 | 0 | 0 |
| 0 | 0 | 0 | 0 |
| 0 | 0 | 0 | 0 |
| 0 | 0 | 0 | 0 |
| 0 | 0 | 0 | 0 |

| | | | |
|---|---|---|---|
| 0 | 0 | 0 | 0 |
| 0 | 0 | 0 | 0 |
| 0 | 0 | 0 | 0 |
| 0 | 0 | 0 | 0 |
| 0 | 0 | 0 | 0 |
| 0 | 0 | 0 | 0 |
| 0 | 0 | 0 | 0 |
| 1 | 1 | 0 | 0 |
| 0 | 0 | 0 | 0 |
| 0 | 0 | 0 | 0 |
| 0 | 0 | 0 | 0 |
| 0 | 0 | 0 | 1 |
| 0 | 0 | 0 | 0 |
| 0 | 0 | 0 | 0 |
| 0 | 0 | 0 | 0 |
| 0 | 0 | 0 | 0 |
| 0 | 0 | 0 | 0 |
| 0 | 0 | 0 | 0 |
| 0 | 0 | 0 | 0 |
| 0 | 0 | 0 | 0 |
| 0 | 0 | 0 | 0 |
| 0 | 0 | 0 | 0 |
| 0 | 0 | 0 | 0 |
| 0 | 0 | 0 | 0 |
| 0 | 0 | 0 | 0 |
| 0 | 0 | 0 | 0 |
| 0 | 0 | 0 | 0 |
| 0 | 0 | 0 | 0 |
| 0 | 0 | 0 | 0 |
| 0 | 1 | 0 | 0 |
| 0 | 1 | 0 | 0 |
| 0 | 0 | 0 | 0 |
| 0 | 0 | 0 | 0 |
| 0 | 0 | 0 | 0 |
| 0 | 0 | 0 | 0 |
| 0 | 1 | 0 | 0 |
| 0 | 0 | 0 | 0 |
| 0 | 0 | 0 | 0 |
| 0 | 0 | 0 | 0 |
| 0 | 0 | 0 | 0 |
| 0 | 0 | 0 | 0 |
| 1 | 1 | 0 | 0 |
| 1 | 0 | 0 | 0 |
| 0 | 0 | 0 | 0 |
| 0 | 0 | 0 | 0 |
| 0 | 0 | 0 | 0 |
| 0 | 0 | 0 | 0 |
| 0 | 0 | 0 | 0 |
| 0 | 0 | 0 | 0 |
| 0 | 0 | 0 | 0 |
| 0 | 0 | 0 | 0 |
| 0 | 0 | 0 | 0 |
| 0 | 0 | 0 | 0 |
| 0 | 0 | 0 | 0 |

| | | | |
|---|---|---|---|
| 0 | 0 | 0 | 0 |
| 0 | 0 | 0 | 0 |
| 0 | 0 | 0 | 0 |
| 0 | 0 | 0 | 0 |
| 0 | 0 | 0 | 0 |
| 0 | 0 | 0 | 0 |
| 0 | 0 | 0 | 0 |
| 0 | 0 | 0 | 0 |
| 1 | 0 | 0 | 0 |
| 1 | 0 | 0 | 0 |
| 0 | 0 | 0 | 0 |
| 0 | 0 | 0 | 0 |
| 0 | 0 | 0 | 0 |
| 1 | 0 | 0 | 0 |
| 0 | 0 | 0 | 0 |
| 0 | 0 | 0 | 0 |
| 0 | 0 | 0 | 0 |
| 0 | 0 | 0 | 0 |
| 0 | 0 | 0 | 0 |
| 1 | 1 | 0 | 0 |
| 0 | 0 | 1 | 0 |
| 0 | 0 | 0 | 0 |
| 0 | 0 | 0 | 0 |
| 0 | 0 | 0 | 0 |
| 0 | 0 | 0 | 0 |
| 0 | 0 | 0 | 0 |
| 0 | 0 | 0 | 0 |
| 0 | 0 | 0 | 0 |
| 0 | 0 | 0 | 0 |
| 0 | 0 | 0 | 0 |
| 0 | 0 | 0 | 0 |
| 0 | 0 | 0 | 0 |
| 0 | 0 | 0 | 0 |
| 0 | 0 | 0 | 0 |
| 0 | 0 | 0 | 0 |
| 0 | 0 | 0 | 0 |
| 0 | 0 | 0 | 0 |
| 0 | 0 | 0 | 0 |
| 0 | 0 | 0 | 0 |
| 0 | 0 | 0 | 0 |
| 0 | 0 | 0 | 0 |
| 0 | 0 | 0 | 0 |
| 0 | 0 | 0 | 0 |
| 0 | 0 | 0 | 0 |
| 0 | 0 | 0 | 0 |
| 0 | 0 | 0 | 0 |
| 1 | 0 | 1 | 0 |
| 1 | 0 | 0 | 0 |
| 0 | 0 | 0 | 0 |
| 0 | 0 | 0 | 0 |
| 0 | 0 | 1 | 0 |
| 0 | 0 | 0 | 0 |
| 0 | 0 | 0 | 0 |
| 0 | 0 | 0 | 0 |

| | | | |
|---|---|---|---|
| 0 | 0 | 0 | 0 |
| 0 | 0 | 0 | 0 |
| 0 | 0 | 0 | 0 |
| 0 | 0 | 0 | 0 |
| 0 | 0 | 0 | 0 |
| 0 | 0 | 0 | 0 |
| 0 | 0 | 0 | 0 |
| 0 | 0 | 0 | 0 |
| 0 | 0 | 0 | 0 |
| 0 | 0 | 0 | 0 |
| 0 | 0 | 0 | 0 |
| 0 | 0 | 0 | 0 |
| 0 | 0 | 0 | 0 |
| 0 | 0 | 0 | 0 |
| 0 | 0 | 0 | 0 |
| 0 | 0 | 0 | 0 |
| 0 | 0 | 0 | 0 |
| 0 | 0 | 0 | 0 |
| 0 | 0 | 0 | 0 |
| 1 | 0 | 0 | 0 |
| 0 | 0 | 0 | 0 |
| 0 | 0 | 0 | 0 |
| 0 | 0 | 0 | 0 |
| 0 | 0 | 0 | 0 |
| 0 | 0 | 0 | 0 |
| 0 | 0 | 0 | 0 |
| 0 | 0 | 0 | 0 |
| 0 | 0 | 0 | 0 |
| 0 | 0 | 0 | 0 |
| 0 | 0 | 0 | 0 |
| 0 | 0 | 0 | 0 |
| 0 | 0 | 0 | 0 |
| 0 | 0 | 0 | 0 |
| 0 | 0 | 0 | 0 |
| 0 | 0 | 0 | 0 |
| 0 | 0 | 0 | 0 |
| 0 | 0 | 0 | 0 |
| 0 | 0 | 0 | 0 |
| 1 | 0 | 0 | 1 |
| 0 | 0 | 0 | 0 |
| 0 | 0 | 0 | 0 |
| 0 | 0 | 0 | 0 |
| 0 | 0 | 0 | 1 |
| 0 | 0 | 0 | 1 |
| 0 | 0 | 0 | 1 |
| 0 | 0 | 0 | 0 |
| 0 | 0 | 0 | 0 |
| 0 | 0 | 0 | 0 |
| 0 | 0 | 0 | 0 |
| 0 | 0 | 0 | 0 |
| 0 | 0 | 0 | 0 |
| 0 | 0 | 0 | 0 |
| 0 | 0 | 0 | 0 |
| 0 | 0 | 0 | 0 |

| | | | |
|---|---|---|---|
| 0 | 0 | 0 | 0 |
| 0 | 0 | 0 | 0 |
| 0 | 0 | 0 | 0 |
| 0 | 0 | 0 | 0 |
| 0 | 0 | 0 | 0 |
| 0 | 0 | 0 | 0 |
| 0 | 0 | 0 | 0 |
| 0 | 0 | 0 | 0 |
| 0 | 0 | 0 | 0 |
| 0 | 0 | 0 | 0 |
| 0 | 0 | 0 | 0 |
| 0 | 0 | 0 | 0 |
| 0 | 0 | 0 | 0 |
| 0 | 0 | 0 | 0 |
| 0 | 0 | 0 | 0 |
| 0 | 0 | 0 | 0 |
| 0 | 0 | 0 | 0 |
| 0 | 0 | 0 | 0 |
| 0 | 0 | 0 | 0 |
| 0 | 0 | 0 | 0 |
| 0 | 0 | 0 | 0 |
| 0 | 0 | 0 | 0 |
| 0 | 0 | 0 | 0 |
| 0 | 0 | 0 | 0 |
| 0 | 0 | 0 | 0 |
| 0 | 0 | 0 | 0 |
| 0 | 0 | 0 | 0 |
| 0 | 0 | 0 | 0 |
| 0 | 0 | 0 | 0 |
| 0 | 0 | 0 | 0 |
| 0 | 0 | 0 | 0 |
| 0 | 0 | 0 | 0 |
| 1 | 1 | 0 | 1 |
| 1 | 1 | 0 | 1 |
| 1 | 0 | 0 | 0 |
| 0 | 0 | 0 | 0 |
| 0 | 0 | 0 | 0 |
| 0 | 0 | 0 | 0 |
| 0 | 0 | 0 | 0 |
| 0 | 0 | 0 | 0 |
| 0 | 0 | 0 | 0 |
| 0 | 0 | 0 | 0 |
| 1 | 1 | 0 | 0 |
| 0 | 0 | 0 | 0 |
| 1 | 0 | 0 | 0 |
| 0 | 0 | 0 | 0 |
| 0 | 0 | 0 | 0 |
| 0 | 0 | 0 | 0 |
| 0 | 0 | 0 | 0 |
| 0 | 0 | 0 | 0 |
| 1 | 0 | 0 | 0 |
| 0 | 0 | 0 | 0 |
| 0 | 0 | 0 | 0 |
| 0 | 0 | 0 | 0 |

| | | | |
|---|---|---|---|
| 0 | 0 | 0 | 0 |
| 0 | 0 | 0 | 0 |
| 0 | 0 | 0 | 0 |
| 0 | 0 | 0 | 0 |
| 0 | 0 | 0 | 0 |
| 0 | 0 | 0 | 0 |
| 0 | 0 | 0 | 0 |
| 0 | 0 | 0 | 0 |
| 0 | 0 | 0 | 0 |
| 0 | 0 | 0 | 0 |
| 0 | 0 | 0 | 0 |
| 0 | 0 | 0 | 0 |
| 0 | 0 | 0 | 0 |
| 0 | 0 | 0 | 0 |
| 0 | 0 | 0 | 0 |
| 0 | 0 | 0 | 0 |
| 0 | 0 | 0 | 0 |
| 0 | 0 | 0 | 0 |
| 0 | 0 | 0 | 0 |
| 0 | 0 | 0 | 1 |
| 0 | 0 | 0 | 0 |
| 0 | 0 | 0 | 1 |
| 0 | 0 | 0 | 1 |
| 0 | 0 | 0 | 1 |
| 0 | 0 | 0 | 0 |
| 0 | 0 | 0 | 0 |
| 0 | 0 | 0 | 0 |
| 0 | 0 | 0 | 0 |
| 0 | 0 | 0 | 0 |
| 0 | 0 | 0 | 0 |
| 0 | 0 | 0 | 0 |
| 0 | 0 | 0 | 0 |
| 0 | 0 | 0 | 0 |
| 0 | 0 | 0 | 0 |
| 0 | 0 | 0 | 0 |
| 0 | 1 | 0 | 0 |
| 0 | 0 | 0 | 0 |
| 0 | 0 | 0 | 0 |
| 0 | 1 | 0 | 0 |
| 0 | 0 | 0 | 0 |
| 0 | 0 | 0 | 0 |
| 0 | 0 | 0 | 0 |
| 0 | 0 | 0 | 0 |
| 0 | 0 | 0 | 0 |
| 0 | 0 | 0 | 0 |
| 0 | 0 | 0 | 0 |
| 0 | 0 | 0 | 0 |
| 0 | 0 | 0 | 0 |
| 0 | 0 | 0 | 0 |
| 0 | 0 | 0 | 0 |
| 0 | 0 | 0 | 0 |
| 0 | 0 | 0 | 0 |
| 0 | 0 | 0 | 0 |
| 0 | 0 | 0 | 0 |

| | | | |
|---|---|---|---|
| 0 | 0 | 0 | 0 |
| 0 | 0 | 0 | 0 |
| 0 | 0 | 0 | 0 |
| 0 | 0 | 0 | 0 |
| 0 | 0 | 0 | 0 |
| 0 | 0 | 0 | 0 |
| 0 | 0 | 0 | 0 |
| 0 | 0 | 0 | 0 |
| 0 | 0 | 1 | 0 |
| 0 | 0 | 1 | 0 |
| 0 | 0 | 0 | 0 |
| 0 | 0 | 0 | 0 |
| 0 | 0 | 0 | 0 |
| 0 | 0 | 0 | 0 |
| 0 | 0 | 0 | 0 |
| 0 | 1 | 0 | 0 |
| 0 | 0 | 0 | 0 |
| 0 | 0 | 0 | 0 |
| 1 | 1 | 0 | 0 |
| 0 | 0 | 0 | 0 |
| 0 | 0 | 0 | 0 |
| 0 | 0 | 0 | 0 |
| 0 | 0 | 0 | 0 |
| 0 | 0 | 0 | 0 |
| 0 | 0 | 0 | 0 |
| 0 | 0 | 0 | 0 |
| 0 | 0 | 0 | 0 |
| 0 | 0 | 0 | 0 |
| 0 | 0 | 0 | 0 |
| 0 | 0 | 0 | 0 |
| 0 | 0 | 0 | 0 |
| 0 | 0 | 0 | 0 |
| 0 | 0 | 0 | 0 |
| 0 | 0 | 0 | 0 |
| 0 | 0 | 0 | 0 |
| 0 | 0 | 0 | 0 |
| 0 | 0 | 0 | 0 |
| 0 | 0 | 0 | 0 |
| 0 | 0 | 0 | 0 |
| 0 | 0 | 0 | 0 |
| 0 | 0 | 0 | 0 |
| 0 | 0 | 0 | 0 |
| 0 | 0 | 0 | 0 |
| 0 | 0 | 0 | 0 |
| 0 | 0 | 0 | 0 |
| 0 | 0 | 0 | 0 |
| 0 | 0 | 0 | 0 |
| 0 | 0 | 0 | 0 |
| 0 | 0 | 0 | 0 |
| 0 | 0 | 0 | 0 |
| 0 | 0 | 0 | 0 |
| 0 | 0 | 0 | 0 |
| 0 | 0 | 0 | 0 |
| 1 | 0 | 0 | 0 |

| | | | |
|---|---|---|---|
| 0 | 0 | 0 | 0 |
| 0 | 0 | 0 | 0 |
| 0 | 0 | 0 | 0 |
| 0 | 0 | 0 | 0 |
| 0 | 0 | 0 | 0 |
| 0 | 0 | 0 | 0 |
| 0 | 0 | 0 | 0 |
| 0 | 0 | 0 | 0 |
| 0 | 0 | 0 | 0 |
| 0 | 0 | 0 | 0 |
| 1 | 0 | 0 | 0 |
| 0 | 0 | 0 | 0 |
| 0 | 0 | 0 | 0 |
| 0 | 0 | 0 | 0 |
| 1 | 1 | 0 | 1 |
| 0 | 0 | 0 | 0 |
| 1 | 0 | 0 | 0 |
| 1 | 1 | 0 | 0 |
| 0 | 0 | 0 | 0 |
| 0 | 0 | 0 | 0 |
| 0 | 0 | 0 | 0 |
| 0 | 0 | 0 | 0 |
| 0 | 0 | 0 | 0 |
| 0 | 0 | 0 | 1 |
| 0 | 0 | 0 | 0 |
| 0 | 0 | 0 | 0 |
| 0 | 0 | 0 | 0 |
| 0 | 0 | 0 | 1 |
| 0 | 0 | 0 | 0 |
| 0 | 0 | 0 | 0 |
| 0 | 0 | 0 | 0 |
| 0 | 0 | 0 | 0 |
| 0 | 0 | 0 | 0 |
| 0 | 0 | 0 | 0 |
| 0 | 0 | 0 | 0 |
| 0 | 0 | 0 | 0 |
| 0 | 0 | 0 | 0 |
| 0 | 0 | 0 | 0 |
| 0 | 0 | 0 | 0 |
| 0 | 0 | 0 | 0 |
| 0 | 0 | 0 | 0 |
| 0 | 0 | 0 | 0 |
| 0 | 0 | 0 | 0 |
| 0 | 0 | 0 | 0 |
| 0 | 0 | 1 | 0 |
| 0 | 0 | 0 | 0 |
| 0 | 0 | 0 | 0 |
| 0 | 0 | 0 | 0 |
| 0 | 0 | 0 | 0 |
| 0 | 0 | 0 | 0 |
| 0 | 0 | 0 | 0 |
| 0 | 0 | 0 | 0 |
| 0 | 0 | 0 | 0 |
| 0 | 0 | 0 | 0 |

| | | | |
|---|---|---|---|
| 0 | 0 | 0 | 0 |
| 0 | 0 | 0 | 0 |
| 0 | 0 | 0 | 0 |
| 0 | 0 | 0 | 0 |
| 0 | 0 | 0 | 0 |
| 0 | 0 | 0 | 0 |
| 1 | 0 | 0 | 1 |
| 1 | 0 | 0 | 0 |
| 0 | 0 | 0 | 0 |
| 0 | 0 | 0 | 0 |
| 0 | 0 | 0 | 0 |
| 0 | 0 | 0 | 0 |
| 0 | 0 | 0 | 0 |
| 0 | 0 | 0 | 0 |
| 0 | 0 | 0 | 0 |
| 0 | 0 | 0 | 0 |
| 0 | 0 | 0 | 0 |
| 0 | 0 | 0 | 0 |
| 0 | 0 | 0 | 0 |
| 0 | 0 | 0 | 0 |
| 1 | 0 | 0 | 0 |
| 0 | 0 | 0 | 0 |
| 0 | 0 | 0 | 0 |
| 0 | 0 | 0 | 0 |
| 0 | 0 | 0 | 0 |
| 0 | 0 | 0 | 0 |
| 0 | 0 | 0 | 0 |
| 0 | 0 | 0 | 0 |
| 0 | 0 | 0 | 0 |
| 0 | 0 | 0 | 0 |
| 0 | 0 | 0 | 0 |
| 0 | 0 | 0 | 0 |
| 0 | 0 | 0 | 0 |
| 0 | 0 | 0 | 0 |
| 0 | 0 | 0 | 0 |
| 0 | 0 | 0 | 0 |
| 0 | 0 | 0 | 0 |
| 0 | 0 | 0 | 0 |
| 0 | 0 | 0 | 0 |
| 0 | 0 | 0 | 0 |
| 0 | 0 | 0 | 0 |
| 0 | 0 | 0 | 0 |
| 0 | 0 | 0 | 0 |
| 0 | 0 | 0 | 0 |
| 0 | 0 | 0 | 0 |
| 0 | 0 | 0 | 0 |
| 0 | 0 | 0 | 0 |
| 0 | 0 | 0 | 0 |
| 0 | 0 | 0 | 0 |
| 0 | 0 | 0 | 0 |
| 0 | 0 | 0 | 0 |
| 0 | 0 | 0 | 0 |
| 0 | 0 | 0 | 0 |
| 0 | 0 | 0 | 0 |

| | | | |
|---|---|---|---|
| 0 | 0 | 0 | 0 |
| 0 | 0 | 0 | 0 |
| 0 | 0 | 0 | 0 |
| 1 | 0 | 0 | 0 |
| 0 | 0 | 0 | 0 |
| 1 | 0 | 0 | 0 |
| 0 | 0 | 0 | 0 |
| 0 | 0 | 0 | 0 |
| 1 | 0 | 0 | 0 |
| 0 | 0 | 0 | 0 |
| 0 | 0 | 0 | 0 |
| 0 | 0 | 0 | 0 |
| 0 | 0 | 0 | 0 |
| 0 | 0 | 0 | 0 |
| 0 | 0 | 0 | 0 |
| 0 | 0 | 0 | 0 |
| 0 | 0 | 0 | 0 |
| 0 | 0 | 0 | 0 |
| 1 | 0 | 0 | 0 |
| 0 | 0 | 0 | 0 |
| 0 | 0 | 0 | 0 |
| 0 | 0 | 0 | 0 |
| 0 | 0 | 0 | 0 |
| 0 | 0 | 0 | 0 |
| 0 | 0 | 0 | 0 |
| 0 | 0 | 0 | 0 |
| 0 | 0 | 0 | 0 |
| 0 | 0 | 0 | 0 |
| 0 | 1 | 0 | 0 |
| 0 | 0 | 0 | 0 |
| 0 | 0 | 0 | 0 |
| 0 | 0 | 0 | 0 |
| 0 | 1 | 0 | 0 |
| 1 | 1 | 0 | 0 |
| 1 | 0 | 0 | 0 |
| 0 | 0 | 0 | 0 |
| 0 | 0 | 0 | 0 |
| 0 | 0 | 0 | 0 |
| 0 | 0 | 0 | 0 |
| 0 | 0 | 0 | 0 |
| 1 | 0 | 0 | 0 |
| 0 | 0 | 0 | 0 |
| 0 | 0 | 0 | 0 |
| 0 | 0 | 0 | 0 |
| 0 | 0 | 0 | 0 |
| 0 | 0 | 0 | 0 |
| 0 | 0 | 0 | 0 |
| 0 | 0 | 0 | 0 |
| 0 | 0 | 0 | 0 |
| 0 | 0 | 0 | 0 |
| 0 | 0 | 0 | 0 |
| 0 | 0 | 0 | 0 |
| 0 | 0 | 0 | 0 |
| 1 | 0 | 0 | 0 |

| | | | |
|---|---|---|---|
| 1 | 0 | 0 | 1 |
| 0 | 0 | 0 | 0 |
| 0 | 0 | 0 | 0 |
| 0 | 0 | 0 | 0 |
| 0 | 0 | 0 | 0 |
| 0 | 0 | 0 | 0 |
| 0 | 0 | 0 | 0 |
| 0 | 0 | 0 | 0 |
| 0 | 0 | 0 | 0 |
| 0 | 0 | 0 | 0 |
| 0 | 0 | 0 | 0 |
| 0 | 0 | 0 | 0 |
| 0 | 0 | 0 | 0 |
| 0 | 0 | 0 | 0 |
| 0 | 0 | 0 | 0 |
| 0 | 0 | 0 | 0 |
| 0 | 0 | 0 | 0 |
| 0 | 0 | 0 | 0 |
| 0 | 0 | 0 | 0 |
| 0 | 0 | 0 | 0 |
| 0 | 0 | 0 | 0 |
| 0 | 0 | 0 | 0 |
| 0 | 1 | 0 | 0 |
| 0 | 0 | 0 | 0 |
| 0 | 0 | 0 | 0 |
| 0 | 0 | 0 | 0 |
| 0 | 0 | 0 | 0 |
| 0 | 0 | 0 | 0 |
| 0 | 0 | 0 | 0 |
| 0 | 0 | 0 | 0 |
| 0 | 0 | 0 | 0 |
| 0 | 0 | 0 | 0 |
| 0 | 0 | 0 | 0 |
| 0 | 0 | 0 | 0 |
| 0 | 0 | 0 | 0 |
| 0 | 0 | 0 | 0 |
| 0 | 0 | 0 | 0 |
| 0 | 0 | 0 | 0 |
| 0 | 0 | 0 | 0 |
| 0 | 0 | 0 | 0 |
| 0 | 0 | 0 | 0 |
| 0 | 0 | 0 | 0 |
| 0 | 0 | 0 | 0 |
| 0 | 0 | 0 | 0 |
| 0 | 0 | 0 | 0 |
| 0 | 0 | 0 | 0 |
| 0 | 0 | 0 | 0 |
| 0 | 0 | 0 | 0 |
| 0 | 0 | 0 | 0 |
| 0 | 0 | 0 | 0 |
| 0 | 0 | 0 | 0 |
| 0 | 0 | 0 | 0 |
| 0 | 0 | 0 | 0 |
| 0 | 0 | 0 | 0 |

| | | | |
|---|---|---|---|
| 0 | 0 | 0 | 0 |
| 0 | 0 | 0 | 0 |
| 0 | 0 | 0 | 0 |
| 0 | 0 | 0 | 0 |
| 0 | 0 | 0 | 0 |
| 0 | 0 | 0 | 0 |
| 0 | 1 | 0 | 0 |
| 0 | 1 | 0 | 0 |
| 0 | 0 | 0 | 0 |
| 0 | 0 | 0 | 0 |
| 0 | 0 | 0 | 0 |
| 0 | 0 | 0 | 0 |
| 0 | 0 | 0 | 0 |
| 0 | 0 | 0 | 0 |
| 0 | 0 | 0 | 0 |
| 0 | 0 | 0 | 0 |
| 1 | 0 | 0 | 0 |
| 0 | 0 | 0 | 0 |
| 1 | 0 | 0 | 0 |
| 1 | 0 | 0 | 0 |
| 1 | 0 | 0 | 0 |
| 1 | 0 | 0 | 0 |
| 0 | 0 | 0 | 0 |
| 0 | 0 | 0 | 0 |
| 0 | 0 | 0 | 0 |
| 0 | 0 | 0 | 0 |
| 0 | 0 | 0 | 0 |
| 0 | 0 | 0 | 0 |
| 0 | 0 | 0 | 0 |
| 0 | 0 | 0 | 0 |
| 0 | 0 | 0 | 0 |
| 0 | 0 | 0 | 0 |
| 0 | 0 | 0 | 0 |
| 0 | 0 | 0 | 0 |
| 0 | 0 | 0 | 0 |
| 0 | 0 | 1 | 0 |
| 1 | 0 | 0 | 0 |
| 1 | 0 | 0 | 0 |
| 1 | 0 | 0 | 0 |
| 0 | 0 | 0 | 0 |
| 0 | 0 | 0 | 0 |
| 0 | 0 | 0 | 0 |
| 0 | 0 | 0 | 0 |
| 1 | 0 | 0 | 0 |
| 1 | 0 | 0 | 0 |
| 1 | 0 | 0 | 0 |
| 0 | 0 | 0 | 0 |
| 0 | 0 | 0 | 0 |
| 1 | 0 | 0 | 0 |
| 0 | 0 | 0 | 0 |
| 0 | 0 | 0 | 0 |
| 1 | 0 | 0 | 0 |
| 0 | 0 | 0 | 0 |
| 0 | 0 | 0 | 0 |

| | | | |
|---|---|---|---|
| 0 | 0 | 0 | 0 |
| 0 | 0 | 0 | 0 |
| 0 | 0 | 0 | 0 |
| 0 | 0 | 0 | 0 |
| 0 | 0 | 0 | 0 |
| 0 | 0 | 0 | 0 |
| 0 | 0 | 0 | 0 |
| 0 | 0 | 0 | 0 |
| 0 | 0 | 0 | 0 |
| 0 | 0 | 0 | 0 |
| 0 | 0 | 0 | 0 |
| 0 | 0 | 0 | 0 |
| 0 | 0 | 0 | 0 |
| 0 | 0 | 0 | 0 |
| 0 | 0 | 0 | 0 |
| 0 | 0 | 0 | 0 |
| 0 | 0 | 0 | 0 |
| 0 | 0 | 0 | 0 |
| 0 | 0 | 0 | 0 |
| 0 | 0 | 0 | 0 |
| 0 | 0 | 0 | 0 |
| 0 | 0 | 0 | 0 |
| 1 | 1 | 0 | 0 |
| 0 | 0 | 0 | 0 |
| 0 | 1 | 0 | 0 |
| 0 | 0 | 0 | 0 |
| 0 | 0 | 0 | 0 |
| 0 | 0 | 0 | 0 |
| 0 | 0 | 0 | 0 |
| 0 | 0 | 0 | 0 |
| 0 | 0 | 0 | 0 |
| 0 | 0 | 0 | 0 |
| 0 | 0 | 0 | 0 |
| 0 | 0 | 0 | 0 |
| 0 | 0 | 0 | 0 |
| 0 | 0 | 0 | 0 |
| 0 | 0 | 0 | 0 |
| 0 | 0 | 0 | 0 |
| 0 | 0 | 0 | 0 |
| 0 | 0 | 0 | 0 |
| 0 | 0 | 0 | 0 |
| 0 | 0 | 0 | 0 |
| 0 | 0 | 0 | 0 |
| 0 | 0 | 0 | 0 |
| 0 | 0 | 0 | 0 |
| 1 | 0 | 0 | 1 |
| 0 | 0 | 0 | 0 |
| 0 | 0 | 0 | 0 |
| 0 | 0 | 0 | 0 |
| 1 | 0 | 0 | 0 |
| 1 | 0 | 0 | 0 |
| 1 | 0 | 0 | 0 |
| 0 | 0 | 0 | 0 |
| 0 | 0 | 0 | 0 |

| | | | |
|---|---|---|---|
| 0 | 0 | 0 | 0 |
| 0 | 0 | 0 | 0 |
| 0 | 0 | 0 | 0 |
| 0 | 0 | 0 | 0 |
| 0 | 0 | 0 | 0 |
| 0 | 0 | 0 | 0 |
| 0 | 0 | 0 | 0 |
| 0 | 0 | 0 | 0 |
| 0 | 0 | 0 | 0 |
| 0 | 0 | 0 | 0 |
| 0 | 0 | 0 | 1 |
| 0 | 0 | 0 | 0 |
| 0 | 0 | 0 | 0 |
| 0 | 0 | 0 | 0 |
| 0 | 0 | 0 | 0 |
| 0 | 0 | 0 | 0 |
| 0 | 0 | 0 | 0 |
| 0 | 0 | 0 | 0 |
| 0 | 0 | 0 | 0 |
| 0 | 0 | 0 | 0 |
| 0 | 0 | 0 | 0 |
| 0 | 0 | 0 | 0 |
| 0 | 0 | 0 | 0 |
| 0 | 0 | 0 | 0 |
| 0 | 0 | 0 | 0 |
| 0 | 0 | 0 | 0 |
| 0 | 0 | 0 | 0 |
| 0 | 0 | 0 | 0 |
| 0 | 0 | 0 | 0 |
| 0 | 0 | 0 | 0 |
| 0 | 1 | 0 | 0 |
| 0 | 0 | 0 | 0 |
| 1 | 1 | 0 | 0 |
| 1 | 1 | 0 | 0 |
| 0 | 0 | 0 | 0 |
| 0 | 0 | 0 | 0 |
| 0 | 0 | 0 | 0 |
| 0 | 0 | 0 | 0 |
| 0 | 0 | 0 | 0 |
| 0 | 0 | 0 | 0 |
| 0 | 0 | 0 | 0 |
| 0 | 1 | 0 | 0 |
| 0 | 0 | 0 | 0 |
| 0 | 0 | 0 | 0 |
| 0 | 0 | 0 | 0 |
| 1 | 0 | 0 | 0 |
| 1 | 0 | 0 | 0 |
| 1 | 0 | 0 | 0 |
| 1 | 1 | 1 | 0 |
| 0 | 0 | 0 | 0 |
| 1 | 1 | 1 | 1 |
| 0 | 0 | 0 | 0 |
| 0 | 0 | 0 | 0 |
| 0 | 0 | 0 | 0 |

| | | | |
|---|---|---|---|
| 0 | 0 | 0 | 0 |
| 0 | 0 | 0 | 0 |
| 0 | 0 | 0 | 0 |
| 0 | 0 | 0 | 0 |
| 1 | 0 | 0 | 0 |
| 1 | 0 | 0 | 0 |
| 0 | 0 | 0 | 0 |
| 0 | 0 | 0 | 0 |
| 0 | 0 | 0 | 0 |
| 0 | 0 | 0 | 0 |
| 0 | 0 | 0 | 0 |
| 0 | 0 | 0 | 0 |
| 0 | 0 | 0 | 0 |
| 0 | 0 | 0 | 0 |
| 0 | 0 | 0 | 0 |
| 0 | 0 | 0 | 0 |
| 0 | 0 | 0 | 0 |
| 0 | 0 | 0 | 0 |
| 0 | 0 | 0 | 0 |
| 0 | 0 | 0 | 0 |
| 0 | 0 | 0 | 0 |
| 0 | 0 | 0 | 0 |
| 0 | 0 | 0 | 0 |
| 0 | 0 | 0 | 0 |
| 0 | 0 | 0 | 0 |
| 0 | 0 | 0 | 0 |
| 0 | 0 | 0 | 0 |
| 0 | 0 | 0 | 0 |
| 0 | 0 | 0 | 0 |
| 0 | 0 | 0 | 0 |
| 0 | 0 | 0 | 0 |
| 0 | 0 | 0 | 0 |
| 0 | 0 | 0 | 0 |
| 0 | 0 | 0 | 0 |
| 0 | 0 | 0 | 0 |
| 0 | 0 | 0 | 0 |
| 0 | 0 | 0 | 0 |
| 0 | 0 | 0 | 0 |
| 0 | 0 | 0 | 0 |
| 0 | 0 | 0 | 0 |
| 0 | 0 | 0 | 0 |
| 0 | 0 | 0 | 0 |
| 0 | 0 | 0 | 0 |
| 0 | 0 | 0 | 0 |
| 0 | 0 | 0 | 0 |
| 0 | 0 | 0 | 0 |
| 0 | 0 | 0 | 0 |
| 0 | 0 | 0 | 0 |
| 0 | 0 | 0 | 0 |
| 0 | 0 | 0 | 0 |
| 0 | 0 | 0 | 0 |
| 0 | 0 | 0 | 0 |
| 0 | 0 | 0 | 0 |
| 0 | 1 | 0 | 0 |

| | | | |
|---|---|---|---|
| 0 | 0 | 0 | 0 |
| 0 | 0 | 0 | 0 |
| 0 | 0 | 0 | 0 |
| 0 | 0 | 0 | 0 |
| 0 | 0 | 0 | 0 |
| 0 | 0 | 0 | 0 |
| 0 | 0 | 0 | 0 |
| 0 | 0 | 0 | 0 |
| 0 | 0 | 0 | 0 |
| 0 | 0 | 0 | 0 |
| 0 | 0 | 0 | 0 |
| 0 | 0 | 0 | 0 |
| 0 | 0 | 0 | 0 |
| 0 | 0 | 0 | 0 |
| 0 | 0 | 0 | 0 |
| 0 | 0 | 0 | 0 |
| 0 | 0 | 0 | 0 |
| 0 | 0 | 0 | 0 |
| 0 | 0 | 0 | 0 |
| 0 | 0 | 0 | 0 |
| 0 | 0 | 0 | 0 |
| 0 | 0 | 0 | 0 |
| 0 | 0 | 0 | 0 |
| 0 | 0 | 0 | 0 |
| 0 | 0 | 0 | 0 |
| 0 | 0 | 0 | 0 |
| 0 | 0 | 0 | 0 |
| 0 | 0 | 0 | 0 |
| 0 | 0 | 0 | 0 |
| 0 | 0 | 0 | 0 |
| 0 | 0 | 0 | 0 |
| 0 | 0 | 0 | 0 |
| 0 | 0 | 0 | 0 |
| 0 | 0 | 0 | 0 |
| 0 | 0 | 0 | 0 |
| 0 | 0 | 0 | 0 |
| 0 | 0 | 0 | 0 |
| 0 | 0 | 0 | 0 |
| 0 | 0 | 0 | 0 |
| 0 | 0 | 0 | 0 |
| 0 | 0 | 0 | 0 |
| 0 | 0 | 0 | 0 |
| 0 | 0 | 0 | 0 |
| 0 | 0 | 0 | 0 |
| 0 | 0 | 0 | 0 |
| 0 | 0 | 0 | 0 |
| 0 | 0 | 0 | 0 |
| 0 | 0 | 0 | 0 |
| 0 | 0 | 0 | 0 |
| 0 | 0 | 0 | 0 |
| 0 | 0 | 0 | 0 |
| 0 | 0 | 0 | 0 |
| 0 | 0 | 0 | 0 |
| 0 | 0 | 0 | 0 |

| | | | |
|---|---|---|---|
| 0 | 0 | 0 | 0 |
| 0 | 0 | 0 | 0 |
| 0 | 0 | 0 | 0 |
| 0 | 0 | 0 | 0 |
| 0 | 0 | 0 | 0 |
| 0 | 0 | 0 | 0 |
| 0 | 0 | 0 | 0 |
| 0 | 0 | 0 | 0 |
| 0 | 0 | 0 | 0 |
| 0 | 0 | 0 | 0 |
| 0 | 0 | 0 | 0 |
| 0 | 0 | 1 | 0 |
| 0 | 0 | 0 | 0 |
| 0 | 0 | 1 | 0 |
| 0 | 0 | 1 | 0 |
| 0 | 0 | 1 | 0 |
| 0 | 0 | 1 | 0 |
| 0 | 0 | 0 | 0 |
| 0 | 0 | 0 | 0 |
| 0 | 0 | 0 | 0 |
| 1 | 0 | 0 | 0 |

| glycan_binding_site | water_binding_site | water_only_binding_site | non_binding_site |
|---|---|---|---|
| 0 | 0 | 0 | 0 |
| 0 | 0 | 0 | 0 |
| 0 | 0 | 0 | 1 |
| 0 | 0 | 0 | 0 |
| 0 | 0 | 0 | 0 |
| 0 | 0 | 0 | 0 |
| 0 | 0 | 0 | 1 |
| 0 | 0 | 0 | 1 |
| 0 | 0 | 0 | 0 |
| 0 | 0 | 0 | 0 |
| 0 | 0 | 0 | 0 |
| 0 | 0 | 0 | 0 |
| 0 | 0 | 0 | 1 |
| 1 | 0 | 0 | 0 |
| 0 | 0 | 0 | 0 |
| 0 | 0 | 0 | 0 |
| 0 | 0 | 0 | 0 |
| 0 | 0 | 0 | 0 |
| 0 | 0 | 0 | 0 |
| 0 | 0 | 0 | 0 |
| 0 | 1 | 0 | 0 |
| 1 | 0 | 0 | 0 |
| 1 | 1 | 0 | 0 |
| 0 | 1 | 0 | 0 |
| 0 | 0 | 0 | 0 |
| 0 | 0 | 0 | 1 |
| 0 | 0 | 0 | 0 |
| 0 | 0 | 0 | 1 |
| 0 | 0 | 0 | 0 |
| 0 | 0 | 0 | 1 |
| 0 | 0 | 0 | 1 |
| 0 | 0 | 0 | 0 |
| 0 | 0 | 0 | 0 |
| 0 | 0 | 0 | 0 |
| 0 | 0 | 0 | 0 |
| 0 | 0 | 0 | 1 |
| 0 | 0 | 0 | 0 |
| 0 | 0 | 0 | 0 |
| 0 | 0 | 0 | 0 |
| 0 | 0 | 0 | 0 |
| 0 | 0 | 0 | 0 |
| 0 | 0 | 0 | 0 |
| 0 | 0 | 0 | 0 |
| 0 | 0 | 0 | 0 |
| 0 | 0 | 0 | 0 |
| 0 | 0 | 0 | 0 |
| 0 | 0 | 0 | 0 |
| 0 | 0 | 0 | 0 |
| 0 | 0 | 0 | 0 |
| 0 | 0 | 0 | 0 |
| 0 | 0 | 0 | 0 |
| 0 | 0 | 0 | 0 |
| 0 | 0 | 0 | 0 |

| | | | |
|---|---|---|---|
| 0 | 0 | 0 | 0 |
| 0 | 0 | 0 | 0 |
| 0 | 0 | 0 | 0 |
| 0 | 0 | 0 | 0 |
| 0 | 0 | 0 | 0 |
| 0 | 0 | 0 | 0 |
| 0 | 0 | 0 | 0 |
| 0 | 0 | 0 | 0 |
| 0 | 0 | 0 | 0 |
| 0 | 0 | 0 | 0 |
| 0 | 0 | 0 | 0 |
| 0 | 0 | 0 | 0 |
| 0 | 0 | 0 | 0 |
| 0 | 0 | 0 | 0 |
| 0 | 0 | 0 | 0 |
| 0 | 0 | 0 | 0 |
| 0 | 0 | 0 | 0 |
| 0 | 0 | 0 | 0 |
| 0 | 0 | 0 | 0 |
| 0 | 0 | 0 | 0 |
| 0 | 0 | 0 | 0 |
| 0 | 0 | 0 | 0 |
| 0 | 0 | 0 | 0 |
| 0 | 0 | 0 | 0 |
| 0 | 0 | 0 | 0 |
| 0 | 0 | 0 | 0 |
| 0 | 0 | 0 | 0 |
| 0 | 0 | 0 | 0 |
| 0 | 0 | 0 | 0 |
| 0 | 0 | 0 | 0 |
| 0 | 0 | 0 | 0 |
| 0 | 0 | 0 | 0 |
| 0 | 0 | 0 | 0 |
| 0 | 0 | 0 | 0 |
| 0 | 0 | 0 | 0 |
| 0 | 0 | 0 | 0 |
| 0 | 0 | 0 | 0 |
| 0 | 0 | 0 | 0 |
| 0 | 0 | 0 | 0 |
| 0 | 0 | 0 | 0 |
| 1 | 0 | 0 | 0 |
| 0 | 0 | 0 | 0 |
| 1 | 0 | 0 | 0 |
| 1 | 0 | 0 | 0 |
| 0 | 0 | 0 | 0 |
| 0 | 0 | 0 | 1 |
| 0 | 0 | 0 | 1 |
| 0 | 0 | 0 | 0 |
| 1 | 0 | 0 | 0 |
| 0 | 0 | 0 | 0 |
| 0 | 0 | 0 | 0 |
| 0 | 0 | 0 | 1 |
| 0 | 0 | 0 | 1 |
| 0 | 0 | 0 | 0 |

| | | | |
|---|---|---|---|
| 0 | 0 | 0 | 1 |
| 0 | 0 | 0 | 1 |
| 0 | 0 | 0 | 0 |
| 0 | 0 | 0 | 0 |
| 0 | 0 | 0 | 0 |
| 0 | 0 | 0 | 0 |
| 0 | 0 | 0 | 1 |
| 0 | 0 | 0 | 0 |
| 0 | 0 | 0 | 0 |
| 0 | 0 | 0 | 0 |
| 0 | 0 | 0 | 0 |
| 0 | 0 | 0 | 0 |
| 0 | 0 | 0 | 0 |
| 1 | 0 | 0 | 0 |
| 1 | 0 | 0 | 0 |
| 0 | 0 | 0 | 1 |
| 0 | 0 | 0 | 0 |
| 0 | 0 | 0 | 0 |
| 0 | 0 | 0 | 0 |
| 0 | 0 | 0 | 0 |
| 0 | 0 | 0 | 0 |
| 0 | 0 | 0 | 0 |
| 0 | 0 | 0 | 0 |
| 0 | 0 | 0 | 0 |
| 0 | 0 | 0 | 0 |
| 0 | 0 | 0 | 0 |
| 0 | 0 | 0 | 0 |
| 0 | 0 | 0 | 0 |
| 0 | 0 | 0 | 0 |
| 0 | 0 | 0 | 0 |
| 0 | 0 | 0 | 0 |
| 0 | 0 | 0 | 0 |
| 0 | 0 | 0 | 0 |
| 0 | 0 | 0 | 0 |
| 0 | 0 | 0 | 0 |
| 0 | 0 | 0 | 0 |
| 0 | 0 | 0 | 0 |
| 0 | 0 | 0 | 0 |
| 0 | 0 | 0 | 0 |
| 0 | 0 | 0 | 0 |
| 0 | 0 | 0 | 0 |
| 0 | 0 | 0 | 0 |
| 0 | 0 | 0 | 0 |
| 0 | 0 | 0 | 0 |
| 0 | 0 | 0 | 0 |
| 0 | 0 | 0 | 0 |
| 1 | 0 | 0 | 0 |
| 0 | 0 | 0 | 0 |
| 0 | 0 | 0 | 0 |
| 0 | 0 | 0 | 0 |
| 0 | 0 | 0 | 0 |
| 0 | 0 | 0 | 0 |
| 0 | 0 | 0 | 0 |
| 0 | 1 | 1 | 1 |

| | | | |
|---|---|---|---|
| 0 | 1 | 1 | 1 |
| 0 | 1 | 1 | 1 |
| 0 | 0 | 0 | 1 |
| 0 | 0 | 0 | 0 |
| 0 | 0 | 0 | 0 |
| 0 | 1 | 1 | 1 |
| 0 | 0 | 0 | 0 |
| 0 | 1 | 1 | 1 |
| 0 | 1 | 1 | 1 |
| 0 | 0 | 0 | 1 |
| 0 | 0 | 0 | 1 |
| 0 | 1 | 0 | 0 |
| 0 | 1 | 1 | 1 |
| 0 | 0 | 0 | 0 |
| 0 | 1 | 1 | 1 |
| 0 | 0 | 0 | 1 |
| 0 | 0 | 0 | 1 |
| 0 | 1 | 1 | 1 |
| 0 | 1 | 1 | 1 |
| 0 | 0 | 0 | 0 |
| 0 | 1 | 0 | 0 |
| 0 | 0 | 0 | 1 |
| 0 | 0 | 0 | 0 |
| 0 | 0 | 0 | 0 |
| 0 | 0 | 0 | 0 |
| 0 | 0 | 0 | 1 |
| 0 | 0 | 0 | 0 |
| 0 | 0 | 0 | 0 |
| 0 | 0 | 0 | 1 |
| 0 | 0 | 0 | 1 |
| 0 | 0 | 0 | 1 |
| 0 | 0 | 0 | 1 |
| 0 | 0 | 0 | 1 |
| 0 | 0 | 0 | 1 |
| 0 | 0 | 0 | 0 |
| 0 | 0 | 0 | 1 |
| 1 | 0 | 0 | 0 |
| 1 | 0 | 0 | 0 |
| 0 | 1 | 1 | 1 |
| 0 | 1 | 1 | 1 |
| 0 | 0 | 0 | 1 |
| 0 | 1 | 0 | 0 |
| 0 | 1 | 0 | 0 |
| 0 | 0 | 0 | 1 |
| 0 | 0 | 0 | 1 |
| 0 | 0 | 0 | 1 |
| 0 | 0 | 0 | 0 |
| 0 | 0 | 0 | 0 |
| 0 | 0 | 0 | 0 |
| 0 | 0 | 0 | 0 |
| 0 | 0 | 0 | 0 |
| 0 | 0 | 0 | 1 |
| 0 | 0 | 0 | 1 |
| 0 | 0 | 0 | 0 |

| | | | |
|---|---|---|---|
| 0 | 0 | 0 | 0 |
| 0 | 0 | 0 | 0 |
| 0 | 0 | 0 | 0 |
| 0 | 0 | 0 | 1 |
| 0 | 0 | 0 | 1 |
| 0 | 0 | 0 | 1 |
| 0 | 0 | 0 | 0 |
| 0 | 0 | 0 | 0 |
| 0 | 0 | 0 | 1 |
| 0 | 0 | 0 | 1 |
| 1 | 0 | 0 | 0 |
| 0 | 0 | 0 | 1 |
| 1 | 0 | 0 | 0 |
| 0 | 0 | 0 | 1 |
| 0 | 0 | 0 | 1 |
| 0 | 0 | 0 | 1 |
| 0 | 0 | 0 | 1 |
| 0 | 0 | 0 | 0 |
| 0 | 0 | 0 | 0 |
| 0 | 0 | 0 | 0 |
| 0 | 0 | 0 | 0 |
| 0 | 0 | 0 | 1 |
| 0 | 0 | 0 | 0 |
| 0 | 0 | 0 | 1 |
| 0 | 0 | 0 | 1 |
| 0 | 0 | 0 | 0 |
| 0 | 0 | 0 | 0 |
| 0 | 0 | 0 | 0 |
| 0 | 0 | 0 | 0 |
| 0 | 0 | 0 | 0 |
| 0 | 0 | 0 | 0 |
| 0 | 0 | 0 | 0 |
| 0 | 0 | 0 | 1 |
| 0 | 0 | 0 | 1 |
| 0 | 0 | 0 | 0 |
| 0 | 0 | 0 | 1 |
| 0 | 0 | 0 | 1 |
| 0 | 0 | 0 | 0 |
| 0 | 0 | 0 | 1 |
| 0 | 0 | 0 | 1 |
| 0 | 0 | 0 | 0 |
| 0 | 0 | 0 | 1 |
| 0 | 0 | 0 | 1 |
| 0 | 0 | 0 | 1 |
| 0 | 0 | 0 | 0 |
| 0 | 0 | 0 | 0 |
| 0 | 0 | 0 | 0 |
| 0 | 0 | 0 | 0 |
| 0 | 0 | 0 | 0 |
| 0 | 0 | 0 | 0 |
| 0 | 0 | 0 | 0 |
| 0 | 0 | 0 | 0 |
| 0 | 0 | 0 | 0 |
| 0 | 0 | 0 | 0 |

| | | | |
|---|---|---|---|
| 0 | 0 | 0 | 0 |
| 0 | 0 | 0 | 0 |
| 0 | 0 | 0 | 0 |
| 0 | 0 | 0 | 0 |
| 0 | 0 | 0 | 0 |
| 0 | 0 | 0 | 0 |
| 0 | 0 | 0 | 1 |
| 0 | 0 | 0 | 1 |
| 0 | 0 | 0 | 0 |
| 0 | 0 | 0 | 0 |
| 0 | 0 | 0 | 0 |
| 0 | 0 | 0 | 0 |
| 0 | 0 | 0 | 0 |
| 0 | 0 | 0 | 1 |
| 0 | 0 | 0 | 1 |
| 0 | 0 | 0 | 0 |
| 0 | 0 | 0 | 1 |
| 0 | 0 | 0 | 0 |
| 0 | 0 | 0 | 1 |
| 0 | 0 | 0 | 0 |
| 0 | 0 | 0 | 1 |
| 0 | 0 | 0 | 1 |
| 0 | 0 | 0 | 1 |
| 0 | 0 | 0 | 0 |
| 0 | 0 | 0 | 0 |
| 0 | 0 | 0 | 0 |
| 0 | 0 | 0 | 0 |
| 0 | 0 | 0 | 0 |
| 0 | 0 | 0 | 0 |
| 0 | 0 | 0 | 0 |
| 0 | 0 | 0 | 1 |
| 0 | 0 | 0 | 1 |
| 0 | 0 | 0 | 0 |
| 0 | 0 | 0 | 0 |
| 0 | 0 | 0 | 0 |
| 0 | 0 | 0 | 1 |
| 0 | 0 | 0 | 0 |
| 0 | 0 | 0 | 0 |
| 0 | 0 | 0 | 0 |
| 0 | 0 | 0 | 0 |
| 0 | 0 | 0 | 1 |
| 0 | 0 | 0 | 1 |
| 0 | 0 | 0 | 0 |
| 0 | 0 | 0 | 1 |
| 0 | 0 | 0 | 0 |
| 0 | 0 | 0 | 0 |
| 0 | 0 | 0 | 0 |
| 0 | 0 | 0 | 1 |
| 0 | 0 | 0 | 1 |
| 0 | 0 | 0 | 0 |
| 0 | 0 | 0 | 1 |
| 0 | 0 | 0 | 1 |
| 0 | 0 | 0 | 0 |
| 0 | 0 | 0 | 1 |

| | | | |
|---|---|---|---|
| 0 | 0 | 0 | 0 |
| 0 | 0 | 0 | 1 |
| 0 | 0 | 0 | 1 |
| 0 | 0 | 0 | 0 |
| 0 | 0 | 0 | 0 |
| 0 | 0 | 0 | 0 |
| 0 | 0 | 0 | 1 |
| 0 | 0 | 0 | 0 |
| 0 | 0 | 0 | 0 |
| 0 | 0 | 0 | 1 |
| 0 | 0 | 0 | 1 |
| 0 | 0 | 0 | 1 |
| 0 | 0 | 0 | 1 |
| 0 | 0 | 0 | 1 |
| 0 | 0 | 0 | 0 |
| 0 | 0 | 0 | 1 |
| 0 | 0 | 0 | 0 |
| 0 | 0 | 0 | 0 |
| 0 | 0 | 0 | 1 |
| 0 | 0 | 0 | 0 |
| 0 | 0 | 0 | 0 |
| 0 | 0 | 0 | 0 |
| 0 | 0 | 0 | 0 |
| 0 | 0 | 0 | 0 |
| 0 | 0 | 0 | 1 |
| 0 | 0 | 0 | 1 |
| 0 | 0 | 0 | 0 |
| 0 | 0 | 0 | 0 |
| 0 | 0 | 0 | 0 |
| 0 | 0 | 0 | 0 |
| 0 | 0 | 0 | 0 |
| 0 | 0 | 0 | 0 |
| 0 | 0 | 0 | 0 |
| 0 | 0 | 0 | 0 |
| 0 | 0 | 0 | 1 |
| 0 | 0 | 0 | 1 |
| 0 | 0 | 0 | 0 |
| 0 | 0 | 0 | 0 |
| 0 | 0 | 0 | 0 |
| 0 | 0 | 0 | 0 |
| 0 | 0 | 0 | 0 |
| 0 | 0 | 0 | 0 |
| 0 | 0 | 0 | 1 |
| 0 | 0 | 0 | 0 |
| 0 | 0 | 0 | 0 |
| 0 | 0 | 0 | 0 |
| 0 | 0 | 0 | 0 |
| 0 | 0 | 0 | 0 |
| 1 | 1 | 0 | 0 |
| 0 | 0 | 0 | 1 |
| 0 | 0 | 0 | 0 |
| 0 | 0 | 0 | 0 |
| 0 | 0 | 0 | 0 |
| 0 | 0 | 0 | 0 |

| | | | |
|---|---|---|---|
| 0 | 0 | 0 | 0 |
| 0 | 0 | 0 | 0 |
| 0 | 0 | 0 | 0 |
| 0 | 0 | 0 | 0 |
| 0 | 0 | 0 | 1 |
| 0 | 0 | 0 | 0 |
| 0 | 0 | 0 | 0 |
| 0 | 0 | 0 | 0 |
| 0 | 0 | 0 | 0 |
| 0 | 0 | 0 | 0 |
| 0 | 0 | 0 | 0 |
| 0 | 0 | 0 | 0 |
| 0 | 0 | 0 | 0 |
| 0 | 0 | 0 | 0 |
| 0 | 0 | 0 | 0 |
| 0 | 0 | 0 | 0 |
| 0 | 0 | 0 | 0 |
| 0 | 0 | 0 | 0 |
| 0 | 0 | 0 | 1 |
| 0 | 0 | 0 | 0 |
| 0 | 0 | 0 | 0 |
| 0 | 0 | 0 | 0 |
| 0 | 0 | 0 | 0 |
| 0 | 0 | 0 | 1 |
| 0 | 0 | 0 | 0 |
| 0 | 0 | 0 | 0 |
| 0 | 0 | 0 | 0 |
| 0 | 0 | 0 | 1 |
| 0 | 0 | 0 | 1 |
| 0 | 0 | 0 | 1 |
| 0 | 0 | 0 | 0 |
| 0 | 0 | 0 | 0 |
| 0 | 0 | 0 | 0 |
| 0 | 0 | 0 | 0 |
| 0 | 0 | 0 | 0 |
| 0 | 0 | 0 | 1 |
| 0 | 0 | 0 | 1 |
| 0 | 0 | 0 | 0 |
| 0 | 0 | 0 | 0 |
| 0 | 0 | 0 | 1 |
| 0 | 0 | 0 | 1 |
| 0 | 0 | 0 | 1 |
| 0 | 0 | 0 | 1 |
| 0 | 0 | 0 | 1 |
| 0 | 0 | 0 | 0 |
| 0 | 0 | 0 | 0 |
| 0 | 0 | 0 | 0 |
| 0 | 0 | 0 | 0 |
| 0 | 0 | 0 | 0 |
| 0 | 0 | 0 | 1 |
| 0 | 0 | 0 | 1 |
| 0 | 0 | 0 | 0 |
| 0 | 0 | 0 | 1 |
| 0 | 0 | 0 | 0 |

| | | | |
|---|---|---|---|
| 0 | 0 | 0 | 1 |
| 0 | 0 | 0 | 1 |
| 0 | 0 | 0 | 0 |
| 0 | 0 | 0 | 1 |
| 0 | 0 | 0 | 1 |
| 0 | 0 | 0 | 1 |
| 0 | 0 | 0 | 1 |
| 0 | 0 | 0 | 0 |
| 0 | 0 | 0 | 1 |
| 0 | 0 | 0 | 1 |
| 0 | 0 | 0 | 0 |
| 0 | 0 | 0 | 1 |
| 0 | 0 | 0 | 1 |
| 0 | 0 | 0 | 1 |
| 0 | 0 | 0 | 0 |
| 0 | 0 | 0 | 0 |
| 0 | 0 | 0 | 1 |
| 0 | 0 | 0 | 1 |
| 0 | 0 | 0 | 0 |
| 0 | 0 | 0 | 0 |
| 0 | 0 | 0 | 0 |
| 0 | 0 | 0 | 0 |
| 0 | 0 | 0 | 0 |
| 0 | 0 | 0 | 0 |
| 0 | 0 | 0 | 0 |
| 0 | 0 | 0 | 1 |
| 0 | 0 | 0 | 0 |
| 0 | 0 | 0 | 0 |
| 0 | 0 | 0 | 0 |
| 0 | 0 | 0 | 1 |
| 0 | 0 | 0 | 0 |
| 0 | 0 | 0 | 0 |
| 0 | 0 | 0 | 1 |
| 0 | 0 | 0 | 0 |
| 0 | 0 | 0 | 0 |
| 0 | 0 | 0 | 0 |
| 0 | 0 | 0 | 0 |
| 0 | 0 | 0 | 0 |
| 0 | 0 | 0 | 0 |
| 0 | 0 | 0 | 0 |
| 0 | 0 | 0 | 1 |
| 0 | 0 | 0 | 0 |
| 0 | 0 | 0 | 1 |
| 0 | 0 | 0 | 0 |
| 0 | 0 | 0 | 0 |
| 1 | 0 | 0 | 0 |
| 0 | 0 | 0 | 0 |
| 1 | 0 | 0 | 0 |
| 0 | 0 | 0 | 0 |
| 0 | 0 | 0 | 0 |
| 0 | 0 | 0 | 1 |
| 0 | 0 | 0 | 0 |
| 0 | 0 | 0 | 0 |
| 0 | 0 | 0 | 0 |

| | | | |
|---|---|---|---|
| 0 | 0 | 0 | 1 |
| 0 | 0 | 0 | 0 |
| 0 | 0 | 0 | 1 |
| 0 | 0 | 0 | 1 |
| 1 | 0 | 0 | 0 |
| 0 | 0 | 0 | 0 |
| 1 | 0 | 0 | 0 |
| 1 | 0 | 0 | 0 |
| 0 | 0 | 0 | 0 |
| 0 | 0 | 0 | 1 |
| 0 | 0 | 0 | 1 |
| 0 | 0 | 0 | 1 |
| 0 | 0 | 0 | 1 |
| 1 | 0 | 0 | 0 |
| 0 | 0 | 0 | 1 |
| 0 | 0 | 0 | 0 |
| 0 | 0 | 0 | 0 |
| 0 | 0 | 0 | 0 |
| 1 | 0 | 0 | 0 |
| 0 | 0 | 0 | 1 |
| 1 | 0 | 0 | 0 |
| 0 | 0 | 0 | 1 |
| 0 | 0 | 0 | 0 |
| 0 | 0 | 0 | 1 |
| 0 | 0 | 0 | 1 |
| 0 | 0 | 0 | 1 |
| 0 | 0 | 0 | 0 |
| 0 | 0 | 0 | 1 |
| 0 | 0 | 0 | 1 |
| 0 | 0 | 0 | 1 |
| 0 | 0 | 0 | 0 |
| 0 | 0 | 0 | 0 |
| 0 | 0 | 0 | 1 |
| 0 | 0 | 0 | 1 |
| 0 | 0 | 0 | 0 |
| 0 | 0 | 0 | 0 |
| 0 | 0 | 0 | 0 |
| 1 | 0 | 0 | 0 |
| 0 | 0 | 0 | 0 |
| 0 | 0 | 0 | 1 |
| 0 | 0 | 0 | 0 |
| 0 | 0 | 0 | 0 |
| 0 | 0 | 0 | 0 |
| 0 | 0 | 0 | 0 |
| 0 | 0 | 0 | 1 |
| 1 | 0 | 0 | 0 |
| 0 | 0 | 0 | 0 |
| 0 | 0 | 0 | 0 |
| 0 | 0 | 0 | 0 |
| 1 | 0 | 0 | 0 |
| 0 | 0 | 0 | 0 |
| 0 | 0 | 0 | 0 |
| 0 | 0 | 0 | 1 |
| 0 | 0 | 0 | 0 |

| | | | |
|---|---|---|---|
| 0 | 0 | 0 | 0 |
| 0 | 0 | 0 | 0 |
| 0 | 0 | 0 | 1 |
| 0 | 0 | 0 | 1 |
| 0 | 0 | 0 | 1 |
| 0 | 0 | 0 | 0 |
| 0 | 0 | 0 | 1 |
| 0 | 0 | 0 | 1 |
| 0 | 0 | 0 | 0 |
| 0 | 0 | 0 | 0 |
| 1 | 0 | 0 | 0 |
| 1 | 0 | 0 | 0 |
| 0 | 0 | 0 | 1 |
| 1 | 0 | 0 | 0 |
| 0 | 0 | 0 | 0 |
| 0 | 0 | 0 | 0 |
| 0 | 1 | 0 | 0 |
| 0 | 0 | 0 | 0 |
| 0 | 0 | 0 | 0 |
| 0 | 0 | 0 | 1 |
| 0 | 0 | 0 | 0 |
| 0 | 0 | 0 | 0 |
| 0 | 0 | 0 | 0 |
| 0 | 0 | 0 | 0 |
| 0 | 0 | 0 | 0 |
| 0 | 0 | 0 | 1 |
| 0 | 0 | 0 | 1 |
| 0 | 0 | 0 | 1 |
| 0 | 0 | 0 | 0 |
| 0 | 0 | 0 | 0 |
| 0 | 0 | 0 | 0 |
| 0 | 0 | 0 | 0 |
| 0 | 0 | 0 | 0 |
| 0 | 0 | 0 | 0 |
| 0 | 0 | 0 | 0 |
| 0 | 0 | 0 | 1 |
| 0 | 0 | 0 | 0 |
| 0 | 0 | 0 | 0 |
| 0 | 0 | 0 | 1 |
| 0 | 0 | 0 | 1 |
| 0 | 0 | 0 | 1 |
| 0 | 0 | 0 | 0 |
| 0 | 0 | 0 | 0 |
| 0 | 0 | 0 | 0 |
| 0 | 0 | 0 | 0 |
| 0 | 0 | 0 | 0 |
| 0 | 0 | 0 | 0 |
| 0 | 0 | 0 | 1 |
| 0 | 0 | 0 | 1 |
| 0 | 0 | 0 | 0 |
| 0 | 0 | 0 | 0 |
| 0 | 0 | 0 | 0 |
| 0 | 0 | 0 | 0 |
| 0 | 0 | 0 | 1 |

| | | | |
|---|---|---|---|
| 0 | 0 | 0 | 1 |
| 0 | 0 | 0 | 0 |
| 0 | 0 | 0 | 1 |
| 0 | 0 | 0 | 0 |
| 0 | 0 | 0 | 1 |
| 0 | 0 | 0 | 1 |
| 0 | 0 | 0 | 0 |
| 0 | 0 | 0 | 1 |
| 0 | 0 | 0 | 1 |
| 0 | 0 | 0 | 1 |
| 0 | 0 | 0 | 1 |
| 0 | 0 | 0 | 1 |
| 0 | 0 | 0 | 1 |
| 0 | 0 | 0 | 1 |
| 0 | 0 | 0 | 1 |
| 0 | 0 | 0 | 1 |
| 0 | 0 | 0 | 0 |
| 0 | 0 | 0 | 0 |
| 0 | 0 | 0 | 1 |
| 0 | 0 | 0 | 0 |
| 0 | 0 | 0 | 0 |
| 0 | 0 | 0 | 0 |
| 0 | 0 | 0 | 1 |
| 0 | 0 | 0 | 0 |
| 0 | 0 | 0 | 0 |
| 0 | 0 | 0 | 1 |
| 0 | 0 | 0 | 1 |
| 0 | 0 | 0 | 1 |
| 0 | 1 | 0 | 0 |
| 0 | 0 | 0 | 0 |
| 0 | 0 | 0 | 1 |
| 0 | 0 | 0 | 0 |
| 0 | 0 | 0 | 0 |
| 0 | 0 | 0 | 0 |
| 0 | 0 | 0 | 0 |
| 0 | 0 | 0 | 0 |
| 0 | 0 | 0 | 1 |
| 0 | 0 | 0 | 1 |
| 0 | 0 | 0 | 1 |
| 0 | 0 | 0 | 1 |
| 0 | 0 | 0 | 0 |
| 0 | 0 | 0 | 1 |
| 0 | 0 | 0 | 0 |
| 0 | 0 | 0 | 1 |
| 1 | 0 | 0 | 0 |
| 0 | 0 | 0 | 1 |
| 0 | 0 | 0 | 0 |
| 0 | 0 | 0 | 0 |
| 1 | 0 | 0 | 0 |
| 0 | 0 | 0 | 0 |
| 0 | 0 | 0 | 0 |
| 0 | 0 | 0 | 0 |
| 0 | 0 | 0 | 0 |
| 0 | 0 | 0 | 0 |

| | | | |
|---|---|---|---|
| 0 | 0 | 0 | 0 |
| 0 | 0 | 0 | 0 |
| 0 | 0 | 0 | 0 |
| 0 | 0 | 0 | 0 |
| 0 | 0 | 0 | 0 |
| 0 | 0 | 0 | 0 |
| 0 | 0 | 0 | 0 |
| 0 | 0 | 0 | 1 |
| 0 | 0 | 0 | 0 |
| 0 | 0 | 0 | 0 |
| 0 | 0 | 0 | 0 |
| 0 | 0 | 0 | 1 |
| 0 | 0 | 0 | 0 |
| 0 | 0 | 0 | 0 |
| 0 | 0 | 0 | 0 |
| 0 | 0 | 0 | 0 |
| 0 | 0 | 0 | 0 |
| 0 | 0 | 0 | 0 |
| 0 | 0 | 0 | 0 |
| 0 | 0 | 0 | 0 |
| 0 | 0 | 0 | 0 |
| 0 | 0 | 0 | 0 |
| 0 | 0 | 0 | 0 |
| 0 | 0 | 0 | 0 |
| 0 | 0 | 0 | 0 |
| 0 | 0 | 0 | 0 |
| 0 | 0 | 0 | 0 |
| 0 | 0 | 0 | 0 |
| 0 | 0 | 0 | 0 |
| 0 | 0 | 0 | 0 |
| 0 | 0 | 0 | 0 |
| 0 | 0 | 0 | 0 |
| 0 | 0 | 0 | 0 |
| 0 | 0 | 0 | 0 |
| 0 | 0 | 0 | 0 |
| 0 | 0 | 0 | 0 |
| 0 | 0 | 0 | 0 |
| 0 | 0 | 0 | 0 |
| 0 | 0 | 0 | 0 |
| 0 | 0 | 0 | 0 |
| 0 | 0 | 0 | 1 |
| 0 | 0 | 0 | 0 |
| 0 | 0 | 0 | 0 |
| 0 | 0 | 0 | 0 |
| 0 | 0 | 0 | 0 |
| 0 | 0 | 0 | 0 |
| 0 | 0 | 0 | 0 |
| 0 | 0 | 0 | 0 |
| 0 | 0 | 0 | 0 |
| 0 | 0 | 0 | 1 |
| 0 | 0 | 0 | 1 |
| 0 | 0 | 0 | 1 |
| 1 | 0 | 0 | 0 |
| 0 | 0 | 0 | 0 |

| | | | |
|---|---|---|---|
| 0 | 0 | 0 | 0 |
| 0 | 0 | 0 | 0 |
| 1 | 0 | 0 | 0 |
| 1 | 0 | 0 | 0 |
| 0 | 0 | 0 | 0 |
| 0 | 0 | 0 | 0 |
| 0 | 0 | 0 | 0 |
| 0 | 0 | 0 | 1 |
| 0 | 0 | 0 | 1 |
| 0 | 0 | 0 | 0 |
| 0 | 0 | 0 | 0 |
| 0 | 0 | 0 | 1 |
| 0 | 0 | 0 | 0 |
| 0 | 0 | 0 | 0 |
| 0 | 0 | 0 | 0 |
| 0 | 0 | 0 | 0 |
| 0 | 0 | 0 | 0 |
| 0 | 0 | 0 | 1 |
| 0 | 0 | 0 | 0 |
| 0 | 0 | 0 | 1 |
| 0 | 0 | 0 | 0 |
| 0 | 0 | 0 | 0 |
| 0 | 0 | 0 | 1 |
| 0 | 0 | 0 | 0 |
| 0 | 0 | 0 | 0 |
| 0 | 0 | 0 | 1 |
| 0 | 0 | 0 | 0 |
| 0 | 0 | 0 | 0 |
| 0 | 0 | 0 | 0 |
| 0 | 0 | 0 | 0 |
| 0 | 0 | 0 | 0 |
| 0 | 0 | 0 | 0 |
| 0 | 0 | 0 | 1 |
| 0 | 0 | 0 | 0 |
| 0 | 0 | 0 | 0 |
| 0 | 0 | 0 | 0 |
| 0 | 0 | 0 | 1 |
| 0 | 0 | 0 | 0 |
| 0 | 0 | 0 | 0 |
| 0 | 0 | 0 | 0 |
| 0 | 0 | 0 | 0 |
| 0 | 1 | 0 | 0 |
| 0 | 1 | 0 | 0 |
| 0 | 1 | 0 | 0 |
| 0 | 0 | 0 | 0 |
| 0 | 0 | 0 | 1 |
| 0 | 0 | 0 | 0 |
| 0 | 0 | 0 | 0 |
| 0 | 0 | 0 | 0 |
| 0 | 0 | 0 | 0 |
| 0 | 0 | 0 | 0 |
| 0 | 0 | 0 | 0 |
| 0 | 0 | 0 | 1 |
| 0 | 0 | 0 | 1 |

| | | | |
|---|---|---|---|
| 0 | 0 | 0 | 1 |
| 0 | 0 | 0 | 1 |
| 0 | 0 | 0 | 0 |
| 0 | 0 | 0 | 0 |
| 0 | 0 | 0 | 0 |
| 0 | 1 | 0 | 0 |
| 0 | 0 | 0 | 0 |
| 0 | 0 | 0 | 1 |
| 0 | 0 | 0 | 0 |
| 0 | 0 | 0 | 1 |
| 0 | 0 | 0 | 1 |
| 0 | 0 | 0 | 1 |
| 0 | 0 | 0 | 0 |
| 0 | 0 | 0 | 0 |
| 0 | 0 | 0 | 0 |
| 0 | 0 | 0 | 0 |
| 0 | 0 | 0 | 0 |
| 1 | 0 | 0 | 0 |
| 0 | 0 | 0 | 1 |
| 0 | 0 | 0 | 1 |
| 0 | 0 | 0 | 1 |
| 0 | 0 | 0 | 0 |
| 0 | 0 | 0 | 1 |
| 0 | 0 | 0 | 0 |
| 0 | 0 | 0 | 0 |
| 0 | 0 | 0 | 0 |
| 0 | 0 | 0 | 0 |
| 0 | 0 | 0 | 0 |
| 0 | 0 | 0 | 0 |
| 0 | 0 | 0 | 0 |
| 0 | 0 | 0 | 0 |
| 0 | 0 | 0 | 0 |
| 0 | 0 | 0 | 0 |
| 0 | 0 | 0 | 0 |
| 0 | 0 | 0 | 0 |
| 0 | 0 | 0 | 0 |
| 0 | 0 | 0 | 0 |
| 0 | 0 | 0 | 0 |
| 1 | 0 | 0 | 0 |
| 0 | 0 | 0 | 0 |
| 0 | 0 | 0 | 0 |
| 0 | 0 | 0 | 1 |
| 0 | 0 | 0 | 0 |
| 0 | 0 | 0 | 0 |
| 0 | 0 | 0 | 0 |
| 0 | 0 | 0 | 0 |
| 0 | 0 | 0 | 1 |
| 0 | 0 | 0 | 0 |
| 0 | 0 | 0 | 0 |
| 0 | 0 | 0 | 0 |
| 0 | 0 | 0 | 0 |
| 0 | 0 | 0 | 0 |
| 0 | 0 | 0 | 1 |
| 0 | 0 | 0 | 1 |

| | | | |
|---|---|---|---|
| 0 | 0 | 0 | 0 |
| 0 | 0 | 0 | 0 |
| 0 | 0 | 0 | 1 |
| 0 | 0 | 0 | 1 |
| 0 | 0 | 0 | 1 |
| 0 | 0 | 0 | 0 |
| 0 | 0 | 0 | 1 |
| 0 | 0 | 0 | 1 |
| 0 | 0 | 0 | 0 |
| 0 | 0 | 0 | 0 |
| 0 | 0 | 0 | 0 |
| 0 | 0 | 0 | 0 |
| 0 | 0 | 0 | 0 |
| 0 | 0 | 0 | 1 |
| 0 | 0 | 0 | 1 |
| 0 | 0 | 0 | 1 |
| 0 | 0 | 0 | 1 |
| 0 | 0 | 0 | 1 |
| 0 | 0 | 0 | 0 |
| 0 | 0 | 0 | 0 |
| 0 | 0 | 0 | 1 |
| 0 | 0 | 0 | 1 |
| 0 | 0 | 0 | 0 |
| 0 | 0 | 0 | 0 |
| 0 | 0 | 0 | 1 |
| 0 | 0 | 0 | 1 |
| 0 | 0 | 0 | 0 |
| 0 | 0 | 0 | 1 |
| 0 | 0 | 0 | 0 |
| 0 | 0 | 0 | 0 |
| 0 | 0 | 0 | 0 |
| 0 | 0 | 0 | 0 |
| 0 | 0 | 0 | 0 |
| 1 | 0 | 0 | 0 |
| 1 | 0 | 0 | 0 |
| 0 | 0 | 0 | 0 |
| 0 | 0 | 0 | 0 |
| 0 | 0 | 0 | 0 |
| 0 | 0 | 0 | 1 |
| 0 | 0 | 0 | 0 |
| 0 | 0 | 0 | 1 |
| 0 | 0 | 0 | 1 |
| 0 | 0 | 0 | 1 |
| 0 | 0 | 0 | 1 |
| 0 | 0 | 0 | 1 |
| 0 | 0 | 0 | 0 |
| 0 | 0 | 0 | 0 |
| 0 | 0 | 0 | 0 |
| 0 | 0 | 0 | 0 |
| 0 | 0 | 0 | 0 |
| 0 | 0 | 0 | 0 |
| 0 | 0 | 0 | 0 |
| 0 | 0 | 0 | 0 |
| 0 | 0 | 0 | 1 |

| | | | |
|---|---|---|---|
| 0 | 0 | 0 | 1 |
| 0 | 0 | 0 | 0 |
| 0 | 0 | 0 | 0 |
| 0 | 0 | 0 | 0 |
| 0 | 0 | 0 | 1 |
| 0 | 0 | 0 | 0 |
| 0 | 0 | 0 | 0 |
| 0 | 0 | 0 | 0 |
| 0 | 0 | 0 | 1 |
| 0 | 0 | 0 | 1 |
| 0 | 0 | 0 | 0 |
| 0 | 0 | 0 | 0 |
| 0 | 0 | 0 | 0 |
| 0 | 0 | 0 | 1 |
| 0 | 0 | 0 | 0 |
| 0 | 0 | 0 | 0 |
| 0 | 0 | 0 | 0 |
| 0 | 0 | 0 | 0 |
| 0 | 0 | 0 | 0 |
| 0 | 0 | 0 | 0 |
| 0 | 0 | 0 | 0 |
| 0 | 0 | 0 | 1 |
| 0 | 0 | 0 | 0 |
| 0 | 0 | 0 | 1 |
| 0 | 0 | 0 | 0 |
| 0 | 0 | 0 | 0 |
| 0 | 0 | 0 | 1 |
| 0 | 0 | 0 | 0 |
| 0 | 0 | 0 | 0 |
| 0 | 0 | 0 | 0 |
| 0 | 0 | 0 | 0 |
| 0 | 0 | 0 | 0 |
| 0 | 0 | 0 | 0 |
| 0 | 0 | 0 | 0 |
| 0 | 0 | 0 | 1 |
| 0 | 0 | 0 | 0 |
| 0 | 0 | 0 | 0 |
| 0 | 0 | 0 | 0 |
| 0 | 0 | 0 | 0 |
| 0 | 0 | 0 | 0 |
| 0 | 0 | 0 | 0 |
| 0 | 0 | 0 | 0 |
| 0 | 0 | 0 | 0 |
| 0 | 0 | 0 | 0 |
| 0 | 0 | 0 | 0 |
| 0 | 0 | 0 | 0 |
| 0 | 0 | 0 | 0 |
| 0 | 0 | 0 | 1 |
| 0 | 0 | 0 | 0 |
| 0 | 0 | 0 | 1 |
| 0 | 0 | 0 | 0 |
| 0 | 0 | 0 | 0 |
| 0 | 0 | 0 | 0 |
| 0 | 0 | 0 | 0 |

| | | | |
|---|---|---|---|
| 0 | 0 | 0 | 0 |
| 0 | 0 | 0 | 0 |
| 0 | 0 | 0 | 1 |
| 0 | 0 | 0 | 0 |
| 0 | 0 | 0 | 0 |
| 0 | 0 | 0 | 1 |
| 0 | 0 | 0 | 1 |
| 0 | 0 | 0 | 1 |
| 0 | 0 | 0 | 0 |
| 0 | 0 | 0 | 1 |
| 0 | 0 | 0 | 0 |
| 0 | 0 | 0 | 0 |
| 0 | 0 | 0 | 0 |
| 0 | 0 | 0 | 1 |
| 0 | 0 | 0 | 0 |
| 0 | 0 | 0 | 1 |
| 0 | 0 | 0 | 0 |
| 0 | 0 | 0 | 0 |
| 1 | 0 | 0 | 0 |
| 0 | 0 | 0 | 1 |
| 0 | 0 | 0 | 1 |
| 0 | 0 | 0 | 0 |
| 0 | 0 | 0 | 1 |
| 0 | 0 | 0 | 0 |
| 0 | 0 | 0 | 0 |
| 0 | 0 | 0 | 1 |
| 0 | 0 | 0 | 1 |
| 0 | 0 | 0 | 0 |
| 0 | 0 | 0 | 1 |
| 1 | 0 | 0 | 0 |
| 0 | 0 | 0 | 1 |
| 0 | 0 | 0 | 1 |
| 0 | 0 | 0 | 1 |
| 0 | 0 | 0 | 1 |
| 0 | 0 | 0 | 1 |
| 0 | 0 | 0 | 1 |
| 0 | 0 | 0 | 1 |
| 0 | 0 | 0 | 1 |
| 0 | 0 | 0 | 1 |
| 0 | 0 | 0 | 1 |
| 0 | 0 | 0 | 1 |
| 0 | 0 | 0 | 1 |
| 0 | 0 | 0 | 1 |
| 0 | 0 | 0 | 1 |
| 0 | 0 | 0 | 1 |
| 0 | 0 | 0 | 1 |
| 0 | 0 | 0 | 1 |
| 0 | 0 | 0 | 1 |
| 0 | 0 | 0 | 1 |
| 0 | 0 | 0 | 1 |
| 0 | 0 | 0 | 1 |
| 0 | 0 | 0 | 1 |
| 0 | 0 | 0 | 1 |
| 0 | 0 | 0 | 1 |

| | | | |
|---|---|---|---|
| 0 | 0 | 0 | 1 |
| 0 | 0 | 0 | 1 |
| 0 | 0 | 0 | 1 |
| 0 | 0 | 0 | 1 |
| 0 | 0 | 0 | 1 |
| 0 | 0 | 0 | 1 |
| 0 | 0 | 0 | 1 |
| 0 | 0 | 0 | 1 |
| 0 | 0 | 0 | 1 |
| 0 | 0 | 0 | 1 |
| 0 | 0 | 0 | 1 |
| 0 | 0 | 0 | 0 |
| 0 | 0 | 0 | 0 |
| 0 | 0 | 0 | 0 |
| 0 | 0 | 0 | 1 |
| 0 | 0 | 0 | 0 |
| 0 | 0 | 0 | 1 |
| 0 | 0 | 0 | 0 |
| 0 | 0 | 0 | 1 |
| 0 | 0 | 0 | 1 |
| 0 | 0 | 0 | 0 |
| 0 | 0 | 0 | 1 |
| 0 | 0 | 0 | 1 |
| 0 | 0 | 0 | 0 |
| 0 | 0 | 0 | 0 |
| 0 | 0 | 0 | 1 |
| 0 | 0 | 0 | 1 |
| 0 | 0 | 0 | 0 |
| 0 | 0 | 0 | 0 |
| 0 | 0 | 0 | 0 |
| 0 | 0 | 0 | 0 |
| 0 | 0 | 0 | 0 |
| 0 | 0 | 0 | 1 |
| 0 | 0 | 0 | 1 |
| 0 | 0 | 0 | 1 |
| 0 | 0 | 0 | 1 |
| 0 | 0 | 0 | 1 |
| 0 | 0 | 0 | 1 |
| 0 | 0 | 0 | 1 |
| 0 | 0 | 0 | 0 |
| 0 | 0 | 0 | 0 |
| 0 | 0 | 0 | 1 |
| 0 | 0 | 0 | 0 |
| 0 | 0 | 0 | 0 |
| 0 | 0 | 0 | 0 |
| 0 | 0 | 0 | 1 |
| 0 | 0 | 0 | 0 |
| 0 | 0 | 0 | 1 |
| 0 | 0 | 0 | 1 |
| 0 | 0 | 0 | 0 |
| 0 | 0 | 0 | 0 |
| 0 | 0 | 0 | 0 |
| 0 | 0 | 0 | 1 |
| 0 | 0 | 0 | 0 |

| | | | |
|---|---|---|---|
| 0 | 0 | 0 | 0 |
| 0 | 0 | 0 | 1 |
| 0 | 0 | 0 | 1 |
| 0 | 0 | 0 | 1 |
| 0 | 0 | 0 | 0 |
| 0 | 0 | 0 | 1 |
| 0 | 0 | 0 | 0 |
| 0 | 0 | 0 | 1 |
| 0 | 0 | 0 | 1 |
| 0 | 0 | 0 | 0 |
| 0 | 1 | 1 | 1 |
| 0 | 0 | 0 | 1 |
| 0 | 0 | 0 | 1 |
| 0 | 0 | 0 | 0 |
| 0 | 0 | 0 | 1 |
| 0 | 0 | 0 | 1 |
| 0 | 0 | 0 | 0 |
| 0 | 0 | 0 | 1 |
| 0 | 0 | 0 | 0 |
| 0 | 0 | 0 | 0 |
| 0 | 0 | 0 | 0 |
| 0 | 0 | 0 | 0 |
| 0 | 0 | 0 | 0 |
| 0 | 0 | 0 | 0 |
| 0 | 0 | 0 | 0 |
| 0 | 0 | 0 | 0 |
| 0 | 0 | 0 | 0 |
| 0 | 0 | 0 | 0 |
| 0 | 0 | 0 | 0 |
| 0 | 0 | 0 | 0 |
| 0 | 0 | 0 | 0 |
| 0 | 0 | 0 | 0 |
| 0 | 0 | 0 | 0 |
| 0 | 0 | 0 | 0 |
| 0 | 0 | 0 | 0 |
| 0 | 0 | 0 | 0 |
| 0 | 0 | 0 | 1 |
| 0 | 0 | 0 | 0 |
| 0 | 0 | 0 | 0 |
| 0 | 0 | 0 | 1 |
| 0 | 0 | 0 | 1 |
| 0 | 0 | 0 | 0 |
| 0 | 0 | 0 | 0 |
| 0 | 0 | 0 | 0 |
| 0 | 0 | 0 | 0 |
| 0 | 0 | 0 | 0 |
| 0 | 0 | 0 | 0 |
| 0 | 0 | 0 | 0 |
| 0 | 0 | 0 | 1 |
| 0 | 0 | 0 | 1 |
| 0 | 0 | 0 | 0 |
| 0 | 0 | 0 | 1 |
| 0 | 0 | 0 | 0 |
| 0 | 0 | 0 | 0 |

| | | | |
|---|---|---|---|
| 0 | 0 | 0 | 0 |
| 0 | 0 | 0 | 0 |
| 0 | 0 | 0 | 0 |
| 0 | 0 | 0 | 0 |
| 0 | 0 | 0 | 0 |
| 0 | 0 | 0 | 0 |
| 0 | 0 | 0 | 0 |
| 0 | 0 | 0 | 0 |
| 0 | 0 | 0 | 0 |
| 0 | 0 | 0 | 0 |
| 0 | 0 | 0 | 0 |
| 0 | 0 | 0 | 0 |
| 0 | 0 | 0 | 0 |
| 0 | 0 | 0 | 0 |
| 0 | 0 | 0 | 1 |
| 0 | 0 | 0 | 0 |
| 0 | 0 | 0 | 0 |
| 0 | 0 | 0 | 0 |
| 0 | 0 | 0 | 1 |
| 0 | 0 | 0 | 0 |
| 0 | 0 | 0 | 0 |
| 0 | 1 | 0 | 0 |
| 0 | 0 | 0 | 0 |
| 0 | 0 | 0 | 0 |
| 0 | 0 | 0 | 0 |
| 0 | 0 | 0 | 1 |
| 0 | 0 | 0 | 0 |
| 0 | 0 | 0 | 0 |
| 0 | 0 | 0 | 0 |
| 0 | 0 | 0 | 0 |
| 0 | 0 | 0 | 1 |
| 0 | 0 | 0 | 0 |
| 0 | 0 | 0 | 1 |
| 0 | 0 | 0 | 1 |
| 1 | 0 | 0 | 0 |
| 0 | 0 | 0 | 0 |
| 0 | 0 | 0 | 1 |
| 0 | 0 | 0 | 1 |
| 0 | 0 | 0 | 1 |
| 0 | 0 | 0 | 1 |
| 0 | 0 | 0 | 0 |
| 0 | 0 | 0 | 0 |
| 0 | 0 | 0 | 0 |
| 0 | 0 | 0 | 0 |
| 0 | 0 | 0 | 0 |
| 0 | 0 | 0 | 0 |
| 0 | 0 | 0 | 0 |
| 0 | 0 | 0 | 0 |
| 0 | 0 | 0 | 1 |
| 0 | 0 | 0 | 1 |
| 0 | 0 | 0 | 0 |
| 0 | 0 | 0 | 0 |
| 0 | 0 | 0 | 0 |
| 0 | 0 | 0 | 1 |

| | | | |
|---|---|---|---|
| 0 | 0 | 0 | 1 |
| 0 | 0 | 0 | 1 |
| 0 | 0 | 0 | 1 |
| 0 | 0 | 0 | 0 |
| 0 | 0 | 0 | 1 |
| 0 | 0 | 0 | 1 |
| 0 | 0 | 0 | 0 |
| 0 | 0 | 0 | 1 |
| 0 | 0 | 0 | 0 |
| 0 | 0 | 0 | 0 |
| 0 | 0 | 0 | 0 |
| 0 | 0 | 0 | 0 |
| 0 | 0 | 0 | 0 |
| 0 | 0 | 0 | 0 |
| 0 | 0 | 0 | 0 |
| 0 | 0 | 0 | 0 |
| 0 | 0 | 0 | 1 |
| 0 | 0 | 0 | 0 |
| 0 | 0 | 0 | 0 |
| 0 | 0 | 0 | 1 |
| 0 | 0 | 0 | 0 |
| 0 | 0 | 0 | 0 |
| 0 | 0 | 0 | 1 |
| 0 | 0 | 0 | 0 |
| 0 | 0 | 0 | 1 |
| 0 | 0 | 0 | 1 |
| 0 | 0 | 0 | 1 |
| 0 | 0 | 0 | 0 |
| 0 | 0 | 0 | 0 |
| 0 | 0 | 0 | 0 |
| 0 | 0 | 0 | 1 |
| 0 | 0 | 0 | 0 |
| 0 | 0 | 0 | 0 |
| 0 | 0 | 0 | 0 |
| 0 | 0 | 0 | 1 |
| 0 | 0 | 0 | 1 |
| 0 | 0 | 0 | 0 |
| 0 | 0 | 0 | 0 |
| 0 | 0 | 0 | 0 |
| 0 | 0 | 0 | 1 |
| 0 | 0 | 0 | 1 |
| 0 | 0 | 0 | 1 |
| 0 | 0 | 0 | 1 |
| 0 | 0 | 0 | 0 |
| 0 | 0 | 0 | 1 |
| 0 | 0 | 0 | 0 |
| 0 | 0 | 0 | 1 |
| 0 | 0 | 0 | 0 |
| 0 | 0 | 0 | 0 |
| 0 | 0 | 0 | 1 |
| 0 | 0 | 0 | 1 |
| 0 | 0 | 0 | 0 |
| 0 | 0 | 0 | 0 |
| 0 | 0 | 0 | 1 |

| | | | |
|---|---|---|---|
| 0 | 0 | 0 | 1 |
| 0 | 0 | 0 | 1 |
| 0 | 0 | 0 | 1 |
| 0 | 0 | 0 | 1 |
| 0 | 0 | 0 | 0 |
| 0 | 0 | 0 | 1 |
| 0 | 0 | 0 | 1 |
| 0 | 0 | 0 | 0 |
| 0 | 0 | 0 | 0 |
| 0 | 0 | 0 | 1 |
| 0 | 0 | 0 | 0 |
| 0 | 0 | 0 | 0 |
| 1 | 0 | 0 | 0 |
| 1 | 0 | 0 | 0 |
| 0 | 0 | 0 | 0 |
| 1 | 0 | 0 | 0 |
| 1 | 0 | 0 | 0 |
| 1 | 0 | 0 | 0 |
| 1 | 0 | 0 | 0 |
| 0 | 0 | 0 | 1 |
| 0 | 0 | 0 | 1 |
| 0 | 0 | 0 | 1 |
| 0 | 0 | 0 | 1 |
| 0 | 0 | 0 | 1 |
| 0 | 0 | 0 | 0 |
| 0 | 0 | 0 | 1 |
| 1 | 0 | 0 | 0 |
| 1 | 0 | 0 | 0 |
| 0 | 0 | 0 | 1 |
| 0 | 0 | 0 | 1 |
| 0 | 0 | 0 | 1 |
| 0 | 0 | 0 | 1 |
| 0 | 0 | 0 | 1 |
| 0 | 0 | 0 | 0 |
| 0 | 0 | 0 | 0 |
| 0 | 0 | 0 | 1 |
| 0 | 0 | 0 | 0 |
| 0 | 0 | 0 | 1 |
| 0 | 0 | 0 | 0 |
| 0 | 0 | 0 | 0 |
| 0 | 0 | 0 | 0 |
| 0 | 0 | 0 | 0 |
| 0 | 0 | 0 | 1 |
| 0 | 0 | 0 | 0 |
| 0 | 0 | 0 | 0 |
| 1 | 0 | 0 | 0 |
| 1 | 0 | 0 | 0 |
| 1 | 0 | 0 | 0 |
| 0 | 0 | 0 | 0 |
| 0 | 0 | 0 | 0 |
| 0 | 0 | 0 | 0 |
| 1 | 0 | 0 | 0 |
| 1 | 0 | 0 | 0 |
| 0 | 0 | 0 | 1 |

| | | | |
|---|---|---|---|
| 0 | 0 | 0 | 0 |
| 0 | 0 | 0 | 1 |
| 0 | 0 | 0 | 0 |
| 0 | 0 | 0 | 0 |
| 0 | 0 | 0 | 0 |
| 0 | 0 | 0 | 1 |
| 0 | 0 | 0 | 0 |
| 0 | 0 | 0 | 1 |
| 0 | 0 | 0 | 1 |
| 0 | 0 | 0 | 0 |
| 0 | 0 | 0 | 1 |
| 0 | 0 | 0 | 1 |
| 0 | 0 | 0 | 1 |
| 0 | 0 | 0 | 1 |
| 0 | 0 | 0 | 1 |
| 0 | 0 | 0 | 0 |
| 0 | 0 | 0 | 0 |
| 0 | 0 | 0 | 1 |
| 0 | 0 | 0 | 0 |
| 0 | 0 | 0 | 1 |
| 0 | 0 | 0 | 0 |
| 0 | 0 | 0 | 0 |
| 0 | 0 | 0 | 0 |
| 0 | 0 | 0 | 0 |
| 0 | 0 | 0 | 0 |
| 0 | 0 | 0 | 0 |
| 0 | 0 | 0 | 0 |
| 0 | 0 | 0 | 0 |
| 0 | 0 | 0 | 0 |
| 0 | 0 | 0 | 0 |
| 0 | 0 | 0 | 0 |
| 0 | 0 | 0 | 0 |
| 0 | 0 | 0 | 1 |
| 0 | 0 | 0 | 1 |
| 0 | 0 | 0 | 0 |
| 0 | 0 | 0 | 0 |
| 0 | 0 | 0 | 1 |
| 0 | 0 | 0 | 0 |
| 0 | 0 | 0 | 1 |
| 0 | 0 | 0 | 0 |
| 0 | 0 | 0 | 0 |
| 0 | 0 | 0 | 0 |
| 0 | 0 | 0 | 1 |
| 0 | 0 | 0 | 1 |
| 0 | 0 | 0 | 0 |
| 0 | 0 | 0 | 0 |
| 0 | 0 | 0 | 0 |
| 0 | 0 | 0 | 0 |
| 0 | 0 | 0 | 0 |
| 0 | 0 | 0 | 0 |
| 0 | 0 | 0 | 1 |
| 0 | 0 | 0 | 0 |
| 0 | 0 | 0 | 0 |
| 0 | 0 | 0 | 1 |

| | | | |
|---|---|---|---|
| 0 | 0 | 0 | 0 |
| 0 | 0 | 0 | 1 |
| 0 | 0 | 0 | 0 |
| 0 | 0 | 0 | 0 |
| 0 | 0 | 0 | 0 |
| 0 | 0 | 0 | 0 |
| 0 | 0 | 0 | 0 |
| 0 | 0 | 0 | 0 |
| 0 | 0 | 0 | 0 |
| 0 | 0 | 0 | 0 |
| 0 | 0 | 0 | 0 |
| 0 | 0 | 0 | 1 |
| 0 | 0 | 0 | 0 |
| 0 | 0 | 0 | 1 |
| 0 | 0 | 0 | 1 |
| 0 | 0 | 0 | 1 |
| 0 | 0 | 0 | 0 |
| 0 | 0 | 0 | 0 |
| 0 | 1 | 1 | 1 |
| 0 | 0 | 0 | 0 |
| 0 | 0 | 0 | 1 |
| 0 | 0 | 0 | 0 |
| 0 | 0 | 0 | 1 |
| 0 | 0 | 0 | 0 |
| 0 | 0 | 0 | 1 |
| 0 | 0 | 0 | 1 |
| 0 | 0 | 0 | 1 |
| 0 | 0 | 0 | 0 |
| 0 | 0 | 0 | 1 |
| 0 | 0 | 0 | 1 |
| 0 | 0 | 0 | 1 |
| 1 | 0 | 0 | 0 |
| 0 | 0 | 0 | 1 |
| 0 | 0 | 0 | 0 |
| 0 | 0 | 0 | 0 |
| 0 | 0 | 0 | 0 |
| 0 | 0 | 0 | 1 |
| 0 | 0 | 0 | 1 |
| 0 | 0 | 0 | 0 |
| 0 | 0 | 0 | 0 |
| 0 | 0 | 0 | 0 |
| 0 | 0 | 0 | 0 |
| 0 | 0 | 0 | 0 |
| 0 | 0 | 0 | 0 |
| 0 | 0 | 0 | 0 |
| 0 | 0 | 0 | 0 |
| 0 | 0 | 0 | 1 |
| 0 | 0 | 0 | 1 |
| 0 | 0 | 0 | 0 |
| 0 | 0 | 0 | 0 |
| 0 | 0 | 0 | 0 |
| 0 | 0 | 0 | 1 |
| 0 | 0 | 0 | 1 |
| 0 | 0 | 0 | 0 |

| | | | |
|---|---|---|---|
| 0 | 0 | 0 | 0 |
| 0 | 0 | 0 | 1 |
| 0 | 0 | 0 | 1 |
| 0 | 0 | 0 | 1 |
| 0 | 0 | 0 | 0 |
| 0 | 0 | 0 | 0 |
| 0 | 0 | 0 | 0 |
| 0 | 0 | 0 | 0 |
| 0 | 0 | 0 | 0 |
| 0 | 0 | 0 | 0 |
| 0 | 0 | 0 | 1 |
| 0 | 0 | 0 | 0 |
| 0 | 0 | 0 | 0 |
| 0 | 0 | 0 | 0 |
| 0 | 0 | 0 | 1 |
| 0 | 0 | 0 | 1 |
| 0 | 0 | 0 | 0 |
| 0 | 0 | 0 | 1 |
| 0 | 0 | 0 | 0 |
| 0 | 0 | 0 | 0 |
| 0 | 0 | 0 | 1 |
| 0 | 0 | 0 | 0 |
| 0 | 0 | 0 | 0 |
| 0 | 0 | 0 | 1 |
| 0 | 0 | 0 | 0 |
| 0 | 0 | 0 | 0 |
| 0 | 0 | 0 | 0 |
| 0 | 0 | 0 | 1 |
| 0 | 0 | 0 | 0 |
| 0 | 0 | 0 | 0 |
| 0 | 0 | 0 | 0 |
| 0 | 0 | 0 | 1 |
| 0 | 0 | 0 | 1 |
| 0 | 0 | 0 | 1 |
| 0 | 0 | 0 | 1 |
| 0 | 0 | 0 | 1 |
| 0 | 0 | 0 | 1 |
| 0 | 0 | 0 | 1 |
| 0 | 0 | 0 | 1 |
| 0 | 0 | 0 | 1 |
| 0 | 0 | 0 | 1 |
| 0 | 0 | 0 | 1 |
| 1 | 0 | 0 | 0 |
| 0 | 0 | 0 | 1 |
| 0 | 0 | 0 | 1 |
| 0 | 0 | 0 | 1 |
| 0 | 0 | 0 | 1 |
| 0 | 0 | 0 | 1 |
| 0 | 0 | 0 | 1 |
| 0 | 0 | 0 | 1 |
| 0 | 0 | 0 | 1 |
| 0 | 0 | 0 | 1 |
| 0 | 0 | 0 | 1 |
| 0 | 0 | 0 | 1 |

| | | | |
|---|---|---|---|
| 0 | 0 | 0 | 1 |
| 0 | 0 | 0 | 1 |
| 0 | 0 | 0 | 1 |
| 0 | 0 | 0 | 1 |
| 0 | 0 | 0 | 1 |
| 0 | 0 | 0 | 1 |
| 0 | 0 | 0 | 1 |
| 0 | 0 | 0 | 1 |
| 0 | 0 | 0 | 0 |
| 0 | 0 | 0 | 0 |
| 0 | 0 | 0 | 0 |
| 0 | 0 | 0 | 0 |
| 0 | 0 | 0 | 0 |
| 0 | 0 | 0 | 0 |
| 0 | 0 | 0 | 0 |
| 0 | 0 | 0 | 0 |
| 0 | 0 | 0 | 0 |
| 0 | 0 | 0 | 0 |
| 0 | 0 | 0 | 0 |
| 0 | 0 | 0 | 0 |
| 1 | 0 | 0 | 0 |
| 1 | 0 | 0 | 0 |
| 0 | 0 | 0 | 0 |
| 0 | 0 | 0 | 0 |
| 0 | 0 | 0 | 0 |
| 0 | 0 | 0 | 1 |
| 0 | 0 | 0 | 1 |
| 0 | 0 | 0 | 0 |
| 0 | 0 | 0 | 0 |
| 1 | 0 | 0 | 0 |
| 1 | 0 | 0 | 0 |
| 0 | 0 | 0 | 0 |
| 1 | 0 | 0 | 0 |
| 1 | 0 | 0 | 0 |
| 0 | 0 | 0 | 0 |
| 0 | 0 | 0 | 0 |
| 0 | 0 | 0 | 0 |
| 0 | 0 | 0 | 0 |
| 0 | 0 | 0 | 0 |
| 0 | 0 | 0 | 0 |
| 0 | 0 | 0 | 0 |
| 0 | 0 | 0 | 0 |
| 1 | 0 | 0 | 0 |
| 1 | 0 | 0 | 0 |
| 1 | 0 | 0 | 0 |
| 0 | 0 | 0 | 0 |
| 0 | 0 | 0 | 1 |
| 0 | 0 | 0 | 1 |
| 0 | 0 | 0 | 1 |
| 0 | 0 | 0 | 1 |
| 0 | 0 | 0 | 1 |
| 0 | 0 | 0 | 1 |
| 0 | 0 | 0 | 0 |
| 0 | 0 | 0 | 0 |

| | | | |
|---|---|---|---|
| 0 | 0 | 0 | 0 |
| 0 | 0 | 0 | 0 |
| 0 | 0 | 0 | 0 |
| 0 | 0 | 0 | 0 |
| 0 | 0 | 0 | 1 |
| 0 | 0 | 0 | 0 |
| 0 | 0 | 0 | 0 |
| 0 | 0 | 0 | 0 |
| 0 | 0 | 0 | 1 |
| 0 | 0 | 0 | 0 |
| 1 | 0 | 0 | 0 |
| 0 | 0 | 0 | 1 |
| 0 | 0 | 0 | 0 |
| 0 | 0 | 0 | 0 |
| 0 | 0 | 0 | 0 |
| 0 | 0 | 0 | 0 |
| 0 | 0 | 0 | 1 |
| 0 | 0 | 0 | 0 |
| 0 | 0 | 0 | 0 |
| 0 | 0 | 0 | 1 |
| 0 | 0 | 0 | 0 |
| 0 | 0 | 0 | 0 |
| 0 | 0 | 0 | 0 |
| 0 | 0 | 0 | 0 |
| 0 | 0 | 0 | 0 |
| 0 | 0 | 0 | 1 |
| 0 | 0 | 0 | 1 |
| 0 | 0 | 0 | 1 |
| 0 | 0 | 0 | 1 |
| 0 | 0 | 0 | 1 |
| 0 | 0 | 0 | 1 |
| 0 | 0 | 0 | 1 |
| 0 | 0 | 0 | 1 |
| 0 | 0 | 0 | 1 |
| 0 | 0 | 0 | 0 |
| 0 | 0 | 0 | 0 |
| 0 | 0 | 0 | 0 |
| 0 | 0 | 0 | 0 |
| 0 | 0 | 0 | 0 |
| 0 | 0 | 0 | 0 |
| 0 | 0 | 0 | 1 |
| 0 | 0 | 0 | 1 |
| 0 | 0 | 0 | 1 |
| 0 | 0 | 0 | 0 |
| 0 | 0 | 0 | 0 |
| 0 | 0 | 0 | 0 |
| 0 | 0 | 0 | 1 |
| 0 | 0 | 0 | 1 |
| 0 | 0 | 0 | 0 |
| 0 | 0 | 0 | 0 |
| 0 | 0 | 0 | 1 |
| 0 | 0 | 0 | 1 |
| 0 | 0 | 0 | 1 |
| 0 | 0 | 0 | 1 |

| | | | |
|---|---|---|---|
| 0 | 0 | 0 | 1 |
| 0 | 0 | 0 | 1 |
| 0 | 0 | 0 | 1 |
| 0 | 0 | 0 | 0 |
| 0 | 0 | 0 | 0 |
| 0 | 0 | 0 | 1 |
| 0 | 0 | 0 | 1 |
| 0 | 0 | 0 | 1 |
| 0 | 0 | 0 | 1 |
| 0 | 0 | 0 | 1 |
| 0 | 0 | 0 | 0 |
| 0 | 0 | 0 | 1 |
| 0 | 0 | 0 | 0 |
| 0 | 0 | 0 | 0 |
| 0 | 0 | 0 | 0 |
| 0 | 0 | 0 | 1 |
| 0 | 0 | 0 | 1 |
| 0 | 0 | 0 | 1 |
| 0 | 0 | 0 | 1 |
| 0 | 0 | 0 | 1 |
| 0 | 0 | 0 | 0 |
| 0 | 0 | 0 | 1 |
| 0 | 0 | 0 | 1 |
| 0 | 0 | 0 | 0 |
| 0 | 0 | 0 | 1 |
| 0 | 0 | 0 | 0 |
| 0 | 0 | 0 | 0 |
| 0 | 0 | 0 | 1 |
| 0 | 0 | 0 | 1 |
| 0 | 0 | 0 | 1 |
| 0 | 0 | 0 | 1 |
| 0 | 0 | 0 | 1 |
| 0 | 0 | 0 | 1 |
| 0 | 0 | 0 | 1 |
| 0 | 0 | 0 | 1 |
| 0 | 0 | 0 | 0 |
| 0 | 0 | 0 | 1 |
| 0 | 0 | 0 | 1 |
| 0 | 0 | 0 | 0 |
| 0 | 0 | 0 | 0 |
| 0 | 0 | 0 | 1 |
| 0 | 0 | 0 | 1 |
| 1 | 0 | 0 | 0 |
| 1 | 0 | 0 | 0 |
| 0 | 0 | 0 | 0 |
| 0 | 0 | 0 | 0 |
| 1 | 0 | 0 | 0 |
| 1 | 0 | 0 | 0 |
| 0 | 0 | 0 | 1 |
| 1 | 0 | 0 | 0 |
| 1 | 0 | 0 | 0 |
| 1 | 0 | 0 | 0 |
| 1 | 0 | 0 | 0 |
| 1 | 0 | 0 | 0 |

| | | | |
|---|---|---|---|
| 0 | 0 | 0 | 0 |
| 0 | 0 | 0 | 0 |
| 1 | 0 | 0 | 0 |
| 1 | 0 | 0 | 0 |
| 1 | 0 | 0 | 0 |
| 1 | 0 | 0 | 0 |
| 0 | 0 | 0 | 0 |
| 0 | 0 | 0 | 0 |
| 0 | 0 | 0 | 0 |
| 0 | 0 | 0 | 1 |
| 0 | 0 | 0 | 1 |
| 0 | 0 | 0 | 0 |
| 0 | 0 | 0 | 1 |
| 0 | 0 | 0 | 0 |
| 0 | 0 | 0 | 0 |
| 0 | 0 | 0 | 1 |
| 0 | 0 | 0 | 0 |
| 0 | 0 | 0 | 1 |
| 0 | 0 | 0 | 1 |
| 0 | 0 | 0 | 1 |
| 0 | 0 | 0 | 1 |
| 0 | 0 | 0 | 1 |
| 0 | 0 | 0 | 1 |
| 0 | 0 | 0 | 0 |
| 0 | 0 | 0 | 0 |
| 0 | 0 | 0 | 1 |
| 0 | 0 | 0 | 1 |
| 0 | 0 | 0 | 1 |
| 0 | 0 | 0 | 0 |
| 0 | 0 | 0 | 0 |
| 0 | 0 | 0 | 1 |
| 0 | 0 | 0 | 1 |
| 0 | 0 | 0 | 1 |
| 0 | 0 | 0 | 0 |
| 0 | 0 | 0 | 0 |
| 0 | 0 | 0 | 1 |
| 0 | 0 | 0 | 0 |
| 0 | 0 | 0 | 1 |
| 0 | 0 | 0 | 1 |
| 0 | 0 | 0 | 0 |
| 0 | 0 | 0 | 0 |
| 0 | 0 | 0 | 0 |
| 0 | 0 | 0 | 0 |
| 0 | 0 | 0 | 0 |
| 0 | 0 | 0 | 0 |
| 0 | 0 | 0 | 1 |
| 0 | 0 | 0 | 1 |
| 0 | 0 | 0 | 0 |
| 0 | 0 | 0 | 0 |
| 0 | 0 | 0 | 0 |
| 0 | 0 | 0 | 1 |
| 0 | 0 | 0 | 1 |
| 0 | 0 | 0 | 0 |
| 0 | 0 | 0 | 0 |

| | | | |
|---|---|---|---|
| 0 | 0 | 0 | 1 |
| 0 | 0 | 0 | 1 |
| 0 | 0 | 0 | 1 |
| 0 | 0 | 0 | 1 |
| 0 | 1 | 0 | 0 |
| 0 | 0 | 0 | 0 |
| 0 | 0 | 0 | 1 |
| 0 | 0 | 0 | 0 |
| 0 | 0 | 0 | 1 |
| 0 | 0 | 0 | 0 |
| 0 | 0 | 0 | 1 |
| 0 | 0 | 0 | 1 |
| 0 | 0 | 0 | 1 |
| 0 | 0 | 0 | 1 |
| 0 | 0 | 0 | 1 |
| 0 | 0 | 0 | 1 |
| 0 | 0 | 0 | 0 |
| 0 | 0 | 0 | 1 |
| 0 | 0 | 0 | 1 |
| 0 | 0 | 0 | 0 |
| 0 | 0 | 0 | 0 |
| 0 | 0 | 0 | 1 |
| 0 | 0 | 0 | 1 |
| 0 | 0 | 0 | 0 |
| 0 | 0 | 0 | 1 |
| 0 | 0 | 0 | 0 |
| 0 | 0 | 0 | 0 |
| 0 | 0 | 0 | 0 |
| 0 | 0 | 0 | 0 |
| 0 | 0 | 0 | 0 |
| 0 | 0 | 0 | 0 |
| 0 | 0 | 0 | 0 |
| 0 | 0 | 0 | 1 |
| 0 | 0 | 0 | 0 |
| 0 | 0 | 0 | 1 |
| 0 | 0 | 0 | 1 |
| 0 | 0 | 0 | 1 |
| 0 | 0 | 0 | 1 |
| 0 | 0 | 0 | 0 |
| 0 | 0 | 0 | 1 |
| 0 | 0 | 0 | 1 |
| 0 | 0 | 0 | 1 |
| 0 | 0 | 0 | 1 |
| 0 | 0 | 0 | 0 |
| 0 | 0 | 0 | 1 |
| 0 | 0 | 0 | 1 |
| 0 | 0 | 0 | 1 |
| 0 | 0 | 0 | 1 |
| 0 | 0 | 0 | 1 |
| 0 | 0 | 0 | 1 |
| 0 | 0 | 0 | 0 |
| 0 | 0 | 0 | 0 |
| 0 | 0 | 0 | 0 |
| 0 | 0 | 0 | 1 |

| | | | |
|---|---|---|---|
| 0 | 0 | 0 | 0 |
| 0 | 0 | 0 | 0 |
| 0 | 0 | 0 | 0 |
| 0 | 0 | 0 | 0 |
| 1 | 0 | 0 | 0 |
| 1 | 0 | 0 | 0 |
| 0 | 0 | 0 | 0 |
| 0 | 0 | 0 | 0 |
| 0 | 0 | 0 | 1 |
| 0 | 0 | 0 | 1 |
| 0 | 0 | 0 | 0 |
| 0 | 0 | 0 | 1 |
| 0 | 0 | 0 | 1 |
| 0 | 0 | 0 | 0 |
| 0 | 0 | 0 | 0 |
| 0 | 0 | 0 | 0 |
| 0 | 0 | 0 | 0 |
| 0 | 0 | 0 | 0 |
| 0 | 0 | 0 | 0 |
| 0 | 0 | 0 | 1 |
| 0 | 0 | 0 | 1 |
| 0 | 0 | 0 | 0 |
| 0 | 0 | 0 | 1 |
| 0 | 0 | 0 | 0 |
| 0 | 0 | 0 | 1 |
| 0 | 0 | 0 | 1 |
| 0 | 0 | 0 | 1 |
| 0 | 0 | 0 | 1 |
| 0 | 0 | 0 | 1 |
| 0 | 0 | 0 | 0 |
| 0 | 0 | 0 | 1 |
| 0 | 0 | 0 | 1 |
| 0 | 0 | 0 | 0 |
| 0 | 0 | 0 | 1 |
| 0 | 0 | 0 | 0 |
| 0 | 0 | 0 | 0 |
| 0 | 0 | 0 | 0 |
| 0 | 0 | 0 | 0 |
| 0 | 0 | 0 | 0 |
| 0 | 0 | 0 | 0 |
| 0 | 0 | 0 | 0 |
| 0 | 0 | 0 | 0 |
| 0 | 0 | 0 | 1 |
| 0 | 0 | 0 | 0 |
| 0 | 0 | 0 | 1 |
| 0 | 0 | 0 | 0 |
| 0 | 0 | 0 | 1 |
| 0 | 0 | 0 | 1 |
| 0 | 0 | 0 | 1 |
| 0 | 0 | 0 | 0 |
| 0 | 0 | 0 | 1 |
| 0 | 0 | 0 | 1 |
| 0 | 0 | 0 | 1 |
| 0 | 0 | 0 | 1 |

| | | | |
|---|---|---|---|
| 1 | 0 | 0 | 0 |
| 0 | 0 | 0 | 1 |
| 0 | 0 | 0 | 1 |
| 0 | 0 | 0 | 1 |
| 0 | 0 | 0 | 1 |
| 0 | 0 | 0 | 1 |
| 0 | 0 | 0 | 1 |
| 0 | 0 | 0 | 1 |
| 0 | 0 | 0 | 1 |
| 0 | 0 | 0 | 1 |
| 0 | 0 | 0 | 0 |
| 0 | 0 | 0 | 1 |
| 0 | 0 | 0 | 1 |
| 0 | 0 | 0 | 1 |
| 0 | 0 | 0 | 0 |
| 0 | 0 | 0 | 0 |
| 0 | 0 | 0 | 0 |
| 0 | 0 | 0 | 0 |
| 0 | 0 | 0 | 0 |
| 0 | 0 | 0 | 1 |
| 0 | 0 | 0 | 1 |
| 0 | 0 | 0 | 1 |
| 0 | 0 | 0 | 0 |
| 0 | 0 | 0 | 1 |
| 0 | 0 | 0 | 1 |
| 0 | 0 | 0 | 1 |
| 0 | 0 | 0 | 1 |
| 0 | 0 | 0 | 1 |
| 0 | 0 | 0 | 0 |
| 0 | 0 | 0 | 0 |
| 0 | 0 | 0 | 0 |
| 0 | 0 | 0 | 0 |
| 0 | 0 | 0 | 0 |
| 0 | 0 | 0 | 0 |
| 0 | 0 | 0 | 1 |
| 0 | 0 | 0 | 1 |
| 0 | 0 | 0 | 0 |
| 0 | 0 | 0 | 0 |
| 0 | 0 | 0 | 0 |
| 0 | 0 | 0 | 0 |
| 0 | 0 | 0 | 0 |
| 0 | 0 | 0 | 0 |
| 0 | 0 | 0 | 1 |
| 0 | 0 | 0 | 1 |
| 0 | 0 | 0 | 0 |
| 0 | 0 | 0 | 1 |
| 0 | 0 | 0 | 1 |
| 0 | 0 | 0 | 1 |
| 0 | 0 | 0 | 1 |
| 0 | 0 | 0 | 1 |
| 0 | 0 | 0 | 1 |
| 0 | 0 | 0 | 1 |
| 0 | 0 | 0 | 1 |
| 0 | 0 | 0 | 1 |

| | | | |
|---|---|---|---|
| 0 | 0 | 0 | 0 |
| 0 | 0 | 0 | 1 |
| 0 | 0 | 0 | 0 |
| 0 | 0 | 0 | 1 |
| 0 | 0 | 0 | 1 |
| 0 | 0 | 0 | 1 |
| 0 | 0 | 0 | 0 |
| 0 | 0 | 0 | 0 |
| 0 | 0 | 0 | 1 |
| 0 | 0 | 0 | 1 |
| 0 | 0 | 0 | 1 |
| 0 | 0 | 0 | 1 |
| 0 | 0 | 0 | 1 |
| 0 | 0 | 0 | 1 |
| 0 | 0 | 0 | 0 |
| 0 | 0 | 0 | 0 |
| 0 | 0 | 0 | 0 |
| 0 | 0 | 0 | 0 |
| 0 | 0 | 0 | 1 |
| 0 | 0 | 0 | 0 |
| 0 | 0 | 0 | 0 |
| 0 | 0 | 0 | 0 |
| 0 | 0 | 0 | 0 |
| 0 | 0 | 0 | 0 |
| 0 | 0 | 0 | 1 |
| 0 | 0 | 0 | 0 |
| 0 | 0 | 0 | 1 |
| 0 | 0 | 0 | 1 |
| 0 | 0 | 0 | 1 |
| 0 | 0 | 0 | 0 |
| 0 | 0 | 0 | 1 |
| 0 | 0 | 0 | 1 |
| 0 | 0 | 0 | 0 |
| 0 | 0 | 0 | 0 |
| 0 | 0 | 0 | 1 |
| 0 | 0 | 0 | 1 |
| 0 | 0 | 0 | 1 |
| 0 | 0 | 0 | 1 |
| 0 | 0 | 0 | 1 |
| 0 | 0 | 0 | 0 |
| 0 | 0 | 0 | 0 |
| 0 | 0 | 0 | 0 |
| 0 | 0 | 0 | 0 |
| 0 | 0 | 0 | 0 |
| 0 | 0 | 0 | 0 |
| 0 | 0 | 0 | 1 |
| 0 | 0 | 0 | 1 |
| 0 | 0 | 0 | 0 |
| 0 | 0 | 0 | 0 |
| 0 | 0 | 0 | 1 |
| 0 | 0 | 0 | 0 |
| 0 | 0 | 0 | 0 |
| 0 | 0 | 0 | 1 |
| 0 | 0 | 0 | 1 |

| | | | |
|---|---|---|---|
| 0 | 0 | 0 | 1 |
| 0 | 0 | 0 | 0 |
| 0 | 0 | 0 | 0 |
| 0 | 0 | 0 | 1 |
| 0 | 0 | 0 | 1 |
| 0 | 0 | 0 | 0 |
| 0 | 0 | 0 | 0 |
| 0 | 0 | 0 | 0 |
| 0 | 0 | 0 | 0 |
| 0 | 0 | 0 | 0 |
| 0 | 0 | 0 | 0 |
| 0 | 0 | 0 | 0 |
| 0 | 0 | 0 | 0 |
| 0 | 0 | 0 | 0 |
| 0 | 0 | 0 | 0 |
| 0 | 0 | 0 | 1 |
| 0 | 0 | 0 | 0 |
| 0 | 0 | 0 | 1 |
| 0 | 0 | 0 | 1 |
| 0 | 0 | 0 | 1 |
| 0 | 0 | 0 | 0 |
| 0 | 0 | 0 | 1 |
| 0 | 0 | 0 | 0 |
| 0 | 0 | 0 | 0 |
| 0 | 0 | 0 | 1 |
| 0 | 0 | 0 | 0 |
| 0 | 0 | 0 | 1 |
| 0 | 0 | 0 | 1 |
| 0 | 0 | 0 | 1 |
| 0 | 0 | 0 | 0 |
| 0 | 0 | 0 | 0 |
| 0 | 0 | 0 | 0 |
| 0 | 0 | 0 | 0 |
| 0 | 0 | 0 | 0 |
| 0 | 0 | 0 | 0 |
| 0 | 0 | 0 | 0 |
| 0 | 0 | 0 | 0 |
| 0 | 0 | 0 | 1 |
| 0 | 0 | 0 | 1 |
| 0 | 0 | 0 | 0 |
| 0 | 0 | 0 | 0 |
| 0 | 0 | 0 | 0 |
| 0 | 0 | 0 | 0 |
| 0 | 0 | 0 | 0 |
| 0 | 0 | 0 | 0 |
| 0 | 0 | 0 | 0 |
| 0 | 0 | 0 | 1 |
| 0 | 0 | 0 | 0 |
| 0 | 0 | 0 | 0 |
| 0 | 0 | 0 | 0 |
| 0 | 0 | 0 | 0 |
| 0 | 0 | 0 | 1 |
| 0 | 0 | 0 | 0 |
| 0 | 0 | 0 | 0 |

| | | | |
|---|---|---|---|
| 0 | 0 | 0 | 0 |
| 0 | 0 | 0 | 0 |
| 0 | 0 | 0 | 0 |
| 0 | 0 | 0 | 0 |
| 0 | 0 | 0 | 0 |
| 0 | 0 | 0 | 0 |
| 0 | 0 | 0 | 0 |
| 0 | 0 | 0 | 0 |
| 1 | 0 | 0 | 0 |
| 0 | 0 | 0 | 0 |
| 0 | 0 | 0 | 1 |
| 0 | 0 | 0 | 0 |
| 0 | 0 | 0 | 1 |
| 0 | 0 | 0 | 0 |
| 0 | 0 | 0 | 1 |
| 0 | 0 | 0 | 1 |
| 0 | 0 | 0 | 1 |
| 0 | 0 | 0 | 1 |
| 0 | 0 | 0 | 0 |
| 0 | 0 | 0 | 0 |
| 0 | 0 | 0 | 1 |
| 0 | 0 | 0 | 0 |
| 0 | 0 | 0 | 0 |
| 0 | 0 | 0 | 1 |
| 0 | 0 | 0 | 0 |
| 0 | 0 | 0 | 0 |
| 0 | 0 | 0 | 0 |
| 0 | 0 | 0 | 1 |
| 0 | 0 | 0 | 0 |
| 0 | 0 | 0 | 0 |
| 0 | 0 | 0 | 1 |
| 1 | 0 | 0 | 0 |
| 0 | 0 | 0 | 1 |
| 0 | 0 | 0 | 1 |
| 1 | 0 | 0 | 0 |
| 1 | 0 | 0 | 0 |
| 1 | 0 | 0 | 0 |
| 0 | 0 | 0 | 1 |
| 0 | 0 | 0 | 1 |
| 0 | 0 | 0 | 0 |
| 1 | 0 | 0 | 0 |
| 0 | 0 | 0 | 0 |
| 0 | 0 | 0 | 0 |
| 1 | 0 | 0 | 0 |
| 1 | 0 | 0 | 0 |
| 0 | 0 | 0 | 0 |
| 0 | 0 | 0 | 1 |
| 0 | 0 | 0 | 1 |
| 0 | 0 | 0 | 1 |
| 0 | 0 | 0 | 1 |
| 1 | 0 | 0 | 0 |
| 0 | 0 | 0 | 0 |
| 0 | 0 | 0 | 0 |
| 0 | 0 | 0 | 0 |

| | | | |
|---|---|---|---|
| 0 | 0 | 0 | 0 |
| 0 | 0 | 0 | 1 |
| 0 | 0 | 0 | 0 |
| 0 | 0 | 0 | 0 |
| 0 | 0 | 0 | 0 |
| 0 | 0 | 0 | 1 |
| 0 | 0 | 0 | 0 |
| 0 | 0 | 0 | 0 |
| 0 | 0 | 0 | 1 |
| 0 | 0 | 0 | 0 |
| 0 | 0 | 0 | 1 |
| 0 | 0 | 0 | 1 |
| 0 | 0 | 0 | 1 |
| 0 | 0 | 0 | 0 |
| 0 | 0 | 0 | 1 |
| 0 | 0 | 0 | 0 |
| 0 | 0 | 0 | 1 |
| 0 | 0 | 0 | 1 |
| 0 | 0 | 0 | 0 |
| 0 | 0 | 0 | 0 |
| 0 | 0 | 0 | 0 |
| 0 | 0 | 0 | 0 |
| 0 | 0 | 0 | 0 |
| 0 | 0 | 0 | 0 |
| 0 | 0 | 0 | 0 |
| 0 | 0 | 0 | 1 |
| 0 | 0 | 0 | 1 |
| 0 | 0 | 0 | 1 |
| 0 | 0 | 0 | 1 |
| 0 | 0 | 0 | 1 |
| 0 | 0 | 0 | 1 |
| 0 | 0 | 0 | 1 |
| 0 | 0 | 0 | 0 |
| 0 | 0 | 0 | 0 |
| 0 | 0 | 0 | 0 |
| 0 | 0 | 0 | 0 |
| 0 | 0 | 0 | 0 |
| 0 | 0 | 0 | 0 |
| 0 | 0 | 0 | 1 |
| 0 | 0 | 0 | 1 |
| 0 | 0 | 0 | 1 |
| 0 | 0 | 0 | 1 |
| 0 | 0 | 0 | 1 |
| 0 | 0 | 0 | 1 |
| 0 | 1 | 0 | 0 |
| 0 | 0 | 0 | 0 |
| 0 | 0 | 0 | 1 |
| 0 | 0 | 0 | 0 |
| 0 | 0 | 0 | 0 |
| 0 | 0 | 0 | 0 |
| 0 | 0 | 0 | 0 |
| 0 | 0 | 0 | 0 |
| 0 | 0 | 0 | 0 |
| 0 | 0 | 0 | 0 |

| | | | |
|---|---|---|---|
| 0 | 0 | 0 | 1 |
| 0 | 0 | 0 | 1 |
| 1 | 0 | 0 | 0 |
| 1 | 0 | 0 | 0 |
| 0 | 0 | 0 | 1 |
| 0 | 0 | 0 | 0 |
| 0 | 0 | 0 | 0 |
| 0 | 0 | 0 | 1 |
| 0 | 0 | 0 | 1 |
| 0 | 0 | 0 | 0 |
| 0 | 0 | 0 | 1 |
| 0 | 0 | 0 | 1 |
| 0 | 0 | 0 | 0 |
| 0 | 0 | 0 | 0 |
| 0 | 0 | 0 | 0 |
| 0 | 0 | 0 | 0 |
| 0 | 0 | 0 | 1 |
| 0 | 0 | 0 | 1 |
| 0 | 0 | 0 | 1 |
| 0 | 0 | 0 | 1 |
| 0 | 0 | 0 | 1 |
| 0 | 0 | 0 | 1 |
| 0 | 0 | 0 | 1 |
| 0 | 0 | 0 | 1 |
| 0 | 0 | 0 | 1 |
| 0 | 0 | 0 | 0 |
| 0 | 0 | 0 | 1 |
| 0 | 0 | 0 | 1 |
| 0 | 0 | 0 | 1 |
| 0 | 0 | 0 | 1 |
| 0 | 0 | 0 | 1 |
| 0 | 0 | 0 | 1 |
| 0 | 0 | 0 | 1 |
| 0 | 0 | 0 | 1 |
| 0 | 0 | 0 | 1 |
| 0 | 0 | 0 | 1 |
| 0 | 0 | 0 | 1 |
| 0 | 0 | 0 | 0 |
| 0 | 0 | 0 | 1 |
| 0 | 0 | 0 | 1 |
| 0 | 0 | 0 | 0 |
| 0 | 0 | 0 | 1 |
| 0 | 0 | 0 | 1 |
| 0 | 0 | 0 | 0 |
| 0 | 0 | 0 | 0 |
| 0 | 0 | 0 | 1 |
| 0 | 0 | 0 | 1 |
| 0 | 0 | 0 | 1 |
| 0 | 0 | 0 | 1 |
| 0 | 0 | 0 | 1 |
| 0 | 0 | 0 | 1 |
| 0 | 0 | 0 | 0 |
| 0 | 0 | 0 | 0 |
| 0 | 0 | 0 | 0 |

| | | | |
|---|---|---|---|
| 0 | 0 | 0 | 0 |
| 0 | 0 | 0 | 1 |
| 0 | 0 | 0 | 1 |
| 0 | 0 | 0 | 1 |
| 0 | 0 | 0 | 1 |
| 0 | 0 | 0 | 1 |
| 0 | 0 | 0 | 1 |
| 0 | 0 | 0 | 1 |
| 0 | 0 | 0 | 0 |
| 0 | 0 | 0 | 1 |
| 0 | 0 | 0 | 1 |
| 0 | 0 | 0 | 1 |
| 0 | 0 | 0 | 1 |
| 0 | 0 | 0 | 1 |
| 0 | 0 | 0 | 1 |
| 0 | 0 | 0 | 1 |
| 0 | 0 | 0 | 1 |
| 0 | 0 | 0 | 1 |
| 0 | 0 | 0 | 0 |
| 0 | 0 | 0 | 0 |
| 0 | 0 | 0 | 0 |
| 0 | 0 | 0 | 0 |
| 0 | 0 | 0 | 1 |
| 0 | 0 | 0 | 0 |
| 0 | 0 | 0 | 0 |
| 0 | 0 | 0 | 1 |
| 0 | 0 | 0 | 1 |
| 0 | 0 | 0 | 1 |
| 0 | 0 | 0 | 1 |
| 0 | 0 | 0 | 1 |
| 0 | 0 | 0 | 1 |
| 0 | 0 | 0 | 1 |
| 0 | 0 | 0 | 0 |
| 0 | 0 | 0 | 1 |
| 0 | 0 | 0 | 1 |
| 0 | 0 | 0 | 1 |
| 0 | 0 | 0 | 0 |
| 0 | 0 | 0 | 1 |
| 0 | 0 | 0 | 0 |
| 0 | 0 | 0 | 0 |
| 0 | 0 | 0 | 0 |
| 0 | 0 | 0 | 0 |
| 0 | 0 | 0 | 0 |
| 0 | 0 | 0 | 1 |
| 0 | 0 | 0 | 1 |
| 0 | 0 | 0 | 1 |
| 0 | 0 | 0 | 0 |
| 0 | 0 | 0 | 0 |
| 0 | 0 | 0 | 0 |
| 0 | 0 | 0 | 0 |
| 0 | 0 | 0 | 0 |
| 0 | 0 | 0 | 0 |
| 0 | 0 | 0 | 0 |
| 0 | 0 | 0 | 1 |

| | | | |
|---|---|---|---|
| 0 | 0 | 0 | 0 |
| 0 | 0 | 0 | 0 |
| 0 | 0 | 0 | 1 |
| 1 | 0 | 0 | 0 |
| 1 | 0 | 0 | 0 |
| 0 | 0 | 0 | 1 |
| 0 | 0 | 0 | 1 |
| 0 | 0 | 0 | 1 |
| 0 | 0 | 0 | 1 |
| 0 | 0 | 0 | 1 |
| 0 | 0 | 0 | 1 |
| 0 | 0 | 0 | 0 |
| 0 | 0 | 0 | 1 |
| 0 | 0 | 0 | 1 |
| 0 | 0 | 0 | 0 |
| 0 | 0 | 0 | 1 |
| 0 | 0 | 0 | 0 |
| 0 | 0 | 0 | 1 |
| 0 | 0 | 0 | 0 |
| 1 | 0 | 0 | 0 |
| 0 | 0 | 0 | 1 |
| 0 | 0 | 0 | 1 |
| 0 | 0 | 0 | 0 |
| 0 | 0 | 0 | 1 |
| 0 | 0 | 0 | 0 |
| 0 | 0 | 0 | 1 |
| 0 | 0 | 0 | 1 |
| 0 | 0 | 0 | 0 |
| 0 | 0 | 0 | 1 |
| 0 | 0 | 0 | 0 |
| 0 | 0 | 0 | 1 |
| 0 | 0 | 0 | 0 |
| 0 | 0 | 0 | 0 |
| 0 | 0 | 0 | 0 |
| 0 | 0 | 0 | 1 |
| 0 | 0 | 0 | 0 |
| 0 | 0 | 0 | 1 |
| 0 | 0 | 0 | 1 |
| 0 | 0 | 0 | 0 |
| 0 | 0 | 0 | 1 |
| 0 | 0 | 0 | 1 |
| 0 | 0 | 0 | 1 |
| 0 | 0 | 0 | 1 |
| 0 | 0 | 0 | 0 |
| 0 | 0 | 0 | 0 |
| 0 | 0 | 0 | 1 |
| 0 | 0 | 0 | 1 |
| 0 | 0 | 0 | 1 |
| 0 | 0 | 0 | 0 |
| 0 | 0 | 0 | 1 |
| 0 | 0 | 0 | 0 |
| 0 | 0 | 0 | 1 |
| 0 | 0 | 0 | 1 |
| 0 | 0 | 0 | 0 |

| | | | |
|---|---|---|---|
| 0 | 0 | 0 | 1 |
| 0 | 0 | 0 | 1 |
| 0 | 0 | 0 | 1 |
| 0 | 0 | 0 | 1 |
| 0 | 0 | 0 | 1 |
| 0 | 0 | 0 | 1 |
| 0 | 0 | 0 | 1 |
| 0 | 0 | 0 | 0 |
| 0 | 0 | 0 | 1 |
| 0 | 0 | 0 | 1 |
| 0 | 0 | 0 | 1 |
| 0 | 0 | 0 | 1 |
| 0 | 0 | 0 | 0 |
| 0 | 0 | 0 | 1 |
| 0 | 0 | 0 | 0 |
| 0 | 0 | 0 | 0 |
| 0 | 0 | 0 | 1 |
| 0 | 0 | 0 | 1 |
| 0 | 0 | 0 | 1 |
| 0 | 0 | 0 | 1 |
| 0 | 0 | 0 | 0 |
| 0 | 0 | 0 | 0 |
| 0 | 0 | 0 | 1 |
| 0 | 0 | 0 | 1 |
| 0 | 0 | 0 | 0 |
| 0 | 0 | 0 | 1 |
| 0 | 0 | 0 | 0 |
| 0 | 0 | 0 | 0 |
| 0 | 0 | 0 | 0 |
| 0 | 0 | 0 | 0 |
| 0 | 0 | 0 | 1 |
| 0 | 0 | 0 | 1 |
| 0 | 0 | 0 | 0 |
| 0 | 0 | 0 | 0 |
| 0 | 0 | 0 | 0 |
| 0 | 0 | 0 | 0 |
| 0 | 0 | 0 | 0 |
| 0 | 0 | 0 | 0 |
| 0 | 0 | 0 | 0 |
| 0 | 0 | 0 | 1 |
| 0 | 0 | 0 | 1 |
| 0 | 0 | 0 | 1 |
| 0 | 0 | 0 | 1 |
| 0 | 0 | 0 | 1 |
| 0 | 0 | 0 | 1 |
| 0 | 0 | 0 | 0 |
| 0 | 0 | 0 | 1 |
| 0 | 0 | 0 | 1 |
| 0 | 0 | 0 | 1 |
| 0 | 0 | 0 | 0 |
| 0 | 0 | 0 | 0 |
| 0 | 0 | 0 | 1 |
| 0 | 0 | 0 | 1 |
| 0 | 0 | 0 | 1 |

| | | | |
|---|---|---|---|
| 0 | 0 | 0 | 1 |
| 0 | 0 | 0 | 1 |
| 0 | 0 | 0 | 1 |
| 0 | 0 | 0 | 1 |
| 0 | 0 | 0 | 0 |
| 0 | 0 | 0 | 1 |
| 0 | 0 | 0 | 0 |
| 0 | 0 | 0 | 0 |
| 0 | 0 | 0 | 0 |
| 0 | 0 | 0 | 0 |
| 0 | 0 | 0 | 0 |
| 0 | 0 | 0 | 1 |
| 0 | 0 | 0 | 0 |
| 0 | 0 | 0 | 0 |
| 0 | 0 | 0 | 1 |
| 0 | 0 | 0 | 1 |
| 0 | 0 | 0 | 1 |
| 0 | 0 | 0 | 1 |
| 0 | 0 | 0 | 1 |
| 0 | 0 | 0 | 1 |
| 0 | 0 | 0 | 1 |
| 0 | 0 | 0 | 1 |
| 0 | 0 | 0 | 1 |
| 0 | 0 | 0 | 0 |
| 0 | 0 | 0 | 1 |
| 0 | 0 | 0 | 1 |
| 0 | 0 | 0 | 1 |
| 0 | 0 | 0 | 0 |
| 0 | 0 | 0 | 1 |
| 0 | 0 | 0 | 1 |
| 0 | 0 | 0 | 0 |
| 0 | 0 | 0 | 0 |
| 0 | 0 | 0 | 1 |
| 0 | 0 | 0 | 0 |
| 0 | 0 | 0 | 1 |
| 0 | 0 | 0 | 1 |
| 0 | 0 | 0 | 1 |
| 0 | 0 | 0 | 1 |
| 0 | 0 | 0 | 1 |
| 0 | 0 | 0 | 1 |
| 0 | 0 | 0 | 1 |
| 0 | 0 | 0 | 1 |
| 0 | 0 | 0 | 1 |
| 0 | 0 | 0 | 1 |
| 0 | 0 | 0 | 1 |
| 0 | 0 | 0 | 1 |
| 0 | 0 | 0 | 1 |
| 0 | 0 | 0 | 1 |
| 0 | 0 | 0 | 0 |
| 0 | 0 | 0 | 1 |
| 0 | 0 | 0 | 1 |
| 0 | 0 | 0 | 1 |
| 0 | 0 | 0 | 1 |
| 0 | 0 | 0 | 1 |

| | | | |
|---|---|---|---|
| 0 | 0 | 0 | 1 |
| 0 | 0 | 0 | 1 |
| 0 | 0 | 0 | 1 |
| 0 | 0 | 0 | 1 |
| 0 | 0 | 0 | 1 |
| 0 | 0 | 0 | 1 |
| 0 | 0 | 0 | 1 |
| 0 | 0 | 0 | 1 |
| 0 | 0 | 0 | 1 |
| 0 | 0 | 0 | 1 |
| 0 | 0 | 0 | 0 |
| 0 | 0 | 0 | 0 |
| 0 | 0 | 0 | 0 |
| 1 | 0 | 0 | 0 |
| 1 | 0 | 0 | 0 |
| 0 | 0 | 0 | 1 |
| 1 | 0 | 0 | 0 |
| 0 | 0 | 0 | 0 |
| 0 | 0 | 0 | 0 |
| 0 | 0 | 0 | 0 |
| 0 | 0 | 0 | 0 |
| 1 | 0 | 0 | 0 |
| 0 | 0 | 0 | 1 |
| 0 | 0 | 0 | 0 |
| 1 | 0 | 0 | 0 |
| 1 | 0 | 0 | 0 |
| 0 | 0 | 0 | 1 |
| 0 | 0 | 0 | 0 |
| 0 | 0 | 0 | 1 |
| 0 | 0 | 0 | 1 |
| 0 | 0 | 0 | 1 |
| 0 | 0 | 0 | 0 |
| 0 | 0 | 0 | 1 |
| 0 | 0 | 0 | 1 |
| 0 | 0 | 0 | 0 |
| 0 | 0 | 0 | 0 |
| 0 | 0 | 0 | 0 |
| 0 | 0 | 0 | 0 |
| 0 | 0 | 0 | 0 |
| 0 | 0 | 0 | 1 |
| 0 | 0 | 0 | 1 |
| 0 | 0 | 0 | 0 |
| 0 | 0 | 0 | 1 |
| 0 | 0 | 0 | 0 |
| 0 | 0 | 0 | 1 |
| 0 | 0 | 0 | 1 |
| 0 | 0 | 0 | 1 |
| 0 | 0 | 0 | 0 |
| 0 | 0 | 0 | 0 |
| 0 | 0 | 0 | 0 |
| 0 | 0 | 0 | 0 |
| 0 | 0 | 0 | 1 |
| 0 | 0 | 0 | 0 |
| 0 | 0 | 0 | 1 |

| | | | |
|---|---|---|---|
| 0 | 0 | 0 | 1 |
| 0 | 0 | 0 | 1 |
| 0 | 0 | 0 | 1 |
| 0 | 0 | 0 | 1 |
| 0 | 0 | 0 | 1 |
| 0 | 0 | 0 | 0 |
| 0 | 0 | 0 | 0 |
| 0 | 0 | 0 | 0 |
| 0 | 0 | 0 | 0 |
| 0 | 0 | 0 | 1 |
| 1 | 0 | 0 | 0 |
| 1 | 0 | 0 | 0 |
| 0 | 0 | 0 | 0 |
| 0 | 0 | 0 | 0 |
| 0 | 0 | 0 | 0 |
| 0 | 0 | 0 | 0 |
| 0 | 0 | 0 | 1 |
| 0 | 0 | 0 | 1 |
| 0 | 0 | 0 | 1 |
| 0 | 0 | 0 | 1 |
| 0 | 0 | 0 | 1 |
| 0 | 0 | 0 | 1 |
| 0 | 0 | 0 | 1 |
| 0 | 0 | 0 | 1 |
| 0 | 0 | 0 | 1 |
| 0 | 0 | 0 | 1 |
| 0 | 0 | 0 | 1 |
| 0 | 0 | 0 | 1 |
| 0 | 0 | 0 | 0 |
| 0 | 0 | 0 | 1 |
| 0 | 0 | 0 | 1 |
| 0 | 0 | 0 | 1 |
| 0 | 0 | 0 | 1 |
| 0 | 0 | 0 | 0 |
| 0 | 0 | 0 | 1 |
| 0 | 0 | 0 | 0 |
| 0 | 0 | 0 | 1 |
| 0 | 0 | 0 | 0 |
| 0 | 0 | 0 | 1 |
| 0 | 0 | 0 | 0 |
| 0 | 0 | 0 | 1 |
| 0 | 0 | 0 | 1 |
| 0 | 0 | 0 | 1 |
| 0 | 0 | 0 | 0 |
| 0 | 0 | 0 | 1 |
| 0 | 0 | 0 | 1 |
| 0 | 0 | 0 | 1 |
| 0 | 0 | 0 | 1 |
| 0 | 0 | 0 | 1 |
| 0 | 0 | 0 | 1 |
| 0 | 0 | 0 | 1 |
| 0 | 0 | 0 | 1 |
| 0 | 0 | 0 | 0 |
| 0 | 0 | 0 | 0 |

| | | | |
|---|---|---|---|
| 0 | 0 | 0 | 1 |
| 0 | 0 | 0 | 0 |
| 0 | 0 | 0 | 1 |
| 0 | 0 | 0 | 1 |
| 0 | 0 | 0 | 0 |
| 0 | 0 | 0 | 0 |
| 0 | 0 | 0 | 0 |
| 0 | 0 | 0 | 1 |
| 0 | 0 | 0 | 1 |
| 0 | 0 | 0 | 0 |
| 0 | 0 | 0 | 0 |
| 0 | 0 | 0 | 0 |
| 0 | 0 | 0 | 0 |
| 0 | 0 | 0 | 0 |
| 0 | 0 | 0 | 1 |
| 0 | 0 | 0 | 0 |
| 0 | 0 | 0 | 0 |
| 0 | 0 | 0 | 0 |
| 0 | 0 | 0 | 1 |
| 0 | 0 | 0 | 1 |
| 0 | 0 | 0 | 1 |
| 0 | 0 | 0 | 1 |
| 0 | 0 | 0 | 1 |
| 0 | 0 | 0 | 0 |
| 0 | 0 | 0 | 1 |
| 0 | 0 | 0 | 1 |
| 0 | 0 | 0 | 0 |
| 0 | 0 | 0 | 0 |
| 0 | 0 | 0 | 1 |
| 0 | 0 | 0 | 1 |
| 0 | 0 | 0 | 0 |
| 0 | 0 | 0 | 1 |
| 0 | 0 | 0 | 1 |
| 0 | 0 | 0 | 1 |
| 0 | 0 | 0 | 1 |
| 0 | 0 | 0 | 0 |
| 0 | 0 | 0 | 1 |
| 0 | 0 | 0 | 0 |
| 0 | 0 | 0 | 0 |
| 0 | 0 | 0 | 1 |
| 0 | 0 | 0 | 1 |
| 0 | 0 | 0 | 0 |
| 0 | 0 | 0 | 0 |
| 0 | 0 | 0 | 1 |
| 0 | 0 | 0 | 0 |
| 0 | 0 | 0 | 0 |
| 0 | 0 | 0 | 0 |
| 0 | 0 | 0 | 1 |
| 0 | 0 | 0 | 0 |
| 0 | 0 | 0 | 0 |
| 0 | 0 | 0 | 0 |
| 0 | 0 | 0 | 1 |
| 0 | 0 | 0 | 0 |
| 0 | 0 | 0 | 1 |

| | | | |
|---|---|---|---|
| 0 | 0 | 0 | 1 |
| 0 | 0 | 0 | 1 |
| 0 | 0 | 0 | 1 |
| 0 | 0 | 0 | 1 |
| 0 | 0 | 0 | 0 |
| 0 | 0 | 0 | 0 |
| 0 | 0 | 0 | 1 |
| 0 | 0 | 0 | 1 |
| 0 | 0 | 0 | 1 |
| 0 | 0 | 0 | 0 |
| 0 | 0 | 0 | 1 |
| 0 | 0 | 0 | 1 |
| 0 | 0 | 0 | 0 |
| 0 | 0 | 0 | 1 |
| 0 | 0 | 0 | 1 |
| 0 | 0 | 0 | 1 |
| 0 | 0 | 0 | 1 |
| 0 | 0 | 0 | 1 |
| 0 | 0 | 0 | 1 |
| 0 | 0 | 0 | 1 |
| 0 | 0 | 0 | 1 |
| 0 | 0 | 0 | 1 |
| 0 | 0 | 0 | 1 |
| 0 | 0 | 0 | 0 |
| 0 | 0 | 0 | 1 |
| 1 | 0 | 0 | 0 |
| 0 | 0 | 0 | 1 |
| 0 | 0 | 0 | 0 |
| 0 | 0 | 0 | 1 |
| 0 | 0 | 0 | 0 |
| 0 | 0 | 0 | 0 |
| 0 | 0 | 0 | 0 |
| 0 | 0 | 0 | 0 |
| 0 | 0 | 0 | 0 |
| 0 | 0 | 0 | 1 |
| 0 | 0 | 0 | 0 |
| 0 | 0 | 0 | 1 |
| 0 | 0 | 0 | 0 |
| 0 | 0 | 0 | 1 |
| 0 | 0 | 0 | 1 |
| 0 | 0 | 0 | 0 |
| 0 | 0 | 0 | 0 |
| 0 | 0 | 0 | 0 |
| 0 | 0 | 0 | 0 |
| 0 | 0 | 0 | 0 |
| 0 | 0 | 0 | 0 |
| 0 | 0 | 0 | 0 |
| 0 | 0 | 0 | 0 |
| 0 | 0 | 0 | 1 |
| 0 | 0 | 0 | 0 |
| 0 | 0 | 0 | 1 |
| 0 | 0 | 0 | 1 |
| 0 | 0 | 0 | 1 |
| 0 | 0 | 0 | 1 |

| | | | |
|---|---|---|---|
| 0 | 0 | 0 | 1 |
| 0 | 0 | 0 | 1 |
| 0 | 0 | 0 | 1 |
| 0 | 0 | 0 | 1 |
| 0 | 0 | 0 | 1 |
| 0 | 0 | 0 | 1 |
| 0 | 0 | 0 | 0 |
| 0 | 0 | 0 | 0 |
| 0 | 0 | 0 | 0 |
| 0 | 0 | 0 | 0 |
| 0 | 0 | 0 | 0 |
| 0 | 0 | 0 | 0 |
| 0 | 0 | 0 | 0 |
| 0 | 0 | 0 | 0 |
| 0 | 0 | 0 | 0 |
| 0 | 0 | 0 | 0 |
| 1 | 0 | 0 | 0 |
| 0 | 0 | 0 | 0 |
| 0 | 0 | 0 | 1 |
| 0 | 0 | 0 | 0 |
| 0 | 0 | 0 | 1 |
| 0 | 0 | 0 | 1 |
| 0 | 0 | 0 | 1 |
| 0 | 0 | 0 | 1 |
| 0 | 0 | 0 | 1 |
| 0 | 0 | 0 | 0 |
| 0 | 0 | 0 | 0 |
| 0 | 0 | 0 | 0 |
| 0 | 0 | 0 | 0 |
| 0 | 0 | 0 | 0 |
| 0 | 0 | 0 | 0 |
| 1 | 0 | 0 | 0 |
| 0 | 0 | 0 | 0 |
| 1 | 0 | 0 | 0 |
| 0 | 0 | 0 | 1 |
| 1 | 0 | 0 | 0 |
| 1 | 0 | 0 | 0 |
| 0 | 0 | 0 | 0 |
| 1 | 0 | 0 | 0 |
| 0 | 0 | 0 | 1 |
| 0 | 0 | 0 | 0 |
| 0 | 0 | 0 | 0 |
| 0 | 0 | 0 | 1 |
| 0 | 0 | 0 | 1 |
| 0 | 0 | 0 | 1 |
| 0 | 0 | 0 | 1 |
| 0 | 0 | 0 | 1 |
| 0 | 0 | 0 | 0 |
| 0 | 0 | 0 | 0 |
| 0 | 0 | 0 | 0 |
| 0 | 0 | 0 | 0 |
| 0 | 0 | 0 | 1 |
| 0 | 0 | 0 | 0 |
| 0 | 0 | 0 | 1 |

| | | | |
|---|---|---|---|
| 0 | 0 | 0 | 1 |
| 0 | 0 | 0 | 1 |
| 0 | 0 | 0 | 1 |
| 0 | 0 | 0 | 1 |
| 0 | 0 | 0 | 1 |
| 0 | 0 | 0 | 0 |
| 0 | 0 | 0 | 1 |
| 0 | 0 | 0 | 1 |
| 0 | 0 | 0 | 1 |
| 0 | 0 | 0 | 1 |
| 0 | 0 | 0 | 0 |
| 0 | 0 | 0 | 0 |
| 0 | 0 | 0 | 0 |
| 0 | 0 | 0 | 1 |
| 0 | 0 | 0 | 1 |
| 0 | 0 | 0 | 1 |
| 0 | 0 | 0 | 1 |
| 0 | 0 | 0 | 0 |
| 0 | 0 | 0 | 1 |
| 0 | 0 | 0 | 1 |
| 0 | 0 | 0 | 1 |
| 0 | 0 | 0 | 1 |
| 0 | 0 | 0 | 0 |
| 0 | 0 | 0 | 1 |
| 0 | 0 | 0 | 0 |
| 0 | 0 | 0 | 0 |
| 0 | 0 | 0 | 0 |
| 0 | 0 | 0 | 0 |
| 0 | 0 | 0 | 1 |
| 0 | 0 | 0 | 0 |
| 0 | 0 | 0 | 1 |
| 0 | 0 | 0 | 1 |
| 0 | 0 | 0 | 1 |
| 0 | 0 | 0 | 1 |
| 0 | 0 | 0 | 1 |
| 0 | 0 | 0 | 1 |
| 0 | 0 | 0 | 1 |
| 0 | 0 | 0 | 0 |
| 0 | 0 | 0 | 0 |
| 0 | 0 | 0 | 0 |
| 0 | 0 | 0 | 0 |
| 0 | 0 | 0 | 0 |
| 0 | 0 | 0 | 1 |
| 0 | 0 | 0 | 0 |
| 0 | 0 | 0 | 0 |
| 0 | 0 | 0 | 1 |
| 0 | 0 | 0 | 0 |
| 0 | 0 | 0 | 0 |
| 0 | 0 | 0 | 0 |
| 1 | 0 | 0 | 0 |
| 0 | 0 | 0 | 1 |
| 0 | 0 | 0 | 1 |
| 0 | 0 | 0 | 1 |
| 0 | 0 | 0 | 0 |

| | | | |
|---|---|---|---|
| 0 | 0 | 0 | 0 |
| 0 | 0 | 0 | 1 |
| 0 | 0 | 0 | 0 |
| 0 | 0 | 0 | 1 |
| 0 | 0 | 0 | 1 |
| 0 | 0 | 0 | 1 |
| 0 | 0 | 0 | 1 |
| 0 | 0 | 0 | 1 |
| 0 | 0 | 0 | 0 |
| 0 | 0 | 0 | 1 |
| 0 | 0 | 0 | 0 |
| 0 | 0 | 0 | 0 |
| 0 | 0 | 0 | 0 |
| 0 | 0 | 0 | 0 |
| 0 | 0 | 0 | 1 |
| 0 | 0 | 0 | 1 |
| 0 | 0 | 0 | 0 |
| 0 | 0 | 0 | 0 |
| 0 | 0 | 0 | 1 |
| 0 | 0 | 0 | 1 |
| 0 | 0 | 0 | 0 |
| 0 | 0 | 0 | 1 |
| 0 | 0 | 0 | 1 |
| 0 | 0 | 0 | 1 |
| 0 | 0 | 0 | 0 |
| 0 | 0 | 0 | 1 |
| 0 | 0 | 0 | 0 |
| 0 | 0 | 0 | 0 |
| 0 | 0 | 0 | 0 |
| 0 | 0 | 0 | 1 |
| 0 | 0 | 0 | 0 |
| 0 | 0 | 0 | 0 |
| 0 | 0 | 0 | 0 |
| 0 | 0 | 0 | 1 |
| 0 | 0 | 0 | 0 |
| 0 | 0 | 0 | 1 |
| 0 | 0 | 0 | 0 |
| 0 | 0 | 0 | 0 |
| 0 | 0 | 0 | 0 |
| 0 | 0 | 0 | 0 |
| 0 | 0 | 0 | 0 |
| 0 | 0 | 0 | 1 |
| 0 | 0 | 0 | 0 |
| 0 | 0 | 0 | 0 |
| 0 | 0 | 0 | 0 |
| 1 | 0 | 0 | 0 |
| 0 | 0 | 0 | 0 |
| 1 | 0 | 0 | 0 |
| 1 | 0 | 0 | 0 |
| 1 | 0 | 0 | 0 |
| 0 | 0 | 0 | 0 |
| 0 | 0 | 0 | 0 |
| 0 | 0 | 0 | 1 |
| 0 | 0 | 0 | 0 |

| | | | |
|---|---|---|---|
| 0 | 0 | 0 | 1 |
| 0 | 0 | 0 | 1 |
| 0 | 0 | 0 | 1 |
| 0 | 0 | 0 | 1 |
| 0 | 0 | 0 | 1 |
| 0 | 0 | 0 | 1 |
| 0 | 0 | 0 | 1 |
| 0 | 0 | 0 | 1 |
| 0 | 0 | 0 | 1 |
| 0 | 0 | 0 | 0 |
| 0 | 0 | 0 | 0 |
| 0 | 0 | 0 | 1 |
| 0 | 0 | 0 | 1 |
| 0 | 0 | 0 | 1 |
| 0 | 0 | 0 | 0 |
| 0 | 0 | 0 | 1 |
| 0 | 0 | 0 | 1 |
| 0 | 0 | 0 | 0 |
| 0 | 0 | 0 | 1 |
| 0 | 0 | 0 | 1 |
| 0 | 0 | 0 | 1 |
| 0 | 0 | 0 | 0 |
| 0 | 0 | 0 | 0 |
| 0 | 0 | 0 | 0 |
| 0 | 0 | 0 | 0 |
| 0 | 0 | 0 | 0 |
| 0 | 0 | 0 | 0 |
| 0 | 0 | 0 | 0 |
| 0 | 0 | 0 | 1 |
| 0 | 0 | 0 | 0 |
| 1 | 0 | 0 | 0 |
| 0 | 0 | 0 | 1 |
| 0 | 0 | 0 | 1 |
| 0 | 0 | 0 | 1 |
| 0 | 0 | 0 | 1 |
| 0 | 0 | 0 | 1 |
| 0 | 0 | 0 | 1 |
| 0 | 0 | 0 | 1 |
| 0 | 0 | 0 | 1 |
| 0 | 0 | 0 | 1 |
| 0 | 0 | 0 | 1 |
| 0 | 0 | 0 | 1 |
| 0 | 0 | 0 | 1 |
| 0 | 0 | 0 | 0 |
| 0 | 0 | 0 | 1 |
| 0 | 0 | 0 | 0 |
| 0 | 0 | 0 | 1 |
| 0 | 0 | 0 | 1 |
| 0 | 0 | 0 | 1 |
| 0 | 0 | 0 | 1 |
| 0 | 0 | 0 | 0 |
| 0 | 0 | 0 | 1 |
| 0 | 0 | 0 | 1 |
| 0 | 0 | 0 | 0 |

| | | | |
|---|---|---|---|
| 0 | 0 | 0 | 0 |
| 0 | 0 | 0 | 0 |
| 0 | 0 | 0 | 0 |
| 0 | 0 | 0 | 0 |
| 0 | 0 | 0 | 1 |
| 0 | 0 | 0 | 1 |
| 0 | 0 | 0 | 1 |
| 0 | 0 | 0 | 0 |
| 0 | 0 | 0 | 0 |
| 0 | 0 | 0 | 0 |
| 0 | 0 | 0 | 0 |
| 0 | 0 | 0 | 1 |
| 0 | 0 | 0 | 1 |
| 0 | 0 | 0 | 1 |
| 0 | 0 | 0 | 1 |
| 0 | 0 | 0 | 0 |
| 0 | 0 | 0 | 0 |
| 0 | 0 | 0 | 1 |
| 0 | 0 | 0 | 1 |
| 0 | 0 | 0 | 1 |
| 0 | 0 | 0 | 1 |
| 0 | 0 | 0 | 0 |
| 0 | 0 | 0 | 1 |
| 0 | 0 | 0 | 1 |
| 0 | 0 | 0 | 0 |
| 0 | 0 | 0 | 0 |
| 0 | 0 | 0 | 0 |
| 0 | 0 | 0 | 0 |
| 0 | 0 | 0 | 1 |
| 0 | 0 | 0 | 1 |
| 0 | 0 | 0 | 1 |
| 0 | 0 | 0 | 0 |
| 0 | 0 | 0 | 1 |
| 0 | 0 | 0 | 1 |
| 0 | 0 | 0 | 1 |
| 0 | 0 | 0 | 1 |
| 0 | 0 | 0 | 1 |
| 0 | 0 | 0 | 1 |
| 0 | 0 | 0 | 1 |
| 0 | 0 | 0 | 1 |
| 0 | 0 | 0 | 1 |
| 0 | 0 | 0 | 1 |
| 0 | 0 | 0 | 0 |
| 0 | 0 | 0 | 1 |
| 0 | 0 | 0 | 1 |
| 0 | 0 | 0 | 1 |
| 0 | 0 | 0 | 1 |
| 0 | 0 | 0 | 1 |
| 0 | 0 | 0 | 1 |
| 0 | 0 | 0 | 0 |
| 0 | 0 | 0 | 1 |
| 0 | 0 | 0 | 1 |
| 0 | 0 | 0 | 1 |
| 0 | 0 | 0 | 1 |

| | | | |
|---|---|---|---|
| 0 | 0 | 0 | 0 |
| 0 | 0 | 0 | 1 |
| 0 | 0 | 0 | 1 |
| 0 | 0 | 0 | 1 |
| 0 | 0 | 0 | 1 |
| 0 | 0 | 0 | 1 |
| 0 | 0 | 0 | 1 |
| 0 | 0 | 0 | 0 |
| 0 | 0 | 0 | 1 |
| 0 | 0 | 0 | 1 |
| 0 | 0 | 0 | 0 |
| 0 | 0 | 0 | 0 |
| 0 | 0 | 0 | 0 |
| 0 | 0 | 0 | 1 |
| 0 | 0 | 0 | 0 |
| 0 | 0 | 0 | 0 |
| 0 | 0 | 0 | 0 |
| 0 | 0 | 0 | 0 |
| 0 | 0 | 0 | 1 |
| 0 | 0 | 0 | 0 |
| 0 | 0 | 0 | 0 |
| 0 | 0 | 0 | 0 |
| 0 | 0 | 0 | 0 |
| 0 | 0 | 0 | 0 |
| 0 | 0 | 0 | 0 |
| 0 | 0 | 0 | 0 |
| 0 | 0 | 0 | 0 |
| 0 | 0 | 0 | 0 |
| 0 | 0 | 0 | 0 |
| 0 | 0 | 0 | 0 |
| 0 | 0 | 0 | 0 |
| 0 | 0 | 0 | 0 |
| 0 | 0 | 0 | 1 |
| 0 | 0 | 0 | 0 |
| 0 | 0 | 0 | 1 |
| 0 | 0 | 0 | 0 |
| 0 | 0 | 0 | 0 |
| 0 | 0 | 0 | 1 |
| 0 | 0 | 0 | 1 |
| 0 | 0 | 0 | 1 |
| 0 | 0 | 0 | 1 |
| 0 | 0 | 0 | 0 |
| 0 | 0 | 0 | 0 |
| 0 | 0 | 0 | 0 |
| 0 | 0 | 0 | 1 |
| 0 | 0 | 0 | 0 |
| 0 | 0 | 0 | 1 |
| 0 | 0 | 0 | 0 |
| 0 | 0 | 0 | 1 |
| 0 | 0 | 0 | 1 |
| 0 | 0 | 0 | 1 |
| 0 | 0 | 0 | 1 |
| 0 | 0 | 0 | 1 |
| 0 | 0 | 0 | 1 |

| | | | |
|---|---|---|---|
| 0 | 0 | 0 | 1 |
| 0 | 0 | 0 | 1 |
| 0 | 0 | 0 | 1 |
| 0 | 0 | 0 | 1 |
| 0 | 0 | 0 | 1 |
| 0 | 0 | 0 | 0 |
| 0 | 0 | 0 | 0 |
| 0 | 0 | 0 | 1 |
| 0 | 0 | 0 | 0 |
| 0 | 0 | 0 | 0 |
| 0 | 0 | 0 | 1 |
| 0 | 0 | 0 | 1 |
| 0 | 0 | 0 | 1 |
| 0 | 0 | 0 | 1 |
| 0 | 0 | 0 | 1 |
| 0 | 0 | 0 | 1 |
| 0 | 0 | 0 | 1 |
| 0 | 0 | 0 | 1 |
| 0 | 0 | 0 | 0 |
| 0 | 0 | 0 | 0 |
| 0 | 0 | 0 | 1 |
| 0 | 0 | 0 | 1 |
| 0 | 0 | 0 | 1 |
| 0 | 0 | 0 | 0 |
| 0 | 0 | 0 | 1 |
| 0 | 0 | 0 | 0 |
| 0 | 0 | 0 | 0 |
| 0 | 0 | 0 | 1 |
| 0 | 0 | 0 | 1 |
| 0 | 0 | 0 | 1 |
| 0 | 0 | 0 | 0 |
| 0 | 0 | 0 | 1 |
| 0 | 0 | 0 | 0 |
| 0 | 0 | 0 | 0 |
| 0 | 0 | 0 | 0 |
| 0 | 0 | 0 | 0 |
| 0 | 0 | 0 | 0 |
| 0 | 0 | 0 | 1 |
| 0 | 0 | 0 | 1 |
| 0 | 0 | 0 | 1 |
| 0 | 0 | 0 | 0 |
| 0 | 0 | 0 | 0 |
| 0 | 0 | 0 | 0 |
| 0 | 0 | 0 | 0 |
| 0 | 0 | 0 | 1 |
| 0 | 0 | 0 | 1 |
| 0 | 0 | 0 | 1 |
| 0 | 0 | 0 | 0 |
| 0 | 0 | 0 | 0 |
| 0 | 0 | 0 | 0 |
| 0 | 0 | 0 | 1 |
| 0 | 0 | 0 | 1 |
| 0 | 0 | 0 | 1 |
| 0 | 0 | 0 | 1 |

| | | | |
|---|---|---|---|
| 0 | 0 | 0 | 1 |
| 0 | 0 | 0 | 1 |
| 0 | 0 | 0 | 1 |
| 0 | 0 | 0 | 0 |
| 0 | 0 | 0 | 0 |
| 0 | 0 | 0 | 1 |
| 0 | 0 | 0 | 0 |
| 0 | 0 | 0 | 1 |
| 0 | 0 | 0 | 1 |
| 0 | 0 | 0 | 1 |
| 0 | 0 | 0 | 1 |
| 0 | 0 | 0 | 0 |
| 0 | 0 | 0 | 1 |
| 0 | 0 | 0 | 0 |
| 0 | 0 | 0 | 1 |
| 0 | 0 | 0 | 0 |
| 0 | 0 | 0 | 1 |
| 0 | 0 | 0 | 1 |
| 0 | 0 | 0 | 1 |
| 0 | 0 | 0 | 0 |
| 0 | 0 | 0 | 0 |
| 0 | 0 | 0 | 0 |
| 0 | 0 | 0 | 0 |
| 0 | 0 | 0 | 0 |
| 0 | 0 | 0 | 0 |
| 0 | 0 | 0 | 1 |
| 0 | 0 | 0 | 0 |
| 0 | 0 | 0 | 1 |
| 0 | 0 | 0 | 1 |
| 0 | 0 | 0 | 1 |
| 0 | 0 | 0 | 1 |
| 0 | 0 | 0 | 1 |
| 0 | 0 | 0 | 0 |
| 0 | 0 | 0 | 0 |
| 0 | 0 | 0 | 1 |
| 0 | 0 | 0 | 1 |
| 0 | 0 | 0 | 0 |
| 0 | 0 | 0 | 0 |
| 0 | 0 | 0 | 0 |
| 0 | 0 | 0 | 1 |
| 0 | 0 | 0 | 0 |
| 0 | 0 | 0 | 0 |
| 0 | 0 | 0 | 1 |
| 0 | 0 | 0 | 0 |
| 0 | 0 | 0 | 1 |
| 0 | 0 | 0 | 1 |
| 0 | 0 | 0 | 1 |
| 0 | 0 | 0 | 1 |
| 0 | 0 | 0 | 1 |
| 1 | 0 | 0 | 0 |
| 0 | 0 | 0 | 1 |
| 0 | 0 | 0 | 1 |
| 0 | 0 | 0 | 1 |
| 0 | 0 | 0 | 1 |

| | | | |
|---|---|---|---|
| 0 | 0 | 0 | 1 |
| 0 | 0 | 0 | 1 |
| 0 | 0 | 0 | 1 |
| 0 | 0 | 0 | 0 |
| 0 | 0 | 0 | 1 |
| 0 | 0 | 0 | 1 |
| 0 | 0 | 0 | 1 |
| 0 | 0 | 0 | 0 |
| 0 | 0 | 0 | 1 |
| 0 | 0 | 0 | 1 |
| 0 | 0 | 0 | 1 |
| 0 | 0 | 0 | 1 |
| 0 | 0 | 0 | 1 |
| 0 | 0 | 0 | 1 |
| 0 | 0 | 0 | 1 |
| 0 | 0 | 0 | 1 |
| 0 | 0 | 0 | 0 |
| 0 | 0 | 0 | 0 |
| 0 | 0 | 0 | 1 |
| 0 | 0 | 0 | 1 |
| 0 | 0 | 0 | 1 |
| 0 | 0 | 0 | 1 |
| 0 | 0 | 0 | 1 |
| 0 | 0 | 0 | 1 |
| 0 | 0 | 0 | 1 |
| 0 | 0 | 0 | 1 |
| 0 | 0 | 0 | 1 |
| 0 | 0 | 0 | 0 |
| 0 | 0 | 0 | 0 |
| 0 | 0 | 0 | 1 |
| 0 | 0 | 0 | 1 |
| 0 | 0 | 0 | 0 |
| 0 | 0 | 0 | 1 |
| 0 | 0 | 0 | 0 |
| 1 | 0 | 0 | 0 |
| 1 | 0 | 0 | 0 |
| 1 | 0 | 0 | 0 |
| 1 | 0 | 0 | 0 |
| 0 | 0 | 0 | 1 |
| 0 | 0 | 0 | 1 |
| 0 | 0 | 0 | 1 |
| 0 | 0 | 0 | 1 |
| 0 | 0 | 0 | 1 |
| 0 | 0 | 0 | 1 |
| 0 | 0 | 0 | 0 |
| 0 | 0 | 0 | 0 |
| 0 | 0 | 0 | 0 |
| 0 | 0 | 0 | 1 |
| 0 | 0 | 0 | 1 |
| 0 | 0 | 0 | 1 |
| 0 | 0 | 0 | 1 |
| 0 | 0 | 0 | 0 |
| 0 | 0 | 0 | 0 |
| 0 | 0 | 0 | 0 |

| | | | |
|---|---|---|---|
| 0 | 0 | 0 | 0 |
| 0 | 0 | 0 | 0 |
| 0 | 0 | 0 | 1 |
| 0 | 0 | 0 | 0 |
| 0 | 0 | 0 | 0 |
| 0 | 0 | 0 | 0 |
| 0 | 0 | 0 | 0 |
| 0 | 0 | 0 | 1 |
| 0 | 0 | 0 | 1 |
| 0 | 0 | 0 | 1 |
| 0 | 0 | 0 | 1 |
| 0 | 0 | 0 | 1 |
| 0 | 0 | 0 | 1 |
| 0 | 0 | 0 | 1 |
| 0 | 0 | 0 | 1 |
| 0 | 0 | 0 | 1 |
| 0 | 0 | 0 | 1 |
| 0 | 0 | 0 | 1 |
| 0 | 0 | 0 | 1 |
| 0 | 0 | 0 | 1 |
| 0 | 0 | 0 | 0 |
| 0 | 0 | 0 | 0 |
| 0 | 0 | 0 | 0 |
| 0 | 0 | 0 | 0 |
| 0 | 0 | 0 | 0 |
| 0 | 0 | 0 | 1 |
| 0 | 0 | 0 | 1 |
| 0 | 0 | 0 | 0 |
| 0 | 0 | 0 | 1 |
| 0 | 0 | 0 | 1 |
| 0 | 0 | 0 | 0 |
| 0 | 0 | 0 | 0 |
| 0 | 0 | 0 | 0 |
| 0 | 0 | 0 | 1 |
| 0 | 0 | 0 | 0 |
| 0 | 0 | 0 | 1 |
| 0 | 0 | 0 | 1 |
| 0 | 0 | 0 | 0 |
| 0 | 0 | 0 | 1 |
| 0 | 0 | 0 | 1 |
| 0 | 0 | 0 | 1 |
| 0 | 0 | 0 | 1 |
| 0 | 0 | 0 | 1 |
| 0 | 0 | 0 | 1 |
| 0 | 0 | 0 | 1 |
| 0 | 0 | 0 | 1 |
| 0 | 0 | 0 | 1 |
| 0 | 0 | 0 | 1 |
| 0 | 0 | 0 | 0 |
| 0 | 0 | 0 | 1 |
| 1 | 0 | 0 | 0 |
| 1 | 0 | 0 | 0 |
| 0 | 0 | 0 | 1 |
| 0 | 0 | 0 | 1 |

| | | | |
|---|---|---|---|
| 1 | 0 | 0 | 0 |
| 0 | 0 | 0 | 1 |
| 0 | 0 | 0 | 1 |
| 0 | 0 | 0 | 1 |
| 0 | 0 | 0 | 1 |
| 1 | 0 | 0 | 0 |
| 1 | 0 | 0 | 0 |
| 0 | 0 | 0 | 1 |
| 0 | 0 | 0 | 1 |
| 0 | 0 | 0 | 1 |
| 0 | 0 | 0 | 1 |
| 0 | 0 | 0 | 0 |
| 0 | 0 | 0 | 1 |
| 0 | 0 | 0 | 1 |
| 0 | 0 | 0 | 1 |
| 0 | 0 | 0 | 1 |
| 0 | 0 | 0 | 0 |
| 0 | 0 | 0 | 0 |
| 0 | 0 | 0 | 1 |
| 0 | 0 | 0 | 1 |
| 0 | 0 | 0 | 1 |
| 0 | 0 | 0 | 1 |
| 0 | 0 | 0 | 1 |
| 0 | 0 | 0 | 1 |
| 0 | 0 | 0 | 0 |
| 0 | 0 | 0 | 1 |
| 0 | 0 | 0 | 1 |
| 0 | 0 | 0 | 1 |
| 0 | 0 | 0 | 1 |
| 0 | 0 | 0 | 1 |
| 0 | 0 | 0 | 1 |
| 0 | 0 | 0 | 0 |
| 0 | 0 | 0 | 0 |
| 0 | 0 | 0 | 1 |
| 0 | 0 | 0 | 1 |
| 0 | 0 | 0 | 0 |
| 1 | 0 | 0 | 0 |
| 0 | 0 | 0 | 0 |
| 1 | 0 | 0 | 0 |
| 0 | 0 | 0 | 0 |
| 0 | 0 | 0 | 0 |
| 0 | 0 | 0 | 0 |
| 0 | 0 | 0 | 0 |
| 0 | 0 | 0 | 0 |
| 0 | 0 | 0 | 0 |
| 0 | 0 | 0 | 0 |
| 0 | 0 | 0 | 0 |
| 1 | 0 | 0 | 0 |
| 0 | 0 | 0 | 1 |
| 0 | 0 | 0 | 1 |
| 0 | 0 | 0 | 1 |
| 0 | 0 | 0 | 1 |
| 0 | 0 | 0 | 0 |
| 0 | 0 | 0 | 1 |

| | | | |
|---|---|---|---|
| 0 | 0 | 0 | 1 |
| 0 | 0 | 0 | 1 |
| 0 | 0 | 0 | 1 |
| 0 | 0 | 0 | 1 |
| 0 | 0 | 0 | 1 |
| 0 | 0 | 0 | 1 |
| 0 | 0 | 0 | 1 |
| 0 | 0 | 0 | 0 |
| 0 | 0 | 0 | 0 |
| 0 | 0 | 0 | 1 |
| 0 | 0 | 0 | 1 |
| 0 | 0 | 0 | 1 |
| 0 | 0 | 0 | 1 |
| 0 | 0 | 0 | 0 |
| 0 | 0 | 0 | 1 |
| 0 | 0 | 0 | 1 |
| 0 | 0 | 0 | 0 |
| 0 | 0 | 0 | 1 |
| 1 | 0 | 0 | 0 |
| 0 | 0 | 0 | 1 |
| 0 | 0 | 0 | 1 |
| 0 | 0 | 0 | 1 |
| 0 | 0 | 0 | 1 |
| 0 | 0 | 0 | 0 |
| 0 | 0 | 0 | 1 |
| 0 | 0 | 0 | 1 |
| 0 | 0 | 0 | 1 |
| 0 | 0 | 0 | 1 |
| 0 | 0 | 0 | 1 |
| 0 | 0 | 0 | 1 |
| 0 | 0 | 0 | 1 |
| 1 | 0 | 0 | 0 |
| 0 | 0 | 0 | 1 |
| 0 | 0 | 0 | 1 |
| 0 | 0 | 0 | 1 |
| 0 | 0 | 0 | 1 |
| 0 | 0 | 0 | 1 |
| 0 | 0 | 0 | 1 |
| 1 | 0 | 0 | 0 |
| 0 | 0 | 0 | 1 |
| 0 | 0 | 0 | 1 |
| 0 | 0 | 0 | 1 |
| 0 | 0 | 0 | 1 |
| 0 | 0 | 0 | 1 |
| 0 | 0 | 0 | 1 |
| 0 | 0 | 0 | 0 |
| 0 | 0 | 0 | 1 |
| 0 | 0 | 0 | 1 |
| 0 | 0 | 0 | 1 |
| 0 | 0 | 0 | 1 |
| 0 | 0 | 0 | 1 |
| 0 | 0 | 0 | 1 |
| 0 | 0 | 0 | 1 |
| 0 | 0 | 0 | 1 |

| | | | |
|---|---|---|---|
| 0 | 0 | 0 | 1 |
| 0 | 0 | 0 | 1 |
| 0 | 0 | 0 | 1 |
| 0 | 0 | 0 | 1 |
| 0 | 0 | 0 | 1 |
| 0 | 0 | 0 | 1 |
| 0 | 0 | 0 | 1 |
| 0 | 0 | 0 | 1 |
| 0 | 0 | 0 | 1 |
| 0 | 0 | 0 | 1 |
| 0 | 0 | 0 | 1 |
| 0 | 0 | 0 | 1 |
| 0 | 0 | 0 | 1 |
| 0 | 0 | 0 | 1 |
| 0 | 0 | 0 | 1 |
| 1 | 0 | 0 | 0 |
| 0 | 0 | 0 | 1 |
| 0 | 0 | 0 | 0 |
| 0 | 0 | 0 | 1 |
| 0 | 0 | 0 | 1 |
| 1 | 0 | 0 | 0 |
| 0 | 0 | 0 | 1 |
| 0 | 0 | 0 | 1 |
| 0 | 0 | 0 | 1 |
| 0 | 0 | 0 | 0 |
| 0 | 0 | 0 | 1 |
| 0 | 0 | 0 | 0 |
| 0 | 0 | 0 | 1 |
| 0 | 0 | 0 | 1 |
| 0 | 0 | 0 | 1 |
| 0 | 0 | 0 | 1 |
| 0 | 0 | 0 | 1 |
| 0 | 0 | 0 | 1 |
| 0 | 0 | 0 | 1 |
| 0 | 0 | 0 | 1 |
| 0 | 0 | 0 | 1 |
| 0 | 0 | 0 | 1 |
| 0 | 0 | 0 | 1 |
| 0 | 0 | 0 | 1 |
| 0 | 0 | 0 | 1 |
| 0 | 0 | 0 | 1 |
| 0 | 0 | 0 | 1 |
| 0 | 0 | 0 | 1 |
| 0 | 0 | 0 | 0 |
| 0 | 0 | 0 | 1 |
| 0 | 0 | 0 | 1 |
| 0 | 0 | 0 | 0 |
| 0 | 0 | 0 | 0 |
| 0 | 0 | 0 | 1 |
| 0 | 0 | 0 | 0 |
| 0 | 0 | 0 | 0 |
| 0 | 0 | 0 | 0 |
| 0 | 0 | 0 | 1 |
| 0 | 0 | 0 | 1 |

| | | | |
|---|---|---|---|
| 0 | 0 | 0 | 1 |
| 0 | 0 | 0 | 1 |
| 0 | 0 | 0 | 1 |
| 0 | 0 | 0 | 1 |
| 0 | 0 | 0 | 1 |
| 0 | 0 | 0 | 1 |
| 0 | 0 | 0 | 1 |
| 1 | 0 | 0 | 0 |
| 0 | 0 | 0 | 1 |
| 0 | 0 | 0 | 1 |
| 0 | 0 | 0 | 1 |
| 0 | 0 | 0 | 1 |
| 0 | 0 | 0 | 1 |
| 0 | 0 | 0 | 1 |
| 0 | 0 | 0 | 0 |
| 0 | 0 | 0 | 0 |
| 0 | 0 | 0 | 1 |
| 0 | 0 | 0 | 0 |
| 0 | 0 | 0 | 1 |
| 0 | 0 | 0 | 1 |
| 0 | 0 | 0 | 0 |
| 0 | 0 | 0 | 1 |
| 0 | 0 | 0 | 1 |
| 0 | 0 | 0 | 1 |
| 0 | 0 | 0 | 0 |
| 0 | 0 | 0 | 1 |
| 0 | 0 | 0 | 1 |
| 1 | 0 | 0 | 0 |
| 0 | 0 | 0 | 0 |
| 1 | 0 | 0 | 0 |
| 1 | 0 | 0 | 0 |
| 0 | 0 | 0 | 1 |
| 0 | 0 | 0 | 1 |
| 0 | 0 | 0 | 1 |
| 0 | 0 | 0 | 1 |
| 0 | 0 | 0 | 1 |
| 0 | 0 | 0 | 0 |
| 0 | 0 | 0 | 0 |
| 0 | 0 | 0 | 1 |
| 0 | 0 | 0 | 1 |
| 0 | 0 | 0 | 1 |
| 0 | 0 | 0 | 1 |
| 0 | 0 | 0 | 0 |
| 0 | 0 | 0 | 1 |
| 0 | 0 | 0 | 1 |
| 0 | 0 | 0 | 1 |
| 0 | 0 | 0 | 1 |
| 0 | 0 | 0 | 1 |
| 0 | 0 | 0 | 0 |
| 0 | 0 | 0 | 0 |
| 0 | 0 | 0 | 0 |
| 0 | 0 | 0 | 0 |
| 0 | 0 | 0 | 0 |
| 0 | 0 | 0 | 1 |

| | | | |
|---|---|---|---|
| 0 | 0 | 0 | 0 |
| 0 | 0 | 0 | 0 |
| 0 | 0 | 0 | 1 |
| 1 | 0 | 0 | 0 |
| 0 | 0 | 0 | 1 |
| 0 | 0 | 0 | 1 |
| 0 | 0 | 0 | 1 |
| 0 | 0 | 0 | 1 |
| 0 | 0 | 0 | 1 |
| 1 | 0 | 0 | 0 |
| 0 | 0 | 0 | 1 |
| 0 | 0 | 0 | 1 |
| 0 | 0 | 0 | 1 |
| 0 | 0 | 0 | 1 |
| 0 | 0 | 0 | 1 |
| 0 | 0 | 0 | 1 |
| 0 | 0 | 0 | 1 |
| 0 | 0 | 0 | 1 |
| 0 | 0 | 0 | 1 |
| 0 | 0 | 0 | 1 |
| 0 | 0 | 0 | 1 |
| 0 | 0 | 0 | 1 |
| 0 | 0 | 0 | 1 |
| 0 | 0 | 0 | 1 |
| 0 | 0 | 0 | 1 |
| 0 | 0 | 0 | 1 |
| 0 | 0 | 0 | 1 |
| 0 | 0 | 0 | 1 |
| 0 | 0 | 0 | 1 |
| 0 | 0 | 0 | 1 |
| 0 | 0 | 0 | 1 |
| 0 | 0 | 0 | 1 |
| 1 | 0 | 0 | 0 |
| 0 | 0 | 0 | 1 |
| 0 | 0 | 0 | 1 |
| 0 | 0 | 0 | 1 |
| 0 | 0 | 0 | 1 |
| 0 | 0 | 0 | 1 |
| 0 | 0 | 0 | 1 |
| 0 | 0 | 0 | 1 |
| 0 | 0 | 0 | 1 |
| 0 | 0 | 0 | 1 |
| 0 | 0 | 0 | 1 |
| 0 | 0 | 0 | 1 |
| 0 | 0 | 0 | 1 |
| 0 | 0 | 0 | 1 |
| 0 | 0 | 0 | 1 |
| 0 | 0 | 0 | 0 |
| 0 | 0 | 0 | 0 |
| 0 | 0 | 0 | 0 |
| 0 | 0 | 0 | 0 |
| 0 | 0 | 0 | 0 |
| 0 | 0 | 0 | 1 |
| 0 | 0 | 0 | 0 |

| | | | |
|---|---|---|---|
| 0 | 0 | 0 | 1 |
| 0 | 0 | 0 | 0 |
| 0 | 0 | 0 | 1 |
| 0 | 0 | 0 | 1 |
| 0 | 0 | 0 | 1 |
| 1 | 0 | 0 | 0 |
| 0 | 0 | 0 | 0 |
| 0 | 0 | 0 | 0 |
| 1 | 0 | 0 | 0 |
| 0 | 0 | 0 | 0 |
| 0 | 0 | 0 | 1 |
| 0 | 0 | 0 | 0 |
| 0 | 0 | 0 | 1 |
| 0 | 0 | 0 | 1 |
| 0 | 0 | 0 | 1 |
| 1 | 0 | 0 | 0 |
| 0 | 0 | 0 | 1 |
| 0 | 0 | 0 | 1 |
| 0 | 0 | 0 | 0 |
| 0 | 0 | 0 | 0 |
| 1 | 0 | 0 | 0 |
| 1 | 0 | 0 | 0 |
| 0 | 0 | 0 | 0 |
| 0 | 0 | 0 | 0 |
| 0 | 0 | 0 | 1 |
| 0 | 0 | 0 | 0 |
| 0 | 0 | 0 | 0 |
| 0 | 0 | 0 | 0 |
| 0 | 0 | 0 | 0 |
| 0 | 0 | 0 | 0 |
| 0 | 0 | 0 | 0 |
| 0 | 0 | 0 | 1 |
| 0 | 0 | 0 | 1 |
| 0 | 0 | 0 | 0 |
| 0 | 0 | 0 | 0 |
| 0 | 0 | 0 | 1 |
| 0 | 0 | 0 | 0 |
| 0 | 0 | 0 | 0 |
| 0 | 0 | 0 | 0 |
| 0 | 0 | 0 | 0 |
| 0 | 0 | 0 | 0 |
| 0 | 0 | 0 | 1 |
| 0 | 0 | 0 | 1 |
| 0 | 0 | 0 | 1 |
| 0 | 0 | 0 | 0 |
| 0 | 0 | 0 | 0 |
| 0 | 0 | 0 | 0 |
| 0 | 0 | 0 | 1 |
| 0 | 0 | 0 | 1 |
| 0 | 0 | 0 | 1 |
| 0 | 0 | 0 | 1 |
| 0 | 0 | 0 | 1 |
| 0 | 0 | 0 | 1 |
| 0 | 0 | 0 | 0 |

| | | | |
|---|---|---|---|
| 0 | 0 | 0 | 1 |
| 0 | 0 | 0 | 1 |
| 0 | 0 | 0 | 1 |
| 0 | 0 | 0 | 1 |
| 1 | 0 | 0 | 0 |
| 0 | 0 | 0 | 0 |
| 0 | 0 | 0 | 1 |
| 0 | 0 | 0 | 1 |
| 0 | 0 | 0 | 1 |
| 0 | 0 | 0 | 1 |
| 0 | 0 | 0 | 1 |
| 0 | 0 | 0 | 0 |
| 0 | 0 | 0 | 1 |
| 0 | 0 | 0 | 0 |
| 0 | 0 | 0 | 0 |
| 0 | 0 | 0 | 0 |
| 0 | 0 | 0 | 0 |
| 0 | 0 | 0 | 0 |
| 0 | 0 | 0 | 0 |
| 0 | 0 | 0 | 0 |
| 0 | 0 | 0 | 0 |
| 0 | 0 | 0 | 0 |
| 0 | 0 | 0 | 0 |
| 0 | 0 | 0 | 1 |
| 0 | 0 | 0 | 1 |
| 0 | 0 | 0 | 1 |
| 0 | 0 | 0 | 1 |
| 0 | 0 | 0 | 1 |
| 0 | 0 | 0 | 1 |
| 0 | 0 | 0 | 1 |
| 0 | 0 | 0 | 1 |
| 0 | 0 | 0 | 0 |
| 0 | 0 | 0 | 1 |
| 0 | 0 | 0 | 0 |
| 0 | 0 | 0 | 0 |
| 0 | 0 | 0 | 0 |
| 0 | 0 | 0 | 0 |
| 0 | 0 | 0 | 1 |
| 0 | 0 | 0 | 1 |
| 0 | 0 | 0 | 1 |
| 0 | 0 | 0 | 1 |
| 0 | 0 | 0 | 1 |
| 0 | 0 | 0 | 1 |
| 0 | 0 | 0 | 1 |
| 0 | 0 | 0 | 1 |
| 1 | 0 | 0 | 0 |
| 0 | 0 | 0 | 1 |
| 0 | 0 | 0 | 1 |
| 0 | 0 | 0 | 1 |
| 1 | 0 | 0 | 0 |
| 1 | 0 | 0 | 0 |
| 0 | 0 | 0 | 0 |
| 0 | 0 | 0 | 1 |
| 0 | 0 | 0 | 1 |

| | | | |
|---|---|---|---|
| 0 | 0 | 0 | 1 |
| 0 | 0 | 0 | 1 |
| 0 | 0 | 0 | 1 |
| 0 | 0 | 0 | 0 |
| 0 | 0 | 0 | 1 |
| 0 | 0 | 0 | 1 |
| 0 | 0 | 0 | 0 |
| 0 | 0 | 0 | 1 |
| 0 | 0 | 0 | 1 |
| 0 | 0 | 0 | 1 |
| 0 | 0 | 0 | 0 |
| 0 | 0 | 0 | 0 |
| 0 | 0 | 0 | 1 |
| 0 | 0 | 0 | 1 |
| 0 | 0 | 0 | 1 |
| 0 | 0 | 0 | 1 |
| 0 | 0 | 0 | 0 |
| 0 | 0 | 0 | 0 |
| 0 | 0 | 0 | 1 |
| 0 | 0 | 0 | 0 |
| 0 | 0 | 0 | 0 |
| 0 | 0 | 0 | 0 |
| 0 | 0 | 0 | 1 |
| 0 | 0 | 0 | 0 |
| 0 | 0 | 0 | 0 |
| 0 | 0 | 0 | 1 |
| 0 | 0 | 0 | 0 |
| 0 | 0 | 0 | 1 |
| 0 | 0 | 0 | 1 |
| 0 | 0 | 0 | 0 |
| 0 | 0 | 0 | 1 |
| 0 | 0 | 0 | 1 |
| 0 | 0 | 0 | 0 |
| 0 | 0 | 0 | 1 |
| 0 | 0 | 0 | 1 |
| 0 | 0 | 0 | 0 |
| 0 | 0 | 0 | 1 |
| 0 | 0 | 0 | 0 |
| 0 | 0 | 0 | 1 |
| 0 | 0 | 0 | 0 |
| 0 | 0 | 0 | 0 |
| 0 | 0 | 0 | 1 |
| 0 | 0 | 0 | 0 |
| 0 | 0 | 0 | 0 |
| 0 | 0 | 0 | 1 |
| 0 | 0 | 0 | 1 |
| 0 | 0 | 0 | 1 |
| 0 | 0 | 0 | 1 |
| 0 | 0 | 0 | 1 |
| 0 | 0 | 0 | 1 |
| 0 | 0 | 0 | 1 |
| 0 | 0 | 0 | 1 |
| 0 | 0 | 0 | 1 |
| 0 | 0 | 0 | 1 |

| | | | |
|---|---|---|---|
| 0 | 0 | 0 | 1 |
| 0 | 0 | 0 | 1 |
| 0 | 0 | 0 | 1 |
| 0 | 0 | 0 | 0 |
| 0 | 0 | 0 | 1 |
| 0 | 0 | 0 | 0 |
| 0 | 0 | 0 | 1 |
| 0 | 0 | 0 | 1 |
| 0 | 0 | 0 | 0 |
| 0 | 0 | 0 | 0 |
| 0 | 0 | 0 | 1 |
| 0 | 0 | 0 | 0 |
| 0 | 0 | 0 | 0 |
| 0 | 0 | 0 | 0 |
| 0 | 0 | 0 | 1 |
| 0 | 0 | 0 | 0 |
| 0 | 0 | 0 | 0 |
| 0 | 0 | 0 | 1 |
| 1 | 0 | 0 | 0 |
| 1 | 0 | 0 | 0 |
| 1 | 0 | 0 | 0 |
| 1 | 0 | 0 | 0 |
| 0 | 0 | 0 | 1 |
| 0 | 0 | 0 | 0 |
| 0 | 0 | 0 | 0 |
| 0 | 0 | 0 | 0 |
| 0 | 0 | 0 | 0 |
| 0 | 0 | 0 | 0 |
| 0 | 0 | 0 | 0 |
| 0 | 0 | 0 | 0 |
| 0 | 0 | 0 | 0 |
| 0 | 0 | 0 | 0 |
| 0 | 0 | 0 | 0 |
| 0 | 0 | 0 | 1 |
| 0 | 0 | 0 | 1 |
| 0 | 0 | 0 | 0 |
| 0 | 0 | 0 | 1 |
| 0 | 0 | 0 | 0 |
| 0 | 0 | 0 | 0 |
| 0 | 0 | 0 | 0 |
| 0 | 0 | 0 | 0 |
| 0 | 0 | 0 | 1 |
| 0 | 0 | 0 | 0 |
| 0 | 0 | 0 | 0 |
| 0 | 0 | 0 | 1 |
| 0 | 0 | 0 | 1 |
| 0 | 0 | 0 | 1 |
| 0 | 0 | 0 | 1 |
| 0 | 0 | 0 | 1 |
| 0 | 0 | 0 | 1 |
| 0 | 0 | 0 | 1 |
| 0 | 0 | 0 | 1 |
| 0 | 0 | 0 | 1 |
| 0 | 0 | 0 | 1 |

| | | | |
|---|---|---|---|
| 0 | 0 | 0 | 1 |
| 0 | 0 | 0 | 0 |
| 0 | 0 | 0 | 1 |
| 0 | 0 | 0 | 1 |
| 0 | 0 | 0 | 1 |
| 0 | 0 | 0 | 1 |
| 0 | 0 | 0 | 1 |
| 0 | 0 | 0 | 1 |
| 0 | 0 | 0 | 1 |
| 0 | 0 | 0 | 1 |
| 0 | 0 | 0 | 1 |
| 0 | 0 | 0 | 0 |
| 0 | 0 | 0 | 1 |
| 0 | 0 | 0 | 1 |
| 0 | 0 | 0 | 0 |
| 0 | 0 | 0 | 1 |
| 0 | 0 | 0 | 1 |
| 0 | 0 | 0 | 1 |
| 0 | 0 | 0 | 1 |
| 0 | 0 | 0 | 1 |
| 1 | 0 | 0 | 0 |
| 0 | 0 | 0 | 1 |
| 1 | 0 | 0 | 0 |
| 1 | 0 | 0 | 0 |
| 0 | 0 | 0 | 1 |
| 0 | 0 | 0 | 0 |
| 0 | 0 | 0 | 1 |
| 0 | 0 | 0 | 1 |
| 0 | 0 | 0 | 1 |
| 0 | 0 | 0 | 1 |
| 0 | 0 | 0 | 1 |
| 0 | 0 | 0 | 0 |
| 0 | 0 | 0 | 1 |
| 0 | 0 | 0 | 0 |
| 0 | 0 | 0 | 1 |
| 0 | 0 | 0 | 1 |
| 0 | 0 | 0 | 1 |
| 0 | 0 | 0 | 1 |
| 0 | 0 | 0 | 1 |
| 0 | 0 | 0 | 0 |
| 0 | 0 | 0 | 1 |
| 0 | 0 | 0 | 1 |
| 0 | 0 | 0 | 1 |
| 0 | 0 | 0 | 0 |
| 0 | 0 | 0 | 1 |
| 0 | 0 | 0 | 1 |
| 0 | 0 | 0 | 1 |
| 1 | 0 | 0 | 0 |
| 0 | 0 | 0 | 0 |
| 0 | 0 | 0 | 1 |
| 0 | 0 | 0 | 1 |
| 0 | 0 | 0 | 0 |
| 1 | 0 | 0 | 0 |
| 0 | 0 | 0 | 0 |

| | | | |
|---|---|---|---|
| 1 | 0 | 0 | 0 |
| 0 | 0 | 0 | 1 |
| 0 | 0 | 0 | 0 |
| 0 | 0 | 0 | 0 |
| 0 | 0 | 0 | 0 |
| 0 | 0 | 0 | 1 |
| 0 | 0 | 0 | 1 |
| 0 | 0 | 0 | 1 |
| 0 | 0 | 0 | 1 |
| 0 | 0 | 0 | 0 |
| 0 | 0 | 0 | 1 |
| 0 | 0 | 0 | 1 |
| 0 | 0 | 0 | 1 |
| 0 | 0 | 0 | 1 |
| 0 | 0 | 0 | 0 |
| 0 | 0 | 0 | 1 |
| 0 | 0 | 0 | 1 |
| 0 | 0 | 0 | 0 |
| 0 | 0 | 0 | 1 |
| 0 | 0 | 0 | 0 |
| 0 | 0 | 0 | 0 |
| 0 | 0 | 0 | 1 |
| 0 | 0 | 0 | 0 |
| 0 | 0 | 0 | 1 |
| 0 | 0 | 0 | 0 |
| 0 | 0 | 0 | 0 |
| 0 | 0 | 0 | 1 |
| 0 | 0 | 0 | 1 |
| 1 | 0 | 0 | 0 |
| 0 | 0 | 0 | 1 |
| 0 | 0 | 0 | 1 |
| 0 | 0 | 0 | 1 |
| 0 | 0 | 0 | 1 |
| 0 | 0 | 0 | 1 |
| 0 | 0 | 0 | 0 |
| 0 | 0 | 0 | 0 |
| 0 | 0 | 0 | 1 |
| 0 | 0 | 0 | 0 |
| 0 | 0 | 0 | 0 |
| 0 | 0 | 0 | 0 |
| 0 | 0 | 0 | 0 |
| 0 | 0 | 0 | 0 |
| 0 | 0 | 0 | 0 |
| 0 | 0 | 0 | 1 |
| 0 | 0 | 0 | 0 |
| 0 | 0 | 0 | 1 |
| 0 | 0 | 0 | 1 |
| 0 | 0 | 0 | 0 |
| 0 | 0 | 0 | 0 |
| 0 | 0 | 0 | 0 |
| 0 | 0 | 0 | 0 |
| 0 | 0 | 0 | 0 |
| 0 | 0 | 0 | 1 |
| 0 | 0 | 0 | 1 |

| | | | |
|---|---|---|---|
| 0 | 0 | 0 | 1 |
| 0 | 0 | 0 | 0 |
| 0 | 0 | 0 | 0 |
| 0 | 0 | 0 | 0 |
| 0 | 0 | 0 | 0 |
| 0 | 0 | 0 | 0 |
| 0 | 0 | 0 | 0 |
| 0 | 0 | 0 | 1 |
| 0 | 0 | 0 | 1 |
| 0 | 0 | 0 | 1 |
| 0 | 0 | 0 | 0 |
| 0 | 0 | 0 | 0 |
| 0 | 0 | 0 | 1 |
| 0 | 0 | 0 | 0 |
| 0 | 0 | 0 | 1 |
| 0 | 0 | 0 | 0 |
| 0 | 0 | 0 | 1 |
| 0 | 0 | 0 | 1 |
| 0 | 0 | 0 | 1 |
| 0 | 0 | 0 | 1 |
| 0 | 0 | 0 | 1 |
| 0 | 0 | 0 | 1 |
| 0 | 0 | 0 | 1 |
| 1 | 0 | 0 | 0 |
| 0 | 0 | 0 | 0 |
| 0 | 0 | 0 | 1 |
| 0 | 0 | 0 | 1 |
| 0 | 0 | 0 | 1 |
| 0 | 0 | 0 | 1 |
| 0 | 0 | 0 | 0 |
| 0 | 0 | 0 | 0 |
| 0 | 0 | 0 | 0 |
| 0 | 0 | 0 | 0 |
| 0 | 0 | 0 | 1 |
| 0 | 0 | 0 | 0 |
| 0 | 0 | 0 | 0 |
| 0 | 0 | 0 | 0 |
| 0 | 0 | 0 | 0 |
| 0 | 0 | 0 | 0 |
| 0 | 0 | 0 | 1 |
| 0 | 0 | 0 | 1 |
| 0 | 0 | 0 | 0 |
| 0 | 0 | 0 | 1 |
| 0 | 0 | 0 | 1 |
| 0 | 0 | 0 | 0 |
| 0 | 0 | 0 | 1 |
| 0 | 0 | 0 | 1 |
| 0 | 0 | 0 | 1 |
| 0 | 0 | 0 | 1 |
| 0 | 0 | 0 | 1 |
| 0 | 0 | 0 | 0 |
| 0 | 0 | 0 | 1 |
| 0 | 0 | 0 | 0 |
| 0 | 0 | 0 | 0 |

| | | | |
|---|---|---|---|
| 0 | 0 | 0 | 1 |
| 0 | 0 | 0 | 1 |
| 0 | 0 | 0 | 0 |
| 0 | 0 | 0 | 0 |
| 0 | 0 | 0 | 1 |
| 0 | 0 | 0 | 1 |
| 0 | 0 | 0 | 1 |
| 0 | 0 | 0 | 1 |
| 0 | 0 | 0 | 1 |
| 0 | 0 | 0 | 1 |
| 0 | 0 | 0 | 1 |
| 0 | 0 | 0 | 1 |
| 0 | 0 | 0 | 0 |
| 0 | 0 | 0 | 1 |
| 0 | 0 | 0 | 1 |
| 0 | 0 | 0 | 1 |
| 0 | 0 | 0 | 1 |
| 0 | 0 | 0 | 1 |
| 0 | 0 | 0 | 1 |
| 0 | 0 | 0 | 1 |
| 0 | 0 | 0 | 1 |
| 0 | 0 | 0 | 1 |
| 0 | 0 | 0 | 1 |
| 0 | 0 | 0 | 1 |
| 0 | 0 | 0 | 1 |
| 0 | 0 | 0 | 1 |
| 0 | 0 | 0 | 1 |
| 0 | 0 | 0 | 1 |
| 0 | 0 | 0 | 1 |
| 0 | 0 | 0 | 1 |
| 0 | 0 | 0 | 1 |
| 0 | 0 | 0 | 0 |
| 0 | 0 | 0 | 1 |
| 0 | 0 | 0 | 1 |
| 0 | 0 | 0 | 1 |
| 0 | 0 | 0 | 1 |
| 0 | 0 | 0 | 1 |
| 0 | 0 | 0 | 1 |
| 0 | 0 | 0 | 1 |
| 0 | 0 | 0 | 1 |
| 0 | 0 | 0 | 0 |
| 0 | 0 | 0 | 0 |
| 0 | 0 | 0 | 1 |
| 0 | 0 | 0 | 1 |
| 0 | 0 | 0 | 1 |
| 0 | 0 | 0 | 1 |
| 0 | 0 | 0 | 1 |
| 0 | 0 | 0 | 1 |
| 0 | 0 | 0 | 0 |
| 0 | 0 | 0 | 0 |
| 0 | 0 | 0 | 1 |
| 0 | 0 | 0 | 1 |
| 0 | 0 | 0 | 0 |
| 0 | 0 | 0 | 1 |

| | | | |
|---|---|---|---|
| 0 | 0 | 0 | 0 |
| 0 | 0 | 0 | 0 |
| 0 | 0 | 0 | 0 |
| 0 | 0 | 0 | 1 |
| 0 | 0 | 0 | 1 |
| 0 | 0 | 0 | 1 |
| 0 | 0 | 0 | 1 |
| 0 | 0 | 0 | 0 |
| 0 | 0 | 0 | 0 |
| 0 | 0 | 0 | 0 |
| 0 | 0 | 0 | 0 |
| 0 | 0 | 0 | 1 |
| 0 | 0 | 0 | 0 |
| 0 | 0 | 0 | 1 |
| 0 | 0 | 0 | 1 |
| 0 | 0 | 0 | 1 |
| 0 | 0 | 0 | 1 |
| 0 | 0 | 0 | 0 |
| 0 | 0 | 0 | 1 |
| 0 | 0 | 0 | 1 |
| 0 | 0 | 0 | 1 |
| 0 | 0 | 0 | 1 |
| 0 | 0 | 0 | 1 |
| 0 | 0 | 0 | 1 |
| 0 | 0 | 0 | 1 |
| 0 | 0 | 0 | 1 |
| 0 | 0 | 0 | 1 |
| 0 | 0 | 0 | 1 |
| 0 | 0 | 0 | 1 |
| 0 | 0 | 0 | 1 |
| 0 | 0 | 0 | 0 |
| 0 | 0 | 0 | 0 |
| 0 | 0 | 0 | 1 |
| 0 | 0 | 0 | 1 |
| 0 | 0 | 0 | 1 |
| 0 | 0 | 0 | 1 |
| 0 | 0 | 0 | 1 |
| 0 | 0 | 0 | 1 |
| 0 | 0 | 0 | 1 |
| 0 | 0 | 0 | 1 |
| 0 | 0 | 0 | 1 |
| 0 | 0 | 0 | 0 |
| 0 | 0 | 0 | 1 |
| 0 | 0 | 0 | 1 |
| 0 | 0 | 0 | 0 |
| 0 | 0 | 0 | 0 |
| 0 | 0 | 0 | 1 |
| 0 | 0 | 0 | 1 |
| 0 | 0 | 0 | 0 |
| 0 | 0 | 0 | 0 |
| 0 | 0 | 0 | 0 |
| 0 | 0 | 0 | 0 |
| 0 | 0 | 0 | 0 |
| 0 | 0 | 0 | 0 |

| | | | |
|---|---|---|---|
| 0 | 0 | 0 | 1 |
| 0 | 0 | 0 | 1 |
| 0 | 0 | 0 | 1 |
| 0 | 0 | 0 | 1 |
| 0 | 0 | 0 | 1 |
| 0 | 0 | 0 | 0 |
| 0 | 0 | 0 | 0 |
| 0 | 0 | 0 | 0 |
| 0 | 0 | 0 | 0 |
| 0 | 0 | 0 | 0 |
| 0 | 0 | 0 | 1 |
| 0 | 0 | 0 | 0 |
| 0 | 0 | 0 | 0 |
| 0 | 0 | 0 | 0 |
| 0 | 0 | 0 | 1 |
| 0 | 0 | 0 | 0 |
| 0 | 0 | 0 | 1 |
| 0 | 0 | 0 | 1 |
| 1 | 0 | 0 | 0 |
| 1 | 0 | 0 | 0 |
| 0 | 0 | 0 | 0 |
| 0 | 0 | 0 | 0 |
| 0 | 0 | 0 | 1 |
| 0 | 0 | 0 | 0 |
| 0 | 0 | 0 | 0 |
| 0 | 0 | 0 | 0 |
| 0 | 0 | 0 | 0 |
| 0 | 0 | 0 | 1 |
| 0 | 0 | 0 | 1 |
| 0 | 0 | 0 | 1 |
| 0 | 0 | 0 | 1 |
| 0 | 0 | 0 | 1 |
| 0 | 0 | 0 | 0 |
| 0 | 0 | 0 | 1 |
| 0 | 0 | 0 | 0 |
| 0 | 0 | 0 | 0 |
| 0 | 0 | 0 | 0 |
| 0 | 0 | 0 | 0 |
| 0 | 0 | 0 | 0 |
| 0 | 0 | 0 | 0 |
| 0 | 0 | 0 | 0 |
| 1 | 0 | 0 | 0 |
| 0 | 0 | 0 | 0 |
| 0 | 0 | 0 | 1 |
| 0 | 0 | 0 | 1 |
| 0 | 0 | 0 | 1 |
| 0 | 0 | 0 | 1 |
| 0 | 0 | 0 | 0 |
| 0 | 0 | 0 | 0 |
| 0 | 0 | 0 | 0 |
| 0 | 0 | 0 | 1 |
| 0 | 0 | 0 | 1 |
| 0 | 0 | 0 | 0 |
| 0 | 0 | 0 | 1 |

| | | | |
|---|---|---|---|
| 0 | 0 | 0 | 1 |
| 0 | 0 | 0 | 0 |
| 0 | 0 | 0 | 1 |
| 0 | 0 | 0 | 1 |
| 0 | 0 | 0 | 1 |
| 0 | 0 | 0 | 1 |
| 0 | 0 | 0 | 1 |
| 0 | 0 | 0 | 1 |
| 0 | 0 | 0 | 1 |
| 0 | 0 | 0 | 1 |
| 0 | 0 | 0 | 1 |
| 0 | 0 | 0 | 1 |
| 0 | 0 | 0 | 0 |
| 0 | 0 | 0 | 0 |
| 0 | 0 | 0 | 0 |
| 0 | 0 | 0 | 0 |
| 0 | 0 | 0 | 0 |
| 0 | 0 | 0 | 0 |
| 0 | 0 | 0 | 0 |
| 0 | 0 | 0 | 1 |
| 1 | 1 | 0 | 0 |
| 0 | 0 | 0 | 0 |
| 0 | 0 | 0 | 0 |
| 0 | 0 | 0 | 0 |
| 0 | 0 | 0 | 0 |
| 0 | 0 | 0 | 1 |
| 0 | 0 | 0 | 1 |
| 0 | 0 | 0 | 0 |
| 0 | 0 | 0 | 1 |
| 0 | 0 | 0 | 0 |
| 0 | 0 | 0 | 0 |
| 0 | 0 | 0 | 1 |
| 0 | 0 | 0 | 0 |
| 0 | 0 | 0 | 1 |
| 0 | 0 | 0 | 1 |
| 0 | 0 | 0 | 1 |
| 0 | 0 | 0 | 1 |
| 0 | 0 | 0 | 1 |
| 0 | 0 | 0 | 1 |
| 0 | 0 | 0 | 1 |
| 0 | 0 | 0 | 0 |
| 0 | 0 | 0 | 1 |
| 0 | 0 | 0 | 1 |
| 0 | 0 | 0 | 0 |
| 0 | 0 | 0 | 0 |
| 0 | 0 | 0 | 1 |
| 0 | 0 | 0 | 0 |
| 0 | 0 | 0 | 0 |
| 0 | 0 | 0 | 1 |
| 0 | 0 | 0 | 1 |
| 0 | 0 | 0 | 0 |
| 0 | 0 | 0 | 1 |
| 0 | 0 | 0 | 1 |
| 0 | 0 | 0 | 1 |

| | | | |
|---|---|---|---|
| 0 | 0 | 0 | 1 |
| 0 | 0 | 0 | 1 |
| 0 | 0 | 0 | 1 |
| 0 | 0 | 0 | 1 |
| 0 | 0 | 0 | 0 |
| 0 | 0 | 0 | 0 |
| 0 | 0 | 0 | 1 |
| 0 | 0 | 0 | 1 |
| 0 | 0 | 0 | 1 |
| 0 | 0 | 0 | 1 |
| 0 | 0 | 0 | 1 |
| 0 | 0 | 0 | 0 |
| 0 | 0 | 0 | 0 |
| 0 | 0 | 0 | 1 |
| 0 | 0 | 0 | 1 |
| 0 | 0 | 0 | 0 |
| 0 | 0 | 0 | 1 |
| 0 | 0 | 0 | 1 |
| 0 | 0 | 0 | 0 |
| 0 | 0 | 0 | 1 |
| 0 | 0 | 0 | 1 |
| 0 | 0 | 0 | 1 |
| 0 | 0 | 0 | 1 |
| 0 | 0 | 0 | 0 |
| 0 | 0 | 0 | 1 |
| 0 | 0 | 0 | 0 |
| 0 | 0 | 0 | 1 |
| 0 | 0 | 0 | 1 |
| 1 | 0 | 0 | 0 |
| 1 | 0 | 0 | 0 |
| 0 | 0 | 0 | 0 |
| 0 | 0 | 0 | 0 |
| 0 | 0 | 0 | 0 |
| 0 | 0 | 0 | 0 |
| 0 | 0 | 0 | 1 |
| 0 | 0 | 0 | 1 |
| 0 | 0 | 0 | 1 |
| 1 | 0 | 0 | 0 |
| 0 | 0 | 0 | 1 |
| 0 | 0 | 0 | 1 |
| 0 | 0 | 0 | 0 |
| 0 | 0 | 0 | 0 |
| 0 | 0 | 0 | 0 |
| 0 | 0 | 0 | 1 |
| 0 | 0 | 0 | 0 |
| 0 | 0 | 0 | 1 |
| 0 | 0 | 0 | 1 |
| 0 | 0 | 0 | 0 |
| 0 | 0 | 0 | 1 |
| 0 | 0 | 0 | 1 |
| 0 | 0 | 0 | 1 |
| 0 | 0 | 0 | 0 |
| 0 | 0 | 0 | 1 |
| 0 | 0 | 0 | 0 |

| | | | |
|---|---|---|---|
| 0 | 0 | 0 | 1 |
| 0 | 0 | 0 | 0 |
| 0 | 0 | 0 | 0 |
| 0 | 0 | 0 | 1 |
| 0 | 0 | 0 | 1 |
| 0 | 0 | 0 | 1 |
| 0 | 0 | 0 | 0 |
| 0 | 0 | 0 | 1 |
| 0 | 0 | 0 | 1 |
| 0 | 0 | 0 | 1 |
| 0 | 0 | 0 | 1 |
| 0 | 0 | 0 | 1 |
| 0 | 0 | 0 | 1 |
| 0 | 0 | 0 | 1 |
| 0 | 0 | 0 | 1 |
| 0 | 0 | 0 | 1 |
| 0 | 0 | 0 | 1 |
| 0 | 0 | 0 | 1 |
| 0 | 0 | 0 | 1 |
| 0 | 0 | 0 | 1 |
| 0 | 0 | 0 | 1 |
| 0 | 0 | 0 | 0 |
| 0 | 0 | 0 | 1 |
| 0 | 0 | 0 | 1 |
| 0 | 0 | 0 | 1 |
| 0 | 0 | 0 | 1 |
| 0 | 0 | 0 | 0 |
| 0 | 0 | 0 | 1 |
| 0 | 0 | 0 | 0 |
| 0 | 0 | 0 | 1 |
| 0 | 0 | 0 | 1 |
| 0 | 0 | 0 | 0 |
| 0 | 0 | 0 | 1 |
| 0 | 0 | 0 | 1 |
| 0 | 0 | 0 | 1 |
| 0 | 0 | 0 | 1 |
| 0 | 0 | 0 | 1 |
| 0 | 0 | 0 | 1 |
| 0 | 0 | 0 | 0 |
| 0 | 0 | 0 | 0 |
| 0 | 0 | 0 | 0 |
| 0 | 0 | 0 | 1 |
| 0 | 0 | 0 | 1 |
| 0 | 0 | 0 | 1 |
| 0 | 0 | 0 | 1 |
| 0 | 0 | 0 | 0 |
| 0 | 0 | 0 | 1 |
| 0 | 0 | 0 | 0 |
| 0 | 0 | 0 | 1 |
| 0 | 0 | 0 | 0 |
| 0 | 0 | 0 | 0 |
| 0 | 0 | 0 | 0 |
| 0 | 0 | 0 | 1 |
| 0 | 0 | 0 | 1 |

| | | | |
|---|---|---|---|
| 0 | 0 | 0 | 0 |
| 0 | 0 | 0 | 0 |
| 0 | 0 | 0 | 1 |
| 0 | 0 | 0 | 1 |
| 0 | 0 | 0 | 1 |
| 0 | 0 | 0 | 1 |
| 0 | 0 | 0 | 1 |
| 0 | 0 | 0 | 1 |
| 0 | 0 | 0 | 1 |
| 0 | 0 | 0 | 1 |
| 0 | 0 | 0 | 1 |
| 0 | 0 | 0 | 1 |
| 0 | 0 | 0 | 1 |
| 0 | 0 | 0 | 1 |
| 0 | 0 | 0 | 1 |
| 0 | 0 | 0 | 0 |
| 0 | 0 | 0 | 0 |
| 0 | 0 | 0 | 0 |
| 0 | 0 | 0 | 1 |
| 0 | 0 | 0 | 1 |
| 0 | 0 | 0 | 1 |
| 0 | 0 | 0 | 0 |
| 0 | 0 | 0 | 0 |
| 0 | 0 | 0 | 0 |
| 0 | 0 | 0 | 0 |
| 0 | 0 | 0 | 1 |
| 0 | 0 | 0 | 1 |
| 0 | 0 | 0 | 1 |
| 0 | 0 | 0 | 1 |
| 0 | 0 | 0 | 1 |
| 0 | 0 | 0 | 0 |
| 0 | 0 | 0 | 1 |
| 0 | 0 | 0 | 0 |
| 0 | 0 | 0 | 1 |
| 0 | 0 | 0 | 1 |
| 0 | 0 | 0 | 1 |
| 0 | 0 | 0 | 0 |
| 0 | 0 | 0 | 0 |
| 0 | 0 | 0 | 1 |
| 0 | 0 | 0 | 1 |
| 0 | 0 | 0 | 1 |
| 0 | 0 | 0 | 0 |
| 0 | 0 | 0 | 1 |
| 0 | 0 | 0 | 1 |
| 0 | 0 | 0 | 0 |
| 0 | 0 | 0 | 1 |
| 0 | 0 | 0 | 0 |
| 0 | 0 | 0 | 0 |
| 0 | 0 | 0 | 1 |
| 0 | 0 | 0 | 1 |
| 0 | 0 | 0 | 0 |
| 0 | 0 | 0 | 0 |
| 0 | 0 | 0 | 1 |
| 0 | 0 | 0 | 0 |

| | | | |
|---|---|---|---|
| 0 | 0 | 0 | 1 |
| 0 | 0 | 0 | 0 |
| 0 | 0 | 0 | 1 |
| 0 | 0 | 0 | 0 |
| 0 | 0 | 0 | 1 |
| 0 | 0 | 0 | 1 |
| 0 | 0 | 0 | 1 |
| 0 | 0 | 0 | 1 |
| 0 | 0 | 0 | 0 |
| 0 | 0 | 0 | 1 |
| 0 | 0 | 0 | 1 |
| 0 | 0 | 0 | 1 |
| 0 | 0 | 0 | 0 |
| 0 | 0 | 0 | 0 |
| 0 | 0 | 0 | 0 |
| 0 | 0 | 0 | 1 |
| 0 | 0 | 0 | 0 |
| 0 | 0 | 0 | 1 |
| 0 | 1 | 0 | 0 |
| 0 | 0 | 0 | 0 |
| 0 | 0 | 0 | 0 |
| 0 | 0 | 0 | 0 |
| 0 | 0 | 0 | 0 |
| 1 | 0 | 0 | 0 |
| 0 | 0 | 0 | 1 |
| 0 | 0 | 0 | 1 |
| 0 | 0 | 0 | 0 |
| 0 | 0 | 0 | 0 |
| 0 | 0 | 0 | 0 |
| 0 | 0 | 0 | 0 |
| 0 | 0 | 0 | 1 |
| 0 | 0 | 0 | 1 |
| 0 | 0 | 0 | 1 |
| 0 | 0 | 0 | 1 |
| 0 | 0 | 0 | 1 |
| 1 | 0 | 0 | 0 |
| 1 | 0 | 0 | 0 |
| 0 | 0 | 0 | 1 |
| 0 | 0 | 0 | 1 |
| 0 | 0 | 0 | 1 |
| 0 | 0 | 0 | 1 |
| 0 | 0 | 0 | 1 |
| 0 | 0 | 0 | 1 |
| 0 | 0 | 0 | 0 |
| 0 | 0 | 0 | 0 |
| 0 | 0 | 0 | 0 |
| 0 | 0 | 0 | 1 |
| 0 | 0 | 0 | 0 |
| 0 | 0 | 0 | 0 |
| 0 | 0 | 0 | 0 |
| 0 | 0 | 0 | 1 |
| 0 | 0 | 0 | 1 |
| 0 | 0 | 0 | 1 |
| 0 | 0 | 0 | 1 |

| | | | |
|---|---|---|---|
| 0 | 0 | 0 | 1 |
| 0 | 0 | 0 | 1 |
| 0 | 0 | 0 | 1 |
| 0 | 0 | 0 | 1 |
| 0 | 0 | 0 | 0 |
| 0 | 0 | 0 | 1 |
| 0 | 0 | 0 | 0 |
| 0 | 0 | 0 | 0 |
| 0 | 0 | 0 | 0 |
| 0 | 0 | 0 | 0 |
| 0 | 0 | 0 | 0 |
| 0 | 0 | 0 | 0 |
| 0 | 0 | 0 | 1 |
| 0 | 0 | 0 | 0 |
| 0 | 0 | 0 | 0 |
| 0 | 0 | 0 | 1 |
| 0 | 0 | 0 | 1 |
| 0 | 0 | 0 | 1 |
| 0 | 0 | 0 | 0 |
| 0 | 0 | 0 | 0 |
| 0 | 0 | 0 | 0 |
| 0 | 0 | 0 | 0 |
| 0 | 0 | 0 | 1 |
| 0 | 0 | 0 | 1 |
| 0 | 0 | 0 | 1 |
| 0 | 0 | 0 | 1 |
| 0 | 0 | 0 | 1 |
| 0 | 0 | 0 | 0 |
| 0 | 0 | 0 | 0 |
| 0 | 0 | 0 | 0 |
| 0 | 0 | 0 | 0 |
| 0 | 0 | 0 | 0 |
| 0 | 0 | 0 | 0 |
| 0 | 0 | 0 | 1 |
| 0 | 0 | 0 | 1 |
| 0 | 0 | 0 | 0 |
| 0 | 0 | 0 | 1 |
| 0 | 0 | 0 | 1 |
| 0 | 0 | 0 | 1 |
| 0 | 0 | 0 | 1 |
| 0 | 0 | 0 | 0 |
| 0 | 0 | 0 | 1 |
| 0 | 0 | 0 | 0 |
| 0 | 0 | 0 | 1 |
| 0 | 0 | 0 | 1 |
| 0 | 0 | 0 | 1 |
| 0 | 0 | 0 | 1 |
| 0 | 0 | 0 | 0 |
| 0 | 0 | 0 | 1 |
| 0 | 0 | 0 | 1 |
| 0 | 0 | 0 | 1 |
| 0 | 0 | 0 | 1 |
| 0 | 0 | 0 | 1 |
| 0 | 0 | 0 | 1 |

| | | | |
|---|---|---|---|
| 0 | 0 | 0 | 1 |
| 0 | 0 | 0 | 1 |
| 0 | 0 | 0 | 1 |
| 0 | 0 | 0 | 1 |
| 0 | 0 | 0 | 1 |
| 0 | 0 | 0 | 1 |
| 0 | 0 | 0 | 1 |
| 0 | 0 | 0 | 1 |
| 0 | 0 | 0 | 1 |
| 0 | 0 | 0 | 1 |
| 0 | 0 | 0 | 1 |
| 0 | 0 | 0 | 0 |
| 0 | 0 | 0 | 1 |
| 0 | 0 | 0 | 1 |
| 0 | 0 | 0 | 1 |
| 0 | 0 | 0 | 1 |
| 0 | 0 | 0 | 1 |
| 0 | 0 | 0 | 0 |
| 0 | 0 | 0 | 0 |
| 0 | 0 | 0 | 0 |
| 0 | 0 | 0 | 0 |
| 0 | 0 | 0 | 1 |
| 1 | 0 | 0 | 0 |
| 0 | 0 | 0 | 1 |
| 0 | 0 | 0 | 1 |
| 0 | 0 | 0 | 1 |
| 0 | 0 | 0 | 1 |
| 0 | 0 | 0 | 1 |
| 0 | 0 | 0 | 0 |
| 0 | 0 | 0 | 0 |
| 0 | 0 | 0 | 1 |
| 0 | 0 | 0 | 1 |
| 1 | 0 | 0 | 0 |
| 0 | 0 | 0 | 1 |
| 0 | 0 | 0 | 0 |
| 1 | 0 | 0 | 0 |
| 1 | 0 | 0 | 0 |
| 0 | 0 | 0 | 1 |
| 1 | 0 | 0 | 0 |
| 0 | 0 | 0 | 1 |
| 1 | 0 | 0 | 0 |
| 0 | 0 | 0 | 1 |
| 0 | 0 | 0 | 0 |
| 0 | 0 | 0 | 1 |
| 0 | 0 | 0 | 1 |
| 0 | 0 | 0 | 1 |
| 0 | 0 | 0 | 1 |
| 0 | 0 | 0 | 1 |
| 0 | 0 | 0 | 1 |
| 0 | 0 | 0 | 0 |
| 0 | 0 | 0 | 1 |
| 1 | 0 | 0 | 0 |
| 0 | 0 | 0 | 1 |
| 0 | 0 | 0 | 1 |

| | | | |
|---|---|---|---|
| 0 | 0 | 0 | 0 |
| 0 | 0 | 0 | 0 |
| 0 | 0 | 0 | 0 |
| 0 | 0 | 0 | 0 |
| 0 | 0 | 0 | 0 |
| 0 | 0 | 0 | 1 |
| 0 | 0 | 0 | 0 |
| 0 | 0 | 0 | 1 |
| 0 | 0 | 0 | 1 |
| 0 | 0 | 0 | 0 |
| 0 | 0 | 0 | 0 |
| 0 | 0 | 0 | 0 |
| 0 | 0 | 0 | 1 |
| 0 | 0 | 0 | 1 |
| 0 | 0 | 0 | 1 |
| 0 | 0 | 0 | 1 |
| 0 | 0 | 0 | 1 |
| 0 | 0 | 0 | 1 |
| 0 | 0 | 0 | 1 |
| 0 | 0 | 0 | 1 |
| 0 | 0 | 0 | 1 |
| 0 | 0 | 0 | 1 |
| 0 | 0 | 0 | 1 |
| 0 | 0 | 0 | 1 |
| 0 | 0 | 0 | 1 |
| 0 | 0 | 0 | 1 |
| 0 | 0 | 0 | 1 |
| 0 | 0 | 0 | 1 |
| 0 | 0 | 0 | 1 |
| 0 | 0 | 0 | 1 |
| 0 | 0 | 0 | 1 |
| 0 | 0 | 0 | 1 |
| 0 | 0 | 0 | 1 |
| 0 | 0 | 0 | 1 |
| 0 | 0 | 0 | 1 |
| 0 | 0 | 0 | 1 |
| 0 | 0 | 0 | 1 |
| 0 | 0 | 0 | 1 |
| 0 | 0 | 0 | 1 |
| 0 | 0 | 0 | 1 |
| 0 | 0 | 0 | 1 |
| 0 | 0 | 0 | 1 |
| 0 | 0 | 0 | 1 |
| 0 | 0 | 0 | 1 |
| 0 | 0 | 0 | 1 |
| 0 | 0 | 0 | 1 |
| 0 | 0 | 0 | 0 |
| 0 | 0 | 0 | 1 |
| 0 | 0 | 0 | 1 |
| 0 | 0 | 0 | 1 |
| 0 | 0 | 0 | 1 |
| 0 | 0 | 0 | 0 |
| 0 | 0 | 0 | 1 |
| 0 | 0 | 0 | 1 |
| 0 | 0 | 0 | 1 |

| | | | |
|---|---|---|---|
| 0 | 0 | 0 | 1 |
| 0 | 0 | 0 | 1 |
| 0 | 0 | 0 | 1 |
| 0 | 0 | 0 | 1 |
| 0 | 0 | 0 | 1 |
| 0 | 0 | 0 | 1 |
| 0 | 0 | 0 | 1 |
| 0 | 0 | 0 | 0 |
| 0 | 0 | 0 | 0 |
| 0 | 0 | 0 | 1 |
| 0 | 0 | 0 | 1 |
| 0 | 0 | 0 | 1 |
| 0 | 0 | 0 | 1 |
| 0 | 0 | 0 | 1 |
| 0 | 0 | 0 | 0 |
| 0 | 0 | 0 | 0 |
| 0 | 0 | 0 | 0 |
| 0 | 0 | 0 | 1 |
| 0 | 0 | 0 | 1 |
| 0 | 0 | 0 | 1 |
| 0 | 0 | 0 | 0 |
| 0 | 0 | 0 | 0 |
| 0 | 0 | 0 | 0 |
| 0 | 0 | 0 | 0 |
| 0 | 0 | 0 | 0 |
| 0 | 0 | 0 | 0 |
| 0 | 0 | 0 | 1 |
| 0 | 0 | 0 | 1 |
| 0 | 0 | 0 | 0 |
| 0 | 0 | 0 | 1 |
| 0 | 0 | 0 | 1 |
| 0 | 0 | 0 | 1 |
| 0 | 0 | 0 | 0 |
| 0 | 0 | 0 | 1 |
| 0 | 0 | 0 | 1 |
| 0 | 0 | 0 | 1 |
| 0 | 0 | 0 | 0 |
| 0 | 0 | 0 | 0 |
| 0 | 0 | 0 | 0 |
| 0 | 0 | 0 | 1 |
| 0 | 0 | 0 | 1 |
| 0 | 0 | 0 | 1 |
| 0 | 0 | 0 | 1 |
| 0 | 0 | 0 | 1 |
| 0 | 0 | 0 | 1 |
| 0 | 0 | 0 | 0 |
| 0 | 0 | 0 | 1 |
| 0 | 0 | 0 | 1 |
| 0 | 0 | 0 | 1 |
| 0 | 0 | 0 | 1 |
| 0 | 0 | 0 | 1 |
| 0 | 0 | 0 | 1 |
| 0 | 0 | 0 | 1 |
| 0 | 0 | 0 | 1 |

| | | | |
|---|---|---|---|
| 0 | 0 | 0 | 1 |
| 0 | 0 | 0 | 1 |
| 0 | 0 | 0 | 1 |
| 0 | 0 | 0 | 0 |
| 0 | 0 | 0 | 1 |
| 0 | 0 | 0 | 1 |
| 0 | 0 | 0 | 1 |
| 0 | 0 | 0 | 0 |
| 0 | 0 | 0 | 0 |
| 0 | 0 | 0 | 1 |
| 0 | 0 | 0 | 1 |
| 0 | 0 | 0 | 1 |
| 0 | 0 | 0 | 0 |
| 0 | 0 | 0 | 1 |
| 0 | 0 | 0 | 1 |
| 0 | 0 | 0 | 1 |
| 0 | 0 | 0 | 0 |
| 0 | 0 | 0 | 1 |
| 0 | 0 | 0 | 1 |
| 0 | 0 | 0 | 1 |
| 0 | 0 | 0 | 1 |
| 0 | 0 | 0 | 1 |
| 0 | 0 | 0 | 1 |
| 0 | 0 | 0 | 1 |
| 0 | 0 | 0 | 1 |
| 0 | 0 | 0 | 1 |
| 0 | 0 | 0 | 1 |
| 0 | 0 | 0 | 0 |
| 0 | 0 | 0 | 0 |
| 0 | 0 | 0 | 0 |
| 0 | 0 | 0 | 0 |
| 0 | 0 | 0 | 0 |
| 0 | 0 | 0 | 0 |
| 0 | 0 | 0 | 1 |
| 0 | 0 | 0 | 0 |
| 0 | 0 | 0 | 0 |
| 0 | 0 | 0 | 0 |
| 0 | 0 | 0 | 1 |
| 0 | 0 | 0 | 1 |
| 0 | 0 | 0 | 1 |
| 0 | 0 | 0 | 1 |
| 0 | 0 | 0 | 1 |
| 0 | 0 | 0 | 0 |
| 0 | 0 | 0 | 0 |
| 0 | 0 | 0 | 1 |
| 0 | 0 | 0 | 1 |
| 0 | 0 | 0 | 0 |
| 0 | 0 | 0 | 1 |
| 0 | 0 | 0 | 1 |
| 0 | 0 | 0 | 1 |
| 0 | 0 | 0 | 1 |
| 0 | 0 | 0 | 1 |
| 0 | 0 | 0 | 0 |
| 0 | 0 | 0 | 0 |

| | | | |
|---|---|---|---|
| 0 | 0 | 0 | 0 |
| 0 | 0 | 0 | 0 |
| 0 | 0 | 0 | 0 |
| 0 | 0 | 0 | 0 |
| 0 | 0 | 0 | 0 |
| 0 | 0 | 0 | 0 |
| 0 | 0 | 0 | 0 |
| 0 | 0 | 0 | 0 |
| 0 | 0 | 0 | 1 |
| 0 | 0 | 0 | 0 |
| 0 | 0 | 0 | 1 |
| 0 | 0 | 0 | 1 |
| 0 | 0 | 0 | 1 |
| 0 | 0 | 0 | 1 |
| 0 | 0 | 0 | 0 |
| 0 | 0 | 0 | 1 |
| 0 | 0 | 0 | 1 |
| 0 | 0 | 0 | 1 |
| 0 | 0 | 0 | 1 |
| 0 | 0 | 0 | 0 |
| 0 | 0 | 0 | 0 |
| 0 | 0 | 0 | 1 |
| 0 | 0 | 0 | 0 |
| 0 | 0 | 0 | 0 |
| 0 | 0 | 0 | 0 |
| 0 | 0 | 0 | 0 |
| 0 | 0 | 0 | 0 |
| 0 | 0 | 0 | 1 |
| 0 | 0 | 0 | 1 |
| 0 | 0 | 0 | 1 |
| 0 | 0 | 0 | 0 |
| 0 | 0 | 0 | 1 |
| 0 | 0 | 0 | 1 |
| 0 | 0 | 0 | 1 |
| 0 | 0 | 0 | 1 |
| 0 | 0 | 0 | 1 |
| 0 | 0 | 0 | 1 |
| 0 | 0 | 0 | 1 |
| 0 | 0 | 0 | 1 |
| 0 | 0 | 0 | 1 |
| 0 | 0 | 0 | 1 |
| 0 | 0 | 0 | 0 |
| 0 | 0 | 0 | 1 |
| 0 | 0 | 0 | 1 |
| 0 | 0 | 0 | 1 |
| 0 | 0 | 0 | 1 |
| 0 | 0 | 0 | 1 |
| 0 | 0 | 0 | 1 |
| 0 | 0 | 0 | 0 |
| 0 | 0 | 0 | 1 |
| 0 | 0 | 0 | 1 |
| 0 | 0 | 0 | 1 |
| 0 | 0 | 0 | 0 |
| 0 | 0 | 0 | 0 |

| | | | |
|---|---|---|---|
| 0 | 0 | 0 | 1 |
| 0 | 0 | 0 | 1 |
| 0 | 0 | 0 | 1 |
| 0 | 0 | 0 | 1 |
| 0 | 0 | 0 | 0 |
| 1 | 0 | 0 | 0 |
| 0 | 0 | 0 | 1 |
| 0 | 0 | 0 | 0 |
| 0 | 0 | 0 | 1 |
| 0 | 0 | 0 | 1 |
| 0 | 0 | 0 | 0 |
| 0 | 0 | 0 | 1 |
| 0 | 0 | 0 | 0 |
| 0 | 0 | 0 | 0 |
| 0 | 0 | 0 | 0 |
| 0 | 0 | 0 | 0 |
| 0 | 0 | 0 | 1 |
| 0 | 0 | 0 | 1 |
| 0 | 0 | 0 | 0 |
| 0 | 0 | 0 | 0 |
| 0 | 0 | 0 | 0 |
| 0 | 0 | 0 | 0 |
| 0 | 0 | 0 | 0 |
| 0 | 0 | 0 | 1 |
| 0 | 0 | 0 | 1 |
| 0 | 0 | 0 | 0 |
| 0 | 0 | 0 | 1 |
| 0 | 0 | 0 | 0 |
| 0 | 0 | 0 | 0 |
| 0 | 0 | 0 | 0 |
| 0 | 0 | 0 | 1 |
| 1 | 0 | 0 | 0 |
| 1 | 0 | 0 | 0 |
| 0 | 0 | 0 | 1 |
| 0 | 0 | 0 | 1 |
| 0 | 0 | 0 | 0 |
| 0 | 0 | 0 | 0 |
| 0 | 0 | 0 | 1 |
| 0 | 0 | 0 | 0 |
| 0 | 0 | 0 | 0 |
| 0 | 0 | 0 | 0 |
| 0 | 0 | 0 | 0 |
| 0 | 0 | 0 | 1 |
| 0 | 0 | 0 | 1 |
| 0 | 0 | 0 | 0 |
| 0 | 0 | 0 | 1 |
| 0 | 0 | 0 | 0 |
| 0 | 0 | 0 | 1 |
| 0 | 0 | 0 | 1 |
| 0 | 0 | 0 | 1 |
| 0 | 0 | 0 | 1 |
| 0 | 0 | 0 | 1 |
| 0 | 0 | 0 | 1 |
| 0 | 0 | 0 | 1 |

| | | | |
|---|---|---|---|
| 0 | 0 | 0 | 1 |
| 0 | 0 | 0 | 1 |
| 0 | 0 | 0 | 1 |
| 0 | 0 | 0 | 1 |
| 0 | 0 | 0 | 1 |
| 0 | 0 | 0 | 1 |
| 0 | 0 | 0 | 1 |
| 0 | 0 | 0 | 0 |
| 0 | 0 | 0 | 1 |
| 1 | 0 | 0 | 0 |
| 0 | 0 | 0 | 0 |
| 0 | 0 | 0 | 0 |
| 0 | 0 | 0 | 1 |
| 0 | 0 | 0 | 1 |
| 0 | 0 | 0 | 0 |
| 0 | 0 | 0 | 1 |
| 0 | 0 | 0 | 1 |
| 0 | 0 | 0 | 1 |
| 0 | 0 | 0 | 1 |
| 0 | 0 | 0 | 1 |
| 0 | 0 | 0 | 1 |
| 0 | 0 | 0 | 1 |
| 0 | 0 | 0 | 1 |
| 0 | 0 | 0 | 1 |
| 0 | 0 | 0 | 1 |
| 0 | 0 | 0 | 1 |
| 0 | 0 | 0 | 1 |
| 0 | 0 | 0 | 1 |
| 0 | 0 | 0 | 1 |
| 0 | 0 | 0 | 0 |
| 0 | 0 | 0 | 0 |
| 0 | 0 | 0 | 1 |
| 0 | 0 | 0 | 1 |
| 0 | 0 | 0 | 1 |
| 0 | 0 | 0 | 1 |
| 0 | 0 | 0 | 0 |
| 0 | 0 | 0 | 1 |
| 0 | 0 | 0 | 1 |
| 0 | 0 | 0 | 1 |
| 0 | 0 | 0 | 1 |
| 0 | 0 | 0 | 1 |
| 0 | 0 | 0 | 0 |
| 0 | 0 | 0 | 0 |
| 0 | 0 | 0 | 0 |
| 0 | 0 | 0 | 1 |
| 0 | 0 | 0 | 1 |
| 0 | 0 | 0 | 1 |
| 0 | 0 | 0 | 1 |
| 0 | 0 | 0 | 1 |
| 0 | 0 | 0 | 1 |
| 0 | 0 | 0 | 1 |
| 0 | 0 | 0 | 1 |
| 0 | 0 | 0 | 1 |
| 0 | 0 | 0 | 0 |

| | | | |
|---|---|---|---|
| 0 | 0 | 0 | 1 |
| 0 | 0 | 0 | 1 |
| 0 | 0 | 0 | 1 |
| 0 | 0 | 0 | 1 |
| 0 | 0 | 0 | 1 |
| 0 | 0 | 0 | 1 |
| 0 | 0 | 0 | 1 |
| 0 | 0 | 0 | 1 |
| 0 | 0 | 0 | 0 |
| 0 | 0 | 0 | 0 |
| 0 | 0 | 0 | 1 |
| 0 | 0 | 0 | 1 |
| 0 | 0 | 0 | 1 |
| 0 | 0 | 0 | 0 |
| 0 | 0 | 0 | 1 |
| 0 | 0 | 0 | 1 |
| 0 | 0 | 0 | 1 |
| 1 | 0 | 0 | 0 |
| 1 | 0 | 0 | 0 |
| 0 | 0 | 0 | 0 |
| 0 | 0 | 0 | 0 |
| 0 | 0 | 0 | 1 |
| 0 | 0 | 0 | 1 |
| 0 | 0 | 0 | 1 |
| 0 | 0 | 0 | 1 |
| 1 | 0 | 0 | 0 |
| 0 | 0 | 0 | 1 |
| 0 | 0 | 0 | 1 |
| 0 | 0 | 0 | 1 |
| 0 | 0 | 0 | 1 |
| 0 | 0 | 0 | 1 |
| 0 | 0 | 0 | 1 |
| 0 | 0 | 0 | 1 |
| 0 | 0 | 0 | 1 |
| 0 | 0 | 0 | 1 |
| 0 | 0 | 0 | 1 |
| 0 | 0 | 0 | 1 |
| 0 | 0 | 0 | 1 |
| 0 | 0 | 0 | 1 |
| 0 | 0 | 0 | 0 |
| 0 | 0 | 0 | 1 |
| 0 | 0 | 0 | 1 |
| 0 | 0 | 0 | 0 |
| 0 | 0 | 0 | 1 |
| 0 | 0 | 0 | 1 |
| 0 | 0 | 0 | 1 |
| 0 | 0 | 0 | 0 |
| 0 | 0 | 0 | 0 |
| 0 | 0 | 0 | 1 |
| 0 | 0 | 0 | 1 |
| 0 | 0 | 0 | 0 |
| 0 | 0 | 0 | 1 |
| 0 | 0 | 0 | 1 |
| 0 | 0 | 0 | 1 |

| | | | |
|---|---|---|---|
| 0 | 0 | 0 | 1 |
| 0 | 0 | 0 | 1 |
| 0 | 0 | 0 | 1 |
| 0 | 0 | 0 | 1 |
| 0 | 0 | 0 | 1 |
| 0 | 0 | 0 | 1 |
| 0 | 0 | 0 | 1 |
| 0 | 0 | 0 | 1 |
| 0 | 0 | 0 | 1 |
| 0 | 0 | 0 | 1 |
| 0 | 0 | 0 | 1 |
| 0 | 0 | 0 | 1 |
| 0 | 0 | 0 | 1 |
| 0 | 0 | 0 | 0 |
| 0 | 0 | 0 | 0 |
| 0 | 0 | 0 | 0 |
| 0 | 0 | 0 | 0 |
| 0 | 0 | 0 | 1 |
| 0 | 0 | 0 | 1 |
| 0 | 0 | 0 | 0 |
| 0 | 0 | 0 | 1 |
| 0 | 0 | 0 | 1 |
| 0 | 0 | 0 | 1 |
| 0 | 0 | 0 | 1 |
| 0 | 0 | 0 | 1 |
| 0 | 0 | 0 | 0 |
| 0 | 0 | 0 | 1 |
| 0 | 0 | 0 | 1 |
| 0 | 0 | 0 | 1 |
| 0 | 0 | 0 | 0 |
| 0 | 0 | 0 | 0 |
| 0 | 0 | 0 | 1 |
| 0 | 0 | 0 | 1 |
| 0 | 0 | 0 | 1 |
| 0 | 0 | 0 | 1 |
| 0 | 0 | 0 | 1 |
| 0 | 0 | 0 | 1 |
| 0 | 0 | 0 | 1 |
| 0 | 0 | 0 | 0 |
| 0 | 0 | 0 | 1 |
| 0 | 0 | 0 | 1 |
| 0 | 0 | 0 | 0 |
| 0 | 0 | 0 | 0 |
| 0 | 0 | 0 | 0 |
| 0 | 0 | 0 | 0 |
| 0 | 0 | 0 | 1 |
| 0 | 0 | 0 | 0 |
| 0 | 0 | 0 | 1 |
| 0 | 0 | 0 | 1 |
| 0 | 0 | 0 | 1 |
| 0 | 0 | 0 | 1 |
| 0 | 0 | 0 | 1 |
| 0 | 0 | 0 | 1 |
| 0 | 0 | 0 | 1 |

| | | | |
|---|---|---|---|
| 0 | 0 | 0 | 1 |
| 0 | 0 | 0 | 1 |
| 0 | 0 | 0 | 0 |
| 0 | 0 | 0 | 1 |
| 0 | 0 | 0 | 1 |
| 0 | 0 | 0 | 1 |
| 0 | 0 | 0 | 1 |
| 0 | 0 | 0 | 1 |
| 0 | 0 | 0 | 1 |
| 0 | 0 | 0 | 1 |
| 0 | 0 | 0 | 1 |
| 0 | 0 | 0 | 1 |
| 0 | 0 | 0 | 1 |
| 0 | 0 | 0 | 1 |
| 0 | 0 | 0 | 1 |
| 0 | 0 | 0 | 1 |
| 0 | 0 | 0 | 1 |
| 0 | 0 | 0 | 1 |
| 0 | 0 | 0 | 1 |
| 0 | 0 | 0 | 1 |
| 0 | 0 | 0 | 1 |
| 0 | 0 | 0 | 1 |
| 0 | 0 | 0 | 1 |
| 0 | 0 | 0 | 1 |
| 0 | 0 | 0 | 1 |
| 0 | 0 | 0 | 1 |
| 0 | 0 | 0 | 1 |
| 0 | 0 | 0 | 1 |
| 0 | 0 | 0 | 1 |
| 0 | 0 | 0 | 1 |
| 1 | 0 | 0 | 0 |
| 0 | 0 | 0 | 1 |
| 0 | 0 | 0 | 0 |
| 0 | 0 | 0 | 0 |
| 0 | 0 | 0 | 0 |
| 0 | 0 | 0 | 0 |
| 0 | 0 | 0 | 1 |
| 0 | 0 | 0 | 0 |
| 0 | 0 | 0 | 1 |
| 0 | 0 | 0 | 1 |
| 1 | 0 | 0 | 0 |
| 1 | 0 | 0 | 0 |
| 0 | 0 | 0 | 0 |
| 0 | 0 | 0 | 1 |
| 0 | 0 | 0 | 0 |
| 0 | 0 | 0 | 0 |
| 0 | 0 | 0 | 1 |
| 0 | 0 | 0 | 1 |
| 0 | 0 | 0 | 1 |
| 0 | 0 | 0 | 1 |
| 0 | 0 | 0 | 0 |
| 0 | 0 | 0 | 1 |
| 0 | 0 | 0 | 1 |
| 0 | 0 | 0 | 1 |

| | | | |
|---|---|---|---|
| 0 | 0 | 0 | 1 |
| 0 | 0 | 0 | 1 |
| 0 | 0 | 0 | 1 |
| 0 | 0 | 0 | 1 |
| 0 | 0 | 0 | 1 |
| 1 | 0 | 0 | 0 |
| 0 | 0 | 0 | 1 |
| 0 | 0 | 0 | 0 |
| 0 | 0 | 0 | 0 |
| 0 | 0 | 0 | 0 |
| 0 | 0 | 0 | 1 |
| 0 | 0 | 0 | 0 |
| 0 | 0 | 0 | 1 |
| 0 | 0 | 0 | 1 |
| 0 | 0 | 0 | 1 |
| 0 | 0 | 0 | 1 |
| 0 | 0 | 0 | 1 |
| 0 | 0 | 0 | 0 |
| 0 | 0 | 0 | 1 |
| 0 | 0 | 0 | 0 |
| 0 | 0 | 0 | 1 |
| 0 | 0 | 0 | 0 |
| 0 | 0 | 0 | 0 |
| 0 | 0 | 0 | 0 |
| 0 | 0 | 0 | 1 |
| 0 | 0 | 0 | 1 |
| 0 | 0 | 0 | 1 |
| 0 | 0 | 0 | 1 |
| 0 | 0 | 0 | 0 |
| 0 | 0 | 0 | 1 |
| 0 | 0 | 0 | 1 |
| 0 | 0 | 0 | 1 |
| 0 | 0 | 0 | 1 |
| 0 | 0 | 0 | 1 |
| 0 | 0 | 0 | 0 |
| 0 | 0 | 0 | 0 |
| 0 | 0 | 0 | 1 |
| 0 | 0 | 0 | 0 |
| 0 | 0 | 0 | 0 |
| 0 | 0 | 0 | 1 |
| 0 | 0 | 0 | 0 |
| 0 | 0 | 0 | 0 |
| 0 | 0 | 0 | 1 |
| 0 | 0 | 0 | 1 |
| 0 | 0 | 0 | 1 |
| 0 | 0 | 0 | 1 |
| 0 | 0 | 0 | 1 |
| 0 | 0 | 0 | 1 |
| 0 | 0 | 0 | 1 |
| 0 | 0 | 0 | 1 |
| 0 | 0 | 0 | 1 |
| 0 | 0 | 0 | 1 |
| 0 | 0 | 0 | 1 |
| 0 | 0 | 0 | 1 |

| | | | |
|---|---|---|---|
| 0 | 0 | 0 | 1 |
| 0 | 0 | 0 | 0 |
| 0 | 0 | 0 | 1 |
| 0 | 0 | 0 | 1 |
| 0 | 0 | 0 | 0 |
| 0 | 0 | 0 | 1 |
| 0 | 0 | 0 | 1 |
| 0 | 0 | 0 | 1 |
| 0 | 0 | 0 | 0 |
| 0 | 0 | 0 | 0 |
| 0 | 0 | 0 | 1 |
| 0 | 0 | 0 | 0 |
| 0 | 0 | 0 | 0 |
| 0 | 0 | 0 | 1 |
| 0 | 0 | 0 | 1 |
| 0 | 0 | 0 | 0 |
| 0 | 0 | 0 | 1 |
| 0 | 0 | 0 | 0 |
| 0 | 0 | 0 | 0 |
| 0 | 0 | 0 | 1 |
| 0 | 0 | 0 | 0 |
| 0 | 0 | 0 | 0 |
| 1 | 0 | 0 | 0 |
| 0 | 0 | 0 | 1 |
| 0 | 0 | 0 | 1 |
| 0 | 0 | 0 | 0 |
| 0 | 0 | 0 | 1 |
| 0 | 0 | 0 | 1 |
| 0 | 0 | 0 | 1 |
| 0 | 0 | 0 | 1 |
| 0 | 0 | 0 | 0 |
| 0 | 0 | 0 | 1 |
| 0 | 0 | 0 | 0 |
| 0 | 0 | 0 | 1 |
| 0 | 0 | 0 | 0 |
| 0 | 0 | 0 | 0 |
| 0 | 0 | 0 | 1 |
| 0 | 0 | 0 | 1 |
| 0 | 0 | 0 | 1 |
| 0 | 0 | 0 | 1 |
| 0 | 0 | 0 | 1 |
| 0 | 0 | 0 | 1 |
| 0 | 0 | 0 | 1 |
| 0 | 0 | 0 | 1 |
| 0 | 0 | 0 | 1 |
| 0 | 0 | 0 | 1 |
| 0 | 0 | 0 | 1 |
| 0 | 0 | 0 | 1 |
| 0 | 0 | 0 | 1 |
| 0 | 0 | 0 | 1 |
| 0 | 0 | 0 | 1 |
| 0 | 0 | 0 | 1 |
| 0 | 0 | 0 | 1 |
| 0 | 0 | 0 | 0 |

| | | | |
|---|---|---|---|
| 0 | 0 | 0 | 1 |
| 0 | 0 | 0 | 1 |
| 0 | 0 | 0 | 1 |
| 0 | 0 | 0 | 1 |
| 0 | 0 | 0 | 0 |
| 0 | 0 | 0 | 1 |
| 0 | 0 | 0 | 1 |
| 0 | 0 | 0 | 1 |
| 0 | 0 | 0 | 1 |
| 0 | 0 | 0 | 1 |
| 0 | 0 | 0 | 0 |
| 0 | 0 | 0 | 1 |
| 0 | 0 | 0 | 1 |
| 0 | 0 | 0 | 1 |
| 0 | 0 | 0 | 0 |
| 0 | 0 | 0 | 0 |
| 0 | 0 | 0 | 0 |
| 0 | 0 | 0 | 0 |
| 0 | 0 | 0 | 1 |
| 0 | 0 | 0 | 0 |
| 0 | 0 | 0 | 0 |
| 1 | 0 | 0 | 0 |
| 0 | 0 | 0 | 1 |
| 0 | 0 | 0 | 0 |
| 1 | 0 | 0 | 0 |
| 0 | 0 | 0 | 1 |
| 1 | 0 | 0 | 0 |
| 0 | 0 | 0 | 0 |
| 1 | 0 | 0 | 0 |
| 1 | 0 | 0 | 0 |
| 1 | 0 | 0 | 0 |
| 0 | 0 | 0 | 1 |
| 0 | 0 | 0 | 1 |
| 0 | 0 | 0 | 1 |
| 0 | 0 | 0 | 1 |
| 0 | 0 | 0 | 1 |
| 0 | 0 | 0 | 1 |
| 0 | 0 | 0 | 0 |
| 0 | 0 | 0 | 0 |
| 0 | 0 | 0 | 1 |
| 0 | 0 | 0 | 1 |
| 0 | 0 | 0 | 1 |
| 0 | 0 | 0 | 0 |
| 0 | 0 | 0 | 1 |
| 0 | 0 | 0 | 0 |
| 0 | 0 | 0 | 1 |
| 0 | 0 | 0 | 1 |
| 0 | 0 | 0 | 1 |
| 0 | 1 | 0 | 0 |
| 0 | 1 | 0 | 0 |
| 0 | 1 | 0 | 0 |
| 0 | 0 | 0 | 1 |
| 0 | 0 | 0 | 1 |
| 0 | 0 | 0 | 0 |

| | | | |
|---|---|---|---|
| 0 | 0 | 0 | 1 |
| 0 | 0 | 0 | 0 |
| 0 | 0 | 0 | 1 |
| 0 | 1 | 0 | 0 |
| 0 | 1 | 0 | 0 |
| 0 | 0 | 0 | 1 |
| 0 | 1 | 0 | 0 |
| 0 | 1 | 0 | 0 |
| 0 | 0 | 0 | 1 |
| 0 | 0 | 0 | 1 |
| 0 | 0 | 0 | 1 |
| 0 | 0 | 0 | 1 |
| 0 | 0 | 0 | 1 |
| 0 | 0 | 0 | 1 |
| 0 | 0 | 0 | 1 |
| 0 | 0 | 0 | 1 |
| 0 | 0 | 0 | 1 |
| 0 | 0 | 0 | 1 |
| 0 | 0 | 0 | 1 |
| 0 | 0 | 0 | 1 |
| 1 | 0 | 0 | 0 |
| 0 | 0 | 0 | 1 |
| 0 | 0 | 0 | 1 |
| 0 | 0 | 0 | 1 |
| 0 | 0 | 0 | 1 |
| 0 | 0 | 0 | 0 |
| 0 | 0 | 0 | 0 |
| 0 | 0 | 0 | 1 |
| 0 | 0 | 0 | 1 |
| 0 | 0 | 0 | 1 |
| 0 | 0 | 0 | 1 |
| 0 | 0 | 0 | 1 |
| 0 | 0 | 0 | 1 |
| 0 | 0 | 0 | 1 |
| 0 | 0 | 0 | 1 |
| 0 | 0 | 0 | 1 |
| 0 | 0 | 0 | 1 |
| 0 | 0 | 0 | 1 |
| 0 | 0 | 0 | 1 |
| 0 | 0 | 0 | 1 |
| 0 | 0 | 0 | 1 |
| 0 | 0 | 0 | 1 |
| 0 | 0 | 0 | 1 |
| 0 | 0 | 0 | 0 |
| 0 | 0 | 0 | 1 |
| 0 | 0 | 0 | 0 |
| 0 | 0 | 0 | 0 |
| 0 | 0 | 0 | 0 |
| 0 | 0 | 0 | 0 |
| 0 | 0 | 0 | 0 |
| 0 | 0 | 0 | 0 |
| 0 | 0 | 0 | 1 |
| 0 | 0 | 0 | 1 |
| 0 | 0 | 0 | 1 |

| | | | |
|---|---|---|---|
| 0 | 0 | 0 | 1 |
| 0 | 0 | 0 | 1 |
| 0 | 0 | 0 | 1 |
| 0 | 0 | 0 | 0 |
| 0 | 0 | 0 | 1 |
| 0 | 0 | 0 | 0 |
| 0 | 0 | 0 | 1 |
| 0 | 0 | 0 | 1 |
| 0 | 0 | 0 | 0 |
| 0 | 0 | 0 | 1 |
| 0 | 0 | 0 | 1 |
| 0 | 0 | 0 | 1 |
| 0 | 0 | 0 | 1 |
| 0 | 0 | 0 | 1 |
| 0 | 0 | 0 | 1 |
| 0 | 0 | 0 | 0 |
| 0 | 0 | 0 | 0 |
| 0 | 0 | 0 | 0 |
| 0 | 0 | 0 | 0 |
| 0 | 0 | 0 | 0 |
| 0 | 0 | 0 | 1 |
| 0 | 0 | 0 | 1 |
| 0 | 0 | 0 | 1 |
| 0 | 0 | 0 | 1 |
| 0 | 0 | 0 | 0 |
| 0 | 0 | 0 | 1 |
| 0 | 0 | 0 | 1 |
| 0 | 0 | 0 | 1 |
| 0 | 0 | 0 | 0 |
| 0 | 0 | 0 | 1 |
| 0 | 0 | 0 | 0 |
| 0 | 0 | 0 | 0 |
| 0 | 0 | 0 | 0 |
| 0 | 0 | 0 | 0 |
| 0 | 0 | 0 | 0 |
| 0 | 0 | 0 | 1 |
| 0 | 0 | 0 | 0 |
| 0 | 0 | 0 | 0 |
| 0 | 0 | 0 | 0 |
| 0 | 0 | 0 | 0 |
| 0 | 0 | 0 | 0 |
| 0 | 0 | 0 | 0 |
| 0 | 0 | 0 | 0 |
| 0 | 0 | 0 | 0 |
| 0 | 0 | 0 | 0 |
| 0 | 0 | 0 | 0 |
| 0 | 0 | 0 | 0 |
| 0 | 0 | 0 | 0 |
| 0 | 0 | 0 | 1 |
| 0 | 0 | 0 | 0 |
| 0 | 0 | 0 | 1 |
| 0 | 0 | 0 | 1 |
| 0 | 0 | 0 | 1 |
| 0 | 0 | 0 | 0 |

| | | | |
|---|---|---|---|
| 0 | 0 | 0 | 0 |
| 0 | 0 | 0 | 1 |
| 0 | 0 | 0 | 1 |
| 0 | 0 | 0 | 1 |
| 0 | 0 | 0 | 1 |
| 0 | 0 | 0 | 0 |
| 0 | 0 | 0 | 1 |
| 0 | 0 | 0 | 1 |
| 0 | 0 | 0 | 1 |
| 0 | 0 | 0 | 1 |
| 0 | 0 | 0 | 1 |
| 0 | 0 | 0 | 1 |
| 0 | 0 | 0 | 1 |
| 1 | 0 | 0 | 0 |
| 0 | 0 | 0 | 1 |
| 0 | 0 | 0 | 1 |
| 0 | 0 | 0 | 1 |
| 0 | 0 | 0 | 0 |
| 0 | 0 | 0 | 1 |
| 0 | 0 | 0 | 1 |
| 0 | 0 | 0 | 1 |
| 0 | 0 | 0 | 1 |
| 0 | 0 | 0 | 0 |
| 0 | 0 | 0 | 1 |
| 0 | 0 | 0 | 1 |
| 0 | 0 | 0 | 0 |
| 0 | 0 | 0 | 0 |
| 0 | 0 | 0 | 0 |
| 0 | 0 | 0 | 1 |
| 0 | 0 | 0 | 1 |
| 0 | 0 | 0 | 1 |
| 0 | 0 | 0 | 1 |
| 0 | 0 | 0 | 1 |
| 0 | 0 | 0 | 1 |
| 0 | 0 | 0 | 1 |
| 0 | 0 | 0 | 1 |
| 0 | 0 | 0 | 1 |
| 0 | 0 | 0 | 1 |
| 0 | 0 | 0 | 1 |
| 0 | 0 | 0 | 1 |
| 0 | 0 | 0 | 1 |
| 0 | 0 | 0 | 1 |
| 0 | 0 | 0 | 1 |
| 0 | 0 | 0 | 1 |
| 0 | 0 | 0 | 1 |
| 0 | 0 | 0 | 1 |
| 0 | 0 | 0 | 1 |
| 0 | 0 | 0 | 1 |
| 0 | 0 | 0 | 1 |
| 0 | 0 | 0 | 1 |
| 0 | 0 | 0 | 1 |
| 0 | 0 | 0 | 1 |
| 0 | 0 | 0 | 1 |

| | | | |
|---|---|---|---|
| 0 | 0 | 0 | 1 |
| 0 | 0 | 0 | 0 |
| 0 | 0 | 0 | 1 |
| 0 | 0 | 0 | 1 |
| 0 | 0 | 0 | 1 |
| 0 | 0 | 0 | 0 |
| 0 | 0 | 0 | 0 |
| 0 | 0 | 0 | 0 |
| 0 | 0 | 0 | 1 |
| 0 | 0 | 0 | 1 |
| 0 | 0 | 0 | 1 |
| 0 | 0 | 0 | 1 |
| 0 | 0 | 0 | 1 |
| 0 | 0 | 0 | 1 |
| 0 | 0 | 0 | 0 |
| 0 | 0 | 0 | 0 |
| 0 | 0 | 0 | 0 |
| 0 | 0 | 0 | 1 |
| 0 | 0 | 0 | 0 |
| 0 | 0 | 0 | 0 |
| 0 | 0 | 0 | 0 |
| 0 | 0 | 0 | 0 |
| 0 | 0 | 0 | 0 |
| 0 | 0 | 0 | 0 |
| 0 | 0 | 0 | 1 |
| 0 | 0 | 0 | 1 |
| 0 | 0 | 0 | 1 |
| 0 | 0 | 0 | 1 |
| 0 | 0 | 0 | 1 |
| 0 | 0 | 0 | 1 |
| 0 | 0 | 0 | 1 |
| 0 | 0 | 0 | 1 |
| 0 | 0 | 0 | 0 |
| 0 | 0 | 0 | 1 |
| 0 | 0 | 0 | 1 |
| 0 | 0 | 0 | 0 |
| 0 | 0 | 0 | 0 |
| 0 | 0 | 0 | 0 |
| 0 | 0 | 0 | 0 |
| 0 | 0 | 0 | 1 |
| 0 | 0 | 0 | 1 |
| 0 | 0 | 0 | 1 |
| 0 | 0 | 0 | 1 |
| 0 | 0 | 0 | 0 |
| 0 | 0 | 0 | 0 |
| 0 | 0 | 0 | 0 |
| 0 | 0 | 0 | 1 |
| 0 | 0 | 0 | 1 |
| 0 | 0 | 0 | 0 |
| 0 | 0 | 0 | 1 |
| 0 | 0 | 0 | 1 |
| 0 | 0 | 0 | 0 |
| 0 | 0 | 0 | 1 |
| 0 | 0 | 0 | 1 |

| | | | |
|---|---|---|---|
| 0 | 0 | 0 | 1 |
| 0 | 0 | 0 | 1 |
| 0 | 0 | 0 | 1 |
| 0 | 0 | 0 | 1 |
| 0 | 0 | 0 | 1 |
| 0 | 0 | 0 | 1 |
| 0 | 0 | 0 | 1 |
| 0 | 0 | 0 | 1 |
| 0 | 0 | 0 | 0 |
| 0 | 0 | 0 | 0 |
| 0 | 0 | 0 | 1 |
| 0 | 0 | 0 | 0 |
| 0 | 0 | 0 | 0 |
| 0 | 0 | 0 | 0 |
| 0 | 0 | 0 | 1 |
| 0 | 0 | 0 | 1 |
| 0 | 0 | 0 | 0 |
| 0 | 0 | 0 | 0 |
| 0 | 0 | 0 | 0 |
| 0 | 0 | 0 | 1 |
| 0 | 0 | 0 | 1 |
| 0 | 0 | 0 | 1 |
| 0 | 0 | 0 | 0 |
| 0 | 0 | 0 | 1 |
| 0 | 0 | 0 | 0 |
| 0 | 0 | 0 | 1 |
| 0 | 0 | 0 | 1 |
| 0 | 0 | 0 | 1 |
| 0 | 0 | 0 | 1 |
| 0 | 0 | 0 | 0 |
| 0 | 0 | 0 | 0 |
| 0 | 0 | 0 | 0 |
| 0 | 0 | 0 | 1 |
| 0 | 0 | 0 | 1 |
| 0 | 0 | 0 | 1 |
| 0 | 0 | 0 | 1 |
| 0 | 0 | 0 | 1 |
| 0 | 0 | 0 | 1 |
| 0 | 0 | 0 | 1 |
| 0 | 0 | 0 | 1 |
| 0 | 0 | 0 | 0 |
| 0 | 0 | 0 | 1 |
| 0 | 0 | 0 | 1 |
| 0 | 0 | 0 | 1 |
| 0 | 0 | 0 | 1 |
| 0 | 0 | 0 | 0 |
| 0 | 0 | 0 | 1 |
| 0 | 0 | 0 | 0 |
| 0 | 0 | 0 | 1 |
| 0 | 0 | 0 | 0 |
| 0 | 0 | 0 | 0 |
| 0 | 0 | 0 | 0 |
| 0 | 0 | 0 | 1 |
| 0 | 0 | 0 | 1 |

| | | | |
|---|---|---|---|
| 0 | 0 | 0 | 1 |
| 0 | 0 | 0 | 1 |
| 0 | 0 | 0 | 1 |
| 0 | 0 | 0 | 1 |
| 0 | 0 | 0 | 1 |
| 0 | 0 | 0 | 1 |
| 0 | 0 | 0 | 1 |
| 0 | 0 | 0 | 0 |
| 0 | 0 | 0 | 0 |
| 0 | 0 | 0 | 1 |
| 0 | 0 | 0 | 0 |
| 0 | 0 | 0 | 1 |
| 0 | 0 | 0 | 1 |
| 0 | 0 | 0 | 1 |
| 0 | 0 | 0 | 1 |
| 0 | 0 | 0 | 1 |
| 0 | 0 | 0 | 1 |
| 0 | 0 | 0 | 1 |
| 0 | 0 | 0 | 1 |
| 0 | 0 | 0 | 1 |
| 0 | 0 | 0 | 1 |
| 0 | 0 | 0 | 1 |
| 0 | 0 | 0 | 1 |
| 0 | 0 | 0 | 1 |
| 0 | 0 | 0 | 1 |
| 0 | 0 | 0 | 1 |
| 0 | 0 | 0 | 1 |
| 0 | 0 | 0 | 1 |
| 0 | 0 | 0 | 0 |
| 0 | 0 | 0 | 1 |
| 0 | 0 | 0 | 0 |
| 0 | 0 | 0 | 1 |
| 0 | 0 | 0 | 0 |
| 0 | 0 | 0 | 0 |
| 0 | 0 | 0 | 1 |
| 0 | 0 | 0 | 0 |
| 0 | 0 | 0 | 0 |
| 0 | 0 | 0 | 1 |
| 0 | 0 | 0 | 1 |
| 0 | 0 | 0 | 1 |
| 0 | 0 | 0 | 1 |
| 0 | 0 | 0 | 0 |
| 0 | 0 | 0 | 0 |
| 0 | 0 | 0 | 1 |
| 0 | 0 | 0 | 1 |
| 0 | 0 | 0 | 0 |
| 0 | 0 | 0 | 0 |
| 0 | 0 | 0 | 0 |
| 0 | 0 | 0 | 0 |
| 0 | 0 | 0 | 1 |
| 0 | 0 | 0 | 0 |
| 0 | 0 | 0 | 0 |
| 0 | 0 | 0 | 1 |
| 0 | 0 | 0 | 1 |

| | | | |
|---|---|---|---|
| 0 | 0 | 0 | 1 |
| 0 | 0 | 0 | 1 |
| 0 | 0 | 0 | 1 |
| 0 | 0 | 0 | 1 |
| 0 | 0 | 0 | 0 |
| 0 | 0 | 0 | 0 |
| 0 | 0 | 0 | 0 |
| 0 | 0 | 0 | 0 |
| 0 | 0 | 0 | 1 |
| 0 | 0 | 0 | 0 |
| 0 | 0 | 0 | 0 |
| 0 | 0 | 0 | 0 |
| 0 | 0 | 0 | 0 |
| 0 | 0 | 0 | 1 |
| 0 | 0 | 0 | 1 |
| 0 | 0 | 0 | 1 |
| 0 | 0 | 0 | 1 |
| 0 | 0 | 0 | 1 |
| 0 | 0 | 0 | 1 |
| 0 | 0 | 0 | 1 |
| 0 | 0 | 0 | 1 |
| 0 | 0 | 0 | 1 |
| 0 | 0 | 0 | 1 |
| 0 | 0 | 0 | 1 |
| 0 | 0 | 0 | 1 |
| 0 | 0 | 0 | 0 |
| 0 | 0 | 0 | 0 |
| 0 | 0 | 0 | 0 |
| 0 | 0 | 0 | 0 |
| 0 | 0 | 0 | 1 |
| 0 | 0 | 0 | 1 |
| 0 | 0 | 0 | 1 |
| 0 | 0 | 0 | 1 |
| 0 | 0 | 0 | 1 |
| 0 | 0 | 0 | 1 |
| 0 | 0 | 0 | 1 |
| 0 | 0 | 0 | 1 |
| 0 | 0 | 0 | 1 |
| 0 | 0 | 0 | 1 |
| 0 | 0 | 0 | 1 |
| 0 | 0 | 0 | 1 |
| 0 | 0 | 0 | 1 |
| 0 | 0 | 0 | 1 |
| 0 | 0 | 0 | 1 |
| 0 | 0 | 0 | 1 |
| 0 | 0 | 0 | 1 |
| 0 | 0 | 0 | 1 |
| 0 | 0 | 0 | 1 |
| 0 | 0 | 0 | 1 |
| 0 | 0 | 0 | 1 |
| 0 | 0 | 0 | 1 |
| 0 | 0 | 0 | 1 |
| 0 | 0 | 0 | 1 |
| 0 | 0 | 0 | 0 |

| | | | |
|---|---|---|---|
| 0 | 0 | 0 | 1 |
| 0 | 0 | 0 | 1 |
| 0 | 0 | 0 | 1 |
| 0 | 0 | 0 | 1 |
| 0 | 0 | 0 | 1 |
| 0 | 0 | 0 | 1 |
| 0 | 0 | 0 | 1 |
| 0 | 0 | 0 | 1 |
| 0 | 0 | 0 | 1 |
| 0 | 0 | 0 | 1 |
| 0 | 0 | 0 | 1 |
| 0 | 0 | 0 | 1 |
| 0 | 0 | 0 | 1 |
| 0 | 0 | 0 | 1 |
| 0 | 0 | 0 | 1 |
| 0 | 0 | 0 | 1 |
| 0 | 0 | 0 | 1 |
| 0 | 0 | 0 | 1 |
| 0 | 0 | 0 | 1 |
| 0 | 0 | 0 | 1 |
| 0 | 0 | 0 | 1 |
| 0 | 0 | 0 | 1 |
| 0 | 0 | 0 | 1 |
| 0 | 0 | 0 | 1 |
| 0 | 0 | 0 | 1 |
| 0 | 0 | 0 | 1 |
| 0 | 0 | 0 | 1 |
| 0 | 0 | 0 | 1 |
| 0 | 0 | 0 | 1 |
| 0 | 0 | 0 | 1 |
| 0 | 0 | 0 | 1 |
| 0 | 0 | 0 | 1 |
| 0 | 0 | 0 | 1 |
| 0 | 0 | 0 | 1 |
| 0 | 0 | 0 | 1 |
| 0 | 0 | 0 | 1 |
| 0 | 0 | 0 | 1 |
| 0 | 0 | 0 | 1 |
| 0 | 0 | 0 | 1 |
| 0 | 0 | 0 | 1 |
| 0 | 0 | 0 | 1 |
| 0 | 0 | 0 | 1 |
| 0 | 0 | 0 | 1 |
| 0 | 0 | 0 | 1 |
| 0 | 0 | 0 | 1 |
| 0 | 0 | 0 | 1 |
| 0 | 0 | 0 | 0 |
| 0 | 0 | 0 | 1 |
| 0 | 0 | 0 | 1 |
| 0 | 0 | 0 | 1 |
| 0 | 0 | 0 | 1 |
| 0 | 0 | 0 | 1 |
| 0 | 0 | 0 | 1 |
| 0 | 0 | 0 | 1 |

| | | | |
|---|---|---|---|
| 0 | 0 | 0 | 1 |
| 0 | 0 | 0 | 1 |
| 0 | 0 | 0 | 0 |
| 0 | 0 | 0 | 1 |
| 0 | 0 | 0 | 1 |
| 0 | 0 | 0 | 1 |
| 0 | 0 | 0 | 0 |
| 0 | 0 | 0 | 1 |
| 0 | 0 | 0 | 1 |
| 0 | 0 | 0 | 1 |
| 0 | 0 | 0 | 1 |
| 0 | 0 | 0 | 0 |
| 0 | 0 | 0 | 1 |
| 0 | 0 | 0 | 0 |
| 0 | 0 | 0 | 0 |
| 0 | 0 | 0 | 0 |
| 0 | 0 | 0 | 0 |
| 0 | 0 | 0 | 0 |
| 0 | 0 | 0 | 0 |
| 0 | 0 | 0 | 0 |
| 0 | 0 | 0 | 0 |

| surface_non_binding_site | non_binding_site_single_water | surface_non_binding_site_single_water |
|---|---|---|
| 0 | 0 | 0 |
| 0 | 0 | 0 |
| 0 | 0 | 0 |
| 0 | 0 | 0 |
| 0 | 0 | 0 |
| 0 | 0 | 0 |
| 1 | 1 | 1 |
| 1 | 1 | 1 |
| 0 | 0 | 0 |
| 0 | 0 | 0 |
| 0 | 0 | 0 |
| 0 | 0 | 0 |
| 0 | 0 | 0 |
| 0 | 0 | 0 |
| 0 | 0 | 0 |
| 0 | 0 | 0 |
| 0 | 0 | 0 |
| 0 | 0 | 0 |
| 0 | 0 | 0 |
| 0 | 0 | 0 |
| 0 | 0 | 0 |
| 0 | 0 | 0 |
| 0 | 0 | 0 |
| 0 | 0 | 0 |
| 0 | 0 | 0 |
| 1 | 0 | 0 |
| 0 | 0 | 0 |
| 1 | 1 | 1 |
| 0 | 0 | 0 |
| 0 | 1 | 0 |
| 0 | 1 | 0 |
| 0 | 0 | 0 |
| 0 | 0 | 0 |
| 0 | 0 | 0 |
| 0 | 0 | 0 |
| 1 | 1 | 1 |
| 0 | 0 | 0 |
| 0 | 0 | 0 |
| 0 | 0 | 0 |
| 0 | 0 | 0 |
| 0 | 0 | 0 |
| 0 | 0 | 0 |
| 0 | 0 | 0 |
| 0 | 0 | 0 |
| 0 | 0 | 0 |
| 0 | 0 | 0 |
| 0 | 0 | 0 |
| 0 | 0 | 0 |
| 0 | 0 | 0 |
| 0 | 0 | 0 |
| 0 | 0 | 0 |
| 0 | 0 | 0 |
| 0 | 0 | 0 |

| | | |
|---|---|---|
| 0 | 0 | 0 |
| 0 | 0 | 0 |
| 0 | 0 | 0 |
| 0 | 0 | 0 |
| 0 | 0 | 0 |
| 0 | 0 | 0 |
| 0 | 0 | 0 |
| 0 | 0 | 0 |
| 0 | 0 | 0 |
| 0 | 0 | 0 |
| 0 | 0 | 0 |
| 0 | 0 | 0 |
| 0 | 0 | 0 |
| 0 | 0 | 0 |
| 0 | 0 | 0 |
| 0 | 0 | 0 |
| 0 | 0 | 0 |
| 0 | 0 | 0 |
| 0 | 0 | 0 |
| 0 | 0 | 0 |
| 0 | 0 | 0 |
| 0 | 0 | 0 |
| 0 | 0 | 0 |
| 0 | 0 | 0 |
| 0 | 0 | 0 |
| 0 | 0 | 0 |
| 0 | 0 | 0 |
| 0 | 0 | 0 |
| 0 | 0 | 0 |
| 0 | 0 | 0 |
| 0 | 0 | 0 |
| 0 | 0 | 0 |
| 0 | 0 | 0 |
| 0 | 0 | 0 |
| 0 | 0 | 0 |
| 0 | 0 | 0 |
| 0 | 0 | 0 |
| 0 | 0 | 0 |
| 0 | 0 | 0 |
| 0 | 0 | 0 |
| 0 | 0 | 0 |
| 0 | 0 | 0 |
| 0 | 0 | 0 |
| 0 | 0 | 0 |
| 0 | 0 | 0 |
| 1 | 1 | 1 |
| 0 | 1 | 0 |
| 0 | 0 | 0 |
| 0 | 0 | 0 |
| 0 | 0 | 0 |
| 0 | 0 | 0 |
| 1 | 1 | 1 |
| 1 | 1 | 1 |
| 0 | 0 | 0 |

| | | |
|---|---|---|
| 0 | 1 | 0 |
| 0 | 1 | 0 |
| 0 | 0 | 0 |
| 0 | 0 | 0 |
| 0 | 0 | 0 |
| 0 | 0 | 0 |
| 0 | 1 | 0 |
| 0 | 0 | 0 |
| 0 | 0 | 0 |
| 0 | 0 | 0 |
| 0 | 0 | 0 |
| 0 | 0 | 0 |
| 0 | 0 | 0 |
| 0 | 0 | 0 |
| 0 | 0 | 0 |
| 1 | 1 | 1 |
| 0 | 0 | 0 |
| 0 | 0 | 0 |
| 0 | 0 | 0 |
| 0 | 0 | 0 |
| 0 | 0 | 0 |
| 0 | 0 | 0 |
| 0 | 0 | 0 |
| 0 | 0 | 0 |
| 0 | 0 | 0 |
| 0 | 0 | 0 |
| 0 | 0 | 0 |
| 0 | 0 | 0 |
| 0 | 0 | 0 |
| 0 | 0 | 0 |
| 0 | 0 | 0 |
| 0 | 0 | 0 |
| 0 | 0 | 0 |
| 0 | 0 | 0 |
| 0 | 0 | 0 |
| 0 | 0 | 0 |
| 0 | 0 | 0 |
| 0 | 0 | 0 |
| 0 | 0 | 0 |
| 0 | 0 | 0 |
| 0 | 0 | 0 |
| 0 | 0 | 0 |
| 0 | 0 | 0 |
| 0 | 0 | 0 |
| 0 | 0 | 0 |
| 0 | 0 | 0 |
| 0 | 0 | 0 |
| 0 | 0 | 0 |
| 0 | 0 | 0 |
| 0 | 0 | 0 |
| 0 | 0 | 0 |
| 0 | 0 | 0 |
| 0 | 0 | 0 |
| 0 | 0 | 0 |

| | | |
|---|---|---|
| 1 | 0 | 0 |
| 1 | 0 | 0 |
| 1 | 0 | 0 |
| 0 | 0 | 0 |
| 0 | 0 | 0 |
| 0 | 0 | 0 |
| 0 | 0 | 0 |
| 1 | 0 | 0 |
| 0 | 0 | 0 |
| 1 | 0 | 0 |
| 1 | 0 | 0 |
| 0 | 0 | 0 |
| 1 | 0 | 0 |
| 0 | 0 | 0 |
| 0 | 0 | 0 |
| 1 | 0 | 0 |
| 1 | 0 | 0 |
| 1 | 0 | 0 |
| 1 | 0 | 0 |
| 0 | 0 | 0 |
| 0 | 0 | 0 |
| 0 | 1 | 0 |
| 0 | 0 | 0 |
| 0 | 0 | 0 |
| 0 | 0 | 0 |
| 1 | 0 | 0 |
| 0 | 0 | 0 |
| 0 | 0 | 0 |
| 1 | 0 | 0 |
| 1 | 0 | 0 |
| 1 | 1 | 1 |
| 0 | 0 | 0 |
| 0 | 0 | 0 |
| 0 | 1 | 0 |
| 0 | 0 | 0 |
| 0 | 0 | 0 |
| 0 | 0 | 0 |
| 0 | 0 | 0 |
| 0 | 0 | 0 |
| 1 | 0 | 0 |
| 1 | 0 | 0 |
| 0 | 0 | 0 |
| 0 | 0 | 0 |
| 1 | 1 | 1 |
| 1 | 0 | 0 |
| 1 | 0 | 0 |
| 0 | 0 | 0 |
| 0 | 0 | 0 |
| 0 | 0 | 0 |
| 0 | 0 | 0 |
| 0 | 0 | 0 |
| 0 | 1 | 0 |
| 1 | 0 | 0 |
| 0 | 0 | 0 |

| | | |
|---|---|---|
| 0 | 0 | 0 |
| 0 | 0 | 0 |
| 0 | 0 | 0 |
| 1 | 1 | 1 |
| 1 | 1 | 1 |
| 1 | 0 | 0 |
| 0 | 0 | 0 |
| 0 | 0 | 0 |
| 1 | 0 | 0 |
| 1 | 0 | 0 |
| 0 | 0 | 0 |
| 0 | 1 | 0 |
| 0 | 0 | 0 |
| 1 | 0 | 0 |
| 0 | 1 | 0 |
| 1 | 1 | 1 |
| 1 | 1 | 1 |
| 0 | 0 | 0 |
| 0 | 0 | 0 |
| 0 | 0 | 0 |
| 0 | 0 | 0 |
| 1 | 0 | 0 |
| 0 | 0 | 0 |
| 0 | 1 | 0 |
| 1 | 1 | 1 |
| 0 | 0 | 0 |
| 0 | 0 | 0 |
| 0 | 0 | 0 |
| 0 | 0 | 0 |
| 0 | 0 | 0 |
| 0 | 0 | 0 |
| 0 | 0 | 0 |
| 1 | 0 | 0 |
| 1 | 0 | 0 |
| 0 | 0 | 0 |
| 1 | 0 | 0 |
| 0 | 0 | 0 |
| 0 | 0 | 0 |
| 1 | 0 | 0 |
| 0 | 1 | 0 |
| 0 | 0 | 0 |
| 1 | 0 | 0 |
| 1 | 0 | 0 |
| 1 | 0 | 0 |
| 0 | 0 | 0 |
| 0 | 0 | 0 |
| 0 | 0 | 0 |
| 0 | 0 | 0 |
| 0 | 0 | 0 |
| 0 | 0 | 0 |
| 0 | 0 | 0 |
| 0 | 0 | 0 |
| 0 | 0 | 0 |
| 0 | 0 | 0 |

| | | |
|---|---|---|
| 0 | 0 | 0 |
| 0 | 0 | 0 |
| 0 | 0 | 0 |
| 0 | 0 | 0 |
| 0 | 0 | 0 |
| 0 | 0 | 0 |
| 0 | 1 | 0 |
| 0 | 1 | 0 |
| 0 | 0 | 0 |
| 0 | 0 | 0 |
| 0 | 0 | 0 |
| 0 | 0 | 0 |
| 0 | 0 | 0 |
| 0 | 1 | 0 |
| 0 | 1 | 0 |
| 0 | 0 | 0 |
| 0 | 1 | 0 |
| 0 | 0 | 0 |
| 1 | 1 | 1 |
| 0 | 0 | 0 |
| 1 | 1 | 1 |
| 1 | 0 | 0 |
| 1 | 1 | 1 |
| 0 | 0 | 0 |
| 0 | 0 | 0 |
| 0 | 0 | 0 |
| 0 | 0 | 0 |
| 0 | 0 | 0 |
| 0 | 0 | 0 |
| 0 | 0 | 0 |
| 0 | 0 | 0 |
| 1 | 0 | 0 |
| 0 | 0 | 0 |
| 0 | 0 | 0 |
| 0 | 0 | 0 |
| 0 | 0 | 0 |
| 0 | 0 | 0 |
| 0 | 0 | 0 |
| 0 | 0 | 0 |
| 0 | 0 | 0 |
| 0 | 1 | 0 |
| 0 | 1 | 0 |
| 0 | 0 | 0 |
| 0 | 1 | 0 |
| 0 | 0 | 0 |
| 0 | 0 | 0 |
| 0 | 0 | 0 |
| 1 | 1 | 1 |
| 1 | 1 | 1 |
| 0 | 0 | 0 |
| 0 | 1 | 0 |
| 1 | 1 | 1 |
| 0 | 0 | 0 |
| 0 | 1 | 0 |

| | | |
|---|---|---|
| 0 | 0 | 0 |
| 1 | 1 | 1 |
| 0 | 0 | 0 |
| 0 | 0 | 0 |
| 0 | 0 | 0 |
| 0 | 0 | 0 |
| 0 | 1 | 0 |
| 0 | 0 | 0 |
| 0 | 0 | 0 |
| 1 | 1 | 1 |
| 1 | 1 | 1 |
| 1 | 1 | 1 |
| 1 | 1 | 1 |
| 1 | 1 | 1 |
| 0 | 0 | 0 |
| 1 | 1 | 1 |
| 0 | 0 | 0 |
| 0 | 0 | 0 |
| 1 | 1 | 1 |
| 0 | 0 | 0 |
| 0 | 0 | 0 |
| 0 | 0 | 0 |
| 0 | 0 | 0 |
| 0 | 0 | 0 |
| 1 | 1 | 1 |
| 1 | 0 | 0 |
| 0 | 0 | 0 |
| 0 | 0 | 0 |
| 0 | 0 | 0 |
| 0 | 0 | 0 |
| 0 | 0 | 0 |
| 0 | 0 | 0 |
| 0 | 0 | 0 |
| 0 | 0 | 0 |
| 1 | 1 | 1 |
| 1 | 1 | 1 |
| 0 | 0 | 0 |
| 0 | 0 | 0 |
| 0 | 0 | 0 |
| 0 | 0 | 0 |
| 0 | 0 | 0 |
| 0 | 0 | 0 |
| 0 | 1 | 0 |
| 0 | 0 | 0 |
| 0 | 0 | 0 |
| 0 | 0 | 0 |
| 0 | 0 | 0 |
| 0 | 0 | 0 |
| 0 | 0 | 0 |
| 1 | 1 | 1 |
| 0 | 0 | 0 |
| 0 | 0 | 0 |
| 0 | 0 | 0 |
| 0 | 0 | 0 |

| | | |
|---|---|---|
| 0 | 0 | 0 |
| 0 | 0 | 0 |
| 0 | 0 | 0 |
| 0 | 0 | 0 |
| 1 | 1 | 1 |
| 0 | 0 | 0 |
| 0 | 0 | 0 |
| 0 | 0 | 0 |
| 0 | 0 | 0 |
| 0 | 0 | 0 |
| 0 | 0 | 0 |
| 0 | 0 | 0 |
| 0 | 0 | 0 |
| 0 | 0 | 0 |
| 0 | 0 | 0 |
| 0 | 0 | 0 |
| 0 | 0 | 0 |
| 0 | 0 | 0 |
| 1 | 1 | 1 |
| 0 | 0 | 0 |
| 0 | 0 | 0 |
| 0 | 0 | 0 |
| 0 | 0 | 0 |
| 1 | 0 | 0 |
| 0 | 0 | 0 |
| 0 | 0 | 0 |
| 0 | 0 | 0 |
| 1 | 0 | 0 |
| 1 | 0 | 0 |
| 1 | 0 | 0 |
| 0 | 0 | 0 |
| 0 | 0 | 0 |
| 0 | 0 | 0 |
| 0 | 0 | 0 |
| 0 | 0 | 0 |
| 1 | 0 | 0 |
| 1 | 0 | 0 |
| 0 | 0 | 0 |
| 0 | 0 | 0 |
| 1 | 0 | 0 |
| 1 | 0 | 0 |
| 1 | 0 | 0 |
| 0 | 1 | 0 |
| 1 | 1 | 1 |
| 0 | 0 | 0 |
| 0 | 0 | 0 |
| 0 | 0 | 0 |
| 0 | 0 | 0 |
| 0 | 0 | 0 |
| 0 | 1 | 0 |
| 1 | 1 | 1 |
| 0 | 0 | 0 |
| 1 | 1 | 1 |
| 0 | 0 | 0 |

| | | |
|---|---|---|
| 1 | 1 | 1 |
| 1 | 1 | 1 |
| 0 | 0 | 0 |
| 1 | 1 | 1 |
| 1 | 1 | 1 |
| 1 | 1 | 1 |
| 1 | 1 | 1 |
| 0 | 0 | 0 |
| 1 | 0 | 0 |
| 1 | 1 | 1 |
| 0 | 0 | 0 |
| 1 | 1 | 1 |
| 1 | 1 | 1 |
| 0 | 1 | 0 |
| 0 | 0 | 0 |
| 0 | 0 | 0 |
| 1 | 1 | 1 |
| 1 | 1 | 1 |
| 0 | 0 | 0 |
| 0 | 0 | 0 |
| 0 | 0 | 0 |
| 0 | 0 | 0 |
| 0 | 0 | 0 |
| 0 | 0 | 0 |
| 0 | 0 | 0 |
| 1 | 1 | 1 |
| 0 | 0 | 0 |
| 0 | 0 | 0 |
| 0 | 0 | 0 |
| 1 | 0 | 0 |
| 0 | 0 | 0 |
| 0 | 0 | 0 |
| 0 | 0 | 0 |
| 0 | 0 | 0 |
| 0 | 0 | 0 |
| 0 | 0 | 0 |
| 0 | 0 | 0 |
| 0 | 0 | 0 |
| 0 | 0 | 0 |
| 0 | 0 | 0 |
| 0 | 1 | 0 |
| 0 | 0 | 0 |
| 1 | 1 | 1 |
| 0 | 0 | 0 |
| 0 | 0 | 0 |
| 0 | 0 | 0 |
| 0 | 0 | 0 |
| 0 | 0 | 0 |
| 0 | 0 | 0 |
| 0 | 0 | 0 |
| 0 | 1 | 0 |
| 0 | 0 | 0 |
| 0 | 0 | 0 |
| 0 | 0 | 0 |

| | | |
|---|---|---|
| 1 | 1 | 1 |
| 0 | 0 | 0 |
| 0 | 1 | 0 |
| 0 | 1 | 0 |
| 0 | 0 | 0 |
| 0 | 0 | 0 |
| 0 | 0 | 0 |
| 0 | 0 | 0 |
| 0 | 0 | 0 |
| 0 | 1 | 0 |
| 1 | 1 | 1 |
| 1 | 1 | 1 |
| 1 | 1 | 1 |
| 0 | 0 | 0 |
| 1 | 1 | 1 |
| 0 | 0 | 0 |
| 0 | 0 | 0 |
| 0 | 0 | 0 |
| 0 | 0 | 0 |
| 1 | 0 | 0 |
| 0 | 0 | 0 |
| 1 | 1 | 1 |
| 0 | 0 | 0 |
| 0 | 1 | 0 |
| 1 | 1 | 1 |
| 1 | 1 | 1 |
| 0 | 0 | 0 |
| 1 | 1 | 1 |
| 1 | 1 | 1 |
| 1 | 1 | 1 |
| 0 | 0 | 0 |
| 0 | 0 | 0 |
| 1 | 1 | 1 |
| 1 | 1 | 1 |
| 0 | 0 | 0 |
| 0 | 0 | 0 |
| 0 | 0 | 0 |
| 0 | 0 | 0 |
| 0 | 0 | 0 |
| 1 | 1 | 1 |
| 0 | 0 | 0 |
| 0 | 0 | 0 |
| 0 | 0 | 0 |
| 0 | 0 | 0 |
| 1 | 0 | 0 |
| 0 | 0 | 0 |
| 0 | 0 | 0 |
| 0 | 0 | 0 |
| 0 | 0 | 0 |
| 0 | 0 | 0 |
| 0 | 0 | 0 |
| 0 | 0 | 0 |
| 1 | 0 | 0 |
| 0 | 0 | 0 |

| | | |
|---|---|---|
| 0 | 0 | 0 |
| 0 | 0 | 0 |
| 0 | 0 | 0 |
| 0 | 0 | 0 |
| 0 | 0 | 0 |
| 0 | 0 | 0 |
| 0 | 0 | 0 |
| 1 | 0 | 0 |
| 0 | 0 | 0 |
| 0 | 0 | 0 |
| 0 | 0 | 0 |
| 0 | 0 | 0 |
| 1 | 1 | 1 |
| 0 | 0 | 0 |
| 0 | 0 | 0 |
| 0 | 0 | 0 |
| 0 | 0 | 0 |
| 0 | 0 | 0 |
| 0 | 0 | 0 |
| 1 | 1 | 1 |
| 0 | 0 | 0 |
| 0 | 0 | 0 |
| 0 | 0 | 0 |
| 0 | 0 | 0 |
| 0 | 0 | 0 |
| 1 | 0 | 0 |
| 1 | 1 | 1 |
| 0 | 1 | 0 |
| 0 | 0 | 0 |
| 0 | 0 | 0 |
| 0 | 0 | 0 |
| 0 | 0 | 0 |
| 0 | 0 | 0 |
| 0 | 0 | 0 |
| 0 | 0 | 0 |
| 1 | 0 | 0 |
| 0 | 0 | 0 |
| 0 | 0 | 0 |
| 1 | 1 | 1 |
| 1 | 0 | 0 |
| 0 | 1 | 0 |
| 0 | 0 | 0 |
| 0 | 0 | 0 |
| 0 | 0 | 0 |
| 0 | 0 | 0 |
| 0 | 0 | 0 |
| 0 | 0 | 0 |
| 1 | 1 | 1 |
| 1 | 1 | 1 |
| 0 | 0 | 0 |
| 0 | 0 | 0 |
| 0 | 0 | 0 |
| 0 | 0 | 0 |
| 1 | 1 | 1 |

| | | |
|---|---|---|
| 1 | 1 | 1 |
| 0 | 0 | 0 |
| 0 | 1 | 0 |
| 0 | 0 | 0 |
| 1 | 0 | 0 |
| 1 | 0 | 0 |
| 0 | 0 | 0 |
| 1 | 0 | 0 |
| 1 | 0 | 0 |
| 1 | 0 | 0 |
| 1 | 0 | 0 |
| 1 | 0 | 0 |
| 1 | 1 | 1 |
| 1 | 1 | 1 |
| 1 | 1 | 1 |
| 1 | 1 | 1 |
| 0 | 0 | 0 |
| 0 | 0 | 0 |
| 1 | 0 | 0 |
| 0 | 0 | 0 |
| 0 | 0 | 0 |
| 0 | 0 | 0 |
| 1 | 0 | 0 |
| 0 | 0 | 0 |
| 0 | 0 | 0 |
| 1 | 1 | 1 |
| 1 | 0 | 0 |
| 1 | 0 | 0 |
| 0 | 0 | 0 |
| 0 | 0 | 0 |
| 0 | 0 | 0 |
| 0 | 0 | 0 |
| 0 | 0 | 0 |
| 0 | 0 | 0 |
| 0 | 0 | 0 |
| 0 | 0 | 0 |
| 1 | 0 | 0 |
| 1 | 1 | 1 |
| 1 | 1 | 1 |
| 1 | 1 | 1 |
| 0 | 0 | 0 |
| 1 | 1 | 1 |
| 0 | 0 | 0 |
| 1 | 1 | 1 |
| 0 | 0 | 0 |
| 0 | 1 | 0 |
| 0 | 0 | 0 |
| 0 | 0 | 0 |
| 0 | 0 | 0 |
| 0 | 0 | 0 |
| 0 | 0 | 0 |
| 0 | 0 | 0 |
| 0 | 0 | 0 |
| 0 | 0 | 0 |

| | | |
|---|---|---|
| 0 | 0 | 0 |
| 0 | 0 | 0 |
| 0 | 0 | 0 |
| 0 | 0 | 0 |
| 0 | 0 | 0 |
| 0 | 0 | 0 |
| 0 | 0 | 0 |
| 1 | 1 | 1 |
| 0 | 0 | 0 |
| 0 | 0 | 0 |
| 0 | 0 | 0 |
| 1 | 0 | 0 |
| 0 | 0 | 0 |
| 0 | 0 | 0 |
| 0 | 0 | 0 |
| 0 | 0 | 0 |
| 0 | 0 | 0 |
| 0 | 0 | 0 |
| 0 | 0 | 0 |
| 0 | 0 | 0 |
| 0 | 0 | 0 |
| 0 | 0 | 0 |
| 0 | 0 | 0 |
| 0 | 0 | 0 |
| 0 | 0 | 0 |
| 0 | 0 | 0 |
| 0 | 0 | 0 |
| 0 | 0 | 0 |
| 0 | 0 | 0 |
| 0 | 0 | 0 |
| 0 | 0 | 0 |
| 0 | 0 | 0 |
| 0 | 0 | 0 |
| 0 | 0 | 0 |
| 0 | 0 | 0 |
| 0 | 0 | 0 |
| 0 | 0 | 0 |
| 0 | 0 | 0 |
| 0 | 0 | 0 |
| 0 | 0 | 0 |
| 1 | 0 | 0 |
| 0 | 0 | 0 |
| 0 | 0 | 0 |
| 0 | 0 | 0 |
| 0 | 0 | 0 |
| 0 | 0 | 0 |
| 0 | 0 | 0 |
| 0 | 0 | 0 |
| 0 | 0 | 0 |
| 0 | 1 | 0 |
| 0 | 1 | 0 |
| 1 | 0 | 0 |
| 0 | 0 | 0 |
| 0 | 0 | 0 |

| | | |
|---|---|---|
| 0 | 0 | 0 |
| 0 | 0 | 0 |
| 0 | 0 | 0 |
| 0 | 0 | 0 |
| 0 | 0 | 0 |
| 0 | 0 | 0 |
| 0 | 0 | 0 |
| 1 | 0 | 0 |
| 1 | 0 | 0 |
| 0 | 0 | 0 |
| 0 | 0 | 0 |
| 1 | 0 | 0 |
| 0 | 0 | 0 |
| 0 | 0 | 0 |
| 0 | 0 | 0 |
| 0 | 0 | 0 |
| 0 | 0 | 0 |
| 1 | 0 | 0 |
| 0 | 0 | 0 |
| 1 | 0 | 0 |
| 0 | 0 | 0 |
| 0 | 0 | 0 |
| 1 | 1 | 1 |
| 0 | 0 | 0 |
| 0 | 0 | 0 |
| 1 | 0 | 0 |
| 0 | 0 | 0 |
| 0 | 0 | 0 |
| 0 | 0 | 0 |
| 0 | 0 | 0 |
| 0 | 0 | 0 |
| 0 | 0 | 0 |
| 1 | 0 | 0 |
| 0 | 0 | 0 |
| 0 | 0 | 0 |
| 0 | 0 | 0 |
| 1 | 1 | 1 |
| 0 | 0 | 0 |
| 0 | 0 | 0 |
| 0 | 0 | 0 |
| 0 | 0 | 0 |
| 0 | 0 | 0 |
| 0 | 0 | 0 |
| 0 | 0 | 0 |
| 0 | 0 | 0 |
| 0 | 0 | 0 |
| 0 | 0 | 0 |
| 0 | 0 | 0 |
| 0 | 0 | 0 |
| 0 | 0 | 0 |
| 0 | 0 | 0 |
| 0 | 0 | 0 |
| 0 | 0 | 0 |
| 0 | 0 | 0 |

| | | |
|---|---|---|
| 1 | 0 | 0 |
| 0 | 1 | 0 |
| 0 | 0 | 0 |
| 0 | 0 | 0 |
| 0 | 0 | 0 |
| 0 | 0 | 0 |
| 0 | 0 | 0 |
| 0 | 0 | 0 |
| 0 | 0 | 0 |
| 1 | 0 | 0 |
| 0 | 1 | 0 |
| 1 | 1 | 1 |
| 0 | 0 | 0 |
| 0 | 0 | 0 |
| 0 | 0 | 0 |
| 0 | 0 | 0 |
| 0 | 0 | 0 |
| 0 | 0 | 0 |
| 1 | 1 | 1 |
| 1 | 0 | 0 |
| 1 | 0 | 0 |
| 0 | 0 | 0 |
| 0 | 1 | 0 |
| 0 | 0 | 0 |
| 0 | 0 | 0 |
| 0 | 0 | 0 |
| 0 | 0 | 0 |
| 0 | 0 | 0 |
| 0 | 0 | 0 |
| 0 | 0 | 0 |
| 0 | 0 | 0 |
| 0 | 0 | 0 |
| 0 | 0 | 0 |
| 0 | 0 | 0 |
| 0 | 0 | 0 |
| 0 | 0 | 0 |
| 0 | 0 | 0 |
| 0 | 0 | 0 |
| 0 | 0 | 0 |
| 0 | 0 | 0 |
| 0 | 0 | 0 |
| 1 | 0 | 0 |
| 0 | 0 | 0 |
| 0 | 0 | 0 |
| 0 | 0 | 0 |
| 0 | 0 | 0 |
| 1 | 0 | 0 |
| 0 | 0 | 0 |
| 0 | 0 | 0 |
| 0 | 0 | 0 |
| 0 | 0 | 0 |
| 0 | 0 | 0 |
| 1 | 0 | 0 |
| 0 | 1 | 0 |

| | | |
|---|---|---|
| 0 | 0 | 0 |
| 0 | 0 | 0 |
| 1 | 0 | 0 |
| 1 | 0 | 0 |
| 1 | 0 | 0 |
| 0 | 0 | 0 |
| 1 | 0 | 0 |
| 0 | 0 | 0 |
| 0 | 0 | 0 |
| 0 | 0 | 0 |
| 0 | 0 | 0 |
| 0 | 0 | 0 |
| 0 | 0 | 0 |
| 1 | 0 | 0 |
| 1 | 0 | 0 |
| 1 | 0 | 0 |
| 0 | 0 | 0 |
| 1 | 1 | 1 |
| 0 | 0 | 0 |
| 0 | 0 | 0 |
| 1 | 0 | 0 |
| 1 | 0 | 0 |
| 0 | 0 | 0 |
| 0 | 0 | 0 |
| 1 | 0 | 0 |
| 0 | 0 | 0 |
| 0 | 0 | 0 |
| 1 | 0 | 0 |
| 0 | 0 | 0 |
| 0 | 0 | 0 |
| 0 | 0 | 0 |
| 0 | 0 | 0 |
| 0 | 0 | 0 |
| 0 | 0 | 0 |
| 0 | 0 | 0 |
| 0 | 0 | 0 |
| 0 | 0 | 0 |
| 0 | 0 | 0 |
| 0 | 0 | 0 |
| 0 | 0 | 0 |
| 1 | 0 | 0 |
| 1 | 0 | 0 |
| 1 | 0 | 0 |
| 1 | 1 | 1 |
| 1 | 1 | 1 |
| 0 | 0 | 0 |
| 0 | 0 | 0 |
| 0 | 0 | 0 |
| 0 | 0 | 0 |
| 0 | 0 | 0 |
| 0 | 0 | 0 |
| 0 | 0 | 0 |
| 0 | 0 | 0 |
| 1 | 0 | 0 |

| | | |
|---|---|---|
| 1 | 0 | 0 |
| 0 | 0 | 0 |
| 0 | 0 | 0 |
| 0 | 0 | 0 |
| 1 | 0 | 0 |
| 0 | 0 | 0 |
| 0 | 0 | 0 |
| 0 | 0 | 0 |
| 1 | 0 | 0 |
| 1 | 0 | 0 |
| 0 | 0 | 0 |
| 0 | 0 | 0 |
| 0 | 0 | 0 |
| 1 | 0 | 0 |
| 0 | 0 | 0 |
| 0 | 0 | 0 |
| 0 | 0 | 0 |
| 0 | 0 | 0 |
| 0 | 0 | 0 |
| 0 | 0 | 0 |
| 0 | 0 | 0 |
| 1 | 1 | 1 |
| 0 | 0 | 0 |
| 0 | 1 | 0 |
| 0 | 0 | 0 |
| 0 | 0 | 0 |
| 0 | 0 | 0 |
| 0 | 0 | 0 |
| 0 | 0 | 0 |
| 0 | 0 | 0 |
| 0 | 0 | 0 |
| 0 | 0 | 0 |
| 0 | 0 | 0 |
| 0 | 0 | 0 |
| 0 | 1 | 0 |
| 0 | 0 | 0 |
| 0 | 0 | 0 |
| 0 | 0 | 0 |
| 0 | 0 | 0 |
| 0 | 0 | 0 |
| 0 | 0 | 0 |
| 0 | 0 | 0 |
| 0 | 0 | 0 |
| 0 | 0 | 0 |
| 0 | 0 | 0 |
| 0 | 0 | 0 |
| 0 | 0 | 0 |
| 1 | 1 | 1 |
| 0 | 0 | 0 |
| 1 | 1 | 1 |
| 0 | 0 | 0 |
| 0 | 0 | 0 |
| 0 | 0 | 0 |
| 0 | 0 | 0 |

| | | |
|---|---|---|
| 0 | 0 | 0 |
| 0 | 0 | 0 |
| 1 | 1 | 1 |
| 0 | 0 | 0 |
| 0 | 0 | 0 |
| 0 | 1 | 0 |
| 0 | 1 | 0 |
| 0 | 1 | 0 |
| 0 | 0 | 0 |
| 0 | 1 | 0 |
| 0 | 0 | 0 |
| 0 | 0 | 0 |
| 0 | 0 | 0 |
| 1 | 1 | 1 |
| 0 | 0 | 0 |
| 0 | 1 | 0 |
| 0 | 0 | 0 |
| 0 | 0 | 0 |
| 0 | 0 | 0 |
| 1 | 1 | 1 |
| 1 | 1 | 1 |
| 0 | 0 | 0 |
| 1 | 1 | 1 |
| 0 | 0 | 0 |
| 0 | 0 | 0 |
| 1 | 1 | 1 |
| 1 | 1 | 1 |
| 0 | 0 | 0 |
| 1 | 1 | 1 |
| 0 | 0 | 0 |
| 0 | 1 | 0 |
| 0 | 1 | 0 |
| 1 | 1 | 1 |
| 0 | 1 | 0 |
| 1 | 1 | 1 |
| 1 | 1 | 1 |
| 1 | 1 | 1 |
| 1 | 1 | 1 |
| 1 | 1 | 1 |
| 1 | 1 | 1 |
| 1 | 1 | 1 |
| 1 | 1 | 1 |
| 0 | 1 | 0 |
| 0 | 1 | 0 |
| 0 | 1 | 0 |
| 0 | 1 | 0 |
| 0 | 1 | 0 |
| 0 | 1 | 0 |
| 0 | 1 | 0 |
| 0 | 1 | 0 |
| 0 | 1 | 0 |
| 0 | 1 | 0 |
| 0 | 1 | 0 |
| 0 | 1 | 0 |

| | | |
|---|---|---|
| 0 | 1 | 0 |
| 1 | 1 | 1 |
| 1 | 1 | 1 |
| 1 | 1 | 1 |
| 1 | 1 | 1 |
| 1 | 1 | 1 |
| 1 | 1 | 1 |
| 0 | 1 | 0 |
| 0 | 1 | 0 |
| 0 | 1 | 0 |
| 0 | 1 | 0 |
| 0 | 0 | 0 |
| 0 | 0 | 0 |
| 0 | 0 | 0 |
| 1 | 0 | 0 |
| 0 | 0 | 0 |
| 1 | 0 | 0 |
| 0 | 0 | 0 |
| 1 | 0 | 0 |
| 1 | 1 | 1 |
| 0 | 0 | 0 |
| 1 | 1 | 1 |
| 1 | 1 | 1 |
| 0 | 0 | 0 |
| 0 | 0 | 0 |
| 1 | 1 | 1 |
| 1 | 1 | 1 |
| 0 | 0 | 0 |
| 0 | 0 | 0 |
| 0 | 0 | 0 |
| 0 | 0 | 0 |
| 0 | 0 | 0 |
| 0 | 1 | 0 |
| 0 | 1 | 0 |
| 1 | 1 | 1 |
| 1 | 1 | 1 |
| 1 | 1 | 1 |
| 1 | 1 | 1 |
| 1 | 1 | 1 |
| 0 | 0 | 0 |
| 0 | 0 | 0 |
| 0 | 1 | 0 |
| 0 | 0 | 0 |
| 0 | 0 | 0 |
| 0 | 0 | 0 |
| 1 | 0 | 0 |
| 0 | 0 | 0 |
| 1 | 0 | 0 |
| 1 | 0 | 0 |
| 0 | 0 | 0 |
| 0 | 0 | 0 |
| 0 | 0 | 0 |
| 0 | 0 | 0 |
| 0 | 0 | 0 |

| | | |
|---|---|---|
| 0 | 0 | 0 |
| 0 | 1 | 0 |
| 1 | 1 | 1 |
| 1 | 1 | 1 |
| 0 | 0 | 0 |
| 1 | 1 | 1 |
| 0 | 0 | 0 |
| 1 | 1 | 1 |
| 1 | 1 | 1 |
| 0 | 0 | 0 |
| 1 | 0 | 0 |
| 1 | 1 | 1 |
| 1 | 1 | 1 |
| 0 | 0 | 0 |
| 1 | 0 | 0 |
| 1 | 0 | 0 |
| 0 | 0 | 0 |
| 1 | 0 | 0 |
| 0 | 0 | 0 |
| 0 | 0 | 0 |
| 0 | 0 | 0 |
| 0 | 0 | 0 |
| 0 | 0 | 0 |
| 0 | 0 | 0 |
| 0 | 0 | 0 |
| 0 | 0 | 0 |
| 0 | 0 | 0 |
| 0 | 0 | 0 |
| 0 | 0 | 0 |
| 0 | 0 | 0 |
| 0 | 0 | 0 |
| 0 | 0 | 0 |
| 0 | 0 | 0 |
| 0 | 0 | 0 |
| 0 | 0 | 0 |
| 0 | 0 | 0 |
| 1 | 0 | 0 |
| 0 | 0 | 0 |
| 0 | 0 | 0 |
| 0 | 0 | 0 |
| 0 | 0 | 0 |
| 0 | 0 | 0 |
| 0 | 0 | 0 |
| 0 | 0 | 0 |
| 0 | 0 | 0 |
| 0 | 0 | 0 |
| 0 | 0 | 0 |
| 0 | 0 | 0 |
| 1 | 0 | 0 |
| 1 | 0 | 0 |
| 0 | 0 | 0 |
| 1 | 1 | 1 |
| 0 | 0 | 0 |
| 0 | 0 | 0 |

| | | |
|---|---|---|
| 0 | 0 | 0 |
| 0 | 0 | 0 |
| 0 | 0 | 0 |
| 0 | 0 | 0 |
| 0 | 0 | 0 |
| 0 | 0 | 0 |
| 0 | 0 | 0 |
| 0 | 0 | 0 |
| 0 | 0 | 0 |
| 0 | 0 | 0 |
| 0 | 0 | 0 |
| 0 | 0 | 0 |
| 0 | 0 | 0 |
| 0 | 0 | 0 |
| 0 | 1 | 0 |
| 0 | 0 | 0 |
| 0 | 0 | 0 |
| 0 | 0 | 0 |
| 1 | 0 | 0 |
| 0 | 0 | 0 |
| 0 | 0 | 0 |
| 0 | 0 | 0 |
| 0 | 0 | 0 |
| 0 | 0 | 0 |
| 0 | 0 | 0 |
| 1 | 0 | 0 |
| 0 | 0 | 0 |
| 0 | 0 | 0 |
| 0 | 0 | 0 |
| 0 | 0 | 0 |
| 1 | 1 | 1 |
| 0 | 0 | 0 |
| 1 | 1 | 1 |
| 1 | 0 | 0 |
| 0 | 0 | 0 |
| 0 | 0 | 0 |
| 1 | 1 | 1 |
| 1 | 1 | 1 |
| 1 | 1 | 1 |
| 1 | 0 | 0 |
| 0 | 0 | 0 |
| 0 | 0 | 0 |
| 0 | 0 | 0 |
| 0 | 0 | 0 |
| 0 | 0 | 0 |
| 0 | 0 | 0 |
| 0 | 0 | 0 |
| 0 | 0 | 0 |
| 1 | 1 | 1 |
| 1 | 1 | 1 |
| 0 | 0 | 0 |
| 0 | 0 | 0 |
| 0 | 0 | 0 |
| 0 | 1 | 0 |

| | | |
|---|---|---|
| 1 | 1 | 1 |
| 1 | 1 | 1 |
| 1 | 0 | 0 |
| 0 | 0 | 0 |
| 1 | 0 | 0 |
| 1 | 1 | 1 |
| 0 | 0 | 0 |
| 1 | 1 | 1 |
| 0 | 0 | 0 |
| 0 | 0 | 0 |
| 0 | 0 | 0 |
| 0 | 0 | 0 |
| 0 | 0 | 0 |
| 0 | 0 | 0 |
| 0 | 0 | 0 |
| 0 | 0 | 0 |
| 0 | 1 | 0 |
| 0 | 0 | 0 |
| 0 | 0 | 0 |
| 1 | 0 | 0 |
| 0 | 0 | 0 |
| 0 | 0 | 0 |
| 1 | 1 | 1 |
| 0 | 0 | 0 |
| 1 | 1 | 1 |
| 1 | 1 | 1 |
| 0 | 1 | 0 |
| 0 | 0 | 0 |
| 0 | 0 | 0 |
| 0 | 0 | 0 |
| 1 | 1 | 1 |
| 0 | 0 | 0 |
| 0 | 0 | 0 |
| 0 | 0 | 0 |
| 0 | 1 | 0 |
| 0 | 1 | 0 |
| 0 | 0 | 0 |
| 0 | 0 | 0 |
| 0 | 0 | 0 |
| 1 | 1 | 1 |
| 1 | 1 | 1 |
| 0 | 1 | 0 |
| 0 | 1 | 0 |
| 0 | 0 | 0 |
| 1 | 1 | 1 |
| 0 | 0 | 0 |
| 1 | 1 | 1 |
| 0 | 0 | 0 |
| 0 | 0 | 0 |
| 1 | 1 | 1 |
| 0 | 1 | 0 |
| 0 | 0 | 0 |
| 0 | 0 | 0 |
| 1 | 1 | 1 |

| | | |
|---|---|---|
| 1 | 1 | 1 |
| 1 | 1 | 1 |
| 0 | 1 | 0 |
| 0 | 0 | 0 |
| 0 | 0 | 0 |
| 1 | 0 | 0 |
| 1 | 0 | 0 |
| 0 | 0 | 0 |
| 0 | 0 | 0 |
| 1 | 1 | 1 |
| 0 | 0 | 0 |
| 0 | 0 | 0 |
| 0 | 0 | 0 |
| 0 | 0 | 0 |
| 0 | 0 | 0 |
| 0 | 0 | 0 |
| 0 | 0 | 0 |
| 0 | 0 | 0 |
| 0 | 0 | 0 |
| 0 | 1 | 0 |
| 1 | 1 | 1 |
| 1 | 1 | 1 |
| 1 | 1 | 1 |
| 1 | 1 | 1 |
| 0 | 0 | 0 |
| 0 | 1 | 0 |
| 0 | 0 | 0 |
| 0 | 0 | 0 |
| 1 | 1 | 1 |
| 1 | 1 | 1 |
| 1 | 1 | 1 |
| 1 | 1 | 1 |
| 1 | 1 | 1 |
| 0 | 0 | 0 |
| 0 | 0 | 0 |
| 1 | 1 | 1 |
| 0 | 0 | 0 |
| 0 | 1 | 0 |
| 0 | 0 | 0 |
| 0 | 0 | 0 |
| 0 | 0 | 0 |
| 0 | 0 | 0 |
| 0 | 1 | 0 |
| 0 | 0 | 0 |
| 0 | 0 | 0 |
| 0 | 0 | 0 |
| 0 | 0 | 0 |
| 0 | 0 | 0 |
| 0 | 0 | 0 |
| 0 | 0 | 0 |
| 0 | 0 | 0 |
| 0 | 0 | 0 |
| 0 | 0 | 0 |
| 0 | 1 | 0 |

| | | |
|---|---|---|
| 0 | 0 | 0 |
| 0 | 1 | 0 |
| 0 | 0 | 0 |
| 0 | 0 | 0 |
| 0 | 0 | 0 |
| 1 | 1 | 1 |
| 0 | 0 | 0 |
| 1 | 1 | 1 |
| 0 | 1 | 0 |
| 0 | 0 | 0 |
| 0 | 1 | 0 |
| 0 | 1 | 0 |
| 0 | 1 | 0 |
| 0 | 1 | 0 |
| 0 | 1 | 0 |
| 0 | 0 | 0 |
| 0 | 0 | 0 |
| 0 | 1 | 0 |
| 0 | 0 | 0 |
| 1 | 1 | 1 |
| 0 | 0 | 0 |
| 0 | 0 | 0 |
| 0 | 0 | 0 |
| 0 | 0 | 0 |
| 0 | 0 | 0 |
| 0 | 0 | 0 |
| 0 | 0 | 0 |
| 0 | 0 | 0 |
| 0 | 0 | 0 |
| 0 | 0 | 0 |
| 0 | 0 | 0 |
| 0 | 0 | 0 |
| 1 | 0 | 0 |
| 1 | 0 | 0 |
| 0 | 0 | 0 |
| 0 | 0 | 0 |
| 0 | 0 | 0 |
| 0 | 0 | 0 |
| 1 | 0 | 0 |
| 0 | 0 | 0 |
| 0 | 0 | 0 |
| 0 | 0 | 0 |
| 1 | 0 | 0 |
| 0 | 1 | 0 |
| 0 | 0 | 0 |
| 0 | 0 | 0 |
| 0 | 0 | 0 |
| 0 | 0 | 0 |
| 0 | 0 | 0 |
| 0 | 0 | 0 |
| 1 | 0 | 0 |
| 0 | 0 | 0 |
| 0 | 0 | 0 |
| 1 | 0 | 0 |

| | | |
|---|---|---|
| 0 | 0 | 0 |
| 1 | 0 | 0 |
| 0 | 0 | 0 |
| 0 | 0 | 0 |
| 0 | 0 | 0 |
| 0 | 0 | 0 |
| 0 | 0 | 0 |
| 0 | 0 | 0 |
| 0 | 0 | 0 |
| 0 | 0 | 0 |
| 0 | 0 | 0 |
| 1 | 1 | 1 |
| 0 | 0 | 0 |
| 1 | 1 | 1 |
| 1 | 0 | 0 |
| 1 | 0 | 0 |
| 0 | 0 | 0 |
| 0 | 0 | 0 |
| 0 | 0 | 0 |
| 0 | 0 | 0 |
| 1 | 0 | 0 |
| 0 | 0 | 0 |
| 0 | 1 | 0 |
| 0 | 0 | 0 |
| 1 | 1 | 1 |
| 1 | 1 | 1 |
| 1 | 1 | 1 |
| 0 | 0 | 0 |
| 1 | 1 | 1 |
| 1 | 1 | 1 |
| 1 | 1 | 1 |
| 0 | 0 | 0 |
| 1 | 1 | 1 |
| 0 | 0 | 0 |
| 0 | 0 | 0 |
| 0 | 0 | 0 |
| 0 | 1 | 0 |
| 1 | 1 | 1 |
| 0 | 0 | 0 |
| 0 | 0 | 0 |
| 0 | 0 | 0 |
| 0 | 0 | 0 |
| 0 | 0 | 0 |
| 0 | 0 | 0 |
| 0 | 0 | 0 |
| 0 | 0 | 0 |
| 1 | 1 | 1 |
| 1 | 1 | 1 |
| 0 | 0 | 0 |
| 0 | 0 | 0 |
| 0 | 0 | 0 |
| 1 | 1 | 1 |
| 1 | 1 | 1 |
| 0 | 0 | 0 |

| | | |
|---|---|---|
| 0 | 0 | 0 |
| 0 | 1 | 0 |
| 0 | 1 | 0 |
| 1 | 1 | 1 |
| 0 | 0 | 0 |
| 0 | 0 | 0 |
| 0 | 0 | 0 |
| 0 | 0 | 0 |
| 0 | 0 | 0 |
| 0 | 0 | 0 |
| 1 | 1 | 1 |
| 0 | 0 | 0 |
| 0 | 0 | 0 |
| 0 | 0 | 0 |
| 1 | 0 | 0 |
| 1 | 1 | 1 |
| 0 | 0 | 0 |
| 1 | 0 | 0 |
| 0 | 0 | 0 |
| 0 | 0 | 0 |
| 1 | 0 | 0 |
| 0 | 0 | 0 |
| 0 | 0 | 0 |
| 1 | 0 | 0 |
| 0 | 0 | 0 |
| 0 | 0 | 0 |
| 0 | 0 | 0 |
| 1 | 0 | 0 |
| 0 | 0 | 0 |
| 0 | 0 | 0 |
| 0 | 0 | 0 |
| 1 | 1 | 1 |
| 1 | 1 | 1 |
| 1 | 1 | 1 |
| 0 | 1 | 0 |
| 0 | 1 | 0 |
| 0 | 1 | 0 |
| 1 | 1 | 1 |
| 1 | 1 | 1 |
| 1 | 1 | 1 |
| 0 | 1 | 0 |
| 1 | 1 | 1 |
| 0 | 0 | 0 |
| 0 | 1 | 0 |
| 1 | 1 | 1 |
| 1 | 1 | 1 |
| 0 | 1 | 0 |
| 1 | 1 | 1 |
| 1 | 1 | 1 |
| 1 | 1 | 1 |
| 1 | 1 | 1 |
| 1 | 1 | 1 |
| 1 | 1 | 1 |
| 1 | 1 | 1 |

| | | |
|---|---|---|
| 0 | 1 | 0 |
| 0 | 1 | 0 |
| 1 | 1 | 1 |
| 1 | 1 | 1 |
| 1 | 1 | 1 |
| 1 | 1 | 1 |
| 1 | 1 | 1 |
| 1 | 1 | 1 |
| 0 | 0 | 0 |
| 0 | 0 | 0 |
| 0 | 0 | 0 |
| 0 | 0 | 0 |
| 0 | 0 | 0 |
| 0 | 0 | 0 |
| 0 | 0 | 0 |
| 0 | 0 | 0 |
| 0 | 0 | 0 |
| 0 | 0 | 0 |
| 0 | 0 | 0 |
| 0 | 0 | 0 |
| 0 | 0 | 0 |
| 0 | 0 | 0 |
| 0 | 0 | 0 |
| 0 | 0 | 0 |
| 0 | 0 | 0 |
| 1 | 1 | 1 |
| 1 | 1 | 1 |
| 0 | 0 | 0 |
| 0 | 0 | 0 |
| 0 | 0 | 0 |
| 0 | 0 | 0 |
| 0 | 0 | 0 |
| 0 | 0 | 0 |
| 0 | 0 | 0 |
| 0 | 0 | 0 |
| 0 | 0 | 0 |
| 0 | 0 | 0 |
| 0 | 0 | 0 |
| 0 | 0 | 0 |
| 0 | 0 | 0 |
| 0 | 0 | 0 |
| 0 | 0 | 0 |
| 0 | 0 | 0 |
| 0 | 0 | 0 |
| 0 | 0 | 0 |
| 0 | 0 | 0 |
| 1 | 1 | 1 |
| 0 | 1 | 0 |
| 1 | 1 | 1 |
| 1 | 1 | 1 |
| 1 | 1 | 1 |
| 1 | 1 | 1 |
| 0 | 0 | 0 |
| 0 | 0 | 0 |

| | | |
|---|---|---|
| 0 | 0 | 0 |
| 0 | 0 | 0 |
| 0 | 0 | 0 |
| 0 | 0 | 0 |
| 0 | 1 | 0 |
| 0 | 0 | 0 |
| 0 | 0 | 0 |
| 0 | 0 | 0 |
| 1 | 1 | 1 |
| 0 | 0 | 0 |
| 0 | 0 | 0 |
| 0 | 1 | 0 |
| 0 | 0 | 0 |
| 0 | 0 | 0 |
| 0 | 0 | 0 |
| 0 | 0 | 0 |
| 1 | 1 | 1 |
| 0 | 0 | 0 |
| 0 | 0 | 0 |
| 1 | 1 | 1 |
| 0 | 0 | 0 |
| 0 | 0 | 0 |
| 0 | 0 | 0 |
| 0 | 0 | 0 |
| 0 | 0 | 0 |
| 0 | 1 | 0 |
| 0 | 1 | 0 |
| 1 | 1 | 1 |
| 1 | 1 | 1 |
| 0 | 1 | 0 |
| 1 | 1 | 1 |
| 1 | 1 | 1 |
| 1 | 0 | 0 |
| 1 | 0 | 0 |
| 0 | 0 | 0 |
| 0 | 0 | 0 |
| 0 | 0 | 0 |
| 0 | 0 | 0 |
| 0 | 0 | 0 |
| 0 | 0 | 0 |
| 1 | 1 | 1 |
| 1 | 1 | 1 |
| 1 | 1 | 1 |
| 0 | 0 | 0 |
| 0 | 0 | 0 |
| 0 | 0 | 0 |
| 1 | 1 | 1 |
| 0 | 1 | 0 |
| 0 | 0 | 0 |
| 0 | 0 | 0 |
| 1 | 1 | 1 |
| 1 | 1 | 1 |
| 1 | 1 | 1 |
| 1 | 1 | 1 |

| | | |
|---|---|---|
| 0 | 1 | 0 |
| 1 | 1 | 1 |
| 0 | 1 | 0 |
| 0 | 0 | 0 |
| 0 | 0 | 0 |
| 1 | 1 | 1 |
| 1 | 1 | 1 |
| 1 | 1 | 1 |
| 1 | 1 | 1 |
| 1 | 1 | 1 |
| 0 | 0 | 0 |
| 0 | 1 | 0 |
| 0 | 0 | 0 |
| 0 | 0 | 0 |
| 0 | 0 | 0 |
| 1 | 1 | 1 |
| 1 | 1 | 1 |
| 0 | 1 | 0 |
| 1 | 1 | 1 |
| 1 | 1 | 1 |
| 0 | 0 | 0 |
| 1 | 1 | 1 |
| 1 | 1 | 1 |
| 0 | 0 | 0 |
| 1 | 0 | 0 |
| 0 | 0 | 0 |
| 0 | 0 | 0 |
| 1 | 1 | 1 |
| 1 | 1 | 1 |
| 1 | 1 | 1 |
| 1 | 1 | 1 |
| 1 | 1 | 1 |
| 1 | 1 | 1 |
| 1 | 1 | 1 |
| 1 | 1 | 1 |
| 0 | 0 | 0 |
| 1 | 1 | 1 |
| 1 | 1 | 1 |
| 0 | 0 | 0 |
| 0 | 0 | 0 |
| 1 | 1 | 1 |
| 0 | 1 | 0 |
| 0 | 0 | 0 |
| 0 | 0 | 0 |
| 0 | 0 | 0 |
| 0 | 0 | 0 |
| 0 | 0 | 0 |
| 0 | 0 | 0 |
| 1 | 1 | 1 |
| 0 | 0 | 0 |
| 0 | 0 | 0 |
| 0 | 0 | 0 |
| 0 | 0 | 0 |
| 0 | 0 | 0 |

| | | |
|---|---|---|
| 0 | 0 | 0 |
| 0 | 0 | 0 |
| 0 | 0 | 0 |
| 0 | 0 | 0 |
| 0 | 0 | 0 |
| 0 | 0 | 0 |
| 0 | 0 | 0 |
| 0 | 0 | 0 |
| 0 | 0 | 0 |
| 0 | 1 | 0 |
| 0 | 1 | 0 |
| 0 | 0 | 0 |
| 1 | 1 | 1 |
| 0 | 0 | 0 |
| 0 | 0 | 0 |
| 0 | 1 | 0 |
| 0 | 0 | 0 |
| 1 | 0 | 0 |
| 1 | 1 | 1 |
| 1 | 1 | 1 |
| 0 | 0 | 0 |
| 0 | 1 | 0 |
| 0 | 1 | 0 |
| 0 | 0 | 0 |
| 0 | 0 | 0 |
| 0 | 1 | 0 |
| 0 | 1 | 0 |
| 1 | 1 | 1 |
| 0 | 0 | 0 |
| 0 | 0 | 0 |
| 1 | 1 | 1 |
| 1 | 1 | 1 |
| 0 | 0 | 0 |
| 0 | 0 | 0 |
| 0 | 0 | 0 |
| 0 | 1 | 0 |
| 0 | 0 | 0 |
| 0 | 1 | 0 |
| 0 | 1 | 0 |
| 0 | 0 | 0 |
| 0 | 0 | 0 |
| 0 | 0 | 0 |
| 0 | 0 | 0 |
| 0 | 0 | 0 |
| 0 | 0 | 0 |
| 1 | 1 | 1 |
| 0 | 1 | 0 |
| 0 | 0 | 0 |
| 0 | 0 | 0 |
| 0 | 0 | 0 |
| 1 | 1 | 1 |
| 1 | 1 | 1 |
| 0 | 0 | 0 |
| 0 | 0 | 0 |

| | | |
|---|---|---|
| 1 | 1 | 1 |
| 1 | 1 | 1 |
| 1 | 1 | 1 |
| 1 | 1 | 1 |
| 0 | 0 | 0 |
| 0 | 0 | 0 |
| 1 | 0 | 0 |
| 0 | 0 | 0 |
| 1 | 0 | 0 |
| 0 | 0 | 0 |
| 1 | 1 | 1 |
| 1 | 1 | 1 |
| 1 | 1 | 1 |
| 1 | 1 | 1 |
| 1 | 1 | 1 |
| 1 | 1 | 1 |
| 0 | 0 | 0 |
| 1 | 1 | 1 |
| 1 | 1 | 1 |
| 0 | 0 | 0 |
| 0 | 0 | 0 |
| 0 | 1 | 0 |
| 0 | 1 | 0 |
| 0 | 0 | 0 |
| 0 | 1 | 0 |
| 0 | 0 | 0 |
| 0 | 0 | 0 |
| 0 | 0 | 0 |
| 0 | 0 | 0 |
| 0 | 0 | 0 |
| 0 | 0 | 0 |
| 0 | 0 | 0 |
| 0 | 1 | 0 |
| 0 | 0 | 0 |
| 1 | 1 | 1 |
| 1 | 1 | 1 |
| 1 | 1 | 1 |
| 1 | 1 | 1 |
| 0 | 0 | 0 |
| 1 | 1 | 1 |
| 1 | 1 | 1 |
| 1 | 1 | 1 |
| 1 | 1 | 1 |
| 0 | 0 | 0 |
| 1 | 1 | 1 |
| 1 | 1 | 1 |
| 1 | 1 | 1 |
| 1 | 1 | 1 |
| 1 | 1 | 1 |
| 1 | 1 | 1 |
| 0 | 0 | 0 |
| 0 | 0 | 0 |
| 0 | 0 | 0 |
| 0 | 1 | 0 |

| | | |
|---|---|---|
| 0 | 0 | 0 |
| 0 | 0 | 0 |
| 0 | 0 | 0 |
| 0 | 0 | 0 |
| 0 | 0 | 0 |
| 0 | 0 | 0 |
| 0 | 0 | 0 |
| 0 | 0 | 0 |
| 1 | 1 | 1 |
| 1 | 1 | 1 |
| 0 | 0 | 0 |
| 1 | 1 | 1 |
| 1 | 1 | 1 |
| 0 | 0 | 0 |
| 0 | 0 | 0 |
| 0 | 0 | 0 |
| 0 | 0 | 0 |
| 0 | 0 | 0 |
| 0 | 0 | 0 |
| 1 | 1 | 1 |
| 1 | 1 | 1 |
| 0 | 0 | 0 |
| 1 | 1 | 1 |
| 0 | 0 | 0 |
| 1 | 1 | 1 |
| 1 | 1 | 1 |
| 1 | 1 | 1 |
| 0 | 1 | 0 |
| 0 | 1 | 0 |
| 0 | 0 | 0 |
| 0 | 1 | 0 |
| 1 | 1 | 1 |
| 0 | 0 | 0 |
| 1 | 1 | 1 |
| 0 | 0 | 0 |
| 0 | 0 | 0 |
| 0 | 0 | 0 |
| 0 | 0 | 0 |
| 0 | 0 | 0 |
| 0 | 0 | 0 |
| 0 | 0 | 0 |
| 0 | 0 | 0 |
| 1 | 1 | 1 |
| 0 | 0 | 0 |
| 1 | 1 | 1 |
| 0 | 0 | 0 |
| 1 | 0 | 0 |
| 1 | 1 | 1 |
| 1 | 1 | 1 |
| 0 | 0 | 0 |
| 1 | 1 | 1 |
| 1 | 1 | 1 |
| 0 | 1 | 0 |
| 1 | 1 | 1 |

| | | |
|---|---|---|
| 0 | 0 | 0 |
| 0 | 1 | 0 |
| 1 | 1 | 1 |
| 0 | 1 | 0 |
| 0 | 1 | 0 |
| 0 | 1 | 0 |
| 0 | 1 | 0 |
| 1 | 1 | 1 |
| 1 | 1 | 1 |
| 1 | 1 | 1 |
| 0 | 0 | 0 |
| 1 | 1 | 1 |
| 1 | 1 | 1 |
| 0 | 1 | 0 |
| 0 | 0 | 0 |
| 0 | 0 | 0 |
| 0 | 0 | 0 |
| 0 | 0 | 0 |
| 0 | 0 | 0 |
| 0 | 1 | 0 |
| 0 | 1 | 0 |
| 1 | 1 | 1 |
| 0 | 0 | 0 |
| 0 | 1 | 0 |
| 0 | 1 | 0 |
| 0 | 1 | 0 |
| 1 | 1 | 1 |
| 1 | 1 | 1 |
| 0 | 0 | 0 |
| 0 | 0 | 0 |
| 0 | 0 | 0 |
| 0 | 0 | 0 |
| 0 | 0 | 0 |
| 0 | 0 | 0 |
| 1 | 1 | 1 |
| 1 | 1 | 1 |
| 0 | 0 | 0 |
| 0 | 0 | 0 |
| 0 | 0 | 0 |
| 0 | 0 | 0 |
| 0 | 0 | 0 |
| 0 | 0 | 0 |
| 1 | 1 | 1 |
| 1 | 1 | 1 |
| 0 | 0 | 0 |
| 1 | 1 | 1 |
| 1 | 1 | 1 |
| 1 | 1 | 1 |
| 1 | 1 | 1 |
| 1 | 1 | 1 |
| 1 | 0 | 0 |
| 0 | 1 | 0 |
| 0 | 1 | 0 |
| 1 | 0 | 0 |

| | | |
|---|---|---|
| 0 | 0 | 0 |
| 1 | 1 | 1 |
| 0 | 0 | 0 |
| 0 | 1 | 0 |
| 1 | 0 | 0 |
| 0 | 0 | 0 |
| 0 | 0 | 0 |
| 0 | 0 | 0 |
| 1 | 0 | 0 |
| 1 | 0 | 0 |
| 0 | 0 | 0 |
| 1 | 1 | 1 |
| 1 | 0 | 0 |
| 1 | 0 | 0 |
| 0 | 0 | 0 |
| 0 | 0 | 0 |
| 0 | 0 | 0 |
| 0 | 0 | 0 |
| 1 | 1 | 1 |
| 0 | 0 | 0 |
| 0 | 0 | 0 |
| 0 | 0 | 0 |
| 0 | 0 | 0 |
| 0 | 0 | 0 |
| 1 | 1 | 1 |
| 0 | 0 | 0 |
| 1 | 1 | 1 |
| 0 | 1 | 0 |
| 1 | 1 | 1 |
| 0 | 0 | 0 |
| 1 | 1 | 1 |
| 1 | 1 | 1 |
| 0 | 0 | 0 |
| 0 | 0 | 0 |
| 1 | 1 | 1 |
| 1 | 1 | 1 |
| 1 | 1 | 1 |
| 0 | 1 | 0 |
| 1 | 0 | 0 |
| 0 | 0 | 0 |
| 0 | 0 | 0 |
| 0 | 0 | 0 |
| 0 | 0 | 0 |
| 0 | 0 | 0 |
| 0 | 0 | 0 |
| 0 | 1 | 0 |
| 0 | 1 | 0 |
| 0 | 0 | 0 |
| 0 | 0 | 0 |
| 1 | 1 | 1 |
| 0 | 0 | 0 |
| 0 | 0 | 0 |
| 0 | 1 | 0 |
| 1 | 1 | 1 |

| | | |
|---|---|---|
| 1 | 1 | 1 |
| 0 | 0 | 0 |
| 0 | 0 | 0 |
| 1 | 0 | 0 |
| 1 | 1 | 1 |
| 0 | 0 | 0 |
| 0 | 0 | 0 |
| 0 | 0 | 0 |
| 0 | 0 | 0 |
| 0 | 0 | 0 |
| 0 | 0 | 0 |
| 0 | 0 | 0 |
| 0 | 0 | 0 |
| 0 | 0 | 0 |
| 0 | 0 | 0 |
| 0 | 1 | 0 |
| 0 | 0 | 0 |
| 1 | 1 | 1 |
| 1 | 0 | 0 |
| 1 | 1 | 1 |
| 0 | 0 | 0 |
| 1 | 1 | 1 |
| 0 | 0 | 0 |
| 0 | 0 | 0 |
| 1 | 1 | 1 |
| 0 | 0 | 0 |
| 0 | 1 | 0 |
| 0 | 1 | 0 |
| 0 | 1 | 0 |
| 0 | 0 | 0 |
| 0 | 0 | 0 |
| 0 | 0 | 0 |
| 0 | 0 | 0 |
| 0 | 0 | 0 |
| 0 | 0 | 0 |
| 0 | 0 | 0 |
| 0 | 0 | 0 |
| 1 | 1 | 1 |
| 1 | 0 | 0 |
| 0 | 0 | 0 |
| 0 | 0 | 0 |
| 0 | 0 | 0 |
| 0 | 0 | 0 |
| 0 | 0 | 0 |
| 0 | 0 | 0 |
| 0 | 0 | 0 |
| 1 | 1 | 1 |
| 0 | 0 | 0 |
| 0 | 0 | 0 |
| 0 | 0 | 0 |
| 0 | 0 | 0 |
| 0 | 1 | 0 |
| 0 | 0 | 0 |
| 0 | 0 | 0 |

| | | |
|---|---|---|
| 0 | 0 | 0 |
| 0 | 0 | 0 |
| 0 | 0 | 0 |
| 0 | 0 | 0 |
| 0 | 0 | 0 |
| 0 | 0 | 0 |
| 0 | 0 | 0 |
| 0 | 0 | 0 |
| 0 | 0 | 0 |
| 0 | 0 | 0 |
| 1 | 1 | 1 |
| 0 | 0 | 0 |
| 1 | 0 | 0 |
| 0 | 0 | 0 |
| 0 | 1 | 0 |
| 1 | 1 | 1 |
| 1 | 1 | 1 |
| 1 | 1 | 1 |
| 0 | 0 | 0 |
| 0 | 0 | 0 |
| 1 | 1 | 1 |
| 0 | 0 | 0 |
| 0 | 0 | 0 |
| 1 | 1 | 1 |
| 0 | 0 | 0 |
| 0 | 0 | 0 |
| 0 | 0 | 0 |
| 1 | 0 | 0 |
| 0 | 0 | 0 |
| 0 | 0 | 0 |
| 1 | 0 | 0 |
| 0 | 0 | 0 |
| 1 | 0 | 0 |
| 1 | 0 | 0 |
| 0 | 0 | 0 |
| 0 | 0 | 0 |
| 0 | 0 | 0 |
| 1 | 0 | 0 |
| 1 | 0 | 0 |
| 0 | 0 | 0 |
| 0 | 0 | 0 |
| 0 | 0 | 0 |
| 0 | 0 | 0 |
| 0 | 0 | 0 |
| 0 | 0 | 0 |
| 0 | 0 | 0 |
| 0 | 0 | 0 |
| 0 | 0 | 0 |
| 0 | 0 | 0 |
| 0 | 0 | 0 |
| 0 | 0 | 0 |
| 0 | 0 | 0 |
| 0 | 0 | 0 |
| 0 | 0 | 0 |

| | | |
|---|---|---|
| 0 | 0 | 0 |
| 1 | 0 | 0 |
| 0 | 0 | 0 |
| 0 | 0 | 0 |
| 0 | 0 | 0 |
| 1 | 0 | 0 |
| 0 | 0 | 0 |
| 0 | 0 | 0 |
| 1 | 1 | 1 |
| 0 | 0 | 0 |
| 0 | 1 | 0 |
| 1 | 1 | 1 |
| 1 | 1 | 1 |
| 0 | 0 | 0 |
| 1 | 1 | 1 |
| 0 | 0 | 0 |
| 1 | 1 | 1 |
| 1 | 1 | 1 |
| 0 | 0 | 0 |
| 0 | 0 | 0 |
| 0 | 0 | 0 |
| 0 | 0 | 0 |
| 0 | 0 | 0 |
| 0 | 0 | 0 |
| 0 | 0 | 0 |
| 0 | 1 | 0 |
| 1 | 1 | 1 |
| 1 | 1 | 1 |
| 1 | 0 | 0 |
| 1 | 1 | 1 |
| 1 | 1 | 1 |
| 0 | 1 | 0 |
| 0 | 0 | 0 |
| 0 | 0 | 0 |
| 0 | 0 | 0 |
| 0 | 0 | 0 |
| 0 | 0 | 0 |
| 0 | 0 | 0 |
| 0 | 1 | 0 |
| 0 | 1 | 0 |
| 0 | 1 | 0 |
| 0 | 1 | 0 |
| 0 | 1 | 0 |
| 0 | 1 | 0 |
| 0 | 0 | 0 |
| 0 | 0 | 0 |
| 0 | 1 | 0 |
| 0 | 0 | 0 |
| 0 | 0 | 0 |
| 0 | 0 | 0 |
| 0 | 0 | 0 |
| 0 | 0 | 0 |
| 0 | 0 | 0 |
| 0 | 0 | 0 |

| | | |
|---|---|---|
| 1 | 1 | 1 |
| 1 | 1 | 1 |
| 0 | 0 | 0 |
| 0 | 0 | 0 |
| 1 | 1 | 1 |
| 0 | 0 | 0 |
| 0 | 0 | 0 |
| 1 | 1 | 1 |
| 1 | 1 | 1 |
| 0 | 0 | 0 |
| 1 | 1 | 1 |
| 1 | 1 | 1 |
| 0 | 0 | 0 |
| 0 | 0 | 0 |
| 0 | 0 | 0 |
| 0 | 0 | 0 |
| 0 | 1 | 0 |
| 0 | 1 | 0 |
| 1 | 1 | 1 |
| 1 | 1 | 1 |
| 1 | 1 | 1 |
| 0 | 1 | 0 |
| 1 | 1 | 1 |
| 1 | 1 | 1 |
| 1 | 1 | 1 |
| 0 | 0 | 0 |
| 1 | 1 | 1 |
| 1 | 1 | 1 |
| 1 | 1 | 1 |
| 0 | 1 | 0 |
| 0 | 1 | 0 |
| 1 | 1 | 1 |
| 1 | 1 | 1 |
| 1 | 1 | 1 |
| 1 | 1 | 1 |
| 1 | 1 | 1 |
| 1 | 1 | 1 |
| 0 | 0 | 0 |
| 0 | 0 | 0 |
| 1 | 0 | 0 |
| 0 | 0 | 0 |
| 1 | 1 | 1 |
| 1 | 1 | 1 |
| 0 | 0 | 0 |
| 0 | 0 | 0 |
| 0 | 1 | 0 |
| 0 | 1 | 0 |
| 0 | 1 | 0 |
| 1 | 1 | 1 |
| 1 | 1 | 1 |
| 1 | 1 | 1 |
| 0 | 0 | 0 |
| 0 | 0 | 0 |
| 0 | 0 | 0 |

| | | |
|---|---|---|
| 0 | 0 | 0 |
| 1 | 1 | 1 |
| 1 | 1 | 1 |
| 1 | 1 | 1 |
| 1 | 1 | 1 |
| 1 | 1 | 1 |
| 1 | 1 | 1 |
| 1 | 1 | 1 |
| 0 | 0 | 0 |
| 1 | 1 | 1 |
| 1 | 1 | 1 |
| 1 | 1 | 1 |
| 0 | 1 | 0 |
| 0 | 1 | 0 |
| 1 | 1 | 1 |
| 1 | 1 | 1 |
| 0 | 1 | 0 |
| 0 | 1 | 0 |
| 0 | 0 | 0 |
| 0 | 0 | 0 |
| 0 | 0 | 0 |
| 0 | 0 | 0 |
| 0 | 1 | 0 |
| 0 | 0 | 0 |
| 0 | 0 | 0 |
| 1 | 1 | 1 |
| 1 | 1 | 1 |
| 1 | 1 | 1 |
| 1 | 1 | 1 |
| 1 | 1 | 1 |
| 1 | 1 | 1 |
| 1 | 1 | 1 |
| 0 | 0 | 0 |
| 1 | 1 | 1 |
| 1 | 1 | 1 |
| 0 | 1 | 0 |
| 0 | 0 | 0 |
| 1 | 1 | 1 |
| 0 | 0 | 0 |
| 0 | 0 | 0 |
| 0 | 0 | 0 |
| 0 | 0 | 0 |
| 0 | 0 | 0 |
| 1 | 1 | 1 |
| 1 | 1 | 1 |
| 1 | 1 | 1 |
| 0 | 0 | 0 |
| 0 | 0 | 0 |
| 0 | 0 | 0 |
| 0 | 0 | 0 |
| 0 | 0 | 0 |
| 0 | 0 | 0 |
| 0 | 0 | 0 |
| 0 | 1 | 0 |

| | | |
|---|---|---|
| 0 | 0 | 0 |
| 0 | 0 | 0 |
| 0 | 1 | 0 |
| 0 | 0 | 0 |
| 0 | 0 | 0 |
| 1 | 1 | 1 |
| 1 | 1 | 1 |
| 1 | 0 | 0 |
| 1 | 1 | 1 |
| 1 | 1 | 1 |
| 1 | 1 | 1 |
| 0 | 0 | 0 |
| 1 | 1 | 1 |
| 1 | 1 | 1 |
| 0 | 0 | 0 |
| 1 | 1 | 1 |
| 0 | 0 | 0 |
| 1 | 1 | 1 |
| 0 | 0 | 0 |
| 0 | 0 | 0 |
| 1 | 1 | 1 |
| 1 | 1 | 1 |
| 0 | 0 | 0 |
| 1 | 1 | 1 |
| 0 | 0 | 0 |
| 1 | 1 | 1 |
| 1 | 1 | 1 |
| 0 | 0 | 0 |
| 1 | 1 | 1 |
| 0 | 0 | 0 |
| 0 | 1 | 0 |
| 0 | 0 | 0 |
| 0 | 0 | 0 |
| 0 | 0 | 0 |
| 1 | 1 | 1 |
| 0 | 0 | 0 |
| 1 | 1 | 1 |
| 1 | 1 | 1 |
| 0 | 0 | 0 |
| 1 | 1 | 1 |
| 1 | 1 | 1 |
| 1 | 1 | 1 |
| 1 | 1 | 1 |
| 0 | 0 | 0 |
| 0 | 0 | 0 |
| 1 | 1 | 1 |
| 1 | 1 | 1 |
| 1 | 0 | 0 |
| 0 | 0 | 0 |
| 1 | 0 | 0 |
| 0 | 0 | 0 |
| 1 | 1 | 1 |
| 1 | 1 | 1 |
| 0 | 0 | 0 |

| | | |
|---|---|---|
| 1 | 1 | 1 |
| 1 | 1 | 1 |
| 1 | 1 | 1 |
| 1 | 1 | 1 |
| 0 | 1 | 0 |
| 1 | 1 | 1 |
| 1 | 1 | 1 |
| 0 | 0 | 0 |
| 1 | 1 | 1 |
| 1 | 1 | 1 |
| 1 | 1 | 1 |
| 1 | 1 | 1 |
| 0 | 0 | 0 |
| 1 | 1 | 1 |
| 0 | 0 | 0 |
| 0 | 0 | 0 |
| 1 | 1 | 1 |
| 1 | 1 | 1 |
| 1 | 1 | 1 |
| 1 | 1 | 1 |
| 0 | 0 | 0 |
| 0 | 0 | 0 |
| 1 | 1 | 1 |
| 1 | 1 | 1 |
| 0 | 0 | 0 |
| 1 | 1 | 1 |
| 0 | 0 | 0 |
| 0 | 0 | 0 |
| 0 | 0 | 0 |
| 0 | 0 | 0 |
| 0 | 1 | 0 |
| 0 | 1 | 0 |
| 0 | 0 | 0 |
| 0 | 0 | 0 |
| 0 | 0 | 0 |
| 0 | 0 | 0 |
| 0 | 0 | 0 |
| 0 | 0 | 0 |
| 0 | 0 | 0 |
| 1 | 0 | 0 |
| 1 | 1 | 1 |
| 0 | 1 | 0 |
| 0 | 1 | 0 |
| 0 | 1 | 0 |
| 0 | 1 | 0 |
| 0 | 0 | 0 |
| 0 | 1 | 0 |
| 0 | 1 | 0 |
| 1 | 1 | 1 |
| 0 | 0 | 0 |
| 0 | 0 | 0 |
| 1 | 1 | 1 |
| 1 | 1 | 1 |
| 1 | 1 | 1 |

| | | |
|---|---|---|
| 1 | 1 | 1 |
| 1 | 1 | 1 |
| 1 | 1 | 1 |
| 1 | 1 | 1 |
| 0 | 0 | 0 |
| 1 | 1 | 1 |
| 0 | 0 | 0 |
| 0 | 0 | 0 |
| 0 | 0 | 0 |
| 0 | 0 | 0 |
| 0 | 0 | 0 |
| 1 | 1 | 1 |
| 0 | 0 | 0 |
| 0 | 0 | 0 |
| 1 | 1 | 1 |
| 1 | 1 | 1 |
| 1 | 1 | 1 |
| 1 | 1 | 1 |
| 1 | 1 | 1 |
| 1 | 1 | 1 |
| 1 | 1 | 1 |
| 1 | 1 | 1 |
| 1 | 1 | 1 |
| 0 | 0 | 0 |
| 1 | 1 | 1 |
| 1 | 1 | 1 |
| 1 | 1 | 1 |
| 0 | 0 | 0 |
| 1 | 0 | 0 |
| 1 | 0 | 0 |
| 0 | 0 | 0 |
| 0 | 0 | 0 |
| 1 | 1 | 1 |
| 0 | 0 | 0 |
| 0 | 1 | 0 |
| 1 | 1 | 1 |
| 1 | 1 | 1 |
| 1 | 1 | 1 |
| 1 | 1 | 1 |
| 1 | 1 | 1 |
| 1 | 1 | 1 |
| 1 | 1 | 1 |
| 1 | 1 | 1 |
| 1 | 1 | 1 |
| 1 | 1 | 1 |
| 1 | 1 | 1 |
| 1 | 1 | 1 |
| 1 | 1 | 1 |
| 0 | 0 | 0 |
| 0 | 1 | 0 |
| 1 | 1 | 1 |
| 1 | 0 | 0 |
| 1 | 0 | 0 |
| 1 | 1 | 1 |

| | | |
|---|---|---|
| 1 | 1 | 1 |
| 0 | 1 | 0 |
| 1 | 1 | 1 |
| 0 | 1 | 0 |
| 1 | 0 | 0 |
| 1 | 1 | 1 |
| 1 | 1 | 1 |
| 1 | 1 | 1 |
| 1 | 1 | 1 |
| 1 | 1 | 1 |
| 0 | 0 | 0 |
| 0 | 0 | 0 |
| 0 | 0 | 0 |
| 0 | 0 | 0 |
| 0 | 0 | 0 |
| 1 | 0 | 0 |
| 0 | 0 | 0 |
| 0 | 0 | 0 |
| 0 | 0 | 0 |
| 0 | 0 | 0 |
| 0 | 0 | 0 |
| 0 | 0 | 0 |
| 0 | 1 | 0 |
| 0 | 0 | 0 |
| 0 | 0 | 0 |
| 0 | 0 | 0 |
| 1 | 1 | 1 |
| 0 | 0 | 0 |
| 0 | 1 | 0 |
| 0 | 1 | 0 |
| 0 | 1 | 0 |
| 0 | 0 | 0 |
| 1 | 1 | 1 |
| 1 | 1 | 1 |
| 0 | 0 | 0 |
| 0 | 0 | 0 |
| 0 | 0 | 0 |
| 0 | 0 | 0 |
| 0 | 0 | 0 |
| 1 | 0 | 0 |
| 1 | 0 | 0 |
| 0 | 0 | 0 |
| 1 | 0 | 0 |
| 0 | 0 | 0 |
| 1 | 1 | 1 |
| 1 | 0 | 0 |
| 1 | 1 | 1 |
| 0 | 0 | 0 |
| 0 | 0 | 0 |
| 0 | 0 | 0 |
| 0 | 0 | 0 |
| 1 | 1 | 1 |
| 0 | 0 | 0 |
| 1 | 1 | 1 |

| | | |
|---|---|---|
| 1 | 1 | 1 |
| 1 | 1 | 1 |
| 1 | 1 | 1 |
| 1 | 1 | 1 |
| 1 | 1 | 1 |
| 0 | 0 | 0 |
| 0 | 0 | 0 |
| 0 | 0 | 0 |
| 0 | 0 | 0 |
| 0 | 1 | 0 |
| 0 | 0 | 0 |
| 0 | 0 | 0 |
| 0 | 0 | 0 |
| 0 | 0 | 0 |
| 0 | 0 | 0 |
| 0 | 0 | 0 |
| 1 | 1 | 1 |
| 0 | 1 | 0 |
| 1 | 1 | 1 |
| 1 | 1 | 1 |
| 1 | 1 | 1 |
| 1 | 1 | 1 |
| 1 | 1 | 1 |
| 1 | 1 | 1 |
| 1 | 1 | 1 |
| 1 | 1 | 1 |
| 1 | 1 | 1 |
| 1 | 1 | 1 |
| 0 | 0 | 0 |
| 0 | 1 | 0 |
| 0 | 1 | 0 |
| 1 | 0 | 0 |
| 1 | 0 | 0 |
| 0 | 0 | 0 |
| 1 | 1 | 1 |
| 0 | 0 | 0 |
| 1 | 0 | 0 |
| 0 | 0 | 0 |
| 1 | 1 | 1 |
| 0 | 0 | 0 |
| 1 | 1 | 1 |
| 1 | 1 | 1 |
| 0 | 1 | 0 |
| 0 | 0 | 0 |
| 1 | 1 | 1 |
| 1 | 1 | 1 |
| 1 | 1 | 1 |
| 1 | 1 | 1 |
| 1 | 1 | 1 |
| 1 | 1 | 1 |
| 1 | 1 | 1 |
| 1 | 0 | 0 |
| 0 | 0 | 0 |
| 0 | 0 | 0 |

| | | |
|---|---|---|
| 1 | 1 | 1 |
| 0 | 0 | 0 |
| 1 | 1 | 1 |
| 1 | 1 | 1 |
| 0 | 0 | 0 |
| 0 | 0 | 0 |
| 0 | 0 | 0 |
| 1 | 1 | 1 |
| 1 | 1 | 1 |
| 0 | 0 | 0 |
| 0 | 0 | 0 |
| 0 | 0 | 0 |
| 0 | 0 | 0 |
| 0 | 0 | 0 |
| 1 | 1 | 1 |
| 0 | 0 | 0 |
| 0 | 0 | 0 |
| 0 | 0 | 0 |
| 1 | 1 | 1 |
| 1 | 1 | 1 |
| 1 | 1 | 1 |
| 1 | 1 | 1 |
| 1 | 1 | 1 |
| 0 | 0 | 0 |
| 0 | 1 | 0 |
| 0 | 1 | 0 |
| 0 | 0 | 0 |
| 0 | 0 | 0 |
| 1 | 1 | 1 |
| 1 | 1 | 1 |
| 0 | 0 | 0 |
| 1 | 1 | 1 |
| 1 | 1 | 1 |
| 0 | 1 | 0 |
| 1 | 1 | 1 |
| 0 | 0 | 0 |
| 0 | 1 | 0 |
| 0 | 0 | 0 |
| 0 | 0 | 0 |
| 0 | 1 | 0 |
| 0 | 1 | 0 |
| 0 | 0 | 0 |
| 0 | 0 | 0 |
| 0 | 1 | 0 |
| 0 | 0 | 0 |
| 0 | 0 | 0 |
| 0 | 0 | 0 |
| 0 | 1 | 0 |
| 0 | 0 | 0 |
| 0 | 0 | 0 |
| 0 | 0 | 0 |
| 1 | 1 | 1 |
| 0 | 0 | 0 |
| 1 | 1 | 1 |

| | | |
|---|---|---|
| 1 | 1 | 1 |
| 1 | 1 | 1 |
| 1 | 1 | 1 |
| 1 | 1 | 1 |
| 0 | 0 | 0 |
| 0 | 0 | 0 |
| 1 | 1 | 1 |
| 1 | 0 | 0 |
| 1 | 1 | 1 |
| 0 | 0 | 0 |
| 1 | 0 | 0 |
| 1 | 0 | 0 |
| 0 | 0 | 0 |
| 1 | 1 | 1 |
| 1 | 1 | 1 |
| 1 | 1 | 1 |
| 1 | 1 | 1 |
| 1 | 1 | 1 |
| 1 | 1 | 1 |
| 1 | 1 | 1 |
| 1 | 1 | 1 |
| 0 | 1 | 0 |
| 1 | 1 | 1 |
| 0 | 0 | 0 |
| 1 | 1 | 1 |
| 0 | 0 | 0 |
| 1 | 1 | 1 |
| 0 | 0 | 0 |
| 1 | 1 | 1 |
| 0 | 0 | 0 |
| 0 | 0 | 0 |
| 0 | 0 | 0 |
| 0 | 0 | 0 |
| 0 | 0 | 0 |
| 0 | 1 | 0 |
| 0 | 0 | 0 |
| 1 | 1 | 1 |
| 0 | 0 | 0 |
| 1 | 1 | 1 |
| 1 | 1 | 1 |
| 0 | 0 | 0 |
| 0 | 0 | 0 |
| 0 | 0 | 0 |
| 0 | 0 | 0 |
| 0 | 0 | 0 |
| 0 | 0 | 0 |
| 0 | 0 | 0 |
| 0 | 0 | 0 |
| 1 | 1 | 1 |
| 0 | 0 | 0 |
| 1 | 0 | 0 |
| 1 | 0 | 0 |
| 1 | 1 | 1 |
| 1 | 1 | 1 |

| | | |
|---|---|---|
| 1 | 1 | 1 |
| 1 | 1 | 1 |
| 1 | 1 | 1 |
| 0 | 1 | 0 |
| 0 | 1 | 0 |
| 0 | 1 | 0 |
| 0 | 0 | 0 |
| 0 | 0 | 0 |
| 0 | 0 | 0 |
| 0 | 0 | 0 |
| 0 | 0 | 0 |
| 0 | 0 | 0 |
| 0 | 0 | 0 |
| 0 | 0 | 0 |
| 0 | 0 | 0 |
| 0 | 0 | 0 |
| 0 | 0 | 0 |
| 0 | 0 | 0 |
| 1 | 1 | 1 |
| 0 | 0 | 0 |
| 0 | 1 | 0 |
| 0 | 1 | 0 |
| 0 | 1 | 0 |
| 1 | 1 | 1 |
| 0 | 1 | 0 |
| 0 | 0 | 0 |
| 0 | 0 | 0 |
| 0 | 0 | 0 |
| 0 | 0 | 0 |
| 0 | 0 | 0 |
| 0 | 0 | 0 |
| 0 | 0 | 0 |
| 0 | 0 | 0 |
| 0 | 0 | 0 |
| 1 | 1 | 1 |
| 0 | 0 | 0 |
| 0 | 0 | 0 |
| 0 | 0 | 0 |
| 0 | 0 | 0 |
| 0 | 1 | 0 |
| 0 | 0 | 0 |
| 0 | 0 | 0 |
| 1 | 1 | 1 |
| 0 | 1 | 0 |
| 1 | 1 | 1 |
| 0 | 1 | 0 |
| 0 | 1 | 0 |
| 0 | 0 | 0 |
| 0 | 0 | 0 |
| 0 | 0 | 0 |
| 0 | 0 | 0 |
| 0 | 1 | 0 |
| 0 | 0 | 0 |
| 1 | 1 | 1 |

| | | |
|---|---|---|
| 0 | 1 | 0 |
| 1 | 1 | 1 |
| 1 | 1 | 1 |
| 1 | 1 | 1 |
| 1 | 1 | 1 |
| 0 | 0 | 0 |
| 0 | 1 | 0 |
| 1 | 1 | 1 |
| 1 | 1 | 1 |
| 1 | 1 | 1 |
| 0 | 0 | 0 |
| 0 | 0 | 0 |
| 0 | 0 | 0 |
| 0 | 0 | 0 |
| 0 | 0 | 0 |
| 0 | 1 | 0 |
| 1 | 0 | 0 |
| 0 | 0 | 0 |
| 1 | 1 | 1 |
| 1 | 1 | 1 |
| 1 | 1 | 1 |
| 1 | 1 | 1 |
| 0 | 0 | 0 |
| 1 | 1 | 1 |
| 0 | 0 | 0 |
| 0 | 0 | 0 |
| 0 | 0 | 0 |
| 0 | 0 | 0 |
| 1 | 1 | 1 |
| 0 | 0 | 0 |
| 1 | 1 | 1 |
| 0 | 1 | 0 |
| 0 | 1 | 0 |
| 0 | 1 | 0 |
| 1 | 1 | 1 |
| 1 | 1 | 1 |
| 0 | 1 | 0 |
| 0 | 0 | 0 |
| 0 | 0 | 0 |
| 0 | 0 | 0 |
| 0 | 0 | 0 |
| 0 | 0 | 0 |
| 1 | 1 | 1 |
| 0 | 0 | 0 |
| 0 | 0 | 0 |
| 1 | 1 | 1 |
| 0 | 0 | 0 |
| 0 | 0 | 0 |
| 0 | 0 | 0 |
| 0 | 0 | 0 |
| 1 | 1 | 1 |
| 1 | 1 | 1 |
| 1 | 1 | 1 |
| 0 | 0 | 0 |

| | | |
|---|---|---|
| 0 | 0 | 0 |
| 1 | 1 | 1 |
| 0 | 0 | 0 |
| 0 | 1 | 0 |
| 1 | 0 | 0 |
| 1 | 0 | 0 |
| 1 | 0 | 0 |
| 1 | 1 | 1 |
| 0 | 0 | 0 |
| 1 | 1 | 1 |
| 0 | 0 | 0 |
| 0 | 0 | 0 |
| 0 | 0 | 0 |
| 0 | 0 | 0 |
| 1 | 1 | 1 |
| 1 | 1 | 1 |
| 0 | 0 | 0 |
| 0 | 0 | 0 |
| 0 | 1 | 0 |
| 1 | 1 | 1 |
| 0 | 0 | 0 |
| 1 | 0 | 0 |
| 1 | 0 | 0 |
| 1 | 1 | 1 |
| 0 | 0 | 0 |
| 1 | 0 | 0 |
| 0 | 0 | 0 |
| 0 | 0 | 0 |
| 0 | 0 | 0 |
| 1 | 1 | 1 |
| 0 | 0 | 0 |
| 0 | 0 | 0 |
| 0 | 0 | 0 |
| 1 | 1 | 1 |
| 0 | 0 | 0 |
| 1 | 1 | 1 |
| 0 | 0 | 0 |
| 0 | 0 | 0 |
| 0 | 0 | 0 |
| 0 | 0 | 0 |
| 0 | 0 | 0 |
| 1 | 1 | 1 |
| 0 | 0 | 0 |
| 0 | 0 | 0 |
| 0 | 0 | 0 |
| 0 | 0 | 0 |
| 0 | 0 | 0 |
| 0 | 0 | 0 |
| 0 | 0 | 0 |
| 0 | 0 | 0 |
| 0 | 0 | 0 |
| 0 | 0 | 0 |
| 1 | 1 | 1 |
| 0 | 0 | 0 |

| | | |
|---|---|---|
| 0 | 1 | 0 |
| 1 | 1 | 1 |
| 1 | 1 | 1 |
| 1 | 1 | 1 |
| 1 | 1 | 1 |
| 1 | 1 | 1 |
| 1 | 1 | 1 |
| 1 | 1 | 1 |
| 1 | 1 | 1 |
| 0 | 0 | 0 |
| 0 | 0 | 0 |
| 1 | 1 | 1 |
| 1 | 1 | 1 |
| 1 | 1 | 1 |
| 0 | 0 | 0 |
| 1 | 1 | 1 |
| 1 | 1 | 1 |
| 0 | 0 | 0 |
| 0 | 1 | 0 |
| 0 | 1 | 0 |
| 0 | 1 | 0 |
| 0 | 0 | 0 |
| 0 | 0 | 0 |
| 0 | 0 | 0 |
| 0 | 0 | 0 |
| 0 | 0 | 0 |
| 0 | 0 | 0 |
| 0 | 0 | 0 |
| 1 | 1 | 1 |
| 0 | 0 | 0 |
| 0 | 0 | 0 |
| 1 | 1 | 1 |
| 1 | 1 | 1 |
| 1 | 1 | 1 |
| 1 | 1 | 1 |
| 1 | 1 | 1 |
| 1 | 1 | 1 |
| 1 | 1 | 1 |
| 1 | 1 | 1 |
| 1 | 1 | 1 |
| 1 | 1 | 1 |
| 1 | 1 | 1 |
| 1 | 1 | 1 |
| 0 | 0 | 0 |
| 1 | 1 | 1 |
| 0 | 0 | 0 |
| 1 | 1 | 1 |
| 1 | 1 | 1 |
| 1 | 1 | 1 |
| 1 | 1 | 1 |
| 0 | 0 | 0 |
| 1 | 1 | 1 |
| 1 | 1 | 1 |
| 0 | 0 | 0 |

| | | |
|---|---|---|
| 0 | 0 | 0 |
| 0 | 0 | 0 |
| 0 | 0 | 0 |
| 0 | 0 | 0 |
| 1 | 1 | 1 |
| 0 | 1 | 0 |
| 1 | 1 | 1 |
| 0 | 0 | 0 |
| 0 | 0 | 0 |
| 0 | 0 | 0 |
| 0 | 0 | 0 |
| 1 | 1 | 1 |
| 1 | 1 | 1 |
| 1 | 1 | 1 |
| 1 | 1 | 1 |
| 0 | 0 | 0 |
| 0 | 0 | 0 |
| 0 | 1 | 0 |
| 0 | 1 | 0 |
| 1 | 1 | 1 |
| 1 | 1 | 1 |
| 0 | 0 | 0 |
| 1 | 1 | 1 |
| 0 | 1 | 0 |
| 0 | 0 | 0 |
| 0 | 0 | 0 |
| 0 | 0 | 0 |
| 0 | 0 | 0 |
| 1 | 1 | 1 |
| 1 | 1 | 1 |
| 1 | 1 | 1 |
| 0 | 0 | 0 |
| 1 | 1 | 1 |
| 1 | 1 | 1 |
| 1 | 1 | 1 |
| 1 | 1 | 1 |
| 1 | 1 | 1 |
| 1 | 1 | 1 |
| 1 | 1 | 1 |
| 1 | 1 | 1 |
| 1 | 1 | 1 |
| 1 | 1 | 1 |
| 0 | 0 | 0 |
| 0 | 1 | 0 |
| 0 | 1 | 0 |
| 1 | 1 | 1 |
| 0 | 1 | 0 |
| 1 | 1 | 1 |
| 1 | 1 | 1 |
| 0 | 0 | 0 |
| 1 | 1 | 1 |
| 1 | 1 | 1 |
| 1 | 1 | 1 |
| 1 | 1 | 1 |

| | | |
|---|---|---|
| 0 | 0 | 0 |
| 1 | 1 | 1 |
| 0 | 0 | 0 |
| 0 | 1 | 0 |
| 1 | 1 | 1 |
| 1 | 1 | 1 |
| 1 | 1 | 1 |
| 0 | 0 | 0 |
| 1 | 1 | 1 |
| 1 | 1 | 1 |
| 0 | 0 | 0 |
| 0 | 0 | 0 |
| 0 | 0 | 0 |
| 1 | 1 | 1 |
| 0 | 0 | 0 |
| 0 | 0 | 0 |
| 0 | 0 | 0 |
| 0 | 0 | 0 |
| 0 | 1 | 0 |
| 0 | 0 | 0 |
| 0 | 0 | 0 |
| 0 | 0 | 0 |
| 0 | 0 | 0 |
| 0 | 0 | 0 |
| 0 | 0 | 0 |
| 0 | 0 | 0 |
| 0 | 0 | 0 |
| 0 | 0 | 0 |
| 0 | 0 | 0 |
| 0 | 0 | 0 |
| 0 | 0 | 0 |
| 0 | 0 | 0 |
| 1 | 1 | 1 |
| 0 | 0 | 0 |
| 1 | 1 | 1 |
| 0 | 0 | 0 |
| 0 | 0 | 0 |
| 1 | 1 | 1 |
| 1 | 1 | 1 |
| 0 | 1 | 0 |
| 0 | 1 | 0 |
| 0 | 0 | 0 |
| 0 | 0 | 0 |
| 0 | 0 | 0 |
| 1 | 1 | 1 |
| 0 | 0 | 0 |
| 1 | 1 | 1 |
| 0 | 0 | 0 |
| 0 | 1 | 0 |
| 0 | 1 | 0 |
| 0 | 1 | 0 |
| 0 | 1 | 0 |
| 0 | 1 | 0 |
| 1 | 1 | 1 |

| | | |
|---|---|---|
| 1 | 1 | 1 |
| 1 | 1 | 1 |
| 0 | 1 | 0 |
| 1 | 1 | 1 |
| 1 | 1 | 1 |
| 0 | 0 | 0 |
| 0 | 0 | 0 |
| 1 | 1 | 1 |
| 0 | 0 | 0 |
| 0 | 0 | 0 |
| 1 | 1 | 1 |
| 0 | 1 | 0 |
| 1 | 1 | 1 |
| 1 | 1 | 1 |
| 1 | 1 | 1 |
| 0 | 0 | 0 |
| 1 | 1 | 1 |
| 1 | 1 | 1 |
| 0 | 0 | 0 |
| 0 | 0 | 0 |
| 0 | 0 | 0 |
| 1 | 1 | 1 |
| 1 | 1 | 1 |
| 0 | 0 | 0 |
| 1 | 1 | 1 |
| 0 | 0 | 0 |
| 0 | 0 | 0 |
| 1 | 1 | 1 |
| 1 | 1 | 1 |
| 1 | 1 | 1 |
| 0 | 0 | 0 |
| 1 | 0 | 0 |
| 0 | 0 | 0 |
| 0 | 0 | 0 |
| 0 | 0 | 0 |
| 0 | 0 | 0 |
| 0 | 0 | 0 |
| 1 | 1 | 1 |
| 0 | 0 | 0 |
| 0 | 1 | 0 |
| 0 | 0 | 0 |
| 0 | 0 | 0 |
| 0 | 0 | 0 |
| 0 | 0 | 0 |
| 1 | 1 | 1 |
| 0 | 1 | 0 |
| 0 | 1 | 0 |
| 0 | 0 | 0 |
| 0 | 0 | 0 |
| 0 | 0 | 0 |
| 1 | 1 | 1 |
| 0 | 1 | 0 |
| 1 | 1 | 1 |
| 1 | 1 | 1 |

| | | |
|---|---|---|
| 1 | 1 | 1 |
| 0 | 1 | 0 |
| 0 | 1 | 0 |
| 0 | 0 | 0 |
| 0 | 0 | 0 |
| 1 | 1 | 1 |
| 0 | 0 | 0 |
| 1 | 1 | 1 |
| 1 | 1 | 1 |
| 1 | 1 | 1 |
| 1 | 1 | 1 |
| 0 | 0 | 0 |
| 1 | 0 | 0 |
| 0 | 0 | 0 |
| 1 | 1 | 1 |
| 0 | 0 | 0 |
| 0 | 0 | 0 |
| 0 | 0 | 0 |
| 1 | 1 | 1 |
| 0 | 0 | 0 |
| 0 | 0 | 0 |
| 0 | 0 | 0 |
| 0 | 0 | 0 |
| 0 | 0 | 0 |
| 0 | 0 | 0 |
| 1 | 1 | 1 |
| 0 | 0 | 0 |
| 0 | 1 | 0 |
| 0 | 1 | 0 |
| 0 | 1 | 0 |
| 0 | 1 | 0 |
| 0 | 1 | 0 |
| 0 | 0 | 0 |
| 0 | 0 | 0 |
| 1 | 1 | 1 |
| 1 | 1 | 1 |
| 0 | 0 | 0 |
| 0 | 0 | 0 |
| 0 | 0 | 0 |
| 1 | 0 | 0 |
| 0 | 0 | 0 |
| 0 | 0 | 0 |
| 1 | 1 | 1 |
| 0 | 0 | 0 |
| 1 | 1 | 1 |
| 1 | 1 | 1 |
| 1 | 1 | 1 |
| 1 | 1 | 1 |
| 1 | 1 | 1 |
| 0 | 0 | 0 |
| 1 | 1 | 1 |
| 1 | 1 | 1 |
| 1 | 1 | 1 |
| 0 | 1 | 0 |

| | | |
|---|---|---|
| 1 | 1 | 1 |
| 1 | 1 | 1 |
| 1 | 1 | 1 |
| 0 | 0 | 0 |
| 1 | 1 | 1 |
| 1 | 1 | 1 |
| 0 | 1 | 0 |
| 0 | 0 | 0 |
| 1 | 1 | 1 |
| 1 | 1 | 1 |
| 1 | 1 | 1 |
| 1 | 1 | 1 |
| 1 | 1 | 1 |
| 0 | 1 | 0 |
| 0 | 1 | 0 |
| 0 | 1 | 0 |
| 0 | 0 | 0 |
| 0 | 0 | 0 |
| 0 | 1 | 0 |
| 1 | 1 | 1 |
| 0 | 1 | 0 |
| 1 | 1 | 1 |
| 0 | 1 | 0 |
| 1 | 1 | 1 |
| 1 | 1 | 1 |
| 1 | 1 | 1 |
| 1 | 1 | 1 |
| 0 | 0 | 0 |
| 0 | 0 | 0 |
| 1 | 1 | 1 |
| 1 | 1 | 1 |
| 0 | 0 | 0 |
| 0 | 1 | 0 |
| 0 | 0 | 0 |
| 0 | 0 | 0 |
| 0 | 0 | 0 |
| 0 | 0 | 0 |
| 0 | 0 | 0 |
| 1 | 1 | 1 |
| 1 | 1 | 1 |
| 1 | 1 | 1 |
| 0 | 1 | 0 |
| 1 | 1 | 1 |
| 1 | 1 | 1 |
| 0 | 0 | 0 |
| 0 | 0 | 0 |
| 0 | 0 | 0 |
| 1 | 1 | 1 |
| 1 | 1 | 1 |
| 1 | 1 | 1 |
| 1 | 1 | 1 |
| 0 | 0 | 0 |
| 0 | 0 | 0 |
| 0 | 0 | 0 |

| | | |
|---|---|---|
| 0 | 0 | 0 |
| 0 | 0 | 0 |
| 1 | 1 | 1 |
| 0 | 0 | 0 |
| 0 | 0 | 0 |
| 0 | 0 | 0 |
| 0 | 0 | 0 |
| 1 | 1 | 1 |
| 1 | 1 | 1 |
| 1 | 1 | 1 |
| 1 | 1 | 1 |
| 1 | 1 | 1 |
| 0 | 1 | 0 |
| 0 | 1 | 0 |
| 1 | 1 | 1 |
| 1 | 1 | 1 |
| 1 | 1 | 1 |
| 1 | 1 | 1 |
| 1 | 1 | 1 |
| 1 | 1 | 1 |
| 0 | 0 | 0 |
| 0 | 0 | 0 |
| 0 | 0 | 0 |
| 0 | 0 | 0 |
| 0 | 0 | 0 |
| 1 | 1 | 1 |
| 1 | 1 | 1 |
| 0 | 0 | 0 |
| 0 | 1 | 0 |
| 0 | 1 | 0 |
| 0 | 0 | 0 |
| 0 | 0 | 0 |
| 0 | 0 | 0 |
| 1 | 1 | 1 |
| 0 | 0 | 0 |
| 1 | 1 | 1 |
| 0 | 1 | 0 |
| 0 | 0 | 0 |
| 1 | 1 | 1 |
| 1 | 1 | 1 |
| 0 | 1 | 0 |
| 0 | 1 | 0 |
| 1 | 1 | 1 |
| 1 | 1 | 1 |
| 1 | 1 | 1 |
| 1 | 1 | 1 |
| 1 | 1 | 1 |
| 1 | 1 | 1 |
| 0 | 0 | 0 |
| 1 | 1 | 1 |
| 0 | 0 | 0 |
| 0 | 0 | 0 |
| 1 | 1 | 1 |
| 0 | 1 | 0 |

| | | |
|---|---|---|
| 0 | 0 | 0 |
| 1 | 1 | 1 |
| 1 | 1 | 1 |
| 1 | 1 | 1 |
| 1 | 1 | 1 |
| 0 | 0 | 0 |
| 0 | 0 | 0 |
| 1 | 1 | 1 |
| 0 | 1 | 0 |
| 1 | 1 | 1 |
| 1 | 1 | 1 |
| 0 | 0 | 0 |
| 0 | 1 | 0 |
| 1 | 1 | 1 |
| 0 | 1 | 0 |
| 1 | 1 | 1 |
| 0 | 0 | 0 |
| 0 | 0 | 0 |
| 1 | 1 | 1 |
| 1 | 1 | 1 |
| 1 | 1 | 1 |
| 1 | 1 | 1 |
| 1 | 1 | 1 |
| 1 | 1 | 1 |
| 0 | 0 | 0 |
| 1 | 1 | 1 |
| 1 | 1 | 1 |
| 1 | 1 | 1 |
| 1 | 1 | 1 |
| 1 | 1 | 1 |
| 0 | 1 | 0 |
| 0 | 0 | 0 |
| 0 | 0 | 0 |
| 0 | 1 | 0 |
| 1 | 1 | 1 |
| 0 | 0 | 0 |
| 0 | 0 | 0 |
| 0 | 0 | 0 |
| 0 | 0 | 0 |
| 0 | 0 | 0 |
| 0 | 0 | 0 |
| 0 | 0 | 0 |
| 0 | 0 | 0 |
| 0 | 0 | 0 |
| 0 | 0 | 0 |
| 0 | 0 | 0 |
| 0 | 0 | 0 |
| 0 | 0 | 0 |
| 1 | 0 | 0 |
| 1 | 1 | 1 |
| 1 | 0 | 0 |
| 1 | 1 | 1 |
| 0 | 0 | 0 |
| 1 | 1 | 1 |

| | | |
|---|---|---|
| 1 | 1 | 1 |
| 1 | 1 | 1 |
| 1 | 1 | 1 |
| 1 | 1 | 1 |
| 1 | 1 | 1 |
| 1 | 1 | 1 |
| 1 | 1 | 1 |
| 0 | 0 | 0 |
| 0 | 0 | 0 |
| 1 | 0 | 0 |
| 1 | 1 | 1 |
| 1 | 1 | 1 |
| 1 | 1 | 1 |
| 0 | 0 | 0 |
| 1 | 1 | 1 |
| 1 | 1 | 1 |
| 0 | 0 | 0 |
| 1 | 1 | 1 |
| 0 | 0 | 0 |
| 0 | 1 | 0 |
| 1 | 1 | 1 |
| 1 | 1 | 1 |
| 1 | 1 | 1 |
| 0 | 0 | 0 |
| 1 | 1 | 1 |
| 1 | 1 | 1 |
| 0 | 1 | 0 |
| 0 | 1 | 0 |
| 0 | 1 | 0 |
| 0 | 1 | 0 |
| 1 | 1 | 1 |
| 0 | 0 | 0 |
| 1 | 1 | 1 |
| 0 | 1 | 0 |
| 1 | 1 | 1 |
| 1 | 1 | 1 |
| 1 | 1 | 1 |
| 1 | 1 | 1 |
| 0 | 0 | 0 |
| 1 | 1 | 1 |
| 1 | 1 | 1 |
| 1 | 1 | 1 |
| 0 | 1 | 0 |
| 1 | 1 | 1 |
| 1 | 1 | 1 |
| 0 | 0 | 0 |
| 0 | 1 | 0 |
| 0 | 1 | 0 |
| 1 | 0 | 0 |
| 1 | 0 | 0 |
| 0 | 1 | 0 |
| 1 | 0 | 0 |
| 0 | 0 | 0 |
| 1 | 0 | 0 |

| | | |
|---|---|---|
| 1 | 0 | 0 |
| 1 | 0 | 0 |
| 1 | 0 | 0 |
| 1 | 0 | 0 |
| 0 | 0 | 0 |
| 0 | 0 | 0 |
| 0 | 0 | 0 |
| 0 | 0 | 0 |
| 1 | 0 | 0 |
| 1 | 0 | 0 |
| 1 | 0 | 0 |
| 1 | 0 | 0 |
| 1 | 0 | 0 |
| 0 | 0 | 0 |
| 1 | 0 | 0 |
| 0 | 0 | 0 |
| 1 | 0 | 0 |
| 0 | 0 | 0 |
| 1 | 0 | 0 |
| 0 | 0 | 0 |
| 0 | 0 | 0 |
| 1 | 0 | 0 |
| 1 | 0 | 0 |
| 0 | 0 | 0 |
| 0 | 0 | 0 |
| 1 | 0 | 0 |
| 0 | 0 | 0 |
| 0 | 0 | 0 |
| 0 | 0 | 0 |
| 1 | 0 | 0 |
| 1 | 0 | 0 |
| 0 | 0 | 0 |
| 1 | 0 | 0 |
| 1 | 0 | 0 |
| 1 | 0 | 0 |
| 1 | 0 | 0 |
| 1 | 0 | 0 |
| 1 | 0 | 0 |
| 1 | 0 | 0 |
| 1 | 0 | 0 |
| 1 | 0 | 0 |
| 1 | 0 | 0 |
| 1 | 0 | 0 |
| 0 | 0 | 0 |
| 1 | 1 | 1 |
| 1 | 1 | 1 |
| 0 | 0 | 0 |
| 0 | 0 | 0 |
| 0 | 1 | 0 |
| 0 | 0 | 0 |
| 0 | 0 | 0 |
| 0 | 0 | 0 |
| 1 | 1 | 1 |
| 1 | 1 | 1 |

| | | |
|---|---|---|
| 1 | 1 | 1 |
| 1 | 1 | 1 |
| 1 | 1 | 1 |
| 1 | 1 | 1 |
| 1 | 1 | 1 |
| 1 | 1 | 1 |
| 1 | 1 | 1 |
| 0 | 0 | 0 |
| 1 | 1 | 1 |
| 1 | 1 | 1 |
| 0 | 1 | 0 |
| 0 | 1 | 0 |
| 1 | 0 | 0 |
| 1 | 1 | 1 |
| 0 | 0 | 0 |
| 0 | 0 | 0 |
| 0 | 1 | 0 |
| 0 | 0 | 0 |
| 1 | 1 | 1 |
| 1 | 1 | 1 |
| 0 | 0 | 0 |
| 0 | 1 | 0 |
| 0 | 1 | 0 |
| 1 | 1 | 1 |
| 0 | 0 | 0 |
| 1 | 1 | 1 |
| 0 | 1 | 0 |
| 0 | 0 | 0 |
| 0 | 0 | 0 |
| 0 | 0 | 0 |
| 0 | 0 | 0 |
| 1 | 1 | 1 |
| 1 | 1 | 1 |
| 0 | 1 | 0 |
| 1 | 1 | 1 |
| 1 | 1 | 1 |
| 0 | 0 | 0 |
| 0 | 0 | 0 |
| 1 | 1 | 1 |
| 1 | 1 | 1 |
| 1 | 1 | 1 |
| 1 | 1 | 1 |
| 0 | 0 | 0 |
| 0 | 1 | 0 |
| 1 | 1 | 1 |
| 1 | 1 | 1 |
| 1 | 1 | 1 |
| 1 | 1 | 1 |
| 0 | 0 | 0 |
| 0 | 0 | 0 |
| 0 | 0 | 0 |
| 0 | 0 | 0 |
| 0 | 0 | 0 |
| 1 | 1 | 1 |

| | | |
|---|---|---|
| 0 | 0 | 0 |
| 0 | 0 | 0 |
| 0 | 1 | 0 |
| 0 | 0 | 0 |
| 1 | 1 | 1 |
| 0 | 1 | 0 |
| 0 | 1 | 0 |
| 1 | 1 | 1 |
| 1 | 1 | 1 |
| 0 | 0 | 0 |
| 1 | 1 | 1 |
| 0 | 1 | 0 |
| 1 | 1 | 1 |
| 0 | 1 | 0 |
| 0 | 1 | 0 |
| 0 | 1 | 0 |
| 0 | 1 | 0 |
| 1 | 1 | 1 |
| 1 | 1 | 1 |
| 0 | 1 | 0 |
| 0 | 1 | 0 |
| 0 | 1 | 0 |
| 0 | 1 | 0 |
| 0 | 1 | 0 |
| 0 | 1 | 0 |
| 0 | 1 | 0 |
| 0 | 1 | 0 |
| 0 | 1 | 0 |
| 0 | 1 | 0 |
| 0 | 1 | 0 |
| 1 | 1 | 1 |
| 1 | 1 | 1 |
| 0 | 0 | 0 |
| 1 | 1 | 1 |
| 1 | 1 | 1 |
| 1 | 1 | 1 |
| 1 | 1 | 1 |
| 0 | 1 | 0 |
| 1 | 1 | 1 |
| 1 | 1 | 1 |
| 1 | 1 | 1 |
| 1 | 1 | 1 |
| 1 | 1 | 1 |
| 0 | 1 | 0 |
| 0 | 1 | 0 |
| 1 | 1 | 1 |
| 1 | 1 | 1 |
| 0 | 0 | 0 |
| 0 | 0 | 0 |
| 0 | 0 | 0 |
| 0 | 0 | 0 |
| 0 | 0 | 0 |
| 1 | 1 | 1 |
| 0 | 0 | 0 |

| | | |
|---|---|---|
| 1 | 1 | 1 |
| 0 | 0 | 0 |
| 1 | 1 | 1 |
| 0 | 1 | 0 |
| 0 | 1 | 0 |
| 0 | 0 | 0 |
| 0 | 0 | 0 |
| 0 | 0 | 0 |
| 0 | 0 | 0 |
| 0 | 0 | 0 |
| 0 | 1 | 0 |
| 0 | 0 | 0 |
| 0 | 1 | 0 |
| 1 | 1 | 1 |
| 0 | 1 | 0 |
| 0 | 0 | 0 |
| 1 | 1 | 1 |
| 1 | 1 | 1 |
| 0 | 0 | 0 |
| 0 | 0 | 0 |
| 0 | 0 | 0 |
| 0 | 0 | 0 |
| 0 | 0 | 0 |
| 0 | 0 | 0 |
| 0 | 1 | 0 |
| 0 | 0 | 0 |
| 0 | 0 | 0 |
| 0 | 0 | 0 |
| 0 | 0 | 0 |
| 0 | 0 | 0 |
| 0 | 0 | 0 |
| 1 | 1 | 1 |
| 1 | 1 | 1 |
| 0 | 0 | 0 |
| 0 | 0 | 0 |
| 1 | 0 | 0 |
| 0 | 0 | 0 |
| 0 | 0 | 0 |
| 0 | 0 | 0 |
| 0 | 0 | 0 |
| 0 | 0 | 0 |
| 1 | 0 | 0 |
| 1 | 1 | 1 |
| 1 | 1 | 1 |
| 0 | 0 | 0 |
| 0 | 0 | 0 |
| 0 | 0 | 0 |
| 1 | 1 | 1 |
| 1 | 1 | 1 |
| 1 | 1 | 1 |
| 1 | 1 | 1 |
| 1 | 1 | 1 |
| 1 | 1 | 1 |
| 0 | 0 | 0 |

| | | |
|---|---|---|
| 1 | 1 | 1 |
| 1 | 1 | 1 |
| 1 | 1 | 1 |
| 1 | 1 | 1 |
| 0 | 0 | 0 |
| 0 | 0 | 0 |
| 1 | 0 | 0 |
| 1 | 0 | 0 |
| 0 | 1 | 0 |
| 1 | 1 | 1 |
| 1 | 1 | 1 |
| 0 | 0 | 0 |
| 1 | 1 | 1 |
| 0 | 0 | 0 |
| 0 | 0 | 0 |
| 0 | 0 | 0 |
| 0 | 0 | 0 |
| 0 | 0 | 0 |
| 0 | 0 | 0 |
| 0 | 0 | 0 |
| 0 | 0 | 0 |
| 0 | 0 | 0 |
| 0 | 0 | 0 |
| 0 | 1 | 0 |
| 0 | 1 | 0 |
| 0 | 1 | 0 |
| 1 | 1 | 1 |
| 1 | 1 | 1 |
| 1 | 1 | 1 |
| 1 | 1 | 1 |
| 1 | 1 | 1 |
| 0 | 0 | 0 |
| 1 | 1 | 1 |
| 0 | 0 | 0 |
| 0 | 0 | 0 |
| 0 | 0 | 0 |
| 0 | 0 | 0 |
| 1 | 1 | 1 |
| 1 | 1 | 1 |
| 0 | 1 | 0 |
| 1 | 1 | 1 |
| 1 | 1 | 1 |
| 1 | 1 | 1 |
| 0 | 1 | 0 |
| 1 | 1 | 1 |
| 0 | 0 | 0 |
| 1 | 1 | 1 |
| 1 | 1 | 1 |
| 1 | 1 | 1 |
| 0 | 0 | 0 |
| 0 | 0 | 0 |
| 0 | 0 | 0 |
| 0 | 1 | 0 |
| 0 | 1 | 0 |

| | | |
|---|---|---|
| 0 | 1 | 0 |
| 0 | 1 | 0 |
| 0 | 1 | 0 |
| 0 | 0 | 0 |
| 0 | 1 | 0 |
| 1 | 1 | 1 |
| 0 | 0 | 0 |
| 0 | 1 | 0 |
| 1 | 1 | 1 |
| 1 | 1 | 1 |
| 0 | 0 | 0 |
| 0 | 0 | 0 |
| 1 | 1 | 1 |
| 1 | 1 | 1 |
| 0 | 1 | 0 |
| 0 | 1 | 0 |
| 0 | 0 | 0 |
| 0 | 0 | 0 |
| 1 | 1 | 1 |
| 0 | 0 | 0 |
| 0 | 0 | 0 |
| 0 | 0 | 0 |
| 0 | 1 | 0 |
| 0 | 0 | 0 |
| 0 | 0 | 0 |
| 1 | 1 | 1 |
| 0 | 0 | 0 |
| 0 | 1 | 0 |
| 1 | 1 | 1 |
| 0 | 0 | 0 |
| 0 | 1 | 0 |
| 1 | 1 | 1 |
| 0 | 0 | 0 |
| 0 | 1 | 0 |
| 0 | 1 | 0 |
| 0 | 0 | 0 |
| 1 | 1 | 1 |
| 0 | 0 | 0 |
| 1 | 1 | 1 |
| 0 | 0 | 0 |
| 0 | 0 | 0 |
| 1 | 0 | 0 |
| 0 | 0 | 0 |
| 0 | 0 | 0 |
| 1 | 1 | 1 |
| 1 | 0 | 0 |
| 0 | 1 | 0 |
| 1 | 1 | 1 |
| 1 | 1 | 1 |
| 1 | 1 | 1 |
| 1 | 1 | 1 |
| 0 | 1 | 0 |
| 0 | 1 | 0 |
| 0 | 1 | 0 |

| | | |
|---|---|---|
| 0 | 1 | 0 |
| 0 | 1 | 0 |
| 0 | 1 | 0 |
| 0 | 0 | 0 |
| 1 | 0 | 0 |
| 0 | 0 | 0 |
| 1 | 1 | 1 |
| 1 | 0 | 0 |
| 0 | 0 | 0 |
| 0 | 0 | 0 |
| 1 | 1 | 1 |
| 0 | 0 | 0 |
| 0 | 0 | 0 |
| 0 | 0 | 0 |
| 1 | 0 | 0 |
| 0 | 0 | 0 |
| 0 | 0 | 0 |
| 0 | 1 | 0 |
| 0 | 0 | 0 |
| 0 | 0 | 0 |
| 0 | 0 | 0 |
| 0 | 0 | 0 |
| 1 | 1 | 1 |
| 0 | 0 | 0 |
| 0 | 0 | 0 |
| 0 | 0 | 0 |
| 0 | 0 | 0 |
| 0 | 0 | 0 |
| 0 | 0 | 0 |
| 0 | 0 | 0 |
| 0 | 0 | 0 |
| 0 | 0 | 0 |
| 0 | 0 | 0 |
| 0 | 1 | 0 |
| 0 | 1 | 0 |
| 0 | 0 | 0 |
| 1 | 1 | 1 |
| 0 | 0 | 0 |
| 0 | 0 | 0 |
| 0 | 0 | 0 |
| 0 | 0 | 0 |
| 1 | 1 | 1 |
| 0 | 0 | 0 |
| 0 | 0 | 0 |
| 1 | 1 | 1 |
| 0 | 1 | 0 |
| 1 | 1 | 1 |
| 1 | 1 | 1 |
| 1 | 1 | 1 |
| 0 | 1 | 0 |
| 1 | 1 | 1 |
| 1 | 1 | 1 |
| 1 | 1 | 1 |
| 1 | 1 | 1 |

| | | |
|---|---|---|
| 1 | 1 | 1 |
| 0 | 0 | 0 |
| 1 | 1 | 1 |
| 1 | 1 | 1 |
| 1 | 1 | 1 |
| 1 | 1 | 1 |
| 1 | 1 | 1 |
| 1 | 1 | 1 |
| 1 | 1 | 1 |
| 0 | 1 | 0 |
| 1 | 1 | 1 |
| 0 | 0 | 0 |
| 1 | 0 | 0 |
| 0 | 1 | 0 |
| 0 | 0 | 0 |
| 0 | 1 | 0 |
| 0 | 1 | 0 |
| 1 | 1 | 1 |
| 1 | 1 | 1 |
| 1 | 1 | 1 |
| 0 | 0 | 0 |
| 0 | 1 | 0 |
| 0 | 0 | 0 |
| 0 | 0 | 0 |
| 1 | 1 | 1 |
| 0 | 0 | 0 |
| 1 | 1 | 1 |
| 1 | 1 | 1 |
| 1 | 1 | 1 |
| 1 | 1 | 1 |
| 1 | 1 | 1 |
| 0 | 0 | 0 |
| 1 | 1 | 1 |
| 0 | 0 | 0 |
| 1 | 1 | 1 |
| 0 | 1 | 0 |
| 0 | 1 | 0 |
| 0 | 1 | 0 |
| 1 | 1 | 1 |
| 0 | 0 | 0 |
| 1 | 1 | 1 |
| 1 | 1 | 1 |
| 0 | 1 | 0 |
| 0 | 0 | 0 |
| 1 | 1 | 1 |
| 1 | 1 | 1 |
| 1 | 1 | 1 |
| 0 | 0 | 0 |
| 0 | 0 | 0 |
| 1 | 1 | 1 |
| 1 | 1 | 1 |
| 0 | 0 | 0 |
| 0 | 0 | 0 |
| 0 | 0 | 0 |

| | | |
|---|---|---|
| 0 | 0 | 0 |
| 1 | 1 | 1 |
| 0 | 0 | 0 |
| 0 | 0 | 0 |
| 0 | 0 | 0 |
| 1 | 1 | 1 |
| 1 | 1 | 1 |
| 0 | 1 | 0 |
| 1 | 1 | 1 |
| 0 | 0 | 0 |
| 1 | 1 | 1 |
| 0 | 1 | 0 |
| 1 | 1 | 1 |
| 1 | 1 | 1 |
| 0 | 0 | 0 |
| 0 | 1 | 0 |
| 1 | 1 | 1 |
| 0 | 0 | 0 |
| 1 | 1 | 1 |
| 0 | 0 | 0 |
| 0 | 0 | 0 |
| 1 | 1 | 1 |
| 0 | 0 | 0 |
| 1 | 1 | 1 |
| 0 | 0 | 0 |
| 0 | 0 | 0 |
| 1 | 1 | 1 |
| 0 | 1 | 0 |
| 0 | 0 | 0 |
| 1 | 1 | 1 |
| 1 | 1 | 1 |
| 1 | 1 | 1 |
| 1 | 1 | 1 |
| 0 | 1 | 0 |
| 0 | 0 | 0 |
| 0 | 0 | 0 |
| 1 | 1 | 1 |
| 0 | 0 | 0 |
| 0 | 0 | 0 |
| 0 | 0 | 0 |
| 0 | 0 | 0 |
| 0 | 0 | 0 |
| 0 | 0 | 0 |
| 0 | 1 | 0 |
| 0 | 0 | 0 |
| 1 | 1 | 1 |
| 0 | 1 | 0 |
| 0 | 0 | 0 |
| 0 | 0 | 0 |
| 0 | 0 | 0 |
| 0 | 0 | 0 |
| 0 | 0 | 0 |
| 1 | 1 | 1 |
| 1 | 1 | 1 |

| | | |
|---|---|---|
| 1 | 1 | 1 |
| 0 | 0 | 0 |
| 0 | 0 | 0 |
| 0 | 0 | 0 |
| 0 | 0 | 0 |
| 0 | 0 | 0 |
| 0 | 0 | 0 |
| 1 | 0 | 0 |
| 1 | 0 | 0 |
| 1 | 1 | 1 |
| 0 | 0 | 0 |
| 0 | 0 | 0 |
| 1 | 1 | 1 |
| 0 | 0 | 0 |
| 1 | 1 | 1 |
| 0 | 0 | 0 |
| 1 | 1 | 1 |
| 1 | 1 | 1 |
| 0 | 1 | 0 |
| 0 | 1 | 0 |
| 0 | 1 | 0 |
| 1 | 1 | 1 |
| 0 | 1 | 0 |
| 0 | 0 | 0 |
| 0 | 0 | 0 |
| 1 | 1 | 1 |
| 1 | 1 | 1 |
| 1 | 1 | 1 |
| 1 | 1 | 1 |
| 0 | 0 | 0 |
| 0 | 0 | 0 |
| 0 | 0 | 0 |
| 0 | 0 | 0 |
| 1 | 1 | 1 |
| 0 | 0 | 0 |
| 0 | 0 | 0 |
| 0 | 0 | 0 |
| 0 | 0 | 0 |
| 0 | 0 | 0 |
| 1 | 1 | 1 |
| 1 | 1 | 1 |
| 0 | 0 | 0 |
| 1 | 1 | 1 |
| 1 | 1 | 1 |
| 0 | 0 | 0 |
| 1 | 0 | 0 |
| 1 | 1 | 1 |
| 0 | 1 | 0 |
| 1 | 0 | 0 |
| 0 | 1 | 0 |
| 0 | 0 | 0 |
| 1 | 1 | 1 |
| 0 | 0 | 0 |
| 0 | 0 | 0 |

| | | |
|---|---|---|
| 0 | 1 | 0 |
| 1 | 1 | 1 |
| 0 | 0 | 0 |
| 0 | 0 | 0 |
| 1 | 1 | 1 |
| 1 | 1 | 1 |
| 1 | 1 | 1 |
| 1 | 1 | 1 |
| 1 | 1 | 1 |
| 1 | 1 | 1 |
| 1 | 1 | 1 |
| 1 | 1 | 1 |
| 0 | 0 | 0 |
| 1 | 1 | 1 |
| 1 | 1 | 1 |
| 1 | 1 | 1 |
| 1 | 1 | 1 |
| 1 | 1 | 1 |
| 1 | 1 | 1 |
| 1 | 1 | 1 |
| 1 | 1 | 1 |
| 1 | 1 | 1 |
| 1 | 1 | 1 |
| 1 | 1 | 1 |
| 1 | 1 | 1 |
| 1 | 1 | 1 |
| 0 | 1 | 0 |
| 1 | 1 | 1 |
| 1 | 1 | 1 |
| 1 | 1 | 1 |
| 1 | 1 | 1 |
| 0 | 0 | 0 |
| 1 | 1 | 1 |
| 0 | 1 | 0 |
| 1 | 1 | 1 |
| 1 | 1 | 1 |
| 0 | 1 | 0 |
| 1 | 1 | 1 |
| 1 | 1 | 1 |
| 1 | 1 | 1 |
| 0 | 0 | 0 |
| 0 | 0 | 0 |
| 1 | 1 | 1 |
| 1 | 1 | 1 |
| 1 | 1 | 1 |
| 1 | 0 | 0 |
| 1 | 0 | 0 |
| 1 | 1 | 1 |
| 0 | 0 | 0 |
| 0 | 0 | 0 |
| 1 | 1 | 1 |
| 0 | 1 | 0 |
| 0 | 0 | 0 |
| 1 | 1 | 1 |

| | | |
|---|---|---|
| 0 | 0 | 0 |
| 0 | 0 | 0 |
| 0 | 0 | 0 |
| 1 | 1 | 1 |
| 1 | 1 | 1 |
| 1 | 1 | 1 |
| 1 | 1 | 1 |
| 0 | 0 | 0 |
| 0 | 0 | 0 |
| 0 | 0 | 0 |
| 0 | 0 | 0 |
| 1 | 1 | 1 |
| 0 | 0 | 0 |
| 1 | 1 | 1 |
| 1 | 1 | 1 |
| 1 | 1 | 1 |
| 1 | 1 | 1 |
| 0 | 0 | 0 |
| 1 | 1 | 1 |
| 1 | 1 | 1 |
| 1 | 1 | 1 |
| 1 | 1 | 1 |
| 0 | 1 | 0 |
| 1 | 1 | 1 |
| 1 | 1 | 1 |
| 0 | 1 | 0 |
| 1 | 1 | 1 |
| 1 | 1 | 1 |
| 1 | 1 | 1 |
| 1 | 1 | 1 |
| 0 | 0 | 0 |
| 0 | 0 | 0 |
| 1 | 1 | 1 |
| 1 | 1 | 1 |
| 1 | 1 | 1 |
| 1 | 1 | 1 |
| 1 | 1 | 1 |
| 1 | 1 | 1 |
| 1 | 1 | 1 |
| 1 | 1 | 1 |
| 1 | 1 | 1 |
| 0 | 0 | 0 |
| 1 | 1 | 1 |
| 1 | 1 | 1 |
| 0 | 0 | 0 |
| 0 | 0 | 0 |
| 1 | 1 | 1 |
| 1 | 1 | 1 |
| 0 | 0 | 0 |
| 0 | 0 | 0 |
| 0 | 0 | 0 |
| 0 | 0 | 0 |
| 0 | 0 | 0 |
| 0 | 0 | 0 |

| | | |
|---|---|---|
| 1 | 1 | 1 |
| 1 | 1 | 1 |
| 1 | 1 | 1 |
| 1 | 1 | 1 |
| 1 | 1 | 1 |
| 0 | 0 | 0 |
| 0 | 0 | 0 |
| 0 | 0 | 0 |
| 0 | 0 | 0 |
| 0 | 0 | 0 |
| 0 | 1 | 0 |
| 0 | 0 | 0 |
| 0 | 0 | 0 |
| 0 | 0 | 0 |
| 0 | 1 | 0 |
| 0 | 0 | 0 |
| 0 | 1 | 0 |
| 0 | 1 | 0 |
| 0 | 0 | 0 |
| 0 | 0 | 0 |
| 0 | 0 | 0 |
| 0 | 0 | 0 |
| 1 | 0 | 0 |
| 0 | 0 | 0 |
| 0 | 0 | 0 |
| 0 | 0 | 0 |
| 0 | 0 | 0 |
| 1 | 1 | 1 |
| 1 | 1 | 1 |
| 1 | 1 | 1 |
| 1 | 1 | 1 |
| 1 | 1 | 1 |
| 0 | 0 | 0 |
| 0 | 1 | 0 |
| 0 | 0 | 0 |
| 0 | 0 | 0 |
| 0 | 0 | 0 |
| 0 | 0 | 0 |
| 0 | 0 | 0 |
| 0 | 0 | 0 |
| 0 | 0 | 0 |
| 0 | 0 | 0 |
| 0 | 0 | 0 |
| 1 | 1 | 1 |
| 1 | 1 | 1 |
| 1 | 1 | 1 |
| 1 | 1 | 1 |
| 0 | 0 | 0 |
| 0 | 0 | 0 |
| 0 | 0 | 0 |
| 1 | 1 | 1 |
| 1 | 1 | 1 |
| 0 | 0 | 0 |
| 1 | 1 | 1 |

| | | |
|---|---|---|
| 1 | 1 | 1 |
| 0 | 0 | 0 |
| 0 | 1 | 0 |
| 0 | 1 | 0 |
| 1 | 1 | 1 |
| 1 | 1 | 1 |
| 0 | 1 | 0 |
| 1 | 1 | 1 |
| 0 | 1 | 0 |
| 1 | 1 | 1 |
| 1 | 1 | 1 |
| 1 | 1 | 1 |
| 0 | 0 | 0 |
| 0 | 0 | 0 |
| 0 | 0 | 0 |
| 0 | 0 | 0 |
| 0 | 0 | 0 |
| 0 | 0 | 0 |
| 0 | 0 | 0 |
| 0 | 1 | 0 |
| 0 | 0 | 0 |
| 0 | 0 | 0 |
| 0 | 0 | 0 |
| 0 | 0 | 0 |
| 0 | 0 | 0 |
| 1 | 1 | 1 |
| 1 | 1 | 1 |
| 0 | 0 | 0 |
| 0 | 1 | 0 |
| 0 | 0 | 0 |
| 0 | 0 | 0 |
| 1 | 1 | 1 |
| 0 | 0 | 0 |
| 1 | 1 | 1 |
| 1 | 1 | 1 |
| 1 | 1 | 1 |
| 1 | 0 | 0 |
| 1 | 1 | 1 |
| 1 | 1 | 1 |
| 1 | 1 | 1 |
| 0 | 0 | 0 |
| 1 | 1 | 1 |
| 1 | 1 | 1 |
| 0 | 0 | 0 |
| 0 | 0 | 0 |
| 1 | 1 | 1 |
| 0 | 0 | 0 |
| 0 | 0 | 0 |
| 1 | 1 | 1 |
| 1 | 1 | 1 |
| 0 | 0 | 0 |
| 1 | 1 | 1 |
| 1 | 1 | 1 |
| 1 | 1 | 1 |

| | | |
|---|---|---|
| 1 | 1 | 1 |
| 1 | 1 | 1 |
| 1 | 1 | 1 |
| 1 | 1 | 1 |
| 0 | 0 | 0 |
| 0 | 0 | 0 |
| 1 | 1 | 1 |
| 1 | 1 | 1 |
| 1 | 1 | 1 |
| 0 | 1 | 0 |
| 1 | 1 | 1 |
| 0 | 0 | 0 |
| 0 | 0 | 0 |
| 1 | 1 | 1 |
| 1 | 1 | 1 |
| 0 | 0 | 0 |
| 0 | 1 | 0 |
| 1 | 1 | 1 |
| 0 | 0 | 0 |
| 1 | 1 | 1 |
| 1 | 1 | 1 |
| 1 | 1 | 1 |
| 1 | 1 | 1 |
| 0 | 0 | 0 |
| 1 | 1 | 1 |
| 0 | 0 | 0 |
| 1 | 1 | 1 |
| 1 | 1 | 1 |
| 0 | 0 | 0 |
| 0 | 0 | 0 |
| 0 | 0 | 0 |
| 0 | 0 | 0 |
| 0 | 0 | 0 |
| 0 | 0 | 0 |
| 1 | 1 | 1 |
| 1 | 1 | 1 |
| 1 | 1 | 1 |
| 0 | 0 | 0 |
| 1 | 1 | 1 |
| 1 | 1 | 1 |
| 0 | 0 | 0 |
| 0 | 0 | 0 |
| 0 | 0 | 0 |
| 1 | 1 | 1 |
| 0 | 0 | 0 |
| 0 | 1 | 0 |
| 0 | 1 | 0 |
| 0 | 0 | 0 |
| 1 | 1 | 1 |
| 1 | 1 | 1 |
| 1 | 1 | 1 |
| 0 | 0 | 0 |
| 1 | 1 | 1 |
| 0 | 0 | 0 |

| | | |
|---|---|---|
| 1 | 1 | 1 |
| 0 | 0 | 0 |
| 0 | 0 | 0 |
| 0 | 1 | 0 |
| 1 | 1 | 1 |
| 1 | 1 | 1 |
| 0 | 0 | 0 |
| 1 | 1 | 1 |
| 0 | 1 | 0 |
| 0 | 1 | 0 |
| 1 | 1 | 1 |
| 1 | 1 | 1 |
| 1 | 1 | 1 |
| 1 | 1 | 1 |
| 1 | 1 | 1 |
| 1 | 1 | 1 |
| 1 | 1 | 1 |
| 1 | 1 | 1 |
| 1 | 1 | 1 |
| 1 | 1 | 1 |
| 1 | 1 | 1 |
| 0 | 0 | 0 |
| 0 | 1 | 0 |
| 0 | 1 | 0 |
| 1 | 1 | 1 |
| 1 | 1 | 1 |
| 0 | 0 | 0 |
| 1 | 1 | 1 |
| 0 | 0 | 0 |
| 0 | 1 | 0 |
| 1 | 1 | 1 |
| 0 | 0 | 0 |
| 0 | 0 | 0 |
| 1 | 0 | 0 |
| 1 | 1 | 1 |
| 1 | 1 | 1 |
| 1 | 1 | 1 |
| 1 | 1 | 1 |
| 0 | 0 | 0 |
| 0 | 0 | 0 |
| 0 | 0 | 0 |
| 1 | 1 | 1 |
| 1 | 1 | 1 |
| 0 | 1 | 0 |
| 0 | 1 | 0 |
| 0 | 0 | 0 |
| 1 | 1 | 1 |
| 0 | 0 | 0 |
| 1 | 1 | 1 |
| 0 | 0 | 0 |
| 0 | 0 | 0 |
| 0 | 0 | 0 |
| 1 | 1 | 1 |
| 1 | 1 | 1 |

| | | |
|---|---|---|
| 0 | 0 | 0 |
| 0 | 0 | 0 |
| 1 | 1 | 1 |
| 1 | 1 | 1 |
| 1 | 1 | 1 |
| 1 | 1 | 1 |
| 1 | 1 | 1 |
| 1 | 1 | 1 |
| 1 | 1 | 1 |
| 1 | 1 | 1 |
| 1 | 1 | 1 |
| 1 | 1 | 1 |
| 1 | 1 | 1 |
| 1 | 1 | 1 |
| 1 | 1 | 1 |
| 0 | 0 | 0 |
| 0 | 0 | 0 |
| 0 | 0 | 0 |
| 1 | 1 | 1 |
| 1 | 1 | 1 |
| 0 | 1 | 0 |
| 0 | 0 | 0 |
| 0 | 0 | 0 |
| 0 | 0 | 0 |
| 0 | 0 | 0 |
| 1 | 1 | 1 |
| 1 | 1 | 1 |
| 1 | 1 | 1 |
| 1 | 1 | 1 |
| 1 | 1 | 1 |
| 0 | 0 | 0 |
| 0 | 1 | 0 |
| 0 | 0 | 0 |
| 1 | 1 | 1 |
| 1 | 0 | 0 |
| 1 | 0 | 0 |
| 0 | 0 | 0 |
| 0 | 0 | 0 |
| 0 | 1 | 0 |
| 1 | 1 | 1 |
| 1 | 0 | 0 |
| 0 | 0 | 0 |
| 1 | 1 | 1 |
| 0 | 1 | 0 |
| 0 | 0 | 0 |
| 1 | 0 | 0 |
| 0 | 0 | 0 |
| 0 | 0 | 0 |
| 1 | 1 | 1 |
| 0 | 1 | 0 |
| 0 | 0 | 0 |
| 0 | 0 | 0 |
| 1 | 1 | 1 |
| 0 | 0 | 0 |

| | | |
|---|---|---|
| 0 | 1 | 0 |
| 0 | 0 | 0 |
| 1 | 0 | 0 |
| 0 | 0 | 0 |
| 1 | 1 | 1 |
| 1 | 1 | 1 |
| 1 | 1 | 1 |
| 0 | 1 | 0 |
| 0 | 0 | 0 |
| 1 | 1 | 1 |
| 0 | 1 | 0 |
| 0 | 1 | 0 |
| 0 | 0 | 0 |
| 0 | 0 | 0 |
| 0 | 0 | 0 |
| 1 | 1 | 1 |
| 0 | 0 | 0 |
| 0 | 1 | 0 |
| 0 | 0 | 0 |
| 0 | 0 | 0 |
| 0 | 0 | 0 |
| 0 | 0 | 0 |
| 0 | 0 | 0 |
| 0 | 0 | 0 |
| 0 | 1 | 0 |
| 0 | 1 | 0 |
| 0 | 0 | 0 |
| 0 | 0 | 0 |
| 0 | 0 | 0 |
| 0 | 0 | 0 |
| 1 | 1 | 1 |
| 1 | 1 | 1 |
| 1 | 1 | 1 |
| 1 | 1 | 1 |
| 1 | 1 | 1 |
| 0 | 0 | 0 |
| 0 | 0 | 0 |
| 0 | 1 | 0 |
| 1 | 1 | 1 |
| 1 | 1 | 1 |
| 0 | 1 | 0 |
| 1 | 1 | 1 |
| 1 | 1 | 1 |
| 0 | 0 | 0 |
| 0 | 0 | 0 |
| 0 | 0 | 0 |
| 1 | 1 | 1 |
| 0 | 0 | 0 |
| 0 | 0 | 0 |
| 0 | 0 | 0 |
| 1 | 1 | 1 |
| 1 | 1 | 1 |
| 0 | 1 | 0 |
| 0 | 1 | 0 |

| | | |
|---|---|---|
| 0 | 1 | 0 |
| 1 | 1 | 1 |
| 1 | 0 | 0 |
| 1 | 1 | 1 |
| 0 | 0 | 0 |
| 1 | 0 | 0 |
| 0 | 0 | 0 |
| 0 | 0 | 0 |
| 0 | 0 | 0 |
| 0 | 0 | 0 |
| 0 | 0 | 0 |
| 0 | 0 | 0 |
| 0 | 1 | 0 |
| 0 | 0 | 0 |
| 0 | 0 | 0 |
| 1 | 1 | 1 |
| 1 | 1 | 1 |
| 1 | 1 | 1 |
| 0 | 0 | 0 |
| 0 | 0 | 0 |
| 0 | 0 | 0 |
| 0 | 0 | 0 |
| 0 | 1 | 0 |
| 0 | 1 | 0 |
| 1 | 1 | 1 |
| 1 | 1 | 1 |
| 1 | 1 | 1 |
| 0 | 0 | 0 |
| 0 | 0 | 0 |
| 0 | 0 | 0 |
| 0 | 0 | 0 |
| 0 | 0 | 0 |
| 0 | 0 | 0 |
| 0 | 1 | 0 |
| 1 | 1 | 1 |
| 0 | 0 | 0 |
| 1 | 1 | 1 |
| 1 | 1 | 1 |
| 1 | 1 | 1 |
| 1 | 1 | 1 |
| 0 | 0 | 0 |
| 1 | 1 | 1 |
| 0 | 0 | 0 |
| 0 | 1 | 0 |
| 1 | 1 | 1 |
| 1 | 1 | 1 |
| 1 | 1 | 1 |
| 0 | 0 | 0 |
| 1 | 1 | 1 |
| 1 | 1 | 1 |
| 1 | 1 | 1 |
| 1 | 1 | 1 |
| 1 | 1 | 1 |
| 0 | 1 | 0 |

| | | |
|---|---|---|
| 0 | 1 | 0 |
| 1 | 1 | 1 |
| 1 | 1 | 1 |
| 0 | 1 | 0 |
| 1 | 1 | 1 |
| 0 | 1 | 0 |
| 0 | 1 | 0 |
| 0 | 1 | 0 |
| 0 | 1 | 0 |
| 0 | 1 | 0 |
| 0 | 1 | 0 |
| 0 | 0 | 0 |
| 0 | 1 | 0 |
| 0 | 1 | 0 |
| 0 | 1 | 0 |
| 0 | 1 | 0 |
| 0 | 1 | 0 |
| 0 | 0 | 0 |
| 0 | 0 | 0 |
| 0 | 0 | 0 |
| 0 | 0 | 0 |
| 0 | 1 | 0 |
| 0 | 0 | 0 |
| 1 | 1 | 1 |
| 1 | 1 | 1 |
| 1 | 1 | 1 |
| 1 | 1 | 1 |
| 0 | 1 | 0 |
| 0 | 0 | 0 |
| 0 | 0 | 0 |
| 1 | 1 | 1 |
| 1 | 1 | 1 |
| 0 | 0 | 0 |
| 1 | 1 | 1 |
| 0 | 0 | 0 |
| 0 | 0 | 0 |
| 0 | 0 | 0 |
| 1 | 1 | 1 |
| 0 | 0 | 0 |
| 1 | 1 | 1 |
| 0 | 0 | 0 |
| 1 | 1 | 1 |
| 0 | 0 | 0 |
| 1 | 0 | 0 |
| 1 | 0 | 0 |
| 1 | 0 | 0 |
| 1 | 0 | 0 |
| 0 | 1 | 0 |
| 0 | 1 | 0 |
| 0 | 0 | 0 |
| 1 | 0 | 0 |
| 0 | 0 | 0 |
| 1 | 0 | 0 |
| 1 | 1 | 1 |

| | | |
|---|---|---|
| 0 | 0 | 0 |
| 0 | 0 | 0 |
| 0 | 0 | 0 |
| 0 | 0 | 0 |
| 0 | 0 | 0 |
| 1 | 1 | 1 |
| 0 | 0 | 0 |
| 0 | 0 | 0 |
| 1 | 1 | 1 |
| 0 | 0 | 0 |
| 0 | 0 | 0 |
| 0 | 0 | 0 |
| 0 | 1 | 0 |
| 0 | 1 | 0 |
| 0 | 1 | 0 |
| 0 | 1 | 0 |
| 0 | 1 | 0 |
| 0 | 1 | 0 |
| 0 | 1 | 0 |
| 0 | 1 | 0 |
| 0 | 1 | 0 |
| 0 | 1 | 0 |
| 0 | 1 | 0 |
| 0 | 1 | 0 |
| 0 | 1 | 0 |
| 0 | 1 | 0 |
| 0 | 1 | 0 |
| 0 | 1 | 0 |
| 0 | 1 | 0 |
| 0 | 1 | 0 |
| 0 | 1 | 0 |
| 0 | 1 | 0 |
| 0 | 1 | 0 |
| 0 | 1 | 0 |
| 0 | 1 | 0 |
| 0 | 1 | 0 |
| 0 | 1 | 0 |
| 0 | 1 | 0 |
| 0 | 1 | 0 |
| 0 | 1 | 0 |
| 0 | 1 | 0 |
| 0 | 1 | 0 |
| 0 | 1 | 0 |
| 0 | 1 | 0 |
| 0 | 1 | 0 |
| 0 | 0 | 0 |
| 1 | 1 | 1 |
| 1 | 1 | 1 |
| 1 | 1 | 1 |
| 1 | 1 | 1 |
| 0 | 0 | 0 |
| 1 | 1 | 1 |
| 1 | 1 | 1 |
| 0 | 1 | 0 |

| | | |
|---|---|---|
| 1 | 1 | 1 |
| 1 | 1 | 1 |
| 1 | 1 | 1 |
| 1 | 1 | 1 |
| 0 | 1 | 0 |
| 0 | 1 | 0 |
| 0 | 1 | 0 |
| 0 | 0 | 0 |
| 0 | 0 | 0 |
| 1 | 1 | 1 |
| 0 | 1 | 0 |
| 1 | 1 | 1 |
| 1 | 1 | 1 |
| 1 | 1 | 1 |
| 0 | 0 | 0 |
| 0 | 0 | 0 |
| 0 | 0 | 0 |
| 1 | 1 | 1 |
| 1 | 0 | 0 |
| 1 | 1 | 1 |
| 0 | 0 | 0 |
| 0 | 0 | 0 |
| 0 | 0 | 0 |
| 0 | 0 | 0 |
| 0 | 0 | 0 |
| 0 | 0 | 0 |
| 0 | 1 | 0 |
| 1 | 1 | 1 |
| 0 | 0 | 0 |
| 1 | 1 | 1 |
| 1 | 1 | 1 |
| 1 | 1 | 1 |
| 0 | 0 | 0 |
| 1 | 1 | 1 |
| 1 | 1 | 1 |
| 1 | 1 | 1 |
| 0 | 0 | 0 |
| 0 | 0 | 0 |
| 0 | 0 | 0 |
| 1 | 1 | 1 |
| 1 | 1 | 1 |
| 1 | 1 | 1 |
| 1 | 1 | 1 |
| 1 | 1 | 1 |
| 0 | 1 | 0 |
| 0 | 0 | 0 |
| 1 | 1 | 1 |
| 0 | 1 | 0 |
| 1 | 1 | 1 |
| 0 | 1 | 0 |
| 1 | 1 | 1 |
| 1 | 1 | 1 |
| 0 | 1 | 0 |
| 1 | 1 | 1 |

| | | |
|---|---|---|
| 1 | 1 | 1 |
| 0 | 1 | 0 |
| 1 | 1 | 1 |
| 0 | 0 | 0 |
| 1 | 1 | 1 |
| 1 | 1 | 1 |
| 1 | 1 | 1 |
| 0 | 0 | 0 |
| 0 | 0 | 0 |
| 1 | 1 | 1 |
| 1 | 1 | 1 |
| 1 | 1 | 1 |
| 0 | 0 | 0 |
| 0 | 1 | 0 |
| 0 | 1 | 0 |
| 0 | 1 | 0 |
| 0 | 0 | 0 |
| 1 | 1 | 1 |
| 1 | 1 | 1 |
| 1 | 1 | 1 |
| 1 | 1 | 1 |
| 1 | 1 | 1 |
| 1 | 1 | 1 |
| 1 | 1 | 1 |
| 0 | 1 | 0 |
| 1 | 1 | 1 |
| 0 | 1 | 0 |
| 0 | 0 | 0 |
| 0 | 0 | 0 |
| 0 | 0 | 0 |
| 0 | 0 | 0 |
| 0 | 0 | 0 |
| 0 | 0 | 0 |
| 0 | 1 | 0 |
| 0 | 0 | 0 |
| 0 | 0 | 0 |
| 0 | 0 | 0 |
| 1 | 1 | 1 |
| 1 | 1 | 1 |
| 1 | 1 | 1 |
| 1 | 1 | 1 |
| 0 | 1 | 0 |
| 0 | 0 | 0 |
| 0 | 0 | 0 |
| 1 | 1 | 1 |
| 1 | 1 | 1 |
| 0 | 0 | 0 |
| 1 | 1 | 1 |
| 1 | 1 | 1 |
| 1 | 1 | 1 |
| 1 | 1 | 1 |
| 1 | 1 | 1 |
| 0 | 0 | 0 |
| 0 | 0 | 0 |

| | | |
|---|---|---|
| 0 | 0 | 0 |
| 0 | 0 | 0 |
| 0 | 0 | 0 |
| 0 | 0 | 0 |
| 0 | 0 | 0 |
| 0 | 0 | 0 |
| 0 | 0 | 0 |
| 0 | 0 | 0 |
| 1 | 1 | 1 |
| 0 | 0 | 0 |
| 1 | 1 | 1 |
| 1 | 1 | 1 |
| 1 | 1 | 1 |
| 1 | 1 | 1 |
| 0 | 0 | 0 |
| 1 | 1 | 1 |
| 1 | 1 | 1 |
| 1 | 1 | 1 |
| 1 | 1 | 1 |
| 0 | 0 | 0 |
| 0 | 0 | 0 |
| 1 | 1 | 1 |
| 0 | 0 | 0 |
| 0 | 0 | 0 |
| 0 | 0 | 0 |
| 0 | 0 | 0 |
| 0 | 0 | 0 |
| 0 | 1 | 0 |
| 0 | 1 | 0 |
| 0 | 1 | 0 |
| 0 | 0 | 0 |
| 1 | 1 | 1 |
| 1 | 1 | 1 |
| 1 | 1 | 1 |
| 1 | 1 | 1 |
| 1 | 1 | 1 |
| 1 | 1 | 1 |
| 1 | 1 | 1 |
| 1 | 1 | 1 |
| 1 | 1 | 1 |
| 0 | 1 | 0 |
| 0 | 0 | 0 |
| 1 | 1 | 1 |
| 1 | 1 | 1 |
| 1 | 1 | 1 |
| 0 | 1 | 0 |
| 1 | 1 | 1 |
| 1 | 1 | 1 |
| 0 | 0 | 0 |
| 0 | 1 | 0 |
| 1 | 1 | 1 |
| 1 | 1 | 1 |
| 0 | 0 | 0 |
| 0 | 0 | 0 |

| | | |
|---|---|---|
| 1 | 1 | 1 |
| 1 | 1 | 1 |
| 0 | 1 | 0 |
| 1 | 0 | 0 |
| 0 | 0 | 0 |
| 0 | 0 | 0 |
| 0 | 1 | 0 |
| 0 | 0 | 0 |
| 1 | 1 | 1 |
| 1 | 1 | 1 |
| 0 | 0 | 0 |
| 1 | 0 | 0 |
| 0 | 0 | 0 |
| 0 | 0 | 0 |
| 0 | 0 | 0 |
| 0 | 0 | 0 |
| 1 | 1 | 1 |
| 1 | 1 | 1 |
| 0 | 0 | 0 |
| 0 | 0 | 0 |
| 0 | 0 | 0 |
| 0 | 0 | 0 |
| 0 | 0 | 0 |
| 1 | 1 | 1 |
| 1 | 1 | 1 |
| 0 | 0 | 0 |
| 1 | 1 | 1 |
| 0 | 0 | 0 |
| 0 | 0 | 0 |
| 0 | 0 | 0 |
| 0 | 1 | 0 |
| 0 | 0 | 0 |
| 0 | 0 | 0 |
| 1 | 1 | 1 |
| 1 | 1 | 1 |
| 0 | 0 | 0 |
| 0 | 0 | 0 |
| 1 | 0 | 0 |
| 0 | 0 | 0 |
| 0 | 0 | 0 |
| 0 | 0 | 0 |
| 0 | 0 | 0 |
| 0 | 1 | 0 |
| 1 | 1 | 1 |
| 0 | 0 | 0 |
| 0 | 1 | 0 |
| 0 | 0 | 0 |
| 1 | 1 | 1 |
| 1 | 1 | 1 |
| 1 | 1 | 1 |
| 1 | 1 | 1 |
| 1 | 1 | 1 |
| 0 | 1 | 0 |
| 1 | 1 | 1 |

| | | |
|---|---|---|
| 1 | 1 | 1 |
| 1 | 1 | 1 |
| 1 | 1 | 1 |
| 0 | 1 | 0 |
| 1 | 1 | 1 |
| 1 | 1 | 1 |
| 0 | 1 | 0 |
| 0 | 0 | 0 |
| 1 | 1 | 1 |
| 0 | 0 | 0 |
| 0 | 0 | 0 |
| 0 | 0 | 0 |
| 0 | 1 | 0 |
| 1 | 1 | 1 |
| 0 | 0 | 0 |
| 0 | 1 | 0 |
| 0 | 1 | 0 |
| 0 | 1 | 0 |
| 0 | 1 | 0 |
| 0 | 1 | 0 |
| 1 | 1 | 1 |
| 1 | 1 | 1 |
| 1 | 1 | 1 |
| 0 | 1 | 0 |
| 1 | 1 | 1 |
| 1 | 1 | 1 |
| 1 | 1 | 1 |
| 1 | 0 | 0 |
| 1 | 1 | 1 |
| 0 | 0 | 0 |
| 0 | 0 | 0 |
| 1 | 1 | 1 |
| 1 | 1 | 1 |
| 0 | 1 | 0 |
| 0 | 1 | 0 |
| 0 | 0 | 0 |
| 1 | 1 | 1 |
| 1 | 1 | 1 |
| 1 | 1 | 1 |
| 1 | 1 | 1 |
| 1 | 1 | 1 |
| 0 | 0 | 0 |
| 0 | 0 | 0 |
| 0 | 0 | 0 |
| 1 | 1 | 1 |
| 1 | 1 | 1 |
| 1 | 1 | 1 |
| 1 | 1 | 1 |
| 1 | 1 | 1 |
| 1 | 1 | 1 |
| 1 | 1 | 1 |
| 1 | 1 | 1 |
| 1 | 1 | 1 |
| 0 | 0 | 0 |

| | | |
|---|---|---|
| 1 | 1 | 1 |
| 1 | 1 | 1 |
| 1 | 1 | 1 |
| 1 | 1 | 1 |
| 1 | 1 | 1 |
| 1 | 1 | 1 |
| 1 | 1 | 1 |
| 0 | 1 | 0 |
| 0 | 0 | 0 |
| 0 | 0 | 0 |
| 1 | 1 | 1 |
| 0 | 1 | 0 |
| 0 | 1 | 0 |
| 0 | 0 | 0 |
| 1 | 1 | 1 |
| 1 | 1 | 1 |
| 1 | 1 | 1 |
| 0 | 0 | 0 |
| 0 | 0 | 0 |
| 0 | 0 | 0 |
| 0 | 0 | 0 |
| 0 | 1 | 0 |
| 1 | 1 | 1 |
| 1 | 1 | 1 |
| 1 | 1 | 1 |
| 0 | 0 | 0 |
| 1 | 1 | 1 |
| 0 | 1 | 0 |
| 1 | 1 | 1 |
| 1 | 1 | 1 |
| 1 | 1 | 1 |
| 1 | 1 | 1 |
| 1 | 1 | 1 |
| 1 | 1 | 1 |
| 1 | 1 | 1 |
| 1 | 1 | 1 |
| 1 | 1 | 1 |
| 1 | 1 | 1 |
| 1 | 1 | 1 |
| 0 | 0 | 0 |
| 1 | 1 | 1 |
| 1 | 1 | 1 |
| 0 | 0 | 0 |
| 1 | 1 | 1 |
| 1 | 1 | 1 |
| 1 | 1 | 1 |
| 0 | 0 | 0 |
| 0 | 0 | 0 |
| 1 | 1 | 1 |
| 1 | 1 | 1 |
| 0 | 0 | 0 |
| 1 | 1 | 1 |
| 0 | 1 | 0 |
| 1 | 1 | 1 |

| | | |
|---|---|---|
| 1 | 1 | 1 |
| 1 | 1 | 1 |
| 1 | 1 | 1 |
| 1 | 1 | 1 |
| 1 | 1 | 1 |
| 1 | 1 | 1 |
| 1 | 1 | 1 |
| 0 | 1 | 0 |
| 1 | 1 | 1 |
| 1 | 1 | 1 |
| 1 | 1 | 1 |
| 1 | 1 | 1 |
| 1 | 1 | 1 |
| 0 | 0 | 0 |
| 0 | 0 | 0 |
| 0 | 0 | 0 |
| 0 | 0 | 0 |
| 0 | 1 | 0 |
| 1 | 1 | 1 |
| 0 | 0 | 0 |
| 1 | 1 | 1 |
| 1 | 0 | 0 |
| 1 | 1 | 1 |
| 1 | 1 | 1 |
| 1 | 1 | 1 |
| 0 | 0 | 0 |
| 0 | 1 | 0 |
| 1 | 1 | 1 |
| 1 | 1 | 1 |
| 0 | 0 | 0 |
| 0 | 0 | 0 |
| 1 | 1 | 1 |
| 1 | 1 | 1 |
| 1 | 1 | 1 |
| 1 | 1 | 1 |
| 1 | 1 | 1 |
| 1 | 1 | 1 |
| 1 | 1 | 1 |
| 0 | 0 | 0 |
| 1 | 1 | 1 |
| 1 | 1 | 1 |
| 0 | 0 | 0 |
| 0 | 0 | 0 |
| 0 | 0 | 0 |
| 0 | 0 | 0 |
| 1 | 1 | 1 |
| 0 | 0 | 0 |
| 1 | 1 | 1 |
| 1 | 1 | 1 |
| 1 | 1 | 1 |
| 0 | 1 | 0 |
| 1 | 0 | 0 |
| 1 | 0 | 0 |
| 1 | 0 | 0 |

| | | |
|---|---|---|
| 1 | 1 | 1 |
| 1 | 1 | 1 |
| 0 | 0 | 0 |
| 1 | 1 | 1 |
| 1 | 1 | 1 |
| 1 | 1 | 1 |
| 1 | 1 | 1 |
| 0 | 1 | 0 |
| 1 | 1 | 1 |
| 1 | 1 | 1 |
| 1 | 1 | 1 |
| 0 | 1 | 0 |
| 0 | 1 | 0 |
| 0 | 1 | 0 |
| 0 | 1 | 0 |
| 0 | 1 | 0 |
| 0 | 1 | 0 |
| 0 | 1 | 0 |
| 0 | 1 | 0 |
| 0 | 1 | 0 |
| 0 | 1 | 0 |
| 0 | 1 | 0 |
| 0 | 1 | 0 |
| 0 | 1 | 0 |
| 0 | 1 | 0 |
| 0 | 1 | 0 |
| 0 | 1 | 0 |
| 0 | 1 | 0 |
| 1 | 1 | 1 |
| 1 | 1 | 1 |
| 0 | 0 | 0 |
| 1 | 1 | 1 |
| 0 | 0 | 0 |
| 0 | 0 | 0 |
| 0 | 0 | 0 |
| 0 | 0 | 0 |
| 1 | 1 | 1 |
| 0 | 0 | 0 |
| 1 | 1 | 1 |
| 1 | 1 | 1 |
| 0 | 0 | 0 |
| 0 | 0 | 0 |
| 0 | 0 | 0 |
| 1 | 1 | 1 |
| 0 | 0 | 0 |
| 0 | 0 | 0 |
| 1 | 1 | 1 |
| 1 | 1 | 1 |
| 1 | 1 | 1 |
| 1 | 1 | 1 |
| 0 | 0 | 0 |
| 1 | 1 | 1 |
| 1 | 1 | 1 |
| 1 | 1 | 1 |

| | | |
|---|---|---|
| 1 | 1 | 1 |
| 1 | 1 | 1 |
| 1 | 1 | 1 |
| 1 | 1 | 1 |
| 1 | 1 | 1 |
| 0 | 0 | 0 |
| 1 | 1 | 1 |
| 0 | 0 | 0 |
| 0 | 0 | 0 |
| 0 | 0 | 0 |
| 1 | 1 | 1 |
| 0 | 0 | 0 |
| 1 | 1 | 1 |
| 1 | 1 | 1 |
| 1 | 1 | 1 |
| 1 | 0 | 0 |
| 1 | 1 | 1 |
| 0 | 0 | 0 |
| 1 | 1 | 1 |
| 0 | 0 | 0 |
| 1 | 1 | 1 |
| 0 | 0 | 0 |
| 0 | 0 | 0 |
| 0 | 0 | 0 |
| 1 | 1 | 1 |
| 0 | 1 | 0 |
| 1 | 1 | 1 |
| 1 | 1 | 1 |
| 0 | 0 | 0 |
| 1 | 1 | 1 |
| 1 | 1 | 1 |
| 1 | 1 | 1 |
| 1 | 1 | 1 |
| 1 | 1 | 1 |
| 0 | 0 | 0 |
| 0 | 0 | 0 |
| 1 | 1 | 1 |
| 0 | 0 | 0 |
| 0 | 0 | 0 |
| 0 | 1 | 0 |
| 0 | 0 | 0 |
| 0 | 0 | 0 |
| 0 | 1 | 0 |
| 0 | 1 | 0 |
| 1 | 1 | 1 |
| 0 | 1 | 0 |
| 1 | 1 | 1 |
| 1 | 1 | 1 |
| 1 | 1 | 1 |
| 1 | 1 | 1 |
| 1 | 1 | 1 |
| 1 | 1 | 1 |
| 1 | 1 | 1 |
| 1 | 1 | 1 |

| | | |
|---|---|---|
| 1 | 1 | 1 |
| 0 | 0 | 0 |
| 0 | 1 | 0 |
| 1 | 1 | 1 |
| 0 | 0 | 0 |
| 1 | 1 | 1 |
| 1 | 1 | 1 |
| 1 | 1 | 1 |
| 0 | 0 | 0 |
| 0 | 0 | 0 |
| 1 | 1 | 1 |
| 0 | 0 | 0 |
| 0 | 0 | 0 |
| 1 | 1 | 1 |
| 1 | 1 | 1 |
| 0 | 0 | 0 |
| 0 | 1 | 0 |
| 0 | 0 | 0 |
| 0 | 0 | 0 |
| 1 | 1 | 1 |
| 0 | 0 | 0 |
| 0 | 0 | 0 |
| 0 | 0 | 0 |
| 1 | 1 | 1 |
| 0 | 1 | 0 |
| 0 | 0 | 0 |
| 1 | 1 | 1 |
| 1 | 1 | 1 |
| 1 | 1 | 1 |
| 1 | 1 | 1 |
| 0 | 0 | 0 |
| 1 | 1 | 1 |
| 0 | 0 | 0 |
| 1 | 1 | 1 |
| 0 | 0 | 0 |
| 0 | 0 | 0 |
| 1 | 1 | 1 |
| 1 | 1 | 1 |
| 1 | 1 | 1 |
| 1 | 1 | 1 |
| 1 | 1 | 1 |
| 1 | 1 | 1 |
| 1 | 1 | 1 |
| 1 | 1 | 1 |
| 1 | 1 | 1 |
| 1 | 1 | 1 |
| 1 | 0 | 0 |
| 1 | 1 | 1 |
| 1 | 1 | 1 |
| 1 | 1 | 1 |
| 1 | 1 | 1 |
| 1 | 1 | 1 |
| 1 | 1 | 1 |
| 0 | 0 | 0 |

| | | |
|---|---|---|
| 1 | 1 | 1 |
| 1 | 1 | 1 |
| 1 | 1 | 1 |
| 1 | 1 | 1 |
| 0 | 0 | 0 |
| 1 | 1 | 1 |
| 0 | 1 | 0 |
| 0 | 1 | 0 |
| 0 | 1 | 0 |
| 1 | 1 | 1 |
| 0 | 0 | 0 |
| 1 | 1 | 1 |
| 1 | 1 | 1 |
| 1 | 1 | 1 |
| 0 | 0 | 0 |
| 0 | 0 | 0 |
| 0 | 0 | 0 |
| 0 | 0 | 0 |
| 0 | 1 | 0 |
| 0 | 0 | 0 |
| 0 | 0 | 0 |
| 0 | 0 | 0 |
| 1 | 1 | 1 |
| 0 | 0 | 0 |
| 0 | 0 | 0 |
| 1 | 1 | 1 |
| 0 | 0 | 0 |
| 0 | 0 | 0 |
| 0 | 0 | 0 |
| 0 | 0 | 0 |
| 0 | 0 | 0 |
| 1 | 1 | 1 |
| 1 | 1 | 1 |
| 0 | 1 | 0 |
| 1 | 1 | 1 |
| 1 | 1 | 1 |
| 1 | 1 | 1 |
| 0 | 0 | 0 |
| 0 | 0 | 0 |
| 0 | 1 | 0 |
| 1 | 1 | 1 |
| 0 | 0 | 0 |
| 0 | 0 | 0 |
| 0 | 1 | 0 |
| 0 | 0 | 0 |
| 1 | 1 | 1 |
| 1 | 1 | 1 |
| 1 | 1 | 1 |
| 0 | 0 | 0 |
| 0 | 0 | 0 |
| 0 | 0 | 0 |
| 1 | 1 | 1 |
| 0 | 1 | 0 |
| 0 | 0 | 0 |

| | | |
|---|---|---|
| 1 | 1 | 1 |
| 0 | 0 | 0 |
| 1 | 1 | 1 |
| 0 | 0 | 0 |
| 0 | 0 | 0 |
| 0 | 1 | 0 |
| 0 | 0 | 0 |
| 0 | 0 | 0 |
| 1 | 1 | 1 |
| 1 | 1 | 1 |
| 1 | 1 | 1 |
| 1 | 1 | 1 |
| 1 | 1 | 1 |
| 1 | 1 | 1 |
| 1 | 1 | 1 |
| 1 | 1 | 1 |
| 1 | 1 | 1 |
| 0 | 1 | 0 |
| 1 | 1 | 1 |
| 1 | 1 | 1 |
| 0 | 0 | 0 |
| 0 | 1 | 0 |
| 0 | 1 | 0 |
| 0 | 1 | 0 |
| 0 | 1 | 0 |
| 0 | 0 | 0 |
| 0 | 0 | 0 |
| 1 | 1 | 1 |
| 1 | 1 | 1 |
| 1 | 1 | 1 |
| 1 | 1 | 1 |
| 1 | 1 | 1 |
| 1 | 1 | 1 |
| 0 | 1 | 0 |
| 1 | 1 | 1 |
| 1 | 1 | 1 |
| 1 | 1 | 1 |
| 1 | 1 | 1 |
| 0 | 1 | 0 |
| 1 | 1 | 1 |
| 1 | 1 | 1 |
| 0 | 1 | 0 |
| 1 | 1 | 1 |
| 0 | 0 | 0 |
| 1 | 1 | 1 |
| 0 | 0 | 0 |
| 0 | 0 | 0 |
| 0 | 0 | 0 |
| 0 | 0 | 0 |
| 0 | 0 | 0 |
| 0 | 0 | 0 |
| 0 | 1 | 0 |
| 1 | 1 | 1 |
| 1 | 1 | 1 |

| | | |
|---|---|---|
| 1 | 1 | 1 |
| 1 | 1 | 1 |
| 1 | 1 | 1 |
| 0 | 0 | 0 |
| 1 | 1 | 1 |
| 0 | 0 | 0 |
| 1 | 1 | 1 |
| 1 | 1 | 1 |
| 0 | 0 | 0 |
| 1 | 1 | 1 |
| 1 | 1 | 1 |
| 0 | 1 | 0 |
| 1 | 1 | 1 |
| 1 | 1 | 1 |
| 1 | 1 | 1 |
| 0 | 0 | 0 |
| 0 | 0 | 0 |
| 0 | 0 | 0 |
| 0 | 0 | 0 |
| 0 | 0 | 0 |
| 1 | 1 | 1 |
| 1 | 1 | 1 |
| 1 | 1 | 1 |
| 1 | 1 | 1 |
| 0 | 0 | 0 |
| 1 | 1 | 1 |
| 1 | 1 | 1 |
| 1 | 1 | 1 |
| 0 | 0 | 0 |
| 1 | 1 | 1 |
| 0 | 0 | 0 |
| 0 | 0 | 0 |
| 0 | 0 | 0 |
| 0 | 0 | 0 |
| 0 | 0 | 0 |
| 1 | 1 | 1 |
| 0 | 0 | 0 |
| 0 | 0 | 0 |
| 0 | 0 | 0 |
| 0 | 0 | 0 |
| 0 | 0 | 0 |
| 0 | 0 | 0 |
| 0 | 0 | 0 |
| 0 | 0 | 0 |
| 0 | 0 | 0 |
| 0 | 0 | 0 |
| 0 | 0 | 0 |
| 0 | 0 | 0 |
| 1 | 1 | 1 |
| 0 | 0 | 0 |
| 1 | 1 | 1 |
| 1 | 1 | 1 |
| 0 | 1 | 0 |
| 0 | 0 | 0 |

| | | |
|---|---|---|
| 0 | 0 | 0 |
| 0 | 1 | 0 |
| 1 | 1 | 1 |
| 1 | 1 | 1 |
| 1 | 1 | 1 |
| 0 | 0 | 0 |
| 1 | 1 | 1 |
| 0 | 1 | 0 |
| 1 | 1 | 1 |
| 0 | 1 | 0 |
| 0 | 1 | 0 |
| 0 | 1 | 0 |
| 1 | 1 | 1 |
| 0 | 0 | 0 |
| 0 | 1 | 0 |
| 1 | 1 | 1 |
| 1 | 1 | 1 |
| 0 | 0 | 0 |
| 1 | 1 | 1 |
| 1 | 1 | 1 |
| 1 | 1 | 1 |
| 1 | 1 | 1 |
| 0 | 0 | 0 |
| 1 | 1 | 1 |
| 1 | 1 | 1 |
| 0 | 0 | 0 |
| 0 | 0 | 0 |
| 0 | 0 | 0 |
| 1 | 1 | 1 |
| 0 | 1 | 0 |
| 1 | 1 | 1 |
| 1 | 1 | 1 |
| 1 | 1 | 1 |
| 1 | 1 | 1 |
| 1 | 1 | 1 |
| 1 | 1 | 1 |
| 1 | 1 | 1 |
| 1 | 1 | 1 |
| 1 | 1 | 1 |
| 1 | 1 | 1 |
| 1 | 1 | 1 |
| 1 | 1 | 1 |
| 1 | 1 | 1 |
| 1 | 1 | 1 |
| 1 | 1 | 1 |
| 1 | 1 | 1 |
| 1 | 1 | 1 |
| 1 | 1 | 1 |
| 1 | 0 | 0 |
| 1 | 1 | 1 |
| 1 | 1 | 1 |
| 1 | 1 | 1 |
| 1 | 1 | 1 |
| 1 | 1 | 1 |

| | | |
|---|---|---|
| 1 | 1 | 1 |
| 0 | 0 | 0 |
| 1 | 1 | 1 |
| 1 | 1 | 1 |
| 1 | 1 | 1 |
| 0 | 0 | 0 |
| 0 | 0 | 0 |
| 0 | 0 | 0 |
| 1 | 1 | 1 |
| 1 | 1 | 1 |
| 0 | 1 | 0 |
| 1 | 1 | 1 |
| 1 | 1 | 1 |
| 1 | 1 | 1 |
| 0 | 0 | 0 |
| 0 | 0 | 0 |
| 0 | 0 | 0 |
| 1 | 1 | 1 |
| 0 | 0 | 0 |
| 0 | 0 | 0 |
| 0 | 0 | 0 |
| 0 | 0 | 0 |
| 0 | 0 | 0 |
| 0 | 0 | 0 |
| 0 | 1 | 0 |
| 1 | 1 | 1 |
| 1 | 1 | 1 |
| 0 | 1 | 0 |
| 0 | 1 | 0 |
| 1 | 1 | 1 |
| 1 | 1 | 1 |
| 1 | 1 | 1 |
| 0 | 0 | 0 |
| 1 | 1 | 1 |
| 0 | 1 | 0 |
| 0 | 0 | 0 |
| 0 | 0 | 0 |
| 0 | 0 | 0 |
| 0 | 0 | 0 |
| 1 | 1 | 1 |
| 0 | 1 | 0 |
| 1 | 1 | 1 |
| 1 | 1 | 1 |
| 0 | 0 | 0 |
| 0 | 0 | 0 |
| 0 | 0 | 0 |
| 0 | 1 | 0 |
| 1 | 1 | 1 |
| 0 | 0 | 0 |
| 1 | 1 | 1 |
| 1 | 1 | 1 |
| 0 | 0 | 0 |
| 1 | 1 | 1 |
| 0 | 1 | 0 |

| | | |
|---|---|---|
| 1 | 1 | 1 |
| 0 | 1 | 0 |
| 1 | 1 | 1 |
| 1 | 1 | 1 |
| 1 | 1 | 1 |
| 1 | 1 | 1 |
| 0 | 1 | 0 |
| 1 | 1 | 1 |
| 0 | 0 | 0 |
| 0 | 0 | 0 |
| 1 | 1 | 1 |
| 0 | 0 | 0 |
| 0 | 0 | 0 |
| 0 | 0 | 0 |
| 1 | 1 | 1 |
| 1 | 1 | 1 |
| 0 | 0 | 0 |
| 0 | 0 | 0 |
| 0 | 0 | 0 |
| 1 | 1 | 1 |
| 1 | 1 | 1 |
| 1 | 1 | 1 |
| 0 | 0 | 0 |
| 1 | 1 | 1 |
| 0 | 0 | 0 |
| 1 | 1 | 1 |
| 1 | 1 | 1 |
| 1 | 1 | 1 |
| 1 | 1 | 1 |
| 0 | 0 | 0 |
| 0 | 0 | 0 |
| 0 | 0 | 0 |
| 1 | 1 | 1 |
| 0 | 1 | 0 |
| 1 | 1 | 1 |
| 1 | 1 | 1 |
| 1 | 1 | 1 |
| 0 | 1 | 0 |
| 0 | 1 | 0 |
| 0 | 1 | 0 |
| 0 | 0 | 0 |
| 1 | 1 | 1 |
| 1 | 1 | 1 |
| 1 | 1 | 1 |
| 1 | 1 | 1 |
| 0 | 0 | 0 |
| 1 | 0 | 0 |
| 0 | 0 | 0 |
| 1 | 1 | 1 |
| 0 | 0 | 0 |
| 0 | 0 | 0 |
| 0 | 0 | 0 |
| 0 | 1 | 0 |
| 1 | 0 | 0 |

| | | |
|---|---|---|
| 1 | 0 | 0 |
| 0 | 0 | 0 |
| 0 | 1 | 0 |
| 1 | 1 | 1 |
| 0 | 1 | 0 |
| 0 | 1 | 0 |
| 0 | 1 | 0 |
| 0 | 0 | 0 |
| 0 | 0 | 0 |
| 0 | 1 | 0 |
| 0 | 0 | 0 |
| 1 | 1 | 1 |
| 1 | 1 | 1 |
| 1 | 1 | 1 |
| 0 | 1 | 0 |
| 1 | 1 | 1 |
| 1 | 1 | 1 |
| 1 | 1 | 1 |
| 1 | 1 | 1 |
| 0 | 1 | 0 |
| 1 | 1 | 1 |
| 1 | 1 | 1 |
| 1 | 1 | 1 |
| 1 | 1 | 1 |
| 0 | 1 | 0 |
| 0 | 1 | 0 |
| 0 | 1 | 0 |
| 0 | 1 | 0 |
| 0 | 0 | 0 |
| 0 | 1 | 0 |
| 0 | 0 | 0 |
| 1 | 1 | 1 |
| 0 | 0 | 0 |
| 0 | 0 | 0 |
| 0 | 1 | 0 |
| 0 | 0 | 0 |
| 0 | 0 | 0 |
| 1 | 1 | 1 |
| 0 | 1 | 0 |
| 0 | 1 | 0 |
| 1 | 1 | 1 |
| 0 | 0 | 0 |
| 0 | 0 | 0 |
| 1 | 1 | 1 |
| 1 | 1 | 1 |
| 0 | 0 | 0 |
| 0 | 0 | 0 |
| 0 | 0 | 0 |
| 0 | 0 | 0 |
| 0 | 1 | 0 |
| 0 | 0 | 0 |
| 0 | 0 | 0 |
| 1 | 1 | 1 |
| 1 | 1 | 1 |

| | | |
|---|---|---|
| 1 | 1 | 1 |
| 1 | 1 | 1 |
| 1 | 1 | 1 |
| 1 | 1 | 1 |
| 0 | 0 | 0 |
| 0 | 0 | 0 |
| 0 | 0 | 0 |
| 0 | 0 | 0 |
| 1 | 1 | 1 |
| 0 | 0 | 0 |
| 0 | 0 | 0 |
| 0 | 0 | 0 |
| 0 | 0 | 0 |
| 1 | 1 | 1 |
| 1 | 0 | 0 |
| 0 | 1 | 0 |
| 1 | 1 | 1 |
| 1 | 1 | 1 |
| 1 | 1 | 1 |
| 0 | 1 | 0 |
| 1 | 1 | 1 |
| 1 | 1 | 1 |
| 1 | 1 | 1 |
| 1 | 1 | 1 |
| 1 | 1 | 1 |
| 0 | 0 | 0 |
| 0 | 0 | 0 |
| 0 | 0 | 0 |
| 0 | 0 | 0 |
| 0 | 1 | 0 |
| 0 | 1 | 0 |
| 0 | 1 | 0 |
| 1 | 1 | 1 |
| 0 | 1 | 0 |
| 0 | 1 | 0 |
| 1 | 1 | 1 |
| 1 | 1 | 1 |
| 1 | 1 | 1 |
| 1 | 1 | 1 |
| 1 | 1 | 1 |
| 1 | 1 | 1 |
| 1 | 1 | 1 |
| 1 | 1 | 1 |
| 1 | 1 | 1 |
| 1 | 1 | 1 |
| 1 | 1 | 1 |
| 1 | 1 | 1 |
| 1 | 1 | 1 |
| 1 | 1 | 1 |
| 1 | 1 | 1 |
| 1 | 1 | 1 |
| 1 | 1 | 1 |
| 1 | 1 | 1 |
| 0 | 0 | 0 |

| | | |
|---|---|---|
| 1 | 1 | 1 |
| 1 | 1 | 1 |
| 1 | 1 | 1 |
| 0 | 1 | 0 |
| 0 | 1 | 0 |
| 0 | 1 | 0 |
| 0 | 1 | 0 |
| 0 | 1 | 0 |
| 1 | 1 | 1 |
| 1 | 1 | 1 |
| 1 | 1 | 1 |
| 1 | 1 | 1 |
| 1 | 1 | 1 |
| 1 | 1 | 1 |
| 1 | 1 | 1 |
| 1 | 1 | 1 |
| 1 | 1 | 1 |
| 1 | 1 | 1 |
| 1 | 1 | 1 |
| 1 | 1 | 1 |
| 1 | 1 | 1 |
| 1 | 1 | 1 |
| 1 | 1 | 1 |
| 1 | 1 | 1 |
| 1 | 1 | 1 |
| 1 | 1 | 1 |
| 1 | 1 | 1 |
| 1 | 1 | 1 |
| 1 | 1 | 1 |
| 1 | 1 | 1 |
| 1 | 1 | 1 |
| 1 | 1 | 1 |
| 1 | 1 | 1 |
| 1 | 1 | 1 |
| 0 | 1 | 0 |
| 1 | 1 | 1 |
| 1 | 1 | 1 |
| 1 | 1 | 1 |
| 1 | 1 | 1 |
| 1 | 1 | 1 |
| 1 | 1 | 1 |
| 0 | 1 | 0 |
| 1 | 1 | 1 |
| 1 | 1 | 1 |
| 1 | 1 | 1 |
| 0 | 1 | 0 |
| 1 | 0 | 0 |
| 0 | 0 | 0 |
| 1 | 1 | 1 |
| 0 | 1 | 0 |
| 1 | 1 | 1 |
| 1 | 1 | 1 |
| 0 | 1 | 0 |
| 0 | 1 | 0 |
| 0 | 1 | 0 |

| | | |
|---|---|---|
| 0 | 1 | 0 |
| 1 | 1 | 1 |
| 0 | 0 | 0 |
| 1 | 1 | 1 |
| 1 | 1 | 1 |
| 1 | 1 | 1 |
| 0 | 0 | 0 |
| 1 | 1 | 1 |
| 1 | 1 | 1 |
| 0 | 1 | 0 |
| 0 | 1 | 0 |
| 0 | 0 | 0 |
| 1 | 1 | 1 |
| 0 | 0 | 0 |
| 0 | 0 | 0 |
| 0 | 0 | 0 |
| 0 | 0 | 0 |
| 0 | 0 | 0 |
| 0 | 0 | 0 |
| 0 | 0 | 0 |
| 0 | 0 | 0 |

| organism_scientific | evo_cons | cluster_id | pdb_id | chain_id | resi | resn | change | uniprot_id |
|---|---|---|---|---|---|---|---|---|
| HOMO SAPIENS | 5 | 87 | 3uzw | A | 106 | LEU | p.Leu106Phe | P51857 |
| HOMO SAPIENS | 3 | 87 | 3uzw | A | 133 | PRO | p.Pro133Arg | P51857 |
| HOMO SAPIENS | 0 | 87 | 3uzw | A | 198 | PRO | p.Pro198Leu | P51857 |
| HOMO SAPIENS | 7 | 87 | 3uzw | A | 261 | ARG | p.Arg261Cys | P51857 |
| HOMO SAPIENS | 0 | 4693 | 3mk1 | A | 3 | PRO | p.Pro25Leu | P05187 |
| HOMO SAPIENS | 8 | 4830 | 2h64 | B | 101 | CYS | p.Cys124Arg | P36894 |
| HOMO SAPIENS | 2 | 4830 | 2h64 | B | 116 | PRO | p.Pro139Ser | P36894 |
| HOMO SAPIENS | 0 | 6937 | 3dar | A | 186 | MET | p.Met186Thr | P21802 |
| HOMO SAPIENS | 3 | 5 | 1aut | C | 30 | GLN | p.Gln226His | P04070 |
| HOMO SAPIENS | 1 | 5 | 1aut | C | 68 | LEU | p.Leu265Phe | P04070 |
| HOMO SAPIENS | 0 | 5 | 1aut | C | 75 | ARG | p.Arg272Cys | P04070 |
| HOMO SAPIENS | 6 | 5 | 1aut | C | 92 | PRO | p.Pro289Leu | P04070 |
| HOMO SAPIENS | 0 | 5 | 1aut | C | 104 | ALA | p.Ala301Val | P04070 |
| HOMO SAPIENS | 1 | 5 | 1aut | C | 112 | ALA | p.Ala309Thr | P04070 |
| HOMO SAPIENS | 2 | 5 | 1aut | C | 115 | SER | p.Ser312Leu | P04070 |
| HOMO SAPIENS | 1 | 5 | 1aut | C | 133 | GLY | p.Gly334Ser | P04070 |
| HOMO SAPIENS | 0 | 5 | 1aut | C | 147 | ARG | p.Arg348Ter | P04070 |
| HOMO SAPIENS | 8 | 5 | 1aut | C | 194 | ASP | p.Asp401Asn | P04070 |
| HOMO SAPIENS | 7 | 5 | 1aut | C | 237 | TRP | p.Trp444Cys | P04070 |
| HOMO SAPIENS | 0 | 5 | 1aut | C | 238 | ILE | p.Ile445Met | P04070 |
| HOMO SAPIENS | 8 | 43 | 1aiy | F | 10 | HIS | p.His34Asp | P01308 |
| HOMO SAPIENS | 2 | 43 | 1aiy | F | 22 | ARG | p.Arg46Gln | P01308 |
| HOMO SAPIENS | 4 | 43 | 1aiy | F | 23 | GLY | p.Gly47Val | P01308 |
| HOMO SAPIENS | 7 | 43 | 1aiy | F | 24 | PHE | p.Phe48Ser | P01308 |
| HOMO SAPIENS | 5 | 24 | 2fg8 | A | 33 | THR | p.Thr30Ile | P02792 |
| HOMO SAPIENS | 4 | 24 | 2fg8 | A | 99 | ALA | p.Ala96Thr | P02792 |
| HOMO SAPIENS | 5 | 24 | 2fg8 | A | 107 | GLU | p.Glu104Ter | P02792 |
| HOMO SAPIENS | 3 | 5409 | 5bnq | A | 199 | MET | p.Met199Lys | O14788 |
| HOMO SAPIENS | -1 | 839 | 3swc | A | 959 | GLN | p.Gln990His | Q9H9B1 |
| HOMO SAPIENS | -1 | 839 | 3swc | A | 968 | ALA | p.Ala999Val | Q9H9B1 |
| HOMO SAPIENS | -1 | 839 | 3swc | A | 973 | SER | p.Ser1004Ile | Q9H9B1 |
| HOMO SAPIENS | 8 | 17 | 1eou | A | 18 | LYS | p.Lys18Glu | P00918 |
| HOMO SAPIENS | 2 | 17 | 1eou | A | 107 | HIS | p.His107Tyr | P00918 |
| HOMO SAPIENS | 1 | 17 | 1eou | A | 236 | PRO | p.Pro236His | P00918 |
| HOMO SAPIENS | 6 | 17 | 1eou | A | 252 | ASN | p.Asn252Asp | P00918 |
| HOMO SAPIENS | 2 | 9140 | 5c4v | A | 445 | ARG | p.Arg445Ter | Q13485 |
| HOMO SAPIENS | 7 | 9140 | 5c4v | A | 496 | ARG | p.Arg496Cys | Q13485 |
| HOMO SAPIENS | -1 | 42 | 4pwi | A | 6 | GLY | p.Gly26Ser | P02766 |
| HOMO SAPIENS | 6 | 42 | 4pwi | A | 10 | CYS | p.Cys30Arg | P02766 |
| HOMO SAPIENS | 8 | 42 | 4pwi | A | 12 | LEU | p.Leu32Pro | P02766 |
| HOMO SAPIENS | 9 | 42 | 4pwi | A | 18 | ASP | p.Asp38Gly | P02766 |
| HOMO SAPIENS | 8 | 42 | 4pwi | A | 20 | VAL | p.Val40Ile | P02766 |
| HOMO SAPIENS | 4 | 42 | 4pwi | A | 30 | MET | p.Val50Ala | P02766 |
| HOMO SAPIENS | 4 | 42 | 4pwi | A | 30 | MET | p.Val50Leu | P02766 |
| HOMO SAPIENS | 6 | 42 | 4pwi | A | 33 | PHE | p.Phe53Ile | P02766 |
| HOMO SAPIENS | 7 | 42 | 4pwi | A | 36 | ALA | p.Ala56Pro | P02766 |
| HOMO SAPIENS | 7 | 42 | 4pwi | A | 42 | GLU | p.Glu62Gly | P02766 |
| HOMO SAPIENS | 8 | 42 | 4pwi | A | 44 | PHE | p.Phe64Ser | P02766 |
| HOMO SAPIENS | 4 | 42 | 4pwi | A | 47 | GLY | p.Gly67Ala | P02766 |
| HOMO SAPIENS | 6 | 42 | 4pwi | A | 49 | THR | p.Thr69Ala | P02766 |
| HOMO SAPIENS | 6 | 42 | 4pwi | A | 50 | SER | p.Ser70Ile | P02766 |
| HOMO SAPIENS | 6 | 42 | 4pwi | A | 50 | SER | p.Ser70Arg | P02766 |
| HOMO SAPIENS | 0 | 42 | 4pwi | A | 53 | GLY | p.Gly73Glu | P02766 |

| | | | | | | | | |
|---|---|---|---|---|---|---|---|---|
| HOMO SAPIENS | 7 | 42 | 4pwi | A | 55 | LEU | p.Leu75Pro | P02766 |
| HOMO SAPIENS | 6 | 42 | 4pwi | A | 58 | LEU | p.Leu78His | P02766 |
| HOMO SAPIENS | 7 | 42 | 4pwi | A | 61 | GLU | p.Glu81Lys | P02766 |
| HOMO SAPIENS | 8 | 42 | 4pwi | A | 64 | PHE | p.Phe84Leu | P02766 |
| HOMO SAPIENS | 8 | 42 | 4pwi | A | 68 | ILE | p.Ile88Leu | P02766 |
| HOMO SAPIENS | 8 | 42 | 4pwi | A | 69 | TYR | p.Tyr89His | P02766 |
| HOMO SAPIENS | 4 | 42 | 4pwi | A | 71 | VAL | p.Val91Ala | P02766 |
| HOMO SAPIENS | 6 | 42 | 4pwi | A | 77 | SER | p.Ser97Tyr | P02766 |
| HOMO SAPIENS | 4 | 42 | 4pwi | A | 84 | ILE | p.Ile104Ser | P02766 |
| HOMO SAPIENS | 8 | 42 | 4pwi | A | 89 | GLU | p.Glu109Gln | P02766 |
| HOMO SAPIENS | 8 | 42 | 4pwi | A | 97 | ALA | p.Ala117Gly | P02766 |
| HOMO SAPIENS | 7 | 42 | 4pwi | A | 99 | ASP | p.Asp119Asn | P02766 |
| HOMO SAPIENS | 9 | 42 | 4pwi | A | 107 | ILE | p.Ile127Val | P02766 |
| HOMO SAPIENS | 9 | 42 | 4pwi | A | 111 | LEU | p.Leu131Met | P02766 |
| HOMO SAPIENS | 7 | 42 | 4pwi | A | 114 | TYR | p.Tyr134Cys | P02766 |
| HOMO SAPIENS | 7 | 42 | 4pwi | A | 114 | TYR | p.Tyr134His | P02766 |
| HOMO SAPIENS | 7 | 42 | 4pwi | A | 122 | VAL | p.Val142Ile | P02766 |
| HOMO SAPIENS | -1 | 42 | 4pwi | A | 126 | LYS | p.Lys146Arg | P02766 |
| HOMO SAPIENS | -1 | 42 | 4pwi | A | 127 | GLU | p.Glu147Ter | P02766 |
| HOMO SAPIENS | -1 | 1312 | 4dce | A | 1275 | ARG | p.Arg1275Gln | Q9UM73 |
| HOMO SAPIENS | -1 | 1312 | 4dce | A | 1280 | ALA | p.Ala1280Val | Q9UM73 |
| HOMO SAPIENS | 1 | 20 | 1y0x | X | 234 | ALA | p.Ala234Thr | P10828 |
| HOMO SAPIENS | 0 | 20 | 1y0x | X | 243 | ARG | p.Arg243Trp | P10828 |
| HOMO SAPIENS | 3 | 20 | 1y0x | X | 316 | ARG | p.Arg316His | P10828 |
| HOMO SAPIENS | 1 | 20 | 1y0x | X | 317 | ALA | p.Ala317Thr | P10828 |
| HOMO SAPIENS | 2 | 20 | 1y0x | X | 320 | ARG | p.Arg320Cys | P10828 |
| HOMO SAPIENS | 2 | 20 | 1y0x | X | 320 | ARG | p.Arg320His | P10828 |
| HOMO SAPIENS | 0 | 20 | 1y0x | X | 332 | GLY | p.Gly332Arg | P10828 |
| HOMO SAPIENS | 1 | 20 | 1y0x | X | 338 | ARG | p.Arg338Trp | P10828 |
| HOMO SAPIENS | 0 | 20 | 1y0x | X | 340 | GLN | p.Gln340His | P10828 |
| HOMO SAPIENS | 0 | 20 | 1y0x | X | 345 | GLY | p.Gly345Val | P10828 |
| HOMO SAPIENS | 0 | 20 | 1y0x | X | 345 | GLY | p.Gly345Ser | P10828 |
| HOMO SAPIENS | 0 | 20 | 1y0x | X | 347 | GLY | p.Gly347Glu | P10828 |
| HOMO SAPIENS | 1 | 20 | 1y0x | X | 438 | ARG | p.Arg438His | P10828 |
| HOMO SAPIENS | 0 | 20 | 1y0x | X | 442 | MET | p.Met442Val | P10828 |
| HOMO SAPIENS | 0 | 20 | 1y0x | X | 443 | LYS | p.Lys443Glu | P10828 |
| HOMO SAPIENS | 0 | 20 | 1y0x | X | 446 | CYS | p.Cys446Arg | P10828 |
| HOMO SAPIENS | 2 | 20 | 1y0x | X | 453 | PRO | p.Pro453His | P10828 |
| HOMO SAPIENS | 2 | 20 | 1y0x | X | 453 | PRO | p.Pro453Thr | P10828 |
| HOMO SAPIENS | 3 | 20 | 1y0x | X | 459 | PHE | p.Phe459Cys | P10828 |
| HOMO SAPIENS | 0 | 4409 | 3qb7 | D | 87 | GLU | p.Glu109Lys | P31785 |
| HOMO SAPIENS | 1 | 4409 | 3qb7 | D | 92 | GLY | p.Gly114Asp | P31785 |
| HOMO SAPIENS | 1 | 4409 | 3qb7 | D | 93 | CYS | p.Cys115Arg | P31785 |
| HOMO SAPIENS | 0 | 4409 | 3qb7 | D | 131 | ILE | p.Ile153Asn | P31785 |
| HOMO SAPIENS | 9 | 4409 | 3qb7 | D | 200 | ARG | p.Arg222Cys | P31785 |
| HOMO SAPIENS | 0 | 3891 | 3ije | B | 33 | LEU | p.Leu59Pro | P05106 |
| HOMO SAPIENS | 0 | 3891 | 3ije | B | 117 | LEU | p.Leu143Trp | P05106 |
| HOMO SAPIENS | 3 | 3891 | 3ije | B | 119 | ASP | p.Asp145Tyr | P05106 |
| HOMO SAPIENS | 2 | 3891 | 3ije | B | 143 | ARG | p.Arg169Gln | P05106 |
| HOMO SAPIENS | 2 | 3891 | 3ije | B | 214 | ARG | p.Arg240Gln | P05106 |
| HOMO SAPIENS | 2 | 3891 | 3ije | B | 214 | ARG | p.Arg240Trp | P05106 |
| HOMO SAPIENS | 2 | 3891 | 3ije | B | 374 | CYS | p.Cys400Tyr | P05106 |
| HOMO SAPIENS | 1 | 3891 | 3ije | B | 407 | PRO | p.Pro433Ala | P05106 |
| HOMO SAPIENS | 0 | 3891 | 3ije | B | 489 | ARG | p.Arg515Gln | P05106 |

| | | | | | | | | |
|---|---|---|---|---|---|---|---|---|
| HOMO SAPIENS | -1 | 9613 | 5mke | B | 28 | ARG | p.Arg28Pro | Q13563 |
| HOMO SAPIENS | -1 | 9613 | 5mke | B | 190 | ALA | p.Ala190Thr | Q13563 |
| HOMO SAPIENS | 4 | 9613 | 5mke | B | 262 | VAL | p.Val262Met | Q13563 |
| HOMO SAPIENS | 0 | 9613 | 5mke | B | 482 | PHE | p.Phe482Cys | Q13563 |
| HOMO SAPIENS | 4 | 9613 | 5mke | B | 511 | ASP | p.Asp511Val | Q13563 |
| HOMO SAPIENS | 5 | 9613 | 5mke | B | 516 | VAL | p.Val516Leu | Q13563 |
| HOMO SAPIENS | -1 | 9613 | 5mke | B | 807 | ARG | p.Arg807Gln | Q13563 |
| HOMO SAPIENS | 4 | 2428 | 1olh | C | 337 | ARG | p.Arg337Cys | P04637 |
| HOMO SAPIENS | 4 | 2428 | 1olh | C | 337 | ARG | p.Arg337His | P04637 |
| HOMO SAPIENS | 8 | 2428 | 1olh | C | 338 | PHE | p.Phe338Leu | P04637 |
| HOMO SAPIENS | 8 | 2428 | 1olh | C | 339 | GLU | p.Glu339Lys | P04637 |
| HOMO SAPIENS | 7 | 2428 | 1olh | C | 344 | LEU | p.Leu344Pro | P04637 |
| HOMO SAPIENS | 0 | 2428 | 1olh | C | 360 | GLY | p.Gly360Ala | P04637 |
| HOMO SAPIENS | 7 | 4867 | 1mox | B | 404 | GLY | p.Gly428Asp | P00533 |
| HOMO SAPIENS | 5 | 4867 | 1mox | B | 497 | ARG | p.Arg521Lys | P00533 |
| HOMO SAPIENS | 1 | 4630 | 1fu1 | A | 43 | TRP | p.Trp43Arg | Q13426 |
| HOMO SAPIENS | 2 | 4630 | 1fu1 | A | 161 | ARG | p.Arg161Ter | Q13426 |
| HOMO SAPIENS | 2 | 15 | 1lfy | B | 2 | HIS | p.His3Gln | P68871 |
| HOMO SAPIENS | 2 | 15 | 1lfy | B | 6 | GLU | p.Glu7Ala | P68871 |
| HOMO SAPIENS | 2 | 15 | 1lfy | B | 6 | GLU | p.Glu7Lys | P68871 |
| HOMO SAPIENS | 4 | 15 | 1lfy | B | 9 | SER | p.Ser10Cys | P68871 |
| HOMO SAPIENS | 6 | 15 | 1lfy | B | 15 | TRP | p.Trp16Ter | P68871 |
| HOMO SAPIENS | 3 | 15 | 1lfy | B | 17 | LYS | p.Lys18Glu | P68871 |
| HOMO SAPIENS | 6 | 15 | 1lfy | B | 19 | ASN | p.Asn20Ser | P68871 |
| HOMO SAPIENS | 6 | 15 | 1lfy | B | 22 | GLU | p.Glu23Ala | P68871 |
| HOMO SAPIENS | 6 | 15 | 1lfy | B | 22 | GLU | p.Glu23Lys | P68871 |
| HOMO SAPIENS | 8 | 15 | 1lfy | B | 26 | GLU | p.Glu27Lys | P68871 |
| HOMO SAPIENS | 8 | 15 | 1lfy | B | 28 | LEU | p.Leu29Pro | P68871 |
| HOMO SAPIENS | 8 | 15 | 1lfy | B | 30 | ARG | p.Arg31Ser | P68871 |
| HOMO SAPIENS | 8 | 15 | 1lfy | B | 30 | ARG | p.Arg31Lys | P68871 |
| HOMO SAPIENS | 6 | 15 | 1lfy | B | 39 | GLN | p.Gln40Ter | P68871 |
| HOMO SAPIENS | 2 | 15 | 1lfy | B | 43 | GLU | p.Glu44Gln | P68871 |
| HOMO SAPIENS | 6 | 15 | 1lfy | B | 57 | ASN | p.Asn58Lys | P68871 |
| HOMO SAPIENS | 6 | 15 | 1lfy | B | 59 | LYS | p.Lys60Asn | P68871 |
| HOMO SAPIENS | 8 | 15 | 1lfy | B | 61 | LYS | p.Lys62Glu | P68871 |
| HOMO SAPIENS | 4 | 15 | 1lfy | B | 73 | ASP | p.Asp74Tyr | P68871 |
| HOMO SAPIENS | 4 | 15 | 1lfy | B | 87 | THR | p.Thr88Ile | P68871 |
| HOMO SAPIENS | 4 | 15 | 1lfy | B | 87 | THR | p.Thr88Pro | P68871 |
| HOMO SAPIENS | 5 | 15 | 1lfy | B | 90 | GLU | p.Glu91Lys | P68871 |
| HOMO SAPIENS | 6 | 15 | 1lfy | B | 91 | LEU | p.Leu92Pro | P68871 |
| HOMO SAPIENS | 5 | 15 | 1lfy | B | 97 | HIS | p.His98Gln | P68871 |
| HOMO SAPIENS | 6 | 15 | 1lfy | B | 110 | LEU | p.Leu111Pro | P68871 |
| HOMO SAPIENS | 6 | 15 | 1lfy | B | 113 | VAL | p.Val114Glu | P68871 |
| HOMO SAPIENS | 2 | 15 | 1lfy | B | 119 | GLY | p.Gly120Ala | P68871 |
| HOMO SAPIENS | 4 | 15 | 1lfy | B | 121 | GLU | p.Glu122Gln | P68871 |
| HOMO SAPIENS | 5 | 15 | 1lfy | B | 134 | VAL | p.Val135Glu | P68871 |
| HOMO SAPIENS | 3 | 15 | 1lfy | B | 143 | HIS | p.His144Gln | P68871 |
| HOMO SAPIENS | 4 | 91 | 5d9k | A | 75 | GLY | p.Gly75Val | P51812 |
| HOMO SAPIENS | 5 | 91 | 5d9k | A | 82 | VAL | p.Val82Phe | P51812 |
| HOMO SAPIENS | 0 | 91 | 5d9k | A | 114 | ARG | p.Arg114Trp | P51812 |
| HOMO SAPIENS | 4 | 91 | 5d9k | A | 189 | ILE | p.Ile189Lys | P51812 |
| HOMO SAPIENS | -1 | 91 | 5d9k | A | 227 | SER | p.Ser227Ala | P51812 |
| HOMO SAPIENS | 1 | 91 | 5d9k | A | 268 | PHE | p.Phe268Ser | P51812 |
| HOMO SAPIENS | 0 | 954 | 2vt0 | A | 15 | VAL | p.Val54Leu | P04062 |

| | | | | | | | | |
|---|---|---|---|---|---|---|---|---|
| HOMO SAPIENS | 5 | 954 | 2vt0 | A | 46 | GLY | p.Gly85Glu | P04062 |
| HOMO SAPIENS | 5 | 954 | 2vt0 | A | 48 | ARG | p.Arg87Trp | P04062 |
| HOMO SAPIENS | 6 | 954 | 2vt0 | A | 79 | LYS | p.Lys118Asn | P04062 |
| HOMO SAPIENS | 0 | 954 | 2vt0 | A | 120 | ARG | p.Arg159Gln | P04062 |
| HOMO SAPIENS | 0 | 954 | 2vt0 | A | 120 | ARG | p.Arg159Trp | P04062 |
| HOMO SAPIENS | 0 | 954 | 2vt0 | A | 122 | PRO | p.Pro161Ser | P04062 |
| HOMO SAPIENS | 5 | 954 | 2vt0 | A | 131 | ARG | p.Arg170Leu | P04062 |
| HOMO SAPIENS | 5 | 954 | 2vt0 | A | 135 | TYR | p.Tyr174Cys | P04062 |
| HOMO SAPIENS | 0 | 954 | 2vt0 | A | 157 | LYS | p.Lys196Gln | P04062 |
| HOMO SAPIENS | 5 | 954 | 2vt0 | A | 163 | ARG | p.Arg202Gln | P04062 |
| HOMO SAPIENS | 5 | 954 | 2vt0 | A | 170 | ARG | p.Arg209Cys | P04062 |
| HOMO SAPIENS | 5 | 954 | 2vt0 | A | 188 | ASN | p.Asn227Ser | P04062 |
| HOMO SAPIENS | 3 | 954 | 2vt0 | A | 196 | SER | p.Ser235Pro | P04062 |
| HOMO SAPIENS | 5 | 954 | 2vt0 | A | 202 | GLY | p.Gly241Arg | P04062 |
| HOMO SAPIENS | 0 | 954 | 2vt0 | A | 212 | TYR | p.Tyr251His | P04062 |
| HOMO SAPIENS | 0 | 954 | 2vt0 | A | 213 | PHE | p.Phe252Ile | P04062 |
| HOMO SAPIENS | 5 | 954 | 2vt0 | A | 216 | PHE | p.Phe255Tyr | P04062 |
| HOMO SAPIENS | 6 | 954 | 2vt0 | A | 251 | PHE | p.Phe290Leu | P04062 |
| HOMO SAPIENS | 5 | 954 | 2vt0 | A | 255 | HIS | p.His294Gln | P04062 |
| HOMO SAPIENS | 6 | 954 | 2vt0 | A | 257 | ARG | p.Arg296Gln | P04062 |
| HOMO SAPIENS | 5 | 954 | 2vt0 | A | 289 | PRO | p.Pro328Leu | P04062 |
| HOMO SAPIENS | 0 | 954 | 2vt0 | A | 309 | ALA | p.Ala348Val | P04062 |
| HOMO SAPIENS | 0 | 954 | 2vt0 | A | 311 | HIS | p.His350Arg | P04062 |
| HOMO SAPIENS | 6 | 954 | 2vt0 | A | 312 | TRP | p.Trp351Cys | P04062 |
| HOMO SAPIENS | 5 | 954 | 2vt0 | A | 323 | THR | p.Thr362Ile | P04062 |
| HOMO SAPIENS | 5 | 954 | 2vt0 | A | 325 | GLY | p.Gly364Arg | P04062 |
| HOMO SAPIENS | 6 | 954 | 2vt0 | A | 342 | CYS | p.Cys381Gly | P04062 |
| HOMO SAPIENS | 5 | 954 | 2vt0 | A | 353 | ARG | p.Arg392Gly | P04062 |
| HOMO SAPIENS | 3 | 954 | 2vt0 | A | 364 | SER | p.Ser403Thr | P04062 |
| HOMO SAPIENS | 5 | 954 | 2vt0 | A | 369 | THR | p.Thr408Met | P04062 |
| HOMO SAPIENS | 3 | 954 | 2vt0 | A | 370 | ASN | p.Asn409Ser | P04062 |
| HOMO SAPIENS | 0 | 954 | 2vt0 | A | 371 | LEU | p.Leu410Val | P04062 |
| HOMO SAPIENS | 0 | 954 | 2vt0 | A | 375 | VAL | p.Val414Leu | P04062 |
| HOMO SAPIENS | 0 | 954 | 2vt0 | A | 377 | GLY | p.Gly416Ser | P04062 |
| HOMO SAPIENS | 8 | 954 | 2vt0 | A | 398 | VAL | p.Val437Phe | P04062 |
| HOMO SAPIENS | 0 | 954 | 2vt0 | A | 401 | PRO | p.Pro440Leu | P04062 |
| HOMO SAPIENS | 3 | 954 | 2vt0 | A | 409 | ASP | p.Asp448His | P04062 |
| HOMO SAPIENS | 3 | 954 | 2vt0 | A | 409 | ASP | p.Asp448Val | P04062 |
| HOMO SAPIENS | 0 | 954 | 2vt0 | A | 415 | PRO | p.Pro454Arg | P04062 |
| HOMO SAPIENS | 5 | 954 | 2vt0 | A | 444 | LEU | p.Leu483Arg | P04062 |
| HOMO SAPIENS | 5 | 954 | 2vt0 | A | 456 | ALA | p.Ala495Pro | P04062 |
| HOMO SAPIENS | 5 | 954 | 2vt0 | A | 463 | ARG | p.Arg502Cys | P04062 |
| HOMO SAPIENS | 5 | 954 | 2vt0 | A | 463 | ARG | p.Arg502His | P04062 |
| HOMO SAPIENS | 5 | 954 | 2vt0 | A | 478 | GLY | p.Gly517Ser | P04062 |
| HOMO SAPIENS | 5 | 954 | 2vt0 | A | 496 | ARG | p.Arg535Cys | P04062 |
| HOMO SAPIENS | 5 | 954 | 2vt0 | A | 496 | ARG | p.Arg535His | P04062 |
| HOMO SAPIENS | 1 | 9037 | 2rqb | A | 946 | ALA | p.Ala946Thr | Q9BYX4 |
| HOMO SAPIENS | 0 | 253 | 5E+092 | A | 308 | LEU | p.Leu308Pro | P37173 |
| HOMO SAPIENS | 0 | 253 | 5E+092 | A | 315 | THR | p.Thr315Met | P37173 |
| HOMO SAPIENS | 0 | 253 | 5E+092 | A | 328 | HIS | p.His328Gln | P37173 |
| HOMO SAPIENS | 0 | 253 | 5E+092 | A | 336 | TYR | p.Tyr336Asn | P37173 |
| HOMO SAPIENS | 0 | 253 | 5E+092 | A | 348 | ARG | p.Arg348Cys | P37173 |
| HOMO SAPIENS | 0 | 253 | 5E+092 | A | 355 | ALA | p.Ala355Pro | P37173 |
| HOMO SAPIENS | 1 | 253 | 5E+092 | A | 356 | ARG | p.Arg356Gly | P37173 |

| | | | | | | | | |
|---|---|---|---|---|---|---|---|---|
| HOMO SAPIENS | 0 | 253 | 5E+092 | A | 357 | GLY | p.Gly357Trp | P37173 |
| HOMO SAPIENS | 1 | 253 | 5E+092 | A | 403 | ARG | p.Arg403His | P37173 |
| HOMO SAPIENS | 5 | 253 | 5E+092 | A | 449 | SER | p.Ser449Phe | P37173 |
| HOMO SAPIENS | 0 | 253 | 5E+092 | A | 460 | ARG | p.Arg460Cys | P37173 |
| HOMO SAPIENS | 0 | 253 | 5E+092 | A | 460 | ARG | p.Arg460His | P37173 |
| HOMO SAPIENS | 2 | 253 | 5E+092 | A | 526 | GLU | p.Glu526Gln | P37173 |
| HOMO SAPIENS | 5 | 253 | 5E+092 | A | 528 | ARG | p.Arg528Cys | P37173 |
| HOMO SAPIENS | 5 | 253 | 5E+092 | A | 528 | ARG | p.Arg528His | P37173 |
| HOMO SAPIENS | 6 | 158 | 5amc | A | 231 | ARG | p.Arg260Leu | P12821 |
| HOMO SAPIENS | 7 | 158 | 5amc | A | 237 | TYR | p.Tyr266Ter | P12821 |
| HOMO SAPIENS | 6 | 158 | 5amc | A | 325 | GLY | p.Gly354Arg | P12821 |
| HOMO SAPIENS | 0 | 158 | 5amc | A | 467 | ARG | p.Arg496Ter | P12821 |
| HOMO SAPIENS | 4 | 158 | 5amc | A | 479 | ARG | p.Arg508Ter | P12821 |
| HOMO SAPIENS | 0 | 158 | 5amc | A | 563 | ASP | p.Asp592Gly | P12821 |
| HOMO SAPIENS | -1 | 33062 | 1gr3 | A | 545 | GLY | p.Gly545Arg | Q03692 |
| HOMO SAPIENS | 5 | 33062 | 1gr3 | A | 591 | CYS | p.Cys591Arg | Q03692 |
| HOMO SAPIENS | 2 | 33062 | 1gr3 | A | 595 | GLY | p.Gly595Glu | Q03692 |
| HOMO SAPIENS | 0 | 33062 | 1gr3 | A | 597 | TYR | p.Tyr597Cys | Q03692 |
| HOMO SAPIENS | 5 | 33062 | 1gr3 | A | 598 | TYR | p.Tyr598Asp | Q03692 |
| HOMO SAPIENS | 7 | 33062 | 1gr3 | A | 600 | SER | p.Ser600Pro | Q03692 |
| HOMO SAPIENS | 1 | 33062 | 1gr3 | A | 614 | LEU | p.Leu614Pro | Q03692 |
| HOMO SAPIENS | 6 | 33062 | 1gr3 | A | 651 | TRP | p.Trp651Arg | Q03692 |
| HOMO SAPIENS | 7 | 33062 | 1gr3 | A | 671 | SER | p.Ser671Pro | Q03692 |
| HOMO SAPIENS | -1 | 38257 | 2l2i | B | 87 | PRO | p.Pro87Ala | Q13351 |
| HOMO SAPIENS | 2 | 2036 | 1ghq | A | 231 | GLY | p.Gly1224Asp | P01024 |
| HOMO SAPIENS | 5 | 3594 | 4cde | F | 388 | TYR | p.Tyr412Cys | P53634 |
| HOMO SAPIENS | 9 | 3594 | 4cde | F | 405 | TRP | p.Trp429Cys | P53634 |
| HOMO SAPIENS | 0 | 3594 | 4cde | F | 429 | ILE | p.Ile453Val | P53634 |
| HOMO SAPIENS | 7 | 18 | 1jkb | A | 34 | TRP | p.Trp52Ter | P61626 |
| HOMO SAPIENS | 9 | 18 | 1jkb | A | 70 | THR | p.Thr88Asn | P61626 |
| HOMO SAPIENS | 8 | 18 | 1jkb | A | 121 | VAL | p.Val139Ile | P61626 |
| HOMO SAPIENS | 2 | 414 | 3fj6 | A | 136 | ARG | p.Arg135Cys | Q02127 |
| HOMO SAPIENS | 1 | 414 | 3fj6 | A | 153 | GLY | p.Gly152Arg | Q02127 |
| HOMO SAPIENS | 0 | 414 | 3fj6 | A | 200 | ARG | p.Arg199Cys | Q02127 |
| HOMO SAPIENS | 0 | 414 | 3fj6 | A | 203 | GLY | p.Gly202Ala | Q02127 |
| HOMO SAPIENS | 0 | 414 | 3fj6 | A | 245 | ARG | p.Arg244Trp | Q02127 |
| HOMO SAPIENS | 0 | 414 | 3fj6 | A | 342 | ALA | p.Ala341Val | Q02127 |
| HOMO SAPIENS | 1 | 414 | 3fj6 | A | 347 | ARG | p.Arg346Trp | Q02127 |
| HOMO SAPIENS | 6 | 103 | 1u3w | A | 77 | GLY | p.Gly78Ter | P00326 |
| HOMO SAPIENS | -1 | 8369 | 4ux8 | F | 16 | ARG | p.Arg93Trp | P39905 |
| HOMO SAPIENS | 0 | 366 | 2zjc | A | 32 | GLY | p.Arg108Trp | P01375 |
| HOMO SAPIENS | 0 | 1457 | 3v9i | A | 352 | LEU | p.Ser352Leu | P30038 |
| HOMO SAPIENS | 0 | 1457 | 3v9i | A | 470 | VAL | p.Val470Ile | P30038 |
| HOMO SAPIENS | 0 | 1457 | 3v9i | A | 473 | THR | p.Thr473Ala | P30038 |
| HOMO SAPIENS | 1 | 2 | 4wo5 | A | 462 | ARG | p.Arg462Ile | P15056 |
| HOMO SAPIENS | 3 | 2 | 4wo5 | A | 463 | ILE | p.Ile463Ser | P15056 |
| HOMO SAPIENS | 1 | 2 | 4wo5 | A | 466 | GLY | p.Gly466Val | P15056 |
| HOMO SAPIENS | 1 | 2 | 4wo5 | A | 468 | PHE | p.Phe468Ser | P15056 |
| HOMO SAPIENS | 2 | 2 | 4wo5 | A | 469 | GLY | p.Gly469Ala | P15056 |
| HOMO SAPIENS | 2 | 2 | 4wo5 | A | 469 | GLY | p.Gly469Arg | P15056 |
| HOMO SAPIENS | 2 | 2 | 4wo5 | A | 485 | LEU | p.Leu485Phe | P15056 |
| HOMO SAPIENS | 0 | 2 | 4wo5 | A | 501 | GLU | p.Glu501Gly | P15056 |
| HOMO SAPIENS | 0 | 2 | 4wo5 | A | 501 | GLU | p.Glu501Lys | P15056 |
| HOMO SAPIENS | 7 | 2 | 4wo5 | A | 581 | ASN | p.Asn581Asp | P15056 |

| | | | | | | | | |
|---|---|---|---|---|---|---|---|---|
| HOMO SAPIENS | 8 | 2 | 4wo5 | A | 594 | ASP | p.Asp594Gly | P15056 |
| HOMO SAPIENS | 0 | 2 | 4wo5 | A | 597 | LEU | p.Leu597Arg | P15056 |
| HOMO SAPIENS | 0 | 2 | 4wo5 | A | 597 | LEU | p.Leu597Val | P15056 |
| HOMO SAPIENS | 0 | 2 | 4wo5 | A | 600 | VAL | p.Val600Glu | P15056 |
| HOMO SAPIENS | 2 | 2 | 4wo5 | A | 601 | LYS | p.Lys601Glu | P15056 |
| HOMO SAPIENS | 8 | 2 | 4wo5 | A | 638 | ASP | p.Asp638Glu | P15056 |
| HOMO SAPIENS | -1 | 30366 | 5bqd | B | 19 | ALA | p.Ala19Ser | Q99593 |
| HOMO SAPIENS | -1 | 30366 | 5bqd | B | 23 | PRO | p.Pro23His | Q99593 |
| HOMO SAPIENS | 1 | 30366 | 5bqd | B | 49 | GLN | p.Gln49Lys | Q99593 |
| HOMO SAPIENS | 2 | 30366 | 5bqd | B | 54 | ILE | p.Ile54Thr | Q99593 |
| HOMO SAPIENS | 8 | 30366 | 5bqd | B | 80 | GLY | p.Gly80Arg | Q99593 |
| HOMO SAPIENS | 6 | 30366 | 5bqd | B | 106 | ILE | p.Ile106Val | Q99593 |
| HOMO SAPIENS | 7 | 30366 | 5bqd | B | 111 | ASP | p.Asp111Tyr | Q99593 |
| HOMO SAPIENS | -1 | 30366 | 5bqd | B | 237 | ARG | p.Arg237Gln | Q99593 |
| HOMO SAPIENS | -1 | 30366 | 5bqd | B | 237 | ARG | p.Arg237Trp | Q99593 |
| HOMO SAPIENS | 5 | 4789 | 1n4m | A | 567 | SER | p.Ser567Leu | P06400 |
| HOMO SAPIENS | 0 | 4789 | 1n4m | A | 658 | ALA | p.Ala658Asp | P06400 |
| HOMO SAPIENS | 7 | 4789 | 1n4m | A | 661 | ARG | p.Arg661Trp | P06400 |
| HOMO SAPIENS | 6 | 4789 | 1n4m | A | 668 | ARG | p.Arg668His | P06400 |
| HOMO SAPIENS | 8 | 4789 | 1n4m | A | 697 | ASP | p.Asp697Glu | P06400 |
| HOMO SAPIENS | 6 | 4789 | 1n4m | A | 712 | CYS | p.Cys712Arg | P06400 |
| HOMO SAPIENS | 1 | 1798 | 2r5g | A | 101 | HIS | p.His101Gln | O15247 |
| HOMO SAPIENS | 0 | 1798 | 2r5g | A | 160 | PRO | p.Pro160Ala | O15247 |
| HOMO SAPIENS | 2 | 22 | 4x0t | A | 82 | ARG | p.Arg110Ter | P49419 |
| HOMO SAPIENS | 0 | 22 | 4x0t | A | 149 | ALA | p.Ala177Pro | P49419 |
| HOMO SAPIENS | 4 | 22 | 4x0t | A | 164 | THR | p.Thr192Met | P49419 |
| HOMO SAPIENS | 9 | 22 | 4x0t | A | 167 | ASN | p.Asn195Ser | P49419 |
| HOMO SAPIENS | 0 | 22 | 4x0t | A | 175 | TRP | p.Trp203Gly | P49419 |
| HOMO SAPIENS | 0 | 22 | 4x0t | A | 194 | THR | p.Thr222Ala | P49419 |
| HOMO SAPIENS | 0 | 22 | 4x0t | A | 273 | ASN | p.Asn301Ile | P49419 |
| HOMO SAPIENS | 0 | 22 | 4x0t | A | 307 | ARG | p.Arg335Gln | P49419 |
| HOMO SAPIENS | 3 | 22 | 4x0t | A | 313 | SER | p.Ser341Asn | P49419 |
| HOMO SAPIENS | 0 | 22 | 4x0t | A | 329 | ILE | p.Ile357Val | P49419 |
| HOMO SAPIENS | 1 | 22 | 4x0t | A | 380 | TYR | p.Tyr408Ter | P49419 |
| HOMO SAPIENS | 9 | 22 | 4x0t | A | 399 | GLU | p.Glu427Gln | P49419 |
| HOMO SAPIENS | 0 | 22 | 4x0t | A | 406 | TYR | p.Tyr434Cys | P49419 |
| HOMO SAPIENS | 0 | 22 | 4x0t | A | 431 | ILE | p.Ile459Phe | P49419 |
| HOMO SAPIENS | 4 | 22 | 4x0t | A | 477 | GLY | p.Gly505Arg | P49419 |
| HOMO SAPIENS | 0 | 22 | 4x0t | A | 495 | THR | p.Thr523Ala | P49419 |
| HOMO SAPIENS | 8 | 11042 | 1y6b | A | 1145 | PRO | p.Pro1147Ser | P35968 |
| HOMO SAPIENS | -1 | 3867 | 2xij | A | 18 | GLN | p.Gln18Ter | P22033 |
| HOMO SAPIENS | -1 | 3867 | 2xij | A | 31 | ARG | p.Arg31Ter | P22033 |
| HOMO SAPIENS | 2 | 3867 | 2xij | A | 93 | ARG | p.Arg93His | P22033 |
| HOMO SAPIENS | 0 | 3867 | 2xij | A | 94 | GLY | p.Gly94Arg | P22033 |
| HOMO SAPIENS | 2 | 3867 | 2xij | A | 95 | PRO | p.Pro95Leu | P22033 |
| HOMO SAPIENS | 2 | 3867 | 2xij | A | 105 | TRP | p.Trp105Arg | P22033 |
| HOMO SAPIENS | 2 | 3867 | 2xij | A | 108 | ARG | p.Arg108Cys | P22033 |
| HOMO SAPIENS | 3 | 3867 | 2xij | A | 154 | ARG | p.Arg154His | P22033 |
| HOMO SAPIENS | 3 | 3867 | 2xij | A | 156 | ASP | p.Asp156Val | P22033 |
| HOMO SAPIENS | 0 | 3867 | 2xij | A | 189 | ASN | p.Asn189Ile | P22033 |
| HOMO SAPIENS | 0 | 3867 | 2xij | A | 191 | ALA | p.Ala191Glu | P22033 |
| HOMO SAPIENS | 0 | 3867 | 2xij | A | 203 | GLY | p.Gly203Arg | P22033 |
| HOMO SAPIENS | 2 | 3867 | 2xij | A | 215 | GLY | p.Gly215Ser | P22033 |
| HOMO SAPIENS | 0 | 3867 | 2xij | A | 219 | ASN | p.Asn219Tyr | P22033 |

| | | | | | | | | |
|---|---|---|---|---|---|---|---|---|
| HOMO SAPIENS | 3 | 3867 | 2xij | A | 228 | ARG | p.Arg228Ter | P22033 |
| HOMO SAPIENS | 1 | 3867 | 2xij | A | 297 | THR | p.Thr297Ile | P22033 |
| HOMO SAPIENS | 0 | 3867 | 2xij | A | 326 | ARG | p.Arg326Lys | P22033 |
| HOMO SAPIENS | 3 | 3867 | 2xij | A | 369 | ARG | p.Arg369His | P22033 |
| HOMO SAPIENS | 3 | 3867 | 2xij | A | 369 | ARG | p.Arg369Cys | P22033 |
| HOMO SAPIENS | 0 | 3867 | 2xij | A | 370 | THR | p.Thr370Pro | P22033 |
| HOMO SAPIENS | 0 | 3867 | 2xij | A | 377 | ALA | p.Ala377Glu | P22033 |
| HOMO SAPIENS | 2 | 3867 | 2xij | A | 426 | GLY | p.Gly426Ala | P22033 |
| HOMO SAPIENS | 2 | 3867 | 2xij | A | 427 | GLY | p.Gly427Asp | P22033 |
| HOMO SAPIENS | 1 | 3867 | 2xij | A | 499 | THR | p.Ala499Thr | P22033 |
| HOMO SAPIENS | 2 | 3867 | 2xij | A | 532 | ARG | p.Arg532His | P22033 |
| HOMO SAPIENS | 3 | 3867 | 2xij | A | 555 | ALA | p.Ala555Thr | P22033 |
| HOMO SAPIENS | 0 | 3867 | 2xij | A | 573 | PHE | p.Phe573Ser | P22033 |
| HOMO SAPIENS | 3 | 3867 | 2xij | A | 623 | GLY | p.Gly623Arg | P22033 |
| HOMO SAPIENS | 5 | 3867 | 2xij | A | 671 | VAL | p.Ile671Val | P22033 |
| HOMO SAPIENS | 4 | 3867 | 2xij | A | 694 | ARG | p.Arg694Trp | P22033 |
| HOMO SAPIENS | 1 | 3867 | 2xij | A | 700 | MET | p.Met700Lys | P22033 |
| HOMO SAPIENS | 8 | 3867 | 2xij | A | 703 | GLY | p.Gly703Arg | P22033 |
| HOMO SAPIENS | 0 | 3867 | 2xij | A | 717 | GLY | p.Gly717Val | P22033 |
| HOMO SAPIENS | 1 | 3867 | 2xij | A | 727 | ARG | p.Arg727Ter | P22033 |
| HOMO SAPIENS | -1 | 52 | 3k91 | B | 72 | GLY | p.Gly73Ser | P00441 |
| HOMO SAPIENS | 6 | 52 | 3k91 | B | 96 | ASP | p.Asp97Asn | P00441 |
| HOMO SAPIENS | 5 | 52 | 3k91 | B | 112 | ILE | p.Ile113Thr | P00441 |
| HOMO SAPIENS | -1 | 52 | 3k91 | B | 134 | SER | p.Ser135Asn | P00441 |
| HOMO SAPIENS | 0 | 3950 | 4fom | A | 296 | ASN | p.Asn296His | Q9NQS3 |
| HOMO SAPIENS | 1 | 10 | 1de4 | A | 31 | VAL | p.Val53Met | Q30201 |
| HOMO SAPIENS | 0 | 10 | 1de4 | A | 37 | VAL | p.Val59Met | Q30201 |
| HOMO SAPIENS | 0 | 10 | 1de4 | A | 71 | GLY | p.Gly93Arg | Q30201 |
| HOMO SAPIENS | 0 | 10 | 1de4 | A | 83 | ILE | p.Ile105Thr | Q30201 |
| HOMO SAPIENS | 0 | 10 | 1de4 | A | 105 | GLN | p.Gln127His | Q30201 |
| HOMO SAPIENS | 1 | 10 | 1de4 | A | 146 | GLU | p.Glu168Ter | Q30201 |
| HOMO SAPIENS | 2 | 10 | 1de4 | A | 261 | GLN | p.Gln283Pro | Q30201 |
| HOMO SAPIENS | 0 | 8038 | 4x7n | A | 586 | ILE | p.Ile586Leu | Q9NZJ5 |
| HOMO SAPIENS | 0 | 8038 | 4x7n | A | 588 | ARG | p.Arg588Gln | Q9NZJ5 |
| HOMO SAPIENS | 0 | 35175 | 5ugw | A | 1062 | ALA | p.Ala1062Thr | O14746 |
| HOMO SAPIENS | 2 | 35175 | 5ugw | A | 1090 | VAL | p.Val1090Met | O14746 |
| HOMO SAPIENS | 0 | 1722 | 4z3k | C | 147 | ARG | p.Arg150Gly | P35270 |
| HOMO SAPIENS | 0 | 1722 | 4z3k | C | 160 | PRO | p.Pro163Leu | P35270 |
| HOMO SAPIENS | 0 | 1722 | 4z3k | C | 207 | ARG | p.Arg210Trp | P35270 |
| HOMO SAPIENS | 0 | 1722 | 4z3k | C | 216 | ARG | p.Arg219Ter | P35270 |
| HOMO SAPIENS | 0 | 1722 | 4z3k | C | 248 | LYS | p.Lys251Ter | P35270 |
| HOMO SAPIENS | 1 | 2832 | 4x8v | L | 115 | HIS | p.His175Gln | P08709 |
| HOMO SAPIENS | 0 | 825 | 1emh | A | 242 | PHE | p.Phe251Ser | P13051 |
| HOMO SAPIENS | 0 | 122 | 5uwk | A | 109 | ARG | p.Arg109Ter | P45452 |
| HOMO SAPIENS | 1 | 122 | 5uwk | A | 170 | LYS | p.Lys170Thr | P45452 |
| HOMO SAPIENS | 9 | 122 | 5uwk | A | 232 | HIS | p.His232Asn | P45452 |
| HOMO SAPIENS | 5 | 122 | 5uwk | A | 257 | ASP | p.Asp257Val | P45452 |
| HOMO SAPIENS | 9 | 1074 | 3p5i | A | 264 | TRP | p.Trp264Arg | Q9UJ71 |
| HOMO SAPIENS | 0 | 146 | 3r3g | A | 14C | GLU | p.Glu352Lys | P00734 |
| HOMO SAPIENS | 3 | 9522 | 1wyq | A | 83 | LEU | p.Leu253Pro | O15020 |
| HOMO SAPIENS | 8 | 67 | 2vgb | A | 130 | SER | p.Ser130Tyr | P30613 |
| HOMO SAPIENS | 8 | 67 | 2vgb | A | 163 | ARG | p.Arg163Cys | P30613 |
| HOMO SAPIENS | 4 | 67 | 2vgb | A | 241 | GLU | p.Glu241Ter | P30613 |
| HOMO SAPIENS | 9 | 67 | 2vgb | A | 384 | THR | p.Thr384Met | P30613 |

| | | | | | | | | |
|---|---|---|---|---|---|---|---|---|
| HOMO SAPIENS | 0 | 67 | 2vgb | A | 421 | GLN | p.Gln421Lys | P30613 |
| HOMO SAPIENS | 6 | 67 | 2vgb | A | 479 | ARG | p.Arg479His | P30613 |
| HOMO SAPIENS | 8 | 67 | 2vgb | A | 510 | ARG | p.Arg510Gln | P30613 |
| HOMO SAPIENS | 4 | 67 | 2vgb | A | 569 | ARG | p.Arg569Trp | P30613 |
| HOMO SAPIENS | 0 | 9088 | 5v44 | A | 232 | ASN | p.Asn232Lys | Q9NZJ4 |
| HOMO SAPIENS | 4 | 16 | 3lbn | A | 12 | GLY | p.Gly12Val | P01112 |
| HOMO SAPIENS | 4 | 16 | 3lbn | A | 12 | GLY | p.Gly12Ser | P01112 |
| HOMO SAPIENS | 4 | 16 | 3lbn | A | 13 | GLY | p.Gly13Cys | P01112 |
| HOMO SAPIENS | 4 | 16 | 3lbn | A | 13 | GLY | p.Gly13Asp | P01112 |
| HOMO SAPIENS | 5 | 16 | 3lbn | A | 22 | GLN | p.Gln22Lys | P01112 |
| HOMO SAPIENS | 5 | 16 | 3lbn | A | 58 | THR | p.Thr58Ile | P01112 |
| HOMO SAPIENS | 3 | 16 | 3lbn | A | 61 | GLN | p.Gln61Lys | P01112 |
| HOMO SAPIENS | 1 | 16 | 3lbn | A | 63 | GLU | p.Glu63Lys | P01112 |
| HOMO SAPIENS | 9 | 16 | 3lbn | A | 117 | LYS | p.Lys117Arg | P01112 |
| HOMO SAPIENS | 6 | 16 | 3lbn | A | 123 | ARG | p.Arg123Cys | P01112 |
| HOMO SAPIENS | 1 | 16 | 3lbn | A | 138 | GLY | p.Gly138Ser | P01112 |
| HOMO SAPIENS | 9 | 16 | 3lbn | A | 146 | ALA | p.Ala146Thr | P01112 |
| HOMO SAPIENS | 9 | 16 | 3lbn | A | 146 | ALA | p.Ala146Val | P01112 |
| HOMO SAPIENS | 1 | 359 | 5nvy | C | 63 | LEU | p.Leu63Pro | P40337 |
| HOMO SAPIENS | 1 | 359 | 5nvy | C | 64 | ARG | p.Arg64Pro | P40337 |
| HOMO SAPIENS | 1 | 359 | 5nvy | C | 70 | GLU | p.Glu70Lys | P40337 |
| HOMO SAPIENS | 0 | 359 | 5nvy | C | 84 | VAL | p.Val84Leu | P40337 |
| HOMO SAPIENS | 1 | 359 | 5nvy | C | 88 | TRP | p.Trp88Ser | P40337 |
| HOMO SAPIENS | 1 | 359 | 5nvy | C | 93 | GLY | p.Gly93Cys | P40337 |
| HOMO SAPIENS | 1 | 359 | 5nvy | C | 98 | TYR | p.Tyr98His | P40337 |
| HOMO SAPIENS | 1 | 359 | 5nvy | C | 111 | SER | p.Ser111Arg | P40337 |
| HOMO SAPIENS | 1 | 359 | 5nvy | C | 112 | TYR | p.Tyr112His | P40337 |
| HOMO SAPIENS | 1 | 359 | 5nvy | C | 126 | ASP | p.Asp126Tyr | P40337 |
| HOMO SAPIENS | 1 | 359 | 5nvy | C | 130 | VAL | p.Val130Leu | P40337 |
| HOMO SAPIENS | 1 | 359 | 5nvy | C | 135 | LEU | p.Leu135Phe | P40337 |
| HOMO SAPIENS | 9 | 359 | 5nvy | C | 161 | ARG | p.Arg161Gly | P40337 |
| HOMO SAPIENS | 7 | 359 | 5nvy | C | 162 | CYS | p.Cys162Phe | P40337 |
| HOMO SAPIENS | 4 | 359 | 5nvy | C | 163 | LEU | p.Leu163Pro | P40337 |
| HOMO SAPIENS | 3 | 359 | 5nvy | C | 164 | GLN | p.Gln164Arg | P40337 |
| HOMO SAPIENS | 9 | 359 | 5nvy | C | 166 | VAL | p.Val166Phe | P40337 |
| HOMO SAPIENS | 7 | 359 | 5nvy | C | 167 | ARG | p.Arg167Gly | P40337 |
| HOMO SAPIENS | 7 | 359 | 5nvy | C | 167 | ARG | p.Arg167Gln | P40337 |
| HOMO SAPIENS | 1 | 359 | 5nvy | C | 183 | SER | p.Ser183Ter | P40337 |
| HOMO SAPIENS | 2 | 359 | 5nvy | C | 188 | LEU | p.Leu188Val | P40337 |
| HOMO SAPIENS | 1 | 359 | 5nvy | C | 191 | HIS | p.His191Asp | P40337 |
| HOMO SAPIENS | 1 | 359 | 5nvy | C | 192 | PRO | p.Pro192Ser | P40337 |
| HOMO SAPIENS | 1 | 359 | 5nvy | C | 200 | ARG | p.Arg200Trp | P40337 |
| HOMO SAPIENS | -1 | 359 | 5nvy | C | 211 | MET | p.Met211Leu | P40337 |
| HOMO SAPIENS | 0 | 14099 | 4btm | A | 282 | ARG | p.Arg258Gln | Q5TCY1 |
| HOMO SAPIENS | 0 | 696 | 5h1b | A | 150 | ARG | p.Arg150Gln | Q06609 |
| HOMO SAPIENS | -1 | 208 | 2bxl | A | 1 | ASP | p.Asp25Val | P02768 |
| HOMO SAPIENS | 0 | 208 | 2bxl | A | 269 | ASP | p.Asp293Gly | P02768 |
| HOMO SAPIENS | 0 | 208 | 2bxl | A | 570 | GLU | p.Glu594Lys | P02768 |
| HOMO SAPIENS | 0 | 208 | 2bxl | A | 573 | LYS | p.Lys597Glu | P02768 |
| HOMO SAPIENS | -1 | 12813 | 2uzx | B | 35 | MET | p.Met35Leu | P08581 |
| HOMO SAPIENS | 1 | 12813 | 2uzx | B | 156 | SER | p.Ser156Leu | P08581 |
| HOMO SAPIENS | 0 | 12813 | 2uzx | B | 168 | GLU | p.Glu168Asp | P08581 |
| HOMO SAPIENS | 2 | 12813 | 2uzx | B | 211 | LEU | p.Leu211Trp | P08581 |
| HOMO SAPIENS | 2 | 12813 | 2uzx | B | 218 | ARG | p.Arg218Ser | P08581 |

| | | | | | | | | |
|---|---|---|---|---|---|---|---|---|
| HOMO SAPIENS | 1 | 12813 | 2uzx | B | 238 | LEU | p.Leu238Ser | P08581 |
| HOMO SAPIENS | 4 | 12813 | 2uzx | B | 320 | ALA | p.Ala320Val | P08581 |
| HOMO SAPIENS | 1 | 12813 | 2uzx | B | 375 | ASN | p.Asn375Ser | P08581 |
| HOMO SAPIENS | 0 | 12813 | 2uzx | B | 575 | LEU | p.Leu575Ile | P08581 |
| HOMO SAPIENS | 1 | 10780 | 4ejx | A | 348 | ARG | p.Arg367Cys | P00450 |
| HOMO SAPIENS | 2 | 10780 | 4ejx | A | 525 | ASP | p.Asp544Glu | P00450 |
| HOMO SAPIENS | 1 | 1407 | 5jn5 | A | 88 | ILE | p.Ile88Val | P36871 |
| HOMO SAPIENS | 7 | 1407 | 5jn5 | A | 115 | THR | p.Thr115Ala | P36871 |
| HOMO SAPIENS | 3 | 1407 | 5jn5 | A | 121 | GLY | p.Gly121Arg | P36871 |
| HOMO SAPIENS | 2 | 1407 | 5jn5 | A | 221 | ARG | p.Arg221Cys | P36871 |
| HOMO SAPIENS | 6 | 1407 | 5jn5 | A | 263 | TYR | p.Asp263Tyr | P36871 |
| HOMO SAPIENS | 3 | 1407 | 5jn5 | A | 420 | TYR | p.Tyr420His | P36871 |
| HOMO SAPIENS | 0 | 1407 | 5jn5 | A | 501 | VAL | p.Val501Ile | P36871 |
| HOMO SAPIENS | -1 | 1878 | 1yow | A | 255 | ARG | p.Arg255Leu | Q13285 |
| HOMO SAPIENS | 6 | 1878 | 1yow | A | 293 | ASP | p.Asp293Asn | Q13285 |
| HOMO SAPIENS | 1 | 1878 | 1yow | A | 437 | LEU | p.Leu437Gln | Q13285 |
| HOMO SAPIENS | 0 | 3071 | 2wpm | E | 145 | ARG | p.Arg191His | P00740 |
| HOMO SAPIENS | 0 | 3071 | 2wpm | E | 145 | ARG | p.Arg191Cys | P00740 |
| HOMO SAPIENS | 3 | 589 | 3ggc | B | 15 | GLY | p.Gly16Ser | P00492 |
| HOMO SAPIENS | 4 | 589 | 3ggc | B | 40 | LEU | p.Leu41Pro | P00492 |
| HOMO SAPIENS | 2 | 589 | 3ggc | B | 50 | ARG | p.Arg51Gly | P00492 |
| HOMO SAPIENS | 5 | 589 | 3ggc | B | 51 | ASP | p.Asp52Gly | P00492 |
| HOMO SAPIENS | 0 | 589 | 3ggc | B | 57 | GLY | p.Gly58Arg | P00492 |
| HOMO SAPIENS | 6 | 589 | 3ggc | B | 69 | GLY | p.Gly70Glu | P00492 |
| HOMO SAPIENS | 1 | 589 | 3ggc | B | 73 | PHE | p.Phe74Leu | P00492 |
| HOMO SAPIENS | 0 | 589 | 3ggc | B | 77 | LEU | p.Leu78Val | P00492 |
| HOMO SAPIENS | 3 | 589 | 3ggc | B | 79 | ASP | p.Asp80Val | P00492 |
| HOMO SAPIENS | 0 | 589 | 3ggc | B | 103 | SER | p.Ser104Arg | P00492 |
| HOMO SAPIENS | 0 | 589 | 3ggc | B | 109 | SER | p.Ser110Leu | P00492 |
| HOMO SAPIENS | 1 | 589 | 3ggc | B | 129 | VAL | p.Val130Asp | P00492 |
| HOMO SAPIENS | 4 | 589 | 3ggc | B | 131 | ILE | p.Ile132Met | P00492 |
| HOMO SAPIENS | 7 | 589 | 3ggc | B | 142 | MET | p.Met143Lys | P00492 |
| HOMO SAPIENS | 0 | 589 | 3ggc | B | 160 | ALA | p.Ala161Ser | P00492 |
| HOMO SAPIENS | 6 | 589 | 3ggc | B | 167 | THR | p.Thr168Ile | P00492 |
| HOMO SAPIENS | 0 | 589 | 3ggc | B | 169 | ARG | p.Arg170Ter | P00492 |
| HOMO SAPIENS | 5 | 589 | 3ggc | B | 175 | PRO | p.Pro176Leu | P00492 |
| HOMO SAPIENS | 8 | 589 | 3ggc | B | 193 | ASP | p.Asp194Glu | P00492 |
| HOMO SAPIENS | 8 | 589 | 3ggc | B | 193 | ASP | p.Asp194Asn | P00492 |
| HOMO SAPIENS | 8 | 589 | 3ggc | B | 198 | PHE | p.Phe199Val | P00492 |
| HOMO SAPIENS | 8 | 589 | 3ggc | B | 200 | ASP | p.Asp201Gly | P00492 |
| HOMO SAPIENS | 0 | 4269 | 2pla | B | 174 | GLU | p.Glu174Lys | Q8N335 |
| HOMO SAPIENS | 1 | 4269 | 2pla | B | 220 | ARG | p.Arg220His | Q8N335 |
| HOMO SAPIENS | 4 | 4269 | 2pla | B | 280 | ALA | p.Ala280Val | Q8N335 |
| HOMO SAPIENS | 0 | 2712 | 1ii4 | E | 252 | TRP | p.Ser252Leu | P21802 |
| HOMO SAPIENS | 0 | 2712 | 1ii4 | E | 253 | PRO | p.Pro253Arg | P21802 |
| HOMO SAPIENS | 5 | 2712 | 1ii4 | E | 267 | SER | p.Ser267Pro | P21802 |
| HOMO SAPIENS | 0 | 2712 | 1ii4 | E | 278 | CYS | p.Cys278Phe | P21802 |
| HOMO SAPIENS | 0 | 2712 | 1ii4 | E | 289 | GLN | p.Gln289Pro | P21802 |
| HOMO SAPIENS | 0 | 2712 | 1ii4 | E | 290 | TRP | p.Trp290Cys | P21802 |
| HOMO SAPIENS | 0 | 2712 | 1ii4 | E | 290 | TRP | p.Trp290Arg | P21802 |
| HOMO SAPIENS | 0 | 2712 | 1ii4 | E | 292 | LYS | p.Lys292Glu | P21802 |
| HOMO SAPIENS | 0 | 2712 | 1ii4 | E | 315 | ALA | p.Ala315Ser | P21802 |
| HOMO SAPIENS | 0 | 2712 | 1ii4 | E | 321 | ASP | p.Asp321Ala | P21802 |
| HOMO SAPIENS | 1 | 2712 | 1ii4 | E | 328 | TYR | p.Tyr328Cys | P21802 |

| | | | | | | | | |
|---|---|---|---|---|---|---|---|---|
| HOMO SAPIENS | 9 | 2712 | 1ii4 | E | 340 | TYR | p.Tyr340His | P21802 |
| HOMO SAPIENS | 1 | 2712 | 1ii4 | E | 341 | THR | p.Thr341Pro | P21802 |
| HOMO SAPIENS | 0 | 2712 | 1ii4 | E | 342 | CYS | p.Cys342Arg | P21802 |
| HOMO SAPIENS | 0 | 2712 | 1ii4 | E | 342 | CYS | p.Cys342Trp | P21802 |
| HOMO SAPIENS | 0 | 2712 | 1ii4 | E | 344 | ALA | p.Ala344Gly | P21802 |
| HOMO SAPIENS | 0 | 2712 | 1ii4 | E | 347 | SER | p.Ser347Cys | P21802 |
| HOMO SAPIENS | 0 | 2712 | 1ii4 | E | 351 | SER | p.Ser351Cys | P21802 |
| HOMO SAPIENS | 1 | 2712 | 1ii4 | E | 354 | SER | p.Ser354Cys | P21802 |
| HOMO SAPIENS | 2 | 53 | 4pj1 | H | 5 | ASP | p.Asp29Gly | P10809 |
| HOMO SAPIENS | 0 | 53 | 4pj1 | H | 74 | VAL | p.Val98Ile | P10809 |
| HOMO SAPIENS | 0 | 29968 | 2fdb | M | 71 | LYS | p.Lys89Glu | P55075 |
| HOMO SAPIENS | 2 | 29968 | 2fdb | M | 98 | ARG | p.Arg116Gly | P55075 |
| HOMO SAPIENS | 2 | 29968 | 2fdb | M | 122 | GLY | p.Gly140Ser | P55075 |
| HOMO SAPIENS | 8 | 5061 | 5f85 | B | 60 | GLY | p.Gly106Ser | P00740 |
| HOMO SAPIENS | 0 | 35944 | 1h6h | A | 85 | THR | p.Thr85Asn | Q15080 |
| HOMO SAPIENS | 8 | 35944 | 1h6h | A | 105 | ARG | p.Arg105Gln | Q15080 |
| HOMO SAPIENS | 7 | 12016 | 1hj7 | A | 295 | ASN | p.Asn316Ser | P01130 |
| HOMO SAPIENS | 5 | 12016 | 1hj7 | A | 302 | GLY | p.Gly323Ser | P01130 |
| HOMO SAPIENS | 6 | 12016 | 1hj7 | A | 319 | CYS | p.Cys340Trp | P01130 |
| HOMO SAPIENS | 7 | 12016 | 1hj7 | A | 329 | ARG | p.Arg350Ter | P01130 |
| HOMO SAPIENS | 5 | 12016 | 1hj7 | A | 331 | CYS | p.Cys352Ter | P01130 |
| HOMO SAPIENS | 2 | 12016 | 1hj7 | A | 345 | GLN | p.Gln366Arg | P01130 |
| HOMO SAPIENS | 2 | 1481 | 2i07 | A | 239 | TYR | p.Tyr261Ter | P01024 |
| HOMO SAPIENS | -1 | 1481 | 2i07 | A | 292 | LEU | p.Pro314Leu | P01024 |
| HOMO SAPIENS | 2 | 1481 | 2i07 | A | 326 | ILE | p.Ile348Val | P01024 |
| HOMO SAPIENS | 2 | 1481 | 2i07 | A | 413 | GLU | p.Glu435Lys | P01024 |
| HOMO SAPIENS | 4 | 1481 | 2i07 | A | 447 | GLU | p.Glu469Asp | P01024 |
| HOMO SAPIENS | 2 | 1481 | 2i07 | A | 597 | VAL | p.Val619Met | P01024 |
| HOMO SAPIENS | 5 | 1481 | 2i07 | A | 603 | ILE | p.Ile625Phe | P01024 |
| HOMO SAPIENS | 0 | 1481 | 2i07 | A | 611 | LYS | p.Lys633Arg | P01024 |
| HOMO SAPIENS | 0 | 11123 | 3c5n | A | 315 | GLY | p.Gly368Trp | O00294 |
| HOMO SAPIENS | 8 | 11123 | 3c5n | A | 329 | PHE | p.Phe382Ser | O00294 |
| HOMO SAPIENS | 5 | 11123 | 3c5n | A | 429 | ARG | p.Arg482Trp | O00294 |
| HOMO SAPIENS | 8 | 11123 | 3c5n | A | 438 | PHE | p.Phe491Leu | O00294 |
| HOMO SAPIENS | 3 | 4398 | 3fx0 | B | 1288 | ALA | p.Ala288Gly | Q9Y6K9 |
| HOMO SAPIENS | 7 | 4398 | 3fx0 | B | 1315 | GLU | p.Glu315Ala | Q9Y6K9 |
| HOMO SAPIENS | 8 | 4398 | 3fx0 | B | 1319 | ARG | p.Arg319Gln | Q9Y6K9 |
| HOMO SAPIENS | 6 | 3989 | 3qd6 | S | 83 | CYS | p.Cys83Arg | P25942 |
| HOMO SAPIENS | 1 | 3989 | 3qd6 | S | 124 | SER | p.Ser124Leu | P25942 |
| HOMO SAPIENS | 2 | 24003 | 2m0v | A | 225 | GLU | p.Glu225Lys | O14745 |
| HOMO SAPIENS | 2 | 46 | 4aqd | A | 70 | ASP | p.Asp98Gly | P06276 |
| HOMO SAPIENS | 4 | 46 | 4aqd | A | 115 | GLY | p.Gly143Asp | P06276 |
| HOMO SAPIENS | 3 | 46 | 4aqd | A | 184 | ALA | p.Ala212Val | P06276 |
| HOMO SAPIENS | 0 | 46 | 4aqd | A | 225 | GLY | p.Gly253Ter | P06276 |
| HOMO SAPIENS | 2 | 46 | 4aqd | A | 243 | THR | p.Thr271Met | P06276 |
| HOMO SAPIENS | 2 | 46 | 4aqd | A | 255 | GLU | p.Glu283Asp | P06276 |
| HOMO SAPIENS | 5 | 46 | 4aqd | A | 267 | LYS | p.Lys295Arg | P06276 |
| HOMO SAPIENS | 3 | 46 | 4aqd | A | 271 | GLU | p.Glu299Ter | P06276 |
| HOMO SAPIENS | 2 | 46 | 4aqd | A | 307 | LEU | p.Leu335Pro | P06276 |
| HOMO SAPIENS | 1 | 46 | 4aqd | A | 330 | LEU | p.Leu358Ile | P06276 |
| HOMO SAPIENS | 0 | 46 | 4aqd | A | 497 | GLU | p.Glu525Val | P06276 |
| HOMO SAPIENS | 3 | 786 | 1ols | B | 51 | GLY | p.Gly101Asp | P21953 |
| HOMO SAPIENS | 2 | 786 | 1ols | B | 87 | ALA | p.Ala137Val | P21953 |
| HOMO SAPIENS | 0 | 786 | 1ols | B | 110 | ILE | p.Ile160Ser | P21953 |

| | | | | | | | | |
|---|---|---|---|---|---|---|---|---|
| HOMO SAPIENS | 4 | 786 | 1ols | B | 120 | ARG | p.Arg170Cys | P21953 |
| HOMO SAPIENS | 2 | 786 | 1ols | B | 126 | ASN | p.Asn176Tyr | P21953 |
| HOMO SAPIENS | 0 | 786 | 1ols | B | 133 | ARG | p.Arg183Gly | P21953 |
| HOMO SAPIENS | 0 | 786 | 1ols | B | 133 | ARG | p.Arg183Pro | P21953 |
| HOMO SAPIENS | 0 | 786 | 1ols | B | 135 | PRO | p.Pro185Leu | P21953 |
| HOMO SAPIENS | 3 | 786 | 1ols | B | 156 | HIS | p.His206Tyr | P21953 |
| HOMO SAPIENS | 0 | 786 | 1ols | B | 228 | GLY | p.Gly278Ser | P21953 |
| HOMO SAPIENS | 1 | 786 | 1ols | B | 235 | ARG | p.Arg285Ter | P21953 |
| HOMO SAPIENS | 6 | 786 | 1ols | B | 274 | ARG | p.Arg324Ter | P21953 |
| HOMO SAPIENS | 1 | 831 | 4zt1 | B | 48 | VAL | p.Val202Ile | P12830 |
| HOMO SAPIENS | 6 | 831 | 4zt1 | B | 70 | ARG | p.Arg224Cys | P12830 |
| HOMO SAPIENS | 5 | 831 | 4zt1 | B | 79 | HIS | p.His233Gln | P12830 |
| HOMO SAPIENS | 4 | 831 | 4zt1 | B | 120 | GLY | p.Gly274Ser | P12830 |
| HOMO SAPIENS | 7 | 831 | 4zt1 | B | 187 | TYR | p.Tyr341Ter | P12830 |
| HOMO SAPIENS | 0 | 8497 | 3sla | A | 287 | ASN | p.Asn287Ser | P35222 |
| HOMO SAPIENS | 7 | 23986 | 2ke1 | A | 301 | VAL | p.Val301Met | O43918 |
| HOMO SAPIENS | 0 | 4967 | 1rj7 | B | 334 | ARG | p.Arg334His | Q92838 |
| HOMO SAPIENS | 0 | 4967 | 1rj7 | B | 338 | THR | p.Thr338Met | Q92838 |
| HOMO SAPIENS | 3 | 4967 | 1rj7 | B | 349 | ALA | p.Ala349Thr | Q92838 |
| HOMO SAPIENS | 2 | 4967 | 1rj7 | B | 358 | GLN | p.Gln358Glu | Q92838 |
| HOMO SAPIENS | 1 | 4967 | 1rj7 | B | 375 | LYS | p.Lys375Glu | Q92838 |
| HOMO SAPIENS | 9 | 7623 | 1d4t | A | 32 | ARG | p.Arg32Thr | O60880 |
| HOMO SAPIENS | 6 | 7623 | 1d4t | A | 55 | ARG | p.Arg55Leu | O60880 |
| HOMO SAPIENS | 0 | 7623 | 1d4t | A | 68 | THR | p.Thr68Ile | O60880 |
| HOMO SAPIENS | 6 | 705 | 3idh | A | 59 | PRO | p.Pro59Ser | P35557 |
| HOMO SAPIENS | 5 | 705 | 3idh | A | 124 | ASP | p.Asp124Asn | P35557 |
| HOMO SAPIENS | 4 | 705 | 3idh | A | 131 | SER | p.Ser131Pro | P35557 |
| HOMO SAPIENS | 0 | 705 | 3idh | A | 147 | GLY | p.Gly147Asp | P35557 |
| HOMO SAPIENS | 5 | 705 | 3idh | A | 186 | ARG | p.Arg186Ter | P35557 |
| HOMO SAPIENS | 0 | 705 | 3idh | A | 202 | MET | p.Met202Thr | P35557 |
| HOMO SAPIENS | 2 | 705 | 3idh | A | 210 | MET | p.Met210Lys | P35557 |
| HOMO SAPIENS | 1 | 705 | 3idh | A | 215 | TYR | p.Tyr215Ter | P35557 |
| HOMO SAPIENS | 0 | 705 | 3idh | A | 226 | VAL | p.Val226Met | P35557 |
| HOMO SAPIENS | 9 | 705 | 3idh | A | 228 | THR | p.Thr228Met | P35557 |
| HOMO SAPIENS | 9 | 705 | 3idh | A | 256 | GLU | p.Glu256Lys | P35557 |
| HOMO SAPIENS | 2 | 705 | 3idh | A | 261 | GLY | p.Gly261Arg | P35557 |
| HOMO SAPIENS | 6 | 705 | 3idh | A | 279 | GLU | p.Glu279Gln | P35557 |
| HOMO SAPIENS | 7 | 705 | 3idh | A | 299 | GLY | p.Gly299Arg | P35557 |
| HOMO SAPIENS | 2 | 705 | 3idh | A | 318 | GLY | p.Gly318Trp | P35557 |
| HOMO SAPIENS | 4 | 705 | 3idh | A | 378 | ALA | p.Ala378Thr | P35557 |
| HOMO SAPIENS | 0 | 705 | 3idh | A | 387 | ALA | p.Ala387Glu | P35557 |
| HOMO SAPIENS | 5 | 705 | 3idh | A | 414 | LYS | p.Lys414Glu | P35557 |
| HOMO SAPIENS | 0 | 705 | 3idh | A | 455 | VAL | p.Val455Met | P35557 |
| HOMO SAPIENS | -1 | 705 | 3idh | A | 462 | MET | p.Met462Ile | P35557 |
| HOMO SAPIENS | 1 | 509 | 4nef | A | 22 | LEU | p.Leu22Val | P41181 |
| HOMO SAPIENS | 4 | 509 | 4nef | A | 57 | GLN | p.Gln57Pro | P41181 |
| HOMO SAPIENS | 8 | 509 | 4nef | A | 64 | GLY | p.Gly64Arg | P41181 |
| HOMO SAPIENS | 7 | 509 | 4nef | A | 68 | ASN | p.Asn68Ser | P41181 |
| HOMO SAPIENS | 1 | 509 | 4nef | A | 100 | GLY | p.Gly100Val | P41181 |
| HOMO SAPIENS | 0 | 509 | 4nef | A | 125 | THR | p.Thr125Met | P41181 |
| HOMO SAPIENS | 0 | 509 | 4nef | A | 126 | THR | p.Thr126Met | P41181 |
| HOMO SAPIENS | 4 | 509 | 4nef | A | 146 | PHE | p.Phe146Leu | P41181 |
| HOMO SAPIENS | 0 | 509 | 4nef | A | 175 | GLY | p.Gly175Arg | P41181 |
| HOMO SAPIENS | 5 | 509 | 4nef | A | 181 | CYS | p.Cys181Trp | P41181 |

| | | | | | | | | | |
|---|---|---|---|---|---|---|---|---|---|
| HOMO SAPIENS | 9 | 509 | 4nef | A | | 187 | ARG | p.Arg187Cys | P41181 |
| HOMO SAPIENS | 0 | 509 | 4nef | A | | 190 | ALA | p.Ala190Thr | P41181 |
| HOMO SAPIENS | 0 | 509 | 4nef | A | | 216 | SER | p.Ser216Pro | P41181 |
| HOMO SAPIENS | 4 | 2242 | 1z70 | X | | 1155 | SER | p.Ser155Pro | Q8NBK3 |
| HOMO SAPIENS | 3 | 2242 | 1z70 | X | | 1218 | CYS | p.Cys218Tyr | Q8NBK3 |
| HOMO SAPIENS | 0 | 2242 | 1z70 | X | | 1279 | ALA | p.Ala279Val | Q8NBK3 |
| HOMO SAPIENS | -1 | 2242 | 1z70 | X | | 1336 | OCS | p.Cys336Arg | Q8NBK3 |
| HOMO SAPIENS | 4 | 2242 | 1z70 | X | | 1345 | ARG | p.Arg345Cys | Q8NBK3 |
| HOMO SAPIENS | 2 | 2242 | 1z70 | X | | 1348 | ALA | p.Ala348Pro | Q8NBK3 |
| HOMO SAPIENS | 5 | 2242 | 1z70 | X | | 1349 | ARG | p.Arg349Gln | Q8NBK3 |
| HOMO SAPIENS | 5 | 2242 | 1z70 | X | | 1349 | ARG | p.Arg349Trp | Q8NBK3 |
| HOMO SAPIENS | 3 | 2242 | 1z70 | X | | 1359 | SER | p.Ser359Ter | Q8NBK3 |
| HOMO SAPIENS | 5 | 8269 | 5wrr | B | | 136 | ARG | p.Arg136Ter | Q96MK3 |
| HOMO SAPIENS | 1 | 8269 | 5wrr | B | | 332 | LYS | p.Asn332Lys | Q96MK3 |
| HOMO SAPIENS | 3 | 8269 | 5wrr | B | | 478 | ARG | p.Arg478Ter | Q96MK3 |
| HOMO SAPIENS | 5 | 8269 | 5wrr | B | | 504 | THR | p.Thr504Ile | Q96MK3 |
| HOMO SAPIENS | 5 | 1411 | 4h1s | B | | 221 | SER | p.Ser221Ter | P21589 |
| HOMO SAPIENS | 1 | 1411 | 4h1s | B | | 358 | CYS | p.Cys358Tyr | P21589 |
| HOMO SAPIENS | 2 | 23 | 5vvd | B | | 298 | GLU | p.Glu298Asp | P29474 |
| HOMO SAPIENS | 0 | 19 | 5jlh | E | | 88 | THR | p.Thr89Ile | P63261 |
| HOMO SAPIENS | 6 | 19 | 5jlh | E | | 117 | LYS | p.Lys118Met | P63261 |
| HOMO SAPIENS | 5 | 19 | 5jlh | E | | 263 | PRO | p.Pro264Leu | P63261 |
| HOMO SAPIENS | 1 | 19 | 5jlh | E | | 277 | THR | p.Thr278Ile | P63261 |
| HOMO SAPIENS | 6 | 19 | 5jlh | E | | 331 | PRO | p.Pro332Ala | P63261 |
| HOMO SAPIENS | 5 | 19 | 5jlh | E | | 369 | VAL | p.Val370Ala | P63261 |
| HOMO SAPIENS | 0 | 14306 | 4w4o | C | | 92 | ARG | p.Arg92Ter | P12314 |
| HOMO SAPIENS | 3 | 603 | 1nuh | A | | 4 | THR | p.Thr5Ile | P06744 |
| HOMO SAPIENS | 2 | 603 | 1nuh | A | | 19 | HIS | p.His20Pro | P06744 |
| HOMO SAPIENS | 8 | 603 | 1nuh | A | | 158 | GLY | p.Gly159Ser | P06744 |
| HOMO SAPIENS | 6 | 603 | 1nuh | A | | 223 | THR | p.Thr224Met | P06744 |
| HOMO SAPIENS | 0 | 603 | 1nuh | A | | 338 | LEU | p.Leu339Pro | P06744 |
| HOMO SAPIENS | 6 | 603 | 1nuh | A | | 342 | GLN | p.Gln343Arg | P06744 |
| HOMO SAPIENS | 6 | 603 | 1nuh | A | | 374 | THR | p.Thr375Arg | P06744 |
| HOMO SAPIENS | 4 | 603 | 1nuh | A | | 524 | ILE | p.Ile525Thr | P06744 |
| HOMO SAPIENS | 4 | 603 | 1nuh | A | | 538 | ASP | p.Asp539Asn | P06744 |
| HOMO SAPIENS | 1 | 3107 | 3kzj | A | | 1197 | VAL | p.Val1197Ala | P08603 |
| HOMO SAPIENS | 9 | 1748 | 2noh | A | | 46 | ARG | p.Arg46Gln | O15527 |
| HOMO SAPIENS | 2 | 11750 | 3s4y | A | | 33 | ARG | p.Arg33His | Q9H3S4 |
| HOMO SAPIENS | 2 | 11750 | 3s4y | A | | 40 | LEU | p.Leu40Pro | Q9H3S4 |
| HOMO SAPIENS | 4 | 11750 | 3s4y | A | | 142 | LEU | p.Leu142Phe | Q9H3S4 |
| HOMO SAPIENS | 5 | 11750 | 3s4y | A | | 219 | ASN | p.Asn219Ser | Q9H3S4 |
| HOMO SAPIENS | 1 | 2591 | 5vnj | A | | 382 | PHE | p.Phe382Leu | Q15436 |
| HOMO SAPIENS | 2 | 2591 | 5vnj | A | | 702 | MET | p.Met702Val | Q15436 |
| HOMO SAPIENS | 1 | 2426 | 5m8t | A | | 87 | ARG | p.Arg87Trp | P17643 |
| HOMO SAPIENS | 1 | 2426 | 5m8t | A | | 166 | SER | p.Ser166Ter | P17643 |
| HOMO SAPIENS | 0 | 2426 | 5m8t | A | | 356 | ARG | p.Arg356Gln | P17643 |
| HOMO SAPIENS | 2 | 2426 | 5m8t | A | | 374 | SER | p.Arg374Ter | P17643 |
| HOMO SAPIENS | 6 | 2426 | 5m8t | A | | 378 | ASN | p.Asn378Ser | P17643 |
| HOMO SAPIENS | 6 | 2426 | 5m8t | A | | 382 | LEU | p.Leu382Pro | P17643 |
| HOMO SAPIENS | 7 | 10097 | 4n6v | | 1 | 46 | GLN | p.Gln46Ter | P04080 |
| HOMO SAPIENS | 3 | 10097 | 4n6v | | 1 | 65 | VAL | p.Val65Ile | P04080 |
| HOMO SAPIENS | 8 | 10097 | 4n6v | | 1 | 68 | ARG | p.Arg68Ter | P04080 |
| HOMO SAPIENS | 2 | 859 | 4z9a | A | | 105 | GLN | p.Gln106Arg | P24666 |
| HOMO SAPIENS | 8 | 1420 | 5nan | F | | 95 | SER | p.Ser95Leu | Q96PD4 |

| | | | | | | | | |
|---|---|---|---|---|---|---|---|---|
| HOMO SAPIENS | 6 | 1420 | 5nan | F | 126 | GLU | p.Glu126Gly | Q96PD4 |
| HOMO SAPIENS | 8 | 1420 | 5nan | F | 155 | VAL | p.Val155Ile | Q96PD4 |
| HOMO SAPIENS | 6 | 1420 | 5nan | F | 157 | PRO | p.Pro157Ser | Q96PD4 |
| HOMO SAPIENS | -1 | 1420 | 5nan | F | 161 | HIS | p.His161Arg | Q96PD4 |
| HOMO SAPIENS | 3 | 288 | 5i96 | B | 140 | GLN | p.Arg140Gly | P48735 |
| HOMO SAPIENS | 3 | 288 | 5i96 | B | 140 | GLN | p.Arg140Gln | P48735 |
| HOMO SAPIENS | 2 | 288 | 5i96 | B | 261 | ARG | p.Arg261His | P48735 |
| HOMO SAPIENS | 2 | 288 | 5i96 | B | 435 | THR | p.Thr435Met | P48735 |
| HOMO SAPIENS | 9 | 120 | 1tup | B | 107 | TYR | p.Tyr107His | P04637 |
| HOMO SAPIENS | 8 | 120 | 1tup | B | 110 | ARG | p.Arg110Leu | P04637 |
| HOMO SAPIENS | 5 | 120 | 1tup | B | 120 | LYS | p.Lys120Glu | P04637 |
| HOMO SAPIENS | 4 | 120 | 1tup | B | 133 | MET | p.Met133Thr | P04637 |
| HOMO SAPIENS | 3 | 120 | 1tup | B | 134 | PHE | p.Phe134Cys | P04637 |
| HOMO SAPIENS | 8 | 120 | 1tup | B | 136 | GLN | p.Gln136His | P04637 |
| HOMO SAPIENS | 9 | 120 | 1tup | B | 138 | ALA | p.Ala138Pro | P04637 |
| HOMO SAPIENS | 9 | 120 | 1tup | B | 151 | PRO | p.Pro151Ser | P04637 |
| HOMO SAPIENS | 9 | 120 | 1tup | B | 152 | PRO | p.Pro152Leu | P04637 |
| HOMO SAPIENS | 9 | 120 | 1tup | B | 158 | ARG | p.Arg158Cys | P04637 |
| HOMO SAPIENS | 9 | 120 | 1tup | B | 158 | ARG | p.Arg158His | P04637 |
| HOMO SAPIENS | 9 | 120 | 1tup | B | 175 | ARG | p.Arg175Cys | P04637 |
| HOMO SAPIENS | 9 | 120 | 1tup | B | 175 | ARG | p.Arg175Leu | P04637 |
| HOMO SAPIENS | 8 | 120 | 1tup | B | 181 | ARG | p.Arg181Leu | P04637 |
| HOMO SAPIENS | 9 | 120 | 1tup | B | 189 | ALA | p.Ala189Val | P04637 |
| HOMO SAPIENS | 9 | 120 | 1tup | B | 196 | ARG | p.Arg196Gly | P04637 |
| HOMO SAPIENS | 9 | 120 | 1tup | B | 213 | ARG | p.Arg213Gln | P04637 |
| HOMO SAPIENS | 9 | 120 | 1tup | B | 213 | ARG | p.Arg213Ter | P04637 |
| HOMO SAPIENS | 9 | 120 | 1tup | B | 214 | HIS | p.His214Gln | P04637 |
| HOMO SAPIENS | 9 | 120 | 1tup | B | 220 | TYR | p.Tyr220Cys | P04637 |
| HOMO SAPIENS | 0 | 120 | 1tup | B | 234 | TYR | p.Tyr234Cys | P04637 |
| HOMO SAPIENS | 9 | 120 | 1tup | B | 237 | MET | p.Met237Ile | P04637 |
| HOMO SAPIENS | 0 | 120 | 1tup | B | 238 | CYS | p.Cys238Tyr | P04637 |
| HOMO SAPIENS | 9 | 120 | 1tup | B | 241 | SER | p.Ser241Phe | P04637 |
| HOMO SAPIENS | 9 | 120 | 1tup | B | 242 | CYS | p.Cys242Ser | P04637 |
| HOMO SAPIENS | 9 | 120 | 1tup | B | 245 | GLY | p.Gly245Cys | P04637 |
| HOMO SAPIENS | 9 | 120 | 1tup | B | 245 | GLY | p.Gly245Asp | P04637 |
| HOMO SAPIENS | 9 | 120 | 1tup | B | 248 | ARG | p.Arg248Leu | P04637 |
| HOMO SAPIENS | 9 | 120 | 1tup | B | 248 | ARG | p.Arg248Trp | P04637 |
| HOMO SAPIENS | 9 | 120 | 1tup | B | 251 | ILE | p.Ile251Ser | P04637 |
| HOMO SAPIENS | 9 | 120 | 1tup | B | 252 | LEU | p.Leu252Pro | P04637 |
| HOMO SAPIENS | 9 | 120 | 1tup | B | 258 | GLU | p.Glu258Lys | P04637 |
| HOMO SAPIENS | 8 | 120 | 1tup | B | 272 | VAL | p.Val272Leu | P04637 |
| HOMO SAPIENS | 9 | 120 | 1tup | B | 273 | ARG | p.Arg273Cys | P04637 |
| HOMO SAPIENS | 9 | 120 | 1tup | B | 273 | ARG | p.Arg273His | P04637 |
| HOMO SAPIENS | 9 | 120 | 1tup | B | 277 | CYS | p.Cys277Phe | P04637 |
| HOMO SAPIENS | 9 | 120 | 1tup | B | 280 | ARG | p.Arg280Ile | P04637 |
| HOMO SAPIENS | 9 | 120 | 1tup | B | 282 | ARG | p.Arg282Trp | P04637 |
| HOMO SAPIENS | 9 | 120 | 1tup | B | 283 | ARG | p.Arg283His | P04637 |
| HOMO SAPIENS | 9 | 120 | 1tup | B | 285 | GLU | p.Glu285Val | P04637 |
| HOMO SAPIENS | -1 | 120 | 1tup | B | 292 | LYS | p.Lys292Ile | P04637 |
| HOMO SAPIENS | -1 | 120 | 1tup | B | 298 | GLU | p.Glu298Lys | P04637 |
| HOMO SAPIENS | -1 | 2791 | 4ygb | A | 39 | VAL | p.Val39Ala | P49257 |
| HOMO SAPIENS | 2 | 106 | 4x6j | A | 32 | GLY | p.Gly146Arg | P43235 |
| HOMO SAPIENS | 0 | 106 | 4x6j | A | 76 | GLN | p.Gln190Ter | P43235 |
| HOMO SAPIENS | 7 | 106 | 4x6j | A | 198 | ARG | p.Arg312Ter | P43235 |

| | | | | | | | | |
|---|---|---|---|---|---|---|---|---|
| HOMO SAPIENS | 5 | 490 | 2hlo | B | 172 | LEU | p.Leu202Gln | P02675 |
| HOMO SAPIENS | 0 | 490 | 2hlo | B | 353 | LEU | p.Leu383Arg | P02675 |
| HOMO SAPIENS | 4 | 490 | 2hlo | B | 400 | GLY | p.Gly430Asp | P02675 |
| HOMO SAPIENS | 7 | 490 | 2hlo | B | 448 | ARG | p.Arg478Lys | P02675 |
| HOMO SAPIENS | 1 | 289 | 4p13 | B | 18 | GLU | p.Glu43Lys | P11310 |
| HOMO SAPIENS | 2 | 289 | 4p13 | B | 20 | GLN | p.Gln45Arg | P11310 |
| HOMO SAPIENS | 5 | 289 | 4p13 | B | 28 | ARG | p.Arg53Cys | P11310 |
| HOMO SAPIENS | 1 | 289 | 4p13 | B | 42 | TYR | p.Tyr67His | P11310 |
| HOMO SAPIENS | 1 | 289 | 4p13 | B | 53 | ILE | p.Ile78Thr | P11310 |
| HOMO SAPIENS | 1 | 289 | 4p13 | B | 59 | LEU | p.Leu84Phe | P11310 |
| HOMO SAPIENS | 2 | 289 | 4p13 | B | 96 | THR | p.Thr121Ile | P11310 |
| HOMO SAPIENS | 0 | 289 | 4p13 | B | 124 | MET | p.Met149Ile | P11310 |
| HOMO SAPIENS | 5 | 289 | 4p13 | B | 143 | ASP | p.Asp168Ala | P11310 |
| HOMO SAPIENS | 7 | 289 | 4p13 | B | 168 | THR | p.Thr193Ala | P11310 |
| HOMO SAPIENS | 1 | 289 | 4p13 | B | 170 | GLY | p.Gly195Arg | P11310 |
| HOMO SAPIENS | 2 | 289 | 4p13 | B | 181 | ARG | p.Arg206Cys | P11310 |
| HOMO SAPIENS | 2 | 289 | 4p13 | B | 181 | ARG | p.Arg206His | P11310 |
| HOMO SAPIENS | 8 | 289 | 4p13 | B | 203 | THR | p.Thr228Ile | P11310 |
| HOMO SAPIENS | 2 | 289 | 4p13 | B | 208 | ILE | p.Ile233Asn | P11310 |
| HOMO SAPIENS | 1 | 289 | 4p13 | B | 219 | CYS | p.Cys244Arg | P11310 |
| HOMO SAPIENS | 6 | 289 | 4p13 | B | 220 | SER | p.Ser245Leu | P11310 |
| HOMO SAPIENS | 5 | 289 | 4p13 | B | 228 | GLU | p.Glu253Lys | P11310 |
| HOMO SAPIENS | 2 | 289 | 4p13 | B | 241 | ASP | p.Asp266Gly | P11310 |
| HOMO SAPIENS | 1 | 289 | 4p13 | B | 242 | GLY | p.Gly267Arg | P11310 |
| HOMO SAPIENS | 7 | 289 | 4p13 | B | 256 | ARG | p.Arg281Thr | P11310 |
| HOMO SAPIENS | 4 | 289 | 4p13 | B | 269 | ARG | p.Arg294Thr | P11310 |
| HOMO SAPIENS | 4 | 289 | 4p13 | B | 304 | GLU | p.Lys329Glu | P11310 |
| HOMO SAPIENS | 1 | 289 | 4p13 | B | 324 | ARG | p.Arg349Ter | P11310 |
| HOMO SAPIENS | 3 | 289 | 4p13 | B | 326 | THR | p.Thr351Ile | P11310 |
| HOMO SAPIENS | 7 | 289 | 4p13 | B | 350 | ILE | p.Ile375Thr | P11310 |
| HOMO SAPIENS | 3 | 289 | 4p13 | B | 359 | GLU | p.Glu384Ter | P11310 |
| HOMO SAPIENS | 1 | 4563 | 3i2b | A | 16 | ARG | p.Arg16Cys | Q03393 |
| HOMO SAPIENS | 5 | 4563 | 3i2b | A | 25 | ARG | p.Arg25Gln | Q03393 |
| HOMO SAPIENS | 1 | 4563 | 3i2b | A | 47 | ASN | p.Asn47Asp | Q03393 |
| HOMO SAPIENS | 8 | 4563 | 3i2b | A | 52 | ASN | p.Asn52Ser | Q03393 |
| HOMO SAPIENS | 1 | 4563 | 3i2b | A | 56 | VAL | p.Val56Met | Q03393 |
| HOMO SAPIENS | 1 | 4563 | 3i2b | A | 87 | PRO | p.Pro87Ser | Q03393 |
| HOMO SAPIENS | 1 | 4563 | 3i2b | A | 96 | ASP | p.Asp96Asn | Q03393 |
| HOMO SAPIENS | 0 | 4563 | 3i2b | A | 99 | TYR | p.Tyr99Ter | Q03393 |
| HOMO SAPIENS | 2 | 4563 | 3i2b | A | 116 | ASP | p.Asp116Gly | Q03393 |
| HOMO SAPIENS | 2 | 41 | 3exx | A | 1 | GLY | p.Gly90Cys | P01308 |
| HOMO SAPIENS | 5 | 41 | 3exx | A | 3 | VAL | p.Val92Leu | P01308 |
| HOMO SAPIENS | 4 | 41 | 3exx | A | 7 | CYS | p.Cys96Ser | P01308 |
| HOMO SAPIENS | 9 | 41 | 3exx | A | 19 | TYR | p.Tyr108Cys | P01308 |
| HOMO SAPIENS | 9 | 131 | 4mjo | D | 118 | ASP | p.Asp119Asn | P09467 |
| HOMO SAPIENS | 0 | 131 | 4mjo | D | 163 | GLY | p.Gly164Ser | P09467 |
| HOMO SAPIENS | 0 | 131 | 4mjo | D | 176 | ALA | p.Ala177Asp | P09467 |
| HOMO SAPIENS | 5 | 131 | 4mjo | D | 193 | PHE | p.Phe194Ser | P09467 |
| HOMO SAPIENS | 4 | 131 | 4mjo | D | 217 | ARG | p.Arg218Lys | P09467 |
| HOMO SAPIENS | 4 | 131 | 4mjo | D | 223 | VAL | p.Val224Ile | P09467 |
| HOMO SAPIENS | 4 | 131 | 4mjo | D | 232 | PHE | p.Phe233Ile | P09467 |
| HOMO SAPIENS | 5 | 131 | 4mjo | D | 243 | ARG | p.Arg244Trp | P09467 |
| HOMO SAPIENS | 0 | 131 | 4mjo | D | 283 | PRO | p.Pro284Arg | P09467 |
| HOMO SAPIENS | 0 | 131 | 4mjo | D | 288 | MET | p.Met289Leu | P09467 |

| | | | | | | | | |
|---|---|---|---|---|---|---|---|---|
| HOMO SAPIENS | 0 | 131 | 4mjo | D | 323 | ASP | p.Asp324Asn | P09467 |
| HOMO SAPIENS | -1 | 13122 | 3hfc | A | 583 | ARG | p.Arg583Cys | P51787 |
| HOMO SAPIENS | 1 | 13122 | 3hfc | A | 587 | THR | p.Thr587Met | P51787 |
| HOMO SAPIENS | 0 | 13122 | 3hfc | A | 589 | GLY | p.Gly589Asp | P51787 |
| HOMO SAPIENS | 4 | 3002 | 3nbq | B | 94 | ARG | p.Arg94Cys | Q16831 |
| HOMO SAPIENS | 1 | 3616 | 3dbp | A | 97 | ILE | p.Ile286Asn | P11172 |
| HOMO SAPIENS | 0 | 3616 | 3dbp | A | 240 | GLY | p.Gly429Arg | P11172 |
| HOMO SAPIENS | 0 | 3616 | 3dbp | A | 257 | ILE | p.Ile446Val | P11172 |
| HOMO SAPIENS | 1 | 584 | 1drf | A | 79 | LEU | p.Leu80Phe | P00374 |
| HOMO SAPIENS | 3 | 1609 | 3qa2 | B | 49 | VAL | p.Val49Ile | P50053 |
| HOMO SAPIENS | 0 | 20560 | 5mug | A | 101 | HIS | p.His101Gln | P49638 |
| HOMO SAPIENS | 2 | 20560 | 5mug | A | 120 | ALA | p.Ala120Thr | P49638 |
| HOMO SAPIENS | 3 | 20560 | 5mug | A | 192 | ARG | p.Arg192His | P49638 |
| HOMO SAPIENS | 3 | 20560 | 5mug | A | 221 | ARG | p.Arg221Trp | P49638 |
| HOMO SAPIENS | 4 | 256 | 3vtv | A | 65 | THR | p.Glu50Lys | Q9GZQ8 |
| HOMO SAPIENS | 4 | 256 | 3vtv | A | 113 | VAL | p.Met98Lys | Q9GZQ8 |
| HOMO SAPIENS | 4 | 5397 | 1m61 | A | 192 | ARG | p.Arg192Trp | P43403 |
| HOMO SAPIENS | 5 | 39881 | 1aut | L | 107 | HIS | p.His149Pro | P04070 |
| HOMO SAPIENS | 5 | 4010 | 4uf8 | I | 33 | ALA | p.Ala33Val | Q13501 |
| HOMO SAPIENS | 3 | 313 | 5w49 | A | 98 | PRO | p.Pro98Leu | P23526 |
| HOMO SAPIENS | 2 | 313 | 5w49 | A | 143 | TYR | p.Tyr143Cys | P23526 |
| HOMO SAPIENS | -1 | 1643 | 2m8t | A | 102 | PRO | p.Pro102Leu | P04156 |
| HOMO SAPIENS | -1 | 1643 | 2m8t | A | 105 | PRO | p.Pro105Leu | P04156 |
| HOMO SAPIENS | -1 | 1643 | 2m8t | A | 117 | ALA | p.Ala117Val | P04156 |
| HOMO SAPIENS | 5 | 1643 | 2m8t | A | 129 | MET | p.Met129Val | P04156 |
| HOMO SAPIENS | 5 | 1643 | 2m8t | A | 131 | GLY | p.Gly131Val | P04156 |
| HOMO SAPIENS | 2 | 1643 | 2m8t | A | 142 | GLY | p.Gly142Ser | P04156 |
| HOMO SAPIENS | 3 | 1643 | 2m8t | A | 171 | ASN | p.Asn171Ser | P04156 |
| HOMO SAPIENS | 9 | 1643 | 2m8t | A | 178 | ASP | p.Asp178Asn | P04156 |
| HOMO SAPIENS | 9 | 1643 | 2m8t | A | 180 | VAL | p.Val180Ile | P04156 |
| HOMO SAPIENS | 9 | 1643 | 2m8t | A | 183 | THR | p.Thr183Ala | P04156 |
| HOMO SAPIENS | 9 | 1643 | 2m8t | A | 187 | HIS | p.His187Arg | P04156 |
| HOMO SAPIENS | 8 | 1643 | 2m8t | A | 198 | PHE | p.Phe198Ser | P04156 |
| HOMO SAPIENS | 9 | 1643 | 2m8t | A | 200 | GLU | p.Glu200Lys | P04156 |
| HOMO SAPIENS | 9 | 1643 | 2m8t | A | 208 | ARG | p.Arg208His | P04156 |
| HOMO SAPIENS | 3 | 1643 | 2m8t | A | 210 | VAL | p.Val210Ile | P04156 |
| HOMO SAPIENS | 6 | 1643 | 2m8t | A | 217 | GLN | p.Gln217Arg | P04156 |
| HOMO SAPIENS | 1 | 29 | 5kos | A | 151 | GLY | p.Gly217Arg | P00797 |
| HOMO SAPIENS | 1 | 29 | 5kos | A | 164 | ARG | p.Arg230Lys | P00797 |
| HOMO SAPIENS | 3 | 504 | 4zkv | B | 37 | ARG | p.Arg37Pro | P49773 |
| HOMO SAPIENS | 2 | 504 | 4zkv | B | 62 | GLN | p.Gln62Ter | P49773 |
| HOMO SAPIENS | 2 | 504 | 4zkv | B | 84 | CYS | p.Cys84Arg | P49773 |
| HOMO SAPIENS | 4 | 504 | 4zkv | B | 93 | GLY | p.Gly93Asp | P49773 |
| HOMO SAPIENS | 9 | 504 | 4zkv | B | 112 | HIS | p.His112Asn | P49773 |
| HOMO SAPIENS | 0 | 74 | 5vwo | C | 54 | ASN | p.Asn54Lys | P04181 |
| HOMO SAPIENS | 0 | 74 | 5vwo | C | 55 | TYR | p.Tyr55His | P04181 |
| HOMO SAPIENS | 2 | 74 | 5vwo | C | 90 | GLN | p.Gln90Glu | P04181 |
| HOMO SAPIENS | 6 | 74 | 5vwo | C | 91 | GLY | p.Gly91Glu | P04181 |
| HOMO SAPIENS | 2 | 74 | 5vwo | C | 93 | CYS | p.Cys93Phe | P04181 |
| HOMO SAPIENS | 1 | 74 | 5vwo | C | 154 | ARG | p.Arg154Leu | P04181 |
| HOMO SAPIENS | 0 | 74 | 5vwo | C | 180 | ARG | p.Arg180Thr | P04181 |
| HOMO SAPIENS | 0 | 74 | 5vwo | C | 199 | PRO | p.Pro199Gln | P04181 |
| HOMO SAPIENS | 4 | 74 | 5vwo | C | 209 | TYR | p.Tyr209Ter | P04181 |
| HOMO SAPIENS | 0 | 74 | 5vwo | C | 226 | ALA | p.Ala226Val | P04181 |

| | | | | | | | | |
|---|---|---|---|---|---|---|---|---|
| HOMO SAPIENS | 5 | 74 | 5vwo | C | 233 | GLN | p.Gln233Arg | P04181 |
| HOMO SAPIENS | 5 | 74 | 5vwo | C | 241 | PRO | p.Pro241Leu | P04181 |
| HOMO SAPIENS | 2 | 74 | 5vwo | C | 245 | TYR | p.Tyr245Cys | P04181 |
| HOMO SAPIENS | 6 | 74 | 5vwo | C | 250 | ARG | p.Arg250Pro | P04181 |
| HOMO SAPIENS | 6 | 74 | 5vwo | C | 250 | ARG | p.Arg250Ter | P04181 |
| HOMO SAPIENS | 8 | 74 | 5vwo | C | 267 | THR | p.Thr267Ile | P04181 |
| HOMO SAPIENS | 0 | 74 | 5vwo | C | 270 | ALA | p.Ala270Pro | P04181 |
| HOMO SAPIENS | 0 | 74 | 5vwo | C | 271 | ARG | p.Arg271Lys | P04181 |
| HOMO SAPIENS | 1 | 74 | 5vwo | C | 299 | TYR | p.Tyr299Ter | P04181 |
| HOMO SAPIENS | 0 | 74 | 5vwo | C | 318 | GLU | p.Glu318Lys | P04181 |
| HOMO SAPIENS | 1 | 74 | 5vwo | C | 319 | HIS | p.His319Tyr | P04181 |
| HOMO SAPIENS | 1 | 74 | 5vwo | C | 331 | ARG | p.Arg331Ter | P04181 |
| HOMO SAPIENS | 0 | 74 | 5vwo | C | 332 | VAL | p.Val332Met | P04181 |
| HOMO SAPIENS | 1 | 74 | 5vwo | C | 353 | GLY | p.Gly353Asp | P04181 |
| HOMO SAPIENS | 2 | 74 | 5vwo | C | 375 | GLY | p.Gly375Ala | P04181 |
| HOMO SAPIENS | 0 | 74 | 5vwo | C | 394 | CYS | p.Cys394Arg | P04181 |
| HOMO SAPIENS | 0 | 74 | 5vwo | C | 402 | LEU | p.Leu402Pro | P04181 |
| HOMO SAPIENS | 0 | 74 | 5vwo | C | 436 | ILE | p.Ile436Asn | P04181 |
| HOMO SAPIENS | 7 | 24356 | 4az0 | A | 104 | VAL | p.Val132Met | P10619 |
| HOMO SAPIENS | 4 | 24356 | 4az0 | A | 221 | TYR | p.Tyr249Asn | P10619 |
| HOMO SAPIENS | 3 | 245 | 1rsz | A | 24 | ARG | p.Arg24Ter | P00491 |
| HOMO SAPIENS | 3 | 245 | 1rsz | A | 51 | SER | p.Gly51Ser | P00491 |
| HOMO SAPIENS | 1 | 245 | 1rsz | A | 58 | ARG | p.Arg58Ter | P00491 |
| HOMO SAPIENS | 4 | 245 | 1rsz | A | 89 | GLU | p.Glu89Lys | P00491 |
| HOMO SAPIENS | 6 | 245 | 1rsz | A | 128 | ASP | p.Asp128Gly | P00491 |
| HOMO SAPIENS | 0 | 245 | 1rsz | A | 174 | ALA | p.Ala174Pro | P00491 |
| HOMO SAPIENS | 8 | 245 | 1rsz | A | 192 | TYR | p.Tyr192Cys | P00491 |
| HOMO SAPIENS | 8 | 245 | 1rsz | A | 234 | ARG | p.Arg234Pro | P00491 |
| HOMO SAPIENS | 0 | 37 | 5tby | E | 22 | GLU | p.Glu22Lys | P10916 |
| HOMO SAPIENS | 1 | 37 | 5tby | E | 86 | PHE | p.Phe86Ser | P10916 |
| HOMO SAPIENS | 9 | 71 | 5m9j | A | 12 | GLN | p.Gln36Leu | P03950 |
| HOMO SAPIENS | 3 | 71 | 5m9j | A | 17 | LYS | p.Lys41Glu | P03950 |
| HOMO SAPIENS | 3 | 71 | 5m9j | A | 17 | LYS | p.Lys41Ile | P03950 |
| HOMO SAPIENS | 4 | 71 | 5m9j | A | 28 | SER | p.Ser52Asn | P03950 |
| HOMO SAPIENS | 7 | 71 | 5m9j | A | 31 | ARG | p.Arg55Lys | P03950 |
| HOMO SAPIENS | 9 | 71 | 5m9j | A | 39 | CYS | p.Cys63Trp | P03950 |
| HOMO SAPIENS | 9 | 71 | 5m9j | A | 40 | LYS | p.Lys64Ile | P03950 |
| HOMO SAPIENS | 2 | 71 | 5m9j | A | 60 | GLU | p.Lys84Glu | P03950 |
| HOMO SAPIENS | 0 | 71 | 5m9j | A | 112 | PRO | p.Pro136Leu | P03950 |
| HOMO SAPIENS | 9 | 71 | 5m9j | A | 113 | VAL | p.Val137Ile | P03950 |
| HOMO SAPIENS | 3 | 326 | 5dqi | A | 209 | GLY | p.Gly209Val | Q9Y253 |
| HOMO SAPIENS | 1 | 326 | 5dqi | A | 242 | SER | p.Ser242Ter | Q9Y253 |
| HOMO SAPIENS | 1 | 326 | 5dqi | A | 303 | ARG | p.Arg303Ter | Q9Y253 |
| HOMO SAPIENS | 1 | 8203 | 3iu3 | K | 132 | VAL | p.Val153Ile | P01589 |
| HOMO SAPIENS | 0 | 4995 | 5ffo | H | 234 | THR | p.Thr263Ile | P01137 |
| HOMO SAPIENS | 5 | 38609 | 2ruk | A | 49 | ASP | p.Asp49His | P04637 |
| HOMO SAPIENS | 5 | 38609 | 2ruk | A | 58 | PRO | p.Pro58Arg | P04637 |
| HOMO SAPIENS | 0 | 1607 | 6pah | A | 117 | THR | p.Thr117Ile | P00439 |
| HOMO SAPIENS | 3 | 1607 | 6pah | A | 145 | ASP | p.Asp145Val | P00439 |
| HOMO SAPIENS | 5 | 1607 | 6pah | A | 158 | ARG | p.Arg158Gln | P00439 |
| HOMO SAPIENS | 5 | 1607 | 6pah | A | 158 | ARG | p.Arg158Trp | P00439 |
| HOMO SAPIENS | 5 | 1607 | 6pah | A | 169 | ARG | p.Arg169His | P00439 |
| HOMO SAPIENS | 5 | 1607 | 6pah | A | 169 | ARG | p.Arg169Cys | P00439 |
| HOMO SAPIENS | 3 | 1607 | 6pah | A | 176 | ARG | p.Arg176Leu | P00439 |

| | | | | | | | | |
|---|---|---|---|---|---|---|---|---|
| HOMO SAPIENS | 3 | 1607 | 6pah | A | 177 | VAL | p.Val177Leu | P00439 |
| HOMO SAPIENS | 1 | 1607 | 6pah | A | 194 | LEU | p.Leu194Pro | P00439 |
| HOMO SAPIENS | 1 | 1607 | 6pah | A | 204 | TYR | p.Tyr204Cys | P00439 |
| HOMO SAPIENS | 1 | 1607 | 6pah | A | 204 | TYR | p.Tyr204Ter | P00439 |
| HOMO SAPIENS | 2 | 1607 | 6pah | A | 230 | VAL | p.Val230Ile | P00439 |
| HOMO SAPIENS | 4 | 1607 | 6pah | A | 232 | GLN | p.Gln232Ter | P00439 |
| HOMO SAPIENS | 4 | 1607 | 6pah | A | 241 | ARG | p.Arg241Cys | P00439 |
| HOMO SAPIENS | 4 | 1607 | 6pah | A | 241 | ARG | p.Arg241His | P00439 |
| HOMO SAPIENS | 5 | 1607 | 6pah | A | 243 | ARG | p.Arg243Gln | P00439 |
| HOMO SAPIENS | 5 | 1607 | 6pah | A | 243 | ARG | p.Arg243Ter | P00439 |
| HOMO SAPIENS | 7 | 1607 | 6pah | A | 249 | LEU | p.Leu249Phe | P00439 |
| HOMO SAPIENS | 5 | 1607 | 6pah | A | 252 | ARG | p.Arg252Gly | P00439 |
| HOMO SAPIENS | 5 | 1607 | 6pah | A | 261 | ARG | p.Arg261Ter | P00439 |
| HOMO SAPIENS | 4 | 1607 | 6pah | A | 267 | GLN | p.Gln267Leu | P00439 |
| HOMO SAPIENS | 1 | 1607 | 6pah | A | 272 | GLY | p.Gly272Ter | P00439 |
| HOMO SAPIENS | 2 | 1607 | 6pah | A | 277 | TYR | p.Tyr277Asp | P00439 |
| HOMO SAPIENS | 7 | 1607 | 6pah | A | 280 | GLU | p.Glu280Lys | P00439 |
| HOMO SAPIENS | 8 | 1607 | 6pah | A | 281 | PRO | p.Pro281Leu | P00439 |
| HOMO SAPIENS | 0 | 1607 | 6pah | A | 299 | PHE | p.Phe299Cys | P00439 |
| HOMO SAPIENS | 5 | 1607 | 6pah | A | 300 | ALA | p.Ala300Ser | P00439 |
| HOMO SAPIENS | 5 | 1607 | 6pah | A | 309 | ALA | p.Ala309Asp | P00439 |
| HOMO SAPIENS | 4 | 1607 | 6pah | A | 320 | LYS | p.Lys320Asn | P00439 |
| HOMO SAPIENS | 0 | 1607 | 6pah | A | 345 | ALA | p.Ala345Ser | P00439 |
| HOMO SAPIENS | 0 | 1607 | 6pah | A | 348 | LEU | p.Leu348Val | P00439 |
| HOMO SAPIENS | 6 | 1607 | 6pah | A | 356 | TYR | p.Tyr356Ter | P00439 |
| HOMO SAPIENS | 2 | 1607 | 6pah | A | 380 | THR | p.Thr380Met | P00439 |
| HOMO SAPIENS | 0 | 1607 | 6pah | A | 386 | TYR | p.Tyr386Cys | P00439 |
| HOMO SAPIENS | 5 | 1607 | 6pah | A | 390 | GLU | p.Glu390Gly | P00439 |
| HOMO SAPIENS | 5 | 1607 | 6pah | A | 403 | ALA | p.Ala403Val | P00439 |
| HOMO SAPIENS | 4 | 1607 | 6pah | A | 408 | ARG | p.Arg408Trp | P00439 |
| HOMO SAPIENS | 3 | 1607 | 6pah | A | 413 | ARG | p.Arg413Pro | P00439 |
| HOMO SAPIENS | 4 | 1607 | 6pah | A | 414 | TYR | p.Tyr414Cys | P00439 |
| HOMO SAPIENS | 4 | 1607 | 6pah | A | 414 | TYR | p.Tyr414His | P00439 |
| HOMO SAPIENS | 4 | 1607 | 6pah | A | 415 | ASP | p.Asp415Asn | P00439 |
| HOMO SAPIENS | -1 | 676 | 5dc5 | A | 7 | PRO | p.Pro7Leu | Q9BY41 |
| HOMO SAPIENS | 0 | 676 | 5dc5 | A | 311 | THR | p.Thr311Met | Q9BY41 |
| HOMO SAPIENS | 1 | 124 | 5ftk | E | 95 | ARG | p.Arg95Gly | P55072 |
| HOMO SAPIENS | 1 | 124 | 5ftk | E | 155 | ARG | p.Arg155Cys | P55072 |
| HOMO SAPIENS | 1 | 124 | 5ftk | E | 155 | ARG | p.Arg155His | P55072 |
| HOMO SAPIENS | 1 | 124 | 5ftk | E | 159 | ARG | p.Arg159His | P55072 |
| HOMO SAPIENS | 1 | 124 | 5ftk | E | 159 | ARG | p.Arg159Gly | P55072 |
| HOMO SAPIENS | 0 | 124 | 5ftk | E | 191 | ARG | p.Arg191Gln | P55072 |
| HOMO SAPIENS | 0 | 124 | 5ftk | E | 232 | ALA | p.Ala232Glu | P55072 |
| HOMO SAPIENS | 5 | 5442 | 1fgg | B | 223 | GLY | p.Gly223Ser | O94766 |
| HOMO SAPIENS | 0 | 5442 | 1fgg | B | 277 | ARG | p.Arg277Gln | O94766 |
| HOMO SAPIENS | 0 | 9072 | 2gys | B | 225 | GLU | p.Glu249Gln | P32927 |
| HOMO SAPIENS | 0 | 9072 | 2gys | B | 247 | SER | p.Ser271Ter | P32927 |
| HOMO SAPIENS | 0 | 9072 | 2gys | B | 294 | ASP | p.Asp312Asn | P32927 |
| HOMO SAPIENS | 0 | 1484 | 4zrj | A | 62 | PHE | p.Phe62Ser | P35240 |
| HOMO SAPIENS | 3 | 1484 | 4zrj | A | 205 | MET | p.Met205Val | P35240 |
| HOMO SAPIENS | 3 | 1484 | 4zrj | A | 301 | ILE | p.Ile301Met | P35240 |
| HOMO SAPIENS | 4 | 40923 | 2enq | A | 68 | ILE | p.Ile391Met | P42336 |
| HOMO SAPIENS | 1 | 6179 | 5ofv | A | 125 | THR | p.Thr125Met | O43175 |
| HOMO SAPIENS | 0 | 6179 | 5ofv | A | 135 | ARG | p.Arg135Trp | O43175 |

| | | | | | | | | |
|---|---|---|---|---|---|---|---|---|
| HOMO SAPIENS | 1 | 6179 | 5ofv | A | 163 | ARG | p.Arg163Gln | O43175 |
| HOMO SAPIENS | 9 | 6179 | 5ofv | A | 261 | VAL | p.Val261Met | O43175 |
| HOMO SAPIENS | 7 | 34118 | 2i5o | A | 24 | MET | p.Met647Leu | Q9Y253 |
| HOMO SAPIENS | 8 | 3234 | 5c84 | A | 311 | LYS | p.Lys311Asn | P98170 |
| HOMO SAPIENS | 3 | 3234 | 5c84 | A | 321 | ALA | p.Ala321Gly | P98170 |
| HOMO SAPIENS | -1 | 15129 | 1zy8 | K | 1 | GLY | p.Gly54Ser | O00330 |
| HOMO SAPIENS | -1 | 15129 | 1zy8 | K | 48 | THR | p.Thr101Ala | O00330 |
| HOMO SAPIENS | -1 | 15129 | 1zy8 | K | 105 | GLU | p.Glu158Asp | O00330 |
| HOMO SAPIENS | 8 | 15129 | 1zy8 | K | 136 | ARG | p.Arg189His | O00330 |
| HOMO SAPIENS | -1 | 15129 | 1zy8 | K | 195 | GLN | p.Gln248Ter | O00330 |
| HOMO SAPIENS | 8 | 202 | 4bzb | A | 123 | HIS | p.His123Pro | Q9Y3Z3 |
| HOMO SAPIENS | 4 | 202 | 4bzb | A | 145 | ARG | p.Arg145Ter | Q9Y3Z3 |
| HOMO SAPIENS | 6 | 202 | 4bzb | A | 164 | ARG | p.Arg164Ter | Q9Y3Z3 |
| HOMO SAPIENS | 3 | 202 | 4bzb | A | 201 | ILE | p.Ile201Asn | Q9Y3Z3 |
| HOMO SAPIENS | 0 | 202 | 4bzb | A | 209 | GLY | p.Gly209Ser | Q9Y3Z3 |
| HOMO SAPIENS | 0 | 202 | 4bzb | A | 254 | MET | p.Met254Val | Q9Y3Z3 |
| HOMO SAPIENS | 5 | 202 | 4bzb | A | 531 | ARG | p.Arg531Ser | Q9Y3Z3 |
| HOMO SAPIENS | 6 | 202 | 4bzb | A | 548 | GLN | p.Gln548Ter | Q9Y3Z3 |
| HOMO SAPIENS | 3 | 30947 | 5lgd | A | 90 | PRO | p.Pro90Ser | P16671 |
| HOMO SAPIENS | 6 | 30947 | 5lgd | A | 254 | PHE | p.Phe254Leu | P16671 |
| HOMO SAPIENS | 0 | 30947 | 5lgd | A | 413 | ILE | p.Ile413Leu | P16671 |
| HOMO SAPIENS | 2 | 3906 | 5jtc | A | 735 | ARG | p.Arg735Trp | Q12797 |
| HOMO SAPIENS | 1 | 45 | 5uyx | C | 146 | GLU | p.Glu146Val | P48643 |
| HOMO SAPIENS | 1 | 45 | 5uyx | C | 147 | HIS | p.His147Arg | P48643 |
| HOMO SAPIENS | 0 | 45 | 5uyx | C | 362 | ILE | p.Ile362Met | P48643 |
| HOMO SAPIENS | 8 | 39596 | 1mhn | A | 95 | GLY | p.Gly95Arg | Q16637 |
| HOMO SAPIENS | 4 | 39596 | 1mhn | A | 111 | ALA | p.Ala111Gly | Q16637 |
| HOMO SAPIENS | 9 | 39596 | 1mhn | A | 116 | ILE | p.Ile116Phe | Q16637 |
| HOMO SAPIENS | 1 | 39596 | 1mhn | A | 136 | GLN | p.Gln136Glu | Q16637 |
| HOMO SAPIENS | 1 | 6408 | 5xjy | A | 85 | PRO | p.Pro85Leu | O95477 |
| HOMO SAPIENS | 0 | 6408 | 5xjy | A | 219 | ARG | p.Arg219Lys | O95477 |
| HOMO SAPIENS | 0 | 6408 | 5xjy | A | 230 | ARG | p.Arg230Cys | O95477 |
| HOMO SAPIENS | 1 | 6408 | 5xjy | A | 281 | MET | p.Met281Thr | O95477 |
| HOMO SAPIENS | -1 | 6408 | 5xjy | A | 343 | ALA | p.Ala343Gly | O95477 |
| HOMO SAPIENS | 1 | 6408 | 5xjy | A | 399 | VAL | p.Val399Ala | O95477 |
| HOMO SAPIENS | 1 | 6408 | 5xjy | A | 496 | ARG | p.Arg496Trp | O95477 |
| HOMO SAPIENS | 1 | 6408 | 5xjy | A | 590 | TRP | p.Trp590Ser | O95477 |
| HOMO SAPIENS | 1 | 6408 | 5xjy | A | 638 | ARG | p.Arg638Gln | O95477 |
| HOMO SAPIENS | 4 | 6408 | 5xjy | A | 697 | ALA | p.Ala697Thr | O95477 |
| HOMO SAPIENS | 1 | 6408 | 5xjy | A | 771 | VAL | p.Val771Met | O95477 |
| HOMO SAPIENS | 1 | 6408 | 5xjy | A | 774 | THR | p.Thr774Pro | O95477 |
| HOMO SAPIENS | 1 | 6408 | 5xjy | A | 776 | LYS | p.Lys776Asn | O95477 |
| HOMO SAPIENS | 0 | 6408 | 5xjy | A | 825 | VAL | p.Val825Ile | O95477 |
| HOMO SAPIENS | -1 | 6408 | 5xjy | A | 868 | GLU | p.Glu868Lys | O95477 |
| HOMO SAPIENS | -1 | 6408 | 5xjy | A | 883 | ILE | p.Ile883Met | O95477 |
| HOMO SAPIENS | 0 | 6408 | 5xjy | A | 935 | ASN | p.Asn935Ser | O95477 |
| HOMO SAPIENS | 0 | 6408 | 5xjy | A | 935 | ASN | p.Asn935His | O95477 |
| HOMO SAPIENS | 0 | 6408 | 5xjy | A | 937 | ALA | p.Ala937Val | O95477 |
| HOMO SAPIENS | 0 | 6408 | 5xjy | A | 1018 | ASP | p.Asp1018Gly | O95477 |
| HOMO SAPIENS | 0 | 6408 | 5xjy | A | 1096 | VAL | p.Val1096Ile | O95477 |
| HOMO SAPIENS | 0 | 6408 | 5xjy | A | 1099 | ASP | p.Asp1099Tyr | O95477 |
| HOMO SAPIENS | 0 | 6408 | 5xjy | A | 1172 | GLU | p.Glu1172Asp | O95477 |
| HOMO SAPIENS | 0 | 6408 | 5xjy | A | 1181 | SER | p.Ser1181Phe | O95477 |
| HOMO SAPIENS | 0 | 6408 | 5xjy | A | 1182 | ALA | p.Ala1182Thr | O95477 |

| | | | | | | | | |
|---|---|---|---|---|---|---|---|---|
| HOMO SAPIENS | -1 | 6408 | 5xjy | A | 1289 | ASP | p.Asp1289Asn | O95477 |
| HOMO SAPIENS | 0 | 6408 | 5xjy | A | 1399 | THR | p.Thr1399Ser | O95477 |
| HOMO SAPIENS | 1 | 6408 | 5xjy | A | 1408 | LEU | p.Leu1408Phe | O95477 |
| HOMO SAPIENS | 0 | 6408 | 5xjy | A | 1587 | LYS | p.Lys1587Arg | O95477 |
| HOMO SAPIENS | 1 | 6408 | 5xjy | A | 1674 | VAL | p.Val1674Ile | O95477 |
| HOMO SAPIENS | 1 | 6408 | 5xjy | A | 1680 | ARG | p.Arg1680Trp | O95477 |
| HOMO SAPIENS | 1 | 6408 | 5xjy | A | 1680 | ARG | p.Arg1680Gln | O95477 |
| HOMO SAPIENS | 0 | 6408 | 5xjy | A | 1800 | ASN | p.Asn1800His | O95477 |
| HOMO SAPIENS | -1 | 6408 | 5xjy | A | 1925 | ARG | p.Arg1925Gln | O95477 |
| HOMO SAPIENS | 0 | 6408 | 5xjy | A | 2009 | PHE | p.Phe2009Ser | O95477 |
| HOMO SAPIENS | -1 | 6408 | 5xjy | A | 2243 | ASP | p.Asp2243Glu | O95477 |
| HOMO SAPIENS | 4 | 23760 | 2mut | A | 231 | LEU | p.Phe231Leu | P07992 |
| HOMO SAPIENS | 6 | 1460 | 2hre | B | 96 | ARG | p.Arg96Gln | P22830 |
| HOMO SAPIENS | 8 | 1460 | 2hre | B | 129 | GLY | p.Gly129Arg | P22830 |
| HOMO SAPIENS | 9 | 1460 | 2hre | B | 172 | GLU | p.Glu172Lys | P22830 |
| HOMO SAPIENS | 4 | 1460 | 2hre | B | 285 | GLN | p.Gln285Arg | P22830 |
| HOMO SAPIENS | 5 | 1460 | 2hre | B | 362 | VAL | p.Val362Gly | P22830 |
| HOMO SAPIENS | 6 | 1460 | 2hre | B | 411 | CYS | p.Cys411Gly | P22830 |
| HOMO SAPIENS | 6 | 1460 | 2hre | B | 417 | PHE | p.Phe417Ser | P22830 |
| HOMO SAPIENS | 0 | 2210 | 5bug | A | 19 | ASP | p.Asp19Asn | P60484 |
| HOMO SAPIENS | 0 | 2210 | 5bug | A | 35 | MET | p.Met35Arg | P60484 |
| HOMO SAPIENS | 1 | 2210 | 5bug | A | 61 | HIS | p.His61Asp | P60484 |
| HOMO SAPIENS | 0 | 2210 | 5bug | A | 68 | TYR | p.Tyr68His | P60484 |
| HOMO SAPIENS | 8 | 2210 | 5bug | A | 70 | LEU | p.Leu70Pro | P60484 |
| HOMO SAPIENS | 6 | 2210 | 5bug | A | 93 | HIS | p.His93Arg | P60484 |
| HOMO SAPIENS | 1 | 2210 | 5bug | A | 112 | LEU | p.Leu112Pro | P60484 |
| HOMO SAPIENS | 0 | 2210 | 5bug | A | 121 | ALA | p.Ala121Gly | P60484 |
| HOMO SAPIENS | 3 | 2210 | 5bug | A | 123 | HIS | p.His123Arg | P60484 |
| HOMO SAPIENS | 7 | 2210 | 5bug | A | 124 | CYS | p.Cys124Arg | P60484 |
| HOMO SAPIENS | 9 | 2210 | 5bug | A | 129 | GLY | p.Gly129Glu | P60484 |
| HOMO SAPIENS | 9 | 2210 | 5bug | A | 130 | ARG | p.Arg130Gln | P60484 |
| HOMO SAPIENS | 9 | 2210 | 5bug | A | 130 | ARG | p.Arg130Gly | P60484 |
| HOMO SAPIENS | 0 | 2210 | 5bug | A | 132 | GLY | p.Gly132Val | P60484 |
| HOMO SAPIENS | 0 | 2210 | 5bug | A | 170 | SER | p.Ser170Arg | P60484 |
| HOMO SAPIENS | 1 | 2210 | 5bug | A | 173 | ARG | p.Arg173His | P60484 |
| HOMO SAPIENS | 0 | 2210 | 5bug | A | 217 | VAL | p.Val217Ile | P60484 |
| HOMO SAPIENS | 0 | 2210 | 5bug | A | 241 | PHE | p.Phe241Ser | P60484 |
| HOMO SAPIENS | 0 | 2210 | 5bug | A | 250 | CYS | p.Cys250Ter | P60484 |
| HOMO SAPIENS | 1 | 2210 | 5bug | A | 252 | ASP | p.Asp252Gly | P60484 |
| HOMO SAPIENS | 3 | 123 | 2ibu | A | 93 | ASN | p.Asn93Ser | P24752 |
| HOMO SAPIENS | 0 | 123 | 2ibu | A | 145 | GLN | p.Gln145Glu | P24752 |
| HOMO SAPIENS | 0 | 123 | 2ibu | A | 152 | GLY | p.Gly152Ala | P24752 |
| HOMO SAPIENS | 2 | 123 | 2ibu | A | 158 | ASN | p.Asn158Asp | P24752 |
| HOMO SAPIENS | 2 | 123 | 2ibu | A | 158 | ASN | p.Asn158Ser | P24752 |
| HOMO SAPIENS | 5 | 123 | 2ibu | A | 183 | GLY | p.Gly183Arg | P24752 |
| HOMO SAPIENS | 3 | 123 | 2ibu | A | 272 | GLN | p.Gln272Ter | P24752 |
| HOMO SAPIENS | 3 | 123 | 2ibu | A | 312 | ILE | p.Ile312Thr | P24752 |
| HOMO SAPIENS | 1 | 123 | 2ibu | A | 333 | ALA | p.Ala333Pro | P24752 |
| HOMO SAPIENS | 5 | 123 | 2ibu | A | 379 | GLY | p.Gly379Val | P24752 |
| HOMO SAPIENS | 0 | 123 | 2ibu | A | 380 | ALA | p.Ala380Thr | P24752 |
| HOMO SAPIENS | 2 | 369 | 5nrw | A | 1175 | TYR | p.Tyr1175Cys | Q92793 |
| HOMO SAPIENS | 2 | 830 | 2vc2 | A | 116 | LEU | p.Leu147Val | P08514 |
| HOMO SAPIENS | 0 | 830 | 2vc2 | A | 183 | LEU | p.Leu214Pro | P08514 |
| HOMO SAPIENS | 4 | 830 | 2vc2 | A | 242 | GLY | p.Gly273Asp | P08514 |

| | | | | | | | | |
|---|---|---|---|---|---|---|---|---|
| HOMO SAPIENS | 6 | 830 | 2vc2 | A | 324 | GLU | p.Glu355Lys | P08514 |
| HOMO SAPIENS | -1 | 10353 | 5ejc | E | 989 | ALA | p.Ala989Ser | Q92574 |
| HOMO SAPIENS | 0 | 15448 | 3lgg | A | 21 | GLY | p.Gly47Arg | Q9NZK5 |
| HOMO SAPIENS | 0 | 15448 | 3lgg | A | 21 | GLY | p.Gly47Val | Q9NZK5 |
| HOMO SAPIENS | 9 | 15448 | 3lgg | A | 86 | HIS | p.His112Gln | Q9NZK5 |
| HOMO SAPIENS | 0 | 15448 | 3lgg | A | 143 | ARG | p.Arg169Gln | Q9NZK5 |
| HOMO SAPIENS | 3 | 15448 | 3lgg | A | 221 | ALA | p.Ala247Val | Q9NZK5 |
| HOMO SAPIENS | 0 | 15448 | 3lgg | A | 225 | PRO | p.Pro251Leu | Q9NZK5 |
| HOMO SAPIENS | 0 | 15448 | 3lgg | A | 283 | MET | p.Met309Ile | Q9NZK5 |
| HOMO SAPIENS | 2 | 15448 | 3lgg | A | 334 | THR | p.Thr360Ala | Q9NZK5 |
| HOMO SAPIENS | 7 | 15448 | 3lgg | A | 357 | GLY | p.Gly383Ser | Q9NZK5 |
| HOMO SAPIENS | 0 | 15448 | 3lgg | A | 427 | TYR | p.Tyr453Cys | Q9NZK5 |
| HOMO SAPIENS | 0 | 872 | 2y3i | A | 87 | LEU | p.Leu88Pro | P00558 |
| HOMO SAPIENS | 1 | 872 | 2y3i | A | 163 | ASP | p.Asp164Val | P00558 |
| HOMO SAPIENS | 4 | 872 | 2y3i | A | 205 | ARG | p.Arg206Pro | P00558 |
| HOMO SAPIENS | 4 | 872 | 2y3i | A | 267 | ASP | p.Asp268Asn | P00558 |
| HOMO SAPIENS | 8 | 872 | 2y3i | A | 284 | ASP | p.Asp285Val | P00558 |
| HOMO SAPIENS | 4 | 8 | 5jco | B | 188 | ILE | p.Ile188Leu | Q71U36 |
| HOMO SAPIENS | 7 | 8 | 5jco | B | 263 | PRO | p.Pro263Thr | Q71U36 |
| HOMO SAPIENS | 6 | 8 | 5jco | B | 264 | ARG | p.Arg264Cys | Q71U36 |
| HOMO SAPIENS | 7 | 8 | 5jco | B | 402 | ARG | p.Arg402His | Q71U36 |
| HOMO SAPIENS | 7 | 8 | 5jco | B | 419 | SER | p.Ser419Leu | Q71U36 |
| HOMO SAPIENS | -1 | 880 | 4nwf | A | 2 | THR | p.Thr2Ile | Q06124 |
| HOMO SAPIENS | 2 | 880 | 4nwf | A | 58 | ASN | p.Asn58Ser | Q06124 |
| HOMO SAPIENS | 4 | 880 | 4nwf | A | 61 | ASP | p.Asp61Gly | Q06124 |
| HOMO SAPIENS | 2 | 880 | 4nwf | A | 62 | TYR | p.Tyr62Asp | Q06124 |
| HOMO SAPIENS | 3 | 880 | 4nwf | A | 63 | TYR | p.Tyr63Cys | Q06124 |
| HOMO SAPIENS | 2 | 880 | 4nwf | A | 69 | GLU | p.Glu69Gln | Q06124 |
| HOMO SAPIENS | 2 | 880 | 4nwf | A | 72 | ALA | p.Ala72Gly | Q06124 |
| HOMO SAPIENS | 2 | 880 | 4nwf | A | 72 | ALA | p.Ala72Ser | Q06124 |
| HOMO SAPIENS | 3 | 880 | 4nwf | A | 73 | THR | p.Thr73Ile | Q06124 |
| HOMO SAPIENS | 5 | 880 | 4nwf | A | 76 | GLU | p.Glu76Ala | Q06124 |
| HOMO SAPIENS | 5 | 880 | 4nwf | A | 76 | GLU | p.Glu76Lys | Q06124 |
| HOMO SAPIENS | 3 | 880 | 4nwf | A | 79 | GLN | p.Gln79Arg | Q06124 |
| HOMO SAPIENS | 5 | 880 | 4nwf | A | 152 | ARG | p.Arg152His | Q06124 |
| HOMO SAPIENS | 2 | 880 | 4nwf | A | 268 | GLY | p.Gly268Cys | Q06124 |
| HOMO SAPIENS | 9 | 880 | 4nwf | A | 276 | LYS | p.Lys276Arg | Q06124 |
| HOMO SAPIENS | 9 | 880 | 4nwf | A | 279 | TYR | p.Tyr279Cys | Q06124 |
| HOMO SAPIENS | 5 | 880 | 4nwf | A | 285 | PHE | p.Phe285Ser | Q06124 |
| HOMO SAPIENS | 0 | 880 | 4nwf | A | 308 | ASP | p.Asn308Ser | Q06124 |
| HOMO SAPIENS | 0 | 880 | 4nwf | A | 308 | ASP | p.Asn308Asp | Q06124 |
| HOMO SAPIENS | 5 | 880 | 4nwf | A | 392 | ALA | p.Ala392Thr | Q06124 |
| HOMO SAPIENS | 1 | 880 | 4nwf | A | 411 | THR | p.Thr415Met | Q06124 |
| HOMO SAPIENS | 0 | 880 | 4nwf | A | 461 | ALA | p.Ala465Thr | Q06124 |
| HOMO SAPIENS | 0 | 880 | 4nwf | A | 464 | GLY | p.Gly468Ala | Q06124 |
| HOMO SAPIENS | 1 | 880 | 4nwf | A | 491 | PRO | p.Pro495Ser | Q06124 |
| HOMO SAPIENS | 1 | 880 | 4nwf | A | 502 | SER | p.Ser506Thr | Q06124 |
| HOMO SAPIENS | 8 | 880 | 4nwf | A | 510 | GLN | p.Gln514Pro | Q06124 |
| HOMO SAPIENS | 4 | 1872 | 3wf1 | D | 109 | ARG | p.Arg109Trp | P16278 |
| HOMO SAPIENS | 0 | 1872 | 3wf1 | D | 521 | ARG | p.Arg521Cys | P16278 |
| HOMO SAPIENS | 5 | 910 | 5xte | A | 45 | SER | p.Ser45Phe | O14949 |
| HOMO SAPIENS | 2 | 9017 | 5f0k | A | 382 | LYS | p.Lys382Arg | Q96QK1 |
| HOMO SAPIENS | 1 | 710 | 4lsj | A | 559 | ILE | p.Ile559Asn | P04150 |
| HOMO SAPIENS | 5 | 710 | 4lsj | A | 571 | VAL | p.Val571Ala | P04150 |

| | | | | | | | | |
|---|---|---|---|---|---|---|---|---|
| HOMO SAPIENS | 0 | 710 | 4lsj | A | 641 | ASP | p.Asp641Val | P04150 |
| HOMO SAPIENS | 1 | 710 | 4lsj | A | 678 | ASP | p.Asp678Glu | P04150 |
| HOMO SAPIENS | 2 | 710 | 4lsj | A | 679 | GLY | p.Gly679Ser | P04150 |
| HOMO SAPIENS | 1 | 710 | 4lsj | A | 737 | PHE | p.Phe737Leu | P04150 |
| HOMO SAPIENS | 1 | 710 | 4lsj | A | 747 | ILE | p.Ile747Met | P04150 |
| HOMO SAPIENS | 1 | 710 | 4lsj | A | 753 | LEU | p.Leu753Phe | P04150 |
| HOMO SAPIENS | 1 | 710 | 4lsj | A | 773 | LEU | p.Leu773Pro | P04150 |
| HOMO SAPIENS | 1 | 2145 | 1av1 | B | 71 | PHE | p.Phe95Tyr | P02647 |
| HOMO SAPIENS | 1 | 2145 | 1av1 | B | 128 | GLU | p.Glu152Lys | P02647 |
| HOMO SAPIENS | 1 | 2145 | 1av1 | B | 136 | GLU | p.Glu160Lys | P02647 |
| HOMO SAPIENS | 2 | 2145 | 1av1 | B | 143 | PRO | p.Pro167Arg | P02647 |
| HOMO SAPIENS | 6 | 2145 | 1av1 | B | 164 | ALA | p.Ala188Ser | P02647 |
| HOMO SAPIENS | 7 | 2145 | 1av1 | B | 173 | ARG | p.Arg197Cys | P02647 |
| HOMO SAPIENS | 2 | 2145 | 1av1 | B | 198 | GLU | p.Glu222Lys | P02647 |
| HOMO SAPIENS | -1 | 3432 | 2beu | A | 1 | SER | p.Ser46Ter | P12694 |
| HOMO SAPIENS | 5 | 3432 | 2beu | A | 79 | ARG | p.Arg124Trp | P12694 |
| HOMO SAPIENS | 7 | 3432 | 2beu | A | 114 | ARG | p.Arg159Gln | P12694 |
| HOMO SAPIENS | 6 | 3432 | 2beu | A | 166 | THR | p.Thr211Met | P12694 |
| HOMO SAPIENS | 4 | 3432 | 2beu | A | 175 | ALA | p.Ala220Val | P12694 |
| HOMO SAPIENS | 5 | 3432 | 2beu | A | 204 | GLY | p.Gly249Ser | P12694 |
| HOMO SAPIENS | 4 | 3432 | 2beu | A | 209 | ALA | p.Ala254Asp | P12694 |
| HOMO SAPIENS | 6 | 3432 | 2beu | A | 220 | ARG | p.Arg265Trp | P12694 |
| HOMO SAPIENS | 6 | 3432 | 2beu | A | 234 | TYR | p.Tyr279Cys | P12694 |
| HOMO SAPIENS | 5 | 3432 | 2beu | A | 240 | ALA | p.Ala285Pro | P12694 |
| HOMO SAPIENS | 4 | 3432 | 2beu | A | 245 | GLY | p.Gly290Arg | P12694 |
| HOMO SAPIENS | 5 | 3432 | 2beu | A | 269 | ARG | p.Arg314Ter | P12694 |
| HOMO SAPIENS | 3 | 3432 | 2beu | A | 301 | ARG | p.Arg346Cys | P12694 |
| HOMO SAPIENS | 5 | 3432 | 2beu | A | 359 | ASN | p.Asn404Ser | P12694 |
| HOMO SAPIENS | 4 | 3432 | 2beu | A | 393 | TYR | p.Tyr438Asn | P12694 |
| HOMO SAPIENS | 8 | 4541 | 1rr8 | C | 533 | ASP | p.Asp533Gly | P11387 |
| HOMO SAPIENS | 2 | 3247 | 1tzn | d | 1105 | GLY | p.Gly105Asp | P58335 |
| HOMO SAPIENS | 3 | 3247 | 1tzn | d | 1111 | ARG | p.Arg111Cys | P58335 |
| HOMO SAPIENS | 2 | 3247 | 1tzn | d | 1189 | ILE | p.Ile189Thr | P58335 |
| HOMO SAPIENS | 9 | 467 | 1suv | C | 58 | ASP | p.Asp77Asn | P02787 |
| HOMO SAPIENS | 2 | 467 | 1suv | C | 277 | ASP | p.Asp296Gly | P02787 |
| HOMO SAPIENS | 9 | 6485 | 1xwd | B | 51 | CYS | p.Cys69Gly | P01225 |
| HOMO SAPIENS | 1 | 694 | 1wuu | A | 28 | PRO | p.Pro28Thr | P51570 |
| HOMO SAPIENS | 0 | 694 | 1wuu | A | 32 | VAL | p.Val32Met | P51570 |
| HOMO SAPIENS | 0 | 694 | 1wuu | A | 198 | ALA | p.Ala198Val | P51570 |
| HOMO SAPIENS | 0 | 143 | 4nkx | A | 93 | PHE | p.Phe93Cys | P05093 |
| HOMO SAPIENS | 3 | 143 | 4nkx | A | 96 | ARG | p.Arg96Trp | P05093 |
| HOMO SAPIENS | 3 | 143 | 4nkx | A | 96 | ARG | p.Arg96Gln | P05093 |
| HOMO SAPIENS | 0 | 143 | 4nkx | A | 106 | SER | p.Ser106Pro | P05093 |
| HOMO SAPIENS | 1 | 143 | 4nkx | A | 114 | PHE | p.Phe114Val | P05093 |
| HOMO SAPIENS | 1 | 143 | 4nkx | A | 116 | ASP | p.Asp116Val | P05093 |
| HOMO SAPIENS | 3 | 143 | 4nkx | A | 125 | ARG | p.Arg125Gln | P05093 |
| HOMO SAPIENS | 0 | 143 | 4nkx | A | 201 | TYR | p.Tyr201Asp | P05093 |
| HOMO SAPIENS | 0 | 143 | 4nkx | A | 239 | ARG | p.Arg239Ter | P05093 |
| HOMO SAPIENS | 0 | 143 | 4nkx | A | 329 | TYR | p.Tyr329Asp | P05093 |
| HOMO SAPIENS | 1 | 143 | 4nkx | A | 342 | PRO | p.Pro342Thr | P05093 |
| HOMO SAPIENS | 2 | 143 | 4nkx | A | 347 | ARG | p.Arg347His | P05093 |
| HOMO SAPIENS | 2 | 143 | 4nkx | A | 347 | ARG | p.Arg347Cys | P05093 |
| HOMO SAPIENS | 1 | 143 | 4nkx | A | 358 | ARG | p.Arg358Gln | P05093 |
| HOMO SAPIENS | 0 | 143 | 4nkx | A | 362 | ARG | p.Arg362Cys | P05093 |

| | | | | | | | | |
|---|---|---|---|---|---|---|---|---|
| HOMO SAPIENS | 1 | 143 | 4nkx | A | 406 | TRP | p.Trp406Arg | P05093 |
| HOMO SAPIENS | 0 | 143 | 4nkx | A | 428 | PRO | p.Pro428Leu | P05093 |
| HOMO SAPIENS | 6 | 1441 | 1uuh | A | 46 | ARG | p.Arg46Gly | P16070 |
| HOMO SAPIENS | 3 | 82 | 4wlv | D | 37 | GLY | p.Gly37Arg | P40926 |
| HOMO SAPIENS | 0 | 82 | 4wlv | D | 133 | PRO | p.Pro133Leu | P40926 |
| HOMO SAPIENS | 2 | 392 | 5xtd | A | 56 | SER | p.Ser56Pro | P49821 |
| HOMO SAPIENS | 2 | 392 | 5xtd | A | 59 | ARG | p.Arg59Ter | P49821 |
| HOMO SAPIENS | 2 | 392 | 5xtd | A | 74 | ASP | p.Asp74Asn | P49821 |
| HOMO SAPIENS | 4 | 392 | 5xtd | A | 116 | ASN | p.Asn116Lys | P49821 |
| HOMO SAPIENS | 4 | 392 | 5xtd | A | 117 | ALA | p.Ala117Thr | P49821 |
| HOMO SAPIENS | 5 | 392 | 5xtd | A | 122 | PRO | p.Pro122Leu | P49821 |
| HOMO SAPIENS | 3 | 392 | 5xtd | A | 160 | GLY | p.Gly160Glu | P49821 |
| HOMO SAPIENS | 4 | 392 | 5xtd | A | 214 | GLU | p.Glu214Lys | P49821 |
| HOMO SAPIENS | 2 | 392 | 5xtd | A | 357 | MET | p.Met357Thr | P49821 |
| HOMO SAPIENS | 4 | 392 | 5xtd | A | 423 | THR | p.Thr423Met | P49821 |
| HOMO SAPIENS | 0 | 22593 | 3pdf | A | 8 | TYR | p.Tyr32Ter | P53634 |
| HOMO SAPIENS | 0 | 22593 | 3pdf | A | 15 | TRP | p.Trp39Ser | P53634 |
| HOMO SAPIENS | 0 | 22593 | 3pdf | A | 103 | HIS | p.His127Pro | P53634 |
| HOMO SAPIENS | 9 | 22593 | 3pdf | A | 228 | GLN | p.Gln252Leu | P53634 |
| HOMO SAPIENS | 3 | 22593 | 3pdf | A | 262 | GLN | p.Gln286Arg | P53634 |
| HOMO SAPIENS | 4 | 22593 | 3pdf | A | 277 | GLY | p.Gly301Ser | P53634 |
| HOMO SAPIENS | 0 | 22593 | 3pdf | A | 323 | TYR | p.Tyr347Cys | P53634 |
| HOMO SAPIENS | 5 | 3461 | 4c2a | A | 1306 | GLN | p.Arg1306Trp | P04275 |
| HOMO SAPIENS | 5 | 3461 | 4c2a | A | 1374 | ARG | p.Arg1374Cys | P04275 |
| HOMO SAPIENS | 5 | 3461 | 4c2a | A | 1379 | ARG | p.Arg1379Cys | P04275 |
| HOMO SAPIENS | 4 | 3461 | 4c2a | A | 1380 | ILE | p.Ile1380Val | P04275 |
| HOMO SAPIENS | 0 | 3461 | 4c2a | A | 1381 | ALA | p.Thr1381Ala | P04275 |
| HOMO SAPIENS | 5 | 3461 | 4c2a | A | 1399 | ARG | p.Arg1399His | P04275 |
| HOMO SAPIENS | 5 | 3461 | 4c2a | A | 1435 | ASN | p.Asn1435Ser | P04275 |
| HOMO SAPIENS | 0 | 4533 | 5uj7 | C | 174 | TYR | p.Tyr174Cys | O43929 |
| HOMO SAPIENS | 0 | 4533 | 5uj7 | C | 202 | LEU | p.Leu202Val | O43929 |
| HOMO SAPIENS | 0 | 1374 | 3j07 | O | 39 | PRO | p.Pro39Leu | P02511 |
| HOMO SAPIENS | 0 | 1374 | 3j07 | O | 107 | ARG | p.Arg107His | P02511 |
| HOMO SAPIENS | 0 | 1374 | 3j07 | O | 157 | ARG | p.Arg157His | P02511 |
| HOMO SAPIENS | -1 | 942 | 1s9i | B | 57 | PHE | p.Phe57Cys | P36507 |
| HOMO SAPIENS | -1 | 942 | 1s9i | B | 57 | PHE | p.Phe57Val | P36507 |
| HOMO SAPIENS | 2 | 942 | 1s9i | B | 128 | PRO | p.Pro128Gln | P36507 |
| HOMO SAPIENS | 1 | 942 | 1s9i | B | 134 | TYR | p.Tyr134His | P36507 |
| HOMO SAPIENS | 1 | 942 | 1s9i | B | 188 | HIS | p.His188Gln | P36507 |
| HOMO SAPIENS | 0 | 942 | 1s9i | B | 262 | VAL | p.Val262Phe | P36507 |
| HOMO SAPIENS | 0 | 942 | 1s9i | B | 263 | GLY | p.Gly263Arg | P36507 |
| HOMO SAPIENS | -1 | 942 | 1s9i | B | 298 | PRO | p.Pro298Leu | P36507 |
| HOMO SAPIENS | -1 | 942 | 1s9i | B | 305 | VAL | p.Val305Ile | P36507 |
| HOMO SAPIENS | 0 | 3245 | 1fbv | A | 367 | GLN | p.Gln367Pro | P22681 |
| HOMO SAPIENS | 0 | 3245 | 1fbv | A | 371 | TYR | p.Tyr371His | P22681 |
| HOMO SAPIENS | 1 | 3245 | 1fbv | A | 382 | LYS | p.Lys382Glu | P22681 |
| HOMO SAPIENS | 2 | 3245 | 1fbv | A | 390 | ASP | p.Asp390Tyr | P22681 |
| HOMO SAPIENS | 9 | 3245 | 1fbv | A | 396 | CYS | p.Cys396Arg | P22681 |
| HOMO SAPIENS | 7 | 3245 | 1fbv | A | 420 | ARG | p.Arg420Gln | P22681 |
| HOMO SAPIENS | 1 | 3245 | 1fbv | A | 429 | ILE | p.Ile429Met | P22681 |
| HOMO SAPIENS | 0 | 2604 | 2pbd | P | 111 | LEU | p.Leu112Met | P07737 |
| HOMO SAPIENS | 3 | 2604 | 2pbd | P | 116 | GLU | p.Glu117Gly | P07737 |
| HOMO SAPIENS | 4 | 2013 | 3b7w | A | 472 | ARG | p.Arg472Gly | Q08AH3 |
| HOMO SAPIENS | 0 | 24346 | 3sh5 | A | 85 | ARG | p.Arg4281His | P98160 |

| | | | | | | | | |
|---|---|---|---|---|---|---|---|---|
| HOMO SAPIENS | 0 | 24346 | 3sh5 | A | 122 | ARG | p.Arg4318Trp | P98160 |
| HOMO SAPIENS | 0 | 24346 | 3sh5 | A | 135 | SER | p.Ser4331Asn | P98160 |
| HOMO SAPIENS | 0 | 24346 | 3sh5 | A | 136 | VAL | p.Val4332Ile | P98160 |
| HOMO SAPIENS | 0 | 707 | 4p2g | A | 26 | MET | p.Met26Ile | Q99497 |
| HOMO SAPIENS | 5 | 707 | 4p2g | A | 39 | ALA | p.Ala39Ser | Q99497 |
| HOMO SAPIENS | 5 | 707 | 4p2g | A | 64 | GLU | p.Glu64Asp | Q99497 |
| HOMO SAPIENS | 4 | 707 | 4p2g | A | 149 | ASP | p.Asp149Ala | Q99497 |
| HOMO SAPIENS | 6 | 707 | 4p2g | A | 163 | GLU | p.Glu163Lys | Q99497 |
| HOMO SAPIENS | 6 | 707 | 4p2g | A | 166 | LEU | p.Leu166Pro | Q99497 |
| HOMO SAPIENS | 3 | 5121 | 4xss | E | 31 | GLY | p.Gly58Arg | P06213 |
| HOMO SAPIENS | 2 | 5121 | 4xss | E | 119 | ILE | p.Ile146Met | P06213 |
| HOMO SAPIENS | 8 | 5121 | 4xss | E | 209 | HIS | p.His236Arg | P06213 |
| HOMO SAPIENS | 2 | 5121 | 4xss | E | 234 | ASP | p.Asp261Glu | P06213 |
| HOMO SAPIENS | 1 | 5121 | 4xss | E | 276 | GLN | p.Gln303His | P06213 |
| HOMO SAPIENS | 1 | 27790 | 5erp | B | 140 | THR | p.Thr275Met | Q02487 |
| HOMO SAPIENS | 0 | 27790 | 5erp | B | 150 | ILE | p.Ile285Thr | Q02487 |
| HOMO SAPIENS | 1 | 27790 | 5erp | B | 205 | THR | p.Thr340Ala | Q02487 |
| HOMO SAPIENS | 1 | 27790 | 5erp | B | 223 | THR | p.Thr358Ile | Q02487 |
| HOMO SAPIENS | 0 | 27790 | 5erp | B | 301 | GLY | p.Gly436Ala | Q02487 |
| HOMO SAPIENS | -1 | 27790 | 5erp | B | 315 | ARG | p.Arg450Ser | Q02487 |
| HOMO SAPIENS | 0 | 27790 | 5erp | B | 369 | SER | p.Ser504Gly | Q02487 |
| HOMO SAPIENS | 0 | 27790 | 5erp | B | 385 | ILE | p.Ile520Thr | Q02487 |
| HOMO SAPIENS | 0 | 27790 | 5erp | B | 461 | ALA | p.Ala596Val | Q02487 |
| HOMO SAPIENS | 0 | 27790 | 5erp | B | 503 | GLN | p.Gln638His | Q02487 |
| HOMO SAPIENS | 5 | 6599 | 4nsc | A | 129 | ARG | p.Arg129Leu | Q9BPX6 |
| HOMO SAPIENS | -1 | 6599 | 4nsc | A | 183 | GLN | p.Gln183Ter | Q9BPX6 |
| HOMO SAPIENS | 0 | 20420 | 3ghm | A | 398 | ARG | p.Arg398His | Q76LX8 |
| HOMO SAPIENS | 0 | 20420 | 3ghm | A | 448 | GLN | p.Gln448Glu | Q76LX8 |
| HOMO SAPIENS | 1 | 20420 | 3ghm | A | 449 | GLN | p.Gln449Ter | Q76LX8 |
| HOMO SAPIENS | 0 | 20420 | 3ghm | A | 456 | GLN | p.Gln456His | Q76LX8 |
| HOMO SAPIENS | 1 | 20420 | 3ghm | A | 528 | ARG | p.Arg528Gly | Q76LX8 |
| HOMO SAPIENS | 0 | 20420 | 3ghm | A | 618 | PRO | p.Pro618Ala | Q76LX8 |
| HOMO SAPIENS | 0 | 20420 | 3ghm | A | 625 | ARG | p.Arg625His | Q76LX8 |
| HOMO SAPIENS | 0 | 275 | 4rmz | A | 293 | GLN | p.Gln293Ter | Q9NWZ3 |
| HOMO SAPIENS | 0 | 275 | 4rmz | A | 428 | ALA | p.Ala428Thr | Q9NWZ3 |
| HOMO SAPIENS | 2 | 10458 | 3puf | B | 138 | LEU | p.Leu138Phe | Q5TBB1 |
| HOMO SAPIENS | 6 | 10458 | 3puf | B | 177 | ALA | p.Ala177Thr | Q5TBB1 |
| HOMO SAPIENS | -1 | 10458 | 3puf | B | 185 | VAL | p.Val185Gly | Q5TBB1 |
| HOMO SAPIENS | 0 | 26648 | 2n90 | A | 36 | ARG | p.Arg444Gln | P04629 |
| HOMO SAPIENS | 2 | 977 | 1dnw | D | 166 | ALA | p.Ala332Val | P05164 |
| HOMO SAPIENS | 4 | 977 | 1dnw | D | 333 | ARG | p.Arg499Cys | P05164 |
| HOMO SAPIENS | 1 | 977 | 1dnw | D | 403 | ARG | p.Arg569Trp | P05164 |
| HOMO SAPIENS | 0 | 25704 | 3ajb | A | 82 | GLN | p.Gln82Arg | P56589 |
| HOMO SAPIENS | 3 | 255 | 3t1p | A | 38 | TYR | p.Tyr62Ter | P01009 |
| HOMO SAPIENS | 3 | 255 | 3t1p | A | 39 | ARG | p.Arg63Cys | P01009 |
| HOMO SAPIENS | 0 | 255 | 3t1p | A | 101 | ARG | p.Arg125His | P01009 |
| HOMO SAPIENS | 0 | 255 | 3t1p | A | 212 | GLN | p.Gln236Arg | P01009 |
| HOMO SAPIENS | 0 | 255 | 3t1p | A | 213 | VAL | p.Val237Ala | P01009 |
| HOMO SAPIENS | 1 | 255 | 3t1p | A | 269 | HIS | p.His293Gln | P01009 |
| HOMO SAPIENS | 8 | 255 | 3t1p | A | 341 | ASP | p.Asp365Asn | P01009 |
| HOMO SAPIENS | 9 | 255 | 3t1p | A | 342 | GLU | p.Glu366Lys | P01009 |
| HOMO SAPIENS | 0 | 255 | 3t1p | A | 369 | PRO | p.Pro393Ser | P01009 |
| HOMO SAPIENS | 0 | 255 | 3t1p | A | 376 | GLU | p.Glu400Asp | P01009 |
| HOMO SAPIENS | -1 | 5175 | 3zxi | A | 35 | ALA | p.Ala35Asp | Q9Y2Z4 |

| | | | | | | | | | |
|---|---|---|---|---|---|---|---|---|---|
| HOMO SAPIENS | 0 | 5175 | 3zxi | A | | 52 | PHE | p.Phe52Leu | Q9Y2Z4 |
| HOMO SAPIENS | -1 | 5175 | 3zxi | A | | 68 | GLY | p.Gly68Ser | Q9Y2Z4 |
| HOMO SAPIENS | 0 | 5175 | 3zxi | A | | 179 | LYS | p.Lys179Gln | Q9Y2Z4 |
| HOMO SAPIENS | 0 | 5175 | 3zxi | A | | 191 | GLY | p.Gly191Val | Q9Y2Z4 |
| HOMO SAPIENS | 0 | 5175 | 3zxi | A | | 251 | ILE | p.Ile251Val | Q9Y2Z4 |
| HOMO SAPIENS | 2 | 5175 | 3zxi | A | | 295 | ASP | p.Asp295Asn | Q9Y2Z4 |
| HOMO SAPIENS | 0 | 5175 | 3zxi | A | | 311 | ASP | p.Asp311Glu | Q9Y2Z4 |
| HOMO SAPIENS | 0 | 5175 | 3zxi | A | | 360 | ARG | p.Arg360Ter | Q9Y2Z4 |
| HOMO SAPIENS | 1 | 3016 | 3exg | | 3 | 4 | ASP | p.Asp33His | P08559 |
| HOMO SAPIENS | 1 | 3016 | 3exg | | 3 | 176 | PHE | p.Phe205Leu | P08559 |
| HOMO SAPIENS | 1 | 3016 | 3exg | | 3 | 214 | TYR | p.Tyr243Asn | P08559 |
| HOMO SAPIENS | 1 | 3016 | 3exg | | 3 | 229 | ASP | p.Asp258Ala | P08559 |
| HOMO SAPIENS | 0 | 3016 | 3exg | | 3 | 233 | VAL | p.Val262Ile | P08559 |
| HOMO SAPIENS | 1 | 3016 | 3exg | | 3 | 234 | ARG | p.Arg263Gly | P08559 |
| HOMO SAPIENS | 0 | 3016 | 3exg | | 3 | 253 | MET | p.Met282Leu | P08559 |
| HOMO SAPIENS | 7 | 3016 | 3exg | | 3 | 259 | ARG | p.Arg288His | P08559 |
| HOMO SAPIENS | -1 | 3016 | 3exg | | 3 | 273 | ARG | p.Arg302Cys | P08559 |
| HOMO SAPIENS | 2 | 3016 | 3exg | | 3 | 286 | ASP | p.Asp315Asn | P08559 |
| HOMO SAPIENS | 1 | 3016 | 3exg | | 3 | 349 | ARG | p.Arg378His | P08559 |
| HOMO SAPIENS | 5 | 6729 | 4a0l | E | | 572 | ARG | p.Arg572Cys | Q13620 |
| HOMO SAPIENS | 0 | 1067 | 4kyo | A | | 9 | THR | p.Thr9Asn | P21549 |
| HOMO SAPIENS | 0 | 1067 | 4kyo | A | | 11 | PRO | p.Pro11Leu | P21549 |
| HOMO SAPIENS | 0 | 1067 | 4kyo | A | | 22 | ASN | p.Asn22Ser | P21549 |
| HOMO SAPIENS | 4 | 1067 | 4kyo | A | | 25 | LEU | p.Leu25Arg | P21549 |
| HOMO SAPIENS | 7 | 1067 | 4kyo | A | | 36 | ARG | p.Arg36His | P21549 |
| HOMO SAPIENS | 3 | 1067 | 4kyo | A | | 41 | GLY | p.Gly41Arg | P21549 |
| HOMO SAPIENS | 0 | 1067 | 4kyo | A | | 42 | GLY | p.Gly42Glu | P21549 |
| HOMO SAPIENS | 3 | 1067 | 4kyo | A | | 56 | ILE | p.Ile56Asn | P21549 |
| HOMO SAPIENS | 3 | 1067 | 4kyo | A | | 59 | GLU | p.Glu59Lys | P21549 |
| HOMO SAPIENS | 3 | 1067 | 4kyo | A | | 66 | TYR | p.Tyr66Ter | P21549 |
| HOMO SAPIENS | 3 | 1067 | 4kyo | A | | 111 | ARG | p.Arg111Leu | P21549 |
| HOMO SAPIENS | 7 | 1067 | 4kyo | A | | 118 | ARG | p.Arg118His | P21549 |
| HOMO SAPIENS | 3 | 1067 | 4kyo | A | | 122 | ARG | p.Arg122Ter | P21549 |
| HOMO SAPIENS | 3 | 1067 | 4kyo | A | | 124 | HIS | p.His124Pro | P21549 |
| HOMO SAPIENS | 0 | 1067 | 4kyo | A | | 152 | PHE | p.Phe152Ile | P21549 |
| HOMO SAPIENS | 1 | 1067 | 4kyo | A | | 156 | GLY | p.Gly156Arg | P21549 |
| HOMO SAPIENS | 0 | 1067 | 4kyo | A | | 161 | GLY | p.Gly161Arg | P21549 |
| HOMO SAPIENS | 3 | 1067 | 4kyo | A | | 166 | LEU | p.Leu166Pro | P21549 |
| HOMO SAPIENS | 5 | 1067 | 4kyo | A | | 170 | GLY | p.Gly170Arg | P21549 |
| HOMO SAPIENS | 5 | 1067 | 4kyo | A | | 186 | ALA | p.Ala186Val | P21549 |
| HOMO SAPIENS | 3 | 1067 | 4kyo | A | | 187 | SER | p.Ser187Phe | P21549 |
| HOMO SAPIENS | 3 | 1067 | 4kyo | A | | 197 | ARG | p.Arg197Pro | P21549 |
| HOMO SAPIENS | 5 | 1067 | 4kyo | A | | 201 | ASP | p.Asp201Glu | P21549 |
| HOMO SAPIENS | 0 | 1067 | 4kyo | A | | 202 | ILE | p.Ile202Asn | P21549 |
| HOMO SAPIENS | 1 | 1067 | 4kyo | A | | 205 | SER | p.Ser205Pro | P21549 |
| HOMO SAPIENS | 1 | 1067 | 4kyo | A | | 205 | SER | p.Ser205Ter | P21549 |
| HOMO SAPIENS | 2 | 1067 | 4kyo | A | | 216 | GLY | p.Gly216Arg | P21549 |
| HOMO SAPIENS | 5 | 1067 | 4kyo | A | | 233 | ARG | p.Arg233Cys | P21549 |
| HOMO SAPIENS | 5 | 1067 | 4kyo | A | | 244 | ILE | p.Ile244Thr | P21549 |
| HOMO SAPIENS | 5 | 1067 | 4kyo | A | | 244 | ILE | p.Ile244Met | P21549 |
| HOMO SAPIENS | 4 | 1067 | 4kyo | A | | 253 | CYS | p.Cys253Arg | P21549 |
| HOMO SAPIENS | 7 | 1067 | 4kyo | A | | 274 | GLU | p.Glu274Asp | P21549 |
| HOMO SAPIENS | 7 | 1067 | 4kyo | A | | 279 | ILE | p.Ile279Thr | P21549 |
| HOMO SAPIENS | 6 | 1067 | 4kyo | A | | 295 | ALA | p.Ala295Thr | P21549 |

| | | | | | | | | |
|---|---|---|---|---|---|---|---|---|
| HOMO SAPIENS | 5 | 1067 | 4kyo | A | 340 | ILE | p.Ile340Met | P21549 |
| HOMO SAPIENS | 1 | 790 | 2ric | C | 270 | SER | p.Ser290Thr | P35247 |
| HOMO SAPIENS | 0 | 790 | 2ric | C | 289 | GLU | p.Glu309Lys | P35247 |
| HOMO SAPIENS | 1 | 212 | 5sxe | A | 545 | GLU | p.Glu545Lys | P42336 |
| HOMO SAPIENS | 3 | 212 | 5sxe | A | 546 | GLN | p.Gln546Glu | P42336 |
| HOMO SAPIENS | 2 | 212 | 5sxe | A | 1047 | HIS | p.His1047Leu | P42336 |
| HOMO SAPIENS | 7 | 152 | 2zqq | F | 197 | ARG | p.Arg197Ter | Q13825 |
| HOMO SAPIENS | 0 | 152 | 2zqq | F | 248 | ALA | p.Ala248Thr | Q13825 |
| HOMO SAPIENS | 0 | 152 | 2zqq | F | 275 | ALA | p.Ala275Val | Q13825 |
| HOMO SAPIENS | 5 | 152 | 2zqq | F | 331 | LYS | p.Lys331Ter | Q13825 |
| HOMO SAPIENS | 6 | 17801 | 2j8l | A | 283 | PHE | p.Phe301Leu | P03951 |
| HOMO SAPIENS | 3 | 17801 | 2j8l | A | 308 | ARG | p.Arg326Cys | P03951 |
| HOMO SAPIENS | 0 | 17801 | 2j8l | A | 321 | CYS | p.Cys339Phe | P03951 |
| HOMO SAPIENS | 3 | 17801 | 2j8l | A | 351 | TYR | p.Tyr369Ter | P03951 |
| HOMO SAPIENS | 7 | 1802 | 5db0 | A | 22 | LEU | p.Leu22Arg | O00255 |
| HOMO SAPIENS | 7 | 1802 | 5db0 | A | 26 | GLU | p.Glu26Lys | O00255 |
| HOMO SAPIENS | 7 | 1802 | 5db0 | A | 139 | HIS | p.His139Asp | O00255 |
| HOMO SAPIENS | 0 | 1802 | 5db0 | A | 171 | ARG | p.Arg176Gln | O00255 |
| HOMO SAPIENS | 0 | 1802 | 5db0 | A | 184 | VAL | p.Val189Glu | O00255 |
| HOMO SAPIENS | 7 | 1802 | 5db0 | A | 255 | GLU | p.Glu260Lys | O00255 |
| HOMO SAPIENS | 4 | 1802 | 5db0 | A | 418 | ASP | p.Asp423Asn | O00255 |
| HOMO SAPIENS | 0 | 1802 | 5db0 | A | 436 | TRP | p.Trp441Arg | O00255 |
| HOMO SAPIENS | -1 | 1802 | 5db0 | A | 541 | ALA | p.Thr546Ala | O00255 |
| HOMO SAPIENS | 4 | 1802 | 5db0 | A | 552 | THR | p.Thr557Ser | O00255 |
| HOMO SAPIENS | 0 | 3450 | 2yuz | A | 36 | PRO | p.Pro461Gln | Q00872 |
| HOMO SAPIENS | 1 | 3450 | 2yuz | A | 56 | GLN | p.His481Gln | Q00872 |
| HOMO SAPIENS | 2 | 3450 | 2yuz | A | 68 | ALA | p.Ala493Val | Q00872 |
| HOMO SAPIENS | 0 | 3211 | 3rqg | A | 72 | VAL | p.Val72Leu | Q9BUL8 |
| HOMO SAPIENS | 2 | 4168 | 4q4f | A | 121 | ARG | p.Arg121Ter | Q14108 |
| HOMO SAPIENS | 2 | 4168 | 4q4f | A | 121 | ARG | p.Arg121Gln | Q14108 |
| HOMO SAPIENS | 2 | 4168 | 4q4f | A | 159 | MET | p.Met159Val | Q14108 |
| HOMO SAPIENS | 4 | 4168 | 4q4f | A | 363 | HIS | p.His363Asn | Q14108 |
| HOMO SAPIENS | 2 | 4168 | 4q4f | A | 396 | VAL | p.Val396Ile | Q14108 |
| HOMO SAPIENS | 0 | 1448 | 1y97 | A | 87 | ARG | p.Arg130Ter | Q9BQ50 |
| HOMO SAPIENS | 0 | 3444 | 4iwp | A | 151 | SER | p.Ser151Phe | Q9UHD2 |
| HOMO SAPIENS | 0 | 3444 | 4iwp | A | 401 | LYS | p.Lys401Glu | Q9UHD2 |
| HOMO SAPIENS | 0 | 116 | 1zy8 | B | 194 | GLY | p.Gly229Cys | P09622 |
| HOMO SAPIENS | 0 | 116 | 1zy8 | B | 255 | ASP | p.Asp290Gly | P09622 |
| HOMO SAPIENS | 2 | 116 | 1zy8 | B | 326 | MET | p.Met361Val | P09622 |
| HOMO SAPIENS | 0 | 116 | 1zy8 | B | 340 | GLU | p.Glu375Lys | P09622 |
| HOMO SAPIENS | 2 | 116 | 1zy8 | B | 358 | ILE | p.Ile393Thr | P09622 |
| HOMO SAPIENS | 1 | 116 | 1zy8 | B | 453 | PRO | p.Pro488Leu | P09622 |
| HOMO SAPIENS | 7 | 127 | 4qo7 | B | 160 | SER | p.Ser161Arg | P00338 |
| HOMO SAPIENS | 0 | 2025 | 1wrl | A | 8 | ALA | p.Ala8Val | P63316 |
| HOMO SAPIENS | 8 | 2025 | 1wrl | A | 24 | PHE | p.Phe24Leu | P63316 |
| HOMO SAPIENS | 0 | 2025 | 1wrl | A | 84 | SER | p.Cys84Tyr | P63316 |
| HOMO SAPIENS | 8 | 3791 | 1mhd | B | 93 | ARG | p.Arg93Ter | P84022 |
| HOMO SAPIENS | 0 | 1575 | 4phc | A | 205 | GLY | p.Gly205Asp | P12081 |
| HOMO SAPIENS | 0 | 1575 | 4phc | A | 454 | TYR | p.Tyr454Ser | P12081 |
| HOMO SAPIENS | 0 | 39620 | 1ozs | B | 135 | ARG | p.Arg136Gln | P19429 |
| HOMO SAPIENS | 5 | 39620 | 1ozs | B | 140 | ARG | p.Arg141Trp | P19429 |
| HOMO SAPIENS | 5 | 39620 | 1ozs | B | 142 | THR | p.Thr143Asn | P19429 |
| HOMO SAPIENS | 5 | 39620 | 1ozs | B | 143 | LEU | p.Leu144Gln | P19429 |
| HOMO SAPIENS | 5 | 39620 | 1ozs | B | 144 | ARG | p.Arg145Gly | P19429 |

| | | | | | | | | |
|---|---|---|---|---|---|---|---|---|
| HOMO SAPIENS | 5 | 39620 | 1ozs | B | 144 | ARG | p.Arg145Gln | P19429 |
| HOMO SAPIENS | -1 | 20934 | 4nuv | D | 42 | GLY | p.Gly42Asp | Q16570 |
| HOMO SAPIENS | -1 | 40565 | 5lcw | S | 4 | VAL | p.Val4Met | O60566 |
| HOMO SAPIENS | 0 | 40565 | 5lcw | S | 40 | THR | p.Thr40Met | O60566 |
| HOMO SAPIENS | 4 | 40565 | 5lcw | S | 198 | ALA | p.Glu390Asp | O60566 |
| HOMO SAPIENS | 4 | 758 | 4oiu | A | 726 | ARG | p.Arg727Leu | P10275 |
| HOMO SAPIENS | 4 | 758 | 4oiu | A | 730 | VAL | p.Val731Met | P10275 |
| HOMO SAPIENS | 2 | 758 | 4oiu | A | 798 | GLN | p.Gln799Glu | P10275 |
| HOMO SAPIENS | 3 | 758 | 4oiu | A | 855 | ARG | p.Arg856His | P10275 |
| HOMO SAPIENS | 1 | 758 | 4oiu | A | 886 | MET | p.Met887Val | P10275 |
| HOMO SAPIENS | 0 | 3303 | 4ie0 | A | 134 | ALA | p.Ala134Thr | Q9C0B1 |
| HOMO SAPIENS | 9 | 3303 | 4ie0 | A | 316 | ARG | p.Arg316Gln | Q9C0B1 |
| HOMO SAPIENS | 1 | 41502 | 3fgh | A | 136 | PRO | p.Pro178Leu | Q00059 |
| HOMO SAPIENS | 5 | 175 | 3ceh | B | 58 | THR | p.Thr59Lys | P06737 |
| HOMO SAPIENS | 8 | 175 | 3ceh | B | 93 | ARG | p.Arg94Ter | P06737 |
| HOMO SAPIENS | 5 | 175 | 3ceh | B | 221 | VAL | p.Val222Ile | P06737 |
| HOMO SAPIENS | 0 | 175 | 3ceh | B | 338 | ASN | p.Asn339Ser | P06737 |
| HOMO SAPIENS | 1 | 175 | 3ceh | B | 372 | PHE | p.Phe373Leu | P06737 |
| HOMO SAPIENS | 9 | 175 | 3ceh | B | 376 | ASN | p.Asn377Lys | P06737 |
| HOMO SAPIENS | 8 | 175 | 3ceh | B | 398 | ARG | p.Arg399Ter | P06737 |
| HOMO SAPIENS | 5 | 175 | 3ceh | B | 424 | ARG | p.Arg425Pro | P06737 |
| HOMO SAPIENS | 9 | 175 | 3ceh | B | 455 | VAL | p.Val456Met | P06737 |
| HOMO SAPIENS | 8 | 175 | 3ceh | B | 490 | ARG | p.Arg491Cys | P06737 |
| HOMO SAPIENS | 8 | 175 | 3ceh | B | 631 | ASN | p.Asn632Ile | P06737 |
| HOMO SAPIENS | 9 | 175 | 3ceh | B | 648 | TYR | p.Tyr649Ter | P06737 |
| HOMO SAPIENS | 9 | 175 | 3ceh | B | 672 | GLU | p.Glu673Lys | P06737 |
| HOMO SAPIENS | 9 | 175 | 3ceh | B | 674 | SER | p.Ser675Leu | P06737 |
| HOMO SAPIENS | 0 | 175 | 3ceh | B | 805 | ILE | p.Ile806Leu | P06737 |
| HOMO SAPIENS | 0 | 175 | 3ceh | B | 820 | TYR | p.Tyr821His | P06737 |
| HOMO SAPIENS | 1 | 6750 | 4fdj | A | 47 | GLY | p.Gly47Arg | P34059 |
| HOMO SAPIENS | 9 | 6750 | 4fdj | A | 60 | ASP | p.Asp60Asn | P34059 |
| HOMO SAPIENS | 5 | 6750 | 4fdj | A | 67 | LEU | p.Leu67Met | P34059 |
| HOMO SAPIENS | 0 | 6750 | 4fdj | A | 69 | PHE | p.Phe69Val | P34059 |
| HOMO SAPIENS | 3 | 6750 | 4fdj | A | 94 | ARG | p.Arg94Gly | P34059 |
| HOMO SAPIENS | 0 | 6750 | 4fdj | A | 113 | ILE | p.Ile113Phe | P34059 |
| HOMO SAPIENS | 0 | 6750 | 4fdj | A | 138 | VAL | p.Val138Ala | P34059 |
| HOMO SAPIENS | 0 | 6750 | 4fdj | A | 139 | GLY | p.Gly139Ser | P34059 |
| HOMO SAPIENS | 0 | 6750 | 4fdj | A | 151 | PRO | p.Pro151Thr | P34059 |
| HOMO SAPIENS | 0 | 6750 | 4fdj | A | 155 | GLY | p.Gly155Arg | P34059 |
| HOMO SAPIENS | 0 | 6750 | 4fdj | A | 159 | TRP | p.Trp159Ter | P34059 |
| HOMO SAPIENS | 0 | 6750 | 4fdj | A | 162 | SER | p.Ser162Phe | P34059 |
| HOMO SAPIENS | 0 | 6750 | 4fdj | A | 200 | THR | p.Thr200Met | P34059 |
| HOMO SAPIENS | 0 | 6750 | 4fdj | A | 204 | ASN | p.Asn204Lys | P34059 |
| HOMO SAPIENS | 0 | 6750 | 4fdj | A | 231 | ALA | p.Ala231Gly | P34059 |
| HOMO SAPIENS | 2 | 6750 | 4fdj | A | 235 | THR | p.Thr235Lys | P34059 |
| HOMO SAPIENS | 1 | 6750 | 4fdj | A | 259 | ARG | p.Arg259Gln | P34059 |
| HOMO SAPIENS | 0 | 6750 | 4fdj | A | 287 | SER | p.Ser287Leu | P34059 |
| HOMO SAPIENS | 0 | 6750 | 4fdj | A | 291 | ALA | p.Ala291Thr | P34059 |
| HOMO SAPIENS | 3 | 6750 | 4fdj | A | 301 | GLY | p.Gly301Cys | P34059 |
| HOMO SAPIENS | 0 | 6750 | 4fdj | A | 312 | THR | p.Thr312Ser | P34059 |
| HOMO SAPIENS | 0 | 6750 | 4fdj | A | 318 | MET | p.Met318Arg | P34059 |
| HOMO SAPIENS | 0 | 6750 | 4fdj | A | 340 | GLY | p.Gly340Asp | P34059 |
| HOMO SAPIENS | 0 | 6750 | 4fdj | A | 386 | ARG | p.Arg386Cys | P34059 |
| HOMO SAPIENS | 0 | 6750 | 4fdj | A | 391 | MET | p.Met391Val | P34059 |

| | | | | | | | | |
|---|---|---|---|---|---|---|---|---|
| HOMO SAPIENS | 0 | 6750 | 4fdj | A | 392 | ALA | p.Ala392Val | P34059 |
| HOMO SAPIENS | 0 | 6750 | 4fdj | A | 393 | ALA | p.Ala393Ser | P34059 |
| HOMO SAPIENS | 0 | 6750 | 4fdj | A | 459 | ALA | p.Ala459Val | P34059 |
| HOMO SAPIENS | 0 | 6750 | 4fdj | A | 473 | GLN | p.Gln473Ter | P34059 |
| HOMO SAPIENS | 2 | 6750 | 4fdj | A | 480 | VAL | p.Val480Phe | P34059 |
| HOMO SAPIENS | 0 | 6750 | 4fdj | A | 487 | ASN | p.Asn487Ser | P34059 |
| HOMO SAPIENS | 0 | 6750 | 4fdj | A | 488 | VAL | p.Val488Met | P34059 |
| HOMO SAPIENS | 0 | 6750 | 4fdj | A | 520 | TRP | p.Trp520Ter | P34059 |
| HOMO SAPIENS | 4 | 1088 | 2p3u | A | 102 | GLU | p.Glu142Lys | P00742 |
| HOMO SAPIENS | 6 | 370 | 1nr1 | D | 73 | ASP | p.Asp126His | P00367 |
| HOMO SAPIENS | 3 | 370 | 1nr1 | D | 221 | ARG | p.Arg274Cys | P00367 |
| HOMO SAPIENS | 2 | 370 | 1nr1 | D | 265 | ARG | p.Arg318Lys | P00367 |
| HOMO SAPIENS | 1 | 370 | 1nr1 | D | 269 | ARG | p.Arg322His | P00367 |
| HOMO SAPIENS | 1 | 370 | 1nr1 | D | 296 | GLU | p.Glu349Ala | P00367 |
| HOMO SAPIENS | 1 | 370 | 1nr1 | D | 445 | SER | p.Ser498Leu | P00367 |
| HOMO SAPIENS | 1 | 370 | 1nr1 | D | 446 | GLY | p.Gly499Asp | P00367 |
| HOMO SAPIENS | 1 | 370 | 1nr1 | D | 446 | GLY | p.Gly499Ser | P00367 |
| HOMO SAPIENS | 1 | 370 | 1nr1 | D | 448 | SER | p.Ser501Pro | P00367 |
| HOMO SAPIENS | 2 | 370 | 1nr1 | D | 454 | HIS | p.His507Tyr | P00367 |
| HOMO SAPIENS | 1 | 7720 | 2he0 | A | 147 | HIS | p.His2018Gln | P46531 |
| HOMO SAPIENS | 2 | 11844 | 5l0v | A | 218 | ARG | p.Arg218Ter | Q8NBL1 |
| HOMO SAPIENS | 2 | 11844 | 5l0v | A | 279 | ARG | p.Arg279Trp | Q8NBL1 |
| HOMO SAPIENS | 9 | 1485 | 5va0 | A | 477 | ARG | p.Arg477His | P04150 |
| HOMO SAPIENS | 0 | 235 | 2fun | B | 2248 | ARG | p.Arg248Trp | Q14790 |
| HOMO SAPIENS | 4 | 235 | 2fun | B | 2285 | HIS | p.Asp285His | Q14790 |
| HOMO SAPIENS | 0 | 4822 | 5y3n | A | 307 | ARG | p.Arg307Gly | Q12931 |
| HOMO SAPIENS | 8 | 32928 | 4m6w | A | 1886 | HIS | p.His1886Tyr | Q8IYD8 |
| HOMO SAPIENS | 0 | 2711 | 4y5v | F | 22 | GLY | p.Gly46Glu | P19235 |
| HOMO SAPIENS | -1 | 2711 | 4y5v | F | 166 | ALA | p.Ala190Ser | P19235 |
| HOMO SAPIENS | 4 | 5161 | 5bpu | A | 42 | HIS | p.His42Arg | Q00604 |
| HOMO SAPIENS | 0 | 5161 | 5bpu | A | 44 | TYR | p.Tyr44Cys | Q00604 |
| HOMO SAPIENS | 0 | 5161 | 5bpu | A | 45 | VAL | p.Val45Glu | Q00604 |
| HOMO SAPIENS | 0 | 5161 | 5bpu | A | 60 | VAL | p.Val60Glu | Q00604 |
| HOMO SAPIENS | 0 | 5161 | 5bpu | A | 61 | LEU | p.Leu61Phe | Q00604 |
| HOMO SAPIENS | 7 | 5161 | 5bpu | A | 69 | CYS | p.Cys69Ser | Q00604 |
| HOMO SAPIENS | 0 | 5161 | 5bpu | A | 75 | SER | p.Ser75Cys | Q00604 |
| HOMO SAPIENS | 0 | 5161 | 5bpu | A | 90 | ARG | p.Arg90Pro | Q00604 |
| HOMO SAPIENS | 9 | 5161 | 5bpu | A | 96 | CYS | p.Cys96Tyr | Q00604 |
| HOMO SAPIENS | 9 | 5161 | 5bpu | A | 96 | CYS | p.Cys96Trp | Q00604 |
| HOMO SAPIENS | 0 | 5161 | 5bpu | A | 101 | SER | p.Ser101Phe | Q00604 |
| HOMO SAPIENS | 0 | 5161 | 5bpu | A | 105 | ALA | p.Ala105Thr | Q00604 |
| HOMO SAPIENS | 0 | 5161 | 5bpu | A | 110 | CYS | p.Cys110Gly | Q00604 |
| HOMO SAPIENS | 0 | 5161 | 5bpu | A | 121 | ARG | p.Arg121Trp | Q00604 |
| HOMO SAPIENS | 0 | 5161 | 5bpu | A | 121 | ARG | p.Arg121Leu | Q00604 |
| HOMO SAPIENS | 0 | 5161 | 5bpu | A | 124 | LEU | p.Leu124Phe | Q00604 |
| HOMO SAPIENS | 8 | 5161 | 5bpu | A | 128 | CYS | p.Cys128Ter | Q00604 |
| HOMO SAPIENS | 1 | 13944 | 5xqh | A | 238 | ARG | p.Arg238His | Q9GZN7 |
| HOMO SAPIENS | -1 | 7508 | 5jl6 | A | 39 | TRP | p.Trp39Arg | P11511 |
| HOMO SAPIENS | 0 | 7508 | 5jl6 | A | 201 | THR | p.Thr201Met | P11511 |
| HOMO SAPIENS | 0 | 7508 | 5jl6 | A | 210 | GLU | p.Glu210Lys | P11511 |
| HOMO SAPIENS | 1 | 7508 | 5jl6 | A | 264 | ARG | p.Arg264Cys | P11511 |
| HOMO SAPIENS | 2 | 7508 | 5jl6 | A | 365 | ARG | p.Arg365Gln | P11511 |
| HOMO SAPIENS | 7 | 7508 | 5jl6 | A | 375 | ARG | p.Arg375Cys | P11511 |
| HOMO SAPIENS | 8 | 7508 | 5jl6 | A | 435 | ARG | p.Arg435Cys | P11511 |

| | | | | | | | | |
|---|---|---|---|---|---|---|---|---|
| HOMO SAPIENS | 9 | 7508 | 5jl6 | A | 437 | CYS | p.Cys437Tyr | P11511 |
| HOMO SAPIENS | -1 | 30206 | 5l95 | B | 246 | ARG | p.Arg246Ter | Q9GZZ9 |
| HOMO SAPIENS | 4 | 875 | 3efh | B | 43 | GLU | p.Glu43Asp | P60891 |
| HOMO SAPIENS | 5 | 875 | 3efh | B | 65 | ASP | p.Asp65Asn | P60891 |
| HOMO SAPIENS | 0 | 875 | 3efh | B | 87 | ALA | p.Ala87Thr | P60891 |
| HOMO SAPIENS | 2 | 875 | 3efh | B | 115 | MET | p.Met115Thr | P60891 |
| HOMO SAPIENS | 8 | 875 | 3efh | B | 133 | GLN | p.Gln133Pro | P60891 |
| HOMO SAPIENS | 8 | 875 | 3efh | B | 152 | LEU | p.Leu152Pro | P60891 |
| HOMO SAPIENS | 7 | 875 | 3efh | B | 290 | ILE | p.Ile290Thr | P60891 |
| HOMO SAPIENS | 6 | 875 | 3efh | B | 306 | GLY | p.Gly306Arg | P60891 |
| HOMO SAPIENS | 0 | 7978 | 5edm | A | 122 | MET | p.Thr165Met | P00734 |
| HOMO SAPIENS | -1 | 7978 | 5edm | A | 257 | ARG | p.Arg314Cys | P00734 |
| HOMO SAPIENS | 0 | 1716 | 1b98 | M | 126 | ARG | p.Arg206Gln | P34130 |
| HOMO SAPIENS | 0 | 1716 | 1b98 | M | 126 | ARG | p.Arg206Trp | P34130 |
| HOMO SAPIENS | 9 | 3960 | 2qyp | B | 28 | ILE | p.Ile338Val | P07602 |
| HOMO SAPIENS | 1 | 3960 | 2qyp | B | 39 | LEU | p.Leu349Pro | P07602 |
| HOMO SAPIENS | 0 | 31414 | 5u1d | A | 333 | ILE | p.Ile393Val | Q03518 |
| HOMO SAPIENS | 0 | 31414 | 5u1d | A | 637 | ASP | p.Asp697Gly | Q03518 |
| HOMO SAPIENS | 0 | 31414 | 5u1d | A | 659 | ARG | p.Arg719Gln | Q03518 |
| HOMO SAPIENS | 3 | 119 | 2pho | A | 11 | ILE | p.Ile11Thr | P05089 |
| HOMO SAPIENS | 8 | 119 | 2pho | A | 21 | ARG | p.Arg21Ter | P05089 |
| HOMO SAPIENS | 0 | 119 | 2pho | A | 122 | TRP | p.Trp122Ter | P05089 |
| HOMO SAPIENS | 6 | 119 | 2pho | A | 138 | GLY | p.Gly138Val | P05089 |
| HOMO SAPIENS | 5 | 119 | 2pho | A | 290 | THR | p.Thr290Ser | P05089 |
| HOMO SAPIENS | 5 | 119 | 2pho | A | 291 | ARG | p.Arg291Ter | P05089 |
| HOMO SAPIENS | 0 | 3732 | 2i99 | A | 247 | TYR | p.Tyr247Ter | Q14894 |
| HOMO SAPIENS | -1 | 1382 | 4ffx | A | 2 | ALA | p.Ala2Val | P30566 |
| HOMO SAPIENS | 1 | 1382 | 4ffx | A | 72 | ILE | p.Ile72Met | P30566 |
| HOMO SAPIENS | 1 | 1382 | 4ffx | A | 100 | PRO | p.Pro100Ala | P30566 |
| HOMO SAPIENS | 0 | 1382 | 4ffx | A | 114 | TYR | p.Tyr114His | P30566 |
| HOMO SAPIENS | 1 | 1382 | 4ffx | A | 141 | ARG | p.Arg141Trp | P30566 |
| HOMO SAPIENS | 0 | 1382 | 4ffx | A | 147 | LYS | p.Lys147Met | P30566 |
| HOMO SAPIENS | 3 | 1382 | 4ffx | A | 190 | ARG | p.Arg190Gln | P30566 |
| HOMO SAPIENS | 3 | 1382 | 4ffx | A | 190 | ARG | p.Arg190Ter | P30566 |
| HOMO SAPIENS | 0 | 1382 | 4ffx | A | 217 | HIS | p.His217Asp | P30566 |
| HOMO SAPIENS | 2 | 1382 | 4ffx | A | 246 | LYS | p.Lys246Glu | P30566 |
| HOMO SAPIENS | 5 | 1382 | 4ffx | A | 303 | ARG | p.Arg303Cys | P30566 |
| HOMO SAPIENS | 1 | 1382 | 4ffx | A | 318 | PRO | p.Pro318Leu | P30566 |
| HOMO SAPIENS | 6 | 1382 | 4ffx | A | 332 | ASP | p.Asp332His | P30566 |
| HOMO SAPIENS | 6 | 1382 | 4ffx | A | 337 | ARG | p.Arg337Ter | P30566 |
| HOMO SAPIENS | 1 | 1382 | 4ffx | A | 396 | ARG | p.Arg396His | P30566 |
| HOMO SAPIENS | 1 | 1382 | 4ffx | A | 426 | ARG | p.Arg426His | P30566 |
| HOMO SAPIENS | 0 | 1382 | 4ffx | A | 438 | SER | p.Ser438Pro | P30566 |
| HOMO SAPIENS | 1 | 1382 | 4ffx | A | 447 | SER | p.Ser447Pro | P30566 |
| HOMO SAPIENS | 2 | 1382 | 4ffx | A | 448 | SER | p.Ser448Pro | P30566 |
| HOMO SAPIENS | 2 | 1382 | 4ffx | A | 450 | THR | p.Thr450Ser | P30566 |
| HOMO SAPIENS | 0 | 6212 | 1ao3 | A | 46 | SER | p.Ser1731Thr | P04275 |
| HOMO SAPIENS | 0 | 1711 | 3f9c | A | 279 | VAL | p.Val279Phe | Q13093 |
| HOMO SAPIENS | 0 | 6112 | 4fvl | A | 312 | HIS | p.His312Arg | P45452 |
| HOMO SAPIENS | 0 | 6112 | 4fvl | A | 323 | THR | p.Thr323Met | P45452 |
| HOMO SAPIENS | 0 | 6112 | 4fvl | A | 390 | ASP | p.Asp390Gly | P45452 |
| HOMO SAPIENS | 0 | 6112 | 4fvl | A | 458 | ARG | p.Arg458Cys | P45452 |
| HOMO SAPIENS | 2 | 6180 | 5ggn | A | 97 | ARG | p.Arg97Trp | Q8WZA1 |
| HOMO SAPIENS | 2 | 6180 | 5ggn | A | 215 | ARG | p.Arg215Ter | Q8WZA1 |

| | | | | | | | | |
|---|---|---|---|---|---|---|---|---|
| HOMO SAPIENS | -1 | 5526 | 4ak3 | A | 11 | ASP | p.Asp1232Asn | P02461 |
| HOMO SAPIENS | 6 | 5526 | 4ak3 | A | 38 | ALA | p.Ala1259Thr | P02461 |
| HOMO SAPIENS | 0 | 5526 | 4ak3 | A | 92 | LYS | p.Lys1313Arg | P02461 |
| HOMO SAPIENS | 0 | 5526 | 4ak3 | A | 132 | GLN | p.His1353Gln | P02461 |
| HOMO SAPIENS | 4 | 16616 | 1hci | B | 433 | SER | p.Ser433Leu | P35609 |
| HOMO SAPIENS | 1 | 16616 | 1hci | B | 475 | ASP | p.Asp475Asn | P35609 |
| HOMO SAPIENS | 1 | 16616 | 1hci | B | 476 | ALA | p.Ala476Ser | P35609 |
| HOMO SAPIENS | 2 | 16616 | 1hci | B | 507 | MET | p.Met507Val | P35609 |
| HOMO SAPIENS | 1 | 16616 | 1hci | B | 604 | MET | p.Met604Val | P35609 |
| HOMO SAPIENS | 2 | 16616 | 1hci | B | 640 | ARG | p.Arg640His | P35609 |
| HOMO SAPIENS | 1 | 16616 | 1hci | B | 721 | ARG | p.Arg721Cys | P35609 |
| HOMO SAPIENS | -1 | 3402 | 5iaa | B | 57 | MET | p.Met57Val | Q9GZZ9 |
| HOMO SAPIENS | 7 | 3250 | 3irq | B | 193 | PRO | p.Pro193Ala | P55265 |
| HOMO SAPIENS | 2 | 15200 | 2wns | A | 30 | SER | p.Ser30Gly | P11172 |
| HOMO SAPIENS | 0 | 15200 | 2wns | A | 96 | ARG | p.Arg96Gly | P11172 |
| HOMO SAPIENS | 0 | 15200 | 2wns | A | 109 | VAL | p.Val109Gly | P11172 |
| HOMO SAPIENS | 2 | 15759 | 4x4w | B | 140 | LEU | p.Leu166Ser | Q96Q11 |
| HOMO SAPIENS | 0 | 15759 | 4x4w | B | 197 | ILE | p.Ile223Thr | Q96Q11 |
| HOMO SAPIENS | 0 | 3156 | 3w03 | B | 14 | ALA | p.Ala14Thr | Q9H9Q4 |
| HOMO SAPIENS | 0 | 3156 | 3w03 | B | 57 | ARG | p.Arg57Gly | Q9H9Q4 |
| HOMO SAPIENS | 0 | 3156 | 3w03 | B | 123 | CYS | p.Cys123Arg | Q9H9Q4 |
| HOMO SAPIENS | 0 | 11693 | 3ojy | A | 63 | GLN | p.Gln93Lys | P07357 |
| HOMO SAPIENS | 0 | 1421 | 2qc8 | E | 275 | GLU | p.Glu275Asp | P15104 |
| HOMO SAPIENS | 9 | 1421 | 2qc8 | E | 324 | ARG | p.Arg324Cys | P15104 |
| HOMO SAPIENS | 6 | 1421 | 2qc8 | E | 341 | ARG | p.Arg341Cys | P15104 |
| HOMO SAPIENS | 6 | 2850 | 3qvb | A | 83 | THR | p.Thr83Met | P46976 |
| HOMO SAPIENS | 8 | 2850 | 3qvb | A | 102 | ASP | p.Asp102His | P46976 |
| HOMO SAPIENS | 1 | 24314 | 4m9p | A | 480 | PRO | p.Pro480Leu | P21333 |
| HOMO SAPIENS | 0 | 24314 | 4m9p | A | 528 | VAL | p.Val528Met | P21333 |
| HOMO SAPIENS | 0 | 24314 | 4m9p | A | 590 | VAL | p.Val590Ile | P21333 |
| HOMO SAPIENS | 0 | 24314 | 4m9p | A | 605 | ASP | p.Asp605Asn | P21333 |
| HOMO SAPIENS | 1 | 24314 | 4m9p | A | 637 | PRO | p.Pro637Gln | P21333 |
| HOMO SAPIENS | 0 | 24314 | 4m9p | A | 656 | LEU | p.Leu656Phe | P21333 |
| HOMO SAPIENS | 2 | 24314 | 4m9p | A | 711 | VAL | p.Val711Asp | P21333 |
| HOMO SAPIENS | 0 | 4951 | 5joe | A | 40 | THR | p.Thr9621Ser | Q8WZ42 |
| HOMO SAPIENS | 1 | 4951 | 5joe | A | 41 | GLU | p.Glu9622Gln | Q8WZ42 |
| HOMO SAPIENS | 0 | 4951 | 5joe | A | 82 | ALA | p.Ala9663Ser | Q8WZ42 |
| HOMO SAPIENS | 4 | 4951 | 5joe | A | 89 | ILE | p.Ile9670Asn | Q8WZ42 |
| HOMO SAPIENS | 0 | 9664 | 3c6m | A | 56 | GLY | p.Gly56Ser | P52788 |
| HOMO SAPIENS | 0 | 9664 | 3c6m | A | 132 | VAL | p.Val132Gly | P52788 |
| HOMO SAPIENS | 1 | 9664 | 3c6m | A | 190 | THR | p.Thr190Ile | P52788 |
| HOMO SAPIENS | -1 | 12118 | 5hcd | B | 19 | GLN | p.Gln19Ter | P01031 |
| HOMO SAPIENS | 0 | 1583 | 2b4x | L | 7 | ILE | p.Ile39Asn | P01008 |
| HOMO SAPIENS | 0 | 1583 | 2b4x | L | 24 | ARG | p.Arg56Cys | P01008 |
| HOMO SAPIENS | 0 | 1583 | 2b4x | L | 47 | ARG | p.Arg79Cys | P01008 |
| HOMO SAPIENS | 0 | 1583 | 2b4x | L | 47 | ARG | p.Arg79His | P01008 |
| HOMO SAPIENS | 1 | 1583 | 2b4x | L | 95 | CYS | p.Cys127Arg | P01008 |
| HOMO SAPIENS | 1 | 1583 | 2b4x | L | 99 | LEU | p.Leu131Phe | P01008 |
| HOMO SAPIENS | 0 | 1583 | 2b4x | L | 116 | SER | p.Ser148Pro | P01008 |
| HOMO SAPIENS | 0 | 1583 | 2b4x | L | 129 | ARG | p.Arg161Gln | P01008 |
| HOMO SAPIENS | 0 | 1583 | 2b4x | L | 135 | ASN | p.Asn167Thr | P01008 |
| HOMO SAPIENS | 2 | 1583 | 2b4x | L | 187 | ASN | p.Asn219Asp | P01008 |
| HOMO SAPIENS | 0 | 1583 | 2b4x | L | 191 | SER | p.Ser223Pro | P01008 |
| HOMO SAPIENS | 2 | 1583 | 2b4x | L | 349 | SER | p.Ser381Pro | P01008 |

| | | | | | | | | |
|---|---|---|---|---|---|---|---|---|
| HOMO SAPIENS | 0 | 1583 | 2b4x | L | 382 | ALA | p.Ala414Thr | P01008 |
| HOMO SAPIENS | 0 | 1583 | 2b4x | L | 387 | ALA | p.Ala419Val | P01008 |
| HOMO SAPIENS | 0 | 1583 | 2b4x | L | 392 | GLY | p.Gly424Asp | P01008 |
| HOMO SAPIENS | 0 | 1583 | 2b4x | L | 393 | ARG | p.Arg425His | P01008 |
| HOMO SAPIENS | 0 | 1583 | 2b4x | L | 393 | ARG | p.Arg425Cys | P01008 |
| HOMO SAPIENS | 0 | 1583 | 2b4x | L | 394 | SER | p.Ser426Leu | P01008 |
| HOMO SAPIENS | 0 | 1583 | 2b4x | L | 404 | ALA | p.Ala436Thr | P01008 |
| HOMO SAPIENS | 2 | 1583 | 2b4x | L | 407 | PRO | p.Pro439Leu | P01008 |
| HOMO SAPIENS | 2 | 1583 | 2b4x | L | 429 | PRO | p.Pro461Leu | P01008 |
| HOMO SAPIENS | 0 | 11647 | 2a1t | S | 78 | ALA | p.Ala78Ser | P38117 |
| HOMO SAPIENS | 0 | 11647 | 2a1t | S | 98 | ARG | p.Arg98Cys | P38117 |
| HOMO SAPIENS | 8 | 11647 | 2a1t | S | 128 | ASP | p.Asp128Asn | P38117 |
| HOMO SAPIENS | 5 | 11647 | 2a1t | S | 154 | THR | p.Thr154Met | P38117 |
| HOMO SAPIENS | 6 | 11647 | 2a1t | S | 164 | ARG | p.Arg164Gln | P38117 |
| HOMO SAPIENS | 5 | 11647 | 2a1t | S | 174 | ARG | p.Arg174His | P38117 |
| HOMO SAPIENS | -1 | 11200 | 4tx5 | A | 4 | ILE | p.Ile59Val | Q9NR28 |
| HOMO SAPIENS | 0 | 11200 | 4tx5 | A | 169 | GLY | p.Gly224Arg | Q9NR28 |
| HOMO SAPIENS | 3 | 417 | 2vr2 | A | 6 | ARG | p.Arg6Gln | Q14117 |
| HOMO SAPIENS | 1 | 417 | 2vr2 | A | 334 | GLN | p.Gln334Arg | Q14117 |
| HOMO SAPIENS | 2 | 417 | 2vr2 | A | 360 | TRP | p.Trp360Arg | Q14117 |
| HOMO SAPIENS | 0 | 417 | 2vr2 | A | 379 | SER | p.Ser379Arg | Q14117 |
| HOMO SAPIENS | 0 | 417 | 2vr2 | A | 435 | GLY | p.Gly435Arg | Q14117 |
| HOMO SAPIENS | 0 | 12498 | 3eo3 | B | 423 | ILE | p.Ile423Val | Q9Y223 |
| HOMO SAPIENS | 5 | 12498 | 3eo3 | B | 572 | VAL | p.Val572Leu | Q9Y223 |
| HOMO SAPIENS | 0 | 12498 | 3eo3 | B | 576 | GLY | p.Gly576Glu | Q9Y223 |
| HOMO SAPIENS | 0 | 12498 | 3eo3 | B | 587 | ILE | p.Ile587Thr | Q9Y223 |
| HOMO SAPIENS | 0 | 12498 | 3eo3 | B | 615 | SER | p.Ser615Ter | Q9Y223 |
| HOMO SAPIENS | 1 | 12498 | 3eo3 | B | 631 | ALA | p.Ala631Val | Q9Y223 |
| HOMO SAPIENS | 1 | 12498 | 3eo3 | B | 669 | GLY | p.Gly669Arg | Q9Y223 |
| HOMO SAPIENS | 1 | 12498 | 3eo3 | B | 696 | VAL | p.Val696Met | Q9Y223 |
| HOMO SAPIENS | 2 | 12498 | 3eo3 | B | 712 | MET | p.Met712Thr | Q9Y223 |
| HOMO SAPIENS | 0 | 1568 | 1r74 | A | 49 | LEU | p.Leu50Pro | Q14749 |
| HOMO SAPIENS | 0 | 1568 | 1r74 | A | 176 | HIS | p.His177Asn | Q14749 |
| HOMO SAPIENS | 3 | 147 | 5eig | A | 67 | THR | p.Thr67Ile | P32929 |
| HOMO SAPIENS | 3 | 147 | 5eig | A | 240 | GLN | p.Gln240Glu | P32929 |
| HOMO SAPIENS | -1 | 147 | 5eig | A | 403 | SER | p.Ser403Ile | P32929 |
| HOMO SAPIENS | 8 | 20036 | 2lsr | A | 31 | ARG | p.Arg31Ter | Q9Y6N9 |
| HOMO SAPIENS | -1 | 41382 | 4uis | B | 431 | UNK | p.Ala431Glu | P49768 |
| HOMO SAPIENS | -1 | 8536 | 3i6w | B | 85 | PRO | p.Pro85Leu | O96017 |
| HOMO SAPIENS | 0 | 8536 | 3i6w | B | 93 | TRP | p.Trp93Ter | O96017 |
| HOMO SAPIENS | 0 | 8536 | 3i6w | B | 95 | ARG | p.Arg95Ter | O96017 |
| HOMO SAPIENS | 0 | 8536 | 3i6w | B | 117 | ARG | p.Arg117Gly | O96017 |
| HOMO SAPIENS | 0 | 8536 | 3i6w | B | 119 | LYS | p.Lys119Arg | O96017 |
| HOMO SAPIENS | 0 | 8536 | 3i6w | B | 137 | ARG | p.Arg137Gln | O96017 |
| HOMO SAPIENS | 0 | 8536 | 3i6w | B | 137 | ARG | p.Arg137Ter | O96017 |
| HOMO SAPIENS | 0 | 8536 | 3i6w | B | 145 | ARG | p.Arg145Trp | O96017 |
| HOMO SAPIENS | 0 | 8536 | 3i6w | B | 160 | ILE | p.Ile160Met | O96017 |
| HOMO SAPIENS | 0 | 41322 | 4bxl | C | 46 | GLU | p.Glu46Lys | P37840 |
| HOMO SAPIENS | 5 | 41322 | 4bxl | C | 53 | ALA | p.Ala53Thr | P37840 |
| HOMO SAPIENS | 2 | 24966 | 3hve | A | 15 | ARG | p.Arg15Ser | Q9H2C0 |
| HOMO SAPIENS | 1 | 24966 | 3hve | A | 138 | ARG | p.Arg138His | Q9H2C0 |
| HOMO SAPIENS | 0 | 24966 | 3hve | A | 201 | ARG | p.Arg201Ter | Q9H2C0 |
| HOMO SAPIENS | 0 | 11680 | 5an9 | J | 8 | ASN | p.Asn8Lys | Q9Y3A5 |
| HOMO SAPIENS | 1 | 11680 | 5an9 | J | 33 | LYS | p.Lys33Arg | Q9Y3A5 |

| | | | | | | | | |
|---|---|---|---|---|---|---|---|---|
| HOMO SAPIENS | 1 | 11680 | 5an9 | J | 126 | ARG | p.Arg126Thr | Q9Y3A5 |
| HOMO SAPIENS | 1 | 11680 | 5an9 | J | 130 | VAL | p.Val130Met | Q9Y3A5 |
| HOMO SAPIENS | 1 | 11680 | 5an9 | J | 169 | ARG | p.Arg169Cys | Q9Y3A5 |
| HOMO SAPIENS | 1 | 11680 | 5an9 | J | 218 | ARG | p.Arg218Ter | Q9Y3A5 |
| HOMO SAPIENS | 4 | 978 | 3zs0 | B | 7 | TYR | p.Tyr173Cys | P05164 |
| HOMO SAPIENS | 1 | 978 | 3zs0 | B | 85 | MET | p.Met251Thr | P05164 |
| HOMO SAPIENS | 4 | 5140 | 2l4r | A | 52 | CYS | p.Cys52Ter | Q12809 |
| HOMO SAPIENS | 4 | 5140 | 2l4r | A | 63 | PRO | p.Pro63His | Q12809 |
| HOMO SAPIENS | 0 | 5140 | 2l4r | A | 100 | ARG | p.Arg100Gly | Q12809 |
| HOMO SAPIENS | 4 | 5140 | 2l4r | A | 129 | PHE | p.Phe129Leu | Q12809 |
| HOMO SAPIENS | 0 | 25836 | 2gy7 | B | 103 | GLU | p.Glu103Asp | Q02763 |
| HOMO SAPIENS | 0 | 25836 | 2gy7 | B | 148 | ILE | p.Ile148Thr | Q02763 |
| HOMO SAPIENS | 0 | 25836 | 2gy7 | B | 226 | ALA | p.Ala226Val | Q02763 |
| HOMO SAPIENS | 0 | 25836 | 2gy7 | B | 391 | THR | p.Thr391Ile | Q02763 |
| HOMO SAPIENS | 4 | 12563 | 5kwy | D | 20 | GLU | p.Glu20Ter | P61916 |
| HOMO SAPIENS | 2 | 12563 | 5kwy | D | 30 | VAL | p.Val30Met | P61916 |
| HOMO SAPIENS | 4 | 12563 | 5kwy | D | 39 | VAL | p.Val39Met | P61916 |
| HOMO SAPIENS | 8 | 12563 | 5kwy | D | 67 | SER | p.Ser67Pro | P61916 |
| HOMO SAPIENS | 8 | 12563 | 5kwy | D | 99 | CYS | p.Cys99Arg | P61916 |
| HOMO SAPIENS | 2 | 12563 | 5kwy | D | 118 | GLU | p.Glu118Ter | P61916 |
| HOMO SAPIENS | 2 | 12563 | 5kwy | D | 120 | PRO | p.Pro120Ser | P61916 |
| HOMO SAPIENS | -1 | 1679 | 1pmk | A | -2 | VAL | p.Val374Phe | P00747 |
| HOMO SAPIENS | -1 | 5315 | 2w2n | P | 53 | ALA | p.Ala53Val | Q8NBP7 |
| HOMO SAPIENS | 8 | 13496 | 2yad | A | 138 | THR | p.Asn138Thr | P11686 |
| HOMO SAPIENS | -1 | 13496 | 2yad | A | 173 | ARG | p.Arg167Gln | P11686 |
| HOMO SAPIENS | 2 | 13496 | 2yad | A | 186 | SER | p.Asn186Ser | P11686 |
| HOMO SAPIENS | 0 | 4426 | 3qd6 | C | 123 | ALA | p.Ala123Glu | P29965 |
| HOMO SAPIENS | 1 | 4426 | 3qd6 | C | 140 | TRP | p.Trp140Gly | P29965 |
| HOMO SAPIENS | 0 | 4426 | 3qd6 | C | 155 | LEU | p.Leu155Pro | P29965 |
| HOMO SAPIENS | 6 | 4426 | 3qd6 | C | 227 | GLY | p.Gly227Val | P29965 |
| HOMO SAPIENS | 0 | 4426 | 3qd6 | C | 235 | ALA | p.Ala235Pro | P29965 |
| HOMO SAPIENS | 0 | 10734 | 3dyd | B | 45 | PRO | p.Pro45Ser | P17735 |
| HOMO SAPIENS | -1 | 10734 | 3dyd | B | 57 | ARG | p.Arg57Ter | P17735 |
| HOMO SAPIENS | 8 | 10734 | 3dyd | B | 417 | ARG | p.Arg417Ter | P17735 |
| HOMO SAPIENS | 0 | 6956 | 4pzs | A | 1600 | LEU | p.Leu1600Phe | Q09472 |
| HOMO SAPIENS | 0 | 9370 | 3hfv | A | 21 | SER | p.Ser57Cys | O95363 |
| HOMO SAPIENS | 1 | 9370 | 3hfv | A | 25 | ASP | p.Asp61Glu | O95363 |
| HOMO SAPIENS | 1 | 9370 | 3hfv | A | 130 | ALA | p.Ala166Thr | O95363 |
| HOMO SAPIENS | 5 | 9370 | 3hfv | A | 148 | ASP | p.Asp184Asn | O95363 |
| HOMO SAPIENS | 0 | 9370 | 3hfv | A | 187 | ARG | p.Arg223Cys | O95363 |
| HOMO SAPIENS | 0 | 9370 | 3hfv | A | 210 | THR | p.Thr246Met | O95363 |
| HOMO SAPIENS | 1 | 9370 | 3hfv | A | 244 | ASN | p.Asn280Ser | O95363 |
| HOMO SAPIENS | 2 | 9370 | 3hfv | A | 288 | TYR | p.Tyr324Cys | O95363 |
| HOMO SAPIENS | 5 | 9370 | 3hfv | A | 293 | ILE | p.Ile329Thr | O95363 |
| HOMO SAPIENS | 1 | 9370 | 3hfv | A | 355 | ASP | p.Asp391Val | O95363 |
| HOMO SAPIENS | 1 | 9370 | 3hfv | A | 371 | THR | p.Thr407Met | O95363 |
| HOMO SAPIENS | 1 | 9370 | 3hfv | A | 383 | ARG | p.Arg419Cys | O95363 |
| HOMO SAPIENS | 1 | 9370 | 3hfv | A | 387 | ARG | p.Arg423Gln | O95363 |
| HOMO SAPIENS | 0 | 14401 | 1q3x | A | 371 | ASP | p.Asp371Tyr | O00187 |
| HOMO SAPIENS | 0 | 14401 | 1q3x | A | 377 | VAL | p.Val377Ala | O00187 |
| HOMO SAPIENS | 0 | 14401 | 1q3x | A | 439 | ARG | p.Arg439His | O00187 |
| HOMO SAPIENS | 0 | 14401 | 1q3x | A | 577 | GLN | p.Gln577His | O00187 |
| HOMO SAPIENS | 2 | 18270 | 3teq | A | 429 | ARG | p.Arg429Cys | Q13586 |
| HOMO SAPIENS | -1 | 16081 | 4mqt | A | 231 | VAL | p.Val231Ile | P08172 |

| | | | | | | | | |
|---|---|---|---|---|---|---|---|---|
| HOMO SAPIENS | 9 | 6607 | 3whu | B | 81 | ASP | p.Asp81His | Q8NI22 |
| HOMO SAPIENS | 9 | 6607 | 3whu | B | 129 | ASP | p.Asp129Glu | Q8NI22 |
| HOMO SAPIENS | 9 | 6607 | 3whu | B | 136 | ILE | p.Ile136Thr | Q8NI22 |
| HOMO SAPIENS | 0 | 7208 | 2e9w | D | 11 | ASN | p.Asn36Ser | P21583 |
| HOMO SAPIENS | 0 | 24566 | 4bdx | A | 10 | LEU | p.Leu140Val | P00748 |
| HOMO SAPIENS | 5 | 24566 | 4bdx | A | 77 | PRO | p.Ala207Pro | P00748 |
| HOMO SAPIENS | 7 | 606 | 3qnf | A | 319 | MET | p.Met319Val | Q9NZ08 |
| HOMO SAPIENS | 8 | 4097 | 1ydl | A | 21 | LEU | p.Leu21Pro | Q6ZYL4 |
| HOMO SAPIENS | 6 | 4097 | 1ydl | A | 56 | ARG | p.Arg56Ter | Q6ZYL4 |
| HOMO SAPIENS | 8 | 36691 | 1dqb | A | 42 | ARG | p.Arg403Lys | P07204 |
| HOMO SAPIENS | 0 | 5642 | 3j7y | X | 59 | ARG | p.Arg59Pro | Q13084 |
| HOMO SAPIENS | 1 | 5642 | 3j7y | X | 204 | VAL | p.Val204Met | Q13084 |
| HOMO SAPIENS | 1 | 1119 | 3nwu | B | 252 | ALA | p.Ala252Thr | Q92743 |
| HOMO SAPIENS | 3 | 1119 | 3nwu | B | 274 | ARG | p.Arg274Gln | Q92743 |
| HOMO SAPIENS | 9 | 1119 | 3nwu | B | 297 | VAL | p.Val297Met | Q92743 |
| HOMO SAPIENS | -1 | 1119 | 3nwu | B | 302 | ARG | p.Arg302Ter | Q92743 |
| HOMO SAPIENS | 9 | 1119 | 3nwu | B | 321 | ALA | p.Ala321Thr | Q92743 |
| HOMO SAPIENS | -1 | 1119 | 3nwu | B | 370 | ARG | p.Arg370Ter | Q92743 |
| HOMO SAPIENS | 0 | 416 | 4ycu | B | 105 | LEU | p.Leu105His | Q15046 |
| HOMO SAPIENS | 0 | 416 | 4ycu | B | 145 | TYR | p.Tyr145His | Q15046 |
| HOMO SAPIENS | 0 | 416 | 4ycu | B | 200 | PRO | p.Pro200Leu | Q15046 |
| HOMO SAPIENS | 0 | 416 | 4ycu | B | 201 | TYR | p.Tyr201His | Q15046 |
| HOMO SAPIENS | 0 | 416 | 4ycu | B | 420 | ARG | p.Arg420Cys | Q15046 |
| HOMO SAPIENS | 9 | 3297 | 4fpb | A | 680 | THR | p.Thr680Ser | P19367 |
| HOMO SAPIENS | 0 | 6771 | 2zt6 | A | 201 | ALA | p.Ala255Val | P41250 |
| HOMO SAPIENS | 0 | 6771 | 2zt6 | A | 202 | ASP | p.Asp256His | P41250 |
| HOMO SAPIENS | 0 | 6771 | 2zt6 | A | 334 | ARG | p.Arg388Gln | P41250 |
| HOMO SAPIENS | -1 | 6771 | 2zt6 | A | 439 | ASN | p.Asn493Ser | P41250 |
| HOMO SAPIENS | -1 | 6771 | 2zt6 | A | 500 | ASP | p.Asp554Asn | P41250 |
| HOMO SAPIENS | 1 | 6771 | 2zt6 | A | 581 | SER | p.Ser635Leu | P41250 |
| HOMO SAPIENS | -1 | 6771 | 2zt6 | A | 684 | GLU | p.Glu738Lys | P41250 |
| HOMO SAPIENS | 0 | 2687 | 4grw | B | 276 | VAL | p.Val298Phe | P29460 |
| HOMO SAPIENS | 6 | 913 | 1m75 | B | 45 | ASP | p.Asp57Glu | Q16836 |
| HOMO SAPIENS | 2 | 913 | 1m75 | B | 74 | PRO | p.Leu86Pro | Q16836 |
| HOMO SAPIENS | 3 | 913 | 1m75 | B | 214 | TYR | p.Tyr226His | Q16836 |
| HOMO SAPIENS | 3 | 913 | 1m75 | B | 224 | ARG | p.Arg236Ter | Q16836 |
| HOMO SAPIENS | 4 | 913 | 1m75 | B | 246 | PRO | p.Pro258Leu | Q16836 |
| HOMO SAPIENS | 2 | 4713 | 4djn | B | 40 | ARG | p.Arg40Cys | Q7LG56 |
| HOMO SAPIENS | 1 | 4713 | 4djn | B | 41 | ARG | p.Arg41Pro | Q7LG56 |
| HOMO SAPIENS | 2 | 4713 | 4djn | B | 110 | ARG | p.Arg110His | Q7LG56 |
| HOMO SAPIENS | 2 | 4713 | 4djn | B | 121 | ARG | p.Arg121His | Q7LG56 |
| HOMO SAPIENS | 9 | 4713 | 4djn | B | 131 | GLU | p.Glu131Lys | Q7LG56 |
| HOMO SAPIENS | 2 | 4713 | 4djn | B | 182 | THR | p.Thr182Ala | Q7LG56 |
| HOMO SAPIENS | 6 | 4713 | 4djn | B | 194 | GLU | p.Glu194Lys | Q7LG56 |
| HOMO SAPIENS | 2 | 4713 | 4djn | B | 211 | ARG | p.Arg211Lys | Q7LG56 |
| HOMO SAPIENS | 0 | 4713 | 4djn | B | 224 | ILE | p.Ile224Ser | Q7LG56 |
| HOMO SAPIENS | 3 | 4713 | 4djn | B | 229 | GLY | p.Gly229Val | Q7LG56 |
| HOMO SAPIENS | 2 | 4713 | 4djn | B | 236 | CYS | p.Cys236Phe | Q7LG56 |
| HOMO SAPIENS | 0 | 4713 | 4djn | B | 273 | GLY | p.Gly273Ser | Q7LG56 |
| HOMO SAPIENS | -1 | 4713 | 4djn | B | 282 | MSE | p.Met282Ile | Q7LG56 |
| HOMO SAPIENS | 3 | 4713 | 4djn | B | 284 | GLN | p.Gln284Ter | Q7LG56 |
| HOMO SAPIENS | 3 | 4713 | 4djn | B | 305 | ALA | p.Ala305Ser | Q7LG56 |
| HOMO SAPIENS | -1 | 8974 | 5n7i | A | 500 | ARG | p.Arg500Ter | Q8N4S9 |
| HOMO SAPIENS | 0 | 17072 | 3fxi | B | 299 | ASP | p.Asp299Gly | O00206 |

| | | | | | | | | |
|---|---|---|---|---|---|---|---|---|
| HOMO SAPIENS | 1 | 14355 | 2wwk | T | 55 | ILE | p.Ile34306Asn | Q8WZ42 |
| HOMO SAPIENS | 0 | 14355 | 2wwk | T | 64 | LEU | p.Leu34315Pro | Q8WZ42 |
| HOMO SAPIENS | 2 | 17797 | 1xu1 | S | 72 | ARG | p.Arg72His | O14836 |
| HOMO SAPIENS | -1 | 10836 | 4ztu | A | 39 | GLN | p.Gln39Arg | P54098 |
| HOMO SAPIENS | -1 | 10836 | 4ztu | A | 42 | ARG | p.Arg42Gln | P54098 |
| HOMO SAPIENS | -1 | 10836 | 4ztu | A | 45 | GLN | p.Gln45Arg | P54098 |
| HOMO SAPIENS | -1 | 10836 | 4ztu | A | 52 | GLN | p.Gln52Glu | P54098 |
| HOMO SAPIENS | 0 | 10836 | 4ztu | A | 88 | PHE | p.Phe88Leu | P54098 |
| HOMO SAPIENS | 1 | 10836 | 4ztu | A | 194 | ALA | p.Ala194Val | P54098 |
| HOMO SAPIENS | 1 | 10836 | 4ztu | A | 227 | ARG | p.Arg227Trp | P54098 |
| HOMO SAPIENS | 0 | 10836 | 4ztu | A | 232 | ARG | p.Arg232His | P54098 |
| HOMO SAPIENS | 1 | 10836 | 4ztu | A | 285 | GLN | p.Gln285His | P54098 |
| HOMO SAPIENS | 1 | 10836 | 4ztu | A | 434 | TYR | p.Tyr434His | P54098 |
| HOMO SAPIENS | -1 | 10836 | 4ztu | A | 511 | SER | p.Ser511Asn | P54098 |
| HOMO SAPIENS | 0 | 10836 | 4ztu | A | 533 | CYS | p.Cys533Trp | P54098 |
| HOMO SAPIENS | 1 | 10836 | 4ztu | A | 546 | ARG | p.Arg546Cys | P54098 |
| HOMO SAPIENS | 1 | 10836 | 4ztu | A | 587 | PRO | p.Pro587Leu | P54098 |
| HOMO SAPIENS | 0 | 10836 | 4ztu | A | 603 | MET | p.Met603Thr | P54098 |
| HOMO SAPIENS | 0 | 10836 | 4ztu | A | 623 | LEU | p.Leu623Trp | P54098 |
| HOMO SAPIENS | -1 | 10836 | 4ztu | A | 627 | ARG | p.Arg627Trp | P54098 |
| HOMO SAPIENS | -1 | 10836 | 4ztu | A | 628 | ARG | p.Arg628Pro | P54098 |
| HOMO SAPIENS | 0 | 10836 | 4ztu | A | 635 | PRO | p.Pro635Leu | P54098 |
| HOMO SAPIENS | 0 | 10836 | 4ztu | A | 662 | GLU | p.Glu662Lys | P54098 |
| HOMO SAPIENS | -1 | 10836 | 4ztu | A | 674 | GLY | p.Gly674Asp | P54098 |
| HOMO SAPIENS | -1 | 10836 | 4ztu | A | 722 | ARG | p.Arg722His | P54098 |
| HOMO SAPIENS | -1 | 10836 | 4ztu | A | 737 | GLY | p.Gly737Arg | P54098 |
| HOMO SAPIENS | 0 | 10836 | 4ztu | A | 748 | TRP | p.Trp748Ser | P54098 |
| HOMO SAPIENS | 0 | 10836 | 4ztu | A | 749 | PHE | p.Phe749Ser | P54098 |
| HOMO SAPIENS | 1 | 10836 | 4ztu | A | 807 | ARG | p.Arg807Cys | P54098 |
| HOMO SAPIENS | 1 | 10836 | 4ztu | A | 831 | TYR | p.Tyr831Cys | P54098 |
| HOMO SAPIENS | 8 | 10836 | 4ztu | A | 848 | GLY | p.Gly848Ser | P54098 |
| HOMO SAPIENS | 1 | 10836 | 4ztu | A | 852 | ARG | p.Arg852Cys | P54098 |
| HOMO SAPIENS | 9 | 10836 | 4ztu | A | 853 | ARG | p.Arg853Trp | P54098 |
| HOMO SAPIENS | 1 | 10836 | 4ztu | A | 864 | ASN | p.Asn864Ser | P54098 |
| HOMO SAPIENS | 1 | 10836 | 4ztu | A | 879 | GLN | p.Gln879Arg | P54098 |
| HOMO SAPIENS | 1 | 10836 | 4ztu | A | 914 | THR | p.Thr914Pro | P54098 |
| HOMO SAPIENS | 6 | 10836 | 4ztu | A | 932 | HIS | p.His932Tyr | P54098 |
| HOMO SAPIENS | 2 | 10836 | 4ztu | A | 944 | GLU | p.Glu944Lys | P54098 |
| HOMO SAPIENS | 6 | 10836 | 4ztu | A | 955 | TYR | p.Tyr955Cys | P54098 |
| HOMO SAPIENS | 1 | 10836 | 4ztu | A | 957 | ALA | p.Ala957Ser | P54098 |
| HOMO SAPIENS | 1 | 10836 | 4ztu | A | 957 | ALA | p.Ala957Val | P54098 |
| HOMO SAPIENS | 1 | 10836 | 4ztu | A | 964 | ARG | p.Arg964Cys | P54098 |
| HOMO SAPIENS | 1 | 10836 | 4ztu | A | 966 | LEU | p.Leu966Arg | P54098 |
| HOMO SAPIENS | 0 | 10836 | 4ztu | A | 1051 | GLY | p.Gly1051Arg | P54098 |
| HOMO SAPIENS | 1 | 10836 | 4ztu | A | 1073 | PRO | p.Pro1073Leu | P54098 |
| HOMO SAPIENS | 1 | 10836 | 4ztu | A | 1096 | ARG | p.Arg1096Cys | P54098 |
| HOMO SAPIENS | 1 | 10836 | 4ztu | A | 1096 | ARG | p.Arg1096His | P54098 |
| HOMO SAPIENS | 0 | 10836 | 4ztu | A | 1116 | MET | p.Met1116Thr | P54098 |
| HOMO SAPIENS | 1 | 10836 | 4ztu | A | 1142 | ARG | p.Arg1142Trp | P54098 |
| HOMO SAPIENS | 1 | 10836 | 4ztu | A | 1146 | ARG | p.Arg1146Cys | P54098 |
| HOMO SAPIENS | 1 | 10836 | 4ztu | A | 1157 | ASN | p.Asn1157Ser | P54098 |
| HOMO SAPIENS | -1 | 10836 | 4ztu | A | 1234 | ARG | p.Arg1234Gln | P54098 |
| HOMO SAPIENS | -1 | 10836 | 4ztu | A | 1236 | GLN | p.Gln1236His | P54098 |
| HOMO SAPIENS | 5 | 2666 | 1jph | A | 62 | PRO | p.Pro62Leu | P06132 |

| | | | | | | | | |
|---|---|---|---|---|---|---|---|---|
| HOMO SAPIENS | 5 | 2666 | 1jph | A | 116 | GLN | p.Gln116Ter | P06132 |
| HOMO SAPIENS | 0 | 2666 | 1jph | A | 165 | MET | p.Met165Arg | P06132 |
| HOMO SAPIENS | 6 | 2666 | 1jph | A | 167 | GLU | p.Glu167Lys | P06132 |
| HOMO SAPIENS | 0 | 2666 | 1jph | A | 195 | LEU | p.Leu195Phe | P06132 |
| HOMO SAPIENS | 7 | 2666 | 1jph | A | 253 | LEU | p.Leu253Gln | P06132 |
| HOMO SAPIENS | 0 | 2666 | 1jph | A | 281 | GLY | p.Gly281Glu | P06132 |
| HOMO SAPIENS | 9 | 2666 | 1jph | A | 304 | ASN | p.Asn304Lys | P06132 |
| HOMO SAPIENS | 8 | 2666 | 1jph | A | 311 | TYR | p.Tyr311Cys | P06132 |
| HOMO SAPIENS | 5 | 2666 | 1jph | A | 332 | ARG | p.Arg332His | P06132 |
| HOMO SAPIENS | 1 | 889 | 1m51 | A | 184 | LEU | p.Val184Leu | P35558 |
| HOMO SAPIENS | 0 | 889 | 1m51 | A | 267 | VAL | p.Ile267Val | P35558 |
| HOMO SAPIENS | 2 | 889 | 1m51 | A | 276 | GLU | p.Glu276Lys | P35558 |
| HOMO SAPIENS | 6 | 889 | 1m51 | A | 368 | VAL | p.Val368Ile | P35558 |
| HOMO SAPIENS | 0 | 889 | 1m51 | A | 579 | SER | p.Ser579Arg | P35558 |
| HOMO SAPIENS | 1 | 4084 | 2gjx | A | 28 | GLN | p.Gln28Ter | P06865 |
| HOMO SAPIENS | 0 | 4084 | 2gjx | A | 39 | LEU | p.Leu39Arg | P06865 |
| HOMO SAPIENS | 0 | 4084 | 2gjx | A | 127 | LEU | p.Leu127Arg | P06865 |
| HOMO SAPIENS | 6 | 4084 | 2gjx | A | 137 | ARG | p.Arg137Ter | P06865 |
| HOMO SAPIENS | 6 | 4084 | 2gjx | A | 170 | ARG | p.Arg170Gln | P06865 |
| HOMO SAPIENS | 9 | 4084 | 2gjx | A | 178 | ARG | p.Arg178Cys | P06865 |
| HOMO SAPIENS | 9 | 4084 | 2gjx | A | 178 | ARG | p.Arg178His | P06865 |
| HOMO SAPIENS | 5 | 4084 | 2gjx | A | 180 | TYR | p.Tyr180His | P06865 |
| HOMO SAPIENS | 5 | 4084 | 2gjx | A | 180 | TYR | p.Tyr180Ter | P06865 |
| HOMO SAPIENS | 1 | 4084 | 2gjx | A | 197 | LYS | p.Lys197Thr | P06865 |
| HOMO SAPIENS | 0 | 4084 | 2gjx | A | 200 | VAL | p.Val200Met | P06865 |
| HOMO SAPIENS | 9 | 4084 | 2gjx | A | 204 | HIS | p.His204Arg | P06865 |
| HOMO SAPIENS | 5 | 4084 | 2gjx | A | 210 | SER | p.Ser210Phe | P06865 |
| HOMO SAPIENS | 0 | 4084 | 2gjx | A | 211 | PHE | p.Phe211Ser | P06865 |
| HOMO SAPIENS | 2 | 4084 | 2gjx | A | 250 | GLY | p.Gly250Asp | P06865 |
| HOMO SAPIENS | 9 | 4084 | 2gjx | A | 258 | ASP | p.Asp258His | P06865 |
| HOMO SAPIENS | 0 | 4084 | 2gjx | A | 269 | GLY | p.Gly269Ser | P06865 |
| HOMO SAPIENS | 0 | 4084 | 2gjx | A | 301 | MET | p.Met301Arg | P06865 |
| HOMO SAPIENS | 3 | 4084 | 2gjx | A | 393 | ARG | p.Arg393Ter | P06865 |
| HOMO SAPIENS | 3 | 4084 | 2gjx | A | 393 | ARG | p.Arg393Pro | P06865 |
| HOMO SAPIENS | 0 | 4084 | 2gjx | A | 399 | ASN | p.Asn399Asp | P06865 |
| HOMO SAPIENS | 2 | 4084 | 2gjx | A | 420 | TRP | p.Trp420Cys | P06865 |
| HOMO SAPIENS | 1 | 4084 | 2gjx | A | 434 | PHE | p.Phe434Leu | P06865 |
| HOMO SAPIENS | 0 | 4084 | 2gjx | A | 454 | GLY | p.Gly454Ser | P06865 |
| HOMO SAPIENS | 1 | 4084 | 2gjx | A | 474 | TRP | p.Trp474Cys | P06865 |
| HOMO SAPIENS | 0 | 4084 | 2gjx | A | 479 | ALA | p.Ala479Thr | P06865 |
| HOMO SAPIENS | 5 | 4084 | 2gjx | A | 485 | TRP | p.Trp485Arg | P06865 |
| HOMO SAPIENS | 4 | 4084 | 2gjx | A | 499 | ARG | p.Arg499Cys | P06865 |
| HOMO SAPIENS | 4 | 4084 | 2gjx | A | 499 | ARG | p.Arg499His | P06865 |
| HOMO SAPIENS | 2 | 4084 | 2gjx | A | 504 | ARG | p.Arg504Cys | P06865 |
| HOMO SAPIENS | 2 | 4084 | 2gjx | A | 504 | ARG | p.Arg504His | P06865 |
| HOMO SAPIENS | -1 | 1943 | 1xmj | A | 417 | ASN | p.Asn417Lys | P13569 |
| HOMO SAPIENS | -1 | 1943 | 1xmj | A | 422 | SER | p.Ser422Phe | P13569 |
| HOMO SAPIENS | 1 | 1943 | 1xmj | A | 438 | THR | p.Thr438Ala | P13569 |
| HOMO SAPIENS | 0 | 1943 | 1xmj | A | 455 | ALA | p.Ala455Glu | P13569 |
| HOMO SAPIENS | 4 | 1943 | 1xmj | A | 456 | VAL | p.Val456Ala | P13569 |
| HOMO SAPIENS | 9 | 1943 | 1xmj | A | 458 | GLY | p.Gly458Val | P13569 |
| HOMO SAPIENS | 8 | 1943 | 1xmj | A | 466 | SER | p.Ser466Ter | P13569 |
| HOMO SAPIENS | 0 | 1943 | 1xmj | A | 467 | LEU | p.Leu467Pro | P13569 |
| HOMO SAPIENS | 1 | 1943 | 1xmj | A | 470 | MET | p.Val470Met | P13569 |

| | | | | | | | | |
|---|---|---|---|---|---|---|---|---|
| HOMO SAPIENS | 5 | 1943 | 1xmj | A | 480 | GLY | p.Gly480Cys | P13569 |
| HOMO SAPIENS | 3 | 1943 | 1xmj | A | 492 | SER | p.Ser492Phe | P13569 |
| HOMO SAPIENS | 8 | 1943 | 1xmj | A | 493 | GLN | p.Gln493Ter | P13569 |
| HOMO SAPIENS | 5 | 1943 | 1xmj | A | 506 | ILE | p.Ile506Met | P13569 |
| HOMO SAPIENS | 3 | 1943 | 1xmj | A | 520 | VAL | p.Val520Phe | P13569 |
| HOMO SAPIENS | 1 | 1943 | 1xmj | A | 528 | GLU | p.Glu528Asp | P13569 |
| HOMO SAPIENS | 1 | 1943 | 1xmj | A | 534 | ALA | p.Ala534Glu | P13569 |
| HOMO SAPIENS | 4 | 1943 | 1xmj | A | 542 | GLY | p.Gly542Ter | P13569 |
| HOMO SAPIENS | 6 | 1943 | 1xmj | A | 549 | SER | p.Ser549Asn | P13569 |
| HOMO SAPIENS | 6 | 1943 | 1xmj | A | 549 | SER | p.Ser549Arg | P13569 |
| HOMO SAPIENS | 6 | 1943 | 1xmj | A | 549 | SER | p.Ser549Arg | P13569 |
| HOMO SAPIENS | 6 | 1943 | 1xmj | A | 551 | GLY | p.Gly551Asp | P13569 |
| HOMO SAPIENS | 6 | 1943 | 1xmj | A | 551 | GLY | p.Gly551Ser | P13569 |
| HOMO SAPIENS | 6 | 1943 | 1xmj | A | 553 | GLN | p.Arg553Gln | P13569 |
| HOMO SAPIENS | 6 | 1943 | 1xmj | A | 553 | GLN | p.Arg553Ter | P13569 |
| HOMO SAPIENS | 0 | 1943 | 1xmj | A | 559 | ALA | p.Ala559Thr | P13569 |
| HOMO SAPIENS | 7 | 1943 | 1xmj | A | 560 | ARG | p.Arg560Lys | P13569 |
| HOMO SAPIENS | 4 | 1943 | 1xmj | A | 562 | VAL | p.Val562Ile | P13569 |
| HOMO SAPIENS | 3 | 1943 | 1xmj | A | 563 | TYR | p.Tyr563Asn | P13569 |
| HOMO SAPIENS | 3 | 1943 | 1xmj | A | 569 | TYR | p.Tyr569Asp | P13569 |
| HOMO SAPIENS | 3 | 1943 | 1xmj | A | 585 | GLU | p.Glu585Ter | P13569 |
| HOMO SAPIENS | 1 | 1943 | 1xmj | A | 618 | ILE | p.Ile618Thr | P13569 |
| HOMO SAPIENS | 9 | 1943 | 1xmj | A | 622 | GLY | p.Gly622Asp | P13569 |
| HOMO SAPIENS | -1 | 1943 | 1xmj | A | 648 | ASP | p.Asp648Val | P13569 |
| HOMO SAPIENS | 0 | 1943 | 1xmj | A | 673 | GLY | p.Gly673Ter | P13569 |
| HOMO SAPIENS | 5 | 5111 | 4rec | A | 749 | ARG | p.Arg749Ter | Q9Y2M0 |
| HOMO SAPIENS | -1 | 4580 | 4l0d | B | 2 | GLY | p.Pro2Leu | P35520 |
| HOMO SAPIENS | -1 | 4580 | 4l0d | B | 12 | PRO | p.Pro12Ser | P35520 |
| HOMO SAPIENS | -1 | 4580 | 4l0d | B | 18 | ARG | p.Arg18Cys | P35520 |
| HOMO SAPIENS | 1 | 4580 | 4l0d | B | 49 | PRO | p.Pro49Leu | P35520 |
| HOMO SAPIENS | 2 | 4580 | 4l0d | B | 70 | PRO | p.Pro70Leu | P35520 |
| HOMO SAPIENS | 1 | 4580 | 4l0d | B | 102 | LYS | p.Lys102Gln | P35520 |
| HOMO SAPIENS | 7 | 4580 | 4l0d | B | 114 | ALA | p.Ala114Val | P35520 |
| HOMO SAPIENS | 5 | 4580 | 4l0d | B | 116 | GLY | p.Gly116Arg | P35520 |
| HOMO SAPIENS | 8 | 4580 | 4l0d | B | 121 | ARG | p.Arg121Cys | P35520 |
| HOMO SAPIENS | 8 | 4580 | 4l0d | B | 121 | ARG | p.Arg121His | P35520 |
| HOMO SAPIENS | 5 | 4580 | 4l0d | B | 125 | ARG | p.Arg125Gln | P35520 |
| HOMO SAPIENS | 4 | 4580 | 4l0d | B | 134 | GLY | p.Gly134Arg | P35520 |
| HOMO SAPIENS | 4 | 4580 | 4l0d | B | 139 | GLY | p.Gly139Arg | P35520 |
| HOMO SAPIENS | 0 | 4580 | 4l0d | B | 145 | PRO | p.Pro145Leu | P35520 |
| HOMO SAPIENS | 0 | 4580 | 4l0d | B | 148 | GLY | p.Gly148Arg | P35520 |
| HOMO SAPIENS | 0 | 4580 | 4l0d | B | 168 | VAL | p.Val168Met | P35520 |
| HOMO SAPIENS | 2 | 4580 | 4l0d | B | 191 | THR | p.Thr191Met | P35520 |
| HOMO SAPIENS | 5 | 4580 | 4l0d | B | 234 | ASP | p.Asp234Asn | P35520 |
| HOMO SAPIENS | 0 | 4580 | 4l0d | B | 257 | THR | p.Thr257Met | P35520 |
| HOMO SAPIENS | 4 | 4580 | 4l0d | B | 262 | THR | p.Thr262Met | P35520 |
| HOMO SAPIENS | 0 | 4580 | 4l0d | B | 278 | ILE | p.Ile278Thr | P35520 |
| HOMO SAPIENS | 2 | 4580 | 4l0d | B | 307 | GLY | p.Gly307Ser | P35520 |
| HOMO SAPIENS | 4 | 4580 | 4l0d | B | 320 | VAL | p.Val320Ala | P35520 |
| HOMO SAPIENS | 2 | 4580 | 4l0d | B | 336 | ARG | p.Arg336Cys | P35520 |
| HOMO SAPIENS | 2 | 4580 | 4l0d | B | 336 | ARG | p.Arg336His | P35520 |
| HOMO SAPIENS | 0 | 4580 | 4l0d | B | 351 | GLY | p.Gly351Arg | P35520 |
| HOMO SAPIENS | 0 | 4580 | 4l0d | B | 353 | THR | p.Thr353Met | P35520 |
| HOMO SAPIENS | 0 | 4580 | 4l0d | B | 371 | VAL | p.Val371Met | P35520 |

| | | | | | | | | |
|---|---|---|---|---|---|---|---|---|
| HOMO SAPIENS | 2 | 4580 | 4l0d | B | 379 | ARG | p.Arg379Gln | P35520 |
| HOMO SAPIENS | 2 | 4580 | 4l0d | B | 379 | ARG | p.Arg379Trp | P35520 |
| HOMO SAPIENS | 2 | 4580 | 4l0d | B | 384 | LYS | p.Lys384Glu | P35520 |
| HOMO SAPIENS | 0 | 4580 | 4l0d | B | 422 | PRO | p.Pro422Leu | P35520 |
| HOMO SAPIENS | 0 | 4580 | 4l0d | B | 444 | ASP | p.Asp444Asn | P35520 |
| HOMO SAPIENS | 0 | 4580 | 4l0d | B | 466 | SER | p.Ser466Leu | P35520 |
| HOMO SAPIENS | 0 | 4580 | 4l0d | B | 539 | LEU | p.Leu539Ser | P35520 |
| HOMO SAPIENS | 0 | 4580 | 4l0d | B | 548 | ARG | p.Arg548Gln | P35520 |
| HOMO SAPIENS | 1 | 6536 | 3ulu | A | 412 | LEU | p.Leu412Phe | O15455 |
| HOMO SAPIENS | 1 | 28493 | 3ii6 | X | 814 | ARG | p.Arg814Ter | P49917 |
| HOMO SAPIENS | 1 | 28493 | 3ii6 | X | 857 | ALA | p.Ala857Thr | P49917 |
| HOMO SAPIENS | 9 | 3543 | 2q51 | A | 24 | GLU | p.Glu24Gly | P45381 |
| HOMO SAPIENS | 1 | 3543 | 2q51 | A | 27 | GLY | p.Gly27Arg | P45381 |
| HOMO SAPIENS | 9 | 3543 | 2q51 | A | 71 | ARG | p.Arg71His | P45381 |
| HOMO SAPIENS | -1 | 3543 | 2q51 | A | 82 | MSE | p.Met82Thr | P45381 |
| HOMO SAPIENS | 0 | 3543 | 2q51 | A | 143 | ILE | p.Ile143Phe | P45381 |
| HOMO SAPIENS | 1 | 3543 | 2q51 | A | 152 | CYS | p.Cys152Arg | P45381 |
| HOMO SAPIENS | 2 | 3543 | 2q51 | A | 218 | CYS | p.Cys218Ter | P45381 |
| HOMO SAPIENS | 1 | 3543 | 2q51 | A | 231 | TYR | p.Tyr231Cys | P45381 |
| HOMO SAPIENS | 1 | 3543 | 2q51 | A | 231 | TYR | p.Tyr231Ter | P45381 |
| HOMO SAPIENS | 2 | 3543 | 2q51 | A | 274 | GLY | p.Gly274Arg | P45381 |
| HOMO SAPIENS | 6 | 3543 | 2q51 | A | 285 | GLU | p.Glu285Ala | P45381 |
| HOMO SAPIENS | 5 | 3543 | 2q51 | A | 287 | ALA | p.Ala287Thr | P45381 |
| HOMO SAPIENS | 2 | 3543 | 2q51 | A | 305 | ALA | p.Ala305Glu | P45381 |
| HOMO SAPIENS | 2 | 24687 | 1hfi | A | 25 | SER | p.Ser890Ile | P08603 |
| HOMO SAPIENS | 0 | 1508 | 3wn5 | C | 45 | LEU | p.Leu66Arg | P08637 |
| HOMO SAPIENS | 0 | 1508 | 3wn5 | C | 155 | VAL | p.Phe176Val | P08637 |
| HOMO SAPIENS | 0 | 750 | 5tpa | A | 54 | VAL | p.Val452Met | Q12879 |
| HOMO SAPIENS | 9 | 750 | 5tpa | A | 120 | ARG | p.Arg518Cys | Q12879 |
| HOMO SAPIENS | 0 | 750 | 5tpa | A | 199 | ALA | p.Ala716Thr | Q12879 |
| HOMO SAPIENS | 4 | 1363 | 3hg2 | B | 40 | PRO | p.Pro40Ser | P06280 |
| HOMO SAPIENS | 3 | 1363 | 3hg2 | B | 65 | SER | p.Ser65Thr | P06280 |
| HOMO SAPIENS | 4 | 1363 | 3hg2 | B | 66 | GLU | p.Glu66Gln | P06280 |
| HOMO SAPIENS | 2 | 1363 | 3hg2 | B | 80 | GLY | p.Gly80Asp | P06280 |
| HOMO SAPIENS | 3 | 1363 | 3hg2 | B | 112 | ARG | p.Arg112Cys | P06280 |
| HOMO SAPIENS | 3 | 1363 | 3hg2 | B | 112 | ARG | p.Arg112His | P06280 |
| HOMO SAPIENS | 0 | 1363 | 3hg2 | B | 139 | ASN | p.Asn139Ser | P06280 |
| HOMO SAPIENS | 7 | 1363 | 3hg2 | B | 156 | ALA | p.Ala156Thr | P06280 |
| HOMO SAPIENS | 5 | 1363 | 3hg2 | B | 162 | TRP | p.Trp162Arg | P06280 |
| HOMO SAPIENS | 5 | 1363 | 3hg2 | B | 207 | TYR | p.Tyr207His | P06280 |
| HOMO SAPIENS | 0 | 1363 | 3hg2 | B | 215 | ASN | p.Asn215Ser | P06280 |
| HOMO SAPIENS | 0 | 1363 | 3hg2 | B | 264 | ASP | p.Asp264Val | P06280 |
| HOMO SAPIENS | 5 | 1363 | 3hg2 | B | 266 | ASP | p.Asp266Val | P06280 |
| HOMO SAPIENS | 2 | 1363 | 3hg2 | B | 269 | VAL | p.Val269Ala | P06280 |
| HOMO SAPIENS | 3 | 1363 | 3hg2 | B | 272 | ASN | p.Asn272Ser | P06280 |
| HOMO SAPIENS | 3 | 1363 | 3hg2 | B | 279 | GLN | p.Gln279Glu | P06280 |
| HOMO SAPIENS | 0 | 1363 | 3hg2 | B | 289 | ILE | p.Ile289Val | P06280 |
| HOMO SAPIENS | 0 | 1363 | 3hg2 | B | 296 | MET | p.Met296Val | P06280 |
| HOMO SAPIENS | 0 | 1363 | 3hg2 | B | 296 | MET | p.Met296Ile | P06280 |
| HOMO SAPIENS | 0 | 1363 | 3hg2 | B | 297 | SER | p.Ser297Phe | P06280 |
| HOMO SAPIENS | 2 | 1363 | 3hg2 | B | 301 | ARG | p.Arg301Gln | P06280 |
| HOMO SAPIENS | 3 | 1363 | 3hg2 | B | 327 | GLN | p.Gln327Lys | P06280 |
| HOMO SAPIENS | 0 | 1363 | 3hg2 | B | 328 | GLY | p.Gly328Arg | P06280 |
| HOMO SAPIENS | 0 | 1363 | 3hg2 | B | 328 | GLY | p.Gly328Ala | P06280 |

| | | | | | | | | |
|---|---|---|---|---|---|---|---|---|
| HOMO SAPIENS | 2 | 1363 | 3hg2 | B | 331 | LEU | p.Leu331Phe | P06280 |
| HOMO SAPIENS | 0 | 1363 | 3hg2 | B | 342 | ARG | p.Arg342Gln | P06280 |
| HOMO SAPIENS | 1 | 1363 | 3hg2 | B | 361 | GLY | p.Gly361Arg | P06280 |
| HOMO SAPIENS | 1 | 1363 | 3hg2 | B | 363 | ARG | p.Arg363His | P06280 |
| HOMO SAPIENS | 1 | 1363 | 3hg2 | B | 385 | THR | p.Thr385Ala | P06280 |
| HOMO SAPIENS | 1 | 1363 | 3hg2 | B | 399 | TRP | p.Trp399Ser | P06280 |
| HOMO SAPIENS | 1 | 1363 | 3hg2 | B | 410 | THR | p.Thr410Ala | P06280 |
| HOMO SAPIENS | 2 | 3904 | 3caa | B | 385 | MET | p.Met414Val | P01011 |
| HOMO SAPIENS | 0 | 558 | 5eub | A | 56 | VAL | p.Val56Ile | Q13126 |
| HOMO SAPIENS | 0 | 558 | 5eub | A | 189 | TRP | p.Trp189Leu | Q13126 |
| HOMO SAPIENS | 0 | 558 | 5eub | A | 218 | ALA | p.Ala218Gly | Q13126 |
| HOMO SAPIENS | 1 | 3865 | 1cb5 | B | 443 | ILE | p.Ile443Val | Q13867 |
| HOMO SAPIENS | 2 | 14329 | 4m92 | A | 24 | ILE | p.Ile65Val | Q13454 |
| HOMO SAPIENS | 1 | 7381 | 5kz5 | k | 50 | GLY | p.Gly50Glu | Q9H1K1 |
| HOMO SAPIENS | 0 | 1102 | 4fdh | E | 86 | PRO | p.Pro86Ala | P19099 |
| HOMO SAPIENS | 0 | 1102 | 4fdh | E | 118 | ALA | p.Ala118Thr | P19099 |
| HOMO SAPIENS | 0 | 1102 | 4fdh | E | 159 | PRO | p.Pro159Leu | P19099 |
| HOMO SAPIENS | 0 | 1102 | 4fdh | E | 173 | LYS | p.Lys173Arg | P19099 |
| HOMO SAPIENS | 0 | 1102 | 4fdh | E | 181 | ARG | p.Arg181Trp | P19099 |
| HOMO SAPIENS | 0 | 1102 | 4fdh | E | 185 | THR | p.Thr185Ile | P19099 |
| HOMO SAPIENS | 0 | 1102 | 4fdh | E | 198 | GLU | p.Glu198Asp | P19099 |
| HOMO SAPIENS | 0 | 1102 | 4fdh | E | 214 | HIS | p.His214Asp | P19099 |
| HOMO SAPIENS | 0 | 1102 | 4fdh | E | 251 | LYS | p.Lys251Arg | P19099 |
| HOMO SAPIENS | 0 | 1102 | 4fdh | E | 255 | GLU | p.Glu255Ter | P19099 |
| HOMO SAPIENS | 0 | 1102 | 4fdh | E | 281 | ASN | p.Asn281Ser | P19099 |
| HOMO SAPIENS | 0 | 1102 | 4fdh | E | 320 | ALA | p.Ala320Val | P19099 |
| HOMO SAPIENS | 0 | 1102 | 4fdh | E | 339 | ILE | p.Ile339Thr | P19099 |
| HOMO SAPIENS | 0 | 1102 | 4fdh | E | 341 | ARG | p.Arg341Cys | P19099 |
| HOMO SAPIENS | 2 | 1102 | 4fdh | E | 374 | ARG | p.Arg374Trp | P19099 |
| HOMO SAPIENS | 0 | 1102 | 4fdh | E | 386 | VAL | p.Val386Ala | P19099 |
| HOMO SAPIENS | 0 | 1102 | 4fdh | E | 390 | LEU | p.Leu390Phe | P19099 |
| HOMO SAPIENS | -1 | 1102 | 4fdh | E | 435 | GLY | p.Gly435Ser | P19099 |
| HOMO SAPIENS | 0 | 1102 | 4fdh | E | 461 | LEU | p.Leu461Pro | P19099 |
| HOMO SAPIENS | 0 | 1102 | 4fdh | E | 498 | THR | p.Thr498Ala | P19099 |
| HOMO SAPIENS | 6 | 10292 | 2of5 | F | 128 | GLY | p.Gly128Arg | P78560 |
| HOMO SAPIENS | 6 | 10292 | 2of5 | F | 170 | ARG | p.Arg170Cys | P78560 |
| HOMO SAPIENS | -1 | 18965 | 3ghg | A | 16 | ARG | p.Arg35Cys | P02671 |
| HOMO SAPIENS | -1 | 18965 | 3ghg | A | 16 | ARG | p.Arg35His | P02671 |
| HOMO SAPIENS | -1 | 18965 | 3ghg | A | 283 | PRO | p.Pro302Ala | P02671 |
| HOMO SAPIENS | -1 | 18965 | 3ghg | A | 312 | THR | p.Thr331Ala | P02671 |
| HOMO SAPIENS | -1 | 18965 | 3ghg | A | 381 | SER | p.Ser400Phe | P02671 |
| HOMO SAPIENS | -1 | 18965 | 3ghg | A | 463 | VAL | p.Val482Met | P02671 |
| HOMO SAPIENS | -1 | 18965 | 3ghg | A | 526 | GLU | p.Glu545Val | P02671 |
| HOMO SAPIENS | -1 | 18965 | 3ghg | A | 554 | ARG | p.Arg573Leu | P02671 |
| HOMO SAPIENS | 8 | 3023 | 4ttb | A | 101 | ARG | p.Arg101Trp | Q6PHW0 |
| HOMO SAPIENS | 2 | 3023 | 4ttb | A | 116 | ILE | p.Ile116Thr | Q6PHW0 |
| HOMO SAPIENS | 0 | 3023 | 4ttb | A | 220 | ALA | p.Ala220Thr | Q6PHW0 |
| HOMO SAPIENS | 2 | 3023 | 4ttb | A | 271 | GLU | p.Glu271Lys | Q6PHW0 |
| HOMO SAPIENS | 3 | 33226 | 1ya5 | T | 11 | SER | p.Ser11Ter | O15273 |
| HOMO SAPIENS | 6 | 33226 | 1ya5 | T | 64 | SER | p.Ser64Leu | O15273 |
| HOMO SAPIENS | 6 | 33226 | 1ya5 | T | 70 | ARG | p.Arg70Gln | O15273 |
| HOMO SAPIENS | 3 | 33226 | 1ya5 | T | 75 | GLY | p.Gly75Ser | O15273 |
| HOMO SAPIENS | 0 | 33226 | 1ya5 | T | 87 | ARG | p.Arg87Trp | O15273 |
| HOMO SAPIENS | 2 | 5667 | 3dax | A | 398 | PRO | p.Pro398Ala | P22680 |

| | | | | | | | | |
|---|---|---|---|---|---|---|---|---|
| HOMO SAPIENS | 0 | 12861 | 5mya | A | 464 | ASN | p.Asn464His | Q02763 |
| HOMO SAPIENS | 2 | 12861 | 5mya | A | 486 | VAL | p.Val486Ile | Q02763 |
| HOMO SAPIENS | 0 | 12861 | 5mya | A | 522 | ARG | p.Arg522His | Q02763 |
| HOMO SAPIENS | 2 | 12861 | 5mya | A | 600 | VAL | p.Val600Leu | Q02763 |
| HOMO SAPIENS | 0 | 12861 | 5mya | A | 634 | LEU | p.Leu634Phe | Q02763 |
| HOMO SAPIENS | 0 | 12861 | 5mya | A | 684 | HIS | p.His684Gln | Q02763 |
| HOMO SAPIENS | 0 | 12861 | 5mya | A | 724 | ALA | p.Ala724Thr | Q02763 |
| HOMO SAPIENS | 0 | 7743 | 3qzw | G | 90 | GLY | p.Gly111Ser | P01732 |
| HOMO SAPIENS | 2 | 9498 | 3kci | A | 4267 | ASP | p.Asp4267Glu | O95714 |
| HOMO SAPIENS | 5 | 201 | 3ips | B | 415 | ARG | p.Arg415Gln | Q13133 |
| HOMO SAPIENS | 3 | 229 | 2q50 | A | 68 | LEU | p.Leu68Pro | Q9UBQ7 |
| HOMO SAPIENS | 2 | 229 | 2q50 | A | 99 | ARG | p.Arg99Ter | Q9UBQ7 |
| HOMO SAPIENS | 3 | 229 | 2q50 | A | 160 | GLY | p.Gly160Arg | Q9UBQ7 |
| HOMO SAPIENS | 5 | 229 | 2q50 | A | 165 | GLY | p.Gly165Asp | Q9UBQ7 |
| HOMO SAPIENS | 4 | 229 | 2q50 | A | 302 | ARG | p.Arg302Cys | Q9UBQ7 |
| HOMO SAPIENS | 4 | 229 | 2q50 | A | 302 | ARG | p.Arg302His | Q9UBQ7 |
| HOMO SAPIENS | -1 | 229 | 2q50 | A | 322 | MSE | p.Met322Arg | Q9UBQ7 |
| HOMO SAPIENS | -1 | 4519 | 5clw | A | 40 | PRO | p.Pro40Thr | Q04446 |
| HOMO SAPIENS | 1 | 4519 | 5clw | A | 59 | ILE | p.Ile59Thr | Q04446 |
| HOMO SAPIENS | 0 | 4519 | 5clw | A | 190 | ARG | p.Arg190Gly | Q04446 |
| HOMO SAPIENS | 1 | 4519 | 5clw | A | 224 | LEU | p.Leu224Pro | Q04446 |
| HOMO SAPIENS | 0 | 4519 | 5clw | A | 236 | GLN | p.Gln236His | Q04446 |
| HOMO SAPIENS | 1 | 4519 | 5clw | A | 254 | THR | p.Thr254Ala | Q04446 |
| HOMO SAPIENS | 0 | 4519 | 5clw | A | 257 | PHE | p.Phe257Leu | Q04446 |
| HOMO SAPIENS | 1 | 4519 | 5clw | A | 262 | ARG | p.Arg262Cys | Q04446 |
| HOMO SAPIENS | 3 | 4519 | 5clw | A | 329 | TYR | p.Tyr329Ser | Q04446 |
| HOMO SAPIENS | 0 | 4519 | 5clw | A | 507 | THR | p.Thr507Ala | Q04446 |
| HOMO SAPIENS | 0 | 4519 | 5clw | A | 515 | ARG | p.Arg515Cys | Q04446 |
| HOMO SAPIENS | 0 | 4519 | 5clw | A | 515 | ARG | p.Arg515His | Q04446 |
| HOMO SAPIENS | 0 | 4519 | 5clw | A | 524 | ARG | p.Arg524Gln | Q04446 |
| HOMO SAPIENS | 0 | 4519 | 5clw | A | 524 | ARG | p.Arg524Ter | Q04446 |
| HOMO SAPIENS | 0 | 4519 | 5clw | A | 545 | HIS | p.His545Arg | Q04446 |
| HOMO SAPIENS | 2 | 4519 | 5clw | A | 548 | TRP | p.Trp548Ter | Q04446 |
| HOMO SAPIENS | 0 | 4519 | 5clw | A | 592 | GLU | p.Glu592Ter | Q04446 |
| HOMO SAPIENS | 0 | 4519 | 5clw | A | 628 | HIS | p.His628Arg | Q04446 |
| HOMO SAPIENS | 0 | 4519 | 5clw | A | 637 | ARG | p.Arg637Ter | Q04446 |
| HOMO SAPIENS | 0 | 4519 | 5clw | A | 673 | ALA | p.Ala673Thr | Q04446 |
| HOMO SAPIENS | 1 | 10405 | 3bg3 | D | 507 | VAL | p.Val507Ile | P11498 |
| HOMO SAPIENS | 0 | 10405 | 3bg3 | D | 583 | ARG | p.Arg583Leu | P11498 |
| HOMO SAPIENS | 1 | 10405 | 3bg3 | D | 589 | LYS | p.Lys589Asn | P11498 |
| HOMO SAPIENS | 6 | 10405 | 3bg3 | D | 610 | ALA | p.Ala610Thr | P11498 |
| HOMO SAPIENS | 2 | 10405 | 3bg3 | D | 631 | ARG | p.Arg631Gln | P11498 |
| HOMO SAPIENS | 1 | 10405 | 3bg3 | D | 732 | ARG | p.Arg732Gln | P11498 |
| HOMO SAPIENS | 6 | 10405 | 3bg3 | D | 743 | MET | p.Met743Ile | P11498 |
| HOMO SAPIENS | 0 | 10405 | 3bg3 | D | 750 | THR | p.Thr750Met | P11498 |
| HOMO SAPIENS | -1 | 13751 | 2o8d | B | 350 | ALA | p.Ala350Val | P52701 |
| HOMO SAPIENS | -1 | 13751 | 2o8d | B | 352 | VAL | p.Val352Ile | P52701 |
| HOMO SAPIENS | -1 | 13751 | 2o8d | B | 355 | GLY | p.Gly355Ser | P52701 |
| HOMO SAPIENS | 0 | 13751 | 2o8d | B | 385 | ARG | p.Arg385Trp | P52701 |
| HOMO SAPIENS | 0 | 13751 | 2o8d | B | 390 | ASP | p.Asp390Glu | P52701 |
| HOMO SAPIENS | 0 | 13751 | 2o8d | B | 396 | LEU | p.Leu396Val | P52701 |
| HOMO SAPIENS | 0 | 13751 | 2o8d | B | 449 | LEU | p.Leu449Pro | P52701 |
| HOMO SAPIENS | 0 | 13751 | 2o8d | B | 468 | ARG | p.Arg468His | P52701 |
| HOMO SAPIENS | 0 | 13751 | 2o8d | B | 503 | SER | p.Ser503Cys | P52701 |

| | | | | | | | | |
|---|---|---|---|---|---|---|---|---|
| HOMO SAPIENS | 1 | 13751 | 2o8d | B | 509 | VAL | p.Val509Ala | P52701 |
| HOMO SAPIENS | 0 | 13751 | 2o8d | B | 675 | GLU | p.Glu675Asp | P52701 |
| HOMO SAPIENS | 0 | 13751 | 2o8d | B | 795 | ILE | p.Ile795Thr | P52701 |
| HOMO SAPIENS | 0 | 13751 | 2o8d | B | 850 | TYR | p.Tyr850Ter | P52701 |
| HOMO SAPIENS | 0 | 13751 | 2o8d | B | 878 | VAL | p.Val878Ala | P52701 |
| HOMO SAPIENS | 0 | 13751 | 2o8d | B | 911 | ARG | p.Arg911Ter | P52701 |
| HOMO SAPIENS | 0 | 13751 | 2o8d | B | 922 | ARG | p.Arg922Ter | P52701 |
| HOMO SAPIENS | 0 | 13751 | 2o8d | B | 969 | TYR | p.Tyr969Cys | P52701 |
| HOMO SAPIENS | 0 | 13751 | 2o8d | B | 1005 | ARG | p.Arg1005Ter | P52701 |
| HOMO SAPIENS | 0 | 13751 | 2o8d | B | 1035 | ARG | p.Arg1035Ter | P52701 |
| HOMO SAPIENS | 1 | 13751 | 2o8d | B | 1048 | GLN | p.Gln1048Ter | P52701 |
| HOMO SAPIENS | 1 | 13751 | 2o8d | B | 1051 | VAL | p.Val1051Ile | P52701 |
| HOMO SAPIENS | 0 | 13751 | 2o8d | B | 1054 | ILE | p.Ile1054Phe | P52701 |
| HOMO SAPIENS | 0 | 13751 | 2o8d | B | 1054 | ILE | p.Ile1054Met | P52701 |
| HOMO SAPIENS | 0 | 13751 | 2o8d | B | 1163 | GLU | p.Glu1163Val | P52701 |
| HOMO SAPIENS | 0 | 13751 | 2o8d | B | 1237 | GLU | p.Glu1237Asp | P52701 |
| HOMO SAPIENS | 0 | 13751 | 2o8d | B | 1242 | ARG | p.Arg1242His | P52701 |
| HOMO SAPIENS | -1 | 13751 | 2o8d | B | 1278 | PRO | p.Pro1278Thr | P52701 |
| HOMO SAPIENS | 0 | 13751 | 2o8d | B | 1304 | ARG | p.Arg1304Lys | P52701 |
| HOMO SAPIENS | 0 | 13751 | 2o8d | B | 1317 | HIS | p.His1317Gln | P52701 |
| HOMO SAPIENS | 0 | 13751 | 2o8d | B | 1329 | SER | p.Ser1329Leu | P52701 |
| HOMO SAPIENS | 3 | 13273 | 4g8k | A | 265 | GLU | p.Glu265Ter | Q05823 |
| HOMO SAPIENS | -1 | 11088 | 5uj7 | A | 666 | ARG | p.Arg666Trp | Q13415 |
| HOMO SAPIENS | 4 | 11088 | 5uj7 | A | 720 | ARG | p.Arg720Gln | Q13415 |
| HOMO SAPIENS | 0 | 11088 | 5uj7 | A | 816 | MET | p.Met816Thr | Q13415 |
| HOMO SAPIENS | 1 | 4352 | 5va1 | A | 421 | THR | p.Thr421Met | Q12809 |
| HOMO SAPIENS | 0 | 4352 | 5va1 | A | 451 | PRO | p.Pro451Leu | Q12809 |
| HOMO SAPIENS | 0 | 4352 | 5va1 | A | 470 | ASN | p.Asn470Asp | Q12809 |
| HOMO SAPIENS | 0 | 4352 | 5va1 | A | 490 | ALA | p.Ala490Thr | Q12809 |
| HOMO SAPIENS | 0 | 4352 | 5va1 | A | 491 | VAL | p.Val491Ile | Q12809 |
| HOMO SAPIENS | 1 | 4352 | 5va1 | A | 533 | VAL | p.Val533Gly | Q12809 |
| HOMO SAPIENS | 0 | 4352 | 5va1 | A | 552 | LEU | p.Leu552Ser | Q12809 |
| HOMO SAPIENS | 0 | 4352 | 5va1 | A | 558 | ALA | p.Ala558Pro | Q12809 |
| HOMO SAPIENS | 1 | 4352 | 5va1 | A | 561 | ALA | p.Ala561Val | Q12809 |
| HOMO SAPIENS | 0 | 4352 | 5va1 | A | 572 | GLY | p.Gly572Arg | Q12809 |
| HOMO SAPIENS | -1 | 4352 | 5va1 | A | 582 | ARG | p.Arg582Cys | Q12809 |
| HOMO SAPIENS | 0 | 4352 | 5va1 | A | 584 | GLY | p.Gly584Ser | Q12809 |
| HOMO SAPIENS | 0 | 4352 | 5va1 | A | 588 | ASN | p.Asn588Lys | Q12809 |
| HOMO SAPIENS | 0 | 4352 | 5va1 | A | 593 | ILE | p.Ile593Arg | Q12809 |
| HOMO SAPIENS | 1 | 4352 | 5va1 | A | 627 | PHE | p.Phe627Leu | Q12809 |
| HOMO SAPIENS | 0 | 4352 | 5va1 | A | 630 | VAL | p.Val630Leu | Q12809 |
| HOMO SAPIENS | 0 | 4352 | 5va1 | A | 652 | TYR | p.Tyr652Ter | Q12809 |
| HOMO SAPIENS | 0 | 4352 | 5va1 | A | 711 | ILE | p.Ile711Val | Q12809 |
| HOMO SAPIENS | 0 | 4352 | 5va1 | A | 735 | SER | p.Ser735Leu | Q12809 |
| HOMO SAPIENS | 0 | 4352 | 5va1 | A | 744 | ARG | p.Arg744Ter | Q12809 |
| HOMO SAPIENS | 0 | 4352 | 5va1 | A | 784 | ARG | p.Arg784Trp | Q12809 |
| HOMO SAPIENS | 1 | 4352 | 5va1 | A | 820 | GLY | p.Gly820Arg | Q12809 |
| HOMO SAPIENS | 5 | 4352 | 5va1 | A | 822 | VAL | p.Val822Met | Q12809 |
| HOMO SAPIENS | 1 | 4352 | 5va1 | A | 863 | ARG | p.Arg863Ter | Q12809 |
| HOMO SAPIENS | 5 | 303 | 2obv | A | 55 | ALA | p.Ala55Asp | Q00266 |
| HOMO SAPIENS | 7 | 303 | 2obv | A | 259 | ALA | p.Ala259Gly | Q00266 |
| HOMO SAPIENS | 9 | 303 | 2obv | A | 264 | ARG | p.Arg264His | Q00266 |
| HOMO SAPIENS | 9 | 303 | 2obv | A | 264 | ARG | p.Arg264Cys | Q00266 |
| HOMO SAPIENS | 0 | 303 | 2obv | A | 305 | LEU | p.Leu305Pro | Q00266 |

| | | | | | | | | |
|---|---|---|---|---|---|---|---|---|
| HOMO SAPIENS | 7 | 303 | 2obv | A | 322 | ILE | p.Ile322Met | Q00266 |
| HOMO SAPIENS | 0 | 303 | 2obv | A | 336 | GLY | p.Gly336Arg | Q00266 |
| HOMO SAPIENS | 0 | 303 | 2obv | A | 357 | PRO | p.Pro357Leu | Q00266 |
| HOMO SAPIENS | 0 | 2077 | 4zxb | E | 462 | ASN | p.Asn489Ser | P06213 |
| HOMO SAPIENS | -1 | 2077 | 4zxb | E | 519 | ASP | p.Asp546Glu | P06213 |
| HOMO SAPIENS | 2 | 2077 | 4zxb | E | 819 | THR | p.Thr858Ala | P06213 |
| HOMO SAPIENS | 0 | 2077 | 4zxb | E | 907 | ASP | p.Asp946Glu | P06213 |
| HOMO SAPIENS | 7 | 3274 | 1wsv | B | 14 | HIS | p.His42Arg | P48728 |
| HOMO SAPIENS | 0 | 3274 | 1wsv | B | 19 | GLY | p.Gly47Arg | P48728 |
| HOMO SAPIENS | 4 | 3274 | 1wsv | B | 45 | ARG | p.Arg73Cys | P48728 |
| HOMO SAPIENS | 1 | 3274 | 1wsv | B | 49 | SER | p.Ser77Leu | P48728 |
| HOMO SAPIENS | 3 | 3274 | 1wsv | B | 241 | GLY | p.Gly269Asp | P48728 |
| HOMO SAPIENS | 7 | 3274 | 1wsv | B | 248 | ASP | p.Asp276His | P48728 |
| HOMO SAPIENS | 0 | 3274 | 1wsv | B | 272 | MET | p.Met300Val | P48728 |
| HOMO SAPIENS | 4 | 3274 | 1wsv | B | 292 | ARG | p.Arg320His | P48728 |
| HOMO SAPIENS | 0 | 3274 | 1wsv | B | 354 | ARG | p.Arg382Gln | P48728 |
| HOMO SAPIENS | -1 | 435 | 3uyt | A | 44 | THR | p.Thr44Ala | P48730 |
| HOMO SAPIENS | 1 | 4698 | 5ife | C | 666 | LYS | p.Lys666Asn | O75533 |
| HOMO SAPIENS | 1 | 16917 | 1apy | C | 37 | GLY | p.Gly60Asp | P20933 |
| HOMO SAPIENS | 0 | 16917 | 1apy | C | 49 | SER | p.Ser72Pro | P20933 |
| HOMO SAPIENS | 0 | 16917 | 1apy | C | 84 | ARG | p.Arg107Ter | P20933 |
| HOMO SAPIENS | 0 | 16917 | 1apy | C | 126 | SER | p.Thr149Ser | P20933 |
| HOMO SAPIENS | 0 | 16917 | 1apy | C | 135 | TRP | p.Trp158Ter | P20933 |
| HOMO SAPIENS | 0 | 16917 | 1apy | C | 140 | CYS | p.Cys163Ser | P20933 |
| HOMO SAPIENS | 0 | 16917 | 1apy | C | 145 | TRP | p.Trp168Ter | P20933 |
| HOMO SAPIENS | 0 | 10413 | 4frw | A | 185 | THR | p.Thr185Met | Q96NY8 |
| HOMO SAPIENS | 0 | 5217 | 5n6w | G | 96 | TRP | p.Trp96Arg | O15537 |
| HOMO SAPIENS | 1 | 5217 | 5n6w | G | 102 | ARG | p.Arg102Trp | O15537 |
| HOMO SAPIENS | 0 | 5217 | 5n6w | G | 109 | GLY | p.Gly109Arg | O15537 |
| HOMO SAPIENS | -1 | 5217 | 5n6w | G | 223 | CYS | p.Cys223Arg | O15537 |
| HOMO SAPIENS | 0 | 9485 | 4z6y | E | 138 | ALA | p.Ala138Asp | Q9P0N9 |
| HOMO SAPIENS | 3 | 4504 | 4mwt | B | 411 | GLY | p.Gly439Ser | P10619 |
| HOMO SAPIENS | 3 | 4504 | 4mwt | B | 412 | PHE | p.Phe440Val | P10619 |
| HOMO SAPIENS | 6 | 6391 | 4kxy | A | 318 | ARG | p.Arg318Cys | P29401 |
| HOMO SAPIENS | 6 | 19546 | 2bx6 | A | 87 | THR | p.Thr87Ile | O75695 |
| HOMO SAPIENS | 1 | 19546 | 2bx6 | A | 118 | ARG | p.Arg118His | O75695 |
| HOMO SAPIENS | 1 | 19546 | 2bx6 | A | 201 | ILE | p.Ile201Val | O75695 |
| HOMO SAPIENS | 1 | 19546 | 2bx6 | A | 282 | ARG | p.Arg282Trp | O75695 |
| HOMO SAPIENS | 1 | 5265 | 5usr | H | 68 | ARG | p.Arg68Leu | Q9HD34 |
| HOMO SAPIENS | 2 | 1012 | 4nyi | S | 569 | LEU | p.Leu569Val | Q07889 |
| HOMO SAPIENS | 2 | 1012 | 4nyi | S | 655 | PRO | p.Pro655Leu | Q07889 |
| HOMO SAPIENS | 4 | 1012 | 4nyi | S | 663 | ILE | p.Ile663Met | Q07889 |
| HOMO SAPIENS | 2 | 1012 | 4nyi | S | 702 | TYR | p.Tyr702Cys | Q07889 |
| HOMO SAPIENS | 2 | 1012 | 4nyi | S | 708 | ALA | p.Ala708Thr | Q07889 |
| HOMO SAPIENS | 4 | 1012 | 4nyi | S | 733 | ILE | p.Ile733Phe | Q07889 |
| HOMO SAPIENS | 1 | 1848 | 3g1n | A | 4013 | ARG | p.Arg4013Trp | Q7Z6Z7 |
| HOMO SAPIENS | 1 | 1848 | 3g1n | A | 4187 | ARG | p.Arg4187Cys | Q7Z6Z7 |
| HOMO SAPIENS | 1 | 77 | 4pod | A | 41 | CYS | p.Cys79Tyr | P60174 |
| HOMO SAPIENS | 9 | 77 | 4pod | A | 104 | GLU | p.Glu142Asp | P60174 |
| HOMO SAPIENS | 1 | 77 | 4pod | A | 122 | GLY | p.Gly160Arg | P60174 |
| HOMO SAPIENS | -1 | 8672 | 3thz | A | 8 | THR | p.Thr8Met | P43246 |
| HOMO SAPIENS | 0 | 8672 | 3thz | A | 19 | PHE | p.Phe19Leu | P43246 |
| HOMO SAPIENS | 0 | 8672 | 3thz | A | 23 | PHE | p.Phe23Leu | P43246 |
| HOMO SAPIENS | 0 | 8672 | 3thz | A | 96 | ARG | p.Arg96His | P43246 |

| | | | | | | | | |
|---|---|---|---|---|---|---|---|---|
| HOMO SAPIENS | 0 | 8672 | 3thz | A | 102 | VAL | p.Val102Ile | P43246 |
| HOMO SAPIENS | 0 | 8672 | 3thz | A | 106 | ARG | p.Arg106Lys | P43246 |
| HOMO SAPIENS | 0 | 8672 | 3thz | A | 127 | ASN | p.Asn127Ser | P43246 |
| HOMO SAPIENS | 0 | 8672 | 3thz | A | 128 | LEU | p.Leu128Val | P43246 |
| HOMO SAPIENS | 0 | 8672 | 3thz | A | 163 | VAL | p.Val163Gly | P43246 |
| HOMO SAPIENS | 0 | 8672 | 3thz | A | 169 | ILE | p.Ile169Val | P43246 |
| HOMO SAPIENS | 0 | 8672 | 3thz | A | 315 | GLY | p.Gly315Asp | P43246 |
| HOMO SAPIENS | -1 | 8672 | 3thz | A | 322 | GLY | p.Gly322Asp | P43246 |
| HOMO SAPIENS | 0 | 8672 | 3thz | A | 331 | ASN | p.Asn331Asp | P43246 |
| HOMO SAPIENS | 1 | 8672 | 3thz | A | 390 | LEU | p.Leu390Phe | P43246 |
| HOMO SAPIENS | 0 | 8672 | 3thz | A | 419 | GLN | p.Gln419Lys | P43246 |
| HOMO SAPIENS | 0 | 8672 | 3thz | A | 447 | PHE | p.Phe447Leu | P43246 |
| HOMO SAPIENS | 0 | 8672 | 3thz | A | 524 | ARG | p.Arg524Pro | P43246 |
| HOMO SAPIENS | 0 | 8672 | 3thz | A | 539 | PHE | p.Phe539Leu | P43246 |
| HOMO SAPIENS | -1 | 8672 | 3thz | A | 546 | LYS | p.Lys546Asn | P43246 |
| HOMO SAPIENS | 0 | 8672 | 3thz | A | 564 | THR | p.Thr564Ala | P43246 |
| HOMO SAPIENS | 0 | 8672 | 3thz | A | 577 | ILE | p.Ile577Thr | P43246 |
| HOMO SAPIENS | 0 | 8672 | 3thz | A | 588 | TYR | p.Tyr588Ter | P43246 |
| HOMO SAPIENS | 0 | 8672 | 3thz | A | 622 | PRO | p.Pro622Leu | P43246 |
| HOMO SAPIENS | 0 | 8672 | 3thz | A | 629 | GLN | p.Gln629Arg | P43246 |
| HOMO SAPIENS | 0 | 8672 | 3thz | A | 636 | ALA | p.Ala636Pro | P43246 |
| HOMO SAPIENS | 2 | 8672 | 3thz | A | 642 | VAL | p.Val642Ile | P43246 |
| HOMO SAPIENS | 1 | 8672 | 3thz | A | 654 | ASP | p.Asp654Glu | P43246 |
| HOMO SAPIENS | 0 | 8672 | 3thz | A | 662 | GLN | p.Gln662His | P43246 |
| HOMO SAPIENS | 2 | 8672 | 3thz | A | 681 | GLN | p.Gln681His | P43246 |
| HOMO SAPIENS | 0 | 8672 | 3thz | A | 707 | CYS | p.Cys707Ser | P43246 |
| HOMO SAPIENS | 0 | 8672 | 3thz | A | 809 | GLU | p.Glu809Lys | P43246 |
| HOMO SAPIENS | 0 | 8672 | 3thz | A | 813 | MET | p.Met813Val | P43246 |
| HOMO SAPIENS | 1 | 8672 | 3thz | A | 834 | ALA | p.Ala834Thr | P43246 |
| HOMO SAPIENS | 0 | 8672 | 3thz | A | 849 | LEU | p.Leu849Val | P43246 |
| HOMO SAPIENS | -1 | 8672 | 3thz | A | 860 | SER | p.Ser860Leu | P43246 |
| HOMO SAPIENS | -1 | 8672 | 3thz | A | 934 | THR | p.Thr934Lys | P43246 |
| HOMO SAPIENS | 2 | 292 | 1i3k | A | 34 | ASN | p.Asn34Ser | Q14376 |
| HOMO SAPIENS | 7 | 292 | 1i3k | A | 90 | GLY | p.Gly90Glu | Q14376 |
| HOMO SAPIENS | 2 | 292 | 1i3k | A | 103 | ASP | p.Asp103Gly | Q14376 |
| HOMO SAPIENS | 2 | 292 | 1i3k | A | 136 | TYR | p.Tyr136Ter | Q14376 |
| HOMO SAPIENS | 0 | 292 | 1i3k | A | 169 | ARG | p.Arg169Trp | Q14376 |
| HOMO SAPIENS | 0 | 292 | 1i3k | A | 183 | LEU | p.Leu183Pro | Q14376 |
| HOMO SAPIENS | 0 | 292 | 1i3k | A | 302 | GLY | p.Gly302Asp | Q14376 |
| HOMO SAPIENS | 0 | 292 | 1i3k | A | 313 | LEU | p.Leu313Met | Q14376 |
| HOMO SAPIENS | 1 | 2084 | 5vc8 | A | 295 | GLN | p.Gln295Leu | O96028 |
| HOMO SAPIENS | -1 | 10157 | 5wrw | A | 7 | GLY | p.Gly7Ala | O76094 |
| HOMO SAPIENS | -1 | 10157 | 5wrw | A | 7 | GLY | p.Gly7Arg | O76094 |
| HOMO SAPIENS | -1 | 10157 | 5wrw | A | 8 | GLY | p.Gly8Ala | O76094 |
| HOMO SAPIENS | 0 | 3113 | 3in5 | A | 29 | GLU | p.Glu29Lys | Q9UBT6 |
| HOMO SAPIENS | 5 | 4546 | 5in3 | B | 44 | VAL | p.Val44Leu | P07902 |
| HOMO SAPIENS | -1 | 4546 | 5in3 | B | 51 | ARG | p.Arg51Leu | P07902 |
| HOMO SAPIENS | -1 | 4546 | 5in3 | B | 62 | LEU | p.Leu62Met | P07902 |
| HOMO SAPIENS | 4 | 4546 | 5in3 | B | 66 | PRO | p.Pro66His | P07902 |
| HOMO SAPIENS | 2 | 4546 | 5in3 | B | 74 | LEU | p.Leu74Pro | P07902 |
| HOMO SAPIENS | 5 | 4546 | 5in3 | B | 97 | ASN | p.Asn97Ser | P07902 |
| HOMO SAPIENS | 1 | 4546 | 5in3 | B | 135 | SER | p.Ser135Leu | P07902 |
| HOMO SAPIENS | 5 | 4546 | 5in3 | B | 138 | THR | p.Thr138Met | P07902 |
| HOMO SAPIENS | 7 | 4546 | 5in3 | B | 139 | LEU | p.Leu139Pro | P07902 |

| | | | | | | | | |
|---|---|---|---|---|---|---|---|---|
| HOMO SAPIENS | 2 | 4546 | 5in3 | B | 142 | MET | p.Met142Lys | P07902 |
| HOMO SAPIENS | 1 | 4546 | 5in3 | B | 143 | SER | p.Ser143Leu | P07902 |
| HOMO SAPIENS | 2 | 4546 | 5in3 | B | 148 | ARG | p.Arg148Gly | P07902 |
| HOMO SAPIENS | 2 | 4546 | 5in3 | B | 148 | ARG | p.Arg148Gln | P07902 |
| HOMO SAPIENS | 5 | 4546 | 5in3 | B | 183 | PRO | p.Pro183Thr | P07902 |
| HOMO SAPIENS | 5 | 4546 | 5in3 | B | 185 | PRO | p.Pro185His | P07902 |
| HOMO SAPIENS | 5 | 4546 | 5in3 | B | 186 | HIS | p.His186Tyr | P07902 |
| HOMO SAPIENS | 7 | 4546 | 5in3 | B | 188 | GLN | p.Gln188Arg | P07902 |
| HOMO SAPIENS | 5 | 4546 | 5in3 | B | 195 | LEU | p.Leu195Pro | P07902 |
| HOMO SAPIENS | 4 | 4546 | 5in3 | B | 201 | ARG | p.Arg201His | P07902 |
| HOMO SAPIENS | 4 | 4546 | 5in3 | B | 201 | ARG | p.Arg201Cys | P07902 |
| HOMO SAPIENS | 4 | 4546 | 5in3 | B | 203 | GLU | p.Glu203Lys | P07902 |
| HOMO SAPIENS | 2 | 4546 | 5in3 | B | 209 | TYR | p.Tyr209Cys | P07902 |
| HOMO SAPIENS | 5 | 4546 | 5in3 | B | 223 | ARG | p.Arg223Ser | P07902 |
| HOMO SAPIENS | 5 | 4546 | 5in3 | B | 226 | LEU | p.Leu226Pro | P07902 |
| HOMO SAPIENS | 5 | 4546 | 5in3 | B | 226 | LEU | p.Leu226Val | P07902 |
| HOMO SAPIENS | 2 | 4546 | 5in3 | B | 227 | LEU | p.Leu227Pro | P07902 |
| HOMO SAPIENS | 8 | 4546 | 5in3 | B | 258 | ARG | p.Arg258Cys | P07902 |
| HOMO SAPIENS | 1 | 4546 | 5in3 | B | 265 | PRO | p.Pro265Ala | P07902 |
| HOMO SAPIENS | 8 | 4546 | 5in3 | B | 268 | THR | p.Thr268Asn | P07902 |
| HOMO SAPIENS | 7 | 4546 | 5in3 | B | 278 | ILE | p.Ile278Asn | P07902 |
| HOMO SAPIENS | 5 | 4546 | 5in3 | B | 285 | LYS | p.Lys285Asn | P07902 |
| HOMO SAPIENS | 1 | 4546 | 5in3 | B | 319 | HIS | p.His319Gln | P07902 |
| HOMO SAPIENS | 4 | 4546 | 5in3 | B | 321 | HIS | p.His321Tyr | P07902 |
| HOMO SAPIENS | 4 | 4546 | 5in3 | B | 325 | PRO | p.Pro325Leu | P07902 |
| HOMO SAPIENS | 2 | 4546 | 5in3 | B | 328 | ARG | p.Arg328His | P07902 |
| HOMO SAPIENS | 2 | 4546 | 5in3 | B | 329 | SER | p.Ser329Phe | P07902 |
| HOMO SAPIENS | 5 | 4546 | 5in3 | B | 334 | LYS | p.Lys334Arg | P07902 |
| HOMO SAPIENS | 6 | 1290 | 5uav | C | 119 | ARG | p.Arg119Gly | P32322 |
| HOMO SAPIENS | 6 | 1290 | 5uav | C | 119 | ARG | p.Arg119His | P32322 |
| HOMO SAPIENS | 3 | 1290 | 5uav | C | 251 | ARG | p.Arg251His | P32322 |
| HOMO SAPIENS | 7 | 1290 | 5uav | C | 266 | ARG | p.Arg266Gln | P32322 |
| HOMO SAPIENS | 0 | 9618 | 2r7e | A | 22 | GLY | p.Gly41Cys | P00451 |
| HOMO SAPIENS | 2 | 9618 | 2r7e | A | 113 | GLU | p.Glu132Asp | P00451 |
| HOMO SAPIENS | -1 | 9618 | 2r7e | A | 336 | ARG | p.Arg355Ter | P00451 |
| HOMO SAPIENS | 0 | 9618 | 2r7e | A | 372 | ARG | p.Arg391His | P00451 |
| HOMO SAPIENS | 0 | 9618 | 2r7e | A | 373 | SER | p.Ser392Leu | P00451 |
| HOMO SAPIENS | 0 | 9618 | 2r7e | A | 373 | SER | p.Ser392Pro | P00451 |
| HOMO SAPIENS | 3 | 9618 | 2r7e | A | 386 | ILE | p.Ile405Ser | P00451 |
| HOMO SAPIENS | 6 | 9618 | 2r7e | A | 390 | GLU | p.Glu409Gly | P00451 |
| HOMO SAPIENS | 2 | 9618 | 2r7e | A | 412 | LEU | p.Leu431Phe | P00451 |
| HOMO SAPIENS | 5 | 9618 | 2r7e | A | 425 | LYS | p.Lys444Arg | P00451 |
| HOMO SAPIENS | 0 | 9618 | 2r7e | A | 484 | ARG | p.Arg503His | P00451 |
| HOMO SAPIENS | 2 | 9618 | 2r7e | A | 531 | ARG | p.Arg550His | P00451 |
| HOMO SAPIENS | 3 | 9618 | 2r7e | A | 535 | SER | p.Ser554Gly | P00451 |
| HOMO SAPIENS | 2 | 9618 | 2r7e | A | 593 | ARG | p.Arg612Cys | P00451 |
| HOMO SAPIENS | 0 | 9618 | 2r7e | A | 704 | ALA | p.Ala723Thr | P00451 |
| HOMO SAPIENS | 0 | 9618 | 2r7e | A | 720 | GLU | p.Glu739Lys | P00451 |
| HOMO SAPIENS | 1 | 13085 | 3ud1 | A | 1075 | ILE | p.Thr1075Ile | P16157 |
| HOMO SAPIENS | -1 | 3289 | 2fy4 | A | 2 | ALA | p.Ala120Thr | P28329 |
| HOMO SAPIENS | 7 | 3289 | 2fy4 | A | 18 | VAL | p.Val136Met | P28329 |
| HOMO SAPIENS | 5 | 3289 | 2fy4 | A | 92 | LEU | p.Leu210Pro | P28329 |
| HOMO SAPIENS | 2 | 3289 | 2fy4 | A | 93 | PRO | p.Pro211Ala | P28329 |
| HOMO SAPIENS | 4 | 3289 | 2fy4 | A | 131 | LEU | p.Leu249Val | P28329 |

| | | | | | | | | |
|---|---|---|---|---|---|---|---|---|
| HOMO SAPIENS | 2 | 3289 | 2fy4 | A | 181 | PRO | p.Pro299Leu | P28329 |
| HOMO SAPIENS | 0 | 3289 | 2fy4 | A | 218 | ILE | p.Ile336Thr | P28329 |
| HOMO SAPIENS | 2 | 3289 | 2fy4 | A | 261 | ASP | p.Asp379His | P28329 |
| HOMO SAPIENS | 0 | 3289 | 2fy4 | A | 302 | ARG | p.Arg420Cys | P28329 |
| HOMO SAPIENS | 5 | 3289 | 2fy4 | A | 323 | GLU | p.Glu441Lys | P28329 |
| HOMO SAPIENS | 0 | 3289 | 2fy4 | A | 343 | MET | p.Val461Met | P28329 |
| HOMO SAPIENS | 2 | 3289 | 2fy4 | A | 364 | ARG | p.Arg482Gly | P28329 |
| HOMO SAPIENS | 3 | 3289 | 2fy4 | A | 380 | SER | p.Ser498Leu | P28329 |
| HOMO SAPIENS | 6 | 3289 | 2fy4 | A | 388 | VAL | p.Val506Leu | P28329 |
| HOMO SAPIENS | 7 | 3289 | 2fy4 | A | 429 | HIS | p.His547Gln | P28329 |
| HOMO SAPIENS | 8 | 3289 | 2fy4 | A | 442 | ARG | p.Arg560His | P28329 |
| HOMO SAPIENS | 3 | 3289 | 2fy4 | A | 443 | ARG | p.Arg561Gln | P28329 |
| HOMO SAPIENS | 0 | 4137 | 3hhr | B | 38 | CYS | p.Cys56Ter | P10912 |
| HOMO SAPIENS | 0 | 4137 | 3hhr | B | 43 | ARG | p.Arg61Ter | P10912 |
| HOMO SAPIENS | 0 | 4137 | 3hhr | B | 44 | GLU | p.Glu62Lys | P10912 |
| HOMO SAPIENS | 0 | 4137 | 3hhr | B | 96 | PHE | p.Phe114Ser | P10912 |
| HOMO SAPIENS | 5 | 4137 | 3hhr | B | 131 | PRO | p.Pro149Gln | P10912 |
| HOMO SAPIENS | 0 | 4137 | 3hhr | B | 144 | VAL | p.Val162Ile | P10912 |
| HOMO SAPIENS | 0 | 4137 | 3hhr | B | 150 | HIS | p.His168Gln | P10912 |
| HOMO SAPIENS | 0 | 4137 | 3hhr | B | 152 | ASP | p.Asp170His | P10912 |
| HOMO SAPIENS | 0 | 4137 | 3hhr | B | 153 | ILE | p.Ile171Thr | P10912 |
| HOMO SAPIENS | 0 | 4137 | 3hhr | B | 154 | GLN | p.Gln172Pro | P10912 |
| HOMO SAPIENS | 0 | 4137 | 3hhr | B | 155 | VAL | p.Val173Gly | P10912 |
| HOMO SAPIENS | 0 | 4137 | 3hhr | B | 217 | ARG | p.Arg235Ter | P10912 |
| HOMO SAPIENS | 5 | 4137 | 3hhr | B | 224 | GLU | p.Glu242Asp | P10912 |
| HOMO SAPIENS | 5 | 4137 | 3hhr | B | 224 | GLU | p.Glu242Ter | P10912 |
| HOMO SAPIENS | 6 | 2493 | 5bpb | A | 69 | HIS | p.His69Tyr | Q9ULV1 |
| HOMO SAPIENS | 6 | 2493 | 5bpb | A | 105 | MET | p.Met105Val | Q9ULV1 |
| HOMO SAPIENS | -1 | 2493 | 5bpb | A | 168 | PRO | p.Pro168Ser | Q9ULV1 |
| HOMO SAPIENS | 0 | 32123 | 4ccz | A | 314 | ASP | p.Asp314Asn | Q99707 |
| HOMO SAPIENS | 0 | 32123 | 4ccz | A | 345 | VAL | p.Val345Phe | Q99707 |
| HOMO SAPIENS | 1 | 32123 | 4ccz | A | 585 | ARG | p.Arg585Ter | Q99707 |
| HOMO SAPIENS | 4 | 4618 | 2ag4 | A | 26 | PRO | p.Pro55Leu | P17900 |
| HOMO SAPIENS | 4 | 4618 | 2ag4 | A | 30 | VAL | p.Ile59Val | P17900 |
| HOMO SAPIENS | 2 | 4618 | 2ag4 | A | 40 | VAL | p.Met69Val | P17900 |
| HOMO SAPIENS | 2 | 4618 | 2ag4 | A | 124 | VAL | p.Val153Ala | P17900 |
| HOMO SAPIENS | 8 | 4618 | 2ag4 | A | 140 | ARG | p.Arg169Pro | P17900 |
| HOMO SAPIENS | 5 | 1082 | 1hk0 | X | 14 | ARG | p.Arg15Cys | P07320 |
| HOMO SAPIENS | 7 | 1082 | 1hk0 | X | 23 | PRO | p.Pro24Thr | P07320 |
| HOMO SAPIENS | 6 | 1082 | 1hk0 | X | 36 | ARG | p.Arg37Ser | P07320 |
| HOMO SAPIENS | 5 | 1082 | 1hk0 | X | 55 | TYR | p.Tyr56Ter | P07320 |
| HOMO SAPIENS | 3 | 1082 | 1hk0 | X | 58 | ARG | p.Arg59His | P07320 |
| HOMO SAPIENS | 7 | 1082 | 1hk0 | X | 88 | HIS | p.His88Gln | P07320 |
| HOMO SAPIENS | 6 | 1082 | 1hk0 | X | 95 | ARG | p.Arg95Ser | P07320 |
| HOMO SAPIENS | 5 | 1082 | 1hk0 | X | 126 | VAL | p.Val126Met | P07320 |
| HOMO SAPIENS | 8 | 1082 | 1hk0 | X | 157 | TRP | p.Trp157Ter | P07320 |
| HOMO SAPIENS | 0 | 22879 | 3j0a | B | 644 | ILE | p.Ile644Phe | O60602 |
| HOMO SAPIENS | 8 | 6 | 2o23 | B | 122 | LEU | p.Leu122Val | Q99714 |
| HOMO SAPIENS | 1 | 6 | 2o23 | B | 130 | ARG | p.Arg130Cys | Q99714 |
| HOMO SAPIENS | 1 | 6 | 2o23 | B | 247 | ASN | p.Asn247Ser | Q99714 |
| HOMO SAPIENS | 0 | 6 | 2o23 | B | 249 | GLU | p.Glu249Gln | Q99714 |
| HOMO SAPIENS | 1 | 12910 | 4hl4 | A | 67 | ARG | p.Arg67Ter | Q96BZ9 |
| HOMO SAPIENS | 8 | 16105 | 1lmj | A | 7 | GLU | p.Glu1073Lys | P35555 |
| HOMO SAPIENS | 8 | 16105 | 1lmj | A | 8 | CYS | p.Cys1074Arg | P35555 |

| | | | | | | | | |
|---|---|---|---|---|---|---|---|---|
| HOMO SAPIENS | 9 | 16105 | 1lmj | A | 51 | CYS | p.Cys1117Tyr | P35555 |
| HOMO SAPIENS | 9 | 16105 | 1lmj | A | 63 | CYS | p.Cys1129Tyr | P35555 |
| HOMO SAPIENS | 9 | 16105 | 1lmj | A | 65 | ASN | p.Asn1131Tyr | P35555 |
| HOMO SAPIENS | 4 | 16105 | 1lmj | A | 71 | ARG | p.Arg1137Pro | P35555 |
| HOMO SAPIENS | 1 | 16105 | 1lmj | A | 82 | PRO | p.Pro1148Ala | P35555 |
| HOMO SAPIENS | 0 | 4795 | 3s5f | A | 106 | LEU | p.Leu106Ser | Q16595 |
| HOMO SAPIENS | 0 | 4795 | 3s5f | A | 130 | GLY | p.Gly130Val | Q16595 |
| HOMO SAPIENS | 0 | 4795 | 3s5f | A | 154 | ILE | p.Ile154Phe | Q16595 |
| HOMO SAPIENS | -1 | 4795 | 3s5f | A | 209 | ASP | p.Asp209Gly | Q16595 |
| HOMO SAPIENS | -1 | 2884 | 4lli | A | 1655 | ASP | p.Asp1655Glu | Q9Y4I1 |
| HOMO SAPIENS | 0 | 1786 | 7pck | C | 56P | TYR | p.Tyr71Ter | P43235 |
| HOMO SAPIENS | 0 | 1786 | 7pck | C | 64P | GLY | p.Gly79Glu | P43235 |
| HOMO SAPIENS | 7 | 38286 | 2l53 | B | 1914 | ARG | p.Arg1914Gly | Q14524 |
| HOMO SAPIENS | 1 | 38286 | 2l53 | B | 1924 | ALA | p.Ala1924Thr | Q14524 |
| HOMO SAPIENS | 0 | 31747 | 3ksy | A | 25 | PRO | p.Pro25Ser | Q07889 |
| HOMO SAPIENS | 3 | 31747 | 3ksy | A | 37 | THR | p.Thr37Ala | Q07889 |
| HOMO SAPIENS | 0 | 31747 | 3ksy | A | 54 | GLN | p.Gln54His | Q07889 |
| HOMO SAPIENS | -1 | 31747 | 3ksy | A | 190 | ASP | p.Asp190Glu | Q07889 |
| HOMO SAPIENS | 2 | 31747 | 3ksy | A | 266 | THR | p.Thr266Lys | Q07889 |
| HOMO SAPIENS | 1 | 31747 | 3ksy | A | 269 | MET | p.Met269Arg | Q07889 |
| HOMO SAPIENS | 1 | 31747 | 3ksy | A | 340 | PRO | p.Pro340Ser | Q07889 |
| HOMO SAPIENS | 1 | 31747 | 3ksy | A | 423 | ASN | p.Asn423Lys | Q07889 |
| HOMO SAPIENS | 1 | 31747 | 3ksy | A | 424 | GLU | p.Glu424Lys | Q07889 |
| HOMO SAPIENS | 1 | 31747 | 3ksy | A | 432 | TRP | p.Trp432Arg | Q07889 |
| HOMO SAPIENS | 1 | 31747 | 3ksy | A | 552 | ARG | p.Arg552Gly | Q07889 |
| HOMO SAPIENS | 1 | 31747 | 3ksy | A | 552 | ARG | p.Arg552Ser | Q07889 |
| HOMO SAPIENS | 3 | 7529 | 6jdw | A | 93 | ILE | p.Ile93Met | P50440 |
| HOMO SAPIENS | 2 | 7529 | 6jdw | A | 110 | GLN | p.Gln110His | P50440 |
| HOMO SAPIENS | 5 | 7529 | 6jdw | A | 149 | TRP | p.Trp149Ter | P50440 |
| HOMO SAPIENS | 0 | 7529 | 6jdw | A | 169 | ARG | p.Arg169Ter | P50440 |
| HOMO SAPIENS | 0 | 7529 | 6jdw | A | 260 | ARG | p.Arg260Ter | P50440 |
| HOMO SAPIENS | 3 | 7529 | 6jdw | A | 344 | ILE | p.Ile344Val | P50440 |
| HOMO SAPIENS | 5 | 7529 | 6jdw | A | 418 | LEU | p.Leu418Ile | P50440 |
| HOMO SAPIENS | 1 | 14788 | 1eak | A | 101 | ARG | p.Arg101His | P08253 |
| HOMO SAPIENS | 0 | 14788 | 1eak | A | 383 | ASP | p.Asp383Glu | P08253 |
| HOMO SAPIENS | 7 | 14788 | 1eak | A | 404 | GLN | p.Glu404Lys | P08253 |
| HOMO SAPIENS | 2 | 14788 | 1eak | A | 447 | ALA | p.Ala447Val | P08253 |
| HOMO SAPIENS | 0 | 1255 | 5xtc | p | 8 | PRO | p.Pro8His | Q9Y6M9 |
| HOMO SAPIENS | 0 | 1255 | 5xtc | p | 146 | PRO | p.Pro146Ser | Q9Y6M9 |
| HOMO SAPIENS | 3 | 6721 | 1a22 | A | 25 | PHE | p.Phe51Tyr | P01241 |
| HOMO SAPIENS | 2 | 6721 | 1a22 | A | 95 | SER | p.Ser121Arg | P01241 |
| HOMO SAPIENS | 2 | 6721 | 1a22 | A | 110 | VAL | p.Val136Ile | P01241 |
| HOMO SAPIENS | 0 | 1380 | 5i6z | A | 605 | LYS | p.Lys605Asn | P31645 |
| HOMO SAPIENS | 0 | 941 | 1qki | D | 48 | ILE | p.Ile48Thr | P11413 |
| HOMO SAPIENS | 3 | 941 | 1qki | D | 68 | VAL | p.Val68Met | P11413 |
| HOMO SAPIENS | 2 | 941 | 1qki | D | 126 | ASN | p.Asn126Asp | P11413 |
| HOMO SAPIENS | 8 | 941 | 1qki | D | 163 | GLY | p.Gly163Ser | P11413 |
| HOMO SAPIENS | 6 | 941 | 1qki | D | 165 | ASN | p.Asn165Asp | P11413 |
| HOMO SAPIENS | 4 | 941 | 1qki | D | 181 | ASP | p.Asp181Val | P11413 |
| HOMO SAPIENS | 4 | 941 | 1qki | D | 182 | ARG | p.Arg182Trp | P11413 |
| HOMO SAPIENS | 5 | 941 | 1qki | D | 188 | SER | p.Ser188Phe | P11413 |
| HOMO SAPIENS | 9 | 941 | 1qki | D | 198 | ARG | p.Arg198Cys | P11413 |
| HOMO SAPIENS | 8 | 941 | 1qki | D | 213 | VAL | p.Val213Leu | P11413 |
| HOMO SAPIENS | 5 | 941 | 1qki | D | 282 | ASP | p.Asp282His | P11413 |

| | | | | | | | | |
|---|---|---|---|---|---|---|---|---|
| HOMO SAPIENS | 7 | 941 | 1qki | D | 285 | ARG | p.Arg285His | P11413 |
| HOMO SAPIENS | 5 | 941 | 1qki | D | 291 | VAL | p.Val291Met | P11413 |
| HOMO SAPIENS | 8 | 941 | 1qki | D | 323 | LEU | p.Leu323Pro | P11413 |
| HOMO SAPIENS | 3 | 941 | 1qki | D | 335 | ALA | p.Ala335Thr | P11413 |
| HOMO SAPIENS | 6 | 941 | 1qki | D | 350 | ASP | p.Asp350His | P11413 |
| HOMO SAPIENS | 5 | 941 | 1qki | D | 393 | ARG | p.Arg393His | P11413 |
| HOMO SAPIENS | 5 | 941 | 1qki | D | 454 | ARG | p.Arg454Cys | P11413 |
| HOMO SAPIENS | 2 | 941 | 1qki | D | 459 | LEU | p.Arg459Leu | P11413 |
| HOMO SAPIENS | 4 | 941 | 1qki | D | 463 | ARG | p.Arg463His | P11413 |
| HOMO SAPIENS | 4 | 2483 | 1tij | A | 68 | LEU | p.Leu94Gln | P01034 |
| HOMO SAPIENS | 2 | 20632 | 3v5b | A | 331 | ARG | p.Arg331Pro | P02545 |
| HOMO SAPIENS | 2 | 20632 | 3v5b | A | 374 | HIS | p.His374Gln | P02545 |
| HOMO SAPIENS | 4 | 13929 | 5xtd | J | 75 | ARG | p.Arg75His | Q16795 |
| HOMO SAPIENS | 3 | 13929 | 5xtd | J | 243 | VAL | p.Val243Ala | Q16795 |
| HOMO SAPIENS | 1 | 13929 | 5xtd | J | 294 | PRO | p.Pro294Leu | Q16795 |
| HOMO SAPIENS | 2 | 13929 | 5xtd | J | 321 | ARG | p.Arg321Pro | Q16795 |
| HOMO SAPIENS | 0 | 9411 | 5thl | A | 41 | ARG | p.Gly41Arg | P54577 |
| HOMO SAPIENS | 0 | 9411 | 5thl | A | 196 | GLU | p.Glu196Lys | P54577 |
| HOMO SAPIENS | 0 | 9411 | 5thl | A | 316 | ALA | p.Ala316Ser | P54577 |
| HOMO SAPIENS | 0 | 9411 | 5thl | A | 339 | ALA | p.Ala339Thr | P54577 |
| HOMO SAPIENS | 4 | 16273 | 2e8n | A | 46 | THR | p.Thr940Ile | P29317 |
| HOMO SAPIENS | 7 | 16273 | 2e8n | A | 54 | GLY | p.Gly948Trp | P29317 |
| HOMO SAPIENS | 6 | 16273 | 2e8n | A | 65 | ALA | p.Ala959Thr | P29317 |
| HOMO SAPIENS | 7 | 30387 | 2k9y | A | 541 | VAL | p.Val541Met | P29317 |
| HOMO SAPIENS | 0 | 6961 | 4nha | A | 843 | ILE | p.Ile853Val | Q9UPY3 |
| HOMO SAPIENS | -1 | 6961 | 4nha | A | 862 | ALA | p.Ala872Thr | Q9UPY3 |
| HOMO SAPIENS | 3 | 15708 | 4pzg | A | 16 | ARG | p.Arg16Ter | Q9Y5X0 |
| HOMO SAPIENS | 4 | 15708 | 4pzg | A | 51 | ARG | p.Arg51Gln | Q9Y5X0 |
| HOMO SAPIENS | 3 | 30166 | 4o00 | A | 44 | ARG | p.Arg14257Ter | Q8WZ42 |
| HOMO SAPIENS | 2 | 30166 | 4o00 | A | 54 | ARG | p.Arg14267His | Q8WZ42 |
| HOMO SAPIENS | 8 | 11158 | 4hca | A | 275 | TRP | p.Trp274Arg | P23771 |
| HOMO SAPIENS | 2 | 4328 | 4e0s | A | 1476 | ARG | p.Arg1476Ter | P01031 |
| HOMO SAPIENS | 0 | 11462 | 1dx5 | I | 455 | ALA | p.Ala473Val | P07204 |
| HOMO SAPIENS | -1 | 11150 | 2fic | A | 45 | LYS | p.Lys35Asn | O00499 |
| HOMO SAPIENS | 6 | 11150 | 2fic | A | 161 | ASP | p.Asp151Asn | O00499 |
| HOMO SAPIENS | 8 | 27382 | 4mqv | B | 17 | ARG | p.Arg17Ter | Q9NZC9 |
| HOMO SAPIENS | -1 | 37991 | 1q0p | A | 231 | GLN | p.Gln256Ter | P00751 |
| HOMO SAPIENS | 1 | 443 | 5xte | U | 215 | LEU | p.Leu215Phe | P08574 |
| HOMO SAPIENS | 8 | 21471 | 5y4e | A | 1461 | ASN | p.Thr692Ala | Q01484 |
| HOMO SAPIENS | -1 | 35443 | 3um3 | B | 129 | ASP | p.Asp129Val | Q9H444 |
| HOMO SAPIENS | -1 | 35443 | 3um3 | B | 161 | GLU | p.Glu161Lys | Q9H444 |
| HOMO SAPIENS | 6 | 28781 | 2n2a | B | 111 | ILE | p.Ile654Val | P04626 |
| HOMO SAPIENS | 7 | 28781 | 2n2a | B | 112 | ILE | p.Ile655Val | P04626 |
| HOMO SAPIENS | -1 | 3346 | 4ztz | C | 61 | GLY | p.Gly61Arg | Q9UHN1 |
| HOMO SAPIENS | 4 | 3346 | 4ztz | C | 103 | GLY | p.Gly103Ser | Q9UHN1 |
| HOMO SAPIENS | 4 | 3346 | 4ztz | C | 133 | HIS | p.His133Tyr | Q9UHN1 |
| HOMO SAPIENS | -1 | 3346 | 4ztz | C | 153 | LEU | p.Leu153Val | Q9UHN1 |
| HOMO SAPIENS | -1 | 3346 | 4ztz | C | 169 | ALA | p.Ala169Thr | Q9UHN1 |
| HOMO SAPIENS | 7 | 3346 | 4ztz | C | 369 | ARG | p.Arg369Gly | Q9UHN1 |
| HOMO SAPIENS | 4 | 3346 | 4ztz | C | 386 | ASP | p.Asp386Glu | Q9UHN1 |
| HOMO SAPIENS | 4 | 3346 | 4ztz | C | 451 | GLY | p.Gly451Glu | Q9UHN1 |
| HOMO SAPIENS | 4 | 2131 | 5fo9 | E | 1549 | LEU | p.Leu1549Met | P01024 |
| HOMO SAPIENS | 2 | 2131 | 5fo9 | E | 1619 | SER | p.Ser1619Arg | P01024 |
| HOMO SAPIENS | 3 | 12662 | 4jhp | C | 75 | ILE | p.Ile75Val | Q92834 |

| | | | | | | | | |
|---|---|---|---|---|---|---|---|---|
| HOMO SAPIENS | 1 | 12662 | 4jhp | C | 173 | GLY | p.Gly173Arg | Q92834 |
| HOMO SAPIENS | 0 | 12662 | 4jhp | C | 184 | GLN | p.Gln184His | Q92834 |
| HOMO SAPIENS | 3 | 12662 | 4jhp | C | 262 | ALA | p.Ala262Gly | Q92834 |
| HOMO SAPIENS | 3 | 12662 | 4jhp | C | 302 | CYS | p.Cys302Tyr | Q92834 |
| HOMO SAPIENS | 5 | 1093 | 2jg9 | A | 186 | GLN | p.Gln208Ter | P02745 |
| HOMO SAPIENS | 0 | 13181 | 4r4l | A | 12 | MET | p.Met13Ile | Q13976 |
| HOMO SAPIENS | 8 | 25939 | 4iyp | A | 106 | ARG | p.Arg106Gln | P78318 |
| HOMO SAPIENS | 1 | 217 | 2xsx | A | 71 | SER | p.Asn71Ser | P13929 |
| HOMO SAPIENS | 0 | 217 | 2xsx | A | 76 | ALA | p.Ala76Pro | P13929 |
| HOMO SAPIENS | 1 | 217 | 2xsx | A | 85 | ALA | p.Val85Ala | P13929 |
| HOMO SAPIENS | 8 | 217 | 2xsx | A | 156 | GLY | p.Gly156Asp | P13929 |
| HOMO SAPIENS | 5 | 217 | 2xsx | A | 332 | VAL | p.Val332Ile | P13929 |
| HOMO SAPIENS | 7 | 217 | 2xsx | A | 374 | GLY | p.Gly374Glu | P13929 |
| HOMO SAPIENS | 1 | 5508 | 3szm | A | 682 | THR | p.Thr682Asn | Q8NEM0 |
| HOMO SAPIENS | 0 | 5508 | 3szm | A | 715 | TRP | p.Trp715Ter | Q8NEM0 |
| HOMO SAPIENS | 1 | 5508 | 3szm | A | 761 | VAL | p.Ala761Val | Q8NEM0 |
| HOMO SAPIENS | 1 | 5508 | 3szm | A | 828 | PRO | p.Pro828Ser | Q8NEM0 |
| HOMO SAPIENS | 4 | 5508 | 3szm | A | 829 | GLU | p.Glu829Asp | Q8NEM0 |
| HOMO SAPIENS | 2 | 16353 | 2q0z | X | 171 | GLY | p.Gly1978Ala | O75643 |
| HOMO SAPIENS | 2 | 16353 | 2q0z | X | 188 | ALA | p.Ala1995Thr | O75643 |
| HOMO SAPIENS | 0 | 4127 | 3swr | A | 860 | GLY | p.Gly860Arg | P26358 |
| HOMO SAPIENS | -1 | 4127 | 3swr | A | 1122 | SER | p.Ser1122Cys | P26358 |
| HOMO SAPIENS | 0 | 4127 | 3swr | A | 1216 | ASP | p.Asp1216Glu | P26358 |
| HOMO SAPIENS | 0 | 6382 | 3o96 | A | 17 | GLU | p.Glu17Lys | P31749 |
| HOMO SAPIENS | 6 | 14314 | 5xtc | Q | 71 | PRO | p.Pro71Leu | O75306 |
| HOMO SAPIENS | 2 | 5886 | 4fao | E | 22 | ARG | p.Arg40His | Q13705 |
| HOMO SAPIENS | 1 | 2534 | 3mp4 | A | 37 | GLU | p.Glu37Ter | P35914 |
| HOMO SAPIENS | 3 | 2534 | 3mp4 | A | 41 | MET | p.Arg41Gln | P35914 |
| HOMO SAPIENS | 1 | 2534 | 3mp4 | A | 70 | VAL | p.Val70Leu | P35914 |
| HOMO SAPIENS | 2 | 2196 | 3ifq | A | 142 | ARG | p.Arg142His | P14923 |
| HOMO SAPIENS | 2 | 2196 | 3ifq | A | 153 | ASP | p.Asp153Glu | P14923 |
| HOMO SAPIENS | 3 | 2196 | 3ifq | A | 176 | ARG | p.Arg176Gln | P14923 |
| HOMO SAPIENS | 3 | 2196 | 3ifq | A | 265 | ARG | p.Arg265His | P14923 |
| HOMO SAPIENS | 2 | 2196 | 3ifq | A | 301 | GLU | p.Glu301Gly | P14923 |
| HOMO SAPIENS | 0 | 2196 | 3ifq | A | 456 | VAL | p.Val456Ile | P14923 |
| HOMO SAPIENS | 3 | 2196 | 3ifq | A | 603 | VAL | p.Val603Leu | P14923 |
| HOMO SAPIENS | 0 | 2196 | 3ifq | A | 648 | VAL | p.Val648Ile | P14923 |
| HOMO SAPIENS | 1 | 8746 | 1cgj | I | 32 | PRO | p.Pro55Ser | P00995 |
| HOMO SAPIENS | 1 | 8746 | 1cgj | I | 44 | ARG | p.Arg67His | P00995 |
| HOMO SAPIENS | 1 | 8746 | 1cgj | I | 44 | ARG | p.Arg67Cys | P00995 |
| HOMO SAPIENS | 3 | 3212 | 1syq | A | 105 | ARG | p.Arg105Ter | P18206 |
| HOMO SAPIENS | 3 | 3212 | 1syq | A | 126 | PHE | p.Phe126Leu | P18206 |
| HOMO SAPIENS | 4 | 10028 | 3eto | A | 1609 | ARG | p.Arg1608His | P46531 |
| HOMO SAPIENS | 1 | 10028 | 3eto | A | 1672 | VAL | p.Val1671Ile | P46531 |
| HOMO SAPIENS | 5 | 10028 | 3eto | A | 1692 | GLN | p.Gln1691His | P46531 |
| HOMO SAPIENS | 3 | 3749 | 5xtc | Z | 70 | GLY | p.Gly70Ter | O43676 |
| HOMO SAPIENS | 2 | 17039 | 3brv | B | 57 | GLU | p.Glu57Lys | Q9Y6K9 |
| HOMO SAPIENS | 8 | 3589 | 2fh3 | A | 433 | VAL | p.Val460Leu | P06396 |
| HOMO SAPIENS | 1 | 3589 | 2fh3 | A | 454 | ARG | p.Arg481Cys | P06396 |
| HOMO SAPIENS | 4 | 3589 | 2fh3 | A | 498 | ARG | p.Arg525His | P06396 |
| HOMO SAPIENS | 2 | 3589 | 2fh3 | A | 520 | TYR | p.Tyr547Cys | P06396 |
| HOMO SAPIENS | 1 | 3589 | 2fh3 | A | 529 | GLY | p.Gly556Arg | P06396 |
| HOMO SAPIENS | 2 | 3589 | 2fh3 | A | 536 | THR | p.Thr563Ser | P06396 |
| HOMO SAPIENS | 2 | 3589 | 2fh3 | A | 579 | VAL | p.Val606Met | P06396 |

| | | | | | | | | |
|---|---|---|---|---|---|---|---|---|
| HOMO SAPIENS | 1 | 3589 | 2fh3 | A | 589 | THR | p.Thr616Met | P06396 |
| HOMO SAPIENS | 5 | 3589 | 2fh3 | A | 668 | THR | p.Thr695Met | P06396 |
| HOMO SAPIENS | 0 | 148 | 1lcl | A | 130 | ILE | p.Ile130Ser | Q05315 |
| HOMO SAPIENS | -1 | 24727 | 4fj3 | P | 237 | ALA | p.Ala237Thr | P04049 |
| HOMO SAPIENS | 0 | 24727 | 4fj3 | P | 257 | SER | p.Ser257Leu | P04049 |
| HOMO SAPIENS | 0 | 24727 | 4fj3 | P | 261 | PRO | p.Pro261Ala | P04049 |
| HOMO SAPIENS | 0 | 2949 | 5xtb | B | 77 | ARG | p.Arg77Gly | O00217 |
| HOMO SAPIENS | 0 | 2949 | 5xtb | B | 79 | PRO | p.Pro79Leu | O00217 |
| HOMO SAPIENS | 0 | 2949 | 5xtb | B | 100 | ALA | p.Ala100Val | O00217 |
| HOMO SAPIENS | 1 | 2949 | 5xtb | B | 115 | LYS | p.Lys115Glu | O00217 |
| HOMO SAPIENS | 0 | 2949 | 5xtb | B | 138 | ARG | p.Arg138His | O00217 |
| HOMO SAPIENS | 1 | 3959 | 5k5s | B | 25 | ARG | p.Arg25Ter | P41180 |
| HOMO SAPIENS | 4 | 3959 | 5k5s | B | 39 | PRO | p.Pro39Ala | P41180 |
| HOMO SAPIENS | 0 | 3959 | 5k5s | B | 47 | LYS | p.Lys47Asn | P41180 |
| HOMO SAPIENS | 2 | 3959 | 5k5s | B | 62 | ARG | p.Arg62Met | P41180 |
| HOMO SAPIENS | 2 | 3959 | 5k5s | B | 66 | ARG | p.Arg66Cys | P41180 |
| HOMO SAPIENS | 4 | 3959 | 5k5s | B | 90 | ASN | p.Asn90Thr | P41180 |
| HOMO SAPIENS | 1 | 3959 | 5k5s | B | 116 | ALA | p.Ala116Thr | P41180 |
| HOMO SAPIENS | 1 | 3959 | 5k5s | B | 118 | ASN | p.Asn118Lys | P41180 |
| HOMO SAPIENS | -1 | 3959 | 5k5s | B | 125 | LEU | p.Leu125Pro | P41180 |
| HOMO SAPIENS | -1 | 3959 | 5k5s | B | 127 | GLU | p.Glu127Ala | P41180 |
| HOMO SAPIENS | -1 | 3959 | 5k5s | B | 128 | PHE | p.Phe128Leu | P41180 |
| HOMO SAPIENS | -1 | 3959 | 5k5s | B | 131 | CYS | p.Cys131Trp | P41180 |
| HOMO SAPIENS | 1 | 3959 | 5k5s | B | 138 | THR | p.Thr138Met | P41180 |
| HOMO SAPIENS | 0 | 3959 | 5k5s | B | 143 | GLY | p.Gly143Glu | P41180 |
| HOMO SAPIENS | 5 | 3959 | 5k5s | B | 151 | THR | p.Thr151Met | P41180 |
| HOMO SAPIENS | 4 | 3959 | 5k5s | B | 180 | PHE | p.Phe180Cys | P41180 |
| HOMO SAPIENS | 0 | 3959 | 5k5s | B | 185 | ARG | p.Arg185Gln | P41180 |
| HOMO SAPIENS | 0 | 3959 | 5k5s | B | 185 | ARG | p.Arg185Ter | P41180 |
| HOMO SAPIENS | 2 | 3959 | 5k5s | B | 191 | GLU | p.Glu191Lys | P41180 |
| HOMO SAPIENS | 2 | 3959 | 5k5s | B | 227 | ARG | p.Arg227Leu | P41180 |
| HOMO SAPIENS | 2 | 3959 | 5k5s | B | 250 | GLU | p.Glu250Lys | P41180 |
| HOMO SAPIENS | 6 | 3959 | 5k5s | B | 297 | GLU | p.Glu297Lys | P41180 |
| HOMO SAPIENS | 1 | 3959 | 5k5s | B | 336 | LYS | p.Lys336Asn | P41180 |
| HOMO SAPIENS | 6 | 3959 | 5k5s | B | 429 | HIS | p.His429Tyr | P41180 |
| HOMO SAPIENS | 0 | 3959 | 5k5s | B | 445 | THR | p.Thr445Ala | P41180 |
| HOMO SAPIENS | 3 | 3959 | 5k5s | B | 509 | GLY | p.Gly509Arg | P41180 |
| HOMO SAPIENS | 1 | 3959 | 5k5s | B | 553 | GLY | p.Gly553Arg | P41180 |
| HOMO SAPIENS | 3 | 3959 | 5k5s | B | 592 | ASN | p.Asn592Ser | P41180 |
| HOMO SAPIENS | -1 | 3959 | 5k5s | B | 604 | GLU | p.Glu604Lys | P41180 |
| HOMO SAPIENS | 4 | 2564 | 5l0c | C | 975 | ARG | p.Arg975Trp | P18206 |
| HOMO SAPIENS | 4 | 2564 | 5l0c | C | 975 | ARG | p.Arg975Leu | P18206 |
| HOMO SAPIENS | 1 | 5695 | 5i8r | A | 294 | ARG | p.Arg294Gln | P17405 |
| HOMO SAPIENS | 0 | 5695 | 5i8r | A | 485 | ALA | p.Ala485Val | P17405 |
| HOMO SAPIENS | 1 | 5695 | 5i8r | A | 506 | GLY | p.Gly506Arg | P17405 |
| HOMO SAPIENS | -1 | 1978 | 3avs | A | 1060 | HIS | p.His1060Leu | O15550 |
| HOMO SAPIENS | 0 | 11173 | 4v0c | B | 362 | LYS | p.Lys362Arg | P51787 |
| HOMO SAPIENS | 3 | 11173 | 4v0c | B | 381 | CYS | p.Cys381Tyr | P51787 |
| HOMO SAPIENS | 7 | 11173 | 4v0c | B | 518 | ARG | p.Arg518Gly | P51787 |
| HOMO SAPIENS | 2 | 11173 | 4v0c | B | 525 | ALA | p.Ala525Thr | P51787 |
| HOMO SAPIENS | 0 | 11173 | 4v0c | B | 530 | GLN | p.Gln530Ter | P51787 |
| HOMO SAPIENS | -1 | 11173 | 4v0c | B | 536 | TYR | p.Tyr536Ter | P51787 |
| HOMO SAPIENS | 2 | 1113 | 5xtd | N | 60 | ARG | p.Arg60Ter | Q9UI09 |
| HOMO SAPIENS | 3 | 6061 | 2mmz | A | 148 | LEU | p.Leu148Ser | Q86SX6 |

| | | | | | | | | |
|---|---|---|---|---|---|---|---|---|
| HOMO SAPIENS | -1 | 26081 | 3eu7 | A | 837 | GLU | p.Glu837Ter | Q86YC2 |
| HOMO SAPIENS | 0 | 26081 | 3eu7 | A | 864 | PRO | p.Pro864Ser | Q86YC2 |
| HOMO SAPIENS | 5 | 26081 | 3eu7 | A | 891 | CYS | p.Cys891Trp | Q86YC2 |
| HOMO SAPIENS | 0 | 26081 | 3eu7 | A | 932 | VAL | p.Val932Met | Q86YC2 |
| HOMO SAPIENS | 5 | 26081 | 3eu7 | A | 961 | LEU | p.Leu961Met | Q86YC2 |
| HOMO SAPIENS | 5 | 26081 | 3eu7 | A | 1038 | TRP | p.Trp1038Ter | Q86YC2 |
| HOMO SAPIENS | -1 | 26081 | 3eu7 | A | 1086 | ARG | p.Arg1086Ter | Q86YC2 |
| HOMO SAPIENS | 0 | 26081 | 3eu7 | A | 1122 | ASP | p.Asp1122Glu | Q86YC2 |
| HOMO SAPIENS | 5 | 26081 | 3eu7 | A | 1158 | HIS | p.His1158Gln | Q86YC2 |
| HOMO SAPIENS | 5 | 26081 | 3eu7 | A | 1183 | TYR | p.Tyr1183Ter | Q86YC2 |
| HOMO SAPIENS | 4 | 199 | 1qo5 | C | 45 | ARG | p.Arg46Trp | P05062 |
| HOMO SAPIENS | 5 | 199 | 1qo5 | C | 147 | TRP | p.Trp148Arg | P05062 |
| HOMO SAPIENS | 0 | 199 | 1qo5 | C | 149 | ALA | p.Ala150Pro | P05062 |
| HOMO SAPIENS | 0 | 199 | 1qo5 | C | 174 | ALA | p.Ala175Asp | P05062 |
| HOMO SAPIENS | 0 | 199 | 1qo5 | C | 334 | ASN | p.Asn335Lys | P05062 |
| HOMO SAPIENS | 0 | 199 | 1qo5 | C | 337 | ALA | p.Ala338Val | P05062 |
| HOMO SAPIENS | 5 | 199 | 1qo5 | C | 342 | TYR | p.Tyr343His | P05062 |
| HOMO SAPIENS | 0 | 6910 | 4zht | B | 59 | ARG | p.Arg59Ter | Q9Y223 |
| HOMO SAPIENS | 4 | 6910 | 4zht | B | 129 | ARG | p.Arg129Ter | Q9Y223 |
| HOMO SAPIENS | 4 | 6910 | 4zht | B | 129 | ARG | p.Arg129Gln | Q9Y223 |
| HOMO SAPIENS | 1 | 6910 | 4zht | B | 162 | ARG | p.Arg162Cys | Q9Y223 |
| HOMO SAPIENS | 0 | 6910 | 4zht | B | 176 | ASP | p.Asp176Val | Q9Y223 |
| HOMO SAPIENS | 0 | 6910 | 4zht | B | 216 | VAL | p.Val216Ala | Q9Y223 |
| HOMO SAPIENS | 0 | 6910 | 4zht | B | 246 | ARG | p.Arg246Trp | Q9Y223 |
| HOMO SAPIENS | 0 | 6910 | 4zht | B | 263 | ARG | p.Arg263Leu | Q9Y223 |
| HOMO SAPIENS | 0 | 6910 | 4zht | B | 266 | ARG | p.Arg266Gln | Q9Y223 |
| HOMO SAPIENS | 0 | 6910 | 4zht | B | 266 | ARG | p.Arg266Trp | Q9Y223 |
| HOMO SAPIENS | 0 | 6910 | 4zht | B | 277 | ARG | p.Arg277Cys | Q9Y223 |
| HOMO SAPIENS | 0 | 6910 | 4zht | B | 378 | ASP | p.Asp378Tyr | Q9Y223 |
| HOMO SAPIENS | 3 | 211 | 5kxi | C | 262 | VAL | p.Val287Leu | P17787 |
| HOMO SAPIENS | 0 | 211 | 5kxi | C | 369 | PHE | p.Phe478Leu | P17787 |
| HOMO SAPIENS | 0 | 2454 | 5iy9 | V | null | PHE | p.Phe99Ser | P19447 |
| HOMO SAPIENS | 0 | 2454 | 5iy9 | V | null | ARG | p.Arg109Ter | P19447 |
| HOMO SAPIENS | 0 | 2454 | 5iy9 | V | null | THR | p.Thr119Pro | P19447 |
| HOMO SAPIENS | 0 | 2454 | 5iy9 | V | 425 | ARG | p.Arg425Ter | P19447 |
| HOMO SAPIENS | 1 | 2454 | 5iy9 | V | 545 | GLN | p.Gln545Ter | P19447 |
| HOMO SAPIENS | -1 | 3805 | 3rbq | A | 57 | LYS | p.Lys57Ter | Q13432 |
| HOMO SAPIENS | 2 | 2649 | 4ido | A | 43 | ASP | p.Asp43Glu | Q8WXF7 |
| HOMO SAPIENS | 2 | 2649 | 4ido | A | 157 | LEU | p.Leu157Trp | Q8WXF7 |
| HOMO SAPIENS | 2 | 2649 | 4ido | A | 193 | PHE | p.Phe193Cys | Q8WXF7 |
| HOMO SAPIENS | 7 | 2649 | 4ido | A | 217 | ARG | p.Arg217Gln | Q8WXF7 |
| HOMO SAPIENS | 4 | 2649 | 4ido | A | 239 | ARG | p.Arg239Cys | Q8WXF7 |
| HOMO SAPIENS | 3 | 2649 | 4ido | A | 258 | HIS | p.His258Arg | Q8WXF7 |
| HOMO SAPIENS | 4 | 2649 | 4ido | A | 259 | SER | p.Ser259Tyr | Q8WXF7 |
| HOMO SAPIENS | 4 | 2649 | 4ido | A | 408 | MET | p.Met408Val | Q8WXF7 |
| HOMO SAPIENS | 2 | 2649 | 4ido | A | 415 | ARG | p.Arg415Trp | Q8WXF7 |
| HOMO SAPIENS | 3 | 2649 | 4ido | A | 416 | ARG | p.Arg416Cys | Q8WXF7 |
| HOMO SAPIENS | 5 | 2203 | 3wff | A | 810 | LEU | p.Ser810Leu | P08235 |
| HOMO SAPIENS | 4 | 12648 | 5ep6 | B | 696 | GLU | p.Glu696Lys | Q9UHD2 |
| HOMO SAPIENS | 0 | 30622 | 2pe4 | A | 90 | GLU | p.Glu90Asp | Q12794 |
| HOMO SAPIENS | 0 | 30622 | 2pe4 | A | 256 | GLY | p.Gly256Arg | Q12794 |
| HOMO SAPIENS | 1 | 30622 | 2pe4 | A | 268 | GLU | p.Glu268Lys | Q12794 |
| HOMO SAPIENS | 0 | 30622 | 2pe4 | A | 392 | THR | p.Thr392Met | Q12794 |
| HOMO SAPIENS | 5 | 5411 | 1lya | A | 93 | SER | p.Ser157Leu | P07339 |

| | | | | | | | | |
|---|---|---|---|---|---|---|---|---|
| HOMO SAPIENS | 3 | 5743 | 2uzp | A | 189 | ASN | p.Asn189Lys | P05129 |
| HOMO SAPIENS | 1 | 12529 | 3kfd | J | 21 | ASN | p.Asn45Ser | P36897 |
| HOMO SAPIENS | 1 | 1063 | 2e1q | B | 149 | ARG | p.Arg149Cys | P47989 |
| HOMO SAPIENS | -1 | 1063 | 2e1q | B | 172 | GLY | p.Gly172Arg | P47989 |
| HOMO SAPIENS | 4 | 1063 | 2e1q | B | 228 | ARG | p.Arg228Ter | P47989 |
| HOMO SAPIENS | 5 | 1063 | 2e1q | B | 395 | LYS | p.Lys395Met | P47989 |
| HOMO SAPIENS | 2 | 1063 | 2e1q | B | 646 | ILE | p.Ile646Val | P47989 |
| HOMO SAPIENS | 4 | 1063 | 2e1q | B | 703 | ILE | p.Ile703Val | P47989 |
| HOMO SAPIENS | 8 | 2525 | 1kqn | F | 18 | ASN | p.Asn18Ser | Q9HAN9 |
| HOMO SAPIENS | 0 | 2525 | 1kqn | F | 151 | VAL | p.Val151Phe | Q9HAN9 |
| HOMO SAPIENS | 0 | 2525 | 1kqn | F | 153 | LEU | p.Leu153Val | Q9HAN9 |
| HOMO SAPIENS | 0 | 2525 | 1kqn | F | 169 | TRP | p.Trp169Ter | Q9HAN9 |
| HOMO SAPIENS | 0 | 2525 | 1kqn | F | 207 | ARG | p.Arg207Trp | Q9HAN9 |
| HOMO SAPIENS | 0 | 2525 | 1kqn | F | 273 | ASN | p.Asn273Asp | Q9HAN9 |
| HOMO SAPIENS | 0 | 12299 | 5duq | A | 432 | VAL | p.Val454Glu | P05155 |
| HOMO SAPIENS | 0 | 12299 | 5duq | A | 436 | ALA | p.Ala458Thr | P05155 |
| HOMO SAPIENS | 0 | 12299 | 5duq | A | 443 | ALA | p.Ala465Val | P05155 |
| HOMO SAPIENS | 0 | 12299 | 5duq | A | 444 | ARG | p.Arg466His | P05155 |
| HOMO SAPIENS | 0 | 12299 | 5duq | A | 458 | MET | p.Val480Met | P05155 |
| HOMO SAPIENS | 0 | 327 | 1uhp | A | 36 | ASP | p.Asp274Asn | Q9UPQ7 |
| HOMO SAPIENS | 0 | 64 | 2mo5 | A | 54 | ALA | p.Ala55Thr | P12104 |
| HOMO SAPIENS | 2 | 35969 | 1jrh | I | 60 | CYS | p.Cys77Tyr | P15260 |
| HOMO SAPIENS | 4 | 35969 | 1jrh | I | 70 | ILE | p.Ile87Thr | P15260 |
| HOMO SAPIENS | 2 | 1068 | 1mu7 | A | 304 | ARG | p.Arg304Gln | Q9NUW8 |
| HOMO SAPIENS | 8 | 1068 | 1mu7 | A | 493 | HIS | p.His493Arg | Q9NUW8 |
| HOMO SAPIENS | 0 | 32360 | 2kqp | A | 31 | ARG | p.Arg55Cys | P01308 |
| HOMO SAPIENS | 0 | 32360 | 2kqp | A | 65 | ARG | p.Arg89His | P01308 |
| HOMO SAPIENS | 0 | 32360 | 2kqp | A | 65 | ARG | p.Arg89Cys | P01308 |
| HOMO SAPIENS | 6 | 10805 | 3nks | A | 12 | ILE | p.Ile12Thr | P50336 |
| HOMO SAPIENS | 0 | 10805 | 3nks | A | 20 | HIS | p.His20Pro | P50336 |
| HOMO SAPIENS | 0 | 10805 | 3nks | A | 59 | ARG | p.Arg59Trp | P50336 |
| HOMO SAPIENS | 0 | 10805 | 3nks | A | 168 | ARG | p.Arg168Cys | P50336 |
| HOMO SAPIENS | 0 | 10805 | 3nks | A | 168 | ARG | p.Arg168His | P50336 |
| HOMO SAPIENS | 2 | 10805 | 3nks | A | 232 | GLY | p.Gly232Arg | P50336 |
| HOMO SAPIENS | 3 | 10805 | 3nks | A | 256 | PRO | p.Pro256Arg | P50336 |
| HOMO SAPIENS | 0 | 10805 | 3nks | A | 304 | ARG | p.Arg304His | P50336 |
| HOMO SAPIENS | 0 | 10805 | 3nks | A | 349 | ASP | p.Asp349Ala | P50336 |
| HOMO SAPIENS | 9 | 7008 | 3u2l | A | 125 | GLN | p.Gln125Ter | P55789 |
| HOMO SAPIENS | 4 | 7008 | 3u2l | A | 172 | HIS | p.His172Asn | P55789 |
| HOMO SAPIENS | 4 | 7008 | 3u2l | A | 194 | ARG | p.Arg194His | P55789 |
| HOMO SAPIENS | 3 | 7008 | 3u2l | A | 198 | GLY | p.Gly198Ser | P55789 |
| HOMO SAPIENS | 0 | 20821 | 3e0g | A | 65 | HIS | p.His116Tyr | P42702 |
| HOMO SAPIENS | 3 | 1151 | 5l9v | A | 278 | ASP | p.Asp278Asn | Q9GZT9 |
| HOMO SAPIENS | 4 | 1151 | 5l9v | A | 317 | CYS | p.Pro317Arg | Q9GZT9 |
| HOMO SAPIENS | 5 | 1151 | 5l9v | A | 371 | ARG | p.Arg371His | Q9GZT9 |
| HOMO SAPIENS | 2 | 3935 | 1gwb | A | 36 | ARG | p.Arg36Gly | P07359 |
| HOMO SAPIENS | 1 | 3935 | 1gwb | A | 73 | LEU | p.Leu73Phe | P07359 |
| HOMO SAPIENS | 1 | 3935 | 1gwb | A | 172 | ALA | p.Ala172Val | P07359 |
| HOMO SAPIENS | 0 | 3935 | 1gwb | A | 249 | GLY | p.Gly249Val | P07359 |
| HOMO SAPIENS | 0 | 3935 | 1gwb | A | 255 | MET | p.Met255Val | P07359 |
| HOMO SAPIENS | 2 | 14541 | 5cuz | A | 138 | GLU | p.Glu138Lys | Q9H3L0 |
| HOMO SAPIENS | 2 | 14541 | 5cuz | A | 143 | SER | p.Ser143Ile | Q9H3L0 |
| HOMO SAPIENS | 4 | 14541 | 5cuz | A | 182 | THR | p.Thr182Asn | Q9H3L0 |
| HOMO SAPIENS | 6 | 14541 | 5cuz | A | 206 | ASN | p.Asn206Ile | Q9H3L0 |

| | | | | | | | | |
|---|---|---|---|---|---|---|---|---|
| HOMO SAPIENS | 0 | 14541 | 5cuz | A | 249 | TYR | p.Tyr249Cys | Q9H3L0 |
| HOMO SAPIENS | 4 | 14541 | 5cuz | A | 250 | ARG | p.Arg250Ter | Q9H3L0 |
| HOMO SAPIENS | 2 | 14541 | 5cuz | A | 259 | LEU | p.Leu259Pro | Q9H3L0 |
| HOMO SAPIENS | 0 | 4206 | 4fao | D | 30 | PRO | p.Pro30Ser | P37023 |
| HOMO SAPIENS | 1 | 4206 | 4fao | D | 50 | TRP | p.Trp50Cys | P37023 |
| HOMO SAPIENS | 0 | 16553 | 3v30 | A | 251 | GLN | p.Gln251Glu | O14593 |
| HOMO SAPIENS | 0 | 5012 | 5hbf | A | 442 | ALA | p.Ala442Gly | Q13231 |
| HOMO SAPIENS | 5 | 4844 | 5tdc | A | 136 | HIS | p.His136Arg | Q8IWV7 |
| HOMO SAPIENS | 0 | 15435 | 4cgz | A | 643 | ARG | p.Arg643His | P54132 |
| HOMO SAPIENS | 1 | 15435 | 4cgz | A | 672 | GLN | p.Gln672Arg | P54132 |
| HOMO SAPIENS | 1 | 15435 | 4cgz | A | 700 | GLN | p.Gln700Ter | P54132 |
| HOMO SAPIENS | 0 | 15435 | 4cgz | A | 707 | PRO | p.Pro707Ser | P54132 |
| HOMO SAPIENS | 0 | 15435 | 4cgz | A | 755 | LYS | p.Lys755Glu | P54132 |
| HOMO SAPIENS | 0 | 15435 | 4cgz | A | 881 | TRP | p.Trp881Ter | P54132 |
| HOMO SAPIENS | 2 | 15435 | 4cgz | A | 899 | ARG | p.Arg899Ter | P54132 |
| HOMO SAPIENS | 0 | 15435 | 4cgz | A | 1036 | CYS | p.Cys1036Phe | P54132 |
| HOMO SAPIENS | 0 | 15435 | 4cgz | A | 1055 | CYS | p.Cys1055Ser | P54132 |
| HOMO SAPIENS | 0 | 15435 | 4cgz | A | 1139 | ARG | p.Arg1139Ter | P54132 |
| HOMO SAPIENS | -1 | 15435 | 4cgz | A | 1198 | VAL | p.Val1198Met | P54132 |
| HOMO SAPIENS | -1 | 15435 | 4cgz | A | 1205 | VAL | p.Val1205Ile | P54132 |
| HOMO SAPIENS | 0 | 15435 | 4cgz | A | 1251 | GLU | p.Glu1251Gln | P54132 |
| HOMO SAPIENS | 0 | 11185 | 1kb4 | A | 30 | ARG | p.Arg30Ter | P11473 |
| HOMO SAPIENS | 5 | 11185 | 1kb4 | A | 33 | GLY | p.Gly33Asp | P11473 |
| HOMO SAPIENS | 5 | 11185 | 1kb4 | A | 46 | GLY | p.Gly46Asp | P11473 |
| HOMO SAPIENS | 6 | 11185 | 1kb4 | A | 50 | ARG | p.Arg50Gln | P11473 |
| HOMO SAPIENS | 9 | 11185 | 1kb4 | A | 73 | ARG | p.Arg73Gln | P11473 |
| HOMO SAPIENS | 9 | 11185 | 1kb4 | A | 80 | ARG | p.Arg80Gln | P11473 |
| HOMO SAPIENS | 0 | 5382 | 3fl7 | A | 41 | LEU | p.Leu41Val | P29317 |
| HOMO SAPIENS | 2 | 5382 | 3fl7 | A | 175 | ARG | p.Arg175Cys | P29317 |
| HOMO SAPIENS | 1 | 5382 | 3fl7 | A | 195 | ARG | p.Arg195Cys | P29317 |
| HOMO SAPIENS | 0 | 5382 | 3fl7 | A | 232 | ASP | p.Asp232Gly | P29317 |
| HOMO SAPIENS | 0 | 5382 | 3fl7 | A | 277 | SER | p.Ser277Leu | P29317 |
| HOMO SAPIENS | 0 | 5382 | 3fl7 | A | 315 | ARG | p.Arg315Gln | P29317 |
| HOMO SAPIENS | 7 | 5382 | 3fl7 | A | 350 | PRO | p.Pro350Thr | P29317 |
| HOMO SAPIENS | 0 | 5382 | 3fl7 | A | 391 | GLY | p.Gly391Arg | P29317 |
| HOMO SAPIENS | 0 | 5382 | 3fl7 | A | 418 | VAL | p.Val418Ile | P29317 |
| HOMO SAPIENS | 0 | 5382 | 3fl7 | A | 511 | THR | p.Thr511Met | P29317 |
| HOMO SAPIENS | 2 | 29168 | 4p22 | B | 144 | VAL | p.Val144Ile | P22314 |
| HOMO SAPIENS | 5 | 4348 | 1bf5 | A | 241 | ARG | p.Arg241Gln | P42224 |
| HOMO SAPIENS | 2 | 4348 | 1bf5 | A | 266 | VAL | p.Val266Ile | P42224 |
| HOMO SAPIENS | 4 | 4348 | 1bf5 | A | 320 | GLU | p.Glu320Gln | P42224 |
| HOMO SAPIENS | 1 | 4348 | 1bf5 | A | 372 | ASN | p.Asn372Lys | P42224 |
| HOMO SAPIENS | 1 | 4348 | 1bf5 | A | 447 | ASP | p.Asp447Glu | P42224 |
| HOMO SAPIENS | 2 | 4348 | 1bf5 | A | 463 | GLN | p.Gln463His | P42224 |
| HOMO SAPIENS | 4 | 4348 | 1bf5 | A | 600 | LEU | p.Leu600Pro | P42224 |
| HOMO SAPIENS | 0 | 4348 | 1bf5 | A | 706 | LEU | p.Leu706Ser | P42224 |
| HOMO SAPIENS | 4 | 4991 | 4bej | B | 141 | ASN | p.Asn141Ser | O00429 |
| HOMO SAPIENS | 0 | 4991 | 4bej | B | 395 | ALA | p.Ala395Asp | O00429 |
| HOMO SAPIENS | 2 | 11792 | 1fvo | A | 40 | ARG | p.Arg40Cys | P00480 |
| HOMO SAPIENS | 2 | 11792 | 1fvo | A | 40 | ARG | p.Arg40His | P00480 |
| HOMO SAPIENS | 0 | 11792 | 1fvo | A | 45 | LEU | p.Leu45Pro | P00480 |
| HOMO SAPIENS | 1 | 11792 | 1fvo | A | 47 | ASN | p.Asn47Ile | P00480 |
| HOMO SAPIENS | 0 | 11792 | 1fvo | A | 50 | GLY | p.Gly50Arg | P00480 |
| HOMO SAPIENS | 5 | 11792 | 1fvo | A | 87 | GLU | p.Glu87Lys | P00480 |

| | | | | | | | | |
|---|---|---|---|---|---|---|---|---|
| HOMO SAPIENS | 3 | 11792 | 1fvo | A | 111 | LEU | p.Leu111Pro | P00480 |
| HOMO SAPIENS | 1 | 11792 | 1fvo | A | 125 | THR | p.Thr125Met | P00480 |
| HOMO SAPIENS | 3 | 11792 | 1fvo | A | 129 | ARG | p.Arg129His | P00480 |
| HOMO SAPIENS | 9 | 11792 | 1fvo | A | 141 | ARG | p.Arg141Pro | P00480 |
| HOMO SAPIENS | 0 | 11792 | 1fvo | A | 206 | MET | p.Met206Arg | P00480 |
| HOMO SAPIENS | 0 | 11792 | 1fvo | A | 207 | SER | p.Ser207Arg | P00480 |
| HOMO SAPIENS | 0 | 11792 | 1fvo | A | 216 | GLN | p.Gln216Glu | P00480 |
| HOMO SAPIENS | 1 | 11792 | 1fvo | A | 225 | PRO | p.Pro225Leu | P00480 |
| HOMO SAPIENS | 0 | 11792 | 1fvo | A | 270 | ARG | p.Gln270Arg | P00480 |
| HOMO SAPIENS | 0 | 11792 | 1fvo | A | 270 | ARG | p.Gln270Ter | P00480 |
| HOMO SAPIENS | 2 | 11792 | 1fvo | A | 277 | ARG | p.Arg277Trp | P00480 |
| HOMO SAPIENS | 0 | 3721 | 4r3z | B | 2 | ASP | p.Asp2Gly | P54136 |
| HOMO SAPIENS | 0 | 3721 | 4r3z | B | 456 | SER | p.Ser456Leu | P54136 |
| HOMO SAPIENS | 5 | 3721 | 4r3z | B | 512 | ARG | p.Arg512Gln | P54136 |
| HOMO SAPIENS | 3 | 5749 | 3hl2 | A | 239 | ALA | p.Ala239Thr | Q9HD40 |
| HOMO SAPIENS | 1 | 5749 | 3hl2 | A | 334 | TYR | p.Tyr334Cys | Q9HD40 |
| HOMO SAPIENS | 1 | 5749 | 3hl2 | A | 452 | LYS | p.Lys452Asn | Q9HD40 |
| HOMO SAPIENS | -1 | 5749 | 3hl2 | A | 465 | ARG | p.Arg465Leu | Q9HD40 |
| HOMO SAPIENS | -1 | 5749 | 3hl2 | A | 489 | ASP | p.Asp489Gly | Q9HD40 |
| HOMO SAPIENS | 5 | 1721 | 3k15 | A | 1652 | MET | p.Met1652Ile | P38398 |
| HOMO SAPIENS | 5 | 1721 | 3k15 | A | 1652 | MET | p.Met1652Lys | P38398 |
| HOMO SAPIENS | 5 | 1721 | 3k15 | A | 1662 | PHE | p.Phe1662Cys | P38398 |
| HOMO SAPIENS | 5 | 1721 | 3k15 | A | 1664 | LEU | p.Leu1664Pro | P38398 |
| HOMO SAPIENS | 0 | 1721 | 3k15 | A | 1665 | VAL | p.Val1665Met | P38398 |
| HOMO SAPIENS | 5 | 1721 | 3k15 | A | 1690 | LYS | p.Lys1690Gln | P38398 |
| HOMO SAPIENS | 5 | 1721 | 3k15 | A | 1692 | ASP | p.Asp1692Asn | P38398 |
| HOMO SAPIENS | 5 | 1721 | 3k15 | A | 1699 | ARG | p.Arg1699Gln | P38398 |
| HOMO SAPIENS | 5 | 1721 | 3k15 | A | 1699 | ARG | p.Arg1699Trp | P38398 |
| HOMO SAPIENS | 3 | 1721 | 3k15 | A | 1706 | GLY | p.Gly1706Glu | P38398 |
| HOMO SAPIENS | 5 | 1721 | 3k15 | A | 1720 | THR | p.Thr1720Ala | P38398 |
| HOMO SAPIENS | 5 | 1721 | 3k15 | A | 1751 | ARG | p.Arg1751Pro | P38398 |
| HOMO SAPIENS | 4 | 1721 | 3k15 | A | 1775 | MET | p.Met1775Arg | P38398 |
| HOMO SAPIENS | 2 | 1721 | 3k15 | A | 1788 | GLY | p.Gly1788Val | P38398 |
| HOMO SAPIENS | 5 | 1721 | 3k15 | A | 1794 | GLU | p.Glu1794Asp | P38398 |
| HOMO SAPIENS | 5 | 1721 | 3k15 | A | 1804 | VAL | p.Val1804Asp | P38398 |
| HOMO SAPIENS | 3 | 1721 | 3k15 | A | 1815 | TRP | p.Trp1815Ter | P38398 |
| HOMO SAPIENS | 1 | 1721 | 3k15 | A | 1819 | ASN | p.Asn1819Ser | P38398 |
| HOMO SAPIENS | 5 | 1721 | 3k15 | A | 1856 | PRO | p.Pro1856Ser | P38398 |
| HOMO SAPIENS | 5 | 1721 | 3k15 | A | 1858 | ILE | p.Ile1858Leu | P38398 |
| HOMO SAPIENS | 5 | 1721 | 3k15 | A | 1858 | ILE | p.Ile1858Thr | P38398 |
| HOMO SAPIENS | 5 | 1721 | 3k15 | A | 1859 | PRO | p.Pro1859Arg | P38398 |
| HOMO SAPIENS | 2 | 22086 | 3kq4 | B | 1297 | PRO | p.Pro1297Leu | O60494 |
| HOMO SAPIENS | 1 | 12135 | 2k9j | B | 724 | ARG | p.Arg750Ter | P05106 |
| HOMO SAPIENS | -1 | 24027 | 2qtl | A | 175 | LEU | p.Ser202Leu | Q9UBK8 |
| HOMO SAPIENS | 0 | 24027 | 2qtl | A | 257 | SER | p.Ser284Thr | Q9UBK8 |
| HOMO SAPIENS | 0 | 24027 | 2qtl | A | 333 | LEU | p.Leu360Val | Q9UBK8 |
| HOMO SAPIENS | 0 | 24027 | 2qtl | A | 350 | LYS | p.Lys377Arg | Q9UBK8 |
| HOMO SAPIENS | 0 | 24027 | 2qtl | A | 415 | ARG | p.Arg442Cys | Q9UBK8 |
| HOMO SAPIENS | 2 | 24027 | 2qtl | A | 450 | PRO | p.Pro477Arg | Q9UBK8 |
| HOMO SAPIENS | 4 | 24027 | 2qtl | A | 487 | GLY | p.Gly514Arg | Q9UBK8 |
| HOMO SAPIENS | 1 | 24027 | 2qtl | A | 595 | HIS | p.His622Tyr | Q9UBK8 |
| HOMO SAPIENS | 0 | 5758 | 3di2 | B | 46 | THR | p.Thr66Ile | P16871 |
| HOMO SAPIENS | 1 | 5758 | 3di2 | B | 118 | VAL | p.Ile138Val | P16871 |
| HOMO SAPIENS | 4 | 848 | 5oa3 | Y | 79 | LEU | p.Leu79Val | P62847 |

| | | | | | | | | |
|---|---|---|---|---|---|---|---|---|
| HOMO SAPIENS | 2 | 848 | 5oa3 | Y | 124 | ASN | p.Asn124Ser | P62847 |
| HOMO SAPIENS | 0 | 81 | 3s5o | A | 36 | ALA | p.Ala36Glu | Q86XE5 |
| HOMO SAPIENS | 0 | 81 | 3s5o | A | 39 | TYR | p.Tyr39Ter | Q86XE5 |
| HOMO SAPIENS | 0 | 81 | 3s5o | A | 45 | PRO | p.Pro45Leu | Q86XE5 |
| HOMO SAPIENS | 0 | 81 | 3s5o | A | 70 | ARG | p.Arg70Ter | Q86XE5 |
| HOMO SAPIENS | 0 | 81 | 3s5o | A | 70 | ARG | p.Arg70Pro | Q86XE5 |
| HOMO SAPIENS | 1 | 81 | 3s5o | A | 97 | ARG | p.Arg97Cys | Q86XE5 |
| HOMO SAPIENS | 4 | 81 | 3s5o | A | 113 | GLU | p.Glu113Lys | Q86XE5 |
| HOMO SAPIENS | 1 | 81 | 3s5o | A | 190 | PRO | p.Pro190Leu | Q86XE5 |
| HOMO SAPIENS | 0 | 81 | 3s5o | A | 245 | VAL | p.Val245Phe | Q86XE5 |
| HOMO SAPIENS | 0 | 81 | 3s5o | A | 257 | CYS | p.Cys257Gly | Q86XE5 |
| HOMO SAPIENS | 0 | 81 | 3s5o | A | 287 | GLY | p.Gly287Val | Q86XE5 |
| HOMO SAPIENS | 7 | 81 | 3s5o | A | 303 | ARG | p.Arg303Cys | Q86XE5 |
| HOMO SAPIENS | 0 | 81 | 3s5o | A | 325 | GLY | p.Gly325Ser | Q86XE5 |
| HOMO SAPIENS | -1 | 622 | 5era | A | 12 | GLY | p.Gly12Arg | P29033 |
| HOMO SAPIENS | -1 | 622 | 5era | A | 12 | GLY | p.Gly12Asp | P29033 |
| HOMO SAPIENS | -1 | 622 | 5era | A | 17 | SER | p.Ser17Phe | P29033 |
| HOMO SAPIENS | 5 | 622 | 5era | A | 24 | TRP | p.Trp24Ter | P29033 |
| HOMO SAPIENS | 0 | 622 | 5era | A | 32 | ARG | p.Arg32Cys | P29033 |
| HOMO SAPIENS | 5 | 622 | 5era | A | 34 | MET | p.Met34Thr | P29033 |
| HOMO SAPIENS | 5 | 622 | 5era | A | 36 | LEU | p.Leu36Pro | P29033 |
| HOMO SAPIENS | 5 | 622 | 5era | A | 37 | VAL | p.Val37Ile | P29033 |
| HOMO SAPIENS | 5 | 622 | 5era | A | 40 | ALA | p.Ala40Glu | P29033 |
| HOMO SAPIENS | 5 | 622 | 5era | A | 44 | TRP | p.Trp44Cys | P29033 |
| HOMO SAPIENS | 5 | 622 | 5era | A | 45 | GLY | p.Gly45Glu | P29033 |
| HOMO SAPIENS | 5 | 622 | 5era | A | 47 | GLU | p.Glu47Ter | P29033 |
| HOMO SAPIENS | 5 | 622 | 5era | A | 50 | ASP | p.Asp50Asn | P29033 |
| HOMO SAPIENS | 5 | 622 | 5era | A | 54 | ASN | p.Asn54Lys | P29033 |
| HOMO SAPIENS | 5 | 622 | 5era | A | 57 | GLN | p.Gln57Ter | P29033 |
| HOMO SAPIENS | 5 | 622 | 5era | A | 59 | GLY | p.Gly59Ala | P29033 |
| HOMO SAPIENS | 5 | 622 | 5era | A | 59 | GLY | p.Gly59Ser | P29033 |
| HOMO SAPIENS | 5 | 622 | 5era | A | 66 | ASP | p.Asp66His | P29033 |
| HOMO SAPIENS | 5 | 622 | 5era | A | 68 | TYR | p.Tyr68Cys | P29033 |
| HOMO SAPIENS | 5 | 622 | 5era | A | 70 | PRO | p.Pro70Ala | P29033 |
| HOMO SAPIENS | 5 | 622 | 5era | A | 73 | HIS | p.His73Arg | P29033 |
| HOMO SAPIENS | 5 | 622 | 5era | A | 75 | ARG | p.Arg75Trp | P29033 |
| HOMO SAPIENS | 5 | 622 | 5era | A | 75 | ARG | p.Arg75Gln | P29033 |
| HOMO SAPIENS | 2 | 622 | 5era | A | 77 | TRP | p.Trp77Arg | P29033 |
| HOMO SAPIENS | 2 | 622 | 5era | A | 77 | TRP | p.Trp77Ter | P29033 |
| HOMO SAPIENS | 0 | 622 | 5era | A | 80 | GLN | p.Gln80Ter | P29033 |
| HOMO SAPIENS | 0 | 622 | 5era | A | 80 | GLN | p.Gln80Pro | P29033 |
| HOMO SAPIENS | 5 | 622 | 5era | A | 81 | LEU | p.Leu81Val | P29033 |
| HOMO SAPIENS | 5 | 622 | 5era | A | 82 | ILE | p.Ile82Met | P29033 |
| HOMO SAPIENS | 5 | 622 | 5era | A | 83 | PHE | p.Phe83Leu | P29033 |
| HOMO SAPIENS | 5 | 622 | 5era | A | 84 | VAL | p.Val84Leu | P29033 |
| HOMO SAPIENS | 5 | 622 | 5era | A | 90 | LEU | p.Leu90Pro | P29033 |
| HOMO SAPIENS | 5 | 622 | 5era | A | 95 | VAL | p.Val95Met | P29033 |
| HOMO SAPIENS | -1 | 622 | 5era | A | 100 | HIS | p.His100Tyr | P29033 |
| HOMO SAPIENS | -1 | 622 | 5era | A | 113 | SER | p.Ser113Arg | P29033 |
| HOMO SAPIENS | -1 | 622 | 5era | A | 119 | GLU | p.Glu119Lys | P29033 |
| HOMO SAPIENS | -1 | 622 | 5era | A | 122 | LYS | p.Lys122Ile | P29033 |
| HOMO SAPIENS | -1 | 622 | 5era | A | 123 | THR | p.Thr123Asn | P29033 |
| HOMO SAPIENS | -1 | 622 | 5era | A | 124 | GLN | p.Gln124Ter | P29033 |
| HOMO SAPIENS | 5 | 622 | 5era | A | 139 | SER | p.Ser139Asn | P29033 |

| | | | | | | | | |
|---|---|---|---|---|---|---|---|---|
| HOMO SAPIENS | 5 | 622 | 5era | A | 143 | ARG | p.Arg143Trp | P29033 |
| HOMO SAPIENS | 5 | 622 | 5era | A | 143 | ARG | p.Arg143Gln | P29033 |
| HOMO SAPIENS | 0 | 622 | 5era | A | 147 | GLU | p.Glu147Lys | P29033 |
| HOMO SAPIENS | 7 | 622 | 5era | A | 155 | TYR | p.Tyr155Ter | P29033 |
| HOMO SAPIENS | 2 | 622 | 5era | A | 159 | ASP | p.Asp159Val | P29033 |
| HOMO SAPIENS | 5 | 622 | 5era | A | 163 | MET | p.Met163Val | P29033 |
| HOMO SAPIENS | 5 | 622 | 5era | A | 165 | ARG | p.Arg165Trp | P29033 |
| HOMO SAPIENS | 5 | 622 | 5era | A | 167 | VAL | p.Val167Met | P29033 |
| HOMO SAPIENS | 5 | 622 | 5era | A | 169 | CYS | p.Cys169Tyr | P29033 |
| HOMO SAPIENS | 5 | 622 | 5era | A | 179 | ASP | p.Asp179Asn | P29033 |
| HOMO SAPIENS | 0 | 622 | 5era | A | 199 | SER | p.Ser199Phe | P29033 |
| HOMO SAPIENS | 0 | 622 | 5era | A | 202 | CYS | p.Cys202Phe | P29033 |
| HOMO SAPIENS | 5 | 622 | 5era | A | 203 | ILE | p.Ile203Thr | P29033 |
| HOMO SAPIENS | 5 | 622 | 5era | A | 206 | ASN | p.Asn206Ser | P29033 |
| HOMO SAPIENS | 0 | 8103 | 5dot | A | 163 | GLY | p.Gly163Ter | P31327 |
| HOMO SAPIENS | 0 | 8103 | 5dot | A | 337 | HIS | p.His337Arg | P31327 |
| HOMO SAPIENS | 3 | 8103 | 5dot | A | 344 | THR | p.Thr344Ala | P31327 |
| HOMO SAPIENS | 1 | 8103 | 5dot | A | 405 | THR | p.Thr405Asn | P31327 |
| HOMO SAPIENS | 5 | 8103 | 5dot | A | 544 | THR | p.Thr544Met | P31327 |
| HOMO SAPIENS | 5 | 8103 | 5dot | A | 716 | ASN | p.Asn716Lys | P31327 |
| HOMO SAPIENS | -1 | 8103 | 5dot | A | 780 | ARG | p.Arg780His | P31327 |
| HOMO SAPIENS | 0 | 8103 | 5dot | A | 982 | GLY | p.Gly982Asp | P31327 |
| HOMO SAPIENS | 3 | 8103 | 5dot | A | 1119 | ALA | p.Ala1119Ser | P31327 |
| HOMO SAPIENS | 1 | 8103 | 5dot | A | 1376 | GLY | p.Gly1376Ser | P31327 |
| HOMO SAPIENS | 4 | 6833 | 1l8l | B | 27 | ARG | p.Arg27Ser | P78330 |
| HOMO SAPIENS | 3 | 6833 | 1l8l | B | 32 | ASP | p.Asp32Gly | P78330 |
| HOMO SAPIENS | 3 | 6833 | 1l8l | B | 32 | ASP | p.Asp32Asn | P78330 |
| HOMO SAPIENS | 1 | 6833 | 1l8l | B | 49 | ARG | p.Arg49Trp | P78330 |
| HOMO SAPIENS | 0 | 6833 | 1l8l | B | 52 | MET | p.Met52Thr | P78330 |
| HOMO SAPIENS | 0 | 6833 | 1l8l | B | 83 | GLN | p.Gln83His | P78330 |
| HOMO SAPIENS | 6 | 810 | 2iw2 | A | 185 | ARG | p.Arg184Gln | P12955 |
| HOMO SAPIENS | 0 | 810 | 2iw2 | A | 213 | ALA | p.Ala212Pro | P12955 |
| HOMO SAPIENS | 1 | 810 | 2iw2 | A | 266 | ARG | p.Arg265Ter | P12955 |
| HOMO SAPIENS | 9 | 810 | 2iw2 | A | 277 | ASP | p.Asp276Asn | P12955 |
| HOMO SAPIENS | 0 | 810 | 2iw2 | A | 279 | GLY | p.Gly278Asp | P12955 |
| HOMO SAPIENS | 9 | 810 | 2iw2 | A | 378 | HIS | p.His377Gln | P12955 |
| HOMO SAPIENS | 0 | 810 | 2iw2 | A | 389 | ARG | p.Arg388His | P12955 |
| HOMO SAPIENS | 0 | 810 | 2iw2 | A | 436 | PHE | p.Leu435Phe | P12955 |
| HOMO SAPIENS | 0 | 810 | 2iw2 | A | 449 | GLY | p.Gly448Arg | P12955 |
| HOMO SAPIENS | -1 | 1530 | 5fwj | A | 388 | ALA | p.Ala388Pro | P41229 |
| HOMO SAPIENS | 4 | 1530 | 5fwj | A | 451 | SER | p.Ser451Arg | P41229 |
| HOMO SAPIENS | 3 | 1530 | 5fwj | A | 731 | LEU | p.Leu731Phe | P41229 |
| HOMO SAPIENS | 3 | 578 | 5net | B | 123 | ALA | p.Ala143Thr | P18564 |
| HOMO SAPIENS | 6 | 578 | 5net | B | 176 | PRO | p.Pro196Thr | P18564 |
| HOMO SAPIENS | 3 | 578 | 5net | B | 280 | GLU | p.Glu300Lys | P18564 |
| HOMO SAPIENS | -1 | 5247 | 4j15 | A | 274 | ALA | p.Ala274Val | P14868 |
| HOMO SAPIENS | -1 | 5247 | 4j15 | A | 280 | HIS | p.His280Leu | P14868 |
| HOMO SAPIENS | 2 | 5247 | 4j15 | A | 460 | ARG | p.Arg460His | P14868 |
| HOMO SAPIENS | 5 | 5247 | 4j15 | A | 494 | ARG | p.Arg494Cys | P14868 |
| HOMO SAPIENS | 7 | 2566 | 1b55 | A | 28 | ARG | p.Arg28His | Q06187 |
| HOMO SAPIENS | 5 | 2566 | 1b55 | A | 33 | THR | p.Thr33Pro | Q06187 |
| HOMO SAPIENS | 0 | 2566 | 1b55 | A | 113 | VAL | p.Val113Asp | Q06187 |
| HOMO SAPIENS | 0 | 356 | 5a2s | A | 786 | ALA | p.Ala786Thr | P56524 |
| HOMO SAPIENS | 5 | 2960 | 4cak | A | 553 | ARG | p.Arg584Ter | P08514 |

| | | | | | | | | |
|---|---|---|---|---|---|---|---|---|
| HOMO SAPIENS | 5 | 2960 | 4cak | A | 565 | ILE | p.Ile596Thr | P08514 |
| HOMO SAPIENS | -1 | 2960 | 4cak | A | 841 | LEU | p.Leu872Met | P08514 |
| HOMO SAPIENS | -1 | 2960 | 4cak | A | 843 | ILE | p.Ile874Ser | P08514 |
| HOMO SAPIENS | 0 | 5054 | 4k8r | B | 216 | PHE | p.Phe216Leu | Q8NBP7 |
| HOMO SAPIENS | 1 | 5054 | 4k8r | B | 417 | HIS | p.His417Gln | Q8NBP7 |
| HOMO SAPIENS | 1 | 5054 | 4k8r | B | 425 | ASN | p.Asn425Ser | Q8NBP7 |
| HOMO SAPIENS | 1 | 5054 | 4k8r | B | 443 | ALA | p.Ala443Thr | Q8NBP7 |
| HOMO SAPIENS | 0 | 27283 | 3n5n | X | 71 | HIS | p.His82Arg | Q9UIF7 |
| HOMO SAPIENS | 1 | 27283 | 3n5n | X | 83 | ARG | p.Arg94Ter | Q9UIF7 |
| HOMO SAPIENS | 8 | 27283 | 3n5n | X | 89 | TRP | p.Trp100Ter | Q9UIF7 |
| HOMO SAPIENS | 8 | 27283 | 3n5n | X | 90 | TYR | p.Tyr101Ter | Q9UIF7 |
| HOMO SAPIENS | 7 | 27283 | 3n5n | X | 95 | ARG | p.Arg106Trp | Q9UIF7 |
| HOMO SAPIENS | 7 | 27283 | 3n5n | X | 117 | TRP | p.Trp128Arg | Q9UIF7 |
| HOMO SAPIENS | 0 | 27283 | 3n5n | X | 137 | GLY | p.Gly148Arg | Q9UIF7 |
| HOMO SAPIENS | 5 | 27283 | 3n5n | X | 142 | TRP | p.Trp153Ter | Q9UIF7 |
| HOMO SAPIENS | 0 | 27283 | 3n5n | X | 165 | TYR | p.Tyr176Cys | Q9UIF7 |
| HOMO SAPIENS | 0 | 27283 | 3n5n | X | 168 | ARG | p.Arg179His | Q9UIF7 |
| HOMO SAPIENS | 0 | 27283 | 3n5n | X | 168 | ARG | p.Arg179Cys | Q9UIF7 |
| HOMO SAPIENS | 0 | 27283 | 3n5n | X | 196 | GLN | p.Gln207Ter | Q9UIF7 |
| HOMO SAPIENS | 8 | 27283 | 3n5n | X | 201 | VAL | p.Val212Met | Q9UIF7 |
| HOMO SAPIENS | 9 | 27283 | 3n5n | X | 227 | ARG | p.Arg238Trp | Q9UIF7 |
| HOMO SAPIENS | 9 | 27283 | 3n5n | X | 231 | ARG | p.Arg242Cys | Q9UIF7 |
| HOMO SAPIENS | 9 | 27283 | 3n5n | X | 231 | ARG | p.Arg242His | Q9UIF7 |
| HOMO SAPIENS | 3 | 27283 | 3n5n | X | 232 | VAL | p.Val243Phe | Q9UIF7 |
| HOMO SAPIENS | 3 | 27283 | 3n5n | X | 246 | GLN | p.Gln257Ter | Q9UIF7 |
| HOMO SAPIENS | 9 | 27283 | 3n5n | X | 272 | GLY | p.Gly283Glu | Q9UIF7 |
| HOMO SAPIENS | 9 | 27283 | 3n5n | X | 281 | PRO | p.Pro292Leu | Q9UIF7 |
| HOMO SAPIENS | -1 | 27283 | 3n5n | X | 314 | ASP | p.Asp325Glu | Q9UIF7 |
| HOMO SAPIENS | 0 | 27283 | 3n5n | X | 324 | GLN | p.Gln335His | Q9UIF7 |
| HOMO SAPIENS | 0 | 27283 | 3n5n | X | 324 | GLN | p.Gln335Arg | Q9UIF7 |
| HOMO SAPIENS | 0 | 27283 | 3n5n | X | 324 | GLN | p.Gln335Ter | Q9UIF7 |
| HOMO SAPIENS | -1 | 20852 | 4p60 | B | 243 | ARG | p.Arg243Lys | O75746 |
| HOMO SAPIENS | -1 | 1875 | 5dd2 | G | 152 | VAL | p.Val179Leu | P06396 |
| HOMO SAPIENS | 9 | 2126 | 3n1p | B | 46 | PRO | p.Pro46Leu | Q14623 |
| HOMO SAPIENS | 8 | 2126 | 3n1p | B | 76 | GLU | p.Glu76Ala | Q14623 |
| HOMO SAPIENS | 9 | 2126 | 3n1p | B | 77 | ARG | p.Arg77Ser | Q14623 |
| HOMO SAPIENS | 6 | 2126 | 3n1p | B | 95 | GLU | p.Glu95Lys | Q14623 |
| HOMO SAPIENS | 9 | 2126 | 3n1p | B | 100 | ASP | p.Asp100Glu | Q14623 |
| HOMO SAPIENS | 9 | 2126 | 3n1p | B | 131 | GLU | p.Glu131Lys | Q14623 |
| HOMO SAPIENS | 0 | 2126 | 3n1p | B | 190 | VAL | p.Val190Ala | Q14623 |
| HOMO SAPIENS | 5 | 40410 | 3ioz | B | 70 | GLN | p.Gln70Leu | P20963 |
| HOMO SAPIENS | 1 | 5754 | 5lkz | A | 1108 | THR | p.Tyr1191Ter | Q09472 |
| HOMO SAPIENS | 3 | 8159 | 1tiu | A | 3 | GLU | p.Lys12679Glu | Q8WZ42 |
| HOMO SAPIENS | 4 | 3275 | 4ldq | A | 210 | ARG | p.Arg210Gln | Q9BV20 |
| HOMO SAPIENS | 0 | 8143 | 4miv | B | 66 | SER | p.Ser66Trp | P51688 |
| HOMO SAPIENS | 1 | 8143 | 4miv | B | 74 | ARG | p.Arg74Cys | P51688 |
| HOMO SAPIENS | 0 | 8143 | 4miv | B | 128 | PRO | p.Pro128Leu | P51688 |
| HOMO SAPIENS | 1 | 8143 | 4miv | B | 150 | ARG | p.Arg150Gln | P51688 |
| HOMO SAPIENS | 0 | 8143 | 4miv | B | 191 | GLY | p.Gly191Arg | P51688 |
| HOMO SAPIENS | 0 | 8143 | 4miv | B | 206 | ARG | p.Arg206Pro | P51688 |
| HOMO SAPIENS | 0 | 8143 | 4miv | B | 225 | PHE | p.Phe225Leu | P51688 |
| HOMO SAPIENS | 0 | 8143 | 4miv | B | 233 | ARG | p.Arg233Ter | P51688 |
| HOMO SAPIENS | 0 | 8143 | 4miv | B | 234 | ALA | p.Ala234Gly | P51688 |
| HOMO SAPIENS | 0 | 8143 | 4miv | B | 245 | ARG | p.Arg245His | P51688 |

| | | | | | | | | |
|---|---|---|---|---|---|---|---|---|
| HOMO SAPIENS | 0 | 8143 | 4miv | B | 293 | PRO | p.Pro293Ser | P51688 |
| HOMO SAPIENS | 1 | 8143 | 4miv | B | 298 | SER | p.Ser298Pro | P51688 |
| HOMO SAPIENS | 0 | 8143 | 4miv | B | 361 | VAL | p.Val361Ile | P51688 |
| HOMO SAPIENS | 0 | 8143 | 4miv | B | 369 | GLU | p.Glu369Lys | P51688 |
| HOMO SAPIENS | 0 | 8143 | 4miv | B | 372 | MET | p.Met372Ile | P51688 |
| HOMO SAPIENS | 1 | 8143 | 4miv | B | 380 | GLN | p.Gln380Arg | P51688 |
| HOMO SAPIENS | 0 | 8143 | 4miv | B | 387 | VAL | p.Val387Met | P51688 |
| HOMO SAPIENS | 0 | 8143 | 4miv | B | 394 | MET | p.Met394Ile | P51688 |
| HOMO SAPIENS | 0 | 8143 | 4miv | B | 433 | ARG | p.Arg433Gln | P51688 |
| HOMO SAPIENS | 1 | 8143 | 4miv | B | 446 | HIS | p.His446Arg | P51688 |
| HOMO SAPIENS | 5 | 8143 | 4miv | B | 447 | GLU | p.Glu447Lys | P51688 |
| HOMO SAPIENS | 2 | 8143 | 4miv | B | 456 | ARG | p.Arg456His | P51688 |
| HOMO SAPIENS | 1 | 21880 | 2oo9 | B | 857 | LEU | p.Leu857Phe | P22681 |
| HOMO SAPIENS | 0 | 9467 | 1s1d | A | 154 | LEU | p.Leu224Pro | Q8WVQ1 |
| HOMO SAPIENS | 0 | 9467 | 1s1d | A | 156 | VAL | p.Val226Met | Q8WVQ1 |
| HOMO SAPIENS | 0 | 9467 | 1s1d | A | 229 | PRO | p.Pro299Leu | Q8WVQ1 |
| HOMO SAPIENS | 4 | 9467 | 1s1d | A | 230 | ARG | p.Arg300Cys | Q8WVQ1 |
| HOMO SAPIENS | 4 | 9467 | 1s1d | A | 230 | ARG | p.Arg300His | Q8WVQ1 |
| HOMO SAPIENS | 0 | 9467 | 1s1d | A | 253 | ALA | p.Ala323Thr | Q8WVQ1 |
| HOMO SAPIENS | 2 | 9467 | 1s1d | A | 305 | MET | p.Met375Leu | Q8WVQ1 |
| HOMO SAPIENS | -1 | 2361 | 5llm | A | 5 | ASN | p.Asn5Lys | P43003 |
| HOMO SAPIENS | 0 | 2361 | 5llm | A | 71 | TYR | p.Tyr71Cys | P43003 |
| HOMO SAPIENS | -1 | 2361 | 5llm | A | 211 | LYS | p.Glu219Asp | Q15758 |
| HOMO SAPIENS | 8 | 2361 | 5llm | A | 270 | PRO | p.Pro290Arg | P43003 |
| HOMO SAPIENS | 1 | 2361 | 5llm | A | 321 | THR | p.Ala329Thr | Q15758 |
| HOMO SAPIENS | 1 | 2361 | 5llm | A | 365 | ARG | p.Arg385His | P43003 |
| HOMO SAPIENS | 1 | 2361 | 5llm | A | 479 | ARG | p.Arg499Gln | P43003 |
| HOMO SAPIENS | 3 | 5412 | 4obz | D | 165 | PHE | p.Phe229Ile | P07339 |
| HOMO SAPIENS | 2 | 5412 | 4obz | D | 187 | ASP | p.Asp251Asn | P07339 |
| HOMO SAPIENS | 0 | 5412 | 4obz | D | 218 | GLY | p.Gly282Arg | P07339 |
| HOMO SAPIENS | 0 | 5412 | 4obz | D | 245 | ARG | p.Arg309His | P07339 |
| HOMO SAPIENS | 2 | 5412 | 4obz | D | 319 | TRP | p.Trp383Cys | P07339 |
| HOMO SAPIENS | 0 | 3544 | 5h64 | A | 277 | MET | p.Met277Ile | P42345 |
| HOMO SAPIENS | 0 | 14230 | 2kbx | A | 145 | ARG | p.Arg145Ser | Q13418 |
| HOMO SAPIENS | 2 | 34970 | 4xhz | A | 834 | ARG | p.Arg855Trp | Q96QU1 |
| HOMO SAPIENS | 4 | 34970 | 4xhz | A | 854 | SER | p.Ser875Ter | Q96QU1 |
| HOMO SAPIENS | 1 | 34970 | 4xhz | A | 908 | ARG | p.Arg929Gln | Q96QU1 |
| HOMO SAPIENS | 1 | 34970 | 4xhz | A | 941 | ARG | p.Arg962Cys | Q96QU1 |
| HOMO SAPIENS | 0 | 34970 | 4xhz | A | 970 | ARG | p.Arg991Ter | Q96QU1 |
| HOMO SAPIENS | 0 | 34970 | 4xhz | A | 976 | GLU | p.Glu997Gly | Q96QU1 |
| HOMO SAPIENS | 0 | 25734 | 4asc | A | 345 | SER | p.Asn345Ser | Q2TBA0 |
| HOMO SAPIENS | 0 | 25734 | 4asc | A | 514 | VAL | p.Val514Met | Q2TBA0 |
| HOMO SAPIENS | 6 | 25734 | 4asc | A | 528 | GLU | p.Glu528Lys | Q2TBA0 |
| HOMO SAPIENS | 0 | 25734 | 4asc | A | 580 | LEU | p.Leu580Ile | Q2TBA0 |
| HOMO SAPIENS | 8 | 2356 | 3kse | D | 22 | LYS | p.Lys22Ter | P01040 |
| HOMO SAPIENS | 1 | 2356 | 3kse | D | 58 | ARG | p.Arg58Ter | P01040 |
| HOMO SAPIENS | 2 | 6000 | 1jqd | A | 60 | GLY | p.Gly60Asp | P50135 |
| HOMO SAPIENS | 9 | 6000 | 1jqd | A | 208 | LEU | p.Leu208Pro | P50135 |
| HOMO SAPIENS | 0 | 14741 | 5eh1 | A | 58 | THR | p.Thr58Arg | P38484 |
| HOMO SAPIENS | 0 | 14741 | 5eh1 | A | 168 | THR | p.Thr168Asn | P38484 |
| HOMO SAPIENS | 1 | 4814 | 4kxk | D | 427 | ARG | p.Arg427Ter | P50542 |
| HOMO SAPIENS | 0 | 188 | 2xt3 | A | 49 | ASP | p.Asp52Asn | Q2M1P5 |
| HOMO SAPIENS | 3 | 188 | 2xt3 | A | 142 | TYR | p.Tyr145Ser | Q2M1P5 |
| HOMO SAPIENS | 9 | 8681 | 5tby | D | 57 | ALA | p.Ala57Asp | P08590 |

| | | | | | | | | |
|---|---|---|---|---|---|---|---|---|
| HOMO SAPIENS | 3 | 8681 | 5tby | D | 62 | ASP | p.Asp62Asn | P08590 |
| HOMO SAPIENS | 0 | 8681 | 5tby | D | 65 | PRO | p.Pro65Arg | P08590 |
| HOMO SAPIENS | 7 | 8681 | 5tby | D | 79 | VAL | p.Val79Ile | P08590 |
| HOMO SAPIENS | 0 | 8681 | 5tby | D | 143 | GLU | p.Glu143Lys | P08590 |
| HOMO SAPIENS | 0 | 8681 | 5tby | D | 149 | MET | p.Met149Val | P08590 |
| HOMO SAPIENS | 3 | 8681 | 5tby | D | 154 | ARG | p.Arg154His | P08590 |
| HOMO SAPIENS | 6 | 30984 | 4xwh | A | 140 | TYR | p.Tyr140Cys | P54802 |
| HOMO SAPIENS | 4 | 30984 | 4xwh | A | 141 | SER | p.Ser141Thr | P54802 |
| HOMO SAPIENS | 2 | 30984 | 4xwh | A | 234 | ARG | p.Arg234Cys | P54802 |
| HOMO SAPIENS | 2 | 30984 | 4xwh | A | 336 | GLU | p.Glu336Ter | P54802 |
| HOMO SAPIENS | 6 | 30984 | 4xwh | A | 482 | ARG | p.Arg482Trp | P54802 |
| HOMO SAPIENS | 2 | 30984 | 4xwh | A | 521 | PRO | p.Pro521Leu | P54802 |
| HOMO SAPIENS | 0 | 30984 | 4xwh | A | 565 | ARG | p.Arg565Gln | P54802 |
| HOMO SAPIENS | 0 | 30984 | 4xwh | A | 565 | ARG | p.Arg565Trp | P54802 |
| HOMO SAPIENS | 6 | 30984 | 4xwh | A | 626 | ARG | p.Arg626Ter | P54802 |
| HOMO SAPIENS | 6 | 30984 | 4xwh | A | 643 | ARG | p.Arg643His | P54802 |
| HOMO SAPIENS | 6 | 30984 | 4xwh | A | 643 | ARG | p.Arg643Cys | P54802 |
| HOMO SAPIENS | 2 | 30984 | 4xwh | A | 674 | ARG | p.Arg674His | P54802 |
| HOMO SAPIENS | 2 | 30984 | 4xwh | A | 737 | ARG | p.Arg737Gly | P54802 |
| HOMO SAPIENS | 5 | 10545 | 1oqe | N | 6 | PRO | p.Pro21Arg | Q96RJ3 |
| HOMO SAPIENS | 1 | 16995 | 5d3a | B | 12 | MET | p.Met947Arg | Q7Z4S6 |
| HOMO SAPIENS | 1 | 16995 | 5d3a | B | 12 | MET | p.Met947Val | Q7Z4S6 |
| HOMO SAPIENS | 2 | 16995 | 5d3a | B | 19 | ARG | p.Arg954Gln | Q7Z4S6 |
| HOMO SAPIENS | 2 | 16995 | 5d3a | B | 19 | ARG | p.Arg954Trp | Q7Z4S6 |
| HOMO SAPIENS | 0 | 16995 | 5d3a | B | 75 | ILE | p.Ile1010Thr | Q7Z4S6 |
| HOMO SAPIENS | 0 | 17091 | 2yin | B | 1324 | GLN | p.Gln1324Ter | Q92608 |
| HOMO SAPIENS | 5 | 27606 | 1lm7 | B | 2366 | ARG | p.Arg2366Cys | P15924 |
| HOMO SAPIENS | 2 | 15731 | 1bi7 | B | 24 | ARG | p.Arg24Pro | P42771 |
| HOMO SAPIENS | 0 | 15731 | 1bi7 | B | 28 | VAL | p.Val28Gly | P42771 |
| HOMO SAPIENS | 0 | 15731 | 1bi7 | B | 49 | ILE | p.Ile49Thr | P42771 |
| HOMO SAPIENS | 0 | 15731 | 1bi7 | B | 53 | MET | p.Met53Ile | P42771 |
| HOMO SAPIENS | 2 | 15731 | 1bi7 | B | 56 | SER | p.Ser56Ile | P42771 |
| HOMO SAPIENS | 1 | 15731 | 1bi7 | B | 57 | ALA | p.Ala57Val | P42771 |
| HOMO SAPIENS | 4 | 15731 | 1bi7 | B | 59 | VAL | p.Val59Gly | P42771 |
| HOMO SAPIENS | 9 | 15731 | 1bi7 | B | 89 | GLY | p.Gly89Asp | P42771 |
| HOMO SAPIENS | 0 | 15731 | 1bi7 | B | 101 | GLY | p.Gly101Trp | P42771 |
| HOMO SAPIENS | 0 | 15731 | 1bi7 | B | 114 | PRO | p.Pro114Ser | P42771 |
| HOMO SAPIENS | 9 | 15731 | 1bi7 | B | 123 | HIS | p.His123Gln | P42771 |
| HOMO SAPIENS | 8 | 15731 | 1bi7 | B | 126 | VAL | p.Val126Asp | P42771 |
| HOMO SAPIENS | 7 | 15731 | 1bi7 | B | 127 | ALA | p.Ala127Ser | P42771 |
| HOMO SAPIENS | -1 | 15731 | 1bi7 | B | 148 | ALA | p.Ala148Thr | P42771 |
| HOMO SAPIENS | -1 | 19858 | 3jbk | M | 866 | ALA | p.Ala934Val | P18206 |
| HOMO SAPIENS | 0 | 19484 | 3ee6 | B | 62 | SER | p.Ser62Leu | O14773 |
| HOMO SAPIENS | 1 | 19484 | 3ee6 | B | 66 | GLN | p.Gln66Ter | O14773 |
| HOMO SAPIENS | 0 | 19484 | 3ee6 | B | 100 | GLN | p.Gln100Arg | O14773 |
| HOMO SAPIENS | 0 | 19484 | 3ee6 | B | 104 | LEU | p.Leu104Ter | O14773 |
| HOMO SAPIENS | 0 | 19484 | 3ee6 | B | 127 | ARG | p.Arg127Gln | O14773 |
| HOMO SAPIENS | 0 | 19484 | 3ee6 | B | 127 | ARG | p.Arg127Ter | O14773 |
| HOMO SAPIENS | -1 | 19484 | 3ee6 | B | 185 | ARG | p.Arg185Cys | O14773 |
| HOMO SAPIENS | 0 | 19484 | 3ee6 | B | 206 | ARG | p.Arg206Cys | O14773 |
| HOMO SAPIENS | 0 | 19484 | 3ee6 | B | 208 | ARG | p.Arg208Ter | O14773 |
| HOMO SAPIENS | 0 | 19484 | 3ee6 | B | 214 | GLN | p.Gln214Ter | O14773 |
| HOMO SAPIENS | 1 | 19484 | 3ee6 | B | 276 | ASP | p.Asp276Val | O14773 |
| HOMO SAPIENS | 0 | 19484 | 3ee6 | B | 284 | GLY | p.Gly284Val | O14773 |

| | | | | | | | | |
|---|---|---|---|---|---|---|---|---|
| HOMO SAPIENS | 0 | 19484 | 3ee6 | B | 286 | ASN | p.Asn286Ser | O14773 |
| HOMO SAPIENS | 0 | 19484 | 3ee6 | B | 339 | ARG | p.Arg339Gly | O14773 |
| HOMO SAPIENS | 2 | 19484 | 3ee6 | B | 345 | MET | p.Met345Leu | O14773 |
| HOMO SAPIENS | 0 | 19484 | 3ee6 | B | 365 | CYS | p.Cys365Arg | O14773 |
| HOMO SAPIENS | 0 | 19484 | 3ee6 | B | 365 | CYS | p.Cys365Tyr | O14773 |
| HOMO SAPIENS | 0 | 19484 | 3ee6 | B | 373 | GLN | p.Gln373Glu | O14773 |
| HOMO SAPIENS | 6 | 19484 | 3ee6 | B | 418 | ARG | p.Arg418Gln | O14773 |
| HOMO SAPIENS | 7 | 19484 | 3ee6 | B | 422 | GLN | p.Gln422His | O14773 |
| HOMO SAPIENS | 0 | 19484 | 3ee6 | B | 447 | ARG | p.Arg447His | O14773 |
| HOMO SAPIENS | 0 | 19484 | 3ee6 | B | 466 | VAL | p.Val466Met | O14773 |
| HOMO SAPIENS | 0 | 4654 | 5t5w | A | 47 | LYS | p.Lys47Glu | Q08334 |
| HOMO SAPIENS | 0 | 4654 | 5t5w | A | 141 | GLU | p.Glu141Ter | Q08334 |
| HOMO SAPIENS | -1 | 18815 | 4wj7 | A | 53 | VAL | p.Val53Ile | Q9BSQ5 |
| HOMO SAPIENS | 0 | 18815 | 4wj7 | A | 120 | VAL | p.Val120Ile | Q9BSQ5 |
| HOMO SAPIENS | 0 | 18815 | 4wj7 | A | 198 | LEU | p.Leu198Arg | Q9BSQ5 |
| HOMO SAPIENS | 0 | 4602 | 1mjd | A | 47 | SER | p.Ser47Arg | O43602 |
| HOMO SAPIENS | 6 | 4602 | 1mjd | A | 59 | ARG | p.Arg59Leu | O43602 |
| HOMO SAPIENS | 7 | 4602 | 1mjd | A | 62 | ASP | p.Asp62Asn | O43602 |
| HOMO SAPIENS | 7 | 4602 | 1mjd | A | 71 | ALA | p.Ala71Ser | O43602 |
| HOMO SAPIENS | 7 | 4602 | 1mjd | A | 78 | ARG | p.Arg78Leu | O43602 |
| HOMO SAPIENS | 7 | 4602 | 1mjd | A | 89 | ARG | p.Arg89Gly | O43602 |
| HOMO SAPIENS | 8 | 4602 | 1mjd | A | 104 | ILE | p.Ile104Phe | O43602 |
| HOMO SAPIENS | 7 | 4602 | 1mjd | A | 125 | TYR | p.Tyr125His | O43602 |
| HOMO SAPIENS | 5 | 34467 | 1n0w | B | 1529 | GLY | p.Gly1529Arg | P51587 |
| HOMO SAPIENS | 8 | 5738 | 1u31 | A | 130 | ASN | p.Asn1009Lys | Q13423 |
| HOMO SAPIENS | 0 | 5760 | 2i6s | A | 298 | GLU | p.Glu318Asp | P06681 |
| HOMO SAPIENS | 0 | 5760 | 2i6s | A | 573 | ARG | p.Arg593Gln | P06681 |
| HOMO SAPIENS | 0 | 5760 | 2i6s | A | 592 | SER | p.Ser612Asn | P06681 |
| HOMO SAPIENS | 0 | 5760 | 2i6s | A | 621 | VAL | p.Val641Ala | P06681 |
| HOMO SAPIENS | 1 | 10977 | 2fby | A | 114 | VAL | p.Val114Ile | Q14191 |
| HOMO SAPIENS | 0 | 10977 | 2fby | A | 135 | LYS | p.Lys135Glu | Q14191 |
| HOMO SAPIENS | -1 | 1175 | 1qrk | A | 34 | VAL | p.Val35Leu | P00488 |
| HOMO SAPIENS | 4 | 1175 | 1qrk | A | 564 | PRO | p.Pro565Leu | P00488 |
| HOMO SAPIENS | 0 | 1175 | 1qrk | A | 650 | VAL | p.Val651Ile | P00488 |
| HOMO SAPIENS | 0 | 1175 | 1qrk | A | 651 | GLU | p.Gln652Glu | P00488 |
| HOMO SAPIENS | 3 | 909 | 4h0w | A | 375 | GLU | p.Glu394Lys | P02787 |
| HOMO SAPIENS | 0 | 909 | 4h0w | A | 458 | ALA | p.Ala477Pro | P02787 |
| HOMO SAPIENS | 6 | 909 | 4h0w | A | 515 | TYR | p.Tyr534Ter | P02787 |
| HOMO SAPIENS | 7 | 909 | 4h0w | A | 570 | PRO | p.Pro589Ser | P02787 |
| HOMO SAPIENS | 3 | 909 | 4h0w | A | 652 | GLY | p.Gly671Glu | P02787 |
| HOMO SAPIENS | 5 | 1964 | 2car | A | 151 | TRP | p.Trp151Ter | Q9BY32 |
| HOMO SAPIENS | 7 | 1964 | 2car | A | 178 | ARG | p.Arg178Cys | Q9BY32 |
| HOMO SAPIENS | 0 | 144 | 2n0a | J | 30 | ALA | p.Ala30Pro | P37840 |
| HOMO SAPIENS | 1 | 12288 | 4q58 | D | 1225 | ARG | p.Arg1225His | P16144 |
| HOMO SAPIENS | 2 | 12288 | 4q58 | D | 1281 | ARG | p.Arg1281Trp | P16144 |
| HOMO SAPIENS | 2 | 12288 | 4q58 | D | 1287 | ASN | p.Asn1287Lys | P16144 |
| HOMO SAPIENS | 0 | 5085 | 3s94 | A | 360 | ARG | p.Arg360His | O75581 |
| HOMO SAPIENS | 0 | 5085 | 3s94 | A | 433 | ASN | p.Asn433Ser | O75581 |
| HOMO SAPIENS | 2 | 5085 | 3s94 | A | 611 | ARG | p.Arg611Cys | O75581 |
| HOMO SAPIENS | 0 | 15452 | 4dur | A | 19 | LYS | p.Lys38Glu | P00747 |
| HOMO SAPIENS | 2 | 15452 | 4dur | A | 216 | ARG | p.Arg235His | P00747 |
| HOMO SAPIENS | 7 | 8406 | 4wo1 | D | 28 | VAL | p.Val55Leu | O43914 |
| HOMO SAPIENS | 2 | 895 | 5xte | K | 183 | ARG | p.Arg183Trp | P22695 |
| HOMO SAPIENS | 2 | 4191 | 4wnd | A | 120 | ARG | p.Arg127Ter | P81274 |

| | | | | | | | | |
|---|---|---|---|---|---|---|---|---|
| HOMO SAPIENS | 2 | 4191 | 4wnd | A | 120 | ARG | p.Arg127Gln | P81274 |
| HOMO SAPIENS | 1 | 4191 | 4wnd | A | 349 | GLY | p.Gly356Arg | P81274 |
| HOMO SAPIENS | 0 | 41734 | 2kbi | A | 1777 | VAL | p.Val1777Met | Q14524 |
| HOMO SAPIENS | 1 | 41734 | 2kbi | A | 1784 | GLU | p.Glu1784Lys | Q14524 |
| HOMO SAPIENS | 5 | 41734 | 2kbi | A | 1795 | TYR | p.Tyr1795Cys | Q14524 |
| HOMO SAPIENS | 5 | 41734 | 2kbi | A | 1795 | TYR | p.Tyr1795His | Q14524 |
| HOMO SAPIENS | 7 | 41734 | 2kbi | A | 1826 | ARG | p.Arg1826Cys | Q14524 |
| HOMO SAPIENS | 3 | 2119 | 1pl0 | A | 426 | LYS | p.Lys426Arg | P31939 |
| HOMO SAPIENS | 5 | 3475 | 5u62 | C | 50 | LEU | p.Leu50Ser | Q15910 |
| HOMO SAPIENS | 3 | 379 | 4uez | A | 200 | TYR | p.Tyr200Ter | P15085 |
| HOMO SAPIENS | 0 | 8195 | 5xaw | B | 67 | ARG | p.Arg67Trp | Q8IWT1 |
| HOMO SAPIENS | 0 | 8195 | 5xaw | B | 94 | THR | p.Thr94Met | Q8IWT1 |
| HOMO SAPIENS | 1 | 5711 | 1fo3 | A | 265 | ARG | p.Arg265His | Q9UKM7 |
| HOMO SAPIENS | 4 | 5711 | 1fo3 | A | 334 | ARG | p.Arg334Cys | Q9UKM7 |
| HOMO SAPIENS | 1 | 5711 | 1fo3 | A | 478 | LYS | p.Lys478Arg | Q9UKM7 |
| HOMO SAPIENS | 1 | 5711 | 1fo3 | A | 682 | ASP | p.Asp682Glu | Q9UKM7 |
| HOMO SAPIENS | 0 | 170 | 2n3y | A | 41 | GLY | p.Gly42Ser | P99999 |
| HOMO SAPIENS | 0 | 4122 | 3w81 | A | 33 | HIS | p.His33Gln | P35475 |
| HOMO SAPIENS | 0 | 4122 | 3w81 | A | 64 | TYR | p.Tyr64Ter | P35475 |
| HOMO SAPIENS | 0 | 4122 | 3w81 | A | 70 | GLN | p.Gln70Ter | P35475 |
| HOMO SAPIENS | 0 | 4122 | 3w81 | A | 75 | ALA | p.Ala75Thr | P35475 |
| HOMO SAPIENS | 0 | 4122 | 3w81 | A | 79 | ALA | p.Ala79Val | P35475 |
| HOMO SAPIENS | 0 | 4122 | 3w81 | A | 79 | ALA | p.Ala79Thr | P35475 |
| HOMO SAPIENS | 0 | 4122 | 3w81 | A | 89 | ARG | p.Arg89Gln | P35475 |
| HOMO SAPIENS | 0 | 4122 | 3w81 | A | 99 | THR | p.Thr99Ile | P35475 |
| HOMO SAPIENS | 0 | 4122 | 3w81 | A | 105 | ARG | p.Arg105Gln | P35475 |
| HOMO SAPIENS | 0 | 4122 | 3w81 | A | 238 | LEU | p.Leu238Gln | P35475 |
| HOMO SAPIENS | 0 | 4122 | 3w81 | A | 300 | ALA | p.Ala300Thr | P35475 |
| HOMO SAPIENS | 0 | 4122 | 3w81 | A | 322 | VAL | p.Val322Glu | P35475 |
| HOMO SAPIENS | 0 | 4122 | 3w81 | A | 327 | ALA | p.Ala327Pro | P35475 |
| HOMO SAPIENS | 0 | 4122 | 3w81 | A | 346 | LEU | p.Leu346Arg | P35475 |
| HOMO SAPIENS | 0 | 4122 | 3w81 | A | 349 | ASP | p.Asp349Asn | P35475 |
| HOMO SAPIENS | 0 | 4122 | 3w81 | A | 361 | ALA | p.Ala361Thr | P35475 |
| HOMO SAPIENS | 0 | 4122 | 3w81 | A | 366 | THR | p.Thr366Pro | P35475 |
| HOMO SAPIENS | 0 | 4122 | 3w81 | A | 402 | TRP | p.Trp402Ter | P35475 |
| HOMO SAPIENS | 0 | 4122 | 3w81 | A | 409 | GLY | p.Gly409Arg | P35475 |
| HOMO SAPIENS | 0 | 4122 | 3w81 | A | 454 | VAL | p.Val454Ile | P35475 |
| HOMO SAPIENS | 1 | 4122 | 3w81 | A | 490 | LEU | p.Leu490Pro | P35475 |
| HOMO SAPIENS | 1 | 4122 | 3w81 | A | 492 | ARG | p.Arg492Pro | P35475 |
| HOMO SAPIENS | 0 | 4122 | 3w81 | A | 533 | PRO | p.Pro533Arg | P35475 |
| HOMO SAPIENS | 0 | 4122 | 3w81 | A | 591 | ALA | p.Ala591Thr | P35475 |
| HOMO SAPIENS | 7 | 4122 | 3w81 | A | 619 | ARG | p.Arg619Gly | P35475 |
| HOMO SAPIENS | 0 | 15325 | 3wvz | B | 54 | VAL | p.Val54Leu | Q53FT3 |
| HOMO SAPIENS | 3 | 3228 | 5iz8 | A | 421 | TRP | p.Trp421Ter | P25054 |
| HOMO SAPIENS | 2 | 3228 | 5iz8 | A | 437 | PRO | p.Pro437Arg | P25054 |
| HOMO SAPIENS | 3 | 3228 | 5iz8 | A | 499 | ARG | p.Arg499Ter | P25054 |
| HOMO SAPIENS | -1 | 29229 | 5buo | A | 539 | SER | p.Ser614Gly | P05067 |
| HOMO SAPIENS | -1 | 34931 | 4n7v | C | 54 | SER | p.Ser54Leu | O94986 |
| HOMO SAPIENS | 0 | 2450 | 5kzx | A | 123 | GLY | p.Gly123Glu | P10253 |
| HOMO SAPIENS | 1 | 2450 | 5kzx | A | 190 | ARG | p.Arg190His | P10253 |
| HOMO SAPIENS | -1 | 2450 | 5kzx | A | 199 | HIS | p.His199Arg | P10253 |
| HOMO SAPIENS | 0 | 2450 | 5kzx | A | 219 | GLY | p.Gly219Arg | P10253 |
| HOMO SAPIENS | 0 | 2450 | 5kzx | A | 220 | VAL | p.Val220Leu | P10253 |
| HOMO SAPIENS | 0 | 2450 | 5kzx | A | 222 | VAL | p.Val222Met | P10253 |

| | | | | | | | | |
|---|---|---|---|---|---|---|---|---|
| HOMO SAPIENS | 0 | 2450 | 5kzx | A | 237 | ALA | p.Ala237Val | P10253 |
| HOMO SAPIENS | 0 | 2450 | 5kzx | A | 251 | SER | p.Ser251Leu | P10253 |
| HOMO SAPIENS | 0 | 2450 | 5kzx | A | 254 | SER | p.Ser254Leu | P10253 |
| HOMO SAPIENS | 0 | 2450 | 5kzx | A | 262 | GLU | p.Glu262Lys | P10253 |
| HOMO SAPIENS | 0 | 2450 | 5kzx | A | 293 | GLY | p.Gly293Arg | P10253 |
| HOMO SAPIENS | 0 | 2450 | 5kzx | A | 299 | LEU | p.Leu299Arg | P10253 |
| HOMO SAPIENS | 0 | 2450 | 5kzx | A | 309 | GLY | p.Gly309Arg | P10253 |
| HOMO SAPIENS | 0 | 2450 | 5kzx | A | 318 | MET | p.Met318Thr | P10253 |
| HOMO SAPIENS | 0 | 2450 | 5kzx | A | 355 | LEU | p.Leu355Pro | P10253 |
| HOMO SAPIENS | 1 | 2450 | 5kzx | A | 361 | PRO | p.Pro361Leu | P10253 |
| HOMO SAPIENS | 4 | 2450 | 5kzx | A | 408 | MET | p.Met408Val | P10253 |
| HOMO SAPIENS | 1 | 2450 | 5kzx | A | 429 | GLN | p.Gln429Glu | P10253 |
| HOMO SAPIENS | 4 | 2450 | 5kzx | A | 437 | ARG | p.Arg437Cys | P10253 |
| HOMO SAPIENS | 0 | 2450 | 5kzx | A | 439 | MET | p.Met439Lys | P10253 |
| HOMO SAPIENS | 1 | 2450 | 5kzx | A | 448 | SER | p.Ser448Thr | P10253 |
| HOMO SAPIENS | 0 | 2450 | 5kzx | A | 451 | PRO | p.Pro451Arg | P10253 |
| HOMO SAPIENS | 1 | 2450 | 5kzx | A | 459 | ASP | p.Asp459His | P10253 |
| HOMO SAPIENS | 8 | 2450 | 5kzx | A | 481 | TRP | p.Trp481Arg | P10253 |
| HOMO SAPIENS | 1 | 2450 | 5kzx | A | 489 | ASP | p.Asp489Asn | P10253 |
| HOMO SAPIENS | 0 | 2450 | 5kzx | A | 521 | GLU | p.Glu521Lys | P10253 |
| HOMO SAPIENS | 0 | 2450 | 5kzx | A | 529 | SER | p.Ser529Val | P10253 |
| HOMO SAPIENS | 0 | 2450 | 5kzx | A | 545 | PRO | p.Pro545Leu | P10253 |
| HOMO SAPIENS | 0 | 2450 | 5kzx | A | 552 | LEU | p.Leu552Pro | P10253 |
| HOMO SAPIENS | 0 | 2450 | 5kzx | A | 576 | GLY | p.Gly576Ser | P10253 |
| HOMO SAPIENS | 9 | 2450 | 5kzx | A | 600 | ARG | p.Arg600His | P10253 |
| HOMO SAPIENS | 8 | 2450 | 5kzx | A | 615 | GLY | p.Gly615Arg | P10253 |
| HOMO SAPIENS | 3 | 2450 | 5kzx | A | 619 | SER | p.Ser619Asn | P10253 |
| HOMO SAPIENS | 0 | 2450 | 5kzx | A | 638 | GLY | p.Gly638Trp | P10253 |
| HOMO SAPIENS | 0 | 2450 | 5kzx | A | 643 | GLY | p.Gly643Arg | P10253 |
| HOMO SAPIENS | 9 | 2450 | 5kzx | A | 645 | ASP | p.Asp645Glu | P10253 |
| HOMO SAPIENS | 9 | 2450 | 5kzx | A | 645 | ASP | p.Asp645His | P10253 |
| HOMO SAPIENS | 0 | 2450 | 5kzx | A | 648 | GLY | p.Gly648Ser | P10253 |
| HOMO SAPIENS | 5 | 2450 | 5kzx | A | 660 | ARG | p.Arg660His | P10253 |
| HOMO SAPIENS | 8 | 2450 | 5kzx | A | 672 | ARG | p.Arg672Gln | P10253 |
| HOMO SAPIENS | 0 | 2450 | 5kzx | A | 702 | ARG | p.Arg702Leu | P10253 |
| HOMO SAPIENS | 3 | 2450 | 5kzx | A | 725 | ARG | p.Arg725Trp | P10253 |
| HOMO SAPIENS | 0 | 2450 | 5kzx | A | 746 | TRP | p.Trp746Cys | P10253 |
| HOMO SAPIENS | 0 | 2450 | 5kzx | A | 780 | ILE | p.Val780Ile | P10253 |
| HOMO SAPIENS | 1 | 2450 | 5kzx | A | 816 | VAL | p.Val816Ile | P10253 |
| HOMO SAPIENS | 0 | 2450 | 5kzx | A | 838 | GLN | p.Gln838Ter | P10253 |
| HOMO SAPIENS | 2 | 2450 | 5kzx | A | 854 | ARG | p.Arg854Ter | P10253 |
| HOMO SAPIENS | 3 | 2450 | 5kzx | A | 870 | ARG | p.Arg870Ter | P10253 |
| HOMO SAPIENS | 0 | 2450 | 5kzx | A | 890 | VAL | p.Val890Leu | P10253 |
| HOMO SAPIENS | 6 | 1882 | 4km6 | A | 175 | CYS | p.Cys175Ter | P15328 |
| HOMO SAPIENS | 3 | 14767 | 1lm5 | A | 2628 | LEU | p.Leu2628Pro | P15924 |
| HOMO SAPIENS | 5 | 14767 | 1lm5 | A | 2655 | ALA | p.Ala2655Gly | P15924 |
| HOMO SAPIENS | 3 | 14767 | 1lm5 | A | 2731 | TYR | p.Tyr2731His | P15924 |
| HOMO SAPIENS | 1 | 14767 | 1lm5 | A | 2767 | THR | p.Thr2767Asn | P15924 |
| HOMO SAPIENS | -1 | 14767 | 1lm5 | A | 2819 | MET | p.Met2819Leu | P15924 |
| HOMO SAPIENS | -1 | 11246 | 4y5o | A | 289 | SER | p.Ser289Asn | Q9BSQ5 |
| HOMO SAPIENS | 0 | 7380 | 5kz5 | I | 60 | ARG | p.Arg60His | Q16595 |
| HOMO SAPIENS | 0 | 7380 | 5kz5 | I | 76 | MET | p.Met76Val | Q16595 |
| HOMO SAPIENS | 2 | 22187 | 3fby | A | 328 | CYS | p.Cys328Arg | P49747 |
| HOMO SAPIENS | 2 | 22187 | 3fby | A | 342 | ASP | p.Asp342Tyr | P49747 |

| | | | | | | | | |
|---|---|---|---|---|---|---|---|---|
| HOMO SAPIENS | 2 | 22187 | 3fby | A | 348 | CYS | p.Cys348Arg | P49747 |
| HOMO SAPIENS | 2 | 22187 | 3fby | A | 386 | ASN | p.Asn386Asp | P49747 |
| HOMO SAPIENS | 2 | 22187 | 3fby | A | 453 | ASN | p.Asn453Ser | P49747 |
| HOMO SAPIENS | 2 | 22187 | 3fby | A | 468 | CYS | p.Cys468Tyr | P49747 |
| HOMO SAPIENS | 3 | 22187 | 3fby | A | 472 | ASP | p.Asp472Tyr | P49747 |
| HOMO SAPIENS | 3 | 22187 | 3fby | A | 473 | ASP | p.Asp473Gly | P49747 |
| HOMO SAPIENS | 2 | 22187 | 3fby | A | 523 | ASN | p.Asn523Lys | P49747 |
| HOMO SAPIENS | 1 | 22187 | 3fby | A | 718 | ARG | p.Arg718Trp | P49747 |
| HOMO SAPIENS | 1 | 22187 | 3fby | A | 719 | GLY | p.Gly719Asp | P49747 |
| HOMO SAPIENS | 0 | 22187 | 3fby | A | 756 | GLN | p.Gln756Arg | P49747 |
| HOMO SAPIENS | -1 | 2784 | 5kvh | A | 88 | MET | p.Met88Val | O95831 |
| HOMO SAPIENS | -1 | 2784 | 5kvh | A | 96 | ASN | p.Asn96Ser | O95831 |
| HOMO SAPIENS | 0 | 2784 | 5kvh | A | 282 | THR | p.Thr282Met | O95831 |
| HOMO SAPIENS | 1 | 2784 | 5kvh | A | 366 | ASN | p.Asn366Ser | O95831 |
| HOMO SAPIENS | 3 | 2784 | 5kvh | A | 463 | ARG | p.Arg463Ile | O95831 |
| HOMO SAPIENS | 0 | 2784 | 5kvh | A | 498 | VAL | p.Val498Met | O95831 |
| HOMO SAPIENS | -1 | 2784 | 5kvh | A | 549 | ALA | p.Ala549Val | O95831 |
| HOMO SAPIENS | 0 | 38316 | 2lnk | C | 1933 | ARG | p.Arg1933Ter | P35579 |
| HOMO SAPIENS | 0 | 33808 | 2miu | A | 29 | VAL | p.Val29Met | Q14192 |
| HOMO SAPIENS | 1 | 33808 | 2miu | A | 37 | ALA | p.Ala37Ser | Q14192 |
| HOMO SAPIENS | 2 | 20994 | 2hey | R | 65 | ARG | p.Arg65Cys | P43489 |
| HOMO SAPIENS | -1 | 5687 | 2xwj | I | 7 | ARG | p.Arg32Gln | P00751 |
| HOMO SAPIENS | -1 | 5687 | 2xwj | I | 7 | ARG | p.Arg32Trp | P00751 |
| HOMO SAPIENS | 0 | 5687 | 2xwj | I | 49 | ARG | p.Arg74His | P00751 |
| HOMO SAPIENS | 0 | 5687 | 2xwj | I | 177 | ARG | p.Arg202Gly | P00751 |
| HOMO SAPIENS | 0 | 5687 | 2xwj | I | 217 | ILE | p.Ile242Leu | P00751 |
| HOMO SAPIENS | -1 | 5687 | 2xwj | I | 227 | GLY | p.Gly252Ser | P00751 |
| HOMO SAPIENS | 0 | 5687 | 2xwj | I | 483 | HIS | p.His508Gln | P00751 |
| HOMO SAPIENS | 0 | 5687 | 2xwj | I | 508 | LYS | p.Lys533Arg | P00751 |
| HOMO SAPIENS | 0 | 5687 | 2xwj | I | 540 | LYS | p.Lys565Glu | P00751 |
| HOMO SAPIENS | 0 | 5687 | 2xwj | I | 541 | GLU | p.Glu566Ala | P00751 |
| HOMO SAPIENS | 2 | 953 | 3d4j | A | 249 | PHE | p.Phe249Ser | P53602 |
| HOMO SAPIENS | 1 | 953 | 3d4j | A | 292 | ASN | p.Asn292Ser | P53602 |
| HOMO SAPIENS | 0 | 3300 | 1s9c | A | 140 | ASN | p.Asn457Tyr | P51659 |
| HOMO SAPIENS | 2 | 3300 | 1s9c | A | 167 | ILE | p.Ile484Val | P51659 |
| HOMO SAPIENS | 0 | 3300 | 1s9c | A | 174 | ALA | p.Ala491Thr | P51659 |
| HOMO SAPIENS | 8 | 3300 | 1s9c | A | 193 | ASP | p.Asp510Asn | P51659 |
| HOMO SAPIENS | 3 | 3300 | 1s9c | A | 194 | TRP | p.Trp511Arg | P51659 |
| HOMO SAPIENS | 0 | 3300 | 1s9c | A | 242 | VAL | p.Ile559Val | P51659 |
| HOMO SAPIENS | 0 | 10810 | 5i25 | A | 5 | GLN | p.Gln23Ter | P03951 |
| HOMO SAPIENS | 0 | 10810 | 5i25 | A | 55 | TRP | p.Trp73Ter | P03951 |
| HOMO SAPIENS | 0 | 10810 | 5i25 | A | 91 | ALA | p.Ala109Thr | P03951 |
| HOMO SAPIENS | 0 | 10810 | 5i25 | A | 117 | GLU | p.Glu135Ter | P03951 |
| HOMO SAPIENS | 0 | 10810 | 5i25 | A | 118 | CYS | p.Cys136Ter | P03951 |
| HOMO SAPIENS | 0 | 10810 | 5i25 | A | 128 | CYS | p.Cys146Ter | P03951 |
| HOMO SAPIENS | 0 | 10810 | 5i25 | A | 210 | ARG | p.Arg228Ter | P03951 |
| HOMO SAPIENS | 0 | 10810 | 5i25 | A | 226 | GLN | p.Gln244Ter | P03951 |
| HOMO SAPIENS | 0 | 10810 | 5i25 | A | 233 | GLN | p.Gln251Ter | P03951 |
| HOMO SAPIENS | 0 | 16668 | 3lbx | A | 28 | ARG | p.Arg28Leu | P02549 |
| HOMO SAPIENS | 0 | 16668 | 3lbx | A | 28 | ARG | p.Arg28Cys | P02549 |
| HOMO SAPIENS | 1 | 16668 | 3lbx | A | 41 | ARG | p.Arg41Trp | P02549 |
| HOMO SAPIENS | 1 | 16668 | 3lbx | A | 45 | ARG | p.Arg45Ser | P02549 |
| HOMO SAPIENS | 2 | 16668 | 3lbx | A | 48 | LYS | p.Lys48Arg | P02549 |
| HOMO SAPIENS | 1 | 16668 | 3lbx | A | 152 | ASP | p.Asp152Asn | P02549 |

| | | | | | | | | |
|---|---|---|---|---|---|---|---|---|
| HOMO SAPIENS | 0 | 226 | 2ble | A | 256 | ILE | p.Phe256Ile | P36959 |
| HOMO SAPIENS | 2 | 36808 | 4c7n | A | 3 | SER | p.Ser357Pro | O75030 |
| HOMO SAPIENS | 0 | 4092 | 5jnx | A | 113 | CYS | p.Cys113Arg | O15118 |
| HOMO SAPIENS | 4 | 4092 | 5jnx | A | 215 | HIS | p.His215Arg | O15118 |
| HOMO SAPIENS | 6 | 4092 | 5jnx | A | 237 | PRO | p.Pro237Ser | O15118 |
| HOMO SAPIENS | 0 | 4092 | 5jnx | A | 363 | ALA | p.Ala363Val | O15118 |
| HOMO SAPIENS | 0 | 4092 | 5jnx | A | 378 | VAL | p.Val378Ala | O15118 |
| HOMO SAPIENS | 8 | 4092 | 5jnx | A | 404 | ARG | p.Arg404Gln | O15118 |
| HOMO SAPIENS | 8 | 4092 | 5jnx | A | 411 | ARG | p.Arg411Gln | O15118 |
| HOMO SAPIENS | 2 | 4092 | 5jnx | A | 474 | PRO | p.Pro474Leu | O15118 |
| HOMO SAPIENS | 8 | 4092 | 5jnx | A | 511 | THR | p.Thr511Met | O15118 |
| HOMO SAPIENS | 0 | 4092 | 5jnx | A | 642 | MET | p.Met642Ile | O15118 |
| HOMO SAPIENS | 2 | 4092 | 5jnx | A | 664 | VAL | p.Val664Met | O15118 |
| HOMO SAPIENS | 0 | 4092 | 5jnx | A | 734 | SER | p.Ser734Ile | O15118 |
| HOMO SAPIENS | -1 | 4092 | 5jnx | A | 775 | GLN | p.Gln775Pro | O15118 |
| HOMO SAPIENS | 0 | 4092 | 5jnx | A | 858 | ILE | p.Ile858Val | O15118 |
| HOMO SAPIENS | 0 | 4092 | 5jnx | A | 889 | VAL | p.Val889Met | O15118 |
| HOMO SAPIENS | 0 | 4092 | 5jnx | A | 910 | GLY | p.Gly910Ser | O15118 |
| HOMO SAPIENS | 0 | 4092 | 5jnx | A | 911 | GLY | p.Gly911Ser | O15118 |
| HOMO SAPIENS | 0 | 4092 | 5jnx | A | 928 | GLN | p.Gln928Pro | O15118 |
| HOMO SAPIENS | 0 | 4092 | 5jnx | A | 931 | ASN | p.Asn931Lys | O15118 |
| HOMO SAPIENS | 0 | 4092 | 5jnx | A | 940 | SER | p.Ser940Leu | O15118 |
| HOMO SAPIENS | 0 | 4092 | 5jnx | A | 950 | VAL | p.Val950Met | O15118 |
| HOMO SAPIENS | 0 | 4092 | 5jnx | A | 954 | SER | p.Ser954Leu | O15118 |
| HOMO SAPIENS | 0 | 4092 | 5jnx | A | 958 | ARG | p.Arg958Gln | O15118 |
| HOMO SAPIENS | 0 | 4092 | 5jnx | A | 958 | ARG | p.Arg958Ter | O15118 |
| HOMO SAPIENS | -1 | 4092 | 5jnx | A | 978 | ARG | p.Arg978Cys | O15118 |
| HOMO SAPIENS | 0 | 4092 | 5jnx | A | 992 | GLY | p.Gly992Trp | O15118 |
| HOMO SAPIENS | 0 | 4092 | 5jnx | A | 1007 | PRO | p.Pro1007Ala | O15118 |
| HOMO SAPIENS | 0 | 4092 | 5jnx | A | 1019 | TYR | p.Tyr1019Cys | O15118 |
| HOMO SAPIENS | 2 | 4092 | 5jnx | A | 1035 | ALA | p.Ala1035Val | O15118 |
| HOMO SAPIENS | 2 | 4092 | 5jnx | A | 1054 | ALA | p.Ala1054Thr | O15118 |
| HOMO SAPIENS | 2 | 4092 | 5jnx | A | 1061 | ILE | p.Ile1061Thr | O15118 |
| HOMO SAPIENS | 0 | 4092 | 5jnx | A | 1073 | GLY | p.Gly1073Ser | O15118 |
| HOMO SAPIENS | 2 | 4092 | 5jnx | A | 1088 | TYR | p.Tyr1088Cys | O15118 |
| HOMO SAPIENS | 2 | 4092 | 5jnx | A | 1142 | MET | p.Met1142Thr | O15118 |
| HOMO SAPIENS | 2 | 4092 | 5jnx | A | 1156 | ASN | p.Asn1156Ser | O15118 |
| HOMO SAPIENS | 0 | 4092 | 5jnx | A | 1179 | MET | p.Met1179Val | O15118 |
| HOMO SAPIENS | 0 | 4092 | 5jnx | A | 1186 | ARG | p.Arg1186His | O15118 |
| HOMO SAPIENS | 0 | 4092 | 5jnx | A | 1187 | ALA | p.Ala1187Gly | O15118 |
| HOMO SAPIENS | 2 | 4092 | 5jnx | A | 1189 | GLU | p.Glu1189Gly | O15118 |
| HOMO SAPIENS | 2 | 4092 | 5jnx | A | 1200 | SER | p.Ser1200Gly | O15118 |
| HOMO SAPIENS | 2 | 4092 | 5jnx | A | 1205 | THR | p.Thr1205Lys | O15118 |
| HOMO SAPIENS | 0 | 4092 | 5jnx | A | 1213 | LEU | p.Leu1213Phe | O15118 |
| HOMO SAPIENS | -1 | 4092 | 5jnx | A | 1266 | ARG | p.Arg1266Gln | O15118 |
| HOMO SAPIENS | 9 | 16498 | 4i6o | A | 64 | ARG | p.Arg735Trp | P01024 |
| HOMO SAPIENS | 1 | 21902 | 3un9 | C | 751 | ARG | p.Arg751His | Q86UT6 |
| HOMO SAPIENS | 7 | 20391 | 2h32 | B | 105 | PRO | p.Pro142Leu | P15814 |
| HOMO SAPIENS | 6 | 1635 | 4x45 | A | 65 | ASP | p.Asp65Val | P07741 |
| HOMO SAPIENS | 1 | 1635 | 4x45 | A | 98 | TRP | p.Trp98Ter | P07741 |
| HOMO SAPIENS | 0 | 1635 | 4x45 | A | 110 | LEU | p.Leu110Pro | P07741 |
| HOMO SAPIENS | 7 | 1635 | 4x45 | A | 136 | MET | p.Met136Thr | P07741 |
| HOMO SAPIENS | 1 | 1635 | 4x45 | A | 150 | VAL | p.Val150Phe | P07741 |
| HOMO SAPIENS | 4 | 34810 | 2rpc | A | 16 | CYS | p.Cys253Ser | O60481 |

| | | | | | | | | |
|---|---|---|---|---|---|---|---|---|
| HOMO SAPIENS | 1 | 34810 | 2rpc | A | 18 | TRP | p.Trp255Gly | O60481 |
| HOMO SAPIENS | 1 | 14359 | 4hxi | A | 78 | MET | p.Met78Val | Q9UH77 |
| HOMO SAPIENS | 0 | 5342 | 4d9t | A | 628 | GLU | p.Glu628Asp | P51812 |
| HOMO SAPIENS | -1 | 5342 | 4d9t | A | 723 | ARG | p.Arg723His | P51812 |
| HOMO SAPIENS | -1 | 5342 | 4d9t | A | 729 | ARG | p.Arg729Gln | P51812 |
| HOMO SAPIENS | 1 | 9619 | 3j2s | B | 1781 | ARG | p.Arg1800Cys | P00451 |
| HOMO SAPIENS | 0 | 9619 | 3j2s | B | 1823 | MET | p.Met1842Ile | P00451 |
| HOMO SAPIENS | 1 | 9619 | 3j2s | B | 1825 | PRO | p.Pro1844Ser | P00451 |
| HOMO SAPIENS | 0 | 9619 | 3j2s | B | 1826 | THR | p.Thr1845Pro | P00451 |
| HOMO SAPIENS | 0 | 9619 | 3j2s | B | 1834 | ALA | p.Ala1853Val | P00451 |
| HOMO SAPIENS | 0 | 9619 | 3j2s | B | 1846 | ASP | p.Asp1865Asn | P00451 |
| HOMO SAPIENS | 1 | 9619 | 3j2s | B | 1848 | HIS | p.His1867Arg | P00451 |
| HOMO SAPIENS | 0 | 9619 | 3j2s | B | 1854 | PRO | p.Pro1873Arg | P00451 |
| HOMO SAPIENS | 1 | 9619 | 3j2s | B | 1885 | GLU | p.Glu1904Lys | P00451 |
| HOMO SAPIENS | 0 | 9619 | 3j2s | B | 1922 | ASN | p.Asn1941Ser | P00451 |
| HOMO SAPIENS | 1 | 9619 | 3j2s | B | 1941 | ARG | p.Arg1960Gln | P00451 |
| HOMO SAPIENS | 4 | 9619 | 3j2s | B | 2105 | TYR | p.Tyr2124Cys | P00451 |
| HOMO SAPIENS | 6 | 9619 | 3j2s | B | 2150 | ARG | p.Arg2169His | P00451 |
| HOMO SAPIENS | 0 | 9619 | 3j2s | B | 2159 | ARG | p.Arg2178Cys | P00451 |
| HOMO SAPIENS | 0 | 9619 | 3j2s | B | 2159 | ARG | p.Arg2178His | P00451 |
| HOMO SAPIENS | 2 | 9619 | 3j2s | B | 2238 | MET | p.Met2257Val | P00451 |
| HOMO SAPIENS | 2 | 9619 | 3j2s | B | 2300 | PRO | p.Pro2319Ser | P00451 |
| HOMO SAPIENS | 1 | 667 | 1z2m | A | 127 | GLU | p.Glu127Ter | P05161 |
| HOMO SAPIENS | 0 | 15717 | 2nn6 | G | 30 | GLY | p.Gly31Ala | Q9NQT5 |
| HOMO SAPIENS | -1 | 15717 | 2nn6 | G | 55 | ASN | p.Asn56His | Q9NQT5 |
| HOMO SAPIENS | 7 | 15717 | 2nn6 | G | 131 | ASP | p.Asp132Ala | Q9NQT5 |
| HOMO SAPIENS | 7 | 784 | 3rkq | A | 161 | ARG | p.Arg161Pro | P52952 |
| HOMO SAPIENS | 6 | 784 | 3rkq | A | 178 | THR | p.Thr178Met | P52952 |
| HOMO SAPIENS | 7 | 784 | 3rkq | A | 183 | LYS | p.Lys183Glu | P52952 |
| HOMO SAPIENS | 8 | 784 | 3rkq | A | 190 | ARG | p.Arg190Cys | P52952 |
| HOMO SAPIENS | 6 | 385 | 30000 | C | 70 | ALA | p.Ala70Thr | P07954 |
| HOMO SAPIENS | 2 | 385 | 30000 | C | 101 | ARG | p.Arg101Pro | P07954 |
| HOMO SAPIENS | 2 | 385 | 30000 | C | 174 | PRO | p.Pro174Arg | P07954 |
| HOMO SAPIENS | 6 | 385 | 30000 | C | 187 | SER | p.Ser187Ter | P07954 |
| HOMO SAPIENS | 9 | 385 | 30000 | C | 233 | ARG | p.Arg233His | P07954 |
| HOMO SAPIENS | 3 | 385 | 30000 | C | 273 | ALA | p.Ala273Thr | P07954 |
| HOMO SAPIENS | 4 | 385 | 30000 | C | 334 | SER | p.Ser334Arg | P07954 |
| HOMO SAPIENS | 6 | 385 | 30000 | C | 340 | ASN | p.Asn340Lys | P07954 |
| HOMO SAPIENS | 8 | 385 | 30000 | C | 343 | ARG | p.Arg343Ter | P07954 |
| HOMO SAPIENS | 3 | 385 | 30000 | C | 350 | ARG | p.Arg350Trp | P07954 |
| HOMO SAPIENS | 4 | 385 | 30000 | C | 376 | GLN | p.Gln376Pro | P07954 |
| HOMO SAPIENS | 6 | 385 | 30000 | C | 434 | CYS | p.Cys434Ter | P07954 |
| HOMO SAPIENS | 0 | 3316 | 4f91 | B | 934 | THR | p.Thr934Ala | O75643 |
| HOMO SAPIENS | 0 | 3316 | 4f91 | B | 1002 | ASN | p.Asn1002Ser | O75643 |
| HOMO SAPIENS | 0 | 3316 | 4f91 | B | 1087 | SER | p.Ser1087Leu | O75643 |
| HOMO SAPIENS | 4 | 3316 | 4f91 | B | 1389 | VAL | p.Val1389Ile | O75643 |
| HOMO SAPIENS | 0 | 619 | 5hbt | B | 195 | ASP | p.Asp240His | P02708 |
| HOMO SAPIENS | 1 | 619 | 5hbt | B | 209 | ARG | p.Arg254Leu | P02708 |
| HOMO SAPIENS | 0 | 18171 | 2y25 | B | 1408 | ASP | p.Asp1408Asn | P52179 |
| HOMO SAPIENS | 0 | 18171 | 2y25 | B | 1453 | MET | p.Met1453Thr | P52179 |
| HOMO SAPIENS | 0 | 18171 | 2y25 | B | 1459 | ILE | p.Ile1459Lys | P52179 |
| HOMO SAPIENS | 0 | 18171 | 2y25 | B | 1558 | PHE | p.Phe1558Leu | P52179 |
| HOMO SAPIENS | 0 | 18171 | 2y25 | B | 1573 | ARG | p.Arg1573Gln | P52179 |
| HOMO SAPIENS | 7 | 18171 | 2y25 | B | 1663 | VAL | p.Val1663Met | P52179 |

| | | | | | | | | |
|---|---|---|---|---|---|---|---|---|
| HOMO SAPIENS | 8 | 14262 | 2fcw | B | 117 | SER | p.Ser138Ter | P01130 |
| HOMO SAPIENS | 8 | 14262 | 2fcw | B | 118 | ASP | p.Asp139Glu | P01130 |
| HOMO SAPIENS | 9 | 14262 | 2fcw | B | 119 | GLU | p.Glu140Lys | P01130 |
| HOMO SAPIENS | 7 | 14262 | 2fcw | B | 122 | CYS | p.Cys143Ter | P01130 |
| HOMO SAPIENS | 6 | 14262 | 2fcw | B | 134 | CYS | p.Cys155Ter | P01130 |
| HOMO SAPIENS | 7 | 14262 | 2fcw | B | 147 | ASP | p.Asp168Glu | P01130 |
| HOMO SAPIENS | 4 | 2624 | 3b7b | A | 855 | VAL | p.Val886Met | Q9H9B1 |
| HOMO SAPIENS | 7 | 2624 | 3b7b | A | 868 | ILE | p.Ile899Val | Q9H9B1 |
| HOMO SAPIENS | 0 | 2624 | 3b7b | A | 892 | GLU | p.Glu923Lys | Q9H9B1 |
| HOMO SAPIENS | 0 | 4791 | 5ctd | B | 68 | LEU | p.Pro549Ala | Q14055 |
| HOMO SAPIENS | 5 | 460 | 4ui1 | D | 80 | CYS | p.Cys80Arg | Q6ZVN8 |
| HOMO SAPIENS | 3 | 460 | 4ui1 | D | 101 | LEU | p.Leu101Pro | Q6ZVN8 |
| HOMO SAPIENS | 0 | 40782 | 1sy6 | A | 47 | LYS | p.Lys69Ter | P09693 |
| HOMO SAPIENS | -1 | 1092 | 2idx | A | 62 | THR | p.Thr62Met | Q96EY8 |
| HOMO SAPIENS | 6 | 1092 | 2idx | A | 183 | HIS | p.His183Leu | Q96EY8 |
| HOMO SAPIENS | 7 | 1092 | 2idx | A | 186 | ARG | p.Arg186Trp | Q96EY8 |
| HOMO SAPIENS | 6 | 1092 | 2idx | A | 190 | ARG | p.Arg190Cys | Q96EY8 |
| HOMO SAPIENS | 6 | 1092 | 2idx | A | 190 | ARG | p.Arg190His | Q96EY8 |
| HOMO SAPIENS | 7 | 1092 | 2idx | A | 191 | ARG | p.Arg191Trp | Q96EY8 |
| HOMO SAPIENS | 7 | 1092 | 2idx | A | 191 | ARG | p.Arg191Gln | Q96EY8 |
| HOMO SAPIENS | 6 | 1092 | 2idx | A | 195 | ARG | p.Arg195His | Q96EY8 |
| HOMO SAPIENS | 1 | 1092 | 2idx | A | 234 | GLN | p.Gln234Ter | Q96EY8 |
| HOMO SAPIENS | 0 | 1092 | 2idx | A | 239 | MET | p.Met239Lys | Q96EY8 |
| HOMO SAPIENS | -1 | 6418 | 5fur | G | 11 | THR | p.Thr11Ile | P21675 |
| HOMO SAPIENS | -1 | 6418 | 5fur | G | 290 | LEU | p.Leu269Val | P21675 |
| HOMO SAPIENS | 0 | 6100 | 1wiq | A | 240 | ARG | p.Arg265Trp | P01730 |
| HOMO SAPIENS | 0 | 13757 | 4ye6 | A | 45 | GLY | p.Gly45Val | P47897 |
| HOMO SAPIENS | 1 | 13757 | 4ye6 | A | 57 | TYR | p.Tyr57His | P47897 |
| HOMO SAPIENS | 8 | 13757 | 4ye6 | A | 403 | ARG | p.Arg403Trp | P47897 |
| HOMO SAPIENS | 7 | 13757 | 4ye6 | A | 524 | ARG | p.Arg524Trp | P47897 |
| HOMO SAPIENS | 6 | 13757 | 4ye6 | A | 690 | ARG | p.Arg690Cys | P47897 |
| HOMO SAPIENS | 0 | 26958 | 2lzl | A | 370 | GLY | p.Gly370Cys | P22607 |
| HOMO SAPIENS | 0 | 26958 | 2lzl | A | 371 | SER | p.Ser371Cys | P22607 |
| HOMO SAPIENS | 0 | 26958 | 2lzl | A | 373 | TYR | p.Tyr373Cys | P22607 |
| HOMO SAPIENS | 0 | 26958 | 2lzl | A | 380 | GLY | p.Gly380Arg | P22607 |
| HOMO SAPIENS | 0 | 26958 | 2lzl | A | 384 | PHE | p.Phe384Leu | P22607 |
| HOMO SAPIENS | 5 | 26958 | 2lzl | A | 391 | ALA | p.Ala391Glu | P22607 |
| HOMO SAPIENS | 6 | 11325 | 2wa6 | A | 121 | TYR | p.Tyr121Phe | O75369 |
| HOMO SAPIENS | 2 | 11325 | 2wa6 | A | 161 | PHE | p.Phe161Cys | O75369 |
| HOMO SAPIENS | 0 | 11325 | 2wa6 | A | 171 | LEU | p.Leu171Arg | O75369 |
| HOMO SAPIENS | 2 | 11325 | 2wa6 | A | 173 | ALA | p.Ala173Val | O75369 |
| HOMO SAPIENS | 0 | 11325 | 2wa6 | A | 202 | MET | p.Met202Val | O75369 |
| HOMO SAPIENS | 3 | 11325 | 2wa6 | A | 227 | GLU | p.Glu227Lys | O75369 |
| HOMO SAPIENS | 0 | 11325 | 2wa6 | A | 234 | LEU | p.Leu234Val | O75369 |
| HOMO SAPIENS | 2 | 11325 | 2wa6 | A | 235 | SER | p.Ser235Pro | O75369 |
| HOMO SAPIENS | 9 | 7240 | 3tgx | A | 137 | TYR | p.Tyr156Cys | Q9HBE5 |
| HOMO SAPIENS | 0 | 35479 | 4elj | A | 307 | THR | p.Thr307Ile | P06400 |
| HOMO SAPIENS | 0 | 35479 | 4elj | A | 310 | GLY | p.Gly310Glu | P06400 |
| HOMO SAPIENS | 0 | 35479 | 4elj | A | 377 | THR | p.Thr377Ser | P06400 |
| HOMO SAPIENS | 9 | 36706 | 1he7 | A | 359 | TYR | p.Tyr359Cys | P04629 |
| HOMO SAPIENS | 4 | 6538 | 4d0e | A | 504 | ARG | p.Arg504His | P46531 |
| HOMO SAPIENS | 1 | 26518 | 5fm5 | M | 600 | GLU | p.Glu600Val | P52179 |
| HOMO SAPIENS | -1 | 26518 | 5fm5 | M | 651 | ARG | p.Arg651Gln | P52179 |
| HOMO SAPIENS | -1 | 26518 | 5fm5 | M | 660 | PRO | p.Pro660Ala | P52179 |

| | | | | | | | | |
|---|---|---|---|---|---|---|---|---|
| HOMO SAPIENS | -1 | 26518 | 5fm5 | M | 688 | THR | p.Thr688Ser | P52179 |
| HOMO SAPIENS | -1 | 26518 | 5fm5 | M | 704 | GLU | p.Glu704Lys | P52179 |
| HOMO SAPIENS | -1 | 26518 | 5fm5 | M | 727 | THR | p.Thr727Ala | P52179 |
| HOMO SAPIENS | 2 | 27054 | 4rhp | B | 209 | LEU | p.Leu209Val | O75208 |
| HOMO SAPIENS | -1 | 27054 | 4rhp | B | 288 | LYS | p.Lys288Asn | O75208 |
| HOMO SAPIENS | 5 | 2384 | 5fqd | B | 419 | ARG | p.Arg419Ter | Q96SW2 |
| HOMO SAPIENS | 4 | 5154 | 2hl3 | A | 56 | LYS | p.Lys56Arg | Q14203 |
| HOMO SAPIENS | 0 | 5154 | 2hl3 | A | 59 | GLY | p.Gly59Ser | Q14203 |
| HOMO SAPIENS | 0 | 5154 | 2hl3 | A | 71 | GLY | p.Gly71Ala | Q14203 |
| HOMO SAPIENS | 0 | 5154 | 2hl3 | A | 71 | GLY | p.Gly71Arg | Q14203 |
| HOMO SAPIENS | 7 | 5154 | 2hl3 | A | 72 | THR | p.Thr72Pro | Q14203 |
| HOMO SAPIENS | 2 | 5154 | 2hl3 | A | 74 | GLN | p.Gln74Pro | Q14203 |
| HOMO SAPIENS | 2 | 20039 | 2l1g | A | 9 | GLY | p.Gly9Cys | Q9NVV9 |
| HOMO SAPIENS | 5 | 20039 | 2l1g | A | 81 | PHE | p.Phe81Leu | Q9NVV9 |
| HOMO SAPIENS | -1 | 97 | 5uz7 | A | 201 | ARG | p.Arg201His | P63092 |
| HOMO SAPIENS | -1 | 97 | 5uz7 | A | 201 | ARG | p.Arg201Cys | P63092 |
| HOMO SAPIENS | 0 | 3257 | 4b7w | B | 278 | GLU | p.Glu278Asp | P41235 |
| HOMO SAPIENS | 1 | 3257 | 4b7w | B | 331 | ARG | p.Arg331Cys | P41235 |
| HOMO SAPIENS | 0 | 2814 | 4qgk | B | 10 | GLN | p.Gln10Ter | P51648 |
| HOMO SAPIENS | 0 | 2814 | 4qgk | B | 177 | ARG | p.Arg177Ter | P51648 |
| HOMO SAPIENS | 9 | 2814 | 4qgk | B | 184 | THR | p.Thr184Met | P51648 |
| HOMO SAPIENS | 1 | 2814 | 4qgk | B | 188 | ALA | p.Ala188Val | P51648 |
| HOMO SAPIENS | 0 | 2814 | 4qgk | B | 214 | CYS | p.Cys214Tyr | P51648 |
| HOMO SAPIENS | 0 | 2814 | 4qgk | B | 245 | ASP | p.Asp245Asn | P51648 |
| HOMO SAPIENS | 0 | 2814 | 4qgk | B | 266 | LYS | p.Lys266Asn | P51648 |
| HOMO SAPIENS | 0 | 2814 | 4qgk | B | 315 | PRO | p.Pro315Ser | P51648 |
| HOMO SAPIENS | 0 | 2814 | 4qgk | B | 386 | ASN | p.Asn386Ser | P51648 |
| HOMO SAPIENS | -1 | 5486 | 5u6o | B | 47 | GLY | p.Gly47Val | O60741 |
| HOMO SAPIENS | 1 | 21475 | 3pdy | A | 679 | THR | p.Thr789Ile | Q15149 |
| HOMO SAPIENS | 0 | 32890 | 4djc | B | 1501 | LEU | p.Leu1501Val | Q14524 |
| HOMO SAPIENS | -1 | 32890 | 4djc | B | 1512 | ARG | p.Arg1512Trp | Q14524 |
| HOMO SAPIENS | 0 | 26683 | 2xze | B | 38 | ARG | p.Arg38Cys | O95630 |
| HOMO SAPIENS | 0 | 26683 | 2xze | B | 100 | PHE | p.Phe100Tyr | O95630 |
| HOMO SAPIENS | -1 | 23175 | 5dka | A | 163 | SER | p.Ser163Leu | Q96EQ8 |
| HOMO SAPIENS | -1 | 23175 | 5dka | A | 174 | ARG | p.Arg174Cys | Q96EQ8 |
| HOMO SAPIENS | 6 | 21177 | 3k1r | A | 155 | ARG | p.Arg155Ter | Q9Y6N9 |
| HOMO SAPIENS | 0 | 1237 | 2wwm | O | 40 | PRO | p.Pro40Arg | O75147 |
| HOMO SAPIENS | 1 | 2543 | 5ewl | B | 64 | VAL | p.Val64Leu | Q13224 |
| HOMO SAPIENS | 2 | 2543 | 5ewl | B | 271 | ALA | p.Ala271Glu | Q13224 |
| HOMO SAPIENS | 3 | 34262 | 2mng | A | 613 | PHE | p.Phe613Leu | Q9Y3Q4 |
| HOMO SAPIENS | 7 | 34262 | 2mng | A | 672 | SER | p.Ser672Arg | Q9Y3Q4 |
| HOMO SAPIENS | 4 | 1667 | 1f05 | A | 321 | LYS | p.Lys321Met | P37837 |
| HOMO SAPIENS | 0 | 25179 | 2knc | B | 752 | SER | p.Ser778Pro | P05106 |
| HOMO SAPIENS | 0 | 2185 | 2ig7 | A | 50 | TYR | p.Tyr50Cys | Q9Y259 |
| HOMO SAPIENS | 0 | 2185 | 2ig7 | A | 224 | ASN | p.Asn224His | Q9Y259 |
| HOMO SAPIENS | 0 | 2185 | 2ig7 | A | 270 | TYR | p.Tyr270Ter | Q9Y259 |
| HOMO SAPIENS | 0 | 2185 | 2ig7 | A | 328 | GLN | p.Gln328Arg | Q9Y259 |
| HOMO SAPIENS | 3 | 2201 | 2r0o | A | 179 | PRO | p.Pro179Leu | O43707 |
| HOMO SAPIENS | 3 | 2201 | 2r0o | A | 255 | GLU | p.Lys255Glu | O43707 |
| HOMO SAPIENS | 3 | 2201 | 2r0o | A | 259 | THR | p.Thr259Ile | O43707 |
| HOMO SAPIENS | 4 | 2201 | 2r0o | A | 262 | SER | p.Ser262Pro | O43707 |
| HOMO SAPIENS | -1 | 31221 | 2wvr | C | 44 | THR | p.Thr44Pro | Q9H211 |
| HOMO SAPIENS | -1 | 31221 | 2wvr | C | 135 | ALA | p.Ala135Val | Q9H211 |
| HOMO SAPIENS | -1 | 31221 | 2wvr | C | 456 | GLU | p.Glu456Ala | Q9H211 |

| | | | | | | | | |
|---|---|---|---|---|---|---|---|---|
| HOMO SAPIENS | -1 | 31221 | 2wvr | C | 462 | ARG | p.Arg462Gln | Q9H211 |
| HOMO SAPIENS | -1 | 31221 | 2wvr | C | 468 | GLU | p.Glu468Lys | Q9H211 |
| HOMO SAPIENS | -1 | 31221 | 2wvr | C | 537 | ALA | p.Ala537Val | Q9H211 |
| HOMO SAPIENS | 5 | 2576 | 5o3l | C | 363 | VAL | p.Val680Ile | P10636 |
| HOMO SAPIENS | 5 | 1345 | 1czt | A | 37 | ARG | p.Arg2102Cys | P12259 |
| HOMO SAPIENS | 1 | 1345 | 1czt | A | 83 | MET | p.Met2148Thr | P12259 |
| HOMO SAPIENS | 1 | 1345 | 1czt | A | 157 | ASP | p.Asp2222Gly | P12259 |
| HOMO SAPIENS | 8 | 35985 | 1lo1 | A | 110 | ALA | p.Ala110Val | O95718 |
| HOMO SAPIENS | 1 | 617 | 2ocp | B | 44 | GLU | p.Glu44Lys | Q16854 |
| HOMO SAPIENS | 5 | 617 | 2ocp | B | 46 | ASN | p.Asn46Ser | Q16854 |
| HOMO SAPIENS | 0 | 617 | 2ocp | B | 59 | THR | p.Thr59Met | Q16854 |
| HOMO SAPIENS | 7 | 617 | 2ocp | B | 142 | ARG | p.Arg142Lys | Q16854 |
| HOMO SAPIENS | 3 | 617 | 2ocp | B | 154 | ASN | p.Asn154Lys | Q16854 |
| HOMO SAPIENS | 3 | 617 | 2ocp | B | 170 | GLN | p.Gln170Arg | Q16854 |
| HOMO SAPIENS | 3 | 617 | 2ocp | B | 227 | GLU | p.Glu227Lys | Q16854 |
| HOMO SAPIENS | 3 | 617 | 2ocp | B | 233 | LYS | p.Lys233Glu | Q16854 |
| HOMO SAPIENS | 3 | 617 | 2ocp | B | 255 | ASP | p.Asp255Tyr | Q16854 |
| HOMO SAPIENS | -1 | 38975 | 5itz | D | 421 | GLN | p.Gln421His | Q9HC77 |
| HOMO SAPIENS | 2 | 6235 | 2f8v | A | 15 | VAL | p.Val15Ile | Q8WZ42 |
| HOMO SAPIENS | 0 | 6235 | 2f8v | A | 54 | VAL | p.Val54Met | Q8WZ42 |
| HOMO SAPIENS | 0 | 6235 | 2f8v | A | 59 | SER | p.Ser59Arg | Q8WZ42 |
| HOMO SAPIENS | 0 | 6235 | 2f8v | A | 60 | ASP | p.Asp60Tyr | Q8WZ42 |
| HOMO SAPIENS | 0 | 12630 | 1ext | B | 12 | ASP | p.Asp41Glu | P19438 |
| HOMO SAPIENS | 3 | 12630 | 1ext | B | 30 | CYS | p.Cys59Arg | P19438 |
| HOMO SAPIENS | 3 | 12630 | 1ext | B | 30 | CYS | p.Cys59Ser | P19438 |
| HOMO SAPIENS | 5 | 12630 | 1ext | B | 33 | CYS | p.Cys62Tyr | P19438 |
| HOMO SAPIENS | 5 | 12630 | 1ext | B | 33 | CYS | p.Cys62Gly | P19438 |
| HOMO SAPIENS | 1 | 12630 | 1ext | B | 46 | PRO | p.Pro75Leu | P19438 |
| HOMO SAPIENS | 3 | 12630 | 1ext | B | 50 | THR | p.Thr79Met | P19438 |
| HOMO SAPIENS | 6 | 12630 | 1ext | B | 52 | CYS | p.Cys81Phe | P19438 |
| HOMO SAPIENS | 6 | 12630 | 1ext | B | 70 | CYS | p.Cys99Ser | P19438 |
| HOMO SAPIENS | 5 | 12630 | 1ext | B | 88 | CYS | p.Cys117Arg | P19438 |
| HOMO SAPIENS | 5 | 12630 | 1ext | B | 88 | CYS | p.Cys117Tyr | P19438 |
| HOMO SAPIENS | -1 | 12630 | 1ext | B | 170 | ILE | p.Ile199Asn | P19438 |
| HOMO SAPIENS | 0 | 994 | 1aos | B | 95 | ARG | p.Arg95Cys | P04424 |
| HOMO SAPIENS | 0 | 994 | 1aos | B | 111 | ARG | p.Arg111Gln | P04424 |
| HOMO SAPIENS | 0 | 994 | 1aos | B | 131 | THR | p.Thr131Met | P04424 |
| HOMO SAPIENS | 6 | 994 | 1aos | B | 178 | VAL | p.Val178Met | P04424 |
| HOMO SAPIENS | 9 | 994 | 1aos | B | 182 | ARG | p.Arg182Gln | P04424 |
| HOMO SAPIENS | 9 | 994 | 1aos | B | 186 | ARG | p.Arg186Trp | P04424 |
| HOMO SAPIENS | 8 | 994 | 1aos | B | 193 | ARG | p.Arg193Gln | P04424 |
| HOMO SAPIENS | 5 | 994 | 1aos | B | 217 | ARG | p.Arg217Ter | P04424 |
| HOMO SAPIENS | 0 | 994 | 1aos | B | 286 | GLN | p.Gln286Arg | P04424 |
| HOMO SAPIENS | 5 | 994 | 1aos | B | 379 | ARG | p.Arg379Cys | P04424 |
| HOMO SAPIENS | 4 | 16066 | 5mv9 | A | 1719 | TYR | p.Tyr1719Cys | Q13402 |
| HOMO SAPIENS | -1 | 16066 | 5mv9 | A | 1739 | ARG | p.Arg1739Ter | Q13402 |
| HOMO SAPIENS | 4 | 16066 | 5mv9 | A | 1775 | ILE | p.Ile1775Thr | Q13402 |
| HOMO SAPIENS | 2 | 16066 | 5mv9 | A | 1786 | SER | p.Ser1786Thr | Q13402 |
| HOMO SAPIENS | 4 | 16066 | 5mv9 | A | 1798 | GLN | p.Gln1798Ter | Q13402 |
| HOMO SAPIENS | 0 | 16066 | 5mv9 | A | 1858 | LEU | p.Leu1858Pro | Q13402 |
| HOMO SAPIENS | 6 | 16066 | 5mv9 | A | 1873 | ARG | p.Arg1873Trp | Q13402 |
| HOMO SAPIENS | 6 | 16066 | 5mv9 | A | 1873 | ARG | p.Arg1873Leu | Q13402 |
| HOMO SAPIENS | 6 | 16066 | 5mv9 | A | 1883 | ARG | p.Arg1883Gln | Q13402 |
| HOMO SAPIENS | 4 | 16066 | 5mv9 | A | 1887 | PRO | p.Pro1887Leu | Q13402 |

| | | | | | | | | |
|---|---|---|---|---|---|---|---|---|
| HOMO SAPIENS | 5 | 16066 | 5mv9 | A | 1942 | GLY | p.Gly1942Ter | Q13402 |
| HOMO SAPIENS | 5 | 16066 | 5mv9 | A | 1954 | LEU | p.Leu1954Ile | Q13402 |
| HOMO SAPIENS | 5 | 16066 | 5mv9 | A | 1956 | VAL | p.Val1956Ile | Q13402 |
| HOMO SAPIENS | 5 | 16066 | 5mv9 | A | 1967 | ARG | p.Arg1967Ter | Q13402 |
| HOMO SAPIENS | 5 | 16066 | 5mv9 | A | 1990 | GLN | p.Gln1990Ter | Q13402 |
| HOMO SAPIENS | 2 | 16066 | 5mv9 | A | 2005 | LYS | p.Lys2005Glu | Q13402 |
| HOMO SAPIENS | 1 | 16066 | 5mv9 | A | 2072 | VAL | p.Val2072Ile | Q13402 |
| HOMO SAPIENS | 4 | 16066 | 5mv9 | A | 2109 | THR | p.Thr2109Ile | Q13402 |
| HOMO SAPIENS | 4 | 16066 | 5mv9 | A | 2142 | ASP | p.Asp2142Asn | Q13402 |
| HOMO SAPIENS | -1 | 16066 | 5mv9 | A | 2214 | GLY | p.Gly2214Ser | Q13402 |
| HOMO SAPIENS | 7 | 27942 | 1deb | A | 24 | ARG | p.Arg24Ter | P25054 |
| HOMO SAPIENS | 0 | 876 | 5hnr | B | 12 | PHE | p.Phe12Leu | P13716 |
| HOMO SAPIENS | 0 | 876 | 5hnr | B | 59 | LYS | p.Lys59Asn | P13716 |
| HOMO SAPIENS | -1 | 876 | 5hnr | B | 133 | GLY | p.Gly133Arg | P13716 |
| HOMO SAPIENS | 2 | 876 | 5hnr | B | 240 | ARG | p.Arg240Trp | P13716 |
| HOMO SAPIENS | 0 | 876 | 5hnr | B | 274 | ALA | p.Ala274Thr | P13716 |
| HOMO SAPIENS | 0 | 876 | 5hnr | B | 275 | VAL | p.Val275Met | P13716 |
| HOMO SAPIENS | 0 | 1142 | 3qe2 | A | 503 | VAL | p.Ala500Val | P16435 |
| HOMO SAPIENS | 8 | 14358 | 4dkd | C | 32 | VAL | p.Val32Gly | P07333 |
| HOMO SAPIENS | 0 | 14358 | 4dkd | C | 60 | LEU | p.Leu60Val | P07333 |
| HOMO SAPIENS | 7 | 14358 | 4dkd | C | 75 | THR | p.Thr75Ile | P07333 |
| HOMO SAPIENS | -1 | 14358 | 4dkd | C | 90 | PRO | p.Pro90Thr | P07333 |
| HOMO SAPIENS | 0 | 14358 | 4dkd | C | 106 | ARG | p.Arg106Trp | P07333 |
| HOMO SAPIENS | 0 | 14358 | 4dkd | C | 241 | ASN | p.Asn241Asp | P07333 |
| HOMO SAPIENS | 0 | 14358 | 4dkd | C | 245 | ALA | p.Ala245Ser | P07333 |
| HOMO SAPIENS | 0 | 14358 | 4dkd | C | 255 | ASN | p.Asn255Ile | P07333 |
| HOMO SAPIENS | 0 | 14358 | 4dkd | C | 268 | VAL | p.Val268Ile | P07333 |
| HOMO SAPIENS | 0 | 14358 | 4dkd | C | 279 | VAL | p.Val279Met | P07333 |
| HOMO SAPIENS | 0 | 14358 | 4dkd | C | 294 | ARG | p.Arg294Gln | P07333 |
| HOMO SAPIENS | 5 | 8968 | 3ppy | A | 1565 | VAL | p.Val1565Leu | P04275 |
| HOMO SAPIENS | 4 | 8968 | 3ppy | A | 1584 | TYR | p.Tyr1584Cys | P04275 |
| HOMO SAPIENS | 5 | 5341 | 3k3n | A | 279 | PHE | p.Phe315Ser | Q9UPP1 |
| HOMO SAPIENS | 0 | 29463 | 1q7l | A | 197 | ARG | p.Arg197Trp | Q03154 |
| HOMO SAPIENS | 0 | 33761 | 2leh | B | 30 | ASP | p.Asp30Asn | Q16637 |
| HOMO SAPIENS | 6 | 33761 | 2leh | B | 44 | ASP | p.Asp44Val | Q16637 |
| HOMO SAPIENS | -1 | 12249 | 2pnr | C | 224 | GLU | p.Glu310Lys | P10515 |
| HOMO SAPIENS | -1 | 12249 | 2pnr | C | 232 | VAL | p.Val318Ala | P10515 |
| HOMO SAPIENS | -1 | 7 | 5whk | B | -9 | ALA | p.Ala11Pro | P61769 |
| HOMO SAPIENS | 8 | 20298 | 4euw | A | 154 | PHE | p.Phe154Leu | P48436 |
| HOMO SAPIENS | 9 | 20298 | 4euw | A | 158 | ALA | p.Ala158Thr | P48436 |
| HOMO SAPIENS | 5 | 20298 | 4euw | A | 165 | HIS | p.His165Tyr | P48436 |
| HOMO SAPIENS | 3 | 20298 | 4euw | A | 169 | HIS | p.His169Gln | P48436 |
| HOMO SAPIENS | 0 | 20298 | 4euw | A | 173 | LYS | p.Lys173Glu | P48436 |
| HOMO SAPIENS | 1 | 16465 | 2jgx | A | 402 | TYR | p.Tyr402His | P08603 |
| HOMO SAPIENS | 0 | 8993 | 3v8o | A | 659 | LEU | p.Leu659Arg | Q14315 |
| HOMO SAPIENS | 6 | 8993 | 3v8o | A | 693 | ASP | p.Asp693Ala | Q14315 |
| HOMO SAPIENS | 0 | 8993 | 3v8o | A | 709 | ALA | p.Ala709Thr | Q14315 |
| HOMO SAPIENS | 2 | 3852 | 5es4 | B | 106 | ASP | p.Asp128Asn | P05107 |
| HOMO SAPIENS | 2 | 3852 | 5es4 | B | 116 | SER | p.Ser138Pro | P05107 |
| HOMO SAPIENS | 1 | 3852 | 5es4 | B | 127 | LEU | p.Leu149Pro | P05107 |
| HOMO SAPIENS | 0 | 3852 | 5es4 | B | 147 | GLY | p.Gly169Arg | P05107 |
| HOMO SAPIENS | 3 | 3852 | 5es4 | B | 156 | PRO | p.Pro178Leu | P05107 |
| HOMO SAPIENS | 0 | 3852 | 5es4 | B | 174 | LYS | p.Lys196Thr | P05107 |
| HOMO SAPIENS | 2 | 3852 | 5es4 | B | 251 | GLY | p.Gly273Arg | P05107 |

| | | | | | | | | |
|---|---|---|---|---|---|---|---|---|
| HOMO SAPIENS | 2 | 3852 | 5es4 | B | 329 | ASN | p.Asn351Ser | P05107 |
| HOMO SAPIENS | 2 | 3852 | 5es4 | B | 571 | ARG | p.Arg593Cys | P05107 |
| HOMO SAPIENS | 7 | 5962 | 5fn2 | B | 146 | MET | p.Met146Leu | P49768 |
| HOMO SAPIENS | 3 | 5962 | 5fn2 | B | 166 | LEU | p.Leu166Pro | P49768 |
| HOMO SAPIENS | 1 | 5962 | 5fn2 | B | 233 | MET | p.Met233Leu | P49768 |
| HOMO SAPIENS | 1 | 5962 | 5fn2 | B | 261 | VAL | p.Val261Gly | P49768 |
| HOMO SAPIENS | 1 | 5962 | 5fn2 | B | 266 | GLY | p.Gly266Ser | P49768 |
| HOMO SAPIENS | -1 | 5962 | 5fn2 | B | 333 | ASP | p.Asp333Gly | P49768 |
| HOMO SAPIENS | 6 | 1918 | 3dlx | A | 58 | THR | p.Thr58Met | P55809 |
| HOMO SAPIENS | 5 | 1918 | 3dlx | A | 133 | VAL | p.Val133Glu | P55809 |
| HOMO SAPIENS | 5 | 1918 | 3dlx | A | 219 | GLY | p.Gly219Glu | P55809 |
| HOMO SAPIENS | 0 | 1918 | 3dlx | A | 221 | VAL | p.Val221Met | P55809 |
| HOMO SAPIENS | 5 | 1918 | 3dlx | A | 283 | SER | p.Ser283Ter | P55809 |
| HOMO SAPIENS | 4 | 1918 | 3dlx | A | 324 | GLY | p.Gly324Glu | P55809 |
| HOMO SAPIENS | 0 | 1918 | 3dlx | A | 456 | CYS | p.Cys456Phe | P55809 |
| HOMO SAPIENS | 0 | 9010 | 2wfr | A | 162 | LEU | p.Leu162Pro | O43323 |
| HOMO SAPIENS | 5 | 3770 | 5n38 | A | 56 | VAL | p.Val56Glu | O60260 |
| HOMO SAPIENS | 1 | 3770 | 5n38 | A | 161 | LYS | p.Lys161Asn | O60260 |
| HOMO SAPIENS | 1 | 3770 | 5n38 | A | 167 | SER | p.Ser167Asn | O60260 |
| HOMO SAPIENS | 1 | 3770 | 5n38 | A | 211 | LYS | p.Lys211Asn | O60260 |
| HOMO SAPIENS | 1 | 3770 | 5n38 | A | 212 | CYS | p.Cys212Tyr | O60260 |
| HOMO SAPIENS | 1 | 3770 | 5n38 | A | 275 | ARG | p.Arg275Trp | O60260 |
| HOMO SAPIENS | 2 | 3770 | 5n38 | A | 334 | ARG | p.Arg334Cys | O60260 |
| HOMO SAPIENS | 1 | 3770 | 5n38 | A | 366 | ARG | p.Arg366Trp | O60260 |
| HOMO SAPIENS | -1 | 3770 | 5n38 | A | 380 | VAL | p.Val380Leu | O60260 |
| HOMO SAPIENS | 1 | 3770 | 5n38 | A | 394 | ASP | p.Asp394Asn | O60260 |
| HOMO SAPIENS | 1 | 3770 | 5n38 | A | 430 | GLY | p.Gly430Asp | O60260 |
| HOMO SAPIENS | -1 | 2467 | 4xx1 | A | 10 | PRO | p.Pro34Leu | P04180 |
| HOMO SAPIENS | 4 | 2467 | 4xx1 | A | 83 | TYR | p.Tyr107Ter | P04180 |
| HOMO SAPIENS | 4 | 2467 | 4xx1 | A | 93 | ALA | p.Ala117Thr | P04180 |
| HOMO SAPIENS | 4 | 2467 | 4xx1 | A | 123 | THR | p.Thr147Ile | P04180 |
| HOMO SAPIENS | 4 | 2467 | 4xx1 | A | 135 | ARG | p.Arg159Trp | P04180 |
| HOMO SAPIENS | 3 | 2467 | 4xx1 | A | 158 | ARG | p.Arg182Cys | P04180 |
| HOMO SAPIENS | 0 | 2467 | 4xx1 | A | 209 | LEU | p.Leu233Pro | P04180 |
| HOMO SAPIENS | 1 | 2467 | 4xx1 | A | 228 | ASN | p.Asn252Lys | P04180 |
| HOMO SAPIENS | 4 | 2467 | 4xx1 | A | 252 | MET | p.Met276Lys | P04180 |
| HOMO SAPIENS | 3 | 2467 | 4xx1 | A | 293 | MET | p.Met317Ile | P04180 |
| HOMO SAPIENS | 5 | 2467 | 4xx1 | A | 321 | THR | p.Thr345Met | P04180 |
| HOMO SAPIENS | 9 | 2467 | 4xx1 | A | 347 | THR | p.Thr371Met | P04180 |
| HOMO SAPIENS | 1 | 5834 | 3qrf | G | 371 | PHE | p.Phe371Cys | Q9BZS1 |
| HOMO SAPIENS | 1 | 5834 | 3qrf | G | 373 | PHE | p.Phe373Ala | Q9BZS1 |
| HOMO SAPIENS | 9 | 5834 | 3qrf | G | 384 | ALA | p.Ala384Thr | Q9BZS1 |
| HOMO SAPIENS | 6 | 5834 | 3qrf | G | 397 | ARG | p.Arg397Trp | Q9BZS1 |
| HOMO SAPIENS | -1 | 11869 | 4km8 | A | 15 | PRO | p.Pro15Leu | Q9UMX1 |
| HOMO SAPIENS | 5 | 262 | 4kbo | A | 126 | ARG | p.Arg126Trp | P38646 |
| HOMO SAPIENS | 3 | 262 | 4kbo | A | 128 | TYR | p.Tyr128Cys | P38646 |
| HOMO SAPIENS | -1 | 11347 | 4uf9 | D | 117 | ALA | p.Ala117Val | Q13501 |
| HOMO SAPIENS | 0 | 2173 | 4ah6 | A | 45 | SER | p.Ser45Gly | Q6PI48 |
| HOMO SAPIENS | 2 | 2173 | 4ah6 | A | 152 | CYS | p.Cys152Phe | Q6PI48 |
| HOMO SAPIENS | 4 | 2173 | 4ah6 | A | 179 | ARG | p.Arg179His | Q6PI48 |
| HOMO SAPIENS | 0 | 2173 | 4ah6 | A | 196 | LYS | p.Lys196Arg | Q6PI48 |
| HOMO SAPIENS | 8 | 2173 | 4ah6 | A | 263 | ARG | p.Arg263Gln | Q6PI48 |
| HOMO SAPIENS | 1 | 2173 | 4ah6 | A | 338 | GLY | p.Gly338Glu | Q6PI48 |
| HOMO SAPIENS | 0 | 2173 | 4ah6 | A | 425 | GLU | p.Glu425Ter | Q6PI48 |

| | | | | | | | | |
|---|---|---|---|---|---|---|---|---|
| HOMO SAPIENS | 1 | 2173 | 4ah6 | A | 548 | LEU | p.Leu548Met | Q6PI48 |
| HOMO SAPIENS | 0 | 2173 | 4ah6 | A | 609 | ARG | p.Arg609Trp | Q6PI48 |
| HOMO SAPIENS | 2 | 2173 | 4ah6 | A | 613 | LEU | p.Leu613Phe | Q6PI48 |
| HOMO SAPIENS | 0 | 2173 | 4ah6 | A | 626 | LEU | p.Leu626Gln | Q6PI48 |
| HOMO SAPIENS | 0 | 2173 | 4ah6 | A | 626 | LEU | p.Leu626Val | Q6PI48 |
| HOMO SAPIENS | 7 | 3203 | 2cn1 | A | 87 | ASP | p.Asp137Val | Q9H0P0 |
| HOMO SAPIENS | 8 | 3203 | 2cn1 | A | 166 | GLN | p.Gln216Ter | Q9H0P0 |
| HOMO SAPIENS | 8 | 3203 | 2cn1 | A | 179 | ASN | p.Asn229Ser | Q9H0P0 |
| HOMO SAPIENS | 7 | 3203 | 2cn1 | A | 230 | GLY | p.Gly280Arg | Q9H0P0 |
| HOMO SAPIENS | 2 | 3424 | 2a1t | R | 116 | GLY | p.Gly116Arg | P13804 |
| HOMO SAPIENS | 2 | 3424 | 2a1t | R | 157 | VAL | p.Val157Gly | P13804 |
| HOMO SAPIENS | 2 | 3424 | 2a1t | R | 171 | THR | p.Thr171Ile | P13804 |
| HOMO SAPIENS | 0 | 3424 | 2a1t | R | 178 | THR | p.Thr178Arg | P13804 |
| HOMO SAPIENS | 0 | 3424 | 2a1t | R | 194 | VAL | p.Val194Met | P13804 |
| HOMO SAPIENS | 1 | 3424 | 2a1t | R | 209 | ARG | p.Arg209Ter | P13804 |
| HOMO SAPIENS | 6 | 3424 | 2a1t | R | 266 | THR | p.Thr266Met | P13804 |
| HOMO SAPIENS | 0 | 27598 | 1g1c | A | 26 | ARG | p.Arg2098Gly | Q8WZ42 |
| HOMO SAPIENS | 1 | 27598 | 1g1c | A | 46 | ILE | p.Ile2118Thr | Q8WZ42 |
| HOMO SAPIENS | 0 | 32885 | 4bsk | A | 149 | ASN | p.Asn149Asp | P35916 |
| HOMO SAPIENS | -1 | 8074 | 3ghg | B | 17 | ARG | p.Arg47Ter | P02675 |
| HOMO SAPIENS | -1 | 447 | 5g4p | A | 190 | ALA | p.Ala190Thr | P84074 |
| HOMO SAPIENS | 0 | 10761 | 5u1d | B | 379 | VAL | p.Val379Ile | Q03519 |
| HOMO SAPIENS | 0 | 10761 | 5u1d | B | 665 | THR | p.Thr665Ala | Q03519 |
| HOMO SAPIENS | 5 | 29251 | 5fn7 | A | 79 | ILE | p.Ile294Leu | P08575 |
| HOMO SAPIENS | -1 | 27727 | 4ky9 | A | 56 | LYS | p.Lys56Glu | P02730 |
| HOMO SAPIENS | 2 | 27727 | 4ky9 | A | 72 | GLU | p.Glu72Asp | P02730 |
| HOMO SAPIENS | 2 | 27727 | 4ky9 | A | 90 | GLU | p.Glu90Lys | P02730 |
| HOMO SAPIENS | 1 | 27727 | 4ky9 | A | 130 | GLY | p.Gly130Arg | P02730 |
| HOMO SAPIENS | 2 | 27727 | 4ky9 | A | 150 | ARG | p.Arg150Ter | P02730 |
| HOMO SAPIENS | 0 | 27727 | 4ky9 | A | 327 | PRO | p.Pro327Arg | P02730 |
| HOMO SAPIENS | 2 | 13202 | 2fxo | B | 870 | ARG | p.Arg870His | P12883 |
| HOMO SAPIENS | 2 | 13202 | 2fxo | B | 870 | ARG | p.Arg870Cys | P12883 |
| HOMO SAPIENS | 4 | 13202 | 2fxo | B | 904 | ARG | p.Arg904Cys | P12883 |
| HOMO SAPIENS | 4 | 13202 | 2fxo | B | 906 | ASP | p.Asp906Gly | P12883 |
| HOMO SAPIENS | 2 | 13202 | 2fxo | B | 908 | LEU | p.Leu908Val | P12883 |
| HOMO SAPIENS | 2 | 13202 | 2fxo | B | 923 | ASN | p.Asn923Lys | P12883 |
| HOMO SAPIENS | 7 | 13202 | 2fxo | B | 935 | GLU | p.Glu935Lys | P12883 |
| HOMO SAPIENS | 7 | 13202 | 2fxo | B | 935 | GLU | p.Glu935Val | P12883 |
| HOMO SAPIENS | 8 | 13202 | 2fxo | B | 949 | GLU | p.Glu949Lys | P12883 |
| HOMO SAPIENS | 0 | 13200 | 2a1s | A | 349 | ARG | p.Arg349Trp | O95453 |
| HOMO SAPIENS | 0 | 13200 | 2a1s | A | 421 | LYS | p.Lys421Arg | O95453 |
| HOMO SAPIENS | 6 | 3196 | 5xtb | T | 115 | CYS | p.Cys115Tyr | O75380 |
| HOMO SAPIENS | 0 | 1108 | 5ukf | A | 89 | ARG | p.Arg89Gln | Q99986 |
| HOMO SAPIENS | 0 | 1108 | 5ukf | A | 119 | HIS | p.His119Arg | Q99986 |
| HOMO SAPIENS | 0 | 1108 | 5ukf | A | 133 | ARG | p.Arg133Cys | Q99986 |
| HOMO SAPIENS | 0 | 1108 | 5ukf | A | 236 | VAL | p.Val236Leu | Q99986 |
| HOMO SAPIENS | 0 | 1108 | 5ukf | A | 321 | ARG | p.Arg321Cys | Q99986 |
| HOMO SAPIENS | -1 | 1108 | 5ukf | A | 358 | ARG | p.Arg358Ter | Q99986 |
| HOMO SAPIENS | 0 | 5842 | 4yli | C | 204 | GLY | p.Gly204Ser | Q9BWP8 |
| HOMO SAPIENS | -1 | 31362 | 4wqo | A | 25 | PRO | p.Pro25Leu | P40337 |
| HOMO SAPIENS | 2 | 10107 | 2j8h | A | 34 | VAL | p.Val31886Ile | Q8WZ42 |
| HOMO SAPIENS | 8 | 20190 | 1wm5 | A | 77 | ARG | p.Arg77Gln | P19878 |
| HOMO SAPIENS | 1 | 20190 | 1wm5 | A | 100 | GLN | p.Gln100Ter | P19878 |
| HOMO SAPIENS | 0 | 20190 | 1wm5 | A | 102 | ARG | p.Arg102Ter | P19878 |

| | | | | | | | | | |
|---|---|---|---|---|---|---|---|---|---|
| HOMO SAPIENS | 0 | 20190 | 1wm5 | A | | 128 | ALA | p.Ala128Val | P19878 |
| HOMO SAPIENS | 0 | 20190 | 1wm5 | A | | 181 | LYS | p.Lys181Arg | P19878 |
| HOMO SAPIENS | 7 | 1278 | 5a2q | a | | 33 | ASP | p.Asp33Asn | P62854 |
| HOMO SAPIENS | 3 | 19515 | 2obd | A | | 11 | ILE | p.Ile28Met | P11597 |
| HOMO SAPIENS | 3 | 19515 | 2obd | A | | 89 | VAL | p.Val106Met | P11597 |
| HOMO SAPIENS | 3 | 19515 | 2obd | A | | 137 | ARG | p.Arg154Trp | P11597 |
| HOMO SAPIENS | 3 | 19515 | 2obd | A | | 139 | ASP | p.Asp156Asn | P11597 |
| HOMO SAPIENS | 3 | 19515 | 2obd | A | | 204 | SER | p.Ser221Arg | P11597 |
| HOMO SAPIENS | 0 | 19515 | 2obd | A | | 297 | GLU | p.Glu314Lys | P11597 |
| HOMO SAPIENS | 0 | 19515 | 2obd | A | | 314 | GLY | p.Gly331Ser | P11597 |
| HOMO SAPIENS | 0 | 19515 | 2obd | A | | 332 | SER | p.Ser349Tyr | P11597 |
| HOMO SAPIENS | 0 | 19515 | 2obd | A | | 368 | VAL | p.Val385Met | P11597 |
| HOMO SAPIENS | 0 | 19515 | 2obd | A | | 373 | ALA | p.Ala390Pro | P11597 |
| HOMO SAPIENS | 3 | 19515 | 2obd | A | | 405 | ILE | p.Val422Ile | P11597 |
| HOMO SAPIENS | 0 | 19515 | 2obd | A | | 451 | ARG | p.Arg468Gln | P11597 |
| HOMO SAPIENS | 0 | 19515 | 2obd | A | | 451 | ARG | p.Arg468Ter | P11597 |
| HOMO SAPIENS | 0 | 6626 | 3hop | A | | 39 | ALA | p.Ala39Gly | P21333 |
| HOMO SAPIENS | 1 | 6626 | 3hop | A | | 82 | GLU | p.Glu82Val | P21333 |
| HOMO SAPIENS | 1 | 6626 | 3hop | A | | 128 | ALA | p.Ala128Val | P21333 |
| HOMO SAPIENS | 2 | 6626 | 3hop | A | | 203 | ASP | p.Asp203Tyr | P21333 |
| HOMO SAPIENS | 1 | 6626 | 3hop | A | | 207 | PRO | p.Pro207Leu | P21333 |
| HOMO SAPIENS | 2 | 6626 | 3hop | A | | 210 | CYS | p.Cys210Phe | P21333 |
| HOMO SAPIENS | 7 | 6626 | 3hop | A | | 254 | GLU | p.Glu254Lys | P21333 |
| HOMO SAPIENS | 4 | 4210 | 2avg | A | | 8 | VAL | p.Val158Met | Q14896 |
| HOMO SAPIENS | 6 | 4210 | 2avg | A | | 15 | GLU | p.Glu165Asp | Q14896 |
| HOMO SAPIENS | 4 | 4210 | 2avg | A | | 27 | ARG | p.Arg177His | Q14896 |
| HOMO SAPIENS | 0 | 4210 | 2avg | A | | 28 | VAL | p.Val178Met | Q14896 |
| HOMO SAPIENS | 6 | 4210 | 2avg | A | | 39 | VAL | p.Val189Ile | Q14896 |
| HOMO SAPIENS | 2 | 4210 | 2avg | A | | 52 | LYS | p.Lys202Gln | Q14896 |
| HOMO SAPIENS | 4 | 4210 | 2avg | A | | 67 | SER | p.Ser217Gly | Q14896 |
| HOMO SAPIENS | 6 | 4210 | 2avg | A | | 69 | VAL | p.Val219Leu | Q14896 |
| HOMO SAPIENS | 2 | 4210 | 2avg | A | | 86 | GLY | p.Ser236Gly | Q14896 |
| HOMO SAPIENS | 4 | 4210 | 2avg | A | | 108 | GLU | p.Glu258Lys | Q14896 |
| HOMO SAPIENS | 0 | 22237 | 1w6k | A | | 588 | GLY | p.Gly588Ser | P48449 |
| HOMO SAPIENS | 0 | 10765 | 4a5w | B | | 98 | GLU | p.Glu98Ala | P13671 |
| HOMO SAPIENS | 0 | 10765 | 4a5w | B | | 596 | ARG | p.Arg596Ter | P13671 |
| HOMO SAPIENS | -1 | 15515 | 3hx3 | A | | 11 | MSE | p.Met10Lys | P12271 |
| HOMO SAPIENS | 8 | 15515 | 3hx3 | A | | 64 | ARG | p.Arg64Gln | P12271 |
| HOMO SAPIENS | 0 | 15515 | 3hx3 | A | | 182 | PHE | p.Phe182Cys | P12271 |
| HOMO SAPIENS | 0 | 15515 | 3hx3 | A | | 216 | ARG | p.Arg216Gln | P12271 |
| HOMO SAPIENS | -1 | 15515 | 3hx3 | A | | 226 | MSE | p.Met226Lys | P12271 |
| HOMO SAPIENS | 0 | 15515 | 3hx3 | A | | 234 | TRP | p.Arg234Trp | P12271 |
| HOMO SAPIENS | 0 | 15515 | 3hx3 | A | | 273 | LEU | p.Leu273Phe | P12271 |
| HOMO SAPIENS | 0 | 15515 | 3hx3 | A | | 292 | THR | p.Thr292Met | P12271 |
| HOMO SAPIENS | 0 | 5152 | 2www | D | | 95 | GLN | p.Gln95Ter | Q8IVH4 |
| HOMO SAPIENS | 0 | 5152 | 2www | D | | 133 | GLN | p.Gln133Ter | Q8IVH4 |
| HOMO SAPIENS | 0 | 5152 | 2www | D | | 145 | ARG | p.Arg145Ter | Q8IVH4 |
| HOMO SAPIENS | 0 | 5152 | 2www | D | | 207 | TYR | p.Tyr207Cys | Q8IVH4 |
| HOMO SAPIENS | 8 | 5152 | 2www | D | | 330 | ARG | p.Arg330Ter | Q8IVH4 |
| HOMO SAPIENS | 0 | 5152 | 2www | D | | 363 | GLN | p.Gln363His | Q8IVH4 |
| HOMO SAPIENS | 1 | 1626 | 1ilr | | 1 | 105 | SER | p.Ser130Arg | P18510 |
| HOMO SAPIENS | -1 | 929 | 5loh | A | | 12 | GLY | p.Gly12Arg | Q96GX5 |
| HOMO SAPIENS | 9 | 3206 | 1x63 | A | | 21 | CYS | p.Cys104Arg | Q13642 |
| HOMO SAPIENS | 8 | 3206 | 1x63 | A | | 39 | TRP | p.Trp122Ser | Q13642 |

| | | | | | | | | |
|---|---|---|---|---|---|---|---|---|
| HOMO SAPIENS | 9 | 3206 | 1x63 | A | 40 | HIS | p.His123Tyr | Q13642 |
| HOMO SAPIENS | 9 | 3206 | 1x63 | A | 40 | HIS | p.His123Leu | Q13642 |
| HOMO SAPIENS | 9 | 3206 | 1x63 | A | 40 | HIS | p.His123Gln | Q13642 |
| HOMO SAPIENS | 3 | 3206 | 1x63 | A | 49 | CYS | p.Cys132Phe | Q13642 |
| HOMO SAPIENS | 3 | 3206 | 1x63 | A | 67 | CYS | p.Cys150Tyr | Q13642 |
| HOMO SAPIENS | 3 | 3206 | 1x63 | A | 70 | CYS | p.Cys153Arg | Q13642 |
| HOMO SAPIENS | 3 | 3206 | 1x63 | A | 70 | CYS | p.Cys153Tyr | Q13642 |
| HOMO SAPIENS | 5 | 1052 | 3ecs | E | 208 | ASN | p.Asn208Tyr | Q14232 |
| HOMO SAPIENS | 2 | 1052 | 3ecs | E | 275 | TYR | p.Tyr275Cys | Q14232 |
| HOMO SAPIENS | 0 | 24253 | 5i35 | A | 271 | ARG | p.Arg271Cys | Q8NI60 |
| HOMO SAPIENS | 0 | 24253 | 5i35 | A | 272 | GLY | p.Gly272Asp | Q8NI60 |
| HOMO SAPIENS | 0 | 24253 | 5i35 | A | 299 | ARG | p.Arg299Trp | Q8NI60 |
| HOMO SAPIENS | 1 | 24253 | 5i35 | A | 339 | ALA | p.Ala339Thr | Q8NI60 |
| HOMO SAPIENS | 0 | 24253 | 5i35 | A | 348 | ARG | p.Arg348Ter | Q8NI60 |
| HOMO SAPIENS | 0 | 24253 | 5i35 | A | 395 | ILE | p.Ile395Met | Q8NI60 |
| HOMO SAPIENS | 0 | 24253 | 5i35 | A | 396 | ASP | p.Asp396Glu | Q8NI60 |
| HOMO SAPIENS | 0 | 24253 | 5i35 | A | 480 | PHE | p.Phe480Leu | Q8NI60 |
| HOMO SAPIENS | 2 | 24253 | 5i35 | A | 511 | THR | p.Thr511Met | Q8NI60 |
| HOMO SAPIENS | 0 | 24253 | 5i35 | A | 514 | TYR | p.Tyr514Cys | Q8NI60 |
| HOMO SAPIENS | 0 | 24253 | 5i35 | A | 538 | ARG | p.Arg538Gln | Q8NI60 |
| HOMO SAPIENS | 0 | 24253 | 5i35 | A | 549 | GLY | p.Gly549Ser | Q8NI60 |
| HOMO SAPIENS | 0 | 24253 | 5i35 | A | 551 | GLU | p.Glu551Lys | Q8NI60 |
| HOMO SAPIENS | -1 | 24253 | 5i35 | A | 555 | MSE | p.Met555Ile | Q8NI60 |
| HOMO SAPIENS | 0 | 24253 | 5i35 | A | 568 | GLU | p.Glu568Ter | Q8NI60 |
| HOMO SAPIENS | 0 | 24253 | 5i35 | A | 602 | PRO | p.Pro602Arg | Q8NI60 |
| HOMO SAPIENS | -1 | 15505 | 4goj | C | 22 | GLY | p.Gly22Val | Q13432 |
| HOMO SAPIENS | 5 | 23902 | 2lrr | A | 765 | HIS | p.His765Gln | P38935 |
| HOMO SAPIENS | 1 | 29650 | 2vj2 | A | 234 | CYS | p.Cys234Tyr | P78504 |
| HOMO SAPIENS | 0 | 29650 | 2vj2 | A | 255 | TYR | p.Tyr255Ter | P78504 |
| HOMO SAPIENS | 3 | 29650 | 2vj2 | A | 274 | GLY | p.Gly274Asp | P78504 |
| HOMO SAPIENS | 1 | 105 | 5he0 | B | 77 | GLY | p.Gly77Ser | P62873 |
| HOMO SAPIENS | 2 | 105 | 5he0 | B | 80 | ILE | p.Ile80Thr | P62873 |
| HOMO SAPIENS | 3 | 4587 | 3rgh | A | 159 | ASP | p.Asp1159Ala | P21333 |
| HOMO SAPIENS | 3 | 4587 | 3rgh | A | 184 | ASP | p.Asp1184Glu | P21333 |
| HOMO SAPIENS | 2 | 4587 | 3rgh | A | 186 | SER | p.Ser1186Leu | P21333 |
| HOMO SAPIENS | 4 | 4587 | 3rgh | A | 188 | ALA | p.Ala1188Thr | P21333 |
| HOMO SAPIENS | 2 | 4587 | 3rgh | A | 199 | SER | p.Ser1199Leu | P21333 |
| HOMO SAPIENS | 0 | 20283 | 4n3w | A | 1278 | GLU | p.Glu1278Lys | Q92793 |
| HOMO SAPIENS | 1 | 14294 | 2d9q | B | 208 | MET | p.Met231Thr | Q99062 |
| HOMO SAPIENS | 6 | 14294 | 2d9q | B | 297 | ASP | p.Asp320Asn | Q99062 |
| HOMO SAPIENS | 0 | 11642 | 3bpt | A | 66 | ARG | p.Arg66Trp | Q6NVY1 |
| HOMO SAPIENS | 0 | 11642 | 3bpt | A | 122 | TYR | p.Tyr122Cys | Q6NVY1 |
| HOMO SAPIENS | 0 | 11642 | 3bpt | A | 345 | GLY | p.Gly345Ser | Q6NVY1 |
| HOMO SAPIENS | 1 | 3942 | 3sby | A | 60 | ASP | p.Asp60His | Q9Y4U1 |
| HOMO SAPIENS | 0 | 3942 | 3sby | A | 111 | ARG | p.Arg111Ter | Q9Y4U1 |
| HOMO SAPIENS | 1 | 3942 | 3sby | A | 116 | LEU | p.Leu116Pro | Q9Y4U1 |
| HOMO SAPIENS | 1 | 3942 | 3sby | A | 130 | TYR | p.Tyr130Cys | Q9Y4U1 |
| HOMO SAPIENS | 1 | 3942 | 3sby | A | 132 | ARG | p.Arg132Ter | Q9Y4U1 |
| HOMO SAPIENS | 1 | 3942 | 3sby | A | 147 | GLY | p.Gly147Ala | Q9Y4U1 |
| HOMO SAPIENS | 1 | 3942 | 3sby | A | 161 | ARG | p.Arg161Gln | Q9Y4U1 |
| HOMO SAPIENS | 1 | 3942 | 3sby | A | 161 | ARG | p.Arg161Ter | Q9Y4U1 |
| HOMO SAPIENS | 1 | 3942 | 3sby | A | 203 | TRP | p.Trp203Ter | Q9Y4U1 |
| HOMO SAPIENS | 1 | 3942 | 3sby | A | 203 | TRP | p.Trp203Ter | Q9Y4U1 |
| HOMO SAPIENS | 1 | 3942 | 3sby | A | 205 | TYR | p.Tyr205Ter | Q9Y4U1 |

| | | | | | | | | | |
|---|---|---|---|---|---|---|---|---|---|
| HOMO SAPIENS | 0 | 38340 | 2mbh | B | | 39 | MET | p.Met39Leu | Q13351 |
| HOMO SAPIENS | 2 | 7360 | 4k30 | A | | 430 | LEU | p.Leu430Pro | Q8N159 |
| HOMO SAPIENS | 2 | 7360 | 4k30 | A | | 433 | GLU | p.Glu433Asp | Q8N159 |
| HOMO SAPIENS | 7 | 7360 | 4k30 | A | | 484 | TRP | p.Trp484Arg | Q8N159 |
| HOMO SAPIENS | 1 | 41903 | 2xs6 | A | | 151 | PRO | p.Pro151Ser | O00459 |
| HOMO SAPIENS | 0 | 172 | 5glh | A | | 183 | ALA | p.Ala183Gly | P24530 |
| HOMO SAPIENS | 0 | 172 | 5glh | A | | 201 | ARG | p.Arg201Ter | P24530 |
| HOMO SAPIENS | 0 | 172 | 5glh | A | | 244 | THR | p.Thr244Met | P24530 |
| HOMO SAPIENS | 0 | 172 | 5glh | A | | 253 | ARG | p.Arg253Ter | P24530 |
| HOMO SAPIENS | 9 | 172 | 5glh | A | | 1017 | ILE | p.Leu17Phe | P00720 |
| HOMO SAPIENS | -1 | 3342 | 1ljm | A | | 1058 | HIS | p.His58Asn | Q01196 |
| HOMO SAPIENS | 4 | 3342 | 1ljm | A | | 1174 | ARG | p.Arg174Gln | Q01196 |
| HOMO SAPIENS | 5 | 25607 | 1p22 | C | | 32 | ASP | p.Asp32Ala | P35222 |
| HOMO SAPIENS | 5 | 25607 | 1p22 | C | | 32 | ASP | p.Asp32Tyr | P35222 |
| HOMO SAPIENS | -1 | 25607 | 1p22 | C | | 33 | SEP | p.Ser33Phe | P35222 |
| HOMO SAPIENS | 5 | 25607 | 1p22 | C | | 34 | GLY | p.Gly34Glu | P35222 |
| HOMO SAPIENS | -1 | 25607 | 1p22 | C | | 37 | SEP | p.Ser37Cys | P35222 |
| HOMO SAPIENS | -1 | 25607 | 1p22 | C | | 41 | THR | p.Thr41Ile | P35222 |
| HOMO SAPIENS | 1 | 48 | 4uow | | 9 | 60 | GLN | p.Arg34111His | Q8WZ42 |
| HOMO SAPIENS | 0 | 48 | 4uow | | 9 | 64 | LEU | p.Val34115Ala | Q8WZ42 |
| HOMO SAPIENS | 0 | 1127 | 5c01 | A | | 638 | ARG | p.Arg638Ter | P29597 |
| HOMO SAPIENS | 0 | 1127 | 5c01 | A | | 684 | ILE | p.Ile684Ser | P29597 |
| HOMO SAPIENS | 0 | 1127 | 5c01 | A | | 703 | ARG | p.Arg703Trp | P29597 |
| HOMO SAPIENS | 5 | 16389 | 2aq0 | B | | 128 | ALA | p.Ala860Asp | Q92889 |
| HOMO SAPIENS | 2 | 16389 | 2aq0 | B | | 141 | ILE | p.Ile873Val | Q92889 |
| HOMO SAPIENS | 5 | 16389 | 2aq0 | B | | 143 | GLU | p.Glu875Gly | Q92889 |
| HOMO SAPIENS | 0 | 16389 | 2aq0 | B | | 180 | GLY | p.Gly912Arg | Q92889 |
| HOMO SAPIENS | 0 | 15816 | 2xrc | A | | 89 | THR | p.Thr107Ala | P05156 |
| HOMO SAPIENS | 0 | 15816 | 2xrc | A | | 133 | ASN | p.Asn151Ser | P05156 |
| HOMO SAPIENS | 0 | 15816 | 2xrc | A | | 185 | THR | p.Thr203Ile | P05156 |
| HOMO SAPIENS | 0 | 15816 | 2xrc | A | | 225 | GLY | p.Gly243Asp | P05156 |
| HOMO SAPIENS | 0 | 15816 | 2xrc | A | | 243 | GLY | p.Gly261Ala | P05156 |
| HOMO SAPIENS | 0 | 15816 | 2xrc | A | | 269 | GLY | p.Gly287Arg | P05156 |
| HOMO SAPIENS | -1 | 15816 | 2xrc | A | | 288 | ILE | p.Ile306Val | P05156 |
| HOMO SAPIENS | -1 | 15816 | 2xrc | A | | 388 | ARG | p.Arg406His | P05156 |
| HOMO SAPIENS | 7 | 15816 | 2xrc | A | | 398 | ILE | p.Ile416Leu | P05156 |
| HOMO SAPIENS | 9 | 15816 | 2xrc | A | | 400 | HIS | p.His418Leu | P05156 |
| HOMO SAPIENS | 0 | 15816 | 2xrc | A | | 423 | LYS | p.Lys441Arg | P05156 |
| HOMO SAPIENS | -1 | 15816 | 2xrc | A | | 530 | GLU | p.Glu548Gln | P05156 |
| HOMO SAPIENS | 0 | 15816 | 2xrc | A | | 535 | PRO | p.Pro553Ala | P05156 |
| HOMO SAPIENS | 0 | 15816 | 2xrc | A | | 536 | GLU | p.Glu554Gly | P05156 |
| HOMO SAPIENS | 1 | 12385 | 4zlp | B | | 1450 | ALA | p.Ala1450Pro | Q9UM47 |
| HOMO SAPIENS | 7 | 12385 | 4zlp | B | | 1482 | GLY | p.Gly1482Arg | Q9UM47 |
| HOMO SAPIENS | 2 | 12385 | 4zlp | B | | 1518 | LEU | p.Leu1518Met | Q9UM47 |
| HOMO SAPIENS | 0 | 12385 | 4zlp | B | | 1560 | ARG | p.Arg1560Pro | Q9UM47 |
| HOMO SAPIENS | 0 | 1227 | 4a04 | A | | 148 | PHE | p.Phe148Tyr | O43435 |
| HOMO SAPIENS | 4 | 1227 | 4a04 | A | | 194 | HIS | p.His194Gln | O43435 |
| HOMO SAPIENS | 2 | 8017 | 3ei4 | B | | 244 | LYS | p.Lys244Glu | Q92466 |
| HOMO SAPIENS | 1 | 8017 | 3ei4 | B | | 273 | ARG | p.Arg273His | Q92466 |
| HOMO SAPIENS | 3 | 36112 | 2k8f | B | | 9 | SER | p.Ser9Arg | P04637 |
| HOMO SAPIENS | 2 | 12880 | 5ggf | A | | 265 | ARG | p.Arg265His | Q8WZA1 |
| HOMO SAPIENS | 2 | 12880 | 5ggf | A | | 287 | ILE | p.Ile287Ser | Q8WZA1 |
| HOMO SAPIENS | 2 | 12880 | 5ggf | A | | 311 | ARG | p.Arg311Gln | Q8WZA1 |
| HOMO SAPIENS | 2 | 12880 | 5ggf | A | | 425 | TRP | p.Trp425Ser | Q8WZA1 |

| | | | | | | | | |
|---|---|---|---|---|---|---|---|---|
| HOMO SAPIENS | 2 | 12880 | 5ggf | A | 442 | ARG | p.Arg442Cys | Q8WZA1 |
| HOMO SAPIENS | 2 | 12880 | 5ggf | A | 448 | GLY | p.Gly448Arg | Q8WZA1 |
| HOMO SAPIENS | 0 | 12880 | 5ggf | A | 490 | CYS | p.Cys490Tyr | Q8WZA1 |
| HOMO SAPIENS | 0 | 12880 | 5ggf | A | 493 | PRO | p.Pro493Arg | Q8WZA1 |
| HOMO SAPIENS | 3 | 12880 | 5ggf | A | 550 | SER | p.Ser550Asn | Q8WZA1 |
| HOMO SAPIENS | 1 | 12880 | 5ggf | A | 556 | ASP | p.Asp556Asn | Q8WZA1 |
| HOMO SAPIENS | 0 | 12880 | 5ggf | A | 580 | ARG | p.Arg580Ter | Q8WZA1 |
| HOMO SAPIENS | 2 | 12880 | 5ggf | A | 605 | ARG | p.Arg605Pro | Q8WZA1 |
| HOMO SAPIENS | 0 | 12880 | 5ggf | A | 623 | VAL | p.Met623Val | Q8WZA1 |
| HOMO SAPIENS | 1 | 10187 | 4qw2 | A | 145 | ALA | p.Ala145Ser | Q06787 |
| HOMO SAPIENS | 1 | 10187 | 4qw2 | A | 174 | ASN | p.Asn174Lys | Q06787 |
| HOMO SAPIENS | 0 | 24002 | 2m0t | A | 110 | LEU | p.Leu110Val | O14745 |
| HOMO SAPIENS | 9 | 3126 | 4mjh | A | 99 | LEU | p.Leu99Met | P04792 |
| HOMO SAPIENS | 1 | 3126 | 4mjh | A | 127 | ARG | p.Arg127Trp | P04792 |
| HOMO SAPIENS | 1 | 3126 | 4mjh | A | 127 | ARG | p.Arg127Leu | P04792 |
| HOMO SAPIENS | 1 | 3126 | 4mjh | A | 135 | SER | p.Ser135Phe | P04792 |
| HOMO SAPIENS | 2 | 3126 | 4mjh | A | 136 | ARG | p.Arg136Trp | P04792 |
| HOMO SAPIENS | 1 | 3126 | 4mjh | A | 140 | ARG | p.Arg140Gly | P04792 |
| HOMO SAPIENS | 0 | 3126 | 4mjh | A | 151 | THR | p.Thr151Ile | P04792 |
| HOMO SAPIENS | -1 | 15780 | 2jlp | A | 213 | ARG | p.Arg231Gly | P08294 |
| HOMO SAPIENS | 8 | 3578 | 1apz | D | 203 | GLY | p.Gly226Asp | P20933 |
| HOMO SAPIENS | 1 | 3578 | 1apz | D | 279 | GLY | p.Gly302Arg | P20933 |
| HOMO SAPIENS | 0 | 3578 | 1apz | D | 283 | CYS | p.Cys306Arg | P20933 |
| HOMO SAPIENS | 5 | 1335 | 5xtd | C | 115 | ARG | p.Arg115Cys | O75251 |
| HOMO SAPIENS | 0 | 1335 | 5xtd | C | 122 | VAL | p.Val122Met | O75251 |
| HOMO SAPIENS | 2 | 5063 | 2le4 | A | 37 | ARG | p.Arg74Pro | P48431 |
| HOMO SAPIENS | 1 | 25870 | 2o10 | A | 44 | LEU | p.Leu44Pro | P50461 |
| HOMO SAPIENS | 4 | 25870 | 2o10 | A | 58 | CYS | p.Cys58Gly | P50461 |
| HOMO SAPIENS | 0 | 18748 | 5fii | C | 39 | PHE | p.Phe39Leu | P00439 |
| HOMO SAPIENS | 6 | 18748 | 5fii | C | 48 | LEU | p.Leu48Ser | P00439 |
| HOMO SAPIENS | 2 | 18748 | 5fii | C | 65 | ILE | p.Ile65Asn | P00439 |
| HOMO SAPIENS | 3 | 18748 | 5fii | C | 95 | ILE | p.Ile95Phe | P00439 |
| HOMO SAPIENS | 1 | 14261 | 2vay | B | 1539 | ARG | p.Arg1539Cys | Q13698 |
| HOMO SAPIENS | 0 | 2494 | 4dx1 | A | 183 | GLU | p.Glu183Asp | Q9HBA0 |
| HOMO SAPIENS | 0 | 2494 | 4dx1 | A | 217 | ALA | p.Ala217Ser | Q9HBA0 |
| HOMO SAPIENS | 0 | 2494 | 4dx1 | A | 217 | ALA | p.Ala217Val | Q9HBA0 |
| HOMO SAPIENS | 4 | 2494 | 4dx1 | A | 224 | ARG | p.Arg224Trp | Q9HBA0 |
| HOMO SAPIENS | 4 | 2494 | 4dx1 | A | 254 | VAL | p.Val254Leu | Q9HBA0 |
| HOMO SAPIENS | 0 | 2494 | 4dx1 | A | 269 | ARG | p.Arg269Cys | Q9HBA0 |
| HOMO SAPIENS | 0 | 2494 | 4dx1 | A | 269 | ARG | p.Arg269His | Q9HBA0 |
| HOMO SAPIENS | 0 | 2494 | 4dx1 | A | 278 | GLU | p.Glu278Lys | Q9HBA0 |
| HOMO SAPIENS | 0 | 2494 | 4dx1 | A | 315 | ARG | p.Arg315Trp | Q9HBA0 |
| HOMO SAPIENS | 0 | 2494 | 4dx1 | A | 316 | ARG | p.Arg316Cys | Q9HBA0 |
| HOMO SAPIENS | 0 | 2494 | 4dx1 | A | 331 | ILE | p.Ile331Phe | Q9HBA0 |
| HOMO SAPIENS | 0 | 2494 | 4dx1 | A | 333 | ASP | p.Asp333Gly | Q9HBA0 |
| HOMO SAPIENS | 1 | 2494 | 4dx1 | A | 365 | VAL | p.Val365Met | Q9HBA0 |
| HOMO SAPIENS | 0 | 23998 | 2l1r | B | 161 | ARG | p.Arg162Trp | P19429 |
| HOMO SAPIENS | 0 | 8354 | 4f0x | D | 187 | SER | p.Ser187Ter | O95822 |
| HOMO SAPIENS | 2 | 8354 | 4f0x | D | 259 | GLY | p.Gly259Ala | O95822 |
| HOMO SAPIENS | 2 | 8354 | 4f0x | D | 284 | THR | p.Thr284Ala | O95822 |
| HOMO SAPIENS | 3 | 8354 | 4f0x | D | 309 | LYS | p.Lys309Gln | O95822 |
| HOMO SAPIENS | 0 | 21182 | 4apo | A | 228 | GLN | p.Gln228Lys | O00170 |
| HOMO SAPIENS | 1 | 21182 | 4apo | A | 307 | GLN | p.Gln307Arg | O00170 |
| HOMO SAPIENS | 0 | 2287 | 5of4 | B | 112 | ARG | p.Arg112His | P18074 |

| | | | | | | | | |
|---|---|---|---|---|---|---|---|---|
| HOMO SAPIENS | 0 | 2287 | 5of4 | B | 312 | ASP | p.Asp312Asn | P18074 |
| HOMO SAPIENS | 3 | 2287 | 5of4 | B | 461 | LEU | p.Leu461Val | P18074 |
| HOMO SAPIENS | 0 | 2287 | 5of4 | B | 485 | LEU | p.Leu485Pro | P18074 |
| HOMO SAPIENS | 1 | 2287 | 5of4 | B | 541 | SER | p.Ser541Arg | P18074 |
| HOMO SAPIENS | 0 | 2287 | 5of4 | B | 616 | ARG | p.Arg616Pro | P18074 |
| HOMO SAPIENS | 0 | 2287 | 5of4 | B | 616 | ARG | p.Arg616Trp | P18074 |
| HOMO SAPIENS | 1 | 2287 | 5of4 | B | 658 | ARG | p.Arg658Cys | P18074 |
| HOMO SAPIENS | 1 | 2287 | 5of4 | B | 681 | ASP | p.Asp681Asn | P18074 |
| HOMO SAPIENS | 1 | 2287 | 5of4 | B | 683 | ARG | p.Arg683Trp | P18074 |
| HOMO SAPIENS | 0 | 2287 | 5of4 | B | 713 | GLY | p.Gly713Arg | P18074 |
| HOMO SAPIENS | 0 | 2287 | 5of4 | B | 717 | ALA | p.Ala717Gly | P18074 |
| HOMO SAPIENS | 0 | 2287 | 5of4 | B | 722 | ARG | p.Arg722Trp | P18074 |
| HOMO SAPIENS | 0 | 2287 | 5of4 | B | 725 | ALA | p.Ala725Pro | P18074 |
| HOMO SAPIENS | 4 | 12188 | 4its | A | 122 | ARG | p.Arg122Gly | Q9Y606 |
| HOMO SAPIENS | 4 | 12188 | 4its | A | 133 | ASP | p.Asp133Asn | Q9Y606 |
| HOMO SAPIENS | 4 | 12188 | 4its | A | 144 | ARG | p.Arg144Trp | Q9Y606 |
| HOMO SAPIENS | 0 | 31529 | 1kgd | A | 761 | ARG | p.Arg766Gln | O14936 |
| HOMO SAPIENS | 1 | 31529 | 1kgd | A | 824 | GLN | p.Gln829Arg | O14936 |
| HOMO SAPIENS | 0 | 10906 | 2h63 | C | 18 | ARG | p.Arg18Ter | P53004 |
| HOMO SAPIENS | 9 | 27767 | 4wyq | A | 280 | LEU | p.Leu278Phe | Q9UPY3 |
| HOMO SAPIENS | 0 | 27767 | 4wyq | A | 297 | SER | p.Ser295Cys | Q9UPY3 |
| HOMO SAPIENS | 1 | 27767 | 4wyq | A | 334 | TYR | p.Tyr332Ter | Q9UPY3 |
| HOMO SAPIENS | 7 | 7775 | 5sv7 | C | 136 | SER | p.Ser136Cys | Q9NZJ5 |
| HOMO SAPIENS | 2 | 7775 | 5sv7 | C | 166 | GLN | p.Gln166Arg | Q9NZJ5 |
| HOMO SAPIENS | 2 | 7775 | 5sv7 | C | 332 | GLU | p.Glu332Ter | Q9NZJ5 |
| HOMO SAPIENS | 0 | 21449 | 2e9w | B | 84 | THR | p.Thr84Met | P10721 |
| HOMO SAPIENS | 0 | 21449 | 2e9w | B | 168 | ALA | p.Ala168Glu | P10721 |
| HOMO SAPIENS | 0 | 21449 | 2e9w | B | 178 | ALA | p.Ala178Thr | P10721 |
| HOMO SAPIENS | 0 | 21449 | 2e9w | B | 293 | ASN | p.Asn293Ser | P10721 |
| HOMO SAPIENS | -1 | 21449 | 2e9w | B | 318 | MET | p.Met318Leu | P10721 |
| HOMO SAPIENS | 0 | 21449 | 2e9w | B | 374 | VAL | p.Val374Ile | P10721 |
| HOMO SAPIENS | 6 | 3010 | 3flg | A | 66 | SER | p.Ser270Pro | Q9UBC3 |
| HOMO SAPIENS | 1 | 33918 | 2xzz | A | 760 | ARG | p.Arg760Ter | P22735 |
| HOMO SAPIENS | 2 | 1254 | 5xtc | w | 65 | ASN | p.Asn65Ser | O95299 |
| HOMO SAPIENS | 1 | 1254 | 5xtc | w | 238 | GLU | p.Glu238Lys | O95299 |
| HOMO SAPIENS | 3 | 10245 | 3siv | B | 194 | ALA | p.Ala194Glu | Q8WWY3 |
| HOMO SAPIENS | 4 | 10245 | 3siv | B | 216 | ALA | p.Ala216Pro | Q8WWY3 |
| HOMO SAPIENS | 0 | 2670 | 3dwb | A | 341 | THR | p.Thr341Ile | P42892 |
| HOMO SAPIENS | 1 | 2670 | 3dwb | A | 462 | SER | p.Ser462Asn | P42892 |
| HOMO SAPIENS | 1 | 2670 | 3dwb | A | 754 | ARG | p.Arg754Cys | P42892 |
| HOMO SAPIENS | 1 | 2961 | 5xtb | O | 40 | VAL | p.Val40Met | P19404 |
| HOMO SAPIENS | 5 | 2961 | 5xtb | O | 172 | ILE | p.Ile172Thr | P19404 |
| HOMO SAPIENS | 5 | 2961 | 5xtb | O | 190 | ASP | p.Asp190Gly | P19404 |
| HOMO SAPIENS | 5 | 29655 | 2xdg | A | 57 | ALA | p.Ala57Thr | Q02643 |
| HOMO SAPIENS | 4 | 29655 | 2xdg | A | 72 | GLU | p.Glu72Ter | Q02643 |
| HOMO SAPIENS | 0 | 29655 | 2xdg | A | 121 | GLU | p.Glu121Asp | Q02643 |
| HOMO SAPIENS | -1 | 17935 | 1hyn | P | 38 | ASP | p.Asp38Ala | P02730 |
| HOMO SAPIENS | 7 | 12825 | 1g82 | D | 99 | SER | p.Ser99Asn | P31371 |
| HOMO SAPIENS | 6 | 24927 | 1jm7 | B | 85 | VAL | p.Val85Leu | Q99728 |
| HOMO SAPIENS | 1 | 24927 | 1jm7 | B | 112 | ARG | p.Arg112Ter | Q99728 |
| HOMO SAPIENS | 0 | 24927 | 1jm7 | B | 116 | HIS | p.His116Tyr | Q99728 |
| HOMO SAPIENS | 7 | 92 | 5hde | A | 61 | LYS | p.Lys61Arg | Q05209 |
| HOMO SAPIENS | 0 | 17246 | 1szb | A | 84 | ARG | p.Arg99Gln | O00187 |
| HOMO SAPIENS | 1 | 17246 | 1szb | A | 105 | ASP | p.Asp120Gly | O00187 |

| | | | | | | | | |
|---|---|---|---|---|---|---|---|---|
| HOMO SAPIENS | 0 | 17246 | 1szb | A | 111 | PRO | p.Pro126Leu | O00187 |
| HOMO SAPIENS | 9 | 17246 | 1szb | A | 141 | CYS | p.Cys156Tyr | O00187 |
| HOMO SAPIENS | 0 | 32473 | 1b72 | A | 207 | ARG | p.Arg207Cys | P14653 |
| HOMO SAPIENS | 0 | 1920 | 4yzf | B | 488 | VAL | p.Val488Met | P02730 |
| HOMO SAPIENS | 0 | 1920 | 4yzf | B | 589 | ARG | p.Arg589Cys | P02730 |
| HOMO SAPIENS | 0 | 1920 | 4yzf | B | 589 | ARG | p.Arg589His | P02730 |
| HOMO SAPIENS | 0 | 1920 | 4yzf | B | 602 | ARG | p.Arg602Pro | P02730 |
| HOMO SAPIENS | 0 | 1920 | 4yzf | B | 613 | SER | p.Ser613Phe | P02730 |
| HOMO SAPIENS | -1 | 1920 | 4yzf | B | 646 | ARG | p.Arg646Gln | P02730 |
| HOMO SAPIENS | -1 | 1920 | 4yzf | B | 646 | ARG | p.Arg646Trp | P02730 |
| HOMO SAPIENS | 0 | 1920 | 4yzf | B | 701 | GLY | p.Gly701Asp | P02730 |
| HOMO SAPIENS | 0 | 1920 | 4yzf | B | 760 | ARG | p.Arg760Gln | P02730 |
| HOMO SAPIENS | 0 | 1920 | 4yzf | B | 771 | GLY | p.Gly771Asp | P02730 |
| HOMO SAPIENS | 0 | 1920 | 4yzf | B | 773 | SER | p.Ser773Pro | P02730 |
| HOMO SAPIENS | 0 | 1920 | 4yzf | B | 837 | THR | p.Thr837Ala | P02730 |
| HOMO SAPIENS | 0 | 1920 | 4yzf | B | 854 | PRO | p.Pro854Leu | P02730 |
| HOMO SAPIENS | 0 | 1920 | 4yzf | B | 858 | ALA | p.Ala858Asp | P02730 |
| HOMO SAPIENS | 0 | 1920 | 4yzf | B | 862 | VAL | p.Val862Ile | P02730 |
| HOMO SAPIENS | 0 | 1920 | 4yzf | B | 868 | PRO | p.Pro868Leu | P02730 |
| HOMO SAPIENS | 0 | 1920 | 4yzf | B | 870 | ARG | p.Arg870Trp | P02730 |
| HOMO SAPIENS | 0 | 9863 | 2juj | A | 54 | VAL | p.Val904Ile | P22681 |
| HOMO SAPIENS | 0 | 21934 | 2na7 | A | 184 | ILE | p.Ile184Val | P25445 |
| HOMO SAPIENS | -1 | 6746 | 1lw3 | A | 3 | THR | p.Lys3Thr | Q13614 |
| HOMO SAPIENS | -1 | 6746 | 1lw3 | A | 44 | VAL | p.Val44Ile | Q13614 |
| HOMO SAPIENS | 3 | 6746 | 1lw3 | A | 183 | ALA | p.Ala183Ser | Q13614 |
| HOMO SAPIENS | 4 | 6746 | 1lw3 | A | 482 | GLN | p.Gln482Ter | Q13614 |
| HOMO SAPIENS | 4 | 6746 | 1lw3 | A | 502 | GLU | p.Glu502Gln | Q13614 |
| HOMO SAPIENS | 3 | 6746 | 1lw3 | A | 545 | ASN | p.Asn545Ser | Q13614 |
| HOMO SAPIENS | 0 | 6746 | 1lw3 | A | 586 | ARG | p.Arg586Trp | Q13614 |
| HOMO SAPIENS | -1 | 6746 | 1lw3 | A | 602 | ALA | p.Ala602Gly | Q13614 |
| HOMO SAPIENS | 1 | 319 | 4ch9 | B | 336 | ARG | p.Arg336Ile | Q9UH77 |
| HOMO SAPIENS | 3 | 319 | 4ch9 | B | 340 | ALA | p.Ala340Val | Q9UH77 |
| HOMO SAPIENS | 0 | 319 | 4ch9 | B | 384 | ARG | p.Arg384Gln | Q9UH77 |
| HOMO SAPIENS | 5 | 319 | 4ch9 | B | 426 | PRO | p.Pro426Leu | Q9UH77 |
| HOMO SAPIENS | 3 | 319 | 4ch9 | B | 431 | ARG | p.Arg431Gln | Q9UH77 |
| HOMO SAPIENS | 2 | 319 | 4ch9 | B | 575 | ARG | p.Arg575Trp | Q9UH77 |
| HOMO SAPIENS | 0 | 574 | 2qkk | A | 142 | VAL | p.Val142Ile | O60930 |
| HOMO SAPIENS | 0 | 24021 | 2n8y | A | 752 | ARG | p.Arg752Gln | P12814 |
| HOMO SAPIENS | 0 | 18693 | 2y7j | B | 44 | ARG | p.Arg44Ter | P15735 |
| HOMO SAPIENS | 0 | 18693 | 2y7j | B | 106 | VAL | p.Val106Glu | P15735 |
| HOMO SAPIENS | 3 | 18693 | 2y7j | B | 189 | GLY | p.Gly189Glu | P15735 |
| HOMO SAPIENS | 0 | 17087 | 2ifg | A | 237 | THR | p.Thr237Met | P04629 |
| HOMO SAPIENS | 0 | 7980 | 1qub | A | 247 | VAL | p.Val266Leu | P02749 |
| HOMO SAPIENS | -1 | 2205 | 1h7u | A | 18 | ILE | p.Ile18Val | P54278 |
| HOMO SAPIENS | -1 | 2205 | 1h7u | A | 20 | ARG | p.Arg20Gln | P54278 |
| HOMO SAPIENS | 9 | 2205 | 1h7u | A | 46 | SER | p.Ser46Ile | P54278 |
| HOMO SAPIENS | 0 | 2205 | 1h7u | A | 60 | ASP | p.Asp60Glu | P54278 |
| HOMO SAPIENS | 0 | 2205 | 1h7u | A | 80 | PHE | p.Phe80Leu | P54278 |
| HOMO SAPIENS | 0 | 2205 | 1h7u | A | 118 | SER | p.Ser118Arg | P54278 |
| HOMO SAPIENS | 0 | 2205 | 1h7u | A | 126 | HIS | p.His126Gln | P54278 |
| HOMO SAPIENS | 0 | 2205 | 1h7u | A | 128 | SER | p.Ser128Leu | P54278 |
| HOMO SAPIENS | 1 | 2205 | 1h7u | A | 134 | ARG | p.Arg134Ter | P54278 |
| HOMO SAPIENS | 0 | 2205 | 1h7u | A | 166 | LEU | p.Leu166Pro | P54278 |
| HOMO SAPIENS | 1 | 2205 | 1h7u | A | 191 | TYR | p.Tyr191Cys | P54278 |

| | | | | | | | | |
|---|---|---|---|---|---|---|---|---|
| HOMO SAPIENS | 0 | 2205 | 1h7u | A | 211 | ARG | p.Arg211Ter | P54278 |
| HOMO SAPIENS | 0 | 2205 | 1h7u | A | 233 | GLN | p.Gln233Ter | P54278 |
| HOMO SAPIENS | 0 | 2205 | 1h7u | A | 255 | TYR | p.Tyr255Ter | P54278 |
| HOMO SAPIENS | 0 | 2205 | 1h7u | A | 277 | THR | p.Thr277Lys | P54278 |
| HOMO SAPIENS | 0 | 2205 | 1h7u | A | 301 | LYS | p.Lys301Asn | P54278 |
| HOMO SAPIENS | 0 | 2205 | 1h7u | A | 312 | MET | p.Met312Ile | P54278 |
| HOMO SAPIENS | 0 | 2205 | 1h7u | A | 312 | MET | p.Met312Thr | P54278 |
| HOMO SAPIENS | 0 | 2205 | 1h7u | A | 315 | ARG | p.Arg315Ter | P54278 |
| HOMO SAPIENS | 0 | 2205 | 1h7u | A | 318 | TYR | p.Tyr318Cys | P54278 |
| HOMO SAPIENS | 0 | 2205 | 1h7u | A | 335 | ASN | p.Asn335Ile | P54278 |
| HOMO SAPIENS | 0 | 2205 | 1h7u | A | 360 | ILE | p.Ile360Met | P54278 |
| HOMO SAPIENS | 6 | 20754 | 5mpo | B | 7 | VAL | p.Val7Phe | O96033 |
| HOMO SAPIENS | 4 | 20754 | 5mpo | B | 64 | VAL | p.Val64Ile | O96033 |
| HOMO SAPIENS | 5 | 34944 | 4qam | B | 1114 | ASP | p.Asp1114Gly | Q96KN7 |
| HOMO SAPIENS | 5 | 34944 | 4qam | B | 1240 | GLY | p.Gly1240Glu | Q96KN7 |
| HOMO SAPIENS | 9 | 40496 | 3vnn | A | 278 | ARG | p.Arg278His | P49917 |
| HOMO SAPIENS | 6 | 30131 | 4dt7 | E | 40 | PRO | p.Pro210Leu | P04070 |
| HOMO SAPIENS | -1 | 30131 | 4dt7 | E | 50 | ARG | p.Arg220Trp | P04070 |
| HOMO SAPIENS | 7 | 15311 | 2pqt | A | 64 | ARG | p.Arg64Trp | P18440 |
| HOMO SAPIENS | 2 | 15311 | 2pqt | A | 149 | VAL | p.Val149Ile | P18440 |
| HOMO SAPIENS | -1 | 19581 | 3hy8 | A | 33 | ASP | p.Asp33Val | Q9NVS9 |
| HOMO SAPIENS | 5 | 19581 | 3hy8 | A | 116 | ARG | p.Arg116Gln | Q9NVS9 |
| HOMO SAPIENS | 4 | 19581 | 3hy8 | A | 225 | ARG | p.Arg225His | Q9NVS9 |
| HOMO SAPIENS | 2 | 19581 | 3hy8 | A | 229 | TRP | p.Arg229Trp | Q9NVS9 |
| HOMO SAPIENS | 2 | 19581 | 3hy8 | A | 229 | TRP | p.Arg229Gln | Q9NVS9 |
| HOMO SAPIENS | 0 | 10266 | 3t1i | A | 41 | ASP | p.Asp41His | P49959 |
| HOMO SAPIENS | 1 | 10266 | 3t1i | A | 305 | ARG | p.Arg305Trp | P49959 |
| HOMO SAPIENS | 0 | 10266 | 3t1i | A | 308 | PHE | p.Phe308Leu | P49959 |
| HOMO SAPIENS | 0 | 10266 | 3t1i | A | 334 | SER | p.Ser334Arg | P49959 |
| HOMO SAPIENS | 1 | 1600 | 3rbf | A | 61 | GLU | p.Glu61Asp | P20711 |
| HOMO SAPIENS | 0 | 1600 | 3rbf | A | 91 | ALA | p.Ala91Val | P20711 |
| HOMO SAPIENS | -1 | 1600 | 3rbf | A | 102 | GLY | p.Gly102Ser | P20711 |
| HOMO SAPIENS | 3 | 1600 | 3rbf | A | 147 | SER | p.Ser147Arg | P20711 |
| HOMO SAPIENS | 1 | 1600 | 3rbf | A | 210 | PRO | p.Pro210Leu | P20711 |
| HOMO SAPIENS | 0 | 1600 | 3rbf | A | 217 | MET | p.Met217Val | P20711 |
| HOMO SAPIENS | 4 | 1600 | 3rbf | A | 250 | SER | p.Ser250Phe | P20711 |
| HOMO SAPIENS | 2 | 1600 | 3rbf | A | 275 | ALA | p.Ala275Thr | P20711 |
| HOMO SAPIENS | 1 | 1600 | 3rbf | A | 309 | PHE | p.Phe309Leu | P20711 |
| HOMO SAPIENS | -1 | 1600 | 3rbf | A | 347 | ARG | p.Arg347Gln | P20711 |
| HOMO SAPIENS | 0 | 1600 | 3rbf | A | 462 | ARG | p.Arg462Gln | P20711 |
| HOMO SAPIENS | 6 | 20830 | 5y4f | A | 714 | THR | p.Thr715Ile | Q01484 |
| HOMO SAPIENS | 7 | 20830 | 5y4f | A | 719 | ASP | p.Asp720Glu | Q01484 |
| HOMO SAPIENS | 0 | 20830 | 5y4f | A | 814 | VAL | p.Val815Ile | Q01484 |
| HOMO SAPIENS | -1 | 302 | 2c9y | A | 9 | GLU | p.Glu9Ter | P54819 |
| HOMO SAPIENS | 0 | 302 | 2c9y | A | 103 | ARG | p.Arg103Trp | P54819 |
| HOMO SAPIENS | 3 | 302 | 2c9y | A | 165 | ASP | p.Asp165Gly | P54819 |
| HOMO SAPIENS | 0 | 302 | 2c9y | A | 182 | ALA | p.Ala182Asp | P54819 |
| HOMO SAPIENS | 2 | 302 | 2c9y | A | 186 | ARG | p.Arg186Cys | P54819 |
| HOMO SAPIENS | 0 | 302 | 2c9y | A | 209 | ALA | p.Ala209Thr | P54819 |
| HOMO SAPIENS | -1 | 16920 | 1jlj | B | 10 | ASN | p.Asn10Tyr | Q9NQX3 |
| HOMO SAPIENS | 9 | 8462 | 1zbq | B | 16 | GLY | p.Gly16Ser | P51659 |
| HOMO SAPIENS | 5 | 8462 | 1zbq | B | 19 | ALA | p.Ala19Gly | P51659 |
| HOMO SAPIENS | 1 | 8462 | 1zbq | B | 34 | ALA | p.Ala34Val | P51659 |
| HOMO SAPIENS | 1 | 8462 | 1zbq | B | 106 | ARG | p.Arg106His | P51659 |

| | | | | | | | | |
|---|---|---|---|---|---|---|---|---|
| HOMO SAPIENS | 0 | 8462 | 1zbq | B | 140 | LYS | p.Lys140Asn | P51659 |
| HOMO SAPIENS | 0 | 8462 | 1zbq | B | 292 | THR | p.Thr292Ser | P51659 |
| HOMO SAPIENS | 0 | 4375 | 4ndi | A | 199 | ARG | p.Arg213His | Q7Z2E3 |
| HOMO SAPIENS | 3 | 4375 | 4ndi | A | 201 | HIS | p.His215Arg | Q7Z2E3 |
| HOMO SAPIENS | 0 | 4375 | 4ndi | A | 206 | PRO | p.Pro220Leu | Q7Z2E3 |
| HOMO SAPIENS | 2 | 4375 | 4ndi | A | 223 | LEU | p.Leu237Phe | Q7Z2E3 |
| HOMO SAPIENS | 0 | 4375 | 4ndi | A | 245 | ARG | p.Arg259His | Q7Z2E3 |
| HOMO SAPIENS | 0 | 4375 | 4ndi | A | 263 | VAL | p.Val277Gly | Q7Z2E3 |
| HOMO SAPIENS | 0 | 4375 | 4ndi | A | 279 | TRP | p.Trp293Ter | Q7Z2E3 |
| HOMO SAPIENS | 0 | 27004 | 4ccg | Y | 321 | ASP | p.Asp321Glu | Q9NW38 |
| HOMO SAPIENS | -1 | 27004 | 4ccg | Y | 372 | GLY | p.Gly372Ala | Q9NW38 |
| HOMO SAPIENS | -1 | 6383 | 5fai | A | 34 | ALA | p.Ala34Gly | Q92979 |
| HOMO SAPIENS | 8 | 6383 | 5fai | A | 86 | ASP | p.Asp86Gly | Q92979 |
| HOMO SAPIENS | 2 | 24585 | 4mi5 | A | 684 | ARG | p.Arg679Cys | Q15910 |
| HOMO SAPIENS | 0 | 15967 | 3nhb | A | 723 | ARG | p.Arg723Gln | Q9NP58 |
| HOMO SAPIENS | 5 | 18152 | 1oey | A | 389 | HIS | p.His389Gln | P19878 |
| HOMO SAPIENS | 5 | 18152 | 1oey | A | 395 | ARG | p.Arg395Trp | P19878 |
| HOMO SAPIENS | -1 | 1044 | 1li1 | F | 2 | VAL | p.Val1486Ile | P08572 |
| HOMO SAPIENS | 6 | 1044 | 1li1 | F | 37 | GLU | p.Glu1521Lys | P08572 |
| HOMO SAPIENS | 5 | 1044 | 1li1 | F | 157 | SER | p.Ser1639Leu | P08572 |
| HOMO SAPIENS | 5 | 1044 | 1li1 | F | 179 | ARG | p.Arg1662His | P08572 |
| HOMO SAPIENS | 5 | 1044 | 1li1 | F | 180 | GLY | p.Gly1663Arg | P08572 |
| HOMO SAPIENS | 6 | 20076 | 4uyi | A | 671 | LEU | p.Leu671Ser | Q8IY92 |
| HOMO SAPIENS | 0 | 19588 | 1jmj | A | 189 | ARG | p.Arg208His | P05546 |
| HOMO SAPIENS | -1 | 38023 | 1ry7 | B | 65 | GLY | p.Gly65Arg | P22607 |
| HOMO SAPIENS | -1 | 38023 | 1ry7 | B | 139 | ASP | p.Asp139Glu | P22607 |
| HOMO SAPIENS | 0 | 38023 | 1ry7 | B | 248 | ARG | p.Arg248Cys | P22607 |
| HOMO SAPIENS | 0 | 38023 | 1ry7 | B | 249 | SER | p.Ser249Cys | P22607 |
| HOMO SAPIENS | 0 | 38023 | 1ry7 | B | 250 | PRO | p.Pro250Arg | P22607 |
| HOMO SAPIENS | 0 | 38023 | 1ry7 | B | 322 | GLU | p.Glu322Lys | P22607 |
| HOMO SAPIENS | 5 | 16884 | 5eli | A | 33 | GLN | p.Gln33Ter | Q9NZC2 |
| HOMO SAPIENS | 0 | 16884 | 5eli | A | 44 | TRP | p.Trp44Ter | Q9NZC2 |
| HOMO SAPIENS | 0 | 16884 | 5eli | A | 47 | ARG | p.Arg47His | Q9NZC2 |
| HOMO SAPIENS | 0 | 16884 | 5eli | A | 66 | THR | p.Thr66Met | Q9NZC2 |
| HOMO SAPIENS | 0 | 16884 | 5eli | A | 78 | TRP | p.Trp78Ter | Q9NZC2 |
| HOMO SAPIENS | 1 | 16884 | 5eli | A | 96 | THR | p.Thr96Lys | Q9NZC2 |
| HOMO SAPIENS | 9 | 16884 | 5eli | A | 126 | VAL | p.Val126Gly | Q9NZC2 |
| HOMO SAPIENS | -1 | 12267 | 3w5o | A | 3 | ALA | p.Ala3Val | P49917 |
| HOMO SAPIENS | 1 | 12267 | 3w5o | A | 9 | THR | p.Thr9Ile | P49917 |
| HOMO SAPIENS | 2 | 12267 | 3w5o | A | 188 | ARG | p.Arg188Gln | P49917 |
| HOMO SAPIENS | 0 | 12267 | 3w5o | A | 469 | GLY | p.Gly469Glu | P49917 |
| HOMO SAPIENS | 1 | 12267 | 3w5o | A | 580 | ARG | p.Arg580Ter | P49917 |
| HOMO SAPIENS | 0 | 12267 | 3w5o | A | 600 | GLU | p.Glu600Lys | P49917 |
| HOMO SAPIENS | 9 | 2839 | 2hw5 | B | 54 | ARG | p.Arg54Cys | P30084 |
| HOMO SAPIENS | 9 | 2839 | 2hw5 | B | 59 | ASN | p.Asn59Ser | P30084 |
| HOMO SAPIENS | 0 | 2839 | 2hw5 | B | 273 | LYS | p.Lys273Glu | P30084 |
| HOMO SAPIENS | 1 | 12266 | 2o2k | B | 939 | GLY | p.Gly939Cys | Q99707 |
| HOMO SAPIENS | 5 | 12266 | 2o2k | B | 1164 | ARG | p.Arg1164His | Q99707 |
| HOMO SAPIENS | 1 | 12266 | 2o2k | B | 1166 | LEU | p.Leu1166Met | Q99707 |
| HOMO SAPIENS | 5 | 12266 | 2o2k | B | 1173 | PRO | p.Pro1173Leu | Q99707 |
| HOMO SAPIENS | 5 | 12266 | 2o2k | B | 1222 | ASN | p.Asn1222Ser | Q99707 |
| HOMO SAPIENS | 0 | 30891 | 3ch4 | B | 138 | ARG | p.Arg138Ter | Q15126 |
| HOMO SAPIENS | -1 | 4433 | 1qlz | A | 54 | GLY | p.Gly54Ser | P04156 |
| HOMO SAPIENS | 0 | 22568 | 3cxl | A | 20 | LEU | p.Leu20Phe | P15882 |

| | | | | | | | | |
|---|---|---|---|---|---|---|---|---|
| HOMO SAPIENS | 0 | 22568 | 3cxl | A | 126 | ILE | p.Ile126Met | P15882 |
| HOMO SAPIENS | 0 | 22568 | 3cxl | A | 141 | PRO | p.Pro141Leu | P15882 |
| HOMO SAPIENS | 0 | 22568 | 3cxl | A | 143 | TYR | p.Tyr143His | P15882 |
| HOMO SAPIENS | 0 | 22568 | 3cxl | A | 223 | ALA | p.Ala223Val | P15882 |
| HOMO SAPIENS | 0 | 22568 | 3cxl | A | 228 | GLY | p.Gly228Ser | P15882 |
| HOMO SAPIENS | 0 | 22568 | 3cxl | A | 252 | PRO | p.Pro252Gln | P15882 |
| HOMO SAPIENS | 0 | 22568 | 3cxl | A | 313 | GLU | p.Glu313Lys | P15882 |
| HOMO SAPIENS | 3 | 2668 | 5eqg | A | 34 | ASN | p.Asn34Ile | P11166 |
| HOMO SAPIENS | 3 | 2668 | 5eqg | A | 77 | MET | p.Met77Val | P11166 |
| HOMO SAPIENS | 6 | 2668 | 5eqg | A | 91 | GLY | p.Gly91Asp | P11166 |
| HOMO SAPIENS | 6 | 2668 | 5eqg | A | 92 | ARG | p.Arg92Trp | P11166 |
| HOMO SAPIENS | 6 | 2668 | 5eqg | A | 93 | ARG | p.Arg93Trp | P11166 |
| HOMO SAPIENS | 0 | 2668 | 5eqg | A | 95 | SER | p.Ser95Ile | P11166 |
| HOMO SAPIENS | 3 | 2668 | 5eqg | A | 126 | ARG | p.Arg126His | P11166 |
| HOMO SAPIENS | 3 | 2668 | 5eqg | A | 126 | ARG | p.Arg126Cys | P11166 |
| HOMO SAPIENS | 4 | 2668 | 5eqg | A | 232 | ARG | p.Arg232Cys | P11166 |
| HOMO SAPIENS | 3 | 2668 | 5eqg | A | 249 | ARG | p.Arg249Gln | P11166 |
| HOMO SAPIENS | 3 | 2668 | 5eqg | A | 255 | LYS | p.Lys255Arg | P11166 |
| HOMO SAPIENS | 0 | 2668 | 5eqg | A | 275 | ALA | p.Ala275Thr | P11166 |
| HOMO SAPIENS | 4 | 2668 | 5eqg | A | 299 | GLU | p.Glu299Lys | P11166 |
| HOMO SAPIENS | 6 | 2668 | 5eqg | A | 314 | GLY | p.Gly314Ser | P11166 |
| HOMO SAPIENS | 7 | 2668 | 5eqg | A | 315 | ILE | p.Ile315Met | P11166 |
| HOMO SAPIENS | -1 | 2668 | 5eqg | A | 458 | ARG | p.Arg458Trp | P11166 |
| HOMO SAPIENS | -1 | 2668 | 5eqg | A | 468 | ARG | p.Arg468Trp | P11166 |
| HOMO SAPIENS | 3 | 2291 | 2amy | A | 31 | PHE | p.Phe31Leu | O15305 |
| HOMO SAPIENS | 7 | 2291 | 2amy | A | 32 | LEU | p.Leu32Arg | O15305 |
| HOMO SAPIENS | 7 | 2291 | 2amy | A | 65 | ASP | p.Asp65Tyr | O15305 |
| HOMO SAPIENS | 1 | 2291 | 2amy | A | 108 | ALA | p.Ala108Val | O15305 |
| HOMO SAPIENS | 3 | 2291 | 2amy | A | 113 | PRO | p.Pro113Leu | O15305 |
| HOMO SAPIENS | 3 | 2291 | 2amy | A | 113 | PRO | p.Pro113Ser | O15305 |
| HOMO SAPIENS | 6 | 2291 | 2amy | A | 117 | GLY | p.Gly117Arg | O15305 |
| HOMO SAPIENS | 6 | 2291 | 2amy | A | 119 | PHE | p.Phe119Leu | O15305 |
| HOMO SAPIENS | 6 | 2291 | 2amy | A | 123 | ARG | p.Arg123Gln | O15305 |
| HOMO SAPIENS | 0 | 2291 | 2amy | A | 129 | VAL | p.Val129Met | O15305 |
| HOMO SAPIENS | 7 | 2291 | 2amy | A | 139 | GLU | p.Glu139Lys | O15305 |
| HOMO SAPIENS | 5 | 2291 | 2amy | A | 141 | ARG | p.Arg141His | O15305 |
| HOMO SAPIENS | 7 | 2291 | 2amy | A | 144 | PHE | p.Phe144Leu | O15305 |
| HOMO SAPIENS | 6 | 2291 | 2amy | A | 148 | ASP | p.Asp148Asn | O15305 |
| HOMO SAPIENS | 6 | 2291 | 2amy | A | 153 | ILE | p.Ile153Thr | O15305 |
| HOMO SAPIENS | 2 | 2291 | 2amy | A | 157 | PHE | p.Phe157Ser | O15305 |
| HOMO SAPIENS | 1 | 2291 | 2amy | A | 162 | ARG | p.Arg162Trp | O15305 |
| HOMO SAPIENS | 6 | 2291 | 2amy | A | 188 | ASP | p.Asp188Gly | O15305 |
| HOMO SAPIENS | 5 | 2291 | 2amy | A | 194 | ARG | p.Arg194Ter | O15305 |
| HOMO SAPIENS | 0 | 2291 | 2amy | A | 207 | PHE | p.Phe207Ser | O15305 |
| HOMO SAPIENS | 5 | 2291 | 2amy | A | 208 | GLY | p.Gly208Ala | O15305 |
| HOMO SAPIENS | 1 | 2291 | 2amy | A | 216 | ASN | p.Asn216Ile | O15305 |
| HOMO SAPIENS | 5 | 2291 | 2amy | A | 218 | HIS | p.His218Leu | O15305 |
| HOMO SAPIENS | 6 | 2291 | 2amy | A | 223 | ASP | p.Asp223Glu | O15305 |
| HOMO SAPIENS | 6 | 2291 | 2amy | A | 226 | THR | p.Thr226Ser | O15305 |
| HOMO SAPIENS | 5 | 2291 | 2amy | A | 231 | VAL | p.Val231Met | O15305 |
| HOMO SAPIENS | 1 | 2291 | 2amy | A | 237 | THR | p.Thr237Met | O15305 |
| HOMO SAPIENS | 1 | 2291 | 2amy | A | 241 | CYS | p.Cys241Ser | O15305 |
| HOMO SAPIENS | 5 | 35907 | 1d6g | A | 21 | GLY | p.Gly21Arg | P32238 |
| HOMO SAPIENS | -1 | 33209 | 1x9d | A | 197 | PRO | p.Pro197Leu | Q9UKM7 |

| | | | | | | | | |
|---|---|---|---|---|---|---|---|---|
| HOMO SAPIENS | -1 | 33209 | 1x9d | A | 201 | PRO | p.Pro201Leu | Q9UKM7 |
| HOMO SAPIENS | 6 | 21305 | 4yep | B | 1326 | ARG | p.Arg1326Ter | P24043 |
| HOMO SAPIENS | 6 | 21305 | 4yep | B | 1350 | ARG | p.Arg1350Ter | P24043 |
| HOMO SAPIENS | 0 | 21553 | 3qyb | A | 1120 | VAL | p.Val1119Ala | O60343 |
| HOMO SAPIENS | 2 | 21553 | 3qyb | A | 1148 | THR | p.Thr1147Met | O60343 |
| HOMO SAPIENS | -1 | 30645 | 3m0c | C | -17 | TRP | p.Trp4Ter | P01130 |
| HOMO SAPIENS | -1 | 30645 | 3m0c | C | -17 | TRP | p.Trp4Ter | P01130 |
| HOMO SAPIENS | -1 | 30645 | 3m0c | C | 12 | GLN | p.Gln33Ter | P01130 |
| HOMO SAPIENS | -1 | 30645 | 3m0c | C | 15 | ASP | p.Asp36Glu | P01130 |
| HOMO SAPIENS | -1 | 30645 | 3m0c | C | 28 | SER | p.Ser49Arg | P01130 |
| HOMO SAPIENS | -1 | 30645 | 3m0c | C | 41 | THR | p.Thr62Met | P01130 |
| HOMO SAPIENS | -1 | 30645 | 3m0c | C | 69 | ASP | p.Asp90Asn | P01130 |
| HOMO SAPIENS | -1 | 30645 | 3m0c | C | 76 | ASN | p.Asn97Lys | P01130 |
| HOMO SAPIENS | -1 | 30645 | 3m0c | C | 78 | SER | p.Ser99Ter | P01130 |
| HOMO SAPIENS | -1 | 30645 | 3m0c | C | 81 | GLN | p.Gln102Ter | P01130 |
| HOMO SAPIENS | -1 | 30645 | 3m0c | C | 84 | PRO | p.Pro105Ser | P01130 |
| HOMO SAPIENS | -1 | 30645 | 3m0c | C | 194 | ARG | p.Arg215Cys | P01130 |
| HOMO SAPIENS | -1 | 30645 | 3m0c | C | 198 | GLY | p.Gly219Asp | P01130 |
| HOMO SAPIENS | -1 | 30645 | 3m0c | C | 201 | CYS | p.Cys222Tyr | P01130 |
| HOMO SAPIENS | -1 | 30645 | 3m0c | C | 201 | CYS | p.Cys222Ter | P01130 |
| HOMO SAPIENS | -1 | 30645 | 3m0c | C | 206 | ASP | p.Asp227Glu | P01130 |
| HOMO SAPIENS | -1 | 30645 | 3m0c | C | 249 | CYS | p.Cys270Ter | P01130 |
| HOMO SAPIENS | -1 | 30645 | 3m0c | C | 250 | VAL | p.Val271Ile | P01130 |
| HOMO SAPIENS | 0 | 30645 | 3m0c | C | 255 | CYS | p.Cys276Tyr | P01130 |
| HOMO SAPIENS | 0 | 30645 | 3m0c | C | 255 | CYS | p.Cys276Ter | P01130 |
| HOMO SAPIENS | 0 | 30645 | 3m0c | C | 256 | GLU | p.Glu277Lys | P01130 |
| HOMO SAPIENS | 0 | 30645 | 3m0c | C | 265 | SER | p.Ser286Arg | P01130 |
| HOMO SAPIENS | 0 | 30645 | 3m0c | C | 284 | TRP | p.Trp305Ter | P01130 |
| HOMO SAPIENS | 0 | 30645 | 3m0c | C | 292 | CYS | p.Cys313Ter | P01130 |
| HOMO SAPIENS | 1 | 30645 | 3m0c | C | 385 | ARG | p.Arg406Trp | P01130 |
| HOMO SAPIENS | 0 | 30645 | 3m0c | C | 395 | ARG | p.Arg416Gln | P01130 |
| HOMO SAPIENS | 0 | 30645 | 3m0c | C | 398 | TYR | p.Tyr419Ter | P01130 |
| HOMO SAPIENS | 1 | 30645 | 3m0c | C | 415 | VAL | p.Val436Ala | P01130 |
| HOMO SAPIENS | 1 | 30645 | 3m0c | C | 424 | ASP | p.Asp445Glu | P01130 |
| HOMO SAPIENS | 1 | 30645 | 3m0c | C | 425 | LEU | p.Leu446Val | P01130 |
| HOMO SAPIENS | -1 | 30645 | 3m0c | C | 440 | GLY | p.Gly461Cys | P01130 |
| HOMO SAPIENS | -1 | 30645 | 3m0c | C | 441 | VAL | p.Val462Ile | P01130 |
| HOMO SAPIENS | 1 | 30645 | 3m0c | C | 447 | VAL | p.Val468Ile | P01130 |
| HOMO SAPIENS | 1 | 30645 | 3m0c | C | 489 | THR | p.Thr510Met | P01130 |
| HOMO SAPIENS | 0 | 30645 | 3m0c | C | 524 | GLY | p.Gly545Glu | P01130 |
| HOMO SAPIENS | 1 | 30645 | 3m0c | C | 615 | GLY | p.Gly636Ser | P01130 |
| HOMO SAPIENS | 1 | 30645 | 3m0c | C | 625 | LEU | p.Leu646Ile | P01130 |
| HOMO SAPIENS | 1 | 30645 | 3m0c | C | 630 | ASP | p.Asp651Gly | P01130 |
| HOMO SAPIENS | 3 | 30645 | 3m0c | C | 646 | CYS | p.Cys667Tyr | P01130 |
| HOMO SAPIENS | 4 | 30645 | 3m0c | C | 654 | GLY | p.Gly675Ser | P01130 |
| HOMO SAPIENS | 6 | 30645 | 3m0c | C | 660 | CYS | p.Cys681Ter | P01130 |
| HOMO SAPIENS | 5 | 30645 | 3m0c | C | 676 | ALA | p.Ala697Pro | P01130 |
| HOMO SAPIENS | -1 | 30645 | 3m0c | C | 749 | GLN | p.Gln770Ter | P01130 |
| HOMO SAPIENS | -1 | 41323 | 4bxo | A | 1812 | PRO | p.Pro1812Ala | Q8IYD8 |
| HOMO SAPIENS | 5 | 8335 | 3h0t | C | 12 | GLY | p.Gly71Asp | P81172 |
| HOMO SAPIENS | 6 | 609 | 1bhg | B | 148 | PRO | p.Pro148Ser | P08236 |
| HOMO SAPIENS | 1 | 609 | 1bhg | B | 176 | LEU | p.Leu176Phe | P08236 |
| HOMO SAPIENS | 6 | 609 | 1bhg | B | 216 | ARG | p.Arg216Trp | P08236 |
| HOMO SAPIENS | 0 | 609 | 1bhg | B | 350 | LYS | p.Lys350Asn | P08236 |

| | | | | | | | | |
|---|---|---|---|---|---|---|---|---|
| HOMO SAPIENS | 1 | 609 | 1bhg | B | 354 | ALA | p.Ala354Val | P08236 |
| HOMO SAPIENS | 1 | 609 | 1bhg | B | 357 | ARG | p.Arg357Ter | P08236 |
| HOMO SAPIENS | 0 | 609 | 1bhg | B | 382 | ARG | p.Arg382Cys | P08236 |
| HOMO SAPIENS | 0 | 609 | 1bhg | B | 495 | TYR | p.Tyr495Cys | P08236 |
| HOMO SAPIENS | 2 | 609 | 1bhg | B | 507 | TRP | p.Trp507Ter | P08236 |
| HOMO SAPIENS | 0 | 609 | 1bhg | B | 577 | ARG | p.Arg577Leu | P08236 |
| HOMO SAPIENS | 1 | 609 | 1bhg | B | 611 | ARG | p.Arg611Trp | P08236 |
| HOMO SAPIENS | 0 | 609 | 1bhg | B | 619 | ALA | p.Ala619Val | P08236 |
| HOMO SAPIENS | 0 | 609 | 1bhg | B | 627 | TRP | p.Trp627Cys | P08236 |
| HOMO SAPIENS | 5 | 3422 | 1x9n | A | 566 | GLU | p.Glu566Lys | P18858 |
| HOMO SAPIENS | 2 | 3422 | 1x9n | A | 771 | ARG | p.Arg771Trp | P18858 |
| HOMO SAPIENS | 0 | 3279 | 5tby | B | 797 | ALA | p.Ala797Thr | P12883 |
| HOMO SAPIENS | -1 | 3279 | 5tby | B | 982 | MET | p.Met982Thr | P12883 |
| HOMO SAPIENS | -1 | 3279 | 5tby | B | 992 | LEU | p.Leu992Met | P12883 |
| HOMO SAPIENS | -1 | 3279 | 5tby | B | 1004 | GLN | p.Gln1004Glu | P12883 |
| HOMO SAPIENS | -1 | 3279 | 5tby | B | 1013 | GLU | p.Glu1013Lys | P12883 |
| HOMO SAPIENS | -1 | 3279 | 5tby | B | 1032 | ASP | p.Asp1032Asn | P12883 |
| HOMO SAPIENS | -1 | 3279 | 5tby | B | 1053 | ARG | p.Arg1053Gln | P12883 |
| HOMO SAPIENS | -1 | 3279 | 5tby | B | 1117 | GLU | p.Glu1117Ter | P12883 |
| HOMO SAPIENS | -1 | 3279 | 5tby | B | 1193 | ARG | p.Arg1193His | P12883 |
| HOMO SAPIENS | -1 | 3279 | 5tby | B | 1205 | GLU | p.Glu1205Lys | P12883 |
| HOMO SAPIENS | -1 | 3279 | 5tby | B | 1207 | ILE | p.Ile1207Met | P12883 |
| HOMO SAPIENS | -1 | 3279 | 5tby | B | 1250 | ARG | p.Arg1250Pro | P12883 |
| HOMO SAPIENS | -1 | 3279 | 5tby | B | 1259 | HIS | p.His1259Gln | P12883 |
| HOMO SAPIENS | -1 | 3279 | 5tby | B | 1272 | ASP | p.Asp1272Asn | P12883 |
| HOMO SAPIENS | -1 | 3279 | 5tby | B | 1295 | GLU | p.Glu1295Lys | P12883 |
| HOMO SAPIENS | -1 | 3279 | 5tby | B | 1335 | SER | p.Ser1335Leu | P12883 |
| HOMO SAPIENS | -1 | 3279 | 5tby | B | 1399 | GLU | p.Glu1399Asp | P12883 |
| HOMO SAPIENS | -1 | 3279 | 5tby | B | 1413 | SER | p.Ser1413Ter | P12883 |
| HOMO SAPIENS | -1 | 3279 | 5tby | B | 1459 | LYS | p.Lys1459Asn | P12883 |
| HOMO SAPIENS | -1 | 3279 | 5tby | B | 1467 | LEU | p.Leu1467Val | P12883 |
| HOMO SAPIENS | -1 | 3279 | 5tby | B | 1491 | SER | p.Ser1491Cys | P12883 |
| HOMO SAPIENS | -1 | 3279 | 5tby | B | 1496 | GLU | p.Glu1496Ala | P12883 |
| HOMO SAPIENS | -1 | 3279 | 5tby | B | 1500 | ARG | p.Arg1500Pro | P12883 |
| HOMO SAPIENS | -1 | 3279 | 5tby | B | 1606 | ARG | p.Arg1606His | P12883 |
| HOMO SAPIENS | -1 | 3279 | 5tby | B | 1664 | ASN | p.Asn1664Lys | P12883 |
| HOMO SAPIENS | -1 | 3279 | 5tby | B | 1712 | ARG | p.Arg1712Trp | P12883 |
| HOMO SAPIENS | -1 | 3279 | 5tby | B | 1769 | LEU | p.Leu1769Met | P12883 |
| HOMO SAPIENS | -1 | 3279 | 5tby | B | 1808 | GLY | p.Gly1808Ser | P12883 |
| HOMO SAPIENS | -1 | 3279 | 5tby | B | 1818 | ARG | p.Arg1818Trp | P12883 |
| HOMO SAPIENS | -1 | 3279 | 5tby | B | 1820 | ARG | p.Arg1820Trp | P12883 |
| HOMO SAPIENS | -1 | 3279 | 5tby | B | 1844 | GLU | p.Glu1844Lys | P12883 |
| HOMO SAPIENS | -1 | 3279 | 5tby | B | 1845 | ARG | p.Arg1845Trp | P12883 |
| HOMO SAPIENS | -1 | 3279 | 5tby | B | 1854 | THR | p.Thr1854Met | P12883 |
| HOMO SAPIENS | -1 | 3279 | 5tby | B | 1909 | ARG | p.Arg1909Trp | P12883 |
| HOMO SAPIENS | 2 | 2260 | 4kl1 | A | 553 | ASP | p.Asp553Asn | Q9Y3Q4 |
| HOMO SAPIENS | 0 | 40570 | 5lst | A | 522 | ARG | p.Arg522Cys | O94761 |
| HOMO SAPIENS | 0 | 40570 | 5lst | A | 522 | ARG | p.Arg522His | O94761 |
| HOMO SAPIENS | 0 | 40570 | 5lst | A | 591 | PRO | p.Pro591Leu | O94761 |
| HOMO SAPIENS | 0 | 40570 | 5lst | A | 793 | PRO | p.Pro793Ser | O94761 |
| HOMO SAPIENS | 4 | 40570 | 5lst | A | 799 | VAL | p.Val799Met | O94761 |
| HOMO SAPIENS | 0 | 40570 | 5lst | A | 1004 | ARG | p.Arg1004Trp | O94761 |
| HOMO SAPIENS | 0 | 40570 | 5lst | A | 1005 | ARG | p.Arg1005Gln | O94761 |
| HOMO SAPIENS | 0 | 40570 | 5lst | A | 1021 | ARG | p.Arg1021Gln | O94761 |

| | | | | | | | | |
|---|---|---|---|---|---|---|---|---|
| HOMO SAPIENS | 0 | 40570 | 5lst | A | 1043 | LEU | p.Leu1043Pro | O94761 |
| HOMO SAPIENS | 0 | 40570 | 5lst | A | 1045 | ALA | p.Ala1045Thr | O94761 |
| HOMO SAPIENS | 0 | 40570 | 5lst | A | 1105 | GLY | p.Gly1105Asp | O94761 |
| HOMO SAPIENS | 0 | 40570 | 5lst | A | 1105 | GLY | p.Gly1105Ser | O94761 |
| HOMO SAPIENS | -1 | 40570 | 5lst | A | 1113 | GLY | p.Gly1113Arg | O94761 |
| HOMO SAPIENS | -1 | 18865 | 1gxc | A | 64 | GLU | p.Glu64Lys | O96017 |
| HOMO SAPIENS | -1 | 11673 | 3dzu | D | 85 | PRO | p.Pro113Gln | P37231 |
| HOMO SAPIENS | 0 | 19383 | 3b96 | A | 63 | LYS | p.Lys103Arg | P49748 |
| HOMO SAPIENS | 4 | 19383 | 3b96 | A | 140 | ALA | p.Ala180Thr | P49748 |
| HOMO SAPIENS | 1 | 19383 | 3b96 | A | 145 | GLY | p.Gly185Ser | P49748 |
| HOMO SAPIENS | 0 | 19383 | 3b96 | A | 220 | THR | p.Thr260Met | P49748 |
| HOMO SAPIENS | 5 | 19383 | 3b96 | A | 243 | VAL | p.Val283Ala | P49748 |
| HOMO SAPIENS | 0 | 19383 | 3b96 | A | 249 | GLY | p.Gly289Arg | P49748 |
| HOMO SAPIENS | 0 | 19383 | 3b96 | A | 254 | GLY | p.Gly294Glu | P49748 |
| HOMO SAPIENS | 7 | 19383 | 3b96 | A | 326 | ARG | p.Arg366Cys | P49748 |
| HOMO SAPIENS | 7 | 19383 | 3b96 | A | 326 | ARG | p.Arg366His | P49748 |
| HOMO SAPIENS | 5 | 19383 | 3b96 | A | 342 | LYS | p.Lys382Gln | P49748 |
| HOMO SAPIENS | 0 | 19383 | 3b96 | A | 369 | THR | p.Thr409Met | P49748 |
| HOMO SAPIENS | 5 | 19383 | 3b96 | A | 376 | ALA | p.Ala416Ser | P49748 |
| HOMO SAPIENS | 0 | 19383 | 3b96 | A | 385 | ALA | p.Ala425Thr | P49748 |
| HOMO SAPIENS | 5 | 19383 | 3b96 | A | 399 | GLY | p.Gly439Asp | P49748 |
| HOMO SAPIENS | 7 | 19383 | 3b96 | A | 401 | GLY | p.Gly441Asp | P49748 |
| HOMO SAPIENS | 9 | 19383 | 3b96 | A | 413 | ARG | p.Arg453Gln | P49748 |
| HOMO SAPIENS | 2 | 19383 | 3b96 | A | 418 | PHE | p.Phe458Leu | P49748 |
| HOMO SAPIENS | 6 | 19383 | 3b96 | A | 419 | ARG | p.Arg459Trp | P49748 |
| HOMO SAPIENS | 6 | 19383 | 3b96 | A | 419 | ARG | p.Arg459Gln | P49748 |
| HOMO SAPIENS | -1 | 19383 | 3b96 | A | 471 | ARG | p.Arg511Gln | P49748 |
| HOMO SAPIENS | 0 | 19383 | 3b96 | A | 573 | ARG | p.Arg613Trp | P49748 |
| HOMO SAPIENS | 0 | 19383 | 3b96 | A | 575 | ARG | p.Arg615Gln | P49748 |
| HOMO SAPIENS | 0 | 22286 | 3k2s | A | 3 | PRO | p.Pro27His | P02647 |
| HOMO SAPIENS | 0 | 22286 | 3k2s | A | 4 | PRO | p.Pro28Arg | P02647 |
| HOMO SAPIENS | 0 | 22286 | 3k2s | A | 10 | ARG | p.Arg34Leu | P02647 |
| HOMO SAPIENS | 0 | 22286 | 3k2s | A | 26 | GLY | p.Gly50Arg | P02647 |
| HOMO SAPIENS | 0 | 22286 | 3k2s | A | 37 | ALA | p.Ala61Thr | P02647 |
| HOMO SAPIENS | 1 | 1622 | 2qrv | D | 631 | ARG | p.Arg635Trp | Q9Y6K1 |
| HOMO SAPIENS | 1 | 1622 | 2qrv | D | 900 | PRO | p.Pro904Leu | Q9Y6K1 |
| HOMO SAPIENS | 6 | 40796 | 1v18 | B | 1509 | LEU | p.Leu1509Val | P25054 |
| HOMO SAPIENS | 1 | 22375 | 4pki | G | 1214 | LYS | p.Asp214Asn | P28289 |
| HOMO SAPIENS | 1 | 16307 | 5mv8 | B | 483 | ALA | p.Ala483Ser | Q9Y6N9 |
| HOMO SAPIENS | 6 | 16307 | 5mv8 | B | 506 | ASN | p.Asn506Lys | Q9Y6N9 |
| HOMO SAPIENS | 4 | 16307 | 5mv8 | B | 519 | ASP | p.Glu519Asp | Q9Y6N9 |
| HOMO SAPIENS | 0 | 16307 | 5mv8 | B | 530 | GLY | p.Gly530Arg | Q9Y6N9 |
| HOMO SAPIENS | 4 | 36221 | 4cvo | A | 127 | VAL | p.Val127Ile | Q03468 |
| HOMO SAPIENS | 7 | 36221 | 4cvo | A | 134 | ARG | p.Arg134Trp | Q03468 |
| HOMO SAPIENS | 0 | 36221 | 4cvo | A | 156 | GLN | p.Gln156Ter | Q03468 |
| HOMO SAPIENS | 0 | 10829 | 4a11 | B | 13 | GLU | p.Glu13Ter | Q13216 |
| HOMO SAPIENS | 1 | 10829 | 4a11 | B | 100 | TYR | p.Tyr100Ter | Q13216 |
| HOMO SAPIENS | 0 | 10829 | 4a11 | B | 160 | ALA | p.Ala160Val | Q13216 |
| HOMO SAPIENS | 1 | 6759 | 5lev | A | 29 | ILE | p.Ile29Phe | Q9H3H5 |
| HOMO SAPIENS | 0 | 6759 | 5lev | A | 108 | MET | p.Met108Ile | Q9H3H5 |
| HOMO SAPIENS | 2 | 6759 | 5lev | A | 117 | VAL | p.Val117Ile | Q9H3H5 |
| HOMO SAPIENS | 1 | 6759 | 5lev | A | 170 | TYR | p.Tyr170Cys | Q9H3H5 |
| HOMO SAPIENS | 0 | 6759 | 5lev | A | 264 | GLY | p.Val264Gly | Q9H3H5 |
| HOMO SAPIENS | 0 | 6759 | 5lev | A | 332 | PHE | p.Phe332Val | Q9H3H5 |

| | | | | | | | | |
|---|---|---|---|---|---|---|---|---|
| HOMO SAPIENS | 7 | 6759 | 5lev | A | 393 | ILE | p.Ile393Val | Q9H3H5 |
| HOMO SAPIENS | -1 | 33473 | 5m5e | C | 9 | ASN | p.Asn31Ser | P31785 |
| HOMO SAPIENS | 0 | 26155 | 1ic8 | B | 98 | ALA | p.Ala98Val | P20823 |
| HOMO SAPIENS | 0 | 15329 | 4iqr | B | 127 | ARG | p.Arg136Trp | P41235 |
| HOMO SAPIENS | 0 | 20725 | 4bsj | A | 350 | GLU | p.Glu350Val | P35916 |
| HOMO SAPIENS | 0 | 20725 | 4bsj | A | 494 | THR | p.Thr494Ala | P35916 |
| HOMO SAPIENS | 0 | 20725 | 4bsj | A | 527 | ASN | p.Asn527Ser | P35916 |
| HOMO SAPIENS | 6 | 29959 | 1xiw | B | 47 | ARG | p.Arg68Ter | P04234 |
| HOMO SAPIENS | 2 | 29959 | 1xiw | B | 72 | CYS | p.Cys93Ter | P04234 |
| HOMO SAPIENS | 3 | 20242 | 5tjc | A | 102 | ARG | p.Arg102Ter | Q9GZU1 |
| HOMO SAPIENS | 3 | 20242 | 5tjc | A | 232 | THR | p.Thr232Pro | Q9GZU1 |
| HOMO SAPIENS | 7 | 9590 | 4ap8 | A | 50 | THR | p.Thr50Ala | O96007 |
| HOMO SAPIENS | 4 | 9590 | 4ap8 | A | 168 | GLU | p.Glu168Lys | O96007 |
| HOMO SAPIENS | -1 | 4297 | 5do7 | B | 19 | ASP | p.Asp19His | Q9H221 |
| HOMO SAPIENS | -1 | 4297 | 5do7 | B | 54 | TYR | p.Tyr54Cys | Q9H221 |
| HOMO SAPIENS | -1 | 4297 | 5do7 | B | 80 | CYS | p.Cys80Tyr | Q9H221 |
| HOMO SAPIENS | 5 | 4297 | 5do7 | B | 107 | SER | p.Ser107Ter | Q9H221 |
| HOMO SAPIENS | 1 | 4297 | 5do7 | B | 164 | ARG | p.Arg164Pro | Q9H221 |
| HOMO SAPIENS | 0 | 4297 | 5do7 | B | 210 | VAL | p.Val210Met | Q9H221 |
| HOMO SAPIENS | 0 | 4297 | 5do7 | B | 231 | PRO | p.Pro231Thr | Q9H221 |
| HOMO SAPIENS | 0 | 4297 | 5do7 | B | 263 | ARG | p.Arg263Gln | Q9H221 |
| HOMO SAPIENS | 0 | 4297 | 5do7 | B | 361 | TRP | p.Trp361Ter | Q9H221 |
| HOMO SAPIENS | 0 | 4297 | 5do7 | B | 400 | THR | p.Thr400Lys | Q9H221 |
| HOMO SAPIENS | 0 | 4297 | 5do7 | B | 549 | ALA | p.Ala549Pro | Q9H221 |
| HOMO SAPIENS | 0 | 4297 | 5do7 | B | 596 | LEU | p.Leu596Arg | Q9H221 |
| HOMO SAPIENS | 0 | 4297 | 5do7 | B | 632 | VAL | p.Val632Ala | Q9H221 |
| HOMO SAPIENS | 0 | 4297 | 5do7 | B | 658 | TYR | p.Tyr658Ter | Q9H221 |
| HOMO SAPIENS | 2 | 9356 | 4igg | B | 206 | GLN | p.Gln206His | P35221 |
| HOMO SAPIENS | 6 | 2179 | 2wk6 | A | 44 | ARG | p.Arg44Gln | P19971 |
| HOMO SAPIENS | 6 | 2179 | 2wk6 | A | 81 | ARG | p.Arg81Gln | P19971 |
| HOMO SAPIENS | 6 | 2179 | 2wk6 | A | 145 | GLY | p.Gly145Arg | P19971 |
| HOMO SAPIENS | 7 | 2179 | 2wk6 | A | 146 | ARG | p.Arg146His | P19971 |
| HOMO SAPIENS | 7 | 2179 | 2wk6 | A | 153 | GLY | p.Gly153Ser | P19971 |
| HOMO SAPIENS | 0 | 2179 | 2wk6 | A | 173 | MET | p.Met173Ile | P19971 |
| HOMO SAPIENS | 6 | 2179 | 2wk6 | A | 202 | ARG | p.Arg202Lys | P19971 |
| HOMO SAPIENS | 6 | 2179 | 2wk6 | A | 208 | VAL | p.Val208Met | P19971 |
| HOMO SAPIENS | 5 | 2179 | 2wk6 | A | 222 | LYS | p.Lys222Arg | P19971 |
| HOMO SAPIENS | 5 | 2179 | 2wk6 | A | 289 | GLU | p.Glu289Ala | P19971 |
| HOMO SAPIENS | 2 | 2179 | 2wk6 | A | 311 | GLY | p.Gly311Arg | P19971 |
| HOMO SAPIENS | 2 | 2179 | 2wk6 | A | 376 | ARG | p.Arg376Pro | P19971 |
| HOMO SAPIENS | 2 | 2179 | 2wk6 | A | 465 | ALA | p.Ala465Thr | P19971 |
| HOMO SAPIENS | -1 | 11856 | 3orh | A | 3 | ALA | p.Ala3Ser | Q14353 |
| HOMO SAPIENS | 1 | 11856 | 3orh | A | 20 | TRP | p.Trp20Ser | Q14353 |
| HOMO SAPIENS | 1 | 11856 | 3orh | A | 50 | MET | p.Met50Leu | Q14353 |
| HOMO SAPIENS | 1 | 11856 | 3orh | A | 110 | VAL | p.Val110Phe | Q14353 |
| HOMO SAPIENS | 8 | 11856 | 3orh | A | 164 | GLY | p.Gly164Asp | Q14353 |
| HOMO SAPIENS | 1 | 11856 | 3orh | A | 169 | CYS | p.Cys169Tyr | Q14353 |
| HOMO SAPIENS | 1 | 11856 | 3orh | A | 174 | TRP | p.Trp174Ter | Q14353 |
| HOMO SAPIENS | 2 | 11856 | 3orh | A | 196 | ALA | p.Ala196Val | Q14353 |
| HOMO SAPIENS | 1 | 11856 | 3orh | A | 209 | THR | p.Thr209Met | Q14353 |
| HOMO SAPIENS | 1 | 11856 | 3orh | A | 219 | ASP | p.Asp219Asn | Q14353 |
| HOMO SAPIENS | 1 | 11856 | 3orh | A | 224 | ALA | p.Ala224Thr | Q14353 |
| HOMO SAPIENS | 1 | 13244 | 2m6i | D | 218 | ARG | p.Arg246Leu | P23415 |
| HOMO SAPIENS | 1 | 13244 | 2m6i | D | 244 | ILE | p.Ile272Asn | P23415 |

| | | | | | | | | |
|---|---|---|---|---|---|---|---|---|
| HOMO SAPIENS | 5 | 13244 | 2m6i | D | 250 | PRO | p.Pro278Thr | P23415 |
| HOMO SAPIENS | 6 | 13244 | 2m6i | D | 266 | GLN | p.Gln294His | P23415 |
| HOMO SAPIENS | 1 | 13244 | 2m6i | D | 271 | ARG | p.Arg299Leu | P23415 |
| HOMO SAPIENS | 1 | 13244 | 2m6i | D | 276 | LYS | p.Lys304Glu | P23415 |
| HOMO SAPIENS | 1 | 13244 | 2m6i | D | 279 | TYR | p.Tyr307Cys | P23415 |
| HOMO SAPIENS | 0 | 13244 | 2m6i | D | 316 | GLY | p.Gly378Ser | P23415 |
| HOMO SAPIENS | 1 | 13244 | 2m6i | D | 400 | ARG | p.Arg428His | P23415 |
| HOMO SAPIENS | 3 | 1967 | 2i6b | A | 301 | ALA | p.Ala318Glu | P55263 |
| HOMO SAPIENS | 4 | 11096 | 1h8b | A | 61 | ALA | p.Ala882Thr | P35609 |
| HOMO SAPIENS | 0 | 15423 | 5ngz | A | 2 | GLN | p.Gln2Glu | Q9NPD8 |
| HOMO SAPIENS | 3 | 30475 | 3lpw | A | 21 | SER | p.Ser22894Tyr | Q8WZ42 |
| HOMO SAPIENS | 6 | 32687 | 2arf | A | 1041 | ARG | p.Arg1041Trp | P35670 |
| HOMO SAPIENS | 8 | 32687 | 2arf | A | 1061 | GLY | p.Gly1061Glu | P35670 |
| HOMO SAPIENS | 6 | 32687 | 2arf | A | 1064 | GLU | p.Glu1064Ala | P35670 |
| HOMO SAPIENS | 2 | 32687 | 2arf | A | 1069 | HIS | p.His1069Gln | P35670 |
| HOMO SAPIENS | 2 | 32687 | 2arf | A | 1088 | LEU | p.Leu1088Ter | P35670 |
| HOMO SAPIENS | 0 | 32687 | 2arf | A | 1099 | GLY | p.Gly1099Ser | P35670 |
| HOMO SAPIENS | 4 | 32687 | 2arf | A | 1106 | VAL | p.Val1106Asp | P35670 |
| HOMO SAPIENS | 2 | 32687 | 2arf | A | 1142 | GLN | p.Gln1142His | P35670 |
| HOMO SAPIENS | 8 | 32687 | 2arf | A | 1148 | ILE | p.Ile1148Thr | P35670 |
| HOMO SAPIENS | 6 | 32687 | 2arf | A | 1151 | ARG | p.Arg1151Cys | P35670 |
| HOMO SAPIENS | 4 | 32687 | 2arf | A | 1169 | MET | p.Met1169Val | P35670 |
| HOMO SAPIENS | 4 | 32687 | 2arf | A | 1173 | GLU | p.Glu1173Lys | P35670 |
| HOMO SAPIENS | 2 | 32687 | 2arf | A | 1176 | GLY | p.Gly1176Arg | P35670 |
| HOMO SAPIENS | 3 | 15544 | 5uq3 | C | 575 | PRO | p.Pro69Leu | P46527 |
| HOMO SAPIENS | 6 | 15544 | 5uq3 | C | 592 | GLU | p.Glu86Asp | P46527 |
| HOMO SAPIENS | 6 | 15544 | 5uq3 | C | 599 | ARG | p.Arg93Trp | P46527 |
| HOMO SAPIENS | -1 | 15544 | 5uq3 | C | 615 | VAL | p.Val109Gly | P46527 |
| HOMO SAPIENS | -1 | 15544 | 5uq3 | C | 631 | SER | p.Ser125Pro | P46527 |
| HOMO SAPIENS | -1 | 15544 | 5uq3 | C | 639 | PRO | p.Pro133Ser | P46527 |
| HOMO SAPIENS | 2 | 2703 | 4wxq | A | 245 | CYS | p.Cys245Tyr | Q99972 |
| HOMO SAPIENS | 0 | 2703 | 4wxq | A | 252 | GLY | p.Gly252Arg | Q99972 |
| HOMO SAPIENS | 1 | 2703 | 4wxq | A | 337 | GLN | p.Gln337Arg | Q99972 |
| HOMO SAPIENS | 5 | 2703 | 4wxq | A | 347 | TYR | p.Tyr347Ter | Q99972 |
| HOMO SAPIENS | 3 | 2703 | 4wxq | A | 364 | GLY | p.Gly364Val | Q99972 |
| HOMO SAPIENS | 2 | 2703 | 4wxq | A | 367 | GLY | p.Gly367Arg | Q99972 |
| HOMO SAPIENS | 2 | 2703 | 4wxq | A | 368 | GLN | p.Gln368Ter | Q99972 |
| HOMO SAPIENS | 6 | 2703 | 4wxq | A | 370 | PRO | p.Pro370Leu | Q99972 |
| HOMO SAPIENS | 1 | 2703 | 4wxq | A | 380 | ASP | p.Asp380His | Q99972 |
| HOMO SAPIENS | 8 | 2703 | 4wxq | A | 399 | GLY | p.Gly399Val | Q99972 |
| HOMO SAPIENS | 0 | 2703 | 4wxq | A | 423 | LYS | p.Lys423Glu | Q99972 |
| HOMO SAPIENS | 3 | 2703 | 4wxq | A | 433 | CYS | p.Cys433Arg | Q99972 |
| HOMO SAPIENS | 3 | 2703 | 4wxq | A | 437 | TYR | p.Tyr437His | Q99972 |
| HOMO SAPIENS | 2 | 2703 | 4wxq | A | 477 | ILE | p.Ile477Ser | Q99972 |
| HOMO SAPIENS | 3 | 2703 | 4wxq | A | 480 | ASN | p.Asn480Lys | Q99972 |
| HOMO SAPIENS | -1 | 34100 | 2fot | C | 1171 | GLN | p.Gln1173Pro | Q13813 |
| HOMO SAPIENS | 0 | 8773 | 1kw2 | A | 432 | ASP | p.Asp432Glu | P02774 |
| HOMO SAPIENS | 0 | 8773 | 1kw2 | A | 436 | LYS | p.Lys436Thr | P02774 |
| HOMO SAPIENS | 0 | 8402 | 4dzo | B | 649 | TYR | p.Tyr649Ter | Q9Y6D9 |
| HOMO SAPIENS | 0 | 39962 | 1nst | A | 611 | GLY | p.Gly611Ser | P52848 |
| HOMO SAPIENS | 2 | 22578 | 3dof | B | 45 | MET | p.Met45Arg | Q9Y2Y0 |
| HOMO SAPIENS | 1 | 23290 | 5j1s | A | 292 | ILE | p.Ile292Thr | O14656 |
| HOMO SAPIENS | 4 | 9505 | 4j1y | A | 534 | ARG | p.Arg534Trp | P09871 |
| HOMO SAPIENS | 9 | 260 | 2h57 | A | 31 | THR | p.Thr31Met | Q9H0F7 |

| | | | | | | | | |
|---|---|---|---|---|---|---|---|---|
| HOMO SAPIENS | 0 | 260 | 2h57 | A | 91 | ILE | p.Ile91Thr | Q9H0F7 |
| HOMO SAPIENS | 0 | 260 | 2h57 | A | 121 | ARG | p.Arg121His | Q9H0F7 |
| HOMO SAPIENS | 0 | 260 | 2h57 | A | 122 | ARG | p.Arg122Ter | Q9H0F7 |
| HOMO SAPIENS | 2 | 260 | 2h57 | A | 169 | GLY | p.Gly169Ala | Q9H0F7 |
| HOMO SAPIENS | 4 | 260 | 2h57 | A | 170 | LEU | p.Leu170Trp | Q9H0F7 |
| HOMO SAPIENS | 4 | 34023 | 2cr3 | A | 18 | GLY | p.Gly48Ser | P11362 |
| HOMO SAPIENS | 0 | 37667 | 3ii7 | A | 420 | ARG | p.Arg420Cys | Q8IXQ5 |
| HOMO SAPIENS | 0 | 41720 | 2k9u | A | 68 | ARG | p.Arg2364His | Q14315 |
| HOMO SAPIENS | 1 | 29250 | 5fmv | A | 568 | HIS | p.His568Gln | P08575 |
| HOMO SAPIENS | 1 | 1280 | 1fb1 | A | 69 | PRO | p.Pro69Leu | P30793 |
| HOMO SAPIENS | 3 | 1280 | 1fb1 | A | 88 | ARG | p.Arg88Trp | P30793 |
| HOMO SAPIENS | 2 | 1280 | 1fb1 | A | 108 | GLY | p.Gly108Asp | P30793 |
| HOMO SAPIENS | 4 | 1280 | 1fb1 | A | 134 | ASP | p.Asp134Val | P30793 |
| HOMO SAPIENS | 6 | 1280 | 1fb1 | A | 135 | ILE | p.Ile135Lys | P30793 |
| HOMO SAPIENS | 9 | 1280 | 1fb1 | A | 144 | HIS | p.His144Pro | P30793 |
| HOMO SAPIENS | 9 | 1280 | 1fb1 | A | 184 | ARG | p.Arg184His | P30793 |
| HOMO SAPIENS | 1 | 1280 | 1fb1 | A | 196 | ALA | p.Ala196Ser | P30793 |
| HOMO SAPIENS | 3 | 1280 | 1fb1 | A | 201 | GLY | p.Gly201Glu | P30793 |
| HOMO SAPIENS | 2 | 1280 | 1fb1 | A | 204 | VAL | p.Val204Ile | P30793 |
| HOMO SAPIENS | 2 | 1280 | 1fb1 | A | 211 | MET | p.Met211Ile | P30793 |
| HOMO SAPIENS | 0 | 1280 | 1fb1 | A | 221 | MET | p.Met221Thr | P30793 |
| HOMO SAPIENS | 3 | 1280 | 1fb1 | A | 224 | LYS | p.Lys224Arg | P30793 |
| HOMO SAPIENS | 2 | 1280 | 1fb1 | A | 249 | ARG | p.Arg249Ser | P30793 |
| HOMO SAPIENS | 8 | 41352 | 4k12 | A | 47 | ILE | p.Ile551Thr | P08603 |
| HOMO SAPIENS | 9 | 20703 | 5nir | A | 57 | CYS | p.Cys57Tyr | P02458 |
| HOMO SAPIENS | 9 | 20703 | 5nir | A | 64 | CYS | p.Cys64Ter | P02458 |
| HOMO SAPIENS | 0 | 20087 | 4cmn | A | 299 | MET | p.Met299Ile | Q01968 |
| HOMO SAPIENS | 1 | 20819 | 3aaf | A | 979 | ILE | p.Ile979Met | Q14191 |
| HOMO SAPIENS | 1 | 20819 | 3aaf | A | 984 | LEU | p.Leu984Ile | Q14191 |
| HOMO SAPIENS | 3 | 20819 | 3aaf | A | 987 | ARG | p.Arg987Ter | Q14191 |
| HOMO SAPIENS | -1 | 20819 | 3aaf | A | 1074 | PHE | p.Leu1074Phe | Q14191 |
| HOMO SAPIENS | 0 | 3619 | 3dem | A | 206 | ASN | p.Asn225Ser | P48740 |
| HOMO SAPIENS | 1 | 3619 | 3dem | A | 269 | ARG | p.Arg288Leu | P48740 |
| HOMO SAPIENS | 0 | 3105 | 3ooi | A | 164 | TYR | p.Tyr2015His | Q96L73 |
| HOMO SAPIENS | 6 | 15828 | 2lkx | A | 16 | LEU | p.Leu100Gln | Q99697 |
| HOMO SAPIENS | 8 | 15828 | 2lkx | A | 24 | ARG | p.Arg108His | Q99697 |
| HOMO SAPIENS | 2 | 15828 | 2lkx | A | 30 | THR | p.Thr114Pro | Q99697 |
| HOMO SAPIENS | 8 | 15828 | 2lkx | A | 45 | VAL | p.Val129Leu | Q99697 |
| HOMO SAPIENS | 8 | 15828 | 2lkx | A | 46 | ARG | p.Arg130Trp | Q99697 |
| HOMO SAPIENS | 9 | 15828 | 2lkx | A | 53 | ARG | p.Arg137Pro | Q99697 |
| HOMO SAPIENS | 9 | 13017 | 4irq | A | 141 | ARG | p.Arg141Trp | Q9UBV7 |
| HOMO SAPIENS | 0 | 13017 | 4irq | A | 186 | ALA | p.Ala186Asp | Q9UBV7 |
| HOMO SAPIENS | 0 | 13017 | 4irq | A | 206 | LEU | p.Leu206Pro | Q9UBV7 |
| HOMO SAPIENS | 1 | 13017 | 4irq | A | 214 | CYS | p.Cys214Tyr | Q9UBV7 |
| HOMO SAPIENS | 0 | 13017 | 4irq | A | 270 | ARG | p.Arg270Cys | Q9UBV7 |
| HOMO SAPIENS | -1 | 13053 | 4mqw | X | 307 | ALA | p.Ala307Thr | P23945 |
| HOMO SAPIENS | 0 | 19406 | 4b3g | A | 43 | ARG | p.Arg43Ter | P38935 |
| HOMO SAPIENS | 0 | 19406 | 4b3g | A | 46 | CYS | p.Cys46Ter | P38935 |
| HOMO SAPIENS | 0 | 19406 | 4b3g | A | 75 | ALA | p.Ala75Thr | P38935 |
| HOMO SAPIENS | 0 | 19406 | 4b3g | A | 201 | LEU | p.Leu201Ser | P38935 |
| HOMO SAPIENS | 3 | 19406 | 4b3g | A | 213 | HIS | p.His213Arg | P38935 |
| HOMO SAPIENS | 0 | 19406 | 4b3g | A | 275 | ILE | p.Ile275Val | P38935 |
| HOMO SAPIENS | -1 | 19406 | 4b3g | A | 320 | ARG | p.Arg320Ter | P38935 |
| HOMO SAPIENS | 0 | 19406 | 4b3g | A | 361 | LEU | p.Leu361Pro | P38935 |

| | | | | | | | | |
|---|---|---|---|---|---|---|---|---|
| HOMO SAPIENS | 0 | 19406 | 4b3g | A | 369 | PHE | p.Phe369Leu | P38935 |
| HOMO SAPIENS | 0 | 19406 | 4b3g | A | 493 | THR | p.Thr493Ile | P38935 |
| HOMO SAPIENS | 0 | 19406 | 4b3g | A | 496 | CYS | p.Cys496Ter | P38935 |
| HOMO SAPIENS | 0 | 19406 | 4b3g | A | 514 | GLU | p.Glu514Lys | P38935 |
| HOMO SAPIENS | 0 | 19406 | 4b3g | A | 528 | ALA | p.Ala528Thr | P38935 |
| HOMO SAPIENS | 0 | 19406 | 4b3g | A | 531 | PRO | p.Pro531Thr | P38935 |
| HOMO SAPIENS | 0 | 19406 | 4b3g | A | 580 | VAL | p.Val580Ile | P38935 |
| HOMO SAPIENS | 6 | 19406 | 4b3g | A | 603 | ARG | p.Arg603His | P38935 |
| HOMO SAPIENS | 0 | 21501 | 5jde | A | 40 | SER | p.Ser2831Asn | Q8WZ42 |
| HOMO SAPIENS | 5 | 21501 | 5jde | A | 189 | ALA | p.Ala2980Thr | Q8WZ42 |
| HOMO SAPIENS | 6 | 15008 | 3mop | H | 12 | ARG | p.Arg12Cys | Q9NWZ3 |
| HOMO SAPIENS | 8 | 17455 | 3jtc | C | 15 | ARG | p.Arg57Trp | P04070 |
| HOMO SAPIENS | -1 | 17455 | 3jtc | C | 20 | CGU | p.Glu62Ala | P04070 |
| HOMO SAPIENS | 0 | 27795 | 5gox | B | 625 | TYR | p.Tyr625Ter | Q92878 |
| HOMO SAPIENS | 0 | 27795 | 5gox | B | 653 | SER | p.Ser653Ter | Q92878 |
| HOMO SAPIENS | -1 | 6558 | 5nug | A | 137 | VAL | p.Val137Ile | Q14204 |
| HOMO SAPIENS | -1 | 6558 | 5nug | A | 567 | ARG | p.Arg567His | Q14204 |
| HOMO SAPIENS | -1 | 6558 | 5nug | A | 1140 | THR | p.Thr1140Met | Q14204 |
| HOMO SAPIENS | -1 | 6558 | 5nug | A | 1991 | ASP | p.Asp1991Asn | Q14204 |
| HOMO SAPIENS | -1 | 6558 | 5nug | A | 2401 | LYS | p.Lys2401Asn | Q14204 |
| HOMO SAPIENS | 0 | 6558 | 5nug | A | 2488 | ARG | p.Arg2488His | Q14204 |
| HOMO SAPIENS | 0 | 6558 | 5nug | A | 3907 | HIS | p.His3907Gln | Q14204 |
| HOMO SAPIENS | 0 | 6558 | 5nug | A | 3981 | THR | p.Thr3981Arg | Q14204 |
| HOMO SAPIENS | 0 | 6558 | 5nug | A | 4394 | THR | p.Thr4394Met | Q14204 |
| HOMO SAPIENS | 0 | 6558 | 5nug | A | 4573 | ASN | p.Asn4573Ser | Q14204 |
| HOMO SAPIENS | 0 | 6558 | 5nug | A | 4603 | SER | p.Ser4603Ile | Q14204 |
| HOMO SAPIENS | 1 | 4244 | 1umk | A | 57 | ARG | p.Arg58Gln | P00387 |
| HOMO SAPIENS | 1 | 4244 | 1umk | A | 72 | LEU | p.Leu73Pro | P00387 |
| HOMO SAPIENS | 1 | 4244 | 1umk | A | 105 | VAL | p.Val106Met | P00387 |
| HOMO SAPIENS | 0 | 4244 | 1umk | A | 116 | THR | p.Thr117Ser | P00387 |
| HOMO SAPIENS | 8 | 4244 | 1umk | A | 127 | SER | p.Ser128Pro | P00387 |
| HOMO SAPIENS | 0 | 4244 | 1umk | A | 148 | LEU | p.Leu149Pro | P00387 |
| HOMO SAPIENS | 0 | 4244 | 1umk | A | 159 | ARG | p.Arg160Ter | P00387 |
| HOMO SAPIENS | 0 | 4244 | 1umk | A | 203 | CYS | p.Cys204Arg | P00387 |
| HOMO SAPIENS | 0 | 4244 | 1umk | A | 203 | CYS | p.Cys204Tyr | P00387 |
| HOMO SAPIENS | 2 | 4244 | 1umk | A | 239 | ASP | p.Asp240Gly | P00387 |
| HOMO SAPIENS | 0 | 4244 | 1umk | A | 291 | GLY | p.Gly292Asp | P00387 |
| HOMO SAPIENS | 4 | 34495 | 1s35 | A | 1151 | ASP | p.Asn1151Asp | P11277 |
| HOMO SAPIENS | 3 | 30110 | 3zqs | A | 224 | THR | p.Thr224Ala | Q9NW38 |
| HOMO SAPIENS | 3 | 17588 | 5dse | B | 53 | LEU | p.Leu53Pro | Q9BYI3 |
| HOMO SAPIENS | 0 | 1564 | 4yqf | B | 72 | HIS | p.His347Gln | Q9UHD8 |
| HOMO SAPIENS | 0 | 17673 | 4jnw | A | 32 | ARG | p.Arg32186His | Q8WZ42 |
| HOMO SAPIENS | 3 | 17673 | 4jnw | A | 140 | ILE | p.Ile32294Val | Q8WZ42 |
| HOMO SAPIENS | 5 | 17673 | 4jnw | A | 269 | ALA | p.Ala32423Thr | Q8WZ42 |
| HOMO SAPIENS | 3 | 12858 | 3s97 | C | 302 | LEU | p.Leu302Pro | Q12860 |
| HOMO SAPIENS | 0 | 6147 | 4nn0 | A | 114 | ARG | p.Arg114Gln | Q9BXJ0 |
| HOMO SAPIENS | 0 | 7549 | 1tr2 | A | 277 | LEU | p.Leu277Met | P18206 |
| HOMO SAPIENS | 1 | 7549 | 1tr2 | A | 413 | ALA | p.Ala413Thr | P18206 |
| HOMO SAPIENS | 1 | 7549 | 1tr2 | A | 519 | ILE | p.Ile519Leu | P18206 |
| HOMO SAPIENS | 5 | 7549 | 1tr2 | A | 547 | ARG | p.Arg547Ter | P18206 |
| HOMO SAPIENS | 2 | 7549 | 1tr2 | A | 636 | HIS | p.His636Arg | P18206 |
| HOMO SAPIENS | 2 | 7549 | 1tr2 | A | 824 | ILE | p.Ile824Met | P18206 |
| HOMO SAPIENS | 2 | 13900 | 3gro | A | 45 | CYS | p.Cys45Tyr | P50897 |
| HOMO SAPIENS | 2 | 13900 | 3gro | A | 75 | THR | p.Thr75Pro | P50897 |

| | | | | | | | | |
|---|---|---|---|---|---|---|---|---|
| HOMO SAPIENS | 2 | 13900 | 3gro | A | 79 | ASP | p.Asp79Gly | P50897 |
| HOMO SAPIENS | 5 | 13900 | 3gro | A | 91 | GLN | p.Gln91Pro | P50897 |
| HOMO SAPIENS | 5 | 13900 | 3gro | A | 96 | CYS | p.Cys96Tyr | P50897 |
| HOMO SAPIENS | 0 | 13900 | 3gro | A | 104 | LYS | p.Lys104Ter | P50897 |
| HOMO SAPIENS | 2 | 13900 | 3gro | A | 108 | GLY | p.Gly108Arg | P50897 |
| HOMO SAPIENS | 2 | 13900 | 3gro | A | 109 | TYR | p.Tyr109Asp | P50897 |
| HOMO SAPIENS | 0 | 13900 | 3gro | A | 122 | ARG | p.Arg122Trp | P50897 |
| HOMO SAPIENS | 0 | 13900 | 3gro | A | 138 | SER | p.Ser138Leu | P50897 |
| HOMO SAPIENS | 5 | 13900 | 3gro | A | 151 | ARG | p.Arg151Ter | P50897 |
| HOMO SAPIENS | 7 | 13900 | 3gro | A | 164 | ARG | p.Arg164Ter | P50897 |
| HOMO SAPIENS | 0 | 13900 | 3gro | A | 177 | GLN | p.Gln177Glu | P50897 |
| HOMO SAPIENS | 0 | 13900 | 3gro | A | 181 | VAL | p.Val181Leu | P50897 |
| HOMO SAPIENS | 0 | 13900 | 3gro | A | 184 | GLU | p.Glu184Lys | P50897 |
| HOMO SAPIENS | 5 | 13900 | 3gro | A | 186 | TRP | p.Trp186Ter | P50897 |
| HOMO SAPIENS | 0 | 13900 | 3gro | A | 219 | LEU | p.Leu219Gln | P50897 |
| HOMO SAPIENS | 0 | 13900 | 3gro | A | 222 | LEU | p.Leu222Pro | P50897 |
| HOMO SAPIENS | 5 | 13900 | 3gro | A | 296 | TRP | p.Trp296Ter | P50897 |
| HOMO SAPIENS | 6 | 35028 | 5mwb | A | 444 | CYS | p.Cys444Tyr | Q04721 |
| HOMO SAPIENS | 0 | 35028 | 5mwb | A | 514 | ASN | p.Asn514Ser | Q04721 |
| HOMO SAPIENS | 5 | 1483 | 4gl2 | A | 822 | ARG | p.Arg822Gln | Q9BYX4 |
| HOMO SAPIENS | 0 | 1483 | 4gl2 | A | 843 | ARG | p.His843Arg | Q9BYX4 |
| HOMO SAPIENS | -1 | 28838 | 3cl3 | D | 153 | LEU | p.Leu153Arg | Q9Y6K9 |
| HOMO SAPIENS | 2 | 22563 | 4zel | B | 79 | ARG | p.Arg79Trp | P09172 |
| HOMO SAPIENS | 2 | 22563 | 4zel | B | 101 | VAL | p.Val101Met | P09172 |
| HOMO SAPIENS | 2 | 22563 | 4zel | B | 114 | ASP | p.Asp114Glu | P09172 |
| HOMO SAPIENS | 0 | 22563 | 4zel | B | 201 | ASN | p.Asn201Ser | P09172 |
| HOMO SAPIENS | 2 | 22563 | 4zel | B | 211 | ALA | p.Ala211Thr | P09172 |
| HOMO SAPIENS | -1 | 22563 | 4zel | B | 290 | ASP | p.Asp290Asn | P09172 |
| HOMO SAPIENS | 2 | 22563 | 4zel | B | 318 | ALA | p.Ala318Ser | P09172 |
| HOMO SAPIENS | 2 | 22563 | 4zel | B | 345 | ASP | p.Asp345Asn | P09172 |
| HOMO SAPIENS | 2 | 22563 | 4zel | B | 362 | ALA | p.Ala362Glu | P09172 |
| HOMO SAPIENS | 2 | 22563 | 4zel | B | 549 | ARG | p.Arg549Cys | P09172 |
| HOMO SAPIENS | 9 | 9780 | 1y6k | R | 36 | TYR | p.Tyr57Cys | Q13651 |
| HOMO SAPIENS | 1 | 9780 | 1y6k | R | 40 | LEU | p.Leu61Val | Q13651 |
| HOMO SAPIENS | 1 | 9780 | 1y6k | R | 63 | THR | p.Thr84Ile | Q13651 |
| HOMO SAPIENS | 9 | 9780 | 1y6k | R | 80 | ARG | p.Arg101Trp | Q13651 |
| HOMO SAPIENS | 0 | 9780 | 1y6k | R | 120 | GLY | p.Gly141Arg | Q13651 |
| HOMO SAPIENS | 0 | 9780 | 1y6k | R | 138 | SER | p.Ser159Gly | Q13651 |
| HOMO SAPIENS | 0 | 9780 | 1y6k | R | 203 | ILE | p.Ile224Val | Q13651 |
| HOMO SAPIENS | 0 | 16019 | 3rep | A | 211 | ARG | p.Arg211Cys | Q13418 |
| HOMO SAPIENS | 3 | 7879 | 5ec3 | A | 33 | ALA | p.Ala33Thr | P32754 |
| HOMO SAPIENS | 4 | 7879 | 5ec3 | A | 113 | ARG | p.Arg113Gln | P32754 |
| HOMO SAPIENS | 3 | 7879 | 5ec3 | A | 340 | VAL | p.Val340Leu | P32754 |
| HOMO SAPIENS | 0 | 8904 | 1qk9 | A | 6 | LYS | p.Lys82Arg | P51608 |
| HOMO SAPIENS | 4 | 8904 | 1qk9 | A | 24 | LEU | p.Leu100Val | P51608 |
| HOMO SAPIENS | 7 | 8904 | 1qk9 | A | 30 | ARG | p.Arg106Trp | P51608 |
| HOMO SAPIENS | 6 | 8904 | 1qk9 | A | 57 | ARG | p.Arg133Cys | P51608 |
| HOMO SAPIENS | 3 | 8904 | 1qk9 | A | 61 | GLU | p.Glu137Gly | P51608 |
| HOMO SAPIENS | 2 | 8904 | 1qk9 | A | 64 | ALA | p.Ala140Val | P51608 |
| HOMO SAPIENS | 5 | 8904 | 1qk9 | A | 79 | PHE | p.Phe155Ser | P51608 |
| HOMO SAPIENS | 3 | 8904 | 1qk9 | A | 82 | THR | p.Thr158Met | P51608 |
| HOMO SAPIENS | 1 | 23121 | 3e9k | A | 198 | THR | p.Thr198Ala | Q16719 |
| HOMO SAPIENS | -1 | 26090 | 3tje | F | 19 | LEU | p.Leu35Met | P25445 |
| HOMO SAPIENS | 8 | 26090 | 3tje | F | 106 | THR | p.Thr122Ile | P25445 |

| | | | | | | | | |
|---|---|---|---|---|---|---|---|---|
| HOMO SAPIENS | 2 | 36404 | 5cqr | A | 765 | GLY | p.Gly765Glu | O95163 |
| HOMO SAPIENS | 2 | 36404 | 5cqr | A | 816 | ILE | p.Ile816Leu | O95163 |
| HOMO SAPIENS | 2 | 36404 | 5cqr | A | 830 | ILE | p.Ile830Met | O95163 |
| HOMO SAPIENS | 5 | 36404 | 5cqr | A | 939 | TYR | p.Tyr939Ter | O95163 |
| HOMO SAPIENS | 2 | 36404 | 5cqr | A | 952 | LYS | p.Lys952Ile | O95163 |
| HOMO SAPIENS | 5 | 36404 | 5cqr | A | 1013 | GLY | p.Gly1013Ser | O95163 |
| HOMO SAPIENS | 5 | 36404 | 5cqr | A | 1072 | CYS | p.Cys1072Ser | O95163 |
| HOMO SAPIENS | -1 | 36404 | 5cqr | A | 1158 | PRO | p.Pro1158Leu | O95163 |
| HOMO SAPIENS | 1 | 34149 | 2k40 | A | 18 | ASN | p.Asn125Ser | Q9UBX0 |
| HOMO SAPIENS | 0 | 34149 | 2k40 | A | 42 | LEU | p.Glu149Lys | Q9UBX0 |
| HOMO SAPIENS | 9 | 34149 | 2k40 | A | 53 | ARG | p.Arg160Cys | Q9UBX0 |
| HOMO SAPIENS | 0 | 34149 | 2k40 | A | 63 | SER | p.Ser170Leu | Q9UBX0 |
| HOMO SAPIENS | 2 | 2468 | 3vp6 | A | 484 | LEU | p.Leu484Met | Q99259 |
| HOMO SAPIENS | 0 | 2468 | 3vp6 | A | 532 | ARG | p.Arg532Gln | Q99259 |
| HOMO SAPIENS | 5 | 1253 | 5xtd | S | 8 | GLY | p.Gly8Arg | O15239 |
| HOMO SAPIENS | 7 | 1253 | 5xtd | S | 32 | GLY | p.Gly32Arg | O15239 |
| HOMO SAPIENS | 7 | 1253 | 5xtd | S | 37 | ARG | p.Arg37Ser | O15239 |
| HOMO SAPIENS | -1 | 13292 | 3u1k | A | 121 | VAL | p.Ile121Val | Q8TCS8 |
| HOMO SAPIENS | 2 | 13292 | 3u1k | A | 165 | PRO | p.Pro165Ser | Q8TCS8 |
| HOMO SAPIENS | 2 | 13292 | 3u1k | A | 192 | ARG | p.Arg192Gln | Q8TCS8 |
| HOMO SAPIENS | 1 | 13292 | 3u1k | A | 315 | LYS | p.Lys315Arg | Q8TCS8 |
| HOMO SAPIENS | 4 | 13292 | 3u1k | A | 509 | VAL | p.Val509Ile | Q8TCS8 |
| HOMO SAPIENS | 7 | 13292 | 3u1k | A | 531 | THR | p.Thr531Arg | Q8TCS8 |
| HOMO SAPIENS | 0 | 13292 | 3u1k | A | 590 | ASN | p.Asn590Asp | Q8TCS8 |
| HOMO SAPIENS | 0 | 27048 | 4p5w | B | 184 | ARG | p.Arg184Ter | Q9UJS0 |
| HOMO SAPIENS | 0 | 27048 | 4p5w | B | 225 | SER | p.Ser225Ter | Q9UJS0 |
| HOMO SAPIENS | -1 | 27048 | 4p5w | B | 631 | ALA | p.Arg360Ter | Q9UJS0 |
| HOMO SAPIENS | 0 | 24240 | 4mdr | A | 318 | ARG | p.Arg318Ter | O00189 |
| HOMO SAPIENS | 0 | 24240 | 4mdr | A | 338 | ARG | p.Arg338Ter | O00189 |
| HOMO SAPIENS | 0 | 13429 | 4cg4 | A | 479 | PHE | p.Phe479Leu | O15553 |
| HOMO SAPIENS | 0 | 13429 | 4cg4 | A | 487 | VAL | p.Val487Leu | O15553 |
| HOMO SAPIENS | 3 | 13429 | 4cg4 | A | 653 | ARG | p.Arg653His | O15553 |
| HOMO SAPIENS | 1 | 13429 | 4cg4 | A | 680 | MET | p.Met680Ile | O15553 |
| HOMO SAPIENS | 1 | 13429 | 4cg4 | A | 688 | TYR | p.Tyr688Ter | O15553 |
| HOMO SAPIENS | 0 | 13429 | 4cg4 | A | 694 | MET | p.Met694Ile | O15553 |
| HOMO SAPIENS | 0 | 13429 | 4cg4 | A | 694 | MET | p.Met694Val | O15553 |
| HOMO SAPIENS | 1 | 13429 | 4cg4 | A | 726 | VAL | p.Val726Ala | O15553 |
| HOMO SAPIENS | 0 | 13429 | 4cg4 | A | 761 | ARG | p.Arg761His | O15553 |
| HOMO SAPIENS | 0 | 2079 | 4okh | A | 691 | GLY | p.Gly691Arg | P20807 |
| HOMO SAPIENS | 2 | 2079 | 4okh | A | 748 | ARG | p.Arg748Ter | P20807 |
| HOMO SAPIENS | 0 | 2079 | 4okh | A | 763 | TYR | p.Tyr763Cys | P20807 |
| HOMO SAPIENS | 1 | 2079 | 4okh | A | 769 | ARG | p.Arg769Gln | P20807 |
| HOMO SAPIENS | 9 | 2079 | 4okh | A | 777 | ILE | p.Ile777Val | P20807 |
| HOMO SAPIENS | 8 | 2079 | 4okh | A | 778 | ASP | p.Asp778His | P20807 |
| HOMO SAPIENS | 5 | 2079 | 4okh | A | 780 | ASP | p.Asp780His | P20807 |
| HOMO SAPIENS | 1 | 1200 | 2gi7 | A | 142 | ARG | p.Arg162Trp | Q9HCN6 |
| HOMO SAPIENS | 4 | 9199 | 3ezq | C | 260 | ASP | p.Asp260Val | P25445 |
| HOMO SAPIENS | 0 | 36522 | 5ms9 | A | 122 | ARG | p.Arg122Cys | P35555 |
| HOMO SAPIENS | 0 | 36522 | 5ms9 | A | 240 | ARG | p.Arg240Cys | P35555 |
| HOMO SAPIENS | 0 | 17448 | 3g2f | A | 225 | ARG | p.Arg225His | Q13873 |
| HOMO SAPIENS | 0 | 17448 | 3g2f | A | 347 | CYS | p.Cys347Tyr | Q13873 |
| HOMO SAPIENS | 3 | 17448 | 3g2f | A | 485 | ASP | p.Asp485Gly | Q13873 |
| HOMO SAPIENS | 7 | 17448 | 3g2f | A | 491 | ARG | p.Arg491Gln | Q13873 |
| HOMO SAPIENS | 7 | 17448 | 3g2f | A | 491 | ARG | p.Arg491Trp | Q13873 |

| | | | | | | | | |
|---|---|---|---|---|---|---|---|---|
| HOMO SAPIENS | 0 | 4861 | 4ux8 | A | 56 | LEU | p.Leu56Met | P07949 |
| HOMO SAPIENS | 1 | 4861 | 4ux8 | A | 88 | ILE | p.Ile88Leu | P07949 |
| HOMO SAPIENS | 0 | 4861 | 4ux8 | A | 114 | ARG | p.Arg114His | P07949 |
| HOMO SAPIENS | 5 | 4861 | 4ux8 | A | 180 | ARG | p.Arg180Gln | P07949 |
| HOMO SAPIENS | 0 | 4861 | 4ux8 | A | 208 | GLU | p.Glu208Asp | P07949 |
| HOMO SAPIENS | 0 | 4861 | 4ux8 | A | 373 | ALA | p.Ala373Val | P07949 |
| HOMO SAPIENS | 0 | 4861 | 4ux8 | A | 375 | LEU | p.Leu375Gln | P07949 |
| HOMO SAPIENS | -1 | 4861 | 4ux8 | A | 386 | ALA | p.Ala386Val | P07949 |
| HOMO SAPIENS | 0 | 4861 | 4ux8 | A | 446 | GLY | p.Gly446Arg | P07949 |
| HOMO SAPIENS | 0 | 4861 | 4ux8 | A | 480 | GLU | p.Glu480Lys | P07949 |
| HOMO SAPIENS | 0 | 4861 | 4ux8 | A | 489 | ASP | p.Asp489Asn | P07949 |
| HOMO SAPIENS | -1 | 4861 | 4ux8 | A | 510 | ALA | p.Ala510Val | P07949 |
| HOMO SAPIENS | -1 | 4861 | 4ux8 | A | 533 | GLY | p.Gly533Cys | P07949 |
| HOMO SAPIENS | -1 | 4861 | 4ux8 | A | 567 | ASP | p.Asp567Asn | P07949 |
| HOMO SAPIENS | -1 | 4861 | 4ux8 | A | 609 | CYS | p.Cys609Tyr | P07949 |
| HOMO SAPIENS | -1 | 4861 | 4ux8 | A | 609 | CYS | p.Cys609Gly | P07949 |
| HOMO SAPIENS | -1 | 4861 | 4ux8 | A | 611 | CYS | p.Cys611Trp | P07949 |
| HOMO SAPIENS | -1 | 4861 | 4ux8 | A | 618 | CYS | p.Cys618Gly | P07949 |
| HOMO SAPIENS | -1 | 4861 | 4ux8 | A | 618 | CYS | p.Cys618Phe | P07949 |
| HOMO SAPIENS | -1 | 4861 | 4ux8 | A | 620 | CYS | p.Cys620Gly | P07949 |
| HOMO SAPIENS | -1 | 4861 | 4ux8 | A | 620 | CYS | p.Cys620Ser | P07949 |
| HOMO SAPIENS | -1 | 4861 | 4ux8 | A | 620 | CYS | p.Cys620Trp | P07949 |
| HOMO SAPIENS | -1 | 4861 | 4ux8 | A | 623 | GLU | p.Glu623Lys | P07949 |
| HOMO SAPIENS | -1 | 4861 | 4ux8 | A | 631 | ASP | p.Asp631Gly | P07949 |
| HOMO SAPIENS | -1 | 4861 | 4ux8 | A | 631 | ASP | p.Asp631Glu | P07949 |
| HOMO SAPIENS | -1 | 4861 | 4ux8 | A | 634 | CYS | p.Cys634Gly | P07949 |
| HOMO SAPIENS | -1 | 4861 | 4ux8 | A | 634 | CYS | p.Cys634Phe | P07949 |
| HOMO SAPIENS | -1 | 4861 | 4ux8 | A | 634 | CYS | p.Cys634Trp | P07949 |
| HOMO SAPIENS | 0 | 36308 | 4nkq | B | 123 | PRO | p.Pro142Leu | P15509 |
| HOMO SAPIENS | 0 | 36308 | 4nkq | B | 145 | ARG | p.Arg164Gln | P15509 |
| HOMO SAPIENS | -1 | 36308 | 4nkq | B | 177 | GLY | p.Gly196Arg | P15509 |
| HOMO SAPIENS | 0 | 36006 | 1oqa | A | 107 | HIS | p.His1860Pro | P38398 |
| HOMO SAPIENS | 7 | 15394 | 2ozo | A | 465 | ARG | p.Arg465His | P43403 |
| HOMO SAPIENS | 7 | 15394 | 2ozo | A | 465 | ARG | p.Arg465Cys | P43403 |
| HOMO SAPIENS | 4 | 15394 | 2ozo | A | 518 | SER | p.Ser518Arg | P43403 |
| HOMO SAPIENS | 1 | 20813 | 3o5z | A | 19 | ARG | p.Arg19Trp | O00459 |
| HOMO SAPIENS | 2 | 4329 | 2bb5 | A | 5 | ILE | p.Ile23Val | P20062 |
| HOMO SAPIENS | 4 | 4329 | 2bb5 | A | 59 | LYS | p.Lys77Met | P20062 |
| HOMO SAPIENS | 4 | 4329 | 2bb5 | A | 197 | ARG | p.Arg215Trp | P20062 |
| HOMO SAPIENS | 1 | 4329 | 2bb5 | A | 241 | PRO | p.Arg259Pro | P20062 |
| HOMO SAPIENS | 2 | 4329 | 2bb5 | A | 315 | THR | p.Thr333Lys | P20062 |
| HOMO SAPIENS | 5 | 4329 | 2bb5 | A | 330 | SER | p.Ser348Phe | P20062 |
| HOMO SAPIENS | 4 | 4329 | 2bb5 | A | 358 | LEU | p.Leu376Ser | P20062 |
| HOMO SAPIENS | 5 | 4329 | 2bb5 | A | 381 | ARG | p.Arg399Gln | P20062 |
| HOMO SAPIENS | 4 | 18294 | 4r30 | A | 163 | ASN | p.Asn163Asp | O95278 |
| HOMO SAPIENS | 6 | 18294 | 4r30 | A | 227 | ALA | p.Ala227Val | O95278 |
| HOMO SAPIENS | 4 | 18294 | 4r30 | A | 241 | ARG | p.Arg241Ter | O95278 |
| HOMO SAPIENS | 4 | 18294 | 4r30 | A | 325 | ARG | p.Arg325His | O95278 |
| HOMO SAPIENS | 0 | 2632 | 4xem | A | 22 | GLU | p.Glu22Lys | P49588 |
| HOMO SAPIENS | 0 | 2632 | 4xem | A | 275 | GLY | p.Gly275Asp | P49588 |
| HOMO SAPIENS | 1 | 2632 | 4xem | A | 329 | ARG | p.Arg329His | P49588 |
| HOMO SAPIENS | 0 | 11431 | 5t4n | A | 284 | GLN | p.Gln305Glu | Q96QU1 |
| HOMO SAPIENS | 0 | 11431 | 5t4n | A | 315 | ARG | p.Arg336Ter | Q96QU1 |
| HOMO SAPIENS | 0 | 11431 | 5t4n | A | 322 | ARG | p.Arg343Lys | Q96QU1 |

| | | | | | | | | |
|---|---|---|---|---|---|---|---|---|
| HOMO SAPIENS | 1 | 11431 | 5t4n | A | 326 | LEU | p.Leu347Phe | Q96QU1 |
| HOMO SAPIENS | 0 | 11431 | 5t4n | A | 373 | TYR | p.Tyr394Cys | Q96QU1 |
| HOMO SAPIENS | 0 | 11431 | 5t4n | A | 398 | ASN | p.Asn419Ser | Q96QU1 |
| HOMO SAPIENS | 0 | 11431 | 5t4n | A | 414 | ALA | p.Asp435Ala | Q96QU1 |
| HOMO SAPIENS | 0 | 11431 | 5t4n | A | 433 | VAL | p.Val454Phe | Q96QU1 |
| HOMO SAPIENS | 1 | 11431 | 5t4n | A | 507 | VAL | p.Val528Asp | Q96QU1 |
| HOMO SAPIENS | 0 | 11431 | 5t4n | A | 547 | ALA | p.Ala568Ser | Q96QU1 |
| HOMO SAPIENS | 3 | 3194 | 5xtd | L | 84 | ARG | p.Arg84His | O43181 |
| HOMO SAPIENS | 1 | 3194 | 5xtd | L | 130 | THR | p.Thr130Ser | O43181 |
| HOMO SAPIENS | 8 | 3194 | 5xtd | L | 136 | SER | p.Ser136Ala | O43181 |
| HOMO SAPIENS | 0 | 40749 | 1nun | A | 106 | CYS | p.Cys106Phe | O15520 |
| HOMO SAPIENS | 9 | 40749 | 1nun | A | 156 | ILE | p.Ile156Arg | O15520 |
| HOMO SAPIENS | 1 | 40749 | 1nun | A | 193 | ARG | p.Arg193Ter | O15520 |
| HOMO SAPIENS | 0 | 40749 | 1nun | A | 207 | HIS | p.His207Pro | O15520 |
| HOMO SAPIENS | 0 | 7026 | 3n1f | D | 875 | SER | p.Ser875Gly | Q4KMG0 |
| HOMO SAPIENS | 6 | 23802 | 3rzv | A | 424 | ARG | p.Arg424Ter | O95630 |
| HOMO SAPIENS | 0 | 36720 | 1jm7 | A | 12 | GLN | p.Gln12Ter | P38398 |
| HOMO SAPIENS | 3 | 36720 | 1jm7 | A | 23 | GLU | p.Glu23Asp | P38398 |
| HOMO SAPIENS | 1 | 36720 | 1jm7 | A | 45 | LYS | p.Lys45Gln | P38398 |
| HOMO SAPIENS | 8 | 36720 | 1jm7 | A | 47 | CYS | p.Cys47Ter | P38398 |
| HOMO SAPIENS | 0 | 36720 | 1jm7 | A | 59 | SER | p.Ser59Ter | P38398 |
| HOMO SAPIENS | 8 | 36720 | 1jm7 | A | 61 | CYS | p.Cys61Gly | P38398 |
| HOMO SAPIENS | 8 | 36720 | 1jm7 | A | 63 | LEU | p.Leu63Ter | P38398 |
| HOMO SAPIENS | 7 | 36720 | 1jm7 | A | 64 | CYS | p.Cys64Gly | P38398 |
| HOMO SAPIENS | 0 | 36720 | 1jm7 | A | 101 | TYR | p.Tyr101Ter | P38398 |
| HOMO SAPIENS | 0 | 36720 | 1jm7 | A | null | TYR | p.Tyr105Cys | P38398 |
| HOMO SAPIENS | 0 | 28500 | 3kuz | A | 1217 | CYS | p.Cys1217Arg | O60486 |
| HOMO SAPIENS | 5 | 8097 | 3p56 | C | 69 | ARG | p.Arg69Trp | Q8TDP1 |
| HOMO SAPIENS | 5 | 8097 | 3p56 | C | 143 | LYS | p.Lys143Ile | Q8TDP1 |
| HOMO SAPIENS | 2 | 30073 | 3nmz | A | 380 | ARG | p.Arg380Gln | P25054 |
| HOMO SAPIENS | 1 | 22573 | 1cjy | A | 111 | SER | p.Ser111Pro | P47712 |
| HOMO SAPIENS | 0 | 22573 | 1cjy | A | 485 | ARG | p.Arg485His | P47712 |
| HOMO SAPIENS | 2 | 965 | 3E+077 | A | 99 | ALA | p.Ala99Val | Q9Y617 |
| HOMO SAPIENS | 2 | 965 | 3E+077 | A | 100 | ASP | p.Asp100Ala | Q9Y617 |
| HOMO SAPIENS | 4 | 1503 | 3kxc | A | 62 | ARG | p.Arg62Trp | O43617 |
| HOMO SAPIENS | 6 | 9418 | 1hdr | A | 22 | GLY | p.Gly23Asp | P09417 |
| HOMO SAPIENS | 2 | 9418 | 1hdr | A | 35 | TRP | p.Trp36Arg | P09417 |
| HOMO SAPIENS | 2 | 9418 | 1hdr | A | 107 | TRP | p.Trp108Gly | P09417 |
| HOMO SAPIENS | 6 | 9418 | 1hdr | A | 149 | TYR | p.Tyr150Cys | P09417 |
| HOMO SAPIENS | -1 | 12079 | 3tnu | B | 373 | GLY | p.Gly373Ala | P13647 |
| HOMO SAPIENS | 1 | 12079 | 3tnu | B | 463 | LEU | p.Leu463Pro | P13647 |
| HOMO SAPIENS | 1 | 12079 | 3tnu | B | 471 | ARG | p.Arg471Cys | P13647 |
| HOMO SAPIENS | 0 | 8135 | 3rbn | B | 89 | SER | p.Ser556Ile | P40692 |
| HOMO SAPIENS | 0 | 8135 | 3rbn | B | 111 | GLU | p.Glu578Gly | P40692 |
| HOMO SAPIENS | 2 | 8135 | 3rbn | B | 114 | PRO | p.Pro581Leu | P40692 |
| HOMO SAPIENS | 0 | 8135 | 3rbn | B | 133 | GLU | p.Glu600Gly | P40692 |
| HOMO SAPIENS | 2 | 8135 | 3rbn | B | 136 | PRO | p.Pro603Arg | P40692 |
| HOMO SAPIENS | 2 | 8135 | 3rbn | B | 136 | PRO | p.Pro603Leu | P40692 |
| HOMO SAPIENS | 7 | 8135 | 3rbn | B | 140 | LEU | p.Leu607His | P40692 |
| HOMO SAPIENS | 2 | 8135 | 3rbn | B | 151 | LYS | p.Lys618Ala | P40692 |
| HOMO SAPIENS | 2 | 8135 | 3rbn | B | 151 | LYS | p.Lys618Thr | P40692 |
| HOMO SAPIENS | 5 | 8135 | 3rbn | B | 181 | PRO | p.Pro648Ser | P40692 |
| HOMO SAPIENS | 2 | 8135 | 3rbn | B | 188 | ILE | p.Ile655Val | P40692 |
| HOMO SAPIENS | 2 | 8135 | 3rbn | B | 214 | ALA | p.Ala681Thr | P40692 |

| | | | | | | | | |
|---|---|---|---|---|---|---|---|---|
| HOMO SAPIENS | 2 | 8135 | 3rbn | B | 220 | ARG | p.Arg687Trp | P40692 |
| HOMO SAPIENS | 2 | 8135 | 3rbn | B | 222 | GLN | p.Gln689Arg | P40692 |
| HOMO SAPIENS | -1 | 8135 | 3rbn | B | 236 | GLN | p.Gln701Ter | P40692 |
| HOMO SAPIENS | 7 | 8135 | 3rbn | B | 251 | HIS | p.His718Tyr | P40692 |
| HOMO SAPIENS | 0 | 8135 | 3rbn | B | 284 | LYS | p.Lys751Arg | P40692 |
| HOMO SAPIENS | 1 | 21998 | 5d68 | A | 355 | LEU | p.Leu355Phe | O00522 |
| HOMO SAPIENS | 3 | 21998 | 5d68 | A | 380 | THR | p.Thr380Met | O00522 |
| HOMO SAPIENS | 0 | 21998 | 5d68 | A | 415 | ILE | p.Ile415Met | O00522 |
| HOMO SAPIENS | 0 | 33789 | 2m0c | A | 13 | ARG | p.Arg218Gln | Q9H161 |
| HOMO SAPIENS | 7 | 33789 | 2m0c | A | 52 | ARG | p.Arg257Thr | Q9H161 |
| HOMO SAPIENS | 0 | 33789 | 2m0c | A | 67 | ARG | p.Arg272Pro | Q9H161 |
| HOMO SAPIENS | 6 | 8263 | 2osg | A | 51 | GLY | p.Gly355Arg | Q9UDY2 |
| HOMO SAPIENS | 5 | 8263 | 2osg | A | 60 | THR | p.Thr364Met | Q9UDY2 |
| HOMO SAPIENS | 1 | 2475 | 5E+026 | B | 234 | THR | p.Thr234Ala | Q9BZ23 |
| HOMO SAPIENS | 2 | 2475 | 5E+026 | B | 264 | ARG | p.Arg264Trp | Q9BZ23 |
| HOMO SAPIENS | 4 | 2475 | 5E+026 | B | 286 | ARG | p.Arg286Cys | Q9BZ23 |
| HOMO SAPIENS | 0 | 2475 | 5E+026 | B | 357 | ARG | p.Arg357Trp | Q9BZ23 |
| HOMO SAPIENS | 2 | 2475 | 5E+026 | B | 481 | ARG | p.Arg481Ter | Q9BZ23 |
| HOMO SAPIENS | 8 | 2475 | 5E+026 | B | 521 | GLY | p.Gly521Arg | Q9BZ23 |
| HOMO SAPIENS | 2 | 2475 | 5E+026 | B | 528 | THR | p.Thr528Met | Q9BZ23 |
| HOMO SAPIENS | 1 | 3742 | 5xtd | E | 108 | HIS | p.His134Tyr | P56556 |
| HOMO SAPIENS | 1 | 16725 | 3g04 | C | 36 | ASP | p.Asp36His | P16473 |
| HOMO SAPIENS | 3 | 16725 | 3g04 | C | 52 | PRO | p.Pro52Thr | P16473 |
| HOMO SAPIENS | 1 | 16725 | 3g04 | C | 162 | PRO | p.Pro162Ala | P16473 |
| HOMO SAPIENS | -1 | 12366 | 5d2d | C | 750 | PRO | p.Pro750Leu | P13569 |
| HOMO SAPIENS | 0 | 12366 | 5d2d | C | 764 | ARG | p.Arg764Ter | P13569 |
| HOMO SAPIENS | 0 | 34845 | 2y23 | A | 1151 | ASP | p.Asp1151Glu | P52179 |
| HOMO SAPIENS | 0 | 34845 | 2y23 | A | 1180 | ASP | p.Asp1180Ala | P52179 |
| HOMO SAPIENS | 6 | 36740 | 3lbx | B | 2053 | ALA | p.Ala2053Pro | P11277 |
| HOMO SAPIENS | 0 | 19289 | 3ihj | A | 404 | ARG | p.Arg404Ter | Q8TD30 |
| HOMO SAPIENS | 5 | 1034 | 5xtd | Q | 228 | ARG | p.Arg228Gln | O75306 |
| HOMO SAPIENS | 5 | 1034 | 5xtd | Q | 229 | PRO | p.Pro229Gln | O75306 |
| HOMO SAPIENS | 4 | 1034 | 5xtd | Q | 323 | ARG | p.Arg323Gln | O75306 |
| HOMO SAPIENS | 4 | 1034 | 5xtd | Q | 352 | PRO | p.Pro352Ala | O75306 |
| HOMO SAPIENS | 5 | 1034 | 5xtd | Q | 413 | SER | p.Ser413Pro | O75306 |
| HOMO SAPIENS | 1 | 2772 | 2vaf | A | 1047 | THR | p.Thr66Ala | O14958 |
| HOMO SAPIENS | 5 | 2772 | 2vaf | A | 1057 | VAL | p.Val76Met | O14958 |
| HOMO SAPIENS | 0 | 2772 | 2vaf | A | 1148 | LEU | p.Leu167His | O14958 |
| HOMO SAPIENS | 1 | 2772 | 2vaf | A | 1149 | ILE | p.Ile168Thr | O14958 |
| HOMO SAPIENS | 8 | 2772 | 2vaf | A | 1225 | HIS | p.His244Arg | O14958 |
| HOMO SAPIENS | 8 | 2772 | 2vaf | A | 1225 | HIS | p.His244Tyr | O14958 |
| HOMO SAPIENS | 7 | 2772 | 2vaf | A | 1251 | ILE | p.Ile270Thr | O14958 |
| HOMO SAPIENS | 3 | 2772 | 2vaf | A | 1288 | ASP | p.Asp307His | O14958 |
| HOMO SAPIENS | 5 | 2772 | 2vaf | A | 1289 | PRO | p.Pro308Leu | O14958 |
| HOMO SAPIENS | 1 | 31885 | 2aex | A | 132 | ALA | p.Ala132Val | P36551 |
| HOMO SAPIENS | 1 | 31885 | 2aex | A | 272 | ASN | p.Asn272His | P36551 |
| HOMO SAPIENS | 8 | 31885 | 2aex | A | 279 | GLY | p.Gly279Arg | P36551 |
| HOMO SAPIENS | 5 | 31885 | 2aex | A | 294 | VAL | p.Val294Ile | P36551 |
| HOMO SAPIENS | 5 | 31885 | 2aex | A | 295 | HIS | p.His295Asp | P36551 |
| HOMO SAPIENS | 1 | 31885 | 2aex | A | 331 | ARG | p.Arg331Trp | P36551 |
| HOMO SAPIENS | 3 | 31885 | 2aex | A | 352 | ARG | p.Arg352Cys | P36551 |
| HOMO SAPIENS | 1 | 31885 | 2aex | A | 404 | LYS | p.Lys404Glu | P36551 |
| HOMO SAPIENS | 2 | 31885 | 2aex | A | 447 | ARG | p.Arg447Cys | P36551 |
| HOMO SAPIENS | 5 | 11721 | 2r3v | A | 20 | HIS | p.His20Pro | Q03426 |

| | | | | | | | | |
|---|---|---|---|---|---|---|---|---|
| HOMO SAPIENS | 1 | 11721 | 2r3v | A | 116 | TYR | p.Tyr116His | Q03426 |
| HOMO SAPIENS | 1 | 11721 | 2r3v | A | 148 | ALA | p.Ala148Thr | Q03426 |
| HOMO SAPIENS | 1 | 11721 | 2r3v | A | 167 | PRO | p.Pro167Leu | Q03426 |
| HOMO SAPIENS | 6 | 11721 | 2r3v | A | 203 | VAL | p.Val203Ala | Q03426 |
| HOMO SAPIENS | 2 | 11721 | 2r3v | A | 215 | ARG | p.Arg215Ter | Q03426 |
| HOMO SAPIENS | 2 | 11721 | 2r3v | A | 237 | THR | p.Thr237Ser | Q03426 |
| HOMO SAPIENS | 0 | 11721 | 2r3v | A | 268 | ILE | p.Ile268Thr | Q03426 |
| HOMO SAPIENS | 0 | 11721 | 2r3v | A | 268 | ILE | p.Ile268Val | Q03426 |
| HOMO SAPIENS | 0 | 11721 | 2r3v | A | 301 | ASN | p.Asn301Ser | Q03426 |
| HOMO SAPIENS | 6 | 11721 | 2r3v | A | 310 | VAL | p.Val310Met | Q03426 |
| HOMO SAPIENS | 5 | 11721 | 2r3v | A | 334 | ALA | p.Ala334Thr | Q03426 |
| HOMO SAPIENS | 2 | 11721 | 2r3v | A | 377 | VAL | p.Val377Ile | Q03426 |
| HOMO SAPIENS | 1 | 11721 | 2r3v | A | 386 | ASP | p.Asp386Asn | Q03426 |
| HOMO SAPIENS | 1 | 11721 | 2r3v | A | 388 | ARG | p.Arg388Ter | Q03426 |
| HOMO SAPIENS | 8 | 5010 | 2hgs | A | 125 | ARG | p.Arg125Cys | P48637 |
| HOMO SAPIENS | 1 | 5010 | 2hgs | A | 164 | ARG | p.Arg164Gln | P48637 |
| HOMO SAPIENS | 0 | 5010 | 2hgs | A | 219 | ASP | p.Asp219Gly | P48637 |
| HOMO SAPIENS | 7 | 5010 | 2hgs | A | 267 | ARG | p.Arg267Trp | P48637 |
| HOMO SAPIENS | 0 | 5010 | 2hgs | A | 283 | ARG | p.Arg283Cys | P48637 |
| HOMO SAPIENS | 0 | 6892 | 5erd | B | 7 | VAL | p.Val56Met | Q14126 |
| HOMO SAPIENS | 1 | 6892 | 5erd | B | 25 | HIS | p.His74Arg | Q14126 |
| HOMO SAPIENS | 0 | 6892 | 5erd | B | 40 | TYR | p.Tyr89Cys | Q14126 |
| HOMO SAPIENS | 0 | 6892 | 5erd | B | 97 | ARG | p.Arg146Leu | Q14126 |
| HOMO SAPIENS | 7 | 6892 | 5erd | B | 217 | ASN | p.Asn266Ser | Q14126 |
| HOMO SAPIENS | 0 | 6892 | 5erd | B | 243 | ARG | p.Arg292His | Q14126 |
| HOMO SAPIENS | 0 | 6892 | 5erd | B | 244 | ILE | p.Ile293Val | Q14126 |
| HOMO SAPIENS | 0 | 6892 | 5erd | B | 343 | VAL | p.Val392Ile | Q14126 |
| HOMO SAPIENS | 4 | 6892 | 5erd | B | 350 | GLU | p.Glu399Lys | Q14126 |
| HOMO SAPIENS | 0 | 6892 | 5erd | B | 410 | TYR | p.Tyr459Cys | Q14126 |
| HOMO SAPIENS | 0 | 6892 | 5erd | B | 466 | VAL | p.Val515Ile | Q14126 |
| HOMO SAPIENS | 0 | 6892 | 5erd | B | 482 | PHE | p.Phe531Cys | Q14126 |
| HOMO SAPIENS | 0 | 6892 | 5erd | B | 484 | VAL | p.Val533Ile | Q14126 |
| HOMO SAPIENS | 0 | 6892 | 5erd | B | 545 | LEU | p.Leu594Pro | Q14126 |
| HOMO SAPIENS | 0 | 1338 | 2ax4 | A | 58 | THR | p.Thr48Arg | O95340 |
| HOMO SAPIENS | 7 | 8675 | 2zkm | X | 385 | LYS | p.Lys385Arg | Q00722 |
| HOMO SAPIENS | 1 | 11769 | 2wya | B | 54 | VAL | p.Val54Met | P54868 |
| HOMO SAPIENS | 1 | 11769 | 2wya | B | 116 | ARG | p.Arg116Cys | P54868 |
| HOMO SAPIENS | 5 | 11769 | 2wya | B | 167 | TYR | p.Tyr167Cys | P54868 |
| HOMO SAPIENS | 1 | 11769 | 2wya | B | 174 | PHE | p.Phe174Leu | P54868 |
| HOMO SAPIENS | 0 | 11769 | 2wya | B | 212 | GLY | p.Gly212Arg | P54868 |
| HOMO SAPIENS | 1 | 11769 | 2wya | B | 288 | ARG | p.Arg288Ter | P54868 |
| HOMO SAPIENS | 0 | 11769 | 2wya | B | 375 | SER | p.Ser375Cys | P54868 |
| HOMO SAPIENS | 4 | 11769 | 2wya | B | 500 | ARG | p.Arg500His | P54868 |
| HOMO SAPIENS | 0 | 3102 | 2iwo | A | 1880 | ARG | p.Arg1908Lys | O75970 |
| HOMO SAPIENS | 6 | 14296 | 2m9w | A | 38 | GLY | p.Gly296Ser | P43694 |
| HOMO SAPIENS | 0 | 14296 | 2m9w | A | 58 | GLN | p.Gln316Glu | P43694 |
| HOMO SAPIENS | 2 | 704 | 5ekd | B | 239 | ARG | p.Arg239Ter | Q9UGM6 |
| HOMO SAPIENS | 2 | 704 | 5ekd | B | 264 | TYR | p.Tyr264Cys | Q9UGM6 |
| HOMO SAPIENS | 6 | 2228 | 1ey2 | A | 47 | SER | p.Ser47Ter | Q93099 |
| HOMO SAPIENS | 6 | 2228 | 1ey2 | A | 48 | ALA | p.Ala48Ser | Q93099 |
| HOMO SAPIENS | 6 | 2228 | 1ey2 | A | 53 | ARG | p.Arg53Gln | Q93099 |
| HOMO SAPIENS | 3 | 2228 | 1ey2 | A | 80 | HIS | p.Gln80His | Q93099 |
| HOMO SAPIENS | 0 | 2228 | 1ey2 | A | 120 | CYS | p.Cys120Trp | Q93099 |
| HOMO SAPIENS | 3 | 2228 | 1ey2 | A | 122 | ALA | p.Ala122Val | Q93099 |

| | | | | | | | | |
|---|---|---|---|---|---|---|---|---|
| HOMO SAPIENS | 3 | 2228 | 1ey2 | A | 161 | GLY | p.Gly161Arg | Q93099 |
| HOMO SAPIENS | 6 | 2228 | 1ey2 | A | 225 | ARG | p.Arg225His | Q93099 |
| HOMO SAPIENS | 0 | 2228 | 1ey2 | A | 230 | PRO | p.Pro230Ser | Q93099 |
| HOMO SAPIENS | 6 | 2228 | 1ey2 | A | 270 | GLY | p.Gly270Arg | Q93099 |
| HOMO SAPIENS | 6 | 2228 | 1ey2 | A | 330 | ARG | p.Arg330Ser | Q93099 |
| HOMO SAPIENS | -1 | 2228 | 1ey2 | A | 368 | MSE | p.Met368Val | Q93099 |
| HOMO SAPIENS | 6 | 2228 | 1ey2 | A | 371 | HIS | p.His371Arg | Q93099 |
| HOMO SAPIENS | 6 | 2228 | 1ey2 | A | 401 | GLU | p.Glu401Gln | Q93099 |
| HOMO SAPIENS | 5 | 34648 | 2k27 | A | 31 | ARG | p.Arg31His | Q06710 |
| HOMO SAPIENS | 7 | 34648 | 2k27 | A | 40 | GLN | p.Gln40Pro | Q06710 |
| HOMO SAPIENS | 5 | 34648 | 2k27 | A | 57 | CYS | p.Cys57Tyr | Q06710 |
| HOMO SAPIENS | 5 | 34648 | 2k27 | A | 62 | LEU | p.Leu62Arg | Q06710 |
| HOMO SAPIENS | 1 | 21399 | 3edu | A | 1819 | GLU | p.Glu1819Ter | P11277 |
| HOMO SAPIENS | 1 | 18153 | 1oey | J | 245 | TYR | p.Tyr245Ter | Q15080 |
| HOMO SAPIENS | 2 | 41567 | 3sw0 | X | 1050 | ASN | p.Asn1050Tyr | P08603 |
| HOMO SAPIENS | 5 | 41567 | 3sw0 | X | 1059 | ILE | p.Ile1059Thr | P08603 |
| HOMO SAPIENS | 0 | 36362 | 4wrm | A | 362 | HIS | p.His362Arg | P07333 |
| HOMO SAPIENS | -1 | 36362 | 4wrm | A | 413 | GLY | p.Gly413Ser | P07333 |
| HOMO SAPIENS | 0 | 31255 | 2w2j | A | 100 | SER | p.Ser100Pro | P35219 |
| HOMO SAPIENS | 0 | 31255 | 2w2j | A | 100 | SER | p.Ser100Leu | P35219 |
| HOMO SAPIENS | 5 | 31255 | 2w2j | A | 237 | ARG | p.Arg237Gln | P35219 |
| HOMO SAPIENS | -1 | 26450 | 1th1 | C | 1395 | SER | p.Ser1395Cys | P25054 |
| HOMO SAPIENS | -1 | 26450 | 1th1 | C | 1413 | MET | p.Met1413Val | P25054 |
| HOMO SAPIENS | -1 | 26450 | 1th1 | C | 1446 | ALA | p.Ala1446Thr | P25054 |
| HOMO SAPIENS | -1 | 26450 | 1th1 | C | 1454 | LYS | p.Lys1454Glu | P25054 |
| HOMO SAPIENS | 0 | 26450 | 1th1 | C | 1474 | ALA | p.Ala1474Thr | P25054 |
| HOMO SAPIENS | 1 | 981 | 5xtb | F | 14 | LEU | p.Leu14Val | O43678 |
| HOMO SAPIENS | 1 | 11069 | 3kq4 | A | 46 | SER | p.Ser46Leu | P27352 |
| HOMO SAPIENS | 1 | 11069 | 3kq4 | A | 97 | MET | p.Met97Thr | P27352 |
| HOMO SAPIENS | 1 | 11069 | 3kq4 | A | 255 | ASN | p.Asn255Ser | P27352 |
| HOMO SAPIENS | 1 | 11069 | 3kq4 | A | 330 | ASN | p.Asn330Lys | P27352 |
| HOMO SAPIENS | 0 | 41489 | 3erb | A | 53 | PRO | p.Pro73Leu | P06681 |
| HOMO SAPIENS | 6 | 8715 | 2bq8 | X | 23 | THR | p.Thr44Met | P13686 |
| HOMO SAPIENS | 0 | 8715 | 2bq8 | X | 88 | GLY | p.Gly109Arg | P13686 |
| HOMO SAPIENS | 6 | 8715 | 2bq8 | X | 127 | VAL | p.Val148Met | P13686 |
| HOMO SAPIENS | 2 | 8715 | 2bq8 | X | 179 | VAL | p.Val200Met | P13686 |
| HOMO SAPIENS | 6 | 8715 | 2bq8 | X | 251 | ARG | p.Arg272Cys | P13686 |
| HOMO SAPIENS | 7 | 8112 | 3p56 | A | 37 | GLY | p.Gly37Ser | O75792 |
| HOMO SAPIENS | -1 | 8112 | 3p56 | A | 202 | LEU | p.Leu202Ser | O75792 |
| HOMO SAPIENS | 1 | 8112 | 3p56 | A | 205 | ASP | p.Asp205Glu | O75792 |
| HOMO SAPIENS | 4 | 8112 | 3p56 | A | 212 | ASN | p.Asn212Ile | O75792 |
| HOMO SAPIENS | 0 | 8112 | 3p56 | A | 235 | ARG | p.Arg235Gln | O75792 |
| HOMO SAPIENS | 4 | 8112 | 3p56 | A | 240 | THR | p.Thr240Met | O75792 |
| HOMO SAPIENS | 0 | 8112 | 3p56 | A | 291 | ARG | p.Arg291His | O75792 |
| HOMO SAPIENS | 0 | 37603 | 2yqg | A | 10 | ARG | p.Arg49His | Q14126 |
| HOMO SAPIENS | 0 | 21178 | 3k1r | B | 458 | ASP | p.Asp458Val | Q495M9 |
| HOMO SAPIENS | -1 | 36828 | 4fq3 | B | 507 | GLY | p.Gly507Asp | P35637 |
| HOMO SAPIENS | 5 | 36828 | 4fq3 | B | 517 | HIS | p.His517Gln | P35637 |
| HOMO SAPIENS | 8 | 36828 | 4fq3 | B | 518 | ARG | p.Arg518Lys | P35637 |
| HOMO SAPIENS | 6 | 36828 | 4fq3 | B | 521 | ARG | p.Arg521Cys | P35637 |
| HOMO SAPIENS | 6 | 36828 | 4fq3 | B | 521 | ARG | p.Arg521His | P35637 |
| HOMO SAPIENS | 0 | 6363 | 3j7y | c | null | HIS | p.His13Arg | Q9H9J2 |
| HOMO SAPIENS | 2 | 6363 | 3j7y | c | 105 | ARG | p.Arg105Cys | Q9H9J2 |
| HOMO SAPIENS | 2 | 6363 | 3j7y | c | 138 | THR | p.Thr138Ile | Q9H9J2 |

| | | | | | | | | |
|---|---|---|---|---|---|---|---|---|
| HOMO SAPIENS | 1 | 6363 | 3j7y | c | 156 | LEU | p.Leu156Arg | Q9H9J2 |
| HOMO SAPIENS | 1 | 6782 | 5l46 | A | 109 | HIS | p.His109Arg | Q9UI17 |
| HOMO SAPIENS | 0 | 23951 | 5t76 | A | 175 | GLY | p.Gly931Ser | P49588 |
| HOMO SAPIENS | 8 | 12604 | 3p0l | A | 182 | ARG | p.Arg182Leu | P49675 |
| HOMO SAPIENS | 2 | 12604 | 3p0l | A | 187 | VAL | p.Val187Met | P49675 |
| HOMO SAPIENS | 2 | 12604 | 3p0l | A | 188 | ARG | p.Arg188Cys | P49675 |
| HOMO SAPIENS | 0 | 12604 | 3p0l | A | 193 | ARG | p.Arg193Ter | P49675 |
| HOMO SAPIENS | 9 | 12604 | 3p0l | A | 217 | ARG | p.Arg217Thr | P49675 |
| HOMO SAPIENS | 4 | 12604 | 3p0l | A | 218 | ALA | p.Ala218Val | P49675 |
| HOMO SAPIENS | 0 | 12604 | 3p0l | A | 250 | TRP | p.Trp250Ter | P49675 |
| HOMO SAPIENS | 1 | 12604 | 3p0l | A | 258 | GLN | p.Gln258Ter | P49675 |
| HOMO SAPIENS | -1 | 27765 | 4wkr | A | 4 | GLU | p.Glu4Lys | Q4G0J3 |
| HOMO SAPIENS | 2 | 3587 | 4kit | C | 2301 | PRO | p.Pro2301Thr | Q6P2Q9 |
| HOMO SAPIENS | 3 | 3587 | 4kit | C | 2304 | PHE | p.Phe2304Leu | Q6P2Q9 |
| HOMO SAPIENS | 3 | 3587 | 4kit | C | 2309 | HIS | p.His2309Pro | Q6P2Q9 |
| HOMO SAPIENS | 3 | 3587 | 4kit | C | 2310 | ARG | p.Arg2310Lys | Q6P2Q9 |
| HOMO SAPIENS | 0 | 9073 | 1n69 | B | 21 | ASN | p.Asn215His | P07602 |
| HOMO SAPIENS | 0 | 9073 | 1n69 | B | 23 | THR | p.Thr217Ile | P07602 |
| HOMO SAPIENS | 0 | 9073 | 1n69 | B | 47 | CYS | p.Cys241Ser | P07602 |
| HOMO SAPIENS | 0 | 475 | 2vd5 | A | 225 | THR | p.Thr225Met | Q09013 |
| HOMO SAPIENS | 1 | 9066 | 4mzv | A | 31 | ASN | p.Asn31Lys | P16422 |
| HOMO SAPIENS | 1 | 9066 | 4mzv | A | 47 | GLN | p.Gln47His | P16422 |
| HOMO SAPIENS | 3 | 9066 | 4mzv | A | 66 | CYS | p.Cys66Tyr | P16422 |
| HOMO SAPIENS | 1 | 9066 | 4mzv | A | 102 | SER | p.Ser102Gly | P16422 |
| HOMO SAPIENS | 5 | 9066 | 4mzv | A | 115 | THR | p.Met115Thr | P16422 |
| HOMO SAPIENS | 1 | 9066 | 4mzv | A | 138 | ARG | p.Arg138Ter | P16422 |
| HOMO SAPIENS | 1 | 9066 | 4mzv | A | 153 | ARG | p.Arg153Thr | P16422 |
| HOMO SAPIENS | 1 | 9066 | 4mzv | A | 165 | ALA | p.Ala165Thr | P16422 |
| HOMO SAPIENS | 8 | 1827 | 2nz2 | A | 14 | GLY | p.Gly14Ser | P00966 |
| HOMO SAPIENS | 5 | 1827 | 2nz2 | A | 18 | SER | p.Ser18Leu | P00966 |
| HOMO SAPIENS | 3 | 1827 | 2nz2 | A | 86 | ARG | p.Arg86Cys | P00966 |
| HOMO SAPIENS | 3 | 1827 | 2nz2 | A | 86 | ARG | p.Arg86His | P00966 |
| HOMO SAPIENS | 1 | 1827 | 2nz2 | A | 179 | TRP | p.Trp179Arg | P00966 |
| HOMO SAPIENS | 7 | 1827 | 2nz2 | A | 180 | SER | p.Ser180Asn | P00966 |
| HOMO SAPIENS | 7 | 1827 | 2nz2 | A | 191 | GLU | p.Glu191Lys | P00966 |
| HOMO SAPIENS | 4 | 1827 | 2nz2 | A | 256 | GLU | p.Glu256Lys | P00966 |
| HOMO SAPIENS | 4 | 1827 | 2nz2 | A | 263 | VAL | p.Val263Met | P00966 |
| HOMO SAPIENS | 5 | 1827 | 2nz2 | A | 265 | ARG | p.Arg265Cys | P00966 |
| HOMO SAPIENS | 5 | 1827 | 2nz2 | A | 268 | ILE | p.Ile268Met | P00966 |
| HOMO SAPIENS | 6 | 1827 | 2nz2 | A | 269 | VAL | p.Val269Met | P00966 |
| HOMO SAPIENS | 7 | 1827 | 2nz2 | A | 272 | ARG | p.Arg272Cys | P00966 |
| HOMO SAPIENS | 3 | 1827 | 2nz2 | A | 304 | ARG | p.Arg304Trp | P00966 |
| HOMO SAPIENS | 3 | 1827 | 2nz2 | A | 307 | ARG | p.Arg307Cys | P00966 |
| HOMO SAPIENS | 3 | 1827 | 2nz2 | A | 310 | LYS | p.Lys310Gln | P00966 |
| HOMO SAPIENS | 2 | 1827 | 2nz2 | A | 362 | GLY | p.Gly362Val | P00966 |
| HOMO SAPIENS | 2 | 1827 | 2nz2 | A | 363 | ARG | p.Arg363Trp | P00966 |
| HOMO SAPIENS | 2 | 1827 | 2nz2 | A | 363 | ARG | p.Arg363Gln | P00966 |
| HOMO SAPIENS | 1 | 1827 | 2nz2 | A | 390 | GLY | p.Gly390Arg | P00966 |
| HOMO SAPIENS | 0 | 29524 | 5vbn | B | 2165 | ARG | p.Arg2165His | Q07864 |
| HOMO SAPIENS | 2 | 29524 | 5vbn | B | 2255 | ILE | p.Ile2255Phe | Q07864 |
| HOMO SAPIENS | 2 | 29524 | 5vbn | B | 2274 | LEU | p.Leu2274Val | Q07864 |
| HOMO SAPIENS | 0 | 31897 | 3t0o | A | 184 | CYS | p.Cys184Arg | O00584 |
| HOMO SAPIENS | -1 | 31897 | 3t0o | A | 215 | GLU | p.Glu215Lys | O00584 |
| HOMO SAPIENS | 0 | 31897 | 3t0o | A | 236 | ARG | p.Arg236Trp | O00584 |

| | | | | | | | | |
|---|---|---|---|---|---|---|---|---|
| HOMO SAPIENS | 7 | 18490 | 4pcw | B | 31 | GLU | p.Glu32Ter | P20930 |
| HOMO SAPIENS | -1 | 39865 | 2h9e | L | 152 | GLY | p.Gly192Arg | P00742 |
| HOMO SAPIENS | 2 | 4309 | 2x0b | F | 174 | THR | p.Thr207Met | P01019 |
| HOMO SAPIENS | 3 | 4309 | 2x0b | F | 342 | ARG | p.Arg375Gln | P01019 |
| HOMO SAPIENS | 2 | 3094 | 1zll | B | 9 | ARG | p.Arg9Cys | P26678 |
| HOMO SAPIENS | 9 | 3094 | 1zll | B | 39 | LEU | p.Leu39Ter | P26678 |
| HOMO SAPIENS | 5 | 3094 | 1zll | B | 49 | VAL | p.Val49Met | P26678 |
| HOMO SAPIENS | -1 | 41498 | 3ffd | P | 8 | LEU | p.Leu44Pro | P12272 |
| HOMO SAPIENS | 1 | 20697 | 1igr | A | 407 | ARG | p.Arg437His | P08069 |
| HOMO SAPIENS | 0 | 20697 | 1igr | A | 416 | MET | p.Met446Val | P08069 |
| HOMO SAPIENS | -1 | 20697 | 1igr | A | 462 | VAL | p.Val492Ile | P08069 |
| HOMO SAPIENS | 0 | 3973 | 4cbz | A | 37 | LEU | p.Leu37Ser | P78504 |
| HOMO SAPIENS | 0 | 3973 | 4cbz | A | 45 | VAL | p.Val45Leu | P78504 |
| HOMO SAPIENS | 0 | 3973 | 4cbz | A | 134 | LEU | p.Leu134Met | P78504 |
| HOMO SAPIENS | -1 | 3973 | 4cbz | A | 146 | VAL | p.Val146Ile | P78504 |
| HOMO SAPIENS | 0 | 3973 | 4cbz | A | 176 | VAL | p.Val176Ile | P78504 |
| HOMO SAPIENS | 0 | 3973 | 4cbz | A | 184 | ARG | p.Arg184Cys | P78504 |
| HOMO SAPIENS | 6 | 1697 | 4hmy | S | 33 | ARG | p.Arg33Trp | Q96PC3 |
| HOMO SAPIENS | 2 | 23752 | 5j4o | A | 13 | ARG | p.Arg1611Ser | P02549 |
| HOMO SAPIENS | 0 | 23752 | 5j4o | A | 42 | ALA | p.Ala1640Gly | P02549 |
| HOMO SAPIENS | 0 | 23752 | 5j4o | A | 108 | HIS | p.His1706Gln | P02549 |
| HOMO SAPIENS | 6 | 23752 | 5j4o | A | 206 | LEU | p.Leu1804Phe | P02549 |
| HOMO SAPIENS | 8 | 531 | 2qz4 | A | 349 | GLY | p.Gly349Ser | Q9UQ90 |
| HOMO SAPIENS | 0 | 531 | 2qz4 | A | 486 | ARG | p.Arg486Gln | Q9UQ90 |
| HOMO SAPIENS | 1 | 531 | 2qz4 | A | 510 | ALA | p.Ala510Val | Q9UQ90 |
| HOMO SAPIENS | 0 | 531 | 2qz4 | A | 529 | ALA | p.Ala529Val | Q9UQ90 |
| HOMO SAPIENS | 0 | 531 | 2qz4 | A | 558 | LYS | p.Lys558Ter | Q9UQ90 |
| HOMO SAPIENS | 1 | 531 | 2qz4 | A | 559 | LYS | p.Lys559Ter | Q9UQ90 |
| HOMO SAPIENS | 2 | 7736 | 1fnh | A | 114 | TRP | p.Trp1834Arg | P02751 |
| HOMO SAPIENS | 0 | 7736 | 1fnh | A | 163 | LEU | p.Leu1883Arg | P02751 |
| HOMO SAPIENS | 3 | 20337 | 5wy5 | B | 264 | LEU | p.Leu264Phe | Q96MG7 |
| HOMO SAPIENS | 8 | 36096 | 2jku | A | 643 | GLY | p.Gly668Arg | P05165 |
| HOMO SAPIENS | 6 | 36096 | 2jku | A | 663 | CYS | p.Cys688Arg | P05165 |
| HOMO SAPIENS | 0 | 9850 | 3kbt | A | 1606 | TYR | p.Tyr1606Ter | P11277 |
| HOMO SAPIENS | 4 | 6876 | 4jkq | A | 217 | ILE | p.Ile217Val | Q92736 |
| HOMO SAPIENS | 5 | 6876 | 4jkq | A | 420 | ARG | p.Arg420Trp | Q92736 |
| HOMO SAPIENS | 2 | 27603 | 1itv | B | 62 | PRO | p.Arg574Pro | P14780 |
| HOMO SAPIENS | 4 | 27603 | 1itv | B | 156 | ARG | p.Arg668Gln | P14780 |
| HOMO SAPIENS | 2 | 22954 | 4f52 | E | 48 | ILE | p.Ile48Met | Q92990 |
| HOMO SAPIENS | 0 | 22954 | 4f52 | E | 91 | ASP | p.Asp91Asn | Q92990 |
| HOMO SAPIENS | 0 | 22954 | 4f52 | E | 146 | ALA | p.Ala146Thr | Q92990 |
| HOMO SAPIENS | 0 | 22954 | 4f52 | E | 181 | ILE | p.Ile181Lys | Q92990 |
| HOMO SAPIENS | 0 | 22954 | 4f52 | E | 194 | ASN | p.Asn194Ser | Q92990 |
| HOMO SAPIENS | 2 | 22954 | 4f52 | E | 254 | PRO | p.Pro254Arg | Q92990 |
| HOMO SAPIENS | 2 | 22954 | 4f52 | E | 300 | ILE | p.Ile300Thr | Q92990 |
| HOMO SAPIENS | 2 | 22954 | 4f52 | E | 336 | LEU | p.Leu336Ser | Q92990 |
| HOMO SAPIENS | 0 | 22954 | 4f52 | E | 353 | TYR | p.Tyr353His | Q92990 |
| HOMO SAPIENS | 0 | 22954 | 4f52 | E | 387 | MET | p.Met387Val | Q92990 |
| HOMO SAPIENS | 4 | 2946 | 5gl7 | A | 344 | TYR | p.Tyr344Cys | P56192 |
| HOMO SAPIENS | 0 | 2946 | 5gl7 | A | 370 | PHE | p.Phe370Leu | P56192 |
| HOMO SAPIENS | 0 | 2946 | 5gl7 | A | 393 | ALA | p.Ala393Thr | P56192 |
| HOMO SAPIENS | 0 | 2946 | 5gl7 | A | 523 | ILE | p.Ile523Thr | P56192 |
| HOMO SAPIENS | 0 | 2946 | 5gl7 | A | 567 | SER | p.Ser567Leu | P56192 |
| HOMO SAPIENS | 0 | 2946 | 5gl7 | A | 800 | PRO | p.Pro800Thr | P56192 |

| | | | | | | | | |
|---|---|---|---|---|---|---|---|---|
| HOMO SAPIENS | 3 | 34961 | 4wy4 | C | 44 | TYR | p.Tyr44His | O95721 |
| HOMO SAPIENS | 8 | 34961 | 4wy4 | C | 89 | GLU | p.Glu89Lys | O95721 |
| HOMO SAPIENS | 0 | 9427 | 4chl | B | 21 | ALA | p.Ala21Ser | O95571 |
| HOMO SAPIENS | 2 | 9427 | 4chl | B | 62 | ALA | p.Ala62Thr | O95571 |
| HOMO SAPIENS | 0 | 9427 | 4chl | B | 114 | ILE | p.Ile114Phe | O95571 |
| HOMO SAPIENS | 2 | 9427 | 4chl | B | 159 | ARG | p.Arg159His | O95571 |
| HOMO SAPIENS | 8 | 9427 | 4chl | B | 163 | ARG | p.Arg163Trp | O95571 |
| HOMO SAPIENS | 8 | 9427 | 4chl | B | 163 | ARG | p.Arg163Gln | O95571 |
| HOMO SAPIENS | 2 | 9427 | 4chl | B | 185 | LEU | p.Leu185Arg | O95571 |
| HOMO SAPIENS | 0 | 40917 | 2e8o | A | 11 | PRO | p.Pro26Leu | Q9Y3Z3 |
| HOMO SAPIENS | -1 | 39190 | 4gow | A | 556 | TYR | p.Tyr556Ter | P56696 |
| HOMO SAPIENS | 0 | 2399 | 2mkv | A | 8 | GLY | p.Gly10Ser | P54710 |
| HOMO SAPIENS | 1 | 2399 | 2mkv | A | 24 | VAL | p.Val26Ile | P54710 |
| HOMO SAPIENS | 4 | 2399 | 2mkv | A | 39 | GLY | p.Gly41Arg | P54710 |
| HOMO SAPIENS | 0 | 20801 | 2zej | B | 1441 | ARG | p.Arg1441Cys | Q5S007 |
| HOMO SAPIENS | 0 | 20801 | 2zej | B | 1441 | ARG | p.Arg1441His | Q5S007 |
| HOMO SAPIENS | 5 | 1019 | 5xtb | M | 62 | ARG | p.Arg62Gln | P28331 |
| HOMO SAPIENS | 1 | 1019 | 5xtb | M | 85 | ALA | p.Ala85Thr | P28331 |
| HOMO SAPIENS | 5 | 1019 | 5xtb | M | 231 | LEU | p.Leu231Val | P28331 |
| HOMO SAPIENS | 2 | 1019 | 5xtb | M | 241 | ARG | p.Arg241Trp | P28331 |
| HOMO SAPIENS | 1 | 1019 | 5xtb | M | 252 | ASP | p.Asp252Gly | P28331 |
| HOMO SAPIENS | 1 | 1019 | 5xtb | M | 408 | ARG | p.Arg408Cys | P28331 |
| HOMO SAPIENS | 2 | 1019 | 5xtb | M | 417 | ARG | p.Arg417Ser | P28331 |
| HOMO SAPIENS | 1 | 1019 | 5xtb | M | 557 | ARG | p.Arg557Ter | P28331 |
| HOMO SAPIENS | 0 | 1019 | 5xtb | M | 567 | ILE | p.Ile567Val | P28331 |
| HOMO SAPIENS | 1 | 1019 | 5xtb | M | 595 | THR | p.Thr595Ala | P28331 |
| HOMO SAPIENS | 1 | 1019 | 5xtb | M | 703 | ALA | p.Ala703Thr | P28331 |
| HOMO SAPIENS | -1 | 3437 | 5hyn | A | 185 | ASP | p.Asp185His | Q15910 |
| HOMO SAPIENS | -1 | 3437 | 5hyn | A | 482 | ALA | p.Ala482Thr | Q15910 |
| HOMO SAPIENS | 0 | 8201 | 2wfd | A | 373 | TYR | p.Tyr373Cys | Q9P2J5 |
| HOMO SAPIENS | 1 | 3590 | 3edi | A | 90 | VAL | p.Val238Ala | O43897 |
| HOMO SAPIENS | 6 | 4555 | 1ytq | A | 64 | ALA | p.Ala65Ser | P43320 |
| HOMO SAPIENS | 0 | 4555 | 1ytq | A | 194 | TRP | p.Trp195Gly | P43320 |
| HOMO SAPIENS | 0 | 1013 | 2nn6 | C | 272 | SER | p.Ser272Thr | Q96B26 |
| HOMO SAPIENS | -1 | 29026 | 5m73 | D | 530 | ILE | p.Ile530Asn | O76094 |
| HOMO SAPIENS | 8 | 2252 | 1d5v | A | 57 | SER | p.Ser125Leu | Q99958 |
| HOMO SAPIENS | 1 | 4337 | 3ecr | A | 29 | ARG | p.Arg26His | P08397 |
| HOMO SAPIENS | 2 | 4337 | 3ecr | A | 34 | ALA | p.Ala31Thr | P08397 |
| HOMO SAPIENS | 2 | 4337 | 3ecr | A | 37 | GLN | p.Gln34Lys | P08397 |
| HOMO SAPIENS | 1 | 4337 | 3ecr | A | 58 | ALA | p.Ala55Ser | P08397 |
| HOMO SAPIENS | 0 | 4337 | 3ecr | A | 84 | LEU | p.Leu81Pro | P08397 |
| HOMO SAPIENS | 0 | 4337 | 3ecr | A | 114 | GLY | p.Gly111Arg | P08397 |
| HOMO SAPIENS | 1 | 4337 | 3ecr | A | 119 | ARG | p.Arg116Trp | P08397 |
| HOMO SAPIENS | 1 | 4337 | 3ecr | A | 152 | ARG | p.Arg149Gln | P08397 |
| HOMO SAPIENS | 1 | 4337 | 3ecr | A | 170 | ARG | p.Arg167Trp | P08397 |
| HOMO SAPIENS | 1 | 4337 | 3ecr | A | 176 | ARG | p.Arg173Gln | P08397 |
| HOMO SAPIENS | 0 | 4337 | 3ecr | A | 180 | LEU | p.Leu177Arg | P08397 |
| HOMO SAPIENS | 0 | 4337 | 3ecr | A | 181 | ASP | p.Asp178Asn | P08397 |
| HOMO SAPIENS | 1 | 4337 | 3ecr | A | 198 | ARG | p.Arg195Cys | P08397 |
| HOMO SAPIENS | 0 | 4337 | 3ecr | A | 201 | TRP | p.Trp198Ter | P08397 |
| HOMO SAPIENS | 1 | 4337 | 3ecr | A | 204 | ARG | p.Arg201Trp | P08397 |
| HOMO SAPIENS | 1 | 4337 | 3ecr | A | 219 | GLY | p.Gly216Asp | P08397 |
| HOMO SAPIENS | 1 | 4337 | 3ecr | A | 226 | GLU | p.Glu223Lys | P08397 |
| HOMO SAPIENS | 0 | 4337 | 3ecr | A | 248 | LEU | p.Leu245Arg | P08397 |

| | | | | | | | | |
|---|---|---|---|---|---|---|---|---|
| HOMO SAPIENS | 1 | 4337 | 3ecr | A | 250 | CYS | p.Cys247Arg | P08397 |
| HOMO SAPIENS | 0 | 4337 | 3ecr | A | 253 | GLU | p.Glu250Lys | P08397 |
| HOMO SAPIENS | 0 | 4337 | 3ecr | A | 255 | ALA | p.Ala252Thr | P08397 |
| HOMO SAPIENS | 0 | 4337 | 3ecr | A | 255 | ALA | p.Ala252Val | P08397 |
| HOMO SAPIENS | 0 | 4337 | 3ecr | A | 259 | HIS | p.His256Asn | P08397 |
| HOMO SAPIENS | 0 | 4337 | 3ecr | A | 324 | ARG | p.Arg321His | P08397 |
| HOMO SAPIENS | -1 | 4337 | 3ecr | A | 362 | ASP | p.Asp359Asn | P08397 |
| HOMO SAPIENS | 1 | 1204 | 5lwv | A | 854 | ILE | p.Ala864Thr | O15294 |
| HOMO SAPIENS | 1 | 25016 | 1m4u | A | 35 | PRO | p.Pro35Arg | Q13253 |
| HOMO SAPIENS | 1 | 25016 | 1m4u | A | 35 | PRO | p.Pro35Ser | Q13253 |
| HOMO SAPIENS | 1 | 25016 | 1m4u | A | 167 | ARG | p.Arg167Gly | Q13253 |
| HOMO SAPIENS | 1 | 25016 | 1m4u | A | 184 | CYS | p.Cys184Tyr | Q13253 |
| HOMO SAPIENS | 0 | 25016 | 1m4u | A | 189 | GLY | p.Gly189Cys | Q13253 |
| HOMO SAPIENS | 1 | 25016 | 1m4u | A | 204 | ARG | p.Arg204Leu | Q13253 |
| HOMO SAPIENS | 1 | 25016 | 1m4u | A | 205 | TRP | p.Trp205Cys | Q13253 |
| HOMO SAPIENS | 1 | 25016 | 1m4u | A | 217 | TRP | p.Trp217Gly | Q13253 |
| HOMO SAPIENS | 0 | 25016 | 1m4u | A | 222 | TYR | p.Tyr222Cys | Q13253 |
| HOMO SAPIENS | 0 | 25016 | 1m4u | A | 222 | TYR | p.Tyr222Asp | Q13253 |
| HOMO SAPIENS | 1 | 25016 | 1m4u | A | 223 | PRO | p.Pro223Leu | Q13253 |
| HOMO SAPIENS | 0 | 1394 | 1um7 | A | 95 | ALA | p.Ala463Val | Q92796 |
| HOMO SAPIENS | 0 | 21294 | 4jpz | B | 1788 | GLU | p.Glu1788Asp | Q99250 |
| HOMO SAPIENS | 9 | 24068 | 2db6 | A | 47 | ARG | p.Arg119Cys | Q96MF2 |
| HOMO SAPIENS | 5 | 1869 | 4omt | A | 39 | ARG | p.Arg39Leu | P08237 |
| HOMO SAPIENS | 2 | 1869 | 4omt | A | 100 | ARG | p.Arg100Gln | P08237 |
| HOMO SAPIENS | 7 | 1869 | 4omt | A | 543 | ASP | p.Asp543Ala | P08237 |
| HOMO SAPIENS | 1 | 1869 | 4omt | A | 686 | TRP | p.Trp686Cys | P08237 |
| HOMO SAPIENS | 1 | 1869 | 4omt | A | 696 | ARG | p.Arg696His | P08237 |
| HOMO SAPIENS | 0 | 12163 | 4wxx | A | 495 | TYR | p.Tyr495Cys | P26358 |
| HOMO SAPIENS | 0 | 12163 | 4wxx | A | 528 | ILE | p.Ile528Met | P26358 |
| HOMO SAPIENS | 0 | 12391 | 1j1d | C | 36 | LYS | p.Lys36Gln | P19429 |
| HOMO SAPIENS | 0 | 12391 | 1j1d | C | 82 | PRO | p.Pro82Ser | P19429 |
| HOMO SAPIENS | 7 | 12391 | 1j1d | C | 102 | ALA | p.Ala102Thr | P19429 |
| HOMO SAPIENS | 1 | 5000 | 4cca | A | 17 | GLY | p.Gly17Arg | Q15833 |
| HOMO SAPIENS | 0 | 5000 | 4cca | A | 205 | VAL | p.Val205Ile | Q15833 |
| HOMO SAPIENS | 0 | 5000 | 4cca | A | 209 | LEU | p.Leu209Pro | Q15833 |
| HOMO SAPIENS | 1 | 5000 | 4cca | A | 459 | ARG | p.Arg459Trp | Q15833 |
| HOMO SAPIENS | 7 | 5000 | 4cca | A | 477 | PRO | p.Pro477Leu | Q15833 |
| HOMO SAPIENS | -1 | 5000 | 4cca | A | 526 | ILE | p.Ile526Val | Q15833 |
| HOMO SAPIENS | 0 | 5000 | 4cca | A | 541 | GLY | p.Gly541Ser | Q15833 |
| HOMO SAPIENS | 0 | 5000 | 4cca | A | 555 | ARG | p.Arg555Gly | Q15833 |
| HOMO SAPIENS | 5 | 28551 | 3vcm | P | 26 | ARG | p.Arg49Ter | P00797 |
| HOMO SAPIENS | 6 | 36453 | 5dzz | A | 2070 | ASP | p.Asp2070His | P15924 |
| HOMO SAPIENS | 0 | 36453 | 5dzz | A | 2166 | ARG | p.Arg2166Ter | P15924 |
| HOMO SAPIENS | 0 | 20268 | 2da6 | A | 96 | ASN | p.Asn321Ser | P35680 |
| HOMO SAPIENS | 1 | 1405 | 3iar | A | 8 | ASP | p.Asp8Asn | P00813 |
| HOMO SAPIENS | 8 | 1405 | 3iar | A | 15 | HIS | p.His15Asp | P00813 |
| HOMO SAPIENS | 6 | 1405 | 3iar | A | 74 | GLY | p.Gly74Val | P00813 |
| HOMO SAPIENS | 6 | 1405 | 3iar | A | 80 | LYS | p.Lys80Arg | P00813 |
| HOMO SAPIENS | 5 | 1405 | 3iar | A | 97 | TYR | p.Tyr97Cys | P00813 |
| HOMO SAPIENS | 5 | 1405 | 3iar | A | 101 | ARG | p.Arg101Leu | P00813 |
| HOMO SAPIENS | 5 | 1405 | 3iar | A | 101 | ARG | p.Arg101Trp | P00813 |
| HOMO SAPIENS | 5 | 1405 | 3iar | A | 106 | LEU | p.Leu106Val | P00813 |
| HOMO SAPIENS | 0 | 1405 | 3iar | A | 107 | LEU | p.Leu107Pro | P00813 |
| HOMO SAPIENS | 9 | 1405 | 3iar | A | 156 | ARG | p.Arg156Cys | P00813 |

| | | | | | | | | |
|---|---|---|---|---|---|---|---|---|
| HOMO SAPIENS | 6 | 1405 | 3iar | A | 211 | ARG | p.Arg211His | P00813 |
| HOMO SAPIENS | 0 | 1405 | 3iar | A | 216 | GLY | p.Gly216Arg | P00813 |
| HOMO SAPIENS | 0 | 1405 | 3iar | A | 291 | SER | p.Ser291Leu | P00813 |
| HOMO SAPIENS | 6 | 1405 | 3iar | A | 304 | LEU | p.Leu304Arg | P00813 |
| HOMO SAPIENS | 0 | 1405 | 3iar | A | 329 | ALA | p.Ala329Val | P00813 |
| HOMO SAPIENS | -1 | 31340 | 4atu | I | 186 | ARG | p.Arg186His | O43602 |
| HOMO SAPIENS | -1 | 31340 | 4atu | I | 192 | ARG | p.Arg192Trp | O43602 |
| HOMO SAPIENS | -1 | 31340 | 4atu | I | 196 | ARG | p.Arg196His | O43602 |
| HOMO SAPIENS | -1 | 31340 | 4atu | I | 203 | THR | p.Thr203Arg | O43602 |
| HOMO SAPIENS | -1 | 31340 | 4atu | I | 286 | ALA | p.Ala286Ser | O43602 |
| HOMO SAPIENS | 4 | 30734 | 1fsu | A | 95 | ARG | p.Arg95Gln | P15848 |
| HOMO SAPIENS | 0 | 30734 | 1fsu | A | 117 | CYS | p.Cys117Arg | P15848 |
| HOMO SAPIENS | 0 | 30734 | 1fsu | A | 137 | GLY | p.Gly137Val | P15848 |
| HOMO SAPIENS | 0 | 30734 | 1fsu | A | 210 | TYR | p.Tyr210Cys | P15848 |
| HOMO SAPIENS | 0 | 30734 | 1fsu | A | 236 | LEU | p.Leu236Pro | P15848 |
| HOMO SAPIENS | 1 | 30734 | 1fsu | A | 315 | ARG | p.Arg315Gln | P15848 |
| HOMO SAPIENS | 0 | 30734 | 1fsu | A | 358 | VAL | p.Val358Met | P15848 |
| HOMO SAPIENS | 2 | 30734 | 1fsu | A | 376 | VAL | p.Val376Met | P15848 |
| HOMO SAPIENS | 0 | 30734 | 1fsu | A | 384 | SER | p.Ser384Asn | P15848 |
| HOMO SAPIENS | 0 | 30734 | 1fsu | A | 393 | HIS | p.His393Pro | P15848 |
| HOMO SAPIENS | 0 | 30734 | 1fsu | A | 405 | CYS | p.Cys405Tyr | P15848 |
| HOMO SAPIENS | 0 | 30734 | 1fsu | A | 458 | ASN | p.Asn458Ser | P15848 |
| HOMO SAPIENS | 1 | 30734 | 1fsu | A | 484 | ARG | p.Arg484Gly | P15848 |
| HOMO SAPIENS | 0 | 38072 | 1wmv | A | 51 | PRO | p.Pro98Leu | Q9NZC7 |
| HOMO SAPIENS | -1 | 5674 | 1j1e | C | 185 | ASN | p.Asn185Lys | P19429 |
| HOMO SAPIENS | -1 | 5674 | 1j1e | C | 192 | ARG | p.Arg192His | P19429 |
| HOMO SAPIENS | -1 | 5674 | 1j1e | C | 196 | ASP | p.Asp196Asn | P19429 |
| HOMO SAPIENS | -1 | 5674 | 1j1e | C | 206 | LYS | p.Lys206Gln | P19429 |
| HOMO SAPIENS | 0 | 38067 | 1wlp | A | 13 | PRO | p.Pro156Gln | P13498 |
| HOMO SAPIENS | -1 | 20441 | 3vfd | A | 289 | HIS | p.His289Tyr | Q9UBP0 |
| HOMO SAPIENS | 2 | 20441 | 3vfd | A | 344 | ILE | p.Ile344Lys | Q9UBP0 |
| HOMO SAPIENS | 0 | 20441 | 3vfd | A | 362 | SER | p.Ser362Cys | Q9UBP0 |
| HOMO SAPIENS | 1 | 20441 | 3vfd | A | 386 | ASN | p.Asn386Ser | Q9UBP0 |
| HOMO SAPIENS | 9 | 20441 | 3vfd | A | 441 | ASP | p.Asp441Gly | Q9UBP0 |
| HOMO SAPIENS | -1 | 20441 | 3vfd | A | 448 | CYS | p.Cys448Tyr | Q9UBP0 |
| HOMO SAPIENS | 2 | 20441 | 3vfd | A | 500 | ARG | p.Arg499Cys | Q9UBP0 |
| HOMO SAPIENS | 1 | 20441 | 3vfd | A | 563 | ARG | p.Arg562Gly | Q9UBP0 |
| HOMO SAPIENS | 3 | 18471 | 4dx8 | J | 26 | ARG | p.Arg26Gln | O00522 |
| HOMO SAPIENS | -1 | 18471 | 4dx8 | J | 101 | GLY | p.Gly101Val | O00522 |
| HOMO SAPIENS | -1 | 34735 | 2ly4 | B | 83 | ALA | p.Ala83Glu | P04637 |
| HOMO SAPIENS | 0 | 31128 | 4cvh | A | 116 | ARG | p.Arg116Cys | A4D126 |
| HOMO SAPIENS | 8 | 31128 | 4cvh | A | 215 | GLN | p.Gln215Ter | A4D126 |
| HOMO SAPIENS | 0 | 31128 | 4cvh | A | 268 | ARG | p.Arg268Ter | A4D126 |
| HOMO SAPIENS | 0 | 31128 | 4cvh | A | 316 | THR | p.Thr316Lys | A4D126 |
| HOMO SAPIENS | 1 | 7164 | 3mt5 | A | 369 | ASP | p.Asp434Gly | Q12791 |
| HOMO SAPIENS | 2 | 26812 | 2wtk | F | 49 | TYR | p.Tyr49Asp | Q15831 |
| HOMO SAPIENS | 0 | 26812 | 2wtk | F | 67 | LEU | p.Leu67Pro | Q15831 |
| HOMO SAPIENS | 4 | 26812 | 2wtk | F | 104 | ARG | p.Arg104Ser | Q15831 |
| HOMO SAPIENS | 0 | 26812 | 2wtk | F | 135 | GLY | p.Gly135Arg | Q15831 |
| HOMO SAPIENS | 0 | 26812 | 2wtk | F | 163 | GLY | p.Gly163Asp | Q15831 |
| HOMO SAPIENS | 2 | 26812 | 2wtk | F | 194 | ALA | p.Asp194Asn | Q15831 |
| HOMO SAPIENS | 2 | 1357 | 5xtd | W | 57 | ARG | p.Arg57His | Q9P0J0 |
| HOMO SAPIENS | 0 | 28086 | 4nn2 | A | 229 | HIS | p.His229Arg | Q8IWS0 |
| HOMO SAPIENS | 0 | 28086 | 4nn2 | A | 234 | LYS | p.Lys234Glu | Q8IWS0 |

| | | | | | | | | |
|---|---|---|---|---|---|---|---|---|
| HOMO SAPIENS | 0 | 28086 | 4nn2 | A | 257 | ARG | p.Arg257Gly | Q8IWS0 |
| HOMO SAPIENS | 5 | 31182 | 2a1x | A | 176 | GLN | p.Gln176Lys | O14832 |
| HOMO SAPIENS | 1 | 31182 | 2a1x | A | 204 | GLY | p.Gly204Ser | O14832 |
| HOMO SAPIENS | 0 | 31182 | 2a1x | A | 269 | ASN | p.Asn269His | O14832 |
| HOMO SAPIENS | 3 | 31182 | 2a1x | A | 275 | ARG | p.Arg275Gln | O14832 |
| HOMO SAPIENS | 3 | 31182 | 2a1x | A | 275 | ARG | p.Arg275Trp | O14832 |
| HOMO SAPIENS | 4 | 2823 | 5xtb | P | 145 | THR | p.Thr145Ile | O75489 |
| HOMO SAPIENS | 5 | 2823 | 5xtb | P | 199 | ARG | p.Arg199Trp | O75489 |
| HOMO SAPIENS | 0 | 8645 | 2h5g | A | 372 | SER | p.Ser372Tyr | P54886 |
| HOMO SAPIENS | 3 | 8645 | 2h5g | A | 715 | ASP | p.Asp715His | P54886 |
| HOMO SAPIENS | 0 | 8645 | 2h5g | A | 765 | ARG | p.Arg765Gln | P54886 |
| HOMO SAPIENS | 0 | 8645 | 2h5g | A | 782 | TYR | p.Tyr782Cys | P54886 |
| HOMO SAPIENS | 0 | 8645 | 2h5g | A | 784 | HIS | p.His784Tyr | P54886 |
| HOMO SAPIENS | 1 | 13992 | 2cue | A | 39 | ARG | p.Arg242Thr | P26367 |
| HOMO SAPIENS | 9 | 13992 | 2cue | A | 55 | PHE | p.Phe258Ser | P26367 |
| HOMO SAPIENS | 0 | 30418 | 2xsw | A | 303 | VAL | p.Val303Met | Q9NRR6 |
| HOMO SAPIENS | 0 | 30418 | 2xsw | A | 315 | PRO | p.Pro315Leu | Q9NRR6 |
| HOMO SAPIENS | 0 | 30418 | 2xsw | A | 341 | GLY | p.Gly341Ser | Q9NRR6 |
| HOMO SAPIENS | 0 | 30418 | 2xsw | A | 378 | ARG | p.Arg378Cys | Q9NRR6 |
| HOMO SAPIENS | 0 | 30418 | 2xsw | A | 435 | ARG | p.Arg435Gln | Q9NRR6 |
| HOMO SAPIENS | 0 | 30418 | 2xsw | A | 515 | ARG | p.Arg515Trp | Q9NRR6 |
| HOMO SAPIENS | 1 | 30418 | 2xsw | A | 526 | PRO | p.Pro526Leu | Q9NRR6 |
| HOMO SAPIENS | 0 | 30418 | 2xsw | A | 563 | ARG | p.Arg563His | Q9NRR6 |
| HOMO SAPIENS | 2 | 30418 | 2xsw | A | 585 | ARG | p.Arg585His | Q9NRR6 |
| HOMO SAPIENS | 2 | 11167 | 4y99 | B | 205 | ARG | p.Arg215Leu | P45379 |
| HOMO SAPIENS | 5 | 11167 | 4y99 | B | 244 | GLU | p.Glu254Asp | P45379 |
| HOMO SAPIENS | 1 | 11167 | 4y99 | B | 253 | LYS | p.Lys263Arg | P45379 |
| HOMO SAPIENS | 8 | 11167 | 4y99 | B | 265 | ARG | p.Arg275Gln | P45379 |
| HOMO SAPIENS | 3 | 11167 | 4y99 | B | 270 | ASP | p.Asp280Asn | P45379 |
| HOMO SAPIENS | -1 | 11167 | 4y99 | B | 278 | ARG | p.Arg288Pro | P45379 |
| HOMO SAPIENS | -1 | 11167 | 4y99 | B | 286 | ARG | p.Arg296Cys | P45379 |
| HOMO SAPIENS | -1 | 11167 | 4y99 | B | 286 | ARG | p.Arg296His | P45379 |
| HOMO SAPIENS | 0 | 15993 | 4po6 | A | 197 | ARG | p.Arg197His | P29597 |
| HOMO SAPIENS | -1 | 15993 | 4po6 | A | 362 | VAL | p.Val362Phe | P29597 |
| HOMO SAPIENS | -1 | 15993 | 4po6 | A | 363 | GLY | p.Gly363Ser | P29597 |
| HOMO SAPIENS | 0 | 5646 | 5uak | A | 7 | GLU | p.Glu7Ter | P13569 |
| HOMO SAPIENS | 0 | 5646 | 5uak | A | 27 | GLY | p.Gly27Ter | P13569 |
| HOMO SAPIENS | 0 | 5646 | 5uak | A | 46 | ALA | p.Ala46Asp | P13569 |
| HOMO SAPIENS | 0 | 5646 | 5uak | A | 60 | GLU | p.Glu60Ter | P13569 |
| HOMO SAPIENS | 0 | 5646 | 5uak | A | 67 | PRO | p.Pro67Leu | P13569 |
| HOMO SAPIENS | 0 | 5646 | 5uak | A | 74 | ARG | p.Arg74Trp | P13569 |
| HOMO SAPIENS | 0 | 5646 | 5uak | A | 75 | ARG | p.Arg75Ter | P13569 |
| HOMO SAPIENS | 0 | 5646 | 5uak | A | 85 | GLY | p.Gly85Glu | P13569 |
| HOMO SAPIENS | 0 | 5646 | 5uak | A | 91 | GLY | p.Gly91Arg | P13569 |
| HOMO SAPIENS | 0 | 5646 | 5uak | A | 92 | GLU | p.Glu92Lys | P13569 |
| HOMO SAPIENS | 0 | 5646 | 5uak | A | 98 | GLN | p.Gln98Arg | P13569 |
| HOMO SAPIENS | 0 | 5646 | 5uak | A | 98 | GLN | p.Gln98Ter | P13569 |
| HOMO SAPIENS | 0 | 5646 | 5uak | A | 109 | TYR | p.Tyr109Cys | P13569 |
| HOMO SAPIENS | 0 | 5646 | 5uak | A | 110 | ASP | p.Asp110His | P13569 |
| HOMO SAPIENS | 0 | 5646 | 5uak | A | 117 | ARG | p.Arg117Cys | P13569 |
| HOMO SAPIENS | 0 | 5646 | 5uak | A | 117 | ARG | p.Arg117His | P13569 |
| HOMO SAPIENS | 0 | 5646 | 5uak | A | 125 | ILE | p.Ile125Thr | P13569 |
| HOMO SAPIENS | 0 | 5646 | 5uak | A | 148 | ILE | p.Ile148Thr | P13569 |
| HOMO SAPIENS | 0 | 5646 | 5uak | A | 170 | ARG | p.Arg170His | P13569 |

| | | | | | | | | |
|---|---|---|---|---|---|---|---|---|
| HOMO SAPIENS | 0 | 5646 | 5uak | A | 178 | GLY | p.Gly178Arg | P13569 |
| HOMO SAPIENS | 0 | 5646 | 5uak | A | 206 | LEU | p.Leu206Trp | P13569 |
| HOMO SAPIENS | 0 | 5646 | 5uak | A | 373 | ASP | p.Asp373Asn | P13569 |
| HOMO SAPIENS | -1 | 5646 | 5uak | A | 709 | ARG | p.Arg709Ter | P13569 |
| HOMO SAPIENS | -1 | 5646 | 5uak | A | 732 | LEU | p.Leu732Ter | P13569 |
| HOMO SAPIENS | -1 | 5646 | 5uak | A | 785 | ARG | p.Arg785Ter | P13569 |
| HOMO SAPIENS | -1 | 5646 | 5uak | A | 831 | GLU | p.Glu831Ter | P13569 |
| HOMO SAPIENS | -1 | 5646 | 5uak | A | 836 | ASP | p.Asp836Tyr | P13569 |
| HOMO SAPIENS | 0 | 5646 | 5uak | A | 851 | ARG | p.Arg851Ter | P13569 |
| HOMO SAPIENS | 0 | 5646 | 5uak | A | 913 | TYR | p.Tyr913Cys | P13569 |
| HOMO SAPIENS | 0 | 5646 | 5uak | A | 913 | TYR | p.Tyr913Ter | P13569 |
| HOMO SAPIENS | 0 | 5646 | 5uak | A | 945 | SER | p.Ser945Leu | P13569 |
| HOMO SAPIENS | 0 | 5646 | 5uak | A | 949 | HIS | p.His949Tyr | P13569 |
| HOMO SAPIENS | 0 | 5646 | 5uak | A | 970 | GLY | p.Gly970Asp | P13569 |
| HOMO SAPIENS | 0 | 5646 | 5uak | A | 1027 | ILE | p.Ile1027Thr | P13569 |
| HOMO SAPIENS | 0 | 5646 | 5uak | A | 1032 | TYR | p.Tyr1032Cys | P13569 |
| HOMO SAPIENS | 0 | 5646 | 5uak | A | 1061 | GLY | p.Gly1061Arg | P13569 |
| HOMO SAPIENS | 1 | 5646 | 5uak | A | 1065 | LEU | p.Leu1065Pro | P13569 |
| HOMO SAPIENS | 1 | 5646 | 5uak | A | 1066 | ARG | p.Arg1066Cys | P13569 |
| HOMO SAPIENS | 1 | 5646 | 5uak | A | 1066 | ARG | p.Arg1066His | P13569 |
| HOMO SAPIENS | 1 | 5646 | 5uak | A | 1067 | ALA | p.Ala1067Thr | P13569 |
| HOMO SAPIENS | 0 | 5646 | 5uak | A | 1069 | GLY | p.Gly1069Arg | P13569 |
| HOMO SAPIENS | 0 | 5646 | 5uak | A | 1070 | ARG | p.Arg1070Trp | P13569 |
| HOMO SAPIENS | 1 | 5646 | 5uak | A | 1071 | GLN | p.Gln1071Pro | P13569 |
| HOMO SAPIENS | 0 | 5646 | 5uak | A | 1074 | PHE | p.Phe1074Leu | P13569 |
| HOMO SAPIENS | 0 | 5646 | 5uak | A | 1077 | LEU | p.Leu1077Pro | P13569 |
| HOMO SAPIENS | 0 | 5646 | 5uak | A | 1089 | TRP | p.Trp1089Ter | P13569 |
| HOMO SAPIENS | 0 | 5646 | 5uak | A | 1092 | TYR | p.Tyr1092Ter | P13569 |
| HOMO SAPIENS | 0 | 5646 | 5uak | A | 1101 | MET | p.Met1101Lys | P13569 |
| HOMO SAPIENS | 0 | 5646 | 5uak | A | 1156 | LEU | p.Leu1156Phe | P13569 |
| HOMO SAPIENS | 0 | 5646 | 5uak | A | 1158 | ARG | p.Arg1158Ter | P13569 |
| HOMO SAPIENS | 0 | 5646 | 5uak | A | 1162 | ARG | p.Arg1162Leu | P13569 |
| HOMO SAPIENS | 0 | 5646 | 5uak | A | 1162 | ARG | p.Arg1162Ter | P13569 |
| HOMO SAPIENS | -1 | 5646 | 5uak | A | 1476 | GLN | p.Gln1476Ter | P13569 |
| HOMO SAPIENS | 0 | 22763 | 1imv | A | 52 | MET | p.Thr72Met | P36955 |
| HOMO SAPIENS | 0 | 22763 | 1imv | A | 301 | TYR | p.Tyr321Ter | P36955 |
| HOMO SAPIENS | 0 | 23179 | 5fql | A | 160 | PRO | p.Pro160Arg | P22304 |
| HOMO SAPIENS | 0 | 23179 | 5fql | A | 172 | ARG | p.Arg172Ter | P22304 |
| HOMO SAPIENS | 0 | 23179 | 5fql | A | 214 | THR | p.Thr214Met | P22304 |
| HOMO SAPIENS | 0 | 23179 | 5fql | A | 309 | THR | p.Thr309Ala | P22304 |
| HOMO SAPIENS | 0 | 23179 | 5fql | A | 333 | SER | p.Ser333Leu | P22304 |
| HOMO SAPIENS | 0 | 23179 | 5fql | A | 443 | ARG | p.Arg443Ter | P22304 |
| HOMO SAPIENS | 0 | 23179 | 5fql | A | 468 | ARG | p.Arg468Leu | P22304 |
| HOMO SAPIENS | 0 | 23179 | 5fql | A | 468 | ARG | p.Arg468Trp | P22304 |
| HOMO SAPIENS | 0 | 23179 | 5fql | A | 502 | TRP | p.Trp502Ser | P22304 |
| HOMO SAPIENS | 9 | 20989 | 1nfi | F | 185 | THR | p.Thr185Met | P25963 |
| HOMO SAPIENS | 4 | 20989 | 1nfi | F | 194 | GLY | p.Gly194Ala | P25963 |
| HOMO SAPIENS | 0 | 11538 | 3eab | A | 162 | VAL | p.Val162Ile | Q9UBP0 |
| HOMO SAPIENS | 0 | 33399 | 2p39 | A | 71 | SER | p.Ser71Gly | Q9GZV9 |
| HOMO SAPIENS | 3 | 33399 | 2p39 | A | 96 | MET | p.Met96Thr | Q9GZV9 |
| HOMO SAPIENS | 0 | 33399 | 2p39 | A | 129 | SER | p.Ser129Phe | Q9GZV9 |
| HOMO SAPIENS | -1 | 33399 | 2p39 | A | 172 | PRO | p.Pro172Leu | Q9GZV9 |
| HOMO SAPIENS | -1 | 33399 | 2p39 | A | 176 | ARG | p.Arg176Gln | Q9GZV9 |
| HOMO SAPIENS | -1 | 33399 | 2p39 | A | 179 | ARG | p.Arg179Trp | Q9GZV9 |

| | | | | | | | | |
|---|---|---|---|---|---|---|---|---|
| HOMO SAPIENS | -1 | 33399 | 2p39 | A | 179 | ARG | p.Arg179Gln | Q9GZV9 |
| HOMO SAPIENS | 0 | 16217 | 2dne | A | 79 | ILE | p.Ile163Leu | P10515 |
| HOMO SAPIENS | 8 | 7666 | 4kbq | A | 65 | ASN | p.Asn65Ser | Q9UNE7 |
| HOMO SAPIENS | 1 | 6407 | 1ikt | A | 71 | THR | p.Thr687Ile | P51659 |
| HOMO SAPIENS | 0 | 6407 | 1ikt | A | 112 | MET | p.Met728Val | P51659 |
| HOMO SAPIENS | 0 | 5849 | 4ihb | A | 67 | VAL | p.Val67Asp | O75923 |
| HOMO SAPIENS | 6 | 17703 | 5fcn | A | 20 | HIS | p.His99Gln | Q66GS9 |
| HOMO SAPIENS | 8 | 17703 | 5fcn | A | 33 | ARG | p.Arg112His | Q66GS9 |
| HOMO SAPIENS | 0 | 34095 | 2E+073 | A | 101 | HIS | p.His101Tyr | P05129 |
| HOMO SAPIENS | -1 | 41358 | 4liq | E | 507 | THR | p.Thr507Met | P07333 |
| HOMO SAPIENS | -1 | 21290 | 1fch | A | 235 | MET | p.Met272Arg | P50542 |
| HOMO SAPIENS | 5 | 23431 | 6pax | A | 23 | ARG | p.Arg26Gly | P26367 |
| HOMO SAPIENS | 1 | 23431 | 6pax | A | 61 | GLY | p.Gly64Val | P26367 |
| HOMO SAPIENS | 1 | 23431 | 6pax | A | 65 | PRO | p.Pro68Ser | P26367 |
| HOMO SAPIENS | 1 | 23431 | 6pax | A | 116 | SER | p.Ser119Arg | P26367 |
| HOMO SAPIENS | 9 | 23431 | 6pax | A | 123 | VAL | p.Val126Asp | P26367 |
| HOMO SAPIENS | 9 | 23431 | 6pax | A | 125 | ARG | p.Arg128Cys | P26367 |
| HOMO SAPIENS | 9 | 33003 | 4zrj | B | 535 | LEU | p.Leu535Pro | P35240 |
| HOMO SAPIENS | 7 | 33003 | 4zrj | B | 538 | GLN | p.Gln538Pro | P35240 |
| HOMO SAPIENS | 0 | 33003 | 4zrj | B | 585 | TRP | p.Ala585Thr | P35240 |
| HOMO SAPIENS | 0 | 3800 | 2nzj | A | 193 | VAL | p.Val193Met | O75628 |

| gene_name | max_cons | max_bs_id | max_pdb_id | max_chain_id |
|---|---|---|---|---|
| AKR1D1 | None | None | None | None |
| AKR1D1 | None | None | None | None |
| AKR1D1 | None | None | None | None |
| AKR1D1 | None | None | None | None |
| ALPP | None | None | None | None |
| BMPR1A | None | None | None | None |
| BMPR1A | None | None | None | None |
| FGFR2 | None | None | None | None |
| PROC | None | None | None | None |
| PROC | None | None | None | None |
| PROC | None | None | None | None |
| PROC | None | None | None | None |
| PROC | None | None | None | None |
| PROC | None | None | None | None |
| PROC | None | None | None | None |
| PROC | None | None | None | None |
| PROC | None | None | None | None |
| PROC | None | None | None | None |
| PROC | None | None | None | None |
| PROC | None | None | None | None |
| INS | 1 | 2 | 3zs2 | J |
| INS | None | None | None | None |
| INS | 1 | 1 | 3zs2 | J |
| INS | 1 | 1 | 3zs2 | J |
| FTL | None | None | None | None |
| FTL | None | None | None | None |
| FTL | None | None | None | None |
| TNFSF11 | None | None | None | None |
| EHMT1 | None | None | None | None |
| EHMT1 | None | None | None | None |
| EHMT1 | None | None | None | None |
| CA2 | None | None | None | None |
| CA2 | None | None | None | None |
| CA2 | None | None | None | None |
| CA2 | None | None | None | None |
| SMAD4 | None | None | None | None |
| SMAD4 | None | None | None | None |
| TTR | None | None | None | None |
| TTR | None | None | None | None |
| TTR | None | None | None | None |
| TTR | None | None | None | None |
| TTR | None | None | None | None |
| TTR | None | None | None | None |
| TTR | None | None | None | None |
| TTR | None | None | None | None |
| TTR | None | None | None | None |
| TTR | None | None | None | None |
| TTR | None | None | None | None |
| TTR | None | None | None | None |
| TTR | None | None | None | None |
| TTR | None | None | None | None |
| TTR | None | None | None | None |
| TTR | None | None | None | None |

| | | | | |
|---|---|---|---|---|
| TTR | None | None | None | None |
| TTR | None | None | None | None |
| TTR | None | None | None | None |
| TTR | None | None | None | None |
| TTR | None | None | None | None |
| TTR | None | None | None | None |
| TTR | None | None | None | None |
| TTR | None | None | None | None |
| TTR | None | None | None | None |
| TTR | None | None | None | None |
| TTR | None | None | None | None |
| TTR | None | None | None | None |
| TTR | None | None | None | None |
| TTR | None | None | None | None |
| TTR | None | None | None | None |
| TTR | None | None | None | None |
| TTR | None | None | None | None |
| TTR | None | None | None | None |
| TTR | None | None | None | None |
| ALK | None | None | None | None |
| ALK | None | None | None | None |
| THRB | None | None | None | None |
| THRB | None | None | None | None |
| THRB | None | None | None | None |
| THRB | None | None | None | None |
| THRB | None | None | None | None |
| THRB | None | None | None | None |
| THRB | None | None | None | None |
| THRB | None | None | None | None |
| THRB | None | None | None | None |
| THRB | None | None | None | None |
| THRB | None | None | None | None |
| THRB | None | None | None | None |
| THRB | None | None | None | None |
| THRB | None | None | None | None |
| THRB | None | None | None | None |
| THRB | None | None | None | None |
| THRB | None | None | None | None |
| THRB | None | None | None | None |
| THRB | None | None | None | None |
| IL2RG | None | None | None | None |
| IL2RG | None | None | None | None |
| IL2RG | None | None | None | None |
| IL2RG | None | None | None | None |
| IL2RG | None | None | None | None |
| ITGB3 | None | None | None | None |
| ITGB3 | None | None | None | None |
| ITGB3 | None | None | None | None |
| ITGB3 | None | None | None | None |
| ITGB3 | None | None | None | None |
| ITGB3 | None | None | None | None |
| ITGB3 | None | None | None | None |
| ITGB3 | None | None | None | None |
| ITGB3 | None | None | None | None |

| | | | | |
|---|---|---|---|---|
| PKD2 | None | None | None | None |
| PKD2 | None | None | None | None |
| PKD2 | None | None | None | None |
| PKD2 | None | None | None | None |
| PKD2 | None | None | None | None |
| PKD2 | None | None | None | None |
| PKD2 | None | None | None | None |
| TP53 | None | None | None | None |
| TP53 | None | None | None | None |
| TP53 | None | None | None | None |
| TP53 | None | None | None | None |
| TP53 | None | None | None | None |
| TP53 | None | None | None | None |
| EGFR | None | None | None | None |
| EGFR | None | None | None | None |
| XRCC4 | None | None | None | None |
| XRCC4 | None | None | None | None |
| HBB | None | None | None | None |
| HBB | None | None | None | None |
| HBB | None | None | None | None |
| HBB | None | None | None | None |
| HBB | None | None | None | None |
| HBB | None | None | None | None |
| HBB | None | None | None | None |
| HBB | None | None | None | None |
| HBB | None | None | None | None |
| HBB | None | None | None | None |
| HBB | None | None | None | None |
| HBB | None | None | None | None |
| HBB | None | None | None | None |
| HBB | None | None | None | None |
| HBB | None | None | None | None |
| HBB | None | None | None | None |
| HBB | None | None | None | None |
| HBB | None | None | None | None |
| HBB | None | None | None | None |
| HBB | None | None | None | None |
| HBB | None | None | None | None |
| HBB | None | None | None | None |
| HBB | None | None | None | None |
| HBB | None | None | None | None |
| HBB | None | None | None | None |
| HBB | None | None | None | None |
| HBB | None | None | None | None |
| HBB | None | None | None | None |
| HBB | None | None | None | None |
| HBB | None | None | None | None |
| RPS6KA3 | None | None | None | None |
| RPS6KA3 | None | None | None | None |
| RPS6KA3 | None | None | None | None |
| RPS6KA3 | None | None | None | None |
| RPS6KA3 | None | None | None | None |
| RPS6KA3 | None | None | None | None |
| GBA | 1 | 15 | 3gxd | B |

| | | | | | |
|---|---|---|---|---|---|
| GBA | | 1 | 55 | 3gxd | B |
| GBA | | 1 | 15 | 3gxd | B |
| GBA | None | None | | None | None |
| GBA | None | None | | None | None |
| GBA | None | None | | None | None |
| GBA | | 1 | 12 | 1ogs | A |
| GBA | None | None | | None | None |
| GBA | | 1 | 2 | 1ogs | A |
| GBA | | 1 | 11 | 1ogs | A |
| GBA | None | None | | None | None |
| GBA | None | None | | None | None |
| GBA | | 1 | 18 | 1ogs | A |
| GBA | | 1 | 3 | 1ogs | A |
| GBA | None | None | | None | None |
| GBA | | 1 | 2 | 1ogs | A |
| GBA | None | None | | None | None |
| GBA | None | None | | None | None |
| GBA | | 1 | 3 | 1ogs | A |
| GBA | | 1 | 3 | 1ogs | A |
| GBA | None | None | | None | None |
| GBA | | 1 | 28 | 1ogs | A |
| GBA | None | None | | None | None |
| GBA | None | None | | None | None |
| GBA | None | None | | None | None |
| GBA | None | None | | None | None |
| GBA | None | None | | None | None |
| GBA | None | None | | None | None |
| GBA | None | None | | None | None |
| GBA | None | None | | None | None |
| GBA | None | None | | None | None |
| GBA | None | None | | None | None |
| GBA | None | None | | None | None |
| GBA | None | None | | None | None |
| GBA | None | None | | None | None |
| GBA | None | None | | None | None |
| GBA | None | None | | None | None |
| GBA | None | None | | None | None |
| GBA | None | None | | None | None |
| GBA | | 1 | 15 | 3gxd | B |
| GBA | | 1 | 44 | 3gxd | B |
| GBA | None | None | | None | None |
| GBA | | 1 | 44 | 3gxd | B |
| GBA | | 1 | 44 | 3gxd | B |
| GBA | None | None | | None | None |
| GBA | None | None | | None | None |
| GBA | None | None | | None | None |
| IFIH1 | None | None | | None | None |
| TGFBR2 | None | None | | None | None |
| TGFBR2 | None | None | | None | None |
| TGFBR2 | None | None | | None | None |
| TGFBR2 | None | None | | None | None |
| TGFBR2 | None | None | | None | None |
| TGFBR2 | None | None | | None | None |
| TGFBR2 | None | None | | None | None |

| | | | | |
|---|---|---|---|---|
| TGFBR2 | None | None | None | None |
| TGFBR2 | None | None | None | None |
| TGFBR2 | None | None | None | None |
| TGFBR2 | None | None | None | None |
| TGFBR2 | None | None | None | None |
| TGFBR2 | None | None | None | None |
| TGFBR2 | None | None | None | None |
| TGFBR2 | None | None | None | None |
| ACE | None | None | None | None |
| ACE | None | None | None | None |
| ACE | None | None | None | None |
| ACE | None | None | None | None |
| ACE | None | None | None | None |
| ACE | None | None | None | None |
| COL10A1 | None | None | None | None |
| COL10A1 | None | None | None | None |
| COL10A1 | None | None | None | None |
| COL10A1 | None | None | None | None |
| COL10A1 | None | None | None | None |
| COL10A1 | None | None | None | None |
| COL10A1 | None | None | None | None |
| COL10A1 | None | None | None | None |
| COL10A1 | None | None | None | None |
| KLF1 | None | None | None | None |
| C3 | None | None | None | None |
| CTSC | None | None | None | None |
| CTSC | None | None | None | None |
| CTSC | None | None | None | None |
| LYZ | None | None | None | None |
| LYZ | None | None | None | None |
| LYZ | None | None | None | None |
| DHODH | None | None | None | None |
| DHODH | None | None | None | None |
| DHODH | None | None | None | None |
| DHODH | None | None | None | None |
| DHODH | None | None | None | None |
| DHODH | None | None | None | None |
| DHODH | None | None | None | None |
| ADH1C | None | None | None | None |
| GDNF | None | None | None | None |
| TNF | None | None | None | None |
| ALDH4A1 | None | None | None | None |
| ALDH4A1 | None | None | None | None |
| ALDH4A1 | None | None | None | None |
| BRAF | None | None | None | None |
| BRAF | None | None | None | None |
| BRAF | None | None | None | None |
| BRAF | None | None | None | None |
| BRAF | None | None | None | None |
| BRAF | None | None | None | None |
| BRAF | None | None | None | None |
| BRAF | None | None | None | None |
| BRAF | None | None | None | None |
| BRAF | None | None | None | None |

| | | | | |
|---|---|---|---|---|
| BRAF | None | None | None | None |
| BRAF | None | None | None | None |
| BRAF | None | None | None | None |
| BRAF | None | None | None | None |
| BRAF | None | None | None | None |
| BRAF | None | None | None | None |
| TBX5 | None | None | None | None |
| TBX5 | None | None | None | None |
| TBX5 | None | None | None | None |
| TBX5 | None | None | None | None |
| TBX5 | None | None | None | None |
| TBX5 | None | None | None | None |
| TBX5 | None | None | None | None |
| TBX5 | None | None | None | None |
| TBX5 | None | None | None | None |
| RB1 | None | None | None | None |
| RB1 | None | None | None | None |
| RB1 | None | None | None | None |
| RB1 | None | None | None | None |
| RB1 | None | None | None | None |
| RB1 | None | None | None | None |
| CLIC2 | None | None | None | None |
| CLIC2 | None | None | None | None |
| ALDH7A1 | None | None | None | None |
| ALDH7A1 | None | None | None | None |
| ALDH7A1 | None | None | None | None |
| ALDH7A1 | None | None | None | None |
| ALDH7A1 | None | None | None | None |
| ALDH7A1 | None | None | None | None |
| ALDH7A1 | None | None | None | None |
| ALDH7A1 | None | None | None | None |
| ALDH7A1 | None | None | None | None |
| ALDH7A1 | None | None | None | None |
| ALDH7A1 | None | None | None | None |
| ALDH7A1 | None | None | None | None |
| ALDH7A1 | None | None | None | None |
| ALDH7A1 | None | None | None | None |
| ALDH7A1 | None | None | None | None |
| ALDH7A1 | None | None | None | None |
| KDR | None | None | None | None |
| MUT | None | None | None | None |
| MUT | None | None | None | None |
| MUT | None | None | None | None |
| MUT | None | None | None | None |
| MUT | None | None | None | None |
| MUT | None | None | None | None |
| MUT | None | None | None | None |
| MUT | None | None | None | None |
| MUT | None | None | None | None |
| MUT | None | None | None | None |
| MUT | None | None | None | None |
| MUT | None | None | None | None |
| MUT | None | None | None | None |
| MUT | None | None | None | None |

| | | | | |
|---|---|---|---|---|
| MUT | None | None | None | None |
| MUT | None | None | None | None |
| MUT | None | None | None | None |
| MUT | None | None | None | None |
| MUT | None | None | None | None |
| MUT | None | None | None | None |
| MUT | None | None | None | None |
| MUT | None | None | None | None |
| MUT | None | None | None | None |
| MUT | None | None | None | None |
| MUT | None | None | None | None |
| MUT | None | None | None | None |
| MUT | None | None | None | None |
| MUT | None | None | None | None |
| MUT | None | None | None | None |
| MUT | None | None | None | None |
| MUT | None | None | None | None |
| MUT | None | None | None | None |
| MUT | None | None | None | None |
| MUT | None | None | None | None |
| SOD1 | None | None | None | None |
| SOD1 | None | None | None | None |
| SOD1 | None | None | None | None |
| SOD1 | None | None | None | None |
| NECTIN3 | None | None | None | None |
| HFE | None | None | None | None |
| HFE | None | None | None | None |
| HFE | None | None | None | None |
| HFE | None | None | None | None |
| HFE | None | None | None | None |
| HFE | None | None | None | None |
| HFE | None | None | None | None |
| EIF2AK3 | None | None | None | None |
| EIF2AK3 | None | None | None | None |
| TERT | None | None | None | None |
| TERT | None | None | None | None |
| SPR | None | None | None | None |
| SPR | None | None | None | None |
| SPR | None | None | None | None |
| SPR | None | None | None | None |
| SPR | None | None | None | None |
| F7 | None | None | None | None |
| UNG | None | None | None | None |
| MMP13 | None | None | None | None |
| MMP13 | None | None | None | None |
| MMP13 | None | None | None | None |
| MMP13 | None | None | None | None |
| CD207 | None | None | None | None |
| F2 | 1 | 2 | 1nrq | L |
| SPTBN2 | None | None | None | None |
| PKLR | None | None | None | None |
| PKLR | None | None | None | None |
| PKLR | None | None | None | None |
| PKLR | None | None | None | None |

| | | | | |
|---|---|---|---|---|
| PKLR | None | None | None | None |
| PKLR | None | None | None | None |
| PKLR | None | None | None | None |
| PKLR | None | None | None | None |
| SACS | None | None | None | None |
| HRAS | None | None | None | None |
| HRAS | None | None | None | None |
| HRAS | None | None | None | None |
| HRAS | None | None | None | None |
| HRAS | None | None | None | None |
| HRAS | None | None | None | None |
| HRAS | None | None | None | None |
| HRAS | None | None | None | None |
| HRAS | None | None | None | None |
| HRAS | None | None | None | None |
| HRAS | None | None | None | None |
| HRAS | None | None | None | None |
| HRAS | None | None | None | None |
| VHL | None | None | None | None |
| VHL | None | None | None | None |
| VHL | None | None | None | None |
| VHL | None | None | None | None |
| VHL | None | None | None | None |
| VHL | None | None | None | None |
| VHL | None | None | None | None |
| VHL | None | None | None | None |
| VHL | None | None | None | None |
| VHL | None | None | None | None |
| VHL | None | None | None | None |
| VHL | None | None | None | None |
| VHL | None | None | None | None |
| VHL | None | None | None | None |
| VHL | None | None | None | None |
| VHL | None | None | None | None |
| VHL | None | None | None | None |
| VHL | None | None | None | None |
| VHL | None | None | None | None |
| VHL | None | None | None | None |
| VHL | None | None | None | None |
| VHL | None | None | None | None |
| VHL | None | None | None | None |
| VHL | None | None | None | None |
| VHL | None | None | None | None |
| TTBK1 | None | None | None | None |
| RAD51 | None | None | None | None |
| ALB | None | None | None | None |
| ALB | None | None | None | None |
| ALB | None | None | None | None |
| ALB | None | None | None | None |
| MET | None | None | None | None |
| MET | None | None | None | None |
| MET | None | None | None | None |
| MET | None | None | None | None |
| MET | None | None | None | None |

| | | | | |
|---|---|---|---|---|
| MET | None | None | None | None |
| MET | None | None | None | None |
| MET | None | None | None | None |
| MET | None | None | None | None |
| CP | None | None | None | None |
| CP | None | None | None | None |
| PGM1 | None | None | None | None |
| PGM1 | None | None | None | None |
| PGM1 | None | None | None | None |
| PGM1 | None | None | None | None |
| PGM1 | None | None | None | None |
| PGM1 | None | None | None | None |
| PGM1 | None | None | None | None |
| NR5A1 | None | None | None | None |
| NR5A1 | None | None | None | None |
| NR5A1 | None | None | None | None |
| F9 | None | None | None | None |
| F9 | None | None | None | None |
| HPRT1 | None | None | None | None |
| HPRT1 | None | None | None | None |
| HPRT1 | None | None | None | None |
| HPRT1 | None | None | None | None |
| HPRT1 | None | None | None | None |
| HPRT1 | None | None | None | None |
| HPRT1 | None | None | None | None |
| HPRT1 | None | None | None | None |
| HPRT1 | None | None | None | None |
| HPRT1 | None | None | None | None |
| HPRT1 | None | None | None | None |
| HPRT1 | None | None | None | None |
| HPRT1 | None | None | None | None |
| HPRT1 | None | None | None | None |
| HPRT1 | None | None | None | None |
| HPRT1 | None | None | None | None |
| HPRT1 | None | None | None | None |
| HPRT1 | None | None | None | None |
| HPRT1 | None | None | None | None |
| HPRT1 | None | None | None | None |
| HPRT1 | None | None | None | None |
| HPRT1 | None | None | None | None |
| GPD1L | None | None | None | None |
| GPD1L | None | None | None | None |
| GPD1L | None | None | None | None |
| FGFR2 | None | None | None | None |
| FGFR2 | None | None | None | None |
| FGFR2 | None | None | None | None |
| FGFR2 | None | None | None | None |
| FGFR2 | None | None | None | None |
| FGFR2 | None | None | None | None |
| FGFR2 | None | None | None | None |
| FGFR2 | None | None | None | None |
| FGFR2 | None | None | None | None |
| FGFR2 | None | None | None | None |
| FGFR2 | None | None | None | None |

| | | | | |
|---|---|---|---|---|
| FGFR2 | None | None | None | None |
| FGFR2 | None | None | None | None |
| FGFR2 | None | None | None | None |
| FGFR2 | None | None | None | None |
| FGFR2 | None | None | None | None |
| FGFR2 | None | None | None | None |
| FGFR2 | None | None | None | None |
| FGFR2 | None | None | None | None |
| HSPD1 | None | None | None | None |
| HSPD1 | None | None | None | None |
| FGF8 | None | None | None | None |
| FGF8 | None | None | None | None |
| FGF8 | None | None | None | None |
| F9 | None | None | None | None |
| NCF4 | None | None | None | None |
| NCF4 | None | None | None | None |
| LDLR | None | None | None | None |
| LDLR | None | None | None | None |
| LDLR | None | None | None | None |
| LDLR | None | None | None | None |
| LDLR | None | None | None | None |
| LDLR | None | None | None | None |
| C3 | None | None | None | None |
| C3 | None | None | None | None |
| C3 | None | None | None | None |
| C3 | None | None | None | None |
| C3 | None | None | None | None |
| C3 | None | None | None | None |
| C3 | None | None | None | None |
| C3 | None | None | None | None |
| TULP1 | None | None | None | None |
| TULP1 | None | None | None | None |
| TULP1 | None | None | None | None |
| TULP1 | None | None | None | None |
| IKBKG | None | None | None | None |
| IKBKG | None | None | None | None |
| IKBKG | None | None | None | None |
| CD40 | None | None | None | None |
| CD40 | None | None | None | None |
| SLC9A3R1 | None | None | None | None |
| BCHE | None | None | None | None |
| BCHE | None | None | None | None |
| BCHE | None | None | None | None |
| BCHE | None | None | None | None |
| BCHE | None | None | None | None |
| BCHE | None | None | None | None |
| BCHE | None | None | None | None |
| BCHE | None | None | None | None |
| BCHE | None | None | None | None |
| BCHE | None | None | None | None |
| BCHE | None | None | None | None |
| BCKDHB | None | None | None | None |
| BCKDHB | None | None | None | None |
| BCKDHB | None | None | None | None |

| | | | | |
|---|---|---|---|---|
| BCKDHB | None | None | None | None |
| BCKDHB | None | None | None | None |
| BCKDHB | None | None | None | None |
| BCKDHB | None | None | None | None |
| BCKDHB | None | None | None | None |
| BCKDHB | None | None | None | None |
| BCKDHB | None | None | None | None |
| BCKDHB | None | None | None | None |
| BCKDHB | None | None | None | None |
| CDH1 | None | None | None | None |
| CDH1 | None | None | None | None |
| CDH1 | None | None | None | None |
| CDH1 | None | None | None | None |
| CDH1 | None | None | None | None |
| CTNNB1 | None | None | None | None |
| AIRE | None | None | None | None |
| EDA | 1 | 1 | 1rj8 | A |
| EDA | None | None | None | None |
| EDA | None | None | None | None |
| EDA | None | None | None | None |
| EDA | None | None | None | None |
| SH2D1A | None | None | None | None |
| SH2D1A | None | None | None | None |
| SH2D1A | None | None | None | None |
| GCK | None | None | None | None |
| GCK | None | None | None | None |
| GCK | None | None | None | None |
| GCK | None | None | None | None |
| GCK | None | None | None | None |
| GCK | None | None | None | None |
| GCK | None | None | None | None |
| GCK | None | None | None | None |
| GCK | None | None | None | None |
| GCK | None | None | None | None |
| GCK | None | None | None | None |
| GCK | None | None | None | None |
| GCK | None | None | None | None |
| GCK | None | None | None | None |
| GCK | None | None | None | None |
| GCK | None | None | None | None |
| GCK | None | None | None | None |
| GCK | None | None | None | None |
| GCK | None | None | None | None |
| GCK | None | None | None | None |
| AQP2 | None | None | None | None |
| AQP2 | None | None | None | None |
| AQP2 | None | None | None | None |
| AQP2 | None | None | None | None |
| AQP2 | None | None | None | None |
| AQP2 | None | None | None | None |
| AQP2 | None | None | None | None |
| AQP2 | None | None | None | None |
| AQP2 | None | None | None | None |
| AQP2 | None | None | None | None |

| | | | | |
|---|---|---|---|---|
| AQP2 | None | None | None | None |
| AQP2 | None | None | None | None |
| AQP2 | None | None | None | None |
| SUMF1 | None | None | None | None |
| SUMF1 | None | None | None | None |
| SUMF1 | None | None | None | None |
| SUMF1 | None | None | None | None |
| SUMF1 | None | None | None | None |
| SUMF1 | None | None | None | None |
| SUMF1 | None | None | None | None |
| SUMF1 | None | None | None | None |
| SUMF1 | None | None | None | None |
| FAM20A | None | None | None | None |
| FAM20A | None | None | None | None |
| FAM20A | None | None | None | None |
| FAM20A | None | None | None | None |
| NT5E | None | None | None | None |
| NT5E | None | None | None | None |
| NOS3 | None | None | None | None |
| ACTG1 | None | None | None | None |
| ACTG1 | None | None | None | None |
| ACTG1 | None | None | None | None |
| ACTG1 | None | None | None | None |
| ACTG1 | None | None | None | None |
| ACTG1 | None | None | None | None |
| FCGR1A | None | None | None | None |
| GPI | None | None | None | None |
| GPI | None | None | None | None |
| GPI | 1 | 3 | 1jiq | A |
| GPI | None | None | None | None |
| GPI | None | None | None | None |
| GPI | None | None | None | None |
| GPI | None | None | None | None |
| GPI | None | None | None | None |
| GPI | None | None | None | None |
| CFH | None | None | None | None |
| OGG1 | None | None | None | None |
| TPK1 | None | None | None | None |
| TPK1 | None | None | None | None |
| TPK1 | None | None | None | None |
| TPK1 | None | None | None | None |
| SEC23A | None | None | None | None |
| SEC23A | None | None | None | None |
| TYRP1 | None | None | None | None |
| TYRP1 | None | None | None | None |
| TYRP1 | None | None | None | None |
| TYRP1 | None | None | None | None |
| TYRP1 | None | None | None | None |
| TYRP1 | None | None | None | None |
| CSTB | None | None | None | None |
| CSTB | None | None | None | None |
| CSTB | None | None | None | None |
| ACP1 | None | None | None | None |
| IL17F | None | None | None | None |

| | | | | |
|---|---|---|---|---|
| IL17F | None | None | None | None |
| IL17F | None | None | None | None |
| IL17F | None | None | None | None |
| IL17F | None | None | None | None |
| IDH2 | None | None | None | None |
| IDH2 | None | None | None | None |
| IDH2 | None | None | None | None |
| IDH2 | None | None | None | None |
| TP53 | None | None | None | None |
| TP53 | None | None | None | None |
| TP53 | None | None | None | None |
| TP53 | None | None | None | None |
| TP53 | None | None | None | None |
| TP53 | None | None | None | None |
| TP53 | None | None | None | None |
| TP53 | None | None | None | None |
| TP53 | None | None | None | None |
| TP53 | None | None | None | None |
| TP53 | None | None | None | None |
| TP53 | None | None | None | None |
| TP53 | None | None | None | None |
| TP53 | None | None | None | None |
| TP53 | None | None | None | None |
| TP53 | None | None | None | None |
| TP53 | None | None | None | None |
| TP53 | None | None | None | None |
| TP53 | None | None | None | None |
| TP53 | None | None | None | None |
| TP53 | None | None | None | None |
| TP53 | None | None | None | None |
| TP53 | None | None | None | None |
| TP53 | None | None | None | None |
| TP53 | None | None | None | None |
| TP53 | None | None | None | None |
| TP53 | None | None | None | None |
| TP53 | None | None | None | None |
| TP53 | None | None | None | None |
| TP53 | None | None | None | None |
| TP53 | None | None | None | None |
| TP53 | None | None | None | None |
| TP53 | None | None | None | None |
| TP53 | None | None | None | None |
| TP53 | None | None | None | None |
| TP53 | None | None | None | None |
| TP53 | None | None | None | None |
| TP53 | None | None | None | None |
| TP53 | None | None | None | None |
| TP53 | None | None | None | None |
| TP53 | None | None | None | None |
| TP53 | None | None | None | None |
| TP53 | None | None | None | None |
| TP53 | None | None | None | None |
| LMAN1 | None | None | None | None |
| CTSK | None | None | None | None |
| CTSK | None | None | None | None |
| CTSK | None | None | None | None |

| | | | | |
|---|---|---|---|---|
| FGB | None | None | None | None |
| FGB | None | None | None | None |
| FGB | None | None | None | None |
| FGB | None | None | None | None |
| ACADM | None | None | None | None |
| ACADM | None | None | None | None |
| ACADM | None | None | None | None |
| ACADM | None | None | None | None |
| ACADM | None | None | None | None |
| ACADM | None | None | None | None |
| ACADM | None | None | None | None |
| ACADM | None | None | None | None |
| ACADM | None | None | None | None |
| ACADM | None | None | None | None |
| ACADM | None | None | None | None |
| ACADM | None | None | None | None |
| ACADM | None | None | None | None |
| ACADM | None | None | None | None |
| ACADM | None | None | None | None |
| ACADM | None | None | None | None |
| ACADM | None | None | None | None |
| ACADM | None | None | None | None |
| ACADM | None | None | None | None |
| ACADM | None | None | None | None |
| ACADM | None | None | None | None |
| ACADM | None | None | None | None |
| ACADM | None | None | None | None |
| ACADM | None | None | None | None |
| ACADM | None | None | None | None |
| ACADM | None | None | None | None |
| ACADM | None | None | None | None |
| PTS | None | None | None | None |
| PTS | None | None | None | None |
| PTS | None | None | None | None |
| PTS | None | None | None | None |
| PTS | None | None | None | None |
| PTS | None | None | None | None |
| PTS | None | None | None | None |
| PTS | None | None | None | None |
| PTS | None | None | None | None |
| INS | None | None | None | None |
| INS | 1 | 5 | 2l1z | A |
| INS | 1 | 5 | 2l1z | A |
| INS | 1 | 1 | 1his | A |
| FBP1 | None | None | None | None |
| FBP1 | None | None | None | None |
| FBP1 | None | None | None | None |
| FBP1 | None | None | None | None |
| FBP1 | None | None | None | None |
| FBP1 | None | None | None | None |
| FBP1 | None | None | None | None |
| FBP1 | None | None | None | None |
| FBP1 | None | None | None | None |
| FBP1 | None | None | None | None |

| | | | | |
|---|---|---|---|---|
| FBP1 | None | None | None | None |
| KCNQ1 | None | None | None | None |
| KCNQ1 | None | None | None | None |
| KCNQ1 | None | None | None | None |
| UPP1 | None | None | None | None |
| UMPS | 1 | 3 | 2qcl | A |
| UMPS | None | None | None | None |
| UMPS | None | None | None | None |
| DHFR | None | None | None | None |
| KHK | None | None | None | None |
| TTPA | None | None | None | None |
| TTPA | None | None | None | None |
| TTPA | None | None | None | None |
| TTPA | None | None | None | None |
| MAP1LC3B | None | None | None | None |
| MAP1LC3B | None | None | None | None |
| ZAP70 | None | None | None | None |
| PROC | None | None | None | None |
| SQSTM1 | None | None | None | None |
| AHCY | None | None | None | None |
| AHCY | None | None | None | None |
| PRNP | None | None | None | None |
| PRNP | None | None | None | None |
| PRNP | None | None | None | None |
| PRNP | None | None | None | None |
| PRNP | None | None | None | None |
| PRNP | None | None | None | None |
| PRNP | None | None | None | None |
| PRNP | None | None | None | None |
| PRNP | None | None | None | None |
| PRNP | None | None | None | None |
| PRNP | None | None | None | None |
| PRNP | None | None | None | None |
| PRNP | None | None | None | None |
| PRNP | None | None | None | None |
| PRNP | None | None | None | None |
| PRNP | None | None | None | None |
| REN | None | None | None | None |
| REN | None | None | None | None |
| HINT1 | None | None | None | None |
| HINT1 | None | None | None | None |
| HINT1 | None | None | None | None |
| HINT1 | None | None | None | None |
| HINT1 | None | None | None | None |
| OAT | None | None | None | None |
| OAT | None | None | None | None |
| OAT | None | None | None | None |
| OAT | None | None | None | None |
| OAT | None | None | None | None |
| OAT | None | None | None | None |
| OAT | None | None | None | None |
| OAT | None | None | None | None |
| OAT | None | None | None | None |
| OAT | None | None | None | None |

| | | | | |
|---|---|---|---|---|
| OAT | None | None | None | None |
| OAT | None | None | None | None |
| OAT | None | None | None | None |
| OAT | None | None | None | None |
| OAT | None | None | None | None |
| OAT | None | None | None | None |
| OAT | None | None | None | None |
| OAT | None | None | None | None |
| OAT | None | None | None | None |
| OAT | None | None | None | None |
| OAT | None | None | None | None |
| OAT | None | None | None | None |
| OAT | None | None | None | None |
| OAT | None | None | None | None |
| OAT | None | None | None | None |
| OAT | None | None | None | None |
| OAT | None | None | None | None |
| OAT | None | None | None | None |
| CTSA | None | None | None | None |
| CTSA | None | None | None | None |
| PNP | None | None | None | None |
| PNP | None | None | None | None |
| PNP | None | None | None | None |
| PNP | None | None | None | None |
| PNP | None | None | None | None |
| PNP | None | None | None | None |
| PNP | None | None | None | None |
| PNP | None | None | None | None |
| MYL2 | None | None | None | None |
| MYL2 | None | None | None | None |
| ANG | None | None | None | None |
| ANG | None | None | None | None |
| ANG | None | None | None | None |
| ANG | None | None | None | None |
| ANG | None | None | None | None |
| ANG | None | None | None | None |
| ANG | None | None | None | None |
| ANG | None | None | None | None |
| ANG | None | None | None | None |
| ANG | None | None | None | None |
| POLH | None | None | None | None |
| POLH | None | None | None | None |
| POLH | None | None | None | None |
| IL2RA | None | None | None | None |
| TGFB1 | None | None | None | None |
| TP53 | None | None | None | None |
| TP53 | None | None | None | None |
| PAH | None | None | None | None |
| PAH | None | None | None | None |
| PAH | None | None | None | None |
| PAH | None | None | None | None |
| PAH | None | None | None | None |
| PAH | None | None | None | None |
| PAH | None | None | None | None |

| | | | | |
|---|---|---|---|---|
| PAH | None | None | None | None |
| PAH | None | None | None | None |
| PAH | None | None | None | None |
| PAH | None | None | None | None |
| PAH | None | None | None | None |
| PAH | None | None | None | None |
| PAH | None | None | None | None |
| PAH | None | None | None | None |
| PAH | None | None | None | None |
| PAH | None | None | None | None |
| PAH | None | None | None | None |
| PAH | None | None | None | None |
| PAH | None | None | None | None |
| PAH | None | None | None | None |
| PAH | None | None | None | None |
| PAH | None | None | None | None |
| PAH | None | None | None | None |
| PAH | None | None | None | None |
| PAH | None | None | None | None |
| PAH | None | None | None | None |
| PAH | None | None | None | None |
| PAH | None | None | None | None |
| PAH | None | None | None | None |
| PAH | None | None | None | None |
| PAH | None | None | None | None |
| PAH | None | None | None | None |
| PAH | None | None | None | None |
| PAH | None | None | None | None |
| PAH | None | None | None | None |
| PAH | None | None | None | None |
| PAH | None | None | None | None |
| PAH | None | None | None | None |
| PAH | None | None | None | None |
| PAH | None | None | None | None |
| HDAC8 | None | None | None | None |
| HDAC8 | None | None | None | None |
| VCP | None | None | None | None |
| VCP | None | None | None | None |
| VCP | None | None | None | None |
| VCP | None | None | None | None |
| VCP | None | None | None | None |
| VCP | None | None | None | None |
| VCP | None | None | None | None |
| B3GAT3 | None | None | None | None |
| B3GAT3 | None | None | None | None |
| CSF2RB | None | None | None | None |
| CSF2RB | None | None | None | None |
| CSF2RB | None | None | None | None |
| NF2 | None | None | None | None |
| NF2 | None | None | None | None |
| NF2 | None | None | None | None |
| PIK3CA | None | None | None | None |
| PHGDH | None | None | None | None |
| PHGDH | None | None | None | None |

| | | | | |
|---|---|---|---|---|
| PHGDH | None | None | None | None |
| PHGDH | None | None | None | None |
| POLH | None | None | None | None |
| XIAP | None | None | None | None |
| XIAP | None | None | None | None |
| PDHX | None | None | None | None |
| PDHX | None | None | None | None |
| PDHX | None | None | None | None |
| PDHX | None | None | None | None |
| PDHX | None | None | None | None |
| SAMHD1 | None | None | None | None |
| SAMHD1 | None | None | None | None |
| SAMHD1 | None | None | None | None |
| SAMHD1 | None | None | None | None |
| SAMHD1 | None | None | None | None |
| SAMHD1 | None | None | None | None |
| SAMHD1 | None | None | None | None |
| SAMHD1 | None | None | None | None |
| CD36 | None | None | None | None |
| CD36 | None | None | None | None |
| CD36 | None | None | None | None |
| ASPH | None | None | None | None |
| CCT5 | None | None | None | None |
| CCT5 | None | None | None | None |
| CCT5 | None | None | None | None |
| SMN1 | None | None | None | None |
| SMN1 | None | None | None | None |
| SMN1 | None | None | None | None |
| SMN1 | None | None | None | None |
| ABCA1 | None | None | None | None |
| ABCA1 | None | None | None | None |
| ABCA1 | None | None | None | None |
| ABCA1 | None | None | None | None |
| ABCA1 | None | None | None | None |
| ABCA1 | None | None | None | None |
| ABCA1 | None | None | None | None |
| ABCA1 | None | None | None | None |
| ABCA1 | None | None | None | None |
| ABCA1 | None | None | None | None |
| ABCA1 | None | None | None | None |
| ABCA1 | None | None | None | None |
| ABCA1 | None | None | None | None |
| ABCA1 | None | None | None | None |
| ABCA1 | None | None | None | None |
| ABCA1 | None | None | None | None |
| ABCA1 | None | None | None | None |
| ABCA1 | None | None | None | None |
| ABCA1 | None | None | None | None |
| ABCA1 | None | None | None | None |
| ABCA1 | None | None | None | None |
| ABCA1 | None | None | None | None |
| ABCA1 | None | None | None | None |
| ABCA1 | None | None | None | None |

| | | | | |
|---|---|---|---|---|
| ABCA1 | None | None | None | None |
| ABCA1 | None | None | None | None |
| ABCA1 | None | None | None | None |
| ABCA1 | None | None | None | None |
| ABCA1 | None | None | None | None |
| ABCA1 | None | None | None | None |
| ABCA1 | None | None | None | None |
| ABCA1 | None | None | None | None |
| ABCA1 | None | None | None | None |
| ABCA1 | None | None | None | None |
| ABCA1 | None | None | None | None |
| ERCC1 | None | None | None | None |
| FECH | None | None | None | None |
| FECH | None | None | None | None |
| FECH | None | None | None | None |
| FECH | None | None | None | None |
| FECH | None | None | None | None |
| FECH | None | None | None | None |
| FECH | None | None | None | None |
| PTEN | None | None | None | None |
| PTEN | None | None | None | None |
| PTEN | None | None | None | None |
| PTEN | None | None | None | None |
| PTEN | None | None | None | None |
| PTEN | None | None | None | None |
| PTEN | None | None | None | None |
| PTEN | None | None | None | None |
| PTEN | None | None | None | None |
| PTEN | None | None | None | None |
| PTEN | None | None | None | None |
| PTEN | None | None | None | None |
| PTEN | None | None | None | None |
| PTEN | None | None | None | None |
| PTEN | None | None | None | None |
| PTEN | None | None | None | None |
| PTEN | None | None | None | None |
| PTEN | None | None | None | None |
| PTEN | None | None | None | None |
| PTEN | None | None | None | None |
| ACAT1 | None | None | None | None |
| ACAT1 | None | None | None | None |
| ACAT1 | None | None | None | None |
| ACAT1 | None | None | None | None |
| ACAT1 | None | None | None | None |
| ACAT1 | None | None | None | None |
| ACAT1 | None | None | None | None |
| ACAT1 | None | None | None | None |
| ACAT1 | None | None | None | None |
| ACAT1 | None | None | None | None |
| ACAT1 | None | None | None | None |
| CREBBP | None | None | None | None |
| ITGA2B | None | None | None | None |
| ITGA2B | None | None | None | None |
| ITGA2B | None | None | None | None |

| | | | | |
|---|---|---|---|---|
| ITGA2B | None | None | None | None |
| TSC1 | None | None | None | None |
| ADA2 | None | None | None | None |
| ADA2 | None | None | None | None |
| ADA2 | None | None | None | None |
| ADA2 | None | None | None | None |
| ADA2 | None | None | None | None |
| ADA2 | None | None | None | None |
| ADA2 | None | None | None | None |
| ADA2 | None | None | None | None |
| ADA2 | | 1 | 1 3lgd | B |
| ADA2 | None | None | None | None |
| PGK1 | None | None | None | None |
| PGK1 | None | None | None | None |
| PGK1 | None | None | None | None |
| PGK1 | None | None | None | None |
| PGK1 | None | None | None | None |
| TUBA1A | None | None | None | None |
| TUBA1A | None | None | None | None |
| TUBA1A | None | None | None | None |
| TUBA1A | None | None | None | None |
| TUBA1A | None | None | None | None |
| PTPN11 | None | None | None | None |
| PTPN11 | None | None | None | None |
| PTPN11 | None | None | None | None |
| PTPN11 | None | None | None | None |
| PTPN11 | None | None | None | None |
| PTPN11 | None | None | None | None |
| PTPN11 | None | None | None | None |
| PTPN11 | None | None | None | None |
| PTPN11 | None | None | None | None |
| PTPN11 | None | None | None | None |
| PTPN11 | None | None | None | None |
| PTPN11 | None | None | None | None |
| PTPN11 | None | None | None | None |
| PTPN11 | None | None | None | None |
| PTPN11 | None | None | None | None |
| PTPN11 | None | None | None | None |
| PTPN11 | None | None | None | None |
| PTPN11 | None | None | None | None |
| PTPN11 | None | None | None | None |
| PTPN11 | None | None | None | None |
| PTPN11 | None | None | None | None |
| PTPN11 | None | None | None | None |
| PTPN11 | None | None | None | None |
| PTPN11 | None | None | None | None |
| PTPN11 | None | None | None | None |
| PTPN11 | None | None | None | None |
| GLB1 | None | None | None | None |
| GLB1 | None | None | None | None |
| UQCRQ | None | None | None | None |
| VPS35 | None | None | None | None |
| NR3C1 | None | None | None | None |
| NR3C1 | None | None | None | None |

| | | | | |
|---|---|---|---|---|
| NR3C1 | None | None | None | None |
| NR3C1 | None | None | None | None |
| NR3C1 | None | None | None | None |
| NR3C1 | None | None | None | None |
| NR3C1 | None | None | None | None |
| NR3C1 | None | None | None | None |
| NR3C1 | None | None | None | None |
| APOA1 | None | None | None | None |
| APOA1 | None | None | None | None |
| APOA1 | None | None | None | None |
| APOA1 | None | None | None | None |
| APOA1 | None | None | None | None |
| APOA1 | None | None | None | None |
| APOA1 | None | None | None | None |
| BCKDHA | None | None | None | None |
| BCKDHA | None | None | None | None |
| BCKDHA | None | None | None | None |
| BCKDHA | None | None | None | None |
| BCKDHA | None | None | None | None |
| BCKDHA | None | None | None | None |
| BCKDHA | None | None | None | None |
| BCKDHA | 1 | 1 | 2beu | A |
| BCKDHA | None | None | None | None |
| BCKDHA | None | None | None | None |
| BCKDHA | None | None | None | None |
| BCKDHA | None | None | None | None |
| BCKDHA | None | None | None | None |
| BCKDHA | None | None | None | None |
| BCKDHA | None | None | None | None |
| TOP1 | None | None | None | None |
| ANTXR2 | None | None | None | None |
| ANTXR2 | None | None | None | None |
| ANTXR2 | None | None | None | None |
| TF | None | None | None | None |
| TF | None | None | None | None |
| FSHB | None | None | None | None |
| GALK1 | None | None | None | None |
| GALK1 | None | None | None | None |
| GALK1 | None | None | None | None |
| CYP17A1 | None | None | None | None |
| CYP17A1 | None | None | None | None |
| CYP17A1 | None | None | None | None |
| CYP17A1 | None | None | None | None |
| CYP17A1 | None | None | None | None |
| CYP17A1 | None | None | None | None |
| CYP17A1 | None | None | None | None |
| CYP17A1 | None | None | None | None |
| CYP17A1 | None | None | None | None |
| CYP17A1 | None | None | None | None |
| CYP17A1 | None | None | None | None |
| CYP17A1 | None | None | None | None |
| CYP17A1 | None | None | None | None |
| CYP17A1 | None | None | None | None |
| CYP17A1 | None | None | None | None |

| | | | | |
|---|---|---|---|---|
| CYP17A1 | None | None | None | None |
| CYP17A1 | None | None | None | None |
| CD44 | None | None | None | None |
| MDH2 | None | None | None | None |
| MDH2 | None | None | None | None |
| NDUFV1 | None | None | None | None |
| NDUFV1 | None | None | None | None |
| NDUFV1 | None | None | None | None |
| NDUFV1 | None | None | None | None |
| NDUFV1 | None | None | None | None |
| NDUFV1 | None | None | None | None |
| NDUFV1 | None | None | None | None |
| NDUFV1 | None | None | None | None |
| NDUFV1 | None | None | None | None |
| NDUFV1 | None | None | None | None |
| CTSC | None | None | None | None |
| CTSC | None | None | None | None |
| CTSC | None | None | None | None |
| CTSC | None | None | None | None |
| CTSC | None | None | None | None |
| CTSC | None | None | None | None |
| CTSC | None | None | None | None |
| VWF | None | None | None | None |
| VWF | None | None | None | None |
| VWF | None | None | None | None |
| VWF | None | None | None | None |
| VWF | None | None | None | None |
| VWF | None | None | None | None |
| VWF | None | None | None | None |
| ORC4 | None | None | None | None |
| ORC4 | None | None | None | None |
| CRYAB | None | None | None | None |
| CRYAB | None | None | None | None |
| CRYAB | None | None | None | None |
| MAP2K2 | None | None | None | None |
| MAP2K2 | None | None | None | None |
| MAP2K2 | None | None | None | None |
| MAP2K2 | None | None | None | None |
| MAP2K2 | None | None | None | None |
| MAP2K2 | None | None | None | None |
| MAP2K2 | None | None | None | None |
| MAP2K2 | None | None | None | None |
| MAP2K2 | None | None | None | None |
| CBL | None | None | None | None |
| CBL | None | None | None | None |
| CBL | None | None | None | None |
| CBL | None | None | None | None |
| CBL | None | None | None | None |
| CBL | None | None | None | None |
| CBL | None | None | None | None |
| PFN1 | None | None | None | None |
| PFN1 | None | None | None | None |
| ACSM2A | None | None | None | None |
| HSPG2 | None | None | None | None |

| | | | | |
|---|---|---|---|---|
| HSPG2 | None | None | None | None |
| HSPG2 | None | None | None | None |
| HSPG2 | None | None | None | None |
| PARK7 | None | None | None | None |
| PARK7 | None | None | None | None |
| PARK7 | None | None | None | None |
| PARK7 | None | None | None | None |
| PARK7 | None | None | None | None |
| PARK7 | None | None | None | None |
| INSR | None | None | None | None |
| INSR | None | None | None | None |
| INSR | None | None | None | None |
| INSR | None | None | None | None |
| INSR | None | None | None | None |
| DSC2 | None | None | None | None |
| DSC2 | None | None | None | None |
| DSC2 | None | None | None | None |
| DSC2 | None | None | None | None |
| DSC2 | None | None | None | None |
| DSC2 | None | None | None | None |
| DSC2 | None | None | None | None |
| DSC2 | None | None | None | None |
| DSC2 | None | None | None | None |
| DSC2 | None | None | None | None |
| MICU1 | None | None | None | None |
| MICU1 | None | None | None | None |
| ADAMTS13 | None | None | None | None |
| ADAMTS13 | None | None | None | None |
| ADAMTS13 | None | None | None | None |
| ADAMTS13 | None | None | None | None |
| ADAMTS13 | None | None | None | None |
| ADAMTS13 | None | None | None | None |
| ADAMTS13 | None | None | None | None |
| IRAK4 | None | None | None | None |
| IRAK4 | None | None | None | None |
| RNASEH2B | None | None | None | None |
| RNASEH2B | None | None | None | None |
| RNASEH2B | None | None | None | None |
| NTRK1 | None | None | None | None |
| MPO | None | None | None | None |
| MPO | None | None | None | None |
| MPO | None | None | None | None |
| PEX3 | None | None | None | None |
| SERPINA1 | None | None | None | None |
| SERPINA1 | None | None | None | None |
| SERPINA1 | None | None | None | None |
| SERPINA1 | None | None | None | None |
| SERPINA1 | None | None | None | None |
| SERPINA1 | None | None | None | None |
| SERPINA1 | None | None | None | None |
| SERPINA1 | None | None | None | None |
| SERPINA1 | None | None | None | None |
| SERPINA1 | None | None | None | None |
| YARS2 | None | None | None | None |

| | | | | |
|---|---|---|---|---|
| YARS2 | None | None | None | None |
| YARS2 | None | None | None | None |
| YARS2 | None | None | None | None |
| YARS2 | None | None | None | None |
| YARS2 | None | None | None | None |
| YARS2 | None | None | None | None |
| YARS2 | None | None | None | None |
| YARS2 | None | None | None | None |
| PDHA1 | None | None | None | None |
| PDHA1 | None | None | None | None |
| PDHA1 | None | None | None | None |
| PDHA1 | None | None | None | None |
| PDHA1 | None | None | None | None |
| PDHA1 | None | None | None | None |
| PDHA1 | None | None | None | None |
| PDHA1 | None | None | None | None |
| PDHA1 | None | None | None | None |
| PDHA1 | None | None | None | None |
| PDHA1 | None | None | None | None |
| CUL4B | None | None | None | None |
| AGXT | None | None | None | None |
| AGXT | None | None | None | None |
| AGXT | None | None | None | None |
| AGXT | None | None | None | None |
| AGXT | None | None | None | None |
| AGXT | None | None | None | None |
| AGXT | None | None | None | None |
| AGXT | None | None | None | None |
| AGXT | None | None | None | None |
| AGXT | None | None | None | None |
| AGXT | None | None | None | None |
| AGXT | None | None | None | None |
| AGXT | None | None | None | None |
| AGXT | None | None | None | None |
| AGXT | None | None | None | None |
| AGXT | None | None | None | None |
| AGXT | None | None | None | None |
| AGXT | None | None | None | None |
| AGXT | None | None | None | None |
| AGXT | None | None | None | None |
| AGXT | None | None | None | None |
| AGXT | None | None | None | None |
| AGXT | None | None | None | None |
| AGXT | None | None | None | None |
| AGXT | None | None | None | None |
| AGXT | None | None | None | None |
| AGXT | None | None | None | None |
| AGXT | None | None | None | None |
| AGXT | None | None | None | None |
| AGXT | None | None | None | None |
| AGXT | None | None | None | None |
| AGXT | None | None | None | None |
| AGXT | None | None | None | None |
| AGXT | None | None | None | None |

| | | | | |
|---|---|---|---|---|
| AGXT | None | None | None | None |
| SFTPD | None | None | None | None |
| SFTPD | None | None | None | None |
| PIK3CA | None | None | None | None |
| PIK3CA | None | None | None | None |
| PIK3CA | None | None | None | None |
| AUH | None | None | None | None |
| AUH | None | None | None | None |
| AUH | None | None | None | None |
| AUH | None | None | None | None |
| F11 | None | None | None | None |
| F11 | None | None | None | None |
| F11 | None | None | None | None |
| F11 | None | None | None | None |
| MEN1 | None | None | None | None |
| MEN1 | None | None | None | None |
| MEN1 | None | None | None | None |
| MEN1 | None | None | None | None |
| MEN1 | 1 | 2 | 4i80 | A |
| MEN1 | None | None | None | None |
| MEN1 | None | None | None | None |
| MEN1 | None | None | None | None |
| MEN1 | None | None | None | None |
| MEN1 | None | None | None | None |
| MYBPC1 | None | None | None | None |
| MYBPC1 | None | None | None | None |
| MYBPC1 | None | None | None | None |
| PDCD10 | None | None | None | None |
| SCARB2 | None | None | None | None |
| SCARB2 | None | None | None | None |
| SCARB2 | None | None | None | None |
| SCARB2 | None | None | None | None |
| SCARB2 | None | None | None | None |
| TREX2 | None | None | None | None |
| TBK1 | None | None | None | None |
| TBK1 | None | None | None | None |
| DLD | None | None | None | None |
| DLD | None | None | None | None |
| DLD | None | None | None | None |
| DLD | None | None | None | None |
| DLD | None | None | None | None |
| DLD | None | None | None | None |
| LDHA | None | None | None | None |
| TNNC1 | None | None | None | None |
| TNNC1 | None | None | None | None |
| TNNC1 | None | None | None | None |
| SMAD3 | None | None | None | None |
| HARS | None | None | None | None |
| HARS | None | None | None | None |
| TNNI3 | None | None | None | None |
| TNNI3 | None | None | None | None |
| TNNI3 | None | None | None | None |
| TNNI3 | None | None | None | None |
| TNNI3 | None | None | None | None |

| | | | | |
|---|---|---|---|---|
| TNNI3 | None | None | None | None |
| ACKR1 | None | None | None | None |
| BUB1B | None | None | None | None |
| BUB1B | None | None | None | None |
| BUB1B | None | None | None | None |
| AR | None | None | None | None |
| AR | None | None | None | None |
| AR | None | None | None | None |
| AR | None | None | None | None |
| AR | None | None | None | None |
| FTO | None | None | None | None |
| FTO | None | None | None | None |
| TFAM | None | None | None | None |
| PYGL | None | None | None | None |
| PYGL | None | None | None | None |
| PYGL | None | None | None | None |
| PYGL | None | None | None | None |
| PYGL | None | None | None | None |
| PYGL | None | None | None | None |
| PYGL | None | None | None | None |
| PYGL | None | None | None | None |
| PYGL | None | None | None | None |
| PYGL | None | None | None | None |
| PYGL | None | None | None | None |
| PYGL | None | None | None | None |
| PYGL | None | None | None | None |
| PYGL | None | None | None | None |
| PYGL | None | None | None | None |
| PYGL | None | None | None | None |
| GALNS | None | None | None | None |
| GALNS | None | None | None | None |
| GALNS | None | None | None | None |
| GALNS | None | None | None | None |
| GALNS | None | None | None | None |
| GALNS | None | None | None | None |
| GALNS | None | None | None | None |
| GALNS | None | None | None | None |
| GALNS | None | None | None | None |
| GALNS | None | None | None | None |
| GALNS | None | None | None | None |
| GALNS | None | None | None | None |
| GALNS | None | None | None | None |
| GALNS | None | None | None | None |
| GALNS | None | None | None | None |
| GALNS | None | None | None | None |
| GALNS | None | None | None | None |
| GALNS | None | None | None | None |
| GALNS | None | None | None | None |
| GALNS | None | None | None | None |
| GALNS | None | None | None | None |
| GALNS | None | None | None | None |
| GALNS | None | None | None | None |
| GALNS | None | None | None | None |
| GALNS | None | None | None | None |

| | | | | |
|---|---|---|---|---|
| GALNS | None | None | None | None |
| GALNS | None | None | None | None |
| GALNS | None | None | None | None |
| GALNS | None | None | None | None |
| GALNS | None | None | None | None |
| GALNS | None | None | None | None |
| GALNS | None | None | None | None |
| GALNS | None | None | None | None |
| F10 | None | None | None | None |
| GLUD1 | None | None | None | None |
| GLUD1 | None | None | None | None |
| GLUD1 | None | None | None | None |
| GLUD1 | None | None | None | None |
| GLUD1 | None | None | None | None |
| GLUD1 | None | None | None | None |
| GLUD1 | None | None | None | None |
| GLUD1 | None | None | None | None |
| GLUD1 | None | None | None | None |
| GLUD1 | None | None | None | None |
| NOTCH1 | None | None | None | None |
| POGLUT1 | None | None | None | None |
| POGLUT1 | None | None | None | None |
| NR3C1 | None | None | None | None |
| CASP8 | None | None | None | None |
| CASP8 | None | None | None | None |
| TRAP1 | None | None | None | None |
| FANCM | None | None | None | None |
| EPOR | None | None | None | None |
| EPOR | None | None | None | None |
| NDP | None | None | None | None |
| NDP | None | None | None | None |
| NDP | None | None | None | None |
| NDP | None | None | None | None |
| NDP | None | None | None | None |
| NDP | None | None | None | None |
| NDP | None | None | None | None |
| NDP | None | None | None | None |
| NDP | None | None | None | None |
| NDP | None | None | None | None |
| NDP | None | None | None | None |
| NDP | None | None | None | None |
| NDP | None | None | None | None |
| NDP | None | None | None | None |
| NDP | None | None | None | None |
| NDP | None | None | None | None |
| NDP | None | None | None | None |
| ROGDI | None | None | None | None |
| CYP19A1 | None | None | None | None |
| CYP19A1 | None | None | None | None |
| CYP19A1 | None | None | None | None |
| CYP19A1 | None | None | None | None |
| CYP19A1 | None | None | None | None |
| CYP19A1 | None | None | None | None |
| CYP19A1 | None | None | None | None |

| | | | | |
|---|---|---|---|---|
| CYP19A1 | None | None | None | None |
| UBA5 | None | None | None | None |
| PRPS1 | None | None | None | None |
| PRPS1 | None | None | None | None |
| PRPS1 | None | None | None | None |
| PRPS1 | None | None | None | None |
| PRPS1 | None | None | None | None |
| PRPS1 | None | None | None | None |
| PRPS1 | None | None | None | None |
| PRPS1 | None | None | None | None |
| F2 | None | None | None | None |
| F2 | None | None | None | None |
| NTF4 | None | None | None | None |
| NTF4 | None | None | None | None |
| PSAP | None | None | None | None |
| PSAP | None | None | None | None |
| TAP1 | None | None | None | None |
| TAP1 | None | None | None | None |
| TAP1 | None | None | None | None |
| ARG1 | None | None | None | None |
| ARG1 | None | None | None | None |
| ARG1 | None | None | None | None |
| ARG1 | None | None | None | None |
| ARG1 | None | None | None | None |
| ARG1 | None | None | None | None |
| CRYM | None | None | None | None |
| ADSL | None | None | None | None |
| ADSL | None | None | None | None |
| ADSL | None | None | None | None |
| ADSL | None | None | None | None |
| ADSL | None | None | None | None |
| ADSL | None | None | None | None |
| ADSL | None | None | None | None |
| ADSL | None | None | None | None |
| ADSL | None | None | None | None |
| ADSL | None | None | None | None |
| ADSL | None | None | None | None |
| ADSL | None | None | None | None |
| ADSL | None | None | None | None |
| ADSL | None | None | None | None |
| ADSL | None | None | None | None |
| ADSL | None | None | None | None |
| ADSL | None | None | None | None |
| ADSL | None | None | None | None |
| ADSL | None | None | None | None |
| ADSL | None | None | None | None |
| VWF | None | None | None | None |
| PLA2G7 | None | None | None | None |
| MMP13 | None | None | None | None |
| MMP13 | None | None | None | None |
| MMP13 | None | None | None | None |
| MMP13 | None | None | None | None |
| POMGNT1 | None | None | None | None |
| POMGNT1 | None | None | None | None |

| | | | | |
|---|---|---|---|---|
| COL3A1 | None | None | None | None |
| COL3A1 | None | None | None | None |
| COL3A1 | None | None | None | None |
| COL3A1 | None | None | None | None |
| ACTN2 | None | None | None | None |
| ACTN2 | None | None | None | None |
| ACTN2 | None | None | None | None |
| ACTN2 | None | None | None | None |
| ACTN2 | None | None | None | None |
| ACTN2 | None | None | None | None |
| ACTN2 | None | None | None | None |
| UBA5 | None | None | None | None |
| ADAR | None | None | None | None |
| UMPS | None | None | None | None |
| UMPS | None | None | None | None |
| UMPS | None | None | None | None |
| TRNT1 | None | None | None | None |
| TRNT1 | None | None | None | None |
| NHEJ1 | None | None | None | None |
| NHEJ1 | None | None | None | None |
| NHEJ1 | None | None | None | None |
| C8A | None | None | None | None |
| GLUL | None | None | None | None |
| GLUL | None | None | None | None |
| GLUL | None | None | None | None |
| GYG1 | None | None | None | None |
| GYG1 | None | None | None | None |
| FLNA | None | None | None | None |
| FLNA | None | None | None | None |
| FLNA | None | None | None | None |
| FLNA | None | None | None | None |
| FLNA | None | None | None | None |
| FLNA | None | None | None | None |
| FLNA | None | None | None | None |
| TTN | None | None | None | None |
| TTN | None | None | None | None |
| TTN | None | None | None | None |
| TTN | None | None | None | None |
| SMS | None | None | None | None |
| SMS | None | None | None | None |
| SMS | None | None | None | None |
| C5 | None | None | None | None |
| SERPINC1 | None | None | None | None |
| SERPINC1 | None | None | None | None |
| SERPINC1 | None | None | None | None |
| SERPINC1 | None | None | None | None |
| SERPINC1 | None | None | None | None |
| SERPINC1 | None | None | None | None |
| SERPINC1 | None | None | None | None |
| SERPINC1 | None | None | None | None |
| SERPINC1 | None | None | None | None |
| SERPINC1 | None | None | None | None |
| SERPINC1 | None | None | None | None |
| SERPINC1 | None | None | None | None |

| | | | | |
|---|---|---|---|---|
| SERPINC1 | None | None | None | None |
| SERPINC1 | None | None | None | None |
| SERPINC1 | None | None | None | None |
| SERPINC1 | None | None | None | None |
| SERPINC1 | None | None | None | None |
| SERPINC1 | None | None | None | None |
| SERPINC1 | None | None | None | None |
| SERPINC1 | None | None | None | None |
| SERPINC1 | None | None | None | None |
| ETFB | None | None | None | None |
| ETFB | None | None | None | None |
| ETFB | None | None | None | None |
| ETFB | None | None | None | None |
| ETFB | None | None | None | None |
| ETFB | None | None | None | None |
| DIABLO | None | None | None | None |
| DIABLO | None | None | None | None |
| DPYS | None | None | None | None |
| DPYS | None | None | None | None |
| DPYS | None | None | None | None |
| DPYS | None | None | None | None |
| DPYS | None | None | None | None |
| GNE | None | None | None | None |
| GNE | None | None | None | None |
| GNE | None | None | None | None |
| GNE | None | None | None | None |
| GNE | None | None | None | None |
| GNE | None | None | None | None |
| GNE | None | None | None | None |
| GNE | None | None | None | None |
| GNE | None | None | None | None |
| GNMT | None | None | None | None |
| GNMT | None | None | None | None |
| CTH | None | None | None | None |
| CTH | None | None | None | None |
| CTH | None | None | None | None |
| USH1C | None | None | None | None |
| PSEN1 | None | None | None | None |
| CHEK2 | None | None | None | None |
| CHEK2 | None | None | None | None |
| CHEK2 | None | None | None | None |
| CHEK2 | None | None | None | None |
| CHEK2 | None | None | None | None |
| CHEK2 | None | None | None | None |
| CHEK2 | None | None | None | None |
| CHEK2 | None | None | None | None |
| CHEK2 | None | None | None | None |
| SNCA | None | None | None | None |
| SNCA | None | None | None | None |
| GAN | None | None | None | None |
| GAN | None | None | None | None |
| GAN | None | None | None | None |
| SBDS | None | None | None | None |
| SBDS | None | None | None | None |

| | | | | |
|---|---|---|---|---|
| SBDS | None | None | None | None |
| SBDS | None | None | None | None |
| SBDS | None | None | None | None |
| SBDS | None | None | None | None |
| MPO | | 1 | 3 | 3f9p | A |
| MPO | None | None | None | None |
| KCNH2 | None | None | None | None |
| KCNH2 | None | None | None | None |
| KCNH2 | None | None | None | None |
| KCNH2 | None | None | None | None |
| TEK | None | None | None | None |
| TEK | None | None | None | None |
| TEK | None | None | None | None |
| TEK | None | None | None | None |
| NPC2 | None | None | None | None |
| NPC2 | None | None | None | None |
| NPC2 | None | None | None | None |
| NPC2 | None | None | None | None |
| NPC2 | None | None | None | None |
| NPC2 | None | None | None | None |
| NPC2 | None | None | None | None |
| PLG | None | None | None | None |
| PCSK9 | None | None | None | None |
| SFTPC | None | None | None | None |
| SFTPC | None | None | None | None |
| SFTPC | None | None | None | None |
| CD40LG | None | None | None | None |
| CD40LG | None | None | None | None |
| CD40LG | None | None | None | None |
| CD40LG | None | None | None | None |
| CD40LG | None | None | None | None |
| TAT | None | None | None | None |
| TAT | None | None | None | None |
| TAT | None | None | None | None |
| EP300 | None | None | None | None |
| FARS2 | None | None | None | None |
| FARS2 | None | None | None | None |
| FARS2 | None | None | None | None |
| FARS2 | None | None | None | None |
| FARS2 | None | None | None | None |
| FARS2 | None | None | None | None |
| FARS2 | None | None | None | None |
| FARS2 | None | None | None | None |
| FARS2 | None | None | None | None |
| FARS2 | None | None | None | None |
| FARS2 | None | None | None | None |
| FARS2 | None | None | None | None |
| FARS2 | None | None | None | None |
| MASP2 | None | None | None | None |
| MASP2 | None | None | None | None |
| MASP2 | None | None | None | None |
| MASP2 | None | None | None | None |
| STIM1 | None | None | None | None |
| CHRM2 | None | None | None | None |

| | | | | |
|---|---|---|---|---|
| MCFD2 | None | None | None | None |
| MCFD2 | None | None | None | None |
| MCFD2 | None | None | None | None |
| KITLG | None | None | None | None |
| F12 | None | None | None | None |
| F12 | None | None | None | None |
| ERAP1 | None | None | None | None |
| GTF2H5 | None | None | None | None |
| GTF2H5 | None | None | None | None |
| THBD | None | None | None | None |
| MRPL28 | None | None | None | None |
| MRPL28 | None | None | None | None |
| HTRA1 | None | None | None | None |
| HTRA1 | None | None | None | None |
| HTRA1 | None | None | None | None |
| HTRA1 | None | None | None | None |
| HTRA1 | None | None | None | None |
| HTRA1 | None | None | None | None |
| KARS | None | None | None | None |
| KARS | None | None | None | None |
| KARS | None | None | None | None |
| KARS | None | None | None | None |
| KARS | None | None | None | None |
| HK1 | None | None | None | None |
| GARS | None | None | None | None |
| GARS | None | None | None | None |
| GARS | None | None | None | None |
| GARS | None | None | None | None |
| GARS | None | None | None | None |
| GARS | None | None | None | None |
| GARS | None | None | None | None |
| IL12B | None | None | None | None |
| HADH | None | None | None | None |
| HADH | None | None | None | None |
| HADH | None | None | None | None |
| HADH | None | None | None | None |
| HADH | None | None | None | None |
| RRM2B | None | None | None | None |
| RRM2B | None | None | None | None |
| RRM2B | None | None | None | None |
| RRM2B | None | None | None | None |
| RRM2B | None | None | None | None |
| RRM2B | None | None | None | None |
| RRM2B | None | None | None | None |
| RRM2B | None | None | None | None |
| RRM2B | None | None | None | None |
| RRM2B | None | None | None | None |
| RRM2B | None | None | None | None |
| RRM2B | None | None | None | None |
| RRM2B | None | None | None | None |
| RRM2B | None | None | None | None |
| RRM2B | None | None | None | None |
| MARVELD2 | None | None | None | None |
| TLR4 | None | None | None | None |

| | | | | |
|---|---|---|---|---|
| TTN | None | None | None | None |
| TTN | None | None | None | None |
| TNFRSF13B | None | None | None | None |
| POLG | None | None | None | None |
| POLG | None | None | None | None |
| POLG | None | None | None | None |
| POLG | None | None | None | None |
| POLG | None | None | None | None |
| POLG | None | None | None | None |
| POLG | None | None | None | None |
| POLG | None | None | None | None |
| POLG | None | None | None | None |
| POLG | None | None | None | None |
| POLG | None | None | None | None |
| POLG | None | None | None | None |
| POLG | None | None | None | None |
| POLG | None | None | None | None |
| POLG | None | None | None | None |
| POLG | None | None | None | None |
| POLG | None | None | None | None |
| POLG | None | None | None | None |
| POLG | None | None | None | None |
| POLG | None | None | None | None |
| POLG | None | None | None | None |
| POLG | None | None | None | None |
| POLG | None | None | None | None |
| POLG | None | None | None | None |
| POLG | None | None | None | None |
| POLG | None | None | None | None |
| POLG | None | None | None | None |
| POLG | None | None | None | None |
| POLG | None | None | None | None |
| POLG | None | None | None | None |
| POLG | None | None | None | None |
| POLG | None | None | None | None |
| POLG | None | None | None | None |
| POLG | None | None | None | None |
| POLG | None | None | None | None |
| POLG | None | None | None | None |
| POLG | None | None | None | None |
| POLG | None | None | None | None |
| POLG | None | None | None | None |
| POLG | None | None | None | None |
| POLG | None | None | None | None |
| POLG | None | None | None | None |
| POLG | None | None | None | None |
| POLG | None | None | None | None |
| POLG | None | None | None | None |
| POLG | None | None | None | None |
| POLG | None | None | None | None |
| POLG | None | None | None | None |
| POLG | None | None | None | None |
| POLG | None | None | None | None |
| POLG | None | None | None | None |
| UROD | None | None | None | None |

| | | | | |
|---|---|---|---|---|
| UROD | None | None | None | None |
| UROD | None | None | None | None |
| UROD | None | None | None | None |
| UROD | None | None | None | None |
| UROD | None | None | None | None |
| UROD | None | None | None | None |
| UROD | None | None | None | None |
| UROD | None | None | None | None |
| UROD | None | None | None | None |
| PCK1 | None | None | None | None |
| PCK1 | None | None | None | None |
| PCK1 | None | None | None | None |
| PCK1 | None | None | None | None |
| PCK1 | None | None | None | None |
| HEXA | None | None | None | None |
| HEXA | None | None | None | None |
| HEXA | None | None | None | None |
| HEXA | None | None | None | None |
| HEXA | None | None | None | None |
| HEXA | None | None | None | None |
| HEXA | None | None | None | None |
| HEXA | None | None | None | None |
| HEXA | None | None | None | None |
| HEXA | None | None | None | None |
| HEXA | None | None | None | None |
| HEXA | None | None | None | None |
| HEXA | None | None | None | None |
| HEXA | None | None | None | None |
| HEXA | None | None | None | None |
| HEXA | None | None | None | None |
| HEXA | None | None | None | None |
| HEXA | None | None | None | None |
| HEXA | None | None | None | None |
| HEXA | None | None | None | None |
| HEXA | None | None | None | None |
| HEXA | None | None | None | None |
| HEXA | None | None | None | None |
| HEXA | None | None | None | None |
| HEXA | None | None | None | None |
| HEXA | None | None | None | None |
| HEXA | None | None | None | None |
| HEXA | None | None | None | None |
| HEXA | None | None | None | None |
| HEXA | None | None | None | None |
| HEXA | None | None | None | None |
| CFTR | None | None | None | None |
| CFTR | None | None | None | None |
| CFTR | None | None | None | None |
| CFTR | None | None | None | None |
| CFTR | None | None | None | None |
| CFTR | None | None | None | None |
| CFTR | None | None | None | None |
| CFTR | None | None | None | None |
| CFTR | None | None | None | None |

| | | | | |
|---|---|---|---|---|
| CFTR | None | None | None | None |
| CFTR | None | None | None | None |
| CFTR | None | None | None | None |
| CFTR | None | None | None | None |
| CFTR | None | None | None | None |
| CFTR | None | None | None | None |
| CFTR | None | None | None | None |
| CFTR | None | None | None | None |
| CFTR | None | None | None | None |
| CFTR | None | None | None | None |
| CFTR | None | None | None | None |
| CFTR | None | None | None | None |
| CFTR | None | None | None | None |
| CFTR | None | None | None | None |
| CFTR | None | None | None | None |
| CFTR | None | None | None | None |
| CFTR | None | None | None | None |
| CFTR | None | None | None | None |
| CFTR | None | None | None | None |
| CFTR | None | None | None | None |
| CFTR | None | None | None | None |
| CFTR | None | None | None | None |
| CFTR | None | None | None | None |
| CFTR | None | None | None | None |
| CFTR | None | None | None | None |
| FAN1 | None | None | None | None |
| CBS | None | None | None | None |
| CBS | None | None | None | None |
| CBS | None | None | None | None |
| CBS | None | None | None | None |
| CBS | None | None | None | None |
| CBS | None | None | None | None |
| CBS | None | None | None | None |
| CBS | None | None | None | None |
| CBS | None | None | None | None |
| CBS | None | None | None | None |
| CBS | None | None | None | None |
| CBS | None | None | None | None |
| CBS | None | None | None | None |
| CBS | None | None | None | None |
| CBS | None | None | None | None |
| CBS | None | None | None | None |
| CBS | None | None | None | None |
| CBS | None | None | None | None |
| CBS | None | None | None | None |
| CBS | None | None | None | None |
| CBS | None | None | None | None |
| CBS | None | None | None | None |
| CBS | None | None | None | None |
| CBS | None | None | None | None |
| CBS | None | None | None | None |
| CBS | None | None | None | None |
| CBS | None | None | None | None |
| CBS | None | None | None | None |

| | | | | |
|---|---|---|---|---|
| CBS | None | None | None | None |
| CBS | None | None | None | None |
| CBS | None | None | None | None |
| CBS | None | None | None | None |
| CBS | None | None | None | None |
| CBS | None | None | None | None |
| CBS | None | None | None | None |
| CBS | None | None | None | None |
| TLR3 | None | None | None | None |
| LIG4 | None | None | None | None |
| LIG4 | None | None | None | None |
| ASPA | None | None | None | None |
| ASPA | None | None | None | None |
| ASPA | None | None | None | None |
| ASPA | None | None | None | None |
| ASPA | None | None | None | None |
| ASPA | None | None | None | None |
| ASPA | None | None | None | None |
| ASPA | None | None | None | None |
| ASPA | None | None | None | None |
| ASPA | None | None | None | None |
| ASPA | None | None | None | None |
| ASPA | None | None | None | None |
| ASPA | None | None | None | None |
| CFH | None | None | None | None |
| FCGR3A | None | None | None | None |
| FCGR3A | None | None | None | None |
| GRIN2A | None | None | None | None |
| GRIN2A | None | None | None | None |
| GRIN2A | None | None | None | None |
| GLA | None | None | None | None |
| GLA | None | None | None | None |
| GLA | None | None | None | None |
| GLA | None | None | None | None |
| GLA | None | None | None | None |
| GLA | None | None | None | None |
| GLA | None | None | None | None |
| GLA | None | None | None | None |
| GLA | None | None | None | None |
| GLA | None | None | None | None |
| GLA | None | None | None | None |
| GLA | None | None | None | None |
| GLA | None | None | None | None |
| GLA | None | None | None | None |
| GLA | None | None | None | None |
| GLA | None | None | None | None |
| GLA | None | None | None | None |
| GLA | None | None | None | None |
| GLA | None | None | None | None |
| GLA | None | None | None | None |
| GLA | None | None | None | None |
| GLA | None | None | None | None |
| GLA | None | None | None | None |
| GLA | None | None | None | None |

| | | | | |
|---|---|---|---|---|
| GLA | None | None | None | None |
| GLA | None | None | None | None |
| GLA | None | None | None | None |
| GLA | None | None | None | None |
| GLA | None | None | None | None |
| GLA | None | None | None | None |
| GLA | None | None | None | None |
| SERPINA3 | None | None | None | None |
| MTAP | None | None | None | None |
| MTAP | None | None | None | None |
| MTAP | None | None | None | None |
| BLMH | None | None | None | None |
| TUSC3 | None | None | None | None |
| ISCU | None | None | None | None |
| CYP11B2 | None | None | None | None |
| CYP11B2 | None | None | None | None |
| CYP11B2 | None | None | None | None |
| CYP11B2 | None | None | None | None |
| CYP11B2 | None | None | None | None |
| CYP11B2 | None | None | None | None |
| CYP11B2 | None | None | None | None |
| CYP11B2 | None | None | None | None |
| CYP11B2 | None | None | None | None |
| CYP11B2 | None | None | None | None |
| CYP11B2 | None | None | None | None |
| CYP11B2 | None | None | None | None |
| CYP11B2 | None | None | None | None |
| CYP11B2 | None | None | None | None |
| CYP11B2 | None | None | None | None |
| CYP11B2 | None | None | None | None |
| CYP11B2 | None | None | None | None |
| CYP11B2 | None | None | None | None |
| CYP11B2 | None | None | None | None |
| CYP11B2 | None | None | None | None |
| CRADD | None | None | None | None |
| CRADD | None | None | None | None |
| FGA | None | None | None | None |
| FGA | None | None | None | None |
| FGA | None | None | None | None |
| FGA | None | None | None | None |
| FGA | None | None | None | None |
| FGA | None | None | None | None |
| FGA | None | None | None | None |
| FGA | None | None | None | None |
| IYD | | 1 | 1 4ttb | A |
| IYD | None | None | None | None |
| IYD | None | None | None | None |
| IYD | None | None | None | None |
| TCAP | None | None | None | None |
| TCAP | None | None | None | None |
| TCAP | None | None | None | None |
| TCAP | None | None | None | None |
| TCAP | None | None | None | None |
| CYP7A1 | None | None | None | None |

| | | | | |
|---|---|---|---|---|
| TEK | None | None | None | None |
| TEK | None | None | None | None |
| TEK | None | None | None | None |
| TEK | None | None | None | None |
| TEK | None | None | None | None |
| TEK | None | None | None | None |
| TEK | None | None | None | None |
| CD8A | None | None | None | None |
| HERC2 | None | None | None | None |
| NR1H3 | None | None | None | None |
| GRHPR | None | None | None | None |
| GRHPR | None | None | None | None |
| GRHPR | None | None | None | None |
| GRHPR | None | None | None | None |
| GRHPR | None | None | None | None |
| GRHPR | None | None | None | None |
| GRHPR | None | None | None | None |
| GBE1 | None | None | None | None |
| GBE1 | None | None | None | None |
| GBE1 | None | None | None | None |
| GBE1 | None | None | None | None |
| GBE1 | None | None | None | None |
| GBE1 | None | None | None | None |
| GBE1 | None | None | None | None |
| GBE1 | None | None | None | None |
| GBE1 | None | None | None | None |
| GBE1 | None | None | None | None |
| GBE1 | None | None | None | None |
| GBE1 | None | None | None | None |
| GBE1 | None | None | None | None |
| GBE1 | None | None | None | None |
| GBE1 | None | None | None | None |
| GBE1 | None | None | None | None |
| GBE1 | None | None | None | None |
| GBE1 | None | None | None | None |
| GBE1 | None | None | None | None |
| PC | None | None | None | None |
| PC | None | None | None | None |
| PC | None | None | None | None |
| PC | None | None | None | None |
| PC | None | None | None | None |
| PC | None | None | None | None |
| PC | None | None | None | None |
| PC | None | None | None | None |
| MSH6 | None | None | None | None |
| MSH6 | None | None | None | None |
| MSH6 | None | None | None | None |
| MSH6 | None | None | None | None |
| MSH6 | None | None | None | None |
| MSH6 | None | None | None | None |
| MSH6 | None | None | None | None |
| MSH6 | None | None | None | None |
| MSH6 | None | None | None | None |

| | | | | |
|---|---|---|---|---|
| MSH6 | None | None | None | None |
| MSH6 | None | None | None | None |
| MSH6 | None | None | None | None |
| MSH6 | None | None | None | None |
| MSH6 | None | None | None | None |
| MSH6 | None | None | None | None |
| MSH6 | None | None | None | None |
| MSH6 | None | None | None | None |
| MSH6 | None | None | None | None |
| MSH6 | None | None | None | None |
| MSH6 | None | None | None | None |
| MSH6 | None | None | None | None |
| MSH6 | None | None | None | None |
| MSH6 | None | None | None | None |
| MSH6 | None | None | None | None |
| MSH6 | None | None | None | None |
| MSH6 | None | None | None | None |
| MSH6 | None | None | None | None |
| MSH6 | None | None | None | None |
| MSH6 | None | None | None | None |
| MSH6 | None | None | None | None |
| RNASEL | None | None | None | None |
| ORC1 | None | None | None | None |
| ORC1 | None | None | None | None |
| ORC1 | None | None | None | None |
| KCNH2 | None | None | None | None |
| KCNH2 | None | None | None | None |
| KCNH2 | None | None | None | None |
| KCNH2 | None | None | None | None |
| KCNH2 | None | None | None | None |
| KCNH2 | None | None | None | None |
| KCNH2 | None | None | None | None |
| KCNH2 | None | None | None | None |
| KCNH2 | None | None | None | None |
| KCNH2 | None | None | None | None |
| KCNH2 | None | None | None | None |
| KCNH2 | None | None | None | None |
| KCNH2 | None | None | None | None |
| KCNH2 | None | None | None | None |
| KCNH2 | None | None | None | None |
| KCNH2 | None | None | None | None |
| KCNH2 | None | None | None | None |
| KCNH2 | None | None | None | None |
| KCNH2 | None | None | None | None |
| KCNH2 | None | None | None | None |
| KCNH2 | None | None | None | None |
| KCNH2 | None | None | None | None |
| KCNH2 | None | None | None | None |
| KCNH2 | None | None | None | None |
| MAT1A | None | None | None | None |
| MAT1A | None | None | None | None |
| MAT1A | None | None | None | None |
| MAT1A | None | None | None | None |
| MAT1A | None | None | None | None |

| | | | | |
|---|---|---|---|---|
| MAT1A | None | None | None | None |
| MAT1A | None | None | None | None |
| MAT1A | None | None | None | None |
| INSR | None | None | None | None |
| INSR | None | None | None | None |
| INSR | None | None | None | None |
| INSR | None | None | None | None |
| AMT | None | None | None | None |
| AMT | None | None | None | None |
| AMT | None | None | None | None |
| AMT | None | None | None | None |
| AMT | None | None | None | None |
| AMT | None | None | None | None |
| AMT | None | None | None | None |
| AMT | None | None | None | None |
| AMT | None | None | None | None |
| CSNK1D | None | None | None | None |
| SF3B1 | None | None | None | None |
| AGA | None | None | None | None |
| AGA | None | None | None | None |
| AGA | None | None | None | None |
| AGA | None | None | None | None |
| AGA | None | None | None | None |
| AGA | None | None | None | None |
| AGA | None | None | None | None |
| NECTIN4 | None | None | None | None |
| RS1 | None | None | None | None |
| RS1 | None | None | None | None |
| RS1 | None | None | None | None |
| RS1 | None | None | None | None |
| TBC1D7 | None | None | None | None |
| CTSA | None | None | None | None |
| CTSA | None | None | None | None |
| TKT | None | None | None | None |
| RP2 | None | None | None | None |
| RP2 | None | None | None | None |
| RP2 | None | None | None | None |
| RP2 | None | None | None | None |
| LYRM4 | None | None | None | None |
| SOS1 | None | None | None | None |
| SOS1 | None | None | None | None |
| SOS1 | None | None | None | None |
| SOS1 | None | None | None | None |
| SOS1 | None | None | None | None |
| SOS1 | None | None | None | None |
| HUWE1 | None | None | None | None |
| HUWE1 | None | None | None | None |
| TPI1 | None | None | None | None |
| TPI1 | None | None | None | None |
| TPI1 | None | None | None | None |
| MSH2 | None | None | None | None |
| MSH2 | None | None | None | None |
| MSH2 | None | None | None | None |
| MSH2 | None | None | None | None |

| | | | | |
|---|---|---|---|---|
| MSH2 | None | None | None | None |
| MSH2 | None | None | None | None |
| MSH2 | None | None | None | None |
| MSH2 | None | None | None | None |
| MSH2 | None | None | None | None |
| MSH2 | None | None | None | None |
| MSH2 | None | None | None | None |
| MSH2 | None | None | None | None |
| MSH2 | None | None | None | None |
| MSH2 | None | None | None | None |
| MSH2 | None | None | None | None |
| MSH2 | None | None | None | None |
| MSH2 | None | None | None | None |
| MSH2 | None | None | None | None |
| MSH2 | None | None | None | None |
| MSH2 | None | None | None | None |
| MSH2 | None | None | None | None |
| MSH2 | None | None | None | None |
| MSH2 | None | None | None | None |
| MSH2 | None | None | None | None |
| MSH2 | None | None | None | None |
| MSH2 | None | None | None | None |
| MSH2 | None | None | None | None |
| MSH2 | None | None | None | None |
| MSH2 | None | None | None | None |
| MSH2 | None | None | None | None |
| MSH2 | None | None | None | None |
| MSH2 | None | None | None | None |
| MSH2 | None | None | None | None |
| MSH2 | None | None | None | None |
| MSH2 | None | None | None | None |
| MSH2 | None | None | None | None |
| GALE | None | None | None | None |
| GALE | None | None | None | None |
| GALE | None | None | None | None |
| GALE | None | None | None | None |
| GALE | None | None | None | None |
| GALE | None | None | None | None |
| GALE | None | None | None | None |
| GALE | None | None | None | None |
| NSD2 | None | None | None | None |
| SRP72 | None | None | None | None |
| SRP72 | None | None | None | None |
| SRP72 | None | None | None | None |
| POLK | None | None | None | None |
| GALT | None | None | None | None |
| GALT | None | None | None | None |
| GALT | None | None | None | None |
| GALT | None | None | None | None |
| GALT | None | None | None | None |
| GALT | None | None | None | None |
| GALT | None | None | None | None |
| GALT | None | None | None | None |
| GALT | None | None | None | None |

| | | | | |
|---|---|---|---|---|
| GALT | None | None | None | None |
| GALT | None | None | None | None |
| GALT | None | None | None | None |
| GALT | None | None | None | None |
| GALT | None | None | None | None |
| GALT | None | None | None | None |
| GALT | None | None | None | None |
| GALT | None | None | None | None |
| GALT | None | None | None | None |
| GALT | None | None | None | None |
| GALT | None | None | None | None |
| GALT | None | None | None | None |
| GALT | None | None | None | None |
| GALT | None | None | None | None |
| GALT | None | None | None | None |
| GALT | None | None | None | None |
| GALT | None | None | None | None |
| GALT | None | None | None | None |
| GALT | None | None | None | None |
| GALT | None | None | None | None |
| GALT | None | None | None | None |
| GALT | None | None | None | None |
| GALT | None | None | None | None |
| GALT | None | None | None | None |
| GALT | None | None | None | None |
| GALT | None | None | None | None |
| GALT | None | None | None | None |
| GALT | None | None | None | None |
| PYCR1 | None | None | None | None |
| PYCR1 | None | None | None | None |
| PYCR1 | None | None | None | None |
| PYCR1 | None | None | None | None |
| F8 | None | None | None | None |
| F8 | None | None | None | None |
| F8 | None | None | None | None |
| F8 | None | None | None | None |
| F8 | None | None | None | None |
| F8 | None | None | None | None |
| F8 | None | None | None | None |
| F8 | None | None | None | None |
| F8 | None | None | None | None |
| F8 | None | None | None | None |
| F8 | None | None | None | None |
| F8 | None | None | None | None |
| F8 | None | None | None | None |
| F8 | None | None | None | None |
| F8 | None | None | None | None |
| F8 | None | None | None | None |
| ANK1 | None | None | None | None |
| CHAT | None | None | None | None |
| CHAT | None | None | None | None |
| CHAT | None | None | None | None |
| CHAT | None | None | None | None |
| CHAT | None | None | None | None |

| | | | | |
|---|---|---|---|---|
| CHAT | None | None | None | None |
| CHAT | None | None | None | None |
| CHAT | None | None | None | None |
| CHAT | None | None | None | None |
| CHAT | None | None | None | None |
| CHAT | None | None | None | None |
| CHAT | None | None | None | None |
| CHAT | None | None | None | None |
| CHAT | None | None | None | None |
| CHAT | None | None | None | None |
| CHAT | None | None | None | None |
| CHAT | None | None | None | None |
| GHR | None | None | None | None |
| GHR | None | None | None | None |
| GHR | None | None | None | None |
| GHR | None | None | None | None |
| GHR | None | None | None | None |
| GHR | None | None | None | None |
| GHR | None | None | None | None |
| GHR | None | None | None | None |
| GHR | None | None | None | None |
| GHR | None | None | None | None |
| GHR | None | None | None | None |
| GHR | None | None | None | None |
| GHR | None | None | None | None |
| GHR | None | None | None | None |
| FZD4 | None | None | None | None |
| FZD4 | None | None | None | None |
| FZD4 | None | None | None | None |
| MTR | None | None | None | None |
| MTR | None | None | None | None |
| MTR | None | None | None | None |
| GM2A | None | None | None | None |
| GM2A | None | None | None | None |
| GM2A | None | None | None | None |
| GM2A | None | None | None | None |
| GM2A | None | None | None | None |
| CRYGD | None | None | None | None |
| CRYGD | None | None | None | None |
| CRYGD | None | None | None | None |
| CRYGD | None | None | None | None |
| CRYGD | None | None | None | None |
| CRYGD | None | None | None | None |
| CRYGD | None | None | None | None |
| CRYGD | None | None | None | None |
| CRYGD | None | None | None | None |
| TLR5 | None | None | None | None |
| HSD17B10 | None | None | None | None |
| HSD17B10 | None | None | None | None |
| HSD17B10 | None | None | None | None |
| HSD17B10 | None | None | None | None |
| TBC1D20 | None | None | None | None |
| FBN1 | None | None | None | None |
| FBN1 | None | None | None | None |

| | | | | |
|---|---|---|---|---|
| FBN1 | None | None | None | None |
| FBN1 | None | None | None | None |
| FBN1 | None | None | None | None |
| FBN1 | None | None | None | None |
| FBN1 | None | None | None | None |
| FXN | None | None | None | None |
| FXN | None | None | None | None |
| FXN | None | None | None | None |
| FXN | None | None | None | None |
| MYO5A | None | None | None | None |
| CTSK | None | None | None | None |
| CTSK | None | None | None | None |
| SCN5A | None | None | None | None |
| SCN5A | None | None | None | None |
| SOS1 | None | None | None | None |
| SOS1 | None | None | None | None |
| SOS1 | None | None | None | None |
| SOS1 | None | None | None | None |
| SOS1 | None | None | None | None |
| SOS1 | None | None | None | None |
| SOS1 | None | None | None | None |
| SOS1 | None | None | None | None |
| SOS1 | None | None | None | None |
| SOS1 | None | None | None | None |
| SOS1 | None | None | None | None |
| SOS1 | None | None | None | None |
| GATM | None | None | None | None |
| GATM | None | None | None | None |
| GATM | None | None | None | None |
| GATM | None | None | None | None |
| GATM | None | None | None | None |
| GATM | None | None | None | None |
| GATM | None | None | None | None |
| MMP2 | None | None | None | None |
| MMP2 | None | None | None | None |
| MMP2 | None | None | None | None |
| MMP2 | None | None | None | None |
| NDUFB9 | None | None | None | None |
| NDUFB9 | None | None | None | None |
| GH1 | None | None | None | None |
| GH1 | None | None | None | None |
| GH1 | None | None | None | None |
| SLC6A4 | None | None | None | None |
| G6PD | None | None | None | None |
| G6PD | None | None | None | None |
| G6PD | None | None | None | None |
| G6PD | None | None | None | None |
| G6PD | None | None | None | None |
| G6PD | None | None | None | None |
| G6PD | None | None | None | None |
| G6PD | None | None | None | None |
| G6PD | None | None | None | None |
| G6PD | None | None | None | None |
| G6PD | None | None | None | None |

| | | | | |
|---|---|---|---|---|
| G6PD | None | None | None | None |
| G6PD | None | None | None | None |
| G6PD | None | None | None | None |
| G6PD | None | None | None | None |
| G6PD | None | None | None | None |
| G6PD | None | None | None | None |
| G6PD | None | None | None | None |
| G6PD | None | None | None | None |
| G6PD | None | None | None | None |
| CST3 | None | None | None | None |
| LMNA | None | None | None | None |
| LMNA | None | None | None | None |
| NDUFA9 | None | None | None | None |
| NDUFA9 | None | None | None | None |
| NDUFA9 | None | None | None | None |
| NDUFA9 | None | None | None | None |
| YARS | None | None | None | None |
| YARS | None | None | None | None |
| YARS | None | None | None | None |
| YARS | None | None | None | None |
| EPHA2 | None | None | None | None |
| EPHA2 | None | None | None | None |
| EPHA2 | None | None | None | None |
| EPHA2 | None | None | None | None |
| DICER1 | None | None | None | None |
| DICER1 | None | None | None | None |
| SNX10 | None | None | None | None |
| SNX10 | None | None | None | None |
| TTN | None | None | None | None |
| TTN | None | None | None | None |
| GATA3 | None | None | None | None |
| C5 | None | None | None | None |
| THBD | None | None | None | None |
| BIN1 | None | None | None | None |
| BIN1 | None | None | None | None |
| SMARCAL1 | None | None | None | None |
| CFB | None | None | None | None |
| CYC1 | None | None | None | None |
| ANK2 | None | None | None | None |
| CHMP4B | None | None | None | None |
| CHMP4B | None | None | None | None |
| ERBB2 | None | None | None | None |
| ERBB2 | None | None | None | None |
| POLG2 | None | None | None | None |
| POLG2 | None | None | None | None |
| POLG2 | None | None | None | None |
| POLG2 | None | None | None | None |
| POLG2 | None | None | None | None |
| POLG2 | None | None | None | None |
| POLG2 | None | None | None | None |
| POLG2 | None | None | None | None |
| C3 | None | None | None | None |
| C3 | None | None | None | None |
| RPGR | None | None | None | None |

| | | | | |
|---|---|---|---|---|
| RPGR | None | None | None | None |
| RPGR | None | None | None | None |
| RPGR | None | None | None | None |
| RPGR | None | None | None | None |
| C1QA | None | None | None | None |
| PRKG1 | None | None | None | None |
| IGBP1 | None | None | None | None |
| ENO3 | None | None | None | None |
| ENO3 | None | None | None | None |
| ENO3 | None | None | None | None |
| ENO3 | None | None | None | None |
| ENO3 | None | None | None | None |
| ENO3 | None | None | None | None |
| MCPH1 | None | None | None | None |
| MCPH1 | None | None | None | None |
| MCPH1 | None | None | None | None |
| MCPH1 | None | None | None | None |
| MCPH1 | None | None | None | None |
| SNRNP200 | None | None | None | None |
| SNRNP200 | None | None | None | None |
| DNMT1 | None | None | None | None |
| DNMT1 | None | None | None | None |
| DNMT1 | None | None | None | None |
| AKT1 | None | None | None | None |
| NDUFS2 | None | None | None | None |
| ACVR2B | None | None | None | None |
| HMGCL | None | None | None | None |
| HMGCL | None | None | None | None |
| HMGCL | None | None | None | None |
| JUP | None | None | None | None |
| JUP | None | None | None | None |
| JUP | None | None | None | None |
| JUP | None | None | None | None |
| JUP | None | None | None | None |
| JUP | None | None | None | None |
| JUP | None | None | None | None |
| JUP | None | None | None | None |
| SPINK1 | None | None | None | None |
| SPINK1 | None | None | None | None |
| SPINK1 | None | None | None | None |
| VCL | None | None | None | None |
| VCL | None | None | None | None |
| NOTCH1 | None | None | None | None |
| NOTCH1 | None | None | None | None |
| NOTCH1 | None | None | None | None |
| NDUFB3 | None | None | None | None |
| IKBKG | None | None | None | None |
| GSN | None | None | None | None |
| GSN | None | None | None | None |
| GSN | None | None | None | None |
| GSN | None | None | None | None |
| GSN | None | None | None | None |
| GSN | None | None | None | None |
| GSN | None | None | None | None |

| | | | | |
|---|---|---|---|---|
| GSN | None | None | None | None |
| GSN | None | None | None | None |
| CLC | None | None | None | None |
| RAF1 | None | None | None | None |
| RAF1 | None | None | None | None |
| RAF1 | None | None | None | None |
| NDUFS8 | None | None | None | None |
| NDUFS8 | None | None | None | None |
| NDUFS8 | None | None | None | None |
| NDUFS8 | None | None | None | None |
| NDUFS8 | None | None | None | None |
| CASR | None | None | None | None |
| CASR | None | None | None | None |
| CASR | None | None | None | None |
| CASR | None | None | None | None |
| CASR | None | None | None | None |
| CASR | None | None | None | None |
| CASR | None | None | None | None |
| CASR | None | None | None | None |
| CASR | None | None | None | None |
| CASR | None | None | None | None |
| CASR | None | None | None | None |
| CASR | None | None | None | None |
| CASR | None | None | None | None |
| CASR | None | None | None | None |
| CASR | None | None | None | None |
| CASR | None | None | None | None |
| CASR | None | None | None | None |
| CASR | None | None | None | None |
| CASR | None | None | None | None |
| CASR | None | None | None | None |
| CASR | None | None | None | None |
| CASR | None | None | None | None |
| CASR | None | None | None | None |
| CASR | None | None | None | None |
| CASR | None | None | None | None |
| CASR | None | None | None | None |
| CASR | None | None | None | None |
| CASR | None | None | None | None |
| CASR | None | None | None | None |
| VCL | None | None | None | None |
| VCL | None | None | None | None |
| SMPD1 | None | None | None | None |
| SMPD1 | None | None | None | None |
| SMPD1 | None | None | None | None |
| KDM6A | None | None | None | None |
| KCNQ1 | None | None | None | None |
| KCNQ1 | None | None | None | None |
| KCNQ1 | None | None | None | None |
| KCNQ1 | None | None | None | None |
| KCNQ1 | None | None | None | None |
| KCNQ1 | None | None | None | None |
| NDUFA12 | None | None | None | None |
| GLRX5 | None | None | None | None |

| | | | | |
|---|---|---|---|---|
| PALB2 | None | None | None | None |
| PALB2 | None | None | None | None |
| PALB2 | None | None | None | None |
| PALB2 | None | None | None | None |
| PALB2 | None | None | None | None |
| PALB2 | None | None | None | None |
| PALB2 | None | None | None | None |
| PALB2 | None | None | None | None |
| PALB2 | None | None | None | None |
| PALB2 | None | None | None | None |
| ALDOB | None | None | None | None |
| ALDOB | None | None | None | None |
| ALDOB | None | None | None | None |
| ALDOB | None | None | None | None |
| ALDOB | None | None | None | None |
| ALDOB | None | None | None | None |
| ALDOB | None | None | None | None |
| GNE | None | None | None | None |
| GNE | None | None | None | None |
| GNE | None | None | None | None |
| GNE | None | None | None | None |
| GNE | None | None | None | None |
| GNE | None | None | None | None |
| GNE | None | None | None | None |
| GNE | None | None | None | None |
| GNE | None | None | None | None |
| GNE | None | None | None | None |
| GNE | None | None | None | None |
| GNE | None | None | None | None |
| CHRNB2 | None | None | None | None |
| CHRNB2 | None | None | None | None |
| ERCC3 | None | None | None | None |
| ERCC3 | None | None | None | None |
| ERCC3 | None | None | None | None |
| ERCC3 | None | None | None | None |
| ERCC3 | None | None | None | None |
| UNC119 | None | None | None | None |
| ATL1 | None | None | None | None |
| ATL1 | None | None | None | None |
| ATL1 | None | None | None | None |
| ATL1 | None | None | None | None |
| ATL1 | None | None | None | None |
| ATL1 | None | None | None | None |
| ATL1 | None | None | None | None |
| ATL1 | None | None | None | None |
| ATL1 | None | None | None | None |
| ATL1 | None | None | None | None |
| NR3C2 | None | None | None | None |
| TBK1 | None | None | None | None |
| HYAL1 | None | None | None | None |
| HYAL1 | None | None | None | None |
| HYAL1 | None | None | None | None |
| HYAL1 | None | None | None | None |
| CTSD | None | None | None | None |

| | | | | |
|---|---|---|---|---|
| PRKCG | None | None | None | None |
| TGFBR1 | None | None | None | None |
| XDH | None | None | None | None |
| XDH | None | None | None | None |
| XDH | None | None | None | None |
| XDH | None | None | None | None |
| XDH | None | None | None | None |
| XDH | None | None | None | None |
| NMNAT1 | None | None | None | None |
| NMNAT1 | None | None | None | None |
| NMNAT1 | None | None | None | None |
| NMNAT1 | None | None | None | None |
| NMNAT1 | None | None | None | None |
| NMNAT1 | None | None | None | None |
| SERPING1 | None | None | None | None |
| SERPING1 | None | None | None | None |
| SERPING1 | None | None | None | None |
| SERPING1 | None | None | None | None |
| SERPING1 | None | None | None | None |
| PDZRN3 | None | None | None | None |
| FABP2 | None | None | None | None |
| IFNGR1 | None | None | None | None |
| IFNGR1 | None | None | None | None |
| TDP1 | None | None | None | None |
| TDP1 | None | None | None | None |
| INS | None | None | None | None |
| INS | None | None | None | None |
| INS | None | None | None | None |
| PPOX | None | None | None | None |
| PPOX | None | None | None | None |
| PPOX | None | None | None | None |
| PPOX | None | None | None | None |
| PPOX | None | None | None | None |
| PPOX | None | None | None | None |
| PPOX | None | None | None | None |
| PPOX | None | None | None | None |
| PPOX | None | None | None | None |
| GFER | None | None | None | None |
| GFER | None | None | None | None |
| GFER | None | None | None | None |
| GFER | None | None | None | None |
| LIFR | None | None | None | None |
| EGLN1 | None | None | None | None |
| EGLN1 | None | None | None | None |
| EGLN1 | None | None | None | None |
| GP1BA | None | None | None | None |
| GP1BA | None | None | None | None |
| GP1BA | None | None | None | None |
| GP1BA | None | None | None | None |
| GP1BA | None | None | None | None |
| MMADHC | None | None | None | None |
| MMADHC | None | None | None | None |
| MMADHC | None | None | None | None |
| MMADHC | None | None | None | None |

| | | | | |
|---|---|---|---|---|
| MMADHC | None | None | None | None |
| MMADHC | None | None | None | None |
| MMADHC | None | None | None | None |
| ACVRL1 | None | None | None | None |
| ACVRL1 | None | None | None | None |
| RFXANK | None | None | None | None |
| CHIT1 | None | None | None | None |
| UBR1 | None | None | None | None |
| BLM | None | None | None | None |
| BLM | None | None | None | None |
| BLM | None | None | None | None |
| BLM | None | None | None | None |
| BLM | None | None | None | None |
| BLM | None | None | None | None |
| BLM | None | None | None | None |
| BLM | None | None | None | None |
| BLM | None | None | None | None |
| BLM | None | None | None | None |
| BLM | None | None | None | None |
| BLM | None | None | None | None |
| BLM | None | None | None | None |
| VDR | None | None | None | None |
| VDR | None | None | None | None |
| VDR | None | None | None | None |
| VDR | None | None | None | None |
| VDR | None | None | None | None |
| VDR | None | None | None | None |
| EPHA2 | None | None | None | None |
| EPHA2 | None | None | None | None |
| EPHA2 | None | None | None | None |
| EPHA2 | None | None | None | None |
| EPHA2 | None | None | None | None |
| EPHA2 | None | None | None | None |
| EPHA2 | None | None | None | None |
| EPHA2 | None | None | None | None |
| EPHA2 | None | None | None | None |
| EPHA2 | None | None | None | None |
| UBA1 | None | None | None | None |
| STAT1 | None | None | None | None |
| STAT1 | None | None | None | None |
| STAT1 | None | None | None | None |
| STAT1 | None | None | None | None |
| STAT1 | None | None | None | None |
| STAT1 | None | None | None | None |
| STAT1 | None | None | None | None |
| STAT1 | None | None | None | None |
| DNM1L | None | None | None | None |
| DNM1L | None | None | None | None |
| OTC | None | None | None | None |
| OTC | None | None | None | None |
| OTC | None | None | None | None |
| OTC | None | None | None | None |
| OTC | None | None | None | None |
| OTC | None | None | None | None |

| | | | | |
|---|---|---|---|---|
| OTC | None | None | None | None |
| OTC | None | None | None | None |
| OTC | None | None | None | None |
| OTC | None | None | None | None |
| OTC | None | None | None | None |
| OTC | None | None | None | None |
| OTC | None | None | None | None |
| OTC | None | None | None | None |
| OTC | None | None | None | None |
| OTC | None | None | None | None |
| OTC | None | None | None | None |
| RARS | None | None | None | None |
| RARS | None | None | None | None |
| RARS | None | None | None | None |
| SEPSECS | None | None | None | None |
| SEPSECS | None | None | None | None |
| SEPSECS | None | None | None | None |
| SEPSECS | None | None | None | None |
| SEPSECS | None | None | None | None |
| BRCA1 | None | None | None | None |
| BRCA1 | None | None | None | None |
| BRCA1 | None | None | None | None |
| BRCA1 | None | None | None | None |
| BRCA1 | None | None | None | None |
| BRCA1 | None | None | None | None |
| BRCA1 | None | None | None | None |
| BRCA1 | None | None | None | None |
| BRCA1 | None | None | None | None |
| BRCA1 | None | None | None | None |
| BRCA1 | None | None | None | None |
| BRCA1 | None | None | None | None |
| BRCA1 | None | None | None | None |
| BRCA1 | None | None | None | None |
| BRCA1 | None | None | None | None |
| BRCA1 | None | None | None | None |
| BRCA1 | None | None | None | None |
| BRCA1 | None | None | None | None |
| BRCA1 | None | None | None | None |
| BRCA1 | None | None | None | None |
| BRCA1 | None | None | None | None |
| BRCA1 | None | None | None | None |
| CUBN | None | None | None | None |
| ITGB3 | None | None | None | None |
| MTRR | None | None | None | None |
| MTRR | None | None | None | None |
| MTRR | None | None | None | None |
| MTRR | None | None | None | None |
| MTRR | None | None | None | None |
| MTRR | None | None | None | None |
| MTRR | None | None | None | None |
| MTRR | None | None | None | None |
| IL7R | None | None | None | None |
| IL7R | None | None | None | None |
| RPS24 | None | None | None | None |

| | | | | |
|---|---|---|---|---|
| RPS24 | None | None | None | None |
| HOGA1 | None | None | None | None |
| HOGA1 | None | None | None | None |
| HOGA1 | None | None | None | None |
| HOGA1 | None | None | None | None |
| HOGA1 | None | None | None | None |
| HOGA1 | None | None | None | None |
| HOGA1 | None | None | None | None |
| HOGA1 | None | None | None | None |
| HOGA1 | None | None | None | None |
| HOGA1 | None | None | None | None |
| HOGA1 | None | None | None | None |
| HOGA1 | None | None | None | None |
| HOGA1 | None | None | None | None |
| GJB2 | None | None | None | None |
| GJB2 | None | None | None | None |
| GJB2 | None | None | None | None |
| GJB2 | None | None | None | None |
| GJB2 | None | None | None | None |
| GJB2 | None | None | None | None |
| GJB2 | None | None | None | None |
| GJB2 | None | None | None | None |
| GJB2 | None | None | None | None |
| GJB2 | None | None | None | None |
| GJB2 | None | None | None | None |
| GJB2 | None | None | None | None |
| GJB2 | None | None | None | None |
| GJB2 | None | None | None | None |
| GJB2 | None | None | None | None |
| GJB2 | None | None | None | None |
| GJB2 | None | None | None | None |
| GJB2 | None | None | None | None |
| GJB2 | None | None | None | None |
| GJB2 | None | None | None | None |
| GJB2 | None | None | None | None |
| GJB2 | None | None | None | None |
| GJB2 | None | None | None | None |
| GJB2 | None | None | None | None |
| GJB2 | None | None | None | None |
| GJB2 | None | None | None | None |
| GJB2 | None | None | None | None |
| GJB2 | None | None | None | None |
| GJB2 | None | None | None | None |
| GJB2 | None | None | None | None |
| GJB2 | None | None | None | None |
| GJB2 | None | None | None | None |
| GJB2 | None | None | None | None |
| GJB2 | None | None | None | None |
| GJB2 | None | None | None | None |
| GJB2 | None | None | None | None |
| GJB2 | None | None | None | None |
| GJB2 | None | None | None | None |
| GJB2 | None | None | None | None |
| GJB2 | None | None | None | None |

| | | | | |
|---|---|---|---|---|
| GJB2 | None | None | None | None |
| GJB2 | None | None | None | None |
| GJB2 | None | None | None | None |
| GJB2 | None | None | None | None |
| GJB2 | None | None | None | None |
| GJB2 | None | None | None | None |
| GJB2 | None | None | None | None |
| GJB2 | None | None | None | None |
| GJB2 | None | None | None | None |
| GJB2 | None | None | None | None |
| GJB2 | None | None | None | None |
| GJB2 | None | None | None | None |
| GJB2 | None | None | None | None |
| GJB2 | None | None | None | None |
| CPS1 | None | None | None | None |
| CPS1 | None | None | None | None |
| CPS1 | None | None | None | None |
| CPS1 | None | None | None | None |
| CPS1 | None | None | None | None |
| CPS1 | None | None | None | None |
| CPS1 | None | None | None | None |
| CPS1 | None | None | None | None |
| CPS1 | None | None | None | None |
| CPS1 | None | None | None | None |
| PSPH | None | None | None | None |
| PSPH | None | None | None | None |
| PSPH | None | None | None | None |
| PSPH | None | None | None | None |
| PSPH | None | None | None | None |
| PSPH | None | None | None | None |
| PEPD | None | None | None | None |
| PEPD | None | None | None | None |
| PEPD | None | None | None | None |
| PEPD | None | None | None | None |
| PEPD | None | None | None | None |
| PEPD | None | None | None | None |
| PEPD | None | None | None | None |
| PEPD | None | None | None | None |
| PEPD | None | None | None | None |
| KDM5C | None | None | None | None |
| KDM5C | None | None | None | None |
| KDM5C | None | None | None | None |
| ITGB6 | None | None | None | None |
| ITGB6 | None | None | None | None |
| ITGB6 | None | None | None | None |
| DARS | None | None | None | None |
| DARS | None | None | None | None |
| DARS | None | None | None | None |
| DARS | None | None | None | None |
| BTK | None | None | None | None |
| BTK | None | None | None | None |
| BTK | None | None | None | None |
| HDAC4 | None | None | None | None |
| ITGA2B | None | None | None | None |

| | | | | |
|---|---|---|---|---|
| ITGA2B | None | None | None | None |
| ITGA2B | None | None | None | None |
| ITGA2B | None | None | None | None |
| PCSK9 | None | None | None | None |
| PCSK9 | None | None | None | None |
| PCSK9 | None | None | None | None |
| PCSK9 | None | None | None | None |
| MUTYH | None | None | None | None |
| MUTYH | None | None | None | None |
| MUTYH | None | None | None | None |
| MUTYH | None | None | None | None |
| MUTYH | None | None | None | None |
| MUTYH | None | None | None | None |
| MUTYH | None | None | None | None |
| MUTYH | None | None | None | None |
| MUTYH | None | None | None | None |
| MUTYH | None | None | None | None |
| MUTYH | None | None | None | None |
| MUTYH | None | None | None | None |
| MUTYH | None | None | None | None |
| MUTYH | None | None | None | None |
| MUTYH | None | None | None | None |
| MUTYH | None | None | None | None |
| MUTYH | None | None | None | None |
| MUTYH | None | None | None | None |
| MUTYH | None | None | None | None |
| MUTYH | None | None | None | None |
| MUTYH | None | None | None | None |
| MUTYH | None | None | None | None |
| MUTYH | None | None | None | None |
| MUTYH | None | None | None | None |
| SLC25A12 | None | None | None | None |
| GSN | None | None | None | None |
| IHH | None | None | None | None |
| IHH | None | None | None | None |
| IHH | None | None | None | None |
| IHH | None | None | None | None |
| IHH | None | None | None | None |
| IHH | None | None | None | None |
| IHH | None | None | None | None |
| CD247 | None | None | None | None |
| EP300 | None | None | None | None |
| TTN | None | None | None | None |
| MRI1 | None | None | None | None |
| SGSH | None | None | None | None |
| SGSH | None | None | None | None |
| SGSH | None | None | None | None |
| SGSH | None | None | None | None |
| SGSH | None | None | None | None |
| SGSH | None | None | None | None |
| SGSH | None | None | None | None |
| SGSH | None | None | None | None |
| SGSH | None | None | None | None |
| SGSH | None | None | None | None |

| | | | | |
|---|---|---|---|---|
| SGSH | None | None | None | None |
| SGSH | None | None | None | None |
| SGSH | None | None | None | None |
| SGSH | None | None | None | None |
| SGSH | None | None | None | None |
| SGSH | None | None | None | None |
| SGSH | None | None | None | None |
| SGSH | None | None | None | None |
| SGSH | None | None | None | None |
| SGSH | None | None | None | None |
| SGSH | None | None | None | None |
| SGSH | None | None | None | None |
| CBL | None | None | None | None |
| CANT1 | None | None | None | None |
| CANT1 | None | None | None | None |
| CANT1 | None | None | None | None |
| CANT1 | None | None | None | None |
| CANT1 | None | None | None | None |
| CANT1 | None | None | None | None |
| CANT1 | None | None | None | None |
| SLC1A3 | None | None | None | None |
| SLC1A3 | None | None | None | None |
| SLC1A5 | None | None | None | None |
| SLC1A3 | None | None | None | None |
| SLC1A5 | None | None | None | None |
| SLC1A3 | None | None | None | None |
| SLC1A3 | None | None | None | None |
| CTSD | None | None | None | None |
| CTSD | None | None | None | None |
| CTSD | None | None | None | None |
| CTSD | None | None | None | None |
| CTSD | None | None | None | None |
| MTOR | None | None | None | None |
| ILK | None | None | None | None |
| PCDH15 | None | None | None | None |
| PCDH15 | None | None | None | None |
| PCDH15 | None | None | None | None |
| PCDH15 | None | None | None | None |
| PCDH15 | None | None | None | None |
| PCDH15 | None | None | None | None |
| KLHL40 | None | None | None | None |
| KLHL40 | None | None | None | None |
| KLHL40 | None | None | None | None |
| KLHL40 | None | None | None | None |
| CSTA | None | None | None | None |
| CSTA | None | None | None | None |
| HNMT | None | None | None | None |
| HNMT | None | None | None | None |
| IFNGR2 | None | None | None | None |
| IFNGR2 | None | None | None | None |
| PEX5 | None | None | None | None |
| KIF7 | None | None | None | None |
| KIF7 | None | None | None | None |
| MYL3 | None | None | None | None |

| | | | | |
|---|---|---|---|---|
| MYL3 | None | None | None | None |
| MYL3 | None | None | None | None |
| MYL3 | None | None | None | None |
| MYL3 | None | None | None | None |
| MYL3 | None | None | None | None |
| MYL3 | None | None | None | None |
| NAGLU | None | None | None | None |
| NAGLU | None | None | None | None |
| NAGLU | None | None | None | None |
| NAGLU | None | None | None | None |
| NAGLU | None | None | None | None |
| NAGLU | None | None | None | None |
| NAGLU | None | None | None | None |
| NAGLU | None | None | None | None |
| NAGLU | None | None | None | None |
| NAGLU | None | None | None | None |
| NAGLU | None | None | None | None |
| NAGLU | None | None | None | None |
| NAGLU | None | None | None | None |
| TNFRSF13C | None | None | None | None |
| KIF21A | None | None | None | None |
| KIF21A | None | None | None | None |
| KIF21A | None | None | None | None |
| KIF21A | None | None | None | None |
| KIF21A | None | None | None | None |
| DOCK2 | None | None | None | None |
| DSP | None | None | None | None |
| CDKN2A | None | None | None | None |
| CDKN2A | None | None | None | None |
| CDKN2A | None | None | None | None |
| CDKN2A | None | None | None | None |
| CDKN2A | None | None | None | None |
| CDKN2A | None | None | None | None |
| CDKN2A | None | None | None | None |
| CDKN2A | None | None | None | None |
| CDKN2A | None | None | None | None |
| CDKN2A | None | None | None | None |
| CDKN2A | None | None | None | None |
| CDKN2A | None | None | None | None |
| CDKN2A | None | None | None | None |
| CDKN2A | None | None | None | None |
| VCL | None | None | None | None |
| TPP1 | None | None | None | None |
| TPP1 | None | None | None | None |
| TPP1 | None | None | None | None |
| TPP1 | None | None | None | None |
| TPP1 | None | None | None | None |
| TPP1 | None | None | None | None |
| TPP1 | None | None | None | None |
| TPP1 | None | None | None | None |
| TPP1 | None | None | None | None |
| TPP1 | None | None | None | None |
| TPP1 | None | None | None | None |
| TPP1 | None | None | None | None |

| | | | | |
|---|---|---|---|---|
| TPP1 | None | None | None | None |
| TPP1 | None | None | None | None |
| TPP1 | None | None | None | None |
| TPP1 | None | None | None | None |
| TPP1 | None | None | None | None |
| TPP1 | None | None | None | None |
| TPP1 | None | None | None | None |
| TPP1 | None | None | None | None |
| TPP1 | None | None | None | None |
| TPP1 | None | None | None | None |
| IL10RB | None | None | None | None |
| IL10RB | None | None | None | None |
| CCM2 | None | None | None | None |
| CCM2 | None | None | None | None |
| CCM2 | None | None | None | None |
| DCX | None | None | None | None |
| DCX | None | None | None | None |
| DCX | None | None | None | None |
| DCX | None | None | None | None |
| DCX | None | None | None | None |
| DCX | None | None | None | None |
| DCX | None | None | None | None |
| DCX | None | None | None | None |
| BRCA2 | None | None | None | None |
| NNT | None | None | None | None |
| C2 | None | None | None | None |
| C2 | None | None | None | None |
| C2 | None | None | None | None |
| C2 | None | None | None | None |
| WRN | None | None | None | None |
| WRN | None | None | None | None |
| F13A1 | None | None | None | None |
| F13A1 | None | None | None | None |
| F13A1 | None | None | None | None |
| F13A1 | None | None | None | None |
| TF | None | None | None | None |
| TF | None | None | None | None |
| TF | None | None | None | None |
| TF | None | None | None | None |
| TF | None | None | None | None |
| ITPA | None | None | None | None |
| ITPA | None | None | None | None |
| SNCA | None | None | None | None |
| ITGB4 | None | None | None | None |
| ITGB4 | None | None | None | None |
| ITGB4 | None | None | None | None |
| LRP6 | None | None | None | None |
| LRP6 | None | None | None | None |
| LRP6 | None | None | None | None |
| PLG | None | None | None | None |
| PLG | None | None | None | None |
| TYROBP | None | None | None | None |
| UQCRC2 | None | None | None | None |
| GPSM2 | None | None | None | None |

| | | | | |
|---|---|---|---|---|
| GPSM2 | None | None | None | None |
| GPSM2 | None | None | None | None |
| SCN5A | None | None | None | None |
| SCN5A | None | None | None | None |
| SCN5A | None | None | None | None |
| SCN5A | None | None | None | None |
| SCN5A | None | None | None | None |
| ATIC | None | None | None | None |
| EZH2 | None | None | None | None |
| CPA1 | None | None | None | None |
| SCN4B | None | None | None | None |
| SCN4B | None | None | None | None |
| MAN1B1 | None | None | None | None |
| MAN1B1 | None | None | None | None |
| MAN1B1 | None | None | None | None |
| MAN1B1 | None | None | None | None |
| CYCS | None | None | None | None |
| IDUA | None | None | None | None |
| IDUA | None | None | None | None |
| IDUA | None | None | None | None |
| IDUA | None | None | None | None |
| IDUA | None | None | None | None |
| IDUA | None | None | None | None |
| IDUA | None | None | None | None |
| IDUA | None | None | None | None |
| IDUA | None | None | None | None |
| IDUA | None | None | None | None |
| IDUA | None | None | None | None |
| IDUA | None | None | None | None |
| IDUA | None | None | None | None |
| IDUA | None | None | None | None |
| IDUA | None | None | None | None |
| IDUA | None | None | None | None |
| IDUA | None | None | None | None |
| IDUA | None | None | None | None |
| IDUA | None | None | None | None |
| IDUA | None | None | None | None |
| IDUA | None | None | None | None |
| IDUA | None | None | None | None |
| IDUA | None | None | None | None |
| IDUA | None | None | None | None |
| IDUA | None | None | None | None |
| HIKESHI | None | None | None | None |
| APC | None | None | None | None |
| APC | None | None | None | None |
| APC | None | None | None | None |
| APP | None | None | None | None |
| CEP152 | None | None | None | None |
| GAA | None | None | None | None |
| GAA | None | None | None | None |
| GAA | None | None | None | None |
| GAA | None | None | None | None |
| GAA | None | None | None | None |
| GAA | None | None | None | None |

| | | | | |
|---|---|---|---|---|
| GAA | None | None | None | None |
| GAA | None | None | None | None |
| GAA | None | None | None | None |
| GAA | None | None | None | None |
| GAA | None | None | None | None |
| GAA | None | None | None | None |
| GAA | None | None | None | None |
| GAA | None | None | None | None |
| GAA | None | None | None | None |
| GAA | None | None | None | None |
| GAA | None | None | None | None |
| GAA | None | None | None | None |
| GAA | None | None | None | None |
| GAA | None | None | None | None |
| GAA | None | None | None | None |
| GAA | None | None | None | None |
| GAA | None | None | None | None |
| GAA | None | None | None | None |
| GAA | None | None | None | None |
| GAA | None | None | None | None |
| GAA | None | None | None | None |
| GAA | None | None | None | None |
| GAA | None | None | None | None |
| GAA | None | None | None | None |
| GAA | None | None | None | None |
| GAA | None | None | None | None |
| GAA | None | None | None | None |
| GAA | None | None | None | None |
| GAA | None | None | None | None |
| GAA | None | None | None | None |
| GAA | None | None | None | None |
| GAA | None | None | None | None |
| GAA | None | None | None | None |
| GAA | None | None | None | None |
| GAA | None | None | None | None |
| GAA | None | None | None | None |
| GAA | None | None | None | None |
| GAA | None | None | None | None |
| GAA | None | None | None | None |
| GAA | None | None | None | None |
| GAA | None | None | None | None |
| GAA | None | None | None | None |
| GAA | None | None | None | None |
| FOLR1 | None | None | None | None |
| DSP | None | None | None | None |
| DSP | None | None | None | None |
| DSP | None | None | None | None |
| DSP | None | None | None | None |
| DSP | None | None | None | None |
| CCM2 | None | None | None | None |
| FXN | None | None | None | None |
| FXN | None | None | None | None |
| COMP | None | None | None | None |
| COMP | None | None | None | None |

| | | | | |
|---|---|---|---|---|
| COMP | None | None | None | None |
| COMP | None | None | None | None |
| COMP | None | None | None | None |
| COMP | None | None | None | None |
| COMP | None | None | None | None |
| COMP | None | None | None | None |
| COMP | None | None | None | None |
| COMP | None | None | None | None |
| COMP | None | None | None | None |
| COMP | None | None | None | None |
| AIFM1 | None | None | None | None |
| AIFM1 | None | None | None | None |
| AIFM1 | None | None | None | None |
| AIFM1 | None | None | None | None |
| AIFM1 | None | None | None | None |
| AIFM1 | None | None | None | None |
| AIFM1 | None | None | None | None |
| MYH9 | None | None | None | None |
| FHL2 | None | None | None | None |
| FHL2 | None | None | None | None |
| TNFRSF4 | None | None | None | None |
| CFB | None | None | None | None |
| CFB | None | None | None | None |
| CFB | None | None | None | None |
| CFB | None | None | None | None |
| CFB | None | None | None | None |
| CFB | None | None | None | None |
| CFB | None | None | None | None |
| CFB | None | None | None | None |
| CFB | None | None | None | None |
| CFB | None | None | None | None |
| MVD | None | None | None | None |
| MVD | None | None | None | None |
| HSD17B4 | None | None | None | None |
| HSD17B4 | None | None | None | None |
| HSD17B4 | None | None | None | None |
| HSD17B4 | None | None | None | None |
| HSD17B4 | None | None | None | None |
| HSD17B4 | None | None | None | None |
| F11 | None | None | None | None |
| F11 | None | None | None | None |
| F11 | None | None | None | None |
| F11 | None | None | None | None |
| F11 | None | None | None | None |
| F11 | None | None | None | None |
| F11 | None | None | None | None |
| F11 | None | None | None | None |
| F11 | None | None | None | None |
| SPTA1 | None | None | None | None |
| SPTA1 | None | None | None | None |
| SPTA1 | None | None | None | None |
| SPTA1 | None | None | None | None |
| SPTA1 | None | None | None | None |
| SPTA1 | None | None | None | None |

| | | | | |
|---|---|---|---|---|
| GMPR | None | None | None | None |
| MITF | None | None | None | None |
| NPC1 | None | None | None | None |
| NPC1 | None | None | None | None |
| NPC1 | None | None | None | None |
| NPC1 | None | None | None | None |
| NPC1 | None | None | None | None |
| NPC1 | None | None | None | None |
| NPC1 | None | None | None | None |
| NPC1 | None | None | None | None |
| NPC1 | None | None | None | None |
| NPC1 | None | None | None | None |
| NPC1 | None | None | None | None |
| NPC1 | None | None | None | None |
| NPC1 | None | None | None | None |
| NPC1 | None | None | None | None |
| NPC1 | None | None | None | None |
| NPC1 | None | None | None | None |
| NPC1 | None | None | None | None |
| NPC1 | None | None | None | None |
| NPC1 | None | None | None | None |
| NPC1 | None | None | None | None |
| NPC1 | None | None | None | None |
| NPC1 | None | None | None | None |
| NPC1 | None | None | None | None |
| NPC1 | None | None | None | None |
| NPC1 | None | None | None | None |
| NPC1 | None | None | None | None |
| NPC1 | None | None | None | None |
| NPC1 | None | None | None | None |
| NPC1 | None | None | None | None |
| NPC1 | None | None | None | None |
| NPC1 | None | None | None | None |
| NPC1 | None | None | None | None |
| NPC1 | None | None | None | None |
| NPC1 | None | None | None | None |
| NPC1 | None | None | None | None |
| NPC1 | None | None | None | None |
| NPC1 | None | None | None | None |
| NPC1 | None | None | None | None |
| NPC1 | None | None | None | None |
| NPC1 | None | None | None | None |
| NPC1 | None | None | None | None |
| NPC1 | None | None | None | None |
| NPC1 | None | None | None | None |
| C3 | None | None | None | None |
| NLRX1 | None | None | None | None |
| IGLL1 | None | None | None | None |
| APRT | None | None | None | None |
| APRT | None | None | None | None |
| APRT | None | None | None | None |
| APRT | None | None | None | None |
| APRT | None | None | None | None |
| ZIC3 | None | None | None | None |

| | | | | |
|---|---|---|---|---|
| ZIC3 | None | None | None | None |
| KLHL3 | None | None | None | None |
| RPS6KA3 | None | None | None | None |
| RPS6KA3 | None | None | None | None |
| RPS6KA3 | None | None | None | None |
| F8 | None | None | None | None |
| F8 | None | None | None | None |
| F8 | None | None | None | None |
| F8 | None | None | None | None |
| F8 | None | None | None | None |
| F8 | None | None | None | None |
| F8 | None | None | None | None |
| F8 | None | None | None | None |
| F8 | None | None | None | None |
| F8 | None | None | None | None |
| F8 | None | None | None | None |
| F8 | None | None | None | None |
| F8 | None | None | None | None |
| F8 | None | None | None | None |
| F8 | None | None | None | None |
| F8 | None | None | None | None |
| F8 | None | None | None | None |
| ISG15 | None | None | None | None |
| EXOSC3 | None | None | None | None |
| EXOSC3 | None | None | None | None |
| EXOSC3 | None | None | None | None |
| NKX2-5 | None | None | None | None |
| NKX2-5 | None | None | None | None |
| NKX2-5 | None | None | None | None |
| NKX2-5 | None | None | None | None |
| FH | None | None | None | None |
| FH | None | None | None | None |
| FH | None | None | None | None |
| FH | None | None | None | None |
| FH | None | None | None | None |
| FH | None | None | None | None |
| FH | None | None | None | None |
| FH | None | None | None | None |
| FH | None | None | None | None |
| FH | None | None | None | None |
| FH | None | None | None | None |
| FH | None | None | None | None |
| SNRNP200 | None | None | None | None |
| SNRNP200 | None | None | None | None |
| SNRNP200 | None | None | None | None |
| SNRNP200 | None | None | None | None |
| CHRNA1 | None | None | None | None |
| CHRNA1 | None | None | None | None |
| MYOM1 | None | None | None | None |
| MYOM1 | None | None | None | None |
| MYOM1 | None | None | None | None |
| MYOM1 | None | None | None | None |
| MYOM1 | None | None | None | None |
| MYOM1 | None | None | None | None |

| | | | | |
|---|---|---|---|---|
| LDLR | None | None | None | None |
| LDLR | None | None | None | None |
| LDLR | None | None | None | None |
| LDLR | None | None | None | None |
| LDLR | None | None | None | None |
| LDLR | None | None | None | None |
| EHMT1 | None | None | None | None |
| EHMT1 | None | None | None | None |
| EHMT1 | None | None | None | None |
| COL9A2 | None | None | None | None |
| HFE2 | None | None | None | None |
| HFE2 | None | None | None | None |
| CD3G | None | None | None | None |
| MMAB | None | None | None | None |
| MMAB | None | None | None | None |
| MMAB | None | None | None | None |
| MMAB | None | None | None | None |
| MMAB | None | None | None | None |
| MMAB | None | None | None | None |
| MMAB | None | None | None | None |
| MMAB | None | None | None | None |
| MMAB | None | None | None | None |
| MMAB | None | None | None | None |
| TAF1 | None | None | None | None |
| TAF1 | None | None | None | None |
| CD4 | None | None | None | None |
| QARS | None | None | None | None |
| QARS | None | None | None | None |
| QARS | None | None | None | None |
| QARS | None | None | None | None |
| QARS | None | None | None | None |
| FGFR3 | None | None | None | None |
| FGFR3 | None | None | None | None |
| FGFR3 | None | None | None | None |
| FGFR3 | None | None | None | None |
| FGFR3 | None | None | None | None |
| FGFR3 | None | None | None | None |
| FLNB | None | None | None | None |
| FLNB | None | None | None | None |
| FLNB | None | None | None | None |
| FLNB | None | None | None | None |
| FLNB | None | None | None | None |
| FLNB | None | None | None | None |
| FLNB | None | None | None | None |
| FLNB | None | None | None | None |
| IL21R | None | None | None | None |
| RB1 | None | None | None | None |
| RB1 | None | None | None | None |
| RB1 | None | None | None | None |
| NTRK1 | None | None | None | None |
| NOTCH1 | None | None | None | None |
| MYOM1 | None | None | None | None |
| MYOM1 | None | None | None | None |
| MYOM1 | None | None | None | None |

| | | | | |
|---|---|---|---|---|
| MYOM1 | None | None | None | None |
| MYOM1 | None | None | None | None |
| MYOM1 | None | None | None | None |
| COQ9 | None | None | None | None |
| COQ9 | None | None | None | None |
| CRBN | None | None | None | None |
| DCTN1 | None | None | None | None |
| DCTN1 | None | None | None | None |
| DCTN1 | None | None | None | None |
| DCTN1 | None | None | None | None |
| DCTN1 | None | None | None | None |
| DCTN1 | None | None | None | None |
| THAP1 | None | None | None | None |
| THAP1 | None | None | None | None |
| GNAS | None | None | None | None |
| GNAS | None | None | None | None |
| HNF4A | None | None | None | None |
| HNF4A | None | None | None | None |
| ALDH3A2 | None | None | None | None |
| ALDH3A2 | None | None | None | None |
| ALDH3A2 | None | None | None | None |
| ALDH3A2 | None | None | None | None |
| ALDH3A2 | None | None | None | None |
| ALDH3A2 | None | None | None | None |
| ALDH3A2 | None | None | None | None |
| ALDH3A2 | None | None | None | None |
| ALDH3A2 | None | None | None | None |
| HCN1 | None | None | None | None |
| PLEC | None | None | None | None |
| SCN5A | None | None | None | None |
| SCN5A | None | None | None | None |
| STAMBP | None | None | None | None |
| STAMBP | None | None | None | None |
| RNF125 | None | None | None | None |
| RNF125 | None | None | None | None |
| USH1C | None | None | None | None |
| OBSL1 | None | None | None | None |
| GRIN2B | None | None | None | None |
| GRIN2B | None | None | None | None |
| HCN4 | None | None | None | None |
| HCN4 | None | None | None | None |
| TALDO1 | None | None | None | None |
| ITGB3 | None | None | None | None |
| CHKB | None | None | None | None |
| CHKB | None | None | None | None |
| CHKB | None | None | None | None |
| CHKB | None | None | None | None |
| ACTN4 | None | None | None | None |
| ACTN4 | None | None | None | None |
| ACTN4 | None | None | None | None |
| ACTN4 | None | None | None | None |
| CDT1 | None | None | None | None |
| CDT1 | None | None | None | None |
| CDT1 | None | None | None | None |

| | | | | |
|---|---|---|---|---|
| CDT1 | None | None | None | None |
| CDT1 | None | None | None | None |
| CDT1 | None | None | None | None |
| MAPT | None | None | None | None |
| F5 | None | None | None | None |
| F5 | None | None | None | None |
| F5 | None | None | None | None |
| ESRRB | None | None | None | None |
| DGUOK | None | None | None | None |
| DGUOK | None | None | None | None |
| DGUOK | None | None | None | None |
| DGUOK | None | None | None | None |
| DGUOK | None | None | None | None |
| DGUOK | None | None | None | None |
| DGUOK | None | None | None | None |
| DGUOK | None | None | None | None |
| DGUOK | None | None | None | None |
| CENPJ | None | None | None | None |
| TTN | None | None | None | None |
| TTN | None | None | None | None |
| TTN | None | None | None | None |
| TTN | None | None | None | None |
| TNFRSF1A | None | None | None | None |
| TNFRSF1A | None | None | None | None |
| TNFRSF1A | None | None | None | None |
| TNFRSF1A | None | None | None | None |
| TNFRSF1A | None | None | None | None |
| TNFRSF1A | None | None | None | None |
| TNFRSF1A | None | None | None | None |
| TNFRSF1A | None | None | None | None |
| TNFRSF1A | None | None | None | None |
| TNFRSF1A | None | None | None | None |
| TNFRSF1A | None | None | None | None |
| TNFRSF1A | None | None | None | None |
| ASL | None | None | None | None |
| ASL | None | None | None | None |
| ASL | None | None | None | None |
| ASL | None | None | None | None |
| ASL | None | None | None | None |
| ASL | None | None | None | None |
| ASL | None | None | None | None |
| ASL | None | None | None | None |
| ASL | None | None | None | None |
| ASL | None | None | None | None |
| MYO7A | None | None | None | None |
| MYO7A | None | None | None | None |
| MYO7A | None | None | None | None |
| MYO7A | None | None | None | None |
| MYO7A | None | None | None | None |
| MYO7A | None | None | None | None |
| MYO7A | None | None | None | None |
| MYO7A | None | None | None | None |
| MYO7A | None | None | None | None |
| MYO7A | None | None | None | None |

| | | | | |
|---|---|---|---|---|
| MYO7A | None | None | None | None |
| MYO7A | None | None | None | None |
| MYO7A | None | None | None | None |
| MYO7A | None | None | None | None |
| MYO7A | None | None | None | None |
| MYO7A | None | None | None | None |
| MYO7A | None | None | None | None |
| MYO7A | None | None | None | None |
| MYO7A | None | None | None | None |
| MYO7A | None | None | None | None |
| APC | None | None | None | None |
| ALAD | None | None | None | None |
| ALAD | None | None | None | None |
| ALAD | None | None | None | None |
| ALAD | None | None | None | None |
| ALAD | None | None | None | None |
| ALAD | None | None | None | None |
| POR | None | None | None | None |
| CSF1R | None | None | None | None |
| CSF1R | None | None | None | None |
| CSF1R | None | None | None | None |
| CSF1R | None | None | None | None |
| CSF1R | None | None | None | None |
| CSF1R | None | None | None | None |
| CSF1R | None | None | None | None |
| CSF1R | None | None | None | None |
| CSF1R | None | None | None | None |
| CSF1R | None | None | None | None |
| CSF1R | None | None | None | None |
| VWF | None | None | None | None |
| VWF | None | None | None | None |
| PHF8 | None | None | None | None |
| ACY1 | None | None | None | None |
| SMN1 | None | None | None | None |
| SMN1 | None | None | None | None |
| DLAT | None | None | None | None |
| DLAT | None | None | None | None |
| B2M | None | None | None | None |
| SOX9 | None | None | None | None |
| SOX9 | None | None | None | None |
| SOX9 | None | None | None | None |
| SOX9 | None | None | None | None |
| SOX9 | None | None | None | None |
| CFH | None | None | None | None |
| FLNC | None | None | None | None |
| FLNC | None | None | None | None |
| FLNC | None | None | None | None |
| ITGB2 | None | None | None | None |
| ITGB2 | None | None | None | None |
| ITGB2 | None | None | None | None |
| ITGB2 | None | None | None | None |
| ITGB2 | None | None | None | None |
| ITGB2 | None | None | None | None |
| ITGB2 | None | None | None | None |

| | | | | |
|---|---|---|---|---|
| ITGB2 | None | None | None | None |
| ITGB2 | None | None | None | None |
| PSEN1 | None | None | None | None |
| PSEN1 | None | None | None | None |
| PSEN1 | None | None | None | None |
| PSEN1 | None | None | None | None |
| PSEN1 | None | None | None | None |
| PSEN1 | None | None | None | None |
| OXCT1 | None | None | None | None |
| OXCT1 | None | None | None | None |
| OXCT1 | None | None | None | None |
| OXCT1 | None | None | None | None |
| OXCT1 | None | None | None | None |
| OXCT1 | None | None | None | None |
| OXCT1 | None | None | None | None |
| DHH | None | None | None | None |
| PRKN | None | None | None | None |
| PRKN | None | None | None | None |
| PRKN | None | None | None | None |
| PRKN | None | None | None | None |
| PRKN | None | None | None | None |
| PRKN | None | None | None | None |
| PRKN | None | None | None | None |
| PRKN | None | None | None | None |
| PRKN | None | None | None | None |
| PRKN | None | None | None | None |
| PRKN | None | None | None | None |
| LCAT | None | None | None | None |
| LCAT | None | None | None | None |
| LCAT | None | None | None | None |
| LCAT | None | None | None | None |
| LCAT | None | None | None | None |
| LCAT | None | None | None | None |
| LCAT | None | None | None | None |
| LCAT | None | None | None | None |
| LCAT | None | None | None | None |
| LCAT | None | None | None | None |
| LCAT | None | None | None | None |
| LCAT | None | None | None | None |
| FOXP3 | None | None | None | None |
| FOXP3 | None | None | None | None |
| FOXP3 | None | None | None | None |
| FOXP3 | None | None | None | None |
| SUFU | None | None | None | None |
| HSPA9 | None | None | None | None |
| HSPA9 | None | None | None | None |
| SQSTM1 | None | None | None | None |
| DARS2 | None | None | None | None |
| DARS2 | None | None | None | None |
| DARS2 | None | None | None | None |
| DARS2 | None | None | None | None |
| DARS2 | None | None | None | None |
| DARS2 | None | None | None | None |
| DARS2 | None | None | None | None |

| | | | | |
|---|---|---|---|---|
| DARS2 | None | None | None | None |
| DARS2 | None | None | None | None |
| DARS2 | None | None | None | None |
| DARS2 | None | None | None | None |
| DARS2 | None | None | None | None |
| NT5C3A | None | None | None | None |
| NT5C3A | None | None | None | None |
| NT5C3A | None | None | None | None |
| NT5C3A | None | None | None | None |
| ETFA | None | None | None | None |
| ETFA | None | None | None | None |
| ETFA | None | None | None | None |
| ETFA | None | None | None | None |
| ETFA | None | None | None | None |
| ETFA | None | None | None | None |
| ETFA | None | None | None | None |
| TTN | None | None | None | None |
| TTN | None | None | None | None |
| FLT4 | None | None | None | None |
| FGB | None | None | None | None |
| HPCA | None | None | None | None |
| TAP2 | None | None | None | None |
| TAP2 | None | None | None | None |
| PTPRC | None | None | None | None |
| SLC4A1 | None | None | None | None |
| SLC4A1 | None | None | None | None |
| SLC4A1 | None | None | None | None |
| SLC4A1 | None | None | None | None |
| SLC4A1 | None | None | None | None |
| SLC4A1 | None | None | None | None |
| MYH7 | None | None | None | None |
| MYH7 | None | None | None | None |
| MYH7 | None | None | None | None |
| MYH7 | None | None | None | None |
| MYH7 | None | None | None | None |
| MYH7 | None | None | None | None |
| MYH7 | None | None | None | None |
| MYH7 | None | None | None | None |
| MYH7 | None | None | None | None |
| PARN | None | None | None | None |
| PARN | None | None | None | None |
| NDUFS6 | None | None | None | None |
| VRK1 | None | None | None | None |
| VRK1 | None | None | None | None |
| VRK1 | None | None | None | None |
| VRK1 | None | None | None | None |
| VRK1 | None | None | None | None |
| VRK1 | None | None | None | None |
| COLEC11 | None | None | None | None |
| VHL | None | None | None | None |
| TTN | None | None | None | None |
| NCF2 | None | None | None | None |
| NCF2 | None | None | None | None |
| NCF2 | None | None | None | None |

| | | | | |
|---|---|---|---|---|
| NCF2 | None | None | None | None |
| NCF2 | None | None | None | None |
| RPS26 | None | None | None | None |
| CETP | None | None | None | None |
| CETP | None | None | None | None |
| CETP | None | None | None | None |
| CETP | None | None | None | None |
| CETP | None | None | None | None |
| CETP | None | None | None | None |
| CETP | None | None | None | None |
| CETP | None | None | None | None |
| CETP | None | None | None | None |
| CETP | None | None | None | None |
| CETP | None | None | None | None |
| CETP | None | None | None | None |
| CETP | None | None | None | None |
| FLNA | None | None | None | None |
| FLNA | None | None | None | None |
| FLNA | None | None | None | None |
| FLNA | None | None | None | None |
| FLNA | None | None | None | None |
| FLNA | None | None | None | None |
| FLNA | None | None | None | None |
| MYBPC3 | None | None | None | None |
| MYBPC3 | None | None | None | None |
| MYBPC3 | None | None | None | None |
| MYBPC3 | None | None | None | None |
| MYBPC3 | None | None | None | None |
| MYBPC3 | None | None | None | None |
| MYBPC3 | None | None | None | None |
| MYBPC3 | None | None | None | None |
| MYBPC3 | None | None | None | None |
| MYBPC3 | None | None | None | None |
| LSS | None | None | None | None |
| C6 | None | None | None | None |
| C6 | None | None | None | None |
| RLBP1 | None | None | None | None |
| RLBP1 | None | None | None | None |
| RLBP1 | None | None | None | None |
| RLBP1 | None | None | None | None |
| RLBP1 | None | None | None | None |
| RLBP1 | None | None | None | None |
| RLBP1 | None | None | None | None |
| RLBP1 | None | None | None | None |
| MMAA | None | None | None | None |
| MMAA | None | None | None | None |
| MMAA | None | None | None | None |
| MMAA | None | None | None | None |
| MMAA | None | None | None | None |
| MMAA | None | None | None | None |
| IL1RN | None | None | None | None |
| MASTL | None | None | None | None |
| FHL1 | None | None | None | None |
| FHL1 | None | None | None | None |

| | | | | |
|---|---|---|---|---|
| FHL1 | None | None | None | None |
| FHL1 | None | None | None | None |
| FHL1 | None | None | None | None |
| FHL1 | None | None | None | None |
| FHL1 | None | None | None | None |
| FHL1 | None | None | None | None |
| FHL1 | None | None | None | None |
| EIF2B1 | None | None | None | None |
| EIF2B1 | None | None | None | None |
| COQ8A | None | None | None | None |
| COQ8A | None | None | None | None |
| COQ8A | None | None | None | None |
| COQ8A | None | None | None | None |
| COQ8A | None | None | None | None |
| COQ8A | None | None | None | None |
| COQ8A | None | None | None | None |
| COQ8A | None | None | None | None |
| COQ8A | None | None | None | None |
| COQ8A | None | None | None | None |
| COQ8A | None | None | None | None |
| COQ8A | None | None | None | None |
| COQ8A | None | None | None | None |
| COQ8A | None | None | None | None |
| COQ8A | None | None | None | None |
| COQ8A | None | None | None | None |
| UNC119 | None | None | None | None |
| IGHMBP2 | None | None | None | None |
| JAG1 | None | None | None | None |
| JAG1 | None | None | None | None |
| JAG1 | None | None | None | None |
| GNB1 | None | None | None | None |
| GNB1 | None | None | None | None |
| FLNA | None | None | None | None |
| FLNA | None | None | None | None |
| FLNA | None | None | None | None |
| FLNA | None | None | None | None |
| FLNA | None | None | None | None |
| CREBBP | None | None | None | None |
| CSF3R | None | None | None | None |
| CSF3R | None | None | None | None |
| HIBCH | None | None | None | None |
| HIBCH | None | None | None | None |
| HIBCH | None | None | None | None |
| MMACHC | None | None | None | None |
| MMACHC | None | None | None | None |
| MMACHC | None | None | None | None |
| MMACHC | None | None | None | None |
| MMACHC | None | None | None | None |
| MMACHC | None | None | None | None |
| MMACHC | None | None | None | None |
| MMACHC | None | None | None | None |
| MMACHC | None | None | None | None |
| MMACHC | None | None | None | None |
| MMACHC | None | None | None | None |

| | | | | |
|---|---|---|---|---|
| KLF1 | None | None | None | None |
| NAGS | None | None | None | None |
| NAGS | None | None | None | None |
| NAGS | None | None | None | None |
| PIK3R2 | None | None | None | None |
| EDNRB | None | None | None | None |
| EDNRB | None | None | None | None |
| EDNRB | None | None | None | None |
| EDNRB | None | None | None | None |
| E | None | None | None | None |
| RUNX1 | None | None | None | None |
| RUNX1 | None | None | None | None |
| CTNNB1 | None | None | None | None |
| CTNNB1 | None | None | None | None |
| CTNNB1 | None | None | None | None |
| CTNNB1 | None | None | None | None |
| CTNNB1 | None | None | None | None |
| CTNNB1 | None | None | None | None |
| TTN | None | None | None | None |
| TTN | None | None | None | None |
| TYK2 | None | None | None | None |
| TYK2 | None | None | None | None |
| TYK2 | None | None | None | None |
| ERCC4 | None | None | None | None |
| ERCC4 | None | None | None | None |
| ERCC4 | None | None | None | None |
| ERCC4 | None | None | None | None |
| CFI | None | None | None | None |
| CFI | None | None | None | None |
| CFI | None | None | None | None |
| CFI | None | None | None | None |
| CFI | None | None | None | None |
| CFI | None | None | None | None |
| CFI | None | None | None | None |
| CFI | None | None | None | None |
| CFI | None | None | None | None |
| CFI | None | None | None | None |
| CFI | None | None | None | None |
| CFI | None | None | None | None |
| CFI | None | None | None | None |
| CFI | None | None | None | None |
| NOTCH3 | None | None | None | None |
| NOTCH3 | None | None | None | None |
| NOTCH3 | None | None | None | None |
| NOTCH3 | None | None | None | None |
| TBX1 | None | None | None | None |
| TBX1 | None | None | None | None |
| DDB2 | None | None | None | None |
| DDB2 | None | None | None | None |
| TP53 | None | None | None | None |
| POMGNT1 | None | None | None | None |
| POMGNT1 | None | None | None | None |
| POMGNT1 | None | None | None | None |
| POMGNT1 | None | None | None | None |

| | | | | |
|---|---|---|---|---|
| POMGNT1 | None | None | None | None |
| POMGNT1 | None | None | None | None |
| POMGNT1 | None | None | None | None |
| POMGNT1 | None | None | None | None |
| POMGNT1 | None | None | None | None |
| POMGNT1 | None | None | None | None |
| POMGNT1 | None | None | None | None |
| POMGNT1 | None | None | None | None |
| POMGNT1 | None | None | None | None |
| FMR1 | None | None | None | None |
| FMR1 | None | None | None | None |
| SLC9A3R1 | None | None | None | None |
| HSPB1 | None | None | None | None |
| HSPB1 | None | None | None | None |
| HSPB1 | None | None | None | None |
| HSPB1 | None | None | None | None |
| HSPB1 | None | None | None | None |
| HSPB1 | None | None | None | None |
| HSPB1 | None | None | None | None |
| SOD3 | None | None | None | None |
| AGA | | 1 | 2 1apz | D |
| AGA | None | None | None | None |
| AGA | None | None | None | None |
| NDUFS7 | None | None | None | None |
| NDUFS7 | None | None | None | None |
| SOX2 | None | None | None | None |
| CSRP3 | None | None | None | None |
| CSRP3 | None | None | None | None |
| PAH | None | None | None | None |
| PAH | None | None | None | None |
| PAH | None | None | None | None |
| PAH | None | None | None | None |
| CACNA1S | None | None | None | None |
| TRPV4 | None | None | None | None |
| TRPV4 | None | None | None | None |
| TRPV4 | None | None | None | None |
| TRPV4 | None | None | None | None |
| TRPV4 | None | None | None | None |
| TRPV4 | None | None | None | None |
| TRPV4 | None | None | None | None |
| TRPV4 | None | None | None | None |
| TRPV4 | None | None | None | None |
| TRPV4 | None | None | None | None |
| TRPV4 | None | None | None | None |
| TRPV4 | None | None | None | None |
| TRPV4 | None | None | None | None |
| TNNI3 | None | None | None | None |
| MLYCD | None | None | None | None |
| MLYCD | None | None | None | None |
| MLYCD | None | None | None | None |
| MLYCD | None | None | None | None |
| AIP | None | None | None | None |
| AIP | None | None | None | None |
| ERCC2 | None | None | None | None |

| | | | | |
|---|---|---|---|---|
| ERCC2 | None | None | None | None |
| ERCC2 | None | None | None | None |
| ERCC2 | None | None | None | None |
| ERCC2 | None | None | None | None |
| ERCC2 | None | None | None | None |
| ERCC2 | None | None | None | None |
| ERCC2 | None | None | None | None |
| ERCC2 | None | None | None | None |
| ERCC2 | None | None | None | None |
| ERCC2 | None | None | None | None |
| ERCC2 | None | None | None | None |
| ERCC2 | None | None | None | None |
| ERCC2 | None | None | None | None |
| PUS1 | None | None | None | None |
| PUS1 | None | None | None | None |
| PUS1 | None | None | None | None |
| CASK | None | None | None | None |
| CASK | None | None | None | None |
| BLVRA | None | None | None | None |
| DICER1 | None | None | None | None |
| DICER1 | None | None | None | None |
| DICER1 | None | None | None | None |
| EIF2AK3 | None | None | None | None |
| EIF2AK3 | None | None | None | None |
| EIF2AK3 | None | None | None | None |
| KIT | None | None | None | None |
| KIT | None | None | None | None |
| KIT | None | None | None | None |
| KIT | None | None | None | None |
| KIT | None | None | None | None |
| KIT | None | None | None | None |
| DNMT3B | None | None | None | None |
| TGM1 | None | None | None | None |
| NDUFA10 | None | None | None | None |
| NDUFA10 | None | None | None | None |
| PRPF31 | None | None | None | None |
| PRPF31 | None | None | None | None |
| ECE1 | None | None | None | None |
| ECE1 | None | None | None | None |
| ECE1 | None | None | None | None |
| NDUFV2 | None | None | None | None |
| NDUFV2 | None | None | None | None |
| NDUFV2 | None | None | None | None |
| GHRHR | None | None | None | None |
| GHRHR | None | None | None | None |
| GHRHR | None | None | None | None |
| SLC4A1 | None | None | None | None |
| FGF9 | None | None | None | None |
| BARD1 | None | None | None | None |
| BARD1 | None | None | None | None |
| BARD1 | None | None | None | None |
| PTPN12 | None | None | None | None |
| MASP2 | None | None | None | None |
| MASP2 | None | None | None | None |

| | | | | |
|---|---|---|---|---|
| MASP2 | None | None | None | None |
| MASP2 | None | None | None | None |
| HOXB1 | None | None | None | None |
| SLC4A1 | None | None | None | None |
| SLC4A1 | None | None | None | None |
| SLC4A1 | None | None | None | None |
| SLC4A1 | None | None | None | None |
| SLC4A1 | None | None | None | None |
| SLC4A1 | None | None | None | None |
| SLC4A1 | None | None | None | None |
| SLC4A1 | None | None | None | None |
| SLC4A1 | None | None | None | None |
| SLC4A1 | None | None | None | None |
| SLC4A1 | None | None | None | None |
| SLC4A1 | None | None | None | None |
| SLC4A1 | None | None | None | None |
| SLC4A1 | None | None | None | None |
| SLC4A1 | None | None | None | None |
| SLC4A1 | None | None | None | None |
| SLC4A1 | None | None | None | None |
| CBL | None | None | None | None |
| FAS | None | None | None | None |
| MTMR2 | None | None | None | None |
| MTMR2 | None | None | None | None |
| MTMR2 | None | None | None | None |
| MTMR2 | None | None | None | None |
| MTMR2 | None | None | None | None |
| MTMR2 | None | None | None | None |
| MTMR2 | None | None | None | None |
| MTMR2 | None | None | None | None |
| KLHL3 | None | None | None | None |
| KLHL3 | None | None | None | None |
| KLHL3 | None | None | None | None |
| KLHL3 | None | None | None | None |
| KLHL3 | None | None | None | None |
| KLHL3 | None | None | None | None |
| RNASEH1 | None | None | None | None |
| ACTN1 | None | None | None | None |
| PHKG2 | None | None | None | None |
| PHKG2 | None | None | None | None |
| PHKG2 | None | None | None | None |
| NTRK1 | None | None | None | None |
| APOH | None | None | None | None |
| PMS2 | None | None | None | None |
| PMS2 | None | None | None | None |
| PMS2 | None | None | None | None |
| PMS2 | None | None | None | None |
| PMS2 | None | None | None | None |
| PMS2 | None | None | None | None |
| PMS2 | None | None | None | None |
| PMS2 | None | None | None | None |
| PMS2 | None | None | None | None |
| PMS2 | None | None | None | None |
| PMS2 | None | None | None | None |

| | | | | |
|---|---|---|---|---|
| PMS2 | None | None | None | None |
| PMS2 | None | None | None | None |
| PMS2 | None | None | None | None |
| PMS2 | None | None | None | None |
| PMS2 | None | None | None | None |
| PMS2 | None | None | None | None |
| PMS2 | None | None | None | None |
| PMS2 | None | None | None | None |
| PMS2 | None | None | None | None |
| PMS2 | None | None | None | None |
| PMS2 | None | None | None | None |
| MOCS2 | None | None | None | None |
| MOCS2 | None | None | None | None |
| RPGRIP1 | None | None | None | None |
| RPGRIP1 | None | None | None | None |
| LIG4 | None | None | None | None |
| PROC | None | None | None | None |
| PROC | None | None | None | None |
| NAT1 | None | None | None | None |
| NAT1 | None | None | None | None |
| PNPO | None | None | None | None |
| PNPO | None | None | None | None |
| PNPO | None | None | None | None |
| PNPO | None | None | None | None |
| PNPO | None | None | None | None |
| MRE11 | None | None | None | None |
| MRE11 | None | None | None | None |
| MRE11 | None | None | None | None |
| MRE11 | None | None | None | None |
| DDC | None | None | None | None |
| DDC | None | None | None | None |
| DDC | None | None | None | None |
| DDC | None | None | None | None |
| DDC | None | None | None | None |
| DDC | None | None | None | None |
| DDC | None | None | None | None |
| DDC | None | None | None | None |
| DDC | None | None | None | None |
| DDC | None | None | None | None |
| DDC | None | None | None | None |
| ANK2 | None | None | None | None |
| ANK2 | None | None | None | None |
| ANK2 | None | None | None | None |
| AK2 | None | None | None | None |
| AK2 | None | None | None | None |
| AK2 | None | None | None | None |
| AK2 | None | None | None | None |
| AK2 | None | None | None | None |
| AK2 | None | None | None | None |
| GPHN | None | None | None | None |
| HSD17B4 | None | None | None | None |
| HSD17B4 | None | None | None | None |
| HSD17B4 | None | None | None | None |
| HSD17B4 | None | None | None | None |

| | | | | |
|---|---|---|---|---|
| HSD17B4 | None | None | None | None |
| HSD17B4 | None | None | None | None |
| APTX | None | None | None | None |
| APTX | None | None | None | None |
| APTX | None | None | None | None |
| APTX | None | None | None | None |
| APTX | None | None | None | None |
| APTX | None | None | None | None |
| APTX | None | None | None | None |
| FANCL | None | None | None | None |
| FANCL | None | None | None | None |
| EMG1 | None | None | None | None |
| EMG1 | None | None | None | None |
| EZH2 | None | None | None | None |
| ABCB6 | None | None | None | None |
| NCF2 | None | None | None | None |
| NCF2 | None | None | None | None |
| COL4A2 | None | None | None | None |
| COL4A2 | 1 | 1 | 1li1 | F |
| COL4A2 | None | None | None | None |
| COL4A2 | None | None | None | None |
| COL4A2 | None | None | None | None |
| SLX4 | None | None | None | None |
| SERPIND1 | None | None | None | None |
| FGFR3 | None | None | None | None |
| FGFR3 | None | None | None | None |
| FGFR3 | None | None | None | None |
| FGFR3 | None | None | None | None |
| FGFR3 | None | None | None | None |
| FGFR3 | None | None | None | None |
| TREM2 | None | None | None | None |
| TREM2 | None | None | None | None |
| TREM2 | None | None | None | None |
| TREM2 | None | None | None | None |
| TREM2 | None | None | None | None |
| TREM2 | None | None | None | None |
| TREM2 | None | None | None | None |
| LIG4 | None | None | None | None |
| LIG4 | None | None | None | None |
| LIG4 | None | None | None | None |
| LIG4 | None | None | None | None |
| LIG4 | None | None | None | None |
| LIG4 | None | None | None | None |
| ECHS1 | None | None | None | None |
| ECHS1 | None | None | None | None |
| ECHS1 | None | None | None | None |
| MTR | None | None | None | None |
| MTR | None | None | None | None |
| MTR | None | None | None | None |
| MTR | None | None | None | None |
| MTR | None | None | None | None |
| PMVK | None | None | None | None |
| PRNP | None | None | None | None |
| CHN1 | None | None | None | None |

| | | | | |
|---|---|---|---|---|
| CHN1 | None | None | None | None |
| CHN1 | None | None | None | None |
| CHN1 | None | None | None | None |
| CHN1 | None | None | None | None |
| CHN1 | None | None | None | None |
| CHN1 | None | None | None | None |
| CHN1 | None | None | None | None |
| SLC2A1 | None | None | None | None |
| SLC2A1 | None | None | None | None |
| SLC2A1 | None | None | None | None |
| SLC2A1 | None | None | None | None |
| SLC2A1 | None | None | None | None |
| SLC2A1 | None | None | None | None |
| SLC2A1 | None | None | None | None |
| SLC2A1 | None | None | None | None |
| SLC2A1 | None | None | None | None |
| SLC2A1 | None | None | None | None |
| SLC2A1 | None | None | None | None |
| SLC2A1 | None | None | None | None |
| SLC2A1 | None | None | None | None |
| SLC2A1 | None | None | None | None |
| SLC2A1 | None | None | None | None |
| SLC2A1 | None | None | None | None |
| SLC2A1 | None | None | None | None |
| PMM2 | None | None | None | None |
| PMM2 | None | None | None | None |
| PMM2 | None | None | None | None |
| PMM2 | None | None | None | None |
| PMM2 | None | None | None | None |
| PMM2 | None | None | None | None |
| PMM2 | None | None | None | None |
| PMM2 | None | None | None | None |
| PMM2 | None | None | None | None |
| PMM2 | None | None | None | None |
| PMM2 | None | None | None | None |
| PMM2 | None | None | None | None |
| PMM2 | None | None | None | None |
| PMM2 | None | None | None | None |
| PMM2 | None | None | None | None |
| PMM2 | None | None | None | None |
| PMM2 | None | None | None | None |
| PMM2 | None | None | None | None |
| PMM2 | None | None | None | None |
| PMM2 | None | None | None | None |
| PMM2 | None | None | None | None |
| PMM2 | None | None | None | None |
| PMM2 | None | None | None | None |
| PMM2 | None | None | None | None |
| PMM2 | None | None | None | None |
| PMM2 | None | None | None | None |
| PMM2 | None | None | None | None |
| PMM2 | None | None | None | None |
| CCKAR | None | None | None | None |
| MAN1B1 | None | None | None | None |

| | | | | |
|---|---|---|---|---|
| MAN1B1 | None | None | None | None |
| LAMA2 | None | None | None | None |
| LAMA2 | None | None | None | None |
| TBC1D4 | None | None | None | None |
| TBC1D4 | None | None | None | None |
| LDLR | None | None | None | None |
| LDLR | None | None | None | None |
| LDLR | None | None | None | None |
| LDLR | None | None | None | None |
| LDLR | None | None | None | None |
| LDLR | None | None | None | None |
| LDLR | None | None | None | None |
| LDLR | None | None | None | None |
| LDLR | None | None | None | None |
| LDLR | None | None | None | None |
| LDLR | None | None | None | None |
| LDLR | None | None | None | None |
| LDLR | None | None | None | None |
| LDLR | None | None | None | None |
| LDLR | None | None | None | None |
| LDLR | None | None | None | None |
| LDLR | None | None | None | None |
| LDLR | None | None | None | None |
| LDLR | None | None | None | None |
| LDLR | None | None | None | None |
| LDLR | None | None | None | None |
| LDLR | None | None | None | None |
| LDLR | None | None | None | None |
| LDLR | None | None | None | None |
| LDLR | None | None | None | None |
| LDLR | None | None | None | None |
| LDLR | None | None | None | None |
| LDLR | None | None | None | None |
| LDLR | None | None | None | None |
| LDLR | None | None | None | None |
| LDLR | None | None | None | None |
| LDLR | None | None | None | None |
| LDLR | None | None | None | None |
| LDLR | None | None | None | None |
| LDLR | None | None | None | None |
| LDLR | None | None | None | None |
| LDLR | None | None | None | None |
| LDLR | None | None | None | None |
| LDLR | None | None | None | None |
| LDLR | None | None | None | None |
| LDLR | None | None | None | None |
| LDLR | None | None | None | None |
| LDLR | None | None | None | None |
| FANCM | None | None | None | None |
| HAMP | None | None | None | None |
| GUSB | None | None | None | None |
| GUSB | None | None | None | None |
| GUSB | None | None | None | None |
| GUSB | None | None | None | None |

| | | | | |
|---|---|---|---|---|
| GUSB | None | None | None | None |
| GUSB | None | None | None | None |
| GUSB | None | None | None | None |
| GUSB | None | None | None | None |
| GUSB | None | None | None | None |
| GUSB | None | None | None | None |
| GUSB | None | None | None | None |
| GUSB | None | None | None | None |
| GUSB | None | None | None | None |
| LIG1 | None | None | None | None |
| LIG1 | None | None | None | None |
| MYH7 | None | None | None | None |
| MYH7 | None | None | None | None |
| MYH7 | None | None | None | None |
| MYH7 | None | None | None | None |
| MYH7 | None | None | None | None |
| MYH7 | None | None | None | None |
| MYH7 | None | None | None | None |
| MYH7 | None | None | None | None |
| MYH7 | None | None | None | None |
| MYH7 | None | None | None | None |
| MYH7 | None | None | None | None |
| MYH7 | None | None | None | None |
| MYH7 | None | None | None | None |
| MYH7 | None | None | None | None |
| MYH7 | None | None | None | None |
| MYH7 | None | None | None | None |
| MYH7 | None | None | None | None |
| MYH7 | None | None | None | None |
| MYH7 | None | None | None | None |
| MYH7 | None | None | None | None |
| MYH7 | None | None | None | None |
| MYH7 | None | None | None | None |
| MYH7 | None | None | None | None |
| MYH7 | None | None | None | None |
| MYH7 | None | None | None | None |
| MYH7 | None | None | None | None |
| MYH7 | None | None | None | None |
| MYH7 | None | None | None | None |
| MYH7 | None | None | None | None |
| MYH7 | None | None | None | None |
| MYH7 | None | None | None | None |
| MYH7 | None | None | None | None |
| MYH7 | None | None | None | None |
| MYH7 | None | None | None | None |
| HCN4 | None | None | None | None |
| RECQL4 | None | None | None | None |
| RECQL4 | None | None | None | None |
| RECQL4 | None | None | None | None |
| RECQL4 | None | None | None | None |
| RECQL4 | None | None | None | None |
| RECQL4 | None | None | None | None |
| RECQL4 | None | None | None | None |
| RECQL4 | None | None | None | None |

| | | | | |
|---|---|---|---|---|
| RECQL4 | None | None | None | None |
| RECQL4 | None | None | None | None |
| RECQL4 | None | None | None | None |
| RECQL4 | None | None | None | None |
| RECQL4 | None | None | None | None |
| CHEK2 | None | None | None | None |
| PPARG | None | None | None | None |
| ACADVL | None | None | None | None |
| ACADVL | None | None | None | None |
| ACADVL | None | None | None | None |
| ACADVL | None | None | None | None |
| ACADVL | None | None | None | None |
| ACADVL | None | None | None | None |
| ACADVL | None | None | None | None |
| ACADVL | None | None | None | None |
| ACADVL | None | None | None | None |
| ACADVL | None | None | None | None |
| ACADVL | None | None | None | None |
| ACADVL | None | None | None | None |
| ACADVL | None | None | None | None |
| ACADVL | None | None | None | None |
| ACADVL | None | None | None | None |
| ACADVL | None | None | None | None |
| ACADVL | None | None | None | None |
| ACADVL | None | None | None | None |
| ACADVL | None | None | None | None |
| ACADVL | None | None | None | None |
| ACADVL | None | None | None | None |
| ACADVL | None | None | None | None |
| APOA1 | None | None | None | None |
| APOA1 | None | None | None | None |
| APOA1 | None | None | None | None |
| APOA1 | None | None | None | None |
| APOA1 | None | None | None | None |
| DNMT3A | None | None | None | None |
| DNMT3A | None | None | None | None |
| APC | None | None | None | None |
| TMOD1 | None | None | None | None |
| USH1C | None | None | None | None |
| USH1C | None | None | None | None |
| USH1C | None | None | None | None |
| USH1C | None | None | None | None |
| ERCC6 | None | None | None | None |
| ERCC6 | None | None | None | None |
| ERCC6 | None | None | None | None |
| ERCC8 | None | None | None | None |
| ERCC8 | None | None | None | None |
| ERCC8 | None | None | None | None |
| DPAGT1 | None | None | None | None |
| DPAGT1 | None | None | None | None |
| DPAGT1 | None | None | None | None |
| DPAGT1 | None | None | None | None |
| DPAGT1 | None | None | None | None |
| DPAGT1 | None | None | None | None |

| | | | | |
|---|---|---|---|---|
| DPAGT1 | None | None | None | None |
| IL2RG | None | None | None | None |
| HNF1A | None | None | None | None |
| HNF4A | None | None | None | None |
| FLT4 | None | None | None | None |
| FLT4 | None | None | None | None |
| FLT4 | None | None | None | None |
| CD3D | None | None | None | None |
| CD3D | None | None | None | None |
| MCOLN1 | None | None | None | None |
| MCOLN1 | None | None | None | None |
| MOCS2 | None | None | None | None |
| MOCS2 | None | None | None | None |
| ABCG8 | None | None | None | None |
| ABCG8 | None | None | None | None |
| ABCG8 | None | None | None | None |
| ABCG8 | None | None | None | None |
| ABCG8 | None | None | None | None |
| ABCG8 | None | None | None | None |
| ABCG8 | None | None | None | None |
| ABCG8 | None | None | None | None |
| ABCG8 | None | None | None | None |
| ABCG8 | None | None | None | None |
| ABCG8 | None | None | None | None |
| ABCG8 | None | None | None | None |
| ABCG8 | None | None | None | None |
| ABCG8 | None | None | None | None |
| CTNNA1 | None | None | None | None |
| TYMP | None | None | None | None |
| TYMP | None | None | None | None |
| TYMP | None | None | None | None |
| TYMP | None | None | None | None |
| TYMP | None | None | None | None |
| TYMP | None | None | None | None |
| TYMP | None | None | None | None |
| TYMP | None | None | None | None |
| TYMP | None | None | None | None |
| TYMP | None | None | None | None |
| TYMP | None | None | None | None |
| TYMP | None | None | None | None |
| TYMP | None | None | None | None |
| GAMT | None | None | None | None |
| GAMT | None | None | None | None |
| GAMT | None | None | None | None |
| GAMT | None | None | None | None |
| GAMT | None | None | None | None |
| GAMT | None | None | None | None |
| GAMT | None | None | None | None |
| GAMT | None | None | None | None |
| GAMT | None | None | None | None |
| GAMT | None | None | None | None |
| GAMT | None | None | None | None |
| GLRA1 | None | None | None | None |
| GLRA1 | None | None | None | None |

| | | | | |
|---|---|---|---|---|
| GLRA1 | None | None | None | None |
| GLRA1 | None | None | None | None |
| GLRA1 | None | None | None | None |
| GLRA1 | None | None | None | None |
| GLRA1 | None | None | None | None |
| GLRA1 | None | None | None | None |
| GLRA1 | None | None | None | None |
| ADK | None | None | None | None |
| ACTN2 | None | None | None | None |
| UBE2T | None | None | None | None |
| TTN | None | None | None | None |
| ATP7B | None | None | None | None |
| ATP7B | None | None | None | None |
| ATP7B | None | None | None | None |
| ATP7B | None | None | None | None |
| ATP7B | None | None | None | None |
| ATP7B | None | None | None | None |
| ATP7B | None | None | None | None |
| ATP7B | None | None | None | None |
| ATP7B | None | None | None | None |
| ATP7B | None | None | None | None |
| ATP7B | None | None | None | None |
| ATP7B | None | None | None | None |
| ATP7B | None | None | None | None |
| CDKN1B | None | None | None | None |
| CDKN1B | None | None | None | None |
| CDKN1B | None | None | None | None |
| CDKN1B | None | None | None | None |
| CDKN1B | None | None | None | None |
| CDKN1B | None | None | None | None |
| MYOC | None | None | None | None |
| MYOC | None | None | None | None |
| MYOC | None | None | None | None |
| MYOC | None | None | None | None |
| MYOC | None | None | None | None |
| MYOC | None | None | None | None |
| MYOC | None | None | None | None |
| MYOC | None | None | None | None |
| MYOC | None | None | None | None |
| MYOC | None | None | None | None |
| MYOC | None | None | None | None |
| MYOC | None | None | None | None |
| MYOC | None | None | None | None |
| MYOC | None | None | None | None |
| MYOC | None | None | None | None |
| SPTAN1 | None | None | None | None |
| GC | None | None | None | None |
| GC | None | None | None | None |
| MAD1L1 | None | None | None | None |
| NDST1 | None | None | None | None |
| ARL2BP | None | None | None | None |
| TOR1A | None | None | None | None |
| C1S | None | None | None | None |
| ARL6 | None | None | None | None |

| | | | | |
|---|---|---|---|---|
| ARL6 | None | None | None | None |
| ARL6 | None | None | None | None |
| ARL6 | None | None | None | None |
| ARL6 | None | None | None | None |
| ARL6 | None | None | None | None |
| FGFR1 | None | None | None | None |
| KLHL7 | None | None | None | None |
| FLNC | None | None | None | None |
| PTPRC | None | None | None | None |
| GCH1 | None | None | None | None |
| GCH1 | None | None | None | None |
| GCH1 | None | None | None | None |
| GCH1 | None | None | None | None |
| GCH1 | None | None | None | None |
| GCH1 | None | None | None | None |
| GCH1 | None | None | None | None |
| GCH1 | None | None | None | None |
| GCH1 | None | None | None | None |
| GCH1 | None | None | None | None |
| GCH1 | None | None | None | None |
| GCH1 | None | None | None | None |
| GCH1 | None | None | None | None |
| GCH1 | None | None | None | None |
| CFH | None | None | None | None |
| COL2A1 | None | None | None | None |
| COL2A1 | None | None | None | None |
| OCRL | None | None | None | None |
| WRN | None | None | None | None |
| WRN | None | None | None | None |
| WRN | None | None | None | None |
| WRN | None | None | None | None |
| MASP1 | None | None | None | None |
| MASP1 | None | None | None | None |
| NSD1 | None | None | None | None |
| PITX2 | None | None | None | None |
| PITX2 | None | None | None | None |
| PITX2 | None | None | None | None |
| PITX2 | None | None | None | None |
| PITX2 | None | None | None | None |
| PITX2 | None | None | None | None |
| B4GALT7 | None | None | None | None |
| B4GALT7 | None | None | None | None |
| B4GALT7 | None | None | None | None |
| B4GALT7 | None | None | None | None |
| B4GALT7 | None | None | None | None |
| FSHR | None | None | None | None |
| IGHMBP2 | None | None | None | None |
| IGHMBP2 | None | None | None | None |
| IGHMBP2 | None | None | None | None |
| IGHMBP2 | None | None | None | None |
| IGHMBP2 | None | None | None | None |
| IGHMBP2 | None | None | None | None |
| IGHMBP2 | None | None | None | None |
| IGHMBP2 | None | None | None | None |

| | | | | |
|---|---|---|---|---|
| IGHMBP2 | None | None | None | None |
| IGHMBP2 | None | None | None | None |
| IGHMBP2 | None | None | None | None |
| IGHMBP2 | None | None | None | None |
| IGHMBP2 | None | None | None | None |
| IGHMBP2 | None | None | None | None |
| IGHMBP2 | None | None | None | None |
| IGHMBP2 | None | None | None | None |
| TTN | None | None | None | None |
| TTN | None | None | None | None |
| IRAK4 | None | None | None | None |
| PROC | None | None | None | None |
| PROC | None | None | None | None |
| RAD50 | None | None | None | None |
| RAD50 | None | None | None | None |
| DYNC1H1 | None | None | None | None |
| DYNC1H1 | None | None | None | None |
| DYNC1H1 | None | None | None | None |
| DYNC1H1 | None | None | None | None |
| DYNC1H1 | None | None | None | None |
| DYNC1H1 | None | None | None | None |
| DYNC1H1 | None | None | None | None |
| DYNC1H1 | None | None | None | None |
| DYNC1H1 | None | None | None | None |
| DYNC1H1 | None | None | None | None |
| DYNC1H1 | None | None | None | None |
| CYB5R3 | None | None | None | None |
| CYB5R3 | None | None | None | None |
| CYB5R3 | None | None | None | None |
| CYB5R3 | None | None | None | None |
| CYB5R3 | None | None | None | None |
| CYB5R3 | None | None | None | None |
| CYB5R3 | None | None | None | None |
| CYB5R3 | None | None | None | None |
| CYB5R3 | None | None | None | None |
| CYB5R3 | None | None | None | None |
| CYB5R3 | None | None | None | None |
| SPTB | None | None | None | None |
| FANCL | None | None | None | None |
| FAM126A | None | None | None | None |
| SEPT9 | None | None | None | None |
| TTN | None | None | None | None |
| TTN | None | None | None | None |
| TTN | None | None | None | None |
| CNTN1 | None | None | None | None |
| C1QTNF5 | None | None | None | None |
| VCL | None | None | None | None |
| VCL | None | None | None | None |
| VCL | None | None | None | None |
| VCL | None | None | None | None |
| VCL | None | None | None | None |
| VCL | None | None | None | None |
| PPT1 | None | None | None | None |
| PPT1 | None | None | None | None |

| | | | | |
|---|---|---|---|---|
| PPT1 | None | None | None | None |
| PPT1 | None | None | None | None |
| PPT1 | None | None | None | None |
| PPT1 | None | None | None | None |
| PPT1 | None | None | None | None |
| PPT1 | None | None | None | None |
| PPT1 | None | None | None | None |
| PPT1 | None | None | None | None |
| PPT1 | None | None | None | None |
| PPT1 | None | None | None | None |
| PPT1 | None | None | None | None |
| PPT1 | None | None | None | None |
| PPT1 | None | None | None | None |
| PPT1 | None | None | None | None |
| PPT1 | None | None | None | None |
| PPT1 | None | None | None | None |
| PPT1 | None | None | None | None |
| NOTCH2 | None | None | None | None |
| NOTCH2 | None | None | None | None |
| IFIH1 | None | None | None | None |
| IFIH1 | None | None | None | None |
| IKBKG | None | None | None | None |
| DBH | None | None | None | None |
| DBH | None | None | None | None |
| DBH | None | None | None | None |
| DBH | None | None | None | None |
| DBH | None | None | None | None |
| DBH | None | None | None | None |
| DBH | None | None | None | None |
| DBH | None | None | None | None |
| DBH | None | None | None | None |
| DBH | None | None | None | None |
| IL10RA | None | None | None | None |
| IL10RA | None | None | None | None |
| IL10RA | None | None | None | None |
| IL10RA | None | None | None | None |
| IL10RA | None | None | None | None |
| IL10RA | None | None | None | None |
| IL10RA | None | None | None | None |
| ILK | None | None | None | None |
| HPD | None | None | None | None |
| HPD | None | None | None | None |
| HPD | None | None | None | None |
| MECP2 | None | None | None | None |
| MECP2 | None | None | None | None |
| MECP2 | None | None | None | None |
| MECP2 | None | None | None | None |
| MECP2 | None | None | None | None |
| MECP2 | None | None | None | None |
| MECP2 | None | None | None | None |
| MECP2 | None | None | None | None |
| KYNU | None | None | None | None |
| FAS | None | None | None | None |
| FAS | None | None | None | None |

| | | | | |
|---|---|---|---|---|
| ELP1 | None | None | None | None |
| ELP1 | None | None | None | None |
| ELP1 | None | None | None | None |
| ELP1 | None | None | None | None |
| ELP1 | None | None | None | None |
| ELP1 | None | None | None | None |
| ELP1 | None | None | None | None |
| ELP1 | None | None | None | None |
| HESX1 | None | None | None | None |
| HESX1 | None | None | None | None |
| HESX1 | None | None | None | None |
| HESX1 | None | None | None | None |
| GAD1 | None | None | None | None |
| GAD1 | None | None | None | None |
| NDUFA1 | None | None | None | None |
| NDUFA1 | None | None | None | None |
| NDUFA1 | None | None | None | None |
| PNPT1 | None | None | None | None |
| PNPT1 | None | None | None | None |
| PNPT1 | None | None | None | None |
| PNPT1 | None | None | None | None |
| PNPT1 | None | None | None | None |
| PNPT1 | None | None | None | None |
| PNPT1 | None | None | None | None |
| SLC25A13 | None | None | None | None |
| SLC25A13 | None | None | None | None |
| SLC25A13 | None | None | None | None |
| AP4M1 | None | None | None | None |
| AP4M1 | None | None | None | None |
| MEFV | None | None | None | None |
| MEFV | None | None | None | None |
| MEFV | None | None | None | None |
| MEFV | None | None | None | None |
| MEFV | None | None | None | None |
| MEFV | None | None | None | None |
| MEFV | None | None | None | None |
| MEFV | None | None | None | None |
| MEFV | None | None | None | None |
| CAPN3 | None | None | None | None |
| CAPN3 | None | None | None | None |
| CAPN3 | None | None | None | None |
| CAPN3 | None | None | None | None |
| CAPN3 | None | None | None | None |
| CAPN3 | None | None | None | None |
| CAPN3 | None | None | None | None |
| GP6 | None | None | None | None |
| FAS | None | None | None | None |
| FBN1 | None | None | None | None |
| FBN1 | None | None | None | None |
| BMPR2 | None | None | None | None |
| BMPR2 | None | None | None | None |
| BMPR2 | None | None | None | None |
| BMPR2 | None | None | None | None |
| BMPR2 | None | None | None | None |

| | | | | |
|---|---|---|---|---|
| RET | None | None | None | None |
| RET | None | None | None | None |
| RET | None | None | None | None |
| RET | None | None | None | None |
| RET | None | None | None | None |
| RET | None | None | None | None |
| RET | None | None | None | None |
| RET | None | None | None | None |
| RET | None | None | None | None |
| RET | None | None | None | None |
| RET | None | None | None | None |
| RET | None | None | None | None |
| RET | None | None | None | None |
| RET | None | None | None | None |
| RET | None | None | None | None |
| RET | None | None | None | None |
| RET | None | None | None | None |
| RET | None | None | None | None |
| RET | None | None | None | None |
| RET | None | None | None | None |
| RET | None | None | None | None |
| RET | None | None | None | None |
| RET | None | None | None | None |
| RET | None | None | None | None |
| RET | None | None | None | None |
| RET | None | None | None | None |
| RET | None | None | None | None |
| RET | None | None | None | None |
| CSF2RA | None | None | None | None |
| CSF2RA | None | None | None | None |
| CSF2RA | None | None | None | None |
| BRCA1 | None | None | None | None |
| ZAP70 | None | None | None | None |
| ZAP70 | None | None | None | None |
| ZAP70 | None | None | None | None |
| PIK3R2 | None | None | None | None |
| TCN2 | None | None | None | None |
| TCN2 | None | None | None | None |
| TCN2 | None | None | None | None |
| TCN2 | None | None | None | None |
| TCN2 | None | None | None | None |
| TCN2 | None | None | None | None |
| TCN2 | None | None | None | None |
| TCN2 | None | None | None | None |
| EPM2A | None | None | None | None |
| EPM2A | None | None | None | None |
| EPM2A | None | None | None | None |
| EPM2A | None | None | None | None |
| AARS | None | None | None | None |
| AARS | None | None | None | None |
| AARS | None | None | None | None |
| PCDH15 | None | None | None | None |
| PCDH15 | None | None | None | None |
| PCDH15 | None | None | None | None |

| | | | | |
|---|---|---|---|---|
| PCDH15 | None | None | None | None |
| PCDH15 | None | None | None | None |
| PCDH15 | None | None | None | None |
| PCDH15 | None | None | None | None |
| PCDH15 | None | None | None | None |
| PCDH15 | None | None | None | None |
| PCDH15 | None | None | None | None |
| NDUFS4 | None | None | None | None |
| NDUFS4 | None | None | None | None |
| NDUFS4 | None | None | None | None |
| FGF10 | None | None | None | None |
| FGF10 | None | None | None | None |
| FGF10 | None | None | None | None |
| FGF10 | None | None | None | None |
| CDON | None | None | None | None |
| STAMBP | None | None | None | None |
| BRCA1 | None | None | None | None |
| BRCA1 | None | None | None | None |
| BRCA1 | None | None | None | None |
| BRCA1 | None | None | None | None |
| BRCA1 | None | None | None | None |
| BRCA1 | None | None | None | None |
| BRCA1 | None | None | None | None |
| BRCA1 | None | None | None | None |
| BRCA1 | None | None | None | None |
| BRCA1 | None | None | None | None |
| PLXNC1 | None | None | None | None |
| RNASEH2C | None | None | None | None |
| RNASEH2C | None | None | None | None |
| APC | None | None | None | None |
| PLA2G4A | None | None | None | None |
| PLA2G4A | None | None | None | None |
| PSAT1 | None | None | None | None |
| PSAT1 | None | None | None | None |
| TRAPPC3 | None | None | None | None |
| QDPR | None | None | None | None |
| QDPR | None | None | None | None |
| QDPR | None | None | None | None |
| QDPR | None | None | None | None |
| KRT5 | None | None | None | None |
| KRT5 | None | None | None | None |
| KRT5 | None | None | None | None |
| MLH1 | None | None | None | None |
| MLH1 | None | None | None | None |
| MLH1 | None | None | None | None |
| MLH1 | None | None | None | None |
| MLH1 | None | None | None | None |
| MLH1 | None | None | None | None |
| MLH1 | None | None | None | None |
| MLH1 | None | None | None | None |
| MLH1 | None | None | None | None |
| MLH1 | None | None | None | None |
| MLH1 | None | None | None | None |
| MLH1 | None | None | None | None |

| | | | | |
|---|---|---|---|---|
| MLH1 | None | None | None | None |
| MLH1 | None | None | None | None |
| MLH1 | None | None | None | None |
| MLH1 | None | None | None | None |
| MLH1 | None | None | None | None |
| KRIT1 | None | None | None | None |
| KRIT1 | None | None | None | None |
| KRIT1 | None | None | None | None |
| ALX4 | None | None | None | None |
| ALX4 | None | None | None | None |
| ALX4 | None | None | None | None |
| TJP2 | None | None | None | None |
| TJP2 | None | None | None | None |
| PANK2 | None | None | None | None |
| PANK2 | None | None | None | None |
| PANK2 | None | None | None | None |
| PANK2 | None | None | None | None |
| PANK2 | None | None | None | None |
| PANK2 | None | None | None | None |
| PANK2 | None | None | None | None |
| NDUFA6 | None | None | None | None |
| TSHR | None | None | None | None |
| TSHR | None | None | None | None |
| TSHR | None | None | None | None |
| CFTR | None | None | None | None |
| CFTR | None | None | None | None |
| MYOM1 | None | None | None | None |
| MYOM1 | None | None | None | None |
| SPTB | None | None | None | None |
| GPT2 | None | None | None | None |
| NDUFS2 | None | None | None | None |
| NDUFS2 | None | None | None | None |
| NDUFS2 | None | None | None | None |
| NDUFS2 | None | None | None | None |
| NDUFS2 | None | None | None | None |
| CASQ2 | None | None | None | None |
| CASQ2 | None | None | None | None |
| CASQ2 | None | None | None | None |
| CASQ2 | None | None | None | None |
| CASQ2 | None | None | None | None |
| CASQ2 | None | None | None | None |
| CASQ2 | None | None | None | None |
| CASQ2 | None | None | None | None |
| CASQ2 | None | None | None | None |
| CPOX | None | None | None | None |
| CPOX | None | None | None | None |
| CPOX | None | None | None | None |
| CPOX | None | None | None | None |
| CPOX | None | None | None | None |
| CPOX | None | None | None | None |
| CPOX | None | None | None | None |
| CPOX | None | None | None | None |
| CPOX | None | None | None | None |
| MVK | None | None | None | None |

| | | | | |
|---|---|---|---|---|
| MVK | None | None | None | None |
| MVK | None | None | None | None |
| MVK | None | None | None | None |
| MVK | None | None | None | None |
| MVK | None | None | None | None |
| MVK | None | None | None | None |
| MVK | None | None | None | None |
| MVK | None | None | None | None |
| MVK | None | None | None | None |
| MVK | None | None | None | None |
| MVK | None | None | None | None |
| MVK | None | None | None | None |
| MVK | None | None | None | None |
| MVK | None | None | None | None |
| GSS | None | None | None | None |
| GSS | None | None | None | None |
| GSS | None | None | None | None |
| GSS | None | None | None | None |
| GSS | None | None | None | None |
| DSG2 | None | None | None | None |
| DSG2 | None | None | None | None |
| DSG2 | None | None | None | None |
| DSG2 | None | None | None | None |
| DSG2 | None | None | None | None |
| DSG2 | None | None | None | None |
| DSG2 | None | None | None | None |
| DSG2 | None | None | None | None |
| DSG2 | None | None | None | None |
| DSG2 | None | None | None | None |
| DSG2 | None | None | None | None |
| DSG2 | None | None | None | None |
| DSG2 | None | None | None | None |
| DSG2 | None | None | None | None |
| PAPSS2 | None | None | None | None |
| PLCB2 | None | None | None | None |
| HMGCS2 | None | None | None | None |
| HMGCS2 | None | None | None | None |
| HMGCS2 | None | None | None | None |
| HMGCS2 | None | None | None | None |
| HMGCS2 | None | None | None | None |
| HMGCS2 | None | None | None | None |
| HMGCS2 | None | None | None | None |
| HMGCS2 | None | None | None | None |
| MPDZ | None | None | None | None |
| GATA4 | None | None | None | None |
| GATA4 | None | None | None | None |
| WARS2 | None | None | None | None |
| WARS2 | None | None | None | None |
| HGD | 1 | 47 | 1eyb | A |
| HGD | 1 | 47 | 1eyb | A |
| HGD | 1 | 70 | 1eyb | A |
| HGD | None | None | None | None |
| HGD | None | None | None | None |
| HGD | None | None | None | None |

| | | | | |
|---|---|---|---|---|
| HGD | None | None | None | None |
| HGD | None | None | None | None |
| HGD | None | None | None | None |
| HGD | 1 | 1 | 1ey2 | A |
| HGD | 1 | 9 | 1ey2 | A |
| HGD | None | None | None | None |
| HGD | 1 | 10 | 1eyb | A |
| HGD | 1 | 7 | 1ey2 | A |
| PAX8 | None | None | None | None |
| PAX8 | None | None | None | None |
| PAX8 | None | None | None | None |
| PAX8 | None | None | None | None |
| SPTB | None | None | None | None |
| NCF4 | None | None | None | None |
| CFH | None | None | None | None |
| CFH | None | None | None | None |
| CSF1R | None | None | None | None |
| CSF1R | None | None | None | None |
| CA8 | None | None | None | None |
| CA8 | None | None | None | None |
| CA8 | None | None | None | None |
| APC | None | None | None | None |
| APC | None | None | None | None |
| APC | None | None | None | None |
| APC | None | None | None | None |
| APC | None | None | None | None |
| NDUFA2 | None | None | None | None |
| GIF | None | None | None | None |
| GIF | None | None | None | None |
| GIF | None | None | None | None |
| GIF | None | None | None | None |
| C2 | None | None | None | None |
| ACP5 | None | None | None | None |
| ACP5 | None | None | None | None |
| ACP5 | None | None | None | None |
| ACP5 | None | None | None | None |
| ACP5 | None | None | None | None |
| RNASEH2A | None | None | None | None |
| RNASEH2A | None | None | None | None |
| RNASEH2A | None | None | None | None |
| RNASEH2A | None | None | None | None |
| RNASEH2A | None | None | None | None |
| RNASEH2A | None | None | None | None |
| RNASEH2A | None | None | None | None |
| DSG2 | None | None | None | None |
| USH1G | None | None | None | None |
| FUS | None | None | None | None |
| FUS | None | None | None | None |
| FUS | None | None | None | None |
| FUS | None | None | None | None |
| FUS | None | None | None | None |
| MRPL44 | None | None | None | None |
| MRPL44 | None | None | None | None |
| MRPL44 | None | None | None | None |

| | | | | |
|---|---|---|---|---|
| MRPL44 | None | None | None | None |
| DMGDH | None | None | None | None |
| AARS | None | None | None | None |
| STAR | None | None | None | None |
| STAR | None | None | None | None |
| STAR | None | None | None | None |
| STAR | None | None | None | None |
| STAR | None | None | None | None |
| STAR | None | None | None | None |
| STAR | None | None | None | None |
| STAR | None | None | None | None |
| LARP7 | None | None | None | None |
| PRPF8 | None | None | None | None |
| PRPF8 | None | None | None | None |
| PRPF8 | None | None | None | None |
| PRPF8 | None | None | None | None |
| PSAP | None | None | None | None |
| PSAP | None | None | None | None |
| PSAP | None | None | None | None |
| DMPK | None | None | None | None |
| EPCAM | None | None | None | None |
| EPCAM | None | None | None | None |
| EPCAM | None | None | None | None |
| EPCAM | None | None | None | None |
| EPCAM | None | None | None | None |
| EPCAM | None | None | None | None |
| EPCAM | None | None | None | None |
| EPCAM | None | None | None | None |
| ASS1 | None | None | None | None |
| ASS1 | None | None | None | None |
| ASS1 | None | None | None | None |
| ASS1 | None | None | None | None |
| ASS1 | None | None | None | None |
| ASS1 | None | None | None | None |
| ASS1 | None | None | None | None |
| ASS1 | None | None | None | None |
| ASS1 | None | None | None | None |
| ASS1 | None | None | None | None |
| ASS1 | None | None | None | None |
| ASS1 | None | None | None | None |
| ASS1 | None | None | None | None |
| ASS1 | None | None | None | None |
| ASS1 | None | None | None | None |
| ASS1 | None | None | None | None |
| ASS1 | None | None | None | None |
| ASS1 | None | None | None | None |
| ASS1 | None | None | None | None |
| ASS1 | None | None | None | None |
| POLE | None | None | None | None |
| POLE | None | None | None | None |
| POLE | None | None | None | None |
| RNASET2 | None | None | None | None |
| RNASET2 | None | None | None | None |
| RNASET2 | None | None | None | None |

| | | | | |
|---|---|---|---|---|
| FLG | None | None | None | None |
| F10 | None | None | None | None |
| AGT | None | None | None | None |
| AGT | None | None | None | None |
| PLN | None | None | None | None |
| PLN | None | None | None | None |
| PLN | None | None | None | None |
| PTHLH | None | None | None | None |
| IGF1R | None | None | None | None |
| IGF1R | None | None | None | None |
| IGF1R | None | None | None | None |
| JAG1 | None | None | None | None |
| JAG1 | None | None | None | None |
| JAG1 | None | None | None | None |
| JAG1 | None | None | None | None |
| JAG1 | None | None | None | None |
| JAG1 | None | None | None | None |
| AP1S3 | None | None | None | None |
| SPTA1 | None | None | None | None |
| SPTA1 | None | None | None | None |
| SPTA1 | None | None | None | None |
| SPTA1 | None | None | None | None |
| SPG7 | None | None | None | None |
| SPG7 | None | None | None | None |
| SPG7 | None | None | None | None |
| SPG7 | None | None | None | None |
| SPG7 | None | None | None | None |
| SPG7 | None | None | None | None |
| FN1 | None | None | None | None |
| FN1 | None | None | None | None |
| NSMCE3 | None | None | None | None |
| PCCA | None | None | None | None |
| PCCA | None | None | None | None |
| SPTB | None | None | None | None |
| RYR2 | None | None | None | None |
| RYR2 | None | None | None | None |
| MMP9 | None | None | None | None |
| MMP9 | None | None | None | None |
| GLMN | None | None | None | None |
| GLMN | None | None | None | None |
| GLMN | None | None | None | None |
| GLMN | None | None | None | None |
| GLMN | None | None | None | None |
| GLMN | None | None | None | None |
| GLMN | None | None | None | None |
| GLMN | None | None | None | None |
| GLMN | None | None | None | None |
| GLMN | None | None | None | None |
| MARS | None | None | None | None |
| MARS | None | None | None | None |
| MARS | None | None | None | None |
| MARS | None | None | None | None |
| MARS | None | None | None | None |
| MARS | None | None | None | None |

| | | | | |
|---|---|---|---|---|
| SNAP29 | None | None | None | None |
| SNAP29 | None | None | None | None |
| ETHE1 | None | None | None | None |
| ETHE1 | None | None | None | None |
| ETHE1 | None | None | None | None |
| ETHE1 | None | None | None | None |
| ETHE1 | None | None | None | None |
| ETHE1 | None | None | None | None |
| ETHE1 | None | None | None | None |
| SAMHD1 | None | None | None | None |
| KCNQ4 | None | None | None | None |
| FXYD2 | None | None | None | None |
| FXYD2 | None | None | None | None |
| FXYD2 | None | None | None | None |
| LRRK2 | None | None | None | None |
| LRRK2 | None | None | None | None |
| NDUFS1 | None | None | None | None |
| NDUFS1 | None | None | None | None |
| NDUFS1 | None | None | None | None |
| NDUFS1 | None | None | None | None |
| NDUFS1 | None | None | None | None |
| NDUFS1 | None | None | None | None |
| NDUFS1 | None | None | None | None |
| NDUFS1 | None | None | None | None |
| NDUFS1 | None | None | None | None |
| NDUFS1 | None | None | None | None |
| NDUFS1 | None | None | None | None |
| EZH2 | None | None | None | None |
| EZH2 | None | None | None | None |
| LARS | None | None | None | None |
| TLL1 | None | None | None | None |
| CRYBB2 | None | None | None | None |
| CRYBB2 | None | None | None | None |
| EXOSC8 | None | None | None | None |
| SRP72 | None | None | None | None |
| FOXC2 | None | None | None | None |
| HMBS | None | None | None | None |
| HMBS | None | None | None | None |
| HMBS | None | None | None | None |
| HMBS | None | None | None | None |
| HMBS | None | None | None | None |
| HMBS | None | None | None | None |
| HMBS | None | None | None | None |
| HMBS | None | None | None | None |
| HMBS | None | None | None | None |
| HMBS | None | None | None | None |
| HMBS | None | None | None | None |
| HMBS | None | None | None | None |
| HMBS | None | None | None | None |
| HMBS | None | None | None | None |
| HMBS | None | None | None | None |
| HMBS | None | None | None | None |
| HMBS | None | None | None | None |
| HMBS | None | None | None | None |

| | | | | |
|---|---|---|---|---|
| HMBS | None | None | None | None |
| HMBS | None | None | None | None |
| HMBS | None | None | None | None |
| HMBS | None | None | None | None |
| HMBS | None | None | None | None |
| HMBS | None | None | None | None |
| HMBS | None | None | None | None |
| OGT | None | None | None | None |
| NOG | None | None | None | None |
| NOG | None | None | None | None |
| NOG | None | None | None | None |
| NOG | None | None | None | None |
| NOG | None | None | None | None |
| NOG | None | None | None | None |
| NOG | None | None | None | None |
| NOG | None | None | None | None |
| NOG | None | None | None | None |
| NOG | None | None | None | None |
| NOG | None | None | None | None |
| DLG3 | None | None | None | None |
| SCN2A | None | None | None | None |
| STAC3 | None | None | None | None |
| PFKM | None | None | None | None |
| PFKM | None | None | None | None |
| PFKM | None | None | None | None |
| PFKM | None | None | None | None |
| PFKM | None | None | None | None |
| DNMT1 | None | None | None | None |
| DNMT1 | None | None | None | None |
| TNNI3 | None | None | None | None |
| TNNI3 | None | None | None | None |
| TNNI3 | None | None | None | None |
| STXBP2 | None | None | None | None |
| STXBP2 | None | None | None | None |
| STXBP2 | None | None | None | None |
| STXBP2 | None | None | None | None |
| STXBP2 | None | None | None | None |
| STXBP2 | None | None | None | None |
| STXBP2 | None | None | None | None |
| STXBP2 | None | None | None | None |
| REN | None | None | None | None |
| DSP | None | None | None | None |
| DSP | None | None | None | None |
| HNF1B | None | None | None | None |
| ADA | None | None | None | None |
| ADA | None | None | None | None |
| ADA | None | None | None | None |
| ADA | None | None | None | None |
| ADA | None | None | None | None |
| ADA | None | None | None | None |
| ADA | None | None | None | None |
| ADA | None | None | None | None |
| ADA | None | None | None | None |
| ADA | None | None | None | None |

| | | | | |
|---|---|---|---|---|
| ADA | None | None | None | None |
| ADA | None | None | None | None |
| ADA | None | None | None | None |
| ADA | None | None | None | None |
| ADA | None | None | None | None |
| DCX | None | None | None | None |
| DCX | None | None | None | None |
| DCX | None | None | None | None |
| DCX | None | None | None | None |
| DCX | None | None | None | None |
| ARSB | None | None | None | None |
| ARSB | None | None | None | None |
| ARSB | None | None | None | None |
| ARSB | None | None | None | None |
| ARSB | None | None | None | None |
| ARSB | None | None | None | None |
| ARSB | None | None | None | None |
| ARSB | None | None | None | None |
| ARSB | None | None | None | None |
| ARSB | None | None | None | None |
| ARSB | None | None | None | None |
| ARSB | None | None | None | None |
| ARSB | None | None | None | None |
| WWOX | None | None | None | None |
| TNNI3 | None | None | None | None |
| TNNI3 | None | None | None | None |
| TNNI3 | None | None | None | None |
| TNNI3 | None | None | None | None |
| CYBA | None | None | None | None |
| SPAST | None | None | None | None |
| SPAST | None | None | None | None |
| SPAST | None | None | None | None |
| SPAST | None | None | None | None |
| SPAST | None | None | None | None |
| SPAST | None | None | None | None |
| SPAST | None | None | None | None |
| SPAST | None | None | None | None |
| KRIT1 | None | None | None | None |
| KRIT1 | None | None | None | None |
| TP53 | None | None | None | None |
| ISPD | None | None | None | None |
| ISPD | None | None | None | None |
| ISPD | None | None | None | None |
| ISPD | None | None | None | None |
| KCNMA1 | None | None | None | None |
| STK11 | None | None | None | None |
| STK11 | None | None | None | None |
| STK11 | None | None | None | None |
| STK11 | None | None | None | None |
| STK11 | None | None | None | None |
| STK11 | None | None | None | None |
| NDUFA13 | None | None | None | None |
| PHF6 | None | None | None | None |
| PHF6 | None | None | None | None |

| | | | | |
|---|---|---|---|---|
| PHF6 | None | None | None | None |
| PHYH | None | None | None | None |
| PHYH | None | None | None | None |
| PHYH | None | None | None | None |
| PHYH | None | None | None | None |
| PHYH | None | None | None | None |
| NDUFS3 | None | None | None | None |
| NDUFS3 | None | None | None | None |
| ALDH18A1 | None | None | None | None |
| ALDH18A1 | None | None | None | None |
| ALDH18A1 | None | None | None | None |
| ALDH18A1 | None | None | None | None |
| ALDH18A1 | None | None | None | None |
| PAX6 | None | None | None | None |
| PAX6 | None | None | None | None |
| INPP5E | None | None | None | None |
| INPP5E | None | None | None | None |
| INPP5E | None | None | None | None |
| INPP5E | None | None | None | None |
| INPP5E | None | None | None | None |
| INPP5E | None | None | None | None |
| INPP5E | None | None | None | None |
| INPP5E | None | None | None | None |
| INPP5E | None | None | None | None |
| TNNT2 | None | None | None | None |
| TNNT2 | None | None | None | None |
| TNNT2 | None | None | None | None |
| TNNT2 | None | None | None | None |
| TNNT2 | None | None | None | None |
| TNNT2 | None | None | None | None |
| TNNT2 | None | None | None | None |
| TNNT2 | None | None | None | None |
| TYK2 | None | None | None | None |
| TYK2 | None | None | None | None |
| TYK2 | None | None | None | None |
| CFTR | None | None | None | None |
| CFTR | None | None | None | None |
| CFTR | None | None | None | None |
| CFTR | None | None | None | None |
| CFTR | None | None | None | None |
| CFTR | None | None | None | None |
| CFTR | None | None | None | None |
| CFTR | None | None | None | None |
| CFTR | None | None | None | None |
| CFTR | None | None | None | None |
| CFTR | None | None | None | None |
| CFTR | None | None | None | None |
| CFTR | None | None | None | None |
| CFTR | None | None | None | None |
| CFTR | None | None | None | None |
| CFTR | None | None | None | None |
| CFTR | None | None | None | None |
| CFTR | None | None | None | None |
| CFTR | None | None | None | None |

| | | | | |
|---|---|---|---|---|
| CFTR | None | None | None | None |
| CFTR | None | None | None | None |
| CFTR | None | None | None | None |
| CFTR | None | None | None | None |
| CFTR | None | None | None | None |
| CFTR | None | None | None | None |
| CFTR | None | None | None | None |
| CFTR | None | None | None | None |
| CFTR | None | None | None | None |
| CFTR | None | None | None | None |
| CFTR | None | None | None | None |
| CFTR | None | None | None | None |
| CFTR | None | None | None | None |
| CFTR | None | None | None | None |
| CFTR | None | None | None | None |
| CFTR | None | None | None | None |
| CFTR | None | None | None | None |
| CFTR | None | None | None | None |
| CFTR | None | None | None | None |
| CFTR | None | None | None | None |
| CFTR | None | None | None | None |
| CFTR | None | None | None | None |
| CFTR | None | None | None | None |
| CFTR | None | None | None | None |
| CFTR | None | None | None | None |
| CFTR | None | None | None | None |
| CFTR | None | None | None | None |
| CFTR | None | None | None | None |
| CFTR | None | None | None | None |
| CFTR | None | None | None | None |
| CFTR | None | None | None | None |
| CFTR | None | None | None | None |
| CFTR | None | None | None | None |
| CFTR | None | None | None | None |
| SERPINF1 | None | None | None | None |
| SERPINF1 | None | None | None | None |
| IDS | None | None | None | None |
| IDS | None | None | None | None |
| IDS | None | None | None | None |
| IDS | None | None | None | None |
| IDS | None | None | None | None |
| IDS | None | None | None | None |
| IDS | None | None | None | None |
| IDS | None | None | None | None |
| IDS | None | None | None | None |
| NFKBIA | None | None | None | None |
| NFKBIA | None | None | None | None |
| SPAST | None | None | None | None |
| FGF23 | None | None | None | None |
| FGF23 | None | None | None | None |
| FGF23 | None | None | None | None |
| FGF23 | None | None | None | None |
| FGF23 | None | None | None | None |
| FGF23 | None | None | None | None |

| | | | | |
|---|---|---|---|---|
| FGF23 | None | None | None | None |
| DLAT | None | None | None | None |
| STUB1 | None | None | None | None |
| HSD17B4 | None | None | None | None |
| HSD17B4 | None | None | None | None |
| DYSF | None | None | None | None |
| CEP135 | None | None | None | None |
| CEP135 | None | None | None | None |
| PRKCG | None | None | None | None |
| CSF1R | None | None | None | None |
| PEX5 | None | None | None | None |
| PAX6 | None | None | None | None |
| PAX6 | None | None | None | None |
| PAX6 | None | None | None | None |
| PAX6 | None | None | None | None |
| PAX6 | None | None | None | None |
| PAX6 | None | None | None | None |
| NF2 | None | None | None | None |
| NF2 | None | None | None | None |
| NF2 | None | None | None | None |
| REM1 | None | None | None | None |
